# A targeted next generation sequencing approach to develop robust, genotype-specific mutation profiles and uncover novel variants in *Saccharomyces cerevisiae*

**DOI:** 10.1101/2020.06.25.171595

**Authors:** Natalie A. Lamb, Jonathan Bard, Michael J. Buck, Jennifer A. Surtees

## Abstract

Distinct mutation signatures arise from environmental exposures and/or from defects in metabolic pathways that promote genome stability. The presence of a particular mutation signature in a cell or a tumor can therefore predict the underlying mechanism of mutagenesis, which, in practice, may be clinically important. These insults to the genome often alter dNTP pools, which itself impacts replication fidelity. Therefore, the impact of altered dNTP pools should be considered when making mechanistic predictions based on mutation signatures. We developed a targeted deep-sequencing approach on the *CAN1* gene in *Saccharomyces cerevisiae* to define information-rich mutational profiles associated with distinct *rnr1* backgrounds that alter replication fidelity by elevating dNTP levels.. The mutation spectra of *rnr1Y285F* and *rnr1Y285A* alleles were characterized previously; our analysis was consistent with this prior work but the sequencing depth achieved in our study allowed a significantly more robust and nuanced computational analysis of the variants observed, generating profiles that integrated information about mutation spectra, position effects, and sequence context. This approach revealed novel, genotype-specific mutation profiles in the presence of even modest changes in dNTP pools. Furthermore, we identified broader sequence contexts and specific nucleotide motifs that influenced variant profiles in different *rnr1* backgrounds, which allowed us to make specific mechanistic predictions about the impact of altered dNTP pools on replication fidelity.

## INTRODUCTION

Specific environmental exposures and/or genetic backgrounds generate distinct mutation signatures (CHAN *et al.* 2012; CHAN *et al.* 2015; SAINI *et al.* 2020). Therefore, characterizing mutations provides insight into the underlying molecular mechanisms of mutagenesis in the absence of information about genotype or exposure. For example, C→T mutations have long been associated with UV exposure (HOWARD AND TESSMAN 1964). This is the basis for the Catalog of Somatic Mutations in Cancer (COSMIC), which has curated somatic mutation signatures in cancers to infer mutagenic mechanisms potentially active in generating the cancer (TATE *et al.* 2018). Our goal in this study was to develop a broader analytic pipeline that allowed us to build genotype-specific mutation profiles from the ground up, using both canonical and signature mutations (BRASH 2015) as well as sequence context.

In *Saccharomyces cerevisiae*, mutation spectra are frequently determined via one of two general strategies. The first requires selection of mutants, most often at the *CAN1* locus (XU *et al.* 2008; KUMAR *et al.* 2011; BUCKLAND *et al.* 2014), which encodes an arginine permease that also imports the toxic arginine analog, L-canavanine, leading to cell death (FANTES AND CREANOR 1984; HOFFMANN 1985). Inactivating mutations in *CAN1* block canavanine uptake and toxicity, allowing selection for mutations in *CAN1*. The *can1* gene from individual resistant colonies is then amplified by PCR and subjected to Sanger sequencing. However, this approach is relatively low throughput (CHABES *et al.* 2003; XU *et al.* 2008; KUMAR *et al.* 2011) and will miss low frequency variants that might exist within a resistant colony. The second approach has been mutation accumulation (MA) experiments followed by WGS, which avoids potential bias associated with focusing on a single genetic locus (i.e. *CAN1*). However, the depth of sequencing is limited because the entire genome is sequenced, complicating statistically significant comparisons among genotypes (LUJAN *et al.* 2014; ZHU *et al.* 2014; RENTOFT *et al.* 2016). Increased depth requires increased numbers of MA lines and many passages are required to accumulate sufficient mutations and, again, low frequency and potentially diagnostic mutations will be missed.

We developed a high-throughput sequencing/bioinformatic pipeline that allowed characterization and comparison of robust mutation profiles from different genetic backgrounds. We focused on three *rnr1* alleles (*rnr1D57N*, *rnr1Y285F*, *rnr1Y285A*) that decrease replication fidelity by altering dNTP pools (CHABES *et al.* 2003; XU *et al.* 2008; KUMAR *et al.* 2010). We chose these *rnr1* alleles because 1) altered ribonucleotide expression levels have been correlated with many cancers (AYE *et al.* 2015), 2) skewed dNTP pools have been observed in some cancer cell lines (MATHEWS 2015) and 3) there was Sanger and MA sequencing data available for comparison and validation of our analytic approach(XU *et al.* 2008; KUMAR *et al.* 2011; BUCKLAND *et al.* 2014; WATT *et al.* 2016). We paired selection for mutations at *CAN1* with next-generation sequencing (NGS) to define mutation profiles for *RNR1*, *rnr1D57N*, *rnr1Y285F* and *rnr1Y285A*. Importantly, *CAN1* has been found to be representative of mutations occurring genome-wide when Sanger and WGS approaches were compared (KUMAR *et al.* 2011; BUCKLAND *et al.* 2014; WATT *et al.* 2016). This approach significantly increased sequencing depth compared with previous studies and allowed us to innovate in our analysis.

*RNR1* encodes the large subunit of ribonucleotide reductase (RNR), the enzyme that catalyzes the rate-limiting step in dNTP synthesis. RNR expression and activity is tightly regulated to promote replication fidelity and is misregulated in many different types of tumors and cancer cells (AYE *et al.* 2015; MATHEWS 2015; MATHEWS 2018). Altered RNR activity is associated with cancer and nucleobase analogs are frequently used as chemotherapeutics, commonly targeting RNR (JORDHEIM *et al.* 2013; KOHNKEN *et al.* 2015). Overexpression of *Rrm2*, encoding the small subunit of RNR, induced lung neoplasms in a mouse model, a result of an elevated mutator phenotype (XU *et al.* 2008). Similarly, mutations in *SAMHD1*, which encodes a dNTP triphosphohydrolase that modulates dNTP pools in mammalian systems, led to oncogenesis likely driven by increases in dNTP pool levels (RENTOFT *et al.* 2016; MAUNEY AND HOLLIS 2018). Mutations that affect the allosteric regulation of RNR (*rnr1D57N*, *rnr1Y285F*, *rnr1Y285A*) increase and/or skew dNTP pools and increase mutation rates to varying degrees (CHABES *et al.* 2003; XU *et al.* 2008; KUMAR *et al.* 2010; KUMAR *et al.* 2011; BUCKLAND *et al.* 2014; WATT *et al.* 2016). Altered dNTP pools compromise replication fidelity by: 1) increasing the frequency of misinsertion and misalignment by replicative DNA polymerases and 2) biasing those polymerases toward synthesis at the expense of proofreading, promoting extension beyond a mispair (PHEAR *et al.* 1987; KUNKEL AND SONI 1988; KUMAR *et al.* 2010).

In this study we defined and characterized mutation profiles in wildtype yeast and in isogenic strains bearing the *rnr1D57N*, *rnr1Y285F* and *rnr1Y285A* alleles (CHABES *et al.* 2003; XU *et al.* 2008; KUMAR *et al.* 2010). We built comprehensive mutation profiles from first principles, which consisted of mutation spectrum analysis of the specific *types* of variants observed *and* their position, the trinucleotide context in which the variants occurred and the broader sequence context of the variants, identifying motifs that were specifically enriched or depleted in distinct *rnr1* backgrounds. Importantly, and in contrast to previous work (CHABES *et al.* 2003; XU *et al.* 2008; KUMAR *et al.* 2010; KUMAR *et al.* 2011; BUCKLAND *et al.* 2014; WATT *et al.* 2016), our analysis defined novel, distinct profiles in all three *rnr1* backgrounds, with *rnr1Y285A* exhibiting the most unique profile. We noted many of the same *rnr1Y285A* mutation motifs determined by WGS (WATT *et al.* 2016), while also revealing new high and low frequency variants for *rnr1Y285A* and unique profiles in *rnr1Y285F* and *rnr1D57N*, indicating that even small changes in dNTP pools contribute to mutagenesis.

## MATERIALS AND METHODS

### Strains and plasmids

All strains in this study were derived from the W303 *RAD5+* background (**Table S1**). Strains containing *rnr1Y28F/A* in the *pGAL-RNR1* background and integration plasmids to re-create these strains (KUMAR *et al.* 2010) were kindly provided by Andrei Chabes. To integrate *rnr1* alleles at the endogenous *RNR1* locus in the absence of *pGAL-RNR1*, we created a new set of *rnr1* integration plasmids. First, we amplified pRS416 (SIKORSKI AND HIETER 1989) with SO261 and SO262, which each contain an *AatII* recognition site (**Table S2)**. The resulting PCR product, which consists of pRS416 without the *ARS/CEN* region, was digested with *AatII* and ligated to generate the yeast integration plasmid, pEM3, which carries a counter-selectable *URA3* marker (**Table S3**). *RNR1* was amplified from JSY13 with SO263 (**Table S2**; encodes a *Xho*I site) and SO265 (**Table S2**; encodes a *Spe*I site) to amplify a product that extends ~1 kb upstream of the endogenous *RNR1* start site to ~3kb downstream of the *RNR1* start site, to capture the full *RNR1* gene. This fragment was digested with *Xho*I and *Spe*I and ligated into pEM3 to generate the *RNR1* integration plasmid pNL1 (**Table S3**). To generate a truncated version of *RNR1* in the same plasmid, *RNR1* was amplified from JSY13 with SO263 (encodes a *Xho*I site) and SO264 (**Table S2**; encodes a *Spe*I site) to amplify a product that extends ~1 kb upstream of the endogenous *RNR1* start site to ~2kb into the *RNR1* open reading frame. The PCR product was digested with *Xho*I and *Spe*I and ligated into pEM3 to generate pNL2 (**Table S3**).

We generated versions of pNL1 and pNL2 that encode *rnr1D57N, rnr1Y285F* and *rnr1Y285A* using the Q5 site-directed mutagenesis kit (New England Biolabs) (**Table S3**). Oligonucleotides used for the mutagenesis are described in **Table S2**. Plasmids were sequenced to confirm the presence of the desired mutations and absence of secondary mutations. To integrate each *rnr1* allele into the endogenous *RNR1* locus, both the full-length and truncated integration plasmid for a given allele were linearized with Sph1 and used to co-transform JSY13 using the standard lithium acetate approach (GIETZ *et al.* 1992). Transformants were selected on synthetic complete medium lacking uracil (SC-URA). Single colonies were patched onto YPD to allow for homologous recombination and transplacement of one copy of *rnr1.* After 2 days growth at 30°C, cells from these patches were struck out to obtain single colonies on plates containing 5-FOA (0.1%), selecting for a loss of *URA3*. Plasmid integration was confirmed by PCR using SO266 (**Table S2**; specific to the integration plasmid) and SO230 (**Table S2**; specific to the genomic locus). Sanger sequencing was used to confirm retention of a mutant *rnr1* allele and a loss of *RNR1. rnr1* was amplified in two fragments for sequencing, using the SO32/SO21 and SO25/SO22 primer pairs (**Table S2**).

### Mutation Rates at *can1*

Mutation rates were determined through canavanine resistance assays as described previously (XU *et al.* 2008). Strains were struck out to single colonies on complete media. individual 2 mm colonies were carefully measured and selected to assay. Colonies were suspended in 100 μl in TE (10 mM Tris-HCl, pH 7.5; 1 mM EDTA); 75 μl of undiluted colony suspension was plated on SC-ARG +canavanine plates. The cell suspension was diluted 1:10,000 and 20 μl was plated under permissive conditions on SC-ARG. At least two independent isolates of 11 colonies were assayed for each genotype. Isolates were first analyzed independently before grouping data to calculate rates. Mutation rates and confidence intervals were calculated utilizing FluCalc fluctuation analysis software [Radchenko 2018].

### Pooling Canavanine resistant colonies

Strains were patched on SC-ARG plates and grown at 30°C for 3 days, ~30-35 generations. A quarter of the patch was used to inoculate a 25 mL SC-ARG liquid culture. A patch was used to inoculate the culture instead of a single colony to reduce the potential effects of jackpot mutations in the cultures. Cells were grown for approximately 3-4 additional generations and then plated on SC-ARG + canavanine plates, to select for approximately 2,000 canavanine resistant (Can^R^) colonies. It required another ~30 generations to generate colonies, for a total of ~65-70 generations of accumulated mutations, although mutations that confer canavanine resistance should occur in the first 35-40 generations. Canavanine resistant colonies were selected in at least four independent experiments per genotype. For each genotype, at least two independent isolates were used (**Table S4**).

Colonies were counted, and collected by adding TE (100 mM Tris-HCl, pH 7.4, 10 mM EDTA) to the plate and using a sterile glass spreader to scrape off the cells. Colonies from multiple plates were pooled and resuspended in TE (pH 7.4) to a final volume of 10-12 mL. One mL of the colony suspension was used to extract genomic DNA (gDNA). Briefly, cells were lysed by vortexing in 200 μl 1:1 phenol: chloroform, 200 μl chromosome preparation buffer (10 mM Tris,-HCl, pH 8.0, 100 mM NaCl, 1 mM EDTA, 1% SDS, 2% Triton X-100) and 0.3 grams acid-washed glass beads (Sigma; 425-600 microns). 200 μl TE pH 8.0 was added and reactions were centrifuged for 5 minutes at 16,000 xg. The resulting supernatant was collected. Three additional phenol:chloroform extractions were performed to increase DNA purity. gDNA was precipitated by addition of ammonium acetate to a final concentration of 100 mM followed by 2 volumes of 95% ethanol. The gDNA was collected by centrifugation at 16,000 xg for 10 minutes, followed by a wash with 70% ethanol. The gDNA pellets were resuspended in 50 μl nuclease free water with RNase A (final concentration of 50 μg/ml) and incubated at room temperature for at least 1 hour after which gDNA was stored at −20°C.

### Pooling unselected samples

In addition to canavanine resistant colonies, we also pooled ~2000 unselected colonies from *RNR1*, *rnr1D57N*, *rnr1Y285F* and *rnr1Y285A* backgrounds as permissive controls. These colonies were grown as described above, except that in the final step, cells were grown on SC-ARG in the absence of canavanine. Colonies were pooled and genomic DNA extracted as described above.

### Library Preparation and Sequencing

*CAN1/can1* was amplified from gDNA in 6 overlapping, 349-350 base pair fragments using primers listed in **Table S2**. KAPA HiFi ReadyMix (Roche) was used to amplify these fragments in 25 μL reactions for each *can1* region for each of the 150 samples (total of 906 reactions).

Two-μL of DNA was added to each reaction, in the range of 50-500 ng. Two-μL of each PCR reaction was electrophoresed on a 1% agarose gel to confirm amplification. For each sample, 20 μL of PCR product from each of the 6 regions were pooled in a 96-well plate and purified using the Zymo ZR-96 DNA Clean-up Kit. A total of 65 pooled sample sets (**Table S4**) were generated for paired-end sequencing (2×300), including technical replicates. PCR products from the same genomic preparation of pooled samples were independently amplified and sequenced.

### Library barcoding and QC

Nextera barcode adapters were added to *can1* amplicons and were then minimally PCR amplified (8 cycles) for attachment of Illumina Nextera XT index primers set A (Illumina). After PCR, excess adapters were removed using Ampure XP beads (Beckman Coulter) and samples were eluted into EB buffer. Barcoded amplicons were checked for quality using an Advanced Analytical Fragment Analyzer and Qubit Fluorescence (Invitrogen). Amplicons were pooled to 10 nM in EB buffer and the final concentration was determined using the Illumina Universal qPCR Amplification kit from Kapa Biosystems. All pooled samples were diluted to 4 nM, denatured using NaOH and loaded onto an Illumina MiSeq sequencing platform (PE300, V3) with 20% PhiX control. The sequencing was performed in two separate runs to increase coverage and as a check for reproducibility.

### Upstream sequencing analysis

Reads were trimmed using a variable length trimmer (CutAdapt version 1.14) specifying a quality score of Q30. Trimmed reads were further processed using CLC Genomics Workbench Version 11. Paired-end reads were merged, primer locations were trimmed, and processed reads were aligned to the SacCer3 reference genome. Variants were then called using the CLC low frequency variant caller with required significance of 0.01%. Variant files were exported from CLC as VCF files and downstream analysis was performed in RStudio (version 1.2.1335), paired with custom python scripting (**Fig. 1**).

**Figure 1.**
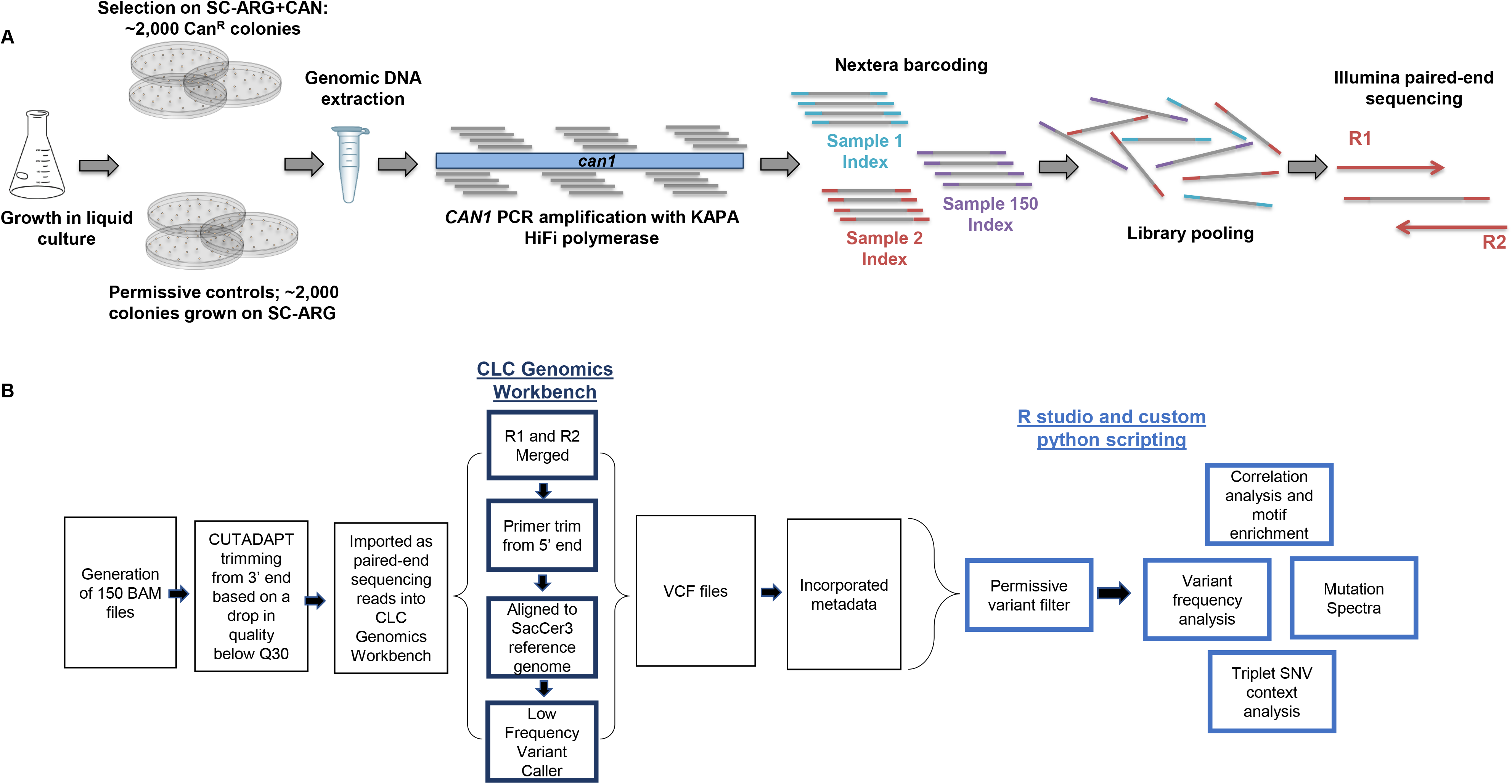
**A schematic of (A)** experimental design and **(B)** pipeline for data analysis.

The variant classes included 6 possible single nucleotide variants (SNVs), single base A/T or G/C insertions and deletions, complex insertions and deletions, as well as mononucleotide variants (MNVs) and replacements (Replac.). MNVs are dinucleotide SNVs, where two neighboring nucleotides are both mutated, ex: CC> AT. Replacements are complex insertions or deletions, where the deleted or replaced base is a variant. Two examples include AAC > G and C > AT. Both MNVs and replacements are extremely low frequency events and rarely occurred in our data set; neither had a significant impact on clustering. Our initial analysis assessed the frequency of each variant type as a function of genotype.

### Mutation Spectra Visualization

Variants across biological replicates were analyzed in two different ways. The first, was a more conservative approach based on presence or absence of a variant at a particular position within *can1* for a given genotype, referred to as “unique counts”. Each position-specific, unique variant was counted only once per replicate and scored based on how many biological replicates it occurred in. In our NGS approach, we cannot easily distinguish between a mutation becoming fixed early in the growth of the culture and a mutation occurring independently multiple times.

By not considering the variant frequency in this analysis, we eliminated this concern. It also mitigated the effects of any “jackpot” mutations that might skew variant frequencies. The second approach incorporated the frequency of each unique variant across *CAN1* (sum of frequencies). This approach added the sum of the variant frequencies if a particular variant occurred in multiple biological replicates. This analysis generated a genotype-specific mutation profile that incorporates variant type, frequency and position. Overall, the results from both types of analysis were consistent when mutation spectra were visualized.

### Permissive variant analysis and filtering

We applied a filter based on our permissive samples (*RNR1*, *rnr1D57N*, *rnr1Y285F*, *rnr1Y285A* grown in the absence of selection) to remove background mutations from Can^R^ samples. The permissive filter removes any variant that occurs below the average permissive sample unique variant frequency of 0.109% (**Fig. 3C)**. We customized the permissive filter because we also observed position-specific variants at a frequency higher than the overall average permissive variant frequency of 0.109%. Thus, these systematically higher frequency variants were removed from the selected samples if they occurred at a frequency below the highest frequency variant that occurred at the same position in the permissive data set. **Fig. 3E-G** shows an example of the permissive filter applied. This is a conservative filter, and undoubtedly low frequency events that are biologically relevant may be removed but are below the sensitivity of this assay. The filtering parameters can be adjusted accordingly, for other applications of this targeted sequencing approach.

### Determining SNV in trinucleotide context

The trinucleotide context surrounding the SNV was determined by taking a 3 bp window surrounding the reference position within *CAN1*. We cannot determine which strand incurred a mutation and therefore all SNVs were categorized as C or T changes for this analysis. There are a total of 96 different possible SNV changes in unique sequence context (ALEXANDROV *et al.* 2015). For a given sample, the number of SNVs in each of these 96 contexts was totaled. This analysis does not take frequency into account and only scores the presence or absence of a particular type of SNV. The frequency plotted is calculated based on unique counts of an SNV that occurred, divided by the total number of times that SNV in tri-context occurred. The data was further condensed by taking the average of each of the 96 different contexts for all the biological replicates in one genotype.

The number of trinucleotide sequence contexts in *CAN1* was calculated using a sliding window approach utilizing python scripting. For each of the 96 different SNV changes in triplet context, the average number of SNVs in a genotype was divided by the number of times the triplet sequence context occurs in *CAN1*. This dataset was imported into R-studio and plotted via the barplot() function. The average number of times an SNV is seen in a genotype out of the total number of times the sequence context occurs across *CAN1* is plotted (**Fig. 5)**.

### Hierarchical cluster analysis and motif enrichment

To identify genotype-specific variants and mutation profiles and to eliminate frequency bias from variants that occurred early on in the growth of a particular sample, we condensed unique variants based on the number of biological replicates sequenced for that genotype. While we were hesitant to include variant frequency in this analysis, we reasoned that observing a variant in multiple replicates increased the probability that it was specific to that genotype. If a variant occurred in 4 out of 4 biological replicates it was represented as 1, if it occurred in 3 out of 6 replicates it was represented as 0.5. This strategy provided an unbiased way to assess the probability that a given variant was genotype-specific. These data were clustered on rows (or unique variants), after applying a row sum cutoff of > 2 to eliminate low frequency variants that are less likely to be driving the observed differences in mutation spectra. Clustering the data based on unique variants allows us to identify different *types* of variants in specific sequence contexts that are potentially diagnostic for a particular genotype. We performed motif enrichment on the different variant classes (i.e. G/C deletion, CG>AT SNVs) independently, with a 12 base window surrounding the variant. Heatmaps were plotted using the pheatmap library in RStudio and motif enrichment was performed using Berkely web logos (CROOKS *et al.* 2004).

All data and reagents are available upon request. All variant sequence are provided in **Table S5**.

## RESULTS

### Rates of canavanine resistance in different genetic backgrounds

To characterize and compare mutation profiles between different isogenetic backgrounds, we genetically altered dNTP pools using previously characterized mutations in *RNR1*, the large subunit of RNR. We chose RNR because alterations in dNTP pool levels generate distinct mutation profiles and RNR is well-associated with cancer(XU *et al.* 2008; KUMAR *et al.* 2010; KUMAR *et al.* 2011; BUCKLAND *et al.* 2014; AYE *et al.* 2015; MATHEWS 2015; WATT *et al.* 2016; MATHEWS 2018). The *rnr1D57N* mutation, in the activity site, leads to a balanced two-fold increase in each of the four dNTPs (CHABES *et al.* 2003). Two mutations in the RNR specificity site, *rnr1Y285F* and *rnr1Y285A*, lead to three- and twenty-fold skewed increases in dCTP and dTTP, respectively (KUMAR *et al.* 2011). The previously characterized *rnr1Y285F* and *rnr1Y285A* strains also encoded a wild-type copy of *RNR1* under control of the inducible *pGAL1* promoter (*pGAL-RNR1*) (KUMAR *et al.* 2010). Therefore, we constructed strains that encoded only *rnr1Y285F* and *rnr1Y285A* for comparison and for use in this study (**Table S1**).

We determined mutation rates of all strains at the *CAN1* locus using a canavanine resistance assay (see Materials and Methods). The *rnr1D57N* and *rnr1Y285F* strains exhibited low mutation rates (2-4 fold increase), while the *rnr1Y285A* strain exhibited a larger ~10-fold increase in mutation rate(KUMAR *et al.* 2010) (**Table S5**), consistent with previous work (CHABES *et al.* 2003; XU *et al.* 2008; KUMAR *et al.* 2010; KUMAR *et al.* 2011). The *pGAL-RNR1* construct did not affect mutation rates in *rnr1Y285F*, but did modulate levels of mutagenesis in the *rnr1Y285A* background (**Table S5**).

### Selection at *CAN1* paired with next generation sequencing (NGS) to define mutation profiles

To generate mutation sequence data in *RNR1*/*rnr1* backgrounds, we selected ~ 2,000 canavanine resistant colonies (“selected samples”), each of which sustained at least one mutation within *CAN1*. Resistant colonies were pooled, genomic DNA was extracted and *can1* was amplified in 6 overlapping regions. To amplify the 5’ end of *CAN1/can1*, which includes a highly repetitive promoter element, we used a primer that annealed to that element, to avoid replication slippage during PCR or sequencing (**Table S2**: CAN1 reg1 Forward_anchored). Amplicons from each pooled sample, representing a single replicate for a given genotype, were barcoded, combined and sequenced using 2×300 paired-end sequencing (see Materials and Methods) (**Fig. 1A**).

Identifying genotype-specific trends in mutation profiles is complicated by the stochastic nature of mutations. To help account for this, we sequenced at least four independent samples for each genotype, using at least two independently generated isolates (**Table S1 & S4**). In parallel, we sequenced pooled samples (~1,000 colonies each) grown in the *absence* of canavanine selection, which we dubbed “permissive” samples (**Figure 1A)**. These permissive controls allowed us to develop a threshold for background mutations at each position along *CAN1* that are a result of low frequency, stochastic mutations and sequencing and/or PCR polymerase bias (see below).

We developed a custom bioinformatic pipeline for analysis. Upstream analysis was performed in CLC Genomics Workbench 11, determining variants utilizing a low frequency variant caller, and downstream analysis was performed in R Studio and through custom python scripts to determine the effect of sequence context and to compare genotypes (**Fig. 1B**; see Materials and Methods and below for details). All sequence data were scored for: 1) single nucleotide variants (SNVs), 2) single base (A/T or G/C) insertions or deletions, 3) complex (>1 base) insertions or deletions, 4) dinucleotide SNVs at adjacent nucleotides, i.e. mononucleotide variants (MNVs), and 5) complex replacements (see Materials and Methods). We could not distinguish which strand incurred a mutation; a C to A transition could also be a G to T change and was represented as such, i.e., CG> AT. NGS allowed deep sequencing of pooled samples at each position along *can1*, providing: 1) large sample sizes for each pooled group, 2) sequencing depth sufficient to uncover low frequency variants and 3) novel insight into mutation profiles and positional effects on mutations, all of which would be unattainable via a whole genome approach because the sequencing depth/coverage is insufficient for this type of analysis.

On average *CAN1* was sequenced at a depth of 16,000x coverage per sample. The total variant frequency for each sample was calculated by taking the sum of the number of variants and dividing by the total number of reads sequenced for that sample. The variant frequencies for all biological replicates within a genotype were averaged. The total variant frequency for all selected samples ranged from ~80% to >100%, with an average of 99.35% (**Fig. 2A**). Frequencies above 100% indicated more than 1 mutation within *can1* in a Can^R^ colony, and were observed in *rnr1Y285A* strains, which had higher mutation rates (**Table S5**).

**Figure 2.**
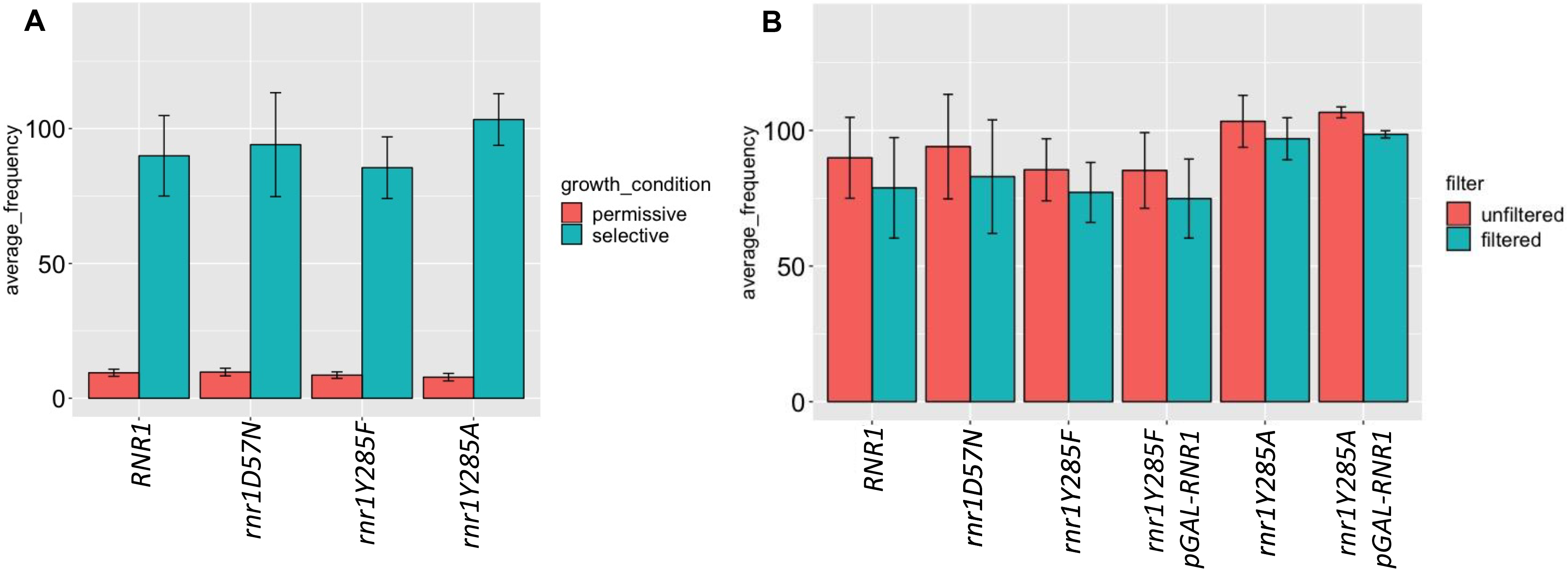
Absolute variant frequency is consistent with selection at *CAN1.* The average absolute variant frequency was calculated for each genotype by dividing the total number of variants by the total number of reads sequenced. The average was then taken for all biological replicates sequenced in a genotype. Error bars represent the standard deviation between biological replicates within each genotype. The number of total biological replicates sequenced varied by genotype; numbers are displayed in **Table S4**. **(A)** The average total variant frequency differed significantly between selected and permissive samples, regardless of genotype. (**B**) The average variant frequency decreased following application of the permissive variant filter.

### Permissive sample filtering removes background mutations

Under permissive conditions (no canavanine selection), the average variant frequency for all four genetic backgrounds (wildtype*, rnr1D57N, rnr1Y285F* and *rnr1Y285A*) was 8.7% compared to >90% in the samples selected in the presence of canavanine (**Fig. 2A**). While the lower variant frequencies in permissive controls were expected, it was higher than predicted given the known mutation rates of replicative polymerases, (BEBENEK *et al.* 1992; KUNKEL 1992; LEE *et al.* 2016) indicating that these background mutations were being introduced as part of the PCR/sequencing pipeline. Consistent with this prediction, the variant distribution in permissive controls from wildtype*, rnr1D57N, rnr1Y285F* and *rnr1Y285A* were virtually identical, despite differences in mutation rates and mutation spectra in selected samples (CHABES *et al.* 2003; XU *et al.* 2008) (**Table S5, Fig. 3A & 3B)**. Of the 296 unique variants observed in all permissive samples, 82 (27.7%) were observed in all 4 genotypes; over half were observed in at least 2 genetic backgrounds. Moreover, the single nucleotide variant (SNV) spectra from permissive samples, independent of genotype, (**Fig. 3A**) closely resembled the SNV spectrum observed for KAPA HiFi polymerase (OYOLA *et al.* 2012; POTAPOV AND ONG 2017), with a large bias toward CG>TA changes.

**Figure 3.**
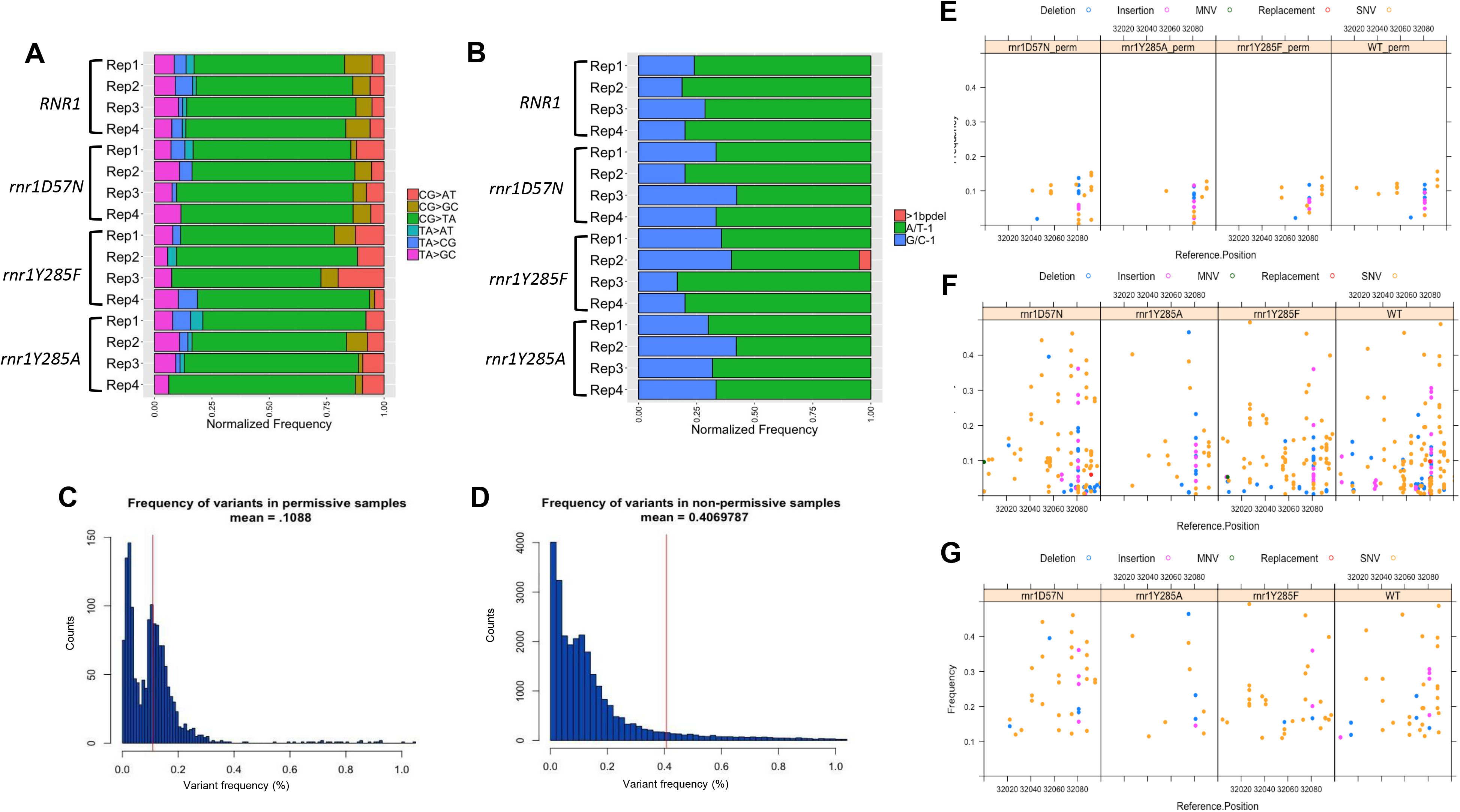
Permissive controls show consistent mutation spectra and are filtered from our data set. **(A)** The SNV spectra for all permissive samples. Plotted are the number of unique variants in a sample normalized out of 100% for comparison purposes. Individual biological replicates for each genotype are shown. **(B)** The deletion spectra for each permissive sample. Shown are single G/C (blue), A/T (green) base deletions and deletions greater than 1 base (pink). Individual biological replicates for each genotype are shown. **(C)** A histogram plotting the number of variants (y-axis) and the frequency (x-axis) at which they occur in both permissive and (**D)** selected samples. The red line represents the average variant frequency in permissive and selected samples respectively. **(E)** A 100 bp window from 32,000 to 32,100 displaying the different types of variants and the frequency at which they occur in 4 different permissive genotypes. **(F)** The same window displaying the variants that occur in the same four genotypes where mutants were selected in the presence of canavanine. **(G)** Variants that remain in selected samples post-permissive variant filter.

To correct for these effects, we developed a permissive variant filter to remove potentially artefactual variants as part of the analytical pipeline. We determined the average frequency of each variant at each position along *CAN1/can1* observed in permissive samples, which was 0.109% compared to 0.407% in the selected samples (**Fig. 3C & 3D).** Any variant that occurred below the average permissive variant frequency of 0.109% was removed by the permissive filter, which represents ~1 variant called for every 1,000 reads sequenced. Some position-specific variants systematically occurred at average frequencies above 0.109%, in multiple biological replicates and genotypes. We incorporated this into the permissive filter, setting the threshold for each position-specific variant to its highest observed frequency in a permissive sample.

This conservative approach ensured that our analysis of mutation profiles in selected samples (see below) was not driven by background variants. We set a blanket cutoff at 0.109%, rather than making the filter *exclusively* position-specific, because we sequenced significantly fewer permissive samples (18) than selected (47). Therefore, the permissive filter likely underestimates stochastic mutations. The higher the number of samples sequenced, the greater the probability that low frequency mutations due to “noise” will be sequenced. **Fig. 3E** illustrates the application of this filter, where the filter cutoff is 0.109%, except for positions 31658 and 31659, where variants occurred at position-specific higher average frequencies in the permissive samples. This reduced the noise in the selected data (**Fig. 3F, G**), decreased variant frequency in selected samples by an average of 7.5% (**Fig. 2B**) and resulted in only minor changes to mutation spectra overall (**Fig. S1**).

### Alterations in dNTP pools change mutation profiles

We determined the mutation profiles for each individual replicate (**Fig. S2**) and for each genotype by combining biological replicates (**Fig. 4**). We compared the relative levels of transitions, transversions, in/dels and other variants in our datasets with similar data from previous analyses of *rnr1D57N*, *rnr1Y285F pGAL-RNR1* and *rnr1Y285A pGAL-RNR1* (XU *et al.* 2008; KUMAR *et al.* 2011) (**Fig. S3**). The distributions for the same genotypes were quite similar, but because of the depth of sequencing in our data, we were able to perform a more nuanced analysis (see below). The most pronounced differences were observed in the *rnr1D57N* spectra, likely a result of the small sample size (n = 16 *can1* colonies) used in the published results [Xu 2008].

**Figure 4.**
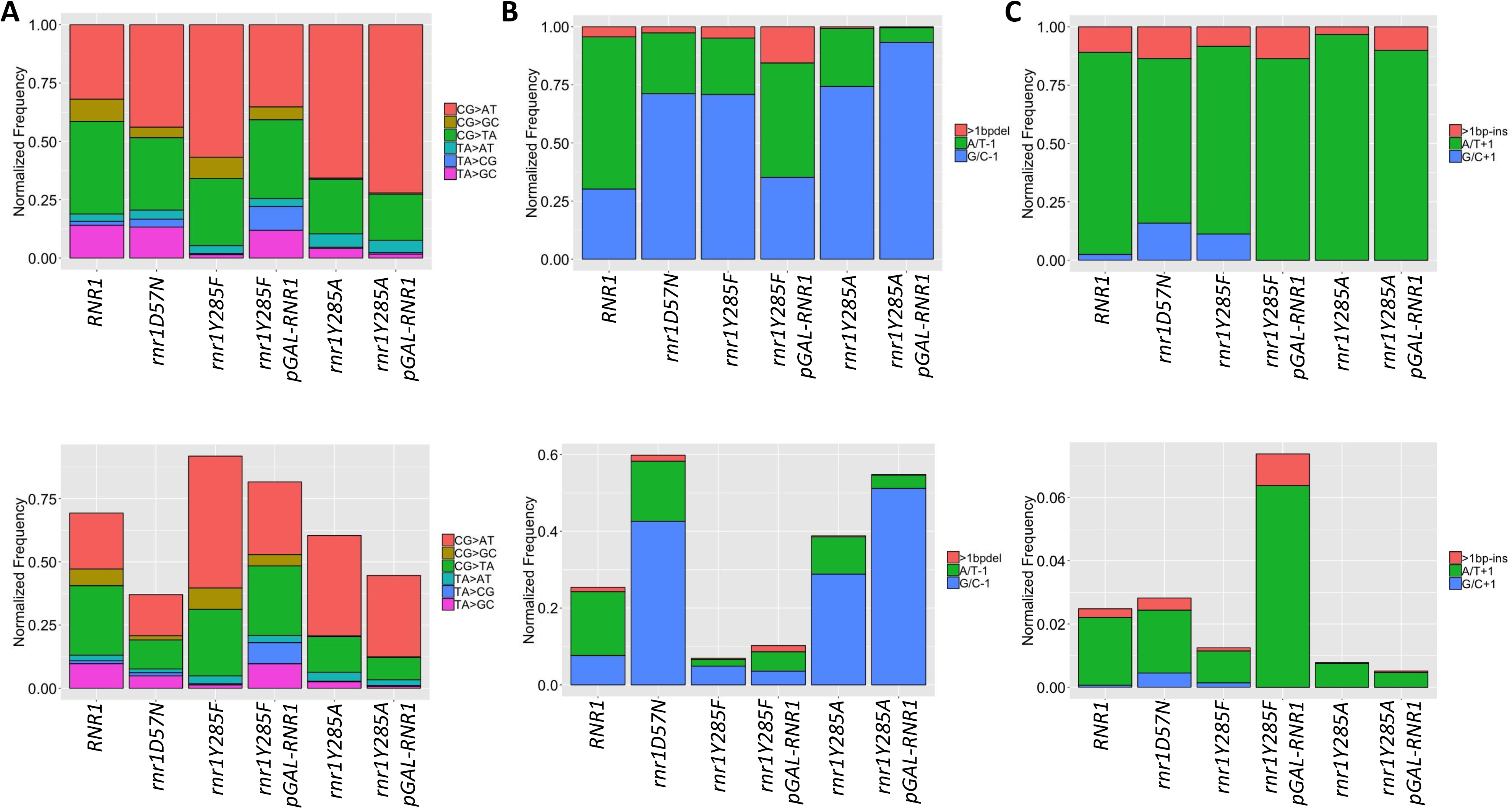
Mutation spectra vary by *rnr1* allele. (**A)** The SNV spectra normalized to total SNVs (upper panel) and normalized out of total variants (lower panel). (**B)** The deletion spectra normalized to total deletions (upper) and normalized to total variants (lower). (**C)** The insertion spectra normalized to total deletions (upper) and normalized out of total variants (lower).

After the initial analysis, we characterized mutation events in two ways: 1) the presence of a particular variant *at a unique position*, i.e., the number of different C>A changes within *can1* (“unique variants”) (Counts in **Table S6**) and 2) the frequency at which each of these unique variants occurred, i.e., the combined frequency of all C>A observed changes along *can1* (“sum of frequencies”) (Freq. in **Table S6**) (see Materials and Methods). The latter essentially provides mutation spectra but is structured to incorporate position information. The former prevented potential “jackpot” mutations from dominating and skewing the mutation spectra.

These analyses allowed us to determine whether different *types* of mutations occurred in a genotype-dependent manner, independent of frequency, and whether variant frequencies were altered in a significant way (Counts/Freq. in **Table S6**). For example, a decreased number for “unique variants” combined with unchanged or increased “sum of frequencies” indicated that variant type is more localized, possibly indicating a mutational hotspot (**Table S6**). Independent biological replicates with the same genotype closely resembled each other (**Fig. S2**); variant frequencies were more variable than unique counts among biological replicates of the same genotype (**Fig. S2**), consistent with the stochastic nature of mutation accumulation during the growth of a culture. Distinct mutagenic events were observed in multiple, *independent*, experiments, consistent with systematic, genotype-specific changes in mutation profiles.

The relative frequency of SNVs, insertions and deletions, normalized by the number of sequence reads, varied significantly by *RNR1*/*rnr1* allele (**Table S6**, **Fig. 4**), as did the absolute variant frequencies (**Fig. 4B**). In *rnr1D57N*, deletions increased substantially compared to *RNR1*, while insertions remained unchanged. The SNV profile of *rnr1D57N* was very similar to wildtype, although the overall SNV frequency was reduced (**Table S6**, **Fig. 4A, B**), consistent with more stochastic rather than systematic mutation events. In contrast, SNVs dominated the *rnr1Y285F* profile. Despite its lower mutation rate (**Table S5**), the normalized variant profiles of *rnr1Y285F* were very similar to those of *rnr1Y285A*, although CG>GC changes were essentially eliminated in *rnr1Y285A*. The proportion of variant type was skewed toward SNVs in *rnr1Y285F* and deletions in *rnr1Y285A*. While the *rnr1Y285A* and *rnr1Y285A pGAL-RNR1* strains resulted in almost indistinguishable mutation profiles, *rnr1Y285F* and *rnr1Y285F pGAL-RNR1* showed more variation compared to one another (**Table S6**, **Fig. S2**), likely because there is a higher proportion of stochastic versus systematic mutations when mutation rates are lower. All three *rnr1* backgrounds exhibited a significant increase in G/C −1 bp deletions compared to wild-type, although the absolute frequency varied (**Table S6**, **Fig. 4C, D**). Overall, very few insertions were observed, but we noted that the G/C +1 bp insertions were extremely rare in *rnr1Y285A* cells compared to other genotypes (**Table S6**, **Fig. 4E, F**). The depth of sequencing coverage in the current study revealed more distinct and detailed mutation profiles than previously identified in *rnr1D57N* and *rnr1Y285F* (XU *et al.* 2008; KUMAR *et al.* 2010; KUMAR *et al.* 2011; BUCKLAND *et al.* 2014; WATT *et al.* 2016), with clear shifts in the types and frequency of mutations that accumulate in the presence of balanced versus skewed elevations in dNTP levels.

### Unique variants occur within *CAN1* in a genotype-specific manner

Mutations occurred across the 1,772 bp *CAN1* in all genotypes tested. For each genotype, we identified unique variants in each replicate and then calculated the average variant frequency of each unique variant (**Fig. 5**, **Fig. S4**). Combined, we identified 860 unique variants in all genotypes tested (**Fig. 5)**; 288 in *rnr1Y285A* genotypes, 570 in *rnr1Y285F* genotypes, 452 in wildtype and 322 in *rrn1D57N* (**Fig. S4**). On average this is over 5 times greater than the number of unique mutational events picked up by previous Sanger sequencing approaches.

**Figure 5.**
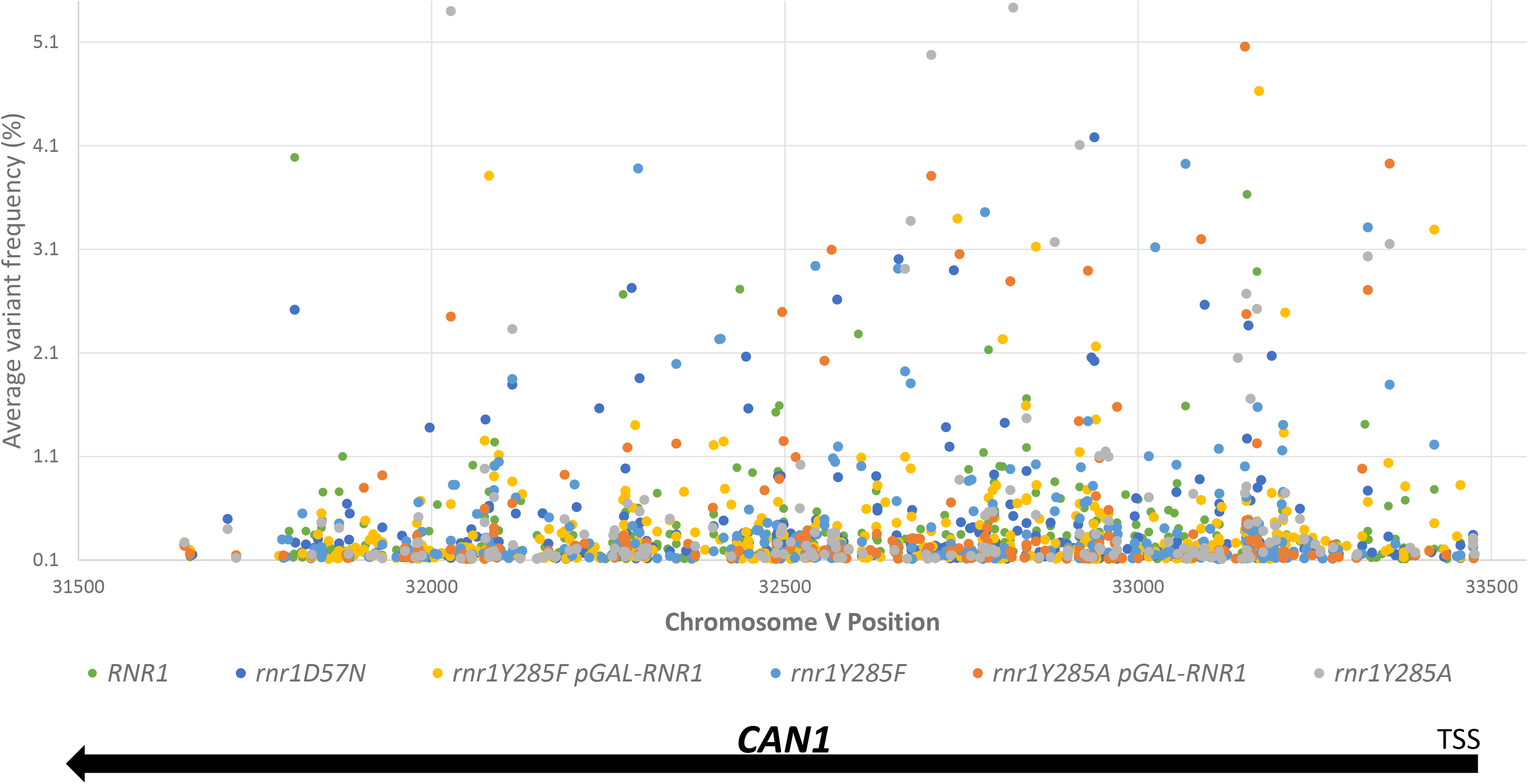
Variant spread across *CAN1*. The average variant frequency of a unique low frequency (<5%, y-axis) variants within a genotype is plotted. High frequency variants are plotted in Figure S5. *CAN1* is annotated by position on Chromosome V on the (−) strand. The transcriptional start site (TSS) and arrow show the direction of transcription from the start 5’ to the end of the gene, 3’.

Many unique variants were observed in a single isolate at low frequency, while others occurred in multiple biological replicates at increased variant frequency. The majority of average variant frequencies were below 5%; 44/860 variants occurred at an average frequency greater than 5% (**Fig. S5**). When a unique variant occurred at a frequency above 25% in only one biological replicate, we defined it as a “jackpot” mutation; a mutation that arose after two generations of growth would result in >25% of the cells (colonies) harboring that mutation. This is distinct from high frequency unique variants, which had an average frequency above 5%, which we analyzed to identify systematic, genotype-specific variants.

We compared the average variant frequency of high frequency variants (>5% average frequency) that were enriched within a particular genotype (**Fig. 6**). In *RNR1* (9 biological replicates), we identified 15 unique high frequency variants, 9 of which occurred in more than one *RNR1* biological replicate (**Fig. 6A**). The majority occurred in a small proportion of biological replicates (**Fig. 6A**). When a high frequency variant was present in multiple *RNR1* replicates, it typically occurred at variant frequencies of less than 1% for the remaining replicates. Most of the *RNR1* high frequency variants were not observed in any of the *rnr1* backgrounds. The exception of the SNV at position 32114 which was observed in all four *rnr1Y285F/A* genotypes. This was also the most significantly mutated position in *RNR1*; it was in over half the biological replicates.

**Figure 6.**
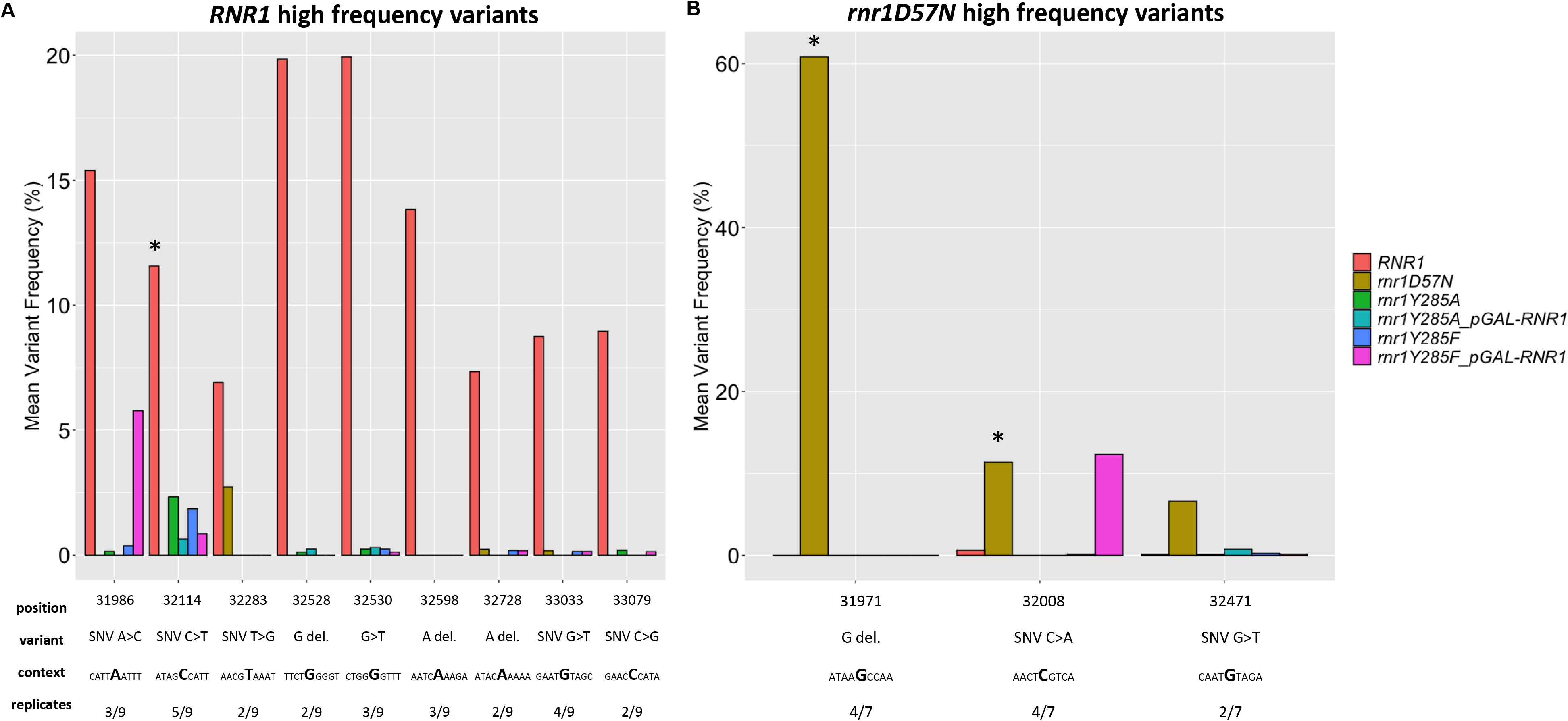
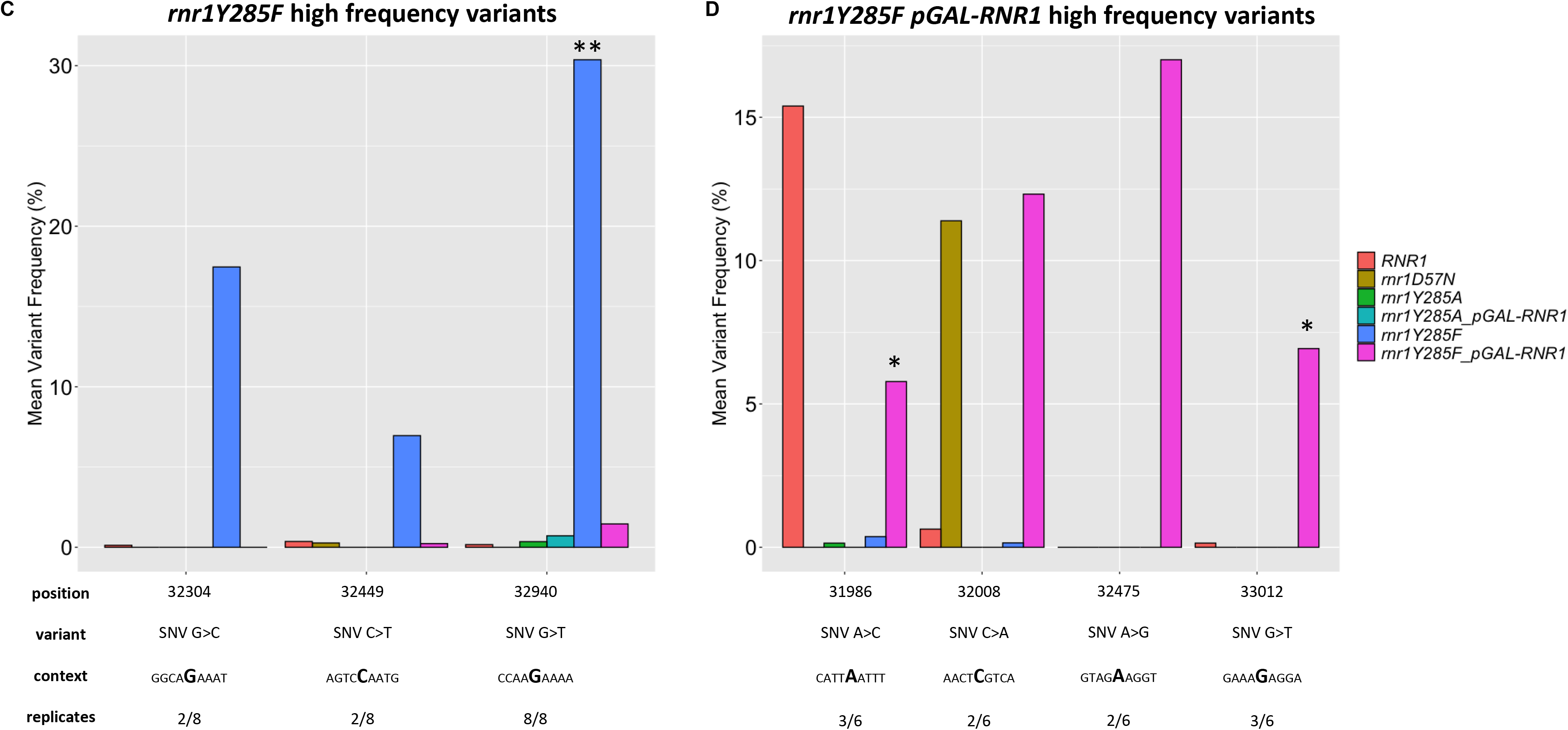
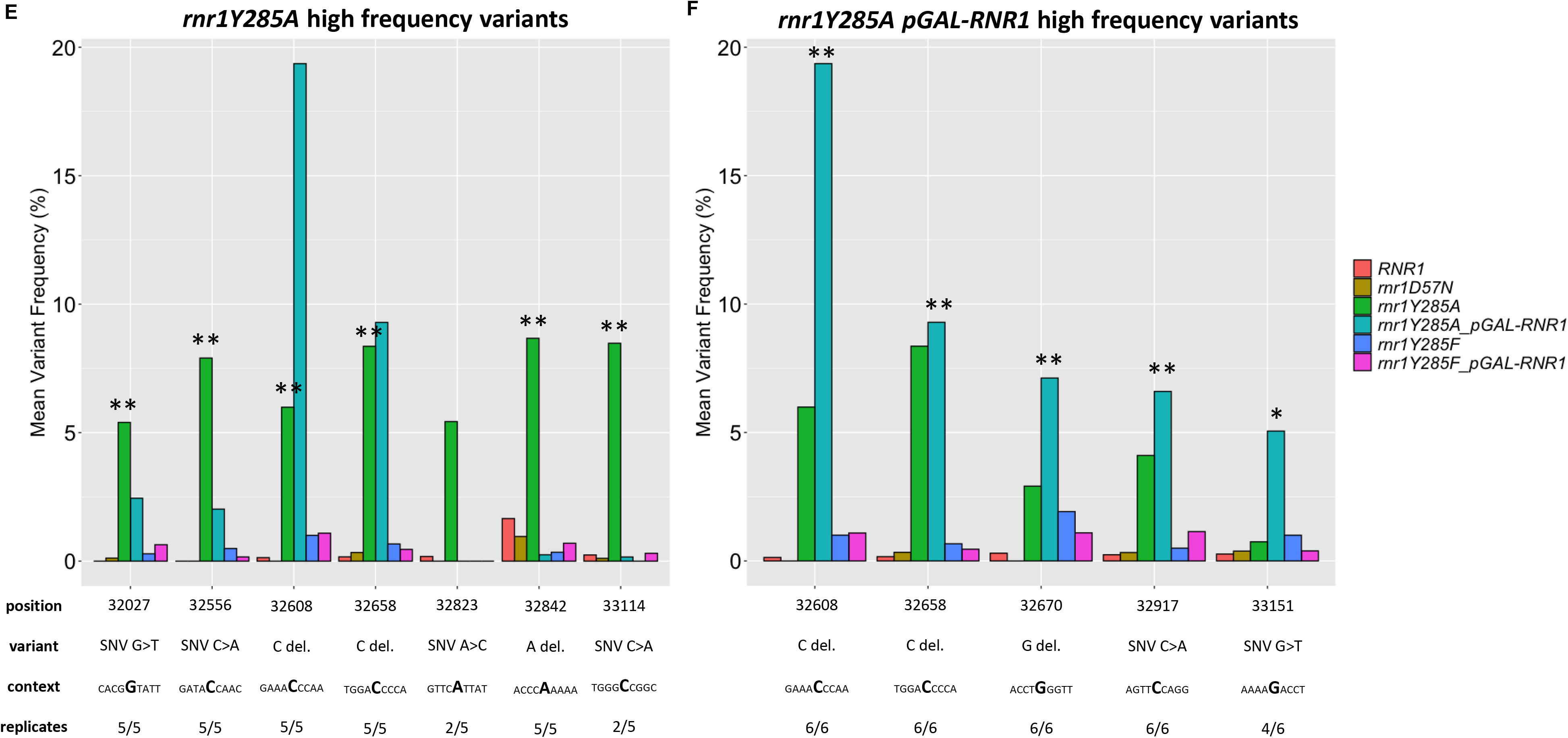
High frequency variants occur systematically in *rnr1Y285F/A* genotypes. Unique variants with an average variant frequency > 5% for (**A**) wildtype, (**B**) *rnr1D57N*, (**C**) *rnr1Y285F-pGAL RNR1*, (**D**) *rnr1Y285F*, (**E**) *rnr1Y285A-pGAL-RNR1*, (**F**) *rnr1Y285A.* For comparison, the average frequency of each variant in other genotypes is also plotted. Below each plot, the fraction of biological replicates in which the variant occurred is observed and the surrounding sequence context are indicated. *, unique variant occurred in ≥ 50% of biological replicates for the genotype analyzed;**, unique variant occurred in 100% of biological replicates for the genotype analyzed.

In *rnr1D57N*, 3 high frequency unique variants were systematically mutated in multiple *rnr1D57N* biological replicates, and were specific to *rnr1D57N* (**Fig. 6B**, orange bars). For example, the G deletion at position 31971 occurred in 4/7 biological replicates (**Fig. 6B**) and drives the observed overall mutation spectrum for this genotype (**Fig. 4**). Similarly, *rnr1Y285F* (periwinkle) and *rnr1Y285F pGAL-RNR1* (pink bars) exhibited very few high frequency unique variants. There was little overlap between these positions (**Fig. 6C & 6D**) between the two *rnr1Y285F* strains – or with any other genetic backgrounds. In contrast, we observed overlapping, systematic high frequency unique variants in multiple replicates of *rnr1Y285A* (green) and *rnr1Y285A pGAL-RNR1* (turquoise) (**Fig. 6E & 6F**), including CG>AT and GC>TA SNVs as well as G/C single base deletions. Notably, several of these high frequency unique variants were also observed in both *rnr1Y285F* genotypes (periwinkle and pink bars) at lower frequencies, but in multiple biological replicates. This indicates that these types of mutations occur in the same sequence contexts when dCTP and dTTP levels are skewed, whether by a modest 3-fold or the more significant 20-fold increase. Variants at these positions, commonly G/C −1 deletions and CG>AT changes, are increasingly probable when dNTP pools are further elevated and skewed.

The high frequency unique variants that we observed in *rnr1Y285A-pGAL-RNR1* and *rnr1Y285A* overlapped significantly with previously defined “hotspots” in *rnr1Y285A pGAL-RNR1* by Sanger sequencing (KUMAR *et al.* 2011; BUCKLAND *et al.* 2014). Our targeted sequencing approach allowed us to identify additional, previously unobserved positions in *CAN1* that were highly susceptible to mutation in the *rrn1Y285A* backgrounds, including the unique G/C deletion at position 32658 and the GC>TA SNV at position 32027 and the CG>AT SNV at position 32917 (**Fig. 6E & 6F)**. These three new high frequency unique variants each occurred in all *rnr1Y285A* biological replicates and may be diagnostic of this genotype.

### Low frequency variant analysis identifies new*rnr1Y285A* hotspots

The sequencing depth across *can1* achieved in our targeted NGS approach also allowed us to analyze low frequency variants for systematic, genotype-specific changes. For this analysis, the inclusion of multiple biological replicates was critical. We analyzed low frequency variants, defined as less than 5% average variant frequency enriched in a genotype-specific manner. *RNR1*, *rnr1D57N* and *rnr1Y285F* mutation profiles exhibited a large number of low frequency variants (56-123) found in only one replicate and/or at low variant frequency, consistent with a high proportion of stochastic mutations. In contrast, both *rnr1Y285A* genotypes exhibited a much smaller number of unique low frequency events (13 deletions, 16 SNVs; **Fig. 7**). The concentration of low frequency events in *rnr1Y285A* backgrounds indicates a more systematic, rather than stochastic, pattern of mutagenesis. Notably, there were many G/C deletions (**Fig. 7A)** and CG>AT SNVs (**Fig. 7B**) unique to, and therefore diagnostic of, *rnr1Y285A* backgrounds. Furthermore, these genotype-specific mutations appeared to drive the mutation profiles of *rnr1Y285A* backgrounds (**Fig. 4**). We further investigated these variants by systematically analyzing sequence context.

**Figure 7.**
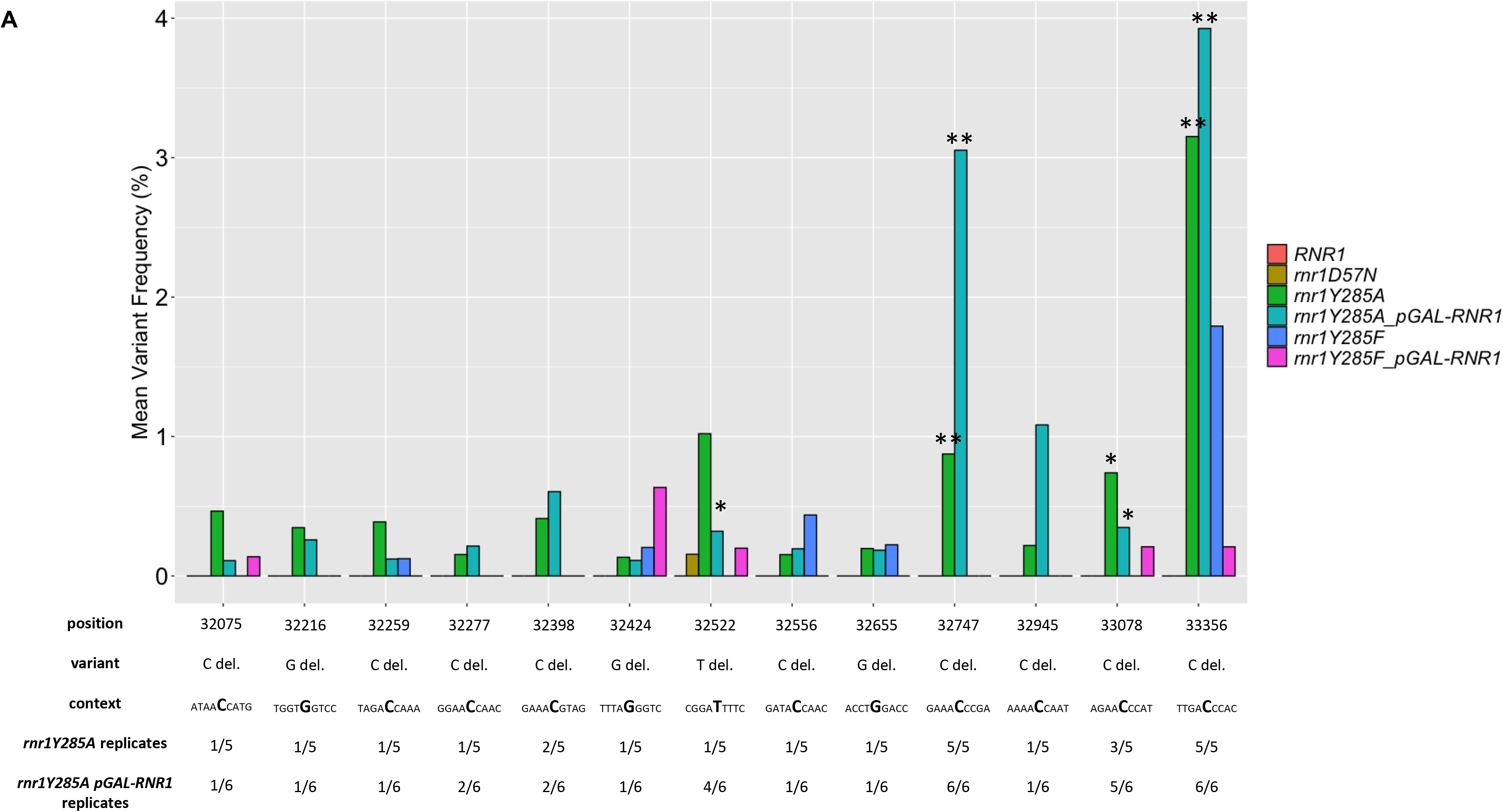
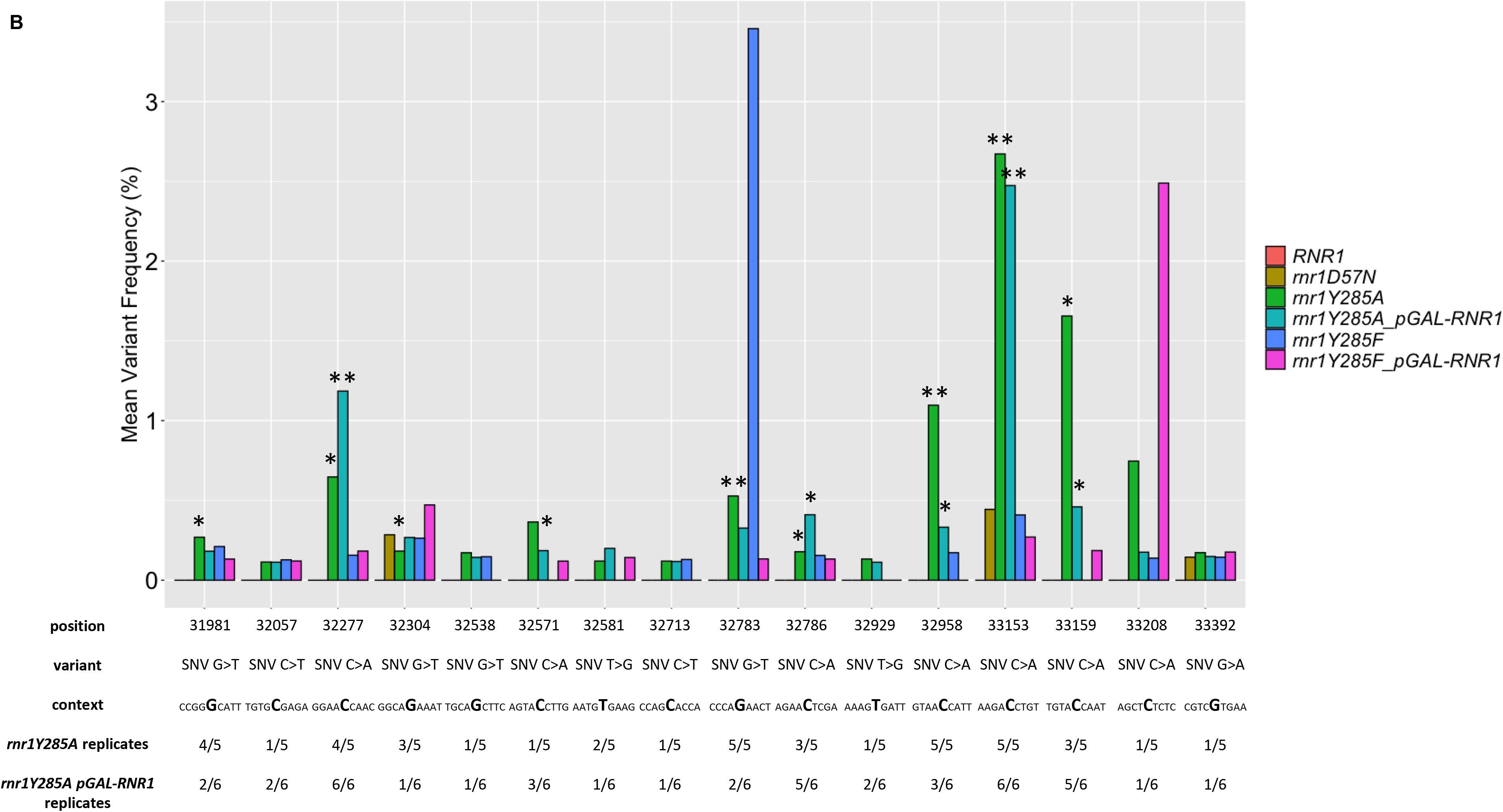
Low frequency variants specific to *rnr1Y285A* genotypes. Low frequency deletions (**A**) and SNVS (**B**) in *rnr1Y285A pGAL-RNR1* (turquoise) and *rnr1Y285A* (green) genotypes. None of these variants were observed in wildtype, but some occurred at low frequencies in *rnr1D57N* (brown) and *rnr1Y285F* (blue and pink) genotypes. *, variant occurred in ≥ 50% of *rnr1Y285A* or *rnr1Y285A pGAL-RNR1* biological replicates;**, variant occurred in 100% of the *rnr1Y285A* or *rnr1Y285A pGAL-RNR1* biological replicates analyzed.

### SNVs in trinucleotide context reveal unique mutation signatures

The analysis of both high and low frequency variants above indicated specific positions within *can1* that were susceptible to mutation in a genotype-specific manner, but the analysis is cumbersome and somewhat biased by frequency. The significant sequencing depth achieved in our study provided the opportunity to analyze genotype-specific variants and sequence context more systematically. We took two distinct approaches, which have not been previously applied to these *rnr1* alleles: 1) trinucleotide context analyses and 2) hierarchical cluster analyses paired with motif enrichment.

Assessing the trinucleotide context of mutations is an increasingly common approach to extract mutation signatures, especially in human cancers (ALEXANDROV *et al.* 2016; HARADHVALA *et al.* 2018). For our analysis, each unique SNV was categorized with respect to the nucleotide immediately 5’ and 3’ to the variant, with 96 possible triplet contexts. We determined the average number of times an SNV was observed in a particular triplet context per genotype, normalized to the number of times the triplet context occurs in *CAN1* (**Fig. 8)**. Pearson correlation coefficients of each SNV in unique trinucleotide context were calculated to evaluate patterns of SNV mutagenesis (**Fig S6 & Table S7**). Notably, in all genotypes, C→T changes (red bars, **Fig. 8**), particularly in G**C**C and G**C**G sequence contexts, dominated. The proportion of G**C**C and G**C**G changes increased with altered dNTPs, most dramatically in *rnr1Y285F* and *rnr1Y285A* samples, which were highly correlated within a genotype (*rnr1Y285A:rnr1Y285A pGAL-RNR1* r_s_ = 0.930, *rnr1Y285F:rnr1Y285F pGAL-RNR1* r_s_ = 0.947) and between *rnr1Y285F* and *rnr1Y285A* genotypes (*rnr1Y285F:rnr1Y285A* r_s_ = 0.924, *rnr1Y285F pGAL-RNR1:rnr1Y285A pGAL-RNR1* r_s_ = 0.909). *RNR1* and *rnr1Y285A* genotypes showed the weakest correlations (*RNR1:rnr1Y285A* r_s_ =0.693, *RNR1:rnr1Y285A pGAL-RNR1* r_s_ = 0.658), with many variants absent in *rnr1Y285A* (C>G (black), T>A (gray), T>C (green) and T>G (pink) variants. For example, C>G errors in A**C** G, A**C** T, C**C** G, G**C** A, G**C** G, G**C** T, T**C** C and T**C** T contexts were completely absent in both *rnr1Y285A* genotypes. Many of these missing variants occurred in repetitive sequences, while errors in different repetitive contexts dominated in *rnr1Y285A*.

**Figure 8.**
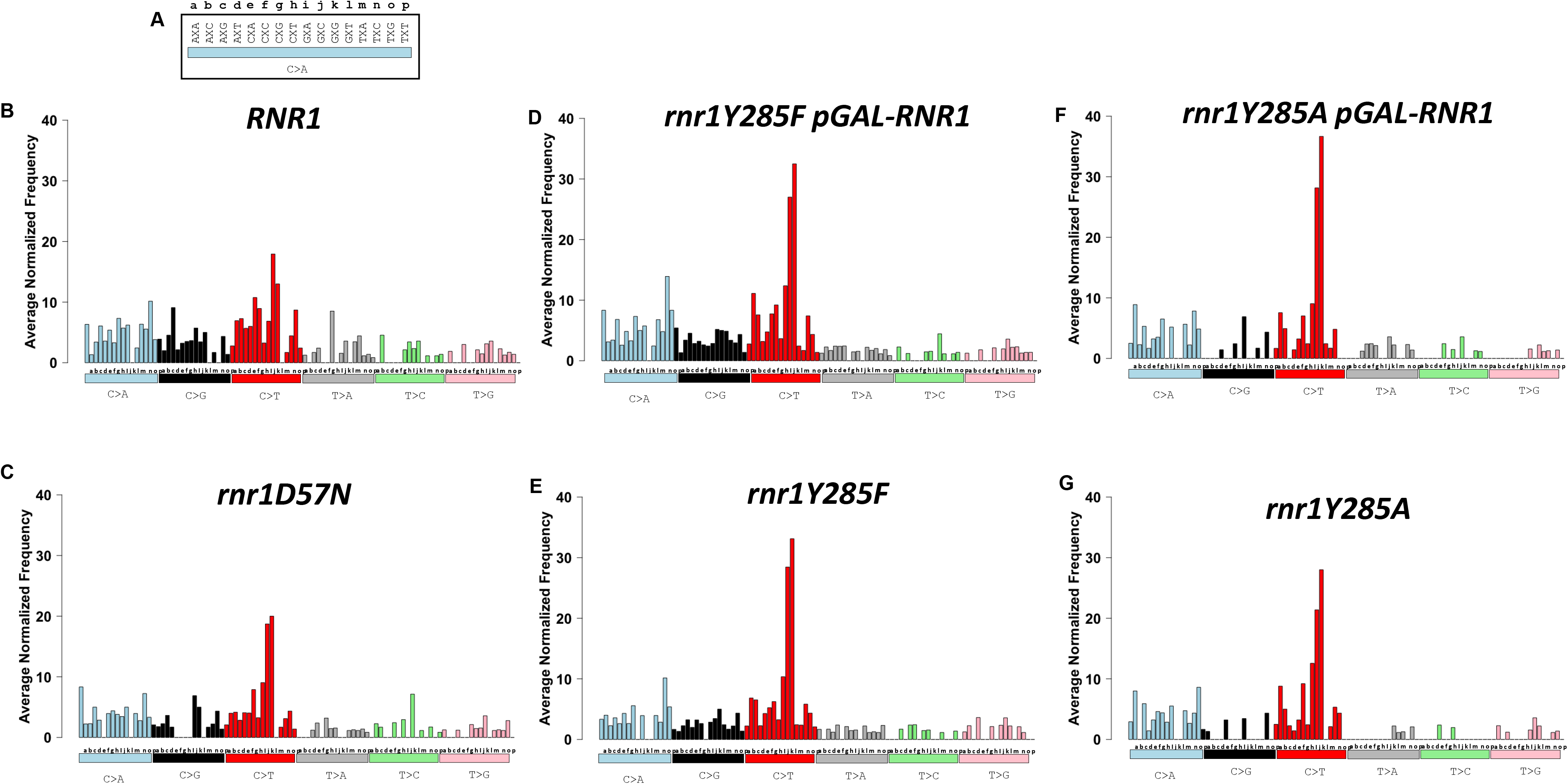
The average number of each SNV as it occurs in unique triplet nucleotide context is distinct between *rnr1* alleles. Bars are colored according to the six different types of SNVs. (**A**) The 16 different triplet contexts are lettered for display purposes. The variant change (C>A, turquoise bar) occurs at the middle nucleotide marked X in each triplet context for (**B**) *RNR1*, (**C**) *rnr1D57N*, (**D**) *rnr1Y285F pGAL-RNR1*, (**E**) *rnr1Y285F*, (**F**) *rnr1Y285A pGAL-RNR1* and (**G**) *rnr1Y285A*.

*RNR1* and *rnr1D57N* or *rnr1Y285F* were more strongly correlated (*RNR1:rnr1D57N* r_s_ =0.774, *RNR1:rnr1Y285F* r_s_ = 0.736), consistent with more subtle differences between SNV spectra in trinucleotide context (**Fig. 8**). In *rnr1D57N*, the most apparent difference is a loss of C>G variants in C**C** A, C**C** C, C**C** G and C**C** T; both *rnr1D57N* and *rnr1Y285F* also exhibited some variability in T>A (grey), T>C (green) and T>G (pink) variants.

### Motif enrichment reveals CC dinucleotides are commonly mutated in *rnr1Y285F/A* genotypes

We performed hierarchical cluster analysis of the unique variants in our dataset to determine differences between mutation profiles and to determine which variants in *CAN1/can1* were driving these differences. This analysis was paired with motif enrichment to determine broader sequence contexts prone to mutagenesis in a particular genetic background. We confirmed that *rnr1Y285A* and *rnr1Y285A pGAL-RNR1* shared distinct features as these genotypes clustered together (**Fig. 9**), as did the *rnr1Y285F* genotypes. Both *rnr1Y285F* and *rnr1Y285A* clustered away from *rnr1D57N* and *RNR1*, which were more closely related in this analysis. We observed three main clusters of unique variants, labeled I, II and III.

**Figure 9.**
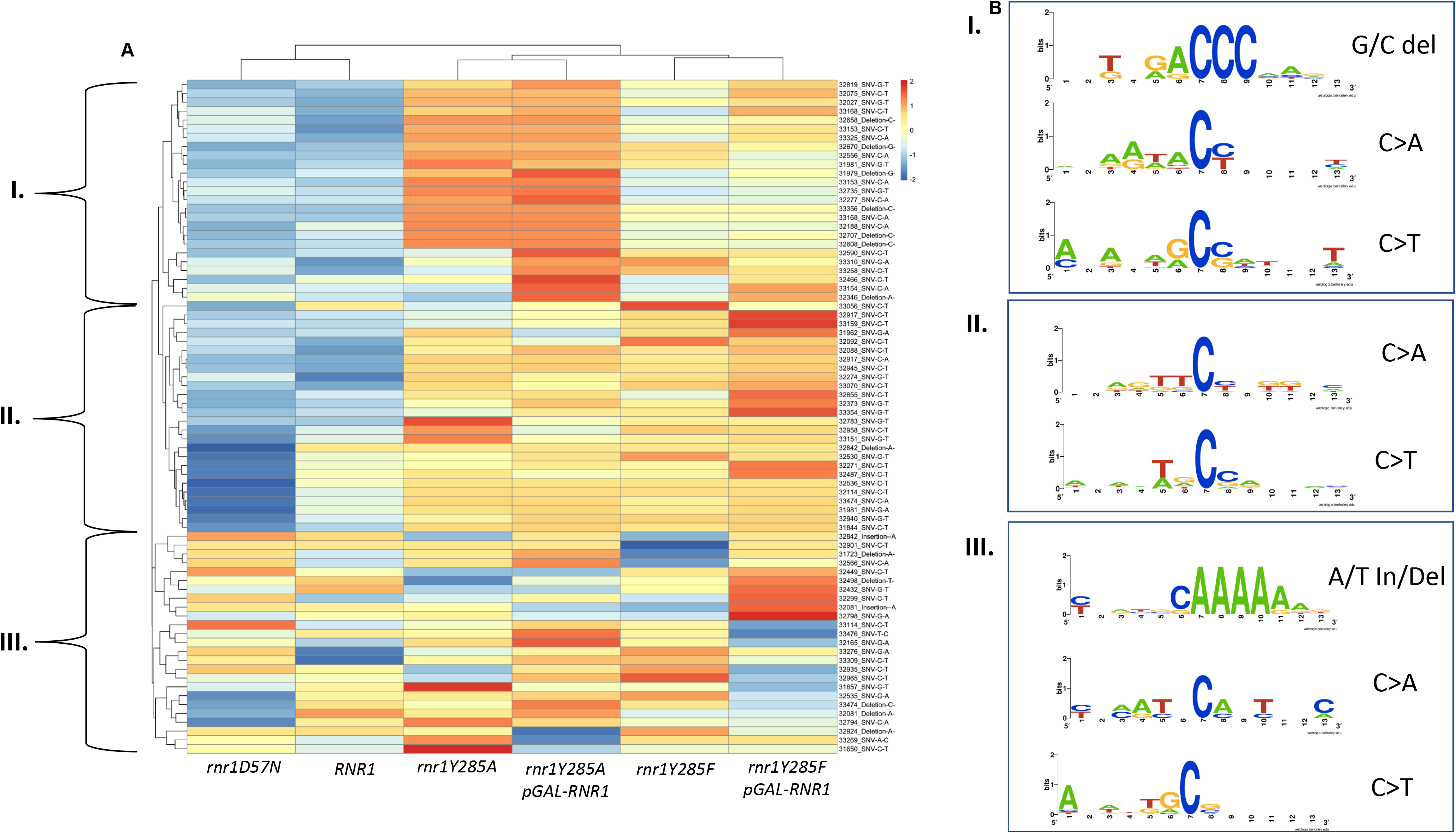
Unique variants in enriched motifs form distinct clusters. **(A)** Hierarchical cluster analysis of the highest occurring variants reveals distinct clusters, annotated I, II and III. **(B)** The different classes of variants enriched in each cluster were subset, and 12 basepairs surrounding the variant was used to perform motif enrichment. The corresponding motifs from the different variant classes in each boxed cluster are displayed on the right and labeled accordingly (box I, II and III).

In the first two clusters (**Fig. 9**, I and II) *RNR1* and *rnr1D57N* were very similar, while *rnr1Y285F/A* exhibited distinct differential enrichment profiles. The variants in cluster I were more significantly overrepresented in *rnr1Y285A* backgrounds, relative to *rnr1Y285F.* The reverse was true of variants in cluster II. Both clusters I and II were underrepresented in *RNR1* and *rnr1D57N*. Within cluster II, there were several variants that were more significantly underrepresented in *rnr1D57N* than in *RNR1*. In contrast, the third cluster (III) did not exhibit clear trends, with different variants under- or over-represented in different genotypes. Combined, these data indicate genotype-specific mutagenesis.

From each of these three clusters, we performed motif enrichment on the sequence surrounding the variants and identified unique contexts differentially enriched in *rnr1Y285F/A* backgrounds, consistent with the trinucleotide context data (**Fig. 8)** as well as high frequency (**Fig. 6**) and low frequency (**Fig. 7**) variant analysis. C>A and C>T changes in cluster I occurred at CC dinucleotides, while those in cluster III, which were underrepresented in *rnr1Y285F/A*, did not. This is consistent with the prediction that repetitive GC sequences are more prone to mutation in the presence of skewed elevations in dCTP and dTTP. Similarly, G/C deletions in repetitive G/C context were differentially enriched in *rnr1Y285A* genotypes (**Fig. 9, cluster II**), while A/T insertions and deletions were enriched across all samples (**Fig. 9, cluster III**).

## DISCUSSION

To build mutation profiles from first principles, we developed a *CAN1* selection-based next generation sequencing approach to determine robust, information-rich, genotype-specific mutation profiles. We focused on RNR alleles; altered RNR activity is clearly implicated in reduced replication fidelity and carcinogenesis (MATHEWS 2015; MATHEWS 2018). While *CAN1* is a widely used reporter gene in yeast, the application of deep-sequencing allowed streamlined sample preparation and more accurate determination of mutation spectra, by increasing both the number of colonies and biological replicates analyzed at one time. This approach allowed us to 1) identify low frequency sequence variants, 2) develop mutational fingerprints that resulted from compromised replication fidelity as a result of altered dNTP pools, and 3) identify broader sequence contexts significantly enriched in specific genotypes. We expanded our understanding of the impact of altered dNTP pools on mutagenesis and developed an approach that can be applied to study other genetic backgrounds or environmental exposures.

Our sequencing/analytic approach allowed us to define genotype-specific mutation profiles, including trinucleotide sequence context for mutations and broader sequence motifs surrounding mutations. Deep sequencing and analysis of only *can1* mutations identified mutation sequence motifs similar to those generated through WGS (WATT *et al.* 2016) and revealed new details for mutation profiles in genotypes with lower mutations rates. From this information, we can infer mechanisms of mutagenesis (see below). Importantly, the motifs identified previously in *rnr1Y285A* lines also carried a *msh2Δ*, effectively eliminating MMR, which might alter the diagnostic sequence contexts in which mutations are specifically present or absent. We identified sequence motifs in *rnr1Y285A* similar to those in *rnr1Y285A msh2Δ* from WGS (WATT *et al.* 2016), indicating that the errors generated when dNTPs are elevated and skewed are substrates for mismatch repair.

### Mechanisms of mutagenesis from distinct elevations in dNTP levels

Despite a 2-fold increase in mutation rate above wildtype, the *rnr1D57N* mutation spectrum closely resembled wildtype (**Fig. 4**), consistent with previous studies (XU *et al.* 2008; KUMAR *et al.* 2011). This indicated that the 2-fold balanced increase in dNTPs causes errors to accumulate in a stochastic manner, similar to wildtype. Given the low mutation rate in *rnr1D57N* (**Table S5**), the majority of these errors are likely corrected by the mismatch repair (MMR) system, which identifies replication errors and targets them for repair(XU *et al.* 2008; KUNKEL AND ERIE 2015). Nonetheless, we observed a significant increase in G/C single base deletions in this genetic background (**Fig. 4B**) and were able to identify new sequence contexts specific to mutations in *rnr1D57N*. This includes CG>TA, CG>AT changes, A/T deletions in repetitive A/T sequences (**Figs. 4, 6B, 9**), which appeared diagnostic of *rnr1D57N* mutagenesis. We predict that these regions are inherently more prone to mutation; the two-fold increase in dNTP levels exacerbates this by favoring replicative polymerase synthesis over proofreading activity.

The high frequency of G/C deletions observed in *rnr1D57N* was driven by a single base G deletion at position 31971 that was mutated in 4 out of 7 *rnr1D57N* replicates and occurs at a high frequency, on average ~60% (**Figs. 4B & 6B, S2D**– compare unique counts vs. frequency of variants). The high frequency indicated the mutation arose early in the growth of the cultures, but the fact that it occurred and was enriched in multiple biological replicates indicated that this position is uniquely susceptible to deletion, *specifically* in *rnr1D57N*. This variant was bordered by repeats on both the 5’ side (repetitive A/T motif) and 3’ side (CC dinucleotide) (ATAA**G**CCA). Therefore, both the excess in dTTP incorporated opposite the AA dinucleotide and the excess of dCTP incorporated on the opposite strand could result in transient misalignment and misincorporation at this position and it is therefore likely this variant occurred during both leading and lagging strand replication. Increased dNTP levels alter replication fork dynamics(DAVIDSON *et al.* 2012; POLI *et al.* 2012), potentially enhancing the next nucleotide effect and resulting in increased frequencies at this position in *rnr1D57N*.

The *rnr1Y285F* allele exhibited a modest effect on mutation rate, but a distinct mutation profile (**Fig. 4**). While the profile shift was more subtle than in *rnr1Y285A*, meta-analysis of the types and positions of variants in *rnr1Y285F* significantly overlapped with that of *rnr1Y285A* (**Figs. 6, 8, 9**), despite the differences in mutation rate (**Table S5**) and dCTP/dTTP levels (KUMAR *et al.* 2010). This was particularly noticeable with G/C variants, i.e. G/C deletions and CG>AT variants. Therefore, even modestly increased dCTP and dTTP pools resulted in distinct error accumulation. We predict an increased probability that replicative polymerases incorporated the excess nucleotides (dCTP, dTTP) during synthesis and more efficiently extended mismatches at the expense of proofreading (KUMAR *et al.* 2011; WATT *et al.* 2016).

We noted a substantial increase in G/C deletions specific to *rnr1Y285A* backgrounds (**Fig. 6E & 6F**), which may be explained by limiting levels of dGTP in *rnr1Y285A*. dGTP levels are already limiting in both yeast and mammalian cells(CHABES *et al.* 2003; HÅKANSSON *et al.* 2006);in *rnr1Y285A*, the proportion of dGTP relative to the total dNTP pool is extremely small, reducing the probability of its incorporation. We predict that this led to this distinct pattern of deletion in *rnr1Y285F/A* (MATHEWS 2015), within specific sequence contexts, i.e. CC and CCC runs (**Fig. 9**). Notably, *rnr1* alleles that dramatically increased dGTP levels were also severely mutagenic (*rnr1K243E* and *rnr1I262V, N291D)*, increasing SNVs and frame shift mutations in repetitive contexts(SCHMIDT *et al.* 2019). This highlights the importance of proper absolute and relative abundance of dGTP.

Our data indicated that the mutation profiles generated using our sequencing and analytical approach were diagnostic of the balanced or unbalanced nature of the dNTP pools, i.e. which dNTPs were elevated. While the mutation rate and variant frequencies reflected the absolute levels of dNTPs; far more in/dels than SNVs were observed for *rnr1Y285A* while the reverse was true for *rnr1Y285F*, but the proportions were similar. This hypothesis could be tested by comparing the *rnr1D57N* mutation profile with that of *RNR1* galactose-induced overexpression, which similarly leads to balanced dNTP pools, but elevated ~10-fold above wildtype levels (CHABES AND STILLMAN 2007), as well as other *rnr1* alleles.

### Implications for understanding mutation signatures in human cancers

Mutation signatures of human tumors are used to identify molecular drivers of carcinogenesis (ALEXANDROV *et al.* 2013; NIK-ZAINAL *et al.* 2016; HARADHVALA *et al.* 2018; ALEXANDROV *et al.* 2020), which have clear implications for diagnosis, prognosis and treatment options for patients (VAN HOECK *et al.* 2019). Tumor mutation signatures are typically extracted from a large sampling of human tumor information via mathematical approaches; implicated molecular pathways are inferred. This has led to spurious associations between mutation profiles and driver mutations, e.g. RefSig 3 and *BRCA1/BRCA2* (DEGASPERI *et al.* 2020). In contrast, we used the *S. cerevisiae* model system to develop information-rich mutation profiles from first principles, using multiple biological replicates of defined genetic backgrounds alongside wild-type controls.

Importantly, elevated dNTP levels (balanced or skewed) have not been considered when evaluating mutation signatures from human tumors, although they almost certainly contribute to mutagenesis in cancer (AYE *et al.* 2015; MATHEWS 2015; MATHEWS 2017; PAI AND KEARSEY 2017; DEGASPERI *et al.* 2020). We noted distinct similarities between *rnr1* SNV triplet mutation profiles (**Fig. 8)** and specific COSMIC signatures, most notably signatures 6 and 15. Signature 6 occurs most commonly in colorectal and uterine cancers and is associated with defective MMR. We previously noted synergistic effects on mutation rate between *rnr1D57N* and MMR deletions(XU *et al.* 2008). The contribution of elevated dNTP pool levels to mutagenesis in combination with MMR is intriguing.

While elevated dNTP levels have been implicated in supporting rapidly proliferating cancer cells, it remains to be determined whether these are skewed or balanced increases, which could be tumor-specific (WILSON *et al.* 2011; KOHNKEN *et al.*; MATHEWS; PURHONEN *et al.* 2020). Quantification of dNTP levels in HeLa cell lines demonstrated elevated levels of all four dNTPs with an approximate 2-fold increase in dATP and dCTP and a 4-fold increase in dTTP and dGTP (MATHEWS 2015). Different skewed elevations result in distinct mutation spectra (SCHMIDT *et al.* 2019) and thus more studies are necessary to determine what dNTP imbalances are relevant to different types of cancers. In the meantime, certain indicator mutations, such as a high number of G/C single base deletions, can point towards specific dNTP imbalances *i.e.* high dCTP and dTTP levels seen in *rnr1Y285A*.

## ACKNOWLEDGEMENTS

We are grateful to Dr. Andrei Chabes for providing strains and plasmids. We are grateful to Dr. Don Yergeau for his advice in next generation sequencing. We are grateful to members of the Surtees lab (past and present) for helpful discussions and input. We are grateful to Dr. Mark Sutton and Dr. John Panepinto for input and helpful discussions.

