## Supplemental figures and tables for "A targeted next generation sequencing approach to develop robust, genotype-specific mutation profiles and uncover novel variants in *Saccharomyces cerevisiae*"

**Supplemental Figure Legends:**

**Figure S1. Mutation spectra become more distinct after permissive filtering. (A)** Total number of unique variants for all biological replicates before (Pre-filter; upper panel) and after (Post-filter; lower panel) application of the permissive filter. **(B)** The sum of all variant frequencies for all biological replicate before (Pre-filter; upper panel) and after (Post-filter; lower panel) application of the permissive filter. Both plots exhibited changes post-permissive filtering, but they are more pronounced in unique variants (**A**).

**Figure S2. Mutation spectra for all biological replicates.** (**A**) The SNV mutation spectra based on the total of unique variants and (**B**) the sum of frequencies from each of the 6 classes observed in each biological replicate. (**C**) The deletion spectra based on the unique variants and (**D**) the sum of frequencies. (**E**) Insertion spectra based on unique variants and (**F**) the sum of frequencies.

**Figure S3. A comparison to previous studies which utilized Sanger sequencing.** Variant counts determined using Sanger sequencing from previous studies {xu, 2008; kumar, 2011} were compared with the normalized sum of frequencies from genotypes in our study.

**Figure S4. The average variant frequency across *CAN1* by genotype.** Low variant frequencies (<5%) and distribution for (**A**) *RNR1*, (**B**) *rnr1D57N*, (**C**) *rnr1Y285F pGAL-RNR1*, (**D**) *rnr1Y285F, (***E***)* *rnr1Y285A pGAL-RNR1*, **(F)** and *rnr1Y285A*. The *rnr1Y285A* genotypes exhibited a lower unique variant frequency

**Figure S5. High frequency variants across *CAN1*.** Variants that occurred at greater than 5% average frequency within a genotype are plotted across *CAN1*.

**Figure S6. Correlation analysis on SNVs in triplet context.** Spearman correlation performed on the average number of each SNV as it occurs in unique triplet nucleotide context. Correlation values are displayed in **Table S8**.

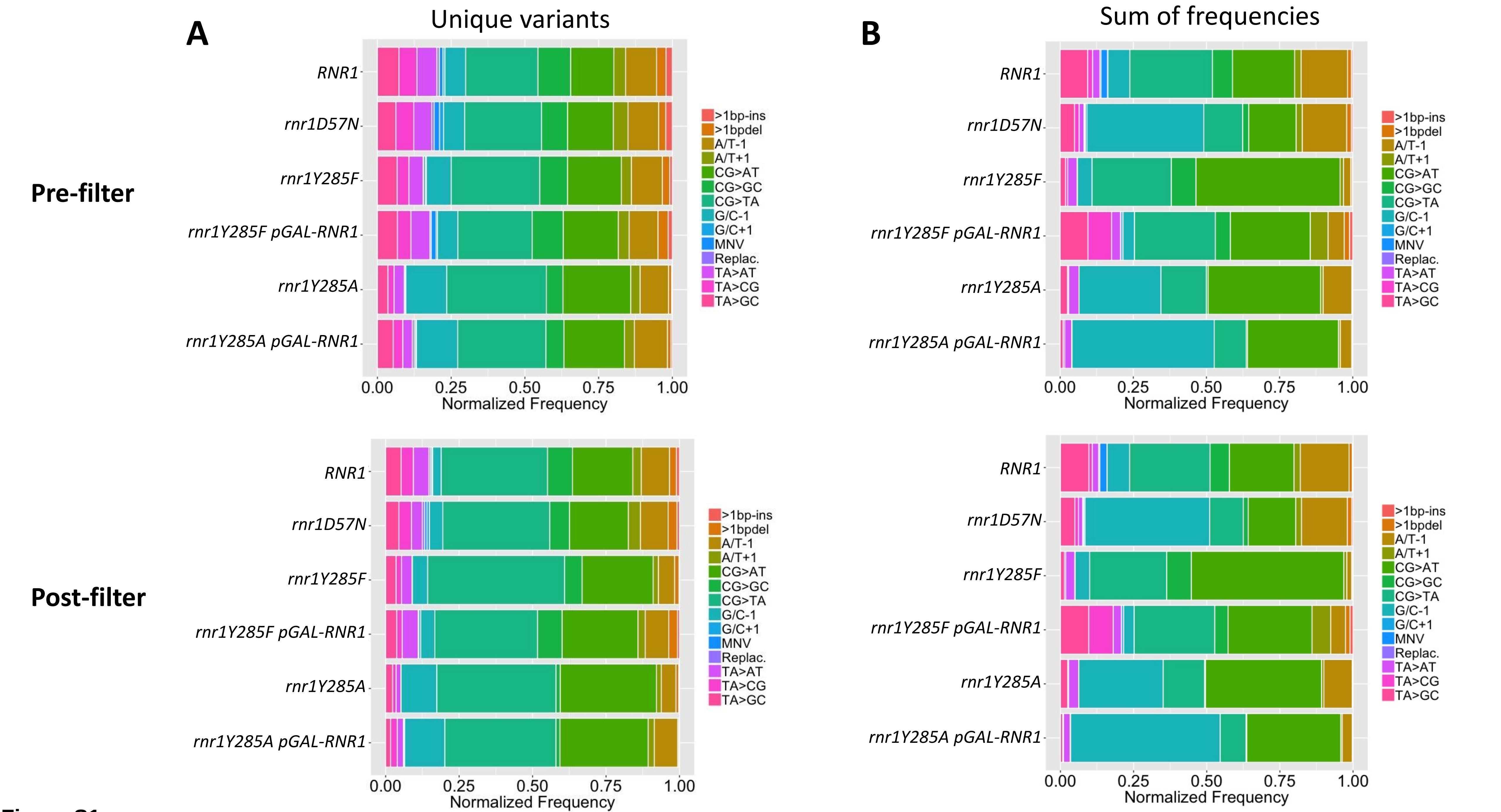
A
Unique variants
B
Sum of frequencies
Pre-filter
Post-filter
RNR1
rnr1D57N
rnr1Y285F
rnr1Y285F pGAL-RNR1
rnr1Y285A
rnr1Y285A pGAL-RNR1
0.00
0.25
0.50
0.75
1.00
Normalized Frequency
>1bp-ins
>1bpdel
A/T-1
A/T+1
CG>AT
CG>GC
CG>TA
G/C-1
G/C+1
MNV
Replac.
TA>AT
TA>CG
TA>GC

Figure S1.

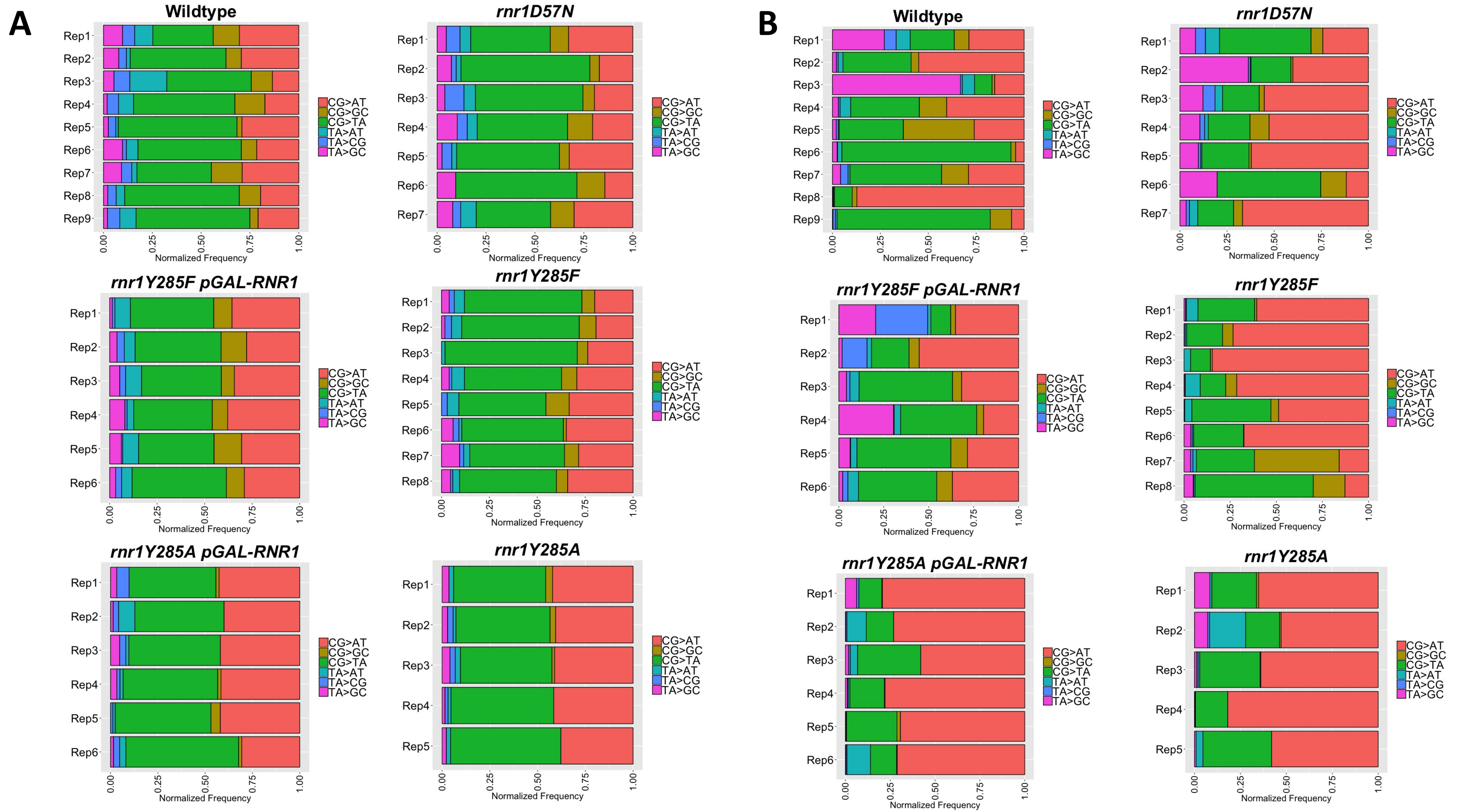

Unique counts

Sum of frequencies

Figure S2.

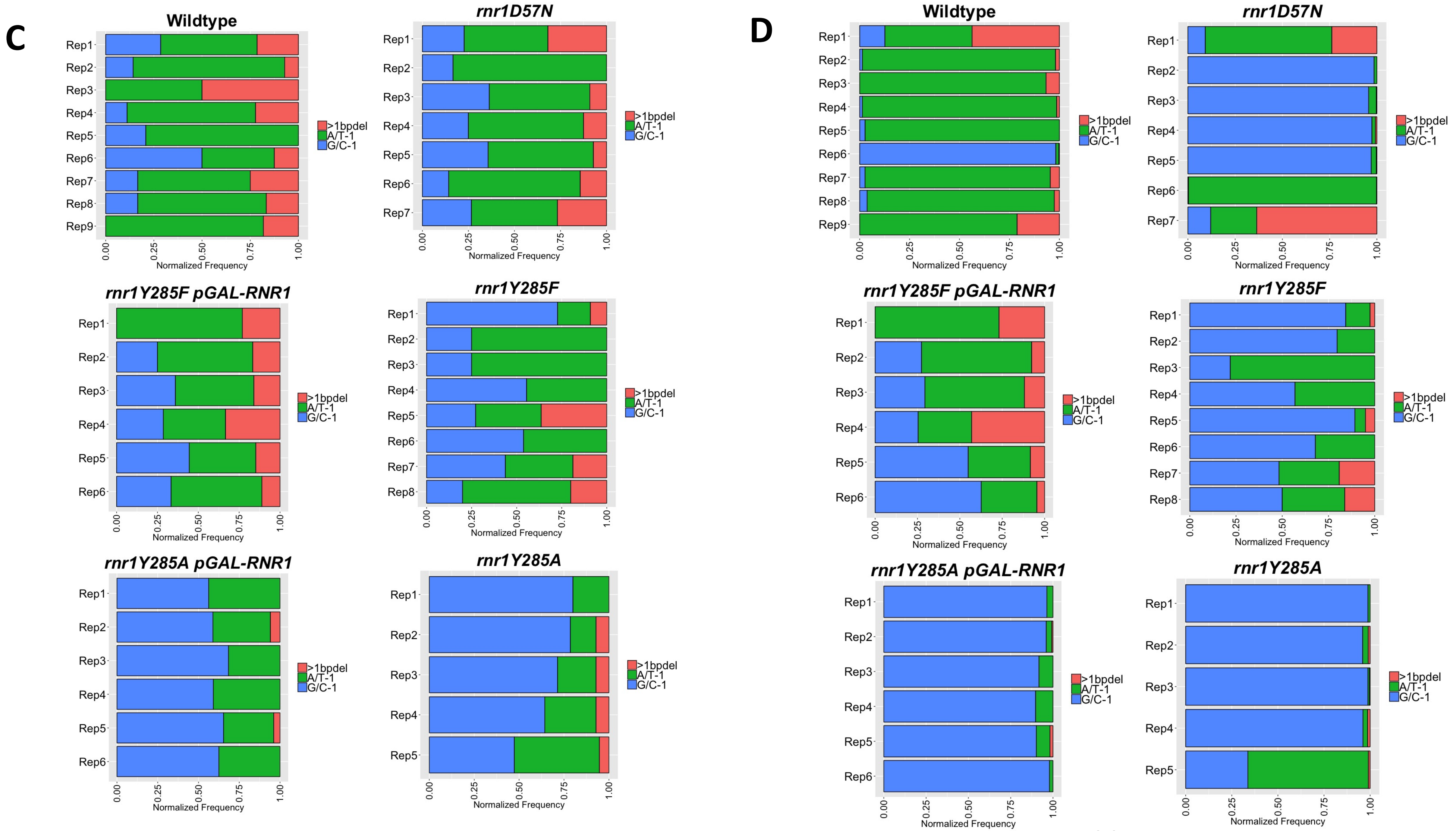

**Figure S2 (cont'd).**

**Unique counts**

**Sum of frequencies**

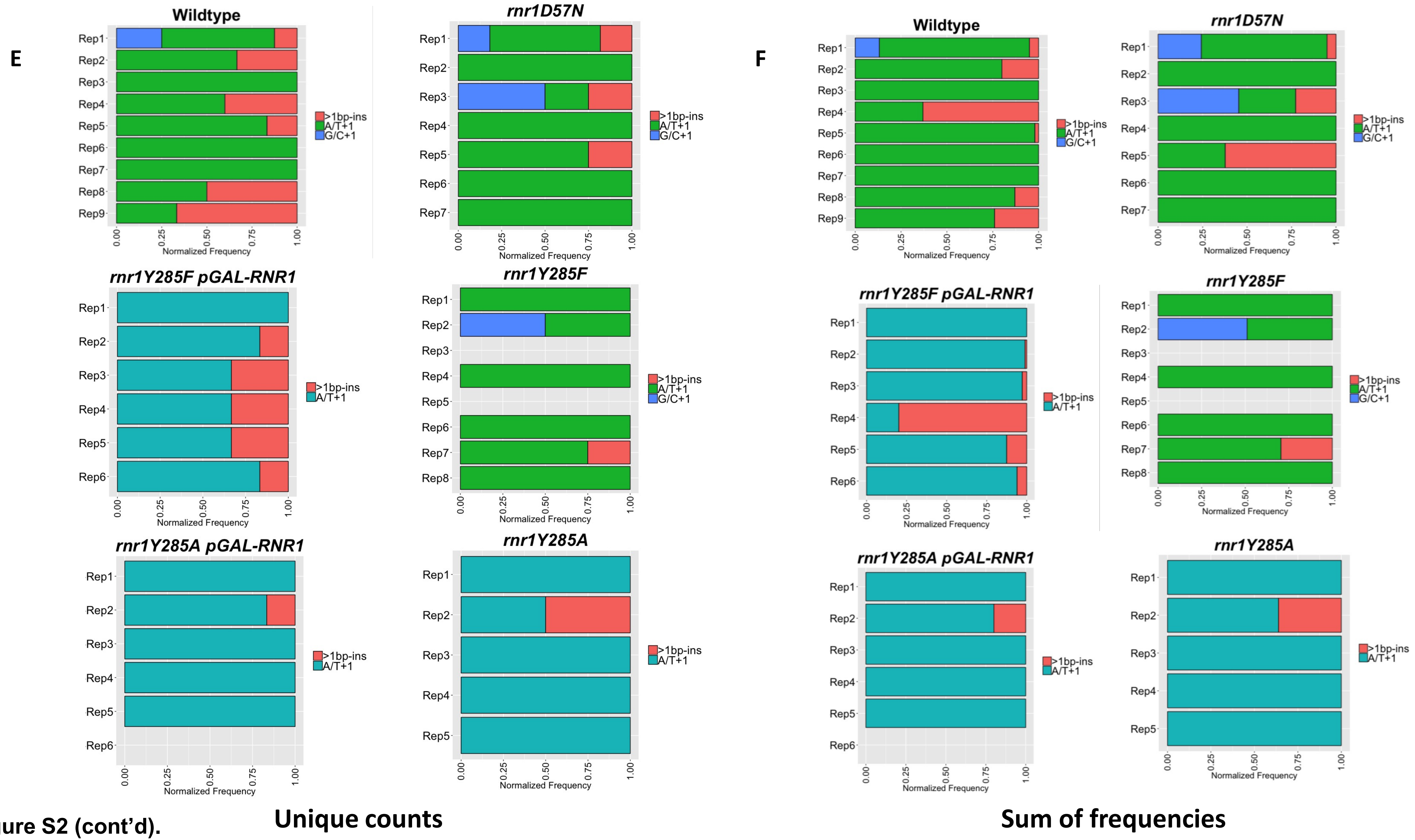

**Figure S2 (cont'd).**

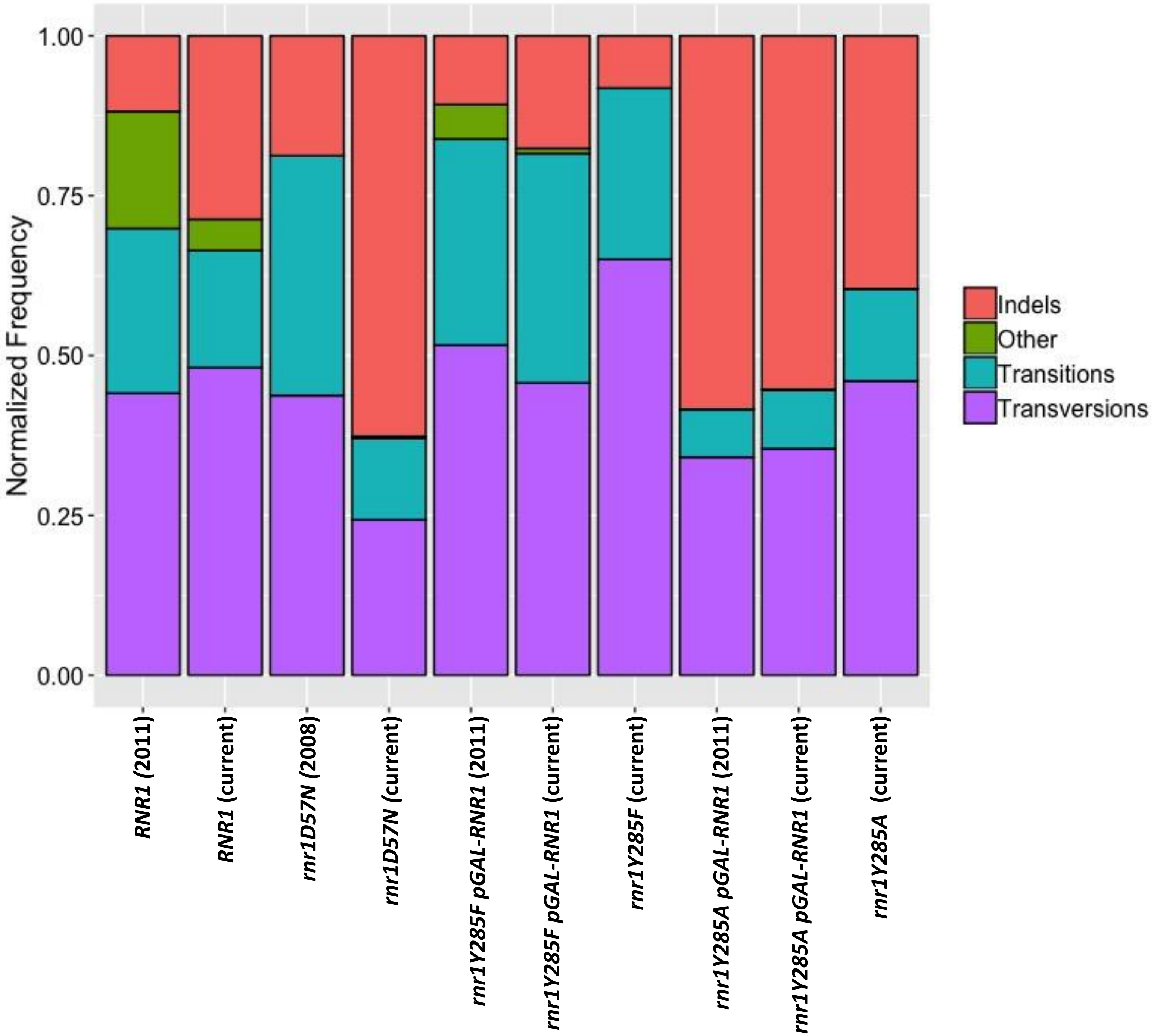
Normalized Frequency
1.00
0.75
0.50
0.25
0.00
Indels
Other
Transitions
Transversions
RNR1 (2011)
RNR1 (current)
rnr1D57N (2008)
rnr1D57N (current)
rnr1Y285F pGAL-RNR1 (2011)
rnr1Y285F pGAL-RNR1 (current)
rnr1Y285F (current)
rnr1Y285A pGAL-RNR1 (2011)
rnr1Y285A pGAL-RNR1 (current)
rnr1Y285A (current)

**Figure S3.**

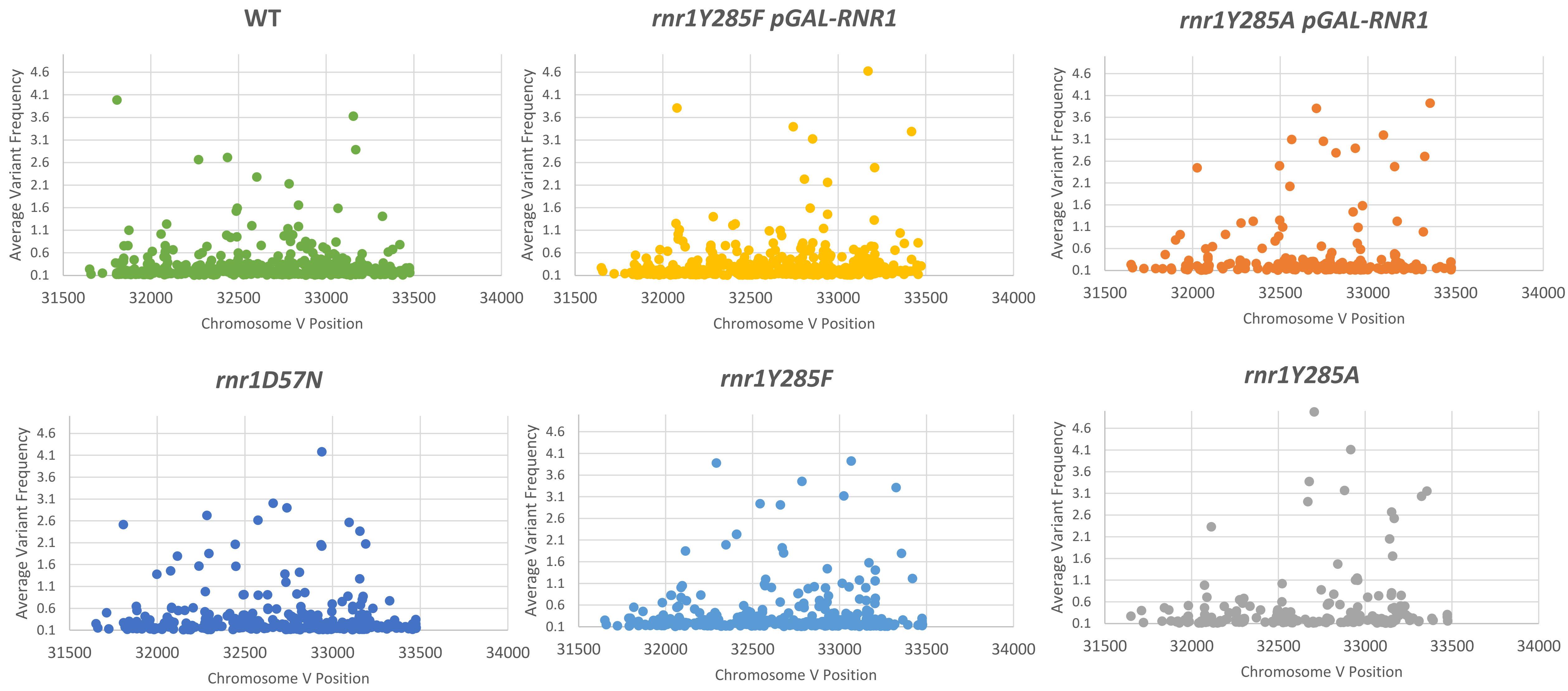
WT
rnr1Y285F pGAL-RNR1
rnr1Y285A pGAL-RNR1
rnr1D57N
rnr1Y285F
rnr1Y285A
Average Variant Frequency
4.6
4.1
3.6
3.1
2.6
2.1
1.6
1.1
0.6
0.1
31500
32000
32500
33000
33500
34000
Chromosome V Position

**Figure S4.**

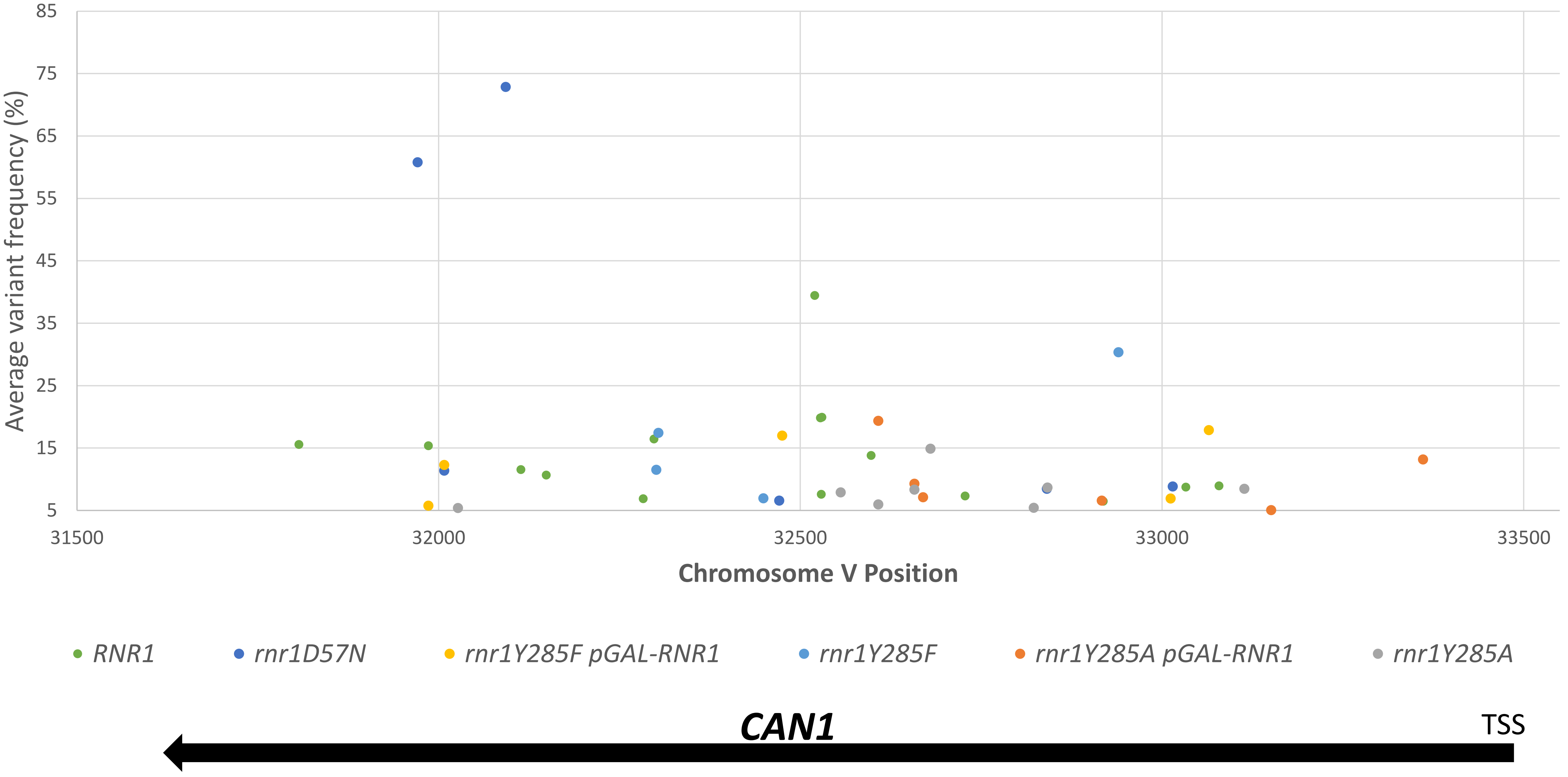

**Figure S5. Dominant mutations across *CAN1***

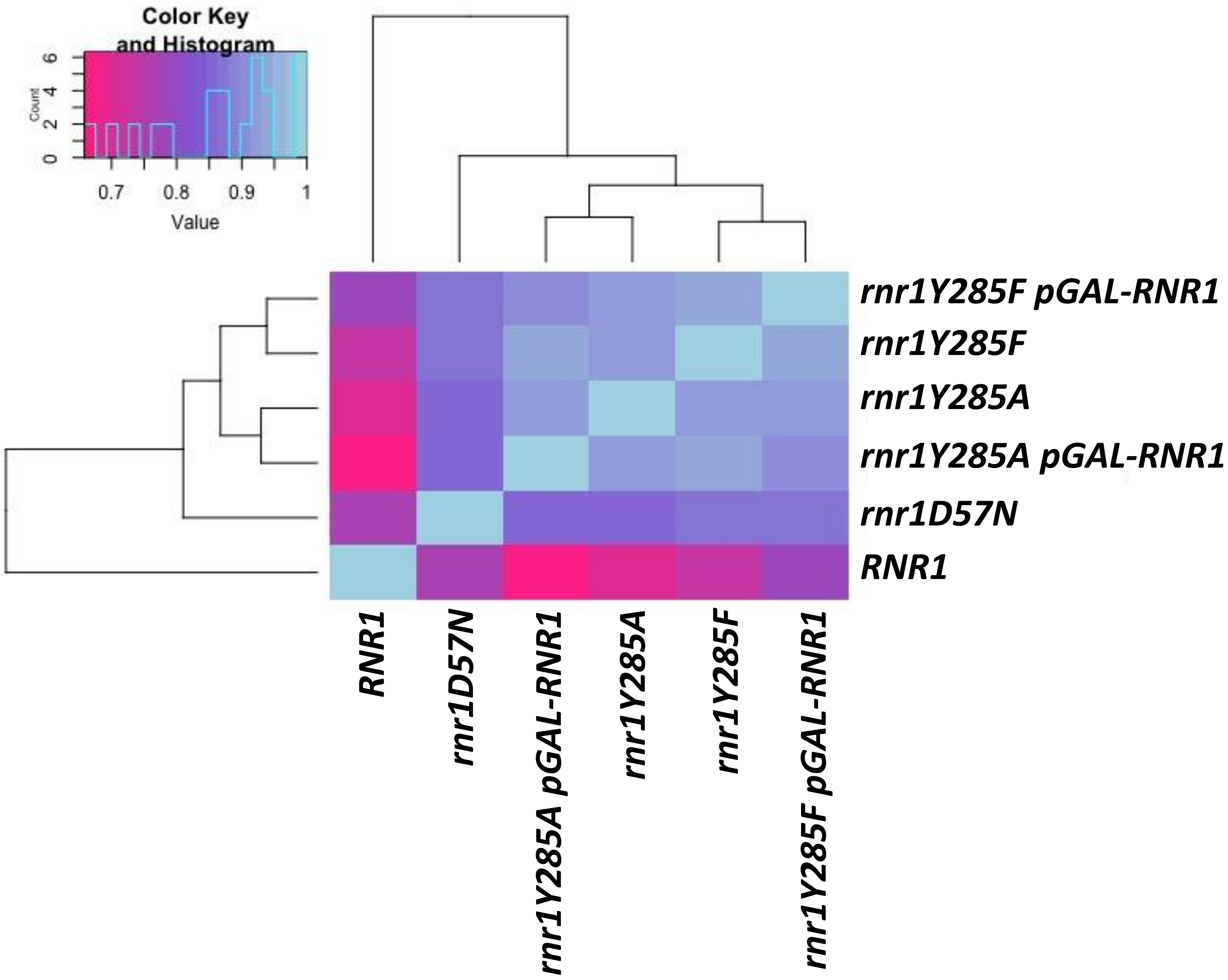

Color Key
and Histogram
Count
0
2
4
6
0.7
0.8
0.9
1
Value
rnr1Y285F pGAL-RNR1
rnr1Y285F
rnr1Y285A
rnr1Y285A pGAL-RNR1
rnr1D57N
RNR1
RNR1
rnr1D57N
rnr1Y285A pGAL-RNR1
rnr1Y285A
rnr1Y285F
rnr1Y285F pGAL-RNR1

**Figure S6.**

**Supplementary Table 1.** Strains used in this study
All strains are in the W303 background:
*CAN1, ade2-1, his3-11,15, leu2-3, 112, trp1-1, ura3-1, RAD5+ or adc2-11 tris3-11,15 leu 2-3 112 trp1-1 URA 3 RAD5+*

| JSY# | Other name | Relevant Genotype | dNTP levels | Source |
|---|---|---|---|---|
| JSY13 | EAY1995/4C | *RNR1, MATa, CAN1, ade2-1, his3-11,15, leu2-3, 112, trp1-1, ura3-1,* | Wildtype | Chabes et al., 2003<br>Xu et al., 2008 |
| JSY2424 | AC402 | *RNR1-ade2-11 tris3-11,15 leu 2-3 112 trp1-1 URA 3 RAD5+* | Wildtype | Chabes et al., 2003 |
| JSY14 | EAY1995/8C | *rnr1D57N* | 2-fold balanced elevation in dNTPs | Chabes et al., 2003 |
| JSY2420, JSY2421 | DK3A, DK3C | *rnr1Y285F -URA3-pGal-RNR1* | 3-fold increase in dCTP and dTTP | Kumar et al., 2010 |
| JSY2422, JSY2423 | DK8A, DK8E | *rnr1Y285A -URA3-pGal-RNR1* | 20-fold increase in dCTP and dTTP | Kumar et al., 2010 |
| JSY3811,JSY3865,JSY3866 | JSY3811,JSY3865,JSY3866 | *rnr1Y285F* | 3-fold increase in dCTP and dTTP | This Study |
| JSY3868, 3869 | JSY3868, 3869 | *rnr1Y285A* | 20-fold increase in dCTP and dTTP | This Study |

**Supplementary Table 2.** Oligonucleotides used in this study

| SO# | Name | Sequence- 5'-3' | Description |
|---|---|---|---|
| 261 | PRS414 FWD + 0_AatII | GAACTAGACGTCGTTTCGGTGATGACGGTGAA | To generate pEM3 integration plasmid |
| 262 | PRS414 REV+4200+AatII | TCCATAGACGTCGGCACTTTTCGGGGAAATGT | To generate pEM3 |
| 263 | RNR1_-1kb_XhoI_fwd | CTAAGACTCGAGGTGTATAGTAGATCTGCAGCT | To amplify *RNR1/rnr1* alleles for ligation into pEM3 |
| 264 | RNR1_2KB_SpeI_rev | CATCTTACTAGTACGTTTGGTAAGCCTTGAATG | To amplify *RNR1/rnr1* alleles for ligation into pEM3 |
| 265 | RNR1_3kb_SpeI_REV | CATCTTACTAGTTATCATGGCCTTCTTACAAGG | To amplify *RNR1/rnr1* alleles for ligation into pEM3 |
| 266 | PRS416_1.9KB_FWD | CGTAATACGACTCACTATAGG | To amplify *RNR1/rnr1* alleles for ligation into pEM3 |
| 230 | RNR1_rev_3 | AAGTTGCGTTCCTTTATAGAACAATG | sequencing primer in *RNR1* |
| 32 | RNR1_16 | GGCGTTACCATAGAAATGTACTG | Upstream *RNR1* seq primer @ NT -683 |
| 21 | RNR1rev12 | GGAAGTCTTACCATCCTCAGAAG | *RNR1* reverse primer at Nt. 1389 (opposite RNR1_4) |
| 25 | RNR1_3 | CTCTCCAACATCAGCTCCTGG | *RNR1* seq primer @ NT 1053 |
| 22 | RNR1rev13 | GAATCTGTATCATGGCCTTCTTAC | *RNR1* reverse primer - 233 NT. downstream of STOP |
| 744 | rnr1D57N_Q5_F | AATCGAACTAAACAACTTAGC | rnr1D57N Q5 forward |
| 745 | rnr1D57N_Q5_R | GTTGTGACACCTTCATAG | rnr1D57N Q5 reverse |
| 746 | rnr1Y285A_Q5_F | CACTGCCCGTGCTGTTGACCAG | rnr1Y285A Q5 forward |
| 747 | rnr1Y285A_Q5_R | TTATTGAAAACACGAATCATAGG | rnr1Y285A Q5 reverse |
| 748 | rnr1Y285F_Q5_F | ACTGCCCGTTTTGTTGACCAG | rnr1Y285F Q5 forward |
| 749 | rnr1Y285F_Q5_R | GTTATTGAAAACACGAATCATAGG | rnr1Y285F Q5 reverse |
| 807 | CAN1 reg1 Forward_anchored | TCGTCGGCAGCGTCAGATGTGTATAAGAGACAGCTCCTGTAAAAACAAAAAAAAA | Sequencing primers with nextera adapters: CAN1-region1 |
| 714 | CAN1 reg1 Reverse | GTCTCGTGGGCTCGGAGATGTGTATAAGAGACAGTAGTACCACCAAGGGCAATC | Sequencing primers with nextera adapters: CAN1-region1 |
| 715 | CAN1 reg2 Forward | TCGTCGGCAGCGTCAGATGTGTATAAGAGACAGAAGCAAAGACATATTGGTAT | Sequencing primers with nextera adapters: CAN1-region2 |
| 716 | CAN1 reg2 Reverse | GTCTCGTGGGCTCGGAGATGTGTATAAGAGACAGATCCATGCGCGCAGTGGAAC | Sequencing primers with nextera adapters: CAN1-region2 |
| 717 | CAN1 reg3 Forward | TCGTCGGCAGCGTCAGATGTGTATAAGAGACAGTTCAATTTTGGACGTACAAA | Sequencing primers with nextera adapters: CAN1-region3 |
| 718 | CAN1 reg3 Reverse | GTCTCGTGGGCTCGGAGATGTGTATAAGAGACAGACCAGCAGTGATACCAACTA | Sequencing primers with nextera adapters: CAN1-region3 |
| 719 | CAN1 reg4 Forward | TCGTCGGCAGCGTCAGATGTGTATAAGAGACAGCACATTTCAAGGTACTGAAC | Sequencing primers with nextera adapters: CAN1-region4 |
| 720 | CAN1 reg4 Reverse | GTCTCGTGGGCTCGGAGATGTGTATAAGAGACAGTAGGAGCCAACTTGTTCTTT | Sequencing primers with nextera adapters: CAN1-region4 |
| 721 | CAN1 reg5 Forward | TCGTCGGCAGCGTCAGATGTGTATAAGAGACAGCGTATTTTATTTGGTCTATC | Sequencing primers with nextera adapters: CAN1-region5 |
| 722 | CAN1 reg5 Reverse | GTCTCGTGGGCTCGGAGATGTGTATAAGAGACAGAAACCTTGAATAATGATAAT | Sequencing primers with nextera adapters: CAN1-region5 |
| 723 | CAN1 reg6 Forward | TCGTCGGCAGCGTCAGATGTGTATAAGAGACAGCGGCCACATTTATGACGATC | Sequencing primers with nextera adapters: CAN1-region6 |
| 724 | CAN1 reg6 Reverse | GTCTCGTGGGCTCGGAGATGTGTATAAGAGACAGCTTTCTTTTCGGTGTATGAC | Sequencing primers with nextera adapters: CAN1-region6 |

**Supplementary Table 3.** Plasmids used in this study

| **Plasmid** | **Composition** | **Reference** |
|---|---|---|
| pRS416 | pBluescript, *URA3*, *CEN6*, *ARSH4* | Sikorski and Hieter (1989) |
| pEM3 | pBluescript, *URA3* | This study |
| pNL1 | pBluescript, *URA3* , *RNR1* | This study |
| pNL2 | pBluescript, *URA3* , *RNR1* -truncated | This study |
| pNL3 | pBluescript, *URA3* , *rnr1D57N* | This study |
| pNL4 | pBluescript, *URA3* , *rnr1Y285A* | This study |
| pNL5 | pBluescript, *URA3* , *rnr1Y285F* | This study |
| pNL6 | pBluescript, *URA3* , *rnr1D57N*- truncated | This study |
| pNL7 | pBluescript, *URA3* , *rnr1Y285F*- truncated | This study |

**Supplementary Table 4.** Samples used in this study

| Sample_Name | Group | Group | Bio_Replicate_Number | GrpRep | Permissive | Technical_Replicate | Tech_Rep_Sample | Parent_Strain | pGAL | Growth_Condition | BioRepNumber | GAL |
|---|---|---|---|---|---|---|---|---|---|---|---|---|
| 12_v.txt | rnr1D57N | rnr1D57N | 1 | rnr1D57N-Rep1 | FALSE | FALSE | | JSY14 | FALSE | CAN | 7 | FALSE |
| 13_v.txt | rnr1D57N | rnr1D57N | 2 | rnr1D57N-Rep2 | FALSE | FALSE | 172 | JSY14 | FALSE | CAN | 7 | FALSE |
| 14_v.txt | rnr1D57N | rnr1D57N | 3 | rnr1D57N-Rep3 | FALSE | FALSE | 173 | JSY14 | FALSE | CAN | 7 | FALSE |
| 15_v.txt | rnr1D57N | rnr1D57N | 4 | rnr1D57N-Rep4 | FALSE | FALSE | | JSY14 | FALSE | CAN | 7 | FALSE |
| 171_v.txt | rnr1D57N | rnr1D57N | 1A | rnr1D57N-Rep1A | FALSE | TRUE | | JSY14 | FALSE | CAN | 7 | FALSE |
| 172_v.txt | rnr1D57N | rnr1D57N | 2A | rnr1D57N-Rep2A | FALSE | TRUE | 13 | JSY14 | FALSE | CAN | 7 | FALSE |
| 173_v.txt | rnr1D57N | rnr1D57N | 3A | rnr1D57N-Rep3A | FALSE | TRUE | 14 | JSY14 | FALSE | CAN | 7 | FALSE |
| 186_v.txt | rnr1D57N | rnr1D57N | 5 | rnr1D57N-Rep5 | FALSE | FALSE | | JSY14 | FALSE | CAN | 7 | FALSE |
| NL3_v.txt | rnr1D57N | rnr1D57N | 6 | rnr1D57N-Rep6 | FALSE | FALSE | | JSY3602 | FALSE | CAN | 7 | FALSE |
| NL4_v.txt | rnr1D57N | rnr1D57N | 7 | rnr1D57N-Rep7 | FALSE | FALSE | | JSY3602 | FALSE | CAN | 7 | FALSE |
| 8_v.txt | rnr1D57N_perm | rnr1D57N | 1 | rnr1D57N_perm-Rep1 | TRUE | FALSE | | JSY14 | FALSE | ARG | 4 | FALSE |
| 9_v.txt | rnr1D57N_perm | rnr1D57N | 2 | rnr1D57N_perm-Rep2 | TRUE | FALSE | | JSY14 | FALSE | ARG | 4 | FALSE |
| 10_v.txt | rnr1D57N_perm | rnr1D57N | 3 | rnr1D57N_perm-Rep3 | TRUE | FALSE | | JSY14 | FALSE | ARG | 4 | FALSE |
| 11_v.txt | rnr1D57N_perm | rnr1D57N | 4 | rnr1D57N_perm-Rep4 | TRUE | FALSE | | JSY14 | FALSE | ARG | 4 | FALSE |
| 62_v.txt | rnr1Y285A | rnr1Y285A | 1 | rnr1Y285A-Rep1 | FALSE | FALSE | | JSY3868 | FALSE | CAN | 5 | FALSE |
| 63_v.txt | rnr1Y285A | rnr1Y285A | 2 | rnr1Y285A-Rep2 | FALSE | FALSE | | JSY3868 | FALSE | CAN | 5 | FALSE |
| 64_v.txt | rnr1Y285A | rnr1Y285A | 3 | rnr1Y285A-Rep3 | FALSE | FALSE | | JSY3869 | FALSE | CAN | 5 | FALSE |
| 65_v.txt | rnr1Y285A | rnr1Y285A | 4 | rnr1Y285A-Rep4 | FALSE | FALSE | | JSY3869 | FALSE | CAN | 5 | FALSE |
| NL9_v.txt | rnr1Y285A | rnr1Y285A | 5 | rnr1Y285A-Rep5 | FALSE | FALSE | | JSY3868 | FALSE | CAN | 5 | FALSE |
| 58_v.txt | rnr1Y285A_perm | rnr1Y285A | 1 | rnr1Y285A_perm-Rep1 | TRUE | FALSE | | JSY3868 | FALSE | ARG | 4 | FALSE |
| 59_v.txt | rnr1Y285A_perm | rnr1Y285A | 2 | rnr1Y285A_perm-Rep2 | TRUE | FALSE | | JSY3868 | FALSE | ARG | 4 | FALSE |
| 60_v.txt | rnr1Y285A_perm | rnr1Y285A | 3 | rnr1Y285A_perm-Rep3 | TRUE | FALSE | | JSY3868 | FALSE | ARG | 4 | FALSE |
| 61_v.txt | rnr1Y285A_perm | rnr1Y285A | 4 | rnr1Y285A_perm-Rep4 | TRUE | FALSE | | JSY3868 | FALSE | ARG | 4 | FALSE |
| 137_v.txt | rnr1Y285A_pGAI | rnr1Y285A | 1 | rnr1Y285A_pGAL-Rep1 | FALSE | FALSE | | JSY2422 | TRUE | CAN | 6 | FALSE |
| 138_v.txt | rnr1Y285A_pGAI | rnr1Y285A | 2 | rnr1Y285A_pGAL-Rep2 | FALSE | FALSE | | JSY2422 | TRUE | CAN | 6 | FALSE |
| 139_v.txt | rnr1Y285A_pGAI | rnr1Y285A | 3 | rnr1Y285A_pGAL-Rep3 | FALSE | FALSE | | JSY2423 | TRUE | CAN | 6 | FALSE |
| 140_v.txt | rnr1Y285A_pGAI | rnr1Y285A | 4 | rnr1Y285A_pGAL-Rep4 | FALSE | FALSE | | JSY2423 | TRUE | CAN | 6 | FALSE |
| 176_v.txt | rnr1Y285A_pGAI | rnr1Y285A | 5 | rnr1Y285A_pGAL-Rep5 | FALSE | FALSE | | JSY2422 | TRUE | CAN | 6 | FALSE |
| 177_v.txt | rnr1Y285A_pGAI | rnr1Y285A | 6 | rnr1Y285A_pGAL-Rep6 | FALSE | FALSE | | JSY2423 | TRUE | CAN | 6 | FALSE |
| 42_v.txt | rnr1Y285F | rnr1Y285F | 1 | rnr1Y285F-Rep1 | FALSE | FALSE | | JSY3811 | FALSE | CAN | 8 | FALSE |
| 43_v.txt | rnr1Y285F | rnr1Y285F | 2 | rnr1Y285F-Rep2 | FALSE | FALSE | | JSY3811 | FALSE | CAN | 8 | FALSE |
| 44_v.txt | rnr1Y285F | rnr1Y285F | 3 | rnr1Y285F-Rep3 | FALSE | FALSE | | JSY3811 | FALSE | CAN | 8 | FALSE |
| 45_v.txt | rnr1Y285F | rnr1Y285F | 4 | rnr1Y285F-Rep4 | FALSE | FALSE | | JSY3811 | FALSE | CAN | 8 | FALSE |
| 161_v.txt | rnr1Y285F | rnr1Y285F | 5 | rnr1Y285F-Rep5 | FALSE | FALSE | | JSY3865 | FALSE | CAN | 8 | FALSE |
| 162_v.txt | rnr1Y285F | rnr1Y285F | 6 | rnr1Y285F-Rep6 | FALSE | FALSE | | JSY3865 | FALSE | CAN | 8 | FALSE |
| 163_v.txt | rnr1Y285F | rnr1Y285F | 7 | rnr1Y285F-Rep7 | FALSE | FALSE | | JSY3866 | FALSE | CAN | 8 | FALSE |
| 164_v.txt | rnr1Y285F | rnr1Y285F | 8 | rnr1Y285F-Rep8 | FALSE | FALSE | | JSY3866 | FALSE | CAN | 8 | FALSE |
| 40_v.txt | rnr1Y285F_perm | rnr1Y285F | 1 | rnr1Y285F_perm-Rep1 | TRUE | FALSE | | JSY3811 | FALSE | ARG | 4 | FALSE |
| 41_v.txt | rnr1Y285F_perm | rnr1Y285F | 2 | rnr1Y285F_perm-Rep2 | TRUE | FALSE | | JSY3811 | FALSE | ARG | 4 | FALSE |
| 165_v.txt | rnr1Y285F_perm | rnr1Y285F | 3 | rnr1Y285F_perm-Rep3 | TRUE | FALSE | | JSY3865 | FALSE | ARG | 4 | FALSE |
| 166_v.txt | rnr1Y285F_perm | rnr1Y285F | 4 | rnr1Y285F_perm-Rep4 | TRUE | FALSE | | JSY3866 | FALSE | ARG | 4 | FALSE |
| 133_v.txt | rnr1Y285F_pGAL | rnr1Y285F | 1 | rnr1Y285F_pGAL-Rep1 | FALSE | FALSE | | JSY2420 | TRUE | CAN | 6 | FALSE |
| 134_v.txt | rnr1Y285F_pGAL | rnr1Y285F | 2 | rnr1Y285F_pGAL-Rep2 | FALSE | FALSE | | JSY2420 | TRUE | CAN | 6 | FALSE |
| 135_v.txt | rnr1Y285F_pGAL | rnr1Y285F | 3 | rnr1Y285F_pGAL-Rep3 | FALSE | FALSE | | JSY2421 | TRUE | CAN | 6 | FALSE |
| 136_v.txt | rnr1Y285F_pGAL | rnr1Y285F | 4 | rnr1Y285F_pGAL-Rep4 | FALSE | FALSE | | JSY2421 | TRUE | CAN | 6 | FALSE |
| 174_v.txt | rnr1Y285F_pGAL | rnr1Y285F | 5 | rnr1Y285F_pGAL-Rep5 | FALSE | FALSE | | JSY2420 | TRUE | CAN | 6 | FALSE |
| 175_v.txt | rnr1Y285F_pGAL | rnr1Y285F | 6 | rnr1Y285F_pGAL-Rep6 | FALSE | FALSE | | JSY2421 | TRUE | CAN | 6 | FALSE |
| 5_v.txt | WT | WT | 1 | WT-Rep1 | FALSE | FALSE | | JSY13 | FALSE | CAN | 9 | FALSE |
| 6_v.txt | WT | WT | 2 | WT-Rep2 | FALSE | FALSE | | JSY13 | FALSE | CAN | 9 | FALSE |
| 7_v.txt | WT | WT | 3 | WT-Rep3 | FALSE | FALSE | | JSY13 | FALSE | CAN | 9 | FALSE |
| 168_v.txt | WT | WT | 1A | WT-Rep1A | FALSE | TRUE | | JSY13 | FALSE | CAN | 9 | FALSE |
| 169_v.txt | WT | WT | 4 | WT-Rep4 | FALSE | FALSE | | JSY13 | FALSE | CAN | 9 | FALSE |
| 170_v.txt | WT | WT | 2A | WT-Rep2A | FALSE | TRUE | | JSY13 | FALSE | CAN | 9 | FALSE |
| 187_v.txt | WT | WT | 3A | WT-Rep3A | FALSE | TRUE | | JSY13 | FALSE | CAN | 9 | FALSE |
| 188_v.txt | WT | WT | 5 | WT-Rep5 | FALSE | FALSE | | JSY13 | FALSE | CAN | 9 | FALSE |
| 129_v.txt | WT | WT | 6 | WT-Rep6 | FALSE | FALSE | | JSY2424 | FALSE | CAN | 9 | FALSE |
| 130_v.txt | WT | WT | 7 | WT-Rep7 | FALSE | FALSE | | JSY2424 | FALSE | CAN | 9 | FALSE |
| 131_v.txt | WT | WT | 8 | WT-Rep8 | FALSE | FALSE | | JSY2424 | FALSE | CAN | 9 | FALSE |
| 132_v.txt | WT | WT | 9 | WT-Rep9 | FALSE | FALSE | | JSY2424 | FALSE | CAN | 9 | FALSE |
| 127_v.txt | WT_perm | WT | 1 | WT_perm-Rep1 | TRUE | FALSE | | JSY2424 | FALSE | ARG | 6 | FALSE |
| 128_v.txt | WT_perm | WT | 2 | WT_perm-Rep2 | TRUE | FALSE | | JSY2424 | FALSE | ARG | 6 | FALSE |
| 1_v.txt | WT_perm | WT | 3 | WT_perm-Rep3 | TRUE | FALSE | | JSY13 | FALSE | ARG | 6 | FALSE |
| 2_v.txt | WT_perm | WT | 4 | WT_perm-Rep4 | TRUE | FALSE | | JSY13 | FALSE | ARG | 6 | FALSE |
| 3_v.txt | WT_perm | WT | 5 | WT_perm-Rep5 | TRUE | FALSE | | JSY13 | FALSE | ARG | 6 | FALSE |
| 4_v.txt | WT_perm | WT | 6 | WT_perm-Rep6 | TRUE | FALSE | | JSY13 | FALSE | ARG | 6 | FALSE |

**Supplementary Table 5 - All variants**

| Source | Reference.Pc | SNP.ID | Type | Length | Reference | Allele | Count | Coverage | Frequency | Average.qual | Group | pGAL | Growth.Conc | Parent.Strain | Bio.Rep | GrpRep | CONTEXT | SNV.Split | Ins.Split | Del.Split | SNV.Split | Clust.I.D. | Context long | Context left | Comp conte | triContext | compofref | compofvariar | SNVtrichange | SNV TriContext | Extended context | Genotype reps | Tech rep | GAL status |
|---|---|---|---|---|---|---|---|---|---|---|---|---|---|---|---|---|---|---|---|---|---|---|---|---|---|---|---|---|---|---|---|---|---|---|
| 12_v.txt | 31723 | 31723-SNV-A | SNV | 1 | A | C | 9 | 11167 | 0.08059461 | 47.7790936 | rnr1D57N | FALSE | CAN | JSY14 | 1 | rnr1D57N-Re | CAA | A>C | G/C+1 | A/T-1 | TA>GC | TA>GC | TGTCCCAAA | AAAAAGT | GTT | GTT | T | G | T>G | T>G_GTT | TA>GC_AAAAAGT | 7 | FALSE | FALSE |
| 12_v.txt | 31840 | 31840-SNV-A | SNV | 1 | A | C | 2 | 11167 | 0.01790991 | 64 | rnr1D57N | FALSE | CAN | JSY14 | 1 | rnr1D57N-Re | AAG | A>C | G/C+1 | A/T-1 | TA>GC | TA>GC | GAAATAAGA | AGATCCA | TTC | TTC | T | G | T>G | T>G_TTC | TA>GC_AGATCCA | 7 | FALSE | FALSE |
| 12_v.txt | 31902 | 31902-SNV-A | SNV | 1 | A | C | 7 | 11162 | 0.06271278 | 38 | rnr1D57N | FALSE | CAN | JSY14 | 1 | rnr1D57N-Re | GAA | A>C | G/C+1 | A/T-1 | TA>GC | TA>GC | CCATTGAAT | AATTTTG | CTT | CTT | T | G | T>G | T>G_CTT | TA>GC_AATTTTG | 7 | FALSE | FALSE |
| 12_v.txt | 31913 | 31913-SNV-A | SNV | 1 | A | C | 1 | 11162 | 8.96E-03 | 26.0000059 | rnr1D57N | FALSE | CAN | JSY14 | 1 | rnr1D57N-Re | CAA | A>C | G/C+1 | A/T-1 | TA>GC | TA>GC | TGGTGCAAA | AAAAGCC | GTT | GTT | T | G | T>G | T>G_GTT | TA>GC_AAAAGCC | 7 | FALSE | FALSE |
| 12_v.txt | 31950 | 31950-SNV-A | SNV | 1 | A | C | 16 | 16625 | 0.0962406 | 29.1875 | rnr1D57N | FALSE | CAN | JSY14 | 1 | rnr1D57N-Re | CAT | A>C | G/C+1 | A/T-1 | TA>GC | TA>GC | ATCGTCATA | ATAAATG | GTA | GTA | T | G | T>G | T>G_GTA | TA>GC_ATAAATG | 7 | FALSE | FALSE |
| 12_v.txt | 32005 | 32005-SNV-A | SNV | 1 | A | C | 17 | 16625 | 0.10225564 | 35.1176471 | rnr1D57N | FALSE | CAN | JSY14 | 1 | rnr1D57N-Re | AAC | A>C | G/C+1 | A/T-1 | TA>GC | TA>GC | ATGGTAACT | ACTCGTC | TTG | TTG | T | G | T>G | T>G_TTG | TA>GC_ACTCGTC | 7 | FALSE | FALSE |
| 12_v.txt | 32086 | 32086-SNV-A | SNV | 1 | A | C | 1 | 16624 | 6.02E-03 | 35.9999939 | rnr1D57N | FALSE | CAN | JSY14 | 1 | rnr1D57N-Re | AAG | A>C | G/C+1 | A/T-1 | TA>GC | TA>GC | CAAAAAAGC | AGCCTGC | TTC | TTC | T | G | T>G | T>G_TTC | TA>GC_AGCCTGC | 7 | FALSE | FALSE |
| 12_v.txt | 32103 | 32103-SNV-A | SNV | 1 | A | C | 7 | 16625 | 0.04210526 | 63.9999547 | rnr1D57N | FALSE | CAN | JSY14 | 1 | rnr1D57N-Re | GAT | A>C | G/C+1 | A/T-1 | TA>GC | TA>GC | CCAGTGATA | ATATTTA | CTA | CTA | T | G | T>G | T>G_CTA | TA>GC_ATATTTA | 7 | FALSE | FALSE |
| 12_v.txt | 32118 | 32118-SNV-T | SNV | 1 | T | G | 1 | 16625 | 6.02E-03 | 63.9999916 | rnr1D57N | FALSE | CAN | JSY14 | 1 | rnr1D57N-Re | TTC | T>G | G/C+1 | A/T-1 | TA>GC | TA>GC | AGCCATTCG | TCGAAAA | AAG | TTC | T | C | T>G | T>G_TTC | TA>GC_TCGAAAA | 7 | FALSE | FALSE |
| 12_v.txt | 32152 | 32152-SNV-T | SNV | 1 | T | G | 13 | 16625 | 0.07819549 | 46.6150131 | rnr1D57N | FALSE | CAN | JSY14 | 1 | rnr1D57N-Re | ATG | T>G | G/C+1 | A/T-1 | TA>GC | TA>GC | TCTCCATGTA | TGTAAGC | TAC | ATG | T | C | T>G | T>G_ATG | TA>GC_TGTAAGC | 7 | FALSE | FALSE |
| 12_v.txt | 32284 | 32284-SNV-A | SNV | 1 | A | C | 25 | 29225 | 0.0855432 | 37.7988099 | rnr1D57N | FALSE | CAN | JSY14 | 1 | rnr1D57N-Re | TAA | A>C | G/C+1 | A/T-1 | TA>GC | TA>GC | CAACGTAAA | AAATATT | ATT | ATT | T | G | T>G | T>G_ATT | TA>GC_AAATATT | 7 | FALSE | FALSE |
| 12_v.txt | 32294 | 32294-SNV-A | SNV | 1 | A | C | 542 | 29225 | 1.85457656 | 53.501193 | rnr1D57N | FALSE | CAN | JSY14 | 1 | rnr1D57N-Re | AAT | A>C | G/C+1 | A/T-1 | TA>GC | TA>GC | ATTTGAATTT | ATTTGCG | TTA | TTA | T | G | T>G | T>G_TTA | TA>GC_ATTTGCG | 7 | FALSE | FALSE |
| 12_v.txt | 32319 | 32319-SNV-A | SNV | 1 | A | C | 27 | 29225 | 0.09238666 | 59.6296296 | rnr1D57N | FALSE | CAN | JSY14 | 1 | rnr1D57N-Re | TAA | A>C | G/C+1 | A/T-1 | TA>GC | TA>GC | GTTGTTAAG | AAGATAA | ATT | ATT | T | G | T>G | T>G_ATT | TA>GC_AAGATAA | 7 | FALSE | FALSE |
| 12_v.txt | 32360 | 32360-SNV-A | SNV | 1 | A | C | 1 | 29224 | 3.42E-03 | 63.9978829 | rnr1D57N | FALSE | CAN | JSY14 | 1 | rnr1D57N-Re | CAG | A>C | G/C+1 | A/T-1 | TA>GC | TA>GC | TGTACCAGA | AGAGTTC | GTC | GTC | T | G | T>G | T>G_GTC | TA>GC_AGAGTTC | 7 | FALSE | FALSE |
| 12_v.txt | 32441 | 32441-SNV-T | SNV | 1 | T | G | 6 | 29225 | 0.02053037 | 51.8333333 | rnr1D57N | FALSE | CAN | JSY14 | 1 | rnr1D57N-Re | CTA | T>G | G/C+1 | A/T-1 | TA>GC | TA>GC | TGGAACTAA | TAAAAGT | GAT | CTA | T | C | T>G | T>G_CTA | TA>GC_TAAAAGT | 7 | FALSE | FALSE |
| 12_v.txt | 32493 | 32493-SNV-A | SNV | 1 | A | C | 8 | 29225 | 0.02737382 | 36.1260689 | rnr1D57N | FALSE | CAN | JSY14 | 1 | rnr1D57N-Re | AAC | A>C | G/C+1 | A/T-1 | TA>GC | TA>GC | CGGAAAACA | ACAACTT | TTG | TTG | T | G | T>G | T>G_TTG | TA>GC_ACAACTT | 7 | FALSE | FALSE |
| 12_v.txt | 32574 | 32574-SNV-T | SNV | 1 | T | G | 681 | 26053 | 2.61390243 | 37.0660078 | rnr1D57N | FALSE | CAN | JSY14 | 1 | rnr1D57N-Re | TTG | T>G | G/C+1 | A/T-1 | TA>GC | TA>GC | GTACCTTGA | TGAAATG | AAC | TTG | T | C | T>G | T>G_TTG | TA>GC_TGAAATG | 7 | FALSE | FALSE |
| 12_v.txt | 32631 | 32631-SNV-T | SNV | 1 | T | G | 2 | 26053 | 7.68E-03 | 53.5 | rnr1D57N | FALSE | CAN | JSY14 | 1 | rnr1D57N-Re | GTT | T>G | G/C+1 | A/T-1 | TA>GC | TA>GC | CCTTCGTTTT | TTTTTAT | CAA | GTT | T | C | T>G | T>G_GTT | TA>GC_TTTTTAT | 7 | FALSE | FALSE |
| 12_v.txt | 32680 | 32680-SNV-A | SNV | 1 | A | C | 9 | 26053 | 0.03454497 | 63.2222222 | rnr1D57N | FALSE | CAN | JSY14 | 1 | rnr1D57N-Re | CAA | A>C | G/C+1 | A/T-1 | TA>GC | TA>GC | TTCTCCAATA | AATAACG | GTT | GTT | T | G | T>G | T>G_GTT | TA>GC_AATAACG | 7 | FALSE | FALSE |
| 12_v.txt | 32739 | 32739-SNV-A | SNV | 1 | A | C | 26 | 26053 | 0.09979657 | 57.6538462 | rnr1D57N | FALSE | CAN | JSY14 | 1 | rnr1D57N-Re | TAT | A>C | G/C+1 | A/T-1 | TA>GC | TA>GC | CAGTATATTA | ATTAGAA | ATA | ATA | T | G | T>G | T>G_ATA | TA>GC_ATTAGAA | 7 | FALSE | FALSE |
| 12_v.txt | 32760 | 32760-SNV-A | SNV | 1 | A | C | 31 | 26053 | 0.11898822 | 53.5159442 | rnr1D57N | FALSE | CAN | JSY14 | 1 | rnr1D57N-Re | TAA | A>C | G/C+1 | A/T-1 | TA>GC | TA>GC | ATGGCTAAA | AAAACTT | ATT | ATT | T | G | T>G | T>G_ATT | TA>GC_AAAACTT | 7 | FALSE | FALSE |
| 12_v.txt | 32785 | 32785-SNV-A | SNV | 1 | A | C | 97 | 26053 | 0.37231797 | 44.5670103 | rnr1D57N | FALSE | CAN | JSY14 | 1 | rnr1D57N-Re | AAC | A>C | G/C+1 | A/T-1 | TA>GC | TA>GC | CCCAGAACT | ACTCGAA | TTG | TTG | T | G | T>G | T>G_TTG | TA>GC_ACTCGAA | 7 | FALSE | FALSE |
| 12_v.txt | 32787 | 32787-SNV-T | SNV | 1 | T | G | 4 | 26053 | 0.01535332 | 36.0000637 | rnr1D57N | FALSE | CAN | JSY14 | 1 | rnr1D57N-Re | CTC | T>G | G/C+1 | A/T-1 | TA>GC | TA>GC | CAGAACTCG | TCGAATT | GAG | CTC | T | C | T>G | T>G_CTC | TA>GC_TCGAATT | 7 | FALSE | FALSE |
| 12_v.txt | 32801 | 32801-SNV-A | SNV | 1 | A | C | 23 | 26053 | 0.08828158 | 49.4347354 | rnr1D57N | FALSE | CAN | JSY14 | 1 | rnr1D57N-Re | AAT | A>C | G/C+1 | A/T-1 | TA>GC | TA>GC | ACCGTAATA | ATATTTG | TTA | TTA | T | G | T>G | T>G_TTA | TA>GC_ATATTTG | 7 | FALSE | FALSE |
| 12_v.txt | 32842 | 32842-SNV-A | SNV | 1 | A | C | 2 | 26053 | 7.68E-03 | 38.000132 | rnr1D57N | FALSE | CAN | JSY14 | 1 | rnr1D57N-Re | CAA | A>C | G/C+1 | A/T-1 | TA>GC | TA>GC | TTACCCAAA | AAAAAAT | GTT | GTT | T | G | T>G | T>G_GTT | TA>GC_AAAAAAT | 7 | FALSE | FALSE |
| 12_v.txt | 32884 | 32884-SNV-A | SNV | 1 | A | C | 2 | 21315 | 9.38E-03 | 37.5000075 | rnr1D57N | FALSE | CAN | JSY14 | 1 | rnr1D57N-Re | CAA | A>C | G/C+1 | A/T-1 | TA>GC | TA>GC | ACGTCCAAA | AAAATTG | GTT | GTT | T | G | T>G | T>G_GTT | TA>GC_AAAATTG | 7 | FALSE | FALSE |
| 12_v.txt | 32915 | 32915-SNV-T | SNV | 1 | T | G | 44 | 21315 | 0.2064274 | 43.7535917 | rnr1D57N | FALSE | CAN | JSY14 | 1 | rnr1D57N-Re | GTT | T>G | G/C+1 | A/T-1 | TA>GC | TA>GC | ACTAAGTTC | TTCCAGG | CAA | GTT | T | C | T>G | T>G_GTT | TA>GC_TTCCAGG | 7 | FALSE | FALSE |
| 12_v.txt | 32953 | 32953-SNV-T | SNV | 1 | T | G | 7 | 21315 | 0.03284072 | 54.4285714 | rnr1D57N | FALSE | CAN | JSY14 | 1 | rnr1D57N-Re | ATG | T>G | G/C+1 | A/T-1 | TA>GC | TA>GC | AATACATGT | TGTAACC | TAC | ATG | T | C | T>G | T>G_ATG | TA>GC_TGTAACC | 7 | FALSE | FALSE |
| 12_v.txt | 32981 | 32981-SNV-A | SNV | 1 | A | C | 9 | 21315 | 0.04222379 | 64 | rnr1D57N | FALSE | CAN | JSY14 | 1 | rnr1D57N-Re | GAG | A>C | G/C+1 | A/T-1 | TA>GC | TA>GC | TGCTGGAGA | AGAAAGG | CTC | CTC | T | G | T>G | T>G_CTC | TA>GC_AGAAAGG | 7 | FALSE | FALSE |
| 12_v.txt | 33054 | 33054-SNV-T | SNV | 1 | T | G | 161 | 21315 | 0.75533662 | 58.9314969 | rnr1D57N | FALSE | CAN | JSY14 | 1 | rnr1D57N-Re | CTG | T>G | G/C+1 | A/T-1 | TA>GC | TA>GC | AAGGACTGC | TGCGTGA | GAC | CTG | T | C | T>G | T>G_CTG | TA>GC_TGCGTGA | 7 | FALSE | FALSE |
| 12_v.txt | 33192 | 33192-SNV-T | SNV | 1 | T | G | 14 | 20300 | 0.06896552 | 31.7857143 | rnr1D57N | FALSE | CAN | JSY14 | 1 | rnr1D57N-Re | ATG | T>G | G/C+1 | A/T-1 | TA>GC | TA>GC | CCAATATGT | TGTCTTT | TAC | ATG | T | C | T>G | T>G_ATG | TA>GC_TGTCTTT | 7 | FALSE | FALSE |
| 12_v.txt | 33204 | 33204-SNV-A | SNV | 1 | A | C | 8 | 20300 | 0.03940887 | 29 | rnr1D57N | FALSE | CAN | JSY14 | 1 | rnr1D57N-Re | AAG | A>C | G/C+1 | A/T-1 | TA>GC | TA>GC | TGCTTAAGC | AGCTCTC | TTC | TTC | T | G | T>G | T>G_TTC | TA>GC_AGCTCTC | 7 | FALSE | FALSE |
| 12_v.txt | 33269 | 33269-SNV-A | SNV | 1 | A | C | 141 | 20312 | 0.69417093 | 15.6595745 | rnr1D57N | FALSE | CAN | JSY14 | 1 | rnr1D57N-Re | TAG | A>C | G/C+1 | A/T-1 | TA>GC | TA>GC | CTCCATAGA | AGAGAAC | ATC | ATC | T | G | T>G | T>G_ATC | TA>GC_AGAGAAC | 7 | FALSE | FALSE |
| 12_v.txt | 33476 | 33476-SNV-T | SNV | 1 | T | G | 2 | 20311 | 9.85E-03 | 36.999036 | rnr1D57N | FALSE | CAN | JSY14 | 1 | rnr1D57N-Re | CTT | T>G | G/C+1 | A/T-1 | TA>GC | TA>GC | TATGCCTTTT | TTTTTTT | GAA | CTT | T | C | T>G | T>G_CTT | TA>GC_TTTTTTT | 7 | FALSE | FALSE |
| 12_v.txt | 33477 | 33477-SNV-T | SNV | 1 | T | G | 7 | 20308 | 0.03446918 | 36.8571362 | rnr1D57N | FALSE | CAN | JSY14 | 1 | rnr1D57N-Re | TTT | T>G | G/C+1 | A/T-1 | TA>GC | TA>GC | ATGCCTTTTT | TTTTTTT | AAA | TTT | T | C | T>G | T>G_TTT | TA>GC_TTTTTTT | 7 | FALSE | FALSE |
| 12_v.txt | 31723 | 31723-SNV-A | SNV | 1 | A | G | 5 | 11167 | 0.04477478 | 43.4015478 | rnr1D57N | FALSE | CAN | JSY14 | 1 | rnr1D57N-Re | CAA | A>G | G/C+1 | A/T-1 | TA>CG | TA>CG | TGTCCCAAA | AAAAAGT | GTT | GTT | T | C | T>C | T>C_GTT | TA>CG_AAAAAGT | 7 | FALSE | FALSE |
| 12_v.txt | 31822 | 31822-SNV-T | SNV | 1 | T | C | 6 | 11167 | 0.05372974 | 50.6666667 | rnr1D57N | FALSE | CAN | JSY14 | 1 | rnr1D57N-Re | CTG | T>C | G/C+1 | A/T-1 | TA>CG | TA>CG | TGCATCTGA | TGAATAT | GAC | CTG | T | G | T>C | T>C_CTG | TA>CG_TGAATAT | 7 | FALSE | FALSE |
| 12_v.txt | 31876 | 31876-SNV-A | SNV | 1 | A | G | 6 | 11167 | 0.05372974 | 33.000522 | rnr1D57N | FALSE | CAN | JSY14 | 1 | rnr1D57N-Re | TAG | A>G | G/C+1 | A/T-1 | TA>CG | TA>CG | AGATATAGG | AGGCGGC | ATC | ATC | T | C | T>C | T>C_ATC | TA>CG_AGGCGGC | 7 | FALSE | FALSE |
| 12_v.txt | 32086 | 32086-SNV-A | SNV | 1 | A | G | 12 | 16624 | 0.07218479 | 61.8333237 | rnr1D57N | FALSE | CAN | JSY14 | 1 | rnr1D57N-Re | AAG | A>G | G/C+1 | A/T-1 | TA>CG | TA>CG | CAAAAAAGC | AGCCTGC | TTC | TTC | T | C | T>C | T>C_TTC | TA>CG_AGCCTGC | 7 | FALSE | FALSE |
| 12_v.txt | 32123 | 32123-SNV-A | SNV | 1 | A | G | 7 | 16622 | 0.04211286 | 27.9997702 | rnr1D57N | FALSE | CAN | JSY14 | 1 | rnr1D57N-Re | AAA | A>G | G/C+1 | A/T-1 | TA>CG | TA>CG | TTCGAAAAC | AACTTTG | TTT | TTT | T | C | T>C | T>C_TTT | TA>CG_AACTTTG | 7 | FALSE | FALSE |
| 12_v.txt | 32126 | 32126-SNV-T | SNV | 1 | T | C | 3 | 16625 | 0.01804511 | 52.3319265 | rnr1D57N | FALSE | CAN | JSY14 | 1 | rnr1D57N-Re | CTT | T>C | G/C+1 | A/T-1 | TA>CG | TA>CG | GAAAACTTT | TTTGTCA | GAA | CTT | T | G | T>C | T>C_CTT | TA>CG_TTTGTCA | 7 | FALSE | FALSE |
| 12_v.txt | 32132 | 32132-SNV-A | SNV | 1 | A | G | 6 | 16625 | 0.03609023 | 38.6666481 | rnr1D57N | FALSE | CAN | JSY14 | 1 | rnr1D57N-Re | CAC | A>G | G/C+1 | A/T-1 | TA>CG | TA>CG | TTTGTCACCA | ACCACCA | GTG | GTG | T | C | T>C | T>C_GTG | TA>CG_ACCACCA | 7 | FALSE | FALSE |
| 12_v.txt | 32275 | 32275-SNV-A | SNV | 1 | A | G | 105 | 29226 | 0.35926914 | 35.5332844 | rnr1D57N | FALSE | CAN | JSY14 | 1 | rnr1D57N-Re | GAA | A>G | G/C+1 | A/T-1 | TA>CG | TA>CG | TACGGGAAC | AACCAAC | CTT | CTT | T | C | T>C | T>C_CTT | TA>CG_AACCAAC | 7 | FALSE | FALSE |
| 12_v.txt | 32293 | 32293-SNV-A | SNV | 1 | A | G | 52 | 29224 | 0.17793594 | 53.7062906 | rnr1D57N | FALSE | CAN | JSY14 | 1 | rnr1D57N-Re | GAA | A>G | G/C+1 | A/T-1 | TA>CG | TA>CG | TATTTGAATT | AATTTGC | CTT | CTT | T | C | T>C | T>C_CTT | TA>CG_AATTTGC | 7 | FALSE | FALSE |
| 12_v.txt | 32346 | 32346-SNV-A | SNV | 1 | A | G | 7 | 29223 | 0.02395374 | 60.2853732 | rnr1D57N | FALSE | CAN | JSY14 | 1 | rnr1D57N-Re | CAA | A>G | G/C+1 | A/T-1 | TA>CG | TA>CG | TGTGGCAAA | AAAACCT | GTT | GTT | T | C | T>C | T>C_GTT | TA>CG_AAAACCT | 7 | FALSE | FALSE |
| 12_v.txt | 32375 | 32375-SNV-A | SNV | 1 | A | G | 10 | 29224 | 0.03421845 | 50.8000001 | rnr1D57N | FALSE | CAN | JSY14 | 1 | rnr1D57N-Re | CAA | A>G | G/C+1 | A/T-1 | TA>CG | TA>CG | AATAGCAAT | AATAATA | GTT | GTT | T | C | T>C | T>C_GTT | TA>CG_AATAATA | 7 | FALSE | FALSE |
| 12_v.txt | 32376 | 32376-SNV-A | SNV | 1 | A | G | 5 | 29225 | 0.01710864 | 51.1996624 | rnr1D57N | FALSE | CAN | JSY14 | 1 | rnr1D57N-Re | AAT | A>G | G/C+1 | A/T-1 | TA>CG | TA>CG | ATAGCAATA | ATAATAA | TTA | TTA | T | C | T>C | T>C_TTA | TA>CG_ATAATAA | 7 | FALSE | FALSE |
| 12_v.txt | 32377 | 32377-SNV-T | SNV | 1 | T | C | 10 | 29224 | 0.03421845 | 43.4994831 | rnr1D57N | FALSE | CAN | JSY14 | 1 | rnr1D57N-Re | ATA | T>C | G/C+1 | A/T-1 | TA>CG | TA>CG | TAGCAATAA | TAATAAA | TAT | ATA | T | G | T>C | T>C_ATA | TA>CG_TAATAAA | 7 | FALSE | FALSE |
| 12_v.txt | 32445 | 32445-SNV-A | SNV | 1 | A | G | 603 | 29225 | 2.06330197 | 53.4742952 | rnr1D57N | FALSE | CAN | JSY14 | 1 | rnr1D57N-Re | AAG | A>G | G/C+1 | A/T-1 | TA>CG | TA>CG | ACTAAAAGT | AGTCCAA | TTC | TTC | T | C | T>C | T>C_TTC | TA>CG_AGTCCAA | 7 | FALSE | FALSE |
| 12_v.txt | 32543 | 32543-SNV-A | SNV | 1 | A | G | 14 | 29224 | 0.04790583 | 20.785539 | rnr1D57N | FALSE | CAN | JSY14 | 1 | rnr1D57N-Re | CAC | A>G | G/C+1 | A/T-1 | TA>CG | TA>CG | AGCTTCACC | ACCAGCA | GTG | GTG | T | C | T>C | T>C_GTG | TA>CG_ACCAGCA | 7 | FALSE | FALSE |
| 12_v.txt | 32631 | 32631-SNV-T | SNV | 1 | T | C | 9 | 26053 | 0.03454497 | 46.3333333 | rnr1D57N | FALSE | CAN | JSY14 | 1 | rnr1D57N-Re | GTT | T>C | G/C+1 | A/T-1 | TA>CG | TA>CG | CCTTCGTTTT | TTTTTAT | CAA | GTT | T | G | T>C | T>C_GTT | TA>CG_TTTTTAT | 7 | FALSE | FALSE |
| 12_v.txt | 32728 | 32728-SNV-A | SNV | 1 | A | G | 14 | 26053 | 0.05373661 | 56.9990563 | rnr1D57N | FALSE | CAN | JSY14 | 1 | rnr1D57N-Re | CAA | A>G | G/C+1 | A/T-1 | TA>CG | TA>CG | CCATACAAA | AAAAACA | GTT | GTT | T | C | T>C | T>C_GTT | TA>CG_AAAAACA | 7 | FALSE | FALSE |
| 12_v.txt | 32734 | 32734-SNV-A | SNV | 1 | A | G | 30 | 26053 | 0.11514989 | 60.1331514 | rnr1D57N | FALSE | CAN | JSY14 | 1 | rnr1D57N-Re | CAG | A>G | G/C+1 | A/T-1 | TA>CG | TA>CG | AAAAACAGT | AGTATAT | GTC | GTC | T | C | T>C | T>C_GTC | TA>CG_AGTATAT | 7 | FALSE | FALSE |
| 12_v.txt | 32759 | 32759-SNV-T | SNV | 1 | T | C | 5 | 26053 | 0.01919165 | 54.5998529 | rnr1D57N | FALSE | CAN | JSY14 | 1 | rnr1D57N-Re | CTA | T>C | G/C+1 | A/T-1 | TA>CG | TA>CG | AATGGCTAA | TAAAACT | GAT | CTA | T | G | T>C | T>C_CTA | TA>CG_TAAAACT | 7 | FALSE | FALSE |
| 12_v.txt | 32787 | 32787-SNV-T | SNV | 1 | T | C | 14 | 26053 | 0.05373661 | 42.9284081 | rnr1D57N | FALSE | CAN | JSY14 | 1 | rnr1D57N-Re | CTC | T>C | G/C+1 | A/T-1 | TA>CG | TA>CG | CAGAACTCG | TCGAATT | GAG | CTC | T | G | T>C | T>C_CTC | TA>CG_TCGAATT | 7 | FALSE | FALSE |
| 12_v.txt | 32814 | 32814-SNV-A | SNV | 1 | A | G | 14 | 26053 | 0.05373661 | 33.8570651 | rnr1D57N | FALSE | CAN | JSY14 | 1 | rnr1D57N-Re | GAA | A>G | G/C+1 | A/T-1 | TA>CG | TA>CG | ACAGGGAAC | AACAAGT | CTT | CTT | T | C | T>C | T>C_CTT | TA>CG_AACAAGT | 7 | FALSE | FALSE |
| 12_v.txt | 32903 | 32903-SNV-A | SNV | 1 | A | G | 5 | 21315 | 0.02345766 | 34.7802111 | rnr1D57N | FALSE | CAN | JSY14 | 1 | rnr1D57N-Re | CAA | A>G | G/C+1 | A/T-1 | TA>CG | TA>CG | TTGGCCAAC | AACTACA | GTT | GTT | T | C | T>C | T>C_GTT | TA>CG_AACTACA | 7 | FALSE | FALSE |
| 12_v.txt | 32915 | 32915-SNV-T | SNV | 1 | T | C | 8 | 21315 | 0.03753225 | 31.4873336 | rnr1D57N | FALSE | CAN | JSY14 | 1 | rnr1D57N-Re | GTT | T>C | G/C+1 | A/T-1 | TA>CG | TA>CG | ACTAAGTTC | TTCCAGG | CAA | GTT | T | G | T>C | T>C_GTT | TA>CG_TTCCAGG | 7 | FALSE | FALSE |
| 12_v.txt | 32924 | 32924-SNV-A | SNV | 1 | A | G | 4 | 21315 | 0.01876613 | 53.4986884 | rnr1D57N | FALSE | CAN | JSY14 | 1 | rnr1D57N-Re | CAA | A>G | G/C+1 | A/T-1 | TA>CG | TA>CG | CAGGGCAAA | AAAAGTG | GTT | GTT | T | C | T>C | T>C_GTT | TA>CG_AAAAGTG | 7 | FALSE | FALSE |
| 12_v.txt | 32947 | 32947-SNV-A | SNV | 1 | A | G | 42 | 21315 | 0.19704434 | 60.2619048 | rnr1D57N | FALSE | CAN | JSY14 | 1 | rnr1D57N-Re | CAA | A>G | G/C+1 | A/T-1 | TA>CG | TA>CG | AAAACCAAT | AATACAT | GTT | GTT | T | C | T>C | T>C_GTT | TA>CG_AATACAT | 7 | FALSE | FALSE |
| 12_v.txt | 32985 | 32985-SNV-A | SNV | 1 | A | G | 44 | 21315 | 0.2064274 | 60.1136364 | rnr1D57N | FALSE | CAN | JSY14 | 1 | rnr1D57N-Re | AAG | A>G | G/C+1 | A/T-1 | TA>CG | TA>CG | GGAGAAAGC | AGGAATC | TTC | TTC | T | C | T>C | T>C_TTC | TA>CG_AGGAATC | 7 | FALSE | FALSE |
| 12_v.txt | 33072 | 33072-SNV-A | SNV | 1 | A | G | 8 | 21315 | 0.03753225 | 57.6244693 | rnr1D57N | FALSE | CAN | JSY14 | 1 | rnr1D57N-Re | CAA | A>G | G/C+1 | A/T-1 | TA>CG | TA>CG | TATGCCAAA | AAAGAAC | GTT | GTT | T | C | T>C | T>C_GTT | TA>CG_AAAGAAC | 7 | FALSE | FALSE |
| 12_v.txt | 33126 | 33126-SNV-A | SNV | 1 | A | G | 54 | 21315 | 0.25334272 | 41.962963 | rnr1D57N | FALSE | CAN | JSY14 | 1 | rnr1D57N-Re | CAG | A>G | G/C+1 | A/T-1 | TA>CG | TA>CG | TTGGTCAGA | AGAGGTG | GTC | GTC | T | C | T>C | T>C_GTC | TA>CG_AGAGGTG | 7 | FALSE | FALSE |
| 12_v.txt | 33192 | 33192-SNV-T | SNV | 1 | T | C | 46 | 20300 | 0.22660099 | 35.2608696 | rnr1D57N | FALSE | CAN | JSY14 | 1 | rnr1D57N-Re | ATG | T>C | G/C+1 | A/T-1 | TA>CG | TA>CG | CCAATATGT | TGTCTTT | TAC | ATG | T | G | T>C | T>C_ATG | TA>CG_TGTCTTT | 7 | FALSE | FALSE |
| 12_v.txt | 33452 | 33452-SNV-T | SNV | 1 | T | C | 9 | 20312 | 0.04430878 | 42.7777778 | rnr1D57N | FALSE | CAN | JSY14 | 1 | rnr1D57N-Re | CTT | T>C | G/C+1 | A/T-1 | TA>CG | TA>CG | GTCTTCTTTT | TTTTGAA | GAA | CTT | T | G | T>C | T>C_CTT | TA>CG_TTTTGAA | 7 | FALSE | FALSE |
| 12_v.txt | 33476 | 33476-SNV-T | SNV | 1 | T | C | 31 | 20311 | 0.15262666 | 31.6774103 | rnr1D57N | FALSE | CAN | JSY14 | 1 | rnr1D57N-Re | CTT | T>C | G/C+1 | A/T-1 | TA>CG | TA>CG | TATGCCTTTT | TTTTTTT | GAA | CTT | T | G | T>C | T>C_CTT | TA>CG_TTTTTTT | 7 | FALSE | FALSE |
| 12_v.txt | 31808 | 31808-SNV-A | SNV | 1 | A | T | 1 | 11167 | 8.96E-03 | 31.998115 | rnr1D57N | FALSE | CAN | JSY14 | 1 | rnr1D57N-Re | AAA | A>T | A/T+1 | A/T-1 | TA>AT | TA>AT | CTTCCAAATA | AATAAAT | TTT | TTT | T | A | T>A | T>A_TTT | TA>AT_AATAAAT | 7 | FALSE | FALSE |
| 12_v.txt | 31809 | 31809-SNV-A | SNV | 1 | A | T | 1 | 11167 | 8.96E-03 | 47.9965568 | rnr1D57N | FALSE | CAN | JSY14 | 1 | rnr1D57N-Re | AAT | A>T | A/T+1 | A/T-1 | TA>AT | TA>AT | TTCCAAATA | ATAAATC | TTA | TTA | T | A | T>A | T>A_TTA | TA>AT_ATAAATC | 7 | FALSE | FALSE |
| 12_v.txt | 31816 | 31816-SNV-T | SNV | 1 | T | A | 17 | 11167 | 0.15223426 | 61.2941177 | rnr1D57N | FALSE | CAN | JSY14 | 1 | rnr1D57N-Re | CTG | T>A | A/T+1 | A/T-1 | TA>AT | TA>AT | TAAATCTGC | TGCATCT | GAC | CTG | T | T | T>A | T>A_CTG | TA>AT_TGCATCT | 7 | FALSE | FALSE |
| 12_v.txt | 31822 | 31822-SNV-T | SNV | 1 | T | A | 28 | 11167 | 0.25073878 | 55.8214286 | rnr1D57N | FALSE | CAN | JSY14 | 1 | rnr1D57N-Re | CTG | T>A | A/T+1 | A/T-1 | TA>AT | TA>AT | TGCATCTGA | TGAATAT | GAC | CTG | T | T | T>A | T>A_CTG | TA>AT_TGAATAT | 7 | FALSE | FALSE |
| 12_v.txt | 31840 | 31840-SNV-A | SNV | 1 | A | T | 6 | 11167 | 0.05372974 | 59.6666667 | rnr1D57N | FALSE | CAN | JSY14 | 1 | rnr1D57N-Re | AAG | A>T | A/T+1 | A/T-1 | TA>AT | TA>AT | GAAATAAGA | AGATCCA | TTC | TTC | T | A | T>A | T>A_TTC | TA>AT_AGATCCA | 7 | FALSE | FALSE |
| 12_v.txt | 31903 | 31903-SNV-A | SNV | 1 | A | T | 11 | 11162 | 0.09854865 | 46.4545455 | rnr1D57N | FALSE | CAN | JSY14 | 1 | rnr1D57N-Re | AAT | A>T | A/T+1 | A/T-1 | TA>AT | TA>AT | CATTGAATTT | ATTTTGG | TTA | TTA | T | A | T>A | T>A_TTA | TA>AT_ATTTTGG | 7 | FALSE | FALSE |
| 12_v.txt | 31996 | 31996-SNV-T | SNV | 1 | T | A | 25 | 16625 | 0.15037594 | 52.36 | rnr1D57N | FALSE | CAN | JSY14 | 1 | rnr1D57N-Re | TTA | T>A | A/T+1 | A/T-1 | TA>AT | TA>AT | TAGCTTTAA | TAAATGG | AAT | TTA | T | T | T>A | T>A_TTA | TA>AT_TAAATGG | 7 | FALSE | FALSE |
| 12_v.txt | 32032 | 32032-SNV-T | SNV | 1 | T | A | 17 | 16625 | 0.10225564 | 58.7058824 | rnr1D57N | FALSE | CAN | JSY14 | 1 | rnr1D57N-Re | TTC | T>A | A/T+1 | A/T-1 | TA>AT | TA>AT | GGTATTTCA | TCAAAGC | AAG | TTC | T | T | T>A | T>A_TTC | TA>AT_TCAAAGC | 7 | FALSE | FALSE |
| 12_v.txt | 32050 | 32050-SNV-T | SNV | 1 | T | A | 57 | 16625 | 0.34285714 | 61.9473684 | rnr1D57N | FALSE | CAN | JSY14 | 1 | rnr1D57N-Re | CTG | T>A | A/T+1 | A/T-1 | TA>AT | TA>AT | TAAATCTGA | TGATGTG | GAC | CTG | T | T | T>A | T>A_CTG | TA>AT_TGATGTG | 7 | FALSE | FALSE |
| 12_v.txt | 32055 | 32055-SNV-T | SNV | 1 | T | A | 12 | 16625 | 0.07218045 | 59.8333333 | rnr1D57N | FALSE | CAN | JSY14 | 1 | rnr1D57N-Re | GTG | T>A | A/T+1 | A/T-1 | TA>AT | TA>AT | CTGATGTGC | TGCGAGA | CAC | GTG | T | T | T>A | T>A_GTG | TA>AT_TGCGAGA | 7 | FALSE | FALSE |
| 12_v.txt | 32101 | 32101-SNV-T | SNV | 1 | T | A | 3 | 16624 | 0.0180462 | 63.9999999 | rnr1D57N | FALSE | CAN | JSY14 | 1 | rnr1D57N-Re | GTG | T>A | A/T+1 | A/T-1 | TA>AT | TA>AT | CACCAGTGA | TGATATT | CAC | GTG | T | T | T>A | T>A_GTG | TA>AT_TGATATT | 7 | FALSE | FALSE |
| 12_v.txt | 32117 | 32117-SNV-T | SNV | 1 | T | A | 3 | 16625 | 0.01804511 | 46.9968601 | rnr1D57N | FALSE | CAN | JSY14 | 1 | rnr1D57N-Re | ATT | T>A | A/T+1 | A/T-1 | TA>AT | TA>AT | TAGCCATTC | TTCGAAA | TAA | ATT | T | T | T>A | T>A_ATT | TA>AT_TTCGAAA | 7 | FALSE | FALSE |
| 12_v.txt | 32123 | 32123-SNV-A | SNV | 1 | A | T | 10 | 16622 | 0.06016123 | 10.599798 | rnr1D57N | FALSE | CAN | JSY14 | 1 | rnr1D57N-Re | AAA | A>T | A/T+1 | A/T-1 | TA>AT | TA>AT | TTCGAAAAC | AACTTTG | TTT | TTT | T | A | T>A | T>A_TTT | TA>AT_AACTTTG | 7 | FALSE | FALSE |
| 12_v.txt | 32124 | 32124-SNV-A | SNV | 1 | A | T | 2 | 16625 | 0.01203008 | 32.4995796 | rnr1D57N | FALSE | CAN | JSY14 | 1 | rnr1D57N-Re | AAC | A>T | A/T+1 | A/T-1 | TA>AT | TA>AT | TCGAAAACT | ACTTTGT | TTG | TTG | T | A | T>A | T>A_TTG | TA>AT_ACTTTGT | 7 | FALSE | FALSE |
| 12_v.txt | 32130 | 32130-SNV-T | SNV | 1 | T | A | 6 | 16625 | 0.03609023 | 44.3332705 | rnr1D57N | FALSE | CAN | JSY14 | 1 | rnr1D57N-Re | GTC | T>A | A/T+1 | A/T-1 | TA>AT | TA>AT | ACTTTGTCAC | TCACCAC | CAG | GTC | T | T | T>A | T>A_GTC | TA>AT_TCACCAC | 7 | FALSE | FALSE |
| 12_v.txt | 32152 | 32152-SNV-T | SNV | 1 | T | A | 2 | 16625 | 0.01203008 | 50.4972226 | rnr1D57N | FALSE | CAN | JSY14 | 1 | rnr1D57N-Re | ATG | T>A | A/T+1 | A/T-1 | TA>AT | TA>AT | TCTCCATGTA | TGTAAGC | TAC | ATG | T | T | T>A | T>A_ATG | TA>AT_TGTAAGC | 7 | FALSE | FALSE |
| 12_v.txt | 32275 | 32275-SNV-A | SNV | 1 | A | T | 3 | 29226 | 0.01026483 | 28.3331925 | rnr1D57N | FALSE | CAN | JSY14 | 1 | rnr1D57N-Re | GAA | A>T | A/T+1 | A/T-1 | TA>AT | TA>AT | TACGGGAAC | AACCAAC | CTT | CTT | T | A | T>A | T>A_CTT | TA>AT_AACCAAC | 7 | FALSE | FALSE |
| 12_v.txt | 32283 | 32283-SNV-T | SNV | 1 | T | A | 30 | 29225 | 0.10265184 | 30.1377531 | rnr1D57N | FALSE | CAN | JSY14 | 1 | rnr1D57N-Re | GTA | T>A | A/T+1 | A/T-1 | TA>AT | TA>AT | CCAACGTAA | TAAATAT | CAT | GTA | T | T | T>A | T>A_GTA | TA>AT_TAAATAT | 7 | FALSE | FALSE |
| 12_v.txt | 32287 | 32287-SNV-T | SNV | 1 | T | A | 21 | 29225 | 0.07185629 | 46.4761905 | rnr1D57N | FALSE | CAN | JSY14 | 1 | rnr1D57N-Re | ATA | T>A | A/T+1 | A/T-1 | TA>AT | TA>AT | CGTAAATAT | TATTTGA | TAT | ATA | T | T | T>A | T>A_ATA | TA>AT_TATTTGA | 7 | FALSE | FALSE |
| 12_v.txt | 32294 | 32294-SNV-A | SNV | 1 | A | T | 12 | 29225 | 0.04106074 | 45.3291065 | rnr1D57N | FALSE | CAN | JSY14 | 1 | rnr1D57N-Re | AAT | A>T | A/T+1 | A/T-1 | TA>AT | TA>AT | ATTTGAATTT | ATTTGCG | TTA | TTA | T | A | T>A | T>A_TTA | TA>AT_ATTTGCG | 7 | FALSE | FALSE |
| 12_v.txt | 32376 | 32376-SNV-A | SNV | 1 | A | T | 1 | 29225 | 3.42E-03 | 48.9992631 | rnr1D57N | FALSE | CAN | JSY14 | 1 | rnr1D57N-Re | AAT | A>T | A/T+1 | A/T-1 | TA>AT | TA>AT | ATAGCAATA | ATAATAA | TTA | TTA | T | A | T>A | T>A_TTA | TA>AT_ATAATAA | 7 | FALSE | FALSE |
| 12_v.txt | 32377 | 32377-SNV-T | SNV | 1 | T | A | 2 | 29224 | 6.84E-03 | 63.9885936 | rnr1D57N | FALSE | CAN | JSY14 | 1 | rnr1D57N-Re | ATA | T>A | A/T+1 | A/T-1 | TA>AT | TA>AT | TAGCAATAA | TAATAAA | TAT | ATA | T | T | T>A | T>A_ATA | TA>AT_TAATAAA | 7 | FALSE | FALSE |
| 12_v.txt | 32475 | 32475-SNV-A | SNV | 1 | A | T | 28 | 29225 | 0.09580838 | 45.7142857 | rnr1D57N | FALSE | CAN | JSY14 | 1 | rnr1D57N-Re | GAA | A>T | A/T+1 | A/T-1 | TA>AT | TA>AT | ATGTAGAAG | AAGGTTA | CTT | CTT | T | A | T>A | T>A_CTT | TA>AT_AAGGTTA | 7 | FALSE | FALSE |
| 12_v.txt | 32498 | 32498-SNV-T | SNV | 1 | T | A | 6 | 29225 | 0.02053037 | 42.165727 | rnr1D57N | FALSE | CAN | JSY14 | 1 | rnr1D57N-Re | CTT | T>A | A/T+1 | A/T-1 | TA>AT | TA>AT | AACAACTTTT | TTTTTTG | GAA | CTT | T | T | T>A | T>A_CTT | TA>AT_TTTTTTG | 7 | FALSE | FALSE |
| 12_v.txt | 32541 | 32541-SNV-T | SNV | 1 | T | A | 4 | 29225 | 0.01368691 | 37.7496349 | rnr1D57N | FALSE | CAN | JSY14 | 1 | rnr1D57N-Re | TTC | T>A | A/T+1 | A/T-1 | TA>AT | TA>AT | GCAGCTTCAC | TCACCAG | AAG | TTC | T | T | T>A | T>A_TTC | TA>AT_TCACCAG | 7 | FALSE | FALSE |
| 12_v.txt | 32543 | 32543-SNV-A | SNV | 1 | A | T | 20 | 29224 | 0.0684369 | 15.0993255 | rnr1D57N | FALSE | CAN | JSY14 | 1 | rnr1D57N-Re | CAC | A>T | A/T+1 | A/T-1 | TA>AT | TA>AT | AGCTTCACC | ACCAGCA | GTG | GTG | T | A | T>A | T>A_GTG | TA>AT_ACCAGCA | 7 | FALSE | FALSE |
| 12_v.txt | 32574 | 32574-SNV-T | SNV | 1 | T | A | 8 | 26053 | 0.03070664 | 37.2499727 | rnr1D57N | FALSE | CAN | JSY14 | 1 | rnr1D57N-Re | TTG | T>A | A/T+1 | A/T-1 | TA>AT | TA>AT | GTACCTTGA | TGAAATG | AAC | TTG | T | T | T>A | T>A_TTG | TA>AT_TGAAATG | 7 | FALSE | FALSE |
| 12_v.txt | 32581 | 32581-SNV-T | SNV | 1 | T | A | 3 | 26053 | 0.01151499 | 33.3304652 | rnr1D57N | FALSE | CAN | JSY14 | 1 | rnr1D57N-Re | GTG | T>A | A/T+1 | A/T-1 | TA>AT | TA>AT | GAAATGTGA | TGAAGGC | CAC | GTG | T | T | T>A | T>A_GTG | TA>AT_TGAAGGC | 7 | FALSE | FALSE |
| 12_v.txt | 32631 | 32631-SNV-T | SNV | 1 | T | A | 6 | 26053 | 0.02302998 | 59.5 | rnr1D57N | FALSE | CAN | JSY14 | 1 | rnr1D57N-Re | GTT | T>A | A/T+1 | A/T-1 | TA>AT | TA>AT | CCTTCGTTTT | TTTTTAT | CAA | GTT | T | T | T>A | T>A_GTT | TA>AT_TTTTTAT | 7 | FALSE | FALSE |
| 12_v.txt | 32734 | 32734-SNV-A | SNV | 1 | A | T | 4 | 26053 | 0.01535332 | 50.2495256 | rnr1D57N | FALSE | CAN | JSY14 | 1 | rnr1D57N-Re | CAG | A>T | A/T+1 | A/T-1 | TA>AT | TA>AT | AAAAACAGT | AGTATAT | GTC | GTC | T | A | T>A | T>A_GTC | TA>AT_AGTATAT | 7 | FALSE | FALSE |
| 12_v.txt | 32759 | 32759-SNV-T | SNV | 1 | T | A | 4 | 26053 | 0.01535332 | 21.2497307 | rnr1D57N | FALSE | CAN | JSY14 | 1 | rnr1D57N-Re | CTA | T>A | A/T+1 | A/T-1 | TA>AT | TA>AT | AATGGCTAA | TAAAACT | GAT | CTA | T | T | T>A | T>A_CTA | TA>AT_TAAAACT | 7 | FALSE | FALSE |
| 12_v.txt | 32803 | 32803-SNV-A | SNV | 1 | A | T | 13 | 26053 | 0.04989828 | 46.3076923 | rnr1D57N | FALSE | CAN | JSY14 | 1 | rnr1D57N-Re | TAT | A>T | A/T+1 | A/T-1 | TA>AT | TA>AT | CGTAATATTT | ATTTGAC | ATA | ATA | T | A | T>A | T>A_ATA | TA>AT_ATTTGAC | 7 | FALSE | FALSE |
| 12_v.txt | 32806 | 32806-SNV-T | SNV | 1 | T | A | 42 | 26053 | 0.16120984 | 44.4047619 | rnr1D57N | FALSE | CAN | JSY14 | 1 | rnr1D57N-Re | TTG | T>A | A/T+1 | A/T-1 | TA>AT | TA>AT | AATATTTGA | TGACAGG | AAC | TTG | T | T | T>A | T>A_TTG | TA>AT_TGACAGG | 7 | FALSE | FALSE |
| 12_v.txt | 32814 | 32814-SNV-A | SNV | 1 | A | T | 5 | 26053 | 0.01919165 | 18.3999795 | rnr1D57N | FALSE | CAN | JSY14 | 1 | rnr1D57N-Re | GAA | A>T | A/T+1 | A/T-1 | TA>AT | TA>AT | ACAGGGAAC | AACAAGT | CTT | CTT | T | A | T>A | T>A_CTT | TA>AT_AACAAGT | 7 | FALSE | FALSE |
| 12_v.txt | 32938 | 32938-SNV-A | SNV | 1 | A | T | 891 | 21315 | 4.18015482 | 56.977415 | rnr1D57N | FALSE | CAN | JSY14 | 1 | rnr1D57N-Re | CAA | A>T | A/T+1 | A/T-1 | TA>AT | TA>AT | TTGCCCAAG | AAGAAAA | GTT | GTT | T | A | T>A | T>A_GTT | TA>AT_AAGAAAA | 7 | FALSE | FALSE |
| 12_v.txt | 32992 | 32992-SNV-T | SNV | 1 | T | A | 25 | 21315 | 0.1172883 | 61.6 | rnr1D57N | FALSE | CAN | JSY14 | 1 | rnr1D57N-Re | CTT | T>A | A/T+1 | A/T-1 | TA>AT | TA>AT | GGAATCTTT | TTTGTGA | GAA | CTT | T | T | T>A | T>A_CTT | TA>AT_TTTGTGA | 7 | FALSE | FALSE |
| 12_v.txt | 33054 | 33054-SNV-T | SNV | 1 | T | A | 4 | 21315 | 0.01876613 | 50.4990827 | rnr1D57N | FALSE | CAN | JSY14 | 1 | rnr1D57N-Re | CTG | T>A | A/T+1 | A/T-1 | TA>AT | TA>AT | AAGGACTGC | TGCGTGA | GAC | CTG | T | T | T>A | T>A_CTG | TA>AT_TGCGTGA | 7 | FALSE | FALSE |
| 12_v.txt | 33066 | 33066-SNV-T | SNV | 1 | T | A | 5 | 21314 | 0.02345876 | 43.7986449 | rnr1D57N | FALSE | CAN | JSY14 | 1 | rnr1D57N-Re | ATA | T>A | A/T+1 | A/T-1 | TA>AT | TA>AT | ACAGAATAT | TATGCCA | TAT | ATA | T | T | T>A | T>A_ATA | TA>AT_TATGCCA | 7 | FALSE | FALSE |
| 12_v.txt | 33090 | 33090-SNV-T | SNV | 1 | T | A | 2 | 21315 | 9.38E-03 | 50.9909227 | rnr1D57N | FALSE | CAN | JSY14 | 1 | rnr1D57N-Re | ATA | T>A | A/T+1 | A/T-1 | TA>AT | TA>AT | AATAAATAT | TATGATA | TAT | ATA | T | T | T>A | T>A_ATA | TA>AT_TATGATA | 7 | FALSE | FALSE |
| 12_v.txt | 33094 | 33094-SNV-A | SNV | 1 | A | T | 5 | 21315 | 0.02345766 | 40.1997349 | rnr1D57N | FALSE | CAN | JSY14 | 1 | rnr1D57N-Re | GAT | A>T | A/T+1 | A/T-1 | TA>AT | TA>AT | AATATGATA | ATATAAG | CTA | CTA | T | A | T>A | T>A_CTA | TA>AT_ATATAAG | 7 | FALSE | FALSE |
| 12_v.txt | 33295 | 33295-SNV-T | SNV | 1 | T | A | 15 | 20312 | 0.07384797 | 50.9333333 | rnr1D57N | FALSE | CAN | JSY14 | 1 | rnr1D57N-Re | CTC | T>A | A/T+1 | A/T-1 | TA>AT | TA>AT | TTACTCTCGT | TCGTCGG | GAG | CTC | T | T | T>A | T>A_CTC | TA>AT_TCGTCGG | 7 | FALSE | FALSE |
| 12_v.txt | 33321 | 33321-SNV-A | SNV | 1 | A | T | 21 | 20312 | 0.10338716 | 61.7142857 | rnr1D57N | FALSE | CAN | JSY14 | 1 | rnr1D57N-Re | CAA | A>T | A/T+1 | A/T-1 | TA>AT | TA>AT | GGATACAAT | AATTCTT | GTT | GTT | T | A | T>A | T>A_GTT | TA>AT_AATTCTT | 7 | FALSE | FALSE |
| 12_v.txt | 31650 | 31650-SNV-C | SNV | 1 | C | T | 34 | 11167 | 0.30446852 | 36.7941177 | rnr1D57N | FALSE | CAN | JSY14 | 1 | rnr1D57N-Re | GCG | C>T | A/T+1 | G/C-1 | CG>TA | CG>TA | AGAATGCGA | CGAAATG | CGC | GCG | C | A | C>T | C>T_GCG | CG>TA_CGAAATG | 7 | FALSE | FALSE |
| 12_v.txt | 31806 | 31806-SNV-C | SNV | 1 | C | T | 281 | 11167 | 2.5163428 | 58.2213327 | rnr1D57N | FALSE | CAN | JSY14 | 1 | rnr1D57N-Re | CCA | C>T | A/T+1 | G/C-1 | CG>TA | CG>TA | ATCTTCCAAA | CAAATAA | GGT | CCA | C | A | C>T | C>T_CCA | CG>TA_CAAATAA | 7 | FALSE | FALSE |
| 12_v.txt | 31881 | 31881-SNV-G | SNV | 1 | G | A | 14 | 11167 | 0.12536939 | 40.24627 | rnr1D57N | FALSE | CAN | JSY14 | 1 | rnr1D57N-Re | GGC | G>A | A/T+1 | G/C-1 | CG>TA | CG>TA | TAGGCGGCA | GCAGCAA | CCG | CCG | C | T | C>T | C>T_CCG | CG>TA_GCAGCAA | 7 | FALSE | FALSE |
| 12_v.txt | 31882 | 31882-SNV-C | SNV | 1 | C | T | 17 | 11167 | 0.15223426 | 46.1130914 | rnr1D57N | FALSE | CAN | JSY14 | 1 | rnr1D57N-Re | GCA | C>T | A/T+1 | G/C-1 | CG>TA | CG>TA | AGGCGGCAG | CAGCAAA | CGT | GCA | C | A | C>T | C>T_GCA | CG>TA_CAGCAAA | 7 | FALSE | FALSE |
| 12_v.txt | 31956 | 31956-SNV-G | SNV | 1 | G | A | 2 | 16625 | 0.01203008 | 37.9994595 | rnr1D57N | FALSE | CAN | JSY14 | 1 | rnr1D57N-Re | TGT | G>A | A/T+1 | G/C-1 | CG>TA | CG>TA | ATAAATGTG | GTGGCCG | ACA | ACA | C | T | C>T | C>T_ACA | CG>TA_GTGGCCG | 7 | FALSE | FALSE |
| 12_v.txt | 31962 | 31962-SNV-G | SNV | 1 | G | A | 34 | 16625 | 0.20451128 | 37.2941177 | rnr1D57N | FALSE | CAN | JSY14 | 1 | rnr1D57N-Re | CGC | G>A | A/T+1 | G/C-1 | CG>TA | CG>TA | GTGGCCGCA | GCATAAT | GCG | GCG | C | T | C>T | C>T_GCG | CG>TA_GCATAAT | 7 | FALSE | FALSE |
| 12_v.txt | 31972 | 31972-SNV-C | SNV | 1 | C | T | 14 | 16625 | 0.08421053 | 37.2142857 | rnr1D57N | FALSE | CAN | JSY14 | 1 | rnr1D57N-Re | GCC | C>T | A/T+1 | G/C-1 | CG>TA | CG>TA | AATAAGCCA | CCAAGCC | CGG | GCC | C | A | C>T | C>T_GCC | CG>TA_CCAAGCC | 7 | FALSE | FALSE |

| | | | | | | | | | | | | | | | | | | | | | | | | | | | | | | | | | | |
|---|---|---|---|---|---|---|---|---|---|---|---|---|---|---|---|---|---|---|---|---|---|---|---|---|---|---|---|---|---|---|---|---|---|---|
| 12_v.txt | 31981 | 31981-SNV-( | SNV | 1 | G | A | 19 | 16625 | 0.11428571 | 51.7894737 | rnr1D57N | FALSE | CAN | JSY14 | 1 | rnr1D57N-Re | GGC | G>A | A/T+1 | G/C-1 | CG>TA | CG>TA | AGCCGGGCA | GCATTAA | CCG | CCG | C | T | C>T | C>T_CCG | CG>TA_GCATTAA | 7 | FALSE | FALSE |
| 12_v.txt | 32041 | 32041-SNV-( | SNV | 1 | G | A | 36 | 16625 | 0.21654135 | 60.6944444 | rnr1D57N | FALSE | CAN | JSY14 | 1 | rnr1D57N-Re | TGC | G>A | A/T+1 | G/C-1 | CG>TA | CG>TA | AAGCTTGCA | GCATAAA | ACG | ACG | C | T | C>T | C>T_ACG | CG>TA_GCATAAA | 7 | FALSE | FALSE |
| 12_v.txt | 32057 | 32057-SNV-( | SNV | 1 | C | T | 16 | 16625 | 0.0962406 | 62.875 | rnr1D57N | FALSE | CAN | JSY14 | 1 | rnr1D57N-Re | GCG | C>T | A/T+1 | G/C-1 | CG>TA | CG>TA | GATGTGCGA | CGAGATT | CGC | GCG | C | A | C>T | C>T_GCG | CG>TA_CGAGATT | 7 | FALSE | FALSE |
| 12_v.txt | 32075 | 32075-SNV-( | SNV | 1 | C | T | 15 | 16625 | 0.09022556 | 59.6 | rnr1D57N | FALSE | CAN | JSY14 | 1 | rnr1D57N-Re | ACC | C>T | A/T+1 | G/C-1 | CG>TA | CG>TA | AAATAACCA | CCATGCA | TGG | ACC | C | A | C>T | C>T_ACC | CG>TA_CCATGCA | 7 | FALSE | FALSE |
| 12_v.txt | 32087 | 32087-SNV-( | SNV | 1 | G | A | 4 | 16625 | 0.02406015 | 58.2495241 | rnr1D57N | FALSE | CAN | JSY14 | 1 | rnr1D57N-Re | AGC | G>A | A/T+1 | G/C-1 | CG>TA | CG>TA | AAAAAAGCC | GCCTGCA | TCG | TCG | C | T | C>T | C>T_TCG | CG>TA_GCCTGCA | 7 | FALSE | FALSE |
| 12_v.txt | 32088 | 32088-SNV-( | SNV | 1 | C | T | 64 | 16625 | 0.38496241 | 59.9688783 | rnr1D57N | FALSE | CAN | JSY14 | 1 | rnr1D57N-Re | GCC | C>T | A/T+1 | G/C-1 | CG>TA | CG>TA | AAAAAGCCT | CCTGCAA | CGG | GCC | C | A | C>T | C>T_GCC | CG>TA_CCTGCAA | 7 | FALSE | FALSE |
| 12_v.txt | 32091 | 32091-SNV-( | SNV | 1 | G | A | 13 | 16624 | 0.07820019 | 61.6923077 | rnr1D57N | FALSE | CAN | JSY14 | 1 | rnr1D57N-Re | TGC | G>A | A/T+1 | G/C-1 | CG>TA | CG>TA | AAGCCTGCA | GCAACAC | ACG | ACG | C | T | C>T | C>T_ACG | CG>TA_GCAACAC | 7 | FALSE | FALSE |
| 12_v.txt | 32092 | 32092-SNV-( | SNV | 1 | C | T | 21 | 16624 | 0.12632339 | 62.3317695 | rnr1D57N | FALSE | CAN | JSY14 | 1 | rnr1D57N-Re | GCA | C>T | A/T+1 | G/C-1 | CG>TA | CG>TA | AGCCTGCAA | CAACACC | CGT | GCA | C | A | C>T | C>T_GCA | CG>TA_CAACACC | 7 | FALSE | FALSE |
| 12_v.txt | 32097 | 32097-SNV-( | SNV | 1 | C | T | 14 | 16625 | 0.08421053 | 62.5714286 | rnr1D57N | FALSE | CAN | JSY14 | 1 | rnr1D57N-Re | ACC | C>T | A/T+1 | G/C-1 | CG>TA | CG>TA | GCAACACCA | CCAGTGA | TGG | ACC | C | A | C>T | C>T_ACC | CG>TA_CCAGTGA | 7 | FALSE | FALSE |
| 12_v.txt | 32100 | 32100-SNV-( | SNV | 1 | G | A | 6 | 16625 | 0.03609023 | 63.9999999 | rnr1D57N | FALSE | CAN | JSY14 | 1 | rnr1D57N-Re | AGT | G>A | A/T+1 | G/C-1 | CG>TA | CG>TA | ACACCAGTG | GTGATAT | TCA | TCA | C | T | C>T | C>T_TCA | CG>TA_GTGATAT | 7 | FALSE | FALSE |
| 12_v.txt | 32114 | 32114-SNV-( | SNV | 1 | C | T | 11 | 16625 | 0.06616541 | 53.1817297 | rnr1D57N | FALSE | CAN | JSY14 | 1 | rnr1D57N-Re | GCC | C>T | A/T+1 | G/C-1 | CG>TA | CG>TA | TAATAGCCA | CCATTCG | CGG | GCC | C | A | C>T | C>T_GCC | CG>TA_CCATTCG | 7 | FALSE | FALSE |
| 12_v.txt | 32115 | 32115-SNV-( | SNV | 1 | C | T | 4 | 16625 | 0.02406015 | 39.2488028 | rnr1D57N | FALSE | CAN | JSY14 | 1 | rnr1D57N-Re | CCA | C>T | A/T+1 | G/C-1 | CG>TA | CG>TA | AATAGCCAT | CATTCGA | GGT | CCA | C | A | C>T | C>T_CCA | CG>TA_CATTCGA | 7 | FALSE | FALSE |
| 12_v.txt | 32165 | 32165-SNV-( | SNV | 1 | G | A | 20 | 16625 | 0.12030075 | 43.8491068 | rnr1D57N | FALSE | CAN | JSY14 | 1 | rnr1D57N-Re | CGC | G>A | A/T+1 | G/C-1 | CG>TA | CG>TA | CAAAGCGCC | GCCAAAT | GCG | GCG | C | T | C>T | C>T_GCG | CG>TA_GCCAAAT | 7 | FALSE | FALSE |
| 12_v.txt | 32271 | 32271-SNV-( | SNV | 1 | C | T | 43 | 29226 | 0.14712927 | 31.5346217 | rnr1D57N | FALSE | CAN | JSY14 | 1 | rnr1D57N-Re | ACG | C>T | A/T+1 | G/C-1 | CG>TA | CG>TA | AAAATACGG | CGGGAAC | TGC | ACG | C | A | C>T | C>T_ACG | CG>TA_CGGGAAC | 7 | FALSE | FALSE |
| 12_v.txt | 32272 | 32272-SNV-( | SNV | 1 | G | A | 152 | 29226 | 0.52008486 | 37.4013659 | rnr1D57N | FALSE | CAN | JSY14 | 1 | rnr1D57N-Re | CGG | G>A | A/T+1 | G/C-1 | CG>TA | CG>TA | AAATACGGG | GGGAACC | GCC | GCC | C | T | C>T | C>T_GCC | CG>TA_GGGAACC | 7 | FALSE | FALSE |
| 12_v.txt | 32274 | 32274-SNV-( | SNV | 1 | G | A | 60 | 29226 | 0.20529665 | 36.8500196 | rnr1D57N | FALSE | CAN | JSY14 | 1 | rnr1D57N-Re | GGA | G>A | A/T+1 | G/C-1 | CG>TA | CG>TA | ATACGGGAA | GAACCAA | CCT | CCT | C | T | C>T | C>T_CCT | CG>TA_GAACCAA | 7 | FALSE | FALSE |
| 12_v.txt | 32299 | 32299-SNV-( | SNV | 1 | C | T | 46 | 29225 | 0.15739949 | 50.6956522 | rnr1D57N | FALSE | CAN | JSY14 | 1 | rnr1D57N-Re | GCG | C>T | A/T+1 | G/C-1 | CG>TA | CG>TA | AATTTGCGG | CGGCAGA | CGC | GCG | C | A | C>T | C>T_GCG | CG>TA_CGGCAGA | 7 | FALSE | FALSE |
| 12_v.txt | 32301 | 32301-SNV-( | SNV | 1 | G | A | 21 | 29225 | 0.07185629 | 59.0952381 | rnr1D57N | FALSE | CAN | JSY14 | 1 | rnr1D57N-Re | GGC | G>A | A/T+1 | G/C-1 | CG>TA | CG>TA | TTTGCGGCA | GCAGAAA | CCG | CCG | C | T | C>T | C>T_CCG | CG>TA_GCAGAAA | 7 | FALSE | FALSE |
| 12_v.txt | 32313 | 32313-SNV-( | SNV | 1 | G | A | 53 | 29225 | 0.18135158 | 58.7735849 | rnr1D57N | FALSE | CAN | JSY14 | 1 | rnr1D57N-Re | GGT | G>A | A/T+1 | G/C-1 | CG>TA | CG>TA | ATAATGGTT | GTTGTTA | CCA | CCA | C | T | C>T | C>T_CCA | CG>TA_GTTGTTA | 7 | FALSE | FALSE |
| 12_v.txt | 32344 | 32344-SNV-( | SNV | 1 | G | A | 23 | 29225 | 0.07869974 | 62.3913044 | rnr1D57N | FALSE | CAN | JSY14 | 1 | rnr1D57N-Re | GGC | G>A | A/T+1 | G/C-1 | CG>TA | CG>TA | TATGTGGCA | GCAAAAC | CCG | CCG | C | T | C>T | C>T_CCG | CG>TA_GCAAAAC | 7 | FALSE | FALSE |
| 12_v.txt | 32361 | 32361-SNV-( | SNV | 1 | G | A | 3 | 29224 | 0.01026554 | 63.9994702 | rnr1D57N | FALSE | CAN | JSY14 | 1 | rnr1D57N-Re | AGA | G>A | A/T+1 | G/C-1 | CG>TA | CG>TA | GTACCAGAG | GAGTTCT | TCT | TCT | C | T | C>T | C>T_TCT | CG>TA_GAGTTCT | 7 | FALSE | FALSE |
| 12_v.txt | 32449 | 32449-SNV-( | SNV | 1 | C | T | 71 | 29225 | 0.24294269 | 50.3802817 | rnr1D57N | FALSE | CAN | JSY14 | 1 | rnr1D57N-Re | CCA | C>T | A/T+1 | G/C-1 | CG>TA | CG>TA | AAAGTCCAA | CAATGAA | GGT | CCA | C | A | C>T | C>T_CCA | CG>TA_CAATGAA | 7 | FALSE | FALSE |
| 12_v.txt | 32466 | 32466-SNV-( | SNV | 1 | C | T | 41 | 29225 | 0.14029085 | 49.7804878 | rnr1D57N | FALSE | CAN | JSY14 | 1 | rnr1D57N-Re | GCC | C>T | A/T+1 | G/C-1 | CG>TA | CG>TA | AGAGAGCCA | CCAATGT | CGG | GCC | C | A | C>T | C>T_GCC | CG>TA_CCAATGT | 7 | FALSE | FALSE |
| 12_v.txt | 32487 | 32487-SNV-( | SNV | 1 | C | T | 29 | 29225 | 0.09923011 | 41.3452298 | rnr1D57N | FALSE | CAN | JSY14 | 1 | rnr1D57N-Re | ACG | C>T | A/T+1 | G/C-1 | CG>TA | CG>TA | AAGATACGG | CGGAAAA | TGC | ACG | C | A | C>T | C>T_ACG | CG>TA_CGGAAAA | 7 | FALSE | FALSE |
| 12_v.txt | 32488 | 32488-SNV-( | SNV | 1 | G | A | 53 | 29225 | 0.18135158 | 46.7934283 | rnr1D57N | FALSE | CAN | JSY14 | 1 | rnr1D57N-Re | CGG | G>A | A/T+1 | G/C-1 | CG>TA | CG>TA | AGATACGGA | GGAAAAC | GCC | GCC | C | T | C>T | C>T_GCC | CG>TA_GGAAAAC | 7 | FALSE | FALSE |
| 12_v.txt | 32535 | 32535-SNV-( | SNV | 1 | G | A | 34 | 29225 | 0.11633875 | 36.8231992 | rnr1D57N | FALSE | CAN | JSY14 | 1 | rnr1D57N-Re | TGC | G>A | A/T+1 | G/C-1 | CG>TA | CG>TA | GGGTTTGCA | GCAGCTT | ACG | ACG | C | T | C>T | C>T_ACG | CG>TA_GCAGCTT | 7 | FALSE | FALSE |
| 12_v.txt | 32536 | 32536-SNV-( | SNV | 1 | C | T | 39 | 29225 | 0.13344739 | 37.0512821 | rnr1D57N | FALSE | CAN | JSY14 | 1 | rnr1D57N-Re | GCA | C>T | A/T+1 | G/C-1 | CG>TA | CG>TA | GGTTTGCAG | CAGCTTC | CGT | GCA | C | A | C>T | C>T_GCA | CG>TA_CAGCTTC | 7 | FALSE | FALSE |
| 12_v.txt | 32542 | 32542-SNV-( | SNV | 1 | C | T | 41 | 29225 | 0.14029085 | 37.1463391 | rnr1D57N | FALSE | CAN | JSY14 | 1 | rnr1D57N-Re | TCA | C>T | A/T+1 | G/C-1 | CG>TA | CG>TA | CAGCTTCAC | CACCAGC | AGT | TCA | C | A | C>T | C>T_TCA | CG>TA_CACCAGC | 7 | FALSE | FALSE |
| 12_v.txt | 32544 | 32544-SNV-( | SNV | 1 | C | T | 142 | 29224 | 0.485902 | 37.4857399 | rnr1D57N | FALSE | CAN | JSY14 | 1 | rnr1D57N-Re | ACC | C>T | A/T+1 | G/C-1 | CG>TA | CG>TA | GCTTCACCA | CCAGCAG | TGG | ACC | C | A | C>T | C>T_ACC | CG>TA_CCAGCAG | 7 | FALSE | FALSE |
| 12_v.txt | 32545 | 32545-SNV-( | SNV | 1 | C | T | 17 | 29224 | 0.05817137 | 37.1671041 | rnr1D57N | FALSE | CAN | JSY14 | 1 | rnr1D57N-Re | CCA | C>T | A/T+1 | G/C-1 | CG>TA | CG>TA | CTTCACCAG | CAGCAGT | GGT | CCA | C | A | C>T | C>T_CCA | CG>TA_CAGCAGT | 7 | FALSE | FALSE |
| 12_v.txt | 32580 | 32580-SNV-( | SNV | 1 | G | A | 10 | 26053 | 0.0383833 | 28.2975228 | rnr1D57N | FALSE | CAN | JSY14 | 1 | rnr1D57N-Re | TGT | G>A | A/T+1 | G/C-1 | CG>TA | CG>TA | TGAAATGTG | GTGAAGG | ACA | ACA | C | T | C>T | C>T_ACA | CG>TA_GTGAAGG | 7 | FALSE | FALSE |
| 12_v.txt | 32590 | 32590-SNV-( | SNV | 1 | C | T | 41 | 26053 | 0.15737151 | 37.2926829 | rnr1D57N | FALSE | CAN | JSY14 | 1 | rnr1D57N-Re | GCG | C>T | A/T+1 | G/C-1 | CG>TA | CG>TA | AGGCAGCGT | CGTTAAT | CGC | GCG | C | A | C>T | C>T_GCG | CG>TA_CGTTAAT | 7 | FALSE | FALSE |
| 12_v.txt | 32609 | 32609-SNV-( | SNV | 1 | C | T | 47 | 26053 | 0.18040149 | 47.6368055 | rnr1D57N | FALSE | CAN | JSY14 | 1 | rnr1D57N-Re | CCC | C>T | A/T+1 | G/C-1 | CG>TA | CG>TA | GGAAACCCA | CCAACCT | GGG | CCC | C | A | C>T | C>T_CCC | CG>TA_CCAACCT | 7 | FALSE | FALSE |
| 12_v.txt | 32610 | 32610-SNV-( | SNV | 1 | C | T | 31 | 26051 | 0.11899735 | 50.6060144 | rnr1D57N | FALSE | CAN | JSY14 | 1 | rnr1D57N-Re | CCA | C>T | A/T+1 | G/C-1 | CG>TA | CG>TA | GAAACCCAA | CAACCTA | GGT | CCA | C | A | C>T | C>T_CCA | CG>TA_CAACCTA | 7 | FALSE | FALSE |
| 12_v.txt | 32678 | 32678-SNV-( | SNV | 1 | C | T | 66 | 26053 | 0.25332975 | 56.0151515 | rnr1D57N | FALSE | CAN | JSY14 | 1 | rnr1D57N-Re | TCC | C>T | A/T+1 | G/C-1 | CG>TA | CG>TA | GTTTCTCCA | CCAATAA | AGG | TCC | C | A | C>T | C>T_TCC | CG>TA_CCAATAA | 7 | FALSE | FALSE |
| 12_v.txt | 32700 | 32700-SNV-( | SNV | 1 | C | T | 26 | 26053 | 0.09979657 | 59.4230769 | rnr1D57N | FALSE | CAN | JSY14 | 1 | rnr1D57N-Re | GCC | C>T | A/T+1 | G/C-1 | CG>TA | CG>TA | ACTGGGCCG | CCGGTAA | CGG | GCC | C | A | C>T | C>T_GCC | CG>TA_CCGGTAA | 7 | FALSE | FALSE |
| 12_v.txt | 32702 | 32702-SNV-( | SNV | 1 | G | A | 22 | 26053 | 0.08444325 | 56.0909091 | rnr1D57N | FALSE | CAN | JSY14 | 1 | rnr1D57N-Re | CGG | G>A | A/T+1 | G/C-1 | CG>TA | CG>TA | TGGGCCGGT | GGTAACC | GCC | GCC | C | T | C>T | C>T_GCC | CG>TA_GGTAACC | 7 | FALSE | FALSE |
| 12_v.txt | 32749 | 32749-SNV-( | SNV | 1 | C | T | 164 | 26053 | 0.62948605 | 54.9573171 | rnr1D57N | FALSE | CAN | JSY14 | 1 | rnr1D57N-Re | CCG | C>T | A/T+1 | G/C-1 | CG>TA | CG>TA | GAAACCCGA | CGATAAT | GGC | CCG | C | A | C>T | C>T_CCG | CG>TA_CGATAAT | 7 | FALSE | FALSE |
| 12_v.txt | 32775 | 32775-SNV-( | SNV | 1 | G | A | 95 | 26053 | 0.36464131 | 51.2842105 | rnr1D57N | FALSE | CAN | JSY14 | 1 | rnr1D57N-Re | AGC | G>A | A/T+1 | G/C-1 | CG>TA | CG>TA | ATGGAAGCG | GCGACCC | TCG | TCG | C | T | C>T | C>T_TCG | CG>TA_GCGACCC | 7 | FALSE | FALSE |
| 12_v.txt | 32776 | 32776-SNV-( | SNV | 1 | C | T | 23 | 26052 | 0.08828497 | 55.4346059 | rnr1D57N | FALSE | CAN | JSY14 | 1 | rnr1D57N-Re | GCG | C>T | A/T+1 | G/C-1 | CG>TA | CG>TA | TGGAAGCGA | CGACCCA | CGC | GCG | C | A | C>T | C>T_GCG | CG>TA_CGACCCA | 7 | FALSE | FALSE |
| 12_v.txt | 32780 | 32780-SNV-( | SNV | 1 | C | T | 63 | 26053 | 0.24181476 | 44.2347126 | rnr1D57N | FALSE | CAN | JSY14 | 1 | rnr1D57N-Re | CCC | C>T | A/T+1 | G/C-1 | CG>TA | CG>TA | AGCGACCCA | CCAGAAC | GGG | CCC | C | A | C>T | C>T_CCC | CG>TA_CCAGAAC | 7 | FALSE | FALSE |
| 12_v.txt | 32788 | 32788-SNV-( | SNV | 1 | C | T | 18 | 26052 | 0.06909258 | 44.3888888 | rnr1D57N | FALSE | CAN | JSY14 | 1 | rnr1D57N-Re | TCG | C>T | A/T+1 | G/C-1 | CG>TA | CG>TA | AGAACTCGA | CGAATTC | AGC | TCG | C | A | C>T | C>T_TCG | CG>TA_CGAATTC | 7 | FALSE | FALSE |
| 12_v.txt | 32794 | 32794-SNV-( | SNV | 1 | C | T | 25 | 26053 | 0.09595824 | 49 | rnr1D57N | FALSE | CAN | JSY14 | 1 | rnr1D57N-Re | TCA | C>T | A/T+1 | G/C-1 | CG>TA | CG>TA | CGAATTCAC | CACCGTA | AGT | TCA | C | A | C>T | C>T_TCA | CG>TA_CACCGTA | 7 | FALSE | FALSE |
| 12_v.txt | 32797 | 32797-SNV-( | SNV | 1 | C | T | 31 | 26053 | 0.11898822 | 44.9032227 | rnr1D57N | FALSE | CAN | JSY14 | 1 | rnr1D57N-Re | CCG | C>T | A/T+1 | G/C-1 | CG>TA | CG>TA | ATTCACCGT | CGTAATA | GGC | CCG | C | A | C>T | C>T_CCG | CG>TA_CGTAATA | 7 | FALSE | FALSE |
| 12_v.txt | 32798 | 32798-SNV-( | SNV | 1 | G | A | 32 | 26053 | 0.12282655 | 45.0932316 | rnr1D57N | FALSE | CAN | JSY14 | 1 | rnr1D57N-Re | CGT | G>A | A/T+1 | G/C-1 | CG>TA | CG>TA | TTCACCGTA | GTAATAT | GCA | GCA | C | T | C>T | C>T_GCA | CG>TA_GTAATAT | 7 | FALSE | FALSE |
| 12_v.txt | 32841 | 32841-SNV-( | SNV | 1 | C | T | 2209 | 26053 | 8.47887 | 42.80602 | rnr1D57N | FALSE | CAN | JSY14 | 1 | rnr1D57N-Re | CCA | C>T | A/T+1 | G/C-1 | CG>TA | CG>TA | ATTACCCAA | CAAAAAA | GGT | CCA | C | A | C>T | C>T_CCA | CG>TA_CAAAAAA | 7 | FALSE | FALSE |
| 12_v.txt | 32855 | 32855-SNV-( | SNV | 1 | C | T | 150 | 26052 | 0.57577153 | 35.5 | rnr1D57N | FALSE | CAN | JSY14 | 1 | rnr1D57N-Re | TCC | C>T | A/T+1 | G/C-1 | CG>TA | CG>TA | ACTAATCCA | CCATGCC | AGG | TCC | C | A | C>T | C>T_TCC | CG>TA_CCATGCC | 7 | FALSE | FALSE |
| 12_v.txt | 32860 | 32860-SNV-( | SNV | 1 | C | T | 28 | 26051 | 0.10748148 | 37.8214286 | rnr1D57N | FALSE | CAN | JSY14 | 1 | rnr1D57N-Re | GCC | C>T | A/T+1 | G/C-1 | CG>TA | CG>TA | TCCATGCCG | CCGCCAG | CGG | GCC | C | A | C>T | C>T_GCC | CG>TA_CCGCCAG | 7 | FALSE | FALSE |
| 12_v.txt | 32883 | 32883-SNV-( | SNV | 1 | C | T | 29 | 21315 | 0.13605442 | 37.0684161 | rnr1D57N | FALSE | CAN | JSY14 | 1 | rnr1D57N-Re | CCA | C>T | A/T+1 | G/C-1 | CG>TA | CG>TA | TACGTCCAA | CAAAATT | GGT | CCA | C | A | C>T | C>T_CCA | CG>TA_CAAAATT | 7 | FALSE | FALSE |
| 12_v.txt | 32900 | 32900-SNV-( | SNV | 1 | G | A | 22 | 21315 | 0.1032137 | 35.7727776 | rnr1D57N | FALSE | CAN | JSY14 | 1 | rnr1D57N-Re | GGC | G>A | A/T+1 | G/C-1 | CG>TA | CG>TA | GACTTGGCC | GCCAACT | CCG | CCG | C | T | C>T | C>T_CCG | CG>TA_GCCAACT | 7 | FALSE | FALSE |
| 12_v.txt | 32901 | 32901-SNV-( | SNV | 1 | C | T | 42 | 21315 | 0.19704434 | 31.8097195 | rnr1D57N | FALSE | CAN | JSY14 | 1 | rnr1D57N-Re | GCC | C>T | A/T+1 | G/C-1 | CG>TA | CG>TA | ACTTGGCCA | CCAACTA | CGG | GCC | C | A | C>T | C>T_GCC | CG>TA_CCAACTA | 7 | FALSE | FALSE |
| 12_v.txt | 32917 | 32917-SNV-( | SNV | 1 | C | T | 58 | 21315 | 0.27210884 | 51.6724138 | rnr1D57N | FALSE | CAN | JSY14 | 1 | rnr1D57N-Re | TCC | C>T | A/T+1 | G/C-1 | CG>TA | CG>TA | TAAGTTCCA | CCAGGGC | AGG | TCC | C | A | C>T | C>T_TCC | CG>TA_CCAGGGC | 7 | FALSE | FALSE |
| 12_v.txt | 32935 | 32935-SNV-( | SNV | 1 | C | T | 29 | 21315 | 0.13605442 | 50 | rnr1D57N | FALSE | CAN | JSY14 | 1 | rnr1D57N-Re | GCC | C>T | A/T+1 | G/C-1 | CG>TA | CG>TA | TGATTGCCC | CCCAAGA | CGG | GCC | C | A | C>T | C>T_GCC | CG>TA_CCCAAGA | 7 | FALSE | FALSE |
| 12_v.txt | 32945 | 32945-SNV-( | SNV | 1 | C | T | 43 | 21315 | 0.20173587 | 55.7906977 | rnr1D57N | FALSE | CAN | JSY14 | 1 | rnr1D57N-Re | ACC | C>T | A/T+1 | G/C-1 | CG>TA | CG>TA | AGAAAACCA | CCAATAC | TGG | ACC | C | A | C>T | C>T_ACC | CG>TA_CCAATAC | 7 | FALSE | FALSE |
| 12_v.txt | 32958 | 32958-SNV-( | SNV | 1 | C | T | 29 | 21315 | 0.13605442 | 50.2391521 | rnr1D57N | FALSE | CAN | JSY14 | 1 | rnr1D57N-Re | ACC | C>T | A/T+1 | G/C-1 | CG>TA | CG>TA | ATGTAACCA | CCATTGG | TGG | ACC | C | A | C>T | C>T_ACC | CG>TA_CCATTGG | 7 | FALSE | FALSE |
| 12_v.txt | 32959 | 32959-SNV-( | SNV | 1 | C | T | 4 | 21315 | 0.01876613 | 40.7496081 | rnr1D57N | FALSE | CAN | JSY14 | 1 | rnr1D57N-Re | CCA | C>T | A/T+1 | G/C-1 | CG>TA | CG>TA | TGTAACCAT | CATTGGC | GGT | CCA | C | A | C>T | C>T_CCA | CG>TA_CATTGGC | 7 | FALSE | FALSE |
| 12_v.txt | 32965 | 32965-SNV-( | SNV | 1 | C | T | 25 | 21315 | 0.1172883 | 54.4 | rnr1D57N | FALSE | CAN | JSY14 | 1 | rnr1D57N-Re | GCC | C>T | A/T+1 | G/C-1 | CG>TA | CG>TA | CATTGGCCG | CCGCACC | CGG | GCC | C | A | C>T | C>T_GCC | CG>TA_CCGCACC | 7 | FALSE | FALSE |
| 12_v.txt | 32967 | 32967-SNV-( | SNV | 1 | G | A | 57 | 21315 | 0.26741731 | 59.3859649 | rnr1D57N | FALSE | CAN | JSY14 | 1 | rnr1D57N-Re | CGC | G>A | A/T+1 | G/C-1 | CG>TA | CG>TA | TTGGCCGCA | GCACCAA | GCG | GCG | C | T | C>T | C>T_GCG | CG>TA_GCACCAA | 7 | FALSE | FALSE |
| 12_v.txt | 32970 | 32970-SNV-( | SNV | 1 | C | T | 27 | 21314 | 0.1266773 | 58.5555556 | rnr1D57N | FALSE | CAN | JSY14 | 1 | rnr1D57N-Re | ACC | C>T | A/T+1 | G/C-1 | CG>TA | CG>TA | GCCGCACCA | CCAAATG | TGG | ACC | C | A | C>T | C>T_ACC | CG>TA_CCAAATG | 7 | FALSE | FALSE |
| 12_v.txt | 33015 | 33015-SNV-( | SNV | 1 | G | A | 1888 | 21315 | 8.85761201 | 61.0058263 | rnr1D57N | FALSE | CAN | JSY14 | 1 | rnr1D57N-Re | GGA | G>A | A/T+1 | G/C-1 | CG>TA | CG>TA | AAAGAGGAT | GATGTAA | CCT | CCT | C | T | C>T | C>T_CCT | CG>TA_GATGTAA | 7 | FALSE | FALSE |
| 12_v.txt | 33055 | 33055-SNV-( | SNV | 1 | G | A | 59 | 21315 | 0.27680038 | 55.3047657 | rnr1D57N | FALSE | CAN | JSY14 | 1 | rnr1D57N-Re | TGC | G>A | A/T+1 | G/C-1 | CG>TA | CG>TA | AGGACTGCG | GCGTGAC | ACG | ACG | C | T | C>T | C>T_ACG | CG>TA_GCGTGAC | 7 | FALSE | FALSE |
| 12_v.txt | 33056 | 33056-SNV-( | SNV | 1 | C | T | 33 | 21314 | 0.15482781 | 52.9396623 | rnr1D57N | FALSE | CAN | JSY14 | 1 | rnr1D57N-Re | GCG | C>T | A/T+1 | G/C-1 | CG>TA | CG>TA | GGACTGCGT | CGTGACA | CGC | GCG | C | A | C>T | C>T_GCG | CG>TA_CGTGACA | 7 | FALSE | FALSE |
| 12_v.txt | 33069 | 33069-SNV-( | SNV | 1 | G | A | 19 | 21315 | 0.0891391 | 47.9484645 | rnr1D57N | FALSE | CAN | JSY14 | 1 | rnr1D57N-Re | TGC | G>A | A/T+1 | G/C-1 | CG>TA | CG>TA | GAATATGCC | GCCAAAG | ACG | ACG | C | T | C>T | C>T_ACG | CG>TA_GCCAAAG | 7 | FALSE | FALSE |
| 12_v.txt | 33070 | 33070-SNV-( | SNV | 1 | C | T | 21 | 21315 | 0.09852217 | 59.3785985 | rnr1D57N | FALSE | CAN | JSY14 | 1 | rnr1D57N-Re | GCC | C>T | A/T+1 | G/C-1 | CG>TA | CG>TA | AATATGCCA | CCAAAGA | CGG | GCC | C | A | C>T | C>T_GCC | CG>TA_CCAAAGA | 7 | FALSE | FALSE |
| 12_v.txt | 33075 | 33075-SNV-( | SNV | 1 | G | A | 22 | 21315 | 0.1032137 | 48.5454546 | rnr1D57N | FALSE | CAN | JSY14 | 1 | rnr1D57N-Re | AGA | G>A | A/T+1 | G/C-1 | CG>TA | CG>TA | GCCAAAGAA | GAACCCA | TCT | TCT | C | T | C>T | C>T_TCT | CG>TA_GAACCCA | 7 | FALSE | FALSE |
| 12_v.txt | 33114 | 33114-SNV-( | SNV | 1 | C | T | 173 | 21315 | 0.811635 | 43.5720546 | rnr1D57N | FALSE | CAN | JSY14 | 1 | rnr1D57N-Re | GCC | C>T | A/T+1 | G/C-1 | CG>TA | CG>TA | ACTGGGCCG | CCGGCGT | CGG | GCC | C | A | C>T | C>T_GCC | CG>TA_CCGGCGT | 7 | FALSE | FALSE |
| 12_v.txt | 33115 | 33115-SNV-( | SNV | 1 | C | T | 137 | 21315 | 0.64273986 | 44 | rnr1D57N | FALSE | CAN | JSY14 | 1 | rnr1D57N-Re | CCG | C>T | A/T+1 | G/C-1 | CG>TA | CG>TA | CTGGGCCGG | CGGCGTT | GGC | CCG | C | A | C>T | C>T_CCG | CG>TA_CGGCGTT | 7 | FALSE | FALSE |
| 12_v.txt | 33153 | 33153-SNV-( | SNV | 1 | C | T | 56 | 21315 | 0.26272578 | 36.5892857 | rnr1D57N | FALSE | CAN | JSY14 | 1 | rnr1D57N-Re | ACC | C>T | A/T+1 | G/C-1 | CG>TA | CG>TA | AAAAGACCT | CCTGTAC | TGG | ACC | C | A | C>T | C>T_ACC | CG>TA_CCTGTAC | 7 | FALSE | FALSE |
| 12_v.txt | 33156 | 33156-SNV-( | SNV | 1 | G | A | 504 | 21315 | 2.36453202 | 37.3869048 | rnr1D57N | FALSE | CAN | JSY14 | 1 | rnr1D57N-Re | TGT | G>A | A/T+1 | G/C-1 | CG>TA | CG>TA | AGACCTGTA | GTACCAA | ACA | ACA | C | T | C>T | C>T_ACA | CG>TA_GTACCAA | 7 | FALSE | FALSE |
| 12_v.txt | 33159 | 33159-SNV-( | SNV | 1 | C | T | 144 | 21315 | 0.67558058 | 36.8316212 | rnr1D57N | FALSE | CAN | JSY14 | 1 | rnr1D57N-Re | ACC | C>T | A/T+1 | G/C-1 | CG>TA | CG>TA | CCTGTACCA | CCAATAG | TGG | ACC | C | A | C>T | C>T_ACC | CG>TA_CCAATAG | 7 | FALSE | FALSE |
| 12_v.txt | 33168 | 33168-SNV-( | SNV | 1 | C | T | 59 | 21296 | 0.27704733 | 37.779661 | rnr1D57N | FALSE | CAN | JSY14 | 1 | rnr1D57N-Re | ACC | C>T | A/T+1 | G/C-1 | CG>TA | CG>TA | ATAGTACCA | CCACCAA | TGG | ACC | C | A | C>T | C>T_ACC | CG>TA_CCACCAA | 7 | FALSE | FALSE |
| 12_v.txt | 33177 | 33177-SNV-( | SNV | 1 | G | A | 28 | 21290 | 0.13151714 | 37.8214286 | rnr1D57N | FALSE | CAN | JSY14 | 1 | rnr1D57N-Re | GGC | G>A | A/T+1 | G/C-1 | CG>TA | CG>TA | CCAAGGGCA | GCAATCA | CCG | CCG | C | T | C>T | C>T_CCG | CG>TA_GCAATCA | 7 | FALSE | FALSE |
| 12_v.txt | 33199 | 33199-SNV-( | SNV | 1 | G | A | 22 | 20300 | 0.10837438 | 35.6363636 | rnr1D57N | FALSE | CAN | JSY14 | 1 | rnr1D57N-Re | TGC | G>A | A/T+1 | G/C-1 | CG>TA | CG>TA | GTCTTTGCT | GCTTAAG | ACG | ACG | C | T | C>T | C>T_ACG | CG>TA_GCTTAAG | 7 | FALSE | FALSE |
| 12_v.txt | 33205 | 33205-SNV-( | SNV | 1 | G | A | 11 | 20300 | 0.05418719 | 39.2708664 | rnr1D57N | FALSE | CAN | JSY14 | 1 | rnr1D57N-Re | AGC | G>A | A/T+1 | G/C-1 | CG>TA | CG>TA | GCTTAAGCT | GCTCTCT | TCG | TCG | C | T | C>T | C>T_TCG | CG>TA_GCTCTCT | 7 | FALSE | FALSE |
| 12_v.txt | 33229 | 33229-SNV-( | SNV | 1 | G | A | 121 | 20312 | 0.59570697 | 39.0743802 | rnr1D57N | FALSE | CAN | JSY14 | 1 | rnr1D57N-Re | TGT | G>A | A/T+1 | G/C-1 | CG>TA | CG>TA | CGTTCTGTA | GTACTTC | ACA | ACA | C | T | C>T | C>T_ACA | CG>TA_GTACTTC | 7 | FALSE | FALSE |
| 12_v.txt | 33258 | 33258-SNV-( | SNV | 1 | C | T | 34 | 20312 | 0.16738874 | 43.7058824 | rnr1D57N | FALSE | CAN | JSY14 | 1 | rnr1D57N-Re | GCC | C>T | A/T+1 | G/C-1 | CG>TA | CG>TA | CCTATGCCA | CCATCCT | CGG | GCC | C | A | C>T | C>T_GCC | CG>TA_CCATCCT | 7 | FALSE | FALSE |
| 12_v.txt | 33276 | 33276-SNV-( | SNV | 1 | G | A | 35 | 20312 | 0.17231193 | 48.3428571 | rnr1D57N | FALSE | CAN | JSY14 | 1 | rnr1D57N-Re | CGT | G>A | A/T+1 | G/C-1 | CG>TA | CG>TA | GAGAACGTA | GTATCCT | GCA | GCA | C | T | C>T | C>T_GCA | CG>TA_GTATCCT | 7 | FALSE | FALSE |
| 12_v.txt | 33284 | 33284-SNV-( | SNV | 1 | G | A | 32 | 20312 | 0.15754234 | 44 | rnr1D57N | FALSE | CAN | JSY14 | 1 | rnr1D57N-Re | CGC | G>A | A/T+1 | G/C-1 | CG>TA | CG>TA | ATCCTCGCC | GCCATTT | GCG | GCG | C | T | C>T | C>T_GCG | CG>TA_GCCATTT | 7 | FALSE | FALSE |
| 12_v.txt | 33309 | 33309-SNV-( | SNV | 1 | C | T | 29 | 20312 | 0.14277275 | 57.0656322 | rnr1D57N | FALSE | CAN | JSY14 | 1 | rnr1D57N-Re | GCG | C>T | A/T+1 | G/C-1 | CG>TA | CG>TA | AAAGAGCGC | CGCAATG | CGC | GCG | C | A | C>T | C>T_GCG | CG>TA_CGCAATG | 7 | FALSE | FALSE |
| 12_v.txt | 33310 | 33310-SNV-( | SNV | 1 | G | A | 22 | 20307 | 0.10833703 | 57.0820247 | rnr1D57N | FALSE | CAN | JSY14 | 1 | rnr1D57N-Re | CGC | G>A | A/T+1 | G/C-1 | CG>TA | CG>TA | AAGAGCGCA | GCAATGG | GCG | GCG | C | T | C>T | C>T_GCG | CG>TA_GCAATGG | 7 | FALSE | FALSE |
| 12_v.txt | 33382 | 33382-SNV-( | SNV | 1 | C | T | 20 | 20312 | 0.09846396 | 56.95 | rnr1D57N | FALSE | CAN | JSY14 | 1 | rnr1D57N-Re | GCT | C>T | A/T+1 | G/C-1 | CG>TA | CG>TA | GTGAAGCTT | CTTCAAC | CGA | GCT | C | A | C>T | C>T_GCT | CG>TA_CTTCAAC | 7 | FALSE | FALSE |
| 12_v.txt | 33392 | 33392-SNV-( | SNV | 1 | G | A | 26 | 20312 | 0.12800315 | 48.0384615 | rnr1D57N | FALSE | CAN | JSY14 | 1 | rnr1D57N-Re | CGT | G>A | A/T+1 | G/C-1 | CG>TA | CG>TA | AACGTCGTG | GTGAAAG | GCA | GCA | C | T | C>T | C>T_GCA | CG>TA_GTGAAAG | 7 | FALSE | FALSE |
| 12_v.txt | 33474 | 33474-SNV-( | SNV | 1 | C | T | 25 | 20311 | 0.12308601 | 34.9997067 | rnr1D57N | FALSE | CAN | JSY14 | 1 | rnr1D57N-Re | GCC | C>T | A/T+1 | G/C-1 | CG>TA | CG>TA | GCTATGCCT | CCTTTTT | CGG | GCC | C | A | C>T | C>T_GCC | CG>TA_CCTTTTT | 7 | FALSE | FALSE |
| 12_v.txt | 31917 | 31917-SNV-( | SNV | 1 | G | C | 10 | 11162 | 0.08958968 | 37.6 | rnr1D57N | FALSE | CAN | JSY14 | 1 | rnr1D57N-Re | AGC | G>C | G/C+1 | G/C-1 | CG>GC | CG>GC | GCAAAAGCC | GCCGTGA | TCG | TCG | C | G | C>G | C>G_TCG | CG>GC_GCCGTGA | 7 | FALSE | FALSE |
| 12_v.txt | 32064 | 32064-SNV-( | SNV | 1 | G | C | 48 | 16625 | 0.28872181 | 61.3541667 | rnr1D57N | FALSE | CAN | JSY14 | 1 | rnr1D57N-Re | TGA | G>C | G/C+1 | G/C-1 | CG>GC | CG>GC | GAGATTGAG | GAGATAA | ACT | ACT | C | G | C>G | C>G_ACT | CG>GC_GAGATAA | 7 | FALSE | FALSE |
| 12_v.txt | 32092 | 32092-SNV-( | SNV | 1 | C | G | 2 | 16624 | 0.0120308 | 50.9983362 | rnr1D57N | FALSE | CAN | JSY14 | 1 | rnr1D57N-Re | GCA | C>G | G/C+1 | G/C-1 | CG>GC | CG>GC | AGCCTGCAA | CAACACC | CGT | GCA | C | C | C>G | C>G_GCA | CG>GC_CAACACC | 7 | FALSE | FALSE |
| 12_v.txt | 32150 | 32150-SNV-( | SNV | 1 | C | G | 1 | 16625 | 6.02E-03 | 38.0018519 | rnr1D57N | FALSE | CAN | JSY14 | 1 | rnr1D57N-Re | CCA | C>G | G/C+1 | G/C-1 | CG>GC | CG>GC | TGTCTCCAT | CATGTAA | GGT | CCA | C | C | C>G | C>G_CCA | CG>GC_CATGTAA | 7 | FALSE | FALSE |
| 12_v.txt | 32223 | 32223-SNV-( | SNV | 1 | G | C | 6 | 16624 | 0.0360924 | 36.1666667 | rnr1D57N | FALSE | CAN | JSY14 | 1 | rnr1D57N-Re | TGA | G>C | G/C+1 | G/C-1 | CG>GC | CG>GC | GTCCTTGAC | GACAGGA | ACT | ACT | C | G | C>G | C>G_ACT | CG>GC_GACAGGA | 7 | FALSE | FALSE |
| 12_v.txt | 32272 | 32272-SNV-( | SNV | 1 | G | C | 42 | 29226 | 0.14370766 | 37.0476459 | rnr1D57N | FALSE | CAN | JSY14 | 1 | rnr1D57N-Re | CGG | G>C | G/C+1 | G/C-1 | CG>GC | CG>GC | AAATACGGG | GGGAACC | GCC | GCC | C | G | C>G | C>G_GCC | CG>GC_GGGAACC | 7 | FALSE | FALSE |
| 12_v.txt | 32278 | 32278-SNV-( | SNV | 1 | C | G | 52 | 29226 | 0.17792377 | 34.7115385 | rnr1D57N | FALSE | CAN | JSY14 | 1 | rnr1D57N-Re | CCA | C>G | G/C+1 | G/C-1 | CG>GC | CG>GC | GGGAACCAA | CAACGTA | GGT | CCA | C | C | C>G | C>G_CCA | CG>GC_CAACGTA | 7 | FALSE | FALSE |
| 12_v.txt | 32292 | 32292-SNV-( | SNV | 1 | G | C | 7 | 29224 | 0.02395292 | 55.8555944 | rnr1D57N | FALSE | CAN | JSY14 | 1 | rnr1D57N-Re | TGA | G>C | G/C+1 | G/C-1 | CG>GC | CG>GC | ATATTTGAA | GAATTTG | ACT | ACT | C | G | C>G | C>G_ACT | CG>GC_GAATTTG | 7 | FALSE | FALSE |
| 12_v.txt | 32399 | 32399-SNV-( | SNV | 1 | G | C | 17 | 29225 | 0.05816938 | 58.7647059 | rnr1D57N | FALSE | CAN | JSY14 | 1 | rnr1D57N-Re | CGT | G>C | G/C+1 | G/C-1 | CG>GC | CG>GC | AGAAACGTA | GTAGGAA | GCA | GCA | C | G | C>G | C>G_GCA | CG>GC_GTAGGAA | 7 | FALSE | FALSE |
| 12_v.txt | 32432 | 32432-SNV-( | SNV | 1 | G | C | 83 | 29225 | 0.28400342 | 57.1566265 | rnr1D57N | FALSE | CAN | JSY14 | 1 | rnr1D57N-Re | TGT | G>C | G/C+1 | G/C-1 | CG>GC | CG>GC | GTCATTGTA | GTATGGA | ACA | ACA | C | G | C>G | C>G_ACA | CG>GC_GTATGGA | 7 | FALSE | FALSE |
| 12_v.txt | 32449 | 32449-SNV-( | SNV | 1 | C | G | 18 | 29225 | 0.0615911 | 55.0555556 | rnr1D57N | FALSE | CAN | JSY14 | 1 | rnr1D57N-Re | CCA | C>G | G/C+1 | G/C-1 | CG>GC | CG>GC | AAAGTCCAA | CAATGAA | GGT | CCA | C | C | C>G | C>G_CCA | CG>GC_CAATGAA | 7 | FALSE | FALSE |
| 12_v.txt | 32471 | 32471-SNV-( | SNV | 1 | G | C | 51 | 29225 | 0.17450813 | 47.8235294 | rnr1D57N | FALSE | CAN | JSY14 | 1 | rnr1D57N-Re | TGT | G>C | G/C+1 | G/C-1 | CG>GC | CG>GC | GCCAATGTA | GTAGAAG | ACA | ACA | C | G | C>G | C>G_ACA | CG>GC_GTAGAAG | 7 | FALSE | FALSE |
| 12_v.txt | 32487 | 32487-SNV-( | SNV | 1 | C | G | 14 | 29225 | 0.04790419 | 47.7155484 | rnr1D57N | FALSE | CAN | JSY14 | 1 | rnr1D57N-Re | ACG | C>G | G/C+1 | G/C-1 | CG>GC | CG>GC | AAGATACGG | CGGAAAA | TGC | ACG | C | C | C>G | C>G_ACG | CG>GC_CGGAAAA | 7 | FALSE | FALSE |
| 12_v.txt | 32536 | 32536-SNV-( | SNV | 1 | C | G | 3 | 29225 | 0.01026518 | 29 | rnr1D57N | FALSE | CAN | JSY14 | 1 | rnr1D57N-Re | GCA | C>G | G/C+1 | G/C-1 | CG>GC | CG>GC | GGTTTGCAG | CAGCTTC | CGT | GCA | C | C | C>G | C>G_GCA | CG>GC_CAGCTTC | 7 | FALSE | FALSE |
| 12_v.txt | 32539 | 32539-SNV-( | SNV | 1 | C | G | 22 | 29225 | 0.07527802 | 37.6363636 | rnr1D57N | FALSE | CAN | JSY14 | 1 | rnr1D57N-Re | GCT | C>G | G/C+1 | G/C-1 | CG>GC | CG>GC | TTGCAGCTT | CTTCACC | CGA | GCT | C | C | C>G | C>G_GCT | CG>GC_CTTCACC | 7 | FALSE | FALSE |
| 12_v.txt | 32544 | 32544-SNV-( | SNV | 1 | C | G | 8 | 29224 | 0.02737476 | 37.8748988 | rnr1D57N | FALSE | CAN | JSY14 | 1 | rnr1D57N-Re | ACC | C>G | G/C+1 | G/C-1 | CG>GC | CG>GC | GCTTCACCA | CCAGCAG | TGG | ACC | C | C | C>G | C>G_ACC | CG>GC_CCAGCAG | 7 | FALSE | FALSE |
| 12_v.txt | 32557 | 32557-SNV-( | SNV | 1 | C | G | 8 | 29207 | 0.02739069 | 37.999426 | rnr1D57N | FALSE | CAN | JSY14 | 1 | rnr1D57N-Re | CCA | C>G | G/C+1 | G/C-1 | CG>GC | CG>GC | TGATACCAA | CAACTAG | GGT | CCA | C | C | C>G | C>G_CCA | CG>GC_CAACTAG | 7 | FALSE | FALSE |
| 12_v.txt | 32743 | 32743-SNV-( | SNV | 1 | G | C | 31 | 26053 | 0.11898822 | 54.2580645 | rnr1D57N | FALSE | CAN | JSY14 | 1 | rnr1D57N-Re | AGA | G>C | G/C+1 | G/C-1 | CG>GC | CG>GC | ATATTAGAA | GAAACCC | TCT | TCT | C | G | C>G | C>G_TCT | CG>GC_GAAACCC | 7 | FALSE | FALSE |
| 12_v.txt | 32758 | 32758-SNV-( | SNV | 1 | C | G | 10 | 26053 | 0.0383833 | 49.4922793 | rnr1D57N | FALSE | CAN | JSY14 | 1 | rnr1D57N-Re | GCT | C>G | G/C+1 | G/C-1 | CG>GC | CG>GC | TAATGGCTA | CTAAAAC | CGA | GCT | C | C | C>G | C>G_GCT | CG>GC_CTAAAAC | 7 | FALSE | FALSE |
| 12_v.txt | 32776 | 32776-SNV-( | SNV | 1 | C | G | 48 | 26052 | 0.18424689 | 47.7531585 | rnr1D57N | FALSE | CAN | JSY14 | 1 | rnr1D57N-Re | GCG | C>G | G/C+1 | G/C-1 | CG>GC | CG>GC | TGGAAGCGA | CGACCCA | CGC | GCG | C | C | C>G | C>G_GCG | CG>GC_CGACCCA | 7 | FALSE | FALSE |
| 12_v.txt | 32798 | 32798-SNV-( | SNV | 1 | G | C | 4 | 26053 | 0.01535332 | 41.4994027 | rnr1D57N | FALSE | CAN | JSY14 | 1 | rnr1D57N-Re | CGT | G>C | G/C+1 | G/C-1 | CG>GC | CG>GC | TTCACCGTA | GTAATAT | GCA | GCA | C | G | C>G | C>G_GCA | CG>GC_GTAATAT | 7 | FALSE | FALSE |
| 12_v.txt | 32809 | 32809-SNV-( | SNV | 1 | C | G | 19 | 26053 | 0.07292826 | 41.7894737 | rnr1D57N | FALSE | CAN | JSY14 | 1 | rnr1D57N-Re | ACA | C>G | G/C+1 | G/C-1 | CG>GC | CG>GC | ATTTGACAG | CAGGGAA | TGT | ACA | C | C | C>G | C>G_ACA | CG>GC_CAGGGAA | 7 | FALSE | FALSE |
| 12_v.txt | 32811 | 32811-SNV-( | SNV | 1 | G | C | 371 | 26053 | 1.42402027 | 44.1132076 | rnr1D57N | FALSE | CAN | JSY14 | 1 | rnr1D57N-Re | AGG | G>C | G/C+1 | G/C-1 | CG>GC | CG>GC | TTGACAGGG | GGGAACA | TCC | TCC | C | G | C>G | C>G_TCC | CG>GC_GGGAACA | 7 | FALSE | FALSE |
| 12_v.txt | 32876 | 32876-SNV-( | SNV | 1 | G | C | 8 | 21306 | 0.03754811 | 34.5 | rnr1D57N | FALSE | CAN | JSY14 | 1 | rnr1D57N-Re | TGT | G>C | G/C+1 | G/C-1 | CG>GC | CG>GC | AACTTTGTA | GTACGTC | ACA | ACA | C | G | C>G | C>G_ACA | CG>GC_GTACGTC | 7 | FALSE | FALSE |
| 12_v.txt | 32914 | 32914-SNV-( | SNV | 1 | G | C | 1 | 21315 | 4.69E-03 | 16.0012485 | rnr1D57N | FALSE | CAN | JSY14 | 1 | rnr1D57N-Re | AGT | G>C | G/C+1 | G/C-1 | CG>GC | CG>GC | CACTAAGTT | GTTCCAG | TCA | TCA | C | G | C>G | C>G_TCA | CG>GC_GTTCCAG | 7 | FALSE | FALSE |
| 12_v.txt | 32937 | 32937-SNV-( | SNV | 1 | C | G | 16 | 21315 | 0.07506451 | 55.6145571 | rnr1D57N | FALSE | CAN | JSY14 | 1 | rnr1D57N-Re | CCA | C>G | G/C+1 | G/C-1 | CG>GC | CG>GC | ATTGCCCAA | CAAGAAA | GGT | CCA | C | C | C>G | C>G_CCA | CG>GC_CAAGAAA | 7 | FALSE | FALSE |
| 12_v.txt | 32959 | 32959-SNV-( | SNV | 1 | C | G | 32 | 21315 | 0.15012902 | 60.9960676 | rnr1D57N | FALSE | CAN | JSY14 | 1 | rnr1D57N-Re | CCA | C>G | G/C+1 | G/C-1 | CG>GC | CG>GC | TGTAACCAT | CATTGGC | GGT | CCA | C | C | C>G | C>G_CCA | CG>GC_CATTGGC | 7 | FALSE | FALSE |
| 12_v.txt | 32970 | 32970-SNV-( | SNV | 1 | C | G | 2 | 21314 | 9.38E-03 | 63 | rnr1D57N | FALSE | CAN | JSY14 | 1 | rnr1D57N-Re | ACC | C>G | G/C+1 | G/C-1 | CG>GC | CG>GC | GCCGCACCA | CCAAATG | TGG | ACC | C | C | C>G | C>G_ACC | CG>GC_CCAAATG | 7 | FALSE | FALSE |
| 12_v.txt | 32971 | 32971-SNV-( | SNV | 1 | C | G | 10 | 21315 | 0.04691532 | 57.5900761 | rnr1D57N | FALSE | CAN | JSY14 | 1 | rnr1D57N-Re | CCA | C>G | G/C+1 | G/C-1 | CG>GC | CG>GC | CCGCACCAA | CAAATGC | GGT | CCA | C | C | C>G | C>G_CCA | CG>GC_CAAATGC | 7 | FALSE | FALSE |
| 12_v.txt | 32977 | 32977-SNV-( | SNV | 1 | C | G | 9 | 21315 | 0.04222379 | 62.3333333 | rnr1D57N | FALSE | CAN | JSY14 | 1 | rnr1D57N-Re | GCT | C>G | G/C+1 | G/C-1 | CG>GC | CG>GC | CAAATGCTG | CTGGAGA | CGA | GCT | C | C | C>G | C>G_GCT | CG>GC_CTGGAGA | 7 | FALSE | FALSE |
| 12_v.txt | 32997 | 32997-SNV-( | SNV | 1 | G | C | 236 | 21315 | 1.1072015 | 61 | rnr1D57N | FALSE | CAN | JSY14 | 1 | rnr1D57N-Re | TGA | G>C | G/C+1 | G/C-1 | CG>GC | CG>GC | CTTTGTGAG | GAGAAAA | ACT | ACT | C | G | C>G | C>G_ACT | CG>GC_GAGAAAA | 7 | FALSE | FALSE |
| 12_v.txt | 33018 | 33018-SNV-( | SNV | 1 | G | C | 16 | 21315 | 0.07506451 | 62.6875 | rnr1D57N | FALSE | CAN | JSY14 | 1 | rnr1D57N-Re | TGT | G>C | G/C+1 | G/C-1 | CG>GC | CG>GC | GAGGATGTA | GTAACAG | ACA | ACA | C | G | C>G | C>G_ACA | CG>GC_GTAACAG | 7 | FALSE | FALSE |
| 12_v.txt | 33037 | 33037-SNV-( | SNV | 1 | C | G | 12 | 21315 | 0.05629838 | 59.0833333 | rnr1D57N | FALSE | CAN | JSY14 | 1 | rnr1D57N-Re | GCC | C>G | G/C+1 | G/C-1 | CG>GC | CG>GC | ATGTAGCCA | CCATTTC | CGG | GCC | C | C | C>G | C>G_GCC | CG>GC_CCATTTC | 7 | FALSE | FALSE |
| 12_v.txt | 33069 | 33069-SNV-( | SNV | 1 | G | C | 6 | 21315 | 0.02814919 | 55.1677763 | rnr1D57N | FALSE | CAN | JSY14 | 1 | rnr1D57N-Re | TGC | G>C | G/C+1 | G/C-1 | CG>GC | CG>GC | GAATATGCC | GCCAAAG | ACG | ACG | C | G | C>G | C>G_ACG | CG>GC_GCCAAAG | 7 | FALSE | FALSE |
| 12_v.txt | 33070 | 33070-SNV-( | SNV | 1 | C | G | 41 | 21315 | 0.1923528 | 54.5834453 | rnr1D57N | FALSE | CAN | JSY14 | 1 | rnr1D57N-Re | GCC | C>G | G/C+1 | G/C-1 | CG>GC | CG>GC | AATATGCCA | CCAAAGA | CGG | GCC | C | C | C>G | C>G_GCC | CG>GC_CCAAAGA | 7 | FALSE | FALSE |
| 12_v.txt | 33115 | 33115-SNV-( | SNV | 1 | C | G | 17 | 21315 | 0.07975604 | 37.1764706 | rnr1D57N | FALSE | CAN | JSY14 | 1 | rnr1D57N-Re | CCG | C>G | G/C+1 | G/C-1 | CG>GC | CG>GC | CTGGGCCGG | CGGCGTT | GGC | CCG | C | C | C>G | C>G_CCG | CG>GC_CGGCGTT | 7 | FALSE | FALSE |

| | | | | | | | | | | | | | | | | | | | | | | | | | | | | | | | | | | |
|---|---|---|---|---|---|---|---|---|---|---|---|---|---|---|---|---|---|---|---|---|---|---|---|---|---|---|---|---|---|---|---|---|---|---|
| 12_v.txt | 33154 | 33154-SNV-( | SNV | 1 | C | G | 15 | 21315 | 0.07037298 | 37.9333394 | rnr1D57N | FALSE | CAN | JSY14 | 1 | rnr1D57N-Re | CCT | C>G | G/C+1 | G/C-1 | CG>GC | CG>GC | AAAGACCTG | CTGTACC | GGA | CCT | C | C | C>G | C>G_CCT | CG>GC_CTGTACC | 7 | FALSE | FALSE |
| 12_v.txt | 33205 | 33205-SNV-( | SNV | 1 | G | C | 8 | 20300 | 0.03940887 | 41.6215151 | rnr1D57N | FALSE | CAN | JSY14 | 1 | rnr1D57N-Re | AGC | G>C | G/C+1 | G/C-1 | CG>GC | CG>GC | GCTTAAGCT | GCTCTCT | TCG | TCG | C | G | C>G | C>G_TCG | CG>GC_GCTCTCT | 7 | FALSE | FALSE |
| 12_v.txt | 33310 | 33310-SNV-( | SNV | 1 | G | C | 3 | 20307 | 0.01477323 | 20.3326873 | rnr1D57N | FALSE | CAN | JSY14 | 1 | rnr1D57N-Re | CGC | G>C | G/C+1 | G/C-1 | CG>GC | CG>GC | AAGAGCGCA | GCAATGG | GCG | GCG | C | G | C>G | C>G_GCG | CG>GC_GCAATGG | 7 | FALSE | FALSE |
| 12_v.txt | 33378 | 33378-SNV-( | SNV | 1 | G | C | 3 | 20312 | 0.01476959 | 64 | rnr1D57N | FALSE | CAN | JSY14 | 1 | rnr1D57N-Re | TGA | G>C | G/C+1 | G/C-1 | CG>GC | CG>GC | GTTTGTGAA | GAAGCTT | ACT | ACT | C | G | C>G | C>G_ACT | CG>GC_GAAGCTT | 7 | FALSE | FALSE |
| 12_v.txt | 33456 | 33456-SNV-( | SNV | 1 | G | C | 49 | 20312 | 0.24123671 | 43.2244898 | rnr1D57N | FALSE | CAN | JSY14 | 1 | rnr1D57N-Re | TGA | G>C | G/C+1 | G/C-1 | CG>GC | CG>GC | TCTTTTGAAT | GAATTTG | ACT | ACT | C | G | C>G | C>G_ACT | CG>GC_GAATTTG | 7 | FALSE | FALSE |
| 12_v.txt | 31657 | 31657-SNV-( | SNV | 1 | G | T | 96 | 11167 | 0.85967583 | 18.0416667 | rnr1D57N | FALSE | CAN | JSY14 | 1 | rnr1D57N-Re | GGC | G>T | A/T+1 | G/C-1 | CG>AT | CG>AT | GAAATGGCG | GCGTGGA | CCG | CCG | C | A | C>A | C>A_CCG | CG>AT_GCGTGGA | 7 | FALSE | FALSE |
| 12_v.txt | 31869 | 31869-SNV-( | SNV | 1 | G | T | 29 | 11167 | 0.25969374 | 43.7586207 | rnr1D57N | FALSE | CAN | JSY14 | 1 | rnr1D57N-Re | AGA | G>T | A/T+1 | G/C-1 | CG>AT | CG>AT | AAAATAGAG | GAGATAT | TCT | TCT | C | A | C>A | C>A_TCT | CG>AT_GAGATAT | 7 | FALSE | FALSE |
| 12_v.txt | 32088 | 32088-SNV-( | SNV | 1 | C | A | 5 | 16625 | 0.03007519 | 53.8027108 | rnr1D57N | FALSE | CAN | JSY14 | 1 | rnr1D57N-Re | GCC | C>A | A/T+1 | G/C-1 | CG>AT | CG>AT | AAAAAGCCT | CCTGCAA | CGG | GCC | C | T | C>A | C>A_GCC | CG>AT_CCTGCAA | 7 | FALSE | FALSE |
| 12_v.txt | 32095 | 32095-SNV-( | SNV | 1 | C | A | 46 | 16625 | 0.27669173 | 61.9565217 | rnr1D57N | FALSE | CAN | JSY14 | 1 | rnr1D57N-Re | ACA | C>A | A/T+1 | G/C-1 | CG>AT | CG>AT | CTGCAACAC | CACCAGT | TGT | ACA | C | T | C>A | C>A_ACA | CG>AT_CACCAGT | 7 | FALSE | FALSE |
| 12_v.txt | 32102 | 32102-SNV-( | SNV | 1 | G | T | 2 | 16625 | 0.01203008 | 63.9994997 | rnr1D57N | FALSE | CAN | JSY14 | 1 | rnr1D57N-Re | TGA | G>T | A/T+1 | G/C-1 | CG>AT | CG>AT | ACCAGTGAT | GATATTT | ACT | ACT | C | A | C>A | C>A_ACT | CG>AT_GATATTT | 7 | FALSE | FALSE |
| 12_v.txt | 32114 | 32114-SNV-( | SNV | 1 | C | A | 3 | 16625 | 0.01804511 | 37.6666132 | rnr1D57N | FALSE | CAN | JSY14 | 1 | rnr1D57N-Re | GCC | C>A | A/T+1 | G/C-1 | CG>AT | CG>AT | TAATAGCCA | CCATTCG | CGG | GCC | C | T | C>A | C>A_GCC | CG>AT_CCATTCG | 7 | FALSE | FALSE |
| 12_v.txt | 32119 | 32119-SNV-( | SNV | 1 | C | A | 69 | 16625 | 0.41503759 | 57.8286642 | rnr1D57N | FALSE | CAN | JSY14 | 1 | rnr1D57N-Re | TCG | C>A | A/T+1 | G/C-1 | CG>AT | CG>AT | GCCATTCGA | CGAAAAC | AGC | TCG | C | T | C>A | C>A_TCG | CG>AT_CGAAAAC | 7 | FALSE | FALSE |
| 12_v.txt | 32120 | 32120-SNV-( | SNV | 1 | G | T | 4 | 16625 | 0.02406015 | 50.7481769 | rnr1D57N | FALSE | CAN | JSY14 | 1 | rnr1D57N-Re | CGA | G>T | A/T+1 | G/C-1 | CG>AT | CG>AT | CCATTCGAA | GAAAACT | GCT | GCT | C | A | C>A | C>A_GCT | CG>AT_GAAAACT | 7 | FALSE | FALSE |
| 12_v.txt | 32149 | 32149-SNV-( | SNV | 1 | C | A | 15 | 16625 | 0.09022556 | 48.5996881 | rnr1D57N | FALSE | CAN | JSY14 | 1 | rnr1D57N-Re | TCC | C>A | A/T+1 | G/C-1 | CG>AT | CG>AT | ATGTCTCCAT | CCATGTA | AGG | TCC | C | T | C>A | C>A_TCC | CG>AT_CCATGTA | 7 | FALSE | FALSE |
| 12_v.txt | 32150 | 32150-SNV-( | SNV | 1 | C | A | 17 | 16625 | 0.10225564 | 41.4132324 | rnr1D57N | FALSE | CAN | JSY14 | 1 | rnr1D57N-Re | CCA | C>A | A/T+1 | G/C-1 | CG>AT | CG>AT | TGTCTCCATG | CATGTAA | GGT | CCA | C | T | C>A | C>A_CCA | CG>AT_CATGTAA | 7 | FALSE | FALSE |
| 12_v.txt | 32153 | 32153-SNV-( | SNV | 1 | G | T | 13 | 16625 | 0.07819549 | 53.6155587 | rnr1D57N | FALSE | CAN | JSY14 | 1 | rnr1D57N-Re | TGT | G>T | A/T+1 | G/C-1 | CG>AT | CG>AT | CTCCATGTA | GTAAGCC | ACA | ACA | C | A | C>A | C>A_ACA | CG>AT_GTAAGCC | 7 | FALSE | FALSE |
| 12_v.txt | 32195 | 32195-SNV-( | SNV | 1 | G | T | 22 | 16625 | 0.13233083 | 43.6363636 | rnr1D57N | FALSE | CAN | JSY14 | 1 | rnr1D57N-Re | TGT | G>T | A/T+1 | G/C-1 | CG>AT | CG>AT | TGCAATGTA | GTATGGA | ACA | ACA | C | A | C>A | C>A_ACA | CG>AT_GTATGGA | 7 | FALSE | FALSE |
| 12_v.txt | 32272 | 32272-SNV-( | SNV | 1 | G | T | 49 | 29226 | 0.16765893 | 34.285886 | rnr1D57N | FALSE | CAN | JSY14 | 1 | rnr1D57N-Re | CGG | G>T | A/T+1 | G/C-1 | CG>AT | CG>AT | AAATACGGG | GGGAACC | GCC | GCC | C | A | C>A | C>A_GCC | CG>AT_GGGAACC | 7 | FALSE | FALSE |
| 12_v.txt | 32274 | 32274-SNV-( | SNV | 1 | G | T | 342 | 29226 | 1.17019093 | 36.0707527 | rnr1D57N | FALSE | CAN | JSY14 | 1 | rnr1D57N-Re | GGA | G>T | A/T+1 | G/C-1 | CG>AT | CG>AT | ATACGGGAA | GAACCAA | CCT | CCT | C | A | C>A | C>A_CCT | CG>AT_GAACCAA | 7 | FALSE | FALSE |
| 12_v.txt | 32292 | 32292-SNV-( | SNV | 1 | G | T | 39 | 29224 | 0.13345196 | 51.1793264 | rnr1D57N | FALSE | CAN | JSY14 | 1 | rnr1D57N-Re | TGA | G>T | A/T+1 | G/C-1 | CG>AT | CG>AT | ATATTTGAA | GAATTTG | ACT | ACT | C | A | C>A | C>A_ACT | CG>AT_GAATTTG | 7 | FALSE | FALSE |
| 12_v.txt | 32301 | 32301-SNV-( | SNV | 1 | G | T | 23 | 29225 | 0.07869974 | 43.6521739 | rnr1D57N | FALSE | CAN | JSY14 | 1 | rnr1D57N-Re | GGC | G>T | A/T+1 | G/C-1 | CG>AT | CG>AT | TTTGCGGCA | GCAGAAA | CCG | CCG | C | A | C>A | C>A_CCG | CG>AT_GCAGAAA | 7 | FALSE | FALSE |
| 12_v.txt | 32368 | 32368-SNV-( | SNV | 1 | C | A | 48 | 29225 | 0.16424294 | 58.25 | rnr1D57N | FALSE | CAN | JSY14 | 1 | rnr1D57N-Re | TCA | C>A | A/T+1 | G/C-1 | CG>AT | CG>AT | AGTTCTCAAT | CAATAGC | AGT | TCA | C | T | C>A | C>A_TCA | CG>AT_CAATAGC | 7 | FALSE | FALSE |
| 12_v.txt | 32373 | 32373-SNV-( | SNV | 1 | G | T | 26 | 29225 | 0.08896493 | 56.7307692 | rnr1D57N | FALSE | CAN | JSY14 | 1 | rnr1D57N-Re | AGC | G>T | A/T+1 | G/C-1 | CG>AT | CG>AT | TCAATAGCA | GCAATAA | TCG | TCG | C | A | C>A | C>A_TCG | CG>AT_GCAATAA | 7 | FALSE | FALSE |
| 12_v.txt | 32399 | 32399-SNV-( | SNV | 1 | G | T | 151 | 29225 | 0.51668092 | 57.5298013 | rnr1D57N | FALSE | CAN | JSY14 | 1 | rnr1D57N-Re | CGT | G>T | A/T+1 | G/C-1 | CG>AT | CG>AT | AGAAACGTA | GTAGGAA | GCA | GCA | C | A | C>A | C>A_GCA | CG>AT_GTAGGAA | 7 | FALSE | FALSE |
| 12_v.txt | 32432 | 32432-SNV-( | SNV | 1 | G | T | 34 | 29225 | 0.11633875 | 58.0588235 | rnr1D57N | FALSE | CAN | JSY14 | 1 | rnr1D57N-Re | TGT | G>T | A/T+1 | G/C-1 | CG>AT | CG>AT | GTCATTGTAT | GTATGGA | ACA | ACA | C | A | C>A | C>A_ACA | CG>AT_GTATGGA | 7 | FALSE | FALSE |
| 12_v.txt | 32449 | 32449-SNV-( | SNV | 1 | C | A | 30 | 29225 | 0.10265184 | 51.4 | rnr1D57N | FALSE | CAN | JSY14 | 1 | rnr1D57N-Re | CCA | C>A | A/T+1 | G/C-1 | CG>AT | CG>AT | AAAGTCCAA | CAATGAA | GGT | CCA | C | T | C>A | C>A_CCA | CG>AT_CAATGAA | 7 | FALSE | FALSE |
| 12_v.txt | 32463 | 32463-SNV-( | SNV | 1 | G | T | 23 | 29225 | 0.07869974 | 46.173913 | rnr1D57N | FALSE | CAN | JSY14 | 1 | rnr1D57N-Re | AGA | G>T | A/T+1 | G/C-1 | CG>AT | CG>AT | AATAGAGAG | GAGCCAA | TCT | TCT | C | A | C>A | C>A_TCT | CG>AT_GAGCCAA | 7 | FALSE | FALSE |
| 12_v.txt | 32471 | 32471-SNV-( | SNV | 1 | G | T | 70 | 29225 | 0.23952096 | 46.9714286 | rnr1D57N | FALSE | CAN | JSY14 | 1 | rnr1D57N-Re | TGT | G>T | A/T+1 | G/C-1 | CG>AT | CG>AT | GCCAATGTA | GTAGAAG | ACA | ACA | C | A | C>A | C>A_ACA | CG>AT_GTAGAAG | 7 | FALSE | FALSE |
| 12_v.txt | 32494 | 32494-SNV-( | SNV | 1 | C | A | 267 | 29224 | 0.91363263 | 47.5240588 | rnr1D57N | FALSE | CAN | JSY14 | 1 | rnr1D57N-Re | ACA | C>A | A/T+1 | G/C-1 | CG>AT | CG>AT | GGAAAACAA | CAACTTT | TGT | ACA | C | T | C>A | C>A_ACA | CG>AT_CAACTTT | 7 | FALSE | FALSE |
| 12_v.txt | 32508 | 32508-SNV-( | SNV | 1 | G | T | 104 | 29225 | 0.35585971 | 43.5769231 | rnr1D57N | FALSE | CAN | JSY14 | 1 | rnr1D57N-Re | GGC | G>T | A/T+1 | G/C-1 | CG>AT | CG>AT | TTGATGGCT | GCTCTTG | CCG | CCG | C | A | C>A | C>A_CCG | CG>AT_GCTCTTG | 7 | FALSE | FALSE |
| 12_v.txt | 32536 | 32536-SNV-( | SNV | 1 | C | A | 9 | 29225 | 0.03079555 | 30.5555556 | rnr1D57N | FALSE | CAN | JSY14 | 1 | rnr1D57N-Re | GCA | C>A | A/T+1 | G/C-1 | CG>AT | CG>AT | GGTTTGCAG | CAGCTTC | CGT | GCA | C | T | C>A | C>A_GCA | CG>AT_CAGCTTC | 7 | FALSE | FALSE |
| 12_v.txt | 32542 | 32542-SNV-( | SNV | 1 | C | A | 25 | 29225 | 0.0855432 | 35.9599984 | rnr1D57N | FALSE | CAN | JSY14 | 1 | rnr1D57N-Re | TCA | C>A | A/T+1 | G/C-1 | CG>AT | CG>AT | CAGCTTCAC | CACCAGC | AGT | TCA | C | T | C>A | C>A_TCA | CG>AT_CACCAGC | 7 | FALSE | FALSE |
| 12_v.txt | 32545 | 32545-SNV-( | SNV | 1 | C | A | 53 | 29224 | 0.18135779 | 36.824479 | rnr1D57N | FALSE | CAN | JSY14 | 1 | rnr1D57N-Re | CCA | C>A | A/T+1 | G/C-1 | CG>AT | CG>AT | CTTCACCAG | CAGCAGT | GGT | CCA | C | T | C>A | C>A_CCA | CG>AT_CAGCAGT | 7 | FALSE | FALSE |
| 12_v.txt | 32556 | 32556-SNV-( | SNV | 1 | C | A | 20 | 29210 | 0.0684697 | 35.2499832 | rnr1D57N | FALSE | CAN | JSY14 | 1 | rnr1D57N-Re | ACC | C>A | A/T+1 | G/C-1 | CG>AT | CG>AT | GTGATACCA | CCAACTA | TGG | ACC | C | T | C>A | C>A_ACC | CG>AT_CCAACTA | 7 | FALSE | FALSE |
| 12_v.txt | 32566 | 32566-SNV-( | SNV | 1 | C | A | 47 | 26044 | 0.18046383 | 37.4893617 | rnr1D57N | FALSE | CAN | JSY14 | 1 | rnr1D57N-Re | TCA | C>A | A/T+1 | G/C-1 | CG>AT | CG>AT | CTAGTTCAGT | CAGTACC | AGT | TCA | C | T | C>A | C>A_TCA | CG>AT_CAGTACC | 7 | FALSE | FALSE |
| 12_v.txt | 32575 | 32575-SNV-( | SNV | 1 | G | T | 38 | 26053 | 0.14585652 | 35.31059 | rnr1D57N | FALSE | CAN | JSY14 | 1 | rnr1D57N-Re | TGA | G>T | A/T+1 | G/C-1 | CG>AT | CG>AT | TACCTTGAA | GAAATGT | ACT | ACT | C | A | C>A | C>A_ACT | CG>AT_GAAATGT | 7 | FALSE | FALSE |
| 12_v.txt | 32580 | 32580-SNV-( | SNV | 1 | G | T | 26 | 26053 | 0.09979657 | 36.5726447 | rnr1D57N | FALSE | CAN | JSY14 | 1 | rnr1D57N-Re | TGT | G>T | A/T+1 | G/C-1 | CG>AT | CG>AT | TGAAATGTG | GTGAAGG | ACA | ACA | C | A | C>A | C>A_ACA | CG>AT_GTGAAGG | 7 | FALSE | FALSE |
| 12_v.txt | 32629 | 32629-SNV-( | SNV | 1 | C | A | 428 | 26053 | 1.64280505 | 55.6962617 | rnr1D57N | FALSE | CAN | JSY14 | 1 | rnr1D57N-Re | TCG | C>A | A/T+1 | G/C-1 | CG>AT | CG>AT | TCCCTTCGTT | CGTTTTT | AGC | TCG | C | T | C>A | C>A_TCG | CG>AT_CGTTTTT | 7 | FALSE | FALSE |
| 12_v.txt | 32658 | 32658-SNV-( | SNV | 1 | C | A | 18 | 26053 | 0.06908993 | 25.780858 | rnr1D57N | FALSE | CAN | JSY14 | 1 | rnr1D57N-Re | ACC | C>A | A/T+1 | G/C-1 | CG>AT | CG>AT | CCTGGACCC | CCCCAGG | TGG | ACC | C | T | C>A | C>A_ACC | CG>AT_CCCCAGG | 7 | FALSE | FALSE |
| 12_v.txt | 32678 | 32678-SNV-( | SNV | 1 | C | A | 13 | 26053 | 0.04989828 | 25.4615385 | rnr1D57N | FALSE | CAN | JSY14 | 1 | rnr1D57N-Re | TCC | C>A | A/T+1 | G/C-1 | CG>AT | CG>AT | GTTTCTCCAA | CCAATAA | AGG | TCC | C | T | C>A | C>A_TCC | CG>AT_CCAATAA | 7 | FALSE | FALSE |
| 12_v.txt | 32692 | 32692-SNV-( | SNV | 1 | C | A | 63 | 26053 | 0.24181476 | 57.8571429 | rnr1D57N | FALSE | CAN | JSY14 | 1 | rnr1D57N-Re | CCA | C>A | A/T+1 | G/C-1 | CG>AT | CG>AT | GGAATCCAA | CAACTGG | GGT | CCA | C | T | C>A | C>A_CCA | CG>AT_CAACTGG | 7 | FALSE | FALSE |
| 12_v.txt | 32733 | 32733-SNV-( | SNV | 1 | C | A | 311 | 26053 | 1.19372049 | 59.6607106 | rnr1D57N | FALSE | CAN | JSY14 | 1 | rnr1D57N-Re | ACA | C>A | A/T+1 | G/C-1 | CG>AT | CG>AT | CAAAAACAG | CAGTATA | TGT | ACA | C | T | C>A | C>A_ACA | CG>AT_CAGTATA | 7 | FALSE | FALSE |
| 12_v.txt | 32735 | 32735-SNV-( | SNV | 1 | G | T | 30 | 26053 | 0.11514989 | 56.3983157 | rnr1D57N | FALSE | CAN | JSY14 | 1 | rnr1D57N-Re | AGT | G>T | A/T+1 | G/C-1 | CG>AT | CG>AT | AAAACAGTA | GTATATT | TCA | TCA | C | A | C>A | C>A_TCA | CG>AT_GTATATT | 7 | FALSE | FALSE |
| 12_v.txt | 32780 | 32780-SNV-( | SNV | 1 | C | A | 10 | 26053 | 0.0383833 | 30.8884262 | rnr1D57N | FALSE | CAN | JSY14 | 1 | rnr1D57N-Re | CCC | C>A | A/T+1 | G/C-1 | CG>AT | CG>AT | AGCGACCCA | CCAGAAC | GGG | CCC | C | T | C>A | C>A_CCC | CG>AT_CCAGAAC | 7 | FALSE | FALSE |
| 12_v.txt | 32788 | 32788-SNV-( | SNV | 1 | C | A | 76 | 26052 | 0.29172424 | 48.0526316 | rnr1D57N | FALSE | CAN | JSY14 | 1 | rnr1D57N-Re | TCG | C>A | A/T+1 | G/C-1 | CG>AT | CG>AT | AGAACTCGA | CGAATTC | AGC | TCG | C | T | C>A | C>A_TCG | CG>AT_CGAATTC | 7 | FALSE | FALSE |
| 12_v.txt | 32794 | 32794-SNV-( | SNV | 1 | C | A | 58 | 26053 | 0.22262311 | 41.9827586 | rnr1D57N | FALSE | CAN | JSY14 | 1 | rnr1D57N-Re | TCA | C>A | A/T+1 | G/C-1 | CG>AT | CG>AT | CGAATTCAC | CACCGTA | AGT | TCA | C | T | C>A | C>A_TCA | CG>AT_CACCGTA | 7 | FALSE | FALSE |
| 12_v.txt | 32797 | 32797-SNV-( | SNV | 1 | C | A | 5 | 26053 | 0.01919165 | 42.9999981 | rnr1D57N | FALSE | CAN | JSY14 | 1 | rnr1D57N-Re | CCG | C>A | A/T+1 | G/C-1 | CG>AT | CG>AT | ATTCACCGTA | CGTAATA | GGC | CCG | C | T | C>A | C>A_CCG | CG>AT_CGTAATA | 7 | FALSE | FALSE |
| 12_v.txt | 32809 | 32809-SNV-( | SNV | 1 | C | A | 14 | 26053 | 0.05373661 | 37.4285714 | rnr1D57N | FALSE | CAN | JSY14 | 1 | rnr1D57N-Re | ACA | C>A | A/T+1 | G/C-1 | CG>AT | CG>AT | ATTTGACAG | CAGGGAA | TGT | ACA | C | T | C>A | C>A_ACA | CG>AT_CAGGGAA | 7 | FALSE | FALSE |
| 12_v.txt | 32819 | 32819-SNV-( | SNV | 1 | G | T | 17 | 26053 | 0.0652516 | 37.0588235 | rnr1D57N | FALSE | CAN | JSY14 | 1 | rnr1D57N-Re | AGT | G>T | A/T+1 | G/C-1 | CG>AT | CG>AT | GAACAAGTT | GTTCATT | TCA | TCA | C | A | C>A | C>A_TCA | CG>AT_GTTCATT | 7 | FALSE | FALSE |
| 12_v.txt | 32841 | 32841-SNV-( | SNV | 1 | C | A | 6 | 26053 | 0.02302998 | 46.662951 | rnr1D57N | FALSE | CAN | JSY14 | 1 | rnr1D57N-Re | CCA | C>A | A/T+1 | G/C-1 | CG>AT | CG>AT | ATTACCCAA | CAAAAAA | GGT | CCA | C | T | C>A | C>A_CCA | CG>AT_CAAAAAA | 7 | FALSE | FALSE |
| 12_v.txt | 32876 | 32876-SNV-( | SNV | 1 | G | T | 98 | 21306 | 0.45996433 | 35.6530612 | rnr1D57N | FALSE | CAN | JSY14 | 1 | rnr1D57N-Re | TGT | G>T | A/T+1 | G/C-1 | CG>AT | CG>AT | AACTTTGTA | GTACGTC | ACA | ACA | C | A | C>A | C>A_ACA | CG>AT_GTACGTC | 7 | FALSE | FALSE |
| 12_v.txt | 32900 | 32900-SNV-( | SNV | 1 | G | T | 17 | 21315 | 0.07975604 | 19.9430464 | rnr1D57N | FALSE | CAN | JSY14 | 1 | rnr1D57N-Re | GGC | G>T | A/T+1 | G/C-1 | CG>AT | CG>AT | GACTTGGCC | GCCAACT | CCG | CCG | C | A | C>A | C>A_CCG | CG>AT_GCCAACT | 7 | FALSE | FALSE |
| 12_v.txt | 32901 | 32901-SNV-( | SNV | 1 | C | A | 6 | 21315 | 0.02814919 | 38 | rnr1D57N | FALSE | CAN | JSY14 | 1 | rnr1D57N-Re | GCC | C>A | A/T+1 | G/C-1 | CG>AT | CG>AT | ACTTGGCCA | CCAACTA | CGG | GCC | C | T | C>A | C>A_GCC | CG>AT_CCAACTA | 7 | FALSE | FALSE |
| 12_v.txt | 32902 | 32902-SNV-( | SNV | 1 | C | A | 3 | 21315 | 0.0140746 | 20.6668028 | rnr1D57N | FALSE | CAN | JSY14 | 1 | rnr1D57N-Re | CCA | C>A | A/T+1 | G/C-1 | CG>AT | CG>AT | CTTGGCCAA | CAACTAC | GGT | CCA | C | T | C>A | C>A_CCA | CG>AT_CAACTAC | 7 | FALSE | FALSE |
| 12_v.txt | 32908 | 32908-SNV-( | SNV | 1 | C | A | 46 | 21315 | 0.21581046 | 48 | rnr1D57N | FALSE | CAN | JSY14 | 1 | rnr1D57N-Re | ACA | C>A | A/T+1 | G/C-1 | CG>AT | CG>AT | CAACTACACT | CACTAAG | TGT | ACA | C | T | C>A | C>A_ACA | CG>AT_CACTAAG | 7 | FALSE | FALSE |
| 12_v.txt | 32917 | 32917-SNV-( | SNV | 1 | C | A | 93 | 21315 | 0.43631246 | 51.7419355 | rnr1D57N | FALSE | CAN | JSY14 | 1 | rnr1D57N-Re | TCC | C>A | A/T+1 | G/C-1 | CG>AT | CG>AT | TAAGTTCCA | CCAGGGC | AGG | TCC | C | T | C>A | C>A_TCC | CG>AT_CCAGGGC | 7 | FALSE | FALSE |
| 12_v.txt | 32928 | 32928-SNV-( | SNV | 1 | G | T | 25 | 21315 | 0.1172883 | 60.28 | rnr1D57N | FALSE | CAN | JSY14 | 1 | rnr1D57N-Re | AGT | G>T | A/T+1 | G/C-1 | CG>AT | CG>AT | GCAAAAGTG | GTGATTG | TCA | TCA | C | A | C>A | C>A_TCA | CG>AT_GTGATTG | 7 | FALSE | FALSE |
| 12_v.txt | 32934 | 32934-SNV-( | SNV | 1 | G | T | 438 | 21315 | 2.05489092 | 52.1064128 | rnr1D57N | FALSE | CAN | JSY14 | 1 | rnr1D57N-Re | TGC | G>T | A/T+1 | G/C-1 | CG>AT | CG>AT | GTGATTGCC | GCCCAAG | ACG | ACG | C | A | C>A | C>A_ACG | CG>AT_GCCCAAG | 7 | FALSE | FALSE |
| 12_v.txt | 32935 | 32935-SNV-( | SNV | 1 | C | A | 3 | 21315 | 0.0140746 | 63.6666667 | rnr1D57N | FALSE | CAN | JSY14 | 1 | rnr1D57N-Re | GCC | C>A | A/T+1 | G/C-1 | CG>AT | CG>AT | TGATTGCCC | CCCAAGA | CGG | GCC | C | T | C>A | C>A_GCC | CG>AT_CCCAAGA | 7 | FALSE | FALSE |
| 12_v.txt | 32937 | 32937-SNV-( | SNV | 1 | C | A | 13 | 21315 | 0.06098991 | 52.9063552 | rnr1D57N | FALSE | CAN | JSY14 | 1 | rnr1D57N-Re | CCA | C>A | A/T+1 | G/C-1 | CG>AT | CG>AT | ATTGCCCAA | CAAGAAA | GGT | CCA | C | T | C>A | C>A_CCA | CG>AT_CAAGAAA | 7 | FALSE | FALSE |
| 12_v.txt | 32940 | 32940-SNV-( | SNV | 1 | G | T | 18 | 21315 | 0.08444757 | 38.7222222 | rnr1D57N | FALSE | CAN | JSY14 | 1 | rnr1D57N-Re | AGA | G>T | A/T+1 | G/C-1 | CG>AT | CG>AT | GCCCAAGAA | GAAAACC | TCT | TCT | C | A | C>A | C>A_TCT | CG>AT_GAAAACC | 7 | FALSE | FALSE |
| 12_v.txt | 32970 | 32970-SNV-( | SNV | 1 | C | A | 24 | 21314 | 0.11260205 | 59.625 | rnr1D57N | FALSE | CAN | JSY14 | 1 | rnr1D57N-Re | ACC | C>A | A/T+1 | G/C-1 | CG>AT | CG>AT | GCCGCACCA | CCAAATG | TGG | ACC | C | T | C>A | C>A_ACC | CG>AT_CCAAATG | 7 | FALSE | FALSE |
| 12_v.txt | 33012 | 33012-SNV-( | SNV | 1 | G | T | 11 | 21315 | 0.05160685 | 61.3636364 | rnr1D57N | FALSE | CAN | JSY14 | 1 | rnr1D57N-Re | AGA | G>T | A/T+1 | G/C-1 | CG>AT | CG>AT | GTGAAAGAG | GAGGATG | TCT | TCT | C | A | C>A | C>A_TCT | CG>AT_GAGGATG | 7 | FALSE | FALSE |
| 12_v.txt | 33025 | 33025-SNV-( | SNV | 1 | G | T | 14 | 21315 | 0.06568145 | 56.8571429 | rnr1D57N | FALSE | CAN | JSY14 | 1 | rnr1D57N-Re | GGG | G>T | A/T+1 | G/C-1 | CG>AT | CG>AT | TAACAGGGA | GGATGAA | CCC | CCC | C | A | C>A | C>A_CCC | CG>AT_GGATGAA | 7 | FALSE | FALSE |
| 12_v.txt | 33033 | 33033-SNV-( | SNV | 1 | G | T | 38 | 21315 | 0.17827821 | 59.1842105 | rnr1D57N | FALSE | CAN | JSY14 | 1 | rnr1D57N-Re | TGT | G>T | A/T+1 | G/C-1 | CG>AT | CG>AT | ATGAATGTA | GTAGCCA | ACA | ACA | C | A | C>A | C>A_ACA | CG>AT_GTAGCCA | 7 | FALSE | FALSE |
| 12_v.txt | 33043 | 33043-SNV-( | SNV | 1 | C | A | 37 | 21315 | 0.17358668 | 63.0810811 | rnr1D57N | FALSE | CAN | JSY14 | 1 | rnr1D57N-Re | TCA | C>A | A/T+1 | G/C-1 | CG>AT | CG>AT | CCATTTCACC | CACCCAA | AGT | TCA | C | T | C>A | C>A_TCA | CG>AT_CACCCAA | 7 | FALSE | FALSE |
| 12_v.txt | 33051 | 33051-SNV-( | SNV | 1 | G | T | 39 | 21315 | 0.18296974 | 59.9230769 | rnr1D57N | FALSE | CAN | JSY14 | 1 | rnr1D57N-Re | GGA | G>T | A/T+1 | G/C-1 | CG>AT | CG>AT | CCCAAGGAC | GACTGCG | CCT | CCT | C | A | C>A | C>A_CCT | CG>AT_GACTGCG | 7 | FALSE | FALSE |
| 12_v.txt | 33056 | 33056-SNV-( | SNV | 1 | C | A | 7 | 21314 | 0.03284226 | 40.2914586 | rnr1D57N | FALSE | CAN | JSY14 | 1 | rnr1D57N-Re | GCG | C>A | A/T+1 | G/C-1 | CG>AT | CG>AT | GGACTGCGT | CGTGACA | CGC | GCG | C | T | C>A | C>A_GCG | CG>AT_CGTGACA | 7 | FALSE | FALSE |
| 12_v.txt | 33114 | 33114-SNV-( | SNV | 1 | C | A | 5 | 21315 | 0.02345766 | 47.9975324 | rnr1D57N | FALSE | CAN | JSY14 | 1 | rnr1D57N-Re | GCC | C>A | A/T+1 | G/C-1 | CG>AT | CG>AT | ACTGGGCCG | CCGGCGT | CGG | GCC | C | T | C>A | C>A_GCC | CG>AT_CCGGCGT | 7 | FALSE | FALSE |
| 12_v.txt | 33115 | 33115-SNV-( | SNV | 1 | C | A | 121 | 21315 | 0.56767535 | 46.5619835 | rnr1D57N | FALSE | CAN | JSY14 | 1 | rnr1D57N-Re | CCG | C>A | A/T+1 | G/C-1 | CG>AT | CG>AT | CTGGGCCGG | CGGCGTT | GGC | CCG | C | T | C>A | C>A_CCG | CG>AT_CGGCGTT | 7 | FALSE | FALSE |
| 12_v.txt | 33118 | 33118-SNV-( | SNV | 1 | C | A | 25 | 21315 | 0.1172883 | 41.4 | rnr1D57N | FALSE | CAN | JSY14 | 1 | rnr1D57N-Re | GCG | C>A | A/T+1 | G/C-1 | CG>AT | CG>AT | GGCCGGCGT | CGTTGGT | CGC | GCG | C | T | C>A | C>A_GCG | CG>AT_CGTTGGT | 7 | FALSE | FALSE |
| 12_v.txt | 33151 | 33151-SNV-( | SNV | 1 | G | T | 27 | 21315 | 0.12667136 | 38.8888889 | rnr1D57N | FALSE | CAN | JSY14 | 1 | rnr1D57N-Re | AGA | G>T | A/T+1 | G/C-1 | CG>AT | CG>AT | TGAAAAGAC | GACCTGT | TCT | TCT | C | A | C>A | C>A_TCT | CG>AT_GACCTGT | 7 | FALSE | FALSE |
| 12_v.txt | 33153 | 33153-SNV-( | SNV | 1 | C | A | 12 | 21315 | 0.05629838 | 37.5 | rnr1D57N | FALSE | CAN | JSY14 | 1 | rnr1D57N-Re | ACC | C>A | A/T+1 | G/C-1 | CG>AT | CG>AT | AAAAGACCT | CCTGTAC | TGG | ACC | C | T | C>A | C>A_ACC | CG>AT_CCTGTAC | 7 | FALSE | FALSE |
| 12_v.txt | 33154 | 33154-SNV-( | SNV | 1 | C | A | 24 | 21315 | 0.11259676 | 35.8751499 | rnr1D57N | FALSE | CAN | JSY14 | 1 | rnr1D57N-Re | CCT | C>A | A/T+1 | G/C-1 | CG>AT | CG>AT | AAAGACCTG | CTGTACC | GGA | CCT | C | T | C>A | C>A_CCT | CG>AT_CTGTACC | 7 | FALSE | FALSE |
| 12_v.txt | 33159 | 33159-SNV-( | SNV | 1 | C | A | 10 | 21315 | 0.04691532 | 35.2943098 | rnr1D57N | FALSE | CAN | JSY14 | 1 | rnr1D57N-Re | ACC | C>A | A/T+1 | G/C-1 | CG>AT | CG>AT | CCTGTACCA | CCAATAG | TGG | ACC | C | T | C>A | C>A_ACC | CG>AT_CCAATAG | 7 | FALSE | FALSE |
| 12_v.txt | 33160 | 33160-SNV-( | SNV | 1 | C | A | 16 | 21315 | 0.07506451 | 37.3121485 | rnr1D57N | FALSE | CAN | JSY14 | 1 | rnr1D57N-Re | CCA | C>A | A/T+1 | G/C-1 | CG>AT | CG>AT | CTGTACCAAT | CAATAGT | GGT | CCA | C | T | C>A | C>A_CCA | CG>AT_CAATAGT | 7 | FALSE | FALSE |
| 12_v.txt | 33168 | 33168-SNV-( | SNV | 1 | C | A | 28 | 21296 | 0.13148009 | 37.9642857 | rnr1D57N | FALSE | CAN | JSY14 | 1 | rnr1D57N-Re | ACC | C>A | A/T+1 | G/C-1 | CG>AT | CG>AT | ATAGTACCA | CCACCAA | TGG | ACC | C | T | C>A | C>A_ACC | CG>AT_CCACCAA | 7 | FALSE | FALSE |
| 12_v.txt | 33169 | 33169-SNV-( | SNV | 1 | C | A | 170 | 21296 | 0.79827198 | 37.7056057 | rnr1D57N | FALSE | CAN | JSY14 | 1 | rnr1D57N-Re | CCA | C>A | A/T+1 | G/C-1 | CG>AT | CG>AT | TAGTACCAC | CACCAAG | GGT | CCA | C | T | C>A | C>A_CCA | CG>AT_CACCAAG | 7 | FALSE | FALSE |
| 12_v.txt | 33177 | 33177-SNV-( | SNV | 1 | G | T | 11 | 21290 | 0.05166745 | 37.8181818 | rnr1D57N | FALSE | CAN | JSY14 | 1 | rnr1D57N-Re | GGC | G>T | A/T+1 | G/C-1 | CG>AT | CG>AT | CCAAGGGCA | GCAATCA | CCG | CCG | C | A | C>A | C>A_CCG | CG>AT_GCAATCA | 7 | FALSE | FALSE |
| 12_v.txt | 33210 | 33210-SNV-( | SNV | 1 | C | A | 18 | 20300 | 0.08866995 | 42.7222222 | rnr1D57N | FALSE | CAN | JSY14 | 1 | rnr1D57N-Re | TCT | C>A | A/T+1 | G/C-1 | CG>AT | CG>AT | AGCTCTCTCT | CTCTTCA | AGA | TCT | C | T | C>A | C>A_TCT | CG>AT_CTCTTCA | 7 | FALSE | FALSE |
| 12_v.txt | 33220 | 33220-SNV-( | SNV | 1 | C | A | 33 | 20312 | 0.16246554 | 41.4545455 | rnr1D57N | FALSE | CAN | JSY14 | 1 | rnr1D57N-Re | TCA | C>A | A/T+1 | G/C-1 | CG>AT | CG>AT | TCACTTCAGC | CAGCGTT | AGT | TCA | C | T | C>A | C>A_TCA | CG>AT_CAGCGTT | 7 | FALSE | FALSE |
| 12_v.txt | 33247 | 33247-SNV-( | SNV | 1 | C | A | 22 | 20312 | 0.10831036 | 41.7272727 | rnr1D57N | FALSE | CAN | JSY14 | 1 | rnr1D57N-Re | TCA | C>A | A/T+1 | G/C-1 | CG>AT | CG>AT | CATCTTCATC | CATCACC | AGT | TCA | C | T | C>A | C>A_TCA | CG>AT_CATCACC | 7 | FALSE | FALSE |
| 12_v.txt | 33325 | 33325-SNV-( | SNV | 1 | C | A | 360 | 20312 | 1.77235132 | 59.7194444 | rnr1D57N | FALSE | CAN | JSY14 | 1 | rnr1D57N-Re | TCT | C>A | A/T+1 | G/C-1 | CG>AT | CG>AT | ACAATTCTTT | CTTTACT | AGA | TCT | C | T | C>A | C>A_TCT | CG>AT_CTTTACT | 7 | FALSE | FALSE |
| 12_v.txt | 33354 | 33354-SNV-( | SNV | 1 | G | T | 163 | 20305 | 0.80275794 | 29.8682994 | rnr1D57N | FALSE | CAN | JSY14 | 1 | rnr1D57N-Re | TGA | G>T | A/T+1 | G/C-1 | CG>AT | CG>AT | GGTATTGAC | GACCCAC | ACT | ACT | C | A | C>A | C>A_ACT | CG>AT_GACCCAC | 7 | FALSE | FALSE |
| 12_v.txt | 33385 | 33385-SNV-( | SNV | 1 | C | A | 21 | 20312 | 0.10338716 | 54.1428571 | rnr1D57N | FALSE | CAN | JSY14 | 1 | rnr1D57N-Re | TCA | C>A | A/T+1 | G/C-1 | CG>AT | CG>AT | AAGCTTCAA | CAACGTC | AGT | TCA | C | T | C>A | C>A_TCA | CG>AT_CAACGTC | 7 | FALSE | FALSE |
| 12_v.txt | 33415 | 33415-SNV-( | SNV | 1 | C | A | 36 | 20312 | 0.17723513 | 34.6388889 | rnr1D57N | FALSE | CAN | JSY14 | 1 | rnr1D57N-Re | TCA | C>A | A/T+1 | G/C-1 | CG>AT | CG>AT | CCGGCTCATT | CATTGTA | AGT | TCA | C | T | C>A | C>A_TCA | CG>AT_CATTGTA | 7 | FALSE | FALSE |
| 12_v.txt | 33433 | 33433-SNV-( | SNV | 1 | C | A | 27 | 20312 | 0.13292635 | 41.7407407 | rnr1D57N | FALSE | CAN | JSY14 | 1 | rnr1D57N-Re | TCC | C>A | A/T+1 | G/C-1 | CG>AT | CG>AT | GCTTCTCCTC | CCTCTAT | AGG | TCC | C | T | C>A | C>A_TCC | CG>AT_CCTCTAT | 7 | FALSE | FALSE |
| 12_v.txt | 33436 | 33436-SNV-( | SNV | 1 | C | A | 34 | 20312 | 0.16738874 | 47.3529412 | rnr1D57N | FALSE | CAN | JSY14 | 1 | rnr1D57N-Re | TCT | C>A | A/T+1 | G/C-1 | CG>AT | CG>AT | TCTCCTCTAT | CTATGTC | AGA | TCT | C | T | C>A | C>A_TCT | CG>AT_CTATGTC | 7 | FALSE | FALSE |
| 12_v.txt | 33456 | 33456-SNV-( | SNV | 1 | G | T | 33 | 20312 | 0.16246554 | 25.6060606 | rnr1D57N | FALSE | CAN | JSY14 | 1 | rnr1D57N-Re | TGA | G>T | A/T+1 | G/C-1 | CG>AT | CG>AT | TCTTTTGAAT | GAATTTG | ACT | ACT | C | A | C>A | C>A_ACT | CG>AT_GAATTTG | 7 | FALSE | FALSE |
| 12_v.txt | 33474 | 33474-SNV-( | SNV | 1 | C | A | 42 | 20311 | 0.2067845 | 33.730436 | rnr1D57N | FALSE | CAN | JSY14 | 1 | rnr1D57N-Re | GCC | C>A | A/T+1 | G/C-1 | CG>AT | CG>AT | GCTATGCCTT | CCTTTTT | CGG | GCC | C | T | C>A | C>A_GCC | CG>AT_CCTTTTT | 7 | FALSE | FALSE |
| 12_v.txt | 33475 | 33475-SNV-( | SNV | 1 | C | A | 19 | 20311 | 0.09354537 | 37.0011018 | rnr1D57N | FALSE | CAN | JSY14 | 1 | rnr1D57N-Re | CCT | C>A | A/T+1 | G/C-1 | CG>AT | CG>AT | CTATGCCTTT | CTTTTTT | GGA | CCT | C | T | C>A | C>A_CCT | CG>AT_CTTTTTT | 7 | FALSE | FALSE |
| 12_v.txt | 31806 | 31806-Repla | Replacement | 2 | C | AA | 1 | 11167 | 8.96E-03 | 38.0006072 | rnr1D57N | FALSE | CAN | JSY14 | 1 | rnr1D57N-Re | CCA | nope | >1bp-ins | G/C-1 | none | Replac. | ATCTTCCAA | CAAATAA | GGT | CCA | C | none | nope | nope_CCA | Replac._CAAATAA | 7 | FALSE | FALSE |
| 12_v.txt | 32164 | 32164-Repla | Replacement | 2 | CG | A | 13 | 16625 | 0.07819549 | 53.7692308 | rnr1D57N | FALSE | CAN | JSY14 | 1 | rnr1D57N-Re | GCG | nope | A/T+1 | >1bpdel | none | Replac. | CCAAAGCGC | CGCCAAA | CGC | GCG | CG | T | NA | NA_GCG | Replac._CGCCAAA | 7 | FALSE | FALSE |
| 12_v.txt | 32494 | 32494-Repla | Replacement | 2 | C | TA | 1 | 29225 | 3.42E-03 | 35.5 | rnr1D57N | FALSE | CAN | JSY14 | 1 | rnr1D57N-Re | ACA | nope | >1bp-ins | G/C-1 | none | Replac. | GGAAAACAA | CAACTTT | TGT | ACA | C | none | nope | nope_ACA | Replac._CAACTTT | 7 | FALSE | FALSE |
| 12_v.txt | 32814 | 32814-Repla | Replacement | 3 | AAC | G | 29 | 26053 | 0.11131156 | 42.9655172 | rnr1D57N | FALSE | CAN | JSY14 | 1 | rnr1D57N-Re | GAA | nope | G/C+1 | >1bpdel | none | Replac. | ACAGGGAAC | AACAAGT | CTT | GAA | AAC | C | NA | NA_GAA | Replac._AACAAGT | 7 | FALSE | FALSE |
| 12_v.txt | 32841 | 32841-Repla | Replacement | 2 | C | TA | 1 | 26053 | 3.84E-03 | 64 | rnr1D57N | FALSE | CAN | JSY14 | 1 | rnr1D57N-Re | CCA | nope | >1bp-ins | G/C-1 | none | Replac. | ATTACCCAA | CAAAAAA | GGT | CCA | C | none | nope | nope_CCA | Replac._CAAAAAA | 7 | FALSE | FALSE |
| 12_v.txt | 32937 | 32937-Repla | Replacement | 2 | C | TG | 1 | 21315 | 4.69E-03 | 22.5 | rnr1D57N | FALSE | CAN | JSY14 | 1 | rnr1D57N-Re | CCA | nope | >1bp-ins | G/C-1 | none | Replac. | ATTGCCCAA | CAAGAAA | GGT | CCA | C | none | nope | nope_CCA | Replac._CAAGAAA | 7 | FALSE | FALSE |
| 12_v.txt | 31875 | 31875-MNV- | MNV | 2 | TA | GG | 8 | 11167 | 0.07163965 | 47.4374983 | rnr1D57N | FALSE | CAN | JSY14 | 1 | rnr1D57N-Re | ATA | nope | >1bp-ins | >1bpdel | none | MNV | GAGATATAG | TAGGCGG | TAT | ATA | TA | none | NA | NA_ATA | MNV_TAGGCGG | 7 | FALSE | FALSE |
| 12_v.txt | 31956 | 31956-MNV- | MNV | 2 | GT | TA | 23 | 16625 | 0.13834587 | 36.8478015 | rnr1D57N | FALSE | CAN | JSY14 | 1 | rnr1D57N-Re | TGT | nope | >1bp-ins | >1bpdel | none | MNV | ATAAATGTG | GTGGCCG | ACA | TGT | GT | none | NA | NA_TGT | MNV_GTGGCCG | 7 | FALSE | FALSE |
| 12_v.txt | 32100 | 32100-MNV- | MNV | 2 | GT | TG | 6 | 16625 | 0.03609023 | 63.75 | rnr1D57N | FALSE | CAN | JSY14 | 1 | rnr1D57N-Re | AGT | nope | >1bp-ins | >1bpdel | none | MNV | ACACCAGTG | GTGATAT | TCA | AGT | GT | none | NA | NA_AGT | MNV_GTGATAT | 7 | FALSE | FALSE |
| 12_v.txt | 32292 | 32292-MNV- | MNV | 3 | GAA | ATT | 3 | 29225 | 0.01026518 | 37.3333334 | rnr1D57N | FALSE | CAN | JSY14 | 1 | rnr1D57N-Re | TGA | nope | >1bp-ins | >1bpdel | none | MNV | ATATTTGAA | GAATTTG | ACT | TGA | GAA | none | NA | NA_TGA | MNV_GAATTTG | 7 | FALSE | FALSE |
| 12_v.txt | 32293 | 32293-MNV- | MNV | 2 | AA | GC | 1 | 29225 | 3.42E-03 | 63.9999839 | rnr1D57N | FALSE | CAN | JSY14 | 1 | rnr1D57N-Re | GAA | nope | >1bp-ins | >1bpdel | none | MNV | TATTTGAAT | AATTTGC | CTT | GAA | AA | none | NA | NA_GAA | MNV_AATTTGC | 7 | FALSE | FALSE |
| 12_v.txt | 32541 | 32541-MNV- | MNV | 5 | TCACC | GATAA | 1 | 29225 | 3.42E-03 | 17.4 | rnr1D57N | FALSE | CAN | JSY14 | 1 | rnr1D57N-Re | TTC | nope | >1bp-ins | >1bpdel | none | MNV | GCAGCTTCA | TCACCAG | AAG | TTC | TCACC | none | NA | NA_TTC | MNV_TCACCAG | 7 | FALSE | FALSE |
| 12_v.txt | 32544 | 32544-MNV- | MNV | 2 | CC | AA | 3 | 29224 | 0.01026554 | 32.6666619 | rnr1D57N | FALSE | CAN | JSY14 | 1 | rnr1D57N-Re | ACC | nope | >1bp-ins | >1bpdel | none | MNV | GCTTCACCA | CCAGCAG | TGG | ACC | CC | none | NA | NA_ACC | MNV_CCAGCAG | 7 | FALSE | FALSE |
| 12_v.txt | 32787 | 32787-MNV- | MNV | 2 | TC | CT | 17 | 26053 | 0.0652516 | 47.1176469 | rnr1D57N | FALSE | CAN | JSY14 | 1 | rnr1D57N-Re | CTC | nope | >1bp-ins | >1bpdel | none | MNV | CAGAACTCG | TCGAATT | GAG | CTC | TC | none | NA | NA_CTC | MNV_TCGAATT | 7 | FALSE | FALSE |
| 12_v.txt | 32797 | 32797-MNV- | MNV | 2 | CG | TT | 51 | 26053 | 0.19575481 | 46.3621979 | rnr1D57N | FALSE | CAN | JSY14 | 1 | rnr1D57N-Re | CCG | nope | >1bp-ins | >1bpdel | none | MNV | ATTCACCGTA | CGTAATA | GGC | CCG | CG | none | NA | NA_CCG | MNV_CGTAATA | 7 | FALSE | FALSE |
| 12_v.txt | 32914 | 32914-MNV- | MNV | 2 | GT | AC | 45 | 21315 | 0.21111893 | 47.8945052 | rnr1D57N | FALSE | CAN | JSY14 | 1 | rnr1D57N-Re | AGT | nope | >1bp-ins | >1bpdel | none | MNV | CACTAAGTT | GTTCCAG | TCA | AGT | GT | none | NA | NA_AGT | MNV_GTTCCAG | 7 | FALSE | FALSE |
| 12_v.txt | 32958 | 32958-MNV- | MNV | 2 | CC | TG | 1 | 21315 | 4.69E-03 | 63.9999994 | rnr1D57N | FALSE | CAN | JSY14 | 1 | rnr1D57N-Re | ACC | nope | >1bp-ins | >1bpdel | none | MNV | ATGTAACCA | CCATTGG | TGG | ACC | CC | none | NA | NA_ACC | MNV_CCATTGG | 7 | FALSE | FALSE |
| 12_v.txt | 33114 | 33114-MNV- | MNV | 2 | CC | AA | 1 | 21315 | 4.69E-03 | 35.5 | rnr1D57N | FALSE | CAN | JSY14 | 1 | rnr1D57N-Re | GCC | nope | >1bp-ins | >1bpdel | none | MNV | ACTGGGCCG | CCGGCGT | CGG | GCC | CC | none | NA | NA_GCC | MNV_CCGGCGT | 7 | FALSE | FALSE |
| 12_v.txt | 33114 | 33114-MNV- | MNV | 2 | CC | TT | 1 | 21315 | 4.69E-03 | 38 | rnr1D57N | FALSE | CAN | JSY14 | 1 | rnr1D57N-Re | GCC | nope | >1bp-ins | >1bpdel | none | MNV | ACTGGGCCG | CCGGCGT | CGG | GCC | CC | none | NA | NA_GCC | MNV_CCGGCGT | 7 | FALSE | FALSE |
| 12_v.txt | 33153 | 33153-MNV- | MNV | 2 | CC | AA | 1 | 21315 | 4.69E-03 | 36 | rnr1D57N | FALSE | CAN | JSY14 | 1 | rnr1D57N-Re | ACC | nope | >1bp-ins | >1bpdel | none | MNV | AAAAGACCT | CCTGTAC | TGG | ACC | CC | none | NA | NA_ACC | MNV_CCTGTAC | 7 | FALSE | FALSE |
| 12_v.txt | 33159 | 33159-MNV- | MNV | 2 | CC | AA | 1 | 21315 | 4.69E-03 | 34 | rnr1D57N | FALSE | CAN | JSY14 | 1 | rnr1D57N-Re | ACC | nope | >1bp-ins | >1bpdel | none | MNV | CCTGTACCA | CCAATAG | TGG | ACC | CC | none | NA | NA_ACC | MNV_CCAATAG | 7 | FALSE | FALSE |
| 12_v.txt | 33159 | 33159-MNV- | MNV | 2 | CC | TT | 2 | 21315 | 9.38E-03 | 37.4999783 | rnr1D57N | FALSE | CAN | JSY14 | 1 | rnr1D57N-Re | ACC | nope | >1bp-ins | >1bpdel | none | MNV | CCTGTACCA | CCAATAG | TGG | ACC | CC | none | NA | NA_ACC | MNV_CCAATAG | 7 | FALSE | FALSE |
| 12_v.txt | 33168 | 33168-MNV- | MNV | 2 | CC | AT | 5 | 21296 | 0.02347859 | 37.999992 | rnr1D57N | FALSE | CAN | JSY14 | 1 | rnr1D57N-Re | ACC | nope | >1bp-ins | >1bpdel | none | MNV | ATAGTACCA | CCACCAA | TGG | ACC | CC | none | NA | NA_ACC | MNV_CCACCAA | 7 | FALSE | FALSE |
| 12_v.txt | 31723 | 31723-Insert | Insertion | 1 | - | A | 3 | 11167 | 0.02686487 | 64 | rnr1D57N | FALSE | CAN | JSY14 | 1 | rnr1D57N-Re | CAA | nope | A/T+1 | >1bpdel | none | A/T+1 | TGTCCCAAA | AAAAAGT | GTT | CAA | - | T | NA | NA_CAA | A/T+1_AAAAAGT | 7 | FALSE | FALSE |
| 12_v.txt | 31881 | 31881-Insert | Insertion | 2 | - | CA | 1 | 11167 | 8.96E-03 | 37 | rnr1D57N | FALSE | CAN | JSY14 | 1 | rnr1D57N-Re | GGC | nope | >1bp-ins | >1bpdel | none | >1bp-ins | TAGGCGGCA | GCAGCAA | CCG | GGC | - | none | NA | NA_GGC | >1bp-ins_GCAGCAA | 7 | FALSE | FALSE |
| 12_v.txt | 31883 | 31883-Insert | Insertion | 2 | - | AA | 1 | 11167 | 8.96E-03 | 38 | rnr1D57N | FALSE | CAN | JSY14 | 1 | rnr1D57N-Re | CAG | nope | >1bp-ins | >1bpdel | none | >1bp-ins | GGCGGCAGC | AGCAAAG | GTC | CAG | - | none | NA | NA_CAG | >1bp-ins_AGCAAAG | 7 | FALSE | FALSE |
| 12_v.txt | 31883 | 31883-Insert | Insertion | 5 | - | AGCAA | 33 | 11167 | 0.29551357 | 44.4484849 | rnr1D57N | FALSE | CAN | JSY14 | 1 | rnr1D57N-Re | CAG | nope | >1bp-ins | >1bpdel | none | >1bp-ins | GGCGGCAGC | AGCAAAG | GTC | CAG | - | none | NA | NA_CAG | >1bp-ins_AGCAAAG | 7 | FALSE | FALSE |
| 12_v.txt | 31902 | 31902-Insert | Insertion | 1 | - | C | 11 | 11162 | 0.09854865 | 44.9090909 | rnr1D57N | FALSE | CAN | JSY14 | 1 | rnr1D57N-Re | GAA | nope | G/C+1 | >1bpdel | none | G/C+1 | CCATTGAATT | AATTTTG | CTT | GAA | - | G | NA | NA_GAA | G/C+1_AATTTTG | 7 | FALSE | FALSE |

| | | | | | | | | | | | | | | | | | | | | | | | | | | | | | | | | | | |
|---|---|---|---|---|---|---|---|---|---|---|---|---|---|---|---|---|---|---|---|---|---|---|---|---|---|---|---|---|---|---|---|---|---|---|
| 12_v.txt | 31913 | 31913-Insert | Insertion | 1 | - | A | 10 | 11162 | 0.08958968 | 38 | rnr1D57N | FALSE | CAN | JSY14 | 1 | rnr1D57N-Re | CAA | nope | A/T+1 | >1bpdel | none | A/T+1 | TGGTGCAAA | AAAAGCC | GTT | CAA | - | T | NA | NA_CAA | A/T+1_AAAAGCC | 7 | FALSE | FALSE |
| 12_v.txt | 31969 | 31969-Insert | Insertion | 1 | - | A | 3 | 16625 | 0.01804511 | 36.6666667 | rnr1D57N | FALSE | CAN | JSY14 | 1 | rnr1D57N-Re | TAA | nope | A/T+1 | >1bpdel | none | A/T+1 | CATAATAAG | AAGCCAA | ATT | TAA | - | T | NA | NA_TAA | A/T+1_AAGCCAA | 7 | FALSE | FALSE |
| 12_v.txt | 32067 | 32067-Insert | Insertion | 2 | - | AA | 10 | 16625 | 0.06015038 | 63.05 | rnr1D57N | FALSE | CAN | JSY14 | 1 | rnr1D57N-Re | GAT | nope | >1bp-ins | >1bpdel | none | >1bp-ins | ATTGAGATA | ATAAATA | CTA | GAT | - | none | NA | NA_GAT | >1bp-ins_ATAAATA | 7 | FALSE | FALSE |
| 12_v.txt | 32081 | 32081-Insert | Insertion | 1 | - | A | 246 | 16625 | 1.47969925 | 63.7926829 | rnr1D57N | FALSE | CAN | JSY14 | 1 | rnr1D57N-Re | CAA | nope | A/T+1 | >1bpdel | none | A/T+1 | CCATGCAAA | AAAAAAG | GTT | CAA | - | T | NA | NA_CAA | A/T+1_AAAAAAG | 7 | FALSE | FALSE |
| 12_v.txt | 32081 | 32081-Insert | Insertion | 2 | - | AA | 7 | 16625 | 0.04210526 | 63.6428571 | rnr1D57N | FALSE | CAN | JSY14 | 1 | rnr1D57N-Re | CAA | nope | >1bp-ins | >1bpdel | none | >1bp-ins | CCATGCAAA | AAAAAAG | GTT | CAA | - | none | NA | NA_CAA | >1bp-ins_AAAAAAG | 7 | FALSE | FALSE |
| 12_v.txt | 32087 | 32087-Insert | Insertion | 1 | - | G | 1 | 16625 | 6.02E-03 | 64 | rnr1D57N | FALSE | CAN | JSY14 | 1 | rnr1D57N-Re | AGC | nope | G/C+1 | >1bpdel | none | G/C+1 | AAAAAAGCC | GCCTGCA | TCG | AGC | - | C | NA | NA_AGC | G/C+1_GCCTGCA | 7 | FALSE | FALSE |
| 12_v.txt | 32166 | 32166-Insert | Insertion | 1 | - | C | 3 | 16625 | 0.01804511 | 46.6666667 | rnr1D57N | FALSE | CAN | JSY14 | 1 | rnr1D57N-Re | GCC | nope | G/C+1 | >1bpdel | none | G/C+1 | AAAGCGCCA | CCAAATG | CGG | GCC | - | G | NA | NA_GCC | G/C+1_CCAAATG | 7 | FALSE | FALSE |
| 12_v.txt | 32261 | 32261-Insert | Insertion | 1 | - | A | 5 | 29225 | 0.01710864 | 36.6 | rnr1D57N | FALSE | CAN | JSY14 | 1 | rnr1D57N-Re | CAA | nope | A/T+1 | >1bpdel | none | A/T+1 | TAGACCAAA | AAATAAA | GTT | CAA | - | T | NA | NA_CAA | A/T+1_AAATAAA | 7 | FALSE | FALSE |
| 12_v.txt | 32289 | 32289-Insert | Insertion | 1 | - | T | 34 | 29225 | 0.11633875 | 55.3823529 | rnr1D57N | FALSE | CAN | JSY14 | 1 | rnr1D57N-Re | ATT | nope | A/T+1 | >1bpdel | none | A/T+1 | TAAATATTTG | TTTGAAT | TAA | ATT | - | A | NA | NA_ATT | A/T+1_TTTGAAT | 7 | FALSE | FALSE |
| 12_v.txt | 32322 | 32322-Insert | Insertion | 1 | - | A | 23 | 29225 | 0.07869974 | 62.8695652 | rnr1D57N | FALSE | CAN | JSY14 | 1 | rnr1D57N-Re | GAT | nope | A/T+1 | >1bpdel | none | A/T+1 | GTTAAGATA | ATAACAG | CTA | GAT | - | T | NA | NA_GAT | A/T+1_ATAACAG | 7 | FALSE | FALSE |
| 12_v.txt | 32346 | 32346-Insert | Insertion | 1 | - | A | 3 | 29225 | 0.01026518 | 64 | rnr1D57N | FALSE | CAN | JSY14 | 1 | rnr1D57N-Re | CAA | nope | A/T+1 | >1bpdel | none | A/T+1 | TGTGGCAAA | AAAACCT | GTT | CAA | - | T | NA | NA_CAA | A/T+1_AAAACCT | 7 | FALSE | FALSE |
| 12_v.txt | 32352 | 32352-Insert | Insertion | 1 | - | T | 9 | 29225 | 0.03079555 | 61.5555556 | rnr1D57N | FALSE | CAN | JSY14 | 1 | rnr1D57N-Re | CTT | nope | A/T+1 | >1bpdel | none | A/T+1 | AAAACCTTTG | TTTGTAC | GAA | CTT | - | A | NA | NA_CTT | A/T+1_TTTGTAC | 7 | FALSE | FALSE |
| 12_v.txt | 32473 | 32473-Insert | Insertion | 3 | - | AGA | 46 | 29225 | 0.15739949 | 43.9130435 | rnr1D57N | FALSE | CAN | JSY14 | 1 | rnr1D57N-Re | TAG | nope | >1bp-ins | >1bpdel | none | >1bp-ins | CAATGTAGA | AGAAGGT | ATC | TAG | - | none | NA | NA_TAG | >1bp-ins_AGAAGGT | 7 | FALSE | FALSE |
| 12_v.txt | 32488 | 32488-Insert | Insertion | 1 | - | G | 1 | 29225 | 3.42E-03 | 64 | rnr1D57N | FALSE | CAN | JSY14 | 1 | rnr1D57N-Re | CGG | nope | G/C+1 | >1bpdel | none | G/C+1 | AGATACGGA | GGAAAAC | GCC | CGG | - | C | NA | NA_CGG | G/C+1_GGAAAAC | 7 | FALSE | FALSE |
| 12_v.txt | 32490 | 32490-Insert | Insertion | 1 | - | A | 265 | 29225 | 0.90675791 | 47.9735849 | rnr1D57N | FALSE | CAN | JSY14 | 1 | rnr1D57N-Re | GAA | nope | A/T+1 | >1bpdel | none | A/T+1 | ATACGGAAA | AAAACAA | CTT | GAA | - | T | NA | NA_GAA | A/T+1_AAAACAA | 7 | FALSE | FALSE |
| 12_v.txt | 32498 | 32498-Insert | Insertion | 1 | - | T | 23 | 29225 | 0.07869974 | 38.7826087 | rnr1D57N | FALSE | CAN | JSY14 | 1 | rnr1D57N-Re | CTT | nope | A/T+1 | >1bpdel | none | A/T+1 | AACAACTTTT | TTTTTTG | GAA | CTT | - | A | NA | NA_CTT | A/T+1_TTTTTTG | 7 | FALSE | FALSE |
| 12_v.txt | 32528 | 32528-Insert | Insertion | 1 | - | G | 21 | 29225 | 0.07185629 | 38.9047619 | rnr1D57N | FALSE | CAN | JSY14 | 1 | rnr1D57N-Re | TGG | nope | G/C+1 | >1bpdel | none | G/C+1 | TTTTCTGGGG | GGGGTTT | ACC | TGG | - | C | NA | NA_TGG | G/C+1_GGGGTTT | 7 | FALSE | FALSE |
| 12_v.txt | 32598 | 32598-Insert | Insertion | 2 | - | AA | 5 | 26053 | 0.01919165 | 46.5 | rnr1D57N | FALSE | CAN | JSY14 | 1 | rnr1D57N-Re | CAA | nope | >1bp-ins | >1bpdel | none | >1bp-ins | TTAATCAAAC | AAAGAGG | GTT | CAA | - | none | NA | NA_CAA | >1bp-ins_AAAGAGG | 7 | FALSE | FALSE |
| 12_v.txt | 32631 | 32631-Insert | Insertion | 1 | - | T | 157 | 26053 | 0.60261774 | 53.3121019 | rnr1D57N | FALSE | CAN | JSY14 | 1 | rnr1D57N-Re | GTT | nope | A/T+1 | >1bpdel | none | A/T+1 | CCTTCGTTTT | TTTTTAT | CAA | GTT | - | A | NA | NA_GTT | A/T+1_TTTTTAT | 7 | FALSE | FALSE |
| 12_v.txt | 32658 | 32658-Insert | Insertion | 1 | - | C | 1 | 26053 | 3.84E-03 | 64 | rnr1D57N | FALSE | CAN | JSY14 | 1 | rnr1D57N-Re | ACC | nope | G/C+1 | >1bpdel | none | G/C+1 | CCTGGACCCC | CCCCAGG | TGG | ACC | - | G | NA | NA_ACC | G/C+1_CCCCAGG | 7 | FALSE | FALSE |
| 12_v.txt | 32665 | 32665-Insert | Insertion | 27 | - | CACCTGGGT | 7 | 26053 | 0.02686831 | 52.6137566 | rnr1D57N | FALSE | CAN | JSY14 | 1 | rnr1D57N-Re | GCA | nope | >1bp-ins | >1bpdel | none | >1bp-ins | CCCAGGCAC | CACCTGG | CGT | GCA | - | none | NA | NA_GCA | >1bp-ins_CACCTGG | 7 | FALSE | FALSE |
| 12_v.txt | 32678 | 32678-Insert | Insertion | 37 | - | CCAATAACG | 28 | 26053 | 0.10747323 | 59.3996139 | rnr1D57N | FALSE | CAN | JSY14 | 1 | rnr1D57N-Re | TCC | nope | >1bp-ins | >1bpdel | none | >1bp-ins | GTTTCTCCAA | CCAATAA | AGG | TCC | - | none | NA | NA_TCC | >1bp-ins_CCAATAA | 7 | FALSE | FALSE |
| 12_v.txt | 32728 | 32728-Insert | Insertion | 1 | - | A | 360 | 26053 | 1.38179864 | 61.5444444 | rnr1D57N | FALSE | CAN | JSY14 | 1 | rnr1D57N-Re | CAA | nope | A/T+1 | >1bpdel | none | A/T+1 | CCATACAAA | AAAAACA | GTT | CAA | - | T | NA | NA_CAA | A/T+1_AAAAACA | 7 | FALSE | FALSE |
| 12_v.txt | 32760 | 32760-Insert | Insertion | 22 | - | AAAACTTTG | 16 | 26053 | 0.06141327 | 54.0909091 | rnr1D57N | FALSE | CAN | JSY14 | 1 | rnr1D57N-Re | TAA | nope | >1bp-ins | >1bpdel | none | >1bp-ins | ATGGCTAAA | AAAACTT | ATT | TAA | - | none | NA | NA_TAA | >1bp-ins_AAAACTT | 7 | FALSE | FALSE |
| 12_v.txt | 32802 | 32802-Insert | Insertion | 1 | - | T | 11 | 26053 | 0.04222163 | 49.3636364 | rnr1D57N | FALSE | CAN | JSY14 | 1 | rnr1D57N-Re | ATA | nope | A/T+1 | >1bpdel | none | A/T+1 | CCGTAATATT | TATTTGA | TAT | ATA | - | A | NA | NA_ATA | A/T+1_TATTTGA | 7 | FALSE | FALSE |
| 12_v.txt | 32839 | 32839-Insert | Insertion | 1 | - | C | 1 | 26053 | 3.84E-03 | 38 | rnr1D57N | FALSE | CAN | JSY14 | 1 | rnr1D57N-Re | ACC | nope | G/C+1 | >1bpdel | none | G/C+1 | TAATTACCCA | CCCAAAA | TGG | ACC | - | G | NA | NA_ACC | G/C+1_CCCAAAA | 7 | FALSE | FALSE |
| 12_v.txt | 32840 | 32840-Insert | Insertion | 1 | - | A | 9 | 26053 | 0.03454497 | 40.8888889 | rnr1D57N | FALSE | CAN | JSY14 | 1 | rnr1D57N-Re | CCC | nope | A/T+1 | >1bpdel | none | A/T+1 | AATTACCCA | CCAAAAA | GGG | CCC | - | T | NA | NA_CCC | A/T+1_CCAAAAA | 7 | FALSE | FALSE |
| 12_v.txt | 32842 | 32842-Insert | Insertion | 1 | - | A | 434 | 26053 | 1.66583503 | 43.7096774 | rnr1D57N | FALSE | CAN | JSY14 | 1 | rnr1D57N-Re | CAA | nope | A/T+1 | >1bpdel | none | A/T+1 | TTACCCAAA | AAAAAAT | GTT | CAA | - | T | NA | NA_CAA | A/T+1_AAAAAAT | 7 | FALSE | FALSE |
| 12_v.txt | 32842 | 32842-Insert | Insertion | 2 | - | AA | 2 | 26053 | 7.68E-03 | 38 | rnr1D57N | FALSE | CAN | JSY14 | 1 | rnr1D57N-Re | CAA | nope | >1bp-ins | >1bpdel | none | >1bp-ins | TTACCCAAA | AAAAAAT | GTT | CAA | - | none | NA | NA_CAA | >1bp-ins_AAAAAAT | 7 | FALSE | FALSE |
| 12_v.txt | 32842 | 32842-Insert | Insertion | 1 | - | G | 40 | 26053 | 0.15353318 | 41.25 | rnr1D57N | FALSE | CAN | JSY14 | 1 | rnr1D57N-Re | CAA | nope | G/C+1 | >1bpdel | none | G/C+1 | TTACCCAAA | AAAAAAT | GTT | CAA | - | C | NA | NA_CAA | G/C+1_AAAAAAT | 7 | FALSE | FALSE |
| 12_v.txt | 32842 | 32842-Insert | Insertion | 1 | - | T | 1 | 26053 | 3.84E-03 | 37 | rnr1D57N | FALSE | CAN | JSY14 | 1 | rnr1D57N-Re | CAA | nope | A/T+1 | >1bpdel | none | A/T+1 | TTACCCAAA | AAAAAAT | GTT | CAA | - | A | NA | NA_CAA | A/T+1_AAAAAAT | 7 | FALSE | FALSE |
| 12_v.txt | 32884 | 32884-Insert | Insertion | 1 | - | A | 1 | 21315 | 4.69E-03 | 38 | rnr1D57N | FALSE | CAN | JSY14 | 1 | rnr1D57N-Re | CAA | nope | A/T+1 | >1bpdel | none | A/T+1 | ACGTCCAAA | AAAATTG | GTT | CAA | - | T | NA | NA_CAA | A/T+1_AAAATTG | 7 | FALSE | FALSE |
| 12_v.txt | 32924 | 32924-Insert | Insertion | 2 | - | AA | 5 | 21315 | 0.02345766 | 61.3 | rnr1D57N | FALSE | CAN | JSY14 | 1 | rnr1D57N-Re | CAA | nope | >1bp-ins | >1bpdel | none | >1bp-ins | CAGGGCAAA | AAAAGTG | GTT | CAA | - | none | NA | NA_CAA | >1bp-ins_AAAAGTG | 7 | FALSE | FALSE |
| 12_v.txt | 32936 | 32936-Insert | Insertion | 1 | - | A | 1 | 21315 | 4.69E-03 | 11 | rnr1D57N | FALSE | CAN | JSY14 | 1 | rnr1D57N-Re | CCC | nope | A/T+1 | >1bpdel | none | A/T+1 | GATTGCCCA | CCAAGAA | GGG | CCC | - | T | NA | NA_CCC | A/T+1_CCAAGAA | 7 | FALSE | FALSE |
| 12_v.txt | 32938 | 32938-Insert | Insertion | 1 | - | G | 431 | 21315 | 2.0220502 | 57.9698376 | rnr1D57N | FALSE | CAN | JSY14 | 1 | rnr1D57N-Re | CAA | nope | G/C+1 | >1bpdel | none | G/C+1 | TTGCCCAAG | AAGAAAA | GTT | CAA | - | C | NA | NA_CAA | G/C+1_AAGAAAA | 7 | FALSE | FALSE |
| 12_v.txt | 32938 | 32938-Insert | Insertion | 1 | - | T | 1 | 21315 | 4.69E-03 | 64 | rnr1D57N | FALSE | CAN | JSY14 | 1 | rnr1D57N-Re | CAA | nope | A/T+1 | >1bpdel | none | A/T+1 | TTGCCCAAG | AAGAAAA | GTT | CAA | - | A | NA | NA_CAA | A/T+1_AAGAAAA | 7 | FALSE | FALSE |
| 12_v.txt | 33000 | 33000-Insert | Insertion | 1 | - | A | 11 | 21315 | 0.05160685 | 64 | rnr1D57N | FALSE | CAN | JSY14 | 1 | rnr1D57N-Re | GAA | nope | A/T+1 | >1bpdel | none | A/T+1 | TGTGAGAAA | AAAACTG | CTT | GAA | - | T | NA | NA_GAA | A/T+1_AAAACTG | 7 | FALSE | FALSE |
| 12_v.txt | 33072 | 33072-Insert | Insertion | 1 | - | A | 18 | 21315 | 0.08444757 | 53.7222222 | rnr1D57N | FALSE | CAN | JSY14 | 1 | rnr1D57N-Re | CAA | nope | A/T+1 | >1bpdel | none | A/T+1 | TATGCCAAA | AAAGAAC | GTT | CAA | - | T | NA | NA_CAA | A/T+1_AAAGAAC | 7 | FALSE | FALSE |
| 12_v.txt | 33072 | 33072-Insert | Insertion | 2 | - | AA | 5 | 21315 | 0.02345766 | 58.8 | rnr1D57N | FALSE | CAN | JSY14 | 1 | rnr1D57N-Re | CAA | nope | >1bp-ins | >1bpdel | none | >1bp-ins | TATGCCAAA | AAAGAAC | GTT | CAA | - | none | NA | NA_CAA | >1bp-ins_AAAGAAC | 7 | FALSE | FALSE |
| 12_v.txt | 33331 | 33331-Insert | Insertion | 1 | - | T | 35 | 20312 | 0.17231193 | 60.9142857 | rnr1D57N | FALSE | CAN | JSY14 | 1 | rnr1D57N-Re | CTT | nope | A/T+1 | >1bpdel | none | A/T+1 | CTTTACTTTT | TTTTCTC | GAA | CTT | - | A | NA | NA_CTT | A/T+1_TTTTCTC | 7 | FALSE | FALSE |
| 12_v.txt | 33355 | 33355-Insert | Insertion | 4 | - | ACCC | 6 | 20312 | 0.02953919 | 57.7083333 | rnr1D57N | FALSE | CAN | JSY14 | 1 | rnr1D57N-Re | GAC | nope | >1bp-ins | >1bpdel | none | >1bp-ins | GTATTGACCC | ACCCACG | CTG | GAC | - | none | NA | NA_GAC | >1bp-ins_ACCCACG | 7 | FALSE | FALSE |
| 12_v.txt | 33452 | 33452-Insert | Insertion | 1 | - | T | 4 | 20312 | 0.01969279 | 37.5 | rnr1D57N | FALSE | CAN | JSY14 | 1 | rnr1D57N-Re | CTT | nope | A/T+1 | >1bpdel | none | A/T+1 | GTCTTCTTTT | TTTTGAA | GAA | CTT | - | A | NA | NA_CTT | A/T+1_TTTTGAA | 7 | FALSE | FALSE |
| 12_v.txt | 31807 | 31807-Deleti | Deletion | 4 | AAAT | - | 30 | 11167 | 0.2686487 | 57.6 | rnr1D57N | FALSE | CAN | JSY14 | 1 | rnr1D57N-Re | CAA | nope | >1bp-ins | >1bpdel | none | >1bpdel | TCTTCCAAAT | AAATAAA | GTT | CAA | AAAT | none | NA | NA_CAA | >1bpdel_AAATAAA | 7 | FALSE | FALSE |
| 12_v.txt | 32086 | 32086-Deleti | Deletion | 3 | AGC | - | 4 | 16625 | 0.02406015 | 64 | rnr1D57N | FALSE | CAN | JSY14 | 1 | rnr1D57N-Re | AAG | nope | >1bp-ins | >1bpdel | none | >1bpdel | CAAAAAAGC | AGCCTGC | TTC | AAG | AGC | none | NA | NA_AAG | >1bpdel_AGCCTGC | 7 | FALSE | FALSE |
| 12_v.txt | 32091 | 32091-Deleti | Deletion | 2 | GC | - | 4 | 16625 | 0.02406015 | 64 | rnr1D57N | FALSE | CAN | JSY14 | 1 | rnr1D57N-Re | TGC | nope | >1bp-ins | >1bpdel | none | >1bpdel | AAGCCTGCA | GCAACAC | ACG | TGC | GC | none | NA | NA_TGC | >1bpdel_GCAACAC | 7 | FALSE | FALSE |
| 12_v.txt | 32100 | 32100-Deleti | Deletion | 4 | GTGA | - | 4 | 16625 | 0.02406015 | 64 | rnr1D57N | FALSE | CAN | JSY14 | 1 | rnr1D57N-Re | AGT | nope | >1bp-ins | >1bpdel | none | >1bpdel | ACACCAGTG | GTGATAT | TCA | AGT | GTGA | none | NA | NA_AGT | >1bpdel_GTGATAT | 7 | FALSE | FALSE |
| 12_v.txt | 32114 | 32114-Deleti | Deletion | 19 | CCATTCGAA | - | 298 | 16625 | 1.7924812 | 59.1979866 | rnr1D57N | FALSE | CAN | JSY14 | 1 | rnr1D57N-Re | GCC | nope | >1bp-ins | >1bpdel | none | >1bpdel | TAATAGCCAT | CCATTCG | CGG | GCC | CCATTCGAA | none | NA | NA_GCC | >1bpdel_CCATTCG | 7 | FALSE | FALSE |
| 12_v.txt | 32360 | 32360-Deleti | Deletion | 2 | AG | - | 55 | 29225 | 0.18819504 | 59.6181818 | rnr1D57N | FALSE | CAN | JSY14 | 1 | rnr1D57N-Re | CAG | nope | >1bp-ins | >1bpdel | none | >1bpdel | TGTACCAGA | AGAGTTC | GTC | CAG | AG | none | NA | NA_CAG | >1bpdel_AGAGTTC | 7 | FALSE | FALSE |
| 12_v.txt | 32375 | 32375-Deleti | Deletion | 3 | AAT | - | 17 | 29225 | 0.05816938 | 59.6470588 | rnr1D57N | FALSE | CAN | JSY14 | 1 | rnr1D57N-Re | CAA | nope | >1bp-ins | >1bpdel | none | >1bpdel | AATAGCAAT | AATAATA | GTT | CAA | AAT | none | NA | NA_CAA | >1bpdel_AATAATA | 7 | FALSE | FALSE |
| 12_v.txt | 32541 | 32541-Deleti | Deletion | 3 | TCA | - | 46 | 29225 | 0.15739949 | 35.5 | rnr1D57N | FALSE | CAN | JSY14 | 1 | rnr1D57N-Re | TTC | nope | >1bp-ins | >1bpdel | none | >1bpdel | GCAGCTTCA | TCACCAG | AAG | TTC | TCA | none | NA | NA_TTC | >1bpdel_TCACCAG | 7 | FALSE | FALSE |
| 12_v.txt | 32550 | 32550-Deleti | Deletion | 3 | GTG | - | 8 | 29220 | 0.02737851 | 37.5 | rnr1D57N | FALSE | CAN | JSY14 | 1 | rnr1D57N-Re | AGT | nope | >1bp-ins | >1bpdel | none | >1bpdel | CCAGCAGTG | GTGATAC | TCA | AGT | GTG | none | NA | NA_AGT | >1bpdel_GTGATAC | 7 | FALSE | FALSE |
| 12_v.txt | 32902 | 32902-Deleti | Deletion | 2 | CA | - | 45 | 21315 | 0.21111893 | 46.0444444 | rnr1D57N | FALSE | CAN | JSY14 | 1 | rnr1D57N-Re | CCA | nope | >1bp-ins | >1bpdel | none | >1bpdel | CTTGGCCAA | CAACTAC | GGT | CCA | CA | none | NA | NA_CCA | >1bpdel_CAACTAC | 7 | FALSE | FALSE |
| 12_v.txt | 33065 | 33065-Deleti | Deletion | 2 | AT | - | 26 | 21315 | 0.12197983 | 57.9615385 | rnr1D57N | FALSE | CAN | JSY14 | 1 | rnr1D57N-Re | AAT | nope | >1bp-ins | >1bpdel | none | >1bpdel | GACAGAATA | ATATGCC | TTA | AAT | AT | none | NA | NA_AAT | >1bpdel_ATATGCC | 7 | FALSE | FALSE |
| 12_v.txt | 33089 | 33089-Deleti | Deletion | 2 | AT | - | 53 | 21315 | 0.24865119 | 55.9622642 | rnr1D57N | FALSE | CAN | JSY14 | 1 | rnr1D57N-Re | AAT | nope | >1bp-ins | >1bpdel | none | >1bpdel | AAATAAATA | ATATGAT | TTA | AAT | AT | none | NA | NA_AAT | >1bpdel_ATATGAT | 7 | FALSE | FALSE |
| 12_v.txt | 33094 | 33094-Deleti | Deletion | 2 | AT | - | 17 | 21315 | 0.07975604 | 57.5294118 | rnr1D57N | FALSE | CAN | JSY14 | 1 | rnr1D57N-Re | GAT | nope | >1bp-ins | >1bpdel | none | >1bpdel | AATATGATA | ATATAAG | CTA | GAT | AT | none | NA | NA_GAT | >1bpdel_ATATAAG | 7 | FALSE | FALSE |
| 12_v.txt | 33474 | 33474-Deleti | Deletion | 2 | CC | - | 1 | 20311 | 4.92E-03 | 10 | rnr1D57N | FALSE | CAN | JSY14 | 1 | rnr1D57N-Re | GCC | nope | >1bp-ins | >1bpdel | none | >1bpdel | GCTATGCCTT | CCTTTTT | CGG | GCC | CC | none | NA | NA_GCC | >1bpdel_CCTTTTT | 7 | FALSE | FALSE |
| 12_v.txt | 31723 | 31723-Deleti | Deletion | 1 | A | - | 14 | 11167 | 0.12536939 | 58.2142934 | rnr1D57N | FALSE | CAN | JSY14 | 1 | rnr1D57N-Re | CAA | nope | >1bp-ins | A/T-1 | none | A/T-1 | TGTCCCAAA | AAAAAGT | GTT | GTT | T | none | T>none | T>none_GTT | A/T-1_AAAAAGT | 7 | FALSE | FALSE |
| 12_v.txt | 31863 | 31863-Deleti | Deletion | 1 | A | - | 5 | 11167 | 0.04477478 | 39.2 | rnr1D57N | FALSE | CAN | JSY14 | 1 | rnr1D57N-Re | GAA | nope | >1bp-ins | A/T-1 | none | A/T-1 | AACAGGAAA | AAAATAG | CTT | CTT | T | none | T>none | T>none_CTT | A/T-1_AAAATAG | 7 | FALSE | FALSE |
| 12_v.txt | 31913 | 31913-Deleti | Deletion | 1 | A | - | 6 | 11162 | 0.05375381 | 42.1666655 | rnr1D57N | FALSE | CAN | JSY14 | 1 | rnr1D57N-Re | CAA | nope | >1bp-ins | A/T-1 | none | A/T-1 | TGGTGCAAA | AAAAGCC | GTT | GTT | T | none | T>none | T>none_GTT | A/T-1_AAAAGCC | 7 | FALSE | FALSE |
| 12_v.txt | 31926 | 31926-Deleti | Deletion | 1 | C | - | 27 | 11161 | 0.24191381 | 37.7037037 | rnr1D57N | FALSE | CAN | JSY14 | 1 | rnr1D57N-Re | ACC | nope | >1bp-ins | G/C-1 | none | G/C-1 | GTGAAACCT | CCTTGAA | TGG | ACC | C | none | nope | nope_ACC | G/C-1_CCTTGAA | 7 | FALSE | FALSE |
| 12_v.txt | 31957 | 31957-Deleti | Deletion | 1 | T | - | 1 | 16625 | 6.02E-03 | 38 | rnr1D57N | FALSE | CAN | JSY14 | 1 | rnr1D57N-Re | GTG | nope | >1bp-ins | A/T-1 | none | A/T-1 | TAAATGTGG | TGGCCGC | CAC | GTG | T | none | nope | nope_GTG | A/T-1_TGGCCGC | 7 | FALSE | FALSE |
| 12_v.txt | 31972 | 31972-Deleti | Deletion | 1 | C | - | 7 | 16625 | 0.04210526 | 31.2857143 | rnr1D57N | FALSE | CAN | JSY14 | 1 | rnr1D57N-Re | GCC | nope | >1bp-ins | G/C-1 | none | G/C-1 | AATAAGCCA | CCAAGCC | CGG | GCC | C | none | nope | nope_GCC | G/C-1_CCAAGCC | 7 | FALSE | FALSE |
| 12_v.txt | 31976 | 31976-Deleti | Deletion | 1 | G | - | 9 | 16625 | 0.05413534 | 46.6666667 | rnr1D57N | FALSE | CAN | JSY14 | 1 | rnr1D57N-Re | AGC | nope | >1bp-ins | G/C-1 | none | G/C-1 | AGCCAAGCC | GCCGGGC | TCG | TCG | C | none | C>none | C>none_TCG | G/C-1_GCCGGGC | 7 | FALSE | FALSE |
| 12_v.txt | 32075 | 32075-Deleti | Deletion | 1 | C | - | 5 | 16625 | 0.03007519 | 64 | rnr1D57N | FALSE | CAN | JSY14 | 1 | rnr1D57N-Re | ACC | nope | >1bp-ins | G/C-1 | none | G/C-1 | AAATAACCA | CCATGCA | TGG | ACC | C | none | nope | nope_ACC | G/C-1_CCATGCA | 7 | FALSE | FALSE |
| 12_v.txt | 32079 | 32079-Deleti | Deletion | 1 | G | - | 4 | 16625 | 0.02406015 | 64 | rnr1D57N | FALSE | CAN | JSY14 | 1 | rnr1D57N-Re | TGC | nope | >1bp-ins | G/C-1 | none | G/C-1 | AACCATGCA | GCAAAAA | ACG | ACG | C | none | C>none | C>none_ACG | G/C-1_GCAAAAA | 7 | FALSE | FALSE |
| 12_v.txt | 32081 | 32081-Deleti | Deletion | 1 | A | - | 32 | 16624 | 0.19249278 | 63.9062501 | rnr1D57N | FALSE | CAN | JSY14 | 1 | rnr1D57N-Re | CAA | nope | >1bp-ins | A/T-1 | none | A/T-1 | CCATGCAAA | AAAAAAG | GTT | GTT | T | none | T>none | T>none_GTT | A/T-1_AAAAAAG | 7 | FALSE | FALSE |
| 12_v.txt | 32095 | 32095-Deleti | Deletion | 1 | C | - | 4 | 16625 | 0.02406015 | 64 | rnr1D57N | FALSE | CAN | JSY14 | 1 | rnr1D57N-Re | ACA | nope | >1bp-ins | G/C-1 | none | G/C-1 | CTGCAACAC | CACCAGT | TGT | ACA | C | none | nope | nope_ACA | G/C-1_CACCAGT | 7 | FALSE | FALSE |
| 12_v.txt | 32097 | 32097-Deleti | Deletion | 1 | C | - | 5 | 16625 | 0.03007519 | 58.8 | rnr1D57N | FALSE | CAN | JSY14 | 1 | rnr1D57N-Re | ACC | nope | >1bp-ins | G/C-1 | none | G/C-1 | GCAACACCA | CCAGTGA | TGG | ACC | C | none | nope | nope_ACC | G/C-1_CCAGTGA | 7 | FALSE | FALSE |
| 12_v.txt | 32148 | 32148-Deleti | Deletion | 1 | T | - | 12 | 16625 | 0.07218045 | 50.6666682 | rnr1D57N | FALSE | CAN | JSY14 | 1 | rnr1D57N-Re | CTC | nope | >1bp-ins | A/T-1 | none | A/T-1 | GATGTCTCCA | TCCATGT | GAG | CTC | T | none | nope | nope_CTC | A/T-1_TCCATGT | 7 | FALSE | FALSE |
| 12_v.txt | 32149 | 32149-Deleti | Deletion | 1 | C | - | 2 | 16625 | 0.01203008 | 63.999992 | rnr1D57N | FALSE | CAN | JSY14 | 1 | rnr1D57N-Re | TCC | nope | >1bp-ins | G/C-1 | none | G/C-1 | ATGTCTCCAT | CCATGTA | AGG | TCC | C | none | nope | nope_TCC | G/C-1_CCATGTA | 7 | FALSE | FALSE |
| 12_v.txt | 32165 | 32165-Deleti | Deletion | 1 | G | - | 43 | 16625 | 0.25864662 | 47.2325483 | rnr1D57N | FALSE | CAN | JSY14 | 1 | rnr1D57N-Re | CGC | nope | >1bp-ins | G/C-1 | none | G/C-1 | CAAAGCGCC | GCCAAAT | GCG | GCG | C | none | C>none | C>none_GCG | G/C-1_GCCAAAT | 7 | FALSE | FALSE |
| 12_v.txt | 32237 | 32237-Deleti | Deletion | 1 | A | - | 260 | 16623 | 1.56409794 | 37.2730769 | rnr1D57N | FALSE | CAN | JSY14 | 1 | rnr1D57N-Re | GAG | nope | >1bp-ins | A/T-1 | none | A/T-1 | TTTAGGAGC | AGCCAAC | CTC | CTC | T | none | T>none | T>none_CTC | A/T-1_AGCCAAC | 7 | FALSE | FALSE |
| 12_v.txt | 32259 | 32259-Deleti | Deletion | 1 | C | - | 14 | 29224 | 0.04790583 | 37.7857143 | rnr1D57N | FALSE | CAN | JSY14 | 1 | rnr1D57N-Re | ACC | nope | >1bp-ins | G/C-1 | none | G/C-1 | GATAGACCA | CCAAATA | TGG | ACC | C | none | nope | nope_ACC | G/C-1_CCAAATA | 7 | FALSE | FALSE |
| 12_v.txt | 32284 | 32284-Deleti | Deletion | 1 | A | - | 5 | 29225 | 0.01710864 | 53.5999965 | rnr1D57N | FALSE | CAN | JSY14 | 1 | rnr1D57N-Re | TAA | nope | >1bp-ins | A/T-1 | none | A/T-1 | CAACGTAAA | AAATATT | ATT | ATT | T | none | T>none | T>none_ATT | A/T-1_AAATATT | 7 | FALSE | FALSE |
| 12_v.txt | 32313 | 32313-Deleti | Deletion | 1 | G | - | 2 | 29225 | 6.84E-03 | 64 | rnr1D57N | FALSE | CAN | JSY14 | 1 | rnr1D57N-Re | GGT | nope | >1bp-ins | G/C-1 | none | G/C-1 | ATAATGGTT | GTTGTTA | CCA | CCA | C | none | C>none | C>none_CCA | G/C-1_GTTGTTA | 7 | FALSE | FALSE |
| 12_v.txt | 32346 | 32346-Deleti | Deletion | 1 | A | - | 86 | 29223 | 0.29428875 | 61.1976743 | rnr1D57N | FALSE | CAN | JSY14 | 1 | rnr1D57N-Re | CAA | nope | >1bp-ins | A/T-1 | none | A/T-1 | TGTGGCAAA | AAAACCT | GTT | GTT | T | none | T>none | T>none_GTT | A/T-1_AAAACCT | 7 | FALSE | FALSE |
| 12_v.txt | 32494 | 32494-Deleti | Deletion | 1 | C | - | 1 | 29224 | 3.42E-03 | 63.9999842 | rnr1D57N | FALSE | CAN | JSY14 | 1 | rnr1D57N-Re | ACA | nope | >1bp-ins | G/C-1 | none | G/C-1 | GGAAAACAA | CAACTTT | TGT | ACA | C | none | nope | nope_ACA | G/C-1_CAACTTT | 7 | FALSE | FALSE |
| 12_v.txt | 32498 | 32498-Deleti | Deletion | 1 | T | - | 138 | 29225 | 0.47219846 | 44.8478238 | rnr1D57N | FALSE | CAN | JSY14 | 1 | rnr1D57N-Re | CTT | nope | >1bp-ins | A/T-1 | none | A/T-1 | AACAACTTTT | TTTTTTG | GAA | CTT | T | none | nope | nope_CTT | A/T-1_TTTTTTG | 7 | FALSE | FALSE |
| 12_v.txt | 32544 | 32544-Deleti | Deletion | 1 | C | - | 3 | 29224 | 0.01026554 | 37.3333333 | rnr1D57N | FALSE | CAN | JSY14 | 1 | rnr1D57N-Re | ACC | nope | >1bp-ins | G/C-1 | none | G/C-1 | GCTTCACCA | CCAGCAG | TGG | ACC | C | none | nope | nope_ACC | G/C-1_CCAGCAG | 7 | FALSE | FALSE |
| 12_v.txt | 32581 | 32581-Deleti | Deletion | 1 | T | - | 15 | 26053 | 0.05757494 | 37.3999988 | rnr1D57N | FALSE | CAN | JSY14 | 1 | rnr1D57N-Re | GTG | nope | >1bp-ins | A/T-1 | none | A/T-1 | GAAATGTGA | TGAAGGC | CAC | GTG | T | none | nope | nope_GTG | A/T-1_TGAAGGC | 7 | FALSE | FALSE |
| 12_v.txt | 32631 | 32631-Deleti | Deletion | 1 | T | - | 203 | 26053 | 0.7791809 | 51.817734 | rnr1D57N | FALSE | CAN | JSY14 | 1 | rnr1D57N-Re | GTT | nope | >1bp-ins | A/T-1 | none | A/T-1 | CCTTCGTTTT | TTTTTAT | CAA | GTT | T | none | nope | nope_GTT | A/T-1_TTTTTAT | 7 | FALSE | FALSE |
| 12_v.txt | 32657 | 32657-Deleti | Deletion | 1 | A | - | 45 | 26053 | 0.17272483 | 56.2444548 | rnr1D57N | FALSE | CAN | JSY14 | 1 | rnr1D57N-Re | GAC | nope | >1bp-ins | A/T-1 | none | A/T-1 | ACCTGGACC | ACCCCAG | CTG | CTG | T | none | T>none | T>none_CTG | A/T-1_ACCCCAG | 7 | FALSE | FALSE |
| 12_v.txt | 32658 | 32658-Deleti | Deletion | 1 | C | - | 110 | 26053 | 0.42221625 | 62.3363609 | rnr1D57N | FALSE | CAN | JSY14 | 1 | rnr1D57N-Re | ACC | nope | >1bp-ins | G/C-1 | none | G/C-1 | CCTGGACCC | CCCCAGG | TGG | ACC | C | none | nope | nope_ACC | G/C-1_CCCCAGG | 7 | FALSE | FALSE |
| 12_v.txt | 32678 | 32678-Deleti | Deletion | 1 | C | - | 2 | 26053 | 7.68E-03 | 62 | rnr1D57N | FALSE | CAN | JSY14 | 1 | rnr1D57N-Re | TCC | nope | >1bp-ins | G/C-1 | none | G/C-1 | GTTTCTCCAA | CCAATAA | AGG | TCC | C | none | nope | nope_TCC | G/C-1_CCAATAA | 7 | FALSE | FALSE |
| 12_v.txt | 32682 | 32682-Deleti | Deletion | 1 | T | - | 14 | 26053 | 0.05373661 | 62.7142857 | rnr1D57N | FALSE | CAN | JSY14 | 1 | rnr1D57N-Re | ATA | nope | >1bp-ins | A/T-1 | none | A/T-1 | CTCCAATAAC | TAACGGA | TAT | ATA | T | none | nope | nope_ATA | A/T-1_TAACGGA | 7 | FALSE | FALSE |
| 12_v.txt | 32728 | 32728-Deleti | Deletion | 1 | A | - | 52 | 26053 | 0.19959314 | 60.6346078 | rnr1D57N | FALSE | CAN | JSY14 | 1 | rnr1D57N-Re | CAA | nope | >1bp-ins | A/T-1 | none | A/T-1 | CCATACAAA | AAAAACA | GTT | GTT | T | none | T>none | T>none_GTT | A/T-1_AAAAACA | 7 | FALSE | FALSE |
| 12_v.txt | 32758 | 32758-Deleti | Deletion | 1 | C | - | 6 | 26053 | 0.02302998 | 49.5000065 | rnr1D57N | FALSE | CAN | JSY14 | 1 | rnr1D57N-Re | GCT | nope | >1bp-ins | G/C-1 | none | G/C-1 | TAATGGCTA | CTAAAAC | CGA | GCT | C | none | nope | nope_GCT | G/C-1_CTAAAAC | 7 | FALSE | FALSE |
| 12_v.txt | 32759 | 32759-Deleti | Deletion | 1 | T | - | 23 | 26053 | 0.08828158 | 51.8695646 | rnr1D57N | FALSE | CAN | JSY14 | 1 | rnr1D57N-Re | CTA | nope | >1bp-ins | A/T-1 | none | A/T-1 | AATGGCTAA | TAAAACT | GAT | CTA | T | none | nope | nope_CTA | A/T-1_TAAAACT | 7 | FALSE | FALSE |
| 12_v.txt | 32760 | 32760-Deleti | Deletion | 1 | A | - | 2 | 26053 | 7.68E-03 | 64 | rnr1D57N | FALSE | CAN | JSY14 | 1 | rnr1D57N-Re | TAA | nope | >1bp-ins | A/T-1 | none | A/T-1 | ATGGCTAAA | AAAACTT | ATT | ATT | T | none | T>none | T>none_ATT | A/T-1_AAAACTT | 7 | FALSE | FALSE |
| 12_v.txt | 32779 | 32779-Deleti | Deletion | 1 | C | - | 7 | 26053 | 0.02686831 | 56.5714239 | rnr1D57N | FALSE | CAN | JSY14 | 1 | rnr1D57N-Re | ACC | nope | >1bp-ins | G/C-1 | none | G/C-1 | AAGCGACCC | CCCAGAA | TGG | ACC | C | none | nope | nope_ACC | G/C-1_CCCAGAA | 7 | FALSE | FALSE |
| 12_v.txt | 32839 | 32839-Deleti | Deletion | 1 | C | - | 9 | 26053 | 0.03454497 | 39.888886 | rnr1D57N | FALSE | CAN | JSY14 | 1 | rnr1D57N-Re | ACC | nope | >1bp-ins | G/C-1 | none | G/C-1 | TAATTACCCA | CCCAAAA | TGG | ACC | C | none | nope | nope_ACC | G/C-1_CCCAAAA | 7 | FALSE | FALSE |
| 12_v.txt | 32842 | 32842-Deleti | Deletion | 1 | A | - | 1113 | 26053 | 4.2720608 | 41.7160881 | rnr1D57N | FALSE | CAN | JSY14 | 1 | rnr1D57N-Re | CAA | nope | >1bp-ins | A/T-1 | none | A/T-1 | TTACCCAAA | AAAAAAT | GTT | GTT | T | none | T>none | T>none_GTT | A/T-1_AAAAAAT | 7 | FALSE | FALSE |
| 12_v.txt | 32884 | 32884-Deleti | Deletion | 1 | A | - | 9 | 21315 | 0.04222379 | 37.2222225 | rnr1D57N | FALSE | CAN | JSY14 | 1 | rnr1D57N-Re | CAA | nope | >1bp-ins | A/T-1 | none | A/T-1 | ACGTCCAAA | AAAATTG | GTT | GTT | T | none | T>none | T>none_GTT | A/T-1_AAAATTG | 7 | FALSE | FALSE |
| 12_v.txt | 32901 | 32901-Deleti | Deletion | 1 | C | - | 1 | 21315 | 4.69E-03 | 38 | rnr1D57N | FALSE | CAN | JSY14 | 1 | rnr1D57N-Re | GCC | nope | >1bp-ins | G/C-1 | none | G/C-1 | ACTTGGCCA | CCAACTA | CGG | GCC | C | none | nope | nope_GCC | G/C-1_CCAACTA | 7 | FALSE | FALSE |
| 12_v.txt | 32924 | 32924-Deleti | Deletion | 1 | A | - | 18 | 21315 | 0.08444757 | 58.4999963 | rnr1D57N | FALSE | CAN | JSY14 | 1 | rnr1D57N-Re | CAA | nope | >1bp-ins | A/T-1 | none | A/T-1 | CAGGGCAAA | AAAAGTG | GTT | GTT | T | none | T>none | T>none_GTT | A/T-1_AAAAGTG | 7 | FALSE | FALSE |
| 12_v.txt | 32935 | 32935-Deleti | Deletion | 1 | C | - | 21 | 21315 | 0.09852217 | 51.1904762 | rnr1D57N | FALSE | CAN | JSY14 | 1 | rnr1D57N-Re | GCC | nope | >1bp-ins | G/C-1 | none | G/C-1 | TGATTGCCCA | CCCAAGA | CGG | GCC | C | none | nope | nope_GCC | G/C-1_CCCAAGA | 7 | FALSE | FALSE |
| 12_v.txt | 32958 | 32958-Deleti | Deletion | 1 | C | - | 2 | 21315 | 9.38E-03 | 64 | rnr1D57N | FALSE | CAN | JSY14 | 1 | rnr1D57N-Re | ACC | nope | >1bp-ins | G/C-1 | none | G/C-1 | ATGTAACCAT | CCATTGG | TGG | ACC | C | none | nope | nope_ACC | G/C-1_CCATTGG | 7 | FALSE | FALSE |
| 12_v.txt | 32970 | 32970-Deleti | Deletion | 1 | C | - | 3 | 21314 | 0.01407526 | 52 | rnr1D57N | FALSE | CAN | JSY14 | 1 | rnr1D57N-Re | ACC | nope | >1bp-ins | G/C-1 | none | G/C-1 | GCCGCACCA | CCAAATG | TGG | ACC | C | none | nope | nope_ACC | G/C-1_CCAAATG | 7 | FALSE | FALSE |
| 12_v.txt | 32972 | 32972-Deleti | Deletion | 1 | A | - | 13 | 21314 | 0.06099278 | 62.6923037 | rnr1D57N | FALSE | CAN | JSY14 | 1 | rnr1D57N-Re | CAA | nope | >1bp-ins | A/T-1 | none | A/T-1 | CGCACCAAAT | AAATGCT | GTT | GTT | T | none | T>none | T>none_GTT | A/T-1_AAATGCT | 7 | FALSE | FALSE |
| 12_v.txt | 33000 | 33000-Deleti | Deletion | 1 | A | - | 68 | 21315 | 0.31902416 | 62.1764622 | rnr1D57N | FALSE | CAN | JSY14 | 1 | rnr1D57N-Re | GAA | nope | >1bp-ins | A/T-1 | none | A/T-1 | TGTGAGAAA | AAAACTG | CTT | CTT | T | none | T>none | T>none_CTT | A/T-1_AAAACTG | 7 | FALSE | FALSE |
| 12_v.txt | 33072 | 33072-Deleti | Deletion | 1 | A | - | 6 | 21315 | 0.02814919 | 59.333328 | rnr1D57N | FALSE | CAN | JSY14 | 1 | rnr1D57N-Re | CAA | nope | >1bp-ins | A/T-1 | none | A/T-1 | TATGCCAAA | AAAGAAC | GTT | GTT | T | none | T>none | T>none_GTT | A/T-1_AAAGAAC | 7 | FALSE | FALSE |
| 12_v.txt | 33147 | 33147-Deleti | Deletion | 1 | A | - | 7 | 21315 | 0.03284072 | 38 | rnr1D57N | FALSE | CAN | JSY14 | 1 | rnr1D57N-Re | GAA | nope | >1bp-ins | A/T-1 | none | A/T-1 | CCAATGAAA | AAAAGAC | CTT | CTT | T | none | T>none | T>none_CTT | A/T-1_AAAAGAC | 7 | FALSE | FALSE |
| 12_v.txt | 33168 | 33168-Deleti | Deletion | 1 | C | - | 2 | 21296 | 9.39E-03 | 38 | rnr1D57N | FALSE | CAN | JSY14 | 1 | rnr1D57N-Re | ACC | nope | >1bp-ins | G/C-1 | none | G/C-1 | ATAGTACCA | CCACCAA | TGG | ACC | C | none | nope | nope_ACC | G/C-1_CCACCAA | 7 | FALSE | FALSE |
| 12_v.txt | 33178 | 33178-Deleti | Deletion | 1 | C | - | 25 | 21290 | 0.11742602 | 37.9999998 | rnr1D57N | FALSE | CAN | JSY14 | 1 | rnr1D57N-Re | GCA | nope | >1bp-ins | G/C-1 | none | G/C-1 | CAAGGGCAA | CAATCAT | CGT | GCA | C | none | nope | nope_GCA | G/C-1_CAATCAT | 7 | FALSE | FALSE |
| 12_v.txt | 33273 | 33273-Deleti | Deletion | 1 | A | - | 22 | 20312 | 0.10831036 | 50.0909091 | rnr1D57N | FALSE | CAN | JSY14 | 1 | rnr1D57N-Re | GAA | nope | >1bp-ins | A/T-1 | none | A/T-1 | ATAGAGAAC | AACGTAT | CTT | CTT | T | none | T>none | T>none_CTT | A/T-1_AACGTAT | 7 | FALSE | FALSE |
| 12_v.txt | 33300 | 33300-Deleti | Deletion | 1 | G | - | 13 | 20312 | 0.06400158 | 59.5384615 | rnr1D57N | FALSE | CAN | JSY14 | 1 | rnr1D57N-Re | CGG | nope | >1bp-ins | G/C-1 | none | G/C-1 | CTCGTCGGG | GGGAAAG | GCC | GCC | C | none | C>none | C>none_GCC | G/C-1_GGGAAAG | 7 | FALSE | FALSE |
| 12_v.txt | 33378 | 33378-Deleti | Deletion | 1 | G | - | 6 | 20312 | 0.02953919 | 59.5 | rnr1D57N | FALSE | CAN | JSY14 | 1 | rnr1D57N-Re | TGA | nope | >1bp-ins | G/C-1 | none | G/C-1 | GTTTGTGAA | GAAGCTT | ACT | ACT | C | none | C>none | C>none_ACT | G/C-1_GAAGCTT | 7 | FALSE | FALSE |
| 12_v.txt | 33452 | 33452-Deleti | Deletion | 1 | T | - | 6 | 20312 | 0.02953919 | 44.1666667 | rnr1D57N | FALSE | CAN | JSY14 | 1 | rnr1D57N-Re | CTT | nope | >1bp-ins | A/T-1 | none | A/T-1 | GTCTTCTTTT | TTTTGAA | GAA | CTT | T | none | nope | nope_CTT | A/T-1_TTTTGAA | 7 | FALSE | FALSE |
| 12_v.txt | 33456 | 33456-Deleti | Deletion | 1 | G | - | 24 | 20312 | 0.11815676 | 44 | rnr1D57N | FALSE | CAN | JSY14 | 1 | rnr1D57N-Re | TGA | nope | >1bp-ins | G/C-1 | none | G/C-1 | TCTTTTGAAT | GAATTTG | ACT | ACT | C | none | C>none | C>none_ACT | G/C-1_GAATTTG | 7 | FALSE | FALSE |
| 12_v.txt | 33474 | 33474-Deleti | Deletion | 1 | C | - | 38 | 20311 | 0.18709074 | 31.473684 | rnr1D57N | FALSE | CAN | JSY14 | 1 | rnr1D57N-Re | GCC | nope | >1bp-ins | G/C-1 | none | G/C-1 | GCTATGCCTT | CCTTTTT | CGG | GCC | C | none | nope | nope_GCC | G/C-1_CCTTTTT | 7 | FALSE | FALSE |
| 13_v.txt | 31723 | 31723-SNV-A | SNV | 1 | A | C | 4 | 5942 | 0.0673174 | 43.75 | rnr1D57N | FALSE | CAN | JSY14 | 2 | rnr1D57N-Re | CAA | A>C | G/C+1 | A/T-1 | TA>GC | TA>GC | TGTCCCAAA | AAAAAGT | GTT | GTT | T | G | T>G | T>G_GTT | TA>GC_AAAAAGT | 7 | FALSE | FALSE |
| 13_v.txt | 31950 | 31950-SNV-A | SNV | 1 | A | C | 9 | 12930 | 0.06960557 | 38 | rnr1D57N | FALSE | CAN | JSY14 | 2 | rnr1D57N-Re | CAT | A>C | G/C+1 | A/T-1 | TA>GC | TA>GC | ATCGTCATA | ATAAATG | GTA | GTA | T | G | T>G | T>G_GTA | TA>GC_ATAAATG | 7 | FALSE | FALSE |
| 13_v.txt | 32283 | 32283-SNV-T | SNV | 1 | T | G | 587 | 13617 | 4.31078799 | 53.8262351 | rnr1D57N | FALSE | CAN | JSY14 | 2 | rnr1D57N-Re | GTA | T>G | G/C+1 | A/T-1 | TA>GC | TA>GC | CCAACGTAA | TAAATAT | CAT | GTA | T | C | T>G | T>G_GTA | TA>GC_TAAATAT | 7 | FALSE | FALSE |
| 13_v.txt | 32574 | 32574-SNV-T | SNV | 1 | T | G | 14 | 16747 | 0.08359706 | 33.7126531 | rnr1D57N | FALSE | CAN | JSY14 | 2 | rnr1D57N-Re | TTG | T>G | G/C+1 | A/T-1 | TA>GC | TA>GC | GTACCTTGA | TGAAATG | AAC | TTG | T | C | T>G | T>G_TTG | TA>GC_TGAAATG | 7 | FALSE | FALSE |
| 13_v.txt | 32739 | 32739-SNV-A | SNV | 1 | A | C | 1329 | 16747 | 7.93574969 | 55.6606471 | rnr1D57N | FALSE | CAN | JSY14 | 2 | rnr1D57N-Re | TAT | A>C | G/C+1 | A/T-1 | TA>GC | TA>GC | CAGTATATTA | ATTAGAA | ATA | ATA | T | G | T>G | T>G_ATA | TA>GC_ATTAGAA | 7 | FALSE | FALSE |
| 13_v.txt | 32799 | 32799-SNV-T | SNV | 1 | T | G | 6 | 16747 | 0.03582731 | 38.0022917 | rnr1D57N | FALSE | CAN | JSY14 | 2 | rnr1D57N-Re | GTA | T>G | G/C+1 | A/T-1 | TA>GC | TA>GC | TCACCGTAAT | TAATATT | CAT | GTA | T | C | T>G | T>G_GTA | TA>GC_TAATATT | 7 | FALSE | FALSE |
| 13_v.txt | 32817 | 32817-SNV-A | SNV | 1 | A | C | 26 | 16747 | 0.15525169 | 42.9615385 | rnr1D57N | FALSE | CAN | JSY14 | 2 | rnr1D57N-Re | CAA | A>C | G/C+1 | A/T-1 | TA>GC | TA>GC | GGGAACAAG | AAGTTCA | GTT | GTT | T | G | T>G | T>G_GTT | TA>GC_AAGTTCA | 7 | FALSE | FALSE |
| 13_v.txt | 32842 | 32842-SNV-A | SNV | 1 | A | C | 3 | 16747 | 0.01791366 | 38 | rnr1D57N | FALSE | CAN | JSY14 | 2 | rnr1D57N-Re | CAA | A>C | G/C+1 | A/T-1 | TA>GC | TA>GC | TTACCCAAA | AAAAAAT | GTT | GTT | T | G | T>G | T>G_GTT | TA>GC_AAAAAAT | 7 | FALSE | FALSE |
| 13_v.txt | 33157 | 33157-SNV-T | SNV | 1 | T | G | 10 | 18234 | 0.0548426 | 36.6 | rnr1D57N | FALSE | CAN | JSY14 | 2 | rnr1D57N-Re | GTA | T>G | G/C+1 | A/T-1 | TA>GC | TA>GC | GACCTGTAC | TACCAAT | CAT | GTA | T | C | T>G | T>G_GTA | TA>GC_TACCAAT | 7 | FALSE | FALSE |

| | | | | | | | | | | | | | | | | | | | | | | | | | | | | | | | | | | |
|---|---|---|---|---|---|---|---|---|---|---|---|---|---|---|---|---|---|---|---|---|---|---|---|---|---|---|---|---|---|---|---|---|---|---|
| 13_v.txt | 33269 | 33269-SNV-A | SNV | 1 | A | C | 71 | 11356 | 0.62522015 | 17.1690141 | rnr1D57N | FALSE | CAN | JSY14 | 2 | rnr1D57N-Re | TAG | A>C | G/C+1 | A/T-1 | TA>GC | TA>GC | CTCCATAGA | AGAGAAC | ATC | ATC | T | G | T>G | T>G_ATC | TA>GC_AGAGAAC | 7 | FALSE | FALSE |
| 13_v.txt | 32583 | 32583-SNV-A | SNV | 1 | A | G | 58 | 16747 | 0.34633069 | 37.1206897 | rnr1D57N | FALSE | CAN | JSY14 | 2 | rnr1D57N-Re | GAA | A>G | G/C+1 | A/T-1 | TA>CG | TA>CG | AATGTGAAG | AAGGCAG | CTT | CTT | T | C | T>C | T>C_CTT | TA>CG_AAGGCAG | 7 | FALSE | FALSE |
| 13_v.txt | 32842 | 32842-SNV-A | SNV | 1 | A | G | 5 | 16747 | 0.02985609 | 41 | rnr1D57N | FALSE | CAN | JSY14 | 2 | rnr1D57N-Re | CAA | A>G | G/C+1 | A/T-1 | TA>CG | TA>CG | TTACCCAAA | AAAAAAT | GTT | GTT | T | C | T>C | T>C_GTT | TA>CG_AAAAAAT | 7 | FALSE | FALSE |
| 13_v.txt | 32498 | 32498-SNV-T | SNV | 1 | T | A | 2 | 13616 | 0.0146886 | 50.9972984 | rnr1D57N | FALSE | CAN | JSY14 | 2 | rnr1D57N-Re | CTT | T>A | A/T+1 | A/T-1 | TA>AT | TA>AT | AACAACTTTT | TTTTTTG | GAA | CTT | T | T | T>A | T>A_CTT | TA>AT_TTTTTTG | 7 | FALSE | FALSE |
| 13_v.txt | 32916 | 32916-SNV-T | SNV | 1 | T | A | 23 | 18234 | 0.12613798 | 55.5217391 | rnr1D57N | FALSE | CAN | JSY14 | 2 | rnr1D57N-Re | TTC | T>A | A/T+1 | A/T-1 | TA>AT | TA>AT | CTAAGTTCC | TCCAGGG | AAG | TTC | T | T | T>A | T>A_TTC | TA>AT_TCCAGGG | 7 | FALSE | FALSE |
| 13_v.txt | 33251 | 33251-SNV-A | SNV | 1 | A | T | 8 | 11356 | 0.07044734 | 23.875 | rnr1D57N | FALSE | CAN | JSY14 | 2 | rnr1D57N-Re | CAC | A>T | A/T+1 | A/T-1 | TA>AT | TA>AT | TTCATCACCT | ACCTATG | GTG | GTG | T | A | T>A | T>A_GTG | TA>AT_ACCTATG | 7 | FALSE | FALSE |
| 13_v.txt | 31650 | 31650-SNV-C | SNV | 1 | C | T | 12 | 5942 | 0.20195221 | 36.1666667 | rnr1D57N | FALSE | CAN | JSY14 | 2 | rnr1D57N-Re | GCG | C>T | A/T+1 | G/C-1 | CG>TA | CG>TA | AGAATGCGA | CGAAATG | CGC | GCG | C | A | C>T | C>T_GCG | CG>TA_CGAAATG | 7 | FALSE | FALSE |
| 13_v.txt | 31834 | 31834-SNV-G | SNV | 1 | G | A | 13 | 5942 | 0.21878156 | 50.1538462 | rnr1D57N | FALSE | CAN | JSY14 | 2 | rnr1D57N-Re | TGA | G>A | A/T+1 | G/C-1 | CG>TA | CG>TA | TGCATTGAA | GAAATAA | ACT | ACT | C | T | C>T | C>T_ACT | CG>TA_GAAATAA | 7 | FALSE | FALSE |
| 13_v.txt | 31881 | 31881-SNV-G | SNV | 1 | G | A | 10 | 5942 | 0.1682935 | 39.8 | rnr1D57N | FALSE | CAN | JSY14 | 2 | rnr1D57N-Re | GGC | G>A | A/T+1 | G/C-1 | CG>TA | CG>TA | TAGGCGGCA | GCAGCAA | CCG | CCG | C | T | C>T | C>T_CCG | CG>TA_GCAGCAA | 7 | FALSE | FALSE |
| 13_v.txt | 32022 | 32022-SNV-C | SNV | 1 | C | T | 21 | 12931 | 0.16240043 | 49.6190476 | rnr1D57N | FALSE | CAN | JSY14 | 2 | rnr1D57N-Re | GCC | C>T | A/T+1 | G/C-1 | CG>TA | CG>TA | GAGATGCCA | CCACGGT | CGG | GCC | C | A | C>T | C>T_GCC | CG>TA_CCACGGT | 7 | FALSE | FALSE |
| 13_v.txt | 32075 | 32075-SNV-C | SNV | 1 | C | T | 23 | 12928 | 0.17790842 | 60.2600855 | rnr1D57N | FALSE | CAN | JSY14 | 2 | rnr1D57N-Re | ACC | C>T | A/T+1 | G/C-1 | CG>TA | CG>TA | AAATAACCA | CCATGCA | TGG | ACC | C | A | C>T | C>T_ACC | CG>TA_CCATGCA | 7 | FALSE | FALSE |
| 13_v.txt | 32076 | 32076-SNV-C | SNV | 1 | C | T | 317 | 12929 | 2.45185243 | 61.9666562 | rnr1D57N | FALSE | CAN | JSY14 | 2 | rnr1D57N-Re | CCA | C>T | A/T+1 | G/C-1 | CG>TA | CG>TA | AATAACCAT | CATGCAA | GGT | CCA | C | A | C>T | C>T_CCA | CG>TA_CATGCAA | 7 | FALSE | FALSE |
| 13_v.txt | 32088 | 32088-SNV-C | SNV | 1 | C | T | 36 | 12931 | 0.27840074 | 60.9722222 | rnr1D57N | FALSE | CAN | JSY14 | 2 | rnr1D57N-Re | GCC | C>T | A/T+1 | G/C-1 | CG>TA | CG>TA | AAAAAGCCT | CCTGCAA | CGG | GCC | C | A | C>T | C>T_GCC | CG>TA_CCTGCAA | 7 | FALSE | FALSE |
| 13_v.txt | 32092 | 32092-SNV-C | SNV | 1 | C | T | 16 | 12931 | 0.12373366 | 58.8125 | rnr1D57N | FALSE | CAN | JSY14 | 2 | rnr1D57N-Re | GCA | C>T | A/T+1 | G/C-1 | CG>TA | CG>TA | AGCCTGCAA | CAACACC | CGT | GCA | C | A | C>T | C>T_GCA | CG>TA_CAACACC | 7 | FALSE | FALSE |
| 13_v.txt | 32165 | 32165-SNV-G | SNV | 1 | G | A | 23 | 12931 | 0.17786714 | 45.3913044 | rnr1D57N | FALSE | CAN | JSY14 | 2 | rnr1D57N-Re | CGC | G>A | A/T+1 | G/C-1 | CG>TA | CG>TA | CAAAGCGCC | GCCAAAT | GCG | GCG | C | T | C>T | C>T_GCG | CG>TA_GCCAAAT | 7 | FALSE | FALSE |
| 13_v.txt | 32173 | 32173-SNV-C | SNV | 1 | C | T | 17 | 12931 | 0.13146702 | 45.4705882 | rnr1D57N | FALSE | CAN | JSY14 | 2 | rnr1D57N-Re | GCA | C>T | A/T+1 | G/C-1 | CG>TA | CG>TA | CAAATGCAG | CAGCAGT | CGT | GCA | C | A | C>T | C>T_GCA | CG>TA_CAGCAGT | 7 | FALSE | FALSE |
| 13_v.txt | 32176 | 32176-SNV-C | SNV | 1 | C | T | 15 | 12931 | 0.11600031 | 48.0666667 | rnr1D57N | FALSE | CAN | JSY14 | 2 | rnr1D57N-Re | GCA | C>T | A/T+1 | G/C-1 | CG>TA | CG>TA | ATGCAGCAG | CAGTAAC | CGT | GCA | C | A | C>T | C>T_GCA | CG>TA_CAGTAAC | 7 | FALSE | FALSE |
| 13_v.txt | 32271 | 32271-SNV-C | SNV | 1 | C | T | 24 | 13617 | 0.17625028 | 32.4166667 | rnr1D57N | FALSE | CAN | JSY14 | 2 | rnr1D57N-Re | ACG | C>T | A/T+1 | G/C-1 | CG>TA | CG>TA | AAAATACGG | CGGGAAC | TGC | ACG | C | A | C>T | C>T_ACG | CG>TA_CGGGAAC | 7 | FALSE | FALSE |
| 13_v.txt | 32299 | 32299-SNV-C | SNV | 1 | C | T | 29 | 13617 | 0.21296908 | 58.7931035 | rnr1D57N | FALSE | CAN | JSY14 | 2 | rnr1D57N-Re | GCG | C>T | A/T+1 | G/C-1 | CG>TA | CG>TA | AATTTGCGG | CGGCAGA | CGC | GCG | C | A | C>T | C>T_GCG | CG>TA_CGGCAGA | 7 | FALSE | FALSE |
| 13_v.txt | 32449 | 32449-SNV-C | SNV | 1 | C | T | 26 | 13616 | 0.19095182 | 51.6153846 | rnr1D57N | FALSE | CAN | JSY14 | 2 | rnr1D57N-Re | CCA | C>T | A/T+1 | G/C-1 | CG>TA | CG>TA | AAAGTCCAA | CAATGAA | GGT | CCA | C | A | C>T | C>T_CCA | CG>TA_CAATGAA | 7 | FALSE | FALSE |
| 13_v.txt | 32466 | 32466-SNV-C | SNV | 1 | C | T | 23 | 13616 | 0.16891892 | 45.4782609 | rnr1D57N | FALSE | CAN | JSY14 | 2 | rnr1D57N-Re | GCC | C>T | A/T+1 | G/C-1 | CG>TA | CG>TA | AGAGAGCCA | CCAATGT | CGG | GCC | C | A | C>T | C>T_GCC | CG>TA_CCAATGT | 7 | FALSE | FALSE |
| 13_v.txt | 32488 | 32488-SNV-G | SNV | 1 | G | A | 68 | 13616 | 0.49941246 | 47.4558824 | rnr1D57N | FALSE | CAN | JSY14 | 2 | rnr1D57N-Re | CGG | G>A | A/T+1 | G/C-1 | CG>TA | CG>TA | AGATACGGA | GGAAAAC | GCC | GCC | C | T | C>T | C>T_GCC | CG>TA_GGAAAAC | 7 | FALSE | FALSE |
| 13_v.txt | 32535 | 32535-SNV-G | SNV | 1 | G | A | 18 | 13616 | 0.13219742 | 36.5 | rnr1D57N | FALSE | CAN | JSY14 | 2 | rnr1D57N-Re | TGC | G>A | A/T+1 | G/C-1 | CG>TA | CG>TA | GGGTTTGCA | GCAGCTT | ACG | ACG | C | T | C>T | C>T_ACG | CG>TA_GCAGCTT | 7 | FALSE | FALSE |
| 13_v.txt | 32575 | 32575-SNV-G | SNV | 1 | G | A | 39 | 16747 | 0.23287753 | 37.4871795 | rnr1D57N | FALSE | CAN | JSY14 | 2 | rnr1D57N-Re | TGA | G>A | A/T+1 | G/C-1 | CG>TA | CG>TA | TACCTTGAA | GAAATGT | ACT | ACT | C | T | C>T | C>T_ACT | CG>TA_GAAATGT | 7 | FALSE | FALSE |
| 13_v.txt | 32589 | 32589-SNV-G | SNV | 1 | G | A | 3 | 16747 | 0.01791366 | 37.6663094 | rnr1D57N | FALSE | CAN | JSY14 | 2 | rnr1D57N-Re | AGC | G>A | A/T+1 | G/C-1 | CG>TA | CG>TA | AAGGCAGCG | GCGTTAA | TCG | TCG | C | T | C>T | C>T_TCG | CG>TA_GCGTTAA | 7 | FALSE | FALSE |
| 13_v.txt | 32590 | 32590-SNV-C | SNV | 1 | C | T | 30 | 16747 | 0.17913656 | 36.0994092 | rnr1D57N | FALSE | CAN | JSY14 | 2 | rnr1D57N-Re | GCG | C>T | A/T+1 | G/C-1 | CG>TA | CG>TA | AGGCAGCGT | CGTTAAT | CGC | GCG | C | A | C>T | C>T_GCG | CG>TA_CGTTAAT | 7 | FALSE | FALSE |
| 13_v.txt | 32700 | 32700-SNV-C | SNV | 1 | C | T | 27 | 16747 | 0.16122291 | 57.2592593 | rnr1D57N | FALSE | CAN | JSY14 | 2 | rnr1D57N-Re | GCC | C>T | A/T+1 | G/C-1 | CG>TA | CG>TA | ACTGGGCCG | CCGGTAA | CGG | GCC | C | A | C>T | C>T_GCC | CG>TA_CCGGTAA | 7 | FALSE | FALSE |
| 13_v.txt | 32776 | 32776-SNV-C | SNV | 1 | C | T | 19 | 16747 | 0.11345316 | 46.1052632 | rnr1D57N | FALSE | CAN | JSY14 | 2 | rnr1D57N-Re | GCG | C>T | A/T+1 | G/C-1 | CG>TA | CG>TA | TGGAAGCGA | CGACCCA | CGC | GCG | C | A | C>T | C>T_GCG | CG>TA_CGACCCA | 7 | FALSE | FALSE |
| 13_v.txt | 32798 | 32798-SNV-G | SNV | 1 | G | A | 27 | 16747 | 0.16122291 | 50.7030865 | rnr1D57N | FALSE | CAN | JSY14 | 2 | rnr1D57N-Re | CGT | G>A | A/T+1 | G/C-1 | CG>TA | CG>TA | TTCACCGTA | GTAATAT | GCA | GCA | C | T | C>T | C>T_GCA | CG>TA_GTAATAT | 7 | FALSE | FALSE |
| 13_v.txt | 32811 | 32811-SNV-G | SNV | 1 | G | A | 16 | 16747 | 0.0955395 | 48.3130223 | rnr1D57N | FALSE | CAN | JSY14 | 2 | rnr1D57N-Re | AGG | G>A | A/T+1 | G/C-1 | CG>TA | CG>TA | TTGACAGGG | GGGAACA | TCC | TCC | C | T | C>T | C>T_TCC | CG>TA_GGGAACA | 7 | FALSE | FALSE |
| 13_v.txt | 32900 | 32900-SNV-G | SNV | 1 | G | A | 28 | 18234 | 0.15355929 | 36.9239463 | rnr1D57N | FALSE | CAN | JSY14 | 2 | rnr1D57N-Re | GGC | G>A | A/T+1 | G/C-1 | CG>TA | CG>TA | GACTTGGCC | GCCAACT | CCG | CCG | C | T | C>T | C>T_CCG | CG>TA_GCCAACT | 7 | FALSE | FALSE |
| 13_v.txt | 32901 | 32901-SNV-C | SNV | 1 | C | T | 45 | 18234 | 0.24679171 | 30.933796 | rnr1D57N | FALSE | CAN | JSY14 | 2 | rnr1D57N-Re | GCC | C>T | A/T+1 | G/C-1 | CG>TA | CG>TA | ACTTGGCCA | CCAACTA | CGG | GCC | C | A | C>T | C>T_GCC | CG>TA_CCAACTA | 7 | FALSE | FALSE |
| 13_v.txt | 32965 | 32965-SNV-C | SNV | 1 | C | T | 23 | 18234 | 0.12613798 | 62 | rnr1D57N | FALSE | CAN | JSY14 | 2 | rnr1D57N-Re | GCC | C>T | A/T+1 | G/C-1 | CG>TA | CG>TA | CATTGGCCG | CCGCACC | CGG | GCC | C | A | C>T | C>T_GCC | CG>TA_CCGCACC | 7 | FALSE | FALSE |
| 13_v.txt | 32967 | 32967-SNV-G | SNV | 1 | G | A | 44 | 18234 | 0.24130745 | 60.8181818 | rnr1D57N | FALSE | CAN | JSY14 | 2 | rnr1D57N-Re | CGC | G>A | A/T+1 | G/C-1 | CG>TA | CG>TA | TTGGCCGCA | GCACCAA | GCG | GCG | C | T | C>T | C>T_GCG | CG>TA_GCACCAA | 7 | FALSE | FALSE |
| 13_v.txt | 33056 | 33056-SNV-C | SNV | 1 | C | T | 22 | 18234 | 0.12065372 | 50.9090909 | rnr1D57N | FALSE | CAN | JSY14 | 2 | rnr1D57N-Re | GCG | C>T | A/T+1 | G/C-1 | CG>TA | CG>TA | GGACTGCGT | CGTGACA | CGC | GCG | C | A | C>T | C>T_GCG | CG>TA_CGTGACA | 7 | FALSE | FALSE |
| 13_v.txt | 33070 | 33070-SNV-C | SNV | 1 | C | T | 14 | 18234 | 0.07677964 | 58.3571429 | rnr1D57N | FALSE | CAN | JSY14 | 2 | rnr1D57N-Re | GCC | C>T | A/T+1 | G/C-1 | CG>TA | CG>TA | AATATGCCA | CCAAAGA | CGG | GCC | C | A | C>T | C>T_GCC | CG>TA_CCAAAGA | 7 | FALSE | FALSE |
| 13_v.txt | 33104 | 33104-SNV-G | SNV | 1 | G | A | 27 | 18234 | 0.14807503 | 42.4074074 | rnr1D57N | FALSE | CAN | JSY14 | 2 | rnr1D57N-Re | CGC | G>A | A/T+1 | G/C-1 | CG>TA | CG>TA | AAGAGCGCC | GCCCACT | GCG | GCG | C | T | C>T | C>T_GCG | CG>TA_GCCCACT | 7 | FALSE | FALSE |
| 13_v.txt | 33111 | 33111-SNV-G | SNV | 1 | G | A | 21 | 18234 | 0.11516946 | 40.6190476 | rnr1D57N | FALSE | CAN | JSY14 | 2 | rnr1D57N-Re | TGG | G>A | A/T+1 | G/C-1 | CG>TA | CG>TA | CCCACTGGG | GGGCCGG | ACC | ACC | C | T | C>T | C>T_ACC | CG>TA_GGGCCGG | 7 | FALSE | FALSE |
| 13_v.txt | 33114 | 33114-SNV-C | SNV | 1 | C | T | 28 | 18234 | 0.15355929 | 43.1428571 | rnr1D57N | FALSE | CAN | JSY14 | 2 | rnr1D57N-Re | GCC | C>T | A/T+1 | G/C-1 | CG>TA | CG>TA | ACTGGGCCG | CCGGCGT | CGG | GCC | C | A | C>T | C>T_GCC | CG>TA_CCGGCGT | 7 | FALSE | FALSE |
| 13_v.txt | 33276 | 33276-SNV-G | SNV | 1 | G | A | 21 | 11356 | 0.18492427 | 49 | rnr1D57N | FALSE | CAN | JSY14 | 2 | rnr1D57N-Re | CGT | G>A | A/T+1 | G/C-1 | CG>TA | CG>TA | GAGAACGTA | GTATCCT | GCA | GCA | C | T | C>T | C>T_GCA | CG>TA_GTATCCT | 7 | FALSE | FALSE |
| 13_v.txt | 33310 | 33310-SNV-G | SNV | 1 | G | A | 17 | 11356 | 0.1497006 | 53.3529412 | rnr1D57N | FALSE | CAN | JSY14 | 2 | rnr1D57N-Re | CGC | G>A | A/T+1 | G/C-1 | CG>TA | CG>TA | AAGAGCGCA | GCAATGG | GCG | GCG | C | T | C>T | C>T_GCG | CG>TA_GCAATGG | 7 | FALSE | FALSE |
| 13_v.txt | 33412 | 33412-SNV-G | SNV | 1 | G | A | 18 | 11356 | 0.15850652 | 46.3888889 | rnr1D57N | FALSE | CAN | JSY14 | 2 | rnr1D57N-Re | GGC | G>A | A/T+1 | G/C-1 | CG>TA | CG>TA | TGACCGGCT | GCTCATT | CCG | CCG | C | T | C>T | C>T_CCG | CG>TA_GCTCATT | 7 | FALSE | FALSE |
| 13_v.txt | 33474 | 33474-SNV-C | SNV | 1 | C | T | 15 | 11356 | 0.13208876 | 35 | rnr1D57N | FALSE | CAN | JSY14 | 2 | rnr1D57N-Re | GCC | C>T | A/T+1 | G/C-1 | CG>TA | CG>TA | GCTATGCCTT | CCTTTTT | CGG | GCC | C | A | C>T | C>T_GCC | CG>TA_CCTTTTT | 7 | FALSE | FALSE |
| 13_v.txt | 32271 | 32271-SNV-C | SNV | 1 | C | G | 23 | 13617 | 0.16890651 | 36.2173913 | rnr1D57N | FALSE | CAN | JSY14 | 2 | rnr1D57N-Re | ACG | C>G | G/C+1 | G/C-1 | CG>GC | CG>GC | AAAATACGG | CGGGAAC | TGC | ACG | C | C | C>G | C>G_ACG | CG>GC_CGGGAAC | 7 | FALSE | FALSE |
| 13_v.txt | 32278 | 32278-SNV-C | SNV | 1 | C | G | 7 | 13617 | 0.05140633 | 37.8571429 | rnr1D57N | FALSE | CAN | JSY14 | 2 | rnr1D57N-Re | CCA | C>G | G/C+1 | G/C-1 | CG>GC | CG>GC | GGGAACCAA | CAACGTA | GGT | CCA | C | C | C>G | C>G_CCA | CG>GC_CAACGTA | 7 | FALSE | FALSE |
| 13_v.txt | 32783 | 32783-SNV-G | SNV | 1 | G | C | 3 | 16747 | 0.01791366 | 37.6666667 | rnr1D57N | FALSE | CAN | JSY14 | 2 | rnr1D57N-Re | AGA | G>C | G/C+1 | G/C-1 | CG>GC | CG>GC | GACCCAGAA | GAACTCG | TCT | TCT | C | G | C>G | C>G_TCT | CG>GC_GAACTCG | 7 | FALSE | FALSE |
| 13_v.txt | 33070 | 33070-SNV-C | SNV | 1 | C | G | 13 | 18234 | 0.07129538 | 55.4615385 | rnr1D57N | FALSE | CAN | JSY14 | 2 | rnr1D57N-Re | GCC | C>G | G/C+1 | G/C-1 | CG>GC | CG>GC | AATATGCCA | CCAAAGA | CGG | GCC | C | C | C>G | C>G_GCC | CG>GC_CCAAAGA | 7 | FALSE | FALSE |
| 13_v.txt | 33172 | 33172-SNV-C | SNV | 1 | C | G | 32 | 18233 | 0.17550595 | 37.96875 | rnr1D57N | FALSE | CAN | JSY14 | 2 | rnr1D57N-Re | CCA | C>G | G/C+1 | G/C-1 | CG>GC | CG>GC | TACCACCAA | CAAGGGC | GGT | CCA | C | C | C>G | C>G_CCA | CG>GC_CAAGGGC | 7 | FALSE | FALSE |
| 13_v.txt | 31657 | 31657-SNV-G | SNV | 1 | G | T | 58 | 5942 | 0.97610232 | 19.4655172 | rnr1D57N | FALSE | CAN | JSY14 | 2 | rnr1D57N-Re | GGC | G>T | A/T+1 | G/C-1 | CG>AT | CG>AT | GAAATGGCG | GCGTGGA | CCG | CCG | C | A | C>A | C>A_CCG | CG>AT_GCGTGGA | 7 | FALSE | FALSE |
| 13_v.txt | 32008 | 32008-SNV-C | SNV | 1 | C | A | 1398 | 12931 | 10.8112288 | 57.7360515 | rnr1D57N | FALSE | CAN | JSY14 | 2 | rnr1D57N-Re | TCG | C>A | A/T+1 | G/C-1 | CG>AT | CG>AT | GTAACTCGT | CGTCACG | AGC | TCG | C | T | C>A | C>A_TCG | CG>AT_CGTCACG | 7 | FALSE | FALSE |
| 13_v.txt | 32119 | 32119-SNV-C | SNV | 1 | C | A | 145 | 12931 | 1.12133632 | 58.6137931 | rnr1D57N | FALSE | CAN | JSY14 | 2 | rnr1D57N-Re | TCG | C>A | A/T+1 | G/C-1 | CG>AT | CG>AT | GCCATTCGA | CGAAAAC | AGC | TCG | C | T | C>A | C>A_TCG | CG>AT_CGAAAAC | 7 | FALSE | FALSE |
| 13_v.txt | 32205 | 32205-SNV-C | SNV | 1 | C | A | 79 | 12931 | 0.61093496 | 39 | rnr1D57N | FALSE | CAN | JSY14 | 2 | rnr1D57N-Re | ACC | C>A | A/T+1 | G/C-1 | CG>AT | CG>AT | GGAACACCA | CCACCTT | TGG | ACC | C | T | C>A | C>A_ACC | CG>AT_CCACCTT | 7 | FALSE | FALSE |
| 13_v.txt | 32399 | 32399-SNV-G | SNV | 1 | G | T | 8 | 13617 | 0.05875009 | 63.5 | rnr1D57N | FALSE | CAN | JSY14 | 2 | rnr1D57N-Re | CGT | G>T | A/T+1 | G/C-1 | CG>AT | CG>AT | AGAAACGTA | GTAGGAA | GCA | GCA | C | A | C>A | C>A_GCA | CG>AT_GTAGGAA | 7 | FALSE | FALSE |
| 13_v.txt | 32575 | 32575-SNV-G | SNV | 1 | G | T | 8 | 16747 | 0.04776975 | 30 | rnr1D57N | FALSE | CAN | JSY14 | 2 | rnr1D57N-Re | TGA | G>T | A/T+1 | G/C-1 | CG>AT | CG>AT | TACCTTGAA | GAAATGT | ACT | ACT | C | A | C>A | C>A_ACT | CG>AT_GAAATGT | 7 | FALSE | FALSE |
| 13_v.txt | 32783 | 32783-SNV-G | SNV | 1 | G | T | 7 | 16747 | 0.04179853 | 49.2857143 | rnr1D57N | FALSE | CAN | JSY14 | 2 | rnr1D57N-Re | AGA | G>T | A/T+1 | G/C-1 | CG>AT | CG>AT | GACCCAGAA | GAACTCG | TCT | TCT | C | A | C>A | C>A_TCT | CG>AT_GAACTCG | 7 | FALSE | FALSE |
| 13_v.txt | 32811 | 32811-SNV-G | SNV | 1 | G | T | 5 | 16747 | 0.02985609 | 37.6008453 | rnr1D57N | FALSE | CAN | JSY14 | 2 | rnr1D57N-Re | AGG | G>T | A/T+1 | G/C-1 | CG>AT | CG>AT | TTGACAGGG | GGGAACA | TCC | TCC | C | A | C>A | C>A_TCC | CG>AT_GGGAACA | 7 | FALSE | FALSE |
| 13_v.txt | 33006 | 33006-SNV-G | SNV | 1 | G | T | 10 | 18234 | 0.0548426 | 63.9 | rnr1D57N | FALSE | CAN | JSY14 | 2 | rnr1D57N-Re | TGT | G>T | A/T+1 | G/C-1 | CG>AT | CG>AT | AAAACTGTG | GTGAAAG | ACA | ACA | C | A | C>A | C>A_ACA | CG>AT_GTGAAAG | 7 | FALSE | FALSE |
| 13_v.txt | 33033 | 33033-SNV-G | SNV | 1 | G | T | 15 | 18234 | 0.0822639 | 59.6 | rnr1D57N | FALSE | CAN | JSY14 | 2 | rnr1D57N-Re | TGT | G>T | A/T+1 | G/C-1 | CG>AT | CG>AT | ATGAATGTA | GTAGCCA | ACA | ACA | C | A | C>A | C>A_ACA | CG>AT_GTAGCCA | 7 | FALSE | FALSE |
| 13_v.txt | 33043 | 33043-SNV-C | SNV | 1 | C | A | 50 | 18234 | 0.27421301 | 57.82 | rnr1D57N | FALSE | CAN | JSY14 | 2 | rnr1D57N-Re | TCA | C>A | A/T+1 | G/C-1 | CG>AT | CG>AT | CCATTTCACC | CACCCAA | AGT | TCA | C | T | C>A | C>A_TCA | CG>AT_CACCCAA | 7 | FALSE | FALSE |
| 13_v.txt | 33153 | 33153-SNV-C | SNV | 1 | C | A | 81 | 18234 | 0.44422507 | 41 | rnr1D57N | FALSE | CAN | JSY14 | 2 | rnr1D57N-Re | ACC | C>A | A/T+1 | G/C-1 | CG>AT | CG>AT | AAAAGACCT | CCTGTAC | TGG | ACC | C | T | C>A | C>A_ACC | CG>AT_CCTGTAC | 7 | FALSE | FALSE |
| 13_v.txt | 33159 | 33159-SNV-C | SNV | 1 | C | A | 16 | 18234 | 0.08774816 | 37.75 | rnr1D57N | FALSE | CAN | JSY14 | 2 | rnr1D57N-Re | ACC | C>A | A/T+1 | G/C-1 | CG>AT | CG>AT | CCTGTACCA | CCAATAG | TGG | ACC | C | T | C>A | C>A_ACC | CG>AT_CCAATAG | 7 | FALSE | FALSE |
| 13_v.txt | 33474 | 33474-SNV-C | SNV | 1 | C | A | 34 | 11356 | 0.2994012 | 35.2058824 | rnr1D57N | FALSE | CAN | JSY14 | 2 | rnr1D57N-Re | GCC | C>A | A/T+1 | G/C-1 | CG>AT | CG>AT | GCTATGCCTT | CCTTTTT | CGG | GCC | C | T | C>A | C>A_GCC | CG>AT_CCTTTTT | 7 | FALSE | FALSE |
| 13_v.txt | 33475 | 33475-SNV-C | SNV | 1 | C | A | 13 | 11356 | 0.11447693 | 35.3846154 | rnr1D57N | FALSE | CAN | JSY14 | 2 | rnr1D57N-Re | CCT | C>A | A/T+1 | G/C-1 | CG>AT | CG>AT | CTATGCCTTT | CTTTTTT | GGA | CCT | C | T | C>A | C>A_CCT | CG>AT_CTTTTTT | 7 | FALSE | FALSE |
| 13_v.txt | 32811 | 32811-Repla | Replacement | 3 | GGG | CA | 26 | 16747 | 0.15525169 | 44.8333334 | rnr1D57N | FALSE | CAN | JSY14 | 2 | rnr1D57N-Re | AGG | nope | >1bp-ins | >1bpdel | none | Replac. | TTGACAGGG | GGGAACA | TCC | AGG | GGG | none | NA | NA_AGG | Replac._GGGAACA | 7 | FALSE | FALSE |
| 13_v.txt | 32081 | 32081-Insert | Insertion | 1 | - | A | 3 | 12931 | 0.02320006 | 54.6666667 | rnr1D57N | FALSE | CAN | JSY14 | 2 | rnr1D57N-Re | CAA | nope | A/T+1 | >1bpdel | none | A/T+1 | CCATGCAAA | AAAAAAG | GTT | CAA | - | T | NA | NA_CAA | A/T+1_AAAAAAG | 7 | FALSE | FALSE |
| 13_v.txt | 32498 | 32498-Insert | Insertion | 1 | - | T | 2 | 13616 | 0.0146886 | 60 | rnr1D57N | FALSE | CAN | JSY14 | 2 | rnr1D57N-Re | CTT | nope | A/T+1 | >1bpdel | none | A/T+1 | AACAACTTTT | TTTTTTG | GAA | CTT | - | A | NA | NA_CTT | A/T+1_TTTTTTG | 7 | FALSE | FALSE |
| 13_v.txt | 32842 | 32842-Insert | Insertion | 1 | - | A | 29 | 16747 | 0.17316534 | 44 | rnr1D57N | FALSE | CAN | JSY14 | 2 | rnr1D57N-Re | CAA | nope | A/T+1 | >1bpdel | none | A/T+1 | TTACCCAAA | AAAAAAT | GTT | CAA | - | T | NA | NA_CAA | A/T+1_AAAAAAT | 7 | FALSE | FALSE |
| 13_v.txt | 32924 | 32924-Insert | Insertion | 1 | - | A | 2 | 18234 | 0.01096852 | 51 | rnr1D57N | FALSE | CAN | JSY14 | 2 | rnr1D57N-Re | CAA | nope | A/T+1 | >1bpdel | none | A/T+1 | CAGGGCAAA | AAAAGTG | GTT | CAA | - | T | NA | NA_CAA | A/T+1_AAAAGTG | 7 | FALSE | FALSE |
| 13_v.txt | 32589 | 32589-Deleti | Deletion | 2 | GC | - | 5 | 16747 | 0.02985609 | 38 | rnr1D57N | FALSE | CAN | JSY14 | 2 | rnr1D57N-Re | AGC | nope | >1bp-ins | >1bpdel | none | >1bpdel | AAGGCAGCG | GCGTTAA | TCG | AGC | GC | none | NA | NA_AGC | >1bpdel_GCGTTAA | 7 | FALSE | FALSE |
| 13_v.txt | 31663 | 31663-Deleti | Deletion | 1 | A | - | 6 | 5942 | 0.1009761 | 33.8333333 | rnr1D57N | FALSE | CAN | JSY14 | 2 | rnr1D57N-Re | GAA | nope | >1bp-ins | A/T-1 | none | A/T-1 | GCGTGGAAA | AAATGTG | CTT | CTT | T | none | T>none | T>none_CTT | A/T-1_AAATGTG | 7 | FALSE | FALSE |
| 13_v.txt | 31723 | 31723-Deleti | Deletion | 1 | A | - | 8 | 5942 | 0.1346348 | 60.25 | rnr1D57N | FALSE | CAN | JSY14 | 2 | rnr1D57N-Re | CAA | nope | >1bp-ins | A/T-1 | none | A/T-1 | TGTCCCAAA | AAAAAGT | GTT | GTT | T | none | T>none | T>none_GTT | A/T-1_AAAAAGT | 7 | FALSE | FALSE |
| 13_v.txt | 31731 | 31731-Deleti | Deletion | 1 | T | - | 4 | 5942 | 0.0673174 | 64 | rnr1D57N | FALSE | CAN | JSY14 | 2 | rnr1D57N-Re | CTT | nope | >1bp-ins | A/T-1 | none | A/T-1 | AAAGTCTTT | TTTGGTT | GAA | CTT | T | none | nope | nope_CTT | A/T-1_TTTGGTT | 7 | FALSE | FALSE |
| 13_v.txt | 31904 | 31904-Deleti | Deletion | 1 | T | - | 5 | 5942 | 0.08414675 | 43.2 | rnr1D57N | FALSE | CAN | JSY14 | 2 | rnr1D57N-Re | ATT | nope | >1bp-ins | A/T-1 | none | A/T-1 | ATTGAATTTT | TTTTGGT | TAA | ATT | T | none | nope | nope_ATT | A/T-1_TTTTGGT | 7 | FALSE | FALSE |
| 13_v.txt | 31913 | 31913-Deleti | Deletion | 1 | A | - | 4 | 5942 | 0.0673174 | 38 | rnr1D57N | FALSE | CAN | JSY14 | 2 | rnr1D57N-Re | CAA | nope | >1bp-ins | A/T-1 | none | A/T-1 | TGGTGCAAA | AAAAGCC | GTT | GTT | T | none | T>none | T>none_GTT | A/T-1_AAAAGCC | 7 | FALSE | FALSE |
| 13_v.txt | 31971 | 31971-Deleti | Deletion | 1 | G | - | 7707 | 12931 | 59.6009589 | 37.3145193 | rnr1D57N | FALSE | CAN | JSY14 | 2 | rnr1D57N-Re | AGC | nope | >1bp-ins | G/C-1 | none | G/C-1 | TAATAAGCC | GCCAAGC | TCG | TCG | C | none | C>none | C>none_TCG | G/C-1_GCCAAGC | 7 | FALSE | FALSE |
| 13_v.txt | 32075 | 32075-Deleti | Deletion | 1 | C | - | 2 | 12928 | 0.0154703 | 64 | rnr1D57N | FALSE | CAN | JSY14 | 2 | rnr1D57N-Re | ACC | nope | >1bp-ins | G/C-1 | none | G/C-1 | AAATAACCA | CCATGCA | TGG | ACC | C | none | nope | nope_ACC | G/C-1_CCATGCA | 7 | FALSE | FALSE |
| 13_v.txt | 32081 | 32081-Deleti | Deletion | 1 | A | - | 13 | 12930 | 0.10054138 | 61.384616 | rnr1D57N | FALSE | CAN | JSY14 | 2 | rnr1D57N-Re | CAA | nope | >1bp-ins | A/T-1 | none | A/T-1 | CCATGCAAA | AAAAAAG | GTT | GTT | T | none | T>none | T>none_GTT | A/T-1_AAAAAAG | 7 | FALSE | FALSE |
| 13_v.txt | 32381 | 32381-Deleti | Deletion | 1 | A | - | 8 | 13617 | 0.05875009 | 56.375 | rnr1D57N | FALSE | CAN | JSY14 | 2 | rnr1D57N-Re | TAA | nope | >1bp-ins | A/T-1 | none | A/T-1 | AATAATAAA | AAAGGGA | ATT | ATT | T | none | T>none | T>none_ATT | A/T-1_AAAGGGA | 7 | FALSE | FALSE |
| 13_v.txt | 32498 | 32498-Deleti | Deletion | 1 | T | - | 17 | 13616 | 0.12485311 | 42.9411748 | rnr1D57N | FALSE | CAN | JSY14 | 2 | rnr1D57N-Re | CTT | nope | >1bp-ins | A/T-1 | none | A/T-1 | AACAACTTTT | TTTTTTG | GAA | CTT | T | none | nope | nope_CTT | A/T-1_TTTTTTG | 7 | FALSE | FALSE |
| 13_v.txt | 32583 | 32583-Deleti | Deletion | 1 | A | - | 2 | 16747 | 0.01194244 | 38 | rnr1D57N | FALSE | CAN | JSY14 | 2 | rnr1D57N-Re | GAA | nope | >1bp-ins | A/T-1 | none | A/T-1 | AATGTGAAG | AAGGCAG | CTT | CTT | T | none | T>none | T>none_CTT | A/T-1_AAGGCAG | 7 | FALSE | FALSE |
| 13_v.txt | 32631 | 32631-Deleti | Deletion | 1 | T | - | 10 | 16747 | 0.05971219 | 47 | rnr1D57N | FALSE | CAN | JSY14 | 2 | rnr1D57N-Re | GTT | nope | >1bp-ins | A/T-1 | none | A/T-1 | CCTTCGTTTT | TTTTTAT | CAA | GTT | T | none | nope | nope_GTT | A/T-1_TTTTTAT | 7 | FALSE | FALSE |
| 13_v.txt | 32658 | 32658-Deleti | Deletion | 1 | C | - | 7 | 16747 | 0.04179853 | 59.1428571 | rnr1D57N | FALSE | CAN | JSY14 | 2 | rnr1D57N-Re | ACC | nope | >1bp-ins | G/C-1 | none | G/C-1 | CCTGGACCC | CCCCAGG | TGG | ACC | C | none | nope | nope_ACC | G/C-1_CCCCAGG | 7 | FALSE | FALSE |
| 13_v.txt | 32722 | 32722-Deleti | Deletion | 1 | C | - | 7 | 16747 | 0.04179853 | 64 | rnr1D57N | FALSE | CAN | JSY14 | 2 | rnr1D57N-Re | ACC | nope | >1bp-ins | G/C-1 | none | G/C-1 | CACAAACCA | CCATACA | TGG | ACC | C | none | nope | nope_ACC | G/C-1_CCATACA | 7 | FALSE | FALSE |
| 13_v.txt | 32744 | 32744-Deleti | Deletion | 1 | A | - | 7 | 16747 | 0.04179853 | 54.8571429 | rnr1D57N | FALSE | CAN | JSY14 | 2 | rnr1D57N-Re | GAA | nope | >1bp-ins | A/T-1 | none | A/T-1 | TATTAGAAA | AAACCCG | CTT | CTT | T | none | T>none | T>none_CTT | A/T-1_AAACCCG | 7 | FALSE | FALSE |
| 13_v.txt | 32783 | 32783-Deleti | Deletion | 1 | G | - | 2 | 16747 | 0.01194244 | 51 | rnr1D57N | FALSE | CAN | JSY14 | 2 | rnr1D57N-Re | AGA | nope | >1bp-ins | G/C-1 | none | G/C-1 | GACCCAGAA | GAACTCG | TCT | TCT | C | none | C>none | C>none_TCT | G/C-1_GAACTCG | 7 | FALSE | FALSE |
| 13_v.txt | 32799 | 32799-Deleti | Deletion | 1 | T | - | 1 | 16747 | 5.97E-03 | 38.0000028 | rnr1D57N | FALSE | CAN | JSY14 | 2 | rnr1D57N-Re | GTA | nope | >1bp-ins | A/T-1 | none | A/T-1 | TCACCGTAAT | TAATATT | CAT | GTA | T | none | nope | nope_GTA | A/T-1_TAATATT | 7 | FALSE | FALSE |
| 13_v.txt | 32842 | 32842-Deleti | Deletion | 1 | A | - | 33 | 16747 | 0.19705022 | 41.9393939 | rnr1D57N | FALSE | CAN | JSY14 | 2 | rnr1D57N-Re | CAA | nope | >1bp-ins | A/T-1 | none | A/T-1 | TTACCCAAA | AAAAAAT | GTT | GTT | T | none | T>none | T>none_GTT | A/T-1_AAAAAAT | 7 | FALSE | FALSE |
| 13_v.txt | 32884 | 32884-Deleti | Deletion | 1 | A | - | 8 | 18236 | 0.04386927 | 36.625 | rnr1D57N | FALSE | CAN | JSY14 | 2 | rnr1D57N-Re | CAA | nope | >1bp-ins | A/T-1 | none | A/T-1 | ACGTCCAAA | AAAATTG | GTT | GTT | T | none | T>none | T>none_GTT | A/T-1_AAAATTG | 7 | FALSE | FALSE |
| 13_v.txt | 32901 | 32901-Deleti | Deletion | 1 | C | - | 1 | 18234 | 5.48E-03 | 38 | rnr1D57N | FALSE | CAN | JSY14 | 2 | rnr1D57N-Re | GCC | nope | >1bp-ins | G/C-1 | none | G/C-1 | ACTTGGCCA | CCAACTA | CGG | GCC | C | none | nope | nope_GCC | G/C-1_CCAACTA | 7 | FALSE | FALSE |
| 13_v.txt | 32924 | 32924-Deleti | Deletion | 1 | A | - | 24 | 18234 | 0.13162224 | 55.2499995 | rnr1D57N | FALSE | CAN | JSY14 | 2 | rnr1D57N-Re | CAA | nope | >1bp-ins | A/T-1 | none | A/T-1 | CAGGGCAAA | AAAAGTG | GTT | GTT | T | none | T>none | T>none_GTT | A/T-1_AAAAGTG | 7 | FALSE | FALSE |
| 13_v.txt | 33000 | 33000-Deleti | Deletion | 1 | A | - | 58 | 18234 | 0.31808709 | 59.3793103 | rnr1D57N | FALSE | CAN | JSY14 | 2 | rnr1D57N-Re | GAA | nope | >1bp-ins | A/T-1 | none | A/T-1 | TGTGAGAAA | AAAACTG | CTT | CTT | T | none | T>none | T>none_CTT | A/T-1_AAAACTG | 7 | FALSE | FALSE |
| 13_v.txt | 33251 | 33251-Deleti | Deletion | 1 | A | - | 5 | 11356 | 0.04402959 | 53.4 | rnr1D57N | FALSE | CAN | JSY14 | 2 | rnr1D57N-Re | CAC | nope | >1bp-ins | A/T-1 | none | A/T-1 | TTCATCACCT | ACCTATG | GTG | GTG | T | none | T>none | T>none_GTG | A/T-1_ACCTATG | 7 | FALSE | FALSE |
| 13_v.txt | 33474 | 33474-Deleti | Deletion | 1 | C | - | 33 | 11356 | 0.29059528 | 26.3333333 | rnr1D57N | FALSE | CAN | JSY14 | 2 | rnr1D57N-Re | GCC | nope | >1bp-ins | G/C-1 | none | G/C-1 | GCTATGCCTT | CCTTTTT | CGG | GCC | C | none | nope | nope_GCC | G/C-1_CCTTTTT | 7 | FALSE | FALSE |
| 14_v.txt | 31723 | 31723-SNV-A | SNV | 1 | A | C | 4 | 10140 | 0.03944773 | 62.25 | rnr1D57N | FALSE | CAN | JSY14 | 3 | rnr1D57N-Re | CAA | A>C | G/C+1 | A/T-1 | TA>GC | TA>GC | TGTCCCAAA | AAAAAGT | GTT | GTT | T | G | T>G | T>G_GTT | TA>GC_AAAAAGT | 7 | FALSE | FALSE |
| 14_v.txt | 32283 | 32283-SNV-T | SNV | 1 | T | G | 722 | 22305 | 3.23694239 | 53.0152355 | rnr1D57N | FALSE | CAN | JSY14 | 3 | rnr1D57N-Re | GTA | T>G | G/C+1 | A/T-1 | TA>GC | TA>GC | CCAACGTAA | TAAATAT | CAT | GTA | T | C | T>G | T>G_GTA | TA>GC_TAAATAT | 7 | FALSE | FALSE |
| 14_v.txt | 32631 | 32631-SNV-T | SNV | 1 | T | G | 4 | 20619 | 0.01939958 | 44.75 | rnr1D57N | FALSE | CAN | JSY14 | 3 | rnr1D57N-Re | GTT | T>G | G/C+1 | A/T-1 | TA>GC | TA>GC | CCTTCGTTTT | TTTTTAT | CAA | GTT | T | C | T>G | T>G_GTT | TA>GC_TTTTTAT | 7 | FALSE | FALSE |
| 14_v.txt | 32739 | 32739-SNV-A | SNV | 1 | A | C | 56 | 20619 | 0.27159416 | 56.1785714 | rnr1D57N | FALSE | CAN | JSY14 | 3 | rnr1D57N-Re | TAT | A>C | G/C+1 | A/T-1 | TA>GC | TA>GC | CAGTATATT | ATTAGAA | ATA | ATA | T | G | T>G | T>G_ATA | TA>GC_ATTAGAA | 7 | FALSE | FALSE |
| 14_v.txt | 32823 | 32823-SNV-A | SNV | 1 | A | C | 12 | 20619 | 0.05819875 | 49.5 | rnr1D57N | FALSE | CAN | JSY14 | 3 | rnr1D57N-Re | CAT | A>C | G/C+1 | A/T-1 | TA>GC | TA>GC | AAGTTCATT | ATTATTG | GTA | GTA | T | G | T>G | T>G_GTA | TA>GC_ATTATTG | 7 | FALSE | FALSE |
| 14_v.txt | 32842 | 32842-SNV-A | SNV | 1 | A | C | 3 | 20619 | 0.01454969 | 41.333313 | rnr1D57N | FALSE | CAN | JSY14 | 3 | rnr1D57N-Re | CAA | A>C | G/C+1 | A/T-1 | TA>GC | TA>GC | TTACCCAAA | AAAAAAT | GTT | GTT | T | G | T>G | T>G_GTT | TA>GC_AAAAAAT | 7 | FALSE | FALSE |
| 14_v.txt | 32884 | 32884-SNV-A | SNV | 1 | A | C | 6 | 22864 | 0.02624213 | 36.1666667 | rnr1D57N | FALSE | CAN | JSY14 | 3 | rnr1D57N-Re | CAA | A>C | G/C+1 | A/T-1 | TA>GC | TA>GC | ACGTCCAAA | AAAATTG | GTT | GTT | T | G | T>G | T>G_GTT | TA>GC_AAAATTG | 7 | FALSE | FALSE |
| 14_v.txt | 33099 | 33099-SNV-A | SNV | 1 | A | C | 10 | 22863 | 0.04373879 | 42.8 | rnr1D57N | FALSE | CAN | JSY14 | 3 | rnr1D57N-Re | AAG | A>C | G/C+1 | A/T-1 | TA>GC | TA>GC | GATATAAGA | AGAGCGC | TTC | TTC | T | G | T>G | T>G_TTC | TA>GC_AGAGCGC | 7 | FALSE | FALSE |
| 14_v.txt | 33269 | 33269-SNV-A | SNV | 1 | A | C | 132 | 16483 | 0.80082509 | 15.6212121 | rnr1D57N | FALSE | CAN | JSY14 | 3 | rnr1D57N-Re | TAG | A>C | G/C+1 | A/T-1 | TA>GC | TA>GC | CTCCATAGA | AGAGAAC | ATC | ATC | T | G | T>G | T>G_ATC | TA>GC_AGAGAAC | 7 | FALSE | FALSE |
| 14_v.txt | 33476 | 33476-SNV-T | SNV | 1 | T | G | 1 | 16482 | 6.07E-03 | 37.9989018 | rnr1D57N | FALSE | CAN | JSY14 | 3 | rnr1D57N-Re | CTT | T>G | G/C+1 | A/T-1 | TA>GC | TA>GC | TATGCCTTTT | TTTTTTT | GAA | CTT | T | C | T>G | T>G_CTT | TA>GC_TTTTTTT | 7 | FALSE | FALSE |
| 14_v.txt | 32055 | 32055-SNV-T | SNV | 1 | T | C | 15 | 14110 | 0.10630758 | 58.2666667 | rnr1D57N | FALSE | CAN | JSY14 | 3 | rnr1D57N-Re | GTG | T>C | G/C+1 | A/T-1 | TA>CG | TA>CG | CTGATGTGC | TGCGAGA | CAC | GTG | T | G | T>C | T>C_GTG | TA>CG_TGCGAGA | 7 | FALSE | FALSE |
| 14_v.txt | 32256 | 32256-SNV-A | SNV | 1 | A | G | 19 | 22297 | 0.08521326 | 36.7368421 | rnr1D57N | FALSE | CAN | JSY14 | 3 | rnr1D57N-Re | TAG | A>G | G/C+1 | A/T-1 | TA>CG | TA>CG | TTTGATAGA | AGACCAA | ATC | ATC | T | C | T>C | T>C_ATC | TA>CG_AGACCAA | 7 | FALSE | FALSE |
| 14_v.txt | 32293 | 32293-SNV-A | SNV | 1 | A | G | 33 | 22305 | 0.14794889 | 54.1818182 | rnr1D57N | FALSE | CAN | JSY14 | 3 | rnr1D57N-Re | GAA | A>G | G/C+1 | A/T-1 | TA>CG | TA>CG | TATTTGAATT | AATTTGC | CTT | CTT | T | C | T>C | T>C_CTT | TA>CG_AATTTGC | 7 | FALSE | FALSE |
| 14_v.txt | 32338 | 32338-SNV-T | SNV | 1 | T | C | 3 | 22305 | 0.0134499 | 54.3331586 | rnr1D57N | FALSE | CAN | JSY14 | 3 | rnr1D57N-Re | ATA | T>C | G/C+1 | A/T-1 | TA>CG | TA>CG | TGAAGATAT | TATGTGG | TAT | ATA | T | G | T>C | T>C_ATA | TA>CG_TATGTGG | 7 | FALSE | FALSE |
| 14_v.txt | 32583 | 32583-SNV-A | SNV | 1 | A | G | 47 | 20619 | 0.2279451 | 37.3191489 | rnr1D57N | FALSE | CAN | JSY14 | 3 | rnr1D57N-Re | GAA | A>G | G/C+1 | A/T-1 | TA>CG | TA>CG | AATGTGAAG | AAGGCAG | CTT | CTT | T | C | T>C | T>C_CTT | TA>CG_AAGGCAG | 7 | FALSE | FALSE |
| 14_v.txt | 32762 | 32762-SNV-A | SNV | 1 | A | G | 84 | 20619 | 0.40739124 | 52.5119048 | rnr1D57N | FALSE | CAN | JSY14 | 3 | rnr1D57N-Re | AAA | A>G | G/C+1 | A/T-1 | TA>CG | TA>CG | GGCTAAAAC | AACTTTG | TTT | TTT | T | C | T>C | T>C_TTT | TA>CG_AACTTTG | 7 | FALSE | FALSE |
| 14_v.txt | 32787 | 32787-SNV-T | SNV | 1 | T | C | 16 | 20619 | 0.07759833 | 45.0625 | rnr1D57N | FALSE | CAN | JSY14 | 3 | rnr1D57N-Re | CTC | T>C | G/C+1 | A/T-1 | TA>CG | TA>CG | CAGAACTCG | TCGAATT | GAG | CTC | T | G | T>C | T>C_CTC | TA>CG_TCGAATT | 7 | FALSE | FALSE |
| 14_v.txt | 33174 | 33174-SNV-A | SNV | 1 | A | G | 199 | 22857 | 0.87063044 | 37.2160804 | rnr1D57N | FALSE | CAN | JSY14 | 3 | rnr1D57N-Re | AAG | A>G | G/C+1 | A/T-1 | TA>CG | TA>CG | CCACCAAGG | AGGGCAA | TTC | TTC | T | C | T>C | T>C_TTC | TA>CG_AGGGCAA | 7 | FALSE | FALSE |
| 14_v.txt | 33476 | 33476-SNV-T | SNV | 1 | T | C | 26 | 16482 | 0.15774785 | 32.6152455 | rnr1D57N | FALSE | CAN | JSY14 | 3 | rnr1D57N-Re | CTT | T>C | G/C+1 | A/T-1 | TA>CG | TA>CG | TATGCCTTTT | TTTTTTT | GAA | CTT | T | G | T>C | T>C_CTT | TA>CG_TTTTTTT | 7 | FALSE | FALSE |
| 14_v.txt | 32338 | 32338-SNV-T | SNV | 1 | T | A | 1 | 22305 | 4.48E-03 | 63.9985005 | rnr1D57N | FALSE | CAN | JSY14 | 3 | rnr1D57N-Re | ATA | T>A | A/T+1 | A/T-1 | TA>AT | TA>AT | TGAAGATAT | TATGTGG | TAT | ATA | T | T | T>A | T>A_ATA | TA>AT_TATGTGG | 7 | FALSE | FALSE |
| 14_v.txt | 32680 | 32680-SNV-A | SNV | 1 | A | T | 120 | 20619 | 0.58198749 | 61.9916667 | rnr1D57N | FALSE | CAN | JSY14 | 3 | rnr1D57N-Re | CAA | A>T | A/T+1 | A/T-1 | TA>AT | TA>AT | TTCTCCAATA | AATAACG | GTT | GTT | T | A | T>A | T>A_GTT | TA>AT_AATAACG | 7 | FALSE | FALSE |
| 14_v.txt | 32766 | 32766-SNV-T | SNV | 1 | T | A | 86 | 20619 | 0.41709103 | 52.5930233 | rnr1D57N | FALSE | CAN | JSY14 | 3 | rnr1D57N-Re | TTT | T>A | A/T+1 | A/T-1 | TA>AT | TA>AT | AAAACTTTG | TTGATGG | AAA | TTT | T | T | T>A | T>A_TTT | TA>AT_TTGATGG | 7 | FALSE | FALSE |
| 14_v.txt | 32787 | 32787-SNV-T | SNV | 1 | T | A | 19 | 20619 | 0.09214802 | 43.2105263 | rnr1D57N | FALSE | CAN | JSY14 | 3 | rnr1D57N-Re | CTC | T>A | A/T+1 | A/T-1 | TA>AT | TA>AT | CAGAACTCG | TCGAATT | GAG | CTC | T | T | T>A | T>A_CTC | TA>AT_TCGAATT | 7 | FALSE | FALSE |
| 14_v.txt | 32823 | 32823-SNV-A | SNV | 1 | A | T | 9 | 20619 | 0.04364906 | 40.7777778 | rnr1D57N | FALSE | CAN | JSY14 | 3 | rnr1D57N-Re | CAT | A>T | A/T+1 | A/T-1 | TA>AT | TA>AT | AAGTTCATT | ATTATTG | GTA | GTA | T | A | T>A | T>A_GTA | TA>AT_ATTATTG | 7 | FALSE | FALSE |
| 14_v.txt | 33072 | 33072-SNV-A | SNV | 1 | A | T | 7 | 22863 | 0.03061715 | 52.4285714 | rnr1D57N | FALSE | CAN | JSY14 | 3 | rnr1D57N-Re | CAA | A>T | A/T+1 | A/T-1 | TA>AT | TA>AT | TATGCCAAA | AAAGAAC | GTT | GTT | T | A | T>A | T>A_GTT | TA>AT_AAAGAAC | 7 | FALSE | FALSE |
| 14_v.txt | 33095 | 33095-SNV-T | SNV | 1 | T | A | 6 | 22863 | 0.02624328 | 41.4988519 | rnr1D57N | FALSE | CAN | JSY14 | 3 | rnr1D57N-Re | ATA | T>A | A/T+1 | A/T-1 | TA>AT | TA>AT | ATATGATAT | TATAAGA | TAT | ATA | T | T | T>A | T>A_ATA | TA>AT_TATAAGA | 7 | FALSE | FALSE |

| | | | | | | | | | | | | | | | | | | | | | | | | | | | | | | | | | | |
|---|---|---|---|---|---|---|---|---|---|---|---|---|---|---|---|---|---|---|---|---|---|---|---|---|---|---|---|---|---|---|---|---|---|---|
| 14_v.txt | 33364 | 33364-SNV-T | SNV | 1 | T | A | 24 | 16483 | 0.14560456 | 52.6666667 | rnr1D57N | FALSE | CAN | JSY14 | 3 | rnr1D57N-Re | CTG | T>A | A/T+1 | A/T-1 | TA>AT | TA>AT | CACGTCTGTC | TGTGGTG | GAC | CTG | T | T | T>A | T>A_CTG | TA>AT_TGTGGTG | 7 | FALSE | FALSE |
| 14_v.txt | 31650 | 31650-SNV-( | SNV | 1 | C | T | 22 | 10140 | 0.21696253 | 34.4090909 | rnr1D57N | FALSE | CAN | JSY14 | 3 | rnr1D57N-Re | GCG | C>T | A/T+1 | G/C-1 | CG>TA | CG>TA | AGAATGCGA | CGAAATG | CGC | GCG | C | A | C>T | C>T_GCG | CG>TA_CGAAATG | 7 | FALSE | FALSE |
| 14_v.txt | 31829 | 31829-SNV-( | SNV | 1 | G | A | 6 | 10140 | 0.0591716 | 59.6666667 | rnr1D57N | FALSE | CAN | JSY14 | 3 | rnr1D57N-Re | TGC | G>A | A/T+1 | G/C-1 | CG>TA | CG>TA | GAATATGCA | GCATTGA | ACG | ACG | C | T | C>T | C>T_ACG | CG>TA_GCATTGA | 7 | FALSE | FALSE |
| 14_v.txt | 31845 | 31845-SNV-( | SNV | 1 | C | T | 17 | 10140 | 0.16765286 | 40.7058824 | rnr1D57N | FALSE | CAN | JSY14 | 3 | rnr1D57N-Re | CCA | C>T | A/T+1 | G/C-1 | CG>TA | CG>TA | AAGATCCAA | CAAACAG | GGT | CCA | C | A | C>T | C>T_CCA | CG>TA_CAAACAG | 7 | FALSE | FALSE |
| 14_v.txt | 31912 | 31912-SNV-( | SNV | 1 | C | T | 13 | 10140 | 0.12820513 | 39.2307692 | rnr1D57N | FALSE | CAN | JSY14 | 3 | rnr1D57N-Re | GCA | C>T | A/T+1 | G/C-1 | CG>TA | CG>TA | TTGGTGCAA | CAAAAGC | CGT | GCA | C | A | C>T | C>T_GCA | CG>TA_CAAAAGC | 7 | FALSE | FALSE |
| 14_v.txt | 31930 | 31930-SNV-( | SNV | 1 | G | A | 42 | 10140 | 0.41420118 | 37.0238095 | rnr1D57N | FALSE | CAN | JSY14 | 3 | rnr1D57N-Re | TGA | G>A | A/T+1 | G/C-1 | CG>TA | CG>TA | AACCTTGAA | GAATAAT | ACT | ACT | C | T | C>T | C>T_ACT | CG>TA_GAATAAT | 7 | FALSE | FALSE |
| 14_v.txt | 31960 | 31960-SNV-( | SNV | 1 | C | T | 18 | 14108 | 0.12758719 | 36.8888889 | rnr1D57N | FALSE | CAN | JSY14 | 3 | rnr1D57N-Re | GCC | C>T | A/T+1 | G/C-1 | CG>TA | CG>TA | ATGTGGCCG | CCGCATA | CGG | GCC | C | A | C>T | C>T_GCC | CG>TA_CCGCATA | 7 | FALSE | FALSE |
| 14_v.txt | 31962 | 31962-SNV-( | SNV | 1 | G | A | 19 | 14108 | 0.13467536 | 34.4736842 | rnr1D57N | FALSE | CAN | JSY14 | 3 | rnr1D57N-Re | CGC | G>A | A/T+1 | G/C-1 | CG>TA | CG>TA | GTGGCCGCA | GCATAAT | GCG | GCG | C | T | C>T | C>T_GCG | CG>TA_GCATAAT | 7 | FALSE | FALSE |
| 14_v.txt | 32057 | 32057-SNV-( | SNV | 1 | C | T | 15 | 14110 | 0.10630758 | 62.2666667 | rnr1D57N | FALSE | CAN | JSY14 | 3 | rnr1D57N-Re | GCG | C>T | A/T+1 | G/C-1 | CG>TA | CG>TA | GATGTGCGA | CGAGATT | CGC | GCG | C | A | C>T | C>T_GCG | CG>TA_CGAGATT | 7 | FALSE | FALSE |
| 14_v.txt | 32075 | 32075-SNV-( | SNV | 1 | C | T | 48 | 14110 | 0.34018427 | 63.3541667 | rnr1D57N | FALSE | CAN | JSY14 | 3 | rnr1D57N-Re | ACC | C>T | A/T+1 | G/C-1 | CG>TA | CG>TA | AAATAACCA | CCATGCA | TGG | ACC | C | A | C>T | C>T_ACC | CG>TA_CCATGCA | 7 | FALSE | FALSE |
| 14_v.txt | 32092 | 32092-SNV-( | SNV | 1 | C | T | 17 | 14110 | 0.12048193 | 60.1764706 | rnr1D57N | FALSE | CAN | JSY14 | 3 | rnr1D57N-Re | GCA | C>T | A/T+1 | G/C-1 | CG>TA | CG>TA | AGCCTGCAA | CAACACC | CGT | GCA | C | A | C>T | C>T_GCA | CG>TA_CAACACC | 7 | FALSE | FALSE |
| 14_v.txt | 32165 | 32165-SNV-( | SNV | 1 | G | A | 21 | 14110 | 0.14883062 | 52 | rnr1D57N | FALSE | CAN | JSY14 | 3 | rnr1D57N-Re | CGC | G>A | A/T+1 | G/C-1 | CG>TA | CG>TA | CAAAGCGCC | GCCAAAT | GCG | GCG | C | T | C>T | C>T_GCG | CG>TA_GCCAAAT | 7 | FALSE | FALSE |
| 14_v.txt | 32173 | 32173-SNV-( | SNV | 1 | C | T | 15 | 14110 | 0.10630758 | 44.6 | rnr1D57N | FALSE | CAN | JSY14 | 3 | rnr1D57N-Re | GCA | C>T | A/T+1 | G/C-1 | CG>TA | CG>TA | CAAATGCAG | CAGCAGT | CGT | GCA | C | A | C>T | C>T_GCA | CG>TA_CAGCAGT | 7 | FALSE | FALSE |
| 14_v.txt | 32271 | 32271-SNV-( | SNV | 1 | C | T | 79 | 22305 | 0.35418068 | 35.2911392 | rnr1D57N | FALSE | CAN | JSY14 | 3 | rnr1D57N-Re | ACG | C>T | A/T+1 | G/C-1 | CG>TA | CG>TA | AAAATACGG | CGGGAAC | TGC | ACG | C | A | C>T | C>T_ACG | CG>TA_CGGGAAC | 7 | FALSE | FALSE |
| 14_v.txt | 32274 | 32274-SNV-( | SNV | 1 | G | A | 22 | 22305 | 0.09863259 | 35.2272727 | rnr1D57N | FALSE | CAN | JSY14 | 3 | rnr1D57N-Re | GGA | G>A | A/T+1 | G/C-1 | CG>TA | CG>TA | ATACGGGAA | GAACCAA | CCT | CCT | C | T | C>T | C>T_CCT | CG>TA_GAACCAA | 7 | FALSE | FALSE |
| 14_v.txt | 32299 | 32299-SNV-( | SNV | 1 | C | T | 47 | 22305 | 0.21071509 | 53 | rnr1D57N | FALSE | CAN | JSY14 | 3 | rnr1D57N-Re | GCG | C>T | A/T+1 | G/C-1 | CG>TA | CG>TA | AATTTGCGG | CGGCAGA | CGC | GCG | C | A | C>T | C>T_GCG | CG>TA_CGGCAGA | 7 | FALSE | FALSE |
| 14_v.txt | 32466 | 32466-SNV-( | SNV | 1 | C | T | 32 | 22305 | 0.14346559 | 44.875 | rnr1D57N | FALSE | CAN | JSY14 | 3 | rnr1D57N-Re | GCC | C>T | A/T+1 | G/C-1 | CG>TA | CG>TA | AGAGAGCCA | CCAATGT | CGG | GCC | C | A | C>T | C>T_GCC | CG>TA_CCAATGT | 7 | FALSE | FALSE |
| 14_v.txt | 32487 | 32487-SNV-( | SNV | 1 | C | T | 36 | 22305 | 0.16139879 | 47.1944444 | rnr1D57N | FALSE | CAN | JSY14 | 3 | rnr1D57N-Re | ACG | C>T | A/T+1 | G/C-1 | CG>TA | CG>TA | AAGATACGG | CGGAAAA | TGC | ACG | C | A | C>T | C>T_ACG | CG>TA_CGGAAAA | 7 | FALSE | FALSE |
| 14_v.txt | 32535 | 32535-SNV-( | SNV | 1 | G | A | 24 | 22304 | 0.10760402 | 36.749569 | rnr1D57N | FALSE | CAN | JSY14 | 3 | rnr1D57N-Re | TGC | G>A | A/T+1 | G/C-1 | CG>TA | CG>TA | GGGTTTGCA | GCAGCTT | ACG | ACG | C | T | C>T | C>T_ACG | CG>TA_GCAGCTT | 7 | FALSE | FALSE |
| 14_v.txt | 32536 | 32536-SNV-( | SNV | 1 | C | T | 34 | 22304 | 0.15243902 | 36.7354196 | rnr1D57N | FALSE | CAN | JSY14 | 3 | rnr1D57N-Re | GCA | C>T | A/T+1 | G/C-1 | CG>TA | CG>TA | GGTTTGCAG | CAGCTTC | CGT | GCA | C | A | C>T | C>T_GCA | CG>TA_CAGCTTC | 7 | FALSE | FALSE |
| 14_v.txt | 32590 | 32590-SNV-( | SNV | 1 | C | T | 36 | 20619 | 0.17459625 | 37.0277778 | rnr1D57N | FALSE | CAN | JSY14 | 3 | rnr1D57N-Re | GCG | C>T | A/T+1 | G/C-1 | CG>TA | CG>TA | AGGCAGCGT | CGTTAAT | CGC | GCG | C | A | C>T | C>T_GCG | CG>TA_CGTTAAT | 7 | FALSE | FALSE |
| 14_v.txt | 32660 | 32660-SNV-( | SNV | 1 | C | T | 34 | 20619 | 0.16489646 | 59.1176471 | rnr1D57N | FALSE | CAN | JSY14 | 3 | rnr1D57N-Re | CCC | C>T | A/T+1 | G/C-1 | CG>TA | CG>TA | TGGACCCCA | CCAGGCA | GGG | CCC | C | A | C>T | C>T_CCC | CG>TA_CCAGGCA | 7 | FALSE | FALSE |
| 14_v.txt | 32700 | 32700-SNV-( | SNV | 1 | C | T | 29 | 20619 | 0.14064698 | 62.6896552 | rnr1D57N | FALSE | CAN | JSY14 | 3 | rnr1D57N-Re | GCC | C>T | A/T+1 | G/C-1 | CG>TA | CG>TA | ACTGGGCCG | CCGGTAA | CGG | GCC | C | A | C>T | C>T_GCC | CG>TA_CCGGTAA | 7 | FALSE | FALSE |
| 14_v.txt | 32702 | 32702-SNV-( | SNV | 1 | G | A | 20 | 20619 | 0.09699792 | 54.8 | rnr1D57N | FALSE | CAN | JSY14 | 3 | rnr1D57N-Re | CGG | G>A | A/T+1 | G/C-1 | CG>TA | CG>TA | TGGGCCGGT | GGTAACC | GCC | GCC | C | T | C>T | C>T_GCC | CG>TA_GGTAACC | 7 | FALSE | FALSE |
| 14_v.txt | 32713 | 32713-SNV-( | SNV | 1 | C | T | 17 | 20619 | 0.08244823 | 59.5294118 | rnr1D57N | FALSE | CAN | JSY14 | 3 | rnr1D57N-Re | GCA | C>T | A/T+1 | G/C-1 | CG>TA | CG>TA | CCCCAGCAC | CACCACA | CGT | GCA | C | A | C>T | C>T_GCA | CG>TA_CACCACA | 7 | FALSE | FALSE |
| 14_v.txt | 32757 | 32757-SNV-( | SNV | 1 | G | A | 22 | 20619 | 0.10669771 | 58.5 | rnr1D57N | FALSE | CAN | JSY14 | 3 | rnr1D57N-Re | GGC | G>A | A/T+1 | G/C-1 | CG>TA | CG>TA | ATAATGGCT | GCTAAAA | CCG | CCG | C | T | C>T | C>T_CCG | CG>TA_GCTAAAA | 7 | FALSE | FALSE |
| 14_v.txt | 32776 | 32776-SNV-( | SNV | 1 | C | T | 26 | 20619 | 0.12609729 | 50.7307692 | rnr1D57N | FALSE | CAN | JSY14 | 3 | rnr1D57N-Re | GCG | C>T | A/T+1 | G/C-1 | CG>TA | CG>TA | TGGAAGCGA | CGACCCA | CGC | GCG | C | A | C>T | C>T_GCG | CG>TA_CGACCCA | 7 | FALSE | FALSE |
| 14_v.txt | 32796 | 32796-SNV-( | SNV | 1 | C | T | 30 | 20619 | 0.14549687 | 42.7666667 | rnr1D57N | FALSE | CAN | JSY14 | 3 | rnr1D57N-Re | ACC | C>T | A/T+1 | G/C-1 | CG>TA | CG>TA | AATTCACCG | CCGTAAT | TGG | ACC | C | A | C>T | C>T_ACC | CG>TA_CCGTAAT | 7 | FALSE | FALSE |
| 14_v.txt | 32798 | 32798-SNV-( | SNV | 1 | G | A | 30 | 20619 | 0.14549687 | 42.7333333 | rnr1D57N | FALSE | CAN | JSY14 | 3 | rnr1D57N-Re | CGT | G>A | A/T+1 | G/C-1 | CG>TA | CG>TA | TTCACCGTA | GTAATAT | GCA | GCA | C | T | C>T | C>T_GCA | CG>TA_GTAATAT | 7 | FALSE | FALSE |
| 14_v.txt | 32855 | 32855-SNV-( | SNV | 1 | C | T | 52 | 20618 | 0.25220681 | 36.1730769 | rnr1D57N | FALSE | CAN | JSY14 | 3 | rnr1D57N-Re | TCC | C>T | A/T+1 | G/C-1 | CG>TA | CG>TA | ACTAATCCAT | CCATGCC | AGG | TCC | C | A | C>T | C>T_TCC | CG>TA_CCATGCC | 7 | FALSE | FALSE |
| 14_v.txt | 32860 | 32860-SNV-( | SNV | 1 | C | T | 22 | 20618 | 0.10670288 | 37.6363636 | rnr1D57N | FALSE | CAN | JSY14 | 3 | rnr1D57N-Re | GCC | C>T | A/T+1 | G/C-1 | CG>TA | CG>TA | TCCATGCCG | CCGCCAG | CGG | GCC | C | A | C>T | C>T_GCC | CG>TA_CCGCCAG | 7 | FALSE | FALSE |
| 14_v.txt | 32901 | 32901-SNV-( | SNV | 1 | C | T | 55 | 22863 | 0.24056336 | 31.2727273 | rnr1D57N | FALSE | CAN | JSY14 | 3 | rnr1D57N-Re | GCC | C>T | A/T+1 | G/C-1 | CG>TA | CG>TA | ACTTGGCCA | CCAACTA | CGG | GCC | C | A | C>T | C>T_GCC | CG>TA_CCAACTA | 7 | FALSE | FALSE |
| 14_v.txt | 32935 | 32935-SNV-( | SNV | 1 | C | T | 29 | 22863 | 0.1268425 | 38.1034483 | rnr1D57N | FALSE | CAN | JSY14 | 3 | rnr1D57N-Re | GCC | C>T | A/T+1 | G/C-1 | CG>TA | CG>TA | TGATTGCCCA | CCCAAGA | CGG | GCC | C | A | C>T | C>T_GCC | CG>TA_CCCAAGA | 7 | FALSE | FALSE |
| 14_v.txt | 32946 | 32946-SNV-( | SNV | 1 | C | T | 39 | 22863 | 0.17058129 | 57.7435897 | rnr1D57N | FALSE | CAN | JSY14 | 3 | rnr1D57N-Re | CCA | C>T | A/T+1 | G/C-1 | CG>TA | CG>TA | GAAAACCAA | CAATACA | GGT | CCA | C | A | C>T | C>T_CCA | CG>TA_CAATACA | 7 | FALSE | FALSE |
| 14_v.txt | 32965 | 32965-SNV-( | SNV | 1 | C | T | 23 | 22863 | 0.10059922 | 58.0869565 | rnr1D57N | FALSE | CAN | JSY14 | 3 | rnr1D57N-Re | GCC | C>T | A/T+1 | G/C-1 | CG>TA | CG>TA | CATTGGCCG | CCGCACC | CGG | GCC | C | A | C>T | C>T_GCC | CG>TA_CCGCACC | 7 | FALSE | FALSE |
| 14_v.txt | 32967 | 32967-SNV-( | SNV | 1 | G | A | 55 | 22863 | 0.24056336 | 59.8909091 | rnr1D57N | FALSE | CAN | JSY14 | 3 | rnr1D57N-Re | CGC | G>A | A/T+1 | G/C-1 | CG>TA | CG>TA | TTGGCCGCA | GCACCAA | GCG | GCG | C | T | C>T | C>T_GCG | CG>TA_GCACCAA | 7 | FALSE | FALSE |
| 14_v.txt | 33012 | 33012-SNV-( | SNV | 1 | G | A | 47 | 22863 | 0.20557232 | 63.5744681 | rnr1D57N | FALSE | CAN | JSY14 | 3 | rnr1D57N-Re | AGA | G>A | A/T+1 | G/C-1 | CG>TA | CG>TA | GTGAAAGAG | GAGGATG | TCT | TCT | C | T | C>T | C>T_TCT | CG>TA_GAGGATG | 7 | FALSE | FALSE |
| 14_v.txt | 33056 | 33056-SNV-( | SNV | 1 | C | T | 21 | 22863 | 0.09185146 | 59 | rnr1D57N | FALSE | CAN | JSY14 | 3 | rnr1D57N-Re | GCG | C>T | A/T+1 | G/C-1 | CG>TA | CG>TA | GGACTGCGT | CGTGACA | CGC | GCG | C | A | C>T | C>T_GCG | CG>TA_CGTGACA | 7 | FALSE | FALSE |
| 14_v.txt | 33069 | 33069-SNV-( | SNV | 1 | G | A | 20 | 22863 | 0.08747758 | 53.5 | rnr1D57N | FALSE | CAN | JSY14 | 3 | rnr1D57N-Re | TGC | G>A | A/T+1 | G/C-1 | CG>TA | CG>TA | GAATATGCA | GCCAAAG | ACG | ACG | C | T | C>T | C>T_ACG | CG>TA_GCCAAAG | 7 | FALSE | FALSE |
| 14_v.txt | 33111 | 33111-SNV-( | SNV | 1 | G | A | 24 | 22863 | 0.1049731 | 40.9166667 | rnr1D57N | FALSE | CAN | JSY14 | 3 | rnr1D57N-Re | TGG | G>A | A/T+1 | G/C-1 | CG>TA | CG>TA | CCCACTGGG | GGGCCGG | ACC | ACC | C | T | C>T | C>T_ACC | CG>TA_GGGCCGG | 7 | FALSE | FALSE |
| 14_v.txt | 33113 | 33113-SNV-( | SNV | 1 | G | A | 23 | 22863 | 0.10059922 | 46.6507879 | rnr1D57N | FALSE | CAN | JSY14 | 3 | rnr1D57N-Re | GGC | G>A | A/T+1 | G/C-1 | CG>TA | CG>TA | CACTGGGCC | GCCGGCG | CCG | CCG | C | T | C>T | C>T_CCG | CG>TA_GCCGGCG | 7 | FALSE | FALSE |
| 14_v.txt | 33114 | 33114-SNV-( | SNV | 1 | C | T | 29 | 22863 | 0.1268425 | 39.7240917 | rnr1D57N | FALSE | CAN | JSY14 | 3 | rnr1D57N-Re | GCC | C>T | A/T+1 | G/C-1 | CG>TA | CG>TA | ACTGGGCCG | CCGGCGT | CGG | GCC | C | A | C>T | C>T_GCC | CG>TA_CCGGCGT | 7 | FALSE | FALSE |
| 14_v.txt | 33118 | 33118-SNV-( | SNV | 1 | C | T | 24 | 22863 | 0.1049731 | 41.4166667 | rnr1D57N | FALSE | CAN | JSY14 | 3 | rnr1D57N-Re | GCG | C>T | A/T+1 | G/C-1 | CG>TA | CG>TA | GGCCGGCGT | CGTTGGT | CGC | GCG | C | A | C>T | C>T_GCG | CG>TA_CGTTGGT | 7 | FALSE | FALSE |
| 14_v.txt | 33153 | 33153-SNV-( | SNV | 1 | C | T | 133 | 22863 | 0.58172593 | 36.3157895 | rnr1D57N | FALSE | CAN | JSY14 | 3 | rnr1D57N-Re | ACC | C>T | A/T+1 | G/C-1 | CG>TA | CG>TA | AAAAGACCT | CCTGTAC | TGG | ACC | C | A | C>T | C>T_ACC | CG>TA_CCTGTAC | 7 | FALSE | FALSE |
| 14_v.txt | 33168 | 33168-SNV-( | SNV | 1 | C | T | 20 | 22862 | 0.08748141 | 35.55 | rnr1D57N | FALSE | CAN | JSY14 | 3 | rnr1D57N-Re | ACC | C>T | A/T+1 | G/C-1 | CG>TA | CG>TA | ATAGTACCA | CCACCAA | TGG | ACC | C | A | C>T | C>T_ACC | CG>TA_CCACCAA | 7 | FALSE | FALSE |
| 14_v.txt | 33258 | 33258-SNV-( | SNV | 1 | C | T | 20 | 16483 | 0.12133714 | 45.35 | rnr1D57N | FALSE | CAN | JSY14 | 3 | rnr1D57N-Re | GCC | C>T | A/T+1 | G/C-1 | CG>TA | CG>TA | CCTATGCCAT | CCATCCT | CGG | GCC | C | A | C>T | C>T_GCC | CG>TA_CCATCCT | 7 | FALSE | FALSE |
| 14_v.txt | 33276 | 33276-SNV-( | SNV | 1 | G | A | 28 | 16483 | 0.16987199 | 46.4642857 | rnr1D57N | FALSE | CAN | JSY14 | 3 | rnr1D57N-Re | CGT | G>A | A/T+1 | G/C-1 | CG>TA | CG>TA | GAGAACGTA | GTATCCT | GCA | GCA | C | T | C>T | C>T_GCA | CG>TA_GTATCCT | 7 | FALSE | FALSE |
| 14_v.txt | 33284 | 33284-SNV-( | SNV | 1 | G | A | 18 | 16483 | 0.10920342 | 49.9444444 | rnr1D57N | FALSE | CAN | JSY14 | 3 | rnr1D57N-Re | CGC | G>A | A/T+1 | G/C-1 | CG>TA | CG>TA | ATCCTCGCCA | GCCATTT | GCG | GCG | C | T | C>T | C>T_GCG | CG>TA_GCCATTT | 7 | FALSE | FALSE |
| 14_v.txt | 33309 | 33309-SNV-( | SNV | 1 | C | T | 20 | 16483 | 0.12133714 | 53.546327 | rnr1D57N | FALSE | CAN | JSY14 | 3 | rnr1D57N-Re | GCG | C>T | A/T+1 | G/C-1 | CG>TA | CG>TA | AAAGAGCGC | CGCAATG | CGC | GCG | C | A | C>T | C>T_GCG | CG>TA_CGCAATG | 7 | FALSE | FALSE |
| 14_v.txt | 33310 | 33310-SNV-( | SNV | 1 | G | A | 27 | 16480 | 0.16383495 | 60.9884415 | rnr1D57N | FALSE | CAN | JSY14 | 3 | rnr1D57N-Re | CGC | G>A | A/T+1 | G/C-1 | CG>TA | CG>TA | AAGAGCGCA | GCAATGG | GCG | GCG | C | T | C>T | C>T_GCG | CG>TA_GCAATGG | 7 | FALSE | FALSE |
| 14_v.txt | 33392 | 33392-SNV-( | SNV | 1 | G | A | 23 | 16483 | 0.13953771 | 50.0434783 | rnr1D57N | FALSE | CAN | JSY14 | 3 | rnr1D57N-Re | CGT | G>A | A/T+1 | G/C-1 | CG>TA | CG>TA | AACGTCGTG | GTGAAAG | GCA | GCA | C | T | C>T | C>T_GCA | CG>TA_GTGAAAG | 7 | FALSE | FALSE |
| 14_v.txt | 33474 | 33474-SNV-( | SNV | 1 | C | T | 23 | 16483 | 0.13953771 | 34.9565217 | rnr1D57N | FALSE | CAN | JSY14 | 3 | rnr1D57N-Re | GCC | C>T | A/T+1 | G/C-1 | CG>TA | CG>TA | GCTATGCCT | CCTTTTT | CGG | GCC | C | A | C>T | C>T_GCC | CG>TA_CCTTTTT | 7 | FALSE | FALSE |
| 14_v.txt | 32200 | 32200-SNV-( | SNV | 1 | G | C | 9 | 14110 | 0.06378455 | 43.5555556 | rnr1D57N | FALSE | CAN | JSY14 | 3 | rnr1D57N-Re | GGA | G>C | G/C+1 | G/C-1 | CG>GC | CG>GC | TGTATGGAA | GAACACC | CCT | CCT | C | G | C>G | C>G_CCT | CG>GC_GAACACC | 7 | FALSE | FALSE |
| 14_v.txt | 32271 | 32271-SNV-( | SNV | 1 | C | G | 7 | 22305 | 0.0313831 | 37 | rnr1D57N | FALSE | CAN | JSY14 | 3 | rnr1D57N-Re | ACG | C>G | G/C+1 | G/C-1 | CG>GC | CG>GC | AAAATACGG | CGGGAAC | TGC | ACG | C | C | C>G | C>G_ACG | CG>GC_CGGGAAC | 7 | FALSE | FALSE |
| 14_v.txt | 32302 | 32302-SNV-( | SNV | 1 | C | G | 7 | 22305 | 0.0313831 | 50.5714286 | rnr1D57N | FALSE | CAN | JSY14 | 3 | rnr1D57N-Re | GCA | C>G | G/C+1 | G/C-1 | CG>GC | CG>GC | TTGCGGCAG | CAGAAAT | CGT | GCA | C | C | C>G | C>G_GCA | CG>GC_CAGAAAT | 7 | FALSE | FALSE |
| 14_v.txt | 32471 | 32471-SNV-( | SNV | 1 | G | C | 7 | 22305 | 0.0313831 | 48.5714286 | rnr1D57N | FALSE | CAN | JSY14 | 3 | rnr1D57N-Re | TGT | G>C | G/C+1 | G/C-1 | CG>GC | CG>GC | GCCAATGTA | GTAGAAG | ACA | ACA | C | G | C>G | C>G_ACA | CG>GC_GTAGAAG | 7 | FALSE | FALSE |
| 14_v.txt | 32536 | 32536-SNV-( | SNV | 1 | C | G | 2 | 22304 | 8.97E-03 | 36.000529 | rnr1D57N | FALSE | CAN | JSY14 | 3 | rnr1D57N-Re | GCA | C>G | G/C+1 | G/C-1 | CG>GC | CG>GC | GGTTTGCAG | CAGCTTC | CGT | GCA | C | C | C>G | C>G_GCA | CG>GC_CAGCTTC | 7 | FALSE | FALSE |
| 14_v.txt | 32749 | 32749-SNV-( | SNV | 1 | C | G | 7 | 20619 | 0.03394927 | 60 | rnr1D57N | FALSE | CAN | JSY14 | 3 | rnr1D57N-Re | CCG | C>G | G/C+1 | G/C-1 | CG>GC | CG>GC | GAAACCCGA | CGATAAT | GGC | CCG | C | C | C>G | C>G_CCG | CG>GC_CGATAAT | 7 | FALSE | FALSE |
| 14_v.txt | 32788 | 32788-SNV-( | SNV | 1 | C | G | 2 | 20619 | 9.70E-03 | 36.5020945 | rnr1D57N | FALSE | CAN | JSY14 | 3 | rnr1D57N-Re | TCG | C>G | G/C+1 | G/C-1 | CG>GC | CG>GC | AGAACTCGA | CGAATTC | AGC | TCG | C | C | C>G | C>G_TCG | CG>GC_CGAATTC | 7 | FALSE | FALSE |
| 14_v.txt | 32829 | 32829-SNV-( | SNV | 1 | G | C | 79 | 20619 | 0.38314176 | 45.1139241 | rnr1D57N | FALSE | CAN | JSY14 | 3 | rnr1D57N-Re | TGT | G>C | G/C+1 | G/C-1 | CG>GC | CG>GC | ATTATTGTGA | GTGATAA | ACA | ACA | C | G | C>G | C>G_ACA | CG>GC_GTGATAA | 7 | FALSE | FALSE |
| 14_v.txt | 33172 | 33172-SNV-( | SNV | 1 | C | G | 46 | 22862 | 0.20120724 | 37.9565217 | rnr1D57N | FALSE | CAN | JSY14 | 3 | rnr1D57N-Re | CCA | C>G | G/C+1 | G/C-1 | CG>GC | CG>GC | TACCACCAA | CAAGGGC | GGT | CCA | C | C | C>G | C>G_CCA | CG>GC_CAAGGGC | 7 | FALSE | FALSE |
| 14_v.txt | 33284 | 33284-SNV-( | SNV | 1 | G | C | 4 | 16483 | 0.02426743 | 41.75 | rnr1D57N | FALSE | CAN | JSY14 | 3 | rnr1D57N-Re | CGC | G>C | G/C+1 | G/C-1 | CG>GC | CG>GC | ATCCTCGCCA | GCCATTT | GCG | GCG | C | G | C>G | C>G_GCG | CG>GC_GCCATTT | 7 | FALSE | FALSE |
| 14_v.txt | 33309 | 33309-SNV-( | SNV | 1 | C | G | 1 | 16483 | 6.07E-03 | 56.9951821 | rnr1D57N | FALSE | CAN | JSY14 | 3 | rnr1D57N-Re | GCG | C>G | G/C+1 | G/C-1 | CG>GC | CG>GC | AAAGAGCGC | CGCAATG | CGC | GCG | C | C | C>G | C>G_GCG | CG>GC_CGCAATG | 7 | FALSE | FALSE |
| 14_v.txt | 33354 | 33354-SNV-( | SNV | 1 | G | C | 30 | 16483 | 0.1820057 | 57.8333333 | rnr1D57N | FALSE | CAN | JSY14 | 3 | rnr1D57N-Re | TGA | G>C | G/C+1 | G/C-1 | CG>GC | CG>GC | GGTATTGAC | GACCCAC | ACT | ACT | C | G | C>G | C>G_ACT | CG>GC_GACCCAC | 7 | FALSE | FALSE |
| 14_v.txt | 33475 | 33475-SNV-( | SNV | 1 | C | G | 2 | 16483 | 0.01213371 | 38 | rnr1D57N | FALSE | CAN | JSY14 | 3 | rnr1D57N-Re | CCT | C>G | G/C+1 | G/C-1 | CG>GC | CG>GC | CTATGCCTT | CTTTTTT | GGA | CCT | C | C | C>G | C>G_CCT | CG>GC_CTTTTTT | 7 | FALSE | FALSE |
| 14_v.txt | 31829 | 31829-SNV-( | SNV | 1 | G | T | 12 | 10140 | 0.1183432 | 56.0833333 | rnr1D57N | FALSE | CAN | JSY14 | 3 | rnr1D57N-Re | TGC | G>T | A/T+1 | G/C-1 | CG>AT | CG>AT | GAATATGCA | GCATTGA | ACG | ACG | C | A | C>A | C>A_ACG | CG>AT_GCATTGA | 7 | FALSE | FALSE |
| 14_v.txt | 31971 | 31971-SNV-( | SNV | 1 | G | T | 14 | 14108 | 0.09923448 | 16.5 | rnr1D57N | FALSE | CAN | JSY14 | 3 | rnr1D57N-Re | AGC | G>T | A/T+1 | G/C-1 | CG>AT | CG>AT | TAATAAGCC | GCCAAGC | TCG | TCG | C | A | C>A | C>A_TCG | CG>AT_GCCAAGC | 7 | FALSE | FALSE |
| 14_v.txt | 32008 | 32008-SNV-( | SNV | 1 | C | A | 1863 | 14108 | 13.2052736 | 57.5990338 | rnr1D57N | FALSE | CAN | JSY14 | 3 | rnr1D57N-Re | TCG | C>A | A/T+1 | G/C-1 | CG>AT | CG>AT | GTAACTCGT | CGTCACG | AGC | TCG | C | T | C>A | C>A_TCG | CG>AT_CGTCACG | 7 | FALSE | FALSE |
| 14_v.txt | 32188 | 32188-SNV-( | SNV | 1 | C | A | 19 | 14110 | 0.13465627 | 41.0526316 | rnr1D57N | FALSE | CAN | JSY14 | 3 | rnr1D57N-Re | ACT | C>A | A/T+1 | G/C-1 | CG>AT | CG>AT | CGAAAACTG | CTGCAAT | TGA | ACT | C | T | C>A | C>A_ACT | CG>AT_CTGCAAT | 7 | FALSE | FALSE |
| 14_v.txt | 32238 | 32238-SNV-( | SNV | 1 | G | T | 9 | 14109 | 0.06378907 | 35 | rnr1D57N | FALSE | CAN | JSY14 | 3 | rnr1D57N-Re | AGC | G>T | A/T+1 | G/C-1 | CG>AT | CG>AT | TTAGGAGCC | GCCAACT | TCG | TCG | C | A | C>A | C>A_TCG | CG>AT_GCCAACT | 7 | FALSE | FALSE |
| 14_v.txt | 32274 | 32274-SNV-( | SNV | 1 | G | T | 10 | 22305 | 0.044833 | 34.3 | rnr1D57N | FALSE | CAN | JSY14 | 3 | rnr1D57N-Re | GGA | G>T | A/T+1 | G/C-1 | CG>AT | CG>AT | ATACGGGAA | GAACCAA | CCT | CCT | C | A | C>A | C>A_CCT | CG>AT_GAACCAA | 7 | FALSE | FALSE |
| 14_v.txt | 32545 | 32545-SNV-( | SNV | 1 | C | A | 12 | 22304 | 0.05380201 | 35.25 | rnr1D57N | FALSE | CAN | JSY14 | 3 | rnr1D57N-Re | CCA | C>A | A/T+1 | G/C-1 | CG>AT | CG>AT | CTTCACCAG | CAGCAGT | GGT | CCA | C | T | C>A | C>A_CCA | CG>AT_CAGCAGT | 7 | FALSE | FALSE |
| 14_v.txt | 32556 | 32556-SNV-( | SNV | 1 | C | A | 19 | 22303 | 0.08519033 | 35.2105263 | rnr1D57N | FALSE | CAN | JSY14 | 3 | rnr1D57N-Re | ACC | C>A | A/T+1 | G/C-1 | CG>AT | CG>AT | GTGATACCA | CCAACTA | TGG | ACC | C | T | C>A | C>A_ACC | CG>AT_CCAACTA | 7 | FALSE | FALSE |
| 14_v.txt | 32572 | 32572-SNV-( | SNV | 1 | C | A | 13 | 20619 | 0.06304865 | 33.0769231 | rnr1D57N | FALSE | CAN | JSY14 | 3 | rnr1D57N-Re | CCT | C>A | A/T+1 | G/C-1 | CG>AT | CG>AT | CAGTACCTTC | CTTGAAA | GGA | CCT | C | T | C>A | C>A_CCT | CG>AT_CTTGAAA | 7 | FALSE | FALSE |
| 14_v.txt | 32575 | 32575-SNV-( | SNV | 1 | G | T | 105 | 20619 | 0.50923905 | 37.0190476 | rnr1D57N | FALSE | CAN | JSY14 | 3 | rnr1D57N-Re | TGA | G>T | A/T+1 | G/C-1 | CG>AT | CG>AT | TACCTTGAA | GAAATGT | ACT | ACT | C | A | C>A | C>A_ACT | CG>AT_GAAATGT | 7 | FALSE | FALSE |
| 14_v.txt | 32692 | 32692-SNV-( | SNV | 1 | C | A | 33 | 20619 | 0.16004656 | 51.9393939 | rnr1D57N | FALSE | CAN | JSY14 | 3 | rnr1D57N-Re | CCA | C>A | A/T+1 | G/C-1 | CG>AT | CG>AT | GGAATCCAA | CAACTGG | GGT | CCA | C | T | C>A | C>A_CCA | CG>AT_CAACTGG | 7 | FALSE | FALSE |
| 14_v.txt | 32713 | 32713-SNV-( | SNV | 1 | C | A | 49 | 20619 | 0.23764489 | 58.6734694 | rnr1D57N | FALSE | CAN | JSY14 | 3 | rnr1D57N-Re | GCA | C>A | A/T+1 | G/C-1 | CG>AT | CG>AT | CCCCAGCAC | CACCACA | CGT | GCA | C | T | C>A | C>A_GCA | CG>AT_CACCACA | 7 | FALSE | FALSE |
| 14_v.txt | 32783 | 32783-SNV-( | SNV | 1 | G | T | 13 | 20619 | 0.06304865 | 45.1538462 | rnr1D57N | FALSE | CAN | JSY14 | 3 | rnr1D57N-Re | AGA | G>T | A/T+1 | G/C-1 | CG>AT | CG>AT | GACCCAGAA | GAACTCG | TCT | TCT | C | A | C>A | C>A_TCT | CG>AT_GAACTCG | 7 | FALSE | FALSE |
| 14_v.txt | 32788 | 32788-SNV-( | SNV | 1 | C | A | 22 | 20619 | 0.10669771 | 46.3193902 | rnr1D57N | FALSE | CAN | JSY14 | 3 | rnr1D57N-Re | TCG | C>A | A/T+1 | G/C-1 | CG>AT | CG>AT | AGAACTCGA | CGAATTC | AGC | TCG | C | T | C>A | C>A_TCG | CG>AT_CGAATTC | 7 | FALSE | FALSE |
| 14_v.txt | 32954 | 32954-SNV-( | SNV | 1 | G | T | 22 | 22863 | 0.09622534 | 51.7727273 | rnr1D57N | FALSE | CAN | JSY14 | 3 | rnr1D57N-Re | TGT | G>T | A/T+1 | G/C-1 | CG>AT | CG>AT | ATACATGTA | GTAACCA | ACA | ACA | C | A | C>A | C>A_ACA | CG>AT_GTAACCA | 7 | FALSE | FALSE |
| 14_v.txt | 32997 | 32997-SNV-( | SNV | 1 | G | T | 29 | 22863 | 0.1268425 | 60.8965517 | rnr1D57N | FALSE | CAN | JSY14 | 3 | rnr1D57N-Re | TGA | G>T | A/T+1 | G/C-1 | CG>AT | CG>AT | CTTTGTGAG | GAGAAAA | ACT | ACT | C | A | C>A | C>A_ACT | CG>AT_GAGAAAA | 7 | FALSE | FALSE |
| 14_v.txt | 33015 | 33015-SNV-( | SNV | 1 | G | T | 15 | 22863 | 0.06560819 | 55.9333333 | rnr1D57N | FALSE | CAN | JSY14 | 3 | rnr1D57N-Re | GGA | G>T | A/T+1 | G/C-1 | CG>AT | CG>AT | AAAGAGGAT | GATGTAA | CCT | CCT | C | A | C>A | C>A_CCT | CG>AT_GATGTAA | 7 | FALSE | FALSE |
| 14_v.txt | 33114 | 33114-SNV-( | SNV | 1 | C | A | 25 | 22863 | 0.10934698 | 42.8779537 | rnr1D57N | FALSE | CAN | JSY14 | 3 | rnr1D57N-Re | GCC | C>A | A/T+1 | G/C-1 | CG>AT | CG>AT | ACTGGGCCG | CCGGCGT | CGG | GCC | C | T | C>A | C>A_GCC | CG>AT_CCGGCGT | 7 | FALSE | FALSE |
| 14_v.txt | 33160 | 33160-SNV-( | SNV | 1 | C | A | 152 | 22863 | 0.66482964 | 37.5592105 | rnr1D57N | FALSE | CAN | JSY14 | 3 | rnr1D57N-Re | CCA | C>A | A/T+1 | G/C-1 | CG>AT | CG>AT | CTGTACCAAT | CAATAGT | GGT | CCA | C | T | C>A | C>A_CCA | CG>AT_CAATAGT | 7 | FALSE | FALSE |
| 14_v.txt | 33195 | 33195-SNV-( | SNV | 1 | C | A | 87 | 16484 | 0.52778452 | 36.3333333 | rnr1D57N | FALSE | CAN | JSY14 | 3 | rnr1D57N-Re | TCT | C>A | A/T+1 | G/C-1 | CG>AT | CG>AT | ATATGTCTTT | CTTTGCT | AGA | TCT | C | T | C>A | C>A_TCT | CG>AT_CTTTGCT | 7 | FALSE | FALSE |
| 14_v.txt | 33474 | 33474-SNV-( | SNV | 1 | C | A | 36 | 16483 | 0.21840684 | 32.3333333 | rnr1D57N | FALSE | CAN | JSY14 | 3 | rnr1D57N-Re | GCC | C>A | A/T+1 | G/C-1 | CG>AT | CG>AT | GCTATGCCT | CCTTTTT | CGG | GCC | C | T | C>A | C>A_GCC | CG>AT_CCTTTTT | 7 | FALSE | FALSE |
| 14_v.txt | 33475 | 33475-SNV-( | SNV | 1 | C | A | 15 | 16483 | 0.09100285 | 36.9333333 | rnr1D57N | FALSE | CAN | JSY14 | 3 | rnr1D57N-Re | CCT | C>A | A/T+1 | G/C-1 | CG>AT | CG>AT | CTATGCCTTT | CTTTTTT | GGA | CCT | C | T | C>A | C>A_CCT | CG>AT_CTTTTTT | 7 | FALSE | FALSE |
| 14_v.txt | 32717 | 32717-Repla | Replacement | 4 | A | GTTC | 42 | 20619 | 0.20369562 | 61.0892857 | rnr1D57N | FALSE | CAN | JSY14 | 3 | rnr1D57N-Re | CAC | nope | >1bp-ins | A/T-1 | none | Replac. | AGCACCACA | ACAAACC | GTG | GTG | T | none | T>none | T>none_GTG | Replac._ACAAACC | 7 | FALSE | FALSE |
| 14_v.txt | 32717 | 32717-Repla | Replacement | 4 | A | GTTT | 1 | 20619 | 4.85E-03 | 59.5 | rnr1D57N | FALSE | CAN | JSY14 | 3 | rnr1D57N-Re | CAC | nope | >1bp-ins | A/T-1 | none | Replac. | AGCACCACA | ACAAACC | GTG | GTG | T | none | T>none | T>none_GTG | Replac._ACAAACC | 7 | FALSE | FALSE |
| 14_v.txt | 33474 | 33474-MNV- | MNV | 2 | CC | TA | 1 | 16483 | 6.07E-03 | 37.5 | rnr1D57N | FALSE | CAN | JSY14 | 3 | rnr1D57N-Re | GCC | nope | >1bp-ins | >1bpdel | none | MNV | GCTATGCCT | CCTTTTT | CGG | GCC | CC | none | NA | NA_GCC | MNV_CCTTTTT | 7 | FALSE | FALSE |
| 14_v.txt | 32079 | 32079-Insert | Insertion | 1 | - | A | 3 | 14110 | 0.02126152 | 64 | rnr1D57N | FALSE | CAN | JSY14 | 3 | rnr1D57N-Re | TGC | nope | A/T+1 | >1bpdel | none | A/T+1 | AACCATGCA | GCAAAAA | ACG | TGC | - | T | NA | NA_TGC | A/T+1_GCAAAAA | 7 | FALSE | FALSE |
| 14_v.txt | 32081 | 32081-Insert | Insertion | 1 | - | A | 14 | 14110 | 0.09922041 | 64 | rnr1D57N | FALSE | CAN | JSY14 | 3 | rnr1D57N-Re | CAA | nope | A/T+1 | >1bpdel | none | A/T+1 | CCATGCAAA | AAAAAAG | GTT | CAA | - | T | NA | NA_CAA | A/T+1_AAAAAAG | 7 | FALSE | FALSE |
| 14_v.txt | 32498 | 32498-Insert | Insertion | 1 | - | T | 19 | 22305 | 0.0851827 | 44.2105263 | rnr1D57N | FALSE | CAN | JSY14 | 3 | rnr1D57N-Re | CTT | nope | A/T+1 | >1bpdel | none | A/T+1 | AACAACTTTT | TTTTTTG | GAA | CTT | - | A | NA | NA_CTT | A/T+1_TTTTTTG | 7 | FALSE | FALSE |
| 14_v.txt | 32597 | 32597-Insert | Insertion | 2 | - | AA | 9 | 20619 | 0.04364906 | 40.8333333 | rnr1D57N | FALSE | CAN | JSY14 | 3 | rnr1D57N-Re | TCA | nope | >1bp-ins | >1bpdel | none | >1bp-ins | GTTAATCAA | CAAAGAG | AGT | TCA | - | none | NA | NA_TCA | >1bp-ins_CAAAGAG | 7 | FALSE | FALSE |
| 14_v.txt | 32715 | 32715-Insert | Insertion | 3 | - | AAT | 44 | 20619 | 0.21339541 | 62.7045455 | rnr1D57N | FALSE | CAN | JSY14 | 3 | rnr1D57N-Re | ACC | nope | >1bp-ins | >1bpdel | none | >1bp-ins | CCAGCACCA | CCACAAA | TGG | ACC | - | none | NA | NA_ACC | >1bp-ins_CCACAAA | 7 | FALSE | FALSE |
| 14_v.txt | 32718 | 32718-Insert | Insertion | 3 | - | TTC | 1 | 20619 | 4.85E-03 | 64 | rnr1D57N | FALSE | CAN | JSY14 | 3 | rnr1D57N-Re | ACA | nope | >1bp-ins | >1bpdel | none | >1bp-ins | GCACCACAA | CAAACCA | TGT | ACA | - | none | NA | NA_ACA | >1bp-ins_CAAACCA | 7 | FALSE | FALSE |
| 14_v.txt | 32721 | 32721-Insert | Insertion | 1 | - | G | 44 | 20619 | 0.21339541 | 59.1590909 | rnr1D57N | FALSE | CAN | JSY14 | 3 | rnr1D57N-Re | AAC | nope | G/C+1 | >1bpdel | none | G/C+1 | CCACAAACC | ACCATAC | TTG | AAC | - | C | NA | NA_AAC | G/C+1_ACCATAC | 7 | FALSE | FALSE |
| 14_v.txt | 32722 | 32722-Insert | Insertion | 1 | - | G | 44 | 20619 | 0.21339541 | 61.0681818 | rnr1D57N | FALSE | CAN | JSY14 | 3 | rnr1D57N-Re | ACC | nope | G/C+1 | >1bpdel | none | G/C+1 | CACAAACCA | CCATACA | TGG | ACC | - | C | NA | NA_ACC | G/C+1_CCATACA | 7 | FALSE | FALSE |
| 14_v.txt | 32842 | 32842-Insert | Insertion | 1 | - | A | 62 | 20619 | 0.30069354 | 40.9354839 | rnr1D57N | FALSE | CAN | JSY14 | 3 | rnr1D57N-Re | CAA | nope | A/T+1 | >1bpdel | none | A/T+1 | TTACCCAAA | AAAAAAT | GTT | CAA | - | T | NA | NA_CAA | A/T+1_AAAAAAT | 7 | FALSE | FALSE |
| 14_v.txt | 32337 | 32337-Deleti | Deletion | 2 | AT | - | 41 | 22305 | 0.18381529 | 62.8536585 | rnr1D57N | FALSE | CAN | JSY14 | 3 | rnr1D57N-Re | GAT | nope | >1bp-ins | >1bpdel | none | >1bpdel | TTGAAGATA | ATATGTG | CTA | GAT | AT | none | NA | NA_GAT | >1bpdel_ATATGTG | 7 | FALSE | FALSE |
| 14_v.txt | 33094 | 33094-Deleti | Deletion | 2 | AT | - | 17 | 22863 | 0.07435595 | 52.8823529 | rnr1D57N | FALSE | CAN | JSY14 | 3 | rnr1D57N-Re | GAT | nope | >1bp-ins | >1bpdel | none | >1bpdel | AATATGATA | ATATAAG | CTA | GAT | AT | none | NA | NA_GAT | >1bpdel_ATATAAG | 7 | FALSE | FALSE |
| 14_v.txt | 31663 | 31663-Deleti | Deletion | 1 | A | - | 6 | 10140 | 0.0591716 | 38.8333333 | rnr1D57N | FALSE | CAN | JSY14 | 3 | rnr1D57N-Re | GAA | nope | >1bp-ins | A/T-1 | none | A/T-1 | GCGTGGAAA | AAATGTG | CTT | CTT | T | none | T>none | T>none_CTT | A/T-1_AAATGTG | 7 | FALSE | FALSE |
| 14_v.txt | 31723 | 31723-Deleti | Deletion | 1 | A | - | 13 | 10140 | 0.12820513 | 60.5384615 | rnr1D57N | FALSE | CAN | JSY14 | 3 | rnr1D57N-Re | CAA | nope | >1bp-ins | A/T-1 | none | A/T-1 | TGTCCCAAA | AAAAAGT | GTT | GTT | T | none | T>none | T>none_GTT | A/T-1_AAAAAGT | 7 | FALSE | FALSE |
| 14_v.txt | 31863 | 31863-Deleti | Deletion | 1 | A | - | 5 | 10140 | 0.04930967 | 38.6 | rnr1D57N | FALSE | CAN | JSY14 | 3 | rnr1D57N-Re | GAA | nope | >1bp-ins | A/T-1 | none | A/T-1 | AACAGGAAA | AAAATAG | CTT | CTT | T | none | T>none | T>none_CTT | A/T-1_AAAATAG | 7 | FALSE | FALSE |
| 14_v.txt | 31904 | 31904-Deleti | Deletion | 1 | T | - | 7 | 10140 | 0.06903353 | 37.4285714 | rnr1D57N | FALSE | CAN | JSY14 | 3 | rnr1D57N-Re | ATT | nope | >1bp-ins | A/T-1 | none | A/T-1 | ATTGAATTTT | TTTTGGT | TAA | ATT | T | none | nope | nope_ATT | A/T-1_TTTTGGT | 7 | FALSE | FALSE |
| 14_v.txt | 31971 | 31971-Deleti | Deletion | 1 | G | - | 8600 | 14108 | 60.9583215 | 37.3912791 | rnr1D57N | FALSE | CAN | JSY14 | 3 | rnr1D57N-Re | AGC | nope | >1bp-ins | G/C-1 | none | G/C-1 | TAATAAGCC | GCCAAGC | TCG | TCG | C | none | C>none | C>none_TCG | G/C-1_GCCAAGC | 7 | FALSE | FALSE |
| 14_v.txt | 32081 | 32081-Deleti | Deletion | 1 | A | - | 201 | 14110 | 1.42452162 | 62.4278618 | rnr1D57N | FALSE | CAN | JSY14 | 3 | rnr1D57N-Re | CAA | nope | >1bp-ins | A/T-1 | none | A/T-1 | CCATGCAAA | AAAAAAG | GTT | GTT | T | none | T>none | T>none_GTT | A/T-1_AAAAAAG | 7 | FALSE | FALSE |
| 14_v.txt | 32442 | 32442-Deleti | Deletion | 1 | A | - | 81 | 22305 | 0.36314728 | 54.4567901 | rnr1D57N | FALSE | CAN | JSY14 | 3 | rnr1D57N-Re | TAA | nope | >1bp-ins | A/T-1 | none | A/T-1 | GGAACTAAA | AAAAGTC | ATT | ATT | T | none | T>none | T>none_ATT | A/T-1_AAAAGTC | 7 | FALSE | FALSE |
| 14_v.txt | 32498 | 32498-Deleti | Deletion | 1 | T | - | 23 | 22305 | 0.10311589 | 41.4347882 | rnr1D57N | FALSE | CAN | JSY14 | 3 | rnr1D57N-Re | CTT | nope | >1bp-ins | A/T-1 | none | A/T-1 | AACAACTTTT | TTTTTTG | GAA | CTT | T | none | nope | nope_CTT | A/T-1_TTTTTTG | 7 | FALSE | FALSE |
| 14_v.txt | 32522 | 32522-Deleti | Deletion | 1 | T | - | 5 | 22305 | 0.0224165 | 38 | rnr1D57N | FALSE | CAN | JSY14 | 3 | rnr1D57N-Re | ATT | nope | >1bp-ins | A/T-1 | none | A/T-1 | AACGGATTTT | TTTTCTG | TAA | ATT | T | none | nope | nope_ATT | A/T-1_TTTTCTG | 7 | FALSE | FALSE |
| 14_v.txt | 32624 | 32624-Deleti | Deletion | 1 | C | - | 30 | 20619 | 0.14549687 | 44.0333333 | rnr1D57N | FALSE | CAN | JSY14 | 3 | rnr1D57N-Re | TCC | nope | >1bp-ins | G/C-1 | none | G/C-1 | GAACCTCCCT | CCCTTCG | AGG | TCC | C | none | nope | nope_TCC | G/C-1_CCCTTCG | 7 | FALSE | FALSE |
| 14_v.txt | 32631 | 32631-Deleti | Deletion | 1 | T | - | 77 | 20619 | 0.37344197 | 49.7142857 | rnr1D57N | FALSE | CAN | JSY14 | 3 | rnr1D57N-Re | GTT | nope | >1bp-ins | A/T-1 | none | A/T-1 | CCTTCGTTTT | TTTTTAT | CAA | GTT | T | none | nope | nope_GTT | A/T-1_TTTTTAT | 7 | FALSE | FALSE |
| 14_v.txt | 32637 | 32637-Deleti | Deletion | 1 | T | - | 22 | 20619 | 0.10669771 | 52.8636364 | rnr1D57N | FALSE | CAN | JSY14 | 3 | rnr1D57N-Re | ATC | nope | >1bp-ins | A/T-1 | none | A/T-1 | TTTTTATCCT | TCCTTAG | TAG | ATC | T | none | nope | nope_ATC | A/T-1_TCCTTAG | 7 | FALSE | FALSE |
| 14_v.txt | 32658 | 32658-Deleti | Deletion | 1 | C | - | 75 | 20619 | 0.36374218 | 61.44 | rnr1D57N | FALSE | CAN | JSY14 | 3 | rnr1D57N-Re | ACC | nope | >1bp-ins | G/C-1 | none | G/C-1 | CCTGGACCCC | CCCCAGG | TGG | ACC | C | none | nope | nope_ACC | G/C-1_CCCCAGG | 7 | FALSE | FALSE |
| 14_v.txt | 32663 | 32663-Deleti | Deletion | 1 | G | - | 7 | 20619 | 0.03394927 | 64 | rnr1D57N | FALSE | CAN | JSY14 | 3 | rnr1D57N-Re | AGG | nope | >1bp-ins | G/C-1 | none | G/C-1 | ACCCCAGGC | GGCACCT | TCC | TCC | C | none | C>none | C>none_TCC | G/C-1_GGCACCT | 7 | FALSE | FALSE |
| 14_v.txt | 32707 | 32707-Deleti | Deletion | 1 | C | - | 5 | 20619 | 0.02424948 | 58.8 | rnr1D57N | FALSE | CAN | JSY14 | 3 | rnr1D57N-Re | ACC | nope | >1bp-ins | G/C-1 | none | G/C-1 | CGGTAACCC | CCCCAGC | TGG | ACC | C | none | nope | nope_ACC | G/C-1_CCCCAGC | 7 | FALSE | FALSE |
| 14_v.txt | 32717 | 32717-Deleti | Deletion | 1 | A | - | 1 | 20619 | 4.85E-03 | 63.9999947 | rnr1D57N | FALSE | CAN | JSY14 | 3 | rnr1D57N-Re | CAC | nope | >1bp-ins | A/T-1 | none | A/T-1 | AGCACCACA | ACAAACC | GTG | GTG | T | none | T>none | T>none_GTG | A/T-1_ACAAACC | 7 | FALSE | FALSE |
| 14_v.txt | 32726 | 32726-Deleti | Deletion | 1 | A | - | 44 | 20619 | 0.21339541 | 59.0454546 | rnr1D57N | FALSE | CAN | JSY14 | 3 | rnr1D57N-Re | TAC | nope | >1bp-ins | A/T-1 | none | A/T-1 | AACCATACA | ACAAAAA | ATG | ATG | T | none | T>none | T>none_ATG | A/T-1_ACAAAAA | 7 | FALSE | FALSE |
| 14_v.txt | 32728 | 32728-Deleti | Deletion | 1 | A | - | 7 | 20619 | 0.03394927 | 59.4285714 | rnr1D57N | FALSE | CAN | JSY14 | 3 | rnr1D57N-Re | CAA | nope | >1bp-ins | A/T-1 | none | A/T-1 | CCATACAAA | AAAAACA | GTT | GTT | T | none | T>none | T>none_GTT | A/T-1_AAAAACA | 7 | FALSE | FALSE |

| | | | | | | | | | | | | | | | | | | | | | | | | | | | | | | | | | | |
|---|---|---|---|---|---|---|---|---|---|---|---|---|---|---|---|---|---|---|---|---|---|---|---|---|---|---|---|---|---|---|---|---|---|---|
| 14_v.txt | 32839 | 32839-Deleti | Deletion | 1 | C | - | 15 | 20619 | 0.07274844 | 44.3333333 | rnr1D57N | FALSE | CAN | JSY14 | 3 | rnr1D57N-Re | ACC | nope | >1bp-ins | G/C-1 | none | G/C-1 | TAATTACCCA | CCCAAAA | TGG | ACC | C | none | nope | nope_ACC | G/C-1_CCCAAAA | 7 | FALSE | FALSE |
| 14_v.txt | 32842 | 32842-Deleti | Deletion | 1 | A | - | 26 | 20619 | 0.12609729 | 40.1538462 | rnr1D57N | FALSE | CAN | JSY14 | 3 | rnr1D57N-Re | CAA | nope | >1bp-ins | A/T-1 | none | A/T-1 | TTACCCAAAA | AAAAAAT | GTT | GTT | T | none | T>none | T>none_GTT | A/T-1_AAAAAAT | 7 | FALSE | FALSE |
| 14_v.txt | 32884 | 32884-Deleti | Deletion | 1 | A | - | 6 | 22864 | 0.02624213 | 38 | rnr1D57N | FALSE | CAN | JSY14 | 3 | rnr1D57N-Re | CAA | nope | >1bp-ins | A/T-1 | none | A/T-1 | ACGTCCAAA | AAAATTG | GTT | GTT | T | none | T>none | T>none_GTT | A/T-1_AAAATTG | 7 | FALSE | FALSE |
| 14_v.txt | 32924 | 32924-Deleti | Deletion | 1 | A | - | 22 | 22863 | 0.09622534 | 55.1363636 | rnr1D57N | FALSE | CAN | JSY14 | 3 | rnr1D57N-Re | CAA | nope | >1bp-ins | A/T-1 | none | A/T-1 | CAGGGCAAA | AAAAGTG | GTT | GTT | T | none | T>none | T>none_GTT | A/T-1_AAAAGTG | 7 | FALSE | FALSE |
| 14_v.txt | 32935 | 32935-Deleti | Deletion | 1 | C | - | 5 | 22863 | 0.0218694 | 36.8 | rnr1D57N | FALSE | CAN | JSY14 | 3 | rnr1D57N-Re | GCC | nope | >1bp-ins | G/C-1 | none | G/C-1 | TGATTGCCCA | CCCAAGA | CGG | GCC | C | none | nope | nope_GCC | G/C-1_CCCAAGA | 7 | FALSE | FALSE |
| 14_v.txt | 32965 | 32965-Deleti | Deletion | 1 | C | - | 3 | 22863 | 0.01312164 | 51.6666667 | rnr1D57N | FALSE | CAN | JSY14 | 3 | rnr1D57N-Re | GCC | nope | >1bp-ins | G/C-1 | none | G/C-1 | CATTGGCCG | CCGCACC | CGG | GCC | C | none | nope | nope_GCC | G/C-1_CCGCACC | 7 | FALSE | FALSE |
| 14_v.txt | 32978 | 32978-Deleti | Deletion | 1 | T | - | 24 | 22863 | 0.1049731 | 62.8333333 | rnr1D57N | FALSE | CAN | JSY14 | 3 | rnr1D57N-Re | CTG | nope | >1bp-ins | A/T-1 | none | A/T-1 | AAATGCTGG | TGGAGAA | GAC | CTG | T | none | nope | nope_CTG | A/T-1_TGGAGAA | 7 | FALSE | FALSE |
| 14_v.txt | 33072 | 33072-Deleti | Deletion | 1 | A | - | 18 | 22863 | 0.07872983 | 58.3888889 | rnr1D57N | FALSE | CAN | JSY14 | 3 | rnr1D57N-Re | CAA | nope | >1bp-ins | A/T-1 | none | A/T-1 | TATGCCAAA | AAAGAAC | GTT | GTT | T | none | T>none | T>none_GTT | A/T-1_AAAGAAC | 7 | FALSE | FALSE |
| 14_v.txt | 33080 | 33080-Deleti | Deletion | 1 | C | - | 5 | 22863 | 0.0218694 | 47 | rnr1D57N | FALSE | CAN | JSY14 | 3 | rnr1D57N-Re | CCA | nope | >1bp-ins | G/C-1 | none | G/C-1 | AGAACCCAT | CATAAAT | GGT | CCA | C | none | nope | nope_CCA | G/C-1_CATAAAT | 7 | FALSE | FALSE |
| 14_v.txt | 33111 | 33111-Deleti | Deletion | 1 | G | - | 28 | 22863 | 0.12246862 | 41.2857143 | rnr1D57N | FALSE | CAN | JSY14 | 3 | rnr1D57N-Re | TGG | nope | >1bp-ins | G/C-1 | none | G/C-1 | CCCACTGGG | GGGCCGG | ACC | ACC | C | none | C>none | C>none_ACC | G/C-1_GGGCCGG | 7 | FALSE | FALSE |
| 14_v.txt | 33168 | 33168-Deleti | Deletion | 1 | C | - | 8 | 22862 | 0.03499256 | 37.875 | rnr1D57N | FALSE | CAN | JSY14 | 3 | rnr1D57N-Re | ACC | nope | >1bp-ins | G/C-1 | none | G/C-1 | ATAGTACCA | CCACCAA | TGG | ACC | C | none | nope | nope_ACC | G/C-1_CCACCAA | 7 | FALSE | FALSE |
| 14_v.txt | 33474 | 33474-Deleti | Deletion | 1 | C | - | 27 | 16483 | 0.16380513 | 27.4074074 | rnr1D57N | FALSE | CAN | JSY14 | 3 | rnr1D57N-Re | GCC | nope | >1bp-ins | G/C-1 | none | G/C-1 | GCTATGCCTT | CCTTTTT | CGG | GCC | C | none | nope | nope_GCC | G/C-1_CCTTTTT | 7 | FALSE | FALSE |
| 15_v.txt | 31723 | 31723-SNV-A | SNV | 1 | A | C | 5 | 25145 | 0.01988467 | 53.4004121 | rnr1D57N | FALSE | CAN | JSY14 | 4 | rnr1D57N-Re | CAA | A>C | G/C+1 | A/T-1 | TA>GC | TA>GC | TGTCCCAAAA | AAAAAGT | GTT | GTT | T | G | T>G | T>G_GTT | TA>GC_AAAAAGT | 7 | FALSE | FALSE |
| 15_v.txt | 32162 | 32162-SNV-A | SNV | 1 | A | C | 8 | 64665 | 0.01237145 | 30.751915 | rnr1D57N | FALSE | CAN | JSY14 | 4 | rnr1D57N-Re | AAG | A>C | G/C+1 | A/T-1 | TA>GC | TA>GC | AGCCAAAGC | AGCGCCA | TTC | TTC | T | G | T>G | T>G_TTC | TA>GC_AGCGCCA | 7 | FALSE | FALSE |
| 15_v.txt | 32187 | 32187-SNV-A | SNV | 1 | A | C | 10 | 64665 | 0.01546432 | 40.5 | rnr1D57N | FALSE | CAN | JSY14 | 4 | rnr1D57N-Re | AAC | A>C | G/C+1 | A/T-1 | TA>GC | TA>GC | ACGAAAACT | ACTGCAA | TTG | TTG | T | G | T>G | T>G_TTG | TA>GC_ACTGCAA | 7 | FALSE | FALSE |
| 15_v.txt | 32262 | 32262-SNV-A | SNV | 1 | A | C | 15 | 72337 | 0.02073628 | 34.4 | rnr1D57N | FALSE | CAN | JSY14 | 4 | rnr1D57N-Re | AAA | A>C | G/C+1 | A/T-1 | TA>GC | TA>GC | AGACCAAAT | AATAAAA | TTT | TTT | T | G | T>G | T>G_TTT | TA>GC_AATAAAA | 7 | FALSE | FALSE |
| 15_v.txt | 32283 | 32283-SNV-T | SNV | 1 | T | G | 252 | 72336 | 0.34837425 | 53.9920635 | rnr1D57N | FALSE | CAN | JSY14 | 4 | rnr1D57N-Re | GTA | T>G | G/C+1 | A/T-1 | TA>GC | TA>GC | CCAACGTAA | TAAATAT | CAT | GTA | T | C | T>G | T>G_GTA | TA>GC_TAAATAT | 7 | FALSE | FALSE |
| 15_v.txt | 32383 | 32383-SNV-A | SNV | 1 | A | C | 7 | 72336 | 9.68E-03 | 54.8571429 | rnr1D57N | FALSE | CAN | JSY14 | 4 | rnr1D57N-Re | AAG | A>C | G/C+1 | A/T-1 | TA>GC | TA>GC | TAATAAAGG | AGGGAGA | TTC | TTC | T | G | T>G | T>G_TTC | TA>GC_AGGGAGA | 7 | FALSE | FALSE |
| 15_v.txt | 32541 | 32541-SNV-T | SNV | 1 | T | G | 248 | 72336 | 0.3428445 | 36.7217742 | rnr1D57N | FALSE | CAN | JSY14 | 4 | rnr1D57N-Re | TTC | T>G | G/C+1 | A/T-1 | TA>GC | TA>GC | GCAGCTTCA | TCACCAG | AAG | TTC | T | C | T>G | T>G_TTC | TA>GC_TCACCAG | 7 | FALSE | FALSE |
| 15_v.txt | 32631 | 32631-SNV-T | SNV | 1 | T | G | 6 | 70822 | 8.47E-03 | 47.6666667 | rnr1D57N | FALSE | CAN | JSY14 | 4 | rnr1D57N-Re | GTT | T>G | G/C+1 | A/T-1 | TA>GC | TA>GC | CCTTCGTTTT | TTTTTAT | CAA | GTT | T | C | T>G | T>G_GTT | TA>GC_TTTTTAT | 7 | FALSE | FALSE |
| 15_v.txt | 32673 | 32673-SNV-T | SNV | 1 | T | G | 31 | 70822 | 0.04377171 | 48.9354839 | rnr1D57N | FALSE | CAN | JSY14 | 4 | rnr1D57N-Re | GTT | T>G | G/C+1 | A/T-1 | TA>GC | TA>GC | CCTGGGTTTC | TTTCTCC | CAA | GTT | T | C | T>G | T>G_GTT | TA>GC_TTTCTCC | 7 | FALSE | FALSE |
| 15_v.txt | 32728 | 32728-SNV-A | SNV | 1 | A | C | 10 | 70820 | 0.01412031 | 13.5003069 | rnr1D57N | FALSE | CAN | JSY14 | 4 | rnr1D57N-Re | CAA | A>C | G/C+1 | A/T-1 | TA>GC | TA>GC | CCATACAAAA | AAAAACA | GTT | GTT | T | G | T>G | T>G_GTT | TA>GC_AAAAACA | 7 | FALSE | FALSE |
| 15_v.txt | 32732 | 32732-SNV-A | SNV | 1 | A | C | 9 | 70822 | 0.01270792 | 48.3333333 | rnr1D57N | FALSE | CAN | JSY14 | 4 | rnr1D57N-Re | AAC | A>C | G/C+1 | A/T-1 | TA>GC | TA>GC | ACAAAAACA | ACAGTAT | TTG | TTG | T | G | T>G | T>G_TTG | TA>GC_ACAGTAT | 7 | FALSE | FALSE |
| 15_v.txt | 32739 | 32739-SNV-A | SNV | 1 | A | C | 340 | 70822 | 0.48007681 | 55.85 | rnr1D57N | FALSE | CAN | JSY14 | 4 | rnr1D57N-Re | TAT | A>C | G/C+1 | A/T-1 | TA>GC | TA>GC | CAGTATATTA | ATTAGAA | ATA | ATA | T | G | T>G | T>G_ATA | TA>GC_ATTAGAA | 7 | FALSE | FALSE |
| 15_v.txt | 32799 | 32799-SNV-T | SNV | 1 | T | G | 25 | 70822 | 0.03529977 | 39.240432 | rnr1D57N | FALSE | CAN | JSY14 | 4 | rnr1D57N-Re | GTA | T>G | G/C+1 | A/T-1 | TA>GC | TA>GC | TCACCGTAAT | TAATATT | CAT | GTA | T | C | T>G | T>G_GTA | TA>GC_TAATATT | 7 | FALSE | FALSE |
| 15_v.txt | 32842 | 32842-SNV-A | SNV | 1 | A | C | 6 | 70822 | 8.47E-03 | 38.0006386 | rnr1D57N | FALSE | CAN | JSY14 | 4 | rnr1D57N-Re | CAA | A>C | G/C+1 | A/T-1 | TA>GC | TA>GC | TTACCCAAAA | AAAAAAT | GTT | GTT | T | G | T>G | T>G_GTT | TA>GC_AAAAAAT | 7 | FALSE | FALSE |
| 15_v.txt | 32884 | 32884-SNV-A | SNV | 1 | A | C | 4 | 75142 | 5.32E-03 | 30.4994554 | rnr1D57N | FALSE | CAN | JSY14 | 4 | rnr1D57N-Re | CAA | A>C | G/C+1 | A/T-1 | TA>GC | TA>GC | ACGTCCAAA | AAAATTG | GTT | GTT | T | G | T>G | T>G_GTT | TA>GC_AAAATTG | 7 | FALSE | FALSE |
| 15_v.txt | 33087 | 33087-SNV-A | SNV | 1 | A | C | 660 | 75142 | 0.87833702 | 52.0469697 | rnr1D57N | FALSE | CAN | JSY14 | 4 | rnr1D57N-Re | TAA | A>C | G/C+1 | A/T-1 | TA>GC | TA>GC | ATAAATAAA | AAATATG | ATT | ATT | T | G | T>G | T>G_ATT | TA>GC_AAATATG | 7 | FALSE | FALSE |
| 15_v.txt | 33204 | 33204-SNV-A | SNV | 1 | A | C | 32 | 61063 | 0.05240489 | 29.0002886 | rnr1D57N | FALSE | CAN | JSY14 | 4 | rnr1D57N-Re | AAG | A>C | G/C+1 | A/T-1 | TA>GC | TA>GC | TGCTTAAGC | AGCTCTC | TTC | TTC | T | G | T>G | T>G_TTC | TA>GC_AGCTCTC | 7 | FALSE | FALSE |
| 15_v.txt | 33269 | 33269-SNV-A | SNV | 1 | A | C | 463 | 61063 | 0.7582333 | 15.7840173 | rnr1D57N | FALSE | CAN | JSY14 | 4 | rnr1D57N-Re | TAG | A>C | G/C+1 | A/T-1 | TA>GC | TA>GC | CTCCATAGA | AGAGAAC | ATC | ATC | T | G | T>G | T>G_ATC | TA>GC_AGAGAAC | 7 | FALSE | FALSE |
| 15_v.txt | 33476 | 33476-SNV-T | SNV | 1 | T | G | 5 | 61062 | 8.19E-03 | 36.2 | rnr1D57N | FALSE | CAN | JSY14 | 4 | rnr1D57N-Re | CTT | T>G | G/C+1 | A/T-1 | TA>GC | TA>GC | TATGCCTTTT | TTTTTTT | GAA | CTT | T | C | T>G | T>G_CTT | TA>GC_TTTTTTT | 7 | FALSE | FALSE |
| 15_v.txt | 31723 | 31723-SNV-A | SNV | 1 | A | G | 8 | 25145 | 0.03181547 | 53.2501268 | rnr1D57N | FALSE | CAN | JSY14 | 4 | rnr1D57N-Re | CAA | A>G | G/C+1 | A/T-1 | TA>CG | TA>CG | TGTCCCAAAA | AAAAAGT | GTT | GTT | T | C | T>C | T>C_GTT | TA>CG_AAAAAGT | 7 | FALSE | FALSE |
| 15_v.txt | 32081 | 32081-SNV-A | SNV | 1 | A | G | 15 | 64727 | 0.02317426 | 61.1999661 | rnr1D57N | FALSE | CAN | JSY14 | 4 | rnr1D57N-Re | CAA | A>G | G/C+1 | A/T-1 | TA>CG | TA>CG | CCATGCAAA | AAAAAAG | GTT | GTT | T | C | T>C | T>C_GTT | TA>CG_AAAAAAG | 7 | FALSE | FALSE |
| 15_v.txt | 32160 | 32160-SNV-A | SNV | 1 | A | G | 16 | 64665 | 0.02474291 | 38.9999469 | rnr1D57N | FALSE | CAN | JSY14 | 4 | rnr1D57N-Re | CAA | A>G | G/C+1 | A/T-1 | TA>CG | TA>CG | TAAGCCAAA | AAAGCGC | GTT | GTT | T | C | T>C | T>C_GTT | TA>CG_AAAGCGC | 7 | FALSE | FALSE |
| 15_v.txt | 32161 | 32161-SNV-A | SNV | 1 | A | G | 17 | 64665 | 0.02628934 | 34.2884334 | rnr1D57N | FALSE | CAN | JSY14 | 4 | rnr1D57N-Re | AAA | A>G | G/C+1 | A/T-1 | TA>CG | TA>CG | AAGCCAAAG | AAGCGCC | TTT | TTT | T | C | T>C | T>C_TTT | TA>CG_AAGCGCC | 7 | FALSE | FALSE |
| 15_v.txt | 32490 | 32490-SNV-A | SNV | 1 | A | G | 26 | 72335 | 0.03594387 | 40.5768764 | rnr1D57N | FALSE | CAN | JSY14 | 4 | rnr1D57N-Re | GAA | A>G | G/C+1 | A/T-1 | TA>CG | TA>CG | ATACGGAAA | AAAACAA | CTT | CTT | T | C | T>C | T>C_CTT | TA>CG_AAAACAA | 7 | FALSE | FALSE |
| 15_v.txt | 32498 | 32498-SNV-T | SNV | 1 | T | C | 6 | 72336 | 8.30E-03 | 29.1666626 | rnr1D57N | FALSE | CAN | JSY14 | 4 | rnr1D57N-Re | CTT | T>C | G/C+1 | A/T-1 | TA>CG | TA>CG | AACAACTTTT | TTTTTTG | GAA | CTT | T | G | T>C | T>C_CTT | TA>CG_TTTTTTG | 7 | FALSE | FALSE |
| 15_v.txt | 32522 | 32522-SNV-T | SNV | 1 | T | C | 8 | 72336 | 0.0110595 | 37.1249707 | rnr1D57N | FALSE | CAN | JSY14 | 4 | rnr1D57N-Re | ATT | T>C | G/C+1 | A/T-1 | TA>CG | TA>CG | AACGGATTTT | TTTTCTG | TAA | ATT | T | G | T>C | T>C_ATT | TA>CG_TTTTCTG | 7 | FALSE | FALSE |
| 15_v.txt | 32583 | 32583-SNV-A | SNV | 1 | A | G | 49 | 70822 | 0.06918754 | 37.5510204 | rnr1D57N | FALSE | CAN | JSY14 | 4 | rnr1D57N-Re | GAA | A>G | G/C+1 | A/T-1 | TA>CG | TA>CG | AATGTGAAG | AAGGCAG | CTT | CTT | T | C | T>C | T>C_CTT | TA>CG_AAGGCAG | 7 | FALSE | FALSE |
| 15_v.txt | 32675 | 32675-SNV-T | SNV | 1 | T | C | 38 | 70822 | 0.05365564 | 54.7894737 | rnr1D57N | FALSE | CAN | JSY14 | 4 | rnr1D57N-Re | TTC | T>C | G/C+1 | A/T-1 | TA>CG | TA>CG | TGGGTTTCTC | TCTCCAA | AAG | TTC | T | G | T>C | T>C_TTC | TA>CG_TCTCCAA | 7 | FALSE | FALSE |
| 15_v.txt | 32677 | 32677-SNV-T | SNV | 1 | T | C | 26 | 70818 | 0.03671383 | 46.3460832 | rnr1D57N | FALSE | CAN | JSY14 | 4 | rnr1D57N-Re | CTC | T>C | G/C+1 | A/T-1 | TA>CG | TA>CG | GGTTTCTCCA | TCCAATA | GAG | CTC | T | G | T>C | T>C_CTC | TA>CG_TCCAATA | 7 | FALSE | FALSE |
| 15_v.txt | 32680 | 32680-SNV-A | SNV | 1 | A | G | 24 | 70822 | 0.03388778 | 61.3749176 | rnr1D57N | FALSE | CAN | JSY14 | 4 | rnr1D57N-Re | CAA | A>G | G/C+1 | A/T-1 | TA>CG | TA>CG | TTCTCCAATA | AATAACG | GTT | GTT | T | C | T>C | T>C_GTT | TA>CG_AATAACG | 7 | FALSE | FALSE |
| 15_v.txt | 32799 | 32799-SNV-T | SNV | 1 | T | C | 22 | 70822 | 0.03106379 | 41.2728836 | rnr1D57N | FALSE | CAN | JSY14 | 4 | rnr1D57N-Re | GTA | T>C | G/C+1 | A/T-1 | TA>CG | TA>CG | TCACCGTAAT | TAATATT | CAT | GTA | T | G | T>C | T>C_GTA | TA>CG_TAATATT | 7 | FALSE | FALSE |
| 15_v.txt | 32800 | 32800-SNV-A | SNV | 1 | A | G | 16 | 70822 | 0.02259185 | 34.1873886 | rnr1D57N | FALSE | CAN | JSY14 | 4 | rnr1D57N-Re | TAA | A>G | G/C+1 | A/T-1 | TA>CG | TA>CG | CACCGTAAT | AATATTT | ATT | ATT | T | C | T>C | T>C_ATT | TA>CG_AATATTT | 7 | FALSE | FALSE |
| 15_v.txt | 32842 | 32842-SNV-A | SNV | 1 | A | G | 16 | 70822 | 0.02259185 | 34.875109 | rnr1D57N | FALSE | CAN | JSY14 | 4 | rnr1D57N-Re | CAA | A>G | G/C+1 | A/T-1 | TA>CG | TA>CG | TTACCCAAAA | AAAAAAT | GTT | GTT | T | C | T>C | T>C_GTT | TA>CG_AAAAAAT | 7 | FALSE | FALSE |
| 15_v.txt | 32929 | 32929-SNV-T | SNV | 1 | T | C | 25 | 75142 | 0.03327034 | 38.7600853 | rnr1D57N | FALSE | CAN | JSY14 | 4 | rnr1D57N-Re | GTG | T>C | G/C+1 | A/T-1 | TA>CG | TA>CG | CAAAAGTGA | TGATTGC | CAC | GTG | T | G | T>C | T>C_GTG | TA>CG_TGATTGC | 7 | FALSE | FALSE |
| 15_v.txt | 33074 | 33074-SNV-A | SNV | 1 | A | G | 61 | 75142 | 0.08117963 | 45.852459 | rnr1D57N | FALSE | CAN | JSY14 | 4 | rnr1D57N-Re | AAG | A>G | G/C+1 | A/T-1 | TA>CG | TA>CG | TGCCAAAGA | AGAACCC | TTC | TTC | T | C | T>C | T>C_TTC | TA>CG_AGAACCC | 7 | FALSE | FALSE |
| 15_v.txt | 33150 | 33150-SNV-A | SNV | 1 | A | G | 50 | 75141 | 0.06654157 | 35.3999092 | rnr1D57N | FALSE | CAN | JSY14 | 4 | rnr1D57N-Re | AAG | A>G | G/C+1 | A/T-1 | TA>CG | TA>CG | ATGAAAAGA | AGACCTG | TTC | TTC | T | C | T>C | T>C_TTC | TA>CG_AGACCTG | 7 | FALSE | FALSE |
| 15_v.txt | 33211 | 33211-SNV-T | SNV | 1 | T | C | 201 | 61063 | 0.32916824 | 37.6766169 | rnr1D57N | FALSE | CAN | JSY14 | 4 | rnr1D57N-Re | CTC | T>C | G/C+1 | A/T-1 | TA>CG | TA>CG | GCTCTCTCTT | TCTTCAC | GAG | CTC | T | G | T>C | T>C_CTC | TA>CG_TCTTCAC | 7 | FALSE | FALSE |
| 15_v.txt | 33326 | 33326-SNV-T | SNV | 1 | T | C | 8 | 61060 | 0.01310187 | 44.4999892 | rnr1D57N | FALSE | CAN | JSY14 | 4 | rnr1D57N-Re | CTT | T>C | G/C+1 | A/T-1 | TA>CG | TA>CG | CAATTCTTTA | TTTACTT | GAA | CTT | T | G | T>C | T>C_CTT | TA>CG_TTTACTT | 7 | FALSE | FALSE |
| 15_v.txt | 33369 | 33369-SNV-T | SNV | 1 | T | C | 8 | 61063 | 0.01310122 | 49.3749741 | rnr1D57N | FALSE | CAN | JSY14 | 4 | rnr1D57N-Re | GTG | T>C | G/C+1 | A/T-1 | TA>CG | TA>CG | CTGTGGTGT | TGTGTTT | CAC | GTG | T | G | T>C | T>C_GTG | TA>CG_TGTGTTT | 7 | FALSE | FALSE |
| 15_v.txt | 33476 | 33476-SNV-T | SNV | 1 | T | C | 95 | 61062 | 0.15557958 | 31.2526316 | rnr1D57N | FALSE | CAN | JSY14 | 4 | rnr1D57N-Re | CTT | T>C | G/C+1 | A/T-1 | TA>CG | TA>CG | TATGCCTTTT | TTTTTTT | GAA | CTT | T | G | T>C | T>C_CTT | TA>CG_TTTTTTT | 7 | FALSE | FALSE |
| 15_v.txt | 31996 | 31996-SNV-T | SNV | 1 | T | A | 79 | 64727 | 0.12205108 | 53.8734177 | rnr1D57N | FALSE | CAN | JSY14 | 4 | rnr1D57N-Re | TTA | T>A | A/T+1 | A/T-1 | TA>AT | TA>AT | TAGCTTTAAA | TAAATGG | AAT | TTA | T | T | T>A | T>A_TTA | TA>AT_TAAATGG | 7 | FALSE | FALSE |
| 15_v.txt | 32081 | 32081-SNV-A | SNV | 1 | A | T | 6 | 64727 | 9.27E-03 | 62.4997281 | rnr1D57N | FALSE | CAN | JSY14 | 4 | rnr1D57N-Re | CAA | A>T | A/T+1 | A/T-1 | TA>AT | TA>AT | CCATGCAAA | AAAAAAG | GTT | GTT | T | A | T>A | T>A_GTT | TA>AT_AAAAAAG | 7 | FALSE | FALSE |
| 15_v.txt | 32160 | 32160-SNV-A | SNV | 1 | A | T | 5 | 64665 | 7.73E-03 | 52.1996389 | rnr1D57N | FALSE | CAN | JSY14 | 4 | rnr1D57N-Re | CAA | A>T | A/T+1 | A/T-1 | TA>AT | TA>AT | TAAGCCAAA | AAAGCGC | GTT | GTT | T | A | T>A | T>A_GTT | TA>AT_AAAGCGC | 7 | FALSE | FALSE |
| 15_v.txt | 32161 | 32161-SNV-A | SNV | 1 | A | T | 4 | 64665 | 6.19E-03 | 63.9985748 | rnr1D57N | FALSE | CAN | JSY14 | 4 | rnr1D57N-Re | AAA | A>T | A/T+1 | A/T-1 | TA>AT | TA>AT | AAGCCAAAG | AAGCGCC | TTT | TTT | T | A | T>A | T>A_TTT | TA>AT_AAGCGCC | 7 | FALSE | FALSE |
| 15_v.txt | 32383 | 32383-SNV-A | SNV | 1 | A | T | 17 | 72336 | 0.02350144 | 61.1764706 | rnr1D57N | FALSE | CAN | JSY14 | 4 | rnr1D57N-Re | AAG | A>T | A/T+1 | A/T-1 | TA>AT | TA>AT | TAATAAAGG | AGGGAGA | TTC | TTC | T | A | T>A | T>A_TTC | TA>AT_AGGGAGA | 7 | FALSE | FALSE |
| 15_v.txt | 32456 | 32456-SNV-T | SNV | 1 | T | A | 11 | 72336 | 0.01520681 | 50.7266771 | rnr1D57N | FALSE | CAN | JSY14 | 4 | rnr1D57N-Re | ATA | T>A | A/T+1 | A/T-1 | TA>AT | TA>AT | AATGAATAA | TAATAGA | TAT | ATA | T | T | T>A | T>A_ATA | TA>AT_TAATAGA | 7 | FALSE | FALSE |
| 15_v.txt | 32459 | 32459-SNV-T | SNV | 1 | T | A | 14 | 72336 | 0.01935413 | 39.7111013 | rnr1D57N | FALSE | CAN | JSY14 | 4 | rnr1D57N-Re | ATA | T>A | A/T+1 | A/T-1 | TA>AT | TA>AT | GAATAATAG | TAGAGAG | TAT | ATA | T | T | T>A | T>A_ATA | TA>AT_TAGAGAG | 7 | FALSE | FALSE |
| 15_v.txt | 32498 | 32498-SNV-T | SNV | 1 | T | A | 7 | 72336 | 9.68E-03 | 55.7140725 | rnr1D57N | FALSE | CAN | JSY14 | 4 | rnr1D57N-Re | CTT | T>A | A/T+1 | A/T-1 | TA>AT | TA>AT | AACAACTTTT | TTTTTTG | GAA | CTT | T | T | T>A | T>A_CTT | TA>AT_TTTTTTG | 7 | FALSE | FALSE |
| 15_v.txt | 32522 | 32522-SNV-T | SNV | 1 | T | A | 56 | 72336 | 0.0774165 | 42.2494007 | rnr1D57N | FALSE | CAN | JSY14 | 4 | rnr1D57N-Re | ATT | T>A | A/T+1 | A/T-1 | TA>AT | TA>AT | AACGGATTTT | TTTTCTG | TAA | ATT | T | T | T>A | T>A_ATT | TA>AT_TTTTCTG | 7 | FALSE | FALSE |
| 15_v.txt | 32677 | 32677-SNV-T | SNV | 1 | T | A | 17 | 70818 | 0.0240052 | 59.9401742 | rnr1D57N | FALSE | CAN | JSY14 | 4 | rnr1D57N-Re | CTC | T>A | A/T+1 | A/T-1 | TA>AT | TA>AT | GGTTTCTCCA | TCCAATA | GAG | CTC | T | T | T>A | T>A_CTC | TA>AT_TCCAATA | 7 | FALSE | FALSE |
| 15_v.txt | 32680 | 32680-SNV-A | SNV | 1 | A | T | 4 | 70822 | 5.65E-03 | 63.4999104 | rnr1D57N | FALSE | CAN | JSY14 | 4 | rnr1D57N-Re | CAA | A>T | A/T+1 | A/T-1 | TA>AT | TA>AT | TTCTCCAATA | AATAACG | GTT | GTT | T | A | T>A | T>A_GTT | TA>AT_AATAACG | 7 | FALSE | FALSE |
| 15_v.txt | 32717 | 32717-SNV-A | SNV | 1 | A | T | 55 | 70822 | 0.07765948 | 54.7454546 | rnr1D57N | FALSE | CAN | JSY14 | 4 | rnr1D57N-Re | CAC | A>T | A/T+1 | A/T-1 | TA>AT | TA>AT | AGCACCACA | ACAAACC | GTG | GTG | T | A | T>A | T>A_GTG | TA>AT_ACAAACC | 7 | FALSE | FALSE |
| 15_v.txt | 32799 | 32799-SNV-T | SNV | 1 | T | A | 11 | 70822 | 0.0155319 | 38.0004354 | rnr1D57N | FALSE | CAN | JSY14 | 4 | rnr1D57N-Re | GTA | T>A | A/T+1 | A/T-1 | TA>AT | TA>AT | TCACCGTAAT | TAATATT | CAT | GTA | T | T | T>A | T>A_GTA | TA>AT_TAATATT | 7 | FALSE | FALSE |
| 15_v.txt | 32800 | 32800-SNV-A | SNV | 1 | A | T | 44 | 70822 | 0.06212759 | 50.6579632 | rnr1D57N | FALSE | CAN | JSY14 | 4 | rnr1D57N-Re | TAA | A>T | A/T+1 | A/T-1 | TA>AT | TA>AT | CACCGTAAT | AATATTT | ATT | ATT | T | A | T>A | T>A_ATT | TA>AT_AATATTT | 7 | FALSE | FALSE |
| 15_v.txt | 32929 | 32929-SNV-T | SNV | 1 | T | A | 17 | 75142 | 0.02262383 | 42.9427002 | rnr1D57N | FALSE | CAN | JSY14 | 4 | rnr1D57N-Re | GTG | T>A | A/T+1 | A/T-1 | TA>AT | TA>AT | CAAAAGTGA | TGATTGC | CAC | GTG | T | T | T>A | T>A_GTG | TA>AT_TGATTGC | 7 | FALSE | FALSE |
| 15_v.txt | 33072 | 33072-SNV-A | SNV | 1 | A | T | 35 | 75142 | 0.04657848 | 50.9428571 | rnr1D57N | FALSE | CAN | JSY14 | 4 | rnr1D57N-Re | CAA | A>T | A/T+1 | A/T-1 | TA>AT | TA>AT | TATGCCAAA | AAAGAAC | GTT | GTT | T | A | T>A | T>A_GTT | TA>AT_AAAGAAC | 7 | FALSE | FALSE |
| 15_v.txt | 33150 | 33150-SNV-A | SNV | 1 | A | T | 5 | 75141 | 6.65E-03 | 31.7997529 | rnr1D57N | FALSE | CAN | JSY14 | 4 | rnr1D57N-Re | AAG | A>T | A/T+1 | A/T-1 | TA>AT | TA>AT | ATGAAAAGA | AGACCTG | TTC | TTC | T | A | T>A | T>A_TTC | TA>AT_AGACCTG | 7 | FALSE | FALSE |
| 15_v.txt | 33317 | 33317-SNV-A | SNV | 1 | A | T | 144 | 61063 | 0.23582202 | 58.3958333 | rnr1D57N | FALSE | CAN | JSY14 | 4 | rnr1D57N-Re | GAT | A>T | A/T+1 | A/T-1 | TA>AT | TA>AT | CAATGGATA | ATACAAT | CTA | CTA | T | A | T>A | T>A_CTA | TA>AT_ATACAAT | 7 | FALSE | FALSE |
| 15_v.txt | 33326 | 33326-SNV-T | SNV | 1 | T | A | 4 | 61060 | 6.55E-03 | 40.7498395 | rnr1D57N | FALSE | CAN | JSY14 | 4 | rnr1D57N-Re | CTT | T>A | A/T+1 | A/T-1 | TA>AT | TA>AT | CAATTCTTTA | TTTACTT | GAA | CTT | T | T | T>A | T>A_CTT | TA>AT_TTTACTT | 7 | FALSE | FALSE |
| 15_v.txt | 33369 | 33369-SNV-T | SNV | 1 | T | A | 9 | 61063 | 0.01473888 | 39.7773783 | rnr1D57N | FALSE | CAN | JSY14 | 4 | rnr1D57N-Re | GTG | T>A | A/T+1 | A/T-1 | TA>AT | TA>AT | CTGTGGTGT | TGTGTTT | CAC | GTG | T | T | T>A | T>A_GTG | TA>AT_TGTGTTT | 7 | FALSE | FALSE |
| 15_v.txt | 33476 | 33476-SNV-T | SNV | 1 | T | A | 9 | 61062 | 0.01473912 | 37.4444444 | rnr1D57N | FALSE | CAN | JSY14 | 4 | rnr1D57N-Re | CTT | T>A | A/T+1 | A/T-1 | TA>AT | TA>AT | TATGCCTTTT | TTTTTTT | GAA | CTT | T | T | T>A | T>A_CTT | TA>AT_TTTTTTT | 7 | FALSE | FALSE |
| 15_v.txt | 31650 | 31650-SNV-C | SNV | 1 | C | T | 40 | 25145 | 0.15907735 | 35.5 | rnr1D57N | FALSE | CAN | JSY14 | 4 | rnr1D57N-Re | GCG | C>T | A/T+1 | G/C-1 | CG>TA | CG>TA | AGAATGCGA | CGAAATG | CGC | GCG | C | A | C>T | C>T_GCG | CG>TA_CGAAATG | 7 | FALSE | FALSE |
| 15_v.txt | 31658 | 31658-SNV-C | SNV | 1 | C | T | 29 | 25145 | 0.11533108 | 35.3103394 | rnr1D57N | FALSE | CAN | JSY14 | 4 | rnr1D57N-Re | GCG | C>T | A/T+1 | G/C-1 | CG>TA | CG>TA | AAATGGCGT | CGTGGAA | CGC | GCG | C | A | C>T | C>T_GCG | CG>TA_CGTGGAA | 7 | FALSE | FALSE |
| 15_v.txt | 31830 | 31830-SNV-C | SNV | 1 | C | T | 21 | 25145 | 0.08351561 | 52.2857143 | rnr1D57N | FALSE | CAN | JSY14 | 4 | rnr1D57N-Re | GCA | C>T | A/T+1 | G/C-1 | CG>TA | CG>TA | AATATGCATT | CATTGAA | CGT | GCA | C | A | C>T | C>T_GCA | CG>TA_CATTGAA | 7 | FALSE | FALSE |
| 15_v.txt | 31834 | 31834-SNV-G | SNV | 1 | G | A | 26 | 25145 | 0.10340028 | 45.4615385 | rnr1D57N | FALSE | CAN | JSY14 | 4 | rnr1D57N-Re | TGA | G>A | A/T+1 | G/C-1 | CG>TA | CG>TA | TGCATTGAA | GAAATAA | ACT | ACT | C | T | C>T | C>T_ACT | CG>TA_GAAATAA | 7 | FALSE | FALSE |
| 15_v.txt | 31879 | 31879-SNV-C | SNV | 1 | C | T | 20 | 25145 | 0.07953868 | 44 | rnr1D57N | FALSE | CAN | JSY14 | 4 | rnr1D57N-Re | GCG | C>T | A/T+1 | G/C-1 | CG>TA | CG>TA | TATAGGCGG | CGGCAGC | CGC | GCG | C | A | C>T | C>T_GCG | CG>TA_CGGCAGC | 7 | FALSE | FALSE |
| 15_v.txt | 31881 | 31881-SNV-G | SNV | 1 | G | A | 21 | 25145 | 0.08351561 | 45.8566986 | rnr1D57N | FALSE | CAN | JSY14 | 4 | rnr1D57N-Re | GGC | G>A | A/T+1 | G/C-1 | CG>TA | CG>TA | TAGGCGGCA | GCAGCAA | CCG | CCG | C | T | C>T | C>T_CCG | CG>TA_GCAGCAA | 7 | FALSE | FALSE |
| 15_v.txt | 31882 | 31882-SNV-C | SNV | 1 | C | T | 24 | 25145 | 0.09544641 | 38.4955489 | rnr1D57N | FALSE | CAN | JSY14 | 4 | rnr1D57N-Re | GCA | C>T | A/T+1 | G/C-1 | CG>TA | CG>TA | AGGCGGCAG | CAGCAAA | CGT | GCA | C | A | C>T | C>T_GCA | CG>TA_CAGCAAA | 7 | FALSE | FALSE |
| 15_v.txt | 31911 | 31911-SNV-G | SNV | 1 | G | A | 21 | 25145 | 0.08351561 | 41.8571429 | rnr1D57N | FALSE | CAN | JSY14 | 4 | rnr1D57N-Re | TGC | G>A | A/T+1 | G/C-1 | CG>TA | CG>TA | TTTGGTGCA | GCAAAAG | ACG | ACG | C | T | C>T | C>T_ACG | CG>TA_GCAAAAG | 7 | FALSE | FALSE |
| 15_v.txt | 31920 | 31920-SNV-G | SNV | 1 | G | A | 21 | 25145 | 0.08351561 | 36.0952381 | rnr1D57N | FALSE | CAN | JSY14 | 4 | rnr1D57N-Re | CGT | G>A | A/T+1 | G/C-1 | CG>TA | CG>TA | AAAGCCGTG | GTGAAAC | GCA | GCA | C | T | C>T | C>T_GCA | CG>TA_GTGAAAC | 7 | FALSE | FALSE |
| 15_v.txt | 31960 | 31960-SNV-C | SNV | 1 | C | T | 71 | 64726 | 0.10969317 | 36.3943662 | rnr1D57N | FALSE | CAN | JSY14 | 4 | rnr1D57N-Re | GCC | C>T | A/T+1 | G/C-1 | CG>TA | CG>TA | ATGTGGCCG | CCGCATA | CGG | GCC | C | A | C>T | C>T_GCC | CG>TA_CCGCATA | 7 | FALSE | FALSE |
| 15_v.txt | 31962 | 31962-SNV-G | SNV | 1 | G | A | 90 | 64726 | 0.13904768 | 36.7111111 | rnr1D57N | FALSE | CAN | JSY14 | 4 | rnr1D57N-Re | CGC | G>A | A/T+1 | G/C-1 | CG>TA | CG>TA | GTGGCCGCA | GCATAAT | GCG | GCG | C | T | C>T | C>T_GCG | CG>TA_GCATAAT | 7 | FALSE | FALSE |
| 15_v.txt | 31971 | 31971-SNV-G | SNV | 1 | G | A | 18 | 64727 | 0.02780911 | 29.8333333 | rnr1D57N | FALSE | CAN | JSY14 | 4 | rnr1D57N-Re | AGC | G>A | A/T+1 | G/C-1 | CG>TA | CG>TA | TAATAAGCC | GCCAAGC | TCG | TCG | C | T | C>T | C>T_TCG | CG>TA_GCCAAGC | 7 | FALSE | FALSE |
| 15_v.txt | 31977 | 31977-SNV-C | SNV | 1 | C | T | 52 | 64727 | 0.08033742 | 40.2884615 | rnr1D57N | FALSE | CAN | JSY14 | 4 | rnr1D57N-Re | GCC | C>T | A/T+1 | G/C-1 | CG>TA | CG>TA | GCCAAGCCG | CCGGGCA | CGG | GCC | C | A | C>T | C>T_GCC | CG>TA_CCGGGCA | 7 | FALSE | FALSE |
| 15_v.txt | 31980 | 31980-SNV-G | SNV | 1 | G | A | 80 | 64725 | 0.12359985 | 46.4458502 | rnr1D57N | FALSE | CAN | JSY14 | 4 | rnr1D57N-Re | GGG | G>A | A/T+1 | G/C-1 | CG>TA | CG>TA | AAGCCGGGC | GGCATTA | CCC | CCC | C | T | C>T | C>T_CCC | CG>TA_GGCATTA | 7 | FALSE | FALSE |
| 15_v.txt | 31981 | 31981-SNV-G | SNV | 1 | G | A | 41 | 64727 | 0.06334296 | 47.3169659 | rnr1D57N | FALSE | CAN | JSY14 | 4 | rnr1D57N-Re | GGC | G>A | A/T+1 | G/C-1 | CG>TA | CG>TA | AGCCGGGCA | GCATTAA | CCG | CCG | C | T | C>T | C>T_CCG | CG>TA_GCATTAA | 7 | FALSE | FALSE |
| 15_v.txt | 32056 | 32056-SNV-G | SNV | 1 | G | A | 7 | 64727 | 0.01081465 | 48.1304172 | rnr1D57N | FALSE | CAN | JSY14 | 4 | rnr1D57N-Re | TGC | G>A | A/T+1 | G/C-1 | CG>TA | CG>TA | TGATGTGCG | GCGAGAT | ACG | ACG | C | T | C>T | C>T_ACG | CG>TA_GCGAGAT | 7 | FALSE | FALSE |
| 15_v.txt | 32057 | 32057-SNV-C | SNV | 1 | C | T | 54 | 64726 | 0.08342861 | 57.5735225 | rnr1D57N | FALSE | CAN | JSY14 | 4 | rnr1D57N-Re | GCG | C>T | A/T+1 | G/C-1 | CG>TA | CG>TA | GATGTGCGA | CGAGATT | CGC | GCG | C | A | C>T | C>T_GCG | CG>TA_CGAGATT | 7 | FALSE | FALSE |
| 15_v.txt | 32088 | 32088-SNV-C | SNV | 1 | C | T | 84 | 64727 | 0.12977583 | 59.3690476 | rnr1D57N | FALSE | CAN | JSY14 | 4 | rnr1D57N-Re | GCC | C>T | A/T+1 | G/C-1 | CG>TA | CG>TA | AAAAAGCCT | CCTGCAA | CGG | GCC | C | A | C>T | C>T_GCC | CG>TA_CCTGCAA | 7 | FALSE | FALSE |
| 15_v.txt | 32092 | 32092-SNV-C | SNV | 1 | C | T | 64 | 64727 | 0.09887682 | 58.890625 | rnr1D57N | FALSE | CAN | JSY14 | 4 | rnr1D57N-Re | GCA | C>T | A/T+1 | G/C-1 | CG>TA | CG>TA | AGCCTGCAA | CAACACC | CGT | GCA | C | A | C>T | C>T_GCA | CG>TA_CAACACC | 7 | FALSE | FALSE |
| 15_v.txt | 32114 | 32114-SNV-C | SNV | 1 | C | T | 52 | 64727 | 0.08033742 | 58.2692308 | rnr1D57N | FALSE | CAN | JSY14 | 4 | rnr1D57N-Re | GCC | C>T | A/T+1 | G/C-1 | CG>TA | CG>TA | TAATAGCCA | CCATTCG | CGG | GCC | C | A | C>T | C>T_GCC | CG>TA_CCATTCG | 7 | FALSE | FALSE |
| 15_v.txt | 32157 | 32157-SNV-G | SNV | 1 | G | A | 30 | 64664 | 0.04639367 | 44.0334466 | rnr1D57N | FALSE | CAN | JSY14 | 4 | rnr1D57N-Re | AGC | G>A | A/T+1 | G/C-1 | CG>TA | CG>TA | ATGTAAGCC | GCCAAAG | TCG | TCG | C | T | C>T | C>T_TCG | CG>TA_GCCAAAG | 7 | FALSE | FALSE |
| 15_v.txt | 32158 | 32158-SNV-C | SNV | 1 | C | T | 20 | 64665 | 0.03092863 | 43.1466219 | rnr1D57N | FALSE | CAN | JSY14 | 4 | rnr1D57N-Re | GCC | C>T | A/T+1 | G/C-1 | CG>TA | CG>TA | TGTAAGCCA | CCAAAGC | CGG | GCC | C | A | C>T | C>T_GCC | CG>TA_CCAAAGC | 7 | FALSE | FALSE |
| 15_v.txt | 32164 | 32164-SNV-C | SNV | 1 | C | T | 22 | 64665 | 0.0340215 | 46.999599 | rnr1D57N | FALSE | CAN | JSY14 | 4 | rnr1D57N-Re | GCG | C>T | A/T+1 | G/C-1 | CG>TA | CG>TA | CCAAAGCGC | CGCCAAA | CGC | GCG | C | A | C>T | C>T_GCG | CG>TA_CGCCAAA | 7 | FALSE | FALSE |
| 15_v.txt | 32165 | 32165-SNV-G | SNV | 1 | G | A | 62 | 64665 | 0.09587876 | 43.4181668 | rnr1D57N | FALSE | CAN | JSY14 | 4 | rnr1D57N-Re | CGC | G>A | A/T+1 | G/C-1 | CG>TA | CG>TA | CAAAGCGCC | GCCAAAT | GCG | GCG | C | T | C>T | C>T_GCG | CG>TA_GCCAAAT | 7 | FALSE | FALSE |
| 15_v.txt | 32166 | 32166-SNV-C | SNV | 1 | C | T | 62 | 64665 | 0.09587876 | 47.177552 | rnr1D57N | FALSE | CAN | JSY14 | 4 | rnr1D57N-Re | GCC | C>T | A/T+1 | G/C-1 | CG>TA | CG>TA | AAAGCGCCA | CCAAATG | CGG | GCC | C | A | C>T | C>T_GCC | CG>TA_CCAAATG | 7 | FALSE | FALSE |
| 15_v.txt | 32173 | 32173-SNV-C | SNV | 1 | C | T | 50 | 64665 | 0.07732158 | 46.62 | rnr1D57N | FALSE | CAN | JSY14 | 4 | rnr1D57N-Re | GCA | C>T | A/T+1 | G/C-1 | CG>TA | CG>TA | CAAATGCAG | CAGCAGT | CGT | GCA | C | A | C>T | C>T_GCA | CG>TA_CAGCAGT | 7 | FALSE | FALSE |
| 15_v.txt | 32176 | 32176-SNV-C | SNV | 1 | C | T | 44 | 64665 | 0.06804299 | 42.9772727 | rnr1D57N | FALSE | CAN | JSY14 | 4 | rnr1D57N-Re | GCA | C>T | A/T+1 | G/C-1 | CG>TA | CG>TA | ATGCAGCAG | CAGTAAC | CGT | GCA | C | A | C>T | C>T_GCA | CG>TA_CAGTAAC | 7 | FALSE | FALSE |
| 15_v.txt | 32271 | 32271-SNV-C | SNV | 1 | C | T | 93 | 72336 | 0.12856669 | 33.7603192 | rnr1D57N | FALSE | CAN | JSY14 | 4 | rnr1D57N-Re | ACG | C>T | A/T+1 | G/C-1 | CG>TA | CG>TA | AAAATACGG | CGGGAAC | TGC | ACG | C | A | C>T | C>T_ACG | CG>TA_CGGGAAC | 7 | FALSE | FALSE |
| 15_v.txt | 32299 | 32299-SNV-C | SNV | 1 | C | T | 180 | 72336 | 0.24883875 | 52.9222222 | rnr1D57N | FALSE | CAN | JSY14 | 4 | rnr1D57N-Re | GCG | C>T | A/T+1 | G/C-1 | CG>TA | CG>TA | AATTTGCGG | CGGCAGA | CGC | GCG | C | A | C>T | C>T_GCG | CG>TA_CGGCAGA | 7 | FALSE | FALSE |
| 15_v.txt | 32301 | 32301-SNV-G | SNV | 1 | G | A | 45 | 72336 | 0.06220969 | 48.6888889 | rnr1D57N | FALSE | CAN | JSY14 | 4 | rnr1D57N-Re | GGC | G>A | A/T+1 | G/C-1 | CG>TA | CG>TA | TTTGCGGCA | GCAGAAA | CCG | CCG | C | T | C>T | C>T_CCG | CG>TA_GCAGAAA | 7 | FALSE | FALSE |
| 15_v.txt | 32413 | 32413-SNV-G | SNV | 1 | G | A | 347 | 72336 | 0.47970582 | 58.037464 | rnr1D57N | FALSE | CAN | JSY14 | 4 | rnr1D57N-Re | TGT | G>A | A/T+1 | G/C-1 | CG>TA | CG>TA | TAGATTGTG | GTGTTAG | ACA | ACA | C | T | C>T | C>T_ACA | CG>TA_GTGTTAG | 7 | FALSE | FALSE |
| 15_v.txt | 32449 | 32449-SNV-C | SNV | 1 | C | T | 394 | 72336 | 0.54468038 | 53.0862944 | rnr1D57N | FALSE | CAN | JSY14 | 4 | rnr1D57N-Re | CCA | C>T | A/T+1 | G/C-1 | CG>TA | CG>TA | AAAGTCCAA | CAATGAA | GGT | CCA | C | A | C>T | C>T_CCA | CG>TA_CAATGAA | 7 | FALSE | FALSE |
| 15_v.txt | 32466 | 32466-SNV-C | SNV | 1 | C | T | 80 | 72336 | 0.110595 | 49.4125 | rnr1D57N | FALSE | CAN | JSY14 | 4 | rnr1D57N-Re | GCC | C>T | A/T+1 | G/C-1 | CG>TA | CG>TA | AGAGAGCCA | CCAATGT | CGG | GCC | C | A | C>T | C>T_GCC | CG>TA_CCAATGT | 7 | FALSE | FALSE |
| 15_v.txt | 32487 | 32487-SNV-C | SNV | 1 | C | T | 52 | 72336 | 0.07188675 | 44.7307692 | rnr1D57N | FALSE | CAN | JSY14 | 4 | rnr1D57N-Re | ACG | C>T | A/T+1 | G/C-1 | CG>TA | CG>TA | AAGATACGG | CGGAAAA | TGC | ACG | C | A | C>T | C>T_ACG | CG>TA_CGGAAAA | 7 | FALSE | FALSE |
| 15_v.txt | 32528 | 32528-SNV-G | SNV | 1 | G | A | 18 | 72336 | 0.02488388 | 29.8843713 | rnr1D57N | FALSE | CAN | JSY14 | 4 | rnr1D57N-Re | TGG | G>A | A/T+1 | G/C-1 | CG>TA | CG>TA | TTTTCTGGG | GGGGTTT | ACC | ACC | C | T | C>T | C>T_ACC | CG>TA_GGGGTTT | 7 | FALSE | FALSE |
| 15_v.txt | 32529 | 32529-SNV-G | SNV | 1 | G | A | 21 | 72336 | 0.02903119 | 37.1428571 | rnr1D57N | FALSE | CAN | JSY14 | 4 | rnr1D57N-Re | GGG | G>A | A/T+1 | G/C-1 | CG>TA | CG>TA | TTTCTGGGG | GGGTTTG | CCC | CCC | C | T | C>T | C>T_CCC | CG>TA_GGGTTTG | 7 | FALSE | FALSE |
| 15_v.txt | 32530 | 32530-SNV-G | SNV | 1 | G | A | 9 | 72336 | 0.01244194 | 25.8880645 | rnr1D57N | FALSE | CAN | JSY14 | 4 | rnr1D57N-Re | GGG | G>A | A/T+1 | G/C-1 | CG>TA | CG>TA | TTCTGGGGT | GGTTTGC | CCC | CCC | C | T | C>T | C>T_CCC | CG>TA_GGTTTGC | 7 | FALSE | FALSE |
| 15_v.txt | 32535 | 32535-SNV-G | SNV | 1 | G | A | 108 | 72336 | 0.14930325 | 36.8418072 | rnr1D57N | FALSE | CAN | JSY14 | 4 | rnr1D57N-Re | TGC | G>A | A/T+1 | G/C-1 | CG>TA | CG>TA | GGGTTTGCA | GCAGCTT | ACG | ACG | C | T | C>T | C>T_ACG | CG>TA_GCAGCTT | 7 | FALSE | FALSE |
| 15_v.txt | 32536 | 32536-SNV-C | SNV | 1 | C | T | 113 | 72336 | 0.15621544 | 36.6902655 | rnr1D57N | FALSE | CAN | JSY14 | 4 | rnr1D57N-Re | GCA | C>T | A/T+1 | G/C-1 | CG>TA | CG>TA | GGTTTGCAG | CAGCTTC | CGT | GCA | C | A | C>T | C>T_GCA | CG>TA_CAGCTTC | 7 | FALSE | FALSE |
| 15_v.txt | 32566 | 32566-SNV-C | SNV | 1 | C | T | 88 | 70794 | 0.12430432 | 36.625 | rnr1D57N | FALSE | CAN | JSY14 | 4 | rnr1D57N-Re | TCA | C>T | A/T+1 | G/C-1 | CG>TA | CG>TA | CTAGTTCAG | CAGTACC | AGT | TCA | C | A | C>T | C>T_TCA | CG>TA_CAGTACC | 7 | FALSE | FALSE |
| 15_v.txt | 32575 | 32575-SNV-G | SNV | 1 | G | A | 282 | 70822 | 0.39818136 | 36.9503546 | rnr1D57N | FALSE | CAN | JSY14 | 4 | rnr1D57N-Re | TGA | G>A | A/T+1 | G/C-1 | CG>TA | CG>TA | TACCTTGAA | GAAATGT | ACT | ACT | C | T | C>T | C>T_ACT | CG>TA_GAAATGT | 7 | FALSE | FALSE |
| 15_v.txt | 32590 | 32590-SNV-C | SNV | 1 | C | T | 57 | 70822 | 0.08048347 | 36.6140351 | rnr1D57N | FALSE | CAN | JSY14 | 4 | rnr1D57N-Re | GCG | C>T | A/T+1 | G/C-1 | CG>TA | CG>TA | AGGCAGCGT | CGTTAAT | CGC | GCG | C | A | C>T | C>T_GCG | CG>TA_CGTTAAT | 7 | FALSE | FALSE |
| 15_v.txt | 32610 | 32610-SNV-C | SNV | 1 | C | T | 72 | 70822 | 0.10166333 | 46.4722222 | rnr1D57N | FALSE | CAN | JSY14 | 4 | rnr1D57N-Re | CCA | C>T | A/T+1 | G/C-1 | CG>TA | CG>TA | GAAACCCAA | CAACCTA | GGT | CCA | C | A | C>T | C>T_CCA | CG>TA_CAACCTA | 7 | FALSE | FALSE |
| 15_v.txt | 32678 | 32678-SNV-C | SNV | 1 | C | T | 23 | 70821 | 0.03247624 | 43.6509191 | rnr1D57N | FALSE | CAN | JSY14 | 4 | rnr1D57N-Re | TCC | C>T | A/T+1 | G/C-1 | CG>TA | CG>TA | GTTTCTCCAA | CCAATAA | AGG | TCC | C | A | C>T | C>T_TCC | CG>TA_CCAATAA | 7 | FALSE | FALSE |
| 15_v.txt | 32679 | 32679-SNV-C | SNV | 1 | C | T | 137 | 70821 | 0.19344545 | 59.7252386 | rnr1D57N | FALSE | CAN | JSY14 | 4 | rnr1D57N-Re | CCA | C>T | A/T+1 | G/C-1 | CG>TA | CG>TA | TTTCTCCAAT | CAATAAC | GGT | CCA | C | A | C>T | C>T_CCA | CG>TA_CAATAAC | 7 | FALSE | FALSE |
| 15_v.txt | 32700 | 32700-SNV-C | SNV | 1 | C | T | 62 | 70822 | 0.08754342 | 57.4354839 | rnr1D57N | FALSE | CAN | JSY14 | 4 | rnr1D57N-Re | GCC | C>T | A/T+1 | G/C-1 | CG>TA | CG>TA | ACTGGGCCG | CCGGTAA | CGG | GCC | C | A | C>T | C>T_GCC | CG>TA_CCGGTAA | 7 | FALSE | FALSE |

| | | | | | | | | | | | | | | | | | | | | | | | | | | | | | | | | | | |
|---|---|---|---|---|---|---|---|---|---|---|---|---|---|---|---|---|---|---|---|---|---|---|---|---|---|---|---|---|---|---|---|---|---|---|
| 15_v.txt | 32776 | 32776-SNV-( | SNV | 1 | C | T | 58 | 70822 | 0.08189546 | 49.7413793 | rnr1D57N | FALSE | CAN | JSY14 | 4 | rnr1D57N-Re | GCG | C>T | A/T+1 | G/C-1 | CG>TA | CG>TA | TGGAAGCGA | CGACCCA | CGC | GCG | C | A | C>T | C>T_GCG | CG>TA_CGACCCA | 7 | FALSE | FALSE |
| 15_v.txt | 32798 | 32798-SNV-( | SNV | 1 | G | A | 83 | 70822 | 0.11719522 | 43.1327124 | rnr1D57N | FALSE | CAN | JSY14 | 4 | rnr1D57N-Re | CGT | G>A | A/T+1 | G/C-1 | CG>TA | CG>TA | TTCACCGTAA | GTAATAT | GCA | GCA | C | T | C>T | C>T_GCA | CG>TA_GTAATAT | 7 | FALSE | FALSE |
| 15_v.txt | 32860 | 32860-SNV-( | SNV | 1 | C | T | 63 | 70816 | 0.08896295 | 34.5714286 | rnr1D57N | FALSE | CAN | JSY14 | 4 | rnr1D57N-Re | GCC | C>T | A/T+1 | G/C-1 | CG>TA | CG>TA | TCCATGCCG | CCGCCAG | CGG | GCC | C | A | C>T | C>T_GCC | CG>TA_CCGCCAG | 7 | FALSE | FALSE |
| 15_v.txt | 32862 | 32862-SNV-( | SNV | 1 | G | A | 57 | 70816 | 0.08049029 | 34.2982456 | rnr1D57N | FALSE | CAN | JSY14 | 4 | rnr1D57N-Re | CGC | G>A | A/T+1 | G/C-1 | CG>TA | CG>TA | CATGCCGCC | GCCAGTG | GCG | GCG | C | T | C>T | C>T_GCG | CG>TA_GCCAGTG | 7 | FALSE | FALSE |
| 15_v.txt | 32879 | 32879-SNV-( | SNV | 1 | C | T | 43 | 75144 | 0.05722346 | 35.1123252 | rnr1D57N | FALSE | CAN | JSY14 | 4 | rnr1D57N-Re | ACG | C>T | A/T+1 | G/C-1 | CG>TA | CG>TA | TTTGTACGTC | CGTCCAA | TGC | ACG | C | A | C>T | C>T_ACG | CG>TA_CGTCCAA | 7 | FALSE | FALSE |
| 15_v.txt | 32880 | 32880-SNV-( | SNV | 1 | G | A | 45 | 75143 | 0.05988582 | 35.9530877 | rnr1D57N | FALSE | CAN | JSY14 | 4 | rnr1D57N-Re | CGT | G>A | A/T+1 | G/C-1 | CG>TA | CG>TA | TTGTACGTC | GTCCAAA | GCA | GCA | C | T | C>T | C>T_GCA | CG>TA_GTCCAAA | 7 | FALSE | FALSE |
| 15_v.txt | 32899 | 32899-SNV-( | SNV | 1 | G | A | 132 | 75142 | 0.1756674 | 34.7760211 | rnr1D57N | FALSE | CAN | JSY14 | 4 | rnr1D57N-Re | TGG | G>A | A/T+1 | G/C-1 | CG>TA | CG>TA | TGACTTGGC | GGCCAAC | ACC | ACC | C | T | C>T | C>T_ACC | CG>TA_GGCCAAC | 7 | FALSE | FALSE |
| 15_v.txt | 32900 | 32900-SNV-( | SNV | 1 | G | A | 56 | 75142 | 0.07452557 | 34.4538595 | rnr1D57N | FALSE | CAN | JSY14 | 4 | rnr1D57N-Re | GGC | G>A | A/T+1 | G/C-1 | CG>TA | CG>TA | GACTTGGCC | GCCAACT | CCG | CCG | C | T | C>T | C>T_CCG | CG>TA_GCCAACT | 7 | FALSE | FALSE |
| 15_v.txt | 32901 | 32901-SNV-( | SNV | 1 | C | T | 161 | 75142 | 0.214261 | 32.8757764 | rnr1D57N | FALSE | CAN | JSY14 | 4 | rnr1D57N-Re | GCC | C>T | A/T+1 | G/C-1 | CG>TA | CG>TA | ACTTGGCCA | CCAACTA | CGG | GCC | C | A | C>T | C>T_GCC | CG>TA_CCAACTA | 7 | FALSE | FALSE |
| 15_v.txt | 32922 | 32922-SNV-( | SNV | 1 | G | A | 38 | 75142 | 0.05057092 | 54.1052632 | rnr1D57N | FALSE | CAN | JSY14 | 4 | rnr1D57N-Re | GGC | G>A | A/T+1 | G/C-1 | CG>TA | CG>TA | TCCAGGGCA | GCAAAAG | CCG | CCG | C | T | C>T | C>T_CCG | CG>TA_GCAAAAG | 7 | FALSE | FALSE |
| 15_v.txt | 32928 | 32928-SNV-( | SNV | 1 | G | A | 7 | 75142 | 9.32E-03 | 55.7112076 | rnr1D57N | FALSE | CAN | JSY14 | 4 | rnr1D57N-Re | AGT | G>A | A/T+1 | G/C-1 | CG>TA | CG>TA | GCAAAAGTG | GTGATTG | TCA | TCA | C | T | C>T | C>T_TCA | CG>TA_GTGATTG | 7 | FALSE | FALSE |
| 15_v.txt | 32935 | 32935-SNV-( | SNV | 1 | C | T | 101 | 75142 | 0.13441218 | 41.4653465 | rnr1D57N | FALSE | CAN | JSY14 | 4 | rnr1D57N-Re | GCC | C>T | A/T+1 | G/C-1 | CG>TA | CG>TA | TGATTGCCC | CCCAAGA | CGG | GCC | C | A | C>T | C>T_GCC | CG>TA_CCCAAGA | 7 | FALSE | FALSE |
| 15_v.txt | 32965 | 32965-SNV-( | SNV | 1 | C | T | 70 | 75142 | 0.09315696 | 59.9571429 | rnr1D57N | FALSE | CAN | JSY14 | 4 | rnr1D57N-Re | GCC | C>T | A/T+1 | G/C-1 | CG>TA | CG>TA | CATTGGCCG | CCGCACC | CGG | GCC | C | A | C>T | C>T_GCC | CG>TA_CCGCACC | 7 | FALSE | FALSE |
| 15_v.txt | 32967 | 32967-SNV-( | SNV | 1 | G | A | 177 | 75142 | 0.23555402 | 60.2485876 | rnr1D57N | FALSE | CAN | JSY14 | 4 | rnr1D57N-Re | CGC | G>A | A/T+1 | G/C-1 | CG>TA | CG>TA | TTGGCCGCA | GCACCAA | GCG | GCG | C | T | C>T | C>T_GCG | CG>TA_GCACCAA | 7 | FALSE | FALSE |
| 15_v.txt | 32977 | 32977-SNV-( | SNV | 1 | C | T | 60 | 75142 | 0.07984882 | 57.85 | rnr1D57N | FALSE | CAN | JSY14 | 4 | rnr1D57N-Re | GCT | C>T | A/T+1 | G/C-1 | CG>TA | CG>TA | CAAATGCTG | CTGGAGA | CGA | GCT | C | A | C>T | C>T_GCT | CG>TA_CTGGAGA | 7 | FALSE | FALSE |
| 15_v.txt | 33012 | 33012-SNV-( | SNV | 1 | G | A | 62 | 75142 | 0.08251045 | 59.0967742 | rnr1D57N | FALSE | CAN | JSY14 | 4 | rnr1D57N-Re | AGA | G>A | A/T+1 | G/C-1 | CG>TA | CG>TA | GTGAAAGAG | GAGGATG | TCT | TCT | C | T | C>T | C>T_TCT | CG>TA_GAGGATG | 7 | FALSE | FALSE |
| 15_v.txt | 33056 | 33056-SNV-( | SNV | 1 | C | T | 80 | 75142 | 0.10646509 | 57.2 | rnr1D57N | FALSE | CAN | JSY14 | 4 | rnr1D57N-Re | GCG | C>T | A/T+1 | G/C-1 | CG>TA | CG>TA | GGACTGCGT | CGTGACA | CGC | GCG | C | A | C>T | C>T_GCG | CG>TA_CGTGACA | 7 | FALSE | FALSE |
| 15_v.txt | 33069 | 33069-SNV-( | SNV | 1 | G | A | 58 | 75142 | 0.07718719 | 46.7881019 | rnr1D57N | FALSE | CAN | JSY14 | 4 | rnr1D57N-Re | TGC | G>A | A/T+1 | G/C-1 | CG>TA | CG>TA | GAATATGCC | GCCAAAG | ACG | ACG | C | T | C>T | C>T_ACG | CG>TA_GCCAAAG | 7 | FALSE | FALSE |
| 15_v.txt | 33070 | 33070-SNV-( | SNV | 1 | C | T | 80 | 75142 | 0.10646509 | 54.2372197 | rnr1D57N | FALSE | CAN | JSY14 | 4 | rnr1D57N-Re | GCC | C>T | A/T+1 | G/C-1 | CG>TA | CG>TA | AATATGCCA | CCAAAGA | CGG | GCC | C | A | C>T | C>T_GCC | CG>TA_CCAAAGA | 7 | FALSE | FALSE |
| 15_v.txt | 33113 | 33113-SNV-( | SNV | 1 | G | A | 51 | 75142 | 0.0678715 | 43.2748794 | rnr1D57N | FALSE | CAN | JSY14 | 4 | rnr1D57N-Re | GGC | G>A | A/T+1 | G/C-1 | CG>TA | CG>TA | CACTGGGCC | GCCGGCG | CCG | CCG | C | T | C>T | C>T_CCG | CG>TA_GCCGGCG | 7 | FALSE | FALSE |
| 15_v.txt | 33114 | 33114-SNV-( | SNV | 1 | C | T | 127 | 75142 | 0.16901334 | 43.882266 | rnr1D57N | FALSE | CAN | JSY14 | 4 | rnr1D57N-Re | GCC | C>T | A/T+1 | G/C-1 | CG>TA | CG>TA | ACTGGGCCG | CCGGCGT | CGG | GCC | C | A | C>T | C>T_GCC | CG>TA_CCGGCGT | 7 | FALSE | FALSE |
| 15_v.txt | 33115 | 33115-SNV-( | SNV | 1 | C | T | 72 | 75142 | 0.09581858 | 42.512568 | rnr1D57N | FALSE | CAN | JSY14 | 4 | rnr1D57N-Re | CCG | C>T | A/T+1 | G/C-1 | CG>TA | CG>TA | CTGGGCCGG | CGGCGTT | GGC | CCG | C | A | C>T | C>T_CCG | CG>TA_CGGCGTT | 7 | FALSE | FALSE |
| 15_v.txt | 33118 | 33118-SNV-( | SNV | 1 | C | T | 54 | 75142 | 0.07186394 | 41.2388598 | rnr1D57N | FALSE | CAN | JSY14 | 4 | rnr1D57N-Re | GCG | C>T | A/T+1 | G/C-1 | CG>TA | CG>TA | GGCCGGCGT | CGTTGGT | CGC | GCG | C | A | C>T | C>T_GCG | CG>TA_CGTTGGT | 7 | FALSE | FALSE |
| 15_v.txt | 33119 | 33119-SNV-( | SNV | 1 | G | A | 62 | 75141 | 0.08251155 | 46.2575939 | rnr1D57N | FALSE | CAN | JSY14 | 4 | rnr1D57N-Re | CGT | G>A | A/T+1 | G/C-1 | CG>TA | CG>TA | GCCGGCGTT | GTTGGTC | GCA | GCA | C | T | C>T | C>T_GCA | CG>TA_GTTGGTC | 7 | FALSE | FALSE |
| 15_v.txt | 33168 | 33168-SNV-( | SNV | 1 | C | T | 493 | 75126 | 0.65623087 | 37.2677485 | rnr1D57N | FALSE | CAN | JSY14 | 4 | rnr1D57N-Re | ACC | C>T | A/T+1 | G/C-1 | CG>TA | CG>TA | ATAGTACCA | CCACCAA | TGG | ACC | C | A | C>T | C>T_ACC | CG>TA_CCACCAA | 7 | FALSE | FALSE |
| 15_v.txt | 33199 | 33199-SNV-( | SNV | 1 | G | A | 35 | 61063 | 0.05731785 | 36.0857143 | rnr1D57N | FALSE | CAN | JSY14 | 4 | rnr1D57N-Re | TGC | G>A | A/T+1 | G/C-1 | CG>TA | CG>TA | GTCTTTGCTT | GCTTAAG | ACG | ACG | C | T | C>T | C>T_ACG | CG>TA_GCTTAAG | 7 | FALSE | FALSE |
| 15_v.txt | 33205 | 33205-SNV-( | SNV | 1 | G | A | 19 | 61063 | 0.03111541 | 40.9467809 | rnr1D57N | FALSE | CAN | JSY14 | 4 | rnr1D57N-Re | AGC | G>A | A/T+1 | G/C-1 | CG>TA | CG>TA | GCTTAAGCT | GCTCTCT | TCG | TCG | C | T | C>T | C>T_TCG | CG>TA_GCTCTCT | 7 | FALSE | FALSE |
| 15_v.txt | 33258 | 33258-SNV-( | SNV | 1 | C | T | 87 | 61063 | 0.1424758 | 42.1034483 | rnr1D57N | FALSE | CAN | JSY14 | 4 | rnr1D57N-Re | GCC | C>T | A/T+1 | G/C-1 | CG>TA | CG>TA | CCTATGCCAT | CCATCCT | CGG | GCC | C | A | C>T | C>T_GCC | CG>TA_CCATCCT | 7 | FALSE | FALSE |
| 15_v.txt | 33276 | 33276-SNV-( | SNV | 1 | G | A | 66 | 61063 | 0.10808509 | 46.469697 | rnr1D57N | FALSE | CAN | JSY14 | 4 | rnr1D57N-Re | CGT | G>A | A/T+1 | G/C-1 | CG>TA | CG>TA | GAGAACGTA | GTATCCT | GCA | GCA | C | T | C>T | C>T_GCA | CG>TA_GTATCCT | 7 | FALSE | FALSE |
| 15_v.txt | 33284 | 33284-SNV-( | SNV | 1 | G | A | 53 | 61063 | 0.08679561 | 44.754717 | rnr1D57N | FALSE | CAN | JSY14 | 4 | rnr1D57N-Re | CGC | G>A | A/T+1 | G/C-1 | CG>TA | CG>TA | ATCCTCGCCA | GCCATTT | GCG | GCG | C | T | C>T | C>T_GCG | CG>TA_GCCATTT | 7 | FALSE | FALSE |
| 15_v.txt | 33309 | 33309-SNV-( | SNV | 1 | C | T | 71 | 61059 | 0.11628097 | 55.911516 | rnr1D57N | FALSE | CAN | JSY14 | 4 | rnr1D57N-Re | GCG | C>T | A/T+1 | G/C-1 | CG>TA | CG>TA | AAAGAGCGC | CGCAATG | CGC | GCG | C | A | C>T | C>T_GCG | CG>TA_CGCAATG | 7 | FALSE | FALSE |
| 15_v.txt | 33310 | 33310-SNV-( | SNV | 1 | G | A | 69 | 61058 | 0.11300731 | 54.0329043 | rnr1D57N | FALSE | CAN | JSY14 | 4 | rnr1D57N-Re | CGC | G>A | A/T+1 | G/C-1 | CG>TA | CG>TA | AAGAGCGCA | GCAATGG | GCG | GCG | C | T | C>T | C>T_GCG | CG>TA_GCAATGG | 7 | FALSE | FALSE |
| 15_v.txt | 33368 | 33368-SNV-( | SNV | 1 | G | A | 6 | 61059 | 9.83E-03 | 50.6656207 | rnr1D57N | FALSE | CAN | JSY14 | 4 | rnr1D57N-Re | GGT | G>A | A/T+1 | G/C-1 | CG>TA | CG>TA | TCTGTGGTGT | GTGTGTT | CCA | CCA | C | T | C>T | C>T_CCA | CG>TA_GTGTGTT | 7 | FALSE | FALSE |
| 15_v.txt | 33392 | 33392-SNV-( | SNV | 1 | G | A | 42 | 61063 | 0.06878142 | 48.6428571 | rnr1D57N | FALSE | CAN | JSY14 | 4 | rnr1D57N-Re | CGT | G>A | A/T+1 | G/C-1 | CG>TA | CG>TA | AACGTCGTG | GTGAAAG | GCA | GCA | C | T | C>T | C>T_GCA | CG>TA_GTGAAAG | 7 | FALSE | FALSE |
| 15_v.txt | 33412 | 33412-SNV-( | SNV | 1 | G | A | 90 | 61063 | 0.14738876 | 46.1777778 | rnr1D57N | FALSE | CAN | JSY14 | 4 | rnr1D57N-Re | GGC | G>A | A/T+1 | G/C-1 | CG>TA | CG>TA | TGACCGGCT | GCTCATT | CCG | CCG | C | T | C>T | C>T_CCG | CG>TA_GCTCATT | 7 | FALSE | FALSE |
| 15_v.txt | 33474 | 33474-SNV-( | SNV | 1 | C | T | 89 | 61063 | 0.14575111 | 33.2696629 | rnr1D57N | FALSE | CAN | JSY14 | 4 | rnr1D57N-Re | GCC | C>T | A/T+1 | G/C-1 | CG>TA | CG>TA | GCTATGCCTT | CCTTTTT | CGG | GCC | C | A | C>T | C>T_GCC | CG>TA_CCTTTTT | 7 | FALSE | FALSE |
| 15_v.txt | 31981 | 31981-SNV-( | SNV | 1 | G | C | 5 | 64727 | 7.73E-03 | 35.3999971 | rnr1D57N | FALSE | CAN | JSY14 | 4 | rnr1D57N-Re | GGC | G>C | G/C+1 | G/C-1 | CG>GC | CG>GC | AGCCGGGCA | GCATTAA | CCG | CCG | C | G | C>G | C>G_CCG | CG>GC_GCATTAA | 7 | FALSE | FALSE |
| 15_v.txt | 32064 | 32064-SNV-( | SNV | 1 | G | C | 356 | 64727 | 0.55000232 | 61.8820225 | rnr1D57N | FALSE | CAN | JSY14 | 4 | rnr1D57N-Re | TGA | G>C | G/C+1 | G/C-1 | CG>GC | CG>GC | GAGATTGAG | GAGATAA | ACT | ACT | C | G | C>G | C>G_ACT | CG>GC_GAGATAA | 7 | FALSE | FALSE |
| 15_v.txt | 32119 | 32119-SNV-( | SNV | 1 | C | G | 5 | 64665 | 7.73E-03 | 64 | rnr1D57N | FALSE | CAN | JSY14 | 4 | rnr1D57N-Re | TCG | C>G | G/C+1 | G/C-1 | CG>GC | CG>GC | GCCATTCGA | CGAAAAC | AGC | TCG | C | C | C>G | C>G_TCG | CG>GC_CGAAAAC | 7 | FALSE | FALSE |
| 15_v.txt | 32157 | 32157-SNV-( | SNV | 1 | G | C | 2 | 64664 | 3.09E-03 | 37.5003924 | rnr1D57N | FALSE | CAN | JSY14 | 4 | rnr1D57N-Re | AGC | G>C | G/C+1 | G/C-1 | CG>GC | CG>GC | ATGTAAGCC | GCCAAAG | TCG | TCG | C | G | C>G | C>G_TCG | CG>GC_GCCAAAG | 7 | FALSE | FALSE |
| 15_v.txt | 32163 | 32163-SNV-( | SNV | 1 | G | C | 2 | 64664 | 3.09E-03 | 42.9995176 | rnr1D57N | FALSE | CAN | JSY14 | 4 | rnr1D57N-Re | AGC | G>C | G/C+1 | G/C-1 | CG>GC | CG>GC | GCCAAAGCG | GCGCCAA | TCG | TCG | C | G | C>G | C>G_TCG | CG>GC_GCGCCAA | 7 | FALSE | FALSE |
| 15_v.txt | 32199 | 32199-SNV-( | SNV | 1 | G | C | 133 | 64665 | 0.2056754 | 43.2706767 | rnr1D57N | FALSE | CAN | JSY14 | 4 | rnr1D57N-Re | TGG | G>C | G/C+1 | G/C-1 | CG>GC | CG>GC | ATGTATGGA | GGAACAC | ACC | ACC | C | G | C>G | C>G_ACC | CG>GC_GGAACAC | 7 | FALSE | FALSE |
| 15_v.txt | 32529 | 32529-SNV-( | SNV | 1 | G | C | 44 | 72336 | 0.06082725 | 42.6590909 | rnr1D57N | FALSE | CAN | JSY14 | 4 | rnr1D57N-Re | GGG | G>C | G/C+1 | G/C-1 | CG>GC | CG>GC | TTTCTGGGG | GGGTTTG | CCC | CCC | C | G | C>G | C>G_CCC | CG>GC_GGGTTTG | 7 | FALSE | FALSE |
| 15_v.txt | 32536 | 32536-SNV-( | SNV | 1 | C | G | 3 | 72336 | 4.15E-03 | 37 | rnr1D57N | FALSE | CAN | JSY14 | 4 | rnr1D57N-Re | GCA | C>G | G/C+1 | G/C-1 | CG>GC | CG>GC | GGTTTGCAG | CAGCTTC | CGT | GCA | C | C | C>G | C>G_GCA | CG>GC_CAGCTTC | 7 | FALSE | FALSE |
| 15_v.txt | 32566 | 32566-SNV-( | SNV | 1 | C | G | 6 | 70794 | 8.48E-03 | 36.1666667 | rnr1D57N | FALSE | CAN | JSY14 | 4 | rnr1D57N-Re | TCA | C>G | G/C+1 | G/C-1 | CG>GC | CG>GC | CTAGTTCAG | CAGTACC | AGT | TCA | C | C | C>G | C>G_TCA | CG>GC_CAGTACC | 7 | FALSE | FALSE |
| 15_v.txt | 32819 | 32819-SNV-( | SNV | 1 | G | C | 451 | 70822 | 0.63680777 | 45.1929047 | rnr1D57N | FALSE | CAN | JSY14 | 4 | rnr1D57N-Re | AGT | G>C | G/C+1 | G/C-1 | CG>GC | CG>GC | GAACAAGTT | GTTCATT | TCA | TCA | C | G | C>G | C>G_TCA | CG>GC_GTTCATT | 7 | FALSE | FALSE |
| 15_v.txt | 32901 | 32901-SNV-( | SNV | 1 | C | G | 5 | 75142 | 6.65E-03 | 43.2 | rnr1D57N | FALSE | CAN | JSY14 | 4 | rnr1D57N-Re | GCC | C>G | G/C+1 | G/C-1 | CG>GC | CG>GC | ACTTGGCCA | CCAACTA | CGG | GCC | C | C | C>G | C>G_GCC | CG>GC_CCAACTA | 7 | FALSE | FALSE |
| 15_v.txt | 32937 | 32937-SNV-( | SNV | 1 | C | G | 30 | 75142 | 0.03992441 | 52 | rnr1D57N | FALSE | CAN | JSY14 | 4 | rnr1D57N-Re | CCA | C>G | G/C+1 | G/C-1 | CG>GC | CG>GC | ATTGCCCAA | CAAGAAA | GGT | CCA | C | C | C>G | C>G_CCA | CG>GC_CAAGAAA | 7 | FALSE | FALSE |
| 15_v.txt | 32997 | 32997-SNV-( | SNV | 1 | G | C | 38 | 75142 | 0.05057092 | 60.2894737 | rnr1D57N | FALSE | CAN | JSY14 | 4 | rnr1D57N-Re | TGA | G>C | G/C+1 | G/C-1 | CG>GC | CG>GC | CTTTGTGAG | GAGAAAA | ACT | ACT | C | G | C>G | C>G_ACT | CG>GC_GAGAAAA | 7 | FALSE | FALSE |
| 15_v.txt | 33113 | 33113-SNV-( | SNV | 1 | G | C | 2 | 75142 | 2.66E-03 | 46.5003023 | rnr1D57N | FALSE | CAN | JSY14 | 4 | rnr1D57N-Re | GGC | G>C | G/C+1 | G/C-1 | CG>GC | CG>GC | CACTGGGCC | GCCGGCG | CCG | CCG | C | G | C>G | C>G_CCG | CG>GC_GCCGGCG | 7 | FALSE | FALSE |
| 15_v.txt | 33114 | 33114-SNV-( | SNV | 1 | C | G | 6 | 75142 | 7.99E-03 | 38.500353 | rnr1D57N | FALSE | CAN | JSY14 | 4 | rnr1D57N-Re | GCC | C>G | G/C+1 | G/C-1 | CG>GC | CG>GC | ACTGGGCCG | CCGGCGT | CGG | GCC | C | C | C>G | C>G_GCC | CG>GC_CCGGCGT | 7 | FALSE | FALSE |
| 15_v.txt | 33115 | 33115-SNV-( | SNV | 1 | C | G | 14 | 75142 | 0.01863139 | 44.8557912 | rnr1D57N | FALSE | CAN | JSY14 | 4 | rnr1D57N-Re | CCG | C>G | G/C+1 | G/C-1 | CG>GC | CG>GC | CTGGGCCGG | CGGCGTT | GGC | CCG | C | C | C>G | C>G_CCG | CG>GC_CGGCGTT | 7 | FALSE | FALSE |
| 15_v.txt | 33172 | 33172-SNV-( | SNV | 1 | C | G | 332 | 75126 | 0.44192423 | 37.5481928 | rnr1D57N | FALSE | CAN | JSY14 | 4 | rnr1D57N-Re | CCA | C>G | G/C+1 | G/C-1 | CG>GC | CG>GC | TACCACCAA | CAAGGGC | GGT | CCA | C | C | C>G | C>G_CCA | CG>GC_CAAGGGC | 7 | FALSE | FALSE |
| 15_v.txt | 33193 | 33193-SNV-( | SNV | 1 | G | C | 74 | 61063 | 0.12118632 | 37.2972973 | rnr1D57N | FALSE | CAN | JSY14 | 4 | rnr1D57N-Re | TGT | G>C | G/C+1 | G/C-1 | CG>GC | CG>GC | CAATATGTCT | GTCTTTG | ACA | ACA | C | G | C>G | C>G_ACA | CG>GC_GTCTTTG | 7 | FALSE | FALSE |
| 15_v.txt | 33205 | 33205-SNV-( | SNV | 1 | G | C | 19 | 61063 | 0.03111541 | 37.419976 | rnr1D57N | FALSE | CAN | JSY14 | 4 | rnr1D57N-Re | AGC | G>C | G/C+1 | G/C-1 | CG>GC | CG>GC | GCTTAAGCT | GCTCTCT | TCG | TCG | C | G | C>G | C>G_TCG | CG>GC_GCTCTCT | 7 | FALSE | FALSE |
| 15_v.txt | 33310 | 33310-SNV-( | SNV | 1 | G | C | 6 | 61058 | 9.83E-03 | 32.1641867 | rnr1D57N | FALSE | CAN | JSY14 | 4 | rnr1D57N-Re | CGC | G>C | G/C+1 | G/C-1 | CG>GC | CG>GC | AAGAGCGCA | GCAATGG | GCG | GCG | C | G | C>G | C>G_GCG | CG>GC_GCAATGG | 7 | FALSE | FALSE |
| 15_v.txt | 33354 | 33354-SNV-( | SNV | 1 | G | C | 15 | 61063 | 0.02456479 | 46.4666667 | rnr1D57N | FALSE | CAN | JSY14 | 4 | rnr1D57N-Re | TGA | G>C | G/C+1 | G/C-1 | CG>GC | CG>GC | GGTATTGAC | GACCCAC | ACT | ACT | C | G | C>G | C>G_ACT | CG>GC_GACCCAC | 7 | FALSE | FALSE |
| 15_v.txt | 33356 | 33356-SNV-( | SNV | 1 | C | G | 2 | 61063 | 3.28E-03 | 55.9896484 | rnr1D57N | FALSE | CAN | JSY14 | 4 | rnr1D57N-Re | ACC | C>G | G/C+1 | G/C-1 | CG>GC | CG>GC | TATTGACCCA | CCCACGT | TGG | ACC | C | C | C>G | C>G_ACC | CG>GC_CCCACGT | 7 | FALSE | FALSE |
| 15_v.txt | 31657 | 31657-SNV-( | SNV | 1 | G | T | 169 | 25145 | 0.67210181 | 18.7193561 | rnr1D57N | FALSE | CAN | JSY14 | 4 | rnr1D57N-Re | GGC | G>T | A/T+1 | G/C-1 | CG>AT | CG>AT | GAAATGGCG | GCGTGGA | CCG | CCG | C | A | C>A | C>A_CCG | CG>AT_GCGTGGA | 7 | FALSE | FALSE |
| 15_v.txt | 31979 | 31979-SNV-( | SNV | 1 | G | T | 2 | 64727 | 3.09E-03 | 30.4998474 | rnr1D57N | FALSE | CAN | JSY14 | 4 | rnr1D57N-Re | CGG | G>T | A/T+1 | G/C-1 | CG>AT | CG>AT | CAAGCCGGG | GGGCATT | GCC | GCC | C | A | C>A | C>A_GCC | CG>AT_GGGCATT | 7 | FALSE | FALSE |
| 15_v.txt | 31981 | 31981-SNV-( | SNV | 1 | G | T | 26 | 64727 | 0.04016871 | 24.730759 | rnr1D57N | FALSE | CAN | JSY14 | 4 | rnr1D57N-Re | GGC | G>T | A/T+1 | G/C-1 | CG>AT | CG>AT | AGCCGGGCA | GCATTAA | CCG | CCG | C | A | C>A | C>A_CCG | CG>AT_GCATTAA | 7 | FALSE | FALSE |
| 15_v.txt | 32008 | 32008-SNV-( | SNV | 1 | C | A | 4488 | 64727 | 6.93373708 | 56.8106061 | rnr1D57N | FALSE | CAN | JSY14 | 4 | rnr1D57N-Re | TCG | C>A | A/T+1 | G/C-1 | CG>AT | CG>AT | GTAACTCGTC | CGTCACG | AGC | TCG | C | T | C>A | C>A_TCG | CG>AT_CGTCACG | 7 | FALSE | FALSE |
| 15_v.txt | 32056 | 32056-SNV-( | SNV | 1 | G | T | 17 | 64727 | 0.02626416 | 22.1745946 | rnr1D57N | FALSE | CAN | JSY14 | 4 | rnr1D57N-Re | TGC | G>T | A/T+1 | G/C-1 | CG>AT | CG>AT | TGATGTGCG | GCGAGAT | ACG | ACG | C | A | C>A | C>A_ACG | CG>AT_GCGAGAT | 7 | FALSE | FALSE |
| 15_v.txt | 32064 | 32064-SNV-( | SNV | 1 | G | T | 113 | 64727 | 0.17457939 | 59.7876106 | rnr1D57N | FALSE | CAN | JSY14 | 4 | rnr1D57N-Re | TGA | G>T | A/T+1 | G/C-1 | CG>AT | CG>AT | GAGATTGAG | GAGATAA | ACT | ACT | C | A | C>A | C>A_ACT | CG>AT_GAGATAA | 7 | FALSE | FALSE |
| 15_v.txt | 32088 | 32088-SNV-( | SNV | 1 | C | A | 55 | 64727 | 0.08497227 | 55.8909091 | rnr1D57N | FALSE | CAN | JSY14 | 4 | rnr1D57N-Re | GCC | C>A | A/T+1 | G/C-1 | CG>AT | CG>AT | AAAAAGCCT | CCTGCAA | CGG | GCC | C | T | C>A | C>A_GCC | CG>AT_CCTGCAA | 7 | FALSE | FALSE |
| 15_v.txt | 32119 | 32119-SNV-( | SNV | 1 | C | A | 62 | 64665 | 0.09587876 | 57.1451613 | rnr1D57N | FALSE | CAN | JSY14 | 4 | rnr1D57N-Re | TCG | C>A | A/T+1 | G/C-1 | CG>AT | CG>AT | GCCATTCGA | CGAAAAC | AGC | TCG | C | T | C>A | C>A_TCG | CG>AT_CGAAAAC | 7 | FALSE | FALSE |
| 15_v.txt | 32158 | 32158-SNV-( | SNV | 1 | C | A | 26 | 64665 | 0.04020722 | 24.5380779 | rnr1D57N | FALSE | CAN | JSY14 | 4 | rnr1D57N-Re | GCC | C>A | A/T+1 | G/C-1 | CG>AT | CG>AT | TGTAAGCCA | CCAAAGC | CGG | GCC | C | T | C>A | C>A_GCC | CG>AT_CCAAAGC | 7 | FALSE | FALSE |
| 15_v.txt | 32159 | 32159-SNV-( | SNV | 1 | C | A | 7 | 64664 | 0.01082519 | 31.5708501 | rnr1D57N | FALSE | CAN | JSY14 | 4 | rnr1D57N-Re | CCA | C>A | A/T+1 | G/C-1 | CG>AT | CG>AT | GTAAGCCAA | CAAAGCG | GGT | CCA | C | T | C>A | C>A_CCA | CG>AT_CAAAGCG | 7 | FALSE | FALSE |
| 15_v.txt | 32166 | 32166-SNV-( | SNV | 1 | C | A | 10 | 64665 | 0.01546432 | 23.3029552 | rnr1D57N | FALSE | CAN | JSY14 | 4 | rnr1D57N-Re | GCC | C>A | A/T+1 | G/C-1 | CG>AT | CG>AT | AAAGCGCCA | CCAAATG | CGG | GCC | C | T | C>A | C>A_GCC | CG>AT_CCAAATG | 7 | FALSE | FALSE |
| 15_v.txt | 32195 | 32195-SNV-( | SNV | 1 | G | T | 47 | 64665 | 0.07268229 | 41.9148936 | rnr1D57N | FALSE | CAN | JSY14 | 4 | rnr1D57N-Re | TGT | G>T | A/T+1 | G/C-1 | CG>AT | CG>AT | TGCAATGTA | GTATGGA | ACA | ACA | C | A | C>A | C>A_ACA | CG>AT_GTATGGA | 7 | FALSE | FALSE |
| 15_v.txt | 32272 | 32272-SNV-( | SNV | 1 | G | T | 68 | 72336 | 0.09400575 | 32.7778054 | rnr1D57N | FALSE | CAN | JSY14 | 4 | rnr1D57N-Re | CGG | G>T | A/T+1 | G/C-1 | CG>AT | CG>AT | AAATACGGG | GGGAACC | GCC | GCC | C | A | C>A | C>A_GCC | CG>AT_GGGAACC | 7 | FALSE | FALSE |
| 15_v.txt | 32274 | 32274-SNV-( | SNV | 1 | G | T | 423 | 72336 | 0.58477107 | 36.0992908 | rnr1D57N | FALSE | CAN | JSY14 | 4 | rnr1D57N-Re | GGA | G>T | A/T+1 | G/C-1 | CG>AT | CG>AT | ATACGGGAA | GAACCAA | CCT | CCT | C | A | C>A | C>A_CCT | CG>AT_GAACCAA | 7 | FALSE | FALSE |
| 15_v.txt | 32292 | 32292-SNV-( | SNV | 1 | G | T | 37 | 72336 | 0.05115019 | 42.3513514 | rnr1D57N | FALSE | CAN | JSY14 | 4 | rnr1D57N-Re | TGA | G>T | A/T+1 | G/C-1 | CG>AT | CG>AT | ATATTTGAA | GAATTTG | ACT | ACT | C | A | C>A | C>A_ACT | CG>AT_GAATTTG | 7 | FALSE | FALSE |
| 15_v.txt | 32304 | 32304-SNV-( | SNV | 1 | G | T | 206 | 72336 | 0.28478213 | 56.2281553 | rnr1D57N | FALSE | CAN | JSY14 | 4 | rnr1D57N-Re | AGA | G>T | A/T+1 | G/C-1 | CG>AT | CG>AT | GCGGCAGAA | GAAATAA | TCT | TCT | C | A | C>A | C>A_TCT | CG>AT_GAAATAA | 7 | FALSE | FALSE |
| 15_v.txt | 32528 | 32528-SNV-( | SNV | 1 | G | T | 48 | 72336 | 0.066357 | 41.9318144 | rnr1D57N | FALSE | CAN | JSY14 | 4 | rnr1D57N-Re | TGG | G>T | A/T+1 | G/C-1 | CG>AT | CG>AT | TTTTCTGGG | GGGGTTT | ACC | ACC | C | A | C>A | C>A_ACC | CG>AT_GGGGTTT | 7 | FALSE | FALSE |
| 15_v.txt | 32529 | 32529-SNV-( | SNV | 1 | G | T | 7 | 72336 | 9.68E-03 | 36.5714286 | rnr1D57N | FALSE | CAN | JSY14 | 4 | rnr1D57N-Re | GGG | G>T | A/T+1 | G/C-1 | CG>AT | CG>AT | TTTCTGGGG | GGGTTTG | CCC | CCC | C | A | C>A | C>A_CCC | CG>AT_GGGTTTG | 7 | FALSE | FALSE |
| 15_v.txt | 32530 | 32530-SNV-( | SNV | 1 | G | T | 53 | 72336 | 0.07326919 | 38.2058721 | rnr1D57N | FALSE | CAN | JSY14 | 4 | rnr1D57N-Re | GGG | G>T | A/T+1 | G/C-1 | CG>AT | CG>AT | TTCTGGGGT | GGTTTGC | CCC | CCC | C | A | C>A | C>A_CCC | CG>AT_GGTTTGC | 7 | FALSE | FALSE |
| 15_v.txt | 32536 | 32536-SNV-( | SNV | 1 | C | A | 8 | 72336 | 0.0110595 | 30.75 | rnr1D57N | FALSE | CAN | JSY14 | 4 | rnr1D57N-Re | GCA | C>A | A/T+1 | G/C-1 | CG>AT | CG>AT | GGTTTGCAG | CAGCTTC | CGT | GCA | C | T | C>A | C>A_GCA | CG>AT_CAGCTTC | 7 | FALSE | FALSE |
| 15_v.txt | 32566 | 32566-SNV-( | SNV | 1 | C | A | 114 | 70794 | 0.1610306 | 37.6403509 | rnr1D57N | FALSE | CAN | JSY14 | 4 | rnr1D57N-Re | TCA | C>A | A/T+1 | G/C-1 | CG>AT | CG>AT | CTAGTTCAG | CAGTACC | AGT | TCA | C | T | C>A | C>A_TCA | CG>AT_CAGTACC | 7 | FALSE | FALSE |
| 15_v.txt | 32575 | 32575-SNV-( | SNV | 1 | G | T | 1224 | 70822 | 1.72827652 | 36.8504902 | rnr1D57N | FALSE | CAN | JSY14 | 4 | rnr1D57N-Re | TGA | G>T | A/T+1 | G/C-1 | CG>AT | CG>AT | TACCTTGAA | GAAATGT | ACT | ACT | C | A | C>A | C>A_ACT | CG>AT_GAAATGT | 7 | FALSE | FALSE |
| 15_v.txt | 32796 | 32796-SNV-( | SNV | 1 | C | A | 27 | 70822 | 0.03812375 | 44.7407407 | rnr1D57N | FALSE | CAN | JSY14 | 4 | rnr1D57N-Re | ACC | C>A | A/T+1 | G/C-1 | CG>AT | CG>AT | AATTCACCG | CCGTAAT | TGG | ACC | C | T | C>A | C>A_ACC | CG>AT_CCGTAAT | 7 | FALSE | FALSE |
| 15_v.txt | 32798 | 32798-SNV-( | SNV | 1 | G | T | 18 | 70822 | 0.02541583 | 44.7218502 | rnr1D57N | FALSE | CAN | JSY14 | 4 | rnr1D57N-Re | CGT | G>T | A/T+1 | G/C-1 | CG>AT | CG>AT | TTCACCGTAA | GTAATAT | GCA | GCA | C | A | C>A | C>A_GCA | CG>AT_GTAATAT | 7 | FALSE | FALSE |
| 15_v.txt | 32876 | 32876-SNV-( | SNV | 1 | G | T | 45 | 75130 | 0.05989618 | 33.4666667 | rnr1D57N | FALSE | CAN | JSY14 | 4 | rnr1D57N-Re | TGT | G>T | A/T+1 | G/C-1 | CG>AT | CG>AT | AACTTTGTAC | GTACGTC | ACA | ACA | C | A | C>A | C>A_ACA | CG>AT_GTACGTC | 7 | FALSE | FALSE |
| 15_v.txt | 32879 | 32879-SNV-( | SNV | 1 | C | A | 14 | 75144 | 0.0186309 | 21.9275651 | rnr1D57N | FALSE | CAN | JSY14 | 4 | rnr1D57N-Re | ACG | C>A | A/T+1 | G/C-1 | CG>AT | CG>AT | TTTGTACGTC | CGTCCAA | TGC | ACG | C | T | C>A | C>A_ACG | CG>AT_CGTCCAA | 7 | FALSE | FALSE |
| 15_v.txt | 32899 | 32899-SNV-( | SNV | 1 | G | T | 173 | 75142 | 0.23023076 | 14.5259406 | rnr1D57N | FALSE | CAN | JSY14 | 4 | rnr1D57N-Re | TGG | G>T | A/T+1 | G/C-1 | CG>AT | CG>AT | TGACTTGGC | GGCCAAC | ACC | ACC | C | A | C>A | C>A_ACC | CG>AT_GGCCAAC | 7 | FALSE | FALSE |
| 15_v.txt | 32901 | 32901-SNV-( | SNV | 1 | C | A | 7 | 75142 | 9.32E-03 | 37.2857143 | rnr1D57N | FALSE | CAN | JSY14 | 4 | rnr1D57N-Re | GCC | C>A | A/T+1 | G/C-1 | CG>AT | CG>AT | ACTTGGCCA | CCAACTA | CGG | GCC | C | T | C>A | C>A_GCC | CG>AT_CCAACTA | 7 | FALSE | FALSE |
| 15_v.txt | 32928 | 32928-SNV-( | SNV | 1 | G | T | 11 | 75142 | 0.01463895 | 35.7254799 | rnr1D57N | FALSE | CAN | JSY14 | 4 | rnr1D57N-Re | AGT | G>T | A/T+1 | G/C-1 | CG>AT | CG>AT | GCAAAAGTG | GTGATTG | TCA | TCA | C | A | C>A | C>A_TCA | CG>AT_GTGATTG | 7 | FALSE | FALSE |
| 15_v.txt | 33006 | 33006-SNV-( | SNV | 1 | G | T | 22 | 75142 | 0.0292779 | 60.2272727 | rnr1D57N | FALSE | CAN | JSY14 | 4 | rnr1D57N-Re | TGT | G>T | A/T+1 | G/C-1 | CG>AT | CG>AT | AAAACTGTG | GTGAAAG | ACA | ACA | C | A | C>A | C>A_ACA | CG>AT_GTGAAAG | 7 | FALSE | FALSE |
| 15_v.txt | 33050 | 33050-SNV-( | SNV | 1 | G | T | 21 | 75142 | 0.02794709 | 48.2857143 | rnr1D57N | FALSE | CAN | JSY14 | 4 | rnr1D57N-Re | AGG | G>T | A/T+1 | G/C-1 | CG>AT | CG>AT | ACCCAAGGA | GGACTGC | TCC | TCC | C | A | C>A | C>A_TCC | CG>AT_GGACTGC | 7 | FALSE | FALSE |
| 15_v.txt | 33070 | 33070-SNV-( | SNV | 1 | C | A | 10 | 75142 | 0.01330814 | 36.5999649 | rnr1D57N | FALSE | CAN | JSY14 | 4 | rnr1D57N-Re | GCC | C>A | A/T+1 | G/C-1 | CG>AT | CG>AT | AATATGCCA | CCAAAGA | CGG | GCC | C | T | C>A | C>A_GCC | CG>AT_CCAAAGA | 7 | FALSE | FALSE |
| 15_v.txt | 33093 | 33093-SNV-( | SNV | 1 | G | T | 129 | 75142 | 0.17167496 | 49.9612403 | rnr1D57N | FALSE | CAN | JSY14 | 4 | rnr1D57N-Re | TGA | G>T | A/T+1 | G/C-1 | CG>AT | CG>AT | AAATATGAT | GATATAA | ACT | ACT | C | A | C>A | C>A_ACT | CG>AT_GATATAA | 7 | FALSE | FALSE |
| 15_v.txt | 33114 | 33114-SNV-( | SNV | 1 | C | A | 64 | 75142 | 0.08517207 | 42.6253128 | rnr1D57N | FALSE | CAN | JSY14 | 4 | rnr1D57N-Re | GCC | C>A | A/T+1 | G/C-1 | CG>AT | CG>AT | ACTGGGCCG | CCGGCGT | CGG | GCC | C | T | C>A | C>A_GCC | CG>AT_CCGGCGT | 7 | FALSE | FALSE |
| 15_v.txt | 33151 | 33151-SNV-( | SNV | 1 | G | T | 58 | 75141 | 0.07718822 | 38.1551618 | rnr1D57N | FALSE | CAN | JSY14 | 4 | rnr1D57N-Re | AGA | G>T | A/T+1 | G/C-1 | CG>AT | CG>AT | TGAAAAGAC | GACCTGT | TCT | TCT | C | A | C>A | C>A_TCT | CG>AT_GACCTGT | 7 | FALSE | FALSE |
| 15_v.txt | 33182 | 33182-SNV-( | SNV | 1 | C | A | 25 | 74051 | 0.03376052 | 37.68 | rnr1D57N | FALSE | CAN | JSY14 | 4 | rnr1D57N-Re | TCA | C>A | A/T+1 | G/C-1 | CG>AT | CG>AT | GGCAATCAT | CATACCA | AGT | TCA | C | T | C>A | C>A_TCA | CG>AT_CATACCA | 7 | FALSE | FALSE |
| 15_v.txt | 33220 | 33220-SNV-( | SNV | 1 | C | A | 39 | 61063 | 0.06386846 | 35.8205128 | rnr1D57N | FALSE | CAN | JSY14 | 4 | rnr1D57N-Re | TCA | C>A | A/T+1 | G/C-1 | CG>AT | CG>AT | TCACTTCAGC | CAGCGTT | AGT | TCA | C | T | C>A | C>A_TCA | CG>AT_CAGCGTT | 7 | FALSE | FALSE |
| 15_v.txt | 33325 | 33325-SNV-( | SNV | 1 | C | A | 69 | 61062 | 0.1129999 | 54.4926693 | rnr1D57N | FALSE | CAN | JSY14 | 4 | rnr1D57N-Re | TCT | C>A | A/T+1 | G/C-1 | CG>AT | CG>AT | ACAATTCTTT | CTTTACT | AGA | TCT | C | T | C>A | C>A_TCT | CG>AT_CTTTACT | 7 | FALSE | FALSE |
| 15_v.txt | 33356 | 33356-SNV-( | SNV | 1 | C | A | 8 | 61063 | 0.01310122 | 35.3754949 | rnr1D57N | FALSE | CAN | JSY14 | 4 | rnr1D57N-Re | ACC | C>A | A/T+1 | G/C-1 | CG>AT | CG>AT | TATTGACCCA | CCCACGT | TGG | ACC | C | T | C>A | C>A_ACC | CG>AT_CCCACGT | 7 | FALSE | FALSE |
| 15_v.txt | 33368 | 33368-SNV-( | SNV | 1 | G | T | 37 | 61059 | 0.06059713 | 12.4594571 | rnr1D57N | FALSE | CAN | JSY14 | 4 | rnr1D57N-Re | GGT | G>T | A/T+1 | G/C-1 | CG>AT | CG>AT | TCTGTGGTGT | GTGTGTT | CCA | CCA | C | A | C>A | C>A_CCA | CG>AT_GTGTGTT | 7 | FALSE | FALSE |
| 15_v.txt | 33474 | 33474-SNV-( | SNV | 1 | C | A | 123 | 61063 | 0.20143131 | 32.9186992 | rnr1D57N | FALSE | CAN | JSY14 | 4 | rnr1D57N-Re | GCC | C>A | A/T+1 | G/C-1 | CG>AT | CG>AT | GCTATGCCTT | CCTTTTT | CGG | GCC | C | T | C>A | C>A_GCC | CG>AT_CCTTTTT | 7 | FALSE | FALSE |
| 15_v.txt | 33475 | 33475-SNV-( | SNV | 1 | C | A | 35 | 61063 | 0.05731785 | 34.7714286 | rnr1D57N | FALSE | CAN | JSY14 | 4 | rnr1D57N-Re | CCT | C>A | A/T+1 | G/C-1 | CG>AT | CG>AT | CTATGCCTTT | CTTTTTT | GGA | CCT | C | T | C>A | C>A_CCT | CG>AT_CTTTTTT | 7 | FALSE | FALSE |
| 15_v.txt | 31979 | 31979-MNV- | MNV | 3 | GGG | TTT | 1 | 64727 | 1.55E-03 | 33.9999934 | rnr1D57N | FALSE | CAN | JSY14 | 4 | rnr1D57N-Re | CGG | nope | >1bp-ins | >1bpdel | none | MNV | CAAGCCGGG | GGGCATT | GCC | CGG | GGG | none | NA | NA_CGG | MNV_GGGCATT | 7 | FALSE | FALSE |
| 15_v.txt | 32677 | 32677-MNV- | MNV | 4 | TCCA | AGGT | 26 | 70822 | 0.03671176 | 59.9923078 | rnr1D57N | FALSE | CAN | JSY14 | 4 | rnr1D57N-Re | CTC | nope | >1bp-ins | >1bpdel | none | MNV | GGTTTCTCCA | TCCAATA | GAG | CTC | TCCA | none | NA | NA_CTC | MNV_TCCAATA | 7 | FALSE | FALSE |
| 15_v.txt | 33368 | 33368-MNV- | MNV | 2 | GT | TG | 2 | 61063 | 3.28E-03 | 19.4999926 | rnr1D57N | FALSE | CAN | JSY14 | 4 | rnr1D57N-Re | GGT | nope | >1bp-ins | >1bpdel | none | MNV | TCTGTGGTGT | GTGTGTT | CCA | GGT | GT | none | NA | NA_GGT | MNV_GTGTGTT | 7 | FALSE | FALSE |
| 15_v.txt | 31723 | 31723-Insert | Insertion | 1 | - | A | 3 | 25145 | 0.0119308 | 64 | rnr1D57N | FALSE | CAN | JSY14 | 4 | rnr1D57N-Re | CAA | nope | A/T+1 | >1bpdel | none | A/T+1 | TGTCCCAAA | AAAAAGT | GTT | CAA | - | T | NA | NA_CAA | A/T+1_AAAAAGT | 7 | FALSE | FALSE |
| 15_v.txt | 31819 | 31819-Insert | Insertion | 23 | - | ATCTGAATAT | 11 | 25145 | 0.04374627 | 55.0434783 | rnr1D57N | FALSE | CAN | JSY14 | 4 | rnr1D57N-Re | CAT | nope | >1bp-ins | >1bpdel | none | >1bp-ins | ATCTGCATCT | ATCTGAA | GTA | CAT | - | none | NA | NA_CAT | >1bp-ins_ATCTGAA | 7 | FALSE | FALSE |
| 15_v.txt | 32081 | 32081-Insert | Insertion | 1 | - | A | 171 | 64727 | 0.26418651 | 63.0467836 | rnr1D57N | FALSE | CAN | JSY14 | 4 | rnr1D57N-Re | CAA | nope | A/T+1 | >1bpdel | none | A/T+1 | CCATGCAAA | AAAAAAG | GTT | CAA | - | T | NA | NA_CAA | A/T+1_AAAAAAG | 7 | FALSE | FALSE |
| 15_v.txt | 32265 | 32265-Insert | Insertion | 1 | - | A | 17 | 72337 | 0.02350111 | 37.4705882 | rnr1D57N | FALSE | CAN | JSY14 | 4 | rnr1D57N-Re | TAA | nope | A/T+1 | >1bpdel | none | A/T+1 | CCAAATAAA | AAAATAC | ATT | TAA | - | T | NA | NA_TAA | A/T+1_AAAATAC | 7 | FALSE | FALSE |
| 15_v.txt | 32346 | 32346-Insert | Insertion | 1 | - | A | 42 | 72336 | 0.05806238 | 63.4285714 | rnr1D57N | FALSE | CAN | JSY14 | 4 | rnr1D57N-Re | CAA | nope | A/T+1 | >1bpdel | none | A/T+1 | TGTGGCAAA | AAAACCT | GTT | CAA | - | T | NA | NA_CAA | A/T+1_AAAACCT | 7 | FALSE | FALSE |
| 15_v.txt | 32457 | 32457-Insert | Insertion | 1 | - | A | 1 | 72336 | 1.38E-03 | 37 | rnr1D57N | FALSE | CAN | JSY14 | 4 | rnr1D57N-Re | TAA | nope | A/T+1 | >1bpdel | none | A/T+1 | ATGAATAAT | AATAGAG | ATT | TAA | - | T | NA | NA_TAA | A/T+1_AATAGAG | 7 | FALSE | FALSE |
| 15_v.txt | 32490 | 32490-Insert | Insertion | 1 | - | A | 14 | 72336 | 0.01935413 | 49.1428571 | rnr1D57N | FALSE | CAN | JSY14 | 4 | rnr1D57N-Re | GAA | nope | A/T+1 | >1bpdel | none | A/T+1 | ATACGGAAA | AAAACAA | CTT | GAA | - | T | NA | NA_GAA | A/T+1_AAAACAA | 7 | FALSE | FALSE |
| 15_v.txt | 32498 | 32498-Insert | Insertion | 1 | - | T | 3 | 72336 | 4.15E-03 | 55 | rnr1D57N | FALSE | CAN | JSY14 | 4 | rnr1D57N-Re | CTT | nope | A/T+1 | >1bpdel | none | A/T+1 | AACAACTTTT | TTTTTTG | GAA | CTT | - | A | NA | NA_CTT | A/T+1_TTTTTTG | 7 | FALSE | FALSE |
| 15_v.txt | 32522 | 32522-Insert | Insertion | 1 | - | T | 2 | 72336 | 2.77E-03 | 38 | rnr1D57N | FALSE | CAN | JSY14 | 4 | rnr1D57N-Re | ATT | nope | A/T+1 | >1bpdel | none | A/T+1 | AACGGATTT | TTTTCTG | TAA | ATT | - | A | NA | NA_ATT | A/T+1_TTTTCTG | 7 | FALSE | FALSE |
| 15_v.txt | 32680 | 32680-Insert | Insertion | 1 | - | A | 62 | 70822 | 0.08754342 | 63.2741915 | rnr1D57N | FALSE | CAN | JSY14 | 4 | rnr1D57N-Re | CAA | nope | A/T+1 | >1bpdel | none | A/T+1 | TTCTCCAATA | AATAACG | GTT | CAA | - | T | NA | NA_CAA | A/T+1_AATAACG | 7 | FALSE | FALSE |
| 15_v.txt | 32728 | 32728-Insert | Insertion | 1 | - | A | 15 | 70822 | 0.02117986 | 60.5333333 | rnr1D57N | FALSE | CAN | JSY14 | 4 | rnr1D57N-Re | CAA | nope | A/T+1 | >1bpdel | none | A/T+1 | CCATACAAA | AAAAACA | GTT | CAA | - | T | NA | NA_CAA | A/T+1_AAAAACA | 7 | FALSE | FALSE |
| 15_v.txt | 32836 | 32836-Insert | Insertion | 1 | - | T | 10 | 70822 | 0.01411991 | 43.1 | rnr1D57N | FALSE | CAN | JSY14 | 4 | rnr1D57N-Re | ATT | nope | A/T+1 | >1bpdel | none | A/T+1 | TGATAATTA | TTACCCA | TAA | ATT | - | A | NA | NA_ATT | A/T+1_TTACCCA | 7 | FALSE | FALSE |
| 15_v.txt | 32842 | 32842-Insert | Insertion | 1 | - | A | 169 | 70822 | 0.23862642 | 42.7633136 | rnr1D57N | FALSE | CAN | JSY14 | 4 | rnr1D57N-Re | CAA | nope | A/T+1 | >1bpdel | none | A/T+1 | TTACCCAAA | AAAAAAT | GTT | CAA | - | T | NA | NA_CAA | A/T+1_AAAAAAT | 7 | FALSE | FALSE |
| 15_v.txt | 32884 | 32884-Insert | Insertion | 1 | - | A | 16 | 75142 | 0.02129302 | 37.9375 | rnr1D57N | FALSE | CAN | JSY14 | 4 | rnr1D57N-Re | CAA | nope | A/T+1 | >1bpdel | none | A/T+1 | ACGTCCAAA | AAAATTG | GTT | CAA | - | T | NA | NA_CAA | A/T+1_AAAATTG | 7 | FALSE | FALSE |
| 15_v.txt | 33206 | 33206-Insert | Insertion | 2 | - | CT | 1 | 61063 | 1.64E-03 | 38 | rnr1D57N | FALSE | CAN | JSY14 | 4 | rnr1D57N-Re | GCT | nope | >1bp-ins | >1bpdel | none | >1bp-ins | CTTAAGCTCT | CTCTCTC | CGA | GCT | - | none | NA | NA_GCT | >1bp-ins_CTCTCTC | 7 | FALSE | FALSE |
| 15_v.txt | 33356 | 33356-Insert | Insertion | 1 | - | C | 4 | 61063 | 6.55E-03 | 54.5 | rnr1D57N | FALSE | CAN | JSY14 | 4 | rnr1D57N-Re | ACC | nope | G/C+1 | >1bpdel | none | G/C+1 | TATTGACCCA | CCCACGT | TGG | ACC | - | G | NA | NA_ACC | G/C+1_CCCACGT | 7 | FALSE | FALSE |

| | | | | | | | | | | | | | | | | | | | | | | | | | | | | | | | | | | |
|---|---|---|---|---|---|---|---|---|---|---|---|---|---|---|---|---|---|---|---|---|---|---|---|---|---|---|---|---|---|---|---|---|---|---|
| 15_v.txt | 32157 | 32157-Deleti | Deletion | 8 | GCCAAAGC | - | 356 | 64665 | 0.55052965 | 51.6797753 | rnr1D57N | FALSE | CAN | JSY14 | 4 | rnr1D57N-Re | AGC | nope | >1bp-ins | >1bpdel | none | >1bpdel | ATGTAAGCC/ | GCCAAAG | TCG | AGC | GCCAAAGC | none | NA | NA_AGC | >1bpdel_GCCAAAG | 7 | FALSE | FALSE |
| 15_v.txt | 32158 | 32158-Deleti | Deletion | 2 | CC | - | 1 | 64665 | 1.55E-03 | 11 | rnr1D57N | FALSE | CAN | JSY14 | 4 | rnr1D57N-Re | GCC | nope | >1bp-ins | >1bpdel | none | >1bpdel | TGTAAGCCA/ | CCAAAGC | CGG | GCC | CC | none | NA | NA_GCC | >1bpdel_CCAAAGC | 7 | FALSE | FALSE |
| 15_v.txt | 32161 | 32161-Deleti | Deletion | 4 | AAGC | - | 1 | 64665 | 1.55E-03 | 38 | rnr1D57N | FALSE | CAN | JSY14 | 4 | rnr1D57N-Re | AAA | nope | >1bp-ins | >1bpdel | none | >1bpdel | AAGCCAAAG | AAGCGCC | TTT | AAA | AAGC | none | NA | NA_AAA | >1bpdel_AAGCGCC | 7 | FALSE | FALSE |
| 15_v.txt | 32455 | 32455-Deleti | Deletion | 2 | AT | - | 48 | 72336 | 0.066357 | 52.5208305 | rnr1D57N | FALSE | CAN | JSY14 | 4 | rnr1D57N-Re | AAT | nope | >1bp-ins | >1bpdel | none | >1bpdel | CAATGAATA/ | ATAATAG | TTA | AAT | AT | none | NA | NA_AAT | >1bpdel_ATAATAG | 7 | FALSE | FALSE |
| 15_v.txt | 32458 | 32458-Deleti | Deletion | 2 | AT | - | 75 | 72336 | 0.10368281 | 53.44 | rnr1D57N | FALSE | CAN | JSY14 | 4 | rnr1D57N-Re | AAT | nope | >1bp-ins | >1bpdel | none | >1bpdel | TGAATAATA( | ATAGAGA | TTA | AAT | AT | none | NA | NA_AAT | >1bpdel_ATAGAGA | 7 | FALSE | FALSE |
| 15_v.txt | 32928 | 32928-Deleti | Deletion | 2 | GT | - | 9 | 75142 | 0.01197732 | 46.1111111 | rnr1D57N | FALSE | CAN | JSY14 | 4 | rnr1D57N-Re | AGT | nope | >1bp-ins | >1bpdel | none | >1bpdel | GCAAAAGTG | GTGATTG | TCA | AGT | GT | none | NA | NA_AGT | >1bpdel_GTGATTG | 7 | FALSE | FALSE |
| 15_v.txt | 33368 | 33368-Deleti | Deletion | 2 | GT | - | 8 | 61063 | 0.01310122 | 60.875 | rnr1D57N | FALSE | CAN | JSY14 | 4 | rnr1D57N-Re | GGT | nope | >1bp-ins | >1bpdel | none | >1bpdel | TCTGTGGTGT | GTGTGTT | CCA | GGT | GT | none | NA | NA_GGT | >1bpdel_GTGTGTT | 7 | FALSE | FALSE |
| 15_v.txt | 31663 | 31663-Deleti | Deletion | 1 | A | - | 6 | 25145 | 0.0238616 | 36.5 | rnr1D57N | FALSE | CAN | JSY14 | 4 | rnr1D57N-Re | GAA | nope | >1bp-ins | A/T-1 | none | A/T-1 | GCGTGGAAA | AAATGTG | CTT | CTT | T | none | T>none | T>none_CTT | A/T-1_AAATGTG | 7 | FALSE | FALSE |
| 15_v.txt | 31723 | 31723-Deleti | Deletion | 1 | A | - | 29 | 25145 | 0.11533108 | 58.7931038 | rnr1D57N | FALSE | CAN | JSY14 | 4 | rnr1D57N-Re | CAA | nope | >1bp-ins | A/T-1 | none | A/T-1 | TGTCCCAAA/ | AAAAAGT | GTT | GTT | T | none | T>none | T>none_GTT | A/T-1_AAAAAGT | 7 | FALSE | FALSE |
| 15_v.txt | 31863 | 31863-Deleti | Deletion | 1 | A | - | 6 | 25145 | 0.0238616 | 42 | rnr1D57N | FALSE | CAN | JSY14 | 4 | rnr1D57N-Re | GAA | nope | >1bp-ins | A/T-1 | none | A/T-1 | AACAGGAAA | AAAATAG | CTT | CTT | T | none | T>none | T>none_CTT | A/T-1_AAAATAG | 7 | FALSE | FALSE |
| 15_v.txt | 31904 | 31904-Deleti | Deletion | 1 | T | - | 11 | 25145 | 0.04374627 | 46.8181818 | rnr1D57N | FALSE | CAN | JSY14 | 4 | rnr1D57N-Re | ATT | nope | >1bp-ins | A/T-1 | none | A/T-1 | ATTGAATTTT | TTTTGGT | TAA | ATT | T | none | nope | nope_ATT | A/T-1_TTTTGGT | 7 | FALSE | FALSE |
| 15_v.txt | 31913 | 31913-Deleti | Deletion | 1 | A | - | 6 | 25145 | 0.0238616 | 38 | rnr1D57N | FALSE | CAN | JSY14 | 4 | rnr1D57N-Re | CAA | nope | >1bp-ins | A/T-1 | none | A/T-1 | TGGTGCAAA. | AAAAGCC | GTT | GTT | T | none | T>none | T>none_GTT | A/T-1_AAAAGCC | 7 | FALSE | FALSE |
| 15_v.txt | 31952 | 31952-Deleti | Deletion | 1 | A | - | 9 | 64726 | 0.01390477 | 33.2222222 | rnr1D57N | FALSE | CAN | JSY14 | 4 | rnr1D57N-Re | TAA | nope | >1bp-ins | A/T-1 | none | A/T-1 | CGTCATAAAT | AAATGTG | ATT | ATT | T | none | T>none | T>none_ATT | A/T-1_AAATGTG | 7 | FALSE | FALSE |
| 15_v.txt | 31971 | 31971-Deleti | Deletion | 1 | G | - | 43490 | 64727 | 67.1898899 | 37.4167395 | rnr1D57N | FALSE | CAN | JSY14 | 4 | rnr1D57N-Re | AGC | nope | >1bp-ins | G/C-1 | none | G/C-1 | TAATAAGCC/ | GCCAAGC | TCG | TCG | C | none | C>none | C>none_TCG | G/C-1_GCCAAGC | 7 | FALSE | FALSE |
| 15_v.txt | 31979 | 31979-Deleti | Deletion | 1 | G | - | 64 | 64727 | 0.09887682 | 43.1406239 | rnr1D57N | FALSE | CAN | JSY14 | 4 | rnr1D57N-Re | CGG | nope | >1bp-ins | G/C-1 | none | G/C-1 | CAAGCCGGG | GGGCATT | GCC | GCC | C | none | C>none | C>none_GCC | G/C-1_GGGCATT | 7 | FALSE | FALSE |
| 15_v.txt | 32056 | 32056-Deleti | Deletion | 1 | G | - | 256 | 64727 | 0.39550728 | 62.6679688 | rnr1D57N | FALSE | CAN | JSY14 | 4 | rnr1D57N-Re | TGC | nope | >1bp-ins | G/C-1 | none | G/C-1 | TGATGTGCG, | GCGAGAT | ACG | ACG | C | none | C>none | C>none_ACG | G/C-1_GCGAGAT | 7 | FALSE | FALSE |
| 15_v.txt | 32069 | 32069-Deleti | Deletion | 1 | A | - | 7 | 64727 | 0.01081465 | 64 | rnr1D57N | FALSE | CAN | JSY14 | 4 | rnr1D57N-Re | TAA | nope | >1bp-ins | A/T-1 | none | A/T-1 | TGAGATAAAT | AAATAAC | ATT | ATT | T | none | T>none | T>none_ATT | A/T-1_AAATAAC | 7 | FALSE | FALSE |
| 15_v.txt | 32081 | 32081-Deleti | Deletion | 1 | A | - | 64 | 64727 | 0.09887682 | 62.59375 | rnr1D57N | FALSE | CAN | JSY14 | 4 | rnr1D57N-Re | CAA | nope | >1bp-ins | A/T-1 | none | A/T-1 | CCATGCAAA/ | AAAAAAG | GTT | GTT | T | none | T>none | T>none_GTT | A/T-1_AAAAAAG | 7 | FALSE | FALSE |
| 15_v.txt | 32133 | 32133-Deleti | Deletion | 1 | C | - | 7 | 64665 | 0.01082502 | 60 | rnr1D57N | FALSE | CAN | JSY14 | 4 | rnr1D57N-Re | ACC | nope | >1bp-ins | G/C-1 | none | G/C-1 | TTGTCACCAC | CCACCAG | TGG | ACC | C | none | nope | nope_ACC | G/C-1_CCACCAG | 7 | FALSE | FALSE |
| 15_v.txt | 32160 | 32160-Deleti | Deletion | 1 | A | - | 13 | 64665 | 0.02010361 | 47.4615385 | rnr1D57N | FALSE | CAN | JSY14 | 4 | rnr1D57N-Re | CAA | nope | >1bp-ins | A/T-1 | none | A/T-1 | TAAGCCAAA( | AAAGCGC | GTT | GTT | T | none | T>none | T>none_GTT | A/T-1_AAAGCGC | 7 | FALSE | FALSE |
| 15_v.txt | 32184 | 32184-Deleti | Deletion | 1 | A | - | 15 | 64665 | 0.02319647 | 48.4666667 | rnr1D57N | FALSE | CAN | JSY14 | 4 | rnr1D57N-Re | GAA | nope | >1bp-ins | A/T-1 | none | A/T-1 | GTAACGAAA | AAAACTG | CTT | CTT | T | none | T>none | T>none_CTT | A/T-1_AAAACTG | 7 | FALSE | FALSE |
| 15_v.txt | 32378 | 32378-Deleti | Deletion | 1 | A | - | 9 | 72336 | 0.01244194 | 63.1111111 | rnr1D57N | FALSE | CAN | JSY14 | 4 | rnr1D57N-Re | TAA | nope | >1bp-ins | A/T-1 | none | A/T-1 | AGCAATAAT, | AATAAAG | ATT | ATT | T | none | T>none | T>none_ATT | A/T-1_AATAAAG | 7 | FALSE | FALSE |
| 15_v.txt | 32416 | 32416-Deleti | Deletion | 1 | T | - | 8 | 72336 | 0.0110595 | 59 | rnr1D57N | FALSE | CAN | JSY14 | 4 | rnr1D57N-Re | GTT | nope | >1bp-ins | A/T-1 | none | A/T-1 | ATTGTGTTA( | TTAGTTT | CAA | GTT | T | none | nope | nope_GTT | A/T-1_TTAGTTT | 7 | FALSE | FALSE |
| 15_v.txt | 32442 | 32442-Deleti | Deletion | 1 | A | - | 12 | 72336 | 0.01658925 | 57.5 | rnr1D57N | FALSE | CAN | JSY14 | 4 | rnr1D57N-Re | TAA | nope | >1bp-ins | A/T-1 | none | A/T-1 | GGAACTAAA | AAAAGTC | ATT | ATT | T | none | T>none | T>none_ATT | A/T-1_AAAAGTC | 7 | FALSE | FALSE |
| 15_v.txt | 32490 | 32490-Deleti | Deletion | 1 | A | - | 441 | 72335 | 0.60966337 | 45.2630375 | rnr1D57N | FALSE | CAN | JSY14 | 4 | rnr1D57N-Re | GAA | nope | >1bp-ins | A/T-1 | none | A/T-1 | ATACGGAAA | AAAACAA | CTT | CTT | T | none | T>none | T>none_CTT | A/T-1_AAAACAA | 7 | FALSE | FALSE |
| 15_v.txt | 32498 | 32498-Deleti | Deletion | 1 | T | - | 82 | 72336 | 0.11335988 | 45.3414634 | rnr1D57N | FALSE | CAN | JSY14 | 4 | rnr1D57N-Re | CTT | nope | >1bp-ins | A/T-1 | none | A/T-1 | AACAACTTTT | TTTTTTG | GAA | CTT | T | none | nope | nope_CTT | A/T-1_TTTTTTG | 7 | FALSE | FALSE |
| 15_v.txt | 32519 | 32519-Deleti | Deletion | 1 | G | - | 46 | 72336 | 0.06359213 | 43.6521739 | rnr1D57N | FALSE | CAN | JSY14 | 4 | rnr1D57N-Re | CGG | nope | >1bp-ins | G/C-1 | none | G/C-1 | TGGAACGGA | GGATTTT | GCC | GCC | C | none | C>none | C>none_GCC | G/C-1_GGATTTT | 7 | FALSE | FALSE |
| 15_v.txt | 32522 | 32522-Deleti | Deletion | 1 | T | - | 30 | 72336 | 0.04147313 | 39.6666665 | rnr1D57N | FALSE | CAN | JSY14 | 4 | rnr1D57N-Re | ATT | nope | >1bp-ins | A/T-1 | none | A/T-1 | AACGGATTTT | TTTTCTG | TAA | ATT | T | none | nope | nope_ATT | A/T-1_TTTTCTG | 7 | FALSE | FALSE |
| 15_v.txt | 32528 | 32528-Deleti | Deletion | 1 | G | - | 13 | 72336 | 0.01797169 | 36.7692308 | rnr1D57N | FALSE | CAN | JSY14 | 4 | rnr1D57N-Re | TGG | nope | >1bp-ins | G/C-1 | none | G/C-1 | TTTTCTGGG( | GGGGTTT | ACC | ACC | C | none | C>none | C>none_ACC | G/C-1_GGGGTTT | 7 | FALSE | FALSE |
| 15_v.txt | 32583 | 32583-Deleti | Deletion | 1 | A | - | 9 | 70822 | 0.01270792 | 37.6666667 | rnr1D57N | FALSE | CAN | JSY14 | 4 | rnr1D57N-Re | GAA | nope | >1bp-ins | A/T-1 | none | A/T-1 | AATGTGAAG | AAGGCAG | CTT | CTT | T | none | T>none | T>none_CTT | A/T-1_AAGGCAG | 7 | FALSE | FALSE |
| 15_v.txt | 32598 | 32598-Deleti | Deletion | 1 | A | - | 12 | 70822 | 0.01694389 | 39.4166667 | rnr1D57N | FALSE | CAN | JSY14 | 4 | rnr1D57N-Re | CAA | nope | >1bp-ins | A/T-1 | none | A/T-1 | TTAATCAAA( | AAAGAGG | GTT | GTT | T | none | T>none | T>none_GTT | A/T-1_AAAGAGG | 7 | FALSE | FALSE |
| 15_v.txt | 32624 | 32624-Deleti | Deletion | 1 | C | - | 18 | 70822 | 0.02541583 | 49.0555556 | rnr1D57N | FALSE | CAN | JSY14 | 4 | rnr1D57N-Re | TCC | nope | >1bp-ins | G/C-1 | none | G/C-1 | GAACCTCCCT | CCCTTCG | AGG | TCC | C | none | nope | nope_TCC | G/C-1_CCCTTCG | 7 | FALSE | FALSE |
| 15_v.txt | 32631 | 32631-Deleti | Deletion | 1 | T | - | 20 | 70822 | 0.02823981 | 46.55 | rnr1D57N | FALSE | CAN | JSY14 | 4 | rnr1D57N-Re | GTT | nope | >1bp-ins | A/T-1 | none | A/T-1 | CCTTCGTTTT | TTTTTAT | CAA | GTT | T | none | nope | nope_GTT | A/T-1_TTTTTAT | 7 | FALSE | FALSE |
| 15_v.txt | 32658 | 32658-Deleti | Deletion | 1 | C | - | 25 | 70822 | 0.03529977 | 54.8 | rnr1D57N | FALSE | CAN | JSY14 | 4 | rnr1D57N-Re | ACC | nope | >1bp-ins | G/C-1 | none | G/C-1 | CCTGGACCC( | CCCCAGG | TGG | ACC | C | none | nope | nope_ACC | G/C-1_CCCCAGG | 7 | FALSE | FALSE |
| 15_v.txt | 32670 | 32670-Deleti | Deletion | 1 | G | - | 28 | 70822 | 0.03953574 | 58.3571429 | rnr1D57N | FALSE | CAN | JSY14 | 4 | rnr1D57N-Re | TGG | nope | >1bp-ins | G/C-1 | none | G/C-1 | GCACCTGGG | GGGTTTC | ACC | ACC | C | none | C>none | C>none_ACC | G/C-1_GGGTTTC | 7 | FALSE | FALSE |
| 15_v.txt | 32678 | 32678-Deleti | Deletion | 1 | C | - | 4 | 70821 | 5.65E-03 | 63.999998 | rnr1D57N | FALSE | CAN | JSY14 | 4 | rnr1D57N-Re | TCC | nope | >1bp-ins | G/C-1 | none | G/C-1 | GTTTCTCCAA | CCAATAA | AGG | TCC | C | none | nope | nope_TCC | G/C-1_CCAATAA | 7 | FALSE | FALSE |
| 15_v.txt | 32680 | 32680-Deleti | Deletion | 1 | A | - | 3 | 70822 | 4.24E-03 | 42 | rnr1D57N | FALSE | CAN | JSY14 | 4 | rnr1D57N-Re | CAA | nope | >1bp-ins | A/T-1 | none | A/T-1 | TTCTCCAATA | AATAACG | GTT | GTT | T | none | T>none | T>none_GTT | A/T-1_AATAACG | 7 | FALSE | FALSE |
| 15_v.txt | 32707 | 32707-Deleti | Deletion | 1 | C | - | 11 | 70822 | 0.0155319 | 61.2727273 | rnr1D57N | FALSE | CAN | JSY14 | 4 | rnr1D57N-Re | ACC | nope | >1bp-ins | G/C-1 | none | G/C-1 | CGGTAACCC( | CCCCAGC | TGG | ACC | C | none | nope | nope_ACC | G/C-1_CCCCAGC | 7 | FALSE | FALSE |
| 15_v.txt | 32728 | 32728-Deleti | Deletion | 1 | A | - | 183 | 70820 | 0.25840158 | 60.759563 | rnr1D57N | FALSE | CAN | JSY14 | 4 | rnr1D57N-Re | CAA | nope | >1bp-ins | A/T-1 | none | A/T-1 | CCATACAAA/ | AAAAACA | GTT | GTT | T | none | T>none | T>none_GTT | A/T-1_AAAAACA | 7 | FALSE | FALSE |
| 15_v.txt | 32760 | 32760-Deleti | Deletion | 1 | A | - | 10 | 70822 | 0.01411991 | 54.6 | rnr1D57N | FALSE | CAN | JSY14 | 4 | rnr1D57N-Re | TAA | nope | >1bp-ins | A/T-1 | none | A/T-1 | ATGGCTAAA. | AAAACTT | ATT | ATT | T | none | T>none | T>none_ATT | A/T-1_AAAACTT | 7 | FALSE | FALSE |
| 15_v.txt | 32779 | 32779-Deleti | Deletion | 1 | C | - | 17 | 70822 | 0.02400384 | 54.2941177 | rnr1D57N | FALSE | CAN | JSY14 | 4 | rnr1D57N-Re | ACC | nope | >1bp-ins | G/C-1 | none | G/C-1 | AAGCGACCC. | CCCAGAA | TGG | ACC | C | none | nope | nope_ACC | G/C-1_CCCAGAA | 7 | FALSE | FALSE |
| 15_v.txt | 32800 | 32800-Deleti | Deletion | 1 | A | - | 3 | 70822 | 4.24E-03 | 38 | rnr1D57N | FALSE | CAN | JSY14 | 4 | rnr1D57N-Re | TAA | nope | >1bp-ins | A/T-1 | none | A/T-1 | CACCGTAAT/ | AATATTT | ATT | ATT | T | none | T>none | T>none_ATT | A/T-1_AATATTT | 7 | FALSE | FALSE |
| 15_v.txt | 32807 | 32807-Deleti | Deletion | 1 | G | - | 45 | 70822 | 0.06353958 | 45.7777778 | rnr1D57N | FALSE | CAN | JSY14 | 4 | rnr1D57N-Re | TGA | nope | >1bp-ins | G/C-1 | none | G/C-1 | ATATTTGACA | GACAGGG | ACT | ACT | C | none | C>none | C>none_ACT | G/C-1_GACAGGG | 7 | FALSE | FALSE |
| 15_v.txt | 32842 | 32842-Deleti | Deletion | 1 | A | - | 130 | 70822 | 0.18355878 | 42.4923085 | rnr1D57N | FALSE | CAN | JSY14 | 4 | rnr1D57N-Re | CAA | nope | >1bp-ins | A/T-1 | none | A/T-1 | TTACCCAAA/ | AAAAAAT | GTT | GTT | T | none | T>none | T>none_GTT | A/T-1_AAAAAAT | 7 | FALSE | FALSE |
| 15_v.txt | 32884 | 32884-Deleti | Deletion | 1 | A | - | 11 | 75142 | 0.01463895 | 37.7272722 | rnr1D57N | FALSE | CAN | JSY14 | 4 | rnr1D57N-Re | CAA | nope | >1bp-ins | A/T-1 | none | A/T-1 | ACGTCCAAA/ | AAAATTG | GTT | GTT | T | none | T>none | T>none_GTT | A/T-1_AAAATTG | 7 | FALSE | FALSE |
| 15_v.txt | 32901 | 32901-Deleti | Deletion | 1 | C | - | 2 | 75142 | 2.66E-03 | 38 | rnr1D57N | FALSE | CAN | JSY14 | 4 | rnr1D57N-Re | GCC | nope | >1bp-ins | G/C-1 | none | G/C-1 | ACTTGGCCA/ | CCAACTA | CGG | GCC | C | none | nope | nope_GCC | G/C-1_CCAACTA | 7 | FALSE | FALSE |
| 15_v.txt | 32924 | 32924-Deleti | Deletion | 1 | A | - | 55 | 75142 | 0.07319475 | 50.5272727 | rnr1D57N | FALSE | CAN | JSY14 | 4 | rnr1D57N-Re | CAA | nope | >1bp-ins | A/T-1 | none | A/T-1 | CAGGGCAAA | AAAAGTG | GTT | GTT | T | none | T>none | T>none_GTT | A/T-1_AAAAGTG | 7 | FALSE | FALSE |
| 15_v.txt | 32935 | 32935-Deleti | Deletion | 1 | C | - | 6 | 75142 | 7.99E-03 | 54 | rnr1D57N | FALSE | CAN | JSY14 | 4 | rnr1D57N-Re | GCC | nope | >1bp-ins | G/C-1 | none | G/C-1 | TGATTGCCCA | CCCAAGA | CGG | GCC | C | none | nope | nope_GCC | G/C-1_CCCAAGA | 7 | FALSE | FALSE |
| 15_v.txt | 33045 | 33045-Deleti | Deletion | 1 | C | - | 10 | 75142 | 0.01330814 | 64 | rnr1D57N | FALSE | CAN | JSY14 | 4 | rnr1D57N-Re | ACC | nope | >1bp-ins | G/C-1 | none | G/C-1 | ATTTCACCCA | CCCAAGG | TGG | ACC | C | none | nope | nope_ACC | G/C-1_CCCAAGG | 7 | FALSE | FALSE |
| 15_v.txt | 33083 | 33083-Deleti | Deletion | 1 | A | - | 11 | 75142 | 0.01463895 | 50.8181818 | rnr1D57N | FALSE | CAN | JSY14 | 4 | rnr1D57N-Re | TAA | nope | >1bp-ins | A/T-1 | none | A/T-1 | ACCCATAAAT | AAATAAA | ATT | ATT | T | none | T>none | T>none_ATT | A/T-1_AAATAAA | 7 | FALSE | FALSE |
| 15_v.txt | 33087 | 33087-Deleti | Deletion | 1 | A | - | 4 | 75142 | 5.32E-03 | 63.75 | rnr1D57N | FALSE | CAN | JSY14 | 4 | rnr1D57N-Re | TAA | nope | >1bp-ins | A/T-1 | none | A/T-1 | ATAAATAAAT | AAATATG | ATT | ATT | T | none | T>none | T>none_ATT | A/T-1_AAATATG | 7 | FALSE | FALSE |
| 15_v.txt | 33111 | 33111-Deleti | Deletion | 1 | G | - | 37 | 75142 | 0.04924011 | 41.027027 | rnr1D57N | FALSE | CAN | JSY14 | 4 | rnr1D57N-Re | TGG | nope | >1bp-ins | G/C-1 | none | G/C-1 | CCCACTGGG( | GGGCCGG | ACC | ACC | C | none | C>none | C>none_ACC | G/C-1_GGGCCGG | 7 | FALSE | FALSE |
| 15_v.txt | 33138 | 33138-Deleti | Deletion | 1 | A | - | 9 | 75141 | 0.01197748 | 43.7777778 | rnr1D57N | FALSE | CAN | JSY14 | 4 | rnr1D57N-Re | TAA | nope | >1bp-ins | A/T-1 | none | A/T-1 | GTGGATAAA | AAACCAA | ATT | ATT | T | none | T>none | T>none_ATT | A/T-1_AAACCAA | 7 | FALSE | FALSE |
| 15_v.txt | 33147 | 33147-Deleti | Deletion | 1 | A | - | 14 | 75141 | 0.01863164 | 39.6428571 | rnr1D57N | FALSE | CAN | JSY14 | 4 | rnr1D57N-Re | GAA | nope | >1bp-ins | A/T-1 | none | A/T-1 | CCAATGAAA. | AAAAGAC | CTT | CTT | T | none | T>none | T>none_CTT | A/T-1_AAAAGAC | 7 | FALSE | FALSE |
| 15_v.txt | 33168 | 33168-Deleti | Deletion | 1 | C | - | 5 | 75126 | 6.66E-03 | 38 | rnr1D57N | FALSE | CAN | JSY14 | 4 | rnr1D57N-Re | ACC | nope | >1bp-ins | G/C-1 | none | G/C-1 | ATAGTACCA( | CCACCAA | TGG | ACC | C | none | nope | nope_ACC | G/C-1_CCACCAA | 7 | FALSE | FALSE |
| 15_v.txt | 33303 | 33303-Deleti | Deletion | 1 | A | - | 10 | 61063 | 0.01637653 | 51.3 | rnr1D57N | FALSE | CAN | JSY14 | 4 | rnr1D57N-Re | GAA | nope | >1bp-ins | A/T-1 | none | A/T-1 | GTCGGGAAA | AAAGAGC | CTT | CTT | T | none | T>none | T>none_CTT | A/T-1_AAAGAGC | 7 | FALSE | FALSE |
| 15_v.txt | 33317 | 33317-Deleti | Deletion | 1 | A | - | 8 | 61063 | 0.01310122 | 59.375 | rnr1D57N | FALSE | CAN | JSY14 | 4 | rnr1D57N-Re | GAT | nope | >1bp-ins | A/T-1 | none | A/T-1 | CAATGGATA( | ATACAAT | CTA | CTA | T | none | T>none | T>none_CTA | A/T-1_ATACAAT | 7 | FALSE | FALSE |
| 15_v.txt | 33326 | 33326-Deleti | Deletion | 1 | T | - | 8 | 61060 | 0.01310187 | 58.625 | rnr1D57N | FALSE | CAN | JSY14 | 4 | rnr1D57N-Re | CTT | nope | >1bp-ins | A/T-1 | none | A/T-1 | CAATTCTTTA | TTTACTT | GAA | CTT | T | none | nope | nope_CTT | A/T-1_TTTACTT | 7 | FALSE | FALSE |
| 15_v.txt | 33356 | 33356-Deleti | Deletion | 1 | C | - | 12 | 61063 | 0.01965184 | 51.8333341 | rnr1D57N | FALSE | CAN | JSY14 | 4 | rnr1D57N-Re | ACC | nope | >1bp-ins | G/C-1 | none | G/C-1 | TATTGACCCA | CCCACGT | TGG | ACC | C | none | nope | nope_ACC | G/C-1_CCCACGT | 7 | FALSE | FALSE |
| 15_v.txt | 33474 | 33474-Deleti | Deletion | 1 | C | - | 148 | 61063 | 0.24237263 | 29.0202703 | rnr1D57N | FALSE | CAN | JSY14 | 4 | rnr1D57N-Re | GCC | nope | >1bp-ins | G/C-1 | none | G/C-1 | GCTATGCCTT | CCTTTTT | CGG | GCC | C | none | nope | nope_GCC | G/C-1_CCTTTTT | 7 | FALSE | FALSE |
| 186_v.txt | 32081 | 32081-SNV-/ | SNV | 1 | A | C | 2 | 12563 | 0.01591976 | 37.5002717 | rnr1D57N | FALSE | CAN | JSY14 | 5 | rnr1D57N-Re | CAA | A>C | G/C+1 | A/T-1 | TA>GC | TA>GC | CCATGCAAA/ | AAAAAAG | GTT | GTT | T | G | T>G | T>G_GTT | TA>GC_AAAAAAG | 7 | FALSE | FALSE |
| 186_v.txt | 32256 | 32256-SNV-/ | SNV | 1 | A | C | 19 | 27637 | 0.06874842 | 24.4736842 | rnr1D57N | FALSE | CAN | JSY14 | 5 | rnr1D57N-Re | TAG | A>C | G/C+1 | A/T-1 | TA>GC | TA>GC | TTTGATAGA( | AGACCAA | ATC | ATC | T | G | T>G | T>G_ATC | TA>GC_AGACCAA | 7 | FALSE | FALSE |
| 186_v.txt | 32265 | 32265-SNV-/ | SNV | 1 | A | C | 25 | 27652 | 0.09040937 | 26.44 | rnr1D57N | FALSE | CAN | JSY14 | 5 | rnr1D57N-Re | TAA | A>C | G/C+1 | A/T-1 | TA>GC | TA>GC | CCAAATAAA. | AAAATAC | ATT | ATT | T | G | T>G | T>G_ATT | TA>GC_AAAATAC | 7 | FALSE | FALSE |
| 186_v.txt | 32283 | 32283-SNV-T | SNV | 1 | T | G | 832 | 27652 | 3.00882396 | 51.3689904 | rnr1D57N | FALSE | CAN | JSY14 | 5 | rnr1D57N-Re | GTA | T>G | G/C+1 | A/T-1 | TA>GC | TA>GC | CCAACGTAA/ | TAAATAT | CAT | GTA | T | C | T>G | T>G_GTA | TA>GC_TAAATAT | 7 | FALSE | FALSE |
| 186_v.txt | 32439 | 32439-SNV-/ | SNV | 1 | A | C | 8 | 27653 | 0.02892995 | 60.75 | rnr1D57N | FALSE | CAN | JSY14 | 5 | rnr1D57N-Re | AAC | A>C | G/C+1 | A/T-1 | TA>GC | TA>GC | TATGGAACT/ | ACTAAAA | TTG | TTG | T | G | T>G | T>G_TTG | TA>GC_ACTAAAA | 7 | FALSE | FALSE |
| 186_v.txt | 32640 | 32640-SNV-T | SNV | 1 | T | G | 4 | 17176 | 0.02328831 | 57.5 | rnr1D57N | FALSE | CAN | JSY14 | 5 | rnr1D57N-Re | CTT | T>G | G/C+1 | A/T-1 | TA>GC | TA>GC | TTATCCTTAG | TTAGATA | GAA | CTT | T | C | T>G | T>G_CTT | TA>GC_TTAGATA | 7 | FALSE | FALSE |
| 186_v.txt | 32842 | 32842-SNV-/ | SNV | 1 | A | C | 2 | 17176 | 0.01164416 | 36.0000349 | rnr1D57N | FALSE | CAN | JSY14 | 5 | rnr1D57N-Re | CAA | A>C | G/C+1 | A/T-1 | TA>GC | TA>GC | TTACCCAAA/ | AAAAAAT | GTT | GTT | T | G | T>G | T>G_GTT | TA>GC_AAAAAAT | 7 | FALSE | FALSE |
| 186_v.txt | 32884 | 32884-SNV-/ | SNV | 1 | A | C | 10 | 29334 | 0.03409013 | 28.2 | rnr1D57N | FALSE | CAN | JSY14 | 5 | rnr1D57N-Re | CAA | A>C | G/C+1 | A/T-1 | TA>GC | TA>GC | ACGTCCAAA/ | AAAATTG | GTT | GTT | T | G | T>G | T>G_GTT | TA>GC_AAAATTG | 7 | FALSE | FALSE |
| 186_v.txt | 32938 | 32938-SNV-/ | SNV | 1 | A | C | 6 | 29334 | 0.02045408 | 31.0008466 | rnr1D57N | FALSE | CAN | JSY14 | 5 | rnr1D57N-Re | CAA | A>C | G/C+1 | A/T-1 | TA>GC | TA>GC | TTGCCCAAG/ | AAGAAAA | GTT | GTT | T | G | T>G | T>G_GTT | TA>GC_AAGAAAA | 7 | FALSE | FALSE |
| 186_v.txt | 33269 | 33269-SNV-/ | SNV | 1 | A | C | 167 | 21090 | 0.79184448 | 16.1736527 | rnr1D57N | FALSE | CAN | JSY14 | 5 | rnr1D57N-Re | TAG | A>C | G/C+1 | A/T-1 | TA>GC | TA>GC | CTCCATAGA( | AGAGAAC | ATC | ATC | T | G | T>G | T>G_ATC | TA>GC_AGAGAAC | 7 | FALSE | FALSE |
| 186_v.txt | 32218 | 32218-SNV-T | SNV | 1 | T | C | 3 | 12564 | 0.02387775 | 32.6656027 | rnr1D57N | FALSE | CAN | JSY14 | 5 | rnr1D57N-Re | GTC | T>C | G/C+1 | A/T-1 | TA>CG | TA>CG | TGGTGGTCCT | TCCTTGA | CAG | GTC | T | G | T>C | T>C_GTC | TA>CG_TCCTTGA | 7 | FALSE | FALSE |
| 186_v.txt | 32256 | 32256-SNV-/ | SNV | 1 | A | G | 16 | 27637 | 0.0578934 | 36.8125 | rnr1D57N | FALSE | CAN | JSY14 | 5 | rnr1D57N-Re | TAG | A>G | G/C+1 | A/T-1 | TA>CG | TA>CG | TTTGATAGA( | AGACCAA | ATC | ATC | T | C | T>C | T>C_ATC | TA>CG_AGACCAA | 7 | FALSE | FALSE |
| 186_v.txt | 32305 | 32305-SNV-/ | SNV | 1 | A | G | 32 | 27653 | 0.11571981 | 59.46875 | rnr1D57N | FALSE | CAN | JSY14 | 5 | rnr1D57N-Re | GAA | A>G | G/C+1 | A/T-1 | TA>CG | TA>CG | CGGCAGAAA | AAATAAT | CTT | CTT | T | C | T>C | T>C_CTT | TA>CG_AAATAAT | 7 | FALSE | FALSE |
| 186_v.txt | 32454 | 32454-SNV-/ | SNV | 1 | A | G | 5 | 27653 | 0.01808122 | 58.7985389 | rnr1D57N | FALSE | CAN | JSY14 | 5 | rnr1D57N-Re | GAA | A>G | G/C+1 | A/T-1 | TA>CG | TA>CG | CCAATGAAT/ | AATAATA | CTT | CTT | T | C | T>C | T>C_CTT | TA>CG_AATAATA | 7 | FALSE | FALSE |
| 186_v.txt | 32455 | 32455-SNV-/ | SNV | 1 | A | G | 4 | 27653 | 0.01446498 | 41.7499109 | rnr1D57N | FALSE | CAN | JSY14 | 5 | rnr1D57N-Re | AAT | A>G | G/C+1 | A/T-1 | TA>CG | TA>CG | CAATGAATA, | ATAATAG | TTA | TTA | T | C | T>C | T>C_TTA | TA>CG_ATAATAG | 7 | FALSE | FALSE |
| 186_v.txt | 32503 | 32503-SNV-T | SNV | 1 | T | C | 23 | 27653 | 0.08317362 | 41.2608696 | rnr1D57N | FALSE | CAN | JSY14 | 5 | rnr1D57N-Re | TTG | T>C | G/C+1 | A/T-1 | TA>CG | TA>CG | CTTTTTTGAT( | TGATGGC | AAC | TTG | T | G | T>C | T>C_TTG | TA>CG_TGATGGC | 7 | FALSE | FALSE |
| 186_v.txt | 32583 | 32583-SNV-/ | SNV | 1 | A | G | 44 | 17176 | 0.2561714 | 36.75 | rnr1D57N | FALSE | CAN | JSY14 | 5 | rnr1D57N-Re | GAA | A>G | G/C+1 | A/T-1 | TA>CG | TA>CG | AATGTGAAG | AAGGCAG | CTT | CTT | T | C | T>C | T>C_CTT | TA>CG_AAGGCAG | 7 | FALSE | FALSE |
| 186_v.txt | 32680 | 32680-SNV-/ | SNV | 1 | A | G | 17 | 17176 | 0.09897531 | 58.8823529 | rnr1D57N | FALSE | CAN | JSY14 | 5 | rnr1D57N-Re | CAA | A>G | G/C+1 | A/T-1 | TA>CG | TA>CG | TTCTCCAATA | AATAACG | GTT | GTT | T | C | T>C | T>C_GTT | TA>CG_AATAACG | 7 | FALSE | FALSE |
| 186_v.txt | 32919 | 32919-SNV-/ | SNV | 1 | A | G | 23 | 29334 | 0.07840731 | 46.3043478 | rnr1D57N | FALSE | CAN | JSY14 | 5 | rnr1D57N-Re | CAG | A>G | G/C+1 | A/T-1 | TA>CG | TA>CG | AGTTCCAGG( | AGGGCAA | GTC | GTC | T | C | T>C | T>C_GTC | TA>CG_AGGGCAA | 7 | FALSE | FALSE |
| 186_v.txt | 32938 | 32938-SNV-/ | SNV | 1 | A | G | 25 | 29334 | 0.08522534 | 55.280019 | rnr1D57N | FALSE | CAN | JSY14 | 5 | rnr1D57N-Re | CAA | A>G | G/C+1 | A/T-1 | TA>CG | TA>CG | TTGCCCAAG/ | AAGAAAA | GTT | GTT | T | C | T>C | T>C_GTT | TA>CG_AAGAAAA | 7 | FALSE | FALSE |
| 186_v.txt | 31816 | 31816-SNV-T | SNV | 1 | T | A | 7 | 3690 | 0.1897019 | 56.4285714 | rnr1D57N | FALSE | CAN | JSY14 | 5 | rnr1D57N-Re | CTG | T>A | A/T+1 | A/T-1 | TA>AT | TA>AT | TAAATCTGC/ | TGCATCT | GAC | CTG | T | T | T>A | T>A_CTG | TA>AT_TGCATCT | 7 | FALSE | FALSE |
| 186_v.txt | 32215 | 32215-SNV-T | SNV | 1 | T | A | 1 | 12564 | 7.96E-03 | 37.9976572 | rnr1D57N | FALSE | CAN | JSY14 | 5 | rnr1D57N-Re | GTG | T>A | A/T+1 | A/T-1 | TA>AT | TA>AT | CTTTGGTGGT | TGGTCCT | CAC | GTG | T | T | T>A | T>A_GTG | TA>AT_TGGTCCT | 7 | FALSE | FALSE |
| 186_v.txt | 32288 | 32288-SNV-/ | SNV | 1 | A | T | 16 | 27652 | 0.057862 | 46.4375 | rnr1D57N | FALSE | CAN | JSY14 | 5 | rnr1D57N-Re | TAT | A>T | A/T+1 | A/T-1 | TA>AT | TA>AT | GTAAATATTT | ATTTGAA | ATA | ATA | T | A | T>A | T>A_ATA | TA>AT_ATTTGAA | 7 | FALSE | FALSE |
| 186_v.txt | 31962 | 31962-SNV-( | SNV | 1 | G | A | 23 | 12564 | 0.18306272 | 37.826087 | rnr1D57N | FALSE | CAN | JSY14 | 5 | rnr1D57N-Re | CGC | G>A | A/T+1 | G/C-1 | CG>TA | CG>TA | GTGGCCGCAT | GCATAAT | GCG | GCG | C | T | C>T | C>T_GCG | CG>TA_GCATAAT | 7 | FALSE | FALSE |
| 186_v.txt | 32076 | 32076-SNV-( | SNV | 1 | C | T | 58 | 12564 | 0.46163642 | 62.1551724 | rnr1D57N | FALSE | CAN | JSY14 | 5 | rnr1D57N-Re | CCA | C>T | A/T+1 | G/C-1 | CG>TA | CG>TA | AATAACCATC | CATGCAA | GGT | CCA | C | A | C>T | C>T_CCA | CG>TA_CATGCAA | 7 | FALSE | FALSE |
| 186_v.txt | 32165 | 32165-SNV-( | SNV | 1 | G | A | 19 | 12564 | 0.15122572 | 53.9473684 | rnr1D57N | FALSE | CAN | JSY14 | 5 | rnr1D57N-Re | CGC | G>A | A/T+1 | G/C-1 | CG>TA | CG>TA | CAAAGCGCC. | GCCAAAT | GCG | GCG | C | T | C>T | C>T_GCG | CG>TA_GCCAAAT | 7 | FALSE | FALSE |
| 186_v.txt | 32190 | 32190-SNV-( | SNV | 1 | G | A | 14 | 12564 | 0.11142948 | 48.1428571 | rnr1D57N | FALSE | CAN | JSY14 | 5 | rnr1D57N-Re | TGC | G>A | A/T+1 | G/C-1 | CG>TA | CG>TA | AAAACTGCA. | GCAATGT | ACG | ACG | C | T | C>T | C>T_ACG | CG>TA_GCAATGT | 7 | FALSE | FALSE |
| 186_v.txt | 32271 | 32271-SNV-( | SNV | 1 | C | T | 35 | 27652 | 0.12657312 | 32.5142857 | rnr1D57N | FALSE | CAN | JSY14 | 5 | rnr1D57N-Re | ACG | C>T | A/T+1 | G/C-1 | CG>TA | CG>TA | AAAATACGG | CGGGAAC | TGC | ACG | C | A | C>T | C>T_ACG | CG>TA_CGGGAAC | 7 | FALSE | FALSE |
| 186_v.txt | 32299 | 32299-SNV-( | SNV | 1 | C | T | 37 | 27653 | 0.13380103 | 50.5135135 | rnr1D57N | FALSE | CAN | JSY14 | 5 | rnr1D57N-Re | GCG | C>T | A/T+1 | G/C-1 | CG>TA | CG>TA | AATTTGCGG( | CGGCAGA | CGC | GCG | C | A | C>T | C>T_GCG | CG>TA_CGGCAGA | 7 | FALSE | FALSE |
| 186_v.txt | 32313 | 32313-SNV-( | SNV | 1 | G | A | 71 | 27653 | 0.25675334 | 59.0985916 | rnr1D57N | FALSE | CAN | JSY14 | 5 | rnr1D57N-Re | GGT | G>A | A/T+1 | G/C-1 | CG>TA | CG>TA | ATAATGGTT( | GTTGTTA | CCA | CCA | C | T | C>T | C>T_CCA | CG>TA_GTTGTTA | 7 | FALSE | FALSE |
| 186_v.txt | 32328 | 32328-SNV-( | SNV | 1 | G | A | 1 | 27653 | 3.62E-03 | 63.9998595 | rnr1D57N | FALSE | CAN | JSY14 | 5 | rnr1D57N-Re | AGC | G>A | A/T+1 | G/C-1 | CG>TA | CG>TA | ATAACAGCGT | GCGTTGA | TCG | TCG | C | T | C>T | C>T_TCG | CG>TA_GCGTTGA | 7 | FALSE | FALSE |
| 186_v.txt | 32449 | 32449-SNV-( | SNV | 1 | C | T | 31 | 27653 | 0.11210357 | 46.0645161 | rnr1D57N | FALSE | CAN | JSY14 | 5 | rnr1D57N-Re | CCA | C>T | A/T+1 | G/C-1 | CG>TA | CG>TA | AAAGTCCAAT | CAATGAA | GGT | CCA | C | A | C>T | C>T_CCA | CG>TA_CAATGAA | 7 | FALSE | FALSE |
| 186_v.txt | 32466 | 32466-SNV-( | SNV | 1 | C | T | 34 | 27653 | 0.1229523 | 45.1764706 | rnr1D57N | FALSE | CAN | JSY14 | 5 | rnr1D57N-Re | GCC | C>T | A/T+1 | G/C-1 | CG>TA | CG>TA | AGAGAGCCA | CCAATGT | CGG | GCC | C | A | C>T | C>T_GCC | CG>TA_CCAATGT | 7 | FALSE | FALSE |
| 186_v.txt | 32535 | 32535-SNV-( | SNV | 1 | G | A | 30 | 27652 | 0.10849125 | 35.2667957 | rnr1D57N | FALSE | CAN | JSY14 | 5 | rnr1D57N-Re | TGC | G>A | A/T+1 | G/C-1 | CG>TA | CG>TA | GGGTTTGCA( | GCAGCTT | ACG | ACG | C | T | C>T | C>T_ACG | CG>TA_GCAGCTT | 7 | FALSE | FALSE |
| 186_v.txt | 32536 | 32536-SNV-( | SNV | 1 | C | T | 42 | 27652 | 0.15188775 | 36.3566836 | rnr1D57N | FALSE | CAN | JSY14 | 5 | rnr1D57N-Re | GCA | C>T | A/T+1 | G/C-1 | CG>TA | CG>TA | GGTTTGCAG( | CAGCTTC | CGT | GCA | C | A | C>T | C>T_GCA | CG>TA_CAGCTTC | 7 | FALSE | FALSE |
| 186_v.txt | 32545 | 32545-SNV-( | SNV | 1 | C | T | 26 | 27651 | 0.09402915 | 37.1923077 | rnr1D57N | FALSE | CAN | JSY14 | 5 | rnr1D57N-Re | CCA | C>T | A/T+1 | G/C-1 | CG>TA | CG>TA | CTTCACCAGC | CAGCAGT | GGT | CCA | C | A | C>T | C>T_CCA | CG>TA_CAGCAGT | 7 | FALSE | FALSE |
| 186_v.txt | 32590 | 32590-SNV-( | SNV | 1 | C | T | 22 | 17176 | 0.1280857 | 35.8636364 | rnr1D57N | FALSE | CAN | JSY14 | 5 | rnr1D57N-Re | GCG | C>T | A/T+1 | G/C-1 | CG>TA | CG>TA | AGGCAGCGT | CGTTAAT | CGC | GCG | C | A | C>T | C>T_GCG | CG>TA_CGTTAAT | 7 | FALSE | FALSE |
| 186_v.txt | 32609 | 32609-SNV-( | SNV | 1 | C | T | 29 | 17176 | 0.16884024 | 48.7897357 | rnr1D57N | FALSE | CAN | JSY14 | 5 | rnr1D57N-Re | CCC | C>T | A/T+1 | G/C-1 | CG>TA | CG>TA | GGAAACCCA | CCAACCT | GGG | CCC | C | A | C>T | C>T_CCC | CG>TA_CCAACCT | 7 | FALSE | FALSE |
| 186_v.txt | 32610 | 32610-SNV-( | SNV | 1 | C | T | 26 | 17176 | 0.15137401 | 52.4167147 | rnr1D57N | FALSE | CAN | JSY14 | 5 | rnr1D57N-Re | CCA | C>T | A/T+1 | G/C-1 | CG>TA | CG>TA | GAAACCCAA( | CAACCTA | GGT | CCA | C | A | C>T | C>T_CCA | CG>TA_CAACCTA | 7 | FALSE | FALSE |
| 186_v.txt | 32661 | 32661-SNV-( | SNV | 1 | C | T | 516 | 17176 | 3.0041919 | 59.003876 | rnr1D57N | FALSE | CAN | JSY14 | 5 | rnr1D57N-Re | CCA | C>T | A/T+1 | G/C-1 | CG>TA | CG>TA | GGACCCCAG( | CAGGCAC | GGT | CCA | C | A | C>T | C>T_CCA | CG>TA_CAGGCAC | 7 | FALSE | FALSE |
| 186_v.txt | 32776 | 32776-SNV-( | SNV | 1 | C | T | 22 | 17176 | 0.1280857 | 48.1363636 | rnr1D57N | FALSE | CAN | JSY14 | 5 | rnr1D57N-Re | GCG | C>T | A/T+1 | G/C-1 | CG>TA | CG>TA | TGGAAGCGA | CGACCCA | CGC | GCG | C | A | C>T | C>T_GCG | CG>TA_CGACCCA | 7 | FALSE | FALSE |
| 186_v.txt | 32798 | 32798-SNV-( | SNV | 1 | G | A | 26 | 17176 | 0.15137401 | 43.6923077 | rnr1D57N | FALSE | CAN | JSY14 | 5 | rnr1D57N-Re | CGT | G>A | A/T+1 | G/C-1 | CG>TA | CG>TA | TTCACCGTA/ | GTAATAT | GCA | GCA | C | T | C>T | C>T_GCA | CG>TA_GTAATAT | 7 | FALSE | FALSE |
| 186_v.txt | 32900 | 32900-SNV-( | SNV | 1 | G | A | 30 | 29334 | 0.1022704 | 35.7982556 | rnr1D57N | FALSE | CAN | JSY14 | 5 | rnr1D57N-Re | GGC | G>A | A/T+1 | G/C-1 | CG>TA | CG>TA | GACTTGGCC/ | GCCAACT | CCG | CCG | C | T | C>T | C>T_CCG | CG>TA_GCCAACT | 7 | FALSE | FALSE |
| 186_v.txt | 32901 | 32901-SNV-( | SNV | 1 | C | T | 48 | 29334 | 0.16363265 | 32.12533 | rnr1D57N | FALSE | CAN | JSY14 | 5 | rnr1D57N-Re | GCC | C>T | A/T+1 | G/C-1 | CG>TA | CG>TA | ACTTGGCCA/ | CCAACTA | CGG | GCC | C | A | C>T | C>T_GCC | CG>TA_CCAACTA | 7 | FALSE | FALSE |
| 186_v.txt | 32917 | 32917-SNV-( | SNV | 1 | C | T | 54 | 29334 | 0.18408673 | 44.6481482 | rnr1D57N | FALSE | CAN | JSY14 | 5 | rnr1D57N-Re | TCC | C>T | A/T+1 | G/C-1 | CG>TA | CG>TA | TAAGTTCCA( | CCAGGGC | AGG | TCC | C | A | C>T | C>T_TCC | CG>TA_CCAGGGC | 7 | FALSE | FALSE |
| 186_v.txt | 32935 | 32935-SNV-( | SNV | 1 | C | T | 34 | 29334 | 0.11590646 | 31.4117647 | rnr1D57N | FALSE | CAN | JSY14 | 5 | rnr1D57N-Re | GCC | C>T | A/T+1 | G/C-1 | CG>TA | CG>TA | TGATTGCCC/ | CCCAAGA | CGG | GCC | C | A | C>T | C>T_GCC | CG>TA_CCCAAGA | 7 | FALSE | FALSE |
| 186_v.txt | 32945 | 32945-SNV-( | SNV | 1 | C | T | 42 | 29334 | 0.14317856 | 52.7591906 | rnr1D57N | FALSE | CAN | JSY14 | 5 | rnr1D57N-Re | ACC | C>T | A/T+1 | G/C-1 | CG>TA | CG>TA | AGAAAACCA | CCAATAC | TGG | ACC | C | A | C>T | C>T_ACC | CG>TA_CCAATAC | 7 | FALSE | FALSE |
| 186_v.txt | 32946 | 32946-SNV-( | SNV | 1 | C | T | 62 | 29334 | 0.21135883 | 57.0631519 | rnr1D57N | FALSE | CAN | JSY14 | 5 | rnr1D57N-Re | CCA | C>T | A/T+1 | G/C-1 | CG>TA | CG>TA | GAAAACCAA | CAATACA | GGT | CCA | C | A | C>T | C>T_CCA | CG>TA_CAATACA | 7 | FALSE | FALSE |
| 186_v.txt | 32967 | 32967-SNV-( | SNV | 1 | G | A | 65 | 29334 | 0.22158587 | 57.4461539 | rnr1D57N | FALSE | CAN | JSY14 | 5 | rnr1D57N-Re | CGC | G>A | A/T+1 | G/C-1 | CG>TA | CG>TA | TTGGCCGCA( | GCACCAA | GCG | GCG | C | T | C>T | C>T_GCG | CG>TA_GCACCAA | 7 | FALSE | FALSE |
| 186_v.txt | 32995 | 32995-SNV-( | SNV | 1 | G | A | 24 | 29334 | 0.08181632 | 62.9166667 | rnr1D57N | FALSE | CAN | JSY14 | 5 | rnr1D57N-Re | TGT | G>A | A/T+1 | G/C-1 | CG>TA | CG>TA | ATCTTTGTGA | GTGAGAA | ACA | ACA | C | T | C>T | C>T_ACA | CG>TA_GTGAGAA | 7 | FALSE | FALSE |
| 186_v.txt | 33056 | 33056-SNV-( | SNV | 1 | C | T | 28 | 29334 | 0.09545238 | 55.7857143 | rnr1D57N | FALSE | CAN | JSY14 | 5 | rnr1D57N-Re | GCG | C>T | A/T+1 | G/C-1 | CG>TA | CG>TA | GGACTGCGT( | CGTGACA | CGC | GCG | C | A | C>T | C>T_GCG | CG>TA_CGTGACA | 7 | FALSE | FALSE |
| 186_v.txt | 33069 | 33069-SNV-( | SNV | 1 | G | A | 24 | 29334 | 0.08181632 | 51.0001414 | rnr1D57N | FALSE | CAN | JSY14 | 5 | rnr1D57N-Re | TGC | G>A | A/T+1 | G/C-1 | CG>TA | CG>TA | GAATATGCC/ | GCCAAAG | ACG | ACG | C | T | C>T | C>T_ACG | CG>TA_GCCAAAG | 7 | FALSE | FALSE |
| 186_v.txt | 33070 | 33070-SNV-( | SNV | 1 | C | T | 35 | 29333 | 0.11931954 | 53.3136986 | rnr1D57N | FALSE | CAN | JSY14 | 5 | rnr1D57N-Re | GCC | C>T | A/T+1 | G/C-1 | CG>TA | CG>TA | AATATGCCA/ | CCAAAGA | CGG | GCC | C | A | C>T | C>T_GCC | CG>TA_CCAAAGA | 7 | FALSE | FALSE |
| 186_v.txt | 33114 | 33114-SNV-( | SNV | 1 | C | T | 26 | 29334 | 0.08863435 | 42.3449368 | rnr1D57N | FALSE | CAN | JSY14 | 5 | rnr1D57N-Re | GCC | C>T | A/T+1 | G/C-1 | CG>TA | CG>TA | ACTGGGCCG( | CCGGCGT | CGG | GCC | C | A | C>T | C>T_GCC | CG>TA_CCGGCGT | 7 | FALSE | FALSE |
| 186_v.txt | 33115 | 33115-SNV-( | SNV | 1 | C | T | 11 | 29334 | 0.03749915 | 37.1818306 | rnr1D57N | FALSE | CAN | JSY14 | 5 | rnr1D57N-Re | CCG | C>T | A/T+1 | G/C-1 | CG>TA | CG>TA | CTGGGCCGG | CGGCGTT | GGC | CCG | C | A | C>T | C>T_CCG | CG>TA_CGGCGTT | 7 | FALSE | FALSE |
| 186_v.txt | 33118 | 33118-SNV-( | SNV | 1 | C | T | 30 | 29334 | 0.1022704 | 41.5666667 | rnr1D57N | FALSE | CAN | JSY14 | 5 | rnr1D57N-Re | GCG | C>T | A/T+1 | G/C-1 | CG>TA | CG>TA | GGCCGGCGT | CGTTGGT | CGC | GCG | C | A | C>T | C>T_GCG | CG>TA_CGTTGGT | 7 | FALSE | FALSE |
| 186_v.txt | 33154 | 33154-SNV-( | SNV | 1 | C | T | 373 | 29334 | 1.27156201 | 37.2600536 | rnr1D57N | FALSE | CAN | JSY14 | 5 | rnr1D57N-Re | CCT | C>T | A/T+1 | G/C-1 | CG>TA | CG>TA | AAAGACCTG( | CTGTACC | GGA | CCT | C | A | C>T | C>T_CCT | CG>TA_CTGTACC | 7 | FALSE | FALSE |
| 186_v.txt | 33276 | 33276-SNV-( | SNV | 1 | G | A | 49 | 21090 | 0.2323376 | 46.0816327 | rnr1D57N | FALSE | CAN | JSY14 | 5 | rnr1D57N-Re | CGT | G>A | A/T+1 | G/C-1 | CG>TA | CG>TA | GAGAACGTA | GTATCCT | GCA | GCA | C | T | C>T | C>T_GCA | CG>TA_GTATCCT | 7 | FALSE | FALSE |
| 186_v.txt | 33284 | 33284-SNV-( | SNV | 1 | G | A | 25 | 21090 | 0.11853959 | 43.84 | rnr1D57N | FALSE | CAN | JSY14 | 5 | rnr1D57N-Re | CGC | G>A | A/T+1 | G/C-1 | CG>TA | CG>TA | ATCCTCGCC/ | GCCATTT | GCG | GCG | C | T | C>T | C>T_GCG | CG>TA_GCCATTT | 7 | FALSE | FALSE |

| | | | | | | | | | | | | | | | | | | | | | | | | | | | | | | | | | | |
|---|---|---|---|---|---|---|---|---|---|---|---|---|---|---|---|---|---|---|---|---|---|---|---|---|---|---|---|---|---|---|---|---|---|---|
| 186_v.txt | 33310 | 33310-SNV- | SNV | 1 | G | A | 33 | 21090 | 0.15647226 | 52.7272727 | rnr1D57N | FALSE | CAN | JSY14 | 5 | rnr1D57N-Re | CGC | G>A | A/T+1 | G/C-1 | CG>TA | CG>TA | AAGAGCGCA | GCAATGG | GCG | GCG | C | T | C>T | C>T_GCG | CG>TA_GCAATGG | 7 | FALSE | FALSE |
| 186_v.txt | 33376 | 33376-SNV- | SNV | 1 | G | A | 41 | 21090 | 0.19440493 | 51.1707317 | rnr1D57N | FALSE | CAN | JSY14 | 5 | rnr1D57N-Re | TGT | G>A | A/T+1 | G/C-1 | CG>TA | CG>TA | GTGTTTGTG | GTGAAGC | ACA | ACA | C | T | C>T | C>T_ACA | CG>TA_GTGAAGC | 7 | FALSE | FALSE |
| 186_v.txt | 33412 | 33412-SNV- | SNV | 1 | G | A | 22 | 21090 | 0.10431484 | 51.0909091 | rnr1D57N | FALSE | CAN | JSY14 | 5 | rnr1D57N-Re | GGC | G>A | A/T+1 | G/C-1 | CG>TA | CG>TA | TGACCGGCT | GCTCATT | CCG | CCG | C | T | C>T | C>T_CCG | CG>TA_GCTCATT | 7 | FALSE | FALSE |
| 186_v.txt | 33474 | 33474-SNV- | SNV | 1 | C | T | 45 | 21089 | 0.21338138 | 32.0444445 | rnr1D57N | FALSE | CAN | JSY14 | 5 | rnr1D57N-Re | GCC | C>T | A/T+1 | G/C-1 | CG>TA | CG>TA | GCTATGCCTT | CCTTTTT | CGG | GCC | C | A | C>T | C>T_GCC | CG>TA_CCTTTTT | 7 | FALSE | FALSE |
| 186_v.txt | 31947 | 31947-SNV- | SNV | 1 | G | C | 5 | 12555 | 0.03982477 | 38 | rnr1D57N | FALSE | CAN | JSY14 | 5 | rnr1D57N-Re | CGT | G>C | G/C+1 | G/C-1 | CG>GC | CG>GC | ATGATCGTC | GTCATAA | GCA | GCA | C | G | C>G | C>G_GCA | CG>GC_GTCATAA | 7 | FALSE | FALSE |
| 186_v.txt | 32057 | 32057-SNV- | SNV | 1 | C | G | 11 | 12564 | 0.08755174 | 58.9090909 | rnr1D57N | FALSE | CAN | JSY14 | 5 | rnr1D57N-Re | GCG | C>G | G/C+1 | G/C-1 | CG>GC | CG>GC | GATGTGCGA | CGAGATT | CGC | GCG | C | C | C>G | C>G_GCG | CG>GC_CGAGATT | 7 | FALSE | FALSE |
| 186_v.txt | 32064 | 32064-SNV- | SNV | 1 | G | C | 7 | 12564 | 0.05571474 | 62 | rnr1D57N | FALSE | CAN | JSY14 | 5 | rnr1D57N-Re | TGA | G>C | G/C+1 | G/C-1 | CG>GC | CG>GC | GAGATTGAG | GAGATAA | ACT | ACT | C | G | C>G | C>G_ACT | CG>GC_GAGATAA | 7 | FALSE | FALSE |
| 186_v.txt | 32271 | 32271-SNV- | SNV | 1 | C | G | 63 | 27652 | 0.22783162 | 35.5396825 | rnr1D57N | FALSE | CAN | JSY14 | 5 | rnr1D57N-Re | ACG | C>G | G/C+1 | G/C-1 | CG>GC | CG>GC | AAAATACGG | CGGGAAC | TGC | ACG | C | C | C>G | C>G_ACG | CG>GC_CGGGAAC | 7 | FALSE | FALSE |
| 186_v.txt | 32328 | 32328-SNV- | SNV | 1 | G | C | 1 | 27653 | 3.62E-03 | 25.9987738 | rnr1D57N | FALSE | CAN | JSY14 | 5 | rnr1D57N-Re | AGC | G>C | G/C+1 | G/C-1 | CG>GC | CG>GC | ATAACAGCG | GCGTTGA | TCG | TCG | C | G | C>G | C>G_TCG | CG>GC_GCGTTGA | 7 | FALSE | FALSE |
| 186_v.txt | 32399 | 32399-SNV- | SNV | 1 | G | C | 18 | 27653 | 0.0650924 | 53.5555556 | rnr1D57N | FALSE | CAN | JSY14 | 5 | rnr1D57N-Re | CGT | G>C | G/C+1 | G/C-1 | CG>GC | CG>GC | AGAAACGTA | GTAGGAA | GCA | GCA | C | G | C>G | C>G_GCA | CG>GC_GTAGGAA | 7 | FALSE | FALSE |
| 186_v.txt | 32829 | 32829-SNV- | SNV | 1 | G | C | 7 | 17176 | 0.04075454 | 42.8571429 | rnr1D57N | FALSE | CAN | JSY14 | 5 | rnr1D57N-Re | TGT | G>C | G/C+1 | G/C-1 | CG>GC | CG>GC | ATTATTGTG | GTGATAA | ACA | ACA | C | G | C>G | C>G_ACA | CG>GC_GTGATAA | 7 | FALSE | FALSE |
| 186_v.txt | 32876 | 32876-SNV- | SNV | 1 | G | C | 46 | 29318 | 0.1569002 | 37.7173913 | rnr1D57N | FALSE | CAN | JSY14 | 5 | rnr1D57N-Re | TGT | G>C | G/C+1 | G/C-1 | CG>GC | CG>GC | AACTTTGTAC | GTACGTC | ACA | ACA | C | G | C>G | C>G_ACA | CG>GC_GTACGTC | 7 | FALSE | FALSE |
| 186_v.txt | 32901 | 32901-SNV- | SNV | 1 | C | G | 5 | 29334 | 0.01704507 | 29.0015763 | rnr1D57N | FALSE | CAN | JSY14 | 5 | rnr1D57N-Re | GCC | C>G | G/C+1 | G/C-1 | CG>GC | CG>GC | ACTTGGCCA | CCAACTA | CGG | GCC | C | C | C>G | C>G_GCC | CG>GC_CCAACTA | 7 | FALSE | FALSE |
| 186_v.txt | 33070 | 33070-SNV- | SNV | 1 | C | G | 9 | 29333 | 0.03068217 | 52.6653394 | rnr1D57N | FALSE | CAN | JSY14 | 5 | rnr1D57N-Re | GCC | C>G | G/C+1 | G/C-1 | CG>GC | CG>GC | AATATGCCA | CCAAAGA | CGG | GCC | C | C | C>G | C>G_GCC | CG>GC_CCAAAGA | 7 | FALSE | FALSE |
| 186_v.txt | 33114 | 33114-SNV- | SNV | 1 | C | G | 1 | 29334 | 3.41E-03 | 30.999896 | rnr1D57N | FALSE | CAN | JSY14 | 5 | rnr1D57N-Re | GCC | C>G | G/C+1 | G/C-1 | CG>GC | CG>GC | ACTGGGCCG | CCGGCGT | CGG | GCC | C | C | C>G | C>G_GCC | CG>GC_CCGGCGT | 7 | FALSE | FALSE |
| 186_v.txt | 33115 | 33115-SNV- | SNV | 1 | C | G | 9 | 29334 | 0.03068112 | 45.5557265 | rnr1D57N | FALSE | CAN | JSY14 | 5 | rnr1D57N-Re | CCG | C>G | G/C+1 | G/C-1 | CG>GC | CG>GC | CTGGGCCGG | CGGCGTT | GGC | CCG | C | C | C>G | C>G_CCG | CG>GC_CGGCGTT | 7 | FALSE | FALSE |
| 186_v.txt | 33118 | 33118-SNV- | SNV | 1 | C | G | 4 | 29334 | 0.01363605 | 42.5 | rnr1D57N | FALSE | CAN | JSY14 | 5 | rnr1D57N-Re | GCG | C>G | G/C+1 | G/C-1 | CG>GC | CG>GC | GGCCGGCGT | CGTTGGT | CGC | GCG | C | C | C>G | C>G_GCG | CG>GC_CGTTGGT | 7 | FALSE | FALSE |
| 186_v.txt | 33154 | 33154-SNV- | SNV | 1 | C | G | 4 | 29334 | 0.01363605 | 38 | rnr1D57N | FALSE | CAN | JSY14 | 5 | rnr1D57N-Re | CCT | C>G | G/C+1 | G/C-1 | CG>GC | CG>GC | AAAGACCTG | CTGTACC | GGA | CCT | C | C | C>G | C>G_CCT | CG>GC_CTGTACC | 7 | FALSE | FALSE |
| 186_v.txt | 33284 | 33284-SNV- | SNV | 1 | G | C | 10 | 21090 | 0.04741584 | 55.4 | rnr1D57N | FALSE | CAN | JSY14 | 5 | rnr1D57N-Re | CGC | G>C | G/C+1 | G/C-1 | CG>GC | CG>GC | ATCCTCGCC | GCCATTT | GCG | GCG | C | G | C>G | C>G_GCG | CG>GC_GCCATTT | 7 | FALSE | FALSE |
| 186_v.txt | 31657 | 31657-SNV- | SNV | 1 | G | T | 33 | 3690 | 0.89430894 | 20.6060606 | rnr1D57N | FALSE | CAN | JSY14 | 5 | rnr1D57N-Re | GGC | G>T | A/T+1 | G/C-1 | CG>AT | CG>AT | GAAATGGCG | GCGTGGA | CCG | CCG | C | A | C>A | C>A_CCG | CG>AT_GCGTGGA | 7 | FALSE | FALSE |
| 186_v.txt | 31980 | 31980-SNV- | SNV | 1 | G | T | 13 | 12564 | 0.10347023 | 37.3076923 | rnr1D57N | FALSE | CAN | JSY14 | 5 | rnr1D57N-Re | GGG | G>T | A/T+1 | G/C-1 | CG>AT | CG>AT | AAGCCGGGC | GGCATTA | CCC | CCC | C | A | C>A | C>A_CCC | CG>AT_GGCATTA | 7 | FALSE | FALSE |
| 186_v.txt | 32008 | 32008-SNV- | SNV | 1 | C | A | 1835 | 12564 | 14.6052213 | 56.6599455 | rnr1D57N | FALSE | CAN | JSY14 | 5 | rnr1D57N-Re | TCG | C>A | A/T+1 | G/C-1 | CG>AT | CG>AT | GTAACTCGT | CGTCACG | AGC | TCG | C | T | C>A | C>A_TCG | CG>AT_CGTCACG | 7 | FALSE | FALSE |
| 186_v.txt | 32027 | 32027-SNV- | SNV | 1 | G | T | 15 | 12564 | 0.11938873 | 44 | rnr1D57N | FALSE | CAN | JSY14 | 5 | rnr1D57N-Re | GGT | G>T | A/T+1 | G/C-1 | CG>AT | CG>AT | GCCACGGTA | GTATTTC | CCA | CCA | C | A | C>A | C>A_CCA | CG>AT_GTATTTC | 7 | FALSE | FALSE |
| 186_v.txt | 32054 | 32054-SNV- | SNV | 1 | G | T | 12 | 12564 | 0.09551098 | 48.0833333 | rnr1D57N | FALSE | CAN | JSY14 | 5 | rnr1D57N-Re | TGT | G>T | A/T+1 | G/C-1 | CG>AT | CG>AT | TCTGATGTG | GTGCGAG | ACA | ACA | C | A | C>A | C>A_ACA | CG>AT_GTGCGAG | 7 | FALSE | FALSE |
| 186_v.txt | 32119 | 32119-SNV- | SNV | 1 | C | A | 14 | 12564 | 0.11142948 | 60.2142857 | rnr1D57N | FALSE | CAN | JSY14 | 5 | rnr1D57N-Re | TCG | C>A | A/T+1 | G/C-1 | CG>AT | CG>AT | GCCATTCGA | CGAAAAC | AGC | TCG | C | T | C>A | C>A_TCG | CG>AT_CGAAAAC | 7 | FALSE | FALSE |
| 186_v.txt | 32149 | 32149-SNV- | SNV | 1 | C | A | 14 | 12564 | 0.11142948 | 42 | rnr1D57N | FALSE | CAN | JSY14 | 5 | rnr1D57N-Re | TCC | C>A | A/T+1 | G/C-1 | CG>AT | CG>AT | ATGTCTCCAT | CCATGTA | AGG | TCC | C | T | C>A | C>A_TCC | CG>AT_CCATGTA | 7 | FALSE | FALSE |
| 186_v.txt | 32220 | 32220-SNV- | SNV | 1 | C | A | 2 | 12564 | 0.0159185 | 51.0030028 | rnr1D57N | FALSE | CAN | JSY14 | 5 | rnr1D57N-Re | CCT | C>A | A/T+1 | G/C-1 | CG>AT | CG>AT | GTGGTCCTT | CTTGACA | GGA | CCT | C | T | C>A | C>A_CCT | CG>AT_CTTGACA | 7 | FALSE | FALSE |
| 186_v.txt | 32274 | 32274-SNV- | SNV | 1 | G | T | 330 | 27652 | 1.19340373 | 36.0333333 | rnr1D57N | FALSE | CAN | JSY14 | 5 | rnr1D57N-Re | GGA | G>T | A/T+1 | G/C-1 | CG>AT | CG>AT | ATACGGGAA | GAACCAA | CCT | CCT | C | A | C>A | C>A_CCT | CG>AT_GAACCAA | 7 | FALSE | FALSE |
| 186_v.txt | 32329 | 32329-SNV- | SNV | 1 | C | A | 13 | 27653 | 0.04701117 | 23.923711 | rnr1D57N | FALSE | CAN | JSY14 | 5 | rnr1D57N-Re | GCG | C>A | A/T+1 | G/C-1 | CG>AT | CG>AT | TAACAGCGT | CGTTGAA | CGC | GCG | C | T | C>A | C>A_GCG | CG>AT_CGTTGAA | 7 | FALSE | FALSE |
| 186_v.txt | 32399 | 32399-SNV- | SNV | 1 | G | T | 23 | 27653 | 0.08317362 | 55.6956522 | rnr1D57N | FALSE | CAN | JSY14 | 5 | rnr1D57N-Re | CGT | G>T | A/T+1 | G/C-1 | CG>AT | CG>AT | AGAAACGTA | GTAGGAA | GCA | GCA | C | A | C>A | C>A_GCA | CG>AT_GTAGGAA | 7 | FALSE | FALSE |
| 186_v.txt | 32471 | 32471-SNV- | SNV | 1 | G | T | 16 | 27653 | 0.05785991 | 52.1875 | rnr1D57N | FALSE | CAN | JSY14 | 5 | rnr1D57N-Re | TGT | G>T | A/T+1 | G/C-1 | CG>AT | CG>AT | GCCAATGTA | GTAGAAG | ACA | ACA | C | A | C>A | C>A_ACA | CG>AT_GTAGAAG | 7 | FALSE | FALSE |
| 186_v.txt | 32474 | 32474-SNV- | SNV | 1 | G | T | 54 | 27653 | 0.19527719 | 47.0740741 | rnr1D57N | FALSE | CAN | JSY14 | 5 | rnr1D57N-Re | AGA | G>T | A/T+1 | G/C-1 | CG>AT | CG>AT | AATGTAGAA | GAAGGTT | TCT | TCT | C | A | C>A | C>A_TCT | CG>AT_GAAGGTT | 7 | FALSE | FALSE |
| 186_v.txt | 32566 | 32566-SNV- | SNV | 1 | C | A | 21 | 17170 | 0.12230635 | 35.1428571 | rnr1D57N | FALSE | CAN | JSY14 | 5 | rnr1D57N-Re | TCA | C>A | A/T+1 | G/C-1 | CG>AT | CG>AT | CTAGTTCAG | CAGTACC | AGT | TCA | C | T | C>A | C>A_TCA | CG>AT_CAGTACC | 7 | FALSE | FALSE |
| 186_v.txt | 32575 | 32575-SNV- | SNV | 1 | G | T | 190 | 17177 | 1.10613029 | 36.7684211 | rnr1D57N | FALSE | CAN | JSY14 | 5 | rnr1D57N-Re | TGA | G>T | A/T+1 | G/C-1 | CG>AT | CG>AT | TACCTTGAA | GAAATGT | ACT | ACT | C | A | C>A | C>A_ACT | CG>AT_GAAATGT | 7 | FALSE | FALSE |
| 186_v.txt | 32783 | 32783-SNV- | SNV | 1 | G | T | 17 | 17176 | 0.09897531 | 41.5882353 | rnr1D57N | FALSE | CAN | JSY14 | 5 | rnr1D57N-Re | AGA | G>T | A/T+1 | G/C-1 | CG>AT | CG>AT | GACCCAGAA | GAACTCG | TCT | TCT | C | A | C>A | C>A_TCT | CG>AT_GAACTCG | 7 | FALSE | FALSE |
| 186_v.txt | 32917 | 32917-SNV- | SNV | 1 | C | A | 11 | 29334 | 0.03749915 | 42 | rnr1D57N | FALSE | CAN | JSY14 | 5 | rnr1D57N-Re | TCC | C>A | A/T+1 | G/C-1 | CG>AT | CG>AT | TAAGTTCCA | CCAGGGC | AGG | TCC | C | T | C>A | C>A_TCC | CG>AT_CCAGGGC | 7 | FALSE | FALSE |
| 186_v.txt | 32937 | 32937-SNV- | SNV | 1 | C | A | 47 | 29334 | 0.16022363 | 49.6923846 | rnr1D57N | FALSE | CAN | JSY14 | 5 | rnr1D57N-Re | CCA | C>A | A/T+1 | G/C-1 | CG>AT | CG>AT | ATTGCCCAA | CAAGAAA | GGT | CCA | C | T | C>A | C>A_CCA | CG>AT_CAAGAAA | 7 | FALSE | FALSE |
| 186_v.txt | 33015 | 33015-SNV- | SNV | 1 | G | T | 21 | 29334 | 0.07158928 | 57.8095238 | rnr1D57N | FALSE | CAN | JSY14 | 5 | rnr1D57N-Re | GGA | G>T | A/T+1 | G/C-1 | CG>AT | CG>AT | AAAGAGGAT | GATGTAA | CCT | CCT | C | A | C>A | C>A_CCT | CG>AT_GATGTAA | 7 | FALSE | FALSE |
| 186_v.txt | 33024 | 33024-SNV- | SNV | 1 | G | T | 26 | 29334 | 0.08863435 | 62.4615385 | rnr1D57N | FALSE | CAN | JSY14 | 5 | rnr1D57N-Re | AGG | G>T | A/T+1 | G/C-1 | CG>AT | CG>AT | GTAACAGGG | GGGATGA | TCC | TCC | C | A | C>A | C>A_TCC | CG>AT_GGGATGA | 7 | FALSE | FALSE |
| 186_v.txt | 33154 | 33154-SNV- | SNV | 1 | C | A | 22 | 29334 | 0.0749983 | 35.3636364 | rnr1D57N | FALSE | CAN | JSY14 | 5 | rnr1D57N-Re | CCT | C>A | A/T+1 | G/C-1 | CG>AT | CG>AT | AAAGACCTG | CTGTACC | GGA | CCT | C | T | C>A | C>A_CCT | CG>AT_CTGTACC | 7 | FALSE | FALSE |
| 186_v.txt | 33156 | 33156-SNV- | SNV | 1 | G | T | 29 | 29334 | 0.09886139 | 36.5862069 | rnr1D57N | FALSE | CAN | JSY14 | 5 | rnr1D57N-Re | TGT | G>T | A/T+1 | G/C-1 | CG>AT | CG>AT | AGACCTGTA | GTACCAA | ACA | ACA | C | A | C>A | C>A_ACA | CG>AT_GTACCAA | 7 | FALSE | FALSE |
| 186_v.txt | 33175 | 33175-SNV- | SNV | 1 | G | T | 10 | 29326 | 0.03409943 | 37.9 | rnr1D57N | FALSE | CAN | JSY14 | 5 | rnr1D57N-Re | AGG | G>T | A/T+1 | G/C-1 | CG>AT | CG>AT | CACCAAGGG | GGGCAAT | TCC | TCC | C | A | C>A | C>A_TCC | CG>AT_GGGCAAT | 7 | FALSE | FALSE |
| 186_v.txt | 33195 | 33195-SNV- | SNV | 1 | C | A | 75 | 21090 | 0.35561878 | 35.48 | rnr1D57N | FALSE | CAN | JSY14 | 5 | rnr1D57N-Re | TCT | C>A | A/T+1 | G/C-1 | CG>AT | CG>AT | ATATGTCTTT | CTTTGCT | AGA | TCT | C | T | C>A | C>A_TCT | CG>AT_CTTTGCT | 7 | FALSE | FALSE |
| 186_v.txt | 33325 | 33325-SNV- | SNV | 1 | C | A | 89 | 21090 | 0.42200095 | 57.1348315 | rnr1D57N | FALSE | CAN | JSY14 | 5 | rnr1D57N-Re | TCT | C>A | A/T+1 | G/C-1 | CG>AT | CG>AT | ACAATTCTTT | CTTTACT | AGA | TCT | C | T | C>A | C>A_TCT | CG>AT_CTTTACT | 7 | FALSE | FALSE |
| 186_v.txt | 33474 | 33474-SNV- | SNV | 1 | C | A | 81 | 21089 | 0.38408649 | 34.3950617 | rnr1D57N | FALSE | CAN | JSY14 | 5 | rnr1D57N-Re | GCC | C>A | A/T+1 | G/C-1 | CG>AT | CG>AT | GCTATGCCTT | CCTTTTT | CGG | GCC | C | T | C>A | C>A_GCC | CG>AT_CCTTTTT | 7 | FALSE | FALSE |
| 186_v.txt | 33475 | 33475-SNV- | SNV | 1 | C | A | 35 | 21089 | 0.1659633 | 35.3151685 | rnr1D57N | FALSE | CAN | JSY14 | 5 | rnr1D57N-Re | CCT | C>A | A/T+1 | G/C-1 | CG>AT | CG>AT | CTATGCCTTT | CTTTTTT | GGA | CCT | C | T | C>A | C>A_CCT | CG>AT_CTTTTTT | 7 | FALSE | FALSE |
| 186_v.txt | 32328 | 32328-MNV- | MNV | 2 | GC | CG | 21 | 27653 | 0.07594113 | 56.904758 | rnr1D57N | FALSE | CAN | JSY14 | 5 | rnr1D57N-Re | AGC | nope | >1bp-ins | >1bpdel | none | MNV | ATAACAGCG | GCGTTGA | TCG | AGC | GC | none | NA | NA_AGC | MNV_GCGTTGA | 7 | FALSE | FALSE |
| 186_v.txt | 33114 | 33114-MNV- | MNV | 2 | CC | AA | 2 | 29334 | 6.82E-03 | 37.7502103 | rnr1D57N | FALSE | CAN | JSY14 | 5 | rnr1D57N-Re | GCC | nope | >1bp-ins | >1bpdel | none | MNV | ACTGGGCCG | CCGGCGT | CGG | GCC | CC | none | NA | NA_GCC | MNV_CCGGCGT | 7 | FALSE | FALSE |
| 186_v.txt | 31926 | 31926-Insert | Insertion | 1 | - | C | 2 | 3690 | 0.05420054 | 38 | rnr1D57N | FALSE | CAN | JSY14 | 5 | rnr1D57N-Re | ACC | nope | G/C+1 | >1bpdel | none | G/C+1 | GTGAAACCT | CCTTGAA | TGG | ACC | - | G | NA | NA_ACC | G/C+1_CCTTGAA | 7 | FALSE | FALSE |
| 186_v.txt | 32081 | 32081-Insert | Insertion | 1 | - | A | 36 | 12564 | 0.28653295 | 63.6944444 | rnr1D57N | FALSE | CAN | JSY14 | 5 | rnr1D57N-Re | CAA | nope | A/T+1 | >1bpdel | none | A/T+1 | CCATGCAAA | AAAAAAG | GTT | CAA | - | T | NA | NA_CAA | A/T+1_AAAAAAG | 7 | FALSE | FALSE |
| 186_v.txt | 32168 | 32168-Insert | Insertion | 1 | - | A | 4 | 12564 | 0.031837 | 51 | rnr1D57N | FALSE | CAN | JSY14 | 5 | rnr1D57N-Re | CAA | nope | A/T+1 | >1bpdel | none | A/T+1 | AGCGCCAAA | AAATGCA | GTT | CAA | - | T | NA | NA_CAA | A/T+1_AAATGCA | 7 | FALSE | FALSE |
| 186_v.txt | 32295 | 32295-Insert | Insertion | 1 | - | T | 128 | 27653 | 0.46287925 | 50.34375 | rnr1D57N | FALSE | CAN | JSY14 | 5 | rnr1D57N-Re | ATT | nope | A/T+1 | >1bpdel | none | A/T+1 | TTTGAATTTG | TTTGCGG | TAA | ATT | - | A | NA | NA_ATT | A/T+1_TTTGCGG | 7 | FALSE | FALSE |
| 186_v.txt | 32402 | 32402-Insert | Insertion | 34 | - | GGAAGTAGA | 7 | 27653 | 0.02531371 | 58.6722689 | rnr1D57N | FALSE | CAN | JSY14 | 5 | rnr1D57N-Re | AGG | nope | >1bp-ins | >1bpdel | none | >1bp-ins | AACGTAGGA | GGAAGTA | TCC | AGG | - | none | NA | NA_AGG | >1bp-ins_GGAAGTA | 7 | FALSE | FALSE |
| 186_v.txt | 32448 | 32448-Insert | Insertion | 18 | - | CCAATGAAT | 1 | 27653 | 3.62E-03 | 60.8333274 | rnr1D57N | FALSE | CAN | JSY14 | 5 | rnr1D57N-Re | TCC | nope | >1bp-ins | >1bpdel | none | >1bp-ins | AAAAGTCCA | CCAATGA | AGG | TCC | - | none | NA | NA_TCC | >1bp-ins_CCAATGA | 7 | FALSE | FALSE |
| 186_v.txt | 32448 | 32448-Insert | Insertion | 18 | - | CCAATGAAT | 432 | 27653 | 1.56221748 | 53.3337192 | rnr1D57N | FALSE | CAN | JSY14 | 5 | rnr1D57N-Re | TCC | nope | >1bp-ins | >1bpdel | none | >1bp-ins | AAAAGTCCA | CCAATGA | AGG | TCC | - | none | NA | NA_TCC | >1bp-ins_CCAATGA | 7 | FALSE | FALSE |
| 186_v.txt | 32448 | 32448-Insert | Insertion | 18 | - | CCAATGTAT | 2 | 27653 | 7.23E-03 | 48.749997 | rnr1D57N | FALSE | CAN | JSY14 | 5 | rnr1D57N-Re | TCC | nope | >1bp-ins | >1bpdel | none | >1bp-ins | AAAAGTCCA | CCAATGA | AGG | TCC | - | none | NA | NA_TCC | >1bp-ins_CCAATGA | 7 | FALSE | FALSE |
| 186_v.txt | 32448 | 32448-Insert | Insertion | 13 | - | GAATAATAG | 1 | 27653 | 3.62E-03 | 37.9230769 | rnr1D57N | FALSE | CAN | JSY14 | 5 | rnr1D57N-Re | TCC | nope | >1bp-ins | >1bpdel | none | >1bp-ins | AAAAGTCCA | CCAATGA | AGG | TCC | - | none | NA | NA_TCC | >1bp-ins_CCAATGA | 7 | FALSE | FALSE |
| 186_v.txt | 32454 | 32454-Insert | Insertion | 18 | - | AATAATAGA | 1 | 27653 | 3.62E-03 | 63.3888889 | rnr1D57N | FALSE | CAN | JSY14 | 5 | rnr1D57N-Re | GAA | nope | >1bp-ins | >1bpdel | none | >1bp-ins | CCAATGAAT | AATAATA | CTT | GAA | - | none | NA | NA_GAA | >1bp-ins_AATAATA | 7 | FALSE | FALSE |
| 186_v.txt | 32498 | 32498-Insert | Insertion | 1 | - | T | 23 | 27653 | 0.08317362 | 46.826087 | rnr1D57N | FALSE | CAN | JSY14 | 5 | rnr1D57N-Re | CTT | nope | A/T+1 | >1bpdel | none | A/T+1 | AACAACTTTT | TTTTTTG | GAA | CTT | - | A | NA | NA_CTT | A/T+1_TTTTTTG | 7 | FALSE | FALSE |
| 186_v.txt | 32593 | 32593-Insert | Insertion | 1 | - | A | 4 | 17176 | 0.02328831 | 50.25 | rnr1D57N | FALSE | CAN | JSY14 | 5 | rnr1D57N-Re | TTA | nope | A/T+1 | >1bpdel | none | A/T+1 | CAGCGTTAA | TAATCAA | AAT | TTA | - | T | NA | NA_TTA | A/T+1_TAATCAA | 7 | FALSE | FALSE |
| 186_v.txt | 32728 | 32728-Insert | Insertion | 1 | - | A | 6 | 17176 | 0.03493246 | 57.3333333 | rnr1D57N | FALSE | CAN | JSY14 | 5 | rnr1D57N-Re | CAA | nope | A/T+1 | >1bpdel | none | A/T+1 | CCATACAAA | AAAAACA | GTT | CAA | - | T | NA | NA_CAA | A/T+1_AAAAACA | 7 | FALSE | FALSE |
| 186_v.txt | 32839 | 32839-Insert | Insertion | 1 | - | C | 2 | 17176 | 0.01164416 | 38 | rnr1D57N | FALSE | CAN | JSY14 | 5 | rnr1D57N-Re | ACC | nope | G/C+1 | >1bpdel | none | G/C+1 | TAATTACCC | CCCAAAA | TGG | ACC | - | G | NA | NA_ACC | G/C+1_CCCAAAA | 7 | FALSE | FALSE |
| 186_v.txt | 32842 | 32842-Insert | Insertion | 1 | - | A | 34 | 17176 | 0.19795063 | 46.3235294 | rnr1D57N | FALSE | CAN | JSY14 | 5 | rnr1D57N-Re | CAA | nope | A/T+1 | >1bpdel | none | A/T+1 | TTACCCAAA | AAAAAAT | GTT | CAA | - | T | NA | NA_CAA | A/T+1_AAAAAAT | 7 | FALSE | FALSE |
| 186_v.txt | 33036 | 33036-Insert | Insertion | 33 | - | GCCATTTCAC | 4 | 29334 | 0.01363605 | 61.030303 | rnr1D57N | FALSE | CAN | JSY14 | 5 | rnr1D57N-Re | AGC | nope | >1bp-ins | >1bpdel | none | >1bp-ins | AATGTAGCC | GCCATTT | TCG | AGC | - | none | NA | NA_AGC | >1bp-ins_GCCATTT | 7 | FALSE | FALSE |
| 186_v.txt | 33219 | 33219-Insert | Insertion | 1 | - | C | 7 | 21090 | 0.03319109 | 37.4285714 | rnr1D57N | FALSE | CAN | JSY14 | 5 | rnr1D57N-Re | TTC | nope | G/C+1 | >1bpdel | none | G/C+1 | TTCACTTCAG | TCAGCGT | AAG | TTC | - | G | NA | NA_TTC | G/C+1_TCAGCGT | 7 | FALSE | FALSE |
| 186_v.txt | 33265 | 33265-Insert | Insertion | 23 | - | CCATAGAGA | 4 | 21090 | 0.01896634 | 44.826087 | rnr1D57N | FALSE | CAN | JSY14 | 5 | rnr1D57N-Re | TCC | nope | >1bp-ins | >1bpdel | none | >1bp-ins | CATCCTCCAT | CCATAGA | AGG | TCC | - | none | NA | NA_TCC | >1bp-ins_CCATAGA | 7 | FALSE | FALSE |
| 186_v.txt | 33288 | 33288-Insert | Insertion | 1 | - | T | 9 | 21090 | 0.04267425 | 54.1111111 | rnr1D57N | FALSE | CAN | JSY14 | 5 | rnr1D57N-Re | ATT | nope | A/T+1 | >1bpdel | none | A/T+1 | TCGCCATTTA | TTTACTC | TAA | ATT | - | A | NA | NA_ATT | A/T+1_TTTACTC | 7 | FALSE | FALSE |
| 186_v.txt | 32215 | 32215-Deleti | Deletion | 6 | TGGTCC | - | 21 | 12564 | 0.16714422 | 41.2857143 | rnr1D57N | FALSE | CAN | JSY14 | 5 | rnr1D57N-Re | GTG | nope | >1bp-ins | >1bpdel | none | >1bpdel | CTTTGGTGG | TGGTCCT | CAC | GTG | TGGTCC | none | NA | NA_GTG | >1bpdel_TGGTCCT | 7 | FALSE | FALSE |
| 186_v.txt | 32454 | 32454-Deleti | Deletion | 3 | AAT | - | 7 | 27653 | 0.02531371 | 56.5714286 | rnr1D57N | FALSE | CAN | JSY14 | 5 | rnr1D57N-Re | GAA | nope | >1bp-ins | >1bpdel | none | >1bpdel | CCAATGAAT | AATAATA | CTT | GAA | AAT | none | NA | NA_GAA | >1bpdel_AATAATA | 7 | FALSE | FALSE |
| 186_v.txt | 32736 | 32736-Deleti | Deletion | 2 | TA | - | 7 | 17176 | 0.04075454 | 60 | rnr1D57N | FALSE | CAN | JSY14 | 5 | rnr1D57N-Re | GTA | nope | >1bp-ins | >1bpdel | none | >1bpdel | AAACAGTAT | TATATTA | CAT | GTA | TA | none | NA | NA_GTA | >1bpdel_TATATTA | 7 | FALSE | FALSE |
| 186_v.txt | 33474 | 33474-Deleti | Deletion | 2 | CC | - | 2 | 21089 | 9.48E-03 | 16 | rnr1D57N | FALSE | CAN | JSY14 | 5 | rnr1D57N-Re | GCC | nope | >1bp-ins | >1bpdel | none | >1bpdel | GCTATGCCTT | CCTTTTT | CGG | GCC | CC | none | NA | NA_GCC | >1bpdel_CCTTTTT | 7 | FALSE | FALSE |
| 186_v.txt | 31723 | 31723-Deleti | Deletion | 1 | A | - | 6 | 3690 | 0.16260163 | 63.5 | rnr1D57N | FALSE | CAN | JSY14 | 5 | rnr1D57N-Re | CAA | nope | >1bp-ins | A/T-1 | none | A/T-1 | TGTCCCAAA | AAAAAGT | GTT | GTT | T | none | T>none | T>none_GTT | A/T-1_AAAAAGT | 7 | FALSE | FALSE |
| 186_v.txt | 31861 | 31861-Deleti | Deletion | 1 | G | - | 5 | 3690 | 0.13550136 | 48.4 | rnr1D57N | FALSE | CAN | JSY14 | 5 | rnr1D57N-Re | AGG | nope | >1bp-ins | G/C-1 | none | G/C-1 | AGAACAGGA | GGAAAAT | TCC | TCC | C | none | C>none | C>none_TCC | G/C-1_GGAAAAT | 7 | FALSE | FALSE |
| 186_v.txt | 31958 | 31958-Deleti | Deletion | 1 | G | - | 42 | 12564 | 0.33428844 | 36.6428571 | rnr1D57N | FALSE | CAN | JSY14 | 5 | rnr1D57N-Re | TGG | nope | >1bp-ins | G/C-1 | none | G/C-1 | AAATGTGGC | GGCCGCA | ACC | ACC | C | none | C>none | C>none_ACC | G/C-1_GGCCGCA | 7 | FALSE | FALSE |
| 186_v.txt | 31971 | 31971-Deleti | Deletion | 1 | G | - | 6972 | 12564 | 55.4918816 | 37.3416523 | rnr1D57N | FALSE | CAN | JSY14 | 5 | rnr1D57N-Re | AGC | nope | >1bp-ins | G/C-1 | none | G/C-1 | TAATAAGCC | GCCAAGC | TCG | TCG | C | none | C>none | C>none_TCG | G/C-1_GCCAAGC | 7 | FALSE | FALSE |
| 186_v.txt | 31988 | 31988-Deleti | Deletion | 1 | T | - | 19 | 12564 | 0.15122572 | 53.4210526 | rnr1D57N | FALSE | CAN | JSY14 | 5 | rnr1D57N-Re | ATT | nope | >1bp-ins | A/T-1 | none | A/T-1 | CATTAATTTA | TTTAGCT | TAA | ATT | T | none | nope | nope_ATT | A/T-1_TTTAGCT | 7 | FALSE | FALSE |
| 186_v.txt | 31997 | 31997-Deleti | Deletion | 1 | A | - | 8 | 12564 | 0.06367399 | 56.625 | rnr1D57N | FALSE | CAN | JSY14 | 5 | rnr1D57N-Re | TAA | nope | >1bp-ins | A/T-1 | none | A/T-1 | AGCTTTAAA | AAATGGT | ATT | ATT | T | none | T>none | T>none_ATT | A/T-1_AAATGGT | 7 | FALSE | FALSE |
| 186_v.txt | 32022 | 32022-Deleti | Deletion | 1 | C | - | 18 | 12564 | 0.14326648 | 62.3333333 | rnr1D57N | FALSE | CAN | JSY14 | 5 | rnr1D57N-Re | GCC | nope | >1bp-ins | G/C-1 | none | G/C-1 | GAGATGCCA | CCACGGT | CGG | GCC | C | none | nope | nope_GCC | G/C-1_CCACGGT | 7 | FALSE | FALSE |
| 186_v.txt | 32081 | 32081-Deleti | Deletion | 1 | A | - | 23 | 12563 | 0.18307729 | 62.3043482 | rnr1D57N | FALSE | CAN | JSY14 | 5 | rnr1D57N-Re | CAA | nope | >1bp-ins | A/T-1 | none | A/T-1 | CCATGCAAA | AAAAAAG | GTT | GTT | T | none | T>none | T>none_GTT | A/T-1_AAAAAAG | 7 | FALSE | FALSE |
| 186_v.txt | 32265 | 32265-Deleti | Deletion | 1 | A | - | 3 | 27652 | 0.01084913 | 37.3333333 | rnr1D57N | FALSE | CAN | JSY14 | 5 | rnr1D57N-Re | TAA | nope | >1bp-ins | A/T-1 | none | A/T-1 | CCAAATAAA | AAAATAC | ATT | ATT | T | none | T>none | T>none_ATT | A/T-1_AAAATAC | 7 | FALSE | FALSE |
| 186_v.txt | 32305 | 32305-Deleti | Deletion | 1 | A | - | 3 | 27653 | 0.01084873 | 45.6666667 | rnr1D57N | FALSE | CAN | JSY14 | 5 | rnr1D57N-Re | GAA | nope | >1bp-ins | A/T-1 | none | A/T-1 | CGGCAGAAA | AAATAAT | CTT | CTT | T | none | T>none | T>none_CTT | A/T-1_AAATAAT | 7 | FALSE | FALSE |
| 186_v.txt | 32346 | 32346-Deleti | Deletion | 1 | A | - | 110 | 27653 | 0.39778686 | 60.3727273 | rnr1D57N | FALSE | CAN | JSY14 | 5 | rnr1D57N-Re | CAA | nope | >1bp-ins | A/T-1 | none | A/T-1 | TGTGGCAAA | AAAACCT | GTT | GTT | T | none | T>none | T>none_GTT | A/T-1_AAAACCT | 7 | FALSE | FALSE |
| 186_v.txt | 32381 | 32381-Deleti | Deletion | 1 | A | - | 6 | 27653 | 0.02169747 | 62.5 | rnr1D57N | FALSE | CAN | JSY14 | 5 | rnr1D57N-Re | TAA | nope | >1bp-ins | A/T-1 | none | A/T-1 | AATAATAAA | AAAGGGA | ATT | ATT | T | none | T>none | T>none_ATT | A/T-1_AAAGGGA | 7 | FALSE | FALSE |
| 186_v.txt | 32420 | 32420-Deleti | Deletion | 1 | T | - | 8 | 27653 | 0.02892995 | 62.25 | rnr1D57N | FALSE | CAN | JSY14 | 5 | rnr1D57N-Re | GTT | nope | >1bp-ins | A/T-1 | none | A/T-1 | TGTTAGTTTA | TTTAGGG | CAA | GTT | T | none | nope | nope_GTT | A/T-1_TTTAGGG | 7 | FALSE | FALSE |
| 186_v.txt | 32442 | 32442-Deleti | Deletion | 1 | A | - | 7 | 27653 | 0.02531371 | 52 | rnr1D57N | FALSE | CAN | JSY14 | 5 | rnr1D57N-Re | TAA | nope | >1bp-ins | A/T-1 | none | A/T-1 | GGAACTAAA | AAAAGTC | ATT | ATT | T | none | T>none | T>none_ATT | A/T-1_AAAAGTC | 7 | FALSE | FALSE |
| 186_v.txt | 32498 | 32498-Deleti | Deletion | 1 | T | - | 42 | 27653 | 0.15188226 | 46.6904594 | rnr1D57N | FALSE | CAN | JSY14 | 5 | rnr1D57N-Re | CTT | nope | >1bp-ins | A/T-1 | none | A/T-1 | AACAACTTTT | TTTTTTG | GAA | CTT | T | none | nope | nope_CTT | A/T-1_TTTTTTG | 7 | FALSE | FALSE |
| 186_v.txt | 32522 | 32522-Deleti | Deletion | 1 | T | - | 43 | 27653 | 0.1554985 | 41.8837209 | rnr1D57N | FALSE | CAN | JSY14 | 5 | rnr1D57N-Re | ATT | nope | >1bp-ins | A/T-1 | none | A/T-1 | AACGGATTT | TTTTCTG | TAA | ATT | T | none | nope | nope_ATT | A/T-1_TTTTCTG | 7 | FALSE | FALSE |
| 186_v.txt | 32528 | 32528-Deleti | Deletion | 1 | G | - | 22 | 27653 | 0.07955737 | 43.9545455 | rnr1D57N | FALSE | CAN | JSY14 | 5 | rnr1D57N-Re | TGG | nope | >1bp-ins | G/C-1 | none | G/C-1 | TTTTCTGGG | GGGGTTT | ACC | ACC | C | none | C>none | C>none_ACC | G/C-1_GGGGTTT | 7 | FALSE | FALSE |
| 186_v.txt | 32532 | 32532-Deleti | Deletion | 1 | T | - | 7 | 27653 | 0.02531371 | 38 | rnr1D57N | FALSE | CAN | JSY14 | 5 | rnr1D57N-Re | GTT | nope | >1bp-ins | A/T-1 | none | A/T-1 | CTGGGGTTT | TTTGCAG | CAA | GTT | T | none | nope | nope_GTT | A/T-1_TTTGCAG | 7 | FALSE | FALSE |
| 186_v.txt | 32583 | 32583-Deleti | Deletion | 1 | A | - | 2 | 17176 | 0.01164416 | 35 | rnr1D57N | FALSE | CAN | JSY14 | 5 | rnr1D57N-Re | GAA | nope | >1bp-ins | A/T-1 | none | A/T-1 | AATGTGAAG | AAGGCAG | CTT | CTT | T | none | T>none | T>none_CTT | A/T-1_AAGGCAG | 7 | FALSE | FALSE |
| 186_v.txt | 32605 | 32605-Deleti | Deletion | 1 | A | - | 9 | 17176 | 0.0523987 | 40.4444444 | rnr1D57N | FALSE | CAN | JSY14 | 5 | rnr1D57N-Re | GAA | nope | >1bp-ins | A/T-1 | none | A/T-1 | AAGAGGAAA | AAACCCA | CTT | CTT | T | none | T>none | T>none_CTT | A/T-1_AAACCCA | 7 | FALSE | FALSE |
| 186_v.txt | 32658 | 32658-Deleti | Deletion | 1 | C | - | 16 | 17176 | 0.09315324 | 60.1875 | rnr1D57N | FALSE | CAN | JSY14 | 5 | rnr1D57N-Re | ACC | nope | >1bp-ins | G/C-1 | none | G/C-1 | CCTGGACCC | CCCCAGG | TGG | ACC | C | none | nope | nope_ACC | G/C-1_CCCCAGG | 7 | FALSE | FALSE |
| 186_v.txt | 32707 | 32707-Deleti | Deletion | 1 | C | - | 5 | 17176 | 0.02911039 | 60.8 | rnr1D57N | FALSE | CAN | JSY14 | 5 | rnr1D57N-Re | ACC | nope | >1bp-ins | G/C-1 | none | G/C-1 | CGGTAACCC | CCCCAGC | TGG | ACC | C | none | nope | nope_ACC | G/C-1_CCCCAGC | 7 | FALSE | FALSE |
| 186_v.txt | 32728 | 32728-Deleti | Deletion | 1 | A | - | 7 | 17175 | 0.04075691 | 55.2857111 | rnr1D57N | FALSE | CAN | JSY14 | 5 | rnr1D57N-Re | CAA | nope | >1bp-ins | A/T-1 | none | A/T-1 | CCATACAAA | AAAAACA | GTT | GTT | T | none | T>none | T>none_GTT | A/T-1_AAAAACA | 7 | FALSE | FALSE |
| 186_v.txt | 32829 | 32829-Deleti | Deletion | 1 | G | - | 2 | 17176 | 0.01164416 | 38 | rnr1D57N | FALSE | CAN | JSY14 | 5 | rnr1D57N-Re | TGT | nope | >1bp-ins | G/C-1 | none | G/C-1 | ATTATTGTG | GTGATAA | ACA | ACA | C | none | C>none | C>none_ACA | G/C-1_GTGATAA | 7 | FALSE | FALSE |
| 186_v.txt | 32842 | 32842-Deleti | Deletion | 1 | A | - | 43 | 17176 | 0.25034933 | 42.2325556 | rnr1D57N | FALSE | CAN | JSY14 | 5 | rnr1D57N-Re | CAA | nope | >1bp-ins | A/T-1 | none | A/T-1 | TTACCCAAA | AAAAAAT | GTT | GTT | T | none | T>none | T>none_GTT | A/T-1_AAAAAAT | 7 | FALSE | FALSE |
| 186_v.txt | 32868 | 32868-Deleti | Deletion | 1 | G | - | 23 | 17174 | 0.13392337 | 37.0869565 | rnr1D57N | FALSE | CAN | JSY14 | 5 | rnr1D57N-Re | TGG | nope | >1bp-ins | G/C-1 | none | G/C-1 | GCCAGTGGA | GGAACTT | ACC | ACC | C | none | C>none | C>none_ACC | G/C-1_GGAACTT | 7 | FALSE | FALSE |
| 186_v.txt | 32884 | 32884-Deleti | Deletion | 1 | A | - | 7 | 29334 | 0.02386309 | 36.8571429 | rnr1D57N | FALSE | CAN | JSY14 | 5 | rnr1D57N-Re | CAA | nope | >1bp-ins | A/T-1 | none | A/T-1 | ACGTCCAAA | AAAATTG | GTT | GTT | T | none | T>none | T>none_GTT | A/T-1_AAAATTG | 7 | FALSE | FALSE |
| 186_v.txt | 32924 | 32924-Deleti | Deletion | 1 | A | - | 33 | 29334 | 0.11249744 | 55.8484849 | rnr1D57N | FALSE | CAN | JSY14 | 5 | rnr1D57N-Re | CAA | nope | >1bp-ins | A/T-1 | none | A/T-1 | CAGGGCAAA | AAAAGTG | GTT | GTT | T | none | T>none | T>none_GTT | A/T-1_AAAAGTG | 7 | FALSE | FALSE |
| 186_v.txt | 32935 | 32935-Deleti | Deletion | 1 | C | - | 6 | 29334 | 0.02045408 | 45 | rnr1D57N | FALSE | CAN | JSY14 | 5 | rnr1D57N-Re | GCC | nope | >1bp-ins | G/C-1 | none | G/C-1 | TGATTGCCC | CCCAAGA | CGG | GCC | C | none | nope | nope_GCC | G/C-1_CCCAAGA | 7 | FALSE | FALSE |
| 186_v.txt | 32945 | 32945-Deleti | Deletion | 1 | C | - | 1 | 29334 | 3.41E-03 | 38.000029 | rnr1D57N | FALSE | CAN | JSY14 | 5 | rnr1D57N-Re | ACC | nope | >1bp-ins | G/C-1 | none | G/C-1 | AGAAAACCA | CCAATAC | TGG | ACC | C | none | nope | nope_ACC | G/C-1_CCAATAC | 7 | FALSE | FALSE |
| 186_v.txt | 33040 | 33040-Deleti | Deletion | 1 | T | - | 7 | 29334 | 0.02386309 | 59.2857143 | rnr1D57N | FALSE | CAN | JSY14 | 5 | rnr1D57N-Re | ATT | nope | >1bp-ins | A/T-1 | none | A/T-1 | TAGCCATTTC | TTTCACC | TAA | ATT | T | none | nope | nope_ATT | A/T-1_TTTCACC | 7 | FALSE | FALSE |
| 186_v.txt | 33070 | 33070-Deleti | Deletion | 1 | C | - | 1 | 29333 | 3.41E-03 | 38.0000291 | rnr1D57N | FALSE | CAN | JSY14 | 5 | rnr1D57N-Re | GCC | nope | >1bp-ins | G/C-1 | none | G/C-1 | AATATGCCA | CCAAAGA | CGG | GCC | C | none | nope | nope_GCC | G/C-1_CCAAAGA | 7 | FALSE | FALSE |
| 186_v.txt | 33114 | 33114-Deleti | Deletion | 1 | C | - | 1 | 29334 | 3.41E-03 | 38 | rnr1D57N | FALSE | CAN | JSY14 | 5 | rnr1D57N-Re | GCC | nope | >1bp-ins | G/C-1 | none | G/C-1 | ACTGGGCCG | CCGGCGT | CGG | GCC | C | none | nope | nope_GCC | G/C-1_CCGGCGT | 7 | FALSE | FALSE |
| 186_v.txt | 33138 | 33138-Deleti | Deletion | 1 | A | - | 6 | 29334 | 0.02045408 | 46.6666667 | rnr1D57N | FALSE | CAN | JSY14 | 5 | rnr1D57N-Re | TAA | nope | >1bp-ins | A/T-1 | none | A/T-1 | GTGGATAAA | AAACCAA | ATT | ATT | T | none | T>none | T>none_ATT | A/T-1_AAACCAA | 7 | FALSE | FALSE |
| 186_v.txt | 33147 | 33147-Deleti | Deletion | 1 | A | - | 5 | 29334 | 0.01704507 | 37.6 | rnr1D57N | FALSE | CAN | JSY14 | 5 | rnr1D57N-Re | GAA | nope | >1bp-ins | A/T-1 | none | A/T-1 | CCAATGAAA | AAAAGAC | CTT | CTT | T | none | T>none | T>none_CTT | A/T-1_AAAAGAC | 7 | FALSE | FALSE |
| 186_v.txt | 33175 | 33175-Deleti | Deletion | 1 | G | - | 2 | 29326 | 6.82E-03 | 38 | rnr1D57N | FALSE | CAN | JSY14 | 5 | rnr1D57N-Re | AGG | nope | >1bp-ins | G/C-1 | none | G/C-1 | CACCAAGGG | GGGCAAT | TCC | TCC | C | none | C>none | C>none_TCC | G/C-1_GGGCAAT | 7 | FALSE | FALSE |
| 186_v.txt | 33459 | 33459-Deleti | Deletion | 1 | T | - | 12 | 21090 | 0.056899 | 35.0833333 | rnr1D57N | FALSE | CAN | JSY14 | 5 | rnr1D57N-Re | ATT | nope | >1bp-ins | A/T-1 | none | A/T-1 | TTTGAATTTG | TTTGTCA | TAA | ATT | T | none | nope | nope_ATT | A/T-1_TTTGTCA | 7 | FALSE | FALSE |
| 186_v.txt | 33474 | 33474-Deleti | Deletion | 1 | C | - | 55 | 21089 | 0.26079947 | 28.6727273 | rnr1D57N | FALSE | CAN | JSY14 | 5 | rnr1D57N-Re | GCC | nope | >1bp-ins | G/C-1 | none | G/C-1 | GCTATGCCTT | CCTTTTT | CGG | GCC | C | none | nope | nope_GCC | G/C-1_CCTTTTT | 7 | FALSE | FALSE |
| NL3_v.txt | 31723 | 31723-SNV- | SNV | 1 | A | C | 3 | 6480 | 0.0462963 | 45.3333333 | rnr1D57N | FALSE | CAN | JSY3602 | 6 | rnr1D57N-Re | CAA | A>C | G/C+1 | A/T-1 | TA>GC | TA>GC | TGTCCCAAA | AAAAAGT | GTT | GTT | T | G | T>G | T>G_GTT | TA>GC_AAAAAGT | 7 | FALSE | FALSE |
| NL3_v.txt | 31840 | 31840-SNV- | SNV | 1 | A | C | 2 | 6480 | 0.0308642 | 64 | rnr1D57N | FALSE | CAN | JSY3602 | 6 | rnr1D57N-Re | AAG | A>C | G/C+1 | A/T-1 | TA>GC | TA>GC | GAAATAAGA | AGATCCA | TTC | TTC | T | G | T>G | T>G_TTC | TA>GC_AGATCCA | 7 | FALSE | FALSE |
| NL3_v.txt | 32445 | 32445-SNV- | SNV | 1 | A | C | 12 | 11034 | 0.10875476 | 51.75 | rnr1D57N | FALSE | CAN | JSY3602 | 6 | rnr1D57N-Re | AAG | A>C | G/C+1 | A/T-1 | TA>GC | TA>GC | ACTAAAAGT | AGTCCAA | TTC | TTC | T | G | T>G | T>G_TTC | TA>GC_AGTCCAA | 7 | FALSE | FALSE |
| NL3_v.txt | 32915 | 32915-SNV-T | SNV | 1 | T | G | 76 | 11230 | 0.67675868 | 36.8947368 | rnr1D57N | FALSE | CAN | JSY3602 | 6 | rnr1D57N-Re | GTT | T>G | G/C+1 | A/T-1 | TA>GC | TA>GC | ACTAAGTTCC | TTCCAGG | CAA | GTT | T | C | T>G | T>G_GTT | TA>GC_TTCCAGG | 7 | FALSE | FALSE |
| NL3_v.txt | 33023 | 33023-SNV- | SNV | 1 | A | C | 1 | 11230 | 8.91E-03 | 64 | rnr1D57N | FALSE | CAN | JSY3602 | 6 | rnr1D57N-Re | CAG | A>C | G/C+1 | A/T-1 | TA>GC | TA>GC | TGTAACAGG | AGGGATG | GTC | GTC | T | G | T>G | T>G_GTC | TA>GC_AGGGATG | 7 | FALSE | FALSE |
| NL3_v.txt | 33166 | 33166-SNV-T | SNV | 1 | T | G | 15 | 11228 | 0.13359459 | 38.8 | rnr1D57N | FALSE | CAN | JSY3602 | 6 | rnr1D57N-Re | GTA | T>G | G/C+1 | A/T-1 | TA>GC | TA>GC | CAATAGTAC | TACCACC | CAT | GTA | T | C | T>G | T>G_GTA | TA>GC_TACCACC | 7 | FALSE | FALSE |
| NL3_v.txt | 32445 | 32445-SNV- | SNV | 1 | A | G | 8 | 11034 | 0.07250317 | 47.875 | rnr1D57N | FALSE | CAN | JSY3602 | 6 | rnr1D57N-Re | AAG | A>G | G/C+1 | A/T-1 | TA>CG | TA>CG | ACTAAAAGT | AGTCCAA | TTC | TTC | T | C | T>C | T>C_TTC | TA>CG_AGTCCAA | 7 | FALSE | FALSE |
| NL3_v.txt | 33023 | 33023-SNV- | SNV | 1 | A | G | 9 | 11230 | 0.08014248 | 59.8888889 | rnr1D57N | FALSE | CAN | JSY3602 | 6 | rnr1D57N-Re | CAG | A>G | G/C+1 | A/T-1 | TA>CG | TA>CG | TGTAACAGG | AGGGATG | GTC | GTC | T | C | T>C | T>C_GTC | TA>CG_AGGGATG | 7 | FALSE | FALSE |
| NL3_v.txt | 31840 | 31840-SNV- | SNV | 1 | A | T | 3 | 6480 | 0.0462963 | 60.6666667 | rnr1D57N | FALSE | CAN | JSY3602 | 6 | rnr1D57N-Re | AAG | A>T | A/T+1 | A/T-1 | TA>AT | TA>AT | GAAATAAGA | AGATCCA | TTC | TTC | T | A | T>A | T>A_TTC | TA>AT_AGATCCA | 7 | FALSE | FALSE |
| NL3_v.txt | 32766 | 32766-SNV-T | SNV | 1 | T | A | 7 | 10674 | 0.06557991 | 56.7142857 | rnr1D57N | FALSE | CAN | JSY3602 | 6 | rnr1D57N-Re | TTT | T>A | A/T+1 | A/T-1 | TA>AT | TA>AT | AAAACTTTG | TTGATGG | AAA | TTT | T | T | T>A | T>A_TTT | TA>AT_TTGATGG | 7 | FALSE | FALSE |

| | | | | | | | | | | | | | | | | | | | | | | | | | | | | | | | | | | |
|---|---|---|---|---|---|---|---|---|---|---|---|---|---|---|---|---|---|---|---|---|---|---|---|---|---|---|---|---|---|---|---|---|---|---|
| NL3_v.txt | 33005 | 33005-SNV-T | SNV | 1 | T | A | 2 | 11230 | 0.01780944 | 58.5000285 | rnr1D57N | FALSE | CAN | JSY3602 | 6 | rnr1D57N-Re | CTG | T>A | A/T+1 | A/T-1 | TA>AT | TA>AT | GAAAACTGT | TGTGAAA | GAC | CTG | T | T | T>A | T>A_CTG | TA>AT_TGTGAAA | 7 | FALSE | FALSE |
| NL3_v.txt | 31650 | 31650-SNV-C | SNV | 1 | C | T | 16 | 6480 | 0.24691358 | 36.875 | rnr1D57N | FALSE | CAN | JSY3602 | 6 | rnr1D57N-Re | GCG | C>T | A/T+1 | G/C-1 | CG>TA | CG>TA | AGAATGCGA | CGAAATG | CGC | GCG | C | A | C>T | C>T_GCG | CG>TA_CGAAATG | 7 | FALSE | FALSE |
| NL3_v.txt | 31882 | 31882-SNV-C | SNV | 1 | C | T | 13 | 6480 | 0.20061728 | 36.6923077 | rnr1D57N | FALSE | CAN | JSY3602 | 6 | rnr1D57N-Re | GCA | C>T | A/T+1 | G/C-1 | CG>TA | CG>TA | AGGCGGCAG | CAGCAAA | CGT | GCA | C | A | C>T | C>T_GCA | CG>TA_CAGCAAA | 7 | FALSE | FALSE |
| NL3_v.txt | 31962 | 31962-SNV-C | SNV | 1 | G | A | 12 | 8348 | 0.14374701 | 34.8333333 | rnr1D57N | FALSE | CAN | JSY3602 | 6 | rnr1D57N-Re | CGC | G>A | A/T+1 | G/C-1 | CG>TA | CG>TA | GTGGCCGCA | GCATAAT | GCG | GCG | C | T | C>T | C>T_GCG | CG>TA_GCATAAT | 7 | FALSE | FALSE |
| NL3_v.txt | 32271 | 32271-SNV-C | SNV | 1 | C | T | 14 | 11034 | 0.12688055 | 37.1428571 | rnr1D57N | FALSE | CAN | JSY3602 | 6 | rnr1D57N-Re | ACG | C>T | A/T+1 | G/C-1 | CG>TA | CG>TA | AAAATACGG | CGGGAAC | TGC | ACG | C | A | C>T | C>T_ACG | CG>TA_CGGGAAC | 7 | FALSE | FALSE |
| NL3_v.txt | 32299 | 32299-SNV-C | SNV | 1 | C | T | 15 | 11034 | 0.13594345 | 36.4 | rnr1D57N | FALSE | CAN | JSY3602 | 6 | rnr1D57N-Re | GCG | C>T | A/T+1 | G/C-1 | CG>TA | CG>TA | AATTTGCGG | CGGCAGA | CGC | GCG | C | A | C>T | C>T_GCG | CG>TA_CGGCAGA | 7 | FALSE | FALSE |
| NL3_v.txt | 32449 | 32449-SNV-C | SNV | 1 | C | T | 28 | 11034 | 0.2537611 | 57.3571429 | rnr1D57N | FALSE | CAN | JSY3602 | 6 | rnr1D57N-Re | CCA | C>T | A/T+1 | G/C-1 | CG>TA | CG>TA | AAAGTCCAA | CAATGAA | GGT | CCA | C | A | C>T | C>T_CCA | CG>TA_CAATGAA | 7 | FALSE | FALSE |
| NL3_v.txt | 32590 | 32590-SNV-C | SNV | 1 | C | T | 15 | 10674 | 0.14052839 | 36.6666667 | rnr1D57N | FALSE | CAN | JSY3602 | 6 | rnr1D57N-Re | GCG | C>T | A/T+1 | G/C-1 | CG>TA | CG>TA | AGGCAGCGT | CGTTAAT | CGC | GCG | C | A | C>T | C>T_GCG | CG>TA_CGTTAAT | 7 | FALSE | FALSE |
| NL3_v.txt | 32700 | 32700-SNV-C | SNV | 1 | C | T | 20 | 10674 | 0.18737118 | 61.05 | rnr1D57N | FALSE | CAN | JSY3602 | 6 | rnr1D57N-Re | GCC | C>T | A/T+1 | G/C-1 | CG>TA | CG>TA | ACTGGGCCG | CCGGTAA | CGG | GCC | C | A | C>T | C>T_GCC | CG>TA_CCGGTAA | 7 | FALSE | FALSE |
| NL3_v.txt | 32797 | 32797-SNV-C | SNV | 1 | C | T | 5 | 10674 | 0.0468428 | 34.4 | rnr1D57N | FALSE | CAN | JSY3602 | 6 | rnr1D57N-Re | CCG | C>T | A/T+1 | G/C-1 | CG>TA | CG>TA | ATTCACCGTA | CGTAATA | GGC | CCG | C | A | C>T | C>T_CCG | CG>TA_CGTAATA | 7 | FALSE | FALSE |
| NL3_v.txt | 32798 | 32798-SNV-C | SNV | 1 | G | A | 13 | 10674 | 0.12179127 | 39 | rnr1D57N | FALSE | CAN | JSY3602 | 6 | rnr1D57N-Re | CGT | G>A | A/T+1 | G/C-1 | CG>TA | CG>TA | TTCACCGTAA | GTAATAT | GCA | GCA | C | T | C>T | C>T_GCA | CG>TA_GTAATAT | 7 | FALSE | FALSE |
| NL3_v.txt | 32855 | 32855-SNV-C | SNV | 1 | C | T | 18 | 10674 | 0.16863406 | 35.1111111 | rnr1D57N | FALSE | CAN | JSY3602 | 6 | rnr1D57N-Re | TCC | C>T | A/T+1 | G/C-1 | CG>TA | CG>TA | ACTAATCCAT | CCATGCC | AGG | TCC | C | A | C>T | C>T_TCC | CG>TA_CCATGCC | 7 | FALSE | FALSE |
| NL3_v.txt | 32860 | 32860-SNV-C | SNV | 1 | C | T | 11 | 10674 | 0.10305415 | 37.4545455 | rnr1D57N | FALSE | CAN | JSY3602 | 6 | rnr1D57N-Re | GCC | C>T | A/T+1 | G/C-1 | CG>TA | CG>TA | TCCATGCCG | CCGCCAG | CGG | GCC | C | A | C>T | C>T_GCC | CG>TA_CCGCCAG | 7 | FALSE | FALSE |
| NL3_v.txt | 32901 | 32901-SNV-C | SNV | 1 | C | T | 22 | 11230 | 0.19590383 | 36.4090909 | rnr1D57N | FALSE | CAN | JSY3602 | 6 | rnr1D57N-Re | GCC | C>T | A/T+1 | G/C-1 | CG>TA | CG>TA | ACTTGGCCA | CCAACTA | CGG | GCC | C | A | C>T | C>T_GCC | CG>TA_CCAACTA | 7 | FALSE | FALSE |
| NL3_v.txt | 32922 | 32922-SNV-C | SNV | 1 | G | A | 13 | 11230 | 0.11576135 | 34.3076923 | rnr1D57N | FALSE | CAN | JSY3602 | 6 | rnr1D57N-Re | GGC | G>A | A/T+1 | G/C-1 | CG>TA | CG>TA | TCCAGGGCA | GCAAAAG | CCG | CCG | C | T | C>T | C>T_CCG | CG>TA_GCAAAAG | 7 | FALSE | FALSE |
| NL3_v.txt | 32967 | 32967-SNV-C | SNV | 1 | G | A | 23 | 11230 | 0.20480855 | 61.3043478 | rnr1D57N | FALSE | CAN | JSY3602 | 6 | rnr1D57N-Re | CGC | G>A | A/T+1 | G/C-1 | CG>TA | CG>TA | TTGGCCGCA | GCACCAA | GCG | GCG | C | T | C>T | C>T_GCG | CG>TA_GCACCAA | 7 | FALSE | FALSE |
| NL3_v.txt | 33006 | 33006-SNV-C | SNV | 1 | G | A | 1 | 11230 | 8.91E-03 | 11.0000007 | rnr1D57N | FALSE | CAN | JSY3602 | 6 | rnr1D57N-Re | TGT | G>A | A/T+1 | G/C-1 | CG>TA | CG>TA | AAAACTGTG | GTGAAAG | ACA | ACA | C | T | C>T | C>T_ACA | CG>TA_GTGAAAG | 7 | FALSE | FALSE |
| NL3_v.txt | 33056 | 33056-SNV-C | SNV | 1 | C | T | 15 | 11230 | 0.13357079 | 53.4666667 | rnr1D57N | FALSE | CAN | JSY3602 | 6 | rnr1D57N-Re | GCG | C>T | A/T+1 | G/C-1 | CG>TA | CG>TA | GGACTGCGT | CGTGACA | CGC | GCG | C | A | C>T | C>T_GCG | CG>TA_CGTGACA | 7 | FALSE | FALSE |
| NL3_v.txt | 33070 | 33070-SNV-C | SNV | 1 | C | T | 18 | 11230 | 0.16028495 | 53.6666667 | rnr1D57N | FALSE | CAN | JSY3602 | 6 | rnr1D57N-Re | GCC | C>T | A/T+1 | G/C-1 | CG>TA | CG>TA | AATATGCCA | CCAAAGA | CGG | GCC | C | A | C>T | C>T_GCC | CG>TA_CCAAAGA | 7 | FALSE | FALSE |
| NL3_v.txt | 33113 | 33113-SNV-C | SNV | 1 | G | A | 15 | 11230 | 0.13357079 | 36.3333333 | rnr1D57N | FALSE | CAN | JSY3602 | 6 | rnr1D57N-Re | GGC | G>A | A/T+1 | G/C-1 | CG>TA | CG>TA | CACTGGGCC | GCCGGCG | CCG | CCG | C | T | C>T | C>T_CCG | CG>TA_GCCGGCG | 7 | FALSE | FALSE |
| NL3_v.txt | 33114 | 33114-SNV-C | SNV | 1 | C | T | 20 | 11230 | 0.17809439 | 37.65 | rnr1D57N | FALSE | CAN | JSY3602 | 6 | rnr1D57N-Re | GCC | C>T | A/T+1 | G/C-1 | CG>TA | CG>TA | ACTGGGCCG | CCGGCGT | CGG | GCC | C | A | C>T | C>T_GCC | CG>TA_CCGGCGT | 7 | FALSE | FALSE |
| NL3_v.txt | 33276 | 33276-SNV-C | SNV | 1 | G | A | 14 | 11357 | 0.12327199 | 53.4285714 | rnr1D57N | FALSE | CAN | JSY3602 | 6 | rnr1D57N-Re | CGT | G>A | A/T+1 | G/C-1 | CG>TA | CG>TA | GAGAACGTA | GTATCCT | GCA | GCA | C | T | C>T | C>T_GCA | CG>TA_GTATCCT | 7 | FALSE | FALSE |
| NL3_v.txt | 33310 | 33310-SNV-C | SNV | 1 | G | A | 15 | 11357 | 0.13207713 | 46.8666667 | rnr1D57N | FALSE | CAN | JSY3602 | 6 | rnr1D57N-Re | CGC | G>A | A/T+1 | G/C-1 | CG>TA | CG>TA | AAGAGCGCA | GCAATGG | GCG | GCG | C | T | C>T | C>T_GCG | CG>TA_GCAATGG | 7 | FALSE | FALSE |
| NL3_v.txt | 33392 | 33392-SNV-C | SNV | 1 | G | A | 17 | 11357 | 0.14968742 | 51.7647059 | rnr1D57N | FALSE | CAN | JSY3602 | 6 | rnr1D57N-Re | CGT | G>A | A/T+1 | G/C-1 | CG>TA | CG>TA | AACGTCGTG | GTGAAAG | GCA | GCA | C | T | C>T | C>T_GCA | CG>TA_GTGAAAG | 7 | FALSE | FALSE |
| NL3_v.txt | 33474 | 33474-SNV-C | SNV | 1 | C | T | 15 | 11355 | 0.1321004 | 34.0666667 | rnr1D57N | FALSE | CAN | JSY3602 | 6 | rnr1D57N-Re | GCC | C>T | A/T+1 | G/C-1 | CG>TA | CG>TA | GCTATGCCTT | CCTTTTT | CGG | GCC | C | A | C>T | C>T_GCC | CG>TA_CCTTTTT | 7 | FALSE | FALSE |
| NL3_v.txt | 31956 | 31956-SNV-C | SNV | 1 | G | C | 6 | 8348 | 0.0718735 | 34.1666667 | rnr1D57N | FALSE | CAN | JSY3602 | 6 | rnr1D57N-Re | TGT | G>C | G/C+1 | G/C-1 | CG>GC | CG>GC | ATAAATGTG | GTGGCCG | ACA | ACA | C | G | C>G | C>G_ACA | CG>GC_GTGGCCG | 7 | FALSE | FALSE |
| NL3_v.txt | 32001 | 32001-SNV-C | SNV | 1 | G | C | 1 | 8348 | 0.01197892 | 38.9995535 | rnr1D57N | FALSE | CAN | JSY3602 | 6 | rnr1D57N-Re | TGG | G>C | G/C+1 | G/C-1 | CG>GC | CG>GC | TTAAATGGT | GGTAACT | ACC | ACC | C | G | C>G | C>G_ACC | CG>GC_GGTAACT | 7 | FALSE | FALSE |
| NL3_v.txt | 32027 | 32027-SNV-C | SNV | 1 | G | C | 4 | 8348 | 0.04791567 | 63.25 | rnr1D57N | FALSE | CAN | JSY3602 | 6 | rnr1D57N-Re | GGT | G>C | G/C+1 | G/C-1 | CG>GC | CG>GC | GCCACGGTA | GTATTTC | CCA | CCA | C | G | C>G | C>G_CCA | CG>GC_GTATTTC | 7 | FALSE | FALSE |
| NL3_v.txt | 32064 | 32064-SNV-C | SNV | 1 | G | C | 11 | 8348 | 0.13176809 | 58.3636364 | rnr1D57N | FALSE | CAN | JSY3602 | 6 | rnr1D57N-Re | TGA | G>C | G/C+1 | G/C-1 | CG>GC | CG>GC | GAGATTGAG | GAGATAA | ACT | ACT | C | G | C>G | C>G_ACT | CG>GC_GAGATAA | 7 | FALSE | FALSE |
| NL3_v.txt | 32449 | 32449-SNV-C | SNV | 1 | C | G | 2 | 11034 | 0.01812579 | 58 | rnr1D57N | FALSE | CAN | JSY3602 | 6 | rnr1D57N-Re | CCA | C>G | G/C+1 | G/C-1 | CG>GC | CG>GC | AAAGTCCAA | CAATGAA | GGT | CCA | C | C | C>G | C>G_CCA | CG>GC_CAATGAA | 7 | FALSE | FALSE |
| NL3_v.txt | 32876 | 32876-SNV-C | SNV | 1 | G | C | 22 | 11223 | 0.19602602 | 21.6818182 | rnr1D57N | FALSE | CAN | JSY3602 | 6 | rnr1D57N-Re | TGT | G>C | G/C+1 | G/C-1 | CG>GC | CG>GC | AACTTTGTAC | GTACGTC | ACA | ACA | C | G | C>G | C>G_ACA | CG>GC_GTACGTC | 7 | FALSE | FALSE |
| NL3_v.txt | 32945 | 32945-SNV-C | SNV | 1 | C | G | 13 | 11230 | 0.11576135 | 34.7692308 | rnr1D57N | FALSE | CAN | JSY3602 | 6 | rnr1D57N-Re | ACC | C>G | G/C+1 | G/C-1 | CG>GC | CG>GC | AGAAAACCA | CCAATAC | TGG | ACC | C | C | C>G | C>G_ACC | CG>GC_CCAATAC | 7 | FALSE | FALSE |
| NL3_v.txt | 32997 | 32997-SNV-C | SNV | 1 | G | C | 4 | 11230 | 0.03561888 | 64 | rnr1D57N | FALSE | CAN | JSY3602 | 6 | rnr1D57N-Re | TGA | G>C | G/C+1 | G/C-1 | CG>GC | CG>GC | CTTTGTGAG | GAGAAAA | ACT | ACT | C | G | C>G | C>G_ACT | CG>GC_GAGAAAA | 7 | FALSE | FALSE |
| NL3_v.txt | 33018 | 33018-SNV-C | SNV | 1 | G | C | 7 | 11230 | 0.06233304 | 64 | rnr1D57N | FALSE | CAN | JSY3602 | 6 | rnr1D57N-Re | TGT | G>C | G/C+1 | G/C-1 | CG>GC | CG>GC | GAGGATGTA | GTAACAG | ACA | ACA | C | G | C>G | C>G_ACA | CG>GC_GTAACAG | 7 | FALSE | FALSE |
| NL3_v.txt | 33169 | 33169-SNV-C | SNV | 1 | C | G | 5 | 11227 | 0.0445355 | 38.8 | rnr1D57N | FALSE | CAN | JSY3602 | 6 | rnr1D57N-Re | CCA | C>G | G/C+1 | G/C-1 | CG>GC | CG>GC | TAGTACCAC | CACCAAG | GGT | CCA | C | C | C>G | C>G_CCA | CG>GC_CACCAAG | 7 | FALSE | FALSE |
| NL3_v.txt | 33205 | 33205-SNV-C | SNV | 1 | G | C | 35 | 11357 | 0.30817998 | 37.6571429 | rnr1D57N | FALSE | CAN | JSY3602 | 6 | rnr1D57N-Re | AGC | G>C | G/C+1 | G/C-1 | CG>GC | CG>GC | GCTTAAGCT | GCTCTCT | TCG | TCG | C | G | C>G | C>G_TCG | CG>GC_GCTCTCT | 7 | FALSE | FALSE |
| NL3_v.txt | 33456 | 33456-SNV-C | SNV | 1 | G | C | 4 | 11355 | 0.03522677 | 39 | rnr1D57N | FALSE | CAN | JSY3602 | 6 | rnr1D57N-Re | TGA | G>C | G/C+1 | G/C-1 | CG>GC | CG>GC | TCTTTTGAAT | GAATTTG | ACT | ACT | C | G | C>G | C>G_ACT | CG>GC_GAATTTG | 7 | FALSE | FALSE |
| NL3_v.txt | 32368 | 32368-SNV-C | SNV | 1 | C | A | 23 | 11034 | 0.20844662 | 63.5652174 | rnr1D57N | FALSE | CAN | JSY3602 | 6 | rnr1D57N-Re | TCA | C>A | A/T+1 | G/C-1 | CG>AT | CG>AT | AGTTCTCAAT | CAATAGC | AGT | TCA | C | T | C>A | C>A_TCA | CG>AT_CAATAGC | 7 | FALSE | FALSE |
| NL3_v.txt | 32399 | 32399-SNV-C | SNV | 1 | G | T | 10 | 11034 | 0.09062897 | 64 | rnr1D57N | FALSE | CAN | JSY3602 | 6 | rnr1D57N-Re | CGT | G>T | A/T+1 | G/C-1 | CG>AT | CG>AT | AGAAACGTA | GTAGGAA | GCA | GCA | C | A | C>A | C>A_GCA | CG>AT_GTAGGAA | 7 | FALSE | FALSE |
| NL3_v.txt | 32432 | 32432-SNV-C | SNV | 1 | G | T | 11 | 11034 | 0.09969186 | 61.7272727 | rnr1D57N | FALSE | CAN | JSY3602 | 6 | rnr1D57N-Re | TGT | G>T | A/T+1 | G/C-1 | CG>AT | CG>AT | GTCATTGTAT | GTATGGA | ACA | ACA | C | A | C>A | C>A_ACA | CG>AT_GTATGGA | 7 | FALSE | FALSE |
| NL3_v.txt | 32629 | 32629-SNV-C | SNV | 1 | C | A | 7 | 10674 | 0.06557991 | 35.4286044 | rnr1D57N | FALSE | CAN | JSY3602 | 6 | rnr1D57N-Re | TCG | C>A | A/T+1 | G/C-1 | CG>AT | CG>AT | TCCCTTCGTT | CGTTTTT | AGC | TCG | C | T | C>A | C>A_TCG | CG>AT_CGTTTTT | 7 | FALSE | FALSE |
| NL3_v.txt | 32735 | 32735-SNV-C | SNV | 1 | G | T | 14 | 10674 | 0.13115983 | 61.5 | rnr1D57N | FALSE | CAN | JSY3602 | 6 | rnr1D57N-Re | AGT | G>T | A/T+1 | G/C-1 | CG>AT | CG>AT | AAAACAGTA | GTATATT | TCA | TCA | C | A | C>A | C>A_TCA | CG>AT_GTATATT | 7 | FALSE | FALSE |
| NL3_v.txt | 32797 | 32797-SNV-C | SNV | 1 | C | A | 10 | 10674 | 0.09368559 | 39 | rnr1D57N | FALSE | CAN | JSY3602 | 6 | rnr1D57N-Re | CCG | C>A | A/T+1 | G/C-1 | CG>AT | CG>AT | ATTCACCGTA | CGTAATA | GGC | CCG | C | T | C>A | C>A_CCG | CG>AT_CGTAATA | 7 | FALSE | FALSE |
| NL3_v.txt | 32798 | 32798-SNV-C | SNV | 1 | G | T | 4 | 10674 | 0.03747424 | 38.75 | rnr1D57N | FALSE | CAN | JSY3602 | 6 | rnr1D57N-Re | CGT | G>T | A/T+1 | G/C-1 | CG>AT | CG>AT | TTCACCGTAA | GTAATAT | GCA | GCA | C | A | C>A | C>A_GCA | CG>AT_GTAATAT | 7 | FALSE | FALSE |
| NL3_v.txt | 32997 | 32997-SNV-C | SNV | 1 | G | T | 6 | 11230 | 0.05342832 | 64 | rnr1D57N | FALSE | CAN | JSY3602 | 6 | rnr1D57N-Re | TGA | G>T | A/T+1 | G/C-1 | CG>AT | CG>AT | CTTTGTGAG | GAGAAAA | ACT | ACT | C | A | C>A | C>A_ACT | CG>AT_GAGAAAA | 7 | FALSE | FALSE |
| NL3_v.txt | 33024 | 33024-SNV-C | SNV | 1 | G | T | 16 | 11230 | 0.14247551 | 61.5625536 | rnr1D57N | FALSE | CAN | JSY3602 | 6 | rnr1D57N-Re | AGG | G>T | A/T+1 | G/C-1 | CG>AT | CG>AT | GTAACAGGG | GGGATGA | TCC | TCC | C | A | C>A | C>A_TCC | CG>AT_GGGATGA | 7 | FALSE | FALSE |
| NL3_v.txt | 33151 | 33151-SNV-C | SNV | 1 | G | T | 11 | 11230 | 0.09795192 | 35.1818182 | rnr1D57N | FALSE | CAN | JSY3602 | 6 | rnr1D57N-Re | AGA | G>T | A/T+1 | G/C-1 | CG>AT | CG>AT | TGAAAAGAC | GACCTGT | TCT | TCT | C | A | C>A | C>A_TCT | CG>AT_GACCTGT | 7 | FALSE | FALSE |
| NL3_v.txt | 33186 | 33186-SNV-C | SNV | 1 | C | A | 57 | 11350 | 0.50220264 | 27.7894737 | rnr1D57N | FALSE | CAN | JSY3602 | 6 | rnr1D57N-Re | ACC | C>A | A/T+1 | G/C-1 | CG>AT | CG>AT | ATCATACCA | CCAATAT | TGG | ACC | C | T | C>A | C>A_ACC | CG>AT_CCAATAT | 7 | FALSE | FALSE |
| NL3_v.txt | 33474 | 33474-SNV-C | SNV | 1 | C | A | 13 | 11355 | 0.11448701 | 36.6153846 | rnr1D57N | FALSE | CAN | JSY3602 | 6 | rnr1D57N-Re | GCC | C>A | A/T+1 | G/C-1 | CG>AT | CG>AT | GCTATGCCTT | CCTTTTT | CGG | GCC | C | T | C>A | C>A_GCC | CG>AT_CCTTTTT | 7 | FALSE | FALSE |
| NL3_v.txt | 32092 | 32092-Repla | Replacement | 2 | CA | T | 5 | 8348 | 0.05989459 | 57.5999997 | rnr1D57N | FALSE | CAN | JSY3602 | 6 | rnr1D57N-Re | GCA | nope | A/T+1 | >1bpdel | none | Replac. | AGCCTGCAA | CAACACC | CGT | GCA | CA | A | NA | NA_GCA | Replac._CAACACC | 7 | FALSE | FALSE |
| NL3_v.txt | 32001 | 32001-MNV- | MNV | 2 | GG | TT | 8 | 8348 | 0.09583134 | 38.8124845 | rnr1D57N | FALSE | CAN | JSY3602 | 6 | rnr1D57N-Re | TGG | nope | >1bp-ins | >1bpdel | none | MNV | TTAAATGGT | GGTAACT | ACC | TGG | GG | none | NA | NA_TGG | MNV_GGTAACT | 7 | FALSE | FALSE |
| NL3_v.txt | 32641 | 32641-MNV- | MNV | 2 | TA | AT | 10 | 10674 | 0.09368559 | 39.4919265 | rnr1D57N | FALSE | CAN | JSY3602 | 6 | rnr1D57N-Re | TTA | nope | >1bp-ins | >1bpdel | none | MNV | TATCCTTAGA | TAGATAT | AAT | TTA | TA | none | NA | NA_TTA | MNV_TAGATAT | 7 | FALSE | FALSE |
| NL3_v.txt | 31997 | 31997-Insert | Insertion | 1 | - | A | 115 | 8348 | 1.37757547 | 38.4608696 | rnr1D57N | FALSE | CAN | JSY3602 | 6 | rnr1D57N-Re | TAA | nope | A/T+1 | >1bpdel | none | A/T+1 | AGCTTTAAAT | AAATGGT | ATT | TAA | - | T | NA | NA_TAA | A/T+1_AAATGGT | 7 | FALSE | FALSE |
| NL3_v.txt | 32081 | 32081-Insert | Insertion | 1 | - | A | 60 | 8348 | 0.71873503 | 64 | rnr1D57N | FALSE | CAN | JSY3602 | 6 | rnr1D57N-Re | CAA | nope | A/T+1 | >1bpdel | none | A/T+1 | CCATGCAAA | AAAAAAG | GTT | CAA | - | T | NA | NA_CAA | A/T+1_AAAAAAG | 7 | FALSE | FALSE |
| NL3_v.txt | 32210 | 32210-Insert | Insertion | 1 | - | T | 2 | 8348 | 0.02395783 | 39 | rnr1D57N | FALSE | CAN | JSY3602 | 6 | rnr1D57N-Re | CTT | nope | A/T+1 | >1bpdel | none | A/T+1 | ACCACCTTTG | TTTGGTG | GAA | CTT | - | A | NA | NA_CTT | A/T+1_TTTGGTG | 7 | FALSE | FALSE |
| NL3_v.txt | 32604 | 32604-Insert | Insertion | 1 | - | C | 5 | 10674 | 0.0468428 | 34 | rnr1D57N | FALSE | CAN | JSY3602 | 6 | rnr1D57N-Re | GGA | nope | G/C+1 | >1bpdel | none | G/C+1 | AAAGAGGAA | GAAACCC | CCT | GGA | - | G | NA | NA_GGA | G/C+1_GAAACCC | 7 | FALSE | FALSE |
| NL3_v.txt | 32631 | 32631-Insert | Insertion | 1 | - | T | 2 | 10674 | 0.01873712 | 38.5 | rnr1D57N | FALSE | CAN | JSY3602 | 6 | rnr1D57N-Re | GTT | nope | A/T+1 | >1bpdel | none | A/T+1 | CCTTCGTTTT | TTTTTAT | CAA | GTT | - | A | NA | NA_GTT | A/T+1_TTTTTAT | 7 | FALSE | FALSE |
| NL3_v.txt | 32842 | 32842-Insert | Insertion | 1 | - | A | 24 | 10674 | 0.22484542 | 37.7083333 | rnr1D57N | FALSE | CAN | JSY3602 | 6 | rnr1D57N-Re | CAA | nope | A/T+1 | >1bpdel | none | A/T+1 | TTACCCAAA | AAAAAAT | GTT | CAA | - | T | NA | NA_CAA | A/T+1_AAAAAAT | 7 | FALSE | FALSE |
| NL3_v.txt | 32991 | 32991-Insert | Insertion | 1 | - | A | 3 | 11230 | 0.02671416 | 60 | rnr1D57N | FALSE | CAN | JSY3602 | 6 | rnr1D57N-Re | TCT | nope | A/T+1 | >1bpdel | none | A/T+1 | AGGAATCTT | CTTTGTG | AGA | TCT | - | T | NA | NA_TCT | A/T+1_CTTTGTG | 7 | FALSE | FALSE |
| NL3_v.txt | 33161 | 33161-Insert | Insertion | 1 | - | A | 3 | 11229 | 0.02671654 | 38.6666667 | rnr1D57N | FALSE | CAN | JSY3602 | 6 | rnr1D57N-Re | CAA | nope | A/T+1 | >1bpdel | none | A/T+1 | TGTACCAATA | AATAGTA | GTT | CAA | - | T | NA | NA_CAA | A/T+1_AATAGTA | 7 | FALSE | FALSE |
| NL3_v.txt | 32629 | 32629-Deleti | Deletion | 2 | CG | - | 10 | 10674 | 0.09368559 | 36.3 | rnr1D57N | FALSE | CAN | JSY3602 | 6 | rnr1D57N-Re | TCG | nope | >1bp-ins | >1bpdel | none | >1bpdel | TCCCTTCGTT | CGTTTTT | AGC | TCG | CG | none | NA | NA_TCG | >1bpdel_CGTTTTT | 7 | FALSE | FALSE |
| NL3_v.txt | 33005 | 33005-Deleti | Deletion | 2 | TG | - | 16 | 11230 | 0.14247551 | 62.8125 | rnr1D57N | FALSE | CAN | JSY3602 | 6 | rnr1D57N-Re | CTG | nope | >1bp-ins | >1bpdel | none | >1bpdel | GAAAACTGT | TGTGAAA | GAC | CTG | TG | none | NA | NA_CTG | >1bpdel_TGTGAAA | 7 | FALSE | FALSE |
| NL3_v.txt | 31711 | 31711-Deleti | Deletion | 1 | A | - | 51 | 6480 | 0.78703704 | 32.3921569 | rnr1D57N | FALSE | CAN | JSY3602 | 6 | rnr1D57N-Re | CAA | nope | >1bp-ins | A/T-1 | none | A/T-1 | CATTCCAAA | AAAATTT | GTT | GTT | T | none | T>none | T>none_GTT | A/T-1_AAAATTT | 7 | FALSE | FALSE |
| NL3_v.txt | 31723 | 31723-Deleti | Deletion | 1 | A | - | 5 | 6480 | 0.07716049 | 64 | rnr1D57N | FALSE | CAN | JSY3602 | 6 | rnr1D57N-Re | CAA | nope | >1bp-ins | A/T-1 | none | A/T-1 | TGTCCCAAA | AAAAAGT | GTT | GTT | T | none | T>none | T>none_GTT | A/T-1_AAAAAGT | 7 | FALSE | FALSE |
| NL3_v.txt | 32001 | 32001-Deleti | Deletion | 1 | G | - | 1 | 8348 | 0.01197892 | 38 | rnr1D57N | FALSE | CAN | JSY3602 | 6 | rnr1D57N-Re | TGG | nope | >1bp-ins | G/C-1 | none | G/C-1 | TTAAATGGT | GGTAACT | ACC | ACC | C | none | C>none | C>none_ACC | G/C-1_GGTAACT | 7 | FALSE | FALSE |
| NL3_v.txt | 32081 | 32081-Deleti | Deletion | 1 | A | - | 6 | 8348 | 0.0718735 | 64 | rnr1D57N | FALSE | CAN | JSY3602 | 6 | rnr1D57N-Re | CAA | nope | >1bp-ins | A/T-1 | none | A/T-1 | CCATGCAAA | AAAAAAG | GTT | GTT | T | none | T>none | T>none_GTT | A/T-1_AAAAAAG | 7 | FALSE | FALSE |
| NL3_v.txt | 32093 | 32093-Deleti | Deletion | 1 | A | - | 6084 | 8348 | 72.8797317 | 61.5026299 | rnr1D57N | FALSE | CAN | JSY3602 | 6 | rnr1D57N-Re | CAA | nope | >1bp-ins | A/T-1 | none | A/T-1 | GCCTGCAAC | AACACCA | GTT | GTT | T | none | T>none | T>none_GTT | A/T-1_AACACCA | 7 | FALSE | FALSE |
| NL3_v.txt | 32490 | 32490-Deleti | Deletion | 1 | A | - | 5 | 11034 | 0.04531448 | 38.6 | rnr1D57N | FALSE | CAN | JSY3602 | 6 | rnr1D57N-Re | GAA | nope | >1bp-ins | A/T-1 | none | A/T-1 | ATACGGAAA | AAAACAA | CTT | CTT | T | none | T>none | T>none_CTT | A/T-1_AAAACAA | 7 | FALSE | FALSE |
| NL3_v.txt | 32498 | 32498-Deleti | Deletion | 1 | T | - | 12 | 11034 | 0.10875476 | 37.75 | rnr1D57N | FALSE | CAN | JSY3602 | 6 | rnr1D57N-Re | CTT | nope | >1bp-ins | A/T-1 | none | A/T-1 | AACAACTTTT | TTTTTTG | GAA | CTT | T | none | nope | nope_CTT | A/T-1_TTTTTTG | 7 | FALSE | FALSE |
| NL3_v.txt | 32658 | 32658-Deleti | Deletion | 1 | C | - | 3 | 10674 | 0.02810568 | 64 | rnr1D57N | FALSE | CAN | JSY3602 | 6 | rnr1D57N-Re | ACC | nope | >1bp-ins | G/C-1 | none | G/C-1 | CCTGGACCC | CCCCAGG | TGG | ACC | C | none | nope | nope_ACC | G/C-1_CCCCAGG | 7 | FALSE | FALSE |
| NL3_v.txt | 32707 | 32707-Deleti | Deletion | 1 | C | - | 4 | 10674 | 0.03747424 | 64 | rnr1D57N | FALSE | CAN | JSY3602 | 6 | rnr1D57N-Re | ACC | nope | >1bp-ins | G/C-1 | none | G/C-1 | CGGTAACCC | CCCCAGC | TGG | ACC | C | none | nope | nope_ACC | G/C-1_CCCCAGC | 7 | FALSE | FALSE |
| NL3_v.txt | 32728 | 32728-Deleti | Deletion | 1 | A | - | 6 | 10674 | 0.05621136 | 49.1666667 | rnr1D57N | FALSE | CAN | JSY3602 | 6 | rnr1D57N-Re | CAA | nope | >1bp-ins | A/T-1 | none | A/T-1 | CCATACAAA | AAAAACA | GTT | GTT | T | none | T>none | T>none_GTT | A/T-1_AAAAACA | 7 | FALSE | FALSE |
| NL3_v.txt | 32796 | 32796-Deleti | Deletion | 1 | C | - | 4 | 10674 | 0.03747424 | 37.75 | rnr1D57N | FALSE | CAN | JSY3602 | 6 | rnr1D57N-Re | ACC | nope | >1bp-ins | G/C-1 | none | G/C-1 | AATTCACCGT | CCGTAAT | TGG | ACC | C | none | nope | nope_ACC | G/C-1_CCGTAAT | 7 | FALSE | FALSE |
| NL3_v.txt | 32842 | 32842-Deleti | Deletion | 1 | A | - | 9 | 10674 | 0.08431703 | 37.4444444 | rnr1D57N | FALSE | CAN | JSY3602 | 6 | rnr1D57N-Re | CAA | nope | >1bp-ins | A/T-1 | none | A/T-1 | TTACCCAAA | AAAAAAT | GTT | GTT | T | none | T>none | T>none_GTT | A/T-1_AAAAAAT | 7 | FALSE | FALSE |
| NL3_v.txt | 32896 | 32896-Deleti | Deletion | 1 | C | - | 11 | 11230 | 0.09795192 | 37 | rnr1D57N | FALSE | CAN | JSY3602 | 6 | rnr1D57N-Re | ACT | nope | >1bp-ins | G/C-1 | none | G/C-1 | GAATGACTT | CTTGGCC | TGA | ACT | C | none | nope | nope_ACT | G/C-1_CTTGGCC | 7 | FALSE | FALSE |
| NL3_v.txt | 32924 | 32924-Deleti | Deletion | 1 | A | - | 14 | 11230 | 0.12466607 | 37.5 | rnr1D57N | FALSE | CAN | JSY3602 | 6 | rnr1D57N-Re | CAA | nope | >1bp-ins | A/T-1 | none | A/T-1 | CAGGGCAAA | AAAAGTG | GTT | GTT | T | none | T>none | T>none_GTT | A/T-1_AAAAGTG | 7 | FALSE | FALSE |
| NL3_v.txt | 32941 | 32941-Deleti | Deletion | 1 | A | - | 15 | 11230 | 0.13357079 | 33.7333333 | rnr1D57N | FALSE | CAN | JSY3602 | 6 | rnr1D57N-Re | GAA | nope | >1bp-ins | A/T-1 | none | A/T-1 | CCCAAGAAA | AAAACCA | CTT | CTT | T | none | T>none | T>none_CTT | A/T-1_AAAACCA | 7 | FALSE | FALSE |
| NL3_v.txt | 33147 | 33147-Deleti | Deletion | 1 | A | - | 6 | 11230 | 0.05342832 | 39 | rnr1D57N | FALSE | CAN | JSY3602 | 6 | rnr1D57N-Re | GAA | nope | >1bp-ins | A/T-1 | none | A/T-1 | CCAATGAAA | AAAAGAC | CTT | CTT | T | none | T>none | T>none_CTT | A/T-1_AAAAGAC | 7 | FALSE | FALSE |
| NL3_v.txt | 33251 | 33251-Deleti | Deletion | 1 | A | - | 32 | 11357 | 0.28176455 | 61.75 | rnr1D57N | FALSE | CAN | JSY3602 | 6 | rnr1D57N-Re | CAC | nope | >1bp-ins | A/T-1 | none | A/T-1 | TTCATCACCT | ACCTATG | GTG | GTG | T | none | T>none | T>none_GTG | A/T-1_ACCTATG | 7 | FALSE | FALSE |
| NL3_v.txt | 33445 | 33445-Deleti | Deletion | 1 | C | - | 9 | 11357 | 0.07924628 | 35.3333333 | rnr1D57N | FALSE | CAN | JSY3602 | 6 | rnr1D57N-Re | GCG | nope | >1bp-ins | G/C-1 | none | G/C-1 | TGTCGGCGT | CGTCTTC | CGC | GCG | C | none | nope | nope_GCG | G/C-1_CGTCTTC | 7 | FALSE | FALSE |
| NL3_v.txt | 33474 | 33474-Deleti | Deletion | 1 | C | - | 27 | 11355 | 0.23778071 | 32.6666667 | rnr1D57N | FALSE | CAN | JSY3602 | 6 | rnr1D57N-Re | GCC | nope | >1bp-ins | G/C-1 | none | G/C-1 | GCTATGCCTT | CCTTTTT | CGG | GCC | C | none | nope | nope_GCC | G/C-1_CCTTTTT | 7 | FALSE | FALSE |
| NL4_v.txt | 31723 | 31723-SNV-A | SNV | 1 | A | C | 1 | 5292 | 0.01889645 | 64 | rnr1D57N | FALSE | CAN | JSY3602 | 7 | rnr1D57N-Re | CAA | A>C | G/C+1 | A/T-1 | TA>GC | TA>GC | TGTCCCAAA | AAAAAGT | GTT | GTT | T | G | T>G | T>G_GTT | TA>GC_AAAAAGT | 7 | FALSE | FALSE |
| NL4_v.txt | 32256 | 32256-SNV-A | SNV | 1 | A | C | 50 | 13146 | 0.38034383 | 18.98 | rnr1D57N | FALSE | CAN | JSY3602 | 7 | rnr1D57N-Re | TAG | A>C | G/C+1 | A/T-1 | TA>GC | TA>GC | TTTGATAGA | AGACCAA | ATC | ATC | T | G | T>G | T>G_ATC | TA>GC_AGACCAA | 7 | FALSE | FALSE |
| NL4_v.txt | 32442 | 32442-SNV-A | SNV | 1 | A | C | 21 | 13151 | 0.15968367 | 62.1904762 | rnr1D57N | FALSE | CAN | JSY3602 | 7 | rnr1D57N-Re | TAA | A>C | G/C+1 | A/T-1 | TA>GC | TA>GC | GGAACTAAA | AAAAGTC | ATT | ATT | T | G | T>G | T>G_ATT | TA>GC_AAAAGTC | 7 | FALSE | FALSE |
| NL4_v.txt | 32472 | 32472-SNV-T | SNV | 1 | T | G | 1 | 13151 | 7.60E-03 | 39 | rnr1D57N | FALSE | CAN | JSY3602 | 7 | rnr1D57N-Re | GTA | T>G | G/C+1 | A/T-1 | TA>GC | TA>GC | CCAATGTAG | TAGAAGG | CAT | GTA | T | C | T>G | T>G_GTA | TA>GC_TAGAAGG | 7 | FALSE | FALSE |
| NL4_v.txt | 32481 | 32481-SNV-A | SNV | 1 | A | C | 8 | 13151 | 0.06083188 | 51.5 | rnr1D57N | FALSE | CAN | JSY3602 | 7 | rnr1D57N-Re | TAA | A>C | G/C+1 | A/T-1 | TA>GC | TA>GC | AAGGTTAAG | AAGATAC | ATT | ATT | T | G | T>G | T>G_ATT | TA>GC_AAGATAC | 7 | FALSE | FALSE |
| NL4_v.txt | 32490 | 32490-SNV-A | SNV | 1 | A | C | 58 | 13151 | 0.4410311 | 37.8103448 | rnr1D57N | FALSE | CAN | JSY3602 | 7 | rnr1D57N-Re | GAA | A>C | G/C+1 | A/T-1 | TA>GC | TA>GC | ATACGGAAA | AAAACAA | CTT | CTT | T | G | T>G | T>G_CTT | TA>GC_AAAACAA | 7 | FALSE | FALSE |
| NL4_v.txt | 33095 | 33095-SNV-T | SNV | 1 | T | G | 7 | 5930 | 0.11804385 | 22.1428572 | rnr1D57N | FALSE | CAN | JSY3602 | 7 | rnr1D57N-Re | ATA | T>G | G/C+1 | A/T-1 | TA>GC | TA>GC | ATATGATAT | TATAAGA | TAT | ATA | T | C | T>G | T>G_ATA | TA>GC_TATAAGA | 7 | FALSE | FALSE |
| NL4_v.txt | 33166 | 33166-SNV-T | SNV | 1 | T | G | 9 | 5928 | 0.15182186 | 35.6666667 | rnr1D57N | FALSE | CAN | JSY3602 | 7 | rnr1D57N-Re | GTA | T>G | G/C+1 | A/T-1 | TA>GC | TA>GC | CAATAGTAC | TACCACC | CAT | GTA | T | C | T>G | T>G_GTA | TA>GC_TACCACC | 7 | FALSE | FALSE |
| NL4_v.txt | 33194 | 33194-SNV-T | SNV | 1 | T | G | 15 | 10091 | 0.14864731 | 38.2 | rnr1D57N | FALSE | CAN | JSY3602 | 7 | rnr1D57N-Re | GTC | T>G | G/C+1 | A/T-1 | TA>GC | TA>GC | AATATGTCTT | TCTTTGC | CAG | GTC | T | C | T>G | T>G_GTC | TA>GC_TCTTTGC | 7 | FALSE | FALSE |
| NL4_v.txt | 32256 | 32256-SNV-A | SNV | 1 | A | G | 20 | 13146 | 0.15213753 | 37.7 | rnr1D57N | FALSE | CAN | JSY3602 | 7 | rnr1D57N-Re | TAG | A>G | G/C+1 | A/T-1 | TA>CG | TA>CG | TTTGATAGA | AGACCAA | ATC | ATC | T | C | T>C | T>C_ATC | TA>CG_AGACCAA | 7 | FALSE | FALSE |
| NL4_v.txt | 32293 | 32293-SNV-A | SNV | 1 | A | G | 10 | 13151 | 0.07603985 | 38.5954233 | rnr1D57N | FALSE | CAN | JSY3602 | 7 | rnr1D57N-Re | GAA | A>G | G/C+1 | A/T-1 | TA>CG | TA>CG | TATTTGAATT | AATTTGC | CTT | CTT | T | C | T>C | T>C_CTT | TA>CG_AATTTGC | 7 | FALSE | FALSE |
| NL4_v.txt | 32472 | 32472-SNV-T | SNV | 1 | T | C | 10 | 13151 | 0.07603985 | 62.7 | rnr1D57N | FALSE | CAN | JSY3602 | 7 | rnr1D57N-Re | GTA | T>C | G/C+1 | A/T-1 | TA>CG | TA>CG | CCAATGTAG | TAGAAGG | CAT | GTA | T | G | T>C | T>C_GTA | TA>CG_TAGAAGG | 7 | FALSE | FALSE |
| NL4_v.txt | 32473 | 32473-SNV-A | SNV | 1 | A | G | 6 | 13151 | 0.04562391 | 48.6666667 | rnr1D57N | FALSE | CAN | JSY3602 | 7 | rnr1D57N-Re | TAG | A>G | G/C+1 | A/T-1 | TA>CG | TA>CG | CAATGTAGA | AGAAGGT | ATC | ATC | T | C | T>C | T>C_ATC | TA>CG_AGAAGGT | 7 | FALSE | FALSE |
| NL4_v.txt | 32490 | 32490-SNV-A | SNV | 1 | A | G | 13 | 13151 | 0.0988518 | 37.9230769 | rnr1D57N | FALSE | CAN | JSY3602 | 7 | rnr1D57N-Re | GAA | A>G | G/C+1 | A/T-1 | TA>CG | TA>CG | ATACGGAAA | AAAACAA | CTT | CTT | T | C | T>C | T>C_CTT | TA>CG_AAAACAA | 7 | FALSE | FALSE |
| NL4_v.txt | 32500 | 32500-SNV-T | SNV | 1 | T | C | 2 | 13151 | 0.01520797 | 38.9996935 | rnr1D57N | FALSE | CAN | JSY3602 | 7 | rnr1D57N-Re | TTT | T>C | G/C+1 | A/T-1 | TA>CG | TA>CG | CAACTTTTT | TTTTGAT | AAA | TTT | T | G | T>C | T>C_TTT | TA>CG_TTTTGAT | 7 | FALSE | FALSE |
| NL4_v.txt | 33095 | 33095-SNV-T | SNV | 1 | T | C | 2 | 5930 | 0.03372681 | 64 | rnr1D57N | FALSE | CAN | JSY3602 | 7 | rnr1D57N-Re | ATA | T>C | G/C+1 | A/T-1 | TA>CG | TA>CG | ATATGATAT | TATAAGA | TAT | ATA | T | G | T>C | T>C_ATA | TA>CG_TATAAGA | 7 | FALSE | FALSE |
| NL4_v.txt | 33190 | 33190-SNV-T | SNV | 1 | T | C | 2 | 10092 | 0.01981768 | 37.9975854 | rnr1D57N | FALSE | CAN | JSY3602 | 7 | rnr1D57N-Re | ATA | T>C | G/C+1 | A/T-1 | TA>CG | TA>CG | TACCAATATG | TATGTCT | TAT | ATA | T | G | T>C | T>C_ATA | TA>CG_TATGTCT | 7 | FALSE | FALSE |
| NL4_v.txt | 33211 | 33211-SNV-T | SNV | 1 | T | C | 31 | 10091 | 0.30720444 | 37.9677419 | rnr1D57N | FALSE | CAN | JSY3602 | 7 | rnr1D57N-Re | CTC | T>C | G/C+1 | A/T-1 | TA>CG | TA>CG | GCTCTCTCTT | TCTTCAC | GAG | CTC | T | G | T>C | T>C_CTC | TA>CG_TCTTCAC | 7 | FALSE | FALSE |
| NL4_v.txt | 31839 | 31839-SNV-A | SNV | 1 | A | T | 16 | 5292 | 0.30234316 | 60.8125 | rnr1D57N | FALSE | CAN | JSY3602 | 7 | rnr1D57N-Re | TAA | A>T | A/T+1 | A/T-1 | TA>AT | TA>AT | TGAAATAAG | AAGATCC | ATT | ATT | T | A | T>A | T>A_ATT | TA>AT_AAGATCC | 7 | FALSE | FALSE |
| NL4_v.txt | 31886 | 31886-SNV-A | SNV | 1 | A | T | 8 | 5292 | 0.15117158 | 36.1193128 | rnr1D57N | FALSE | CAN | JSY3602 | 7 | rnr1D57N-Re | CAA | A>T | A/T+1 | A/T-1 | TA>AT | TA>AT | GGCAGCAAA | AAAGCTA | GTT | GTT | T | A | T>A | T>A_GTT | TA>AT_AAAGCTA | 7 | FALSE | FALSE |
| NL4_v.txt | 32050 | 32050-SNV-T | SNV | 1 | T | A | 4 | 1937 | 0.20650491 | 64 | rnr1D57N | FALSE | CAN | JSY3602 | 7 | rnr1D57N-Re | CTG | T>A | A/T+1 | A/T-1 | TA>AT | TA>AT | TAAATCTGAT | TGATGTG | GAC | CTG | T | T | T>A | T>A_CTG | TA>AT_TGATGTG | 7 | FALSE | FALSE |
| NL4_v.txt | 32473 | 32473-SNV-A | SNV | 1 | A | T | 12 | 13151 | 0.09124781 | 54.6666667 | rnr1D57N | FALSE | CAN | JSY3602 | 7 | rnr1D57N-Re | TAG | A>T | A/T+1 | A/T-1 | TA>AT | TA>AT | CAATGTAGA | AGAAGGT | ATC | ATC | T | A | T>A | T>A_ATC | TA>AT_AGAAGGT | 7 | FALSE | FALSE |
| NL4_v.txt | 32499 | 32499-SNV-T | SNV | 1 | T | A | 1 | 13151 | 7.60E-03 | 38.9992631 | rnr1D57N | FALSE | CAN | JSY3602 | 7 | rnr1D57N-Re | TTT | T>A | A/T+1 | A/T-1 | TA>AT | TA>AT | ACAACTTTTT | TTTTTGA | AAA | TTT | T | T | T>A | T>A_TTT | TA>AT_TTTTTGA | 7 | FALSE | FALSE |
| NL4_v.txt | 32823 | 32823-SNV-A | SNV | 1 | A | T | 21 | 3997 | 0.52539405 | 37.5238095 | rnr1D57N | FALSE | CAN | JSY3602 | 7 | rnr1D57N-Re | CAT | A>T | A/T+1 | A/T-1 | TA>AT | TA>AT | AAGTTCATTA | ATTATTG | GTA | GTA | T | A | T>A | T>A_GTA | TA>AT_ATTATTG | 7 | FALSE | FALSE |
| NL4_v.txt | 33108 | 33108-SNV-A | SNV | 1 | A | T | 2 | 5930 | 0.03372681 | 37.5 | rnr1D57N | FALSE | CAN | JSY3602 | 7 | rnr1D57N-Re | CAC | A>T | A/T+1 | A/T-1 | TA>AT | TA>AT | GCGCCCACT | ACTGGGC | GTG | GTG | T | A | T>A | T>A_GTG | TA>AT_ACTGGGC | 7 | FALSE | FALSE |
| NL4_v.txt | 33236 | 33236-SNV-T | SNV | 1 | T | A | 2 | 10091 | 0.01981964 | 38.489686 | rnr1D57N | FALSE | CAN | JSY3602 | 7 | rnr1D57N-Re | CTC | T>A | A/T+1 | A/T-1 | TA>AT | TA>AT | TACTTCTCCT | TCCTTCA | GAG | CTC | T | T | T>A | T>A_CTC | TA>AT_TCCTTCA | 7 | FALSE | FALSE |
| NL4_v.txt | 31650 | 31650-SNV-C | SNV | 1 | C | T | 12 | 5291 | 0.22680023 | 35 | rnr1D57N | FALSE | CAN | JSY3602 | 7 | rnr1D57N-Re | GCG | C>T | A/T+1 | G/C-1 | CG>TA | CG>TA | AGAATGCGA | CGAAATG | CGC | GCG | C | A | C>T | C>T_GCG | CG>TA_CGAAATG | 7 | FALSE | FALSE |
| NL4_v.txt | 31880 | 31880-SNV-C | SNV | 1 | G | A | 34 | 5292 | 0.64247921 | 35.6764706 | rnr1D57N | FALSE | CAN | JSY3602 | 7 | rnr1D57N-Re | CGG | G>A | A/T+1 | G/C-1 | CG>TA | CG>TA | ATAGGCGGC | GGCAGCA | GCC | GCC | C | T | C>T | C>T_GCC | CG>TA_GGCAGCA | 7 | FALSE | FALSE |
| NL4_v.txt | 31884 | 31884-SNV-C | SNV | 1 | G | A | 7 | 5292 | 0.13227513 | 35.5696343 | rnr1D57N | FALSE | CAN | JSY3602 | 7 | rnr1D57N-Re | AGC | G>A | A/T+1 | G/C-1 | CG>TA | CG>TA | GCGGCAGCA | GCAAAGC | TCG | TCG | C | T | C>T | C>T_TCG | CG>TA_GCAAAGC | 7 | FALSE | FALSE |
| NL4_v.txt | 32041 | 32041-SNV-C | SNV | 1 | G | A | 6 | 1937 | 0.30975736 | 61 | rnr1D57N | FALSE | CAN | JSY3602 | 7 | rnr1D57N-Re | TGC | G>A | A/T+1 | G/C-1 | CG>TA | CG>TA | AAGCTTGCA | GCATAAA | ACG | ACG | C | T | C>T | C>T_ACG | CG>TA_GCATAAA | 7 | FALSE | FALSE |
| NL4_v.txt | 32075 | 32075-SNV-C | SNV | 1 | C | T | 8 | 1937 | 0.41300981 | 64 | rnr1D57N | FALSE | CAN | JSY3602 | 7 | rnr1D57N-Re | ACC | C>T | A/T+1 | G/C-1 | CG>TA | CG>TA | AAATAACCA | CCATGCA | TGG | ACC | C | A | C>T | C>T_ACC | CG>TA_CCATGCA | 7 | FALSE | FALSE |
| NL4_v.txt | 32271 | 32271-SNV-C | SNV | 1 | C | T | 15 | 13151 | 0.11405977 | 37.2666667 | rnr1D57N | FALSE | CAN | JSY3602 | 7 | rnr1D57N-Re | ACG | C>T | A/T+1 | G/C-1 | CG>TA | CG>TA | AAAATACGG | CGGGAAC | TGC | ACG | C | A | C>T | C>T_ACG | CG>TA_CGGGAAC | 7 | FALSE | FALSE |
| NL4_v.txt | 32299 | 32299-SNV-C | SNV | 1 | C | T | 31 | 13151 | 0.23572352 | 36.3548387 | rnr1D57N | FALSE | CAN | JSY3602 | 7 | rnr1D57N-Re | GCG | C>T | A/T+1 | G/C-1 | CG>TA | CG>TA | AATTTGCGG | CGGCAGA | CGC | GCG | C | A | C>T | C>T_GCG | CG>TA_CGGCAGA | 7 | FALSE | FALSE |
| NL4_v.txt | 32466 | 32466-SNV-C | SNV | 1 | C | T | 13 | 13151 | 0.0988518 | 62.2307692 | rnr1D57N | FALSE | CAN | JSY3602 | 7 | rnr1D57N-Re | GCC | C>T | A/T+1 | G/C-1 | CG>TA | CG>TA | AGAGAGCCA | CCAATGT | CGG | GCC | C | A | C>T | C>T_GCC | CG>TA_CCAATGT | 7 | FALSE | FALSE |
| NL4_v.txt | 32487 | 32487-SNV-C | SNV | 1 | C | T | 8 | 13151 | 0.06083188 | 36.125 | rnr1D57N | FALSE | CAN | JSY3602 | 7 | rnr1D57N-Re | ACG | C>T | A/T+1 | G/C-1 | CG>TA | CG>TA | AAGATACGG | CGGAAAA | TGC | ACG | C | A | C>T | C>T_ACG | CG>TA_CGGAAAA | 7 | FALSE | FALSE |
| NL4_v.txt | 32535 | 32535-SNV-C | SNV | 1 | G | A | 15 | 13147 | 0.11409447 | 38.5333333 | rnr1D57N | FALSE | CAN | JSY3602 | 7 | rnr1D57N-Re | TGC | G>A | A/T+1 | G/C-1 | CG>TA | CG>TA | GGGTTTGCA | GCAGCTT | ACG | ACG | C | T | C>T | C>T_ACG | CG>TA_GCAGCTT | 7 | FALSE | FALSE |
| NL4_v.txt | 32572 | 32572-SNV-C | SNV | 1 | C | T | 12 | 3968 | 0.30241936 | 37.25 | rnr1D57N | FALSE | CAN | JSY3602 | 7 | rnr1D57N-Re | CCT | C>T | A/T+1 | G/C-1 | CG>TA | CG>TA | CAGTACCTTG | CTTGAAA | GGA | CCT | C | A | C>T | C>T_CCT | CG>TA_CTTGAAA | 7 | FALSE | FALSE |
| NL4_v.txt | 32575 | 32575-SNV-C | SNV | 1 | G | A | 3 | 3968 | 0.07560484 | 38.9809174 | rnr1D57N | FALSE | CAN | JSY3602 | 7 | rnr1D57N-Re | TGA | G>A | A/T+1 | G/C-1 | CG>TA | CG>TA | TACCTTGAA | GAAATGT | ACT | ACT | C | T | C>T | C>T_ACT | CG>TA_GAAATGT | 7 | FALSE | FALSE |

| | | | | | | | | | | | | | | | | | | | | | | | | | | | | | | | | | | |
|---|---|---|---|---|---|---|---|---|---|---|---|---|---|---|---|---|---|---|---|---|---|---|---|---|---|---|---|---|---|---|---|---|---|---|
| NL4_v.txt | 32749 | 32749-SNV- | SNV | 1 | C | T | 12 | 3997 | 0.30022517 | 61.75 | rnr1D57N | FALSE | CAN | JSY3602 | 7 | rnr1D57N-Re | CCG | C>T | A/T+1 | G/C-1 | CG>TA | CG>TA | GAAACCCGA | CGATAAT | GGC | CCG | C | A | C>T | C>T_CCG | CG>TA_CGATAAT | 7 | FALSE | FALSE |
| NL4_v.txt | 32797 | 32797-SNV- | SNV | 1 | C | T | 15 | 3997 | 0.37528146 | 36.9333333 | rnr1D57N | FALSE | CAN | JSY3602 | 7 | rnr1D57N-Re | CCG | C>T | A/T+1 | G/C-1 | CG>TA | CG>TA | ATTCACCGTA | CGTAATA | GGC | CCG | C | A | C>T | C>T_CCG | CG>TA_CGTAATA | 7 | FALSE | FALSE |
| NL4_v.txt | 32798 | 32798-SNV- | SNV | 1 | G | A | 4 | 3997 | 0.10007506 | 38.75 | rnr1D57N | FALSE | CAN | JSY3602 | 7 | rnr1D57N-Re | CGT | G>A | A/T+1 | G/C-1 | CG>TA | CG>TA | TTCACCGTAA | GTAATAT | GCA | GCA | C | T | C>T | C>T_GCA | CG>TA_GTAATAT | 7 | FALSE | FALSE |
| NL4_v.txt | 32882 | 32882-SNV- | SNV | 1 | C | T | 12 | 5930 | 0.20236088 | 36.6570223 | rnr1D57N | FALSE | CAN | JSY3602 | 7 | rnr1D57N-Re | TCC | C>T | A/T+1 | G/C-1 | CG>TA | CG>TA | GTACGTCCA | CCAAAAT | AGG | TCC | C | A | C>T | C>T_TCC | CG>TA_CCAAAAT | 7 | FALSE | FALSE |
| NL4_v.txt | 32883 | 32883-SNV- | SNV | 1 | C | T | 10 | 5930 | 0.16863406 | 36.3983112 | rnr1D57N | FALSE | CAN | JSY3602 | 7 | rnr1D57N-Re | CCA | C>T | A/T+1 | G/C-1 | CG>TA | CG>TA | TACGTCCAA | CAAAATT | GGT | CCA | C | A | C>T | C>T_CCA | CG>TA_CAAAATT | 7 | FALSE | FALSE |
| NL4_v.txt | 32890 | 32890-SNV- | SNV | 1 | G | A | 16 | 5930 | 0.2698145 | 38.875 | rnr1D57N | FALSE | CAN | JSY3602 | 7 | rnr1D57N-Re | TGA | G>A | A/T+1 | G/C-1 | CG>TA | CG>TA | AAAATTGAA | GAATGAC | ACT | ACT | C | T | C>T | C>T_ACT | CG>TA_GAATGAC | 7 | FALSE | FALSE |
| NL4_v.txt | 32899 | 32899-SNV- | SNV | 1 | G | A | 20 | 5930 | 0.33726813 | 38.85 | rnr1D57N | FALSE | CAN | JSY3602 | 7 | rnr1D57N-Re | TGG | G>A | A/T+1 | G/C-1 | CG>TA | CG>TA | TGACTTGGC | GGCCAAC | ACC | ACC | C | T | C>T | C>T_ACC | CG>TA_GGCCAAC | 7 | FALSE | FALSE |
| NL4_v.txt | 32901 | 32901-SNV- | SNV | 1 | C | T | 10 | 5930 | 0.16863406 | 39 | rnr1D57N | FALSE | CAN | JSY3602 | 7 | rnr1D57N-Re | GCC | C>T | A/T+1 | G/C-1 | CG>TA | CG>TA | ACTTGGCCA | CCAACTA | CGG | GCC | C | A | C>T | C>T_GCC | CG>TA_CCAACTA | 7 | FALSE | FALSE |
| NL4_v.txt | 32995 | 32995-SNV- | SNV | 1 | G | A | 31 | 5930 | 0.5227656 | 63.4516129 | rnr1D57N | FALSE | CAN | JSY3602 | 7 | rnr1D57N-Re | TGT | G>A | A/T+1 | G/C-1 | CG>TA | CG>TA | ATCTTTGTGA | GTGAGAA | ACA | ACA | C | T | C>T | C>T_ACA | CG>TA_GTGAGAA | 7 | FALSE | FALSE |
| NL4_v.txt | 33043 | 33043-SNV- | SNV | 1 | C | T | 8 | 5930 | 0.13490725 | 62.25 | rnr1D57N | FALSE | CAN | JSY3602 | 7 | rnr1D57N-Re | TCA | C>T | A/T+1 | G/C-1 | CG>TA | CG>TA | CCATTTCACC | CACCCAA | AGT | TCA | C | A | C>T | C>T_TCA | CG>TA_CACCCAA | 7 | FALSE | FALSE |
| NL4_v.txt | 33070 | 33070-SNV- | SNV | 1 | C | T | 4 | 5930 | 0.06745363 | 64 | rnr1D57N | FALSE | CAN | JSY3602 | 7 | rnr1D57N-Re | GCC | C>T | A/T+1 | G/C-1 | CG>TA | CG>TA | AATATGCCA | CCAAAGA | CGG | GCC | C | A | C>T | C>T_GCC | CG>TA_CCAAAGA | 7 | FALSE | FALSE |
| NL4_v.txt | 33114 | 33114-SNV- | SNV | 1 | C | T | 5 | 5930 | 0.08431703 | 37.4 | rnr1D57N | FALSE | CAN | JSY3602 | 7 | rnr1D57N-Re | GCC | C>T | A/T+1 | G/C-1 | CG>TA | CG>TA | ACTGGGCCG | CCGGCGT | CGG | GCC | C | A | C>T | C>T_GCC | CG>TA_CCGGCGT | 7 | FALSE | FALSE |
| NL4_v.txt | 33153 | 33153-SNV- | SNV | 1 | C | T | 7 | 5930 | 0.11804385 | 38.8567157 | rnr1D57N | FALSE | CAN | JSY3602 | 7 | rnr1D57N-Re | ACC | C>T | A/T+1 | G/C-1 | CG>TA | CG>TA | AAAAGACCT | CCTGTAC | TGG | ACC | C | A | C>T | C>T_ACC | CG>TA_CCTGTAC | 7 | FALSE | FALSE |
| NL4_v.txt | 33168 | 33168-SNV- | SNV | 1 | C | T | 6 | 5928 | 0.10121458 | 35.3333333 | rnr1D57N | FALSE | CAN | JSY3602 | 7 | rnr1D57N-Re | ACC | C>T | A/T+1 | G/C-1 | CG>TA | CG>TA | ATAGTACCA | CCACCAA | TGG | ACC | C | A | C>T | C>T_ACC | CG>TA_CCACCAA | 7 | FALSE | FALSE |
| NL4_v.txt | 33258 | 33258-SNV- | SNV | 1 | C | T | 14 | 10091 | 0.13873749 | 54.3571429 | rnr1D57N | FALSE | CAN | JSY3602 | 7 | rnr1D57N-Re | GCC | C>T | A/T+1 | G/C-1 | CG>TA | CG>TA | CCTATGCCAT | CCATCCT | CGG | GCC | C | A | C>T | C>T_GCC | CG>TA_CCATCCT | 7 | FALSE | FALSE |
| NL4_v.txt | 33365 | 33365-SNV- | SNV | 1 | G | A | 2 | 10089 | 0.01982357 | 56.4999989 | rnr1D57N | FALSE | CAN | JSY3602 | 7 | rnr1D57N-Re | TGT | G>A | A/T+1 | G/C-1 | CG>TA | CG>TA | ACGTCTGTG | GTGGTGT | ACA | ACA | C | T | C>T | C>T_ACA | CG>TA_GTGGTGT | 7 | FALSE | FALSE |
| NL4_v.txt | 33392 | 33392-SNV- | SNV | 1 | G | A | 9 | 10091 | 0.08918839 | 58.5555556 | rnr1D57N | FALSE | CAN | JSY3602 | 7 | rnr1D57N-Re | CGT | G>A | A/T+1 | G/C-1 | CG>TA | CG>TA | AACGTCGTG | GTGAAAG | GCA | GCA | C | T | C>T | C>T_GCA | CG>TA_GTGAAAG | 7 | FALSE | FALSE |
| NL4_v.txt | 33412 | 33412-SNV- | SNV | 1 | G | A | 13 | 10091 | 0.12882767 | 55.4615385 | rnr1D57N | FALSE | CAN | JSY3602 | 7 | rnr1D57N-Re | GGC | G>A | A/T+1 | G/C-1 | CG>TA | CG>TA | TGACCGGCT | GCTCATT | CCG | CCG | C | T | C>T | C>T_CCG | CG>TA_GCTCATT | 7 | FALSE | FALSE |
| NL4_v.txt | 33474 | 33474-SNV- | SNV | 1 | C | T | 11 | 10090 | 0.10901883 | 31 | rnr1D57N | FALSE | CAN | JSY3602 | 7 | rnr1D57N-Re | GCC | C>T | A/T+1 | G/C-1 | CG>TA | CG>TA | GCTATGCCTT | CCTTTTT | CGG | GCC | C | A | C>T | C>T_GCC | CG>TA_CCTTTTT | 7 | FALSE | FALSE |
| NL4_v.txt | 31661 | 31661-SNV- | SNV | 1 | G | C | 8 | 5292 | 0.15117158 | 34 | rnr1D57N | FALSE | CAN | JSY3602 | 7 | rnr1D57N-Re | TGG | G>C | G/C+1 | G/C-1 | CG>GC | CG>GC | TGGCGTGGA | GGAAATG | ACC | ACC | C | G | C>G | C>G_ACC | CG>GC_GGAAATG | 7 | FALSE | FALSE |
| NL4_v.txt | 32271 | 32271-SNV- | SNV | 1 | C | G | 9 | 13151 | 0.06843586 | 37.7777778 | rnr1D57N | FALSE | CAN | JSY3602 | 7 | rnr1D57N-Re | ACG | C>G | G/C+1 | G/C-1 | CG>GC | CG>GC | AAAATACGG | CGGGAAC | TGC | ACG | C | C | C>G | C>G_ACG | CG>GC_CGGGAAC | 7 | FALSE | FALSE |
| NL4_v.txt | 32292 | 32292-SNV- | SNV | 1 | G | C | 52 | 13151 | 0.39540719 | 38.5384811 | rnr1D57N | FALSE | CAN | JSY3602 | 7 | rnr1D57N-Re | TGA | G>C | G/C+1 | G/C-1 | CG>GC | CG>GC | ATATTTGAAT | GAATTTG | ACT | ACT | C | G | C>G | C>G_ACT | CG>GC_GAATTTG | 7 | FALSE | FALSE |
| NL4_v.txt | 32432 | 32432-SNV- | SNV | 1 | G | C | 8 | 13151 | 0.06083188 | 58.5 | rnr1D57N | FALSE | CAN | JSY3602 | 7 | rnr1D57N-Re | TGT | G>C | G/C+1 | G/C-1 | CG>GC | CG>GC | GTCATTGTAT | GTATGGA | ACA | ACA | C | G | C>G | C>G_ACA | CG>GC_GTATGGA | 7 | FALSE | FALSE |
| NL4_v.txt | 32474 | 32474-SNV- | SNV | 1 | G | C | 4 | 13151 | 0.03041594 | 54.9958793 | rnr1D57N | FALSE | CAN | JSY3602 | 7 | rnr1D57N-Re | AGA | G>C | G/C+1 | G/C-1 | CG>GC | CG>GC | AATGTAGAA | GAAGGTT | TCT | TCT | C | G | C>G | C>G_TCT | CG>GC_GAAGGTT | 7 | FALSE | FALSE |
| NL4_v.txt | 32487 | 32487-SNV- | SNV | 1 | C | G | 6 | 13151 | 0.04562391 | 39 | rnr1D57N | FALSE | CAN | JSY3602 | 7 | rnr1D57N-Re | ACG | C>G | G/C+1 | G/C-1 | CG>GC | CG>GC | AAGATACGG | CGGAAAA | TGC | ACG | C | C | C>G | C>G_ACG | CG>GC_CGGAAAA | 7 | FALSE | FALSE |
| NL4_v.txt | 32776 | 32776-SNV- | SNV | 1 | C | G | 5 | 3997 | 0.12509382 | 50.6 | rnr1D57N | FALSE | CAN | JSY3602 | 7 | rnr1D57N-Re | GCG | C>G | G/C+1 | G/C-1 | CG>GC | CG>GC | TGGAAGCGA | CGACCCA | CGC | GCG | C | C | C>G | C>G_GCG | CG>GC_CGACCCA | 7 | FALSE | FALSE |
| NL4_v.txt | 32796 | 32796-SNV- | SNV | 1 | C | G | 1 | 3997 | 0.02501876 | 38 | rnr1D57N | FALSE | CAN | JSY3602 | 7 | rnr1D57N-Re | ACC | C>G | G/C+1 | G/C-1 | CG>GC | CG>GC | AATTCACCGT | CCGTAAT | TGG | ACC | C | C | C>G | C>G_ACC | CG>GC_CCGTAAT | 7 | FALSE | FALSE |
| NL4_v.txt | 32917 | 32917-SNV- | SNV | 1 | C | G | 3 | 5930 | 0.05059022 | 39 | rnr1D57N | FALSE | CAN | JSY3602 | 7 | rnr1D57N-Re | TCC | C>G | G/C+1 | G/C-1 | CG>GC | CG>GC | TAAGTTCCA | CCAGGGC | AGG | TCC | C | C | C>G | C>G_TCC | CG>GC_CCAGGGC | 7 | FALSE | FALSE |
| NL4_v.txt | 32937 | 32937-SNV- | SNV | 1 | C | G | 12 | 5930 | 0.20236088 | 37.4166667 | rnr1D57N | FALSE | CAN | JSY3602 | 7 | rnr1D57N-Re | CCA | C>G | G/C+1 | G/C-1 | CG>GC | CG>GC | ATTGCCCAA | CAAGAAA | GGT | CCA | C | C | C>G | C>G_CCA | CG>GC_CAAGAAA | 7 | FALSE | FALSE |
| NL4_v.txt | 32997 | 32997-SNV- | SNV | 1 | G | C | 17 | 5930 | 0.28667791 | 64 | rnr1D57N | FALSE | CAN | JSY3602 | 7 | rnr1D57N-Re | TGA | G>C | G/C+1 | G/C-1 | CG>GC | CG>GC | CTTTGTGAG | GAGAAAA | ACT | ACT | C | G | C>G | C>G_ACT | CG>GC_GAGAAAA | 7 | FALSE | FALSE |
| NL4_v.txt | 33111 | 33111-SNV- | SNV | 1 | G | C | 4 | 5930 | 0.06745363 | 38 | rnr1D57N | FALSE | CAN | JSY3602 | 7 | rnr1D57N-Re | TGG | G>C | G/C+1 | G/C-1 | CG>GC | CG>GC | CCCACTGGG | GGGCCGG | ACC | ACC | C | G | C>G | C>G_ACC | CG>GC_GGGCCGG | 7 | FALSE | FALSE |
| NL4_v.txt | 33114 | 33114-SNV- | SNV | 1 | C | G | 3 | 5930 | 0.05059022 | 38 | rnr1D57N | FALSE | CAN | JSY3602 | 7 | rnr1D57N-Re | GCC | C>G | G/C+1 | G/C-1 | CG>GC | CG>GC | ACTGGGCCG | CCGGCGT | CGG | GCC | C | C | C>G | C>G_GCC | CG>GC_CCGGCGT | 7 | FALSE | FALSE |
| NL4_v.txt | 33153 | 33153-SNV- | SNV | 1 | C | G | 7 | 5930 | 0.11804385 | 35.1418387 | rnr1D57N | FALSE | CAN | JSY3602 | 7 | rnr1D57N-Re | ACC | C>G | G/C+1 | G/C-1 | CG>GC | CG>GC | AAAAGACCT | CCTGTAC | TGG | ACC | C | C | C>G | C>G_ACC | CG>GC_CCTGTAC | 7 | FALSE | FALSE |
| NL4_v.txt | 33168 | 33168-SNV- | SNV | 1 | C | G | 5 | 5928 | 0.08434548 | 39 | rnr1D57N | FALSE | CAN | JSY3602 | 7 | rnr1D57N-Re | ACC | C>G | G/C+1 | G/C-1 | CG>GC | CG>GC | ATAGTACCA | CCACCAA | TGG | ACC | C | C | C>G | C>G_ACC | CG>GC_CCACCAA | 7 | FALSE | FALSE |
| NL4_v.txt | 33275 | 33275-SNV- | SNV | 1 | C | G | 4 | 10091 | 0.03963928 | 64 | rnr1D57N | FALSE | CAN | JSY3602 | 7 | rnr1D57N-Re | ACG | C>G | G/C+1 | G/C-1 | CG>GC | CG>GC | AGAGAACGT | CGTATCC | TGC | ACG | C | C | C>G | C>G_ACG | CG>GC_CGTATCC | 7 | FALSE | FALSE |
| NL4_v.txt | 33367 | 33367-SNV- | SNV | 1 | G | C | 1 | 10091 | 9.91E-03 | 63.9999979 | rnr1D57N | FALSE | CAN | JSY3602 | 7 | rnr1D57N-Re | TGG | G>C | G/C+1 | G/C-1 | CG>GC | CG>GC | GTCTGTGGT | GGTGTGT | ACC | ACC | C | G | C>G | C>G_ACC | CG>GC_GGTGTGT | 7 | FALSE | FALSE |
| NL4_v.txt | 32292 | 32292-SNV- | SNV | 1 | G | T | 5 | 13151 | 0.03801992 | 35.2000337 | rnr1D57N | FALSE | CAN | JSY3602 | 7 | rnr1D57N-Re | TGA | G>T | A/T+1 | G/C-1 | CG>AT | CG>AT | ATATTTGAAT | GAATTTG | ACT | ACT | C | A | C>A | C>A_ACT | CG>AT_GAATTTG | 7 | FALSE | FALSE |
| NL4_v.txt | 32301 | 32301-SNV- | SNV | 1 | G | T | 13 | 13151 | 0.0988518 | 35.5384615 | rnr1D57N | FALSE | CAN | JSY3602 | 7 | rnr1D57N-Re | GGC | G>T | A/T+1 | G/C-1 | CG>AT | CG>AT | TTTGCGGCA | GCAGAAA | CCG | CCG | C | A | C>A | C>A_CCG | CG>AT_GCAGAAA | 7 | FALSE | FALSE |
| NL4_v.txt | 32373 | 32373-SNV- | SNV | 1 | G | T | 9 | 13151 | 0.06843586 | 64 | rnr1D57N | FALSE | CAN | JSY3602 | 7 | rnr1D57N-Re | AGC | G>T | A/T+1 | G/C-1 | CG>AT | CG>AT | TCAATAGCA | GCAATAA | TCG | TCG | C | A | C>A | C>A_TCG | CG>AT_GCAATAA | 7 | FALSE | FALSE |
| NL4_v.txt | 32399 | 32399-SNV- | SNV | 1 | G | T | 44 | 13151 | 0.33457532 | 62.1363636 | rnr1D57N | FALSE | CAN | JSY3602 | 7 | rnr1D57N-Re | CGT | G>T | A/T+1 | G/C-1 | CG>AT | CG>AT | AGAAACGTA | GTAGGAA | GCA | GCA | C | A | C>A | C>A_GCA | CG>AT_GTAGGAA | 7 | FALSE | FALSE |
| NL4_v.txt | 32432 | 32432-SNV- | SNV | 1 | G | T | 6 | 13151 | 0.04562391 | 59.5 | rnr1D57N | FALSE | CAN | JSY3602 | 7 | rnr1D57N-Re | TGT | G>T | A/T+1 | G/C-1 | CG>AT | CG>AT | GTCATTGTAT | GTATGGA | ACA | ACA | C | A | C>A | C>A_ACA | CG>AT_GTATGGA | 7 | FALSE | FALSE |
| NL4_v.txt | 32471 | 32471-SNV- | SNV | 1 | G | T | 1705 | 13151 | 12.9647936 | 60.9584267 | rnr1D57N | FALSE | CAN | JSY3602 | 7 | rnr1D57N-Re | TGT | G>T | A/T+1 | G/C-1 | CG>AT | CG>AT | GCCAATGTA | GTAGAAG | ACA | ACA | C | A | C>A | C>A_ACA | CG>AT_GTAGAAG | 7 | FALSE | FALSE |
| NL4_v.txt | 32474 | 32474-SNV- | SNV | 1 | G | T | 4 | 13151 | 0.03041594 | 55.2448806 | rnr1D57N | FALSE | CAN | JSY3602 | 7 | rnr1D57N-Re | AGA | G>T | A/T+1 | G/C-1 | CG>AT | CG>AT | AATGTAGAA | GAAGGTT | TCT | TCT | C | A | C>A | C>A_TCT | CG>AT_GAAGGTT | 7 | FALSE | FALSE |
| NL4_v.txt | 32494 | 32494-SNV- | SNV | 1 | C | A | 6 | 13151 | 0.04562391 | 37.8333333 | rnr1D57N | FALSE | CAN | JSY3602 | 7 | rnr1D57N-Re | ACA | C>A | A/T+1 | G/C-1 | CG>AT | CG>AT | GGAAAACAA | CAACTTT | TGT | ACA | C | T | C>A | C>A_ACA | CG>AT_CAACTTT | 7 | FALSE | FALSE |
| NL4_v.txt | 32575 | 32575-SNV- | SNV | 1 | G | T | 40 | 3968 | 1.00806452 | 37.1707929 | rnr1D57N | FALSE | CAN | JSY3602 | 7 | rnr1D57N-Re | TGA | G>T | A/T+1 | G/C-1 | CG>AT | CG>AT | TACCTTGAA | GAAATGT | ACT | ACT | C | A | C>A | C>A_ACT | CG>AT_GAAATGT | 7 | FALSE | FALSE |
| NL4_v.txt | 32629 | 32629-SNV- | SNV | 1 | C | A | 7 | 3968 | 0.17641129 | 39 | rnr1D57N | FALSE | CAN | JSY3602 | 7 | rnr1D57N-Re | TCG | C>A | A/T+1 | G/C-1 | CG>AT | CG>AT | TCCCTTCGTT | CGTTTTT | AGC | TCG | C | T | C>A | C>A_TCG | CG>AT_CGTTTTT | 7 | FALSE | FALSE |
| NL4_v.txt | 32796 | 32796-SNV- | SNV | 1 | C | A | 37 | 3997 | 0.92569427 | 37.7297297 | rnr1D57N | FALSE | CAN | JSY3602 | 7 | rnr1D57N-Re | ACC | C>A | A/T+1 | G/C-1 | CG>AT | CG>AT | AATTCACCGT | CCGTAAT | TGG | ACC | C | T | C>A | C>A_ACC | CG>AT_CCGTAAT | 7 | FALSE | FALSE |
| NL4_v.txt | 32797 | 32797-SNV- | SNV | 1 | C | A | 5 | 3997 | 0.12509382 | 38.6 | rnr1D57N | FALSE | CAN | JSY3602 | 7 | rnr1D57N-Re | CCG | C>A | A/T+1 | G/C-1 | CG>AT | CG>AT | ATTCACCGTA | CGTAATA | GGC | CCG | C | T | C>A | C>A_CCG | CG>AT_CGTAATA | 7 | FALSE | FALSE |
| NL4_v.txt | 32798 | 32798-SNV- | SNV | 1 | G | T | 11 | 3997 | 0.27520641 | 35.9090909 | rnr1D57N | FALSE | CAN | JSY3602 | 7 | rnr1D57N-Re | CGT | G>T | A/T+1 | G/C-1 | CG>AT | CG>AT | TTCACCGTAA | GTAATAT | GCA | GCA | C | A | C>A | C>A_GCA | CG>AT_GTAATAT | 7 | FALSE | FALSE |
| NL4_v.txt | 32917 | 32917-SNV- | SNV | 1 | C | A | 13 | 5930 | 0.21922428 | 37.0769231 | rnr1D57N | FALSE | CAN | JSY3602 | 7 | rnr1D57N-Re | TCC | C>A | A/T+1 | G/C-1 | CG>AT | CG>AT | TAAGTTCCA | CCAGGGC | AGG | TCC | C | T | C>A | C>A_TCC | CG>AT_CCAGGGC | 7 | FALSE | FALSE |
| NL4_v.txt | 33043 | 33043-SNV- | SNV | 1 | C | A | 8 | 5930 | 0.13490725 | 58.375 | rnr1D57N | FALSE | CAN | JSY3602 | 7 | rnr1D57N-Re | TCA | C>A | A/T+1 | G/C-1 | CG>AT | CG>AT | CCATTTCACC | CACCCAA | AGT | TCA | C | T | C>A | C>A_TCA | CG>AT_CACCCAA | 7 | FALSE | FALSE |
| NL4_v.txt | 33051 | 33051-SNV- | SNV | 1 | G | T | 14 | 5930 | 0.23608769 | 63.8571429 | rnr1D57N | FALSE | CAN | JSY3602 | 7 | rnr1D57N-Re | GGA | G>T | A/T+1 | G/C-1 | CG>AT | CG>AT | CCCAAGGAC | GACTGCG | CCT | CCT | C | A | C>A | C>A_CCT | CG>AT_GACTGCG | 7 | FALSE | FALSE |
| NL4_v.txt | 33151 | 33151-SNV- | SNV | 1 | G | T | 38 | 5930 | 0.64080944 | 36.5263158 | rnr1D57N | FALSE | CAN | JSY3602 | 7 | rnr1D57N-Re | AGA | G>T | A/T+1 | G/C-1 | CG>AT | CG>AT | TGAAAAGAC | GACCTGT | TCT | TCT | C | A | C>A | C>A_TCT | CG>AT_GACCTGT | 7 | FALSE | FALSE |
| NL4_v.txt | 33154 | 33154-SNV- | SNV | 1 | C | A | 9 | 5928 | 0.15182186 | 30.106386 | rnr1D57N | FALSE | CAN | JSY3602 | 7 | rnr1D57N-Re | CCT | C>A | A/T+1 | G/C-1 | CG>AT | CG>AT | AAAGACCTG | CTGTACC | GGA | CCT | C | T | C>A | C>A_CCT | CG>AT_CTGTACC | 7 | FALSE | FALSE |
| NL4_v.txt | 33160 | 33160-SNV- | SNV | 1 | C | A | 13 | 5928 | 0.21929825 | 38.7692308 | rnr1D57N | FALSE | CAN | JSY3602 | 7 | rnr1D57N-Re | CCA | C>A | A/T+1 | G/C-1 | CG>AT | CG>AT | CTGTACCAAT | CAATAGT | GGT | CCA | C | T | C>A | C>A_CCA | CG>AT_CAATAGT | 7 | FALSE | FALSE |
| NL4_v.txt | 33168 | 33168-SNV- | SNV | 1 | C | A | 6 | 5928 | 0.10121458 | 38.1666667 | rnr1D57N | FALSE | CAN | JSY3602 | 7 | rnr1D57N-Re | ACC | C>A | A/T+1 | G/C-1 | CG>AT | CG>AT | ATAGTACCA | CCACCAA | TGG | ACC | C | T | C>A | C>A_ACC | CG>AT_CCACCAA | 7 | FALSE | FALSE |
| NL4_v.txt | 33235 | 33235-SNV- | SNV | 1 | C | A | 21 | 10091 | 0.20810623 | 38.9523821 | rnr1D57N | FALSE | CAN | JSY3602 | 7 | rnr1D57N-Re | TCT | C>A | A/T+1 | G/C-1 | CG>AT | CG>AT | GTACTTCTCC | CTCCTTC | AGA | TCT | C | T | C>A | C>A_TCT | CG>AT_CTCCTTC | 7 | FALSE | FALSE |
| NL4_v.txt | 33354 | 33354-SNV- | SNV | 1 | G | T | 33 | 10091 | 0.32702408 | 57.2727273 | rnr1D57N | FALSE | CAN | JSY3602 | 7 | rnr1D57N-Re | TGA | G>T | A/T+1 | G/C-1 | CG>AT | CG>AT | GGTATTGAC | GACCCAC | ACT | ACT | C | A | C>A | C>A_ACT | CG>AT_GACCCAC | 7 | FALSE | FALSE |
| NL4_v.txt | 33365 | 33365-SNV- | SNV | 1 | G | T | 6 | 10089 | 0.05947071 | 32.9559604 | rnr1D57N | FALSE | CAN | JSY3602 | 7 | rnr1D57N-Re | TGT | G>T | A/T+1 | G/C-1 | CG>AT | CG>AT | ACGTCTGTG | GTGGTGT | ACA | ACA | C | A | C>A | C>A_ACA | CG>AT_GTGGTGT | 7 | FALSE | FALSE |
| NL4_v.txt | 33367 | 33367-SNV- | SNV | 1 | G | T | 4 | 10091 | 0.03963928 | 48.33519 | rnr1D57N | FALSE | CAN | JSY3602 | 7 | rnr1D57N-Re | TGG | G>T | A/T+1 | G/C-1 | CG>AT | CG>AT | GTCTGTGGT | GGTGTGT | ACC | ACC | C | A | C>A | C>A_ACC | CG>AT_GGTGTGT | 7 | FALSE | FALSE |
| NL4_v.txt | 33370 | 33370-SNV- | SNV | 1 | G | T | 1 | 10087 | 9.91E-03 | 13.0000003 | rnr1D57N | FALSE | CAN | JSY3602 | 7 | rnr1D57N-Re | TGT | G>T | A/T+1 | G/C-1 | CG>AT | CG>AT | TGTGGTGTG | GTGTTTG | ACA | ACA | C | A | C>A | C>A_ACA | CG>AT_GTGTTTG | 7 | FALSE | FALSE |
| NL4_v.txt | 33372 | 33372-SNV- | SNV | 1 | G | T | 6 | 10091 | 0.05945892 | 20.9997523 | rnr1D57N | FALSE | CAN | JSY3602 | 7 | rnr1D57N-Re | TGT | G>T | A/T+1 | G/C-1 | CG>AT | CG>AT | TGGTGTGTT | GTTTGTG | ACA | ACA | C | A | C>A | C>A_ACA | CG>AT_GTTTGTG | 7 | FALSE | FALSE |
| NL4_v.txt | 33456 | 33456-SNV- | SNV | 1 | G | T | 19 | 10090 | 0.18830525 | 28.3684211 | rnr1D57N | FALSE | CAN | JSY3602 | 7 | rnr1D57N-Re | TGA | G>T | A/T+1 | G/C-1 | CG>AT | CG>AT | TCTTTTGAAT | GAATTTG | ACT | ACT | C | A | C>A | C>A_ACT | CG>AT_GAATTTG | 7 | FALSE | FALSE |
| NL4_v.txt | 33474 | 33474-SNV- | SNV | 1 | C | A | 13 | 10090 | 0.12884044 | 36.1538462 | rnr1D57N | FALSE | CAN | JSY3602 | 7 | rnr1D57N-Re | GCC | C>A | A/T+1 | G/C-1 | CG>AT | CG>AT | GCTATGCCTT | CCTTTTT | CGG | GCC | C | T | C>A | C>A_GCC | CG>AT_CCTTTTT | 7 | FALSE | FALSE |
| NL4_v.txt | 31884 | 31884-Repla | Replacement | 2 | GC | A | 29 | 5292 | 0.54799698 | 38.6724138 | rnr1D57N | FALSE | CAN | JSY3602 | 7 | rnr1D57N-Re | AGC | nope | A/T+1 | >1bpdel | none | Replac. | GCGGCAGCA | GCAAAGC | TCG | AGC | GC | T | NA | NA_AGC | Replac._GCAAAGC | 7 | FALSE | FALSE |
| NL4_v.txt | 32575 | 32575-Repla | Replacement | 2 | G | CA | 3 | 3968 | 0.07560484 | 39 | rnr1D57N | FALSE | CAN | JSY3602 | 7 | rnr1D57N-Re | TGA | nope | >1bp-ins | G/C-1 | none | Replac. | TACCTTGAA | GAAATGT | ACT | ACT | C | none | C>none | C>none_ACT | Replac._GAAATGT | 7 | FALSE | FALSE |
| NL4_v.txt | 31884 | 31884-MNV- | MNV | 2 | GC | AA | 1 | 5292 | 0.01889645 | 27.99985 | rnr1D57N | FALSE | CAN | JSY3602 | 7 | rnr1D57N-Re | AGC | nope | >1bp-ins | >1bpdel | none | MNV | GCGGCAGCA | GCAAAGC | TCG | AGC | GC | none | NA | NA_AGC | MNV_GCAAAGC | 7 | FALSE | FALSE |
| NL4_v.txt | 32498 | 32498-MNV- | MNV | 3 | TTT | ACC | 58 | 13151 | 0.4410311 | 37.3793103 | rnr1D57N | FALSE | CAN | JSY3602 | 7 | rnr1D57N-Re | CTT | nope | >1bp-ins | >1bpdel | none | MNV | AACAACTTTT | TTTTTTG | GAA | CTT | TTT | none | NA | NA_CTT | MNV_TTTTTTG | 7 | FALSE | FALSE |
| NL4_v.txt | 33153 | 33153-MNV- | MNV | 2 | CC | AA | 2 | 5930 | 0.03372681 | 37.7496295 | rnr1D57N | FALSE | CAN | JSY3602 | 7 | rnr1D57N-Re | ACC | nope | >1bp-ins | >1bpdel | none | MNV | AAAAGACCT | CCTGTAC | TGG | ACC | CC | none | NA | NA_ACC | MNV_CCTGTAC | 7 | FALSE | FALSE |
| NL4_v.txt | 31886 | 31886-Insert | Insertion | 1 | - | A | 1 | 5292 | 0.01889645 | 39 | rnr1D57N | FALSE | CAN | JSY3602 | 7 | rnr1D57N-Re | CAA | nope | A/T+1 | >1bpdel | none | A/T+1 | GGCAGCAAA | AAAGCTA | GTT | CAA | - | T | NA | NA_CAA | A/T+1_AAAGCTA | 7 | FALSE | FALSE |
| NL4_v.txt | 31913 | 31913-Insert | Insertion | 1 | - | A | 3 | 5283 | 0.05678592 | 38.6666667 | rnr1D57N | FALSE | CAN | JSY3602 | 7 | rnr1D57N-Re | CAA | nope | A/T+1 | >1bpdel | none | A/T+1 | TGGTGCAAA | AAAAGCC | GTT | CAA | - | T | NA | NA_CAA | A/T+1_AAAAGCC | 7 | FALSE | FALSE |
| NL4_v.txt | 31977 | 31977-Insert | Insertion | 1 | - | C | 2 | 1937 | 0.10325245 | 37.5 | rnr1D57N | FALSE | CAN | JSY3602 | 7 | rnr1D57N-Re | GCC | nope | G/C+1 | >1bpdel | none | G/C+1 | GCCAAGCCG | CCGGGCA | CGG | GCC | - | G | NA | NA_GCC | G/C+1_CCGGGCA | 7 | FALSE | FALSE |
| NL4_v.txt | 32081 | 32081-Insert | Insertion | 1 | - | A | 7 | 1937 | 0.36138358 | 64 | rnr1D57N | FALSE | CAN | JSY3602 | 7 | rnr1D57N-Re | CAA | nope | A/T+1 | >1bpdel | none | A/T+1 | CCATGCAAA | AAAAAAG | GTT | CAA | - | T | NA | NA_CAA | A/T+1_AAAAAAG | 7 | FALSE | FALSE |
| NL4_v.txt | 32352 | 32352-Insert | Insertion | 1 | - | T | 3 | 13151 | 0.02281195 | 64 | rnr1D57N | FALSE | CAN | JSY3602 | 7 | rnr1D57N-Re | CTT | nope | A/T+1 | >1bpdel | none | A/T+1 | AAAACCTTTG | TTTGTAC | GAA | CTT | - | A | NA | NA_CTT | A/T+1_TTTGTAC | 7 | FALSE | FALSE |
| NL4_v.txt | 32402 | 32402-Insert | Insertion | 34 | - | GGAAGTAGA | 5 | 13151 | 0.03801992 | 63.6411765 | rnr1D57N | FALSE | CAN | JSY3602 | 7 | rnr1D57N-Re | AGG | nope | >1bp-ins | >1bpdel | none | >1bp-ins | AACGTAGGA | GGAAGTA | TCC | AGG | - | none | NA | NA_AGG | >1bp-ins_GGAAGTA | 7 | FALSE | FALSE |
| NL4_v.txt | 32498 | 32498-Insert | Insertion | 1 | - | T | 5 | 13151 | 0.03801992 | 32 | rnr1D57N | FALSE | CAN | JSY3602 | 7 | rnr1D57N-Re | CTT | nope | A/T+1 | >1bpdel | none | A/T+1 | AACAACTTTT | TTTTTTG | GAA | CTT | - | A | NA | NA_CTT | A/T+1_TTTTTTG | 7 | FALSE | FALSE |
| NL4_v.txt | 32842 | 32842-Insert | Insertion | 1 | - | A | 16 | 3997 | 0.40030023 | 37.375 | rnr1D57N | FALSE | CAN | JSY3602 | 7 | rnr1D57N-Re | CAA | nope | A/T+1 | >1bpdel | none | A/T+1 | TTACCCAAA | AAAAAAT | GTT | CAA | - | T | NA | NA_CAA | A/T+1_AAAAAAT | 7 | FALSE | FALSE |
| NL4_v.txt | 33087 | 33087-Insert | Insertion | 1 | - | A | 13 | 5930 | 0.21922428 | 62.0769231 | rnr1D57N | FALSE | CAN | JSY3602 | 7 | rnr1D57N-Re | TAA | nope | A/T+1 | >1bpdel | none | A/T+1 | ATAAATAAA | AAATATG | ATT | TAA | - | T | NA | NA_TAA | A/T+1_AAATATG | 7 | FALSE | FALSE |
| NL4_v.txt | 33173 | 33173-Insert | Insertion | 1 | - | A | 6 | 5927 | 0.10123165 | 39 | rnr1D57N | FALSE | CAN | JSY3602 | 7 | rnr1D57N-Re | CAA | nope | A/T+1 | >1bpdel | none | A/T+1 | ACCACCAAG | AAGGGCA | GTT | CAA | - | T | NA | NA_CAA | A/T+1_AAGGGCA | 7 | FALSE | FALSE |
| NL4_v.txt | 33206 | 33206-Insert | Insertion | 2 | - | CT | 4 | 10091 | 0.03963928 | 38.875 | rnr1D57N | FALSE | CAN | JSY3602 | 7 | rnr1D57N-Re | GCT | nope | >1bp-ins | >1bpdel | none | >1bp-ins | CTTAAGCTCT | CTCTCTC | CGA | GCT | - | none | NA | NA_GCT | >1bp-ins_CTCTCTC | 7 | FALSE | FALSE |
| NL4_v.txt | 33244 | 33244-Insert | Insertion | 2 | - | CC | 8 | 10091 | 0.07927857 | 38.8125 | rnr1D57N | FALSE | CAN | JSY3602 | 7 | rnr1D57N-Re | TCT | nope | >1bp-ins | >1bpdel | none | >1bp-ins | CTTCATCTTC | CTTCATC | AGA | TCT | - | none | NA | NA_TCT | >1bp-ins_CTTCATC | 7 | FALSE | FALSE |
| NL4_v.txt | 33356 | 33356-Insert | Insertion | 1 | - | C | 9 | 10091 | 0.08918839 | 54.8888889 | rnr1D57N | FALSE | CAN | JSY3602 | 7 | rnr1D57N-Re | ACC | nope | G/C+1 | >1bpdel | none | G/C+1 | TATTGACCCA | CCCACGT | TGG | ACC | - | G | NA | NA_ACC | G/C+1_CCCACGT | 7 | FALSE | FALSE |
| NL4_v.txt | 32492 | 32492-Deleti | Deletion | 3 | AAC | - | 34 | 13151 | 0.25853547 | 38.4411765 | rnr1D57N | FALSE | CAN | JSY3602 | 7 | rnr1D57N-Re | AAA | nope | >1bp-ins | >1bpdel | none | >1bpdel | ACGGAAAAC | AACAACT | TTT | AAA | AAC | none | NA | NA_AAA | >1bpdel_AACAACT | 7 | FALSE | FALSE |
| NL4_v.txt | 33094 | 33094-Deleti | Deletion | 2 | AT | - | 152 | 5930 | 2.56323777 | 60.5328947 | rnr1D57N | FALSE | CAN | JSY3602 | 7 | rnr1D57N-Re | GAT | nope | >1bp-ins | >1bpdel | none | >1bpdel | AATATGATA | ATATAAG | CTA | GAT | AT | none | NA | NA_GAT | >1bpdel_ATATAAG | 7 | FALSE | FALSE |
| NL4_v.txt | 33189 | 33189-Deleti | Deletion | 2 | AT | - | 209 | 10092 | 2.07094729 | 38.3971292 | rnr1D57N | FALSE | CAN | JSY3602 | 7 | rnr1D57N-Re | AAT | nope | >1bp-ins | >1bpdel | none | >1bpdel | ATACCAATA | ATATGTC | TTA | AAT | AT | none | NA | NA_AAT | >1bpdel_ATATGTC | 7 | FALSE | FALSE |
| NL4_v.txt | 33206 | 33206-Deleti | Deletion | 2 | CT | - | 9 | 10091 | 0.08918839 | 38.4444444 | rnr1D57N | FALSE | CAN | JSY3602 | 7 | rnr1D57N-Re | GCT | nope | >1bp-ins | >1bpdel | none | >1bpdel | CTTAAGCTCT | CTCTCTC | CGA | GCT | CT | none | NA | NA_GCT | >1bpdel_CTCTCTC | 7 | FALSE | FALSE |
| NL4_v.txt | 33364 | 33364-Deleti | Deletion | 11 | TGTGGTGTG | - | 16 | 10091 | 0.15855713 | 62.3125 | rnr1D57N | FALSE | CAN | JSY3602 | 7 | rnr1D57N-Re | CTG | nope | >1bp-ins | >1bpdel | none | >1bpdel | CACGTCTGTG | TGTGGTG | GAC | CTG | TGTGGTGTG | none | NA | NA_CTG | >1bpdel_TGTGGTG | 7 | FALSE | FALSE |
| NL4_v.txt | 31711 | 31711-Deleti | Deletion | 1 | A | - | 11 | 5292 | 0.20786092 | 33.6363636 | rnr1D57N | FALSE | CAN | JSY3602 | 7 | rnr1D57N-Re | CAA | nope | >1bp-ins | A/T-1 | none | A/T-1 | CATTCCAAA | AAAATTT | GTT | GTT | T | none | T>none | T>none_GTT | A/T-1_AAAATTT | 7 | FALSE | FALSE |
| NL4_v.txt | 31723 | 31723-Deleti | Deletion | 1 | A | - | 6 | 5292 | 0.11337869 | 64 | rnr1D57N | FALSE | CAN | JSY3602 | 7 | rnr1D57N-Re | CAA | nope | >1bp-ins | A/T-1 | none | A/T-1 | TGTCCCAAA | AAAAAGT | GTT | GTT | T | none | T>none | T>none_GTT | A/T-1_AAAAAGT | 7 | FALSE | FALSE |
| NL4_v.txt | 32442 | 32442-Deleti | Deletion | 1 | A | - | 11 | 13151 | 0.08364383 | 61.6363636 | rnr1D57N | FALSE | CAN | JSY3602 | 7 | rnr1D57N-Re | TAA | nope | >1bp-ins | A/T-1 | none | A/T-1 | GGAACTAAA | AAAAGTC | ATT | ATT | T | none | T>none | T>none_ATT | A/T-1_AAAAGTC | 7 | FALSE | FALSE |
| NL4_v.txt | 32490 | 32490-Deleti | Deletion | 1 | A | - | 17 | 13151 | 0.12926774 | 38.4705882 | rnr1D57N | FALSE | CAN | JSY3602 | 7 | rnr1D57N-Re | GAA | nope | >1bp-ins | A/T-1 | none | A/T-1 | ATACGGAAA | AAAACAA | CTT | CTT | T | none | T>none | T>none_CTT | A/T-1_AAAACAA | 7 | FALSE | FALSE |
| NL4_v.txt | 32494 | 32494-Deleti | Deletion | 1 | C | - | 58 | 13151 | 0.4410311 | 37.3103448 | rnr1D57N | FALSE | CAN | JSY3602 | 7 | rnr1D57N-Re | ACA | nope | >1bp-ins | G/C-1 | none | G/C-1 | GGAAAACAA | CAACTTT | TGT | ACA | C | none | nope | nope_ACA | G/C-1_CAACTTT | 7 | FALSE | FALSE |
| NL4_v.txt | 32498 | 32498-Deleti | Deletion | 1 | T | - | 60 | 13151 | 0.45623907 | 37.4333292 | rnr1D57N | FALSE | CAN | JSY3602 | 7 | rnr1D57N-Re | CTT | nope | >1bp-ins | A/T-1 | none | A/T-1 | AACAACTTTT | TTTTTTG | GAA | CTT | T | none | nope | nope_CTT | A/T-1_TTTTTTG | 7 | FALSE | FALSE |
| NL4_v.txt | 32528 | 32528-Deleti | Deletion | 1 | G | - | 5 | 13150 | 0.03802281 | 38 | rnr1D57N | FALSE | CAN | JSY3602 | 7 | rnr1D57N-Re | TGG | nope | >1bp-ins | G/C-1 | none | G/C-1 | TTTTCTGGG | GGGGTTT | ACC | ACC | C | none | C>none | C>none_ACC | G/C-1_GGGGTTT | 7 | FALSE | FALSE |
| NL4_v.txt | 32658 | 32658-Deleti | Deletion | 1 | C | - | 9 | 3970 | 0.22670025 | 62.6666667 | rnr1D57N | FALSE | CAN | JSY3602 | 7 | rnr1D57N-Re | ACC | nope | >1bp-ins | G/C-1 | none | G/C-1 | CCTGGACCC | CCCCAGG | TGG | ACC | C | none | nope | nope_ACC | G/C-1_CCCCAGG | 7 | FALSE | FALSE |
| NL4_v.txt | 32707 | 32707-Deleti | Deletion | 1 | C | - | 3 | 3997 | 0.07505629 | 60 | rnr1D57N | FALSE | CAN | JSY3602 | 7 | rnr1D57N-Re | ACC | nope | >1bp-ins | G/C-1 | none | G/C-1 | CGGTAACCC | CCCCAGC | TGG | ACC | C | none | nope | nope_ACC | G/C-1_CCCCAGC | 7 | FALSE | FALSE |
| NL4_v.txt | 32842 | 32842-Deleti | Deletion | 1 | A | - | 29 | 3997 | 0.72554416 | 37.5862069 | rnr1D57N | FALSE | CAN | JSY3602 | 7 | rnr1D57N-Re | CAA | nope | >1bp-ins | A/T-1 | none | A/T-1 | TTACCCAAA | AAAAAAT | GTT | GTT | T | none | T>none | T>none_GTT | A/T-1_AAAAAAT | 7 | FALSE | FALSE |
| NL4_v.txt | 32860 | 32860-Deleti | Deletion | 1 | C | - | 7 | 3997 | 0.17513135 | 38 | rnr1D57N | FALSE | CAN | JSY3602 | 7 | rnr1D57N-Re | GCC | nope | >1bp-ins | G/C-1 | none | G/C-1 | TCCATGCCG | CCGCCAG | CGG | GCC | C | none | nope | nope_GCC | G/C-1_CCGCCAG | 7 | FALSE | FALSE |
| NL4_v.txt | 32924 | 32924-Deleti | Deletion | 1 | A | - | 5 | 5930 | 0.08431703 | 38 | rnr1D57N | FALSE | CAN | JSY3602 | 7 | rnr1D57N-Re | CAA | nope | >1bp-ins | A/T-1 | none | A/T-1 | CAGGGCAAA | AAAAGTG | GTT | GTT | T | none | T>none | T>none_GTT | A/T-1_AAAAGTG | 7 | FALSE | FALSE |
| NL4_v.txt | 33024 | 33024-Deleti | Deletion | 1 | G | - | 4 | 5930 | 0.06745363 | 64 | rnr1D57N | FALSE | CAN | JSY3602 | 7 | rnr1D57N-Re | AGG | nope | >1bp-ins | G/C-1 | none | G/C-1 | GTAACAGGG | GGGATGA | TCC | TCC | C | none | C>none | C>none_TCC | G/C-1_GGGATGA | 7 | FALSE | FALSE |
| NL4_v.txt | 33070 | 33070-Deleti | Deletion | 1 | C | - | 6 | 5930 | 0.10118044 | 55.1666667 | rnr1D57N | FALSE | CAN | JSY3602 | 7 | rnr1D57N-Re | GCC | nope | >1bp-ins | G/C-1 | none | G/C-1 | AATATGCCA | CCAAAGA | CGG | GCC | C | none | nope | nope_GCC | G/C-1_CCAAAGA | 7 | FALSE | FALSE |
| NL4_v.txt | 33108 | 33108-Deleti | Deletion | 1 | A | - | 9 | 5930 | 0.15177066 | 37.2222222 | rnr1D57N | FALSE | CAN | JSY3602 | 7 | rnr1D57N-Re | CAC | nope | >1bp-ins | A/T-1 | none | A/T-1 | GCGCCCACT | ACTGGGC | GTG | GTG | T | none | T>none | T>none_GTG | A/T-1_ACTGGGC | 7 | FALSE | FALSE |
| NL4_v.txt | 33153 | 33153-Deleti | Deletion | 1 | C | - | 1 | 5930 | 0.01686341 | 38.9999999 | rnr1D57N | FALSE | CAN | JSY3602 | 7 | rnr1D57N-Re | ACC | nope | >1bp-ins | G/C-1 | none | G/C-1 | AAAAGACCT | CCTGTAC | TGG | ACC | C | none | nope | nope_ACC | G/C-1_CCTGTAC | 7 | FALSE | FALSE |
| NL4_v.txt | 33168 | 33168-Deleti | Deletion | 1 | C | - | 7 | 5928 | 0.11808367 | 37.8571429 | rnr1D57N | FALSE | CAN | JSY3602 | 7 | rnr1D57N-Re | ACC | nope | >1bp-ins | G/C-1 | none | G/C-1 | ATAGTACCA | CCACCAA | TGG | ACC | C | none | nope | nope_ACC | G/C-1_CCACCAA | 7 | FALSE | FALSE |
| NL4_v.txt | 33236 | 33236-Deleti | Deletion | 1 | T | - | 7 | 10091 | 0.06936874 | 36.8571429 | rnr1D57N | FALSE | CAN | JSY3602 | 7 | rnr1D57N-Re | CTC | nope | >1bp-ins | A/T-1 | none | A/T-1 | TACTTCTCCT | TCCTTCA | GAG | CTC | T | none | nope | nope_CTC | A/T-1_TCCTTCA | 7 | FALSE | FALSE |
| NL4_v.txt | 33251 | 33251-Deleti | Deletion | 1 | A | - | 15 | 10091 | 0.14864731 | 60.8666667 | rnr1D57N | FALSE | CAN | JSY3602 | 7 | rnr1D57N-Re | CAC | nope | >1bp-ins | A/T-1 | none | A/T-1 | TTCATCACCT | ACCTATG | GTG | GTG | T | none | T>none | T>none_GTG | A/T-1_ACCTATG | 7 | FALSE | FALSE |
| NL4_v.txt | 33258 | 33258-Deleti | Deletion | 1 | C | - | 2 | 10091 | 0.01981964 | 51 | rnr1D57N | FALSE | CAN | JSY3602 | 7 | rnr1D57N-Re | GCC | nope | >1bp-ins | G/C-1 | none | G/C-1 | CCTATGCCAT | CCATCCT | CGG | GCC | C | none | nope | nope_GCC | G/C-1_CCATCCT | 7 | FALSE | FALSE |
| NL4_v.txt | 33408 | 33408-Deleti | Deletion | 1 | A | - | 9 | 10091 | 0.08918839 | 53 | rnr1D57N | FALSE | CAN | JSY3602 | 7 | rnr1D57N-Re | GAC | nope | >1bp-ins | A/T-1 | none | A/T-1 | GTTGTGACC | ACCGGCT | CTG | CTG | T | none | T>none | T>none_CTG | A/T-1_ACCGGCT | 7 | FALSE | FALSE |
| NL4_v.txt | 33474 | 33474-Deleti | Deletion | 1 | C | - | 19 | 10090 | 0.18830525 | 31 | rnr1D57N | FALSE | CAN | JSY3602 | 7 | rnr1D57N-Re | GCC | nope | >1bp-ins | G/C-1 | none | G/C-1 | GCTATGCCTT | CCTTTTT | CGG | GCC | C | none | nope | nope_GCC | G/C-1_CCTTTTT | 7 | FALSE | FALSE |
| 171_v.txt | 31723 | 31723-SNV- | SNV | 1 | A | C | 3 | 10179 | 0.02947244 | 62.6666667 | rnr1D57N | FALSE | CAN | JSY14 | 1A | rnr1D57N-Re | CAA | A>C | G/C+1 | A/T-1 | TA>GC | TA>GC | TGTCCCAAA | AAAAAGT | GTT | GTT | T | G | T>G | T>G_GTT | TA>GC_AAAAAGT | 7 | TRUE | FALSE |
| 171_v.txt | 31902 | 31902-SNV- | SNV | 1 | A | C | 5 | 10171 | 0.04915938 | 40.5999589 | rnr1D57N | FALSE | CAN | JSY14 | 1A | rnr1D57N-Re | GAA | A>C | G/C+1 | A/T-1 | TA>GC | TA>GC | CCATTGAATT | AATTTTG | CTT | CTT | T | G | T>G | T>G_CTT | TA>GC_AATTTTG | 7 | TRUE | FALSE |
| 171_v.txt | 31950 | 31950-SNV- | SNV | 1 | A | C | 16 | 24201 | 0.06611297 | 31.6875 | rnr1D57N | FALSE | CAN | JSY14 | 1A | rnr1D57N-Re | CAT | A>C | G/C+1 | A/T-1 | TA>GC | TA>GC | ATCGTCATA | ATAAATG | GTA | GTA | T | G | T>G | T>G_GTA | TA>GC_ATAAATG | 7 | TRUE | FALSE |
| 171_v.txt | 32005 | 32005-SNV- | SNV | 1 | A | C | 15 | 24202 | 0.06197835 | 37.8 | rnr1D57N | FALSE | CAN | JSY14 | 1A | rnr1D57N-Re | AAC | A>C | G/C+1 | A/T-1 | TA>GC | TA>GC | ATGGTAACT | ACTCGTC | TTG | TTG | T | G | T>G | T>G_TTG | TA>GC_ACTCGTC | 7 | TRUE | FALSE |
| 171_v.txt | 32101 | 32101-SNV-T | SNV | 1 | T | G | 9 | 24202 | 0.03718701 | 53.7777778 | rnr1D57N | FALSE | CAN | JSY14 | 1A | rnr1D57N-Re | GTG | T>G | G/C+1 | A/T-1 | TA>GC | TA>GC | CACCAGTGA | TGATATT | CAC | GTG | T | C | T>G | T>G_GTG | TA>GC_TGATATT | 7 | TRUE | FALSE |
| 171_v.txt | 32103 | 32103-SNV- | SNV | 1 | A | C | 10 | 24202 | 0.0413189 | 53.1 | rnr1D57N | FALSE | CAN | JSY14 | 1A | rnr1D57N-Re | GAT | A>C | G/C+1 | A/T-1 | TA>GC | TA>GC | CCAGTGATA | ATATTTA | CTA | CTA | T | G | T>G | T>G_CTA | TA>GC_ATATTTA | 7 | TRUE | FALSE |
| 171_v.txt | 32109 | 32109-SNV- | SNV | 1 | A | C | 6 | 24202 | 0.02479134 | 59.6666667 | rnr1D57N | FALSE | CAN | JSY14 | 1A | rnr1D57N-Re | TAA | A>C | G/C+1 | A/T-1 | TA>GC | TA>GC | ATATTTAATA | AATAGCC | ATT | ATT | T | G | T>G | T>G_ATT | TA>GC_AATAGCC | 7 | TRUE | FALSE |
| 171_v.txt | 32123 | 32123-SNV- | SNV | 1 | A | C | 5 | 24200 | 0.02066116 | 13.8001831 | rnr1D57N | FALSE | CAN | JSY14 | 1A | rnr1D57N-Re | AAA | A>C | G/C+1 | A/T-1 | TA>GC | TA>GC | TTCGAAAAC | AACTTTG | TTT | TTT | T | G | T>G | T>G_TTT | TA>GC_AACTTTG | 7 | TRUE | FALSE |
| 171_v.txt | 32124 | 32124-SNV- | SNV | 1 | A | C | 2 | 24202 | 8.26E-03 | 52.9967384 | rnr1D57N | FALSE | CAN | JSY14 | 1A | rnr1D57N-Re | AAC | A>C | G/C+1 | A/T-1 | TA>GC | TA>GC | TCGAAAACT | ACTTTGT | TTG | TTG | T | G | T>G | T>G_TTG | TA>GC_ACTTTGT | 7 | TRUE | FALSE |
| 171_v.txt | 32132 | 32132-SNV- | SNV | 1 | A | C | 1 | 24202 | 4.13E-03 | 44.9983861 | rnr1D57N | FALSE | CAN | JSY14 | 1A | rnr1D57N-Re | CAC | A>C | G/C+1 | A/T-1 | TA>GC | TA>GC | TTTGTCACCA | ACCACCA | GTG | GTG | T | G | T>G | T>G_GTG | TA>GC_ACCACCA | 7 | TRUE | FALSE |

| | | | | | | | | | | | | | | | | | | | | | | | | | | | | | | | | | | |
|---|---|---|---|---|---|---|---|---|---|---|---|---|---|---|---|---|---|---|---|---|---|---|---|---|---|---|---|---|---|---|---|---|---|---|
| 171_v.txt | 32152 | 32152-SNV-T | SNV | 1 | T | G | 21 | 24202 | 0.08676969 | 50.1427635 | rnr1D57N | FALSE | CAN | JSY14 | 1A | rnr1D57N-Re | ATG | T>G | G/C+1 | A/T-1 | TA>GC | TA>GC | TCTCCATGTA | TGTAAGC | TAC | ATG | T | C | T>G | T>G_ATG | TA>GC_TGTAAGC | 7 | TRUE | FALSE |
| 171_v.txt | 32283 | 32283-SNV-T | SNV | 1 | T | G | 1 | 31383 | 3.19E-03 | 37.999828 | rnr1D57N | FALSE | CAN | JSY14 | 1A | rnr1D57N-Re | GTA | T>G | G/C+1 | A/T-1 | TA>GC | TA>GC | CCAACGTAA | TAAATAT | CAT | GTA | T | C | T>G | T>G_GTA | TA>GC_TAAATAT | 7 | TRUE | FALSE |
| 171_v.txt | 32284 | 32284-SNV-A | SNV | 1 | A | C | 34 | 31383 | 0.10833891 | 27.3528027 | rnr1D57N | FALSE | CAN | JSY14 | 1A | rnr1D57N-Re | TAA | A>C | G/C+1 | A/T-1 | TA>GC | TA>GC | CAACGTAAA | AAATATT | ATT | ATT | T | G | T>G | T>G_ATT | TA>GC_AAATATT | 7 | TRUE | FALSE |
| 171_v.txt | 32294 | 32294-SNV-A | SNV | 1 | A | C | 613 | 31383 | 1.95328681 | 51.2217012 | rnr1D57N | FALSE | CAN | JSY14 | 1A | rnr1D57N-Re | AAT | A>C | G/C+1 | A/T-1 | TA>GC | TA>GC | ATTTGAATTT | ATTTGCG | TTA | TTA | T | G | T>G | T>G_TTA | TA>GC_ATTTGCG | 7 | TRUE | FALSE |
| 171_v.txt | 32319 | 32319-SNV-A | SNV | 1 | A | C | 26 | 31383 | 0.0828474 | 60.6923077 | rnr1D57N | FALSE | CAN | JSY14 | 1A | rnr1D57N-Re | TAA | A>C | G/C+1 | A/T-1 | TA>GC | TA>GC | GTTGTTAAG | AAGATAA | ATT | ATT | T | G | T>G | T>G_ATT | TA>GC_AAGATAA | 7 | TRUE | FALSE |
| 171_v.txt | 32376 | 32376-SNV-A | SNV | 1 | A | C | 1 | 31383 | 3.19E-03 | 59.9965354 | rnr1D57N | FALSE | CAN | JSY14 | 1A | rnr1D57N-Re | AAT | A>C | G/C+1 | A/T-1 | TA>GC | TA>GC | ATAGCAATA | ATAATAA | TTA | TTA | T | G | T>G | T>G_TTA | TA>GC_ATAATAA | 7 | TRUE | FALSE |
| 171_v.txt | 32441 | 32441-SNV-T | SNV | 1 | T | G | 15 | 31383 | 0.04779658 | 50.7990231 | rnr1D57N | FALSE | CAN | JSY14 | 1A | rnr1D57N-Re | CTA | T>G | G/C+1 | A/T-1 | TA>GC | TA>GC | TGGAACTAA | TAAAAGT | GAT | CTA | T | C | T>G | T>G_CTA | TA>GC_TAAAAGT | 7 | TRUE | FALSE |
| 171_v.txt | 32574 | 32574-SNV-T | SNV | 1 | T | G | 741 | 30555 | 2.425135 | 36.7581279 | rnr1D57N | FALSE | CAN | JSY14 | 1A | rnr1D57N-Re | TTG | T>G | G/C+1 | A/T-1 | TA>GC | TA>GC | GTACCTTGA | TGAAATG | AAC | TTG | T | C | T>G | T>G_TTG | TA>GC_TGAAATG | 7 | TRUE | FALSE |
| 171_v.txt | 32631 | 32631-SNV-T | SNV | 1 | T | G | 2 | 30555 | 6.55E-03 | 35.4989067 | rnr1D57N | FALSE | CAN | JSY14 | 1A | rnr1D57N-Re | GTT | T>G | G/C+1 | A/T-1 | TA>GC | TA>GC | CCTTCGTTTT | TTTTTAT | CAA | GTT | T | C | T>G | T>G_GTT | TA>GC_TTTTTAT | 7 | TRUE | FALSE |
| 171_v.txt | 32657 | 32657-SNV-A | SNV | 1 | A | C | 1 | 30555 | 3.27E-03 | 63.9972521 | rnr1D57N | FALSE | CAN | JSY14 | 1A | rnr1D57N-Re | GAC | A>C | G/C+1 | A/T-1 | TA>GC | TA>GC | ACCTGGACC | ACCCCAG | CTG | CTG | T | G | T>G | T>G_CTG | TA>GC_ACCCCAG | 7 | TRUE | FALSE |
| 171_v.txt | 32680 | 32680-SNV-A | SNV | 1 | A | C | 12 | 30555 | 0.03927344 | 60.25 | rnr1D57N | FALSE | CAN | JSY14 | 1A | rnr1D57N-Re | CAA | A>C | G/C+1 | A/T-1 | TA>GC | TA>GC | TTCTCCAATA | AATAACG | GTT | GTT | T | G | T>G | T>G_GTT | TA>GC_AATAACG | 7 | TRUE | FALSE |
| 171_v.txt | 32739 | 32739-SNV-A | SNV | 1 | A | C | 26 | 30555 | 0.08509246 | 57.3461539 | rnr1D57N | FALSE | CAN | JSY14 | 1A | rnr1D57N-Re | TAT | A>C | G/C+1 | A/T-1 | TA>GC | TA>GC | CAGTATATTA | ATTAGAA | ATA | ATA | T | G | T>G | T>G_ATA | TA>GC_ATTAGAA | 7 | TRUE | FALSE |
| 171_v.txt | 32760 | 32760-SNV-A | SNV | 1 | A | C | 37 | 30555 | 0.12109311 | 49.9722413 | rnr1D57N | FALSE | CAN | JSY14 | 1A | rnr1D57N-Re | TAA | A>C | G/C+1 | A/T-1 | TA>GC | TA>GC | ATGGCTAAA | AAAACTT | ATT | ATT | T | G | T>G | T>G_ATT | TA>GC_AAAACTT | 7 | TRUE | FALSE |
| 171_v.txt | 32785 | 32785-SNV-A | SNV | 1 | A | C | 98 | 30555 | 0.3207331 | 47.5612245 | rnr1D57N | FALSE | CAN | JSY14 | 1A | rnr1D57N-Re | AAC | A>C | G/C+1 | A/T-1 | TA>GC | TA>GC | CCCAGAACT | ACTCGAA | TTG | TTG | T | G | T>G | T>G_TTG | TA>GC_ACTCGAA | 7 | TRUE | FALSE |
| 171_v.txt | 32801 | 32801-SNV-A | SNV | 1 | A | C | 17 | 30555 | 0.05563738 | 43.1147929 | rnr1D57N | FALSE | CAN | JSY14 | 1A | rnr1D57N-Re | AAT | A>C | G/C+1 | A/T-1 | TA>GC | TA>GC | ACCGTAATA | ATATTTG | TTA | TTA | T | G | T>G | T>G_TTA | TA>GC_ATATTTG | 7 | TRUE | FALSE |
| 171_v.txt | 32808 | 32808-SNV-A | SNV | 1 | A | C | 8 | 30555 | 0.02618229 | 42.2497147 | rnr1D57N | FALSE | CAN | JSY14 | 1A | rnr1D57N-Re | GAC | A>C | G/C+1 | A/T-1 | TA>GC | TA>GC | TATTTGACA | ACAGGGA | CTG | CTG | T | G | T>G | T>G_CTG | TA>GC_ACAGGGA | 7 | TRUE | FALSE |
| 171_v.txt | 32842 | 32842-SNV-A | SNV | 1 | A | C | 4 | 30555 | 0.01309115 | 38.0007497 | rnr1D57N | FALSE | CAN | JSY14 | 1A | rnr1D57N-Re | CAA | A>C | G/C+1 | A/T-1 | TA>GC | TA>GC | TTACCCAAA | AAAAAAT | GTT | GTT | T | G | T>G | T>G_GTT | TA>GC_AAAAAAT | 7 | TRUE | FALSE |
| 171_v.txt | 32915 | 32915-SNV-T | SNV | 1 | T | G | 69 | 31348 | 0.22010974 | 46.3276586 | rnr1D57N | FALSE | CAN | JSY14 | 1A | rnr1D57N-Re | GTT | T>G | G/C+1 | A/T-1 | TA>GC | TA>GC | ACTAAGTTC | TTCCAGG | CAA | GTT | T | C | T>G | T>G_GTT | TA>GC_TTCCAGG | 7 | TRUE | FALSE |
| 171_v.txt | 32919 | 32919-SNV-A | SNV | 1 | A | C | 13 | 31348 | 0.04146995 | 28.3846154 | rnr1D57N | FALSE | CAN | JSY14 | 1A | rnr1D57N-Re | CAG | A>C | G/C+1 | A/T-1 | TA>GC | TA>GC | AGTTCCAGG | AGGGCAA | GTC | GTC | T | G | T>G | T>G_GTC | TA>GC_AGGGCAA | 7 | TRUE | FALSE |
| 171_v.txt | 32938 | 32938-SNV-A | SNV | 1 | A | C | 6 | 31348 | 0.01913998 | 24.1677153 | rnr1D57N | FALSE | CAN | JSY14 | 1A | rnr1D57N-Re | CAA | A>C | G/C+1 | A/T-1 | TA>GC | TA>GC | TTGCCCAAG | AAGAAAA | GTT | GTT | T | G | T>G | T>G_GTT | TA>GC_AAGAAAA | 7 | TRUE | FALSE |
| 171_v.txt | 32953 | 32953-SNV-T | SNV | 1 | T | G | 18 | 31348 | 0.05741993 | 49.8888889 | rnr1D57N | FALSE | CAN | JSY14 | 1A | rnr1D57N-Re | ATG | T>G | G/C+1 | A/T-1 | TA>GC | TA>GC | AATACATGT | TGTAACC | TAC | ATG | T | C | T>G | T>G_ATG | TA>GC_TGTAACC | 7 | TRUE | FALSE |
| 171_v.txt | 32981 | 32981-SNV-A | SNV | 1 | A | C | 1 | 31348 | 3.19E-03 | 63.9991055 | rnr1D57N | FALSE | CAN | JSY14 | 1A | rnr1D57N-Re | GAG | A>C | G/C+1 | A/T-1 | TA>GC | TA>GC | TGCTGGAGA | AGAAAGG | CTC | CTC | T | G | T>G | T>G_CTC | TA>GC_AGAAAGG | 7 | TRUE | FALSE |
| 171_v.txt | 33054 | 33054-SNV-T | SNV | 1 | T | G | 251 | 31348 | 0.80068904 | 57.5548024 | rnr1D57N | FALSE | CAN | JSY14 | 1A | rnr1D57N-Re | CTG | T>G | G/C+1 | A/T-1 | TA>GC | TA>GC | AAGGACTGC | TGCGTGA | GAC | CTG | T | C | T>G | T>G_CTG | TA>GC_TGCGTGA | 7 | TRUE | FALSE |
| 171_v.txt | 33192 | 33192-SNV-T | SNV | 1 | T | G | 11 | 13896 | 0.07915947 | 33.4545455 | rnr1D57N | FALSE | CAN | JSY14 | 1A | rnr1D57N-Re | ATG | T>G | G/C+1 | A/T-1 | TA>GC | TA>GC | CCAATATGT | TGTCTTT | TAC | ATG | T | C | T>G | T>G_ATG | TA>GC_TGTCTTT | 7 | TRUE | FALSE |
| 171_v.txt | 33269 | 33269-SNV-A | SNV | 1 | A | C | 83 | 13904 | 0.59695052 | 15.0963855 | rnr1D57N | FALSE | CAN | JSY14 | 1A | rnr1D57N-Re | TAG | A>C | G/C+1 | A/T-1 | TA>GC | TA>GC | CTCCATAGA | AGAGAAC | ATC | ATC | T | G | T>G | T>G_ATC | TA>GC_AGAGAAC | 7 | TRUE | FALSE |
| 171_v.txt | 33476 | 33476-SNV-T | SNV | 1 | T | G | 4 | 13902 | 0.02877284 | 37.5 | rnr1D57N | FALSE | CAN | JSY14 | 1A | rnr1D57N-Re | CTT | T>G | G/C+1 | A/T-1 | TA>GC | TA>GC | TATGCCTTTT | TTTTTTT | GAA | CTT | T | C | T>G | T>G_CTT | TA>GC_TTTTTTT | 7 | TRUE | FALSE |
| 171_v.txt | 31808 | 31808-SNV-A | SNV | 1 | A | G | 3 | 10178 | 0.02947534 | 63.9995384 | rnr1D57N | FALSE | CAN | JSY14 | 1A | rnr1D57N-Re | AAA | A>G | G/C+1 | A/T-1 | TA>CG | TA>CG | CTTCCAAATA | AATAAAT | TTT | TTT | T | C | T>C | T>C_TTT | TA>CG_AATAAAT | 7 | TRUE | FALSE |
| 171_v.txt | 31876 | 31876-SNV-A | SNV | 1 | A | G | 7 | 10179 | 0.06876903 | 41.4285685 | rnr1D57N | FALSE | CAN | JSY14 | 1A | rnr1D57N-Re | TAG | A>G | G/C+1 | A/T-1 | TA>CG | TA>CG | AGATATAGG | AGGCGGC | ATC | ATC | T | C | T>C | T>C_ATC | TA>CG_AGGCGGC | 7 | TRUE | FALSE |
| 171_v.txt | 32117 | 32117-SNV-T | SNV | 1 | T | C | 4 | 24201 | 0.01652824 | 62.7499182 | rnr1D57N | FALSE | CAN | JSY14 | 1A | rnr1D57N-Re | ATT | T>C | G/C+1 | A/T-1 | TA>CG | TA>CG | TAGCCATTC | TTCGAAA | TAA | ATT | T | G | T>C | T>C_ATT | TA>CG_TTCGAAA | 7 | TRUE | FALSE |
| 171_v.txt | 32121 | 32121-SNV-A | SNV | 1 | A | G | 10 | 24202 | 0.0413189 | 42.9013212 | rnr1D57N | FALSE | CAN | JSY14 | 1A | rnr1D57N-Re | GAA | A>G | G/C+1 | A/T-1 | TA>CG | TA>CG | CATTCGAAA | AAAACTT | CTT | CTT | T | C | T>C | T>C_CTT | TA>CG_AAAACTT | 7 | TRUE | FALSE |
| 171_v.txt | 32123 | 32123-SNV-A | SNV | 1 | A | G | 11 | 24200 | 0.04545455 | 32.8104125 | rnr1D57N | FALSE | CAN | JSY14 | 1A | rnr1D57N-Re | AAA | A>G | G/C+1 | A/T-1 | TA>CG | TA>CG | TTCGAAAAC | AACTTTG | TTT | TTT | T | C | T>C | T>C_TTT | TA>CG_AACTTTG | 7 | TRUE | FALSE |
| 171_v.txt | 32148 | 32148-SNV-T | SNV | 1 | T | C | 4 | 24202 | 0.01652756 | 60.2497564 | rnr1D57N | FALSE | CAN | JSY14 | 1A | rnr1D57N-Re | CTC | T>C | G/C+1 | A/T-1 | TA>CG | TA>CG | GATGTCTCC | TCCATGT | GAG | CTC | T | G | T>C | T>C_CTC | TA>CG_TCCATGT | 7 | TRUE | FALSE |
| 171_v.txt | 32256 | 32256-SNV-A | SNV | 1 | A | G | 41 | 31371 | 0.13069395 | 36.0487805 | rnr1D57N | FALSE | CAN | JSY14 | 1A | rnr1D57N-Re | TAG | A>G | G/C+1 | A/T-1 | TA>CG | TA>CG | TTTGATAGA | AGACCAA | ATC | ATC | T | C | T>C | T>C_ATC | TA>CG_AGACCAA | 7 | TRUE | FALSE |
| 171_v.txt | 32275 | 32275-SNV-A | SNV | 1 | A | G | 136 | 31385 | 0.43332802 | 37.0955429 | rnr1D57N | FALSE | CAN | JSY14 | 1A | rnr1D57N-Re | GAA | A>G | G/C+1 | A/T-1 | TA>CG | TA>CG | TACGGGAAC | AACCAAC | CTT | CTT | T | C | T>C | T>C_CTT | TA>CG_AACCAAC | 7 | TRUE | FALSE |
| 171_v.txt | 32293 | 32293-SNV-A | SNV | 1 | A | G | 48 | 31383 | 0.15294905 | 51.7682976 | rnr1D57N | FALSE | CAN | JSY14 | 1A | rnr1D57N-Re | GAA | A>G | G/C+1 | A/T-1 | TA>CG | TA>CG | TATTTGAATT | AATTTGC | CTT | CTT | T | C | T>C | T>C_CTT | TA>CG_AATTTGC | 7 | TRUE | FALSE |
| 171_v.txt | 32445 | 32445-SNV-A | SNV | 1 | A | G | 645 | 31383 | 2.05525284 | 53.4821705 | rnr1D57N | FALSE | CAN | JSY14 | 1A | rnr1D57N-Re | AAG | A>G | G/C+1 | A/T-1 | TA>CG | TA>CG | ACTAAAAGT | AGTCCAA | TTC | TTC | T | C | T>C | T>C_TTC | TA>CG_AGTCCAA | 7 | TRUE | FALSE |
| 171_v.txt | 32541 | 32541-SNV-T | SNV | 1 | T | C | 7 | 31383 | 0.02230507 | 31.8570728 | rnr1D57N | FALSE | CAN | JSY14 | 1A | rnr1D57N-Re | TTC | T>C | G/C+1 | A/T-1 | TA>CG | TA>CG | GCAGCTTCA | TCACCAG | AAG | TTC | T | G | T>C | T>C_TTC | TA>CG_TCACCAG | 7 | TRUE | FALSE |
| 171_v.txt | 32543 | 32543-SNV-A | SNV | 1 | A | G | 17 | 31383 | 0.05416946 | 24.1098065 | rnr1D57N | FALSE | CAN | JSY14 | 1A | rnr1D57N-Re | CAC | A>G | G/C+1 | A/T-1 | TA>CG | TA>CG | AGCTTCACC | ACCAGCA | GTG | GTG | T | C | T>C | T>C_GTG | TA>CG_ACCAGCA | 7 | TRUE | FALSE |
| 171_v.txt | 32581 | 32581-SNV-T | SNV | 1 | T | C | 9 | 30554 | 0.02945605 | 36.3332487 | rnr1D57N | FALSE | CAN | JSY14 | 1A | rnr1D57N-Re | GTG | T>C | G/C+1 | A/T-1 | TA>CG | TA>CG | GAAATGTGA | TGAAGGC | CAC | GTG | T | G | T>C | T>C_GTG | TA>CG_TGAAGGC | 7 | TRUE | FALSE |
| 171_v.txt | 32631 | 32631-SNV-T | SNV | 1 | T | C | 2 | 30555 | 6.55E-03 | 63.9970322 | rnr1D57N | FALSE | CAN | JSY14 | 1A | rnr1D57N-Re | GTT | T>C | G/C+1 | A/T-1 | TA>CG | TA>CG | CCTTCGTTTT | TTTTTAT | CAA | GTT | T | G | T>C | T>C_GTT | TA>CG_TTTTTAT | 7 | TRUE | FALSE |
| 171_v.txt | 32662 | 32662-SNV-A | SNV | 1 | A | G | 21 | 30555 | 0.06872852 | 60.3809524 | rnr1D57N | FALSE | CAN | JSY14 | 1A | rnr1D57N-Re | CAG | A>G | G/C+1 | A/T-1 | TA>CG | TA>CG | GACCCCAGG | AGGCACC | GTC | GTC | T | C | T>C | T>C_GTC | TA>CG_AGGCACC | 7 | TRUE | FALSE |
| 171_v.txt | 32728 | 32728-SNV-A | SNV | 1 | A | G | 15 | 30554 | 0.04909341 | 60.9322661 | rnr1D57N | FALSE | CAN | JSY14 | 1A | rnr1D57N-Re | CAA | A>G | G/C+1 | A/T-1 | TA>CG | TA>CG | CCATACAAA | AAAAACA | GTT | GTT | T | C | T>C | T>C_GTT | TA>CG_AAAAACA | 7 | TRUE | FALSE |
| 171_v.txt | 32734 | 32734-SNV-A | SNV | 1 | A | G | 26 | 30555 | 0.08509246 | 59.5385072 | rnr1D57N | FALSE | CAN | JSY14 | 1A | rnr1D57N-Re | CAG | A>G | G/C+1 | A/T-1 | TA>CG | TA>CG | AAAAACAGT | AGTATAT | GTC | GTC | T | C | T>C | T>C_GTC | TA>CG_AGTATAT | 7 | TRUE | FALSE |
| 171_v.txt | 32759 | 32759-SNV-T | SNV | 1 | T | C | 2 | 30555 | 6.55E-03 | 37.4999091 | rnr1D57N | FALSE | CAN | JSY14 | 1A | rnr1D57N-Re | CTA | T>C | G/C+1 | A/T-1 | TA>CG | TA>CG | AATGGCTAA | TAAAACT | GAT | CTA | T | G | T>C | T>C_CTA | TA>CG_TAAAACT | 7 | TRUE | FALSE |
| 171_v.txt | 32787 | 32787-SNV-T | SNV | 1 | T | C | 12 | 30555 | 0.03927344 | 50.499907 | rnr1D57N | FALSE | CAN | JSY14 | 1A | rnr1D57N-Re | CTC | T>C | G/C+1 | A/T-1 | TA>CG | TA>CG | CAGAACTCG | TCGAATT | GAG | CTC | T | G | T>C | T>C_CTC | TA>CG_TCGAATT | 7 | TRUE | FALSE |
| 171_v.txt | 32814 | 32814-SNV-A | SNV | 1 | A | G | 11 | 30555 | 0.03600066 | 32.452985 | rnr1D57N | FALSE | CAN | JSY14 | 1A | rnr1D57N-Re | GAA | A>G | G/C+1 | A/T-1 | TA>CG | TA>CG | ACAGGGAAC | AACAAGT | CTT | CTT | T | C | T>C | T>C_CTT | TA>CG_AACAAGT | 7 | TRUE | FALSE |
| 171_v.txt | 32919 | 32919-SNV-A | SNV | 1 | A | G | 27 | 31348 | 0.0861299 | 43.962963 | rnr1D57N | FALSE | CAN | JSY14 | 1A | rnr1D57N-Re | CAG | A>G | G/C+1 | A/T-1 | TA>CG | TA>CG | AGTTCCAGG | AGGGCAA | GTC | GTC | T | C | T>C | T>C_GTC | TA>CG_AGGGCAA | 7 | TRUE | FALSE |
| 171_v.txt | 32929 | 32929-SNV-T | SNV | 1 | T | C | 26 | 31348 | 0.0829399 | 52.6891248 | rnr1D57N | FALSE | CAN | JSY14 | 1A | rnr1D57N-Re | GTG | T>C | G/C+1 | A/T-1 | TA>CG | TA>CG | CAAAAGTGA | TGATTGC | CAC | GTG | T | G | T>C | T>C_GTG | TA>CG_TGATTGC | 7 | TRUE | FALSE |
| 171_v.txt | 32947 | 32947-SNV-A | SNV | 1 | A | G | 48 | 31348 | 0.15311982 | 57 | rnr1D57N | FALSE | CAN | JSY14 | 1A | rnr1D57N-Re | CAA | A>G | G/C+1 | A/T-1 | TA>CG | TA>CG | AAAACCAAT | AATACAT | GTT | GTT | T | C | T>C | T>C_GTT | TA>CG_AATACAT | 7 | TRUE | FALSE |
| 171_v.txt | 32985 | 32985-SNV-A | SNV | 1 | A | G | 46 | 31348 | 0.14673982 | 57.2391304 | rnr1D57N | FALSE | CAN | JSY14 | 1A | rnr1D57N-Re | AAG | A>G | G/C+1 | A/T-1 | TA>CG | TA>CG | GGAGAAAGC | AGGAATC | TTC | TTC | T | C | T>C | T>C_TTC | TA>CG_AGGAATC | 7 | TRUE | FALSE |
| 171_v.txt | 33000 | 33000-SNV-A | SNV | 1 | A | G | 6 | 31348 | 0.01913998 | 40.6663479 | rnr1D57N | FALSE | CAN | JSY14 | 1A | rnr1D57N-Re | GAA | A>G | G/C+1 | A/T-1 | TA>CG | TA>CG | TGTGAGAAA | AAAACTG | CTT | CTT | T | C | T>C | T>C_CTT | TA>CG_AAAACTG | 7 | TRUE | FALSE |
| 171_v.txt | 33094 | 33094-SNV-A | SNV | 1 | A | G | 4 | 31348 | 0.01275999 | 30.2499165 | rnr1D57N | FALSE | CAN | JSY14 | 1A | rnr1D57N-Re | GAT | A>G | G/C+1 | A/T-1 | TA>CG | TA>CG | AATATGATA | ATATAAG | CTA | CTA | T | C | T>C | T>C_CTA | TA>CG_ATATAAG | 7 | TRUE | FALSE |
| 171_v.txt | 33126 | 33126-SNV-A | SNV | 1 | A | G | 38 | 31348 | 0.12121986 | 42.1052632 | rnr1D57N | FALSE | CAN | JSY14 | 1A | rnr1D57N-Re | CAG | A>G | G/C+1 | A/T-1 | TA>CG | TA>CG | TTGGTCAGA | AGAGGTG | GTC | GTC | T | C | T>C | T>C_GTC | TA>CG_AGAGGTG | 7 | TRUE | FALSE |
| 171_v.txt | 33147 | 33147-SNV-A | SNV | 1 | A | G | 20 | 31348 | 0.06379992 | 35.2 | rnr1D57N | FALSE | CAN | JSY14 | 1A | rnr1D57N-Re | GAA | A>G | G/C+1 | A/T-1 | TA>CG | TA>CG | CCAATGAAA | AAAAGAC | CTT | CTT | T | C | T>C | T>C_CTT | TA>CG_AAAAGAC | 7 | TRUE | FALSE |
| 171_v.txt | 33192 | 33192-SNV-T | SNV | 1 | T | C | 26 | 13896 | 0.1871042 | 35.5384615 | rnr1D57N | FALSE | CAN | JSY14 | 1A | rnr1D57N-Re | ATG | T>C | G/C+1 | A/T-1 | TA>CG | TA>CG | CCAATATGT | TGTCTTT | TAC | ATG | T | G | T>C | T>C_ATG | TA>CG_TGTCTTT | 7 | TRUE | FALSE |
| 171_v.txt | 33476 | 33476-SNV-T | SNV | 1 | T | C | 24 | 13902 | 0.17263703 | 31.7083333 | rnr1D57N | FALSE | CAN | JSY14 | 1A | rnr1D57N-Re | CTT | T>C | G/C+1 | A/T-1 | TA>CG | TA>CG | TATGCCTTTT | TTTTTTT | GAA | CTT | T | G | T>C | T>C_CTT | TA>CG_TTTTTTT | 7 | TRUE | FALSE |
| 171_v.txt | 31822 | 31822-SNV-T | SNV | 1 | T | A | 16 | 10179 | 0.15718636 | 50.125 | rnr1D57N | FALSE | CAN | JSY14 | 1A | rnr1D57N-Re | CTG | T>A | A/T+1 | A/T-1 | TA>AT | TA>AT | TGCATCTGA | TGAATAT | GAC | CTG | T | T | T>A | T>A_CTG | TA>AT_TGAATAT | 7 | TRUE | FALSE |
| 171_v.txt | 31839 | 31839-SNV-A | SNV | 1 | A | T | 9 | 10179 | 0.08841733 | 55.3333333 | rnr1D57N | FALSE | CAN | JSY14 | 1A | rnr1D57N-Re | TAA | A>T | A/T+1 | A/T-1 | TA>AT | TA>AT | TGAAATAAG | AAGATCC | ATT | ATT | T | A | T>A | T>A_ATT | TA>AT_AAGATCC | 7 | TRUE | FALSE |
| 171_v.txt | 31903 | 31903-SNV-A | SNV | 1 | A | T | 10 | 10171 | 0.09831875 | 41.7001128 | rnr1D57N | FALSE | CAN | JSY14 | 1A | rnr1D57N-Re | AAT | A>T | A/T+1 | A/T-1 | TA>AT | TA>AT | CATTGAATTT | ATTTTGG | TTA | TTA | T | A | T>A | T>A_TTA | TA>AT_ATTTTGG | 7 | TRUE | FALSE |
| 171_v.txt | 31957 | 31957-SNV-T | SNV | 1 | T | A | 2 | 24202 | 8.26E-03 | 30.9997232 | rnr1D57N | FALSE | CAN | JSY14 | 1A | rnr1D57N-Re | GTG | T>A | A/T+1 | A/T-1 | TA>AT | TA>AT | TAAATGTGG | TGGCCGC | CAC | GTG | T | T | T>A | T>A_GTG | TA>AT_TGGCCGC | 7 | TRUE | FALSE |
| 171_v.txt | 31964 | 31964-SNV-A | SNV | 1 | A | T | 20 | 24202 | 0.0826378 | 35.25 | rnr1D57N | FALSE | CAN | JSY14 | 1A | rnr1D57N-Re | CAT | A>T | A/T+1 | A/T-1 | TA>AT | TA>AT | GGCCGCATA | ATAATAA | GTA | GTA | T | A | T>A | T>A_GTA | TA>AT_ATAATAA | 7 | TRUE | FALSE |
| 171_v.txt | 31996 | 31996-SNV-T | SNV | 1 | T | A | 30 | 24202 | 0.1239567 | 55.9333333 | rnr1D57N | FALSE | CAN | JSY14 | 1A | rnr1D57N-Re | TTA | T>A | A/T+1 | A/T-1 | TA>AT | TA>AT | TAGCTTTAA | TAAATGG | AAT | TTA | T | T | T>A | T>A_TTA | TA>AT_TAAATGG | 7 | TRUE | FALSE |
| 171_v.txt | 32032 | 32032-SNV-T | SNV | 1 | T | A | 32 | 24202 | 0.13222048 | 60.25 | rnr1D57N | FALSE | CAN | JSY14 | 1A | rnr1D57N-Re | TTC | T>A | A/T+1 | A/T-1 | TA>AT | TA>AT | GGTATTTCA | TCAAAGC | AAG | TTC | T | T | T>A | T>A_TTC | TA>AT_TCAAAGC | 7 | TRUE | FALSE |
| 171_v.txt | 32050 | 32050-SNV-T | SNV | 1 | T | A | 107 | 24202 | 0.44211222 | 62.8878505 | rnr1D57N | FALSE | CAN | JSY14 | 1A | rnr1D57N-Re | CTG | T>A | A/T+1 | A/T-1 | TA>AT | TA>AT | TAAATCTGA | TGATGTG | GAC | CTG | T | T | T>A | T>A_CTG | TA>AT_TGATGTG | 7 | TRUE | FALSE |
| 171_v.txt | 32055 | 32055-SNV-T | SNV | 1 | T | A | 23 | 24202 | 0.09503347 | 60.7391304 | rnr1D57N | FALSE | CAN | JSY14 | 1A | rnr1D57N-Re | GTG | T>A | A/T+1 | A/T-1 | TA>AT | TA>AT | CTGATGTGC | TGCGAGA | CAC | GTG | T | T | T>A | T>A_GTG | TA>AT_TGCGAGA | 7 | TRUE | FALSE |
| 171_v.txt | 32117 | 32117-SNV-T | SNV | 1 | T | A | 10 | 24201 | 0.04132061 | 43.8995168 | rnr1D57N | FALSE | CAN | JSY14 | 1A | rnr1D57N-Re | ATT | T>A | A/T+1 | A/T-1 | TA>AT | TA>AT | TAGCCATTC | TTCGAAA | TAA | ATT | T | T | T>A | T>A_ATT | TA>AT_TTCGAAA | 7 | TRUE | FALSE |
| 171_v.txt | 32126 | 32126-SNV-T | SNV | 1 | T | A | 2 | 24201 | 8.26E-03 | 37.4996221 | rnr1D57N | FALSE | CAN | JSY14 | 1A | rnr1D57N-Re | CTT | T>A | A/T+1 | A/T-1 | TA>AT | TA>AT | GAAAACTTT | TTTGTCA | GAA | CTT | T | T | T>A | T>A_CTT | TA>AT_TTTGTCA | 7 | TRUE | FALSE |
| 171_v.txt | 32127 | 32127-SNV-T | SNV | 1 | T | A | 4 | 24202 | 0.01652756 | 50.4968767 | rnr1D57N | FALSE | CAN | JSY14 | 1A | rnr1D57N-Re | TTT | T>A | A/T+1 | A/T-1 | TA>AT | TA>AT | AAAACTTTG | TTGTCAC | AAA | TTT | T | T | T>A | T>A_TTT | TA>AT_TTGTCAC | 7 | TRUE | FALSE |
| 171_v.txt | 32130 | 32130-SNV-T | SNV | 1 | T | A | 8 | 24202 | 0.03305512 | 49.2493442 | rnr1D57N | FALSE | CAN | JSY14 | 1A | rnr1D57N-Re | GTC | T>A | A/T+1 | A/T-1 | TA>AT | TA>AT | ACTTTGTCAC | TCACCAC | CAG | GTC | T | T | T>A | T>A_GTC | TA>AT_TCACCAC | 7 | TRUE | FALSE |
| 171_v.txt | 32148 | 32148-SNV-T | SNV | 1 | T | A | 8 | 24202 | 0.03305512 | 37.3743069 | rnr1D57N | FALSE | CAN | JSY14 | 1A | rnr1D57N-Re | CTC | T>A | A/T+1 | A/T-1 | TA>AT | TA>AT | GATGTCTCC | TCCATGT | GAG | CTC | T | T | T>A | T>A_CTC | TA>AT_TCCATGT | 7 | TRUE | FALSE |
| 171_v.txt | 32152 | 32152-SNV-T | SNV | 1 | T | A | 3 | 24202 | 0.01239567 | 61.666519 | rnr1D57N | FALSE | CAN | JSY14 | 1A | rnr1D57N-Re | ATG | T>A | A/T+1 | A/T-1 | TA>AT | TA>AT | TCTCCATGTA | TGTAAGC | TAC | ATG | T | T | T>A | T>A_ATG | TA>AT_TGTAAGC | 7 | TRUE | FALSE |
| 171_v.txt | 32275 | 32275-SNV-A | SNV | 1 | A | T | 6 | 31385 | 0.01911741 | 28.6664113 | rnr1D57N | FALSE | CAN | JSY14 | 1A | rnr1D57N-Re | GAA | A>T | A/T+1 | A/T-1 | TA>AT | TA>AT | TACGGGAAC | AACCAAC | CTT | CTT | T | A | T>A | T>A_CTT | TA>AT_AACCAAC | 7 | TRUE | FALSE |
| 171_v.txt | 32283 | 32283-SNV-T | SNV | 1 | T | A | 28 | 31383 | 0.08922028 | 32.9973397 | rnr1D57N | FALSE | CAN | JSY14 | 1A | rnr1D57N-Re | GTA | T>A | A/T+1 | A/T-1 | TA>AT | TA>AT | CCAACGTAA | TAAATAT | CAT | GTA | T | T | T>A | T>A_GTA | TA>AT_TAAATAT | 7 | TRUE | FALSE |
| 171_v.txt | 32287 | 32287-SNV-T | SNV | 1 | T | A | 20 | 31383 | 0.06372877 | 49.55 | rnr1D57N | FALSE | CAN | JSY14 | 1A | rnr1D57N-Re | ATA | T>A | A/T+1 | A/T-1 | TA>AT | TA>AT | CGTAAATAT | TATTTGA | TAT | ATA | T | T | T>A | T>A_ATA | TA>AT_TATTTGA | 7 | TRUE | FALSE |
| 171_v.txt | 32294 | 32294-SNV-A | SNV | 1 | A | T | 13 | 31383 | 0.0414237 | 50.3064145 | rnr1D57N | FALSE | CAN | JSY14 | 1A | rnr1D57N-Re | AAT | A>T | A/T+1 | A/T-1 | TA>AT | TA>AT | ATTTGAATTT | ATTTGCG | TTA | TTA | T | A | T>A | T>A_TTA | TA>AT_ATTTGCG | 7 | TRUE | FALSE |
| 171_v.txt | 32377 | 32377-SNV-T | SNV | 1 | T | A | 6 | 31375 | 0.01912351 | 52.3231313 | rnr1D57N | FALSE | CAN | JSY14 | 1A | rnr1D57N-Re | ATA | T>A | A/T+1 | A/T-1 | TA>AT | TA>AT | TAGCAATAA | TAATAAA | TAT | ATA | T | T | T>A | T>A_ATA | TA>AT_TAATAAA | 7 | TRUE | FALSE |
| 171_v.txt | 32475 | 32475-SNV-A | SNV | 1 | A | T | 17 | 31383 | 0.05416946 | 45.2352941 | rnr1D57N | FALSE | CAN | JSY14 | 1A | rnr1D57N-Re | GAA | A>T | A/T+1 | A/T-1 | TA>AT | TA>AT | ATGTAGAAG | AAGGTTA | CTT | CTT | T | A | T>A | T>A_CTT | TA>AT_AAGGTTA | 7 | TRUE | FALSE |
| 171_v.txt | 32498 | 32498-SNV-T | SNV | 1 | T | A | 6 | 31383 | 0.01911863 | 42.1655206 | rnr1D57N | FALSE | CAN | JSY14 | 1A | rnr1D57N-Re | CTT | T>A | A/T+1 | A/T-1 | TA>AT | TA>AT | AACAACTTTT | TTTTTTG | GAA | CTT | T | T | T>A | T>A_CTT | TA>AT_TTTTTTG | 7 | TRUE | FALSE |
| 171_v.txt | 32541 | 32541-SNV-T | SNV | 1 | T | A | 1 | 31383 | 3.19E-03 | 37.9997802 | rnr1D57N | FALSE | CAN | JSY14 | 1A | rnr1D57N-Re | TTC | T>A | A/T+1 | A/T-1 | TA>AT | TA>AT | GCAGCTTCA | TCACCAG | AAG | TTC | T | T | T>A | T>A_TTC | TA>AT_TCACCAG | 7 | TRUE | FALSE |
| 171_v.txt | 32551 | 32551-SNV-T | SNV | 1 | T | A | 5 | 31379 | 0.01593422 | 37.9999532 | rnr1D57N | FALSE | CAN | JSY14 | 1A | rnr1D57N-Re | GTG | T>A | A/T+1 | A/T-1 | TA>AT | TA>AT | CAGCAGTGA | TGATACC | CAC | GTG | T | T | T>A | T>A_GTG | TA>AT_TGATACC | 7 | TRUE | FALSE |
| 171_v.txt | 32574 | 32574-SNV-T | SNV | 1 | T | A | 5 | 30555 | 0.01636393 | 37.798673 | rnr1D57N | FALSE | CAN | JSY14 | 1A | rnr1D57N-Re | TTG | T>A | A/T+1 | A/T-1 | TA>AT | TA>AT | GTACCTTGA | TGAAATG | AAC | TTG | T | T | T>A | T>A_TTG | TA>AT_TGAAATG | 7 | TRUE | FALSE |
| 171_v.txt | 32759 | 32759-SNV-T | SNV | 1 | T | A | 2 | 30555 | 6.55E-03 | 36.9995679 | rnr1D57N | FALSE | CAN | JSY14 | 1A | rnr1D57N-Re | CTA | T>A | A/T+1 | A/T-1 | TA>AT | TA>AT | AATGGCTAA | TAAAACT | GAT | CTA | T | T | T>A | T>A_CTA | TA>AT_TAAAACT | 7 | TRUE | FALSE |
| 171_v.txt | 32787 | 32787-SNV-T | SNV | 1 | T | A | 4 | 30555 | 0.01309115 | 37.4998073 | rnr1D57N | FALSE | CAN | JSY14 | 1A | rnr1D57N-Re | CTC | T>A | A/T+1 | A/T-1 | TA>AT | TA>AT | CAGAACTCG | TCGAATT | GAG | CTC | T | T | T>A | T>A_CTC | TA>AT_TCGAATT | 7 | TRUE | FALSE |
| 171_v.txt | 32806 | 32806-SNV-T | SNV | 1 | T | A | 64 | 30555 | 0.20945835 | 46.140625 | rnr1D57N | FALSE | CAN | JSY14 | 1A | rnr1D57N-Re | TTG | T>A | A/T+1 | A/T-1 | TA>AT | TA>AT | AATATTTGAC | TGACAGG | AAC | TTG | T | T | T>A | T>A_TTG | TA>AT_TGACAGG | 7 | TRUE | FALSE |
| 171_v.txt | 32808 | 32808-SNV-A | SNV | 1 | A | T | 5 | 30555 | 0.01636393 | 36.7985487 | rnr1D57N | FALSE | CAN | JSY14 | 1A | rnr1D57N-Re | GAC | A>T | A/T+1 | A/T-1 | TA>AT | TA>AT | TATTTGACA | ACAGGGA | CTG | CTG | T | A | T>A | T>A_CTG | TA>AT_ACAGGGA | 7 | TRUE | FALSE |
| 171_v.txt | 32938 | 32938-SNV-A | SNV | 1 | A | T | 1287 | 31348 | 4.10552507 | 56.0018787 | rnr1D57N | FALSE | CAN | JSY14 | 1A | rnr1D57N-Re | CAA | A>T | A/T+1 | A/T-1 | TA>AT | TA>AT | TTGCCCAAG | AAGAAAA | GTT | GTT | T | A | T>A | T>A_GTT | TA>AT_AAGAAAA | 7 | TRUE | FALSE |
| 171_v.txt | 32992 | 32992-SNV-T | SNV | 1 | T | A | 39 | 31348 | 0.12440985 | 57.1025641 | rnr1D57N | FALSE | CAN | JSY14 | 1A | rnr1D57N-Re | CTT | T>A | A/T+1 | A/T-1 | TA>AT | TA>AT | GGAATCTTT | TTTGTGA | GAA | CTT | T | T | T>A | T>A_CTT | TA>AT_TTTGTGA | 7 | TRUE | FALSE |
| 171_v.txt | 33000 | 33000-SNV-A | SNV | 1 | A | T | 3 | 31348 | 9.57E-03 | 37.9991037 | rnr1D57N | FALSE | CAN | JSY14 | 1A | rnr1D57N-Re | GAA | A>T | A/T+1 | A/T-1 | TA>AT | TA>AT | TGTGAGAAA | AAAACTG | CTT | CTT | T | A | T>A | T>A_CTT | TA>AT_AAAACTG | 7 | TRUE | FALSE |
| 171_v.txt | 33066 | 33066-SNV-T | SNV | 1 | T | A | 2 | 31348 | 6.38E-03 | 63.9946301 | rnr1D57N | FALSE | CAN | JSY14 | 1A | rnr1D57N-Re | ATA | T>A | A/T+1 | A/T-1 | TA>AT | TA>AT | ACAGAATAT | TATGCCA | TAT | ATA | T | T | T>A | T>A_ATA | TA>AT_TATGCCA | 7 | TRUE | FALSE |
| 171_v.txt | 33090 | 33090-SNV-T | SNV | 1 | T | A | 3 | 31348 | 9.57E-03 | 54.9952892 | rnr1D57N | FALSE | CAN | JSY14 | 1A | rnr1D57N-Re | ATA | T>A | A/T+1 | A/T-1 | TA>AT | TA>AT | AATAAATAT | TATGATA | TAT | ATA | T | T | T>A | T>A_ATA | TA>AT_TATGATA | 7 | TRUE | FALSE |
| 171_v.txt | 33095 | 33095-SNV-T | SNV | 1 | T | A | 9 | 31348 | 0.02870997 | 37.6661168 | rnr1D57N | FALSE | CAN | JSY14 | 1A | rnr1D57N-Re | ATA | T>A | A/T+1 | A/T-1 | TA>AT | TA>AT | ATATGATAT | TATAAGA | TAT | ATA | T | T | T>A | T>A_ATA | TA>AT_TATAAGA | 7 | TRUE | FALSE |
| 171_v.txt | 33295 | 33295-SNV-T | SNV | 1 | T | A | 13 | 13904 | 0.09349827 | 53.2307692 | rnr1D57N | FALSE | CAN | JSY14 | 1A | rnr1D57N-Re | CTC | T>A | A/T+1 | A/T-1 | TA>AT | TA>AT | TTACTCTCGT | TCGTCGG | GAG | CTC | T | T | T>A | T>A_CTC | TA>AT_TCGTCGG | 7 | TRUE | FALSE |
| 171_v.txt | 33321 | 33321-SNV-A | SNV | 1 | A | T | 14 | 13904 | 0.10069045 | 53.4285714 | rnr1D57N | FALSE | CAN | JSY14 | 1A | rnr1D57N-Re | CAA | A>T | A/T+1 | A/T-1 | TA>AT | TA>AT | GGATACAAT | AATTCTT | GTT | GTT | T | A | T>A | T>A_GTT | TA>AT_AATTCTT | 7 | TRUE | FALSE |
| 171_v.txt | 31650 | 31650-SNV-C | SNV | 1 | C | T | 25 | 10179 | 0.24560369 | 36.8 | rnr1D57N | FALSE | CAN | JSY14 | 1A | rnr1D57N-Re | GCG | C>T | A/T+1 | G/C-1 | CG>TA | CG>TA | AGAATGCGA | CGAAATG | CGC | GCG | C | A | C>T | C>T_GCG | CG>TA_CGAAATG | 7 | TRUE | FALSE |
| 171_v.txt | 31806 | 31806-SNV-C | SNV | 1 | C | T | 257 | 10178 | 2.52505404 | 59.4427379 | rnr1D57N | FALSE | CAN | JSY14 | 1A | rnr1D57N-Re | CCA | C>T | A/T+1 | G/C-1 | CG>TA | CG>TA | ATCTTCCAAA | CAAATAA | GGT | CCA | C | A | C>T | C>T_CCA | CG>TA_CAAATAA | 7 | TRUE | FALSE |
| 171_v.txt | 31962 | 31962-SNV-G | SNV | 1 | G | A | 45 | 24202 | 0.18593505 | 36.8 | rnr1D57N | FALSE | CAN | JSY14 | 1A | rnr1D57N-Re | CGC | G>A | A/T+1 | G/C-1 | CG>TA | CG>TA | GTGGCCGCA | GCATAAT | GCG | GCG | C | T | C>T | C>T_GCG | CG>TA_GCATAAT | 7 | TRUE | FALSE |
| 171_v.txt | 32041 | 32041-SNV-G | SNV | 1 | G | A | 56 | 24202 | 0.23138584 | 62.1785714 | rnr1D57N | FALSE | CAN | JSY14 | 1A | rnr1D57N-Re | TGC | G>A | A/T+1 | G/C-1 | CG>TA | CG>TA | AAGCTTGCA | GCATAAA | ACG | ACG | C | T | C>T | C>T_ACG | CG>TA_GCATAAA | 7 | TRUE | FALSE |
| 171_v.txt | 32088 | 32088-SNV-C | SNV | 1 | C | T | 84 | 24202 | 0.34707875 | 60.3333333 | rnr1D57N | FALSE | CAN | JSY14 | 1A | rnr1D57N-Re | GCC | C>T | A/T+1 | G/C-1 | CG>TA | CG>TA | AAAAAGCCT | CCTGCAA | CGG | GCC | C | A | C>T | C>T_GCC | CG>TA_CCTGCAA | 7 | TRUE | FALSE |
| 171_v.txt | 32092 | 32092-SNV-C | SNV | 1 | C | T | 28 | 24202 | 0.11569292 | 60.5 | rnr1D57N | FALSE | CAN | JSY14 | 1A | rnr1D57N-Re | GCA | C>T | A/T+1 | G/C-1 | CG>TA | CG>TA | AGCCTGCAA | CAACACC | CGT | GCA | C | A | C>T | C>T_GCA | CG>TA_CAACACC | 7 | TRUE | FALSE |
| 171_v.txt | 32114 | 32114-SNV-C | SNV | 1 | C | T | 22 | 24202 | 0.09090158 | 49.5908792 | rnr1D57N | FALSE | CAN | JSY14 | 1A | rnr1D57N-Re | GCC | C>T | A/T+1 | G/C-1 | CG>TA | CG>TA | TAATAGCCA | CCATTCG | CGG | GCC | C | A | C>T | C>T_GCC | CG>TA_CCATTCG | 7 | TRUE | FALSE |
| 171_v.txt | 32115 | 32115-SNV-C | SNV | 1 | C | T | 9 | 24202 | 0.03718701 | 51.1116928 | rnr1D57N | FALSE | CAN | JSY14 | 1A | rnr1D57N-Re | CCA | C>T | A/T+1 | G/C-1 | CG>TA | CG>TA | AATAGCCAT | CATTCGA | GGT | CCA | C | A | C>T | C>T_CCA | CG>TA_CATTCGA | 7 | TRUE | FALSE |
| 171_v.txt | 32120 | 32120-SNV-G | SNV | 1 | G | A | 5 | 24202 | 0.02065945 | 63.9783258 | rnr1D57N | FALSE | CAN | JSY14 | 1A | rnr1D57N-Re | CGA | G>A | A/T+1 | G/C-1 | CG>TA | CG>TA | CCATTCGAA | GAAAACT | GCT | GCT | C | T | C>T | C>T_GCT | CG>TA_GAAAACT | 7 | TRUE | FALSE |
| 171_v.txt | 32125 | 32125-SNV-C | SNV | 1 | C | T | 1 | 24202 | 4.13E-03 | 63.9999996 | rnr1D57N | FALSE | CAN | JSY14 | 1A | rnr1D57N-Re | ACT | C>T | A/T+1 | G/C-1 | CG>TA | CG>TA | CGAAAACTT | CTTTGTC | TGA | ACT | C | A | C>T | C>T_ACT | CG>TA_CTTTGTC | 7 | TRUE | FALSE |
| 171_v.txt | 32131 | 32131-SNV-C | SNV | 1 | C | T | 7 | 24202 | 0.02892323 | 48.2859187 | rnr1D57N | FALSE | CAN | JSY14 | 1A | rnr1D57N-Re | TCA | C>T | A/T+1 | G/C-1 | CG>TA | CG>TA | CTTTGTCACC | CACCACC | AGT | TCA | C | A | C>T | C>T_TCA | CG>TA_CACCACC | 7 | TRUE | FALSE |
| 171_v.txt | 32164 | 32164-SNV-C | SNV | 1 | C | T | 5 | 24202 | 0.02065945 | 44.7991644 | rnr1D57N | FALSE | CAN | JSY14 | 1A | rnr1D57N-Re | GCG | C>T | A/T+1 | G/C-1 | CG>TA | CG>TA | CCAAAGCGC | CGCCAAA | CGC | GCG | C | A | C>T | C>T_GCG | CG>TA_CGCCAAA | 7 | TRUE | FALSE |
| 171_v.txt | 32165 | 32165-SNV-G | SNV | 1 | G | A | 35 | 24202 | 0.14461615 | 47.1709455 | rnr1D57N | FALSE | CAN | JSY14 | 1A | rnr1D57N-Re | CGC | G>A | A/T+1 | G/C-1 | CG>TA | CG>TA | CAAAGCGCC | GCCAAAT | GCG | GCG | C | T | C>T | C>T_GCG | CG>TA_GCCAAAT | 7 | TRUE | FALSE |
| 171_v.txt | 32271 | 32271-SNV-C | SNV | 1 | C | T | 33 | 31385 | 0.10514577 | 34.0910056 | rnr1D57N | FALSE | CAN | JSY14 | 1A | rnr1D57N-Re | ACG | C>T | A/T+1 | G/C-1 | CG>TA | CG>TA | AAAATACGG | CGGGAAC | TGC | ACG | C | A | C>T | C>T_ACG | CG>TA_CGGGAAC | 7 | TRUE | FALSE |
| 171_v.txt | 32272 | 32272-SNV-G | SNV | 1 | G | A | 148 | 31385 | 0.47156285 | 36.7094595 | rnr1D57N | FALSE | CAN | JSY14 | 1A | rnr1D57N-Re | CGG | G>A | A/T+1 | G/C-1 | CG>TA | CG>TA | AAATACGGG | GGGAACC | GCC | GCC | C | T | C>T | C>T_GCC | CG>TA_GGGAACC | 7 | TRUE | FALSE |
| 171_v.txt | 32274 | 32274-SNV-G | SNV | 1 | G | A | 70 | 31385 | 0.22303648 | 36.8124083 | rnr1D57N | FALSE | CAN | JSY14 | 1A | rnr1D57N-Re | GGA | G>A | A/T+1 | G/C-1 | CG>TA | CG>TA | ATACGGGAA | GAACCAA | CCT | CCT | C | T | C>T | C>T_CCT | CG>TA_GAACCAA | 7 | TRUE | FALSE |
| 171_v.txt | 32299 | 32299-SNV-C | SNV | 1 | C | T | 52 | 31383 | 0.1656948 | 49.5961539 | rnr1D57N | FALSE | CAN | JSY14 | 1A | rnr1D57N-Re | GCG | C>T | A/T+1 | G/C-1 | CG>TA | CG>TA | AATTTGCGG | CGGCAGA | CGC | GCG | C | A | C>T | C>T_GCG | CG>TA_CGGCAGA | 7 | TRUE | FALSE |
| 171_v.txt | 32301 | 32301-SNV-G | SNV | 1 | G | A | 17 | 31383 | 0.05416946 | 50.2941177 | rnr1D57N | FALSE | CAN | JSY14 | 1A | rnr1D57N-Re | GGC | G>A | A/T+1 | G/C-1 | CG>TA | CG>TA | TTTGCGGCA | GCAGAAA | CCG | CCG | C | T | C>T | C>T_CCG | CG>TA_GCAGAAA | 7 | TRUE | FALSE |
| 171_v.txt | 32313 | 32313-SNV-G | SNV | 1 | G | A | 55 | 31383 | 0.17525412 | 60.4909091 | rnr1D57N | FALSE | CAN | JSY14 | 1A | rnr1D57N-Re | GGT | G>A | A/T+1 | G/C-1 | CG>TA | CG>TA | ATAATGGTT | GTTGTTA | CCA | CCA | C | T | C>T | C>T_CCA | CG>TA_GTTGTTA | 7 | TRUE | FALSE |
| 171_v.txt | 32361 | 32361-SNV-G | SNV | 1 | G | A | 4 | 31383 | 0.01274575 | 50.2498255 | rnr1D57N | FALSE | CAN | JSY14 | 1A | rnr1D57N-Re | AGA | G>A | A/T+1 | G/C-1 | CG>TA | CG>TA | GTACCAGAG | GAGTTCT | TCT | TCT | C | T | C>T | C>T_TCT | CG>TA_GAGTTCT | 7 | TRUE | FALSE |
| 171_v.txt | 32449 | 32449-SNV-C | SNV | 1 | C | T | 68 | 31383 | 0.21667782 | 51.4558824 | rnr1D57N | FALSE | CAN | JSY14 | 1A | rnr1D57N-Re | CCA | C>T | A/T+1 | G/C-1 | CG>TA | CG>TA | AAAGTCCAA | CAATGAA | GGT | CCA | C | A | C>T | C>T_CCA | CG>TA_CAATGAA | 7 | TRUE | FALSE |
| 171_v.txt | 32466 | 32466-SNV-C | SNV | 1 | C | T | 33 | 31383 | 0.10515247 | 48.0606061 | rnr1D57N | FALSE | CAN | JSY14 | 1A | rnr1D57N-Re | GCC | C>T | A/T+1 | G/C-1 | CG>TA | CG>TA | AGAGAGCCA | CCAATGT | CGG | GCC | C | A | C>T | C>T_GCC | CG>TA_CCAATGT | 7 | TRUE | FALSE |
| 171_v.txt | 32487 | 32487-SNV-C | SNV | 1 | C | T | 18 | 31383 | 0.05735589 | 44.1661837 | rnr1D57N | FALSE | CAN | JSY14 | 1A | rnr1D57N-Re | ACG | C>T | A/T+1 | G/C-1 | CG>TA | CG>TA | AAGATACGG | CGGAAAA | TGC | ACG | C | A | C>T | C>T_ACG | CG>TA_CGGAAAA | 7 | TRUE | FALSE |
| 171_v.txt | 32488 | 32488-SNV-G | SNV | 1 | G | A | 62 | 31383 | 0.19755919 | 47.6293824 | rnr1D57N | FALSE | CAN | JSY14 | 1A | rnr1D57N-Re | CGG | G>A | A/T+1 | G/C-1 | CG>TA | CG>TA | AGATACGGA | GGAAAAC | GCC | GCC | C | T | C>T | C>T_GCC | CG>TA_GGAAAAC | 7 | TRUE | FALSE |
| 171_v.txt | 32535 | 32535-SNV-G | SNV | 1 | G | A | 32 | 31383 | 0.10196603 | 37.3703908 | rnr1D57N | FALSE | CAN | JSY14 | 1A | rnr1D57N-Re | TGC | G>A | A/T+1 | G/C-1 | CG>TA | CG>TA | GGGTTTGCA | GCAGCTT | ACG | ACG | C | T | C>T | C>T_ACG | CG>TA_GCAGCTT | 7 | TRUE | FALSE |
| 171_v.txt | 32536 | 32536-SNV-C | SNV | 1 | C | T | 51 | 31383 | 0.16250836 | 36.9799044 | rnr1D57N | FALSE | CAN | JSY14 | 1A | rnr1D57N-Re | GCA | C>T | A/T+1 | G/C-1 | CG>TA | CG>TA | GGTTTGCAG | CAGCTTC | CGT | GCA | C | A | C>T | C>T_GCA | CG>TA_CAGCTTC | 7 | TRUE | FALSE |

| | | | | | | | | | | | | | | | | | | | | | | | | | | | | | | | | | | |
|---|---|---|---|---|---|---|---|---|---|---|---|---|---|---|---|---|---|---|---|---|---|---|---|---|---|---|---|---|---|---|---|---|---|---|
| 171_v.txt | 32542 | 32542-SNV-( | SNV | 1 | C | T | 43 | 31383 | 0.13701686 | 37.3486412 | rnr1D57N | FALSE | CAN | JSY14 | 1A | rnr1D57N-Re | TCA | C>T | A/T+1 | G/C-1 | CG>TA | CG>TA | CAGCTTCAC( | CACCAGC | AGT | TCA | C | A | C>T | C>T_TCA | CG>TA_CACCAGC | 7 | TRUE | FALSE |
| 171_v.txt | 32544 | 32544-SNV-( | SNV | 1 | C | T | 116 | 31383 | 0.36962687 | 37.835771 | rnr1D57N | FALSE | CAN | JSY14 | 1A | rnr1D57N-Re | ACC | C>T | A/T+1 | G/C-1 | CG>TA | CG>TA | GCTTCACCA( | CCAGCAG | TGG | ACC | C | A | C>T | C>T_ACC | CG>TA_CCAGCAG | 7 | TRUE | FALSE |
| 171_v.txt | 32545 | 32545-SNV-( | SNV | 1 | C | T | 13 | 31383 | 0.0414237 | 34.7692308 | rnr1D57N | FALSE | CAN | JSY14 | 1A | rnr1D57N-Re | CCA | C>T | A/T+1 | G/C-1 | CG>TA | CG>TA | CTTCACCAG( | CAGCAGT | GGT | CCA | C | A | C>T | C>T_CCA | CG>TA_CAGCAGT | 7 | TRUE | FALSE |
| 171_v.txt | 32550 | 32550-SNV-( | SNV | 1 | G | A | 2 | 31379 | 6.37E-03 | 24.499845 | rnr1D57N | FALSE | CAN | JSY14 | 1A | rnr1D57N-Re | AGT | G>A | A/T+1 | G/C-1 | CG>TA | CG>TA | CCAGCAGTG. | GTGATAC | TCA | TCA | C | T | C>T | C>T_TCA | CG>TA_GTGATAC | 7 | TRUE | FALSE |
| 171_v.txt | 32580 | 32580-SNV-( | SNV | 1 | G | A | 7 | 30554 | 0.02291026 | 29.2833482 | rnr1D57N | FALSE | CAN | JSY14 | 1A | rnr1D57N-Re | TGT | G>A | A/T+1 | G/C-1 | CG>TA | CG>TA | TGAAATGTG. | GTGAAGG | ACA | ACA | C | T | C>T | C>T_ACA | CG>TA_GTGAAGG | 7 | TRUE | FALSE |
| 171_v.txt | 32590 | 32590-SNV-( | SNV | 1 | C | T | 34 | 30554 | 0.11127839 | 34.5 | rnr1D57N | FALSE | CAN | JSY14 | 1A | rnr1D57N-Re | GCG | C>T | A/T+1 | G/C-1 | CG>TA | CG>TA | AGGCAGCGT | CGTTAAT | CGC | GCG | C | A | C>T | C>T_GCG | CG>TA_CGTTAAT | 7 | TRUE | FALSE |
| 171_v.txt | 32609 | 32609-SNV-( | SNV | 1 | C | T | 42 | 30555 | 0.13745705 | 45.737016 | rnr1D57N | FALSE | CAN | JSY14 | 1A | rnr1D57N-Re | CCC | C>T | A/T+1 | G/C-1 | CG>TA | CG>TA | GGAAACCCA | CCAACCT | GGG | CCC | C | A | C>T | C>T_CCC | CG>TA_CCAACCT | 7 | TRUE | FALSE |
| 171_v.txt | 32610 | 32610-SNV-( | SNV | 1 | C | T | 31 | 30555 | 0.10145639 | 45.0204353 | rnr1D57N | FALSE | CAN | JSY14 | 1A | rnr1D57N-Re | CCA | C>T | A/T+1 | G/C-1 | CG>TA | CG>TA | GAAACCCAA | CAACCTA | GGT | CCA | C | A | C>T | C>T_CCA | CG>TA_CAACCTA | 7 | TRUE | FALSE |
| 171_v.txt | 32658 | 32658-SNV-( | SNV | 1 | C | T | 6 | 30555 | 0.01963672 | 51.995839 | rnr1D57N | FALSE | CAN | JSY14 | 1A | rnr1D57N-Re | ACC | C>T | A/T+1 | G/C-1 | CG>TA | CG>TA | CCTGGACCC( | CCCCAGG | TGG | ACC | C | A | C>T | C>T_ACC | CG>TA_CCCCAGG | 7 | TRUE | FALSE |
| 171_v.txt | 32678 | 32678-SNV-( | SNV | 1 | C | T | 85 | 30554 | 0.27819598 | 58.4315724 | rnr1D57N | FALSE | CAN | JSY14 | 1A | rnr1D57N-Re | TCC | C>T | A/T+1 | G/C-1 | CG>TA | CG>TA | GTTTCTCCAA | CCAATAA | AGG | TCC | C | A | C>T | C>T_TCC | CG>TA_CCAATAA | 7 | TRUE | FALSE |
| 171_v.txt | 32700 | 32700-SNV-( | SNV | 1 | C | T | 29 | 30555 | 0.09491082 | 55.6896552 | rnr1D57N | FALSE | CAN | JSY14 | 1A | rnr1D57N-Re | GCC | C>T | A/T+1 | G/C-1 | CG>TA | CG>TA | ACTGGGCCG | CCGGTAA | CGG | GCC | C | A | C>T | C>T_GCC | CG>TA_CCGGTAA | 7 | TRUE | FALSE |
| 171_v.txt | 32749 | 32749-SNV-( | SNV | 1 | C | T | 158 | 30555 | 0.51710031 | 57.3101266 | rnr1D57N | FALSE | CAN | JSY14 | 1A | rnr1D57N-Re | CCG | C>T | A/T+1 | G/C-1 | CG>TA | CG>TA | GAAACCCGA | CGATAAT | GGC | CCG | C | A | C>T | C>T_CCG | CG>TA_CGATAAT | 7 | TRUE | FALSE |
| 171_v.txt | 32775 | 32775-SNV-( | SNV | 1 | G | A | 112 | 30555 | 0.36655212 | 49.6339286 | rnr1D57N | FALSE | CAN | JSY14 | 1A | rnr1D57N-Re | AGC | G>A | A/T+1 | G/C-1 | CG>TA | CG>TA | ATGGAAGCG | GCGACCC | TCG | TCG | C | T | C>T | C>T_TCG | CG>TA_GCGACCC | 7 | TRUE | FALSE |
| 171_v.txt | 32776 | 32776-SNV-( | SNV | 1 | C | T | 18 | 30554 | 0.05891209 | 45.3333333 | rnr1D57N | FALSE | CAN | JSY14 | 1A | rnr1D57N-Re | GCG | C>T | A/T+1 | G/C-1 | CG>TA | CG>TA | TGGAAGCGA | CGACCCA | CGC | GCG | C | A | C>T | C>T_GCG | CG>TA_CGACCCA | 7 | TRUE | FALSE |
| 171_v.txt | 32780 | 32780-SNV-( | SNV | 1 | C | T | 70 | 30555 | 0.22909508 | 48.3428571 | rnr1D57N | FALSE | CAN | JSY14 | 1A | rnr1D57N-Re | CCC | C>T | A/T+1 | G/C-1 | CG>TA | CG>TA | AGCGACCCA( | CCAGAAC | GGG | CCC | C | A | C>T | C>T_CCC | CG>TA_CCAGAAC | 7 | TRUE | FALSE |
| 171_v.txt | 32788 | 32788-SNV-( | SNV | 1 | C | T | 13 | 30555 | 0.04254623 | 40.6153846 | rnr1D57N | FALSE | CAN | JSY14 | 1A | rnr1D57N-Re | TCG | C>T | A/T+1 | G/C-1 | CG>TA | CG>TA | AGAACTCGA | CGAATTC | AGC | TCG | C | A | C>T | C>T_TCG | CG>TA_CGAATTC | 7 | TRUE | FALSE |
| 171_v.txt | 32794 | 32794-SNV-( | SNV | 1 | C | T | 31 | 30555 | 0.10145639 | 36.7741936 | rnr1D57N | FALSE | CAN | JSY14 | 1A | rnr1D57N-Re | TCA | C>T | A/T+1 | G/C-1 | CG>TA | CG>TA | CGAATTCAC( | CACCGTA | AGT | TCA | C | A | C>T | C>T_TCA | CG>TA_CACCGTA | 7 | TRUE | FALSE |
| 171_v.txt | 32797 | 32797-SNV-( | SNV | 1 | C | T | 27 | 30555 | 0.08836524 | 48.6207034 | rnr1D57N | FALSE | CAN | JSY14 | 1A | rnr1D57N-Re | CCG | C>T | A/T+1 | G/C-1 | CG>TA | CG>TA | ATTCACCGTA | CGTAATA | GGC | CCG | C | A | C>T | C>T_CCG | CG>TA_CGTAATA | 7 | TRUE | FALSE |
| 171_v.txt | 32798 | 32798-SNV-( | SNV | 1 | G | A | 40 | 30554 | 0.13091576 | 48.5999694 | rnr1D57N | FALSE | CAN | JSY14 | 1A | rnr1D57N-Re | CGT | G>A | A/T+1 | G/C-1 | CG>TA | CG>TA | TTCACCGTAA | GTAATAT | GCA | GCA | C | T | C>T | C>T_GCA | CG>TA_GTAATAT | 7 | TRUE | FALSE |
| 171_v.txt | 32809 | 32809-SNV-( | SNV | 1 | C | T | 8 | 30555 | 0.02618229 | 53.25 | rnr1D57N | FALSE | CAN | JSY14 | 1A | rnr1D57N-Re | ACA | C>T | A/T+1 | G/C-1 | CG>TA | CG>TA | ATTTGACAG( | CAGGGAA | TGT | ACA | C | A | C>T | C>T_ACA | CG>TA_CAGGGAA | 7 | TRUE | FALSE |
| 171_v.txt | 32841 | 32841-SNV-( | SNV | 1 | C | T | 2484 | 30555 | 8.12960236 | 42.770847 | rnr1D57N | FALSE | CAN | JSY14 | 1A | rnr1D57N-Re | CCA | C>T | A/T+1 | G/C-1 | CG>TA | CG>TA | ATTACCCAA/ | CAAAAAA | GGT | CCA | C | A | C>T | C>T_CCA | CG>TA_CAAAAAA | 7 | TRUE | FALSE |
| 171_v.txt | 32855 | 32855-SNV-( | SNV | 1 | C | T | 191 | 30553 | 0.62514319 | 35.7225131 | rnr1D57N | FALSE | CAN | JSY14 | 1A | rnr1D57N-Re | TCC | C>T | A/T+1 | G/C-1 | CG>TA | CG>TA | ACTAATCCAT | CCATGCC | AGG | TCC | C | A | C>T | C>T_TCC | CG>TA_CCATGCC | 7 | TRUE | FALSE |
| 171_v.txt | 32860 | 32860-SNV-( | SNV | 1 | C | T | 29 | 30552 | 0.09492014 | 36.1034483 | rnr1D57N | FALSE | CAN | JSY14 | 1A | rnr1D57N-Re | GCC | C>T | A/T+1 | G/C-1 | CG>TA | CG>TA | TCCATGCCG( | CCGCCAG | CGG | GCC | C | A | C>T | C>T_GCC | CG>TA_CCGCCAG | 7 | TRUE | FALSE |
| 171_v.txt | 32862 | 32862-SNV-( | SNV | 1 | G | A | 25 | 30552 | 0.0818277 | 36.32 | rnr1D57N | FALSE | CAN | JSY14 | 1A | rnr1D57N-Re | CGC | G>A | A/T+1 | G/C-1 | CG>TA | CG>TA | CATGCCGCC/ | GCCAGTG | GCG | GCG | C | T | C>T | C>T_GCG | CG>TA_GCCAGTG | 7 | TRUE | FALSE |
| 171_v.txt | 32883 | 32883-SNV-( | SNV | 1 | C | T | 36 | 31348 | 0.11483986 | 35.9722222 | rnr1D57N | FALSE | CAN | JSY14 | 1A | rnr1D57N-Re | CCA | C>T | A/T+1 | G/C-1 | CG>TA | CG>TA | TACGTCCAA/ | CAAAATT | GGT | CCA | C | A | C>T | C>T_CCA | CG>TA_CAAAATT | 7 | TRUE | FALSE |
| 171_v.txt | 32900 | 32900-SNV-( | SNV | 1 | G | A | 32 | 31348 | 0.10207988 | 35.2070446 | rnr1D57N | FALSE | CAN | JSY14 | 1A | rnr1D57N-Re | GGC | G>A | A/T+1 | G/C-1 | CG>TA | CG>TA | GACTTGGCC/ | GCCAACT | CCG | CCG | C | T | C>T | C>T_CCG | CG>TA_GCCAACT | 7 | TRUE | FALSE |
| 171_v.txt | 32901 | 32901-SNV-( | SNV | 1 | C | T | 72 | 31348 | 0.22967972 | 32.4532052 | rnr1D57N | FALSE | CAN | JSY14 | 1A | rnr1D57N-Re | GCC | C>T | A/T+1 | G/C-1 | CG>TA | CG>TA | ACTTGGCCA/ | CCAACTA | CGG | GCC | C | A | C>T | C>T_GCC | CG>TA_CCAACTA | 7 | TRUE | FALSE |
| 171_v.txt | 32914 | 32914-SNV-( | SNV | 1 | G | A | 2 | 31348 | 6.38E-03 | 24.4999834 | rnr1D57N | FALSE | CAN | JSY14 | 1A | rnr1D57N-Re | AGT | G>A | A/T+1 | G/C-1 | CG>TA | CG>TA | CACTAAGTT( | GTTCCAG | TCA | TCA | C | T | C>T | C>T_TCA | CG>TA_GTTCCAG | 7 | TRUE | FALSE |
| 171_v.txt | 32917 | 32917-SNV-( | SNV | 1 | C | T | 92 | 31348 | 0.29347965 | 46.4673913 | rnr1D57N | FALSE | CAN | JSY14 | 1A | rnr1D57N-Re | TCC | C>T | A/T+1 | G/C-1 | CG>TA | CG>TA | TAAGTTCCA( | CCAGGGC | AGG | TCC | C | A | C>T | C>T_TCC | CG>TA_CCAGGGC | 7 | TRUE | FALSE |
| 171_v.txt | 32934 | 32934-SNV-( | SNV | 1 | G | A | 18 | 31348 | 0.05741993 | 41.0405856 | rnr1D57N | FALSE | CAN | JSY14 | 1A | rnr1D57N-Re | TGC | G>A | A/T+1 | G/C-1 | CG>TA | CG>TA | GTGATTGCC( | GCCCAAG | ACG | ACG | C | T | C>T | C>T_ACG | CG>TA_GCCCAAG | 7 | TRUE | FALSE |
| 171_v.txt | 32935 | 32935-SNV-( | SNV | 1 | C | T | 30 | 31348 | 0.09569989 | 36.8661234 | rnr1D57N | FALSE | CAN | JSY14 | 1A | rnr1D57N-Re | GCC | C>T | A/T+1 | G/C-1 | CG>TA | CG>TA | TGATTGCCC/ | CCCAAGA | CGG | GCC | C | A | C>T | C>T_GCC | CG>TA_CCCAAGA | 7 | TRUE | FALSE |
| 171_v.txt | 32945 | 32945-SNV-( | SNV | 1 | C | T | 58 | 31348 | 0.18501978 | 55.5517241 | rnr1D57N | FALSE | CAN | JSY14 | 1A | rnr1D57N-Re | ACC | C>T | A/T+1 | G/C-1 | CG>TA | CG>TA | AGAAAACCA | CCAATAC | TGG | ACC | C | A | C>T | C>T_ACC | CG>TA_CCAATAC | 7 | TRUE | FALSE |
| 171_v.txt | 32958 | 32958-SNV-( | SNV | 1 | C | T | 46 | 31348 | 0.14673982 | 44.9131737 | rnr1D57N | FALSE | CAN | JSY14 | 1A | rnr1D57N-Re | ACC | C>T | A/T+1 | G/C-1 | CG>TA | CG>TA | ATGTAACCA' | CCATTGG | TGG | ACC | C | A | C>T | C>T_ACC | CG>TA_CCATTGG | 7 | TRUE | FALSE |
| 171_v.txt | 32965 | 32965-SNV-( | SNV | 1 | C | T | 30 | 31348 | 0.09569989 | 56.2666667 | rnr1D57N | FALSE | CAN | JSY14 | 1A | rnr1D57N-Re | GCC | C>T | A/T+1 | G/C-1 | CG>TA | CG>TA | CATTGGCCG( | CCGCACC | CGG | GCC | C | A | C>T | C>T_GCC | CG>TA_CCGCACC | 7 | TRUE | FALSE |
| 171_v.txt | 32967 | 32967-SNV-( | SNV | 1 | G | A | 68 | 31348 | 0.21691974 | 60.9852941 | rnr1D57N | FALSE | CAN | JSY14 | 1A | rnr1D57N-Re | CGC | G>A | A/T+1 | G/C-1 | CG>TA | CG>TA | TTGGCCGCA( | GCACCAA | GCG | GCG | C | T | C>T | C>T_GCG | CG>TA_GCACCAA | 7 | TRUE | FALSE |
| 171_v.txt | 32970 | 32970-SNV-( | SNV | 1 | C | T | 33 | 31348 | 0.10526987 | 60.7878788 | rnr1D57N | FALSE | CAN | JSY14 | 1A | rnr1D57N-Re | ACC | C>T | A/T+1 | G/C-1 | CG>TA | CG>TA | GCCGCACCA/ | CCAAATG | TGG | ACC | C | A | C>T | C>T_ACC | CG>TA_CCAAATG | 7 | TRUE | FALSE |
| 171_v.txt | 32977 | 32977-SNV-( | SNV | 1 | C | T | 22 | 31348 | 0.07017992 | 56.7272727 | rnr1D57N | FALSE | CAN | JSY14 | 1A | rnr1D57N-Re | GCT | C>T | A/T+1 | G/C-1 | CG>TA | CG>TA | CAAATGCTG( | CTGGAGA | CGA | GCT | C | A | C>T | C>T_GCT | CG>TA_CTGGAGA | 7 | TRUE | FALSE |
| 171_v.txt | 32979 | 32979-SNV-( | SNV | 1 | G | A | 6 | 31348 | 0.01913998 | 52.3333247 | rnr1D57N | FALSE | CAN | JSY14 | 1A | rnr1D57N-Re | TGG | G>A | A/T+1 | G/C-1 | CG>TA | CG>TA | AATGCTGGA | GGAGAAA | ACC | ACC | C | T | C>T | C>T_ACC | CG>TA_GGAGAAA | 7 | TRUE | FALSE |
| 171_v.txt | 32980 | 32980-SNV-( | SNV | 1 | G | A | 6 | 31348 | 0.01913998 | 51.999692 | rnr1D57N | FALSE | CAN | JSY14 | 1A | rnr1D57N-Re | GGA | G>A | A/T+1 | G/C-1 | CG>TA | CG>TA | ATGCTGGAG | GAGAAAG | CCT | CCT | C | T | C>T | C>T_CCT | CG>TA_GAGAAAG | 7 | TRUE | FALSE |
| 171_v.txt | 33015 | 33015-SNV-( | SNV | 1 | G | A | 2751 | 31348 | 8.77567947 | 61.3315158 | rnr1D57N | FALSE | CAN | JSY14 | 1A | rnr1D57N-Re | GGA | G>A | A/T+1 | G/C-1 | CG>TA | CG>TA | AAAGAGGAT | GATGTAA | CCT | CCT | C | T | C>T | C>T_CCT | CG>TA_GATGTAA | 7 | TRUE | FALSE |
| 171_v.txt | 33037 | 33037-SNV-( | SNV | 1 | C | T | 15 | 31348 | 0.04784994 | 58.9333333 | rnr1D57N | FALSE | CAN | JSY14 | 1A | rnr1D57N-Re | GCC | C>T | A/T+1 | G/C-1 | CG>TA | CG>TA | ATGTAGCCA' | CCATTTC | CGG | GCC | C | A | C>T | C>T_GCC | CG>TA_CCATTTC | 7 | TRUE | FALSE |
| 171_v.txt | 33055 | 33055-SNV-( | SNV | 1 | G | A | 69 | 31348 | 0.22010974 | 58.1741901 | rnr1D57N | FALSE | CAN | JSY14 | 1A | rnr1D57N-Re | TGC | G>A | A/T+1 | G/C-1 | CG>TA | CG>TA | AGGACTGCG | GCGTGAC | ACG | ACG | C | T | C>T | C>T_ACG | CG>TA_GCGTGAC | 7 | TRUE | FALSE |
| 171_v.txt | 33056 | 33056-SNV-( | SNV | 1 | C | T | 36 | 31347 | 0.11484353 | 55.3021521 | rnr1D57N | FALSE | CAN | JSY14 | 1A | rnr1D57N-Re | GCG | C>T | A/T+1 | G/C-1 | CG>TA | CG>TA | GGACTGCGT( | CGTGACA | CGC | GCG | C | A | C>T | C>T_GCG | CG>TA_CGTGACA | 7 | TRUE | FALSE |
| 171_v.txt | 33070 | 33070-SNV-( | SNV | 1 | C | T | 30 | 31348 | 0.09569989 | 52.6 | rnr1D57N | FALSE | CAN | JSY14 | 1A | rnr1D57N-Re | GCC | C>T | A/T+1 | G/C-1 | CG>TA | CG>TA | AATATGCCA/ | CCAAAGA | CGG | GCC | C | A | C>T | C>T_GCC | CG>TA_CCAAAGA | 7 | TRUE | FALSE |
| 171_v.txt | 33075 | 33075-SNV-( | SNV | 1 | G | A | 29 | 31348 | 0.09250989 | 46.7586207 | rnr1D57N | FALSE | CAN | JSY14 | 1A | rnr1D57N-Re | AGA | G>A | A/T+1 | G/C-1 | CG>TA | CG>TA | GCCAAAGAA | GAACCCA | TCT | TCT | C | T | C>T | C>T_TCT | CG>TA_GAACCCA | 7 | TRUE | FALSE |
| 171_v.txt | 33114 | 33114-SNV-( | SNV | 1 | C | T | 243 | 31348 | 0.77516907 | 44.2139918 | rnr1D57N | FALSE | CAN | JSY14 | 1A | rnr1D57N-Re | GCC | C>T | A/T+1 | G/C-1 | CG>TA | CG>TA | ACTGGGCCG | CCGGCGT | CGG | GCC | C | A | C>T | C>T_GCC | CG>TA_CCGGCGT | 7 | TRUE | FALSE |
| 171_v.txt | 33115 | 33115-SNV-( | SNV | 1 | C | T | 231 | 31348 | 0.73688912 | 43.961039 | rnr1D57N | FALSE | CAN | JSY14 | 1A | rnr1D57N-Re | CCG | C>T | A/T+1 | G/C-1 | CG>TA | CG>TA | CTGGGCCGG | CGGCGTT | GGC | CCG | C | A | C>T | C>T_CCG | CG>TA_CGGCGTT | 7 | TRUE | FALSE |
| 171_v.txt | 33118 | 33118-SNV-( | SNV | 1 | C | T | 25 | 31348 | 0.0797499 | 40.08 | rnr1D57N | FALSE | CAN | JSY14 | 1A | rnr1D57N-Re | GCG | C>T | A/T+1 | G/C-1 | CG>TA | CG>TA | GGCCGGCGT | CGTTGGT | CGC | GCG | C | A | C>T | C>T_GCG | CG>TA_CGTTGGT | 7 | TRUE | FALSE |
| 171_v.txt | 33153 | 33153-SNV-( | SNV | 1 | C | T | 70 | 31348 | 0.22329973 | 36.3572594 | rnr1D57N | FALSE | CAN | JSY14 | 1A | rnr1D57N-Re | ACC | C>T | A/T+1 | G/C-1 | CG>TA | CG>TA | AAAAGACCT( | CCTGTAC | TGG | ACC | C | A | C>T | C>T_ACC | CG>TA_CCTGTAC | 7 | TRUE | FALSE |
| 171_v.txt | 33156 | 33156-SNV-( | SNV | 1 | G | A | 714 | 31344 | 2.27794793 | 37.3319328 | rnr1D57N | FALSE | CAN | JSY14 | 1A | rnr1D57N-Re | TGT | G>A | A/T+1 | G/C-1 | CG>TA | CG>TA | AGACCTGTA( | GTACCAA | ACA | ACA | C | T | C>T | C>T_ACA | CG>TA_GTACCAA | 7 | TRUE | FALSE |
| 171_v.txt | 33159 | 33159-SNV-( | SNV | 1 | C | T | 207 | 31342 | 0.66045562 | 36.7855397 | rnr1D57N | FALSE | CAN | JSY14 | 1A | rnr1D57N-Re | ACC | C>T | A/T+1 | G/C-1 | CG>TA | CG>TA | CCTGTACCA/ | CCAATAG | TGG | ACC | C | A | C>T | C>T_ACC | CG>TA_CCAATAG | 7 | TRUE | FALSE |
| 171_v.txt | 33168 | 33168-SNV-( | SNV | 1 | C | T | 101 | 31300 | 0.32268371 | 37.2339208 | rnr1D57N | FALSE | CAN | JSY14 | 1A | rnr1D57N-Re | ACC | C>T | A/T+1 | G/C-1 | CG>TA | CG>TA | ATAGTACCA( | CCACCAA | TGG | ACC | C | A | C>T | C>T_ACC | CG>TA_CCACCAA | 7 | TRUE | FALSE |
| 171_v.txt | 33177 | 33177-SNV-( | SNV | 1 | G | A | 36 | 31297 | 0.115027 | 36.1111388 | rnr1D57N | FALSE | CAN | JSY14 | 1A | rnr1D57N-Re | GGC | G>A | A/T+1 | G/C-1 | CG>TA | CG>TA | CCAAGGGCA | GCAATCA | CCG | CCG | C | T | C>T | C>T_CCG | CG>TA_GCAATCA | 7 | TRUE | FALSE |
| 171_v.txt | 33178 | 33178-SNV-( | SNV | 1 | C | T | 15 | 31295 | 0.04793098 | 36.400117 | rnr1D57N | FALSE | CAN | JSY14 | 1A | rnr1D57N-Re | GCA | C>T | A/T+1 | G/C-1 | CG>TA | CG>TA | CAAGGGCAA | CAATCAT | CGT | GCA | C | A | C>T | C>T_GCA | CG>TA_CAATCAT | 7 | TRUE | FALSE |
| 171_v.txt | 33199 | 33199-SNV-( | SNV | 1 | G | A | 17 | 13895 | 0.12234617 | 36.2941177 | rnr1D57N | FALSE | CAN | JSY14 | 1A | rnr1D57N-Re | TGC | G>A | A/T+1 | G/C-1 | CG>TA | CG>TA | GTCTTTGCTT | GCTTAAG | ACG | ACG | C | T | C>T | C>T_ACG | CG>TA_GCTTAAG | 7 | TRUE | FALSE |
| 171_v.txt | 33229 | 33229-SNV-( | SNV | 1 | G | A | 109 | 13904 | 0.78394707 | 35.3119266 | rnr1D57N | FALSE | CAN | JSY14 | 1A | rnr1D57N-Re | TGT | G>A | A/T+1 | G/C-1 | CG>TA | CG>TA | CGTTCTGTAC | GTACTTC | ACA | ACA | C | T | C>T | C>T_ACA | CG>TA_GTACTTC | 7 | TRUE | FALSE |
| 171_v.txt | 33276 | 33276-SNV-( | SNV | 1 | G | A | 25 | 13904 | 0.17980437 | 39.52 | rnr1D57N | FALSE | CAN | JSY14 | 1A | rnr1D57N-Re | CGT | G>A | A/T+1 | G/C-1 | CG>TA | CG>TA | GAGAACGTA | GTATCCT | GCA | GCA | C | T | C>T | C>T_GCA | CG>TA_GTATCCT | 7 | TRUE | FALSE |
| 171_v.txt | 33309 | 33309-SNV-( | SNV | 1 | C | T | 15 | 13904 | 0.10788262 | 54.7292495 | rnr1D57N | FALSE | CAN | JSY14 | 1A | rnr1D57N-Re | GCG | C>T | A/T+1 | G/C-1 | CG>TA | CG>TA | AAAGAGCGC | CGCAATG | CGC | GCG | C | A | C>T | C>T_GCG | CG>TA_CGCAATG | 7 | TRUE | FALSE |
| 171_v.txt | 33310 | 33310-SNV-( | SNV | 1 | G | A | 18 | 13903 | 0.12946846 | 55.7133463 | rnr1D57N | FALSE | CAN | JSY14 | 1A | rnr1D57N-Re | CGC | G>A | A/T+1 | G/C-1 | CG>TA | CG>TA | AAGAGCGCA | GCAATGG | GCG | GCG | C | T | C>T | C>T_GCG | CG>TA_GCAATGG | 7 | TRUE | FALSE |
| 171_v.txt | 33354 | 33354-SNV-( | SNV | 1 | G | A | 4 | 13896 | 0.02878526 | 47.7226568 | rnr1D57N | FALSE | CAN | JSY14 | 1A | rnr1D57N-Re | TGA | G>A | A/T+1 | G/C-1 | CG>TA | CG>TA | GGTATTGAC( | GACCCAC | ACT | ACT | C | T | C>T | C>T_ACT | CG>TA_GACCCAC | 7 | TRUE | FALSE |
| 171_v.txt | 33382 | 33382-SNV-( | SNV | 1 | C | T | 17 | 13904 | 0.12226697 | 56.1764706 | rnr1D57N | FALSE | CAN | JSY14 | 1A | rnr1D57N-Re | GCT | C>T | A/T+1 | G/C-1 | CG>TA | CG>TA | GTGAAGCTT( | CTTCAAC | CGA | GCT | C | A | C>T | C>T_GCT | CG>TA_CTTCAAC | 7 | TRUE | FALSE |
| 171_v.txt | 33412 | 33412-SNV-( | SNV | 1 | G | A | 15 | 13904 | 0.10788262 | 51.6 | rnr1D57N | FALSE | CAN | JSY14 | 1A | rnr1D57N-Re | GGC | G>A | A/T+1 | G/C-1 | CG>TA | CG>TA | TGACCGGCT( | GCTCATT | CCG | CCG | C | T | C>T | C>T_CCG | CG>TA_GCTCATT | 7 | TRUE | FALSE |
| 171_v.txt | 33474 | 33474-SNV-( | SNV | 1 | C | T | 18 | 13903 | 0.12946846 | 36.6111111 | rnr1D57N | FALSE | CAN | JSY14 | 1A | rnr1D57N-Re | GCC | C>T | A/T+1 | G/C-1 | CG>TA | CG>TA | GCTATGCCTT | CCTTTTT | CGG | GCC | C | A | C>T | C>T_GCC | CG>TA_CCTTTTT | 7 | TRUE | FALSE |
| 171_v.txt | 31806 | 31806-SNV-( | SNV | 1 | C | G | 1 | 10178 | 9.83E-03 | 63.9944878 | rnr1D57N | FALSE | CAN | JSY14 | 1A | rnr1D57N-Re | CCA | C>G | G/C+1 | G/C-1 | CG>GC | CG>GC | ATCTTCCAAA | CAAATAA | GGT | CCA | C | C | C>G | C>G_CCA | CG>GC_CAAATAA | 7 | TRUE | FALSE |
| 171_v.txt | 32064 | 32064-SNV-( | SNV | 1 | G | C | 65 | 24202 | 0.26857285 | 62.3384615 | rnr1D57N | FALSE | CAN | JSY14 | 1A | rnr1D57N-Re | TGA | G>C | G/C+1 | G/C-1 | CG>GC | CG>GC | GAGATTGAG | GAGATAA | ACT | ACT | C | G | C>G | C>G_ACT | CG>GC_GAGATAA | 7 | TRUE | FALSE |
| 171_v.txt | 32164 | 32164-SNV-( | SNV | 1 | C | G | 1 | 24202 | 4.13E-03 | 37.9989049 | rnr1D57N | FALSE | CAN | JSY14 | 1A | rnr1D57N-Re | GCG | C>G | G/C+1 | G/C-1 | CG>GC | CG>GC | CCAAAGCGC( | CGCCAAA | CGC | GCG | C | C | C>G | C>G_GCG | CG>GC_CGCCAAA | 7 | TRUE | FALSE |
| 171_v.txt | 32223 | 32223-SNV-( | SNV | 1 | G | C | 6 | 24202 | 0.02479134 | 37.8333333 | rnr1D57N | FALSE | CAN | JSY14 | 1A | rnr1D57N-Re | TGA | G>C | G/C+1 | G/C-1 | CG>GC | CG>GC | GTCCTTGAC/ | GACAGGA | ACT | ACT | C | G | C>G | C>G_ACT | CG>GC_GACAGGA | 7 | TRUE | FALSE |
| 171_v.txt | 32272 | 32272-SNV-( | SNV | 1 | G | C | 46 | 31385 | 0.14656683 | 35.9347826 | rnr1D57N | FALSE | CAN | JSY14 | 1A | rnr1D57N-Re | CGG | G>C | G/C+1 | G/C-1 | CG>GC | CG>GC | AAATACGGG | GGGAACC | GCC | GCC | C | G | C>G | C>G_GCC | CG>GC_GGGAACC | 7 | TRUE | FALSE |
| 171_v.txt | 32278 | 32278-SNV-( | SNV | 1 | C | G | 81 | 31385 | 0.25808507 | 35.8518519 | rnr1D57N | FALSE | CAN | JSY14 | 1A | rnr1D57N-Re | CCA | C>G | G/C+1 | G/C-1 | CG>GC | CG>GC | GGGAACCAA | CAACGTA | GGT | CCA | C | C | C>G | C>G_CCA | CG>GC_CAACGTA | 7 | TRUE | FALSE |
| 171_v.txt | 32292 | 32292-SNV-( | SNV | 1 | G | C | 12 | 31383 | 0.03823726 | 52.9148918 | rnr1D57N | FALSE | CAN | JSY14 | 1A | rnr1D57N-Re | TGA | G>C | G/C+1 | G/C-1 | CG>GC | CG>GC | ATATTTGAAT | GAATTTG | ACT | ACT | C | G | C>G | C>G_ACT | CG>GC_GAATTTG | 7 | TRUE | FALSE |
| 171_v.txt | 32316 | 32316-SNV-( | SNV | 1 | G | C | 8 | 31383 | 0.02549151 | 63.625 | rnr1D57N | FALSE | CAN | JSY14 | 1A | rnr1D57N-Re | TGT | G>C | G/C+1 | G/C-1 | CG>GC | CG>GC | ATGGTTGTT/ | GTTAAGA | ACA | ACA | C | G | C>G | C>G_ACA | CG>GC_GTTAAGA | 7 | TRUE | FALSE |
| 171_v.txt | 32399 | 32399-SNV-( | SNV | 1 | G | C | 13 | 31383 | 0.0414237 | 61.7692308 | rnr1D57N | FALSE | CAN | JSY14 | 1A | rnr1D57N-Re | CGT | G>C | G/C+1 | G/C-1 | CG>GC | CG>GC | AGAAACGTA | GTAGGAA | GCA | GCA | C | G | C>G | C>G_GCA | CG>GC_GTAGGAA | 7 | TRUE | FALSE |
| 171_v.txt | 32432 | 32432-SNV-( | SNV | 1 | G | C | 84 | 31383 | 0.26766084 | 55 | rnr1D57N | FALSE | CAN | JSY14 | 1A | rnr1D57N-Re | TGT | G>C | G/C+1 | G/C-1 | CG>GC | CG>GC | GTCATTGTAT | GTATGGA | ACA | ACA | C | G | C>G | C>G_ACA | CG>GC_GTATGGA | 7 | TRUE | FALSE |
| 171_v.txt | 32449 | 32449-SNV-( | SNV | 1 | C | G | 20 | 31383 | 0.06372877 | 50.5 | rnr1D57N | FALSE | CAN | JSY14 | 1A | rnr1D57N-Re | CCA | C>G | G/C+1 | G/C-1 | CG>GC | CG>GC | AAAGTCCAA' | CAATGAA | GGT | CCA | C | C | C>G | C>G_CCA | CG>GC_CAATGAA | 7 | TRUE | FALSE |
| 171_v.txt | 32471 | 32471-SNV-( | SNV | 1 | G | C | 38 | 31383 | 0.12108466 | 46.8157895 | rnr1D57N | FALSE | CAN | JSY14 | 1A | rnr1D57N-Re | TGT | G>C | G/C+1 | G/C-1 | CG>GC | CG>GC | GCCAATGTA( | GTAGAAG | ACA | ACA | C | G | C>G | C>G_ACA | CG>GC_GTAGAAG | 7 | TRUE | FALSE |
| 171_v.txt | 32487 | 32487-SNV-( | SNV | 1 | C | G | 29 | 31383 | 0.09240672 | 42.4123475 | rnr1D57N | FALSE | CAN | JSY14 | 1A | rnr1D57N-Re | ACG | C>G | G/C+1 | G/C-1 | CG>GC | CG>GC | AAGATACGG | CGGAAAA | TGC | ACG | C | C | C>G | C>G_ACG | CG>GC_CGGAAAA | 7 | TRUE | FALSE |
| 171_v.txt | 32539 | 32539-SNV-( | SNV | 1 | C | G | 39 | 31383 | 0.1242711 | 35.9230769 | rnr1D57N | FALSE | CAN | JSY14 | 1A | rnr1D57N-Re | GCT | C>G | G/C+1 | G/C-1 | CG>GC | CG>GC | TTGCAGCTTC | CTTCACC | CGA | GCT | C | C | C>G | C>G_GCT | CG>GC_CTTCACC | 7 | TRUE | FALSE |
| 171_v.txt | 32544 | 32544-SNV-( | SNV | 1 | C | G | 11 | 31383 | 0.03505082 | 36.6359383 | rnr1D57N | FALSE | CAN | JSY14 | 1A | rnr1D57N-Re | ACC | C>G | G/C+1 | G/C-1 | CG>GC | CG>GC | GCTTCACCA( | CCAGCAG | TGG | ACC | C | C | C>G | C>G_ACC | CG>GC_CCAGCAG | 7 | TRUE | FALSE |
| 171_v.txt | 32545 | 32545-SNV-( | SNV | 1 | C | G | 3 | 31383 | 9.56E-03 | 38 | rnr1D57N | FALSE | CAN | JSY14 | 1A | rnr1D57N-Re | CCA | C>G | G/C+1 | G/C-1 | CG>GC | CG>GC | CTTCACCAG( | CAGCAGT | GGT | CCA | C | C | C>G | C>G_CCA | CG>GC_CAGCAGT | 7 | TRUE | FALSE |
| 171_v.txt | 32552 | 32552-SNV-( | SNV | 1 | G | C | 1 | 31378 | 3.19E-03 | 30.8604291 | rnr1D57N | FALSE | CAN | JSY14 | 1A | rnr1D57N-Re | TGA | G>C | G/C+1 | G/C-1 | CG>GC | CG>GC | AGCAGTGAT. | GATACCA | ACT | ACT | C | G | C>G | C>G_ACT | CG>GC_GATACCA | 7 | TRUE | FALSE |
| 171_v.txt | 32557 | 32557-SNV-( | SNV | 1 | C | G | 10 | 31372 | 0.03187556 | 35.1 | rnr1D57N | FALSE | CAN | JSY14 | 1A | rnr1D57N-Re | CCA | C>G | G/C+1 | G/C-1 | CG>GC | CG>GC | TGATACCAA( | CAACTAG | GGT | CCA | C | C | C>G | C>G_CCA | CG>GC_CAACTAG | 7 | TRUE | FALSE |
| 171_v.txt | 32743 | 32743-SNV-( | SNV | 1 | G | C | 39 | 30555 | 0.12763868 | 55.025641 | rnr1D57N | FALSE | CAN | JSY14 | 1A | rnr1D57N-Re | AGA | G>C | G/C+1 | G/C-1 | CG>GC | CG>GC | ATATTAGAA/ | GAAACCC | TCT | TCT | C | G | C>G | C>G_TCT | CG>GC_GAAACCC | 7 | TRUE | FALSE |
| 171_v.txt | 32758 | 32758-SNV-( | SNV | 1 | C | G | 12 | 30555 | 0.03927344 | 54.5822743 | rnr1D57N | FALSE | CAN | JSY14 | 1A | rnr1D57N-Re | GCT | C>G | G/C+1 | G/C-1 | CG>GC | CG>GC | TAATGGCTA/ | CTAAAAC | CGA | GCT | C | C | C>G | C>G_GCT | CG>GC_CTAAAAC | 7 | TRUE | FALSE |
| 171_v.txt | 32776 | 32776-SNV-( | SNV | 1 | C | G | 54 | 30554 | 0.17673627 | 51.2777778 | rnr1D57N | FALSE | CAN | JSY14 | 1A | rnr1D57N-Re | GCG | C>G | G/C+1 | G/C-1 | CG>GC | CG>GC | TGGAAGCGA | CGACCCA | CGC | GCG | C | C | C>G | C>G_GCG | CG>GC_CGACCCA | 7 | TRUE | FALSE |
| 171_v.txt | 32794 | 32794-SNV-( | SNV | 1 | C | G | 5 | 30555 | 0.01636393 | 40.6 | rnr1D57N | FALSE | CAN | JSY14 | 1A | rnr1D57N-Re | TCA | C>G | G/C+1 | G/C-1 | CG>GC | CG>GC | CGAATTCAC( | CACCGTA | AGT | TCA | C | C | C>G | C>G_TCA | CG>GC_CACCGTA | 7 | TRUE | FALSE |
| 171_v.txt | 32798 | 32798-SNV-( | SNV | 1 | G | C | 8 | 30554 | 0.02618315 | 38.6249956 | rnr1D57N | FALSE | CAN | JSY14 | 1A | rnr1D57N-Re | CGT | G>C | G/C+1 | G/C-1 | CG>GC | CG>GC | TTCACCGTA/ | GTAATAT | GCA | GCA | C | G | C>G | C>G_GCA | CG>GC_GTAATAT | 7 | TRUE | FALSE |
| 171_v.txt | 32809 | 32809-SNV-( | SNV | 1 | C | G | 40 | 30555 | 0.13091147 | 44.625 | rnr1D57N | FALSE | CAN | JSY14 | 1A | rnr1D57N-Re | ACA | C>G | G/C+1 | G/C-1 | CG>GC | CG>GC | ATTTGACAG( | CAGGGAA | TGT | ACA | C | C | C>G | C>G_ACA | CG>GC_CAGGGAA | 7 | TRUE | FALSE |
| 171_v.txt | 32811 | 32811-SNV-( | SNV | 1 | G | C | 393 | 30555 | 1.2862052 | 44.1043257 | rnr1D57N | FALSE | CAN | JSY14 | 1A | rnr1D57N-Re | AGG | G>C | G/C+1 | G/C-1 | CG>GC | CG>GC | TTGACAGGG | GGGAACA | TCC | TCC | C | G | C>G | C>G_TCC | CG>GC_GGGAACA | 7 | TRUE | FALSE |
| 171_v.txt | 32876 | 32876-SNV-( | SNV | 1 | G | C | 14 | 31336 | 0.04467705 | 37.9285714 | rnr1D57N | FALSE | CAN | JSY14 | 1A | rnr1D57N-Re | TGT | G>C | G/C+1 | G/C-1 | CG>GC | CG>GC | AACTTTGTAC | GTACGTC | ACA | ACA | C | G | C>G | C>G_ACA | CG>GC_GTACGTC | 7 | TRUE | FALSE |
| 171_v.txt | 32935 | 32935-SNV-( | SNV | 1 | C | G | 1 | 31348 | 3.19E-03 | 36.9989748 | rnr1D57N | FALSE | CAN | JSY14 | 1A | rnr1D57N-Re | GCC | C>G | G/C+1 | G/C-1 | CG>GC | CG>GC | TGATTGCCC/ | CCCAAGA | CGG | GCC | C | C | C>G | C>G_GCC | CG>GC_CCCAAGA | 7 | TRUE | FALSE |
| 171_v.txt | 32937 | 32937-SNV-( | SNV | 1 | C | G | 21 | 31348 | 0.06698992 | 56.1448597 | rnr1D57N | FALSE | CAN | JSY14 | 1A | rnr1D57N-Re | CCA | C>G | G/C+1 | G/C-1 | CG>GC | CG>GC | ATTGCCCAA( | CAAGAAA | GGT | CCA | C | C | C>G | C>G_CCA | CG>GC_CAAGAAA | 7 | TRUE | FALSE |
| 171_v.txt | 32959 | 32959-SNV-( | SNV | 1 | C | G | 44 | 31347 | 0.14036431 | 57.2142991 | rnr1D57N | FALSE | CAN | JSY14 | 1A | rnr1D57N-Re | CCA | C>G | G/C+1 | G/C-1 | CG>GC | CG>GC | TGTAACCATT | CATTGGC | GGT | CCA | C | C | C>G | C>G_CCA | CG>GC_CATTGGC | 7 | TRUE | FALSE |
| 171_v.txt | 32967 | 32967-SNV-( | SNV | 1 | G | C | 4 | 31348 | 0.01275999 | 51 | rnr1D57N | FALSE | CAN | JSY14 | 1A | rnr1D57N-Re | CGC | G>C | G/C+1 | G/C-1 | CG>GC | CG>GC | TTGGCCGCA( | GCACCAA | GCG | GCG | C | G | C>G | C>G_GCG | CG>GC_GCACCAA | 7 | TRUE | FALSE |
| 171_v.txt | 32971 | 32971-SNV-( | SNV | 1 | C | G | 17 | 31348 | 0.05422994 | 60.4713168 | rnr1D57N | FALSE | CAN | JSY14 | 1A | rnr1D57N-Re | CCA | C>G | G/C+1 | G/C-1 | CG>GC | CG>GC | CCGCACCAA/ | CAAATGC | GGT | CCA | C | C | C>G | C>G_CCA | CG>GC_CAAATGC | 7 | TRUE | FALSE |
| 171_v.txt | 32977 | 32977-SNV-( | SNV | 1 | C | G | 16 | 31348 | 0.05103994 | 61.875 | rnr1D57N | FALSE | CAN | JSY14 | 1A | rnr1D57N-Re | GCT | C>G | G/C+1 | G/C-1 | CG>GC | CG>GC | CAAATGCTG( | CTGGAGA | CGA | GCT | C | C | C>G | C>G_GCT | CG>GC_CTGGAGA | 7 | TRUE | FALSE |
| 171_v.txt | 32979 | 32979-SNV-( | SNV | 1 | G | C | 5 | 31348 | 0.01594998 | 27.2032598 | rnr1D57N | FALSE | CAN | JSY14 | 1A | rnr1D57N-Re | TGG | G>C | G/C+1 | G/C-1 | CG>GC | CG>GC | AATGCTGGA | GGAGAAA | ACC | ACC | C | G | C>G | C>G_ACC | CG>GC_GGAGAAA | 7 | TRUE | FALSE |
| 171_v.txt | 32980 | 32980-SNV-( | SNV | 1 | G | C | 1 | 31348 | 3.19E-03 | 20.9995104 | rnr1D57N | FALSE | CAN | JSY14 | 1A | rnr1D57N-Re | GGA | G>C | G/C+1 | G/C-1 | CG>GC | CG>GC | ATGCTGGAG | GAGAAAG | CCT | CCT | C | G | C>G | C>G_CCT | CG>GC_GAGAAAG | 7 | TRUE | FALSE |
| 171_v.txt | 32997 | 32997-SNV-( | SNV | 1 | G | C | 371 | 31348 | 1.18348858 | 61.4123989 | rnr1D57N | FALSE | CAN | JSY14 | 1A | rnr1D57N-Re | TGA | G>C | G/C+1 | G/C-1 | CG>GC | CG>GC | CTTTGTGAG/ | GAGAAAA | ACT | ACT | C | G | C>G | C>G_ACT | CG>GC_GAGAAAA | 7 | TRUE | FALSE |
| 171_v.txt | 33018 | 33018-SNV-( | SNV | 1 | G | C | 21 | 31348 | 0.06698992 | 61.8571429 | rnr1D57N | FALSE | CAN | JSY14 | 1A | rnr1D57N-Re | TGT | G>C | G/C+1 | G/C-1 | CG>GC | CG>GC | GAGGATGTA | GTAACAG | ACA | ACA | C | G | C>G | C>G_ACA | CG>GC_GTAACAG | 7 | TRUE | FALSE |
| 171_v.txt | 33037 | 33037-SNV-( | SNV | 1 | C | G | 17 | 31348 | 0.05422994 | 58 | rnr1D57N | FALSE | CAN | JSY14 | 1A | rnr1D57N-Re | GCC | C>G | G/C+1 | G/C-1 | CG>GC | CG>GC | ATGTAGCCA' | CCATTTC | CGG | GCC | C | C | C>G | C>G_GCC | CG>GC_CCATTTC | 7 | TRUE | FALSE |
| 171_v.txt | 33056 | 33056-SNV-( | SNV | 1 | C | G | 2 | 31347 | 6.38E-03 | 47.9978004 | rnr1D57N | FALSE | CAN | JSY14 | 1A | rnr1D57N-Re | GCG | C>G | G/C+1 | G/C-1 | CG>GC | CG>GC | GGACTGCGT( | CGTGACA | CGC | GCG | C | C | C>G | C>G_GCG | CG>GC_CGTGACA | 7 | TRUE | FALSE |
| 171_v.txt | 33070 | 33070-SNV-( | SNV | 1 | C | G | 65 | 31348 | 0.20734975 | 52.3230769 | rnr1D57N | FALSE | CAN | JSY14 | 1A | rnr1D57N-Re | GCC | C>G | G/C+1 | G/C-1 | CG>GC | CG>GC | AATATGCCA/ | CCAAAGA | CGG | GCC | C | C | C>G | C>G_GCC | CG>GC_CCAAAGA | 7 | TRUE | FALSE |
| 171_v.txt | 33115 | 33115-SNV-( | SNV | 1 | C | G | 10 | 31348 | 0.03189996 | 39.5 | rnr1D57N | FALSE | CAN | JSY14 | 1A | rnr1D57N-Re | CCG | C>G | G/C+1 | G/C-1 | CG>GC | CG>GC | CTGGGCCGG | CGGCGTT | GGC | CCG | C | C | C>G | C>G_CCG | CG>GC_CGGCGTT | 7 | TRUE | FALSE |
| 171_v.txt | 33154 | 33154-SNV-( | SNV | 1 | C | G | 22 | 31348 | 0.07017992 | 37.5439447 | rnr1D57N | FALSE | CAN | JSY14 | 1A | rnr1D57N-Re | CCT | C>G | G/C+1 | G/C-1 | CG>GC | CG>GC | AAAGACCTG | CTGTACC | GGA | CCT | C | C | C>G | C>G_CCT | CG>GC_CTGTACC | 7 | TRUE | FALSE |
| 171_v.txt | 33169 | 33169-SNV-( | SNV | 1 | C | G | 3 | 31300 | 9.59E-03 | 37.9985618 | rnr1D57N | FALSE | CAN | JSY14 | 1A | rnr1D57N-Re | CCA | C>G | G/C+1 | G/C-1 | CG>GC | CG>GC | TAGTACCAC( | CACCAAG | GGT | CCA | C | C | C>G | C>G_CCA | CG>GC_CACCAAG | 7 | TRUE | FALSE |
| 171_v.txt | 33205 | 33205-SNV-( | SNV | 1 | G | C | 10 | 13895 | 0.07196833 | 36.9 | rnr1D57N | FALSE | CAN | JSY14 | 1A | rnr1D57N-Re | AGC | G>C | G/C+1 | G/C-1 | CG>GC | CG>GC | GCTTAAGCT( | GCTCTCT | TCG | TCG | C | G | C>G | C>G_TCG | CG>GC_GCTCTCT | 7 | TRUE | FALSE |
| 171_v.txt | 33309 | 33309-SNV-( | SNV | 1 | C | G | 1 | 13904 | 7.19E-03 | 63.9888348 | rnr1D57N | FALSE | CAN | JSY14 | 1A | rnr1D57N-Re | GCG | C>G | G/C+1 | G/C-1 | CG>GC | CG>GC | AAAGAGCGC | CGCAATG | CGC | GCG | C | C | C>G | C>G_GCG | CG>GC_CGCAATG | 7 | TRUE | FALSE |
| 171_v.txt | 33310 | 33310-SNV-( | SNV | 1 | G | C | 3 | 13903 | 0.02157808 | 25.334381 | rnr1D57N | FALSE | CAN | JSY14 | 1A | rnr1D57N-Re | CGC | G>C | G/C+1 | G/C-1 | CG>GC | CG>GC | AAGAGCGCA | GCAATGG | GCG | GCG | C | G | C>G | C>G_GCG | CG>GC_GCAATGG | 7 | TRUE | FALSE |
| 171_v.txt | 33456 | 33456-SNV-( | SNV | 1 | G | C | 38 | 13904 | 0.27330265 | 44.3157895 | rnr1D57N | FALSE | CAN | JSY14 | 1A | rnr1D57N-Re | TGA | G>C | G/C+1 | G/C-1 | CG>GC | CG>GC | TCTTTTGAAT | GAATTTG | ACT | ACT | C | G | C>G | C>G_ACT | CG>GC_GAATTTG | 7 | TRUE | FALSE |
| 171_v.txt | 31869 | 31869-SNV-( | SNV | 1 | G | T | 35 | 10179 | 0.34384517 | 44.0571429 | rnr1D57N | FALSE | CAN | JSY14 | 1A | rnr1D57N-Re | AGA | G>T | A/T+1 | G/C-1 | CG>AT | CG>AT | AAAATAGAG | GAGATAT | TCT | TCT | C | A | C>A | C>A_TCT | CG>AT_GAGATAT | 7 | TRUE | FALSE |
| 171_v.txt | 32008 | 32008-SNV-( | SNV | 1 | C | A | 25 | 24202 | 0.10329725 | 44.44 | rnr1D57N | FALSE | CAN | JSY14 | 1A | rnr1D57N-Re | TCG | C>A | A/T+1 | G/C-1 | CG>AT | CG>AT | GTAACTCGT( | CGTCACG | AGC | TCG | C | T | C>A | C>A_TCG | CG>AT_CGTCACG | 7 | TRUE | FALSE |
| 171_v.txt | 32095 | 32095-SNV-( | SNV | 1 | C | A | 65 | 24202 | 0.26857285 | 59.8307692 | rnr1D57N | FALSE | CAN | JSY14 | 1A | rnr1D57N-Re | ACA | C>A | A/T+1 | G/C-1 | CG>AT | CG>AT | CTGCAACAC( | CACCAGT | TGT | ACA | C | T | C>A | C>A_ACA | CG>AT_CACCAGT | 7 | TRUE | FALSE |
| 171_v.txt | 32114 | 32114-SNV-( | SNV | 1 | C | A | 15 | 24202 | 0.06197835 | 39.6062837 | rnr1D57N | FALSE | CAN | JSY14 | 1A | rnr1D57N-Re | GCC | C>A | A/T+1 | G/C-1 | CG>AT | CG>AT | TAATAGCCA' | CCATTCG | CGG | GCC | C | T | C>A | C>A_GCC | CG>AT_CCATTCG | 7 | TRUE | FALSE |
| 171_v.txt | 32119 | 32119-SNV-( | SNV | 1 | C | A | 82 | 24202 | 0.33881497 | 59.0591106 | rnr1D57N | FALSE | CAN | JSY14 | 1A | rnr1D57N-Re | TCG | C>A | A/T+1 | G/C-1 | CG>AT | CG>AT | GCCATTCGA/ | CGAAAAC | AGC | TCG | C | T | C>A | C>A_TCG | CG>AT_CGAAAAC | 7 | TRUE | FALSE |
| 171_v.txt | 32120 | 32120-SNV-( | SNV | 1 | G | T | 3 | 24202 | 0.01239567 | 63.9815659 | rnr1D57N | FALSE | CAN | JSY14 | 1A | rnr1D57N-Re | CGA | G>T | A/T+1 | G/C-1 | CG>AT | CG>AT | CCATTCGAA/ | GAAAACT | GCT | GCT | C | A | C>A | C>A_GCT | CG>AT_GAAAACT | 7 | TRUE | FALSE |
| 171_v.txt | 32149 | 32149-SNV-( | SNV | 1 | C | A | 30 | 24202 | 0.1239567 | 47.5069015 | rnr1D57N | FALSE | CAN | JSY14 | 1A | rnr1D57N-Re | TCC | C>A | A/T+1 | G/C-1 | CG>AT | CG>AT | ATGTCTCCAT | CCATGTA | AGG | TCC | C | T | C>A | C>A_TCC | CG>AT_CCATGTA | 7 | TRUE | FALSE |
| 171_v.txt | 32150 | 32150-SNV-( | SNV | 1 | C | A | 25 | 24202 | 0.10329725 | 47.7251603 | rnr1D57N | FALSE | CAN | JSY14 | 1A | rnr1D57N-Re | CCA | C>A | A/T+1 | G/C-1 | CG>AT | CG>AT | TGTCTCCATG | CATGTAA | GGT | CCA | C | T | C>A | C>A_CCA | CG>AT_CATGTAA | 7 | TRUE | FALSE |
| 171_v.txt | 32153 | 32153-SNV-( | SNV | 1 | G | T | 17 | 24202 | 0.07024213 | 53.4065078 | rnr1D57N | FALSE | CAN | JSY14 | 1A | rnr1D57N-Re | TGT | G>T | A/T+1 | G/C-1 | CG>AT | CG>AT | CTCCATGTA/ | GTAAGCC | ACA | ACA | C | A | C>A | C>A_ACA | CG>AT_GTAAGCC | 7 | TRUE | FALSE |
| 171_v.txt | 32195 | 32195-SNV-( | SNV | 1 | G | T | 30 | 24202 | 0.1239567 | 43.0666667 | rnr1D57N | FALSE | CAN | JSY14 | 1A | rnr1D57N-Re | TGT | G>T | A/T+1 | G/C-1 | CG>AT | CG>AT | TGCAATGTA' | GTATGGA | ACA | ACA | C | A | C>A | C>A_ACA | CG>AT_GTATGGA | 7 | TRUE | FALSE |

| | | | | | | | | | | | | | | | | | | | | | | | | | | | | | | | | | | |
|---|---|---|---|---|---|---|---|---|---|---|---|---|---|---|---|---|---|---|---|---|---|---|---|---|---|---|---|---|---|---|---|---|---|---|
| 171_v.txt | 32272 | 32272-SNV-( | SNV | 1 | G | T | 48 | 31385 | 0.1529393 | 34.125 | rnr1D57N | FALSE | CAN | JSY14 | 1A | rnr1D57N-Re | CGG | G>T | A/T+1 | G/C-1 | CG>AT | CG>AT | AAATACGGG | GGGAACC | GCC | GCC | C | A | C>A | C>A_GCC | CG>AT_GGGAACC | 7 | TRUE | FALSE |
| 171_v.txt | 32274 | 32274-SNV-( | SNV | 1 | G | T | 324 | 31385 | 1.03234029 | 36.1118497 | rnr1D57N | FALSE | CAN | JSY14 | 1A | rnr1D57N-Re | GGA | G>T | A/T+1 | G/C-1 | CG>AT | CG>AT | ATACGGGAA | GAACCAA | CCT | CCT | C | A | C>A | C>A_CCT | CG>AT_GAACCAA | 7 | TRUE | FALSE |
| 171_v.txt | 32292 | 32292-SNV-( | SNV | 1 | G | T | 37 | 31383 | 0.11789823 | 51.0262428 | rnr1D57N | FALSE | CAN | JSY14 | 1A | rnr1D57N-Re | TGA | G>T | A/T+1 | G/C-1 | CG>AT | CG>AT | ATATTTGAAT | GAATTTG | ACT | ACT | C | A | C>A | C>A_ACT | CG>AT_GAATTTG | 7 | TRUE | FALSE |
| 171_v.txt | 32301 | 32301-SNV-( | SNV | 1 | G | T | 29 | 31383 | 0.09240672 | 48.2068966 | rnr1D57N | FALSE | CAN | JSY14 | 1A | rnr1D57N-Re | GGC | G>T | A/T+1 | G/C-1 | CG>AT | CG>AT | TTTGCGGCA | GCAGAAA | CCG | CCG | C | A | C>A | C>A_CCG | CG>AT_GCAGAAA | 7 | TRUE | FALSE |
| 171_v.txt | 32368 | 32368-SNV-( | SNV | 1 | C | A | 40 | 31383 | 0.12745754 | 55.6702428 | rnr1D57N | FALSE | CAN | JSY14 | 1A | rnr1D57N-Re | TCA | C>A | A/T+1 | G/C-1 | CG>AT | CG>AT | AGTTCTCAAT | CAATAGC | AGT | TCA | C | T | C>A | C>A_TCA | CG>AT_CAATAGC | 7 | TRUE | FALSE |
| 171_v.txt | 32373 | 32373-SNV-( | SNV | 1 | G | T | 36 | 31383 | 0.11471179 | 58.9444444 | rnr1D57N | FALSE | CAN | JSY14 | 1A | rnr1D57N-Re | AGC | G>T | A/T+1 | G/C-1 | CG>AT | CG>AT | TCAATAGCA | GCAATAA | TCG | TCG | C | A | C>A | C>A_TCG | CG>AT_GCAATAA | 7 | TRUE | FALSE |
| 171_v.txt | 32399 | 32399-SNV-( | SNV | 1 | G | T | 163 | 31383 | 0.51938948 | 59.0184049 | rnr1D57N | FALSE | CAN | JSY14 | 1A | rnr1D57N-Re | CGT | G>T | A/T+1 | G/C-1 | CG>AT | CG>AT | AGAAACGTA | GTAGGAA | GCA | GCA | C | A | C>A | C>A_GCA | CG>AT_GTAGGAA | 7 | TRUE | FALSE |
| 171_v.txt | 32432 | 32432-SNV-( | SNV | 1 | G | T | 43 | 31383 | 0.13701686 | 52.4883721 | rnr1D57N | FALSE | CAN | JSY14 | 1A | rnr1D57N-Re | TGT | G>T | A/T+1 | G/C-1 | CG>AT | CG>AT | GTCATTGTAT | GTATGGA | ACA | ACA | C | A | C>A | C>A_ACA | CG>AT_GTATGGA | 7 | TRUE | FALSE |
| 171_v.txt | 32449 | 32449-SNV-( | SNV | 1 | C | A | 33 | 31383 | 0.10515247 | 46.8787879 | rnr1D57N | FALSE | CAN | JSY14 | 1A | rnr1D57N-Re | CCA | C>A | A/T+1 | G/C-1 | CG>AT | CG>AT | AAAGTCCAA | CAATGAA | GGT | CCA | C | T | C>A | C>A_CCA | CG>AT_CAATGAA | 7 | TRUE | FALSE |
| 171_v.txt | 32463 | 32463-SNV-( | SNV | 1 | G | T | 33 | 31383 | 0.10515247 | 52.4242424 | rnr1D57N | FALSE | CAN | JSY14 | 1A | rnr1D57N-Re | AGA | G>T | A/T+1 | G/C-1 | CG>AT | CG>AT | AATAGAGAG | GAGCCAA | TCT | TCT | C | A | C>A | C>A_TCT | CG>AT_GAGCCAA | 7 | TRUE | FALSE |
| 171_v.txt | 32471 | 32471-SNV-( | SNV | 1 | G | T | 92 | 31383 | 0.29315234 | 48.9021739 | rnr1D57N | FALSE | CAN | JSY14 | 1A | rnr1D57N-Re | TGT | G>T | A/T+1 | G/C-1 | CG>AT | CG>AT | GCCAATGTA | GTAGAAG | ACA | ACA | C | A | C>A | C>A_ACA | CG>AT_GTAGAAG | 7 | TRUE | FALSE |
| 171_v.txt | 32494 | 32494-SNV-( | SNV | 1 | C | A | 286 | 31383 | 0.91132142 | 45.0559441 | rnr1D57N | FALSE | CAN | JSY14 | 1A | rnr1D57N-Re | ACA | C>A | A/T+1 | G/C-1 | CG>AT | CG>AT | GGAAAACAA | CAACTTT | TGT | ACA | C | T | C>A | C>A_ACA | CG>AT_CAACTTT | 7 | TRUE | FALSE |
| 171_v.txt | 32508 | 32508-SNV-( | SNV | 1 | G | T | 119 | 31383 | 0.37918618 | 41.907563 | rnr1D57N | FALSE | CAN | JSY14 | 1A | rnr1D57N-Re | GGC | G>T | A/T+1 | G/C-1 | CG>AT | CG>AT | TTGATGGCT | GCTCTTG | CCG | CCG | C | A | C>A | C>A_CCG | CG>AT_GCTCTTG | 7 | TRUE | FALSE |
| 171_v.txt | 32542 | 32542-SNV-( | SNV | 1 | C | A | 34 | 31383 | 0.10833891 | 36.2941454 | rnr1D57N | FALSE | CAN | JSY14 | 1A | rnr1D57N-Re | TCA | C>A | A/T+1 | G/C-1 | CG>AT | CG>AT | CAGCTTCAC | CACCAGC | AGT | TCA | C | T | C>A | C>A_TCA | CG>AT_CACCAGC | 7 | TRUE | FALSE |
| 171_v.txt | 32545 | 32545-SNV-( | SNV | 1 | C | A | 52 | 31383 | 0.1656948 | 37.0769231 | rnr1D57N | FALSE | CAN | JSY14 | 1A | rnr1D57N-Re | CCA | C>A | A/T+1 | G/C-1 | CG>AT | CG>AT | CTTCACCAG | CAGCAGT | GGT | CCA | C | T | C>A | C>A_CCA | CG>AT_CAGCAGT | 7 | TRUE | FALSE |
| 171_v.txt | 32566 | 32566-SNV-( | SNV | 1 | C | A | 63 | 30540 | 0.20628684 | 37.0476191 | rnr1D57N | FALSE | CAN | JSY14 | 1A | rnr1D57N-Re | TCA | C>A | A/T+1 | G/C-1 | CG>AT | CG>AT | CTAGTTCAGT | CAGTACC | AGT | TCA | C | T | C>A | C>A_TCA | CG>AT_CAGTACC | 7 | TRUE | FALSE |
| 171_v.txt | 32575 | 32575-SNV-( | SNV | 1 | G | T | 60 | 30555 | 0.19636721 | 35.8326041 | rnr1D57N | FALSE | CAN | JSY14 | 1A | rnr1D57N-Re | TGA | G>T | A/T+1 | G/C-1 | CG>AT | CG>AT | TACCTTGAA | GAAATGT | ACT | ACT | C | A | C>A | C>A_ACT | CG>AT_GAAATGT | 7 | TRUE | FALSE |
| 171_v.txt | 32580 | 32580-SNV-( | SNV | 1 | G | T | 17 | 30554 | 0.0556392 | 35.2906654 | rnr1D57N | FALSE | CAN | JSY14 | 1A | rnr1D57N-Re | TGT | G>T | A/T+1 | G/C-1 | CG>AT | CG>AT | TGAAATGTG. | GTGAAGG | ACA | ACA | C | A | C>A | C>A_ACA | CG>AT_GTGAAGG | 7 | TRUE | FALSE |
| 171_v.txt | 32629 | 32629-SNV-( | SNV | 1 | C | A | 514 | 30555 | 1.6822124 | 54.3560311 | rnr1D57N | FALSE | CAN | JSY14 | 1A | rnr1D57N-Re | TCG | C>A | A/T+1 | G/C-1 | CG>AT | CG>AT | TCCCTTCGTT | CGTTTTT | AGC | TCG | C | T | C>A | C>A_TCG | CG>AT_CGTTTTT | 7 | TRUE | FALSE |
| 171_v.txt | 32658 | 32658-SNV-( | SNV | 1 | C | A | 6 | 30555 | 0.01963672 | 21.8331823 | rnr1D57N | FALSE | CAN | JSY14 | 1A | rnr1D57N-Re | ACC | C>A | A/T+1 | G/C-1 | CG>AT | CG>AT | CCTGGACCC | CCCCAGG | TGG | ACC | C | T | C>A | C>A_ACC | CG>AT_CCCCAGG | 7 | TRUE | FALSE |
| 171_v.txt | 32692 | 32692-SNV-( | SNV | 1 | C | A | 75 | 30555 | 0.24545901 | 60.0266667 | rnr1D57N | FALSE | CAN | JSY14 | 1A | rnr1D57N-Re | CCA | C>A | A/T+1 | G/C-1 | CG>AT | CG>AT | GGAATCCAA | CAACTGG | GGT | CCA | C | T | C>A | C>A_CCA | CG>AT_CAACTGG | 7 | TRUE | FALSE |
| 171_v.txt | 32733 | 32733-SNV-( | SNV | 1 | C | A | 301 | 30555 | 0.98510882 | 60.538294 | rnr1D57N | FALSE | CAN | JSY14 | 1A | rnr1D57N-Re | ACA | C>A | A/T+1 | G/C-1 | CG>AT | CG>AT | CAAAAACAG | CAGTATA | TGT | ACA | C | T | C>A | C>A_ACA | CG>AT_CAGTATA | 7 | TRUE | FALSE |
| 171_v.txt | 32735 | 32735-SNV-( | SNV | 1 | G | T | 61 | 30555 | 0.19963999 | 57.4279236 | rnr1D57N | FALSE | CAN | JSY14 | 1A | rnr1D57N-Re | AGT | G>T | A/T+1 | G/C-1 | CG>AT | CG>AT | AAAACAGTA | GTATATT | TCA | TCA | C | A | C>A | C>A_TCA | CG>AT_GTATATT | 7 | TRUE | FALSE |
| 171_v.txt | 32758 | 32758-SNV-( | SNV | 1 | C | A | 7 | 30555 | 0.02290951 | 37.8576446 | rnr1D57N | FALSE | CAN | JSY14 | 1A | rnr1D57N-Re | GCT | C>A | A/T+1 | G/C-1 | CG>AT | CG>AT | TAATGGCTA | CTAAAAC | CGA | GCT | C | T | C>A | C>A_GCT | CG>AT_CTAAAAC | 7 | TRUE | FALSE |
| 171_v.txt | 32776 | 32776-SNV-( | SNV | 1 | C | A | 6 | 30554 | 0.01963736 | 40.3333333 | rnr1D57N | FALSE | CAN | JSY14 | 1A | rnr1D57N-Re | GCG | C>A | A/T+1 | G/C-1 | CG>AT | CG>AT | TGGAAGCGA | CGACCCA | CGC | GCG | C | T | C>A | C>A_GCG | CG>AT_CGACCCA | 7 | TRUE | FALSE |
| 171_v.txt | 32788 | 32788-SNV-( | SNV | 1 | C | A | 85 | 30555 | 0.27818688 | 44.9882353 | rnr1D57N | FALSE | CAN | JSY14 | 1A | rnr1D57N-Re | TCG | C>A | A/T+1 | G/C-1 | CG>AT | CG>AT | AGAACTCGA. | CGAATTC | AGC | TCG | C | T | C>A | C>A_TCG | CG>AT_CGAATTC | 7 | TRUE | FALSE |
| 171_v.txt | 32794 | 32794-SNV-( | SNV | 1 | C | A | 82 | 30555 | 0.26836852 | 42.0365854 | rnr1D57N | FALSE | CAN | JSY14 | 1A | rnr1D57N-Re | TCA | C>A | A/T+1 | G/C-1 | CG>AT | CG>AT | CGAATTCAC | CACCGTA | AGT | TCA | C | T | C>A | C>A_TCA | CG>AT_CACCGTA | 7 | TRUE | FALSE |
| 171_v.txt | 32797 | 32797-SNV-( | SNV | 1 | C | A | 3 | 30555 | 9.82E-03 | 37.6652549 | rnr1D57N | FALSE | CAN | JSY14 | 1A | rnr1D57N-Re | CCG | C>A | A/T+1 | G/C-1 | CG>AT | CG>AT | ATTCACCGTA | CGTAATA | GGC | CCG | C | T | C>A | C>A_CCG | CG>AT_CGTAATA | 7 | TRUE | FALSE |
| 171_v.txt | 32809 | 32809-SNV-( | SNV | 1 | C | A | 16 | 30555 | 0.05236459 | 37.25 | rnr1D57N | FALSE | CAN | JSY14 | 1A | rnr1D57N-Re | ACA | C>A | A/T+1 | G/C-1 | CG>AT | CG>AT | ATTTGACAG | CAGGGAA | TGT | ACA | C | T | C>A | C>A_ACA | CG>AT_CAGGGAA | 7 | TRUE | FALSE |
| 171_v.txt | 32819 | 32819-SNV-( | SNV | 1 | G | T | 30 | 30555 | 0.0981836 | 40.2333333 | rnr1D57N | FALSE | CAN | JSY14 | 1A | rnr1D57N-Re | AGT | G>T | A/T+1 | G/C-1 | CG>AT | CG>AT | GAACAAGTT | GTTCATT | TCA | TCA | C | A | C>A | C>A_TCA | CG>AT_GTTCATT | 7 | TRUE | FALSE |
| 171_v.txt | 32876 | 32876-SNV-( | SNV | 1 | G | T | 139 | 31336 | 0.44357927 | 36.9280576 | rnr1D57N | FALSE | CAN | JSY14 | 1A | rnr1D57N-Re | TGT | G>T | A/T+1 | G/C-1 | CG>AT | CG>AT | AACTTTGTAC | GTACGTC | ACA | ACA | C | A | C>A | C>A_ACA | CG>AT_GTACGTC | 7 | TRUE | FALSE |
| 171_v.txt | 32901 | 32901-SNV-( | SNV | 1 | C | A | 7 | 31348 | 0.02232997 | 34.1330679 | rnr1D57N | FALSE | CAN | JSY14 | 1A | rnr1D57N-Re | GCC | C>A | A/T+1 | G/C-1 | CG>AT | CG>AT | ACTTGGCCA | CCAACTA | CGG | GCC | C | T | C>A | C>A_GCC | CG>AT_CCAACTA | 7 | TRUE | FALSE |
| 171_v.txt | 32902 | 32902-SNV-( | SNV | 1 | C | A | 5 | 31348 | 0.01594998 | 42.9972319 | rnr1D57N | FALSE | CAN | JSY14 | 1A | rnr1D57N-Re | CCA | C>A | A/T+1 | G/C-1 | CG>AT | CG>AT | CTTGGCCAA | CAACTAC | GGT | CCA | C | T | C>A | C>A_CCA | CG>AT_CAACTAC | 7 | TRUE | FALSE |
| 171_v.txt | 32908 | 32908-SNV-( | SNV | 1 | C | A | 68 | 31348 | 0.21691974 | 46.6470588 | rnr1D57N | FALSE | CAN | JSY14 | 1A | rnr1D57N-Re | ACA | C>A | A/T+1 | G/C-1 | CG>AT | CG>AT | CAACTACACT | CACTAAG | TGT | ACA | C | T | C>A | C>A_ACA | CG>AT_CACTAAG | 7 | TRUE | FALSE |
| 171_v.txt | 32917 | 32917-SNV-( | SNV | 1 | C | A | 158 | 31348 | 0.5040194 | 50.8101266 | rnr1D57N | FALSE | CAN | JSY14 | 1A | rnr1D57N-Re | TCC | C>A | A/T+1 | G/C-1 | CG>AT | CG>AT | TAAGTTCCA | CCAGGGC | AGG | TCC | C | T | C>A | C>A_TCC | CG>AT_CCAGGGC | 7 | TRUE | FALSE |
| 171_v.txt | 32928 | 32928-SNV-( | SNV | 1 | G | T | 54 | 31348 | 0.17225979 | 51.2433509 | rnr1D57N | FALSE | CAN | JSY14 | 1A | rnr1D57N-Re | AGT | G>T | A/T+1 | G/C-1 | CG>AT | CG>AT | GCAAAAGTG | GTGATTG | TCA | TCA | C | A | C>A | C>A_TCA | CG>AT_GTGATTG | 7 | TRUE | FALSE |
| 171_v.txt | 32934 | 32934-SNV-( | SNV | 1 | G | T | 621 | 31348 | 1.98098762 | 50.4890756 | rnr1D57N | FALSE | CAN | JSY14 | 1A | rnr1D57N-Re | TGC | G>T | A/T+1 | G/C-1 | CG>AT | CG>AT | GTGATTGCC | GCCCAAG | ACG | ACG | C | A | C>A | C>A_ACG | CG>AT_GCCCAAG | 7 | TRUE | FALSE |
| 171_v.txt | 32937 | 32937-SNV-( | SNV | 1 | C | A | 24 | 31348 | 0.07655991 | 47.0298676 | rnr1D57N | FALSE | CAN | JSY14 | 1A | rnr1D57N-Re | CCA | C>A | A/T+1 | G/C-1 | CG>AT | CG>AT | ATTGCCCAA | CAAGAAA | GGT | CCA | C | T | C>A | C>A_CCA | CG>AT_CAAGAAA | 7 | TRUE | FALSE |
| 171_v.txt | 32940 | 32940-SNV-( | SNV | 1 | G | T | 33 | 31348 | 0.10526987 | 41 | rnr1D57N | FALSE | CAN | JSY14 | 1A | rnr1D57N-Re | AGA | G>T | A/T+1 | G/C-1 | CG>AT | CG>AT | GCCCAAGAA | GAAAACC | TCT | TCT | C | A | C>A | C>A_TCT | CG>AT_GAAAACC | 7 | TRUE | FALSE |
| 171_v.txt | 32970 | 32970-SNV-( | SNV | 1 | C | A | 33 | 31348 | 0.10526987 | 60.6060606 | rnr1D57N | FALSE | CAN | JSY14 | 1A | rnr1D57N-Re | ACC | C>A | A/T+1 | G/C-1 | CG>AT | CG>AT | GCCGCACCA | CCAAATG | TGG | ACC | C | T | C>A | C>A_ACC | CG>AT_CCAAATG | 7 | TRUE | FALSE |
| 171_v.txt | 33012 | 33012-SNV-( | SNV | 1 | G | T | 20 | 31348 | 0.06379992 | 58.75 | rnr1D57N | FALSE | CAN | JSY14 | 1A | rnr1D57N-Re | AGA | G>T | A/T+1 | G/C-1 | CG>AT | CG>AT | GTGAAAGAG | GAGGATG | TCT | TCT | C | A | C>A | C>A_TCT | CG>AT_GAGGATG | 7 | TRUE | FALSE |
| 171_v.txt | 33033 | 33033-SNV-( | SNV | 1 | G | T | 64 | 31348 | 0.20415976 | 56.8125 | rnr1D57N | FALSE | CAN | JSY14 | 1A | rnr1D57N-Re | TGT | G>T | A/T+1 | G/C-1 | CG>AT | CG>AT | ATGAATGTA | GTAGCCA | ACA | ACA | C | A | C>A | C>A_ACA | CG>AT_GTAGCCA | 7 | TRUE | FALSE |
| 171_v.txt | 33043 | 33043-SNV-( | SNV | 1 | C | A | 37 | 31348 | 0.11802986 | 60.2972973 | rnr1D57N | FALSE | CAN | JSY14 | 1A | rnr1D57N-Re | TCA | C>A | A/T+1 | G/C-1 | CG>AT | CG>AT | CCATTTCACC | CACCCAA | AGT | TCA | C | T | C>A | C>A_TCA | CG>AT_CACCCAA | 7 | TRUE | FALSE |
| 171_v.txt | 33051 | 33051-SNV-( | SNV | 1 | G | T | 63 | 31348 | 0.20096976 | 59.7460318 | rnr1D57N | FALSE | CAN | JSY14 | 1A | rnr1D57N-Re | GGA | G>T | A/T+1 | G/C-1 | CG>AT | CG>AT | CCCAAGGACT | GACTGCG | CCT | CCT | C | A | C>A | C>A_CCT | CG>AT_GACTGCG | 7 | TRUE | FALSE |
| 171_v.txt | 33114 | 33114-SNV-( | SNV | 1 | C | A | 5 | 31348 | 0.01594998 | 41.2 | rnr1D57N | FALSE | CAN | JSY14 | 1A | rnr1D57N-Re | GCC | C>A | A/T+1 | G/C-1 | CG>AT | CG>AT | ACTGGGCCG | CCGGCGT | CGG | GCC | C | T | C>A | C>A_GCC | CG>AT_CCGGCGT | 7 | TRUE | FALSE |
| 171_v.txt | 33115 | 33115-SNV-( | SNV | 1 | C | A | 150 | 31348 | 0.47849943 | 45.1466667 | rnr1D57N | FALSE | CAN | JSY14 | 1A | rnr1D57N-Re | CCG | C>A | A/T+1 | G/C-1 | CG>AT | CG>AT | CTGGGCCGG | CGGCGTT | GGC | CCG | C | T | C>A | C>A_CCG | CG>AT_CGGCGTT | 7 | TRUE | FALSE |
| 171_v.txt | 33118 | 33118-SNV-( | SNV | 1 | C | A | 24 | 31348 | 0.07655991 | 44.5833333 | rnr1D57N | FALSE | CAN | JSY14 | 1A | rnr1D57N-Re | GCG | C>A | A/T+1 | G/C-1 | CG>AT | CG>AT | GGCCGGCGT | CGTTGGT | CGC | GCG | C | T | C>A | C>A_GCG | CG>AT_CGTTGGT | 7 | TRUE | FALSE |
| 171_v.txt | 33151 | 33151-SNV-( | SNV | 1 | G | T | 29 | 31348 | 0.09250989 | 36.0689655 | rnr1D57N | FALSE | CAN | JSY14 | 1A | rnr1D57N-Re | AGA | G>T | A/T+1 | G/C-1 | CG>AT | CG>AT | TGAAAAGAC | GACCTGT | TCT | TCT | C | A | C>A | C>A_TCT | CG>AT_GACCTGT | 7 | TRUE | FALSE |
| 171_v.txt | 33153 | 33153-SNV-( | SNV | 1 | C | A | 21 | 31348 | 0.06698992 | 37.4286889 | rnr1D57N | FALSE | CAN | JSY14 | 1A | rnr1D57N-Re | ACC | C>A | A/T+1 | G/C-1 | CG>AT | CG>AT | AAAAGACCT | CCTGTAC | TGG | ACC | C | T | C>A | C>A_ACC | CG>AT_CCTGTAC | 7 | TRUE | FALSE |
| 171_v.txt | 33154 | 33154-SNV-( | SNV | 1 | C | A | 36 | 31348 | 0.11483986 | 36.0551693 | rnr1D57N | FALSE | CAN | JSY14 | 1A | rnr1D57N-Re | CCT | C>A | A/T+1 | G/C-1 | CG>AT | CG>AT | AAAGACCTG | CTGTACC | GGA | CCT | C | T | C>A | C>A_CCT | CG>AT_CTGTACC | 7 | TRUE | FALSE |
| 171_v.txt | 33159 | 33159-SNV-( | SNV | 1 | C | A | 14 | 31342 | 0.0446685 | 35.9972757 | rnr1D57N | FALSE | CAN | JSY14 | 1A | rnr1D57N-Re | ACC | C>A | A/T+1 | G/C-1 | CG>AT | CG>AT | CCTGTACCA | CCAATAG | TGG | ACC | C | T | C>A | C>A_ACC | CG>AT_CCAATAG | 7 | TRUE | FALSE |
| 171_v.txt | 33160 | 33160-SNV-( | SNV | 1 | C | A | 29 | 31342 | 0.0925276 | 36.8616317 | rnr1D57N | FALSE | CAN | JSY14 | 1A | rnr1D57N-Re | CCA | C>A | A/T+1 | G/C-1 | CG>AT | CG>AT | CTGTACCAAT | CAATAGT | GGT | CCA | C | T | C>A | C>A_CCA | CG>AT_CAATAGT | 7 | TRUE | FALSE |
| 171_v.txt | 33168 | 33168-SNV-( | SNV | 1 | C | A | 56 | 31300 | 0.17891374 | 36.5319019 | rnr1D57N | FALSE | CAN | JSY14 | 1A | rnr1D57N-Re | ACC | C>A | A/T+1 | G/C-1 | CG>AT | CG>AT | ATAGTACCA | CCACCAA | TGG | ACC | C | T | C>A | C>A_ACC | CG>AT_CCACCAA | 7 | TRUE | FALSE |
| 171_v.txt | 33169 | 33169-SNV-( | SNV | 1 | C | A | 231 | 31300 | 0.73801917 | 37.5318597 | rnr1D57N | FALSE | CAN | JSY14 | 1A | rnr1D57N-Re | CCA | C>A | A/T+1 | G/C-1 | CG>AT | CG>AT | TAGTACCAC | CACCAAG | GGT | CCA | C | T | C>A | C>A_CCA | CG>AT_CACCAAG | 7 | TRUE | FALSE |
| 171_v.txt | 33177 | 33177-SNV-( | SNV | 1 | G | T | 21 | 31297 | 0.06709908 | 37.5238871 | rnr1D57N | FALSE | CAN | JSY14 | 1A | rnr1D57N-Re | GGC | G>T | A/T+1 | G/C-1 | CG>AT | CG>AT | CCAAGGGCA | GCAATCA | CCG | CCG | C | A | C>A | C>A_CCG | CG>AT_GCAATCA | 7 | TRUE | FALSE |
| 171_v.txt | 33205 | 33205-SNV-( | SNV | 1 | G | T | 8 | 13895 | 0.05757467 | 39.625 | rnr1D57N | FALSE | CAN | JSY14 | 1A | rnr1D57N-Re | AGC | G>T | A/T+1 | G/C-1 | CG>AT | CG>AT | GCTTAAGCT | GCTCTCT | TCG | TCG | C | A | C>A | C>A_TCG | CG>AT_GCTCTCT | 7 | TRUE | FALSE |
| 171_v.txt | 33220 | 33220-SNV-( | SNV | 1 | C | A | 29 | 13904 | 0.20857307 | 39.2758621 | rnr1D57N | FALSE | CAN | JSY14 | 1A | rnr1D57N-Re | TCA | C>A | A/T+1 | G/C-1 | CG>AT | CG>AT | TCACTTCAGC | CAGCGTT | AGT | TCA | C | T | C>A | C>A_TCA | CG>AT_CAGCGTT | 7 | TRUE | FALSE |
| 171_v.txt | 33241 | 33241-SNV-( | SNV | 1 | C | A | 12 | 13904 | 0.0863061 | 41.1666667 | rnr1D57N | FALSE | CAN | JSY14 | 1A | rnr1D57N-Re | TCA | C>A | A/T+1 | G/C-1 | CG>AT | CG>AT | CTCCTTCATC | CATCTTC | AGT | TCA | C | T | C>A | C>A_TCA | CG>AT_CATCTTC | 7 | TRUE | FALSE |
| 171_v.txt | 33247 | 33247-SNV-( | SNV | 1 | C | A | 16 | 13904 | 0.1150748 | 43.125 | rnr1D57N | FALSE | CAN | JSY14 | 1A | rnr1D57N-Re | TCA | C>A | A/T+1 | G/C-1 | CG>AT | CG>AT | CATCTTCATC | CATCACC | AGT | TCA | C | T | C>A | C>A_TCA | CG>AT_CATCACC | 7 | TRUE | FALSE |
| 171_v.txt | 33325 | 33325-SNV-( | SNV | 1 | C | A | 245 | 13904 | 1.76208285 | 58.3510204 | rnr1D57N | FALSE | CAN | JSY14 | 1A | rnr1D57N-Re | TCT | C>A | A/T+1 | G/C-1 | CG>AT | CG>AT | ACAATTCTTT | CTTTACT | AGA | TCT | C | T | C>A | C>A_TCT | CG>AT_CTTTACT | 7 | TRUE | FALSE |
| 171_v.txt | 33354 | 33354-SNV-( | SNV | 1 | G | T | 110 | 13896 | 0.7915947 | 31.6490192 | rnr1D57N | FALSE | CAN | JSY14 | 1A | rnr1D57N-Re | TGA | G>T | A/T+1 | G/C-1 | CG>AT | CG>AT | GGTATTGAC | GACCCAC | ACT | ACT | C | A | C>A | C>A_ACT | CG>AT_GACCCAC | 7 | TRUE | FALSE |
| 171_v.txt | 33385 | 33385-SNV-( | SNV | 1 | C | A | 17 | 13904 | 0.12226697 | 53.8823529 | rnr1D57N | FALSE | CAN | JSY14 | 1A | rnr1D57N-Re | TCA | C>A | A/T+1 | G/C-1 | CG>AT | CG>AT | AAGCTTCAA | CAACGTC | AGT | TCA | C | T | C>A | C>A_TCA | CG>AT_CAACGTC | 7 | TRUE | FALSE |
| 171_v.txt | 33415 | 33415-SNV-( | SNV | 1 | C | A | 23 | 13904 | 0.16542002 | 34.8695652 | rnr1D57N | FALSE | CAN | JSY14 | 1A | rnr1D57N-Re | TCA | C>A | A/T+1 | G/C-1 | CG>AT | CG>AT | CCGGCTCATT | CATTGTA | AGT | TCA | C | T | C>A | C>A_TCA | CG>AT_CATTGTA | 7 | TRUE | FALSE |
| 171_v.txt | 33436 | 33436-SNV-( | SNV | 1 | C | A | 28 | 13904 | 0.2013809 | 44 | rnr1D57N | FALSE | CAN | JSY14 | 1A | rnr1D57N-Re | TCT | C>A | A/T+1 | G/C-1 | CG>AT | CG>AT | TCTCCTCTAT | CTATGTC | AGA | TCT | C | T | C>A | C>A_TCT | CG>AT_CTATGTC | 7 | TRUE | FALSE |
| 171_v.txt | 33474 | 33474-SNV-( | SNV | 1 | C | A | 32 | 13903 | 0.23016615 | 33.625 | rnr1D57N | FALSE | CAN | JSY14 | 1A | rnr1D57N-Re | GCC | C>A | A/T+1 | G/C-1 | CG>AT | CG>AT | GCTATGCCTT | CCTTTTT | CGG | GCC | C | T | C>A | C>A_GCC | CG>AT_CCTTTTT | 7 | TRUE | FALSE |
| 171_v.txt | 32164 | 32164-Repla | Replacement | 2 | CG | A | 11 | 24202 | 0.04545079 | 46.6363636 | rnr1D57N | FALSE | CAN | JSY14 | 1A | rnr1D57N-Re | GCG | nope | A/T+1 | >1bpdel | none | Replac. | CCAAAGCGC | CGCCAAA | CGC | GCG | CG | T | NA | NA_GCG | Replac._CGCCAAA | 7 | TRUE | FALSE |
| 171_v.txt | 32814 | 32814-Repla | Replacement | 3 | AAC | G | 26 | 30555 | 0.08509246 | 44.9358974 | rnr1D57N | FALSE | CAN | JSY14 | 1A | rnr1D57N-Re | GAA | nope | G/C+1 | >1bpdel | none | Replac. | ACAGGGAAC | AACAAGT | CTT | GAA | AAC | C | NA | NA_GAA | Replac._AACAAGT | 7 | TRUE | FALSE |
| 171_v.txt | 32841 | 32841-Repla | Replacement | 2 | C | TA | 1 | 30555 | 3.27E-03 | 37.9999966 | rnr1D57N | FALSE | CAN | JSY14 | 1A | rnr1D57N-Re | CCA | nope | >1bp-ins | G/C-1 | none | Replac. | ATTACCCAA | CAAAAAA | GGT | CCA | C | none | nope | nope_CCA | Replac._CAAAAAA | 7 | TRUE | FALSE |
| 171_v.txt | 32937 | 32937-Repla | Replacement | 2 | C | TG | 1 | 31348 | 3.19E-03 | 22.4999982 | rnr1D57N | FALSE | CAN | JSY14 | 1A | rnr1D57N-Re | CCA | nope | >1bp-ins | G/C-1 | none | Replac. | ATTGCCCAA | CAAGAAA | GGT | CCA | C | none | nope | nope_CCA | Replac._CAAGAAA | 7 | TRUE | FALSE |
| 171_v.txt | 31875 | 31875-MNV- | MNV | 2 | TA | GG | 5 | 10179 | 0.04912074 | 42.2999913 | rnr1D57N | FALSE | CAN | JSY14 | 1A | rnr1D57N-Re | ATA | nope | >1bp-ins | >1bpdel | none | MNV | GAGATATAG | TAGGCGG | TAT | ATA | TA | none | NA | NA_ATA | MNV_TAGGCGG | 7 | TRUE | FALSE |
| 171_v.txt | 31956 | 31956-MNV- | MNV | 2 | GT | TA | 18 | 24202 | 0.07437402 | 36.8889357 | rnr1D57N | FALSE | CAN | JSY14 | 1A | rnr1D57N-Re | TGT | nope | >1bp-ins | >1bpdel | none | MNV | ATAAATGTG | GTGGCCG | ACA | TGT | GT | none | NA | NA_TGT | MNV_GTGGCCG | 7 | TRUE | FALSE |
| 171_v.txt | 32292 | 32292-MNV- | MNV | 3 | GAA | ATT | 5 | 31383 | 0.01593219 | 56.6666663 | rnr1D57N | FALSE | CAN | JSY14 | 1A | rnr1D57N-Re | TGA | nope | >1bp-ins | >1bpdel | none | MNV | ATATTTGAAT | GAATTTG | ACT | TGA | GAA | none | NA | NA_TGA | MNV_GAATTTG | 7 | TRUE | FALSE |
| 171_v.txt | 32535 | 32535-MNV- | MNV | 2 | GC | AT | 1 | 31383 | 3.19E-03 | 37.9999732 | rnr1D57N | FALSE | CAN | JSY14 | 1A | rnr1D57N-Re | TGC | nope | >1bp-ins | >1bpdel | none | MNV | GGGTTTGCA | GCAGCTT | ACG | TGC | GC | none | NA | NA_TGC | MNV_GCAGCTT | 7 | TRUE | FALSE |
| 171_v.txt | 32544 | 32544-MNV- | MNV | 2 | CC | AA | 4 | 31383 | 0.01274575 | 31.6249995 | rnr1D57N | FALSE | CAN | JSY14 | 1A | rnr1D57N-Re | ACC | nope | >1bp-ins | >1bpdel | none | MNV | GCTTCACCA | CCAGCAG | TGG | ACC | CC | none | NA | NA_ACC | MNV_CCAGCAG | 7 | TRUE | FALSE |
| 171_v.txt | 32787 | 32787-MNV- | MNV | 2 | TC | CT | 36 | 30555 | 0.11782032 | 45.9583333 | rnr1D57N | FALSE | CAN | JSY14 | 1A | rnr1D57N-Re | CTC | nope | >1bp-ins | >1bpdel | none | MNV | CAGAACTCG. | TCGAATT | GAG | CTC | TC | none | NA | NA_CTC | MNV_TCGAATT | 7 | TRUE | FALSE |
| 171_v.txt | 32797 | 32797-MNV- | MNV | 2 | CG | TT | 57 | 30555 | 0.18654885 | 43.9468409 | rnr1D57N | FALSE | CAN | JSY14 | 1A | rnr1D57N-Re | CCG | nope | >1bp-ins | >1bpdel | none | MNV | ATTCACCGTA | CGTAATA | GGC | CCG | CG | none | NA | NA_CCG | MNV_CGTAATA | 7 | TRUE | FALSE |
| 171_v.txt | 32900 | 32900-MNV- | MNV | 2 | GC | AA | 1 | 31348 | 3.19E-03 | 35.4999843 | rnr1D57N | FALSE | CAN | JSY14 | 1A | rnr1D57N-Re | GGC | nope | >1bp-ins | >1bpdel | none | MNV | GACTTGGCC | GCCAACT | CCG | GGC | GC | none | NA | NA_GGC | MNV_GCCAACT | 7 | TRUE | FALSE |
| 171_v.txt | 32914 | 32914-MNV- | MNV | 2 | GT | AC | 66 | 31348 | 0.21053975 | 46.2429002 | rnr1D57N | FALSE | CAN | JSY14 | 1A | rnr1D57N-Re | AGT | nope | >1bp-ins | >1bpdel | none | MNV | CACTAAGTT | GTTCCAG | TCA | AGT | GT | none | NA | NA_AGT | MNV_GTTCCAG | 7 | TRUE | FALSE |
| 171_v.txt | 32979 | 32979-MNV- | MNV | 3 | GGA | TAC | 16 | 31348 | 0.05103994 | 61.4166666 | rnr1D57N | FALSE | CAN | JSY14 | 1A | rnr1D57N-Re | TGG | nope | >1bp-ins | >1bpdel | none | MNV | AATGCTGGA | GGAGAAA | ACC | TGG | GGA | none | NA | NA_TGG | MNV_GGAGAAA | 7 | TRUE | FALSE |
| 171_v.txt | 33153 | 33153-MNV- | MNV | 2 | CC | AA | 2 | 31348 | 6.38E-03 | 35.25 | rnr1D57N | FALSE | CAN | JSY14 | 1A | rnr1D57N-Re | ACC | nope | >1bp-ins | >1bpdel | none | MNV | AAAAGACCT | CCTGTAC | TGG | ACC | CC | none | NA | NA_ACC | MNV_CCTGTAC | 7 | TRUE | FALSE |
| 171_v.txt | 33159 | 33159-MNV- | MNV | 2 | CC | AA | 1 | 31342 | 3.19E-03 | 31.5 | rnr1D57N | FALSE | CAN | JSY14 | 1A | rnr1D57N-Re | ACC | nope | >1bp-ins | >1bpdel | none | MNV | CCTGTACCA | CCAATAG | TGG | ACC | CC | none | NA | NA_ACC | MNV_CCAATAG | 7 | TRUE | FALSE |
| 171_v.txt | 33168 | 33168-MNV- | MNV | 2 | CC | AT | 8 | 31300 | 0.02555911 | 37.9999264 | rnr1D57N | FALSE | CAN | JSY14 | 1A | rnr1D57N-Re | ACC | nope | >1bp-ins | >1bpdel | none | MNV | ATAGTACCA | CCACCAA | TGG | ACC | CC | none | NA | NA_ACC | MNV_CCACCAA | 7 | TRUE | FALSE |
| 171_v.txt | 31883 | 31883-Insert | Insertion | 5 | - | AGCAA | 25 | 10179 | 0.24560369 | 43.4 | rnr1D57N | FALSE | CAN | JSY14 | 1A | rnr1D57N-Re | CAG | nope | >1bp-ins | >1bpdel | none | >1bp-ins | GGCGGCAGC | AGCAAAG | GTC | CAG | - | none | NA | NA_CAG | >1bp-ins_AGCAAAG | 7 | TRUE | FALSE |
| 171_v.txt | 31902 | 31902-Insert | Insertion | 1 | - | C | 10 | 10171 | 0.09831875 | 42 | rnr1D57N | FALSE | CAN | JSY14 | 1A | rnr1D57N-Re | GAA | nope | G/C+1 | >1bpdel | none | G/C+1 | CCATTGAATT | AATTTTG | CTT | GAA | - | G | NA | NA_GAA | G/C+1_AATTTTG | 7 | TRUE | FALSE |
| 171_v.txt | 31913 | 31913-Insert | Insertion | 1 | - | A | 2 | 10171 | 0.01966375 | 38 | rnr1D57N | FALSE | CAN | JSY14 | 1A | rnr1D57N-Re | CAA | nope | A/T+1 | >1bpdel | none | A/T+1 | TGGTGCAAA. | AAAAGCC | GTT | CAA | - | T | NA | NA_CAA | A/T+1_AAAAGCC | 7 | TRUE | FALSE |
| 171_v.txt | 31969 | 31969-Insert | Insertion | 1 | - | A | 4 | 24202 | 0.01652756 | 38 | rnr1D57N | FALSE | CAN | JSY14 | 1A | rnr1D57N-Re | TAA | nope | A/T+1 | >1bpdel | none | A/T+1 | CATAATAAG | AAGCCAA | ATT | TAA | - | T | NA | NA_TAA | A/T+1_AAGCCAA | 7 | TRUE | FALSE |
| 171_v.txt | 32067 | 32067-Insert | Insertion | 2 | - | AA | 11 | 24202 | 0.04545079 | 64 | rnr1D57N | FALSE | CAN | JSY14 | 1A | rnr1D57N-Re | GAT | nope | >1bp-ins | >1bpdel | none | >1bp-ins | ATTGAGATA | ATAAATA | CTA | GAT | - | none | NA | NA_GAT | >1bp-ins_ATAAATA | 7 | TRUE | FALSE |
| 171_v.txt | 32081 | 32081-Insert | Insertion | 1 | - | A | 356 | 24202 | 1.47095281 | 63.5252809 | rnr1D57N | FALSE | CAN | JSY14 | 1A | rnr1D57N-Re | CAA | nope | A/T+1 | >1bpdel | none | A/T+1 | CCATGCAAA | AAAAAAG | GTT | CAA | - | T | NA | NA_CAA | A/T+1_AAAAAAG | 7 | TRUE | FALSE |
| 171_v.txt | 32081 | 32081-Insert | Insertion | 2 | - | AA | 13 | 24202 | 0.05371457 | 64 | rnr1D57N | FALSE | CAN | JSY14 | 1A | rnr1D57N-Re | CAA | nope | >1bp-ins | >1bpdel | none | >1bp-ins | CCATGCAAA | AAAAAAG | GTT | CAA | - | none | NA | NA_CAA | >1bp-ins_AAAAAAG | 7 | TRUE | FALSE |
| 171_v.txt | 32166 | 32166-Insert | Insertion | 1 | - | C | 2 | 24202 | 8.26E-03 | 51 | rnr1D57N | FALSE | CAN | JSY14 | 1A | rnr1D57N-Re | GCC | nope | G/C+1 | >1bpdel | none | G/C+1 | AAAGCGCCA | CCAAATG | CGG | GCC | - | G | NA | NA_GCC | G/C+1_CCAAATG | 7 | TRUE | FALSE |
| 171_v.txt | 32228 | 32228-Insert | Insertion | 1 | - | A | 3 | 24201 | 0.01239618 | 38 | rnr1D57N | FALSE | CAN | JSY14 | 1A | rnr1D57N-Re | GGA | nope | A/T+1 | >1bpdel | none | A/T+1 | TGACAGGAA | GAATTTA | CCT | GGA | - | T | NA | NA_GGA | A/T+1_GAATTTA | 7 | TRUE | FALSE |
| 171_v.txt | 32289 | 32289-Insert | Insertion | 1 | - | T | 23 | 31383 | 0.07328809 | 47.9565217 | rnr1D57N | FALSE | CAN | JSY14 | 1A | rnr1D57N-Re | ATT | nope | A/T+1 | >1bpdel | none | A/T+1 | TAAATATTTG | TTTGAAT | TAA | ATT | - | A | NA | NA_ATT | A/T+1_TTTGAAT | 7 | TRUE | FALSE |
| 171_v.txt | 32322 | 32322-Insert | Insertion | 1 | - | A | 21 | 31383 | 0.06691521 | 62.0952381 | rnr1D57N | FALSE | CAN | JSY14 | 1A | rnr1D57N-Re | GAT | nope | A/T+1 | >1bpdel | none | A/T+1 | GTTAAGATA. | ATAACAG | CTA | GAT | - | T | NA | NA_GAT | A/T+1_ATAACAG | 7 | TRUE | FALSE |
| 171_v.txt | 32352 | 32352-Insert | Insertion | 1 | - | T | 9 | 31383 | 0.02867795 | 63 | rnr1D57N | FALSE | CAN | JSY14 | 1A | rnr1D57N-Re | CTT | nope | A/T+1 | >1bpdel | none | A/T+1 | AAAACCTTTG | TTTGTAC | GAA | CTT | - | A | NA | NA_CTT | A/T+1_TTTGTAC | 7 | TRUE | FALSE |
| 171_v.txt | 32369 | 32369-Insert | Insertion | 1 | - | A | 4 | 31383 | 0.01274575 | 64 | rnr1D57N | FALSE | CAN | JSY14 | 1A | rnr1D57N-Re | CAA | nope | A/T+1 | >1bpdel | none | A/T+1 | GTTCTCAATA | AATAGCA | GTT | CAA | - | T | NA | NA_CAA | A/T+1_AATAGCA | 7 | TRUE | FALSE |
| 171_v.txt | 32442 | 32442-Insert | Insertion | 1 | - | A | 1 | 31383 | 3.19E-03 | 64 | rnr1D57N | FALSE | CAN | JSY14 | 1A | rnr1D57N-Re | TAA | nope | A/T+1 | >1bpdel | none | A/T+1 | GGAACTAAA | AAAAGTC | ATT | TAA | - | T | NA | NA_TAA | A/T+1_AAAAGTC | 7 | TRUE | FALSE |
| 171_v.txt | 32473 | 32473-Insert | Insertion | 3 | - | AGA | 38 | 31383 | 0.12108466 | 52.122807 | rnr1D57N | FALSE | CAN | JSY14 | 1A | rnr1D57N-Re | TAG | nope | >1bp-ins | >1bpdel | none | >1bp-ins | CAATGTAGA. | AGAAGGT | ATC | TAG | - | none | NA | NA_TAG | >1bp-ins_AGAAGGT | 7 | TRUE | FALSE |
| 171_v.txt | 32490 | 32490-Insert | Insertion | 1 | - | A | 283 | 31383 | 0.9017621 | 45.3957597 | rnr1D57N | FALSE | CAN | JSY14 | 1A | rnr1D57N-Re | GAA | nope | A/T+1 | >1bpdel | none | A/T+1 | ATACGGAAA | AAAACAA | CTT | GAA | - | T | NA | NA_GAA | A/T+1_AAAACAA | 7 | TRUE | FALSE |
| 171_v.txt | 32498 | 32498-Insert | Insertion | 1 | - | T | 31 | 31383 | 0.09877959 | 42.0967742 | rnr1D57N | FALSE | CAN | JSY14 | 1A | rnr1D57N-Re | CTT | nope | A/T+1 | >1bpdel | none | A/T+1 | AACAACTTTT | TTTTTTG | GAA | CTT | - | A | NA | NA_CTT | A/T+1_TTTTTTG | 7 | TRUE | FALSE |
| 171_v.txt | 32528 | 32528-Insert | Insertion | 1 | - | G | 17 | 31383 | 0.05416946 | 40.6470588 | rnr1D57N | FALSE | CAN | JSY14 | 1A | rnr1D57N-Re | TGG | nope | G/C+1 | >1bpdel | none | G/C+1 | TTTTCTGGGG | GGGGTTT | ACC | TGG | - | C | NA | NA_TGG | G/C+1_GGGGTTT | 7 | TRUE | FALSE |
| 171_v.txt | 32598 | 32598-Insert | Insertion | 2 | - | AA | 8 | 30554 | 0.02618315 | 43.5625 | rnr1D57N | FALSE | CAN | JSY14 | 1A | rnr1D57N-Re | CAA | nope | >1bp-ins | >1bpdel | none | >1bp-ins | TTAATCAAA | AAAGAGG | GTT | CAA | - | none | NA | NA_CAA | >1bp-ins_AAAGAGG | 7 | TRUE | FALSE |
| 171_v.txt | 32631 | 32631-Insert | Insertion | 1 | - | T | 176 | 30555 | 0.57601047 | 51.0909091 | rnr1D57N | FALSE | CAN | JSY14 | 1A | rnr1D57N-Re | GTT | nope | A/T+1 | >1bpdel | none | A/T+1 | CCTTCGTTTT | TTTTTAT | CAA | GTT | - | A | NA | NA_GTT | A/T+1_TTTTTAT | 7 | TRUE | FALSE |
| 171_v.txt | 32665 | 32665-Insert | Insertion | 27 | - | CACCTGGGT | 5 | 30555 | 0.01636393 | 55.2222222 | rnr1D57N | FALSE | CAN | JSY14 | 1A | rnr1D57N-Re | GCA | nope | >1bp-ins | >1bpdel | none | >1bp-ins | CCCAGGCAC | CACCTGG | CGT | GCA | - | none | NA | NA_GCA | >1bp-ins_CACCTGG | 7 | TRUE | FALSE |
| 171_v.txt | 32678 | 32678-Insert | Insertion | 37 | - | CCAATAACG | 25 | 30555 | 0.08181967 | 55.1005405 | rnr1D57N | FALSE | CAN | JSY14 | 1A | rnr1D57N-Re | TCC | nope | >1bp-ins | >1bpdel | none | >1bp-ins | GTTTCTCCAA | CCAATAA | AGG | TCC | - | none | NA | NA_TCC | >1bp-ins_CCAATAA | 7 | TRUE | FALSE |
| 171_v.txt | 32728 | 32728-Insert | Insertion | 1 | - | A | 349 | 30555 | 1.14220259 | 62.0487106 | rnr1D57N | FALSE | CAN | JSY14 | 1A | rnr1D57N-Re | CAA | nope | A/T+1 | >1bpdel | none | A/T+1 | CCATACAAA | AAAAACA | GTT | CAA | - | T | NA | NA_CAA | A/T+1_AAAAACA | 7 | TRUE | FALSE |
| 171_v.txt | 32728 | 32728-Insert | Insertion | 2 | - | AA | 2 | 30555 | 6.55E-03 | 64 | rnr1D57N | FALSE | CAN | JSY14 | 1A | rnr1D57N-Re | CAA | nope | >1bp-ins | >1bpdel | none | >1bp-ins | CCATACAAA | AAAAACA | GTT | CAA | - | none | NA | NA_CAA | >1bp-ins_AAAAACA | 7 | TRUE | FALSE |
| 171_v.txt | 32760 | 32760-Insert | Insertion | 1 | - | A | 1 | 30555 | 3.27E-03 | 38 | rnr1D57N | FALSE | CAN | JSY14 | 1A | rnr1D57N-Re | TAA | nope | A/T+1 | >1bpdel | none | A/T+1 | ATGGCTAAA. | AAAACTT | ATT | TAA | - | T | NA | NA_TAA | A/T+1_AAAACTT | 7 | TRUE | FALSE |
| 171_v.txt | 32760 | 32760-Insert | Insertion | 22 | - | AAAACTTTG | 15 | 30555 | 0.0490918 | 53.430303 | rnr1D57N | FALSE | CAN | JSY14 | 1A | rnr1D57N-Re | TAA | nope | >1bp-ins | >1bpdel | none | >1bp-ins | ATGGCTAAA. | AAAACTT | ATT | TAA | - | none | NA | NA_TAA | >1bp-ins_AAAACTT | 7 | TRUE | FALSE |
| 171_v.txt | 32802 | 32802-Insert | Insertion | 1 | - | T | 11 | 30555 | 0.03600066 | 41.5454546 | rnr1D57N | FALSE | CAN | JSY14 | 1A | rnr1D57N-Re | ATA | nope | A/T+1 | >1bpdel | none | A/T+1 | CCGTAATATT | TATTTGA | TAT | ATA | - | A | NA | NA_ATA | A/T+1_TATTTGA | 7 | TRUE | FALSE |
| 171_v.txt | 32839 | 32839-Insert | Insertion | 1 | - | C | 3 | 30555 | 9.82E-03 | 27.6666667 | rnr1D57N | FALSE | CAN | JSY14 | 1A | rnr1D57N-Re | ACC | nope | G/C+1 | >1bpdel | none | G/C+1 | TAATTACCCA | CCCAAAA | TGG | ACC | - | G | NA | NA_ACC | G/C+1_CCCAAAA | 7 | TRUE | FALSE |
| 171_v.txt | 32840 | 32840-Insert | Insertion | 1 | - | A | 9 | 30555 | 0.02945508 | 46.5555556 | rnr1D57N | FALSE | CAN | JSY14 | 1A | rnr1D57N-Re | CCC | nope | A/T+1 | >1bpdel | none | A/T+1 | AATTACCCA | CCAAAAA | GGG | CCC | - | T | NA | NA_CCC | A/T+1_CCAAAAA | 7 | TRUE | FALSE |
| 171_v.txt | 32842 | 32842-Insert | Insertion | 1 | - | A | 510 | 30555 | 1.66912126 | 41.9058819 | rnr1D57N | FALSE | CAN | JSY14 | 1A | rnr1D57N-Re | CAA | nope | A/T+1 | >1bpdel | none | A/T+1 | TTACCCAAA | AAAAAAT | GTT | CAA | - | T | NA | NA_CAA | A/T+1_AAAAAAT | 7 | TRUE | FALSE |
| 171_v.txt | 32842 | 32842-Insert | Insertion | 1 | - | G | 61 | 30555 | 0.19963999 | 38.9508196 | rnr1D57N | FALSE | CAN | JSY14 | 1A | rnr1D57N-Re | CAA | nope | G/C+1 | >1bpdel | none | G/C+1 | TTACCCAAA | AAAAAAT | GTT | CAA | - | C | NA | NA_CAA | G/C+1_AAAAAAT | 7 | TRUE | FALSE |
| 171_v.txt | 32924 | 32924-Insert | Insertion | 2 | - | AA | 18 | 31348 | 0.05741993 | 52.8888889 | rnr1D57N | FALSE | CAN | JSY14 | 1A | rnr1D57N-Re | CAA | nope | >1bp-ins | >1bpdel | none | >1bp-ins | CAGGGCAAA | AAAAGTG | GTT | CAA | - | none | NA | NA_CAA | >1bp-ins_AAAAGTG | 7 | TRUE | FALSE |
| 171_v.txt | 32938 | 32938-Insert | Insertion | 1 | - | G | 615 | 31348 | 1.96184765 | 55.895935 | rnr1D57N | FALSE | CAN | JSY14 | 1A | rnr1D57N-Re | CAA | nope | G/C+1 | >1bpdel | none | G/C+1 | TTGCCCAAG | AAGAAAA | GTT | CAA | - | C | NA | NA_CAA | G/C+1_AAGAAAA | 7 | TRUE | FALSE |
| 171_v.txt | 32972 | 32972-Insert | Insertion | 1 | - | A | 1 | 31348 | 3.19E-03 | 64 | rnr1D57N | FALSE | CAN | JSY14 | 1A | rnr1D57N-Re | CAA | nope | A/T+1 | >1bpdel | none | A/T+1 | CGCACCAAA | AAATGCT | GTT | CAA | - | T | NA | NA_CAA | A/T+1_AAATGCT | 7 | TRUE | FALSE |
| 171_v.txt | 33000 | 33000-Insert | Insertion | 1 | - | A | 19 | 31348 | 0.06060993 | 62.4210526 | rnr1D57N | FALSE | CAN | JSY14 | 1A | rnr1D57N-Re | GAA | nope | A/T+1 | >1bpdel | none | A/T+1 | TGTGAGAAA | AAAACTG | CTT | GAA | - | T | NA | NA_GAA | A/T+1_AAAACTG | 7 | TRUE | FALSE |
| 171_v.txt | 33072 | 33072-Insert | Insertion | 1 | - | A | 26 | 31348 | 0.0829399 | 50.8846154 | rnr1D57N | FALSE | CAN | JSY14 | 1A | rnr1D57N-Re | CAA | nope | A/T+1 | >1bpdel | none | A/T+1 | TATGCCAAA | AAAGAAC | GTT | CAA | - | T | NA | NA_CAA | A/T+1_AAAGAAC | 7 | TRUE | FALSE |
| 171_v.txt | 33072 | 33072-Insert | Insertion | 2 | - | AA | 4 | 31348 | 0.01275999 | 50.875 | rnr1D57N | FALSE | CAN | JSY14 | 1A | rnr1D57N-Re | CAA | nope | >1bp-ins | >1bpdel | none | >1bp-ins | TATGCCAAA | AAAGAAC | GTT | CAA | - | none | NA | NA_CAA | >1bp-ins_AAAGAAC | 7 | TRUE | FALSE |

| | | | | | | | | | | | | | | | | | | | | | | | | | | | | | | | | | | |
|---|---|---|---|---|---|---|---|---|---|---|---|---|---|---|---|---|---|---|---|---|---|---|---|---|---|---|---|---|---|---|---|---|---|---|
| 171_v.txt | 33331 | 33331-Insert | Insertion | 1 | - | T | 19 | 13904 | 0.13665132 | 62.4210526 | rnr1D57N | FALSE | CAN | JSY14 | 1A | rnr1D57N-Re | CTT | nope | A/T+1 | >1bpdel | none | A/T+1 | CTTTACTTTT( | TTTTCTC | GAA | CTT | - | A | NA | NA_CTT | A/T+1_TTTTCTC | 7 | TRUE | FALSE |
| 171_v.txt | 33355 | 33355-Insert | Insertion | 4 | - | ACCC | 2 | 13904 | 0.01438435 | 63.875 | rnr1D57N | FALSE | CAN | JSY14 | 1A | rnr1D57N-Re | GAC | nope | >1bp-ins | >1bpdel | none | >1bp-ins | GTATTGACC( | ACCCACG | CTG | GAC | - | none | NA | NA_GAC | >1bp-ins_ACCCACG | 7 | TRUE | FALSE |
| 171_v.txt | 31807 | 31807-Deleti | Deletion | 4 | AAAT | - | 23 | 10179 | 0.2259554 | 59.4347826 | rnr1D57N | FALSE | CAN | JSY14 | 1A | rnr1D57N-Re | CAA | nope | >1bp-ins | >1bpdel | none | >1bpdel | TCTTCCAAAT | AAATAAA | GTT | CAA | AAAT | none | NA | NA_CAA | >1bpdel_AAATAAA | 7 | TRUE | FALSE |
| 171_v.txt | 32114 | 32114-Deleti | Deletion | 19 | CCATTCGAA/ | - | 421 | 24202 | 1.73952566 | 59.1187649 | rnr1D57N | FALSE | CAN | JSY14 | 1A | rnr1D57N-Re | GCC | nope | >1bp-ins | >1bpdel | none | >1bpdel | TAATAGCCA' | CCATTCG | CGG | GCC | CCATTCGAA/ | none | NA | NA_GCC | >1bpdel_CCATTCG | 7 | TRUE | FALSE |
| 171_v.txt | 32360 | 32360-Deleti | Deletion | 2 | AG | - | 68 | 31383 | 0.21667782 | 61.4117648 | rnr1D57N | FALSE | CAN | JSY14 | 1A | rnr1D57N-Re | CAG | nope | >1bp-ins | >1bpdel | none | >1bpdel | TGTACCAGA( | AGAGTTC | GTC | CAG | AG | none | NA | NA_CAG | >1bpdel_AGAGTTC | 7 | TRUE | FALSE |
| 171_v.txt | 32375 | 32375-Deleti | Deletion | 3 | AAT | - | 16 | 31383 | 0.05098302 | 60.125 | rnr1D57N | FALSE | CAN | JSY14 | 1A | rnr1D57N-Re | CAA | nope | >1bp-ins | >1bpdel | none | >1bpdel | AATAGCAAT. | AATAATA | GTT | CAA | AAT | none | NA | NA_CAA | >1bpdel_AATAATA | 7 | TRUE | FALSE |
| 171_v.txt | 32541 | 32541-Deleti | Deletion | 3 | TCA | - | 46 | 31383 | 0.14657617 | 37.5217391 | rnr1D57N | FALSE | CAN | JSY14 | 1A | rnr1D57N-Re | TTC | nope | >1bp-ins | >1bpdel | none | >1bpdel | GCAGCTTCA( | TCACCAG | AAG | TTC | TCA | none | NA | NA_TTC | >1bpdel_TCACCAG | 7 | TRUE | FALSE |
| 171_v.txt | 32550 | 32550-Deleti | Deletion | 3 | GTG | - | 10 | 31379 | 0.03186845 | 38 | rnr1D57N | FALSE | CAN | JSY14 | 1A | rnr1D57N-Re | AGT | nope | >1bp-ins | >1bpdel | none | >1bpdel | CCAGCAGTG. | GTGATAC | TCA | AGT | GTG | none | NA | NA_AGT | >1bpdel_GTGATAC | 7 | TRUE | FALSE |
| 171_v.txt | 32657 | 32657-Deleti | Deletion | 2 | AC | - | 1 | 30555 | 3.27E-03 | 11 | rnr1D57N | FALSE | CAN | JSY14 | 1A | rnr1D57N-Re | GAC | nope | >1bp-ins | >1bpdel | none | >1bpdel | ACCTGGACC( | ACCCCAG | CTG | GAC | AC | none | NA | NA_GAC | >1bpdel_ACCCCAG | 7 | TRUE | FALSE |
| 171_v.txt | 32841 | 32841-Deleti | Deletion | 2 | CA | - | 1 | 30555 | 3.27E-03 | 38 | rnr1D57N | FALSE | CAN | JSY14 | 1A | rnr1D57N-Re | CCA | nope | >1bp-ins | >1bpdel | none | >1bpdel | ATTACCCAA/ | CAAAAAA | GGT | CCA | CA | none | NA | NA_CCA | >1bpdel_CAAAAAA | 7 | TRUE | FALSE |
| 171_v.txt | 32902 | 32902-Deleti | Deletion | 2 | CA | - | 66 | 31348 | 0.21053975 | 43.1515152 | rnr1D57N | FALSE | CAN | JSY14 | 1A | rnr1D57N-Re | CCA | nope | >1bp-ins | >1bpdel | none | >1bpdel | CTTGGCCAA( | CAACTAC | GGT | CCA | CA | none | NA | NA_CCA | >1bpdel_CAACTAC | 7 | TRUE | FALSE |
| 171_v.txt | 33065 | 33065-Deleti | Deletion | 2 | AT | - | 27 | 31348 | 0.0861299 | 56.7777778 | rnr1D57N | FALSE | CAN | JSY14 | 1A | rnr1D57N-Re | AAT | nope | >1bp-ins | >1bpdel | none | >1bpdel | GACAGAATA | ATATGCC | TTA | AAT | AT | none | NA | NA_AAT | >1bpdel_ATATGCC | 7 | TRUE | FALSE |
| 171_v.txt | 33089 | 33089-Deleti | Deletion | 2 | AT | - | 78 | 31348 | 0.2488197 | 54.8846154 | rnr1D57N | FALSE | CAN | JSY14 | 1A | rnr1D57N-Re | AAT | nope | >1bp-ins | >1bpdel | none | >1bpdel | AAATAAATA' | ATATGAT | TTA | AAT | AT | none | NA | NA_AAT | >1bpdel_ATATGAT | 7 | TRUE | FALSE |
| 171_v.txt | 33094 | 33094-Deleti | Deletion | 2 | AT | - | 38 | 31348 | 0.12121986 | 52.6842105 | rnr1D57N | FALSE | CAN | JSY14 | 1A | rnr1D57N-Re | GAT | nope | >1bp-ins | >1bpdel | none | >1bpdel | AATATGATA' | ATATAAG | CTA | GAT | AT | none | NA | NA_GAT | >1bpdel_ATATAAG | 7 | TRUE | FALSE |
| 171_v.txt | 33168 | 33168-Deleti | Deletion | 2 | CC | - | 1 | 31300 | 3.20E-03 | 38 | rnr1D57N | FALSE | CAN | JSY14 | 1A | rnr1D57N-Re | ACC | nope | >1bp-ins | >1bpdel | none | >1bpdel | ATAGTACCA( | CCACCAA | TGG | ACC | CC | none | NA | NA_ACC | >1bpdel_CCACCAA | 7 | TRUE | FALSE |
| 171_v.txt | 31723 | 31723-Deleti | Deletion | 1 | A | - | 12 | 10179 | 0.11788977 | 58.5 | rnr1D57N | FALSE | CAN | JSY14 | 1A | rnr1D57N-Re | CAA | nope | >1bp-ins | A/T-1 | none | A/T-1 | TGTCCCAAA/ | AAAAAGT | GTT | GTT | T | none | T>none | T>none_GTT | A/T-1_AAAAAGT | 7 | TRUE | FALSE |
| 171_v.txt | 31926 | 31926-Deleti | Deletion | 1 | C | - | 21 | 10171 | 0.20646937 | 37.1904762 | rnr1D57N | FALSE | CAN | JSY14 | 1A | rnr1D57N-Re | ACC | nope | >1bp-ins | G/C-1 | none | G/C-1 | GTGAAACCT' | CCTTGAA | TGG | ACC | C | none | nope | nope_ACC | G/C-1_CCTTGAA | 7 | TRUE | FALSE |
| 171_v.txt | 31976 | 31976-Deleti | Deletion | 1 | G | - | 9 | 24202 | 0.03718701 | 45.2222222 | rnr1D57N | FALSE | CAN | JSY14 | 1A | rnr1D57N-Re | AGC | nope | >1bp-ins | G/C-1 | none | G/C-1 | AGCCAAGCC | GCCGGGC | TCG | TCG | C | none | C>none | C>none_TCG | G/C-1_GCCGGGC | 7 | TRUE | FALSE |
| 171_v.txt | 32081 | 32081-Deleti | Deletion | 1 | A | - | 31 | 24202 | 0.12808859 | 60.0967743 | rnr1D57N | FALSE | CAN | JSY14 | 1A | rnr1D57N-Re | CAA | nope | >1bp-ins | A/T-1 | none | A/T-1 | CCATGCAAA/ | AAAAAAG | GTT | GTT | T | none | T>none | T>none_GTT | A/T-1_AAAAAAG | 7 | TRUE | FALSE |
| 171_v.txt | 32088 | 32088-Deleti | Deletion | 1 | C | - | 3 | 24202 | 0.01239567 | 55.3333333 | rnr1D57N | FALSE | CAN | JSY14 | 1A | rnr1D57N-Re | GCC | nope | >1bp-ins | G/C-1 | none | G/C-1 | AAAAAGCCT' | CCTGCAA | CGG | GCC | C | none | nope | nope_GCC | G/C-1_CCTGCAA | 7 | TRUE | FALSE |
| 171_v.txt | 32092 | 32092-Deleti | Deletion | 1 | C | - | 4 | 24202 | 0.01652756 | 57.5 | rnr1D57N | FALSE | CAN | JSY14 | 1A | rnr1D57N-Re | GCA | nope | >1bp-ins | G/C-1 | none | G/C-1 | AGCCTGCAA( | CAACACC | CGT | GCA | C | none | nope | nope_GCA | G/C-1_CAACACC | 7 | TRUE | FALSE |
| 171_v.txt | 32095 | 32095-Deleti | Deletion | 1 | C | - | 3 | 24202 | 0.01239567 | 55.3333333 | rnr1D57N | FALSE | CAN | JSY14 | 1A | rnr1D57N-Re | ACA | nope | >1bp-ins | G/C-1 | none | G/C-1 | CTGCAACAC( | CACCAGT | TGT | ACA | C | none | nope | nope_ACA | G/C-1_CACCAGT | 7 | TRUE | FALSE |
| 171_v.txt | 32101 | 32101-Deleti | Deletion | 1 | T | - | 3 | 24202 | 0.01239567 | 55.3333333 | rnr1D57N | FALSE | CAN | JSY14 | 1A | rnr1D57N-Re | GTG | nope | >1bp-ins | A/T-1 | none | A/T-1 | CACCAGTGA' | TGATATT | CAC | GTG | T | none | nope | nope_GTG | A/T-1_TGATATT | 7 | TRUE | FALSE |
| 171_v.txt | 32103 | 32103-Deleti | Deletion | 1 | A | - | 3 | 24202 | 0.01239567 | 55.3333333 | rnr1D57N | FALSE | CAN | JSY14 | 1A | rnr1D57N-Re | GAT | nope | >1bp-ins | A/T-1 | none | A/T-1 | CCAGTGATA' | ATATTTA | CTA | CTA | T | none | T>none | T>none_CTA | A/T-1_ATATTTA | 7 | TRUE | FALSE |
| 171_v.txt | 32114 | 32114-Deleti | Deletion | 1 | C | - | 2 | 24202 | 8.26E-03 | 62 | rnr1D57N | FALSE | CAN | JSY14 | 1A | rnr1D57N-Re | GCC | nope | >1bp-ins | G/C-1 | none | G/C-1 | TAATAGCCA' | CCATTCG | CGG | GCC | C | none | nope | nope_GCC | G/C-1_CCATTCG | 7 | TRUE | FALSE |
| 171_v.txt | 32148 | 32148-Deleti | Deletion | 1 | T | - | 20 | 24202 | 0.0826378 | 49.1499984 | rnr1D57N | FALSE | CAN | JSY14 | 1A | rnr1D57N-Re | CTC | nope | >1bp-ins | A/T-1 | none | A/T-1 | GATGTCTCC/ | TCCATGT | GAG | CTC | T | none | nope | nope_CTC | A/T-1_TCCATGT | 7 | TRUE | FALSE |
| 171_v.txt | 32165 | 32165-Deleti | Deletion | 1 | G | - | 70 | 24202 | 0.2892323 | 49.7142844 | rnr1D57N | FALSE | CAN | JSY14 | 1A | rnr1D57N-Re | CGC | nope | >1bp-ins | G/C-1 | none | G/C-1 | CAAAGCGCC. | GCCAAAT | GCG | GCG | C | none | C>none | C>none_GCG | G/C-1_GCCAAAT | 7 | TRUE | FALSE |
| 171_v.txt | 32237 | 32237-Deleti | Deletion | 1 | A | - | 382 | 24197 | 1.5787081 | 37.0837696 | rnr1D57N | FALSE | CAN | JSY14 | 1A | rnr1D57N-Re | GAG | nope | >1bp-ins | A/T-1 | none | A/T-1 | TTTAGGAGC | AGCCAAC | CTC | CTC | T | none | T>none | T>none_CTC | A/T-1_AGCCAAC | 7 | TRUE | FALSE |
| 171_v.txt | 32259 | 32259-Deleti | Deletion | 1 | C | - | 12 | 31378 | 0.03824336 | 38 | rnr1D57N | FALSE | CAN | JSY14 | 1A | rnr1D57N-Re | ACC | nope | >1bp-ins | G/C-1 | none | G/C-1 | GATAGACCA. | CCAAATA | TGG | ACC | C | none | nope | nope_ACC | G/C-1_CCAAATA | 7 | TRUE | FALSE |
| 171_v.txt | 32284 | 32284-Deleti | Deletion | 1 | A | - | 2 | 31383 | 6.37E-03 | 50.4999981 | rnr1D57N | FALSE | CAN | JSY14 | 1A | rnr1D57N-Re | TAA | nope | >1bp-ins | A/T-1 | none | A/T-1 | CAACGTAAA' | AAATATT | ATT | ATT | T | none | T>none | T>none_ATT | A/T-1_AAATATT | 7 | TRUE | FALSE |
| 171_v.txt | 32346 | 32346-Deleti | Deletion | 1 | A | - | 115 | 31383 | 0.36644043 | 60.0869565 | rnr1D57N | FALSE | CAN | JSY14 | 1A | rnr1D57N-Re | CAA | nope | >1bp-ins | A/T-1 | none | A/T-1 | TGTGGCAAA. | AAAACCT | GTT | GTT | T | none | T>none | T>none_GTT | A/T-1_AAAACCT | 7 | TRUE | FALSE |
| 171_v.txt | 32442 | 32442-Deleti | Deletion | 1 | A | - | 7 | 31383 | 0.02230507 | 52.8571414 | rnr1D57N | FALSE | CAN | JSY14 | 1A | rnr1D57N-Re | TAA | nope | >1bp-ins | A/T-1 | none | A/T-1 | GGAACTAAA. | AAAAGTC | ATT | ATT | T | none | T>none | T>none_ATT | A/T-1_AAAAGTC | 7 | TRUE | FALSE |
| 171_v.txt | 32498 | 32498-Deleti | Deletion | 1 | T | - | 136 | 31383 | 0.43335564 | 43.8602942 | rnr1D57N | FALSE | CAN | JSY14 | 1A | rnr1D57N-Re | CTT | nope | >1bp-ins | A/T-1 | none | A/T-1 | AACAACTTTT | TTTTTTG | GAA | CTT | T | none | nope | nope_CTT | A/T-1_TTTTTTG | 7 | TRUE | FALSE |
| 171_v.txt | 32581 | 32581-Deleti | Deletion | 1 | T | - | 10 | 30554 | 0.03272894 | 37.2999992 | rnr1D57N | FALSE | CAN | JSY14 | 1A | rnr1D57N-Re | GTG | nope | >1bp-ins | A/T-1 | none | A/T-1 | GAAATGTGA. | TGAAGGC | CAC | GTG | T | none | nope | nope_GTG | A/T-1_TGAAGGC | 7 | TRUE | FALSE |
| 171_v.txt | 32631 | 32631-Deleti | Deletion | 1 | T | - | 264 | 30555 | 0.86401571 | 48.3068175 | rnr1D57N | FALSE | CAN | JSY14 | 1A | rnr1D57N-Re | GTT | nope | >1bp-ins | A/T-1 | none | A/T-1 | CCTTCGTTTT | TTTTTAT | CAA | GTT | T | none | nope | nope_GTT | A/T-1_TTTTTAT | 7 | TRUE | FALSE |
| 171_v.txt | 32657 | 32657-Deleti | Deletion | 1 | A | - | 30 | 30555 | 0.0981836 | 57.3333394 | rnr1D57N | FALSE | CAN | JSY14 | 1A | rnr1D57N-Re | GAC | nope | >1bp-ins | A/T-1 | none | A/T-1 | ACCTGGACC( | ACCCCAG | CTG | CTG | T | none | T>none | T>none_CTG | A/T-1_ACCCCAG | 7 | TRUE | FALSE |
| 171_v.txt | 32658 | 32658-Deleti | Deletion | 1 | C | - | 138 | 30555 | 0.45164458 | 59.9130357 | rnr1D57N | FALSE | CAN | JSY14 | 1A | rnr1D57N-Re | ACC | nope | >1bp-ins | G/C-1 | none | G/C-1 | CCTGGACC( | CCCCAGG | TGG | ACC | C | none | nope | nope_ACC | G/C-1_CCCCAGG | 7 | TRUE | FALSE |
| 171_v.txt | 32682 | 32682-Deleti | Deletion | 1 | T | - | 22 | 30555 | 0.07200131 | 59.7272727 | rnr1D57N | FALSE | CAN | JSY14 | 1A | rnr1D57N-Re | ATA | nope | >1bp-ins | A/T-1 | none | A/T-1 | CTCCAATAA( | TAACGGA | TAT | ATA | T | none | nope | nope_ATA | A/T-1_TAACGGA | 7 | TRUE | FALSE |
| 171_v.txt | 32707 | 32707-Deleti | Deletion | 1 | C | - | 9 | 30555 | 0.02945508 | 64 | rnr1D57N | FALSE | CAN | JSY14 | 1A | rnr1D57N-Re | ACC | nope | >1bp-ins | G/C-1 | none | G/C-1 | CGGTAACCC( | CCCCAGC | TGG | ACC | C | none | nope | nope_ACC | G/C-1_CCCCAGC | 7 | TRUE | FALSE |
| 171_v.txt | 32728 | 32728-Deleti | Deletion | 1 | A | - | 64 | 30554 | 0.20946521 | 59.71875 | rnr1D57N | FALSE | CAN | JSY14 | 1A | rnr1D57N-Re | CAA | nope | >1bp-ins | A/T-1 | none | A/T-1 | CCATACAAA/ | AAAAACA | GTT | GTT | T | none | T>none | T>none_GTT | A/T-1_AAAAACA | 7 | TRUE | FALSE |
| 171_v.txt | 32758 | 32758-Deleti | Deletion | 1 | C | - | 8 | 30555 | 0.02618229 | 57.3750001 | rnr1D57N | FALSE | CAN | JSY14 | 1A | rnr1D57N-Re | GCT | nope | >1bp-ins | G/C-1 | none | G/C-1 | TAATGGCTA/ | CTAAAAC | CGA | GCT | C | none | nope | nope_GCT | G/C-1_CTAAAAC | 7 | TRUE | FALSE |
| 171_v.txt | 32759 | 32759-Deleti | Deletion | 1 | T | - | 21 | 30555 | 0.06872852 | 52.3333312 | rnr1D57N | FALSE | CAN | JSY14 | 1A | rnr1D57N-Re | CTA | nope | >1bp-ins | A/T-1 | none | A/T-1 | AATGGCTAA. | TAAAACT | GAT | CTA | T | none | nope | nope_CTA | A/T-1_TAAAACT | 7 | TRUE | FALSE |
| 171_v.txt | 32760 | 32760-Deleti | Deletion | 1 | A | - | 6 | 30555 | 0.01963672 | 54.9999971 | rnr1D57N | FALSE | CAN | JSY14 | 1A | rnr1D57N-Re | TAA | nope | >1bp-ins | A/T-1 | none | A/T-1 | ATGGCTAAA. | AAAACTT | ATT | ATT | T | none | T>none | T>none_ATT | A/T-1_AAAACTT | 7 | TRUE | FALSE |
| 171_v.txt | 32839 | 32839-Deleti | Deletion | 1 | C | - | 16 | 30555 | 0.05236459 | 48.624994 | rnr1D57N | FALSE | CAN | JSY14 | 1A | rnr1D57N-Re | ACC | nope | >1bp-ins | G/C-1 | none | G/C-1 | TAATTACCCA | CCCAAAA | TGG | ACC | C | none | nope | nope_ACC | G/C-1_CCCAAAA | 7 | TRUE | FALSE |
| 171_v.txt | 32842 | 32842-Deleti | Deletion | 1 | A | - | 1381 | 30555 | 4.51971854 | 41.9804517 | rnr1D57N | FALSE | CAN | JSY14 | 1A | rnr1D57N-Re | CAA | nope | >1bp-ins | A/T-1 | none | A/T-1 | TTACCCAAA/ | AAAAAAT | GTT | GTT | T | none | T>none | T>none_GTT | A/T-1_AAAAAAT | 7 | TRUE | FALSE |
| 171_v.txt | 32876 | 32876-Deleti | Deletion | 1 | G | - | 2 | 31336 | 6.38E-03 | 38 | rnr1D57N | FALSE | CAN | JSY14 | 1A | rnr1D57N-Re | TGT | nope | >1bp-ins | G/C-1 | none | G/C-1 | AACTTTGTAC | GTACGTC | ACA | ACA | C | none | C>none | C>none_ACA | G/C-1_GTACGTC | 7 | TRUE | FALSE |
| 171_v.txt | 32901 | 32901-Deleti | Deletion | 1 | C | - | 2 | 31348 | 6.38E-03 | 37.4999964 | rnr1D57N | FALSE | CAN | JSY14 | 1A | rnr1D57N-Re | GCC | nope | >1bp-ins | G/C-1 | none | G/C-1 | ACTTGGCCA( | CCAACTA | CGG | GCC | C | none | nope | nope_GCC | G/C-1_CCAACTA | 7 | TRUE | FALSE |
| 171_v.txt | 32915 | 32915-Deleti | Deletion | 1 | T | - | 2 | 31348 | 6.38E-03 | 50.4999954 | rnr1D57N | FALSE | CAN | JSY14 | 1A | rnr1D57N-Re | GTT | nope | >1bp-ins | A/T-1 | none | A/T-1 | ACTAAGTTC( | TTCCAGG | CAA | GTT | T | none | nope | nope_GTT | A/T-1_TTCCAGG | 7 | TRUE | FALSE |
| 171_v.txt | 32924 | 32924-Deleti | Deletion | 1 | A | - | 18 | 31348 | 0.05741993 | 53.7777717 | rnr1D57N | FALSE | CAN | JSY14 | 1A | rnr1D57N-Re | CAA | nope | >1bp-ins | A/T-1 | none | A/T-1 | CAGGGCAAA | AAAAGTG | GTT | GTT | T | none | T>none | T>none_GTT | A/T-1_AAAAGTG | 7 | TRUE | FALSE |
| 171_v.txt | 32935 | 32935-Deleti | Deletion | 1 | C | - | 33 | 31348 | 0.10526987 | 50.212121 | rnr1D57N | FALSE | CAN | JSY14 | 1A | rnr1D57N-Re | GCC | nope | >1bp-ins | G/C-1 | none | G/C-1 | TGATTGCCCA | CCCAAGA | CGG | GCC | C | none | nope | nope_GCC | G/C-1_CCCAAGA | 7 | TRUE | FALSE |
| 171_v.txt | 32970 | 32970-Deleti | Deletion | 1 | C | - | 2 | 31348 | 6.38E-03 | 64 | rnr1D57N | FALSE | CAN | JSY14 | 1A | rnr1D57N-Re | ACC | nope | >1bp-ins | G/C-1 | none | G/C-1 | GCCGCACCA. | CCAAATG | TGG | ACC | C | none | nope | nope_ACC | G/C-1_CCAAATG | 7 | TRUE | FALSE |
| 171_v.txt | 32972 | 32972-Deleti | Deletion | 1 | A | - | 18 | 31348 | 0.05741993 | 58.7222187 | rnr1D57N | FALSE | CAN | JSY14 | 1A | rnr1D57N-Re | CAA | nope | >1bp-ins | A/T-1 | none | A/T-1 | CGCACCAAA' | AAATGCT | GTT | GTT | T | none | T>none | T>none_GTT | A/T-1_AAATGCT | 7 | TRUE | FALSE |
| 171_v.txt | 33000 | 33000-Deleti | Deletion | 1 | A | - | 66 | 31348 | 0.21053975 | 60.7272681 | rnr1D57N | FALSE | CAN | JSY14 | 1A | rnr1D57N-Re | GAA | nope | >1bp-ins | A/T-1 | none | A/T-1 | TGTGAGAAA | AAAACTG | CTT | CTT | T | none | T>none | T>none_CTT | A/T-1_AAAACTG | 7 | TRUE | FALSE |
| 171_v.txt | 33037 | 33037-Deleti | Deletion | 1 | C | - | 5 | 31348 | 0.01594998 | 58.6 | rnr1D57N | FALSE | CAN | JSY14 | 1A | rnr1D57N-Re | GCC | nope | >1bp-ins | G/C-1 | none | G/C-1 | ATGTAGCCA' | CCATTTC | CGG | GCC | C | none | nope | nope_GCC | G/C-1_CCATTTC | 7 | TRUE | FALSE |
| 171_v.txt | 33138 | 33138-Deleti | Deletion | 1 | A | - | 12 | 31348 | 0.03827995 | 37.9166667 | rnr1D57N | FALSE | CAN | JSY14 | 1A | rnr1D57N-Re | TAA | nope | >1bp-ins | A/T-1 | none | A/T-1 | GTGGATAAA | AAACCAA | ATT | ATT | T | none | T>none | T>none_ATT | A/T-1_AAACCAA | 7 | TRUE | FALSE |
| 171_v.txt | 33145 | 33145-Deleti | Deletion | 1 | T | - | 15 | 31348 | 0.04784994 | 40.6666667 | rnr1D57N | FALSE | CAN | JSY14 | 1A | rnr1D57N-Re | ATG | nope | >1bp-ins | A/T-1 | none | A/T-1 | AACCAATGA. | TGAAAAG | TAC | ATG | T | none | nope | nope_ATG | A/T-1_TGAAAAG | 7 | TRUE | FALSE |
| 171_v.txt | 33147 | 33147-Deleti | Deletion | 1 | A | - | 7 | 31348 | 0.02232997 | 43.8571429 | rnr1D57N | FALSE | CAN | JSY14 | 1A | rnr1D57N-Re | GAA | nope | >1bp-ins | A/T-1 | none | A/T-1 | CCAATGAAA. | AAAAGAC | CTT | CTT | T | none | T>none | T>none_CTT | A/T-1_AAAAGAC | 7 | TRUE | FALSE |
| 171_v.txt | 33159 | 33159-Deleti | Deletion | 1 | C | - | 2 | 31342 | 6.38E-03 | 36.9999972 | rnr1D57N | FALSE | CAN | JSY14 | 1A | rnr1D57N-Re | ACC | nope | >1bp-ins | G/C-1 | none | G/C-1 | CCTGTACCA/ | CCAATAG | TGG | ACC | C | none | nope | nope_ACC | G/C-1_CCAATAG | 7 | TRUE | FALSE |
| 171_v.txt | 33175 | 33175-Deleti | Deletion | 1 | G | - | 6 | 31297 | 0.01917117 | 37.3333333 | rnr1D57N | FALSE | CAN | JSY14 | 1A | rnr1D57N-Re | AGG | nope | >1bp-ins | G/C-1 | none | G/C-1 | CACCAAGGG | GGGCAAT | TCC | TCC | C | none | C>none | C>none_TCC | G/C-1_GGGCAAT | 7 | TRUE | FALSE |
| 171_v.txt | 33178 | 33178-Deleti | Deletion | 1 | C | - | 32 | 31295 | 0.10225276 | 37.2500003 | rnr1D57N | FALSE | CAN | JSY14 | 1A | rnr1D57N-Re | GCA | nope | >1bp-ins | G/C-1 | none | G/C-1 | CAAGGGCAA | CAATCAT | CGT | GCA | C | none | nope | nope_GCA | G/C-1_CAATCAT | 7 | TRUE | FALSE |
| 171_v.txt | 33273 | 33273-Deleti | Deletion | 1 | A | - | 17 | 13904 | 0.12226697 | 49.2941177 | rnr1D57N | FALSE | CAN | JSY14 | 1A | rnr1D57N-Re | GAA | nope | >1bp-ins | A/T-1 | none | A/T-1 | ATAGAGAAC | AACGTAT | CTT | CTT | T | none | T>none | T>none_CTT | A/T-1_AACGTAT | 7 | TRUE | FALSE |
| 171_v.txt | 33300 | 33300-Deleti | Deletion | 1 | G | - | 8 | 13904 | 0.0575374 | 56.5 | rnr1D57N | FALSE | CAN | JSY14 | 1A | rnr1D57N-Re | CGG | nope | >1bp-ins | G/C-1 | none | G/C-1 | CTCGTCGGG/ | GGGAAAG | GCC | GCC | C | none | C>none | C>none_GCC | G/C-1_GGGAAAG | 7 | TRUE | FALSE |
| 171_v.txt | 33456 | 33456-Deleti | Deletion | 1 | G | - | 13 | 13904 | 0.09349827 | 38.3076923 | rnr1D57N | FALSE | CAN | JSY14 | 1A | rnr1D57N-Re | TGA | nope | >1bp-ins | G/C-1 | none | G/C-1 | TCTTTTGAAT | GAATTTG | ACT | ACT | C | none | C>none | C>none_ACT | G/C-1_GAATTTG | 7 | TRUE | FALSE |
| 171_v.txt | 33474 | 33474-Deleti | Deletion | 1 | C | - | 32 | 13903 | 0.23016615 | 30.1875 | rnr1D57N | FALSE | CAN | JSY14 | 1A | rnr1D57N-Re | GCC | nope | >1bp-ins | G/C-1 | none | G/C-1 | GCTATGCCTT | CCTTTTT | CGG | GCC | C | none | nope | nope_GCC | G/C-1_CCTTTTT | 7 | TRUE | FALSE |
| 172_v.txt | 31723 | 31723-SNV-/ | SNV | 1 | A | C | 5 | 12214 | 0.04093663 | 64 | rnr1D57N | FALSE | CAN | JSY14 | 2A | rnr1D57N-Re | CAA | A>C | G/C+1 | A/T-1 | TA>GC | TA>GC | TGTCCCAAA/ | AAAAAGT | GTT | GTT | T | G | T>G | T>G_GTT | TA>GC_AAAAAGT | 7 | TRUE | FALSE |
| 172_v.txt | 31950 | 31950-SNV-/ | SNV | 1 | A | C | 15 | 18829 | 0.07966435 | 35.7333333 | rnr1D57N | FALSE | CAN | JSY14 | 2A | rnr1D57N-Re | CAT | A>C | G/C+1 | A/T-1 | TA>GC | TA>GC | ATCGTCATA/ | ATAAATG | GTA | GTA | T | G | T>G | T>G_GTA | TA>GC_ATAAATG | 7 | TRUE | FALSE |
| 172_v.txt | 32283 | 32283-SNV-T | SNV | 1 | T | G | 1001 | 26233 | 3.81580452 | 53.108878 | rnr1D57N | FALSE | CAN | JSY14 | 2A | rnr1D57N-Re | GTA | T>G | G/C+1 | A/T-1 | TA>GC | TA>GC | CCAACGTAA. | TAAATAT | CAT | GTA | T | C | T>G | T>G_GTA | TA>GC_TAAATAT | 7 | TRUE | FALSE |
| 172_v.txt | 32573 | 32573-SNV-T | SNV | 1 | T | G | 11 | 41543 | 0.02647859 | 30.0909091 | rnr1D57N | FALSE | CAN | JSY14 | 2A | rnr1D57N-Re | CTT | T>G | G/C+1 | A/T-1 | TA>GC | TA>GC | AGTACCTTG/ | TTGAAAT | GAA | CTT | T | C | T>G | T>G_CTT | TA>GC_TTGAAAT | 7 | TRUE | FALSE |
| 172_v.txt | 32574 | 32574-SNV-T | SNV | 1 | T | G | 44 | 41543 | 0.10591435 | 35.8857138 | rnr1D57N | FALSE | CAN | JSY14 | 2A | rnr1D57N-Re | TTG | T>G | G/C+1 | A/T-1 | TA>GC | TA>GC | GTACCTTGA/ | TGAAATG | AAC | TTG | T | C | T>G | T>G_TTG | TA>GC_TGAAATG | 7 | TRUE | FALSE |
| 172_v.txt | 32739 | 32739-SNV-/ | SNV | 1 | A | C | 3237 | 41541 | 7.79230158 | 57.527958 | rnr1D57N | FALSE | CAN | JSY14 | 2A | rnr1D57N-Re | TAT | A>C | G/C+1 | A/T-1 | TA>GC | TA>GC | CAGTATATT/ | ATTAGAA | ATA | ATA | T | G | T>G | T>G_ATA | TA>GC_ATTAGAA | 7 | TRUE | FALSE |
| 172_v.txt | 32799 | 32799-SNV-T | SNV | 1 | T | G | 19 | 41541 | 0.04573795 | 37.0003651 | rnr1D57N | FALSE | CAN | JSY14 | 2A | rnr1D57N-Re | GTA | T>G | G/C+1 | A/T-1 | TA>GC | TA>GC | TCACCGTAAT | TAATATT | CAT | GTA | T | C | T>G | T>G_GTA | TA>GC_TAATATT | 7 | TRUE | FALSE |
| 172_v.txt | 32817 | 32817-SNV-/ | SNV | 1 | A | C | 46 | 41541 | 0.11073397 | 44.4565217 | rnr1D57N | FALSE | CAN | JSY14 | 2A | rnr1D57N-Re | CAA | A>C | G/C+1 | A/T-1 | TA>GC | TA>GC | GGGAACAAC | AAGTTCA | GTT | GTT | T | G | T>G | T>G_GTT | TA>GC_AAGTTCA | 7 | TRUE | FALSE |
| 172_v.txt | 32842 | 32842-SNV-/ | SNV | 1 | A | C | 8 | 41541 | 0.01925808 | 35.3750888 | rnr1D57N | FALSE | CAN | JSY14 | 2A | rnr1D57N-Re | CAA | A>C | G/C+1 | A/T-1 | TA>GC | TA>GC | TTACCCAAA/ | AAAAAAT | GTT | GTT | T | G | T>G | T>G_GTT | TA>GC_AAAAAAT | 7 | TRUE | FALSE |
| 172_v.txt | 32884 | 32884-SNV-/ | SNV | 1 | A | C | 7 | 26528 | 0.02638721 | 31.8571429 | rnr1D57N | FALSE | CAN | JSY14 | 2A | rnr1D57N-Re | CAA | A>C | G/C+1 | A/T-1 | TA>GC | TA>GC | ACGTCCAAA/ | AAAATTG | GTT | GTT | T | G | T>G | T>G_GTT | TA>GC_AAAATTG | 7 | TRUE | FALSE |
| 172_v.txt | 32919 | 32919-SNV-/ | SNV | 1 | A | C | 10 | 26528 | 0.03769602 | 27 | rnr1D57N | FALSE | CAN | JSY14 | 2A | rnr1D57N-Re | CAG | A>C | G/C+1 | A/T-1 | TA>GC | TA>GC | AGTTCCAGG( | AGGGCAA | GTC | GTC | T | G | T>G | T>G_GTC | TA>GC_AGGGCAA | 7 | TRUE | FALSE |
| 172_v.txt | 33157 | 33157-SNV-T | SNV | 1 | T | G | 10 | 26527 | 0.03769744 | 37.7 | rnr1D57N | FALSE | CAN | JSY14 | 2A | rnr1D57N-Re | GTA | T>G | G/C+1 | A/T-1 | TA>GC | TA>GC | GACCTGTAC( | TACCAAT | CAT | GTA | T | C | T>G | T>G_GTA | TA>GC_TACCAAT | 7 | TRUE | FALSE |
| 172_v.txt | 32081 | 32081-SNV-/ | SNV | 1 | A | G | 5 | 18830 | 0.02655337 | 63.9997247 | rnr1D57N | FALSE | CAN | JSY14 | 2A | rnr1D57N-Re | CAA | A>G | G/C+1 | A/T-1 | TA>CG | TA>CG | CCATGCAAA/ | AAAAAAG | GTT | GTT | T | C | T>C | T>C_GTT | TA>CG_AAAAAAG | 7 | TRUE | FALSE |
| 172_v.txt | 32283 | 32283-SNV-T | SNV | 1 | T | C | 8 | 26233 | 0.03049594 | 19.6246045 | rnr1D57N | FALSE | CAN | JSY14 | 2A | rnr1D57N-Re | GTA | T>C | G/C+1 | A/T-1 | TA>CG | TA>CG | CCAACGTAA. | TAAATAT | CAT | GTA | T | G | T>C | T>C_GTA | TA>CG_TAAATAT | 7 | TRUE | FALSE |
| 172_v.txt | 32284 | 32284-SNV-/ | SNV | 1 | A | G | 4 | 26233 | 0.01524797 | 26.998717 | rnr1D57N | FALSE | CAN | JSY14 | 2A | rnr1D57N-Re | TAA | A>G | G/C+1 | A/T-1 | TA>CG | TA>CG | CAACGTAAA' | AAATATT | ATT | ATT | T | C | T>C | T>C_ATT | TA>CG_AAATATT | 7 | TRUE | FALSE |
| 172_v.txt | 32573 | 32573-SNV-T | SNV | 1 | T | C | 13 | 41543 | 0.03129288 | 34.6923077 | rnr1D57N | FALSE | CAN | JSY14 | 2A | rnr1D57N-Re | CTT | T>C | G/C+1 | A/T-1 | TA>CG | TA>CG | AGTACCTTG/ | TTGAAAT | GAA | CTT | T | G | T>C | T>C_CTT | TA>CG_TTGAAAT | 7 | TRUE | FALSE |
| 172_v.txt | 32574 | 32574-SNV-T | SNV | 1 | T | C | 18 | 41543 | 0.0433286 | 32.4992973 | rnr1D57N | FALSE | CAN | JSY14 | 2A | rnr1D57N-Re | TTG | T>C | G/C+1 | A/T-1 | TA>CG | TA>CG | GTACCTTGA/ | TGAAATG | AAC | TTG | T | G | T>C | T>C_TTG | TA>CG_TGAAATG | 7 | TRUE | FALSE |
| 172_v.txt | 32583 | 32583-SNV-/ | SNV | 1 | A | G | 170 | 41541 | 0.40923425 | 36.8647059 | rnr1D57N | FALSE | CAN | JSY14 | 2A | rnr1D57N-Re | GAA | A>G | G/C+1 | A/T-1 | TA>CG | TA>CG | AATGTGAAG | AAGGCAG | CTT | CTT | T | C | T>C | T>C_CTT | TA>CG_AAGGCAG | 7 | TRUE | FALSE |
| 172_v.txt | 32919 | 32919-SNV-/ | SNV | 1 | A | G | 20 | 26528 | 0.07539204 | 46.15 | rnr1D57N | FALSE | CAN | JSY14 | 2A | rnr1D57N-Re | CAG | A>G | G/C+1 | A/T-1 | TA>CG | TA>CG | AGTTCCAGG( | AGGGCAA | GTC | GTC | T | C | T>C | T>C_GTC | TA>CG_AGGGCAA | 7 | TRUE | FALSE |
| 172_v.txt | 32927 | 32927-SNV-/ | SNV | 1 | A | G | 24 | 26528 | 0.09047045 | 51.1666667 | rnr1D57N | FALSE | CAN | JSY14 | 2A | rnr1D57N-Re | AAG | A>G | G/C+1 | A/T-1 | TA>CG | TA>CG | GGCAAAAGT | AGTGATT | TTC | TTC | T | C | T>C | T>C_TTC | TA>CG_AGTGATT | 7 | TRUE | FALSE |
| 172_v.txt | 33476 | 33476-SNV-T | SNV | 1 | T | C | 21 | 11106 | 0.18908698 | 31.2857143 | rnr1D57N | FALSE | CAN | JSY14 | 2A | rnr1D57N-Re | CTT | T>C | G/C+1 | A/T-1 | TA>CG | TA>CG | TATGCCTTTT | TTTTTTT | GAA | CTT | T | G | T>C | T>C_CTT | TA>CG_TTTTTTT | 7 | TRUE | FALSE |
| 172_v.txt | 32573 | 32573-SNV-T | SNV | 1 | T | A | 14 | 41543 | 0.03370002 | 33.8571429 | rnr1D57N | FALSE | CAN | JSY14 | 2A | rnr1D57N-Re | CTT | T>A | A/T+1 | A/T-1 | TA>AT | TA>AT | AGTACCTTG/ | TTGAAAT | GAA | CTT | T | T | T>A | T>A_CTT | TA>AT_TTGAAAT | 7 | TRUE | FALSE |
| 172_v.txt | 32916 | 32916-SNV-T | SNV | 1 | T | A | 38 | 26528 | 0.14324487 | 51 | rnr1D57N | FALSE | CAN | JSY14 | 2A | rnr1D57N-Re | TTC | T>A | A/T+1 | A/T-1 | TA>AT | TA>AT | CTAAGTTCC/ | TCCAGGG | AAG | TTC | T | T | T>A | T>A_TTC | TA>AT_TCCAGGG | 7 | TRUE | FALSE |
| 172_v.txt | 31650 | 31650-SNV-( | SNV | 1 | C | T | 24 | 12214 | 0.19649583 | 35.3333333 | rnr1D57N | FALSE | CAN | JSY14 | 2A | rnr1D57N-Re | GCG | C>T | A/T+1 | G/C-1 | CG>TA | CG>TA | AGAATGCGA | CGAAATG | CGC | GCG | C | A | C>T | C>T_GCG | CG>TA_CGAAATG | 7 | TRUE | FALSE |
| 172_v.txt | 31658 | 31658-SNV-( | SNV | 1 | C | T | 21 | 12214 | 0.17193385 | 37.5228671 | rnr1D57N | FALSE | CAN | JSY14 | 2A | rnr1D57N-Re | GCG | C>T | A/T+1 | G/C-1 | CG>TA | CG>TA | AAATGGCGT | CGTGGAA | CGC | GCG | C | A | C>T | C>T_GCG | CG>TA_CGTGGAA | 7 | TRUE | FALSE |
| 172_v.txt | 31834 | 31834-SNV-( | SNV | 1 | G | A | 22 | 12214 | 0.18012117 | 50.5 | rnr1D57N | FALSE | CAN | JSY14 | 2A | rnr1D57N-Re | TGA | G>A | A/T+1 | G/C-1 | CG>TA | CG>TA | TGCATTGAA/ | GAAATAA | ACT | ACT | C | T | C>T | C>T_ACT | CG>TA_GAAATAA | 7 | TRUE | FALSE |
| 172_v.txt | 31882 | 31882-SNV-( | SNV | 1 | C | T | 17 | 12214 | 0.13918454 | 43.7647059 | rnr1D57N | FALSE | CAN | JSY14 | 2A | rnr1D57N-Re | GCA | C>T | A/T+1 | G/C-1 | CG>TA | CG>TA | AGGCGGCAC | CAGCAAA | CGT | GCA | C | A | C>T | C>T_GCA | CG>TA_CAGCAAA | 7 | TRUE | FALSE |
| 172_v.txt | 31962 | 31962-SNV-( | SNV | 1 | G | A | 29 | 18831 | 0.15400138 | 34.6896552 | rnr1D57N | FALSE | CAN | JSY14 | 2A | rnr1D57N-Re | CGC | G>A | A/T+1 | G/C-1 | CG>TA | CG>TA | GTGGCCGCA | GCATAAT | GCG | GCG | C | T | C>T | C>T_GCG | CG>TA_GCATAAT | 7 | TRUE | FALSE |
| 172_v.txt | 31971 | 31971-SNV-( | SNV | 1 | G | A | 3 | 18831 | 0.01593118 | 34.3048544 | rnr1D57N | FALSE | CAN | JSY14 | 2A | rnr1D57N-Re | AGC | G>A | A/T+1 | G/C-1 | CG>TA | CG>TA | TAATAAGCC/ | GCCAAGC | TCG | TCG | C | T | C>T | C>T_TCG | CG>TA_GCCAAGC | 7 | TRUE | FALSE |
| 172_v.txt | 32057 | 32057-SNV-( | SNV | 1 | C | T | 19 | 18831 | 0.10089746 | 60.1052632 | rnr1D57N | FALSE | CAN | JSY14 | 2A | rnr1D57N-Re | GCG | C>T | A/T+1 | G/C-1 | CG>TA | CG>TA | GATGTGCGA | CGAGATT | CGC | GCG | C | A | C>T | C>T_GCG | CG>TA_CGAGATT | 7 | TRUE | FALSE |
| 172_v.txt | 32075 | 32075-SNV-( | SNV | 1 | C | T | 23 | 18827 | 0.12216498 | 58.1721591 | rnr1D57N | FALSE | CAN | JSY14 | 2A | rnr1D57N-Re | ACC | C>T | A/T+1 | G/C-1 | CG>TA | CG>TA | AAATAACCA' | CCATGCA | TGG | ACC | C | A | C>T | C>T_ACC | CG>TA_CCATGCA | 7 | TRUE | FALSE |
| 172_v.txt | 32076 | 32076-SNV-( | SNV | 1 | C | T | 487 | 18828 | 2.58657319 | 62.2759905 | rnr1D57N | FALSE | CAN | JSY14 | 2A | rnr1D57N-Re | CCA | C>T | A/T+1 | G/C-1 | CG>TA | CG>TA | AATAACCAT( | CATGCAA | GGT | CCA | C | A | C>T | C>T_CCA | CG>TA_CATGCAA | 7 | TRUE | FALSE |
| 172_v.txt | 32088 | 32088-SNV-( | SNV | 1 | C | T | 58 | 18831 | 0.30800276 | 58.4482759 | rnr1D57N | FALSE | CAN | JSY14 | 2A | rnr1D57N-Re | GCC | C>T | A/T+1 | G/C-1 | CG>TA | CG>TA | AAAAAGCCT( | CCTGCAA | CGG | GCC | C | A | C>T | C>T_GCC | CG>TA_CCTGCAA | 7 | TRUE | FALSE |
| 172_v.txt | 32092 | 32092-SNV-( | SNV | 1 | C | T | 16 | 18831 | 0.08496628 | 63.75 | rnr1D57N | FALSE | CAN | JSY14 | 2A | rnr1D57N-Re | GCA | C>T | A/T+1 | G/C-1 | CG>TA | CG>TA | AGCCTGCAA( | CAACACC | CGT | GCA | C | A | C>T | C>T_GCA | CG>TA_CAACACC | 7 | TRUE | FALSE |
| 172_v.txt | 32165 | 32165-SNV-( | SNV | 1 | G | A | 30 | 18831 | 0.15931177 | 50.1333333 | rnr1D57N | FALSE | CAN | JSY14 | 2A | rnr1D57N-Re | CGC | G>A | A/T+1 | G/C-1 | CG>TA | CG>TA | CAAAGCGCC. | GCCAAAT | GCG | GCG | C | T | C>T | C>T_GCG | CG>TA_GCCAAAT | 7 | TRUE | FALSE |
| 172_v.txt | 32173 | 32173-SNV-( | SNV | 1 | C | T | 20 | 18831 | 0.10620785 | 45.35 | rnr1D57N | FALSE | CAN | JSY14 | 2A | rnr1D57N-Re | GCA | C>T | A/T+1 | G/C-1 | CG>TA | CG>TA | CAAATGCAG | CAGCAGT | CGT | GCA | C | A | C>T | C>T_GCA | CG>TA_CAGCAGT | 7 | TRUE | FALSE |
| 172_v.txt | 32271 | 32271-SNV-( | SNV | 1 | C | T | 39 | 26233 | 0.14866771 | 32.3333333 | rnr1D57N | FALSE | CAN | JSY14 | 2A | rnr1D57N-Re | ACG | C>T | A/T+1 | G/C-1 | CG>TA | CG>TA | AAAATACGG | CGGGAAC | TGC | ACG | C | A | C>T | C>T_ACG | CG>TA_CGGGAAC | 7 | TRUE | FALSE |
| 172_v.txt | 32299 | 32299-SNV-( | SNV | 1 | C | T | 41 | 26233 | 0.15629169 | 52.0731707 | rnr1D57N | FALSE | CAN | JSY14 | 2A | rnr1D57N-Re | GCG | C>T | A/T+1 | G/C-1 | CG>TA | CG>TA | AATTTGCGG( | CGGCAGA | CGC | GCG | C | A | C>T | C>T_GCG | CG>TA_CGGCAGA | 7 | TRUE | FALSE |
| 172_v.txt | 32344 | 32344-SNV-( | SNV | 1 | G | A | 22 | 26233 | 0.08386384 | 61.1818182 | rnr1D57N | FALSE | CAN | JSY14 | 2A | rnr1D57N-Re | GGC | G>A | A/T+1 | G/C-1 | CG>TA | CG>TA | TATGTGGCA/ | GCAAAAC | CCG | CCG | C | T | C>T | C>T_CCG | CG>TA_GCAAAAC | 7 | TRUE | FALSE |
| 172_v.txt | 32449 | 32449-SNV-( | SNV | 1 | C | T | 47 | 26233 | 0.17916365 | 54.3191489 | rnr1D57N | FALSE | CAN | JSY14 | 2A | rnr1D57N-Re | CCA | C>T | A/T+1 | G/C-1 | CG>TA | CG>TA | AAAGTCCAA' | CAATGAA | GGT | CCA | C | A | C>T | C>T_CCA | CG>TA_CAATGAA | 7 | TRUE | FALSE |
| 172_v.txt | 32466 | 32466-SNV-( | SNV | 1 | C | T | 40 | 26233 | 0.1524797 | 51.675 | rnr1D57N | FALSE | CAN | JSY14 | 2A | rnr1D57N-Re | GCC | C>T | A/T+1 | G/C-1 | CG>TA | CG>TA | AGAGAGCCA | CCAATGT | CGG | GCC | C | A | C>T | C>T_GCC | CG>TA_CCAATGT | 7 | TRUE | FALSE |
| 172_v.txt | 32488 | 32488-SNV-( | SNV | 1 | G | A | 131 | 26233 | 0.49937102 | 46.0992366 | rnr1D57N | FALSE | CAN | JSY14 | 2A | rnr1D57N-Re | CGG | G>A | A/T+1 | G/C-1 | CG>TA | CG>TA | AGATACGGA | GGAAAAC | GCC | GCC | C | T | C>T | C>T_GCC | CG>TA_GGAAAAC | 7 | TRUE | FALSE |
| 172_v.txt | 32535 | 32535-SNV-( | SNV | 1 | G | A | 39 | 26233 | 0.14866771 | 36.9744084 | rnr1D57N | FALSE | CAN | JSY14 | 2A | rnr1D57N-Re | TGC | G>A | A/T+1 | G/C-1 | CG>TA | CG>TA | GGGTTTGCA( | GCAGCTT | ACG | ACG | C | T | C>T | C>T_ACG | CG>TA_GCAGCTT | 7 | TRUE | FALSE |
| 172_v.txt | 32536 | 32536-SNV-( | SNV | 1 | C | T | 35 | 26233 | 0.13341974 | 36.1420256 | rnr1D57N | FALSE | CAN | JSY14 | 2A | rnr1D57N-Re | GCA | C>T | A/T+1 | G/C-1 | CG>TA | CG>TA | GGTTTGCAG( | CAGCTTC | CGT | GCA | C | A | C>T | C>T_GCA | CG>TA_CAGCTTC | 7 | TRUE | FALSE |
| 172_v.txt | 32545 | 32545-SNV-( | SNV | 1 | C | T | 25 | 26233 | 0.09529981 | 36.9167243 | rnr1D57N | FALSE | CAN | JSY14 | 2A | rnr1D57N-Re | CCA | C>T | A/T+1 | G/C-1 | CG>TA | CG>TA | CTTCACCAGC | CAGCAGT | GGT | CCA | C | A | C>T | C>T_CCA | CG>TA_CAGCAGT | 7 | TRUE | FALSE |
| 172_v.txt | 32575 | 32575-SNV-( | SNV | 1 | G | A | 112 | 41543 | 0.26960017 | 36.6607271 | rnr1D57N | FALSE | CAN | JSY14 | 2A | rnr1D57N-Re | TGA | G>A | A/T+1 | G/C-1 | CG>TA | CG>TA | TACCTTGAA/ | GAAATGT | ACT | ACT | C | T | C>T | C>T_ACT | CG>TA_GAAATGT | 7 | TRUE | FALSE |
| 172_v.txt | 32590 | 32590-SNV-( | SNV | 1 | C | T | 68 | 41541 | 0.1636937 | 35.999754 | rnr1D57N | FALSE | CAN | JSY14 | 2A | rnr1D57N-Re | GCG | C>T | A/T+1 | G/C-1 | CG>TA | CG>TA | AGGCAGCGT | CGTTAAT | CGC | GCG | C | A | C>T | C>T_GCG | CG>TA_CGTTAAT | 7 | TRUE | FALSE |
| 172_v.txt | 32700 | 32700-SNV-( | SNV | 1 | C | T | 43 | 41541 | 0.10351219 | 58.9767442 | rnr1D57N | FALSE | CAN | JSY14 | 2A | rnr1D57N-Re | GCC | C>T | A/T+1 | G/C-1 | CG>TA | CG>TA | ACTGGGCCG | CCGGTAA | CGG | GCC | C | A | C>T | C>T_GCC | CG>TA_CCGGTAA | 7 | TRUE | FALSE |
| 172_v.txt | 32702 | 32702-SNV-( | SNV | 1 | G | A | 35 | 41541 | 0.08425411 | 54.1428571 | rnr1D57N | FALSE | CAN | JSY14 | 2A | rnr1D57N-Re | CGG | G>A | A/T+1 | G/C-1 | CG>TA | CG>TA | TGGGCCGGT. | GGTAACC | GCC | GCC | C | T | C>T | C>T_GCC | CG>TA_GGTAACC | 7 | TRUE | FALSE |
| 172_v.txt | 32757 | 32757-SNV-( | SNV | 1 | G | A | 30 | 41541 | 0.07221781 | 52.7 | rnr1D57N | FALSE | CAN | JSY14 | 2A | rnr1D57N-Re | GGC | G>A | A/T+1 | G/C-1 | CG>TA | CG>TA | ATAATGGCT/ | GCTAAAA | CCG | CCG | C | T | C>T | C>T_CCG | CG>TA_GCTAAAA | 7 | TRUE | FALSE |
| 172_v.txt | 32776 | 32776-SNV-( | SNV | 1 | C | T | 49 | 41541 | 0.11795576 | 45.1020408 | rnr1D57N | FALSE | CAN | JSY14 | 2A | rnr1D57N-Re | GCG | C>T | A/T+1 | G/C-1 | CG>TA | CG>TA | TGGAAGCGA | CGACCCA | CGC | GCG | C | A | C>T | C>T_GCG | CG>TA_CGACCCA | 7 | TRUE | FALSE |
| 172_v.txt | 32798 | 32798-SNV-( | SNV | 1 | G | A | 68 | 41541 | 0.1636937 | 45.9411765 | rnr1D57N | FALSE | CAN | JSY14 | 2A | rnr1D57N-Re | CGT | G>A | A/T+1 | G/C-1 | CG>TA | CG>TA | TTCACCGTA/ | GTAATAT | GCA | GCA | C | T | C>T | C>T_GCA | CG>TA_GTAATAT | 7 | TRUE | FALSE |
| 172_v.txt | 32811 | 32811-SNV-( | SNV | 1 | G | A | 27 | 41541 | 0.06499603 | 44.407311 | rnr1D57N | FALSE | CAN | JSY14 | 2A | rnr1D57N-Re | AGG | G>A | A/T+1 | G/C-1 | CG>TA | CG>TA | TTGACAGGG | GGGAACA | TCC | TCC | C | T | C>T | C>T_TCC | CG>TA_GGGAACA | 7 | TRUE | FALSE |
| 172_v.txt | 32860 | 32860-SNV-( | SNV | 1 | C | T | 65 | 41535 | 0.15649452 | 36.3076923 | rnr1D57N | FALSE | CAN | JSY14 | 2A | rnr1D57N-Re | GCC | C>T | A/T+1 | G/C-1 | CG>TA | CG>TA | TCCATGCCGC | CCGCCAG | CGG | GCC | C | A | C>T | C>T_GCC | CG>TA_CCGCCAG | 7 | TRUE | FALSE |
| 172_v.txt | 32900 | 32900-SNV-( | SNV | 1 | G | A | 36 | 26528 | 0.13570567 | 34.5812302 | rnr1D57N | FALSE | CAN | JSY14 | 2A | rnr1D57N-Re | GGC | G>A | A/T+1 | G/C-1 | CG>TA | CG>TA | GACTTGGCC | GCCAACT | CCG | CCG | C | T | C>T | C>T_CCG | CG>TA_GCCAACT | 7 | TRUE | FALSE |

| | | | | | | | | | | | | | | | | | | | | | | | | | | | | | | | | | | |
|---|---|---|---|---|---|---|---|---|---|---|---|---|---|---|---|---|---|---|---|---|---|---|---|---|---|---|---|---|---|---|---|---|---|---|
| 172_v.txt | 32901 | 32901-SNV- | SNV | 1 | C | T | 47 | 26528 | 0.17717129 | 31.914875 | rnr1D57N | FALSE | CAN | JSY14 | 2A | rnr1D57N-Re | GCC | C>T | A/T+1 | G/C-1 | CG>TA | CG>TA | ACTTGGCCA | CCAACTA | CGG | GCC | C | A | C>T | C>T_GCC | CG>TA_CCAACTA | 7 | TRUE | FALSE |
| 172_v.txt | 32935 | 32935-SNV- | SNV | 1 | C | T | 41 | 26528 | 0.15455368 | 42.2195122 | rnr1D57N | FALSE | CAN | JSY14 | 2A | rnr1D57N-Re | GCC | C>T | A/T+1 | G/C-1 | CG>TA | CG>TA | TGATTGCCC | CCCAAGA | CGG | GCC | C | A | C>T | C>T_GCC | CG>TA_CCCAAGA | 7 | TRUE | FALSE |
| 172_v.txt | 32967 | 32967-SNV- | SNV | 1 | G | A | 55 | 26528 | 0.20732811 | 61.2181818 | rnr1D57N | FALSE | CAN | JSY14 | 2A | rnr1D57N-Re | CGC | G>A | A/T+1 | G/C-1 | CG>TA | CG>TA | TTGGCCGCA | GCACCAA | GCG | GCG | C | T | C>T | C>T_GCG | CG>TA_GCACCAA | 7 | TRUE | FALSE |
| 172_v.txt | 33056 | 33056-SNV- | SNV | 1 | C | T | 39 | 26528 | 0.14701448 | 55.5128205 | rnr1D57N | FALSE | CAN | JSY14 | 2A | rnr1D57N-Re | GCG | C>T | A/T+1 | G/C-1 | CG>TA | CG>TA | GGACTGCGT | CGTGACA | CGC | GCG | C | A | C>T | C>T_GCG | CG>TA_CGTGACA | 7 | TRUE | FALSE |
| 172_v.txt | 33070 | 33070-SNV- | SNV | 1 | C | T | 27 | 26528 | 0.10177925 | 53.5925926 | rnr1D57N | FALSE | CAN | JSY14 | 2A | rnr1D57N-Re | GCC | C>T | A/T+1 | G/C-1 | CG>TA | CG>TA | AATATGCCA | CCAAAGA | CGG | GCC | C | A | C>T | C>T_GCC | CG>TA_CCAAAGA | 7 | TRUE | FALSE |
| 172_v.txt | 33111 | 33111-SNV- | SNV | 1 | G | A | 27 | 26528 | 0.10177925 | 42.0740741 | rnr1D57N | FALSE | CAN | JSY14 | 2A | rnr1D57N-Re | TGG | G>A | A/T+1 | G/C-1 | CG>TA | CG>TA | CCCACTGGG | GGGCCGG | ACC | ACC | C | T | C>T | C>T_ACC | CG>TA_GGGCCGG | 7 | TRUE | FALSE |
| 172_v.txt | 33114 | 33114-SNV- | SNV | 1 | C | T | 47 | 26528 | 0.17717129 | 44.5528476 | rnr1D57N | FALSE | CAN | JSY14 | 2A | rnr1D57N-Re | GCC | C>T | A/T+1 | G/C-1 | CG>TA | CG>TA | ACTGGGCCG | CCGGCGT | CGG | GCC | C | A | C>T | C>T_GCC | CG>TA_CCGGCGT | 7 | TRUE | FALSE |
| 172_v.txt | 33115 | 33115-SNV- | SNV | 1 | C | T | 24 | 26528 | 0.09047045 | 43.8329339 | rnr1D57N | FALSE | CAN | JSY14 | 2A | rnr1D57N-Re | CCG | C>T | A/T+1 | G/C-1 | CG>TA | CG>TA | CTGGGCCGG | CGGCGTT | GGC | CCG | C | A | C>T | C>T_CCG | CG>TA_CGGCGTT | 7 | TRUE | FALSE |
| 172_v.txt | 33118 | 33118-SNV- | SNV | 1 | C | T | 28 | 26528 | 0.10554885 | 38.1785714 | rnr1D57N | FALSE | CAN | JSY14 | 2A | rnr1D57N-Re | GCG | C>T | A/T+1 | G/C-1 | CG>TA | CG>TA | GGCCGGCGT | CGTTGGT | CGC | GCG | C | A | C>T | C>T_GCG | CG>TA_CGTTGGT | 7 | TRUE | FALSE |
| 172_v.txt | 33258 | 33258-SNV- | SNV | 1 | C | T | 19 | 11109 | 0.1710325 | 41.5263158 | rnr1D57N | FALSE | CAN | JSY14 | 2A | rnr1D57N-Re | GCC | C>T | A/T+1 | G/C-1 | CG>TA | CG>TA | CCTATGCCAT | CCATCCT | CGG | GCC | C | A | C>T | C>T_GCC | CG>TA_CCATCCT | 7 | TRUE | FALSE |
| 172_v.txt | 33276 | 33276-SNV- | SNV | 1 | G | A | 29 | 11109 | 0.2610496 | 45.7241379 | rnr1D57N | FALSE | CAN | JSY14 | 2A | rnr1D57N-Re | CGT | G>A | A/T+1 | G/C-1 | CG>TA | CG>TA | GAGAACGTA | GTATCCT | GCA | GCA | C | T | C>T | C>T_GCA | CG>TA_GTATCCT | 7 | TRUE | FALSE |
| 172_v.txt | 33309 | 33309-SNV- | SNV | 1 | C | T | 15 | 11109 | 0.13502566 | 61.3333333 | rnr1D57N | FALSE | CAN | JSY14 | 2A | rnr1D57N-Re | GCG | C>T | A/T+1 | G/C-1 | CG>TA | CG>TA | AAAGAGCGC | CGCAATG | CGC | GCG | C | A | C>T | C>T_GCG | CG>TA_CGCAATG | 7 | TRUE | FALSE |
| 172_v.txt | 33412 | 33412-SNV- | SNV | 1 | G | A | 16 | 11109 | 0.14402737 | 48.1875 | rnr1D57N | FALSE | CAN | JSY14 | 2A | rnr1D57N-Re | GGC | G>A | A/T+1 | G/C-1 | CG>TA | CG>TA | TGACCGGCT | GCTCATT | CCG | CCG | C | T | C>T | C>T_CCG | CG>TA_GCTCATT | 7 | TRUE | FALSE |
| 172_v.txt | 33474 | 33474-SNV- | SNV | 1 | C | T | 15 | 11107 | 0.13504997 | 36.7333333 | rnr1D57N | FALSE | CAN | JSY14 | 2A | rnr1D57N-Re | GCC | C>T | A/T+1 | G/C-1 | CG>TA | CG>TA | GCTATGCCTT | CCTTTTT | CGG | GCC | C | A | C>T | C>T_GCC | CG>TA_CCTTTTT | 7 | TRUE | FALSE |
| 172_v.txt | 31972 | 31972-SNV- | SNV | 1 | C | G | 1 | 18831 | 5.31E-03 | 37.9905574 | rnr1D57N | FALSE | CAN | JSY14 | 2A | rnr1D57N-Re | GCC | C>G | G/C+1 | G/C-1 | CG>GC | CG>GC | AATAAGCCA | CCAAGCC | CGG | GCC | C | C | C>G | C>G_GCC | CG>GC_CCAAGCC | 7 | TRUE | FALSE |
| 172_v.txt | 32271 | 32271-SNV- | SNV | 1 | C | G | 44 | 26233 | 0.16772767 | 35.6818182 | rnr1D57N | FALSE | CAN | JSY14 | 2A | rnr1D57N-Re | ACG | C>G | G/C+1 | G/C-1 | CG>GC | CG>GC | AAAATACGG | CGGGAAC | TGC | ACG | C | C | C>G | C>G_ACG | CG>GC_CGGGAAC | 7 | TRUE | FALSE |
| 172_v.txt | 32278 | 32278-SNV- | SNV | 1 | C | G | 12 | 26233 | 0.04574391 | 36.5 | rnr1D57N | FALSE | CAN | JSY14 | 2A | rnr1D57N-Re | CCA | C>G | G/C+1 | G/C-1 | CG>GC | CG>GC | GGGAACCAA | CAACGTA | GGT | CCA | C | C | C>G | C>G_CCA | CG>GC_CAACGTA | 7 | TRUE | FALSE |
| 172_v.txt | 32449 | 32449-SNV- | SNV | 1 | C | G | 5 | 26233 | 0.01905996 | 34.6 | rnr1D57N | FALSE | CAN | JSY14 | 2A | rnr1D57N-Re | CCA | C>G | G/C+1 | G/C-1 | CG>GC | CG>GC | AAAGTCCAA | CAATGAA | GGT | CCA | C | C | C>G | C>G_CCA | CG>GC_CAATGAA | 7 | TRUE | FALSE |
| 172_v.txt | 32813 | 32813-SNV- | SNV | 1 | G | C | 4 | 41541 | 9.63E-03 | 50.4972845 | rnr1D57N | FALSE | CAN | JSY14 | 2A | rnr1D57N-Re | GGA | G>C | G/C+1 | G/C-1 | CG>GC | CG>GC | GACAGGGAA | GAACAAG | CCT | CCT | C | G | C>G | C>G_CCT | CG>GC_GAACAAG | 7 | TRUE | FALSE |
| 172_v.txt | 32959 | 32959-SNV- | SNV | 1 | C | G | 12 | 26528 | 0.04523522 | 61.25 | rnr1D57N | FALSE | CAN | JSY14 | 2A | rnr1D57N-Re | CCA | C>G | G/C+1 | G/C-1 | CG>GC | CG>GC | TGTAACCATT | CATTGGC | GGT | CCA | C | C | C>G | C>G_CCA | CG>GC_CATTGGC | 7 | TRUE | FALSE |
| 172_v.txt | 33070 | 33070-SNV- | SNV | 1 | C | G | 23 | 26528 | 0.08670084 | 58.0434783 | rnr1D57N | FALSE | CAN | JSY14 | 2A | rnr1D57N-Re | GCC | C>G | G/C+1 | G/C-1 | CG>GC | CG>GC | AATATGCCA | CCAAAGA | CGG | GCC | C | C | C>G | C>G_GCC | CG>GC_CCAAAGA | 7 | TRUE | FALSE |
| 172_v.txt | 33172 | 33172-SNV- | SNV | 1 | C | G | 54 | 26524 | 0.2035892 | 37.9074074 | rnr1D57N | FALSE | CAN | JSY14 | 2A | rnr1D57N-Re | CCA | C>G | G/C+1 | G/C-1 | CG>GC | CG>GC | TACCACCAA | CAAGGGC | GGT | CCA | C | C | C>G | C>G_CCA | CG>GC_CAAGGGC | 7 | TRUE | FALSE |
| 172_v.txt | 31657 | 31657-SNV- | SNV | 1 | G | T | 86 | 12214 | 0.70411004 | 20.0662934 | rnr1D57N | FALSE | CAN | JSY14 | 2A | rnr1D57N-Re | GGC | G>T | A/T+1 | G/C-1 | CG>AT | CG>AT | GAAATGGCG | GCGTGGA | CCG | CCG | C | A | C>A | C>A_CCG | CG>AT_GCGTGGA | 7 | TRUE | FALSE |
| 172_v.txt | 31971 | 31971-SNV- | SNV | 1 | G | T | 9 | 18831 | 0.04779353 | 16.9982769 | rnr1D57N | FALSE | CAN | JSY14 | 2A | rnr1D57N-Re | AGC | G>T | A/T+1 | G/C-1 | CG>AT | CG>AT | TAATAAGCC | GCCAAGC | TCG | TCG | C | A | C>A | C>A_TCG | CG>AT_GCCAAGC | 7 | TRUE | FALSE |
| 172_v.txt | 32008 | 32008-SNV- | SNV | 1 | C | A | 2206 | 18831 | 11.7147257 | 57.6119674 | rnr1D57N | FALSE | CAN | JSY14 | 2A | rnr1D57N-Re | TCG | C>A | A/T+1 | G/C-1 | CG>AT | CG>AT | GTAACTCGT | CGTCACG | AGC | TCG | C | T | C>A | C>A_TCG | CG>AT_CGTCACG | 7 | TRUE | FALSE |
| 172_v.txt | 32119 | 32119-SNV- | SNV | 1 | C | A | 221 | 18831 | 1.17359673 | 58.5746606 | rnr1D57N | FALSE | CAN | JSY14 | 2A | rnr1D57N-Re | TCG | C>A | A/T+1 | G/C-1 | CG>AT | CG>AT | GCCATTCGA | CGAAAAC | AGC | TCG | C | T | C>A | C>A_TCG | CG>AT_CGAAAAC | 7 | TRUE | FALSE |
| 172_v.txt | 32205 | 32205-SNV- | SNV | 1 | C | A | 74 | 18831 | 0.39296904 | 41.9324324 | rnr1D57N | FALSE | CAN | JSY14 | 2A | rnr1D57N-Re | ACC | C>A | A/T+1 | G/C-1 | CG>AT | CG>AT | GGAACACCA | CCACCTT | TGG | ACC | C | T | C>A | C>A_ACC | CG>AT_CCACCTT | 7 | TRUE | FALSE |
| 172_v.txt | 32488 | 32488-SNV- | SNV | 1 | G | T | 7 | 26233 | 0.02668395 | 48.8571429 | rnr1D57N | FALSE | CAN | JSY14 | 2A | rnr1D57N-Re | CGG | G>T | A/T+1 | G/C-1 | CG>AT | CG>AT | AGATACGGA | GGAAAAC | GCC | GCC | C | A | C>A | C>A_GCC | CG>AT_GGAAAAC | 7 | TRUE | FALSE |
| 172_v.txt | 32575 | 32575-SNV- | SNV | 1 | G | T | 12 | 41543 | 0.02888573 | 33.0838628 | rnr1D57N | FALSE | CAN | JSY14 | 2A | rnr1D57N-Re | TGA | G>T | A/T+1 | G/C-1 | CG>AT | CG>AT | TACCTTGAA | GAAATGT | ACT | ACT | C | A | C>A | C>A_ACT | CG>AT_GAAATGT | 7 | TRUE | FALSE |
| 172_v.txt | 32783 | 32783-SNV- | SNV | 1 | G | T | 17 | 41541 | 0.04092343 | 39.7058824 | rnr1D57N | FALSE | CAN | JSY14 | 2A | rnr1D57N-Re | AGA | G>T | A/T+1 | G/C-1 | CG>AT | CG>AT | GACCCAGAA | GAACTCG | TCT | TCT | C | A | C>A | C>A_TCT | CG>AT_GAACTCG | 7 | TRUE | FALSE |
| 172_v.txt | 32811 | 32811-SNV- | SNV | 1 | G | T | 10 | 41541 | 0.0240726 | 45.5997577 | rnr1D57N | FALSE | CAN | JSY14 | 2A | rnr1D57N-Re | AGG | G>T | A/T+1 | G/C-1 | CG>AT | CG>AT | TTGACAGGG | GGGAACA | TCC | TCC | C | A | C>A | C>A_TCC | CG>AT_GGGAACA | 7 | TRUE | FALSE |
| 172_v.txt | 33006 | 33006-SNV- | SNV | 1 | G | T | 14 | 26528 | 0.05277443 | 59.2142857 | rnr1D57N | FALSE | CAN | JSY14 | 2A | rnr1D57N-Re | TGT | G>T | A/T+1 | G/C-1 | CG>AT | CG>AT | AAAACTGTG | GTGAAAG | ACA | ACA | C | A | C>A | C>A_ACA | CG>AT_GTGAAAG | 7 | TRUE | FALSE |
| 172_v.txt | 33033 | 33033-SNV- | SNV | 1 | G | T | 21 | 26528 | 0.07916164 | 57.2380952 | rnr1D57N | FALSE | CAN | JSY14 | 2A | rnr1D57N-Re | TGT | G>T | A/T+1 | G/C-1 | CG>AT | CG>AT | ATGAATGTA | GTAGCCA | ACA | ACA | C | A | C>A | C>A_ACA | CG>AT_GTAGCCA | 7 | TRUE | FALSE |
| 172_v.txt | 33043 | 33043-SNV- | SNV | 1 | C | A | 49 | 26528 | 0.1847105 | 57.9795918 | rnr1D57N | FALSE | CAN | JSY14 | 2A | rnr1D57N-Re | TCA | C>A | A/T+1 | G/C-1 | CG>AT | CG>AT | CCATTTCACC | CACCCAA | AGT | TCA | C | T | C>A | C>A_TCA | CG>AT_CACCCAA | 7 | TRUE | FALSE |
| 172_v.txt | 33153 | 33153-SNV- | SNV | 1 | C | A | 132 | 26527 | 0.49760621 | 41.8106061 | rnr1D57N | FALSE | CAN | JSY14 | 2A | rnr1D57N-Re | ACC | C>A | A/T+1 | G/C-1 | CG>AT | CG>AT | AAAAGACCT | CCTGTAC | TGG | ACC | C | T | C>A | C>A_ACC | CG>AT_CCTGTAC | 7 | TRUE | FALSE |
| 172_v.txt | 33159 | 33159-SNV- | SNV | 1 | C | A | 18 | 26526 | 0.06785795 | 36.9444444 | rnr1D57N | FALSE | CAN | JSY14 | 2A | rnr1D57N-Re | ACC | C>A | A/T+1 | G/C-1 | CG>AT | CG>AT | CCTGTACCA | CCAATAG | TGG | ACC | C | T | C>A | C>A_ACC | CG>AT_CCAATAG | 7 | TRUE | FALSE |
| 172_v.txt | 33474 | 33474-SNV- | SNV | 1 | C | A | 31 | 11107 | 0.27910327 | 33.0322581 | rnr1D57N | FALSE | CAN | JSY14 | 2A | rnr1D57N-Re | GCC | C>A | A/T+1 | G/C-1 | CG>AT | CG>AT | GCTATGCCTT | CCTTTTT | CGG | GCC | C | T | C>A | C>A_GCC | CG>AT_CCTTTTT | 7 | TRUE | FALSE |
| 172_v.txt | 32811 | 32811-Repla | Replacement | 3 | GGG | CA | 45 | 41541 | 0.10832671 | 44.7037037 | rnr1D57N | FALSE | CAN | JSY14 | 2A | rnr1D57N-Re | AGG | nope | >1bp-ins | >1bpdel | none | Replac. | TTGACAGGG | GGGAACA | TCC | AGG | GGG | none | NA | NA_AGG | Replac._GGGAACA | 7 | TRUE | FALSE |
| 172_v.txt | 32544 | 32544-MNV- | MNV | 2 | CC | AA | 1 | 26233 | 3.81E-03 | 32.0000416 | rnr1D57N | FALSE | CAN | JSY14 | 2A | rnr1D57N-Re | ACC | nope | >1bp-ins | >1bpdel | none | MNV | GCTTCACCA | CCAGCAG | TGG | ACC | CC | none | NA | NA_ACC | MNV_CCAGCAG | 7 | TRUE | FALSE |
| 172_v.txt | 32811 | 32811-MNV- | MNV | 2 | GG | CA | 1 | 41541 | 2.41E-03 | 33.5001848 | rnr1D57N | FALSE | CAN | JSY14 | 2A | rnr1D57N-Re | AGG | nope | >1bp-ins | >1bpdel | none | MNV | TTGACAGGG | GGGAACA | TCC | AGG | GG | none | NA | NA_AGG | MNV_GGGAACA | 7 | TRUE | FALSE |
| 172_v.txt | 33114 | 33114-MNV- | MNV | 2 | CC | AA | 1 | 26528 | 3.77E-03 | 37.5002008 | rnr1D57N | FALSE | CAN | JSY14 | 2A | rnr1D57N-Re | GCC | nope | >1bp-ins | >1bpdel | none | MNV | ACTGGGCCG | CCGGCGT | CGG | GCC | CC | none | NA | NA_GCC | MNV_CCGGCGT | 7 | TRUE | FALSE |
| 172_v.txt | 31886 | 31886-Insert | Insertion | 1 | - | T | 2 | 12214 | 0.01637465 | 38 | rnr1D57N | FALSE | CAN | JSY14 | 2A | rnr1D57N-Re | CAA | nope | A/T+1 | >1bpdel | none | A/T+1 | GGCAGCAAA | AAAGCTA | GTT | CAA | - | A | NA | NA_CAA | A/T+1_AAAGCTA | 7 | TRUE | FALSE |
| 172_v.txt | 32081 | 32081-Insert | Insertion | 1 | - | A | 15 | 18831 | 0.07965589 | 63.8666667 | rnr1D57N | FALSE | CAN | JSY14 | 2A | rnr1D57N-Re | CAA | nope | A/T+1 | >1bpdel | none | A/T+1 | CCATGCAAA | AAAAAAG | GTT | CAA | - | T | NA | NA_CAA | A/T+1_AAAAAAG | 7 | TRUE | FALSE |
| 172_v.txt | 32498 | 32498-Insert | Insertion | 1 | - | T | 5 | 26233 | 0.01905996 | 41 | rnr1D57N | FALSE | CAN | JSY14 | 2A | rnr1D57N-Re | CTT | nope | A/T+1 | >1bpdel | none | A/T+1 | AACAACTTTT | TTTTTTG | GAA | CTT | - | A | NA | NA_CTT | A/T+1_TTTTTTG | 7 | TRUE | FALSE |
| 172_v.txt | 32597 | 32597-Insert | Insertion | 1 | - | A | 8 | 41541 | 0.01925808 | 49.625 | rnr1D57N | FALSE | CAN | JSY14 | 2A | rnr1D57N-Re | TCA | nope | A/T+1 | >1bpdel | none | A/T+1 | GTTAATCAA | CAAAGAG | AGT | TCA | - | T | NA | NA_TCA | A/T+1_CAAAGAG | 7 | TRUE | FALSE |
| 172_v.txt | 32728 | 32728-Insert | Insertion | 1 | - | A | 6 | 41541 | 0.01444356 | 61.5 | rnr1D57N | FALSE | CAN | JSY14 | 2A | rnr1D57N-Re | CAA | nope | A/T+1 | >1bpdel | none | A/T+1 | CCATACAAA | AAAAACA | GTT | CAA | - | T | NA | NA_CAA | A/T+1_AAAAACA | 7 | TRUE | FALSE |
| 172_v.txt | 32814 | 32814-Insert | Insertion | 1 | - | A | 1 | 41541 | 2.41E-03 | 38 | rnr1D57N | FALSE | CAN | JSY14 | 2A | rnr1D57N-Re | GAA | nope | A/T+1 | >1bpdel | none | A/T+1 | ACAGGGAAC | AACAAGT | CTT | GAA | - | T | NA | NA_GAA | A/T+1_AACAAGT | 7 | TRUE | FALSE |
| 172_v.txt | 32842 | 32842-Insert | Insertion | 1 | - | A | 55 | 41541 | 0.13239932 | 46.4363636 | rnr1D57N | FALSE | CAN | JSY14 | 2A | rnr1D57N-Re | CAA | nope | A/T+1 | >1bpdel | none | A/T+1 | TTACCCAAA | AAAAAAT | GTT | CAA | - | T | NA | NA_CAA | A/T+1_AAAAAAT | 7 | TRUE | FALSE |
| 172_v.txt | 32842 | 32842-Insert | Insertion | 1 | - | G | 1 | 41541 | 2.41E-03 | 32 | rnr1D57N | FALSE | CAN | JSY14 | 2A | rnr1D57N-Re | CAA | nope | G/C+1 | >1bpdel | none | G/C+1 | TTACCCAAA | AAAAAAT | GTT | CAA | - | C | NA | NA_CAA | G/C+1_AAAAAAT | 7 | TRUE | FALSE |
| 172_v.txt | 33000 | 33000-Insert | Insertion | 2 | - | AA | 8 | 26528 | 0.03015682 | 64 | rnr1D57N | FALSE | CAN | JSY14 | 2A | rnr1D57N-Re | GAA | nope | >1bp-ins | >1bpdel | none | >1bp-ins | TGTGAGAAA | AAAACTG | CTT | GAA | - | none | NA | NA_GAA | >1bp-ins_AAAACTG | 7 | TRUE | FALSE |
| 172_v.txt | 32589 | 32589-Deleti | Deletion | 2 | GC | - | 10 | 41541 | 0.0240726 | 37.7 | rnr1D57N | FALSE | CAN | JSY14 | 2A | rnr1D57N-Re | AGC | nope | >1bp-ins | >1bpdel | none | >1bpdel | AAGGCAGCG | GCGTTAA | TCG | AGC | GC | none | NA | NA_AGC | >1bpdel_GCGTTAA | 7 | TRUE | FALSE |
| 172_v.txt | 31711 | 31711-Deleti | Deletion | 1 | A | - | 7 | 12214 | 0.05731128 | 61.7142857 | rnr1D57N | FALSE | CAN | JSY14 | 2A | rnr1D57N-Re | CAA | nope | >1bp-ins | A/T-1 | none | A/T-1 | CATTCCAAA | AAAATTT | GTT | GTT | T | none | T>none | T>none_GTT | A/T-1_AAAATTT | 7 | TRUE | FALSE |
| 172_v.txt | 31723 | 31723-Deleti | Deletion | 1 | A | - | 17 | 12214 | 0.13918454 | 55.2352941 | rnr1D57N | FALSE | CAN | JSY14 | 2A | rnr1D57N-Re | CAA | nope | >1bp-ins | A/T-1 | none | A/T-1 | TGTCCCAAA | AAAAAGT | GTT | GTT | T | none | T>none | T>none_GTT | A/T-1_AAAAAGT | 7 | TRUE | FALSE |
| 172_v.txt | 31904 | 31904-Deleti | Deletion | 1 | T | - | 7 | 12214 | 0.05731128 | 44.8571429 | rnr1D57N | FALSE | CAN | JSY14 | 2A | rnr1D57N-Re | ATT | nope | >1bp-ins | A/T-1 | none | A/T-1 | ATTGAATTTT | TTTTGGT | TAA | ATT | T | none | nope | nope_ATT | A/T-1_TTTTGGT | 7 | TRUE | FALSE |
| 172_v.txt | 31913 | 31913-Deleti | Deletion | 1 | A | - | 6 | 12214 | 0.04912396 | 33.3333333 | rnr1D57N | FALSE | CAN | JSY14 | 2A | rnr1D57N-Re | CAA | nope | >1bp-ins | A/T-1 | none | A/T-1 | TGGTGCAAA | AAAAGCC | GTT | GTT | T | none | T>none | T>none_GTT | A/T-1_AAAAGCC | 7 | TRUE | FALSE |
| 172_v.txt | 31971 | 31971-Deleti | Deletion | 1 | G | - | 11065 | 18831 | 58.7594923 | 37.410122 | rnr1D57N | FALSE | CAN | JSY14 | 2A | rnr1D57N-Re | AGC | nope | >1bp-ins | G/C-1 | none | G/C-1 | TAATAAGCC | GCCAAGC | TCG | TCG | C | none | C>none | C>none_TCG | G/C-1_GCCAAGC | 7 | TRUE | FALSE |
| 172_v.txt | 31972 | 31972-Deleti | Deletion | 1 | C | - | 5 | 18831 | 0.02655196 | 36.4000005 | rnr1D57N | FALSE | CAN | JSY14 | 2A | rnr1D57N-Re | GCC | nope | >1bp-ins | G/C-1 | none | G/C-1 | AATAAGCCA | CCAAGCC | CGG | GCC | C | none | nope | nope_GCC | G/C-1_CCAAGCC | 7 | TRUE | FALSE |
| 172_v.txt | 32081 | 32081-Deleti | Deletion | 1 | A | - | 25 | 18830 | 0.13276686 | 63.2000001 | rnr1D57N | FALSE | CAN | JSY14 | 2A | rnr1D57N-Re | CAA | nope | >1bp-ins | A/T-1 | none | A/T-1 | CCATGCAAA | AAAAAAG | GTT | GTT | T | none | T>none | T>none_GTT | A/T-1_AAAAAAG | 7 | TRUE | FALSE |
| 172_v.txt | 32210 | 32210-Deleti | Deletion | 1 | T | - | 7 | 18831 | 0.03717275 | 41.5714286 | rnr1D57N | FALSE | CAN | JSY14 | 2A | rnr1D57N-Re | CTT | nope | >1bp-ins | A/T-1 | none | A/T-1 | ACCACCTTTG | TTTGGTG | GAA | CTT | T | none | nope | nope_CTT | A/T-1_TTTGGTG | 7 | TRUE | FALSE |
| 172_v.txt | 32247 | 32247-Deleti | Deletion | 1 | T | - | 6 | 18825 | 0.03187251 | 38 | rnr1D57N | FALSE | CAN | JSY14 | 2A | rnr1D57N-Re | GTT | nope | >1bp-ins | A/T-1 | none | A/T-1 | AACTTGTTCT | TTCTTTG | CAA | GTT | T | none | nope | nope_GTT | A/T-1_TTCTTTG | 7 | TRUE | FALSE |
| 172_v.txt | 32284 | 32284-Deleti | Deletion | 1 | A | - | 5 | 26233 | 0.01905996 | 48.4 | rnr1D57N | FALSE | CAN | JSY14 | 2A | rnr1D57N-Re | TAA | nope | >1bp-ins | A/T-1 | none | A/T-1 | CAACGTAAA | AAATATT | ATT | ATT | T | none | T>none | T>none_ATT | A/T-1_AAATATT | 7 | TRUE | FALSE |
| 172_v.txt | 32416 | 32416-Deleti | Deletion | 1 | T | - | 6 | 26233 | 0.02287196 | 59.6666667 | rnr1D57N | FALSE | CAN | JSY14 | 2A | rnr1D57N-Re | GTT | nope | >1bp-ins | A/T-1 | none | A/T-1 | ATTGTGTTAC | TTAGTTT | CAA | GTT | T | none | nope | nope_GTT | A/T-1_TTAGTTT | 7 | TRUE | FALSE |
| 172_v.txt | 32498 | 32498-Deleti | Deletion | 1 | T | - | 38 | 26233 | 0.14485572 | 46.394736 | rnr1D57N | FALSE | CAN | JSY14 | 2A | rnr1D57N-Re | CTT | nope | >1bp-ins | A/T-1 | none | A/T-1 | AACAACTTTT | TTTTTTG | GAA | CTT | T | none | nope | nope_CTT | A/T-1_TTTTTTG | 7 | TRUE | FALSE |
| 172_v.txt | 32544 | 32544-Deleti | Deletion | 1 | C | - | 4 | 26233 | 0.01524797 | 37.9999987 | rnr1D57N | FALSE | CAN | JSY14 | 2A | rnr1D57N-Re | ACC | nope | >1bp-ins | G/C-1 | none | G/C-1 | GCTTCACCA | CCAGCAG | TGG | ACC | C | none | nope | nope_ACC | G/C-1_CCAGCAG | 7 | TRUE | FALSE |
| 172_v.txt | 32573 | 32573-Deleti | Deletion | 1 | T | - | 2 | 41543 | 4.81E-03 | 38 | rnr1D57N | FALSE | CAN | JSY14 | 2A | rnr1D57N-Re | CTT | nope | >1bp-ins | A/T-1 | none | A/T-1 | AGTACCTTG | TTGAAAT | GAA | CTT | T | none | nope | nope_CTT | A/T-1_TTGAAAT | 7 | TRUE | FALSE |
| 172_v.txt | 32608 | 32608-Deleti | Deletion | 1 | C | - | 6 | 41541 | 0.01444356 | 39.8333333 | rnr1D57N | FALSE | CAN | JSY14 | 2A | rnr1D57N-Re | ACC | nope | >1bp-ins | G/C-1 | none | G/C-1 | AGGAAACCC | CCCAACC | TGG | ACC | C | none | nope | nope_ACC | G/C-1_CCCAACC | 7 | TRUE | FALSE |
| 172_v.txt | 32658 | 32658-Deleti | Deletion | 1 | C | - | 8 | 41541 | 0.01925808 | 57.375 | rnr1D57N | FALSE | CAN | JSY14 | 2A | rnr1D57N-Re | ACC | nope | >1bp-ins | G/C-1 | none | G/C-1 | CCTGGACCC | CCCCAGG | TGG | ACC | C | none | nope | nope_ACC | G/C-1_CCCCAGG | 7 | TRUE | FALSE |
| 172_v.txt | 32707 | 32707-Deleti | Deletion | 1 | C | - | 8 | 41541 | 0.01925808 | 63.75 | rnr1D57N | FALSE | CAN | JSY14 | 2A | rnr1D57N-Re | ACC | nope | >1bp-ins | G/C-1 | none | G/C-1 | CGGTAACCC | CCCCAGC | TGG | ACC | C | none | nope | nope_ACC | G/C-1_CCCCAGC | 7 | TRUE | FALSE |
| 172_v.txt | 32722 | 32722-Deleti | Deletion | 1 | C | - | 10 | 41541 | 0.0240726 | 62.9 | rnr1D57N | FALSE | CAN | JSY14 | 2A | rnr1D57N-Re | ACC | nope | >1bp-ins | G/C-1 | none | G/C-1 | CACAAACCA | CCATACA | TGG | ACC | C | none | nope | nope_ACC | G/C-1_CCATACA | 7 | TRUE | FALSE |
| 172_v.txt | 32728 | 32728-Deleti | Deletion | 1 | A | - | 15 | 41538 | 0.03611151 | 61.3999956 | rnr1D57N | FALSE | CAN | JSY14 | 2A | rnr1D57N-Re | CAA | nope | >1bp-ins | A/T-1 | none | A/T-1 | CCATACAAA | AAAAACA | GTT | GTT | T | none | T>none | T>none_GTT | A/T-1_AAAAACA | 7 | TRUE | FALSE |
| 172_v.txt | 32744 | 32744-Deleti | Deletion | 1 | A | - | 16 | 41541 | 0.03851617 | 55.25 | rnr1D57N | FALSE | CAN | JSY14 | 2A | rnr1D57N-Re | GAA | nope | >1bp-ins | A/T-1 | none | A/T-1 | TATTAGAAA | AAACCCG | CTT | CTT | T | none | T>none | T>none_CTT | A/T-1_AAACCCG | 7 | TRUE | FALSE |
| 172_v.txt | 32760 | 32760-Deleti | Deletion | 1 | A | - | 11 | 41541 | 0.02647986 | 53.6363636 | rnr1D57N | FALSE | CAN | JSY14 | 2A | rnr1D57N-Re | TAA | nope | >1bp-ins | A/T-1 | none | A/T-1 | ATGGCTAAA | AAAACTT | ATT | ATT | T | none | T>none | T>none_ATT | A/T-1_AAAACTT | 7 | TRUE | FALSE |
| 172_v.txt | 32779 | 32779-Deleti | Deletion | 1 | C | - | 6 | 41541 | 0.01444356 | 51 | rnr1D57N | FALSE | CAN | JSY14 | 2A | rnr1D57N-Re | ACC | nope | >1bp-ins | G/C-1 | none | G/C-1 | AAGCGACCC | CCCAGAA | TGG | ACC | C | none | nope | nope_ACC | G/C-1_CCCAGAA | 7 | TRUE | FALSE |
| 172_v.txt | 32842 | 32842-Deleti | Deletion | 1 | A | - | 76 | 41541 | 0.18295178 | 41.1842101 | rnr1D57N | FALSE | CAN | JSY14 | 2A | rnr1D57N-Re | CAA | nope | >1bp-ins | A/T-1 | none | A/T-1 | TTACCCAAA | AAAAAAT | GTT | GTT | T | none | T>none | T>none_GTT | A/T-1_AAAAAAT | 7 | TRUE | FALSE |
| 172_v.txt | 32884 | 32884-Deleti | Deletion | 1 | A | - | 5 | 26528 | 0.01884801 | 37.8 | rnr1D57N | FALSE | CAN | JSY14 | 2A | rnr1D57N-Re | CAA | nope | >1bp-ins | A/T-1 | none | A/T-1 | ACGTCCAAA | AAAATTG | GTT | GTT | T | none | T>none | T>none_GTT | A/T-1_AAAATTG | 7 | TRUE | FALSE |
| 172_v.txt | 32924 | 32924-Deleti | Deletion | 1 | A | - | 23 | 26528 | 0.08670084 | 54.6521739 | rnr1D57N | FALSE | CAN | JSY14 | 2A | rnr1D57N-Re | CAA | nope | >1bp-ins | A/T-1 | none | A/T-1 | CAGGGCAAA | AAAAGTG | GTT | GTT | T | none | T>none | T>none_GTT | A/T-1_AAAAGTG | 7 | TRUE | FALSE |
| 172_v.txt | 33000 | 33000-Deleti | Deletion | 1 | A | - | 74 | 26528 | 0.27895054 | 61.4324326 | rnr1D57N | FALSE | CAN | JSY14 | 2A | rnr1D57N-Re | GAA | nope | >1bp-ins | A/T-1 | none | A/T-1 | TGTGAGAAA | AAAACTG | CTT | CTT | T | none | T>none | T>none_CTT | A/T-1_AAAACTG | 7 | TRUE | FALSE |
| 172_v.txt | 33083 | 33083-Deleti | Deletion | 1 | A | - | 6 | 26528 | 0.02261761 | 50.3333333 | rnr1D57N | FALSE | CAN | JSY14 | 2A | rnr1D57N-Re | TAA | nope | >1bp-ins | A/T-1 | none | A/T-1 | ACCCATAAA | AAATAAA | ATT | ATT | T | none | T>none | T>none_ATT | A/T-1_AAATAAA | 7 | TRUE | FALSE |
| 172_v.txt | 33138 | 33138-Deleti | Deletion | 1 | A | - | 5 | 26527 | 0.01884872 | 43.2 | rnr1D57N | FALSE | CAN | JSY14 | 2A | rnr1D57N-Re | TAA | nope | >1bp-ins | A/T-1 | none | A/T-1 | GTGGATAAA | AAACCAA | ATT | ATT | T | none | T>none | T>none_ATT | A/T-1_AAACCAA | 7 | TRUE | FALSE |
| 172_v.txt | 33147 | 33147-Deleti | Deletion | 1 | A | - | 15 | 26527 | 0.05654616 | 40.9333333 | rnr1D57N | FALSE | CAN | JSY14 | 2A | rnr1D57N-Re | GAA | nope | >1bp-ins | A/T-1 | none | A/T-1 | CCAATGAAA | AAAAGAC | CTT | CTT | T | none | T>none | T>none_CTT | A/T-1_AAAAGAC | 7 | TRUE | FALSE |
| 172_v.txt | 33474 | 33474-Deleti | Deletion | 1 | C | - | 35 | 11107 | 0.31511659 | 30.2857143 | rnr1D57N | FALSE | CAN | JSY14 | 2A | rnr1D57N-Re | GCC | nope | >1bp-ins | G/C-1 | none | G/C-1 | GCTATGCCTT | CCTTTTT | CGG | GCC | C | none | nope | nope_GCC | G/C-1_CCTTTTT | 7 | TRUE | FALSE |
| 173_v.txt | 31840 | 31840-SNV- | SNV | 1 | A | C | 3 | 4049 | 0.07409237 | 64 | rnr1D57N | FALSE | CAN | JSY14 | 3A | rnr1D57N-Re | AAG | A>C | G/C+1 | A/T-1 | TA>GC | TA>GC | GAAATAAGA | AGATCCA | TTC | TTC | T | G | T>G | T>G_TTC | TA>GC_AGATCCA | 7 | TRUE | FALSE |
| 173_v.txt | 32283 | 32283-SNV-T | SNV | 1 | T | G | 310 | 9772 | 3.1723291 | 51.0451613 | rnr1D57N | FALSE | CAN | JSY14 | 3A | rnr1D57N-Re | GTA | T>G | G/C+1 | A/T-1 | TA>GC | TA>GC | CCAACGTAA | TAAATAT | CAT | GTA | T | C | T>G | T>G_GTA | TA>GC_TAAATAT | 7 | TRUE | FALSE |
| 173_v.txt | 32337 | 32337-SNV- | SNV | 1 | A | C | 1 | 9772 | 0.01023332 | 63.9994869 | rnr1D57N | FALSE | CAN | JSY14 | 3A | rnr1D57N-Re | GAT | A>C | G/C+1 | A/T-1 | TA>GC | TA>GC | TTGAAGATA | ATATGTG | CTA | CTA | T | G | T>G | T>G_CTA | TA>GC_ATATGTG | 7 | TRUE | FALSE |
| 173_v.txt | 32732 | 32732-SNV- | SNV | 1 | A | C | 6 | 13670 | 0.04389173 | 45.8333333 | rnr1D57N | FALSE | CAN | JSY14 | 3A | rnr1D57N-Re | AAC | A>C | G/C+1 | A/T-1 | TA>GC | TA>GC | ACAAAAACA | ACAGTAT | TTG | TTG | T | G | T>G | T>G_TTG | TA>GC_ACAGTAT | 7 | TRUE | FALSE |
| 173_v.txt | 32739 | 32739-SNV- | SNV | 1 | A | C | 26 | 13670 | 0.19019751 | 53.2692308 | rnr1D57N | FALSE | CAN | JSY14 | 3A | rnr1D57N-Re | TAT | A>C | G/C+1 | A/T-1 | TA>GC | TA>GC | CAGTATATTA | ATTAGAA | ATA | ATA | T | G | T>G | T>G_ATA | TA>GC_ATTAGAA | 7 | TRUE | FALSE |
| 173_v.txt | 32823 | 32823-SNV- | SNV | 1 | A | C | 5 | 13670 | 0.03657645 | 42 | rnr1D57N | FALSE | CAN | JSY14 | 3A | rnr1D57N-Re | CAT | A>C | G/C+1 | A/T-1 | TA>GC | TA>GC | AAGTTCATTA | ATTATTG | GTA | GTA | T | G | T>G | T>G_GTA | TA>GC_ATTATTG | 7 | TRUE | FALSE |
| 173_v.txt | 32842 | 32842-SNV- | SNV | 1 | A | C | 3 | 13670 | 0.02194587 | 37.9999855 | rnr1D57N | FALSE | CAN | JSY14 | 3A | rnr1D57N-Re | CAA | A>C | G/C+1 | A/T-1 | TA>GC | TA>GC | TTACCCAAA | AAAAAAT | GTT | GTT | T | G | T>G | T>G_GTT | TA>GC_AAAAAAT | 7 | TRUE | FALSE |
| 173_v.txt | 32884 | 32884-SNV- | SNV | 1 | A | C | 4 | 12003 | 0.033325 | 33 | rnr1D57N | FALSE | CAN | JSY14 | 3A | rnr1D57N-Re | CAA | A>C | G/C+1 | A/T-1 | TA>GC | TA>GC | ACGTCCAAA | AAAATTG | GTT | GTT | T | G | T>G | T>G_GTT | TA>GC_AAAATTG | 7 | TRUE | FALSE |
| 173_v.txt | 33269 | 33269-SNV- | SNV | 1 | A | C | 48 | 7739 | 0.62023517 | 14.8541667 | rnr1D57N | FALSE | CAN | JSY14 | 3A | rnr1D57N-Re | TAG | A>C | G/C+1 | A/T-1 | TA>GC | TA>GC | CTCCATAGA | AGAGAAC | ATC | ATC | T | G | T>G | T>G_ATC | TA>GC_AGAGAAC | 7 | TRUE | FALSE |
| 173_v.txt | 32583 | 32583-SNV- | SNV | 1 | A | G | 34 | 13670 | 0.24871982 | 37 | rnr1D57N | FALSE | CAN | JSY14 | 3A | rnr1D57N-Re | GAA | A>G | G/C+1 | A/T-1 | TA>CG | TA>CG | AATGTGAAG | AAGGCAG | CTT | CTT | T | C | T>C | T>C_CTT | TA>CG_AAGGCAG | 7 | TRUE | FALSE |
| 173_v.txt | 32762 | 32762-SNV- | SNV | 1 | A | G | 53 | 13670 | 0.38771032 | 51.8679245 | rnr1D57N | FALSE | CAN | JSY14 | 3A | rnr1D57N-Re | AAA | A>G | G/C+1 | A/T-1 | TA>CG | TA>CG | GGCTAAAAC | AACTTTG | TTT | TTT | T | C | T>C | T>C_TTT | TA>CG_AACTTTG | 7 | TRUE | FALSE |
| 173_v.txt | 32787 | 32787-SNV-T | SNV | 1 | T | C | 10 | 13670 | 0.07315289 | 46.4 | rnr1D57N | FALSE | CAN | JSY14 | 3A | rnr1D57N-Re | CTC | T>C | G/C+1 | A/T-1 | TA>CG | TA>CG | CAGAACTCG | TCGAATT | GAG | CTC | T | G | T>C | T>C_CTC | TA>CG_TCGAATT | 7 | TRUE | FALSE |
| 173_v.txt | 33174 | 33174-SNV- | SNV | 1 | A | G | 91 | 12000 | 0.75833333 | 37.2967033 | rnr1D57N | FALSE | CAN | JSY14 | 3A | rnr1D57N-Re | AAG | A>G | G/C+1 | A/T-1 | TA>CG | TA>CG | CCACCAAGG | AGGGCAA | TTC | TTC | T | C | T>C | T>C_TTC | TA>CG_AGGGCAA | 7 | TRUE | FALSE |
| 173_v.txt | 32498 | 32498-SNV-T | SNV | 1 | T | A | 3 | 9772 | 0.03069996 | 55.3314568 | rnr1D57N | FALSE | CAN | JSY14 | 3A | rnr1D57N-Re | CTT | T>A | A/T+1 | A/T-1 | TA>AT | TA>AT | AACAACTTTT | TTTTTTG | GAA | CTT | T | T | T>A | T>A_CTT | TA>AT_TTTTTTG | 7 | TRUE | FALSE |
| 173_v.txt | 32680 | 32680-SNV- | SNV | 1 | A | T | 70 | 13670 | 0.51207023 | 60.8857143 | rnr1D57N | FALSE | CAN | JSY14 | 3A | rnr1D57N-Re | CAA | A>T | A/T+1 | A/T-1 | TA>AT | TA>AT | TTCTCCAATA | AATAACG | GTT | GTT | T | A | T>A | T>A_GTT | TA>AT_AATAACG | 7 | TRUE | FALSE |
| 173_v.txt | 32717 | 32717-SNV- | SNV | 1 | A | T | 1 | 13668 | 7.32E-03 | 27.9980688 | rnr1D57N | FALSE | CAN | JSY14 | 3A | rnr1D57N-Re | CAC | A>T | A/T+1 | A/T-1 | TA>AT | TA>AT | AGCACCACA | ACAAACC | GTG | GTG | T | A | T>A | T>A_GTG | TA>AT_ACAAACC | 7 | TRUE | FALSE |
| 173_v.txt | 32766 | 32766-SNV-T | SNV | 1 | T | A | 53 | 13670 | 0.38771032 | 49.6603774 | rnr1D57N | FALSE | CAN | JSY14 | 3A | rnr1D57N-Re | TTT | T>A | A/T+1 | A/T-1 | TA>AT | TA>AT | AAAACTTTG | TTGATGG | AAA | TTT | T | T | T>A | T>A_TTT | TA>AT_TTGATGG | 7 | TRUE | FALSE |
| 173_v.txt | 32787 | 32787-SNV-T | SNV | 1 | T | A | 11 | 13670 | 0.08046818 | 41.6363636 | rnr1D57N | FALSE | CAN | JSY14 | 3A | rnr1D57N-Re | CTC | T>A | A/T+1 | A/T-1 | TA>AT | TA>AT | CAGAACTCG | TCGAATT | GAG | CTC | T | T | T>A | T>A_CTC | TA>AT_TCGAATT | 7 | TRUE | FALSE |
| 173_v.txt | 32823 | 32823-SNV- | SNV | 1 | A | T | 5 | 13670 | 0.03657645 | 48.4 | rnr1D57N | FALSE | CAN | JSY14 | 3A | rnr1D57N-Re | CAT | A>T | A/T+1 | A/T-1 | TA>AT | TA>AT | AAGTTCATTA | ATTATTG | GTA | GTA | T | A | T>A | T>A_GTA | TA>AT_ATTATTG | 7 | TRUE | FALSE |
| 173_v.txt | 33364 | 33364-SNV-T | SNV | 1 | T | A | 11 | 7739 | 0.14213723 | 55.9090909 | rnr1D57N | FALSE | CAN | JSY14 | 3A | rnr1D57N-Re | CTG | T>A | A/T+1 | A/T-1 | TA>AT | TA>AT | CACGTCTGTG | TGTGGTG | GAC | CTG | T | T | T>A | T>A_CTG | TA>AT_TGTGGTG | 7 | TRUE | FALSE |
| 173_v.txt | 31658 | 31658-SNV- | SNV | 1 | C | T | 8 | 4049 | 0.19757965 | 37.625 | rnr1D57N | FALSE | CAN | JSY14 | 3A | rnr1D57N-Re | GCG | C>T | A/T+1 | G/C-1 | CG>TA | CG>TA | AAATGGCGT | CGTGGAA | CGC | GCG | C | A | C>T | C>T_GCG | CG>TA_CGTGGAA | 7 | TRUE | FALSE |
| 173_v.txt | 31930 | 31930-SNV- | SNV | 1 | G | A | 17 | 4049 | 0.41985676 | 37.8235294 | rnr1D57N | FALSE | CAN | JSY14 | 3A | rnr1D57N-Re | TGA | G>A | A/T+1 | G/C-1 | CG>TA | CG>TA | AACCTTGAAT | GAATAAT | ACT | ACT | C | T | C>T | C>T_ACT | CG>TA_GAATAAT | 7 | TRUE | FALSE |
| 173_v.txt | 31962 | 31962-SNV- | SNV | 1 | G | A | 14 | 7041 | 0.19883539 | 36.6428571 | rnr1D57N | FALSE | CAN | JSY14 | 3A | rnr1D57N-Re | CGC | G>A | A/T+1 | G/C-1 | CG>TA | CG>TA | GTGGCCGCA | GCATAAT | GCG | GCG | C | T | C>T | C>T_GCG | CG>TA_GCATAAT | 7 | TRUE | FALSE |
| 173_v.txt | 32075 | 32075-SNV- | SNV | 1 | C | T | 26 | 7041 | 0.36926573 | 61.3076923 | rnr1D57N | FALSE | CAN | JSY14 | 3A | rnr1D57N-Re | ACC | C>T | A/T+1 | G/C-1 | CG>TA | CG>TA | AAATAACCA | CCATGCA | TGG | ACC | C | A | C>T | C>T_ACC | CG>TA_CCATGCA | 7 | TRUE | FALSE |
| 173_v.txt | 32165 | 32165-SNV- | SNV | 1 | G | A | 13 | 7041 | 0.18463287 | 44.9230769 | rnr1D57N | FALSE | CAN | JSY14 | 3A | rnr1D57N-Re | CGC | G>A | A/T+1 | G/C-1 | CG>TA | CG>TA | CAAAGCGCC | GCCAAAT | GCG | GCG | C | T | C>T | C>T_GCG | CG>TA_GCCAAAT | 7 | TRUE | FALSE |
| 173_v.txt | 32271 | 32271-SNV- | SNV | 1 | C | T | 44 | 9772 | 0.45026607 | 35.0909091 | rnr1D57N | FALSE | CAN | JSY14 | 3A | rnr1D57N-Re | ACG | C>T | A/T+1 | G/C-1 | CG>TA | CG>TA | AAAATACGG | CGGGAAC | TGC | ACG | C | A | C>T | C>T_ACG | CG>TA_CGGGAAC | 7 | TRUE | FALSE |
| 173_v.txt | 32299 | 32299-SNV- | SNV | 1 | C | T | 18 | 9772 | 0.18419975 | 54.6111111 | rnr1D57N | FALSE | CAN | JSY14 | 3A | rnr1D57N-Re | GCG | C>T | A/T+1 | G/C-1 | CG>TA | CG>TA | AATTTGCGG | CGGCAGA | CGC | GCG | C | A | C>T | C>T_GCG | CG>TA_CGGCAGA | 7 | TRUE | FALSE |
| 173_v.txt | 32487 | 32487-SNV- | SNV | 1 | C | T | 22 | 9772 | 0.22513303 | 45 | rnr1D57N | FALSE | CAN | JSY14 | 3A | rnr1D57N-Re | ACG | C>T | A/T+1 | G/C-1 | CG>TA | CG>TA | AAGATACGG | CGGAAAA | TGC | ACG | C | A | C>T | C>T_ACG | CG>TA_CGGAAAA | 7 | TRUE | FALSE |
| 173_v.txt | 32535 | 32535-SNV- | SNV | 1 | G | A | 12 | 9772 | 0.12279984 | 38 | rnr1D57N | FALSE | CAN | JSY14 | 3A | rnr1D57N-Re | TGC | G>A | A/T+1 | G/C-1 | CG>TA | CG>TA | GGGTTTGCA | GCAGCTT | ACG | ACG | C | T | C>T | C>T_ACG | CG>TA_GCAGCTT | 7 | TRUE | FALSE |
| 173_v.txt | 32590 | 32590-SNV- | SNV | 1 | C | T | 29 | 13670 | 0.21214338 | 35.137931 | rnr1D57N | FALSE | CAN | JSY14 | 3A | rnr1D57N-Re | GCG | C>T | A/T+1 | G/C-1 | CG>TA | CG>TA | AGGCAGCGT | CGTTAAT | CGC | GCG | C | A | C>T | C>T_GCG | CG>TA_CGTTAAT | 7 | TRUE | FALSE |
| 173_v.txt | 32700 | 32700-SNV- | SNV | 1 | C | T | 20 | 13670 | 0.14630578 | 61.45 | rnr1D57N | FALSE | CAN | JSY14 | 3A | rnr1D57N-Re | GCC | C>T | A/T+1 | G/C-1 | CG>TA | CG>TA | ACTGGGCCG | CCGGTAA | CGG | GCC | C | A | C>T | C>T_GCC | CG>TA_CCGGTAA | 7 | TRUE | FALSE |
| 173_v.txt | 32713 | 32713-SNV- | SNV | 1 | C | T | 10 | 13670 | 0.07315289 | 60.7 | rnr1D57N | FALSE | CAN | JSY14 | 3A | rnr1D57N-Re | GCA | C>T | A/T+1 | G/C-1 | CG>TA | CG>TA | CCCCAGCAC | CACCACA | CGT | GCA | C | A | C>T | C>T_GCA | CG>TA_CACCACA | 7 | TRUE | FALSE |
| 173_v.txt | 32757 | 32757-SNV- | SNV | 1 | G | A | 18 | 13670 | 0.1316752 | 51.2222222 | rnr1D57N | FALSE | CAN | JSY14 | 3A | rnr1D57N-Re | GGC | G>A | A/T+1 | G/C-1 | CG>TA | CG>TA | ATAATGGCT | GCTAAAA | CCG | CCG | C | T | C>T | C>T_CCG | CG>TA_GCTAAAA | 7 | TRUE | FALSE |
| 173_v.txt | 32796 | 32796-SNV- | SNV | 1 | C | T | 20 | 13670 | 0.14630578 | 42.95 | rnr1D57N | FALSE | CAN | JSY14 | 3A | rnr1D57N-Re | ACC | C>T | A/T+1 | G/C-1 | CG>TA | CG>TA | AATTCACCGT | CCGTAAT | TGG | ACC | C | A | C>T | C>T_ACC | CG>TA_CCGTAAT | 7 | TRUE | FALSE |
| 173_v.txt | 32798 | 32798-SNV- | SNV | 1 | G | A | 21 | 13670 | 0.15362107 | 44.4761905 | rnr1D57N | FALSE | CAN | JSY14 | 3A | rnr1D57N-Re | CGT | G>A | A/T+1 | G/C-1 | CG>TA | CG>TA | TTCACCGTA | GTAATAT | GCA | GCA | C | T | C>T | C>T_GCA | CG>TA_GTAATAT | 7 | TRUE | FALSE |
| 173_v.txt | 32855 | 32855-SNV- | SNV | 1 | C | T | 33 | 13670 | 0.24140454 | 35.8787879 | rnr1D57N | FALSE | CAN | JSY14 | 3A | rnr1D57N-Re | TCC | C>T | A/T+1 | G/C-1 | CG>TA | CG>TA | ACTAATCCAT | CCATGCC | AGG | TCC | C | A | C>T | C>T_TCC | CG>TA_CCATGCC | 7 | TRUE | FALSE |
| 173_v.txt | 32860 | 32860-SNV- | SNV | 1 | C | T | 24 | 13670 | 0.17556694 | 37.1666667 | rnr1D57N | FALSE | CAN | JSY14 | 3A | rnr1D57N-Re | GCC | C>T | A/T+1 | G/C-1 | CG>TA | CG>TA | TCCATGCCG | CCGCCAG | CGG | GCC | C | A | C>T | C>T_GCC | CG>TA_CCGCCAG | 7 | TRUE | FALSE |
| 173_v.txt | 32946 | 32946-SNV- | SNV | 1 | C | T | 19 | 12003 | 0.15829376 | 57.8947368 | rnr1D57N | FALSE | CAN | JSY14 | 3A | rnr1D57N-Re | CCA | C>T | A/T+1 | G/C-1 | CG>TA | CG>TA | GAAAACCAA | CAATACA | GGT | CCA | C | A | C>T | C>T_CCA | CG>TA_CAATACA | 7 | TRUE | FALSE |

| | | | | | | | | | | | | | | | | | | | | | | | | | | | | | | | | | | |
|---|---|---|---|---|---|---|---|---|---|---|---|---|---|---|---|---|---|---|---|---|---|---|---|---|---|---|---|---|---|---|---|---|---|---|
| 173_v.txt | 32967 | 32967-SNV- | SNV | 1 | G | A | 33 | 12003 | 0.27493127 | 62.4242424 | rnr1D57N | FALSE | CAN | JSY14 | 3A | rnr1D57N-Re | CGC | G>A | A/T+1 | G/C-1 | CG>TA | CG>TA | TTGGCCGCA | GCACCAA | GCG | GCG | C | T | C>T | C>T_GCG | CG>TA_GCACCAA | 7 | TRUE | FALSE |
| 173_v.txt | 33012 | 33012-SNV- | SNV | 1 | G | A | 27 | 12003 | 0.22494376 | 56.2962963 | rnr1D57N | FALSE | CAN | JSY14 | 3A | rnr1D57N-Re | AGA | G>A | A/T+1 | G/C-1 | CG>TA | CG>TA | GTGAAAGAG | GAGGATG | TCT | TCT | C | T | C>T | C>T_TCT | CG>TA_GAGGATG | 7 | TRUE | FALSE |
| 173_v.txt | 33070 | 33070-SNV- | SNV | 1 | C | T | 17 | 12003 | 0.14163126 | 54.5294118 | rnr1D57N | FALSE | CAN | JSY14 | 3A | rnr1D57N-Re | GCC | C>T | A/T+1 | G/C-1 | CG>TA | CG>TA | AATATGCCA | CCAAAGA | CGG | GCC | C | A | C>T | C>T_GCC | CG>TA_CCAAAGA | 7 | TRUE | FALSE |
| 173_v.txt | 33111 | 33111-SNV- | SNV | 1 | G | A | 18 | 12003 | 0.14996251 | 36.2777778 | rnr1D57N | FALSE | CAN | JSY14 | 3A | rnr1D57N-Re | TGG | G>A | A/T+1 | G/C-1 | CG>TA | CG>TA | CCCACTGGG | GGGCCGG | ACC | ACC | C | T | C>T | C>T_ACC | CG>TA_GGGCCGG | 7 | TRUE | FALSE |
| 173_v.txt | 33118 | 33118-SNV- | SNV | 1 | C | T | 18 | 12003 | 0.14996251 | 43.3333333 | rnr1D57N | FALSE | CAN | JSY14 | 3A | rnr1D57N-Re | GCG | C>T | A/T+1 | G/C-1 | CG>TA | CG>TA | GGCCGGCGT | CGTTGGT | CGC | GCG | C | A | C>T | C>T_GCG | CG>TA_CGTTGGT | 7 | TRUE | FALSE |
| 173_v.txt | 33153 | 33153-SNV- | SNV | 1 | C | T | 66 | 12003 | 0.54986253 | 37.0909091 | rnr1D57N | FALSE | CAN | JSY14 | 3A | rnr1D57N-Re | ACC | C>T | A/T+1 | G/C-1 | CG>TA | CG>TA | AAAAGACCT | CCTGTAC | TGG | ACC | C | A | C>T | C>T_ACC | CG>TA_CCTGTAC | 7 | TRUE | FALSE |
| 173_v.txt | 33258 | 33258-SNV- | SNV | 1 | C | T | 18 | 7739 | 0.23258819 | 40.1666667 | rnr1D57N | FALSE | CAN | JSY14 | 3A | rnr1D57N-Re | GCC | C>T | A/T+1 | G/C-1 | CG>TA | CG>TA | CCTATGCCAT | CCATCCT | CGG | GCC | C | A | C>T | C>T_GCC | CG>TA_CCATCCT | 7 | TRUE | FALSE |
| 173_v.txt | 33276 | 33276-SNV- | SNV | 1 | G | A | 16 | 7739 | 0.20674506 | 48 | rnr1D57N | FALSE | CAN | JSY14 | 3A | rnr1D57N-Re | CGT | G>A | A/T+1 | G/C-1 | CG>TA | CG>TA | GAGAACGTA | GTATCCT | GCA | GCA | C | T | C>T | C>T_GCA | CG>TA_GTATCCT | 7 | TRUE | FALSE |
| 173_v.txt | 33392 | 33392-SNV- | SNV | 1 | G | A | 14 | 7739 | 0.18090193 | 55.1428571 | rnr1D57N | FALSE | CAN | JSY14 | 3A | rnr1D57N-Re | CGT | G>A | A/T+1 | G/C-1 | CG>TA | CG>TA | AACGTCGTG | GTGAAAG | GCA | GCA | C | T | C>T | C>T_GCA | CG>TA_GTGAAAG | 7 | TRUE | FALSE |
| 173_v.txt | 33474 | 33474-SNV- | SNV | 1 | C | T | 7 | 7739 | 0.09045096 | 37 | rnr1D57N | FALSE | CAN | JSY14 | 3A | rnr1D57N-Re | GCC | C>T | A/T+1 | G/C-1 | CG>TA | CG>TA | GCTATGCCT | CCTTTTT | CGG | GCC | C | A | C>T | C>T_GCC | CG>TA_CCTTTTT | 7 | TRUE | FALSE |
| 173_v.txt | 32200 | 32200-SNV- | SNV | 1 | G | C | 4 | 7041 | 0.05681011 | 44.25 | rnr1D57N | FALSE | CAN | JSY14 | 3A | rnr1D57N-Re | GGA | G>C | G/C+1 | G/C-1 | CG>GC | CG>GC | TGTATGGAA | GAACACC | CCT | CCT | C | G | C>G | C>G_CCT | CG>GC_GAACACC | 7 | TRUE | FALSE |
| 173_v.txt | 32271 | 32271-SNV- | SNV | 1 | C | G | 6 | 9772 | 0.06139992 | 31 | rnr1D57N | FALSE | CAN | JSY14 | 3A | rnr1D57N-Re | ACG | C>G | G/C+1 | G/C-1 | CG>GC | CG>GC | AAAATACGG | CGGGAAC | TGC | ACG | C | C | C>G | C>G_ACG | CG>GC_CGGGAAC | 7 | TRUE | FALSE |
| 173_v.txt | 32829 | 32829-SNV- | SNV | 1 | G | C | 45 | 13670 | 0.329188 | 45.6 | rnr1D57N | FALSE | CAN | JSY14 | 3A | rnr1D57N-Re | TGT | G>C | G/C+1 | G/C-1 | CG>GC | CG>GC | ATTATTGTGA | GTGATAA | ACA | ACA | C | G | C>G | C>G_ACA | CG>GC_GTGATAA | 7 | TRUE | FALSE |
| 173_v.txt | 33172 | 33172-SNV- | SNV | 1 | C | G | 19 | 12003 | 0.15829376 | 37.631579 | rnr1D57N | FALSE | CAN | JSY14 | 3A | rnr1D57N-Re | CCA | C>G | G/C+1 | G/C-1 | CG>GC | CG>GC | TACCACCAA | CAAGGGC | GGT | CCA | C | C | C>G | C>G_CCA | CG>GC_CAAGGGC | 7 | TRUE | FALSE |
| 173_v.txt | 33205 | 33205-SNV- | SNV | 1 | G | C | 3 | 7739 | 0.0387647 | 46.3333333 | rnr1D57N | FALSE | CAN | JSY14 | 3A | rnr1D57N-Re | AGC | G>C | G/C+1 | G/C-1 | CG>GC | CG>GC | GCTTAAGCT | GCTCTCT | TCG | TCG | C | G | C>G | C>G_TCG | CG>GC_GCTCTCT | 7 | TRUE | FALSE |
| 173_v.txt | 33354 | 33354-SNV- | SNV | 1 | G | C | 12 | 7739 | 0.15505879 | 61.1666667 | rnr1D57N | FALSE | CAN | JSY14 | 3A | rnr1D57N-Re | TGA | G>C | G/C+1 | G/C-1 | CG>GC | CG>GC | GGTATTGAC | GACCCAC | ACT | ACT | C | G | C>G | C>G_ACT | CG>GC_GACCCAC | 7 | TRUE | FALSE |
| 173_v.txt | 31829 | 31829-SNV- | SNV | 1 | G | T | 6 | 4049 | 0.14818474 | 47.1666667 | rnr1D57N | FALSE | CAN | JSY14 | 3A | rnr1D57N-Re | TGC | G>T | A/T+1 | G/C-1 | CG>AT | CG>AT | GAATATGCA | GCATTGA | ACG | ACG | C | A | C>A | C>A_ACG | CG>AT_GCATTGA | 7 | TRUE | FALSE |
| 173_v.txt | 32008 | 32008-SNV- | SNV | 1 | C | A | 965 | 7041 | 13.7054396 | 57.6 | rnr1D57N | FALSE | CAN | JSY14 | 3A | rnr1D57N-Re | TCG | C>A | A/T+1 | G/C-1 | CG>AT | CG>AT | GTAACTCGT | CGTCACG | AGC | TCG | C | T | C>A | C>A_TCG | CG>AT_CGTCACG | 7 | TRUE | FALSE |
| 173_v.txt | 32575 | 32575-SNV- | SNV | 1 | G | T | 66 | 13670 | 0.48280907 | 37.1212121 | rnr1D57N | FALSE | CAN | JSY14 | 3A | rnr1D57N-Re | TGA | G>T | A/T+1 | G/C-1 | CG>AT | CG>AT | TACCTTGAA | GAAATGT | ACT | ACT | C | A | C>A | C>A_ACT | CG>AT_GAAATGT | 7 | TRUE | FALSE |
| 173_v.txt | 32692 | 32692-SNV- | SNV | 1 | C | A | 9 | 13670 | 0.0658376 | 58.3333333 | rnr1D57N | FALSE | CAN | JSY14 | 3A | rnr1D57N-Re | CCA | C>A | A/T+1 | G/C-1 | CG>AT | CG>AT | GGAATCCAA | CAACTGG | GGT | CCA | C | T | C>A | C>A_CCA | CG>AT_CAACTGG | 7 | TRUE | FALSE |
| 173_v.txt | 32713 | 32713-SNV- | SNV | 1 | C | A | 29 | 13670 | 0.21214338 | 55.6551724 | rnr1D57N | FALSE | CAN | JSY14 | 3A | rnr1D57N-Re | GCA | C>A | A/T+1 | G/C-1 | CG>AT | CG>AT | CCCCAGCAC | CACCACA | CGT | GCA | C | T | C>A | C>A_GCA | CG>AT_CACCACA | 7 | TRUE | FALSE |
| 173_v.txt | 32783 | 32783-SNV- | SNV | 1 | G | T | 8 | 13670 | 0.05852231 | 50.875 | rnr1D57N | FALSE | CAN | JSY14 | 3A | rnr1D57N-Re | AGA | G>T | A/T+1 | G/C-1 | CG>AT | CG>AT | GACCCAGAA | GAACTCG | TCT | TCT | C | A | C>A | C>A_TCT | CG>AT_GAACTCG | 7 | TRUE | FALSE |
| 173_v.txt | 32954 | 32954-SNV- | SNV | 1 | G | T | 13 | 12003 | 0.10830626 | 52 | rnr1D57N | FALSE | CAN | JSY14 | 3A | rnr1D57N-Re | TGT | G>T | A/T+1 | G/C-1 | CG>AT | CG>AT | ATACATGTA | GTAACCA | ACA | ACA | C | A | C>A | C>A_ACA | CG>AT_GTAACCA | 7 | TRUE | FALSE |
| 173_v.txt | 32997 | 32997-SNV- | SNV | 1 | G | T | 24 | 12003 | 0.19995001 | 61.875 | rnr1D57N | FALSE | CAN | JSY14 | 3A | rnr1D57N-Re | TGA | G>T | A/T+1 | G/C-1 | CG>AT | CG>AT | CTTTGTGAG | GAGAAAA | ACT | ACT | C | A | C>A | C>A_ACT | CG>AT_GAGAAAA | 7 | TRUE | FALSE |
| 173_v.txt | 33160 | 33160-SNV- | SNV | 1 | C | A | 89 | 12003 | 0.7414813 | 37.3820225 | rnr1D57N | FALSE | CAN | JSY14 | 3A | rnr1D57N-Re | CCA | C>A | A/T+1 | G/C-1 | CG>AT | CG>AT | CTGTACCAA | CAATAGT | GGT | CCA | C | T | C>A | C>A_CCA | CG>AT_CAATAGT | 7 | TRUE | FALSE |
| 173_v.txt | 33195 | 33195-SNV- | SNV | 1 | C | A | 32 | 7740 | 0.41343669 | 36.71875 | rnr1D57N | FALSE | CAN | JSY14 | 3A | rnr1D57N-Re | TCT | C>A | A/T+1 | G/C-1 | CG>AT | CG>AT | ATATGTCTTT | CTTTGCT | AGA | TCT | C | T | C>A | C>A_TCT | CG>AT_CTTTGCT | 7 | TRUE | FALSE |
| 173_v.txt | 33325 | 33325-SNV- | SNV | 1 | C | A | 11 | 7739 | 0.14213723 | 43.9090909 | rnr1D57N | FALSE | CAN | JSY14 | 3A | rnr1D57N-Re | TCT | C>A | A/T+1 | G/C-1 | CG>AT | CG>AT | ACAATTCTTT | CTTTACT | AGA | TCT | C | T | C>A | C>A_TCT | CG>AT_CTTTACT | 7 | TRUE | FALSE |
| 173_v.txt | 33474 | 33474-SNV- | SNV | 1 | C | A | 22 | 7739 | 0.28427445 | 32.6363636 | rnr1D57N | FALSE | CAN | JSY14 | 3A | rnr1D57N-Re | GCC | C>A | A/T+1 | G/C-1 | CG>AT | CG>AT | GCTATGCCT | CCTTTTT | CGG | GCC | C | T | C>A | C>A_GCC | CG>AT_CCTTTTT | 7 | TRUE | FALSE |
| 173_v.txt | 33475 | 33475-SNV- | SNV | 1 | C | A | 10 | 7739 | 0.12921566 | 37.4 | rnr1D57N | FALSE | CAN | JSY14 | 3A | rnr1D57N-Re | CCT | C>A | A/T+1 | G/C-1 | CG>AT | CG>AT | CTATGCCTTT | CTTTTTT | GGA | CCT | C | T | C>A | C>A_CCT | CG>AT_CTTTTTT | 7 | TRUE | FALSE |
| 173_v.txt | 32717 | 32717-Repla | Replacement | 4 | A | GTTC | 25 | 13670 | 0.18288222 | 61.02 | rnr1D57N | FALSE | CAN | JSY14 | 3A | rnr1D57N-Re | CAC | nope | >1bp-ins | A/T-1 | none | Replac. | AGCACCACA | ACAAACC | GTG | GTG | T | none | T>none | T>none_GTG | Replac._ACAAACC | 7 | TRUE | FALSE |
| 173_v.txt | 32081 | 32081-Insert | Insertion | 1 | - | A | 11 | 7041 | 0.15622781 | 63.7272727 | rnr1D57N | FALSE | CAN | JSY14 | 3A | rnr1D57N-Re | CAA | nope | A/T+1 | >1bpdel | none | A/T+1 | CCATGCAAA | AAAAAAG | GTT | CAA | - | T | NA | NA_CAA | A/T+1_AAAAAAG | 7 | TRUE | FALSE |
| 173_v.txt | 32498 | 32498-Insert | Insertion | 1 | - | T | 6 | 9772 | 0.06139992 | 46.6666667 | rnr1D57N | FALSE | CAN | JSY14 | 3A | rnr1D57N-Re | CTT | nope | A/T+1 | >1bpdel | none | A/T+1 | AACAACTTTT | TTTTTTG | GAA | CTT | - | A | NA | NA_CTT | A/T+1_TTTTTTG | 7 | TRUE | FALSE |
| 173_v.txt | 32597 | 32597-Insert | Insertion | 2 | - | AA | 8 | 13670 | 0.05852231 | 41 | rnr1D57N | FALSE | CAN | JSY14 | 3A | rnr1D57N-Re | TCA | nope | >1bp-ins | >1bpdel | none | >1bp-ins | GTTAATCAA | CAAAGAG | AGT | TCA | - | none | NA | NA_TCA | >1bp-ins_CAAAGAG | 7 | TRUE | FALSE |
| 173_v.txt | 32597 | 32597-Insert | Insertion | 3 | - | AAA | 1 | 13670 | 7.32E-03 | 38 | rnr1D57N | FALSE | CAN | JSY14 | 3A | rnr1D57N-Re | TCA | nope | >1bp-ins | >1bpdel | none | >1bp-ins | GTTAATCAA | CAAAGAG | AGT | TCA | - | none | NA | NA_TCA | >1bp-ins_CAAAGAG | 7 | TRUE | FALSE |
| 173_v.txt | 32715 | 32715-Insert | Insertion | 3 | - | AAT | 25 | 13670 | 0.18288222 | 59.84 | rnr1D57N | FALSE | CAN | JSY14 | 3A | rnr1D57N-Re | ACC | nope | >1bp-ins | >1bpdel | none | >1bp-ins | CCAGCACCA | CCACAAA | TGG | ACC | - | none | NA | NA_ACC | >1bp-ins_CCACAAA | 7 | TRUE | FALSE |
| 173_v.txt | 32721 | 32721-Insert | Insertion | 1 | - | G | 25 | 13670 | 0.18288222 | 61.4 | rnr1D57N | FALSE | CAN | JSY14 | 3A | rnr1D57N-Re | AAC | nope | G/C+1 | >1bpdel | none | G/C+1 | CCACAAACC | ACCATAC | TTG | AAC | - | C | NA | NA_AAC | G/C+1_ACCATAC | 7 | TRUE | FALSE |
| 173_v.txt | 32722 | 32722-Insert | Insertion | 1 | - | G | 25 | 13670 | 0.18288222 | 61.96 | rnr1D57N | FALSE | CAN | JSY14 | 3A | rnr1D57N-Re | ACC | nope | G/C+1 | >1bpdel | none | G/C+1 | CACAAACCA | CCATACA | TGG | ACC | - | C | NA | NA_ACC | G/C+1_CCATACA | 7 | TRUE | FALSE |
| 173_v.txt | 32842 | 32842-Insert | Insertion | 1 | - | A | 38 | 13670 | 0.27798098 | 41.9473684 | rnr1D57N | FALSE | CAN | JSY14 | 3A | rnr1D57N-Re | CAA | nope | A/T+1 | >1bpdel | none | A/T+1 | TTACCCAAA | AAAAAAT | GTT | CAA | - | T | NA | NA_CAA | A/T+1_AAAAAAT | 7 | TRUE | FALSE |
| 173_v.txt | 32337 | 32337-Deleti | Deletion | 2 | AT | - | 13 | 9772 | 0.13303316 | 63.9230769 | rnr1D57N | FALSE | CAN | JSY14 | 3A | rnr1D57N-Re | GAT | nope | >1bp-ins | >1bpdel | none | >1bpdel | TTGAAGATA | ATATGTG | CTA | GAT | AT | none | NA | NA_GAT | >1bpdel_ATATGTG | 7 | TRUE | FALSE |
| 173_v.txt | 33094 | 33094-Deleti | Deletion | 2 | AT | - | 5 | 12003 | 0.04165625 | 48.4 | rnr1D57N | FALSE | CAN | JSY14 | 3A | rnr1D57N-Re | GAT | nope | >1bp-ins | >1bpdel | none | >1bpdel | AATATGATA | ATATAAG | CTA | GAT | AT | none | NA | NA_GAT | >1bpdel_ATATAAG | 7 | TRUE | FALSE |
| 173_v.txt | 31663 | 31663-Deleti | Deletion | 1 | A | - | 3 | 4049 | 0.07409237 | 38 | rnr1D57N | FALSE | CAN | JSY14 | 3A | rnr1D57N-Re | GAA | nope | >1bp-ins | A/T-1 | none | A/T-1 | GCGTGGAAA | AAATGTG | CTT | CTT | T | none | T>none | T>none_CTT | A/T-1_AAATGTG | 7 | TRUE | FALSE |
| 173_v.txt | 31971 | 31971-Deleti | Deletion | 1 | G | - | 4334 | 7041 | 61.5537566 | 37.3382557 | rnr1D57N | FALSE | CAN | JSY14 | 3A | rnr1D57N-Re | AGC | nope | >1bp-ins | G/C-1 | none | G/C-1 | TAATAAGCC | GCCAAGC | TCG | TCG | C | none | C>none | C>none_TCG | G/C-1_GCCAAGC | 7 | TRUE | FALSE |
| 173_v.txt | 32081 | 32081-Deleti | Deletion | 1 | A | - | 96 | 7041 | 1.36344269 | 63.083334 | rnr1D57N | FALSE | CAN | JSY14 | 3A | rnr1D57N-Re | CAA | nope | >1bp-ins | A/T-1 | none | A/T-1 | CCATGCAAA | AAAAAAG | GTT | GTT | T | none | T>none | T>none_GTT | A/T-1_AAAAAAG | 7 | TRUE | FALSE |
| 173_v.txt | 32381 | 32381-Deleti | Deletion | 1 | A | - | 4 | 9772 | 0.04093328 | 63.5 | rnr1D57N | FALSE | CAN | JSY14 | 3A | rnr1D57N-Re | TAA | nope | >1bp-ins | A/T-1 | none | A/T-1 | AATAATAAA | AAAGGGA | ATT | ATT | T | none | T>none | T>none_ATT | A/T-1_AAAGGGA | 7 | TRUE | FALSE |
| 173_v.txt | 32442 | 32442-Deleti | Deletion | 1 | A | - | 37 | 9772 | 0.37863283 | 55.945946 | rnr1D57N | FALSE | CAN | JSY14 | 3A | rnr1D57N-Re | TAA | nope | >1bp-ins | A/T-1 | none | A/T-1 | GGAACTAAA | AAAAGTC | ATT | ATT | T | none | T>none | T>none_ATT | A/T-1_AAAAGTC | 7 | TRUE | FALSE |
| 173_v.txt | 32498 | 32498-Deleti | Deletion | 1 | T | - | 10 | 9772 | 0.1023332 | 41.6 | rnr1D57N | FALSE | CAN | JSY14 | 3A | rnr1D57N-Re | CTT | nope | >1bp-ins | A/T-1 | none | A/T-1 | AACAACTTTT | TTTTTTG | GAA | CTT | T | none | nope | nope_CTT | A/T-1_TTTTTTG | 7 | TRUE | FALSE |
| 173_v.txt | 32624 | 32624-Deleti | Deletion | 1 | C | - | 16 | 13670 | 0.11704462 | 49.875 | rnr1D57N | FALSE | CAN | JSY14 | 3A | rnr1D57N-Re | TCC | nope | >1bp-ins | G/C-1 | none | G/C-1 | GAACCTCCCT | CCCTTCG | AGG | TCC | C | none | nope | nope_TCC | G/C-1_CCCTTCG | 7 | TRUE | FALSE |
| 173_v.txt | 32631 | 32631-Deleti | Deletion | 1 | T | - | 43 | 13670 | 0.31455743 | 52.2790698 | rnr1D57N | FALSE | CAN | JSY14 | 3A | rnr1D57N-Re | GTT | nope | >1bp-ins | A/T-1 | none | A/T-1 | CCTTCGTTTT | TTTTTAT | CAA | GTT | T | none | nope | nope_GTT | A/T-1_TTTTTAT | 7 | TRUE | FALSE |
| 173_v.txt | 32637 | 32637-Deleti | Deletion | 1 | T | - | 23 | 13670 | 0.16825165 | 52.0869565 | rnr1D57N | FALSE | CAN | JSY14 | 3A | rnr1D57N-Re | ATC | nope | >1bp-ins | A/T-1 | none | A/T-1 | TTTTTATCCT | TCCTTAG | TAG | ATC | T | none | nope | nope_ATC | A/T-1_TCCTTAG | 7 | TRUE | FALSE |
| 173_v.txt | 32658 | 32658-Deleti | Deletion | 1 | C | - | 34 | 13670 | 0.24871982 | 58.8235294 | rnr1D57N | FALSE | CAN | JSY14 | 3A | rnr1D57N-Re | ACC | nope | >1bp-ins | G/C-1 | none | G/C-1 | CCTGGACCC | CCCCAGG | TGG | ACC | C | none | nope | nope_ACC | G/C-1_CCCCAGG | 7 | TRUE | FALSE |
| 173_v.txt | 32663 | 32663-Deleti | Deletion | 1 | G | - | 8 | 13670 | 0.05852231 | 60.125 | rnr1D57N | FALSE | CAN | JSY14 | 3A | rnr1D57N-Re | AGG | nope | >1bp-ins | G/C-1 | none | G/C-1 | ACCCCAGGC | GGCACCT | TCC | TCC | C | none | C>none | C>none_TCC | G/C-1_GGCACCT | 7 | TRUE | FALSE |
| 173_v.txt | 32726 | 32726-Deleti | Deletion | 1 | A | - | 25 | 13670 | 0.18288222 | 61.32 | rnr1D57N | FALSE | CAN | JSY14 | 3A | rnr1D57N-Re | TAC | nope | >1bp-ins | A/T-1 | none | A/T-1 | AACCATACA | ACAAAAA | ATG | ATG | T | none | T>none | T>none_ATG | A/T-1_ACAAAAA | 7 | TRUE | FALSE |
| 173_v.txt | 32829 | 32829-Deleti | Deletion | 1 | G | - | 2 | 13670 | 0.01463058 | 38 | rnr1D57N | FALSE | CAN | JSY14 | 3A | rnr1D57N-Re | TGT | nope | >1bp-ins | G/C-1 | none | G/C-1 | ATTATTGTGA | GTGATAA | ACA | ACA | C | none | C>none | C>none_ACA | G/C-1_GTGATAA | 7 | TRUE | FALSE |
| 173_v.txt | 32839 | 32839-Deleti | Deletion | 1 | C | - | 14 | 13670 | 0.10241405 | 41.4285714 | rnr1D57N | FALSE | CAN | JSY14 | 3A | rnr1D57N-Re | ACC | nope | >1bp-ins | G/C-1 | none | G/C-1 | TAATTACCCA | CCCAAAA | TGG | ACC | C | none | nope | nope_ACC | G/C-1_CCCAAAA | 7 | TRUE | FALSE |
| 173_v.txt | 32842 | 32842-Deleti | Deletion | 1 | A | - | 14 | 13670 | 0.10241405 | 44.6428571 | rnr1D57N | FALSE | CAN | JSY14 | 3A | rnr1D57N-Re | CAA | nope | >1bp-ins | A/T-1 | none | A/T-1 | TTACCCAAA | AAAAAAT | GTT | GTT | T | none | T>none | T>none_GTT | A/T-1_AAAAAAT | 7 | TRUE | FALSE |
| 173_v.txt | 32884 | 32884-Deleti | Deletion | 1 | A | - | 7 | 12003 | 0.05831875 | 37.8571429 | rnr1D57N | FALSE | CAN | JSY14 | 3A | rnr1D57N-Re | CAA | nope | >1bp-ins | A/T-1 | none | A/T-1 | ACGTCCAAA | AAAATTG | GTT | GTT | T | none | T>none | T>none_GTT | A/T-1_AAAATTG | 7 | TRUE | FALSE |
| 173_v.txt | 32924 | 32924-Deleti | Deletion | 1 | A | - | 15 | 12003 | 0.12496876 | 49.2 | rnr1D57N | FALSE | CAN | JSY14 | 3A | rnr1D57N-Re | CAA | nope | >1bp-ins | A/T-1 | none | A/T-1 | CAGGGCAAA | AAAAGTG | GTT | GTT | T | none | T>none | T>none_GTT | A/T-1_AAAAGTG | 7 | TRUE | FALSE |
| 173_v.txt | 32941 | 32941-Deleti | Deletion | 1 | A | - | 6 | 12003 | 0.0499875 | 64 | rnr1D57N | FALSE | CAN | JSY14 | 3A | rnr1D57N-Re | GAA | nope | >1bp-ins | A/T-1 | none | A/T-1 | CCCAAGAAA | AAAACCA | CTT | CTT | T | none | T>none | T>none_CTT | A/T-1_AAAACCA | 7 | TRUE | FALSE |
| 173_v.txt | 32978 | 32978-Deleti | Deletion | 1 | T | - | 6 | 12003 | 0.0499875 | 58 | rnr1D57N | FALSE | CAN | JSY14 | 3A | rnr1D57N-Re | CTG | nope | >1bp-ins | A/T-1 | none | A/T-1 | AAATGCTGG | TGGAGAA | GAC | CTG | T | none | nope | nope_CTG | A/T-1_TGGAGAA | 7 | TRUE | FALSE |
| 173_v.txt | 33072 | 33072-Deleti | Deletion | 1 | A | - | 13 | 12003 | 0.10830626 | 52.3076923 | rnr1D57N | FALSE | CAN | JSY14 | 3A | rnr1D57N-Re | CAA | nope | >1bp-ins | A/T-1 | none | A/T-1 | TATGCCAAA | AAAGAAC | GTT | GTT | T | none | T>none | T>none_GTT | A/T-1_AAAGAAC | 7 | TRUE | FALSE |
| 173_v.txt | 33111 | 33111-Deleti | Deletion | 1 | G | - | 9 | 12003 | 0.07498126 | 39 | rnr1D57N | FALSE | CAN | JSY14 | 3A | rnr1D57N-Re | TGG | nope | >1bp-ins | G/C-1 | none | G/C-1 | CCCACTGGG | GGGCCGG | ACC | ACC | C | none | C>none | C>none_ACC | G/C-1_GGGCCGG | 7 | TRUE | FALSE |
| 173_v.txt | 33258 | 33258-Deleti | Deletion | 1 | C | - | 1 | 7739 | 0.01292157 | 38 | rnr1D57N | FALSE | CAN | JSY14 | 3A | rnr1D57N-Re | GCC | nope | >1bp-ins | G/C-1 | none | G/C-1 | CCTATGCCAT | CCATCCT | CGG | GCC | C | none | nope | nope_GCC | G/C-1_CCATCCT | 7 | TRUE | FALSE |
| 173_v.txt | 33474 | 33474-Deleti | Deletion | 1 | C | - | 24 | 7739 | 0.31011759 | 32.625 | rnr1D57N | FALSE | CAN | JSY14 | 3A | rnr1D57N-Re | GCC | nope | >1bp-ins | G/C-1 | none | G/C-1 | GCTATGCCT | CCTTTTT | CGG | GCC | C | none | nope | nope_GCC | G/C-1_CCTTTTT | 7 | TRUE | FALSE |
| 8_v.txt | 31723 | 31723-SNV- | SNV | 1 | A | C | 6 | 17740 | 0.03382187 | 61 | rnr1D57N_per | FALSE | ARG | JSY14 | 1 | rnr1D57N_pe | CAA | A>C | G/C+1 | A/T-1 | TA>GC | TA>GC | TGTCCCAAA | AAAAAGT | GTT | GTT | T | G | T>G | T>G_GTT | TA>GC_AAAAAGT | 4 | FALSE | FALSE |
| 8_v.txt | 31840 | 31840-SNV- | SNV | 1 | A | C | 5 | 17740 | 0.02818489 | 58.8 | rnr1D57N_per | FALSE | ARG | JSY14 | 1 | rnr1D57N_pe | AAG | A>C | G/C+1 | A/T-1 | TA>GC | TA>GC | GAAATAAGA | AGATCCA | TTC | TTC | T | G | T>G | T>G_TTC | TA>GC_AGATCCA | 4 | FALSE | FALSE |
| 8_v.txt | 32669 | 32669-SNV-T | SNV | 1 | T | G | 3 | 22419 | 0.01338151 | 61.9991214 | rnr1D57N_per | FALSE | ARG | JSY14 | 1 | rnr1D57N_pe | CTG | T>G | G/C+1 | A/T-1 | TA>GC | TA>GC | GGCACCTGG | TGGGTTT | GAC | CTG | T | C | T>G | T>G_CTG | TA>GC_TGGGTTT | 4 | FALSE | FALSE |
| 8_v.txt | 32842 | 32842-SNV- | SNV | 1 | A | C | 4 | 22419 | 0.01784201 | 37.5000277 | rnr1D57N_per | FALSE | ARG | JSY14 | 1 | rnr1D57N_pe | CAA | A>C | G/C+1 | A/T-1 | TA>GC | TA>GC | TTACCCAAA | AAAAAAT | GTT | GTT | T | G | T>G | T>G_GTT | TA>GC_AAAAAAT | 4 | FALSE | FALSE |
| 8_v.txt | 33099 | 33099-SNV- | SNV | 1 | A | C | 8 | 38692 | 0.02067611 | 34 | rnr1D57N_per | FALSE | ARG | JSY14 | 1 | rnr1D57N_pe | AAG | A>C | G/C+1 | A/T-1 | TA>GC | TA>GC | GATATAAGA | AGAGCGC | TTC | TTC | T | G | T>G | T>G_TTC | TA>GC_AGAGCGC | 4 | FALSE | FALSE |
| 8_v.txt | 33479 | 33479-SNV-T | SNV | 1 | T | G | 5 | 19048 | 0.02624948 | 38 | rnr1D57N_per | FALSE | ARG | JSY14 | 1 | rnr1D57N_pe | TTT | T>G | G/C+1 | A/T-1 | TA>GC | TA>GC | GCCTTTTTTT | TTTTTTT | AAA | TTT | T | C | T>G | T>G_TTT | TA>GC_TTTTTTT | 4 | FALSE | FALSE |
| 8_v.txt | 31840 | 31840-SNV- | SNV | 1 | A | G | 10 | 17740 | 0.05636979 | 53.4 | rnr1D57N_per | FALSE | ARG | JSY14 | 1 | rnr1D57N_pe | AAG | A>G | G/C+1 | A/T-1 | TA>CG | TA>CG | GAAATAAGA | AGATCCA | TTC | TTC | T | C | T>C | T>C_TTC | TA>CG_AGATCCA | 4 | FALSE | FALSE |
| 8_v.txt | 32081 | 32081-SNV- | SNV | 1 | A | G | 5 | 32081 | 0.01558555 | 45.999929 | rnr1D57N_per | FALSE | ARG | JSY14 | 1 | rnr1D57N_pe | CAA | A>G | G/C+1 | A/T-1 | TA>CG | TA>CG | CCATGCAAA | AAAAAAG | GTT | GTT | T | C | T>C | T>C_GTT | TA>CG_AAAAAAG | 4 | FALSE | FALSE |
| 8_v.txt | 32086 | 32086-SNV- | SNV | 1 | A | G | 28 | 32081 | 0.08727908 | 57.6071429 | rnr1D57N_per | FALSE | ARG | JSY14 | 1 | rnr1D57N_pe | AAG | A>G | G/C+1 | A/T-1 | TA>CG | TA>CG | CAAAAAAGC | AGCCTGC | TTC | TTC | T | C | T>C | T>C_TTC | TA>CG_AGCCTGC | 4 | FALSE | FALSE |
| 8_v.txt | 32927 | 32927-SNV- | SNV | 1 | A | G | 38 | 38692 | 0.09821152 | 50.1578947 | rnr1D57N_per | FALSE | ARG | JSY14 | 1 | rnr1D57N_pe | AAG | A>G | G/C+1 | A/T-1 | TA>CG | TA>CG | GGCAAAAGT | AGTGATT | TTC | TTC | T | C | T>C | T>C_TTC | TA>CG_AGTGATT | 4 | FALSE | FALSE |
| 8_v.txt | 33099 | 33099-SNV- | SNV | 1 | A | G | 25 | 38692 | 0.06461284 | 43 | rnr1D57N_per | FALSE | ARG | JSY14 | 1 | rnr1D57N_pe | AAG | A>G | G/C+1 | A/T-1 | TA>CG | TA>CG | GATATAAGA | AGAGCGC | TTC | TTC | T | C | T>C | T>C_TTC | TA>CG_AGAGCGC | 4 | FALSE | FALSE |
| 8_v.txt | 32081 | 32081-SNV- | SNV | 1 | A | T | 1 | 32081 | 3.12E-03 | 63.9996275 | rnr1D57N_per | FALSE | ARG | JSY14 | 1 | rnr1D57N_pe | CAA | A>T | A/T+1 | A/T-1 | TA>AT | TA>AT | CCATGCAAA | AAAAAAG | GTT | GTT | T | A | T>A | T>A_GTT | TA>AT_AAAAAAG | 4 | FALSE | FALSE |
| 8_v.txt | 32669 | 32669-SNV-T | SNV | 1 | T | A | 10 | 22419 | 0.04460502 | 58.4944456 | rnr1D57N_per | FALSE | ARG | JSY14 | 1 | rnr1D57N_pe | CTG | T>A | A/T+1 | A/T-1 | TA>AT | TA>AT | GGCACCTGG | TGGGTTT | GAC | CTG | T | T | T>A | T>A_CTG | TA>AT_TGGGTTT | 4 | FALSE | FALSE |
| 8_v.txt | 33099 | 33099-SNV- | SNV | 1 | A | T | 11 | 38692 | 0.02842965 | 40 | rnr1D57N_per | FALSE | ARG | JSY14 | 1 | rnr1D57N_pe | AAG | A>T | A/T+1 | A/T-1 | TA>AT | TA>AT | GATATAAGA | AGAGCGC | TTC | TTC | T | A | T>A | T>A_TTC | TA>AT_AGAGCGC | 4 | FALSE | FALSE |
| 8_v.txt | 31650 | 31650-SNV- | SNV | 1 | C | T | 26 | 17740 | 0.14656144 | 36.5769231 | rnr1D57N_per | FALSE | ARG | JSY14 | 1 | rnr1D57N_pe | GCG | C>T | A/T+1 | G/C-1 | CG>TA | CG>TA | AGAATGCGA | CGAAATG | CGC | GCG | C | A | C>T | C>T_GCG | CG>TA_CGAAATG | 4 | FALSE | FALSE |
| 8_v.txt | 31658 | 31658-SNV- | SNV | 1 | C | T | 31 | 17740 | 0.17474634 | 31.1933118 | rnr1D57N_per | FALSE | ARG | JSY14 | 1 | rnr1D57N_pe | GCG | C>T | A/T+1 | G/C-1 | CG>TA | CG>TA | AAATGGCGT | CGTGGAA | CGC | GCG | C | A | C>T | C>T_GCG | CG>TA_CGTGGAA | 4 | FALSE | FALSE |
| 8_v.txt | 31817 | 31817-SNV- | SNV | 1 | G | A | 21 | 17740 | 0.11837655 | 48.2857143 | rnr1D57N_per | FALSE | ARG | JSY14 | 1 | rnr1D57N_pe | TGC | G>A | A/T+1 | G/C-1 | CG>TA | CG>TA | AAATCTGCA | GCATCTG | ACG | ACG | C | T | C>T | C>T_ACG | CG>TA_GCATCTG | 4 | FALSE | FALSE |
| 8_v.txt | 31830 | 31830-SNV- | SNV | 1 | C | T | 25 | 17740 | 0.14092447 | 49.24 | rnr1D57N_per | FALSE | ARG | JSY14 | 1 | rnr1D57N_pe | GCA | C>T | A/T+1 | G/C-1 | CG>TA | CG>TA | AATATGCATT | CATTGAA | CGT | GCA | C | A | C>T | C>T_GCA | CG>TA_CATTGAA | 4 | FALSE | FALSE |
| 8_v.txt | 31879 | 31879-SNV- | SNV | 1 | C | T | 20 | 17740 | 0.11273957 | 43.9 | rnr1D57N_per | FALSE | ARG | JSY14 | 1 | rnr1D57N_pe | GCG | C>T | A/T+1 | G/C-1 | CG>TA | CG>TA | TATAGGCGG | CGGCAGC | CGC | GCG | C | A | C>T | C>T_GCG | CG>TA_CGGCAGC | 4 | FALSE | FALSE |
| 8_v.txt | 31882 | 31882-SNV- | SNV | 1 | C | T | 25 | 17740 | 0.14092447 | 41.8 | rnr1D57N_per | FALSE | ARG | JSY14 | 1 | rnr1D57N_pe | GCA | C>T | A/T+1 | G/C-1 | CG>TA | CG>TA | AGGCGGCAG | CAGCAAA | CGT | GCA | C | A | C>T | C>T_GCA | CG>TA_CAGCAAA | 4 | FALSE | FALSE |
| 8_v.txt | 31884 | 31884-SNV- | SNV | 1 | G | A | 20 | 17740 | 0.11273957 | 37.55 | rnr1D57N_per | FALSE | ARG | JSY14 | 1 | rnr1D57N_pe | AGC | G>A | A/T+1 | G/C-1 | CG>TA | CG>TA | GCGGCAGCA | GCAAAGC | TCG | TCG | C | T | C>T | C>T_TCG | CG>TA_GCAAAGC | 4 | FALSE | FALSE |
| 8_v.txt | 31912 | 31912-SNV- | SNV | 1 | C | T | 17 | 17740 | 0.09582864 | 39.1176471 | rnr1D57N_per | FALSE | ARG | JSY14 | 1 | rnr1D57N_pe | GCA | C>T | A/T+1 | G/C-1 | CG>TA | CG>TA | TTGGTGCAA | CAAAAGC | CGT | GCA | C | A | C>T | C>T_GCA | CG>TA_CAAAAGC | 4 | FALSE | FALSE |
| 8_v.txt | 31960 | 31960-SNV- | SNV | 1 | C | T | 40 | 32083 | 0.12467662 | 35.95 | rnr1D57N_per | FALSE | ARG | JSY14 | 1 | rnr1D57N_pe | GCC | C>T | A/T+1 | G/C-1 | CG>TA | CG>TA | ATGTGGCCG | CCGCATA | CGG | GCC | C | A | C>T | C>T_GCC | CG>TA_CCGCATA | 4 | FALSE | FALSE |
| 8_v.txt | 31962 | 31962-SNV- | SNV | 1 | G | A | 68 | 32083 | 0.21195025 | 36.5 | rnr1D57N_per | FALSE | ARG | JSY14 | 1 | rnr1D57N_pe | CGC | G>A | A/T+1 | G/C-1 | CG>TA | CG>TA | GTGGCCGCA | GCATAAT | GCG | GCG | C | T | C>T | C>T_GCG | CG>TA_GCATAAT | 4 | FALSE | FALSE |
| 8_v.txt | 31972 | 31972-SNV- | SNV | 1 | C | T | 30 | 32081 | 0.0935133 | 37.1333333 | rnr1D57N_per | FALSE | ARG | JSY14 | 1 | rnr1D57N_pe | GCC | C>T | A/T+1 | G/C-1 | CG>TA | CG>TA | AATAAGCCA | CCAAGCC | CGG | GCC | C | A | C>T | C>T_GCC | CG>TA_CCAAGCC | 4 | FALSE | FALSE |
| 8_v.txt | 31977 | 31977-SNV- | SNV | 1 | C | T | 32 | 32081 | 0.09974751 | 43.21875 | rnr1D57N_per | FALSE | ARG | JSY14 | 1 | rnr1D57N_pe | GCC | C>T | A/T+1 | G/C-1 | CG>TA | CG>TA | GCCAAGCCG | CCGGGCA | CGG | GCC | C | A | C>T | C>T_GCC | CG>TA_CCGGGCA | 4 | FALSE | FALSE |
| 8_v.txt | 31981 | 31981-SNV- | SNV | 1 | G | A | 33 | 32081 | 0.10286462 | 51.3030303 | rnr1D57N_per | FALSE | ARG | JSY14 | 1 | rnr1D57N_pe | GGC | G>A | A/T+1 | G/C-1 | CG>TA | CG>TA | AGCCGGGCA | GCATTAA | CCG | CCG | C | T | C>T | C>T_CCG | CG>TA_GCATTAA | 4 | FALSE | FALSE |
| 8_v.txt | 32057 | 32057-SNV- | SNV | 1 | C | T | 30 | 32081 | 0.0935133 | 55.8666667 | rnr1D57N_per | FALSE | ARG | JSY14 | 1 | rnr1D57N_pe | GCG | C>T | A/T+1 | G/C-1 | CG>TA | CG>TA | GATGTGCGA | CGAGATT | CGC | GCG | C | A | C>T | C>T_GCG | CG>TA_CGAGATT | 4 | FALSE | FALSE |
| 8_v.txt | 32079 | 32079-SNV- | SNV | 1 | G | A | 38 | 32081 | 0.11845017 | 62.0263158 | rnr1D57N_per | FALSE | ARG | JSY14 | 1 | rnr1D57N_pe | TGC | G>A | A/T+1 | G/C-1 | CG>TA | CG>TA | AACCATGCA | GCAAAAA | ACG | ACG | C | T | C>T | C>T_ACG | CG>TA_GCAAAAA | 4 | FALSE | FALSE |
| 8_v.txt | 32092 | 32092-SNV- | SNV | 1 | C | T | 49 | 32081 | 0.15273838 | 60.1020408 | rnr1D57N_per | FALSE | ARG | JSY14 | 1 | rnr1D57N_pe | GCA | C>T | A/T+1 | G/C-1 | CG>TA | CG>TA | AGCCTGCAA | CAACACC | CGT | GCA | C | A | C>T | C>T_GCA | CG>TA_CAACACC | 4 | FALSE | FALSE |
| 8_v.txt | 32157 | 32157-SNV- | SNV | 1 | G | A | 27 | 32081 | 0.08416197 | 49.1851852 | rnr1D57N_per | FALSE | ARG | JSY14 | 1 | rnr1D57N_pe | AGC | G>A | A/T+1 | G/C-1 | CG>TA | CG>TA | ATGTAAGCC | GCCAAAG | TCG | TCG | C | T | C>T | C>T_TCG | CG>TA_GCCAAAG | 4 | FALSE | FALSE |
| 8_v.txt | 32165 | 32165-SNV- | SNV | 1 | G | A | 62 | 32081 | 0.19326081 | 46.4354839 | rnr1D57N_per | FALSE | ARG | JSY14 | 1 | rnr1D57N_pe | CGC | G>A | A/T+1 | G/C-1 | CG>TA | CG>TA | CAAAGCGCC | GCCAAAT | GCG | GCG | C | T | C>T | C>T_GCG | CG>TA_GCCAAAT | 4 | FALSE | FALSE |
| 8_v.txt | 32173 | 32173-SNV- | SNV | 1 | C | T | 35 | 32081 | 0.10909884 | 50.6 | rnr1D57N_per | FALSE | ARG | JSY14 | 1 | rnr1D57N_pe | GCA | C>T | A/T+1 | G/C-1 | CG>TA | CG>TA | CAAATGCAG | CAGCAGT | CGT | GCA | C | A | C>T | C>T_GCA | CG>TA_CAGCAGT | 4 | FALSE | FALSE |
| 8_v.txt | 32176 | 32176-SNV- | SNV | 1 | C | T | 33 | 32081 | 0.10286462 | 44.6363636 | rnr1D57N_per | FALSE | ARG | JSY14 | 1 | rnr1D57N_pe | GCA | C>T | A/T+1 | G/C-1 | CG>TA | CG>TA | ATGCAGCAG | CAGTAAC | CGT | GCA | C | A | C>T | C>T_GCA | CG>TA_CAGTAAC | 4 | FALSE | FALSE |
| 8_v.txt | 32191 | 32191-SNV- | SNV | 1 | C | T | 24 | 32081 | 0.07481064 | 46.25 | rnr1D57N_per | FALSE | ARG | JSY14 | 1 | rnr1D57N_pe | GCA | C>T | A/T+1 | G/C-1 | CG>TA | CG>TA | AAACTGCAA | CAATGTA | CGT | GCA | C | A | C>T | C>T_GCA | CG>TA_CAATGTA | 4 | FALSE | FALSE |
| 8_v.txt | 32271 | 32271-SNV- | SNV | 1 | C | T | 52 | 26939 | 0.1930287 | 33.3461539 | rnr1D57N_per | FALSE | ARG | JSY14 | 1 | rnr1D57N_pe | ACG | C>T | A/T+1 | G/C-1 | CG>TA | CG>TA | AAAATACGG | CGGGAAC | TGC | ACG | C | A | C>T | C>T_ACG | CG>TA_CGGGAAC | 4 | FALSE | FALSE |
| 8_v.txt | 32299 | 32299-SNV- | SNV | 1 | C | T | 56 | 26939 | 0.20787706 | 51.2142857 | rnr1D57N_per | FALSE | ARG | JSY14 | 1 | rnr1D57N_pe | GCG | C>T | A/T+1 | G/C-1 | CG>TA | CG>TA | AATTTGCGG | CGGCAGA | CGC | GCG | C | A | C>T | C>T_GCG | CG>TA_CGGCAGA | 4 | FALSE | FALSE |
| 8_v.txt | 32344 | 32344-SNV- | SNV | 1 | G | A | 21 | 26939 | 0.0779539 | 63.5714286 | rnr1D57N_per | FALSE | ARG | JSY14 | 1 | rnr1D57N_pe | GGC | G>A | A/T+1 | G/C-1 | CG>TA | CG>TA | TATGTGGCA | GCAAAAC | CCG | CCG | C | T | C>T | C>T_CCG | CG>TA_GCAAAAC | 4 | FALSE | FALSE |
| 8_v.txt | 32466 | 32466-SNV- | SNV | 1 | C | T | 49 | 26939 | 0.18189242 | 47.3469388 | rnr1D57N_per | FALSE | ARG | JSY14 | 1 | rnr1D57N_pe | GCC | C>T | A/T+1 | G/C-1 | CG>TA | CG>TA | AGAGAGCCA | CCAATGT | CGG | GCC | C | A | C>T | C>T_GCC | CG>TA_CCAATGT | 4 | FALSE | FALSE |
| 8_v.txt | 32487 | 32487-SNV- | SNV | 1 | C | T | 25 | 26939 | 0.09280226 | 44.68 | rnr1D57N_per | FALSE | ARG | JSY14 | 1 | rnr1D57N_pe | ACG | C>T | A/T+1 | G/C-1 | CG>TA | CG>TA | AAGATACGG | CGGAAAA | TGC | ACG | C | A | C>T | C>T_ACG | CG>TA_CGGAAAA | 4 | FALSE | FALSE |
| 8_v.txt | 32535 | 32535-SNV- | SNV | 1 | G | A | 35 | 26939 | 0.12992316 | 37.903647 | rnr1D57N_per | FALSE | ARG | JSY14 | 1 | rnr1D57N_pe | TGC | G>A | A/T+1 | G/C-1 | CG>TA | CG>TA | GGGTTTGCA | GCAGCTT | ACG | ACG | C | T | C>T | C>T_ACG | CG>TA_GCAGCTT | 4 | FALSE | FALSE |
| 8_v.txt | 32536 | 32536-SNV- | SNV | 1 | C | T | 49 | 26939 | 0.18189242 | 37.3460344 | rnr1D57N_per | FALSE | ARG | JSY14 | 1 | rnr1D57N_pe | GCA | C>T | A/T+1 | G/C-1 | CG>TA | CG>TA | GGTTTGCAG | CAGCTTC | CGT | GCA | C | A | C>T | C>T_GCA | CG>TA_CAGCTTC | 4 | FALSE | FALSE |
| 8_v.txt | 32590 | 32590-SNV- | SNV | 1 | C | T | 56 | 22419 | 0.24978813 | 36.625 | rnr1D57N_per | FALSE | ARG | JSY14 | 1 | rnr1D57N_pe | GCG | C>T | A/T+1 | G/C-1 | CG>TA | CG>TA | AGGCAGCGT | CGTTAAT | CGC | GCG | C | A | C>T | C>T_GCG | CG>TA_CGTTAAT | 4 | FALSE | FALSE |
| 8_v.txt | 32700 | 32700-SNV- | SNV | 1 | C | T | 40 | 22419 | 0.17842009 | 56.1 | rnr1D57N_per | FALSE | ARG | JSY14 | 1 | rnr1D57N_pe | GCC | C>T | A/T+1 | G/C-1 | CG>TA | CG>TA | ACTGGGCCG | CCGGTAA | CGG | GCC | C | A | C>T | C>T_GCC | CG>TA_CCGGTAA | 4 | FALSE | FALSE |
| 8_v.txt | 32776 | 32776-SNV- | SNV | 1 | C | T | 34 | 22419 | 0.15165708 | 49.0294118 | rnr1D57N_per | FALSE | ARG | JSY14 | 1 | rnr1D57N_pe | GCG | C>T | A/T+1 | G/C-1 | CG>TA | CG>TA | TGGAAGCGA | CGACCCA | CGC | GCG | C | A | C>T | C>T_GCG | CG>TA_CGACCCA | 4 | FALSE | FALSE |
| 8_v.txt | 32798 | 32798-SNV- | SNV | 1 | G | A | 42 | 22419 | 0.1873411 | 42.452381 | rnr1D57N_per | FALSE | ARG | JSY14 | 1 | rnr1D57N_pe | CGT | G>A | A/T+1 | G/C-1 | CG>TA | CG>TA | TTCACCGTA | GTAATAT | GCA | GCA | C | T | C>T | C>T_GCA | CG>TA_GTAATAT | 4 | FALSE | FALSE |
| 8_v.txt | 32860 | 32860-SNV- | SNV | 1 | C | T | 25 | 22416 | 0.11152748 | 37.4 | rnr1D57N_per | FALSE | ARG | JSY14 | 1 | rnr1D57N_pe | GCC | C>T | A/T+1 | G/C-1 | CG>TA | CG>TA | TCCATGCCG | CCGCCAG | CGG | GCC | C | A | C>T | C>T_GCC | CG>TA_CCGCCAG | 4 | FALSE | FALSE |
| 8_v.txt | 32862 | 32862-SNV- | SNV | 1 | G | A | 18 | 22416 | 0.08029979 | 36.3888889 | rnr1D57N_per | FALSE | ARG | JSY14 | 1 | rnr1D57N_pe | CGC | G>A | A/T+1 | G/C-1 | CG>TA | CG>TA | CATGCCGCC | GCCAGTG | GCG | GCG | C | T | C>T | C>T_GCG | CG>TA_GCCAGTG | 4 | FALSE | FALSE |
| 8_v.txt | 32879 | 32879-SNV- | SNV | 1 | C | T | 35 | 38688 | 0.09046733 | 34.9714286 | rnr1D57N_per | FALSE | ARG | JSY14 | 1 | rnr1D57N_pe | ACG | C>T | A/T+1 | G/C-1 | CG>TA | CG>TA | TTTGTACGT | CGTCCAA | TGC | ACG | C | A | C>T | C>T_ACG | CG>TA_CGTCCAA | 4 | FALSE | FALSE |
| 8_v.txt | 32900 | 32900-SNV- | SNV | 1 | G | A | 44 | 38692 | 0.1137186 | 35.6079854 | rnr1D57N_per | FALSE | ARG | JSY14 | 1 | rnr1D57N_pe | GGC | G>A | A/T+1 | G/C-1 | CG>TA | CG>TA | GACTTGGCC | GCCAACT | CCG | CCG | C | T | C>T | C>T_CCG | CG>TA_GCCAACT | 4 | FALSE | FALSE |
| 8_v.txt | 32901 | 32901-SNV- | SNV | 1 | C | T | 93 | 38692 | 0.24035976 | 29.1406514 | rnr1D57N_per | FALSE | ARG | JSY14 | 1 | rnr1D57N_pe | GCC | C>T | A/T+1 | G/C-1 | CG>TA | CG>TA | ACTTGGCCA | CCAACTA | CGG | GCC | C | A | C>T | C>T_GCC | CG>TA_CCAACTA | 4 | FALSE | FALSE |
| 8_v.txt | 32922 | 32922-SNV- | SNV | 1 | G | A | 30 | 38692 | 0.07753541 | 54.5333333 | rnr1D57N_per | FALSE | ARG | JSY14 | 1 | rnr1D57N_pe | GGC | G>A | A/T+1 | G/C-1 | CG>TA | CG>TA | TCCAGGGCA | GCAAAAG | CCG | CCG | C | T | C>T | C>T_CCG | CG>TA_GCAAAAG | 4 | FALSE | FALSE |
| 8_v.txt | 32965 | 32965-SNV- | SNV | 1 | C | T | 48 | 38692 | 0.12405665 | 56.2916667 | rnr1D57N_per | FALSE | ARG | JSY14 | 1 | rnr1D57N_pe | GCC | C>T | A/T+1 | G/C-1 | CG>TA | CG>TA | CATTGGCCG | CCGCACC | CGG | GCC | C | A | C>T | C>T_GCC | CG>TA_CCGCACC | 4 | FALSE | FALSE |
| 8_v.txt | 32967 | 32967-SNV- | SNV | 1 | G | A | 97 | 38692 | 0.25069782 | 59.8969072 | rnr1D57N_per | FALSE | ARG | JSY14 | 1 | rnr1D57N_pe | CGC | G>A | A/T+1 | G/C-1 | CG>TA | CG>TA | TTGGCCGCA | GCACCAA | GCG | GCG | C | T | C>T | C>T_GCG | CG>TA_GCACCAA | 4 | FALSE | FALSE |
| 8_v.txt | 32977 | 32977-SNV- | SNV | 1 | C | T | 34 | 38692 | 0.08787346 | 58.7941177 | rnr1D57N_per | FALSE | ARG | JSY14 | 1 | rnr1D57N_pe | GCT | C>T | A/T+1 | G/C-1 | CG>TA | CG>TA | CAAATGCTG | CTGGAGA | CGA | GCT | C | A | C>T | C>T_GCT | CG>TA_CTGGAGA | 4 | FALSE | FALSE |
| 8_v.txt | 33056 | 33056-SNV- | SNV | 1 | C | T | 43 | 38692 | 0.11113409 | 60.5813954 | rnr1D57N_per | FALSE | ARG | JSY14 | 1 | rnr1D57N_pe | GCG | C>T | A/T+1 | G/C-1 | CG>TA | CG>TA | GGACTGCGT | CGTGACA | CGC | GCG | C | A | C>T | C>T_GCG | CG>TA_CGTGACA | 4 | FALSE | FALSE |

| | | | | | | | | | | | | | | | | | | | | | | | | | | | | | | | | | | |
|---|---|---|---|---|---|---|---|---|---|---|---|---|---|---|---|---|---|---|---|---|---|---|---|---|---|---|---|---|---|---|---|---|---|---|
| 8_v.txt | 33069 | 33069-SNV- | SNV | 1 | G | A | 44 | 38692 | 0.1137186 | 51.659349 | rnr1D57N_per | FALSE | ARG | JSY14 | 1 | rnr1D57N_pe | TGC | G>A | A/T+1 | G/C-1 | CG>TA | CG>TA | GAATATGCC | GCCAAAG | ACG | ACG | C | T | C>T | C>T_ACG | CG>TA_GCCAAAG | 4 | FALSE | FALSE |
| 8_v.txt | 33070 | 33070-SNV- | SNV | 1 | C | T | 40 | 38692 | 0.10338054 | 54.4 | rnr1D57N_per | FALSE | ARG | JSY14 | 1 | rnr1D57N_pe | GCC | C>T | A/T+1 | G/C-1 | CG>TA | CG>TA | AATATGCCA | CCAAAGA | CGG | GCC | C | A | C>T | C>T_GCC | CG>TA_CCAAAGA | 4 | FALSE | FALSE |
| 8_v.txt | 33104 | 33104-SNV- | SNV | 1 | G | A | 45 | 38692 | 0.11630311 | 37.0888889 | rnr1D57N_per | FALSE | ARG | JSY14 | 1 | rnr1D57N_pe | CGC | G>A | A/T+1 | G/C-1 | CG>TA | CG>TA | AAGAGCGCC | GCCCACT | GCG | GCG | C | T | C>T | C>T_GCG | CG>TA_GCCCACT | 4 | FALSE | FALSE |
| 8_v.txt | 33113 | 33113-SNV- | SNV | 1 | G | A | 36 | 38692 | 0.09304249 | 41.7219889 | rnr1D57N_per | FALSE | ARG | JSY14 | 1 | rnr1D57N_pe | GGC | G>A | A/T+1 | G/C-1 | CG>TA | CG>TA | CACTGGGCC | GCCGGCG | CCG | CCG | C | T | C>T | C>T_CCG | CG>TA_GCCGGCG | 4 | FALSE | FALSE |
| 8_v.txt | 33114 | 33114-SNV- | SNV | 1 | C | T | 41 | 38692 | 0.10596506 | 39.9023686 | rnr1D57N_per | FALSE | ARG | JSY14 | 1 | rnr1D57N_pe | GCC | C>T | A/T+1 | G/C-1 | CG>TA | CG>TA | ACTGGGCCG | CCGGCGT | CGG | GCC | C | A | C>T | C>T_GCC | CG>TA_CCGGCGT | 4 | FALSE | FALSE |
| 8_v.txt | 33118 | 33118-SNV- | SNV | 1 | C | T | 45 | 38692 | 0.11630311 | 40.1777778 | rnr1D57N_per | FALSE | ARG | JSY14 | 1 | rnr1D57N_pe | GCG | C>T | A/T+1 | G/C-1 | CG>TA | CG>TA | GGCCGGCGT | CGTTGGT | CGC | GCG | C | A | C>T | C>T_GCG | CG>TA_CGTTGGT | 4 | FALSE | FALSE |
| 8_v.txt | 33119 | 33119-SNV- | SNV | 1 | G | A | 27 | 38692 | 0.06978187 | 39.8151141 | rnr1D57N_per | FALSE | ARG | JSY14 | 1 | rnr1D57N_pe | CGT | G>A | A/T+1 | G/C-1 | CG>TA | CG>TA | GCCGGCGTT | GTTGGTC | GCA | GCA | C | T | C>T | C>T_GCA | CG>TA_GTTGGTC | 4 | FALSE | FALSE |
| 8_v.txt | 33258 | 33258-SNV- | SNV | 1 | C | T | 34 | 19154 | 0.17750861 | 41.3235294 | rnr1D57N_per | FALSE | ARG | JSY14 | 1 | rnr1D57N_pe | GCC | C>T | A/T+1 | G/C-1 | CG>TA | CG>TA | CCTATGCCAT | CCATCCT | CGG | GCC | C | A | C>T | C>T_GCC | CG>TA_CCATCCT | 4 | FALSE | FALSE |
| 8_v.txt | 33275 | 33275-SNV- | SNV | 1 | C | T | 21 | 19154 | 0.10963767 | 51.5495557 | rnr1D57N_per | FALSE | ARG | JSY14 | 1 | rnr1D57N_pe | ACG | C>T | A/T+1 | G/C-1 | CG>TA | CG>TA | AGAGAACGT | CGTATCC | TGC | ACG | C | A | C>T | C>T_ACG | CG>TA_CGTATCC | 4 | FALSE | FALSE |
| 8_v.txt | 33276 | 33276-SNV- | SNV | 1 | G | A | 48 | 19154 | 0.2506004 | 47.2862084 | rnr1D57N_per | FALSE | ARG | JSY14 | 1 | rnr1D57N_pe | CGT | G>A | A/T+1 | G/C-1 | CG>TA | CG>TA | GAGAACGTA | GTATCCT | GCA | GCA | C | T | C>T | C>T_GCA | CG>TA_GTATCCT | 4 | FALSE | FALSE |
| 8_v.txt | 33309 | 33309-SNV- | SNV | 1 | C | T | 23 | 19152 | 0.1200919 | 57.1692419 | rnr1D57N_per | FALSE | ARG | JSY14 | 1 | rnr1D57N_pe | GCG | C>T | A/T+1 | G/C-1 | CG>TA | CG>TA | AAAGAGCGC | CGCAATG | CGC | GCG | C | A | C>T | C>T_GCG | CG>TA_CGCAATG | 4 | FALSE | FALSE |
| 8_v.txt | 33310 | 33310-SNV- | SNV | 1 | G | A | 31 | 19147 | 0.16190526 | 53.2810079 | rnr1D57N_per | FALSE | ARG | JSY14 | 1 | rnr1D57N_pe | CGC | G>A | A/T+1 | G/C-1 | CG>TA | CG>TA | AAGAGCGCA | GCAATGG | GCG | GCG | C | T | C>T | C>T_GCG | CG>TA_GCAATGG | 4 | FALSE | FALSE |
| 8_v.txt | 33392 | 33392-SNV- | SNV | 1 | G | A | 31 | 19154 | 0.16184609 | 46.2903226 | rnr1D57N_per | FALSE | ARG | JSY14 | 1 | rnr1D57N_pe | CGT | G>A | A/T+1 | G/C-1 | CG>TA | CG>TA | AACGTCGTG | GTGAAAG | GCA | GCA | C | T | C>T | C>T_GCA | CG>TA_GTGAAAG | 4 | FALSE | FALSE |
| 8_v.txt | 33412 | 33412-SNV- | SNV | 1 | G | A | 29 | 19154 | 0.15140441 | 46.7931035 | rnr1D57N_per | FALSE | ARG | JSY14 | 1 | rnr1D57N_pe | GGC | G>A | A/T+1 | G/C-1 | CG>TA | CG>TA | TGACCGGCT | GCTCATT | CCG | CCG | C | T | C>T | C>T_CCG | CG>TA_GCTCATT | 4 | FALSE | FALSE |
| 8_v.txt | 33474 | 33474-SNV- | SNV | 1 | C | T | 25 | 19154 | 0.13052104 | 33.08 | rnr1D57N_per | FALSE | ARG | JSY14 | 1 | rnr1D57N_pe | GCC | C>T | A/T+1 | G/C-1 | CG>TA | CG>TA | GCTATGCCTT | CCTTTTT | CGG | GCC | C | A | C>T | C>T_GCC | CG>TA_CCTTTTT | 4 | FALSE | FALSE |
| 8_v.txt | 33118 | 33118-SNV- | SNV | 1 | C | G | 3 | 38692 | 7.75E-03 | 36.6666667 | rnr1D57N_per | FALSE | ARG | JSY14 | 1 | rnr1D57N_pe | GCG | C>G | G/C+1 | G/C-1 | CG>GC | CG>GC | GGCCGGCGT | CGTTGGT | CGC | GCG | C | C | C>G | C>G_GCG | CG>GC_CGTTGGT | 4 | FALSE | FALSE |
| 8_v.txt | 33310 | 33310-SNV- | SNV | 1 | G | C | 4 | 19147 | 0.020891 | 14.2653746 | rnr1D57N_per | FALSE | ARG | JSY14 | 1 | rnr1D57N_pe | CGC | G>C | G/C+1 | G/C-1 | CG>GC | CG>GC | AAGAGCGCA | GCAATGG | GCG | GCG | C | G | C>G | C>G_GCG | CG>GC_GCAATGG | 4 | FALSE | FALSE |
| 8_v.txt | 31657 | 31657-SNV- | SNV | 1 | G | T | 185 | 17740 | 1.04284104 | 18.3276392 | rnr1D57N_per | FALSE | ARG | JSY14 | 1 | rnr1D57N_pe | GGC | G>T | A/T+1 | G/C-1 | CG>AT | CG>AT | GAAATGGCG | GCGTGGA | CCG | CCG | C | A | C>A | C>A_CCG | CG>AT_GCGTGGA | 4 | FALSE | FALSE |
| 8_v.txt | 32424 | 32424-SNV- | SNV | 1 | G | T | 10 | 26939 | 0.0371209 | 53.5 | rnr1D57N_per | FALSE | ARG | JSY14 | 1 | rnr1D57N_pe | AGG | G>T | A/T+1 | G/C-1 | CG>AT | CG>AT | AGTTTAGGG | GGGTCAT | TCC | TCC | C | A | C>A | C>A_TCC | CG>AT_GGGTCAT | 4 | FALSE | FALSE |
| 8_v.txt | 32900 | 32900-SNV- | SNV | 1 | G | T | 36 | 38692 | 0.09304249 | 21.4438956 | rnr1D57N_per | FALSE | ARG | JSY14 | 1 | rnr1D57N_pe | GGC | G>T | A/T+1 | G/C-1 | CG>AT | CG>AT | GACTTGGCC | GCCAACT | CCG | CCG | C | A | C>A | C>A_CCG | CG>AT_GCCAACT | 4 | FALSE | FALSE |
| 8_v.txt | 33050 | 33050-SNV- | SNV | 1 | G | T | 22 | 38692 | 0.0568593 | 56.1818182 | rnr1D57N_per | FALSE | ARG | JSY14 | 1 | rnr1D57N_pe | AGG | G>T | A/T+1 | G/C-1 | CG>AT | CG>AT | ACCCAAGGA | GGACTGC | TCC | TCC | C | A | C>A | C>A_TCC | CG>AT_GGACTGC | 4 | FALSE | FALSE |
| 8_v.txt | 33070 | 33070-SNV- | SNV | 1 | C | A | 17 | 38692 | 0.04393673 | 45.5882353 | rnr1D57N_per | FALSE | ARG | JSY14 | 1 | rnr1D57N_pe | GCC | C>A | A/T+1 | G/C-1 | CG>AT | CG>AT | AATATGCCA | CCAAAGA | CGG | GCC | C | T | C>A | C>A_GCC | CG>AT_CCAAAGA | 4 | FALSE | FALSE |
| 8_v.txt | 33114 | 33114-SNV- | SNV | 1 | C | A | 7 | 38692 | 0.0180916 | 41.4285239 | rnr1D57N_per | FALSE | ARG | JSY14 | 1 | rnr1D57N_pe | GCC | C>A | A/T+1 | G/C-1 | CG>AT | CG>AT | ACTGGGCCG | CCGGCGT | CGG | GCC | C | T | C>A | C>A_GCC | CG>AT_CCGGCGT | 4 | FALSE | FALSE |
| 8_v.txt | 33118 | 33118-SNV- | SNV | 1 | C | A | 8 | 38692 | 0.02067611 | 33.625 | rnr1D57N_per | FALSE | ARG | JSY14 | 1 | rnr1D57N_pe | GCG | C>A | A/T+1 | G/C-1 | CG>AT | CG>AT | GGCCGGCGT | CGTTGGT | CGC | GCG | C | T | C>A | C>A_GCG | CG>AT_CGTTGGT | 4 | FALSE | FALSE |
| 8_v.txt | 33275 | 33275-SNV- | SNV | 1 | C | A | 5 | 19154 | 0.02610421 | 31.7977618 | rnr1D57N_per | FALSE | ARG | JSY14 | 1 | rnr1D57N_pe | ACG | C>A | A/T+1 | G/C-1 | CG>AT | CG>AT | AGAGAACGT | CGTATCC | TGC | ACG | C | T | C>A | C>A_ACG | CG>AT_CGTATCC | 4 | FALSE | FALSE |
| 8_v.txt | 33474 | 33474-SNV- | SNV | 1 | C | A | 62 | 19154 | 0.32369218 | 33.4193548 | rnr1D57N_per | FALSE | ARG | JSY14 | 1 | rnr1D57N_pe | GCC | C>A | A/T+1 | G/C-1 | CG>AT | CG>AT | GCTATGCCTT | CCTTTTT | CGG | GCC | C | T | C>A | C>A_GCC | CG>AT_CCTTTTT | 4 | FALSE | FALSE |
| 8_v.txt | 33475 | 33475-SNV- | SNV | 1 | C | A | 24 | 19154 | 0.1253002 | 37.2916667 | rnr1D57N_per | FALSE | ARG | JSY14 | 1 | rnr1D57N_pe | CCT | C>A | A/T+1 | G/C-1 | CG>AT | CG>AT | CTATGCCTTT | CTTTTTT | GGA | CCT | C | T | C>A | C>A_CCT | CG>AT_CTTTTTT | 4 | FALSE | FALSE |
| 8_v.txt | 33069 | 33069-MNV- | MNV | 2 | GC | CA | 1 | 38692 | 2.59E-03 | 27.5000229 | rnr1D57N_per | FALSE | ARG | JSY14 | 1 | rnr1D57N_pe | TGC | nope | >1bp-ins | >1bpdel | none | MNV | GAATATGCC | GCCAAAG | ACG | TGC | GC | none | NA | NA_TGC | MNV_GCCAAAG | 4 | FALSE | FALSE |
| 8_v.txt | 33474 | 33474-MNV- | MNV | 2 | CC | AA | 1 | 19154 | 5.22E-03 | 38 | rnr1D57N_per | FALSE | ARG | JSY14 | 1 | rnr1D57N_pe | GCC | nope | >1bp-ins | >1bpdel | none | MNV | GCTATGCCTT | CCTTTTT | CGG | GCC | CC | none | NA | NA_GCC | MNV_CCTTTTT | 4 | FALSE | FALSE |
| 8_v.txt | 31711 | 31711-Insert | Insertion | 1 | - | A | 3 | 17740 | 0.01691094 | 64 | rnr1D57N_per | FALSE | ARG | JSY14 | 1 | rnr1D57N_pe | CAA | nope | A/T+1 | >1bpdel | none | A/T+1 | CATTCCAAA | AAAATTT | GTT | CAA | - | T | NA | NA_CAA | A/T+1_AAAATTT | 4 | FALSE | FALSE |
| 8_v.txt | 32081 | 32081-Insert | Insertion | 1 | - | A | 18 | 32081 | 0.05610798 | 64 | rnr1D57N_per | FALSE | ARG | JSY14 | 1 | rnr1D57N_pe | CAA | nope | A/T+1 | >1bpdel | none | A/T+1 | CCATGCAAA | AAAAAAG | GTT | CAA | - | T | NA | NA_CAA | A/T+1_AAAAAAG | 4 | FALSE | FALSE |
| 8_v.txt | 32842 | 32842-Insert | Insertion | 1 | - | A | 4 | 22419 | 0.01784201 | 38 | rnr1D57N_per | FALSE | ARG | JSY14 | 1 | rnr1D57N_pe | CAA | nope | A/T+1 | >1bpdel | none | A/T+1 | TTACCCAAA | AAAAAAT | GTT | CAA | - | T | NA | NA_CAA | A/T+1_AAAAAAT | 4 | FALSE | FALSE |
| 8_v.txt | 32972 | 32972-Insert | Insertion | 1 | - | A | 3 | 38692 | 7.75E-03 | 63 | rnr1D57N_per | FALSE | ARG | JSY14 | 1 | rnr1D57N_pe | CAA | nope | A/T+1 | >1bpdel | none | A/T+1 | CGCACCAAAT | AAATGCT | GTT | CAA | - | T | NA | NA_CAA | A/T+1_AAATGCT | 4 | FALSE | FALSE |
| 8_v.txt | 31663 | 31663-Deleti | Deletion | 1 | A | - | 10 | 17740 | 0.05636979 | 38.4 | rnr1D57N_per | FALSE | ARG | JSY14 | 1 | rnr1D57N_pe | GAA | nope | >1bp-ins | A/T-1 | none | A/T-1 | GCGTGGAAA | AAATGTG | CTT | CTT | T | none | T>none | T>none_CTT | A/T-1_AAATGTG | 4 | FALSE | FALSE |
| 8_v.txt | 31720 | 31720-Deleti | Deletion | 1 | C | - | 4 | 17740 | 0.02254791 | 57 | rnr1D57N_per | FALSE | ARG | JSY14 | 1 | rnr1D57N_pe | TCC | nope | >1bp-ins | G/C-1 | none | G/C-1 | ATTTGTCCCA | CCCAAAA | AGG | TCC | C | none | nope | nope_TCC | G/C-1_CCCAAAA | 4 | FALSE | FALSE |
| 8_v.txt | 31723 | 31723-Deleti | Deletion | 1 | A | - | 27 | 17740 | 0.15219842 | 59.5925926 | rnr1D57N_per | FALSE | ARG | JSY14 | 1 | rnr1D57N_pe | CAA | nope | >1bp-ins | A/T-1 | none | A/T-1 | TGTCCCAAA | AAAAAGT | GTT | GTT | T | none | T>none | T>none_GTT | A/T-1_AAAAAGT | 4 | FALSE | FALSE |
| 8_v.txt | 31879 | 31879-Deleti | Deletion | 1 | C | - | 2 | 17740 | 0.01127396 | 38 | rnr1D57N_per | FALSE | ARG | JSY14 | 1 | rnr1D57N_pe | GCG | nope | >1bp-ins | G/C-1 | none | G/C-1 | TATAGGCGG | CGGCAGC | CGC | GCG | C | none | nope | nope_GCG | G/C-1_CGGCAGC | 4 | FALSE | FALSE |
| 8_v.txt | 31904 | 31904-Deleti | Deletion | 1 | T | - | 7 | 17740 | 0.03945885 | 48.7142857 | rnr1D57N_per | FALSE | ARG | JSY14 | 1 | rnr1D57N_pe | ATT | nope | >1bp-ins | A/T-1 | none | A/T-1 | ATTGAATTTT | TTTTGGT | TAA | ATT | T | none | nope | nope_ATT | A/T-1_TTTTGGT | 4 | FALSE | FALSE |
| 8_v.txt | 31972 | 31972-Deleti | Deletion | 1 | C | - | 8 | 32081 | 0.02493688 | 35.875 | rnr1D57N_per | FALSE | ARG | JSY14 | 1 | rnr1D57N_pe | GCC | nope | >1bp-ins | G/C-1 | none | G/C-1 | AATAAGCCA | CCAAGCC | CGG | GCC | C | none | nope | nope_GCC | G/C-1_CCAAGCC | 4 | FALSE | FALSE |
| 8_v.txt | 32045 | 32045-Deleti | Deletion | 1 | A | - | 6 | 32081 | 0.01870266 | 64 | rnr1D57N_per | FALSE | ARG | JSY14 | 1 | rnr1D57N_pe | TAA | nope | >1bp-ins | A/T-1 | none | A/T-1 | TTGCATAAAT | AAATCTG | ATT | ATT | T | none | T>none | T>none_ATT | A/T-1_AAATCTG | 4 | FALSE | FALSE |
| 8_v.txt | 32081 | 32081-Deleti | Deletion | 1 | A | - | 44 | 32081 | 0.13715283 | 63.1818182 | rnr1D57N_per | FALSE | ARG | JSY14 | 1 | rnr1D57N_pe | CAA | nope | >1bp-ins | A/T-1 | none | A/T-1 | CCATGCAAA | AAAAAAG | GTT | GTT | T | none | T>none | T>none_GTT | A/T-1_AAAAAAG | 4 | FALSE | FALSE |
| 8_v.txt | 32210 | 32210-Deleti | Deletion | 1 | T | - | 6 | 32081 | 0.01870266 | 46.3333333 | rnr1D57N_per | FALSE | ARG | JSY14 | 1 | rnr1D57N_pe | CTT | nope | >1bp-ins | A/T-1 | none | A/T-1 | ACCACCTTTG | TTTGGTG | GAA | CTT | T | none | nope | nope_CTT | A/T-1_TTTGGTG | 4 | FALSE | FALSE |
| 8_v.txt | 32424 | 32424-Deleti | Deletion | 1 | G | - | 5 | 26939 | 0.01856045 | 53.4 | rnr1D57N_per | FALSE | ARG | JSY14 | 1 | rnr1D57N_pe | AGG | nope | >1bp-ins | G/C-1 | none | G/C-1 | AGTTTAGGG | GGGTCAT | TCC | TCC | C | none | C>none | C>none_TCC | G/C-1_GGGTCAT | 4 | FALSE | FALSE |
| 8_v.txt | 32442 | 32442-Deleti | Deletion | 1 | A | - | 8 | 26939 | 0.02969672 | 60.125 | rnr1D57N_per | FALSE | ARG | JSY14 | 1 | rnr1D57N_pe | TAA | nope | >1bp-ins | A/T-1 | none | A/T-1 | GGAACTAAA | AAAAGTC | ATT | ATT | T | none | T>none | T>none_ATT | A/T-1_AAAAGTC | 4 | FALSE | FALSE |
| 8_v.txt | 32498 | 32498-Deleti | Deletion | 1 | T | - | 47 | 26939 | 0.17446824 | 45.1914894 | rnr1D57N_per | FALSE | ARG | JSY14 | 1 | rnr1D57N_pe | CTT | nope | >1bp-ins | A/T-1 | none | A/T-1 | AACAACTTTT | TTTTTTG | GAA | CTT | T | none | nope | nope_CTT | A/T-1_TTTTTTG | 4 | FALSE | FALSE |
| 8_v.txt | 32522 | 32522-Deleti | Deletion | 1 | T | - | 8 | 26939 | 0.02969672 | 41 | rnr1D57N_per | FALSE | ARG | JSY14 | 1 | rnr1D57N_pe | ATT | nope | >1bp-ins | A/T-1 | none | A/T-1 | AACGGATTTT | TTTTCTG | TAA | ATT | T | none | nope | nope_ATT | A/T-1_TTTTCTG | 4 | FALSE | FALSE |
| 8_v.txt | 32631 | 32631-Deleti | Deletion | 1 | T | - | 7 | 22419 | 0.03122352 | 47.4285714 | rnr1D57N_per | FALSE | ARG | JSY14 | 1 | rnr1D57N_pe | GTT | nope | >1bp-ins | A/T-1 | none | A/T-1 | CCTTCGTTTT | TTTTTAT | CAA | GTT | T | none | nope | nope_GTT | A/T-1_TTTTTAT | 4 | FALSE | FALSE |
| 8_v.txt | 32658 | 32658-Deleti | Deletion | 1 | C | - | 8 | 22419 | 0.03568402 | 53.375 | rnr1D57N_per | FALSE | ARG | JSY14 | 1 | rnr1D57N_pe | ACC | nope | >1bp-ins | G/C-1 | none | G/C-1 | CCTGGACCC | CCCCAGG | TGG | ACC | C | none | nope | nope_ACC | G/C-1_CCCCAGG | 4 | FALSE | FALSE |
| 8_v.txt | 32670 | 32670-Deleti | Deletion | 1 | G | - | 6 | 22419 | 0.02676301 | 53.0000017 | rnr1D57N_per | FALSE | ARG | JSY14 | 1 | rnr1D57N_pe | TGG | nope | >1bp-ins | G/C-1 | none | G/C-1 | GCACCTGGG | GGGTTTC | ACC | ACC | C | none | C>none | C>none_ACC | G/C-1_GGGTTTC | 4 | FALSE | FALSE |
| 8_v.txt | 32779 | 32779-Deleti | Deletion | 1 | C | - | 7 | 22419 | 0.03122352 | 52.8571429 | rnr1D57N_per | FALSE | ARG | JSY14 | 1 | rnr1D57N_pe | ACC | nope | >1bp-ins | G/C-1 | none | G/C-1 | AAGCGACCC | CCCAGAA | TGG | ACC | C | none | nope | nope_ACC | G/C-1_CCCAGAA | 4 | FALSE | FALSE |
| 8_v.txt | 32842 | 32842-Deleti | Deletion | 1 | A | - | 36 | 22419 | 0.16057808 | 43.6111106 | rnr1D57N_per | FALSE | ARG | JSY14 | 1 | rnr1D57N_pe | CAA | nope | >1bp-ins | A/T-1 | none | A/T-1 | TTACCCAAA | AAAAAAT | GTT | GTT | T | none | T>none | T>none_GTT | A/T-1_AAAAAAT | 4 | FALSE | FALSE |
| 8_v.txt | 32924 | 32924-Deleti | Deletion | 1 | A | - | 51 | 38692 | 0.13181019 | 53.2745098 | rnr1D57N_per | FALSE | ARG | JSY14 | 1 | rnr1D57N_pe | CAA | nope | >1bp-ins | A/T-1 | none | A/T-1 | CAGGGCAAA | AAAAGTG | GTT | GTT | T | none | T>none | T>none_GTT | A/T-1_AAAAGTG | 4 | FALSE | FALSE |
| 8_v.txt | 33000 | 33000-Deleti | Deletion | 1 | A | - | 9 | 38692 | 0.02326062 | 63.2222222 | rnr1D57N_per | FALSE | ARG | JSY14 | 1 | rnr1D57N_pe | GAA | nope | >1bp-ins | A/T-1 | none | A/T-1 | TGTGAGAAA | AAAACTG | CTT | CTT | T | none | T>none | T>none_CTT | A/T-1_AAAACTG | 4 | FALSE | FALSE |
| 8_v.txt | 33040 | 33040-Deleti | Deletion | 1 | T | - | 7 | 38692 | 0.0180916 | 60.2857143 | rnr1D57N_per | FALSE | ARG | JSY14 | 1 | rnr1D57N_pe | ATT | nope | >1bp-ins | A/T-1 | none | A/T-1 | TAGCCATTTC | TTTCACC | TAA | ATT | T | none | nope | nope_ATT | A/T-1_TTTCACC | 4 | FALSE | FALSE |
| 8_v.txt | 33147 | 33147-Deleti | Deletion | 1 | A | - | 9 | 38692 | 0.02326062 | 38 | rnr1D57N_per | FALSE | ARG | JSY14 | 1 | rnr1D57N_pe | GAA | nope | >1bp-ins | A/T-1 | none | A/T-1 | CCAATGAAA | AAAAGAC | CTT | CTT | T | none | T>none | T>none_CTT | A/T-1_AAAAGAC | 4 | FALSE | FALSE |
| 8_v.txt | 33288 | 33288-Deleti | Deletion | 1 | T | - | 5 | 19154 | 0.02610421 | 53.4 | rnr1D57N_per | FALSE | ARG | JSY14 | 1 | rnr1D57N_pe | ATT | nope | >1bp-ins | A/T-1 | none | A/T-1 | TCGCCATTTA | TTTACTC | TAA | ATT | T | none | nope | nope_ATT | A/T-1_TTTACTC | 4 | FALSE | FALSE |
| 8_v.txt | 33474 | 33474-Deleti | Deletion | 1 | C | - | 56 | 19154 | 0.29236713 | 28.7321429 | rnr1D57N_per | FALSE | ARG | JSY14 | 1 | rnr1D57N_pe | GCC | nope | >1bp-ins | G/C-1 | none | G/C-1 | GCTATGCCTT | CCTTTTT | CGG | GCC | C | none | nope | nope_GCC | G/C-1_CCTTTTT | 4 | FALSE | FALSE |
| 9_v.txt | 31723 | 31723-SNV- | SNV | 1 | A | C | 7 | 12376 | 0.05656109 | 50.2857143 | rnr1D57N_per | FALSE | ARG | JSY14 | 2 | rnr1D57N_pe | CAA | A>C | G/C+1 | A/T-1 | TA>GC | TA>GC | TGTCCCAAA | AAAAAGT | GTT | GTT | T | G | T>G | T>G_GTT | TA>GC_AAAAAGT | 4 | FALSE | FALSE |
| 9_v.txt | 32627 | 32627-SNV-T | SNV | 1 | T | G | 6 | 24361 | 0.02462953 | 50.6666667 | rnr1D57N_per | FALSE | ARG | JSY14 | 2 | rnr1D57N_pe | CTT | T>G | G/C+1 | A/T-1 | TA>GC | TA>GC | CCTCCCTTCG | TTCGTTT | GAA | CTT | T | C | T>G | T>G_CTT | TA>GC_TTCGTTT | 4 | FALSE | FALSE |
| 9_v.txt | 32842 | 32842-SNV- | SNV | 1 | A | C | 3 | 24361 | 0.01231477 | 38 | rnr1D57N_per | FALSE | ARG | JSY14 | 2 | rnr1D57N_pe | CAA | A>C | G/C+1 | A/T-1 | TA>GC | TA>GC | TTACCCAAA | AAAAAAT | GTT | GTT | T | G | T>G | T>G_GTT | TA>GC_AAAAAAT | 4 | FALSE | FALSE |
| 9_v.txt | 32927 | 32927-SNV- | SNV | 1 | A | C | 13 | 15180 | 0.085639 | 18.1538462 | rnr1D57N_per | FALSE | ARG | JSY14 | 2 | rnr1D57N_pe | AAG | A>C | G/C+1 | A/T-1 | TA>GC | TA>GC | GGCAAAAGT | AGTGATT | TTC | TTC | T | G | T>G | T>G_TTC | TA>GC_AGTGATT | 4 | FALSE | FALSE |
| 9_v.txt | 33269 | 33269-SNV- | SNV | 1 | A | C | 105 | 17670 | 0.5942275 | 17.0571429 | rnr1D57N_per | FALSE | ARG | JSY14 | 2 | rnr1D57N_pe | TAG | A>C | G/C+1 | A/T-1 | TA>GC | TA>GC | CTCCATAGA | AGAGAAC | ATC | ATC | T | G | T>G | T>G_ATC | TA>GC_AGAGAAC | 4 | FALSE | FALSE |
| 9_v.txt | 33476 | 33476-SNV-T | SNV | 1 | T | G | 3 | 17667 | 0.01698081 | 28 | rnr1D57N_per | FALSE | ARG | JSY14 | 2 | rnr1D57N_pe | CTT | T>G | G/C+1 | A/T-1 | TA>GC | TA>GC | TATGCCTTTT | TTTTTTT | GAA | CTT | T | C | T>G | T>G_CTT | TA>GC_TTTTTTT | 4 | FALSE | FALSE |
| 9_v.txt | 32081 | 32081-SNV- | SNV | 1 | A | G | 6 | 18882 | 0.0317763 | 64 | rnr1D57N_per | FALSE | ARG | JSY14 | 2 | rnr1D57N_pe | CAA | A>G | G/C+1 | A/T-1 | TA>CG | TA>CG | CCATGCAAA | AAAAAAG | GTT | GTT | T | C | T>C | T>C_GTT | TA>CG_AAAAAAG | 4 | FALSE | FALSE |
| 9_v.txt | 32927 | 32927-SNV- | SNV | 1 | A | G | 18 | 15180 | 0.11857708 | 52.0555556 | rnr1D57N_per | FALSE | ARG | JSY14 | 2 | rnr1D57N_pe | AAG | A>G | G/C+1 | A/T-1 | TA>CG | TA>CG | GGCAAAAGT | AGTGATT | TTC | TTC | T | C | T>C | T>C_TTC | TA>CG_AGTGATT | 4 | FALSE | FALSE |
| 9_v.txt | 33476 | 33476-SNV-T | SNV | 1 | T | C | 28 | 17667 | 0.15848758 | 31.5714286 | rnr1D57N_per | FALSE | ARG | JSY14 | 2 | rnr1D57N_pe | CTT | T>C | G/C+1 | A/T-1 | TA>CG | TA>CG | TATGCCTTTT | TTTTTTT | GAA | CTT | T | G | T>C | T>C_CTT | TA>CG_TTTTTTT | 4 | FALSE | FALSE |
| 9_v.txt | 31650 | 31650-SNV- | SNV | 1 | C | T | 20 | 12376 | 0.1616031 | 36 | rnr1D57N_per | FALSE | ARG | JSY14 | 2 | rnr1D57N_pe | GCG | C>T | A/T+1 | G/C-1 | CG>TA | CG>TA | AGAATGCGA | CGAAATG | CGC | GCG | C | A | C>T | C>T_GCG | CG>TA_CGAAATG | 4 | FALSE | FALSE |
| 9_v.txt | 31658 | 31658-SNV- | SNV | 1 | C | T | 15 | 12376 | 0.12120233 | 36.5333333 | rnr1D57N_per | FALSE | ARG | JSY14 | 2 | rnr1D57N_pe | GCG | C>T | A/T+1 | G/C-1 | CG>TA | CG>TA | AAATGGCGT | CGTGGAA | CGC | GCG | C | A | C>T | C>T_GCG | CG>TA_CGTGGAA | 4 | FALSE | FALSE |
| 9_v.txt | 31778 | 31778-SNV- | SNV | 1 | G | A | 15 | 12376 | 0.12120233 | 52.6 | rnr1D57N_per | FALSE | ARG | JSY14 | 2 | rnr1D57N_pe | CGG | G>A | A/T+1 | G/C-1 | CG>TA | CG>TA | TCTATCGGA | GGAATCG | GCC | GCC | C | T | C>T | C>T_GCC | CG>TA_GGAATCG | 4 | FALSE | FALSE |
| 9_v.txt | 31830 | 31830-SNV- | SNV | 1 | C | T | 14 | 12376 | 0.11312217 | 56.3571429 | rnr1D57N_per | FALSE | ARG | JSY14 | 2 | rnr1D57N_pe | GCA | C>T | A/T+1 | G/C-1 | CG>TA | CG>TA | AATATGCATT | CATTGAA | CGT | GCA | C | A | C>T | C>T_GCA | CG>TA_CATTGAA | 4 | FALSE | FALSE |
| 9_v.txt | 31881 | 31881-SNV- | SNV | 1 | G | A | 14 | 12376 | 0.11312217 | 44.0714286 | rnr1D57N_per | FALSE | ARG | JSY14 | 2 | rnr1D57N_pe | GGC | G>A | A/T+1 | G/C-1 | CG>TA | CG>TA | TAGGCGGCA | GCAGCAA | CCG | CCG | C | T | C>T | C>T_CCG | CG>TA_GCAGCAA | 4 | FALSE | FALSE |
| 9_v.txt | 31911 | 31911-SNV- | SNV | 1 | G | A | 16 | 12376 | 0.12928248 | 39.4375 | rnr1D57N_per | FALSE | ARG | JSY14 | 2 | rnr1D57N_pe | TGC | G>A | A/T+1 | G/C-1 | CG>TA | CG>TA | TTTGGTGCA | GCAAAAG | ACG | ACG | C | T | C>T | C>T_ACG | CG>TA_GCAAAAG | 4 | FALSE | FALSE |
| 9_v.txt | 31962 | 31962-SNV- | SNV | 1 | G | A | 35 | 18882 | 0.18536172 | 37.1428571 | rnr1D57N_per | FALSE | ARG | JSY14 | 2 | rnr1D57N_pe | CGC | G>A | A/T+1 | G/C-1 | CG>TA | CG>TA | GTGGCCGCAT | GCATAAT | GCG | GCG | C | T | C>T | C>T_GCG | CG>TA_GCATAAT | 4 | FALSE | FALSE |
| 9_v.txt | 31972 | 31972-SNV- | SNV | 1 | C | T | 18 | 18882 | 0.09532889 | 36.7222222 | rnr1D57N_per | FALSE | ARG | JSY14 | 2 | rnr1D57N_pe | GCC | C>T | A/T+1 | G/C-1 | CG>TA | CG>TA | AATAAGCCA | CCAAGCC | CGG | GCC | C | A | C>T | C>T_GCC | CG>TA_CCAAGCC | 4 | FALSE | FALSE |
| 9_v.txt | 32041 | 32041-SNV- | SNV | 1 | G | A | 19 | 18882 | 0.10062493 | 59.6842105 | rnr1D57N_per | FALSE | ARG | JSY14 | 2 | rnr1D57N_pe | TGC | G>A | A/T+1 | G/C-1 | CG>TA | CG>TA | AAGCTTGCAT | GCATAAA | ACG | ACG | C | T | C>T | C>T_ACG | CG>TA_GCATAAA | 4 | FALSE | FALSE |
| 9_v.txt | 32057 | 32057-SNV- | SNV | 1 | C | T | 22 | 18882 | 0.11651308 | 61.9545455 | rnr1D57N_per | FALSE | ARG | JSY14 | 2 | rnr1D57N_pe | GCG | C>T | A/T+1 | G/C-1 | CG>TA | CG>TA | GATGTGCGA | CGAGATT | CGC | GCG | C | A | C>T | C>T_GCG | CG>TA_CGAGATT | 4 | FALSE | FALSE |
| 9_v.txt | 32092 | 32092-SNV- | SNV | 1 | C | T | 21 | 18882 | 0.11121703 | 59.8571429 | rnr1D57N_per | FALSE | ARG | JSY14 | 2 | rnr1D57N_pe | GCA | C>T | A/T+1 | G/C-1 | CG>TA | CG>TA | AGCCTGCAA | CAACACC | CGT | GCA | C | A | C>T | C>T_GCA | CG>TA_CAACACC | 4 | FALSE | FALSE |
| 9_v.txt | 32165 | 32165-SNV- | SNV | 1 | G | A | 18 | 18882 | 0.09532889 | 49.5555556 | rnr1D57N_per | FALSE | ARG | JSY14 | 2 | rnr1D57N_pe | CGC | G>A | A/T+1 | G/C-1 | CG>TA | CG>TA | CAAAGCGCC | GCCAAAT | GCG | GCG | C | T | C>T | C>T_GCG | CG>TA_GCCAAAT | 4 | FALSE | FALSE |
| 9_v.txt | 32173 | 32173-SNV- | SNV | 1 | C | T | 18 | 18882 | 0.09532889 | 42.6111111 | rnr1D57N_per | FALSE | ARG | JSY14 | 2 | rnr1D57N_pe | GCA | C>T | A/T+1 | G/C-1 | CG>TA | CG>TA | CAAATGCAG | CAGCAGT | CGT | GCA | C | A | C>T | C>T_GCA | CG>TA_CAGCAGT | 4 | FALSE | FALSE |
| 9_v.txt | 32176 | 32176-SNV- | SNV | 1 | C | T | 20 | 18882 | 0.10592098 | 44.55 | rnr1D57N_per | FALSE | ARG | JSY14 | 2 | rnr1D57N_pe | GCA | C>T | A/T+1 | G/C-1 | CG>TA | CG>TA | ATGCAGCAG | CAGTAAC | CGT | GCA | C | A | C>T | C>T_GCA | CG>TA_CAGTAAC | 4 | FALSE | FALSE |
| 9_v.txt | 32271 | 32271-SNV- | SNV | 1 | C | T | 36 | 20443 | 0.1760994 | 30.3611111 | rnr1D57N_per | FALSE | ARG | JSY14 | 2 | rnr1D57N_pe | ACG | C>T | A/T+1 | G/C-1 | CG>TA | CG>TA | AAAATACGG | CGGGAAC | TGC | ACG | C | A | C>T | C>T_ACG | CG>TA_CGGGAAC | 4 | FALSE | FALSE |
| 9_v.txt | 32299 | 32299-SNV- | SNV | 1 | C | T | 52 | 20442 | 0.25437824 | 53.5384615 | rnr1D57N_per | FALSE | ARG | JSY14 | 2 | rnr1D57N_pe | GCG | C>T | A/T+1 | G/C-1 | CG>TA | CG>TA | AATTTGCGG | CGGCAGA | CGC | GCG | C | A | C>T | C>T_GCG | CG>TA_CGGCAGA | 4 | FALSE | FALSE |
| 9_v.txt | 32466 | 32466-SNV- | SNV | 1 | C | T | 26 | 20442 | 0.12718912 | 48.0769231 | rnr1D57N_per | FALSE | ARG | JSY14 | 2 | rnr1D57N_pe | GCC | C>T | A/T+1 | G/C-1 | CG>TA | CG>TA | AGAGAGCCA | CCAATGT | CGG | GCC | C | A | C>T | C>T_GCC | CG>TA_CCAATGT | 4 | FALSE | FALSE |
| 9_v.txt | 32535 | 32535-SNV- | SNV | 1 | G | A | 20 | 20442 | 0.09783779 | 37.4482495 | rnr1D57N_per | FALSE | ARG | JSY14 | 2 | rnr1D57N_pe | TGC | G>A | A/T+1 | G/C-1 | CG>TA | CG>TA | GGGTTTGCA | GCAGCTT | ACG | ACG | C | T | C>T | C>T_ACG | CG>TA_GCAGCTT | 4 | FALSE | FALSE |
| 9_v.txt | 32536 | 32536-SNV- | SNV | 1 | C | T | 22 | 20442 | 0.10762156 | 35.4086382 | rnr1D57N_per | FALSE | ARG | JSY14 | 2 | rnr1D57N_pe | GCA | C>T | A/T+1 | G/C-1 | CG>TA | CG>TA | GGTTTGCAG | CAGCTTC | CGT | GCA | C | A | C>T | C>T_GCA | CG>TA_CAGCTTC | 4 | FALSE | FALSE |
| 9_v.txt | 32590 | 32590-SNV- | SNV | 1 | C | T | 46 | 24361 | 0.1888264 | 35.5217391 | rnr1D57N_per | FALSE | ARG | JSY14 | 2 | rnr1D57N_pe | GCG | C>T | A/T+1 | G/C-1 | CG>TA | CG>TA | AGGCAGCGT | CGTTAAT | CGC | GCG | C | A | C>T | C>T_GCG | CG>TA_CGTTAAT | 4 | FALSE | FALSE |
| 9_v.txt | 32700 | 32700-SNV- | SNV | 1 | C | T | 27 | 24361 | 0.11083289 | 60.1111111 | rnr1D57N_per | FALSE | ARG | JSY14 | 2 | rnr1D57N_pe | GCC | C>T | A/T+1 | G/C-1 | CG>TA | CG>TA | ACTGGGCCG | CCGGTAA | CGG | GCC | C | A | C>T | C>T_GCC | CG>TA_CCGGTAA | 4 | FALSE | FALSE |
| 9_v.txt | 32702 | 32702-SNV- | SNV | 1 | G | A | 25 | 24361 | 0.10262305 | 54.52 | rnr1D57N_per | FALSE | ARG | JSY14 | 2 | rnr1D57N_pe | CGG | G>A | A/T+1 | G/C-1 | CG>TA | CG>TA | TGGGCCGGT | GGTAACC | GCC | GCC | C | T | C>T | C>T_GCC | CG>TA_GGTAACC | 4 | FALSE | FALSE |
| 9_v.txt | 32776 | 32776-SNV- | SNV | 1 | C | T | 24 | 24361 | 0.09851812 | 49.7916667 | rnr1D57N_per | FALSE | ARG | JSY14 | 2 | rnr1D57N_pe | GCG | C>T | A/T+1 | G/C-1 | CG>TA | CG>TA | TGGAAGCGA | CGACCCA | CGC | GCG | C | A | C>T | C>T_GCG | CG>TA_CGACCCA | 4 | FALSE | FALSE |
| 9_v.txt | 32798 | 32798-SNV- | SNV | 1 | G | A | 38 | 24361 | 0.15598703 | 43.1052632 | rnr1D57N_per | FALSE | ARG | JSY14 | 2 | rnr1D57N_pe | CGT | G>A | A/T+1 | G/C-1 | CG>TA | CG>TA | TTCACCGTAA | GTAATAT | GCA | GCA | C | T | C>T | C>T_GCA | CG>TA_GTAATAT | 4 | FALSE | FALSE |
| 9_v.txt | 32860 | 32860-SNV- | SNV | 1 | C | T | 33 | 24359 | 0.13547354 | 36.8484849 | rnr1D57N_per | FALSE | ARG | JSY14 | 2 | rnr1D57N_pe | GCC | C>T | A/T+1 | G/C-1 | CG>TA | CG>TA | TCCATGCCG | CCGCCAG | CGG | GCC | C | A | C>T | C>T_GCC | CG>TA_CCGCCAG | 4 | FALSE | FALSE |
| 9_v.txt | 32900 | 32900-SNV- | SNV | 1 | G | A | 20 | 15180 | 0.13175231 | 34.4478473 | rnr1D57N_per | FALSE | ARG | JSY14 | 2 | rnr1D57N_pe | GGC | G>A | A/T+1 | G/C-1 | CG>TA | CG>TA | GACTTGGCC | GCCAACT | CCG | CCG | C | T | C>T | C>T_CCG | CG>TA_GCCAACT | 4 | FALSE | FALSE |
| 9_v.txt | 32901 | 32901-SNV- | SNV | 1 | C | T | 32 | 15180 | 0.21080369 | 30.3621314 | rnr1D57N_per | FALSE | ARG | JSY14 | 2 | rnr1D57N_pe | GCC | C>T | A/T+1 | G/C-1 | CG>TA | CG>TA | ACTTGGCCA | CCAACTA | CGG | GCC | C | A | C>T | C>T_GCC | CG>TA_CCAACTA | 4 | FALSE | FALSE |
| 9_v.txt | 32965 | 32965-SNV- | SNV | 1 | C | T | 20 | 15180 | 0.13175231 | 56.2 | rnr1D57N_per | FALSE | ARG | JSY14 | 2 | rnr1D57N_pe | GCC | C>T | A/T+1 | G/C-1 | CG>TA | CG>TA | CATTGGCCG | CCGCACC | CGG | GCC | C | A | C>T | C>T_GCC | CG>TA_CCGCACC | 4 | FALSE | FALSE |
| 9_v.txt | 32967 | 32967-SNV- | SNV | 1 | G | A | 28 | 15180 | 0.18445323 | 59.5 | rnr1D57N_per | FALSE | ARG | JSY14 | 2 | rnr1D57N_pe | CGC | G>A | A/T+1 | G/C-1 | CG>TA | CG>TA | TTGGCCGCA | GCACCAA | GCG | GCG | C | T | C>T | C>T_GCG | CG>TA_GCACCAA | 4 | FALSE | FALSE |
| 9_v.txt | 33056 | 33056-SNV- | SNV | 1 | C | T | 21 | 15180 | 0.13833992 | 57.6666667 | rnr1D57N_per | FALSE | ARG | JSY14 | 2 | rnr1D57N_pe | GCG | C>T | A/T+1 | G/C-1 | CG>TA | CG>TA | GGACTGCGT | CGTGACA | CGC | GCG | C | A | C>T | C>T_GCG | CG>TA_CGTGACA | 4 | FALSE | FALSE |
| 9_v.txt | 33070 | 33070-SNV- | SNV | 1 | C | T | 22 | 15180 | 0.14492754 | 58.6363636 | rnr1D57N_per | FALSE | ARG | JSY14 | 2 | rnr1D57N_pe | GCC | C>T | A/T+1 | G/C-1 | CG>TA | CG>TA | AATATGCCA | CCAAAGA | CGG | GCC | C | A | C>T | C>T_GCC | CG>TA_CCAAAGA | 4 | FALSE | FALSE |
| 9_v.txt | 33114 | 33114-SNV- | SNV | 1 | C | T | 19 | 15180 | 0.12516469 | 41.1578947 | rnr1D57N_per | FALSE | ARG | JSY14 | 2 | rnr1D57N_pe | GCC | C>T | A/T+1 | G/C-1 | CG>TA | CG>TA | ACTGGGCCG | CCGGCGT | CGG | GCC | C | A | C>T | C>T_GCC | CG>TA_CCGGCGT | 4 | FALSE | FALSE |
| 9_v.txt | 33118 | 33118-SNV- | SNV | 1 | C | T | 17 | 15180 | 0.11198946 | 46.8235294 | rnr1D57N_per | FALSE | ARG | JSY14 | 2 | rnr1D57N_pe | GCG | C>T | A/T+1 | G/C-1 | CG>TA | CG>TA | GGCCGGCGT | CGTTGGT | CGC | GCG | C | A | C>T | C>T_GCG | CG>TA_CGTTGGT | 4 | FALSE | FALSE |
| 9_v.txt | 33258 | 33258-SNV- | SNV | 1 | C | T | 27 | 17670 | 0.15280136 | 38.4814815 | rnr1D57N_per | FALSE | ARG | JSY14 | 2 | rnr1D57N_pe | GCC | C>T | A/T+1 | G/C-1 | CG>TA | CG>TA | CCTATGCCAT | CCATCCT | CGG | GCC | C | A | C>T | C>T_GCC | CG>TA_CCATCCT | 4 | FALSE | FALSE |
| 9_v.txt | 33276 | 33276-SNV- | SNV | 1 | G | A | 36 | 17670 | 0.20373514 | 45.6111111 | rnr1D57N_per | FALSE | ARG | JSY14 | 2 | rnr1D57N_pe | CGT | G>A | A/T+1 | G/C-1 | CG>TA | CG>TA | GAGAACGTA | GTATCCT | GCA | GCA | C | T | C>T | C>T_GCA | CG>TA_GTATCCT | 4 | FALSE | FALSE |
| 9_v.txt | 33309 | 33309-SNV- | SNV | 1 | C | T | 21 | 17670 | 0.1188455 | 56.472136 | rnr1D57N_per | FALSE | ARG | JSY14 | 2 | rnr1D57N_pe | GCG | C>T | A/T+1 | G/C-1 | CG>TA | CG>TA | AAAGAGCGC | CGCAATG | CGC | GCG | C | A | C>T | C>T_GCG | CG>TA_CGCAATG | 4 | FALSE | FALSE |
| 9_v.txt | 33310 | 33310-SNV- | SNV | 1 | G | A | 28 | 17669 | 0.15846964 | 55.050722 | rnr1D57N_per | FALSE | ARG | JSY14 | 2 | rnr1D57N_pe | CGC | G>A | A/T+1 | G/C-1 | CG>TA | CG>TA | AAGAGCGCA | GCAATGG | GCG | GCG | C | T | C>T | C>T_GCG | CG>TA_GCAATGG | 4 | FALSE | FALSE |
| 9_v.txt | 33412 | 33412-SNV- | SNV | 1 | G | A | 33 | 17670 | 0.18675722 | 49.3636364 | rnr1D57N_per | FALSE | ARG | JSY14 | 2 | rnr1D57N_pe | GGC | G>A | A/T+1 | G/C-1 | CG>TA | CG>TA | TGACCGGCT | GCTCATT | CCG | CCG | C | T | C>T | C>T_CCG | CG>TA_GCTCATT | 4 | FALSE | FALSE |
| 9_v.txt | 33474 | 33474-SNV- | SNV | 1 | C | T | 19 | 17668 | 0.10753905 | 34.9885446 | rnr1D57N_per | FALSE | ARG | JSY14 | 2 | rnr1D57N_pe | GCC | C>T | A/T+1 | G/C-1 | CG>TA | CG>TA | GCTATGCCTT | CCTTTTT | CGG | GCC | C | A | C>T | C>T_GCC | CG>TA_CCTTTTT | 4 | FALSE | FALSE |
| 9_v.txt | 32092 | 32092-SNV- | SNV | 1 | C | G | 3 | 18882 | 0.01588815 | 64 | rnr1D57N_per | FALSE | ARG | JSY14 | 2 | rnr1D57N_pe | GCA | C>G | G/C+1 | G/C-1 | CG>GC | CG>GC | AGCCTGCAA | CAACACC | CGT | GCA | C | C | C>G | C>G_GCA | CG>GC_CAACACC | 4 | FALSE | FALSE |
| 9_v.txt | 32900 | 32900-SNV- | SNV | 1 | G | C | 2 | 15180 | 0.01317523 | 36.4966261 | rnr1D57N_per | FALSE | ARG | JSY14 | 2 | rnr1D57N_pe | GGC | G>C | G/C+1 | G/C-1 | CG>GC | CG>GC | GACTTGGCC | GCCAACT | CCG | CCG | C | G | C>G | C>G_CCG | CG>GC_GCCAACT | 4 | FALSE | FALSE |
| 9_v.txt | 33056 | 33056-SNV- | SNV | 1 | C | G | 3 | 15180 | 0.01976285 | 50 | rnr1D57N_per | FALSE | ARG | JSY14 | 2 | rnr1D57N_pe | GCG | C>G | G/C+1 | G/C-1 | CG>GC | CG>GC | GGACTGCGT | CGTGACA | CGC | GCG | C | C | C>G | C>G_GCG | CG>GC_CGTGACA | 4 | FALSE | FALSE |
| 9_v.txt | 33309 | 33309-SNV- | SNV | 1 | C | G | 1 | 17670 | 5.66E-03 | 63.9930984 | rnr1D57N_per | FALSE | ARG | JSY14 | 2 | rnr1D57N_pe | GCG | C>G | G/C+1 | G/C-1 | CG>GC | CG>GC | AAAGAGCGC | CGCAATG | CGC | GCG | C | C | C>G | C>G_GCG | CG>GC_CGCAATG | 4 | FALSE | FALSE |
| 9_v.txt | 31657 | 31657-SNV- | SNV | 1 | G | T | 106 | 12376 | 0.85649645 | 19.0892481 | rnr1D57N_per | FALSE | ARG | JSY14 | 2 | rnr1D57N_pe | GGC | G>T | A/T+1 | G/C-1 | CG>AT | CG>AT | GAAATGGCG | GCGTGGA | CCG | CCG | C | A | C>A | C>A_CCG | CG>AT_GCGTGGA | 4 | FALSE | FALSE |
| 9_v.txt | 33474 | 33474-SNV- | SNV | 1 | C | A | 67 | 17668 | 0.37921666 | 35.3838589 | rnr1D57N_per | FALSE | ARG | JSY14 | 2 | rnr1D57N_pe | GCC | C>A | A/T+1 | G/C-1 | CG>AT | CG>AT | GCTATGCCTT | CCTTTTT | CGG | GCC | C | T | C>A | C>A_GCC | CG>AT_CCTTTTT | 4 | FALSE | FALSE |
| 9_v.txt | 33475 | 33475-SNV- | SNV | 1 | C | A | 17 | 17667 | 0.0962246 | 37.8720534 | rnr1D57N_per | FALSE | ARG | JSY14 | 2 | rnr1D57N_pe | CCT | C>A | A/T+1 | G/C-1 | CG>AT | CG>AT | CTATGCCTTT | CTTTTTT | GGA | CCT | C | T | C>A | C>A_CCT | CG>AT_CTTTTTT | 4 | FALSE | FALSE |
| 9_v.txt | 32081 | 32081-Insert | Insertion | 1 | - | A | 9 | 18882 | 0.04766444 | 64 | rnr1D57N_per | FALSE | ARG | JSY14 | 2 | rnr1D57N_pe | CAA | nope | A/T+1 | >1bpdel | none | A/T+1 | CCATGCAAA | AAAAAAG | GTT | CAA | - | T | NA | NA_CAA | A/T+1_AAAAAAG | 4 | FALSE | FALSE |
| 9_v.txt | 32842 | 32842-Insert | Insertion | 1 | - | A | 9 | 24361 | 0.0369443 | 40.8888889 | rnr1D57N_per | FALSE | ARG | JSY14 | 2 | rnr1D57N_pe | CAA | nope | A/T+1 | >1bpdel | none | A/T+1 | TTACCCAAA | AAAAAAT | GTT | CAA | - | T | NA | NA_CAA | A/T+1_AAAAAAT | 4 | FALSE | FALSE |
| 9_v.txt | 31663 | 31663-Deleti | Deletion | 1 | A | - | 6 | 12376 | 0.04848093 | 45.8333333 | rnr1D57N_per | FALSE | ARG | JSY14 | 2 | rnr1D57N_pe | GAA | nope | >1bp-ins | A/T-1 | none | A/T-1 | GCGTGGAAA | AAATGTG | CTT | CTT | T | none | T>none | T>none_CTT | A/T-1_AAATGTG | 4 | FALSE | FALSE |
| 9_v.txt | 31723 | 31723-Deleti | Deletion | 1 | A | - | 9 | 12376 | 0.0727214 | 53 | rnr1D57N_per | FALSE | ARG | JSY14 | 2 | rnr1D57N_pe | CAA | nope | >1bp-ins | A/T-1 | none | A/T-1 | TGTCCCAAA | AAAAAGT | GTT | GTT | T | none | T>none | T>none_GTT | A/T-1_AAAAAGT | 4 | FALSE | FALSE |
| 9_v.txt | 31863 | 31863-Deleti | Deletion | 1 | A | - | 5 | 12376 | 0.04040078 | 53.6 | rnr1D57N_per | FALSE | ARG | JSY14 | 2 | rnr1D57N_pe | GAA | nope | >1bp-ins | A/T-1 | none | A/T-1 | AACAGGAAA | AAAATAG | CTT | CTT | T | none | T>none | T>none_CTT | A/T-1_AAAATAG | 4 | FALSE | FALSE |
| 9_v.txt | 31904 | 31904-Deleti | Deletion | 1 | T | - | 6 | 12376 | 0.04848093 | 59.6666667 | rnr1D57N_per | FALSE | ARG | JSY14 | 2 | rnr1D57N_pe | ATT | nope | >1bp-ins | A/T-1 | none | A/T-1 | ATTGAATTTT | TTTTGGT | TAA | ATT | T | none | nope | nope_ATT | A/T-1_TTTTGGT | 4 | FALSE | FALSE |
| 9_v.txt | 31913 | 31913-Deleti | Deletion | 1 | A | - | 7 | 12376 | 0.05656109 | 37.8571429 | rnr1D57N_per | FALSE | ARG | JSY14 | 2 | rnr1D57N_pe | CAA | nope | >1bp-ins | A/T-1 | none | A/T-1 | TGGTGCAAA | AAAAGCC | GTT | GTT | T | none | T>none | T>none_GTT | A/T-1_AAAAGCC | 4 | FALSE | FALSE |

| | | | | | | | | | | | | | | | | | | | | | | | | | | | | | | | | | | |
|---|---|---|---|---|---|---|---|---|---|---|---|---|---|---|---|---|---|---|---|---|---|---|---|---|---|---|---|---|---|---|---|---|---|---|
| 9_v.txt | 31952 | 31952-Deleti | Deletion | 1 | A | - | 5 | 18882 | 0.02648025 | 38 | rnr1D57N_per | FALSE | ARG | JSY14 | 2 | rnr1D57N_pe | TAA | nope | >1bp-ins | A/T-1 | none | A/T-1 | CGTCATAAAT | AAATGTG | ATT | ATT | T | none | T>none | T>none_ATT | A/T-1_AAATGTG | 4 | FALSE | FALSE |
| 9_v.txt | 31972 | 31972-Deleti | Deletion | 1 | C | - | 5 | 18882 | 0.02648025 | 36.8 | rnr1D57N_per | FALSE | ARG | JSY14 | 2 | rnr1D57N_pe | GCC | nope | >1bp-ins | G/C-1 | none | G/C-1 | AATAAGCCA | CCAAGCC | CGG | GCC | C | none | nope | nope_GCC | G/C-1_CCAAGCC | 4 | FALSE | FALSE |
| 9_v.txt | 32081 | 32081-Deleti | Deletion | 1 | A | - | 22 | 18882 | 0.11651308 | 62.8636364 | rnr1D57N_per | FALSE | ARG | JSY14 | 2 | rnr1D57N_pe | CAA | nope | >1bp-ins | A/T-1 | none | A/T-1 | CCATGCAAA | AAAAAAG | GTT | GTT | T | none | T>none | T>none_GTT | A/T-1_AAAAAAG | 4 | FALSE | FALSE |
| 9_v.txt | 32210 | 32210-Deleti | Deletion | 1 | T | - | 5 | 18882 | 0.02648025 | 42.4 | rnr1D57N_per | FALSE | ARG | JSY14 | 2 | rnr1D57N_pe | CTT | nope | >1bp-ins | A/T-1 | none | A/T-1 | ACCACCTTTG | TTTGGTG | GAA | CTT | T | none | nope | nope_CTT | A/T-1_TTTGGTG | 4 | FALSE | FALSE |
| 9_v.txt | 32231 | 32231-Deleti | Deletion | 1 | T | - | 5 | 18882 | 0.02648025 | 37 | rnr1D57N_per | FALSE | ARG | JSY14 | 2 | rnr1D57N_pe | ATT | nope | >1bp-ins | A/T-1 | none | A/T-1 | CAGGAATTT | TTTAGGA | TAA | ATT | T | none | nope | nope_ATT | A/T-1_TTTAGGA | 4 | FALSE | FALSE |
| 9_v.txt | 32346 | 32346-Deleti | Deletion | 1 | A | - | 5 | 20442 | 0.02445945 | 53.2 | rnr1D57N_per | FALSE | ARG | JSY14 | 2 | rnr1D57N_pe | CAA | nope | >1bp-ins | A/T-1 | none | A/T-1 | TGTGGCAAA | AAAACCT | GTT | GTT | T | none | T>none | T>none_GTT | A/T-1_AAAACCT | 4 | FALSE | FALSE |
| 9_v.txt | 32381 | 32381-Deleti | Deletion | 1 | A | - | 6 | 20442 | 0.02935134 | 56.3333333 | rnr1D57N_per | FALSE | ARG | JSY14 | 2 | rnr1D57N_pe | TAA | nope | >1bp-ins | A/T-1 | none | A/T-1 | AATAATAAA | AAAGGGA | ATT | ATT | T | none | T>none | T>none_ATT | A/T-1_AAAGGGA | 4 | FALSE | FALSE |
| 9_v.txt | 32498 | 32498-Deleti | Deletion | 1 | T | - | 30 | 20442 | 0.14675668 | 45.7333333 | rnr1D57N_per | FALSE | ARG | JSY14 | 2 | rnr1D57N_pe | CTT | nope | >1bp-ins | A/T-1 | none | A/T-1 | AACAACTTTT | TTTTTTG | GAA | CTT | T | none | nope | nope_CTT | A/T-1_TTTTTTG | 4 | FALSE | FALSE |
| 9_v.txt | 32528 | 32528-Deleti | Deletion | 1 | G | - | 4 | 20442 | 0.01956756 | 37 | rnr1D57N_per | FALSE | ARG | JSY14 | 2 | rnr1D57N_pe | TGG | nope | >1bp-ins | G/C-1 | none | G/C-1 | TTTTCTGGG | GGGGTTT | ACC | ACC | C | none | C>none | C>none_ACC | G/C-1_GGGGTTT | 4 | FALSE | FALSE |
| 9_v.txt | 32627 | 32627-Deleti | Deletion | 1 | T | - | 3 | 24361 | 0.01231477 | 54.6666667 | rnr1D57N_per | FALSE | ARG | JSY14 | 2 | rnr1D57N_pe | CTT | nope | >1bp-ins | A/T-1 | none | A/T-1 | CCTCCCTTCG | TTCGTTT | GAA | CTT | T | none | nope | nope_CTT | A/T-1_TTCGTTT | 4 | FALSE | FALSE |
| 9_v.txt | 32631 | 32631-Deleti | Deletion | 1 | T | - | 6 | 24361 | 0.02462953 | 37.1666667 | rnr1D57N_per | FALSE | ARG | JSY14 | 2 | rnr1D57N_pe | GTT | nope | >1bp-ins | A/T-1 | none | A/T-1 | CCTTCGTTTT | TTTTTAT | CAA | GTT | T | none | nope | nope_GTT | A/T-1_TTTTTAT | 4 | FALSE | FALSE |
| 9_v.txt | 32647 | 32647-Deleti | Deletion | 1 | T | - | 5 | 24361 | 0.02052461 | 63 | rnr1D57N_per | FALSE | ARG | JSY14 | 2 | rnr1D57N_pe | ATT | nope | >1bp-ins | A/T-1 | none | A/T-1 | TAGATATTAT | TTATACC | TAA | ATT | T | none | nope | nope_ATT | A/T-1_TTATACC | 4 | FALSE | FALSE |
| 9_v.txt | 32658 | 32658-Deleti | Deletion | 1 | C | - | 10 | 24361 | 0.04104922 | 52.8 | rnr1D57N_per | FALSE | ARG | JSY14 | 2 | rnr1D57N_pe | ACC | nope | >1bp-ins | G/C-1 | none | G/C-1 | CCTGGACCC | CCCCAGG | TGG | ACC | C | none | nope | nope_ACC | G/C-1_CCCCAGG | 4 | FALSE | FALSE |
| 9_v.txt | 32707 | 32707-Deleti | Deletion | 1 | C | - | 6 | 24361 | 0.02462953 | 61.5 | rnr1D57N_per | FALSE | ARG | JSY14 | 2 | rnr1D57N_pe | ACC | nope | >1bp-ins | G/C-1 | none | G/C-1 | CGGTAACCC | CCCCAGC | TGG | ACC | C | none | nope | nope_ACC | G/C-1_CCCCAGC | 4 | FALSE | FALSE |
| 9_v.txt | 32728 | 32728-Deleti | Deletion | 1 | A | - | 5 | 24361 | 0.02052461 | 58.8 | rnr1D57N_per | FALSE | ARG | JSY14 | 2 | rnr1D57N_pe | CAA | nope | >1bp-ins | A/T-1 | none | A/T-1 | CCATACAAA | AAAAACA | GTT | GTT | T | none | T>none | T>none_GTT | A/T-1_AAAAACA | 4 | FALSE | FALSE |
| 9_v.txt | 32842 | 32842-Deleti | Deletion | 1 | A | - | 31 | 24361 | 0.12725258 | 43.1612903 | rnr1D57N_per | FALSE | ARG | JSY14 | 2 | rnr1D57N_pe | CAA | nope | >1bp-ins | A/T-1 | none | A/T-1 | TTACCCAAA | AAAAAAT | GTT | GTT | T | none | T>none | T>none_GTT | A/T-1_AAAAAAT | 4 | FALSE | FALSE |
| 9_v.txt | 32884 | 32884-Deleti | Deletion | 1 | A | - | 6 | 15180 | 0.03952569 | 37.3333333 | rnr1D57N_per | FALSE | ARG | JSY14 | 2 | rnr1D57N_pe | CAA | nope | >1bp-ins | A/T-1 | none | A/T-1 | ACGTCCAAA | AAAATTG | GTT | GTT | T | none | T>none | T>none_GTT | A/T-1_AAAATTG | 4 | FALSE | FALSE |
| 9_v.txt | 32924 | 32924-Deleti | Deletion | 1 | A | - | 15 | 15180 | 0.09881423 | 56.4 | rnr1D57N_per | FALSE | ARG | JSY14 | 2 | rnr1D57N_pe | CAA | nope | >1bp-ins | A/T-1 | none | A/T-1 | CAGGGCAAA | AAAAGTG | GTT | GTT | T | none | T>none | T>none_GTT | A/T-1_AAAAGTG | 4 | FALSE | FALSE |
| 9_v.txt | 33040 | 33040-Deleti | Deletion | 1 | T | - | 4 | 15180 | 0.02635046 | 64 | rnr1D57N_per | FALSE | ARG | JSY14 | 2 | rnr1D57N_pe | ATT | nope | >1bp-ins | A/T-1 | none | A/T-1 | TAGCCATTTC | TTTCACC | TAA | ATT | T | none | nope | nope_ATT | A/T-1_TTTCACC | 4 | FALSE | FALSE |
| 9_v.txt | 33474 | 33474-Deleti | Deletion | 1 | C | - | 43 | 17668 | 0.24337786 | 28.7906977 | rnr1D57N_per | FALSE | ARG | JSY14 | 2 | rnr1D57N_pe | GCC | nope | >1bp-ins | G/C-1 | none | G/C-1 | GCTATGCCTT | CCTTTTT | CGG | GCC | C | none | nope | nope_GCC | G/C-1_CCTTTTT | 4 | FALSE | FALSE |
| 10_v.txt | 31723 | 31723-SNV-A | SNV | 1 | A | C | 5 | 12936 | 0.03865182 | 57.6 | rnr1D57N_per | FALSE | ARG | JSY14 | 3 | rnr1D57N_pe | CAA | A>C | G/C+1 | A/T-1 | TA>GC | TA>GC | TGTCCCAAA | AAAAAGT | GTT | GTT | T | G | T>G | T>G_GTT | TA>GC_AAAAAGT | 4 | FALSE | FALSE |
| 10_v.txt | 32081 | 32081-SNV-A | SNV | 1 | A | C | 1 | 21511 | 4.65E-03 | 17.0037806 | rnr1D57N_per | FALSE | ARG | JSY14 | 3 | rnr1D57N_pe | CAA | A>C | G/C+1 | A/T-1 | TA>GC | TA>GC | CCATGCAAA | AAAAAAG | GTT | GTT | T | G | T>G | T>G_GTT | TA>GC_AAAAAAG | 4 | FALSE | FALSE |
| 10_v.txt | 32842 | 32842-SNV-A | SNV | 1 | A | C | 2 | 21882 | 9.14E-03 | 37.9966175 | rnr1D57N_per | FALSE | ARG | JSY14 | 3 | rnr1D57N_pe | CAA | A>C | G/C+1 | A/T-1 | TA>GC | TA>GC | TTACCCAAA | AAAAAAT | GTT | GTT | T | G | T>G | T>G_GTT | TA>GC_AAAAAAT | 4 | FALSE | FALSE |
| 10_v.txt | 33269 | 33269-SNV-A | SNV | 1 | A | C | 141 | 19820 | 0.71140262 | 17.1205674 | rnr1D57N_per | FALSE | ARG | JSY14 | 3 | rnr1D57N_pe | TAG | A>C | G/C+1 | A/T-1 | TA>GC | TA>GC | CTCCATAGA | AGAGAAC | ATC | ATC | T | G | T>G | T>G_ATC | TA>GC_AGAGAAC | 4 | FALSE | FALSE |
| 10_v.txt | 33476 | 33476-SNV-T | SNV | 1 | T | C | 31 | 19819 | 0.15641556 | 32.0006552 | rnr1D57N_per | FALSE | ARG | JSY14 | 3 | rnr1D57N_pe | CTT | T>C | G/C+1 | A/T-1 | TA>CG | TA>CG | TATGCCTTTT | TTTTTTT | GAA | CTT | T | G | T>C | T>C_CTT | TA>CG_TTTTTTT | 4 | FALSE | FALSE |
| 10_v.txt | 31650 | 31650-SNV-C | SNV | 1 | C | T | 29 | 12936 | 0.22418058 | 35.5862069 | rnr1D57N_per | FALSE | ARG | JSY14 | 3 | rnr1D57N_pe | GCG | C>T | A/T+1 | G/C-1 | CG>TA | CG>TA | AGAATGCGA | CGAAATG | CGC | GCG | C | A | C>T | C>T_GCG | CG>TA_CGAAATG | 4 | FALSE | FALSE |
| 10_v.txt | 31658 | 31658-SNV-C | SNV | 1 | C | T | 23 | 12936 | 0.17779839 | 32.9130435 | rnr1D57N_per | FALSE | ARG | JSY14 | 3 | rnr1D57N_pe | GCG | C>T | A/T+1 | G/C-1 | CG>TA | CG>TA | AAATGGCGT | CGTGGAA | CGC | GCG | C | A | C>T | C>T_GCG | CG>TA_CGTGGAA | 4 | FALSE | FALSE |
| 10_v.txt | 31790 | 31790-SNV-G | SNV | 1 | G | A | 15 | 12936 | 0.11595547 | 58 | rnr1D57N_per | FALSE | ARG | JSY14 | 3 | rnr1D57N_pe | CGA | G>A | A/T+1 | G/C-1 | CG>TA | CG>TA | GATGTCGAC | GACATCT | GCT | GCT | C | T | C>T | C>T_GCT | CG>TA_GACATCT | 4 | FALSE | FALSE |
| 10_v.txt | 31830 | 31830-SNV-C | SNV | 1 | C | T | 17 | 12936 | 0.1314162 | 56.1764706 | rnr1D57N_per | FALSE | ARG | JSY14 | 3 | rnr1D57N_pe | GCA | C>T | A/T+1 | G/C-1 | CG>TA | CG>TA | AATATGCATT | CATTGAA | CGT | GCA | C | A | C>T | C>T_GCA | CG>TA_CATTGAA | 4 | FALSE | FALSE |
| 10_v.txt | 31911 | 31911-SNV-G | SNV | 1 | G | A | 17 | 12936 | 0.1314162 | 37.1176471 | rnr1D57N_per | FALSE | ARG | JSY14 | 3 | rnr1D57N_pe | TGC | G>A | A/T+1 | G/C-1 | CG>TA | CG>TA | TTTGGTGCA | GCAAAAG | ACG | ACG | C | T | C>T | C>T_ACG | CG>TA_GCAAAAG | 4 | FALSE | FALSE |
| 10_v.txt | 31962 | 31962-SNV-G | SNV | 1 | G | A | 44 | 21512 | 0.204537 | 36.3636364 | rnr1D57N_per | FALSE | ARG | JSY14 | 3 | rnr1D57N_pe | CGC | G>A | A/T+1 | G/C-1 | CG>TA | CG>TA | GTGGCCGCA | GCATAAT | GCG | GCG | C | T | C>T | C>T_GCG | CG>TA_GCATAAT | 4 | FALSE | FALSE |
| 10_v.txt | 31972 | 31972-SNV-C | SNV | 1 | C | T | 19 | 21512 | 0.0883228 | 37.1052632 | rnr1D57N_per | FALSE | ARG | JSY14 | 3 | rnr1D57N_pe | GCC | C>T | A/T+1 | G/C-1 | CG>TA | CG>TA | AATAAGCCA | CCAAGCC | CGG | GCC | C | A | C>T | C>T_GCC | CG>TA_CCAAGCC | 4 | FALSE | FALSE |
| 10_v.txt | 32057 | 32057-SNV-C | SNV | 1 | C | T | 20 | 21512 | 0.09297137 | 61.5 | rnr1D57N_per | FALSE | ARG | JSY14 | 3 | rnr1D57N_pe | GCG | C>T | A/T+1 | G/C-1 | CG>TA | CG>TA | GATGTGCGA | CGAGATT | CGC | GCG | C | A | C>T | C>T_GCG | CG>TA_CGAGATT | 4 | FALSE | FALSE |
| 10_v.txt | 32092 | 32092-SNV-C | SNV | 1 | C | T | 22 | 21512 | 0.1022685 | 60.0454546 | rnr1D57N_per | FALSE | ARG | JSY14 | 3 | rnr1D57N_pe | GCA | C>T | A/T+1 | G/C-1 | CG>TA | CG>TA | AGCCTGCAA | CAACACC | CGT | GCA | C | A | C>T | C>T_GCA | CG>TA_CAACACC | 4 | FALSE | FALSE |
| 10_v.txt | 32165 | 32165-SNV-G | SNV | 1 | G | A | 32 | 21512 | 0.14875418 | 44.15625 | rnr1D57N_per | FALSE | ARG | JSY14 | 3 | rnr1D57N_pe | CGC | G>A | A/T+1 | G/C-1 | CG>TA | CG>TA | CAAAGCGCC | GCCAAAT | GCG | GCG | C | T | C>T | C>T_GCG | CG>TA_GCCAAAT | 4 | FALSE | FALSE |
| 10_v.txt | 32173 | 32173-SNV-C | SNV | 1 | C | T | 20 | 21512 | 0.09297137 | 48.05 | rnr1D57N_per | FALSE | ARG | JSY14 | 3 | rnr1D57N_pe | GCA | C>T | A/T+1 | G/C-1 | CG>TA | CG>TA | CAAATGCAG | CAGCAGT | CGT | GCA | C | A | C>T | C>T_GCA | CG>TA_CAGCAGT | 4 | FALSE | FALSE |
| 10_v.txt | 32271 | 32271-SNV-C | SNV | 1 | C | T | 34 | 23299 | 0.14592901 | 28.2647059 | rnr1D57N_per | FALSE | ARG | JSY14 | 3 | rnr1D57N_pe | ACG | C>T | A/T+1 | G/C-1 | CG>TA | CG>TA | AAAATACGG | CGGGAAC | TGC | ACG | C | A | C>T | C>T_ACG | CG>TA_CGGGAAC | 4 | FALSE | FALSE |
| 10_v.txt | 32299 | 32299-SNV-C | SNV | 1 | C | T | 35 | 23299 | 0.15022104 | 53.1428571 | rnr1D57N_per | FALSE | ARG | JSY14 | 3 | rnr1D57N_pe | GCG | C>T | A/T+1 | G/C-1 | CG>TA | CG>TA | AATTTGCGG | CGGCAGA | CGC | GCG | C | A | C>T | C>T_GCG | CG>TA_CGGCAGA | 4 | FALSE | FALSE |
| 10_v.txt | 32466 | 32466-SNV-C | SNV | 1 | C | T | 26 | 23299 | 0.11159277 | 47.6153846 | rnr1D57N_per | FALSE | ARG | JSY14 | 3 | rnr1D57N_pe | GCC | C>T | A/T+1 | G/C-1 | CG>TA | CG>TA | AGAGAGCCA | CCAATGT | CGG | GCC | C | A | C>T | C>T_GCC | CG>TA_CCAATGT | 4 | FALSE | FALSE |
| 10_v.txt | 32535 | 32535-SNV-G | SNV | 1 | G | A | 31 | 23298 | 0.13305863 | 36.8065346 | rnr1D57N_per | FALSE | ARG | JSY14 | 3 | rnr1D57N_pe | TGC | G>A | A/T+1 | G/C-1 | CG>TA | CG>TA | GGGTTTGCA | GCAGCTT | ACG | ACG | C | T | C>T | C>T_ACG | CG>TA_GCAGCTT | 4 | FALSE | FALSE |
| 10_v.txt | 32536 | 32536-SNV-C | SNV | 1 | C | T | 39 | 23298 | 0.16739634 | 37.3333333 | rnr1D57N_per | FALSE | ARG | JSY14 | 3 | rnr1D57N_pe | GCA | C>T | A/T+1 | G/C-1 | CG>TA | CG>TA | GGTTTGCAG | CAGCTTC | CGT | GCA | C | A | C>T | C>T_GCA | CG>TA_CAGCTTC | 4 | FALSE | FALSE |
| 10_v.txt | 32590 | 32590-SNV-C | SNV | 1 | C | T | 43 | 21882 | 0.19650855 | 35.5116279 | rnr1D57N_per | FALSE | ARG | JSY14 | 3 | rnr1D57N_pe | GCG | C>T | A/T+1 | G/C-1 | CG>TA | CG>TA | AGGCAGCGT | CGTTAAT | CGC | GCG | C | A | C>T | C>T_GCG | CG>TA_CGTTAAT | 4 | FALSE | FALSE |
| 10_v.txt | 32700 | 32700-SNV-C | SNV | 1 | C | T | 30 | 21882 | 0.13709899 | 57.8666667 | rnr1D57N_per | FALSE | ARG | JSY14 | 3 | rnr1D57N_pe | GCC | C>T | A/T+1 | G/C-1 | CG>TA | CG>TA | ACTGGGCCG | CCGGTAA | CGG | GCC | C | A | C>T | C>T_GCC | CG>TA_CCGGTAA | 4 | FALSE | FALSE |
| 10_v.txt | 32757 | 32757-SNV-G | SNV | 1 | G | A | 22 | 21882 | 0.10053926 | 59.0909091 | rnr1D57N_per | FALSE | ARG | JSY14 | 3 | rnr1D57N_pe | GGC | G>A | A/T+1 | G/C-1 | CG>TA | CG>TA | ATAATGGCT | GCTAAAA | CCG | CCG | C | T | C>T | C>T_CCG | CG>TA_GCTAAAA | 4 | FALSE | FALSE |
| 10_v.txt | 32776 | 32776-SNV-C | SNV | 1 | C | T | 32 | 21882 | 0.14623892 | 54.15625 | rnr1D57N_per | FALSE | ARG | JSY14 | 3 | rnr1D57N_pe | GCG | C>T | A/T+1 | G/C-1 | CG>TA | CG>TA | TGGAAGCGA | CGACCCA | CGC | GCG | C | A | C>T | C>T_GCG | CG>TA_CGACCCA | 4 | FALSE | FALSE |
| 10_v.txt | 32798 | 32798-SNV-G | SNV | 1 | G | A | 27 | 21882 | 0.12338909 | 40.2222222 | rnr1D57N_per | FALSE | ARG | JSY14 | 3 | rnr1D57N_pe | CGT | G>A | A/T+1 | G/C-1 | CG>TA | CG>TA | TTCACCGTA | GTAATAT | GCA | GCA | C | T | C>T | C>T_GCA | CG>TA_GTAATAT | 4 | FALSE | FALSE |
| 10_v.txt | 32860 | 32860-SNV-C | SNV | 1 | C | T | 22 | 21880 | 0.10054845 | 37.6363636 | rnr1D57N_per | FALSE | ARG | JSY14 | 3 | rnr1D57N_pe | GCC | C>T | A/T+1 | G/C-1 | CG>TA | CG>TA | TCCATGCCG | CCGCCAG | CGG | GCC | C | A | C>T | C>T_GCC | CG>TA_CCGCCAG | 4 | FALSE | FALSE |
| 10_v.txt | 32880 | 32880-SNV-G | SNV | 1 | G | A | 22 | 19173 | 0.11474469 | 33 | rnr1D57N_per | FALSE | ARG | JSY14 | 3 | rnr1D57N_pe | CGT | G>A | A/T+1 | G/C-1 | CG>TA | CG>TA | TTGTACGTC | GTCCAAA | GCA | GCA | C | T | C>T | C>T_GCA | CG>TA_GTCCAAA | 4 | FALSE | FALSE |
| 10_v.txt | 32901 | 32901-SNV-C | SNV | 1 | C | T | 40 | 19173 | 0.20862672 | 32.175 | rnr1D57N_per | FALSE | ARG | JSY14 | 3 | rnr1D57N_pe | GCC | C>T | A/T+1 | G/C-1 | CG>TA | CG>TA | ACTTGGCCA | CCAACTA | CGG | GCC | C | A | C>T | C>T_GCC | CG>TA_CCAACTA | 4 | FALSE | FALSE |
| 10_v.txt | 32922 | 32922-SNV-G | SNV | 1 | G | A | 21 | 19173 | 0.10952903 | 51.952381 | rnr1D57N_per | FALSE | ARG | JSY14 | 3 | rnr1D57N_pe | GGC | G>A | A/T+1 | G/C-1 | CG>TA | CG>TA | TCCAGGGCA | GCAAAAG | CCG | CCG | C | T | C>T | C>T_CCG | CG>TA_GCAAAAG | 4 | FALSE | FALSE |
| 10_v.txt | 32935 | 32935-SNV-C | SNV | 1 | C | T | 35 | 19173 | 0.18254838 | 37.8 | rnr1D57N_per | FALSE | ARG | JSY14 | 3 | rnr1D57N_pe | GCC | C>T | A/T+1 | G/C-1 | CG>TA | CG>TA | TGATTGCCC | CCCAAGA | CGG | GCC | C | A | C>T | C>T_GCC | CG>TA_CCCAAGA | 4 | FALSE | FALSE |
| 10_v.txt | 32965 | 32965-SNV-C | SNV | 1 | C | T | 17 | 19173 | 0.08866635 | 61.1176471 | rnr1D57N_per | FALSE | ARG | JSY14 | 3 | rnr1D57N_pe | GCC | C>T | A/T+1 | G/C-1 | CG>TA | CG>TA | CATTGGCCG | CCGCACC | CGG | GCC | C | A | C>T | C>T_GCC | CG>TA_CCGCACC | 4 | FALSE | FALSE |
| 10_v.txt | 32967 | 32967-SNV-G | SNV | 1 | G | A | 52 | 19173 | 0.27121473 | 60.2307692 | rnr1D57N_per | FALSE | ARG | JSY14 | 3 | rnr1D57N_pe | CGC | G>A | A/T+1 | G/C-1 | CG>TA | CG>TA | TTGGCCGCA | GCACCAA | GCG | GCG | C | T | C>T | C>T_GCG | CG>TA_GCACCAA | 4 | FALSE | FALSE |
| 10_v.txt | 33056 | 33056-SNV-C | SNV | 1 | C | T | 26 | 19173 | 0.13560737 | 56.2692308 | rnr1D57N_per | FALSE | ARG | JSY14 | 3 | rnr1D57N_pe | GCG | C>T | A/T+1 | G/C-1 | CG>TA | CG>TA | GGACTGCGT | CGTGACA | CGC | GCG | C | A | C>T | C>T_GCG | CG>TA_CGTGACA | 4 | FALSE | FALSE |
| 10_v.txt | 33070 | 33070-SNV-C | SNV | 1 | C | T | 22 | 19173 | 0.11474469 | 53 | rnr1D57N_per | FALSE | ARG | JSY14 | 3 | rnr1D57N_pe | GCC | C>T | A/T+1 | G/C-1 | CG>TA | CG>TA | AATATGCCA | CCAAAGA | CGG | GCC | C | A | C>T | C>T_GCC | CG>TA_CCAAAGA | 4 | FALSE | FALSE |
| 10_v.txt | 33113 | 33113-SNV-G | SNV | 1 | G | A | 21 | 19173 | 0.10952903 | 46.0473334 | rnr1D57N_per | FALSE | ARG | JSY14 | 3 | rnr1D57N_pe | GGC | G>A | A/T+1 | G/C-1 | CG>TA | CG>TA | CACTGGGCC | GCCGGCG | CCG | CCG | C | T | C>T | C>T_CCG | CG>TA_GCCGGCG | 4 | FALSE | FALSE |
| 10_v.txt | 33114 | 33114-SNV-C | SNV | 1 | C | T | 20 | 19173 | 0.10431336 | 39.4007964 | rnr1D57N_per | FALSE | ARG | JSY14 | 3 | rnr1D57N_pe | GCC | C>T | A/T+1 | G/C-1 | CG>TA | CG>TA | ACTGGGCCG | CCGGCGT | CGG | GCC | C | A | C>T | C>T_GCC | CG>TA_CCGGCGT | 4 | FALSE | FALSE |
| 10_v.txt | 33118 | 33118-SNV-C | SNV | 1 | C | T | 25 | 19173 | 0.1303917 | 40.2 | rnr1D57N_per | FALSE | ARG | JSY14 | 3 | rnr1D57N_pe | GCG | C>T | A/T+1 | G/C-1 | CG>TA | CG>TA | GGCCGGCGT | CGTTGGT | CGC | GCG | C | A | C>T | C>T_GCG | CG>TA_CGTTGGT | 4 | FALSE | FALSE |
| 10_v.txt | 33258 | 33258-SNV-C | SNV | 1 | C | T | 24 | 19820 | 0.12108981 | 41.6666667 | rnr1D57N_per | FALSE | ARG | JSY14 | 3 | rnr1D57N_pe | GCC | C>T | A/T+1 | G/C-1 | CG>TA | CG>TA | CCTATGCCAT | CCATCCT | CGG | GCC | C | A | C>T | C>T_GCC | CG>TA_CCATCCT | 4 | FALSE | FALSE |
| 10_v.txt | 33276 | 33276-SNV-G | SNV | 1 | G | A | 37 | 19820 | 0.18668012 | 43.8108108 | rnr1D57N_per | FALSE | ARG | JSY14 | 3 | rnr1D57N_pe | CGT | G>A | A/T+1 | G/C-1 | CG>TA | CG>TA | GAGAACGTA | GTATCCT | GCA | GCA | C | T | C>T | C>T_GCA | CG>TA_GTATCCT | 4 | FALSE | FALSE |
| 10_v.txt | 33309 | 33309-SNV-C | SNV | 1 | C | T | 23 | 19819 | 0.11605026 | 56.2147059 | rnr1D57N_per | FALSE | ARG | JSY14 | 3 | rnr1D57N_pe | GCG | C>T | A/T+1 | G/C-1 | CG>TA | CG>TA | AAAGAGCGC | CGCAATG | CGC | GCG | C | A | C>T | C>T_GCG | CG>TA_CGCAATG | 4 | FALSE | FALSE |
| 10_v.txt | 33310 | 33310-SNV-G | SNV | 1 | G | A | 35 | 19817 | 0.17661604 | 56.5613813 | rnr1D57N_per | FALSE | ARG | JSY14 | 3 | rnr1D57N_pe | CGC | G>A | A/T+1 | G/C-1 | CG>TA | CG>TA | AAGAGCGCA | GCAATGG | GCG | GCG | C | T | C>T | C>T_GCG | CG>TA_GCAATGG | 4 | FALSE | FALSE |
| 10_v.txt | 33392 | 33392-SNV-G | SNV | 1 | G | A | 28 | 19820 | 0.14127144 | 54.3928571 | rnr1D57N_per | FALSE | ARG | JSY14 | 3 | rnr1D57N_pe | CGT | G>A | A/T+1 | G/C-1 | CG>TA | CG>TA | AACGTCGTG | GTGAAAG | GCA | GCA | C | T | C>T | C>T_GCA | CG>TA_GTGAAAG | 4 | FALSE | FALSE |
| 10_v.txt | 33412 | 33412-SNV-G | SNV | 1 | G | A | 31 | 19820 | 0.15640767 | 49.1290323 | rnr1D57N_per | FALSE | ARG | JSY14 | 3 | rnr1D57N_pe | GGC | G>A | A/T+1 | G/C-1 | CG>TA | CG>TA | TGACCGGCT | GCTCATT | CCG | CCG | C | T | C>T | C>T_CCG | CG>TA_GCTCATT | 4 | FALSE | FALSE |
| 10_v.txt | 33474 | 33474-SNV-C | SNV | 1 | C | T | 28 | 19820 | 0.14127144 | 34.6787115 | rnr1D57N_per | FALSE | ARG | JSY14 | 3 | rnr1D57N_pe | GCC | C>T | A/T+1 | G/C-1 | CG>TA | CG>TA | GCTATGCCTT | CCTTTTT | CGG | GCC | C | A | C>T | C>T_GCC | CG>TA_CCTTTTT | 4 | FALSE | FALSE |
| 10_v.txt | 32536 | 32536-SNV-C | SNV | 1 | C | G | 2 | 23298 | 8.58E-03 | 38 | rnr1D57N_per | FALSE | ARG | JSY14 | 3 | rnr1D57N_pe | GCA | C>G | G/C+1 | G/C-1 | CG>GC | CG>GC | GGTTTGCAG | CAGCTTC | CGT | GCA | C | C | C>G | C>G_GCA | CG>GC_CAGCTTC | 4 | FALSE | FALSE |
| 10_v.txt | 33114 | 33114-SNV-C | SNV | 1 | C | G | 2 | 19173 | 0.01043134 | 38.0035127 | rnr1D57N_per | FALSE | ARG | JSY14 | 3 | rnr1D57N_pe | GCC | C>G | G/C+1 | G/C-1 | CG>GC | CG>GC | ACTGGGCCG | CCGGCGT | CGG | GCC | C | C | C>G | C>G_GCC | CG>GC_CCGGCGT | 4 | FALSE | FALSE |
| 10_v.txt | 33309 | 33309-SNV-C | SNV | 1 | C | G | 1 | 19819 | 5.05E-03 | 28.0075669 | rnr1D57N_per | FALSE | ARG | JSY14 | 3 | rnr1D57N_pe | GCG | C>G | G/C+1 | G/C-1 | CG>GC | CG>GC | AAAGAGCGC | CGCAATG | CGC | GCG | C | C | C>G | C>G_GCG | CG>GC_CGCAATG | 4 | FALSE | FALSE |
| 10_v.txt | 31657 | 31657-SNV-G | SNV | 1 | G | T | 118 | 12936 | 0.91218306 | 18.0409495 | rnr1D57N_per | FALSE | ARG | JSY14 | 3 | rnr1D57N_pe | GGC | G>T | A/T+1 | G/C-1 | CG>AT | CG>AT | GAAATGGCG | GCGTGGA | CCG | CCG | C | A | C>A | C>A_CCG | CG>AT_GCGTGGA | 4 | FALSE | FALSE |
| 10_v.txt | 32536 | 32536-SNV-C | SNV | 1 | C | A | 4 | 23298 | 0.01716886 | 37.5 | rnr1D57N_per | FALSE | ARG | JSY14 | 3 | rnr1D57N_pe | GCA | C>A | A/T+1 | G/C-1 | CG>AT | CG>AT | GGTTTGCAG | CAGCTTC | CGT | GCA | C | T | C>A | C>A_GCA | CG>AT_CAGCTTC | 4 | FALSE | FALSE |
| 10_v.txt | 33474 | 33474-SNV-C | SNV | 1 | C | A | 56 | 19820 | 0.28254289 | 33.8230719 | rnr1D57N_per | FALSE | ARG | JSY14 | 3 | rnr1D57N_pe | GCC | C>A | A/T+1 | G/C-1 | CG>AT | CG>AT | GCTATGCCTT | CCTTTTT | CGG | GCC | C | T | C>A | C>A_GCC | CG>AT_CCTTTTT | 4 | FALSE | FALSE |
| 10_v.txt | 33475 | 33475-SNV-C | SNV | 1 | C | A | 17 | 19820 | 0.08577195 | 35.7555819 | rnr1D57N_per | FALSE | ARG | JSY14 | 3 | rnr1D57N_pe | CCT | C>A | A/T+1 | G/C-1 | CG>AT | CG>AT | CTATGCCTTT | CTTTTTT | GGA | CCT | C | T | C>A | C>A_CCT | CG>AT_CTTTTTT | 4 | FALSE | FALSE |
| 10_v.txt | 32081 | 32081-Insert | Insertion | 1 | - | A | 11 | 21512 | 0.05113425 | 64 | rnr1D57N_per | FALSE | ARG | JSY14 | 3 | rnr1D57N_pe | CAA | nope | A/T+1 | >1bpdel | none | A/T+1 | CCATGCAAA | AAAAAAG | GTT | CAA | - | T | NA | NA_CAA | A/T+1_AAAAAAG | 4 | FALSE | FALSE |
| 10_v.txt | 32842 | 32842-Insert | Insertion | 1 | - | A | 8 | 21882 | 0.03655973 | 44.125 | rnr1D57N_per | FALSE | ARG | JSY14 | 3 | rnr1D57N_pe | CAA | nope | A/T+1 | >1bpdel | none | A/T+1 | TTACCCAAA | AAAAAAT | GTT | CAA | - | T | NA | NA_CAA | A/T+1_AAAAAAT | 4 | FALSE | FALSE |
| 10_v.txt | 32884 | 32884-Insert | Insertion | 1 | - | A | 2 | 19173 | 0.01043134 | 38 | rnr1D57N_per | FALSE | ARG | JSY14 | 3 | rnr1D57N_pe | CAA | nope | A/T+1 | >1bpdel | none | A/T+1 | ACGTCCAAA | AAAATTG | GTT | CAA | - | T | NA | NA_CAA | A/T+1_AAAATTG | 4 | FALSE | FALSE |
| 10_v.txt | 31663 | 31663-Deleti | Deletion | 1 | A | - | 5 | 12936 | 0.03865182 | 35.8 | rnr1D57N_per | FALSE | ARG | JSY14 | 3 | rnr1D57N_pe | GAA | nope | >1bp-ins | A/T-1 | none | A/T-1 | GCGTGGAAA | AAATGTG | CTT | CTT | T | none | T>none | T>none_CTT | A/T-1_AAATGTG | 4 | FALSE | FALSE |
| 10_v.txt | 31723 | 31723-Deleti | Deletion | 1 | A | - | 21 | 12936 | 0.16233766 | 62.5238095 | rnr1D57N_per | FALSE | ARG | JSY14 | 3 | rnr1D57N_pe | CAA | nope | >1bp-ins | A/T-1 | none | A/T-1 | TGTCCCAAA | AAAAAGT | GTT | GTT | T | none | T>none | T>none_GTT | A/T-1_AAAAAGT | 4 | FALSE | FALSE |
| 10_v.txt | 31904 | 31904-Deleti | Deletion | 1 | T | - | 12 | 12936 | 0.09276438 | 43.6666667 | rnr1D57N_per | FALSE | ARG | JSY14 | 3 | rnr1D57N_pe | ATT | nope | >1bp-ins | A/T-1 | none | A/T-1 | ATTGAATTTT | TTTTGGT | TAA | ATT | T | none | nope | nope_ATT | A/T-1_TTTTGGT | 4 | FALSE | FALSE |
| 10_v.txt | 31958 | 31958-Deleti | Deletion | 1 | G | - | 5 | 21512 | 0.02324284 | 34.6 | rnr1D57N_per | FALSE | ARG | JSY14 | 3 | rnr1D57N_pe | TGG | nope | >1bp-ins | G/C-1 | none | G/C-1 | AAATGTGGC | GGCCGCA | ACC | ACC | C | none | C>none | C>none_ACC | G/C-1_GGCCGCA | 4 | FALSE | FALSE |
| 10_v.txt | 31972 | 31972-Deleti | Deletion | 1 | C | - | 8 | 21512 | 0.03718855 | 35.5 | rnr1D57N_per | FALSE | ARG | JSY14 | 3 | rnr1D57N_pe | GCC | nope | >1bp-ins | G/C-1 | none | G/C-1 | AATAAGCCA | CCAAGCC | CGG | GCC | C | none | nope | nope_GCC | G/C-1_CCAAGCC | 4 | FALSE | FALSE |
| 10_v.txt | 32081 | 32081-Deleti | Deletion | 1 | A | - | 20 | 21511 | 0.09297569 | 64 | rnr1D57N_per | FALSE | ARG | JSY14 | 3 | rnr1D57N_pe | CAA | nope | >1bp-ins | A/T-1 | none | A/T-1 | CCATGCAAA | AAAAAAG | GTT | GTT | T | none | T>none | T>none_GTT | A/T-1_AAAAAAG | 4 | FALSE | FALSE |
| 10_v.txt | 32210 | 32210-Deleti | Deletion | 1 | T | - | 9 | 21512 | 0.04183711 | 37 | rnr1D57N_per | FALSE | ARG | JSY14 | 3 | rnr1D57N_pe | CTT | nope | >1bp-ins | A/T-1 | none | A/T-1 | ACCACCTTTG | TTTGGTG | GAA | CTT | T | none | nope | nope_CTT | A/T-1_TTTGGTG | 4 | FALSE | FALSE |
| 10_v.txt | 32346 | 32346-Deleti | Deletion | 1 | A | - | 9 | 23299 | 0.03862827 | 52.6666667 | rnr1D57N_per | FALSE | ARG | JSY14 | 3 | rnr1D57N_pe | CAA | nope | >1bp-ins | A/T-1 | none | A/T-1 | TGTGGCAAA | AAAACCT | GTT | GTT | T | none | T>none | T>none_GTT | A/T-1_AAAACCT | 4 | FALSE | FALSE |
| 10_v.txt | 32416 | 32416-Deleti | Deletion | 1 | T | - | 5 | 23299 | 0.02146015 | 53.2 | rnr1D57N_per | FALSE | ARG | JSY14 | 3 | rnr1D57N_pe | GTT | nope | >1bp-ins | A/T-1 | none | A/T-1 | ATTGTGTTAG | TTAGTTT | CAA | GTT | T | none | nope | nope_GTT | A/T-1_TTAGTTT | 4 | FALSE | FALSE |
| 10_v.txt | 32442 | 32442-Deleti | Deletion | 1 | A | - | 10 | 23299 | 0.0429203 | 42.4 | rnr1D57N_per | FALSE | ARG | JSY14 | 3 | rnr1D57N_pe | TAA | nope | >1bp-ins | A/T-1 | none | A/T-1 | GGAACTAAA | AAAAGTC | ATT | ATT | T | none | T>none | T>none_ATT | A/T-1_AAAAGTC | 4 | FALSE | FALSE |
| 10_v.txt | 32498 | 32498-Deleti | Deletion | 1 | T | - | 29 | 23299 | 0.12446886 | 42.8965517 | rnr1D57N_per | FALSE | ARG | JSY14 | 3 | rnr1D57N_pe | CTT | nope | >1bp-ins | A/T-1 | none | A/T-1 | AACAACTTTT | TTTTTTG | GAA | CTT | T | none | nope | nope_CTT | A/T-1_TTTTTTG | 4 | FALSE | FALSE |
| 10_v.txt | 32528 | 32528-Deleti | Deletion | 1 | G | - | 7 | 23299 | 0.03004421 | 44.7142857 | rnr1D57N_per | FALSE | ARG | JSY14 | 3 | rnr1D57N_pe | TGG | nope | >1bp-ins | G/C-1 | none | G/C-1 | TTTTCTGGG | GGGGTTT | ACC | ACC | C | none | C>none | C>none_ACC | G/C-1_GGGGTTT | 4 | FALSE | FALSE |
| 10_v.txt | 32536 | 32536-Deleti | Deletion | 1 | C | - | 1 | 23298 | 4.29E-03 | 38 | rnr1D57N_per | FALSE | ARG | JSY14 | 3 | rnr1D57N_pe | GCA | nope | >1bp-ins | G/C-1 | none | G/C-1 | GGTTTGCAG | CAGCTTC | CGT | GCA | C | none | nope | nope_GCA | G/C-1_CAGCTTC | 4 | FALSE | FALSE |
| 10_v.txt | 32544 | 32544-Deleti | Deletion | 1 | C | - | 7 | 23298 | 0.0300455 | 37.8571429 | rnr1D57N_per | FALSE | ARG | JSY14 | 3 | rnr1D57N_pe | ACC | nope | >1bp-ins | G/C-1 | none | G/C-1 | GCTTCACCA | CCAGCAG | TGG | ACC | C | none | nope | nope_ACC | G/C-1_CCAGCAG | 4 | FALSE | FALSE |
| 10_v.txt | 32631 | 32631-Deleti | Deletion | 1 | T | - | 9 | 21882 | 0.0411297 | 42.7777778 | rnr1D57N_per | FALSE | ARG | JSY14 | 3 | rnr1D57N_pe | GTT | nope | >1bp-ins | A/T-1 | none | A/T-1 | CCTTCGTTTT | TTTTTAT | CAA | GTT | T | none | nope | nope_GTT | A/T-1_TTTTTAT | 4 | FALSE | FALSE |
| 10_v.txt | 32658 | 32658-Deleti | Deletion | 1 | C | - | 9 | 21882 | 0.0411297 | 64 | rnr1D57N_per | FALSE | ARG | JSY14 | 3 | rnr1D57N_pe | ACC | nope | >1bp-ins | G/C-1 | none | G/C-1 | CCTGGACCC | CCCCAGG | TGG | ACC | C | none | nope | nope_ACC | G/C-1_CCCCAGG | 4 | FALSE | FALSE |
| 10_v.txt | 32707 | 32707-Deleti | Deletion | 1 | C | - | 5 | 21882 | 0.02284983 | 64 | rnr1D57N_per | FALSE | ARG | JSY14 | 3 | rnr1D57N_pe | ACC | nope | >1bp-ins | G/C-1 | none | G/C-1 | CGGTAACCC | CCCCAGC | TGG | ACC | C | none | nope | nope_ACC | G/C-1_CCCCAGC | 4 | FALSE | FALSE |
| 10_v.txt | 32842 | 32842-Deleti | Deletion | 1 | A | - | 36 | 21882 | 0.16451878 | 42.5555544 | rnr1D57N_per | FALSE | ARG | JSY14 | 3 | rnr1D57N_pe | CAA | nope | >1bp-ins | A/T-1 | none | A/T-1 | TTACCCAAA | AAAAAAT | GTT | GTT | T | none | T>none | T>none_GTT | A/T-1_AAAAAAT | 4 | FALSE | FALSE |
| 10_v.txt | 32924 | 32924-Deleti | Deletion | 1 | A | - | 21 | 19173 | 0.10952903 | 56.4761905 | rnr1D57N_per | FALSE | ARG | JSY14 | 3 | rnr1D57N_pe | CAA | nope | >1bp-ins | A/T-1 | none | A/T-1 | CAGGGCAAA | AAAAGTG | GTT | GTT | T | none | T>none | T>none_GTT | A/T-1_AAAAGTG | 4 | FALSE | FALSE |
| 10_v.txt | 32935 | 32935-Deleti | Deletion | 1 | C | - | 4 | 19173 | 0.02086267 | 40.25 | rnr1D57N_per | FALSE | ARG | JSY14 | 3 | rnr1D57N_pe | GCC | nope | >1bp-ins | G/C-1 | none | G/C-1 | TGATTGCCC | CCCAAGA | CGG | GCC | C | none | nope | nope_GCC | G/C-1_CCCAAGA | 4 | FALSE | FALSE |
| 10_v.txt | 32992 | 32992-Deleti | Deletion | 1 | T | - | 4 | 19173 | 0.02086267 | 62.25 | rnr1D57N_per | FALSE | ARG | JSY14 | 3 | rnr1D57N_pe | CTT | nope | >1bp-ins | A/T-1 | none | A/T-1 | GGAATCTTTC | TTTGTGA | GAA | CTT | T | none | nope | nope_CTT | A/T-1_TTTGTGA | 4 | FALSE | FALSE |
| 10_v.txt | 33147 | 33147-Deleti | Deletion | 1 | A | - | 6 | 19173 | 0.03129401 | 38 | rnr1D57N_per | FALSE | ARG | JSY14 | 3 | rnr1D57N_pe | GAA | nope | >1bp-ins | A/T-1 | none | A/T-1 | CCAATGAAA | AAAAGAC | CTT | CTT | T | none | T>none | T>none_CTT | A/T-1_AAAAGAC | 4 | FALSE | FALSE |
| 10_v.txt | 33258 | 33258-Deleti | Deletion | 1 | C | - | 2 | 19820 | 0.01009082 | 47 | rnr1D57N_per | FALSE | ARG | JSY14 | 3 | rnr1D57N_pe | GCC | nope | >1bp-ins | G/C-1 | none | G/C-1 | CCTATGCCAT | CCATCCT | CGG | GCC | C | none | nope | nope_GCC | G/C-1_CCATCCT | 4 | FALSE | FALSE |
| 10_v.txt | 33303 | 33303-Deleti | Deletion | 1 | A | - | 6 | 19820 | 0.03027245 | 63 | rnr1D57N_per | FALSE | ARG | JSY14 | 3 | rnr1D57N_pe | GAA | nope | >1bp-ins | A/T-1 | none | A/T-1 | GTCGGGAAA | AAAGAGC | CTT | CTT | T | none | T>none | T>none_CTT | A/T-1_AAAGAGC | 4 | FALSE | FALSE |
| 10_v.txt | 33367 | 33367-Deleti | Deletion | 1 | G | - | 5 | 19820 | 0.02522704 | 58.4 | rnr1D57N_per | FALSE | ARG | JSY14 | 3 | rnr1D57N_pe | TGG | nope | >1bp-ins | G/C-1 | none | G/C-1 | GTCTGTGGT | GGTGTGT | ACC | ACC | C | none | C>none | C>none_ACC | G/C-1_GGTGTGT | 4 | FALSE | FALSE |
| 10_v.txt | 33474 | 33474-Deleti | Deletion | 1 | C | - | 44 | 19820 | 0.22199798 | 29.9318182 | rnr1D57N_per | FALSE | ARG | JSY14 | 3 | rnr1D57N_pe | GCC | nope | >1bp-ins | G/C-1 | none | G/C-1 | GCTATGCCTT | CCTTTTT | CGG | GCC | C | none | nope | nope_GCC | G/C-1_CCTTTTT | 4 | FALSE | FALSE |
| 11_v.txt | 31723 | 31723-SNV-A | SNV | 1 | A | C | 6 | 10862 | 0.05523845 | 63.3333333 | rnr1D57N_per | FALSE | ARG | JSY14 | 4 | rnr1D57N_pe | CAA | A>C | G/C+1 | A/T-1 | TA>GC | TA>GC | TGTCCCAAA | AAAAAGT | GTT | GTT | T | G | T>G | T>G_GTT | TA>GC_AAAAAGT | 4 | FALSE | FALSE |
| 11_v.txt | 31907 | 31907-SNV-T | SNV | 1 | T | G | 5 | 10862 | 0.04603204 | 32.4 | rnr1D57N_per | FALSE | ARG | JSY14 | 4 | rnr1D57N_pe | TTG | T>G | G/C+1 | A/T-1 | TA>GC | TA>GC | GAATTTTGG | TGGTGCA | AAC | TTG | T | C | T>G | T>G_TTG | TA>GC_TGGTGCA | 4 | FALSE | FALSE |
| 11_v.txt | 32631 | 32631-SNV-T | SNV | 1 | T | G | 3 | 19186 | 0.0156364 | 37.6666667 | rnr1D57N_per | FALSE | ARG | JSY14 | 4 | rnr1D57N_pe | GTT | T>G | G/C+1 | A/T-1 | TA>GC | TA>GC | CCTTCGTTTT | TTTTTAT | CAA | GTT | T | C | T>G | T>G_GTT | TA>GC_TTTTTAT | 4 | FALSE | FALSE |
| 11_v.txt | 32842 | 32842-SNV-A | SNV | 1 | A | C | 4 | 19186 | 0.02084854 | 27 | rnr1D57N_per | FALSE | ARG | JSY14 | 4 | rnr1D57N_pe | CAA | A>C | G/C+1 | A/T-1 | TA>GC | TA>GC | TTACCCAAA | AAAAAAT | GTT | GTT | T | G | T>G | T>G_GTT | TA>GC_AAAAAAT | 4 | FALSE | FALSE |
| 11_v.txt | 32884 | 32884-SNV-A | SNV | 1 | A | C | 6 | 18502 | 0.03242893 | 33.1666667 | rnr1D57N_per | FALSE | ARG | JSY14 | 4 | rnr1D57N_pe | CAA | A>C | G/C+1 | A/T-1 | TA>GC | TA>GC | ACGTCCAAA | AAAATTG | GTT | GTT | T | G | T>G | T>G_GTT | TA>GC_AAAATTG | 4 | FALSE | FALSE |
| 11_v.txt | 33269 | 33269-SNV-A | SNV | 1 | A | C | 111 | 14359 | 0.77303433 | 16.4774775 | rnr1D57N_per | FALSE | ARG | JSY14 | 4 | rnr1D57N_pe | TAG | A>C | G/C+1 | A/T-1 | TA>GC | TA>GC | CTCCATAGA | AGAGAAC | ATC | ATC | T | G | T>G | T>G_ATC | TA>GC_AGAGAAC | 4 | FALSE | FALSE |
| 11_v.txt | 31650 | 31650-SNV-C | SNV | 1 | C | T | 30 | 10862 | 0.27619223 | 35.7333333 | rnr1D57N_per | FALSE | ARG | JSY14 | 4 | rnr1D57N_pe | GCG | C>T | A/T+1 | G/C-1 | CG>TA | CG>TA | AGAATGCGA | CGAAATG | CGC | GCG | C | A | C>T | C>T_GCG | CG>TA_CGAAATG | 4 | FALSE | FALSE |
| 11_v.txt | 31830 | 31830-SNV-C | SNV | 1 | C | T | 12 | 10862 | 0.11047689 | 50.9166667 | rnr1D57N_per | FALSE | ARG | JSY14 | 4 | rnr1D57N_pe | GCA | C>T | A/T+1 | G/C-1 | CG>TA | CG>TA | AATATGCATT | CATTGAA | CGT | GCA | C | A | C>T | C>T_GCA | CG>TA_CATTGAA | 4 | FALSE | FALSE |
| 11_v.txt | 31882 | 31882-SNV-C | SNV | 1 | C | T | 17 | 10862 | 0.15650893 | 36.7647059 | rnr1D57N_per | FALSE | ARG | JSY14 | 4 | rnr1D57N_pe | GCA | C>T | A/T+1 | G/C-1 | CG>TA | CG>TA | AGGCGGCAG | CAGCAAA | CGT | GCA | C | A | C>T | C>T_GCA | CG>TA_CAGCAAA | 4 | FALSE | FALSE |
| 11_v.txt | 31947 | 31947-SNV-G | SNV | 1 | G | A | 17 | 15161 | 0.11212981 | 35.7647059 | rnr1D57N_per | FALSE | ARG | JSY14 | 4 | rnr1D57N_pe | CGT | G>A | A/T+1 | G/C-1 | CG>TA | CG>TA | ATGATCGTC | GTCATAA | GCA | GCA | C | T | C>T | C>T_GCA | CG>TA_GTCATAA | 4 | FALSE | FALSE |
| 11_v.txt | 31962 | 31962-SNV-G | SNV | 1 | G | A | 28 | 15168 | 0.18459916 | 37.7857143 | rnr1D57N_per | FALSE | ARG | JSY14 | 4 | rnr1D57N_pe | CGC | G>A | A/T+1 | G/C-1 | CG>TA | CG>TA | GTGGCCGCA | GCATAAT | GCG | GCG | C | T | C>T | C>T_GCG | CG>TA_GCATAAT | 4 | FALSE | FALSE |
| 11_v.txt | 31971 | 31971-SNV-G | SNV | 1 | G | A | 5 | 15168 | 0.03296414 | 37.7760411 | rnr1D57N_per | FALSE | ARG | JSY14 | 4 | rnr1D57N_pe | AGC | G>A | A/T+1 | G/C-1 | CG>TA | CG>TA | TAATAAGCC | GCCAAGC | TCG | TCG | C | T | C>T | C>T_TCG | CG>TA_GCCAAGC | 4 | FALSE | FALSE |
| 11_v.txt | 31972 | 31972-SNV-C | SNV | 1 | C | T | 12 | 15168 | 0.07911392 | 36.0833333 | rnr1D57N_per | FALSE | ARG | JSY14 | 4 | rnr1D57N_pe | GCC | C>T | A/T+1 | G/C-1 | CG>TA | CG>TA | AATAAGCCA | CCAAGCC | CGG | GCC | C | A | C>T | C>T_GCC | CG>TA_CCAAGCC | 4 | FALSE | FALSE |
| 11_v.txt | 31981 | 31981-SNV-G | SNV | 1 | G | A | 16 | 15168 | 0.10548523 | 49.4375 | rnr1D57N_per | FALSE | ARG | JSY14 | 4 | rnr1D57N_pe | GGC | G>A | A/T+1 | G/C-1 | CG>TA | CG>TA | AGCCGGGCA | GCATTAA | CCG | CCG | C | T | C>T | C>T_CCG | CG>TA_GCATTAA | 4 | FALSE | FALSE |
| 11_v.txt | 32057 | 32057-SNV-C | SNV | 1 | C | T | 15 | 15168 | 0.09889241 | 59.8 | rnr1D57N_per | FALSE | ARG | JSY14 | 4 | rnr1D57N_pe | GCG | C>T | A/T+1 | G/C-1 | CG>TA | CG>TA | GATGTGCGA | CGAGATT | CGC | GCG | C | A | C>T | C>T_GCG | CG>TA_CGAGATT | 4 | FALSE | FALSE |
| 11_v.txt | 32092 | 32092-SNV-C | SNV | 1 | C | T | 22 | 15168 | 0.14504219 | 58.0909091 | rnr1D57N_per | FALSE | ARG | JSY14 | 4 | rnr1D57N_pe | GCA | C>T | A/T+1 | G/C-1 | CG>TA | CG>TA | AGCCTGCAA | CAACACC | CGT | GCA | C | A | C>T | C>T_GCA | CG>TA_CAACACC | 4 | FALSE | FALSE |
| 11_v.txt | 32165 | 32165-SNV-G | SNV | 1 | G | A | 27 | 15168 | 0.17800633 | 53.8148148 | rnr1D57N_per | FALSE | ARG | JSY14 | 4 | rnr1D57N_pe | CGC | G>A | A/T+1 | G/C-1 | CG>TA | CG>TA | CAAAGCGCC | GCCAAAT | GCG | GCG | C | T | C>T | C>T_GCG | CG>TA_GCCAAAT | 4 | FALSE | FALSE |
| 11_v.txt | 32173 | 32173-SNV-C | SNV | 1 | C | T | 15 | 15168 | 0.09889241 | 50.1333333 | rnr1D57N_per | FALSE | ARG | JSY14 | 4 | rnr1D57N_pe | GCA | C>T | A/T+1 | G/C-1 | CG>TA | CG>TA | CAAATGCAG | CAGCAGT | CGT | GCA | C | A | C>T | C>T_GCA | CG>TA_CAGCAGT | 4 | FALSE | FALSE |

| | | | | | | | | | | | | | | | | | | | | | | | | | | | | | | | | | | |
|---|---|---|---|---|---|---|---|---|---|---|---|---|---|---|---|---|---|---|---|---|---|---|---|---|---|---|---|---|---|---|---|---|---|---|
| 11_v.txt | 32176 | 32176-SNV-C | SNV | 1 | C | T | 15 | 15168 | 0.09889241 | 44.6666667 | rnr1D57N_per | FALSE | ARG | JSY14 | 4 | rnr1D57N_pe | GCA | C>T | A/T+1 | G/C-1 | CG>TA | CG>TA | ATGCAGCAG | CAGTAAC | CGT | GCA | C | A | C>T | C>T_GCA | CG>TA_CAGTAAC | 4 | FALSE | FALSE |
| 11_v.txt | 32271 | 32271-SNV-C | SNV | 1 | C | T | 39 | 17022 | 0.22911526 | 31.7692308 | rnr1D57N_per | FALSE | ARG | JSY14 | 4 | rnr1D57N_pe | ACG | C>T | A/T+1 | G/C-1 | CG>TA | CG>TA | AAAATACGG | CGGGAAC | TGC | ACG | C | A | C>T | C>T_ACG | CG>TA_CGGGAAC | 4 | FALSE | FALSE |
| 11_v.txt | 32299 | 32299-SNV-C | SNV | 1 | C | T | 36 | 17022 | 0.21149101 | 54.1111111 | rnr1D57N_per | FALSE | ARG | JSY14 | 4 | rnr1D57N_pe | GCG | C>T | A/T+1 | G/C-1 | CG>TA | CG>TA | AATTTGCGG | CGGCAGA | CGC | GCG | C | A | C>T | C>T_GCG | CG>TA_CGGCAGA | 4 | FALSE | FALSE |
| 11_v.txt | 32466 | 32466-SNV-C | SNV | 1 | C | T | 28 | 17022 | 0.16449301 | 48.1071429 | rnr1D57N_per | FALSE | ARG | JSY14 | 4 | rnr1D57N_pe | GCC | C>T | A/T+1 | G/C-1 | CG>TA | CG>TA | AGAGAGCCA | CCAATGT | CGG | GCC | C | A | C>T | C>T_GCC | CG>TA_CCAATGT | 4 | FALSE | FALSE |
| 11_v.txt | 32535 | 32535-SNV-C | SNV | 1 | G | A | 16 | 17022 | 0.09399601 | 35.9876429 | rnr1D57N_per | FALSE | ARG | JSY14 | 4 | rnr1D57N_pe | TGC | G>A | A/T+1 | G/C-1 | CG>TA | CG>TA | GGGTTTGCA | GCAGCTT | ACG | ACG | C | T | C>T | C>T_ACG | CG>TA_GCAGCTT | 4 | FALSE | FALSE |
| 11_v.txt | 32536 | 32536-SNV-C | SNV | 1 | C | T | 26 | 17022 | 0.15274351 | 34.6153846 | rnr1D57N_per | FALSE | ARG | JSY14 | 4 | rnr1D57N_pe | GCA | C>T | A/T+1 | G/C-1 | CG>TA | CG>TA | GGTTTGCAG | CAGCTTC | CGT | GCA | C | A | C>T | C>T_GCA | CG>TA_CAGCTTC | 4 | FALSE | FALSE |
| 11_v.txt | 32590 | 32590-SNV-C | SNV | 1 | C | T | 32 | 19186 | 0.16678828 | 37.1875 | rnr1D57N_per | FALSE | ARG | JSY14 | 4 | rnr1D57N_pe | GCG | C>T | A/T+1 | G/C-1 | CG>TA | CG>TA | AGGCAGCGT | CGTTAAT | CGC | GCG | C | A | C>T | C>T_GCG | CG>TA_CGTTAAT | 4 | FALSE | FALSE |
| 11_v.txt | 32700 | 32700-SNV-C | SNV | 1 | C | T | 31 | 19186 | 0.16157615 | 55.3870968 | rnr1D57N_per | FALSE | ARG | JSY14 | 4 | rnr1D57N_pe | GCC | C>T | A/T+1 | G/C-1 | CG>TA | CG>TA | ACTGGGCCG | CCGGTAA | CGG | GCC | C | A | C>T | C>T_GCC | CG>TA_CCGGTAA | 4 | FALSE | FALSE |
| 11_v.txt | 32702 | 32702-SNV-C | SNV | 1 | G | A | 24 | 19186 | 0.12509121 | 48.8333333 | rnr1D57N_per | FALSE | ARG | JSY14 | 4 | rnr1D57N_pe | CGG | G>A | A/T+1 | G/C-1 | CG>TA | CG>TA | TGGGCCGGT | GGTAACC | GCC | GCC | C | T | C>T | C>T_GCC | CG>TA_GGTAACC | 4 | FALSE | FALSE |
| 11_v.txt | 32757 | 32757-SNV-C | SNV | 1 | G | A | 18 | 19186 | 0.09381841 | 52.8333333 | rnr1D57N_per | FALSE | ARG | JSY14 | 4 | rnr1D57N_pe | GGC | G>A | A/T+1 | G/C-1 | CG>TA | CG>TA | ATAATGGCT | GCTAAAA | CCG | CCG | C | T | C>T | C>T_CCG | CG>TA_GCTAAAA | 4 | FALSE | FALSE |
| 11_v.txt | 32776 | 32776-SNV-C | SNV | 1 | C | T | 22 | 19186 | 0.11466695 | 50.0454546 | rnr1D57N_per | FALSE | ARG | JSY14 | 4 | rnr1D57N_pe | GCG | C>T | A/T+1 | G/C-1 | CG>TA | CG>TA | TGGAAGCGA | CGACCCA | CGC | GCG | C | A | C>T | C>T_GCG | CG>TA_CGACCCA | 4 | FALSE | FALSE |
| 11_v.txt | 32789 | 32789-SNV-C | SNV | 1 | G | A | 17 | 19186 | 0.08860628 | 48.5294118 | rnr1D57N_per | FALSE | ARG | JSY14 | 4 | rnr1D57N_pe | CGA | G>A | A/T+1 | G/C-1 | CG>TA | CG>TA | GAACTCGAA | GAATTCA | GCT | GCT | C | T | C>T | C>T_GCT | CG>TA_GAATTCA | 4 | FALSE | FALSE |
| 11_v.txt | 32798 | 32798-SNV-C | SNV | 1 | G | A | 21 | 19186 | 0.10945481 | 50.5714286 | rnr1D57N_per | FALSE | ARG | JSY14 | 4 | rnr1D57N_pe | CGT | G>A | A/T+1 | G/C-1 | CG>TA | CG>TA | TTCACCGTA | GTAATAT | GCA | GCA | C | T | C>T | C>T_GCA | CG>TA_GTAATAT | 4 | FALSE | FALSE |
| 11_v.txt | 32860 | 32860-SNV-C | SNV | 1 | C | T | 20 | 19184 | 0.10425355 | 36.6 | rnr1D57N_per | FALSE | ARG | JSY14 | 4 | rnr1D57N_pe | GCC | C>T | A/T+1 | G/C-1 | CG>TA | CG>TA | TCCATGCCG | CCGCCAG | CGG | GCC | C | A | C>T | C>T_GCC | CG>TA_CCGCCAG | 4 | FALSE | FALSE |
| 11_v.txt | 32901 | 32901-SNV-C | SNV | 1 | C | T | 32 | 18502 | 0.17295428 | 31.09375 | rnr1D57N_per | FALSE | ARG | JSY14 | 4 | rnr1D57N_pe | GCC | C>T | A/T+1 | G/C-1 | CG>TA | CG>TA | ACTTGGCCA | CCAACTA | CGG | GCC | C | A | C>T | C>T_GCC | CG>TA_CCAACTA | 4 | FALSE | FALSE |
| 11_v.txt | 32935 | 32935-SNV-C | SNV | 1 | C | T | 30 | 18502 | 0.16214463 | 37.5666667 | rnr1D57N_per | FALSE | ARG | JSY14 | 4 | rnr1D57N_pe | GCC | C>T | A/T+1 | G/C-1 | CG>TA | CG>TA | TGATTGCCC | CCCAAGA | CGG | GCC | C | A | C>T | C>T_GCC | CG>TA_CCCAAGA | 4 | FALSE | FALSE |
| 11_v.txt | 32967 | 32967-SNV-C | SNV | 1 | G | A | 46 | 18502 | 0.24862177 | 59.2173913 | rnr1D57N_per | FALSE | ARG | JSY14 | 4 | rnr1D57N_pe | CGC | G>A | A/T+1 | G/C-1 | CG>TA | CG>TA | TTGGCCGCA | GCACCAA | GCG | GCG | C | T | C>T | C>T_GCG | CG>TA_GCACCAA | 4 | FALSE | FALSE |
| 11_v.txt | 33056 | 33056-SNV-C | SNV | 1 | C | T | 32 | 18502 | 0.17295428 | 59.15625 | rnr1D57N_per | FALSE | ARG | JSY14 | 4 | rnr1D57N_pe | GCG | C>T | A/T+1 | G/C-1 | CG>TA | CG>TA | GGACTGCGT | CGTGACA | CGC | GCG | C | A | C>T | C>T_GCG | CG>TA_CGTGACA | 4 | FALSE | FALSE |
| 11_v.txt | 33070 | 33070-SNV-C | SNV | 1 | C | T | 26 | 18502 | 0.14052535 | 52.5769231 | rnr1D57N_per | FALSE | ARG | JSY14 | 4 | rnr1D57N_pe | GCC | C>T | A/T+1 | G/C-1 | CG>TA | CG>TA | AATATGCCA | CCAAAGA | CGG | GCC | C | A | C>T | C>T_GCC | CG>TA_CCAAAGA | 4 | FALSE | FALSE |
| 11_v.txt | 33104 | 33104-SNV-C | SNV | 1 | G | A | 24 | 18502 | 0.12971571 | 43.75 | rnr1D57N_per | FALSE | ARG | JSY14 | 4 | rnr1D57N_pe | CGC | G>A | A/T+1 | G/C-1 | CG>TA | CG>TA | AAGAGCGCC | GCCCACT | GCG | GCG | C | T | C>T | C>T_GCG | CG>TA_GCCCACT | 4 | FALSE | FALSE |
| 11_v.txt | 33114 | 33114-SNV-C | SNV | 1 | C | T | 18 | 18502 | 0.09728678 | 39.5 | rnr1D57N_per | FALSE | ARG | JSY14 | 4 | rnr1D57N_pe | GCC | C>T | A/T+1 | G/C-1 | CG>TA | CG>TA | ACTGGGCCG | CCGGCGT | CGG | GCC | C | A | C>T | C>T_GCC | CG>TA_CCGGCGT | 4 | FALSE | FALSE |
| 11_v.txt | 33258 | 33258-SNV-C | SNV | 1 | C | T | 19 | 14359 | 0.13232119 | 44.4736842 | rnr1D57N_per | FALSE | ARG | JSY14 | 4 | rnr1D57N_pe | GCC | C>T | A/T+1 | G/C-1 | CG>TA | CG>TA | CCTATGCCAT | CCATCCT | CGG | GCC | C | A | C>T | C>T_GCC | CG>TA_CCATCCT | 4 | FALSE | FALSE |
| 11_v.txt | 33276 | 33276-SNV-C | SNV | 1 | G | A | 37 | 14359 | 0.25767811 | 46.8378378 | rnr1D57N_per | FALSE | ARG | JSY14 | 4 | rnr1D57N_pe | CGT | G>A | A/T+1 | G/C-1 | CG>TA | CG>TA | GAGAACGTA | GTATCCT | GCA | GCA | C | T | C>T | C>T_GCA | CG>TA_GTATCCT | 4 | FALSE | FALSE |
| 11_v.txt | 33309 | 33309-SNV-C | SNV | 1 | C | T | 15 | 14359 | 0.1044641 | 60.7958041 | rnr1D57N_per | FALSE | ARG | JSY14 | 4 | rnr1D57N_pe | GCG | C>T | A/T+1 | G/C-1 | CG>TA | CG>TA | AAAGAGCGC | CGCAATG | CGC | GCG | C | A | C>T | C>T_GCG | CG>TA_CGCAATG | 4 | FALSE | FALSE |
| 11_v.txt | 33310 | 33310-SNV-C | SNV | 1 | G | A | 25 | 14358 | 0.17411896 | 52.6078204 | rnr1D57N_per | FALSE | ARG | JSY14 | 4 | rnr1D57N_pe | CGC | G>A | A/T+1 | G/C-1 | CG>TA | CG>TA | AAGAGCGCA | GCAATGG | GCG | GCG | C | T | C>T | C>T_GCG | CG>TA_GCAATGG | 4 | FALSE | FALSE |
| 11_v.txt | 33412 | 33412-SNV-C | SNV | 1 | G | A | 32 | 14359 | 0.22285675 | 50.875 | rnr1D57N_per | FALSE | ARG | JSY14 | 4 | rnr1D57N_pe | GGC | G>A | A/T+1 | G/C-1 | CG>TA | CG>TA | TGACCGGCT | GCTCATT | CCG | CCG | C | T | C>T | C>T_CCG | CG>TA_GCTCATT | 4 | FALSE | FALSE |
| 11_v.txt | 33474 | 33474-SNV-C | SNV | 1 | C | T | 13 | 14359 | 0.09053555 | 35 | rnr1D57N_per | FALSE | ARG | JSY14 | 4 | rnr1D57N_pe | GCC | C>T | A/T+1 | G/C-1 | CG>TA | CG>TA | GCTATGCCTT | CCTTTTT | CGG | GCC | C | A | C>T | C>T_GCC | CG>TA_CCTTTTT | 4 | FALSE | FALSE |
| 11_v.txt | 31830 | 31830-SNV-C | SNV | 1 | C | G | 2 | 10862 | 0.01841282 | 63 | rnr1D57N_per | FALSE | ARG | JSY14 | 4 | rnr1D57N_pe | GCA | C>G | G/C+1 | G/C-1 | CG>GC | CG>GC | AATATGCATT | CATTGAA | CGT | GCA | C | C | C>G | C>G_GCA | CG>GC_CATTGAA | 4 | FALSE | FALSE |
| 11_v.txt | 31972 | 31972-SNV-C | SNV | 1 | C | G | 2 | 15168 | 0.01318565 | 25.5 | rnr1D57N_per | FALSE | ARG | JSY14 | 4 | rnr1D57N_pe | GCC | C>G | G/C+1 | G/C-1 | CG>GC | CG>GC | AATAAGCCA | CCAAGCC | CGG | GCC | C | C | C>G | C>G_GCC | CG>GC_CCAAGCC | 4 | FALSE | FALSE |
| 11_v.txt | 32165 | 32165-SNV-C | SNV | 1 | G | C | 4 | 15168 | 0.02637131 | 37.75 | rnr1D57N_per | FALSE | ARG | JSY14 | 4 | rnr1D57N_pe | CGC | G>C | G/C+1 | G/C-1 | CG>GC | CG>GC | CAAAGCGCC | GCCAAAT | GCG | GCG | C | G | C>G | C>G_GCG | CG>GC_GCCAAAT | 4 | FALSE | FALSE |
| 11_v.txt | 32536 | 32536-SNV-C | SNV | 1 | C | G | 1 | 17022 | 5.88E-03 | 38 | rnr1D57N_per | FALSE | ARG | JSY14 | 4 | rnr1D57N_pe | GCA | C>G | G/C+1 | G/C-1 | CG>GC | CG>GC | GGTTTGCAG | CAGCTTC | CGT | GCA | C | C | C>G | C>G_GCA | CG>GC_CAGCTTC | 4 | FALSE | FALSE |
| 11_v.txt | 32536 | 32536-SNV-C | SNV | 1 | C | A | 4 | 17022 | 0.023499 | 36 | rnr1D57N_per | FALSE | ARG | JSY14 | 4 | rnr1D57N_pe | GCA | C>A | A/T+1 | G/C-1 | CG>AT | CG>AT | GGTTTGCAG | CAGCTTC | CGT | GCA | C | T | C>A | C>A_GCA | CG>AT_CAGCTTC | 4 | FALSE | FALSE |
| 11_v.txt | 33474 | 33474-SNV-C | SNV | 1 | C | A | 44 | 14359 | 0.30642802 | 34.0458289 | rnr1D57N_per | FALSE | ARG | JSY14 | 4 | rnr1D57N_pe | GCC | C>A | A/T+1 | G/C-1 | CG>AT | CG>AT | GCTATGCCTT | CCTTTTT | CGG | GCC | C | T | C>A | C>A_GCC | CG>AT_CCTTTTT | 4 | FALSE | FALSE |
| 11_v.txt | 33475 | 33475-SNV-C | SNV | 1 | C | A | 11 | 14359 | 0.07660701 | 34.8181818 | rnr1D57N_per | FALSE | ARG | JSY14 | 4 | rnr1D57N_pe | CCT | C>A | A/T+1 | G/C-1 | CG>AT | CG>AT | CTATGCCTTT | CTTTTTT | GGA | CCT | C | T | C>A | C>A_CCT | CG>AT_CTTTTTT | 4 | FALSE | FALSE |
| 11_v.txt | 32081 | 32081-Insert | Insertion | 1 | - | A | 9 | 15168 | 0.05933544 | 64 | rnr1D57N_per | FALSE | ARG | JSY14 | 4 | rnr1D57N_pe | CAA | nope | A/T+1 | >1bpdel | none | A/T+1 | CCATGCAAA | AAAAAAG | GTT | CAA | - | T | NA | NA_CAA | A/T+1_AAAAAAG | 4 | FALSE | FALSE |
| 11_v.txt | 31663 | 31663-Deleti | Deletion | 1 | A | - | 9 | 10862 | 0.08285767 | 35 | rnr1D57N_per | FALSE | ARG | JSY14 | 4 | rnr1D57N_pe | GAA | nope | >1bp-ins | A/T-1 | none | A/T-1 | GCGTGGAAA | AAATGTG | CTT | CTT | T | none | T>none | T>none_CTT | A/T-1_AAATGTG | 4 | FALSE | FALSE |
| 11_v.txt | 31723 | 31723-Deleti | Deletion | 1 | A | - | 11 | 10862 | 0.10127048 | 60.2727273 | rnr1D57N_per | FALSE | ARG | JSY14 | 4 | rnr1D57N_pe | CAA | nope | >1bp-ins | A/T-1 | none | A/T-1 | TGTCCCAAA | AAAAAGT | GTT | GTT | T | none | T>none | T>none_GTT | A/T-1_AAAAAGT | 4 | FALSE | FALSE |
| 11_v.txt | 31863 | 31863-Deleti | Deletion | 1 | A | - | 5 | 10862 | 0.04603204 | 52.8 | rnr1D57N_per | FALSE | ARG | JSY14 | 4 | rnr1D57N_pe | GAA | nope | >1bp-ins | A/T-1 | none | A/T-1 | AACAGGAAA | AAAATAG | CTT | CTT | T | none | T>none | T>none_CTT | A/T-1_AAAATAG | 4 | FALSE | FALSE |
| 11_v.txt | 31904 | 31904-Deleti | Deletion | 1 | T | - | 7 | 10862 | 0.06444485 | 38 | rnr1D57N_per | FALSE | ARG | JSY14 | 4 | rnr1D57N_pe | ATT | nope | >1bp-ins | A/T-1 | none | A/T-1 | ATTGAATTTT | TTTTGGT | TAA | ATT | T | none | nope | nope_ATT | A/T-1_TTTTGGT | 4 | FALSE | FALSE |
| 11_v.txt | 31971 | 31971-Deleti | Deletion | 1 | G | - | 6 | 15168 | 0.03955696 | 33.1666667 | rnr1D57N_per | FALSE | ARG | JSY14 | 4 | rnr1D57N_pe | AGC | nope | >1bp-ins | G/C-1 | none | G/C-1 | TAATAAGCC | GCCAAGC | TCG | TCG | C | none | C>none | C>none_TCG | G/C-1_GCCAAGC | 4 | FALSE | FALSE |
| 11_v.txt | 31972 | 31972-Deleti | Deletion | 1 | C | - | 5 | 15168 | 0.03296414 | 37.2 | rnr1D57N_per | FALSE | ARG | JSY14 | 4 | rnr1D57N_pe | GCC | nope | >1bp-ins | G/C-1 | none | G/C-1 | AATAAGCCA | CCAAGCC | CGG | GCC | C | none | nope | nope_GCC | G/C-1_CCAAGCC | 4 | FALSE | FALSE |
| 11_v.txt | 32081 | 32081-Deleti | Deletion | 1 | A | - | 15 | 15168 | 0.09889241 | 62.2666673 | rnr1D57N_per | FALSE | ARG | JSY14 | 4 | rnr1D57N_pe | CAA | nope | >1bp-ins | A/T-1 | none | A/T-1 | CCATGCAAA | AAAAAAG | GTT | GTT | T | none | T>none | T>none_GTT | A/T-1_AAAAAAG | 4 | FALSE | FALSE |
| 11_v.txt | 32498 | 32498-Deleti | Deletion | 1 | T | - | 19 | 17022 | 0.11162026 | 38.1578947 | rnr1D57N_per | FALSE | ARG | JSY14 | 4 | rnr1D57N_pe | CTT | nope | >1bp-ins | A/T-1 | none | A/T-1 | AACAACTTTT | TTTTTTG | GAA | CTT | T | none | nope | nope_CTT | A/T-1_TTTTTTG | 4 | FALSE | FALSE |
| 11_v.txt | 32528 | 32528-Deleti | Deletion | 1 | G | - | 4 | 17022 | 0.023499 | 38 | rnr1D57N_per | FALSE | ARG | JSY14 | 4 | rnr1D57N_pe | TGG | nope | >1bp-ins | G/C-1 | none | G/C-1 | TTTTCTGGG | GGGGTTT | ACC | ACC | C | none | C>none | C>none_ACC | G/C-1_GGGGTTT | 4 | FALSE | FALSE |
| 11_v.txt | 32631 | 32631-Deleti | Deletion | 1 | T | - | 8 | 19186 | 0.04169707 | 50.875 | rnr1D57N_per | FALSE | ARG | JSY14 | 4 | rnr1D57N_pe | GTT | nope | >1bp-ins | A/T-1 | none | A/T-1 | CCTTCGTTTT | TTTTTAT | CAA | GTT | T | none | nope | nope_GTT | A/T-1_TTTTTAT | 4 | FALSE | FALSE |
| 11_v.txt | 32658 | 32658-Deleti | Deletion | 1 | C | - | 6 | 19186 | 0.0312728 | 51.8333333 | rnr1D57N_per | FALSE | ARG | JSY14 | 4 | rnr1D57N_pe | ACC | nope | >1bp-ins | G/C-1 | none | G/C-1 | CCTGGACCC | CCCCAGG | TGG | ACC | C | none | nope | nope_ACC | G/C-1_CCCCAGG | 4 | FALSE | FALSE |
| 11_v.txt | 32728 | 32728-Deleti | Deletion | 1 | A | - | 6 | 19186 | 0.0312728 | 58.3333333 | rnr1D57N_per | FALSE | ARG | JSY14 | 4 | rnr1D57N_pe | CAA | nope | >1bp-ins | A/T-1 | none | A/T-1 | CCATACAAA | AAAAACA | GTT | GTT | T | none | T>none | T>none_GTT | A/T-1_AAAAACA | 4 | FALSE | FALSE |
| 11_v.txt | 32842 | 32842-Deleti | Deletion | 1 | A | - | 31 | 19186 | 0.16157615 | 39.4516129 | rnr1D57N_per | FALSE | ARG | JSY14 | 4 | rnr1D57N_pe | CAA | nope | >1bp-ins | A/T-1 | none | A/T-1 | TTACCCAAA | AAAAAAT | GTT | GTT | T | none | T>none | T>none_GTT | A/T-1_AAAAAAT | 4 | FALSE | FALSE |
| 11_v.txt | 32884 | 32884-Deleti | Deletion | 1 | A | - | 6 | 18502 | 0.03242893 | 36 | rnr1D57N_per | FALSE | ARG | JSY14 | 4 | rnr1D57N_pe | CAA | nope | >1bp-ins | A/T-1 | none | A/T-1 | ACGTCCAAA | AAAATTG | GTT | GTT | T | none | T>none | T>none_GTT | A/T-1_AAAATTG | 4 | FALSE | FALSE |
| 11_v.txt | 32924 | 32924-Deleti | Deletion | 1 | A | - | 18 | 18502 | 0.09728678 | 52.6666667 | rnr1D57N_per | FALSE | ARG | JSY14 | 4 | rnr1D57N_pe | CAA | nope | >1bp-ins | A/T-1 | none | A/T-1 | CAGGGCAAA | AAAAGTG | GTT | GTT | T | none | T>none | T>none_GTT | A/T-1_AAAAGTG | 4 | FALSE | FALSE |
| 11_v.txt | 32935 | 32935-Deleti | Deletion | 1 | C | - | 2 | 18502 | 0.01080964 | 60.5 | rnr1D57N_per | FALSE | ARG | JSY14 | 4 | rnr1D57N_pe | GCC | nope | >1bp-ins | G/C-1 | none | G/C-1 | TGATTGCCC | CCCAAGA | CGG | GCC | C | none | nope | nope_GCC | G/C-1_CCCAAGA | 4 | FALSE | FALSE |
| 11_v.txt | 33147 | 33147-Deleti | Deletion | 1 | A | - | 5 | 18502 | 0.02702411 | 38 | rnr1D57N_per | FALSE | ARG | JSY14 | 4 | rnr1D57N_pe | GAA | nope | >1bp-ins | A/T-1 | none | A/T-1 | CCAATGAAA | AAAAGAC | CTT | CTT | T | none | T>none | T>none_CTT | A/T-1_AAAAGAC | 4 | FALSE | FALSE |
| 11_v.txt | 33474 | 33474-Deleti | Deletion | 1 | C | - | 35 | 14359 | 0.24374957 | 29.4285714 | rnr1D57N_per | FALSE | ARG | JSY14 | 4 | rnr1D57N_pe | GCC | nope | >1bp-ins | G/C-1 | none | G/C-1 | GCTATGCCTT | CCTTTTT | CGG | GCC | C | none | nope | nope_GCC | G/C-1_CCTTTTT | 4 | FALSE | FALSE |
| 62_v.txt | 31723 | 31723-SNV-A | SNV | 1 | A | C | 4 | 7804 | 0.05125577 | 53.25 | rnr1Y285A | FALSE | CAN | JSY3868 | 1 | rnr1Y285A-R | CAA | A>C | G/C+1 | A/T-1 | TA>GC | TA>GC | TGTCCCAAA | AAAAAGT | GTT | GTT | T | G | T>G | T>G_GTT | TA>GC_AAAAAGT | 5 | FALSE | FALSE |
| 62_v.txt | 31986 | 31986-SNV-A | SNV | 1 | A | C | 18 | 12415 | 0.1449859 | 50.5555556 | rnr1Y285A | FALSE | CAN | JSY3868 | 1 | rnr1Y285A-R | TAA | A>C | G/C+1 | A/T-1 | TA>GC | TA>GC | GGCATTAATT | AATTTAG | ATT | ATT | T | G | T>G | T>G_ATT | TA>GC_AATTTAG | 5 | FALSE | FALSE |
| 62_v.txt | 32162 | 32162-SNV-A | SNV | 1 | A | C | 4 | 12415 | 0.03221909 | 50.5 | rnr1Y285A | FALSE | CAN | JSY3868 | 1 | rnr1Y285A-R | AAG | A>C | G/C+1 | A/T-1 | TA>GC | TA>GC | AGCCAAAGC | AGCGCCA | TTC | TTC | T | G | T>G | T>G_TTC | TA>GC_AGCGCCA | 5 | FALSE | FALSE |
| 62_v.txt | 32581 | 32581-SNV-T | SNV | 1 | T | G | 21 | 16394 | 0.12809565 | 35 | rnr1Y285A | FALSE | CAN | JSY3868 | 1 | rnr1Y285A-R | GTG | T>G | G/C+1 | A/T-1 | TA>GC | TA>GC | GAAATGTGA | TGAAGGC | CAC | GTG | T | C | T>G | T>G_GTG | TA>GC_TGAAGGC | 5 | FALSE | FALSE |
| 62_v.txt | 32823 | 32823-SNV-A | SNV | 1 | A | C | 900 | 16394 | 5.48981335 | 43.2788889 | rnr1Y285A | FALSE | CAN | JSY3868 | 1 | rnr1Y285A-R | CAT | A>C | G/C+1 | A/T-1 | TA>GC | TA>GC | AAGTTCATT | ATTATTG | GTA | GTA | T | G | T>G | T>G_GTA | TA>GC_ATTATTG | 5 | FALSE | FALSE |
| 62_v.txt | 32924 | 32924-SNV-A | SNV | 1 | A | C | 3 | 20579 | 0.01457797 | 37 | rnr1Y285A | FALSE | CAN | JSY3868 | 1 | rnr1Y285A-R | CAA | A>C | G/C+1 | A/T-1 | TA>GC | TA>GC | CAGGGCAAA | AAAAGTG | GTT | GTT | T | G | T>G | T>G_GTT | TA>GC_AAAAGTG | 5 | FALSE | FALSE |
| 62_v.txt | 33269 | 33269-SNV-A | SNV | 1 | A | C | 94 | 12457 | 0.75459581 | 16.1595745 | rnr1Y285A | FALSE | CAN | JSY3868 | 1 | rnr1Y285A-R | TAG | A>C | G/C+1 | A/T-1 | TA>GC | TA>GC | CTCCATAGA | AGAGAAC | ATC | ATC | T | G | T>G | T>G_ATC | TA>GC_AGAGAAC | 5 | FALSE | FALSE |
| 62_v.txt | 33209 | 33209-SNV-T | SNV | 1 | T | C | 5 | 12457 | 0.04013808 | 41.8 | rnr1Y285A | FALSE | CAN | JSY3868 | 1 | rnr1Y285A-R | CTC | T>C | G/C+1 | A/T-1 | TA>CG | TA>CG | AAGCTCTCT | TCTCTTC | GAG | CTC | T | G | T>C | T>C_CTC | TA>CG_TCTCTTC | 5 | FALSE | FALSE |
| 62_v.txt | 32496 | 32496-SNV-A | SNV | 1 | A | T | 81 | 14963 | 0.54133529 | 44.2345679 | rnr1Y285A | FALSE | CAN | JSY3868 | 1 | rnr1Y285A-R | AAC | A>T | A/T+1 | A/T-1 | TA>AT | TA>AT | AAAACAACT | ACTTTTT | TTG | TTG | T | A | T>A | T>A_TTG | TA>AT_ACTTTTT | 5 | FALSE | FALSE |
| 62_v.txt | 32498 | 32498-SNV-T | SNV | 1 | T | A | 3 | 14963 | 0.02004946 | 42.6666221 | rnr1Y285A | FALSE | CAN | JSY3868 | 1 | rnr1Y285A-R | CTT | T>A | A/T+1 | A/T-1 | TA>AT | TA>AT | AACAACTTTT | TTTTTTG | GAA | CTT | T | T | T>A | T>A_CTT | TA>AT_TTTTTTG | 5 | FALSE | FALSE |
| 62_v.txt | 32947 | 32947-SNV-A | SNV | 1 | A | T | 17 | 20579 | 0.08260848 | 57.9411765 | rnr1Y285A | FALSE | CAN | JSY3868 | 1 | rnr1Y285A-R | CAA | A>T | A/T+1 | A/T-1 | TA>AT | TA>AT | AAAACCAAT | AATACAT | GTT | GTT | T | A | T>A | T>A_GTT | TA>AT_AATACAT | 5 | FALSE | FALSE |
| 62_v.txt | 33065 | 33065-SNV-A | SNV | 1 | A | T | 44 | 20579 | 0.2138102 | 54.9545455 | rnr1Y285A | FALSE | CAN | JSY3868 | 1 | rnr1Y285A-R | AAT | A>T | A/T+1 | A/T-1 | TA>AT | TA>AT | GACAGAATA | ATATGCC | TTA | TTA | T | A | T>A | T>A_TTA | TA>AT_ATATGCC | 5 | FALSE | FALSE |
| 62_v.txt | 33209 | 33209-SNV-T | SNV | 1 | T | A | 13 | 12457 | 0.104359 | 37.6923077 | rnr1Y285A | FALSE | CAN | JSY3868 | 1 | rnr1Y285A-R | CTC | T>A | A/T+1 | A/T-1 | TA>AT | TA>AT | AAGCTCTCT | TCTCTTC | GAG | CTC | T | T | T>A | T>A_CTC | TA>AT_TCTCTTC | 5 | FALSE | FALSE |
| 62_v.txt | 31650 | 31650-SNV-C | SNV | 1 | C | T | 17 | 7804 | 0.21783701 | 36.1764706 | rnr1Y285A | FALSE | CAN | JSY3868 | 1 | rnr1Y285A-R | GCG | C>T | A/T+1 | G/C-1 | CG>TA | CG>TA | AGAATGCGA | CGAAATG | CGC | GCG | C | A | C>T | C>T_GCG | CG>TA_CGAAATG | 5 | FALSE | FALSE |
| 62_v.txt | 31844 | 31844-SNV-C | SNV | 1 | C | T | 41 | 7804 | 0.5253716 | 46.9268293 | rnr1Y285A | FALSE | CAN | JSY3868 | 1 | rnr1Y285A-R | TCC | C>T | A/T+1 | G/C-1 | CG>TA | CG>TA | TAAGATCCA | CCAAACA | AGG | TCC | C | A | C>T | C>T_TCC | CG>TA_CCAAACA | 5 | FALSE | FALSE |
| 62_v.txt | 31882 | 31882-SNV-C | SNV | 1 | C | T | 13 | 7804 | 0.16658124 | 40.3076923 | rnr1Y285A | FALSE | CAN | JSY3868 | 1 | rnr1Y285A-R | GCA | C>T | A/T+1 | G/C-1 | CG>TA | CG>TA | AGGCGGCAG | CAGCAAA | CGT | GCA | C | A | C>T | C>T_GCA | CG>TA_CAGCAAA | 5 | FALSE | FALSE |
| 62_v.txt | 31962 | 31962-SNV-C | SNV | 1 | G | A | 24 | 12415 | 0.19331454 | 37.0833333 | rnr1Y285A | FALSE | CAN | JSY3868 | 1 | rnr1Y285A-R | CGC | G>A | A/T+1 | G/C-1 | CG>TA | CG>TA | GTGGCCGCA | GCATAAT | GCG | GCG | C | T | C>T | C>T_GCG | CG>TA_GCATAAT | 5 | FALSE | FALSE |
| 62_v.txt | 31981 | 31981-SNV-C | SNV | 1 | G | A | 35 | 12415 | 0.28191704 | 51.6857143 | rnr1Y285A | FALSE | CAN | JSY3868 | 1 | rnr1Y285A-R | GGC | G>A | A/T+1 | G/C-1 | CG>TA | CG>TA | AGCCGGGCA | GCATTAA | CCG | CCG | C | T | C>T | C>T_CCG | CG>TA_GCATTAA | 5 | FALSE | FALSE |
| 62_v.txt | 32075 | 32075-SNV-C | SNV | 1 | C | T | 82 | 12415 | 0.66049134 | 61.5487805 | rnr1Y285A | FALSE | CAN | JSY3868 | 1 | rnr1Y285A-R | ACC | C>T | A/T+1 | G/C-1 | CG>TA | CG>TA | AAATAACCA | CCATGCA | TGG | ACC | C | A | C>T | C>T_ACC | CG>TA_CCATGCA | 5 | FALSE | FALSE |
| 62_v.txt | 32088 | 32088-SNV-C | SNV | 1 | C | T | 70 | 12415 | 0.56383407 | 61.7857143 | rnr1Y285A | FALSE | CAN | JSY3868 | 1 | rnr1Y285A-R | GCC | C>T | A/T+1 | G/C-1 | CG>TA | CG>TA | AAAAAGCCT | CCTGCAA | CGG | GCC | C | A | C>T | C>T_GCC | CG>TA_CCTGCAA | 5 | FALSE | FALSE |
| 62_v.txt | 32092 | 32092-SNV-C | SNV | 1 | C | T | 15 | 12415 | 0.12082159 | 63.6666667 | rnr1Y285A | FALSE | CAN | JSY3868 | 1 | rnr1Y285A-R | GCA | C>T | A/T+1 | G/C-1 | CG>TA | CG>TA | AGCCTGCAA | CAACACC | CGT | GCA | C | A | C>T | C>T_GCA | CG>TA_CAACACC | 5 | FALSE | FALSE |
| 62_v.txt | 32114 | 32114-SNV-C | SNV | 1 | C | T | 96 | 12415 | 0.77325816 | 58.0520833 | rnr1Y285A | FALSE | CAN | JSY3868 | 1 | rnr1Y285A-R | GCC | C>T | A/T+1 | G/C-1 | CG>TA | CG>TA | TAATAGCCAT | CCATTCG | CGG | GCC | C | A | C>T | C>T_GCC | CG>TA_CCATTCG | 5 | FALSE | FALSE |
| 62_v.txt | 32165 | 32165-SNV-C | SNV | 1 | G | A | 22 | 12415 | 0.17720499 | 44.9090909 | rnr1Y285A | FALSE | CAN | JSY3868 | 1 | rnr1Y285A-R | CGC | G>A | A/T+1 | G/C-1 | CG>TA | CG>TA | CAAAGCGCC | GCCAAAT | GCG | GCG | C | T | C>T | C>T_GCG | CG>TA_GCCAAAT | 5 | FALSE | FALSE |
| 62_v.txt | 32173 | 32173-SNV-C | SNV | 1 | C | T | 19 | 12415 | 0.15304068 | 48 | rnr1Y285A | FALSE | CAN | JSY3868 | 1 | rnr1Y285A-R | GCA | C>T | A/T+1 | G/C-1 | CG>TA | CG>TA | CAAATGCAG | CAGCAGT | CGT | GCA | C | A | C>T | C>T_GCA | CG>TA_CAGCAGT | 5 | FALSE | FALSE |
| 62_v.txt | 32292 | 32292-SNV-C | SNV | 1 | G | A | 20 | 14963 | 0.13366304 | 57.65 | rnr1Y285A | FALSE | CAN | JSY3868 | 1 | rnr1Y285A-R | TGA | G>A | A/T+1 | G/C-1 | CG>TA | CG>TA | ATATTTGAAT | GAATTTG | ACT | ACT | C | T | C>T | C>T_ACT | CG>TA_GAATTTG | 5 | FALSE | FALSE |
| 62_v.txt | 32344 | 32344-SNV-C | SNV | 1 | G | A | 13 | 14963 | 0.08688097 | 57.3076923 | rnr1Y285A | FALSE | CAN | JSY3868 | 1 | rnr1Y285A-R | GGC | G>A | A/T+1 | G/C-1 | CG>TA | CG>TA | TATGTGGCA | GCAAAAC | CCG | CCG | C | T | C>T | C>T_CCG | CG>TA_GCAAAAC | 5 | FALSE | FALSE |
| 62_v.txt | 32508 | 32508-SNV-C | SNV | 1 | G | A | 44 | 14963 | 0.29405868 | 45.0227273 | rnr1Y285A | FALSE | CAN | JSY3868 | 1 | rnr1Y285A-R | GGC | G>A | A/T+1 | G/C-1 | CG>TA | CG>TA | TTGATGGCT | GCTCTTG | CCG | CCG | C | T | C>T | C>T_CCG | CG>TA_GCTCTTG | 5 | FALSE | FALSE |
| 62_v.txt | 32535 | 32535-SNV-C | SNV | 1 | G | A | 19 | 14963 | 0.12697988 | 35.8420009 | rnr1Y285A | FALSE | CAN | JSY3868 | 1 | rnr1Y285A-R | TGC | G>A | A/T+1 | G/C-1 | CG>TA | CG>TA | GGGTTTGCA | GCAGCTT | ACG | ACG | C | T | C>T | C>T_ACG | CG>TA_GCAGCTT | 5 | FALSE | FALSE |
| 62_v.txt | 32536 | 32536-SNV-C | SNV | 1 | C | T | 30 | 14963 | 0.20049455 | 36.4 | rnr1Y285A | FALSE | CAN | JSY3868 | 1 | rnr1Y285A-R | GCA | C>T | A/T+1 | G/C-1 | CG>TA | CG>TA | GGTTTGCAG | CAGCTTC | CGT | GCA | C | A | C>T | C>T_GCA | CG>TA_CAGCTTC | 5 | FALSE | FALSE |
| 62_v.txt | 32590 | 32590-SNV-C | SNV | 1 | C | T | 30 | 16394 | 0.18299378 | 35.8333333 | rnr1Y285A | FALSE | CAN | JSY3868 | 1 | rnr1Y285A-R | GCG | C>T | A/T+1 | G/C-1 | CG>TA | CG>TA | AGGCAGCGT | CGTTAAT | CGC | GCG | C | A | C>T | C>T_GCG | CG>TA_CGTTAAT | 5 | FALSE | FALSE |
| 62_v.txt | 32610 | 32610-SNV-C | SNV | 1 | C | T | 27 | 16394 | 0.1646944 | 49.1851852 | rnr1Y285A | FALSE | CAN | JSY3868 | 1 | rnr1Y285A-R | CCA | C>T | A/T+1 | G/C-1 | CG>TA | CG>TA | GAAACCCAA | CAACCTA | GGT | CCA | C | A | C>T | C>T_CCA | CG>TA_CAACCTA | 5 | FALSE | FALSE |
| 62_v.txt | 32658 | 32658-SNV-C | SNV | 1 | C | T | 16 | 16394 | 0.09759668 | 53.625 | rnr1Y285A | FALSE | CAN | JSY3868 | 1 | rnr1Y285A-R | ACC | C>T | A/T+1 | G/C-1 | CG>TA | CG>TA | CCTGGACCC | CCCCAGG | TGG | ACC | C | A | C>T | C>T_ACC | CG>TA_CCCCAGG | 5 | FALSE | FALSE |
| 62_v.txt | 32678 | 32678-SNV-C | SNV | 1 | C | T | 553 | 16394 | 3.37318531 | 59.8227848 | rnr1Y285A | FALSE | CAN | JSY3868 | 1 | rnr1Y285A-R | TCC | C>T | A/T+1 | G/C-1 | CG>TA | CG>TA | GTTTCTCCAA | CCAATAA | AGG | TCC | C | A | C>T | C>T_TCC | CG>TA_CCAATAA | 5 | FALSE | FALSE |
| 62_v.txt | 32776 | 32776-SNV-C | SNV | 1 | C | T | 21 | 16394 | 0.12809565 | 50.6190476 | rnr1Y285A | FALSE | CAN | JSY3868 | 1 | rnr1Y285A-R | GCG | C>T | A/T+1 | G/C-1 | CG>TA | CG>TA | TGGAAGCGA | CGACCCA | CGC | GCG | C | A | C>T | C>T_GCG | CG>TA_CGACCCA | 5 | FALSE | FALSE |
| 62_v.txt | 32780 | 32780-SNV-C | SNV | 1 | C | T | 36 | 16394 | 0.21959253 | 43.1944444 | rnr1Y285A | FALSE | CAN | JSY3868 | 1 | rnr1Y285A-R | CCC | C>T | A/T+1 | G/C-1 | CG>TA | CG>TA | AGCGACCCA | CCAGAAC | GGG | CCC | C | A | C>T | C>T_CCC | CG>TA_CCAGAAC | 5 | FALSE | FALSE |
| 62_v.txt | 32794 | 32794-SNV-C | SNV | 1 | C | T | 31 | 16394 | 0.18909357 | 46.1612903 | rnr1Y285A | FALSE | CAN | JSY3868 | 1 | rnr1Y285A-R | TCA | C>T | A/T+1 | G/C-1 | CG>TA | CG>TA | CGAATTCAC | CACCGTA | AGT | TCA | C | A | C>T | C>T_TCA | CG>TA_CACCGTA | 5 | FALSE | FALSE |
| 62_v.txt | 32796 | 32796-SNV-C | SNV | 1 | C | T | 29 | 16394 | 0.17689399 | 44.3793103 | rnr1Y285A | FALSE | CAN | JSY3868 | 1 | rnr1Y285A-R | ACC | C>T | A/T+1 | G/C-1 | CG>TA | CG>TA | AATTCACCGT | CCGTAAT | TGG | ACC | C | A | C>T | C>T_ACC | CG>TA_CCGTAAT | 5 | FALSE | FALSE |
| 62_v.txt | 32798 | 32798-SNV-C | SNV | 1 | G | A | 31 | 16394 | 0.18909357 | 47.7741936 | rnr1Y285A | FALSE | CAN | JSY3868 | 1 | rnr1Y285A-R | CGT | G>A | A/T+1 | G/C-1 | CG>TA | CG>TA | TTCACCGTA | GTAATAT | GCA | GCA | C | T | C>T | C>T_GCA | CG>TA_GTAATAT | 5 | FALSE | FALSE |
| 62_v.txt | 32855 | 32855-SNV-C | SNV | 1 | C | T | 48 | 16394 | 0.29279005 | 36.6875 | rnr1Y285A | FALSE | CAN | JSY3868 | 1 | rnr1Y285A-R | TCC | C>T | A/T+1 | G/C-1 | CG>TA | CG>TA | ACTAATCCAT | CCATGCC | AGG | TCC | C | A | C>T | C>T_TCC | CG>TA_CCATGCC | 5 | FALSE | FALSE |
| 62_v.txt | 32860 | 32860-SNV-C | SNV | 1 | C | T | 15 | 16394 | 0.09149689 | 38 | rnr1Y285A | FALSE | CAN | JSY3868 | 1 | rnr1Y285A-R | GCC | C>T | A/T+1 | G/C-1 | CG>TA | CG>TA | TCCATGCCG | CCGCCAG | CGG | GCC | C | A | C>T | C>T_GCC | CG>TA_CCGCCAG | 5 | FALSE | FALSE |
| 62_v.txt | 32882 | 32882-SNV-C | SNV | 1 | C | T | 652 | 20579 | 3.16827834 | 36.3711656 | rnr1Y285A | FALSE | CAN | JSY3868 | 1 | rnr1Y285A-R | TCC | C>T | A/T+1 | G/C-1 | CG>TA | CG>TA | GTACGTCCA | CCAAAAT | AGG | TCC | C | A | C>T | C>T_TCC | CG>TA_CCAAAAT | 5 | FALSE | FALSE |
| 62_v.txt | 32901 | 32901-SNV-C | SNV | 1 | C | T | 34 | 20579 | 0.16521697 | 35.4705882 | rnr1Y285A | FALSE | CAN | JSY3868 | 1 | rnr1Y285A-R | GCC | C>T | A/T+1 | G/C-1 | CG>TA | CG>TA | ACTTGGCCA | CCAACTA | CGG | GCC | C | A | C>T | C>T_GCC | CG>TA_CCAACTA | 5 | FALSE | FALSE |
| 62_v.txt | 32917 | 32917-SNV-C | SNV | 1 | C | T | 25 | 20579 | 0.12148307 | 46.16 | rnr1Y285A | FALSE | CAN | JSY3868 | 1 | rnr1Y285A-R | TCC | C>T | A/T+1 | G/C-1 | CG>TA | CG>TA | TAAGTTCCA | CCAGGGC | AGG | TCC | C | A | C>T | C>T_TCC | CG>TA_CCAGGGC | 5 | FALSE | FALSE |
| 62_v.txt | 32945 | 32945-SNV-C | SNV | 1 | C | T | 71 | 20579 | 0.34501191 | 56.0422535 | rnr1Y285A | FALSE | CAN | JSY3868 | 1 | rnr1Y285A-R | ACC | C>T | A/T+1 | G/C-1 | CG>TA | CG>TA | AGAAAACCA | CCAATAC | TGG | ACC | C | A | C>T | C>T_ACC | CG>TA_CCAATAC | 5 | FALSE | FALSE |
| 62_v.txt | 32958 | 32958-SNV-C | SNV | 1 | C | T | 132 | 20579 | 0.64143059 | 57.8409091 | rnr1Y285A | FALSE | CAN | JSY3868 | 1 | rnr1Y285A-R | ACC | C>T | A/T+1 | G/C-1 | CG>TA | CG>TA | ATGTAACCAT | CCATTGG | TGG | ACC | C | A | C>T | C>T_ACC | CG>TA_CCATTGG | 5 | FALSE | FALSE |
| 62_v.txt | 32967 | 32967-SNV-C | SNV | 1 | G | A | 38 | 20579 | 0.18465426 | 59.3421053 | rnr1Y285A | FALSE | CAN | JSY3868 | 1 | rnr1Y285A-R | CGC | G>A | A/T+1 | G/C-1 | CG>TA | CG>TA | TTGGCCGCA | GCACCAA | GCG | GCG | C | T | C>T | C>T_GCG | CG>TA_GCACCAA | 5 | FALSE | FALSE |
| 62_v.txt | 32970 | 32970-SNV-C | SNV | 1 | C | T | 58 | 20579 | 0.28184071 | 57.9655172 | rnr1Y285A | FALSE | CAN | JSY3868 | 1 | rnr1Y285A-R | ACC | C>T | A/T+1 | G/C-1 | CG>TA | CG>TA | GCCGCACCA | CCAAATG | TGG | ACC | C | A | C>T | C>T_ACC | CG>TA_CCAAATG | 5 | FALSE | FALSE |
| 62_v.txt | 33015 | 33015-SNV-C | SNV | 1 | G | A | 37 | 20579 | 0.17979494 | 60.1081081 | rnr1Y285A | FALSE | CAN | JSY3868 | 1 | rnr1Y285A-R | GGA | G>A | A/T+1 | G/C-1 | CG>TA | CG>TA | AAAGAGGAT | GATGTAA | CCT | CCT | C | T | C>T | C>T_CCT | CG>TA_GATGTAA | 5 | FALSE | FALSE |
| 62_v.txt | 33043 | 33043-SNV-C | SNV | 1 | C | T | 56 | 20579 | 0.27212207 | 59.0535714 | rnr1Y285A | FALSE | CAN | JSY3868 | 1 | rnr1Y285A-R | TCA | C>T | A/T+1 | G/C-1 | CG>TA | CG>TA | CCATTTCACC | CACCCAA | AGT | TCA | C | A | C>T | C>T_TCA | CG>TA_CACCCAA | 5 | FALSE | FALSE |
| 62_v.txt | 33056 | 33056-SNV-C | SNV | 1 | C | T | 24 | 20579 | 0.11662374 | 55.1666667 | rnr1Y285A | FALSE | CAN | JSY3868 | 1 | rnr1Y285A-R | GCG | C>T | A/T+1 | G/C-1 | CG>TA | CG>TA | GGACTGCGT | CGTGACA | CGC | GCG | C | A | C>T | C>T_GCG | CG>TA_CGTGACA | 5 | FALSE | FALSE |
| 62_v.txt | 33069 | 33069-SNV-C | SNV | 1 | G | A | 20 | 20579 | 0.09718645 | 54.8994171 | rnr1Y285A | FALSE | CAN | JSY3868 | 1 | rnr1Y285A-R | TGC | G>A | A/T+1 | G/C-1 | CG>TA | CG>TA | GAATATGCC | GCCAAAG | ACG | ACG | C | T | C>T | C>T_ACG | CG>TA_GCCAAAG | 5 | FALSE | FALSE |
| 62_v.txt | 33070 | 33070-SNV-C | SNV | 1 | C | T | 21 | 20579 | 0.10204578 | 53.8571429 | rnr1Y285A | FALSE | CAN | JSY3868 | 1 | rnr1Y285A-R | GCC | C>T | A/T+1 | G/C-1 | CG>TA | CG>TA | AATATGCCA | CCAAAGA | CGG | GCC | C | A | C>T | C>T_GCC | CG>TA_CCAAAGA | 5 | FALSE | FALSE |
| 62_v.txt | 33078 | 33078-SNV-C | SNV | 1 | C | T | 34 | 20579 | 0.16521697 | 50.7352941 | rnr1Y285A | FALSE | CAN | JSY3868 | 1 | rnr1Y285A-R | ACC | C>T | A/T+1 | G/C-1 | CG>TA | CG>TA | AAAGAACCC | CCCATAA | TGG | ACC | C | A | C>T | C>T_ACC | CG>TA_CCCATAA | 5 | FALSE | FALSE |
| 62_v.txt | 33104 | 33104-SNV-C | SNV | 1 | G | A | 22 | 20579 | 0.1069051 | 47.5454546 | rnr1Y285A | FALSE | CAN | JSY3868 | 1 | rnr1Y285A-R | CGC | G>A | A/T+1 | G/C-1 | CG>TA | CG>TA | AAGAGCGCC | GCCCACT | GCG | GCG | C | T | C>T | C>T_GCG | CG>TA_GCCCACT | 5 | FALSE | FALSE |
| 62_v.txt | 33114 | 33114-SNV-C | SNV | 1 | C | T | 31 | 20579 | 0.150639 | 38.9243499 | rnr1Y285A | FALSE | CAN | JSY3868 | 1 | rnr1Y285A-R | GCC | C>T | A/T+1 | G/C-1 | CG>TA | CG>TA | ACTGGGCCG | CCGGCGT | CGG | GCC | C | A | C>T | C>T_GCC | CG>TA_CCGGCGT | 5 | FALSE | FALSE |
| 62_v.txt | 33115 | 33115-SNV-C | SNV | 1 | C | T | 20 | 20579 | 0.09718645 | 46.4 | rnr1Y285A | FALSE | CAN | JSY3868 | 1 | rnr1Y285A-R | CCG | C>T | A/T+1 | G/C-1 | CG>TA | CG>TA | CTGGGCCGG | CGGCGTT | GGC | CCG | C | A | C>T | C>T_CCG | CG>TA_CGGCGTT | 5 | FALSE | FALSE |
| 62_v.txt | 33153 | 33153-SNV-C | SNV | 1 | C | T | 120 | 20579 | 0.58311871 | 36.8083334 | rnr1Y285A | FALSE | CAN | JSY3868 | 1 | rnr1Y285A-R | ACC | C>T | A/T+1 | G/C-1 | CG>TA | CG>TA | AAAAGACCT | CCTGTAC | TGG | ACC | C | A | C>T | C>T_ACC | CG>TA_CCTGTAC | 5 | FALSE | FALSE |
| 62_v.txt | 33159 | 33159-SNV-C | SNV | 1 | C | T | 25 | 20579 | 0.12148307 | 37.92 | rnr1Y285A | FALSE | CAN | JSY3868 | 1 | rnr1Y285A-R | ACC | C>T | A/T+1 | G/C-1 | CG>TA | CG>TA | CCTGTACCA | CCAATAG | TGG | ACC | C | A | C>T | C>T_ACC | CG>TA_CCAATAG | 5 | FALSE | FALSE |
| 62_v.txt | 33168 | 33168-SNV-C | SNV | 1 | C | T | 91 | 20579 | 0.44219836 | 37.5384615 | rnr1Y285A | FALSE | CAN | JSY3868 | 1 | rnr1Y285A-R | ACC | C>T | A/T+1 | G/C-1 | CG>TA | CG>TA | ATAGTACCA | CCACCAA | TGG | ACC | C | A | C>T | C>T_ACC | CG>TA_CCACCAA | 5 | FALSE | FALSE |
| 62_v.txt | 33177 | 33177-SNV-C | SNV | 1 | G | A | 35 | 20576 | 0.17010109 | 37.4571429 | rnr1Y285A | FALSE | CAN | JSY3868 | 1 | rnr1Y285A-R | GGC | G>A | A/T+1 | G/C-1 | CG>TA | CG>TA | CCAAGGGCA | GCAATCA | CCG | CCG | C | T | C>T | C>T_CCG | CG>TA_GCAATCA | 5 | FALSE | FALSE |
| 62_v.txt | 33205 | 33205-SNV-C | SNV | 1 | G | A | 11 | 12457 | 0.08830377 | 32.4545455 | rnr1Y285A | FALSE | CAN | JSY3868 | 1 | rnr1Y285A-R | AGC | G>A | A/T+1 | G/C-1 | CG>TA | CG>TA | GCTTAAGCT | GCTCTCT | TCG | TCG | C | T | C>T | C>T_TCG | CG>TA_GCTCTCT | 5 | FALSE | FALSE |
| 62_v.txt | 33258 | 33258-SNV-C | SNV | 1 | C | T | 25 | 12457 | 0.20069038 | 46.08 | rnr1Y285A | FALSE | CAN | JSY3868 | 1 | rnr1Y285A-R | GCC | C>T | A/T+1 | G/C-1 | CG>TA | CG>TA | CCTATGCCAT | CCATCCT | CGG | GCC | C | A | C>T | C>T_GCC | CG>TA_CCATCCT | 5 | FALSE | FALSE |
| 62_v.txt | 33276 | 33276-SNV-C | SNV | 1 | G | A | 19 | 12457 | 0.15252469 | 42.3684211 | rnr1Y285A | FALSE | CAN | JSY3868 | 1 | rnr1Y285A-R | CGT | G>A | A/T+1 | G/C-1 | CG>TA | CG>TA | GAGAACGTA | GTATCCT | GCA | GCA | C | T | C>T | C>T_GCA | CG>TA_GTATCCT | 5 | FALSE | FALSE |
| 62_v.txt | 33309 | 33309-SNV-C | SNV | 1 | C | T | 20 | 12457 | 0.1605523 | 58 | rnr1Y285A | FALSE | CAN | JSY3868 | 1 | rnr1Y285A-R | GCG | C>T | A/T+1 | G/C-1 | CG>TA | CG>TA | AAAGAGCGC | CGCAATG | CGC | GCG | C | A | C>T | C>T_GCG | CG>TA_CGCAATG | 5 | FALSE | FALSE |
| 62_v.txt | 33310 | 33310-SNV-C | SNV | 1 | G | A | 24 | 12456 | 0.19267823 | 57.2195789 | rnr1Y285A | FALSE | CAN | JSY3868 | 1 | rnr1Y285A-R | CGC | G>A | A/T+1 | G/C-1 | CG>TA | CG>TA | AAGAGCGCA | GCAATGG | GCG | GCG | C | T | C>T | C>T_GCG | CG>TA_GCAATGG | 5 | FALSE | FALSE |
| 62_v.txt | 33392 | 33392-SNV-C | SNV | 1 | G | A | 16 | 12457 | 0.12844184 | 48.25 | rnr1Y285A | FALSE | CAN | JSY3868 | 1 | rnr1Y285A-R | CGT | G>A | A/T+1 | G/C-1 | CG>TA | CG>TA | AACGTCGTG | GTGAAAG | GCA | GCA | C | T | C>T | C>T_GCA | CG>TA_GTGAAAG | 5 | FALSE | FALSE |

| | | | | | | | | | | | | | | | | | | | | | | | | | | | | | | | | | | |
|---|---|---|---|---|---|---|---|---|---|---|---|---|---|---|---|---|---|---|---|---|---|---|---|---|---|---|---|---|---|---|---|---|---|---|
| 62_v.txt | 33412 | 33412-SNV-( | SNV | 1 | G | A | 17 | 12457 | 0.13646946 | 47.0588235 | rnr1Y285A | FALSE | CAN | JSY3868 | 1 | rnr1Y285A-R | GGC | G>A | A/T+1 | G/C-1 | CG>TA | CG>TA | TGACCGGCT( | GCTCATT | CCG | CCG | C | T | C>T | C>T_CCG | CG>TA_GCTCATT | 5 | FALSE | FALSE |
| 62_v.txt | 33474 | 33474-SNV-( | SNV | 1 | C | T | 14 | 12457 | 0.11238661 | 36.188967 | rnr1Y285A | FALSE | CAN | JSY3868 | 1 | rnr1Y285A-R | GCC | C>T | A/T+1 | G/C-1 | CG>TA | CG>TA | GCTATGCCTT | CCTTTTT | CGG | GCC | C | A | C>T | C>T_GCC | CG>TA_CCTTTTT | 5 | FALSE | FALSE |
| 62_v.txt | 31956 | 31956-SNV-( | SNV | 1 | G | C | 7 | 12415 | 0.05638341 | 38 | rnr1Y285A | FALSE | CAN | JSY3868 | 1 | rnr1Y285A-R | TGT | G>C | G/C+1 | G/C-1 | CG>GC | CG>GC | ATAAATGTG( | GTGGCCG | ACA | ACA | C | G | C>G | C>G_ACA | CG>GC_GTGGCCG | 5 | FALSE | FALSE |
| 62_v.txt | 31981 | 31981-SNV-( | SNV | 1 | G | C | 8 | 12415 | 0.06443818 | 43.5 | rnr1Y285A | FALSE | CAN | JSY3868 | 1 | rnr1Y285A-R | GGC | G>C | G/C+1 | G/C-1 | CG>GC | CG>GC | AGCCGGGCA | GCATTAA | CCG | CCG | C | G | C>G | C>G_CCG | CG>GC_GCATTAA | 5 | FALSE | FALSE |
| 62_v.txt | 32344 | 32344-SNV-( | SNV | 1 | G | C | 3 | 14963 | 0.02004946 | 58.6666667 | rnr1Y285A | FALSE | CAN | JSY3868 | 1 | rnr1Y285A-R | GGC | G>C | G/C+1 | G/C-1 | CG>GC | CG>GC | TATGTGGCA/ | GCAAAAC | CCG | CCG | C | G | C>G | C>G_CCG | CG>GC_GCAAAAC | 5 | FALSE | FALSE |
| 62_v.txt | 32536 | 32536-SNV-( | SNV | 1 | C | G | 1 | 14963 | 6.68E-03 | 38 | rnr1Y285A | FALSE | CAN | JSY3868 | 1 | rnr1Y285A-R | GCA | C>G | G/C+1 | G/C-1 | CG>GC | CG>GC | GGTTTGCAG( | CAGCTTC | CGT | GCA | C | C | C>G | C>G_GCA | CG>GC_CAGCTTC | 5 | FALSE | FALSE |
| 62_v.txt | 32542 | 32542-SNV-( | SNV | 1 | C | G | 14 | 14963 | 0.09356413 | 37.7857143 | rnr1Y285A | FALSE | CAN | JSY3868 | 1 | rnr1Y285A-R | TCA | C>G | G/C+1 | G/C-1 | CG>GC | CG>GC | CAGCTTCACC | CACCAGC | AGT | TCA | C | C | C>G | C>G_TCA | CG>GC_CACCAGC | 5 | FALSE | FALSE |
| 62_v.txt | 33070 | 33070-SNV-( | SNV | 1 | C | G | 25 | 20579 | 0.12148307 | 53.12 | rnr1Y285A | FALSE | CAN | JSY3868 | 1 | rnr1Y285A-R | GCC | C>G | G/C+1 | G/C-1 | CG>GC | CG>GC | AATATGCCA/ | CCAAAGA | CGG | GCC | C | C | C>G | C>G_GCC | CG>GC_CCAAAGA | 5 | FALSE | FALSE |
| 62_v.txt | 33153 | 33153-SNV-( | SNV | 1 | C | G | 26 | 20579 | 0.12634239 | 37.5384615 | rnr1Y285A | FALSE | CAN | JSY3868 | 1 | rnr1Y285A-R | ACC | C>G | G/C+1 | G/C-1 | CG>GC | CG>GC | AAAAGACCT( | CCTGTAC | TGG | ACC | C | C | C>G | C>G_ACC | CG>GC_CCTGTAC | 5 | FALSE | FALSE |
| 62_v.txt | 33205 | 33205-SNV-( | SNV | 1 | G | C | 68 | 12457 | 0.54587782 | 39.0735294 | rnr1Y285A | FALSE | CAN | JSY3868 | 1 | rnr1Y285A-R | AGC | G>C | G/C+1 | G/C-1 | CG>GC | CG>GC | GCTTAAGCT( | GCTCTCT | TCG | TCG | C | G | C>G | C>G_TCG | CG>GC_GCTCTCT | 5 | FALSE | FALSE |
| 62_v.txt | 33208 | 33208-SNV-( | SNV | 1 | C | G | 1 | 12457 | 8.03E-03 | 38.0005939 | rnr1Y285A | FALSE | CAN | JSY3868 | 1 | rnr1Y285A-R | TCT | C>G | G/C+1 | G/C-1 | CG>GC | CG>GC | TAAGCTCTCT | CTCTCTT | AGA | TCT | C | C | C>G | C>G_TCT | CG>GC_CTCTCTT | 5 | FALSE | FALSE |
| 62_v.txt | 33309 | 33309-SNV-( | SNV | 1 | C | G | 1 | 12457 | 8.03E-03 | 64 | rnr1Y285A | FALSE | CAN | JSY3868 | 1 | rnr1Y285A-R | GCG | C>G | G/C+1 | G/C-1 | CG>GC | CG>GC | AAAGAGCGC | CGCAATG | CGC | GCG | C | C | C>G | C>G_GCG | CG>GC_CGCAATG | 5 | FALSE | FALSE |
| 62_v.txt | 33419 | 33419-SNV-( | SNV | 1 | G | C | 9 | 12457 | 0.07224854 | 44.2222222 | rnr1Y285A | FALSE | CAN | JSY3868 | 1 | rnr1Y285A-R | TGT | G>C | G/C+1 | G/C-1 | CG>GC | CG>GC | CTCATTGTAC | GTACATA | ACA | ACA | C | G | C>G | C>G_ACA | CG>GC_GTACATA | 5 | FALSE | FALSE |
| 62_v.txt | 31657 | 31657-SNV-( | SNV | 1 | G | T | 63 | 7804 | 0.80727832 | 19.4285714 | rnr1Y285A | FALSE | CAN | JSY3868 | 1 | rnr1Y285A-R | GGC | G>T | A/T+1 | G/C-1 | CG>AT | CG>AT | GAAATGGCG | GCGTGGA | CCG | CCG | C | A | C>A | C>A_CCG | CG>AT_GCGTGGA | 5 | FALSE | FALSE |
| 62_v.txt | 31981 | 31981-SNV-( | SNV | 1 | G | T | 26 | 12415 | 0.20942408 | 38.3461539 | rnr1Y285A | FALSE | CAN | JSY3868 | 1 | rnr1Y285A-R | GGC | G>T | A/T+1 | G/C-1 | CG>AT | CG>AT | AGCCGGGCA | GCATTAA | CCG | CCG | C | A | C>A | C>A_CCG | CG>AT_GCATTAA | 5 | FALSE | FALSE |
| 62_v.txt | 32027 | 32027-SNV-( | SNV | 1 | G | T | 1707 | 12415 | 13.7494966 | 60.3192736 | rnr1Y285A | FALSE | CAN | JSY3868 | 1 | rnr1Y285A-R | GGT | G>T | A/T+1 | G/C-1 | CG>AT | CG>AT | GCCACGGTA' | GTATTTC | CCA | CCA | C | A | C>A | C>A_CCA | CG>AT_GTATTTC | 5 | FALSE | FALSE |
| 62_v.txt | 32088 | 32088-SNV-( | SNV | 1 | C | A | 23 | 12415 | 0.18525977 | 59.826087 | rnr1Y285A | FALSE | CAN | JSY3868 | 1 | rnr1Y285A-R | GCC | C>A | A/T+1 | G/C-1 | CG>AT | CG>AT | AAAAGCCT( | CCTGCAA | CGG | GCC | C | T | C>A | C>A_GCC | CG>AT_CCTGCAA | 5 | FALSE | FALSE |
| 62_v.txt | 32149 | 32149-SNV-( | SNV | 1 | C | A | 14 | 12415 | 0.11276681 | 49.6428571 | rnr1Y285A | FALSE | CAN | JSY3868 | 1 | rnr1Y285A-R | TCC | C>A | A/T+1 | G/C-1 | CG>AT | CG>AT | ATGTCTCCAT | CCATGTA | AGG | TCC | C | T | C>A | C>A_TCC | CG>AT_CCATGTA | 5 | FALSE | FALSE |
| 62_v.txt | 32188 | 32188-SNV-( | SNV | 1 | C | A | 62 | 12415 | 0.49939589 | 47.3225807 | rnr1Y285A | FALSE | CAN | JSY3868 | 1 | rnr1Y285A-R | ACT | C>A | A/T+1 | G/C-1 | CG>AT | CG>AT | CGAAAACTG | CTGCAAT | TGA | ACT | C | T | C>A | C>A_ACT | CG>AT_CTGCAAT | 5 | FALSE | FALSE |
| 62_v.txt | 32195 | 32195-SNV-( | SNV | 1 | G | T | 24 | 12415 | 0.19331454 | 44.75 | rnr1Y285A | FALSE | CAN | JSY3868 | 1 | rnr1Y285A-R | TGT | G>T | A/T+1 | G/C-1 | CG>AT | CG>AT | TGCAATGTA' | GTATGGA | ACA | ACA | C | A | C>A | C>A_ACA | CG>AT_GTATGGA | 5 | FALSE | FALSE |
| 62_v.txt | 32274 | 32274-SNV-( | SNV | 1 | G | T | 27 | 14963 | 0.1804451 | 34.6666667 | rnr1Y285A | FALSE | CAN | JSY3868 | 1 | rnr1Y285A-R | GGA | G>T | A/T+1 | G/C-1 | CG>AT | CG>AT | ATACGGGAA | GAACCAA | CCT | CCT | C | A | C>A | C>A_CCT | CG>AT_GAACCAA | 5 | FALSE | FALSE |
| 62_v.txt | 32277 | 32277-SNV-( | SNV | 1 | C | A | 244 | 14963 | 1.63068903 | 36.3893443 | rnr1Y285A | FALSE | CAN | JSY3868 | 1 | rnr1Y285A-R | ACC | C>A | A/T+1 | G/C-1 | CG>AT | CG>AT | CGGGAACCA | CCAACGT | TGG | ACC | C | T | C>A | C>A_ACC | CG>AT_CCAACGT | 5 | FALSE | FALSE |
| 62_v.txt | 32292 | 32292-SNV-( | SNV | 1 | G | T | 20 | 14963 | 0.13366304 | 41.7 | rnr1Y285A | FALSE | CAN | JSY3868 | 1 | rnr1Y285A-R | TGA | G>T | A/T+1 | G/C-1 | CG>AT | CG>AT | ATATTTGAAT | GAATTTG | ACT | ACT | C | A | C>A | C>A_ACT | CG>AT_GAATTTG | 5 | FALSE | FALSE |
| 62_v.txt | 32304 | 32304-SNV-( | SNV | 1 | G | T | 21 | 14963 | 0.14034619 | 55.2380952 | rnr1Y285A | FALSE | CAN | JSY3868 | 1 | rnr1Y285A-R | AGA | G>T | A/T+1 | G/C-1 | CG>AT | CG>AT | GCGGCAGAA | GAAATAA | TCT | TCT | C | A | C>A | C>A_TCT | CG>AT_GAAATAA | 5 | FALSE | FALSE |
| 62_v.txt | 32373 | 32373-SNV-( | SNV | 1 | G | T | 45 | 14963 | 0.30074183 | 62.2444444 | rnr1Y285A | FALSE | CAN | JSY3868 | 1 | rnr1Y285A-R | AGC | G>T | A/T+1 | G/C-1 | CG>AT | CG>AT | TCAATAGCA/ | GCAATAA | TCG | TCG | C | A | C>A | C>A_TCG | CG>AT_GCAATAA | 5 | FALSE | FALSE |
| 62_v.txt | 32432 | 32432-SNV-( | SNV | 1 | G | T | 13 | 14963 | 0.08688097 | 59.6153846 | rnr1Y285A | FALSE | CAN | JSY3868 | 1 | rnr1Y285A-R | TGT | G>T | A/T+1 | G/C-1 | CG>AT | CG>AT | GTCATTGTAT | GTATGGA | ACA | ACA | C | A | C>A | C>A_ACA | CG>AT_GTATGGA | 5 | FALSE | FALSE |
| 62_v.txt | 32471 | 32471-SNV-( | SNV | 1 | G | T | 18 | 14963 | 0.12029673 | 47.9444444 | rnr1Y285A | FALSE | CAN | JSY3868 | 1 | rnr1Y285A-R | TGT | G>T | A/T+1 | G/C-1 | CG>AT | CG>AT | GCCAATGTA( | GTAGAAG | ACA | ACA | C | A | C>A | C>A_ACA | CG>AT_GTAGAAG | 5 | FALSE | FALSE |
| 62_v.txt | 32530 | 32530-SNV-( | SNV | 1 | G | T | 28 | 14963 | 0.18712825 | 40.6785714 | rnr1Y285A | FALSE | CAN | JSY3868 | 1 | rnr1Y285A-R | GGG | G>T | A/T+1 | G/C-1 | CG>AT | CG>AT | TTCTGGGGT | GGTTTGC | CCC | CCC | C | A | C>A | C>A_CCC | CG>AT_GGTTTGC | 5 | FALSE | FALSE |
| 62_v.txt | 32556 | 32556-SNV-( | SNV | 1 | C | A | 255 | 14961 | 1.70443152 | 37.1882353 | rnr1Y285A | FALSE | CAN | JSY3868 | 1 | rnr1Y285A-R | ACC | C>A | A/T+1 | G/C-1 | CG>AT | CG>AT | GTGATACCA/ | CCAACTA | TGG | ACC | C | T | C>A | C>A_ACC | CG>AT_CCAACTA | 5 | FALSE | FALSE |
| 62_v.txt | 32566 | 32566-SNV-( | SNV | 1 | C | A | 57 | 16389 | 0.34779425 | 37.0175439 | rnr1Y285A | FALSE | CAN | JSY3868 | 1 | rnr1Y285A-R | TCA | C>A | A/T+1 | G/C-1 | CG>AT | CG>AT | CTAGTTCAGT | CAGTACC | AGT | TCA | C | T | C>A | C>A_TCA | CG>AT_CAGTACC | 5 | FALSE | FALSE |
| 62_v.txt | 32575 | 32575-SNV-( | SNV | 1 | G | T | 20 | 16394 | 0.12199585 | 36.5 | rnr1Y285A | FALSE | CAN | JSY3868 | 1 | rnr1Y285A-R | TGA | G>T | A/T+1 | G/C-1 | CG>AT | CG>AT | TACCTTGAA/ | GAAATGT | ACT | ACT | C | A | C>A | C>A_ACT | CG>AT_GAAATGT | 5 | FALSE | FALSE |
| 62_v.txt | 32735 | 32735-SNV-( | SNV | 1 | G | T | 22 | 16394 | 0.13419544 | 54.5 | rnr1Y285A | FALSE | CAN | JSY3868 | 1 | rnr1Y285A-R | AGT | G>T | A/T+1 | G/C-1 | CG>AT | CG>AT | AAAACAGTA | GTATATT | TCA | TCA | C | A | C>A | C>A_TCA | CG>AT_GTATATT | 5 | FALSE | FALSE |
| 62_v.txt | 32783 | 32783-SNV-( | SNV | 1 | G | T | 33 | 16394 | 0.20129316 | 47.7878788 | rnr1Y285A | FALSE | CAN | JSY3868 | 1 | rnr1Y285A-R | AGA | G>T | A/T+1 | G/C-1 | CG>AT | CG>AT | GACCCAGAA | GAACTCG | TCT | TCT | C | A | C>A | C>A_TCT | CG>AT_GAACTCG | 5 | FALSE | FALSE |
| 62_v.txt | 32786 | 32786-SNV-( | SNV | 1 | C | A | 27 | 16394 | 0.1646944 | 44.5555556 | rnr1Y285A | FALSE | CAN | JSY3868 | 1 | rnr1Y285A-R | ACT | C>A | A/T+1 | G/C-1 | CG>AT | CG>AT | CCAGAACTC( | CTCGAAT | TGA | ACT | C | T | C>A | C>A_ACT | CG>AT_CTCGAAT | 5 | FALSE | FALSE |
| 62_v.txt | 32794 | 32794-SNV-( | SNV | 1 | C | A | 31 | 16394 | 0.18909357 | 41.8064516 | rnr1Y285A | FALSE | CAN | JSY3868 | 1 | rnr1Y285A-R | TCA | C>A | A/T+1 | G/C-1 | CG>AT | CG>AT | CGAATTCACC | CACCGTA | AGT | TCA | C | T | C>A | C>A_TCA | CG>AT_CACCGTA | 5 | FALSE | FALSE |
| 62_v.txt | 32798 | 32798-SNV-( | SNV | 1 | G | T | 26 | 16394 | 0.15859461 | 46.1153846 | rnr1Y285A | FALSE | CAN | JSY3868 | 1 | rnr1Y285A-R | CGT | G>T | A/T+1 | G/C-1 | CG>AT | CG>AT | TTCACCGTAA | GTAATAT | GCA | GCA | C | A | C>A | C>A_GCA | CG>AT_GTAATAT | 5 | FALSE | FALSE |
| 62_v.txt | 32819 | 32819-SNV-( | SNV | 1 | G | T | 232 | 16394 | 1.41515189 | 43.9353448 | rnr1Y285A | FALSE | CAN | JSY3868 | 1 | rnr1Y285A-R | AGT | G>T | A/T+1 | G/C-1 | CG>AT | CG>AT | GAACAAGTT | GTTCATT | TCA | TCA | C | A | C>A | C>A_TCA | CG>AT_GTTCATT | 5 | FALSE | FALSE |
| 62_v.txt | 32917 | 32917-SNV-( | SNV | 1 | C | A | 489 | 20579 | 2.37620876 | 53.1717791 | rnr1Y285A | FALSE | CAN | JSY3868 | 1 | rnr1Y285A-R | TCC | C>A | A/T+1 | G/C-1 | CG>AT | CG>AT | TAAGTTCCAC | CCAGGGC | AGG | TCC | C | T | C>A | C>A_TCC | CG>AT_CCAGGGC | 5 | FALSE | FALSE |
| 62_v.txt | 32940 | 32940-SNV-( | SNV | 1 | G | T | 33 | 20579 | 0.16035765 | 54.6969697 | rnr1Y285A | FALSE | CAN | JSY3868 | 1 | rnr1Y285A-R | AGA | G>T | A/T+1 | G/C-1 | CG>AT | CG>AT | GCCCAAGAA | GAAAACC | TCT | TCT | C | A | C>A | C>A_TCT | CG>AT_GAAAACC | 5 | FALSE | FALSE |
| 62_v.txt | 32954 | 32954-SNV-( | SNV | 1 | G | T | 11 | 20579 | 0.05345255 | 50.2727273 | rnr1Y285A | FALSE | CAN | JSY3868 | 1 | rnr1Y285A-R | TGT | G>T | A/T+1 | G/C-1 | CG>AT | CG>AT | ATACATGTA/ | GTAACCA | ACA | ACA | C | A | C>A | C>A_ACA | CG>AT_GTAACCA | 5 | FALSE | FALSE |
| 62_v.txt | 32958 | 32958-SNV-( | SNV | 1 | C | A | 321 | 20579 | 1.55984256 | 56.9813084 | rnr1Y285A | FALSE | CAN | JSY3868 | 1 | rnr1Y285A-R | ACC | C>A | A/T+1 | G/C-1 | CG>AT | CG>AT | ATGTAACCAT | CCATTGG | TGG | ACC | C | T | C>A | C>A_ACC | CG>AT_CCATTGG | 5 | FALSE | FALSE |
| 62_v.txt | 33015 | 33015-SNV-( | SNV | 1 | G | T | 27 | 20579 | 0.13120171 | 58 | rnr1Y285A | FALSE | CAN | JSY3868 | 1 | rnr1Y285A-R | GGA | G>T | A/T+1 | G/C-1 | CG>AT | CG>AT | AAAGAGGAT | GATGTAA | CCT | CCT | C | A | C>A | C>A_CCT | CG>AT_GATGTAA | 5 | FALSE | FALSE |
| 62_v.txt | 33070 | 33070-SNV-( | SNV | 1 | C | A | 6 | 20579 | 0.02915594 | 46.5 | rnr1Y285A | FALSE | CAN | JSY3868 | 1 | rnr1Y285A-R | GCC | C>A | A/T+1 | G/C-1 | CG>AT | CG>AT | AATATGCCA/ | CCAAAGA | CGG | GCC | C | T | C>A | C>A_GCC | CG>AT_CCAAAGA | 5 | FALSE | FALSE |
| 62_v.txt | 33093 | 33093-SNV-( | SNV | 1 | G | T | 11 | 20579 | 0.05345255 | 49.1818182 | rnr1Y285A | FALSE | CAN | JSY3868 | 1 | rnr1Y285A-R | TGA | G>T | A/T+1 | G/C-1 | CG>AT | CG>AT | AAATATGAT/ | GATATAA | ACT | ACT | C | A | C>A | C>A_ACT | CG>AT_GATATAA | 5 | FALSE | FALSE |
| 62_v.txt | 33115 | 33115-SNV-( | SNV | 1 | C | A | 18 | 20579 | 0.08746781 | 49.2222222 | rnr1Y285A | FALSE | CAN | JSY3868 | 1 | rnr1Y285A-R | CCG | C>A | A/T+1 | G/C-1 | CG>AT | CG>AT | CTGGGCCGG | CGGCGTT | GGC | CCG | C | T | C>A | C>A_CCG | CG>AT_CGGCGTT | 5 | FALSE | FALSE |
| 62_v.txt | 33151 | 33151-SNV-( | SNV | 1 | G | T | 229 | 20579 | 1.11278488 | 39.5633188 | rnr1Y285A | FALSE | CAN | JSY3868 | 1 | rnr1Y285A-R | AGA | G>T | A/T+1 | G/C-1 | CG>AT | CG>AT | TGAAAAGAC | GACCTGT | TCT | TCT | C | A | C>A | C>A_TCT | CG>AT_GACCTGT | 5 | FALSE | FALSE |
| 62_v.txt | 33153 | 33153-SNV-( | SNV | 1 | C | A | 1031 | 20579 | 5.00996161 | 40.3773036 | rnr1Y285A | FALSE | CAN | JSY3868 | 1 | rnr1Y285A-R | ACC | C>A | A/T+1 | G/C-1 | CG>AT | CG>AT | AAAAGACCT( | CCTGTAC | TGG | ACC | C | T | C>A | C>A_ACC | CG>AT_CCTGTAC | 5 | FALSE | FALSE |
| 62_v.txt | 33154 | 33154-SNV-( | SNV | 1 | C | A | 19 | 20579 | 0.09232713 | 37.6840359 | rnr1Y285A | FALSE | CAN | JSY3868 | 1 | rnr1Y285A-R | CCT | C>A | A/T+1 | G/C-1 | CG>AT | CG>AT | AAAGACCTG | CTGTACC | GGA | CCT | C | T | C>A | C>A_CCT | CG>AT_CTGTACC | 5 | FALSE | FALSE |
| 62_v.txt | 33156 | 33156-SNV-( | SNV | 1 | G | T | 92 | 20579 | 0.44705768 | 37.2826087 | rnr1Y285A | FALSE | CAN | JSY3868 | 1 | rnr1Y285A-R | TGT | G>T | A/T+1 | G/C-1 | CG>AT | CG>AT | AGACCTGTA( | GTACCAA | ACA | ACA | C | A | C>A | C>A_ACA | CG>AT_GTACCAA | 5 | FALSE | FALSE |
| 62_v.txt | 33159 | 33159-SNV-( | SNV | 1 | C | A | 41 | 20579 | 0.19923223 | 37.2195122 | rnr1Y285A | FALSE | CAN | JSY3868 | 1 | rnr1Y285A-R | ACC | C>A | A/T+1 | G/C-1 | CG>AT | CG>AT | CCTGTACCA/ | CCAATAG | TGG | ACC | C | T | C>A | C>A_ACC | CG>AT_CCAATAG | 5 | FALSE | FALSE |
| 62_v.txt | 33168 | 33168-SNV-( | SNV | 1 | C | A | 150 | 20579 | 0.72889839 | 37.32 | rnr1Y285A | FALSE | CAN | JSY3868 | 1 | rnr1Y285A-R | ACC | C>A | A/T+1 | G/C-1 | CG>AT | CG>AT | ATAGTACCA( | CCACCAA | TGG | ACC | C | T | C>A | C>A_ACC | CG>AT_CCACCAA | 5 | FALSE | FALSE |
| 62_v.txt | 33195 | 33195-SNV-( | SNV | 1 | C | A | 33 | 12457 | 0.2649113 | 35.8484849 | rnr1Y285A | FALSE | CAN | JSY3868 | 1 | rnr1Y285A-R | TCT | C>A | A/T+1 | G/C-1 | CG>AT | CG>AT | ATATGTCTTT | CTTTGCT | AGA | TCT | C | T | C>A | C>A_TCT | CG>AT_CTTTGCT | 5 | FALSE | FALSE |
| 62_v.txt | 33208 | 33208-SNV-( | SNV | 1 | C | A | 9 | 12457 | 0.07224854 | 34.5595649 | rnr1Y285A | FALSE | CAN | JSY3868 | 1 | rnr1Y285A-R | TCT | C>A | A/T+1 | G/C-1 | CG>AT | CG>AT | TAAGCTCTCT | CTCTCTT | AGA | TCT | C | T | C>A | C>A_TCT | CG>AT_CTCTCTT | 5 | FALSE | FALSE |
| 62_v.txt | 33235 | 33235-SNV-( | SNV | 1 | C | A | 24 | 12457 | 0.19266276 | 37.4166667 | rnr1Y285A | FALSE | CAN | JSY3868 | 1 | rnr1Y285A-R | TCT | C>A | A/T+1 | G/C-1 | CG>AT | CG>AT | GTACTTCTCC | CTCCTTC | AGA | TCT | C | T | C>A | C>A_TCT | CG>AT_CTCCTTC | 5 | FALSE | FALSE |
| 62_v.txt | 33325 | 33325-SNV-( | SNV | 1 | C | A | 1249 | 12457 | 10.0264911 | 58.8791033 | rnr1Y285A | FALSE | CAN | JSY3868 | 1 | rnr1Y285A-R | TCT | C>A | A/T+1 | G/C-1 | CG>AT | CG>AT | ACAATTCTTT | CTTTACT | AGA | TCT | C | T | C>A | C>A_TCT | CG>AT_CTTTACT | 5 | FALSE | FALSE |
| 62_v.txt | 33474 | 33474-SNV-( | SNV | 1 | C | A | 50 | 12457 | 0.40138075 | 35.086859 | rnr1Y285A | FALSE | CAN | JSY3868 | 1 | rnr1Y285A-R | GCC | C>A | A/T+1 | G/C-1 | CG>AT | CG>AT | GCTATGCCTT | CCTTTTT | CGG | GCC | C | T | C>A | C>A_GCC | CG>AT_CCTTTTT | 5 | FALSE | FALSE |
| 62_v.txt | 33475 | 33475-SNV-( | SNV | 1 | C | A | 11 | 12457 | 0.08830377 | 37.6363636 | rnr1Y285A | FALSE | CAN | JSY3868 | 1 | rnr1Y285A-R | CCT | C>A | A/T+1 | G/C-1 | CG>AT | CG>AT | CTATGCCTTT | CTTTTTT | GGA | CCT | C | T | C>A | C>A_CCT | CG>AT_CTTTTTT | 5 | FALSE | FALSE |
| 62_v.txt | 33153 | 33153-MNV- | MNV | 2 | CC | GG | 1 | 20579 | 4.86E-03 | 18.5 | rnr1Y285A | FALSE | CAN | JSY3868 | 1 | rnr1Y285A-R | ACC | nope | >1bp-ins | >1bpdel | none | MNV | AAAAGACCT( | CCTGTAC | TGG | ACC | CC | none | NA | NA_ACC | MNV_CCTGTAC | 5 | FALSE | FALSE |
| 62_v.txt | 31984 | 31984-Insert | Insertion | 1 | - | T | 4 | 12415 | 0.03221909 | 44.5 | rnr1Y285A | FALSE | CAN | JSY3868 | 1 | rnr1Y285A-R | ATT | nope | A/T+1 | >1bpdel | none | A/T+1 | CGGGCATTA/ | TTAATTT | TAA | ATT | - | A | NA | NA_ATT | A/T+1_TTAATTT | 5 | FALSE | FALSE |
| 62_v.txt | 32081 | 32081-Insert | Insertion | 1 | - | A | 18 | 12415 | 0.1449859 | 63.1666667 | rnr1Y285A | FALSE | CAN | JSY3868 | 1 | rnr1Y285A-R | CAA | nope | A/T+1 | >1bpdel | none | A/T+1 | CCATGCAAA/ | AAAAAAG | GTT | CAA | - | T | NA | NA_CAA | A/T+1_AAAAAAG | 5 | FALSE | FALSE |
| 62_v.txt | 32498 | 32498-Insert | Insertion | 1 | - | T | 3 | 14963 | 0.02004946 | 37.6666667 | rnr1Y285A | FALSE | CAN | JSY3868 | 1 | rnr1Y285A-R | CTT | nope | A/T+1 | >1bpdel | none | A/T+1 | AACAACTTTT | TTTTTTG | GAA | CTT | - | A | NA | NA_CTT | A/T+1_TTTTTTG | 5 | FALSE | FALSE |
| 62_v.txt | 32631 | 32631-Insert | Insertion | 1 | - | T | 9 | 16394 | 0.05489813 | 46.1111111 | rnr1Y285A | FALSE | CAN | JSY3868 | 1 | rnr1Y285A-R | GTT | nope | A/T+1 | >1bpdel | none | A/T+1 | CCTTCGTTTT | TTTTTAT | CAA | GTT | - | A | NA | NA_GTT | A/T+1_TTTTTAT | 5 | FALSE | FALSE |
| 62_v.txt | 32744 | 32744-Insert | Insertion | 1 | - | A | 12 | 16394 | 0.07319751 | 60.9166667 | rnr1Y285A | FALSE | CAN | JSY3868 | 1 | rnr1Y285A-R | GAA | nope | A/T+1 | >1bpdel | none | A/T+1 | TATTAGAAA( | AAACCCG | CTT | GAA | - | T | NA | NA_GAA | A/T+1_AAACCCG | 5 | FALSE | FALSE |
| 62_v.txt | 32842 | 32842-Insert | Insertion | 1 | - | A | 9 | 16394 | 0.05489813 | 43.5555556 | rnr1Y285A | FALSE | CAN | JSY3868 | 1 | rnr1Y285A-R | CAA | nope | A/T+1 | >1bpdel | none | A/T+1 | TTACCCAAA/ | AAAAAAT | GTT | CAA | - | T | NA | NA_CAA | A/T+1_AAAAAAT | 5 | FALSE | FALSE |
| 62_v.txt | 31723 | 31723-Deleti | Deletion | 1 | A | - | 9 | 7804 | 0.11532547 | 60.6666667 | rnr1Y285A | FALSE | CAN | JSY3868 | 1 | rnr1Y285A-R | CAA | nope | >1bp-ins | A/T-1 | none | A/T-1 | TGTCCCAAA/ | AAAAAGT | GTT | GTT | T | none | T>none | T>none_GTT | A/T-1_AAAAAGT | 5 | FALSE | FALSE |
| 62_v.txt | 31747 | 31747-Deleti | Deletion | 1 | C | - | 4 | 7804 | 0.05125577 | 64 | rnr1Y285A | FALSE | CAN | JSY3868 | 1 | rnr1Y285A-R | TCC | nope | >1bp-ins | G/C-1 | none | G/C-1 | GATCTTCCCA | CCCATAC | AGG | TCC | C | none | nope | nope_TCC | G/C-1_CCCATAC | 5 | FALSE | FALSE |
| 62_v.txt | 31913 | 31913-Deleti | Deletion | 1 | A | - | 5 | 7804 | 0.06406971 | 37.6 | rnr1Y285A | FALSE | CAN | JSY3868 | 1 | rnr1Y285A-R | CAA | nope | >1bp-ins | A/T-1 | none | A/T-1 | TGGTGCAAA | AAAAGCC | GTT | GTT | T | none | T>none | T>none_GTT | A/T-1_AAAAGCC | 5 | FALSE | FALSE |
| 62_v.txt | 32081 | 32081-Deleti | Deletion | 1 | A | - | 14 | 12415 | 0.11276681 | 63.5 | rnr1Y285A | FALSE | CAN | JSY3868 | 1 | rnr1Y285A-R | CAA | nope | >1bp-ins | A/T-1 | none | A/T-1 | CCATGCAAA/ | AAAAAAG | GTT | GTT | T | none | T>none | T>none_GTT | A/T-1_AAAAAAG | 5 | FALSE | FALSE |
| 62_v.txt | 32149 | 32149-Deleti | Deletion | 1 | C | - | 2 | 12415 | 0.01610955 | 64 | rnr1Y285A | FALSE | CAN | JSY3868 | 1 | rnr1Y285A-R | TCC | nope | >1bp-ins | G/C-1 | none | G/C-1 | ATGTCTCCAT | CCATGTA | AGG | TCC | C | none | nope | nope_TCC | G/C-1_CCATGTA | 5 | FALSE | FALSE |
| 62_v.txt | 32216 | 32216-Deleti | Deletion | 1 | G | - | 43 | 12415 | 0.34635522 | 39.5813954 | rnr1Y285A | FALSE | CAN | JSY3868 | 1 | rnr1Y285A-R | TGG | nope | >1bp-ins | G/C-1 | none | G/C-1 | TTTGGTGGT( | GGTCCTT | ACC | ACC | C | none | C>none | C>none_ACC | G/C-1_GGTCCTT | 5 | FALSE | FALSE |
| 62_v.txt | 32259 | 32259-Deleti | Deletion | 1 | C | - | 58 | 14962 | 0.38764871 | 37.6034483 | rnr1Y285A | FALSE | CAN | JSY3868 | 1 | rnr1Y285A-R | ACC | nope | >1bp-ins | G/C-1 | none | G/C-1 | GATAGACCA | CCAAATA | TGG | ACC | C | none | nope | nope_ACC | G/C-1_CCAAATA | 5 | FALSE | FALSE |
| 62_v.txt | 32277 | 32277-Deleti | Deletion | 1 | C | - | 23 | 14963 | 0.15371249 | 37.5652174 | rnr1Y285A | FALSE | CAN | JSY3868 | 1 | rnr1Y285A-R | ACC | nope | >1bp-ins | G/C-1 | none | G/C-1 | CGGGAACCA | CCAACGT | TGG | ACC | C | none | nope | nope_ACC | G/C-1_CCAACGT | 5 | FALSE | FALSE |
| 62_v.txt | 32346 | 32346-Deleti | Deletion | 1 | A | - | 5 | 14963 | 0.03341576 | 63.2 | rnr1Y285A | FALSE | CAN | JSY3868 | 1 | rnr1Y285A-R | CAA | nope | >1bp-ins | A/T-1 | none | A/T-1 | TGTGGCAAA | AAAACCT | GTT | GTT | T | none | T>none | T>none_GTT | A/T-1_AAAACCT | 5 | FALSE | FALSE |
| 62_v.txt | 32381 | 32381-Deleti | Deletion | 1 | A | - | 5 | 14963 | 0.03341576 | 58.6 | rnr1Y285A | FALSE | CAN | JSY3868 | 1 | rnr1Y285A-R | TAA | nope | >1bp-ins | A/T-1 | none | A/T-1 | AATAATAAA( | AAAGGGA | ATT | ATT | T | none | T>none | T>none_ATT | A/T-1_AAAGGGA | 5 | FALSE | FALSE |
| 62_v.txt | 32398 | 32398-Deleti | Deletion | 1 | C | - | 89 | 14963 | 0.59480051 | 59.5842697 | rnr1Y285A | FALSE | CAN | JSY3868 | 1 | rnr1Y285A-R | ACG | nope | >1bp-ins | G/C-1 | none | G/C-1 | TAGAAACGT. | CGTAGGA | TGC | ACG | C | none | nope | nope_ACG | G/C-1_CGTAGGA | 5 | FALSE | FALSE |
| 62_v.txt | 32442 | 32442-Deleti | Deletion | 1 | A | - | 5 | 14963 | 0.03341576 | 53.6 | rnr1Y285A | FALSE | CAN | JSY3868 | 1 | rnr1Y285A-R | TAA | nope | >1bp-ins | A/T-1 | none | A/T-1 | GGAACTAAA | AAAAGTC | ATT | ATT | T | none | T>none | T>none_ATT | A/T-1_AAAAGTC | 5 | FALSE | FALSE |
| 62_v.txt | 32448 | 32448-Deleti | Deletion | 1 | C | - | 8 | 14963 | 0.05346521 | 59.25 | rnr1Y285A | FALSE | CAN | JSY3868 | 1 | rnr1Y285A-R | TCC | nope | >1bp-ins | G/C-1 | none | G/C-1 | AAAAGTCCA. | CCAATGA | AGG | TCC | C | none | nope | nope_TCC | G/C-1_CCAATGA | 5 | FALSE | FALSE |
| 62_v.txt | 32498 | 32498-Deleti | Deletion | 1 | T | - | 9 | 14963 | 0.06014837 | 44.7777778 | rnr1Y285A | FALSE | CAN | JSY3868 | 1 | rnr1Y285A-R | CTT | nope | >1bp-ins | A/T-1 | none | A/T-1 | AACAACTTTT | TTTTTTG | GAA | CTT | T | none | nope | nope_CTT | A/T-1_TTTTTTG | 5 | FALSE | FALSE |
| 62_v.txt | 32528 | 32528-Deleti | Deletion | 1 | G | - | 6 | 14963 | 0.04009891 | 37.1666667 | rnr1Y285A | FALSE | CAN | JSY3868 | 1 | rnr1Y285A-R | TGG | nope | >1bp-ins | G/C-1 | none | G/C-1 | TTTTCTGGGC | GGGGTTT | ACC | ACC | C | none | C>none | C>none_ACC | G/C-1_GGGGTTT | 5 | FALSE | FALSE |
| 62_v.txt | 32556 | 32556-Deleti | Deletion | 1 | C | - | 11 | 14961 | 0.0735245 | 37.8181818 | rnr1Y285A | FALSE | CAN | JSY3868 | 1 | rnr1Y285A-R | ACC | nope | >1bp-ins | G/C-1 | none | G/C-1 | GTGATACCA/ | CCAACTA | TGG | ACC | C | none | nope | nope_ACC | G/C-1_CCAACTA | 5 | FALSE | FALSE |
| 62_v.txt | 32603 | 32603-Deleti | Deletion | 1 | G | - | 15 | 16394 | 0.09149689 | 49.2666667 | rnr1Y285A | FALSE | CAN | JSY3868 | 1 | rnr1Y285A-R | AGG | nope | >1bp-ins | G/C-1 | none | G/C-1 | CAAAGAGGA | GGAAACC | TCC | TCC | C | none | C>none | C>none_TCC | G/C-1_GGAAACC | 5 | FALSE | FALSE |
| 62_v.txt | 32608 | 32608-Deleti | Deletion | 1 | C | - | 574 | 16394 | 3.50128096 | 43.5853659 | rnr1Y285A | FALSE | CAN | JSY3868 | 1 | rnr1Y285A-R | ACC | nope | >1bp-ins | G/C-1 | none | G/C-1 | AGGAAACCC | CCCAACC | TGG | ACC | C | none | nope | nope_ACC | G/C-1_CCCAACC | 5 | FALSE | FALSE |
| 62_v.txt | 32658 | 32658-Deleti | Deletion | 1 | C | - | 1178 | 16394 | 7.18555569 | 60.0933786 | rnr1Y285A | FALSE | CAN | JSY3868 | 1 | rnr1Y285A-R | ACC | nope | >1bp-ins | G/C-1 | none | G/C-1 | CCTGGACCC( | CCCCAGG | TGG | ACC | C | none | nope | nope_ACC | G/C-1_CCCCAGG | 5 | FALSE | FALSE |
| 62_v.txt | 32670 | 32670-Deleti | Deletion | 1 | G | - | 772 | 16394 | 4.70903989 | 59.1502591 | rnr1Y285A | FALSE | CAN | JSY3868 | 1 | rnr1Y285A-R | TGG | nope | >1bp-ins | G/C-1 | none | G/C-1 | GCACCTGGG | GGGTTTC | ACC | ACC | C | none | C>none | C>none_ACC | G/C-1_GGGTTTC | 5 | FALSE | FALSE |
| 62_v.txt | 32707 | 32707-Deleti | Deletion | 1 | C | - | 1407 | 16394 | 8.5824082 | 62.3439943 | rnr1Y285A | FALSE | CAN | JSY3868 | 1 | rnr1Y285A-R | ACC | nope | >1bp-ins | G/C-1 | none | G/C-1 | CGGTAACCC( | CCCCAGC | TGG | ACC | C | none | nope | nope_ACC | G/C-1_CCCCAGC | 5 | FALSE | FALSE |
| 62_v.txt | 32747 | 32747-Deleti | Deletion | 1 | C | - | 41 | 16394 | 0.2500915 | 57.0243902 | rnr1Y285A | FALSE | CAN | JSY3868 | 1 | rnr1Y285A-R | ACC | nope | >1bp-ins | G/C-1 | none | G/C-1 | TAGAAACCC( | CCCGATA | TGG | ACC | C | none | nope | nope_ACC | G/C-1_CCCGATA | 5 | FALSE | FALSE |
| 62_v.txt | 32842 | 32842-Deleti | Deletion | 1 | A | - | 21 | 16394 | 0.12809565 | 48.9047612 | rnr1Y285A | FALSE | CAN | JSY3868 | 1 | rnr1Y285A-R | CAA | nope | >1bp-ins | A/T-1 | none | A/T-1 | TTACCCAAA/ | AAAAAAT | GTT | GTT | T | none | T>none | T>none_GTT | A/T-1_AAAAAAT | 5 | FALSE | FALSE |
| 62_v.txt | 32855 | 32855-Deleti | Deletion | 1 | C | - | 3 | 16394 | 0.01829938 | 37.6666667 | rnr1Y285A | FALSE | CAN | JSY3868 | 1 | rnr1Y285A-R | TCC | nope | >1bp-ins | G/C-1 | none | G/C-1 | ACTAATCCAT | CCATGCC | AGG | TCC | C | none | nope | nope_TCC | G/C-1_CCATGCC | 5 | FALSE | FALSE |
| 62_v.txt | 32860 | 32860-Deleti | Deletion | 1 | C | - | 2 | 16394 | 0.01219959 | 38 | rnr1Y285A | FALSE | CAN | JSY3868 | 1 | rnr1Y285A-R | GCC | nope | >1bp-ins | G/C-1 | none | G/C-1 | TCCATGCCG( | CCGCCAG | CGG | GCC | C | none | nope | nope_GCC | G/C-1_CCGCCAG | 5 | FALSE | FALSE |
| 62_v.txt | 32882 | 32882-Deleti | Deletion | 1 | C | - | 3 | 20579 | 0.01457797 | 38 | rnr1Y285A | FALSE | CAN | JSY3868 | 1 | rnr1Y285A-R | TCC | nope | >1bp-ins | G/C-1 | none | G/C-1 | GTACGTCCA/ | CCAAAAT | AGG | TCC | C | none | nope | nope_TCC | G/C-1_CCAAAAT | 5 | FALSE | FALSE |
| 62_v.txt | 32884 | 32884-Deleti | Deletion | 1 | A | - | 19 | 20579 | 0.09232713 | 36.8947368 | rnr1Y285A | FALSE | CAN | JSY3868 | 1 | rnr1Y285A-R | CAA | nope | >1bp-ins | A/T-1 | none | A/T-1 | ACGTCCAAA/ | AAAATTG | GTT | GTT | T | none | T>none | T>none_GTT | A/T-1_AAAATTG | 5 | FALSE | FALSE |
| 62_v.txt | 32917 | 32917-Deleti | Deletion | 1 | C | - | 19 | 20579 | 0.09232713 | 47.9473684 | rnr1Y285A | FALSE | CAN | JSY3868 | 1 | rnr1Y285A-R | TCC | nope | >1bp-ins | G/C-1 | none | G/C-1 | TAAGTTCCAC | CCAGGGC | AGG | TCC | C | none | nope | nope_TCC | G/C-1_CCAGGGC | 5 | FALSE | FALSE |
| 62_v.txt | 32920 | 32920-Deleti | Deletion | 1 | G | - | 4 | 20579 | 0.01943729 | 57.5 | rnr1Y285A | FALSE | CAN | JSY3868 | 1 | rnr1Y285A-R | AGG | nope | >1bp-ins | G/C-1 | none | G/C-1 | GTTCCAGGG | GGGCAAA | TCC | TCC | C | none | C>none | C>none_TCC | G/C-1_GGGCAAA | 5 | FALSE | FALSE |
| 62_v.txt | 32924 | 32924-Deleti | Deletion | 1 | A | - | 14 | 20579 | 0.06803052 | 50.0714286 | rnr1Y285A | FALSE | CAN | JSY3868 | 1 | rnr1Y285A-R | CAA | nope | >1bp-ins | A/T-1 | none | A/T-1 | CAGGGCAAA | AAAAGTG | GTT | GTT | T | none | T>none | T>none_GTT | A/T-1_AAAAGTG | 5 | FALSE | FALSE |
| 62_v.txt | 32945 | 32945-Deleti | Deletion | 1 | C | - | 8 | 20579 | 0.03887458 | 58.625 | rnr1Y285A | FALSE | CAN | JSY3868 | 1 | rnr1Y285A-R | ACC | nope | >1bp-ins | G/C-1 | none | G/C-1 | AGAAAACCA | CCAATAC | TGG | ACC | C | none | nope | nope_ACC | G/C-1_CCAATAC | 5 | FALSE | FALSE |
| 62_v.txt | 32997 | 32997-Deleti | Deletion | 1 | G | - | 4 | 20579 | 0.01943729 | 64 | rnr1Y285A | FALSE | CAN | JSY3868 | 1 | rnr1Y285A-R | TGA | nope | >1bp-ins | G/C-1 | none | G/C-1 | CTTTGTGAG/ | GAGAAAA | ACT | ACT | C | none | C>none | C>none_ACT | G/C-1_GAGAAAA | 5 | FALSE | FALSE |
| 62_v.txt | 33045 | 33045-Deleti | Deletion | 1 | C | - | 6 | 20579 | 0.02915594 | 54.6666667 | rnr1Y285A | FALSE | CAN | JSY3868 | 1 | rnr1Y285A-R | ACC | nope | >1bp-ins | G/C-1 | none | G/C-1 | ATTTCACCCA | CCCAAGG | TGG | ACC | C | none | nope | nope_ACC | G/C-1_CCCAAGG | 5 | FALSE | FALSE |
| 62_v.txt | 33078 | 33078-Deleti | Deletion | 1 | C | - | 35 | 20579 | 0.17007629 | 50.0571429 | rnr1Y285A | FALSE | CAN | JSY3868 | 1 | rnr1Y285A-R | ACC | nope | >1bp-ins | G/C-1 | none | G/C-1 | AAAGAACCC | CCCATAA | TGG | ACC | C | none | nope | nope_ACC | G/C-1_CCCATAA | 5 | FALSE | FALSE |
| 62_v.txt | 33141 | 33141-Deleti | Deletion | 1 | C | - | 422 | 20579 | 2.05063414 | 39.8412322 | rnr1Y285A | FALSE | CAN | JSY3868 | 1 | rnr1Y285A-R | ACC | nope | >1bp-ins | G/C-1 | none | G/C-1 | GATAAACCA. | CCAATGA | TGG | ACC | C | none | nope | nope_ACC | G/C-1_CCAATGA | 5 | FALSE | FALSE |
| 62_v.txt | 33147 | 33147-Deleti | Deletion | 1 | A | - | 7 | 20579 | 0.03401526 | 45.4285714 | rnr1Y285A | FALSE | CAN | JSY3868 | 1 | rnr1Y285A-R | GAA | nope | >1bp-ins | A/T-1 | none | A/T-1 | CCAATGAAA. | AAAAGAC | CTT | CTT | T | none | T>none | T>none_CTT | A/T-1_AAAAGAC | 5 | FALSE | FALSE |
| 62_v.txt | 33159 | 33159-Deleti | Deletion | 1 | C | - | 3 | 20579 | 0.01457797 | 38 | rnr1Y285A | FALSE | CAN | JSY3868 | 1 | rnr1Y285A-R | ACC | nope | >1bp-ins | G/C-1 | none | G/C-1 | CCTGTACCA/ | CCAATAG | TGG | ACC | C | none | nope | nope_ACC | G/C-1_CCAATAG | 5 | FALSE | FALSE |
| 62_v.txt | 33303 | 33303-Deleti | Deletion | 1 | A | - | 8 | 12457 | 0.06422092 | 49.75 | rnr1Y285A | FALSE | CAN | JSY3868 | 1 | rnr1Y285A-R | GAA | nope | >1bp-ins | A/T-1 | none | A/T-1 | GTCGGGAAA | AAAGAGC | CTT | CTT | T | none | T>none | T>none_CTT | A/T-1_AAAGAGC | 5 | FALSE | FALSE |
| 62_v.txt | 33356 | 33356-Deleti | Deletion | 1 | C | - | 258 | 12457 | 2.07112467 | 53.6395349 | rnr1Y285A | FALSE | CAN | JSY3868 | 1 | rnr1Y285A-R | ACC | nope | >1bp-ins | G/C-1 | none | G/C-1 | TATTGACCCA | CCCACGT | TGG | ACC | C | none | nope | nope_ACC | G/C-1_CCCACGT | 5 | FALSE | FALSE |
| 62_v.txt | 33474 | 33474-Deleti | Deletion | 1 | C | - | 33 | 12457 | 0.2649113 | 27.6969642 | rnr1Y285A | FALSE | CAN | JSY3868 | 1 | rnr1Y285A-R | GCC | nope | >1bp-ins | G/C-1 | none | G/C-1 | GCTATGCCTT | CCTTTTT | CGG | GCC | C | none | nope | nope_GCC | G/C-1_CCTTTTT | 5 | FALSE | FALSE |
| 63_v.txt | 31675 | 31675-SNV-/ | SNV | 1 | A | C | 4 | 11813 | 0.033861 | 44 | rnr1Y285A | FALSE | CAN | JSY3868 | 2 | rnr1Y285A-R | AAG | A>C | G/C+1 | A/T-1 | TA>GC | TA>GC | GATCAAAGG | AGGTAAT | TTC | TTC | T | G | T>G | T>G_TTC | TA>GC_AGGTAAT | 5 | FALSE | FALSE |
| 63_v.txt | 31723 | 31723-SNV-/ | SNV | 1 | A | C | 3 | 11813 | 0.02539575 | 62.6666667 | rnr1Y285A | FALSE | CAN | JSY3868 | 2 | rnr1Y285A-R | CAA | A>C | G/C+1 | A/T-1 | TA>GC | TA>GC | TGTCCCAAA/ | AAAAAGT | GTT | GTT | T | G | T>G | T>G_GTT | TA>GC_AAAAAGT | 5 | FALSE | FALSE |
| 63_v.txt | 31854 | 31854-SNV-/ | SNV | 1 | A | C | 7 | 11813 | 0.05925675 | 48.2857143 | rnr1Y285A | FALSE | CAN | JSY3868 | 2 | rnr1Y285A-R | TAA | A>C | G/C+1 | A/T-1 | TA>GC | TA>GC | ACAGCTAAG | AAGAACA | ATT | ATT | T | G | T>G | T>G_ATT | TA>GC_AAGAACA | 5 | FALSE | FALSE |
| 63_v.txt | 32081 | 32081-SNV-/ | SNV | 1 | A | C | 1 | 21212 | 4.71E-03 | 63.9990542 | rnr1Y285A | FALSE | CAN | JSY3868 | 2 | rnr1Y285A-R | CAA | A>C | G/C+1 | A/T-1 | TA>GC | TA>GC | CCATGCAAA/ | AAAAAAG | GTT | GTT | T | G | T>G | T>G_GTT | TA>GC_AAAAAAG | 5 | FALSE | FALSE |
| 63_v.txt | 32445 | 32445-SNV-/ | SNV | 1 | A | C | 12 | 25364 | 0.04731115 | 55.1666667 | rnr1Y285A | FALSE | CAN | JSY3868 | 2 | rnr1Y285A-R | AAG | A>C | G/C+1 | A/T-1 | TA>GC | TA>GC | ACTAAAAGT( | AGTCCAA | TTC | TTC | T | G | T>G | T>G_TTC | TA>GC_AGTCCAA | 5 | FALSE | FALSE |
| 63_v.txt | 32823 | 32823-SNV-/ | SNV | 1 | A | C | 1098 | 20429 | 5.37471242 | 43.7750455 | rnr1Y285A | FALSE | CAN | JSY3868 | 2 | rnr1Y285A-R | CAT | A>C | G/C+1 | A/T-1 | TA>GC | TA>GC | AAGTTCATT/ | ATTATTG | GTA | GTA | T | G | T>G | T>G_GTA | TA>GC_ATTATTG | 5 | FALSE | FALSE |
| 63_v.txt | 32929 | 32929-SNV-T | SNV | 1 | T | G | 36 | 27083 | 0.13292471 | 51.8055556 | rnr1Y285A | FALSE | CAN | JSY3868 | 2 | rnr1Y285A-R | GTG | T>G | G/C+1 | A/T-1 | TA>GC | TA>GC | CAAAAGTGA | TGATTGC | CAC | GTG | T | C | T>G | T>G_GTG | TA>GC_TGATTGC | 5 | FALSE | FALSE |
| 63_v.txt | 32952 | 32952-SNV-/ | SNV | 1 | A | C | 16 | 27083 | 0.05907765 | 40.375 | rnr1Y285A | FALSE | CAN | JSY3868 | 2 | rnr1Y285A-R | CAT | A>C | G/C+1 | A/T-1 | TA>GC | TA>GC | CAATACATGT | ATGTAAC | GTA | GTA | T | G | T>G | T>G_GTA | TA>GC_ATGTAAC | 5 | FALSE | FALSE |
| 63_v.txt | 33138 | 33138-SNV-/ | SNV | 1 | A | C | 5 | 27083 | 0.01846177 | 43.2 | rnr1Y285A | FALSE | CAN | JSY3868 | 2 | rnr1Y285A-R | TAA | A>C | G/C+1 | A/T-1 | TA>GC | TA>GC | GTGGATAAA | AAACCAA | ATT | ATT | T | G | T>G | T>G_ATT | TA>GC_AAACCAA | 5 | FALSE | FALSE |
| 63_v.txt | 33166 | 33166-SNV-T | SNV | 1 | T | G | 28 | 27082 | 0.10338971 | 36.8214286 | rnr1Y285A | FALSE | CAN | JSY3868 | 2 | rnr1Y285A-R | GTA | T>G | G/C+1 | A/T-1 | TA>GC | TA>GC | CAATAGTAC( | TACCACC | CAT | GTA | T | C | T>G | T>G_GTA | TA>GC_TACCACC | 5 | FALSE | FALSE |
| 63_v.txt | 33269 | 33269-SNV-/ | SNV | 1 | A | C | 79 | 10539 | 0.74959674 | 17.3924051 | rnr1Y285A | FALSE | CAN | JSY3868 | 2 | rnr1Y285A-R | TAG | A>C | G/C+1 | A/T-1 | TA>GC | TA>GC | CTCCATAGA( | AGAGAAC | ATC | ATC | T | G | T>G | T>G_ATC | TA>GC_AGAGAAC | 5 | FALSE | FALSE |
| 63_v.txt | 33476 | 33476-SNV-T | SNV | 1 | T | G | 3 | 10539 | 0.0284657 | 33 | rnr1Y285A | FALSE | CAN | JSY3868 | 2 | rnr1Y285A-R | CTT | T>G | G/C+1 | A/T-1 | TA>GC | TA>GC | TATGCCTTTT | TTTTTTT | GAA | CTT | T | C | T>G | T>G_CTT | TA>GC_TTTTTTT | 5 | FALSE | FALSE |
| 63_v.txt | 32521 | 32521-SNV-/ | SNV | 1 | A | G | 152 | 25364 | 0.59927456 | 40.9473684 | rnr1Y285A | FALSE | CAN | JSY3868 | 2 | rnr1Y285A-R | GAT | A>G | G/C+1 | A/T-1 | TA>CG | TA>CG | GAACGGATT | ATTTTCT | CTA | CTA | T | C | T>C | T>C_CTA | TA>CG_ATTTTCT | 5 | FALSE | FALSE |
| 63_v.txt | 32941 | 32941-SNV-/ | SNV | 1 | A | G | 7 | 27083 | 0.02584647 | 53.5524156 | rnr1Y285A | FALSE | CAN | JSY3868 | 2 | rnr1Y285A-R | GAA | A>G | G/C+1 | A/T-1 | TA>CG | TA>CG | CCCAAGAAA | AAAACCA | CTT | CTT | T | C | T>C | T>C_CTT | TA>CG_AAAACCA | 5 | FALSE | FALSE |
| 63_v.txt | 33052 | 33052-SNV-/ | SNV | 1 | A | G | 27 | 27083 | 0.09969354 | 51.4444444 | rnr1Y285A | FALSE | CAN | JSY3868 | 2 | rnr1Y285A-R | GAC | A>G | G/C+1 | A/T-1 | TA>CG | TA>CG | CCAAGGACT( | ACTGCGT | CTG | CTG | T | C | T>C | T>C_CTG | TA>CG_ACTGCGT | 5 | FALSE | FALSE |

| | | | | | | | | | | | | | | | | | | | | | | | | | | | | | | | | | | |
|---|---|---|---|---|---|---|---|---|---|---|---|---|---|---|---|---|---|---|---|---|---|---|---|---|---|---|---|---|---|---|---|---|---|---|
| 63_v.txt | 33476 | 33476-SNV-T | SNV | 1 | T | C | 18 | 10539 | 0.17079419 | 33.6111111 | rnr1Y285A | FALSE | CAN | JSY3868 | 2 | rnr1Y285A-R | CTT | T>C | G/C+1 | A/T-1 | TA>CG | TA>CG | TATGCCTTTT | TTTTTTT | GAA | CTT | T | G | T>C | T>C_CTT | TA>CG_TTTTTTT | 5 | FALSE | FALSE |
| 63_v.txt | 32496 | 32496-SNV-A | SNV | 1 | A | T | 19 | 25364 | 0.07490932 | 45.6842105 | rnr1Y285A | FALSE | CAN | JSY3868 | 2 | rnr1Y285A-R | AAC | A>T | A/T+1 | A/T-1 | TA>AT | TA>AT | AAAACAACT | ACTTTTT | TTG | TTG | T | A | T>A | T>A_TTG | TA>AT_ACTTTTT | 5 | FALSE | FALSE |
| 63_v.txt | 32498 | 32498-SNV-T | SNV | 1 | T | A | 6 | 25364 | 0.02365558 | 50.3332448 | rnr1Y285A | FALSE | CAN | JSY3868 | 2 | rnr1Y285A-R | CTT | T>A | A/T+1 | A/T-1 | TA>AT | TA>AT | AACAACTTTT | TTTTTTG | GAA | CTT | T | T | T>A | T>A_CTT | TA>AT_TTTTTTG | 5 | FALSE | FALSE |
| 63_v.txt | 32680 | 32680-SNV-A | SNV | 1 | A | T | 3045 | 20429 | 14.9052817 | 61.6735632 | rnr1Y285A | FALSE | CAN | JSY3868 | 2 | rnr1Y285A-R | CAA | A>T | A/T+1 | A/T-1 | TA>AT | TA>AT | TTCTCCAATA | AATAACG | GTT | GTT | T | A | T>A | T>A_GTT | TA>AT_AATAACG | 5 | FALSE | FALSE |
| 63_v.txt | 32848 | 32848-SNV-T | SNV | 1 | T | A | 17 | 20428 | 0.08321911 | 37.8823529 | rnr1Y285A | FALSE | CAN | JSY3868 | 2 | rnr1Y285A-R | ATA | T>A | A/T+1 | A/T-1 | TA>AT | TA>AT | AAAAAATAC | TACTAAT | TAT | ATA | T | T | T>A | T>A_ATA | TA>AT_TACTAAT | 5 | FALSE | FALSE |
| 63_v.txt | 33095 | 33095-SNV-T | SNV | 1 | T | A | 6 | 27083 | 0.02215412 | 50.9975633 | rnr1Y285A | FALSE | CAN | JSY3868 | 2 | rnr1Y285A-R | ATA | T>A | A/T+1 | A/T-1 | TA>AT | TA>AT | ATATGATAT | TATAAGA | TAT | ATA | T | T | T>A | T>A_ATA | TA>AT_TATAAGA | 5 | FALSE | FALSE |
| 63_v.txt | 31650 | 31650-SNV-C | SNV | 1 | C | T | 30 | 11813 | 0.2539575 | 36.8 | rnr1Y285A | FALSE | CAN | JSY3868 | 2 | rnr1Y285A-R | GCG | C>T | A/T+1 | G/C-1 | CG>TA | CG>TA | AGAATGCGA | CGAAATG | CGC | GCG | C | A | C>T | C>T_GCG | CG>TA_CGAAATG | 5 | FALSE | FALSE |
| 63_v.txt | 31830 | 31830-SNV-C | SNV | 1 | C | T | 14 | 11813 | 0.1185135 | 56.5714286 | rnr1Y285A | FALSE | CAN | JSY3868 | 2 | rnr1Y285A-R | GCA | C>T | A/T+1 | G/C-1 | CG>TA | CG>TA | AATATGCAT | CATTGAA | CGT | GCA | C | A | C>T | C>T_GCA | CG>TA_CATTGAA | 5 | FALSE | FALSE |
| 63_v.txt | 31844 | 31844-SNV-C | SNV | 1 | C | T | 78 | 11813 | 0.66028951 | 52.1538462 | rnr1Y285A | FALSE | CAN | JSY3868 | 2 | rnr1Y285A-R | TCC | C>T | A/T+1 | G/C-1 | CG>TA | CG>TA | TAAGATCCA | CCAAACA | AGG | TCC | C | A | C>T | C>T_TCC | CG>TA_CCAAACA | 5 | FALSE | FALSE |
| 63_v.txt | 31920 | 31920-SNV-G | SNV | 1 | G | A | 15 | 11812 | 0.1269895 | 34.3333333 | rnr1Y285A | FALSE | CAN | JSY3868 | 2 | rnr1Y285A-R | CGT | G>A | A/T+1 | G/C-1 | CG>TA | CG>TA | AAAGCCGTG | GTGAAAC | GCA | GCA | C | T | C>T | C>T_GCA | CG>TA_GTGAAAC | 5 | FALSE | FALSE |
| 63_v.txt | 31962 | 31962-SNV-G | SNV | 1 | G | A | 31 | 21214 | 0.14612991 | 37 | rnr1Y285A | FALSE | CAN | JSY3868 | 2 | rnr1Y285A-R | CGC | G>A | A/T+1 | G/C-1 | CG>TA | CG>TA | GTGGCCGCA | GCATAAT | GCG | GCG | C | T | C>T | C>T_GCG | CG>TA_GCATAAT | 5 | FALSE | FALSE |
| 63_v.txt | 31972 | 31972-SNV-C | SNV | 1 | C | T | 21 | 21212 | 0.09900057 | 37.8571429 | rnr1Y285A | FALSE | CAN | JSY3868 | 2 | rnr1Y285A-R | GCC | C>T | A/T+1 | G/C-1 | CG>TA | CG>TA | AATAAGCCA | CCAAGCC | CGG | GCC | C | A | C>T | C>T_GCC | CG>TA_CCAAGCC | 5 | FALSE | FALSE |
| 63_v.txt | 31981 | 31981-SNV-G | SNV | 1 | G | A | 147 | 21212 | 0.69300396 | 51.9931973 | rnr1Y285A | FALSE | CAN | JSY3868 | 2 | rnr1Y285A-R | GGC | G>A | A/T+1 | G/C-1 | CG>TA | CG>TA | AGCCGGGCA | GCATTAA | CCG | CCG | C | T | C>T | C>T_CCG | CG>TA_GCATTAA | 5 | FALSE | FALSE |
| 63_v.txt | 32075 | 32075-SNV-C | SNV | 1 | C | T | 81 | 21212 | 0.38185933 | 62.814859 | rnr1Y285A | FALSE | CAN | JSY3868 | 2 | rnr1Y285A-R | ACC | C>T | A/T+1 | G/C-1 | CG>TA | CG>TA | AAATAACCA | CCATGCA | TGG | ACC | C | A | C>T | C>T_ACC | CG>TA_CCATGCA | 5 | FALSE | FALSE |
| 63_v.txt | 32076 | 32076-SNV-C | SNV | 1 | C | T | 65 | 21212 | 0.30643032 | 61.1540492 | rnr1Y285A | FALSE | CAN | JSY3868 | 2 | rnr1Y285A-R | CCA | C>T | A/T+1 | G/C-1 | CG>TA | CG>TA | AATAACCAT | CATGCAA | GGT | CCA | C | A | C>T | C>T_CCA | CG>TA_CATGCAA | 5 | FALSE | FALSE |
| 63_v.txt | 32088 | 32088-SNV-C | SNV | 1 | C | T | 26 | 21212 | 0.12257213 | 61.8461539 | rnr1Y285A | FALSE | CAN | JSY3868 | 2 | rnr1Y285A-R | GCC | C>T | A/T+1 | G/C-1 | CG>TA | CG>TA | AAAAAGCCT | CCTGCAA | CGG | GCC | C | A | C>T | C>T_GCC | CG>TA_CCTGCAA | 5 | FALSE | FALSE |
| 63_v.txt | 32092 | 32092-SNV-C | SNV | 1 | C | T | 24 | 21212 | 0.1131435 | 62.0833333 | rnr1Y285A | FALSE | CAN | JSY3868 | 2 | rnr1Y285A-R | GCA | C>T | A/T+1 | G/C-1 | CG>TA | CG>TA | AGCCTGCAA | CAACACC | CGT | GCA | C | A | C>T | C>T_GCA | CG>TA_CAACACC | 5 | FALSE | FALSE |
| 63_v.txt | 32114 | 32114-SNV-C | SNV | 1 | C | T | 694 | 21212 | 3.27173298 | 60.2809798 | rnr1Y285A | FALSE | CAN | JSY3868 | 2 | rnr1Y285A-R | GCC | C>T | A/T+1 | G/C-1 | CG>TA | CG>TA | TAATAGCCA | CCATTCG | CGG | GCC | C | A | C>T | C>T_GCC | CG>TA_CCATTCG | 5 | FALSE | FALSE |
| 63_v.txt | 32165 | 32165-SNV-G | SNV | 1 | G | A | 33 | 21212 | 0.15557232 | 48.030303 | rnr1Y285A | FALSE | CAN | JSY3868 | 2 | rnr1Y285A-R | CGC | G>A | A/T+1 | G/C-1 | CG>TA | CG>TA | CAAAGCGCC | GCCAAAT | GCG | GCG | C | T | C>T | C>T_GCG | CG>TA_GCCAAAT | 5 | FALSE | FALSE |
| 63_v.txt | 32205 | 32205-SNV-C | SNV | 1 | C | T | 44 | 21212 | 0.20742976 | 39.2727273 | rnr1Y285A | FALSE | CAN | JSY3868 | 2 | rnr1Y285A-R | ACC | C>T | A/T+1 | G/C-1 | CG>TA | CG>TA | GGAACACCA | CCACCTT | TGG | ACC | C | A | C>T | C>T_ACC | CG>TA_CCACCTT | 5 | FALSE | FALSE |
| 63_v.txt | 32259 | 32259-SNV-C | SNV | 1 | C | T | 51 | 25363 | 0.20108031 | 32.4313726 | rnr1Y285A | FALSE | CAN | JSY3868 | 2 | rnr1Y285A-R | ACC | C>T | A/T+1 | G/C-1 | CG>TA | CG>TA | GATAGACCA | CCAAATA | TGG | ACC | C | A | C>T | C>T_ACC | CG>TA_CCAAATA | 5 | FALSE | FALSE |
| 63_v.txt | 32271 | 32271-SNV-C | SNV | 1 | C | T | 53 | 25363 | 0.20896582 | 34.5849057 | rnr1Y285A | FALSE | CAN | JSY3868 | 2 | rnr1Y285A-R | ACG | C>T | A/T+1 | G/C-1 | CG>TA | CG>TA | AAAATACGG | CGGGAAC | TGC | ACG | C | A | C>T | C>T_ACG | CG>TA_CGGGAAC | 5 | FALSE | FALSE |
| 63_v.txt | 32274 | 32274-SNV-G | SNV | 1 | G | A | 22 | 25363 | 0.08674053 | 36.1363636 | rnr1Y285A | FALSE | CAN | JSY3868 | 2 | rnr1Y285A-R | GGA | G>A | A/T+1 | G/C-1 | CG>TA | CG>TA | ATACGGGAA | GAACCAA | CCT | CCT | C | T | C>T | C>T_CCT | CG>TA_GAACCAA | 5 | FALSE | FALSE |
| 63_v.txt | 32299 | 32299-SNV-C | SNV | 1 | C | T | 38 | 25364 | 0.14981864 | 53.7105263 | rnr1Y285A | FALSE | CAN | JSY3868 | 2 | rnr1Y285A-R | GCG | C>T | A/T+1 | G/C-1 | CG>TA | CG>TA | AATTTGCGG | CGGCAGA | CGC | GCG | C | A | C>T | C>T_GCG | CG>TA_CGGCAGA | 5 | FALSE | FALSE |
| 63_v.txt | 32304 | 32304-SNV-G | SNV | 1 | G | A | 20 | 25364 | 0.07885192 | 57.4 | rnr1Y285A | FALSE | CAN | JSY3868 | 2 | rnr1Y285A-R | AGA | G>A | A/T+1 | G/C-1 | CG>TA | CG>TA | GCGGCAGAA | GAAATAA | TCT | TCT | C | T | C>T | C>T_TCT | CG>TA_GAAATAA | 5 | FALSE | FALSE |
| 63_v.txt | 32344 | 32344-SNV-G | SNV | 1 | G | A | 22 | 25364 | 0.08673711 | 58.8636364 | rnr1Y285A | FALSE | CAN | JSY3868 | 2 | rnr1Y285A-R | GGC | G>A | A/T+1 | G/C-1 | CG>TA | CG>TA | TATGTGGCA | GCAAAAC | CCG | CCG | C | T | C>T | C>T_CCG | CG>TA_GCAAAAC | 5 | FALSE | FALSE |
| 63_v.txt | 32466 | 32466-SNV-C | SNV | 1 | C | T | 25 | 25364 | 0.0985649 | 48.56 | rnr1Y285A | FALSE | CAN | JSY3868 | 2 | rnr1Y285A-R | GCC | C>T | A/T+1 | G/C-1 | CG>TA | CG>TA | AGAGAGCCA | CCAATGT | CGG | GCC | C | A | C>T | C>T_GCC | CG>TA_CCAATGT | 5 | FALSE | FALSE |
| 63_v.txt | 32487 | 32487-SNV-C | SNV | 1 | C | T | 41 | 25364 | 0.16164643 | 47.7073171 | rnr1Y285A | FALSE | CAN | JSY3868 | 2 | rnr1Y285A-R | ACG | C>T | A/T+1 | G/C-1 | CG>TA | CG>TA | AAGATACGG | CGGAAAA | TGC | ACG | C | A | C>T | C>T_ACG | CG>TA_CGGAAAA | 5 | FALSE | FALSE |
| 63_v.txt | 32508 | 32508-SNV-G | SNV | 1 | G | A | 31 | 25364 | 0.12222047 | 35.5483871 | rnr1Y285A | FALSE | CAN | JSY3868 | 2 | rnr1Y285A-R | GGC | G>A | A/T+1 | G/C-1 | CG>TA | CG>TA | TTGATGGCT | GCTCTTG | CCG | CCG | C | T | C>T | C>T_CCG | CG>TA_GCTCTTG | 5 | FALSE | FALSE |
| 63_v.txt | 32530 | 32530-SNV-G | SNV | 1 | G | A | 24 | 25364 | 0.0946223 | 39.25 | rnr1Y285A | FALSE | CAN | JSY3868 | 2 | rnr1Y285A-R | GGG | G>A | A/T+1 | G/C-1 | CG>TA | CG>TA | TTCTGGGGT | GGTTTGC | CCC | CCC | C | T | C>T | C>T_CCC | CG>TA_GGTTTGC | 5 | FALSE | FALSE |
| 63_v.txt | 32535 | 32535-SNV-G | SNV | 1 | G | A | 41 | 25364 | 0.16164643 | 36.4382638 | rnr1Y285A | FALSE | CAN | JSY3868 | 2 | rnr1Y285A-R | TGC | G>A | A/T+1 | G/C-1 | CG>TA | CG>TA | GGGTTTGCA | GCAGCTT | ACG | ACG | C | T | C>T | C>T_ACG | CG>TA_GCAGCTT | 5 | FALSE | FALSE |
| 63_v.txt | 32536 | 32536-SNV-C | SNV | 1 | C | T | 43 | 25364 | 0.16953162 | 36.325364 | rnr1Y285A | FALSE | CAN | JSY3868 | 2 | rnr1Y285A-R | GCA | C>T | A/T+1 | G/C-1 | CG>TA | CG>TA | GGTTTGCAG | CAGCTTC | CGT | GCA | C | A | C>T | C>T_GCA | CG>TA_CAGCTTC | 5 | FALSE | FALSE |
| 63_v.txt | 32571 | 32571-SNV-C | SNV | 1 | C | T | 31 | 20429 | 0.15174507 | 34.966297 | rnr1Y285A | FALSE | CAN | JSY3868 | 2 | rnr1Y285A-R | ACC | C>T | A/T+1 | G/C-1 | CG>TA | CG>TA | TCAGTACCTT | CCTTGAA | TGG | ACC | C | A | C>T | C>T_ACC | CG>TA_CCTTGAA | 5 | FALSE | FALSE |
| 63_v.txt | 32590 | 32590-SNV-C | SNV | 1 | C | T | 32 | 20429 | 0.15664007 | 36.40625 | rnr1Y285A | FALSE | CAN | JSY3868 | 2 | rnr1Y285A-R | GCG | C>T | A/T+1 | G/C-1 | CG>TA | CG>TA | AGGCAGCGT | CGTTAAT | CGC | GCG | C | A | C>T | C>T_GCG | CG>TA_CGTTAAT | 5 | FALSE | FALSE |
| 63_v.txt | 32700 | 32700-SNV-C | SNV | 1 | C | T | 21 | 20429 | 0.10279505 | 63.7619048 | rnr1Y285A | FALSE | CAN | JSY3868 | 2 | rnr1Y285A-R | GCC | C>T | A/T+1 | G/C-1 | CG>TA | CG>TA | ACTGGGCCG | CCGGTAA | CGG | GCC | C | A | C>T | C>T_GCC | CG>TA_CCGGTAA | 5 | FALSE | FALSE |
| 63_v.txt | 32757 | 32757-SNV-G | SNV | 1 | G | A | 21 | 20429 | 0.10279505 | 50.5714286 | rnr1Y285A | FALSE | CAN | JSY3868 | 2 | rnr1Y285A-R | GGC | G>A | A/T+1 | G/C-1 | CG>TA | CG>TA | ATAATGGCT | GCTAAAA | CCG | CCG | C | T | C>T | C>T_CCG | CG>TA_GCTAAAA | 5 | FALSE | FALSE |
| 63_v.txt | 32776 | 32776-SNV-C | SNV | 1 | C | T | 24 | 20429 | 0.11748005 | 48.2916667 | rnr1Y285A | FALSE | CAN | JSY3868 | 2 | rnr1Y285A-R | GCG | C>T | A/T+1 | G/C-1 | CG>TA | CG>TA | TGGAAGCGA | CGACCCA | CGC | GCG | C | A | C>T | C>T_GCG | CG>TA_CGACCCA | 5 | FALSE | FALSE |
| 63_v.txt | 32780 | 32780-SNV-C | SNV | 1 | C | T | 67 | 20429 | 0.32796515 | 47.7164179 | rnr1Y285A | FALSE | CAN | JSY3868 | 2 | rnr1Y285A-R | CCC | C>T | A/T+1 | G/C-1 | CG>TA | CG>TA | AGCGACCCA | CCAGAAC | GGG | CCC | C | A | C>T | C>T_CCC | CG>TA_CCAGAAC | 5 | FALSE | FALSE |
| 63_v.txt | 32794 | 32794-SNV-C | SNV | 1 | C | T | 193 | 20429 | 0.94473543 | 46.1554404 | rnr1Y285A | FALSE | CAN | JSY3868 | 2 | rnr1Y285A-R | TCA | C>T | A/T+1 | G/C-1 | CG>TA | CG>TA | CGAATTCAC | CACCGTA | AGT | TCA | C | A | C>T | C>T_TCA | CG>TA_CACCGTA | 5 | FALSE | FALSE |
| 63_v.txt | 32798 | 32798-SNV-G | SNV | 1 | G | A | 24 | 20429 | 0.11748005 | 43.0833333 | rnr1Y285A | FALSE | CAN | JSY3868 | 2 | rnr1Y285A-R | CGT | G>A | A/T+1 | G/C-1 | CG>TA | CG>TA | TTCACCGTA | GTAATAT | GCA | GCA | C | T | C>T | C>T_GCA | CG>TA_GTAATAT | 5 | FALSE | FALSE |
| 63_v.txt | 32855 | 32855-SNV-C | SNV | 1 | C | T | 59 | 20427 | 0.28883341 | 35.8474576 | rnr1Y285A | FALSE | CAN | JSY3868 | 2 | rnr1Y285A-R | TCC | C>T | A/T+1 | G/C-1 | CG>TA | CG>TA | ACTAATCCA | CCATGCC | AGG | TCC | C | A | C>T | C>T_TCC | CG>TA_CCATGCC | 5 | FALSE | FALSE |
| 63_v.txt | 32882 | 32882-SNV-C | SNV | 1 | C | T | 79 | 27084 | 0.29168513 | 28.835443 | rnr1Y285A | FALSE | CAN | JSY3868 | 2 | rnr1Y285A-R | TCC | C>T | A/T+1 | G/C-1 | CG>TA | CG>TA | GTACGTCCA | CCAAAAT | AGG | TCC | C | A | C>T | C>T_TCC | CG>TA_CCAAAAT | 5 | FALSE | FALSE |
| 63_v.txt | 32900 | 32900-SNV-G | SNV | 1 | G | A | 38 | 27083 | 0.14030942 | 35.524537 | rnr1Y285A | FALSE | CAN | JSY3868 | 2 | rnr1Y285A-R | GGC | G>A | A/T+1 | G/C-1 | CG>TA | CG>TA | GACTTGGCC | GCCAACT | CCG | CCG | C | T | C>T | C>T_CCG | CG>TA_GCCAACT | 5 | FALSE | FALSE |
| 63_v.txt | 32901 | 32901-SNV-C | SNV | 1 | C | T | 61 | 27083 | 0.22523354 | 33.9016393 | rnr1Y285A | FALSE | CAN | JSY3868 | 2 | rnr1Y285A-R | GCC | C>T | A/T+1 | G/C-1 | CG>TA | CG>TA | ACTTGGCCA | CCAACTA | CGG | GCC | C | A | C>T | C>T_GCC | CG>TA_CCAACTA | 5 | FALSE | FALSE |
| 63_v.txt | 32935 | 32935-SNV-C | SNV | 1 | C | T | 34 | 27083 | 0.12554001 | 43.5 | rnr1Y285A | FALSE | CAN | JSY3868 | 2 | rnr1Y285A-R | GCC | C>T | A/T+1 | G/C-1 | CG>TA | CG>TA | TGATTGCCC | CCCAAGA | CGG | GCC | C | A | C>T | C>T_GCC | CG>TA_CCCAAGA | 5 | FALSE | FALSE |
| 63_v.txt | 32945 | 32945-SNV-C | SNV | 1 | C | T | 823 | 27083 | 3.03880663 | 57.291616 | rnr1Y285A | FALSE | CAN | JSY3868 | 2 | rnr1Y285A-R | ACC | C>T | A/T+1 | G/C-1 | CG>TA | CG>TA | AGAAAACCA | CCAATAC | TGG | ACC | C | A | C>T | C>T_ACC | CG>TA_CCAATAC | 5 | FALSE | FALSE |
| 63_v.txt | 32958 | 32958-SNV-C | SNV | 1 | C | T | 37 | 27083 | 0.13661707 | 52.945946 | rnr1Y285A | FALSE | CAN | JSY3868 | 2 | rnr1Y285A-R | ACC | C>T | A/T+1 | G/C-1 | CG>TA | CG>TA | ATGTAACCA | CCATTGG | TGG | ACC | C | A | C>T | C>T_ACC | CG>TA_CCATTGG | 5 | FALSE | FALSE |
| 63_v.txt | 32965 | 32965-SNV-C | SNV | 1 | C | T | 20 | 27083 | 0.07384706 | 61.2 | rnr1Y285A | FALSE | CAN | JSY3868 | 2 | rnr1Y285A-R | GCC | C>T | A/T+1 | G/C-1 | CG>TA | CG>TA | CATTGGCCG | CCGCACC | CGG | GCC | C | A | C>T | C>T_GCC | CG>TA_CCGCACC | 5 | FALSE | FALSE |
| 63_v.txt | 32967 | 32967-SNV-G | SNV | 1 | G | A | 52 | 27083 | 0.19200236 | 61.5576923 | rnr1Y285A | FALSE | CAN | JSY3868 | 2 | rnr1Y285A-R | CGC | G>A | A/T+1 | G/C-1 | CG>TA | CG>TA | TTGGCCGCA | GCACCAA | GCG | GCG | C | T | C>T | C>T_GCG | CG>TA_GCACCAA | 5 | FALSE | FALSE |
| 63_v.txt | 33015 | 33015-SNV-G | SNV | 1 | G | A | 28 | 27083 | 0.10338589 | 57.7857143 | rnr1Y285A | FALSE | CAN | JSY3868 | 2 | rnr1Y285A-R | GGA | G>A | A/T+1 | G/C-1 | CG>TA | CG>TA | AAAGAGGAT | GATGTAA | CCT | CCT | C | T | C>T | C>T_CCT | CG>TA_GATGTAA | 5 | FALSE | FALSE |
| 63_v.txt | 33056 | 33056-SNV-C | SNV | 1 | C | T | 51 | 27083 | 0.18831001 | 56.2352941 | rnr1Y285A | FALSE | CAN | JSY3868 | 2 | rnr1Y285A-R | GCG | C>T | A/T+1 | G/C-1 | CG>TA | CG>TA | GGACTGCGT | CGTGACA | CGC | GCG | C | A | C>T | C>T_GCG | CG>TA_CGTGACA | 5 | FALSE | FALSE |
| 63_v.txt | 33069 | 33069-SNV-G | SNV | 1 | G | A | 31 | 27083 | 0.11446295 | 51.3867347 | rnr1Y285A | FALSE | CAN | JSY3868 | 2 | rnr1Y285A-R | TGC | G>A | A/T+1 | G/C-1 | CG>TA | CG>TA | GAATATGCC | GCCAAAG | ACG | ACG | C | T | C>T | C>T_ACG | CG>TA_GCCAAAG | 5 | FALSE | FALSE |
| 63_v.txt | 33070 | 33070-SNV-C | SNV | 1 | C | T | 31 | 27083 | 0.11446295 | 50.6116702 | rnr1Y285A | FALSE | CAN | JSY3868 | 2 | rnr1Y285A-R | GCC | C>T | A/T+1 | G/C-1 | CG>TA | CG>TA | AATATGCCA | CCAAAGA | CGG | GCC | C | A | C>T | C>T_GCC | CG>TA_CCAAAGA | 5 | FALSE | FALSE |
| 63_v.txt | 33104 | 33104-SNV-G | SNV | 1 | G | A | 34 | 27083 | 0.12554001 | 39.4411765 | rnr1Y285A | FALSE | CAN | JSY3868 | 2 | rnr1Y285A-R | CGC | G>A | A/T+1 | G/C-1 | CG>TA | CG>TA | AAGAGCGCC | GCCCACT | GCG | GCG | C | T | C>T | C>T_GCG | CG>TA_GCCCACT | 5 | FALSE | FALSE |
| 63_v.txt | 33114 | 33114-SNV-C | SNV | 1 | C | T | 34 | 27083 | 0.12554001 | 41.0294118 | rnr1Y285A | FALSE | CAN | JSY3868 | 2 | rnr1Y285A-R | GCC | C>T | A/T+1 | G/C-1 | CG>TA | CG>TA | ACTGGGCCG | CCGGCGT | CGG | GCC | C | A | C>T | C>T_GCC | CG>TA_CCGGCGT | 5 | FALSE | FALSE |
| 63_v.txt | 33119 | 33119-SNV-G | SNV | 1 | G | A | 21 | 27083 | 0.07753942 | 46 | rnr1Y285A | FALSE | CAN | JSY3868 | 2 | rnr1Y285A-R | CGT | G>A | A/T+1 | G/C-1 | CG>TA | CG>TA | GCCGGCGTT | GTTGGTC | GCA | GCA | C | T | C>T | C>T_GCA | CG>TA_GTTGGTC | 5 | FALSE | FALSE |
| 63_v.txt | 33153 | 33153-SNV-C | SNV | 1 | C | T | 85 | 27083 | 0.31385002 | 36.6823529 | rnr1Y285A | FALSE | CAN | JSY3868 | 2 | rnr1Y285A-R | ACC | C>T | A/T+1 | G/C-1 | CG>TA | CG>TA | AAAAGACCT | CCTGTAC | TGG | ACC | C | A | C>T | C>T_ACC | CG>TA_CCTGTAC | 5 | FALSE | FALSE |
| 63_v.txt | 33177 | 33177-SNV-G | SNV | 1 | G | A | 29 | 27063 | 0.10715737 | 36.7241379 | rnr1Y285A | FALSE | CAN | JSY3868 | 2 | rnr1Y285A-R | GGC | G>A | A/T+1 | G/C-1 | CG>TA | CG>TA | CCAAGGGCA | GCAATCA | CCG | CCG | C | T | C>T | C>T_CCG | CG>TA_GCAATCA | 5 | FALSE | FALSE |
| 63_v.txt | 33276 | 33276-SNV-G | SNV | 1 | G | A | 24 | 10539 | 0.22772559 | 44.8333333 | rnr1Y285A | FALSE | CAN | JSY3868 | 2 | rnr1Y285A-R | CGT | G>A | A/T+1 | G/C-1 | CG>TA | CG>TA | GAGAACGTA | GTATCCT | GCA | GCA | C | T | C>T | C>T_GCA | CG>TA_GTATCCT | 5 | FALSE | FALSE |
| 63_v.txt | 33310 | 33310-SNV-G | SNV | 1 | G | A | 18 | 10539 | 0.17079419 | 56.8888889 | rnr1Y285A | FALSE | CAN | JSY3868 | 2 | rnr1Y285A-R | CGC | G>A | A/T+1 | G/C-1 | CG>TA | CG>TA | AAGAGCGCA | GCAATGG | GCG | GCG | C | T | C>T | C>T_GCG | CG>TA_GCAATGG | 5 | FALSE | FALSE |
| 63_v.txt | 33474 | 33474-SNV-C | SNV | 1 | C | T | 16 | 10539 | 0.15181706 | 31.0625 | rnr1Y285A | FALSE | CAN | JSY3868 | 2 | rnr1Y285A-R | GCC | C>T | A/T+1 | G/C-1 | CG>TA | CG>TA | GCTATGCCTT | CCTTTTT | CGG | GCC | C | A | C>T | C>T_GCC | CG>TA_CCTTTTT | 5 | FALSE | FALSE |
| 63_v.txt | 32027 | 32027-SNV-G | SNV | 1 | G | C | 6 | 21212 | 0.02828588 | 60.1666667 | rnr1Y285A | FALSE | CAN | JSY3868 | 2 | rnr1Y285A-R | GGT | G>C | G/C+1 | G/C-1 | CG>GC | CG>GC | GCCACGGTA | GTATTTC | CCA | CCA | C | G | C>G | C>G_CCA | CG>GC_GTATTTC | 5 | FALSE | FALSE |
| 63_v.txt | 32292 | 32292-SNV-G | SNV | 1 | G | C | 8 | 25364 | 0.03154077 | 59.25 | rnr1Y285A | FALSE | CAN | JSY3868 | 2 | rnr1Y285A-R | TGA | G>C | G/C+1 | G/C-1 | CG>GC | CG>GC | ATATTTGAA | GAATTTG | ACT | ACT | C | G | C>G | C>G_ACT | CG>GC_GAATTTG | 5 | FALSE | FALSE |
| 63_v.txt | 32557 | 32557-SNV-C | SNV | 1 | C | G | 21 | 25358 | 0.0828141 | 36.7124178 | rnr1Y285A | FALSE | CAN | JSY3868 | 2 | rnr1Y285A-R | CCA | C>G | G/C+1 | G/C-1 | CG>GC | CG>GC | TGATACCAA | CAACTAG | GGT | CCA | C | C | C>G | C>G_CCA | CG>GC_CAACTAG | 5 | FALSE | FALSE |
| 63_v.txt | 32876 | 32876-SNV-G | SNV | 1 | G | C | 34 | 27072 | 0.12559102 | 37.0882353 | rnr1Y285A | FALSE | CAN | JSY3868 | 2 | rnr1Y285A-R | TGT | G>C | G/C+1 | G/C-1 | CG>GC | CG>GC | AACTTTGTA | GTACGTC | ACA | ACA | C | G | C>G | C>G_ACA | CG>GC_GTACGTC | 5 | FALSE | FALSE |
| 63_v.txt | 32940 | 32940-SNV-G | SNV | 1 | G | C | 8 | 27083 | 0.02953883 | 56.2498203 | rnr1Y285A | FALSE | CAN | JSY3868 | 2 | rnr1Y285A-R | AGA | G>C | G/C+1 | G/C-1 | CG>GC | CG>GC | GCCCAAGAA | GAAAACC | TCT | TCT | C | G | C>G | C>G_TCT | CG>GC_GAAAACC | 5 | FALSE | FALSE |
| 63_v.txt | 32954 | 32954-SNV-G | SNV | 1 | G | C | 17 | 27083 | 0.06277 | 57.5294118 | rnr1Y285A | FALSE | CAN | JSY3868 | 2 | rnr1Y285A-R | TGT | G>C | G/C+1 | G/C-1 | CG>GC | CG>GC | ATACATGTA | GTAACCA | ACA | ACA | C | G | C>G | C>G_ACA | CG>GC_GTAACCA | 5 | FALSE | FALSE |
| 63_v.txt | 32965 | 32965-SNV-C | SNV | 1 | C | G | 5 | 27083 | 0.01846177 | 52.8 | rnr1Y285A | FALSE | CAN | JSY3868 | 2 | rnr1Y285A-R | GCC | C>G | G/C+1 | G/C-1 | CG>GC | CG>GC | CATTGGCCG | CCGCACC | CGG | GCC | C | C | C>G | C>G_GCC | CG>GC_CCGCACC | 5 | FALSE | FALSE |
| 63_v.txt | 33069 | 33069-SNV-G | SNV | 1 | G | C | 2 | 27083 | 7.39E-03 | 40.4996366 | rnr1Y285A | FALSE | CAN | JSY3868 | 2 | rnr1Y285A-R | TGC | G>C | G/C+1 | G/C-1 | CG>GC | CG>GC | GAATATGCC | GCCAAAG | ACG | ACG | C | G | C>G | C>G_ACG | CG>GC_GCCAAAG | 5 | FALSE | FALSE |
| 63_v.txt | 33153 | 33153-SNV-C | SNV | 1 | C | G | 6 | 27083 | 0.02215412 | 38 | rnr1Y285A | FALSE | CAN | JSY3868 | 2 | rnr1Y285A-R | ACC | C>G | G/C+1 | G/C-1 | CG>GC | CG>GC | AAAAGACCT | CCTGTAC | TGG | ACC | C | C | C>G | C>G_ACC | CG>GC_CCTGTAC | 5 | FALSE | FALSE |
| 63_v.txt | 33156 | 33156-SNV-G | SNV | 1 | G | C | 16 | 27082 | 0.05907983 | 38 | rnr1Y285A | FALSE | CAN | JSY3868 | 2 | rnr1Y285A-R | TGT | G>C | G/C+1 | G/C-1 | CG>GC | CG>GC | AGACCTGTA | GTACCAA | ACA | ACA | C | G | C>G | C>G_ACA | CG>GC_GTACCAA | 5 | FALSE | FALSE |
| 63_v.txt | 33205 | 33205-SNV-G | SNV | 1 | G | C | 49 | 10539 | 0.46493975 | 39.0401235 | rnr1Y285A | FALSE | CAN | JSY3868 | 2 | rnr1Y285A-R | AGC | G>C | G/C+1 | G/C-1 | CG>GC | CG>GC | GCTTAAGCT | GCTCTCT | TCG | TCG | C | G | C>G | C>G_TCG | CG>GC_GCTCTCT | 5 | FALSE | FALSE |
| 63_v.txt | 31657 | 31657-SNV-G | SNV | 1 | G | T | 103 | 11813 | 0.87192077 | 18.8446602 | rnr1Y285A | FALSE | CAN | JSY3868 | 2 | rnr1Y285A-R | GGC | G>T | A/T+1 | G/C-1 | CG>AT | CG>AT | GAAATGGCG | GCGTGGA | CCG | CCG | C | A | C>A | C>A_CCG | CG>AT_GCGTGGA | 5 | FALSE | FALSE |
| 63_v.txt | 31869 | 31869-SNV-G | SNV | 1 | G | T | 68 | 11813 | 0.57563701 | 46.6911765 | rnr1Y285A | FALSE | CAN | JSY3868 | 2 | rnr1Y285A-R | AGA | G>T | A/T+1 | G/C-1 | CG>AT | CG>AT | AAAATAGAG | GAGATAT | TCT | TCT | C | A | C>A | C>A_TCT | CG>AT_GAGATAT | 5 | FALSE | FALSE |
| 63_v.txt | 31981 | 31981-SNV-G | SNV | 1 | G | T | 53 | 21212 | 0.24985857 | 41.6415094 | rnr1Y285A | FALSE | CAN | JSY3868 | 2 | rnr1Y285A-R | GGC | G>T | A/T+1 | G/C-1 | CG>AT | CG>AT | AGCCGGGCA | GCATTAA | CCG | CCG | C | A | C>A | C>A_CCG | CG>AT_GCATTAA | 5 | FALSE | FALSE |
| 63_v.txt | 32027 | 32027-SNV-G | SNV | 1 | G | T | 1962 | 21212 | 9.24948143 | 60.5397554 | rnr1Y285A | FALSE | CAN | JSY3868 | 2 | rnr1Y285A-R | GGT | G>T | A/T+1 | G/C-1 | CG>AT | CG>AT | GCCACGGTA | GTATTTC | CCA | CCA | C | A | C>A | C>A_CCA | CG>AT_GTATTTC | 5 | FALSE | FALSE |
| 63_v.txt | 32114 | 32114-SNV-C | SNV | 1 | C | A | 9 | 21212 | 0.04242881 | 48.5555556 | rnr1Y285A | FALSE | CAN | JSY3868 | 2 | rnr1Y285A-R | GCC | C>A | A/T+1 | G/C-1 | CG>AT | CG>AT | TAATAGCCA | CCATTCG | CGG | GCC | C | T | C>A | C>A_GCC | CG>AT_CCATTCG | 5 | FALSE | FALSE |
| 63_v.txt | 32188 | 32188-SNV-C | SNV | 1 | C | A | 174 | 21212 | 0.8202904 | 46.5862069 | rnr1Y285A | FALSE | CAN | JSY3868 | 2 | rnr1Y285A-R | ACT | C>A | A/T+1 | G/C-1 | CG>AT | CG>AT | CGAAAACTG | CTGCAAT | TGA | ACT | C | T | C>A | C>A_ACT | CG>AT_CTGCAAT | 5 | FALSE | FALSE |
| 63_v.txt | 32274 | 32274-SNV-G | SNV | 1 | G | T | 74 | 25363 | 0.29176359 | 36.6891892 | rnr1Y285A | FALSE | CAN | JSY3868 | 2 | rnr1Y285A-R | GGA | G>T | A/T+1 | G/C-1 | CG>AT | CG>AT | ATACGGGAA | GAACCAA | CCT | CCT | C | A | C>A | C>A_CCT | CG>AT_GAACCAA | 5 | FALSE | FALSE |
| 63_v.txt | 32277 | 32277-SNV-C | SNV | 1 | C | A | 42 | 25363 | 0.16559555 | 34.5952381 | rnr1Y285A | FALSE | CAN | JSY3868 | 2 | rnr1Y285A-R | ACC | C>A | A/T+1 | G/C-1 | CG>AT | CG>AT | CGGGAACCA | CCAACGT | TGG | ACC | C | T | C>A | C>A_ACC | CG>AT_CCAACGT | 5 | FALSE | FALSE |
| 63_v.txt | 32304 | 32304-SNV-G | SNV | 1 | G | T | 15 | 25364 | 0.05913894 | 48.7333333 | rnr1Y285A | FALSE | CAN | JSY3868 | 2 | rnr1Y285A-R | AGA | G>T | A/T+1 | G/C-1 | CG>AT | CG>AT | GCGGCAGAA | GAAATAA | TCT | TCT | C | A | C>A | C>A_TCT | CG>AT_GAAATAA | 5 | FALSE | FALSE |
| 63_v.txt | 32373 | 32373-SNV-G | SNV | 1 | G | T | 42 | 25364 | 0.16558902 | 60.3809524 | rnr1Y285A | FALSE | CAN | JSY3868 | 2 | rnr1Y285A-R | AGC | G>T | A/T+1 | G/C-1 | CG>AT | CG>AT | TCAATAGCA | GCAATAA | TCG | TCG | C | A | C>A | C>A_TCG | CG>AT_GCAATAA | 5 | FALSE | FALSE |
| 63_v.txt | 32424 | 32424-SNV-G | SNV | 1 | G | T | 4 | 25364 | 0.01577038 | 60 | rnr1Y285A | FALSE | CAN | JSY3868 | 2 | rnr1Y285A-R | AGG | G>T | A/T+1 | G/C-1 | CG>AT | CG>AT | AGTTTAGGG | GGGTCAT | TCC | TCC | C | A | C>A | C>A_TCC | CG>AT_GGGTCAT | 5 | FALSE | FALSE |
| 63_v.txt | 32508 | 32508-SNV-G | SNV | 1 | G | T | 17 | 25364 | 0.06702413 | 43.5882353 | rnr1Y285A | FALSE | CAN | JSY3868 | 2 | rnr1Y285A-R | GGC | G>T | A/T+1 | G/C-1 | CG>AT | CG>AT | TTGATGGCT | GCTCTTG | CCG | CCG | C | A | C>A | C>A_CCG | CG>AT_GCTCTTG | 5 | FALSE | FALSE |
| 63_v.txt | 32530 | 32530-SNV-G | SNV | 1 | G | T | 73 | 25364 | 0.28780949 | 38.369863 | rnr1Y285A | FALSE | CAN | JSY3868 | 2 | rnr1Y285A-R | GGG | G>T | A/T+1 | G/C-1 | CG>AT | CG>AT | TTCTGGGGT | GGTTTGC | CCC | CCC | C | A | C>A | C>A_CCC | CG>AT_GGTTTGC | 5 | FALSE | FALSE |
| 63_v.txt | 32556 | 32556-SNV-C | SNV | 1 | C | A | 155 | 25359 | 0.61122284 | 37.2389552 | rnr1Y285A | FALSE | CAN | JSY3868 | 2 | rnr1Y285A-R | ACC | C>A | A/T+1 | G/C-1 | CG>AT | CG>AT | GTGATACCA | CCAACTA | TGG | ACC | C | T | C>A | C>A_ACC | CG>AT_CCAACTA | 5 | FALSE | FALSE |
| 63_v.txt | 32557 | 32557-SNV-C | SNV | 1 | C | A | 5 | 25358 | 0.01971764 | 27.6000772 | rnr1Y285A | FALSE | CAN | JSY3868 | 2 | rnr1Y285A-R | CCA | C>A | A/T+1 | G/C-1 | CG>AT | CG>AT | TGATACCAA | CAACTAG | GGT | CCA | C | T | C>A | C>A_CCA | CG>AT_CAACTAG | 5 | FALSE | FALSE |
| 63_v.txt | 32572 | 32572-SNV-C | SNV | 1 | C | A | 45 | 20429 | 0.2202751 | 36.2648008 | rnr1Y285A | FALSE | CAN | JSY3868 | 2 | rnr1Y285A-R | CCT | C>A | A/T+1 | G/C-1 | CG>AT | CG>AT | CAGTACCTT | CTTGAAA | GGA | CCT | C | T | C>A | C>A_CCT | CG>AT_CTTGAAA | 5 | FALSE | FALSE |
| 63_v.txt | 32691 | 32691-SNV-C | SNV | 1 | C | A | 20 | 20429 | 0.09790004 | 49.5 | rnr1Y285A | FALSE | CAN | JSY3868 | 2 | rnr1Y285A-R | TCC | C>A | A/T+1 | G/C-1 | CG>AT | CG>AT | CGGAATCCA | CCAACTG | AGG | TCC | C | T | C>A | C>A_TCC | CG>AT_CCAACTG | 5 | FALSE | FALSE |
| 63_v.txt | 32735 | 32735-SNV-G | SNV | 1 | G | T | 24 | 20429 | 0.11748005 | 61.5416667 | rnr1Y285A | FALSE | CAN | JSY3868 | 2 | rnr1Y285A-R | AGT | G>T | A/T+1 | G/C-1 | CG>AT | CG>AT | AAAACAGTA | GTATATT | TCA | TCA | C | A | C>A | C>A_TCA | CG>AT_GTATATT | 5 | FALSE | FALSE |
| 63_v.txt | 32783 | 32783-SNV-G | SNV | 1 | G | T | 111 | 20429 | 0.54334525 | 48.8468469 | rnr1Y285A | FALSE | CAN | JSY3868 | 2 | rnr1Y285A-R | AGA | G>T | A/T+1 | G/C-1 | CG>AT | CG>AT | GACCCAGAA | GAACTCG | TCT | TCT | C | A | C>A | C>A_TCT | CG>AT_GAACTCG | 5 | FALSE | FALSE |
| 63_v.txt | 32786 | 32786-SNV-C | SNV | 1 | C | A | 43 | 20429 | 0.2104851 | 47.8139535 | rnr1Y285A | FALSE | CAN | JSY3868 | 2 | rnr1Y285A-R | ACT | C>A | A/T+1 | G/C-1 | CG>AT | CG>AT | CCAGAACTC | CTCGAAT | TGA | ACT | C | T | C>A | C>A_ACT | CG>AT_CTCGAAT | 5 | FALSE | FALSE |
| 63_v.txt | 32794 | 32794-SNV-C | SNV | 1 | C | A | 16 | 20429 | 0.07832004 | 37.375 | rnr1Y285A | FALSE | CAN | JSY3868 | 2 | rnr1Y285A-R | TCA | C>A | A/T+1 | G/C-1 | CG>AT | CG>AT | CGAATTCAC | CACCGTA | AGT | TCA | C | T | C>A | C>A_TCA | CG>AT_CACCGTA | 5 | FALSE | FALSE |
| 63_v.txt | 32819 | 32819-SNV-G | SNV | 1 | G | T | 168 | 20429 | 0.82236037 | 44.9345238 | rnr1Y285A | FALSE | CAN | JSY3868 | 2 | rnr1Y285A-R | AGT | G>T | A/T+1 | G/C-1 | CG>AT | CG>AT | GAACAAGTT | GTTCATT | TCA | TCA | C | A | C>A | C>A_TCA | CG>AT_GTTCATT | 5 | FALSE | FALSE |
| 63_v.txt | 32901 | 32901-SNV-C | SNV | 1 | C | A | 3 | 27083 | 0.01107706 | 38 | rnr1Y285A | FALSE | CAN | JSY3868 | 2 | rnr1Y285A-R | GCC | C>A | A/T+1 | G/C-1 | CG>AT | CG>AT | ACTTGGCCA | CCAACTA | CGG | GCC | C | T | C>A | C>A_GCC | CG>AT_CCAACTA | 5 | FALSE | FALSE |
| 63_v.txt | 32917 | 32917-SNV-C | SNV | 1 | C | A | 777 | 27083 | 2.86895839 | 51.6808237 | rnr1Y285A | FALSE | CAN | JSY3868 | 2 | rnr1Y285A-R | TCC | C>A | A/T+1 | G/C-1 | CG>AT | CG>AT | TAAGTTCCA | CCAGGGC | AGG | TCC | C | T | C>A | C>A_TCC | CG>AT_CCAGGGC | 5 | FALSE | FALSE |
| 63_v.txt | 32940 | 32940-SNV-G | SNV | 1 | G | T | 70 | 27083 | 0.25846472 | 48.3570641 | rnr1Y285A | FALSE | CAN | JSY3868 | 2 | rnr1Y285A-R | AGA | G>T | A/T+1 | G/C-1 | CG>AT | CG>AT | GCCCAAGAA | GAAAACC | TCT | TCT | C | A | C>A | C>A_TCT | CG>AT_GAAAACC | 5 | FALSE | FALSE |
| 63_v.txt | 32954 | 32954-SNV-G | SNV | 1 | G | T | 563 | 27083 | 2.07879482 | 55.2095915 | rnr1Y285A | FALSE | CAN | JSY3868 | 2 | rnr1Y285A-R | TGT | G>T | A/T+1 | G/C-1 | CG>AT | CG>AT | ATACATGTA | GTAACCA | ACA | ACA | C | A | C>A | C>A_ACA | CG>AT_GTAACCA | 5 | FALSE | FALSE |
| 63_v.txt | 32958 | 32958-SNV-C | SNV | 1 | C | A | 716 | 27083 | 2.64372485 | 58.1801676 | rnr1Y285A | FALSE | CAN | JSY3868 | 2 | rnr1Y285A-R | ACC | C>A | A/T+1 | G/C-1 | CG>AT | CG>AT | ATGTAACCA | CCATTGG | TGG | ACC | C | T | C>A | C>A_ACC | CG>AT_CCATTGG | 5 | FALSE | FALSE |
| 63_v.txt | 33015 | 33015-SNV-G | SNV | 1 | G | T | 472 | 27083 | 1.74279068 | 58.6885593 | rnr1Y285A | FALSE | CAN | JSY3868 | 2 | rnr1Y285A-R | GGA | G>T | A/T+1 | G/C-1 | CG>AT | CG>AT | AAAGAGGAT | GATGTAA | CCT | CCT | C | A | C>A | C>A_CCT | CG>AT_GATGTAA | 5 | FALSE | FALSE |
| 63_v.txt | 33025 | 33025-SNV-G | SNV | 1 | G | T | 13 | 27083 | 0.04800059 | 58.1538462 | rnr1Y285A | FALSE | CAN | JSY3868 | 2 | rnr1Y285A-R | GGG | G>T | A/T+1 | G/C-1 | CG>AT | CG>AT | TAACAGGGA | GGATGAA | CCC | CCC | C | A | C>A | C>A_CCC | CG>AT_GGATGAA | 5 | FALSE | FALSE |
| 63_v.txt | 33050 | 33050-SNV-G | SNV | 1 | G | T | 14 | 27083 | 0.05169294 | 43.3571429 | rnr1Y285A | FALSE | CAN | JSY3868 | 2 | rnr1Y285A-R | AGG | G>T | A/T+1 | G/C-1 | CG>AT | CG>AT | ACCCAAGGA | GGACTGC | TCC | TCC | C | A | C>A | C>A_TCC | CG>AT_GGACTGC | 5 | FALSE | FALSE |
| 63_v.txt | 33114 | 33114-SNV-C | SNV | 1 | C | A | 10 | 27083 | 0.03692353 | 45.8 | rnr1Y285A | FALSE | CAN | JSY3868 | 2 | rnr1Y285A-R | GCC | C>A | A/T+1 | G/C-1 | CG>AT | CG>AT | ACTGGGCCG | CCGGCGT | CGG | GCC | C | T | C>A | C>A_GCC | CG>AT_CCGGCGT | 5 | FALSE | FALSE |
| 63_v.txt | 33151 | 33151-SNV-G | SNV | 1 | G | T | 62 | 27083 | 0.2289259 | 37.1129032 | rnr1Y285A | FALSE | CAN | JSY3868 | 2 | rnr1Y285A-R | AGA | G>T | A/T+1 | G/C-1 | CG>AT | CG>AT | TGAAAAGAC | GACCTGT | TCT | TCT | C | A | C>A | C>A_TCT | CG>AT_GACCTGT | 5 | FALSE | FALSE |
| 63_v.txt | 33153 | 33153-SNV-C | SNV | 1 | C | A | 1415 | 27083 | 5.22467969 | 40.4720848 | rnr1Y285A | FALSE | CAN | JSY3868 | 2 | rnr1Y285A-R | ACC | C>A | A/T+1 | G/C-1 | CG>AT | CG>AT | AAAAGACCT | CCTGTAC | TGG | ACC | C | T | C>A | C>A_ACC | CG>AT_CCTGTAC | 5 | FALSE | FALSE |
| 63_v.txt | 33156 | 33156-SNV-G | SNV | 1 | G | T | 10 | 27082 | 0.0369249 | 36 | rnr1Y285A | FALSE | CAN | JSY3868 | 2 | rnr1Y285A-R | TGT | G>T | A/T+1 | G/C-1 | CG>AT | CG>AT | AGACCTGTA | GTACCAA | ACA | ACA | C | A | C>A | C>A_ACA | CG>AT_GTACCAA | 5 | FALSE | FALSE |
| 63_v.txt | 33159 | 33159-SNV-C | SNV | 1 | C | A | 1126 | 27082 | 4.15774315 | 37.7690941 | rnr1Y285A | FALSE | CAN | JSY3868 | 2 | rnr1Y285A-R | ACC | C>A | A/T+1 | G/C-1 | CG>AT | CG>AT | CCTGTACCA | CCAATAG | TGG | ACC | C | T | C>A | C>A_ACC | CG>AT_CCAATAG | 5 | FALSE | FALSE |
| 63_v.txt | 33168 | 33168-SNV-C | SNV | 1 | C | A | 966 | 27081 | 3.56707655 | 37.3281574 | rnr1Y285A | FALSE | CAN | JSY3868 | 2 | rnr1Y285A-R | ACC | C>A | A/T+1 | G/C-1 | CG>AT | CG>AT | ATAGTACCA | CCACCAA | TGG | ACC | C | T | C>A | C>A_ACC | CG>AT_CCACCAA | 5 | FALSE | FALSE |
| 63_v.txt | 33177 | 33177-SNV-G | SNV | 1 | G | T | 138 | 27063 | 0.5099213 | 36.442029 | rnr1Y285A | FALSE | CAN | JSY3868 | 2 | rnr1Y285A-R | GGC | G>T | A/T+1 | G/C-1 | CG>AT | CG>AT | CCAAGGGCA | GCAATCA | CCG | CCG | C | A | C>A | C>A_CCG | CG>AT_GCAATCA | 5 | FALSE | FALSE |
| 63_v.txt | 33195 | 33195-SNV-C | SNV | 1 | C | A | 45 | 10539 | 0.42698548 | 36.1777778 | rnr1Y285A | FALSE | CAN | JSY3868 | 2 | rnr1Y285A-R | TCT | C>A | A/T+1 | G/C-1 | CG>AT | CG>AT | ATATGTCTTT | CTTTGCT | AGA | TCT | C | T | C>A | C>A_TCT | CG>AT_CTTTGCT | 5 | FALSE | FALSE |
| 63_v.txt | 33205 | 33205-SNV-G | SNV | 1 | G | T | 5 | 10539 | 0.04744283 | 29.8006345 | rnr1Y285A | FALSE | CAN | JSY3868 | 2 | rnr1Y285A-R | AGC | G>T | A/T+1 | G/C-1 | CG>AT | CG>AT | GCTTAAGCT | GCTCTCT | TCG | TCG | C | A | C>A | C>A_TCG | CG>AT_GCTCTCT | 5 | FALSE | FALSE |
| 63_v.txt | 33325 | 33325-SNV-C | SNV | 1 | C | A | 173 | 10539 | 1.64152197 | 58.7687861 | rnr1Y285A | FALSE | CAN | JSY3868 | 2 | rnr1Y285A-R | TCT | C>A | A/T+1 | G/C-1 | CG>AT | CG>AT | ACAATTCTTT | CTTTACT | AGA | TCT | C | T | C>A | C>A_TCT | CG>AT_CTTTACT | 5 | FALSE | FALSE |
| 63_v.txt | 33354 | 33354-SNV-G | SNV | 1 | G | T | 56 | 10539 | 0.53135971 | 17.5892857 | rnr1Y285A | FALSE | CAN | JSY3868 | 2 | rnr1Y285A-R | TGA | G>T | A/T+1 | G/C-1 | CG>AT | CG>AT | GGTATTGAC | GACCCAC | ACT | ACT | C | A | C>A | C>A_ACT | CG>AT_GACCCAC | 5 | FALSE | FALSE |
| 63_v.txt | 33385 | 33385-SNV-C | SNV | 1 | C | A | 23 | 10539 | 0.21823702 | 54.1304348 | rnr1Y285A | FALSE | CAN | JSY3868 | 2 | rnr1Y285A-R | TCA | C>A | A/T+1 | G/C-1 | CG>AT | CG>AT | AAGCTTCAA | CAACGTC | AGT | TCA | C | T | C>A | C>A_TCA | CG>AT_CAACGTC | 5 | FALSE | FALSE |
| 63_v.txt | 33474 | 33474-SNV-C | SNV | 1 | C | A | 34 | 10539 | 0.32261125 | 34.0294118 | rnr1Y285A | FALSE | CAN | JSY3868 | 2 | rnr1Y285A-R | GCC | C>A | A/T+1 | G/C-1 | CG>AT | CG>AT | GCTATGCCTT | CCTTTTT | CGG | GCC | C | T | C>A | C>A_GCC | CG>AT_CCTTTTT | 5 | FALSE | FALSE |
| 63_v.txt | 32081 | 32081-Insert | Insertion | 1 | - | A | 18 | 21212 | 0.08485763 | 62.4444444 | rnr1Y285A | FALSE | CAN | JSY3868 | 2 | rnr1Y285A-R | CAA | nope | A/T+1 | >1bpdel | none | A/T+1 | CCATGCAAA | AAAAAAG | GTT | CAA | - | T | NA | NA_CAA | A/T+1_AAAAAAG | 5 | FALSE | FALSE |
| 63_v.txt | 32157 | 32157-Insert | Insertion | 8 | - | GCCAAAGC | 26 | 21212 | 0.12257213 | 53.2932692 | rnr1Y285A | FALSE | CAN | JSY3868 | 2 | rnr1Y285A-R | AGC | nope | >1bp-ins | >1bpdel | none | >1bp-ins | ATGTAAGCC | GCCAAAG | TCG | AGC | - | none | NA | NA_AGC | >1bp-ins_GCCAAAG | 5 | FALSE | FALSE |
| 63_v.txt | 32305 | 32305-Insert | Insertion | 1 | - | A | 55 | 25364 | 0.21684277 | 57.6363636 | rnr1Y285A | FALSE | CAN | JSY3868 | 2 | rnr1Y285A-R | GAA | nope | A/T+1 | >1bpdel | none | A/T+1 | CGGCAGAAA | AAATAAT | CTT | GAA | - | T | NA | NA_GAA | A/T+1_AAATAAT | 5 | FALSE | FALSE |
| 63_v.txt | 32498 | 32498-Insert | Insertion | 1 | - | T | 2 | 25364 | 7.89E-03 | 63 | rnr1Y285A | FALSE | CAN | JSY3868 | 2 | rnr1Y285A-R | CTT | nope | A/T+1 | >1bpdel | none | A/T+1 | AACAACTTTT | TTTTTTG | GAA | CTT | - | A | NA | NA_CTT | A/T+1_TTTTTTG | 5 | FALSE | FALSE |

| | | | | | | | | | | | | | | | | | | | | | | | | | | | | | | | | | | |
|---|---|---|---|---|---|---|---|---|---|---|---|---|---|---|---|---|---|---|---|---|---|---|---|---|---|---|---|---|---|---|---|---|---|---|
| 63_v.txt | 32842 | 32842-Insert | Insertion | 1 | - | A | 21 | 20429 | 0.10279505 | 44.1904762 | rnr1Y285A | FALSE | CAN | JSY3868 | 2 | rnr1Y285A-R | CAA | nope | A/T+1 | >1bpdel | none | A/T+1 | TTACCCAAAA | AAAAAAT | GTT | CAA | - | T | NA | NA_CAA | A/T+1_AAAAAAT | 5 | FALSE | FALSE |
| 63_v.txt | 32884 | 32884-Insert | Insertion | 1 | - | A | 10 | 27084 | 0.03692217 | 36.4 | rnr1Y285A | FALSE | CAN | JSY3868 | 2 | rnr1Y285A-R | CAA | nope | A/T+1 | >1bpdel | none | A/T+1 | ACGTCCAAAA | AAAATTG | GTT | CAA | - | T | NA | NA_CAA | A/T+1_AAAATTG | 5 | FALSE | FALSE |
| 63_v.txt | 33206 | 33206-Insert | Insertion | 2 | - | CT | 2 | 10539 | 0.01897713 | 37.75 | rnr1Y285A | FALSE | CAN | JSY3868 | 2 | rnr1Y285A-R | GCT | nope | >1bp-ins | >1bpdel | none | >1bp-ins | CTTAAGCTCT | CTCTCTC | CGA | GCT | - | none | NA | NA_GCT | >1bp-ins_CTCTCTC | 5 | FALSE | FALSE |
| 63_v.txt | 33094 | 33094-Deleti | Deletion | 2 | AT | - | 58 | 27083 | 0.21415648 | 47.7068966 | rnr1Y285A | FALSE | CAN | JSY3868 | 2 | rnr1Y285A-R | GAT | nope | >1bp-ins | >1bpdel | none | >1bpdel | AATATGATAT | ATATAAG | CTA | GAT | AT | none | NA | NA_GAT | >1bpdel_ATATAAG | 5 | FALSE | FALSE |
| 63_v.txt | 31723 | 31723-Deleti | Deletion | 1 | A | - | 13 | 11813 | 0.11004825 | 58.6153846 | rnr1Y285A | FALSE | CAN | JSY3868 | 2 | rnr1Y285A-R | CAA | nope | >1bp-ins | A/T-1 | none | A/T-1 | TGTCCCAAAA | AAAAAGT | GTT | GTT | T | none | T>none | T>none_GTT | A/T-1_AAAAAGT | 5 | FALSE | FALSE |
| 63_v.txt | 31904 | 31904-Deleti | Deletion | 1 | T | - | 5 | 11813 | 0.04232625 | 48 | rnr1Y285A | FALSE | CAN | JSY3868 | 2 | rnr1Y285A-R | ATT | nope | >1bp-ins | A/T-1 | none | A/T-1 | ATTGAATTTT | TTTTGGT | TAA | ATT | T | none | nope | nope_ATT | A/T-1_TTTTGGT | 5 | FALSE | FALSE |
| 63_v.txt | 31913 | 31913-Deleti | Deletion | 1 | A | - | 6 | 11813 | 0.0507915 | 38 | rnr1Y285A | FALSE | CAN | JSY3868 | 2 | rnr1Y285A-R | CAA | nope | >1bp-ins | A/T-1 | none | A/T-1 | TGGTGCAAAA | AAAAGCC | GTT | GTT | T | none | T>none | T>none_GTT | A/T-1_AAAAGCC | 5 | FALSE | FALSE |
| 63_v.txt | 31972 | 31972-Deleti | Deletion | 1 | C | - | 2 | 21212 | 9.43E-03 | 38 | rnr1Y285A | FALSE | CAN | JSY3868 | 2 | rnr1Y285A-R | GCC | nope | >1bp-ins | G/C-1 | none | G/C-1 | AATAAGCCAA | CCAAGCC | CGG | GCC | C | none | nope | nope_GCC | G/C-1_CCAAGCC | 5 | FALSE | FALSE |
| 63_v.txt | 31979 | 31979-Deleti | Deletion | 1 | G | - | 28 | 21212 | 0.13200075 | 37.8928571 | rnr1Y285A | FALSE | CAN | JSY3868 | 2 | rnr1Y285A-R | CGG | nope | >1bp-ins | G/C-1 | none | G/C-1 | CAAGCCGGG | GGGCATT | GCC | GCC | C | none | C>none | C>none_GCC | G/C-1_GGGCATT | 5 | FALSE | FALSE |
| 63_v.txt | 32075 | 32075-Deleti | Deletion | 1 | C | - | 2 | 21212 | 9.43E-03 | 64 | rnr1Y285A | FALSE | CAN | JSY3868 | 2 | rnr1Y285A-R | ACC | nope | >1bp-ins | G/C-1 | none | G/C-1 | AAATAACCAT | CCATGCA | TGG | ACC | C | none | nope | nope_ACC | G/C-1_CCATGCA | 5 | FALSE | FALSE |
| 63_v.txt | 32081 | 32081-Deleti | Deletion | 1 | A | - | 15 | 21212 | 0.07071469 | 58.533334 | rnr1Y285A | FALSE | CAN | JSY3868 | 2 | rnr1Y285A-R | CAA | nope | >1bp-ins | A/T-1 | none | A/T-1 | CCATGCAAAA | AAAAAAG | GTT | GTT | T | none | T>none | T>none_GTT | A/T-1_AAAAAAG | 5 | FALSE | FALSE |
| 63_v.txt | 32210 | 32210-Deleti | Deletion | 1 | T | - | 5 | 21212 | 0.02357156 | 43.2 | rnr1Y285A | FALSE | CAN | JSY3868 | 2 | rnr1Y285A-R | CTT | nope | >1bp-ins | A/T-1 | none | A/T-1 | ACCACCTTTG | TTTGGTG | GAA | CTT | T | none | nope | nope_CTT | A/T-1_TTTGGTG | 5 | FALSE | FALSE |
| 63_v.txt | 32265 | 32265-Deleti | Deletion | 1 | A | - | 6 | 25363 | 0.02365651 | 37.3333333 | rnr1Y285A | FALSE | CAN | JSY3868 | 2 | rnr1Y285A-R | TAA | nope | >1bp-ins | A/T-1 | none | A/T-1 | CCAAATAAAA | AAAATAC | ATT | ATT | T | none | T>none | T>none_ATT | A/T-1_AAAATAC | 5 | FALSE | FALSE |
| 63_v.txt | 32277 | 32277-Deleti | Deletion | 1 | C | - | 18 | 25363 | 0.07096952 | 36.5 | rnr1Y285A | FALSE | CAN | JSY3868 | 2 | rnr1Y285A-R | ACC | nope | >1bp-ins | G/C-1 | none | G/C-1 | CGGGAACCA | CCAACGT | TGG | ACC | C | none | nope | nope_ACC | G/C-1_CCAACGT | 5 | FALSE | FALSE |
| 63_v.txt | 32381 | 32381-Deleti | Deletion | 1 | A | - | 6 | 25364 | 0.02365558 | 64 | rnr1Y285A | FALSE | CAN | JSY3868 | 2 | rnr1Y285A-R | TAA | nope | >1bp-ins | A/T-1 | none | A/T-1 | AATAATAAA | AAAGGGA | ATT | ATT | T | none | T>none | T>none_ATT | A/T-1_AAAGGGA | 5 | FALSE | FALSE |
| 63_v.txt | 32398 | 32398-Deleti | Deletion | 1 | C | - | 58 | 25364 | 0.22867056 | 60.8275862 | rnr1Y285A | FALSE | CAN | JSY3868 | 2 | rnr1Y285A-R | ACG | nope | >1bp-ins | G/C-1 | none | G/C-1 | TAGAAACGT | CGTAGGA | TGC | ACG | C | none | nope | nope_ACG | G/C-1_CGTAGGA | 5 | FALSE | FALSE |
| 63_v.txt | 32424 | 32424-Deleti | Deletion | 1 | G | - | 34 | 25364 | 0.13404826 | 57.5882353 | rnr1Y285A | FALSE | CAN | JSY3868 | 2 | rnr1Y285A-R | AGG | nope | >1bp-ins | G/C-1 | none | G/C-1 | AGTTTAGGG | GGGTCAT | TCC | TCC | C | none | C>none | C>none_TCC | G/C-1_GGGTCAT | 5 | FALSE | FALSE |
| 63_v.txt | 32498 | 32498-Deleti | Deletion | 1 | T | - | 22 | 25364 | 0.08673711 | 46.9999961 | rnr1Y285A | FALSE | CAN | JSY3868 | 2 | rnr1Y285A-R | CTT | nope | >1bp-ins | A/T-1 | none | A/T-1 | AACAACTTTT | TTTTTTG | GAA | CTT | T | none | nope | nope_CTT | A/T-1_TTTTTTG | 5 | FALSE | FALSE |
| 63_v.txt | 32522 | 32522-Deleti | Deletion | 1 | T | - | 14 | 25364 | 0.05519634 | 38.6428568 | rnr1Y285A | FALSE | CAN | JSY3868 | 2 | rnr1Y285A-R | ATT | nope | >1bp-ins | A/T-1 | none | A/T-1 | AACGGATTTT | TTTTCTG | TAA | ATT | T | none | nope | nope_ATT | A/T-1_TTTTCTG | 5 | FALSE | FALSE |
| 63_v.txt | 32528 | 32528-Deleti | Deletion | 1 | G | - | 32 | 25364 | 0.12616307 | 39.6875 | rnr1Y285A | FALSE | CAN | JSY3868 | 2 | rnr1Y285A-R | TGG | nope | >1bp-ins | G/C-1 | none | G/C-1 | TTTTCTGGGG | GGGGTTT | ACC | ACC | C | none | C>none | C>none_ACC | G/C-1_GGGGTTT | 5 | FALSE | FALSE |
| 63_v.txt | 32556 | 32556-Deleti | Deletion | 1 | C | - | 6 | 25359 | 0.02366024 | 37.6666669 | rnr1Y285A | FALSE | CAN | JSY3868 | 2 | rnr1Y285A-R | ACC | nope | >1bp-ins | G/C-1 | none | G/C-1 | GTGATACCAA | CCAACTA | TGG | ACC | C | none | nope | nope_ACC | G/C-1_CCAACTA | 5 | FALSE | FALSE |
| 63_v.txt | 32608 | 32608-Deleti | Deletion | 1 | C | - | 506 | 20429 | 2.47687112 | 43.9229249 | rnr1Y285A | FALSE | CAN | JSY3868 | 2 | rnr1Y285A-R | ACC | nope | >1bp-ins | G/C-1 | none | G/C-1 | AGGAAACCC | CCCAACC | TGG | ACC | C | none | nope | nope_ACC | G/C-1_CCCAACC | 5 | FALSE | FALSE |
| 63_v.txt | 32631 | 32631-Deleti | Deletion | 1 | T | - | 8 | 20429 | 0.03916002 | 53.125 | rnr1Y285A | FALSE | CAN | JSY3868 | 2 | rnr1Y285A-R | GTT | nope | >1bp-ins | A/T-1 | none | A/T-1 | CCTTCGTTTT | TTTTTAT | CAA | GTT | T | none | nope | nope_GTT | A/T-1_TTTTTAT | 5 | FALSE | FALSE |
| 63_v.txt | 32640 | 32640-Deleti | Deletion | 1 | T | - | 5 | 20429 | 0.02447501 | 53 | rnr1Y285A | FALSE | CAN | JSY3868 | 2 | rnr1Y285A-R | CTT | nope | >1bp-ins | A/T-1 | none | A/T-1 | TTATCCTTAG | TTAGATA | GAA | CTT | T | none | nope | nope_CTT | A/T-1_TTAGATA | 5 | FALSE | FALSE |
| 63_v.txt | 32658 | 32658-Deleti | Deletion | 1 | C | - | 1186 | 20429 | 5.80547261 | 60.7740304 | rnr1Y285A | FALSE | CAN | JSY3868 | 2 | rnr1Y285A-R | ACC | nope | >1bp-ins | G/C-1 | none | G/C-1 | CCTGGACCCC | CCCCAGG | TGG | ACC | C | none | nope | nope_ACC | G/C-1_CCCCAGG | 5 | FALSE | FALSE |
| 63_v.txt | 32670 | 32670-Deleti | Deletion | 1 | G | - | 811 | 20429 | 3.96984679 | 59.6991369 | rnr1Y285A | FALSE | CAN | JSY3868 | 2 | rnr1Y285A-R | TGG | nope | >1bp-ins | G/C-1 | none | G/C-1 | GCACCTGGGT | GGGTTTC | ACC | ACC | C | none | C>none | C>none_ACC | G/C-1_GGGTTTC | 5 | FALSE | FALSE |
| 63_v.txt | 32707 | 32707-Deleti | Deletion | 1 | C | - | 371 | 20429 | 1.81604582 | 62.1590297 | rnr1Y285A | FALSE | CAN | JSY3868 | 2 | rnr1Y285A-R | ACC | nope | >1bp-ins | G/C-1 | none | G/C-1 | CGGTAACCCC | CCCCAGC | TGG | ACC | C | none | nope | nope_ACC | G/C-1_CCCCAGC | 5 | FALSE | FALSE |
| 63_v.txt | 32728 | 32728-Deleti | Deletion | 1 | A | - | 9 | 20429 | 0.04405502 | 56.7777778 | rnr1Y285A | FALSE | CAN | JSY3868 | 2 | rnr1Y285A-R | CAA | nope | >1bp-ins | A/T-1 | none | A/T-1 | CCATACAAAA | AAAAACA | GTT | GTT | T | none | T>none | T>none_GTT | A/T-1_AAAAACA | 5 | FALSE | FALSE |
| 63_v.txt | 32747 | 32747-Deleti | Deletion | 1 | C | - | 96 | 20429 | 0.46992021 | 54.6354167 | rnr1Y285A | FALSE | CAN | JSY3868 | 2 | rnr1Y285A-R | ACC | nope | >1bp-ins | G/C-1 | none | G/C-1 | TAGAAACCCC | CCCGATA | TGG | ACC | C | none | nope | nope_ACC | G/C-1_CCCGATA | 5 | FALSE | FALSE |
| 63_v.txt | 32842 | 32842-Deleti | Deletion | 1 | A | - | 105 | 20429 | 0.51397523 | 42.8 | rnr1Y285A | FALSE | CAN | JSY3868 | 2 | rnr1Y285A-R | CAA | nope | >1bp-ins | A/T-1 | none | A/T-1 | TTACCCAAAA | AAAAAAT | GTT | GTT | T | none | T>none | T>none_GTT | A/T-1_AAAAAAT | 5 | FALSE | FALSE |
| 63_v.txt | 32917 | 32917-Deleti | Deletion | 1 | C | - | 4 | 27083 | 0.01476941 | 47.75 | rnr1Y285A | FALSE | CAN | JSY3868 | 2 | rnr1Y285A-R | TCC | nope | >1bp-ins | G/C-1 | none | G/C-1 | TAAGTTCCAC | CCAGGGC | AGG | TCC | C | none | nope | nope_TCC | G/C-1_CCAGGGC | 5 | FALSE | FALSE |
| 63_v.txt | 32924 | 32924-Deleti | Deletion | 1 | A | - | 25 | 27083 | 0.09230883 | 56.2 | rnr1Y285A | FALSE | CAN | JSY3868 | 2 | rnr1Y285A-R | CAA | nope | >1bp-ins | A/T-1 | none | A/T-1 | CAGGGCAAA | AAAAGTG | GTT | GTT | T | none | T>none | T>none_GTT | A/T-1_AAAAGTG | 5 | FALSE | FALSE |
| 63_v.txt | 32935 | 32935-Deleti | Deletion | 1 | C | - | 6 | 27083 | 0.02215412 | 49 | rnr1Y285A | FALSE | CAN | JSY3868 | 2 | rnr1Y285A-R | GCC | nope | >1bp-ins | G/C-1 | none | G/C-1 | TGATTGCCCA | CCCAAGA | CGG | GCC | C | none | nope | nope_GCC | G/C-1_CCCAAGA | 5 | FALSE | FALSE |
| 63_v.txt | 32941 | 32941-Deleti | Deletion | 1 | A | - | 5 | 27083 | 0.01846177 | 58.5999813 | rnr1Y285A | FALSE | CAN | JSY3868 | 2 | rnr1Y285A-R | GAA | nope | >1bp-ins | A/T-1 | none | A/T-1 | CCCAAGAAA | AAAACCA | CTT | CTT | T | none | T>none | T>none_CTT | A/T-1_AAAACCA | 5 | FALSE | FALSE |
| 63_v.txt | 32958 | 32958-Deleti | Deletion | 1 | C | - | 20 | 27083 | 0.07384706 | 56.9 | rnr1Y285A | FALSE | CAN | JSY3868 | 2 | rnr1Y285A-R | ACC | nope | >1bp-ins | G/C-1 | none | G/C-1 | ATGTAACCAT | CCATTGG | TGG | ACC | C | none | nope | nope_ACC | G/C-1_CCATTGG | 5 | FALSE | FALSE |
| 63_v.txt | 33045 | 33045-Deleti | Deletion | 1 | C | - | 6 | 27083 | 0.02215412 | 56.8333333 | rnr1Y285A | FALSE | CAN | JSY3868 | 2 | rnr1Y285A-R | ACC | nope | >1bp-ins | G/C-1 | none | G/C-1 | ATTTCACCCA | CCCAAGG | TGG | ACC | C | none | nope | nope_ACC | G/C-1_CCCAAGG | 5 | FALSE | FALSE |
| 63_v.txt | 33070 | 33070-Deleti | Deletion | 1 | C | - | 1 | 27083 | 3.69E-03 | 25.0000149 | rnr1Y285A | FALSE | CAN | JSY3868 | 2 | rnr1Y285A-R | GCC | nope | >1bp-ins | G/C-1 | none | G/C-1 | AATATGCCAA | CCAAAGA | CGG | GCC | C | none | nope | nope_GCC | G/C-1_CCAAAGA | 5 | FALSE | FALSE |
| 63_v.txt | 33078 | 33078-Deleti | Deletion | 1 | C | - | 131 | 27083 | 0.48369826 | 54.0687023 | rnr1Y285A | FALSE | CAN | JSY3868 | 2 | rnr1Y285A-R | ACC | nope | >1bp-ins | G/C-1 | none | G/C-1 | AAAGAACCC | CCCATAA | TGG | ACC | C | none | nope | nope_ACC | G/C-1_CCCATAA | 5 | FALSE | FALSE |
| 63_v.txt | 33138 | 33138-Deleti | Deletion | 1 | A | - | 4 | 27083 | 0.01476941 | 37.75 | rnr1Y285A | FALSE | CAN | JSY3868 | 2 | rnr1Y285A-R | TAA | nope | >1bp-ins | A/T-1 | none | A/T-1 | GTGGATAAA | AAACCAA | ATT | ATT | T | none | T>none | T>none_ATT | A/T-1_AAACCAA | 5 | FALSE | FALSE |
| 63_v.txt | 33147 | 33147-Deleti | Deletion | 1 | A | - | 9 | 27083 | 0.03323118 | 40.1111111 | rnr1Y285A | FALSE | CAN | JSY3868 | 2 | rnr1Y285A-R | GAA | nope | >1bp-ins | A/T-1 | none | A/T-1 | CCAATGAAA | AAAAGAC | CTT | CTT | T | none | T>none | T>none_CTT | A/T-1_AAAAGAC | 5 | FALSE | FALSE |
| 63_v.txt | 33356 | 33356-Deleti | Deletion | 1 | C | - | 461 | 10539 | 4.37422905 | 54 | rnr1Y285A | FALSE | CAN | JSY3868 | 2 | rnr1Y285A-R | ACC | nope | >1bp-ins | G/C-1 | none | G/C-1 | TATTGACCCA | CCCACGT | TGG | ACC | C | none | nope | nope_ACC | G/C-1_CCCACGT | 5 | FALSE | FALSE |
| 63_v.txt | 33474 | 33474-Deleti | Deletion | 1 | C | - | 21 | 10539 | 0.19925989 | 30.8571429 | rnr1Y285A | FALSE | CAN | JSY3868 | 2 | rnr1Y285A-R | GCC | nope | >1bp-ins | G/C-1 | none | G/C-1 | GCTATGCCTT | CCTTTTT | CGG | GCC | C | none | nope | nope_GCC | G/C-1_CCTTTTT | 5 | FALSE | FALSE |
| 64_v.txt | 31723 | 31723-SNV-A | SNV | 1 | A | C | 3 | 11779 | 0.02546906 | 57.6666667 | rnr1Y285A | FALSE | CAN | JSY3869 | 3 | rnr1Y285A-R | CAA | A>C | G/C+1 | A/T-1 | TA>GC | TA>GC | TGTCCCAAAA | AAAAAGT | GTT | GTT | T | G | T>G | T>G_GTT | TA>GC_AAAAAGT | 5 | FALSE | FALSE |
| 64_v.txt | 32256 | 32256-SNV-A | SNV | 1 | A | C | 32 | 23515 | 0.13608335 | 31.375 | rnr1Y285A | FALSE | CAN | JSY3869 | 3 | rnr1Y285A-R | TAG | A>C | G/C+1 | A/T-1 | TA>GC | TA>GC | TTTGATAGAC | AGACCAA | ATC | ATC | T | G | T>G | T>G_ATC | TA>GC_AGACCAA | 5 | FALSE | FALSE |
| 64_v.txt | 32763 | 32763-SNV-A | SNV | 1 | A | C | 35 | 24916 | 0.14047199 | 49.2571429 | rnr1Y285A | FALSE | CAN | JSY3869 | 3 | rnr1Y285A-R | AAC | A>C | G/C+1 | A/T-1 | TA>GC | TA>GC | GCTAAAACTT | ACTTTGA | TTG | TTG | T | G | T>G | T>G_TTG | TA>GC_ACTTTGA | 5 | FALSE | FALSE |
| 64_v.txt | 32842 | 32842-SNV-A | SNV | 1 | A | C | 8 | 24916 | 0.03210788 | 34.2502714 | rnr1Y285A | FALSE | CAN | JSY3869 | 3 | rnr1Y285A-R | CAA | A>C | G/C+1 | A/T-1 | TA>GC | TA>GC | TTACCCAAAA | AAAAAAT | GTT | GTT | T | G | T>G | T>G_GTT | TA>GC_AAAAAAT | 5 | FALSE | FALSE |
| 64_v.txt | 33194 | 33194-SNV-T | SNV | 1 | T | G | 75 | 17362 | 0.43197788 | 35.7053006 | rnr1Y285A | FALSE | CAN | JSY3869 | 3 | rnr1Y285A-R | GTC | T>G | G/C+1 | A/T-1 | TA>GC | TA>GC | AATATGTCTT | TCTTTGC | CAG | GTC | T | C | T>G | T>G_GTC | TA>GC_TCTTTGC | 5 | FALSE | FALSE |
| 64_v.txt | 33269 | 33269-SNV-A | SNV | 1 | A | C | 134 | 17362 | 0.77180048 | 16.4626866 | rnr1Y285A | FALSE | CAN | JSY3869 | 3 | rnr1Y285A-R | TAG | A>C | G/C+1 | A/T-1 | TA>GC | TA>GC | CTCCATAGAC | AGAGAAC | ATC | ATC | T | G | T>G | T>G_ATC | TA>GC_AGAGAAC | 5 | FALSE | FALSE |
| 64_v.txt | 32081 | 32081-SNV-A | SNV | 1 | A | G | 4 | 19297 | 0.02072861 | 64 | rnr1Y285A | FALSE | CAN | JSY3869 | 3 | rnr1Y285A-R | CAA | A>G | G/C+1 | A/T-1 | TA>CG | TA>CG | CCATGCAAAA | AAAAAAG | GTT | GTT | T | C | T>C | T>C_GTT | TA>CG_AAAAAAG | 5 | FALSE | FALSE |
| 64_v.txt | 32490 | 32490-SNV-A | SNV | 1 | A | G | 61 | 23523 | 0.25932067 | 45.3278689 | rnr1Y285A | FALSE | CAN | JSY3869 | 3 | rnr1Y285A-R | GAA | A>G | G/C+1 | A/T-1 | TA>CG | TA>CG | ATACGGAAA | AAAACAA | CTT | CTT | T | C | T>C | T>C_CTT | TA>CG_AAAACAA | 5 | FALSE | FALSE |
| 64_v.txt | 32498 | 32498-SNV-T | SNV | 1 | T | C | 7 | 23523 | 0.02975811 | 27.4285714 | rnr1Y285A | FALSE | CAN | JSY3869 | 3 | rnr1Y285A-R | CTT | T>C | G/C+1 | A/T-1 | TA>CG | TA>CG | AACAACTTTT | TTTTTTG | GAA | CTT | T | G | T>C | T>C_CTT | TA>CG_TTTTTTG | 5 | FALSE | FALSE |
| 64_v.txt | 32787 | 32787-SNV-T | SNV | 1 | T | C | 19 | 24916 | 0.07625622 | 47.7894737 | rnr1Y285A | FALSE | CAN | JSY3869 | 3 | rnr1Y285A-R | CTC | T>C | G/C+1 | A/T-1 | TA>CG | TA>CG | CAGAACTCGA | TCGAATT | GAG | CTC | T | G | T>C | T>C_CTC | TA>CG_TCGAATT | 5 | FALSE | FALSE |
| 64_v.txt | 33089 | 33089-SNV-A | SNV | 1 | A | G | 2 | 19702 | 0.01015125 | 34.4996219 | rnr1Y285A | FALSE | CAN | JSY3869 | 3 | rnr1Y285A-R | AAT | A>G | G/C+1 | A/T-1 | TA>CG | TA>CG | AAATAAATAT | ATATGAT | TTA | TTA | T | C | T>C | T>C_TTA | TA>CG_ATATGAT | 5 | FALSE | FALSE |
| 64_v.txt | 33194 | 33194-SNV-T | SNV | 1 | T | C | 2 | 17362 | 0.01151941 | 24.4996361 | rnr1Y285A | FALSE | CAN | JSY3869 | 3 | rnr1Y285A-R | GTC | T>C | G/C+1 | A/T-1 | TA>CG | TA>CG | AATATGTCTT | TCTTTGC | CAG | GTC | T | G | T>C | T>C_GTC | TA>CG_TCTTTGC | 5 | FALSE | FALSE |
| 64_v.txt | 33476 | 33476-SNV-T | SNV | 1 | T | C | 25 | 17360 | 0.14400922 | 34.32 | rnr1Y285A | FALSE | CAN | JSY3869 | 3 | rnr1Y285A-R | CTT | T>C | G/C+1 | A/T-1 | TA>CG | TA>CG | TATGCCTTTT | TTTTTTT | GAA | CTT | T | G | T>C | T>C_CTT | TA>CG_TTTTTTT | 5 | FALSE | FALSE |
| 64_v.txt | 32496 | 32496-SNV-A | SNV | 1 | A | T | 66 | 23523 | 0.28057646 | 46.4090909 | rnr1Y285A | FALSE | CAN | JSY3869 | 3 | rnr1Y285A-R | AAC | A>T | A/T+1 | A/T-1 | TA>AT | TA>AT | AAAACAACT | ACTTTTT | TTG | TTG | T | A | T>A | T>A_TTG | TA>AT_ACTTTTT | 5 | FALSE | FALSE |
| 64_v.txt | 32498 | 32498-SNV-T | SNV | 1 | T | A | 5 | 23523 | 0.02125579 | 47.6 | rnr1Y285A | FALSE | CAN | JSY3869 | 3 | rnr1Y285A-R | CTT | T>A | A/T+1 | A/T-1 | TA>AT | TA>AT | AACAACTTTT | TTTTTTG | GAA | CTT | T | T | T>A | T>A_CTT | TA>AT_TTTTTTG | 5 | FALSE | FALSE |
| 64_v.txt | 32785 | 32785-SNV-A | SNV | 1 | A | T | 54 | 24916 | 0.21672821 | 45.6851852 | rnr1Y285A | FALSE | CAN | JSY3869 | 3 | rnr1Y285A-R | AAC | A>T | A/T+1 | A/T-1 | TA>AT | TA>AT | CCCAGAACTC | ACTCGAA | TTG | TTG | T | A | T>A | T>A_TTG | TA>AT_ACTCGAA | 5 | FALSE | FALSE |
| 64_v.txt | 32787 | 32787-SNV-T | SNV | 1 | T | A | 9 | 24916 | 0.03612137 | 49.1111111 | rnr1Y285A | FALSE | CAN | JSY3869 | 3 | rnr1Y285A-R | CTC | T>A | A/T+1 | A/T-1 | TA>AT | TA>AT | CAGAACTCGA | TCGAATT | GAG | CTC | T | T | T>A | T>A_CTC | TA>AT_TCGAATT | 5 | FALSE | FALSE |
| 64_v.txt | 33090 | 33090-SNV-T | SNV | 1 | T | A | 3 | 19702 | 0.01522688 | 37.6588015 | rnr1Y285A | FALSE | CAN | JSY3869 | 3 | rnr1Y285A-R | ATA | T>A | A/T+1 | A/T-1 | TA>AT | TA>AT | AATAAATATC | TATGATA | TAT | ATA | T | T | T>A | T>A_ATA | TA>AT_TATGATA | 5 | FALSE | FALSE |
| 64_v.txt | 31650 | 31650-SNV-C | SNV | 1 | C | T | 25 | 11779 | 0.21224213 | 36.28 | rnr1Y285A | FALSE | CAN | JSY3869 | 3 | rnr1Y285A-R | GCG | C>T | A/T+1 | G/C-1 | CG>TA | CG>TA | AGAATGCGA | CGAAATG | CGC | GCG | C | A | C>T | C>T_GCG | CG>TA_CGAAATG | 5 | FALSE | FALSE |
| 64_v.txt | 31830 | 31830-SNV-C | SNV | 1 | C | T | 18 | 11779 | 0.15281433 | 52.7777778 | rnr1Y285A | FALSE | CAN | JSY3869 | 3 | rnr1Y285A-R | GCA | C>T | A/T+1 | G/C-1 | CG>TA | CG>TA | AATATGCATT | CATTGAA | CGT | GCA | C | A | C>T | C>T_GCA | CG>TA_CATTGAA | 5 | FALSE | FALSE |
| 64_v.txt | 31844 | 31844-SNV-C | SNV | 1 | C | T | 45 | 11779 | 0.38203583 | 45.9333333 | rnr1Y285A | FALSE | CAN | JSY3869 | 3 | rnr1Y285A-R | TCC | C>T | A/T+1 | G/C-1 | CG>TA | CG>TA | TAAGATCCAA | CCAAACA | AGG | TCC | C | A | C>T | C>T_TCC | CG>TA_CCAAACA | 5 | FALSE | FALSE |
| 64_v.txt | 31947 | 31947-SNV-G | SNV | 1 | G | A | 18 | 19289 | 0.09331744 | 37.0555556 | rnr1Y285A | FALSE | CAN | JSY3869 | 3 | rnr1Y285A-R | CGT | G>A | A/T+1 | G/C-1 | CG>TA | CG>TA | ATGATCGTCA | GTCATAA | GCA | GCA | C | T | C>T | C>T_GCA | CG>TA_GTCATAA | 5 | FALSE | FALSE |
| 64_v.txt | 31962 | 31962-SNV-G | SNV | 1 | G | A | 39 | 19297 | 0.20210395 | 36.9230769 | rnr1Y285A | FALSE | CAN | JSY3869 | 3 | rnr1Y285A-R | CGC | G>A | A/T+1 | G/C-1 | CG>TA | CG>TA | GTGGCCGCAT | GCATAAT | GCG | GCG | C | T | C>T | C>T_GCG | CG>TA_GCATAAT | 5 | FALSE | FALSE |
| 64_v.txt | 31981 | 31981-SNV-G | SNV | 1 | G | A | 109 | 19297 | 0.56485464 | 48.7247706 | rnr1Y285A | FALSE | CAN | JSY3869 | 3 | rnr1Y285A-R | GGC | G>A | A/T+1 | G/C-1 | CG>TA | CG>TA | AGCCGGGCA | GCATTAA | CCG | CCG | C | T | C>T | C>T_CCG | CG>TA_GCATTAA | 5 | FALSE | FALSE |
| 64_v.txt | 32041 | 32041-SNV-G | SNV | 1 | G | A | 22 | 19297 | 0.11400736 | 62.8181818 | rnr1Y285A | FALSE | CAN | JSY3869 | 3 | rnr1Y285A-R | TGC | G>A | A/T+1 | G/C-1 | CG>TA | CG>TA | AAGCTTGCAT | GCATAAA | ACG | ACG | C | T | C>T | C>T_ACG | CG>TA_GCATAAA | 5 | FALSE | FALSE |
| 64_v.txt | 32075 | 32075-SNV-C | SNV | 1 | C | T | 320 | 19297 | 1.65828885 | 61.39375 | rnr1Y285A | FALSE | CAN | JSY3869 | 3 | rnr1Y285A-R | ACC | C>T | A/T+1 | G/C-1 | CG>TA | CG>TA | AAATAACCAT | CCATGCA | TGG | ACC | C | A | C>T | C>T_ACC | CG>TA_CCATGCA | 5 | FALSE | FALSE |
| 64_v.txt | 32088 | 32088-SNV-C | SNV | 1 | C | T | 275 | 19297 | 1.42509198 | 61.2072727 | rnr1Y285A | FALSE | CAN | JSY3869 | 3 | rnr1Y285A-R | GCC | C>T | A/T+1 | G/C-1 | CG>TA | CG>TA | AAAAAGCCTG | CCTGCAA | CGG | GCC | C | A | C>T | C>T_GCC | CG>TA_CCTGCAA | 5 | FALSE | FALSE |
| 64_v.txt | 32092 | 32092-SNV-C | SNV | 1 | C | T | 27 | 19297 | 0.13991812 | 60.7777778 | rnr1Y285A | FALSE | CAN | JSY3869 | 3 | rnr1Y285A-R | GCA | C>T | A/T+1 | G/C-1 | CG>TA | CG>TA | AGCCTGCAAC | CAACACC | CGT | GCA | C | A | C>T | C>T_GCA | CG>TA_CAACACC | 5 | FALSE | FALSE |
| 64_v.txt | 32114 | 32114-SNV-C | SNV | 1 | C | T | 1242 | 19295 | 6.43690075 | 59.2157808 | rnr1Y285A | FALSE | CAN | JSY3869 | 3 | rnr1Y285A-R | GCC | C>T | A/T+1 | G/C-1 | CG>TA | CG>TA | TAATAGCCAT | CCATTCG | CGG | GCC | C | A | C>T | C>T_GCC | CG>TA_CCATTCG | 5 | FALSE | FALSE |
| 64_v.txt | 32115 | 32115-SNV-C | SNV | 1 | C | T | 47 | 19297 | 0.24356118 | 58.5908571 | rnr1Y285A | FALSE | CAN | JSY3869 | 3 | rnr1Y285A-R | CCA | C>T | A/T+1 | G/C-1 | CG>TA | CG>TA | AATAGCCATT | CATTCGA | GGT | CCA | C | A | C>T | C>T_CCA | CG>TA_CATTCGA | 5 | FALSE | FALSE |
| 64_v.txt | 32165 | 32165-SNV-G | SNV | 1 | G | A | 32 | 19297 | 0.16582889 | 49.96875 | rnr1Y285A | FALSE | CAN | JSY3869 | 3 | rnr1Y285A-R | CGC | G>A | A/T+1 | G/C-1 | CG>TA | CG>TA | CAAAGCGCCA | GCCAAAT | GCG | GCG | C | T | C>T | C>T_GCG | CG>TA_GCCAAAT | 5 | FALSE | FALSE |
| 64_v.txt | 32176 | 32176-SNV-C | SNV | 1 | C | T | 18 | 19297 | 0.09327875 | 46.1111111 | rnr1Y285A | FALSE | CAN | JSY3869 | 3 | rnr1Y285A-R | GCA | C>T | A/T+1 | G/C-1 | CG>TA | CG>TA | ATGCAGCAG | CAGTAAC | CGT | GCA | C | A | C>T | C>T_GCA | CG>TA_CAGTAAC | 5 | FALSE | FALSE |
| 64_v.txt | 32259 | 32259-SNV-C | SNV | 1 | C | T | 50 | 23523 | 0.21255792 | 31.1 | rnr1Y285A | FALSE | CAN | JSY3869 | 3 | rnr1Y285A-R | ACC | C>T | A/T+1 | G/C-1 | CG>TA | CG>TA | GATAGACCAA | CCAAATA | TGG | ACC | C | A | C>T | C>T_ACC | CG>TA_CCAAATA | 5 | FALSE | FALSE |
| 64_v.txt | 32271 | 32271-SNV-C | SNV | 1 | C | T | 38 | 23523 | 0.16154402 | 34.1825856 | rnr1Y285A | FALSE | CAN | JSY3869 | 3 | rnr1Y285A-R | ACG | C>T | A/T+1 | G/C-1 | CG>TA | CG>TA | AAAATACGG | CGGGAAC | TGC | ACG | C | A | C>T | C>T_ACG | CG>TA_CGGGAAC | 5 | FALSE | FALSE |
| 64_v.txt | 32274 | 32274-SNV-G | SNV | 1 | G | A | 25 | 23523 | 0.10627896 | 35.12 | rnr1Y285A | FALSE | CAN | JSY3869 | 3 | rnr1Y285A-R | GGA | G>A | A/T+1 | G/C-1 | CG>TA | CG>TA | ATACGGGAA | GAACCAA | CCT | CCT | C | T | C>T | C>T_CCT | CG>TA_GAACCAA | 5 | FALSE | FALSE |
| 64_v.txt | 32299 | 32299-SNV-C | SNV | 1 | C | T | 38 | 23523 | 0.16154402 | 56.0789474 | rnr1Y285A | FALSE | CAN | JSY3869 | 3 | rnr1Y285A-R | GCG | C>T | A/T+1 | G/C-1 | CG>TA | CG>TA | AATTTGCGGC | CGGCAGA | CGC | GCG | C | A | C>T | C>T_GCG | CG>TA_CGGCAGA | 5 | FALSE | FALSE |
| 64_v.txt | 32301 | 32301-SNV-G | SNV | 1 | G | A | 35 | 23523 | 0.14879055 | 54.8857143 | rnr1Y285A | FALSE | CAN | JSY3869 | 3 | rnr1Y285A-R | GGC | G>A | A/T+1 | G/C-1 | CG>TA | CG>TA | TTTGCGGCAC | GCAGAAA | CCG | CCG | C | T | C>T | C>T_CCG | CG>TA_GCAGAAA | 5 | FALSE | FALSE |
| 64_v.txt | 32413 | 32413-SNV-G | SNV | 1 | G | A | 21 | 23523 | 0.08927433 | 58.0952381 | rnr1Y285A | FALSE | CAN | JSY3869 | 3 | rnr1Y285A-R | TGT | G>A | A/T+1 | G/C-1 | CG>TA | CG>TA | TAGATTGTGT | GTGTTAG | ACA | ACA | C | T | C>T | C>T_ACA | CG>TA_GTGTTAG | 5 | FALSE | FALSE |
| 64_v.txt | 32466 | 32466-SNV-C | SNV | 1 | C | T | 35 | 23523 | 0.14879055 | 49.7139104 | rnr1Y285A | FALSE | CAN | JSY3869 | 3 | rnr1Y285A-R | GCC | C>T | A/T+1 | G/C-1 | CG>TA | CG>TA | AGAGAGCCA | CCAATGT | CGG | GCC | C | A | C>T | C>T_GCC | CG>TA_CCAATGT | 5 | FALSE | FALSE |
| 64_v.txt | 32467 | 32467-SNV-C | SNV | 1 | C | T | 6 | 23523 | 0.02550695 | 46.499233 | rnr1Y285A | FALSE | CAN | JSY3869 | 3 | rnr1Y285A-R | CCA | C>T | A/T+1 | G/C-1 | CG>TA | CG>TA | GAGAGCCAA | CAATGTA | GGT | CCA | C | A | C>T | C>T_CCA | CG>TA_CAATGTA | 5 | FALSE | FALSE |
| 64_v.txt | 32487 | 32487-SNV-C | SNV | 1 | C | T | 25 | 23523 | 0.10627896 | 39.56 | rnr1Y285A | FALSE | CAN | JSY3869 | 3 | rnr1Y285A-R | ACG | C>T | A/T+1 | G/C-1 | CG>TA | CG>TA | AAGATACGG | CGGAAAA | TGC | ACG | C | A | C>T | C>T_ACG | CG>TA_CGGAAAA | 5 | FALSE | FALSE |
| 64_v.txt | 32508 | 32508-SNV-G | SNV | 1 | G | A | 127 | 23523 | 0.53989712 | 42.8267717 | rnr1Y285A | FALSE | CAN | JSY3869 | 3 | rnr1Y285A-R | GGC | G>A | A/T+1 | G/C-1 | CG>TA | CG>TA | TTGATGGCTC | GCTCTTG | CCG | CCG | C | T | C>T | C>T_CCG | CG>TA_GCTCTTG | 5 | FALSE | FALSE |
| 64_v.txt | 32535 | 32535-SNV-G | SNV | 1 | G | A | 38 | 23523 | 0.16154402 | 37.6578947 | rnr1Y285A | FALSE | CAN | JSY3869 | 3 | rnr1Y285A-R | TGC | G>A | A/T+1 | G/C-1 | CG>TA | CG>TA | GGGTTTGCAC | GCAGCTT | ACG | ACG | C | T | C>T | C>T_ACG | CG>TA_GCAGCTT | 5 | FALSE | FALSE |
| 64_v.txt | 32536 | 32536-SNV-C | SNV | 1 | C | T | 38 | 23523 | 0.16154402 | 36.8674359 | rnr1Y285A | FALSE | CAN | JSY3869 | 3 | rnr1Y285A-R | GCA | C>T | A/T+1 | G/C-1 | CG>TA | CG>TA | GGTTTGCAGC | CAGCTTC | CGT | GCA | C | A | C>T | C>T_GCA | CG>TA_CAGCTTC | 5 | FALSE | FALSE |
| 64_v.txt | 32566 | 32566-SNV-C | SNV | 1 | C | T | 71 | 24910 | 0.28502609 | 36.6760563 | rnr1Y285A | FALSE | CAN | JSY3869 | 3 | rnr1Y285A-R | TCA | C>T | A/T+1 | G/C-1 | CG>TA | CG>TA | CTAGTTCAGT | CAGTACC | AGT | TCA | C | A | C>T | C>T_TCA | CG>TA_CAGTACC | 5 | FALSE | FALSE |
| 64_v.txt | 32571 | 32571-SNV-C | SNV | 1 | C | T | 121 | 24916 | 0.48563172 | 36.0413223 | rnr1Y285A | FALSE | CAN | JSY3869 | 3 | rnr1Y285A-R | ACC | C>T | A/T+1 | G/C-1 | CG>TA | CG>TA | TCAGTACCTT | CCTTGAA | TGG | ACC | C | A | C>T | C>T_ACC | CG>TA_CCTTGAA | 5 | FALSE | FALSE |
| 64_v.txt | 32590 | 32590-SNV-C | SNV | 1 | C | T | 38 | 24916 | 0.15251244 | 36.5 | rnr1Y285A | FALSE | CAN | JSY3869 | 3 | rnr1Y285A-R | GCG | C>T | A/T+1 | G/C-1 | CG>TA | CG>TA | AGGCAGCGT | CGTTAAT | CGC | GCG | C | A | C>T | C>T_GCG | CG>TA_CGTTAAT | 5 | FALSE | FALSE |
| 64_v.txt | 32700 | 32700-SNV-C | SNV | 1 | C | T | 38 | 24916 | 0.15251244 | 62.2368421 | rnr1Y285A | FALSE | CAN | JSY3869 | 3 | rnr1Y285A-R | GCC | C>T | A/T+1 | G/C-1 | CG>TA | CG>TA | ACTGGGCCG | CCGGTAA | CGG | GCC | C | A | C>T | C>T_GCC | CG>TA_CCGGTAA | 5 | FALSE | FALSE |
| 64_v.txt | 32702 | 32702-SNV-G | SNV | 1 | G | A | 23 | 24916 | 0.09231016 | 56.0869565 | rnr1Y285A | FALSE | CAN | JSY3869 | 3 | rnr1Y285A-R | CGG | G>A | A/T+1 | G/C-1 | CG>TA | CG>TA | TGGGCCGGT | GGTAACC | GCC | GCC | C | T | C>T | C>T_GCC | CG>TA_GGTAACC | 5 | FALSE | FALSE |
| 64_v.txt | 32776 | 32776-SNV-C | SNV | 1 | C | T | 27 | 24916 | 0.1083641 | 49.4814815 | rnr1Y285A | FALSE | CAN | JSY3869 | 3 | rnr1Y285A-R | GCG | C>T | A/T+1 | G/C-1 | CG>TA | CG>TA | TGGAAGCGA | CGACCCA | CGC | GCG | C | A | C>T | C>T_GCG | CG>TA_CGACCCA | 5 | FALSE | FALSE |
| 64_v.txt | 32788 | 32788-SNV-C | SNV | 1 | C | T | 140 | 24915 | 0.5619105 | 47.2278089 | rnr1Y285A | FALSE | CAN | JSY3869 | 3 | rnr1Y285A-R | TCG | C>T | A/T+1 | G/C-1 | CG>TA | CG>TA | AGAACTCGA | CGAATTC | AGC | TCG | C | A | C>T | C>T_TCG | CG>TA_CGAATTC | 5 | FALSE | FALSE |
| 64_v.txt | 32798 | 32798-SNV-G | SNV | 1 | G | A | 38 | 24916 | 0.15251244 | 45.0263158 | rnr1Y285A | FALSE | CAN | JSY3869 | 3 | rnr1Y285A-R | CGT | G>A | A/T+1 | G/C-1 | CG>TA | CG>TA | TTCACCGTAA | GTAATAT | GCA | GCA | C | T | C>T | C>T_GCA | CG>TA_GTAATAT | 5 | FALSE | FALSE |
| 64_v.txt | 32860 | 32860-SNV-C | SNV | 1 | C | T | 25 | 24912 | 0.10035324 | 37.08 | rnr1Y285A | FALSE | CAN | JSY3869 | 3 | rnr1Y285A-R | GCC | C>T | A/T+1 | G/C-1 | CG>TA | CG>TA | TCCATGCCGC | CCGCCAG | CGG | GCC | C | A | C>T | C>T_GCC | CG>TA_CCGCCAG | 5 | FALSE | FALSE |
| 64_v.txt | 32882 | 32882-SNV-C | SNV | 1 | C | T | 62 | 19701 | 0.31470484 | 29.6935484 | rnr1Y285A | FALSE | CAN | JSY3869 | 3 | rnr1Y285A-R | TCC | C>T | A/T+1 | G/C-1 | CG>TA | CG>TA | GTACGTCCAA | CCAAAAT | AGG | TCC | C | A | C>T | C>T_TCC | CG>TA_CCAAAAT | 5 | FALSE | FALSE |
| 64_v.txt | 32900 | 32900-SNV-G | SNV | 1 | G | A | 27 | 19701 | 0.13704888 | 33.850364 | rnr1Y285A | FALSE | CAN | JSY3869 | 3 | rnr1Y285A-R | GGC | G>A | A/T+1 | G/C-1 | CG>TA | CG>TA | GACTTGGCCA | GCCAACT | CCG | CCG | C | T | C>T | C>T_CCG | CG>TA_GCCAACT | 5 | FALSE | FALSE |
| 64_v.txt | 32901 | 32901-SNV-C | SNV | 1 | C | T | 38 | 19701 | 0.19288361 | 30.5004472 | rnr1Y285A | FALSE | CAN | JSY3869 | 3 | rnr1Y285A-R | GCC | C>T | A/T+1 | G/C-1 | CG>TA | CG>TA | ACTTGGCCAA | CCAACTA | CGG | GCC | C | A | C>T | C>T_GCC | CG>TA_CCAACTA | 5 | FALSE | FALSE |
| 64_v.txt | 32940 | 32940-SNV-G | SNV | 1 | G | A | 10 | 19702 | 0.05075627 | 57.2 | rnr1Y285A | FALSE | CAN | JSY3869 | 3 | rnr1Y285A-R | AGA | G>A | A/T+1 | G/C-1 | CG>TA | CG>TA | GCCCAAGAA | GAAAACC | TCT | TCT | C | T | C>T | C>T_TCT | CG>TA_GAAAACC | 5 | FALSE | FALSE |
| 64_v.txt | 32945 | 32945-SNV-C | SNV | 1 | C | T | 156 | 19702 | 0.79179779 | 56.3461539 | rnr1Y285A | FALSE | CAN | JSY3869 | 3 | rnr1Y285A-R | ACC | C>T | A/T+1 | G/C-1 | CG>TA | CG>TA | AGAAAACCA | CCAATAC | TGG | ACC | C | A | C>T | C>T_ACC | CG>TA_CCAATAC | 5 | FALSE | FALSE |
| 64_v.txt | 32958 | 32958-SNV-C | SNV | 1 | C | T | 77 | 19702 | 0.39082327 | 52.3116883 | rnr1Y285A | FALSE | CAN | JSY3869 | 3 | rnr1Y285A-R | ACC | C>T | A/T+1 | G/C-1 | CG>TA | CG>TA | ATGTAACCAT | CCATTGG | TGG | ACC | C | A | C>T | C>T_ACC | CG>TA_CCATTGG | 5 | FALSE | FALSE |
| 64_v.txt | 32965 | 32965-SNV-C | SNV | 1 | C | T | 29 | 19702 | 0.14719318 | 54.6896552 | rnr1Y285A | FALSE | CAN | JSY3869 | 3 | rnr1Y285A-R | GCC | C>T | A/T+1 | G/C-1 | CG>TA | CG>TA | CATTGGCCGC | CCGCACC | CGG | GCC | C | A | C>T | C>T_GCC | CG>TA_CCGCACC | 5 | FALSE | FALSE |
| 64_v.txt | 32967 | 32967-SNV-G | SNV | 1 | G | A | 43 | 19702 | 0.21825195 | 59.7441861 | rnr1Y285A | FALSE | CAN | JSY3869 | 3 | rnr1Y285A-R | CGC | G>A | A/T+1 | G/C-1 | CG>TA | CG>TA | TTGGCCGCAC | GCACCAA | GCG | GCG | C | T | C>T | C>T_GCG | CG>TA_GCACCAA | 5 | FALSE | FALSE |
| 64_v.txt | 33056 | 33056-SNV-C | SNV | 1 | C | T | 24 | 19702 | 0.12181504 | 55.7083333 | rnr1Y285A | FALSE | CAN | JSY3869 | 3 | rnr1Y285A-R | GCG | C>T | A/T+1 | G/C-1 | CG>TA | CG>TA | GGACTGCGTC | CGTGACA | CGC | GCG | C | A | C>T | C>T_GCG | CG>TA_CGTGACA | 5 | FALSE | FALSE |
| 64_v.txt | 33070 | 33070-SNV-C | SNV | 1 | C | T | 20 | 19702 | 0.10151254 | 56.05 | rnr1Y285A | FALSE | CAN | JSY3869 | 3 | rnr1Y285A-R | GCC | C>T | A/T+1 | G/C-1 | CG>TA | CG>TA | AATATGCCAA | CCAAAGA | CGG | GCC | C | A | C>T | C>T_GCC | CG>TA_CCAAAGA | 5 | FALSE | FALSE |
| 64_v.txt | 33078 | 33078-SNV-C | SNV | 1 | C | T | 7 | 19702 | 0.03552939 | 51.5712867 | rnr1Y285A | FALSE | CAN | JSY3869 | 3 | rnr1Y285A-R | ACC | C>T | A/T+1 | G/C-1 | CG>TA | CG>TA | AAAGAACCC | CCCATAA | TGG | ACC | C | A | C>T | C>T_ACC | CG>TA_CCCATAA | 5 | FALSE | FALSE |
| 64_v.txt | 33114 | 33114-SNV-C | SNV | 1 | C | T | 29 | 19702 | 0.14719318 | 41.5862069 | rnr1Y285A | FALSE | CAN | JSY3869 | 3 | rnr1Y285A-R | GCC | C>T | A/T+1 | G/C-1 | CG>TA | CG>TA | ACTGGGCCG | CCGGCGT | CGG | GCC | C | A | C>T | C>T_GCC | CG>TA_CCGGCGT | 5 | FALSE | FALSE |
| 64_v.txt | 33153 | 33153-SNV-C | SNV | 1 | C | T | 204 | 19702 | 1.03542788 | 36.9117647 | rnr1Y285A | FALSE | CAN | JSY3869 | 3 | rnr1Y285A-R | ACC | C>T | A/T+1 | G/C-1 | CG>TA | CG>TA | AAAAGACCTC | CCTGTAC | TGG | ACC | C | A | C>T | C>T_ACC | CG>TA_CCTGTAC | 5 | FALSE | FALSE |
| 64_v.txt | 33168 | 33168-SNV-C | SNV | 1 | C | T | 49 | 19701 | 0.24871834 | 35.8979592 | rnr1Y285A | FALSE | CAN | JSY3869 | 3 | rnr1Y285A-R | ACC | C>T | A/T+1 | G/C-1 | CG>TA | CG>TA | ATAGTACCAC | CCACCAA | TGG | ACC | C | A | C>T | C>T_ACC | CG>TA_CCACCAA | 5 | FALSE | FALSE |
| 64_v.txt | 33177 | 33177-SNV-G | SNV | 1 | G | A | 36 | 19700 | 0.18274112 | 37.5833333 | rnr1Y285A | FALSE | CAN | JSY3869 | 3 | rnr1Y285A-R | GGC | G>A | A/T+1 | G/C-1 | CG>TA | CG>TA | CCAAGGGCA | GCAATCA | CCG | CCG | C | T | C>T | C>T_CCG | CG>TA_GCAATCA | 5 | FALSE | FALSE |
| 64_v.txt | 33258 | 33258-SNV-C | SNV | 1 | C | T | 26 | 17362 | 0.14975233 | 44.6538462 | rnr1Y285A | FALSE | CAN | JSY3869 | 3 | rnr1Y285A-R | GCC | C>T | A/T+1 | G/C-1 | CG>TA | CG>TA | CCTATGCCAT | CCATCCT | CGG | GCC | C | A | C>T | C>T_GCC | CG>TA_CCATCCT | 5 | FALSE | FALSE |
| 64_v.txt | 33284 | 33284-SNV-G | SNV | 1 | G | A | 19 | 17362 | 0.1094344 | 48.6842105 | rnr1Y285A | FALSE | CAN | JSY3869 | 3 | rnr1Y285A-R | CGC | G>A | A/T+1 | G/C-1 | CG>TA | CG>TA | ATCCTCGCCA | GCCATTT | GCG | GCG | C | T | C>T | C>T_GCG | CG>TA_GCCATTT | 5 | FALSE | FALSE |
| 64_v.txt | 33309 | 33309-SNV-C | SNV | 1 | C | T | 22 | 17362 | 0.12671351 | 59.3068693 | rnr1Y285A | FALSE | CAN | JSY3869 | 3 | rnr1Y285A-R | GCG | C>T | A/T+1 | G/C-1 | CG>TA | CG>TA | AAAGAGCGC | CGCAATG | CGC | GCG | C | A | C>T | C>T_GCG | CG>TA_CGCAATG | 5 | FALSE | FALSE |
| 64_v.txt | 33310 | 33310-SNV-G | SNV | 1 | G | A | 17 | 17358 | 0.09793755 | 60.788923 | rnr1Y285A | FALSE | CAN | JSY3869 | 3 | rnr1Y285A-R | CGC | G>A | A/T+1 | G/C-1 | CG>TA | CG>TA | AAGAGCGCA | GCAATGG | GCG | GCG | C | T | C>T | C>T_GCG | CG>TA_GCAATGG | 5 | FALSE | FALSE |
| 64_v.txt | 33392 | 33392-SNV-G | SNV | 1 | G | A | 30 | 17362 | 0.17279115 | 45.1 | rnr1Y285A | FALSE | CAN | JSY3869 | 3 | rnr1Y285A-R | CGT | G>A | A/T+1 | G/C-1 | CG>TA | CG>TA | AACGTCGTGA | GTGAAAG | GCA | GCA | C | T | C>T | C>T_GCA | CG>TA_GTGAAAG | 5 | FALSE | FALSE |
| 64_v.txt | 33412 | 33412-SNV-G | SNV | 1 | G | A | 25 | 17362 | 0.14399263 | 50.76 | rnr1Y285A | FALSE | CAN | JSY3869 | 3 | rnr1Y285A-R | GGC | G>A | A/T+1 | G/C-1 | CG>TA | CG>TA | TGACCGGCTC | GCTCATT | CCG | CCG | C | T | C>T | C>T_CCG | CG>TA_GCTCATT | 5 | FALSE | FALSE |
| 64_v.txt | 33474 | 33474-SNV-C | SNV | 1 | C | T | 20 | 17360 | 0.11520737 | 35.7 | rnr1Y285A | FALSE | CAN | JSY3869 | 3 | rnr1Y285A-R | GCC | C>T | A/T+1 | G/C-1 | CG>TA | CG>TA | GCTATGCCTT | CCTTTTT | CGG | GCC | C | A | C>T | C>T_GCC | CG>TA_CCTTTTT | 5 | FALSE | FALSE |
| 64_v.txt | 31947 | 31947-SNV-G | SNV | 1 | G | C | 2 | 19289 | 0.0103686 | 38 | rnr1Y285A | FALSE | CAN | JSY3869 | 3 | rnr1Y285A-R | CGT | G>C | G/C+1 | G/C-1 | CG>GC | CG>GC | ATGATCGTCA | GTCATAA | GCA | GCA | C | G | C>G | C>G_GCA | CG>GC_GTCATAA | 5 | FALSE | FALSE |
| 64_v.txt | 32467 | 32467-SNV-C | SNV | 1 | C | G | 18 | 23523 | 0.07652085 | 46.2206693 | rnr1Y285A | FALSE | CAN | JSY3869 | 3 | rnr1Y285A-R | CCA | C>G | G/C+1 | G/C-1 | CG>GC | CG>GC | GAGAGCCAA | CAATGTA | GGT | CCA | C | C | C>G | C>G_CCA | CG>GC_CAATGTA | 5 | FALSE | FALSE |

| | | | | | | | | | | | | | | | | | | | | | | | | | | | | | | | | | | |
|---|---|---|---|---|---|---|---|---|---|---|---|---|---|---|---|---|---|---|---|---|---|---|---|---|---|---|---|---|---|---|---|---|---|---|
| 64_v.txt | 32529 | 32529-SNV- | SNV | 1 | G | C | 14 | 23523 | 0.05951622 | 43.5681071 | rnr1Y285A | FALSE | CAN | JSY3869 | 3 | rnr1Y285A-R | GGG | G>C | G/C+1 | G/C-1 | CG>GC | CG>GC | TTTCTGGGG | GGGTTTG | CCC | CCC | C | G | C>G | C>G_CCC | CG>GC_GGGTTTG | 5 | FALSE | FALSE |
| 64_v.txt | 32535 | 32535-SNV- | SNV | 1 | G | C | 2 | 23523 | 8.50E-03 | 25 | rnr1Y285A | FALSE | CAN | JSY3869 | 3 | rnr1Y285A-R | TGC | G>C | G/C+1 | G/C-1 | CG>GC | CG>GC | GGGTTTGCA | GCAGCTT | ACG | ACG | C | G | C>G | C>G_ACG | CG>GC_GCAGCTT | 5 | FALSE | FALSE |
| 64_v.txt | 32544 | 32544-SNV- | SNV | 1 | C | G | 7 | 23522 | 0.02975937 | 38 | rnr1Y285A | FALSE | CAN | JSY3869 | 3 | rnr1Y285A-R | ACC | C>G | G/C+1 | G/C-1 | CG>GC | CG>GC | GCTTCACCA | CCAGCAG | TGG | ACC | C | C | C>G | C>G_ACC | CG>GC_CCAGCAG | 5 | FALSE | FALSE |
| 64_v.txt | 32670 | 32670-SNV- | SNV | 1 | G | C | 2 | 24916 | 8.03E-03 | 62 | rnr1Y285A | FALSE | CAN | JSY3869 | 3 | rnr1Y285A-R | TGG | G>C | G/C+1 | G/C-1 | CG>GC | CG>GC | GCACCTGGG | GGGTTTC | ACC | ACC | C | G | C>G | C>G_ACC | CG>GC_GGGTTTC | 5 | FALSE | FALSE |
| 64_v.txt | 32936 | 32936-SNV- | SNV | 1 | C | G | 9 | 19702 | 0.04568064 | 60.7777778 | rnr1Y285A | FALSE | CAN | JSY3869 | 3 | rnr1Y285A-R | CCC | C>G | G/C+1 | G/C-1 | CG>GC | CG>GC | GATTGCCCA | CCAAGAA | GGG | CCC | C | C | C>G | C>G_CCC | CG>GC_CCAAGAA | 5 | FALSE | FALSE |
| 64_v.txt | 32954 | 32954-SNV- | SNV | 1 | G | C | 3 | 19702 | 0.01522688 | 46.6666667 | rnr1Y285A | FALSE | CAN | JSY3869 | 3 | rnr1Y285A-R | TGT | G>C | G/C+1 | G/C-1 | CG>GC | CG>GC | ATACATGTA | GTAACCA | ACA | ACA | C | G | C>G | C>G_ACA | CG>GC_GTAACCA | 5 | FALSE | FALSE |
| 64_v.txt | 33037 | 33037-SNV- | SNV | 1 | C | G | 17 | 19702 | 0.08628566 | 59.8823529 | rnr1Y285A | FALSE | CAN | JSY3869 | 3 | rnr1Y285A-R | GCC | C>G | G/C+1 | G/C-1 | CG>GC | CG>GC | ATGTAGCCA | CCATTTC | CGG | GCC | C | C | C>G | C>G_GCC | CG>GC_CCATTTC | 5 | FALSE | FALSE |
| 64_v.txt | 33070 | 33070-SNV- | SNV | 1 | C | G | 13 | 19702 | 0.06598315 | 57 | rnr1Y285A | FALSE | CAN | JSY3869 | 3 | rnr1Y285A-R | GCC | C>G | G/C+1 | G/C-1 | CG>GC | CG>GC | AATATGCCA | CCAAAGA | CGG | GCC | C | C | C>G | C>G_GCC | CG>GC_CCAAAGA | 5 | FALSE | FALSE |
| 64_v.txt | 33079 | 33079-SNV- | SNV | 1 | C | G | 38 | 19701 | 0.19288361 | 55.4152154 | rnr1Y285A | FALSE | CAN | JSY3869 | 3 | rnr1Y285A-R | CCC | C>G | G/C+1 | G/C-1 | CG>GC | CG>GC | AAGAACCCA | CCATAAA | GGG | CCC | C | C | C>G | C>G_CCC | CG>GC_CCATAAA | 5 | FALSE | FALSE |
| 64_v.txt | 33153 | 33153-SNV- | SNV | 1 | C | G | 6 | 19702 | 0.03045376 | 37.3333333 | rnr1Y285A | FALSE | CAN | JSY3869 | 3 | rnr1Y285A-R | ACC | C>G | G/C+1 | G/C-1 | CG>GC | CG>GC | AAAAGACCT | CCTGTAC | TGG | ACC | C | C | C>G | C>G_ACC | CG>GC_CCTGTAC | 5 | FALSE | FALSE |
| 64_v.txt | 33195 | 33195-SNV- | SNV | 1 | C | G | 1 | 17362 | 5.76E-03 | 37.9944743 | rnr1Y285A | FALSE | CAN | JSY3869 | 3 | rnr1Y285A-R | TCT | C>G | G/C+1 | G/C-1 | CG>GC | CG>GC | ATATGTCTTT | CTTTGCT | AGA | TCT | C | C | C>G | C>G_TCT | CG>GC_CTTTGCT | 5 | FALSE | FALSE |
| 64_v.txt | 33205 | 33205-SNV- | SNV | 1 | G | C | 17 | 17362 | 0.09791499 | 39.2352941 | rnr1Y285A | FALSE | CAN | JSY3869 | 3 | rnr1Y285A-R | AGC | G>C | G/C+1 | G/C-1 | CG>GC | CG>GC | GCTTAAGCT | GCTCTCT | TCG | TCG | C | G | C>G | C>G_TCG | CG>GC_GCTCTCT | 5 | FALSE | FALSE |
| 64_v.txt | 33309 | 33309-SNV- | SNV | 1 | C | G | 1 | 17362 | 5.76E-03 | 63.9904752 | rnr1Y285A | FALSE | CAN | JSY3869 | 3 | rnr1Y285A-R | GCG | C>G | G/C+1 | G/C-1 | CG>GC | CG>GC | AAAGAGCGC | CGCAATG | CGC | GCG | C | C | C>G | C>G_GCG | CG>GC_CGCAATG | 5 | FALSE | FALSE |
| 64_v.txt | 31657 | 31657-SNV- | SNV | 1 | G | T | 88 | 11779 | 0.74709228 | 19.0454546 | rnr1Y285A | FALSE | CAN | JSY3869 | 3 | rnr1Y285A-R | GGC | G>T | A/T+1 | G/C-1 | CG>AT | CG>AT | GAAATGGCG | GCGTGGA | CCG | CCG | C | A | C>A | C>A_CCG | CG>AT_GCGTGGA | 5 | FALSE | FALSE |
| 64_v.txt | 31869 | 31869-SNV- | SNV | 1 | G | T | 29 | 11779 | 0.24620087 | 45.137931 | rnr1Y285A | FALSE | CAN | JSY3869 | 3 | rnr1Y285A-R | AGA | G>T | A/T+1 | G/C-1 | CG>AT | CG>AT | AAAATAGAG | GAGATAT | TCT | TCT | C | A | C>A | C>A_TCT | CG>AT_GAGATAT | 5 | FALSE | FALSE |
| 64_v.txt | 31981 | 31981-SNV- | SNV | 1 | G | T | 80 | 19297 | 0.41457221 | 45.5125 | rnr1Y285A | FALSE | CAN | JSY3869 | 3 | rnr1Y285A-R | GGC | G>T | A/T+1 | G/C-1 | CG>AT | CG>AT | AGCCGGGCA | GCATTAA | CCG | CCG | C | A | C>A | C>A_CCG | CG>AT_GCATTAA | 5 | FALSE | FALSE |
| 64_v.txt | 32027 | 32027-SNV- | SNV | 1 | G | T | 384 | 19297 | 1.98994662 | 60.2526042 | rnr1Y285A | FALSE | CAN | JSY3869 | 3 | rnr1Y285A-R | GGT | G>T | A/T+1 | G/C-1 | CG>AT | CG>AT | GCCACGGTA | GTATTTC | CCA | CCA | C | A | C>A | C>A_CCA | CG>AT_GTATTTC | 5 | FALSE | FALSE |
| 64_v.txt | 32114 | 32114-SNV- | SNV | 1 | C | A | 4 | 19295 | 0.02073076 | 47.25 | rnr1Y285A | FALSE | CAN | JSY3869 | 3 | rnr1Y285A-R | GCC | C>A | A/T+1 | G/C-1 | CG>AT | CG>AT | TAATAGCCA | CCATTCG | CGG | GCC | C | T | C>A | C>A_GCC | CG>AT_CCATTCG | 5 | FALSE | FALSE |
| 64_v.txt | 32188 | 32188-SNV- | SNV | 1 | C | A | 44 | 19297 | 0.22801472 | 46.2045455 | rnr1Y285A | FALSE | CAN | JSY3869 | 3 | rnr1Y285A-R | ACT | C>A | A/T+1 | G/C-1 | CG>AT | CG>AT | CGAAAACTG | CTGCAAT | TGA | ACT | C | T | C>A | C>A_ACT | CG>AT_CTGCAAT | 5 | FALSE | FALSE |
| 64_v.txt | 32274 | 32274-SNV- | SNV | 1 | G | T | 30 | 23523 | 0.12753475 | 35.4333333 | rnr1Y285A | FALSE | CAN | JSY3869 | 3 | rnr1Y285A-R | GGA | G>T | A/T+1 | G/C-1 | CG>AT | CG>AT | ATACGGGAA | GAACCAA | CCT | CCT | C | A | C>A | C>A_CCT | CG>AT_GAACCAA | 5 | FALSE | FALSE |
| 64_v.txt | 32277 | 32277-SNV- | SNV | 1 | C | A | 154 | 23523 | 0.6546784 | 35.9155844 | rnr1Y285A | FALSE | CAN | JSY3869 | 3 | rnr1Y285A-R | ACC | C>A | A/T+1 | G/C-1 | CG>AT | CG>AT | CGGGAACCA | CCAACGT | TGG | ACC | C | T | C>A | C>A_ACC | CG>AT_CCAACGT | 5 | FALSE | FALSE |
| 64_v.txt | 32301 | 32301-SNV- | SNV | 1 | G | T | 160 | 23523 | 0.68018535 | 52.79375 | rnr1Y285A | FALSE | CAN | JSY3869 | 3 | rnr1Y285A-R | GGC | G>T | A/T+1 | G/C-1 | CG>AT | CG>AT | TTTGCGGCA | GCAGAAA | CCG | CCG | C | A | C>A | C>A_CCG | CG>AT_GCAGAAA | 5 | FALSE | FALSE |
| 64_v.txt | 32304 | 32304-SNV- | SNV | 1 | G | T | 36 | 23523 | 0.1530417 | 54.5555556 | rnr1Y285A | FALSE | CAN | JSY3869 | 3 | rnr1Y285A-R | AGA | G>T | A/T+1 | G/C-1 | CG>AT | CG>AT | GCGGCAGAA | GAAATAA | TCT | TCT | C | A | C>A | C>A_TCT | CG>AT_GAAATAA | 5 | FALSE | FALSE |
| 64_v.txt | 32373 | 32373-SNV- | SNV | 1 | G | T | 18 | 23523 | 0.07652085 | 57.8333333 | rnr1Y285A | FALSE | CAN | JSY3869 | 3 | rnr1Y285A-R | AGC | G>T | A/T+1 | G/C-1 | CG>AT | CG>AT | TCAATAGCA | GCAATAA | TCG | TCG | C | A | C>A | C>A_TCG | CG>AT_GCAATAA | 5 | FALSE | FALSE |
| 64_v.txt | 32466 | 32466-SNV- | SNV | 1 | C | A | 5 | 23523 | 0.02125579 | 37.6000933 | rnr1Y285A | FALSE | CAN | JSY3869 | 3 | rnr1Y285A-R | GCC | C>A | A/T+1 | G/C-1 | CG>AT | CG>AT | AGAGAGCCA | CCAATGT | CGG | GCC | C | T | C>A | C>A_GCC | CG>AT_CCAATGT | 5 | FALSE | FALSE |
| 64_v.txt | 32487 | 32487-SNV- | SNV | 1 | C | A | 23 | 23523 | 0.09777664 | 42.6956522 | rnr1Y285A | FALSE | CAN | JSY3869 | 3 | rnr1Y285A-R | ACG | C>A | A/T+1 | G/C-1 | CG>AT | CG>AT | AAGATACGG | CGGAAAA | TGC | ACG | C | T | C>A | C>A_ACG | CG>AT_CGGAAAA | 5 | FALSE | FALSE |
| 64_v.txt | 32494 | 32494-SNV- | SNV | 1 | C | A | 104 | 23523 | 0.44212048 | 47.6346154 | rnr1Y285A | FALSE | CAN | JSY3869 | 3 | rnr1Y285A-R | ACA | C>A | A/T+1 | G/C-1 | CG>AT | CG>AT | GGAAAACAA | CAACTTT | TGT | ACA | C | T | C>A | C>A_ACA | CG>AT_CAACTTT | 5 | FALSE | FALSE |
| 64_v.txt | 32508 | 32508-SNV- | SNV | 1 | G | T | 14 | 23523 | 0.05951622 | 43.5 | rnr1Y285A | FALSE | CAN | JSY3869 | 3 | rnr1Y285A-R | GGC | G>T | A/T+1 | G/C-1 | CG>AT | CG>AT | TTGATGGCT | GCTCTTG | CCG | CCG | C | A | C>A | C>A_CCG | CG>AT_GCTCTTG | 5 | FALSE | FALSE |
| 64_v.txt | 32556 | 32556-SNV- | SNV | 1 | C | A | 659 | 23516 | 2.80234734 | 37.2776935 | rnr1Y285A | FALSE | CAN | JSY3869 | 3 | rnr1Y285A-R | ACC | C>A | A/T+1 | G/C-1 | CG>AT | CG>AT | GTGATACCA | CCAACTA | TGG | ACC | C | T | C>A | C>A_ACC | CG>AT_CCAACTA | 5 | FALSE | FALSE |
| 64_v.txt | 32566 | 32566-SNV- | SNV | 1 | C | A | 26 | 24910 | 0.10437575 | 36.3461539 | rnr1Y285A | FALSE | CAN | JSY3869 | 3 | rnr1Y285A-R | TCA | C>A | A/T+1 | G/C-1 | CG>AT | CG>AT | CTAGTTCAG | CAGTACC | AGT | TCA | C | T | C>A | C>A_TCA | CG>AT_CAGTACC | 5 | FALSE | FALSE |
| 64_v.txt | 32571 | 32571-SNV- | SNV | 1 | C | A | 91 | 24916 | 0.36522716 | 36.1098901 | rnr1Y285A | FALSE | CAN | JSY3869 | 3 | rnr1Y285A-R | ACC | C>A | A/T+1 | G/C-1 | CG>AT | CG>AT | TCAGTACCTT | CCTTGAA | TGG | ACC | C | T | C>A | C>A_ACC | CG>AT_CCTTGAA | 5 | FALSE | FALSE |
| 64_v.txt | 32572 | 32572-SNV- | SNV | 1 | C | A | 15 | 24916 | 0.06020228 | 36.0643183 | rnr1Y285A | FALSE | CAN | JSY3869 | 3 | rnr1Y285A-R | CCT | C>A | A/T+1 | G/C-1 | CG>AT | CG>AT | CAGTACCTT | CTTGAAA | GGA | CCT | C | T | C>A | C>A_CCT | CG>AT_CTTGAAA | 5 | FALSE | FALSE |
| 64_v.txt | 32691 | 32691-SNV- | SNV | 1 | C | A | 24 | 24916 | 0.09632365 | 60.9583333 | rnr1Y285A | FALSE | CAN | JSY3869 | 3 | rnr1Y285A-R | TCC | C>A | A/T+1 | G/C-1 | CG>AT | CG>AT | CGGAATCCA | CCAACTG | AGG | TCC | C | T | C>A | C>A_TCC | CG>AT_CCAACTG | 5 | FALSE | FALSE |
| 64_v.txt | 32735 | 32735-SNV- | SNV | 1 | G | T | 39 | 24916 | 0.15652593 | 56.974359 | rnr1Y285A | FALSE | CAN | JSY3869 | 3 | rnr1Y285A-R | AGT | G>T | A/T+1 | G/C-1 | CG>AT | CG>AT | AAAACAGTA | GTATATT | TCA | TCA | C | A | C>A | C>A_TCA | CG>AT_GTATATT | 5 | FALSE | FALSE |
| 64_v.txt | 32783 | 32783-SNV- | SNV | 1 | G | T | 89 | 24916 | 0.35720019 | 46.6179775 | rnr1Y285A | FALSE | CAN | JSY3869 | 3 | rnr1Y285A-R | AGA | G>T | A/T+1 | G/C-1 | CG>AT | CG>AT | GACCCAGAA | GAACTCG | TCT | TCT | C | A | C>A | C>A_TCT | CG>AT_GAACTCG | 5 | FALSE | FALSE |
| 64_v.txt | 32794 | 32794-SNV- | SNV | 1 | C | A | 63 | 24916 | 0.25284958 | 41.8571429 | rnr1Y285A | FALSE | CAN | JSY3869 | 3 | rnr1Y285A-R | TCA | C>A | A/T+1 | G/C-1 | CG>AT | CG>AT | CGAATTCAC | CACCGTA | AGT | TCA | C | T | C>A | C>A_TCA | CG>AT_CACCGTA | 5 | FALSE | FALSE |
| 64_v.txt | 32811 | 32811-SNV- | SNV | 1 | G | T | 13 | 24916 | 0.05217531 | 46.2307692 | rnr1Y285A | FALSE | CAN | JSY3869 | 3 | rnr1Y285A-R | AGG | G>T | A/T+1 | G/C-1 | CG>AT | CG>AT | TTGACAGGG | GGGAACA | TCC | TCC | C | A | C>A | C>A_TCC | CG>AT_GGGAACA | 5 | FALSE | FALSE |
| 64_v.txt | 32819 | 32819-SNV- | SNV | 1 | G | T | 44 | 24916 | 0.17659335 | 44.8181818 | rnr1Y285A | FALSE | CAN | JSY3869 | 3 | rnr1Y285A-R | AGT | G>T | A/T+1 | G/C-1 | CG>AT | CG>AT | GAACAAGTT | GTTCATT | TCA | TCA | C | A | C>A | C>A_TCA | CG>AT_GTTCATT | 5 | FALSE | FALSE |
| 64_v.txt | 32917 | 32917-SNV- | SNV | 1 | C | A | 1435 | 19702 | 7.28352452 | 51.5365854 | rnr1Y285A | FALSE | CAN | JSY3869 | 3 | rnr1Y285A-R | TCC | C>A | A/T+1 | G/C-1 | CG>AT | CG>AT | TAAGTTCCA | CCAGGGC | AGG | TCC | C | T | C>A | C>A_TCC | CG>AT_CCAGGGC | 5 | FALSE | FALSE |
| 64_v.txt | 32928 | 32928-SNV- | SNV | 1 | G | T | 16 | 19702 | 0.08121003 | 53.5625 | rnr1Y285A | FALSE | CAN | JSY3869 | 3 | rnr1Y285A-R | AGT | G>T | A/T+1 | G/C-1 | CG>AT | CG>AT | GCAAAAGTG | GTGATTG | TCA | TCA | C | A | C>A | C>A_TCA | CG>AT_GTGATTG | 5 | FALSE | FALSE |
| 64_v.txt | 32940 | 32940-SNV- | SNV | 1 | G | T | 77 | 19702 | 0.39082327 | 50.948052 | rnr1Y285A | FALSE | CAN | JSY3869 | 3 | rnr1Y285A-R | AGA | G>T | A/T+1 | G/C-1 | CG>AT | CG>AT | GCCCAAGAA | GAAAACC | TCT | TCT | C | A | C>A | C>A_TCT | CG>AT_GAAAACC | 5 | FALSE | FALSE |
| 64_v.txt | 32954 | 32954-SNV- | SNV | 1 | G | T | 43 | 19702 | 0.21825195 | 55.2325581 | rnr1Y285A | FALSE | CAN | JSY3869 | 3 | rnr1Y285A-R | TGT | G>T | A/T+1 | G/C-1 | CG>AT | CG>AT | ATACATGTA | GTAACCA | ACA | ACA | C | A | C>A | C>A_ACA | CG>AT_GTAACCA | 5 | FALSE | FALSE |
| 64_v.txt | 32958 | 32958-SNV- | SNV | 1 | C | A | 125 | 19702 | 0.63445336 | 57.728 | rnr1Y285A | FALSE | CAN | JSY3869 | 3 | rnr1Y285A-R | ACC | C>A | A/T+1 | G/C-1 | CG>AT | CG>AT | ATGTAACCA | CCATTGG | TGG | ACC | C | T | C>A | C>A_ACC | CG>AT_CCATTGG | 5 | FALSE | FALSE |
| 64_v.txt | 32970 | 32970-SNV- | SNV | 1 | C | A | 39 | 19702 | 0.19794945 | 61.7435897 | rnr1Y285A | FALSE | CAN | JSY3869 | 3 | rnr1Y285A-R | ACC | C>A | A/T+1 | G/C-1 | CG>AT | CG>AT | GCCGCACCA | CCAAATG | TGG | ACC | C | T | C>A | C>A_ACC | CG>AT_CCAAATG | 5 | FALSE | FALSE |
| 64_v.txt | 33093 | 33093-SNV- | SNV | 1 | G | T | 10 | 19702 | 0.05075627 | 52.7 | rnr1Y285A | FALSE | CAN | JSY3869 | 3 | rnr1Y285A-R | TGA | G>T | A/T+1 | G/C-1 | CG>AT | CG>AT | AAATATGAT | GATATAA | ACT | ACT | C | A | C>A | C>A_ACT | CG>AT_GATATAA | 5 | FALSE | FALSE |
| 64_v.txt | 33114 | 33114-SNV- | SNV | 1 | C | A | 2461 | 19702 | 12.4911177 | 43.6802113 | rnr1Y285A | FALSE | CAN | JSY3869 | 3 | rnr1Y285A-R | GCC | C>A | A/T+1 | G/C-1 | CG>AT | CG>AT | ACTGGGCCG | CCGGCGT | CGG | GCC | C | T | C>A | C>A_GCC | CG>AT_CCGGCGT | 5 | FALSE | FALSE |
| 64_v.txt | 33151 | 33151-SNV- | SNV | 1 | G | T | 124 | 19702 | 0.62937773 | 39.0806452 | rnr1Y285A | FALSE | CAN | JSY3869 | 3 | rnr1Y285A-R | AGA | G>T | A/T+1 | G/C-1 | CG>AT | CG>AT | TGAAAAGAC | GACCTGT | TCT | TCT | C | A | C>A | C>A_TCT | CG>AT_GACCTGT | 5 | FALSE | FALSE |
| 64_v.txt | 33153 | 33153-SNV- | SNV | 1 | C | A | 115 | 19702 | 0.58369709 | 40.3826087 | rnr1Y285A | FALSE | CAN | JSY3869 | 3 | rnr1Y285A-R | ACC | C>A | A/T+1 | G/C-1 | CG>AT | CG>AT | AAAAGACCT | CCTGTAC | TGG | ACC | C | T | C>A | C>A_ACC | CG>AT_CCTGTAC | 5 | FALSE | FALSE |
| 64_v.txt | 33154 | 33154-SNV- | SNV | 1 | C | A | 38 | 19702 | 0.19287382 | 37.341022 | rnr1Y285A | FALSE | CAN | JSY3869 | 3 | rnr1Y285A-R | CCT | C>A | A/T+1 | G/C-1 | CG>AT | CG>AT | AAAGACCTG | CTGTACC | GGA | CCT | C | T | C>A | C>A_CCT | CG>AT_CTGTACC | 5 | FALSE | FALSE |
| 64_v.txt | 33160 | 33160-SNV- | SNV | 1 | C | A | 18 | 19703 | 0.09135665 | 34.0555556 | rnr1Y285A | FALSE | CAN | JSY3869 | 3 | rnr1Y285A-R | CCA | C>A | A/T+1 | G/C-1 | CG>AT | CG>AT | CTGTACCAA | CAATAGT | GGT | CCA | C | T | C>A | C>A_CCA | CG>AT_CAATAGT | 5 | FALSE | FALSE |
| 64_v.txt | 33168 | 33168-SNV- | SNV | 1 | C | A | 304 | 19701 | 1.54306888 | 37.0789474 | rnr1Y285A | FALSE | CAN | JSY3869 | 3 | rnr1Y285A-R | ACC | C>A | A/T+1 | G/C-1 | CG>AT | CG>AT | ATAGTACCA | CCACCAA | TGG | ACC | C | T | C>A | C>A_ACC | CG>AT_CCACCAA | 5 | FALSE | FALSE |
| 64_v.txt | 33177 | 33177-SNV- | SNV | 1 | G | T | 17 | 19700 | 0.08629442 | 36.4705882 | rnr1Y285A | FALSE | CAN | JSY3869 | 3 | rnr1Y285A-R | GGC | G>T | A/T+1 | G/C-1 | CG>AT | CG>AT | CCAAGGGCA | GCAATCA | CCG | CCG | C | A | C>A | C>A_CCG | CG>AT_GCAATCA | 5 | FALSE | FALSE |
| 64_v.txt | 33186 | 33186-SNV- | SNV | 1 | C | A | 46 | 17361 | 0.2649617 | 37.6956522 | rnr1Y285A | FALSE | CAN | JSY3869 | 3 | rnr1Y285A-R | ACC | C>A | A/T+1 | G/C-1 | CG>AT | CG>AT | ATCATACCA | CCAATAT | TGG | ACC | C | T | C>A | C>A_ACC | CG>AT_CCAATAT | 5 | FALSE | FALSE |
| 64_v.txt | 33195 | 33195-SNV- | SNV | 1 | C | A | 31 | 17362 | 0.17855086 | 35.6106132 | rnr1Y285A | FALSE | CAN | JSY3869 | 3 | rnr1Y285A-R | TCT | C>A | A/T+1 | G/C-1 | CG>AT | CG>AT | ATATGTCTTT | CTTTGCT | AGA | TCT | C | T | C>A | C>A_TCT | CG>AT_CTTTGCT | 5 | FALSE | FALSE |
| 64_v.txt | 33205 | 33205-SNV- | SNV | 1 | G | T | 17 | 17362 | 0.09791499 | 41.5294118 | rnr1Y285A | FALSE | CAN | JSY3869 | 3 | rnr1Y285A-R | AGC | G>T | A/T+1 | G/C-1 | CG>AT | CG>AT | GCTTAAGCT | GCTCTCT | TCG | TCG | C | A | C>A | C>A_TCG | CG>AT_GCTCTCT | 5 | FALSE | FALSE |
| 64_v.txt | 33235 | 33235-SNV- | SNV | 1 | C | A | 71 | 17362 | 0.40893906 | 38.0422535 | rnr1Y285A | FALSE | CAN | JSY3869 | 3 | rnr1Y285A-R | TCT | C>A | A/T+1 | G/C-1 | CG>AT | CG>AT | GTACTTCTCC | CTCCTTC | AGA | TCT | C | T | C>A | C>A_TCT | CG>AT_CTCCTTC | 5 | FALSE | FALSE |
| 64_v.txt | 33310 | 33310-SNV- | SNV | 1 | G | T | 9 | 17358 | 0.05184929 | 27.2158051 | rnr1Y285A | FALSE | CAN | JSY3869 | 3 | rnr1Y285A-R | CGC | G>T | A/T+1 | G/C-1 | CG>AT | CG>AT | AAGAGCGCA | GCAATGG | GCG | GCG | C | A | C>A | C>A_GCG | CG>AT_GCAATGG | 5 | FALSE | FALSE |
| 64_v.txt | 33325 | 33325-SNV- | SNV | 1 | C | A | 257 | 17362 | 1.48024421 | 60.2062257 | rnr1Y285A | FALSE | CAN | JSY3869 | 3 | rnr1Y285A-R | TCT | C>A | A/T+1 | G/C-1 | CG>AT | CG>AT | ACAATTCTTT | CTTTACT | AGA | TCT | C | T | C>A | C>A_TCT | CG>AT_CTTTACT | 5 | FALSE | FALSE |
| 64_v.txt | 33474 | 33474-SNV- | SNV | 1 | C | A | 49 | 17360 | 0.28225807 | 32.8163265 | rnr1Y285A | FALSE | CAN | JSY3869 | 3 | rnr1Y285A-R | GCC | C>A | A/T+1 | G/C-1 | CG>AT | CG>AT | GCTATGCCTT | CCTTTTT | CGG | GCC | C | T | C>A | C>A_GCC | CG>AT_CCTTTTT | 5 | FALSE | FALSE |
| 64_v.txt | 32114 | 32114-MNV- | MNV | 2 | CC | TT | 1 | 19297 | 5.18E-03 | 63.9999993 | rnr1Y285A | FALSE | CAN | JSY3869 | 3 | rnr1Y285A-R | GCC | nope | >1bp-ins | >1bpdel | none | MNV | TAATAGCCA | CCATTCG | CGG | GCC | CC | none | NA | NA_GCC | MNV_CCATTCG | 5 | FALSE | FALSE |
| 64_v.txt | 32466 | 32466-MNV- | MNV | 2 | CC | AA | 1 | 23523 | 4.25E-03 | 31.499999 | rnr1Y285A | FALSE | CAN | JSY3869 | 3 | rnr1Y285A-R | GCC | nope | >1bp-ins | >1bpdel | none | MNV | AGAGAGCCA | CCAATGT | CGG | GCC | CC | none | NA | NA_GCC | MNV_CCAATGT | 5 | FALSE | FALSE |
| 64_v.txt | 31723 | 31723-Insert | Insertion | 1 | - | A | 5 | 11779 | 0.04244843 | 64 | rnr1Y285A | FALSE | CAN | JSY3869 | 3 | rnr1Y285A-R | CAA | nope | A/T+1 | >1bpdel | none | A/T+1 | TGTCCCAAA | AAAAAGT | GTT | CAA | - | T | NA | NA_CAA | A/T+1_AAAAAGT | 5 | FALSE | FALSE |
| 64_v.txt | 32081 | 32081-Insert | Insertion | 1 | - | A | 8 | 19297 | 0.04145722 | 64 | rnr1Y285A | FALSE | CAN | JSY3869 | 3 | rnr1Y285A-R | CAA | nope | A/T+1 | >1bpdel | none | A/T+1 | CCATGCAAA | AAAAAAG | GTT | CAA | - | T | NA | NA_CAA | A/T+1_AAAAAAG | 5 | FALSE | FALSE |
| 64_v.txt | 32346 | 32346-Insert | Insertion | 1 | - | A | 6 | 23523 | 0.02550695 | 64 | rnr1Y285A | FALSE | CAN | JSY3869 | 3 | rnr1Y285A-R | CAA | nope | A/T+1 | >1bpdel | none | A/T+1 | TGTGGCAAA | AAAACCT | GTT | CAA | - | T | NA | NA_CAA | A/T+1_AAAACCT | 5 | FALSE | FALSE |
| 64_v.txt | 32498 | 32498-Insert | Insertion | 1 | - | T | 2 | 23523 | 8.50E-03 | 37 | rnr1Y285A | FALSE | CAN | JSY3869 | 3 | rnr1Y285A-R | CTT | nope | A/T+1 | >1bpdel | none | A/T+1 | AACAACTTTT | TTTTTTG | GAA | CTT | - | A | NA | NA_CTT | A/T+1_TTTTTTG | 5 | FALSE | FALSE |
| 64_v.txt | 32842 | 32842-Insert | Insertion | 1 | - | A | 102 | 24916 | 0.4093755 | 42.5686275 | rnr1Y285A | FALSE | CAN | JSY3869 | 3 | rnr1Y285A-R | CAA | nope | A/T+1 | >1bpdel | none | A/T+1 | TTACCCAAA | AAAAAAT | GTT | CAA | - | T | NA | NA_CAA | A/T+1_AAAAAAT | 5 | FALSE | FALSE |
| 64_v.txt | 33089 | 33089-Deleti | Deletion | 2 | AT | - | 28 | 19702 | 0.14211755 | 47.5714286 | rnr1Y285A | FALSE | CAN | JSY3869 | 3 | rnr1Y285A-R | AAT | nope | >1bp-ins | >1bpdel | none | >1bpdel | AAATAAATA | ATATGAT | TTA | AAT | AT | none | NA | NA_AAT | >1bpdel_ATATGAT | 5 | FALSE | FALSE |
| 64_v.txt | 31723 | 31723-Deleti | Deletion | 1 | A | - | 16 | 11779 | 0.13583496 | 58.375 | rnr1Y285A | FALSE | CAN | JSY3869 | 3 | rnr1Y285A-R | CAA | nope | >1bp-ins | A/T-1 | none | A/T-1 | TGTCCCAAA | AAAAAGT | GTT | GTT | T | none | T>none | T>none_GTT | A/T-1_AAAAAGT | 5 | FALSE | FALSE |
| 64_v.txt | 31904 | 31904-Deleti | Deletion | 1 | T | - | 9 | 11779 | 0.07640717 | 46.4444444 | rnr1Y285A | FALSE | CAN | JSY3869 | 3 | rnr1Y285A-R | ATT | nope | >1bp-ins | A/T-1 | none | A/T-1 | ATTGAATTTT | TTTTGGT | TAA | ATT | T | none | nope | nope_ATT | A/T-1_TTTTGGT | 5 | FALSE | FALSE |
| 64_v.txt | 31913 | 31913-Deleti | Deletion | 1 | A | - | 5 | 11779 | 0.04244843 | 43 | rnr1Y285A | FALSE | CAN | JSY3869 | 3 | rnr1Y285A-R | CAA | nope | >1bp-ins | A/T-1 | none | A/T-1 | TGGTGCAAA | AAAAGCC | GTT | GTT | T | none | T>none | T>none_GTT | A/T-1_AAAAGCC | 5 | FALSE | FALSE |
| 64_v.txt | 31972 | 31972-Deleti | Deletion | 1 | C | - | 8 | 19297 | 0.04145722 | 34 | rnr1Y285A | FALSE | CAN | JSY3869 | 3 | rnr1Y285A-R | GCC | nope | >1bp-ins | G/C-1 | none | G/C-1 | AATAAGCCA | CCAAGCC | CGG | GCC | C | none | nope | nope_GCC | G/C-1_CCAAGCC | 5 | FALSE | FALSE |
| 64_v.txt | 31979 | 31979-Deleti | Deletion | 1 | G | - | 57 | 19297 | 0.2953827 | 40.3157895 | rnr1Y285A | FALSE | CAN | JSY3869 | 3 | rnr1Y285A-R | CGG | nope | >1bp-ins | G/C-1 | none | G/C-1 | CAAGCCGGG | GGGCATT | GCC | GCC | C | none | C>none | C>none_GCC | G/C-1_GGGCATT | 5 | FALSE | FALSE |
| 64_v.txt | 32069 | 32069-Deleti | Deletion | 1 | A | - | 6 | 19297 | 0.03109292 | 59.6666667 | rnr1Y285A | FALSE | CAN | JSY3869 | 3 | rnr1Y285A-R | TAA | nope | >1bp-ins | A/T-1 | none | A/T-1 | TGAGATAAA | AAATAAC | ATT | ATT | T | none | T>none | T>none_ATT | A/T-1_AAATAAC | 5 | FALSE | FALSE |
| 64_v.txt | 32075 | 32075-Deleti | Deletion | 1 | C | - | 2 | 19297 | 0.01036431 | 64 | rnr1Y285A | FALSE | CAN | JSY3869 | 3 | rnr1Y285A-R | ACC | nope | >1bp-ins | G/C-1 | none | G/C-1 | AAATAACCA | CCATGCA | TGG | ACC | C | none | nope | nope_ACC | G/C-1_CCATGCA | 5 | FALSE | FALSE |
| 64_v.txt | 32081 | 32081-Deleti | Deletion | 1 | A | - | 12 | 19297 | 0.06218583 | 62.1666667 | rnr1Y285A | FALSE | CAN | JSY3869 | 3 | rnr1Y285A-R | CAA | nope | >1bp-ins | A/T-1 | none | A/T-1 | CCATGCAAA | AAAAAAG | GTT | GTT | T | none | T>none | T>none_GTT | A/T-1_AAAAAAG | 5 | FALSE | FALSE |
| 64_v.txt | 32272 | 32272-Deleti | Deletion | 1 | G | - | 21 | 23523 | 0.08927433 | 37.5238096 | rnr1Y285A | FALSE | CAN | JSY3869 | 3 | rnr1Y285A-R | CGG | nope | >1bp-ins | G/C-1 | none | G/C-1 | AAATACGGG | GGGAACC | GCC | GCC | C | none | C>none | C>none_GCC | G/C-1_GGGAACC | 5 | FALSE | FALSE |
| 64_v.txt | 32466 | 32466-Deleti | Deletion | 1 | C | - | 2 | 23523 | 8.50E-03 | 38 | rnr1Y285A | FALSE | CAN | JSY3869 | 3 | rnr1Y285A-R | GCC | nope | >1bp-ins | G/C-1 | none | G/C-1 | AGAGAGCCA | CCAATGT | CGG | GCC | C | none | nope | nope_GCC | G/C-1_CCAATGT | 5 | FALSE | FALSE |
| 64_v.txt | 32498 | 32498-Deleti | Deletion | 1 | T | - | 25 | 23523 | 0.10627896 | 44.48 | rnr1Y285A | FALSE | CAN | JSY3869 | 3 | rnr1Y285A-R | CTT | nope | >1bp-ins | A/T-1 | none | A/T-1 | AACAACTTTT | TTTTTTG | GAA | CTT | T | none | nope | nope_CTT | A/T-1_TTTTTTG | 5 | FALSE | FALSE |
| 64_v.txt | 32522 | 32522-Deleti | Deletion | 1 | T | - | 6 | 23523 | 0.02550695 | 37.8333333 | rnr1Y285A | FALSE | CAN | JSY3869 | 3 | rnr1Y285A-R | ATT | nope | >1bp-ins | A/T-1 | none | A/T-1 | AACGGATTTT | TTTTCTG | TAA | ATT | T | none | nope | nope_ATT | A/T-1_TTTTCTG | 5 | FALSE | FALSE |
| 64_v.txt | 32528 | 32528-Deleti | Deletion | 1 | G | - | 27 | 23523 | 0.11478128 | 41.8148146 | rnr1Y285A | FALSE | CAN | JSY3869 | 3 | rnr1Y285A-R | TGG | nope | >1bp-ins | G/C-1 | none | G/C-1 | TTTTCTGGGG | GGGGTTT | ACC | ACC | C | none | C>none | C>none_ACC | G/C-1_GGGGTTT | 5 | FALSE | FALSE |
| 64_v.txt | 32556 | 32556-Deleti | Deletion | 1 | C | - | 36 | 23516 | 0.15308726 | 37.8333333 | rnr1Y285A | FALSE | CAN | JSY3869 | 3 | rnr1Y285A-R | ACC | nope | >1bp-ins | G/C-1 | none | G/C-1 | GTGATACCA | CCAACTA | TGG | ACC | C | none | nope | nope_ACC | G/C-1_CCAACTA | 5 | FALSE | FALSE |
| 64_v.txt | 32603 | 32603-Deleti | Deletion | 1 | G | - | 6 | 24916 | 0.02408091 | 42 | rnr1Y285A | FALSE | CAN | JSY3869 | 3 | rnr1Y285A-R | AGG | nope | >1bp-ins | G/C-1 | none | G/C-1 | CAAAGAGGA | GGAAACC | TCC | TCC | C | none | C>none | C>none_TCC | G/C-1_GGAAACC | 5 | FALSE | FALSE |
| 64_v.txt | 32608 | 32608-Deleti | Deletion | 1 | C | - | 3096 | 24916 | 12.4257505 | 44.3307494 | rnr1Y285A | FALSE | CAN | JSY3869 | 3 | rnr1Y285A-R | ACC | nope | >1bp-ins | G/C-1 | none | G/C-1 | AGGAAACCC | CCCAACC | TGG | ACC | C | none | nope | nope_ACC | G/C-1_CCCAACC | 5 | FALSE | FALSE |
| 64_v.txt | 32631 | 32631-Deleti | Deletion | 1 | T | - | 11 | 24916 | 0.04414834 | 48.3636364 | rnr1Y285A | FALSE | CAN | JSY3869 | 3 | rnr1Y285A-R | GTT | nope | >1bp-ins | A/T-1 | none | A/T-1 | CCTTCGTTTT | TTTTTAT | CAA | GTT | T | none | nope | nope_GTT | A/T-1_TTTTTAT | 5 | FALSE | FALSE |
| 64_v.txt | 32640 | 32640-Deleti | Deletion | 1 | T | - | 7 | 24916 | 0.0280944 | 49.1428571 | rnr1Y285A | FALSE | CAN | JSY3869 | 3 | rnr1Y285A-R | CTT | nope | >1bp-ins | A/T-1 | none | A/T-1 | TTATCCTTAG | TTAGATA | GAA | CTT | T | none | nope | nope_CTT | A/T-1_TTAGATA | 5 | FALSE | FALSE |
| 64_v.txt | 32658 | 32658-Deleti | Deletion | 1 | C | - | 4451 | 24916 | 17.8640231 | 60.3886767 | rnr1Y285A | FALSE | CAN | JSY3869 | 3 | rnr1Y285A-R | ACC | nope | >1bp-ins | G/C-1 | none | G/C-1 | CCTGGACCC | CCCCAGG | TGG | ACC | C | none | nope | nope_ACC | G/C-1_CCCCAGG | 5 | FALSE | FALSE |
| 64_v.txt | 32670 | 32670-Deleti | Deletion | 1 | G | - | 339 | 24916 | 1.36057152 | 58.8584071 | rnr1Y285A | FALSE | CAN | JSY3869 | 3 | rnr1Y285A-R | TGG | nope | >1bp-ins | G/C-1 | none | G/C-1 | GCACCTGGG | GGGTTTC | ACC | ACC | C | none | C>none | C>none_ACC | G/C-1_GGGTTTC | 5 | FALSE | FALSE |
| 64_v.txt | 32707 | 32707-Deleti | Deletion | 1 | C | - | 1107 | 24916 | 4.44292824 | 62.0379404 | rnr1Y285A | FALSE | CAN | JSY3869 | 3 | rnr1Y285A-R | ACC | nope | >1bp-ins | G/C-1 | none | G/C-1 | CGGTAACCC | CCCCAGC | TGG | ACC | C | none | nope | nope_ACC | G/C-1_CCCCAGC | 5 | FALSE | FALSE |
| 64_v.txt | 32728 | 32728-Deleti | Deletion | 1 | A | - | 9 | 24916 | 0.03612137 | 60.8888889 | rnr1Y285A | FALSE | CAN | JSY3869 | 3 | rnr1Y285A-R | CAA | nope | >1bp-ins | A/T-1 | none | A/T-1 | CCATACAAA | AAAAACA | GTT | GTT | T | none | T>none | T>none_GTT | A/T-1_AAAAACA | 5 | FALSE | FALSE |
| 64_v.txt | 32747 | 32747-Deleti | Deletion | 1 | C | - | 726 | 24916 | 2.91379034 | 55.4876033 | rnr1Y285A | FALSE | CAN | JSY3869 | 3 | rnr1Y285A-R | ACC | nope | >1bp-ins | G/C-1 | none | G/C-1 | TAGAAACCC | CCCGATA | TGG | ACC | C | none | nope | nope_ACC | G/C-1_CCCGATA | 5 | FALSE | FALSE |
| 64_v.txt | 32779 | 32779-Deleti | Deletion | 1 | C | - | 9 | 24916 | 0.03612137 | 55.2222222 | rnr1Y285A | FALSE | CAN | JSY3869 | 3 | rnr1Y285A-R | ACC | nope | >1bp-ins | G/C-1 | none | G/C-1 | AAGCGACCC | CCCAGAA | TGG | ACC | C | none | nope | nope_ACC | G/C-1_CCCAGAA | 5 | FALSE | FALSE |
| 64_v.txt | 32842 | 32842-Deleti | Deletion | 1 | A | - | 30 | 24916 | 0.12040456 | 44.6999989 | rnr1Y285A | FALSE | CAN | JSY3869 | 3 | rnr1Y285A-R | CAA | nope | >1bp-ins | A/T-1 | none | A/T-1 | TTACCCAAA | AAAAAAT | GTT | GTT | T | none | T>none | T>none_GTT | A/T-1_AAAAAAT | 5 | FALSE | FALSE |
| 64_v.txt | 32901 | 32901-Deleti | Deletion | 1 | C | - | 1 | 19701 | 5.08E-03 | 30 | rnr1Y285A | FALSE | CAN | JSY3869 | 3 | rnr1Y285A-R | GCC | nope | >1bp-ins | G/C-1 | none | G/C-1 | ACTTGGCCA | CCAACTA | CGG | GCC | C | none | nope | nope_GCC | G/C-1_CCAACTA | 5 | FALSE | FALSE |
| 64_v.txt | 32924 | 32924-Deleti | Deletion | 1 | A | - | 22 | 19702 | 0.11166379 | 52 | rnr1Y285A | FALSE | CAN | JSY3869 | 3 | rnr1Y285A-R | CAA | nope | >1bp-ins | A/T-1 | none | A/T-1 | CAGGGCAAA | AAAAGTG | GTT | GTT | T | none | T>none | T>none_GTT | A/T-1_AAAAGTG | 5 | FALSE | FALSE |
| 64_v.txt | 32945 | 32945-Deleti | Deletion | 1 | C | - | 43 | 19702 | 0.21825195 | 60.8139535 | rnr1Y285A | FALSE | CAN | JSY3869 | 3 | rnr1Y285A-R | ACC | nope | >1bp-ins | G/C-1 | none | G/C-1 | AGAAAACCA | CCAATAC | TGG | ACC | C | none | nope | nope_ACC | G/C-1_CCAATAC | 5 | FALSE | FALSE |
| 64_v.txt | 32958 | 32958-Deleti | Deletion | 1 | C | - | 3 | 19702 | 0.01522688 | 64 | rnr1Y285A | FALSE | CAN | JSY3869 | 3 | rnr1Y285A-R | ACC | nope | >1bp-ins | G/C-1 | none | G/C-1 | ATGTAACCA | CCATTGG | TGG | ACC | C | none | nope | nope_ACC | G/C-1_CCATTGG | 5 | FALSE | FALSE |
| 64_v.txt | 33078 | 33078-Deleti | Deletion | 1 | C | - | 8 | 19702 | 0.04060502 | 49.3750016 | rnr1Y285A | FALSE | CAN | JSY3869 | 3 | rnr1Y285A-R | ACC | nope | >1bp-ins | G/C-1 | none | G/C-1 | AAAGAACCC | CCCATAA | TGG | ACC | C | none | nope | nope_ACC | G/C-1_CCCATAA | 5 | FALSE | FALSE |
| 64_v.txt | 33168 | 33168-Deleti | Deletion | 1 | C | - | 3 | 19701 | 0.01522765 | 38 | rnr1Y285A | FALSE | CAN | JSY3869 | 3 | rnr1Y285A-R | ACC | nope | >1bp-ins | G/C-1 | none | G/C-1 | ATAGTACCA | CCACCAA | TGG | ACC | C | none | nope | nope_ACC | G/C-1_CCACCAA | 5 | FALSE | FALSE |
| 64_v.txt | 33356 | 33356-Deleti | Deletion | 1 | C | - | 348 | 17362 | 2.00437738 | 56.5718391 | rnr1Y285A | FALSE | CAN | JSY3869 | 3 | rnr1Y285A-R | ACC | nope | >1bp-ins | G/C-1 | none | G/C-1 | TATTGACCCA | CCCACGT | TGG | ACC | C | none | nope | nope_ACC | G/C-1_CCCACGT | 5 | FALSE | FALSE |
| 64_v.txt | 33424 | 33424-Deleti | Deletion | 1 | T | - | 6 | 17362 | 0.03455823 | 48.3333333 | rnr1Y285A | FALSE | CAN | JSY3869 | 3 | rnr1Y285A-R | ATA | nope | >1bp-ins | A/T-1 | none | A/T-1 | TGTACATATG | TATGCTT | TAT | ATA | T | none | nope | nope_ATA | A/T-1_TATGCTT | 5 | FALSE | FALSE |
| 64_v.txt | 33474 | 33474-Deleti | Deletion | 1 | C | - | 49 | 17360 | 0.28225807 | 29.3469388 | rnr1Y285A | FALSE | CAN | JSY3869 | 3 | rnr1Y285A-R | GCC | nope | >1bp-ins | G/C-1 | none | G/C-1 | GCTATGCCTT | CCTTTTT | CGG | GCC | C | none | nope | nope_GCC | G/C-1_CCTTTTT | 5 | FALSE | FALSE |
| 65_v.txt | 32581 | 32581-SNV-T | SNV | 1 | T | G | 25 | 22365 | 0.1117818 | 33.92 | rnr1Y285A | FALSE | CAN | JSY3869 | 4 | rnr1Y285A-R | GTG | T>G | G/C+1 | A/T-1 | TA>GC | TA>GC | GAAATGTGA | TGAAGGC | CAC | GTG | T | C | T>G | T>G_GTG | TA>GC_TGAAGGC | 5 | FALSE | FALSE |
| 65_v.txt | 32884 | 32884-SNV- | SNV | 1 | A | C | 9 | 29325 | 0.03069054 | 32.4444444 | rnr1Y285A | FALSE | CAN | JSY3869 | 4 | rnr1Y285A-R | CAA | A>C | G/C+1 | A/T-1 | TA>GC | TA>GC | ACGTCCAAA | AAAATTG | GTT | GTT | T | G | T>G | T>G_GTT | TA>GC_AAAATTG | 5 | FALSE | FALSE |
| 65_v.txt | 33099 | 33099-SNV- | SNV | 1 | A | C | 9 | 29325 | 0.03069054 | 34.2222222 | rnr1Y285A | FALSE | CAN | JSY3869 | 4 | rnr1Y285A-R | AAG | A>C | G/C+1 | A/T-1 | TA>GC | TA>GC | GATATAAGA | AGAGCGC | TTC | TTC | T | G | T>G | T>G_TTC | TA>GC_AGAGCGC | 5 | FALSE | FALSE |
| 65_v.txt | 33150 | 33150-SNV- | SNV | 1 | A | C | 18 | 29325 | 0.06138107 | 37.4993306 | rnr1Y285A | FALSE | CAN | JSY3869 | 4 | rnr1Y285A-R | AAG | A>C | G/C+1 | A/T-1 | TA>GC | TA>GC | ATGAAAAGA | AGACCTG | TTC | TTC | T | G | T>G | T>G_TTC | TA>GC_AGACCTG | 5 | FALSE | FALSE |
| 65_v.txt | 33269 | 33269-SNV- | SNV | 1 | A | C | 108 | 15273 | 0.70713023 | 15.8148148 | rnr1Y285A | FALSE | CAN | JSY3869 | 4 | rnr1Y285A-R | TAG | A>C | G/C+1 | A/T-1 | TA>GC | TA>GC | CTCCATAGA | AGAGAAC | ATC | ATC | T | G | T>G | T>G_ATC | TA>GC_AGAGAAC | 5 | FALSE | FALSE |
| 65_v.txt | 32338 | 32338-SNV-T | SNV | 1 | T | C | 5 | 36638 | 0.01364703 | 51.9999804 | rnr1Y285A | FALSE | CAN | JSY3869 | 4 | rnr1Y285A-R | ATA | T>C | G/C+1 | A/T-1 | TA>CG | TA>CG | TGAAGATAT | TATGTGG | TAT | ATA | T | G | T>C | T>C_ATA | TA>CG_TATGTGG | 5 | FALSE | FALSE |
| 65_v.txt | 32498 | 32498-SNV-T | SNV | 1 | T | C | 6 | 36638 | 0.01637644 | 37.4999749 | rnr1Y285A | FALSE | CAN | JSY3869 | 4 | rnr1Y285A-R | CTT | T>C | G/C+1 | A/T-1 | TA>CG | TA>CG | AACAACTTTT | TTTTTTG | GAA | CTT | T | G | T>C | T>C_CTT | TA>CG_TTTTTTG | 5 | FALSE | FALSE |
| 65_v.txt | 33099 | 33099-SNV- | SNV | 1 | A | G | 16 | 29325 | 0.05456096 | 45.0625 | rnr1Y285A | FALSE | CAN | JSY3869 | 4 | rnr1Y285A-R | AAG | A>G | G/C+1 | A/T-1 | TA>CG | TA>CG | GATATAAGA | AGAGCGC | TTC | TTC | T | C | T>C | T>C_TTC | TA>CG_AGAGCGC | 5 | FALSE | FALSE |
| 65_v.txt | 33150 | 33150-SNV- | SNV | 1 | A | G | 12 | 29325 | 0.04092072 | 32.916185 | rnr1Y285A | FALSE | CAN | JSY3869 | 4 | rnr1Y285A-R | AAG | A>G | G/C+1 | A/T-1 | TA>CG | TA>CG | ATGAAAAGA | AGACCTG | TTC | TTC | T | C | T>C | T>C_TTC | TA>CG_AGACCTG | 5 | FALSE | FALSE |
| 65_v.txt | 33476 | 33476-SNV-T | SNV | 1 | T | C | 24 | 15273 | 0.15714005 | 35 | rnr1Y285A | FALSE | CAN | JSY3869 | 4 | rnr1Y285A-R | CTT | T>C | G/C+1 | A/T-1 | TA>CG | TA>CG | TATGCCTTTT | TTTTTTT | GAA | CTT | T | G | T>C | T>C_CTT | TA>CG_TTTTTTT | 5 | FALSE | FALSE |
| 65_v.txt | 31816 | 31816-SNV-T | SNV | 1 | T | A | 11 | 11290 | 0.09743136 | 60.7272727 | rnr1Y285A | FALSE | CAN | JSY3869 | 4 | rnr1Y285A-R | CTG | T>A | A/T+1 | A/T-1 | TA>AT | TA>AT | TAAATCTGC | TGCATCT | GAC | CTG | T | T | T>A | T>A_CTG | TA>AT_TGCATCT | 5 | FALSE | FALSE |
| 65_v.txt | 32055 | 32055-SNV-T | SNV | 1 | T | A | 34 | 21944 | 0.15493985 | 61.2352941 | rnr1Y285A | FALSE | CAN | JSY3869 | 4 | rnr1Y285A-R | GTG | T>A | A/T+1 | A/T-1 | TA>AT | TA>AT | CTGATGTGC | TGCGAGA | CAC | GTG | T | T | T>A | T>A_GTG | TA>AT_TGCGAGA | 5 | FALSE | FALSE |
| 65_v.txt | 32160 | 32160-SNV- | SNV | 1 | A | T | 13 | 21944 | 0.05924171 | 52.8461539 | rnr1Y285A | FALSE | CAN | JSY3869 | 4 | rnr1Y285A-R | CAA | A>T | A/T+1 | A/T-1 | TA>AT | TA>AT | TAAGCCAAA | AAAGCGC | GTT | GTT | T | A | T>A | T>A_GTT | TA>AT_AAAGCGC | 5 | FALSE | FALSE |
| 65_v.txt | 32338 | 32338-SNV-T | SNV | 1 | T | A | 4 | 36638 | 0.01091763 | 62.9995706 | rnr1Y285A | FALSE | CAN | JSY3869 | 4 | rnr1Y285A-R | ATA | T>A | A/T+1 | A/T-1 | TA>AT | TA>AT | TGAAGATAT | TATGTGG | TAT | ATA | T | T | T>A | T>A_ATA | TA>AT_TATGTGG | 5 | FALSE | FALSE |
| 65_v.txt | 32498 | 32498-SNV-T | SNV | 1 | T | A | 5 | 36638 | 0.01364703 | 42.9997412 | rnr1Y285A | FALSE | CAN | JSY3869 | 4 | rnr1Y285A-R | CTT | T>A | A/T+1 | A/T-1 | TA>AT | TA>AT | AACAACTTTT | TTTTTTG | GAA | CTT | T | T | T>A | T>A_CTT | TA>AT_TTTTTTG | 5 | FALSE | FALSE |
| 65_v.txt | 33099 | 33099-SNV- | SNV | 1 | A | T | 12 | 29325 | 0.04092072 | 47 | rnr1Y285A | FALSE | CAN | JSY3869 | 4 | rnr1Y285A-R | AAG | A>T | A/T+1 | A/T-1 | TA>AT | TA>AT | GATATAAGA | AGAGCGC | TTC | TTC | T | A | T>A | T>A_TTC | TA>AT_AGAGCGC | 5 | FALSE | FALSE |
| 65_v.txt | 31650 | 31650-SNV- | SNV | 1 | C | T | 31 | 11290 | 0.27457927 | 36.6774194 | rnr1Y285A | FALSE | CAN | JSY3869 | 4 | rnr1Y285A-R | GCG | C>T | A/T+1 | G/C-1 | CG>TA | CG>TA | AGAATGCGA | CGAAATG | CGC | GCG | C | A | C>T | C>T_GCG | CG>TA_CGAAATG | 5 | FALSE | FALSE |
| 65_v.txt | 31844 | 31844-SNV- | SNV | 1 | C | T | 32 | 11290 | 0.28343667 | 47.5 | rnr1Y285A | FALSE | CAN | JSY3869 | 4 | rnr1Y285A-R | TCC | C>T | A/T+1 | G/C-1 | CG>TA | CG>TA | TAAGATCCA | CCAAACA | AGG | TCC | C | A | C>T | C>T_TCC | CG>TA_CCAAACA | 5 | FALSE | FALSE |

| | | | | | | | | | | | | | | | | | | | | | | | | | | | | | | | | | | |
|---|---|---|---|---|---|---|---|---|---|---|---|---|---|---|---|---|---|---|---|---|---|---|---|---|---|---|---|---|---|---|---|---|---|---|
| 65_v.txt | 31882 | 31882-SNV-C | SNV | 1 | C | T | 17 | 11290 | 0.15057573 | 40.6470588 | rnr1Y285A | FALSE | CAN | JSY3869 | 4 | rnr1Y285A-R | GCA | C>T | A/T+1 | G/C-1 | CG>TA | CG>TA | AGGCGGCAG | CAGCAAA | CGT | GCA | C | A | C>T | C>T_GCA | CG>TA_CAGCAAA | 5 | FALSE | FALSE |
| 65_v.txt | 31962 | 31962-SNV-C | SNV | 1 | G | A | 38 | 21944 | 0.17316806 | 37.1052632 | rnr1Y285A | FALSE | CAN | JSY3869 | 4 | rnr1Y285A-R | CGC | G>A | A/T+1 | G/C-1 | CG>TA | CG>TA | GTGGCCGCA | GCATAAT | GCG | GCG | C | T | C>T | C>T_GCG | CG>TA_GCATAAT | 5 | FALSE | FALSE |
| 65_v.txt | 31972 | 31972-SNV-C | SNV | 1 | C | T | 20 | 21944 | 0.09114109 | 36.85 | rnr1Y285A | FALSE | CAN | JSY3869 | 4 | rnr1Y285A-R | GCC | C>T | A/T+1 | G/C-1 | CG>TA | CG>TA | AATAAGCCA | CCAAGCC | CGG | GCC | C | A | C>T | C>T_GCC | CG>TA_CCAAGCC | 5 | FALSE | FALSE |
| 65_v.txt | 31981 | 31981-SNV-C | SNV | 1 | G | A | 113 | 21944 | 0.51494714 | 50.4513274 | rnr1Y285A | FALSE | CAN | JSY3869 | 4 | rnr1Y285A-R | GGC | G>A | A/T+1 | G/C-1 | CG>TA | CG>TA | AGCCGGGCA | GCATTAA | CCG | CCG | C | T | C>T | C>T_CCG | CG>TA_GCATTAA | 5 | FALSE | FALSE |
| 65_v.txt | 32057 | 32057-SNV-C | SNV | 1 | C | T | 25 | 21944 | 0.11392636 | 57.96 | rnr1Y285A | FALSE | CAN | JSY3869 | 4 | rnr1Y285A-R | GCG | C>T | A/T+1 | G/C-1 | CG>TA | CG>TA | GATGTGCGA | CGAGATT | CGC | GCG | C | A | C>T | C>T_GCG | CG>TA_CGAGATT | 5 | FALSE | FALSE |
| 65_v.txt | 32075 | 32075-SNV-C | SNV | 1 | C | T | 344 | 21944 | 1.56762669 | 61.7645349 | rnr1Y285A | FALSE | CAN | JSY3869 | 4 | rnr1Y285A-R | ACC | C>T | A/T+1 | G/C-1 | CG>TA | CG>TA | AAATAACCA | CCATGCA | TGG | ACC | C | A | C>T | C>T_ACC | CG>TA_CCATGCA | 5 | FALSE | FALSE |
| 65_v.txt | 32088 | 32088-SNV-C | SNV | 1 | C | T | 157 | 21944 | 0.71545753 | 61.433121 | rnr1Y285A | FALSE | CAN | JSY3869 | 4 | rnr1Y285A-R | GCC | C>T | A/T+1 | G/C-1 | CG>TA | CG>TA | AAAAAGCCT | CCTGCAA | CGG | GCC | C | A | C>T | C>T_GCC | CG>TA_CCTGCAA | 5 | FALSE | FALSE |
| 65_v.txt | 32092 | 32092-SNV-C | SNV | 1 | C | T | 19 | 21944 | 0.08658403 | 60.1578947 | rnr1Y285A | FALSE | CAN | JSY3869 | 4 | rnr1Y285A-R | GCA | C>T | A/T+1 | G/C-1 | CG>TA | CG>TA | AGCCTGCAA | CAACACC | CGT | GCA | C | A | C>T | C>T_GCA | CG>TA_CAACACC | 5 | FALSE | FALSE |
| 65_v.txt | 32114 | 32114-SNV-C | SNV | 1 | C | T | 182 | 21944 | 0.82938389 | 59.7527473 | rnr1Y285A | FALSE | CAN | JSY3869 | 4 | rnr1Y285A-R | GCC | C>T | A/T+1 | G/C-1 | CG>TA | CG>TA | TAATAGCCA | CCATTCG | CGG | GCC | C | A | C>T | C>T_GCC | CG>TA_CCATTCG | 5 | FALSE | FALSE |
| 65_v.txt | 32165 | 32165-SNV-C | SNV | 1 | G | A | 23 | 21944 | 0.10481225 | 48.0869565 | rnr1Y285A | FALSE | CAN | JSY3869 | 4 | rnr1Y285A-R | CGC | G>A | A/T+1 | G/C-1 | CG>TA | CG>TA | CAAAGCGCC | GCCAAAT | GCG | GCG | C | T | C>T | C>T_GCG | CG>TA_GCCAAAT | 5 | FALSE | FALSE |
| 65_v.txt | 32173 | 32173-SNV-C | SNV | 1 | C | T | 26 | 21944 | 0.11848341 | 46.1923077 | rnr1Y285A | FALSE | CAN | JSY3869 | 4 | rnr1Y285A-R | GCA | C>T | A/T+1 | G/C-1 | CG>TA | CG>TA | CAAATGCAG | CAGCAGT | CGT | GCA | C | A | C>T | C>T_GCA | CG>TA_CAGCAGT | 5 | FALSE | FALSE |
| 65_v.txt | 32271 | 32271-SNV-C | SNV | 1 | C | T | 62 | 36638 | 0.16922321 | 30.4350231 | rnr1Y285A | FALSE | CAN | JSY3869 | 4 | rnr1Y285A-R | ACG | C>T | A/T+1 | G/C-1 | CG>TA | CG>TA | AAAATACGG | CGGGAAC | TGC | ACG | C | A | C>T | C>T_ACG | CG>TA_CGGGAAC | 5 | FALSE | FALSE |
| 65_v.txt | 32274 | 32274-SNV-C | SNV | 1 | G | A | 49 | 36638 | 0.13374093 | 35.7346939 | rnr1Y285A | FALSE | CAN | JSY3869 | 4 | rnr1Y285A-R | GGA | G>A | A/T+1 | G/C-1 | CG>TA | CG>TA | ATACGGGAA | GAACCAA | CCT | CCT | C | T | C>T | C>T_CCT | CG>TA_GAACCAA | 5 | FALSE | FALSE |
| 65_v.txt | 32299 | 32299-SNV-C | SNV | 1 | C | T | 80 | 36638 | 0.21835253 | 50.4875 | rnr1Y285A | FALSE | CAN | JSY3869 | 4 | rnr1Y285A-R | GCG | C>T | A/T+1 | G/C-1 | CG>TA | CG>TA | AATTTGCGG | CGGCAGA | CGC | GCG | C | A | C>T | C>T_GCG | CG>TA_CGGCAGA | 5 | FALSE | FALSE |
| 65_v.txt | 32413 | 32413-SNV-C | SNV | 1 | G | A | 5 | 36637 | 0.01364741 | 57.3992854 | rnr1Y285A | FALSE | CAN | JSY3869 | 4 | rnr1Y285A-R | TGT | G>A | A/T+1 | G/C-1 | CG>TA | CG>TA | TAGATTGTG | GTGTTAG | ACA | ACA | C | T | C>T | C>T_ACA | CG>TA_GTGTTAG | 5 | FALSE | FALSE |
| 65_v.txt | 32466 | 32466-SNV-C | SNV | 1 | C | T | 51 | 36638 | 0.13919974 | 51.6470588 | rnr1Y285A | FALSE | CAN | JSY3869 | 4 | rnr1Y285A-R | GCC | C>T | A/T+1 | G/C-1 | CG>TA | CG>TA | AGAGAGCCA | CCAATGT | CGG | GCC | C | A | C>T | C>T_GCC | CG>TA_CCAATGT | 5 | FALSE | FALSE |
| 65_v.txt | 32487 | 32487-SNV-C | SNV | 1 | C | T | 62 | 36638 | 0.16922321 | 44.8709677 | rnr1Y285A | FALSE | CAN | JSY3869 | 4 | rnr1Y285A-R | ACG | C>T | A/T+1 | G/C-1 | CG>TA | CG>TA | AAGATACGG | CGGAAAA | TGC | ACG | C | A | C>T | C>T_ACG | CG>TA_CGGAAAA | 5 | FALSE | FALSE |
| 65_v.txt | 32508 | 32508-SNV-C | SNV | 1 | G | A | 51 | 36638 | 0.13919974 | 41.0784314 | rnr1Y285A | FALSE | CAN | JSY3869 | 4 | rnr1Y285A-R | GGC | G>A | A/T+1 | G/C-1 | CG>TA | CG>TA | TTGATGGCT | GCTCTTG | CCG | CCG | C | T | C>T | C>T_CCG | CG>TA_GCTCTTG | 5 | FALSE | FALSE |
| 65_v.txt | 32535 | 32535-SNV-C | SNV | 1 | G | A | 35 | 36635 | 0.09553706 | 37.3421546 | rnr1Y285A | FALSE | CAN | JSY3869 | 4 | rnr1Y285A-R | TGC | G>A | A/T+1 | G/C-1 | CG>TA | CG>TA | GGGTTTGCA | GCAGCTT | ACG | ACG | C | T | C>T | C>T_ACG | CG>TA_GCAGCTT | 5 | FALSE | FALSE |
| 65_v.txt | 32536 | 32536-SNV-C | SNV | 1 | C | T | 61 | 36635 | 0.16650744 | 36.1221243 | rnr1Y285A | FALSE | CAN | JSY3869 | 4 | rnr1Y285A-R | GCA | C>T | A/T+1 | G/C-1 | CG>TA | CG>TA | GGTTTGCAG | CAGCTTC | CGT | GCA | C | A | C>T | C>T_GCA | CG>TA_CAGCTTC | 5 | FALSE | FALSE |
| 65_v.txt | 32542 | 32542-SNV-C | SNV | 1 | C | T | 134 | 36635 | 0.36577044 | 36.6791045 | rnr1Y285A | FALSE | CAN | JSY3869 | 4 | rnr1Y285A-R | TCA | C>T | A/T+1 | G/C-1 | CG>TA | CG>TA | CAGCTTCAC | CACCAGC | AGT | TCA | C | A | C>T | C>T_TCA | CG>TA_CACCAGC | 5 | FALSE | FALSE |
| 65_v.txt | 32568 | 32568-SNV-C | SNV | 1 | G | A | 19 | 22365 | 0.08495417 | 37.8421053 | rnr1Y285A | FALSE | CAN | JSY3869 | 4 | rnr1Y285A-R | AGT | G>A | A/T+1 | G/C-1 | CG>TA | CG>TA | AGTTCAGTA | GTACCTT | TCA | TCA | C | T | C>T | C>T_TCA | CG>TA_GTACCTT | 5 | FALSE | FALSE |
| 65_v.txt | 32571 | 32571-SNV-C | SNV | 1 | C | T | 30 | 22365 | 0.13413816 | 36.6666667 | rnr1Y285A | FALSE | CAN | JSY3869 | 4 | rnr1Y285A-R | ACC | C>T | A/T+1 | G/C-1 | CG>TA | CG>TA | TCAGTACCT | CCTTGAA | TGG | ACC | C | A | C>T | C>T_ACC | CG>TA_CCTTGAA | 5 | FALSE | FALSE |
| 65_v.txt | 32575 | 32575-SNV-C | SNV | 1 | G | A | 18 | 22365 | 0.0804829 | 37.1666667 | rnr1Y285A | FALSE | CAN | JSY3869 | 4 | rnr1Y285A-R | TGA | G>A | A/T+1 | G/C-1 | CG>TA | CG>TA | TACCTTGAA | GAAATGT | ACT | ACT | C | T | C>T | C>T_ACT | CG>TA_GAAATGT | 5 | FALSE | FALSE |
| 65_v.txt | 32590 | 32590-SNV-C | SNV | 1 | C | T | 33 | 22365 | 0.14755198 | 36.8484849 | rnr1Y285A | FALSE | CAN | JSY3869 | 4 | rnr1Y285A-R | GCG | C>T | A/T+1 | G/C-1 | CG>TA | CG>TA | AGGCAGCGT | CGTTAAT | CGC | GCG | C | A | C>T | C>T_GCG | CG>TA_CGTTAAT | 5 | FALSE | FALSE |
| 65_v.txt | 32700 | 32700-SNV-C | SNV | 1 | C | T | 28 | 22365 | 0.12519562 | 62.6428571 | rnr1Y285A | FALSE | CAN | JSY3869 | 4 | rnr1Y285A-R | GCC | C>T | A/T+1 | G/C-1 | CG>TA | CG>TA | ACTGGGCCG | CCGGTAA | CGG | GCC | C | A | C>T | C>T_GCC | CG>TA_CCGGTAA | 5 | FALSE | FALSE |
| 65_v.txt | 32702 | 32702-SNV-C | SNV | 1 | G | A | 25 | 22365 | 0.1117818 | 53.64 | rnr1Y285A | FALSE | CAN | JSY3869 | 4 | rnr1Y285A-R | CGG | G>A | A/T+1 | G/C-1 | CG>TA | CG>TA | TGGGCCGGT | GGTAACC | GCC | GCC | C | T | C>T | C>T_GCC | CG>TA_GGTAACC | 5 | FALSE | FALSE |
| 65_v.txt | 32757 | 32757-SNV-C | SNV | 1 | G | A | 22 | 22365 | 0.09836799 | 60.7272727 | rnr1Y285A | FALSE | CAN | JSY3869 | 4 | rnr1Y285A-R | GGC | G>A | A/T+1 | G/C-1 | CG>TA | CG>TA | ATAATGGCT | GCTAAAA | CCG | CCG | C | T | C>T | C>T_CCG | CG>TA_GCTAAAA | 5 | FALSE | FALSE |
| 65_v.txt | 32780 | 32780-SNV-C | SNV | 1 | C | T | 27 | 22365 | 0.12072435 | 50.9877035 | rnr1Y285A | FALSE | CAN | JSY3869 | 4 | rnr1Y285A-R | CCC | C>T | A/T+1 | G/C-1 | CG>TA | CG>TA | AGCGACCCA | CCAGAAC | GGG | CCC | C | A | C>T | C>T_CCC | CG>TA_CCAGAAC | 5 | FALSE | FALSE |
| 65_v.txt | 32798 | 32798-SNV-C | SNV | 1 | G | A | 37 | 22365 | 0.16543707 | 48.972973 | rnr1Y285A | FALSE | CAN | JSY3869 | 4 | rnr1Y285A-R | CGT | G>A | A/T+1 | G/C-1 | CG>TA | CG>TA | TTCACCGTA | GTAATAT | GCA | GCA | C | T | C>T | C>T_GCA | CG>TA_GTAATAT | 5 | FALSE | FALSE |
| 65_v.txt | 32855 | 32855-SNV-C | SNV | 1 | C | T | 229 | 22364 | 1.02396709 | 36.0655022 | rnr1Y285A | FALSE | CAN | JSY3869 | 4 | rnr1Y285A-R | TCC | C>T | A/T+1 | G/C-1 | CG>TA | CG>TA | ACTAATCCA | CCATGCC | AGG | TCC | C | A | C>T | C>T_TCC | CG>TA_CCATGCC | 5 | FALSE | FALSE |
| 65_v.txt | 32860 | 32860-SNV-C | SNV | 1 | C | T | 22 | 22364 | 0.09837238 | 36.3636364 | rnr1Y285A | FALSE | CAN | JSY3869 | 4 | rnr1Y285A-R | GCC | C>T | A/T+1 | G/C-1 | CG>TA | CG>TA | TCCATGCCG | CCGCCAG | CGG | GCC | C | A | C>T | C>T_GCC | CG>TA_CCGCCAG | 5 | FALSE | FALSE |
| 65_v.txt | 32862 | 32862-SNV-C | SNV | 1 | G | A | 17 | 22364 | 0.07601502 | 37.4705882 | rnr1Y285A | FALSE | CAN | JSY3869 | 4 | rnr1Y285A-R | CGC | G>A | A/T+1 | G/C-1 | CG>TA | CG>TA | CATGCCGCC | GCCAGTG | GCG | GCG | C | T | C>T | C>T_GCG | CG>TA_GCCAGTG | 5 | FALSE | FALSE |
| 65_v.txt | 32880 | 32880-SNV-C | SNV | 1 | G | A | 24 | 29324 | 0.08184422 | 36.375 | rnr1Y285A | FALSE | CAN | JSY3869 | 4 | rnr1Y285A-R | CGT | G>A | A/T+1 | G/C-1 | CG>TA | CG>TA | TTGTACGTC | GTCCAAA | GCA | GCA | C | T | C>T | C>T_GCA | CG>TA_GTCCAAA | 5 | FALSE | FALSE |
| 65_v.txt | 32900 | 32900-SNV-C | SNV | 1 | G | A | 27 | 29325 | 0.09207161 | 35.0978761 | rnr1Y285A | FALSE | CAN | JSY3869 | 4 | rnr1Y285A-R | GGC | G>A | A/T+1 | G/C-1 | CG>TA | CG>TA | GACTTGGCC | GCCAACT | CCG | CCG | C | T | C>T | C>T_CCG | CG>TA_GCCAACT | 5 | FALSE | FALSE |
| 65_v.txt | 32901 | 32901-SNV-C | SNV | 1 | C | T | 65 | 29325 | 0.22165388 | 30.8379421 | rnr1Y285A | FALSE | CAN | JSY3869 | 4 | rnr1Y285A-R | GCC | C>T | A/T+1 | G/C-1 | CG>TA | CG>TA | ACTTGGCCA | CCAACTA | CGG | GCC | C | A | C>T | C>T_GCC | CG>TA_CCAACTA | 5 | FALSE | FALSE |
| 65_v.txt | 32917 | 32917-SNV-C | SNV | 1 | C | T | 173 | 29325 | 0.58994032 | 49.6473988 | rnr1Y285A | FALSE | CAN | JSY3869 | 4 | rnr1Y285A-R | TCC | C>T | A/T+1 | G/C-1 | CG>TA | CG>TA | TAAGTTCCA | CCAGGGC | AGG | TCC | C | A | C>T | C>T_TCC | CG>TA_CCAGGGC | 5 | FALSE | FALSE |
| 65_v.txt | 32922 | 32922-SNV-C | SNV | 1 | G | A | 23 | 29325 | 0.07843137 | 53.7826087 | rnr1Y285A | FALSE | CAN | JSY3869 | 4 | rnr1Y285A-R | GGC | G>A | A/T+1 | G/C-1 | CG>TA | CG>TA | TCCAGGGCA | GCAAAAG | CCG | CCG | C | T | C>T | C>T_CCG | CG>TA_GCAAAAG | 5 | FALSE | FALSE |
| 65_v.txt | 32945 | 32945-SNV-C | SNV | 1 | C | T | 69 | 29325 | 0.23529412 | 54.9256483 | rnr1Y285A | FALSE | CAN | JSY3869 | 4 | rnr1Y285A-R | ACC | C>T | A/T+1 | G/C-1 | CG>TA | CG>TA | AGAAAACCA | CCAATAC | TGG | ACC | C | A | C>T | C>T_ACC | CG>TA_CCAATAC | 5 | FALSE | FALSE |
| 65_v.txt | 32946 | 32946-SNV-C | SNV | 1 | C | T | 61 | 29325 | 0.20801364 | 55.621694 | rnr1Y285A | FALSE | CAN | JSY3869 | 4 | rnr1Y285A-R | CCA | C>T | A/T+1 | G/C-1 | CG>TA | CG>TA | GAAAACCAA | CAATACA | GGT | CCA | C | A | C>T | C>T_CCA | CG>TA_CAATACA | 5 | FALSE | FALSE |
| 65_v.txt | 32958 | 32958-SNV-C | SNV | 1 | C | T | 218 | 29325 | 0.74339301 | 57.7798165 | rnr1Y285A | FALSE | CAN | JSY3869 | 4 | rnr1Y285A-R | ACC | C>T | A/T+1 | G/C-1 | CG>TA | CG>TA | ATGTAACCA | CCATTGG | TGG | ACC | C | A | C>T | C>T_ACC | CG>TA_CCATTGG | 5 | FALSE | FALSE |
| 65_v.txt | 32965 | 32965-SNV-C | SNV | 1 | C | T | 30 | 29325 | 0.10230179 | 56.1 | rnr1Y285A | FALSE | CAN | JSY3869 | 4 | rnr1Y285A-R | GCC | C>T | A/T+1 | G/C-1 | CG>TA | CG>TA | CATTGGCCG | CCGCACC | CGG | GCC | C | A | C>T | C>T_GCC | CG>TA_CCGCACC | 5 | FALSE | FALSE |
| 65_v.txt | 32967 | 32967-SNV-C | SNV | 1 | G | A | 63 | 29325 | 0.21483376 | 60.9047619 | rnr1Y285A | FALSE | CAN | JSY3869 | 4 | rnr1Y285A-R | CGC | G>A | A/T+1 | G/C-1 | CG>TA | CG>TA | TTGGCCGCA | GCACCAA | GCG | GCG | C | T | C>T | C>T_GCG | CG>TA_GCACCAA | 5 | FALSE | FALSE |
| 65_v.txt | 32977 | 32977-SNV-C | SNV | 1 | C | T | 23 | 29325 | 0.07843137 | 63.826087 | rnr1Y285A | FALSE | CAN | JSY3869 | 4 | rnr1Y285A-R | GCT | C>T | A/T+1 | G/C-1 | CG>TA | CG>TA | CAAATGCTG | CTGGAGA | CGA | GCT | C | A | C>T | C>T_GCT | CG>TA_CTGGAGA | 5 | FALSE | FALSE |
| 65_v.txt | 33056 | 33056-SNV-C | SNV | 1 | C | T | 39 | 29325 | 0.13299233 | 55.7948718 | rnr1Y285A | FALSE | CAN | JSY3869 | 4 | rnr1Y285A-R | GCG | C>T | A/T+1 | G/C-1 | CG>TA | CG>TA | GGACTGCGT | CGTGACA | CGC | GCG | C | A | C>T | C>T_GCG | CG>TA_CGTGACA | 5 | FALSE | FALSE |
| 65_v.txt | 33069 | 33069-SNV-C | SNV | 1 | G | A | 28 | 29325 | 0.09548167 | 51.3217201 | rnr1Y285A | FALSE | CAN | JSY3869 | 4 | rnr1Y285A-R | TGC | G>A | A/T+1 | G/C-1 | CG>TA | CG>TA | GAATATGCC | GCCAAAG | ACG | ACG | C | T | C>T | C>T_ACG | CG>TA_GCCAAAG | 5 | FALSE | FALSE |
| 65_v.txt | 33070 | 33070-SNV-C | SNV | 1 | C | T | 32 | 29323 | 0.10912935 | 55.0904543 | rnr1Y285A | FALSE | CAN | JSY3869 | 4 | rnr1Y285A-R | GCC | C>T | A/T+1 | G/C-1 | CG>TA | CG>TA | AATATGCCA | CCAAAGA | CGG | GCC | C | A | C>T | C>T_GCC | CG>TA_CCAAAGA | 5 | FALSE | FALSE |
| 65_v.txt | 33114 | 33114-SNV-C | SNV | 1 | C | T | 26 | 29325 | 0.08866155 | 44.8856191 | rnr1Y285A | FALSE | CAN | JSY3869 | 4 | rnr1Y285A-R | GCC | C>T | A/T+1 | G/C-1 | CG>TA | CG>TA | ACTGGGCCG | CCGGCGT | CGG | GCC | C | A | C>T | C>T_GCC | CG>TA_CCGGCGT | 5 | FALSE | FALSE |
| 65_v.txt | 33115 | 33115-SNV-C | SNV | 1 | C | T | 54 | 29325 | 0.18414322 | 42.8445137 | rnr1Y285A | FALSE | CAN | JSY3869 | 4 | rnr1Y285A-R | CCG | C>T | A/T+1 | G/C-1 | CG>TA | CG>TA | CTGGGCCGG | CGGCGTT | GGC | CCG | C | A | C>T | C>T_CCG | CG>TA_CGGCGTT | 5 | FALSE | FALSE |
| 65_v.txt | 33118 | 33118-SNV-C | SNV | 1 | C | T | 30 | 29325 | 0.10230179 | 40.8986215 | rnr1Y285A | FALSE | CAN | JSY3869 | 4 | rnr1Y285A-R | GCG | C>T | A/T+1 | G/C-1 | CG>TA | CG>TA | GGCCGGCGT | CGTTGGT | CGC | GCG | C | A | C>T | C>T_GCG | CG>TA_CGTTGGT | 5 | FALSE | FALSE |
| 65_v.txt | 33119 | 33119-SNV-C | SNV | 1 | G | A | 23 | 29325 | 0.07843137 | 43.0434783 | rnr1Y285A | FALSE | CAN | JSY3869 | 4 | rnr1Y285A-R | CGT | G>A | A/T+1 | G/C-1 | CG>TA | CG>TA | GCCGGCGTT | GTTGGTC | GCA | GCA | C | T | C>T | C>T_GCA | CG>TA_GTTGGTC | 5 | FALSE | FALSE |
| 65_v.txt | 33153 | 33153-SNV-C | SNV | 1 | C | T | 551 | 29325 | 1.87894288 | 37.0798548 | rnr1Y285A | FALSE | CAN | JSY3869 | 4 | rnr1Y285A-R | ACC | C>T | A/T+1 | G/C-1 | CG>TA | CG>TA | AAAAGACCT | CCTGTAC | TGG | ACC | C | A | C>T | C>T_ACC | CG>TA_CCTGTAC | 5 | FALSE | FALSE |
| 65_v.txt | 33168 | 33168-SNV-C | SNV | 1 | C | T | 210 | 29323 | 0.71616138 | 37.4952381 | rnr1Y285A | FALSE | CAN | JSY3869 | 4 | rnr1Y285A-R | ACC | C>T | A/T+1 | G/C-1 | CG>TA | CG>TA | ATAGTACCA | CCACCAA | TGG | ACC | C | A | C>T | C>T_ACC | CG>TA_CCACCAA | 5 | FALSE | FALSE |
| 65_v.txt | 33171 | 33171-SNV-C | SNV | 1 | C | T | 30 | 29323 | 0.10230877 | 37.5 | rnr1Y285A | FALSE | CAN | JSY3869 | 4 | rnr1Y285A-R | ACC | C>T | A/T+1 | G/C-1 | CG>TA | CG>TA | GTACCACCA | CCAAGGG | TGG | ACC | C | A | C>T | C>T_ACC | CG>TA_CCAAGGG | 5 | FALSE | FALSE |
| 65_v.txt | 33177 | 33177-SNV-C | SNV | 1 | G | A | 15 | 29319 | 0.05116136 | 37.6 | rnr1Y285A | FALSE | CAN | JSY3869 | 4 | rnr1Y285A-R | GGC | G>A | A/T+1 | G/C-1 | CG>TA | CG>TA | CCAAGGGCA | GCAATCA | CCG | CCG | C | T | C>T | C>T_CCG | CG>TA_GCAATCA | 5 | FALSE | FALSE |
| 65_v.txt | 33229 | 33229-SNV-C | SNV | 1 | G | A | 76 | 15273 | 0.49761016 | 38.8289474 | rnr1Y285A | FALSE | CAN | JSY3869 | 4 | rnr1Y285A-R | TGT | G>A | A/T+1 | G/C-1 | CG>TA | CG>TA | CGTTCTGTAC | GTACTTC | ACA | ACA | C | T | C>T | C>T_ACA | CG>TA_GTACTTC | 5 | FALSE | FALSE |
| 65_v.txt | 33258 | 33258-SNV-C | SNV | 1 | C | T | 20 | 15273 | 0.13095004 | 43.8 | rnr1Y285A | FALSE | CAN | JSY3869 | 4 | rnr1Y285A-R | GCC | C>T | A/T+1 | G/C-1 | CG>TA | CG>TA | CCTATGCCAT | CCATCCT | CGG | GCC | C | A | C>T | C>T_GCC | CG>TA_CCATCCT | 5 | FALSE | FALSE |
| 65_v.txt | 33276 | 33276-SNV-C | SNV | 1 | G | A | 21 | 15273 | 0.13749755 | 43.9047619 | rnr1Y285A | FALSE | CAN | JSY3869 | 4 | rnr1Y285A-R | CGT | G>A | A/T+1 | G/C-1 | CG>TA | CG>TA | GAGAACGTA | GTATCCT | GCA | GCA | C | T | C>T | C>T_GCA | CG>TA_GTATCCT | 5 | FALSE | FALSE |
| 65_v.txt | 33310 | 33310-SNV-C | SNV | 1 | G | A | 24 | 15273 | 0.15714005 | 58.9166667 | rnr1Y285A | FALSE | CAN | JSY3869 | 4 | rnr1Y285A-R | CGC | G>A | A/T+1 | G/C-1 | CG>TA | CG>TA | AAGAGCGCA | GCAATGG | GCG | GCG | C | T | C>T | C>T_GCG | CG>TA_GCAATGG | 5 | FALSE | FALSE |
| 65_v.txt | 33392 | 33392-SNV-C | SNV | 1 | G | A | 20 | 15273 | 0.13095004 | 52.9 | rnr1Y285A | FALSE | CAN | JSY3869 | 4 | rnr1Y285A-R | CGT | G>A | A/T+1 | G/C-1 | CG>TA | CG>TA | AACGTCGTG | GTGAAAG | GCA | GCA | C | T | C>T | C>T_GCA | CG>TA_GTGAAAG | 5 | FALSE | FALSE |
| 65_v.txt | 33412 | 33412-SNV-C | SNV | 1 | G | A | 20 | 15273 | 0.13095004 | 43.5 | rnr1Y285A | FALSE | CAN | JSY3869 | 4 | rnr1Y285A-R | GGC | G>A | A/T+1 | G/C-1 | CG>TA | CG>TA | TGACCGGCT | GCTCATT | CCG | CCG | C | T | C>T | C>T_CCG | CG>TA_GCTCATT | 5 | FALSE | FALSE |
| 65_v.txt | 33474 | 33474-SNV-C | SNV | 1 | C | T | 24 | 15273 | 0.15714005 | 35.1666667 | rnr1Y285A | FALSE | CAN | JSY3869 | 4 | rnr1Y285A-R | GCC | C>T | A/T+1 | G/C-1 | CG>TA | CG>TA | GCTATGCCTT | CCTTTTT | CGG | GCC | C | A | C>T | C>T_GCC | CG>TA_CCTTTTT | 5 | FALSE | FALSE |
| 65_v.txt | 32092 | 32092-SNV-C | SNV | 1 | C | G | 5 | 21944 | 0.02278527 | 64 | rnr1Y285A | FALSE | CAN | JSY3869 | 4 | rnr1Y285A-R | GCA | C>G | G/C+1 | G/C-1 | CG>GC | CG>GC | AGCCTGCAA | CAACACC | CGT | GCA | C | C | C>G | C>G_GCA | CG>GC_CAACACC | 5 | FALSE | FALSE |
| 65_v.txt | 32271 | 32271-SNV-C | SNV | 1 | C | G | 1 | 36638 | 2.73E-03 | 31.9989297 | rnr1Y285A | FALSE | CAN | JSY3869 | 4 | rnr1Y285A-R | ACG | C>G | G/C+1 | G/C-1 | CG>GC | CG>GC | AAAATACGG | CGGGAAC | TGC | ACG | C | C | C>G | C>G_ACG | CG>GC_CGGGAAC | 5 | FALSE | FALSE |
| 65_v.txt | 32866 | 32866-SNV-C | SNV | 1 | G | C | 7 | 22362 | 0.0313031 | 37.4285714 | rnr1Y285A | FALSE | CAN | JSY3869 | 4 | rnr1Y285A-R | AGT | G>C | G/C+1 | G/C-1 | CG>GC | CG>GC | CCGCCAGTG | GTGGAAC | TCA | TCA | C | G | C>G | C>G_TCA | CG>GC_GTGGAAC | 5 | FALSE | FALSE |
| 65_v.txt | 33118 | 33118-SNV-C | SNV | 1 | C | G | 3 | 29325 | 0.01023018 | 27.6668258 | rnr1Y285A | FALSE | CAN | JSY3869 | 4 | rnr1Y285A-R | GCG | C>G | G/C+1 | G/C-1 | CG>GC | CG>GC | GGCCGGCGT | CGTTGGT | CGC | GCG | C | C | C>G | C>G_GCG | CG>GC_CGTTGGT | 5 | FALSE | FALSE |
| 65_v.txt | 33205 | 33205-SNV-C | SNV | 1 | G | C | 6 | 15273 | 0.03928501 | 37.5 | rnr1Y285A | FALSE | CAN | JSY3869 | 4 | rnr1Y285A-R | AGC | G>C | G/C+1 | G/C-1 | CG>GC | CG>GC | GCTTAAGCT | GCTCTCT | TCG | TCG | C | G | C>G | C>G_TCG | CG>GC_GCTCTCT | 5 | FALSE | FALSE |
| 65_v.txt | 33325 | 33325-SNV-C | SNV | 1 | C | G | 3 | 15273 | 0.01964251 | 63.3333333 | rnr1Y285A | FALSE | CAN | JSY3869 | 4 | rnr1Y285A-R | TCT | C>G | G/C+1 | G/C-1 | CG>GC | CG>GC | ACAATTCTTT | CTTTACT | AGA | TCT | C | C | C>G | C>G_TCT | CG>GC_CTTTACT | 5 | FALSE | FALSE |
| 65_v.txt | 31657 | 31657-SNV-C | SNV | 1 | G | T | 98 | 11290 | 0.8680248 | 19.3367347 | rnr1Y285A | FALSE | CAN | JSY3869 | 4 | rnr1Y285A-R | GGC | G>T | A/T+1 | G/C-1 | CG>AT | CG>AT | GAAATGGCG | GCGTGGA | CCG | CCG | C | A | C>A | C>A_CCG | CG>AT_GCGTGGA | 5 | FALSE | FALSE |
| 65_v.txt | 31930 | 31930-SNV-C | SNV | 1 | G | T | 13 | 11289 | 0.11515635 | 37.9230769 | rnr1Y285A | FALSE | CAN | JSY3869 | 4 | rnr1Y285A-R | TGA | G>T | A/T+1 | G/C-1 | CG>AT | CG>AT | AACCTTGAA | GAATAAT | ACT | ACT | C | A | C>A | C>A_ACT | CG>AT_GAATAAT | 5 | FALSE | FALSE |
| 65_v.txt | 31981 | 31981-SNV-C | SNV | 1 | G | T | 45 | 21944 | 0.20506744 | 42.9777778 | rnr1Y285A | FALSE | CAN | JSY3869 | 4 | rnr1Y285A-R | GGC | G>T | A/T+1 | G/C-1 | CG>AT | CG>AT | AGCCGGGCA | GCATTAA | CCG | CCG | C | A | C>A | C>A_CCG | CG>AT_GCATTAA | 5 | FALSE | FALSE |
| 65_v.txt | 32027 | 32027-SNV-C | SNV | 1 | G | T | 352 | 21944 | 1.60408312 | 61.03125 | rnr1Y285A | FALSE | CAN | JSY3869 | 4 | rnr1Y285A-R | GGT | G>T | A/T+1 | G/C-1 | CG>AT | CG>AT | GCCACGGTA | GTATTTC | CCA | CCA | C | A | C>A | C>A_CCA | CG>AT_GTATTTC | 5 | FALSE | FALSE |
| 65_v.txt | 32188 | 32188-SNV-C | SNV | 1 | C | A | 64 | 21944 | 0.29165148 | 45.453125 | rnr1Y285A | FALSE | CAN | JSY3869 | 4 | rnr1Y285A-R | ACT | C>A | A/T+1 | G/C-1 | CG>AT | CG>AT | CGAAAACTG | CTGCAAT | TGA | ACT | C | T | C>A | C>A_ACT | CG>AT_CTGCAAT | 5 | FALSE | FALSE |
| 65_v.txt | 32203 | 32203-SNV-C | SNV | 1 | C | A | 18 | 21944 | 0.08202698 | 38.6111111 | rnr1Y285A | FALSE | CAN | JSY3869 | 4 | rnr1Y285A-R | ACA | C>A | A/T+1 | G/C-1 | CG>AT | CG>AT | ATGGAACAC | CACCACC | TGT | ACA | C | T | C>A | C>A_ACA | CG>AT_CACCACC | 5 | FALSE | FALSE |
| 65_v.txt | 32257 | 32257-SNV-C | SNV | 1 | G | T | 125 | 36628 | 0.34126898 | 35.984 | rnr1Y285A | FALSE | CAN | JSY3869 | 4 | rnr1Y285A-R | AGA | G>T | A/T+1 | G/C-1 | CG>AT | CG>AT | TTGATAGAC | GACCAAA | TCT | TCT | C | A | C>A | C>A_TCT | CG>AT_GACCAAA | 5 | FALSE | FALSE |
| 65_v.txt | 32274 | 32274-SNV-C | SNV | 1 | G | T | 45 | 36638 | 0.1228233 | 36.4222222 | rnr1Y285A | FALSE | CAN | JSY3869 | 4 | rnr1Y285A-R | GGA | G>T | A/T+1 | G/C-1 | CG>AT | CG>AT | ATACGGGAA | GAACCAA | CCT | CCT | C | A | C>A | C>A_CCT | CG>AT_GAACCAA | 5 | FALSE | FALSE |
| 65_v.txt | 32277 | 32277-SNV-C | SNV | 1 | C | A | 51 | 36638 | 0.13919974 | 31.9019608 | rnr1Y285A | FALSE | CAN | JSY3869 | 4 | rnr1Y285A-R | ACC | C>A | A/T+1 | G/C-1 | CG>AT | CG>AT | CGGGAACCA | CCAACGT | TGG | ACC | C | T | C>A | C>A_ACC | CG>AT_CCAACGT | 5 | FALSE | FALSE |
| 65_v.txt | 32304 | 32304-SNV-C | SNV | 1 | G | T | 93 | 36638 | 0.25383482 | 55.2150538 | rnr1Y285A | FALSE | CAN | JSY3869 | 4 | rnr1Y285A-R | AGA | G>T | A/T+1 | G/C-1 | CG>AT | CG>AT | GCGGCAGAA | GAAATAA | TCT | TCT | C | A | C>A | C>A_TCT | CG>AT_GAAATAA | 5 | FALSE | FALSE |
| 65_v.txt | 32373 | 32373-SNV-C | SNV | 1 | G | T | 24 | 36638 | 0.06550576 | 61.375 | rnr1Y285A | FALSE | CAN | JSY3869 | 4 | rnr1Y285A-R | AGC | G>T | A/T+1 | G/C-1 | CG>AT | CG>AT | TCAATAGCA | GCAATAA | TCG | TCG | C | A | C>A | C>A_TCG | CG>AT_GCAATAA | 5 | FALSE | FALSE |
| 65_v.txt | 32556 | 32556-SNV-C | SNV | 1 | C | A | 12462 | 36621 | 34.0296551 | 37.4776922 | rnr1Y285A | FALSE | CAN | JSY3869 | 4 | rnr1Y285A-R | ACC | C>A | A/T+1 | G/C-1 | CG>AT | CG>AT | GTGATACCA | CCAACTA | TGG | ACC | C | T | C>A | C>A_ACC | CG>AT_CCAACTA | 5 | FALSE | FALSE |
| 65_v.txt | 32566 | 32566-SNV-C | SNV | 1 | C | A | 82 | 22363 | 0.3666771 | 36.9756098 | rnr1Y285A | FALSE | CAN | JSY3869 | 4 | rnr1Y285A-R | TCA | C>A | A/T+1 | G/C-1 | CG>AT | CG>AT | CTAGTTCAGT | CAGTACC | AGT | TCA | C | T | C>A | C>A_TCA | CG>AT_CAGTACC | 5 | FALSE | FALSE |
| 65_v.txt | 32571 | 32571-SNV-C | SNV | 1 | C | A | 18 | 22365 | 0.0804829 | 37.6111111 | rnr1Y285A | FALSE | CAN | JSY3869 | 4 | rnr1Y285A-R | ACC | C>A | A/T+1 | G/C-1 | CG>AT | CG>AT | TCAGTACCTT | CCTTGAA | TGG | ACC | C | T | C>A | C>A_ACC | CG>AT_CCTTGAA | 5 | FALSE | FALSE |
| 65_v.txt | 32735 | 32735-SNV-C | SNV | 1 | G | T | 22 | 22365 | 0.09836799 | 58.6363636 | rnr1Y285A | FALSE | CAN | JSY3869 | 4 | rnr1Y285A-R | AGT | G>T | A/T+1 | G/C-1 | CG>AT | CG>AT | AAAACAGTA | GTATATT | TCA | TCA | C | A | C>A | C>A_TCA | CG>AT_GTATATT | 5 | FALSE | FALSE |
| 65_v.txt | 32780 | 32780-SNV-C | SNV | 1 | C | A | 6 | 22365 | 0.02682763 | 29.997966 | rnr1Y285A | FALSE | CAN | JSY3869 | 4 | rnr1Y285A-R | CCC | C>A | A/T+1 | G/C-1 | CG>AT | CG>AT | AGCGACCCA | CCAGAAC | GGG | CCC | C | T | C>A | C>A_CCC | CG>AT_CCAGAAC | 5 | FALSE | FALSE |
| 65_v.txt | 32783 | 32783-SNV-C | SNV | 1 | G | T | 67 | 22365 | 0.29957523 | 51.2985075 | rnr1Y285A | FALSE | CAN | JSY3869 | 4 | rnr1Y285A-R | AGA | G>T | A/T+1 | G/C-1 | CG>AT | CG>AT | GACCCAGAA | GAACTCG | TCT | TCT | C | A | C>A | C>A_TCT | CG>AT_GAACTCG | 5 | FALSE | FALSE |
| 65_v.txt | 32786 | 32786-SNV-C | SNV | 1 | C | A | 36 | 22365 | 0.1609658 | 44.8333333 | rnr1Y285A | FALSE | CAN | JSY3869 | 4 | rnr1Y285A-R | ACT | C>A | A/T+1 | G/C-1 | CG>AT | CG>AT | CCAGAACTC | CTCGAAT | TGA | ACT | C | T | C>A | C>A_ACT | CG>AT_CTCGAAT | 5 | FALSE | FALSE |
| 65_v.txt | 32794 | 32794-SNV-C | SNV | 1 | C | A | 37 | 22365 | 0.16543707 | 41.2972973 | rnr1Y285A | FALSE | CAN | JSY3869 | 4 | rnr1Y285A-R | TCA | C>A | A/T+1 | G/C-1 | CG>AT | CG>AT | CGAATTCAC | CACCGTA | AGT | TCA | C | T | C>A | C>A_TCA | CG>AT_CACCGTA | 5 | FALSE | FALSE |
| 65_v.txt | 32798 | 32798-SNV-C | SNV | 1 | G | T | 42 | 22365 | 0.18779343 | 46.5952381 | rnr1Y285A | FALSE | CAN | JSY3869 | 4 | rnr1Y285A-R | CGT | G>T | A/T+1 | G/C-1 | CG>AT | CG>AT | TTCACCGTA | GTAATAT | GCA | GCA | C | A | C>A | C>A_GCA | CG>AT_GTAATAT | 5 | FALSE | FALSE |
| 65_v.txt | 32819 | 32819-SNV-C | SNV | 1 | G | T | 153 | 22365 | 0.68410463 | 45.0261438 | rnr1Y285A | FALSE | CAN | JSY3869 | 4 | rnr1Y285A-R | AGT | G>T | A/T+1 | G/C-1 | CG>AT | CG>AT | GAACAAGTT | GTTCATT | TCA | TCA | C | A | C>A | C>A_TCA | CG>AT_GTTCATT | 5 | FALSE | FALSE |
| 65_v.txt | 32917 | 32917-SNV-C | SNV | 1 | C | A | 2197 | 29325 | 7.49190111 | 53.2325899 | rnr1Y285A | FALSE | CAN | JSY3869 | 4 | rnr1Y285A-R | TCC | C>A | A/T+1 | G/C-1 | CG>AT | CG>AT | TAAGTTCCA | CCAGGGC | AGG | TCC | C | T | C>A | C>A_TCC | CG>AT_CCAGGGC | 5 | FALSE | FALSE |
| 65_v.txt | 32958 | 32958-SNV-C | SNV | 1 | C | A | 135 | 29325 | 0.46035806 | 57.0222222 | rnr1Y285A | FALSE | CAN | JSY3869 | 4 | rnr1Y285A-R | ACC | C>A | A/T+1 | G/C-1 | CG>AT | CG>AT | ATGTAACCA | CCATTGG | TGG | ACC | C | T | C>A | C>A_ACC | CG>AT_CCATTGG | 5 | FALSE | FALSE |
| 65_v.txt | 33093 | 33093-SNV-C | SNV | 1 | G | T | 32 | 29325 | 0.10912191 | 48.71875 | rnr1Y285A | FALSE | CAN | JSY3869 | 4 | rnr1Y285A-R | TGA | G>T | A/T+1 | G/C-1 | CG>AT | CG>AT | AAATATGAT | GATATAA | ACT | ACT | C | A | C>A | C>A_ACT | CG>AT_GATATAA | 5 | FALSE | FALSE |
| 65_v.txt | 33114 | 33114-SNV-C | SNV | 1 | C | A | 1310 | 29325 | 4.46717818 | 43.3503078 | rnr1Y285A | FALSE | CAN | JSY3869 | 4 | rnr1Y285A-R | GCC | C>A | A/T+1 | G/C-1 | CG>AT | CG>AT | ACTGGGCCG | CCGGCGT | CGG | GCC | C | T | C>A | C>A_GCC | CG>AT_CCGGCGT | 5 | FALSE | FALSE |
| 65_v.txt | 33151 | 33151-SNV-C | SNV | 1 | G | T | 351 | 29325 | 1.19693095 | 39.5982696 | rnr1Y285A | FALSE | CAN | JSY3869 | 4 | rnr1Y285A-R | AGA | G>T | A/T+1 | G/C-1 | CG>AT | CG>AT | TGAAAAGAC | GACCTGT | TCT | TCT | C | A | C>A | C>A_TCT | CG>AT_GACCTGT | 5 | FALSE | FALSE |
| 65_v.txt | 33153 | 33153-SNV-C | SNV | 1 | C | A | 273 | 29325 | 0.93094629 | 39.7948718 | rnr1Y285A | FALSE | CAN | JSY3869 | 4 | rnr1Y285A-R | ACC | C>A | A/T+1 | G/C-1 | CG>AT | CG>AT | AAAAGACCT | CCTGTAC | TGG | ACC | C | T | C>A | C>A_ACC | CG>AT_CCTGTAC | 5 | FALSE | FALSE |
| 65_v.txt | 33159 | 33159-SNV-C | SNV | 1 | C | A | 179 | 29323 | 0.61044232 | 37.7039106 | rnr1Y285A | FALSE | CAN | JSY3869 | 4 | rnr1Y285A-R | ACC | C>A | A/T+1 | G/C-1 | CG>AT | CG>AT | CCTGTACCA | CCAATAG | TGG | ACC | C | T | C>A | C>A_ACC | CG>AT_CCAATAG | 5 | FALSE | FALSE |
| 65_v.txt | 33168 | 33168-SNV-C | SNV | 1 | C | A | 1738 | 29323 | 5.92708795 | 37.3411968 | rnr1Y285A | FALSE | CAN | JSY3869 | 4 | rnr1Y285A-R | ACC | C>A | A/T+1 | G/C-1 | CG>AT | CG>AT | ATAGTACCA | CCACCAA | TGG | ACC | C | T | C>A | C>A_ACC | CG>AT_CCACCAA | 5 | FALSE | FALSE |
| 65_v.txt | 33177 | 33177-SNV-C | SNV | 1 | G | T | 73 | 29319 | 0.2489853 | 36.1643836 | rnr1Y285A | FALSE | CAN | JSY3869 | 4 | rnr1Y285A-R | GGC | G>T | A/T+1 | G/C-1 | CG>AT | CG>AT | CCAAGGGCA | GCAATCA | CCG | CCG | C | A | C>A | C>A_CCG | CG>AT_GCAATCA | 5 | FALSE | FALSE |
| 65_v.txt | 33205 | 33205-SNV-C | SNV | 1 | G | T | 6 | 15273 | 0.03928501 | 33.3333333 | rnr1Y285A | FALSE | CAN | JSY3869 | 4 | rnr1Y285A-R | AGC | G>T | A/T+1 | G/C-1 | CG>AT | CG>AT | GCTTAAGCT | GCTCTCT | TCG | TCG | C | A | C>A | C>A_TCG | CG>AT_GCTCTCT | 5 | FALSE | FALSE |
| 65_v.txt | 33208 | 33208-SNV-C | SNV | 1 | C | A | 114 | 15273 | 0.74641524 | 39.9122807 | rnr1Y285A | FALSE | CAN | JSY3869 | 4 | rnr1Y285A-R | TCT | C>A | A/T+1 | G/C-1 | CG>AT | CG>AT | TAAGCTCTCT | CTCTCTT | AGA | TCT | C | T | C>A | C>A_TCT | CG>AT_CTCTCTT | 5 | FALSE | FALSE |
| 65_v.txt | 33325 | 33325-SNV-C | SNV | 1 | C | A | 211 | 15273 | 1.38152295 | 58.5402844 | rnr1Y285A | FALSE | CAN | JSY3869 | 4 | rnr1Y285A-R | TCT | C>A | A/T+1 | G/C-1 | CG>AT | CG>AT | ACAATTCTTT | CTTTACT | AGA | TCT | C | T | C>A | C>A_TCT | CG>AT_CTTTACT | 5 | FALSE | FALSE |
| 65_v.txt | 33354 | 33354-SNV-C | SNV | 1 | G | T | 112 | 15273 | 0.73332024 | 27.9375 | rnr1Y285A | FALSE | CAN | JSY3869 | 4 | rnr1Y285A-R | TGA | G>T | A/T+1 | G/C-1 | CG>AT | CG>AT | GGTATTGAC | GACCCAC | ACT | ACT | C | A | C>A | C>A_ACT | CG>AT_GACCCAC | 5 | FALSE | FALSE |
| 65_v.txt | 33436 | 33436-SNV-C | SNV | 1 | C | A | 15 | 15273 | 0.09821253 | 41.5333333 | rnr1Y285A | FALSE | CAN | JSY3869 | 4 | rnr1Y285A-R | TCT | C>A | A/T+1 | G/C-1 | CG>AT | CG>AT | TCTCCTCTAT | CTATGTC | AGA | TCT | C | T | C>A | C>A_TCT | CG>AT_CTATGTC | 5 | FALSE | FALSE |
| 65_v.txt | 33474 | 33474-SNV-C | SNV | 1 | C | A | 43 | 15273 | 0.28154259 | 34.255814 | rnr1Y285A | FALSE | CAN | JSY3869 | 4 | rnr1Y285A-R | GCC | C>A | A/T+1 | G/C-1 | CG>AT | CG>AT | GCTATGCCTT | CCTTTTT | CGG | GCC | C | T | C>A | C>A_GCC | CG>AT_CCTTTTT | 5 | FALSE | FALSE |
| 65_v.txt | 33114 | 33114-MNV- | MNV | 2 | CC | AA | 2 | 29325 | 6.82E-03 | 47.2499942 | rnr1Y285A | FALSE | CAN | JSY3869 | 4 | rnr1Y285A-R | GCC | nope | >1bp-ins | >1bpdel | none | MNV | ACTGGGCCG | CCGGCGT | CGG | GCC | CC | none | NA | NA_GCC | MNV_CCGGCGT | 5 | FALSE | FALSE |
| 65_v.txt | 31966 | 31966-Insert | Insertion | 1 | - | A | 5 | 21944 | 0.02278527 | 38 | rnr1Y285A | FALSE | CAN | JSY3869 | 4 | rnr1Y285A-R | TAA | nope | A/T+1 | >1bpdel | none | A/T+1 | CCGCATAAT | AATAAGC | ATT | TAA | - | T | NA | NA_TAA | A/T+1_AATAAGC | 5 | FALSE | FALSE |
| 65_v.txt | 32081 | 32081-Insert | Insertion | 1 | - | A | 24 | 21944 | 0.1093693 | 64 | rnr1Y285A | FALSE | CAN | JSY3869 | 4 | rnr1Y285A-R | CAA | nope | A/T+1 | >1bpdel | none | A/T+1 | CCATGCAAA | AAAAAAG | GTT | CAA | - | T | NA | NA_CAA | A/T+1_AAAAAAG | 5 | FALSE | FALSE |
| 65_v.txt | 32498 | 32498-Insert | Insertion | 1 | - | T | 3 | 36638 | 8.19E-03 | 46.3333333 | rnr1Y285A | FALSE | CAN | JSY3869 | 4 | rnr1Y285A-R | CTT | nope | A/T+1 | >1bpdel | none | A/T+1 | AACAACTTTT | TTTTTTG | GAA | CTT | - | A | NA | NA_CTT | A/T+1_TTTTTTG | 5 | FALSE | FALSE |
| 65_v.txt | 32842 | 32842-Insert | Insertion | 1 | - | A | 565 | 22365 | 2.52626872 | 42.4920354 | rnr1Y285A | FALSE | CAN | JSY3869 | 4 | rnr1Y285A-R | CAA | nope | A/T+1 | >1bpdel | none | A/T+1 | TTACCCAAA | AAAAAAT | GTT | CAA | - | T | NA | NA_CAA | A/T+1_AAAAAAT | 5 | FALSE | FALSE |
| 65_v.txt | 33013 | 33013-Insert | Insertion | 1 | - | A | 16 | 29325 | 0.05456096 | 63.9375 | rnr1Y285A | FALSE | CAN | JSY3869 | 4 | rnr1Y285A-R | GAG | nope | A/T+1 | >1bpdel | none | A/T+1 | TGAAAGAGG | AGGATGT | CTC | GAG | - | T | NA | NA_GAG | A/T+1_AGGATGT | 5 | FALSE | FALSE |
| 65_v.txt | 32337 | 32337-Deleti | Deletion | 2 | AT | - | 127 | 36638 | 0.34663464 | 61.0472441 | rnr1Y285A | FALSE | CAN | JSY3869 | 4 | rnr1Y285A-R | GAT | nope | >1bp-ins | >1bpdel | none | >1bpdel | TTGAAGATA | ATATGTG | CTA | GAT | AT | none | NA | NA_GAT | >1bpdel_ATATGTG | 5 | FALSE | FALSE |
| 65_v.txt | 32360 | 32360-Deleti | Deletion | 2 | AG | - | 9 | 36638 | 0.02456466 | 60.4444444 | rnr1Y285A | FALSE | CAN | JSY3869 | 4 | rnr1Y285A-R | CAG | nope | >1bp-ins | >1bpdel | none | >1bpdel | TGTACCAGA | AGAGTTC | GTC | CAG | AG | none | NA | NA_CAG | >1bpdel_AGAGTTC | 5 | FALSE | FALSE |
| 65_v.txt | 32412 | 32412-Deleti | Deletion | 2 | TG | - | 21 | 36638 | 0.05731754 | 60.8571429 | rnr1Y285A | FALSE | CAN | JSY3869 | 4 | rnr1Y285A-R | TTG | nope | >1bp-ins | >1bpdel | none | >1bpdel | GTAGATTGT | TGTGTTA | AAC | TTG | TG | none | NA | NA_TTG | >1bpdel_TGTGTTA | 5 | FALSE | FALSE |
| 65_v.txt | 31663 | 31663-Deleti | Deletion | 1 | A | - | 6 | 11290 | 0.05314438 | 38.6666667 | rnr1Y285A | FALSE | CAN | JSY3869 | 4 | rnr1Y285A-R | GAA | nope | >1bp-ins | A/T-1 | none | A/T-1 | GCGTGGAAA | AAATGTG | CTT | CTT | T | none | T>none | T>none_CTT | A/T-1_AAATGTG | 5 | FALSE | FALSE |
| 65_v.txt | 31723 | 31723-Deleti | Deletion | 1 | A | - | 14 | 11290 | 0.12400354 | 58.5714286 | rnr1Y285A | FALSE | CAN | JSY3869 | 4 | rnr1Y285A-R | CAA | nope | >1bp-ins | A/T-1 | none | A/T-1 | TGTCCCAAA | AAAAAGT | GTT | GTT | T | none | T>none | T>none_GTT | A/T-1_AAAAAGT | 5 | FALSE | FALSE |
| 65_v.txt | 31904 | 31904-Deleti | Deletion | 1 | T | - | 12 | 11290 | 0.10628875 | 40.25 | rnr1Y285A | FALSE | CAN | JSY3869 | 4 | rnr1Y285A-R | ATT | nope | >1bp-ins | A/T-1 | none | A/T-1 | ATTGAATTTT | TTTTGGT | TAA | ATT | T | none | nope | nope_ATT | A/T-1_TTTTGGT | 5 | FALSE | FALSE |
| 65_v.txt | 31972 | 31972-Deleti | Deletion | 1 | C | - | 7 | 21944 | 0.03189938 | 36.8571429 | rnr1Y285A | FALSE | CAN | JSY3869 | 4 | rnr1Y285A-R | GCC | nope | >1bp-ins | G/C-1 | none | G/C-1 | AATAAGCCA | CCAAGCC | CGG | GCC | C | none | nope | nope_GCC | G/C-1_CCAAGCC | 5 | FALSE | FALSE |
| 65_v.txt | 31979 | 31979-Deleti | Deletion | 1 | G | - | 15 | 21944 | 0.06835582 | 43.3333333 | rnr1Y285A | FALSE | CAN | JSY3869 | 4 | rnr1Y285A-R | CGG | nope | >1bp-ins | G/C-1 | none | G/C-1 | CAAGCCGGG | GGGCATT | GCC | GCC | C | none | C>none | C>none_GCC | G/C-1_GGGCATT | 5 | FALSE | FALSE |
| 65_v.txt | 32075 | 32075-Deleti | Deletion | 1 | C | - | 102 | 21944 | 0.46481954 | 63.0392157 | rnr1Y285A | FALSE | CAN | JSY3869 | 4 | rnr1Y285A-R | ACC | nope | >1bp-ins | G/C-1 | none | G/C-1 | AAATAACCA | CCATGCA | TGG | ACC | C | none | nope | nope_ACC | G/C-1_CCATGCA | 5 | FALSE | FALSE |

| | | | | | | | | | | | | | | | | | | | | | | | | | | | | | | | | | | |
|---|---|---|---|---|---|---|---|---|---|---|---|---|---|---|---|---|---|---|---|---|---|---|---|---|---|---|---|---|---|---|---|---|---|---|
| 65_v.txt | 32081 | 32081-Deleti | Deletion | 1 | A | - | 36 | 21944 | 0.16405396 | 63.1111112 | rnr1Y285A | FALSE | CAN | JSY3869 | 4 | rnr1Y285A-R | CAA | nope | >1bp-ins | A/T-1 | none | A/T-1 | CCATGCAAA | AAAAAAG | GTT | GTT | T | none | T>none | T>none_GTT | A/T-1_AAAAAAG | 5 | FALSE | FALSE |
| 65_v.txt | 32114 | 32114-Deleti | Deletion | 1 | C | - | 3 | 21944 | 0.01367116 | 63 | rnr1Y285A | FALSE | CAN | JSY3869 | 4 | rnr1Y285A-R | GCC | nope | >1bp-ins | G/C-1 | none | G/C-1 | TAATAGCCA | CCATTCG | CGG | GCC | C | none | nope | nope_GCC | G/C-1_CCATTCG | 5 | FALSE | FALSE |
| 65_v.txt | 32125 | 32125-Deleti | Deletion | 1 | C | - | 25 | 21944 | 0.11392636 | 57.52 | rnr1Y285A | FALSE | CAN | JSY3869 | 4 | rnr1Y285A-R | ACT | nope | >1bp-ins | G/C-1 | none | G/C-1 | CGAAAACTT | CTTTGTC | TGA | ACT | C | none | nope | nope_ACT | G/C-1_CTTTGTC | 5 | FALSE | FALSE |
| 65_v.txt | 32210 | 32210-Deleti | Deletion | 1 | T | - | 9 | 21944 | 0.04101349 | 38.6666667 | rnr1Y285A | FALSE | CAN | JSY3869 | 4 | rnr1Y285A-R | CTT | nope | >1bp-ins | A/T-1 | none | A/T-1 | ACCACCTTTG | TTTGGTG | GAA | CTT | T | none | nope | nope_CTT | A/T-1_TTTGGTG | 5 | FALSE | FALSE |
| 65_v.txt | 32216 | 32216-Deleti | Deletion | 1 | G | - | 10 | 21944 | 0.04557054 | 39.6 | rnr1Y285A | FALSE | CAN | JSY3869 | 4 | rnr1Y285A-R | TGG | nope | >1bp-ins | G/C-1 | none | G/C-1 | TTTGGTGGT | GGTCCTT | ACC | ACC | C | none | C>none | C>none_ACC | G/C-1_GGTCCTT | 5 | FALSE | FALSE |
| 65_v.txt | 32259 | 32259-Deleti | Deletion | 1 | C | - | 6 | 36634 | 0.01637823 | 38 | rnr1Y285A | FALSE | CAN | JSY3869 | 4 | rnr1Y285A-R | ACC | nope | >1bp-ins | G/C-1 | none | G/C-1 | GATAGACCA | CCAAATA | TGG | ACC | C | none | nope | nope_ACC | G/C-1_CCAAATA | 5 | FALSE | FALSE |
| 65_v.txt | 32264 | 32264-Deleti | Deletion | 1 | T | - | 8 | 36638 | 0.02183525 | 37.625 | rnr1Y285A | FALSE | CAN | JSY3869 | 4 | rnr1Y285A-R | ATA | nope | >1bp-ins | A/T-1 | none | A/T-1 | ACCAAATAA | TAAAATA | TAT | ATA | T | none | nope | nope_ATA | A/T-1_TAAAATA | 5 | FALSE | FALSE |
| 65_v.txt | 32272 | 32272-Deleti | Deletion | 1 | G | - | 16 | 36638 | 0.04367051 | 35.8125005 | rnr1Y285A | FALSE | CAN | JSY3869 | 4 | rnr1Y285A-R | CGG | nope | >1bp-ins | G/C-1 | none | G/C-1 | AAATACGGG | GGGAACC | GCC | GCC | C | none | C>none | C>none_GCC | G/C-1_GGGAACC | 5 | FALSE | FALSE |
| 65_v.txt | 32346 | 32346-Deleti | Deletion | 1 | A | - | 23 | 36638 | 0.06277635 | 59.9565217 | rnr1Y285A | FALSE | CAN | JSY3869 | 4 | rnr1Y285A-R | CAA | nope | >1bp-ins | A/T-1 | none | A/T-1 | TGTGGCAAA | AAAACCT | GTT | GTT | T | none | T>none | T>none_GTT | A/T-1_AAAACCT | 5 | FALSE | FALSE |
| 65_v.txt | 32358 | 32358-Deleti | Deletion | 1 | C | - | 14 | 36638 | 0.03821169 | 62.0714286 | rnr1Y285A | FALSE | CAN | JSY3869 | 4 | rnr1Y285A-R | ACC | nope | >1bp-ins | G/C-1 | none | G/C-1 | TTTGTACCAG | CCAGAGT | TGG | ACC | C | none | nope | nope_ACC | G/C-1_CCAGAGT | 5 | FALSE | FALSE |
| 65_v.txt | 32442 | 32442-Deleti | Deletion | 1 | A | - | 6 | 36638 | 0.01637644 | 54.1666667 | rnr1Y285A | FALSE | CAN | JSY3869 | 4 | rnr1Y285A-R | TAA | nope | >1bp-ins | A/T-1 | none | A/T-1 | GGAACTAAA | AAAAGTC | ATT | ATT | T | none | T>none | T>none_ATT | A/T-1_AAAAGTC | 5 | FALSE | FALSE |
| 65_v.txt | 32498 | 32498-Deleti | Deletion | 1 | T | - | 43 | 36638 | 0.11736449 | 47.4418603 | rnr1Y285A | FALSE | CAN | JSY3869 | 4 | rnr1Y285A-R | CTT | nope | >1bp-ins | A/T-1 | none | A/T-1 | AACAACTTTT | TTTTTTG | GAA | CTT | T | none | nope | nope_CTT | A/T-1_TTTTTTG | 5 | FALSE | FALSE |
| 65_v.txt | 32528 | 32528-Deleti | Deletion | 1 | G | - | 8 | 36637 | 0.02183585 | 39.625 | rnr1Y285A | FALSE | CAN | JSY3869 | 4 | rnr1Y285A-R | TGG | nope | >1bp-ins | G/C-1 | none | G/C-1 | TTTTCTGGG | GGGGTTT | ACC | ACC | C | none | C>none | C>none_ACC | G/C-1_GGGGTTT | 5 | FALSE | FALSE |
| 65_v.txt | 32556 | 32556-Deleti | Deletion | 1 | C | - | 27 | 36621 | 0.07372819 | 37.3703704 | rnr1Y285A | FALSE | CAN | JSY3869 | 4 | rnr1Y285A-R | ACC | nope | >1bp-ins | G/C-1 | none | G/C-1 | GTGATACCA | CCAACTA | TGG | ACC | C | none | nope | nope_ACC | G/C-1_CCAACTA | 5 | FALSE | FALSE |
| 65_v.txt | 32598 | 32598-Deleti | Deletion | 1 | A | - | 6 | 22365 | 0.02682763 | 46.6666667 | rnr1Y285A | FALSE | CAN | JSY3869 | 4 | rnr1Y285A-R | CAA | nope | >1bp-ins | A/T-1 | none | A/T-1 | TTAATCAAA | AAAGAGG | GTT | GTT | T | none | T>none | T>none_GTT | A/T-1_AAAGAGG | 5 | FALSE | FALSE |
| 65_v.txt | 32608 | 32608-Deleti | Deletion | 1 | C | - | 772 | 22365 | 3.45182204 | 43.8329016 | rnr1Y285A | FALSE | CAN | JSY3869 | 4 | rnr1Y285A-R | ACC | nope | >1bp-ins | G/C-1 | none | G/C-1 | AGGAAACCC | CCCAACC | TGG | ACC | C | none | nope | nope_ACC | G/C-1_CCCAACC | 5 | FALSE | FALSE |
| 65_v.txt | 32654 | 32654-Deleti | Deletion | 1 | T | - | 13 | 22365 | 0.05812654 | 55.7692308 | rnr1Y285A | FALSE | CAN | JSY3869 | 4 | rnr1Y285A-R | CTG | nope | >1bp-ins | A/T-1 | none | A/T-1 | TATACCTGG | TGGACCC | GAC | CTG | T | none | nope | nope_CTG | A/T-1_TGGACCC | 5 | FALSE | FALSE |
| 65_v.txt | 32658 | 32658-Deleti | Deletion | 1 | C | - | 548 | 22365 | 2.4502571 | 59.8649635 | rnr1Y285A | FALSE | CAN | JSY3869 | 4 | rnr1Y285A-R | ACC | nope | >1bp-ins | G/C-1 | none | G/C-1 | CCTGGACCC | CCCCAGG | TGG | ACC | C | none | nope | nope_ACC | G/C-1_CCCCAGG | 5 | FALSE | FALSE |
| 65_v.txt | 32670 | 32670-Deleti | Deletion | 1 | G | - | 637 | 22365 | 2.84820031 | 59.6452119 | rnr1Y285A | FALSE | CAN | JSY3869 | 4 | rnr1Y285A-R | TGG | nope | >1bp-ins | G/C-1 | none | G/C-1 | GCACCTGGG | GGGTTTC | ACC | ACC | C | none | C>none | C>none_ACC | G/C-1_GGGTTTC | 5 | FALSE | FALSE |
| 65_v.txt | 32697 | 32697-Deleti | Deletion | 1 | G | - | 5 | 22365 | 0.02235636 | 63.8 | rnr1Y285A | FALSE | CAN | JSY3869 | 4 | rnr1Y285A-R | TGG | nope | >1bp-ins | G/C-1 | none | G/C-1 | CCAACTGGG | GGGCCGG | ACC | ACC | C | none | C>none | C>none_ACC | G/C-1_GGGCCGG | 5 | FALSE | FALSE |
| 65_v.txt | 32707 | 32707-Deleti | Deletion | 1 | C | - | 1973 | 22365 | 8.82181981 | 62.2802838 | rnr1Y285A | FALSE | CAN | JSY3869 | 4 | rnr1Y285A-R | ACC | nope | >1bp-ins | G/C-1 | none | G/C-1 | CGGTAACCC | CCCCAGC | TGG | ACC | C | none | nope | nope_ACC | G/C-1_CCCCAGC | 5 | FALSE | FALSE |
| 65_v.txt | 32747 | 32747-Deleti | Deletion | 1 | C | - | 38 | 22365 | 0.16990834 | 56.131579 | rnr1Y285A | FALSE | CAN | JSY3869 | 4 | rnr1Y285A-R | ACC | nope | >1bp-ins | G/C-1 | none | G/C-1 | TAGAAACCC | CCCGATA | TGG | ACC | C | none | nope | nope_ACC | G/C-1_CCCGATA | 5 | FALSE | FALSE |
| 65_v.txt | 32779 | 32779-Deleti | Deletion | 1 | C | - | 20 | 22364 | 0.08942944 | 53.6999967 | rnr1Y285A | FALSE | CAN | JSY3869 | 4 | rnr1Y285A-R | ACC | nope | >1bp-ins | G/C-1 | none | G/C-1 | AAGCGACCC | CCCAGAA | TGG | ACC | C | none | nope | nope_ACC | G/C-1_CCCAGAA | 5 | FALSE | FALSE |
| 65_v.txt | 32842 | 32842-Deleti | Deletion | 1 | A | - | 57 | 22365 | 0.25486251 | 42.2105228 | rnr1Y285A | FALSE | CAN | JSY3869 | 4 | rnr1Y285A-R | CAA | nope | >1bp-ins | A/T-1 | none | A/T-1 | TTACCCAAA | AAAAAAT | GTT | GTT | T | none | T>none | T>none_GTT | A/T-1_AAAAAAT | 5 | FALSE | FALSE |
| 65_v.txt | 32884 | 32884-Deleti | Deletion | 1 | A | - | 4 | 29325 | 0.01364024 | 36 | rnr1Y285A | FALSE | CAN | JSY3869 | 4 | rnr1Y285A-R | CAA | nope | >1bp-ins | A/T-1 | none | A/T-1 | ACGTCCAAA | AAAATTG | GTT | GTT | T | none | T>none | T>none_GTT | A/T-1_AAAATTG | 5 | FALSE | FALSE |
| 65_v.txt | 32924 | 32924-Deleti | Deletion | 1 | A | - | 21 | 29325 | 0.07161125 | 59.0476191 | rnr1Y285A | FALSE | CAN | JSY3869 | 4 | rnr1Y285A-R | CAA | nope | >1bp-ins | A/T-1 | none | A/T-1 | CAGGGCAAA | AAAAGTG | GTT | GTT | T | none | T>none | T>none_GTT | A/T-1_AAAAGTG | 5 | FALSE | FALSE |
| 65_v.txt | 32945 | 32945-Deleti | Deletion | 1 | C | - | 5 | 29325 | 0.0170503 | 63.999999 | rnr1Y285A | FALSE | CAN | JSY3869 | 4 | rnr1Y285A-R | ACC | nope | >1bp-ins | G/C-1 | none | G/C-1 | AGAAAACCA | CCAATAC | TGG | ACC | C | none | nope | nope_ACC | G/C-1_CCAATAC | 5 | FALSE | FALSE |
| 65_v.txt | 32958 | 32958-Deleti | Deletion | 1 | C | - | 5 | 29325 | 0.0170503 | 60.4 | rnr1Y285A | FALSE | CAN | JSY3869 | 4 | rnr1Y285A-R | ACC | nope | >1bp-ins | G/C-1 | none | G/C-1 | ATGTAACCA | CCATTGG | TGG | ACC | C | none | nope | nope_ACC | G/C-1_CCATTGG | 5 | FALSE | FALSE |
| 65_v.txt | 33078 | 33078-Deleti | Deletion | 1 | C | - | 459 | 29325 | 1.56521739 | 54.7864924 | rnr1Y285A | FALSE | CAN | JSY3869 | 4 | rnr1Y285A-R | ACC | nope | >1bp-ins | G/C-1 | none | G/C-1 | AAAGAACCC | CCCATAA | TGG | ACC | C | none | nope | nope_ACC | G/C-1_CCCATAA | 5 | FALSE | FALSE |
| 65_v.txt | 33168 | 33168-Deleti | Deletion | 1 | C | - | 10 | 29323 | 0.03410292 | 37.5 | rnr1Y285A | FALSE | CAN | JSY3869 | 4 | rnr1Y285A-R | ACC | nope | >1bp-ins | G/C-1 | none | G/C-1 | ATAGTACCA | CCACCAA | TGG | ACC | C | none | nope | nope_ACC | G/C-1_CCACCAA | 5 | FALSE | FALSE |
| 65_v.txt | 33252 | 33252-Deleti | Deletion | 1 | C | - | 15 | 15273 | 0.09821253 | 40.4666667 | rnr1Y285A | FALSE | CAN | JSY3869 | 4 | rnr1Y285A-R | ACC | nope | >1bp-ins | G/C-1 | none | G/C-1 | TCATCACCTA | CCTATGC | TGG | ACC | C | none | nope | nope_ACC | G/C-1_CCTATGC | 5 | FALSE | FALSE |
| 65_v.txt | 33356 | 33356-Deleti | Deletion | 1 | C | - | 767 | 15273 | 5.02193413 | 55.5215124 | rnr1Y285A | FALSE | CAN | JSY3869 | 4 | rnr1Y285A-R | ACC | nope | >1bp-ins | G/C-1 | none | G/C-1 | TATTGACCCA | CCCACGT | TGG | ACC | C | none | nope | nope_ACC | G/C-1_CCCACGT | 5 | FALSE | FALSE |
| 65_v.txt | 33403 | 33403-Deleti | Deletion | 1 | T | - | 12 | 15273 | 0.07857003 | 50 | rnr1Y285A | FALSE | CAN | JSY3869 | 4 | rnr1Y285A-R | GTT | nope | >1bp-ins | A/T-1 | none | A/T-1 | AGAGGGTTG | TTGTGAC | CAA | GTT | T | none | nope | nope_GTT | A/T-1_TTGTGAC | 5 | FALSE | FALSE |
| 65_v.txt | 33474 | 33474-Deleti | Deletion | 1 | C | - | 39 | 15273 | 0.25535258 | 28.974359 | rnr1Y285A | FALSE | CAN | JSY3869 | 4 | rnr1Y285A-R | GCC | nope | >1bp-ins | G/C-1 | none | G/C-1 | GCTATGCCTT | CCTTTTT | CGG | GCC | C | none | nope | nope_GCC | G/C-1_CCTTTTT | 5 | FALSE | FALSE |
| NL9_v.txt | 31731 | 31731-SNV-T | SNV | 1 | T | G | 3 | 5008 | 0.05990415 | 64 | rnr1Y285A | FALSE | CAN | JSY3868 | 5 | rnr1Y285A-R | CTT | T>G | G/C+1 | A/T-1 | TA>GC | TA>GC | AAAGTCTTT | TTTGGTT | GAA | CTT | T | C | T>G | T>G_CTT | TA>GC_TTTGGTT | 5 | FALSE | FALSE |
| NL9_v.txt | 32541 | 32541-SNV-T | SNV | 1 | T | G | 10 | 7547 | 0.13250298 | 33.2 | rnr1Y285A | FALSE | CAN | JSY3868 | 5 | rnr1Y285A-R | TTC | T>G | G/C+1 | A/T-1 | TA>GC | TA>GC | GCAGCTTCA | TCACCAG | AAG | TTC | T | C | T>G | T>G_TTC | TA>GC_TCACCAG | 5 | FALSE | FALSE |
| NL9_v.txt | 32884 | 32884-SNV- | SNV | 1 | A | C | 6 | 14556 | 0.04122012 | 29.5 | rnr1Y285A | FALSE | CAN | JSY3868 | 5 | rnr1Y285A-R | CAA | A>C | G/C+1 | A/T-1 | TA>GC | TA>GC | ACGTCCAAA | AAAATTG | GTT | GTT | T | G | T>G | T>G_GTT | TA>GC_AAAATTG | 5 | FALSE | FALSE |
| NL9_v.txt | 32927 | 32927-SNV- | SNV | 1 | A | G | 15 | 14556 | 0.10305029 | 37.0666667 | rnr1Y285A | FALSE | CAN | JSY3868 | 5 | rnr1Y285A-R | AAG | A>G | G/C+1 | A/T-1 | TA>CG | TA>CG | GGCAAAAGT | AGTGATT | TTC | TTC | T | C | T>C | T>C_TTC | TA>CG_AGTGATT | 5 | FALSE | FALSE |
| NL9_v.txt | 31714 | 31714-SNV- | SNV | 1 | A | T | 5 | 5008 | 0.09984026 | 62.4 | rnr1Y285A | FALSE | CAN | JSY3868 | 5 | rnr1Y285A-R | AAT | A>T | A/T+1 | A/T-1 | TA>AT | TA>AT | TCCAAAATTT | ATTTGTC | TTA | TTA | T | A | T>A | T>A_TTA | TA>AT_ATTTGTC | 5 | FALSE | FALSE |
| NL9_v.txt | 31863 | 31863-SNV- | SNV | 1 | A | T | 3 | 5008 | 0.05990415 | 32.3333333 | rnr1Y285A | FALSE | CAN | JSY3868 | 5 | rnr1Y285A-R | GAA | A>T | A/T+1 | A/T-1 | TA>AT | TA>AT | AACAGGAAA | AAAATAG | CTT | CTT | T | A | T>A | T>A_CTT | TA>AT_AAAATAG | 5 | FALSE | FALSE |
| NL9_v.txt | 32065 | 32065-SNV- | SNV | 1 | A | T | 6 | 11192 | 0.05360972 | 64 | rnr1Y285A | FALSE | CAN | JSY3868 | 5 | rnr1Y285A-R | GAG | A>T | A/T+1 | A/T-1 | TA>AT | TA>AT | AGATTGAGA | AGATAAA | CTC | CTC | T | A | T>A | T>A_CTC | TA>AT_AGATAAA | 5 | FALSE | FALSE |
| NL9_v.txt | 32294 | 32294-SNV- | SNV | 1 | A | T | 43 | 7548 | 0.56968733 | 37.4883721 | rnr1Y285A | FALSE | CAN | JSY3868 | 5 | rnr1Y285A-R | AAT | A>T | A/T+1 | A/T-1 | TA>AT | TA>AT | ATTTGAATTT | ATTTGCG | TTA | TTA | T | A | T>A | T>A_TTA | TA>AT_ATTTGCG | 5 | FALSE | FALSE |
| NL9_v.txt | 32541 | 32541-SNV-T | SNV | 1 | T | A | 6 | 7547 | 0.07950179 | 37.5 | rnr1Y285A | FALSE | CAN | JSY3868 | 5 | rnr1Y285A-R | TTC | T>A | A/T+1 | A/T-1 | TA>AT | TA>AT | GCAGCTTCA | TCACCAG | AAG | TTC | T | T | T>A | T>A_TTC | TA>AT_TCACCAG | 5 | FALSE | FALSE |
| NL9_v.txt | 32948 | 32948-SNV- | SNV | 1 | A | T | 11 | 14556 | 0.07557021 | 38.2727273 | rnr1Y285A | FALSE | CAN | JSY3868 | 5 | rnr1Y285A-R | AAT | A>T | A/T+1 | A/T-1 | TA>AT | TA>AT | AAACCAATA | ATACATG | TTA | TTA | T | A | T>A | T>A_TTA | TA>AT_ATACATG | 5 | FALSE | FALSE |
| NL9_v.txt | 31650 | 31650-SNV- | SNV | 1 | C | T | 19 | 5008 | 0.37939297 | 33.2105263 | rnr1Y285A | FALSE | CAN | JSY3868 | 5 | rnr1Y285A-R | GCG | C>T | A/T+1 | G/C-1 | CG>TA | CG>TA | AGAATGCGA | CGAAATG | CGC | GCG | C | A | C>T | C>T_GCG | CG>TA_CGAAATG | 5 | FALSE | FALSE |
| NL9_v.txt | 31920 | 31920-SNV- | SNV | 1 | G | A | 9 | 5007 | 0.17974835 | 38.5555556 | rnr1Y285A | FALSE | CAN | JSY3868 | 5 | rnr1Y285A-R | CGT | G>A | A/T+1 | G/C-1 | CG>TA | CG>TA | AAAGCCGTG | GTGAAAC | GCA | GCA | C | T | C>T | C>T_GCA | CG>TA_GTGAAAC | 5 | FALSE | FALSE |
| NL9_v.txt | 31962 | 31962-SNV- | SNV | 1 | G | A | 21 | 11192 | 0.18763402 | 38.6666667 | rnr1Y285A | FALSE | CAN | JSY3868 | 5 | rnr1Y285A-R | CGC | G>A | A/T+1 | G/C-1 | CG>TA | CG>TA | GTGGCCGCA | GCATAAT | GCG | GCG | C | T | C>T | C>T_GCG | CG>TA_GCATAAT | 5 | FALSE | FALSE |
| NL9_v.txt | 31977 | 31977-SNV- | SNV | 1 | C | T | 14 | 11192 | 0.12508935 | 36.1428571 | rnr1Y285A | FALSE | CAN | JSY3868 | 5 | rnr1Y285A-R | GCC | C>T | A/T+1 | G/C-1 | CG>TA | CG>TA | GCCAAGCCG | CCGGGCA | CGG | GCC | C | A | C>T | C>T_GCC | CG>TA_CCGGGCA | 5 | FALSE | FALSE |
| NL9_v.txt | 32075 | 32075-SNV- | SNV | 1 | C | T | 71 | 11192 | 0.6343817 | 62.0422535 | rnr1Y285A | FALSE | CAN | JSY3868 | 5 | rnr1Y285A-R | ACC | C>T | A/T+1 | G/C-1 | CG>TA | CG>TA | AAATAACCA | CCATGCA | TGG | ACC | C | A | C>T | C>T_ACC | CG>TA_CCATGCA | 5 | FALSE | FALSE |
| NL9_v.txt | 32092 | 32092-SNV- | SNV | 1 | C | T | 17 | 11192 | 0.15189421 | 60.0588235 | rnr1Y285A | FALSE | CAN | JSY3868 | 5 | rnr1Y285A-R | GCA | C>T | A/T+1 | G/C-1 | CG>TA | CG>TA | AGCCTGCAA | CAACACC | CGT | GCA | C | A | C>T | C>T_GCA | CG>TA_CAACACC | 5 | FALSE | FALSE |
| NL9_v.txt | 32114 | 32114-SNV- | SNV | 1 | C | T | 38 | 11192 | 0.33952824 | 61.131579 | rnr1Y285A | FALSE | CAN | JSY3868 | 5 | rnr1Y285A-R | GCC | C>T | A/T+1 | G/C-1 | CG>TA | CG>TA | TAATAGCCA | CCATTCG | CGG | GCC | C | A | C>T | C>T_GCC | CG>TA_CCATTCG | 5 | FALSE | FALSE |
| NL9_v.txt | 32176 | 32176-SNV- | SNV | 1 | C | T | 15 | 11192 | 0.1340243 | 37.1333333 | rnr1Y285A | FALSE | CAN | JSY3868 | 5 | rnr1Y285A-R | GCA | C>T | A/T+1 | G/C-1 | CG>TA | CG>TA | ATGCAGCAG | CAGTAAC | CGT | GCA | C | A | C>T | C>T_GCA | CG>TA_CAGTAAC | 5 | FALSE | FALSE |
| NL9_v.txt | 32259 | 32259-SNV- | SNV | 1 | C | T | 17 | 7548 | 0.22522523 | 38.0588235 | rnr1Y285A | FALSE | CAN | JSY3868 | 5 | rnr1Y285A-R | ACC | C>T | A/T+1 | G/C-1 | CG>TA | CG>TA | GATAGACCA | CCAAATA | TGG | ACC | C | A | C>T | C>T_ACC | CG>TA_CCAAATA | 5 | FALSE | FALSE |
| NL9_v.txt | 32299 | 32299-SNV- | SNV | 1 | C | T | 15 | 7548 | 0.19872814 | 37.0666667 | rnr1Y285A | FALSE | CAN | JSY3868 | 5 | rnr1Y285A-R | GCG | C>T | A/T+1 | G/C-1 | CG>TA | CG>TA | AATTTGCGG | CGGCAGA | CGC | GCG | C | A | C>T | C>T_GCG | CG>TA_CGGCAGA | 5 | FALSE | FALSE |
| NL9_v.txt | 32466 | 32466-SNV- | SNV | 1 | C | T | 11 | 7548 | 0.14573397 | 55.9999996 | rnr1Y285A | FALSE | CAN | JSY3868 | 5 | rnr1Y285A-R | GCC | C>T | A/T+1 | G/C-1 | CG>TA | CG>TA | AGAGAGCCA | CCAATGT | CGG | GCC | C | A | C>T | C>T_GCC | CG>TA_CCAATGT | 5 | FALSE | FALSE |
| NL9_v.txt | 32535 | 32535-SNV- | SNV | 1 | G | A | 11 | 7547 | 0.14575328 | 38.8166673 | rnr1Y285A | FALSE | CAN | JSY3868 | 5 | rnr1Y285A-R | TGC | G>A | A/T+1 | G/C-1 | CG>TA | CG>TA | GGGTTTGCA | GCAGCTT | ACG | ACG | C | T | C>T | C>T_ACG | CG>TA_GCAGCTT | 5 | FALSE | FALSE |
| NL9_v.txt | 32536 | 32536-SNV- | SNV | 1 | C | T | 11 | 7547 | 0.14575328 | 39 | rnr1Y285A | FALSE | CAN | JSY3868 | 5 | rnr1Y285A-R | GCA | C>T | A/T+1 | G/C-1 | CG>TA | CG>TA | GGTTTGCAG | CAGCTTC | CGT | GCA | C | A | C>T | C>T_GCA | CG>TA_CAGCTTC | 5 | FALSE | FALSE |
| NL9_v.txt | 32580 | 32580-SNV- | SNV | 1 | G | A | 18 | 14213 | 0.12664462 | 38.7222222 | rnr1Y285A | FALSE | CAN | JSY3868 | 5 | rnr1Y285A-R | TGT | G>A | A/T+1 | G/C-1 | CG>TA | CG>TA | TGAAATGTG | GTGAAGG | ACA | ACA | C | T | C>T | C>T_ACA | CG>TA_GTGAAGG | 5 | FALSE | FALSE |
| NL9_v.txt | 32590 | 32590-SNV- | SNV | 1 | C | T | 29 | 14213 | 0.20403856 | 37.2068966 | rnr1Y285A | FALSE | CAN | JSY3868 | 5 | rnr1Y285A-R | GCG | C>T | A/T+1 | G/C-1 | CG>TA | CG>TA | AGGCAGCGT | CGTTAAT | CGC | GCG | C | A | C>T | C>T_GCG | CG>TA_CGTTAAT | 5 | FALSE | FALSE |
| NL9_v.txt | 32700 | 32700-SNV- | SNV | 1 | C | T | 17 | 14214 | 0.11960039 | 64 | rnr1Y285A | FALSE | CAN | JSY3868 | 5 | rnr1Y285A-R | GCC | C>T | A/T+1 | G/C-1 | CG>TA | CG>TA | ACTGGGCCG | CCGGTAA | CGG | GCC | C | A | C>T | C>T_GCC | CG>TA_CCGGTAA | 5 | FALSE | FALSE |
| NL9_v.txt | 32713 | 32713-SNV- | SNV | 1 | C | T | 17 | 14214 | 0.11960039 | 64 | rnr1Y285A | FALSE | CAN | JSY3868 | 5 | rnr1Y285A-R | GCA | C>T | A/T+1 | G/C-1 | CG>TA | CG>TA | CCCCAGCAC | CACCACA | CGT | GCA | C | A | C>T | C>T_GCA | CG>TA_CACCACA | 5 | FALSE | FALSE |
| NL9_v.txt | 32798 | 32798-SNV- | SNV | 1 | G | A | 19 | 14214 | 0.13367103 | 36.8947368 | rnr1Y285A | FALSE | CAN | JSY3868 | 5 | rnr1Y285A-R | CGT | G>A | A/T+1 | G/C-1 | CG>TA | CG>TA | TTCACCGTA | GTAATAT | GCA | GCA | C | T | C>T | C>T_GCA | CG>TA_GTAATAT | 5 | FALSE | FALSE |
| NL9_v.txt | 32855 | 32855-SNV- | SNV | 1 | C | T | 77 | 14212 | 0.54179567 | 34.5974026 | rnr1Y285A | FALSE | CAN | JSY3868 | 5 | rnr1Y285A-R | TCC | C>T | A/T+1 | G/C-1 | CG>TA | CG>TA | ACTAATCCAT | CCATGCC | AGG | TCC | C | A | C>T | C>T_TCC | CG>TA_CCATGCC | 5 | FALSE | FALSE |
| NL9_v.txt | 32860 | 32860-SNV- | SNV | 1 | C | T | 18 | 14212 | 0.12665353 | 34.5555556 | rnr1Y285A | FALSE | CAN | JSY3868 | 5 | rnr1Y285A-R | GCC | C>T | A/T+1 | G/C-1 | CG>TA | CG>TA | TCCATGCCG | CCGCCAG | CGG | GCC | C | A | C>T | C>T_GCC | CG>TA_CCGCCAG | 5 | FALSE | FALSE |
| NL9_v.txt | 32901 | 32901-SNV- | SNV | 1 | C | T | 22 | 14556 | 0.15114042 | 38.4090909 | rnr1Y285A | FALSE | CAN | JSY3868 | 5 | rnr1Y285A-R | GCC | C>T | A/T+1 | G/C-1 | CG>TA | CG>TA | ACTTGGCCA | CCAACTA | CGG | GCC | C | A | C>T | C>T_GCC | CG>TA_CCAACTA | 5 | FALSE | FALSE |
| NL9_v.txt | 32935 | 32935-SNV- | SNV | 1 | C | T | 15 | 14556 | 0.10305029 | 38.8666667 | rnr1Y285A | FALSE | CAN | JSY3868 | 5 | rnr1Y285A-R | GCC | C>T | A/T+1 | G/C-1 | CG>TA | CG>TA | TGATTGCCC | CCCAAGA | CGG | GCC | C | A | C>T | C>T_GCC | CG>TA_CCCAAGA | 5 | FALSE | FALSE |
| NL9_v.txt | 32958 | 32958-SNV- | SNV | 1 | C | T | 72 | 14556 | 0.49464139 | 60.5138889 | rnr1Y285A | FALSE | CAN | JSY3868 | 5 | rnr1Y285A-R | ACC | C>T | A/T+1 | G/C-1 | CG>TA | CG>TA | ATGTAACCAT | CCATTGG | TGG | ACC | C | A | C>T | C>T_ACC | CG>TA_CCATTGG | 5 | FALSE | FALSE |
| NL9_v.txt | 32967 | 32967-SNV- | SNV | 1 | G | A | 33 | 14556 | 0.22671064 | 61.8181818 | rnr1Y285A | FALSE | CAN | JSY3868 | 5 | rnr1Y285A-R | CGC | G>A | A/T+1 | G/C-1 | CG>TA | CG>TA | TTGGCCGCA | GCACCAA | GCG | GCG | C | T | C>T | C>T_GCG | CG>TA_GCACCAA | 5 | FALSE | FALSE |
| NL9_v.txt | 33070 | 33070-SNV- | SNV | 1 | C | T | 20 | 14556 | 0.13740039 | 61.6 | rnr1Y285A | FALSE | CAN | JSY3868 | 5 | rnr1Y285A-R | GCC | C>T | A/T+1 | G/C-1 | CG>TA | CG>TA | AATATGCCA | CCAAAGA | CGG | GCC | C | A | C>T | C>T_GCC | CG>TA_CCAAAGA | 5 | FALSE | FALSE |
| NL9_v.txt | 33118 | 33118-SNV- | SNV | 1 | C | T | 16 | 14556 | 0.10992031 | 37.8125 | rnr1Y285A | FALSE | CAN | JSY3868 | 5 | rnr1Y285A-R | GCG | C>T | A/T+1 | G/C-1 | CG>TA | CG>TA | GGCCGGCGT | CGTTGGT | CGC | GCG | C | A | C>T | C>T_GCG | CG>TA_CGTTGGT | 5 | FALSE | FALSE |
| NL9_v.txt | 33153 | 33153-SNV- | SNV | 1 | C | T | 34 | 14556 | 0.23358065 | 36.0882353 | rnr1Y285A | FALSE | CAN | JSY3868 | 5 | rnr1Y285A-R | ACC | C>T | A/T+1 | G/C-1 | CG>TA | CG>TA | AAAAGACCT | CCTGTAC | TGG | ACC | C | A | C>T | C>T_ACC | CG>TA_CCTGTAC | 5 | FALSE | FALSE |
| NL9_v.txt | 33156 | 33156-SNV- | SNV | 1 | G | A | 21 | 14556 | 0.1442704 | 38.8571429 | rnr1Y285A | FALSE | CAN | JSY3868 | 5 | rnr1Y285A-R | TGT | G>A | A/T+1 | G/C-1 | CG>TA | CG>TA | AGACCTGTA | GTACCAA | ACA | ACA | C | T | C>T | C>T_ACA | CG>TA_GTACCAA | 5 | FALSE | FALSE |
| NL9_v.txt | 33258 | 33258-SNV- | SNV | 1 | C | T | 22 | 21078 | 0.10437423 | 56.9090909 | rnr1Y285A | FALSE | CAN | JSY3868 | 5 | rnr1Y285A-R | GCC | C>T | A/T+1 | G/C-1 | CG>TA | CG>TA | CCTATGCCAT | CCATCCT | CGG | GCC | C | A | C>T | C>T_GCC | CG>TA_CCATCCT | 5 | FALSE | FALSE |
| NL9_v.txt | 33276 | 33276-SNV- | SNV | 1 | G | A | 46 | 21078 | 0.21823702 | 58.1521739 | rnr1Y285A | FALSE | CAN | JSY3868 | 5 | rnr1Y285A-R | CGT | G>A | A/T+1 | G/C-1 | CG>TA | CG>TA | GAGAACGTA | GTATCCT | GCA | GCA | C | T | C>T | C>T_GCA | CG>TA_GTATCCT | 5 | FALSE | FALSE |
| NL9_v.txt | 33310 | 33310-SNV- | SNV | 1 | G | A | 24 | 21078 | 0.1138628 | 57.4166667 | rnr1Y285A | FALSE | CAN | JSY3868 | 5 | rnr1Y285A-R | CGC | G>A | A/T+1 | G/C-1 | CG>TA | CG>TA | AAGAGCGCA | GCAATGG | GCG | GCG | C | T | C>T | C>T_GCG | CG>TA_GCAATGG | 5 | FALSE | FALSE |
| NL9_v.txt | 33392 | 33392-SNV- | SNV | 1 | G | A | 26 | 21078 | 0.12335136 | 59.2692308 | rnr1Y285A | FALSE | CAN | JSY3868 | 5 | rnr1Y285A-R | CGT | G>A | A/T+1 | G/C-1 | CG>TA | CG>TA | AACGTCGTG | GTGAAAG | GCA | GCA | C | T | C>T | C>T_GCA | CG>TA_GTGAAAG | 5 | FALSE | FALSE |
| NL9_v.txt | 33412 | 33412-SNV- | SNV | 1 | G | A | 30 | 21078 | 0.14232849 | 47 | rnr1Y285A | FALSE | CAN | JSY3868 | 5 | rnr1Y285A-R | GGC | G>A | A/T+1 | G/C-1 | CG>TA | CG>TA | TGACCGGCT | GCTCATT | CCG | CCG | C | T | C>T | C>T_CCG | CG>TA_GCTCATT | 5 | FALSE | FALSE |
| NL9_v.txt | 33474 | 33474-SNV- | SNV | 1 | C | T | 21 | 21074 | 0.09964886 | 30.1428571 | rnr1Y285A | FALSE | CAN | JSY3868 | 5 | rnr1Y285A-R | GCC | C>T | A/T+1 | G/C-1 | CG>TA | CG>TA | GCTATGCCTT | CCTTTTT | CGG | GCC | C | A | C>T | C>T_GCC | CG>TA_CCTTTTT | 5 | FALSE | FALSE |
| NL9_v.txt | 32058 | 32058-SNV- | SNV | 1 | G | C | 10 | 11192 | 0.08934954 | 62.9 | rnr1Y285A | FALSE | CAN | JSY3868 | 5 | rnr1Y285A-R | CGA | G>C | G/C+1 | G/C-1 | CG>GC | CG>GC | ATGTGCGAG | GAGATTG | GCT | GCT | C | G | C>G | C>G_GCT | CG>GC_GAGATTG | 5 | FALSE | FALSE |
| NL9_v.txt | 32299 | 32299-SNV- | SNV | 1 | C | G | 1 | 7548 | 0.01324854 | 39 | rnr1Y285A | FALSE | CAN | JSY3868 | 5 | rnr1Y285A-R | GCG | C>G | G/C+1 | G/C-1 | CG>GC | CG>GC | AATTTGCGG | CGGCAGA | CGC | GCG | C | C | C>G | C>G_GCG | CG>GC_CGGCAGA | 5 | FALSE | FALSE |
| NL9_v.txt | 32901 | 32901-SNV- | SNV | 1 | C | G | 2 | 14556 | 0.01374004 | 38 | rnr1Y285A | FALSE | CAN | JSY3868 | 5 | rnr1Y285A-R | GCC | C>G | G/C+1 | G/C-1 | CG>GC | CG>GC | ACTTGGCCA | CCAACTA | CGG | GCC | C | C | C>G | C>G_GCC | CG>GC_CCAACTA | 5 | FALSE | FALSE |
| NL9_v.txt | 32027 | 32027-SNV- | SNV | 1 | G | T | 45 | 11192 | 0.40207291 | 62.3333333 | rnr1Y285A | FALSE | CAN | JSY3868 | 5 | rnr1Y285A-R | GGT | G>T | A/T+1 | G/C-1 | CG>AT | CG>AT | GCCACGGTA | GTATTTC | CCA | CCA | C | A | C>A | C>A_CCA | CG>AT_GTATTTC | 5 | FALSE | FALSE |
| NL9_v.txt | 32188 | 32188-SNV- | SNV | 1 | C | A | 71 | 11192 | 0.6343817 | 38.1126761 | rnr1Y285A | FALSE | CAN | JSY3868 | 5 | rnr1Y285A-R | ACT | C>A | A/T+1 | G/C-1 | CG>AT | CG>AT | CGAAAACTG | CTGCAAT | TGA | ACT | C | T | C>A | C>A_ACT | CG>AT_CTGCAAT | 5 | FALSE | FALSE |
| NL9_v.txt | 32465 | 32465-SNV- | SNV | 1 | G | T | 5 | 7548 | 0.06624271 | 35.4006904 | rnr1Y285A | FALSE | CAN | JSY3868 | 5 | rnr1Y285A-R | AGC | G>T | A/T+1 | G/C-1 | CG>AT | CG>AT | TAGAGAGCC | GCCAATG | TCG | TCG | C | A | C>A | C>A_TCG | CG>AT_GCCAATG | 5 | FALSE | FALSE |
| NL9_v.txt | 32494 | 32494-SNV- | SNV | 1 | C | A | 22 | 7548 | 0.29146794 | 38.8636364 | rnr1Y285A | FALSE | CAN | JSY3868 | 5 | rnr1Y285A-R | ACA | C>A | A/T+1 | G/C-1 | CG>AT | CG>AT | GGAAAACAA | CAACTTT | TGT | ACA | C | T | C>A | C>A_ACA | CG>AT_CAACTTT | 5 | FALSE | FALSE |
| NL9_v.txt | 32536 | 32536-SNV- | SNV | 1 | C | A | 3 | 7547 | 0.03975089 | 38.3333333 | rnr1Y285A | FALSE | CAN | JSY3868 | 5 | rnr1Y285A-R | GCA | C>A | A/T+1 | G/C-1 | CG>AT | CG>AT | GGTTTGCAG | CAGCTTC | CGT | GCA | C | T | C>A | C>A_GCA | CG>AT_CAGCTTC | 5 | FALSE | FALSE |
| NL9_v.txt | 32538 | 32538-SNV- | SNV | 1 | G | T | 13 | 7547 | 0.17225388 | 34.4615385 | rnr1Y285A | FALSE | CAN | JSY3868 | 5 | rnr1Y285A-R | AGC | G>T | A/T+1 | G/C-1 | CG>AT | CG>AT | TTTGCAGCTT | GCTTCAC | TCG | TCG | C | A | C>A | C>A_TCG | CG>AT_GCTTCAC | 5 | FALSE | FALSE |
| NL9_v.txt | 32556 | 32556-SNV- | SNV | 1 | C | A | 28 | 7546 | 0.37105751 | 38.7857143 | rnr1Y285A | FALSE | CAN | JSY3868 | 5 | rnr1Y285A-R | ACC | C>A | A/T+1 | G/C-1 | CG>AT | CG>AT | GTGATACCA | CCAACTA | TGG | ACC | C | T | C>A | C>A_ACC | CG>AT_CCAACTA | 5 | FALSE | FALSE |
| NL9_v.txt | 32735 | 32735-SNV- | SNV | 1 | G | T | 25 | 14214 | 0.17588293 | 56.64 | rnr1Y285A | FALSE | CAN | JSY3868 | 5 | rnr1Y285A-R | AGT | G>T | A/T+1 | G/C-1 | CG>AT | CG>AT | AAAACAGTA | GTATATT | TCA | TCA | C | A | C>A | C>A_TCA | CG>AT_GTATATT | 5 | FALSE | FALSE |
| NL9_v.txt | 32783 | 32783-SNV- | SNV | 1 | G | T | 176 | 14214 | 1.23821584 | 54.8522727 | rnr1Y285A | FALSE | CAN | JSY3868 | 5 | rnr1Y285A-R | AGA | G>T | A/T+1 | G/C-1 | CG>AT | CG>AT | GACCCAGAA | GAACTCG | TCT | TCT | C | A | C>A | C>A_TCT | CG>AT_GAACTCG | 5 | FALSE | FALSE |
| NL9_v.txt | 32917 | 32917-SNV- | SNV | 1 | C | A | 75 | 14556 | 0.51525144 | 37.9466667 | rnr1Y285A | FALSE | CAN | JSY3868 | 5 | rnr1Y285A-R | TCC | C>A | A/T+1 | G/C-1 | CG>AT | CG>AT | TAAGTTCCA | CCAGGGC | AGG | TCC | C | T | C>A | C>A_TCC | CG>AT_CCAGGGC | 5 | FALSE | FALSE |
| NL9_v.txt | 32940 | 32940-SNV- | SNV | 1 | G | T | 86 | 14556 | 0.59082165 | 37.6744186 | rnr1Y285A | FALSE | CAN | JSY3868 | 5 | rnr1Y285A-R | AGA | G>T | A/T+1 | G/C-1 | CG>AT | CG>AT | GCCCAAGAA | GAAAACC | TCT | TCT | C | A | C>A | C>A_TCT | CG>AT_GAAAACC | 5 | FALSE | FALSE |
| NL9_v.txt | 32954 | 32954-SNV- | SNV | 1 | G | T | 11 | 14556 | 0.07557021 | 56.4545455 | rnr1Y285A | FALSE | CAN | JSY3868 | 5 | rnr1Y285A-R | TGT | G>T | A/T+1 | G/C-1 | CG>AT | CG>AT | ATACATGTA | GTAACCA | ACA | ACA | C | A | C>A | C>A_ACA | CG>AT_GTAACCA | 5 | FALSE | FALSE |
| NL9_v.txt | 32958 | 32958-SNV- | SNV | 1 | C | A | 27 | 14556 | 0.18549052 | 61.5925926 | rnr1Y285A | FALSE | CAN | JSY3868 | 5 | rnr1Y285A-R | ACC | C>A | A/T+1 | G/C-1 | CG>AT | CG>AT | ATGTAACCAT | CCATTGG | TGG | ACC | C | T | C>A | C>A_ACC | CG>AT_CCATTGG | 5 | FALSE | FALSE |
| NL9_v.txt | 33015 | 33015-SNV- | SNV | 1 | G | T | 35 | 14556 | 0.24045067 | 59.6 | rnr1Y285A | FALSE | CAN | JSY3868 | 5 | rnr1Y285A-R | GGA | G>T | A/T+1 | G/C-1 | CG>AT | CG>AT | AAAGAGGAT | GATGTAA | CCT | CCT | C | A | C>A | C>A_CCT | CG>AT_GATGTAA | 5 | FALSE | FALSE |
| NL9_v.txt | 33093 | 33093-SNV- | SNV | 1 | G | T | 23 | 14556 | 0.15801044 | 57.173913 | rnr1Y285A | FALSE | CAN | JSY3868 | 5 | rnr1Y285A-R | TGA | G>T | A/T+1 | G/C-1 | CG>AT | CG>AT | AAATATGAT | GATATAA | ACT | ACT | C | A | C>A | C>A_ACT | CG>AT_GATATAA | 5 | FALSE | FALSE |
| NL9_v.txt | 33151 | 33151-SNV- | SNV | 1 | G | T | 82 | 14556 | 0.56334158 | 36.8658537 | rnr1Y285A | FALSE | CAN | JSY3868 | 5 | rnr1Y285A-R | AGA | G>T | A/T+1 | G/C-1 | CG>AT | CG>AT | TGAAAAGAC | GACCTGT | TCT | TCT | C | A | C>A | C>A_TCT | CG>AT_GACCTGT | 5 | FALSE | FALSE |
| NL9_v.txt | 33153 | 33153-SNV- | SNV | 1 | C | A | 234 | 14556 | 1.6075845 | 37.1282051 | rnr1Y285A | FALSE | CAN | JSY3868 | 5 | rnr1Y285A-R | ACC | C>A | A/T+1 | G/C-1 | CG>AT | CG>AT | AAAAGACCT | CCTGTAC | TGG | ACC | C | T | C>A | C>A_ACC | CG>AT_CCTGTAC | 5 | FALSE | FALSE |
| NL9_v.txt | 33168 | 33168-SNV- | SNV | 1 | C | A | 124 | 14556 | 0.85188239 | 37.9516129 | rnr1Y285A | FALSE | CAN | JSY3868 | 5 | rnr1Y285A-R | ACC | C>A | A/T+1 | G/C-1 | CG>AT | CG>AT | ATAGTACCA | CCACCAA | TGG | ACC | C | T | C>A | C>A_ACC | CG>AT_CCACCAA | 5 | FALSE | FALSE |
| NL9_v.txt | 33325 | 33325-SNV- | SNV | 1 | C | A | 132 | 21078 | 0.62624537 | 62.0227273 | rnr1Y285A | FALSE | CAN | JSY3868 | 5 | rnr1Y285A-R | TCT | C>A | A/T+1 | G/C-1 | CG>AT | CG>AT | ACAATTCTTT | CTTTACT | AGA | TCT | C | T | C>A | C>A_TCT | CG>AT_CTTTACT | 5 | FALSE | FALSE |
| NL9_v.txt | 33474 | 33474-SNV- | SNV | 1 | C | A | 50 | 21074 | 0.23725918 | 34.18 | rnr1Y285A | FALSE | CAN | JSY3868 | 5 | rnr1Y285A-R | GCC | C>A | A/T+1 | G/C-1 | CG>AT | CG>AT | GCTATGCCTT | CCTTTTT | CGG | GCC | C | T | C>A | C>A_GCC | CG>AT_CCTTTTT | 5 | FALSE | FALSE |
| NL9_v.txt | 32842 | 32842-Repla | Replacement | 2 | AA | C | 1 | 14214 | 7.04E-03 | 37.9924343 | rnr1Y285A | FALSE | CAN | JSY3868 | 5 | rnr1Y285A-R | CAA | nope | G/C+1 | >1bpdel | none | Replac. | TTACCCAAA | AAAAAAT | GTT | CAA | AA | G | NA | NA_CAA | Replac._AAAAAAT | 5 | FALSE | FALSE |
| NL9_v.txt | 32081 | 32081-Insert | Insertion | 1 | - | A | 14 | 11192 | 0.12508935 | 64 | rnr1Y285A | FALSE | CAN | JSY3868 | 5 | rnr1Y285A-R | CAA | nope | A/T+1 | >1bpdel | none | A/T+1 | CCATGCAAA | AAAAAAG | GTT | CAA | - | T | NA | NA_CAA | A/T+1_AAAAAAG | 5 | FALSE | FALSE |
| NL9_v.txt | 32498 | 32498-Insert | Insertion | 1 | - | T | 3 | 7548 | 0.03974563 | 37.3333333 | rnr1Y285A | FALSE | CAN | JSY3868 | 5 | rnr1Y285A-R | CTT | nope | A/T+1 | >1bpdel | none | A/T+1 | AACAACTTTT | TTTTTTG | GAA | CTT | - | A | NA | NA_CTT | A/T+1_TTTTTTG | 5 | FALSE | FALSE |
| NL9_v.txt | 32673 | 32673-Insert | Insertion | 1 | - | T | 16 | 14214 | 0.11256508 | 64 | rnr1Y285A | FALSE | CAN | JSY3868 | 5 | rnr1Y285A-R | GTT | nope | A/T+1 | >1bpdel | none | A/T+1 | CCTGGGTTT | TTTCTCC | CAA | GTT | - | A | NA | NA_GTT | A/T+1_TTTCTCC | 5 | FALSE | FALSE |
| NL9_v.txt | 32842 | 32842-Insert | Insertion | 1 | - | A | 7 | 14214 | 0.04924722 | 38 | rnr1Y285A | FALSE | CAN | JSY3868 | 5 | rnr1Y285A-R | CAA | nope | A/T+1 | >1bpdel | none | A/T+1 | TTACCCAAA | AAAAAAT | GTT | CAA | - | T | NA | NA_CAA | A/T+1_AAAAAAT | 5 | FALSE | FALSE |
| NL9_v.txt | 32337 | 32337-Deleti | Deletion | 2 | AT | - | 49 | 7548 | 0.64917859 | 64 | rnr1Y285A | FALSE | CAN | JSY3868 | 5 | rnr1Y285A-R | GAT | nope | >1bp-ins | >1bpdel | none | >1bpdel | TTGAAGATA | ATATGTG | CTA | GAT | AT | none | NA | NA_GAT | >1bpdel_ATATGTG | 5 | FALSE | FALSE |
| NL9_v.txt | 32465 | 32465-Deleti | Deletion | 3 | GCC | - | 3 | 7548 | 0.03974563 | 55.6666667 | rnr1Y285A | FALSE | CAN | JSY3868 | 5 | rnr1Y285A-R | AGC | nope | >1bp-ins | >1bpdel | none | >1bpdel | TAGAGAGCC | GCCAATG | TCG | AGC | GCC | none | NA | NA_AGC | >1bpdel_GCCAATG | 5 | FALSE | FALSE |
| NL9_v.txt | 32842 | 32842-Deleti | Deletion | 2 | AA | - | 8 | 14214 | 0.05628254 | 38.75 | rnr1Y285A | FALSE | CAN | JSY3868 | 5 | rnr1Y285A-R | CAA | nope | >1bp-ins | >1bpdel | none | >1bpdel | TTACCCAAA | AAAAAAT | GTT | CAA | AA | none | NA | NA_CAA | >1bpdel_AAAAAAT | 5 | FALSE | FALSE |
| NL9_v.txt | 31711 | 31711-Deleti | Deletion | 1 | A | - | 20 | 5008 | 0.39936102 | 34.55 | rnr1Y285A | FALSE | CAN | JSY3868 | 5 | rnr1Y285A-R | CAA | nope | >1bp-ins | A/T-1 | none | A/T-1 | CATTCCAAA | AAAATTT | GTT | GTT | T | none | T>none | T>none_GTT | A/T-1_AAAATTT | 5 | FALSE | FALSE |
| NL9_v.txt | 31723 | 31723-Deleti | Deletion | 1 | A | - | 5 | 5008 | 0.09984026 | 64 | rnr1Y285A | FALSE | CAN | JSY3868 | 5 | rnr1Y285A-R | CAA | nope | >1bp-ins | A/T-1 | none | A/T-1 | TGTCCCAAA | AAAAAGT | GTT | GTT | T | none | T>none | T>none_GTT | A/T-1_AAAAAGT | 5 | FALSE | FALSE |
| NL9_v.txt | 31731 | 31731-Deleti | Deletion | 1 | T | - | 2 | 5008 | 0.0399361 | 64 | rnr1Y285A | FALSE | CAN | JSY3868 | 5 | rnr1Y285A-R | CTT | nope | >1bp-ins | A/T-1 | none | A/T-1 | AAAGTCTTT | TTTGGTT | GAA | CTT | T | none | nope | nope_CTT | A/T-1_TTTGGTT | 5 | FALSE | FALSE |
| NL9_v.txt | 31844 | 31844-Deleti | Deletion | 1 | C | - | 4 | 5008 | 0.07987221 | 61.25 | rnr1Y285A | FALSE | CAN | JSY3868 | 5 | rnr1Y285A-R | TCC | nope | >1bp-ins | G/C-1 | none | G/C-1 | TAAGATCCA | CCAAACA | AGG | TCC | C | none | nope | nope_TCC | G/C-1_CCAAACA | 5 | FALSE | FALSE |
| NL9_v.txt | 31863 | 31863-Deleti | Deletion | 1 | A | - | 4 | 5008 | 0.07987221 | 57.5 | rnr1Y285A | FALSE | CAN | JSY3868 | 5 | rnr1Y285A-R | GAA | nope | >1bp-ins | A/T-1 | none | A/T-1 | AACAGGAAA | AAAATAG | CTT | CTT | T | none | T>none | T>none_CTT | A/T-1_AAAATAG | 5 | FALSE | FALSE |
| NL9_v.txt | 31904 | 31904-Deleti | Deletion | 1 | T | - | 5 | 5008 | 0.09984026 | 38.6 | rnr1Y285A | FALSE | CAN | JSY3868 | 5 | rnr1Y285A-R | ATT | nope | >1bp-ins | A/T-1 | none | A/T-1 | ATTGAATTTT | TTTTGGT | TAA | ATT | T | none | nope | nope_ATT | A/T-1_TTTTGGT | 5 | FALSE | FALSE |
| NL9_v.txt | 31913 | 31913-Deleti | Deletion | 1 | A | - | 7 | 5008 | 0.13977636 | 38.7142857 | rnr1Y285A | FALSE | CAN | JSY3868 | 5 | rnr1Y285A-R | CAA | nope | >1bp-ins | A/T-1 | none | A/T-1 | TGGTGCAAA | AAAAGCC | GTT | GTT | T | none | T>none | T>none_GTT | A/T-1_AAAAGCC | 5 | FALSE | FALSE |
| NL9_v.txt | 31979 | 31979-Deleti | Deletion | 1 | G | - | 41 | 11192 | 0.3663331 | 36.3902439 | rnr1Y285A | FALSE | CAN | JSY3868 | 5 | rnr1Y285A-R | CGG | nope | >1bp-ins | G/C-1 | none | G/C-1 | CAAGCCGGG | GGGCATT | GCC | GCC | C | none | C>none | C>none_GCC | G/C-1_GGGCATT | 5 | FALSE | FALSE |
| NL9_v.txt | 32075 | 32075-Deleti | Deletion | 1 | C | - | 1 | 11192 | 8.94E-03 | 64 | rnr1Y285A | FALSE | CAN | JSY3868 | 5 | rnr1Y285A-R | ACC | nope | >1bp-ins | G/C-1 | none | G/C-1 | AAATAACCA | CCATGCA | TGG | ACC | C | none | nope | nope_ACC | G/C-1_CCATGCA | 5 | FALSE | FALSE |
| NL9_v.txt | 32081 | 32081-Deleti | Deletion | 1 | A | - | 26 | 11192 | 0.23230879 | 64 | rnr1Y285A | FALSE | CAN | JSY3868 | 5 | rnr1Y285A-R | CAA | nope | >1bp-ins | A/T-1 | none | A/T-1 | CCATGCAAA | AAAAAAG | GTT | GTT | T | none | T>none | T>none_GTT | A/T-1_AAAAAAG | 5 | FALSE | FALSE |

| | | | | | | | | | | | | | | | | | | | | | | | | | | | | | | | | | | |
|---|---|---|---|---|---|---|---|---|---|---|---|---|---|---|---|---|---|---|---|---|---|---|---|---|---|---|---|---|---|---|---|---|---|---|
| NL9_v.txt | 32299 | 32299-Deleti | Deletion | 1 | C | - | 1 | 7548 | 0.01324854 | 39 | rnr1Y285A | FALSE | CAN | JSY3868 | 5 | rnr1Y285A-R | GCG | nope | >1bp-ins | G/C-1 | none | G/C-1 | AATTTGCGG | CGGCAGA | CGC | GCG | C | none | nope | nope_GCG | G/C-1_CGGCAGA | 5 | FALSE | FALSE |
| NL9_v.txt | 32346 | 32346-Deleti | Deletion | 1 | A | - | 5 | 7548 | 0.06624271 | 64 | rnr1Y285A | FALSE | CAN | JSY3868 | 5 | rnr1Y285A-R | CAA | nope | >1bp-ins | A/T-1 | none | A/T-1 | TGTGGCAAA | AAAACCT | GTT | GTT | T | none | T>none | T>none_GTT | A/T-1_AAAACCT | 5 | FALSE | FALSE |
| NL9_v.txt | 32461 | 32461-Deleti | Deletion | 1 | G | - | 3 | 7548 | 0.03974563 | 55.6666667 | rnr1Y285A | FALSE | CAN | JSY3868 | 5 | rnr1Y285A-R | AGA | nope | >1bp-ins | G/C-1 | none | G/C-1 | ATAATAGAG | GAGAGCC | TCT | TCT | C | none | C>none | C>none_TCT | G/C-1_GAGAGCC | 5 | FALSE | FALSE |
| NL9_v.txt | 32463 | 32463-Deleti | Deletion | 1 | G | - | 3 | 7548 | 0.03974563 | 55.6666667 | rnr1Y285A | FALSE | CAN | JSY3868 | 5 | rnr1Y285A-R | AGA | nope | >1bp-ins | G/C-1 | none | G/C-1 | AATAGAGAG | GAGCCAA | TCT | TCT | C | none | C>none | C>none_TCT | G/C-1_GAGCCAA | 5 | FALSE | FALSE |
| NL9_v.txt | 32471 | 32471-Deleti | Deletion | 1 | G | - | 3 | 7548 | 0.03974563 | 55.6666667 | rnr1Y285A | FALSE | CAN | JSY3868 | 5 | rnr1Y285A-R | TGT | nope | >1bp-ins | G/C-1 | none | G/C-1 | GCCAATGTA | GTAGAAG | ACA | ACA | C | none | C>none | C>none_ACA | G/C-1_GTAGAAG | 5 | FALSE | FALSE |
| NL9_v.txt | 32477 | 32477-Deleti | Deletion | 1 | G | - | 6 | 7548 | 0.07949126 | 57.8333333 | rnr1Y285A | FALSE | CAN | JSY3868 | 5 | rnr1Y285A-R | AGG | nope | >1bp-ins | G/C-1 | none | G/C-1 | GTAGAAGGT | GGTTAAG | TCC | TCC | C | none | C>none | C>none_TCC | G/C-1_GGTTAAG | 5 | FALSE | FALSE |
| NL9_v.txt | 32498 | 32498-Deleti | Deletion | 1 | T | - | 15 | 7548 | 0.19872814 | 37.0666667 | rnr1Y285A | FALSE | CAN | JSY3868 | 5 | rnr1Y285A-R | CTT | nope | >1bp-ins | A/T-1 | none | A/T-1 | AACAACTTTT | TTTTTTG | GAA | CTT | T | none | nope | nope_CTT | A/T-1_TTTTTTG | 5 | FALSE | FALSE |
| NL9_v.txt | 32522 | 32522-Deleti | Deletion | 1 | T | - | 77 | 7548 | 1.02013779 | 37.012987 | rnr1Y285A | FALSE | CAN | JSY3868 | 5 | rnr1Y285A-R | ATT | nope | >1bp-ins | A/T-1 | none | A/T-1 | AACGGATTTT | TTTTCTG | TAA | ATT | T | none | nope | nope_ATT | A/T-1_TTTTCTG | 5 | FALSE | FALSE |
| NL9_v.txt | 32608 | 32608-Deleti | Deletion | 1 | C | - | 1151 | 14213 | 8.09821994 | 38.3544744 | rnr1Y285A | FALSE | CAN | JSY3868 | 5 | rnr1Y285A-R | ACC | nope | >1bp-ins | G/C-1 | none | G/C-1 | AGGAAACCC | CCCAACC | TGG | ACC | C | none | nope | nope_ACC | G/C-1_CCCAACC | 5 | FALSE | FALSE |
| NL9_v.txt | 32631 | 32631-Deleti | Deletion | 1 | T | - | 12 | 14214 | 0.08442381 | 38.8333333 | rnr1Y285A | FALSE | CAN | JSY3868 | 5 | rnr1Y285A-R | GTT | nope | >1bp-ins | A/T-1 | none | A/T-1 | CCTTCGTTTT | TTTTTAT | CAA | GTT | T | none | nope | nope_GTT | A/T-1_TTTTTAT | 5 | FALSE | FALSE |
| NL9_v.txt | 32640 | 32640-Deleti | Deletion | 1 | T | - | 4 | 14214 | 0.02814127 | 38.25 | rnr1Y285A | FALSE | CAN | JSY3868 | 5 | rnr1Y285A-R | CTT | nope | >1bp-ins | A/T-1 | none | A/T-1 | TTATCCTTAG | TTAGATA | GAA | CTT | T | none | nope | nope_CTT | A/T-1_TTAGATA | 5 | FALSE | FALSE |
| NL9_v.txt | 32655 | 32655-Deleti | Deletion | 1 | G | - | 28 | 14214 | 0.19698888 | 61 | rnr1Y285A | FALSE | CAN | JSY3868 | 5 | rnr1Y285A-R | TGG | nope | >1bp-ins | G/C-1 | none | G/C-1 | ATACCTGGA | GGACCCC | ACC | ACC | C | none | C>none | C>none_ACC | G/C-1_GGACCCC | 5 | FALSE | FALSE |
| NL9_v.txt | 32658 | 32658-Deleti | Deletion | 1 | C | - | 1208 | 14214 | 8.49866329 | 62.4519868 | rnr1Y285A | FALSE | CAN | JSY3868 | 5 | rnr1Y285A-R | ACC | nope | >1bp-ins | G/C-1 | none | G/C-1 | CCTGGACCC | CCCCAGG | TGG | ACC | C | none | nope | nope_ACC | G/C-1_CCCCAGG | 5 | FALSE | FALSE |
| NL9_v.txt | 32670 | 32670-Deleti | Deletion | 1 | G | - | 237 | 14214 | 1.6673702 | 62.1392405 | rnr1Y285A | FALSE | CAN | JSY3868 | 5 | rnr1Y285A-R | TGG | nope | >1bp-ins | G/C-1 | none | G/C-1 | GCACCTGGG | GGGTTTC | ACC | ACC | C | none | C>none | C>none_ACC | G/C-1_GGGTTTC | 5 | FALSE | FALSE |
| NL9_v.txt | 32707 | 32707-Deleti | Deletion | 1 | C | - | 173 | 14214 | 1.21710989 | 62.3526012 | rnr1Y285A | FALSE | CAN | JSY3868 | 5 | rnr1Y285A-R | ACC | nope | >1bp-ins | G/C-1 | none | G/C-1 | CGGTAACCC | CCCCAGC | TGG | ACC | C | none | nope | nope_ACC | G/C-1_CCCCAGC | 5 | FALSE | FALSE |
| NL9_v.txt | 32728 | 32728-Deleti | Deletion | 1 | A | - | 8 | 14214 | 0.05628254 | 61.25 | rnr1Y285A | FALSE | CAN | JSY3868 | 5 | rnr1Y285A-R | CAA | nope | >1bp-ins | A/T-1 | none | A/T-1 | CCATACAAA | AAAAACA | GTT | GTT | T | none | T>none | T>none_GTT | A/T-1_AAAAACA | 5 | FALSE | FALSE |
| NL9_v.txt | 32747 | 32747-Deleti | Deletion | 1 | C | - | 81 | 14214 | 0.5698607 | 60.0864198 | rnr1Y285A | FALSE | CAN | JSY3868 | 5 | rnr1Y285A-R | ACC | nope | >1bp-ins | G/C-1 | none | G/C-1 | TAGAAACCC | CCCGATA | TGG | ACC | C | none | nope | nope_ACC | G/C-1_CCCGATA | 5 | FALSE | FALSE |
| NL9_v.txt | 32839 | 32839-Deleti | Deletion | 1 | C | - | 5 | 14214 | 0.03517659 | 32.2 | rnr1Y285A | FALSE | CAN | JSY3868 | 5 | rnr1Y285A-R | ACC | nope | >1bp-ins | G/C-1 | none | G/C-1 | TAATTACCCA | CCCAAAA | TGG | ACC | C | none | nope | nope_ACC | G/C-1_CCCAAAA | 5 | FALSE | FALSE |
| NL9_v.txt | 32842 | 32842-Deleti | Deletion | 1 | A | - | 6019 | 14214 | 42.3455748 | 36.9991644 | rnr1Y285A | FALSE | CAN | JSY3868 | 5 | rnr1Y285A-R | CAA | nope | >1bp-ins | A/T-1 | none | A/T-1 | TTACCCAAA | AAAAAAT | GTT | GTT | T | none | T>none | T>none_GTT | A/T-1_AAAAAAT | 5 | FALSE | FALSE |
| NL9_v.txt | 32884 | 32884-Deleti | Deletion | 1 | A | - | 9 | 14556 | 0.06183017 | 38 | rnr1Y285A | FALSE | CAN | JSY3868 | 5 | rnr1Y285A-R | CAA | nope | >1bp-ins | A/T-1 | none | A/T-1 | ACGTCCAAA | AAAATTG | GTT | GTT | T | none | T>none | T>none_GTT | A/T-1_AAAATTG | 5 | FALSE | FALSE |
| NL9_v.txt | 32924 | 32924-Deleti | Deletion | 1 | A | - | 30 | 14556 | 0.20610058 | 36.7 | rnr1Y285A | FALSE | CAN | JSY3868 | 5 | rnr1Y285A-R | CAA | nope | >1bp-ins | A/T-1 | none | A/T-1 | CAGGGCAAA | AAAAGTG | GTT | GTT | T | none | T>none | T>none_GTT | A/T-1_AAAAGTG | 5 | FALSE | FALSE |
| NL9_v.txt | 33147 | 33147-Deleti | Deletion | 1 | A | - | 16 | 14556 | 0.10992031 | 39 | rnr1Y285A | FALSE | CAN | JSY3868 | 5 | rnr1Y285A-R | GAA | nope | >1bp-ins | A/T-1 | none | A/T-1 | CCAATGAAA | AAAAGAC | CTT | CTT | T | none | T>none | T>none_CTT | A/T-1_AAAAGAC | 5 | FALSE | FALSE |
| NL9_v.txt | 33175 | 33175-Deleti | Deletion | 1 | G | - | 9 | 14552 | 0.06184717 | 38.7777778 | rnr1Y285A | FALSE | CAN | JSY3868 | 5 | rnr1Y285A-R | AGG | nope | >1bp-ins | G/C-1 | none | G/C-1 | CACCAAGGG | GGGCAAT | TCC | TCC | C | none | C>none | C>none_TCC | G/C-1_GGGCAAT | 5 | FALSE | FALSE |
| NL9_v.txt | 33251 | 33251-Deleti | Deletion | 1 | A | - | 53 | 21078 | 0.25144701 | 60.8301887 | rnr1Y285A | FALSE | CAN | JSY3868 | 5 | rnr1Y285A-R | CAC | nope | >1bp-ins | A/T-1 | none | A/T-1 | TTCATCACCT | ACCTATG | GTG | GTG | T | none | T>none | T>none_GTG | A/T-1_ACCTATG | 5 | FALSE | FALSE |
| NL9_v.txt | 33356 | 33356-Deleti | Deletion | 1 | C | - | 482 | 21078 | 2.28674447 | 59.8775934 | rnr1Y285A | FALSE | CAN | JSY3868 | 5 | rnr1Y285A-R | ACC | nope | >1bp-ins | G/C-1 | none | G/C-1 | TATTGACCCA | CCCACGT | TGG | ACC | C | none | nope | nope_ACC | G/C-1_CCCACGT | 5 | FALSE | FALSE |
| NL9_v.txt | 33474 | 33474-Deleti | Deletion | 1 | C | - | 53 | 21074 | 0.25149473 | 34.6226415 | rnr1Y285A | FALSE | CAN | JSY3868 | 5 | rnr1Y285A-R | GCC | nope | >1bp-ins | G/C-1 | none | G/C-1 | GCTATGCCTT | CCTTTTT | CGG | GCC | C | none | nope | nope_GCC | G/C-1_CCTTTTT | 5 | FALSE | FALSE |
| 58_v.txt | 31928 | 31928-SNV-T | SNV | 1 | T | G | 2 | 7593 | 0.02634005 | 38 | rnr1Y285A_pe | FALSE | ARG | JSY3868 | 1 | rnr1Y285A_p | CTT | T>G | G/C+1 | A/T-1 | TA>GC | TA>GC | GAAACCTTG | TTGAATA | GAA | CTT | T | C | T>G | T>G_CTT | TA>GC_TTGAATA | 4 | FALSE | FALSE |
| 58_v.txt | 32884 | 32884-SNV-A | SNV | 1 | A | C | 6 | 21417 | 0.02801513 | 38 | rnr1Y285A_pe | FALSE | ARG | JSY3868 | 1 | rnr1Y285A_p | CAA | A>C | G/C+1 | A/T-1 | TA>GC | TA>GC | ACGTCCAAA | AAAATTG | GTT | GTT | T | G | T>G | T>G_GTT | TA>GC_AAAATTG | 4 | FALSE | FALSE |
| 58_v.txt | 33269 | 33269-SNV-A | SNV | 1 | A | C | 101 | 13211 | 0.76451442 | 16.3069307 | rnr1Y285A_pe | FALSE | ARG | JSY3868 | 1 | rnr1Y285A_p | TAG | A>C | G/C+1 | A/T-1 | TA>GC | TA>GC | CTCCATAGA | AGAGAAC | ATC | ATC | T | G | T>G | T>G_ATC | TA>GC_AGAGAAC | 4 | FALSE | FALSE |
| 58_v.txt | 31928 | 31928-SNV-T | SNV | 1 | T | C | 7 | 7593 | 0.09219018 | 38 | rnr1Y285A_pe | FALSE | ARG | JSY3868 | 1 | rnr1Y285A_p | CTT | T>C | G/C+1 | A/T-1 | TA>CG | TA>CG | GAAACCTTG | TTGAATA | GAA | CTT | T | G | T>C | T>C_CTT | TA>CG_TTGAATA | 4 | FALSE | FALSE |
| 58_v.txt | 32842 | 32842-SNV-A | SNV | 1 | A | G | 5 | 18166 | 0.02752395 | 48.3994917 | rnr1Y285A_pe | FALSE | ARG | JSY3868 | 1 | rnr1Y285A_p | CAA | A>G | G/C+1 | A/T-1 | TA>CG | TA>CG | TTACCCAAA | AAAAAAT | GTT | GTT | T | C | T>C | T>C_GTT | TA>CG_AAAAAAT | 4 | FALSE | FALSE |
| 58_v.txt | 33476 | 33476-SNV-T | SNV | 1 | T | C | 22 | 13209 | 0.16655311 | 30.4090909 | rnr1Y285A_pe | FALSE | ARG | JSY3868 | 1 | rnr1Y285A_p | CTT | T>C | G/C+1 | A/T-1 | TA>CG | TA>CG | TATGCCTTTT | TTTTTTT | GAA | CTT | T | G | T>C | T>C_CTT | TA>CG_TTTTTTT | 4 | FALSE | FALSE |
| 58_v.txt | 31928 | 31928-SNV-T | SNV | 1 | T | A | 5 | 7593 | 0.06585013 | 38 | rnr1Y285A_pe | FALSE | ARG | JSY3868 | 1 | rnr1Y285A_p | CTT | T>A | A/T+1 | A/T-1 | TA>AT | TA>AT | GAAACCTTG | TTGAATA | GAA | CTT | T | T | T>A | T>A_CTT | TA>AT_TTGAATA | 4 | FALSE | FALSE |
| 58_v.txt | 32498 | 32498-SNV-T | SNV | 1 | T | A | 6 | 25654 | 0.02338817 | 37.9990562 | rnr1Y285A_pe | FALSE | ARG | JSY3868 | 1 | rnr1Y285A_p | CTT | T>A | A/T+1 | A/T-1 | TA>AT | TA>AT | AACAACTTTT | TTTTTTG | GAA | CTT | T | T | T>A | T>A_CTT | TA>AT_TTTTTTG | 4 | FALSE | FALSE |
| 58_v.txt | 31650 | 31650-SNV-C | SNV | 1 | C | T | 17 | 7594 | 0.22386094 | 35.8235294 | rnr1Y285A_pe | FALSE | ARG | JSY3868 | 1 | rnr1Y285A_p | GCG | C>T | A/T+1 | G/C-1 | CG>TA | CG>TA | AGAATGCGA | CGAAATG | CGC | GCG | C | A | C>T | C>T_GCG | CG>TA_CGAAATG | 4 | FALSE | FALSE |
| 58_v.txt | 31912 | 31912-SNV-C | SNV | 1 | C | T | 12 | 7594 | 0.15801949 | 37.5 | rnr1Y285A_pe | FALSE | ARG | JSY3868 | 1 | rnr1Y285A_p | GCA | C>T | A/T+1 | G/C-1 | CG>TA | CG>TA | TTGGTGCAA | CAAAAGC | CGT | GCA | C | A | C>T | C>T_GCA | CG>TA_CAAAAGC | 4 | FALSE | FALSE |
| 58_v.txt | 31962 | 31962-SNV-G | SNV | 1 | G | A | 24 | 14659 | 0.16372195 | 35.5416667 | rnr1Y285A_pe | FALSE | ARG | JSY3868 | 1 | rnr1Y285A_p | CGC | G>A | A/T+1 | G/C-1 | CG>TA | CG>TA | GTGGCCGCA | GCATAAT | GCG | GCG | C | T | C>T | C>T_GCG | CG>TA_GCATAAT | 4 | FALSE | FALSE |
| 58_v.txt | 31972 | 31972-SNV-C | SNV | 1 | C | T | 14 | 14659 | 0.09550447 | 35.8571429 | rnr1Y285A_pe | FALSE | ARG | JSY3868 | 1 | rnr1Y285A_p | GCC | C>T | A/T+1 | G/C-1 | CG>TA | CG>TA | AATAAGCCA | CCAAGCC | CGG | GCC | C | A | C>T | C>T_GCC | CG>TA_CCAAGCC | 4 | FALSE | FALSE |
| 58_v.txt | 32092 | 32092-SNV-C | SNV | 1 | C | T | 16 | 14659 | 0.10914796 | 59.8125 | rnr1Y285A_pe | FALSE | ARG | JSY3868 | 1 | rnr1Y285A_p | GCA | C>T | A/T+1 | G/C-1 | CG>TA | CG>TA | AGCCTGCAA | CAACACC | CGT | GCA | C | A | C>T | C>T_GCA | CG>TA_CAACACC | 4 | FALSE | FALSE |
| 58_v.txt | 32165 | 32165-SNV-G | SNV | 1 | G | A | 17 | 14659 | 0.11596971 | 46.4117647 | rnr1Y285A_pe | FALSE | ARG | JSY3868 | 1 | rnr1Y285A_p | CGC | G>A | A/T+1 | G/C-1 | CG>TA | CG>TA | CAAAGCGCC | GCCAAAT | GCG | GCG | C | T | C>T | C>T_GCG | CG>TA_GCCAAAT | 4 | FALSE | FALSE |
| 58_v.txt | 32271 | 32271-SNV-C | SNV | 1 | C | T | 43 | 25655 | 0.16760865 | 32.3255814 | rnr1Y285A_pe | FALSE | ARG | JSY3868 | 1 | rnr1Y285A_p | ACG | C>T | A/T+1 | G/C-1 | CG>TA | CG>TA | AAAATACGG | CGGGAAC | TGC | ACG | C | A | C>T | C>T_ACG | CG>TA_CGGGAAC | 4 | FALSE | FALSE |
| 58_v.txt | 32299 | 32299-SNV-C | SNV | 1 | C | T | 50 | 25654 | 0.19490138 | 55.38 | rnr1Y285A_pe | FALSE | ARG | JSY3868 | 1 | rnr1Y285A_p | GCG | C>T | A/T+1 | G/C-1 | CG>TA | CG>TA | AATTTGCGG | CGGCAGA | CGC | GCG | C | A | C>T | C>T_GCG | CG>TA_CGGCAGA | 4 | FALSE | FALSE |
| 58_v.txt | 32466 | 32466-SNV-C | SNV | 1 | C | T | 34 | 25654 | 0.13253294 | 45.5 | rnr1Y285A_pe | FALSE | ARG | JSY3868 | 1 | rnr1Y285A_p | GCC | C>T | A/T+1 | G/C-1 | CG>TA | CG>TA | AGAGAGCCA | CCAATGT | CGG | GCC | C | A | C>T | C>T_GCC | CG>TA_CCAATGT | 4 | FALSE | FALSE |
| 58_v.txt | 32487 | 32487-SNV-C | SNV | 1 | C | T | 25 | 25654 | 0.09745069 | 45.52 | rnr1Y285A_pe | FALSE | ARG | JSY3868 | 1 | rnr1Y285A_p | ACG | C>T | A/T+1 | G/C-1 | CG>TA | CG>TA | AAGATACGG | CGGAAAA | TGC | ACG | C | A | C>T | C>T_ACG | CG>TA_CGGAAAA | 4 | FALSE | FALSE |
| 58_v.txt | 32535 | 32535-SNV-G | SNV | 1 | G | A | 40 | 25654 | 0.1559211 | 36.9750563 | rnr1Y285A_pe | FALSE | ARG | JSY3868 | 1 | rnr1Y285A_p | TGC | G>A | A/T+1 | G/C-1 | CG>TA | CG>TA | GGGTTTGCA | GCAGCTT | ACG | ACG | C | T | C>T | C>T_ACG | CG>TA_GCAGCTT | 4 | FALSE | FALSE |
| 58_v.txt | 32536 | 32536-SNV-C | SNV | 1 | C | T | 39 | 25654 | 0.15202308 | 36.5565266 | rnr1Y285A_pe | FALSE | ARG | JSY3868 | 1 | rnr1Y285A_p | GCA | C>T | A/T+1 | G/C-1 | CG>TA | CG>TA | GGTTTGCAG | CAGCTTC | CGT | GCA | C | A | C>T | C>T_GCA | CG>TA_CAGCTTC | 4 | FALSE | FALSE |
| 58_v.txt | 32700 | 32700-SNV-C | SNV | 1 | C | T | 23 | 18166 | 0.12661015 | 59.9130435 | rnr1Y285A_pe | FALSE | ARG | JSY3868 | 1 | rnr1Y285A_p | GCC | C>T | A/T+1 | G/C-1 | CG>TA | CG>TA | ACTGGGCCG | CCGGTAA | CGG | GCC | C | A | C>T | C>T_GCC | CG>TA_CCGGTAA | 4 | FALSE | FALSE |
| 58_v.txt | 32713 | 32713-SNV-C | SNV | 1 | C | T | 18 | 18166 | 0.09908621 | 62.7222222 | rnr1Y285A_pe | FALSE | ARG | JSY3868 | 1 | rnr1Y285A_p | GCA | C>T | A/T+1 | G/C-1 | CG>TA | CG>TA | CCCCAGCAC | CACCACA | CGT | GCA | C | A | C>T | C>T_GCA | CG>TA_CACCACA | 4 | FALSE | FALSE |
| 58_v.txt | 32798 | 32798-SNV-G | SNV | 1 | G | A | 30 | 18166 | 0.16514368 | 43.5 | rnr1Y285A_pe | FALSE | ARG | JSY3868 | 1 | rnr1Y285A_p | CGT | G>A | A/T+1 | G/C-1 | CG>TA | CG>TA | TTCACCGTAA | GTAATAT | GCA | GCA | C | T | C>T | C>T_GCA | CG>TA_GTAATAT | 4 | FALSE | FALSE |
| 58_v.txt | 32862 | 32862-SNV-G | SNV | 1 | G | A | 19 | 18164 | 0.10460251 | 36.3684211 | rnr1Y285A_pe | FALSE | ARG | JSY3868 | 1 | rnr1Y285A_p | CGC | G>A | A/T+1 | G/C-1 | CG>TA | CG>TA | CATGCCGCC | GCCAGTG | GCG | GCG | C | T | C>T | C>T_GCG | CG>TA_GCCAGTG | 4 | FALSE | FALSE |
| 58_v.txt | 32901 | 32901-SNV-C | SNV | 1 | C | T | 42 | 21417 | 0.1961059 | 32.0952381 | rnr1Y285A_pe | FALSE | ARG | JSY3868 | 1 | rnr1Y285A_p | GCC | C>T | A/T+1 | G/C-1 | CG>TA | CG>TA | ACTTGGCCA | CCAACTA | CGG | GCC | C | A | C>T | C>T_GCC | CG>TA_CCAACTA | 4 | FALSE | FALSE |
| 58_v.txt | 32965 | 32965-SNV-C | SNV | 1 | C | T | 20 | 21417 | 0.09338376 | 58.2 | rnr1Y285A_pe | FALSE | ARG | JSY3868 | 1 | rnr1Y285A_p | GCC | C>T | A/T+1 | G/C-1 | CG>TA | CG>TA | CATTGGCCG | CCGCACC | CGG | GCC | C | A | C>T | C>T_GCC | CG>TA_CCGCACC | 4 | FALSE | FALSE |
| 58_v.txt | 32967 | 32967-SNV-G | SNV | 1 | G | A | 66 | 21417 | 0.30816641 | 59.9090909 | rnr1Y285A_pe | FALSE | ARG | JSY3868 | 1 | rnr1Y285A_p | CGC | G>A | A/T+1 | G/C-1 | CG>TA | CG>TA | TTGGCCGCA | GCACCAA | GCG | GCG | C | T | C>T | C>T_GCG | CG>TA_GCACCAA | 4 | FALSE | FALSE |
| 58_v.txt | 33056 | 33056-SNV-C | SNV | 1 | C | T | 25 | 21417 | 0.1167297 | 55.88 | rnr1Y285A_pe | FALSE | ARG | JSY3868 | 1 | rnr1Y285A_p | GCG | C>T | A/T+1 | G/C-1 | CG>TA | CG>TA | GGACTGCGT | CGTGACA | CGC | GCG | C | A | C>T | C>T_GCG | CG>TA_CGTGACA | 4 | FALSE | FALSE |
| 58_v.txt | 33069 | 33069-SNV-G | SNV | 1 | G | A | 21 | 21417 | 0.09805295 | 51.3817955 | rnr1Y285A_pe | FALSE | ARG | JSY3868 | 1 | rnr1Y285A_p | TGC | G>A | A/T+1 | G/C-1 | CG>TA | CG>TA | GAATATGCC | GCCAAAG | ACG | ACG | C | T | C>T | C>T_ACG | CG>TA_GCCAAAG | 4 | FALSE | FALSE |
| 58_v.txt | 33070 | 33070-SNV-C | SNV | 1 | C | T | 28 | 21417 | 0.13073727 | 50.5719334 | rnr1Y285A_pe | FALSE | ARG | JSY3868 | 1 | rnr1Y285A_p | GCC | C>T | A/T+1 | G/C-1 | CG>TA | CG>TA | AATATGCCA | CCAAAGA | CGG | GCC | C | A | C>T | C>T_GCC | CG>TA_CCAAAGA | 4 | FALSE | FALSE |
| 58_v.txt | 33113 | 33113-SNV-G | SNV | 1 | G | A | 21 | 21417 | 0.09805295 | 38.4761905 | rnr1Y285A_pe | FALSE | ARG | JSY3868 | 1 | rnr1Y285A_p | GGC | G>A | A/T+1 | G/C-1 | CG>TA | CG>TA | CACTGGGCC | GCCGGCG | CCG | CCG | C | T | C>T | C>T_CCG | CG>TA_GCCGGCG | 4 | FALSE | FALSE |
| 58_v.txt | 33276 | 33276-SNV-G | SNV | 1 | G | A | 24 | 13211 | 0.18166679 | 44.125 | rnr1Y285A_pe | FALSE | ARG | JSY3868 | 1 | rnr1Y285A_p | CGT | G>A | A/T+1 | G/C-1 | CG>TA | CG>TA | GAGAACGTA | GTATCCT | GCA | GCA | C | T | C>T | C>T_GCA | CG>TA_GTATCCT | 4 | FALSE | FALSE |
| 58_v.txt | 33392 | 33392-SNV-G | SNV | 1 | G | A | 19 | 13211 | 0.14381954 | 43.9473684 | rnr1Y285A_pe | FALSE | ARG | JSY3868 | 1 | rnr1Y285A_p | CGT | G>A | A/T+1 | G/C-1 | CG>TA | CG>TA | AACGTCGTG | GTGAAAG | GCA | GCA | C | T | C>T | C>T_GCA | CG>TA_GTGAAAG | 4 | FALSE | FALSE |
| 58_v.txt | 33412 | 33412-SNV-G | SNV | 1 | G | A | 26 | 13211 | 0.19680569 | 50.4230769 | rnr1Y285A_pe | FALSE | ARG | JSY3868 | 1 | rnr1Y285A_p | GGC | G>A | A/T+1 | G/C-1 | CG>TA | CG>TA | TGACCGGCT | GCTCATT | CCG | CCG | C | T | C>T | C>T_CCG | CG>TA_GCTCATT | 4 | FALSE | FALSE |
| 58_v.txt | 33474 | 33474-SNV-C | SNV | 1 | C | T | 26 | 13210 | 0.19682059 | 33.7692308 | rnr1Y285A_pe | FALSE | ARG | JSY3868 | 1 | rnr1Y285A_p | GCC | C>T | A/T+1 | G/C-1 | CG>TA | CG>TA | GCTATGCCTT | CCTTTTT | CGG | GCC | C | A | C>T | C>T_GCC | CG>TA_CCTTTTT | 4 | FALSE | FALSE |
| 58_v.txt | 31657 | 31657-SNV-G | SNV | 1 | G | T | 70 | 7594 | 0.92178035 | 18.1857143 | rnr1Y285A_pe | FALSE | ARG | JSY3868 | 1 | rnr1Y285A_p | GGC | G>T | A/T+1 | G/C-1 | CG>AT | CG>AT | GAAATGGCG | GCGTGGA | CCG | CCG | C | A | C>A | C>A_CCG | CG>AT_GCGTGGA | 4 | FALSE | FALSE |
| 58_v.txt | 32811 | 32811-SNV-G | SNV | 1 | G | T | 7 | 18166 | 0.03853352 | 49.7142857 | rnr1Y285A_pe | FALSE | ARG | JSY3868 | 1 | rnr1Y285A_p | AGG | G>T | A/T+1 | G/C-1 | CG>AT | CG>AT | TTGACAGGG | GGGAACA | TCC | TCC | C | A | C>A | C>A_TCC | CG>AT_GGGAACA | 4 | FALSE | FALSE |
| 58_v.txt | 33474 | 33474-SNV-C | SNV | 1 | C | A | 47 | 13210 | 0.35579107 | 33.212766 | rnr1Y285A_pe | FALSE | ARG | JSY3868 | 1 | rnr1Y285A_p | GCC | C>A | A/T+1 | G/C-1 | CG>AT | CG>AT | GCTATGCCTT | CCTTTTT | CGG | GCC | C | T | C>A | C>A_GCC | CG>AT_CCTTTTT | 4 | FALSE | FALSE |
| 58_v.txt | 32081 | 32081-Insert | Insertion | 1 | - | A | 3 | 14659 | 0.02046524 | 64 | rnr1Y285A_pe | FALSE | ARG | JSY3868 | 1 | rnr1Y285A_p | CAA | nope | A/T+1 | >1bpdel | none | A/T+1 | CCATGCAAA | AAAAAAG | GTT | CAA | - | T | NA | NA_CAA | A/T+1_AAAAAAG | 4 | FALSE | FALSE |
| 58_v.txt | 32498 | 32498-Insert | Insertion | 1 | - | T | 5 | 25654 | 0.01949014 | 35.2 | rnr1Y285A_pe | FALSE | ARG | JSY3868 | 1 | rnr1Y285A_p | CTT | nope | A/T+1 | >1bpdel | none | A/T+1 | AACAACTTTT | TTTTTTG | GAA | CTT | - | A | NA | NA_CTT | A/T+1_TTTTTTG | 4 | FALSE | FALSE |
| 58_v.txt | 32842 | 32842-Insert | Insertion | 1 | - | A | 9 | 18166 | 0.0495431 | 40.5555556 | rnr1Y285A_pe | FALSE | ARG | JSY3868 | 1 | rnr1Y285A_p | CAA | nope | A/T+1 | >1bpdel | none | A/T+1 | TTACCCAAA | AAAAAAT | GTT | CAA | - | T | NA | NA_CAA | A/T+1_AAAAAAT | 4 | FALSE | FALSE |
| 58_v.txt | 31723 | 31723-Deleti | Deletion | 1 | A | - | 4 | 7594 | 0.05267316 | 63 | rnr1Y285A_pe | FALSE | ARG | JSY3868 | 1 | rnr1Y285A_p | CAA | nope | >1bp-ins | A/T-1 | none | A/T-1 | TGTCCCAAA | AAAAAGT | GTT | GTT | T | none | T>none | T>none_GTT | A/T-1_AAAAAGT | 4 | FALSE | FALSE |
| 58_v.txt | 31904 | 31904-Deleti | Deletion | 1 | T | - | 4 | 7594 | 0.05267316 | 37.75 | rnr1Y285A_pe | FALSE | ARG | JSY3868 | 1 | rnr1Y285A_p | ATT | nope | >1bp-ins | A/T-1 | none | A/T-1 | ATTGAATTTT | TTTTGGT | TAA | ATT | T | none | nope | nope_ATT | A/T-1_TTTTGGT | 4 | FALSE | FALSE |
| 58_v.txt | 31952 | 31952-Deleti | Deletion | 1 | A | - | 4 | 14659 | 0.02728699 | 36.25 | rnr1Y285A_pe | FALSE | ARG | JSY3868 | 1 | rnr1Y285A_p | TAA | nope | >1bp-ins | A/T-1 | none | A/T-1 | CGTCATAAA | AAATGTG | ATT | ATT | T | none | T>none | T>none_ATT | A/T-1_AAATGTG | 4 | FALSE | FALSE |
| 58_v.txt | 31972 | 31972-Deleti | Deletion | 1 | C | - | 4 | 14659 | 0.02728699 | 34.5 | rnr1Y285A_pe | FALSE | ARG | JSY3868 | 1 | rnr1Y285A_p | GCC | nope | >1bp-ins | G/C-1 | none | G/C-1 | AATAAGCCA | CCAAGCC | CGG | GCC | C | none | nope | nope_GCC | G/C-1_CCAAGCC | 4 | FALSE | FALSE |
| 58_v.txt | 31979 | 31979-Deleti | Deletion | 1 | G | - | 3 | 14659 | 0.02046524 | 36.3333333 | rnr1Y285A_pe | FALSE | ARG | JSY3868 | 1 | rnr1Y285A_p | CGG | nope | >1bp-ins | G/C-1 | none | G/C-1 | CAAGCCGGG | GGGCATT | GCC | GCC | C | none | C>none | C>none_GCC | G/C-1_GGGCATT | 4 | FALSE | FALSE |
| 58_v.txt | 32081 | 32081-Deleti | Deletion | 1 | A | - | 13 | 14659 | 0.08868272 | 61.0000011 | rnr1Y285A_pe | FALSE | ARG | JSY3868 | 1 | rnr1Y285A_p | CAA | nope | >1bp-ins | A/T-1 | none | A/T-1 | CCATGCAAA | AAAAAAG | GTT | GTT | T | none | T>none | T>none_GTT | A/T-1_AAAAAAG | 4 | FALSE | FALSE |
| 58_v.txt | 32160 | 32160-Deleti | Deletion | 1 | A | - | 5 | 14659 | 0.03410874 | 58.8 | rnr1Y285A_pe | FALSE | ARG | JSY3868 | 1 | rnr1Y285A_p | CAA | nope | >1bp-ins | A/T-1 | none | A/T-1 | TAAGCCAAA | AAAGCGC | GTT | GTT | T | none | T>none | T>none_GTT | A/T-1_AAAGCGC | 4 | FALSE | FALSE |
| 58_v.txt | 32299 | 32299-Deleti | Deletion | 1 | C | - | 2 | 25654 | 7.80E-03 | 51 | rnr1Y285A_pe | FALSE | ARG | JSY3868 | 1 | rnr1Y285A_p | GCG | nope | >1bp-ins | G/C-1 | none | G/C-1 | AATTTGCGG | CGGCAGA | CGC | GCG | C | none | nope | nope_GCG | G/C-1_CGGCAGA | 4 | FALSE | FALSE |
| 58_v.txt | 32381 | 32381-Deleti | Deletion | 1 | A | - | 9 | 25654 | 0.03508225 | 62.1111111 | rnr1Y285A_pe | FALSE | ARG | JSY3868 | 1 | rnr1Y285A_p | TAA | nope | >1bp-ins | A/T-1 | none | A/T-1 | AATAATAAA | AAAGGGA | ATT | ATT | T | none | T>none | T>none_ATT | A/T-1_AAAGGGA | 4 | FALSE | FALSE |
| 58_v.txt | 32442 | 32442-Deleti | Deletion | 1 | A | - | 7 | 25654 | 0.02728619 | 53.4285714 | rnr1Y285A_pe | FALSE | ARG | JSY3868 | 1 | rnr1Y285A_p | TAA | nope | >1bp-ins | A/T-1 | none | A/T-1 | GGAACTAAA | AAAAGTC | ATT | ATT | T | none | T>none | T>none_ATT | A/T-1_AAAAGTC | 4 | FALSE | FALSE |
| 58_v.txt | 32498 | 32498-Deleti | Deletion | 1 | T | - | 22 | 25654 | 0.08575661 | 42.5909195 | rnr1Y285A_pe | FALSE | ARG | JSY3868 | 1 | rnr1Y285A_p | CTT | nope | >1bp-ins | A/T-1 | none | A/T-1 | AACAACTTTT | TTTTTTG | GAA | CTT | T | none | nope | nope_CTT | A/T-1_TTTTTTG | 4 | FALSE | FALSE |
| 58_v.txt | 32707 | 32707-Deleti | Deletion | 1 | C | - | 5 | 18166 | 0.02752395 | 61.8 | rnr1Y285A_pe | FALSE | ARG | JSY3868 | 1 | rnr1Y285A_p | ACC | nope | >1bp-ins | G/C-1 | none | G/C-1 | CGGTAACCC | CCCCAGC | TGG | ACC | C | none | nope | nope_ACC | G/C-1_CCCCAGC | 4 | FALSE | FALSE |
| 58_v.txt | 32842 | 32842-Deleti | Deletion | 1 | A | - | 23 | 18166 | 0.12661015 | 41.0869565 | rnr1Y285A_pe | FALSE | ARG | JSY3868 | 1 | rnr1Y285A_p | CAA | nope | >1bp-ins | A/T-1 | none | A/T-1 | TTACCCAAA | AAAAAAT | GTT | GTT | T | none | T>none | T>none_GTT | A/T-1_AAAAAAT | 4 | FALSE | FALSE |
| 58_v.txt | 32884 | 32884-Deleti | Deletion | 1 | A | - | 4 | 21417 | 0.01867675 | 37.5 | rnr1Y285A_pe | FALSE | ARG | JSY3868 | 1 | rnr1Y285A_p | CAA | nope | >1bp-ins | A/T-1 | none | A/T-1 | ACGTCCAAA | AAAATTG | GTT | GTT | T | none | T>none | T>none_GTT | A/T-1_AAAATTG | 4 | FALSE | FALSE |
| 58_v.txt | 32901 | 32901-Deleti | Deletion | 1 | C | - | 3 | 21417 | 0.01400756 | 38 | rnr1Y285A_pe | FALSE | ARG | JSY3868 | 1 | rnr1Y285A_p | GCC | nope | >1bp-ins | G/C-1 | none | G/C-1 | ACTTGGCCA | CCAACTA | CGG | GCC | C | none | nope | nope_GCC | G/C-1_CCAACTA | 4 | FALSE | FALSE |
| 58_v.txt | 32924 | 32924-Deleti | Deletion | 1 | A | - | 23 | 21417 | 0.10739133 | 54.7391304 | rnr1Y285A_pe | FALSE | ARG | JSY3868 | 1 | rnr1Y285A_p | CAA | nope | >1bp-ins | A/T-1 | none | A/T-1 | CAGGGCAAA | AAAAGTG | GTT | GTT | T | none | T>none | T>none_GTT | A/T-1_AAAAGTG | 4 | FALSE | FALSE |
| 58_v.txt | 33147 | 33147-Deleti | Deletion | 1 | A | - | 7 | 21416 | 0.03268584 | 41.1428571 | rnr1Y285A_pe | FALSE | ARG | JSY3868 | 1 | rnr1Y285A_p | GAA | nope | >1bp-ins | A/T-1 | none | A/T-1 | CCAATGAAA | AAAAGAC | CTT | CTT | T | none | T>none | T>none_CTT | A/T-1_AAAAGAC | 4 | FALSE | FALSE |
| 58_v.txt | 33288 | 33288-Deleti | Deletion | 1 | T | - | 5 | 13211 | 0.03784725 | 48.4 | rnr1Y285A_pe | FALSE | ARG | JSY3868 | 1 | rnr1Y285A_p | ATT | nope | >1bp-ins | A/T-1 | none | A/T-1 | TCGCCATTTA | TTTACTC | TAA | ATT | T | none | nope | nope_ATT | A/T-1_TTTACTC | 4 | FALSE | FALSE |
| 58_v.txt | 33303 | 33303-Deleti | Deletion | 1 | A | - | 4 | 13211 | 0.0302778 | 63 | rnr1Y285A_pe | FALSE | ARG | JSY3868 | 1 | rnr1Y285A_p | GAA | nope | >1bp-ins | A/T-1 | none | A/T-1 | GTCGGGAAA | AAAGAGC | CTT | CTT | T | none | T>none | T>none_CTT | A/T-1_AAAGAGC | 4 | FALSE | FALSE |
| 58_v.txt | 33474 | 33474-Deleti | Deletion | 1 | C | - | 38 | 13210 | 0.28766086 | 26.0263158 | rnr1Y285A_pe | FALSE | ARG | JSY3868 | 1 | rnr1Y285A_p | GCC | nope | >1bp-ins | G/C-1 | none | G/C-1 | GCTATGCCTT | CCTTTTT | CGG | GCC | C | none | nope | nope_GCC | G/C-1_CCTTTTT | 4 | FALSE | FALSE |
| 59_v.txt | 31723 | 31723-SNV-A | SNV | 1 | A | C | 6 | 8296 | 0.07232401 | 51.3333333 | rnr1Y285A_pe | FALSE | ARG | JSY3868 | 2 | rnr1Y285A_p | CAA | A>C | G/C+1 | A/T-1 | TA>GC | TA>GC | TGTCCCAAA | AAAAAGT | GTT | GTT | T | G | T>G | T>G_GTT | TA>GC_AAAAAGT | 4 | FALSE | FALSE |
| 59_v.txt | 32081 | 32081-SNV-A | SNV | 1 | A | C | 1 | 15088 | 6.63E-03 | 63 | rnr1Y285A_pe | FALSE | ARG | JSY3868 | 2 | rnr1Y285A_p | CAA | A>C | G/C+1 | A/T-1 | TA>GC | TA>GC | CCATGCAAA | AAAAAAG | GTT | GTT | T | G | T>G | T>G_GTT | TA>GC_AAAAAAG | 4 | FALSE | FALSE |
| 59_v.txt | 32842 | 32842-SNV-A | SNV | 1 | A | C | 3 | 20939 | 0.01432733 | 37.9999277 | rnr1Y285A_pe | FALSE | ARG | JSY3868 | 2 | rnr1Y285A_p | CAA | A>C | G/C+1 | A/T-1 | TA>GC | TA>GC | TTACCCAAA | AAAAAAT | GTT | GTT | T | G | T>G | T>G_GTT | TA>GC_AAAAAAT | 4 | FALSE | FALSE |
| 59_v.txt | 32884 | 32884-SNV-A | SNV | 1 | A | C | 7 | 27562 | 0.02539729 | 33.7142857 | rnr1Y285A_pe | FALSE | ARG | JSY3868 | 2 | rnr1Y285A_p | CAA | A>C | G/C+1 | A/T-1 | TA>GC | TA>GC | ACGTCCAAA | AAAATTG | GTT | GTT | T | G | T>G | T>G_GTT | TA>GC_AAAATTG | 4 | FALSE | FALSE |
| 59_v.txt | 33269 | 33269-SNV-A | SNV | 1 | A | C | 109 | 16162 | 0.67442148 | 16.0091743 | rnr1Y285A_pe | FALSE | ARG | JSY3868 | 2 | rnr1Y285A_p | TAG | A>C | G/C+1 | A/T-1 | TA>GC | TA>GC | CTCCATAGA | AGAGAAC | ATC | ATC | T | G | T>G | T>G_ATC | TA>GC_AGAGAAC | 4 | FALSE | FALSE |
| 59_v.txt | 33479 | 33479-SNV-T | SNV | 1 | T | G | 4 | 16085 | 0.02486789 | 36.25 | rnr1Y285A_pe | FALSE | ARG | JSY3868 | 2 | rnr1Y285A_p | TTT | T>G | G/C+1 | A/T-1 | TA>GC | TA>GC | GCCTTTTTTT | TTTTTTT | AAA | TTT | T | C | T>G | T>G_TTT | TA>GC_TTTTTTT | 4 | FALSE | FALSE |
| 59_v.txt | 32081 | 32081-SNV-A | SNV | 1 | A | G | 6 | 15088 | 0.0397667 | 55.1666667 | rnr1Y285A_pe | FALSE | ARG | JSY3868 | 2 | rnr1Y285A_p | CAA | A>G | G/C+1 | A/T-1 | TA>CG | TA>CG | CCATGCAAA | AAAAAAG | GTT | GTT | T | C | T>C | T>C_GTT | TA>CG_AAAAAAG | 4 | FALSE | FALSE |
| 59_v.txt | 33476 | 33476-SNV-T | SNV | 1 | T | C | 22 | 16160 | 0.13613861 | 30.6361405 | rnr1Y285A_pe | FALSE | ARG | JSY3868 | 2 | rnr1Y285A_p | CTT | T>C | G/C+1 | A/T-1 | TA>CG | TA>CG | TATGCCTTTT | TTTTTTT | GAA | CTT | T | G | T>C | T>C_CTT | TA>CG_TTTTTTT | 4 | FALSE | FALSE |
| 59_v.txt | 33476 | 33476-SNV-T | SNV | 1 | T | A | 2 | 16160 | 0.01237624 | 37.9962923 | rnr1Y285A_pe | FALSE | ARG | JSY3868 | 2 | rnr1Y285A_p | CTT | T>A | A/T+1 | A/T-1 | TA>AT | TA>AT | TATGCCTTTT | TTTTTTT | GAA | CTT | T | T | T>A | T>A_CTT | TA>AT_TTTTTTT | 4 | FALSE | FALSE |
| 59_v.txt | 31650 | 31650-SNV-C | SNV | 1 | C | T | 21 | 8296 | 0.25313404 | 36 | rnr1Y285A_pe | FALSE | ARG | JSY3868 | 2 | rnr1Y285A_p | GCG | C>T | A/T+1 | G/C-1 | CG>TA | CG>TA | AGAATGCGA | CGAAATG | CGC | GCG | C | A | C>T | C>T_GCG | CG>TA_CGAAATG | 4 | FALSE | FALSE |
| 59_v.txt | 31658 | 31658-SNV-C | SNV | 1 | C | T | 13 | 8296 | 0.15670203 | 33.6153846 | rnr1Y285A_pe | FALSE | ARG | JSY3868 | 2 | rnr1Y285A_p | GCG | C>T | A/T+1 | G/C-1 | CG>TA | CG>TA | AAATGGCGT | CGTGGAA | CGC | GCG | C | A | C>T | C>T_GCG | CG>TA_CGTGGAA | 4 | FALSE | FALSE |
| 59_v.txt | 31879 | 31879-SNV-C | SNV | 1 | C | T | 15 | 8296 | 0.18081003 | 39.8 | rnr1Y285A_pe | FALSE | ARG | JSY3868 | 2 | rnr1Y285A_p | GCG | C>T | A/T+1 | G/C-1 | CG>TA | CG>TA | TATAGGCGG | CGGCAGC | CGC | GCG | C | A | C>T | C>T_GCG | CG>TA_CGGCAGC | 4 | FALSE | FALSE |
| 59_v.txt | 31960 | 31960-SNV-C | SNV | 1 | C | T | 19 | 15088 | 0.12592789 | 35.0526316 | rnr1Y285A_pe | FALSE | ARG | JSY3868 | 2 | rnr1Y285A_p | GCC | C>T | A/T+1 | G/C-1 | CG>TA | CG>TA | ATGTGGCCG | CCGCATA | CGG | GCC | C | A | C>T | C>T_GCC | CG>TA_CCGCATA | 4 | FALSE | FALSE |
| 59_v.txt | 31962 | 31962-SNV-G | SNV | 1 | G | A | 20 | 15088 | 0.13255567 | 37.55 | rnr1Y285A_pe | FALSE | ARG | JSY3868 | 2 | rnr1Y285A_p | CGC | G>A | A/T+1 | G/C-1 | CG>TA | CG>TA | GTGGCCGCA | GCATAAT | GCG | GCG | C | T | C>T | C>T_GCG | CG>TA_GCATAAT | 4 | FALSE | FALSE |
| 59_v.txt | 31972 | 31972-SNV-C | SNV | 1 | C | T | 19 | 15088 | 0.12592789 | 37.4736842 | rnr1Y285A_pe | FALSE | ARG | JSY3868 | 2 | rnr1Y285A_p | GCC | C>T | A/T+1 | G/C-1 | CG>TA | CG>TA | AATAAGCCA | CCAAGCC | CGG | GCC | C | A | C>T | C>T_GCC | CG>TA_CCAAGCC | 4 | FALSE | FALSE |
| 59_v.txt | 31981 | 31981-SNV-G | SNV | 1 | G | A | 20 | 15088 | 0.13255567 | 44.55 | rnr1Y285A_pe | FALSE | ARG | JSY3868 | 2 | rnr1Y285A_p | GGC | G>A | A/T+1 | G/C-1 | CG>TA | CG>TA | AGCCGGGCA | GCATTAA | CCG | CCG | C | T | C>T | C>T_CCG | CG>TA_GCATTAA | 4 | FALSE | FALSE |
| 59_v.txt | 32092 | 32092-SNV-C | SNV | 1 | C | T | 16 | 15088 | 0.10604454 | 62.875 | rnr1Y285A_pe | FALSE | ARG | JSY3868 | 2 | rnr1Y285A_p | GCA | C>T | A/T+1 | G/C-1 | CG>TA | CG>TA | AGCCTGCAA | CAACACC | CGT | GCA | C | A | C>T | C>T_GCA | CG>TA_CAACACC | 4 | FALSE | FALSE |
| 59_v.txt | 32165 | 32165-SNV-G | SNV | 1 | G | A | 31 | 15088 | 0.20546129 | 50.4193548 | rnr1Y285A_pe | FALSE | ARG | JSY3868 | 2 | rnr1Y285A_p | CGC | G>A | A/T+1 | G/C-1 | CG>TA | CG>TA | CAAAGCGCC | GCCAAAT | GCG | GCG | C | T | C>T | C>T_GCG | CG>TA_GCCAAAT | 4 | FALSE | FALSE |
| 59_v.txt | 32173 | 32173-SNV-C | SNV | 1 | C | T | 17 | 15088 | 0.11267232 | 48.7058824 | rnr1Y285A_pe | FALSE | ARG | JSY3868 | 2 | rnr1Y285A_p | GCA | C>T | A/T+1 | G/C-1 | CG>TA | CG>TA | CAAATGCAG | CAGCAGT | CGT | GCA | C | A | C>T | C>T_GCA | CG>TA_CAGCAGT | 4 | FALSE | FALSE |
| 59_v.txt | 32271 | 32271-SNV-C | SNV | 1 | C | T | 32 | 17039 | 0.18780445 | 32.125 | rnr1Y285A_pe | FALSE | ARG | JSY3868 | 2 | rnr1Y285A_p | ACG | C>T | A/T+1 | G/C-1 | CG>TA | CG>TA | AAAATACGG | CGGGAAC | TGC | ACG | C | A | C>T | C>T_ACG | CG>TA_CGGGAAC | 4 | FALSE | FALSE |
| 59_v.txt | 32299 | 32299-SNV-C | SNV | 1 | C | T | 30 | 17039 | 0.17606667 | 53.3333333 | rnr1Y285A_pe | FALSE | ARG | JSY3868 | 2 | rnr1Y285A_p | GCG | C>T | A/T+1 | G/C-1 | CG>TA | CG>TA | AATTTGCGG | CGGCAGA | CGC | GCG | C | A | C>T | C>T_GCG | CG>TA_CGGCAGA | 4 | FALSE | FALSE |
| 59_v.txt | 32466 | 32466-SNV-C | SNV | 1 | C | T | 23 | 17039 | 0.13498445 | 49.0434783 | rnr1Y285A_pe | FALSE | ARG | JSY3868 | 2 | rnr1Y285A_p | GCC | C>T | A/T+1 | G/C-1 | CG>TA | CG>TA | AGAGAGCCA | CCAATGT | CGG | GCC | C | A | C>T | C>T_GCC | CG>TA_CCAATGT | 4 | FALSE | FALSE |
| 59_v.txt | 32487 | 32487-SNV-C | SNV | 1 | C | T | 16 | 17039 | 0.09390222 | 44 | rnr1Y285A_pe | FALSE | ARG | JSY3868 | 2 | rnr1Y285A_p | ACG | C>T | A/T+1 | G/C-1 | CG>TA | CG>TA | AAGATACGG | CGGAAAA | TGC | ACG | C | A | C>T | C>T_ACG | CG>TA_CGGAAAA | 4 | FALSE | FALSE |
| 59_v.txt | 32535 | 32535-SNV-G | SNV | 1 | G | A | 23 | 17039 | 0.13498445 | 37.3479301 | rnr1Y285A_pe | FALSE | ARG | JSY3868 | 2 | rnr1Y285A_p | TGC | G>A | A/T+1 | G/C-1 | CG>TA | CG>TA | GGGTTTGCA | GCAGCTT | ACG | ACG | C | T | C>T | C>T_ACG | CG>TA_GCAGCTT | 4 | FALSE | FALSE |
| 59_v.txt | 32536 | 32536-SNV-C | SNV | 1 | C | T | 23 | 17039 | 0.13498445 | 37.6077362 | rnr1Y285A_pe | FALSE | ARG | JSY3868 | 2 | rnr1Y285A_p | GCA | C>T | A/T+1 | G/C-1 | CG>TA | CG>TA | GGTTTGCAG | CAGCTTC | CGT | GCA | C | A | C>T | C>T_GCA | CG>TA_CAGCTTC | 4 | FALSE | FALSE |
| 59_v.txt | 32590 | 32590-SNV-C | SNV | 1 | C | T | 36 | 20939 | 0.17192798 | 36.4166667 | rnr1Y285A_pe | FALSE | ARG | JSY3868 | 2 | rnr1Y285A_p | GCG | C>T | A/T+1 | G/C-1 | CG>TA | CG>TA | AGGCAGCGT | CGTTAAT | CGC | GCG | C | A | C>T | C>T_GCG | CG>TA_CGTTAAT | 4 | FALSE | FALSE |
| 59_v.txt | 32700 | 32700-SNV-C | SNV | 1 | C | T | 31 | 20939 | 0.1480491 | 58.0967742 | rnr1Y285A_pe | FALSE | ARG | JSY3868 | 2 | rnr1Y285A_p | GCC | C>T | A/T+1 | G/C-1 | CG>TA | CG>TA | ACTGGGCCG | CCGGTAA | CGG | GCC | C | A | C>T | C>T_GCC | CG>TA_CCGGTAA | 4 | FALSE | FALSE |
| 59_v.txt | 32776 | 32776-SNV-C | SNV | 1 | C | T | 17 | 20939 | 0.08118821 | 52.1176471 | rnr1Y285A_pe | FALSE | ARG | JSY3868 | 2 | rnr1Y285A_p | GCG | C>T | A/T+1 | G/C-1 | CG>TA | CG>TA | TGGAAGCGA | CGACCCA | CGC | GCG | C | A | C>T | C>T_GCG | CG>TA_CGACCCA | 4 | FALSE | FALSE |
| 59_v.txt | 32798 | 32798-SNV-G | SNV | 1 | G | A | 33 | 20939 | 0.15760065 | 43.4545455 | rnr1Y285A_pe | FALSE | ARG | JSY3868 | 2 | rnr1Y285A_p | CGT | G>A | A/T+1 | G/C-1 | CG>TA | CG>TA | TTCACCGTAA | GTAATAT | GCA | GCA | C | T | C>T | C>T_GCA | CG>TA_GTAATAT | 4 | FALSE | FALSE |
| 59_v.txt | 32860 | 32860-SNV-C | SNV | 1 | C | T | 28 | 20938 | 0.13372815 | 34.6428571 | rnr1Y285A_pe | FALSE | ARG | JSY3868 | 2 | rnr1Y285A_p | GCC | C>T | A/T+1 | G/C-1 | CG>TA | CG>TA | TCCATGCCG | CCGCCAG | CGG | GCC | C | A | C>T | C>T_GCC | CG>TA_CCGCCAG | 4 | FALSE | FALSE |
| 59_v.txt | 32900 | 32900-SNV-G | SNV | 1 | G | A | 29 | 27562 | 0.10521733 | 33.9284486 | rnr1Y285A_pe | FALSE | ARG | JSY3868 | 2 | rnr1Y285A_p | GGC | G>A | A/T+1 | G/C-1 | CG>TA | CG>TA | GACTTGGCC | GCCAACT | CCG | CCG | C | T | C>T | C>T_CCG | CG>TA_GCCAACT | 4 | FALSE | FALSE |
| 59_v.txt | 32901 | 32901-SNV-C | SNV | 1 | C | T | 70 | 27562 | 0.25397286 | 32.4290726 | rnr1Y285A_pe | FALSE | ARG | JSY3868 | 2 | rnr1Y285A_p | GCC | C>T | A/T+1 | G/C-1 | CG>TA | CG>TA | ACTTGGCCA | CCAACTA | CGG | GCC | C | A | C>T | C>T_GCC | CG>TA_CCAACTA | 4 | FALSE | FALSE |

| | | | | | | | | | | | | | | | | | | | | | | | | | | | | | | | | | | |
|---|---|---|---|---|---|---|---|---|---|---|---|---|---|---|---|---|---|---|---|---|---|---|---|---|---|---|---|---|---|---|---|---|---|---|
| 59_v.txt | 32935 | 32935-SNV- | SNV | 1 | C | T | 34 | 27562 | 0.12335825 | 35.6176471 | rnr1Y285A_pe | FALSE | ARG | JSY3868 | 2 | rnr1Y285A_p | GCC | C>T | A/T+1 | G/C-1 | CG>TA | CG>TA | TGATTGCCC | CCCAAGA | CGG | GCC | C | A | C>T | C>T_GCC | CG>TA_CCCAAGA | 4 | FALSE | FALSE |
| 59_v.txt | 32965 | 32965-SNV- | SNV | 1 | C | T | 26 | 27562 | 0.09433278 | 57.3846154 | rnr1Y285A_pe | FALSE | ARG | JSY3868 | 2 | rnr1Y285A_p | GCC | C>T | A/T+1 | G/C-1 | CG>TA | CG>TA | CATTGGCCG | CCGCACC | CGG | GCC | C | A | C>T | C>T_GCC | CG>TA_CCGCACC | 4 | FALSE | FALSE |
| 59_v.txt | 32967 | 32967-SNV- | SNV | 1 | G | A | 66 | 27562 | 0.23946013 | 61.1515152 | rnr1Y285A_pe | FALSE | ARG | JSY3868 | 2 | rnr1Y285A_p | CGC | G>A | A/T+1 | G/C-1 | CG>TA | CG>TA | TTGGCCGCA | GCACCAA | GCG | GCG | C | T | C>T | C>T_GCG | CG>TA_GCACCAA | 4 | FALSE | FALSE |
| 59_v.txt | 33056 | 33056-SNV- | SNV | 1 | C | T | 28 | 27562 | 0.10158915 | 55.1071429 | rnr1Y285A_pe | FALSE | ARG | JSY3868 | 2 | rnr1Y285A_p | GCG | C>T | A/T+1 | G/C-1 | CG>TA | CG>TA | GGACTGCGT | CGTGACA | CGC | GCG | C | A | C>T | C>T_GCG | CG>TA_CGTGACA | 4 | FALSE | FALSE |
| 59_v.txt | 33070 | 33070-SNV- | SNV | 1 | C | T | 29 | 27562 | 0.10521733 | 55.4827586 | rnr1Y285A_pe | FALSE | ARG | JSY3868 | 2 | rnr1Y285A_p | GCC | C>T | A/T+1 | G/C-1 | CG>TA | CG>TA | AATATGCCA | CCAAAGA | CGG | GCC | C | A | C>T | C>T_GCC | CG>TA_CCAAAGA | 4 | FALSE | FALSE |
| 59_v.txt | 33104 | 33104-SNV- | SNV | 1 | G | A | 32 | 27562 | 0.11610188 | 39.96875 | rnr1Y285A_pe | FALSE | ARG | JSY3868 | 2 | rnr1Y285A_p | CGC | G>A | A/T+1 | G/C-1 | CG>TA | CG>TA | AAGAGCGCC | GCCCACT | GCG | GCG | C | T | C>T | C>T_GCG | CG>TA_GCCCACT | 4 | FALSE | FALSE |
| 59_v.txt | 33114 | 33114-SNV- | SNV | 1 | C | T | 30 | 27562 | 0.10884551 | 41.5333333 | rnr1Y285A_pe | FALSE | ARG | JSY3868 | 2 | rnr1Y285A_p | GCC | C>T | A/T+1 | G/C-1 | CG>TA | CG>TA | ACTGGGCCG | CCGGCGT | CGG | GCC | C | A | C>T | C>T_GCC | CG>TA_CCGGCGT | 4 | FALSE | FALSE |
| 59_v.txt | 33118 | 33118-SNV- | SNV | 1 | C | T | 28 | 27562 | 0.10158915 | 40.7492964 | rnr1Y285A_pe | FALSE | ARG | JSY3868 | 2 | rnr1Y285A_p | GCG | C>T | A/T+1 | G/C-1 | CG>TA | CG>TA | GGCCGGCGT | CGTTGGT | CGC | GCG | C | A | C>T | C>T_GCG | CG>TA_CGTTGGT | 4 | FALSE | FALSE |
| 59_v.txt | 33119 | 33119-SNV- | SNV | 1 | G | A | 22 | 27562 | 0.07982004 | 40.7269337 | rnr1Y285A_pe | FALSE | ARG | JSY3868 | 2 | rnr1Y285A_p | CGT | G>A | A/T+1 | G/C-1 | CG>TA | CG>TA | GCCGGCGTT | GTTGGTC | GCA | GCA | C | T | C>T | C>T_GCA | CG>TA_GTTGGTC | 4 | FALSE | FALSE |
| 59_v.txt | 33258 | 33258-SNV- | SNV | 1 | C | T | 30 | 16162 | 0.18562059 | 38.5 | rnr1Y285A_pe | FALSE | ARG | JSY3868 | 2 | rnr1Y285A_p | GCC | C>T | A/T+1 | G/C-1 | CG>TA | CG>TA | CCTATGCCAT | CCATCCT | CGG | GCC | C | A | C>T | C>T_GCC | CG>TA_CCATCCT | 4 | FALSE | FALSE |
| 59_v.txt | 33276 | 33276-SNV- | SNV | 1 | G | A | 25 | 16162 | 0.15468383 | 47.92 | rnr1Y285A_pe | FALSE | ARG | JSY3868 | 2 | rnr1Y285A_p | CGT | G>A | A/T+1 | G/C-1 | CG>TA | CG>TA | GAGAACGTA | GTATCCT | GCA | GCA | C | T | C>T | C>T_GCA | CG>TA_GTATCCT | 4 | FALSE | FALSE |
| 59_v.txt | 33310 | 33310-SNV- | SNV | 1 | G | A | 21 | 16162 | 0.12993441 | 57.6190476 | rnr1Y285A_pe | FALSE | ARG | JSY3868 | 2 | rnr1Y285A_p | CGC | G>A | A/T+1 | G/C-1 | CG>TA | CG>TA | AAGAGCGCA | GCAATGG | GCG | GCG | C | T | C>T | C>T_GCG | CG>TA_GCAATGG | 4 | FALSE | FALSE |
| 59_v.txt | 33412 | 33412-SNV- | SNV | 1 | G | A | 24 | 16162 | 0.14849647 | 54.3333333 | rnr1Y285A_pe | FALSE | ARG | JSY3868 | 2 | rnr1Y285A_p | GGC | G>A | A/T+1 | G/C-1 | CG>TA | CG>TA | TGACCGGCT | GCTCATT | CCG | CCG | C | T | C>T | C>T_CCG | CG>TA_GCTCATT | 4 | FALSE | FALSE |
| 59_v.txt | 33474 | 33474-SNV- | SNV | 1 | C | T | 25 | 16162 | 0.15468383 | 35.08 | rnr1Y285A_pe | FALSE | ARG | JSY3868 | 2 | rnr1Y285A_p | GCC | C>T | A/T+1 | G/C-1 | CG>TA | CG>TA | GCTATGCCTT | CCTTTTT | CGG | GCC | C | A | C>T | C>T_GCC | CG>TA_CCTTTTT | 4 | FALSE | FALSE |
| 59_v.txt | 31650 | 31650-SNV- | SNV | 1 | C | G | 2 | 8296 | 0.024108 | 37.5 | rnr1Y285A_pe | FALSE | ARG | JSY3868 | 2 | rnr1Y285A_p | GCG | C>G | G/C+1 | G/C-1 | CG>GC | CG>GC | AGAATGCGA | CGAAATG | CGC | GCG | C | C | C>G | C>G_GCG | CG>GC_CGAAATG | 4 | FALSE | FALSE |
| 59_v.txt | 31657 | 31657-SNV- | SNV | 1 | G | C | 3 | 8296 | 0.03616201 | 19.9997211 | rnr1Y285A_pe | FALSE | ARG | JSY3868 | 2 | rnr1Y285A_p | GGC | G>C | G/C+1 | G/C-1 | CG>GC | CG>GC | GAAATGGCG | GCGTGGA | CCG | CCG | C | G | C>G | C>G_CCG | CG>GC_GCGTGGA | 4 | FALSE | FALSE |
| 59_v.txt | 32901 | 32901-SNV- | SNV | 1 | C | G | 3 | 27562 | 0.01088455 | 36.6669227 | rnr1Y285A_pe | FALSE | ARG | JSY3868 | 2 | rnr1Y285A_p | GCC | C>G | G/C+1 | G/C-1 | CG>GC | CG>GC | ACTTGGCCA | CCAACTA | CGG | GCC | C | C | C>G | C>G_GCC | CG>GC_CCAACTA | 4 | FALSE | FALSE |
| 59_v.txt | 33118 | 33118-SNV- | SNV | 1 | C | G | 3 | 27562 | 0.01088455 | 27.333225 | rnr1Y285A_pe | FALSE | ARG | JSY3868 | 2 | rnr1Y285A_p | GCG | C>G | G/C+1 | G/C-1 | CG>GC | CG>GC | GGCCGGCGT | CGTTGGT | CGC | GCG | C | C | C>G | C>G_GCG | CG>GC_CGTTGGT | 4 | FALSE | FALSE |
| 59_v.txt | 33474 | 33474-SNV- | SNV | 1 | C | G | 1 | 16162 | 6.19E-03 | 27 | rnr1Y285A_pe | FALSE | ARG | JSY3868 | 2 | rnr1Y285A_p | GCC | C>G | G/C+1 | G/C-1 | CG>GC | CG>GC | GCTATGCCTT | CCTTTTT | CGG | GCC | C | C | C>G | C>G_GCC | CG>GC_CCTTTTT | 4 | FALSE | FALSE |
| 59_v.txt | 31657 | 31657-SNV- | SNV | 1 | G | T | 65 | 8296 | 0.78351013 | 18.7846813 | rnr1Y285A_pe | FALSE | ARG | JSY3868 | 2 | rnr1Y285A_p | GGC | G>T | A/T+1 | G/C-1 | CG>AT | CG>AT | GAAATGGCG | GCGTGGA | CCG | CCG | C | A | C>A | C>A_CCG | CG>AT_GCGTGGA | 4 | FALSE | FALSE |
| 59_v.txt | 33310 | 33310-SNV- | SNV | 1 | G | T | 10 | 16162 | 0.06187353 | 24.8 | rnr1Y285A_pe | FALSE | ARG | JSY3868 | 2 | rnr1Y285A_p | CGC | G>T | A/T+1 | G/C-1 | CG>AT | CG>AT | AAGAGCGCA | GCAATGG | GCG | GCG | C | A | C>A | C>A_GCG | CG>AT_GCAATGG | 4 | FALSE | FALSE |
| 59_v.txt | 33474 | 33474-SNV- | SNV | 1 | C | A | 52 | 16162 | 0.32174236 | 34 | rnr1Y285A_pe | FALSE | ARG | JSY3868 | 2 | rnr1Y285A_p | GCC | C>A | A/T+1 | G/C-1 | CG>AT | CG>AT | GCTATGCCTT | CCTTTTT | CGG | GCC | C | T | C>A | C>A_GCC | CG>AT_CCTTTTT | 4 | FALSE | FALSE |
| 59_v.txt | 33475 | 33475-SNV- | SNV | 1 | C | A | 12 | 16161 | 0.07425283 | 33.75 | rnr1Y285A_pe | FALSE | ARG | JSY3868 | 2 | rnr1Y285A_p | CCT | C>A | A/T+1 | G/C-1 | CG>AT | CG>AT | CTATGCCTTT | CTTTTTT | GGA | CCT | C | T | C>A | C>A_CCT | CG>AT_CTTTTTT | 4 | FALSE | FALSE |
| 59_v.txt | 32081 | 32081-Insert | Insertion | 1 | - | A | 8 | 15088 | 0.05302227 | 60.375 | rnr1Y285A_pe | FALSE | ARG | JSY3868 | 2 | rnr1Y285A_p | CAA | nope | A/T+1 | >1bpdel | none | A/T+1 | CCATGCAAA | AAAAAAG | GTT | CAA | - | T | NA | NA_CAA | A/T+1_AAAAAAG | 4 | FALSE | FALSE |
| 59_v.txt | 32842 | 32842-Insert | Insertion | 1 | - | A | 10 | 20939 | 0.04775777 | 45.2 | rnr1Y285A_pe | FALSE | ARG | JSY3868 | 2 | rnr1Y285A_p | CAA | nope | A/T+1 | >1bpdel | none | A/T+1 | TTACCCAAA | AAAAAAT | GTT | CAA | - | T | NA | NA_CAA | A/T+1_AAAAAAT | 4 | FALSE | FALSE |
| 59_v.txt | 33087 | 33087-Insert | Insertion | 1 | - | A | 2 | 27562 | 7.26E-03 | 64 | rnr1Y285A_pe | FALSE | ARG | JSY3868 | 2 | rnr1Y285A_p | TAA | nope | A/T+1 | >1bpdel | none | A/T+1 | ATAAATAAA | AAATATG | ATT | TAA | - | T | NA | NA_TAA | A/T+1_AAATATG | 4 | FALSE | FALSE |
| 59_v.txt | 31723 | 31723-Deleti | Deletion | 1 | A | - | 10 | 8296 | 0.12054002 | 62 | rnr1Y285A_pe | FALSE | ARG | JSY3868 | 2 | rnr1Y285A_p | CAA | nope | >1bp-ins | A/T-1 | none | A/T-1 | TGTCCCAAA | AAAAAGT | GTT | GTT | T | none | T>none | T>none_GTT | A/T-1_AAAAAGT | 4 | FALSE | FALSE |
| 59_v.txt | 31913 | 31913-Deleti | Deletion | 1 | A | - | 4 | 8296 | 0.04821601 | 35 | rnr1Y285A_pe | FALSE | ARG | JSY3868 | 2 | rnr1Y285A_p | CAA | nope | >1bp-ins | A/T-1 | none | A/T-1 | TGGTGCAAA | AAAAGCC | GTT | GTT | T | none | T>none | T>none_GTT | A/T-1_AAAAGCC | 4 | FALSE | FALSE |
| 59_v.txt | 31979 | 31979-Deleti | Deletion | 1 | G | - | 5 | 15088 | 0.03313892 | 34.6 | rnr1Y285A_pe | FALSE | ARG | JSY3868 | 2 | rnr1Y285A_p | CGG | nope | >1bp-ins | G/C-1 | none | G/C-1 | CAAGCCGGG | GGGCATT | GCC | GCC | C | none | C>none | C>none_GCC | G/C-1_GGGCATT | 4 | FALSE | FALSE |
| 59_v.txt | 32081 | 32081-Deleti | Deletion | 1 | A | - | 12 | 15088 | 0.0795334 | 64 | rnr1Y285A_pe | FALSE | ARG | JSY3868 | 2 | rnr1Y285A_p | CAA | nope | >1bp-ins | A/T-1 | none | A/T-1 | CCATGCAAA | AAAAAAG | GTT | GTT | T | none | T>none | T>none_GTT | A/T-1_AAAAAAG | 4 | FALSE | FALSE |
| 59_v.txt | 32346 | 32346-Deleti | Deletion | 1 | A | - | 6 | 17039 | 0.03521333 | 59.6666667 | rnr1Y285A_pe | FALSE | ARG | JSY3868 | 2 | rnr1Y285A_p | CAA | nope | >1bp-ins | A/T-1 | none | A/T-1 | TGTGGCAAA | AAAACCT | GTT | GTT | T | none | T>none | T>none_GTT | A/T-1_AAAACCT | 4 | FALSE | FALSE |
| 59_v.txt | 32381 | 32381-Deleti | Deletion | 1 | A | - | 6 | 17039 | 0.03521333 | 56.6666667 | rnr1Y285A_pe | FALSE | ARG | JSY3868 | 2 | rnr1Y285A_p | TAA | nope | >1bp-ins | A/T-1 | none | A/T-1 | AATAATAAA | AAAGGGA | ATT | ATT | T | none | T>none | T>none_ATT | A/T-1_AAAGGGA | 4 | FALSE | FALSE |
| 59_v.txt | 32498 | 32498-Deleti | Deletion | 1 | T | - | 22 | 17039 | 0.12911556 | 48.5454546 | rnr1Y285A_pe | FALSE | ARG | JSY3868 | 2 | rnr1Y285A_p | CTT | nope | >1bp-ins | A/T-1 | none | A/T-1 | AACAACTTTT | TTTTTTG | GAA | CTT | T | none | nope | nope_CTT | A/T-1_TTTTTTG | 4 | FALSE | FALSE |
| 59_v.txt | 32631 | 32631-Deleti | Deletion | 1 | T | - | 8 | 20939 | 0.03820622 | 48.375 | rnr1Y285A_pe | FALSE | ARG | JSY3868 | 2 | rnr1Y285A_p | GTT | nope | >1bp-ins | A/T-1 | none | A/T-1 | CCTTCGTTTT | TTTTTAT | CAA | GTT | T | none | nope | nope_GTT | A/T-1_TTTTTAT | 4 | FALSE | FALSE |
| 59_v.txt | 32658 | 32658-Deleti | Deletion | 1 | C | - | 8 | 20939 | 0.03820622 | 60.5 | rnr1Y285A_pe | FALSE | ARG | JSY3868 | 2 | rnr1Y285A_p | ACC | nope | >1bp-ins | G/C-1 | none | G/C-1 | CCTGGACCC | CCCCAGG | TGG | ACC | C | none | nope | nope_ACC | G/C-1_CCCCAGG | 4 | FALSE | FALSE |
| 59_v.txt | 32697 | 32697-Deleti | Deletion | 1 | G | - | 4 | 20939 | 0.01910311 | 57.5 | rnr1Y285A_pe | FALSE | ARG | JSY3868 | 2 | rnr1Y285A_p | TGG | nope | >1bp-ins | G/C-1 | none | G/C-1 | CCAACTGGG | GGGCCGG | ACC | ACC | C | none | C>none | C>none_ACC | G/C-1_GGGCCGG | 4 | FALSE | FALSE |
| 59_v.txt | 32707 | 32707-Deleti | Deletion | 1 | C | - | 7 | 20939 | 0.03343044 | 63.1428571 | rnr1Y285A_pe | FALSE | ARG | JSY3868 | 2 | rnr1Y285A_p | ACC | nope | >1bp-ins | G/C-1 | none | G/C-1 | CGGTAACCC | CCCCAGC | TGG | ACC | C | none | nope | nope_ACC | G/C-1_CCCCAGC | 4 | FALSE | FALSE |
| 59_v.txt | 32842 | 32842-Deleti | Deletion | 1 | A | - | 31 | 20939 | 0.1480491 | 40.6129031 | rnr1Y285A_pe | FALSE | ARG | JSY3868 | 2 | rnr1Y285A_p | CAA | nope | >1bp-ins | A/T-1 | none | A/T-1 | TTACCCAAA | AAAAAAT | GTT | GTT | T | none | T>none | T>none_GTT | A/T-1_AAAAAAT | 4 | FALSE | FALSE |
| 59_v.txt | 32884 | 32884-Deleti | Deletion | 1 | A | - | 6 | 27562 | 0.0217691 | 35.6666667 | rnr1Y285A_pe | FALSE | ARG | JSY3868 | 2 | rnr1Y285A_p | CAA | nope | >1bp-ins | A/T-1 | none | A/T-1 | ACGTCCAAA | AAAATTG | GTT | GTT | T | none | T>none | T>none_GTT | A/T-1_AAAATTG | 4 | FALSE | FALSE |
| 59_v.txt | 32901 | 32901-Deleti | Deletion | 1 | C | - | 1 | 27562 | 3.63E-03 | 38 | rnr1Y285A_pe | FALSE | ARG | JSY3868 | 2 | rnr1Y285A_p | GCC | nope | >1bp-ins | G/C-1 | none | G/C-1 | ACTTGGCCA | CCAACTA | CGG | GCC | C | none | nope | nope_GCC | G/C-1_CCAACTA | 4 | FALSE | FALSE |
| 59_v.txt | 32924 | 32924-Deleti | Deletion | 1 | A | - | 28 | 27562 | 0.10158915 | 56.1071429 | rnr1Y285A_pe | FALSE | ARG | JSY3868 | 2 | rnr1Y285A_p | CAA | nope | >1bp-ins | A/T-1 | none | A/T-1 | CAGGGCAAA | AAAAGTG | GTT | GTT | T | none | T>none | T>none_GTT | A/T-1_AAAAGTG | 4 | FALSE | FALSE |
| 59_v.txt | 32935 | 32935-Deleti | Deletion | 1 | C | - | 9 | 27562 | 0.03265365 | 47.1111111 | rnr1Y285A_pe | FALSE | ARG | JSY3868 | 2 | rnr1Y285A_p | GCC | nope | >1bp-ins | G/C-1 | none | G/C-1 | TGATTGCCC | CCCAAGA | CGG | GCC | C | none | nope | nope_GCC | G/C-1_CCCAAGA | 4 | FALSE | FALSE |
| 59_v.txt | 32965 | 32965-Deleti | Deletion | 1 | C | - | 3 | 27562 | 0.01088455 | 54.3333333 | rnr1Y285A_pe | FALSE | ARG | JSY3868 | 2 | rnr1Y285A_p | GCC | nope | >1bp-ins | G/C-1 | none | G/C-1 | CATTGGCCG | CCGCACC | CGG | GCC | C | none | nope | nope_GCC | G/C-1_CCGCACC | 4 | FALSE | FALSE |
| 59_v.txt | 33147 | 33147-Deleti | Deletion | 1 | A | - | 11 | 27562 | 0.03991002 | 37.6363636 | rnr1Y285A_pe | FALSE | ARG | JSY3868 | 2 | rnr1Y285A_p | GAA | nope | >1bp-ins | A/T-1 | none | A/T-1 | CCAATGAAA | AAAAGAC | CTT | CTT | T | none | T>none | T>none_CTT | A/T-1_AAAAGAC | 4 | FALSE | FALSE |
| 59_v.txt | 33474 | 33474-Deleti | Deletion | 1 | C | - | 43 | 16162 | 0.26605618 | 30.3255814 | rnr1Y285A_pe | FALSE | ARG | JSY3868 | 2 | rnr1Y285A_p | GCC | nope | >1bp-ins | G/C-1 | none | G/C-1 | GCTATGCCTT | CCTTTTT | CGG | GCC | C | none | nope | nope_GCC | G/C-1_CCTTTTT | 4 | FALSE | FALSE |
| 60_v.txt | 31723 | 31723-SNV- | SNV | 1 | A | C | 5 | 15246 | 0.03279549 | 62.4 | rnr1Y285A_pe | FALSE | ARG | JSY3868 | 3 | rnr1Y285A_p | CAA | A>C | G/C+1 | A/T-1 | TA>GC | TA>GC | TGTCCCAAA | AAAAAGT | GTT | GTT | T | G | T>G | T>G_GTT | TA>GC_AAAAAGT | 4 | FALSE | FALSE |
| 60_v.txt | 31840 | 31840-SNV- | SNV | 1 | A | C | 4 | 15246 | 0.02623639 | 51 | rnr1Y285A_pe | FALSE | ARG | JSY3868 | 3 | rnr1Y285A_p | AAG | A>C | G/C+1 | A/T-1 | TA>GC | TA>GC | GAAATAAGA | AGATCCA | TTC | TTC | T | G | T>G | T>G_TTC | TA>GC_AGATCCA | 4 | FALSE | FALSE |
| 60_v.txt | 32842 | 32842-SNV- | SNV | 1 | A | C | 5 | 23655 | 0.02113718 | 32.2001154 | rnr1Y285A_pe | FALSE | ARG | JSY3868 | 3 | rnr1Y285A_p | CAA | A>C | G/C+1 | A/T-1 | TA>GC | TA>GC | TTACCCAAA | AAAAAAT | GTT | GTT | T | G | T>G | T>G_GTT | TA>GC_AAAAAAT | 4 | FALSE | FALSE |
| 60_v.txt | 33099 | 33099-SNV- | SNV | 1 | A | C | 10 | 22582 | 0.04428306 | 42.6 | rnr1Y285A_pe | FALSE | ARG | JSY3868 | 3 | rnr1Y285A_p | AAG | A>C | G/C+1 | A/T-1 | TA>GC | TA>GC | GATATAAGA | AGAGCGC | TTC | TTC | T | G | T>G | T>G_TTC | TA>GC_AGAGCGC | 4 | FALSE | FALSE |
| 60_v.txt | 33269 | 33269-SNV- | SNV | 1 | A | C | 134 | 19077 | 0.70241652 | 16.4776119 | rnr1Y285A_pe | FALSE | ARG | JSY3868 | 3 | rnr1Y285A_p | TAG | A>C | G/C+1 | A/T-1 | TA>GC | TA>GC | CTCCATAGA | AGAGAAC | ATC | ATC | T | G | T>G | T>G_ATC | TA>GC_AGAGAAC | 4 | FALSE | FALSE |
| 60_v.txt | 32081 | 32081-SNV- | SNV | 1 | A | G | 6 | 23101 | 0.0259729 | 59.3333333 | rnr1Y285A_pe | FALSE | ARG | JSY3868 | 3 | rnr1Y285A_p | CAA | A>G | G/C+1 | A/T-1 | TA>CG | TA>CG | CCATGCAAA | AAAAAAG | GTT | GTT | T | C | T>C | T>C_GTT | TA>CG_AAAAAAG | 4 | FALSE | FALSE |
| 60_v.txt | 33099 | 33099-SNV- | SNV | 1 | A | T | 6 | 22582 | 0.02656983 | 46 | rnr1Y285A_pe | FALSE | ARG | JSY3868 | 3 | rnr1Y285A_p | AAG | A>T | A/T+1 | A/T-1 | TA>AT | TA>AT | GATATAAGA | AGAGCGC | TTC | TTC | T | A | T>A | T>A_TTC | TA>AT_AGAGCGC | 4 | FALSE | FALSE |
| 60_v.txt | 31650 | 31650-SNV- | SNV | 1 | C | T | 24 | 15246 | 0.15741834 | 37.0833333 | rnr1Y285A_pe | FALSE | ARG | JSY3868 | 3 | rnr1Y285A_p | GCG | C>T | A/T+1 | G/C-1 | CG>TA | CG>TA | AGAATGCGA | CGAAATG | CGC | GCG | C | A | C>T | C>T_GCG | CG>TA_CGAAATG | 4 | FALSE | FALSE |
| 60_v.txt | 31658 | 31658-SNV- | SNV | 1 | C | T | 25 | 15246 | 0.16397744 | 32.519213 | rnr1Y285A_pe | FALSE | ARG | JSY3868 | 3 | rnr1Y285A_p | GCG | C>T | A/T+1 | G/C-1 | CG>TA | CG>TA | AAATGGCGT | CGTGGAA | CGC | GCG | C | A | C>T | C>T_GCG | CG>TA_CGTGGAA | 4 | FALSE | FALSE |
| 60_v.txt | 31830 | 31830-SNV- | SNV | 1 | C | T | 15 | 15246 | 0.09838646 | 51.6666667 | rnr1Y285A_pe | FALSE | ARG | JSY3868 | 3 | rnr1Y285A_p | GCA | C>T | A/T+1 | G/C-1 | CG>TA | CG>TA | AATATGCATT | CATTGAA | CGT | GCA | C | A | C>T | C>T_GCA | CG>TA_CATTGAA | 4 | FALSE | FALSE |
| 60_v.txt | 31882 | 31882-SNV- | SNV | 1 | C | T | 20 | 15246 | 0.13118195 | 39.8 | rnr1Y285A_pe | FALSE | ARG | JSY3868 | 3 | rnr1Y285A_p | GCA | C>T | A/T+1 | G/C-1 | CG>TA | CG>TA | AGGCGGCAG | CAGCAAA | CGT | GCA | C | A | C>T | C>T_GCA | CG>TA_CAGCAAA | 4 | FALSE | FALSE |
| 60_v.txt | 31911 | 31911-SNV- | SNV | 1 | G | A | 14 | 15246 | 0.09182737 | 37.2142857 | rnr1Y285A_pe | FALSE | ARG | JSY3868 | 3 | rnr1Y285A_p | TGC | G>A | A/T+1 | G/C-1 | CG>TA | CG>TA | TTTGGTGCA | GCAAAAG | ACG | ACG | C | T | C>T | C>T_ACG | CG>TA_GCAAAAG | 4 | FALSE | FALSE |
| 60_v.txt | 31962 | 31962-SNV- | SNV | 1 | G | A | 40 | 23100 | 0.17316017 | 35.5 | rnr1Y285A_pe | FALSE | ARG | JSY3868 | 3 | rnr1Y285A_p | CGC | G>A | A/T+1 | G/C-1 | CG>TA | CG>TA | GTGGCCGCA | GCATAAT | GCG | GCG | C | T | C>T | C>T_GCG | CG>TA_GCATAAT | 4 | FALSE | FALSE |
| 60_v.txt | 31972 | 31972-SNV- | SNV | 1 | C | T | 24 | 23100 | 0.1038961 | 32.5 | rnr1Y285A_pe | FALSE | ARG | JSY3868 | 3 | rnr1Y285A_p | GCC | C>T | A/T+1 | G/C-1 | CG>TA | CG>TA | AATAAGCCA | CCAAGCC | CGG | GCC | C | A | C>T | C>T_GCC | CG>TA_CCAAGCC | 4 | FALSE | FALSE |
| 60_v.txt | 31981 | 31981-SNV- | SNV | 1 | G | A | 21 | 23100 | 0.09090909 | 51 | rnr1Y285A_pe | FALSE | ARG | JSY3868 | 3 | rnr1Y285A_p | GGC | G>A | A/T+1 | G/C-1 | CG>TA | CG>TA | AGCCGGGCA | GCATTAA | CCG | CCG | C | T | C>T | C>T_CCG | CG>TA_GCATTAA | 4 | FALSE | FALSE |
| 60_v.txt | 32057 | 32057-SNV- | SNV | 1 | C | T | 23 | 23101 | 0.09956279 | 61.9130435 | rnr1Y285A_pe | FALSE | ARG | JSY3868 | 3 | rnr1Y285A_p | GCG | C>T | A/T+1 | G/C-1 | CG>TA | CG>TA | GATGTGCGA | CGAGATT | CGC | GCG | C | A | C>T | C>T_GCG | CG>TA_CGAGATT | 4 | FALSE | FALSE |
| 60_v.txt | 32088 | 32088-SNV- | SNV | 1 | C | T | 19 | 23101 | 0.08224752 | 60.5263158 | rnr1Y285A_pe | FALSE | ARG | JSY3868 | 3 | rnr1Y285A_p | GCC | C>T | A/T+1 | G/C-1 | CG>TA | CG>TA | AAAAAGCCT | CCTGCAA | CGG | GCC | C | A | C>T | C>T_GCC | CG>TA_CCTGCAA | 4 | FALSE | FALSE |
| 60_v.txt | 32165 | 32165-SNV- | SNV | 1 | G | A | 35 | 23101 | 0.15150859 | 49.4571429 | rnr1Y285A_pe | FALSE | ARG | JSY3868 | 3 | rnr1Y285A_p | CGC | G>A | A/T+1 | G/C-1 | CG>TA | CG>TA | CAAAGCGCC | GCCAAAT | GCG | GCG | C | T | C>T | C>T_GCG | CG>TA_GCCAAAT | 4 | FALSE | FALSE |
| 60_v.txt | 32271 | 32271-SNV- | SNV | 1 | C | T | 49 | 29666 | 0.16517225 | 32.9183674 | rnr1Y285A_pe | FALSE | ARG | JSY3868 | 3 | rnr1Y285A_p | ACG | C>T | A/T+1 | G/C-1 | CG>TA | CG>TA | AAAATACGG | CGGGAAC | TGC | ACG | C | A | C>T | C>T_ACG | CG>TA_CGGGAAC | 4 | FALSE | FALSE |
| 60_v.txt | 32299 | 32299-SNV- | SNV | 1 | C | T | 42 | 29666 | 0.14157622 | 48.6904762 | rnr1Y285A_pe | FALSE | ARG | JSY3868 | 3 | rnr1Y285A_p | GCG | C>T | A/T+1 | G/C-1 | CG>TA | CG>TA | AATTTGCGG | CGGCAGA | CGC | GCG | C | A | C>T | C>T_GCG | CG>TA_CGGCAGA | 4 | FALSE | FALSE |
| 60_v.txt | 32466 | 32466-SNV- | SNV | 1 | C | T | 35 | 29666 | 0.11798018 | 45.8 | rnr1Y285A_pe | FALSE | ARG | JSY3868 | 3 | rnr1Y285A_p | GCC | C>T | A/T+1 | G/C-1 | CG>TA | CG>TA | AGAGAGCCA | CCAATGT | CGG | GCC | C | A | C>T | C>T_GCC | CG>TA_CCAATGT | 4 | FALSE | FALSE |
| 60_v.txt | 32487 | 32487-SNV- | SNV | 1 | C | T | 31 | 29666 | 0.10449673 | 43.0967742 | rnr1Y285A_pe | FALSE | ARG | JSY3868 | 3 | rnr1Y285A_p | ACG | C>T | A/T+1 | G/C-1 | CG>TA | CG>TA | AAGATACGG | CGGAAAA | TGC | ACG | C | A | C>T | C>T_ACG | CG>TA_CGGAAAA | 4 | FALSE | FALSE |
| 60_v.txt | 32535 | 32535-SNV- | SNV | 1 | G | A | 33 | 29666 | 0.11123846 | 37.7172957 | rnr1Y285A_pe | FALSE | ARG | JSY3868 | 3 | rnr1Y285A_p | TGC | G>A | A/T+1 | G/C-1 | CG>TA | CG>TA | GGGTTTGCA | GCAGCTT | ACG | ACG | C | T | C>T | C>T_ACG | CG>TA_GCAGCTT | 4 | FALSE | FALSE |
| 60_v.txt | 32536 | 32536-SNV- | SNV | 1 | C | T | 39 | 29666 | 0.13146363 | 36.7948718 | rnr1Y285A_pe | FALSE | ARG | JSY3868 | 3 | rnr1Y285A_p | GCA | C>T | A/T+1 | G/C-1 | CG>TA | CG>TA | GGTTTGCAG | CAGCTTC | CGT | GCA | C | A | C>T | C>T_GCA | CG>TA_CAGCTTC | 4 | FALSE | FALSE |
| 60_v.txt | 32590 | 32590-SNV- | SNV | 1 | C | T | 31 | 23655 | 0.13105052 | 35.483871 | rnr1Y285A_pe | FALSE | ARG | JSY3868 | 3 | rnr1Y285A_p | GCG | C>T | A/T+1 | G/C-1 | CG>TA | CG>TA | AGGCAGCGT | CGTTAAT | CGC | GCG | C | A | C>T | C>T_GCG | CG>TA_CGTTAAT | 4 | FALSE | FALSE |
| 60_v.txt | 32700 | 32700-SNV- | SNV | 1 | C | T | 32 | 23655 | 0.13527795 | 55.65625 | rnr1Y285A_pe | FALSE | ARG | JSY3868 | 3 | rnr1Y285A_p | GCC | C>T | A/T+1 | G/C-1 | CG>TA | CG>TA | ACTGGGCCG | CCGGTAA | CGG | GCC | C | A | C>T | C>T_GCC | CG>TA_CCGGTAA | 4 | FALSE | FALSE |
| 60_v.txt | 32757 | 32757-SNV- | SNV | 1 | G | A | 23 | 23655 | 0.09723103 | 49.4347826 | rnr1Y285A_pe | FALSE | ARG | JSY3868 | 3 | rnr1Y285A_p | GGC | G>A | A/T+1 | G/C-1 | CG>TA | CG>TA | ATAATGGCT | GCTAAAA | CCG | CCG | C | T | C>T | C>T_CCG | CG>TA_GCTAAAA | 4 | FALSE | FALSE |
| 60_v.txt | 32776 | 32776-SNV- | SNV | 1 | C | T | 19 | 23655 | 0.08032129 | 48.1578947 | rnr1Y285A_pe | FALSE | ARG | JSY3868 | 3 | rnr1Y285A_p | GCG | C>T | A/T+1 | G/C-1 | CG>TA | CG>TA | TGGAAGCGA | CGACCCA | CGC | GCG | C | A | C>T | C>T_GCG | CG>TA_CGACCCA | 4 | FALSE | FALSE |
| 60_v.txt | 32798 | 32798-SNV- | SNV | 1 | G | A | 34 | 23655 | 0.14373283 | 39.3235294 | rnr1Y285A_pe | FALSE | ARG | JSY3868 | 3 | rnr1Y285A_p | CGT | G>A | A/T+1 | G/C-1 | CG>TA | CG>TA | TTCACCGTA | GTAATAT | GCA | GCA | C | T | C>T | C>T_GCA | CG>TA_GTAATAT | 4 | FALSE | FALSE |
| 60_v.txt | 32860 | 32860-SNV- | SNV | 1 | C | T | 30 | 23655 | 0.12682308 | 36.1 | rnr1Y285A_pe | FALSE | ARG | JSY3868 | 3 | rnr1Y285A_p | GCC | C>T | A/T+1 | G/C-1 | CG>TA | CG>TA | TCCATGCCG | CCGCCAG | CGG | GCC | C | A | C>T | C>T_GCC | CG>TA_CCGCCAG | 4 | FALSE | FALSE |
| 60_v.txt | 32862 | 32862-SNV- | SNV | 1 | G | A | 20 | 23655 | 0.08454872 | 35.75 | rnr1Y285A_pe | FALSE | ARG | JSY3868 | 3 | rnr1Y285A_p | CGC | G>A | A/T+1 | G/C-1 | CG>TA | CG>TA | CATGCCGCC | GCCAGTG | GCG | GCG | C | T | C>T | C>T_GCG | CG>TA_GCCAGTG | 4 | FALSE | FALSE |
| 60_v.txt | 32879 | 32879-SNV- | SNV | 1 | C | T | 22 | 22581 | 0.09742704 | 33.8636364 | rnr1Y285A_pe | FALSE | ARG | JSY3868 | 3 | rnr1Y285A_p | ACG | C>T | A/T+1 | G/C-1 | CG>TA | CG>TA | TTTGTACGTC | CGTCCAA | TGC | ACG | C | A | C>T | C>T_ACG | CG>TA_CGTCCAA | 4 | FALSE | FALSE |
| 60_v.txt | 32900 | 32900-SNV- | SNV | 1 | G | A | 30 | 22582 | 0.13284917 | 33.1622191 | rnr1Y285A_pe | FALSE | ARG | JSY3868 | 3 | rnr1Y285A_p | GGC | G>A | A/T+1 | G/C-1 | CG>TA | CG>TA | GACTTGGCC | GCCAACT | CCG | CCG | C | T | C>T | C>T_CCG | CG>TA_GCCAACT | 4 | FALSE | FALSE |
| 60_v.txt | 32901 | 32901-SNV- | SNV | 1 | C | T | 51 | 22582 | 0.22584359 | 31.4905187 | rnr1Y285A_pe | FALSE | ARG | JSY3868 | 3 | rnr1Y285A_p | GCC | C>T | A/T+1 | G/C-1 | CG>TA | CG>TA | ACTTGGCCA | CCAACTA | CGG | GCC | C | A | C>T | C>T_GCC | CG>TA_CCAACTA | 4 | FALSE | FALSE |
| 60_v.txt | 32935 | 32935-SNV- | SNV | 1 | C | T | 27 | 22582 | 0.11956426 | 43.8888889 | rnr1Y285A_pe | FALSE | ARG | JSY3868 | 3 | rnr1Y285A_p | GCC | C>T | A/T+1 | G/C-1 | CG>TA | CG>TA | TGATTGCCC | CCCAAGA | CGG | GCC | C | A | C>T | C>T_GCC | CG>TA_CCCAAGA | 4 | FALSE | FALSE |
| 60_v.txt | 32965 | 32965-SNV- | SNV | 1 | C | T | 26 | 22582 | 0.11513595 | 57 | rnr1Y285A_pe | FALSE | ARG | JSY3868 | 3 | rnr1Y285A_p | GCC | C>T | A/T+1 | G/C-1 | CG>TA | CG>TA | CATTGGCCG | CCGCACC | CGG | GCC | C | A | C>T | C>T_GCC | CG>TA_CCGCACC | 4 | FALSE | FALSE |
| 60_v.txt | 32967 | 32967-SNV- | SNV | 1 | G | A | 54 | 22582 | 0.23912851 | 58.1296296 | rnr1Y285A_pe | FALSE | ARG | JSY3868 | 3 | rnr1Y285A_p | CGC | G>A | A/T+1 | G/C-1 | CG>TA | CG>TA | TTGGCCGCA | GCACCAA | GCG | GCG | C | T | C>T | C>T_GCG | CG>TA_GCACCAA | 4 | FALSE | FALSE |
| 60_v.txt | 33056 | 33056-SNV- | SNV | 1 | C | T | 33 | 22582 | 0.14613409 | 53.7878788 | rnr1Y285A_pe | FALSE | ARG | JSY3868 | 3 | rnr1Y285A_p | GCG | C>T | A/T+1 | G/C-1 | CG>TA | CG>TA | GGACTGCGT | CGTGACA | CGC | GCG | C | A | C>T | C>T_GCG | CG>TA_CGTGACA | 4 | FALSE | FALSE |
| 60_v.txt | 33070 | 33070-SNV- | SNV | 1 | C | T | 23 | 22582 | 0.10185103 | 58.3043478 | rnr1Y285A_pe | FALSE | ARG | JSY3868 | 3 | rnr1Y285A_p | GCC | C>T | A/T+1 | G/C-1 | CG>TA | CG>TA | AATATGCCA | CCAAAGA | CGG | GCC | C | A | C>T | C>T_GCC | CG>TA_CCAAAGA | 4 | FALSE | FALSE |
| 60_v.txt | 33104 | 33104-SNV- | SNV | 1 | G | A | 23 | 22582 | 0.10185103 | 44.2608696 | rnr1Y285A_pe | FALSE | ARG | JSY3868 | 3 | rnr1Y285A_p | CGC | G>A | A/T+1 | G/C-1 | CG>TA | CG>TA | AAGAGCGCC | GCCCACT | GCG | GCG | C | T | C>T | C>T_GCG | CG>TA_GCCCACT | 4 | FALSE | FALSE |
| 60_v.txt | 33118 | 33118-SNV- | SNV | 1 | C | T | 24 | 22582 | 0.10627934 | 40.8333333 | rnr1Y285A_pe | FALSE | ARG | JSY3868 | 3 | rnr1Y285A_p | GCG | C>T | A/T+1 | G/C-1 | CG>TA | CG>TA | GGCCGGCGT | CGTTGGT | CGC | GCG | C | A | C>T | C>T_GCG | CG>TA_CGTTGGT | 4 | FALSE | FALSE |
| 60_v.txt | 33258 | 33258-SNV- | SNV | 1 | C | T | 27 | 19077 | 0.14153169 | 39.037037 | rnr1Y285A_pe | FALSE | ARG | JSY3868 | 3 | rnr1Y285A_p | GCC | C>T | A/T+1 | G/C-1 | CG>TA | CG>TA | CCTATGCCAT | CCATCCT | CGG | GCC | C | A | C>T | C>T_GCC | CG>TA_CCATCCT | 4 | FALSE | FALSE |
| 60_v.txt | 33275 | 33275-SNV- | SNV | 1 | C | T | 21 | 19077 | 0.1100802 | 46.0473557 | rnr1Y285A_pe | FALSE | ARG | JSY3868 | 3 | rnr1Y285A_p | ACG | C>T | A/T+1 | G/C-1 | CG>TA | CG>TA | AGAGAACGT | CGTATCC | TGC | ACG | C | A | C>T | C>T_ACG | CG>TA_CGTATCC | 4 | FALSE | FALSE |
| 60_v.txt | 33276 | 33276-SNV- | SNV | 1 | G | A | 35 | 19077 | 0.183467 | 45.8755496 | rnr1Y285A_pe | FALSE | ARG | JSY3868 | 3 | rnr1Y285A_p | CGT | G>A | A/T+1 | G/C-1 | CG>TA | CG>TA | GAGAACGTA | GTATCCT | GCA | GCA | C | T | C>T | C>T_GCA | CG>TA_GTATCCT | 4 | FALSE | FALSE |
| 60_v.txt | 33309 | 33309-SNV- | SNV | 1 | C | T | 23 | 19076 | 0.12057035 | 56.4347826 | rnr1Y285A_pe | FALSE | ARG | JSY3868 | 3 | rnr1Y285A_p | GCG | C>T | A/T+1 | G/C-1 | CG>TA | CG>TA | AAAGAGCGC | CGCAATG | CGC | GCG | C | A | C>T | C>T_GCG | CG>TA_CGCAATG | 4 | FALSE | FALSE |
| 60_v.txt | 33310 | 33310-SNV- | SNV | 1 | G | A | 21 | 19076 | 0.11008597 | 55.3045586 | rnr1Y285A_pe | FALSE | ARG | JSY3868 | 3 | rnr1Y285A_p | CGC | G>A | A/T+1 | G/C-1 | CG>TA | CG>TA | AAGAGCGCA | GCAATGG | GCG | GCG | C | T | C>T | C>T_GCG | CG>TA_GCAATGG | 4 | FALSE | FALSE |
| 60_v.txt | 33412 | 33412-SNV- | SNV | 1 | G | A | 28 | 19077 | 0.1467736 | 49.8214286 | rnr1Y285A_pe | FALSE | ARG | JSY3868 | 3 | rnr1Y285A_p | GGC | G>A | A/T+1 | G/C-1 | CG>TA | CG>TA | TGACCGGCT | GCTCATT | CCG | CCG | C | T | C>T | C>T_CCG | CG>TA_GCTCATT | 4 | FALSE | FALSE |
| 60_v.txt | 33474 | 33474-SNV- | SNV | 1 | C | T | 23 | 19076 | 0.12057035 | 34.826087 | rnr1Y285A_pe | FALSE | ARG | JSY3868 | 3 | rnr1Y285A_p | GCC | C>T | A/T+1 | G/C-1 | CG>TA | CG>TA | GCTATGCCTT | CCTTTTT | CGG | GCC | C | A | C>T | C>T_GCC | CG>TA_CCTTTTT | 4 | FALSE | FALSE |
| 60_v.txt | 32536 | 32536-SNV- | SNV | 1 | C | G | 3 | 29666 | 0.01011259 | 33.3333333 | rnr1Y285A_pe | FALSE | ARG | JSY3868 | 3 | rnr1Y285A_p | GCA | C>G | G/C+1 | G/C-1 | CG>GC | CG>GC | GGTTTGCAG | CAGCTTC | CGT | GCA | C | C | C>G | C>G_GCA | CG>GC_CAGCTTC | 4 | FALSE | FALSE |
| 60_v.txt | 31657 | 31657-SNV- | SNV | 1 | G | T | 138 | 15246 | 0.90515545 | 20.2632069 | rnr1Y285A_pe | FALSE | ARG | JSY3868 | 3 | rnr1Y285A_p | GGC | G>T | A/T+1 | G/C-1 | CG>AT | CG>AT | GAAATGGCG | GCGTGGA | CCG | CCG | C | A | C>A | C>A_CCG | CG>AT_GCGTGGA | 4 | FALSE | FALSE |
| 60_v.txt | 32384 | 32384-SNV- | SNV | 1 | G | T | 14 | 29666 | 0.04719207 | 54.7142857 | rnr1Y285A_pe | FALSE | ARG | JSY3868 | 3 | rnr1Y285A_p | AGG | G>T | A/T+1 | G/C-1 | CG>AT | CG>AT | AATAAAGGG | GGGAGAA | TCC | TCC | C | A | C>A | C>A_TCC | CG>AT_GGGAGAA | 4 | FALSE | FALSE |
| 60_v.txt | 32535 | 32535-SNV- | SNV | 1 | G | T | 2 | 29666 | 6.74E-03 | 37.9952626 | rnr1Y285A_pe | FALSE | ARG | JSY3868 | 3 | rnr1Y285A_p | TGC | G>T | A/T+1 | G/C-1 | CG>AT | CG>AT | GGGTTTGCA | GCAGCTT | ACG | ACG | C | A | C>A | C>A_ACG | CG>AT_GCAGCTT | 4 | FALSE | FALSE |
| 60_v.txt | 32536 | 32536-SNV- | SNV | 1 | C | A | 4 | 29666 | 0.01348345 | 38 | rnr1Y285A_pe | FALSE | ARG | JSY3868 | 3 | rnr1Y285A_p | GCA | C>A | A/T+1 | G/C-1 | CG>AT | CG>AT | GGTTTGCAG | CAGCTTC | CGT | GCA | C | T | C>A | C>A_GCA | CG>AT_CAGCTTC | 4 | FALSE | FALSE |
| 60_v.txt | 33474 | 33474-SNV- | SNV | 1 | C | A | 52 | 19076 | 0.27259384 | 33.5576923 | rnr1Y285A_pe | FALSE | ARG | JSY3868 | 3 | rnr1Y285A_p | GCC | C>A | A/T+1 | G/C-1 | CG>AT | CG>AT | GCTATGCCTT | CCTTTTT | CGG | GCC | C | T | C>A | C>A_GCC | CG>AT_CCTTTTT | 4 | FALSE | FALSE |
| 60_v.txt | 32081 | 32081-Insert | Insertion | 1 | - | A | 16 | 23101 | 0.06926107 | 64 | rnr1Y285A_pe | FALSE | ARG | JSY3868 | 3 | rnr1Y285A_p | CAA | nope | A/T+1 | >1bpdel | none | A/T+1 | CCATGCAAA | AAAAAAG | GTT | CAA | - | T | NA | NA_CAA | A/T+1_AAAAAAG | 4 | FALSE | FALSE |
| 60_v.txt | 32498 | 32498-Insert | Insertion | 1 | - | T | 2 | 29666 | 6.74E-03 | 51 | rnr1Y285A_pe | FALSE | ARG | JSY3868 | 3 | rnr1Y285A_p | CTT | nope | A/T+1 | >1bpdel | none | A/T+1 | AACAACTTTT | TTTTTTG | GAA | CTT | - | A | NA | NA_CTT | A/T+1_TTTTTTG | 4 | FALSE | FALSE |
| 60_v.txt | 32842 | 32842-Insert | Insertion | 1 | - | A | 4 | 23655 | 0.01690974 | 44.5 | rnr1Y285A_pe | FALSE | ARG | JSY3868 | 3 | rnr1Y285A_p | CAA | nope | A/T+1 | >1bpdel | none | A/T+1 | TTACCCAAA | AAAAAAT | GTT | CAA | - | T | NA | NA_CAA | A/T+1_AAAAAAT | 4 | FALSE | FALSE |
| 60_v.txt | 31682 | 31682-Deleti | Deletion | 1 | A | - | 5 | 15246 | 0.03279549 | 43.2 | rnr1Y285A_pe | FALSE | ARG | JSY3868 | 3 | rnr1Y285A_p | TAA | nope | >1bp-ins | A/T-1 | none | A/T-1 | GGTAATAAA | AAAACGT | ATT | ATT | T | none | T>none | T>none_ATT | A/T-1_AAAACGT | 4 | FALSE | FALSE |
| 60_v.txt | 31711 | 31711-Deleti | Deletion | 1 | A | - | 6 | 15246 | 0.03935459 | 53.1666667 | rnr1Y285A_pe | FALSE | ARG | JSY3868 | 3 | rnr1Y285A_p | CAA | nope | >1bp-ins | A/T-1 | none | A/T-1 | CATTCCAAA | AAAATTT | GTT | GTT | T | none | T>none | T>none_GTT | A/T-1_AAAATTT | 4 | FALSE | FALSE |
| 60_v.txt | 31723 | 31723-Deleti | Deletion | 1 | A | - | 16 | 15246 | 0.10494556 | 62.125 | rnr1Y285A_pe | FALSE | ARG | JSY3868 | 3 | rnr1Y285A_p | CAA | nope | >1bp-ins | A/T-1 | none | A/T-1 | TGTCCCAAA | AAAAAGT | GTT | GTT | T | none | T>none | T>none_GTT | A/T-1_AAAAAGT | 4 | FALSE | FALSE |
| 60_v.txt | 31904 | 31904-Deleti | Deletion | 1 | T | - | 6 | 15246 | 0.03935459 | 41.8333333 | rnr1Y285A_pe | FALSE | ARG | JSY3868 | 3 | rnr1Y285A_p | ATT | nope | >1bp-ins | A/T-1 | none | A/T-1 | ATTGAATTTT | TTTTGGT | TAA | ATT | T | none | nope | nope_ATT | A/T-1_TTTTGGT | 4 | FALSE | FALSE |
| 60_v.txt | 31966 | 31966-Deleti | Deletion | 1 | A | - | 6 | 23100 | 0.02597403 | 37.6666667 | rnr1Y285A_pe | FALSE | ARG | JSY3868 | 3 | rnr1Y285A_p | TAA | nope | >1bp-ins | A/T-1 | none | A/T-1 | CCGCATAAT | AATAAGC | ATT | ATT | T | none | T>none | T>none_ATT | A/T-1_AATAAGC | 4 | FALSE | FALSE |
| 60_v.txt | 31972 | 31972-Deleti | Deletion | 1 | C | - | 10 | 23100 | 0.04329004 | 36 | rnr1Y285A_pe | FALSE | ARG | JSY3868 | 3 | rnr1Y285A_p | GCC | nope | >1bp-ins | G/C-1 | none | G/C-1 | AATAAGCCA | CCAAGCC | CGG | GCC | C | none | nope | nope_GCC | G/C-1_CCAAGCC | 4 | FALSE | FALSE |
| 60_v.txt | 32081 | 32081-Deleti | Deletion | 1 | A | - | 26 | 23101 | 0.11254924 | 62.3076923 | rnr1Y285A_pe | FALSE | ARG | JSY3868 | 3 | rnr1Y285A_p | CAA | nope | >1bp-ins | A/T-1 | none | A/T-1 | CCATGCAAA | AAAAAAG | GTT | GTT | T | none | T>none | T>none_GTT | A/T-1_AAAAAAG | 4 | FALSE | FALSE |
| 60_v.txt | 32265 | 32265-Deleti | Deletion | 1 | A | - | 8 | 29666 | 0.0269669 | 36.875 | rnr1Y285A_pe | FALSE | ARG | JSY3868 | 3 | rnr1Y285A_p | TAA | nope | >1bp-ins | A/T-1 | none | A/T-1 | CCAAATAAA | AAAATAC | ATT | ATT | T | none | T>none | T>none_ATT | A/T-1_AAAATAC | 4 | FALSE | FALSE |
| 60_v.txt | 32384 | 32384-Deleti | Deletion | 1 | G | - | 2 | 29666 | 6.74E-03 | 64 | rnr1Y285A_pe | FALSE | ARG | JSY3868 | 3 | rnr1Y285A_p | AGG | nope | >1bp-ins | G/C-1 | none | G/C-1 | AATAAAGGG | GGGAGAA | TCC | TCC | C | none | C>none | C>none_TCC | G/C-1_GGGAGAA | 4 | FALSE | FALSE |
| 60_v.txt | 32490 | 32490-Deleti | Deletion | 1 | A | - | 6 | 29666 | 0.02022517 | 46.6666667 | rnr1Y285A_pe | FALSE | ARG | JSY3868 | 3 | rnr1Y285A_p | GAA | nope | >1bp-ins | A/T-1 | none | A/T-1 | ATACGGAAA | AAAACAA | CTT | CTT | T | none | T>none | T>none_CTT | A/T-1_AAAACAA | 4 | FALSE | FALSE |
| 60_v.txt | 32498 | 32498-Deleti | Deletion | 1 | T | - | 36 | 29666 | 0.12135104 | 42.1944439 | rnr1Y285A_pe | FALSE | ARG | JSY3868 | 3 | rnr1Y285A_p | CTT | nope | >1bp-ins | A/T-1 | none | A/T-1 | AACAACTTTT | TTTTTTG | GAA | CTT | T | none | nope | nope_CTT | A/T-1_TTTTTTG | 4 | FALSE | FALSE |
| 60_v.txt | 32631 | 32631-Deleti | Deletion | 1 | T | - | 7 | 23655 | 0.02959205 | 42.8571429 | rnr1Y285A_pe | FALSE | ARG | JSY3868 | 3 | rnr1Y285A_p | GTT | nope | >1bp-ins | A/T-1 | none | A/T-1 | CCTTCGTTTT | TTTTTAT | CAA | GTT | T | none | nope | nope_GTT | A/T-1_TTTTTAT | 4 | FALSE | FALSE |
| 60_v.txt | 32658 | 32658-Deleti | Deletion | 1 | C | - | 6 | 23655 | 0.02536462 | 55.3333333 | rnr1Y285A_pe | FALSE | ARG | JSY3868 | 3 | rnr1Y285A_p | ACC | nope | >1bp-ins | G/C-1 | none | G/C-1 | CCTGGACCC | CCCCAGG | TGG | ACC | C | none | nope | nope_ACC | G/C-1_CCCCAGG | 4 | FALSE | FALSE |
| 60_v.txt | 32707 | 32707-Deleti | Deletion | 1 | C | - | 7 | 23655 | 0.02959205 | 61.8571429 | rnr1Y285A_pe | FALSE | ARG | JSY3868 | 3 | rnr1Y285A_p | ACC | nope | >1bp-ins | G/C-1 | none | G/C-1 | CGGTAACCC | CCCCAGC | TGG | ACC | C | none | nope | nope_ACC | G/C-1_CCCCAGC | 4 | FALSE | FALSE |
| 60_v.txt | 32728 | 32728-Deleti | Deletion | 1 | A | - | 10 | 23655 | 0.04227436 | 61.4 | rnr1Y285A_pe | FALSE | ARG | JSY3868 | 3 | rnr1Y285A_p | CAA | nope | >1bp-ins | A/T-1 | none | A/T-1 | CCATACAAA | AAAAACA | GTT | GTT | T | none | T>none | T>none_GTT | A/T-1_AAAAACA | 4 | FALSE | FALSE |
| 60_v.txt | 32842 | 32842-Deleti | Deletion | 1 | A | - | 35 | 23655 | 0.14796026 | 40.7714284 | rnr1Y285A_pe | FALSE | ARG | JSY3868 | 3 | rnr1Y285A_p | CAA | nope | >1bp-ins | A/T-1 | none | A/T-1 | TTACCCAAA | AAAAAAT | GTT | GTT | T | none | T>none | T>none_GTT | A/T-1_AAAAAAT | 4 | FALSE | FALSE |
| 60_v.txt | 32901 | 32901-Deleti | Deletion | 1 | C | - | 3 | 22582 | 0.01328492 | 29.0000049 | rnr1Y285A_pe | FALSE | ARG | JSY3868 | 3 | rnr1Y285A_p | GCC | nope | >1bp-ins | G/C-1 | none | G/C-1 | ACTTGGCCA | CCAACTA | CGG | GCC | C | none | nope | nope_GCC | G/C-1_CCAACTA | 4 | FALSE | FALSE |

| | | | | | | | | | | | | | | | | | | | | | | | | | | | | | | | | | | |
|---|---|---|---|---|---|---|---|---|---|---|---|---|---|---|---|---|---|---|---|---|---|---|---|---|---|---|---|---|---|---|---|---|---|---|
| 60_v.txt | 32924 | 32924-Deleti | Deletion | 1 | A | - | 21 | 22582 | 0.09299442 | 54.7619048 | rnr1Y285A_pe | FALSE | ARG | JSY3868 | 3 | rnr1Y285A_p | CAA | nope | >1bp-ins | A/T-1 | none | A/T-1 | CAGGGCAAA | AAAAGTG | GTT | GTT | T | none | T>none | T>none_GTT | A/T-1_AAAAGTG | 4 | FALSE | FALSE |
| 60_v.txt | 32935 | 32935-Deleti | Deletion | 1 | C | - | 4 | 22582 | 0.01771322 | 34 | rnr1Y285A_pe | FALSE | ARG | JSY3868 | 3 | rnr1Y285A_p | GCC | nope | >1bp-ins | G/C-1 | none | G/C-1 | TGATTGCCC | CCCAAGA | CGG | GCC | C | none | nope | nope_GCC | G/C-1_CCCAAGA | 4 | FALSE | FALSE |
| 60_v.txt | 33147 | 33147-Deleti | Deletion | 1 | A | - | 9 | 22582 | 0.03985475 | 40.7777778 | rnr1Y285A_pe | FALSE | ARG | JSY3868 | 3 | rnr1Y285A_p | GAA | nope | >1bp-ins | A/T-1 | none | A/T-1 | CCAATGAAA | AAAAGAC | CTT | CTT | T | none | T>none | T>none_CTT | A/T-1_AAAAGAC | 4 | FALSE | FALSE |
| 60_v.txt | 33303 | 33303-Deleti | Deletion | 1 | A | - | 7 | 19077 | 0.0366934 | 58.4285714 | rnr1Y285A_pe | FALSE | ARG | JSY3868 | 3 | rnr1Y285A_p | GAA | nope | >1bp-ins | A/T-1 | none | A/T-1 | GTCGGGAAA | AAAGAGC | CTT | CTT | T | none | T>none | T>none_CTT | A/T-1_AAAGAGC | 4 | FALSE | FALSE |
| 60_v.txt | 33474 | 33474-Deleti | Deletion | 1 | C | - | 35 | 19076 | 0.18347662 | 27.3714286 | rnr1Y285A_pe | FALSE | ARG | JSY3868 | 3 | rnr1Y285A_p | GCC | nope | >1bp-ins | G/C-1 | none | G/C-1 | GCTATGCCTT | CCTTTTT | CGG | GCC | C | none | nope | nope_GCC | G/C-1_CCTTTTT | 4 | FALSE | FALSE |
| 61_v.txt | 31723 | 31723-SNV-A | SNV | 1 | A | C | 4 | 5066 | 0.07895776 | 57 | rnr1Y285A_pe | FALSE | ARG | JSY3868 | 4 | rnr1Y285A_p | CAA | A>C | G/C+1 | A/T-1 | TA>GC | TA>GC | TGTCCCAAA | AAAAAGT | GTT | GTT | T | G | T>G | T>G_GTT | TA>GC_AAAAAGT | 4 | FALSE | FALSE |
| 61_v.txt | 32884 | 32884-SNV-A | SNV | 1 | A | C | 5 | 11995 | 0.04168404 | 31.8 | rnr1Y285A_pe | FALSE | ARG | JSY3868 | 4 | rnr1Y285A_p | CAA | A>C | G/C+1 | A/T-1 | TA>GC | TA>GC | ACGTCCAAA | AAAATTG | GTT | GTT | T | G | T>G | T>G_GTT | TA>GC_AAAATTG | 4 | FALSE | FALSE |
| 61_v.txt | 31650 | 31650-SNV-C | SNV | 1 | C | T | 13 | 5066 | 0.25661271 | 34.1538462 | rnr1Y285A_pe | FALSE | ARG | JSY3868 | 4 | rnr1Y285A_p | GCG | C>T | A/T+1 | G/C-1 | CG>TA | CG>TA | AGAATGCGA | CGAAATG | CGC | GCG | C | A | C>T | C>T_GCG | CG>TA_CGAAATG | 4 | FALSE | FALSE |
| 61_v.txt | 31962 | 31962-SNV-C | SNV | 1 | G | A | 16 | 8646 | 0.18505667 | 37.0625 | rnr1Y285A_pe | FALSE | ARG | JSY3868 | 4 | rnr1Y285A_p | CGC | G>A | A/T+1 | G/C-1 | CG>TA | CG>TA | GTGGCCGCA | GCATAAT | GCG | GCG | C | T | C>T | C>T_GCG | CG>TA_GCATAAT | 4 | FALSE | FALSE |
| 61_v.txt | 31972 | 31972-SNV-C | SNV | 1 | C | T | 7 | 8646 | 0.0809623 | 38 | rnr1Y285A_pe | FALSE | ARG | JSY3868 | 4 | rnr1Y285A_p | GCC | C>T | A/T+1 | G/C-1 | CG>TA | CG>TA | AATAAGCCA | CCAAGCC | CGG | GCC | C | A | C>T | C>T_GCC | CG>TA_CCAAGCC | 4 | FALSE | FALSE |
| 61_v.txt | 32092 | 32092-SNV-C | SNV | 1 | C | T | 11 | 8646 | 0.12722646 | 63.5454546 | rnr1Y285A_pe | FALSE | ARG | JSY3868 | 4 | rnr1Y285A_p | GCA | C>T | A/T+1 | G/C-1 | CG>TA | CG>TA | AGCCTGCAA | CAACACC | CGT | GCA | C | A | C>T | C>T_GCA | CG>TA_CAACACC | 4 | FALSE | FALSE |
| 61_v.txt | 32173 | 32173-SNV-C | SNV | 1 | C | T | 14 | 8646 | 0.16192459 | 45.6428571 | rnr1Y285A_pe | FALSE | ARG | JSY3868 | 4 | rnr1Y285A_p | GCA | C>T | A/T+1 | G/C-1 | CG>TA | CG>TA | CAAATGCAG | CAGCAGT | CGT | GCA | C | A | C>T | C>T_GCA | CG>TA_CAGCAGT | 4 | FALSE | FALSE |
| 61_v.txt | 32271 | 32271-SNV-C | SNV | 1 | C | T | 28 | 17933 | 0.15613673 | 32.1071429 | rnr1Y285A_pe | FALSE | ARG | JSY3868 | 4 | rnr1Y285A_p | ACG | C>T | A/T+1 | G/C-1 | CG>TA | CG>TA | AAAATACGG | CGGGAAC | TGC | ACG | C | A | C>T | C>T_ACG | CG>TA_CGGGAAC | 4 | FALSE | FALSE |
| 61_v.txt | 32299 | 32299-SNV-C | SNV | 1 | C | T | 41 | 17933 | 0.22862879 | 57.3414634 | rnr1Y285A_pe | FALSE | ARG | JSY3868 | 4 | rnr1Y285A_p | GCG | C>T | A/T+1 | G/C-1 | CG>TA | CG>TA | AATTTGCGG | CGGCAGA | CGC | GCG | C | A | C>T | C>T_GCG | CG>TA_CGGCAGA | 4 | FALSE | FALSE |
| 61_v.txt | 32466 | 32466-SNV-C | SNV | 1 | C | T | 26 | 17933 | 0.14498411 | 48.3846154 | rnr1Y285A_pe | FALSE | ARG | JSY3868 | 4 | rnr1Y285A_p | GCC | C>T | A/T+1 | G/C-1 | CG>TA | CG>TA | AGAGAGCCA | CCAATGT | CGG | GCC | C | A | C>T | C>T_GCC | CG>TA_CCAATGT | 4 | FALSE | FALSE |
| 61_v.txt | 32535 | 32535-SNV-C | SNV | 1 | G | A | 18 | 17933 | 0.10037361 | 37.221749 | rnr1Y285A_pe | FALSE | ARG | JSY3868 | 4 | rnr1Y285A_p | TGC | G>A | A/T+1 | G/C-1 | CG>TA | CG>TA | GGGTTTGCA | GCAGCTT | ACG | ACG | C | T | C>T | C>T_ACG | CG>TA_GCAGCTT | 4 | FALSE | FALSE |
| 61_v.txt | 32536 | 32536-SNV-C | SNV | 1 | C | T | 31 | 17933 | 0.17286567 | 37.1612903 | rnr1Y285A_pe | FALSE | ARG | JSY3868 | 4 | rnr1Y285A_p | GCA | C>T | A/T+1 | G/C-1 | CG>TA | CG>TA | GGTTTGCAG | CAGCTTC | CGT | GCA | C | A | C>T | C>T_GCA | CG>TA_CAGCTTC | 4 | FALSE | FALSE |
| 61_v.txt | 32700 | 32700-SNV-C | SNV | 1 | C | T | 20 | 14200 | 0.14084507 | 62.7 | rnr1Y285A_pe | FALSE | ARG | JSY3868 | 4 | rnr1Y285A_p | GCC | C>T | A/T+1 | G/C-1 | CG>TA | CG>TA | ACTGGGCCG | CCGGTAA | CGG | GCC | C | A | C>T | C>T_GCC | CG>TA_CCGGTAA | 4 | FALSE | FALSE |
| 61_v.txt | 32776 | 32776-SNV-C | SNV | 1 | C | T | 21 | 14200 | 0.14788732 | 45.7619048 | rnr1Y285A_pe | FALSE | ARG | JSY3868 | 4 | rnr1Y285A_p | GCG | C>T | A/T+1 | G/C-1 | CG>TA | CG>TA | TGGAAGCGA | CGACCCA | CGC | GCG | C | A | C>T | C>T_GCG | CG>TA_CGACCCA | 4 | FALSE | FALSE |
| 61_v.txt | 32798 | 32798-SNV-C | SNV | 1 | G | A | 18 | 14200 | 0.12676056 | 44.6666667 | rnr1Y285A_pe | FALSE | ARG | JSY3868 | 4 | rnr1Y285A_p | CGT | G>A | A/T+1 | G/C-1 | CG>TA | CG>TA | TTCACCGTA | GTAATAT | GCA | GCA | C | T | C>T | C>T_GCA | CG>TA_GTAATAT | 4 | FALSE | FALSE |
| 61_v.txt | 32860 | 32860-SNV-C | SNV | 1 | C | T | 23 | 14199 | 0.16198324 | 37.4347826 | rnr1Y285A_pe | FALSE | ARG | JSY3868 | 4 | rnr1Y285A_p | GCC | C>T | A/T+1 | G/C-1 | CG>TA | CG>TA | TCCATGCCG | CCGCCAG | CGG | GCC | C | A | C>T | C>T_GCC | CG>TA_CCGCCAG | 4 | FALSE | FALSE |
| 61_v.txt | 32900 | 32900-SNV-C | SNV | 1 | G | A | 17 | 11995 | 0.14172572 | 37.0482133 | rnr1Y285A_pe | FALSE | ARG | JSY3868 | 4 | rnr1Y285A_p | GGC | G>A | A/T+1 | G/C-1 | CG>TA | CG>TA | GACTTGGCC | GCCAACT | CCG | CCG | C | T | C>T | C>T_CCG | CG>TA_GCCAACT | 4 | FALSE | FALSE |
| 61_v.txt | 32901 | 32901-SNV-C | SNV | 1 | C | T | 26 | 11995 | 0.21675698 | 28.2307692 | rnr1Y285A_pe | FALSE | ARG | JSY3868 | 4 | rnr1Y285A_p | GCC | C>T | A/T+1 | G/C-1 | CG>TA | CG>TA | ACTTGGCCA | CCAACTA | CGG | GCC | C | A | C>T | C>T_GCC | CG>TA_CCAACTA | 4 | FALSE | FALSE |
| 61_v.txt | 32922 | 32922-SNV-C | SNV | 1 | G | A | 22 | 11995 | 0.18340975 | 47.7272727 | rnr1Y285A_pe | FALSE | ARG | JSY3868 | 4 | rnr1Y285A_p | GGC | G>A | A/T+1 | G/C-1 | CG>TA | CG>TA | TCCAGGGCA | GCAAAAG | CCG | CCG | C | T | C>T | C>T_CCG | CG>TA_GCAAAAG | 4 | FALSE | FALSE |
| 61_v.txt | 32965 | 32965-SNV-C | SNV | 1 | C | T | 16 | 11995 | 0.13338891 | 59.5625 | rnr1Y285A_pe | FALSE | ARG | JSY3868 | 4 | rnr1Y285A_p | GCC | C>T | A/T+1 | G/C-1 | CG>TA | CG>TA | CATTGGCCG | CCGCACC | CGG | GCC | C | A | C>T | C>T_GCC | CG>TA_CCGCACC | 4 | FALSE | FALSE |
| 61_v.txt | 32967 | 32967-SNV-C | SNV | 1 | G | A | 28 | 11995 | 0.2334306 | 57.3571429 | rnr1Y285A_pe | FALSE | ARG | JSY3868 | 4 | rnr1Y285A_p | CGC | G>A | A/T+1 | G/C-1 | CG>TA | CG>TA | TTGGCCGCA | GCACCAA | GCG | GCG | C | T | C>T | C>T_GCG | CG>TA_GCACCAA | 4 | FALSE | FALSE |
| 61_v.txt | 33056 | 33056-SNV-C | SNV | 1 | C | T | 16 | 11995 | 0.13338891 | 53.875 | rnr1Y285A_pe | FALSE | ARG | JSY3868 | 4 | rnr1Y285A_p | GCG | C>T | A/T+1 | G/C-1 | CG>TA | CG>TA | GGACTGCGT | CGTGACA | CGC | GCG | C | A | C>T | C>T_GCG | CG>TA_CGTGACA | 4 | FALSE | FALSE |
| 61_v.txt | 33118 | 33118-SNV-C | SNV | 1 | C | T | 15 | 11995 | 0.12505211 | 40.1333333 | rnr1Y285A_pe | FALSE | ARG | JSY3868 | 4 | rnr1Y285A_p | GCG | C>T | A/T+1 | G/C-1 | CG>TA | CG>TA | GGCCGGCGT | CGTTGGT | CGC | GCG | C | A | C>T | C>T_GCG | CG>TA_CGTTGGT | 4 | FALSE | FALSE |
| 61_v.txt | 33258 | 33258-SNV-C | SNV | 1 | C | T | 13 | 7906 | 0.16443208 | 46.9230769 | rnr1Y285A_pe | FALSE | ARG | JSY3868 | 4 | rnr1Y285A_p | GCC | C>T | A/T+1 | G/C-1 | CG>TA | CG>TA | CCTATGCCAT | CCATCCT | CGG | GCC | C | A | C>T | C>T_GCC | CG>TA_CCATCCT | 4 | FALSE | FALSE |
| 61_v.txt | 33276 | 33276-SNV-C | SNV | 1 | G | A | 13 | 7906 | 0.16443208 | 44.4615385 | rnr1Y285A_pe | FALSE | ARG | JSY3868 | 4 | rnr1Y285A_p | CGT | G>A | A/T+1 | G/C-1 | CG>TA | CG>TA | GAGAACGTA | GTATCCT | GCA | GCA | C | T | C>T | C>T_GCA | CG>TA_GTATCCT | 4 | FALSE | FALSE |
| 61_v.txt | 33309 | 33309-SNV-C | SNV | 1 | C | T | 14 | 7906 | 0.1770807 | 57.7857143 | rnr1Y285A_pe | FALSE | ARG | JSY3868 | 4 | rnr1Y285A_p | GCG | C>T | A/T+1 | G/C-1 | CG>TA | CG>TA | AAAGAGCGC | CGCAATG | CGC | GCG | C | A | C>T | C>T_GCG | CG>TA_CGCAATG | 4 | FALSE | FALSE |
| 61_v.txt | 33412 | 33412-SNV-C | SNV | 1 | G | A | 12 | 7906 | 0.15178346 | 54.1666667 | rnr1Y285A_pe | FALSE | ARG | JSY3868 | 4 | rnr1Y285A_p | GGC | G>A | A/T+1 | G/C-1 | CG>TA | CG>TA | TGACCGGCT | GCTCATT | CCG | CCG | C | T | C>T | C>T_CCG | CG>TA_GCTCATT | 4 | FALSE | FALSE |
| 61_v.txt | 33474 | 33474-SNV-C | SNV | 1 | C | T | 10 | 7906 | 0.12648621 | 36.4 | rnr1Y285A_pe | FALSE | ARG | JSY3868 | 4 | rnr1Y285A_p | GCC | C>T | A/T+1 | G/C-1 | CG>TA | CG>TA | GCTATGCCTT | CCTTTTT | CGG | GCC | C | A | C>T | C>T_GCC | CG>TA_CCTTTTT | 4 | FALSE | FALSE |
| 61_v.txt | 32536 | 32536-SNV-C | SNV | 1 | C | G | 1 | 17933 | 5.58E-03 | 38 | rnr1Y285A_pe | FALSE | ARG | JSY3868 | 4 | rnr1Y285A_p | GCA | C>G | G/C+1 | G/C-1 | CG>GC | CG>GC | GGTTTGCAG | CAGCTTC | CGT | GCA | C | C | C>G | C>G_GCA | CG>GC_CAGCTTC | 4 | FALSE | FALSE |
| 61_v.txt | 32536 | 32536-SNV-C | SNV | 1 | C | A | 4 | 17933 | 0.02230525 | 35.5 | rnr1Y285A_pe | FALSE | ARG | JSY3868 | 4 | rnr1Y285A_p | GCA | C>A | A/T+1 | G/C-1 | CG>AT | CG>AT | GGTTTGCAG | CAGCTTC | CGT | GCA | C | T | C>A | C>A_GCA | CG>AT_CAGCTTC | 4 | FALSE | FALSE |
| 61_v.txt | 32860 | 32860-SNV-C | SNV | 1 | C | A | 5 | 14199 | 0.03521375 | 38 | rnr1Y285A_pe | FALSE | ARG | JSY3868 | 4 | rnr1Y285A_p | GCC | C>A | A/T+1 | G/C-1 | CG>AT | CG>AT | TCCATGCCG | CCGCCAG | CGG | GCC | C | T | C>A | C>A_GCC | CG>AT_CCGCCAG | 4 | FALSE | FALSE |
| 61_v.txt | 33474 | 33474-SNV-C | SNV | 1 | C | A | 29 | 7906 | 0.36681002 | 31.5172414 | rnr1Y285A_pe | FALSE | ARG | JSY3868 | 4 | rnr1Y285A_p | GCC | C>A | A/T+1 | G/C-1 | CG>AT | CG>AT | GCTATGCCTT | CCTTTTT | CGG | GCC | C | T | C>A | C>A_GCC | CG>AT_CCTTTTT | 4 | FALSE | FALSE |
| 61_v.txt | 32081 | 32081-Insert | Insertion | 1 | - | A | 10 | 8646 | 0.11566042 | 64 | rnr1Y285A_pe | FALSE | ARG | JSY3868 | 4 | rnr1Y285A_p | CAA | nope | A/T+1 | >1bpdel | none | A/T+1 | CCATGCAAA | AAAAAAG | GTT | CAA | - | T | NA | NA_CAA | A/T+1_AAAAAAG | 4 | FALSE | FALSE |
| 61_v.txt | 32842 | 32842-Insert | Insertion | 1 | - | A | 5 | 14200 | 0.03521127 | 43.2 | rnr1Y285A_pe | FALSE | ARG | JSY3868 | 4 | rnr1Y285A_p | CAA | nope | A/T+1 | >1bpdel | none | A/T+1 | TTACCCAAA | AAAAAAT | GTT | CAA | - | T | NA | NA_CAA | A/T+1_AAAAAAT | 4 | FALSE | FALSE |
| 61_v.txt | 31723 | 31723-Deleti | Deletion | 1 | A | - | 5 | 5066 | 0.0986972 | 64 | rnr1Y285A_pe | FALSE | ARG | JSY3868 | 4 | rnr1Y285A_p | CAA | nope | >1bp-ins | A/T-1 | none | A/T-1 | TGTCCCAAA | AAAAAGT | GTT | GTT | T | none | T>none | T>none_GTT | A/T-1_AAAAAGT | 4 | FALSE | FALSE |
| 61_v.txt | 31904 | 31904-Deleti | Deletion | 1 | T | - | 4 | 5066 | 0.07895776 | 44.5 | rnr1Y285A_pe | FALSE | ARG | JSY3868 | 4 | rnr1Y285A_p | ATT | nope | >1bp-ins | A/T-1 | none | A/T-1 | ATTGAATTT | TTTTGGT | TAA | ATT | T | none | nope | nope_ATT | A/T-1_TTTTGGT | 4 | FALSE | FALSE |
| 61_v.txt | 31972 | 31972-Deleti | Deletion | 1 | C | - | 6 | 8646 | 0.06939625 | 41.5 | rnr1Y285A_pe | FALSE | ARG | JSY3868 | 4 | rnr1Y285A_p | GCC | nope | >1bp-ins | G/C-1 | none | G/C-1 | AATAAGCCA | CCAAGCC | CGG | GCC | C | none | nope | nope_GCC | G/C-1_CCAAGCC | 4 | FALSE | FALSE |
| 61_v.txt | 32081 | 32081-Deleti | Deletion | 1 | A | - | 8 | 8646 | 0.09252834 | 64 | rnr1Y285A_pe | FALSE | ARG | JSY3868 | 4 | rnr1Y285A_p | CAA | nope | >1bp-ins | A/T-1 | none | A/T-1 | CCATGCAAA | AAAAAAG | GTT | GTT | T | none | T>none | T>none_GTT | A/T-1_AAAAAAG | 4 | FALSE | FALSE |
| 61_v.txt | 32442 | 32442-Deleti | Deletion | 1 | A | - | 6 | 17933 | 0.03345787 | 49.1666667 | rnr1Y285A_pe | FALSE | ARG | JSY3868 | 4 | rnr1Y285A_p | TAA | nope | >1bp-ins | A/T-1 | none | A/T-1 | GGAACTAAA | AAAAGTC | ATT | ATT | T | none | T>none | T>none_ATT | A/T-1_AAAAGTC | 4 | FALSE | FALSE |
| 61_v.txt | 32498 | 32498-Deleti | Deletion | 1 | T | - | 23 | 17933 | 0.12825517 | 44.8695652 | rnr1Y285A_pe | FALSE | ARG | JSY3868 | 4 | rnr1Y285A_p | CTT | nope | >1bp-ins | A/T-1 | none | A/T-1 | AACAACTTT | TTTTTTG | GAA | CTT | T | none | nope | nope_CTT | A/T-1_TTTTTTG | 4 | FALSE | FALSE |
| 61_v.txt | 32528 | 32528-Deleti | Deletion | 1 | G | - | 5 | 17933 | 0.02788156 | 43.2 | rnr1Y285A_pe | FALSE | ARG | JSY3868 | 4 | rnr1Y285A_p | TGG | nope | >1bp-ins | G/C-1 | none | G/C-1 | TTTTCTGGG | GGGGTTT | ACC | ACC | C | none | C>none | C>none_ACC | G/C-1_GGGGTTT | 4 | FALSE | FALSE |
| 61_v.txt | 32842 | 32842-Deleti | Deletion | 1 | A | - | 16 | 14200 | 0.11267606 | 42.9375058 | rnr1Y285A_pe | FALSE | ARG | JSY3868 | 4 | rnr1Y285A_p | CAA | nope | >1bp-ins | A/T-1 | none | A/T-1 | TTACCCAAA | AAAAAAT | GTT | GTT | T | none | T>none | T>none_GTT | A/T-1_AAAAAAT | 4 | FALSE | FALSE |
| 61_v.txt | 32884 | 32884-Deleti | Deletion | 1 | A | - | 2 | 11995 | 0.01667361 | 38 | rnr1Y285A_pe | FALSE | ARG | JSY3868 | 4 | rnr1Y285A_p | CAA | nope | >1bp-ins | A/T-1 | none | A/T-1 | ACGTCCAAA | AAAATTG | GTT | GTT | T | none | T>none | T>none_GTT | A/T-1_AAAATTG | 4 | FALSE | FALSE |
| 61_v.txt | 32924 | 32924-Deleti | Deletion | 1 | A | - | 7 | 11995 | 0.05835765 | 59.8571429 | rnr1Y285A_pe | FALSE | ARG | JSY3868 | 4 | rnr1Y285A_p | CAA | nope | >1bp-ins | A/T-1 | none | A/T-1 | CAGGGCAAA | AAAAGTG | GTT | GTT | T | none | T>none | T>none_GTT | A/T-1_AAAAGTG | 4 | FALSE | FALSE |
| 61_v.txt | 33111 | 33111-Deleti | Deletion | 1 | G | - | 4 | 11995 | 0.03334723 | 43.25 | rnr1Y285A_pe | FALSE | ARG | JSY3868 | 4 | rnr1Y285A_p | TGG | nope | >1bp-ins | G/C-1 | none | G/C-1 | CCCACTGGG | GGGCCGG | ACC | ACC | C | none | C>none | C>none_ACC | G/C-1_GGGCCGG | 4 | FALSE | FALSE |
| 61_v.txt | 33474 | 33474-Deleti | Deletion | 1 | C | - | 23 | 7906 | 0.29091829 | 24.6521739 | rnr1Y285A_pe | FALSE | ARG | JSY3868 | 4 | rnr1Y285A_p | GCC | nope | >1bp-ins | G/C-1 | none | G/C-1 | GCTATGCCTT | CCTTTTT | CGG | GCC | C | none | nope | nope_GCC | G/C-1_CCTTTTT | 4 | FALSE | FALSE |
| 137_v.txt | 31723 | 31723-SNV-A | SNV | 1 | A | C | 4 | 9267 | 0.04316392 | 60 | rnr1Y285A_pG | TRUE | CAN | JSY2422 | 1 | rnr1Y285A_p | CAA | A>C | G/C+1 | A/T-1 | TA>GC | TA>GC | TGTCCCAAA | AAAAAGT | GTT | GTT | T | G | T>G | T>G_GTT | TA>GC_AAAAAGT | 6 | FALSE | FALSE |
| 137_v.txt | 32283 | 32283-SNV-T | SNV | 1 | T | G | 17 | 21218 | 0.08012065 | 53.0588235 | rnr1Y285A_pG | TRUE | CAN | JSY2422 | 1 | rnr1Y285A_p | GTA | T>G | G/C+1 | A/T-1 | TA>GC | TA>GC | CCAACGTAA | TAAATAT | CAT | GTA | T | C | T>G | T>G_GTA | TA>GC_TAAATAT | 6 | FALSE | FALSE |
| 137_v.txt | 32294 | 32294-SNV-A | SNV | 1 | A | C | 13 | 21218 | 0.06126873 | 48.9230769 | rnr1Y285A_pG | TRUE | CAN | JSY2422 | 1 | rnr1Y285A_p | AAT | A>C | G/C+1 | A/T-1 | TA>GC | TA>GC | ATTTGAATTT | ATTTGCG | TTA | TTA | T | G | T>G | T>G_TTA | TA>GC_ATTTGCG | 6 | FALSE | FALSE |
| 137_v.txt | 32581 | 32581-SNV-T | SNV | 1 | T | G | 21 | 20426 | 0.10281014 | 32.1428571 | rnr1Y285A_pG | TRUE | CAN | JSY2422 | 1 | rnr1Y285A_p | GTG | T>G | G/C+1 | A/T-1 | TA>GC | TA>GC | GAAATGTGA | TGAAGGC | CAC | GTG | T | C | T>G | T>G_GTG | TA>GC_TGAAGGC | 6 | FALSE | FALSE |
| 137_v.txt | 32734 | 32734-SNV-A | SNV | 1 | A | C | 3 | 20426 | 0.01468716 | 25.6675643 | rnr1Y285A_pG | TRUE | CAN | JSY2422 | 1 | rnr1Y285A_p | CAG | A>C | G/C+1 | A/T-1 | TA>GC | TA>GC | AAAAACAGT | AGTATAT | GTC | GTC | T | G | T>G | T>G_GTC | TA>GC_AGTATAT | 6 | FALSE | FALSE |
| 137_v.txt | 32803 | 32803-SNV-A | SNV | 1 | A | C | 6 | 20426 | 0.02937433 | 41.3333333 | rnr1Y285A_pG | TRUE | CAN | JSY2422 | 1 | rnr1Y285A_p | TAT | A>C | G/C+1 | A/T-1 | TA>GC | TA>GC | CGTAATATTT | ATTTGAC | ATA | ATA | T | G | T>G | T>G_ATA | TA>GC_ATTTGAC | 6 | FALSE | FALSE |
| 137_v.txt | 32842 | 32842-SNV-A | SNV | 1 | A | C | 5 | 20426 | 0.02447861 | 34.5999933 | rnr1Y285A_pG | TRUE | CAN | JSY2422 | 1 | rnr1Y285A_p | CAA | A>C | G/C+1 | A/T-1 | TA>GC | TA>GC | TTACCCAAA | AAAAAAT | GTT | GTT | T | G | T>G | T>G_GTT | TA>GC_AAAAAAT | 6 | FALSE | FALSE |
| 137_v.txt | 32884 | 32884-SNV-A | SNV | 1 | A | C | 4 | 19511 | 0.02050126 | 37.75 | rnr1Y285A_pG | TRUE | CAN | JSY2422 | 1 | rnr1Y285A_p | CAA | A>C | G/C+1 | A/T-1 | TA>GC | TA>GC | ACGTCCAAA | AAAATTG | GTT | GTT | T | G | T>G | T>G_GTT | TA>GC_AAAATTG | 6 | FALSE | FALSE |
| 137_v.txt | 32929 | 32929-SNV-T | SNV | 1 | T | G | 22 | 19511 | 0.11275691 | 55.6818182 | rnr1Y285A_pG | TRUE | CAN | JSY2422 | 1 | rnr1Y285A_p | GTG | T>G | G/C+1 | A/T-1 | TA>GC | TA>GC | CAAAAGTGA | TGATTGC | CAC | GTG | T | C | T>G | T>G_GTG | TA>GC_TGATTGC | 6 | FALSE | FALSE |
| 137_v.txt | 33089 | 33089-SNV-A | SNV | 1 | A | C | 624 | 19511 | 3.19819589 | 51.5192308 | rnr1Y285A_pG | TRUE | CAN | JSY2422 | 1 | rnr1Y285A_p | AAT | A>C | G/C+1 | A/T-1 | TA>GC | TA>GC | AAATAAATA | ATATGAT | TTA | TTA | T | G | T>G | T>G_TTA | TA>GC_ATATGAT | 6 | FALSE | FALSE |
| 137_v.txt | 32081 | 32081-SNV-A | SNV | 1 | A | G | 4 | 15315 | 0.02611819 | 64 | rnr1Y285A_pG | TRUE | CAN | JSY2422 | 1 | rnr1Y285A_p | CAA | A>G | G/C+1 | A/T-1 | TA>CG | TA>CG | CCATGCAAA | AAAAAAG | GTT | GTT | T | C | T>C | T>C_GTT | TA>CG_AAAAAAG | 6 | FALSE | FALSE |
| 137_v.txt | 32578 | 32578-SNV-A | SNV | 1 | A | G | 29 | 20426 | 0.14197591 | 37.4482759 | rnr1Y285A_pG | TRUE | CAN | JSY2422 | 1 | rnr1Y285A_p | AAT | A>G | G/C+1 | A/T-1 | TA>CG | TA>CG | CTTGAAATG | ATGTGAA | TTA | TTA | T | C | T>C | T>C_TTA | TA>CG_ATGTGAA | 6 | FALSE | FALSE |
| 137_v.txt | 32734 | 32734-SNV-A | SNV | 1 | A | G | 40 | 20426 | 0.19582885 | 57.325146 | rnr1Y285A_pG | TRUE | CAN | JSY2422 | 1 | rnr1Y285A_p | CAG | A>G | G/C+1 | A/T-1 | TA>CG | TA>CG | AAAAACAGT | AGTATAT | GTC | GTC | T | C | T>C | T>C_GTC | TA>CG_AGTATAT | 6 | FALSE | FALSE |
| 137_v.txt | 32916 | 32916-SNV-T | SNV | 1 | T | C | 29 | 19511 | 0.1486341 | 48.79279 | rnr1Y285A_pG | TRUE | CAN | JSY2422 | 1 | rnr1Y285A_p | TTC | T>C | G/C+1 | A/T-1 | TA>CG | TA>CG | CTAAGTTCC | TCCAGGG | AAG | TTC | T | G | T>C | T>C_TTC | TA>CG_TCCAGGG | 6 | FALSE | FALSE |
| 137_v.txt | 32919 | 32919-SNV-A | SNV | 1 | A | G | 20 | 19511 | 0.10250628 | 47.1 | rnr1Y285A_pG | TRUE | CAN | JSY2422 | 1 | rnr1Y285A_p | CAG | A>G | G/C+1 | A/T-1 | TA>CG | TA>CG | AGTTCCAGG | AGGGCAA | GTC | GTC | T | C | T>C | T>C_GTC | TA>CG_AGGGCAA | 6 | FALSE | FALSE |
| 137_v.txt | 33476 | 33476-SNV-T | SNV | 1 | T | C | 27 | 12780 | 0.21126761 | 32.8888889 | rnr1Y285A_pG | TRUE | CAN | JSY2422 | 1 | rnr1Y285A_p | CTT | T>C | G/C+1 | A/T-1 | TA>CG | TA>CG | TATGCCTTTT | TTTTTTT | GAA | CTT | T | G | T>C | T>C_CTT | TA>CG_TTTTTTT | 6 | FALSE | FALSE |
| 137_v.txt | 32631 | 32631-SNV-T | SNV | 1 | T | A | 2 | 20426 | 9.79E-03 | 37.4986769 | rnr1Y285A_pG | TRUE | CAN | JSY2422 | 1 | rnr1Y285A_p | GTT | T>A | A/T+1 | A/T-1 | TA>AT | TA>AT | CCTTCGTTTT | TTTTTAT | CAA | GTT | T | T | T>A | T>A_GTT | TA>AT_TTTTTAT | 6 | FALSE | FALSE |
| 137_v.txt | 32763 | 32763-SNV-A | SNV | 1 | A | T | 15 | 20426 | 0.07343582 | 51.9333333 | rnr1Y285A_pG | TRUE | CAN | JSY2422 | 1 | rnr1Y285A_p | AAC | A>T | A/T+1 | A/T-1 | TA>AT | TA>AT | GCTAAAACT | ACTTTGA | TTG | TTG | T | A | T>A | T>A_TTG | TA>AT_ACTTTGA | 6 | FALSE | FALSE |
| 137_v.txt | 32941 | 32941-SNV-A | SNV | 1 | A | T | 1 | 19511 | 5.13E-03 | 59.9971128 | rnr1Y285A_pG | TRUE | CAN | JSY2422 | 1 | rnr1Y285A_p | GAA | A>T | A/T+1 | A/T-1 | TA>AT | TA>AT | CCCAAGAAA | AAAACCA | CTT | CTT | T | A | T>A | T>A_CTT | TA>AT_AAAACCA | 6 | FALSE | FALSE |
| 137_v.txt | 31844 | 31844-SNV-C | SNV | 1 | C | T | 50 | 9267 | 0.53954894 | 50.42 | rnr1Y285A_pG | TRUE | CAN | JSY2422 | 1 | rnr1Y285A_p | TCC | C>T | A/T+1 | G/C-1 | CG>TA | CG>TA | TAAGATCCA | CCAAACA | AGG | TCC | C | A | C>T | C>T_TCC | CG>TA_CCAAACA | 6 | FALSE | FALSE |
| 137_v.txt | 31879 | 31879-SNV-C | SNV | 1 | C | T | 14 | 9267 | 0.1510737 | 42.4285714 | rnr1Y285A_pG | TRUE | CAN | JSY2422 | 1 | rnr1Y285A_p | GCG | C>T | A/T+1 | G/C-1 | CG>TA | CG>TA | TATAGGCGG | CGGCAGC | CGC | GCG | C | A | C>T | C>T_GCG | CG>TA_CGGCAGC | 6 | FALSE | FALSE |
| 137_v.txt | 31962 | 31962-SNV-C | SNV | 1 | G | A | 25 | 15315 | 0.16323866 | 36.52 | rnr1Y285A_pG | TRUE | CAN | JSY2422 | 1 | rnr1Y285A_p | CGC | G>A | A/T+1 | G/C-1 | CG>TA | CG>TA | GTGGCCGCA | GCATAAT | GCG | GCG | C | T | C>T | C>T_GCG | CG>TA_GCATAAT | 6 | FALSE | FALSE |
| 137_v.txt | 31981 | 31981-SNV-C | SNV | 1 | G | A | 29 | 15315 | 0.18935684 | 52.6896552 | rnr1Y285A_pG | TRUE | CAN | JSY2422 | 1 | rnr1Y285A_p | GGC | G>A | A/T+1 | G/C-1 | CG>TA | CG>TA | AGCCGGGCA | GCATTAA | CCG | CCG | C | T | C>T | C>T_CCG | CG>TA_GCATTAA | 6 | FALSE | FALSE |
| 137_v.txt | 32057 | 32057-SNV-C | SNV | 1 | C | T | 17 | 15315 | 0.11100229 | 57.8823529 | rnr1Y285A_pG | TRUE | CAN | JSY2422 | 1 | rnr1Y285A_p | GCG | C>T | A/T+1 | G/C-1 | CG>TA | CG>TA | GATGTGCGA | CGAGATT | CGC | GCG | C | A | C>T | C>T_GCG | CG>TA_CGAGATT | 6 | FALSE | FALSE |
| 137_v.txt | 32075 | 32075-SNV-C | SNV | 1 | C | T | 25 | 15315 | 0.16323866 | 60.04 | rnr1Y285A_pG | TRUE | CAN | JSY2422 | 1 | rnr1Y285A_p | ACC | C>T | A/T+1 | G/C-1 | CG>TA | CG>TA | AAATAACCA | CCATGCA | TGG | ACC | C | A | C>T | C>T_ACC | CG>TA_CCATGCA | 6 | FALSE | FALSE |
| 137_v.txt | 32088 | 32088-SNV-C | SNV | 1 | C | T | 42 | 15315 | 0.27424094 | 62.165415 | rnr1Y285A_pG | TRUE | CAN | JSY2422 | 1 | rnr1Y285A_p | GCC | C>T | A/T+1 | G/C-1 | CG>TA | CG>TA | AAAAAGCCT | CCTGCAA | CGG | GCC | C | A | C>T | C>T_GCC | CG>TA_CCTGCAA | 6 | FALSE | FALSE |
| 137_v.txt | 32114 | 32114-SNV-C | SNV | 1 | C | T | 57 | 15315 | 0.37218413 | 59.1052632 | rnr1Y285A_pG | TRUE | CAN | JSY2422 | 1 | rnr1Y285A_p | GCC | C>T | A/T+1 | G/C-1 | CG>TA | CG>TA | TAATAGCCA | CCATTCG | CGG | GCC | C | A | C>T | C>T_GCC | CG>TA_CCATTCG | 6 | FALSE | FALSE |
| 137_v.txt | 32165 | 32165-SNV-C | SNV | 1 | G | A | 37 | 15315 | 0.24159321 | 45.2432432 | rnr1Y285A_pG | TRUE | CAN | JSY2422 | 1 | rnr1Y285A_p | CGC | G>A | A/T+1 | G/C-1 | CG>TA | CG>TA | CAAAGCGCC | GCCAAAT | GCG | GCG | C | T | C>T | C>T_GCG | CG>TA_GCCAAAT | 6 | FALSE | FALSE |
| 137_v.txt | 32271 | 32271-SNV-C | SNV | 1 | C | T | 59 | 21218 | 0.27806579 | 35.7627119 | rnr1Y285A_pG | TRUE | CAN | JSY2422 | 1 | rnr1Y285A_p | ACG | C>T | A/T+1 | G/C-1 | CG>TA | CG>TA | AAAATACGG | CGGGAAC | TGC | ACG | C | A | C>T | C>T_ACG | CG>TA_CGGGAAC | 6 | FALSE | FALSE |
| 137_v.txt | 32299 | 32299-SNV-C | SNV | 1 | C | T | 35 | 21218 | 0.16495428 | 46.3142857 | rnr1Y285A_pG | TRUE | CAN | JSY2422 | 1 | rnr1Y285A_p | GCG | C>T | A/T+1 | G/C-1 | CG>TA | CG>TA | AATTTGCGG | CGGCAGA | CGC | GCG | C | A | C>T | C>T_GCG | CG>TA_CGGCAGA | 6 | FALSE | FALSE |
| 137_v.txt | 32301 | 32301-SNV-C | SNV | 1 | G | A | 24 | 21218 | 0.11311151 | 52.2916667 | rnr1Y285A_pG | TRUE | CAN | JSY2422 | 1 | rnr1Y285A_p | GGC | G>A | A/T+1 | G/C-1 | CG>TA | CG>TA | TTTGCGGCA | GCAGAAA | CCG | CCG | C | T | C>T | C>T_CCG | CG>TA_GCAGAAA | 6 | FALSE | FALSE |
| 137_v.txt | 32487 | 32487-SNV-C | SNV | 1 | C | T | 27 | 21218 | 0.12725045 | 43.8888889 | rnr1Y285A_pG | TRUE | CAN | JSY2422 | 1 | rnr1Y285A_p | ACG | C>T | A/T+1 | G/C-1 | CG>TA | CG>TA | AAGATACGG | CGGAAAA | TGC | ACG | C | A | C>T | C>T_ACG | CG>TA_CGGAAAA | 6 | FALSE | FALSE |
| 137_v.txt | 32508 | 32508-SNV-C | SNV | 1 | G | A | 16 | 21218 | 0.07540767 | 39.6875 | rnr1Y285A_pG | TRUE | CAN | JSY2422 | 1 | rnr1Y285A_p | GGC | G>A | A/T+1 | G/C-1 | CG>TA | CG>TA | TTGATGGCT | GCTCTTG | CCG | CCG | C | T | C>T | C>T_CCG | CG>TA_GCTCTTG | 6 | FALSE | FALSE |
| 137_v.txt | 32535 | 32535-SNV-C | SNV | 1 | G | A | 21 | 21217 | 0.09897724 | 36.6667394 | rnr1Y285A_pG | TRUE | CAN | JSY2422 | 1 | rnr1Y285A_p | TGC | G>A | A/T+1 | G/C-1 | CG>TA | CG>TA | GGGTTTGCA | GCAGCTT | ACG | ACG | C | T | C>T | C>T_ACG | CG>TA_GCAGCTT | 6 | FALSE | FALSE |
| 137_v.txt | 32536 | 32536-SNV-C | SNV | 1 | C | T | 31 | 21217 | 0.14610925 | 36.9031228 | rnr1Y285A_pG | TRUE | CAN | JSY2422 | 1 | rnr1Y285A_p | GCA | C>T | A/T+1 | G/C-1 | CG>TA | CG>TA | GGTTTGCAG | CAGCTTC | CGT | GCA | C | A | C>T | C>T_GCA | CG>TA_CAGCTTC | 6 | FALSE | FALSE |
| 137_v.txt | 32572 | 32572-SNV-C | SNV | 1 | C | T | 35 | 20426 | 0.17135024 | 34.7142857 | rnr1Y285A_pG | TRUE | CAN | JSY2422 | 1 | rnr1Y285A_p | CCT | C>T | A/T+1 | G/C-1 | CG>TA | CG>TA | CAGTACCTT | CTTGAAA | GGA | CCT | C | A | C>T | C>T_CCT | CG>TA_CTTGAAA | 6 | FALSE | FALSE |
| 137_v.txt | 32590 | 32590-SNV-C | SNV | 1 | C | T | 40 | 20426 | 0.19582885 | 36.4 | rnr1Y285A_pG | TRUE | CAN | JSY2422 | 1 | rnr1Y285A_p | GCG | C>T | A/T+1 | G/C-1 | CG>TA | CG>TA | AGGCAGCGT | CGTTAAT | CGC | GCG | C | A | C>T | C>T_GCG | CG>TA_CGTTAAT | 6 | FALSE | FALSE |
| 137_v.txt | 32713 | 32713-SNV-C | SNV | 1 | C | T | 24 | 20426 | 0.11749731 | 58.5833333 | rnr1Y285A_pG | TRUE | CAN | JSY2422 | 1 | rnr1Y285A_p | GCA | C>T | A/T+1 | G/C-1 | CG>TA | CG>TA | CCCCAGCAC | CACCACA | CGT | GCA | C | A | C>T | C>T_GCA | CG>TA_CACCACA | 6 | FALSE | FALSE |
| 137_v.txt | 32757 | 32757-SNV-C | SNV | 1 | G | A | 25 | 20426 | 0.12239303 | 50.28 | rnr1Y285A_pG | TRUE | CAN | JSY2422 | 1 | rnr1Y285A_p | GGC | G>A | A/T+1 | G/C-1 | CG>TA | CG>TA | ATAATGGCT | GCTAAAA | CCG | CCG | C | T | C>T | C>T_CCG | CG>TA_GCTAAAA | 6 | FALSE | FALSE |
| 137_v.txt | 32776 | 32776-SNV-C | SNV | 1 | C | T | 24 | 20426 | 0.11749731 | 46.875 | rnr1Y285A_pG | TRUE | CAN | JSY2422 | 1 | rnr1Y285A_p | GCG | C>T | A/T+1 | G/C-1 | CG>TA | CG>TA | TGGAAGCGA | CGACCCA | CGC | GCG | C | A | C>T | C>T_GCG | CG>TA_CGACCCA | 6 | FALSE | FALSE |
| 137_v.txt | 32798 | 32798-SNV-C | SNV | 1 | G | A | 30 | 20426 | 0.14687163 | 45.7666667 | rnr1Y285A_pG | TRUE | CAN | JSY2422 | 1 | rnr1Y285A_p | CGT | G>A | A/T+1 | G/C-1 | CG>TA | CG>TA | TTCACCGTA | GTAATAT | GCA | GCA | C | T | C>T | C>T_GCA | CG>TA_GTAATAT | 6 | FALSE | FALSE |
| 137_v.txt | 32860 | 32860-SNV-C | SNV | 1 | C | T | 21 | 20424 | 0.10282021 | 36.9047619 | rnr1Y285A_pG | TRUE | CAN | JSY2422 | 1 | rnr1Y285A_p | GCC | C>T | A/T+1 | G/C-1 | CG>TA | CG>TA | TCCATGCCG | CCGCCAG | CGG | GCC | C | A | C>T | C>T_GCC | CG>TA_CCGCCAG | 6 | FALSE | FALSE |
| 137_v.txt | 32862 | 32862-SNV-C | SNV | 1 | G | A | 20 | 20423 | 0.09792881 | 35.75 | rnr1Y285A_pG | TRUE | CAN | JSY2422 | 1 | rnr1Y285A_p | CGC | G>A | A/T+1 | G/C-1 | CG>TA | CG>TA | CATGCCGCC | GCCAGTG | GCG | GCG | C | T | C>T | C>T_GCG | CG>TA_GCCAGTG | 6 | FALSE | FALSE |
| 137_v.txt | 32901 | 32901-SNV-C | SNV | 1 | C | T | 40 | 19511 | 0.20501256 | 34 | rnr1Y285A_pG | TRUE | CAN | JSY2422 | 1 | rnr1Y285A_p | GCC | C>T | A/T+1 | G/C-1 | CG>TA | CG>TA | ACTTGGCCA | CCAACTA | CGG | GCC | C | A | C>T | C>T_GCC | CG>TA_CCAACTA | 6 | FALSE | FALSE |
| 137_v.txt | 32945 | 32945-SNV-C | SNV | 1 | C | T | 48 | 19511 | 0.24601507 | 58.8333333 | rnr1Y285A_pG | TRUE | CAN | JSY2422 | 1 | rnr1Y285A_p | ACC | C>T | A/T+1 | G/C-1 | CG>TA | CG>TA | AGAAAACCA | CCAATAC | TGG | ACC | C | A | C>T | C>T_ACC | CG>TA_CCAATAC | 6 | FALSE | FALSE |
| 137_v.txt | 32958 | 32958-SNV-C | SNV | 1 | C | T | 237 | 19511 | 1.2146994 | 58.8185654 | rnr1Y285A_pG | TRUE | CAN | JSY2422 | 1 | rnr1Y285A_p | ACC | C>T | A/T+1 | G/C-1 | CG>TA | CG>TA | ATGTAACCA | CCATTGG | TGG | ACC | C | A | C>T | C>T_ACC | CG>TA_CCATTGG | 6 | FALSE | FALSE |
| 137_v.txt | 32967 | 32967-SNV-C | SNV | 1 | G | A | 50 | 19511 | 0.2562657 | 61.52 | rnr1Y285A_pG | TRUE | CAN | JSY2422 | 1 | rnr1Y285A_p | CGC | G>A | A/T+1 | G/C-1 | CG>TA | CG>TA | TTGGCCGCA | GCACCAA | GCG | GCG | C | T | C>T | C>T_GCG | CG>TA_GCACCAA | 6 | FALSE | FALSE |
| 137_v.txt | 32977 | 32977-SNV-C | SNV | 1 | C | T | 20 | 19511 | 0.10250628 | 54.5 | rnr1Y285A_pG | TRUE | CAN | JSY2422 | 1 | rnr1Y285A_p | GCT | C>T | A/T+1 | G/C-1 | CG>TA | CG>TA | CAAATGCTG | CTGGAGA | CGA | GCT | C | A | C>T | C>T_GCT | CG>TA_CTGGAGA | 6 | FALSE | FALSE |
| 137_v.txt | 33056 | 33056-SNV-C | SNV | 1 | C | T | 31 | 19511 | 0.15888473 | 53.0322581 | rnr1Y285A_pG | TRUE | CAN | JSY2422 | 1 | rnr1Y285A_p | GCG | C>T | A/T+1 | G/C-1 | CG>TA | CG>TA | GGACTGCGT | CGTGACA | CGC | GCG | C | A | C>T | C>T_GCG | CG>TA_CGTGACA | 6 | FALSE | FALSE |
| 137_v.txt | 33114 | 33114-SNV-C | SNV | 1 | C | T | 54 | 19511 | 0.27676695 | 45.0740741 | rnr1Y285A_pG | TRUE | CAN | JSY2422 | 1 | rnr1Y285A_p | GCC | C>T | A/T+1 | G/C-1 | CG>TA | CG>TA | ACTGGGCCG | CCGGCGT | CGG | GCC | C | A | C>T | C>T_GCC | CG>TA_CCGGCGT | 6 | FALSE | FALSE |
| 137_v.txt | 33153 | 33153-SNV-C | SNV | 1 | C | T | 28 | 19511 | 0.14350879 | 35.9642857 | rnr1Y285A_pG | TRUE | CAN | JSY2422 | 1 | rnr1Y285A_p | ACC | C>T | A/T+1 | G/C-1 | CG>TA | CG>TA | AAAAGACCT | CCTGTAC | TGG | ACC | C | A | C>T | C>T_ACC | CG>TA_CCTGTAC | 6 | FALSE | FALSE |
| 137_v.txt | 33159 | 33159-SNV-C | SNV | 1 | C | T | 74 | 19509 | 0.37931211 | 37.2162162 | rnr1Y285A_pG | TRUE | CAN | JSY2422 | 1 | rnr1Y285A_p | ACC | C>T | A/T+1 | G/C-1 | CG>TA | CG>TA | CCTGTACCA | CCAATAG | TGG | ACC | C | A | C>T | C>T_ACC | CG>TA_CCAATAG | 6 | FALSE | FALSE |
| 137_v.txt | 33168 | 33168-SNV-C | SNV | 1 | C | T | 17 | 19507 | 0.0871482 | 37.3529412 | rnr1Y285A_pG | TRUE | CAN | JSY2422 | 1 | rnr1Y285A_p | ACC | C>T | A/T+1 | G/C-1 | CG>TA | CG>TA | ATAGTACCA | CCACCAA | TGG | ACC | C | A | C>T | C>T_ACC | CG>TA_CCACCAA | 6 | FALSE | FALSE |
| 137_v.txt | 33258 | 33258-SNV-C | SNV | 1 | C | T | 24 | 12781 | 0.18777873 | 46.125 | rnr1Y285A_pG | TRUE | CAN | JSY2422 | 1 | rnr1Y285A_p | GCC | C>T | A/T+1 | G/C-1 | CG>TA | CG>TA | CCTATGCCAT | CCATCCT | CGG | GCC | C | A | C>T | C>T_GCC | CG>TA_CCATCCT | 6 | FALSE | FALSE |
| 137_v.txt | 33276 | 33276-SNV-C | SNV | 1 | G | A | 15 | 12781 | 0.11736171 | 47.3333333 | rnr1Y285A_pG | TRUE | CAN | JSY2422 | 1 | rnr1Y285A_p | CGT | G>A | A/T+1 | G/C-1 | CG>TA | CG>TA | GAGAACGTA | GTATCCT | GCA | GCA | C | T | C>T | C>T_GCA | CG>TA_GTATCCT | 6 | FALSE | FALSE |
| 137_v.txt | 33310 | 33310-SNV-C | SNV | 1 | G | A | 23 | 12781 | 0.17995462 | 56.2608696 | rnr1Y285A_pG | TRUE | CAN | JSY2422 | 1 | rnr1Y285A_p | CGC | G>A | A/T+1 | G/C-1 | CG>TA | CG>TA | AAGAGCGCA | GCAATGG | GCG | GCG | C | T | C>T | C>T_GCG | CG>TA_GCAATGG | 6 | FALSE | FALSE |
| 137_v.txt | 33392 | 33392-SNV-C | SNV | 1 | G | A | 19 | 12781 | 0.14865817 | 48.8421053 | rnr1Y285A_pG | TRUE | CAN | JSY2422 | 1 | rnr1Y285A_p | CGT | G>A | A/T+1 | G/C-1 | CG>TA | CG>TA | AACGTCGTG | GTGAAAG | GCA | GCA | C | T | C>T | C>T_GCA | CG>TA_GTGAAAG | 6 | FALSE | FALSE |
| 137_v.txt | 32536 | 32536-SNV-C | SNV | 1 | C | G | 2 | 21217 | 9.43E-03 | 37.9997878 | rnr1Y285A_pG | TRUE | CAN | JSY2422 | 1 | rnr1Y285A_p | GCA | C>G | G/C+1 | G/C-1 | CG>GC | CG>GC | GGTTTGCAG | CAGCTTC | CGT | GCA | C | C | C>G | C>G_GCA | CG>GC_CAGCTTC | 6 | FALSE | FALSE |
| 137_v.txt | 32735 | 32735-SNV-C | SNV | 1 | G | C | 2 | 20426 | 9.79E-03 | 64 | rnr1Y285A_pG | TRUE | CAN | JSY2422 | 1 | rnr1Y285A_p | AGT | G>C | G/C+1 | G/C-1 | CG>GC | CG>GC | AAAACAGTA | GTATATT | TCA | TCA | C | G | C>G | C>G_TCA | CG>GC_GTATATT | 6 | FALSE | FALSE |
| 137_v.txt | 32776 | 32776-SNV-C | SNV | 1 | C | G | 20 | 20426 | 0.09791442 | 46.8 | rnr1Y285A_pG | TRUE | CAN | JSY2422 | 1 | rnr1Y285A_p | GCG | C>G | G/C+1 | G/C-1 | CG>GC | CG>GC | TGGAAGCGA | CGACCCA | CGC | GCG | C | C | C>G | C>G_GCG | CG>GC_CGACCCA | 6 | FALSE | FALSE |
| 137_v.txt | 33205 | 33205-SNV-C | SNV | 1 | G | C | 15 | 12781 | 0.11736171 | 39.4 | rnr1Y285A_pG | TRUE | CAN | JSY2422 | 1 | rnr1Y285A_p | AGC | G>C | G/C+1 | G/C-1 | CG>GC | CG>GC | GCTTAAGCT | GCTCTCT | TCG | TCG | C | G | C>G | C>G_TCG | CG>GC_GCTCTCT | 6 | FALSE | FALSE |
| 137_v.txt | 31657 | 31657-SNV-C | SNV | 1 | G | T | 70 | 9267 | 0.75536851 | 18.5285714 | rnr1Y285A_pG | TRUE | CAN | JSY2422 | 1 | rnr1Y285A_p | GGC | G>T | A/T+1 | G/C-1 | CG>AT | CG>AT | GAAATGGCG | GCGTGGA | CCG | CCG | C | A | C>A | C>A_CCG | CG>AT_GCGTGGA | 6 | FALSE | FALSE |
| 137_v.txt | 31930 | 31930-SNV-C | SNV | 1 | G | T | 85 | 9267 | 0.91723319 | 36.9882353 | rnr1Y285A_pG | TRUE | CAN | JSY2422 | 1 | rnr1Y285A_p | TGA | G>T | A/T+1 | G/C-1 | CG>AT | CG>AT | AACCTTGAA | GAATAAT | ACT | ACT | C | A | C>A | C>A_ACT | CG>AT_GAATAAT | 6 | FALSE | FALSE |
| 137_v.txt | 31981 | 31981-SNV-C | SNV | 1 | G | T | 28 | 15315 | 0.18282729 | 32.5714286 | rnr1Y285A_pG | TRUE | CAN | JSY2422 | 1 | rnr1Y285A_p | GGC | G>T | A/T+1 | G/C-1 | CG>AT | CG>AT | AGCCGGGCA | GCATTAA | CCG | CCG | C | A | C>A | C>A_CCG | CG>AT_GCATTAA | 6 | FALSE | FALSE |
| 137_v.txt | 32027 | 32027-SNV-C | SNV | 1 | G | T | 453 | 15315 | 2.95788443 | 59.5960265 | rnr1Y285A_pG | TRUE | CAN | JSY2422 | 1 | rnr1Y285A_p | GGT | G>T | A/T+1 | G/C-1 | CG>AT | CG>AT | GCCACGGTA | GTATTTC | CCA | CCA | C | A | C>A | C>A_CCA | CG>AT_GTATTTC | 6 | FALSE | FALSE |
| 137_v.txt | 32089 | 32089-SNV-C | SNV | 1 | C | A | 22 | 15315 | 0.14365002 | 55.0506622 | rnr1Y285A_pG | TRUE | CAN | JSY2422 | 1 | rnr1Y285A_p | CCT | C>A | A/T+1 | G/C-1 | CG>AT | CG>AT | AAAAGCCTG | CTGCAAC | GGA | CCT | C | T | C>A | C>A_CCT | CG>AT_CTGCAAC | 6 | FALSE | FALSE |
| 137_v.txt | 32188 | 32188-SNV-C | SNV | 1 | C | A | 49 | 15315 | 0.31994776 | 44.4285714 | rnr1Y285A_pG | TRUE | CAN | JSY2422 | 1 | rnr1Y285A_p | ACT | C>A | A/T+1 | G/C-1 | CG>AT | CG>AT | CGAAAACTG | CTGCAAT | TGA | ACT | C | T | C>A | C>A_ACT | CG>AT_CTGCAAT | 6 | FALSE | FALSE |
| 137_v.txt | 32274 | 32274-SNV-C | SNV | 1 | G | T | 38 | 21218 | 0.17909322 | 37.5263158 | rnr1Y285A_pG | TRUE | CAN | JSY2422 | 1 | rnr1Y285A_p | GGA | G>T | A/T+1 | G/C-1 | CG>AT | CG>AT | ATACGGGAA | GAACCAA | CCT | CCT | C | A | C>A | C>A_CCT | CG>AT_GAACCAA | 6 | FALSE | FALSE |

| | | | | | | | | | | | | | | | | | | | | | | | | | | | | | | | | | | |
|---|---|---|---|---|---|---|---|---|---|---|---|---|---|---|---|---|---|---|---|---|---|---|---|---|---|---|---|---|---|---|---|---|---|---|
| 137_v.txt | 32277 | 32277-SNV- | SNV | 1 | C | A | 289 | 21218 | 1.36205109 | 35.7923875 | rnr1Y285A_p | TRUE | CAN | JSY2422 | 1 | rnr1Y285A_p | ACC | C>A | A/T+1 | G/C-1 | CG>AT | CG>AT | CGGGAACCA | CCAACGT | TGG | ACC | C | T | C>A | C>A_ACC | CG>AT_CCAACGT | 6 | FALSE | FALSE |
| 137_v.txt | 32373 | 32373-SNV- | SNV | 1 | G | T | 39 | 21218 | 0.1838062 | 59.5128205 | rnr1Y285A_p | TRUE | CAN | JSY2422 | 1 | rnr1Y285A_p | AGC | G>T | A/T+1 | G/C-1 | CG>AT | CG>AT | TCAATAGCA | GCAATAA | TCG | TCG | C | A | C>A | C>A_TCG | CG>AT_GCAATAA | 6 | FALSE | FALSE |
| 137_v.txt | 32508 | 32508-SNV- | SNV | 1 | G | T | 23 | 21218 | 0.10839853 | 41.1304348 | rnr1Y285A_p | TRUE | CAN | JSY2422 | 1 | rnr1Y285A_p | GGC | G>T | A/T+1 | G/C-1 | CG>AT | CG>AT | TTGATGGCT | GCTCTTG | CCG | CCG | C | A | C>A | C>A_CCG | CG>AT_GCTCTTG | 6 | FALSE | FALSE |
| 137_v.txt | 32530 | 32530-SNV- | SNV | 1 | G | T | 19 | 21218 | 0.08954661 | 38.1052632 | rnr1Y285A_p | TRUE | CAN | JSY2422 | 1 | rnr1Y285A_p | GGG | G>T | A/T+1 | G/C-1 | CG>AT | CG>AT | TTCTGGGGT | GGTTTGC | CCC | CCC | C | A | C>A | C>A_CCC | CG>AT_GGTTTGC | 6 | FALSE | FALSE |
| 137_v.txt | 32538 | 32538-SNV- | SNV | 1 | G | T | 14 | 21217 | 0.06598482 | 37.9285714 | rnr1Y285A_p | TRUE | CAN | JSY2422 | 1 | rnr1Y285A_p | AGC | G>T | A/T+1 | G/C-1 | CG>AT | CG>AT | TTTGCAGCTT | GCTTCAC | TCG | TCG | C | A | C>A | C>A_TCG | CG>AT_GCTTCAC | 6 | FALSE | FALSE |
| 137_v.txt | 32556 | 32556-SNV- | SNV | 1 | C | A | 118 | 21212 | 0.55628889 | 37.2711864 | rnr1Y285A_p | TRUE | CAN | JSY2422 | 1 | rnr1Y285A_p | ACC | C>A | A/T+1 | G/C-1 | CG>AT | CG>AT | GTGATACCA | CCAACTA | TGG | ACC | C | T | C>A | C>A_ACC | CG>AT_CCAACTA | 6 | FALSE | FALSE |
| 137_v.txt | 32566 | 32566-SNV- | SNV | 1 | C | A | 16 | 20414 | 0.07837758 | 37.6875 | rnr1Y285A_p | TRUE | CAN | JSY2422 | 1 | rnr1Y285A_p | TCA | C>A | A/T+1 | G/C-1 | CG>AT | CG>AT | CTAGTTCAGT | CAGTACC | AGT | TCA | C | T | C>A | C>A_TCA | CG>AT_CAGTACC | 6 | FALSE | FALSE |
| 137_v.txt | 32735 | 32735-SNV- | SNV | 1 | G | T | 49 | 20426 | 0.23989034 | 57.4900634 | rnr1Y285A_p | TRUE | CAN | JSY2422 | 1 | rnr1Y285A_p | AGT | G>T | A/T+1 | G/C-1 | CG>AT | CG>AT | AAAACAGTA | GTATATT | TCA | TCA | C | A | C>A | C>A_TCA | CG>AT_GTATATT | 6 | FALSE | FALSE |
| 137_v.txt | 32783 | 32783-SNV- | SNV | 1 | G | T | 38 | 20426 | 0.1860374 | 44.7105263 | rnr1Y285A_p | TRUE | CAN | JSY2422 | 1 | rnr1Y285A_p | AGA | G>T | A/T+1 | G/C-1 | CG>AT | CG>AT | GACCCAGAA | GAACTCG | TCT | TCT | C | A | C>A | C>A_TCT | CG>AT_GAACTCG | 6 | FALSE | FALSE |
| 137_v.txt | 32786 | 32786-SNV- | SNV | 1 | C | A | 35 | 20426 | 0.17135024 | 45.2857143 | rnr1Y285A_p | TRUE | CAN | JSY2422 | 1 | rnr1Y285A_p | ACT | C>A | A/T+1 | G/C-1 | CG>AT | CG>AT | CCAGAACTC | CTCGAAT | TGA | ACT | C | T | C>A | C>A_ACT | CG>AT_CTCGAAT | 6 | FALSE | FALSE |
| 137_v.txt | 32794 | 32794-SNV- | SNV | 1 | C | A | 167 | 20426 | 0.81758543 | 44.9281437 | rnr1Y285A_p | TRUE | CAN | JSY2422 | 1 | rnr1Y285A_p | TCA | C>A | A/T+1 | G/C-1 | CG>AT | CG>AT | CGAATTCAC | CACCGTA | AGT | TCA | C | T | C>A | C>A_TCA | CG>AT_CACCGTA | 6 | FALSE | FALSE |
| 137_v.txt | 32811 | 32811-SNV- | SNV | 1 | G | T | 10 | 20426 | 0.04895721 | 41.9 | rnr1Y285A_p | TRUE | CAN | JSY2422 | 1 | rnr1Y285A_p | AGG | G>T | A/T+1 | G/C-1 | CG>AT | CG>AT | TTGACAGGG | GGGAACA | TCC | TCC | C | A | C>A | C>A_TCC | CG>AT_GGGAACA | 6 | FALSE | FALSE |
| 137_v.txt | 32819 | 32819-SNV- | SNV | 1 | G | T | 1552 | 20426 | 7.59815921 | 44.3421392 | rnr1Y285A_p | TRUE | CAN | JSY2422 | 1 | rnr1Y285A_p | AGT | G>T | A/T+1 | G/C-1 | CG>AT | CG>AT | GAACAAGTT | GTTCATT | TCA | TCA | C | A | C>A | C>A_TCA | CG>AT_GTTCATT | 6 | FALSE | FALSE |
| 137_v.txt | 32917 | 32917-SNV- | SNV | 1 | C | A | 2141 | 19511 | 10.9732971 | 51.9882115 | rnr1Y285A_p | TRUE | CAN | JSY2422 | 1 | rnr1Y285A_p | TCC | C>A | A/T+1 | G/C-1 | CG>AT | CG>AT | TAAGTTCCA | CCAGGGC | AGG | TCC | C | T | C>A | C>A_TCC | CG>AT_CCAGGGC | 6 | FALSE | FALSE |
| 137_v.txt | 32940 | 32940-SNV- | SNV | 1 | G | T | 33 | 19511 | 0.16913536 | 45.7320151 | rnr1Y285A_p | TRUE | CAN | JSY2422 | 1 | rnr1Y285A_p | AGA | G>T | A/T+1 | G/C-1 | CG>AT | CG>AT | GCCCAAGAA | GAAAACC | TCT | TCT | C | A | C>A | C>A_TCT | CG>AT_GAAAACC | 6 | FALSE | FALSE |
| 137_v.txt | 32958 | 32958-SNV- | SNV | 1 | C | A | 32 | 19511 | 0.16401005 | 52.34375 | rnr1Y285A_p | TRUE | CAN | JSY2422 | 1 | rnr1Y285A_p | ACC | C>A | A/T+1 | G/C-1 | CG>AT | CG>AT | ATGTAACCAT | CCATTGG | TGG | ACC | C | T | C>A | C>A_ACC | CG>AT_CCATTGG | 6 | FALSE | FALSE |
| 137_v.txt | 32970 | 32970-SNV- | SNV | 1 | C | A | 308 | 19511 | 1.57859669 | 61.7337662 | rnr1Y285A_p | TRUE | CAN | JSY2422 | 1 | rnr1Y285A_p | ACC | C>A | A/T+1 | G/C-1 | CG>AT | CG>AT | GCCGCACCA | CCAAATG | TGG | ACC | C | T | C>A | C>A_ACC | CG>AT_CCAAATG | 6 | FALSE | FALSE |
| 137_v.txt | 33093 | 33093-SNV- | SNV | 1 | G | T | 12 | 19511 | 0.06150377 | 56 | rnr1Y285A_p | TRUE | CAN | JSY2422 | 1 | rnr1Y285A_p | TGA | G>T | A/T+1 | G/C-1 | CG>AT | CG>AT | AAATATGAT | GATATAA | ACT | ACT | C | A | C>A | C>A_ACT | CG>AT_GATATAA | 6 | FALSE | FALSE |
| 137_v.txt | 33151 | 33151-SNV- | SNV | 1 | G | T | 19 | 19511 | 0.09738097 | 34.5263158 | rnr1Y285A_p | TRUE | CAN | JSY2422 | 1 | rnr1Y285A_p | AGA | G>T | A/T+1 | G/C-1 | CG>AT | CG>AT | TGAAAAGAC | GACCTGT | TCT | TCT | C | A | C>A | C>A_TCT | CG>AT_GACCTGT | 6 | FALSE | FALSE |
| 137_v.txt | 33153 | 33153-SNV- | SNV | 1 | C | A | 839 | 19511 | 4.30013838 | 40.7973778 | rnr1Y285A_p | TRUE | CAN | JSY2422 | 1 | rnr1Y285A_p | ACC | C>A | A/T+1 | G/C-1 | CG>AT | CG>AT | AAAAGACCT | CCTGTAC | TGG | ACC | C | T | C>A | C>A_ACC | CG>AT_CCTGTAC | 6 | FALSE | FALSE |
| 137_v.txt | 33154 | 33154-SNV- | SNV | 1 | C | A | 96 | 19511 | 0.49203014 | 37.3428071 | rnr1Y285A_p | TRUE | CAN | JSY2422 | 1 | rnr1Y285A_p | CCT | C>A | A/T+1 | G/C-1 | CG>AT | CG>AT | AAAGACCTG | CTGTACC | GGA | CCT | C | T | C>A | C>A_CCT | CG>AT_CTGTACC | 6 | FALSE | FALSE |
| 137_v.txt | 33159 | 33159-SNV- | SNV | 1 | C | A | 43 | 19509 | 0.22041109 | 37.7674419 | rnr1Y285A_p | TRUE | CAN | JSY2422 | 1 | rnr1Y285A_p | ACC | C>A | A/T+1 | G/C-1 | CG>AT | CG>AT | CCTGTACCA | CCAATAG | TGG | ACC | C | T | C>A | C>A_ACC | CG>AT_CCAATAG | 6 | FALSE | FALSE |
| 137_v.txt | 33168 | 33168-SNV- | SNV | 1 | C | A | 106 | 19507 | 0.54339468 | 36.8018868 | rnr1Y285A_p | TRUE | CAN | JSY2422 | 1 | rnr1Y285A_p | ACC | C>A | A/T+1 | G/C-1 | CG>AT | CG>AT | ATAGTACCA | CCACCAA | TGG | ACC | C | T | C>A | C>A_ACC | CG>AT_CCACCAA | 6 | FALSE | FALSE |
| 137_v.txt | 33186 | 33186-SNV- | SNV | 1 | C | A | 51 | 12780 | 0.39906103 | 36.6078431 | rnr1Y285A_p | TRUE | CAN | JSY2422 | 1 | rnr1Y285A_p | ACC | C>A | A/T+1 | G/C-1 | CG>AT | CG>AT | ATCATACCA | CCAATAT | TGG | ACC | C | T | C>A | C>A_ACC | CG>AT_CCAATAT | 6 | FALSE | FALSE |
| 137_v.txt | 33205 | 33205-SNV- | SNV | 1 | G | T | 8 | 12781 | 0.06259291 | 30.875 | rnr1Y285A_p | TRUE | CAN | JSY2422 | 1 | rnr1Y285A_p | AGC | G>T | A/T+1 | G/C-1 | CG>AT | CG>AT | GCTTAAGCT | GCTCTCT | TCG | TCG | C | A | C>A | C>A_TCG | CG>AT_GCTCTCT | 6 | FALSE | FALSE |
| 137_v.txt | 33325 | 33325-SNV- | SNV | 1 | C | A | 919 | 12781 | 7.19036069 | 58.5843308 | rnr1Y285A_p | TRUE | CAN | JSY2422 | 1 | rnr1Y285A_p | TCT | C>A | A/T+1 | G/C-1 | CG>AT | CG>AT | ACAATTCTTT | CTTTACT | AGA | TCT | C | T | C>A | C>A_TCT | CG>AT_CTTTACT | 6 | FALSE | FALSE |
| 137_v.txt | 33354 | 33354-SNV- | SNV | 1 | G | T | 83 | 12781 | 0.64940146 | 20.0481928 | rnr1Y285A_p | TRUE | CAN | JSY2422 | 1 | rnr1Y285A_p | TGA | G>T | A/T+1 | G/C-1 | CG>AT | CG>AT | GGTATTGAC | GACCCAC | ACT | ACT | C | A | C>A | C>A_ACT | CG>AT_GACCCAC | 6 | FALSE | FALSE |
| 137_v.txt | 33436 | 33436-SNV- | SNV | 1 | C | A | 19 | 12781 | 0.14865817 | 39.7894737 | rnr1Y285A_p | TRUE | CAN | JSY2422 | 1 | rnr1Y285A_p | TCT | C>A | A/T+1 | G/C-1 | CG>AT | CG>AT | TCTCCTCTAT | CTATGTC | AGA | TCT | C | T | C>A | C>A_TCT | CG>AT_CTATGTC | 6 | FALSE | FALSE |
| 137_v.txt | 33474 | 33474-SNV- | SNV | 1 | C | A | 29 | 12780 | 0.22691706 | 35.6206897 | rnr1Y285A_p | TRUE | CAN | JSY2422 | 1 | rnr1Y285A_p | GCC | C>A | A/T+1 | G/C-1 | CG>AT | CG>AT | GCTATGCCTT | CCTTTTT | CGG | GCC | C | T | C>A | C>A_GCC | CG>AT_CCTTTTT | 6 | FALSE | FALSE |
| 137_v.txt | 33153 | 33153-MNV- | MNV | 2 | CC | AA | 2 | 19511 | 0.01025063 | 34.25 | rnr1Y285A_p | TRUE | CAN | JSY2422 | 1 | rnr1Y285A_p | ACC | nope | >1bp-ins | >1bpdel | none | MNV | AAAAGACCT | CCTGTAC | TGG | ACC | CC | none | NA | NA_ACC | MNV_CCTGTAC | 6 | FALSE | FALSE |
| 137_v.txt | 32081 | 32081-Insert | Insertion | 1 | - | A | 12 | 15315 | 0.07835455 | 61.8333333 | rnr1Y285A_p | TRUE | CAN | JSY2422 | 1 | rnr1Y285A_p | CAA | nope | A/T+1 | >1bpdel | none | A/T+1 | CCATGCAAA | AAAAAAG | GTT | CAA | - | T | NA | NA_CAA | A/T+1_AAAAAAG | 6 | FALSE | FALSE |
| 137_v.txt | 32375 | 32375-Insert | Insertion | 1 | - | A | 12 | 21218 | 0.05655576 | 64 | rnr1Y285A_p | TRUE | CAN | JSY2422 | 1 | rnr1Y285A_p | CAA | nope | A/T+1 | >1bpdel | none | A/T+1 | AATAGCAAT | AATAATA | GTT | CAA | - | T | NA | NA_CAA | A/T+1_AATAATA | 6 | FALSE | FALSE |
| 137_v.txt | 32532 | 32532-Insert | Insertion | 1 | - | T | 8 | 21218 | 0.03770384 | 41.125 | rnr1Y285A_p | TRUE | CAN | JSY2422 | 1 | rnr1Y285A_p | GTT | nope | A/T+1 | >1bpdel | none | A/T+1 | CTGGGGTTT | TTTGCAG | CAA | GTT | - | A | NA | NA_GTT | A/T+1_TTTGCAG | 6 | FALSE | FALSE |
| 137_v.txt | 32631 | 32631-Insert | Insertion | 1 | - | T | 17 | 20426 | 0.08322726 | 53.0588235 | rnr1Y285A_p | TRUE | CAN | JSY2422 | 1 | rnr1Y285A_p | GTT | nope | A/T+1 | >1bpdel | none | A/T+1 | CCTTCGTTTT | TTTTTAT | CAA | GTT | - | A | NA | NA_GTT | A/T+1_TTTTTAT | 6 | FALSE | FALSE |
| 137_v.txt | 32842 | 32842-Insert | Insertion | 1 | - | A | 28 | 20426 | 0.13708019 | 44.1071429 | rnr1Y285A_p | TRUE | CAN | JSY2422 | 1 | rnr1Y285A_p | CAA | nope | A/T+1 | >1bpdel | none | A/T+1 | TTACCCAAA | AAAAAAT | GTT | CAA | - | T | NA | NA_CAA | A/T+1_AAAAAAT | 6 | FALSE | FALSE |
| 137_v.txt | 32947 | 32947-Insert | Insertion | 1 | - | A | 19 | 19511 | 0.09738097 | 62.0526316 | rnr1Y285A_p | TRUE | CAN | JSY2422 | 1 | rnr1Y285A_p | CAA | nope | A/T+1 | >1bpdel | none | A/T+1 | AAAACCAAT | AATACAT | GTT | CAA | - | T | NA | NA_CAA | A/T+1_AATACAT | 6 | FALSE | FALSE |
| 137_v.txt | 31723 | 31723-Deleti | Deletion | 1 | A | - | 13 | 9267 | 0.14028272 | 61 | rnr1Y285A_p | TRUE | CAN | JSY2422 | 1 | rnr1Y285A_p | CAA | nope | >1bp-ins | A/T-1 | none | A/T-1 | TGTCCCAAA | AAAAAGT | GTT | GTT | T | none | T>none | T>none_GTT | A/T-1_AAAAAGT | 6 | FALSE | FALSE |
| 137_v.txt | 31863 | 31863-Deleti | Deletion | 1 | A | - | 5 | 9267 | 0.05395489 | 47.4 | rnr1Y285A_p | TRUE | CAN | JSY2422 | 1 | rnr1Y285A_p | GAA | nope | >1bp-ins | A/T-1 | none | A/T-1 | AACAGGAAA | AAAATAG | CTT | CTT | T | none | T>none | T>none_CTT | A/T-1_AAAATAG | 6 | FALSE | FALSE |
| 137_v.txt | 31904 | 31904-Deleti | Deletion | 1 | T | - | 53 | 9267 | 0.57192187 | 43.0754717 | rnr1Y285A_p | TRUE | CAN | JSY2422 | 1 | rnr1Y285A_p | ATT | nope | >1bp-ins | A/T-1 | none | A/T-1 | ATTGAATTTT | TTTTGGT | TAA | ATT | T | none | nope | nope_ATT | A/T-1_TTTTGGT | 6 | FALSE | FALSE |
| 137_v.txt | 31972 | 31972-Deleti | Deletion | 1 | C | - | 7 | 15315 | 0.04570682 | 35.4285714 | rnr1Y285A_p | TRUE | CAN | JSY2422 | 1 | rnr1Y285A_p | GCC | nope | >1bp-ins | G/C-1 | none | G/C-1 | AATAAGCCA | CCAAGCC | CGG | GCC | C | none | nope | nope_GCC | G/C-1_CCAAGCC | 6 | FALSE | FALSE |
| 137_v.txt | 31979 | 31979-Deleti | Deletion | 1 | G | - | 59 | 15315 | 0.38524323 | 41.6271186 | rnr1Y285A_p | TRUE | CAN | JSY2422 | 1 | rnr1Y285A_p | CGG | nope | >1bp-ins | G/C-1 | none | G/C-1 | CAAGCCGGG | GGGCATT | GCC | GCC | C | none | C>none | C>none_GCC | G/C-1_GGGCATT | 6 | FALSE | FALSE |
| 137_v.txt | 32057 | 32057-Deleti | Deletion | 1 | C | - | 14 | 15315 | 0.09141365 | 59.7857143 | rnr1Y285A_p | TRUE | CAN | JSY2422 | 1 | rnr1Y285A_p | GCG | nope | >1bp-ins | G/C-1 | none | G/C-1 | GATGTGCGA | CGAGATT | CGC | GCG | C | none | nope | nope_GCG | G/C-1_CGAGATT | 6 | FALSE | FALSE |
| 137_v.txt | 32081 | 32081-Deleti | Deletion | 1 | A | - | 54 | 15315 | 0.3525955 | 63.9444444 | rnr1Y285A_p | TRUE | CAN | JSY2422 | 1 | rnr1Y285A_p | CAA | nope | >1bp-ins | A/T-1 | none | A/T-1 | CCATGCAAA | AAAAAAG | GTT | GTT | T | none | T>none | T>none_GTT | A/T-1_AAAAAAG | 6 | FALSE | FALSE |
| 137_v.txt | 32277 | 32277-Deleti | Deletion | 1 | C | - | 3 | 21218 | 0.01413894 | 36.6666667 | rnr1Y285A_p | TRUE | CAN | JSY2422 | 1 | rnr1Y285A_p | ACC | nope | >1bp-ins | G/C-1 | none | G/C-1 | CGGGAACCA | CCAACGT | TGG | ACC | C | none | nope | nope_ACC | G/C-1_CCAACGT | 6 | FALSE | FALSE |
| 137_v.txt | 32346 | 32346-Deleti | Deletion | 1 | A | - | 24 | 21218 | 0.11311151 | 60.7916667 | rnr1Y285A_p | TRUE | CAN | JSY2422 | 1 | rnr1Y285A_p | CAA | nope | >1bp-ins | A/T-1 | none | A/T-1 | TGTGGCAAA | AAAACCT | GTT | GTT | T | none | T>none | T>none_GTT | A/T-1_AAAACCT | 6 | FALSE | FALSE |
| 137_v.txt | 32424 | 32424-Deleti | Deletion | 1 | G | - | 10 | 21218 | 0.0471298 | 53.4 | rnr1Y285A_p | TRUE | CAN | JSY2422 | 1 | rnr1Y285A_p | AGG | nope | >1bp-ins | G/C-1 | none | G/C-1 | AGTTTAGGG | GGGTCAT | TCC | TCC | C | none | C>none | C>none_TCC | G/C-1_GGGTCAT | 6 | FALSE | FALSE |
| 137_v.txt | 32435 | 32435-Deleti | Deletion | 1 | T | - | 40 | 21218 | 0.18851918 | 54.4 | rnr1Y285A_p | TRUE | CAN | JSY2422 | 1 | rnr1Y285A_p | ATG | nope | >1bp-ins | A/T-1 | none | A/T-1 | ATTGTATGG | TGGAACT | TAC | ATG | T | none | nope | nope_ATG | A/T-1_TGGAACT | 6 | FALSE | FALSE |
| 137_v.txt | 32498 | 32498-Deleti | Deletion | 1 | T | - | 19 | 21218 | 0.08954661 | 42.5789474 | rnr1Y285A_p | TRUE | CAN | JSY2422 | 1 | rnr1Y285A_p | CTT | nope | >1bp-ins | A/T-1 | none | A/T-1 | AACAACTTTT | TTTTTTG | GAA | CTT | T | none | nope | nope_CTT | A/T-1_TTTTTTG | 6 | FALSE | FALSE |
| 137_v.txt | 32522 | 32522-Deleti | Deletion | 1 | T | - | 25 | 21218 | 0.11782449 | 42.04 | rnr1Y285A_p | TRUE | CAN | JSY2422 | 1 | rnr1Y285A_p | ATT | nope | >1bp-ins | A/T-1 | none | A/T-1 | AACGGATTTT | TTTTCTG | TAA | ATT | T | none | nope | nope_ATT | A/T-1_TTTTCTG | 6 | FALSE | FALSE |
| 137_v.txt | 32528 | 32528-Deleti | Deletion | 1 | G | - | 5 | 21218 | 0.0235649 | 43.4 | rnr1Y285A_p | TRUE | CAN | JSY2422 | 1 | rnr1Y285A_p | TGG | nope | >1bp-ins | G/C-1 | none | G/C-1 | TTTTCTGGG | GGGGTTT | ACC | ACC | C | none | C>none | C>none_ACC | G/C-1_GGGGTTT | 6 | FALSE | FALSE |
| 137_v.txt | 32556 | 32556-Deleti | Deletion | 1 | C | - | 2 | 21212 | 9.43E-03 | 38 | rnr1Y285A_p | TRUE | CAN | JSY2422 | 1 | rnr1Y285A_p | ACC | nope | >1bp-ins | G/C-1 | none | G/C-1 | GTGATACCA | CCAACTA | TGG | ACC | C | none | nope | nope_ACC | G/C-1_CCAACTA | 6 | FALSE | FALSE |
| 137_v.txt | 32598 | 32598-Deleti | Deletion | 1 | A | - | 7 | 20426 | 0.03427005 | 41.4285714 | rnr1Y285A_p | TRUE | CAN | JSY2422 | 1 | rnr1Y285A_p | CAA | nope | >1bp-ins | A/T-1 | none | A/T-1 | TTAATCAAA | AAAGAGG | GTT | GTT | T | none | T>none | T>none_GTT | A/T-1_AAAGAGG | 6 | FALSE | FALSE |
| 137_v.txt | 32608 | 32608-Deleti | Deletion | 1 | C | - | 1882 | 20426 | 9.21374719 | 43.7449522 | rnr1Y285A_p | TRUE | CAN | JSY2422 | 1 | rnr1Y285A_p | ACC | nope | >1bp-ins | G/C-1 | none | G/C-1 | AGGAAACCC | CCCAACC | TGG | ACC | C | none | nope | nope_ACC | G/C-1_CCCAACC | 6 | FALSE | FALSE |
| 137_v.txt | 32624 | 32624-Deleti | Deletion | 1 | C | - | 8 | 20426 | 0.03916577 | 41.125 | rnr1Y285A_p | TRUE | CAN | JSY2422 | 1 | rnr1Y285A_p | TCC | nope | >1bp-ins | G/C-1 | none | G/C-1 | GAACCTCCCT | CCCTTCG | AGG | TCC | C | none | nope | nope_TCC | G/C-1_CCCTTCG | 6 | FALSE | FALSE |
| 137_v.txt | 32631 | 32631-Deleti | Deletion | 1 | T | - | 7 | 20426 | 0.03427005 | 40.4285714 | rnr1Y285A_p | TRUE | CAN | JSY2422 | 1 | rnr1Y285A_p | GTT | nope | >1bp-ins | A/T-1 | none | A/T-1 | CCTTCGTTTT | TTTTTAT | CAA | GTT | T | none | nope | nope_GTT | A/T-1_TTTTTAT | 6 | FALSE | FALSE |
| 137_v.txt | 32658 | 32658-Deleti | Deletion | 1 | C | - | 1705 | 20426 | 8.34720454 | 60.5642229 | rnr1Y285A_p | TRUE | CAN | JSY2422 | 1 | rnr1Y285A_p | ACC | nope | >1bp-ins | G/C-1 | none | G/C-1 | CCTGGACCC | CCCCAGG | TGG | ACC | C | none | nope | nope_ACC | G/C-1_CCCCAGG | 6 | FALSE | FALSE |
| 137_v.txt | 32670 | 32670-Deleti | Deletion | 1 | G | - | 2855 | 20426 | 13.9772839 | 59.4322242 | rnr1Y285A_p | TRUE | CAN | JSY2422 | 1 | rnr1Y285A_p | TGG | nope | >1bp-ins | G/C-1 | none | G/C-1 | GCACCTGGG | GGGTTTC | ACC | ACC | C | none | C>none | C>none_ACC | G/C-1_GGGTTTC | 6 | FALSE | FALSE |
| 137_v.txt | 32707 | 32707-Deleti | Deletion | 1 | C | - | 1194 | 20426 | 5.84549104 | 62.237856 | rnr1Y285A_p | TRUE | CAN | JSY2422 | 1 | rnr1Y285A_p | ACC | nope | >1bp-ins | G/C-1 | none | G/C-1 | CGGTAACCC | CCCCAGC | TGG | ACC | C | none | nope | nope_ACC | G/C-1_CCCCAGC | 6 | FALSE | FALSE |
| 137_v.txt | 32719 | 32719-Deleti | Deletion | 1 | A | - | 14 | 20426 | 0.0685401 | 61.0714286 | rnr1Y285A_p | TRUE | CAN | JSY2422 | 1 | rnr1Y285A_p | CAA | nope | >1bp-ins | A/T-1 | none | A/T-1 | CACCACAAA | AAACCAT | GTT | GTT | T | none | T>none | T>none_GTT | A/T-1_AAACCAT | 6 | FALSE | FALSE |
| 137_v.txt | 32747 | 32747-Deleti | Deletion | 1 | C | - | 1103 | 20426 | 5.39998042 | 55.9664551 | rnr1Y285A_p | TRUE | CAN | JSY2422 | 1 | rnr1Y285A_p | ACC | nope | >1bp-ins | G/C-1 | none | G/C-1 | TAGAAACCC | CCCGATA | TGG | ACC | C | none | nope | nope_ACC | G/C-1_CCCGATA | 6 | FALSE | FALSE |
| 137_v.txt | 32760 | 32760-Deleti | Deletion | 1 | A | - | 17 | 20426 | 0.08322726 | 51.0588235 | rnr1Y285A_p | TRUE | CAN | JSY2422 | 1 | rnr1Y285A_p | TAA | nope | >1bp-ins | A/T-1 | none | A/T-1 | ATGGCTAAA | AAAACTT | ATT | ATT | T | none | T>none | T>none_ATT | A/T-1_AAAACTT | 6 | FALSE | FALSE |
| 137_v.txt | 32779 | 32779-Deleti | Deletion | 1 | C | - | 16 | 20426 | 0.07833154 | 53.625 | rnr1Y285A_p | TRUE | CAN | JSY2422 | 1 | rnr1Y285A_p | ACC | nope | >1bp-ins | G/C-1 | none | G/C-1 | AAGCGACCC | CCCAGAA | TGG | ACC | C | none | nope | nope_ACC | G/C-1_CCCAGAA | 6 | FALSE | FALSE |
| 137_v.txt | 32811 | 32811-Deleti | Deletion | 1 | G | - | 2 | 20426 | 9.79E-03 | 38 | rnr1Y285A_p | TRUE | CAN | JSY2422 | 1 | rnr1Y285A_p | AGG | nope | >1bp-ins | G/C-1 | none | G/C-1 | TTGACAGGG | GGGAACA | TCC | TCC | C | none | C>none | C>none_TCC | G/C-1_GGGAACA | 6 | FALSE | FALSE |
| 137_v.txt | 32839 | 32839-Deleti | Deletion | 1 | C | - | 25 | 20426 | 0.12239303 | 46.28 | rnr1Y285A_p | TRUE | CAN | JSY2422 | 1 | rnr1Y285A_p | ACC | nope | >1bp-ins | G/C-1 | none | G/C-1 | TAATTACCCA | CCCAAAA | TGG | ACC | C | none | nope | nope_ACC | G/C-1_CCCAAAA | 6 | FALSE | FALSE |
| 137_v.txt | 32842 | 32842-Deleti | Deletion | 1 | A | - | 40 | 20426 | 0.19582885 | 42.15 | rnr1Y285A_p | TRUE | CAN | JSY2422 | 1 | rnr1Y285A_p | CAA | nope | >1bp-ins | A/T-1 | none | A/T-1 | TTACCCAAA | AAAAAAT | GTT | GTT | T | none | T>none | T>none_GTT | A/T-1_AAAAAAT | 6 | FALSE | FALSE |
| 137_v.txt | 32884 | 32884-Deleti | Deletion | 1 | A | - | 4 | 19511 | 0.02050126 | 37 | rnr1Y285A_p | TRUE | CAN | JSY2422 | 1 | rnr1Y285A_p | CAA | nope | >1bp-ins | A/T-1 | none | A/T-1 | ACGTCCAAA | AAAATTG | GTT | GTT | T | none | T>none | T>none_GTT | A/T-1_AAAATTG | 6 | FALSE | FALSE |
| 137_v.txt | 32917 | 32917-Deleti | Deletion | 1 | C | - | 1 | 19511 | 5.13E-03 | 38.0000067 | rnr1Y285A_p | TRUE | CAN | JSY2422 | 1 | rnr1Y285A_p | TCC | nope | >1bp-ins | G/C-1 | none | G/C-1 | TAAGTTCCA | CCAGGGC | AGG | TCC | C | none | nope | nope_TCC | G/C-1_CCAGGGC | 6 | FALSE | FALSE |
| 137_v.txt | 32924 | 32924-Deleti | Deletion | 1 | A | - | 17 | 19511 | 0.08713034 | 53.7058824 | rnr1Y285A_p | TRUE | CAN | JSY2422 | 1 | rnr1Y285A_p | CAA | nope | >1bp-ins | A/T-1 | none | A/T-1 | CAGGGCAAA | AAAAGTG | GTT | GTT | T | none | T>none | T>none_GTT | A/T-1_AAAAGTG | 6 | FALSE | FALSE |
| 137_v.txt | 32935 | 32935-Deleti | Deletion | 1 | C | - | 7 | 19511 | 0.0358772 | 39 | rnr1Y285A_p | TRUE | CAN | JSY2422 | 1 | rnr1Y285A_p | GCC | nope | >1bp-ins | G/C-1 | none | G/C-1 | TGATTGCCC | CCCAAGA | CGG | GCC | C | none | nope | nope_GCC | G/C-1_CCCAAGA | 6 | FALSE | FALSE |
| 137_v.txt | 32941 | 32941-Deleti | Deletion | 1 | A | - | 6 | 19511 | 0.03075188 | 58.3333335 | rnr1Y285A_p | TRUE | CAN | JSY2422 | 1 | rnr1Y285A_p | GAA | nope | >1bp-ins | A/T-1 | none | A/T-1 | CCCAAGAAA | AAAACCA | CTT | CTT | T | none | T>none | T>none_CTT | A/T-1_AAAACCA | 6 | FALSE | FALSE |
| 137_v.txt | 32945 | 32945-Deleti | Deletion | 1 | C | - | 6 | 19511 | 0.03075188 | 56.5 | rnr1Y285A_p | TRUE | CAN | JSY2422 | 1 | rnr1Y285A_p | ACC | nope | >1bp-ins | G/C-1 | none | G/C-1 | AGAAAACCA | CCAATAC | TGG | ACC | C | none | nope | nope_ACC | G/C-1_CCAATAC | 6 | FALSE | FALSE |
| 137_v.txt | 32956 | 32956-Deleti | Deletion | 1 | A | - | 5 | 19511 | 0.02562657 | 64 | rnr1Y285A_p | TRUE | CAN | JSY2422 | 1 | rnr1Y285A_p | TAA | nope | >1bp-ins | A/T-1 | none | A/T-1 | ACATGTAAC | AACCATT | ATT | ATT | T | none | T>none | T>none_ATT | A/T-1_AACCATT | 6 | FALSE | FALSE |
| 137_v.txt | 32958 | 32958-Deleti | Deletion | 1 | C | - | 15 | 19511 | 0.07687971 | 61.8666667 | rnr1Y285A_p | TRUE | CAN | JSY2422 | 1 | rnr1Y285A_p | ACC | nope | >1bp-ins | G/C-1 | none | G/C-1 | ATGTAACCAT | CCATTGG | TGG | ACC | C | none | nope | nope_ACC | G/C-1_CCATTGG | 6 | FALSE | FALSE |
| 137_v.txt | 33000 | 33000-Deleti | Deletion | 1 | A | - | 9 | 19511 | 0.04612783 | 61.8888889 | rnr1Y285A_p | TRUE | CAN | JSY2422 | 1 | rnr1Y285A_p | GAA | nope | >1bp-ins | A/T-1 | none | A/T-1 | TGTGAGAAA | AAAACTG | CTT | CTT | T | none | T>none | T>none_CTT | A/T-1_AAAACTG | 6 | FALSE | FALSE |
| 137_v.txt | 33078 | 33078-Deleti | Deletion | 1 | C | - | 99 | 19511 | 0.50740608 | 55.8484849 | rnr1Y285A_p | TRUE | CAN | JSY2422 | 1 | rnr1Y285A_p | ACC | nope | >1bp-ins | G/C-1 | none | G/C-1 | AAAGAACCC | CCCATAA | TGG | ACC | C | none | nope | nope_ACC | G/C-1_CCCATAA | 6 | FALSE | FALSE |
| 137_v.txt | 33138 | 33138-Deleti | Deletion | 1 | A | - | 6 | 19511 | 0.03075188 | 40 | rnr1Y285A_p | TRUE | CAN | JSY2422 | 1 | rnr1Y285A_p | TAA | nope | >1bp-ins | A/T-1 | none | A/T-1 | GTGGATAAA | AAACCAA | ATT | ATT | T | none | T>none | T>none_ATT | A/T-1_AAACCAA | 6 | FALSE | FALSE |
| 137_v.txt | 33147 | 33147-Deleti | Deletion | 1 | A | - | 5 | 19511 | 0.02562657 | 43.2 | rnr1Y285A_p | TRUE | CAN | JSY2422 | 1 | rnr1Y285A_p | GAA | nope | >1bp-ins | A/T-1 | none | A/T-1 | CCAATGAAA | AAAAGAC | CTT | CTT | T | none | T>none | T>none_CTT | A/T-1_AAAAGAC | 6 | FALSE | FALSE |
| 137_v.txt | 33168 | 33168-Deleti | Deletion | 1 | C | - | 15 | 19507 | 0.07689547 | 37.8 | rnr1Y285A_p | TRUE | CAN | JSY2422 | 1 | rnr1Y285A_p | ACC | nope | >1bp-ins | G/C-1 | none | G/C-1 | ATAGTACCA | CCACCAA | TGG | ACC | C | none | nope | nope_ACC | G/C-1_CCACCAA | 6 | FALSE | FALSE |
| 137_v.txt | 33356 | 33356-Deleti | Deletion | 1 | C | - | 116 | 12781 | 0.90759722 | 55.7155172 | rnr1Y285A_p | TRUE | CAN | JSY2422 | 1 | rnr1Y285A_p | ACC | nope | >1bp-ins | G/C-1 | none | G/C-1 | TATTGACCCA | CCCACGT | TGG | ACC | C | none | nope | nope_ACC | G/C-1_CCCACGT | 6 | FALSE | FALSE |
| 137_v.txt | 33474 | 33474-Deleti | Deletion | 1 | C | - | 29 | 12780 | 0.22691706 | 29.0689655 | rnr1Y285A_p | TRUE | CAN | JSY2422 | 1 | rnr1Y285A_p | GCC | nope | >1bp-ins | G/C-1 | none | G/C-1 | GCTATGCCTT | CCTTTTT | CGG | GCC | C | none | nope | nope_GCC | G/C-1_CCTTTTT | 6 | FALSE | FALSE |
| 138_v.txt | 31723 | 31723-SNV- | SNV | 1 | A | C | 4 | 9697 | 0.04124987 | 52.5 | rnr1Y285A_p | TRUE | CAN | JSY2422 | 2 | rnr1Y285A_p | CAA | A>C | G/C+1 | A/T-1 | TA>GC | TA>GC | TGTCCCAAA | AAAAAGT | GTT | GTT | T | G | T>G | T>G_GTT | TA>GC_AAAAAGT | 6 | FALSE | FALSE |
| 138_v.txt | 32763 | 32763-SNV- | SNV | 1 | A | C | 21 | 19418 | 0.10814708 | 49.7122163 | rnr1Y285A_p | TRUE | CAN | JSY2422 | 2 | rnr1Y285A_p | AAC | A>C | G/C+1 | A/T-1 | TA>GC | TA>GC | GCTAAAACTT | ACTTTGA | TTG | TTG | T | G | T>G | T>G_TTG | TA>GC_ACTTTGA | 6 | FALSE | FALSE |
| 138_v.txt | 32842 | 32842-SNV- | SNV | 1 | A | C | 2 | 19418 | 0.01029972 | 37.9996128 | rnr1Y285A_p | TRUE | CAN | JSY2422 | 2 | rnr1Y285A_p | CAA | A>C | G/C+1 | A/T-1 | TA>GC | TA>GC | TTACCCAAA | AAAAAAT | GTT | GTT | T | G | T>G | T>G_GTT | TA>GC_AAAAAAT | 6 | FALSE | FALSE |
| 138_v.txt | 32884 | 32884-SNV- | SNV | 1 | A | C | 8 | 24363 | 0.03283668 | 35 | rnr1Y285A_p | TRUE | CAN | JSY2422 | 2 | rnr1Y285A_p | CAA | A>C | G/C+1 | A/T-1 | TA>GC | TA>GC | ACGTCCAAA | AAAATTG | GTT | GTT | T | G | T>G | T>G_GTT | TA>GC_AAAATTG | 6 | FALSE | FALSE |
| 138_v.txt | 32929 | 32929-SNV-T | SNV | 1 | T | G | 22 | 24363 | 0.09030087 | 47.1818182 | rnr1Y285A_p | TRUE | CAN | JSY2422 | 2 | rnr1Y285A_p | GTG | T>G | G/C+1 | A/T-1 | TA>GC | TA>GC | CAAAAGTGA | TGATTGC | CAC | GTG | T | C | T>G | T>G_GTG | TA>GC_TGATTGC | 6 | FALSE | FALSE |
| 138_v.txt | 33089 | 33089-SNV- | SNV | 1 | A | C | 16 | 24363 | 0.06567336 | 51.375 | rnr1Y285A_p | TRUE | CAN | JSY2422 | 2 | rnr1Y285A_p | AAT | A>C | G/C+1 | A/T-1 | TA>GC | TA>GC | AAATAAATA | ATATGAT | TTA | TTA | T | G | T>G | T>G_TTA | TA>GC_ATATGAT | 6 | FALSE | FALSE |
| 138_v.txt | 33162 | 33162-SNV- | SNV | 1 | A | C | 32 | 24362 | 0.13135211 | 34.96875 | rnr1Y285A_p | TRUE | CAN | JSY2422 | 2 | rnr1Y285A_p | AAT | A>C | G/C+1 | A/T-1 | TA>GC | TA>GC | GTACCAATA | ATAGTAC | TTA | TTA | T | G | T>G | T>G_TTA | TA>GC_ATAGTAC | 6 | FALSE | FALSE |
| 138_v.txt | 33269 | 33269-SNV- | SNV | 1 | A | C | 128 | 15623 | 0.81930487 | 15.7109375 | rnr1Y285A_p | TRUE | CAN | JSY2422 | 2 | rnr1Y285A_p | TAG | A>C | G/C+1 | A/T-1 | TA>GC | TA>GC | CTCCATAGA | AGAGAAC | ATC | ATC | T | G | T>G | T>G_ATC | TA>GC_AGAGAAC | 6 | FALSE | FALSE |
| 138_v.txt | 33476 | 33476-SNV-T | SNV | 1 | T | G | 3 | 15620 | 0.01920615 | 37.6667843 | rnr1Y285A_p | TRUE | CAN | JSY2422 | 2 | rnr1Y285A_p | CTT | T>G | G/C+1 | A/T-1 | TA>GC | TA>GC | TATGCCTTTT | TTTTTTT | GAA | CTT | T | C | T>G | T>G_CTT | TA>GC_TTTTTTT | 6 | FALSE | FALSE |
| 138_v.txt | 32498 | 32498-SNV-T | SNV | 1 | T | C | 9 | 24410 | 0.03687014 | 41.3332489 | rnr1Y285A_p | TRUE | CAN | JSY2422 | 2 | rnr1Y285A_p | CTT | T>C | G/C+1 | A/T-1 | TA>CG | TA>CG | AACAACTTTT | TTTTTTG | GAA | CTT | T | G | T>C | T>C_CTT | TA>CG_TTTTTTG | 6 | FALSE | FALSE |
| 138_v.txt | 32694 | 32694-SNV- | SNV | 1 | A | G | 5 | 19418 | 0.02574931 | 53.3997589 | rnr1Y285A_p | TRUE | CAN | JSY2422 | 2 | rnr1Y285A_p | AAC | A>G | G/C+1 | A/T-1 | TA>CG | TA>CG | AATCCAACT | ACTGGGC | TTG | TTG | T | C | T>C | T>C_TTG | TA>CG_ACTGGGC | 6 | FALSE | FALSE |
| 138_v.txt | 32734 | 32734-SNV- | SNV | 1 | A | G | 35 | 19418 | 0.18024513 | 57.7711722 | rnr1Y285A_p | TRUE | CAN | JSY2422 | 2 | rnr1Y285A_p | CAG | A>G | G/C+1 | A/T-1 | TA>CG | TA>CG | AAAAACAGT | AGTATAT | GTC | GTC | T | C | T>C | T>C_GTC | TA>CG_AGTATAT | 6 | FALSE | FALSE |
| 138_v.txt | 32916 | 32916-SNV-T | SNV | 1 | T | C | 6 | 24362 | 0.02462852 | 21.9999729 | rnr1Y285A_p | TRUE | CAN | JSY2422 | 2 | rnr1Y285A_p | TTC | T>C | G/C+1 | A/T-1 | TA>CG | TA>CG | CTAAGTTCCA | TCCAGGG | AAG | TTC | T | G | T>C | T>C_TTC | TA>CG_TCCAGGG | 6 | FALSE | FALSE |
| 138_v.txt | 32929 | 32929-SNV-T | SNV | 1 | T | C | 17 | 24363 | 0.06977794 | 50.2352941 | rnr1Y285A_p | TRUE | CAN | JSY2422 | 2 | rnr1Y285A_p | GTG | T>C | G/C+1 | A/T-1 | TA>CG | TA>CG | CAAAAGTGA | TGATTGC | CAC | GTG | T | G | T>C | T>C_GTG | TA>CG_TGATTGC | 6 | FALSE | FALSE |
| 138_v.txt | 33476 | 33476-SNV-T | SNV | 1 | T | C | 21 | 15620 | 0.13444302 | 30.0483787 | rnr1Y285A_p | TRUE | CAN | JSY2422 | 2 | rnr1Y285A_p | CTT | T>C | G/C+1 | A/T-1 | TA>CG | TA>CG | TATGCCTTTT | TTTTTTT | GAA | CTT | T | G | T>C | T>C_CTT | TA>CG_TTTTTTT | 6 | FALSE | FALSE |
| 138_v.txt | 32050 | 32050-SNV-T | SNV | 1 | T | A | 19 | 17424 | 0.109045 | 60.3157895 | rnr1Y285A_p | TRUE | CAN | JSY2422 | 2 | rnr1Y285A_p | CTG | T>A | A/T+1 | A/T-1 | TA>AT | TA>AT | TAAATCTGAT | TGATGTG | GAC | CTG | T | T | T>A | T>A_CTG | TA>AT_TGATGTG | 6 | FALSE | FALSE |
| 138_v.txt | 32496 | 32496-SNV- | SNV | 1 | A | T | 252 | 24410 | 1.03236379 | 44.9880952 | rnr1Y285A_p | TRUE | CAN | JSY2422 | 2 | rnr1Y285A_p | AAC | A>T | A/T+1 | A/T-1 | TA>AT | TA>AT | AAAACAACT | ACTTTTT | TTG | TTG | T | A | T>A | T>A_TTG | TA>AT_ACTTTTT | 6 | FALSE | FALSE |
| 138_v.txt | 32498 | 32498-SNV-T | SNV | 1 | T | A | 1 | 24410 | 4.10E-03 | 37.9997036 | rnr1Y285A_p | TRUE | CAN | JSY2422 | 2 | rnr1Y285A_p | CTT | T>A | A/T+1 | A/T-1 | TA>AT | TA>AT | AACAACTTTT | TTTTTTG | GAA | CTT | T | T | T>A | T>A_CTT | TA>AT_TTTTTTG | 6 | FALSE | FALSE |
| 138_v.txt | 32641 | 32641-SNV-T | SNV | 1 | T | A | 21 | 19418 | 0.10814708 | 52.6190476 | rnr1Y285A_p | TRUE | CAN | JSY2422 | 2 | rnr1Y285A_p | TTA | T>A | A/T+1 | A/T-1 | TA>AT | TA>AT | TATCCTTAGA | TAGATAT | AAT | TTA | T | T | T>A | T>A_TTA | TA>AT_TAGATAT | 6 | FALSE | FALSE |
| 138_v.txt | 32694 | 32694-SNV- | SNV | 1 | A | T | 2 | 19418 | 0.01029972 | 42.9997358 | rnr1Y285A_p | TRUE | CAN | JSY2422 | 2 | rnr1Y285A_p | AAC | A>T | A/T+1 | A/T-1 | TA>AT | TA>AT | AATCCAACT | ACTGGGC | TTG | TTG | T | A | T>A | T>A_TTG | TA>AT_ACTGGGC | 6 | FALSE | FALSE |
| 138_v.txt | 32763 | 32763-SNV- | SNV | 1 | A | T | 11 | 19418 | 0.05664847 | 57.2517291 | rnr1Y285A_p | TRUE | CAN | JSY2422 | 2 | rnr1Y285A_p | AAC | A>T | A/T+1 | A/T-1 | TA>AT | TA>AT | GCTAAAACTT | ACTTTGA | TTG | TTG | T | A | T>A | T>A_TTG | TA>AT_ACTTTGA | 6 | FALSE | FALSE |
| 138_v.txt | 32821 | 32821-SNV-T | SNV | 1 | T | A | 25 | 19418 | 0.12874652 | 43.88 | rnr1Y285A_p | TRUE | CAN | JSY2422 | 2 | rnr1Y285A_p | TTC | T>A | A/T+1 | A/T-1 | TA>AT | TA>AT | ACAAGTTCAT | TCATTAT | AAG | TTC | T | T | T>A | T>A_TTC | TA>AT_TCATTAT | 6 | FALSE | FALSE |
| 138_v.txt | 32916 | 32916-SNV-T | SNV | 1 | T | A | 351 | 24362 | 1.44076841 | 50.5309895 | rnr1Y285A_p | TRUE | CAN | JSY2422 | 2 | rnr1Y285A_p | TTC | T>A | A/T+1 | A/T-1 | TA>AT | TA>AT | CTAAGTTCCA | TCCAGGG | AAG | TTC | T | T | T>A | T>A_TTC | TA>AT_TCCAGGG | 6 | FALSE | FALSE |
| 138_v.txt | 32929 | 32929-SNV-T | SNV | 1 | T | A | 705 | 24363 | 2.8937323 | 52.0056738 | rnr1Y285A_p | TRUE | CAN | JSY2422 | 2 | rnr1Y285A_p | GTG | T>A | A/T+1 | A/T-1 | TA>AT | TA>AT | CAAAAGTGA | TGATTGC | CAC | GTG | T | T | T>A | T>A_GTG | TA>AT_TGATTGC | 6 | FALSE | FALSE |
| 138_v.txt | 33138 | 33138-SNV- | SNV | 1 | A | T | 28 | 24363 | 0.11492838 | 39.5357143 | rnr1Y285A_p | TRUE | CAN | JSY2422 | 2 | rnr1Y285A_p | TAA | A>T | A/T+1 | A/T-1 | TA>AT | TA>AT | GTGGATAAA | AAACCAA | ATT | ATT | T | A | T>A | T>A_ATT | TA>AT_AAACCAA | 6 | FALSE | FALSE |
| 138_v.txt | 31650 | 31650-SNV- | SNV | 1 | C | T | 22 | 9697 | 0.22687429 | 35.9090909 | rnr1Y285A_p | TRUE | CAN | JSY2422 | 2 | rnr1Y285A_p | GCG | C>T | A/T+1 | G/C-1 | CG>TA | CG>TA | AGAATGCGA | CGAAATG | CGC | GCG | C | A | C>T | C>T_GCG | CG>TA_CGAAATG | 6 | FALSE | FALSE |
| 138_v.txt | 31658 | 31658-SNV- | SNV | 1 | C | T | 15 | 9697 | 0.15468702 | 34.1333333 | rnr1Y285A_p | TRUE | CAN | JSY2422 | 2 | rnr1Y285A_p | GCG | C>T | A/T+1 | G/C-1 | CG>TA | CG>TA | AAATGGCGT | CGTGGAA | CGC | GCG | C | A | C>T | C>T_GCG | CG>TA_CGTGGAA | 6 | FALSE | FALSE |
| 138_v.txt | 31790 | 31790-SNV- | SNV | 1 | G | A | 14 | 9697 | 0.14437455 | 60.2142857 | rnr1Y285A_p | TRUE | CAN | JSY2422 | 2 | rnr1Y285A_p | CGA | G>A | A/T+1 | G/C-1 | CG>TA | CG>TA | GATGTCGAC | GACATCT | GCT | GCT | C | T | C>T | C>T_GCT | CG>TA_GACATCT | 6 | FALSE | FALSE |
| 138_v.txt | 31844 | 31844-SNV- | SNV | 1 | C | T | 26 | 9697 | 0.26812416 | 51.1538462 | rnr1Y285A_p | TRUE | CAN | JSY2422 | 2 | rnr1Y285A_p | TCC | C>T | A/T+1 | G/C-1 | CG>TA | CG>TA | TAAGATCCA | CCAAACA | AGG | TCC | C | A | C>T | C>T_TCC | CG>TA_CCAAACA | 6 | FALSE | FALSE |
| 138_v.txt | 31962 | 31962-SNV- | SNV | 1 | G | A | 38 | 17424 | 0.21808999 | 36.3157895 | rnr1Y285A_p | TRUE | CAN | JSY2422 | 2 | rnr1Y285A_p | CGC | G>A | A/T+1 | G/C-1 | CG>TA | CG>TA | GTGGCCGCA | GCATAAT | GCG | GCG | C | T | C>T | C>T_GCG | CG>TA_GCATAAT | 6 | FALSE | FALSE |
| 138_v.txt | 31972 | 31972-SNV- | SNV | 1 | C | T | 10 | 17424 | 0.0573921 | 38 | rnr1Y285A_p | TRUE | CAN | JSY2422 | 2 | rnr1Y285A_p | GCC | C>T | A/T+1 | G/C-1 | CG>TA | CG>TA | AATAAGCCA | CCAAGCC | CGG | GCC | C | A | C>T | C>T_GCC | CG>TA_CCAAGCC | 6 | FALSE | FALSE |
| 138_v.txt | 31981 | 31981-SNV- | SNV | 1 | G | A | 47 | 17424 | 0.26974288 | 46.2765957 | rnr1Y285A_p | TRUE | CAN | JSY2422 | 2 | rnr1Y285A_p | GGC | G>A | A/T+1 | G/C-1 | CG>TA | CG>TA | AGCCGGGCA | GCATTAA | CCG | CCG | C | T | C>T | C>T_CCG | CG>TA_GCATTAA | 6 | FALSE | FALSE |
| 138_v.txt | 32057 | 32057-SNV- | SNV | 1 | C | T | 20 | 17424 | 0.11478421 | 57.05 | rnr1Y285A_p | TRUE | CAN | JSY2422 | 2 | rnr1Y285A_p | GCG | C>T | A/T+1 | G/C-1 | CG>TA | CG>TA | GATGTGCGA | CGAGATT | CGC | GCG | C | A | C>T | C>T_GCG | CG>TA_CGAGATT | 6 | FALSE | FALSE |
| 138_v.txt | 32075 | 32075-SNV- | SNV | 1 | C | T | 63 | 17424 | 0.36157025 | 61.2698413 | rnr1Y285A_p | TRUE | CAN | JSY2422 | 2 | rnr1Y285A_p | ACC | C>T | A/T+1 | G/C-1 | CG>TA | CG>TA | AAATAACCAT | CCATGCA | TGG | ACC | C | A | C>T | C>T_ACC | CG>TA_CCATGCA | 6 | FALSE | FALSE |
| 138_v.txt | 32088 | 32088-SNV- | SNV | 1 | C | T | 109 | 17424 | 0.62557392 | 61.853211 | rnr1Y285A_p | TRUE | CAN | JSY2422 | 2 | rnr1Y285A_p | GCC | C>T | A/T+1 | G/C-1 | CG>TA | CG>TA | AAAAAGCCT | CCTGCAA | CGG | GCC | C | A | C>T | C>T_GCC | CG>TA_CCTGCAA | 6 | FALSE | FALSE |
| 138_v.txt | 32114 | 32114-SNV- | SNV | 1 | C | T | 193 | 17424 | 1.10766759 | 59.2797928 | rnr1Y285A_p | TRUE | CAN | JSY2422 | 2 | rnr1Y285A_p | GCC | C>T | A/T+1 | G/C-1 | CG>TA | CG>TA | TAATAGCCAT | CCATTCG | CGG | GCC | C | A | C>T | C>T_GCC | CG>TA_CCATTCG | 6 | FALSE | FALSE |
| 138_v.txt | 32165 | 32165-SNV- | SNV | 1 | G | A | 26 | 17424 | 0.14921947 | 50.8461539 | rnr1Y285A_p | TRUE | CAN | JSY2422 | 2 | rnr1Y285A_p | CGC | G>A | A/T+1 | G/C-1 | CG>TA | CG>TA | CAAAGCGCC | GCCAAAT | GCG | GCG | C | T | C>T | C>T_GCG | CG>TA_GCCAAAT | 6 | FALSE | FALSE |
| 138_v.txt | 32271 | 32271-SNV- | SNV | 1 | C | T | 37 | 24410 | 0.15157722 | 33.5945946 | rnr1Y285A_p | TRUE | CAN | JSY2422 | 2 | rnr1Y285A_p | ACG | C>T | A/T+1 | G/C-1 | CG>TA | CG>TA | AAAATACGG | CGGGAAC | TGC | ACG | C | A | C>T | C>T_ACG | CG>TA_CGGGAAC | 6 | FALSE | FALSE |
| 138_v.txt | 32299 | 32299-SNV- | SNV | 1 | C | T | 55 | 24410 | 0.22531749 | 54.4181818 | rnr1Y285A_p | TRUE | CAN | JSY2422 | 2 | rnr1Y285A_p | GCG | C>T | A/T+1 | G/C-1 | CG>TA | CG>TA | AATTTGCGG | CGGCAGA | CGC | GCG | C | A | C>T | C>T_GCG | CG>TA_CGGCAGA | 6 | FALSE | FALSE |

| | | | | | | | | | | | | | | | | | | | | | | | | | | | | | | | | | | |
|---|---|---|---|---|---|---|---|---|---|---|---|---|---|---|---|---|---|---|---|---|---|---|---|---|---|---|---|---|---|---|---|---|---|---|
| 138_v.txt | 32466 | 32466-SNV-( | SNV | 1 | C | T | 33 | 24410 | 0.1351905 | 51.5151515 | rnr1Y285A_p( | TRUE | CAN | JSY2422 | 2 | rnr1Y285A_p | GCC | C>T | A/T+1 | G/C-1 | CG>TA | CG>TA | AGAGAGCCA | CCAATGT | CGG | GCC | C | A | C>T | C>T_GCC | CG>TA_CCAATGT | 6 | FALSE | FALSE |
| 138_v.txt | 32535 | 32535-SNV-( | SNV | 1 | G | A | 36 | 24410 | 0.14748054 | 37.9704691 | rnr1Y285A_p( | TRUE | CAN | JSY2422 | 2 | rnr1Y285A_p | TGC | G>A | A/T+1 | G/C-1 | CG>TA | CG>TA | GGGTTTGCA | GCAGCTT | ACG | ACG | C | T | C>T | C>T_ACG | CG>TA_GCAGCTT | 6 | FALSE | FALSE |
| 138_v.txt | 32536 | 32536-SNV-( | SNV | 1 | C | T | 32 | 24410 | 0.13109381 | 37.1872778 | rnr1Y285A_p( | TRUE | CAN | JSY2422 | 2 | rnr1Y285A_p | GCA | C>T | A/T+1 | G/C-1 | CG>TA | CG>TA | GGTTTGCAG | CAGCTTC | CGT | GCA | C | A | C>T | C>T_GCA | CG>TA_CAGCTTC | 6 | FALSE | FALSE |
| 138_v.txt | 32539 | 32539-SNV-( | SNV | 1 | C | T | 25 | 24409 | 0.10242124 | 37.8394039 | rnr1Y285A_p( | TRUE | CAN | JSY2422 | 2 | rnr1Y285A_p | GCT | C>T | A/T+1 | G/C-1 | CG>TA | CG>TA | TTGCAGCTTC | CTTCACC | CGA | GCT | C | A | C>T | C>T_GCT | CG>TA_CTTCACC | 6 | FALSE | FALSE |
| 138_v.txt | 32544 | 32544-SNV-( | SNV | 1 | C | T | 35 | 24409 | 0.14338973 | 37.4564904 | rnr1Y285A_p( | TRUE | CAN | JSY2422 | 2 | rnr1Y285A_p | ACC | C>T | A/T+1 | G/C-1 | CG>TA | CG>TA | GCTTCACCAG | CCAGCAG | TGG | ACC | C | A | C>T | C>T_ACC | CG>TA_CCAGCAG | 6 | FALSE | FALSE |
| 138_v.txt | 32590 | 32590-SNV-( | SNV | 1 | C | T | 40 | 19418 | 0.20599444 | 36.15 | rnr1Y285A_p( | TRUE | CAN | JSY2422 | 2 | rnr1Y285A_p | GCG | C>T | A/T+1 | G/C-1 | CG>TA | CG>TA | AGGCAGCGT | CGTTAAT | CGC | GCG | C | A | C>T | C>T_GCG | CG>TA_CGTTAAT | 6 | FALSE | FALSE |
| 138_v.txt | 32658 | 32658-SNV-( | SNV | 1 | C | T | 18 | 19418 | 0.0926975 | 56.8888889 | rnr1Y285A_p( | TRUE | CAN | JSY2422 | 2 | rnr1Y285A_p | ACC | C>T | A/T+1 | G/C-1 | CG>TA | CG>TA | CCTGGACCCC | CCCCAGG | TGG | ACC | C | A | C>T | C>T_ACC | CG>TA_CCCCAGG | 6 | FALSE | FALSE |
| 138_v.txt | 32700 | 32700-SNV-( | SNV | 1 | C | T | 20 | 19418 | 0.10299722 | 59.4 | rnr1Y285A_p( | TRUE | CAN | JSY2422 | 2 | rnr1Y285A_p | GCC | C>T | A/T+1 | G/C-1 | CG>TA | CG>TA | ACTGGGCCG | CCGGTAA | CGG | GCC | C | A | C>T | C>T_GCC | CG>TA_CCGGTAA | 6 | FALSE | FALSE |
| 138_v.txt | 32702 | 32702-SNV-( | SNV | 1 | G | A | 21 | 19418 | 0.10814708 | 56.952381 | rnr1Y285A_p( | TRUE | CAN | JSY2422 | 2 | rnr1Y285A_p | CGG | G>A | A/T+1 | G/C-1 | CG>TA | CG>TA | TGGGCCGGT | GGTAACC | GCC | GCC | C | T | C>T | C>T_GCC | CG>TA_GGTAACC | 6 | FALSE | FALSE |
| 138_v.txt | 32776 | 32776-SNV-( | SNV | 1 | C | T | 23 | 19418 | 0.1184468 | 53.2608696 | rnr1Y285A_p( | TRUE | CAN | JSY2422 | 2 | rnr1Y285A_p | GCG | C>T | A/T+1 | G/C-1 | CG>TA | CG>TA | TGGAAGCGA | CGACCCA | CGC | GCG | C | A | C>T | C>T_GCG | CG>TA_CGACCCA | 6 | FALSE | FALSE |
| 138_v.txt | 32798 | 32798-SNV-( | SNV | 1 | G | A | 29 | 19418 | 0.14934597 | 41.8275862 | rnr1Y285A_p( | TRUE | CAN | JSY2422 | 2 | rnr1Y285A_p | CGT | G>A | A/T+1 | G/C-1 | CG>TA | CG>TA | TTCACCGTAA | GTAATAT | GCA | GCA | C | T | C>T | C>T_GCA | CG>TA_GTAATAT | 6 | FALSE | FALSE |
| 138_v.txt | 32855 | 32855-SNV-( | SNV | 1 | C | T | 32 | 19418 | 0.16479555 | 35.375 | rnr1Y285A_p( | TRUE | CAN | JSY2422 | 2 | rnr1Y285A_p | TCC | C>T | A/T+1 | G/C-1 | CG>TA | CG>TA | ACTAATCCAT | CCATGCC | AGG | TCC | C | A | C>T | C>T_TCC | CG>TA_CCATGCC | 6 | FALSE | FALSE |
| 138_v.txt | 32860 | 32860-SNV-( | SNV | 1 | C | T | 25 | 19415 | 0.12876642 | 36.84 | rnr1Y285A_p( | TRUE | CAN | JSY2422 | 2 | rnr1Y285A_p | GCC | C>T | A/T+1 | G/C-1 | CG>TA | CG>TA | TCCATGCCGC | CCGCCAG | CGG | GCC | C | A | C>T | C>T_GCC | CG>TA_CCGCCAG | 6 | FALSE | FALSE |
| 138_v.txt | 32900 | 32900-SNV-( | SNV | 1 | G | A | 37 | 24363 | 0.15186964 | 35.7459872 | rnr1Y285A_p( | TRUE | CAN | JSY2422 | 2 | rnr1Y285A_p | GGC | G>A | A/T+1 | G/C-1 | CG>TA | CG>TA | GACTTGGCC | GCCAACT | CCG | CCG | C | T | C>T | C>T_CCG | CG>TA_GCCAACT | 6 | FALSE | FALSE |
| 138_v.txt | 32901 | 32901-SNV-( | SNV | 1 | C | T | 46 | 24363 | 0.1888109 | 33.517011 | rnr1Y285A_p( | TRUE | CAN | JSY2422 | 2 | rnr1Y285A_p | GCC | C>T | A/T+1 | G/C-1 | CG>TA | CG>TA | ACTTGGCCA | CCAACTA | CGG | GCC | C | A | C>T | C>T_GCC | CG>TA_CCAACTA | 6 | FALSE | FALSE |
| 138_v.txt | 32917 | 32917-SNV-( | SNV | 1 | C | T | 35 | 24363 | 0.14366047 | 47.1147135 | rnr1Y285A_p( | TRUE | CAN | JSY2422 | 2 | rnr1Y285A_p | TCC | C>T | A/T+1 | G/C-1 | CG>TA | CG>TA | TAAGTTCCAC | CCAGGGC | AGG | TCC | C | A | C>T | C>T_TCC | CG>TA_CCAGGGC | 6 | FALSE | FALSE |
| 138_v.txt | 32922 | 32922-SNV-( | SNV | 1 | G | A | 20 | 24363 | 0.0820917 | 53.15 | rnr1Y285A_p( | TRUE | CAN | JSY2422 | 2 | rnr1Y285A_p | GGC | G>A | A/T+1 | G/C-1 | CG>TA | CG>TA | TCCAGGGCA | GCAAAAG | CCG | CCG | C | T | C>T | C>T_CCG | CG>TA_GCAAAAG | 6 | FALSE | FALSE |
| 138_v.txt | 32958 | 32958-SNV-( | SNV | 1 | C | T | 26 | 24363 | 0.10671921 | 46.8461539 | rnr1Y285A_p( | TRUE | CAN | JSY2422 | 2 | rnr1Y285A_p | ACC | C>T | A/T+1 | G/C-1 | CG>TA | CG>TA | ATGTAACCAT | CCATTGG | TGG | ACC | C | A | C>T | C>T_ACC | CG>TA_CCATTGG | 6 | FALSE | FALSE |
| 138_v.txt | 32965 | 32965-SNV-( | SNV | 1 | C | T | 27 | 24363 | 0.11082379 | 56.037037 | rnr1Y285A_p( | TRUE | CAN | JSY2422 | 2 | rnr1Y285A_p | GCC | C>T | A/T+1 | G/C-1 | CG>TA | CG>TA | CATTGGCCGC | CCGCACC | CGG | GCC | C | A | C>T | C>T_GCC | CG>TA_CCGCACC | 6 | FALSE | FALSE |
| 138_v.txt | 32967 | 32967-SNV-( | SNV | 1 | G | A | 66 | 24363 | 0.2709026 | 57.6212121 | rnr1Y285A_p( | TRUE | CAN | JSY2422 | 2 | rnr1Y285A_p | CGC | G>A | A/T+1 | G/C-1 | CG>TA | CG>TA | TTGGCCGCAC | GCACCAA | GCG | GCG | C | T | C>T | C>T_GCG | CG>TA_GCACCAA | 6 | FALSE | FALSE |
| 138_v.txt | 33056 | 33056-SNV-( | SNV | 1 | C | T | 30 | 24363 | 0.12313755 | 58.0333333 | rnr1Y285A_p( | TRUE | CAN | JSY2422 | 2 | rnr1Y285A_p | GCG | C>T | A/T+1 | G/C-1 | CG>TA | CG>TA | GGACTGCGT | CGTGACA | CGC | GCG | C | A | C>T | C>T_GCG | CG>TA_CGTGACA | 6 | FALSE | FALSE |
| 138_v.txt | 33070 | 33070-SNV-( | SNV | 1 | C | T | 25 | 24363 | 0.10261462 | 57.64 | rnr1Y285A_p( | TRUE | CAN | JSY2422 | 2 | rnr1Y285A_p | GCC | C>T | A/T+1 | G/C-1 | CG>TA | CG>TA | AATATGCCA | CCAAAGA | CGG | GCC | C | A | C>T | C>T_GCC | CG>TA_CCAAAGA | 6 | FALSE | FALSE |
| 138_v.txt | 33104 | 33104-SNV-( | SNV | 1 | G | A | 23 | 24363 | 0.09440545 | 40.4782609 | rnr1Y285A_p( | TRUE | CAN | JSY2422 | 2 | rnr1Y285A_p | CGC | G>A | A/T+1 | G/C-1 | CG>TA | CG>TA | AAGAGCGCC | GCCCACT | GCG | GCG | C | T | C>T | C>T_GCG | CG>TA_GCCCACT | 6 | FALSE | FALSE |
| 138_v.txt | 33114 | 33114-SNV-( | SNV | 1 | C | T | 58 | 24363 | 0.23806592 | 40.8965517 | rnr1Y285A_p( | TRUE | CAN | JSY2422 | 2 | rnr1Y285A_p | GCC | C>T | A/T+1 | G/C-1 | CG>TA | CG>TA | ACTGGGCCG | CCGGCGT | CGG | GCC | C | A | C>T | C>T_GCC | CG>TA_CCGGCGT | 6 | FALSE | FALSE |
| 138_v.txt | 33153 | 33153-SNV-( | SNV | 1 | C | T | 236 | 24363 | 0.96868202 | 36.9322069 | rnr1Y285A_p( | TRUE | CAN | JSY2422 | 2 | rnr1Y285A_p | ACC | C>T | A/T+1 | G/C-1 | CG>TA | CG>TA | AAAAGACCT | CCTGTAC | TGG | ACC | C | A | C>T | C>T_ACC | CG>TA_CCTGTAC | 6 | FALSE | FALSE |
| 138_v.txt | 33156 | 33156-SNV-( | SNV | 1 | G | A | 40 | 24363 | 0.16418339 | 37.2 | rnr1Y285A_p( | TRUE | CAN | JSY2422 | 2 | rnr1Y285A_p | TGT | G>A | A/T+1 | G/C-1 | CG>TA | CG>TA | AGACCTGTA | GTACCAA | ACA | ACA | C | T | C>T | C>T_ACA | CG>TA_GTACCAA | 6 | FALSE | FALSE |
| 138_v.txt | 33168 | 33168-SNV-( | SNV | 1 | C | T | 28 | 24359 | 0.11494725 | 34.3214286 | rnr1Y285A_p( | TRUE | CAN | JSY2422 | 2 | rnr1Y285A_p | ACC | C>T | A/T+1 | G/C-1 | CG>TA | CG>TA | ATAGTACCAC | CCACCAA | TGG | ACC | C | A | C>T | C>T_ACC | CG>TA_CCACCAA | 6 | FALSE | FALSE |
| 138_v.txt | 33171 | 33171-SNV-( | SNV | 1 | C | T | 60 | 24359 | 0.24631553 | 35.95 | rnr1Y285A_p( | TRUE | CAN | JSY2422 | 2 | rnr1Y285A_p | ACC | C>T | A/T+1 | G/C-1 | CG>TA | CG>TA | GTACCACCA | CCAAGGG | TGG | ACC | C | A | C>T | C>T_ACC | CG>TA_CCAAGGG | 6 | FALSE | FALSE |
| 138_v.txt | 33177 | 33177-SNV-( | SNV | 1 | G | A | 17 | 24358 | 0.06979227 | 37.6470588 | rnr1Y285A_p( | TRUE | CAN | JSY2422 | 2 | rnr1Y285A_p | GGC | G>A | A/T+1 | G/C-1 | CG>TA | CG>TA | CCAAGGGCA | GCAATCA | CCG | CCG | C | T | C>T | C>T_CCG | CG>TA_GCAATCA | 6 | FALSE | FALSE |
| 138_v.txt | 33258 | 33258-SNV-( | SNV | 1 | C | T | 22 | 15623 | 0.14081803 | 42.5454546 | rnr1Y285A_p( | TRUE | CAN | JSY2422 | 2 | rnr1Y285A_p | GCC | C>T | A/T+1 | G/C-1 | CG>TA | CG>TA | CCTATGCCAT | CCATCCT | CGG | GCC | C | A | C>T | C>T_GCC | CG>TA_CCATCCT | 6 | FALSE | FALSE |
| 138_v.txt | 33276 | 33276-SNV-( | SNV | 1 | G | A | 23 | 15623 | 0.14721884 | 39.3478261 | rnr1Y285A_p( | TRUE | CAN | JSY2422 | 2 | rnr1Y285A_p | CGT | G>A | A/T+1 | G/C-1 | CG>TA | CG>TA | GAGAACGTA | GTATCCT | GCA | GCA | C | T | C>T | C>T_GCA | CG>TA_GTATCCT | 6 | FALSE | FALSE |
| 138_v.txt | 33310 | 33310-SNV-( | SNV | 1 | G | A | 23 | 15623 | 0.14721884 | 56.7391304 | rnr1Y285A_p( | TRUE | CAN | JSY2422 | 2 | rnr1Y285A_p | CGC | G>A | A/T+1 | G/C-1 | CG>TA | CG>TA | AAGAGCGCA | GCAATGG | GCG | GCG | C | T | C>T | C>T_GCG | CG>TA_GCAATGG | 6 | FALSE | FALSE |
| 138_v.txt | 33412 | 33412-SNV-( | SNV | 1 | G | A | 29 | 15623 | 0.18562376 | 47.137931 | rnr1Y285A_p( | TRUE | CAN | JSY2422 | 2 | rnr1Y285A_p | GGC | G>A | A/T+1 | G/C-1 | CG>TA | CG>TA | TGACCGGCTC | GCTCATT | CCG | CCG | C | T | C>T | C>T_CCG | CG>TA_GCTCATT | 6 | FALSE | FALSE |
| 138_v.txt | 33474 | 33474-SNV-( | SNV | 1 | C | T | 21 | 15622 | 0.13442581 | 36.6666667 | rnr1Y285A_p( | TRUE | CAN | JSY2422 | 2 | rnr1Y285A_p | GCC | C>T | A/T+1 | G/C-1 | CG>TA | CG>TA | GCTATGCCTT | CCTTTTT | CGG | GCC | C | A | C>T | C>T_GCC | CG>TA_CCTTTTT | 6 | FALSE | FALSE |
| 138_v.txt | 31658 | 31658-SNV-( | SNV | 1 | C | G | 1 | 9697 | 0.01031247 | 38 | rnr1Y285A_p( | TRUE | CAN | JSY2422 | 2 | rnr1Y285A_p | GCG | C>G | G/C+1 | G/C-1 | CG>GC | CG>GC | AAATGGCGT | CGTGGAA | CGC | GCG | C | C | C>G | C>G_GCG | CG>GC_CGTGGAA | 6 | FALSE | FALSE |
| 138_v.txt | 31981 | 31981-SNV-( | SNV | 1 | G | C | 6 | 17424 | 0.03443526 | 36.3333333 | rnr1Y285A_p( | TRUE | CAN | JSY2422 | 2 | rnr1Y285A_p | GGC | G>C | G/C+1 | G/C-1 | CG>GC | CG>GC | AGCCGGGCA | GCATTAA | CCG | CCG | C | G | C>G | C>G_CCG | CG>GC_GCATTAA | 6 | FALSE | FALSE |
| 138_v.txt | 32114 | 32114-SNV-( | SNV | 1 | C | G | 2 | 17424 | 0.01147842 | 62.5 | rnr1Y285A_p( | TRUE | CAN | JSY2422 | 2 | rnr1Y285A_p | GCC | C>G | G/C+1 | G/C-1 | CG>GC | CG>GC | TAATAGCCAT | CCATTCG | CGG | GCC | C | C | C>G | C>G_GCC | CG>GC_CCATTCG | 6 | FALSE | FALSE |
| 138_v.txt | 32545 | 32545-SNV-( | SNV | 1 | C | G | 2 | 24409 | 8.19E-03 | 36.9999508 | rnr1Y285A_p( | TRUE | CAN | JSY2422 | 2 | rnr1Y285A_p | CCA | C>G | G/C+1 | G/C-1 | CG>GC | CG>GC | CTTCACCAGC | CAGCAGT | GGT | CCA | C | C | C>G | C>G_CCA | CG>GC_CAGCAGT | 6 | FALSE | FALSE |
| 138_v.txt | 32678 | 32678-SNV-( | SNV | 1 | C | G | 3 | 19418 | 0.01544958 | 50.653075 | rnr1Y285A_p( | TRUE | CAN | JSY2422 | 2 | rnr1Y285A_p | TCC | C>G | G/C+1 | G/C-1 | CG>GC | CG>GC | GTTTCTCCAA | CCAATAA | AGG | TCC | C | C | C>G | C>G_TCC | CG>GC_CCAATAA | 6 | FALSE | FALSE |
| 138_v.txt | 33114 | 33114-SNV-( | SNV | 1 | C | G | 3 | 24363 | 0.01231376 | 38 | rnr1Y285A_p( | TRUE | CAN | JSY2422 | 2 | rnr1Y285A_p | GCC | C>G | G/C+1 | G/C-1 | CG>GC | CG>GC | ACTGGGCCG | CCGGCGT | CGG | GCC | C | C | C>G | C>G_GCC | CG>GC_CCGGCGT | 6 | FALSE | FALSE |
| 138_v.txt | 31657 | 31657-SNV-( | SNV | 1 | G | T | 68 | 9697 | 0.70124781 | 18.4476206 | rnr1Y285A_p( | TRUE | CAN | JSY2422 | 2 | rnr1Y285A_p | GGC | G>T | A/T+1 | G/C-1 | CG>AT | CG>AT | GAAATGGCG | GCGTGGA | CCG | CCG | C | A | C>A | C>A_CCG | CG>AT_GCGTGGA | 6 | FALSE | FALSE |
| 138_v.txt | 31981 | 31981-SNV-( | SNV | 1 | G | T | 35 | 17424 | 0.20087236 | 29.8857143 | rnr1Y285A_p( | TRUE | CAN | JSY2422 | 2 | rnr1Y285A_p | GGC | G>T | A/T+1 | G/C-1 | CG>AT | CG>AT | AGCCGGGCA | GCATTAA | CCG | CCG | C | A | C>A | C>A_CCG | CG>AT_GCATTAA | 6 | FALSE | FALSE |
| 138_v.txt | 32027 | 32027-SNV-( | SNV | 1 | G | T | 351 | 17424 | 2.01446281 | 59.4131054 | rnr1Y285A_p( | TRUE | CAN | JSY2422 | 2 | rnr1Y285A_p | GGT | G>T | A/T+1 | G/C-1 | CG>AT | CG>AT | GCCACGGTA | GTATTTC | CCA | CCA | C | A | C>A | C>A_CCA | CG>AT_GTATTTC | 6 | FALSE | FALSE |
| 138_v.txt | 32088 | 32088-SNV-( | SNV | 1 | C | A | 63 | 17424 | 0.36157025 | 61.8253968 | rnr1Y285A_p( | TRUE | CAN | JSY2422 | 2 | rnr1Y285A_p | GCC | C>A | A/T+1 | G/C-1 | CG>AT | CG>AT | AAAAAGCCT | CCTGCAA | CGG | GCC | C | T | C>A | C>A_GCC | CG>AT_CCTGCAA | 6 | FALSE | FALSE |
| 138_v.txt | 32188 | 32188-SNV-( | SNV | 1 | C | A | 67 | 17424 | 0.38452709 | 46.0298508 | rnr1Y285A_p( | TRUE | CAN | JSY2422 | 2 | rnr1Y285A_p | ACT | C>A | A/T+1 | G/C-1 | CG>AT | CG>AT | CGAAAACTG | CTGCAAT | TGA | ACT | C | T | C>A | C>A_ACT | CG>AT_CTGCAAT | 6 | FALSE | FALSE |
| 138_v.txt | 32274 | 32274-SNV-( | SNV | 1 | G | T | 132 | 24410 | 0.54076198 | 35.8484849 | rnr1Y285A_p( | TRUE | CAN | JSY2422 | 2 | rnr1Y285A_p | GGA | G>T | A/T+1 | G/C-1 | CG>AT | CG>AT | ATACGGGAA | GAACCAA | CCT | CCT | C | A | C>A | C>A_CCT | CG>AT_GAACCAA | 6 | FALSE | FALSE |
| 138_v.txt | 32277 | 32277-SNV-( | SNV | 1 | C | A | 711 | 24410 | 2.91274068 | 36.5063291 | rnr1Y285A_p( | TRUE | CAN | JSY2422 | 2 | rnr1Y285A_p | ACC | C>A | A/T+1 | G/C-1 | CG>AT | CG>AT | CGGGAACCA | CCAACGT | TGG | ACC | C | T | C>A | C>A_ACC | CG>AT_CCAACGT | 6 | FALSE | FALSE |
| 138_v.txt | 32373 | 32373-SNV-( | SNV | 1 | G | T | 19 | 24410 | 0.07783695 | 59.3684211 | rnr1Y285A_p( | TRUE | CAN | JSY2422 | 2 | rnr1Y285A_p | AGC | G>T | A/T+1 | G/C-1 | CG>AT | CG>AT | TCAATAGCA | GCAATAA | TCG | TCG | C | A | C>A | C>A_TCG | CG>AT_GCAATAA | 6 | FALSE | FALSE |
| 138_v.txt | 32424 | 32424-SNV-( | SNV | 1 | G | T | 7 | 24410 | 0.02867677 | 50.5714286 | rnr1Y285A_p( | TRUE | CAN | JSY2422 | 2 | rnr1Y285A_p | AGG | G>T | A/T+1 | G/C-1 | CG>AT | CG>AT | AGTTTAGGG | GGGTCAT | TCC | TCC | C | A | C>A | C>A_TCC | CG>AT_GGGTCAT | 6 | FALSE | FALSE |
| 138_v.txt | 32508 | 32508-SNV-( | SNV | 1 | G | T | 44 | 24410 | 0.18025399 | 41.4545455 | rnr1Y285A_p( | TRUE | CAN | JSY2422 | 2 | rnr1Y285A_p | GGC | G>T | A/T+1 | G/C-1 | CG>AT | CG>AT | TTGATGGCTC | GCTCTTG | CCG | CCG | C | A | C>A | C>A_CCG | CG>AT_GCTCTTG | 6 | FALSE | FALSE |
| 138_v.txt | 32515 | 32515-SNV-( | SNV | 1 | G | T | 267 | 24410 | 1.09381401 | 42.8127341 | rnr1Y285A_p( | TRUE | CAN | JSY2422 | 2 | rnr1Y285A_p | GGA | G>T | A/T+1 | G/C-1 | CG>AT | CG>AT | CTCTTGGAAC | GAACGGA | CCT | CCT | C | A | C>A | C>A_CCT | CG>AT_GAACGGA | 6 | FALSE | FALSE |
| 138_v.txt | 32530 | 32530-SNV-( | SNV | 1 | G | T | 91 | 24410 | 0.37279803 | 38.6703297 | rnr1Y285A_p( | TRUE | CAN | JSY2422 | 2 | rnr1Y285A_p | GGG | G>T | A/T+1 | G/C-1 | CG>AT | CG>AT | TTCTGGGGTT | GGTTTGC | CCC | CCC | C | A | C>A | C>A_CCC | CG>AT_GGTTTGC | 6 | FALSE | FALSE |
| 138_v.txt | 32538 | 32538-SNV-( | SNV | 1 | G | T | 35 | 24410 | 0.14338386 | 35.0572394 | rnr1Y285A_p( | TRUE | CAN | JSY2422 | 2 | rnr1Y285A_p | AGC | G>T | A/T+1 | G/C-1 | CG>AT | CG>AT | TTTGCAGCTT | GCTTCAC | TCG | TCG | C | A | C>A | C>A_TCG | CG>AT_GCTTCAC | 6 | FALSE | FALSE |
| 138_v.txt | 32545 | 32545-SNV-( | SNV | 1 | C | A | 10 | 24409 | 0.0409685 | 37.899873 | rnr1Y285A_p( | TRUE | CAN | JSY2422 | 2 | rnr1Y285A_p | CCA | C>A | A/T+1 | G/C-1 | CG>AT | CG>AT | CTTCACCAGC | CAGCAGT | GGT | CCA | C | T | C>A | C>A_CCA | CG>AT_CAGCAGT | 6 | FALSE | FALSE |
| 138_v.txt | 32547 | 32547-SNV-( | SNV | 1 | G | T | 18 | 24408 | 0.07374631 | 34.3333333 | rnr1Y285A_p( | TRUE | CAN | JSY2422 | 2 | rnr1Y285A_p | AGC | G>T | A/T+1 | G/C-1 | CG>AT | CG>AT | TCACCAGCAC | GCAGTGA | TCG | TCG | C | A | C>A | C>A_TCG | CG>AT_GCAGTGA | 6 | FALSE | FALSE |
| 138_v.txt | 32556 | 32556-SNV-( | SNV | 1 | C | A | 282 | 24399 | 1.15578507 | 37.3404255 | rnr1Y285A_p( | TRUE | CAN | JSY2422 | 2 | rnr1Y285A_p | ACC | C>A | A/T+1 | G/C-1 | CG>AT | CG>AT | GTGATACCA | CCAACTA | TGG | ACC | C | T | C>A | C>A_ACC | CG>AT_CCAACTA | 6 | FALSE | FALSE |
| 138_v.txt | 32572 | 32572-SNV-( | SNV | 1 | C | A | 48 | 19418 | 0.24719333 | 36.2083333 | rnr1Y285A_p( | TRUE | CAN | JSY2422 | 2 | rnr1Y285A_p | CCT | C>A | A/T+1 | G/C-1 | CG>AT | CG>AT | CAGTACCTTG | CTTGAAA | GGA | CCT | C | T | C>A | C>A_CCT | CG>AT_CTTGAAA | 6 | FALSE | FALSE |
| 138_v.txt | 32692 | 32692-SNV-( | SNV | 1 | C | A | 9 | 19416 | 0.04635352 | 43.2228282 | rnr1Y285A_p( | TRUE | CAN | JSY2422 | 2 | rnr1Y285A_p | CCA | C>A | A/T+1 | G/C-1 | CG>AT | CG>AT | GGAATCCAA | CAACTGG | GGT | CCA | C | T | C>A | C>A_CCA | CG>AT_CAACTGG | 6 | FALSE | FALSE |
| 138_v.txt | 32735 | 32735-SNV-( | SNV | 1 | G | T | 184 | 19418 | 0.94757442 | 57.7881427 | rnr1Y285A_p( | TRUE | CAN | JSY2422 | 2 | rnr1Y285A_p | AGT | G>T | A/T+1 | G/C-1 | CG>AT | CG>AT | AAAACAGTA | GTATATT | TCA | TCA | C | A | C>A | C>A_TCA | CG>AT_GTATATT | 6 | FALSE | FALSE |
| 138_v.txt | 32783 | 32783-SNV-( | SNV | 1 | G | T | 20 | 19418 | 0.10299722 | 46 | rnr1Y285A_p( | TRUE | CAN | JSY2422 | 2 | rnr1Y285A_p | AGA | G>T | A/T+1 | G/C-1 | CG>AT | CG>AT | GACCCAGAA | GAACTCG | TCT | TCT | C | A | C>A | C>A_TCT | CG>AT_GAACTCG | 6 | FALSE | FALSE |
| 138_v.txt | 32786 | 32786-SNV-( | SNV | 1 | C | A | 16 | 19418 | 0.08239778 | 42.875 | rnr1Y285A_p( | TRUE | CAN | JSY2422 | 2 | rnr1Y285A_p | ACT | C>A | A/T+1 | G/C-1 | CG>AT | CG>AT | CCAGAACTCC | CTCGAAT | TGA | ACT | C | T | C>A | C>A_ACT | CG>AT_CTCGAAT | 6 | FALSE | FALSE |
| 138_v.txt | 32794 | 32794-SNV-( | SNV | 1 | C | A | 42 | 19418 | 0.21629416 | 41 | rnr1Y285A_p( | TRUE | CAN | JSY2422 | 2 | rnr1Y285A_p | TCA | C>A | A/T+1 | G/C-1 | CG>AT | CG>AT | CGAATTCACC | CACCGTA | AGT | TCA | C | T | C>A | C>A_TCA | CG>AT_CACCGTA | 6 | FALSE | FALSE |
| 138_v.txt | 32819 | 32819-SNV-( | SNV | 1 | G | T | 130 | 19418 | 0.66948192 | 43.4846154 | rnr1Y285A_p( | TRUE | CAN | JSY2422 | 2 | rnr1Y285A_p | AGT | G>T | A/T+1 | G/C-1 | CG>AT | CG>AT | GAACAAGTT | GTTCATT | TCA | TCA | C | A | C>A | C>A_TCA | CG>AT_GTTCATT | 6 | FALSE | FALSE |
| 138_v.txt | 32917 | 32917-SNV-( | SNV | 1 | C | A | 336 | 24363 | 1.3791405 | 52.1646383 | rnr1Y285A_p( | TRUE | CAN | JSY2422 | 2 | rnr1Y285A_p | TCC | C>A | A/T+1 | G/C-1 | CG>AT | CG>AT | TAAGTTCCAC | CCAGGGC | AGG | TCC | C | T | C>A | C>A_TCC | CG>AT_CCAGGGC | 6 | FALSE | FALSE |
| 138_v.txt | 32940 | 32940-SNV-( | SNV | 1 | G | T | 534 | 24363 | 2.1918483 | 54.9702382 | rnr1Y285A_p( | TRUE | CAN | JSY2422 | 2 | rnr1Y285A_p | AGA | G>T | A/T+1 | G/C-1 | CG>AT | CG>AT | GCCCAAGAA | GAAAACC | TCT | TCT | C | A | C>A | C>A_TCT | CG>AT_GAAAACC | 6 | FALSE | FALSE |
| 138_v.txt | 32958 | 32958-SNV-( | SNV | 1 | C | A | 155 | 24363 | 0.63621065 | 57.0580645 | rnr1Y285A_p( | TRUE | CAN | JSY2422 | 2 | rnr1Y285A_p | ACC | C>A | A/T+1 | G/C-1 | CG>AT | CG>AT | ATGTAACCAT | CCATTGG | TGG | ACC | C | T | C>A | C>A_ACC | CG>AT_CCATTGG | 6 | FALSE | FALSE |
| 138_v.txt | 33015 | 33015-SNV-( | SNV | 1 | G | T | 44 | 24363 | 0.18060173 | 59.1590909 | rnr1Y285A_p( | TRUE | CAN | JSY2422 | 2 | rnr1Y285A_p | GGA | G>T | A/T+1 | G/C-1 | CG>AT | CG>AT | AAAGAGGAT | GATGTAA | CCT | CCT | C | A | C>A | C>A_CCT | CG>AT_GATGTAA | 6 | FALSE | FALSE |
| 138_v.txt | 33056 | 33056-SNV-( | SNV | 1 | C | A | 9 | 24363 | 0.03694126 | 37.5555556 | rnr1Y285A_p( | TRUE | CAN | JSY2422 | 2 | rnr1Y285A_p | GCG | C>A | A/T+1 | G/C-1 | CG>AT | CG>AT | GGACTGCGT | CGTGACA | CGC | GCG | C | T | C>A | C>A_GCG | CG>AT_CGTGACA | 6 | FALSE | FALSE |
| 138_v.txt | 33112 | 33112-SNV-( | SNV | 1 | G | T | 12 | 24363 | 0.04925502 | 45.4166667 | rnr1Y285A_p( | TRUE | CAN | JSY2422 | 2 | rnr1Y285A_p | GGG | G>T | A/T+1 | G/C-1 | CG>AT | CG>AT | CCACTGGGC | GGCCGGC | CCC | CCC | C | A | C>A | C>A_CCC | CG>AT_GGCCGGC | 6 | FALSE | FALSE |
| 138_v.txt | 33114 | 33114-SNV-( | SNV | 1 | C | A | 12 | 24363 | 0.04925502 | 42.0833333 | rnr1Y285A_p( | TRUE | CAN | JSY2422 | 2 | rnr1Y285A_p | GCC | C>A | A/T+1 | G/C-1 | CG>AT | CG>AT | ACTGGGCCG | CCGGCGT | CGG | GCC | C | T | C>A | C>A_GCC | CG>AT_CCGGCGT | 6 | FALSE | FALSE |
| 138_v.txt | 33151 | 33151-SNV-( | SNV | 1 | G | T | 4746 | 24363 | 19.4803596 | 38.9837758 | rnr1Y285A_p( | TRUE | CAN | JSY2422 | 2 | rnr1Y285A_p | AGA | G>T | A/T+1 | G/C-1 | CG>AT | CG>AT | TGAAAAGAC | GACCTGT | TCT | TCT | C | A | C>A | C>A_TCT | CG>AT_GACCTGT | 6 | FALSE | FALSE |
| 138_v.txt | 33153 | 33153-SNV-( | SNV | 1 | C | A | 101 | 24363 | 0.41456307 | 42.9105853 | rnr1Y285A_p( | TRUE | CAN | JSY2422 | 2 | rnr1Y285A_p | ACC | C>A | A/T+1 | G/C-1 | CG>AT | CG>AT | AAAAGACCT | CCTGTAC | TGG | ACC | C | T | C>A | C>A_ACC | CG>AT_CCTGTAC | 6 | FALSE | FALSE |
| 138_v.txt | 33154 | 33154-SNV-( | SNV | 1 | C | A | 176 | 24363 | 0.72240693 | 37.176204 | rnr1Y285A_p( | TRUE | CAN | JSY2422 | 2 | rnr1Y285A_p | CCT | C>A | A/T+1 | G/C-1 | CG>AT | CG>AT | AAAGACCTG | CTGTACC | GGA | CCT | C | T | C>A | C>A_CCT | CG>AT_CTGTACC | 6 | FALSE | FALSE |
| 138_v.txt | 33159 | 33159-SNV-( | SNV | 1 | C | A | 47 | 24363 | 0.19291549 | 36.6170213 | rnr1Y285A_p( | TRUE | CAN | JSY2422 | 2 | rnr1Y285A_p | ACC | C>A | A/T+1 | G/C-1 | CG>AT | CG>AT | CCTGTACCA | CCAATAG | TGG | ACC | C | T | C>A | C>A_ACC | CG>AT_CCAATAG | 6 | FALSE | FALSE |
| 138_v.txt | 33168 | 33168-SNV-( | SNV | 1 | C | A | 243 | 24359 | 0.9975779 | 37.1563786 | rnr1Y285A_p( | TRUE | CAN | JSY2422 | 2 | rnr1Y285A_p | ACC | C>A | A/T+1 | G/C-1 | CG>AT | CG>AT | ATAGTACCAC | CCACCAA | TGG | ACC | C | T | C>A | C>A_ACC | CG>AT_CCACCAA | 6 | FALSE | FALSE |
| 138_v.txt | 33177 | 33177-SNV-( | SNV | 1 | G | T | 52 | 24358 | 0.21348222 | 37.3076923 | rnr1Y285A_p( | TRUE | CAN | JSY2422 | 2 | rnr1Y285A_p | GGC | G>T | A/T+1 | G/C-1 | CG>AT | CG>AT | CCAAGGGCA | GCAATCA | CCG | CCG | C | A | C>A | C>A_CCG | CG>AT_GCAATCA | 6 | FALSE | FALSE |
| 138_v.txt | 33205 | 33205-SNV-( | SNV | 1 | G | T | 22 | 15623 | 0.14081803 | 38.2272727 | rnr1Y285A_p( | TRUE | CAN | JSY2422 | 2 | rnr1Y285A_p | AGC | G>T | A/T+1 | G/C-1 | CG>AT | CG>AT | GCTTAAGCTC | GCTCTCT | TCG | TCG | C | A | C>A | C>A_TCG | CG>AT_GCTCTCT | 6 | FALSE | FALSE |
| 138_v.txt | 33220 | 33220-SNV-( | SNV | 1 | C | A | 23 | 15623 | 0.14721884 | 37.6521739 | rnr1Y285A_p( | TRUE | CAN | JSY2422 | 2 | rnr1Y285A_p | TCA | C>A | A/T+1 | G/C-1 | CG>AT | CG>AT | TCACTTCAGC | CAGCGTT | AGT | TCA | C | T | C>A | C>A_TCA | CG>AT_CAGCGTT | 6 | FALSE | FALSE |
| 138_v.txt | 33310 | 33310-SNV-( | SNV | 1 | G | T | 11 | 15623 | 0.07040901 | 30.7272727 | rnr1Y285A_p( | TRUE | CAN | JSY2422 | 2 | rnr1Y285A_p | CGC | G>T | A/T+1 | G/C-1 | CG>AT | CG>AT | AAGAGCGCA | GCAATGG | GCG | GCG | C | A | C>A | C>A_GCG | CG>AT_GCAATGG | 6 | FALSE | FALSE |
| 138_v.txt | 33325 | 33325-SNV-( | SNV | 1 | C | A | 90 | 15623 | 0.57607374 | 58.2555556 | rnr1Y285A_p( | TRUE | CAN | JSY2422 | 2 | rnr1Y285A_p | TCT | C>A | A/T+1 | G/C-1 | CG>AT | CG>AT | ACAATTCTTT | CTTTACT | AGA | TCT | C | T | C>A | C>A_TCT | CG>AT_CTTTACT | 6 | FALSE | FALSE |
| 138_v.txt | 33354 | 33354-SNV-( | SNV | 1 | G | T | 134 | 15623 | 0.85770979 | 25.5074627 | rnr1Y285A_p( | TRUE | CAN | JSY2422 | 2 | rnr1Y285A_p | TGA | G>T | A/T+1 | G/C-1 | CG>AT | CG>AT | GGTATTGAC | GACCCAC | ACT | ACT | C | A | C>A | C>A_ACT | CG>AT_GACCCAC | 6 | FALSE | FALSE |
| 138_v.txt | 33474 | 33474-SNV-( | SNV | 1 | C | A | 53 | 15622 | 0.33926514 | 33.3773585 | rnr1Y285A_p( | TRUE | CAN | JSY2422 | 2 | rnr1Y285A_p | GCC | C>A | A/T+1 | G/C-1 | CG>AT | CG>AT | GCTATGCCTT | CCTTTTT | CGG | GCC | C | T | C>A | C>A_GCC | CG>AT_CCTTTTT | 6 | FALSE | FALSE |
| 138_v.txt | 33475 | 33475-SNV-( | SNV | 1 | C | A | 14 | 15622 | 0.08961721 | 37.8571429 | rnr1Y285A_p( | TRUE | CAN | JSY2422 | 2 | rnr1Y285A_p | CCT | C>A | A/T+1 | G/C-1 | CG>AT | CG>AT | CTATGCCTTT | CTTTTTT | GGA | CCT | C | T | C>A | C>A_CCT | CG>AT_CTTTTTT | 6 | FALSE | FALSE |
| 138_v.txt | 32689 | 32689-Repla | Replacement | 7 | A | TGGTTTG | 59 | 19418 | 0.3038418 | 58.1961259 | rnr1Y285A_p( | TRUE | CAN | JSY2422 | 2 | rnr1Y285A_p | AAT | nope | >1bp-ins | A/T-1 | none | Replac. | AACGGAATC | ATCCAAC | TTA | TTA | T | none | T>none | T>none_TTA | Replac._ATCCAAC | 6 | FALSE | FALSE |
| 138_v.txt | 32940 | 32940-Repla | Replacement | 2 | G | TA | 8 | 24363 | 0.03283668 | 59.8749555 | rnr1Y285A_p( | TRUE | CAN | JSY2422 | 2 | rnr1Y285A_p | AGA | nope | >1bp-ins | G/C-1 | none | Replac. | GCCCAAGAA | GAAAACC | TCT | TCT | C | none | C>none | C>none_TCT | Replac._GAAAACC | 6 | FALSE | FALSE |
| 138_v.txt | 32544 | 32544-MNV- | MNV | 2 | CC | AA | 3 | 24409 | 0.01229055 | 31.9999747 | rnr1Y285A_p( | TRUE | CAN | JSY2422 | 2 | rnr1Y285A_p | ACC | nope | >1bp-ins | >1bpdel | none | MNV | GCTTCACCAG | CCAGCAG | TGG | ACC | CC | none | NA | NA_ACC | MNV_CCAGCAG | 6 | FALSE | FALSE |
| 138_v.txt | 32691 | 32691-MNV- | MNV | 4 | CCAA | GGTG | 59 | 19418 | 0.3038418 | 58.9152542 | rnr1Y285A_p( | TRUE | CAN | JSY2422 | 2 | rnr1Y285A_p | TCC | nope | >1bp-ins | >1bpdel | none | MNV | CGGAATCCA | CCAACTG | AGG | TCC | CCAA | none | NA | NA_TCC | MNV_CCAACTG | 6 | FALSE | FALSE |
| 138_v.txt | 33153 | 33153-MNV- | MNV | 2 | CC | AA | 3 | 24363 | 0.01231376 | 28.3333333 | rnr1Y285A_p( | TRUE | CAN | JSY2422 | 2 | rnr1Y285A_p | ACC | nope | >1bp-ins | >1bpdel | none | MNV | AAAAGACCT | CCTGTAC | TGG | ACC | CC | none | NA | NA_ACC | MNV_CCTGTAC | 6 | FALSE | FALSE |
| 138_v.txt | 32081 | 32081-Insert | Insertion | 1 | - | A | 23 | 17424 | 0.13200184 | 64 | rnr1Y285A_p( | TRUE | CAN | JSY2422 | 2 | rnr1Y285A_p | CAA | nope | A/T+1 | >1bpdel | none | A/T+1 | CCATGCAAA | AAAAAAG | GTT | CAA | - | T | NA | NA_CAA | A/T+1_AAAAAAG | 6 | FALSE | FALSE |
| 138_v.txt | 32265 | 32265-Insert | Insertion | 1 | - | A | 2 | 24411 | 8.19E-03 | 38 | rnr1Y285A_p( | TRUE | CAN | JSY2422 | 2 | rnr1Y285A_p | TAA | nope | A/T+1 | >1bpdel | none | A/T+1 | CCAAATAAA | AAAATAC | ATT | TAA | - | T | NA | NA_TAA | A/T+1_AAAATAC | 6 | FALSE | FALSE |
| 138_v.txt | 32498 | 32498-Insert | Insertion | 1 | - | T | 2 | 24410 | 8.19E-03 | 38 | rnr1Y285A_p( | TRUE | CAN | JSY2422 | 2 | rnr1Y285A_p | CTT | nope | A/T+1 | >1bpdel | none | A/T+1 | AACAACTTTT | TTTTTTG | GAA | CTT | - | A | NA | NA_CTT | A/T+1_TTTTTTG | 6 | FALSE | FALSE |
| 138_v.txt | 32532 | 32532-Insert | Insertion | 1 | - | T | 4 | 24410 | 0.01638673 | 38 | rnr1Y285A_p( | TRUE | CAN | JSY2422 | 2 | rnr1Y285A_p | GTT | nope | A/T+1 | >1bpdel | none | A/T+1 | CTGGGGTTTC | TTTGCAG | CAA | GTT | - | A | NA | NA_GTT | A/T+1_TTTGCAG | 6 | FALSE | FALSE |
| 138_v.txt | 32631 | 32631-Insert | Insertion | 1 | - | T | 2 | 19418 | 0.01029972 | 51 | rnr1Y285A_p( | TRUE | CAN | JSY2422 | 2 | rnr1Y285A_p | GTT | nope | A/T+1 | >1bpdel | none | A/T+1 | CCTTCGTTTT | TTTTTAT | CAA | GTT | - | A | NA | NA_GTT | A/T+1_TTTTTAT | 6 | FALSE | FALSE |
| 138_v.txt | 32684 | 32684-Insert | Insertion | 1 | - | T | 59 | 19418 | 0.3038418 | 55.1525424 | rnr1Y285A_p( | TRUE | CAN | JSY2422 | 2 | rnr1Y285A_p | AAC | nope | A/T+1 | >1bpdel | none | A/T+1 | CCAATAACGC | ACGGAAT | TTG | AAC | - | A | NA | NA_AAC | A/T+1_ACGGAAT | 6 | FALSE | FALSE |
| 138_v.txt | 32686 | 32686-Insert | Insertion | 1 | - | T | 59 | 19418 | 0.3038418 | 56.5084746 | rnr1Y285A_p( | TRUE | CAN | JSY2422 | 2 | rnr1Y285A_p | CGG | nope | A/T+1 | >1bpdel | none | A/T+1 | AATAACGGA | GGAATCC | GCC | CGG | - | A | NA | NA_CGG | A/T+1_GGAATCC | 6 | FALSE | FALSE |
| 138_v.txt | 32687 | 32687-Insert | Insertion | 5 | - | TTTTT | 59 | 19418 | 0.3038418 | 58.359322 | rnr1Y285A_p( | TRUE | CAN | JSY2422 | 2 | rnr1Y285A_p | GGA | nope | >1bp-ins | >1bpdel | none | >1bp-ins | ATAACGGAA | GAATCCA | CCT | GGA | - | none | NA | NA_GGA | >1bp-ins_GAATCCA | 6 | FALSE | FALSE |
| 138_v.txt | 32688 | 32688-Insert | Insertion | 1 | - | T | 59 | 19418 | 0.3038418 | 59.8135593 | rnr1Y285A_p( | TRUE | CAN | JSY2422 | 2 | rnr1Y285A_p | GAA | nope | A/T+1 | >1bpdel | none | A/T+1 | TAACGGAAT | AATCCAA | CTT | GAA | - | A | NA | NA_GAA | A/T+1_AATCCAA | 6 | FALSE | FALSE |
| 138_v.txt | 32842 | 32842-Insert | Insertion | 1 | - | A | 34 | 19418 | 0.17509527 | 43.9411765 | rnr1Y285A_p( | TRUE | CAN | JSY2422 | 2 | rnr1Y285A_p | CAA | nope | A/T+1 | >1bpdel | none | A/T+1 | TTACCCAAA | AAAAAAT | GTT | CAA | - | T | NA | NA_CAA | A/T+1_AAAAAAT | 6 | FALSE | FALSE |
| 138_v.txt | 32858 | 32858-Insert | Insertion | 1 | - | T | 10 | 19415 | 0.05150657 | 38 | rnr1Y285A_p( | TRUE | CAN | JSY2422 | 2 | rnr1Y285A_p | ATG | nope | A/T+1 | >1bpdel | none | A/T+1 | AATCCATGC | TGCCGCC | TAC | ATG | - | A | NA | NA_ATG | A/T+1_TGCCGCC | 6 | FALSE | FALSE |
| 138_v.txt | 32678 | 32678-Deleti | Deletion | 2 | CC | - | 59 | 19418 | 0.3038418 | 53.661017 | rnr1Y285A_p( | TRUE | CAN | JSY2422 | 2 | rnr1Y285A_p | TCC | nope | >1bp-ins | >1bpdel | none | >1bpdel | GTTTCTCCAA | CCAATAA | AGG | TCC | CC | none | NA | NA_TCC | >1bpdel_CCAATAA | 6 | FALSE | FALSE |
| 138_v.txt | 31723 | 31723-Deleti | Deletion | 1 | A | - | 11 | 9697 | 0.11343715 | 61.8181818 | rnr1Y285A_p( | TRUE | CAN | JSY2422 | 2 | rnr1Y285A_p | CAA | nope | >1bp-ins | A/T-1 | none | A/T-1 | TGTCCCAAA | AAAAAGT | GTT | GTT | T | none | T>none | T>none_GTT | A/T-1_AAAAAGT | 6 | FALSE | FALSE |
| 138_v.txt | 31904 | 31904-Deleti | Deletion | 1 | T | - | 5 | 9687 | 0.05161557 | 41.4 | rnr1Y285A_p( | TRUE | CAN | JSY2422 | 2 | rnr1Y285A_p | ATT | nope | >1bp-ins | A/T-1 | none | A/T-1 | ATTGAATTTT | TTTTGGT | TAA | ATT | T | none | nope | nope_ATT | A/T-1_TTTTGGT | 6 | FALSE | FALSE |
| 138_v.txt | 31972 | 31972-Deleti | Deletion | 1 | C | - | 6 | 17424 | 0.03443526 | 37 | rnr1Y285A_p( | TRUE | CAN | JSY2422 | 2 | rnr1Y285A_p | GCC | nope | >1bp-ins | G/C-1 | none | G/C-1 | AATAAGCCA | CCAAGCC | CGG | GCC | C | none | nope | nope_GCC | G/C-1_CCAAGCC | 6 | FALSE | FALSE |
| 138_v.txt | 31979 | 31979-Deleti | Deletion | 1 | G | - | 31 | 17424 | 0.17791552 | 37.3548387 | rnr1Y285A_p( | TRUE | CAN | JSY2422 | 2 | rnr1Y285A_p | CGG | nope | >1bp-ins | G/C-1 | none | G/C-1 | CAAGCCGGG | GGGCATT | GCC | GCC | C | none | C>none | C>none_GCC | G/C-1_GGGCATT | 6 | FALSE | FALSE |
| 138_v.txt | 32006 | 32006-Deleti | Deletion | 1 | C | - | 6 | 17424 | 0.03443526 | 54.3333333 | rnr1Y285A_p( | TRUE | CAN | JSY2422 | 2 | rnr1Y285A_p | ACT | nope | >1bp-ins | G/C-1 | none | G/C-1 | TGGTAACTC | CTCGTCA | TGA | ACT | C | none | nope | nope_ACT | G/C-1_CTCGTCA | 6 | FALSE | FALSE |
| 138_v.txt | 32081 | 32081-Deleti | Deletion | 1 | A | - | 22 | 17424 | 0.12626263 | 63.8636364 | rnr1Y285A_p( | TRUE | CAN | JSY2422 | 2 | rnr1Y285A_p | CAA | nope | >1bp-ins | A/T-1 | none | A/T-1 | CCATGCAAA | AAAAAAG | GTT | GTT | T | none | T>none | T>none_GTT | A/T-1_AAAAAAG | 6 | FALSE | FALSE |
| 138_v.txt | 32114 | 32114-Deleti | Deletion | 1 | C | - | 3 | 17424 | 0.01721763 | 55.3333333 | rnr1Y285A_p( | TRUE | CAN | JSY2422 | 2 | rnr1Y285A_p | GCC | nope | >1bp-ins | G/C-1 | none | G/C-1 | TAATAGCCAT | CCATTCG | CGG | GCC | C | none | nope | nope_GCC | G/C-1_CCATTCG | 6 | FALSE | FALSE |
| 138_v.txt | 32216 | 32216-Deleti | Deletion | 1 | G | - | 4 | 17424 | 0.02295684 | 44.5 | rnr1Y285A_p( | TRUE | CAN | JSY2422 | 2 | rnr1Y285A_p | TGG | nope | >1bp-ins | G/C-1 | none | G/C-1 | TTTGGTGGTC | GGTCCTT | ACC | ACC | C | none | C>none | C>none_ACC | G/C-1_GGTCCTT | 6 | FALSE | FALSE |
| 138_v.txt | 32231 | 32231-Deleti | Deletion | 1 | T | - | 4 | 17423 | 0.02295816 | 38 | rnr1Y285A_p( | TRUE | CAN | JSY2422 | 2 | rnr1Y285A_p | ATT | nope | >1bp-ins | A/T-1 | none | A/T-1 | CAGGAATTTA | TTTAGGA | TAA | ATT | T | none | nope | nope_ATT | A/T-1_TTTAGGA | 6 | FALSE | FALSE |
| 138_v.txt | 32259 | 32259-Deleti | Deletion | 1 | C | - | 13 | 24410 | 0.05325686 | 35.6153846 | rnr1Y285A_p( | TRUE | CAN | JSY2422 | 2 | rnr1Y285A_p | ACC | nope | >1bp-ins | G/C-1 | none | G/C-1 | GATAGACCA | CCAAATA | TGG | ACC | C | none | nope | nope_ACC | G/C-1_CCAAATA | 6 | FALSE | FALSE |
| 138_v.txt | 32277 | 32277-Deleti | Deletion | 1 | C | - | 8 | 24410 | 0.03277345 | 36.5 | rnr1Y285A_p( | TRUE | CAN | JSY2422 | 2 | rnr1Y285A_p | ACC | nope | >1bp-ins | G/C-1 | none | G/C-1 | CGGGAACCA | CCAACGT | TGG | ACC | C | none | nope | nope_ACC | G/C-1_CCAACGT | 6 | FALSE | FALSE |
| 138_v.txt | 32346 | 32346-Deleti | Deletion | 1 | A | - | 107 | 24410 | 0.43834494 | 61.4579439 | rnr1Y285A_p( | TRUE | CAN | JSY2422 | 2 | rnr1Y285A_p | CAA | nope | >1bp-ins | A/T-1 | none | A/T-1 | TGTGGCAAA | AAAACCT | GTT | GTT | T | none | T>none | T>none_GTT | A/T-1_AAAACCT | 6 | FALSE | FALSE |
| 138_v.txt | 32358 | 32358-Deleti | Deletion | 1 | C | - | 15 | 24410 | 0.06145023 | 61.2666667 | rnr1Y285A_p( | TRUE | CAN | JSY2422 | 2 | rnr1Y285A_p | ACC | nope | >1bp-ins | G/C-1 | none | G/C-1 | TTTGTACCAG | CCAGAGT | TGG | ACC | C | none | nope | nope_ACC | G/C-1_CCAGAGT | 6 | FALSE | FALSE |
| 138_v.txt | 32398 | 32398-Deleti | Deletion | 1 | C | - | 13 | 24410 | 0.05325686 | 58.6153846 | rnr1Y285A_p( | TRUE | CAN | JSY2422 | 2 | rnr1Y285A_p | ACG | nope | >1bp-ins | G/C-1 | none | G/C-1 | TAGAAACGT | CGTAGGA | TGC | ACG | C | none | nope | nope_ACG | G/C-1_CGTAGGA | 6 | FALSE | FALSE |
| 138_v.txt | 32424 | 32424-Deleti | Deletion | 1 | G | - | 15 | 24410 | 0.06145023 | 64 | rnr1Y285A_p( | TRUE | CAN | JSY2422 | 2 | rnr1Y285A_p | AGG | nope | >1bp-ins | G/C-1 | none | G/C-1 | AGTTTAGGG | GGGTCAT | TCC | TCC | C | none | C>none | C>none_TCC | G/C-1_GGGTCAT | 6 | FALSE | FALSE |
| 138_v.txt | 32498 | 32498-Deleti | Deletion | 1 | T | - | 23 | 24410 | 0.09422368 | 44.999999 | rnr1Y285A_p( | TRUE | CAN | JSY2422 | 2 | rnr1Y285A_p | CTT | nope | >1bp-ins | A/T-1 | none | A/T-1 | AACAACTTTT | TTTTTTG | GAA | CTT | T | none | nope | nope_CTT | A/T-1_TTTTTTG | 6 | FALSE | FALSE |
| 138_v.txt | 32522 | 32522-Deleti | Deletion | 1 | T | - | 47 | 24410 | 0.19254404 | 41.3617021 | rnr1Y285A_p( | TRUE | CAN | JSY2422 | 2 | rnr1Y285A_p | ATT | nope | >1bp-ins | A/T-1 | none | A/T-1 | AACGGATTTT | TTTTCTG | TAA | ATT | T | none | nope | nope_ATT | A/T-1_TTTTCTG | 6 | FALSE | FALSE |
| 138_v.txt | 32544 | 32544-Deleti | Deletion | 1 | C | - | 2 | 24409 | 8.19E-03 | 37.9999997 | rnr1Y285A_p( | TRUE | CAN | JSY2422 | 2 | rnr1Y285A_p | ACC | nope | >1bp-ins | G/C-1 | none | G/C-1 | GCTTCACCAG | CCAGCAG | TGG | ACC | C | none | nope | nope_ACC | G/C-1_CCAGCAG | 6 | FALSE | FALSE |
| 138_v.txt | 32608 | 32608-Deleti | Deletion | 1 | C | - | 2909 | 19418 | 14.9809455 | 43.2310072 | rnr1Y285A_p( | TRUE | CAN | JSY2422 | 2 | rnr1Y285A_p | ACC | nope | >1bp-ins | G/C-1 | none | G/C-1 | AGGAAACCC | CCCAACC | TGG | ACC | C | none | nope | nope_ACC | G/C-1_CCCAACC | 6 | FALSE | FALSE |

| | | | | | | | | | | | | | | | | | | | | | | | | | | | | | | | | | | |
|---|---|---|---|---|---|---|---|---|---|---|---|---|---|---|---|---|---|---|---|---|---|---|---|---|---|---|---|---|---|---|---|---|---|---|
| 138_v.txt | 32631 | 32631-Deleti | Deletion | 1 | T | - | 6 | 19418 | 0.03089917 | 46 | rnr1Y285A_p | TRUE | CAN | JSY2422 | 2 | rnr1Y285A_p | GTT | nope | >1bp-ins | A/T-1 | none | A/T-1 | CCTTCGTTTT | TTTTTAT | CAA | GTT | T | none | nope | nope_GTT | A/T-1_TTTTTAT | 6 | FALSE | FALSE |
| 138_v.txt | 32658 | 32658-Deleti | Deletion | 1 | C | - | 1135 | 19418 | 5.84509218 | 59.8123348 | rnr1Y285A_p | TRUE | CAN | JSY2422 | 2 | rnr1Y285A_p | ACC | nope | >1bp-ins | G/C-1 | none | G/C-1 | CCTGGACCC | CCCCAGG | TGG | ACC | C | none | nope | nope_ACC | G/C-1_CCCCAGG | 6 | FALSE | FALSE |
| 138_v.txt | 32670 | 32670-Deleti | Deletion | 1 | G | - | 968 | 19418 | 4.9850654 | 58.6167355 | rnr1Y285A_p | TRUE | CAN | JSY2422 | 2 | rnr1Y285A_p | TGG | nope | >1bp-ins | G/C-1 | none | G/C-1 | GCACCTGGG | GGGTTTC | ACC | ACC | C | none | C>none | C>none_ACC | G/C-1_GGGTTTC | 6 | FALSE | FALSE |
| 138_v.txt | 32707 | 32707-Deleti | Deletion | 1 | C | - | 409 | 19418 | 2.10629313 | 62.5819071 | rnr1Y285A_p | TRUE | CAN | JSY2422 | 2 | rnr1Y285A_p | ACC | nope | >1bp-ins | G/C-1 | none | G/C-1 | CGGTAACCC | CCCCAGC | TGG | ACC | C | none | nope | nope_ACC | G/C-1_CCCCAGC | 6 | FALSE | FALSE |
| 138_v.txt | 32719 | 32719-Deleti | Deletion | 1 | A | - | 18 | 19418 | 0.0926975 | 57.3333333 | rnr1Y285A_p | TRUE | CAN | JSY2422 | 2 | rnr1Y285A_p | CAA | nope | >1bp-ins | A/T-1 | none | A/T-1 | CACCACAAA | AAACCAT | GTT | GTT | T | none | T>none | T>none_GTT | A/T-1_AAACCAT | 6 | FALSE | FALSE |
| 138_v.txt | 32740 | 32740-Deleti | Deletion | 1 | T | - | 6 | 19418 | 0.03089917 | 54.3333333 | rnr1Y285A_p | TRUE | CAN | JSY2422 | 2 | rnr1Y285A_p | ATT | nope | >1bp-ins | A/T-1 | none | A/T-1 | AGTATATTA | TTAGAAA | TAA | ATT | T | none | nope | nope_ATT | A/T-1_TTAGAAA | 6 | FALSE | FALSE |
| 138_v.txt | 32747 | 32747-Deleti | Deletion | 1 | C | - | 181 | 19418 | 0.93212483 | 56.1436464 | rnr1Y285A_p | TRUE | CAN | JSY2422 | 2 | rnr1Y285A_p | ACC | nope | >1bp-ins | G/C-1 | none | G/C-1 | TAGAAACCC | CCCGATA | TGG | ACC | C | none | nope | nope_ACC | G/C-1_CCCGATA | 6 | FALSE | FALSE |
| 138_v.txt | 32760 | 32760-Deleti | Deletion | 1 | A | - | 41 | 19418 | 0.2111443 | 53.097561 | rnr1Y285A_p | TRUE | CAN | JSY2422 | 2 | rnr1Y285A_p | TAA | nope | >1bp-ins | A/T-1 | none | A/T-1 | ATGGCTAAA | AAAACTT | ATT | ATT | T | none | T>none | T>none_ATT | A/T-1_AAAACTT | 6 | FALSE | FALSE |
| 138_v.txt | 32764 | 32764-Deleti | Deletion | 1 | C | - | 23 | 19418 | 0.1184468 | 55.8695632 | rnr1Y285A_p | TRUE | CAN | JSY2422 | 2 | rnr1Y285A_p | ACT | nope | >1bp-ins | G/C-1 | none | G/C-1 | CTAAAACTT | CTTTGAT | TGA | ACT | C | none | nope | nope_ACT | G/C-1_CTTTGAT | 6 | FALSE | FALSE |
| 138_v.txt | 32779 | 32779-Deleti | Deletion | 1 | C | - | 28 | 19418 | 0.14419611 | 47.0714286 | rnr1Y285A_p | TRUE | CAN | JSY2422 | 2 | rnr1Y285A_p | ACC | nope | >1bp-ins | G/C-1 | none | G/C-1 | AAGCGACCC | CCCAGAA | TGG | ACC | C | none | nope | nope_ACC | G/C-1_CCCAGAA | 6 | FALSE | FALSE |
| 138_v.txt | 32842 | 32842-Deleti | Deletion | 1 | A | - | 69 | 19418 | 0.35534041 | 40.3768116 | rnr1Y285A_p | TRUE | CAN | JSY2422 | 2 | rnr1Y285A_p | CAA | nope | >1bp-ins | A/T-1 | none | A/T-1 | TTACCCAAA | AAAAAAT | GTT | GTT | T | none | T>none | T>none_GTT | A/T-1_AAAAAAT | 6 | FALSE | FALSE |
| 138_v.txt | 32884 | 32884-Deleti | Deletion | 1 | A | - | 4 | 24363 | 0.01641834 | 37.75 | rnr1Y285A_p | TRUE | CAN | JSY2422 | 2 | rnr1Y285A_p | CAA | nope | >1bp-ins | A/T-1 | none | A/T-1 | ACGTCCAAA | AAAATTG | GTT | GTT | T | none | T>none | T>none_GTT | A/T-1_AAAATTG | 6 | FALSE | FALSE |
| 138_v.txt | 32924 | 32924-Deleti | Deletion | 1 | A | - | 21 | 24363 | 0.08619628 | 55.8095238 | rnr1Y285A_p | TRUE | CAN | JSY2422 | 2 | rnr1Y285A_p | CAA | nope | >1bp-ins | A/T-1 | none | A/T-1 | CAGGGCAAA | AAAAGTG | GTT | GTT | T | none | T>none | T>none_GTT | A/T-1_AAAAGTG | 6 | FALSE | FALSE |
| 138_v.txt | 33000 | 33000-Deleti | Deletion | 1 | A | - | 21 | 24363 | 0.08619628 | 60.6666667 | rnr1Y285A_p | TRUE | CAN | JSY2422 | 2 | rnr1Y285A_p | GAA | nope | >1bp-ins | A/T-1 | none | A/T-1 | TGTGAGAAA | AAAACTG | CTT | CTT | T | none | T>none | T>none_CTT | A/T-1_AAAACTG | 6 | FALSE | FALSE |
| 138_v.txt | 33138 | 33138-Deleti | Deletion | 1 | A | - | 4 | 24363 | 0.01641834 | 37.75 | rnr1Y285A_p | TRUE | CAN | JSY2422 | 2 | rnr1Y285A_p | TAA | nope | >1bp-ins | A/T-1 | none | A/T-1 | GTGGATAAA | AAACCAA | ATT | ATT | T | none | T>none | T>none_ATT | A/T-1_AAACCAA | 6 | FALSE | FALSE |
| 138_v.txt | 33356 | 33356-Deleti | Deletion | 1 | C | - | 2002 | 15623 | 12.8144403 | 54.3086913 | rnr1Y285A_p | TRUE | CAN | JSY2422 | 2 | rnr1Y285A_p | ACC | nope | >1bp-ins | G/C-1 | none | G/C-1 | TATTGACCC | CCCACGT | TGG | ACC | C | none | nope | nope_ACC | G/C-1_CCCACGT | 6 | FALSE | FALSE |
| 138_v.txt | 33474 | 33474-Deleti | Deletion | 1 | C | - | 46 | 15622 | 0.29445654 | 30.7826087 | rnr1Y285A_p | TRUE | CAN | JSY2422 | 2 | rnr1Y285A_p | GCC | nope | >1bp-ins | G/C-1 | none | G/C-1 | GCTATGCCT | CCTTTTT | CGG | GCC | C | none | nope | nope_GCC | G/C-1_CCTTTTT | 6 | FALSE | FALSE |
| 139_v.txt | 32262 | 32262-SNV- | SNV | 1 | A | C | 15 | 27298 | 0.05494908 | 27.8 | rnr1Y285A_p | TRUE | CAN | JSY2423 | 3 | rnr1Y285A_p | AAA | A>C | G/C+1 | A/T-1 | TA>GC | TA>GC | AGACCAAAT | AATAAAA | TTT | TTT | T | G | T>G | T>G_TTT | TA>GC_AATAAAA | 6 | FALSE | FALSE |
| 139_v.txt | 32265 | 32265-SNV- | SNV | 1 | A | C | 14 | 27298 | 0.05128581 | 26.2857143 | rnr1Y285A_p | TRUE | CAN | JSY2423 | 3 | rnr1Y285A_p | TAA | A>C | G/C+1 | A/T-1 | TA>GC | TA>GC | CCAAATAAA | AAAATAC | ATT | ATT | T | G | T>G | T>G_ATT | TA>GC_AAAATAC | 6 | FALSE | FALSE |
| 139_v.txt | 32283 | 32283-SNV-T | SNV | 1 | T | G | 27 | 27298 | 0.09890835 | 50.6666667 | rnr1Y285A_p | TRUE | CAN | JSY2423 | 3 | rnr1Y285A_p | GTA | T>G | G/C+1 | A/T-1 | TA>GC | TA>GC | CCAACGTAA | TAAATAT | CAT | GTA | T | C | T>G | T>G_GTA | TA>GC_TAAATAT | 6 | FALSE | FALSE |
| 139_v.txt | 32441 | 32441-SNV-T | SNV | 1 | T | G | 7 | 27298 | 0.0256429 | 52.4298346 | rnr1Y285A_p | TRUE | CAN | JSY2423 | 3 | rnr1Y285A_p | CTA | T>G | G/C+1 | A/T-1 | TA>GC | TA>GC | TGGAACTAA | TAAAAGT | GAT | CTA | T | C | T>G | T>G_CTA | TA>GC_TAAAAGT | 6 | FALSE | FALSE |
| 139_v.txt | 32442 | 32442-SNV- | SNV | 1 | A | C | 1 | 27298 | 3.66E-03 | 63.9997088 | rnr1Y285A_p | TRUE | CAN | JSY2423 | 3 | rnr1Y285A_p | TAA | A>C | G/C+1 | A/T-1 | TA>GC | TA>GC | GGAACTAAA | AAAAGTC | ATT | ATT | T | G | T>G | T>G_ATT | TA>GC_AAAAGTC | 6 | FALSE | FALSE |
| 139_v.txt | 32532 | 32532-SNV-T | SNV | 1 | T | G | 66 | 27298 | 0.24177595 | 37.5 | rnr1Y285A_p | TRUE | CAN | JSY2423 | 3 | rnr1Y285A_p | GTT | T>G | G/C+1 | A/T-1 | TA>GC | TA>GC | CTGGGGTTT | TTTGCAG | CAA | GTT | T | C | T>G | T>G_GTT | TA>GC_TTTGCAG | 6 | FALSE | FALSE |
| 139_v.txt | 32581 | 32581-SNV-T | SNV | 1 | T | G | 6 | 20501 | 0.02926687 | 32.6666667 | rnr1Y285A_p | TRUE | CAN | JSY2423 | 3 | rnr1Y285A_p | GTG | T>G | G/C+1 | A/T-1 | TA>GC | TA>GC | GAAATGTGA | TGAAGGC | CAC | GTG | T | C | T>G | T>G_GTG | TA>GC_TGAAGGC | 6 | FALSE | FALSE |
| 139_v.txt | 32746 | 32746-SNV- | SNV | 1 | A | C | 44 | 20501 | 0.21462368 | 55.3851561 | rnr1Y285A_p | TRUE | CAN | JSY2423 | 3 | rnr1Y285A_p | AAC | A>C | G/C+1 | A/T-1 | TA>GC | TA>GC | TTAGAAACC | ACCCGAT | TTG | TTG | T | G | T>G | T>G_TTG | TA>GC_ACCCGAT | 6 | FALSE | FALSE |
| 139_v.txt | 32763 | 32763-SNV- | SNV | 1 | A | C | 9 | 20501 | 0.0439003 | 58 | rnr1Y285A_p | TRUE | CAN | JSY2423 | 3 | rnr1Y285A_p | AAC | A>C | G/C+1 | A/T-1 | TA>GC | TA>GC | GCTAAAACT | ACTTTGA | TTG | TTG | T | G | T>G | T>G_TTG | TA>GC_ACTTTGA | 6 | FALSE | FALSE |
| 139_v.txt | 32842 | 32842-SNV- | SNV | 1 | A | C | 2 | 20501 | 9.76E-03 | 50.9996828 | rnr1Y285A_p | TRUE | CAN | JSY2423 | 3 | rnr1Y285A_p | CAA | A>C | G/C+1 | A/T-1 | TA>GC | TA>GC | TTACCCAAA | AAAAAAT | GTT | GTT | T | G | T>G | T>G_GTT | TA>GC_AAAAAAT | 6 | FALSE | FALSE |
| 139_v.txt | 32884 | 32884-SNV- | SNV | 1 | A | C | 4 | 23058 | 0.01734756 | 37.9970858 | rnr1Y285A_p | TRUE | CAN | JSY2423 | 3 | rnr1Y285A_p | CAA | A>C | G/C+1 | A/T-1 | TA>GC | TA>GC | ACGTCCAAA | AAAATTG | GTT | GTT | T | G | T>G | T>G_GTT | TA>GC_AAAATTG | 6 | FALSE | FALSE |
| 139_v.txt | 32929 | 32929-SNV-T | SNV | 1 | T | G | 26 | 23058 | 0.11275913 | 55.3846154 | rnr1Y285A_p | TRUE | CAN | JSY2423 | 3 | rnr1Y285A_p | GTG | T>G | G/C+1 | A/T-1 | TA>GC | TA>GC | CAAAAGTGA | TGATTGC | CAC | GTG | T | C | T>G | T>G_GTG | TA>GC_TGATTGC | 6 | FALSE | FALSE |
| 139_v.txt | 33054 | 33054-SNV-T | SNV | 1 | T | G | 9 | 23058 | 0.03903201 | 42.3333333 | rnr1Y285A_p | TRUE | CAN | JSY2423 | 3 | rnr1Y285A_p | CTG | T>G | G/C+1 | A/T-1 | TA>GC | TA>GC | AAGGACTGC | TGCGTGA | GAC | CTG | T | C | T>G | T>G_CTG | TA>GC_TGCGTGA | 6 | FALSE | FALSE |
| 139_v.txt | 33099 | 33099-SNV- | SNV | 1 | A | C | 7 | 23058 | 0.03035823 | 21.7142857 | rnr1Y285A_p | TRUE | CAN | JSY2423 | 3 | rnr1Y285A_p | AAG | A>C | G/C+1 | A/T-1 | TA>GC | TA>GC | GATATAAGA | AGAGCGC | TTC | TTC | T | G | T>G | T>G_TTC | TA>GC_AGAGCGC | 6 | FALSE | FALSE |
| 139_v.txt | 33476 | 33476-SNV-T | SNV | 1 | T | G | 4 | 15617 | 0.02561311 | 36.750223 | rnr1Y285A_p | TRUE | CAN | JSY2423 | 3 | rnr1Y285A_p | CTT | T>G | G/C+1 | A/T-1 | TA>GC | TA>GC | TATGCCTTTT | TTTTTTT | GAA | CTT | T | C | T>G | T>G_CTT | TA>GC_TTTTTTT | 6 | FALSE | FALSE |
| 139_v.txt | 32081 | 32081-SNV- | SNV | 1 | A | G | 6 | 16284 | 0.03684598 | 57.0001382 | rnr1Y285A_p | TRUE | CAN | JSY2423 | 3 | rnr1Y285A_p | CAA | A>G | G/C+1 | A/T-1 | TA>CG | TA>CG | CCATGCAAA | AAAAAAG | GTT | GTT | T | C | T>C | T>C_GTT | TA>CG_AAAAAAG | 6 | FALSE | FALSE |
| 139_v.txt | 32505 | 32505-SNV- | SNV | 1 | A | G | 10 | 27298 | 0.03663272 | 40.1999041 | rnr1Y285A_p | TRUE | CAN | JSY2423 | 3 | rnr1Y285A_p | GAT | A>G | G/C+1 | A/T-1 | TA>CG | TA>CG | TTTTTGATG | ATGGCTC | CTA | CTA | T | C | T>C | T>C_CTA | TA>CG_ATGGCTC | 6 | FALSE | FALSE |
| 139_v.txt | 32581 | 32581-SNV-T | SNV | 1 | T | C | 9 | 20501 | 0.0439003 | 37.8888889 | rnr1Y285A_p | TRUE | CAN | JSY2423 | 3 | rnr1Y285A_p | GTG | T>C | G/C+1 | A/T-1 | TA>CG | TA>CG | GAAATGTGA | TGAAGGC | CAC | GTG | T | G | T>C | T>C_GTG | TA>CG_TGAAGGC | 6 | FALSE | FALSE |
| 139_v.txt | 32746 | 32746-SNV- | SNV | 1 | A | G | 5 | 20501 | 0.02438905 | 47.9998057 | rnr1Y285A_p | TRUE | CAN | JSY2423 | 3 | rnr1Y285A_p | AAC | A>G | G/C+1 | A/T-1 | TA>CG | TA>CG | TTAGAAACC | ACCCGAT | TTG | TTG | T | C | T>C | T>C_TTG | TA>CG_ACCCGAT | 6 | FALSE | FALSE |
| 139_v.txt | 32787 | 32787-SNV-T | SNV | 1 | T | C | 25 | 20501 | 0.12194527 | 38.76 | rnr1Y285A_p | TRUE | CAN | JSY2423 | 3 | rnr1Y285A_p | CTC | T>C | G/C+1 | A/T-1 | TA>CG | TA>CG | CAGAACTCG | TCGAATT | GAG | CTC | T | G | T>C | T>C_CTC | TA>CG_TCGAATT | 6 | FALSE | FALSE |
| 139_v.txt | 32884 | 32884-SNV- | SNV | 1 | A | G | 3 | 23058 | 0.01301067 | 37.3296977 | rnr1Y285A_p | TRUE | CAN | JSY2423 | 3 | rnr1Y285A_p | CAA | A>G | G/C+1 | A/T-1 | TA>CG | TA>CG | ACGTCCAAA | AAAATTG | GTT | GTT | T | C | T>C | T>C_GTT | TA>CG_AAAATTG | 6 | FALSE | FALSE |
| 139_v.txt | 32941 | 32941-SNV- | SNV | 1 | A | G | 4 | 23058 | 0.01734756 | 56.2496205 | rnr1Y285A_p | TRUE | CAN | JSY2423 | 3 | rnr1Y285A_p | GAA | A>G | G/C+1 | A/T-1 | TA>CG | TA>CG | CCCAAGAAA | AAAACCA | CTT | CTT | T | C | T>C | T>C_CTT | TA>CG_AAAACCA | 6 | FALSE | FALSE |
| 139_v.txt | 33099 | 33099-SNV- | SNV | 1 | A | G | 22 | 23058 | 0.09541157 | 45.5 | rnr1Y285A_p | TRUE | CAN | JSY2423 | 3 | rnr1Y285A_p | AAG | A>G | G/C+1 | A/T-1 | TA>CG | TA>CG | GATATAAGA | AGAGCGC | TTC | TTC | T | C | T>C | T>C_TTC | TA>CG_AGAGCGC | 6 | FALSE | FALSE |
| 139_v.txt | 33189 | 33189-SNV- | SNV | 1 | A | G | 2 | 15619 | 0.01280492 | 37.4993255 | rnr1Y285A_p | TRUE | CAN | JSY2423 | 3 | rnr1Y285A_p | AAT | A>G | G/C+1 | A/T-1 | TA>CG | TA>CG | ATACCAATA | ATATGTC | TTA | TTA | T | C | T>C | T>C_TTA | TA>CG_ATATGTC | 6 | FALSE | FALSE |
| 139_v.txt | 33476 | 33476-SNV-T | SNV | 1 | T | C | 27 | 15617 | 0.17288852 | 30.5929198 | rnr1Y285A_p | TRUE | CAN | JSY2423 | 3 | rnr1Y285A_p | CTT | T>C | G/C+1 | A/T-1 | TA>CG | TA>CG | TATGCCTTTT | TTTTTTT | GAA | CTT | T | G | T>C | T>C_CTT | TA>CG_TTTTTTT | 6 | FALSE | FALSE |
| 139_v.txt | 32496 | 32496-SNV- | SNV | 1 | A | T | 348 | 27298 | 1.27481867 | 43.8304598 | rnr1Y285A_p | TRUE | CAN | JSY2423 | 3 | rnr1Y285A_p | AAC | A>T | A/T+1 | A/T-1 | TA>AT | TA>AT | AAAACAACT | ACTTTTT | TTG | TTG | T | A | T>A | T>A_TTG | TA>AT_ACTTTTT | 6 | FALSE | FALSE |
| 139_v.txt | 32502 | 32502-SNV-T | SNV | 1 | T | A | 18 | 27298 | 0.0659389 | 42.3333333 | rnr1Y285A_p | TRUE | CAN | JSY2423 | 3 | rnr1Y285A_p | TTT | T>A | A/T+1 | A/T-1 | TA>AT | TA>AT | ACTTTTTGA | TTGATGG | AAA | TTT | T | T | T>A | T>A_TTT | TA>AT_TTGATGG | 6 | FALSE | FALSE |
| 139_v.txt | 32785 | 32785-SNV- | SNV | 1 | A | T | 20 | 20501 | 0.09755622 | 49.00088 | rnr1Y285A_p | TRUE | CAN | JSY2423 | 3 | rnr1Y285A_p | AAC | A>T | A/T+1 | A/T-1 | TA>AT | TA>AT | CCCAGAACT | ACTCGAA | TTG | TTG | T | A | T>A | T>A_TTG | TA>AT_ACTCGAA | 6 | FALSE | FALSE |
| 139_v.txt | 32941 | 32941-SNV- | SNV | 1 | A | T | 4 | 23058 | 0.01734756 | 29.2492426 | rnr1Y285A_p | TRUE | CAN | JSY2423 | 3 | rnr1Y285A_p | GAA | A>T | A/T+1 | A/T-1 | TA>AT | TA>AT | CCCAAGAAA | AAAACCA | CTT | CTT | T | A | T>A | T>A_CTT | TA>AT_AAAACCA | 6 | FALSE | FALSE |
| 139_v.txt | 33060 | 33060-SNV- | SNV | 1 | A | T | 16 | 23058 | 0.06939023 | 56.75 | rnr1Y285A_p | TRUE | CAN | JSY2423 | 3 | rnr1Y285A_p | GAC | A>T | A/T+1 | A/T-1 | TA>AT | TA>AT | TGCGTGACA | ACAGAAT | CTG | CTG | T | A | T>A | T>A_CTG | TA>AT_ACAGAAT | 6 | FALSE | FALSE |
| 139_v.txt | 31650 | 31650-SNV- | SNV | 1 | C | T | 17 | 8823 | 0.19267823 | 36.8235294 | rnr1Y285A_p | TRUE | CAN | JSY2423 | 3 | rnr1Y285A_p | GCG | C>T | A/T+1 | G/C-1 | CG>TA | CG>TA | AGAATGCGA | CGAAATG | CGC | GCG | C | A | C>T | C>T_GCG | CG>TA_CGAAATG | 6 | FALSE | FALSE |
| 139_v.txt | 31947 | 31947-SNV- | SNV | 1 | G | A | 14 | 16270 | 0.08604794 | 31.5 | rnr1Y285A_p | TRUE | CAN | JSY2423 | 3 | rnr1Y285A_p | CGT | G>A | A/T+1 | G/C-1 | CG>TA | CG>TA | ATGATCGTC | GTCATAA | GCA | GCA | C | T | C>T | C>T_GCA | CG>TA_GTCATAA | 6 | FALSE | FALSE |
| 139_v.txt | 31962 | 31962-SNV- | SNV | 1 | G | A | 21 | 16284 | 0.12896094 | 36.8095238 | rnr1Y285A_p | TRUE | CAN | JSY2423 | 3 | rnr1Y285A_p | CGC | G>A | A/T+1 | G/C-1 | CG>TA | CG>TA | GTGGCCGCA | GCATAAT | GCG | GCG | C | T | C>T | C>T_GCG | CG>TA_GCATAAT | 6 | FALSE | FALSE |
| 139_v.txt | 31972 | 31972-SNV- | SNV | 1 | C | T | 19 | 16284 | 0.11667895 | 36.3157895 | rnr1Y285A_p | TRUE | CAN | JSY2423 | 3 | rnr1Y285A_p | GCC | C>T | A/T+1 | G/C-1 | CG>TA | CG>TA | AATAAGCCA | CCAAGCC | CGG | GCC | C | A | C>T | C>T_GCC | CG>TA_CCAAGCC | 6 | FALSE | FALSE |
| 139_v.txt | 31981 | 31981-SNV- | SNV | 1 | G | A | 26 | 16284 | 0.15966593 | 51.8461539 | rnr1Y285A_p | TRUE | CAN | JSY2423 | 3 | rnr1Y285A_p | GGC | G>A | A/T+1 | G/C-1 | CG>TA | CG>TA | AGCCGGGCA | GCATTAA | CCG | CCG | C | T | C>T | C>T_CCG | CG>TA_GCATTAA | 6 | FALSE | FALSE |
| 139_v.txt | 32026 | 32026-SNV- | SNV | 1 | G | A | 3 | 16284 | 0.01842299 | 54.9996781 | rnr1Y285A_p | TRUE | CAN | JSY2423 | 3 | rnr1Y285A_p | CGG | G>A | A/T+1 | G/C-1 | CG>TA | CG>TA | TGCCACGGT | GGTATTT | GCC | GCC | C | T | C>T | C>T_GCC | CG>TA_GGTATTT | 6 | FALSE | FALSE |
| 139_v.txt | 32075 | 32075-SNV- | SNV | 1 | C | T | 51 | 16284 | 0.31319086 | 60.2745098 | rnr1Y285A_p | TRUE | CAN | JSY2423 | 3 | rnr1Y285A_p | ACC | C>T | A/T+1 | G/C-1 | CG>TA | CG>TA | AAATAACCA | CCATGCA | TGG | ACC | C | A | C>T | C>T_ACC | CG>TA_CCATGCA | 6 | FALSE | FALSE |
| 139_v.txt | 32088 | 32088-SNV- | SNV | 1 | C | T | 29 | 16284 | 0.17808892 | 58.6896552 | rnr1Y285A_p | TRUE | CAN | JSY2423 | 3 | rnr1Y285A_p | GCC | C>T | A/T+1 | G/C-1 | CG>TA | CG>TA | AAAAAGCCT | CCTGCAA | CGG | GCC | C | A | C>T | C>T_GCC | CG>TA_CCTGCAA | 6 | FALSE | FALSE |
| 139_v.txt | 32092 | 32092-SNV- | SNV | 1 | C | T | 22 | 16284 | 0.13510194 | 57.5454546 | rnr1Y285A_p | TRUE | CAN | JSY2423 | 3 | rnr1Y285A_p | GCA | C>T | A/T+1 | G/C-1 | CG>TA | CG>TA | AGCCTGCAA | CAACACC | CGT | GCA | C | A | C>T | C>T_GCA | CG>TA_CAACACC | 6 | FALSE | FALSE |
| 139_v.txt | 32114 | 32114-SNV- | SNV | 1 | C | T | 26 | 16284 | 0.15966593 | 57.3076923 | rnr1Y285A_p | TRUE | CAN | JSY2423 | 3 | rnr1Y285A_p | GCC | C>T | A/T+1 | G/C-1 | CG>TA | CG>TA | TAATAGCCA | CCATTCG | CGG | GCC | C | A | C>T | C>T_GCC | CG>TA_CCATTCG | 6 | FALSE | FALSE |
| 139_v.txt | 32165 | 32165-SNV- | SNV | 1 | G | A | 29 | 16284 | 0.17808892 | 46.4482759 | rnr1Y285A_p | TRUE | CAN | JSY2423 | 3 | rnr1Y285A_p | CGC | G>A | A/T+1 | G/C-1 | CG>TA | CG>TA | CAAAGCGCC | GCCAAAT | GCG | GCG | C | T | C>T | C>T_GCG | CG>TA_GCCAAAT | 6 | FALSE | FALSE |
| 139_v.txt | 32173 | 32173-SNV- | SNV | 1 | C | T | 23 | 16284 | 0.14124294 | 45.0869565 | rnr1Y285A_p | TRUE | CAN | JSY2423 | 3 | rnr1Y285A_p | GCA | C>T | A/T+1 | G/C-1 | CG>TA | CG>TA | CAAATGCAG | CAGCAGT | CGT | GCA | C | A | C>T | C>T_GCA | CG>TA_CAGCAGT | 6 | FALSE | FALSE |
| 139_v.txt | 32271 | 32271-SNV- | SNV | 1 | C | T | 43 | 27298 | 0.1575207 | 32.8139535 | rnr1Y285A_p | TRUE | CAN | JSY2423 | 3 | rnr1Y285A_p | ACG | C>T | A/T+1 | G/C-1 | CG>TA | CG>TA | AAAATACGG | CGGGAAC | TGC | ACG | C | A | C>T | C>T_ACG | CG>TA_CGGGAAC | 6 | FALSE | FALSE |
| 139_v.txt | 32274 | 32274-SNV- | SNV | 1 | G | A | 87 | 27298 | 0.31870467 | 36.7701149 | rnr1Y285A_p | TRUE | CAN | JSY2423 | 3 | rnr1Y285A_p | GGA | G>A | A/T+1 | G/C-1 | CG>TA | CG>TA | ATACGGGAA | GAACCAA | CCT | CCT | C | T | C>T | C>T_CCT | CG>TA_GAACCAA | 6 | FALSE | FALSE |
| 139_v.txt | 32299 | 32299-SNV- | SNV | 1 | C | T | 48 | 27298 | 0.17583706 | 52.3333333 | rnr1Y285A_p | TRUE | CAN | JSY2423 | 3 | rnr1Y285A_p | GCG | C>T | A/T+1 | G/C-1 | CG>TA | CG>TA | AATTTGCGG | CGGCAGA | CGC | GCG | C | A | C>T | C>T_GCG | CG>TA_CGGCAGA | 6 | FALSE | FALSE |
| 139_v.txt | 32466 | 32466-SNV- | SNV | 1 | C | T | 38 | 27298 | 0.13920434 | 46.6578947 | rnr1Y285A_p | TRUE | CAN | JSY2423 | 3 | rnr1Y285A_p | GCC | C>T | A/T+1 | G/C-1 | CG>TA | CG>TA | AGAGAGCCA | CCAATGT | CGG | GCC | C | A | C>T | C>T_GCC | CG>TA_CCAATGT | 6 | FALSE | FALSE |
| 139_v.txt | 32487 | 32487-SNV- | SNV | 1 | C | T | 34 | 27298 | 0.12455125 | 43.1764706 | rnr1Y285A_p | TRUE | CAN | JSY2423 | 3 | rnr1Y285A_p | ACG | C>T | A/T+1 | G/C-1 | CG>TA | CG>TA | AAGATACGG | CGGAAAA | TGC | ACG | C | A | C>T | C>T_ACG | CG>TA_CGGAAAA | 6 | FALSE | FALSE |
| 139_v.txt | 32535 | 32535-SNV- | SNV | 1 | G | A | 34 | 27297 | 0.12455581 | 37.8820059 | rnr1Y285A_p | TRUE | CAN | JSY2423 | 3 | rnr1Y285A_p | TGC | G>A | A/T+1 | G/C-1 | CG>TA | CG>TA | GGGTTTGCA | GCAGCTT | ACG | ACG | C | T | C>T | C>T_ACG | CG>TA_GCAGCTT | 6 | FALSE | FALSE |
| 139_v.txt | 32536 | 32536-SNV- | SNV | 1 | C | T | 50 | 27297 | 0.18317031 | 37.24 | rnr1Y285A_p | TRUE | CAN | JSY2423 | 3 | rnr1Y285A_p | GCA | C>T | A/T+1 | G/C-1 | CG>TA | CG>TA | GGTTTGCAG | CAGCTTC | CGT | GCA | C | A | C>T | C>T_GCA | CG>TA_CAGCTTC | 6 | FALSE | FALSE |
| 139_v.txt | 32566 | 32566-SNV- | SNV | 1 | C | T | 1243 | 20494 | 6.06518981 | 37.1858407 | rnr1Y285A_p | TRUE | CAN | JSY2423 | 3 | rnr1Y285A_p | TCA | C>T | A/T+1 | G/C-1 | CG>TA | CG>TA | CTAGTTCAG | CAGTACC | AGT | TCA | C | A | C>T | C>T_TCA | CG>TA_CAGTACC | 6 | FALSE | FALSE |
| 139_v.txt | 32589 | 32589-SNV- | SNV | 1 | G | A | 4 | 20501 | 0.01951124 | 35.2497393 | rnr1Y285A_p | TRUE | CAN | JSY2423 | 3 | rnr1Y285A_p | AGC | G>A | A/T+1 | G/C-1 | CG>TA | CG>TA | AAGGCAGCG | GCGTTAA | TCG | TCG | C | T | C>T | C>T_TCG | CG>TA_GCGTTAA | 6 | FALSE | FALSE |
| 139_v.txt | 32590 | 32590-SNV- | SNV | 1 | C | T | 37 | 20501 | 0.180479 | 36.5675675 | rnr1Y285A_p | TRUE | CAN | JSY2423 | 3 | rnr1Y285A_p | GCG | C>T | A/T+1 | G/C-1 | CG>TA | CG>TA | AGGCAGCGT | CGTTAAT | CGC | GCG | C | A | C>T | C>T_GCG | CG>TA_CGTTAAT | 6 | FALSE | FALSE |
| 139_v.txt | 32658 | 32658-SNV- | SNV | 1 | C | T | 40 | 20501 | 0.19511243 | 53.125 | rnr1Y285A_p | TRUE | CAN | JSY2423 | 3 | rnr1Y285A_p | ACC | C>T | A/T+1 | G/C-1 | CG>TA | CG>TA | CCTGGACCC | CCCCAGG | TGG | ACC | C | A | C>T | C>T_ACC | CG>TA_CCCCAGG | 6 | FALSE | FALSE |
| 139_v.txt | 32700 | 32700-SNV- | SNV | 1 | C | T | 22 | 20501 | 0.10731184 | 62.5909091 | rnr1Y285A_p | TRUE | CAN | JSY2423 | 3 | rnr1Y285A_p | GCC | C>T | A/T+1 | G/C-1 | CG>TA | CG>TA | ACTGGGCCG | CCGGTAA | CGG | GCC | C | A | C>T | C>T_GCC | CG>TA_CCGGTAA | 6 | FALSE | FALSE |
| 139_v.txt | 32757 | 32757-SNV- | SNV | 1 | G | A | 23 | 20501 | 0.11218965 | 49.3043478 | rnr1Y285A_p | TRUE | CAN | JSY2423 | 3 | rnr1Y285A_p | GGC | G>A | A/T+1 | G/C-1 | CG>TA | CG>TA | ATAATGGCT | GCTAAAA | CCG | CCG | C | T | C>T | C>T_CCG | CG>TA_GCTAAAA | 6 | FALSE | FALSE |
| 139_v.txt | 32798 | 32798-SNV- | SNV | 1 | G | A | 14 | 20501 | 0.06828935 | 45.6428571 | rnr1Y285A_p | TRUE | CAN | JSY2423 | 3 | rnr1Y285A_p | CGT | G>A | A/T+1 | G/C-1 | CG>TA | CG>TA | TTCACCGTA | GTAATAT | GCA | GCA | C | T | C>T | C>T_GCA | CG>TA_GTAATAT | 6 | FALSE | FALSE |
| 139_v.txt | 32855 | 32855-SNV- | SNV | 1 | C | T | 88 | 20501 | 0.42924735 | 36.2272727 | rnr1Y285A_p | TRUE | CAN | JSY2423 | 3 | rnr1Y285A_p | TCC | C>T | A/T+1 | G/C-1 | CG>TA | CG>TA | ACTAATCCA | CCATGCC | AGG | TCC | C | A | C>T | C>T_TCC | CG>TA_CCATGCC | 6 | FALSE | FALSE |
| 139_v.txt | 32860 | 32860-SNV- | SNV | 1 | C | T | 23 | 20499 | 0.1122006 | 37.4347826 | rnr1Y285A_p | TRUE | CAN | JSY2423 | 3 | rnr1Y285A_p | GCC | C>T | A/T+1 | G/C-1 | CG>TA | CG>TA | TCCATGCCG | CCGCCAG | CGG | GCC | C | A | C>T | C>T_GCC | CG>TA_CCGCCAG | 6 | FALSE | FALSE |
| 139_v.txt | 32899 | 32899-SNV- | SNV | 1 | G | A | 26 | 23058 | 0.11275913 | 36.1337672 | rnr1Y285A_p | TRUE | CAN | JSY2423 | 3 | rnr1Y285A_p | TGG | G>A | A/T+1 | G/C-1 | CG>TA | CG>TA | TGACTTGGC | GGCCAAC | ACC | ACC | C | T | C>T | C>T_ACC | CG>TA_GGCCAAC | 6 | FALSE | FALSE |
| 139_v.txt | 32900 | 32900-SNV- | SNV | 1 | G | A | 30 | 23058 | 0.13010669 | 37.0967335 | rnr1Y285A_p | TRUE | CAN | JSY2423 | 3 | rnr1Y285A_p | GGC | G>A | A/T+1 | G/C-1 | CG>TA | CG>TA | GACTTGGCC | GCCAACT | CCG | CCG | C | T | C>T | C>T_CCG | CG>TA_GCCAACT | 6 | FALSE | FALSE |
| 139_v.txt | 32901 | 32901-SNV- | SNV | 1 | C | T | 43 | 23058 | 0.18648625 | 32.4782169 | rnr1Y285A_p | TRUE | CAN | JSY2423 | 3 | rnr1Y285A_p | GCC | C>T | A/T+1 | G/C-1 | CG>TA | CG>TA | ACTTGGCCA | CCAACTA | CGG | GCC | C | A | C>T | C>T_GCC | CG>TA_CCAACTA | 6 | FALSE | FALSE |
| 139_v.txt | 32917 | 32917-SNV- | SNV | 1 | C | T | 54 | 23058 | 0.23419204 | 42.037037 | rnr1Y285A_p | TRUE | CAN | JSY2423 | 3 | rnr1Y285A_p | TCC | C>T | A/T+1 | G/C-1 | CG>TA | CG>TA | TAAGTTCCA | CCAGGGC | AGG | TCC | C | A | C>T | C>T_TCC | CG>TA_CCAGGGC | 6 | FALSE | FALSE |
| 139_v.txt | 32922 | 32922-SNV- | SNV | 1 | G | A | 20 | 23058 | 0.08673779 | 54 | rnr1Y285A_p | TRUE | CAN | JSY2423 | 3 | rnr1Y285A_p | GGC | G>A | A/T+1 | G/C-1 | CG>TA | CG>TA | TCCAGGGCA | GCAAAAG | CCG | CCG | C | T | C>T | C>T_CCG | CG>TA_GCAAAAG | 6 | FALSE | FALSE |
| 139_v.txt | 32945 | 32945-SNV- | SNV | 1 | C | T | 29 | 23058 | 0.1257698 | 59.2758621 | rnr1Y285A_p | TRUE | CAN | JSY2423 | 3 | rnr1Y285A_p | ACC | C>T | A/T+1 | G/C-1 | CG>TA | CG>TA | AGAAAACCA | CCAATAC | TGG | ACC | C | A | C>T | C>T_ACC | CG>TA_CCAATAC | 6 | FALSE | FALSE |
| 139_v.txt | 32965 | 32965-SNV- | SNV | 1 | C | T | 29 | 23058 | 0.1257698 | 57.1724138 | rnr1Y285A_p | TRUE | CAN | JSY2423 | 3 | rnr1Y285A_p | GCC | C>T | A/T+1 | G/C-1 | CG>TA | CG>TA | CATTGGCCG | CCGCACC | CGG | GCC | C | A | C>T | C>T_GCC | CG>TA_CCGCACC | 6 | FALSE | FALSE |
| 139_v.txt | 32967 | 32967-SNV- | SNV | 1 | G | A | 49 | 23058 | 0.21250759 | 62.3877551 | rnr1Y285A_p | TRUE | CAN | JSY2423 | 3 | rnr1Y285A_p | CGC | G>A | A/T+1 | G/C-1 | CG>TA | CG>TA | TTGGCCGCA | GCACCAA | GCG | GCG | C | T | C>T | C>T_GCG | CG>TA_GCACCAA | 6 | FALSE | FALSE |
| 139_v.txt | 33015 | 33015-SNV- | SNV | 1 | G | A | 14 | 23058 | 0.06071645 | 54.0714286 | rnr1Y285A_p | TRUE | CAN | JSY2423 | 3 | rnr1Y285A_p | GGA | G>A | A/T+1 | G/C-1 | CG>TA | CG>TA | AAAGAGGAT | GATGTAA | CCT | CCT | C | T | C>T | C>T_CCT | CG>TA_GATGTAA | 6 | FALSE | FALSE |
| 139_v.txt | 33056 | 33056-SNV- | SNV | 1 | C | T | 23 | 23058 | 0.09974846 | 56 | rnr1Y285A_p | TRUE | CAN | JSY2423 | 3 | rnr1Y285A_p | GCG | C>T | A/T+1 | G/C-1 | CG>TA | CG>TA | GGACTGCGT | CGTGACA | CGC | GCG | C | A | C>T | C>T_GCG | CG>TA_CGTGACA | 6 | FALSE | FALSE |
| 139_v.txt | 33070 | 33070-SNV- | SNV | 1 | C | T | 28 | 23058 | 0.12143291 | 58.7857143 | rnr1Y285A_p | TRUE | CAN | JSY2423 | 3 | rnr1Y285A_p | GCC | C>T | A/T+1 | G/C-1 | CG>TA | CG>TA | AATATGCCA | CCAAAGA | CGG | GCC | C | A | C>T | C>T_GCC | CG>TA_CCAAAGA | 6 | FALSE | FALSE |
| 139_v.txt | 33114 | 33114-SNV- | SNV | 1 | C | T | 32 | 23058 | 0.13878047 | 43.7668788 | rnr1Y285A_p | TRUE | CAN | JSY2423 | 3 | rnr1Y285A_p | GCC | C>T | A/T+1 | G/C-1 | CG>TA | CG>TA | ACTGGGCCG | CCGGCGT | CGG | GCC | C | A | C>T | C>T_GCC | CG>TA_CCGGCGT | 6 | FALSE | FALSE |
| 139_v.txt | 33115 | 33115-SNV- | SNV | 1 | C | T | 12 | 23058 | 0.05204268 | 29.9166667 | rnr1Y285A_p | TRUE | CAN | JSY2423 | 3 | rnr1Y285A_p | CCG | C>T | A/T+1 | G/C-1 | CG>TA | CG>TA | CTGGGCCGG | CGGCGTT | GGC | CCG | C | A | C>T | C>T_CCG | CG>TA_CGGCGTT | 6 | FALSE | FALSE |
| 139_v.txt | 33153 | 33153-SNV- | SNV | 1 | C | T | 48 | 23058 | 0.2081707 | 36.3541667 | rnr1Y285A_p | TRUE | CAN | JSY2423 | 3 | rnr1Y285A_p | ACC | C>T | A/T+1 | G/C-1 | CG>TA | CG>TA | AAAAGACCT | CCTGTAC | TGG | ACC | C | A | C>T | C>T_ACC | CG>TA_CCTGTAC | 6 | FALSE | FALSE |
| 139_v.txt | 33159 | 33159-SNV- | SNV | 1 | C | T | 8 | 23058 | 0.03469512 | 34.500055 | rnr1Y285A_p | TRUE | CAN | JSY2423 | 3 | rnr1Y285A_p | ACC | C>T | A/T+1 | G/C-1 | CG>TA | CG>TA | CCTGTACCA | CCAATAG | TGG | ACC | C | A | C>T | C>T_ACC | CG>TA_CCAATAG | 6 | FALSE | FALSE |
| 139_v.txt | 33258 | 33258-SNV- | SNV | 1 | C | T | 24 | 15619 | 0.15365901 | 44.0833333 | rnr1Y285A_p | TRUE | CAN | JSY2423 | 3 | rnr1Y285A_p | GCC | C>T | A/T+1 | G/C-1 | CG>TA | CG>TA | CCTATGCCAT | CCATCCT | CGG | GCC | C | A | C>T | C>T_GCC | CG>TA_CCATCCT | 6 | FALSE | FALSE |
| 139_v.txt | 33276 | 33276-SNV- | SNV | 1 | G | A | 33 | 15619 | 0.21128113 | 45.030303 | rnr1Y285A_p | TRUE | CAN | JSY2423 | 3 | rnr1Y285A_p | CGT | G>A | A/T+1 | G/C-1 | CG>TA | CG>TA | GAGAACGTA | GTATCCT | GCA | GCA | C | T | C>T | C>T_GCA | CG>TA_GTATCCT | 6 | FALSE | FALSE |
| 139_v.txt | 33309 | 33309-SNV- | SNV | 1 | C | T | 17 | 15618 | 0.10884876 | 52.350812 | rnr1Y285A_p | TRUE | CAN | JSY2423 | 3 | rnr1Y285A_p | GCG | C>T | A/T+1 | G/C-1 | CG>TA | CG>TA | AAAGAGCGC | CGCAATG | CGC | GCG | C | A | C>T | C>T_GCG | CG>TA_CGCAATG | 6 | FALSE | FALSE |
| 139_v.txt | 33310 | 33310-SNV- | SNV | 1 | G | A | 19 | 15618 | 0.1216545 | 54.271637 | rnr1Y285A_p | TRUE | CAN | JSY2423 | 3 | rnr1Y285A_p | CGC | G>A | A/T+1 | G/C-1 | CG>TA | CG>TA | AAGAGCGCA | GCAATGG | GCG | GCG | C | T | C>T | C>T_GCG | CG>TA_GCAATGG | 6 | FALSE | FALSE |
| 139_v.txt | 33412 | 33412-SNV- | SNV | 1 | G | A | 18 | 15619 | 0.11524425 | 47.8333333 | rnr1Y285A_p | TRUE | CAN | JSY2423 | 3 | rnr1Y285A_p | GGC | G>A | A/T+1 | G/C-1 | CG>TA | CG>TA | TGACCGGCT | GCTCATT | CCG | CCG | C | T | C>T | C>T_CCG | CG>TA_GCTCATT | 6 | FALSE | FALSE |
| 139_v.txt | 33474 | 33474-SNV- | SNV | 1 | C | T | 30 | 15617 | 0.19209835 | 31.2 | rnr1Y285A_p | TRUE | CAN | JSY2423 | 3 | rnr1Y285A_p | GCC | C>T | A/T+1 | G/C-1 | CG>TA | CG>TA | GCTATGCCT | CCTTTTT | CGG | GCC | C | A | C>T | C>T_GCC | CG>TA_CCTTTTT | 6 | FALSE | FALSE |
| 139_v.txt | 31947 | 31947-SNV- | SNV | 1 | G | C | 14 | 16270 | 0.08604794 | 36.8571429 | rnr1Y285A_p | TRUE | CAN | JSY2423 | 3 | rnr1Y285A_p | CGT | G>C | G/C+1 | G/C-1 | CG>GC | CG>GC | ATGATCGTC | GTCATAA | GCA | GCA | C | G | C>G | C>G_GCA | CG>GC_GTCATAA | 6 | FALSE | FALSE |
| 139_v.txt | 32203 | 32203-SNV- | SNV | 1 | C | G | 1 | 16284 | 6.14E-03 | 63.9980761 | rnr1Y285A_p | TRUE | CAN | JSY2423 | 3 | rnr1Y285A_p | ACA | C>G | G/C+1 | G/C-1 | CG>GC | CG>GC | ATGGAACAC | CACCACC | TGT | ACA | C | C | C>G | C>G_ACA | CG>GC_CACCACC | 6 | FALSE | FALSE |
| 139_v.txt | 32608 | 32608-SNV- | SNV | 1 | C | G | 2 | 20501 | 9.76E-03 | 43.490297 | rnr1Y285A_p | TRUE | CAN | JSY2423 | 3 | rnr1Y285A_p | ACC | C>G | G/C+1 | G/C-1 | CG>GC | CG>GC | AGGAAACCC | CCCAACC | TGG | ACC | C | C | C>G | C>G_ACC | CG>GC_CCCAACC | 6 | FALSE | FALSE |
| 139_v.txt | 32747 | 32747-SNV- | SNV | 1 | C | G | 2 | 20501 | 9.76E-03 | 37.0000902 | rnr1Y285A_p | TRUE | CAN | JSY2423 | 3 | rnr1Y285A_p | ACC | C>G | G/C+1 | G/C-1 | CG>GC | CG>GC | TAGAAACCC | CCCGATA | TGG | ACC | C | C | C>G | C>G_ACC | CG>GC_CCCGATA | 6 | FALSE | FALSE |
| 139_v.txt | 32786 | 32786-SNV- | SNV | 1 | C | G | 2 | 20501 | 9.76E-03 | 63.9987669 | rnr1Y285A_p | TRUE | CAN | JSY2423 | 3 | rnr1Y285A_p | ACT | C>G | G/C+1 | G/C-1 | CG>GC | CG>GC | CCAGAACTC | CTCGAAT | TGA | ACT | C | C | C>G | C>G_ACT | CG>GC_CTCGAAT | 6 | FALSE | FALSE |
| 139_v.txt | 32819 | 32819-SNV- | SNV | 1 | G | C | 19 | 20501 | 0.09267841 | 45.7894737 | rnr1Y285A_p | TRUE | CAN | JSY2423 | 3 | rnr1Y285A_p | AGT | G>C | G/C+1 | G/C-1 | CG>GC | CG>GC | GAACAAGTT | GTTCATT | TCA | TCA | C | G | C>G | C>G_TCA | CG>GC_GTTCATT | 6 | FALSE | FALSE |
| 139_v.txt | 32901 | 32901-SNV- | SNV | 1 | C | G | 3 | 23058 | 0.01301067 | 34.6653248 | rnr1Y285A_p | TRUE | CAN | JSY2423 | 3 | rnr1Y285A_p | GCC | C>G | G/C+1 | G/C-1 | CG>GC | CG>GC | ACTTGGCCA | CCAACTA | CGG | GCC | C | C | C>G | C>G_GCC | CG>GC_CCAACTA | 6 | FALSE | FALSE |
| 139_v.txt | 32971 | 32971-SNV- | SNV | 1 | C | G | 15 | 23058 | 0.06505334 | 60.9333333 | rnr1Y285A_p | TRUE | CAN | JSY2423 | 3 | rnr1Y285A_p | CCA | C>G | G/C+1 | G/C-1 | CG>GC | CG>GC | CCGCACCAA | CAAATGC | GGT | CCA | C | C | C>G | C>G_CCA | CG>GC_CAAATGC | 6 | FALSE | FALSE |
| 139_v.txt | 31657 | 31657-SNV- | SNV | 1 | G | T | 64 | 8823 | 0.72537686 | 20.53125 | rnr1Y285A_p | TRUE | CAN | JSY2423 | 3 | rnr1Y285A_p | GGC | G>T | A/T+1 | G/C-1 | CG>AT | CG>AT | GAAATGGCG | GCGTGGA | CCG | CCG | C | A | C>A | C>A_CCG | CG>AT_GCGTGGA | 6 | FALSE | FALSE |
| 139_v.txt | 31981 | 31981-SNV- | SNV | 1 | G | T | 13 | 16284 | 0.07983297 | 23 | rnr1Y285A_p | TRUE | CAN | JSY2423 | 3 | rnr1Y285A_p | GGC | G>T | A/T+1 | G/C-1 | CG>AT | CG>AT | AGCCGGGCA | GCATTAA | CCG | CCG | C | A | C>A | C>A_CCG | CG>AT_GCATTAA | 6 | FALSE | FALSE |
| 139_v.txt | 32027 | 32027-SNV- | SNV | 1 | G | T | 364 | 16284 | 2.23532302 | 59.6448236 | rnr1Y285A_p | TRUE | CAN | JSY2423 | 3 | rnr1Y285A_p | GGT | G>T | A/T+1 | G/C-1 | CG>AT | CG>AT | GCCACGGTA | GTATTTC | CCA | CCA | C | A | C>A | C>A_CCA | CG>AT_GTATTTC | 6 | FALSE | FALSE |
| 139_v.txt | 32188 | 32188-SNV- | SNV | 1 | C | A | 42 | 16284 | 0.25792189 | 43.3333333 | rnr1Y285A_p | TRUE | CAN | JSY2423 | 3 | rnr1Y285A_p | ACT | C>A | A/T+1 | G/C-1 | CG>AT | CG>AT | CGAAAACTG | CTGCAAT | TGA | ACT | C | T | C>A | C>A_ACT | CG>AT_CTGCAAT | 6 | FALSE | FALSE |
| 139_v.txt | 32203 | 32203-SNV- | SNV | 1 | C | A | 14 | 16284 | 0.08597396 | 39.4998176 | rnr1Y285A_p | TRUE | CAN | JSY2423 | 3 | rnr1Y285A_p | ACA | C>A | A/T+1 | G/C-1 | CG>AT | CG>AT | ATGGAACAC | CACCACC | TGT | ACA | C | T | C>A | C>A_ACA | CG>AT_CACCACC | 6 | FALSE | FALSE |
| 139_v.txt | 32274 | 32274-SNV- | SNV | 1 | G | T | 31 | 27298 | 0.11356143 | 33.6774194 | rnr1Y285A_p | TRUE | CAN | JSY2423 | 3 | rnr1Y285A_p | GGA | G>T | A/T+1 | G/C-1 | CG>AT | CG>AT | ATACGGGAA | GAACCAA | CCT | CCT | C | A | C>A | C>A_CCT | CG>AT_GAACCAA | 6 | FALSE | FALSE |
| 139_v.txt | 32277 | 32277-SNV- | SNV | 1 | C | A | 56 | 27298 | 0.20514323 | 35.0714286 | rnr1Y285A_p | TRUE | CAN | JSY2423 | 3 | rnr1Y285A_p | ACC | C>A | A/T+1 | G/C-1 | CG>AT | CG>AT | CGGGAACCA | CCAACGT | TGG | ACC | C | T | C>A | C>A_ACC | CG>AT_CCAACGT | 6 | FALSE | FALSE |
| 139_v.txt | 32292 | 32292-SNV- | SNV | 1 | G | T | 42 | 27298 | 0.15385743 | 46.2857143 | rnr1Y285A_p | TRUE | CAN | JSY2423 | 3 | rnr1Y285A_p | TGA | G>T | A/T+1 | G/C-1 | CG>AT | CG>AT | ATATTTGAA | GAATTTG | ACT | ACT | C | A | C>A | C>A_ACT | CG>AT_GAATTTG | 6 | FALSE | FALSE |
| 139_v.txt | 32304 | 32304-SNV- | SNV | 1 | G | T | 27 | 27298 | 0.09890835 | 55.4814815 | rnr1Y285A_p | TRUE | CAN | JSY2423 | 3 | rnr1Y285A_p | AGA | G>T | A/T+1 | G/C-1 | CG>AT | CG>AT | GCGGCAGAA | GAAATAA | TCT | TCT | C | A | C>A | C>A_TCT | CG>AT_GAAATAA | 6 | FALSE | FALSE |
| 139_v.txt | 32373 | 32373-SNV- | SNV | 1 | G | T | 33 | 27298 | 0.12088798 | 58.2424242 | rnr1Y285A_p | TRUE | CAN | JSY2423 | 3 | rnr1Y285A_p | AGC | G>T | A/T+1 | G/C-1 | CG>AT | CG>AT | TCAATAGCA | GCAATAA | TCG | TCG | C | A | C>A | C>A_TCG | CG>AT_GCAATAA | 6 | FALSE | FALSE |
| 139_v.txt | 32471 | 32471-SNV- | SNV | 1 | G | T | 211 | 27298 | 0.7729504 | 47.5545024 | rnr1Y285A_p | TRUE | CAN | JSY2423 | 3 | rnr1Y285A_p | TGT | G>T | A/T+1 | G/C-1 | CG>AT | CG>AT | GCCAATGTA | GTAGAAG | ACA | ACA | C | A | C>A | C>A_ACA | CG>AT_GTAGAAG | 6 | FALSE | FALSE |
| 139_v.txt | 32474 | 32474-SNV- | SNV | 1 | G | T | 25 | 27298 | 0.0915818 | 51.52 | rnr1Y285A_p | TRUE | CAN | JSY2423 | 3 | rnr1Y285A_p | AGA | G>T | A/T+1 | G/C-1 | CG>AT | CG>AT | AATGTAGAA | GAAGGTT | TCT | TCT | C | A | C>A | C>A_TCT | CG>AT_GAAGGTT | 6 | FALSE | FALSE |
| 139_v.txt | 32508 | 32508-SNV- | SNV | 1 | G | T | 84 | 27298 | 0.30771485 | 43.3095238 | rnr1Y285A_p | TRUE | CAN | JSY2423 | 3 | rnr1Y285A_p | GGC | G>T | A/T+1 | G/C-1 | CG>AT | CG>AT | TTGATGGCT | GCTCTTG | CCG | CCG | C | A | C>A | C>A_CCG | CG>AT_GCTCTTG | 6 | FALSE | FALSE |
| 139_v.txt | 32530 | 32530-SNV- | SNV | 1 | G | T | 107 | 27298 | 0.39197011 | 39.2616822 | rnr1Y285A_p | TRUE | CAN | JSY2423 | 3 | rnr1Y285A_p | GGG | G>T | A/T+1 | G/C-1 | CG>AT | CG>AT | TTCTGGGGT | GGTTTGC | CCC | CCC | C | A | C>A | C>A_CCC | CG>AT_GGTTTGC | 6 | FALSE | FALSE |
| 139_v.txt | 32536 | 32536-SNV- | SNV | 1 | C | A | 4 | 27297 | 0.01465363 | 31 | rnr1Y285A_p | TRUE | CAN | JSY2423 | 3 | rnr1Y285A_p | GCA | C>A | A/T+1 | G/C-1 | CG>AT | CG>AT | GGTTTGCAG | CAGCTTC | CGT | GCA | C | T | C>A | C>A_GCA | CG>AT_CAGCTTC | 6 | FALSE | FALSE |

| | | | | | | | | | | | | | | | | | | | | | | | | | | | | | | | | | | |
|---|---|---|---|---|---|---|---|---|---|---|---|---|---|---|---|---|---|---|---|---|---|---|---|---|---|---|---|---|---|---|---|---|---|---|
| 139_v.txt | 32556 | 32556-SNV- | SNV | 1 | C | A | 212 | 27289 | 0.7768698 | 37.0943396 | rnr1Y285A_p | TRUE | CAN | JSY2423 | 3 | rnr1Y285A_p | ACC | C>A | A/T+1 | G/C-1 | CG>AT | CG>AT | GTGATACCA | CCAACTA | TGG | ACC | C | T | C>A | C>A_ACC | CG>AT_CCAACTA | 6 | FALSE | FALSE |
| 139_v.txt | 32572 | 32572-SNV- | SNV | 1 | C | A | 15 | 20501 | 0.07316716 | 37.6666667 | rnr1Y285A_p | TRUE | CAN | JSY2423 | 3 | rnr1Y285A_p | CCT | C>A | A/T+1 | G/C-1 | CG>AT | CG>AT | CAGTACCTT | CTTGAAA | GGA | CCT | C | T | C>A | C>A_CCT | CG>AT_CTTGAAA | 6 | FALSE | FALSE |
| 139_v.txt | 32735 | 32735-SNV- | SNV | 1 | G | T | 140 | 20501 | 0.68289352 | 56.4642857 | rnr1Y285A_p | TRUE | CAN | JSY2423 | 3 | rnr1Y285A_p | AGT | G>T | A/T+1 | G/C-1 | CG>AT | CG>AT | AAAACAGTA | GTATATT | TCA | TCA | C | A | C>A | C>A_TCA | CG>AT_GTATATT | 6 | FALSE | FALSE |
| 139_v.txt | 32786 | 32786-SNV- | SNV | 1 | C | A | 131 | 20501 | 0.63899322 | 47.6713709 | rnr1Y285A_p | TRUE | CAN | JSY2423 | 3 | rnr1Y285A_p | ACT | C>A | A/T+1 | G/C-1 | CG>AT | CG>AT | CCAGAACTC | CTCGAAT | TGA | ACT | C | T | C>A | C>A_ACT | CG>AT_CTCGAAT | 6 | FALSE | FALSE |
| 139_v.txt | 32798 | 32798-SNV- | SNV | 1 | G | T | 19 | 20501 | 0.09267841 | 46.6842105 | rnr1Y285A_p | TRUE | CAN | JSY2423 | 3 | rnr1Y285A_p | CGT | G>T | A/T+1 | G/C-1 | CG>AT | CG>AT | TTCACCGTAA | GTAATAT | GCA | GCA | C | A | C>A | C>A_GCA | CG>AT_GTAATAT | 6 | FALSE | FALSE |
| 139_v.txt | 32819 | 32819-SNV- | SNV | 1 | G | T | 517 | 20501 | 2.5218282 | 44.9651838 | rnr1Y285A_p | TRUE | CAN | JSY2423 | 3 | rnr1Y285A_p | AGT | G>T | A/T+1 | G/C-1 | CG>AT | CG>AT | GAACAAGTT | GTTCATT | TCA | TCA | C | A | C>A | C>A_TCA | CG>AT_GTTCATT | 6 | FALSE | FALSE |
| 139_v.txt | 32917 | 32917-SNV- | SNV | 1 | C | A | 524 | 23058 | 2.27253014 | 52.1183206 | rnr1Y285A_p | TRUE | CAN | JSY2423 | 3 | rnr1Y285A_p | TCC | C>A | A/T+1 | G/C-1 | CG>AT | CG>AT | TAAGTTCCA | CCAGGGC | AGG | TCC | C | T | C>A | C>A_TCC | CG>AT_CCAGGGC | 6 | FALSE | FALSE |
| 139_v.txt | 32940 | 32940-SNV- | SNV | 1 | G | T | 101 | 23057 | 0.43804485 | 51.514624 | rnr1Y285A_p | TRUE | CAN | JSY2423 | 3 | rnr1Y285A_p | AGA | G>T | A/T+1 | G/C-1 | CG>AT | CG>AT | GCCCAAGAA | GAAAACC | TCT | TCT | C | A | C>A | C>A_TCT | CG>AT_GAAAACC | 6 | FALSE | FALSE |
| 139_v.txt | 32958 | 32958-SNV- | SNV | 1 | C | A | 9 | 23058 | 0.03903201 | 52.5555556 | rnr1Y285A_p | TRUE | CAN | JSY2423 | 3 | rnr1Y285A_p | ACC | C>A | A/T+1 | G/C-1 | CG>AT | CG>AT | ATGTAACCA | CCATTGG | TGG | ACC | C | T | C>A | C>A_ACC | CG>AT_CCATTGG | 6 | FALSE | FALSE |
| 139_v.txt | 33015 | 33015-SNV- | SNV | 1 | G | T | 14 | 23058 | 0.06071645 | 49.9285714 | rnr1Y285A_p | TRUE | CAN | JSY2423 | 3 | rnr1Y285A_p | GGA | G>T | A/T+1 | G/C-1 | CG>AT | CG>AT | AAAGAGGAT | GATGTAA | CCT | CCT | C | A | C>A | C>A_CCT | CG>AT_GATGTAA | 6 | FALSE | FALSE |
| 139_v.txt | 33024 | 33024-SNV- | SNV | 1 | G | T | 19 | 23058 | 0.0824009 | 59.7894737 | rnr1Y285A_p | TRUE | CAN | JSY2423 | 3 | rnr1Y285A_p | AGG | G>T | A/T+1 | G/C-1 | CG>AT | CG>AT | GTAACAGGG | GGGATGA | TCC | TCC | C | A | C>A | C>A_TCC | CG>AT_GGGATGA | 6 | FALSE | FALSE |
| 139_v.txt | 33114 | 33114-SNV- | SNV | 1 | C | A | 46 | 23058 | 0.19949692 | 44.7611208 | rnr1Y285A_p | TRUE | CAN | JSY2423 | 3 | rnr1Y285A_p | GCC | C>A | A/T+1 | G/C-1 | CG>AT | CG>AT | ACTGGGCCG | CCGGCGT | CGG | GCC | C | T | C>A | C>A_GCC | CG>AT_CCGGCGT | 6 | FALSE | FALSE |
| 139_v.txt | 33115 | 33115-SNV- | SNV | 1 | C | A | 15 | 23058 | 0.06505334 | 37.7333333 | rnr1Y285A_p | TRUE | CAN | JSY2423 | 3 | rnr1Y285A_p | CCG | C>A | A/T+1 | G/C-1 | CG>AT | CG>AT | CTGGGCCGG | CGGCGTT | GGC | CCG | C | T | C>A | C>A_CCG | CG>AT_CGGCGTT | 6 | FALSE | FALSE |
| 139_v.txt | 33151 | 33151-SNV- | SNV | 1 | G | T | 31 | 23058 | 0.13444358 | 38.3548387 | rnr1Y285A_p | TRUE | CAN | JSY2423 | 3 | rnr1Y285A_p | AGA | G>T | A/T+1 | G/C-1 | CG>AT | CG>AT | TGAAAAGAC | GACCTGT | TCT | TCT | C | A | C>A | C>A_TCT | CG>AT_GACCTGT | 6 | FALSE | FALSE |
| 139_v.txt | 33153 | 33153-SNV- | SNV | 1 | C | A | 235 | 23058 | 1.01916905 | 39.5914894 | rnr1Y285A_p | TRUE | CAN | JSY2423 | 3 | rnr1Y285A_p | ACC | C>A | A/T+1 | G/C-1 | CG>AT | CG>AT | AAAAGACCT | CCTGTAC | TGG | ACC | C | T | C>A | C>A_ACC | CG>AT_CCTGTAC | 6 | FALSE | FALSE |
| 139_v.txt | 33154 | 33154-SNV- | SNV | 1 | C | A | 55 | 23058 | 0.23852893 | 34.9820788 | rnr1Y285A_p | TRUE | CAN | JSY2423 | 3 | rnr1Y285A_p | CCT | C>A | A/T+1 | G/C-1 | CG>AT | CG>AT | AAAGACCTG | CTGTACC | GGA | CCT | C | T | C>A | C>A_CCT | CG>AT_CTGTACC | 6 | FALSE | FALSE |
| 139_v.txt | 33159 | 33159-SNV- | SNV | 1 | C | A | 58 | 23058 | 0.2515396 | 37.3622938 | rnr1Y285A_p | TRUE | CAN | JSY2423 | 3 | rnr1Y285A_p | ACC | C>A | A/T+1 | G/C-1 | CG>AT | CG>AT | CCTGTACCA | CCAATAG | TGG | ACC | C | T | C>A | C>A_ACC | CG>AT_CCAATAG | 6 | FALSE | FALSE |
| 139_v.txt | 33160 | 33160-SNV- | SNV | 1 | C | A | 40 | 23058 | 0.17347558 | 37.1423685 | rnr1Y285A_p | TRUE | CAN | JSY2423 | 3 | rnr1Y285A_p | CCA | C>A | A/T+1 | G/C-1 | CG>AT | CG>AT | CTGTACCAAT | CAATAGT | GGT | CCA | C | T | C>A | C>A_CCA | CG>AT_CAATAGT | 6 | FALSE | FALSE |
| 139_v.txt | 33168 | 33168-SNV- | SNV | 1 | C | A | 200 | 23057 | 0.86741554 | 36.92 | rnr1Y285A_p | TRUE | CAN | JSY2423 | 3 | rnr1Y285A_p | ACC | C>A | A/T+1 | G/C-1 | CG>AT | CG>AT | ATAGTACCA | CCACCAA | TGG | ACC | C | T | C>A | C>A_ACC | CG>AT_CCACCAA | 6 | FALSE | FALSE |
| 139_v.txt | 33175 | 33175-SNV- | SNV | 1 | G | T | 11 | 23052 | 0.0477182 | 37.8181818 | rnr1Y285A_p | TRUE | CAN | JSY2423 | 3 | rnr1Y285A_p | AGG | G>T | A/T+1 | G/C-1 | CG>AT | CG>AT | CACCAAGGG | GGGCAAT | TCC | TCC | C | A | C>A | C>A_TCC | CG>AT_GGGCAAT | 6 | FALSE | FALSE |
| 139_v.txt | 33186 | 33186-SNV- | SNV | 1 | C | A | 22 | 15617 | 0.14087213 | 37.3181818 | rnr1Y285A_p | TRUE | CAN | JSY2423 | 3 | rnr1Y285A_p | ACC | C>A | A/T+1 | G/C-1 | CG>AT | CG>AT | ATCATACCA | CCAATAT | TGG | ACC | C | T | C>A | C>A_ACC | CG>AT_CCAATAT | 6 | FALSE | FALSE |
| 139_v.txt | 33205 | 33205-SNV- | SNV | 1 | G | T | 9 | 15619 | 0.05762213 | 35 | rnr1Y285A_p | TRUE | CAN | JSY2423 | 3 | rnr1Y285A_p | AGC | G>T | A/T+1 | G/C-1 | CG>AT | CG>AT | GCTTAAGCT | GCTCTCT | TCG | TCG | C | A | C>A | C>A_TCG | CG>AT_GCTCTCT | 6 | FALSE | FALSE |
| 139_v.txt | 33247 | 33247-SNV- | SNV | 1 | C | A | 23 | 15619 | 0.14725655 | 34.9565217 | rnr1Y285A_p | TRUE | CAN | JSY2423 | 3 | rnr1Y285A_p | TCA | C>A | A/T+1 | G/C-1 | CG>AT | CG>AT | CATCTTCATC | CATCACC | AGT | TCA | C | T | C>A | C>A_TCA | CG>AT_CATCACC | 6 | FALSE | FALSE |
| 139_v.txt | 33310 | 33310-SNV- | SNV | 1 | G | T | 8 | 15618 | 0.05122295 | 13.108884 | rnr1Y285A_p | TRUE | CAN | JSY2423 | 3 | rnr1Y285A_p | CGC | G>T | A/T+1 | G/C-1 | CG>AT | CG>AT | AAGAGCGCA | GCAATGG | GCG | GCG | C | A | C>A | C>A_GCG | CG>AT_GCAATGG | 6 | FALSE | FALSE |
| 139_v.txt | 33325 | 33325-SNV- | SNV | 1 | C | A | 451 | 15619 | 2.8875088 | 59.3370288 | rnr1Y285A_p | TRUE | CAN | JSY2423 | 3 | rnr1Y285A_p | TCT | C>A | A/T+1 | G/C-1 | CG>AT | CG>AT | ACAATTCTTT | CTTTACT | AGA | TCT | C | T | C>A | C>A_TCT | CG>AT_CTTTACT | 6 | FALSE | FALSE |
| 139_v.txt | 33474 | 33474-SNV- | SNV | 1 | C | A | 40 | 15617 | 0.25613114 | 32.925 | rnr1Y285A_p | TRUE | CAN | JSY2423 | 3 | rnr1Y285A_p | GCC | C>A | A/T+1 | G/C-1 | CG>AT | CG>AT | GCTATGCCTT | CCTTTTT | CGG | GCC | C | T | C>A | C>A_GCC | CG>AT_CCTTTTT | 6 | FALSE | FALSE |
| 139_v.txt | 33475 | 33475-SNV- | SNV | 1 | C | A | 11 | 15617 | 0.07043606 | 35.2727273 | rnr1Y285A_p | TRUE | CAN | JSY2423 | 3 | rnr1Y285A_p | CCT | C>A | A/T+1 | G/C-1 | CG>AT | CG>AT | CTATGCCTTT | CTTTTTT | GGA | CCT | C | T | C>A | C>A_CCT | CG>AT_CTTTTTT | 6 | FALSE | FALSE |
| 139_v.txt | 32608 | 32608-Repla | Replacement | 2 | CC | A | 9 | 20501 | 0.0439003 | 49.3888889 | rnr1Y285A_p | TRUE | CAN | JSY2423 | 3 | rnr1Y285A_p | ACC | nope | A/T+1 | >1bpdel | none | Replac. | AGGAAACCC | CCCAACC | TGG | ACC | CC | T | NA | NA_ACC | Replac._CCCAACC | 6 | FALSE | FALSE |
| 139_v.txt | 32608 | 32608-Repla | Replacement | 2 | CC | T | 3 | 20501 | 0.01463343 | 38 | rnr1Y285A_p | TRUE | CAN | JSY2423 | 3 | rnr1Y285A_p | ACC | nope | A/T+1 | >1bpdel | none | Replac. | AGGAAACCC | CCCAACC | TGG | ACC | CC | A | NA | NA_ACC | Replac._CCCAACC | 6 | FALSE | FALSE |
| 139_v.txt | 32900 | 32900-MNV- | MNV | 2 | GC | CT | 1 | 23058 | 4.34E-03 | 20.9920709 | rnr1Y285A_p | TRUE | CAN | JSY2423 | 3 | rnr1Y285A_p | GGC | nope | >1bp-ins | >1bpdel | none | MNV | GACTTGGCC | GCCAACT | CCG | GGC | GC | none | NA | NA_GGC | MNV_GCCAACT | 6 | FALSE | FALSE |
| 139_v.txt | 33159 | 33159-MNV- | MNV | 2 | CC | AA | 1 | 23058 | 4.34E-03 | 34 | rnr1Y285A_p | TRUE | CAN | JSY2423 | 3 | rnr1Y285A_p | ACC | nope | >1bp-ins | >1bpdel | none | MNV | CCTGTACCA | CCAATAG | TGG | ACC | CC | none | NA | NA_ACC | MNV_CCAATAG | 6 | FALSE | FALSE |
| 139_v.txt | 32081 | 32081-Insert | Insertion | 1 | - | A | 10 | 16284 | 0.06140997 | 64 | rnr1Y285A_p | TRUE | CAN | JSY2423 | 3 | rnr1Y285A_p | CAA | nope | A/T+1 | >1bpdel | none | A/T+1 | CCATGCAAA | AAAAAAG | GTT | CAA | - | T | NA | NA_CAA | A/T+1_AAAAAAG | 6 | FALSE | FALSE |
| 139_v.txt | 32597 | 32597-Insert | Insertion | 1 | - | A | 6 | 20501 | 0.02926687 | 42.3333333 | rnr1Y285A_p | TRUE | CAN | JSY2423 | 3 | rnr1Y285A_p | TCA | nope | A/T+1 | >1bpdel | none | A/T+1 | GTTAATCAA | CAAAGAG | AGT | TCA | - | T | NA | NA_TCA | A/T+1_CAAAGAG | 6 | FALSE | FALSE |
| 139_v.txt | 32842 | 32842-Insert | Insertion | 1 | - | A | 23 | 20501 | 0.11218965 | 42.3043478 | rnr1Y285A_p | TRUE | CAN | JSY2423 | 3 | rnr1Y285A_p | CAA | nope | A/T+1 | >1bpdel | none | A/T+1 | TTACCCAAA | AAAAAAT | GTT | CAA | - | T | NA | NA_CAA | A/T+1_AAAAAAT | 6 | FALSE | FALSE |
| 139_v.txt | 32884 | 32884-Insert | Insertion | 1 | - | A | 2 | 23058 | 8.67E-03 | 38 | rnr1Y285A_p | TRUE | CAN | JSY2423 | 3 | rnr1Y285A_p | CAA | nope | A/T+1 | >1bpdel | none | A/T+1 | ACGTCCAAA | AAAATTG | GTT | CAA | - | T | NA | NA_CAA | A/T+1_AAAATTG | 6 | FALSE | FALSE |
| 139_v.txt | 32201 | 32201-Deleti | Deletion | 3 | AAC | - | 12 | 16284 | 0.07369197 | 41.9166667 | rnr1Y285A_p | TRUE | CAN | JSY2423 | 3 | rnr1Y285A_p | GAA | nope | >1bp-ins | >1bpdel | none | >1bpdel | GTATGGAAC | AACACCA | CTT | GAA | AAC | none | NA | NA_GAA | >1bpdel_AACACCA | 6 | FALSE | FALSE |
| 139_v.txt | 32504 | 32504-Deleti | Deletion | 2 | GA | - | 18 | 27298 | 0.0659389 | 40.8333333 | rnr1Y285A_p | TRUE | CAN | JSY2423 | 3 | rnr1Y285A_p | TGA | nope | >1bp-ins | >1bpdel | none | >1bpdel | TTTTTTGATG | GATGGCT | ACT | TGA | GA | none | NA | NA_TGA | >1bpdel_GATGGCT | 6 | FALSE | FALSE |
| 139_v.txt | 32589 | 32589-Deleti | Deletion | 2 | GC | - | 6 | 20501 | 0.02926687 | 37.5 | rnr1Y285A_p | TRUE | CAN | JSY2423 | 3 | rnr1Y285A_p | AGC | nope | >1bp-ins | >1bpdel | none | >1bpdel | AAGGCAGCG | GCGTTAA | TCG | AGC | GC | none | NA | NA_AGC | >1bpdel_GCGTTAA | 6 | FALSE | FALSE |
| 139_v.txt | 32608 | 32608-Deleti | Deletion | 2 | CC | - | 1 | 20501 | 4.88E-03 | 38 | rnr1Y285A_p | TRUE | CAN | JSY2423 | 3 | rnr1Y285A_p | ACC | nope | >1bp-ins | >1bpdel | none | >1bpdel | AGGAAACCC | CCCAACC | TGG | ACC | CC | none | NA | NA_ACC | >1bpdel_CCCAACC | 6 | FALSE | FALSE |
| 139_v.txt | 33189 | 33189-Deleti | Deletion | 2 | AT | - | 6 | 15619 | 0.03841475 | 36.8333333 | rnr1Y285A_p | TRUE | CAN | JSY2423 | 3 | rnr1Y285A_p | AAT | nope | >1bp-ins | >1bpdel | none | >1bpdel | ATACCAATAT | ATATGTC | TTA | AAT | AT | none | NA | NA_AAT | >1bpdel_ATATGTC | 6 | FALSE | FALSE |
| 139_v.txt | 31723 | 31723-Deleti | Deletion | 1 | A | - | 18 | 8823 | 0.20401224 | 59.7222222 | rnr1Y285A_p | TRUE | CAN | JSY2423 | 3 | rnr1Y285A_p | CAA | nope | >1bp-ins | A/T-1 | none | A/T-1 | TGTCCCAAA | AAAAAGT | GTT | GTT | T | none | T>none | T>none_GTT | A/T-1_AAAAAGT | 6 | FALSE | FALSE |
| 139_v.txt | 31904 | 31904-Deleti | Deletion | 1 | T | - | 8 | 8823 | 0.09067211 | 46.375 | rnr1Y285A_p | TRUE | CAN | JSY2423 | 3 | rnr1Y285A_p | ATT | nope | >1bp-ins | A/T-1 | none | A/T-1 | ATTGAATTTT | TTTTGGT | TAA | ATT | T | none | nope | nope_ATT | A/T-1_TTTTGGT | 6 | FALSE | FALSE |
| 139_v.txt | 31972 | 31972-Deleti | Deletion | 1 | C | - | 8 | 16284 | 0.04912798 | 35 | rnr1Y285A_p | TRUE | CAN | JSY2423 | 3 | rnr1Y285A_p | GCC | nope | >1bp-ins | G/C-1 | none | G/C-1 | AATAAGCCA | CCAAGCC | CGG | GCC | C | none | nope | nope_GCC | G/C-1_CCAAGCC | 6 | FALSE | FALSE |
| 139_v.txt | 31979 | 31979-Deleti | Deletion | 1 | G | - | 46 | 16284 | 0.28248588 | 40.5 | rnr1Y285A_p | TRUE | CAN | JSY2423 | 3 | rnr1Y285A_p | CGG | nope | >1bp-ins | G/C-1 | none | G/C-1 | CAAGCCGGG | GGGCATT | GCC | GCC | C | none | C>none | C>none_GCC | G/C-1_GGGCATT | 6 | FALSE | FALSE |
| 139_v.txt | 31988 | 31988-Deleti | Deletion | 1 | T | - | 12 | 16284 | 0.07369197 | 54.1666667 | rnr1Y285A_p | TRUE | CAN | JSY2423 | 3 | rnr1Y285A_p | ATT | nope | >1bp-ins | A/T-1 | none | A/T-1 | CATTAATTTA | TTTAGCT | TAA | ATT | T | none | nope | nope_ATT | A/T-1_TTTAGCT | 6 | FALSE | FALSE |
| 139_v.txt | 32026 | 32026-Deleti | Deletion | 1 | G | - | 11 | 16284 | 0.06755097 | 56.5454546 | rnr1Y285A_p | TRUE | CAN | JSY2423 | 3 | rnr1Y285A_p | CGG | nope | >1bp-ins | G/C-1 | none | G/C-1 | TGCCACGGT | GGTATTT | GCC | GCC | C | none | C>none | C>none_GCC | G/C-1_GGTATTT | 6 | FALSE | FALSE |
| 139_v.txt | 32045 | 32045-Deleti | Deletion | 1 | A | - | 25 | 16284 | 0.15352493 | 60.56 | rnr1Y285A_p | TRUE | CAN | JSY2423 | 3 | rnr1Y285A_p | TAA | nope | >1bp-ins | A/T-1 | none | A/T-1 | TTGCATAAAT | AAATCTG | ATT | ATT | T | none | T>none | T>none_ATT | A/T-1_AAATCTG | 6 | FALSE | FALSE |
| 139_v.txt | 32075 | 32075-Deleti | Deletion | 1 | C | - | 2 | 16284 | 0.012282 | 64 | rnr1Y285A_p | TRUE | CAN | JSY2423 | 3 | rnr1Y285A_p | ACC | nope | >1bp-ins | G/C-1 | none | G/C-1 | AAATAACCAT | CCATGCA | TGG | ACC | C | none | nope | nope_ACC | G/C-1_CCATGCA | 6 | FALSE | FALSE |
| 139_v.txt | 32081 | 32081-Deleti | Deletion | 1 | A | - | 14 | 16284 | 0.08597396 | 61.8571434 | rnr1Y285A_p | TRUE | CAN | JSY2423 | 3 | rnr1Y285A_p | CAA | nope | >1bp-ins | A/T-1 | none | A/T-1 | CCATGCAAA | AAAAAAG | GTT | GTT | T | none | T>none | T>none_GTT | A/T-1_AAAAAAG | 6 | FALSE | FALSE |
| 139_v.txt | 32106 | 32106-Deleti | Deletion | 1 | T | - | 5 | 16284 | 0.03070499 | 53.6 | rnr1Y285A_p | TRUE | CAN | JSY2423 | 3 | rnr1Y285A_p | ATT | nope | >1bp-ins | A/T-1 | none | A/T-1 | GTGATATTTA | TTTAATA | TAA | ATT | T | none | nope | nope_ATT | A/T-1_TTTAATA | 6 | FALSE | FALSE |
| 139_v.txt | 32114 | 32114-Deleti | Deletion | 1 | C | - | 2 | 16284 | 0.012282 | 51 | rnr1Y285A_p | TRUE | CAN | JSY2423 | 3 | rnr1Y285A_p | GCC | nope | >1bp-ins | G/C-1 | none | G/C-1 | TAATAGCCAT | CCATTCG | CGG | GCC | C | none | nope | nope_GCC | G/C-1_CCATTCG | 6 | FALSE | FALSE |
| 139_v.txt | 32259 | 32259-Deleti | Deletion | 1 | C | - | 13 | 27296 | 0.04762603 | 37.8461539 | rnr1Y285A_p | TRUE | CAN | JSY2423 | 3 | rnr1Y285A_p | ACC | nope | >1bp-ins | G/C-1 | none | G/C-1 | GATAGACCA | CCAAATA | TGG | ACC | C | none | nope | nope_ACC | G/C-1_CCAAATA | 6 | FALSE | FALSE |
| 139_v.txt | 32265 | 32265-Deleti | Deletion | 1 | A | - | 3 | 27298 | 0.01098982 | 38 | rnr1Y285A_p | TRUE | CAN | JSY2423 | 3 | rnr1Y285A_p | TAA | nope | >1bp-ins | A/T-1 | none | A/T-1 | CCAAATAAA | AAAATAC | ATT | ATT | T | none | T>none | T>none_ATT | A/T-1_AAAATAC | 6 | FALSE | FALSE |
| 139_v.txt | 32277 | 32277-Deleti | Deletion | 1 | C | - | 48 | 27298 | 0.17583706 | 37.1666667 | rnr1Y285A_p | TRUE | CAN | JSY2423 | 3 | rnr1Y285A_p | ACC | nope | >1bp-ins | G/C-1 | none | G/C-1 | CGGGAACCA | CCAACGT | TGG | ACC | C | none | nope | nope_ACC | G/C-1_CCAACGT | 6 | FALSE | FALSE |
| 139_v.txt | 32346 | 32346-Deleti | Deletion | 1 | A | - | 46 | 27298 | 0.16851051 | 61.9565217 | rnr1Y285A_p | TRUE | CAN | JSY2423 | 3 | rnr1Y285A_p | CAA | nope | >1bp-ins | A/T-1 | none | A/T-1 | TGTGGCAAA | AAAACCT | GTT | GTT | T | none | T>none | T>none_GTT | A/T-1_AAAACCT | 6 | FALSE | FALSE |
| 139_v.txt | 32381 | 32381-Deleti | Deletion | 1 | A | - | 7 | 27298 | 0.0256429 | 63.8571429 | rnr1Y285A_p | TRUE | CAN | JSY2423 | 3 | rnr1Y285A_p | TAA | nope | >1bp-ins | A/T-1 | none | A/T-1 | AATAATAAA | AAAGGGA | ATT | ATT | T | none | T>none | T>none_ATT | A/T-1_AAAGGGA | 6 | FALSE | FALSE |
| 139_v.txt | 32398 | 32398-Deleti | Deletion | 1 | C | - | 256 | 27298 | 0.93779764 | 60.1132813 | rnr1Y285A_p | TRUE | CAN | JSY2423 | 3 | rnr1Y285A_p | ACG | nope | >1bp-ins | G/C-1 | none | G/C-1 | TAGAAACGT | CGTAGGA | TGC | ACG | C | none | nope | nope_ACG | G/C-1_CGTAGGA | 6 | FALSE | FALSE |
| 139_v.txt | 32442 | 32442-Deleti | Deletion | 1 | A | - | 6 | 27298 | 0.02197963 | 50.4999999 | rnr1Y285A_p | TRUE | CAN | JSY2423 | 3 | rnr1Y285A_p | TAA | nope | >1bp-ins | A/T-1 | none | A/T-1 | GGAACTAAA | AAAAGTC | ATT | ATT | T | none | T>none | T>none_ATT | A/T-1_AAAAGTC | 6 | FALSE | FALSE |
| 139_v.txt | 32498 | 32498-Deleti | Deletion | 1 | T | - | 1263 | 27298 | 4.62671258 | 44.3840063 | rnr1Y285A_p | TRUE | CAN | JSY2423 | 3 | rnr1Y285A_p | CTT | nope | >1bp-ins | A/T-1 | none | A/T-1 | AACAACTTTT | TTTTTTG | GAA | CTT | T | none | nope | nope_CTT | A/T-1_TTTTTTG | 6 | FALSE | FALSE |
| 139_v.txt | 32522 | 32522-Deleti | Deletion | 1 | T | - | 27 | 27298 | 0.09890835 | 41.1481482 | rnr1Y285A_p | TRUE | CAN | JSY2423 | 3 | rnr1Y285A_p | ATT | nope | >1bp-ins | A/T-1 | none | A/T-1 | AACGGATTTT | TTTTCTG | TAA | ATT | T | none | nope | nope_ATT | A/T-1_TTTTCTG | 6 | FALSE | FALSE |
| 139_v.txt | 32532 | 32532-Deleti | Deletion | 1 | T | - | 3 | 27298 | 0.01098982 | 46.6666667 | rnr1Y285A_p | TRUE | CAN | JSY2423 | 3 | rnr1Y285A_p | GTT | nope | >1bp-ins | A/T-1 | none | A/T-1 | CTGGGGTTT | TTTGCAG | CAA | GTT | T | none | nope | nope_GTT | A/T-1_TTTGCAG | 6 | FALSE | FALSE |
| 139_v.txt | 32556 | 32556-Deleti | Deletion | 1 | C | - | 19 | 27289 | 0.06962512 | 37.8947368 | rnr1Y285A_p | TRUE | CAN | JSY2423 | 3 | rnr1Y285A_p | ACC | nope | >1bp-ins | G/C-1 | none | G/C-1 | GTGATACCA | CCAACTA | TGG | ACC | C | none | nope | nope_ACC | G/C-1_CCAACTA | 6 | FALSE | FALSE |
| 139_v.txt | 32608 | 32608-Deleti | Deletion | 1 | C | - | 8908 | 20501 | 43.451539 | 43.4823743 | rnr1Y285A_p | TRUE | CAN | JSY2423 | 3 | rnr1Y285A_p | ACC | nope | >1bp-ins | G/C-1 | none | G/C-1 | AGGAAACCC | CCCAACC | TGG | ACC | C | none | nope | nope_ACC | G/C-1_CCCAACC | 6 | FALSE | FALSE |
| 139_v.txt | 32621 | 32621-Deleti | Deletion | 1 | C | - | 40 | 20501 | 0.19511243 | 41.425 | rnr1Y285A_p | TRUE | CAN | JSY2423 | 3 | rnr1Y285A_p | ACC | nope | >1bp-ins | G/C-1 | none | G/C-1 | TAAGAACCT | CCTCCCT | TGG | ACC | C | none | nope | nope_ACC | G/C-1_CCTCCCT | 6 | FALSE | FALSE |
| 139_v.txt | 32637 | 32637-Deleti | Deletion | 1 | T | - | 7 | 20501 | 0.03414468 | 47.8571429 | rnr1Y285A_p | TRUE | CAN | JSY2423 | 3 | rnr1Y285A_p | ATC | nope | >1bp-ins | A/T-1 | none | A/T-1 | TTTTTATCCT | TCCTTAG | TAG | ATC | T | none | nope | nope_ATC | A/T-1_TCCTTAG | 6 | FALSE | FALSE |
| 139_v.txt | 32658 | 32658-Deleti | Deletion | 1 | C | - | 786 | 20501 | 3.83395932 | 59.8015267 | rnr1Y285A_p | TRUE | CAN | JSY2423 | 3 | rnr1Y285A_p | ACC | nope | >1bp-ins | G/C-1 | none | G/C-1 | CCTGGACCC | CCCCAGG | TGG | ACC | C | none | nope | nope_ACC | G/C-1_CCCCAGG | 6 | FALSE | FALSE |
| 139_v.txt | 32670 | 32670-Deleti | Deletion | 1 | G | - | 256 | 20501 | 1.24871958 | 59.2578125 | rnr1Y285A_p | TRUE | CAN | JSY2423 | 3 | rnr1Y285A_p | TGG | nope | >1bp-ins | G/C-1 | none | G/C-1 | GCACCTGGG | GGGTTTC | ACC | ACC | C | none | C>none | C>none_ACC | G/C-1_GGGTTTC | 6 | FALSE | FALSE |
| 139_v.txt | 32707 | 32707-Deleti | Deletion | 1 | C | - | 308 | 20501 | 1.50236574 | 62.0064935 | rnr1Y285A_p | TRUE | CAN | JSY2423 | 3 | rnr1Y285A_p | ACC | nope | >1bp-ins | G/C-1 | none | G/C-1 | CGGTAACCC | CCCCAGC | TGG | ACC | C | none | nope | nope_ACC | G/C-1_CCCCAGC | 6 | FALSE | FALSE |
| 139_v.txt | 32747 | 32747-Deleti | Deletion | 1 | C | - | 1644 | 20501 | 8.01912102 | 56.472631 | rnr1Y285A_p | TRUE | CAN | JSY2423 | 3 | rnr1Y285A_p | ACC | nope | >1bp-ins | G/C-1 | none | G/C-1 | TAGAAACCC | CCCGATA | TGG | ACC | C | none | nope | nope_ACC | G/C-1_CCCGATA | 6 | FALSE | FALSE |
| 139_v.txt | 32760 | 32760-Deleti | Deletion | 1 | A | - | 46 | 20501 | 0.2243793 | 52.5 | rnr1Y285A_p | TRUE | CAN | JSY2423 | 3 | rnr1Y285A_p | TAA | nope | >1bp-ins | A/T-1 | none | A/T-1 | ATGGCTAAA | AAAACTT | ATT | ATT | T | none | T>none | T>none_ATT | A/T-1_AAAACTT | 6 | FALSE | FALSE |
| 139_v.txt | 32779 | 32779-Deleti | Deletion | 1 | C | - | 54 | 20501 | 0.26340179 | 49.5 | rnr1Y285A_p | TRUE | CAN | JSY2423 | 3 | rnr1Y285A_p | ACC | nope | >1bp-ins | G/C-1 | none | G/C-1 | AAGCGACCC | CCCAGAA | TGG | ACC | C | none | nope | nope_ACC | G/C-1_CCCAGAA | 6 | FALSE | FALSE |
| 139_v.txt | 32839 | 32839-Deleti | Deletion | 1 | C | - | 15 | 20501 | 0.07316716 | 44.8 | rnr1Y285A_p | TRUE | CAN | JSY2423 | 3 | rnr1Y285A_p | ACC | nope | >1bp-ins | G/C-1 | none | G/C-1 | TAATTACCCA | CCCAAAA | TGG | ACC | C | none | nope | nope_ACC | G/C-1_CCCAAAA | 6 | FALSE | FALSE |
| 139_v.txt | 32842 | 32842-Deleti | Deletion | 1 | A | - | 34 | 20501 | 0.16584557 | 41.5882343 | rnr1Y285A_p | TRUE | CAN | JSY2423 | 3 | rnr1Y285A_p | CAA | nope | >1bp-ins | A/T-1 | none | A/T-1 | TTACCCAAA | AAAAAAT | GTT | GTT | T | none | T>none | T>none_GTT | A/T-1_AAAAAAT | 6 | FALSE | FALSE |
| 139_v.txt | 32868 | 32868-Deleti | Deletion | 1 | G | - | 39 | 20497 | 0.19027175 | 37.2307692 | rnr1Y285A_p | TRUE | CAN | JSY2423 | 3 | rnr1Y285A_p | TGG | nope | >1bp-ins | G/C-1 | none | G/C-1 | GCCAGTGGA | GGAACTT | ACC | ACC | C | none | C>none | C>none_ACC | G/C-1_GGAACTT | 6 | FALSE | FALSE |
| 139_v.txt | 32884 | 32884-Deleti | Deletion | 1 | A | - | 8 | 23058 | 0.03469512 | 34.5 | rnr1Y285A_p | TRUE | CAN | JSY2423 | 3 | rnr1Y285A_p | CAA | nope | >1bp-ins | A/T-1 | none | A/T-1 | ACGTCCAAA | AAAATTG | GTT | GTT | T | none | T>none | T>none_GTT | A/T-1_AAAATTG | 6 | FALSE | FALSE |
| 139_v.txt | 32901 | 32901-Deleti | Deletion | 1 | C | - | 1 | 23058 | 4.34E-03 | 38 | rnr1Y285A_p | TRUE | CAN | JSY2423 | 3 | rnr1Y285A_p | GCC | nope | >1bp-ins | G/C-1 | none | G/C-1 | ACTTGGCCA | CCAACTA | CGG | GCC | C | none | nope | nope_GCC | G/C-1_CCAACTA | 6 | FALSE | FALSE |
| 139_v.txt | 32924 | 32924-Deleti | Deletion | 1 | A | - | 24 | 23058 | 0.10408535 | 54.75 | rnr1Y285A_p | TRUE | CAN | JSY2423 | 3 | rnr1Y285A_p | CAA | nope | >1bp-ins | A/T-1 | none | A/T-1 | CAGGGCAAA | AAAAGTG | GTT | GTT | T | none | T>none | T>none_GTT | A/T-1_AAAAGTG | 6 | FALSE | FALSE |
| 139_v.txt | 32941 | 32941-Deleti | Deletion | 1 | A | - | 18 | 23058 | 0.07806401 | 50.7777778 | rnr1Y285A_p | TRUE | CAN | JSY2423 | 3 | rnr1Y285A_p | GAA | nope | >1bp-ins | A/T-1 | none | A/T-1 | CCCAAGAAA | AAAACCA | CTT | CTT | T | none | T>none | T>none_CTT | A/T-1_AAAACCA | 6 | FALSE | FALSE |
| 139_v.txt | 32945 | 32945-Deleti | Deletion | 1 | C | - | 21 | 23058 | 0.09107468 | 60.952381 | rnr1Y285A_p | TRUE | CAN | JSY2423 | 3 | rnr1Y285A_p | ACC | nope | >1bp-ins | G/C-1 | none | G/C-1 | AGAAAACCA | CCAATAC | TGG | ACC | C | none | nope | nope_ACC | G/C-1_CCAATAC | 6 | FALSE | FALSE |
| 139_v.txt | 32958 | 32958-Deleti | Deletion | 1 | C | - | 8 | 23058 | 0.03469512 | 61.625 | rnr1Y285A_p | TRUE | CAN | JSY2423 | 3 | rnr1Y285A_p | ACC | nope | >1bp-ins | G/C-1 | none | G/C-1 | ATGTAACCAT | CCATTGG | TGG | ACC | C | none | nope | nope_ACC | G/C-1_CCATTGG | 6 | FALSE | FALSE |
| 139_v.txt | 33040 | 33040-Deleti | Deletion | 1 | T | - | 5 | 23058 | 0.02168445 | 64 | rnr1Y285A_p | TRUE | CAN | JSY2423 | 3 | rnr1Y285A_p | ATT | nope | >1bp-ins | A/T-1 | none | A/T-1 | TAGCCATTTC | TTTCACC | TAA | ATT | T | none | nope | nope_ATT | A/T-1_TTTCACC | 6 | FALSE | FALSE |
| 139_v.txt | 33078 | 33078-Deleti | Deletion | 1 | C | - | 73 | 23058 | 0.31659294 | 53.3424658 | rnr1Y285A_p | TRUE | CAN | JSY2423 | 3 | rnr1Y285A_p | ACC | nope | >1bp-ins | G/C-1 | none | G/C-1 | AAAGAACCC | CCCATAA | TGG | ACC | C | none | nope | nope_ACC | G/C-1_CCCATAA | 6 | FALSE | FALSE |
| 139_v.txt | 33105 | 33105-Deleti | Deletion | 1 | C | - | 8 | 23058 | 0.03469512 | 37.125 | rnr1Y285A_p | TRUE | CAN | JSY2423 | 3 | rnr1Y285A_p | GCC | nope | >1bp-ins | G/C-1 | none | G/C-1 | AGAGCGCCC | CCCACTG | CGG | GCC | C | none | nope | nope_GCC | G/C-1_CCCACTG | 6 | FALSE | FALSE |
| 139_v.txt | 33141 | 33141-Deleti | Deletion | 1 | C | - | 14 | 23058 | 0.06071645 | 41.0714286 | rnr1Y285A_p | TRUE | CAN | JSY2423 | 3 | rnr1Y285A_p | ACC | nope | >1bp-ins | G/C-1 | none | G/C-1 | GATAAACCA | CCAATGA | TGG | ACC | C | none | nope | nope_ACC | G/C-1_CCAATGA | 6 | FALSE | FALSE |
| 139_v.txt | 33147 | 33147-Deleti | Deletion | 1 | A | - | 7 | 23058 | 0.03035823 | 38 | rnr1Y285A_p | TRUE | CAN | JSY2423 | 3 | rnr1Y285A_p | GAA | nope | >1bp-ins | A/T-1 | none | A/T-1 | CCAATGAAA | AAAAGAC | CTT | CTT | T | none | T>none | T>none_CTT | A/T-1_AAAAGAC | 6 | FALSE | FALSE |
| 139_v.txt | 33153 | 33153-Deleti | Deletion | 1 | C | - | 3 | 23058 | 0.01301067 | 38 | rnr1Y285A_p | TRUE | CAN | JSY2423 | 3 | rnr1Y285A_p | ACC | nope | >1bp-ins | G/C-1 | none | G/C-1 | AAAAGACCT | CCTGTAC | TGG | ACC | C | none | nope | nope_ACC | G/C-1_CCTGTAC | 6 | FALSE | FALSE |
| 139_v.txt | 33159 | 33159-Deleti | Deletion | 1 | C | - | 1 | 23058 | 4.34E-03 | 38 | rnr1Y285A_p | TRUE | CAN | JSY2423 | 3 | rnr1Y285A_p | ACC | nope | >1bp-ins | G/C-1 | none | G/C-1 | CCTGTACCA | CCAATAG | TGG | ACC | C | none | nope | nope_ACC | G/C-1_CCAATAG | 6 | FALSE | FALSE |
| 139_v.txt | 33252 | 33252-Deleti | Deletion | 1 | C | - | 10 | 15619 | 0.06402459 | 39.8 | rnr1Y285A_p | TRUE | CAN | JSY2423 | 3 | rnr1Y285A_p | ACC | nope | >1bp-ins | G/C-1 | none | G/C-1 | TCATCACCTA | CCTATGC | TGG | ACC | C | none | nope | nope_ACC | G/C-1_CCTATGC | 6 | FALSE | FALSE |
| 139_v.txt | 33356 | 33356-Deleti | Deletion | 1 | C | - | 170 | 15619 | 1.08841795 | 53.6117647 | rnr1Y285A_p | TRUE | CAN | JSY2423 | 3 | rnr1Y285A_p | ACC | nope | >1bp-ins | G/C-1 | none | G/C-1 | TATTGACCCA | CCCACGT | TGG | ACC | C | none | nope | nope_ACC | G/C-1_CCCACGT | 6 | FALSE | FALSE |
| 139_v.txt | 33474 | 33474-Deleti | Deletion | 1 | C | - | 59 | 15617 | 0.37779343 | 27.7118644 | rnr1Y285A_p | TRUE | CAN | JSY2423 | 3 | rnr1Y285A_p | GCC | nope | >1bp-ins | G/C-1 | none | G/C-1 | GCTATGCCTT | CCTTTTT | CGG | GCC | C | none | nope | nope_GCC | G/C-1_CCTTTTT | 6 | FALSE | FALSE |
| 140_v.txt | 31964 | 31964-SNV- | SNV | 1 | A | C | 28 | 20031 | 0.13978334 | 36.5346052 | rnr1Y285A_p | TRUE | CAN | JSY2423 | 4 | rnr1Y285A_p | CAT | A>C | G/C+1 | A/T-1 | TA>GC | TA>GC | GGCCGCATA | ATAATAA | GTA | GTA | T | G | T>G | T>G_GTA | TA>GC_ATAATAA | 6 | FALSE | FALSE |
| 140_v.txt | 32532 | 32532-SNV-T | SNV | 1 | T | G | 5 | 28383 | 0.01761618 | 42.9977782 | rnr1Y285A_p | TRUE | CAN | JSY2423 | 4 | rnr1Y285A_p | GTT | T>G | G/C+1 | A/T-1 | TA>GC | TA>GC | CTGGGGTTT | TTTGCAG | CAA | GTT | T | C | T>G | T>G_GTT | TA>GC_TTTGCAG | 6 | FALSE | FALSE |
| 140_v.txt | 32581 | 32581-SNV-T | SNV | 1 | T | G | 41 | 20555 | 0.19946485 | 34.3170732 | rnr1Y285A_p | TRUE | CAN | JSY2423 | 4 | rnr1Y285A_p | GTG | T>G | G/C+1 | A/T-1 | TA>GC | TA>GC | GAAATGTGA | TGAAGGC | CAC | GTG | T | C | T>G | T>G_GTG | TA>GC_TGAAGGC | 6 | FALSE | FALSE |
| 140_v.txt | 32631 | 32631-SNV-T | SNV | 1 | T | G | 3 | 20555 | 0.01459499 | 34.3333333 | rnr1Y285A_p | TRUE | CAN | JSY2423 | 4 | rnr1Y285A_p | GTT | T>G | G/C+1 | A/T-1 | TA>GC | TA>GC | CCTTCGTTTT | TTTTTAT | CAA | GTT | T | C | T>G | T>G_GTT | TA>GC_TTTTTAT | 6 | FALSE | FALSE |
| 140_v.txt | 32746 | 32746-SNV- | SNV | 1 | A | C | 6 | 20555 | 0.02918998 | 59.0002473 | rnr1Y285A_p | TRUE | CAN | JSY2423 | 4 | rnr1Y285A_p | AAC | A>C | G/C+1 | A/T-1 | TA>GC | TA>GC | TTAGAAACC | ACCCGAT | TTG | TTG | T | G | T>G | T>G_TTG | TA>GC_ACCCGAT | 6 | FALSE | FALSE |
| 140_v.txt | 32763 | 32763-SNV- | SNV | 1 | A | C | 8 | 20555 | 0.03891997 | 43.5 | rnr1Y285A_p | TRUE | CAN | JSY2423 | 4 | rnr1Y285A_p | AAC | A>C | G/C+1 | A/T-1 | TA>GC | TA>GC | GCTAAAACTT | ACTTTGA | TTG | TTG | T | G | T>G | T>G_TTG | TA>GC_ACTTTGA | 6 | FALSE | FALSE |
| 140_v.txt | 32842 | 32842-SNV- | SNV | 1 | A | C | 3 | 20555 | 0.01459499 | 42.3341964 | rnr1Y285A_p | TRUE | CAN | JSY2423 | 4 | rnr1Y285A_p | CAA | A>C | G/C+1 | A/T-1 | TA>GC | TA>GC | TTACCCAAA | AAAAAAT | GTT | GTT | T | G | T>G | T>G_GTT | TA>GC_AAAAAAT | 6 | FALSE | FALSE |
| 140_v.txt | 32884 | 32884-SNV- | SNV | 1 | A | C | 7 | 28487 | 0.02457261 | 33.7142857 | rnr1Y285A_p | TRUE | CAN | JSY2423 | 4 | rnr1Y285A_p | CAA | A>C | G/C+1 | A/T-1 | TA>GC | TA>GC | ACGTCCAAA | AAAATTG | GTT | GTT | T | G | T>G | T>G_GTT | TA>GC_AAAATTG | 6 | FALSE | FALSE |
| 140_v.txt | 32913 | 32913-SNV- | SNV | 1 | A | C | 22 | 28487 | 0.07722821 | 34.3181818 | rnr1Y285A_p | TRUE | CAN | JSY2423 | 4 | rnr1Y285A_p | AAG | A>C | G/C+1 | A/T-1 | TA>GC | TA>GC | ACACTAAGTT | AGTTCCA | TTC | TTC | T | G | T>G | T>G_TTC | TA>GC_AGTTCCA | 6 | FALSE | FALSE |
| 140_v.txt | 32929 | 32929-SNV-T | SNV | 1 | T | G | 9 | 28487 | 0.03159336 | 51.8888889 | rnr1Y285A_p | TRUE | CAN | JSY2423 | 4 | rnr1Y285A_p | GTG | T>G | G/C+1 | A/T-1 | TA>GC | TA>GC | CAAAAGTGA | TGATTGC | CAC | GTG | T | C | T>G | T>G_GTG | TA>GC_TGATTGC | 6 | FALSE | FALSE |
| 140_v.txt | 33166 | 33166-SNV-T | SNV | 1 | T | G | 12 | 28484 | 0.04212892 | 37.75 | rnr1Y285A_p | TRUE | CAN | JSY2423 | 4 | rnr1Y285A_p | GTA | T>G | G/C+1 | A/T-1 | TA>GC | TA>GC | CAATAGTAC | TACCACC | CAT | GTA | T | C | T>G | T>G_GTA | TA>GC_TACCACC | 6 | FALSE | FALSE |
| 140_v.txt | 33476 | 33476-SNV-T | SNV | 1 | T | G | 4 | 21081 | 0.01897443 | 29.2514092 | rnr1Y285A_p | TRUE | CAN | JSY2423 | 4 | rnr1Y285A_p | CTT | T>G | G/C+1 | A/T-1 | TA>GC | TA>GC | TATGCCTTTT | TTTTTTT | GAA | CTT | T | C | T>G | T>G_CTT | TA>GC_TTTTTTT | 6 | FALSE | FALSE |
| 140_v.txt | 32081 | 32081-SNV- | SNV | 1 | A | G | 9 | 20031 | 0.04493036 | 48.0003138 | rnr1Y285A_p | TRUE | CAN | JSY2423 | 4 | rnr1Y285A_p | CAA | A>G | G/C+1 | A/T-1 | TA>CG | TA>CG | CCATGCAAA | AAAAAAG | GTT | GTT | T | C | T>C | T>C_GTT | TA>CG_AAAAAAG | 6 | FALSE | FALSE |
| 140_v.txt | 32787 | 32787-SNV-T | SNV | 1 | T | C | 5 | 20555 | 0.02432498 | 42.7999095 | rnr1Y285A_p | TRUE | CAN | JSY2423 | 4 | rnr1Y285A_p | CTC | T>C | G/C+1 | A/T-1 | TA>CG | TA>CG | CAGAACTCG | TCGAATT | GAG | CTC | T | G | T>C | T>C_CTC | TA>CG_TCGAATT | 6 | FALSE | FALSE |
| 140_v.txt | 33089 | 33089-SNV- | SNV | 1 | A | G | 4 | 28487 | 0.01404149 | 46.9999654 | rnr1Y285A_p | TRUE | CAN | JSY2423 | 4 | rnr1Y285A_p | AAT | A>G | G/C+1 | A/T-1 | TA>CG | TA>CG | AAATAAATAT | ATATGAT | TTA | TTA | T | C | T>C | T>C_TTA | TA>CG_ATATGAT | 6 | FALSE | FALSE |
| 140_v.txt | 33476 | 33476-SNV-T | SNV | 1 | T | C | 31 | 21081 | 0.14705185 | 34.4516 | rnr1Y285A_p | TRUE | CAN | JSY2423 | 4 | rnr1Y285A_p | CTT | T>C | G/C+1 | A/T-1 | TA>CG | TA>CG | TATGCCTTTT | TTTTTTT | GAA | CTT | T | G | T>C | T>C_CTT | TA>CG_TTTTTTT | 6 | FALSE | FALSE |
| 140_v.txt | 31663 | 31663-SNV- | SNV | 1 | A | T | 42 | 9956 | 0.42185617 | 19.4285714 | rnr1Y285A_p | TRUE | CAN | JSY2423 | 4 | rnr1Y285A_p | GAA | A>T | A/T+1 | A/T-1 | TA>AT | TA>AT | GCGTGGAAA | AAATGTG | CTT | CTT | T | A | T>A | T>A_CTT | TA>AT_AAATGTG | 6 | FALSE | FALSE |
| 140_v.txt | 32496 | 32496-SNV- | SNV | 1 | A | T | 26 | 28383 | 0.09160413 | 43.4615385 | rnr1Y285A_p | TRUE | CAN | JSY2423 | 4 | rnr1Y285A_p | AAC | A>T | A/T+1 | A/T-1 | TA>AT | TA>AT | AAAACAACT | ACTTTTT | TTG | TTG | T | A | T>A | T>A_TTG | TA>AT_ACTTTTT | 6 | FALSE | FALSE |
| 140_v.txt | 32755 | 32755-SNV-T | SNV | 1 | T | A | 16 | 20555 | 0.07783994 | 57.1875 | rnr1Y285A_p | TRUE | CAN | JSY2423 | 4 | rnr1Y285A_p | ATG | T>A | A/T+1 | A/T-1 | TA>AT | TA>AT | CGATAATGG | TGGCTAA | TAC | ATG | T | T | T>A | T>A_ATG | TA>AT_TGGCTAA | 6 | FALSE | FALSE |
| 140_v.txt | 32787 | 32787-SNV-T | SNV | 1 | T | A | 55 | 20555 | 0.2675748 | 46.0354019 | rnr1Y285A_p | TRUE | CAN | JSY2423 | 4 | rnr1Y285A_p | CTC | T>A | A/T+1 | A/T-1 | TA>AT | TA>AT | CAGAACTCG | TCGAATT | GAG | CTC | T | T | T>A | T>A_CTC | TA>AT_TCGAATT | 6 | FALSE | FALSE |
| 140_v.txt | 32948 | 32948-SNV- | SNV | 1 | A | T | 17 | 28487 | 0.05967634 | 63.4117647 | rnr1Y285A_p | TRUE | CAN | JSY2423 | 4 | rnr1Y285A_p | AAT | A>T | A/T+1 | A/T-1 | TA>AT | TA>AT | AAACCAATA | ATACATG | TTA | TTA | T | A | T>A | T>A_TTA | TA>AT_ATACATG | 6 | FALSE | FALSE |
| 140_v.txt | 33089 | 33089-SNV- | SNV | 1 | A | T | 2 | 28487 | 7.02E-03 | 50.9997844 | rnr1Y285A_p | TRUE | CAN | JSY2423 | 4 | rnr1Y285A_p | AAT | A>T | A/T+1 | A/T-1 | TA>AT | TA>AT | AAATAAATAT | ATATGAT | TTA | TTA | T | A | T>A | T>A_TTA | TA>AT_ATATGAT | 6 | FALSE | FALSE |
| 140_v.txt | 33090 | 33090-SNV-T | SNV | 1 | T | A | 2 | 28487 | 7.02E-03 | 24.4997928 | rnr1Y285A_p | TRUE | CAN | JSY2423 | 4 | rnr1Y285A_p | ATA | T>A | A/T+1 | A/T-1 | TA>AT | TA>AT | AATAAATAT | TATGATA | TAT | ATA | T | T | T>A | T>A_ATA | TA>AT_TATGATA | 6 | FALSE | FALSE |
| 140_v.txt | 31650 | 31650-SNV- | SNV | 1 | C | T | 17 | 9956 | 0.17075131 | 37.0588235 | rnr1Y285A_p | TRUE | CAN | JSY2423 | 4 | rnr1Y285A_p | GCG | C>T | A/T+1 | G/C-1 | CG>TA | CG>TA | AGAATGCGA | CGAAATG | CGC | GCG | C | A | C>T | C>T_GCG | CG>TA_CGAAATG | 6 | FALSE | FALSE |
| 140_v.txt | 31844 | 31844-SNV- | SNV | 1 | C | T | 14 | 9956 | 0.14061872 | 48.4285714 | rnr1Y285A_p | TRUE | CAN | JSY2423 | 4 | rnr1Y285A_p | TCC | C>T | A/T+1 | G/C-1 | CG>TA | CG>TA | TAAGATCCA | CCAAACA | AGG | TCC | C | A | C>T | C>T_TCC | CG>TA_CCAAACA | 6 | FALSE | FALSE |
| 140_v.txt | 31947 | 31947-SNV- | SNV | 1 | G | A | 15 | 20022 | 0.07491759 | 37 | rnr1Y285A_p | TRUE | CAN | JSY2423 | 4 | rnr1Y285A_p | CGT | G>A | A/T+1 | G/C-1 | CG>TA | CG>TA | ATGATCGTC | GTCATAA | GCA | GCA | C | T | C>T | C>T_GCA | CG>TA_GTCATAA | 6 | FALSE | FALSE |
| 140_v.txt | 31960 | 31960-SNV- | SNV | 1 | C | T | 29 | 20031 | 0.1447756 | 33.5862069 | rnr1Y285A_p | TRUE | CAN | JSY2423 | 4 | rnr1Y285A_p | GCC | C>T | A/T+1 | G/C-1 | CG>TA | CG>TA | ATGTGGCCG | CCGCATA | CGG | GCC | C | A | C>T | C>T_GCC | CG>TA_CCGCATA | 6 | FALSE | FALSE |
| 140_v.txt | 31962 | 31962-SNV- | SNV | 1 | G | A | 34 | 20031 | 0.16973691 | 37.6176471 | rnr1Y285A_p | TRUE | CAN | JSY2423 | 4 | rnr1Y285A_p | CGC | G>A | A/T+1 | G/C-1 | CG>TA | CG>TA | GTGGCCGCAT | GCATAAT | GCG | GCG | C | T | C>T | C>T_GCG | CG>TA_GCATAAT | 6 | FALSE | FALSE |
| 140_v.txt | 31972 | 31972-SNV- | SNV | 1 | C | T | 19 | 20031 | 0.09485298 | 35 | rnr1Y285A_p | TRUE | CAN | JSY2423 | 4 | rnr1Y285A_p | GCC | C>T | A/T+1 | G/C-1 | CG>TA | CG>TA | AATAAGCCA | CCAAGCC | CGG | GCC | C | A | C>T | C>T_GCC | CG>TA_CCAAGCC | 6 | FALSE | FALSE |

| | | | | | | | | | | | | | | | | | | | | | | | | | | | | | | | | | | |
|---|---|---|---|---|---|---|---|---|---|---|---|---|---|---|---|---|---|---|---|---|---|---|---|---|---|---|---|---|---|---|---|---|---|---|
| 140_v.txt | 31981 | 31981-SNV- | SNV | 1 | G | A | 27 | 20031 | 0.13479107 | 52.5555556 | rnr1Y285A_p | TRUE | CAN | JSY2423 | 4 | rnr1Y285A_p | GGC | G>A | A/T+1 | G/C-1 | CG>TA | CG>TA | AGCCGGGCA | GCATTAA | CCG | CCG | C | T | C>T | C>T_CCG | CG>TA_GCATTAA | 6 | FALSE | FALSE |
| 140_v.txt | 32037 | 32037-SNV- | SNV | 1 | G | A | 20 | 20031 | 0.09984524 | 58.2 | rnr1Y285A_p | TRUE | CAN | JSY2423 | 4 | rnr1Y285A_p | AGC | G>A | A/T+1 | G/C-1 | CG>TA | CG>TA | TTCAAAGCT | GCTTGCA | TCG | TCG | C | T | C>T | C>T_TCG | CG>TA_GCTTGCA | 6 | FALSE | FALSE |
| 140_v.txt | 32041 | 32041-SNV- | SNV | 1 | G | A | 18 | 20031 | 0.08986072 | 58.8333333 | rnr1Y285A_p | TRUE | CAN | JSY2423 | 4 | rnr1Y285A_p | TGC | G>A | A/T+1 | G/C-1 | CG>TA | CG>TA | AAGCTTGCA | GCATAAA | ACG | ACG | C | T | C>T | C>T_ACG | CG>TA_GCATAAA | 6 | FALSE | FALSE |
| 140_v.txt | 32057 | 32057-SNV- | SNV | 1 | C | T | 19 | 20031 | 0.09485298 | 54.8421053 | rnr1Y285A_p | TRUE | CAN | JSY2423 | 4 | rnr1Y285A_p | GCG | C>T | A/T+1 | G/C-1 | CG>TA | CG>TA | GATGTGCGA | CGAGATT | CGC | GCG | C | A | C>T | C>T_GCG | CG>TA_CGAGATT | 6 | FALSE | FALSE |
| 140_v.txt | 32075 | 32075-SNV- | SNV | 1 | C | T | 97 | 20031 | 0.48424941 | 61.6597938 | rnr1Y285A_p | TRUE | CAN | JSY2423 | 4 | rnr1Y285A_p | ACC | C>T | A/T+1 | G/C-1 | CG>TA | CG>TA | AAATAACCA | CCATGCA | TGG | ACC | C | A | C>T | C>T_ACC | CG>TA_CCATGCA | 6 | FALSE | FALSE |
| 140_v.txt | 32088 | 32088-SNV- | SNV | 1 | C | T | 141 | 20031 | 0.70390894 | 61.070922 | rnr1Y285A_p | TRUE | CAN | JSY2423 | 4 | rnr1Y285A_p | GCC | C>T | A/T+1 | G/C-1 | CG>TA | CG>TA | AAAAAGCCT | CCTGCAA | CGG | GCC | C | A | C>T | C>T_GCC | CG>TA_CCTGCAA | 6 | FALSE | FALSE |
| 140_v.txt | 32092 | 32092-SNV- | SNV | 1 | C | T | 19 | 20031 | 0.09485298 | 64 | rnr1Y285A_p | TRUE | CAN | JSY2423 | 4 | rnr1Y285A_p | GCA | C>T | A/T+1 | G/C-1 | CG>TA | CG>TA | AGCCTGCAA | CAACACC | CGT | GCA | C | A | C>T | C>T_GCA | CG>TA_CAACACC | 6 | FALSE | FALSE |
| 140_v.txt | 32114 | 32114-SNV- | SNV | 1 | C | T | 59 | 20031 | 0.29454346 | 59.8135593 | rnr1Y285A_p | TRUE | CAN | JSY2423 | 4 | rnr1Y285A_p | GCC | C>T | A/T+1 | G/C-1 | CG>TA | CG>TA | TAATAGCCA | CCATTCG | CGG | GCC | C | A | C>T | C>T_GCC | CG>TA_CCATTCG | 6 | FALSE | FALSE |
| 140_v.txt | 32165 | 32165-SNV- | SNV | 1 | G | A | 22 | 20031 | 0.10982976 | 48.7272727 | rnr1Y285A_p | TRUE | CAN | JSY2423 | 4 | rnr1Y285A_p | CGC | G>A | A/T+1 | G/C-1 | CG>TA | CG>TA | CAAAGCGCC | GCCAAAT | GCG | GCG | C | T | C>T | C>T_GCG | CG>TA_GCCAAAT | 6 | FALSE | FALSE |
| 140_v.txt | 32183 | 32183-SNV- | SNV | 1 | G | A | 10 | 20031 | 0.04992262 | 42.9009467 | rnr1Y285A_p | TRUE | CAN | JSY2423 | 4 | rnr1Y285A_p | CGA | G>A | A/T+1 | G/C-1 | CG>TA | CG>TA | AGTAACGAA | GAAAACT | GCT | GCT | C | T | C>T | C>T_GCT | CG>TA_GAAAACT | 6 | FALSE | FALSE |
| 140_v.txt | 32259 | 32259-SNV- | SNV | 1 | C | T | 33 | 28382 | 0.11627088 | 32.4848485 | rnr1Y285A_p | TRUE | CAN | JSY2423 | 4 | rnr1Y285A_p | ACC | C>T | A/T+1 | G/C-1 | CG>TA | CG>TA | GATAGACCA | CCAAATA | TGG | ACC | C | A | C>T | C>T_ACC | CG>TA_CCAAATA | 6 | FALSE | FALSE |
| 140_v.txt | 32271 | 32271-SNV- | SNV | 1 | C | T | 50 | 28383 | 0.17616179 | 34.08 | rnr1Y285A_p | TRUE | CAN | JSY2423 | 4 | rnr1Y285A_p | ACG | C>T | A/T+1 | G/C-1 | CG>TA | CG>TA | AAAATACGG | CGGGAAC | TGC | ACG | C | A | C>T | C>T_ACG | CG>TA_CGGGAAC | 6 | FALSE | FALSE |
| 140_v.txt | 32299 | 32299-SNV- | SNV | 1 | C | T | 50 | 28383 | 0.17616179 | 54.64 | rnr1Y285A_p | TRUE | CAN | JSY2423 | 4 | rnr1Y285A_p | GCG | C>T | A/T+1 | G/C-1 | CG>TA | CG>TA | AATTTGCGG | CGGCAGA | CGC | GCG | C | A | C>T | C>T_GCG | CG>TA_CGGCAGA | 6 | FALSE | FALSE |
| 140_v.txt | 32466 | 32466-SNV- | SNV | 1 | C | T | 44 | 28383 | 0.15502237 | 45.3181818 | rnr1Y285A_p | TRUE | CAN | JSY2423 | 4 | rnr1Y285A_p | GCC | C>T | A/T+1 | G/C-1 | CG>TA | CG>TA | AGAGAGCCA | CCAATGT | CGG | GCC | C | A | C>T | C>T_GCC | CG>TA_CCAATGT | 6 | FALSE | FALSE |
| 140_v.txt | 32487 | 32487-SNV- | SNV | 1 | C | T | 21 | 28383 | 0.07398795 | 47.7595505 | rnr1Y285A_p | TRUE | CAN | JSY2423 | 4 | rnr1Y285A_p | ACG | C>T | A/T+1 | G/C-1 | CG>TA | CG>TA | AAGATACGG | CGGAAAA | TGC | ACG | C | A | C>T | C>T_ACG | CG>TA_CGGAAAA | 6 | FALSE | FALSE |
| 140_v.txt | 32488 | 32488-SNV- | SNV | 1 | G | A | 33 | 28383 | 0.11626678 | 44.1526146 | rnr1Y285A_p | TRUE | CAN | JSY2423 | 4 | rnr1Y285A_p | CGG | G>A | A/T+1 | G/C-1 | CG>TA | CG>TA | AGATACGGA | GGAAAAC | GCC | GCC | C | T | C>T | C>T_GCC | CG>TA_GGAAAAC | 6 | FALSE | FALSE |
| 140_v.txt | 32508 | 32508-SNV- | SNV | 1 | G | A | 18 | 28383 | 0.06341824 | 40.0555556 | rnr1Y285A_p | TRUE | CAN | JSY2423 | 4 | rnr1Y285A_p | GGC | G>A | A/T+1 | G/C-1 | CG>TA | CG>TA | TTGATGGCT | GCTCTTG | CCG | CCG | C | T | C>T | C>T_CCG | CG>TA_GCTCTTG | 6 | FALSE | FALSE |
| 140_v.txt | 32535 | 32535-SNV- | SNV | 1 | G | A | 37 | 28382 | 0.13036432 | 36.6486989 | rnr1Y285A_p | TRUE | CAN | JSY2423 | 4 | rnr1Y285A_p | TGC | G>A | A/T+1 | G/C-1 | CG>TA | CG>TA | GGGTTTGCA | GCAGCTT | ACG | ACG | C | T | C>T | C>T_ACG | CG>TA_GCAGCTT | 6 | FALSE | FALSE |
| 140_v.txt | 32536 | 32536-SNV- | SNV | 1 | C | T | 37 | 28382 | 0.13036432 | 36.9722784 | rnr1Y285A_p | TRUE | CAN | JSY2423 | 4 | rnr1Y285A_p | GCA | C>T | A/T+1 | G/C-1 | CG>TA | CG>TA | GGTTTGCAG | CAGCTTC | CGT | GCA | C | A | C>T | C>T_GCA | CG>TA_CAGCTTC | 6 | FALSE | FALSE |
| 140_v.txt | 32566 | 32566-SNV- | SNV | 1 | C | T | 26 | 20551 | 0.12651453 | 33.9230769 | rnr1Y285A_p | TRUE | CAN | JSY2423 | 4 | rnr1Y285A_p | TCA | C>T | A/T+1 | G/C-1 | CG>TA | CG>TA | CTAGTTCAG | CAGTACC | AGT | TCA | C | A | C>T | C>T_TCA | CG>TA_CAGTACC | 6 | FALSE | FALSE |
| 140_v.txt | 32571 | 32571-SNV- | SNV | 1 | C | T | 31 | 20555 | 0.15081489 | 37.3870968 | rnr1Y285A_p | TRUE | CAN | JSY2423 | 4 | rnr1Y285A_p | ACC | C>T | A/T+1 | G/C-1 | CG>TA | CG>TA | TCAGTACCT | CCTTGAA | TGG | ACC | C | A | C>T | C>T_ACC | CG>TA_CCTTGAA | 6 | FALSE | FALSE |
| 140_v.txt | 32590 | 32590-SNV- | SNV | 1 | C | T | 39 | 20555 | 0.18973486 | 35.4615385 | rnr1Y285A_p | TRUE | CAN | JSY2423 | 4 | rnr1Y285A_p | GCG | C>T | A/T+1 | G/C-1 | CG>TA | CG>TA | AGGCAGCGT | CGTTAAT | CGC | GCG | C | A | C>T | C>T_GCG | CG>TA_CGTTAAT | 6 | FALSE | FALSE |
| 140_v.txt | 32658 | 32658-SNV- | SNV | 1 | C | T | 13 | 20555 | 0.06324495 | 56.3038755 | rnr1Y285A_p | TRUE | CAN | JSY2423 | 4 | rnr1Y285A_p | ACC | C>T | A/T+1 | G/C-1 | CG>TA | CG>TA | CCTGGACCC | CCCCAGG | TGG | ACC | C | A | C>T | C>T_ACC | CG>TA_CCCCAGG | 6 | FALSE | FALSE |
| 140_v.txt | 32700 | 32700-SNV- | SNV | 1 | C | T | 32 | 20555 | 0.15567988 | 61.625 | rnr1Y285A_p | TRUE | CAN | JSY2423 | 4 | rnr1Y285A_p | GCC | C>T | A/T+1 | G/C-1 | CG>TA | CG>TA | ACTGGGCCG | CCGGTAA | CGG | GCC | C | A | C>T | C>T_GCC | CG>TA_CCGGTAA | 6 | FALSE | FALSE |
| 140_v.txt | 32702 | 32702-SNV- | SNV | 1 | G | A | 16 | 20555 | 0.07783994 | 51.3125 | rnr1Y285A_p | TRUE | CAN | JSY2423 | 4 | rnr1Y285A_p | CGG | G>A | A/T+1 | G/C-1 | CG>TA | CG>TA | TGGGCCGGT | GGTAACC | GCC | GCC | C | T | C>T | C>T_GCC | CG>TA_GGTAACC | 6 | FALSE | FALSE |
| 140_v.txt | 32747 | 32747-SNV- | SNV | 1 | C | T | 2 | 20555 | 9.73E-03 | 36.9900522 | rnr1Y285A_p | TRUE | CAN | JSY2423 | 4 | rnr1Y285A_p | ACC | C>T | A/T+1 | G/C-1 | CG>TA | CG>TA | TAGAAACCC | CCCGATA | TGG | ACC | C | A | C>T | C>T_ACC | CG>TA_CCCGATA | 6 | FALSE | FALSE |
| 140_v.txt | 32758 | 32758-SNV- | SNV | 1 | C | T | 15 | 20555 | 0.07297495 | 54 | rnr1Y285A_p | TRUE | CAN | JSY2423 | 4 | rnr1Y285A_p | GCT | C>T | A/T+1 | G/C-1 | CG>TA | CG>TA | TAATGGCTA | CTAAAAC | CGA | GCT | C | A | C>T | C>T_GCT | CG>TA_CTAAAAC | 6 | FALSE | FALSE |
| 140_v.txt | 32776 | 32776-SNV- | SNV | 1 | C | T | 17 | 20555 | 0.08270494 | 51.7058824 | rnr1Y285A_p | TRUE | CAN | JSY2423 | 4 | rnr1Y285A_p | GCG | C>T | A/T+1 | G/C-1 | CG>TA | CG>TA | TGGAAGCGA | CGACCCA | CGC | GCG | C | A | C>T | C>T_GCG | CG>TA_CGACCCA | 6 | FALSE | FALSE |
| 140_v.txt | 32797 | 32797-SNV- | SNV | 1 | C | T | 38 | 20555 | 0.18486986 | 47.3180631 | rnr1Y285A_p | TRUE | CAN | JSY2423 | 4 | rnr1Y285A_p | CCG | C>T | A/T+1 | G/C-1 | CG>TA | CG>TA | ATTCACCGTA | CGTAATA | GGC | CCG | C | A | C>T | C>T_CCG | CG>TA_CGTAATA | 6 | FALSE | FALSE |
| 140_v.txt | 32798 | 32798-SNV- | SNV | 1 | G | A | 27 | 20555 | 0.1313549 | 42.9254566 | rnr1Y285A_p | TRUE | CAN | JSY2423 | 4 | rnr1Y285A_p | CGT | G>A | A/T+1 | G/C-1 | CG>TA | CG>TA | TTCACCGTA | GTAATAT | GCA | GCA | C | T | C>T | C>T_GCA | CG>TA_GTAATAT | 6 | FALSE | FALSE |
| 140_v.txt | 32855 | 32855-SNV- | SNV | 1 | C | T | 33 | 20556 | 0.16053707 | 34.1818182 | rnr1Y285A_p | TRUE | CAN | JSY2423 | 4 | rnr1Y285A_p | TCC | C>T | A/T+1 | G/C-1 | CG>TA | CG>TA | ACTAATCCAT | CCATGCC | AGG | TCC | C | A | C>T | C>T_TCC | CG>TA_CCATGCC | 6 | FALSE | FALSE |
| 140_v.txt | 32860 | 32860-SNV- | SNV | 1 | C | T | 23 | 20554 | 0.11190036 | 37.3913044 | rnr1Y285A_p | TRUE | CAN | JSY2423 | 4 | rnr1Y285A_p | GCC | C>T | A/T+1 | G/C-1 | CG>TA | CG>TA | TCCATGCCG | CCGCCAG | CGG | GCC | C | A | C>T | C>T_GCC | CG>TA_CCGCCAG | 6 | FALSE | FALSE |
| 140_v.txt | 32862 | 32862-SNV- | SNV | 1 | G | A | 17 | 20554 | 0.08270896 | 36.2352941 | rnr1Y285A_p | TRUE | CAN | JSY2423 | 4 | rnr1Y285A_p | CGC | G>A | A/T+1 | G/C-1 | CG>TA | CG>TA | CATGCCGCC | GCCAGTG | GCG | GCG | C | T | C>T | C>T_GCG | CG>TA_GCCAGTG | 6 | FALSE | FALSE |
| 140_v.txt | 32900 | 32900-SNV- | SNV | 1 | G | A | 34 | 28487 | 0.11935269 | 34.6589447 | rnr1Y285A_p | TRUE | CAN | JSY2423 | 4 | rnr1Y285A_p | GGC | G>A | A/T+1 | G/C-1 | CG>TA | CG>TA | GACTTGGCC | GCCAACT | CCG | CCG | C | T | C>T | C>T_CCG | CG>TA_GCCAACT | 6 | FALSE | FALSE |
| 140_v.txt | 32901 | 32901-SNV- | SNV | 1 | C | T | 57 | 28487 | 0.20009127 | 34.5265474 | rnr1Y285A_p | TRUE | CAN | JSY2423 | 4 | rnr1Y285A_p | GCC | C>T | A/T+1 | G/C-1 | CG>TA | CG>TA | ACTTGGCCA | CCAACTA | CGG | GCC | C | A | C>T | C>T_GCC | CG>TA_CCAACTA | 6 | FALSE | FALSE |
| 140_v.txt | 32935 | 32935-SNV- | SNV | 1 | C | T | 39 | 28487 | 0.13690455 | 39.3846154 | rnr1Y285A_p | TRUE | CAN | JSY2423 | 4 | rnr1Y285A_p | GCC | C>T | A/T+1 | G/C-1 | CG>TA | CG>TA | TGATTGCCC | CCCAAGA | CGG | GCC | C | A | C>T | C>T_GCC | CG>TA_CCCAAGA | 6 | FALSE | FALSE |
| 140_v.txt | 32945 | 32945-SNV- | SNV | 1 | C | T | 93 | 28487 | 0.3264647 | 56.8817204 | rnr1Y285A_p | TRUE | CAN | JSY2423 | 4 | rnr1Y285A_p | ACC | C>T | A/T+1 | G/C-1 | CG>TA | CG>TA | AGAAAACCA | CCAATAC | TGG | ACC | C | A | C>T | C>T_ACC | CG>TA_CCAATAC | 6 | FALSE | FALSE |
| 140_v.txt | 32958 | 32958-SNV- | SNV | 1 | C | T | 42 | 28487 | 0.14743567 | 44.8333333 | rnr1Y285A_p | TRUE | CAN | JSY2423 | 4 | rnr1Y285A_p | ACC | C>T | A/T+1 | G/C-1 | CG>TA | CG>TA | ATGTAACCA | CCATTGG | TGG | ACC | C | A | C>T | C>T_ACC | CG>TA_CCATTGG | 6 | FALSE | FALSE |
| 140_v.txt | 32967 | 32967-SNV- | SNV | 1 | G | A | 73 | 28487 | 0.25625724 | 59.7123288 | rnr1Y285A_p | TRUE | CAN | JSY2423 | 4 | rnr1Y285A_p | CGC | G>A | A/T+1 | G/C-1 | CG>TA | CG>TA | TTGGCCGCA | GCACCAA | GCG | GCG | C | T | C>T | C>T_GCG | CG>TA_GCACCAA | 6 | FALSE | FALSE |
| 140_v.txt | 32970 | 32970-SNV- | SNV | 1 | C | T | 26 | 28487 | 0.0912697 | 57.9230769 | rnr1Y285A_p | TRUE | CAN | JSY2423 | 4 | rnr1Y285A_p | ACC | C>T | A/T+1 | G/C-1 | CG>TA | CG>TA | GCCGCACCA | CCAAATG | TGG | ACC | C | A | C>T | C>T_ACC | CG>TA_CCAAATG | 6 | FALSE | FALSE |
| 140_v.txt | 33056 | 33056-SNV- | SNV | 1 | C | T | 40 | 28487 | 0.14041493 | 58.575 | rnr1Y285A_p | TRUE | CAN | JSY2423 | 4 | rnr1Y285A_p | GCG | C>T | A/T+1 | G/C-1 | CG>TA | CG>TA | GGACTGCGT | CGTGACA | CGC | GCG | C | A | C>T | C>T_GCG | CG>TA_CGTGACA | 6 | FALSE | FALSE |
| 140_v.txt | 33069 | 33069-SNV- | SNV | 1 | G | A | 24 | 28487 | 0.08424896 | 56.5827449 | rnr1Y285A_p | TRUE | CAN | JSY2423 | 4 | rnr1Y285A_p | TGC | G>A | A/T+1 | G/C-1 | CG>TA | CG>TA | GAATATGCC | GCCAAAG | ACG | ACG | C | T | C>T | C>T_ACG | CG>TA_GCCAAAG | 6 | FALSE | FALSE |
| 140_v.txt | 33070 | 33070-SNV- | SNV | 1 | C | T | 28 | 28487 | 0.09829045 | 57.0357143 | rnr1Y285A_p | TRUE | CAN | JSY2423 | 4 | rnr1Y285A_p | GCC | C>T | A/T+1 | G/C-1 | CG>TA | CG>TA | AATATGCCA | CCAAAGA | CGG | GCC | C | A | C>T | C>T_GCC | CG>TA_CCAAAGA | 6 | FALSE | FALSE |
| 140_v.txt | 33104 | 33104-SNV- | SNV | 1 | G | A | 33 | 28487 | 0.11584231 | 37.6969697 | rnr1Y285A_p | TRUE | CAN | JSY2423 | 4 | rnr1Y285A_p | CGC | G>A | A/T+1 | G/C-1 | CG>TA | CG>TA | AAGAGCGCC | GCCCACT | GCG | GCG | C | T | C>T | C>T_GCG | CG>TA_GCCCACT | 6 | FALSE | FALSE |
| 140_v.txt | 33113 | 33113-SNV- | SNV | 1 | G | A | 21 | 28487 | 0.07371784 | 50.2380952 | rnr1Y285A_p | TRUE | CAN | JSY2423 | 4 | rnr1Y285A_p | GGC | G>A | A/T+1 | G/C-1 | CG>TA | CG>TA | CACTGGGCC | GCCGGCG | CCG | CCG | C | T | C>T | C>T_CCG | CG>TA_GCCGGCG | 6 | FALSE | FALSE |
| 140_v.txt | 33114 | 33114-SNV- | SNV | 1 | C | T | 26 | 28487 | 0.0912697 | 47 | rnr1Y285A_p | TRUE | CAN | JSY2423 | 4 | rnr1Y285A_p | GCC | C>T | A/T+1 | G/C-1 | CG>TA | CG>TA | ACTGGGCCG | CCGGCGT | CGG | GCC | C | A | C>T | C>T_GCC | CG>TA_CCGGCGT | 6 | FALSE | FALSE |
| 140_v.txt | 33118 | 33118-SNV- | SNV | 1 | C | T | 26 | 28487 | 0.0912697 | 40.2692308 | rnr1Y285A_p | TRUE | CAN | JSY2423 | 4 | rnr1Y285A_p | GCG | C>T | A/T+1 | G/C-1 | CG>TA | CG>TA | GGCCGGCGT | CGTTGGT | CGC | GCG | C | A | C>T | C>T_GCG | CG>TA_CGTTGGT | 6 | FALSE | FALSE |
| 140_v.txt | 33153 | 33153-SNV- | SNV | 1 | C | T | 57 | 28487 | 0.20009127 | 37.3333333 | rnr1Y285A_p | TRUE | CAN | JSY2423 | 4 | rnr1Y285A_p | ACC | C>T | A/T+1 | G/C-1 | CG>TA | CG>TA | AAAAGACCT | CCTGTAC | TGG | ACC | C | A | C>T | C>T_ACC | CG>TA_CCTGTAC | 6 | FALSE | FALSE |
| 140_v.txt | 33159 | 33159-SNV- | SNV | 1 | C | T | 57 | 28486 | 0.20009829 | 36.245614 | rnr1Y285A_p | TRUE | CAN | JSY2423 | 4 | rnr1Y285A_p | ACC | C>T | A/T+1 | G/C-1 | CG>TA | CG>TA | CCTGTACCA | CCAATAG | TGG | ACC | C | A | C>T | C>T_ACC | CG>TA_CCAATAG | 6 | FALSE | FALSE |
| 140_v.txt | 33168 | 33168-SNV- | SNV | 1 | C | T | 45 | 28484 | 0.15798343 | 37.3777778 | rnr1Y285A_p | TRUE | CAN | JSY2423 | 4 | rnr1Y285A_p | ACC | C>T | A/T+1 | G/C-1 | CG>TA | CG>TA | ATAGTACCA | CCACCAA | TGG | ACC | C | A | C>T | C>T_ACC | CG>TA_CCACCAA | 6 | FALSE | FALSE |
| 140_v.txt | 33229 | 33229-SNV- | SNV | 1 | G | A | 21 | 21084 | 0.09960159 | 34.8095238 | rnr1Y285A_p | TRUE | CAN | JSY2423 | 4 | rnr1Y285A_p | TGT | G>A | A/T+1 | G/C-1 | CG>TA | CG>TA | CGTTCTGTAC | GTACTTC | ACA | ACA | C | T | C>T | C>T_ACA | CG>TA_GTACTTC | 6 | FALSE | FALSE |
| 140_v.txt | 33258 | 33258-SNV- | SNV | 1 | C | T | 28 | 21084 | 0.13280213 | 44.2142857 | rnr1Y285A_p | TRUE | CAN | JSY2423 | 4 | rnr1Y285A_p | GCC | C>T | A/T+1 | G/C-1 | CG>TA | CG>TA | CCTATGCCAT | CCATCCT | CGG | GCC | C | A | C>T | C>T_GCC | CG>TA_CCATCCT | 6 | FALSE | FALSE |
| 140_v.txt | 33276 | 33276-SNV- | SNV | 1 | G | A | 32 | 21084 | 0.15177386 | 42.8125 | rnr1Y285A_p | TRUE | CAN | JSY2423 | 4 | rnr1Y285A_p | CGT | G>A | A/T+1 | G/C-1 | CG>TA | CG>TA | GAGAACGTA | GTATCCT | GCA | GCA | C | T | C>T | C>T_GCA | CG>TA_GTATCCT | 6 | FALSE | FALSE |
| 140_v.txt | 33284 | 33284-SNV- | SNV | 1 | G | A | 21 | 21084 | 0.09960159 | 50.8571429 | rnr1Y285A_p | TRUE | CAN | JSY2423 | 4 | rnr1Y285A_p | CGC | G>A | A/T+1 | G/C-1 | CG>TA | CG>TA | ATCCTCGCCA | GCCATTT | GCG | GCG | C | T | C>T | C>T_GCG | CG>TA_GCCATTT | 6 | FALSE | FALSE |
| 140_v.txt | 33309 | 33309-SNV- | SNV | 1 | C | T | 26 | 21084 | 0.12331626 | 59.0384615 | rnr1Y285A_p | TRUE | CAN | JSY2423 | 4 | rnr1Y285A_p | GCG | C>T | A/T+1 | G/C-1 | CG>TA | CG>TA | AAAGAGCGC | CGCAATG | CGC | GCG | C | A | C>T | C>T_GCG | CG>TA_CGCAATG | 6 | FALSE | FALSE |
| 140_v.txt | 33310 | 33310-SNV- | SNV | 1 | G | A | 24 | 21083 | 0.11383579 | 55.2298649 | rnr1Y285A_p | TRUE | CAN | JSY2423 | 4 | rnr1Y285A_p | CGC | G>A | A/T+1 | G/C-1 | CG>TA | CG>TA | AAGAGCGCA | GCAATGG | GCG | GCG | C | T | C>T | C>T_GCG | CG>TA_GCAATGG | 6 | FALSE | FALSE |
| 140_v.txt | 33412 | 33412-SNV- | SNV | 1 | G | A | 30 | 21084 | 0.14228799 | 48.2333333 | rnr1Y285A_p | TRUE | CAN | JSY2423 | 4 | rnr1Y285A_p | GGC | G>A | A/T+1 | G/C-1 | CG>TA | CG>TA | TGACCGGCT | GCTCATT | CCG | CCG | C | T | C>T | C>T_CCG | CG>TA_GCTCATT | 6 | FALSE | FALSE |
| 140_v.txt | 33474 | 33474-SNV- | SNV | 1 | C | T | 23 | 21083 | 0.10909263 | 36.9994131 | rnr1Y285A_p | TRUE | CAN | JSY2423 | 4 | rnr1Y285A_p | GCC | C>T | A/T+1 | G/C-1 | CG>TA | CG>TA | GCTATGCCTT | CCTTTTT | CGG | GCC | C | A | C>T | C>T_GCC | CG>TA_CCTTTTT | 6 | FALSE | FALSE |
| 140_v.txt | 31947 | 31947-SNV- | SNV | 1 | G | C | 12 | 20022 | 0.05993407 | 38 | rnr1Y285A_p | TRUE | CAN | JSY2423 | 4 | rnr1Y285A_p | CGT | G>C | G/C+1 | G/C-1 | CG>GC | CG>GC | ATGATCGTC | GTCATAA | GCA | GCA | C | G | C>G | C>G_GCA | CG>GC_GTCATAA | 6 | FALSE | FALSE |
| 140_v.txt | 32057 | 32057-SNV- | SNV | 1 | C | G | 4 | 20031 | 0.01996905 | 58.75 | rnr1Y285A_p | TRUE | CAN | JSY2423 | 4 | rnr1Y285A_p | GCG | C>G | G/C+1 | G/C-1 | CG>GC | CG>GC | GATGTGCGA | CGAGATT | CGC | GCG | C | C | C>G | C>G_GCG | CG>GC_CGAGATT | 6 | FALSE | FALSE |
| 140_v.txt | 32089 | 32089-SNV- | SNV | 1 | C | G | 3 | 20030 | 0.01497753 | 48.667471 | rnr1Y285A_p | TRUE | CAN | JSY2423 | 4 | rnr1Y285A_p | CCT | C>G | G/C+1 | G/C-1 | CG>GC | CG>GC | AAAAGCCTG | CTGCAAC | GGA | CCT | C | C | C>G | C>G_CCT | CG>GC_CTGCAAC | 6 | FALSE | FALSE |
| 140_v.txt | 32487 | 32487-SNV- | SNV | 1 | C | G | 6 | 28383 | 0.02113941 | 59.3217689 | rnr1Y285A_p | TRUE | CAN | JSY2423 | 4 | rnr1Y285A_p | ACG | C>G | G/C+1 | G/C-1 | CG>GC | CG>GC | AAGATACGG | CGGAAAA | TGC | ACG | C | C | C>G | C>G_ACG | CG>GC_CGGAAAA | 6 | FALSE | FALSE |
| 140_v.txt | 32488 | 32488-SNV- | SNV | 1 | G | C | 1 | 28383 | 3.52E-03 | 53.0007522 | rnr1Y285A_p | TRUE | CAN | JSY2423 | 4 | rnr1Y285A_p | CGG | G>C | G/C+1 | G/C-1 | CG>GC | CG>GC | AGATACGGA | GGAAAAC | GCC | GCC | C | G | C>G | C>G_GCC | CG>GC_GGAAAAC | 6 | FALSE | FALSE |
| 140_v.txt | 32571 | 32571-SNV- | SNV | 1 | C | G | 5 | 20555 | 0.02432498 | 37.2 | rnr1Y285A_p | TRUE | CAN | JSY2423 | 4 | rnr1Y285A_p | ACC | C>G | G/C+1 | G/C-1 | CG>GC | CG>GC | TCAGTACCTT | CCTTGAA | TGG | ACC | C | C | C>G | C>G_ACC | CG>GC_CCTTGAA | 6 | FALSE | FALSE |
| 140_v.txt | 32735 | 32735-SNV- | SNV | 1 | G | C | 4 | 20555 | 0.01945999 | 53.75 | rnr1Y285A_p | TRUE | CAN | JSY2423 | 4 | rnr1Y285A_p | AGT | G>C | G/C+1 | G/C-1 | CG>GC | CG>GC | AAAACAGTA | GTATATT | TCA | TCA | C | G | C>G | C>G_TCA | CG>GC_GTATATT | 6 | FALSE | FALSE |
| 140_v.txt | 32758 | 32758-SNV- | SNV | 1 | C | G | 13 | 20555 | 0.06324495 | 52.5384615 | rnr1Y285A_p | TRUE | CAN | JSY2423 | 4 | rnr1Y285A_p | GCT | C>G | G/C+1 | G/C-1 | CG>GC | CG>GC | TAATGGCTA | CTAAAAC | CGA | GCT | C | C | C>G | C>G_GCT | CG>GC_CTAAAAC | 6 | FALSE | FALSE |
| 140_v.txt | 32937 | 32937-SNV- | SNV | 1 | C | G | 36 | 28487 | 0.12637343 | 57.25 | rnr1Y285A_p | TRUE | CAN | JSY2423 | 4 | rnr1Y285A_p | CCA | C>G | G/C+1 | G/C-1 | CG>GC | CG>GC | ATTGCCCAA | CAAGAAA | GGT | CCA | C | C | C>G | C>G_CCA | CG>GC_CAAGAAA | 6 | FALSE | FALSE |
| 140_v.txt | 33070 | 33070-SNV- | SNV | 1 | C | G | 2 | 28487 | 7.02E-03 | 64 | rnr1Y285A_p | TRUE | CAN | JSY2423 | 4 | rnr1Y285A_p | GCC | C>G | G/C+1 | G/C-1 | CG>GC | CG>GC | AATATGCCA | CCAAAGA | CGG | GCC | C | C | C>G | C>G_GCC | CG>GC_CCAAAGA | 6 | FALSE | FALSE |
| 140_v.txt | 33078 | 33078-SNV- | SNV | 1 | C | G | 3 | 28487 | 0.01053112 | 64 | rnr1Y285A_p | TRUE | CAN | JSY2423 | 4 | rnr1Y285A_p | ACC | C>G | G/C+1 | G/C-1 | CG>GC | CG>GC | AAAGAACCC | CCCATAA | TGG | ACC | C | C | C>G | C>G_ACC | CG>GC_CCCATAA | 6 | FALSE | FALSE |
| 140_v.txt | 33114 | 33114-SNV- | SNV | 1 | C | G | 4 | 28487 | 0.01404149 | 43.75 | rnr1Y285A_p | TRUE | CAN | JSY2423 | 4 | rnr1Y285A_p | GCC | C>G | G/C+1 | G/C-1 | CG>GC | CG>GC | ACTGGGCCG | CCGGCGT | CGG | GCC | C | C | C>G | C>G_GCC | CG>GC_CCGGCGT | 6 | FALSE | FALSE |
| 140_v.txt | 33309 | 33309-SNV- | SNV | 1 | C | G | 2 | 21084 | 9.49E-03 | 64 | rnr1Y285A_p | TRUE | CAN | JSY2423 | 4 | rnr1Y285A_p | GCG | C>G | G/C+1 | G/C-1 | CG>GC | CG>GC | AAAGAGCGC | CGCAATG | CGC | GCG | C | C | C>G | C>G_GCG | CG>GC_CGCAATG | 6 | FALSE | FALSE |
| 140_v.txt | 31979 | 31979-SNV- | SNV | 1 | G | T | 11 | 20031 | 0.05491488 | 22.2728929 | rnr1Y285A_p | TRUE | CAN | JSY2423 | 4 | rnr1Y285A_p | CGG | G>T | A/T+1 | G/C-1 | CG>AT | CG>AT | CAAGCCGGG | GGGCATT | GCC | GCC | C | A | C>A | C>A_GCC | CG>AT_GGGCATT | 6 | FALSE | FALSE |
| 140_v.txt | 31980 | 31980-SNV- | SNV | 1 | G | T | 12 | 20031 | 0.05990714 | 45.7479107 | rnr1Y285A_p | TRUE | CAN | JSY2423 | 4 | rnr1Y285A_p | GGG | G>T | A/T+1 | G/C-1 | CG>AT | CG>AT | AAGCCGGGC | GGCATTA | CCC | CCC | C | A | C>A | C>A_CCC | CG>AT_GGCATTA | 6 | FALSE | FALSE |
| 140_v.txt | 31981 | 31981-SNV- | SNV | 1 | G | T | 27 | 20031 | 0.13479107 | 29.1851852 | rnr1Y285A_p | TRUE | CAN | JSY2423 | 4 | rnr1Y285A_p | GGC | G>T | A/T+1 | G/C-1 | CG>AT | CG>AT | AGCCGGGCA | GCATTAA | CCG | CCG | C | A | C>A | C>A_CCG | CG>AT_GCATTAA | 6 | FALSE | FALSE |
| 140_v.txt | 32027 | 32027-SNV- | SNV | 1 | G | T | 414 | 20031 | 2.06679647 | 60.1038647 | rnr1Y285A_p | TRUE | CAN | JSY2423 | 4 | rnr1Y285A_p | GGT | G>T | A/T+1 | G/C-1 | CG>AT | CG>AT | GCCACGGTA | GTATTTC | CCA | CCA | C | A | C>A | C>A_CCA | CG>AT_GTATTTC | 6 | FALSE | FALSE |
| 140_v.txt | 32075 | 32075-SNV- | SNV | 1 | C | A | 7 | 20031 | 0.03494583 | 48.8571429 | rnr1Y285A_p | TRUE | CAN | JSY2423 | 4 | rnr1Y285A_p | ACC | C>A | A/T+1 | G/C-1 | CG>AT | CG>AT | AAATAACCA | CCATGCA | TGG | ACC | C | T | C>A | C>A_ACC | CG>AT_CCATGCA | 6 | FALSE | FALSE |
| 140_v.txt | 32088 | 32088-SNV- | SNV | 1 | C | A | 12 | 20031 | 0.05990714 | 58.9166667 | rnr1Y285A_p | TRUE | CAN | JSY2423 | 4 | rnr1Y285A_p | GCC | C>A | A/T+1 | G/C-1 | CG>AT | CG>AT | AAAAAGCCT | CCTGCAA | CGG | GCC | C | T | C>A | C>A_GCC | CG>AT_CCTGCAA | 6 | FALSE | FALSE |
| 140_v.txt | 32089 | 32089-SNV- | SNV | 1 | C | A | 15 | 20030 | 0.07488767 | 60.0670173 | rnr1Y285A_p | TRUE | CAN | JSY2423 | 4 | rnr1Y285A_p | CCT | C>A | A/T+1 | G/C-1 | CG>AT | CG>AT | AAAAGCCTG | CTGCAAC | GGA | CCT | C | T | C>A | C>A_CCT | CG>AT_CTGCAAC | 6 | FALSE | FALSE |
| 140_v.txt | 32095 | 32095-SNV- | SNV | 1 | C | A | 19 | 20031 | 0.09485298 | 60.2631579 | rnr1Y285A_p | TRUE | CAN | JSY2423 | 4 | rnr1Y285A_p | ACA | C>A | A/T+1 | G/C-1 | CG>AT | CG>AT | CTGCAACAC | CACCAGT | TGT | ACA | C | T | C>A | C>A_ACA | CG>AT_CACCAGT | 6 | FALSE | FALSE |
| 140_v.txt | 32119 | 32119-SNV- | SNV | 1 | C | A | 12 | 20031 | 0.05990714 | 61.8333333 | rnr1Y285A_p | TRUE | CAN | JSY2423 | 4 | rnr1Y285A_p | TCG | C>A | A/T+1 | G/C-1 | CG>AT | CG>AT | GCCATTCGA | CGAAAAC | AGC | TCG | C | T | C>A | C>A_TCG | CG>AT_CGAAAAC | 6 | FALSE | FALSE |
| 140_v.txt | 32188 | 32188-SNV- | SNV | 1 | C | A | 228 | 20031 | 1.13823574 | 44.0789474 | rnr1Y285A_p | TRUE | CAN | JSY2423 | 4 | rnr1Y285A_p | ACT | C>A | A/T+1 | G/C-1 | CG>AT | CG>AT | CGAAAACTG | CTGCAAT | TGA | ACT | C | T | C>A | C>A_ACT | CG>AT_CTGCAAT | 6 | FALSE | FALSE |
| 140_v.txt | 32271 | 32271-SNV- | SNV | 1 | C | A | 26 | 28383 | 0.09160413 | 37.1538462 | rnr1Y285A_p | TRUE | CAN | JSY2423 | 4 | rnr1Y285A_p | ACG | C>A | A/T+1 | G/C-1 | CG>AT | CG>AT | AAAATACGG | CGGGAAC | TGC | ACG | C | T | C>A | C>A_ACG | CG>AT_CGGGAAC | 6 | FALSE | FALSE |
| 140_v.txt | 32274 | 32274-SNV- | SNV | 1 | G | T | 16 | 28383 | 0.05637177 | 33.9375 | rnr1Y285A_p | TRUE | CAN | JSY2423 | 4 | rnr1Y285A_p | GGA | G>T | A/T+1 | G/C-1 | CG>AT | CG>AT | ATACGGGAA | GAACCAA | CCT | CCT | C | A | C>A | C>A_CCT | CG>AT_GAACCAA | 6 | FALSE | FALSE |
| 140_v.txt | 32277 | 32277-SNV- | SNV | 1 | C | A | 33 | 28383 | 0.11626678 | 32.8484849 | rnr1Y285A_p | TRUE | CAN | JSY2423 | 4 | rnr1Y285A_p | ACC | C>A | A/T+1 | G/C-1 | CG>AT | CG>AT | CGGGAACCA | CCAACGT | TGG | ACC | C | T | C>A | C>A_ACC | CG>AT_CCAACGT | 6 | FALSE | FALSE |
| 140_v.txt | 32363 | 32363-SNV- | SNV | 1 | G | T | 73 | 28383 | 0.25719621 | 57.1643836 | rnr1Y285A_p | TRUE | CAN | JSY2423 | 4 | rnr1Y285A_p | AGT | G>T | A/T+1 | G/C-1 | CG>AT | CG>AT | ACCAGAGTT | GTTCTCA | TCA | TCA | C | A | C>A | C>A_TCA | CG>AT_GTTCTCA | 6 | FALSE | FALSE |
| 140_v.txt | 32373 | 32373-SNV- | SNV | 1 | G | T | 35 | 28383 | 0.12331325 | 59.0857143 | rnr1Y285A_p | TRUE | CAN | JSY2423 | 4 | rnr1Y285A_p | AGC | G>T | A/T+1 | G/C-1 | CG>AT | CG>AT | TCAATAGCA | GCAATAA | TCG | TCG | C | A | C>A | C>A_TCG | CG>AT_GCAATAA | 6 | FALSE | FALSE |
| 140_v.txt | 32424 | 32424-SNV- | SNV | 1 | G | T | 6 | 28383 | 0.02113941 | 55.1666667 | rnr1Y285A_p | TRUE | CAN | JSY2423 | 4 | rnr1Y285A_p | AGG | G>T | A/T+1 | G/C-1 | CG>AT | CG>AT | AGTTTAGGG | GGGTCAT | TCC | TCC | C | A | C>A | C>A_TCC | CG>AT_GGGTCAT | 6 | FALSE | FALSE |
| 140_v.txt | 32463 | 32463-SNV- | SNV | 1 | G | T | 15 | 28383 | 0.05284854 | 53.4666667 | rnr1Y285A_p | TRUE | CAN | JSY2423 | 4 | rnr1Y285A_p | AGA | G>T | A/T+1 | G/C-1 | CG>AT | CG>AT | AATAGAGAG | GAGCCAA | TCT | TCT | C | A | C>A | C>A_TCT | CG>AT_GAGCCAA | 6 | FALSE | FALSE |
| 140_v.txt | 32508 | 32508-SNV- | SNV | 1 | G | T | 70 | 28383 | 0.2466265 | 42.6428571 | rnr1Y285A_p | TRUE | CAN | JSY2423 | 4 | rnr1Y285A_p | GGC | G>T | A/T+1 | G/C-1 | CG>AT | CG>AT | TTGATGGCT | GCTCTTG | CCG | CCG | C | A | C>A | C>A_CCG | CG>AT_GCTCTTG | 6 | FALSE | FALSE |
| 140_v.txt | 32542 | 32542-SNV- | SNV | 1 | C | A | 28 | 28381 | 0.09865755 | 36.25 | rnr1Y285A_p | TRUE | CAN | JSY2423 | 4 | rnr1Y285A_p | TCA | C>A | A/T+1 | G/C-1 | CG>AT | CG>AT | CAGCTTCACC | CACCAGC | AGT | TCA | C | T | C>A | C>A_TCA | CG>AT_CACCAGC | 6 | FALSE | FALSE |
| 140_v.txt | 32556 | 32556-SNV- | SNV | 1 | C | A | 1164 | 28371 | 4.10278101 | 37.4536083 | rnr1Y285A_p | TRUE | CAN | JSY2423 | 4 | rnr1Y285A_p | ACC | C>A | A/T+1 | G/C-1 | CG>AT | CG>AT | GTGATACCA | CCAACTA | TGG | ACC | C | T | C>A | C>A_ACC | CG>AT_CCAACTA | 6 | FALSE | FALSE |
| 140_v.txt | 32566 | 32566-SNV- | SNV | 1 | C | A | 101 | 20551 | 0.49146027 | 37.1386139 | rnr1Y285A_p | TRUE | CAN | JSY2423 | 4 | rnr1Y285A_p | TCA | C>A | A/T+1 | G/C-1 | CG>AT | CG>AT | CTAGTTCAG | CAGTACC | AGT | TCA | C | T | C>A | C>A_TCA | CG>AT_CAGTACC | 6 | FALSE | FALSE |
| 140_v.txt | 32571 | 32571-SNV- | SNV | 1 | C | A | 25 | 20555 | 0.12162491 | 35.72 | rnr1Y285A_p | TRUE | CAN | JSY2423 | 4 | rnr1Y285A_p | ACC | C>A | A/T+1 | G/C-1 | CG>AT | CG>AT | TCAGTACCTT | CCTTGAA | TGG | ACC | C | T | C>A | C>A_ACC | CG>AT_CCTTGAA | 6 | FALSE | FALSE |
| 140_v.txt | 32629 | 32629-SNV- | SNV | 1 | C | A | 25 | 20555 | 0.12162491 | 54.2 | rnr1Y285A_p | TRUE | CAN | JSY2423 | 4 | rnr1Y285A_p | TCG | C>A | A/T+1 | G/C-1 | CG>AT | CG>AT | TCCCTTCGTT | CGTTTTT | AGC | TCG | C | T | C>A | C>A_TCG | CG>AT_CGTTTTT | 6 | FALSE | FALSE |
| 140_v.txt | 32708 | 32708-SNV- | SNV | 1 | C | A | 5 | 20555 | 0.02432498 | 54.8021231 | rnr1Y285A_p | TRUE | CAN | JSY2423 | 4 | rnr1Y285A_p | CCC | C>A | A/T+1 | G/C-1 | CG>AT | CG>AT | GGTAACCCC | CCCAGCA | GGG | CCC | C | T | C>A | C>A_CCC | CG>AT_CCCAGCA | 6 | FALSE | FALSE |
| 140_v.txt | 32722 | 32722-SNV- | SNV | 1 | C | A | 14 | 20555 | 0.06810995 | 55.1428571 | rnr1Y285A_p | TRUE | CAN | JSY2423 | 4 | rnr1Y285A_p | ACC | C>A | A/T+1 | G/C-1 | CG>AT | CG>AT | CACAAACCA | CCATACA | TGG | ACC | C | T | C>A | C>A_ACC | CG>AT_CCATACA | 6 | FALSE | FALSE |
| 140_v.txt | 32735 | 32735-SNV- | SNV | 1 | G | T | 65 | 20555 | 0.31622476 | 54.6153846 | rnr1Y285A_p | TRUE | CAN | JSY2423 | 4 | rnr1Y285A_p | AGT | G>T | A/T+1 | G/C-1 | CG>AT | CG>AT | AAAACAGTA | GTATATT | TCA | TCA | C | A | C>A | C>A_TCA | CG>AT_GTATATT | 6 | FALSE | FALSE |
| 140_v.txt | 32783 | 32783-SNV- | SNV | 1 | G | T | 19 | 20555 | 0.09243493 | 54.1052632 | rnr1Y285A_p | TRUE | CAN | JSY2423 | 4 | rnr1Y285A_p | AGA | G>T | A/T+1 | G/C-1 | CG>AT | CG>AT | GACCCAGAA | GAACTCG | TCT | TCT | C | A | C>A | C>A_TCT | CG>AT_GAACTCG | 6 | FALSE | FALSE |
| 140_v.txt | 32786 | 32786-SNV- | SNV | 1 | C | A | 45 | 20555 | 0.21892484 | 50.9564863 | rnr1Y285A_p | TRUE | CAN | JSY2423 | 4 | rnr1Y285A_p | ACT | C>A | A/T+1 | G/C-1 | CG>AT | CG>AT | CCAGAACTC | CTCGAAT | TGA | ACT | C | T | C>A | C>A_ACT | CG>AT_CTCGAAT | 6 | FALSE | FALSE |
| 140_v.txt | 32794 | 32794-SNV- | SNV | 1 | C | A | 19 | 20555 | 0.09243493 | 46.2631579 | rnr1Y285A_p | TRUE | CAN | JSY2423 | 4 | rnr1Y285A_p | TCA | C>A | A/T+1 | G/C-1 | CG>AT | CG>AT | CGAATTCACC | CACCGTA | AGT | TCA | C | T | C>A | C>A_TCA | CG>AT_CACCGTA | 6 | FALSE | FALSE |
| 140_v.txt | 32811 | 32811-SNV- | SNV | 1 | G | T | 12 | 20555 | 0.05837996 | 43.75 | rnr1Y285A_p | TRUE | CAN | JSY2423 | 4 | rnr1Y285A_p | AGG | G>T | A/T+1 | G/C-1 | CG>AT | CG>AT | TTGACAGGG | GGGAACA | TCC | TCC | C | A | C>A | C>A_TCC | CG>AT_GGGAACA | 6 | FALSE | FALSE |
| 140_v.txt | 32819 | 32819-SNV- | SNV | 1 | G | T | 271 | 20555 | 1.31841401 | 44.8597786 | rnr1Y285A_p | TRUE | CAN | JSY2423 | 4 | rnr1Y285A_p | AGT | G>T | A/T+1 | G/C-1 | CG>AT | CG>AT | GAACAAGTT | GTTCATT | TCA | TCA | C | A | C>A | C>A_TCA | CG>AT_GTTCATT | 6 | FALSE | FALSE |
| 140_v.txt | 32917 | 32917-SNV- | SNV | 1 | C | A | 2107 | 28487 | 7.39635623 | 53.5386806 | rnr1Y285A_p | TRUE | CAN | JSY2423 | 4 | rnr1Y285A_p | TCC | C>A | A/T+1 | G/C-1 | CG>AT | CG>AT | TAAGTTCCA | CCAGGGC | AGG | TCC | C | T | C>A | C>A_TCC | CG>AT_CCAGGGC | 6 | FALSE | FALSE |
| 140_v.txt | 32940 | 32940-SNV- | SNV | 1 | G | T | 66 | 28487 | 0.23168463 | 47.8787879 | rnr1Y285A_p | TRUE | CAN | JSY2423 | 4 | rnr1Y285A_p | AGA | G>T | A/T+1 | G/C-1 | CG>AT | CG>AT | GCCCAAGAA | GAAAACC | TCT | TCT | C | A | C>A | C>A_TCT | CG>AT_GAAAACC | 6 | FALSE | FALSE |
| 140_v.txt | 32958 | 32958-SNV- | SNV | 1 | C | A | 14 | 28487 | 0.04914522 | 49.5714286 | rnr1Y285A_p | TRUE | CAN | JSY2423 | 4 | rnr1Y285A_p | ACC | C>A | A/T+1 | G/C-1 | CG>AT | CG>AT | ATGTAACCA | CCATTGG | TGG | ACC | C | T | C>A | C>A_ACC | CG>AT_CCATTGG | 6 | FALSE | FALSE |
| 140_v.txt | 33051 | 33051-SNV- | SNV | 1 | G | T | 16 | 28487 | 0.05616597 | 51.1875 | rnr1Y285A_p | TRUE | CAN | JSY2423 | 4 | rnr1Y285A_p | GGA | G>T | A/T+1 | G/C-1 | CG>AT | CG>AT | CCCAAGGAC | GACTGCG | CCT | CCT | C | A | C>A | C>A_CCT | CG>AT_GACTGCG | 6 | FALSE | FALSE |
| 140_v.txt | 33070 | 33070-SNV- | SNV | 1 | C | A | 5 | 28487 | 0.01755187 | 37.6 | rnr1Y285A_p | TRUE | CAN | JSY2423 | 4 | rnr1Y285A_p | GCC | C>A | A/T+1 | G/C-1 | CG>AT | CG>AT | AATATGCCA | CCAAAGA | CGG | GCC | C | T | C>A | C>A_GCC | CG>AT_CCAAAGA | 6 | FALSE | FALSE |
| 140_v.txt | 33093 | 33093-SNV- | SNV | 1 | G | T | 39 | 28487 | 0.13690455 | 54.9743046 | rnr1Y285A_p | TRUE | CAN | JSY2423 | 4 | rnr1Y285A_p | TGA | G>T | A/T+1 | G/C-1 | CG>AT | CG>AT | AAATATGAT | GATATAA | ACT | ACT | C | A | C>A | C>A_ACT | CG>AT_GATATAA | 6 | FALSE | FALSE |
| 140_v.txt | 33114 | 33114-SNV- | SNV | 1 | C | A | 35 | 28487 | 0.12286306 | 42.2857143 | rnr1Y285A_p | TRUE | CAN | JSY2423 | 4 | rnr1Y285A_p | GCC | C>A | A/T+1 | G/C-1 | CG>AT | CG>AT | ACTGGGCCG | CCGGCGT | CGG | GCC | C | T | C>A | C>A_GCC | CG>AT_CCGGCGT | 6 | FALSE | FALSE |
| 140_v.txt | 33151 | 33151-SNV- | SNV | 1 | G | T | 25 | 28487 | 0.08775933 | 40.4 | rnr1Y285A_p | TRUE | CAN | JSY2423 | 4 | rnr1Y285A_p | AGA | G>T | A/T+1 | G/C-1 | CG>AT | CG>AT | TGAAAAGAC | GACCTGT | TCT | TCT | C | A | C>A | C>A_TCT | CG>AT_GACCTGT | 6 | FALSE | FALSE |
| 140_v.txt | 33153 | 33153-SNV- | SNV | 1 | C | A | 221 | 28487 | 0.77579247 | 40.561086 | rnr1Y285A_p | TRUE | CAN | JSY2423 | 4 | rnr1Y285A_p | ACC | C>A | A/T+1 | G/C-1 | CG>AT | CG>AT | AAAAGACCT | CCTGTAC | TGG | ACC | C | T | C>A | C>A_ACC | CG>AT_CCTGTAC | 6 | FALSE | FALSE |
| 140_v.txt | 33154 | 33154-SNV- | SNV | 1 | C | A | 76 | 28487 | 0.26678836 | 37.1835339 | rnr1Y285A_p | TRUE | CAN | JSY2423 | 4 | rnr1Y285A_p | CCT | C>A | A/T+1 | G/C-1 | CG>AT | CG>AT | AAAGACCTG | CTGTACC | GGA | CCT | C | T | C>A | C>A_CCT | CG>AT_CTGTACC | 6 | FALSE | FALSE |
| 140_v.txt | 33159 | 33159-SNV- | SNV | 1 | C | A | 98 | 28486 | 0.34402865 | 37.8877551 | rnr1Y285A_p | TRUE | CAN | JSY2423 | 4 | rnr1Y285A_p | ACC | C>A | A/T+1 | G/C-1 | CG>AT | CG>AT | CCTGTACCA | CCAATAG | TGG | ACC | C | T | C>A | C>A_ACC | CG>AT_CCAATAG | 6 | FALSE | FALSE |
| 140_v.txt | 33168 | 33168-SNV- | SNV | 1 | C | A | 334 | 28484 | 1.17258812 | 36.7095808 | rnr1Y285A_p | TRUE | CAN | JSY2423 | 4 | rnr1Y285A_p | ACC | C>A | A/T+1 | G/C-1 | CG>AT | CG>AT | ATAGTACCA | CCACCAA | TGG | ACC | C | T | C>A | C>A_ACC | CG>AT_CCACCAA | 6 | FALSE | FALSE |
| 140_v.txt | 33186 | 33186-SNV- | SNV | 1 | C | A | 35 | 21078 | 0.16604991 | 37.5142857 | rnr1Y285A_p | TRUE | CAN | JSY2423 | 4 | rnr1Y285A_p | ACC | C>A | A/T+1 | G/C-1 | CG>AT | CG>AT | ATCATACCA | CCAATAT | TGG | ACC | C | T | C>A | C>A_ACC | CG>AT_CCAATAT | 6 | FALSE | FALSE |
| 140_v.txt | 33205 | 33205-SNV- | SNV | 1 | G | T | 13 | 21084 | 0.06165813 | 33.7692308 | rnr1Y285A_p | TRUE | CAN | JSY2423 | 4 | rnr1Y285A_p | AGC | G>T | A/T+1 | G/C-1 | CG>AT | CG>AT | GCTTAAGCT | GCTCTCT | TCG | TCG | C | A | C>A | C>A_TCG | CG>AT_GCTCTCT | 6 | FALSE | FALSE |
| 140_v.txt | 33208 | 33208-SNV- | SNV | 1 | C | A | 37 | 21084 | 0.17548852 | 41.5945946 | rnr1Y285A_p | TRUE | CAN | JSY2423 | 4 | rnr1Y285A_p | TCT | C>A | A/T+1 | G/C-1 | CG>AT | CG>AT | TAAGCTCTCT | CTCTCTT | AGA | TCT | C | T | C>A | C>A_TCT | CG>AT_CTCTCTT | 6 | FALSE | FALSE |
| 140_v.txt | 33220 | 33220-SNV- | SNV | 1 | C | A | 13 | 21084 | 0.06165813 | 36.8461539 | rnr1Y285A_p | TRUE | CAN | JSY2423 | 4 | rnr1Y285A_p | TCA | C>A | A/T+1 | G/C-1 | CG>AT | CG>AT | TCACTTCAG | CAGCGTT | AGT | TCA | C | T | C>A | C>A_TCA | CG>AT_CAGCGTT | 6 | FALSE | FALSE |
| 140_v.txt | 33309 | 33309-SNV- | SNV | 1 | C | A | 2 | 21084 | 9.49E-03 | 40 | rnr1Y285A_p | TRUE | CAN | JSY2423 | 4 | rnr1Y285A_p | GCG | C>A | A/T+1 | G/C-1 | CG>AT | CG>AT | AAAGAGCGC | CGCAATG | CGC | GCG | C | T | C>A | C>A_GCG | CG>AT_CGCAATG | 6 | FALSE | FALSE |

| | | | | | | | | | | | | | | | | | | | | | | | | | | | | | | | | | | |
|---|---|---|---|---|---|---|---|---|---|---|---|---|---|---|---|---|---|---|---|---|---|---|---|---|---|---|---|---|---|---|---|---|---|---|
| 140_v.txt | 33325 | 33325-SNV-C | SNV | 1 | C | A | 407 | 21084 | 1.93037374 | 57.6879607 | rnr1Y285A_pC | TRUE | CAN | JSY2423 | 4 | rnr1Y285A_p | TCT | C>A | A/T+1 | G/C-1 | CG>AT | CG>AT | ACAATTCTTT | CTTTACT | AGA | TCT | C | T | C>A | C>A_TCT | CG>AT_CTTTACT | 6 | FALSE | FALSE |
| 140_v.txt | 33474 | 33474-SNV-C | SNV | 1 | C | A | 49 | 21083 | 0.23241474 | 34.6994191 | rnr1Y285A_pC | TRUE | CAN | JSY2423 | 4 | rnr1Y285A_p | GCC | C>A | A/T+1 | G/C-1 | CG>AT | CG>AT | GCTATGCCTT | CCTTTTT | CGG | GCC | C | T | C>A | C>A_GCC | CG>AT_CCTTTTT | 6 | FALSE | FALSE |
| 140_v.txt | 33475 | 33475-SNV-C | SNV | 1 | C | A | 16 | 21082 | 0.07589413 | 36.2964726 | rnr1Y285A_pC | TRUE | CAN | JSY2423 | 4 | rnr1Y285A_p | CCT | C>A | A/T+1 | G/C-1 | CG>AT | CG>AT | CTATGCCTTT | CTTTTTT | GGA | CCT | C | T | C>A | C>A_CCT | CG>AT_CTTTTTT | 6 | FALSE | FALSE |
| 140_v.txt | 32033 | 32033-Repla | Replacement | 2 | C | AA | 14 | 20031 | 0.06989167 | 61.5714613 | rnr1Y285A_pC | TRUE | CAN | JSY2423 | 4 | rnr1Y285A_p | TCA | nope | >1bp-ins | G/C-1 | none | Replac. | GTATTTCAA | CAAAGCT | AGT | TCA | C | none | nope | nope_TCA | Replac._CAAAGCT | 6 | FALSE | FALSE |
| 140_v.txt | 32183 | 32183-Repla | Replacement | 2 | G | TA | 8 | 20031 | 0.0399381 | 57.5 | rnr1Y285A_pC | TRUE | CAN | JSY2423 | 4 | rnr1Y285A_p | CGA | nope | >1bp-ins | G/C-1 | none | Replac. | AGTAACGAA | GAAAACT | GCT | GCT | C | none | C>none | C>none_GCT | Replac._GAAAACT | 6 | FALSE | FALSE |
| 140_v.txt | 31964 | 31964-Insert | Insertion | 2 | - | AT | 5 | 20031 | 0.02496131 | 38 | rnr1Y285A_pC | TRUE | CAN | JSY2423 | 4 | rnr1Y285A_p | CAT | nope | >1bp-ins | >1bpdel | none | >1bp-ins | GGCCGCATA | ATAATAA | GTA | CAT | - | none | NA | NA_CAT | >1bp-ins_ATAATAA | 6 | FALSE | FALSE |
| 140_v.txt | 32081 | 32081-Insert | Insertion | 1 | - | A | 17 | 20031 | 0.08486845 | 63.1176471 | rnr1Y285A_pC | TRUE | CAN | JSY2423 | 4 | rnr1Y285A_p | CAA | nope | A/T+1 | >1bpdel | none | A/T+1 | CCATGCAAA | AAAAAAG | GTT | CAA | - | T | NA | NA_CAA | A/T+1_AAAAAAG | 6 | FALSE | FALSE |
| 140_v.txt | 32337 | 32337-Insert | Insertion | 2 | - | AT | 10 | 28383 | 0.03523236 | 61.8 | rnr1Y285A_pC | TRUE | CAN | JSY2423 | 4 | rnr1Y285A_p | GAT | nope | >1bp-ins | >1bpdel | none | >1bp-ins | TTGAAGATA | ATATGTG | CTA | GAT | - | none | NA | NA_GAT | >1bp-ins_ATATGTG | 6 | FALSE | FALSE |
| 140_v.txt | 32532 | 32532-Insert | Insertion | 1 | - | T | 13 | 28383 | 0.04580207 | 42 | rnr1Y285A_pC | TRUE | CAN | JSY2423 | 4 | rnr1Y285A_p | GTT | nope | A/T+1 | >1bpdel | none | A/T+1 | CTGGGGTTT | TTTGCAG | CAA | GTT | - | A | NA | NA_GTT | A/T+1_TTTGCAG | 6 | FALSE | FALSE |
| 140_v.txt | 32647 | 32647-Insert | Insertion | 1 | - | T | 2 | 20555 | 9.73E-03 | 64 | rnr1Y285A_pC | TRUE | CAN | JSY2423 | 4 | rnr1Y285A_p | ATT | nope | A/T+1 | >1bpdel | none | A/T+1 | TAGATATTA | TTATACC | TAA | ATT | - | A | NA | NA_ATT | A/T+1_TTATACC | 6 | FALSE | FALSE |
| 140_v.txt | 32658 | 32658-Insert | Insertion | 1 | - | C | 2 | 20555 | 9.73E-03 | 64 | rnr1Y285A_pC | TRUE | CAN | JSY2423 | 4 | rnr1Y285A_p | ACC | nope | G/C+1 | >1bpdel | none | G/C+1 | CCTGGACCC | CCCCAGG | TGG | ACC | - | G | NA | NA_ACC | G/C+1_CCCCAGG | 6 | FALSE | FALSE |
| 140_v.txt | 32760 | 32760-Insert | Insertion | 1 | - | A | 13 | 20555 | 0.06324495 | 59.3076923 | rnr1Y285A_pC | TRUE | CAN | JSY2423 | 4 | rnr1Y285A_p | TAA | nope | A/T+1 | >1bpdel | none | A/T+1 | ATGGCTAAA | AAAACTT | ATT | TAA | - | T | NA | NA_TAA | A/T+1_AAAACTT | 6 | FALSE | FALSE |
| 140_v.txt | 32842 | 32842-Insert | Insertion | 1 | - | A | 56 | 20555 | 0.2724398 | 42.4642857 | rnr1Y285A_pC | TRUE | CAN | JSY2423 | 4 | rnr1Y285A_p | CAA | nope | A/T+1 | >1bpdel | none | A/T+1 | TTACCCAAA | AAAAAAT | GTT | CAA | - | T | NA | NA_CAA | A/T+1_AAAAAAT | 6 | FALSE | FALSE |
| 140_v.txt | 32843 | 32843-Insert | Insertion | 1 | - | G | 1 | 20555 | 4.87E-03 | 38 | rnr1Y285A_pC | TRUE | CAN | JSY2423 | 4 | rnr1Y285A_p | AAA | nope | G/C+1 | >1bpdel | none | G/C+1 | TACCCAAAA | AAAAATA | TTT | AAA | - | C | NA | NA_AAA | G/C+1_AAAAATA | 6 | FALSE | FALSE |
| 140_v.txt | 33338 | 33338-Insert | Insertion | 2 | - | AT | 15 | 21084 | 0.071144 | 62.3 | rnr1Y285A_pC | TRUE | CAN | JSY2423 | 4 | rnr1Y285A_p | CAT | nope | >1bp-ins | >1bpdel | none | >1bp-ins | TTTCTCATCT | ATCTTTC | GTA | CAT | - | none | NA | NA_CAT | >1bp-ins_ATCTTTC | 6 | FALSE | FALSE |
| 140_v.txt | 33475 | 33475-Insert | Insertion | 1 | - | T | 1 | 21082 | 4.74E-03 | 37.9999995 | rnr1Y285A_pC | TRUE | CAN | JSY2423 | 4 | rnr1Y285A_p | CCT | nope | A/T+1 | >1bpdel | none | A/T+1 | CTATGCCTTT | CTTTTTT | GGA | CCT | - | A | NA | NA_CCT | A/T+1_CTTTTTT | 6 | FALSE | FALSE |
| 140_v.txt | 32707 | 32707-Deleti | Deletion | 2 | CC | - | 8 | 20555 | 0.03891997 | 64 | rnr1Y285A_pC | TRUE | CAN | JSY2423 | 4 | rnr1Y285A_p | ACC | nope | >1bp-ins | >1bpdel | none | >1bpdel | CGGTAACCC | CCCCAGC | TGG | ACC | CC | none | NA | NA_ACC | >1bpdel_CCCCAGC | 6 | FALSE | FALSE |
| 140_v.txt | 33089 | 33089-Deleti | Deletion | 5 | ATATG | - | 11 | 28487 | 0.03861411 | 51.9090909 | rnr1Y285A_pC | TRUE | CAN | JSY2423 | 4 | rnr1Y285A_p | AAT | nope | >1bp-ins | >1bpdel | none | >1bpdel | AAATAAATA | ATATGAT | TTA | AAT | ATATG | none | NA | NA_AAT | >1bpdel_ATATGAT | 6 | FALSE | FALSE |
| 140_v.txt | 33474 | 33474-Deleti | Deletion | 2 | CC | - | 1 | 21083 | 4.74E-03 | 10 | rnr1Y285A_pC | TRUE | CAN | JSY2423 | 4 | rnr1Y285A_p | GCC | nope | >1bp-ins | >1bpdel | none | >1bpdel | GCTATGCCTT | CCTTTTT | CGG | GCC | CC | none | NA | NA_GCC | >1bpdel_CCTTTTT | 6 | FALSE | FALSE |
| 140_v.txt | 31663 | 31663-Deleti | Deletion | 1 | A | - | 5 | 9956 | 0.05022097 | 33.6 | rnr1Y285A_pC | TRUE | CAN | JSY2423 | 4 | rnr1Y285A_p | GAA | nope | >1bp-ins | A/T-1 | none | A/T-1 | GCGTGGAAA | AAATGTG | CTT | CTT | T | none | T>none | T>none_CTT | A/T-1_AAATGTG | 6 | FALSE | FALSE |
| 140_v.txt | 31673 | 31673-Deleti | Deletion | 1 | A | - | 4 | 9956 | 0.04017678 | 55.25 | rnr1Y285A_pC | TRUE | CAN | JSY2423 | 4 | rnr1Y285A_p | CAA | nope | >1bp-ins | A/T-1 | none | A/T-1 | GTGATCAAA | AAAGGTA | GTT | GTT | T | none | T>none | T>none_GTT | A/T-1_AAAGGTA | 6 | FALSE | FALSE |
| 140_v.txt | 31723 | 31723-Deleti | Deletion | 1 | A | - | 11 | 9956 | 0.11048614 | 63.7272727 | rnr1Y285A_pC | TRUE | CAN | JSY2423 | 4 | rnr1Y285A_p | CAA | nope | >1bp-ins | A/T-1 | none | A/T-1 | TGTCCCAAA | AAAAAGT | GTT | GTT | T | none | T>none | T>none_GTT | A/T-1_AAAAAGT | 6 | FALSE | FALSE |
| 140_v.txt | 31904 | 31904-Deleti | Deletion | 1 | T | - | 15 | 9956 | 0.15066292 | 45.6 | rnr1Y285A_pC | TRUE | CAN | JSY2423 | 4 | rnr1Y285A_p | ATT | nope | >1bp-ins | A/T-1 | none | A/T-1 | ATTGAATTTT | TTTTGGT | TAA | ATT | T | none | nope | nope_ATT | A/T-1_TTTTGGT | 6 | FALSE | FALSE |
| 140_v.txt | 31964 | 31964-Deleti | Deletion | 1 | A | - | 1 | 20031 | 4.99E-03 | 38 | rnr1Y285A_pC | TRUE | CAN | JSY2423 | 4 | rnr1Y285A_p | CAT | nope | >1bp-ins | A/T-1 | none | A/T-1 | GGCCGCATA | ATAATAA | GTA | GTA | T | none | T>none | T>none_GTA | A/T-1_ATAATAA | 6 | FALSE | FALSE |
| 140_v.txt | 31972 | 31972-Deleti | Deletion | 1 | C | - | 8 | 20031 | 0.0399381 | 35 | rnr1Y285A_pC | TRUE | CAN | JSY2423 | 4 | rnr1Y285A_p | GCC | nope | >1bp-ins | G/C-1 | none | G/C-1 | AATAAGCCA | CCAAGCC | CGG | GCC | C | none | nope | nope_GCC | G/C-1_CCAAGCC | 6 | FALSE | FALSE |
| 140_v.txt | 31979 | 31979-Deleti | Deletion | 1 | G | - | 7 | 20031 | 0.03494583 | 45.2857142 | rnr1Y285A_pC | TRUE | CAN | JSY2423 | 4 | rnr1Y285A_p | CGG | nope | >1bp-ins | G/C-1 | none | G/C-1 | CAAGCCGGG | GGGCATT | GCC | GCC | C | none | C>none | C>none_GCC | G/C-1_GGGCATT | 6 | FALSE | FALSE |
| 140_v.txt | 31988 | 31988-Deleti | Deletion | 1 | T | - | 8 | 20031 | 0.0399381 | 44.125 | rnr1Y285A_pC | TRUE | CAN | JSY2423 | 4 | rnr1Y285A_p | ATT | nope | >1bp-ins | A/T-1 | none | A/T-1 | CATTAATTTA | TTTAGCT | TAA | ATT | T | none | nope | nope_ATT | A/T-1_TTTAGCT | 6 | FALSE | FALSE |
| 140_v.txt | 32060 | 32060-Deleti | Deletion | 1 | G | - | 8 | 20031 | 0.0399381 | 60.75 | rnr1Y285A_pC | TRUE | CAN | JSY2423 | 4 | rnr1Y285A_p | AGA | nope | >1bp-ins | G/C-1 | none | G/C-1 | GTGCGAGAT | GATTGAG | TCT | TCT | C | none | C>none | C>none_TCT | G/C-1_GATTGAG | 6 | FALSE | FALSE |
| 140_v.txt | 32075 | 32075-Deleti | Deletion | 1 | C | - | 22 | 20031 | 0.10982976 | 58.7727273 | rnr1Y285A_pC | TRUE | CAN | JSY2423 | 4 | rnr1Y285A_p | ACC | nope | >1bp-ins | G/C-1 | none | G/C-1 | AAATAACCA | CCATGCA | TGG | ACC | C | none | nope | nope_ACC | G/C-1_CCATGCA | 6 | FALSE | FALSE |
| 140_v.txt | 32081 | 32081-Deleti | Deletion | 1 | A | - | 19 | 20031 | 0.09485298 | 62.2631582 | rnr1Y285A_pC | TRUE | CAN | JSY2423 | 4 | rnr1Y285A_p | CAA | nope | >1bp-ins | A/T-1 | none | A/T-1 | CCATGCAAA | AAAAAAG | GTT | GTT | T | none | T>none | T>none_GTT | A/T-1_AAAAAAG | 6 | FALSE | FALSE |
| 140_v.txt | 32121 | 32121-Deleti | Deletion | 1 | A | - | 13 | 20031 | 0.06489941 | 61.3076923 | rnr1Y285A_pC | TRUE | CAN | JSY2423 | 4 | rnr1Y285A_p | GAA | nope | >1bp-ins | A/T-1 | none | A/T-1 | CATTCGAAA | AAAACTT | CTT | CTT | T | none | T>none | T>none_CTT | A/T-1_AAAACTT | 6 | FALSE | FALSE |
| 140_v.txt | 32219 | 32219-Deleti | Deletion | 1 | C | - | 7 | 20031 | 0.03494583 | 37.4285714 | rnr1Y285A_pC | TRUE | CAN | JSY2423 | 4 | rnr1Y285A_p | TCC | nope | >1bp-ins | G/C-1 | none | G/C-1 | GGTGGTCCT | CCTTGAC | AGG | TCC | C | none | nope | nope_TCC | G/C-1_CCTTGAC | 6 | FALSE | FALSE |
| 140_v.txt | 32231 | 32231-Deleti | Deletion | 1 | T | - | 6 | 20031 | 0.02995357 | 37.8333333 | rnr1Y285A_pC | TRUE | CAN | JSY2423 | 4 | rnr1Y285A_p | ATT | nope | >1bp-ins | A/T-1 | none | A/T-1 | CAGGAATTT | TTTAGGA | TAA | ATT | T | none | nope | nope_ATT | A/T-1_TTTAGGA | 6 | FALSE | FALSE |
| 140_v.txt | 32277 | 32277-Deleti | Deletion | 1 | C | - | 15 | 28383 | 0.05284854 | 37.8666667 | rnr1Y285A_pC | TRUE | CAN | JSY2423 | 4 | rnr1Y285A_p | ACC | nope | >1bp-ins | G/C-1 | none | G/C-1 | CGGGAACCA | CCAACGT | TGG | ACC | C | none | nope | nope_ACC | G/C-1_CCAACGT | 6 | FALSE | FALSE |
| 140_v.txt | 32346 | 32346-Deleti | Deletion | 1 | A | - | 1390 | 28383 | 4.89729768 | 61.518705 | rnr1Y285A_pC | TRUE | CAN | JSY2423 | 4 | rnr1Y285A_p | CAA | nope | >1bp-ins | A/T-1 | none | A/T-1 | TGTGGCAAA | AAAACCT | GTT | GTT | T | none | T>none | T>none_GTT | A/T-1_AAAACCT | 6 | FALSE | FALSE |
| 140_v.txt | 32358 | 32358-Deleti | Deletion | 1 | C | - | 10 | 28383 | 0.03523236 | 61.1 | rnr1Y285A_pC | TRUE | CAN | JSY2423 | 4 | rnr1Y285A_p | ACC | nope | >1bp-ins | G/C-1 | none | G/C-1 | TTTGTACCA | CCAGAGT | TGG | ACC | C | none | nope | nope_ACC | G/C-1_CCAGAGT | 6 | FALSE | FALSE |
| 140_v.txt | 32398 | 32398-Deleti | Deletion | 1 | C | - | 77 | 28383 | 0.27128915 | 60.7402597 | rnr1Y285A_pC | TRUE | CAN | JSY2423 | 4 | rnr1Y285A_p | ACG | nope | >1bp-ins | G/C-1 | none | G/C-1 | TAGAAACGT | CGTAGGA | TGC | ACG | C | none | nope | nope_ACG | G/C-1_CGTAGGA | 6 | FALSE | FALSE |
| 140_v.txt | 32420 | 32420-Deleti | Deletion | 1 | T | - | 6 | 28383 | 0.02113941 | 64 | rnr1Y285A_pC | TRUE | CAN | JSY2423 | 4 | rnr1Y285A_p | GTT | nope | >1bp-ins | A/T-1 | none | A/T-1 | TGTTAGTTTA | TTTAGGG | CAA | GTT | T | none | nope | nope_GTT | A/T-1_TTTAGGG | 6 | FALSE | FALSE |
| 140_v.txt | 32424 | 32424-Deleti | Deletion | 1 | G | - | 5 | 28383 | 0.01761618 | 62.8 | rnr1Y285A_pC | TRUE | CAN | JSY2423 | 4 | rnr1Y285A_p | AGG | nope | >1bp-ins | G/C-1 | none | G/C-1 | AGTTTAGGG | GGGTCAT | TCC | TCC | C | none | C>none | C>none_TCC | G/C-1_GGGTCAT | 6 | FALSE | FALSE |
| 140_v.txt | 32488 | 32488-Deleti | Deletion | 1 | G | - | 4 | 28383 | 0.01409294 | 44.5000041 | rnr1Y285A_pC | TRUE | CAN | JSY2423 | 4 | rnr1Y285A_p | CGG | nope | >1bp-ins | G/C-1 | none | G/C-1 | AGATACGGA | GGAAAAC | GCC | GCC | C | none | C>none | C>none_GCC | G/C-1_GGAAAAC | 6 | FALSE | FALSE |
| 140_v.txt | 32498 | 32498-Deleti | Deletion | 1 | T | - | 34 | 28383 | 0.11979002 | 46.0882353 | rnr1Y285A_pC | TRUE | CAN | JSY2423 | 4 | rnr1Y285A_p | CTT | nope | >1bp-ins | A/T-1 | none | A/T-1 | AACAACTTT | TTTTTTG | GAA | CTT | T | none | nope | nope_CTT | A/T-1_TTTTTTG | 6 | FALSE | FALSE |
| 140_v.txt | 32519 | 32519-Deleti | Deletion | 1 | G | - | 14 | 28383 | 0.0493253 | 44.7857143 | rnr1Y285A_pC | TRUE | CAN | JSY2423 | 4 | rnr1Y285A_p | CGG | nope | >1bp-ins | G/C-1 | none | G/C-1 | TGGAACGGA | GGATTTT | GCC | GCC | C | none | C>none | C>none_GCC | G/C-1_GGATTTT | 6 | FALSE | FALSE |
| 140_v.txt | 32522 | 32522-Deleti | Deletion | 1 | T | - | 173 | 28383 | 0.60951978 | 40.0520231 | rnr1Y285A_pC | TRUE | CAN | JSY2423 | 4 | rnr1Y285A_p | ATT | nope | >1bp-ins | A/T-1 | none | A/T-1 | AACGGATTT | TTTTCTG | TAA | ATT | T | none | nope | nope_ATT | A/T-1_TTTTCTG | 6 | FALSE | FALSE |
| 140_v.txt | 32532 | 32532-Deleti | Deletion | 1 | T | - | 3 | 28383 | 0.01056971 | 37.9999998 | rnr1Y285A_pC | TRUE | CAN | JSY2423 | 4 | rnr1Y285A_p | GTT | nope | >1bp-ins | A/T-1 | none | A/T-1 | CTGGGGTTT | TTTGCAG | CAA | GTT | T | none | nope | nope_GTT | A/T-1_TTTGCAG | 6 | FALSE | FALSE |
| 140_v.txt | 32576 | 32576-Deleti | Deletion | 1 | A | - | 25 | 20555 | 0.12162491 | 36.32 | rnr1Y285A_pC | TRUE | CAN | JSY2423 | 4 | rnr1Y285A_p | GAA | nope | >1bp-ins | A/T-1 | none | A/T-1 | ACCTTGAAA | AAATGTG | CTT | CTT | T | none | T>none | T>none_CTT | A/T-1_AAATGTG | 6 | FALSE | FALSE |
| 140_v.txt | 32608 | 32608-Deleti | Deletion | 1 | C | - | 3900 | 20555 | 18.9734858 | 45.7758974 | rnr1Y285A_pC | TRUE | CAN | JSY2423 | 4 | rnr1Y285A_p | ACC | nope | >1bp-ins | G/C-1 | none | G/C-1 | AGGAAACCC | CCCAACC | TGG | ACC | C | none | nope | nope_ACC | G/C-1_CCCAACC | 6 | FALSE | FALSE |
| 140_v.txt | 32631 | 32631-Deleti | Deletion | 1 | T | - | 10 | 20555 | 0.04864996 | 51.6 | rnr1Y285A_pC | TRUE | CAN | JSY2423 | 4 | rnr1Y285A_p | GTT | nope | >1bp-ins | A/T-1 | none | A/T-1 | CCTTCGTTTT | TTTTTAT | CAA | GTT | T | none | nope | nope_GTT | A/T-1_TTTTTAT | 6 | FALSE | FALSE |
| 140_v.txt | 32655 | 32655-Deleti | Deletion | 1 | G | - | 38 | 20555 | 0.18486986 | 58.3684211 | rnr1Y285A_pC | TRUE | CAN | JSY2423 | 4 | rnr1Y285A_p | TGG | nope | >1bp-ins | G/C-1 | none | G/C-1 | ATACCTGGA | GGACCCC | ACC | ACC | C | none | C>none | C>none_ACC | G/C-1_GGACCCC | 6 | FALSE | FALSE |
| 140_v.txt | 32658 | 32658-Deleti | Deletion | 1 | C | - | 1657 | 20555 | 8.06129895 | 61.0126731 | rnr1Y285A_pC | TRUE | CAN | JSY2423 | 4 | rnr1Y285A_p | ACC | nope | >1bp-ins | G/C-1 | none | G/C-1 | CCTGGACCC | CCCCAGG | TGG | ACC | C | none | nope | nope_ACC | G/C-1_CCCCAGG | 6 | FALSE | FALSE |
| 140_v.txt | 32670 | 32670-Deleti | Deletion | 1 | G | - | 1020 | 20555 | 4.96229628 | 60.4019608 | rnr1Y285A_pC | TRUE | CAN | JSY2423 | 4 | rnr1Y285A_p | TGG | nope | >1bp-ins | G/C-1 | none | G/C-1 | GCACCTGGG | GGGTTTC | ACC | ACC | C | none | C>none | C>none_ACC | G/C-1_GGGTTTC | 6 | FALSE | FALSE |
| 140_v.txt | 32702 | 32702-Deleti | Deletion | 1 | G | - | 44 | 20555 | 0.21405984 | 60.4772727 | rnr1Y285A_pC | TRUE | CAN | JSY2423 | 4 | rnr1Y285A_p | CGG | nope | >1bp-ins | G/C-1 | none | G/C-1 | TGGGCCGGT | GGTAACC | GCC | GCC | C | none | C>none | C>none_GCC | G/C-1_GGTAACC | 6 | FALSE | FALSE |
| 140_v.txt | 32707 | 32707-Deleti | Deletion | 1 | C | - | 1088 | 20555 | 5.29311603 | 62.2288612 | rnr1Y285A_pC | TRUE | CAN | JSY2423 | 4 | rnr1Y285A_p | ACC | nope | >1bp-ins | G/C-1 | none | G/C-1 | CGGTAACCC | CCCCAGC | TGG | ACC | C | none | nope | nope_ACC | G/C-1_CCCCAGC | 6 | FALSE | FALSE |
| 140_v.txt | 32728 | 32728-Deleti | Deletion | 1 | A | - | 14 | 20555 | 0.06810995 | 60.2857143 | rnr1Y285A_pC | TRUE | CAN | JSY2423 | 4 | rnr1Y285A_p | CAA | nope | >1bp-ins | A/T-1 | none | A/T-1 | CCATACAAA | AAAAACA | GTT | GTT | T | none | T>none | T>none_GTT | A/T-1_AAAAACA | 6 | FALSE | FALSE |
| 140_v.txt | 32747 | 32747-Deleti | Deletion | 1 | C | - | 269 | 20555 | 1.30868402 | 56.4386637 | rnr1Y285A_pC | TRUE | CAN | JSY2423 | 4 | rnr1Y285A_p | ACC | nope | >1bp-ins | G/C-1 | none | G/C-1 | TAGAAACCC | CCCGATA | TGG | ACC | C | none | nope | nope_ACC | G/C-1_CCCGATA | 6 | FALSE | FALSE |
| 140_v.txt | 32760 | 32760-Deleti | Deletion | 1 | A | - | 68 | 20555 | 0.33081975 | 52.9411728 | rnr1Y285A_pC | TRUE | CAN | JSY2423 | 4 | rnr1Y285A_p | TAA | nope | >1bp-ins | A/T-1 | none | A/T-1 | ATGGCTAAA | AAAACTT | ATT | ATT | T | none | T>none | T>none_ATT | A/T-1_AAAACTT | 6 | FALSE | FALSE |
| 140_v.txt | 32779 | 32779-Deleti | Deletion | 1 | C | - | 5 | 20555 | 0.02432498 | 48.4 | rnr1Y285A_pC | TRUE | CAN | JSY2423 | 4 | rnr1Y285A_p | ACC | nope | >1bp-ins | G/C-1 | none | G/C-1 | AAGCGACCC | CCCAGAA | TGG | ACC | C | none | nope | nope_ACC | G/C-1_CCCAGAA | 6 | FALSE | FALSE |
| 140_v.txt | 32839 | 32839-Deleti | Deletion | 1 | C | - | 14 | 20555 | 0.06810995 | 45 | rnr1Y285A_pC | TRUE | CAN | JSY2423 | 4 | rnr1Y285A_p | ACC | nope | >1bp-ins | G/C-1 | none | G/C-1 | TAATTACCCA | CCCAAAA | TGG | ACC | C | none | nope | nope_ACC | G/C-1_CCCAAAA | 6 | FALSE | FALSE |
| 140_v.txt | 32842 | 32842-Deleti | Deletion | 1 | A | - | 50 | 20555 | 0.24324982 | 41.2400035 | rnr1Y285A_pC | TRUE | CAN | JSY2423 | 4 | rnr1Y285A_p | CAA | nope | >1bp-ins | A/T-1 | none | A/T-1 | TTACCCAAA | AAAAAAT | GTT | GTT | T | none | T>none | T>none_GTT | A/T-1_AAAAAAT | 6 | FALSE | FALSE |
| 140_v.txt | 32884 | 32884-Deleti | Deletion | 1 | A | - | 10 | 28487 | 0.03510373 | 37.5 | rnr1Y285A_pC | TRUE | CAN | JSY2423 | 4 | rnr1Y285A_p | CAA | nope | >1bp-ins | A/T-1 | none | A/T-1 | ACGTCCAAA | AAAATTG | GTT | GTT | T | none | T>none | T>none_GTT | A/T-1_AAAATTG | 6 | FALSE | FALSE |
| 140_v.txt | 32901 | 32901-Deleti | Deletion | 1 | C | - | 2 | 28487 | 7.02E-03 | 34.5000005 | rnr1Y285A_pC | TRUE | CAN | JSY2423 | 4 | rnr1Y285A_p | GCC | nope | >1bp-ins | G/C-1 | none | G/C-1 | ACTTGGCCA | CCAACTA | CGG | GCC | C | none | nope | nope_GCC | G/C-1_CCAACTA | 6 | FALSE | FALSE |
| 140_v.txt | 32924 | 32924-Deleti | Deletion | 1 | A | - | 23 | 28487 | 0.08073858 | 53.173913 | rnr1Y285A_pC | TRUE | CAN | JSY2423 | 4 | rnr1Y285A_p | CAA | nope | >1bp-ins | A/T-1 | none | A/T-1 | CAGGGCAAA | AAAAGTG | GTT | GTT | T | none | T>none | T>none_GTT | A/T-1_AAAAGTG | 6 | FALSE | FALSE |
| 140_v.txt | 32935 | 32935-Deleti | Deletion | 1 | C | - | 8 | 28487 | 0.02808299 | 52.75 | rnr1Y285A_pC | TRUE | CAN | JSY2423 | 4 | rnr1Y285A_p | GCC | nope | >1bp-ins | G/C-1 | none | G/C-1 | TGATTGCCC | CCCAAGA | CGG | GCC | C | none | nope | nope_GCC | G/C-1_CCCAAGA | 6 | FALSE | FALSE |
| 140_v.txt | 32945 | 32945-Deleti | Deletion | 1 | C | - | 9 | 28487 | 0.03159336 | 56.5555556 | rnr1Y285A_pC | TRUE | CAN | JSY2423 | 4 | rnr1Y285A_p | ACC | nope | >1bp-ins | G/C-1 | none | G/C-1 | AGAAAACCA | CCAATAC | TGG | ACC | C | none | nope | nope_ACC | G/C-1_CCAATAC | 6 | FALSE | FALSE |
| 140_v.txt | 32951 | 32951-Deleti | Deletion | 1 | C | - | 16 | 28487 | 0.05616597 | 62.75 | rnr1Y285A_pC | TRUE | CAN | JSY2423 | 4 | rnr1Y285A_p | ACA | nope | >1bp-ins | G/C-1 | none | G/C-1 | CCAATACAT | CATGTAA | TGT | ACA | C | none | nope | nope_ACA | G/C-1_CATGTAA | 6 | FALSE | FALSE |
| 140_v.txt | 32958 | 32958-Deleti | Deletion | 1 | C | - | 32 | 28487 | 0.11233194 | 59.71875 | rnr1Y285A_pC | TRUE | CAN | JSY2423 | 4 | rnr1Y285A_p | ACC | nope | >1bp-ins | G/C-1 | none | G/C-1 | ATGTAACCA | CCATTGG | TGG | ACC | C | none | nope | nope_ACC | G/C-1_CCATTGG | 6 | FALSE | FALSE |
| 140_v.txt | 33000 | 33000-Deleti | Deletion | 1 | A | - | 33 | 28487 | 0.11584231 | 63.0909091 | rnr1Y285A_pC | TRUE | CAN | JSY2423 | 4 | rnr1Y285A_p | GAA | nope | >1bp-ins | A/T-1 | none | A/T-1 | TGTGAGAAA | AAAACTG | CTT | CTT | T | none | T>none | T>none_CTT | A/T-1_AAAACTG | 6 | FALSE | FALSE |
| 140_v.txt | 33004 | 33004-Deleti | Deletion | 1 | C | - | 16 | 28487 | 0.05616597 | 61.125 | rnr1Y285A_pC | TRUE | CAN | JSY2423 | 4 | rnr1Y285A_p | ACT | nope | >1bp-ins | G/C-1 | none | G/C-1 | AGAAAACTG | CTGTGAA | TGA | ACT | C | none | nope | nope_ACT | G/C-1_CTGTGAA | 6 | FALSE | FALSE |
| 140_v.txt | 33078 | 33078-Deleti | Deletion | 1 | C | - | 66 | 28487 | 0.23168463 | 53.2575758 | rnr1Y285A_pC | TRUE | CAN | JSY2423 | 4 | rnr1Y285A_p | ACC | nope | >1bp-ins | G/C-1 | none | G/C-1 | AAAGAACCC | CCCATAA | TGG | ACC | C | none | nope | nope_ACC | G/C-1_CCCATAA | 6 | FALSE | FALSE |
| 140_v.txt | 33114 | 33114-Deleti | Deletion | 1 | C | - | 1 | 28487 | 3.51E-03 | 38 | rnr1Y285A_pC | TRUE | CAN | JSY2423 | 4 | rnr1Y285A_p | GCC | nope | >1bp-ins | G/C-1 | none | G/C-1 | ACTGGGCCG | CCGGCGT | CGG | GCC | C | none | nope | nope_GCC | G/C-1_CCGGCGT | 6 | FALSE | FALSE |
| 140_v.txt | 33153 | 33153-Deleti | Deletion | 1 | C | - | 1 | 28487 | 3.51E-03 | 26 | rnr1Y285A_pC | TRUE | CAN | JSY2423 | 4 | rnr1Y285A_p | ACC | nope | >1bp-ins | G/C-1 | none | G/C-1 | AAAAGACCT | CCTGTAC | TGG | ACC | C | none | nope | nope_ACC | G/C-1_CCTGTAC | 6 | FALSE | FALSE |
| 140_v.txt | 33239 | 33239-Deleti | Deletion | 1 | T | - | 5 | 21084 | 0.02371467 | 43 | rnr1Y285A_pC | TRUE | CAN | JSY2423 | 4 | rnr1Y285A_p | CTT | nope | >1bp-ins | A/T-1 | none | A/T-1 | TTCTCCTTCA | TTCATCT | GAA | CTT | T | none | nope | nope_CTT | A/T-1_TTCATCT | 6 | FALSE | FALSE |
| 140_v.txt | 33356 | 33356-Deleti | Deletion | 1 | C | - | 1233 | 21084 | 5.84803643 | 54.4557989 | rnr1Y285A_pC | TRUE | CAN | JSY2423 | 4 | rnr1Y285A_p | ACC | nope | >1bp-ins | G/C-1 | none | G/C-1 | TATTGACCC | CCCACGT | TGG | ACC | C | none | nope | nope_ACC | G/C-1_CCCACGT | 6 | FALSE | FALSE |
| 140_v.txt | 33361 | 33361-Deleti | Deletion | 1 | G | - | 2780 | 21084 | 13.1853538 | 57.8629496 | rnr1Y285A_pC | TRUE | CAN | JSY2423 | 4 | rnr1Y285A_p | CGT | nope | >1bp-ins | G/C-1 | none | G/C-1 | ACCCACGTC | GTCTGTG | GCA | GCA | C | none | C>none | C>none_GCA | G/C-1_GTCTGTG | 6 | FALSE | FALSE |
| 140_v.txt | 33474 | 33474-Deleti | Deletion | 1 | C | - | 47 | 21083 | 0.22292843 | 26.1914893 | rnr1Y285A_pC | TRUE | CAN | JSY2423 | 4 | rnr1Y285A_p | GCC | nope | >1bp-ins | G/C-1 | none | G/C-1 | GCTATGCCTT | CCTTTTT | CGG | GCC | C | none | nope | nope_GCC | G/C-1_CCTTTTT | 6 | FALSE | FALSE |
| 176_v.txt | 31723 | 31723-SNV-A | SNV | 1 | A | C | 6 | 17332 | 0.03461805 | 60.3333333 | rnr1Y285A_pC | TRUE | CAN | JSY2422 | 5 | rnr1Y285A_p | CAA | A>C | G/C+1 | A/T-1 | TA>GC | TA>GC | TGTCCCAAA | AAAAAGT | GTT | GTT | T | G | T>G | T>G_GTT | TA>GC_AAAAAGT | 6 | FALSE | FALSE |
| 176_v.txt | 31840 | 31840-SNV-A | SNV | 1 | A | C | 3 | 17332 | 0.01730902 | 55.3333333 | rnr1Y285A_pC | TRUE | CAN | JSY2422 | 5 | rnr1Y285A_p | AAG | A>C | G/C+1 | A/T-1 | TA>GC | TA>GC | GAAATAAGA | AGATCCA | TTC | TTC | T | G | T>G | T>G_TTC | TA>GC_AGATCCA | 6 | FALSE | FALSE |
| 176_v.txt | 31913 | 31913-SNV-A | SNV | 1 | A | C | 1 | 17332 | 5.77E-03 | 37.9997145 | rnr1Y285A_pC | TRUE | CAN | JSY2422 | 5 | rnr1Y285A_p | CAA | A>C | G/C+1 | A/T-1 | TA>GC | TA>GC | TGGTGCAAA | AAAAGCC | GTT | GTT | T | G | T>G | T>G_GTT | TA>GC_AAAAGCC | 6 | FALSE | FALSE |
| 176_v.txt | 32383 | 32383-SNV-A | SNV | 1 | A | C | 6 | 40333 | 0.01487616 | 48.5 | rnr1Y285A_pC | TRUE | CAN | JSY2422 | 5 | rnr1Y285A_p | AAG | A>C | G/C+1 | A/T-1 | TA>GC | TA>GC | TAATAAAGG | AGGGAGA | TTC | TTC | T | G | T>G | T>G_TTC | TA>GC_AGGGAGA | 6 | FALSE | FALSE |
| 176_v.txt | 32493 | 32493-SNV-A | SNV | 1 | A | C | 7 | 40333 | 0.01735552 | 40.2848042 | rnr1Y285A_pC | TRUE | CAN | JSY2422 | 5 | rnr1Y285A_p | AAC | A>C | G/C+1 | A/T-1 | TA>GC | TA>GC | CGGAAAACA | ACAACTT | TTG | TTG | T | G | T>G | T>G_TTG | TA>GC_ACAACTT | 6 | FALSE | FALSE |
| 176_v.txt | 32559 | 32559-SNV-A | SNV | 1 | A | C | 29 | 40289 | 0.07197995 | 36.6896552 | rnr1Y285A_pC | TRUE | CAN | JSY2422 | 5 | rnr1Y285A_p | AAC | A>C | G/C+1 | A/T-1 | TA>GC | TA>GC | ATACCAACT | ACTAGTT | TTG | TTG | T | G | T>G | T>G_TTG | TA>GC_ACTAGTT | 6 | FALSE | FALSE |
| 176_v.txt | 32598 | 32598-SNV-A | SNV | 1 | A | C | 25 | 44912 | 0.05566441 | 42.16 | rnr1Y285A_pC | TRUE | CAN | JSY2422 | 5 | rnr1Y285A_p | CAA | A>C | G/C+1 | A/T-1 | TA>GC | TA>GC | TTAATCAAA | AAAGAGG | GTT | GTT | T | G | T>G | T>G_GTT | TA>GC_AAAGAGG | 6 | FALSE | FALSE |
| 176_v.txt | 32763 | 32763-SNV-A | SNV | 1 | A | C | 10 | 44912 | 0.02226576 | 51.8 | rnr1Y285A_pC | TRUE | CAN | JSY2422 | 5 | rnr1Y285A_p | AAC | A>C | G/C+1 | A/T-1 | TA>GC | TA>GC | GCTAAAACT | ACTTTGA | TTG | TTG | T | G | T>G | T>G_TTG | TA>GC_ACTTTGA | 6 | FALSE | FALSE |
| 176_v.txt | 32803 | 32803-SNV-A | SNV | 1 | A | C | 21 | 44912 | 0.04675811 | 46.6190476 | rnr1Y285A_pC | TRUE | CAN | JSY2422 | 5 | rnr1Y285A_p | TAT | A>C | G/C+1 | A/T-1 | TA>GC | TA>GC | CGTAATATT | ATTTGAC | ATA | ATA | T | G | T>G | T>G_ATA | TA>GC_ATTTGAC | 6 | FALSE | FALSE |
| 176_v.txt | 32810 | 32810-SNV-A | SNV | 1 | A | C | 5 | 44912 | 0.01113288 | 42.6 | rnr1Y285A_pC | TRUE | CAN | JSY2422 | 5 | rnr1Y285A_p | CAG | A>C | G/C+1 | A/T-1 | TA>GC | TA>GC | TTTGACAGG | AGGGAAC | GTC | GTC | T | G | T>G | T>G_GTC | TA>GC_AGGGAAC | 6 | FALSE | FALSE |
| 176_v.txt | 32842 | 32842-SNV-A | SNV | 1 | A | C | 6 | 44912 | 0.01335946 | 38.9999107 | rnr1Y285A_pC | TRUE | CAN | JSY2422 | 5 | rnr1Y285A_p | CAA | A>C | G/C+1 | A/T-1 | TA>GC | TA>GC | TTACCCAAA | AAAAAAT | GTT | GTT | T | G | T>G | T>G_GTT | TA>GC_AAAAAAT | 6 | FALSE | FALSE |
| 176_v.txt | 32884 | 32884-SNV-A | SNV | 1 | A | C | 18 | 69133 | 0.02603677 | 33.6111111 | rnr1Y285A_pC | TRUE | CAN | JSY2422 | 5 | rnr1Y285A_p | CAA | A>C | G/C+1 | A/T-1 | TA>GC | TA>GC | ACGTCCAAA | AAAATTG | GTT | GTT | T | G | T>G | T>G_GTT | TA>GC_AAAATTG | 6 | FALSE | FALSE |
| 176_v.txt | 32913 | 32913-SNV-A | SNV | 1 | A | C | 69 | 69133 | 0.09980762 | 32.5797101 | rnr1Y285A_pC | TRUE | CAN | JSY2422 | 5 | rnr1Y285A_p | AAG | A>C | G/C+1 | A/T-1 | TA>GC | TA>GC | ACACTAAGT | AGTTCCA | TTC | TTC | T | G | T>G | T>G_TTC | TA>GC_AGTTCCA | 6 | FALSE | FALSE |
| 176_v.txt | 32915 | 32915-SNV-T | SNV | 1 | T | G | 27 | 69133 | 0.03905516 | 33.9259259 | rnr1Y285A_pC | TRUE | CAN | JSY2422 | 5 | rnr1Y285A_p | GTT | T>G | G/C+1 | A/T-1 | TA>GC | TA>GC | ACTAAGTTC | TTCCAGG | CAA | GTT | T | C | T>G | T>G_GTT | TA>GC_TTCCAGG | 6 | FALSE | FALSE |
| 176_v.txt | 32916 | 32916-SNV-T | SNV | 1 | T | G | 7 | 69133 | 0.01012541 | 19.8571429 | rnr1Y285A_pC | TRUE | CAN | JSY2422 | 5 | rnr1Y285A_p | TTC | T>G | G/C+1 | A/T-1 | TA>GC | TA>GC | CTAAGTTCC | TCCAGGG | AAG | TTC | T | C | T>G | T>G_TTC | TA>GC_TCCAGGG | 6 | FALSE | FALSE |
| 176_v.txt | 32919 | 32919-SNV-A | SNV | 1 | A | C | 18 | 69133 | 0.02603677 | 26.7238404 | rnr1Y285A_pC | TRUE | CAN | JSY2422 | 5 | rnr1Y285A_p | CAG | A>C | G/C+1 | A/T-1 | TA>GC | TA>GC | AGTTCCAGG | AGGGCAA | GTC | GTC | T | G | T>G | T>G_GTC | TA>GC_AGGGCAA | 6 | FALSE | FALSE |
| 176_v.txt | 32941 | 32941-SNV-A | SNV | 1 | A | C | 18 | 69133 | 0.02603677 | 10.6667678 | rnr1Y285A_pC | TRUE | CAN | JSY2422 | 5 | rnr1Y285A_p | GAA | A>C | G/C+1 | A/T-1 | TA>GC | TA>GC | CCCAAGAAA | AAAACCA | CTT | CTT | T | G | T>G | T>G_CTT | TA>GC_AAAACCA | 6 | FALSE | FALSE |
| 176_v.txt | 33099 | 33099-SNV-A | SNV | 1 | A | C | 11 | 69133 | 0.01591136 | 42.1818182 | rnr1Y285A_pC | TRUE | CAN | JSY2422 | 5 | rnr1Y285A_p | AAG | A>C | G/C+1 | A/T-1 | TA>GC | TA>GC | GATATAAGA | AGAGCGC | TTC | TTC | T | G | T>G | T>G_TTC | TA>GC_AGAGCGC | 6 | FALSE | FALSE |
| 176_v.txt | 33269 | 33269-SNV-A | SNV | 1 | A | C | 223 | 33429 | 0.66708547 | 16.1524664 | rnr1Y285A_pC | TRUE | CAN | JSY2422 | 5 | rnr1Y285A_p | TAG | A>C | G/C+1 | A/T-1 | TA>GC | TA>GC | CTCCATAGA | AGAGAAC | ATC | ATC | T | G | T>G | T>G_ATC | TA>GC_AGAGAAC | 6 | FALSE | FALSE |
| 176_v.txt | 33476 | 33476-SNV-T | SNV | 1 | T | G | 6 | 33428 | 0.01794903 | 32.0003601 | rnr1Y285A_pC | TRUE | CAN | JSY2422 | 5 | rnr1Y285A_p | CTT | T>G | G/C+1 | A/T-1 | TA>GC | TA>GC | TATGCCTTTT | TTTTTTT | GAA | CTT | T | C | T>G | T>G_CTT | TA>GC_TTTTTTT | 6 | FALSE | FALSE |
| 176_v.txt | 31840 | 31840-SNV-A | SNV | 1 | A | G | 12 | 17332 | 0.0692361 | 55.5833333 | rnr1Y285A_pC | TRUE | CAN | JSY2422 | 5 | rnr1Y285A_p | AAG | A>G | G/C+1 | A/T-1 | TA>CG | TA>CG | GAAATAAGA | AGATCCA | TTC | TTC | T | C | T>C | T>C_TTC | TA>CG_AGATCCA | 6 | FALSE | FALSE |
| 176_v.txt | 32068 | 32068-SNV-T | SNV | 1 | T | C | 2 | 37085 | 5.39E-03 | 63.9999826 | rnr1Y285A_pC | TRUE | CAN | JSY2422 | 5 | rnr1Y285A_p | ATA | T>C | G/C+1 | A/T-1 | TA>CG | TA>CG | TTGAGATAA | TAAATAA | TAT | ATA | T | G | T>C | T>C_ATA | TA>CG_TAAATAA | 6 | FALSE | FALSE |
| 176_v.txt | 32086 | 32086-SNV-A | SNV | 1 | A | G | 38 | 37086 | 0.10246454 | 58.7368421 | rnr1Y285A_pC | TRUE | CAN | JSY2422 | 5 | rnr1Y285A_p | AAG | A>G | G/C+1 | A/T-1 | TA>CG | TA>CG | CAAAAAAGC | AGCCTGC | TTC | TTC | T | C | T>C | T>C_TTC | TA>CG_AGCCTGC | 6 | FALSE | FALSE |
| 176_v.txt | 32810 | 32810-SNV-A | SNV | 1 | A | G | 35 | 44912 | 0.07793018 | 40.7428571 | rnr1Y285A_pC | TRUE | CAN | JSY2422 | 5 | rnr1Y285A_p | CAG | A>G | G/C+1 | A/T-1 | TA>CG | TA>CG | TTTGACAGG | AGGGAAC | GTC | GTC | T | C | T>C | T>C_GTC | TA>CG_AGGGAAC | 6 | FALSE | FALSE |
| 176_v.txt | 32915 | 32915-SNV-T | SNV | 1 | T | C | 35 | 69133 | 0.05062705 | 25.7428571 | rnr1Y285A_pC | TRUE | CAN | JSY2422 | 5 | rnr1Y285A_p | GTT | T>C | G/C+1 | A/T-1 | TA>CG | TA>CG | ACTAAGTTC | TTCCAGG | CAA | GTT | T | G | T>C | T>C_GTT | TA>CG_TTCCAGG | 6 | FALSE | FALSE |
| 176_v.txt | 32916 | 32916-SNV-T | SNV | 1 | T | C | 15 | 69133 | 0.02169731 | 31.5333333 | rnr1Y285A_pC | TRUE | CAN | JSY2422 | 5 | rnr1Y285A_p | TTC | T>C | G/C+1 | A/T-1 | TA>CG | TA>CG | CTAAGTTCC | TCCAGGG | AAG | TTC | T | G | T>C | T>C_TTC | TA>CG_TCCAGGG | 6 | FALSE | FALSE |
| 176_v.txt | 32919 | 32919-SNV-A | SNV | 1 | A | G | 49 | 69133 | 0.07087787 | 44.3442823 | rnr1Y285A_pC | TRUE | CAN | JSY2422 | 5 | rnr1Y285A_p | CAG | A>G | G/C+1 | A/T-1 | TA>CG | TA>CG | AGTTCCAGG | AGGGCAA | GTC | GTC | T | C | T>C | T>C_GTC | TA>CG_AGGGCAA | 6 | FALSE | FALSE |
| 176_v.txt | 32927 | 32927-SNV-A | SNV | 1 | A | G | 72 | 69133 | 0.10414708 | 49.6666667 | rnr1Y285A_pC | TRUE | CAN | JSY2422 | 5 | rnr1Y285A_p | AAG | A>G | G/C+1 | A/T-1 | TA>CG | TA>CG | GGCAAAAGT | AGTGATT | TTC | TTC | T | C | T>C | T>C_TTC | TA>CG_AGTGATT | 6 | FALSE | FALSE |
| 176_v.txt | 33099 | 33099-SNV-A | SNV | 1 | A | G | 36 | 69133 | 0.05207354 | 49.3333333 | rnr1Y285A_pC | TRUE | CAN | JSY2422 | 5 | rnr1Y285A_p | AAG | A>G | G/C+1 | A/T-1 | TA>CG | TA>CG | GATATAAGA | AGAGCGC | TTC | TTC | T | C | T>C | T>C_TTC | TA>CG_AGAGCGC | 6 | FALSE | FALSE |
| 176_v.txt | 33150 | 33150-SNV-A | SNV | 1 | A | G | 49 | 69133 | 0.07087787 | 36.4490745 | rnr1Y285A_pC | TRUE | CAN | JSY2422 | 5 | rnr1Y285A_p | AAG | A>G | G/C+1 | A/T-1 | TA>CG | TA>CG | ATGAAAAGA | AGACCTG | TTC | TTC | T | C | T>C | T>C_TTC | TA>CG_AGACCTG | 6 | FALSE | FALSE |
| 176_v.txt | 33326 | 33326-SNV-T | SNV | 1 | T | C | 5 | 33427 | 0.01495797 | 63.9997511 | rnr1Y285A_pC | TRUE | CAN | JSY2422 | 5 | rnr1Y285A_p | CTT | T>C | G/C+1 | A/T-1 | TA>CG | TA>CG | CAATTCTTTA | TTTACTT | GAA | CTT | T | G | T>C | T>C_CTT | TA>CG_TTTACTT | 6 | FALSE | FALSE |
| 176_v.txt | 33476 | 33476-SNV-T | SNV | 1 | T | C | 52 | 33428 | 0.15555822 | 30.6539925 | rnr1Y285A_pC | TRUE | CAN | JSY2422 | 5 | rnr1Y285A_p | CTT | T>C | G/C+1 | A/T-1 | TA>CG | TA>CG | TATGCCTTTT | TTTTTTT | GAA | CTT | T | G | T>C | T>C_CTT | TA>CG_TTTTTTT | 6 | FALSE | FALSE |
| 176_v.txt | 31840 | 31840-SNV-A | SNV | 1 | A | T | 7 | 17332 | 0.04038772 | 48.7142857 | rnr1Y285A_pC | TRUE | CAN | JSY2422 | 5 | rnr1Y285A_p | AAG | A>T | A/T+1 | A/T-1 | TA>AT | TA>AT | GAAATAAGA | AGATCCA | TTC | TTC | T | A | T>A | T>A_TTC | TA>AT_AGATCCA | 6 | FALSE | FALSE |
| 176_v.txt | 32068 | 32068-SNV-T | SNV | 1 | T | A | 6 | 37085 | 0.01617905 | 63.9994276 | rnr1Y285A_pC | TRUE | CAN | JSY2422 | 5 | rnr1Y285A_p | ATA | T>A | A/T+1 | A/T-1 | TA>AT | TA>AT | TTGAGATAA | TAAATAA | TAT | ATA | T | T | T>A | T>A_ATA | TA>AT_TAAATAA | 6 | FALSE | FALSE |
| 176_v.txt | 32070 | 32070-SNV-A | SNV | 1 | A | T | 3 | 37085 | 8.09E-03 | 63.9996946 | rnr1Y285A_pC | TRUE | CAN | JSY2422 | 5 | rnr1Y285A_p | AAA | A>T | A/T+1 | A/T-1 | TA>AT | TA>AT | GAGATAAAT | AATAACC | TTT | TTT | T | A | T>A | T>A_TTT | TA>AT_AATAACC | 6 | FALSE | FALSE |
| 176_v.txt | 32383 | 32383-SNV-A | SNV | 1 | A | T | 13 | 40333 | 0.03223167 | 56.9230769 | rnr1Y285A_pC | TRUE | CAN | JSY2422 | 5 | rnr1Y285A_p | AAG | A>T | A/T+1 | A/T-1 | TA>AT | TA>AT | TAATAAAGG | AGGGAGA | TTC | TTC | T | A | T>A | T>A_TTC | TA>AT_AGGGAGA | 6 | FALSE | FALSE |
| 176_v.txt | 32427 | 32427-SNV-T | SNV | 1 | T | A | 59 | 40333 | 0.1462822 | 55.0338983 | rnr1Y285A_pC | TRUE | CAN | JSY2422 | 5 | rnr1Y285A_p | GTC | T>A | A/T+1 | A/T-1 | TA>AT | TA>AT | TTAGGGTCA | TCATTGT | CAG | GTC | T | T | T>A | T>A_GTC | TA>AT_TCATTGT | 6 | FALSE | FALSE |
| 176_v.txt | 32496 | 32496-SNV-A | SNV | 1 | A | T | 24 | 40333 | 0.05950462 | 43.3333333 | rnr1Y285A_pC | TRUE | CAN | JSY2422 | 5 | rnr1Y285A_p | AAC | A>T | A/T+1 | A/T-1 | TA>AT | TA>AT | AAAACAACT | ACTTTTT | TTG | TTG | T | A | T>A | T>A_TTG | TA>AT_ACTTTTT | 6 | FALSE | FALSE |
| 176_v.txt | 32801 | 32801-SNV-A | SNV | 1 | A | T | 32 | 44912 | 0.07125045 | 45.65625 | rnr1Y285A_pC | TRUE | CAN | JSY2422 | 5 | rnr1Y285A_p | AAT | A>T | A/T+1 | A/T-1 | TA>AT | TA>AT | ACCGTAATA | ATATTTG | TTA | TTA | T | A | T>A | T>A_TTA | TA>AT_ATATTTG | 6 | FALSE | FALSE |
| 176_v.txt | 32915 | 32915-SNV-T | SNV | 1 | T | A | 33 | 69133 | 0.04773408 | 16.5454546 | rnr1Y285A_pC | TRUE | CAN | JSY2422 | 5 | rnr1Y285A_p | GTT | T>A | A/T+1 | A/T-1 | TA>AT | TA>AT | ACTAAGTTC | TTCCAGG | CAA | GTT | T | T | T>A | T>A_GTT | TA>AT_TTCCAGG | 6 | FALSE | FALSE |
| 176_v.txt | 32916 | 32916-SNV-T | SNV | 1 | T | A | 67 | 69133 | 0.09691464 | 50.0746269 | rnr1Y285A_pC | TRUE | CAN | JSY2422 | 5 | rnr1Y285A_p | TTC | T>A | A/T+1 | A/T-1 | TA>AT | TA>AT | CTAAGTTCC | TCCAGGG | AAG | TTC | T | T | T>A | T>A_TTC | TA>AT_TCCAGGG | 6 | FALSE | FALSE |
| 176_v.txt | 33094 | 33094-SNV-A | SNV | 1 | A | T | 11 | 69132 | 0.01591159 | 46.1818089 | rnr1Y285A_pC | TRUE | CAN | JSY2422 | 5 | rnr1Y285A_p | GAT | A>T | A/T+1 | A/T-1 | TA>AT | TA>AT | AATATGATA | ATATAAG | CTA | CTA | T | A | T>A | T>A_CTA | TA>AT_ATATAAG | 6 | FALSE | FALSE |
| 176_v.txt | 33095 | 33095-SNV-T | SNV | 1 | T | A | 9 | 69133 | 0.01301839 | 43.664852 | rnr1Y285A_pC | TRUE | CAN | JSY2422 | 5 | rnr1Y285A_p | ATA | T>A | A/T+1 | A/T-1 | TA>AT | TA>AT | ATATGATAT | TATAAGA | TAT | ATA | T | T | T>A | T>A_ATA | TA>AT_TATAAGA | 6 | FALSE | FALSE |
| 176_v.txt | 33180 | 33180-SNV-A | SNV | 1 | A | T | 74 | 69084 | 0.10711598 | 36.4324324 | rnr1Y285A_pC | TRUE | CAN | JSY2422 | 5 | rnr1Y285A_p | AAT | A>T | A/T+1 | A/T-1 | TA>AT | TA>AT | AGGGCAATC | ATCATAC | TTA | TTA | T | A | T>A | T>A_TTA | TA>AT_ATCATAC | 6 | FALSE | FALSE |
| 176_v.txt | 33326 | 33326-SNV-T | SNV | 1 | T | A | 2 | 33427 | 5.98E-03 | 37.4990575 | rnr1Y285A_pC | TRUE | CAN | JSY2422 | 5 | rnr1Y285A_p | CTT | T>A | A/T+1 | A/T-1 | TA>AT | TA>AT | CAATTCTTTA | TTTACTT | GAA | CTT | T | T | T>A | T>A_CTT | TA>AT_TTTACTT | 6 | FALSE | FALSE |

| | | | | | | | | | | | | | | | | | | | | | | | | | | | | | | | | | | |
|---|---|---|---|---|---|---|---|---|---|---|---|---|---|---|---|---|---|---|---|---|---|---|---|---|---|---|---|---|---|---|---|---|---|---|
| 176_v.txt | 31650 | 31650-SNV-( | SNV | 1 | C | T | 30 | 17332 | 0.17309024 | 35.0333333 | rnr1Y285A_p( | TRUE | CAN | JSY2422 | 5 | rnr1Y285A_p | GCG | C>T | A/T+1 | G/C-1 | CG>TA | CG>TA | AGAATGCGA | CGAAATG | CGC | GCG | C | A | C>T | C>T_GCG | CG>TA_CGAAATG | 6 | FALSE | FALSE |
| 176_v.txt | 31658 | 31658-SNV-( | SNV | 1 | C | T | 34 | 17332 | 0.19616894 | 34.970731 | rnr1Y285A_p( | TRUE | CAN | JSY2422 | 5 | rnr1Y285A_p | GCG | C>T | A/T+1 | G/C-1 | CG>TA | CG>TA | AAATGGCGT | CGTGGAA | CGC | GCG | C | A | C>T | C>T_GCG | CG>TA_CGTGGAA | 6 | FALSE | FALSE |
| 176_v.txt | 31830 | 31830-SNV-( | SNV | 1 | C | T | 27 | 17332 | 0.15578121 | 55.7777778 | rnr1Y285A_p( | TRUE | CAN | JSY2422 | 5 | rnr1Y285A_p | GCA | C>T | A/T+1 | G/C-1 | CG>TA | CG>TA | AATATGCATT | CATTGAA | CGT | GCA | C | A | C>T | C>T_GCA | CG>TA_CATTGAA | 6 | FALSE | FALSE |
| 176_v.txt | 31844 | 31844-SNV-( | SNV | 1 | C | T | 211 | 17332 | 1.21740134 | 53.2748815 | rnr1Y285A_p( | TRUE | CAN | JSY2422 | 5 | rnr1Y285A_p | TCC | C>T | A/T+1 | G/C-1 | CG>TA | CG>TA | TAAGATCCA | CCAAACA | AGG | TCC | C | A | C>T | C>T_TCC | CG>TA_CCAAACA | 6 | FALSE | FALSE |
| 176_v.txt | 31879 | 31879-SNV-( | SNV | 1 | C | T | 16 | 17332 | 0.09231479 | 42.75 | rnr1Y285A_p( | TRUE | CAN | JSY2422 | 5 | rnr1Y285A_p | GCG | C>T | A/T+1 | G/C-1 | CG>TA | CG>TA | TATAGGCGG | CGGCAGC | CGC | GCG | C | A | C>T | C>T_GCG | CG>TA_CGGCAGC | 6 | FALSE | FALSE |
| 176_v.txt | 31881 | 31881-SNV-( | SNV | 1 | G | A | 21 | 17332 | 0.12116317 | 49.7140829 | rnr1Y285A_p( | TRUE | CAN | JSY2422 | 5 | rnr1Y285A_p | GGC | G>A | A/T+1 | G/C-1 | CG>TA | CG>TA | TAGGCGGCA | GCAGCAA | CCG | CCG | C | T | C>T | C>T_CCG | CG>TA_GCAGCAA | 6 | FALSE | FALSE |
| 176_v.txt | 31882 | 31882-SNV-( | SNV | 1 | C | T | 23 | 17332 | 0.13270252 | 42.7341557 | rnr1Y285A_p( | TRUE | CAN | JSY2422 | 5 | rnr1Y285A_p | GCA | C>T | A/T+1 | G/C-1 | CG>TA | CG>TA | AGGCGGCAC | CAGCAAA | CGT | GCA | C | A | C>T | C>T_GCA | CG>TA_CAGCAAA | 6 | FALSE | FALSE |
| 176_v.txt | 31884 | 31884-SNV-( | SNV | 1 | G | A | 20 | 17332 | 0.11539349 | 36.15 | rnr1Y285A_p( | TRUE | CAN | JSY2422 | 5 | rnr1Y285A_p | AGC | G>A | A/T+1 | G/C-1 | CG>TA | CG>TA | GCGGCAGCA | GCAAAGC | TCG | TCG | C | T | C>T | C>T_TCG | CG>TA_GCAAAGC | 6 | FALSE | FALSE |
| 176_v.txt | 31912 | 31912-SNV-( | SNV | 1 | C | T | 18 | 17332 | 0.10385414 | 38.7198203 | rnr1Y285A_p( | TRUE | CAN | JSY2422 | 5 | rnr1Y285A_p | GCA | C>T | A/T+1 | G/C-1 | CG>TA | CG>TA | TTGGTGCAA | CAAAAGC | CGT | GCA | C | A | C>T | C>T_GCA | CG>TA_CAAAAGC | 6 | FALSE | FALSE |
| 176_v.txt | 31920 | 31920-SNV-( | SNV | 1 | G | A | 17 | 17329 | 0.09810145 | 37.0588235 | rnr1Y285A_p( | TRUE | CAN | JSY2422 | 5 | rnr1Y285A_p | CGT | G>A | A/T+1 | G/C-1 | CG>TA | CG>TA | AAAGCCGTG | GTGAAAC | GCA | GCA | C | T | C>T | C>T_GCA | CG>TA_GTGAAAC | 6 | FALSE | FALSE |
| 176_v.txt | 31947 | 31947-SNV-( | SNV | 1 | G | A | 36 | 37050 | 0.09716599 | 35.6111111 | rnr1Y285A_p( | TRUE | CAN | JSY2422 | 5 | rnr1Y285A_p | CGT | G>A | A/T+1 | G/C-1 | CG>TA | CG>TA | ATGATCGTC | GTCATAA | GCA | GCA | C | T | C>T | C>T_GCA | CG>TA_GTCATAA | 6 | FALSE | FALSE |
| 176_v.txt | 31962 | 31962-SNV-( | SNV | 1 | G | A | 61 | 37087 | 0.16447812 | 36.6557377 | rnr1Y285A_p( | TRUE | CAN | JSY2422 | 5 | rnr1Y285A_p | CGC | G>A | A/T+1 | G/C-1 | CG>TA | CG>TA | GTGGCCGCA | GCATAAT | GCG | GCG | C | T | C>T | C>T_GCG | CG>TA_GCATAAT | 6 | FALSE | FALSE |
| 176_v.txt | 31972 | 31972-SNV-( | SNV | 1 | C | T | 36 | 37086 | 0.09707167 | 36.6666667 | rnr1Y285A_p( | TRUE | CAN | JSY2422 | 5 | rnr1Y285A_p | GCC | C>T | A/T+1 | G/C-1 | CG>TA | CG>TA | AATAAGCCA | CCAAGCC | CGG | GCC | C | A | C>T | C>T_GCC | CG>TA_CCAAGCC | 6 | FALSE | FALSE |
| 176_v.txt | 31977 | 31977-SNV-( | SNV | 1 | C | T | 37 | 37086 | 0.09976811 | 42.5405405 | rnr1Y285A_p( | TRUE | CAN | JSY2422 | 5 | rnr1Y285A_p | GCC | C>T | A/T+1 | G/C-1 | CG>TA | CG>TA | GCCAAGCCG | CCGGGCA | CGG | GCC | C | A | C>T | C>T_GCC | CG>TA_CCGGGCA | 6 | FALSE | FALSE |
| 176_v.txt | 31981 | 31981-SNV-( | SNV | 1 | G | A | 117 | 37086 | 0.31548293 | 50 | rnr1Y285A_p( | TRUE | CAN | JSY2422 | 5 | rnr1Y285A_p | GGC | G>A | A/T+1 | G/C-1 | CG>TA | CG>TA | AGCCGGGCA | GCATTAA | CCG | CCG | C | T | C>T | C>T_CCG | CG>TA_GCATTAA | 6 | FALSE | FALSE |
| 176_v.txt | 32041 | 32041-SNV-( | SNV | 1 | G | A | 27 | 37086 | 0.07280375 | 58.5185185 | rnr1Y285A_p( | TRUE | CAN | JSY2422 | 5 | rnr1Y285A_p | TGC | G>A | A/T+1 | G/C-1 | CG>TA | CG>TA | AAGCTTGCA | GCATAAA | ACG | ACG | C | T | C>T | C>T_ACG | CG>TA_GCATAAA | 6 | FALSE | FALSE |
| 176_v.txt | 32057 | 32057-SNV-( | SNV | 1 | C | T | 35 | 37086 | 0.09437524 | 60.4571429 | rnr1Y285A_p( | TRUE | CAN | JSY2422 | 5 | rnr1Y285A_p | GCG | C>T | A/T+1 | G/C-1 | CG>TA | CG>TA | GATGTGCGA | CGAGATT | CGC | GCG | C | A | C>T | C>T_GCG | CG>TA_CGAGATT | 6 | FALSE | FALSE |
| 176_v.txt | 32075 | 32075-SNV-( | SNV | 1 | C | T | 627 | 37086 | 1.69066494 | 60.8357257 | rnr1Y285A_p( | TRUE | CAN | JSY2422 | 5 | rnr1Y285A_p | ACC | C>T | A/T+1 | G/C-1 | CG>TA | CG>TA | AAATAACCA | CCATGCA | TGG | ACC | C | A | C>T | C>T_ACC | CG>TA_CCATGCA | 6 | FALSE | FALSE |
| 176_v.txt | 32088 | 32088-SNV-( | SNV | 1 | C | T | 123 | 37082 | 0.33169732 | 60.2671473 | rnr1Y285A_p( | TRUE | CAN | JSY2422 | 5 | rnr1Y285A_p | GCC | C>T | A/T+1 | G/C-1 | CG>TA | CG>TA | AAAAAGCCT | CCTGCAA | CGG | GCC | C | A | C>T | C>T_GCC | CG>TA_CCTGCAA | 6 | FALSE | FALSE |
| 176_v.txt | 32092 | 32092-SNV-( | SNV | 1 | C | T | 49 | 37086 | 0.13212533 | 61.1020408 | rnr1Y285A_p( | TRUE | CAN | JSY2422 | 5 | rnr1Y285A_p | GCA | C>T | A/T+1 | G/C-1 | CG>TA | CG>TA | AGCCTGCAA | CAACACC | CGT | GCA | C | A | C>T | C>T_GCA | CG>TA_CAACACC | 6 | FALSE | FALSE |
| 176_v.txt | 32114 | 32114-SNV-( | SNV | 1 | C | T | 658 | 37086 | 1.77425444 | 60.2218845 | rnr1Y285A_p( | TRUE | CAN | JSY2422 | 5 | rnr1Y285A_p | GCC | C>T | A/T+1 | G/C-1 | CG>TA | CG>TA | TAATAGCCA | CCATTCG | CGG | GCC | C | A | C>T | C>T_GCC | CG>TA_CCATTCG | 6 | FALSE | FALSE |
| 176_v.txt | 32157 | 32157-SNV-( | SNV | 1 | G | A | 31 | 37086 | 0.0835895 | 47.516129 | rnr1Y285A_p( | TRUE | CAN | JSY2422 | 5 | rnr1Y285A_p | AGC | G>A | A/T+1 | G/C-1 | CG>TA | CG>TA | ATGTAAGCC | GCCAAAG | TCG | TCG | C | T | C>T | C>T_TCG | CG>TA_GCCAAAG | 6 | FALSE | FALSE |
| 176_v.txt | 32165 | 32165-SNV-( | SNV | 1 | G | A | 66 | 37086 | 0.17796473 | 51.6363636 | rnr1Y285A_p( | TRUE | CAN | JSY2422 | 5 | rnr1Y285A_p | CGC | G>A | A/T+1 | G/C-1 | CG>TA | CG>TA | CAAAGCGCC | GCCAAAT | GCG | GCG | C | T | C>T | C>T_GCG | CG>TA_GCCAAAT | 6 | FALSE | FALSE |
| 176_v.txt | 32173 | 32173-SNV-( | SNV | 1 | C | T | 42 | 37086 | 0.11325028 | 45.5238095 | rnr1Y285A_p( | TRUE | CAN | JSY2422 | 5 | rnr1Y285A_p | GCA | C>T | A/T+1 | G/C-1 | CG>TA | CG>TA | CAAATGCAG | CAGCAGT | CGT | GCA | C | A | C>T | C>T_GCA | CG>TA_CAGCAGT | 6 | FALSE | FALSE |
| 176_v.txt | 32176 | 32176-SNV-( | SNV | 1 | C | T | 39 | 37086 | 0.10516098 | 46.948718 | rnr1Y285A_p( | TRUE | CAN | JSY2422 | 5 | rnr1Y285A_p | GCA | C>T | A/T+1 | G/C-1 | CG>TA | CG>TA | ATGCAGCAG | CAGTAAC | CGT | GCA | C | A | C>T | C>T_GCA | CG>TA_CAGTAAC | 6 | FALSE | FALSE |
| 176_v.txt | 32190 | 32190-SNV-( | SNV | 1 | G | A | 24 | 37086 | 0.06471445 | 45.3333333 | rnr1Y285A_p( | TRUE | CAN | JSY2422 | 5 | rnr1Y285A_p | TGC | G>A | A/T+1 | G/C-1 | CG>TA | CG>TA | AAAACTGCA | GCAATGT | ACG | ACG | C | T | C>T | C>T_ACG | CG>TA_GCAATGT | 6 | FALSE | FALSE |
| 176_v.txt | 32259 | 32259-SNV-( | SNV | 1 | C | T | 53 | 40331 | 0.13141256 | 30.2264151 | rnr1Y285A_p( | TRUE | CAN | JSY2422 | 5 | rnr1Y285A_p | ACC | C>T | A/T+1 | G/C-1 | CG>TA | CG>TA | GATAGACCA | CCAAATA | TGG | ACC | C | A | C>T | C>T_ACC | CG>TA_CCAAATA | 6 | FALSE | FALSE |
| 176_v.txt | 32271 | 32271-SNV-( | SNV | 1 | C | T | 220 | 40332 | 0.54547258 | 35.2067017 | rnr1Y285A_p( | TRUE | CAN | JSY2422 | 5 | rnr1Y285A_p | ACG | C>T | A/T+1 | G/C-1 | CG>TA | CG>TA | AAAATACGG | CGGGAAC | TGC | ACG | C | A | C>T | C>T_ACG | CG>TA_CGGGAAC | 6 | FALSE | FALSE |
| 176_v.txt | 32274 | 32274-SNV-( | SNV | 1 | G | A | 80 | 40332 | 0.19835367 | 37.25 | rnr1Y285A_p( | TRUE | CAN | JSY2422 | 5 | rnr1Y285A_p | GGA | G>A | A/T+1 | G/C-1 | CG>TA | CG>TA | ATACGGGAA | GAACCAA | CCT | CCT | C | T | C>T | C>T_CCT | CG>TA_GAACCAA | 6 | FALSE | FALSE |
| 176_v.txt | 32299 | 32299-SNV-( | SNV | 1 | C | T | 75 | 40333 | 0.18595195 | 53.1333333 | rnr1Y285A_p( | TRUE | CAN | JSY2422 | 5 | rnr1Y285A_p | GCG | C>T | A/T+1 | G/C-1 | CG>TA | CG>TA | AATTTGCGG | CGGCAGA | CGC | GCG | C | A | C>T | C>T_GCG | CG>TA_CGGCAGA | 6 | FALSE | FALSE |
| 176_v.txt | 32301 | 32301-SNV-( | SNV | 1 | G | A | 69 | 40333 | 0.17107579 | 57.8985507 | rnr1Y285A_p( | TRUE | CAN | JSY2422 | 5 | rnr1Y285A_p | GGC | G>A | A/T+1 | G/C-1 | CG>TA | CG>TA | TTTGCGGCA | GCAGAAA | CCG | CCG | C | T | C>T | C>T_CCG | CG>TA_GCAGAAA | 6 | FALSE | FALSE |
| 176_v.txt | 32344 | 32344-SNV-( | SNV | 1 | G | A | 31 | 40333 | 0.07686014 | 57.7741936 | rnr1Y285A_p( | TRUE | CAN | JSY2422 | 5 | rnr1Y285A_p | GGC | G>A | A/T+1 | G/C-1 | CG>TA | CG>TA | TATGTGGCA | GCAAAAC | CCG | CCG | C | T | C>T | C>T_CCG | CG>TA_GCAAAAC | 6 | FALSE | FALSE |
| 176_v.txt | 32466 | 32466-SNV-( | SNV | 1 | C | T | 60 | 40333 | 0.14876156 | 46.7666667 | rnr1Y285A_p( | TRUE | CAN | JSY2422 | 5 | rnr1Y285A_p | GCC | C>T | A/T+1 | G/C-1 | CG>TA | CG>TA | AGAGAGCCA | CCAATGT | CGG | GCC | C | A | C>T | C>T_GCC | CG>TA_CCAATGT | 6 | FALSE | FALSE |
| 176_v.txt | 32487 | 32487-SNV-( | SNV | 1 | C | T | 23 | 40333 | 0.05702527 | 46.9491823 | rnr1Y285A_p( | TRUE | CAN | JSY2422 | 5 | rnr1Y285A_p | ACG | C>T | A/T+1 | G/C-1 | CG>TA | CG>TA | AAGATACGG | CGGAAAA | TGC | ACG | C | A | C>T | C>T_ACG | CG>TA_CGGAAAA | 6 | FALSE | FALSE |
| 176_v.txt | 32488 | 32488-SNV-( | SNV | 1 | G | A | 43 | 40333 | 0.10661245 | 44.7911729 | rnr1Y285A_p( | TRUE | CAN | JSY2422 | 5 | rnr1Y285A_p | CGG | G>A | A/T+1 | G/C-1 | CG>TA | CG>TA | AGATACGGA | GGAAAAC | GCC | GCC | C | T | C>T | C>T_GCC | CG>TA_GGAAAAC | 6 | FALSE | FALSE |
| 176_v.txt | 32508 | 32508-SNV-( | SNV | 1 | G | A | 30 | 40333 | 0.07438078 | 35.7666667 | rnr1Y285A_p( | TRUE | CAN | JSY2422 | 5 | rnr1Y285A_p | GGC | G>A | A/T+1 | G/C-1 | CG>TA | CG>TA | TTGATGGCT | GCTCTTG | CCG | CCG | C | T | C>T | C>T_CCG | CG>TA_GCTCTTG | 6 | FALSE | FALSE |
| 176_v.txt | 32535 | 32535-SNV-( | SNV | 1 | G | A | 61 | 40333 | 0.15124092 | 36.6016172 | rnr1Y285A_p( | TRUE | CAN | JSY2422 | 5 | rnr1Y285A_p | TGC | G>A | A/T+1 | G/C-1 | CG>TA | CG>TA | GGGTTTGCA | GCAGCTT | ACG | ACG | C | T | C>T | C>T_ACG | CG>TA_GCAGCTT | 6 | FALSE | FALSE |
| 176_v.txt | 32536 | 32536-SNV-( | SNV | 1 | C | T | 76 | 40333 | 0.18843131 | 37.2461733 | rnr1Y285A_p( | TRUE | CAN | JSY2422 | 5 | rnr1Y285A_p | GCA | C>T | A/T+1 | G/C-1 | CG>TA | CG>TA | GGTTTGCAG | CAGCTTC | CGT | GCA | C | A | C>T | C>T_GCA | CG>TA_CAGCTTC | 6 | FALSE | FALSE |
| 176_v.txt | 32539 | 32539-SNV-( | SNV | 1 | C | T | 82 | 40333 | 0.20330747 | 37.2195122 | rnr1Y285A_p( | TRUE | CAN | JSY2422 | 5 | rnr1Y285A_p | GCT | C>T | A/T+1 | G/C-1 | CG>TA | CG>TA | TTGCAGCTT | CTTCACC | CGA | GCT | C | A | C>T | C>T_GCT | CG>TA_CTTCACC | 6 | FALSE | FALSE |
| 176_v.txt | 32571 | 32571-SNV-( | SNV | 1 | C | T | 8 | 44914 | 0.01781182 | 32.2476494 | rnr1Y285A_p( | TRUE | CAN | JSY2422 | 5 | rnr1Y285A_p | ACC | C>T | A/T+1 | G/C-1 | CG>TA | CG>TA | TCAGTACCT | CCTTGAA | TGG | ACC | C | A | C>T | C>T_ACC | CG>TA_CCTTGAA | 6 | FALSE | FALSE |
| 176_v.txt | 32587 | 32587-SNV-( | SNV | 1 | C | T | 35 | 44912 | 0.07793018 | 33.6 | rnr1Y285A_p( | TRUE | CAN | JSY2422 | 5 | rnr1Y285A_p | GCA | C>T | A/T+1 | G/C-1 | CG>TA | CG>TA | TGAAGGCAG | CAGCGTT | CGT | GCA | C | A | C>T | C>T_GCA | CG>TA_CAGCGTT | 6 | FALSE | FALSE |
| 176_v.txt | 32590 | 32590-SNV-( | SNV | 1 | C | T | 91 | 44912 | 0.20261845 | 36.8461539 | rnr1Y285A_p( | TRUE | CAN | JSY2422 | 5 | rnr1Y285A_p | GCG | C>T | A/T+1 | G/C-1 | CG>TA | CG>TA | AGGCAGCGT | CGTTAAT | CGC | GCG | C | A | C>T | C>T_GCG | CG>TA_CGTTAAT | 6 | FALSE | FALSE |
| 176_v.txt | 32658 | 32658-SNV-( | SNV | 1 | C | T | 33 | 44912 | 0.07347702 | 53.7575758 | rnr1Y285A_p( | TRUE | CAN | JSY2422 | 5 | rnr1Y285A_p | ACC | C>T | A/T+1 | G/C-1 | CG>TA | CG>TA | CCTGGACCC | CCCCAGG | TGG | ACC | C | A | C>T | C>T_ACC | CG>TA_CCCCAGG | 6 | FALSE | FALSE |
| 176_v.txt | 32661 | 32661-SNV-( | SNV | 1 | C | T | 100 | 44912 | 0.22265764 | 59.97 | rnr1Y285A_p( | TRUE | CAN | JSY2422 | 5 | rnr1Y285A_p | CCA | C>T | A/T+1 | G/C-1 | CG>TA | CG>TA | GGACCCCAG | CAGGCAC | GGT | CCA | C | A | C>T | C>T_CCA | CG>TA_CAGGCAC | 6 | FALSE | FALSE |
| 176_v.txt | 32679 | 32679-SNV-( | SNV | 1 | C | T | 40 | 44912 | 0.08906306 | 57.675 | rnr1Y285A_p( | TRUE | CAN | JSY2422 | 5 | rnr1Y285A_p | CCA | C>T | A/T+1 | G/C-1 | CG>TA | CG>TA | TTTCTCCAAT | CAATAAC | GGT | CCA | C | A | C>T | C>T_CCA | CG>TA_CAATAAC | 6 | FALSE | FALSE |
| 176_v.txt | 32700 | 32700-SNV-( | SNV | 1 | C | T | 41 | 44912 | 0.09128963 | 58.7804878 | rnr1Y285A_p( | TRUE | CAN | JSY2422 | 5 | rnr1Y285A_p | GCC | C>T | A/T+1 | G/C-1 | CG>TA | CG>TA | ACTGGGCCG | CCGGTAA | CGG | GCC | C | A | C>T | C>T_GCC | CG>TA_CCGGTAA | 6 | FALSE | FALSE |
| 176_v.txt | 32702 | 32702-SNV-( | SNV | 1 | G | A | 39 | 44912 | 0.08683648 | 57.3333333 | rnr1Y285A_p( | TRUE | CAN | JSY2422 | 5 | rnr1Y285A_p | CGG | G>A | A/T+1 | G/C-1 | CG>TA | CG>TA | TGGGCCGGT | GGTAACC | GCC | GCC | C | T | C>T | C>T_GCC | CG>TA_GGTAACC | 6 | FALSE | FALSE |
| 176_v.txt | 32713 | 32713-SNV-( | SNV | 1 | C | T | 29 | 44912 | 0.06457072 | 60.6551724 | rnr1Y285A_p( | TRUE | CAN | JSY2422 | 5 | rnr1Y285A_p | GCA | C>T | A/T+1 | G/C-1 | CG>TA | CG>TA | CCCCAGCAC | CACCACA | CGT | GCA | C | A | C>T | C>T_GCA | CG>TA_CACCACA | 6 | FALSE | FALSE |
| 176_v.txt | 32757 | 32757-SNV-( | SNV | 1 | G | A | 42 | 44912 | 0.09351621 | 45.5 | rnr1Y285A_p( | TRUE | CAN | JSY2422 | 5 | rnr1Y285A_p | GGC | G>A | A/T+1 | G/C-1 | CG>TA | CG>TA | ATAATGGCT | GCTAAAA | CCG | CCG | C | T | C>T | C>T_CCG | CG>TA_GCTAAAA | 6 | FALSE | FALSE |
| 176_v.txt | 32776 | 32776-SNV-( | SNV | 1 | C | T | 59 | 44912 | 0.13136801 | 49.3559322 | rnr1Y285A_p( | TRUE | CAN | JSY2422 | 5 | rnr1Y285A_p | GCG | C>T | A/T+1 | G/C-1 | CG>TA | CG>TA | TGGAAGCGA | CGACCCA | CGC | GCG | C | A | C>T | C>T_GCG | CG>TA_CGACCCA | 6 | FALSE | FALSE |
| 176_v.txt | 32780 | 32780-SNV-( | SNV | 1 | C | T | 46 | 44912 | 0.10242252 | 43.9706453 | rnr1Y285A_p( | TRUE | CAN | JSY2422 | 5 | rnr1Y285A_p | CCC | C>T | A/T+1 | G/C-1 | CG>TA | CG>TA | AGCGACCCA | CCAGAAC | GGG | CCC | C | A | C>T | C>T_CCC | CG>TA_CCAGAAC | 6 | FALSE | FALSE |
| 176_v.txt | 32798 | 32798-SNV-( | SNV | 1 | G | A | 74 | 44912 | 0.16476666 | 44.7027027 | rnr1Y285A_p( | TRUE | CAN | JSY2422 | 5 | rnr1Y285A_p | CGT | G>A | A/T+1 | G/C-1 | CG>TA | CG>TA | TTCACCGTA | GTAATAT | GCA | GCA | C | T | C>T | C>T_GCA | CG>TA_GTAATAT | 6 | FALSE | FALSE |
| 176_v.txt | 32855 | 32855-SNV-( | SNV | 1 | C | T | 237 | 44913 | 0.52768686 | 36.0042194 | rnr1Y285A_p( | TRUE | CAN | JSY2422 | 5 | rnr1Y285A_p | TCC | C>T | A/T+1 | G/C-1 | CG>TA | CG>TA | ACTAATCCAT | CCATGCC | AGG | TCC | C | A | C>T | C>T_TCC | CG>TA_CCATGCC | 6 | FALSE | FALSE |
| 176_v.txt | 32860 | 32860-SNV-( | SNV | 1 | C | T | 62 | 44907 | 0.13806311 | 36.4032258 | rnr1Y285A_p( | TRUE | CAN | JSY2422 | 5 | rnr1Y285A_p | GCC | C>T | A/T+1 | G/C-1 | CG>TA | CG>TA | TCCATGCCG | CCGCCAG | CGG | GCC | C | A | C>T | C>T_GCC | CG>TA_CCGCCAG | 6 | FALSE | FALSE |
| 176_v.txt | 32862 | 32862-SNV-( | SNV | 1 | G | A | 44 | 44907 | 0.09798027 | 34.6590909 | rnr1Y285A_p( | TRUE | CAN | JSY2422 | 5 | rnr1Y285A_p | CGC | G>A | A/T+1 | G/C-1 | CG>TA | CG>TA | CATGCCGCC | GCCAGTG | GCG | GCG | C | T | C>T | C>T_GCG | CG>TA_GCCAGTG | 6 | FALSE | FALSE |
| 176_v.txt | 32879 | 32879-SNV-( | SNV | 1 | C | T | 57 | 69130 | 0.08245335 | 36.6666667 | rnr1Y285A_p( | TRUE | CAN | JSY2422 | 5 | rnr1Y285A_p | ACG | C>T | A/T+1 | G/C-1 | CG>TA | CG>TA | TTTGTACGTC | CGTCCAA | TGC | ACG | C | A | C>T | C>T_ACG | CG>TA_CGTCCAA | 6 | FALSE | FALSE |
| 176_v.txt | 32900 | 32900-SNV-( | SNV | 1 | G | A | 72 | 69133 | 0.10414708 | 35.4700184 | rnr1Y285A_p( | TRUE | CAN | JSY2422 | 5 | rnr1Y285A_p | GGC | G>A | A/T+1 | G/C-1 | CG>TA | CG>TA | GACTTGGCC | GCCAACT | CCG | CCG | C | T | C>T | C>T_CCG | CG>TA_GCCAACT | 6 | FALSE | FALSE |
| 176_v.txt | 32901 | 32901-SNV-( | SNV | 1 | C | T | 149 | 69133 | 0.21552659 | 32.9468002 | rnr1Y285A_p( | TRUE | CAN | JSY2422 | 5 | rnr1Y285A_p | GCC | C>T | A/T+1 | G/C-1 | CG>TA | CG>TA | ACTTGGCCA | CCAACTA | CGG | GCC | C | A | C>T | C>T_GCC | CG>TA_CCAACTA | 6 | FALSE | FALSE |
| 176_v.txt | 32917 | 32917-SNV-( | SNV | 1 | C | T | 144 | 69133 | 0.20829416 | 45.5486111 | rnr1Y285A_p( | TRUE | CAN | JSY2422 | 5 | rnr1Y285A_p | TCC | C>T | A/T+1 | G/C-1 | CG>TA | CG>TA | TAAGTTCCA | CCAGGGC | AGG | TCC | C | A | C>T | C>T_TCC | CG>TA_CCAGGGC | 6 | FALSE | FALSE |
| 176_v.txt | 32920 | 32920-SNV-( | SNV | 1 | G | A | 8 | 69133 | 0.0115719 | 35.6238605 | rnr1Y285A_p( | TRUE | CAN | JSY2422 | 5 | rnr1Y285A_p | AGG | G>A | A/T+1 | G/C-1 | CG>TA | CG>TA | GTTCCAGGG | GGGCAAA | TCC | TCC | C | T | C>T | C>T_TCC | CG>TA_GGGCAAA | 6 | FALSE | FALSE |
| 176_v.txt | 32922 | 32922-SNV-( | SNV | 1 | G | A | 64 | 69133 | 0.09257518 | 54.25 | rnr1Y285A_p( | TRUE | CAN | JSY2422 | 5 | rnr1Y285A_p | GGC | G>A | A/T+1 | G/C-1 | CG>TA | CG>TA | TCCAGGGCA | GCAAAAG | CCG | CCG | C | T | C>T | C>T_CCG | CG>TA_GCAAAAG | 6 | FALSE | FALSE |
| 176_v.txt | 32935 | 32935-SNV-( | SNV | 1 | C | T | 91 | 69131 | 0.13163414 | 36.6158256 | rnr1Y285A_p( | TRUE | CAN | JSY2422 | 5 | rnr1Y285A_p | GCC | C>T | A/T+1 | G/C-1 | CG>TA | CG>TA | TGATTGCCC | CCCAAGA | CGG | GCC | C | A | C>T | C>T_GCC | CG>TA_CCCAAGA | 6 | FALSE | FALSE |
| 176_v.txt | 32940 | 32940-SNV-( | SNV | 1 | G | A | 6 | 69132 | 8.68E-03 | 43.1653294 | rnr1Y285A_p( | TRUE | CAN | JSY2422 | 5 | rnr1Y285A_p | AGA | G>A | A/T+1 | G/C-1 | CG>TA | CG>TA | GCCCAAGAA | GAAAACC | TCT | TCT | C | T | C>T | C>T_TCT | CG>TA_GAAAACC | 6 | FALSE | FALSE |
| 176_v.txt | 32945 | 32945-SNV-( | SNV | 1 | C | T | 293 | 69133 | 0.42382075 | 55.5358362 | rnr1Y285A_p( | TRUE | CAN | JSY2422 | 5 | rnr1Y285A_p | ACC | C>T | A/T+1 | G/C-1 | CG>TA | CG>TA | AGAAAACCA | CCAATAC | TGG | ACC | C | A | C>T | C>T_ACC | CG>TA_CCAATAC | 6 | FALSE | FALSE |
| 176_v.txt | 32958 | 32958-SNV-( | SNV | 1 | C | T | 266 | 69133 | 0.3847656 | 54.9210526 | rnr1Y285A_p( | TRUE | CAN | JSY2422 | 5 | rnr1Y285A_p | ACC | C>T | A/T+1 | G/C-1 | CG>TA | CG>TA | ATGTAACCA | CCATTGG | TGG | ACC | C | A | C>T | C>T_ACC | CG>TA_CCATTGG | 6 | FALSE | FALSE |
| 176_v.txt | 32965 | 32965-SNV-( | SNV | 1 | C | T | 80 | 69133 | 0.11571898 | 58.6875 | rnr1Y285A_p( | TRUE | CAN | JSY2422 | 5 | rnr1Y285A_p | GCC | C>T | A/T+1 | G/C-1 | CG>TA | CG>TA | CATTGGCCG | CCGCACC | CGG | GCC | C | A | C>T | C>T_GCC | CG>TA_CCGCACC | 6 | FALSE | FALSE |
| 176_v.txt | 32967 | 32967-SNV-( | SNV | 1 | G | A | 147 | 69133 | 0.21263362 | 59.5170068 | rnr1Y285A_p( | TRUE | CAN | JSY2422 | 5 | rnr1Y285A_p | CGC | G>A | A/T+1 | G/C-1 | CG>TA | CG>TA | TTGGCCGCA | GCACCAA | GCG | GCG | C | T | C>T | C>T_GCG | CG>TA_GCACCAA | 6 | FALSE | FALSE |
| 176_v.txt | 32977 | 32977-SNV-( | SNV | 1 | C | T | 50 | 69133 | 0.07232436 | 57.6 | rnr1Y285A_p( | TRUE | CAN | JSY2422 | 5 | rnr1Y285A_p | GCT | C>T | A/T+1 | G/C-1 | CG>TA | CG>TA | CAAATGCTG | CTGGAGA | CGA | GCT | C | A | C>T | C>T_GCT | CG>TA_CTGGAGA | 6 | FALSE | FALSE |
| 176_v.txt | 33037 | 33037-SNV-( | SNV | 1 | C | T | 40 | 69133 | 0.05785949 | 59.225 | rnr1Y285A_p( | TRUE | CAN | JSY2422 | 5 | rnr1Y285A_p | GCC | C>T | A/T+1 | G/C-1 | CG>TA | CG>TA | ATGTAGCCA | CCATTTC | CGG | GCC | C | A | C>T | C>T_GCC | CG>TA_CCATTTC | 6 | FALSE | FALSE |
| 176_v.txt | 33056 | 33056-SNV-( | SNV | 1 | C | T | 85 | 69133 | 0.12295141 | 53.9647059 | rnr1Y285A_p( | TRUE | CAN | JSY2422 | 5 | rnr1Y285A_p | GCG | C>T | A/T+1 | G/C-1 | CG>TA | CG>TA | GGACTGCGT | CGTGACA | CGC | GCG | C | A | C>T | C>T_GCG | CG>TA_CGTGACA | 6 | FALSE | FALSE |
| 176_v.txt | 33069 | 33069-SNV-( | SNV | 1 | G | A | 75 | 69133 | 0.10848654 | 51.7731046 | rnr1Y285A_p( | TRUE | CAN | JSY2422 | 5 | rnr1Y285A_p | TGC | G>A | A/T+1 | G/C-1 | CG>TA | CG>TA | GAATATGCC | GCCAAAG | ACG | ACG | C | T | C>T | C>T_ACG | CG>TA_GCCAAAG | 6 | FALSE | FALSE |
| 176_v.txt | 33070 | 33070-SNV-( | SNV | 1 | C | T | 100 | 69129 | 0.14465709 | 52.9490758 | rnr1Y285A_p( | TRUE | CAN | JSY2422 | 5 | rnr1Y285A_p | GCC | C>T | A/T+1 | G/C-1 | CG>TA | CG>TA | AATATGCCA | CCAAAGA | CGG | GCC | C | A | C>T | C>T_GCC | CG>TA_CCAAAGA | 6 | FALSE | FALSE |
| 176_v.txt | 33078 | 33078-SNV-( | SNV | 1 | C | T | 85 | 69133 | 0.12295141 | 51.9058824 | rnr1Y285A_p( | TRUE | CAN | JSY2422 | 5 | rnr1Y285A_p | ACC | C>T | A/T+1 | G/C-1 | CG>TA | CG>TA | AAAGAACCC | CCCATAA | TGG | ACC | C | A | C>T | C>T_ACC | CG>TA_CCCATAA | 6 | FALSE | FALSE |
| 176_v.txt | 33104 | 33104-SNV-( | SNV | 1 | G | A | 91 | 69133 | 0.13163034 | 39.0879121 | rnr1Y285A_p( | TRUE | CAN | JSY2422 | 5 | rnr1Y285A_p | CGC | G>A | A/T+1 | G/C-1 | CG>TA | CG>TA | AAGAGCGCC | GCCCACT | GCG | GCG | C | T | C>T | C>T_GCG | CG>TA_GCCCACT | 6 | FALSE | FALSE |
| 176_v.txt | 33113 | 33113-SNV-( | SNV | 1 | G | A | 57 | 69133 | 0.08244977 | 38.7163213 | rnr1Y285A_p( | TRUE | CAN | JSY2422 | 5 | rnr1Y285A_p | GGC | G>A | A/T+1 | G/C-1 | CG>TA | CG>TA | CACTGGGCC | GCCGGCG | CCG | CCG | C | T | C>T | C>T_CCG | CG>TA_GCCGGCG | 6 | FALSE | FALSE |
| 176_v.txt | 33114 | 33114-SNV-( | SNV | 1 | C | T | 72 | 69133 | 0.10414708 | 41.9720059 | rnr1Y285A_p( | TRUE | CAN | JSY2422 | 5 | rnr1Y285A_p | GCC | C>T | A/T+1 | G/C-1 | CG>TA | CG>TA | ACTGGGCCG | CCGGCGT | CGG | GCC | C | A | C>T | C>T_GCC | CG>TA_CCGGCGT | 6 | FALSE | FALSE |
| 176_v.txt | 33118 | 33118-SNV-( | SNV | 1 | C | T | 72 | 69133 | 0.10414708 | 42.1791715 | rnr1Y285A_p( | TRUE | CAN | JSY2422 | 5 | rnr1Y285A_p | GCG | C>T | A/T+1 | G/C-1 | CG>TA | CG>TA | GGCCGGCGT | CGTTGGT | CGC | GCG | C | A | C>T | C>T_GCG | CG>TA_CGTTGGT | 6 | FALSE | FALSE |
| 176_v.txt | 33119 | 33119-SNV-( | SNV | 1 | G | A | 56 | 69133 | 0.08100328 | 41.6431087 | rnr1Y285A_p( | TRUE | CAN | JSY2422 | 5 | rnr1Y285A_p | CGT | G>A | A/T+1 | G/C-1 | CG>TA | CG>TA | GCCGGCGTT | GTTGGTC | GCA | GCA | C | T | C>T | C>T_GCA | CG>TA_GTTGGTC | 6 | FALSE | FALSE |
| 176_v.txt | 33153 | 33153-SNV-( | SNV | 1 | C | T | 575 | 69133 | 0.83173014 | 36.7322341 | rnr1Y285A_p( | TRUE | CAN | JSY2422 | 5 | rnr1Y285A_p | ACC | C>T | A/T+1 | G/C-1 | CG>TA | CG>TA | AAAAGACCT | CCTGTAC | TGG | ACC | C | A | C>T | C>T_ACC | CG>TA_CCTGTAC | 6 | FALSE | FALSE |
| 176_v.txt | 33154 | 33154-SNV-( | SNV | 1 | C | T | 55 | 69133 | 0.0795568 | 37.1818182 | rnr1Y285A_p( | TRUE | CAN | JSY2422 | 5 | rnr1Y285A_p | CCT | C>T | A/T+1 | G/C-1 | CG>TA | CG>TA | AAAGACCTG | CTGTACC | GGA | CCT | C | A | C>T | C>T_CCT | CG>TA_CTGTACC | 6 | FALSE | FALSE |
| 176_v.txt | 33168 | 33168-SNV-( | SNV | 1 | C | T | 514 | 69130 | 0.74352669 | 37.4299611 | rnr1Y285A_p( | TRUE | CAN | JSY2422 | 5 | rnr1Y285A_p | ACC | C>T | A/T+1 | G/C-1 | CG>TA | CG>TA | ATAGTACCA | CCACCAA | TGG | ACC | C | A | C>T | C>T_ACC | CG>TA_CCACCAA | 6 | FALSE | FALSE |
| 176_v.txt | 33205 | 33205-SNV-( | SNV | 1 | G | A | 22 | 33429 | 0.06581112 | 39.6363636 | rnr1Y285A_p( | TRUE | CAN | JSY2422 | 5 | rnr1Y285A_p | AGC | G>A | A/T+1 | G/C-1 | CG>TA | CG>TA | GCTTAAGCT | GCTCTCT | TCG | TCG | C | T | C>T | C>T_TCG | CG>TA_GCTCTCT | 6 | FALSE | FALSE |
| 176_v.txt | 33258 | 33258-SNV-( | SNV | 1 | C | T | 62 | 33429 | 0.18546771 | 43.8064516 | rnr1Y285A_p( | TRUE | CAN | JSY2422 | 5 | rnr1Y285A_p | GCC | C>T | A/T+1 | G/C-1 | CG>TA | CG>TA | CCTATGCCAT | CCATCCT | CGG | GCC | C | A | C>T | C>T_GCC | CG>TA_CCATCCT | 6 | FALSE | FALSE |
| 176_v.txt | 33275 | 33275-SNV-( | SNV | 1 | C | T | 39 | 33429 | 0.11666517 | 50.9465487 | rnr1Y285A_p( | TRUE | CAN | JSY2422 | 5 | rnr1Y285A_p | ACG | C>T | A/T+1 | G/C-1 | CG>TA | CG>TA | AGAGAACGT | CGTATCC | TGC | ACG | C | A | C>T | C>T_ACG | CG>TA_CGTATCC | 6 | FALSE | FALSE |
| 176_v.txt | 33276 | 33276-SNV-( | SNV | 1 | G | A | 75 | 33424 | 0.22438966 | 46.8472879 | rnr1Y285A_p( | TRUE | CAN | JSY2422 | 5 | rnr1Y285A_p | CGT | G>A | A/T+1 | G/C-1 | CG>TA | CG>TA | GAGAACGTA | GTATCCT | GCA | GCA | C | T | C>T | C>T_GCA | CG>TA_GTATCCT | 6 | FALSE | FALSE |
| 176_v.txt | 33284 | 33284-SNV-( | SNV | 1 | G | A | 43 | 33429 | 0.12863083 | 47.4186047 | rnr1Y285A_p( | TRUE | CAN | JSY2422 | 5 | rnr1Y285A_p | CGC | G>A | A/T+1 | G/C-1 | CG>TA | CG>TA | ATCCTCGCC | GCCATTT | GCG | GCG | C | T | C>T | C>T_GCG | CG>TA_GCCATTT | 6 | FALSE | FALSE |
| 176_v.txt | 33309 | 33309-SNV-( | SNV | 1 | C | T | 31 | 33428 | 0.09273663 | 59.3818865 | rnr1Y285A_p( | TRUE | CAN | JSY2422 | 5 | rnr1Y285A_p | GCG | C>T | A/T+1 | G/C-1 | CG>TA | CG>TA | AAAGAGCGC | CGCAATG | CGC | GCG | C | A | C>T | C>T_GCG | CG>TA_CGCAATG | 6 | FALSE | FALSE |
| 176_v.txt | 33310 | 33310-SNV-( | SNV | 1 | G | A | 52 | 33426 | 0.15556752 | 56.5290693 | rnr1Y285A_p( | TRUE | CAN | JSY2422 | 5 | rnr1Y285A_p | CGC | G>A | A/T+1 | G/C-1 | CG>TA | CG>TA | AAGAGCGCA | GCAATGG | GCG | GCG | C | T | C>T | C>T_GCG | CG>TA_GCAATGG | 6 | FALSE | FALSE |
| 176_v.txt | 33325 | 33325-SNV-( | SNV | 1 | C | T | 3 | 33428 | 8.98E-03 | 53.3288357 | rnr1Y285A_p( | TRUE | CAN | JSY2422 | 5 | rnr1Y285A_p | TCT | C>T | A/T+1 | G/C-1 | CG>TA | CG>TA | ACAATTCTTT | CTTTACT | AGA | TCT | C | A | C>T | C>T_TCT | CG>TA_CTTTACT | 6 | FALSE | FALSE |
| 176_v.txt | 33412 | 33412-SNV-( | SNV | 1 | G | A | 46 | 33429 | 0.13760507 | 52.2608696 | rnr1Y285A_p( | TRUE | CAN | JSY2422 | 5 | rnr1Y285A_p | GGC | G>A | A/T+1 | G/C-1 | CG>TA | CG>TA | TGACCGGCT | GCTCATT | CCG | CCG | C | T | C>T | C>T_CCG | CG>TA_GCTCATT | 6 | FALSE | FALSE |
| 176_v.txt | 33474 | 33474-SNV-( | SNV | 1 | C | T | 56 | 33428 | 0.16752423 | 34.4285714 | rnr1Y285A_p( | TRUE | CAN | JSY2422 | 5 | rnr1Y285A_p | GCC | C>T | A/T+1 | G/C-1 | CG>TA | CG>TA | GCTATGCCT | CCTTTTT | CGG | GCC | C | A | C>T | C>T_GCC | CG>TA_CCTTTTT | 6 | FALSE | FALSE |
| 176_v.txt | 31657 | 31657-SNV-( | SNV | 1 | G | C | 9 | 17332 | 0.05192707 | 15.4443418 | rnr1Y285A_p( | TRUE | CAN | JSY2422 | 5 | rnr1Y285A_p | GGC | G>C | G/C+1 | G/C-1 | CG>GC | CG>GC | GAAATGGCG | GCGTGGA | CCG | CCG | C | G | C>G | C>G_CCG | CG>GC_GCGTGGA | 6 | FALSE | FALSE |
| 176_v.txt | 31658 | 31658-SNV-( | SNV | 1 | C | G | 1 | 17332 | 5.77E-03 | 37.9994175 | rnr1Y285A_p( | TRUE | CAN | JSY2422 | 5 | rnr1Y285A_p | GCG | C>G | G/C+1 | G/C-1 | CG>GC | CG>GC | AAATGGCGT | CGTGGAA | CGC | GCG | C | C | C>G | C>G_GCG | CG>GC_CGTGGAA | 6 | FALSE | FALSE |
| 176_v.txt | 31882 | 31882-SNV-( | SNV | 1 | C | G | 1 | 17332 | 5.77E-03 | 37.9992345 | rnr1Y285A_p( | TRUE | CAN | JSY2422 | 5 | rnr1Y285A_p | GCA | C>G | G/C+1 | G/C-1 | CG>GC | CG>GC | AGGCGGCAC | CAGCAAA | CGT | GCA | C | C | C>G | C>G_GCA | CG>GC_CAGCAAA | 6 | FALSE | FALSE |
| 176_v.txt | 31981 | 31981-SNV-( | SNV | 1 | G | C | 5 | 37086 | 0.01348218 | 31.6 | rnr1Y285A_p( | TRUE | CAN | JSY2422 | 5 | rnr1Y285A_p | GGC | G>C | G/C+1 | G/C-1 | CG>GC | CG>GC | AGCCGGGCA | GCATTAA | CCG | CCG | C | G | C>G | C>G_CCG | CG>GC_GCATTAA | 6 | FALSE | FALSE |
| 176_v.txt | 32027 | 32027-SNV-( | SNV | 1 | G | C | 75 | 37086 | 0.20223265 | 57.7866667 | rnr1Y285A_p( | TRUE | CAN | JSY2422 | 5 | rnr1Y285A_p | GGT | G>C | G/C+1 | G/C-1 | CG>GC | CG>GC | GCCACGGTA | GTATTTC | CCA | CCA | C | G | C>G | C>G_CCA | CG>GC_GTATTTC | 6 | FALSE | FALSE |
| 176_v.txt | 32272 | 32272-SNV-( | SNV | 1 | G | C | 135 | 40332 | 0.33472181 | 36.1691341 | rnr1Y285A_p( | TRUE | CAN | JSY2422 | 5 | rnr1Y285A_p | CGG | G>C | G/C+1 | G/C-1 | CG>GC | CG>GC | AAATACGGG | GGGAACC | GCC | GCC | C | G | C>G | C>G_GCC | CG>GC_GGGAACC | 6 | FALSE | FALSE |
| 176_v.txt | 32488 | 32488-SNV-( | SNV | 1 | G | C | 101 | 40333 | 0.25041529 | 47.634166 | rnr1Y285A_p( | TRUE | CAN | JSY2422 | 5 | rnr1Y285A_p | CGG | G>C | G/C+1 | G/C-1 | CG>GC | CG>GC | AGATACGGA | GGAAAAC | GCC | GCC | C | G | C>G | C>G_GCC | CG>GC_GGAAAAC | 6 | FALSE | FALSE |
| 176_v.txt | 32538 | 32538-SNV-( | SNV | 1 | G | C | 13 | 40333 | 0.03223167 | 35.9961749 | rnr1Y285A_p( | TRUE | CAN | JSY2422 | 5 | rnr1Y285A_p | AGC | G>C | G/C+1 | G/C-1 | CG>GC | CG>GC | TTTGCAGCT | GCTTCAC | TCG | TCG | C | G | C>G | C>G_TCG | CG>GC_GCTTCAC | 6 | FALSE | FALSE |
| 176_v.txt | 32572 | 32572-SNV-( | SNV | 1 | C | G | 39 | 44914 | 0.08683261 | 37.1692244 | rnr1Y285A_p( | TRUE | CAN | JSY2422 | 5 | rnr1Y285A_p | CCT | C>G | G/C+1 | G/C-1 | CG>GC | CG>GC | CAGTACCTC | CTTGAAA | GGA | CCT | C | C | C>G | C>G_CCT | CG>GC_CTTGAAA | 6 | FALSE | FALSE |
| 176_v.txt | 32901 | 32901-SNV-( | SNV | 1 | C | G | 8 | 69133 | 0.0115719 | 37.6246064 | rnr1Y285A_p( | TRUE | CAN | JSY2422 | 5 | rnr1Y285A_p | GCC | C>G | G/C+1 | G/C-1 | CG>GC | CG>GC | ACTTGGCCA | CCAACTA | CGG | GCC | C | C | C>G | C>G_GCC | CG>GC_CCAACTA | 6 | FALSE | FALSE |
| 176_v.txt | 32917 | 32917-SNV-( | SNV | 1 | C | G | 3 | 69133 | 4.34E-03 | 53.6666667 | rnr1Y285A_p( | TRUE | CAN | JSY2422 | 5 | rnr1Y285A_p | TCC | C>G | G/C+1 | G/C-1 | CG>GC | CG>GC | TAAGTTCCA | CCAGGGC | AGG | TCC | C | C | C>G | C>G_TCC | CG>GC_CCAGGGC | 6 | FALSE | FALSE |
| 176_v.txt | 32934 | 32934-SNV-( | SNV | 1 | G | C | 4 | 69133 | 5.79E-03 | 43.4953822 | rnr1Y285A_p( | TRUE | CAN | JSY2422 | 5 | rnr1Y285A_p | TGC | G>C | G/C+1 | G/C-1 | CG>GC | CG>GC | GTGATTGCC | GCCCAAG | ACG | ACG | C | G | C>G | C>G_ACG | CG>GC_GCCCAAG | 6 | FALSE | FALSE |
| 176_v.txt | 32935 | 32935-SNV-( | SNV | 1 | C | G | 12 | 69131 | 0.01735835 | 52.9150429 | rnr1Y285A_p( | TRUE | CAN | JSY2422 | 5 | rnr1Y285A_p | GCC | C>G | G/C+1 | G/C-1 | CG>GC | CG>GC | TGATTGCCC | CCCAAGA | CGG | GCC | C | C | C>G | C>G_GCC | CG>GC_CCCAAGA | 6 | FALSE | FALSE |
| 176_v.txt | 32997 | 32997-SNV-( | SNV | 1 | G | C | 47 | 69133 | 0.0679849 | 62 | rnr1Y285A_p( | TRUE | CAN | JSY2422 | 5 | rnr1Y285A_p | TGA | G>C | G/C+1 | G/C-1 | CG>GC | CG>GC | CTTTGTGAG | GAGAAAA | ACT | ACT | C | G | C>G | C>G_ACT | CG>GC_GAGAAAA | 6 | FALSE | FALSE |
| 176_v.txt | 33056 | 33056-SNV-( | SNV | 1 | C | G | 7 | 69133 | 0.01012541 | 56.2857143 | rnr1Y285A_p( | TRUE | CAN | JSY2422 | 5 | rnr1Y285A_p | GCG | C>G | G/C+1 | G/C-1 | CG>GC | CG>GC | GGACTGCGT | CGTGACA | CGC | GCG | C | C | C>G | C>G_GCG | CG>GC_CGTGACA | 6 | FALSE | FALSE |
| 176_v.txt | 33069 | 33069-SNV-( | SNV | 1 | G | C | 4 | 69133 | 5.79E-03 | 55.4996464 | rnr1Y285A_p( | TRUE | CAN | JSY2422 | 5 | rnr1Y285A_p | TGC | G>C | G/C+1 | G/C-1 | CG>GC | CG>GC | GAATATGCC | GCCAAAG | ACG | ACG | C | G | C>G | C>G_ACG | CG>GC_GCCAAAG | 6 | FALSE | FALSE |
| 176_v.txt | 33070 | 33070-SNV-( | SNV | 1 | C | G | 2 | 69129 | 2.89E-03 | 50.999492 | rnr1Y285A_p( | TRUE | CAN | JSY2422 | 5 | rnr1Y285A_p | GCC | C>G | G/C+1 | G/C-1 | CG>GC | CG>GC | AATATGCCA | CCAAAGA | CGG | GCC | C | C | C>G | C>G_GCC | CG>GC_CCAAAGA | 6 | FALSE | FALSE |
| 176_v.txt | 33093 | 33093-SNV-( | SNV | 1 | G | C | 38 | 69133 | 0.05496651 | 44.892249 | rnr1Y285A_p( | TRUE | CAN | JSY2422 | 5 | rnr1Y285A_p | TGA | G>C | G/C+1 | G/C-1 | CG>GC | CG>GC | AAATATGAT | GATATAA | ACT | ACT | C | G | C>G | C>G_ACT | CG>GC_GATATAA | 6 | FALSE | FALSE |
| 176_v.txt | 33118 | 33118-SNV-( | SNV | 1 | C | G | 2 | 69133 | 2.89E-03 | 37.999092 | rnr1Y285A_p( | TRUE | CAN | JSY2422 | 5 | rnr1Y285A_p | GCG | C>G | G/C+1 | G/C-1 | CG>GC | CG>GC | GGCCGGCGT | CGTTGGT | CGC | GCG | C | C | C>G | C>G_GCG | CG>GC_CGTTGGT | 6 | FALSE | FALSE |
| 176_v.txt | 33154 | 33154-SNV-( | SNV | 1 | C | G | 109 | 69133 | 0.15766711 | 37.3027523 | rnr1Y285A_p( | TRUE | CAN | JSY2422 | 5 | rnr1Y285A_p | CCT | C>G | G/C+1 | G/C-1 | CG>GC | CG>GC | AAAGACCTG | CTGTACC | GGA | CCT | C | C | C>G | C>G_CCT | CG>GC_CTGTACC | 6 | FALSE | FALSE |
| 176_v.txt | 33276 | 33276-SNV-( | SNV | 1 | G | C | 5 | 33424 | 0.01495931 | 33.3957314 | rnr1Y285A_p( | TRUE | CAN | JSY2422 | 5 | rnr1Y285A_p | CGT | G>C | G/C+1 | G/C-1 | CG>GC | CG>GC | GAGAACGTA | GTATCCT | GCA | GCA | C | G | C>G | C>G_GCA | CG>GC_GTATCCT | 6 | FALSE | FALSE |
| 176_v.txt | 33310 | 33310-SNV-( | SNV | 1 | G | C | 4 | 33426 | 0.01196673 | 33.9855164 | rnr1Y285A_p( | TRUE | CAN | JSY2422 | 5 | rnr1Y285A_p | CGC | G>C | G/C+1 | G/C-1 | CG>GC | CG>GC | AAGAGCGCA | GCAATGG | GCG | GCG | C | G | C>G | C>G_GCG | CG>GC_GCAATGG | 6 | FALSE | FALSE |
| 176_v.txt | 31657 | 31657-SNV-( | SNV | 1 | G | T | 112 | 17332 | 0.64620355 | 18.2229567 | rnr1Y285A_p( | TRUE | CAN | JSY2422 | 5 | rnr1Y285A_p | GGC | G>T | A/T+1 | G/C-1 | CG>AT | CG>AT | GAAATGGCG | GCGTGGA | CCG | CCG | C | A | C>A | C>A_CCG | CG>AT_GCGTGGA | 6 | FALSE | FALSE |
| 176_v.txt | 31981 | 31981-SNV-( | SNV | 1 | G | T | 67 | 37086 | 0.18066117 | 39.9104478 | rnr1Y285A_p( | TRUE | CAN | JSY2422 | 5 | rnr1Y285A_p | GGC | G>T | A/T+1 | G/C-1 | CG>AT | CG>AT | AGCCGGGCA | GCATTAA | CCG | CCG | C | A | C>A | C>A_CCG | CG>AT_GCATTAA | 6 | FALSE | FALSE |
| 176_v.txt | 32027 | 32027-SNV-( | SNV | 1 | G | T | 1200 | 37086 | 3.23572238 | 59.9116667 | rnr1Y285A_p( | TRUE | CAN | JSY2422 | 5 | rnr1Y285A_p | GGT | G>T | A/T+1 | G/C-1 | CG>AT | CG>AT | GCCACGGTA | GTATTTC | CCA | CCA | C | A | C>A | C>A_CCA | CG>AT_GTATTTC | 6 | FALSE | FALSE |
| 176_v.txt | 32089 | 32089-SNV-( | SNV | 1 | C | A | 73 | 37082 | 0.19686101 | 57.079761 | rnr1Y285A_p( | TRUE | CAN | JSY2422 | 5 | rnr1Y285A_p | CCT | C>A | A/T+1 | G/C-1 | CG>AT | CG>AT | AAAAGCCTG | CTGCAAC | GGA | CCT | C | T | C>A | C>A_CCT | CG>AT_CTGCAAC | 6 | FALSE | FALSE |
| 176_v.txt | 32095 | 32095-SNV-( | SNV | 1 | C | A | 79 | 37086 | 0.21301839 | 60.7594937 | rnr1Y285A_p( | TRUE | CAN | JSY2422 | 5 | rnr1Y285A_p | ACA | C>A | A/T+1 | G/C-1 | CG>AT | CG>AT | CTGCAACAC | CACCAGT | TGT | ACA | C | T | C>A | C>A_ACA | CG>AT_CACCAGT | 6 | FALSE | FALSE |
| 176_v.txt | 32153 | 32153-SNV-( | SNV | 1 | G | T | 19 | 37086 | 0.05123227 | 51.3684211 | rnr1Y285A_p( | TRUE | CAN | JSY2422 | 5 | rnr1Y285A_p | TGT | G>T | A/T+1 | G/C-1 | CG>AT | CG>AT | CTCCATGTA | GTAAGCC | ACA | ACA | C | A | C>A | C>A_ACA | CG>AT_GTAAGCC | 6 | FALSE | FALSE |

| | | | | | | | | | | | | | | | | | | | | | | | | | | | | | | | | | | |
|---|---|---|---|---|---|---|---|---|---|---|---|---|---|---|---|---|---|---|---|---|---|---|---|---|---|---|---|---|---|---|---|---|---|---|
| 176_v.txt | 32188 | 32188-SNV-C | SNV | 1 | C | A | 855 | 37086 | 2.30545219 | 46.6795322 | rnr1Y285A_pC | TRUE | CAN | JSY2422 | 5 | rnr1Y285A_p | ACT | C>A | A/T+1 | G/C-1 | CG>AT | CG>AT | CGAAAACTG | CTGCAAT | TGA | ACT | C | T | C>A | C>A_ACT | CG>AT_CTGCAAT | 6 | FALSE | FALSE |
| 176_v.txt | 32203 | 32203-SNV-C | SNV | 1 | C | A | 88 | 37086 | 0.23728631 | 40.4431818 | rnr1Y285A_pC | TRUE | CAN | JSY2422 | 5 | rnr1Y285A_p | ACA | C>A | A/T+1 | G/C-1 | CG>AT | CG>AT | ATGGAACAC | CACCACC | TGT | ACA | C | T | C>A | C>A_ACA | CG>AT_CACCACC | 6 | FALSE | FALSE |
| 176_v.txt | 32205 | 32205-SNV-C | SNV | 1 | C | A | 34 | 37086 | 0.0916788 | 35.7647059 | rnr1Y285A_pC | TRUE | CAN | JSY2422 | 5 | rnr1Y285A_p | ACC | C>A | A/T+1 | G/C-1 | CG>AT | CG>AT | GGAACACCA | CCACCTT | TGG | ACC | C | T | C>A | C>A_ACC | CG>AT_CCACCTT | 6 | FALSE | FALSE |
| 176_v.txt | 32271 | 32271-SNV-C | SNV | 1 | C | A | 21 | 40332 | 0.05206784 | 36.1404756 | rnr1Y285A_pC | TRUE | CAN | JSY2422 | 5 | rnr1Y285A_p | ACG | C>A | A/T+1 | G/C-1 | CG>AT | CG>AT | AAAATACGG | CGGGAAC | TGC | ACG | C | T | C>A | C>A_ACG | CG>AT_CGGGAAC | 6 | FALSE | FALSE |
| 176_v.txt | 32272 | 32272-SNV-C | SNV | 1 | G | T | 41 | 40332 | 0.10165625 | 31.1220062 | rnr1Y285A_pC | TRUE | CAN | JSY2422 | 5 | rnr1Y285A_p | CGG | G>T | A/T+1 | G/C-1 | CG>AT | CG>AT | AAATACGGG | GGGAACC | GCC | GCC | C | A | C>A | C>A_GCC | CG>AT_GGGAACC | 6 | FALSE | FALSE |
| 176_v.txt | 32274 | 32274-SNV-C | SNV | 1 | G | T | 173 | 40332 | 0.4289398 | 35.4046243 | rnr1Y285A_pC | TRUE | CAN | JSY2422 | 5 | rnr1Y285A_p | GGA | G>T | A/T+1 | G/C-1 | CG>AT | CG>AT | ATACGGGAA | GAACCAA | CCT | CCT | C | A | C>A | C>A_CCT | CG>AT_GAACCAA | 6 | FALSE | FALSE |
| 176_v.txt | 32277 | 32277-SNV-C | SNV | 1 | C | A | 86 | 40332 | 0.21323019 | 33.7325581 | rnr1Y285A_pC | TRUE | CAN | JSY2422 | 5 | rnr1Y285A_p | ACC | C>A | A/T+1 | G/C-1 | CG>AT | CG>AT | CGGGAACCA | CCAACGT | TGG | ACC | C | T | C>A | C>A_ACC | CG>AT_CCAACGT | 6 | FALSE | FALSE |
| 176_v.txt | 32304 | 32304-SNV-C | SNV | 1 | G | T | 108 | 40333 | 0.26777081 | 58.6666667 | rnr1Y285A_pC | TRUE | CAN | JSY2422 | 5 | rnr1Y285A_p | AGA | G>T | A/T+1 | G/C-1 | CG>AT | CG>AT | GCGGCAGAA | GAAATAA | TCT | TCT | C | A | C>A | C>A_TCT | CG>AT_GAAATAA | 6 | FALSE | FALSE |
| 176_v.txt | 32373 | 32373-SNV-C | SNV | 1 | G | T | 60 | 40333 | 0.14876156 | 60.6166667 | rnr1Y285A_pC | TRUE | CAN | JSY2422 | 5 | rnr1Y285A_p | AGC | G>T | A/T+1 | G/C-1 | CG>AT | CG>AT | TCAATAGCA | GCAATAA | TCG | TCG | C | A | C>A | C>A_TCG | CG>AT_GCAATAA | 6 | FALSE | FALSE |
| 176_v.txt | 32474 | 32474-SNV-C | SNV | 1 | G | T | 35 | 40333 | 0.08677758 | 46.0285714 | rnr1Y285A_pC | TRUE | CAN | JSY2422 | 5 | rnr1Y285A_p | AGA | G>T | A/T+1 | G/C-1 | CG>AT | CG>AT | AATGTAGAA | GAAGGTT | TCT | TCT | C | A | C>A | C>A_TCT | CG>AT_GAAGGTT | 6 | FALSE | FALSE |
| 176_v.txt | 32487 | 32487-SNV-C | SNV | 1 | C | A | 44 | 40333 | 0.10909181 | 45.6993009 | rnr1Y285A_pC | TRUE | CAN | JSY2422 | 5 | rnr1Y285A_p | ACG | C>A | A/T+1 | G/C-1 | CG>AT | CG>AT | AAGATACGG | CGGAAAA | TGC | ACG | C | T | C>A | C>A_ACG | CG>AT_CGGAAAA | 6 | FALSE | FALSE |
| 176_v.txt | 32508 | 32508-SNV-C | SNV | 1 | G | T | 105 | 40333 | 0.26033273 | 44.6571429 | rnr1Y285A_pC | TRUE | CAN | JSY2422 | 5 | rnr1Y285A_p | GGC | G>T | A/T+1 | G/C-1 | CG>AT | CG>AT | TTGATGGCT | GCTCTTG | CCG | CCG | C | A | C>A | C>A_CCG | CG>AT_GCTCTTG | 6 | FALSE | FALSE |
| 176_v.txt | 32530 | 32530-SNV-C | SNV | 1 | G | T | 54 | 40333 | 0.1338854 | 37.537037 | rnr1Y285A_pC | TRUE | CAN | JSY2422 | 5 | rnr1Y285A_p | GGG | G>T | A/T+1 | G/C-1 | CG>AT | CG>AT | TTCTGGGGT | GGTTTGC | CCC | CCC | C | A | C>A | C>A_CCC | CG>AT_GGTTTGC | 6 | FALSE | FALSE |
| 176_v.txt | 32539 | 32539-SNV-C | SNV | 1 | C | A | 9 | 40333 | 0.02231423 | 23 | rnr1Y285A_pC | TRUE | CAN | JSY2422 | 5 | rnr1Y285A_p | GCT | C>A | A/T+1 | G/C-1 | CG>AT | CG>AT | TTGCAGCTT | CTTCACC | CGA | GCT | C | T | C>A | C>A_GCT | CG>AT_CTTCACC | 6 | FALSE | FALSE |
| 176_v.txt | 32556 | 32556-SNV-C | SNV | 1 | C | A | 2018 | 40316 | 5.00545689 | 37.3785927 | rnr1Y285A_pC | TRUE | CAN | JSY2422 | 5 | rnr1Y285A_p | ACC | C>A | A/T+1 | G/C-1 | CG>AT | CG>AT | GTGATACCA | CCAACTA | TGG | ACC | C | T | C>A | C>A_ACC | CG>AT_CCAACTA | 6 | FALSE | FALSE |
| 176_v.txt | 32566 | 32566-SNV-C | SNV | 1 | C | A | 146 | 44897 | 0.32518877 | 36.7876712 | rnr1Y285A_pC | TRUE | CAN | JSY2422 | 5 | rnr1Y285A_p | TCA | C>A | A/T+1 | G/C-1 | CG>AT | CG>AT | CTAGTTCAG | CAGTACC | AGT | TCA | C | T | C>A | C>A_TCA | CG>AT_CAGTACC | 6 | FALSE | FALSE |
| 176_v.txt | 32571 | 32571-SNV-C | SNV | 1 | C | A | 136 | 44914 | 0.30280091 | 37.3934134 | rnr1Y285A_pC | TRUE | CAN | JSY2422 | 5 | rnr1Y285A_p | ACC | C>A | A/T+1 | G/C-1 | CG>AT | CG>AT | TCAGTACCT | CCTTGAA | TGG | ACC | C | T | C>A | C>A_ACC | CG>AT_CCTTGAA | 6 | FALSE | FALSE |
| 176_v.txt | 32629 | 32629-SNV-C | SNV | 1 | C | A | 99 | 44912 | 0.22043107 | 52.2323232 | rnr1Y285A_pC | TRUE | CAN | JSY2422 | 5 | rnr1Y285A_p | TCG | C>A | A/T+1 | G/C-1 | CG>AT | CG>AT | TCCCTTCGT | CGTTTTT | AGC | TCG | C | T | C>A | C>A_TCG | CG>AT_CGTTTTT | 6 | FALSE | FALSE |
| 176_v.txt | 32735 | 32735-SNV-C | SNV | 1 | G | T | 488 | 44912 | 1.08656929 | 58.2561475 | rnr1Y285A_pC | TRUE | CAN | JSY2422 | 5 | rnr1Y285A_p | AGT | G>T | A/T+1 | G/C-1 | CG>AT | CG>AT | AAAACAGTA | GTATATT | TCA | TCA | C | A | C>A | C>A_TCA | CG>AT_GTATATT | 6 | FALSE | FALSE |
| 176_v.txt | 32780 | 32780-SNV-C | SNV | 1 | C | A | 13 | 44912 | 0.02894549 | 23.9219564 | rnr1Y285A_pC | TRUE | CAN | JSY2422 | 5 | rnr1Y285A_p | CCC | C>A | A/T+1 | G/C-1 | CG>AT | CG>AT | AGCGACCCA | CCAGAAC | GGG | CCC | C | T | C>A | C>A_CCC | CG>AT_CCAGAAC | 6 | FALSE | FALSE |
| 176_v.txt | 32783 | 32783-SNV-C | SNV | 1 | G | T | 210 | 44912 | 0.46758105 | 50.0619048 | rnr1Y285A_pC | TRUE | CAN | JSY2422 | 5 | rnr1Y285A_p | AGA | G>T | A/T+1 | G/C-1 | CG>AT | CG>AT | GACCCAGAA | GAACTCG | TCT | TCT | C | A | C>A | C>A_TCT | CG>AT_GAACTCG | 6 | FALSE | FALSE |
| 176_v.txt | 32786 | 32786-SNV-C | SNV | 1 | C | A | 90 | 44912 | 0.20039188 | 46.2555556 | rnr1Y285A_pC | TRUE | CAN | JSY2422 | 5 | rnr1Y285A_p | ACT | C>A | A/T+1 | G/C-1 | CG>AT | CG>AT | CCAGAACTC | CTCGAAT | TGA | ACT | C | T | C>A | C>A_ACT | CG>AT_CTCGAAT | 6 | FALSE | FALSE |
| 176_v.txt | 32794 | 32794-SNV-C | SNV | 1 | C | A | 220 | 44912 | 0.48984681 | 44.2727273 | rnr1Y285A_pC | TRUE | CAN | JSY2422 | 5 | rnr1Y285A_p | TCA | C>A | A/T+1 | G/C-1 | CG>AT | CG>AT | CGAATTCAC | CACCGTA | AGT | TCA | C | T | C>A | C>A_TCA | CG>AT_CACCGTA | 6 | FALSE | FALSE |
| 176_v.txt | 32819 | 32819-SNV-C | SNV | 1 | G | T | 1498 | 44912 | 3.33541147 | 45.246996 | rnr1Y285A_pC | TRUE | CAN | JSY2422 | 5 | rnr1Y285A_p | AGT | G>T | A/T+1 | G/C-1 | CG>AT | CG>AT | GAACAAGTT | GTTCATT | TCA | TCA | C | A | C>A | C>A_TCA | CG>AT_GTTCATT | 6 | FALSE | FALSE |
| 176_v.txt | 32917 | 32917-SNV-C | SNV | 1 | C | A | 3590 | 69133 | 5.19288907 | 52.0013928 | rnr1Y285A_pC | TRUE | CAN | JSY2422 | 5 | rnr1Y285A_p | TCC | C>A | A/T+1 | G/C-1 | CG>AT | CG>AT | TAAGTTCCA | CCAGGGC | AGG | TCC | C | T | C>A | C>A_TCC | CG>AT_CCAGGGC | 6 | FALSE | FALSE |
| 176_v.txt | 32920 | 32920-SNV-C | SNV | 1 | G | T | 16 | 69133 | 0.0231438 | 25.9992807 | rnr1Y285A_pC | TRUE | CAN | JSY2422 | 5 | rnr1Y285A_p | AGG | G>T | A/T+1 | G/C-1 | CG>AT | CG>AT | GTTCCAGGG | GGGCAAA | TCC | TCC | C | A | C>A | C>A_TCC | CG>AT_GGGCAAA | 6 | FALSE | FALSE |
| 176_v.txt | 32940 | 32940-SNV-C | SNV | 1 | G | T | 728 | 69132 | 1.05305792 | 54.9326477 | rnr1Y285A_pC | TRUE | CAN | JSY2422 | 5 | rnr1Y285A_p | AGA | G>T | A/T+1 | G/C-1 | CG>AT | CG>AT | GCCCAAGAA | GAAAACC | TCT | TCT | C | A | C>A | C>A_TCT | CG>AT_GAAAACC | 6 | FALSE | FALSE |
| 176_v.txt | 32958 | 32958-SNV-C | SNV | 1 | C | A | 136 | 69133 | 0.19672226 | 52.4558824 | rnr1Y285A_pC | TRUE | CAN | JSY2422 | 5 | rnr1Y285A_p | ACC | C>A | A/T+1 | G/C-1 | CG>AT | CG>AT | ATGTAACCA | CCATTGG | TGG | ACC | C | T | C>A | C>A_ACC | CG>AT_CCATTGG | 6 | FALSE | FALSE |
| 176_v.txt | 33051 | 33051-SNV-C | SNV | 1 | G | T | 39 | 69133 | 0.056413 | 55.2051282 | rnr1Y285A_pC | TRUE | CAN | JSY2422 | 5 | rnr1Y285A_p | GGA | G>T | A/T+1 | G/C-1 | CG>AT | CG>AT | CCCAAGGAC | GACTGCG | CCT | CCT | C | A | C>A | C>A_CCT | CG>AT_GACTGCG | 6 | FALSE | FALSE |
| 176_v.txt | 33093 | 33093-SNV-C | SNV | 1 | G | T | 38 | 69133 | 0.05496651 | 50.0247456 | rnr1Y285A_pC | TRUE | CAN | JSY2422 | 5 | rnr1Y285A_p | TGA | G>T | A/T+1 | G/C-1 | CG>AT | CG>AT | AAATATGAT | GATATAA | ACT | ACT | C | A | C>A | C>A_ACT | CG>AT_GATATAA | 6 | FALSE | FALSE |
| 176_v.txt | 33114 | 33114-SNV-C | SNV | 1 | C | A | 8 | 69133 | 0.0115719 | 32.1263208 | rnr1Y285A_pC | TRUE | CAN | JSY2422 | 5 | rnr1Y285A_p | GCC | C>A | A/T+1 | G/C-1 | CG>AT | CG>AT | ACTGGGCCG | CCGGCGT | CGG | GCC | C | T | C>A | C>A_GCC | CG>AT_CCGGCGT | 6 | FALSE | FALSE |
| 176_v.txt | 33151 | 33151-SNV-C | SNV | 1 | G | T | 324 | 69133 | 0.46866186 | 38.8085808 | rnr1Y285A_pC | TRUE | CAN | JSY2422 | 5 | rnr1Y285A_p | AGA | G>T | A/T+1 | G/C-1 | CG>AT | CG>AT | TGAAAAGAC | GACCTGT | TCT | TCT | C | A | C>A | C>A_TCT | CG>AT_GACCTGT | 6 | FALSE | FALSE |
| 176_v.txt | 33153 | 33153-SNV-C | SNV | 1 | C | A | 442 | 69133 | 0.63934735 | 40.0426517 | rnr1Y285A_pC | TRUE | CAN | JSY2422 | 5 | rnr1Y285A_p | ACC | C>A | A/T+1 | G/C-1 | CG>AT | CG>AT | AAAAGACCT | CCTGTAC | TGG | ACC | C | T | C>A | C>A_ACC | CG>AT_CCTGTAC | 6 | FALSE | FALSE |
| 176_v.txt | 33154 | 33154-SNV-C | SNV | 1 | C | A | 401 | 69133 | 0.58004137 | 36.872818 | rnr1Y285A_pC | TRUE | CAN | JSY2422 | 5 | rnr1Y285A_p | CCT | C>A | A/T+1 | G/C-1 | CG>AT | CG>AT | AAAGACCTG | CTGTACC | GGA | CCT | C | T | C>A | C>A_CCT | CG>AT_CTGTACC | 6 | FALSE | FALSE |
| 176_v.txt | 33156 | 33156-SNV-C | SNV | 1 | G | T | 71 | 69132 | 0.10270208 | 35.2957747 | rnr1Y285A_pC | TRUE | CAN | JSY2422 | 5 | rnr1Y285A_p | TGT | G>T | A/T+1 | G/C-1 | CG>AT | CG>AT | AGACCTGTA | GTACCAA | ACA | ACA | C | A | C>A | C>A_ACA | CG>AT_GTACCAA | 6 | FALSE | FALSE |
| 176_v.txt | 33159 | 33159-SNV-C | SNV | 1 | C | A | 892 | 69132 | 1.29028525 | 37.5964126 | rnr1Y285A_pC | TRUE | CAN | JSY2422 | 5 | rnr1Y285A_p | ACC | C>A | A/T+1 | G/C-1 | CG>AT | CG>AT | CCTGTACCA | CCAATAG | TGG | ACC | C | T | C>A | C>A_ACC | CG>AT_CCAATAG | 6 | FALSE | FALSE |
| 176_v.txt | 33168 | 33168-SNV-C | SNV | 1 | C | A | 1683 | 69130 | 2.43454361 | 37.1604278 | rnr1Y285A_pC | TRUE | CAN | JSY2422 | 5 | rnr1Y285A_p | ACC | C>A | A/T+1 | G/C-1 | CG>AT | CG>AT | ATAGTACCA | CCACCAA | TGG | ACC | C | T | C>A | C>A_ACC | CG>AT_CCACCAA | 6 | FALSE | FALSE |
| 176_v.txt | 33175 | 33175-SNV-C | SNV | 1 | G | T | 18 | 69115 | 0.02604355 | 36.3333333 | rnr1Y285A_pC | TRUE | CAN | JSY2422 | 5 | rnr1Y285A_p | AGG | G>T | A/T+1 | G/C-1 | CG>AT | CG>AT | CACCAAGGG | GGGCAAT | TCC | TCC | C | A | C>A | C>A_TCC | CG>AT_GGGCAAT | 6 | FALSE | FALSE |
| 176_v.txt | 33186 | 33186-SNV-C | SNV | 1 | C | A | 116 | 33433 | 0.34696258 | 37.112069 | rnr1Y285A_pC | TRUE | CAN | JSY2422 | 5 | rnr1Y285A_p | ACC | C>A | A/T+1 | G/C-1 | CG>AT | CG>AT | ATCATACCA | CCAATAT | TGG | ACC | C | T | C>A | C>A_ACC | CG>AT_CCAATAT | 6 | FALSE | FALSE |
| 176_v.txt | 33195 | 33195-SNV-C | SNV | 1 | C | A | 25 | 33432 | 0.07477866 | 37.52 | rnr1Y285A_pC | TRUE | CAN | JSY2422 | 5 | rnr1Y285A_p | TCT | C>A | A/T+1 | G/C-1 | CG>AT | CG>AT | ATATGTCTT | CTTTGCT | AGA | TCT | C | T | C>A | C>A_TCT | CG>AT_CTTTGCT | 6 | FALSE | FALSE |
| 176_v.txt | 33205 | 33205-SNV-C | SNV | 1 | G | T | 75 | 33429 | 0.2243561 | 38.8933333 | rnr1Y285A_pC | TRUE | CAN | JSY2422 | 5 | rnr1Y285A_p | AGC | G>T | A/T+1 | G/C-1 | CG>AT | CG>AT | GCTTAAGCT | GCTCTCT | TCG | TCG | C | A | C>A | C>A_TCG | CG>AT_GCTCTCT | 6 | FALSE | FALSE |
| 176_v.txt | 33325 | 33325-SNV-C | SNV | 1 | C | A | 540 | 33428 | 1.61541223 | 58.6509679 | rnr1Y285A_pC | TRUE | CAN | JSY2422 | 5 | rnr1Y285A_p | TCT | C>A | A/T+1 | G/C-1 | CG>AT | CG>AT | ACAATTCTT | CTTTACT | AGA | TCT | C | T | C>A | C>A_TCT | CG>AT_CTTTACT | 6 | FALSE | FALSE |
| 176_v.txt | 33474 | 33474-SNV-C | SNV | 1 | C | A | 122 | 33428 | 0.3649635 | 33.9918033 | rnr1Y285A_pC | TRUE | CAN | JSY2422 | 5 | rnr1Y285A_p | GCC | C>A | A/T+1 | G/C-1 | CG>AT | CG>AT | GCTATGCCT | CCTTTTT | CGG | GCC | C | T | C>A | C>A_GCC | CG>AT_CCTTTTT | 6 | FALSE | FALSE |
| 176_v.txt | 33475 | 33475-SNV-C | SNV | 1 | C | A | 25 | 33428 | 0.0747876 | 37.52 | rnr1Y285A_pC | TRUE | CAN | JSY2422 | 5 | rnr1Y285A_p | CCT | C>A | A/T+1 | G/C-1 | CG>AT | CG>AT | CTATGCCTT | CTTTTTT | GGA | CCT | C | T | C>A | C>A_CCT | CG>AT_CTTTTTT | 6 | FALSE | FALSE |
| 176_v.txt | 32940 | 32940-Repla | Replacement | 2 | GA | T | 15 | 69133 | 0.02169731 | 57.4 | rnr1Y285A_pC | TRUE | CAN | JSY2422 | 5 | rnr1Y285A_p | AGA | nope | A/T+1 | >1bpdel | none | Replac. | GCCCAAGAA | GAAAACC | TCT | AGA | GA | A | NA | NA_AGA | Replac._GAAAACC | 6 | FALSE | FALSE |
| 176_v.txt | 31657 | 31657-MNV- | MNV | 2 | GC | TT | 1 | 17332 | 5.77E-03 | 30.4999994 | rnr1Y285A_pC | TRUE | CAN | JSY2422 | 5 | rnr1Y285A_p | GGC | nope | >1bp-ins | >1bpdel | none | MNV | GAAATGGCG | GCGTGGA | CCG | GGC | GC | none | NA | NA_GGC | MNV_GCGTGGA | 6 | FALSE | FALSE |
| 176_v.txt | 32081 | 32081-Insert | Insertion | 1 | - | A | 67 | 37086 | 0.18066117 | 63.5373134 | rnr1Y285A_pC | TRUE | CAN | JSY2422 | 5 | rnr1Y285A_p | CAA | nope | A/T+1 | >1bpdel | none | A/T+1 | CCATGCAAA | AAAAAAG | GTT | CAA | - | T | NA | NA_CAA | A/T+1_AAAAAAG | 6 | FALSE | FALSE |
| 176_v.txt | 32168 | 32168-Insert | Insertion | 1 | - | A | 3 | 37086 | 8.09E-03 | 53 | rnr1Y285A_pC | TRUE | CAN | JSY2422 | 5 | rnr1Y285A_p | CAA | nope | A/T+1 | >1bpdel | none | A/T+1 | AGCGCCAAA | AAATGCA | GTT | CAA | - | T | NA | NA_CAA | A/T+1_AAATGCA | 6 | FALSE | FALSE |
| 176_v.txt | 32498 | 32498-Insert | Insertion | 1 | - | T | 26 | 40333 | 0.06446334 | 41.3846154 | rnr1Y285A_pC | TRUE | CAN | JSY2422 | 5 | rnr1Y285A_p | CTT | nope | A/T+1 | >1bpdel | none | A/T+1 | AACAACTTT | TTTTTTG | GAA | CTT | - | A | NA | NA_CTT | A/T+1_TTTTTTG | 6 | FALSE | FALSE |
| 176_v.txt | 32532 | 32532-Insert | Insertion | 1 | - | T | 156 | 40333 | 0.38678006 | 41.4102564 | rnr1Y285A_pC | TRUE | CAN | JSY2422 | 5 | rnr1Y285A_p | GTT | nope | A/T+1 | >1bpdel | none | A/T+1 | CTGGGGTTT | TTTGCAG | CAA | GTT | - | A | NA | NA_GTT | A/T+1_TTTGCAG | 6 | FALSE | FALSE |
| 176_v.txt | 32605 | 32605-Insert | Insertion | 1 | - | A | 18 | 44912 | 0.04007838 | 40 | rnr1Y285A_pC | TRUE | CAN | JSY2422 | 5 | rnr1Y285A_p | GAA | nope | A/T+1 | >1bpdel | none | A/T+1 | AAGAGGAAA | AAACCCA | CTT | GAA | - | T | NA | NA_GAA | A/T+1_AAACCCA | 6 | FALSE | FALSE |
| 176_v.txt | 32631 | 32631-Insert | Insertion | 1 | - | T | 34 | 44912 | 0.0757036 | 52.7941177 | rnr1Y285A_pC | TRUE | CAN | JSY2422 | 5 | rnr1Y285A_p | GTT | nope | A/T+1 | >1bpdel | none | A/T+1 | CCTTCGTTT | TTTTTAT | CAA | GTT | - | A | NA | NA_GTT | A/T+1_TTTTTAT | 6 | FALSE | FALSE |
| 176_v.txt | 32842 | 32842-Insert | Insertion | 1 | - | A | 185 | 44912 | 0.41191664 | 42 | rnr1Y285A_pC | TRUE | CAN | JSY2422 | 5 | rnr1Y285A_p | CAA | nope | A/T+1 | >1bpdel | none | A/T+1 | TTACCCAAA | AAAAAAT | GTT | CAA | - | T | NA | NA_CAA | A/T+1_AAAAAAT | 6 | FALSE | FALSE |
| 176_v.txt | 33206 | 33206-Insert | Insertion | 2 | - | CT | 26 | 33429 | 0.07777678 | 38.0192308 | rnr1Y285A_pC | TRUE | CAN | JSY2422 | 5 | rnr1Y285A_p | GCT | nope | >1bp-ins | >1bpdel | none | >1bp-ins | CTTAAGCTC | CTCTCTC | CGA | GCT | - | none | NA | NA_GCT | >1bp-ins_CTCTCTC | 6 | FALSE | FALSE |
| 176_v.txt | 32067 | 32067-Deleti | Deletion | 4 | ATAA | - | 26 | 37086 | 0.07010732 | 63 | rnr1Y285A_pC | TRUE | CAN | JSY2422 | 5 | rnr1Y285A_p | GAT | nope | >1bp-ins | >1bpdel | none | >1bpdel | ATTGAGATA | ATAAATA | CTA | GAT | ATAA | none | NA | NA_GAT | >1bpdel_ATAAATA | 6 | FALSE | FALSE |
| 176_v.txt | 32337 | 32337-Deleti | Deletion | 2 | AT | - | 12 | 40333 | 0.02975231 | 63.0833333 | rnr1Y285A_pC | TRUE | CAN | JSY2422 | 5 | rnr1Y285A_p | GAT | nope | >1bp-ins | >1bpdel | none | >1bpdel | TTGAAGATA | ATATGTG | CTA | GAT | AT | none | NA | NA_GAT | >1bpdel_ATATGTG | 6 | FALSE | FALSE |
| 176_v.txt | 32492 | 32492-Deleti | Deletion | 3 | AAC | - | 356 | 40333 | 0.88265192 | 46.0646067 | rnr1Y285A_pC | TRUE | CAN | JSY2422 | 5 | rnr1Y285A_p | AAA | nope | >1bp-ins | >1bpdel | none | >1bpdel | ACGGAAAAC | AACAACT | TTT | AAA | AAC | none | NA | NA_AAA | >1bpdel_AACAACT | 6 | FALSE | FALSE |
| 176_v.txt | 33094 | 33094-Deleti | Deletion | 2 | AT | - | 12 | 69133 | 0.01735785 | 46.5833333 | rnr1Y285A_pC | TRUE | CAN | JSY2422 | 5 | rnr1Y285A_p | GAT | nope | >1bp-ins | >1bpdel | none | >1bpdel | AATATGATA | ATATAAG | CTA | GAT | AT | none | NA | NA_GAT | >1bpdel_ATATAAG | 6 | FALSE | FALSE |
| 176_v.txt | 33474 | 33474-Deleti | Deletion | 2 | CC | - | 1 | 33428 | 2.99E-03 | 10 | rnr1Y285A_pC | TRUE | CAN | JSY2422 | 5 | rnr1Y285A_p | GCC | nope | >1bp-ins | >1bpdel | none | >1bpdel | GCTATGCCT | CCTTTTT | CGG | GCC | CC | none | NA | NA_GCC | >1bpdel_CCTTTTT | 6 | FALSE | FALSE |
| 176_v.txt | 31658 | 31658-Deleti | Deletion | 1 | C | - | 1 | 17332 | 5.77E-03 | 37.9999989 | rnr1Y285A_pC | TRUE | CAN | JSY2422 | 5 | rnr1Y285A_p | GCG | nope | >1bp-ins | G/C-1 | none | G/C-1 | AAATGGCGT | CGTGGAA | CGC | GCG | C | none | nope | nope_GCG | G/C-1_CGTGGAA | 6 | FALSE | FALSE |
| 176_v.txt | 31723 | 31723-Deleti | Deletion | 1 | A | - | 31 | 17332 | 0.17885991 | 58.7096774 | rnr1Y285A_pC | TRUE | CAN | JSY2422 | 5 | rnr1Y285A_p | CAA | nope | >1bp-ins | A/T-1 | none | A/T-1 | TGTCCCAAA | AAAAAGT | GTT | GTT | T | none | T>none | T>none_GTT | A/T-1_AAAAAGT | 6 | FALSE | FALSE |
| 176_v.txt | 31904 | 31904-Deleti | Deletion | 1 | T | - | 394 | 17332 | 2.27325179 | 43.8020305 | rnr1Y285A_pC | TRUE | CAN | JSY2422 | 5 | rnr1Y285A_p | ATT | nope | >1bp-ins | A/T-1 | none | A/T-1 | ATTGAATTT | TTTTGGT | TAA | ATT | T | none | nope | nope_ATT | A/T-1_TTTTGGT | 6 | FALSE | FALSE |
| 176_v.txt | 31913 | 31913-Deleti | Deletion | 1 | A | - | 15 | 17332 | 0.08654512 | 37.8 | rnr1Y285A_pC | TRUE | CAN | JSY2422 | 5 | rnr1Y285A_p | CAA | nope | >1bp-ins | A/T-1 | none | A/T-1 | TGGTGCAAA | AAAAGCC | GTT | GTT | T | none | T>none | T>none_GTT | A/T-1_AAAAGCC | 6 | FALSE | FALSE |
| 176_v.txt | 31920 | 31920-Deleti | Deletion | 1 | G | - | 2 | 17329 | 0.01154135 | 34.5 | rnr1Y285A_pC | TRUE | CAN | JSY2422 | 5 | rnr1Y285A_p | CGT | nope | >1bp-ins | G/C-1 | none | G/C-1 | AAAGCCGTG | GTGAAAC | GCA | GCA | C | none | C>none | C>none_GCA | G/C-1_GTGAAAC | 6 | FALSE | FALSE |
| 176_v.txt | 31958 | 31958-Deleti | Deletion | 1 | G | - | 5 | 37087 | 0.01348181 | 37.6 | rnr1Y285A_pC | TRUE | CAN | JSY2422 | 5 | rnr1Y285A_p | TGG | nope | >1bp-ins | G/C-1 | none | G/C-1 | AAATGTGGC | GGCCGCA | ACC | ACC | C | none | C>none | C>none_ACC | G/C-1_GGCCGCA | 6 | FALSE | FALSE |
| 176_v.txt | 31979 | 31979-Deleti | Deletion | 1 | G | - | 253 | 37086 | 0.68219813 | 42.0909091 | rnr1Y285A_pC | TRUE | CAN | JSY2422 | 5 | rnr1Y285A_p | CGG | nope | >1bp-ins | G/C-1 | none | G/C-1 | CAAGCCGGG | GGGCATT | GCC | GCC | C | none | C>none | C>none_GCC | G/C-1_GGGCATT | 6 | FALSE | FALSE |
| 176_v.txt | 32069 | 32069-Deleti | Deletion | 1 | A | - | 7 | 37084 | 0.01887607 | 60.2857156 | rnr1Y285A_pC | TRUE | CAN | JSY2422 | 5 | rnr1Y285A_p | TAA | nope | >1bp-ins | A/T-1 | none | A/T-1 | TGAGATAAA | AAATAAC | ATT | ATT | T | none | T>none | T>none_ATT | A/T-1_AAATAAC | 6 | FALSE | FALSE |
| 176_v.txt | 32081 | 32081-Deleti | Deletion | 1 | A | - | 41 | 37086 | 0.11055385 | 63.7073172 | rnr1Y285A_pC | TRUE | CAN | JSY2422 | 5 | rnr1Y285A_p | CAA | nope | >1bp-ins | A/T-1 | none | A/T-1 | CCATGCAAA | AAAAAAG | GTT | GTT | T | none | T>none | T>none_GTT | A/T-1_AAAAAAG | 6 | FALSE | FALSE |
| 176_v.txt | 32210 | 32210-Deleti | Deletion | 1 | T | - | 14 | 37086 | 0.03775009 | 40.7857143 | rnr1Y285A_pC | TRUE | CAN | JSY2422 | 5 | rnr1Y285A_p | CTT | nope | >1bp-ins | A/T-1 | none | A/T-1 | ACCACCTTTG | TTTGGTG | GAA | CTT | T | none | nope | nope_CTT | A/T-1_TTTGGTG | 6 | FALSE | FALSE |
| 176_v.txt | 32216 | 32216-Deleti | Deletion | 1 | G | - | 96 | 37086 | 0.25885779 | 41.1145833 | rnr1Y285A_pC | TRUE | CAN | JSY2422 | 5 | rnr1Y285A_p | TGG | nope | >1bp-ins | G/C-1 | none | G/C-1 | TTTGGTGGT | GGTCCTT | ACC | ACC | C | none | C>none | C>none_ACC | G/C-1_GGTCCTT | 6 | FALSE | FALSE |
| 176_v.txt | 32259 | 32259-Deleti | Deletion | 1 | C | - | 49 | 40331 | 0.12149463 | 37.8163265 | rnr1Y285A_pC | TRUE | CAN | JSY2422 | 5 | rnr1Y285A_p | ACC | nope | >1bp-ins | G/C-1 | none | G/C-1 | GATAGACCA | CCAAATA | TGG | ACC | C | none | nope | nope_ACC | G/C-1_CCAAATA | 6 | FALSE | FALSE |
| 176_v.txt | 32277 | 32277-Deleti | Deletion | 1 | C | - | 102 | 40332 | 0.25290092 | 36.5882353 | rnr1Y285A_pC | TRUE | CAN | JSY2422 | 5 | rnr1Y285A_p | ACC | nope | >1bp-ins | G/C-1 | none | G/C-1 | CGGGAACCA | CCAACGT | TGG | ACC | C | none | nope | nope_ACC | G/C-1_CCAACGT | 6 | FALSE | FALSE |
| 176_v.txt | 32346 | 32346-Deleti | Deletion | 1 | A | - | 201 | 40333 | 0.49835123 | 61.0945274 | rnr1Y285A_pC | TRUE | CAN | JSY2422 | 5 | rnr1Y285A_p | CAA | nope | >1bp-ins | A/T-1 | none | A/T-1 | TGTGGCAAA | AAAACCT | GTT | GTT | T | none | T>none | T>none_GTT | A/T-1_AAAACCT | 6 | FALSE | FALSE |
| 176_v.txt | 32381 | 32381-Deleti | Deletion | 1 | A | - | 6 | 40333 | 0.01487616 | 63.5 | rnr1Y285A_pC | TRUE | CAN | JSY2422 | 5 | rnr1Y285A_p | TAA | nope | >1bp-ins | A/T-1 | none | A/T-1 | AATAATAAA | AAAGGGA | ATT | ATT | T | none | T>none | T>none_ATT | A/T-1_AAAGGGA | 6 | FALSE | FALSE |
| 176_v.txt | 32398 | 32398-Deleti | Deletion | 1 | C | - | 8 | 40333 | 0.01983488 | 60 | rnr1Y285A_pC | TRUE | CAN | JSY2422 | 5 | rnr1Y285A_p | ACG | nope | >1bp-ins | G/C-1 | none | G/C-1 | TAGAAACGT | CGTAGGA | TGC | ACG | C | none | nope | nope_ACG | G/C-1_CGTAGGA | 6 | FALSE | FALSE |
| 176_v.txt | 32416 | 32416-Deleti | Deletion | 1 | T | - | 7 | 40333 | 0.01735552 | 60.2857143 | rnr1Y285A_pC | TRUE | CAN | JSY2422 | 5 | rnr1Y285A_p | GTT | nope | >1bp-ins | A/T-1 | none | A/T-1 | ATTGTGTTA | TTAGTTT | CAA | GTT | T | none | nope | nope_GTT | A/T-1_TTAGTTT | 6 | FALSE | FALSE |
| 176_v.txt | 32420 | 32420-Deleti | Deletion | 1 | T | - | 7 | 40333 | 0.01735552 | 61.7142857 | rnr1Y285A_pC | TRUE | CAN | JSY2422 | 5 | rnr1Y285A_p | GTT | nope | >1bp-ins | A/T-1 | none | A/T-1 | TGTTAGTTTA | TTTAGGG | CAA | GTT | T | none | nope | nope_GTT | A/T-1_TTTAGGG | 6 | FALSE | FALSE |
| 176_v.txt | 32424 | 32424-Deleti | Deletion | 1 | G | - | 45 | 40333 | 0.11157117 | 58.1777778 | rnr1Y285A_pC | TRUE | CAN | JSY2422 | 5 | rnr1Y285A_p | AGG | nope | >1bp-ins | G/C-1 | none | G/C-1 | AGTTTAGGG | GGGTCAT | TCC | TCC | C | none | C>none | C>none_TCC | G/C-1_GGGTCAT | 6 | FALSE | FALSE |
| 176_v.txt | 32442 | 32442-Deleti | Deletion | 1 | A | - | 9 | 40333 | 0.02231423 | 60.5555556 | rnr1Y285A_pC | TRUE | CAN | JSY2422 | 5 | rnr1Y285A_p | TAA | nope | >1bp-ins | A/T-1 | none | A/T-1 | GGAACTAAA | AAAAGTC | ATT | ATT | T | none | T>none | T>none_ATT | A/T-1_AAAAGTC | 6 | FALSE | FALSE |
| 176_v.txt | 32496 | 32496-Deleti | Deletion | 1 | A | - | 3 | 40333 | 7.44E-03 | 38 | rnr1Y285A_pC | TRUE | CAN | JSY2422 | 5 | rnr1Y285A_p | AAC | nope | >1bp-ins | A/T-1 | none | A/T-1 | AAAACAACT | ACTTTTT | TTG | TTG | T | none | T>none | T>none_TTG | A/T-1_ACTTTTT | 6 | FALSE | FALSE |
| 176_v.txt | 32498 | 32498-Deleti | Deletion | 1 | T | - | 50 | 40333 | 0.12396797 | 45.2600026 | rnr1Y285A_pC | TRUE | CAN | JSY2422 | 5 | rnr1Y285A_p | CTT | nope | >1bp-ins | A/T-1 | none | A/T-1 | AACAACTTT | TTTTTTG | GAA | CTT | T | none | nope | nope_CTT | A/T-1_TTTTTTG | 6 | FALSE | FALSE |
| 176_v.txt | 32522 | 32522-Deleti | Deletion | 1 | T | - | 146 | 40333 | 0.36198646 | 40.5068493 | rnr1Y285A_pC | TRUE | CAN | JSY2422 | 5 | rnr1Y285A_p | ATT | nope | >1bp-ins | A/T-1 | none | A/T-1 | AACGGATTT | TTTTCTG | TAA | ATT | T | none | nope | nope_ATT | A/T-1_TTTTCTG | 6 | FALSE | FALSE |
| 176_v.txt | 32528 | 32528-Deleti | Deletion | 1 | G | - | 97 | 40333 | 0.24049786 | 42.1237113 | rnr1Y285A_pC | TRUE | CAN | JSY2422 | 5 | rnr1Y285A_p | TGG | nope | >1bp-ins | G/C-1 | none | G/C-1 | TTTTCTGGG | GGGGTTT | ACC | ACC | C | none | C>none | C>none_ACC | G/C-1_GGGGTTT | 6 | FALSE | FALSE |
| 176_v.txt | 32598 | 32598-Deleti | Deletion | 1 | A | - | 6 | 44912 | 0.01335946 | 37.6666667 | rnr1Y285A_pC | TRUE | CAN | JSY2422 | 5 | rnr1Y285A_p | CAA | nope | >1bp-ins | A/T-1 | none | A/T-1 | TTAATCAAA | AAAGAGG | GTT | GTT | T | none | T>none | T>none_GTT | A/T-1_AAAGAGG | 6 | FALSE | FALSE |
| 176_v.txt | 32608 | 32608-Deleti | Deletion | 1 | C | - | 4529 | 44912 | 10.0841646 | 43.2779863 | rnr1Y285A_pC | TRUE | CAN | JSY2422 | 5 | rnr1Y285A_p | ACC | nope | >1bp-ins | G/C-1 | none | G/C-1 | AGGAAACCC | CCCAACC | TGG | ACC | C | none | nope | nope_ACC | G/C-1_CCCAACC | 6 | FALSE | FALSE |
| 176_v.txt | 32621 | 32621-Deleti | Deletion | 1 | C | - | 8 | 44912 | 0.01781261 | 38 | rnr1Y285A_pC | TRUE | CAN | JSY2422 | 5 | rnr1Y285A_p | ACC | nope | >1bp-ins | G/C-1 | none | G/C-1 | TAAGAACCT | CCTCCCT | TGG | ACC | C | none | nope | nope_ACC | G/C-1_CCTCCCT | 6 | FALSE | FALSE |
| 176_v.txt | 32631 | 32631-Deleti | Deletion | 1 | T | - | 17 | 44912 | 0.0378518 | 55.8235274 | rnr1Y285A_pC | TRUE | CAN | JSY2422 | 5 | rnr1Y285A_p | GTT | nope | >1bp-ins | A/T-1 | none | A/T-1 | CCTTCGTTT | TTTTTAT | CAA | GTT | T | none | nope | nope_GTT | A/T-1_TTTTTAT | 6 | FALSE | FALSE |
| 176_v.txt | 32647 | 32647-Deleti | Deletion | 1 | T | - | 7 | 44912 | 0.01558604 | 56.5714286 | rnr1Y285A_pC | TRUE | CAN | JSY2422 | 5 | rnr1Y285A_p | ATT | nope | >1bp-ins | A/T-1 | none | A/T-1 | TAGATATTA | TTATACC | TAA | ATT | T | none | nope | nope_ATT | A/T-1_TTATACC | 6 | FALSE | FALSE |
| 176_v.txt | 32652 | 32652-Deleti | Deletion | 1 | C | - | 128 | 44912 | 0.28500178 | 59.140625 | rnr1Y285A_pC | TRUE | CAN | JSY2422 | 5 | rnr1Y285A_p | ACC | nope | >1bp-ins | G/C-1 | none | G/C-1 | ATTATACCTG | CCTGGAC | TGG | ACC | C | none | nope | nope_ACC | G/C-1_CCTGGAC | 6 | FALSE | FALSE |
| 176_v.txt | 32658 | 32658-Deleti | Deletion | 1 | C | - | 2435 | 44912 | 5.42171357 | 59.8903491 | rnr1Y285A_pC | TRUE | CAN | JSY2422 | 5 | rnr1Y285A_p | ACC | nope | >1bp-ins | G/C-1 | none | G/C-1 | CCTGGACCC | CCCCAGG | TGG | ACC | C | none | nope | nope_ACC | G/C-1_CCCCAGG | 6 | FALSE | FALSE |
| 176_v.txt | 32670 | 32670-Deleti | Deletion | 1 | G | - | 7491 | 44912 | 16.6792839 | 59.4306501 | rnr1Y285A_pC | TRUE | CAN | JSY2422 | 5 | rnr1Y285A_p | TGG | nope | >1bp-ins | G/C-1 | none | G/C-1 | GCACCTGGG | GGGTTTC | ACC | ACC | C | none | C>none | C>none_ACC | G/C-1_GGGTTTC | 6 | FALSE | FALSE |
| 176_v.txt | 32705 | 32705-Deleti | Deletion | 1 | A | - | 100 | 44912 | 0.22265764 | 61.42 | rnr1Y285A_pC | TRUE | CAN | JSY2422 | 5 | rnr1Y285A_p | TAA | nope | >1bp-ins | A/T-1 | none | A/T-1 | GCCGGTAAC | AACCCCA | ATT | ATT | T | none | T>none | T>none_ATT | A/T-1_AACCCCA | 6 | FALSE | FALSE |
| 176_v.txt | 32707 | 32707-Deleti | Deletion | 1 | C | - | 2244 | 44912 | 4.99643748 | 62.4594474 | rnr1Y285A_pC | TRUE | CAN | JSY2422 | 5 | rnr1Y285A_p | ACC | nope | >1bp-ins | G/C-1 | none | G/C-1 | CGGTAACCC | CCCCAGC | TGG | ACC | C | none | nope | nope_ACC | G/C-1_CCCCAGC | 6 | FALSE | FALSE |
| 176_v.txt | 32719 | 32719-Deleti | Deletion | 1 | A | - | 30 | 44912 | 0.06679729 | 62.3 | rnr1Y285A_pC | TRUE | CAN | JSY2422 | 5 | rnr1Y285A_p | CAA | nope | >1bp-ins | A/T-1 | none | A/T-1 | CACCACAAA | AAACCAT | GTT | GTT | T | none | T>none | T>none_GTT | A/T-1_AAACCAT | 6 | FALSE | FALSE |
| 176_v.txt | 32728 | 32728-Deleti | Deletion | 1 | A | - | 12 | 44912 | 0.02671892 | 63.75 | rnr1Y285A_pC | TRUE | CAN | JSY2422 | 5 | rnr1Y285A_p | CAA | nope | >1bp-ins | A/T-1 | none | A/T-1 | CCATACAAA | AAAAACA | GTT | GTT | T | none | T>none | T>none_GTT | A/T-1_AAAAACA | 6 | FALSE | FALSE |
| 176_v.txt | 32747 | 32747-Deleti | Deletion | 1 | C | - | 835 | 44912 | 1.85919131 | 55.9017964 | rnr1Y285A_pC | TRUE | CAN | JSY2422 | 5 | rnr1Y285A_p | ACC | nope | >1bp-ins | G/C-1 | none | G/C-1 | TAGAAACCC | CCCGATA | TGG | ACC | C | none | nope | nope_ACC | G/C-1_CCCGATA | 6 | FALSE | FALSE |
| 176_v.txt | 32760 | 32760-Deleti | Deletion | 1 | A | - | 6 | 44912 | 0.01335946 | 59.6666667 | rnr1Y285A_pC | TRUE | CAN | JSY2422 | 5 | rnr1Y285A_p | TAA | nope | >1bp-ins | A/T-1 | none | A/T-1 | ATGGCTAAA | AAAACTT | ATT | ATT | T | none | T>none | T>none_ATT | A/T-1_AAAACTT | 6 | FALSE | FALSE |
| 176_v.txt | 32779 | 32779-Deleti | Deletion | 1 | C | - | 67 | 44912 | 0.14918062 | 47.8208953 | rnr1Y285A_pC | TRUE | CAN | JSY2422 | 5 | rnr1Y285A_p | ACC | nope | >1bp-ins | G/C-1 | none | G/C-1 | AAGCGACCC | CCCAGAA | TGG | ACC | C | none | nope | nope_ACC | G/C-1_CCCAGAA | 6 | FALSE | FALSE |
| 176_v.txt | 32839 | 32839-Deleti | Deletion | 1 | C | - | 15 | 44912 | 0.03339865 | 44.2666667 | rnr1Y285A_pC | TRUE | CAN | JSY2422 | 5 | rnr1Y285A_p | ACC | nope | >1bp-ins | G/C-1 | none | G/C-1 | TAATTACCCA | CCCAAAA | TGG | ACC | C | none | nope | nope_ACC | G/C-1_CCCAAAA | 6 | FALSE | FALSE |
| 176_v.txt | 32842 | 32842-Deleti | Deletion | 1 | A | - | 91 | 44912 | 0.20261845 | 42.7802195 | rnr1Y285A_pC | TRUE | CAN | JSY2422 | 5 | rnr1Y285A_p | CAA | nope | >1bp-ins | A/T-1 | none | A/T-1 | TTACCCAAA | AAAAAAT | GTT | GTT | T | none | T>none | T>none_GTT | A/T-1_AAAAAAT | 6 | FALSE | FALSE |
| 176_v.txt | 32855 | 32855-Deleti | Deletion | 1 | C | - | 4 | 44913 | 8.91E-03 | 36.5 | rnr1Y285A_pC | TRUE | CAN | JSY2422 | 5 | rnr1Y285A_p | TCC | nope | >1bp-ins | G/C-1 | none | G/C-1 | ACTAATCCAT | CCATGCC | AGG | TCC | C | none | nope | nope_TCC | G/C-1_CCATGCC | 6 | FALSE | FALSE |
| 176_v.txt | 32884 | 32884-Deleti | Deletion | 1 | A | - | 29 | 69133 | 0.04194813 | 37.4137931 | rnr1Y285A_pC | TRUE | CAN | JSY2422 | 5 | rnr1Y285A_p | CAA | nope | >1bp-ins | A/T-1 | none | A/T-1 | ACGTCCAAA | AAAATTG | GTT | GTT | T | none | T>none | T>none_GTT | A/T-1_AAAATTG | 6 | FALSE | FALSE |
| 176_v.txt | 32915 | 32915-Deleti | Deletion | 1 | T | - | 2 | 69133 | 2.89E-03 | 39 | rnr1Y285A_pC | TRUE | CAN | JSY2422 | 5 | rnr1Y285A_p | GTT | nope | >1bp-ins | A/T-1 | none | A/T-1 | ACTAAGTTC | TTCCAGG | CAA | GTT | T | none | nope | nope_GTT | A/T-1_TTCCAGG | 6 | FALSE | FALSE |
| 176_v.txt | 32917 | 32917-Deleti | Deletion | 1 | C | - | 3 | 69133 | 4.34E-03 | 51 | rnr1Y285A_pC | TRUE | CAN | JSY2422 | 5 | rnr1Y285A_p | TCC | nope | >1bp-ins | G/C-1 | none | G/C-1 | TAAGTTCCA | CCAGGGC | AGG | TCC | C | none | nope | nope_TCC | G/C-1_CCAGGGC | 6 | FALSE | FALSE |
| 176_v.txt | 32920 | 32920-Deleti | Deletion | 1 | G | - | 8 | 69133 | 0.0115719 | 53 | rnr1Y285A_pC | TRUE | CAN | JSY2422 | 5 | rnr1Y285A_p | AGG | nope | >1bp-ins | G/C-1 | none | G/C-1 | GTTCCAGGG | GGGCAAA | TCC | TCC | C | none | C>none | C>none_TCC | G/C-1_GGGCAAA | 6 | FALSE | FALSE |
| 176_v.txt | 32924 | 32924-Deleti | Deletion | 1 | A | - | 70 | 69133 | 0.1012541 | 53.0428571 | rnr1Y285A_pC | TRUE | CAN | JSY2422 | 5 | rnr1Y285A_p | CAA | nope | >1bp-ins | A/T-1 | none | A/T-1 | CAGGGCAAA | AAAAGTG | GTT | GTT | T | none | T>none | T>none_GTT | A/T-1_AAAAGTG | 6 | FALSE | FALSE |
| 176_v.txt | 32934 | 32934-Deleti | Deletion | 1 | G | - | 8 | 69133 | 0.0115719 | 46.8749972 | rnr1Y285A_pC | TRUE | CAN | JSY2422 | 5 | rnr1Y285A_p | TGC | nope | >1bp-ins | G/C-1 | none | G/C-1 | GTGATTGCC | GCCCAAG | ACG | ACG | C | none | C>none | C>none_ACG | G/C-1_GCCCAAG | 6 | FALSE | FALSE |
| 176_v.txt | 32935 | 32935-Deleti | Deletion | 1 | C | - | 13 | 69131 | 0.01880488 | 36.9230768 | rnr1Y285A_pC | TRUE | CAN | JSY2422 | 5 | rnr1Y285A_p | GCC | nope | >1bp-ins | G/C-1 | none | G/C-1 | TGATTGCCC | CCCAAGA | CGG | GCC | C | none | nope | nope_GCC | G/C-1_CCCAAGA | 6 | FALSE | FALSE |
| 176_v.txt | 32941 | 32941-Deleti | Deletion | 1 | A | - | 9 | 69133 | 0.01301839 | 55.3333294 | rnr1Y285A_pC | TRUE | CAN | JSY2422 | 5 | rnr1Y285A_p | GAA | nope | >1bp-ins | A/T-1 | none | A/T-1 | CCCAAGAAA | AAAACCA | CTT | CTT | T | none | T>none | T>none_CTT | A/T-1_AAAACCA | 6 | FALSE | FALSE |
| 176_v.txt | 32945 | 32945-Deleti | Deletion | 1 | C | - | 749 | 69133 | 1.08341892 | 58.8664887 | rnr1Y285A_pC | TRUE | CAN | JSY2422 | 5 | rnr1Y285A_p | ACC | nope | >1bp-ins | G/C-1 | none | G/C-1 | AGAAAACCA | CCAATAC | TGG | ACC | C | none | nope | nope_ACC | G/C-1_CCAATAC | 6 | FALSE | FALSE |
| 176_v.txt | 32958 | 32958-Deleti | Deletion | 1 | C | - | 15 | 69133 | 0.02169731 | 52.8666667 | rnr1Y285A_pC | TRUE | CAN | JSY2422 | 5 | rnr1Y285A_p | ACC | nope | >1bp-ins | G/C-1 | none | G/C-1 | ATGTAACCA | CCATTGG | TGG | ACC | C | none | nope | nope_ACC | G/C-1_CCATTGG | 6 | FALSE | FALSE |
| 176_v.txt | 32972 | 32972-Deleti | Deletion | 1 | A | - | 7 | 69133 | 0.01012541 | 64 | rnr1Y285A_pC | TRUE | CAN | JSY2422 | 5 | rnr1Y285A_p | CAA | nope | >1bp-ins | A/T-1 | none | A/T-1 | CGCACCAAA | AAATGCT | GTT | GTT | T | none | T>none | T>none_GTT | A/T-1_AAATGCT | 6 | FALSE | FALSE |
| 176_v.txt | 33000 | 33000-Deleti | Deletion | 1 | A | - | 9 | 69133 | 0.01301839 | 59.8888889 | rnr1Y285A_pC | TRUE | CAN | JSY2422 | 5 | rnr1Y285A_p | GAA | nope | >1bp-ins | A/T-1 | none | A/T-1 | TGTGAGAAA | AAAACTG | CTT | CTT | T | none | T>none | T>none_CTT | A/T-1_AAAACTG | 6 | FALSE | FALSE |
| 176_v.txt | 33045 | 33045-Deleti | Deletion | 1 | C | - | 8 | 69133 | 0.0115719 | 57.375 | rnr1Y285A_pC | TRUE | CAN | JSY2422 | 5 | rnr1Y285A_p | ACC | nope | >1bp-ins | G/C-1 | none | G/C-1 | ATTTCACCCA | CCCAAGG | TGG | ACC | C | none | nope | nope_ACC | G/C-1_CCCAAGG | 6 | FALSE | FALSE |
| 176_v.txt | 33078 | 33078-Deleti | Deletion | 1 | C | - | 278 | 69133 | 0.40212344 | 53.4784173 | rnr1Y285A_pC | TRUE | CAN | JSY2422 | 5 | rnr1Y285A_p | ACC | nope | >1bp-ins | G/C-1 | none | G/C-1 | AAAGAACCC | CCCATAA | TGG | ACC | C | none | nope | nope_ACC | G/C-1_CCCATAA | 6 | FALSE | FALSE |
| 176_v.txt | 33122 | 33122-Deleti | Deletion | 1 | G | - | 132 | 69133 | 0.19093631 | 43.7121212 | rnr1Y285A_pC | TRUE | CAN | JSY2422 | 5 | rnr1Y285A_p | TGG | nope | >1bp-ins | G/C-1 | none | G/C-1 | GGCGTTGGT | GGTCAGA | ACC | ACC | C | none | C>none | C>none_ACC | G/C-1_GGTCAGA | 6 | FALSE | FALSE |
| 176_v.txt | 33141 | 33141-Deleti | Deletion | 1 | C | - | 12 | 69133 | 0.01735785 | 37.9166667 | rnr1Y285A_pC | TRUE | CAN | JSY2422 | 5 | rnr1Y285A_p | ACC | nope | >1bp-ins | G/C-1 | none | G/C-1 | GATAAACCA | CCAATGA | TGG | ACC | C | none | nope | nope_ACC | G/C-1_CCAATGA | 6 | FALSE | FALSE |
| 176_v.txt | 33147 | 33147-Deleti | Deletion | 1 | A | - | 21 | 69133 | 0.03037623 | 37.9047619 | rnr1Y285A_pC | TRUE | CAN | JSY2422 | 5 | rnr1Y285A_p | GAA | nope | >1bp-ins | A/T-1 | none | A/T-1 | CCAATGAAA | AAAAGAC | CTT | CTT | T | none | T>none | T>none_CTT | A/T-1_AAAAGAC | 6 | FALSE | FALSE |
| 176_v.txt | 33168 | 33168-Deleti | Deletion | 1 | C | - | 11 | 69130 | 0.01591205 | 37.9090909 | rnr1Y285A_pC | TRUE | CAN | JSY2422 | 5 | rnr1Y285A_p | ACC | nope | >1bp-ins | G/C-1 | none | G/C-1 | ATAGTACCA | CCACCAA | TGG | ACC | C | none | nope | nope_ACC | G/C-1_CCACCAA | 6 | FALSE | FALSE |

| | | | | | | | | | | | | | | | | | | | | | | | | | | | | | | | | | | |
|---|---|---|---|---|---|---|---|---|---|---|---|---|---|---|---|---|---|---|---|---|---|---|---|---|---|---|---|---|---|---|---|---|---|---|
| 176_v.txt | 33175 | 33175-Deleti | Deletion | 1 | G | - | 7 | 69115 | 0.01012805 | 37.4285714 | rnr1Y285A_p | TRUE | CAN | JSY2422 | 5 | rnr1Y285A_p | AGG | nope | >1bp-ins | G/C-1 | none | G/C-1 | CACCAAGGG | GGGCAAT | TCC | TCC | C | none | C>none | C>none_TCC | G/C-1_GGGCAAT | 6 | FALSE | FALSE |
| 176_v.txt | 33326 | 33326-Deleti | Deletion | 1 | T | - | 8 | 33427 | 0.02393275 | 60.2499999 | rnr1Y285A_p | TRUE | CAN | JSY2422 | 5 | rnr1Y285A_p | CTT | nope | >1bp-ins | A/T-1 | none | A/T-1 | CAATTCTTTA | TTTACTT | GAA | CTT | T | none | nope | nope_CTT | A/T-1_TTTACTT | 6 | FALSE | FALSE |
| 176_v.txt | 33356 | 33356-Deleti | Deletion | 1 | C | - | 557 | 33429 | 1.66621796 | 55.4955117 | rnr1Y285A_p | TRUE | CAN | JSY2422 | 5 | rnr1Y285A_p | ACC | nope | >1bp-ins | G/C-1 | none | G/C-1 | TATTGACCCA | CCCACGT | TGG | ACC | C | none | nope | nope_ACC | G/C-1_CCCACGT | 6 | FALSE | FALSE |
| 176_v.txt | 33474 | 33474-Deleti | Deletion | 1 | C | - | 97 | 33428 | 0.2901759 | 29.8762887 | rnr1Y285A_p | TRUE | CAN | JSY2422 | 5 | rnr1Y285A_p | GCC | nope | >1bp-ins | G/C-1 | none | G/C-1 | GCTATGCCTT | CCTTTTT | CGG | GCC | C | none | nope | nope_GCC | G/C-1_CCTTTTT | 6 | FALSE | FALSE |
| 177_v.txt | 31723 | 31723-SNV- | SNV | 1 | A | C | 6 | 16790 | 0.03573556 | 55.1666667 | rnr1Y285A_p | TRUE | CAN | JSY2423 | 6 | rnr1Y285A_p | CAA | A>C | G/C+1 | A/T-1 | TA>GC | TA>GC | TGTCCCAAA | AAAAAGT | GTT | GTT | T | G | T>G | T>G_GTT | TA>GC_AAAAAGT | 6 | FALSE | FALSE |
| 177_v.txt | 32109 | 32109-SNV- | SNV | 1 | A | C | 7 | 26238 | 0.02667886 | 63.5714286 | rnr1Y285A_p | TRUE | CAN | JSY2423 | 6 | rnr1Y285A_p | TAA | A>C | G/C+1 | A/T-1 | TA>GC | TA>GC | ATATTTAATA | AATAGCC | ATT | ATT | T | G | T>G | T>G_ATT | TA>GC_AATAGCC | 6 | FALSE | FALSE |
| 177_v.txt | 32186 | 32186-SNV- | SNV | 1 | A | C | 13 | 26238 | 0.04954646 | 44.5384615 | rnr1Y285A_p | TRUE | CAN | JSY2423 | 6 | rnr1Y285A_p | AAA | A>C | G/C+1 | A/T-1 | TA>GC | TA>GC | AACGAAAAC | AACTGCA | TTT | TTT | T | G | T>G | T>G_TTT | TA>GC_AACTGCA | 6 | FALSE | FALSE |
| 177_v.txt | 32283 | 32283-SNV- | SNV | 1 | T | G | 10 | 32844 | 0.03044696 | 52.6 | rnr1Y285A_p | TRUE | CAN | JSY2423 | 6 | rnr1Y285A_p | GTA | T>G | G/C+1 | A/T-1 | TA>GC | TA>GC | CCAACGTAA | TAAATAT | CAT | GTA | T | C | T>G | T>G_GTA | TA>GC_TAAATAT | 6 | FALSE | FALSE |
| 177_v.txt | 32532 | 32532-SNV- | SNV | 1 | T | G | 50 | 32843 | 0.15223944 | 39.3789969 | rnr1Y285A_p | TRUE | CAN | JSY2423 | 6 | rnr1Y285A_p | GTT | T>G | G/C+1 | A/T-1 | TA>GC | TA>GC | CTGGGGTTT | TTTGCAG | CAA | GTT | T | C | T>G | T>G_GTT | TA>GC_TTTGCAG | 6 | FALSE | FALSE |
| 177_v.txt | 32631 | 32631-SNV- | SNV | 1 | T | G | 2 | 38633 | 5.18E-03 | 36.5002362 | rnr1Y285A_p | TRUE | CAN | JSY2423 | 6 | rnr1Y285A_p | GTT | T>G | G/C+1 | A/T-1 | TA>GC | TA>GC | CCTTCGTTTT | TTTTTAT | CAA | GTT | T | C | T>G | T>G_GTT | TA>GC_TTTTTAT | 6 | FALSE | FALSE |
| 177_v.txt | 32810 | 32810-SNV- | SNV | 1 | A | C | 7 | 38633 | 0.01811923 | 34.1428571 | rnr1Y285A_p | TRUE | CAN | JSY2423 | 6 | rnr1Y285A_p | CAG | A>C | G/C+1 | A/T-1 | TA>GC | TA>GC | TTTGACAGG | AGGGAAC | GTC | GTC | T | G | T>G | T>G_GTC | TA>GC_AGGGAAC | 6 | FALSE | FALSE |
| 177_v.txt | 32842 | 32842-SNV- | SNV | 1 | A | C | 5 | 38633 | 0.0129423 | 40.5998727 | rnr1Y285A_p | TRUE | CAN | JSY2423 | 6 | rnr1Y285A_p | CAA | A>C | G/C+1 | A/T-1 | TA>GC | TA>GC | TTACCCAAA | AAAAAAT | GTT | GTT | T | G | T>G | T>G_GTT | TA>GC_AAAAAAT | 6 | FALSE | FALSE |
| 177_v.txt | 32884 | 32884-SNV- | SNV | 1 | A | C | 14 | 41638 | 0.03362313 | 32.8571429 | rnr1Y285A_p | TRUE | CAN | JSY2423 | 6 | rnr1Y285A_p | CAA | A>C | G/C+1 | A/T-1 | TA>GC | TA>GC | ACGTCCAAA | AAAATTG | GTT | GTT | T | G | T>G | T>G_GTT | TA>GC_AAAATTG | 6 | FALSE | FALSE |
| 177_v.txt | 32086 | 32086-SNV- | SNV | 1 | A | G | 24 | 26238 | 0.09147039 | 59.8333333 | rnr1Y285A_p | TRUE | CAN | JSY2423 | 6 | rnr1Y285A_p | AAG | A>G | G/C+1 | A/T-1 | TA>CG | TA>CG | CAAAAAAGC | AGCCTGC | TTC | TTC | T | C | T>C | T>C_TTC | TA>CG_AGCCTGC | 6 | FALSE | FALSE |
| 177_v.txt | 32919 | 32919-SNV- | SNV | 1 | A | G | 30 | 41638 | 0.07204957 | 48.5666667 | rnr1Y285A_p | TRUE | CAN | JSY2423 | 6 | rnr1Y285A_p | CAG | A>G | G/C+1 | A/T-1 | TA>CG | TA>CG | AGTTCCAGG | AGGGCAA | GTC | GTC | T | C | T>C | T>C_GTC | TA>CG_AGGGCAA | 6 | FALSE | FALSE |
| 177_v.txt | 32927 | 32927-SNV- | SNV | 1 | A | G | 54 | 41638 | 0.12968923 | 51.4444444 | rnr1Y285A_p | TRUE | CAN | JSY2423 | 6 | rnr1Y285A_p | AAG | A>G | G/C+1 | A/T-1 | TA>CG | TA>CG | GGCAAAAGT | AGTGATT | TTC | TTC | T | C | T>C | T>C_TTC | TA>CG_AGTGATT | 6 | FALSE | FALSE |
| 177_v.txt | 33207 | 33207-SNV- | SNV | 1 | T | C | 10 | 14680 | 0.06811989 | 28.3999535 | rnr1Y285A_p | TRUE | CAN | JSY2423 | 6 | rnr1Y285A_p | CTC | T>C | G/C+1 | A/T-1 | TA>CG | TA>CG | TTAAGCTCTC | TCTCTCT | GAG | CTC | T | G | T>C | T>C_CTC | TA>CG_TCTCTCT | 6 | FALSE | FALSE |
| 177_v.txt | 33476 | 33476-SNV- | SNV | 1 | T | C | 19 | 14678 | 0.12944543 | 33.2633247 | rnr1Y285A_p | TRUE | CAN | JSY2423 | 6 | rnr1Y285A_p | CTT | T>C | G/C+1 | A/T-1 | TA>CG | TA>CG | TATGCCTTTT | TTTTTTT | GAA | CTT | T | G | T>C | T>C_CTT | TA>CG_TTTTTTT | 6 | FALSE | FALSE |
| 177_v.txt | 32109 | 32109-SNV- | SNV | 1 | A | T | 10 | 26238 | 0.03811266 | 63.9 | rnr1Y285A_p | TRUE | CAN | JSY2423 | 6 | rnr1Y285A_p | TAA | A>T | A/T+1 | A/T-1 | TA>AT | TA>AT | ATATTTAATA | AATAGCC | ATT | ATT | T | A | T>A | T>A_ATT | TA>AT_AATAGCC | 6 | FALSE | FALSE |
| 177_v.txt | 32231 | 32231-SNV- | SNV | 1 | T | A | 3 | 26237 | 0.01143423 | 27.6665685 | rnr1Y285A_p | TRUE | CAN | JSY2423 | 6 | rnr1Y285A_p | ATT | T>A | A/T+1 | A/T-1 | TA>AT | TA>AT | CAGGAATTT | TTTAGGA | TAA | ATT | T | T | T>A | T>A_ATT | TA>AT_TTTAGGA | 6 | FALSE | FALSE |
| 177_v.txt | 32265 | 32265-SNV- | SNV | 1 | A | T | 32 | 32844 | 0.09743028 | 36.96875 | rnr1Y285A_p | TRUE | CAN | JSY2423 | 6 | rnr1Y285A_p | TAA | A>T | A/T+1 | A/T-1 | TA>AT | TA>AT | CCAAATAAA | AAAATAC | ATT | ATT | T | A | T>A | T>A_ATT | TA>AT_AAAATAC | 6 | FALSE | FALSE |
| 177_v.txt | 32496 | 32496-SNV- | SNV | 1 | A | T | 1700 | 32844 | 5.17598344 | 45.7770588 | rnr1Y285A_p | TRUE | CAN | JSY2423 | 6 | rnr1Y285A_p | AAC | A>T | A/T+1 | A/T-1 | TA>AT | TA>AT | AAAACAACT | ACTTTTT | TTG | TTG | T | A | T>A | T>A_TTG | TA>AT_ACTTTTT | 6 | FALSE | FALSE |
| 177_v.txt | 32498 | 32498-SNV- | SNV | 1 | T | A | 5 | 32844 | 0.01522348 | 56.9998829 | rnr1Y285A_p | TRUE | CAN | JSY2423 | 6 | rnr1Y285A_p | CTT | T>A | A/T+1 | A/T-1 | TA>AT | TA>AT | AACAACTTTT | TTTTTTG | GAA | CTT | T | T | T>A | T>A_CTT | TA>AT_TTTTTTG | 6 | FALSE | FALSE |
| 177_v.txt | 33317 | 33317-SNV- | SNV | 1 | A | T | 144 | 14680 | 0.98092643 | 58.9722222 | rnr1Y285A_p | TRUE | CAN | JSY2423 | 6 | rnr1Y285A_p | GAT | A>T | A/T+1 | A/T-1 | TA>AT | TA>AT | CAATGGATA | ATACAAT | CTA | CTA | T | A | T>A | T>A_CTA | TA>AT_ATACAAT | 6 | FALSE | FALSE |
| 177_v.txt | 31650 | 31650-SNV- | SNV | 1 | C | T | 41 | 16789 | 0.24420752 | 35.0243902 | rnr1Y285A_p | TRUE | CAN | JSY2423 | 6 | rnr1Y285A_p | GCG | C>T | A/T+1 | G/C-1 | CG>TA | CG>TA | AGAATGCGA | CGAAATG | CGC | GCG | C | A | C>T | C>T_GCG | CG>TA_CGAAATG | 6 | FALSE | FALSE |
| 177_v.txt | 31658 | 31658-SNV- | SNV | 1 | C | T | 23 | 16789 | 0.13699446 | 32.3913044 | rnr1Y285A_p | TRUE | CAN | JSY2423 | 6 | rnr1Y285A_p | GCG | C>T | A/T+1 | G/C-1 | CG>TA | CG>TA | AAATGGCGT | CGTGGAA | CGC | GCG | C | A | C>T | C>T_GCG | CG>TA_CGTGGAA | 6 | FALSE | FALSE |
| 177_v.txt | 31830 | 31830-SNV- | SNV | 1 | C | T | 21 | 16790 | 0.12507445 | 53.3333333 | rnr1Y285A_p | TRUE | CAN | JSY2423 | 6 | rnr1Y285A_p | GCA | C>T | A/T+1 | G/C-1 | CG>TA | CG>TA | AATATGCATT | CATTGAA | CGT | GCA | C | A | C>T | C>T_GCA | CG>TA_CATTGAA | 6 | FALSE | FALSE |
| 177_v.txt | 31844 | 31844-SNV- | SNV | 1 | C | T | 26 | 16790 | 0.15485408 | 53.0384615 | rnr1Y285A_p | TRUE | CAN | JSY2423 | 6 | rnr1Y285A_p | TCC | C>T | A/T+1 | G/C-1 | CG>TA | CG>TA | TAAGATCCA | CCAAACA | AGG | TCC | C | A | C>T | C>T_TCC | CG>TA_CCAAACA | 6 | FALSE | FALSE |
| 177_v.txt | 31881 | 31881-SNV- | SNV | 1 | G | A | 19 | 16790 | 0.1131626 | 41.4206902 | rnr1Y285A_p | TRUE | CAN | JSY2423 | 6 | rnr1Y285A_p | GGC | G>A | A/T+1 | G/C-1 | CG>TA | CG>TA | TAGGCGGCA | GCAGCAA | CCG | CCG | C | T | C>T | C>T_CCG | CG>TA_GCAGCAA | 6 | FALSE | FALSE |
| 177_v.txt | 31882 | 31882-SNV- | SNV | 1 | C | T | 21 | 16790 | 0.12507445 | 43.5714286 | rnr1Y285A_p | TRUE | CAN | JSY2423 | 6 | rnr1Y285A_p | GCA | C>T | A/T+1 | G/C-1 | CG>TA | CG>TA | AGGCGGCAG | CAGCAAA | CGT | GCA | C | A | C>T | C>T_GCA | CG>TA_CAGCAAA | 6 | FALSE | FALSE |
| 177_v.txt | 31911 | 31911-SNV- | SNV | 1 | G | A | 17 | 16790 | 0.10125075 | 43.1176471 | rnr1Y285A_p | TRUE | CAN | JSY2423 | 6 | rnr1Y285A_p | TGC | G>A | A/T+1 | G/C-1 | CG>TA | CG>TA | TTTGGTGCA | GCAAAAG | ACG | ACG | C | T | C>T | C>T_ACG | CG>TA_GCAAAAG | 6 | FALSE | FALSE |
| 177_v.txt | 31960 | 31960-SNV- | SNV | 1 | C | T | 31 | 26238 | 0.11814925 | 35.1935484 | rnr1Y285A_p | TRUE | CAN | JSY2423 | 6 | rnr1Y285A_p | GCC | C>T | A/T+1 | G/C-1 | CG>TA | CG>TA | ATGTGGCCG | CCGCATA | CGG | GCC | C | A | C>T | C>T_GCC | CG>TA_CCGCATA | 6 | FALSE | FALSE |
| 177_v.txt | 31962 | 31962-SNV- | SNV | 1 | G | A | 55 | 26238 | 0.20961964 | 36.6363636 | rnr1Y285A_p | TRUE | CAN | JSY2423 | 6 | rnr1Y285A_p | CGC | G>A | A/T+1 | G/C-1 | CG>TA | CG>TA | GTGGCCGCA | GCATAAT | GCG | GCG | C | T | C>T | C>T_GCG | CG>TA_GCATAAT | 6 | FALSE | FALSE |
| 177_v.txt | 31972 | 31972-SNV- | SNV | 1 | C | T | 25 | 26238 | 0.09528165 | 35.96 | rnr1Y285A_p | TRUE | CAN | JSY2423 | 6 | rnr1Y285A_p | GCC | C>T | A/T+1 | G/C-1 | CG>TA | CG>TA | AATAAGCCA | CCAAGCC | CGG | GCC | C | A | C>T | C>T_GCC | CG>TA_CCAAGCC | 6 | FALSE | FALSE |
| 177_v.txt | 31977 | 31977-SNV- | SNV | 1 | C | T | 27 | 26238 | 0.10290419 | 50.1111111 | rnr1Y285A_p | TRUE | CAN | JSY2423 | 6 | rnr1Y285A_p | GCC | C>T | A/T+1 | G/C-1 | CG>TA | CG>TA | GCCAAGCCG | CCGGGCA | CGG | GCC | C | A | C>T | C>T_GCC | CG>TA_CCGGGCA | 6 | FALSE | FALSE |
| 177_v.txt | 32041 | 32041-SNV- | SNV | 1 | G | A | 33 | 26238 | 0.12577178 | 55.119933 | rnr1Y285A_p | TRUE | CAN | JSY2423 | 6 | rnr1Y285A_p | TGC | G>A | A/T+1 | G/C-1 | CG>TA | CG>TA | AAGCTTGCA | GCATAAA | ACG | ACG | C | T | C>T | C>T_ACG | CG>TA_GCATAAA | 6 | FALSE | FALSE |
| 177_v.txt | 32042 | 32042-SNV- | SNV | 1 | C | T | 27 | 26235 | 0.10291595 | 58.0240161 | rnr1Y285A_p | TRUE | CAN | JSY2423 | 6 | rnr1Y285A_p | GCA | C>T | A/T+1 | G/C-1 | CG>TA | CG>TA | AGCTTGCAT | CATAAAT | CGT | GCA | C | A | C>T | C>T_GCA | CG>TA_CATAAAT | 6 | FALSE | FALSE |
| 177_v.txt | 32057 | 32057-SNV- | SNV | 1 | C | T | 25 | 26238 | 0.09528165 | 64 | rnr1Y285A_p | TRUE | CAN | JSY2423 | 6 | rnr1Y285A_p | GCG | C>T | A/T+1 | G/C-1 | CG>TA | CG>TA | GATGTGCGA | CGAGATT | CGC | GCG | C | A | C>T | C>T_GCG | CG>TA_CGAGATT | 6 | FALSE | FALSE |
| 177_v.txt | 32075 | 32075-SNV- | SNV | 1 | C | T | 147 | 26238 | 0.56025612 | 59.537415 | rnr1Y285A_p | TRUE | CAN | JSY2423 | 6 | rnr1Y285A_p | ACC | C>T | A/T+1 | G/C-1 | CG>TA | CG>TA | AAATAACCA | CCATGCA | TGG | ACC | C | A | C>T | C>T_ACC | CG>TA_CCATGCA | 6 | FALSE | FALSE |
| 177_v.txt | 32079 | 32079-SNV- | SNV | 1 | G | A | 20 | 26238 | 0.07622532 | 62.15 | rnr1Y285A_p | TRUE | CAN | JSY2423 | 6 | rnr1Y285A_p | TGC | G>A | A/T+1 | G/C-1 | CG>TA | CG>TA | AACCATGCA | GCAAAAA | ACG | ACG | C | T | C>T | C>T_ACG | CG>TA_GCAAAAA | 6 | FALSE | FALSE |
| 177_v.txt | 32088 | 32088-SNV- | SNV | 1 | C | T | 93 | 26238 | 0.35444775 | 61.8387097 | rnr1Y285A_p | TRUE | CAN | JSY2423 | 6 | rnr1Y285A_p | GCC | C>T | A/T+1 | G/C-1 | CG>TA | CG>TA | AAAAAGCCT | CCTGCAA | CGG | GCC | C | A | C>T | C>T_GCC | CG>TA_CCTGCAA | 6 | FALSE | FALSE |
| 177_v.txt | 32092 | 32092-SNV- | SNV | 1 | C | T | 32 | 26238 | 0.12196052 | 62 | rnr1Y285A_p | TRUE | CAN | JSY2423 | 6 | rnr1Y285A_p | GCA | C>T | A/T+1 | G/C-1 | CG>TA | CG>TA | AGCCTGCAA | CAACACC | CGT | GCA | C | A | C>T | C>T_GCA | CG>TA_CAACACC | 6 | FALSE | FALSE |
| 177_v.txt | 32114 | 32114-SNV- | SNV | 1 | C | T | 45 | 26238 | 0.17150698 | 59.5111111 | rnr1Y285A_p | TRUE | CAN | JSY2423 | 6 | rnr1Y285A_p | GCC | C>T | A/T+1 | G/C-1 | CG>TA | CG>TA | TAATAGCCA | CCATTCG | CGG | GCC | C | A | C>T | C>T_GCC | CG>TA_CCATTCG | 6 | FALSE | FALSE |
| 177_v.txt | 32157 | 32157-SNV- | SNV | 1 | G | A | 26 | 26238 | 0.09909292 | 50.3461539 | rnr1Y285A_p | TRUE | CAN | JSY2423 | 6 | rnr1Y285A_p | AGC | G>A | A/T+1 | G/C-1 | CG>TA | CG>TA | ATGTAAGCC | GCCAAAG | TCG | TCG | C | T | C>T | C>T_TCG | CG>TA_GCCAAAG | 6 | FALSE | FALSE |
| 177_v.txt | 32165 | 32165-SNV- | SNV | 1 | G | A | 53 | 26238 | 0.2019971 | 48.3962264 | rnr1Y285A_p | TRUE | CAN | JSY2423 | 6 | rnr1Y285A_p | CGC | G>A | A/T+1 | G/C-1 | CG>TA | CG>TA | CAAAGCGCC | GCCAAAT | GCG | GCG | C | T | C>T | C>T_GCG | CG>TA_GCCAAAT | 6 | FALSE | FALSE |
| 177_v.txt | 32190 | 32190-SNV- | SNV | 1 | G | A | 26 | 26238 | 0.09909292 | 43.7307692 | rnr1Y285A_p | TRUE | CAN | JSY2423 | 6 | rnr1Y285A_p | TGC | G>A | A/T+1 | G/C-1 | CG>TA | CG>TA | AAAACTGCA | GCAATGT | ACG | ACG | C | T | C>T | C>T_ACG | CG>TA_GCAATGT | 6 | FALSE | FALSE |
| 177_v.txt | 32271 | 32271-SNV- | SNV | 1 | C | T | 73 | 32844 | 0.22226282 | 35.4657534 | rnr1Y285A_p | TRUE | CAN | JSY2423 | 6 | rnr1Y285A_p | ACG | C>T | A/T+1 | G/C-1 | CG>TA | CG>TA | AAAATACGG | CGGGAAC | TGC | ACG | C | A | C>T | C>T_ACG | CG>TA_CGGGAAC | 6 | FALSE | FALSE |
| 177_v.txt | 32299 | 32299-SNV- | SNV | 1 | C | T | 73 | 32844 | 0.22226282 | 54.037841 | rnr1Y285A_p | TRUE | CAN | JSY2423 | 6 | rnr1Y285A_p | GCG | C>T | A/T+1 | G/C-1 | CG>TA | CG>TA | AATTTGCGG | CGGCAGA | CGC | GCG | C | A | C>T | C>T_GCG | CG>TA_CGGCAGA | 6 | FALSE | FALSE |
| 177_v.txt | 32344 | 32344-SNV- | SNV | 1 | G | A | 25 | 32844 | 0.0761174 | 60.08 | rnr1Y285A_p | TRUE | CAN | JSY2423 | 6 | rnr1Y285A_p | GGC | G>A | A/T+1 | G/C-1 | CG>TA | CG>TA | TATGTGGCA | GCAAAAC | CCG | CCG | C | T | C>T | C>T_CCG | CG>TA_GCAAAAC | 6 | FALSE | FALSE |
| 177_v.txt | 32466 | 32466-SNV- | SNV | 1 | C | T | 56 | 32844 | 0.17050298 | 46.9285714 | rnr1Y285A_p | TRUE | CAN | JSY2423 | 6 | rnr1Y285A_p | GCC | C>T | A/T+1 | G/C-1 | CG>TA | CG>TA | AGAGAGCCA | CCAATGT | CGG | GCC | C | A | C>T | C>T_GCC | CG>TA_CCAATGT | 6 | FALSE | FALSE |
| 177_v.txt | 32508 | 32508-SNV- | SNV | 1 | G | A | 27 | 32844 | 0.0822068 | 38.2962963 | rnr1Y285A_p | TRUE | CAN | JSY2423 | 6 | rnr1Y285A_p | GGC | G>A | A/T+1 | G/C-1 | CG>TA | CG>TA | TTGATGGCT | GCTCTTG | CCG | CCG | C | T | C>T | C>T_CCG | CG>TA_GCTCTTG | 6 | FALSE | FALSE |
| 177_v.txt | 32535 | 32535-SNV- | SNV | 1 | G | A | 36 | 32843 | 0.1096124 | 37.1666717 | rnr1Y285A_p | TRUE | CAN | JSY2423 | 6 | rnr1Y285A_p | TGC | G>A | A/T+1 | G/C-1 | CG>TA | CG>TA | GGGTTTGCA | GCAGCTT | ACG | ACG | C | T | C>T | C>T_ACG | CG>TA_GCAGCTT | 6 | FALSE | FALSE |
| 177_v.txt | 32536 | 32536-SNV- | SNV | 1 | C | T | 50 | 32843 | 0.15223944 | 37.2993822 | rnr1Y285A_p | TRUE | CAN | JSY2423 | 6 | rnr1Y285A_p | GCA | C>T | A/T+1 | G/C-1 | CG>TA | CG>TA | GGTTTGCAG | CAGCTTC | CGT | GCA | C | A | C>T | C>T_GCA | CG>TA_CAGCTTC | 6 | FALSE | FALSE |
| 177_v.txt | 32571 | 32571-SNV- | SNV | 1 | C | T | 17 | 38635 | 0.04400155 | 32.7647059 | rnr1Y285A_p | TRUE | CAN | JSY2423 | 6 | rnr1Y285A_p | ACC | C>T | A/T+1 | G/C-1 | CG>TA | CG>TA | TCAGTACCTT | CCTTGAA | TGG | ACC | C | A | C>T | C>T_ACC | CG>TA_CCTTGAA | 6 | FALSE | FALSE |
| 177_v.txt | 32587 | 32587-SNV- | SNV | 1 | C | T | 45 | 38633 | 0.11648073 | 34.6444444 | rnr1Y285A_p | TRUE | CAN | JSY2423 | 6 | rnr1Y285A_p | GCA | C>T | A/T+1 | G/C-1 | CG>TA | CG>TA | TGAAGGCAG | CAGCGTT | CGT | GCA | C | A | C>T | C>T_GCA | CG>TA_CAGCGTT | 6 | FALSE | FALSE |
| 177_v.txt | 32590 | 32590-SNV- | SNV | 1 | C | T | 76 | 38633 | 0.19672301 | 37.0131579 | rnr1Y285A_p | TRUE | CAN | JSY2423 | 6 | rnr1Y285A_p | GCG | C>T | A/T+1 | G/C-1 | CG>TA | CG>TA | AGGCAGCGT | CGTTAAT | CGC | GCG | C | A | C>T | C>T_GCG | CG>TA_CGTTAAT | 6 | FALSE | FALSE |
| 177_v.txt | 32630 | 32630-SNV- | SNV | 1 | G | A | 7 | 38633 | 0.01811923 | 45.282017 | rnr1Y285A_p | TRUE | CAN | JSY2423 | 6 | rnr1Y285A_p | CGT | G>A | A/T+1 | G/C-1 | CG>TA | CG>TA | CCCTTCGTTT | GTTTTTA | GCA | GCA | C | T | C>T | C>T_GCA | CG>TA_GTTTTTA | 6 | FALSE | FALSE |
| 177_v.txt | 32658 | 32658-SNV- | SNV | 1 | C | T | 18 | 38633 | 0.04659229 | 59.5555556 | rnr1Y285A_p | TRUE | CAN | JSY2423 | 6 | rnr1Y285A_p | ACC | C>T | A/T+1 | G/C-1 | CG>TA | CG>TA | CCTGGACCC | CCCCAGG | TGG | ACC | C | A | C>T | C>T_ACC | CG>TA_CCCCAGG | 6 | FALSE | FALSE |
| 177_v.txt | 32660 | 32660-SNV- | SNV | 1 | C | T | 43 | 38633 | 0.11130381 | 53.7906977 | rnr1Y285A_p | TRUE | CAN | JSY2423 | 6 | rnr1Y285A_p | CCC | C>T | A/T+1 | G/C-1 | CG>TA | CG>TA | TGGACCCCA | CCAGGCA | GGG | CCC | C | A | C>T | C>T_CCC | CG>TA_CCAGGCA | 6 | FALSE | FALSE |
| 177_v.txt | 32700 | 32700-SNV- | SNV | 1 | C | T | 40 | 38633 | 0.10353843 | 55.575 | rnr1Y285A_p | TRUE | CAN | JSY2423 | 6 | rnr1Y285A_p | GCC | C>T | A/T+1 | G/C-1 | CG>TA | CG>TA | ACTGGGCCG | CCGGTAA | CGG | GCC | C | A | C>T | C>T_GCC | CG>TA_CCGGTAA | 6 | FALSE | FALSE |
| 177_v.txt | 32702 | 32702-SNV- | SNV | 1 | G | A | 42 | 38633 | 0.10871535 | 57.5238095 | rnr1Y285A_p | TRUE | CAN | JSY2423 | 6 | rnr1Y285A_p | CGG | G>A | A/T+1 | G/C-1 | CG>TA | CG>TA | TGGGCCGGT | GGTAACC | GCC | GCC | C | T | C>T | C>T_GCC | CG>TA_GGTAACC | 6 | FALSE | FALSE |
| 177_v.txt | 32713 | 32713-SNV- | SNV | 1 | C | T | 32 | 38633 | 0.08283074 | 59.40625 | rnr1Y285A_p | TRUE | CAN | JSY2423 | 6 | rnr1Y285A_p | GCA | C>T | A/T+1 | G/C-1 | CG>TA | CG>TA | CCCCAGCACC | CACCACA | CGT | GCA | C | A | C>T | C>T_GCA | CG>TA_CACCACA | 6 | FALSE | FALSE |
| 177_v.txt | 32757 | 32757-SNV- | SNV | 1 | G | A | 40 | 38633 | 0.10353843 | 55.175 | rnr1Y285A_p | TRUE | CAN | JSY2423 | 6 | rnr1Y285A_p | GGC | G>A | A/T+1 | G/C-1 | CG>TA | CG>TA | ATAATGGCT | GCTAAAA | CCG | CCG | C | T | C>T | C>T_CCG | CG>TA_GCTAAAA | 6 | FALSE | FALSE |
| 177_v.txt | 32776 | 32776-SNV- | SNV | 1 | C | T | 47 | 38633 | 0.12165765 | 45.9148936 | rnr1Y285A_p | TRUE | CAN | JSY2423 | 6 | rnr1Y285A_p | GCG | C>T | A/T+1 | G/C-1 | CG>TA | CG>TA | TGGAAGCGA | CGACCCA | CGC | GCG | C | A | C>T | C>T_GCG | CG>TA_CGACCCA | 6 | FALSE | FALSE |
| 177_v.txt | 32798 | 32798-SNV- | SNV | 1 | G | A | 52 | 38633 | 0.13459995 | 42.3461539 | rnr1Y285A_p | TRUE | CAN | JSY2423 | 6 | rnr1Y285A_p | CGT | G>A | A/T+1 | G/C-1 | CG>TA | CG>TA | TTCACCGTAA | GTAATAT | GCA | GCA | C | T | C>T | C>T_GCA | CG>TA_GTAATAT | 6 | FALSE | FALSE |
| 177_v.txt | 32855 | 32855-SNV- | SNV | 1 | C | T | 46 | 38633 | 0.11906919 | 35 | rnr1Y285A_p | TRUE | CAN | JSY2423 | 6 | rnr1Y285A_p | TCC | C>T | A/T+1 | G/C-1 | CG>TA | CG>TA | ACTAATCCAT | CCATGCC | AGG | TCC | C | A | C>T | C>T_TCC | CG>TA_CCATGCC | 6 | FALSE | FALSE |
| 177_v.txt | 32860 | 32860-SNV- | SNV | 1 | C | T | 46 | 38631 | 0.11907535 | 36.5652174 | rnr1Y285A_p | TRUE | CAN | JSY2423 | 6 | rnr1Y285A_p | GCC | C>T | A/T+1 | G/C-1 | CG>TA | CG>TA | TCCATGCCG | CCGCCAG | CGG | GCC | C | A | C>T | C>T_GCC | CG>TA_CCGCCAG | 6 | FALSE | FALSE |
| 177_v.txt | 32862 | 32862-SNV- | SNV | 1 | G | A | 35 | 38631 | 0.09060081 | 33.1428571 | rnr1Y285A_p | TRUE | CAN | JSY2423 | 6 | rnr1Y285A_p | CGC | G>A | A/T+1 | G/C-1 | CG>TA | CG>TA | CATGCCGCC | GCCAGTG | GCG | GCG | C | T | C>T | C>T_GCG | CG>TA_GCCAGTG | 6 | FALSE | FALSE |
| 177_v.txt | 32879 | 32879-SNV- | SNV | 1 | C | T | 35 | 41635 | 0.08406389 | 36.1428571 | rnr1Y285A_p | TRUE | CAN | JSY2423 | 6 | rnr1Y285A_p | ACG | C>T | A/T+1 | G/C-1 | CG>TA | CG>TA | TTTGTACGTC | CGTCCAA | TGC | ACG | C | A | C>T | C>T_ACG | CG>TA_CGTCCAA | 6 | FALSE | FALSE |
| 177_v.txt | 32882 | 32882-SNV- | SNV | 1 | C | T | 83 | 41638 | 0.19933714 | 28.3855422 | rnr1Y285A_p | TRUE | CAN | JSY2423 | 6 | rnr1Y285A_p | TCC | C>T | A/T+1 | G/C-1 | CG>TA | CG>TA | GTACGTCCA | CCAAAAT | AGG | TCC | C | A | C>T | C>T_TCC | CG>TA_CCAAAAT | 6 | FALSE | FALSE |
| 177_v.txt | 32900 | 32900-SNV- | SNV | 1 | G | A | 46 | 41638 | 0.11047601 | 36.3863315 | rnr1Y285A_p | TRUE | CAN | JSY2423 | 6 | rnr1Y285A_p | GGC | G>A | A/T+1 | G/C-1 | CG>TA | CG>TA | GACTTGGCC | GCCAACT | CCG | CCG | C | T | C>T | C>T_CCG | CG>TA_GCCAACT | 6 | FALSE | FALSE |
| 177_v.txt | 32901 | 32901-SNV- | SNV | 1 | C | T | 97 | 41638 | 0.23296028 | 33.9896907 | rnr1Y285A_p | TRUE | CAN | JSY2423 | 6 | rnr1Y285A_p | GCC | C>T | A/T+1 | G/C-1 | CG>TA | CG>TA | ACTTGGCCA | CCAACTA | CGG | GCC | C | A | C>T | C>T_GCC | CG>TA_CCAACTA | 6 | FALSE | FALSE |
| 177_v.txt | 32922 | 32922-SNV- | SNV | 1 | G | A | 35 | 41638 | 0.08405783 | 54.3428571 | rnr1Y285A_p | TRUE | CAN | JSY2423 | 6 | rnr1Y285A_p | GGC | G>A | A/T+1 | G/C-1 | CG>TA | CG>TA | TCCAGGGCA | GCAAAAG | CCG | CCG | C | T | C>T | C>T_CCG | CG>TA_GCAAAAG | 6 | FALSE | FALSE |
| 177_v.txt | 32934 | 32934-SNV- | SNV | 1 | G | A | 13 | 41638 | 0.03122148 | 47.3806253 | rnr1Y285A_p | TRUE | CAN | JSY2423 | 6 | rnr1Y285A_p | TGC | G>A | A/T+1 | G/C-1 | CG>TA | CG>TA | GTGATTGCC | GCCCAAG | ACG | ACG | C | T | C>T | C>T_ACG | CG>TA_GCCCAAG | 6 | FALSE | FALSE |
| 177_v.txt | 32935 | 32935-SNV- | SNV | 1 | C | T | 58 | 41638 | 0.13929584 | 43 | rnr1Y285A_p | TRUE | CAN | JSY2423 | 6 | rnr1Y285A_p | GCC | C>T | A/T+1 | G/C-1 | CG>TA | CG>TA | TGATTGCCC | CCCAAGA | CGG | GCC | C | A | C>T | C>T_GCC | CG>TA_CCCAAGA | 6 | FALSE | FALSE |
| 177_v.txt | 32945 | 32945-SNV- | SNV | 1 | C | T | 70 | 41638 | 0.16811566 | 56.7 | rnr1Y285A_p | TRUE | CAN | JSY2423 | 6 | rnr1Y285A_p | ACC | C>T | A/T+1 | G/C-1 | CG>TA | CG>TA | AGAAAACCA | CCAATAC | TGG | ACC | C | A | C>T | C>T_ACC | CG>TA_CCAATAC | 6 | FALSE | FALSE |
| 177_v.txt | 32958 | 32958-SNV- | SNV | 1 | C | T | 41 | 41638 | 0.09846775 | 46.9756098 | rnr1Y285A_p | TRUE | CAN | JSY2423 | 6 | rnr1Y285A_p | ACC | C>T | A/T+1 | G/C-1 | CG>TA | CG>TA | ATGTAACCAT | CCATTGG | TGG | ACC | C | A | C>T | C>T_ACC | CG>TA_CCATTGG | 6 | FALSE | FALSE |
| 177_v.txt | 32965 | 32965-SNV- | SNV | 1 | C | T | 45 | 41638 | 0.10807436 | 61.1333333 | rnr1Y285A_p | TRUE | CAN | JSY2423 | 6 | rnr1Y285A_p | GCC | C>T | A/T+1 | G/C-1 | CG>TA | CG>TA | CATTGGCCG | CCGCACC | CGG | GCC | C | A | C>T | C>T_GCC | CG>TA_CCGCACC | 6 | FALSE | FALSE |
| 177_v.txt | 32967 | 32967-SNV- | SNV | 1 | G | A | 90 | 41638 | 0.21614871 | 61.1555556 | rnr1Y285A_p | TRUE | CAN | JSY2423 | 6 | rnr1Y285A_p | CGC | G>A | A/T+1 | G/C-1 | CG>TA | CG>TA | TTGGCCGCA | GCACCAA | GCG | GCG | C | T | C>T | C>T_GCG | CG>TA_GCACCAA | 6 | FALSE | FALSE |
| 177_v.txt | 32977 | 32977-SNV- | SNV | 1 | C | T | 48 | 41638 | 0.11527931 | 58.875 | rnr1Y285A_p | TRUE | CAN | JSY2423 | 6 | rnr1Y285A_p | GCT | C>T | A/T+1 | G/C-1 | CG>TA | CG>TA | CAAATGCTG | CTGGAGA | CGA | GCT | C | A | C>T | C>T_GCT | CG>TA_CTGGAGA | 6 | FALSE | FALSE |
| 177_v.txt | 33056 | 33056-SNV- | SNV | 1 | C | T | 67 | 41638 | 0.16091071 | 57.1492537 | rnr1Y285A_p | TRUE | CAN | JSY2423 | 6 | rnr1Y285A_p | GCG | C>T | A/T+1 | G/C-1 | CG>TA | CG>TA | GGACTGCGT | CGTGACA | CGC | GCG | C | A | C>T | C>T_GCG | CG>TA_CGTGACA | 6 | FALSE | FALSE |
| 177_v.txt | 33069 | 33069-SNV- | SNV | 1 | G | A | 44 | 41638 | 0.1056727 | 51.8415522 | rnr1Y285A_p | TRUE | CAN | JSY2423 | 6 | rnr1Y285A_p | TGC | G>A | A/T+1 | G/C-1 | CG>TA | CG>TA | GAATATGCC | GCCAAAG | ACG | ACG | C | T | C>T | C>T_ACG | CG>TA_GCCAAAG | 6 | FALSE | FALSE |
| 177_v.txt | 33070 | 33070-SNV- | SNV | 1 | C | T | 53 | 41638 | 0.12728757 | 57.5252236 | rnr1Y285A_p | TRUE | CAN | JSY2423 | 6 | rnr1Y285A_p | GCC | C>T | A/T+1 | G/C-1 | CG>TA | CG>TA | AATATGCCA | CCAAAGA | CGG | GCC | C | A | C>T | C>T_GCC | CG>TA_CCAAAGA | 6 | FALSE | FALSE |
| 177_v.txt | 33104 | 33104-SNV- | SNV | 1 | G | A | 44 | 41638 | 0.1056727 | 40.6590909 | rnr1Y285A_p | TRUE | CAN | JSY2423 | 6 | rnr1Y285A_p | CGC | G>A | A/T+1 | G/C-1 | CG>TA | CG>TA | AAGAGCGCC | GCCCACT | GCG | GCG | C | T | C>T | C>T_GCG | CG>TA_GCCCACT | 6 | FALSE | FALSE |
| 177_v.txt | 33113 | 33113-SNV- | SNV | 1 | G | A | 34 | 41638 | 0.08165618 | 45.8756422 | rnr1Y285A_p | TRUE | CAN | JSY2423 | 6 | rnr1Y285A_p | GGC | G>A | A/T+1 | G/C-1 | CG>TA | CG>TA | CACTGGGCC | GCCGGCG | CCG | CCG | C | T | C>T | C>T_CCG | CG>TA_GCCGGCG | 6 | FALSE | FALSE |
| 177_v.txt | 33114 | 33114-SNV- | SNV | 1 | C | T | 47 | 41638 | 0.11287766 | 43.7234043 | rnr1Y285A_p | TRUE | CAN | JSY2423 | 6 | rnr1Y285A_p | GCC | C>T | A/T+1 | G/C-1 | CG>TA | CG>TA | ACTGGGCCG | CCGGCGT | CGG | GCC | C | A | C>T | C>T_GCC | CG>TA_CCGGCGT | 6 | FALSE | FALSE |
| 177_v.txt | 33115 | 33115-SNV- | SNV | 1 | C | T | 13 | 41638 | 0.03122148 | 36.8461539 | rnr1Y285A_p | TRUE | CAN | JSY2423 | 6 | rnr1Y285A_p | CCG | C>T | A/T+1 | G/C-1 | CG>TA | CG>TA | CTGGGCCGG | CGGCGTT | GGC | CCG | C | A | C>T | C>T_CCG | CG>TA_CGGCGTT | 6 | FALSE | FALSE |
| 177_v.txt | 33118 | 33118-SNV- | SNV | 1 | C | T | 39 | 41638 | 0.09366444 | 41.2307692 | rnr1Y285A_p | TRUE | CAN | JSY2423 | 6 | rnr1Y285A_p | GCG | C>T | A/T+1 | G/C-1 | CG>TA | CG>TA | GGCCGGCGT | CGTTGGT | CGC | GCG | C | A | C>T | C>T_GCG | CG>TA_CGTTGGT | 6 | FALSE | FALSE |
| 177_v.txt | 33119 | 33119-SNV- | SNV | 1 | G | A | 30 | 41638 | 0.07204957 | 42.8666667 | rnr1Y285A_p | TRUE | CAN | JSY2423 | 6 | rnr1Y285A_p | CGT | G>A | A/T+1 | G/C-1 | CG>TA | CG>TA | GCCGGCGTT | GTTGGTC | GCA | GCA | C | T | C>T | C>T_GCA | CG>TA_GTTGGTC | 6 | FALSE | FALSE |
| 177_v.txt | 33153 | 33153-SNV- | SNV | 1 | C | T | 224 | 41638 | 0.53797012 | 36.3393548 | rnr1Y285A_p | TRUE | CAN | JSY2423 | 6 | rnr1Y285A_p | ACC | C>T | A/T+1 | G/C-1 | CG>TA | CG>TA | AAAAGACCT | CCTGTAC | TGG | ACC | C | A | C>T | C>T_ACC | CG>TA_CCTGTAC | 6 | FALSE | FALSE |
| 177_v.txt | 33159 | 33159-SNV- | SNV | 1 | C | T | 28 | 41638 | 0.06724627 | 36.6071429 | rnr1Y285A_p | TRUE | CAN | JSY2423 | 6 | rnr1Y285A_p | ACC | C>T | A/T+1 | G/C-1 | CG>TA | CG>TA | CCTGTACCA | CCAATAG | TGG | ACC | C | A | C>T | C>T_ACC | CG>TA_CCAATAG | 6 | FALSE | FALSE |
| 177_v.txt | 33168 | 33168-SNV- | SNV | 1 | C | T | 70 | 41634 | 0.16813182 | 37.5714286 | rnr1Y285A_p | TRUE | CAN | JSY2423 | 6 | rnr1Y285A_p | ACC | C>T | A/T+1 | G/C-1 | CG>TA | CG>TA | ATAGTACCA | CCACCAA | TGG | ACC | C | A | C>T | C>T_ACC | CG>TA_CCACCAA | 6 | FALSE | FALSE |
| 177_v.txt | 33205 | 33205-SNV- | SNV | 1 | G | A | 8 | 14680 | 0.05449591 | 37.875 | rnr1Y285A_p | TRUE | CAN | JSY2423 | 6 | rnr1Y285A_p | AGC | G>A | A/T+1 | G/C-1 | CG>TA | CG>TA | GCTTAAGCT | GCTCTCT | TCG | TCG | C | T | C>T | C>T_TCG | CG>TA_GCTCTCT | 6 | FALSE | FALSE |
| 177_v.txt | 33258 | 33258-SNV- | SNV | 1 | C | T | 31 | 14680 | 0.21117166 | 42.516129 | rnr1Y285A_p | TRUE | CAN | JSY2423 | 6 | rnr1Y285A_p | GCC | C>T | A/T+1 | G/C-1 | CG>TA | CG>TA | CCTATGCCAT | CCATCCT | CGG | GCC | C | A | C>T | C>T_GCC | CG>TA_CCATCCT | 6 | FALSE | FALSE |
| 177_v.txt | 33276 | 33276-SNV- | SNV | 1 | G | A | 44 | 14680 | 0.29972752 | 48.3636364 | rnr1Y285A_p | TRUE | CAN | JSY2423 | 6 | rnr1Y285A_p | CGT | G>A | A/T+1 | G/C-1 | CG>TA | CG>TA | GAGAACGTA | GTATCCT | GCA | GCA | C | T | C>T | C>T_GCA | CG>TA_GTATCCT | 6 | FALSE | FALSE |
| 177_v.txt | 33284 | 33284-SNV- | SNV | 1 | G | A | 18 | 14680 | 0.1226158 | 49 | rnr1Y285A_p | TRUE | CAN | JSY2423 | 6 | rnr1Y285A_p | CGC | G>A | A/T+1 | G/C-1 | CG>TA | CG>TA | ATCCTCGCCA | GCCATTT | GCG | GCG | C | T | C>T | C>T_GCG | CG>TA_GCCATTT | 6 | FALSE | FALSE |
| 177_v.txt | 33309 | 33309-SNV- | SNV | 1 | C | T | 19 | 14679 | 0.12943661 | 59.0669043 | rnr1Y285A_p | TRUE | CAN | JSY2423 | 6 | rnr1Y285A_p | GCG | C>T | A/T+1 | G/C-1 | CG>TA | CG>TA | AAAGAGCGC | CGCAATG | CGC | GCG | C | A | C>T | C>T_GCG | CG>TA_CGCAATG | 6 | FALSE | FALSE |
| 177_v.txt | 33310 | 33310-SNV- | SNV | 1 | G | A | 22 | 14677 | 0.14989439 | 55.9699902 | rnr1Y285A_p | TRUE | CAN | JSY2423 | 6 | rnr1Y285A_p | CGC | G>A | A/T+1 | G/C-1 | CG>TA | CG>TA | AAGAGCGCA | GCAATGG | GCG | GCG | C | T | C>T | C>T_GCG | CG>TA_GCAATGG | 6 | FALSE | FALSE |
| 177_v.txt | 33412 | 33412-SNV- | SNV | 1 | G | A | 23 | 14680 | 0.15667575 | 50.6086957 | rnr1Y285A_p | TRUE | CAN | JSY2423 | 6 | rnr1Y285A_p | GGC | G>A | A/T+1 | G/C-1 | CG>TA | CG>TA | TGACCGGCT | GCTCATT | CCG | CCG | C | T | C>T | C>T_CCG | CG>TA_GCTCATT | 6 | FALSE | FALSE |
| 177_v.txt | 33474 | 33474-SNV- | SNV | 1 | C | T | 30 | 14679 | 0.2043736 | 35.5333333 | rnr1Y285A_p | TRUE | CAN | JSY2423 | 6 | rnr1Y285A_p | GCC | C>T | A/T+1 | G/C-1 | CG>TA | CG>TA | GCTATGCCTT | CCTTTTT | CGG | GCC | C | A | C>T | C>T_GCC | CG>TA_CCTTTTT | 6 | FALSE | FALSE |
| 177_v.txt | 31844 | 31844-SNV- | SNV | 1 | C | G | 3 | 16790 | 0.01786778 | 44.6666667 | rnr1Y285A_p | TRUE | CAN | JSY2423 | 6 | rnr1Y285A_p | TCC | C>G | G/C+1 | G/C-1 | CG>GC | CG>GC | TAAGATCCA | CCAAACA | AGG | TCC | C | C | C>G | C>G_TCC | CG>GC_CCAAACA | 6 | FALSE | FALSE |
| 177_v.txt | 31979 | 31979-SNV- | SNV | 1 | G | C | 3 | 26238 | 0.0114338 | 38 | rnr1Y285A_p | TRUE | CAN | JSY2423 | 6 | rnr1Y285A_p | CGG | G>C | G/C+1 | G/C-1 | CG>GC | CG>GC | CAAGCCGGG | GGGCATT | GCC | GCC | C | G | C>G | C>G_GCC | CG>GC_GGGCATT | 6 | FALSE | FALSE |
| 177_v.txt | 32027 | 32027-SNV- | SNV | 1 | G | C | 9 | 26238 | 0.0343014 | 64 | rnr1Y285A_p | TRUE | CAN | JSY2423 | 6 | rnr1Y285A_p | GGT | G>C | G/C+1 | G/C-1 | CG>GC | CG>GC | GCCACGGTA | GTATTTC | CCA | CCA | C | G | C>G | C>G_CCA | CG>GC_GTATTTC | 6 | FALSE | FALSE |
| 177_v.txt | 32041 | 32041-SNV- | SNV | 1 | G | C | 4 | 26238 | 0.01524506 | 50.4982046 | rnr1Y285A_p | TRUE | CAN | JSY2423 | 6 | rnr1Y285A_p | TGC | G>C | G/C+1 | G/C-1 | CG>GC | CG>GC | AAGCTTGCA | GCATAAA | ACG | ACG | C | G | C>G | C>G_ACG | CG>GC_GCATAAA | 6 | FALSE | FALSE |
| 177_v.txt | 32042 | 32042-SNV- | SNV | 1 | C | G | 2 | 26235 | 7.62E-03 | 63.9884668 | rnr1Y285A_p | TRUE | CAN | JSY2423 | 6 | rnr1Y285A_p | GCA | C>G | G/C+1 | G/C-1 | CG>GC | CG>GC | AGCTTGCAT | CATAAAT | CGT | GCA | C | C | C>G | C>G_GCA | CG>GC_CATAAAT | 6 | FALSE | FALSE |
| 177_v.txt | 32228 | 32228-SNV- | SNV | 1 | G | C | 6 | 26237 | 0.02286847 | 38 | rnr1Y285A_p | TRUE | CAN | JSY2423 | 6 | rnr1Y285A_p | GGA | G>C | G/C+1 | G/C-1 | CG>GC | CG>GC | TGACAGGAA | GAATTTA | CCT | CCT | C | G | C>G | C>G_CCT | CG>GC_GAATTTA | 6 | FALSE | FALSE |
| 177_v.txt | 32299 | 32299-SNV- | SNV | 1 | C | G | 1 | 32844 | 3.05E-03 | 45.9960959 | rnr1Y285A_p | TRUE | CAN | JSY2423 | 6 | rnr1Y285A_p | GCG | C>G | G/C+1 | G/C-1 | CG>GC | CG>GC | AATTTGCGG | CGGCAGA | CGC | GCG | C | C | C>G | C>G_GCG | CG>GC_CGGCAGA | 6 | FALSE | FALSE |
| 177_v.txt | 32536 | 32536-SNV- | SNV | 1 | C | G | 1 | 32843 | 3.05E-03 | 37.9992266 | rnr1Y285A_p | TRUE | CAN | JSY2423 | 6 | rnr1Y285A_p | GCA | C>G | G/C+1 | G/C-1 | CG>GC | CG>GC | GGTTTGCAG | CAGCTTC | CGT | GCA | C | C | C>G | C>G_GCA | CG>GC_CAGCTTC | 6 | FALSE | FALSE |
| 177_v.txt | 32749 | 32749-SNV- | SNV | 1 | C | G | 15 | 38633 | 0.03882691 | 54.8 | rnr1Y285A_p | TRUE | CAN | JSY2423 | 6 | rnr1Y285A_p | CCG | C>G | G/C+1 | G/C-1 | CG>GC | CG>GC | GAAACCCGA | CGATAAT | GGC | CCG | C | C | C>G | C>G_CCG | CG>GC_CGATAAT | 6 | FALSE | FALSE |
| 177_v.txt | 32794 | 32794-SNV- | SNV | 1 | C | G | 5 | 38633 | 0.0129423 | 46.8 | rnr1Y285A_p | TRUE | CAN | JSY2423 | 6 | rnr1Y285A_p | TCA | C>G | G/C+1 | G/C-1 | CG>GC | CG>GC | CGAATTCACC | CACCGTA | AGT | TCA | C | C | C>G | C>G_TCA | CG>GC_CACCGTA | 6 | FALSE | FALSE |
| 177_v.txt | 32819 | 32819-SNV- | SNV | 1 | G | C | 70 | 38633 | 0.18119225 | 44.8 | rnr1Y285A_p | TRUE | CAN | JSY2423 | 6 | rnr1Y285A_p | AGT | G>C | G/C+1 | G/C-1 | CG>GC | CG>GC | GAACAAGTT | GTTCATT | TCA | TCA | C | G | C>G | C>G_TCA | CG>GC_GTTCATT | 6 | FALSE | FALSE |
| 177_v.txt | 32876 | 32876-SNV- | SNV | 1 | G | C | 14 | 41619 | 0.03363848 | 37.5 | rnr1Y285A_p | TRUE | CAN | JSY2423 | 6 | rnr1Y285A_p | TGT | G>C | G/C+1 | G/C-1 | CG>GC | CG>GC | AACTTTGTAC | GTACGTC | ACA | ACA | C | G | C>G | C>G_ACA | CG>GC_GTACGTC | 6 | FALSE | FALSE |
| 177_v.txt | 32935 | 32935-SNV- | SNV | 1 | C | G | 3 | 41638 | 7.21E-03 | 44.3333333 | rnr1Y285A_p | TRUE | CAN | JSY2423 | 6 | rnr1Y285A_p | GCC | C>G | G/C+1 | G/C-1 | CG>GC | CG>GC | TGATTGCCC | CCCAAGA | CGG | GCC | C | C | C>G | C>G_GCC | CG>GC_CCCAAGA | 6 | FALSE | FALSE |
| 177_v.txt | 32954 | 32954-SNV- | SNV | 1 | G | C | 11 | 41638 | 0.02641818 | 57 | rnr1Y285A_p | TRUE | CAN | JSY2423 | 6 | rnr1Y285A_p | TGT | G>C | G/C+1 | G/C-1 | CG>GC | CG>GC | ATACATGTA | GTAACCA | ACA | ACA | C | G | C>G | C>G_ACA | CG>GC_GTAACCA | 6 | FALSE | FALSE |
| 177_v.txt | 33070 | 33070-SNV- | SNV | 1 | C | G | 2 | 41638 | 4.80E-03 | 63.9942736 | rnr1Y285A_p | TRUE | CAN | JSY2423 | 6 | rnr1Y285A_p | GCC | C>G | G/C+1 | G/C-1 | CG>GC | CG>GC | AATATGCCA | CCAAAGA | CGG | GCC | C | C | C>G | C>G_GCC | CG>GC_CCAAAGA | 6 | FALSE | FALSE |
| 177_v.txt | 33078 | 33078-SNV- | SNV | 1 | C | G | 2 | 41637 | 4.80E-03 | 63.9884413 | rnr1Y285A_p | TRUE | CAN | JSY2423 | 6 | rnr1Y285A_p | ACC | C>G | G/C+1 | G/C-1 | CG>GC | CG>GC | AAAGAACCC | CCCATAA | TGG | ACC | C | C | C>G | C>G_ACC | CG>GC_CCCATAA | 6 | FALSE | FALSE |
| 177_v.txt | 33079 | 33079-SNV- | SNV | 1 | C | G | 9 | 41638 | 0.02161487 | 51.1107598 | rnr1Y285A_p | TRUE | CAN | JSY2423 | 6 | rnr1Y285A_p | CCC | C>G | G/C+1 | G/C-1 | CG>GC | CG>GC | AAGAACCCA | CCATAAA | GGG | CCC | C | C | C>G | C>G_CCC | CG>GC_CCATAAA | 6 | FALSE | FALSE |
| 177_v.txt | 33114 | 33114-SNV- | SNV | 1 | C | G | 1 | 41638 | 2.40E-03 | 56 | rnr1Y285A_p | TRUE | CAN | JSY2423 | 6 | rnr1Y285A_p | GCC | C>G | G/C+1 | G/C-1 | CG>GC | CG>GC | ACTGGGCCG | CCGGCGT | CGG | GCC | C | C | C>G | C>G_GCC | CG>GC_CCGGCGT | 6 | FALSE | FALSE |
| 177_v.txt | 33115 | 33115-SNV- | SNV | 1 | C | G | 1 | 41638 | 2.40E-03 | 36 | rnr1Y285A_p | TRUE | CAN | JSY2423 | 6 | rnr1Y285A_p | CCG | C>G | G/C+1 | G/C-1 | CG>GC | CG>GC | CTGGGCCGG | CGGCGTT | GGC | CCG | C | C | C>G | C>G_CCG | CG>GC_CGGCGTT | 6 | FALSE | FALSE |

| | | | | | | | | | | | | | | | | | | | | | | | | | | | | | | | | | | |
|---|---|---|---|---|---|---|---|---|---|---|---|---|---|---|---|---|---|---|---|---|---|---|---|---|---|---|---|---|---|---|---|---|---|---|
| 177_v.txt | 33118 | 33118-SNV-( | SNV | 1 | C | G | 3 | 41638 | 7.21E-03 | 37.6666667 | rnr1Y285A_pC | TRUE | CAN | JSY2423 | 6 | rnr1Y285A_p | GCG | C>G | G/C+1 | G/C-1 | CG>GC | CG>GC | GGCCGGCGT | CGTTGGT | CGC | GCG | C | C | C>G | C>G_GCG | CG>GC_CGTTGGT | 6 | FALSE | FALSE |
| 177_v.txt | 33153 | 33153-SNV-( | SNV | 1 | C | G | 11 | 41638 | 0.02641818 | 32.9092273 | rnr1Y285A_pC | TRUE | CAN | JSY2423 | 6 | rnr1Y285A_p | ACC | C>G | G/C+1 | G/C-1 | CG>GC | CG>GC | AAAAGACCT | CCTGTAC | TGG | ACC | C | C | C>G | C>G_ACC | CG>GC_CCTGTAC | 6 | FALSE | FALSE |
| 177_v.txt | 33205 | 33205-SNV-( | SNV | 1 | G | C | 6 | 14680 | 0.04087194 | 40.8333333 | rnr1Y285A_pC | TRUE | CAN | JSY2423 | 6 | rnr1Y285A_p | AGC | G>C | G/C+1 | G/C-1 | CG>GC | CG>GC | GCTTAAGCT | GCTCTCT | TCG | TCG | C | G | C>G | C>G_TCG | CG>GC_GCTCTCT | 6 | FALSE | FALSE |
| 177_v.txt | 33354 | 33354-SNV-( | SNV | 1 | G | C | 2 | 14680 | 0.01362398 | 63.5 | rnr1Y285A_pC | TRUE | CAN | JSY2423 | 6 | rnr1Y285A_p | TGA | G>C | G/C+1 | G/C-1 | CG>GC | CG>GC | GGTATTGAC | GACCCAC | ACT | ACT | C | G | C>G | C>G_ACT | CG>GC_GACCCAC | 6 | FALSE | FALSE |
| 177_v.txt | 31657 | 31657-SNV-( | SNV | 1 | G | T | 104 | 16789 | 0.61945321 | 19.0736156 | rnr1Y285A_pC | TRUE | CAN | JSY2423 | 6 | rnr1Y285A_p | GGC | G>T | A/T+1 | G/C-1 | CG>AT | CG>AT | GAAATGGCG | GCGTGGA | CCG | CCG | C | A | C>A | C>A_CCG | CG>AT_GCGTGGA | 6 | FALSE | FALSE |
| 177_v.txt | 31882 | 31882-SNV-( | SNV | 1 | C | A | 13 | 16790 | 0.07742704 | 25.2307692 | rnr1Y285A_pC | TRUE | CAN | JSY2423 | 6 | rnr1Y285A_p | GCA | C>A | A/T+1 | G/C-1 | CG>AT | CG>AT | AGGCGGCAG | CAGCAAA | CGT | GCA | C | T | C>A | C>A_GCA | CG>AT_CAGCAAA | 6 | FALSE | FALSE |
| 177_v.txt | 32027 | 32027-SNV-( | SNV | 1 | G | T | 573 | 26238 | 2.18385548 | 60.7050611 | rnr1Y285A_pC | TRUE | CAN | JSY2423 | 6 | rnr1Y285A_p | GGT | G>T | A/T+1 | G/C-1 | CG>AT | CG>AT | GCCACGGTA | GTATTTC | CCA | CCA | C | A | C>A | C>A_CCA | CG>AT_GTATTTC | 6 | FALSE | FALSE |
| 177_v.txt | 32088 | 32088-SNV-( | SNV | 1 | C | A | 23 | 26238 | 0.08765912 | 53.2173913 | rnr1Y285A_pC | TRUE | CAN | JSY2423 | 6 | rnr1Y285A_p | GCC | C>A | A/T+1 | G/C-1 | CG>AT | CG>AT | AAAAAGCCT | CCTGCAA | CGG | GCC | C | T | C>A | C>A_GCC | CG>AT_CCTGCAA | 6 | FALSE | FALSE |
| 177_v.txt | 32188 | 32188-SNV-( | SNV | 1 | C | A | 298 | 26238 | 1.1357573 | 45.7718121 | rnr1Y285A_pC | TRUE | CAN | JSY2423 | 6 | rnr1Y285A_p | ACT | C>A | A/T+1 | G/C-1 | CG>AT | CG>AT | CGAAAACTG | CTGCAAT | TGA | ACT | C | T | C>A | C>A_ACT | CG>AT_CTGCAAT | 6 | FALSE | FALSE |
| 177_v.txt | 32274 | 32274-SNV-( | SNV | 1 | G | T | 33 | 32844 | 0.10047497 | 33.3636364 | rnr1Y285A_pC | TRUE | CAN | JSY2423 | 6 | rnr1Y285A_p | GGA | G>T | A/T+1 | G/C-1 | CG>AT | CG>AT | ATACGGGAA | GAACCAA | CCT | CCT | C | A | C>A | C>A_CCT | CG>AT_GAACCAA | 6 | FALSE | FALSE |
| 177_v.txt | 32277 | 32277-SNV-( | SNV | 1 | C | A | 756 | 32844 | 2.30179028 | 36.5873016 | rnr1Y285A_pC | TRUE | CAN | JSY2423 | 6 | rnr1Y285A_p | ACC | C>A | A/T+1 | G/C-1 | CG>AT | CG>AT | CGGGAACCA | CCAACGT | TGG | ACC | C | T | C>A | C>A_ACC | CG>AT_CCAACGT | 6 | FALSE | FALSE |
| 177_v.txt | 32373 | 32373-SNV-( | SNV | 1 | G | T | 20 | 32844 | 0.06089392 | 61 | rnr1Y285A_pC | TRUE | CAN | JSY2423 | 6 | rnr1Y285A_p | AGC | G>T | A/T+1 | G/C-1 | CG>AT | CG>AT | TCAATAGCA | GCAATAA | TCG | TCG | C | A | C>A | C>A_TCG | CG>AT_GCAATAA | 6 | FALSE | FALSE |
| 177_v.txt | 32508 | 32508-SNV-( | SNV | 1 | G | T | 38 | 32844 | 0.11569845 | 43.5263158 | rnr1Y285A_pC | TRUE | CAN | JSY2423 | 6 | rnr1Y285A_p | GGC | G>T | A/T+1 | G/C-1 | CG>AT | CG>AT | TTGATGGCT | GCTCTTG | CCG | CCG | C | A | C>A | C>A_CCG | CG>AT_GCTCTTG | 6 | FALSE | FALSE |
| 177_v.txt | 32530 | 32530-SNV-( | SNV | 1 | G | T | 22 | 32844 | 0.06698332 | 41 | rnr1Y285A_pC | TRUE | CAN | JSY2423 | 6 | rnr1Y285A_p | GGG | G>T | A/T+1 | G/C-1 | CG>AT | CG>AT | TTCTGGGGT | GGTTTGC | CCC | CCC | C | A | C>A | C>A_CCC | CG>AT_GGTTTGC | 6 | FALSE | FALSE |
| 177_v.txt | 32556 | 32556-SNV-( | SNV | 1 | C | A | 179 | 32831 | 0.54521641 | 37.6201117 | rnr1Y285A_pC | TRUE | CAN | JSY2423 | 6 | rnr1Y285A_p | ACC | C>A | A/T+1 | G/C-1 | CG>AT | CG>AT | GTGATACCA | CCAACTA | TGG | ACC | C | T | C>A | C>A_ACC | CG>AT_CCAACTA | 6 | FALSE | FALSE |
| 177_v.txt | 32566 | 32566-SNV-( | SNV | 1 | C | A | 94 | 38619 | 0.24340351 | 36.6489362 | rnr1Y285A_pC | TRUE | CAN | JSY2423 | 6 | rnr1Y285A_p | TCA | C>A | A/T+1 | G/C-1 | CG>AT | CG>AT | CTAGTTCAG | CAGTACC | AGT | TCA | C | T | C>A | C>A_TCA | CG>AT_CAGTACC | 6 | FALSE | FALSE |
| 177_v.txt | 32571 | 32571-SNV-( | SNV | 1 | C | A | 51 | 38635 | 0.13200466 | 36.8627451 | rnr1Y285A_pC | TRUE | CAN | JSY2423 | 6 | rnr1Y285A_p | ACC | C>A | A/T+1 | G/C-1 | CG>AT | CG>AT | TCAGTACCT | CCTTGAA | TGG | ACC | C | T | C>A | C>A_ACC | CG>AT_CCTTGAA | 6 | FALSE | FALSE |
| 177_v.txt | 32572 | 32572-SNV-( | SNV | 1 | C | A | 26 | 38634 | 0.06729824 | 36.3053753 | rnr1Y285A_pC | TRUE | CAN | JSY2423 | 6 | rnr1Y285A_p | CCT | C>A | A/T+1 | G/C-1 | CG>AT | CG>AT | CAGTACCTT | CTTGAAA | GGA | CCT | C | T | C>A | C>A_CCT | CG>AT_CTTGAAA | 6 | FALSE | FALSE |
| 177_v.txt | 32735 | 32735-SNV-( | SNV | 1 | G | T | 31 | 38633 | 0.08024228 | 54.6451613 | rnr1Y285A_pC | TRUE | CAN | JSY2423 | 6 | rnr1Y285A_p | AGT | G>T | A/T+1 | G/C-1 | CG>AT | CG>AT | AAAACAGTA | GTATATT | TCA | TCA | C | A | C>A | C>A_TCA | CG>AT_GTATATT | 6 | FALSE | FALSE |
| 177_v.txt | 32783 | 32783-SNV-( | SNV | 1 | G | T | 23 | 38633 | 0.0595346 | 47.6086957 | rnr1Y285A_pC | TRUE | CAN | JSY2423 | 6 | rnr1Y285A_p | AGA | G>T | A/T+1 | G/C-1 | CG>AT | CG>AT | GACCCAGAA | GAACTCG | TCT | TCT | C | A | C>A | C>A_TCT | CG>AT_GAACTCG | 6 | FALSE | FALSE |
| 177_v.txt | 32786 | 32786-SNV-( | SNV | 1 | C | A | 318 | 38633 | 0.82313048 | 49.1949686 | rnr1Y285A_pC | TRUE | CAN | JSY2423 | 6 | rnr1Y285A_p | ACT | C>A | A/T+1 | G/C-1 | CG>AT | CG>AT | CCAGAACTC | CTCGAAT | TGA | ACT | C | T | C>A | C>A_ACT | CG>AT_CTCGAAT | 6 | FALSE | FALSE |
| 177_v.txt | 32794 | 32794-SNV-( | SNV | 1 | C | A | 34 | 38633 | 0.08800766 | 39.0588235 | rnr1Y285A_pC | TRUE | CAN | JSY2423 | 6 | rnr1Y285A_p | TCA | C>A | A/T+1 | G/C-1 | CG>AT | CG>AT | CGAATTCAC | CACCGTA | AGT | TCA | C | T | C>A | C>A_TCA | CG>AT_CACCGTA | 6 | FALSE | FALSE |
| 177_v.txt | 32819 | 32819-SNV-( | SNV | 1 | G | T | 504 | 38633 | 1.30458416 | 44.0634921 | rnr1Y285A_pC | TRUE | CAN | JSY2423 | 6 | rnr1Y285A_p | AGT | G>T | A/T+1 | G/C-1 | CG>AT | CG>AT | GAACAAGTT | GTTCATT | TCA | TCA | C | A | C>A | C>A_TCA | CG>AT_GTTCATT | 6 | FALSE | FALSE |
| 177_v.txt | 32900 | 32900-SNV-( | SNV | 1 | G | T | 26 | 41638 | 0.06244296 | 21.0766067 | rnr1Y285A_pC | TRUE | CAN | JSY2423 | 6 | rnr1Y285A_p | GGC | G>T | A/T+1 | G/C-1 | CG>AT | CG>AT | GACTTGGCC | GCCAACT | CCG | CCG | C | A | C>A | C>A_CCG | CG>AT_GCCAACT | 6 | FALSE | FALSE |
| 177_v.txt | 32901 | 32901-SNV-( | SNV | 1 | C | A | 5 | 41638 | 0.01200826 | 38 | rnr1Y285A_pC | TRUE | CAN | JSY2423 | 6 | rnr1Y285A_p | GCC | C>A | A/T+1 | G/C-1 | CG>AT | CG>AT | ACTTGGCCA | CCAACTA | CGG | GCC | C | T | C>A | C>A_GCC | CG>AT_CCAACTA | 6 | FALSE | FALSE |
| 177_v.txt | 32917 | 32917-SNV-( | SNV | 1 | C | A | 5156 | 41638 | 12.3829195 | 52.394104 | rnr1Y285A_pC | TRUE | CAN | JSY2423 | 6 | rnr1Y285A_p | TCC | C>A | A/T+1 | G/C-1 | CG>AT | CG>AT | TAAGTTCCA | CCAGGGC | AGG | TCC | C | T | C>A | C>A_TCC | CG>AT_CCAGGGC | 6 | FALSE | FALSE |
| 177_v.txt | 32935 | 32935-SNV-( | SNV | 1 | C | A | 8 | 41638 | 0.01921322 | 40.75 | rnr1Y285A_pC | TRUE | CAN | JSY2423 | 6 | rnr1Y285A_p | GCC | C>A | A/T+1 | G/C-1 | CG>AT | CG>AT | TGATTGCCC | CCCAAGA | CGG | GCC | C | T | C>A | C>A_GCC | CG>AT_CCCAAGA | 6 | FALSE | FALSE |
| 177_v.txt | 32936 | 32936-SNV-( | SNV | 1 | C | A | 53 | 41637 | 0.12729063 | 36.6398468 | rnr1Y285A_pC | TRUE | CAN | JSY2423 | 6 | rnr1Y285A_p | CCC | C>A | A/T+1 | G/C-1 | CG>AT | CG>AT | GATTGCCCA | CCAAGAA | GGG | CCC | C | T | C>A | C>A_CCC | CG>AT_CCAAGAA | 6 | FALSE | FALSE |
| 177_v.txt | 32940 | 32940-SNV-( | SNV | 1 | G | T | 89 | 41638 | 0.21374706 | 48.6629214 | rnr1Y285A_pC | TRUE | CAN | JSY2423 | 6 | rnr1Y285A_p | AGA | G>T | A/T+1 | G/C-1 | CG>AT | CG>AT | GCCCAAGAA | GAAAACC | TCT | TCT | C | A | C>A | C>A_TCT | CG>AT_GAAAACC | 6 | FALSE | FALSE |
| 177_v.txt | 32958 | 32958-SNV-( | SNV | 1 | C | A | 26 | 41638 | 0.06244296 | 49.9230769 | rnr1Y285A_pC | TRUE | CAN | JSY2423 | 6 | rnr1Y285A_p | ACC | C>A | A/T+1 | G/C-1 | CG>AT | CG>AT | ATGTAACCA | CCATTGG | TGG | ACC | C | T | C>A | C>A_ACC | CG>AT_CCATTGG | 6 | FALSE | FALSE |
| 177_v.txt | 33051 | 33051-SNV-( | SNV | 1 | G | T | 18 | 41638 | 0.04322974 | 61.1111111 | rnr1Y285A_pC | TRUE | CAN | JSY2423 | 6 | rnr1Y285A_p | GGA | G>T | A/T+1 | G/C-1 | CG>AT | CG>AT | CCCAAGGAC | GACTGCG | CCT | CCT | C | A | C>A | C>A_CCT | CG>AT_GACTGCG | 6 | FALSE | FALSE |
| 177_v.txt | 33079 | 33079-SNV-( | SNV | 1 | C | A | 9 | 41638 | 0.02161487 | 48.8886884 | rnr1Y285A_pC | TRUE | CAN | JSY2423 | 6 | rnr1Y285A_p | CCC | C>A | A/T+1 | G/C-1 | CG>AT | CG>AT | AAGAACCCA | CCATAAA | GGG | CCC | C | T | C>A | C>A_CCC | CG>AT_CCATAAA | 6 | FALSE | FALSE |
| 177_v.txt | 33093 | 33093-SNV-( | SNV | 1 | G | T | 15 | 41638 | 0.03602479 | 54.5333333 | rnr1Y285A_pC | TRUE | CAN | JSY2423 | 6 | rnr1Y285A_p | TGA | G>T | A/T+1 | G/C-1 | CG>AT | CG>AT | AAATATGAT | GATATAA | ACT | ACT | C | A | C>A | C>A_ACT | CG>AT_GATATAA | 6 | FALSE | FALSE |
| 177_v.txt | 33113 | 33113-SNV-( | SNV | 1 | G | T | 1 | 41638 | 2.40E-03 | 60.9983449 | rnr1Y285A_pC | TRUE | CAN | JSY2423 | 6 | rnr1Y285A_p | GGC | G>T | A/T+1 | G/C-1 | CG>AT | CG>AT | CACTGGGCC | GCCGGCG | CCG | CCG | C | A | C>A | C>A_CCG | CG>AT_GCCGGCG | 6 | FALSE | FALSE |
| 177_v.txt | 33114 | 33114-SNV-( | SNV | 1 | C | A | 28 | 41638 | 0.06724627 | 42.4285714 | rnr1Y285A_pC | TRUE | CAN | JSY2423 | 6 | rnr1Y285A_p | GCC | C>A | A/T+1 | G/C-1 | CG>AT | CG>AT | ACTGGGCCG | CCGGCGT | CGG | GCC | C | T | C>A | C>A_GCC | CG>AT_CCGGCGT | 6 | FALSE | FALSE |
| 177_v.txt | 33115 | 33115-SNV-( | SNV | 1 | C | A | 44 | 41638 | 0.1056727 | 41.9318182 | rnr1Y285A_pC | TRUE | CAN | JSY2423 | 6 | rnr1Y285A_p | CCG | C>A | A/T+1 | G/C-1 | CG>AT | CG>AT | CTGGGCCGG | CGGCGTT | GGC | CCG | C | T | C>A | C>A_CCG | CG>AT_CGGCGTT | 6 | FALSE | FALSE |
| 177_v.txt | 33118 | 33118-SNV-( | SNV | 1 | C | A | 4 | 41638 | 9.61E-03 | 45.25 | rnr1Y285A_pC | TRUE | CAN | JSY2423 | 6 | rnr1Y285A_p | GCG | C>A | A/T+1 | G/C-1 | CG>AT | CG>AT | GGCCGGCGT | CGTTGGT | CGC | GCG | C | T | C>A | C>A_GCG | CG>AT_CGTTGGT | 6 | FALSE | FALSE |
| 177_v.txt | 33151 | 33151-SNV-( | SNV | 1 | G | T | 58 | 41638 | 0.13929584 | 39.2931035 | rnr1Y285A_pC | TRUE | CAN | JSY2423 | 6 | rnr1Y285A_p | AGA | G>T | A/T+1 | G/C-1 | CG>AT | CG>AT | TGAAAAGAC | GACCTGT | TCT | TCT | C | A | C>A | C>A_TCT | CG>AT_GACCTGT | 6 | FALSE | FALSE |
| 177_v.txt | 33153 | 33153-SNV-( | SNV | 1 | C | A | 3202 | 41638 | 7.69009078 | 40.7022862 | rnr1Y285A_pC | TRUE | CAN | JSY2423 | 6 | rnr1Y285A_p | ACC | C>A | A/T+1 | G/C-1 | CG>AT | CG>AT | AAAAGACCT | CCTGTAC | TGG | ACC | C | T | C>A | C>A_ACC | CG>AT_CCTGTAC | 6 | FALSE | FALSE |
| 177_v.txt | 33154 | 33154-SNV-( | SNV | 1 | C | A | 123 | 41638 | 0.29540324 | 36.4835509 | rnr1Y285A_pC | TRUE | CAN | JSY2423 | 6 | rnr1Y285A_p | CCT | C>A | A/T+1 | G/C-1 | CG>AT | CG>AT | AAAGACCTG | CTGTACC | GGA | CCT | C | T | C>A | C>A_CCT | CG>AT_CTGTACC | 6 | FALSE | FALSE |
| 177_v.txt | 33159 | 33159-SNV-( | SNV | 1 | C | A | 30 | 41638 | 0.07204957 | 35.9333333 | rnr1Y285A_pC | TRUE | CAN | JSY2423 | 6 | rnr1Y285A_p | ACC | C>A | A/T+1 | G/C-1 | CG>AT | CG>AT | CCTGTACCA | CCAATAG | TGG | ACC | C | T | C>A | C>A_ACC | CG>AT_CCAATAG | 6 | FALSE | FALSE |
| 177_v.txt | 33168 | 33168-SNV-( | SNV | 1 | C | A | 560 | 41634 | 1.34505452 | 37.3785714 | rnr1Y285A_pC | TRUE | CAN | JSY2423 | 6 | rnr1Y285A_p | ACC | C>A | A/T+1 | G/C-1 | CG>AT | CG>AT | ATAGTACCA | CCACCAA | TGG | ACC | C | T | C>A | C>A_ACC | CG>AT_CCACCAA | 6 | FALSE | FALSE |
| 177_v.txt | 33186 | 33186-SNV-( | SNV | 1 | C | A | 13 | 14681 | 0.08854983 | 37.6923077 | rnr1Y285A_pC | TRUE | CAN | JSY2423 | 6 | rnr1Y285A_p | ACC | C>A | A/T+1 | G/C-1 | CG>AT | CG>AT | ATCATACCA | CCAATAT | TGG | ACC | C | T | C>A | C>A_ACC | CG>AT_CCAATAT | 6 | FALSE | FALSE |
| 177_v.txt | 33195 | 33195-SNV-( | SNV | 1 | C | A | 10 | 14680 | 0.06811989 | 37.9 | rnr1Y285A_pC | TRUE | CAN | JSY2423 | 6 | rnr1Y285A_p | TCT | C>A | A/T+1 | G/C-1 | CG>AT | CG>AT | ATATGTCTT | CTTTGCT | AGA | TCT | C | T | C>A | C>A_TCT | CG>AT_CTTTGCT | 6 | FALSE | FALSE |
| 177_v.txt | 33205 | 33205-SNV-( | SNV | 1 | G | T | 6 | 14680 | 0.04087194 | 39.6666667 | rnr1Y285A_pC | TRUE | CAN | JSY2423 | 6 | rnr1Y285A_p | AGC | G>T | A/T+1 | G/C-1 | CG>AT | CG>AT | GCTTAAGCT | GCTCTCT | TCG | TCG | C | A | C>A | C>A_TCG | CG>AT_GCTCTCT | 6 | FALSE | FALSE |
| 177_v.txt | 33309 | 33309-SNV-( | SNV | 1 | C | A | 7 | 14679 | 0.04768717 | 27.8546569 | rnr1Y285A_pC | TRUE | CAN | JSY2423 | 6 | rnr1Y285A_p | GCG | C>A | A/T+1 | G/C-1 | CG>AT | CG>AT | AAAGAGCGC | CGCAATG | CGC | GCG | C | T | C>A | C>A_GCG | CG>AT_CGCAATG | 6 | FALSE | FALSE |
| 177_v.txt | 33310 | 33310-SNV-( | SNV | 1 | G | T | 11 | 14677 | 0.0749472 | 19.4514174 | rnr1Y285A_pC | TRUE | CAN | JSY2423 | 6 | rnr1Y285A_p | CGC | G>T | A/T+1 | G/C-1 | CG>AT | CG>AT | AAGAGCGCA | GCAATGG | GCG | GCG | C | A | C>A | C>A_GCG | CG>AT_GCAATGG | 6 | FALSE | FALSE |
| 177_v.txt | 33325 | 33325-SNV-( | SNV | 1 | C | A | 299 | 14680 | 2.03678474 | 60.4080268 | rnr1Y285A_pC | TRUE | CAN | JSY2423 | 6 | rnr1Y285A_p | TCT | C>A | A/T+1 | G/C-1 | CG>AT | CG>AT | ACAATTCTT | CTTTACT | AGA | TCT | C | T | C>A | C>A_TCT | CG>AT_CTTTACT | 6 | FALSE | FALSE |
| 177_v.txt | 33354 | 33354-SNV-( | SNV | 1 | G | T | 81 | 14680 | 0.55177112 | 17.1481482 | rnr1Y285A_pC | TRUE | CAN | JSY2423 | 6 | rnr1Y285A_p | TGA | G>T | A/T+1 | G/C-1 | CG>AT | CG>AT | GGTATTGAC | GACCCAC | ACT | ACT | C | A | C>A | C>A_ACT | CG>AT_GACCCAC | 6 | FALSE | FALSE |
| 177_v.txt | 33474 | 33474-SNV-( | SNV | 1 | C | A | 47 | 14679 | 0.3201853 | 33.5744681 | rnr1Y285A_pC | TRUE | CAN | JSY2423 | 6 | rnr1Y285A_p | GCC | C>A | A/T+1 | G/C-1 | CG>AT | CG>AT | GCTATGCCT | CCTTTTT | CGG | GCC | C | T | C>A | C>A_GCC | CG>AT_CCTTTTT | 6 | FALSE | FALSE |
| 177_v.txt | 33475 | 33475-SNV-( | SNV | 1 | C | A | 17 | 14678 | 0.11581959 | 36.2941177 | rnr1Y285A_pC | TRUE | CAN | JSY2423 | 6 | rnr1Y285A_p | CCT | C>A | A/T+1 | G/C-1 | CG>AT | CG>AT | CTATGCCTT | CTTTTTT | GGA | CCT | C | T | C>A | C>A_CCT | CG>AT_CTTTTTT | 6 | FALSE | FALSE |
| 177_v.txt | 33474 | 33474-Repla | Replacement | 2 | CC | A | 1 | 14679 | 6.81E-03 | 36.5 | rnr1Y285A_pC | TRUE | CAN | JSY2423 | 6 | rnr1Y285A_p | GCC | nope | A/T+1 | >1bpdel | none | Replac. | GCTATGCCT | CCTTTTT | CGG | GCC | CC | T | NA | NA_GCC | Replac._CCTTTTT | 6 | FALSE | FALSE |
| 177_v.txt | 33011 | 33011-MNV- | MNV | 2 | AG | CT | 19 | 41638 | 0.04563139 | 60.4473233 | rnr1Y285A_pC | TRUE | CAN | JSY2423 | 6 | rnr1Y285A_p | AAG | nope | >1bp-ins | >1bpdel | none | MNV | TGTGAAAGA | AGAGGAT | TTC | AAG | AG | none | NA | NA_AAG | MNV_AGAGGAT | 6 | FALSE | FALSE |
| 177_v.txt | 31863 | 31863-Insert | Insertion | 1 | - | A | 2 | 16790 | 0.01191185 | 37.5 | rnr1Y285A_pC | TRUE | CAN | JSY2423 | 6 | rnr1Y285A_p | GAA | nope | A/T+1 | >1bpdel | none | A/T+1 | AACAGGAAA | AAAATAG | CTT | GAA | - | T | NA | NA_GAA | A/T+1_AAAATAG | 6 | FALSE | FALSE |
| 177_v.txt | 31892 | 31892-Insert | Insertion | 1 | - | A | 7 | 16790 | 0.04169148 | 38 | rnr1Y285A_pC | TRUE | CAN | JSY2423 | 6 | rnr1Y285A_p | TAA | nope | A/T+1 | >1bpdel | none | A/T+1 | AAAGCTAAC | AACACCA | ATT | TAA | - | T | NA | NA_TAA | A/T+1_AACACCA | 6 | FALSE | FALSE |
| 177_v.txt | 32081 | 32081-Insert | Insertion | 1 | - | A | 23 | 26238 | 0.08765912 | 64 | rnr1Y285A_pC | TRUE | CAN | JSY2423 | 6 | rnr1Y285A_p | CAA | nope | A/T+1 | >1bpdel | none | A/T+1 | CCATGCAAA | AAAAAAG | GTT | CAA | - | T | NA | NA_CAA | A/T+1_AAAAAAG | 6 | FALSE | FALSE |
| 177_v.txt | 32231 | 32231-Insert | Insertion | 1 | - | T | 6 | 26237 | 0.02286847 | 38 | rnr1Y285A_pC | TRUE | CAN | JSY2423 | 6 | rnr1Y285A_p | ATT | nope | A/T+1 | >1bpdel | none | A/T+1 | CAGGAATTT | TTTAGGA | TAA | ATT | - | A | NA | NA_ATT | A/T+1_TTTAGGA | 6 | FALSE | FALSE |
| 177_v.txt | 32248 | 32248-Insert | Insertion | 1 | - | C | 4 | 26227 | 0.01525146 | 37.25 | rnr1Y285A_pC | TRUE | CAN | JSY2423 | 6 | rnr1Y285A_p | TTC | nope | G/C+1 | >1bpdel | none | G/C+1 | ACTTGTTCT | TCTTTGA | AAG | TTC | - | G | NA | NA_TTC | G/C+1_TCTTTGA | 6 | FALSE | FALSE |
| 177_v.txt | 32299 | 32299-Insert | Insertion | 1 | - | A | 11 | 32844 | 0.03349166 | 58.8181818 | rnr1Y285A_pC | TRUE | CAN | JSY2423 | 6 | rnr1Y285A_p | GCG | nope | A/T+1 | >1bpdel | none | A/T+1 | AATTTGCGG | CGGCAGA | CGC | GCG | - | T | NA | NA_GCG | A/T+1_CGGCAGA | 6 | FALSE | FALSE |
| 177_v.txt | 32480 | 32480-Insert | Insertion | 4 | - | TAAG | 4 | 32844 | 0.01217879 | 49.9375 | rnr1Y285A_pC | TRUE | CAN | JSY2423 | 6 | rnr1Y285A_p | TTA | nope | >1bp-ins | >1bpdel | none | >1bp-ins | GAAGGTTAA | TAAGATA | AAT | TTA | - | none | NA | NA_TTA | >1bp-ins_TAAGATA | 6 | FALSE | FALSE |
| 177_v.txt | 32498 | 32498-Insert | Insertion | 1 | - | T | 19 | 32844 | 0.05784923 | 47.4736842 | rnr1Y285A_pC | TRUE | CAN | JSY2423 | 6 | rnr1Y285A_p | CTT | nope | A/T+1 | >1bpdel | none | A/T+1 | AACAACTTT | TTTTTTG | GAA | CTT | - | A | NA | NA_CTT | A/T+1_TTTTTTG | 6 | FALSE | FALSE |
| 177_v.txt | 32532 | 32532-Insert | Insertion | 1 | - | T | 4 | 32843 | 0.01217916 | 38 | rnr1Y285A_pC | TRUE | CAN | JSY2423 | 6 | rnr1Y285A_p | GTT | nope | A/T+1 | >1bpdel | none | A/T+1 | CTGGGGTTT | TTTGCAG | CAA | GTT | - | A | NA | NA_GTT | A/T+1_TTTGCAG | 6 | FALSE | FALSE |
| 177_v.txt | 32842 | 32842-Insert | Insertion | 1 | - | A | 23 | 38633 | 0.0595346 | 44.5217391 | rnr1Y285A_pC | TRUE | CAN | JSY2423 | 6 | rnr1Y285A_p | CAA | nope | A/T+1 | >1bpdel | none | A/T+1 | TTACCCAAA | AAAAAAT | GTT | CAA | - | T | NA | NA_CAA | A/T+1_AAAAAAT | 6 | FALSE | FALSE |
| 177_v.txt | 32630 | 32630-Deleti | Deletion | 2 | GT | - | 1 | 38633 | 2.59E-03 | 64 | rnr1Y285A_pC | TRUE | CAN | JSY2423 | 6 | rnr1Y285A_p | CGT | nope | >1bp-ins | >1bpdel | none | >1bpdel | CCCTTCGTT | GTTTTTA | GCA | CGT | GT | none | NA | NA_CGT | >1bpdel_GTTTTTA | 6 | FALSE | FALSE |
| 177_v.txt | 33206 | 33206-Deleti | Deletion | 2 | CT | - | 10 | 14680 | 0.06811989 | 37.8 | rnr1Y285A_pC | TRUE | CAN | JSY2423 | 6 | rnr1Y285A_p | GCT | nope | >1bp-ins | >1bpdel | none | >1bpdel | CTTAAGCTC | CTCTCTC | CGA | GCT | CT | none | NA | NA_GCT | >1bpdel_CTCTCTC | 6 | FALSE | FALSE |
| 177_v.txt | 31663 | 31663-Deleti | Deletion | 1 | A | - | 7 | 16790 | 0.04169148 | 40.4285714 | rnr1Y285A_pC | TRUE | CAN | JSY2423 | 6 | rnr1Y285A_p | GAA | nope | >1bp-ins | A/T-1 | none | A/T-1 | GCGTGGAAA | AAATGTG | CTT | CTT | T | none | T>none | T>none_CTT | A/T-1_AAATGTG | 6 | FALSE | FALSE |
| 177_v.txt | 31723 | 31723-Deleti | Deletion | 1 | A | - | 21 | 16790 | 0.12507445 | 60.7142857 | rnr1Y285A_pC | TRUE | CAN | JSY2423 | 6 | rnr1Y285A_p | CAA | nope | >1bp-ins | A/T-1 | none | A/T-1 | TGTCCCAAA | AAAAAGT | GTT | GTT | T | none | T>none | T>none_GTT | A/T-1_AAAAAGT | 6 | FALSE | FALSE |
| 177_v.txt | 31834 | 31834-Deleti | Deletion | 1 | G | - | 5 | 16790 | 0.02977963 | 64 | rnr1Y285A_pC | TRUE | CAN | JSY2423 | 6 | rnr1Y285A_p | TGA | nope | >1bp-ins | G/C-1 | none | G/C-1 | TGCATTGAA | GAAATAA | ACT | ACT | C | none | C>none | C>none_ACT | G/C-1_GAAATAA | 6 | FALSE | FALSE |
| 177_v.txt | 31904 | 31904-Deleti | Deletion | 1 | T | - | 32 | 16790 | 0.19058964 | 44.3125 | rnr1Y285A_pC | TRUE | CAN | JSY2423 | 6 | rnr1Y285A_p | ATT | nope | >1bp-ins | A/T-1 | none | A/T-1 | ATTGAATTT | TTTTGGT | TAA | ATT | T | none | nope | nope_ATT | A/T-1_TTTTGGT | 6 | FALSE | FALSE |
| 177_v.txt | 31913 | 31913-Deleti | Deletion | 1 | A | - | 7 | 16790 | 0.04169148 | 41.2857143 | rnr1Y285A_pC | TRUE | CAN | JSY2423 | 6 | rnr1Y285A_p | CAA | nope | >1bp-ins | A/T-1 | none | A/T-1 | TGGTGCAAA | AAAAGCC | GTT | GTT | T | none | T>none | T>none_GTT | A/T-1_AAAAGCC | 6 | FALSE | FALSE |
| 177_v.txt | 31972 | 31972-Deleti | Deletion | 1 | C | - | 7 | 26238 | 0.02667886 | 36.7142857 | rnr1Y285A_pC | TRUE | CAN | JSY2423 | 6 | rnr1Y285A_p | GCC | nope | >1bp-ins | G/C-1 | none | G/C-1 | AATAAGCCA | CCAAGCC | CGG | GCC | C | none | nope | nope_GCC | G/C-1_CCAAGCC | 6 | FALSE | FALSE |
| 177_v.txt | 31979 | 31979-Deleti | Deletion | 1 | G | - | 33 | 26238 | 0.12577178 | 37.6363636 | rnr1Y285A_pC | TRUE | CAN | JSY2423 | 6 | rnr1Y285A_p | CGG | nope | >1bp-ins | G/C-1 | none | G/C-1 | CAAGCCGGG | GGGCATT | GCC | GCC | C | none | C>none | C>none_GCC | G/C-1_GGGCATT | 6 | FALSE | FALSE |
| 177_v.txt | 32075 | 32075-Deleti | Deletion | 1 | C | - | 22 | 26238 | 0.08384785 | 61.5909091 | rnr1Y285A_pC | TRUE | CAN | JSY2423 | 6 | rnr1Y285A_p | ACC | nope | >1bp-ins | G/C-1 | none | G/C-1 | AAATAACCA | CCATGCA | TGG | ACC | C | none | nope | nope_ACC | G/C-1_CCATGCA | 6 | FALSE | FALSE |
| 177_v.txt | 32081 | 32081-Deleti | Deletion | 1 | A | - | 50 | 26238 | 0.19056331 | 62.4600009 | rnr1Y285A_pC | TRUE | CAN | JSY2423 | 6 | rnr1Y285A_p | CAA | nope | >1bp-ins | A/T-1 | none | A/T-1 | CCATGCAAA | AAAAAAG | GTT | GTT | T | none | T>none | T>none_GTT | A/T-1_AAAAAAG | 6 | FALSE | FALSE |
| 177_v.txt | 32102 | 32102-Deleti | Deletion | 1 | G | - | 8 | 26238 | 0.03049013 | 62.75 | rnr1Y285A_pC | TRUE | CAN | JSY2423 | 6 | rnr1Y285A_p | TGA | nope | >1bp-ins | G/C-1 | none | G/C-1 | ACCAGTGAT | GATATTT | ACT | ACT | C | none | C>none | C>none_ACT | G/C-1_GATATTT | 6 | FALSE | FALSE |
| 177_v.txt | 32184 | 32184-Deleti | Deletion | 1 | A | - | 19 | 26238 | 0.07241406 | 45.8421053 | rnr1Y285A_pC | TRUE | CAN | JSY2423 | 6 | rnr1Y285A_p | GAA | nope | >1bp-ins | A/T-1 | none | A/T-1 | GTAACGAAA | AAAACTG | CTT | CTT | T | none | T>none | T>none_CTT | A/T-1_AAAACTG | 6 | FALSE | FALSE |
| 177_v.txt | 32201 | 32201-Deleti | Deletion | 1 | A | - | 23 | 26238 | 0.08765912 | 45.3913044 | rnr1Y285A_pC | TRUE | CAN | JSY2423 | 6 | rnr1Y285A_p | GAA | nope | >1bp-ins | A/T-1 | none | A/T-1 | GTATGGAAC | AACACCA | CTT | CTT | T | none | T>none | T>none_CTT | A/T-1_AACACCA | 6 | FALSE | FALSE |
| 177_v.txt | 32210 | 32210-Deleti | Deletion | 1 | T | - | 9 | 26238 | 0.0343014 | 45.7777778 | rnr1Y285A_pC | TRUE | CAN | JSY2423 | 6 | rnr1Y285A_p | CTT | nope | >1bp-ins | A/T-1 | none | A/T-1 | ACCACCTTT | TTTGGTG | GAA | CTT | T | none | nope | nope_CTT | A/T-1_TTTGGTG | 6 | FALSE | FALSE |
| 177_v.txt | 32231 | 32231-Deleti | Deletion | 1 | T | - | 7 | 26237 | 0.02667988 | 37.9999998 | rnr1Y285A_pC | TRUE | CAN | JSY2423 | 6 | rnr1Y285A_p | ATT | nope | >1bp-ins | A/T-1 | none | A/T-1 | CAGGAATTT | TTTAGGA | TAA | ATT | T | none | nope | nope_ATT | A/T-1_TTTAGGA | 6 | FALSE | FALSE |
| 177_v.txt | 32265 | 32265-Deleti | Deletion | 1 | A | - | 9 | 32844 | 0.02740227 | 38 | rnr1Y285A_pC | TRUE | CAN | JSY2423 | 6 | rnr1Y285A_p | TAA | nope | >1bp-ins | A/T-1 | none | A/T-1 | CCAAATAAA | AAAATAC | ATT | ATT | T | none | T>none | T>none_ATT | A/T-1_AAAATAC | 6 | FALSE | FALSE |
| 177_v.txt | 32277 | 32277-Deleti | Deletion | 1 | C | - | 12 | 32844 | 0.03653635 | 36.1666667 | rnr1Y285A_pC | TRUE | CAN | JSY2423 | 6 | rnr1Y285A_p | ACC | nope | >1bp-ins | G/C-1 | none | G/C-1 | CGGGAACCA | CCAACGT | TGG | ACC | C | none | nope | nope_ACC | G/C-1_CCAACGT | 6 | FALSE | FALSE |
| 177_v.txt | 32299 | 32299-Deleti | Deletion | 1 | C | - | 1 | 32844 | 3.05E-03 | 63.9999965 | rnr1Y285A_pC | TRUE | CAN | JSY2423 | 6 | rnr1Y285A_p | GCG | nope | >1bp-ins | G/C-1 | none | G/C-1 | AATTTGCGG | CGGCAGA | CGC | GCG | C | none | nope | nope_GCG | G/C-1_CGGCAGA | 6 | FALSE | FALSE |
| 177_v.txt | 32304 | 32304-Deleti | Deletion | 1 | G | - | 14 | 32844 | 0.04262575 | 59.1428571 | rnr1Y285A_pC | TRUE | CAN | JSY2423 | 6 | rnr1Y285A_p | AGA | nope | >1bp-ins | G/C-1 | none | G/C-1 | GCGGCAGAA | GAAATAA | TCT | TCT | C | none | C>none | C>none_TCT | G/C-1_GAAATAA | 6 | FALSE | FALSE |
| 177_v.txt | 32338 | 32338-Deleti | Deletion | 1 | T | - | 6 | 32844 | 0.01826818 | 63.1666667 | rnr1Y285A_pC | TRUE | CAN | JSY2423 | 6 | rnr1Y285A_p | ATA | nope | >1bp-ins | A/T-1 | none | A/T-1 | TGAAGATAT | TATGTGG | TAT | ATA | T | none | nope | nope_ATA | A/T-1_TATGTGG | 6 | FALSE | FALSE |
| 177_v.txt | 32346 | 32346-Deleti | Deletion | 1 | A | - | 23 | 32844 | 0.07002801 | 61.7391304 | rnr1Y285A_pC | TRUE | CAN | JSY2423 | 6 | rnr1Y285A_p | CAA | nope | >1bp-ins | A/T-1 | none | A/T-1 | TGTGGCAAA | AAAACCT | GTT | GTT | T | none | T>none | T>none_GTT | A/T-1_AAAACCT | 6 | FALSE | FALSE |
| 177_v.txt | 32398 | 32398-Deleti | Deletion | 1 | C | - | 11 | 32844 | 0.03349166 | 59 | rnr1Y285A_pC | TRUE | CAN | JSY2423 | 6 | rnr1Y285A_p | ACG | nope | >1bp-ins | G/C-1 | none | G/C-1 | TAGAAACGT | CGTAGGA | TGC | ACG | C | none | nope | nope_ACG | G/C-1_CGTAGGA | 6 | FALSE | FALSE |
| 177_v.txt | 32424 | 32424-Deleti | Deletion | 1 | G | - | 10 | 32844 | 0.03044696 | 64 | rnr1Y285A_pC | TRUE | CAN | JSY2423 | 6 | rnr1Y285A_p | AGG | nope | >1bp-ins | G/C-1 | none | G/C-1 | AGTTTAGGG | GGGTCAT | TCC | TCC | C | none | C>none | C>none_TCC | G/C-1_GGGTCAT | 6 | FALSE | FALSE |
| 177_v.txt | 32442 | 32442-Deleti | Deletion | 1 | A | - | 9 | 32844 | 0.02740227 | 61.1111111 | rnr1Y285A_pC | TRUE | CAN | JSY2423 | 6 | rnr1Y285A_p | TAA | nope | >1bp-ins | A/T-1 | none | A/T-1 | GGAACTAAA | AAAAGTC | ATT | ATT | T | none | T>none | T>none_ATT | A/T-1_AAAAGTC | 6 | FALSE | FALSE |
| 177_v.txt | 32490 | 32490-Deleti | Deletion | 1 | A | - | 20 | 32844 | 0.06089392 | 45.7 | rnr1Y285A_pC | TRUE | CAN | JSY2423 | 6 | rnr1Y285A_p | GAA | nope | >1bp-ins | A/T-1 | none | A/T-1 | ATACGGAAA | AAAACAA | CTT | CTT | T | none | T>none | T>none_CTT | A/T-1_AAAACAA | 6 | FALSE | FALSE |
| 177_v.txt | 32498 | 32498-Deleti | Deletion | 1 | T | - | 41 | 32844 | 0.12483254 | 44.6585366 | rnr1Y285A_pC | TRUE | CAN | JSY2423 | 6 | rnr1Y285A_p | CTT | nope | >1bp-ins | A/T-1 | none | A/T-1 | AACAACTTT | TTTTTTG | GAA | CTT | T | none | nope | nope_CTT | A/T-1_TTTTTTG | 6 | FALSE | FALSE |
| 177_v.txt | 32522 | 32522-Deleti | Deletion | 1 | T | - | 23 | 32844 | 0.07002801 | 38.5217391 | rnr1Y285A_pC | TRUE | CAN | JSY2423 | 6 | rnr1Y285A_p | ATT | nope | >1bp-ins | A/T-1 | none | A/T-1 | AACGGATTT | TTTTCTG | TAA | ATT | T | none | nope | nope_ATT | A/T-1_TTTTCTG | 6 | FALSE | FALSE |
| 177_v.txt | 32528 | 32528-Deleti | Deletion | 1 | G | - | 8 | 32844 | 0.02435757 | 40.375 | rnr1Y285A_pC | TRUE | CAN | JSY2423 | 6 | rnr1Y285A_p | TGG | nope | >1bp-ins | G/C-1 | none | G/C-1 | TTTTCTGGG | GGGGTTT | ACC | ACC | C | none | C>none | C>none_ACC | G/C-1_GGGGTTT | 6 | FALSE | FALSE |
| 177_v.txt | 32532 | 32532-Deleti | Deletion | 1 | T | - | 13 | 32843 | 0.03958226 | 41.3846091 | rnr1Y285A_pC | TRUE | CAN | JSY2423 | 6 | rnr1Y285A_p | GTT | nope | >1bp-ins | A/T-1 | none | A/T-1 | CTGGGGTTT | TTTGCAG | CAA | GTT | T | none | nope | nope_GTT | A/T-1_TTTGCAG | 6 | FALSE | FALSE |
| 177_v.txt | 32556 | 32556-Deleti | Deletion | 1 | C | - | 64 | 32831 | 0.19493771 | 37.8125 | rnr1Y285A_pC | TRUE | CAN | JSY2423 | 6 | rnr1Y285A_p | ACC | nope | >1bp-ins | G/C-1 | none | G/C-1 | GTGATACCA | CCAACTA | TGG | ACC | C | none | nope | nope_ACC | G/C-1_CCAACTA | 6 | FALSE | FALSE |
| 177_v.txt | 32608 | 32608-Deleti | Deletion | 1 | C | - | 7516 | 38633 | 19.4548702 | 44.5730442 | rnr1Y285A_pC | TRUE | CAN | JSY2423 | 6 | rnr1Y285A_p | ACC | nope | >1bp-ins | G/C-1 | none | G/C-1 | AGGAAACCC | CCCAACC | TGG | ACC | C | none | nope | nope_ACC | G/C-1_CCCAACC | 6 | FALSE | FALSE |
| 177_v.txt | 32627 | 32627-Deleti | Deletion | 1 | T | - | 12 | 38633 | 0.03106153 | 50.75 | rnr1Y285A_pC | TRUE | CAN | JSY2423 | 6 | rnr1Y285A_p | CTT | nope | >1bp-ins | A/T-1 | none | A/T-1 | CCTCCCTTC | TTCGTTT | GAA | CTT | T | none | nope | nope_CTT | A/T-1_TTCGTTT | 6 | FALSE | FALSE |
| 177_v.txt | 32630 | 32630-Deleti | Deletion | 1 | G | - | 136 | 38633 | 0.35203065 | 48.9558751 | rnr1Y285A_pC | TRUE | CAN | JSY2423 | 6 | rnr1Y285A_p | CGT | nope | >1bp-ins | G/C-1 | none | G/C-1 | CCCTTCGTT | GTTTTTA | GCA | GCA | C | none | C>none | C>none_GCA | G/C-1_GTTTTTA | 6 | FALSE | FALSE |
| 177_v.txt | 32631 | 32631-Deleti | Deletion | 1 | T | - | 17 | 38633 | 0.04400383 | 47.3529362 | rnr1Y285A_pC | TRUE | CAN | JSY2423 | 6 | rnr1Y285A_p | GTT | nope | >1bp-ins | A/T-1 | none | A/T-1 | CCTTCGTTT | TTTTTAT | CAA | GTT | T | none | nope | nope_GTT | A/T-1_TTTTTAT | 6 | FALSE | FALSE |
| 177_v.txt | 32655 | 32655-Deleti | Deletion | 1 | G | - | 6 | 38633 | 0.01553076 | 52.5 | rnr1Y285A_pC | TRUE | CAN | JSY2423 | 6 | rnr1Y285A_p | TGG | nope | >1bp-ins | G/C-1 | none | G/C-1 | ATACCTGGA | GGACCCC | ACC | ACC | C | none | C>none | C>none_ACC | G/C-1_GGACCCC | 6 | FALSE | FALSE |
| 177_v.txt | 32658 | 32658-Deleti | Deletion | 1 | C | - | 9364 | 38633 | 24.2383455 | 60.365015 | rnr1Y285A_pC | TRUE | CAN | JSY2423 | 6 | rnr1Y285A_p | ACC | nope | >1bp-ins | G/C-1 | none | G/C-1 | CCTGGACCC | CCCCAGG | TGG | ACC | C | none | nope | nope_ACC | G/C-1_CCCCAGG | 6 | FALSE | FALSE |
| 177_v.txt | 32670 | 32670-Deleti | Deletion | 1 | G | - | 334 | 38633 | 0.86454586 | 59.8562874 | rnr1Y285A_pC | TRUE | CAN | JSY2423 | 6 | rnr1Y285A_p | TGG | nope | >1bp-ins | G/C-1 | none | G/C-1 | GCACCTGGG | GGGTTTC | ACC | ACC | C | none | C>none | C>none_ACC | G/C-1_GGGTTTC | 6 | FALSE | FALSE |
| 177_v.txt | 32707 | 32707-Deleti | Deletion | 1 | C | - | 1201 | 38633 | 3.10874123 | 61.9650291 | rnr1Y285A_pC | TRUE | CAN | JSY2423 | 6 | rnr1Y285A_p | ACC | nope | >1bp-ins | G/C-1 | none | G/C-1 | CGGTAACCC | CCCCAGC | TGG | ACC | C | none | nope | nope_ACC | G/C-1_CCCCAGC | 6 | FALSE | FALSE |
| 177_v.txt | 32719 | 32719-Deleti | Deletion | 1 | A | - | 8 | 38633 | 0.02070769 | 57.25 | rnr1Y285A_pC | TRUE | CAN | JSY2423 | 6 | rnr1Y285A_p | CAA | nope | >1bp-ins | A/T-1 | none | A/T-1 | CACCACAAA | AAACCAT | GTT | GTT | T | none | T>none | T>none_GTT | A/T-1_AAACCAT | 6 | FALSE | FALSE |
| 177_v.txt | 32747 | 32747-Deleti | Deletion | 1 | C | - | 309 | 38633 | 0.79983434 | 56.1747573 | rnr1Y285A_pC | TRUE | CAN | JSY2423 | 6 | rnr1Y285A_p | ACC | nope | >1bp-ins | G/C-1 | none | G/C-1 | TAGAAACCC | CCCGATA | TGG | ACC | C | none | nope | nope_ACC | G/C-1_CCCGATA | 6 | FALSE | FALSE |
| 177_v.txt | 32760 | 32760-Deleti | Deletion | 1 | A | - | 13 | 38633 | 0.03364999 | 49.1538462 | rnr1Y285A_pC | TRUE | CAN | JSY2423 | 6 | rnr1Y285A_p | TAA | nope | >1bp-ins | A/T-1 | none | A/T-1 | ATGGCTAAA | AAAACTT | ATT | ATT | T | none | T>none | T>none_ATT | A/T-1_AAAACTT | 6 | FALSE | FALSE |
| 177_v.txt | 32779 | 32779-Deleti | Deletion | 1 | C | - | 15 | 38633 | 0.03882691 | 50.5333333 | rnr1Y285A_pC | TRUE | CAN | JSY2423 | 6 | rnr1Y285A_p | ACC | nope | >1bp-ins | G/C-1 | none | G/C-1 | AAGCGACCC | CCCAGAA | TGG | ACC | C | none | nope | nope_ACC | G/C-1_CCCAGAA | 6 | FALSE | FALSE |
| 177_v.txt | 32839 | 32839-Deleti | Deletion | 1 | C | - | 14 | 38633 | 0.03623845 | 35.8571429 | rnr1Y285A_pC | TRUE | CAN | JSY2423 | 6 | rnr1Y285A_p | ACC | nope | >1bp-ins | G/C-1 | none | G/C-1 | TAATTACCC | CCCAAAA | TGG | ACC | C | none | nope | nope_ACC | G/C-1_CCCAAAA | 6 | FALSE | FALSE |
| 177_v.txt | 32842 | 32842-Deleti | Deletion | 1 | A | - | 126 | 38633 | 0.32614604 | 43.6825391 | rnr1Y285A_pC | TRUE | CAN | JSY2423 | 6 | rnr1Y285A_p | CAA | nope | >1bp-ins | A/T-1 | none | A/T-1 | TTACCCAAA | AAAAAAT | GTT | GTT | T | none | T>none | T>none_GTT | A/T-1_AAAAAAT | 6 | FALSE | FALSE |
| 177_v.txt | 32855 | 32855-Deleti | Deletion | 1 | C | - | 11 | 38633 | 0.02847307 | 37.4545455 | rnr1Y285A_pC | TRUE | CAN | JSY2423 | 6 | rnr1Y285A_p | TCC | nope | >1bp-ins | G/C-1 | none | G/C-1 | ACTAATCCA | CCATGCC | AGG | TCC | C | none | nope | nope_TCC | G/C-1_CCATGCC | 6 | FALSE | FALSE |
| 177_v.txt | 32884 | 32884-Deleti | Deletion | 1 | A | - | 30 | 41638 | 0.07204957 | 37.3666667 | rnr1Y285A_pC | TRUE | CAN | JSY2423 | 6 | rnr1Y285A_p | CAA | nope | >1bp-ins | A/T-1 | none | A/T-1 | ACGTCCAAA | AAAATTG | GTT | GTT | T | none | T>none | T>none_GTT | A/T-1_AAAATTG | 6 | FALSE | FALSE |
| 177_v.txt | 32901 | 32901-Deleti | Deletion | 1 | C | - | 2 | 41638 | 4.80E-03 | 38 | rnr1Y285A_pC | TRUE | CAN | JSY2423 | 6 | rnr1Y285A_p | GCC | nope | >1bp-ins | G/C-1 | none | G/C-1 | ACTTGGCCA | CCAACTA | CGG | GCC | C | none | nope | nope_GCC | G/C-1_CCAACTA | 6 | FALSE | FALSE |
| 177_v.txt | 32917 | 32917-Deleti | Deletion | 1 | C | - | 10 | 41638 | 0.02401652 | 52.1 | rnr1Y285A_pC | TRUE | CAN | JSY2423 | 6 | rnr1Y285A_p | TCC | nope | >1bp-ins | G/C-1 | none | G/C-1 | TAAGTTCCA | CCAGGGC | AGG | TCC | C | none | nope | nope_TCC | G/C-1_CCAGGGC | 6 | FALSE | FALSE |
| 177_v.txt | 32924 | 32924-Deleti | Deletion | 1 | A | - | 48 | 41638 | 0.11527931 | 53.7291667 | rnr1Y285A_pC | TRUE | CAN | JSY2423 | 6 | rnr1Y285A_p | CAA | nope | >1bp-ins | A/T-1 | none | A/T-1 | CAGGGCAAA | AAAAGTG | GTT | GTT | T | none | T>none | T>none_GTT | A/T-1_AAAAGTG | 6 | FALSE | FALSE |
| 177_v.txt | 32934 | 32934-Deleti | Deletion | 1 | G | - | 21 | 41638 | 0.0504347 | 52.7618965 | rnr1Y285A_pC | TRUE | CAN | JSY2423 | 6 | rnr1Y285A_p | TGC | nope | >1bp-ins | G/C-1 | none | G/C-1 | GTGATTGCC | GCCCAAG | ACG | ACG | C | none | C>none | C>none_ACG | G/C-1_GCCCAAG | 6 | FALSE | FALSE |
| 177_v.txt | 32935 | 32935-Deleti | Deletion | 1 | C | - | 5 | 41638 | 0.01200826 | 43 | rnr1Y285A_pC | TRUE | CAN | JSY2423 | 6 | rnr1Y285A_p | GCC | nope | >1bp-ins | G/C-1 | none | G/C-1 | TGATTGCCC | CCCAAGA | CGG | GCC | C | none | nope | nope_GCC | G/C-1_CCCAAGA | 6 | FALSE | FALSE |
| 177_v.txt | 32945 | 32945-Deleti | Deletion | 1 | C | - | 28 | 41638 | 0.06724627 | 54.8214286 | rnr1Y285A_pC | TRUE | CAN | JSY2423 | 6 | rnr1Y285A_p | ACC | nope | >1bp-ins | G/C-1 | none | G/C-1 | AGAAAACCA | CCAATAC | TGG | ACC | C | none | nope | nope_ACC | G/C-1_CCAATAC | 6 | FALSE | FALSE |
| 177_v.txt | 32958 | 32958-Deleti | Deletion | 1 | C | - | 14 | 41638 | 0.03362313 | 57.3571429 | rnr1Y285A_pC | TRUE | CAN | JSY2423 | 6 | rnr1Y285A_p | ACC | nope | >1bp-ins | G/C-1 | none | G/C-1 | ATGTAACCA | CCATTGG | TGG | ACC | C | none | nope | nope_ACC | G/C-1_CCATTGG | 6 | FALSE | FALSE |
| 177_v.txt | 33000 | 33000-Deleti | Deletion | 1 | A | - | 36 | 41638 | 0.08645948 | 60.5277778 | rnr1Y285A_pC | TRUE | CAN | JSY2423 | 6 | rnr1Y285A_p | GAA | nope | >1bp-ins | A/T-1 | none | A/T-1 | TGTGAGAAA | AAAACTG | CTT | CTT | T | none | T>none | T>none_CTT | A/T-1_AAAACTG | 6 | FALSE | FALSE |
| 177_v.txt | 33004 | 33004-Deleti | Deletion | 1 | C | - | 10 | 41638 | 0.02401652 | 64 | rnr1Y285A_pC | TRUE | CAN | JSY2423 | 6 | rnr1Y285A_p | ACT | nope | >1bp-ins | G/C-1 | none | G/C-1 | AGAAAACTG | CTGTGAA | TGA | ACT | C | none | nope | nope_ACT | G/C-1_CTGTGAA | 6 | FALSE | FALSE |

| | | | | | | | | | | | | | | | | | | | | | | | | | | | | | | | | | | |
|---|---|---|---|---|---|---|---|---|---|---|---|---|---|---|---|---|---|---|---|---|---|---|---|---|---|---|---|---|---|---|---|---|---|---|
| 177_v.txt | 33040 | 33040-Deleti | Deletion | 1 | T | - | 9 | 41638 | 0.02161487 | 57.8888889 | rnr1Y285A_pC | TRUE | CAN | JSY2423 | 6 | rnr1Y285A_p | ATT | nope | >1bp-ins | A/T-1 | none | A/T-1 | TAGCCATTTC | TTTCACC | TAA | ATT | T | none | nope | nope_ATT | A/T-1_TTTCACC | 6 | FALSE | FALSE |
| 177_v.txt | 33045 | 33045-Deleti | Deletion | 1 | C | - | 10 | 41638 | 0.02401652 | 58.8 | rnr1Y285A_pC | TRUE | CAN | JSY2423 | 6 | rnr1Y285A_p | ACC | nope | >1bp-ins | G/C-1 | none | G/C-1 | ATTTCACCCA | CCCAAGG | TGG | ACC | C | none | nope | nope_ACC | G/C-1_CCCAAGG | 6 | FALSE | FALSE |
| 177_v.txt | 33078 | 33078-Deleti | Deletion | 1 | C | - | 117 | 41637 | 0.28100007 | 56.0598304 | rnr1Y285A_pC | TRUE | CAN | JSY2423 | 6 | rnr1Y285A_p | ACC | nope | >1bp-ins | G/C-1 | none | G/C-1 | AAAGAACCC | CCCATAA | TGG | ACC | C | none | nope | nope_ACC | G/C-1_CCCATAA | 6 | FALSE | FALSE |
| 177_v.txt | 33114 | 33114-Deleti | Deletion | 1 | C | - | 17 | 41638 | 0.04082809 | 43.1176471 | rnr1Y285A_pC | TRUE | CAN | JSY2423 | 6 | rnr1Y285A_p | GCC | nope | >1bp-ins | G/C-1 | none | G/C-1 | ACTGGGCCG | CCGGCGT | CGG | GCC | C | none | nope | nope_GCC | G/C-1_CCGGCGT | 6 | FALSE | FALSE |
| 177_v.txt | 33147 | 33147-Deleti | Deletion | 1 | A | - | 9 | 41638 | 0.02161487 | 37.8888889 | rnr1Y285A_pC | TRUE | CAN | JSY2423 | 6 | rnr1Y285A_p | GAA | nope | >1bp-ins | A/T-1 | none | A/T-1 | CCAATGAAA | AAAAGAC | CTT | CTT | T | none | T>none | T>none_CTT | A/T-1_AAAAGAC | 6 | FALSE | FALSE |
| 177_v.txt | 33159 | 33159-Deleti | Deletion | 1 | C | - | 3 | 41638 | 7.21E-03 | 38 | rnr1Y285A_pC | TRUE | CAN | JSY2423 | 6 | rnr1Y285A_p | ACC | nope | >1bp-ins | G/C-1 | none | G/C-1 | CCTGTACCAA | CCAATAG | TGG | ACC | C | none | nope | nope_ACC | G/C-1_CCAATAG | 6 | FALSE | FALSE |
| 177_v.txt | 33168 | 33168-Deleti | Deletion | 1 | C | - | 4 | 41634 | 9.61E-03 | 37 | rnr1Y285A_pC | TRUE | CAN | JSY2423 | 6 | rnr1Y285A_p | ACC | nope | >1bp-ins | G/C-1 | none | G/C-1 | ATAGTACCAC | CCACCAA | TGG | ACC | C | none | nope | nope_ACC | G/C-1_CCACCAA | 6 | FALSE | FALSE |
| 177_v.txt | 33354 | 33354-Deleti | Deletion | 1 | G | - | 3 | 14680 | 0.02043597 | 64 | rnr1Y285A_pC | TRUE | CAN | JSY2423 | 6 | rnr1Y285A_p | TGA | nope | >1bp-ins | G/C-1 | none | G/C-1 | GGTATTGACC | GACCCAC | ACT | ACT | C | none | C>none | C>none_ACT | G/C-1_GACCCAC | 6 | FALSE | FALSE |
| 177_v.txt | 33356 | 33356-Deleti | Deletion | 1 | C | - | 181 | 14680 | 1.23297003 | 54.1381216 | rnr1Y285A_pC | TRUE | CAN | JSY2423 | 6 | rnr1Y285A_p | ACC | nope | >1bp-ins | G/C-1 | none | G/C-1 | TATTGACCCA | CCCACGT | TGG | ACC | C | none | nope | nope_ACC | G/C-1_CCCACGT | 6 | FALSE | FALSE |
| 177_v.txt | 33474 | 33474-Deleti | Deletion | 1 | C | - | 38 | 14679 | 0.25887322 | 29.131579 | rnr1Y285A_pC | TRUE | CAN | JSY2423 | 6 | rnr1Y285A_p | GCC | nope | >1bp-ins | G/C-1 | none | G/C-1 | GCTATGCCTT | CCTTTTT | CGG | GCC | C | none | nope | nope_GCC | G/C-1_CCTTTTT | 6 | FALSE | FALSE |
| 42_v.txt | 31723 | 31723-SNV-A | SNV | 1 | A | C | 4 | 11596 | 0.03449465 | 61.75 | rnr1Y285F | FALSE | CAN | JSY3811 | 1 | rnr1Y285F-R | CAA | A>C | G/C+1 | A/T-1 | TA>GC | TA>GC | TGTCCCAAAA | AAAAAGT | GTT | GTT | T | G | T>G | T>G_GTT | TA>GC_AAAAAGT | 8 | FALSE | FALSE |
| 42_v.txt | 31950 | 31950-SNV-A | SNV | 1 | A | C | 27 | 23772 | 0.113579 | 33.962963 | rnr1Y285F | FALSE | CAN | JSY3811 | 1 | rnr1Y285F-R | CAT | A>C | G/C+1 | A/T-1 | TA>GC | TA>GC | ATCGTCATAA | ATAAATG | GTA | GTA | T | G | T>G | T>G_GTA | TA>GC_ATAAATG | 8 | FALSE | FALSE |
| 42_v.txt | 31967 | 31967-SNV-A | SNV | 1 | A | C | 49 | 23773 | 0.20611618 | 32.3469388 | rnr1Y285F | FALSE | CAN | JSY3811 | 1 | rnr1Y285F-R | AAT | A>C | G/C+1 | A/T-1 | TA>GC | TA>GC | CGCATAATA | ATAAGCC | TTA | TTA | T | G | T>G | T>G_TTA | TA>GC_ATAAGCC | 8 | FALSE | FALSE |
| 42_v.txt | 32294 | 32294-SNV-A | SNV | 1 | A | C | 103 | 28272 | 0.36431805 | 48.5048544 | rnr1Y285F | FALSE | CAN | JSY3811 | 1 | rnr1Y285F-R | AAT | A>C | G/C+1 | A/T-1 | TA>GC | TA>GC | ATTTGAATTT | ATTTGCG | TTA | TTA | T | G | T>G | T>G_TTA | TA>GC_ATTTGCG | 8 | FALSE | FALSE |
| 42_v.txt | 32472 | 32472-SNV-T | SNV | 1 | T | G | 13 | 28272 | 0.04598189 | 48.8461539 | rnr1Y285F | FALSE | CAN | JSY3811 | 1 | rnr1Y285F-R | GTA | T>G | G/C+1 | A/T-1 | TA>GC | TA>GC | CCAATGTAGA | TAGAAGG | CAT | GTA | T | C | T>G | T>G_GTA | TA>GC_TAGAAGG | 8 | FALSE | FALSE |
| 42_v.txt | 32493 | 32493-SNV-A | SNV | 1 | A | C | 6 | 28272 | 0.02122241 | 33.5 | rnr1Y285F | FALSE | CAN | JSY3811 | 1 | rnr1Y285F-R | AAC | A>C | G/C+1 | A/T-1 | TA>GC | TA>GC | CGGAAAACA | ACAACTT | TTG | TTG | T | G | T>G | T>G_TTG | TA>GC_ACAACTT | 8 | FALSE | FALSE |
| 42_v.txt | 32581 | 32581-SNV-T | SNV | 1 | T | G | 19 | 23846 | 0.07967793 | 34.1045923 | rnr1Y285F | FALSE | CAN | JSY3811 | 1 | rnr1Y285F-R | GTG | T>G | G/C+1 | A/T-1 | TA>GC | TA>GC | GAAATGTGA | TGAAGGC | CAC | GTG | T | C | T>G | T>G_GTG | TA>GC_TGAAGGC | 8 | FALSE | FALSE |
| 42_v.txt | 32631 | 32631-SNV-T | SNV | 1 | T | G | 2 | 23846 | 8.39E-03 | 64 | rnr1Y285F | FALSE | CAN | JSY3811 | 1 | rnr1Y285F-R | GTT | T>G | G/C+1 | A/T-1 | TA>GC | TA>GC | CCTTCGTTTT | TTTTTAT | CAA | GTT | T | C | T>G | T>G_GTT | TA>GC_TTTTTAT | 8 | FALSE | FALSE |
| 42_v.txt | 32763 | 32763-SNV-A | SNV | 1 | A | C | 18 | 23846 | 0.07548436 | 51 | rnr1Y285F | FALSE | CAN | JSY3811 | 1 | rnr1Y285F-R | AAC | A>C | G/C+1 | A/T-1 | TA>GC | TA>GC | GCTAAAACTT | ACTTTGA | TTG | TTG | T | G | T>G | T>G_TTG | TA>GC_ACTTTGA | 8 | FALSE | FALSE |
| 42_v.txt | 32801 | 32801-SNV-A | SNV | 1 | A | C | 14 | 23846 | 0.05871006 | 42.6428571 | rnr1Y285F | FALSE | CAN | JSY3811 | 1 | rnr1Y285F-R | AAT | A>C | G/C+1 | A/T-1 | TA>GC | TA>GC | ACCGTAATAT | ATATTTG | TTA | TTA | T | G | T>G | T>G_TTA | TA>GC_ATATTTG | 8 | FALSE | FALSE |
| 42_v.txt | 32842 | 32842-SNV-A | SNV | 1 | A | C | 3 | 23846 | 0.01258073 | 46.6661073 | rnr1Y285F | FALSE | CAN | JSY3811 | 1 | rnr1Y285F-R | CAA | A>C | G/C+1 | A/T-1 | TA>GC | TA>GC | TTACCCAAAA | AAAAAAT | GTT | GTT | T | G | T>G | T>G_GTT | TA>GC_AAAAAAT | 8 | FALSE | FALSE |
| 42_v.txt | 32884 | 32884-SNV-A | SNV | 1 | A | C | 4 | 33529 | 0.01192997 | 38 | rnr1Y285F | FALSE | CAN | JSY3811 | 1 | rnr1Y285F-R | CAA | A>C | G/C+1 | A/T-1 | TA>GC | TA>GC | ACGTCCAAAA | AAAATTG | GTT | GTT | T | G | T>G | T>G_GTT | TA>GC_AAAATTG | 8 | FALSE | FALSE |
| 42_v.txt | 32938 | 32938-SNV-A | SNV | 1 | A | C | 4 | 33529 | 0.01192997 | 19.7500005 | rnr1Y285F | FALSE | CAN | JSY3811 | 1 | rnr1Y285F-R | CAA | A>C | G/C+1 | A/T-1 | TA>GC | TA>GC | TTGCCCAAGA | AAGAAAA | GTT | GTT | T | G | T>G | T>G_GTT | TA>GC_AAGAAAA | 8 | FALSE | FALSE |
| 42_v.txt | 33089 | 33089-SNV-A | SNV | 1 | A | C | 21 | 33529 | 0.06263235 | 45.5228122 | rnr1Y285F | FALSE | CAN | JSY3811 | 1 | rnr1Y285F-R | AAT | A>C | G/C+1 | A/T-1 | TA>GC | TA>GC | AAATAAATAT | ATATGAT | TTA | TTA | T | G | T>G | T>G_TTA | TA>GC_ATATGAT | 8 | FALSE | FALSE |
| 42_v.txt | 33269 | 33269-SNV-A | SNV | 1 | A | C | 172 | 23943 | 0.7183728 | 16.2383721 | rnr1Y285F | FALSE | CAN | JSY3811 | 1 | rnr1Y285F-R | TAG | A>C | G/C+1 | A/T-1 | TA>GC | TA>GC | CTCCATAGAC | AGAGAAC | ATC | ATC | T | G | T>G | T>G_ATC | TA>GC_AGAGAAC | 8 | FALSE | FALSE |
| 42_v.txt | 32427 | 32427-SNV-T | SNV | 1 | T | C | 28 | 28272 | 0.09903792 | 53.2857143 | rnr1Y285F | FALSE | CAN | JSY3811 | 1 | rnr1Y285F-R | GTC | T>C | G/C+1 | A/T-1 | TA>CG | TA>CG | TTAGGGTCAT | TCATTGT | CAG | GTC | T | G | T>C | T>C_GTC | TA>CG_TCATTGT | 8 | FALSE | FALSE |
| 42_v.txt | 32476 | 32476-SNV-A | SNV | 1 | A | G | 28 | 28272 | 0.09903792 | 46.3571429 | rnr1Y285F | FALSE | CAN | JSY3811 | 1 | rnr1Y285F-R | AAG | A>G | G/C+1 | A/T-1 | TA>CG | TA>CG | TGTAGAAGG | AGGTTAA | TTC | TTC | T | C | T>C | T>C_TTC | TA>CG_AGGTTAA | 8 | FALSE | FALSE |
| 42_v.txt | 32498 | 32498-SNV-T | SNV | 1 | T | C | 5 | 28272 | 0.01768534 | 38.3999735 | rnr1Y285F | FALSE | CAN | JSY3811 | 1 | rnr1Y285F-R | CTT | T>C | G/C+1 | A/T-1 | TA>CG | TA>CG | AACAACTTTT | TTTTTTG | GAA | CTT | T | G | T>C | T>C_CTT | TA>CG_TTTTTTG | 8 | FALSE | FALSE |
| 42_v.txt | 32521 | 32521-SNV-A | SNV | 1 | A | G | 74 | 28272 | 0.26174307 | 40.8107634 | rnr1Y285F | FALSE | CAN | JSY3811 | 1 | rnr1Y285F-R | GAT | A>G | G/C+1 | A/T-1 | TA>CG | TA>CG | GAACGGATT | ATTTTCT | CTA | CTA | T | C | T>C | T>C_CTA | TA>CG_ATTTTCT | 8 | FALSE | FALSE |
| 42_v.txt | 32690 | 32690-SNV-T | SNV | 1 | T | C | 25 | 23846 | 0.10483939 | 60.7199234 | rnr1Y285F | FALSE | CAN | JSY3811 | 1 | rnr1Y285F-R | ATC | T>C | G/C+1 | A/T-1 | TA>CG | TA>CG | ACGGAATCCA | TCCAACT | TAG | ATC | T | G | T>C | T>C_ATC | TA>CG_TCCAACT | 8 | FALSE | FALSE |
| 42_v.txt | 32734 | 32734-SNV-A | SNV | 1 | A | G | 19 | 23846 | 0.07967793 | 59.4210526 | rnr1Y285F | FALSE | CAN | JSY3811 | 1 | rnr1Y285F-R | CAG | A>G | G/C+1 | A/T-1 | TA>CG | TA>CG | AAAAACAGT | AGTATAT | GTC | GTC | T | C | T>C | T>C_GTC | TA>CG_AGTATAT | 8 | FALSE | FALSE |
| 42_v.txt | 32938 | 32938-SNV-A | SNV | 1 | A | G | 23 | 33529 | 0.06859733 | 54.3912998 | rnr1Y285F | FALSE | CAN | JSY3811 | 1 | rnr1Y285F-R | CAA | A>G | G/C+1 | A/T-1 | TA>CG | TA>CG | TTGCCCAAGA | AAGAAAA | GTT | GTT | T | C | T>C | T>C_GTT | TA>CG_AAGAAAA | 8 | FALSE | FALSE |
| 42_v.txt | 33089 | 33089-SNV-A | SNV | 1 | A | G | 3 | 33529 | 8.95E-03 | 44.9995515 | rnr1Y285F | FALSE | CAN | JSY3811 | 1 | rnr1Y285F-R | AAT | A>G | G/C+1 | A/T-1 | TA>CG | TA>CG | AAATAAATAT | ATATGAT | TTA | TTA | T | C | T>C | T>C_TTA | TA>CG_ATATGAT | 8 | FALSE | FALSE |
| 42_v.txt | 33174 | 33174-SNV-A | SNV | 1 | A | G | 29 | 33518 | 0.08652068 | 35.0676178 | rnr1Y285F | FALSE | CAN | JSY3811 | 1 | rnr1Y285F-R | AAG | A>G | G/C+1 | A/T-1 | TA>CG | TA>CG | CCACCAAGG | AGGGCAA | TTC | TTC | T | C | T>C | T>C_TTC | TA>CG_AGGGCAA | 8 | FALSE | FALSE |
| 42_v.txt | 33476 | 33476-SNV-T | SNV | 1 | T | C | 34 | 23938 | 0.14203359 | 32.0886014 | rnr1Y285F | FALSE | CAN | JSY3811 | 1 | rnr1Y285F-R | CTT | T>C | G/C+1 | A/T-1 | TA>CG | TA>CG | TATGCCTTTT | TTTTTTT | GAA | CTT | T | G | T>C | T>C_CTT | TA>CG_TTTTTTT | 8 | FALSE | FALSE |
| 42_v.txt | 32128 | 32128-SNV-T | SNV | 1 | T | A | 62 | 23773 | 0.26080007 | 57.2903226 | rnr1Y285F | FALSE | CAN | JSY3811 | 1 | rnr1Y285F-R | TTG | T>A | A/T+1 | A/T-1 | TA>AT | TA>AT | AAACTTTGTC | TGTCACC | AAC | TTG | T | T | T>A | T>A_TTG | TA>AT_TGTCACC | 8 | FALSE | FALSE |
| 42_v.txt | 32255 | 32255-SNV-T | SNV | 1 | T | A | 42 | 28250 | 0.14867257 | 36.9047619 | rnr1Y285F | FALSE | CAN | JSY3811 | 1 | rnr1Y285F-R | ATA | T>A | A/T+1 | A/T-1 | TA>AT | TA>AT | CTTTGATAGA | TAGACCA | TAT | ATA | T | T | T>A | T>A_ATA | TA>AT_TAGACCA | 8 | FALSE | FALSE |
| 42_v.txt | 32337 | 32337-SNV-A | SNV | 1 | A | T | 6 | 28271 | 0.02122316 | 51.3309759 | rnr1Y285F | FALSE | CAN | JSY3811 | 1 | rnr1Y285F-R | GAT | A>T | A/T+1 | A/T-1 | TA>AT | TA>AT | TTGAAGATAT | ATATGTG | CTA | CTA | T | A | T>A | T>A_CTA | TA>AT_ATATGTG | 8 | FALSE | FALSE |
| 42_v.txt | 32338 | 32338-SNV-T | SNV | 1 | T | A | 4 | 28272 | 0.01414827 | 63.993936 | rnr1Y285F | FALSE | CAN | JSY3811 | 1 | rnr1Y285F-R | ATA | T>A | A/T+1 | A/T-1 | TA>AT | TA>AT | TGAAGATATC | TATGTGG | TAT | ATA | T | T | T>A | T>A_ATA | TA>AT_TATGTGG | 8 | FALSE | FALSE |
| 42_v.txt | 32498 | 32498-SNV-T | SNV | 1 | T | A | 1 | 28272 | 3.54E-03 | 37.9998399 | rnr1Y285F | FALSE | CAN | JSY3811 | 1 | rnr1Y285F-R | CTT | T>A | A/T+1 | A/T-1 | TA>AT | TA>AT | AACAACTTTT | TTTTTTG | GAA | CTT | T | T | T>A | T>A_CTT | TA>AT_TTTTTTG | 8 | FALSE | FALSE |
| 42_v.txt | 32690 | 32690-SNV-T | SNV | 1 | T | A | 5 | 23846 | 0.02096788 | 49.9990029 | rnr1Y285F | FALSE | CAN | JSY3811 | 1 | rnr1Y285F-R | ATC | T>A | A/T+1 | A/T-1 | TA>AT | TA>AT | ACGGAATCCA | TCCAACT | TAG | ATC | T | T | T>A | T>A_ATC | TA>AT_TCCAACT | 8 | FALSE | FALSE |
| 42_v.txt | 32765 | 32765-SNV-T | SNV | 1 | T | A | 45 | 23846 | 0.1887109 | 54.7333333 | rnr1Y285F | FALSE | CAN | JSY3811 | 1 | rnr1Y285F-R | CTT | T>A | A/T+1 | A/T-1 | TA>AT | TA>AT | TAAAACTTTG | TTTGATG | GAA | CTT | T | T | T>A | T>A_CTT | TA>AT_TTTGATG | 8 | FALSE | FALSE |
| 42_v.txt | 32938 | 32938-SNV-A | SNV | 1 | A | T | 23 | 33529 | 0.06859733 | 53.4782576 | rnr1Y285F | FALSE | CAN | JSY3811 | 1 | rnr1Y285F-R | CAA | A>T | A/T+1 | A/T-1 | TA>AT | TA>AT | TTGCCCAAGA | AAGAAAA | GTT | GTT | T | A | T>A | T>A_GTT | TA>AT_AAGAAAA | 8 | FALSE | FALSE |
| 42_v.txt | 33067 | 33067-SNV-A | SNV | 1 | A | T | 1516 | 33529 | 4.52145904 | 55.7612137 | rnr1Y285F | FALSE | CAN | JSY3811 | 1 | rnr1Y285F-R | TAT | A>T | A/T+1 | A/T-1 | TA>AT | TA>AT | CAGAATATGC | ATGCCAA | ATA | ATA | T | A | T>A | T>A_ATA | TA>AT_ATGCCAA | 8 | FALSE | FALSE |
| 42_v.txt | 33090 | 33090-SNV-T | SNV | 1 | T | A | 8 | 33529 | 0.02385994 | 37.1226895 | rnr1Y285F | FALSE | CAN | JSY3811 | 1 | rnr1Y285F-R | ATA | T>A | A/T+1 | A/T-1 | TA>AT | TA>AT | AATAAATATC | TATGATA | TAT | ATA | T | T | T>A | T>A_ATA | TA>AT_TATGATA | 8 | FALSE | FALSE |
| 42_v.txt | 31650 | 31650-SNV-C | SNV | 1 | C | T | 28 | 11596 | 0.24146257 | 35.8928571 | rnr1Y285F | FALSE | CAN | JSY3811 | 1 | rnr1Y285F-R | GCG | C>T | A/T+1 | G/C-1 | CG>TA | CG>TA | AGAATGCGA | CGAAATG | CGC | GCG | C | A | C>T | C>T_GCG | CG>TA_CGAAATG | 8 | FALSE | FALSE |
| 42_v.txt | 31844 | 31844-SNV-C | SNV | 1 | C | T | 18 | 11596 | 0.15522594 | 48.6525071 | rnr1Y285F | FALSE | CAN | JSY3811 | 1 | rnr1Y285F-R | TCC | C>T | A/T+1 | G/C-1 | CG>TA | CG>TA | TAAGATCCAA | CCAAACA | AGG | TCC | C | A | C>T | C>T_TCC | CG>TA_CCAAACA | 8 | FALSE | FALSE |
| 42_v.txt | 31845 | 31845-SNV-C | SNV | 1 | C | T | 15 | 11596 | 0.12935495 | 53.1049606 | rnr1Y285F | FALSE | CAN | JSY3811 | 1 | rnr1Y285F-R | CCA | C>T | A/T+1 | G/C-1 | CG>TA | CG>TA | AAGATCCAA | CAAACAG | GGT | CCA | C | A | C>T | C>T_CCA | CG>TA_CAAACAG | 8 | FALSE | FALSE |
| 42_v.txt | 31881 | 31881-SNV-G | SNV | 1 | G | A | 12 | 11596 | 0.10348396 | 38.9983157 | rnr1Y285F | FALSE | CAN | JSY3811 | 1 | rnr1Y285F-R | GGC | G>A | A/T+1 | G/C-1 | CG>TA | CG>TA | TAGGCGGCA | GCAGCAA | CCG | CCG | C | T | C>T | C>T_CCG | CG>TA_GCAGCAA | 8 | FALSE | FALSE |
| 42_v.txt | 31882 | 31882-SNV-C | SNV | 1 | C | T | 13 | 11596 | 0.11210762 | 49.5714974 | rnr1Y285F | FALSE | CAN | JSY3811 | 1 | rnr1Y285F-R | GCA | C>T | A/T+1 | G/C-1 | CG>TA | CG>TA | AGGCGGCAG | CAGCAAA | CGT | GCA | C | A | C>T | C>T_GCA | CG>TA_CAGCAAA | 8 | FALSE | FALSE |
| 42_v.txt | 31960 | 31960-SNV-C | SNV | 1 | C | T | 23 | 23773 | 0.09674841 | 36.9130435 | rnr1Y285F | FALSE | CAN | JSY3811 | 1 | rnr1Y285F-R | GCC | C>T | A/T+1 | G/C-1 | CG>TA | CG>TA | ATGTGGCCG | CCGCATA | CGG | GCC | C | A | C>T | C>T_GCC | CG>TA_CCGCATA | 8 | FALSE | FALSE |
| 42_v.txt | 31962 | 31962-SNV-G | SNV | 1 | G | A | 49 | 23773 | 0.20611618 | 36.0204082 | rnr1Y285F | FALSE | CAN | JSY3811 | 1 | rnr1Y285F-R | CGC | G>A | A/T+1 | G/C-1 | CG>TA | CG>TA | GTGGCCGCAT | GCATAAT | GCG | GCG | C | T | C>T | C>T_GCG | CG>TA_GCATAAT | 8 | FALSE | FALSE |
| 42_v.txt | 31981 | 31981-SNV-G | SNV | 1 | G | A | 133 | 23773 | 0.55945821 | 46.593985 | rnr1Y285F | FALSE | CAN | JSY3811 | 1 | rnr1Y285F-R | GGC | G>A | A/T+1 | G/C-1 | CG>TA | CG>TA | AGCCGGGCA | GCATTAA | CCG | CCG | C | T | C>T | C>T_CCG | CG>TA_GCATTAA | 8 | FALSE | FALSE |
| 42_v.txt | 32022 | 32022-SNV-C | SNV | 1 | C | T | 19 | 23773 | 0.0799226 | 55.7894737 | rnr1Y285F | FALSE | CAN | JSY3811 | 1 | rnr1Y285F-R | GCC | C>T | A/T+1 | G/C-1 | CG>TA | CG>TA | GAGATGCCA | CCACGGT | CGG | GCC | C | A | C>T | C>T_GCC | CG>TA_CCACGGT | 8 | FALSE | FALSE |
| 42_v.txt | 32041 | 32041-SNV-G | SNV | 1 | G | A | 49 | 23773 | 0.20611618 | 59.0612245 | rnr1Y285F | FALSE | CAN | JSY3811 | 1 | rnr1Y285F-R | TGC | G>A | A/T+1 | G/C-1 | CG>TA | CG>TA | AAGCTTGCAT | GCATAAA | ACG | ACG | C | T | C>T | C>T_ACG | CG>TA_GCATAAA | 8 | FALSE | FALSE |
| 42_v.txt | 32057 | 32057-SNV-C | SNV | 1 | C | T | 32 | 23773 | 0.13460649 | 57.625 | rnr1Y285F | FALSE | CAN | JSY3811 | 1 | rnr1Y285F-R | GCG | C>T | A/T+1 | G/C-1 | CG>TA | CG>TA | GATGTGCGA | CGAGATT | CGC | GCG | C | A | C>T | C>T_GCG | CG>TA_CGAGATT | 8 | FALSE | FALSE |
| 42_v.txt | 32075 | 32075-SNV-C | SNV | 1 | C | T | 70 | 23773 | 0.29445169 | 62.8142857 | rnr1Y285F | FALSE | CAN | JSY3811 | 1 | rnr1Y285F-R | ACC | C>T | A/T+1 | G/C-1 | CG>TA | CG>TA | AAATAACCAT | CCATGCA | TGG | ACC | C | A | C>T | C>T_ACC | CG>TA_CCATGCA | 8 | FALSE | FALSE |
| 42_v.txt | 32088 | 32088-SNV-C | SNV | 1 | C | T | 329 | 23773 | 1.38392294 | 59.9027356 | rnr1Y285F | FALSE | CAN | JSY3811 | 1 | rnr1Y285F-R | GCC | C>T | A/T+1 | G/C-1 | CG>TA | CG>TA | AAAAAGCCTC | CCTGCAA | CGG | GCC | C | A | C>T | C>T_GCC | CG>TA_CCTGCAA | 8 | FALSE | FALSE |
| 42_v.txt | 32092 | 32092-SNV-C | SNV | 1 | C | T | 32 | 23773 | 0.13460649 | 57.59375 | rnr1Y285F | FALSE | CAN | JSY3811 | 1 | rnr1Y285F-R | GCA | C>T | A/T+1 | G/C-1 | CG>TA | CG>TA | AGCCTGCAAC | CAACACC | CGT | GCA | C | A | C>T | C>T_GCA | CG>TA_CAACACC | 8 | FALSE | FALSE |
| 42_v.txt | 32114 | 32114-SNV-C | SNV | 1 | C | T | 84 | 23773 | 0.35334203 | 59.6666667 | rnr1Y285F | FALSE | CAN | JSY3811 | 1 | rnr1Y285F-R | GCC | C>T | A/T+1 | G/C-1 | CG>TA | CG>TA | TAATAGCCAT | CCATTCG | CGG | GCC | C | A | C>T | C>T_GCC | CG>TA_CCATTCG | 8 | FALSE | FALSE |
| 42_v.txt | 32165 | 32165-SNV-G | SNV | 1 | G | A | 43 | 23773 | 0.18087747 | 46.6733093 | rnr1Y285F | FALSE | CAN | JSY3811 | 1 | rnr1Y285F-R | CGC | G>A | A/T+1 | G/C-1 | CG>TA | CG>TA | CAAAGCGCCA | GCCAAAT | GCG | GCG | C | T | C>T | C>T_GCG | CG>TA_GCCAAAT | 8 | FALSE | FALSE |
| 42_v.txt | 32166 | 32166-SNV-C | SNV | 1 | C | T | 294 | 23773 | 1.23669709 | 46.1058274 | rnr1Y285F | FALSE | CAN | JSY3811 | 1 | rnr1Y285F-R | GCC | C>T | A/T+1 | G/C-1 | CG>TA | CG>TA | AAAGCGCCA | CCAAATG | CGG | GCC | C | A | C>T | C>T_GCC | CG>TA_CCAAATG | 8 | FALSE | FALSE |
| 42_v.txt | 32173 | 32173-SNV-C | SNV | 1 | C | T | 21 | 23773 | 0.08833551 | 50.2857143 | rnr1Y285F | FALSE | CAN | JSY3811 | 1 | rnr1Y285F-R | GCA | C>T | A/T+1 | G/C-1 | CG>TA | CG>TA | CAAATGCAG | CAGCAGT | CGT | GCA | C | A | C>T | C>T_GCA | CG>TA_CAGCAGT | 8 | FALSE | FALSE |
| 42_v.txt | 32176 | 32176-SNV-C | SNV | 1 | C | T | 22 | 23773 | 0.09254196 | 45.3181818 | rnr1Y285F | FALSE | CAN | JSY3811 | 1 | rnr1Y285F-R | GCA | C>T | A/T+1 | G/C-1 | CG>TA | CG>TA | ATGCAGCAG | CAGTAAC | CGT | GCA | C | A | C>T | C>T_GCA | CG>TA_CAGTAAC | 8 | FALSE | FALSE |
| 42_v.txt | 32271 | 32271-SNV-C | SNV | 1 | C | T | 46 | 28272 | 0.16270515 | 30.2164817 | rnr1Y285F | FALSE | CAN | JSY3811 | 1 | rnr1Y285F-R | ACG | C>T | A/T+1 | G/C-1 | CG>TA | CG>TA | AAAATACGG | CGGGAAC | TGC | ACG | C | A | C>T | C>T_ACG | CG>TA_CGGGAAC | 8 | FALSE | FALSE |
| 42_v.txt | 32274 | 32274-SNV-G | SNV | 1 | G | A | 101 | 28272 | 0.35724392 | 35.8316832 | rnr1Y285F | FALSE | CAN | JSY3811 | 1 | rnr1Y285F-R | GGA | G>A | A/T+1 | G/C-1 | CG>TA | CG>TA | ATACGGGAA | GAACCAA | CCT | CCT | C | T | C>T | C>T_CCT | CG>TA_GAACCAA | 8 | FALSE | FALSE |
| 42_v.txt | 32299 | 32299-SNV-C | SNV | 1 | C | T | 70 | 28272 | 0.24759479 | 53.3285714 | rnr1Y285F | FALSE | CAN | JSY3811 | 1 | rnr1Y285F-R | GCG | C>T | A/T+1 | G/C-1 | CG>TA | CG>TA | AATTTGCGGC | CGGCAGA | CGC | GCG | C | A | C>T | C>T_GCG | CG>TA_CGGCAGA | 8 | FALSE | FALSE |
| 42_v.txt | 32449 | 32449-SNV-C | SNV | 1 | C | T | 602 | 28272 | 2.12931522 | 52.5830565 | rnr1Y285F | FALSE | CAN | JSY3811 | 1 | rnr1Y285F-R | CCA | C>T | A/T+1 | G/C-1 | CG>TA | CG>TA | AAAGTCCAAT | CAATGAA | GGT | CCA | C | A | C>T | C>T_CCA | CG>TA_CAATGAA | 8 | FALSE | FALSE |
| 42_v.txt | 32466 | 32466-SNV-C | SNV | 1 | C | T | 53 | 28272 | 0.18746463 | 43.1320755 | rnr1Y285F | FALSE | CAN | JSY3811 | 1 | rnr1Y285F-R | GCC | C>T | A/T+1 | G/C-1 | CG>TA | CG>TA | AGAGAGCCA | CCAATGT | CGG | GCC | C | A | C>T | C>T_GCC | CG>TA_CCAATGT | 8 | FALSE | FALSE |
| 42_v.txt | 32535 | 32535-SNV-G | SNV | 1 | G | A | 33 | 28272 | 0.11672326 | 37.9393975 | rnr1Y285F | FALSE | CAN | JSY3811 | 1 | rnr1Y285F-R | TGC | G>A | A/T+1 | G/C-1 | CG>TA | CG>TA | GGGTTTGCAC | GCAGCTT | ACG | ACG | C | T | C>T | C>T_ACG | CG>TA_GCAGCTT | 8 | FALSE | FALSE |
| 42_v.txt | 32536 | 32536-SNV-C | SNV | 1 | C | T | 37 | 28272 | 0.13087153 | 37.1080278 | rnr1Y285F | FALSE | CAN | JSY3811 | 1 | rnr1Y285F-R | GCA | C>T | A/T+1 | G/C-1 | CG>TA | CG>TA | GGTTTGCAGC | CAGCTTC | CGT | GCA | C | A | C>T | C>T_GCA | CG>TA_CAGCTTC | 8 | FALSE | FALSE |
| 42_v.txt | 32545 | 32545-SNV-C | SNV | 1 | C | T | 160 | 28272 | 0.56593096 | 37.6551544 | rnr1Y285F | FALSE | CAN | JSY3811 | 1 | rnr1Y285F-R | CCA | C>T | A/T+1 | G/C-1 | CG>TA | CG>TA | CTTCACCAGC | CAGCAGT | GGT | CCA | C | A | C>T | C>T_CCA | CG>TA_CAGCAGT | 8 | FALSE | FALSE |
| 42_v.txt | 32568 | 32568-SNV-G | SNV | 1 | G | A | 258 | 23846 | 1.08194246 | 37.5736434 | rnr1Y285F | FALSE | CAN | JSY3811 | 1 | rnr1Y285F-R | AGT | G>A | A/T+1 | G/C-1 | CG>TA | CG>TA | AGTTCAGTAC | GTACCTT | TCA | TCA | C | T | C>T | C>T_TCA | CG>TA_GTACCTT | 8 | FALSE | FALSE |
| 42_v.txt | 32580 | 32580-SNV-G | SNV | 1 | G | A | 59 | 23846 | 0.24742095 | 37.1186441 | rnr1Y285F | FALSE | CAN | JSY3811 | 1 | rnr1Y285F-R | TGT | G>A | A/T+1 | G/C-1 | CG>TA | CG>TA | TGAAATGTGA | GTGAAGG | ACA | ACA | C | T | C>T | C>T_ACA | CG>TA_GTGAAGG | 8 | FALSE | FALSE |
| 42_v.txt | 32590 | 32590-SNV-C | SNV | 1 | C | T | 48 | 23846 | 0.20129162 | 36.8958333 | rnr1Y285F | FALSE | CAN | JSY3811 | 1 | rnr1Y285F-R | GCG | C>T | A/T+1 | G/C-1 | CG>TA | CG>TA | AGGCAGCGT | CGTTAAT | CGC | GCG | C | A | C>T | C>T_GCG | CG>TA_CGTTAAT | 8 | FALSE | FALSE |
| 42_v.txt | 32660 | 32660-SNV-C | SNV | 1 | C | T | 2661 | 23846 | 11.1591043 | 56.6163097 | rnr1Y285F | FALSE | CAN | JSY3811 | 1 | rnr1Y285F-R | CCC | C>T | A/T+1 | G/C-1 | CG>TA | CG>TA | TGGACCCCAC | CCAGGCA | GGG | CCC | C | A | C>T | C>T_CCC | CG>TA_CCAGGCA | 8 | FALSE | FALSE |
| 42_v.txt | 32679 | 32679-SNV-C | SNV | 1 | C | T | 57 | 23846 | 0.2390338 | 57.2982456 | rnr1Y285F | FALSE | CAN | JSY3811 | 1 | rnr1Y285F-R | CCA | C>T | A/T+1 | G/C-1 | CG>TA | CG>TA | TTTCTCCAAT | CAATAAC | GGT | CCA | C | A | C>T | C>T_CCA | CG>TA_CAATAAC | 8 | FALSE | FALSE |
| 42_v.txt | 32691 | 32691-SNV-C | SNV | 1 | C | T | 7 | 23846 | 0.02935503 | 57.5715511 | rnr1Y285F | FALSE | CAN | JSY3811 | 1 | rnr1Y285F-R | TCC | C>T | A/T+1 | G/C-1 | CG>TA | CG>TA | CGGAATCCA | CCAACTG | AGG | TCC | C | A | C>T | C>T_TCC | CG>TA_CCAACTG | 8 | FALSE | FALSE |
| 42_v.txt | 32700 | 32700-SNV-C | SNV | 1 | C | T | 22 | 23846 | 0.09225866 | 60.9090909 | rnr1Y285F | FALSE | CAN | JSY3811 | 1 | rnr1Y285F-R | GCC | C>T | A/T+1 | G/C-1 | CG>TA | CG>TA | ACTGGGCCG | CCGGTAA | CGG | GCC | C | A | C>T | C>T_GCC | CG>TA_CCGGTAA | 8 | FALSE | FALSE |
| 42_v.txt | 32757 | 32757-SNV-G | SNV | 1 | G | A | 27 | 23846 | 0.11322654 | 49 | rnr1Y285F | FALSE | CAN | JSY3811 | 1 | rnr1Y285F-R | GGC | G>A | A/T+1 | G/C-1 | CG>TA | CG>TA | ATAATGGCTA | GCTAAAA | CCG | CCG | C | T | C>T | C>T_CCG | CG>TA_GCTAAAA | 8 | FALSE | FALSE |
| 42_v.txt | 32776 | 32776-SNV-C | SNV | 1 | C | T | 29 | 23846 | 0.12161369 | 46.6896552 | rnr1Y285F | FALSE | CAN | JSY3811 | 1 | rnr1Y285F-R | GCG | C>T | A/T+1 | G/C-1 | CG>TA | CG>TA | TGGAAGCGA | CGACCCA | CGC | GCG | C | A | C>T | C>T_GCG | CG>TA_CGACCCA | 8 | FALSE | FALSE |
| 42_v.txt | 32798 | 32798-SNV-G | SNV | 1 | G | A | 35 | 23846 | 0.14677514 | 43 | rnr1Y285F | FALSE | CAN | JSY3811 | 1 | rnr1Y285F-R | CGT | G>A | A/T+1 | G/C-1 | CG>TA | CG>TA | TTCACCGTAA | GTAATAT | GCA | GCA | C | T | C>T | C>T_GCA | CG>TA_GTAATAT | 8 | FALSE | FALSE |
| 42_v.txt | 32809 | 32809-SNV-C | SNV | 1 | C | T | 16 | 23846 | 0.06709721 | 44.1875 | rnr1Y285F | FALSE | CAN | JSY3811 | 1 | rnr1Y285F-R | ACA | C>T | A/T+1 | G/C-1 | CG>TA | CG>TA | ATTTGACAGC | CAGGGAA | TGT | ACA | C | A | C>T | C>T_ACA | CG>TA_CAGGGAA | 8 | FALSE | FALSE |
| 42_v.txt | 32855 | 32855-SNV-C | SNV | 1 | C | T | 81 | 23846 | 0.33967961 | 35.4074074 | rnr1Y285F | FALSE | CAN | JSY3811 | 1 | rnr1Y285F-R | TCC | C>T | A/T+1 | G/C-1 | CG>TA | CG>TA | ACTAATCCAT | CCATGCC | AGG | TCC | C | A | C>T | C>T_TCC | CG>TA_CCATGCC | 8 | FALSE | FALSE |
| 42_v.txt | 32860 | 32860-SNV-C | SNV | 1 | C | T | 28 | 23844 | 0.11742996 | 37.0357143 | rnr1Y285F | FALSE | CAN | JSY3811 | 1 | rnr1Y285F-R | GCC | C>T | A/T+1 | G/C-1 | CG>TA | CG>TA | TCCATGCCGC | CCGCCAG | CGG | GCC | C | A | C>T | C>T_GCC | CG>TA_CCGCCAG | 8 | FALSE | FALSE |
| 42_v.txt | 32862 | 32862-SNV-G | SNV | 1 | G | A | 25 | 23844 | 0.10484818 | 37.2 | rnr1Y285F | FALSE | CAN | JSY3811 | 1 | rnr1Y285F-R | CGC | G>A | A/T+1 | G/C-1 | CG>TA | CG>TA | CATGCCGCCA | GCCAGTG | GCG | GCG | C | T | C>T | C>T_GCG | CG>TA_GCCAGTG | 8 | FALSE | FALSE |
| 42_v.txt | 32882 | 32882-SNV-C | SNV | 1 | C | T | 50 | 33527 | 0.14913353 | 26.46 | rnr1Y285F | FALSE | CAN | JSY3811 | 1 | rnr1Y285F-R | TCC | C>T | A/T+1 | G/C-1 | CG>TA | CG>TA | GTACGTCCAA | CCAAAAT | AGG | TCC | C | A | C>T | C>T_TCC | CG>TA_CCAAAAT | 8 | FALSE | FALSE |
| 42_v.txt | 32900 | 32900-SNV-G | SNV | 1 | G | A | 47 | 33529 | 0.14017716 | 35.2683788 | rnr1Y285F | FALSE | CAN | JSY3811 | 1 | rnr1Y285F-R | GGC | G>A | A/T+1 | G/C-1 | CG>TA | CG>TA | GACTTGGCCA | GCCAACT | CCG | CCG | C | T | C>T | C>T_CCG | CG>TA_GCCAACT | 8 | FALSE | FALSE |
| 42_v.txt | 32901 | 32901-SNV-C | SNV | 1 | C | T | 69 | 33529 | 0.205792 | 30.0004688 | rnr1Y285F | FALSE | CAN | JSY3811 | 1 | rnr1Y285F-R | GCC | C>T | A/T+1 | G/C-1 | CG>TA | CG>TA | ACTTGGCCAA | CCAACTA | CGG | GCC | C | A | C>T | C>T_GCC | CG>TA_CCAACTA | 8 | FALSE | FALSE |
| 42_v.txt | 32917 | 32917-SNV-C | SNV | 1 | C | T | 29 | 33529 | 0.08649229 | 39.9655172 | rnr1Y285F | FALSE | CAN | JSY3811 | 1 | rnr1Y285F-R | TCC | C>T | A/T+1 | G/C-1 | CG>TA | CG>TA | TAAGTTCCAC | CCAGGGC | AGG | TCC | C | A | C>T | C>T_TCC | CG>TA_CCAGGGC | 8 | FALSE | FALSE |
| 42_v.txt | 32935 | 32935-SNV-C | SNV | 1 | C | T | 53 | 33529 | 0.15807212 | 44.5849057 | rnr1Y285F | FALSE | CAN | JSY3811 | 1 | rnr1Y285F-R | GCC | C>T | A/T+1 | G/C-1 | CG>TA | CG>TA | TGATTGCCCA | CCCAAGA | CGG | GCC | C | A | C>T | C>T_GCC | CG>TA_CCCAAGA | 8 | FALSE | FALSE |
| 42_v.txt | 32937 | 32937-SNV-C | SNV | 1 | C | T | 83 | 33529 | 0.2475469 | 43.1903008 | rnr1Y285F | FALSE | CAN | JSY3811 | 1 | rnr1Y285F-R | CCA | C>T | A/T+1 | G/C-1 | CG>TA | CG>TA | ATTGCCCAAC | CAAGAAA | GGT | CCA | C | A | C>T | C>T_CCA | CG>TA_CAAGAAA | 8 | FALSE | FALSE |
| 42_v.txt | 32945 | 32945-SNV-C | SNV | 1 | C | T | 48 | 33529 | 0.14315965 | 54.6666667 | rnr1Y285F | FALSE | CAN | JSY3811 | 1 | rnr1Y285F-R | ACC | C>T | A/T+1 | G/C-1 | CG>TA | CG>TA | AGAAAACCA | CCAATAC | TGG | ACC | C | A | C>T | C>T_ACC | CG>TA_CCAATAC | 8 | FALSE | FALSE |
| 42_v.txt | 32958 | 32958-SNV-C | SNV | 1 | C | T | 97 | 33529 | 0.2893018 | 52.7010309 | rnr1Y285F | FALSE | CAN | JSY3811 | 1 | rnr1Y285F-R | ACC | C>T | A/T+1 | G/C-1 | CG>TA | CG>TA | ATGTAACCAT | CCATTGG | TGG | ACC | C | A | C>T | C>T_ACC | CG>TA_CCATTGG | 8 | FALSE | FALSE |
| 42_v.txt | 32965 | 32965-SNV-C | SNV | 1 | C | T | 46 | 33529 | 0.13719467 | 54.7173913 | rnr1Y285F | FALSE | CAN | JSY3811 | 1 | rnr1Y285F-R | GCC | C>T | A/T+1 | G/C-1 | CG>TA | CG>TA | CATTGGCCGC | CCGCACC | CGG | GCC | C | A | C>T | C>T_GCC | CG>TA_CCGCACC | 8 | FALSE | FALSE |
| 42_v.txt | 32967 | 32967-SNV-G | SNV | 1 | G | A | 81 | 33529 | 0.24158191 | 59.3950617 | rnr1Y285F | FALSE | CAN | JSY3811 | 1 | rnr1Y285F-R | CGC | G>A | A/T+1 | G/C-1 | CG>TA | CG>TA | TTGGCCGCAC | GCACCAA | GCG | GCG | C | T | C>T | C>T_GCG | CG>TA_GCACCAA | 8 | FALSE | FALSE |
| 42_v.txt | 32995 | 32995-SNV-G | SNV | 1 | G | A | 33 | 33529 | 0.09842226 | 56.8484849 | rnr1Y285F | FALSE | CAN | JSY3811 | 1 | rnr1Y285F-R | TGT | G>A | A/T+1 | G/C-1 | CG>TA | CG>TA | ATCTTTGTGA | GTGAGAA | ACA | ACA | C | T | C>T | C>T_ACA | CG>TA_GTGAGAA | 8 | FALSE | FALSE |
| 42_v.txt | 33056 | 33056-SNV-C | SNV | 1 | C | T | 42 | 33529 | 0.1252647 | 55.1428571 | rnr1Y285F | FALSE | CAN | JSY3811 | 1 | rnr1Y285F-R | GCG | C>T | A/T+1 | G/C-1 | CG>TA | CG>TA | GGACTGCGTC | CGTGACA | CGC | GCG | C | A | C>T | C>T_GCG | CG>TA_CGTGACA | 8 | FALSE | FALSE |
| 42_v.txt | 33070 | 33070-SNV-C | SNV | 1 | C | T | 41 | 33529 | 0.1222822 | 54.8292683 | rnr1Y285F | FALSE | CAN | JSY3811 | 1 | rnr1Y285F-R | GCC | C>T | A/T+1 | G/C-1 | CG>TA | CG>TA | AATATGCCAA | CCAAAGA | CGG | GCC | C | A | C>T | C>T_GCC | CG>TA_CCAAAGA | 8 | FALSE | FALSE |
| 42_v.txt | 33114 | 33114-SNV-C | SNV | 1 | C | T | 41 | 33529 | 0.1222822 | 41.4395715 | rnr1Y285F | FALSE | CAN | JSY3811 | 1 | rnr1Y285F-R | GCC | C>T | A/T+1 | G/C-1 | CG>TA | CG>TA | ACTGGGCCG | CCGGCGT | CGG | GCC | C | A | C>T | C>T_GCC | CG>TA_CCGGCGT | 8 | FALSE | FALSE |
| 42_v.txt | 33115 | 33115-SNV-C | SNV | 1 | C | T | 38 | 33529 | 0.11333473 | 43.4734207 | rnr1Y285F | FALSE | CAN | JSY3811 | 1 | rnr1Y285F-R | CCG | C>T | A/T+1 | G/C-1 | CG>TA | CG>TA | CTGGGCCGG | CGGCGTT | GGC | CCG | C | A | C>T | C>T_CCG | CG>TA_CGGCGTT | 8 | FALSE | FALSE |
| 42_v.txt | 33118 | 33118-SNV-C | SNV | 1 | C | T | 34 | 33529 | 0.10140475 | 41.3797447 | rnr1Y285F | FALSE | CAN | JSY3811 | 1 | rnr1Y285F-R | GCG | C>T | A/T+1 | G/C-1 | CG>TA | CG>TA | GGCCGGCGT | CGTTGGT | CGC | GCG | C | A | C>T | C>T_GCG | CG>TA_CGTTGGT | 8 | FALSE | FALSE |
| 42_v.txt | 33119 | 33119-SNV-G | SNV | 1 | G | A | 24 | 33529 | 0.07157983 | 47.124491 | rnr1Y285F | FALSE | CAN | JSY3811 | 1 | rnr1Y285F-R | CGT | G>A | A/T+1 | G/C-1 | CG>TA | CG>TA | GCCGGCGTTC | GTTGGTC | GCA | GCA | C | T | C>T | C>T_GCA | CG>TA_GTTGGTC | 8 | FALSE | FALSE |
| 42_v.txt | 33153 | 33153-SNV-C | SNV | 1 | C | T | 19 | 33529 | 0.05666736 | 33.5263697 | rnr1Y285F | FALSE | CAN | JSY3811 | 1 | rnr1Y285F-R | ACC | C>T | A/T+1 | G/C-1 | CG>TA | CG>TA | AAAAGACCTC | CCTGTAC | TGG | ACC | C | A | C>T | C>T_ACC | CG>TA_CCTGTAC | 8 | FALSE | FALSE |
| 42_v.txt | 33156 | 33156-SNV-G | SNV | 1 | G | A | 38 | 33528 | 0.11333811 | 36.5263158 | rnr1Y285F | FALSE | CAN | JSY3811 | 1 | rnr1Y285F-R | TGT | G>A | A/T+1 | G/C-1 | CG>TA | CG>TA | AGACCTGTAC | GTACCAA | ACA | ACA | C | T | C>T | C>T_ACA | CG>TA_GTACCAA | 8 | FALSE | FALSE |
| 42_v.txt | 33159 | 33159-SNV-C | SNV | 1 | C | T | 53 | 33527 | 0.15808155 | 36.9811321 | rnr1Y285F | FALSE | CAN | JSY3811 | 1 | rnr1Y285F-R | ACC | C>T | A/T+1 | G/C-1 | CG>TA | CG>TA | CCTGTACCAA | CCAATAG | TGG | ACC | C | A | C>T | C>T_ACC | CG>TA_CCAATAG | 8 | FALSE | FALSE |
| 42_v.txt | 33177 | 33177-SNV-G | SNV | 1 | G | A | 39 | 33518 | 0.11635539 | 36.5641026 | rnr1Y285F | FALSE | CAN | JSY3811 | 1 | rnr1Y285F-R | GGC | G>A | A/T+1 | G/C-1 | CG>TA | CG>TA | CCAAGGGCA | GCAATCA | CCG | CCG | C | T | C>T | C>T_CCG | CG>TA_GCAATCA | 8 | FALSE | FALSE |
| 42_v.txt | 33258 | 33258-SNV-C | SNV | 1 | C | T | 30 | 23943 | 0.12529758 | 44.8333333 | rnr1Y285F | FALSE | CAN | JSY3811 | 1 | rnr1Y285F-R | GCC | C>T | A/T+1 | G/C-1 | CG>TA | CG>TA | CCTATGCCAT | CCATCCT | CGG | GCC | C | A | C>T | C>T_GCC | CG>TA_CCATCCT | 8 | FALSE | FALSE |
| 42_v.txt | 33276 | 33276-SNV-G | SNV | 1 | G | A | 51 | 23943 | 0.21300589 | 44.8235294 | rnr1Y285F | FALSE | CAN | JSY3811 | 1 | rnr1Y285F-R | CGT | G>A | A/T+1 | G/C-1 | CG>TA | CG>TA | GAGAACGTA | GTATCCT | GCA | GCA | C | T | C>T | C>T_GCA | CG>TA_GTATCCT | 8 | FALSE | FALSE |
| 42_v.txt | 33309 | 33309-SNV-C | SNV | 1 | C | T | 29 | 23942 | 0.12112606 | 55.3705666 | rnr1Y285F | FALSE | CAN | JSY3811 | 1 | rnr1Y285F-R | GCG | C>T | A/T+1 | G/C-1 | CG>TA | CG>TA | AAAGAGCGC | CGCAATG | CGC | GCG | C | A | C>T | C>T_GCG | CG>TA_CGCAATG | 8 | FALSE | FALSE |
| 42_v.txt | 33310 | 33310-SNV-G | SNV | 1 | G | A | 32 | 23938 | 0.13367867 | 52.958328 | rnr1Y285F | FALSE | CAN | JSY3811 | 1 | rnr1Y285F-R | CGC | G>A | A/T+1 | G/C-1 | CG>TA | CG>TA | AAGAGCGCA | GCAATGG | GCG | GCG | C | T | C>T | C>T_GCG | CG>TA_GCAATGG | 8 | FALSE | FALSE |
| 42_v.txt | 33412 | 33412-SNV-G | SNV | 1 | G | A | 35 | 23943 | 0.14618051 | 46.6285714 | rnr1Y285F | FALSE | CAN | JSY3811 | 1 | rnr1Y285F-R | GGC | G>A | A/T+1 | G/C-1 | CG>TA | CG>TA | TGACCGGCTC | GCTCATT | CCG | CCG | C | T | C>T | C>T_CCG | CG>TA_GCTCATT | 8 | FALSE | FALSE |
| 42_v.txt | 33474 | 33474-SNV-C | SNV | 1 | C | T | 27 | 23940 | 0.11278196 | 33.3334965 | rnr1Y285F | FALSE | CAN | JSY3811 | 1 | rnr1Y285F-R | GCC | C>T | A/T+1 | G/C-1 | CG>TA | CG>TA | GCTATGCCTT | CCTTTTT | CGG | GCC | C | A | C>T | C>T_GCC | CG>TA_CCTTTTT | 8 | FALSE | FALSE |
| 42_v.txt | 31844 | 31844-SNV-C | SNV | 1 | C | G | 1 | 11596 | 8.62E-03 | 63.9784689 | rnr1Y285F | FALSE | CAN | JSY3811 | 1 | rnr1Y285F-R | TCC | C>G | G/C+1 | G/C-1 | CG>GC | CG>GC | TAAGATCCAA | CCAAACA | AGG | TCC | C | C | C>G | C>G_TCC | CG>GC_CCAAACA | 8 | FALSE | FALSE |
| 42_v.txt | 31881 | 31881-SNV-G | SNV | 1 | G | C | 1 | 11596 | 8.62E-03 | 22.9994442 | rnr1Y285F | FALSE | CAN | JSY3811 | 1 | rnr1Y285F-R | GGC | G>C | G/C+1 | G/C-1 | CG>GC | CG>GC | TAGGCGGCA | GCAGCAA | CCG | CCG | C | G | C>G | C>G_CCG | CG>GC_GCAGCAA | 8 | FALSE | FALSE |
| 42_v.txt | 32165 | 32165-SNV-G | SNV | 1 | G | C | 2 | 23773 | 8.41E-03 | 30.0003611 | rnr1Y285F | FALSE | CAN | JSY3811 | 1 | rnr1Y285F-R | CGC | G>C | G/C+1 | G/C-1 | CG>GC | CG>GC | CAAAGCGCCA | GCCAAAT | GCG | GCG | C | G | C>G | C>G_GCG | CG>GC_GCCAAAT | 8 | FALSE | FALSE |
| 42_v.txt | 32271 | 32271-SNV-C | SNV | 1 | C | G | 83 | 28272 | 0.29357668 | 36.490963 | rnr1Y285F | FALSE | CAN | JSY3811 | 1 | rnr1Y285F-R | ACG | C>G | G/C+1 | G/C-1 | CG>GC | CG>GC | AAAATACGG | CGGGAAC | TGC | ACG | C | C | C>G | C>G_ACG | CG>GC_CGGGAAC | 8 | FALSE | FALSE |
| 42_v.txt | 32272 | 32272-SNV-G | SNV | 1 | G | C | 14 | 28272 | 0.04951896 | 34.5682848 | rnr1Y285F | FALSE | CAN | JSY3811 | 1 | rnr1Y285F-R | CGG | G>C | G/C+1 | G/C-1 | CG>GC | CG>GC | AAATACGGG | GGGAACC | GCC | GCC | C | G | C>G | C>G_GCC | CG>GC_GGGAACC | 8 | FALSE | FALSE |
| 42_v.txt | 32316 | 32316-SNV-G | SNV | 1 | G | C | 7 | 28272 | 0.02475948 | 57.8571429 | rnr1Y285F | FALSE | CAN | JSY3811 | 1 | rnr1Y285F-R | TGT | G>C | G/C+1 | G/C-1 | CG>GC | CG>GC | ATGGTTGTTA | GTTAAGA | ACA | ACA | C | G | C>G | C>G_ACA | CG>GC_GTTAAGA | 8 | FALSE | FALSE |

| | | | | | | | | | | | | | | | | | | | | | | | | | | | | | | | | | | |
|---|---|---|---|---|---|---|---|---|---|---|---|---|---|---|---|---|---|---|---|---|---|---|---|---|---|---|---|---|---|---|---|---|---|---|
| 42_v.txt | 32488 | 32488-SNV- | SNV | 1 | G | C | 13 | 28272 | 0.04598189 | 39.6923077 | rnr1Y285F | FALSE | CAN | JSY3811 | 1 | rnr1Y285F-R | CGG | G>C | G/C+1 | G/C-1 | CG>GC | CG>GC | AGATACGGA | GGAAAAC | GCC | GCC | C | G | C>G | C>G_GCC | CG>GC_GGAAAAC | 8 | FALSE | FALSE |
| 42_v.txt | 32545 | 32545-SNV- | SNV | 1 | C | G | 4 | 28272 | 0.01414827 | 37.4990986 | rnr1Y285F | FALSE | CAN | JSY3811 | 1 | rnr1Y285F-R | CCA | C>G | G/C+1 | G/C-1 | CG>GC | CG>GC | CTTCACCAG | CAGCAGT | GGT | CCA | C | C | C>G | C>G_CCA | CG>GC_CAGCAGT | 8 | FALSE | FALSE |
| 42_v.txt | 32566 | 32566-SNV- | SNV | 1 | C | G | 39 | 23843 | 0.16357002 | 37.3333333 | rnr1Y285F | FALSE | CAN | JSY3811 | 1 | rnr1Y285F-R | TCA | C>G | G/C+1 | G/C-1 | CG>GC | CG>GC | CTAGTTCAG | CAGTACC | AGT | TCA | C | C | C>G | C>G_TCA | CG>GC_CAGTACC | 8 | FALSE | FALSE |
| 42_v.txt | 32776 | 32776-SNV- | SNV | 1 | C | G | 33 | 23846 | 0.13838799 | 46.3939394 | rnr1Y285F | FALSE | CAN | JSY3811 | 1 | rnr1Y285F-R | GCG | C>G | G/C+1 | G/C-1 | CG>GC | CG>GC | TGGAAGCGA | CGACCCA | CGC | GCG | C | C | C>G | C>G_GCG | CG>GC_CGACCCA | 8 | FALSE | FALSE |
| 42_v.txt | 32783 | 32783-SNV- | SNV | 1 | G | C | 11 | 23846 | 0.04612933 | 42.6363636 | rnr1Y285F | FALSE | CAN | JSY3811 | 1 | rnr1Y285F-R | AGA | G>C | G/C+1 | G/C-1 | CG>GC | CG>GC | GACCCAGAA | GAACTCG | TCT | TCT | C | G | C>G | C>G_TCT | CG>GC_GAACTCG | 8 | FALSE | FALSE |
| 42_v.txt | 32876 | 32876-SNV- | SNV | 1 | G | C | 32 | 33518 | 0.09547109 | 37.15625 | rnr1Y285F | FALSE | CAN | JSY3811 | 1 | rnr1Y285F-R | TGT | G>C | G/C+1 | G/C-1 | CG>GC | CG>GC | AACTTTGTA | GTACGTC | ACA | ACA | C | G | C>G | C>G_ACA | CG>GC_GTACGTC | 8 | FALSE | FALSE |
| 42_v.txt | 32937 | 32937-SNV- | SNV | 1 | C | G | 11 | 33529 | 0.03280742 | 45.8155412 | rnr1Y285F | FALSE | CAN | JSY3811 | 1 | rnr1Y285F-R | CCA | C>G | G/C+1 | G/C-1 | CG>GC | CG>GC | ATTGCCCAA | CAAGAAA | GGT | CCA | C | C | C>G | C>G_CCA | CG>GC_CAAGAAA | 8 | FALSE | FALSE |
| 42_v.txt | 32997 | 32997-SNV- | SNV | 1 | G | C | 80 | 33529 | 0.23859942 | 59.3375 | rnr1Y285F | FALSE | CAN | JSY3811 | 1 | rnr1Y285F-R | TGA | G>C | G/C+1 | G/C-1 | CG>GC | CG>GC | CTTTGTGAG | GAGAAAA | ACT | ACT | C | G | C>G | C>G_ACT | CG>GC_GAGAAAA | 8 | FALSE | FALSE |
| 42_v.txt | 33115 | 33115-SNV- | SNV | 1 | C | G | 9 | 33529 | 0.02684244 | 43.1125936 | rnr1Y285F | FALSE | CAN | JSY3811 | 1 | rnr1Y285F-R | CCG | C>G | G/C+1 | G/C-1 | CG>GC | CG>GC | CTGGGCCGG | CGGCGTT | GGC | CCG | C | C | C>G | C>G_CCG | CG>GC_CGGCGTT | 8 | FALSE | FALSE |
| 42_v.txt | 33142 | 33142-SNV- | SNV | 1 | C | G | 24 | 33529 | 0.07157983 | 41.5416667 | rnr1Y285F | FALSE | CAN | JSY3811 | 1 | rnr1Y285F-R | CCA | C>G | G/C+1 | G/C-1 | CG>GC | CG>GC | ATAAACCAA | CAATGAA | GGT | CCA | C | C | C>G | C>G_CCA | CG>GC_CAATGAA | 8 | FALSE | FALSE |
| 42_v.txt | 33156 | 33156-SNV- | SNV | 1 | G | C | 11 | 33528 | 0.0328084 | 37.5454546 | rnr1Y285F | FALSE | CAN | JSY3811 | 1 | rnr1Y285F-R | TGT | G>C | G/C+1 | G/C-1 | CG>GC | CG>GC | AGACCTGTA | GTACCAA | ACA | ACA | C | G | C>G | C>G_ACA | CG>GC_GTACCAA | 8 | FALSE | FALSE |
| 42_v.txt | 33205 | 33205-SNV- | SNV | 1 | G | C | 10 | 23943 | 0.04176586 | 36.4 | rnr1Y285F | FALSE | CAN | JSY3811 | 1 | rnr1Y285F-R | AGC | G>C | G/C+1 | G/C-1 | CG>GC | CG>GC | GCTTAAGCT | GCTCTCT | TCG | TCG | C | G | C>G | C>G_TCG | CG>GC_GCTCTCT | 8 | FALSE | FALSE |
| 42_v.txt | 33210 | 33210-SNV- | SNV | 1 | C | G | 12 | 23943 | 0.05011903 | 40 | rnr1Y285F | FALSE | CAN | JSY3811 | 1 | rnr1Y285F-R | TCT | C>G | G/C+1 | G/C-1 | CG>GC | CG>GC | AGCTCTCTCT | CTCTTCA | AGA | TCT | C | C | C>G | C>G_TCT | CG>GC_CTCTTCA | 8 | FALSE | FALSE |
| 42_v.txt | 33310 | 33310-SNV- | SNV | 1 | G | C | 4 | 23938 | 0.01670983 | 8.50189251 | rnr1Y285F | FALSE | CAN | JSY3811 | 1 | rnr1Y285F-R | CGC | G>C | G/C+1 | G/C-1 | CG>GC | CG>GC | AAGAGCGCA | GCAATGG | GCG | GCG | C | G | C>G | C>G_GCG | CG>GC_GCAATGG | 8 | FALSE | FALSE |
| 42_v.txt | 33378 | 33378-SNV- | SNV | 1 | G | C | 5 | 23943 | 0.02088293 | 58.4 | rnr1Y285F | FALSE | CAN | JSY3811 | 1 | rnr1Y285F-R | TGA | G>C | G/C+1 | G/C-1 | CG>GC | CG>GC | GTTTGTGAA | GAAGCTT | ACT | ACT | C | G | C>G | C>G_ACT | CG>GC_GAAGCTT | 8 | FALSE | FALSE |
| 42_v.txt | 33456 | 33456-SNV- | SNV | 1 | G | C | 27 | 23943 | 0.11276782 | 40.8148148 | rnr1Y285F | FALSE | CAN | JSY3811 | 1 | rnr1Y285F-R | TGA | G>C | G/C+1 | G/C-1 | CG>GC | CG>GC | TCTTTTGAAT | GAATTTG | ACT | ACT | C | G | C>G | C>G_ACT | CG>GC_GAATTTG | 8 | FALSE | FALSE |
| 42_v.txt | 31981 | 31981-SNV- | SNV | 1 | G | T | 66 | 23773 | 0.27762588 | 38.6969697 | rnr1Y285F | FALSE | CAN | JSY3811 | 1 | rnr1Y285F-R | GGC | G>T | A/T+1 | G/C-1 | CG>AT | CG>AT | AGCCGGGCA | GCATTAA | CCG | CCG | C | A | C>A | C>A_CCG | CG>AT_GCATTAA | 8 | FALSE | FALSE |
| 42_v.txt | 32188 | 32188-SNV- | SNV | 1 | C | A | 46 | 23773 | 0.19349682 | 46.9347826 | rnr1Y285F | FALSE | CAN | JSY3811 | 1 | rnr1Y285F-R | ACT | C>A | A/T+1 | G/C-1 | CG>AT | CG>AT | CGAAAACTG | CTGCAAT | TGA | ACT | C | T | C>A | C>A_ACT | CG>AT_CTGCAAT | 8 | FALSE | FALSE |
| 42_v.txt | 32272 | 32272-SNV- | SNV | 1 | G | T | 6 | 28272 | 0.02122241 | 30.8325961 | rnr1Y285F | FALSE | CAN | JSY3811 | 1 | rnr1Y285F-R | CGG | G>T | A/T+1 | G/C-1 | CG>AT | CG>AT | AAATACGGG | GGGAACC | GCC | GCC | C | A | C>A | C>A_GCC | CG>AT_GGGAACC | 8 | FALSE | FALSE |
| 42_v.txt | 32274 | 32274-SNV- | SNV | 1 | G | T | 106 | 28272 | 0.37492926 | 35.8962264 | rnr1Y285F | FALSE | CAN | JSY3811 | 1 | rnr1Y285F-R | GGA | G>T | A/T+1 | G/C-1 | CG>AT | CG>AT | ATACGGGAA | GAACCAA | CCT | CCT | C | A | C>A | C>A_CCT | CG>AT_GAACCAA | 8 | FALSE | FALSE |
| 42_v.txt | 32373 | 32373-SNV- | SNV | 1 | G | T | 26 | 28272 | 0.09196378 | 60.6538462 | rnr1Y285F | FALSE | CAN | JSY3811 | 1 | rnr1Y285F-R | AGC | G>T | A/T+1 | G/C-1 | CG>AT | CG>AT | TCAATAGCA | GCAATAA | TCG | TCG | C | A | C>A | C>A_TCG | CG>AT_GCAATAA | 8 | FALSE | FALSE |
| 42_v.txt | 32399 | 32399-SNV- | SNV | 1 | G | T | 14 | 28272 | 0.04951896 | 53.9285714 | rnr1Y285F | FALSE | CAN | JSY3811 | 1 | rnr1Y285F-R | CGT | G>T | A/T+1 | G/C-1 | CG>AT | CG>AT | AGAAACGTA | GTAGGAA | GCA | GCA | C | A | C>A | C>A_GCA | CG>AT_GTAGGAA | 8 | FALSE | FALSE |
| 42_v.txt | 32432 | 32432-SNV- | SNV | 1 | G | T | 14 | 28272 | 0.04951896 | 51.9285714 | rnr1Y285F | FALSE | CAN | JSY3811 | 1 | rnr1Y285F-R | TGT | G>T | A/T+1 | G/C-1 | CG>AT | CG>AT | GTCATTGTAT | GTATGGA | ACA | ACA | C | A | C>A | C>A_ACA | CG>AT_GTATGGA | 8 | FALSE | FALSE |
| 42_v.txt | 32530 | 32530-SNV- | SNV | 1 | G | T | 68 | 28272 | 0.24052066 | 39.1323529 | rnr1Y285F | FALSE | CAN | JSY3811 | 1 | rnr1Y285F-R | GGG | G>T | A/T+1 | G/C-1 | CG>AT | CG>AT | TTCTGGGGT | GGTTTGC | CCC | CCC | C | A | C>A | C>A_CCC | CG>AT_GGTTTGC | 8 | FALSE | FALSE |
| 42_v.txt | 32544 | 32544-SNV- | SNV | 1 | C | A | 3 | 28272 | 0.01061121 | 37.666588 | rnr1Y285F | FALSE | CAN | JSY3811 | 1 | rnr1Y285F-R | ACC | C>A | A/T+1 | G/C-1 | CG>AT | CG>AT | GCTTCACCAG | CCAGCAG | TGG | ACC | C | T | C>A | C>A_ACC | CG>AT_CCAGCAG | 8 | FALSE | FALSE |
| 42_v.txt | 32556 | 32556-SNV- | SNV | 1 | C | A | 34 | 28268 | 0.12027735 | 36.7647059 | rnr1Y285F | FALSE | CAN | JSY3811 | 1 | rnr1Y285F-R | ACC | C>A | A/T+1 | G/C-1 | CG>AT | CG>AT | GTGATACCA | CCAACTA | TGG | ACC | C | T | C>A | C>A_ACC | CG>AT_CCAACTA | 8 | FALSE | FALSE |
| 42_v.txt | 32566 | 32566-SNV- | SNV | 1 | C | A | 11 | 23843 | 0.04613513 | 30.6363636 | rnr1Y285F | FALSE | CAN | JSY3811 | 1 | rnr1Y285F-R | TCA | C>A | A/T+1 | G/C-1 | CG>AT | CG>AT | CTAGTTCAG | CAGTACC | AGT | TCA | C | T | C>A | C>A_TCA | CG>AT_CAGTACC | 8 | FALSE | FALSE |
| 42_v.txt | 32580 | 32580-SNV- | SNV | 1 | G | T | 7 | 23846 | 0.02935503 | 36.7142857 | rnr1Y285F | FALSE | CAN | JSY3811 | 1 | rnr1Y285F-R | TGT | G>T | A/T+1 | G/C-1 | CG>AT | CG>AT | TGAAATGTG | GTGAAGG | ACA | ACA | C | A | C>A | C>A_ACA | CG>AT_GTGAAGG | 8 | FALSE | FALSE |
| 42_v.txt | 32691 | 32691-SNV- | SNV | 1 | C | A | 10 | 23846 | 0.04193575 | 56.3020217 | rnr1Y285F | FALSE | CAN | JSY3811 | 1 | rnr1Y285F-R | TCC | C>A | A/T+1 | G/C-1 | CG>AT | CG>AT | CGGAATCCA | CCAACTG | AGG | TCC | C | T | C>A | C>A_TCC | CG>AT_CCAACTG | 8 | FALSE | FALSE |
| 42_v.txt | 32783 | 32783-SNV- | SNV | 1 | G | T | 8 | 23846 | 0.0335486 | 37.875 | rnr1Y285F | FALSE | CAN | JSY3811 | 1 | rnr1Y285F-R | AGA | G>T | A/T+1 | G/C-1 | CG>AT | CG>AT | GACCCAGAA | GAACTCG | TCT | TCT | C | A | C>A | C>A_TCT | CG>AT_GAACTCG | 8 | FALSE | FALSE |
| 42_v.txt | 32794 | 32794-SNV- | SNV | 1 | C | A | 78 | 23846 | 0.32709889 | 39.2435897 | rnr1Y285F | FALSE | CAN | JSY3811 | 1 | rnr1Y285F-R | TCA | C>A | A/T+1 | G/C-1 | CG>AT | CG>AT | CGAATTCAC | CACCGTA | AGT | TCA | C | T | C>A | C>A_TCA | CG>AT_CACCGTA | 8 | FALSE | FALSE |
| 42_v.txt | 32809 | 32809-SNV- | SNV | 1 | C | A | 24 | 23846 | 0.10064581 | 45.9166667 | rnr1Y285F | FALSE | CAN | JSY3811 | 1 | rnr1Y285F-R | ACA | C>A | A/T+1 | G/C-1 | CG>AT | CG>AT | ATTTGACAG | CAGGGAA | TGT | ACA | C | T | C>A | C>A_ACA | CG>AT_CAGGGAA | 8 | FALSE | FALSE |
| 42_v.txt | 32819 | 32819-SNV- | SNV | 1 | G | T | 28 | 23846 | 0.11742011 | 46.6071429 | rnr1Y285F | FALSE | CAN | JSY3811 | 1 | rnr1Y285F-R | AGT | G>T | A/T+1 | G/C-1 | CG>AT | CG>AT | GAACAAGTT | GTTCATT | TCA | TCA | C | A | C>A | C>A_TCA | CG>AT_GTTCATT | 8 | FALSE | FALSE |
| 42_v.txt | 32917 | 32917-SNV- | SNV | 1 | C | A | 39 | 33529 | 0.11631722 | 49.7948718 | rnr1Y285F | FALSE | CAN | JSY3811 | 1 | rnr1Y285F-R | TCC | C>A | A/T+1 | G/C-1 | CG>AT | CG>AT | TAAGTTCCAC | CCAGGGC | AGG | TCC | C | T | C>A | C>A_TCC | CG>AT_CCAGGGC | 8 | FALSE | FALSE |
| 42_v.txt | 32940 | 32940-SNV- | SNV | 1 | G | T | 15727 | 33529 | 46.9056638 | 55.4304063 | rnr1Y285F | FALSE | CAN | JSY3811 | 1 | rnr1Y285F-R | AGA | G>T | A/T+1 | G/C-1 | CG>AT | CG>AT | GCCCAAGAA | GAAAACC | TCT | TCT | C | A | C>A | C>A_TCT | CG>AT_GAAAACC | 8 | FALSE | FALSE |
| 42_v.txt | 32958 | 32958-SNV- | SNV | 1 | C | A | 16 | 33529 | 0.04771988 | 40.4375 | rnr1Y285F | FALSE | CAN | JSY3811 | 1 | rnr1Y285F-R | ACC | C>A | A/T+1 | G/C-1 | CG>AT | CG>AT | ATGTAACCAT | CCATTGG | TGG | ACC | C | T | C>A | C>A_ACC | CG>AT_CCATTGG | 8 | FALSE | FALSE |
| 42_v.txt | 33015 | 33015-SNV- | SNV | 1 | G | T | 21 | 33529 | 0.06263235 | 55.6666667 | rnr1Y285F | FALSE | CAN | JSY3811 | 1 | rnr1Y285F-R | GGA | G>T | A/T+1 | G/C-1 | CG>AT | CG>AT | AAAGAGGAT | GATGTAA | CCT | CCT | C | A | C>A | C>A_CCT | CG>AT_GATGTAA | 8 | FALSE | FALSE |
| 42_v.txt | 33033 | 33033-SNV- | SNV | 1 | G | T | 49 | 33529 | 0.14614215 | 58.6326531 | rnr1Y285F | FALSE | CAN | JSY3811 | 1 | rnr1Y285F-R | TGT | G>T | A/T+1 | G/C-1 | CG>AT | CG>AT | ATGAATGTA | GTAGCCA | ACA | ACA | C | A | C>A | C>A_ACA | CG>AT_GTAGCCA | 8 | FALSE | FALSE |
| 42_v.txt | 33043 | 33043-SNV- | SNV | 1 | C | A | 21 | 33529 | 0.06263235 | 58.0952381 | rnr1Y285F | FALSE | CAN | JSY3811 | 1 | rnr1Y285F-R | TCA | C>A | A/T+1 | G/C-1 | CG>AT | CG>AT | CCATTTCACC | CACCCAA | AGT | TCA | C | T | C>A | C>A_TCA | CG>AT_CACCCAA | 8 | FALSE | FALSE |
| 42_v.txt | 33112 | 33112-SNV- | SNV | 1 | G | T | 43 | 33529 | 0.12824719 | 44.6744186 | rnr1Y285F | FALSE | CAN | JSY3811 | 1 | rnr1Y285F-R | GGG | G>T | A/T+1 | G/C-1 | CG>AT | CG>AT | CCACTGGGC | GGCCGGC | CCC | CCC | C | A | C>A | C>A_CCC | CG>AT_GGCCGGC | 8 | FALSE | FALSE |
| 42_v.txt | 33151 | 33151-SNV- | SNV | 1 | G | T | 32 | 33529 | 0.09543977 | 35.03125 | rnr1Y285F | FALSE | CAN | JSY3811 | 1 | rnr1Y285F-R | AGA | G>T | A/T+1 | G/C-1 | CG>AT | CG>AT | TGAAAAGAC | GACCTGT | TCT | TCT | C | A | C>A | C>A_TCT | CG>AT_GACCTGT | 8 | FALSE | FALSE |
| 42_v.txt | 33153 | 33153-SNV- | SNV | 1 | C | A | 64 | 33529 | 0.19087954 | 39.3746598 | rnr1Y285F | FALSE | CAN | JSY3811 | 1 | rnr1Y285F-R | ACC | C>A | A/T+1 | G/C-1 | CG>AT | CG>AT | AAAAGACCT | CCTGTAC | TGG | ACC | C | T | C>A | C>A_ACC | CG>AT_CCTGTAC | 8 | FALSE | FALSE |
| 42_v.txt | 33154 | 33154-SNV- | SNV | 1 | C | A | 26 | 33529 | 0.07754481 | 36.5386267 | rnr1Y285F | FALSE | CAN | JSY3811 | 1 | rnr1Y285F-R | CCT | C>A | A/T+1 | G/C-1 | CG>AT | CG>AT | AAAGACCTG | CTGTACC | GGA | CCT | C | T | C>A | C>A_CCT | CG>AT_CTGTACC | 8 | FALSE | FALSE |
| 42_v.txt | 33168 | 33168-SNV- | SNV | 1 | C | A | 69 | 33524 | 0.20582269 | 36.8115942 | rnr1Y285F | FALSE | CAN | JSY3811 | 1 | rnr1Y285F-R | ACC | C>A | A/T+1 | G/C-1 | CG>AT | CG>AT | ATAGTACCA | CCACCAA | TGG | ACC | C | T | C>A | C>A_ACC | CG>AT_CCACCAA | 8 | FALSE | FALSE |
| 42_v.txt | 33175 | 33175-SNV- | SNV | 1 | G | T | 11 | 33518 | 0.03281819 | 37.9082912 | rnr1Y285F | FALSE | CAN | JSY3811 | 1 | rnr1Y285F-R | AGG | G>T | A/T+1 | G/C-1 | CG>AT | CG>AT | CACCAAGGG | GGGCAAT | TCC | TCC | C | A | C>A | C>A_TCC | CG>AT_GGGCAAT | 8 | FALSE | FALSE |
| 42_v.txt | 33195 | 33195-SNV- | SNV | 1 | C | A | 18 | 23943 | 0.07517855 | 37 | rnr1Y285F | FALSE | CAN | JSY3811 | 1 | rnr1Y285F-R | TCT | C>A | A/T+1 | G/C-1 | CG>AT | CG>AT | ATATGTCTTT | CTTTGCT | AGA | TCT | C | T | C>A | C>A_TCT | CG>AT_CTTTGCT | 8 | FALSE | FALSE |
| 42_v.txt | 33354 | 33354-SNV- | SNV | 1 | G | T | 142 | 23943 | 0.59307522 | 15.8309859 | rnr1Y285F | FALSE | CAN | JSY3811 | 1 | rnr1Y285F-R | TGA | G>T | A/T+1 | G/C-1 | CG>AT | CG>AT | GGTATTGAC | GACCCAC | ACT | ACT | C | A | C>A | C>A_ACT | CG>AT_GACCCAC | 8 | FALSE | FALSE |
| 42_v.txt | 33378 | 33378-SNV- | SNV | 1 | G | T | 13 | 23943 | 0.05429562 | 53.8461539 | rnr1Y285F | FALSE | CAN | JSY3811 | 1 | rnr1Y285F-R | TGA | G>T | A/T+1 | G/C-1 | CG>AT | CG>AT | GTTTGTGAA | GAAGCTT | ACT | ACT | C | A | C>A | C>A_ACT | CG>AT_GAAGCTT | 8 | FALSE | FALSE |
| 42_v.txt | 33474 | 33474-SNV- | SNV | 1 | C | A | 80 | 23940 | 0.33416876 | 34.4257917 | rnr1Y285F | FALSE | CAN | JSY3811 | 1 | rnr1Y285F-R | GCC | C>A | A/T+1 | G/C-1 | CG>AT | CG>AT | GCTATGCCTT | CCTTTTT | CGG | GCC | C | T | C>A | C>A_GCC | CG>AT_CCTTTTT | 8 | FALSE | FALSE |
| 42_v.txt | 33475 | 33475-SNV- | SNV | 1 | C | A | 28 | 23939 | 0.11696395 | 36.6071429 | rnr1Y285F | FALSE | CAN | JSY3811 | 1 | rnr1Y285F-R | CCT | C>A | A/T+1 | G/C-1 | CG>AT | CG>AT | CTATGCCTTT | CTTTTTT | GGA | CCT | C | T | C>A | C>A_CCT | CG>AT_CTTTTTT | 8 | FALSE | FALSE |
| 42_v.txt | 32545 | 32545-Repla | Replacement | 2 | C | AA | 1 | 28272 | 3.54E-03 | 33.9999984 | rnr1Y285F | FALSE | CAN | JSY3811 | 1 | rnr1Y285F-R | CCA | nope | >1bp-ins | G/C-1 | none | Replac. | CTTCACCAG | CAGCAGT | GGT | CCA | C | none | nope | nope_CCA | Replac._CAGCAGT | 8 | FALSE | FALSE |
| 42_v.txt | 32544 | 32544-MNV- | MNV | 2 | CC | AA | 2 | 28272 | 7.07E-03 | 33.3333316 | rnr1Y285F | FALSE | CAN | JSY3811 | 1 | rnr1Y285F-R | ACC | nope | >1bp-ins | >1bpdel | none | MNV | GCTTCACCAG | CCAGCAG | TGG | ACC | CC | none | NA | NA_ACC | MNV_CCAGCAG | 8 | FALSE | FALSE |
| 42_v.txt | 33153 | 33153-MNV- | MNV | 2 | CC | AA | 1 | 33529 | 2.98E-03 | 37.5 | rnr1Y285F | FALSE | CAN | JSY3811 | 1 | rnr1Y285F-R | ACC | nope | >1bp-ins | >1bpdel | none | MNV | AAAAGACCT | CCTGTAC | TGG | ACC | CC | none | NA | NA_ACC | MNV_CCTGTAC | 8 | FALSE | FALSE |
| 42_v.txt | 32081 | 32081-Insert | Insertion | 1 | - | A | 13 | 23773 | 0.05468389 | 61.8461539 | rnr1Y285F | FALSE | CAN | JSY3811 | 1 | rnr1Y285F-R | CAA | nope | A/T+1 | >1bpdel | none | A/T+1 | CCATGCAAA | AAAAAAG | GTT | CAA | - | T | NA | NA_CAA | A/T+1_AAAAAAG | 8 | FALSE | FALSE |
| 42_v.txt | 32146 | 32146-Insert | Insertion | 2 | - | TC | 13 | 23773 | 0.05468389 | 48.9230769 | rnr1Y285F | FALSE | CAN | JSY3811 | 1 | rnr1Y285F-R | GTC | nope | >1bp-ins | >1bpdel | none | >1bp-ins | TAGATGTCT | TCTCCAT | CAG | GTC | - | none | NA | NA_GTC | >1bp-ins_TCTCCAT | 8 | FALSE | FALSE |
| 42_v.txt | 32196 | 32196-Insert | Insertion | 2 | - | TA | 7 | 23773 | 0.02944517 | 40.8571429 | rnr1Y285F | FALSE | CAN | JSY3811 | 1 | rnr1Y285F-R | GTA | nope | >1bp-ins | >1bpdel | none | >1bp-ins | GCAATGTAT | TATGGAA | CAT | GTA | - | none | NA | NA_GTA | >1bp-ins_TATGGAA | 8 | FALSE | FALSE |
| 42_v.txt | 32330 | 32330-Insert | Insertion | 1 | - | T | 5 | 28272 | 0.01768534 | 60.2 | rnr1Y285F | FALSE | CAN | JSY3811 | 1 | rnr1Y285F-R | CGT | nope | A/T+1 | >1bpdel | none | A/T+1 | AACAGCGTT | GTTGAAG | GCA | CGT | - | A | NA | NA_CGT | A/T+1_GTTGAAG | 8 | FALSE | FALSE |
| 42_v.txt | 32498 | 32498-Insert | Insertion | 1 | - | T | 18 | 28272 | 0.06366723 | 41.8333333 | rnr1Y285F | FALSE | CAN | JSY3811 | 1 | rnr1Y285F-R | CTT | nope | A/T+1 | >1bpdel | none | A/T+1 | AACAACTTTT | TTTTTTG | GAA | CTT | - | A | NA | NA_CTT | A/T+1_TTTTTTG | 8 | FALSE | FALSE |
| 42_v.txt | 32635 | 32635-Insert | Insertion | 2 | - | TA | 5 | 23846 | 0.02096788 | 53.1 | rnr1Y285F | FALSE | CAN | JSY3811 | 1 | rnr1Y285F-R | TTA | nope | >1bp-ins | >1bpdel | none | >1bp-ins | CGTTTTTATC | TATCCTT | AAT | TTA | - | none | NA | NA_TTA | >1bp-ins_TATCCTT | 8 | FALSE | FALSE |
| 42_v.txt | 32693 | 32693-Insert | Insertion | 1 | - | A | 23 | 23846 | 0.09645224 | 64 | rnr1Y285F | FALSE | CAN | JSY3811 | 1 | rnr1Y285F-R | CAA | nope | A/T+1 | >1bpdel | none | A/T+1 | GAATCCAAC | AACTGGG | GTT | CAA | - | T | NA | NA_CAA | A/T+1_AACTGGG | 8 | FALSE | FALSE |
| 42_v.txt | 32728 | 32728-Insert | Insertion | 1 | - | A | 27 | 23846 | 0.11322654 | 59.8148148 | rnr1Y285F | FALSE | CAN | JSY3811 | 1 | rnr1Y285F-R | CAA | nope | A/T+1 | >1bpdel | none | A/T+1 | CCATACAAA | AAAAACA | GTT | CAA | - | T | NA | NA_CAA | A/T+1_AAAAACA | 8 | FALSE | FALSE |
| 42_v.txt | 32760 | 32760-Insert | Insertion | 1 | - | A | 45 | 23846 | 0.1887109 | 55.2444444 | rnr1Y285F | FALSE | CAN | JSY3811 | 1 | rnr1Y285F-R | TAA | nope | A/T+1 | >1bpdel | none | A/T+1 | ATGGCTAAA | AAAACTT | ATT | TAA | - | T | NA | NA_TAA | A/T+1_AAAACTT | 8 | FALSE | FALSE |
| 42_v.txt | 32828 | 32828-Insert | Insertion | 2 | - | TG | 12 | 23846 | 0.05032291 | 39.125 | rnr1Y285F | FALSE | CAN | JSY3811 | 1 | rnr1Y285F-R | TTG | nope | >1bp-ins | >1bpdel | none | >1bp-ins | CATTATTGTG | TGTGATA | AAC | TTG | - | none | NA | NA_TTG | >1bp-ins_TGTGATA | 8 | FALSE | FALSE |
| 42_v.txt | 32842 | 32842-Insert | Insertion | 1 | - | A | 20 | 23846 | 0.08387151 | 41.75 | rnr1Y285F | FALSE | CAN | JSY3811 | 1 | rnr1Y285F-R | CAA | nope | A/T+1 | >1bpdel | none | A/T+1 | TTACCCAAA | AAAAAAT | GTT | CAA | - | T | NA | NA_CAA | A/T+1_AAAAAAT | 8 | FALSE | FALSE |
| 42_v.txt | 33064 | 33064-Insert | Insertion | 24 | - | AATATGCCA | 11 | 33529 | 0.03280742 | 55.9962121 | rnr1Y285F | FALSE | CAN | JSY3811 | 1 | rnr1Y285F-R | GAA | nope | >1bp-ins | >1bpdel | none | >1bp-ins | TGACAGAAT | AATATGC | CTT | GAA | - | none | NA | NA_GAA | >1bp-ins_AATATGC | 8 | FALSE | FALSE |
| 42_v.txt | 33161 | 33161-Insert | Insertion | 1 | - | A | 7 | 33524 | 0.02088056 | 37.8571429 | rnr1Y285F | FALSE | CAN | JSY3811 | 1 | rnr1Y285F-R | CAA | nope | A/T+1 | >1bpdel | none | A/T+1 | TGTACCAAT | AATAGTA | GTT | CAA | - | T | NA | NA_CAA | A/T+1_AATAGTA | 8 | FALSE | FALSE |
| 42_v.txt | 32337 | 32337-Deleti | Deletion | 2 | AT | - | 28 | 28272 | 0.09903792 | 62.5357143 | rnr1Y285F | FALSE | CAN | JSY3811 | 1 | rnr1Y285F-R | GAT | nope | >1bp-ins | >1bpdel | none | >1bpdel | TTGAAGATA | ATATGTG | CTA | GAT | AT | none | NA | NA_GAT | >1bpdel_ATATGTG | 8 | FALSE | FALSE |
| 42_v.txt | 32937 | 32937-Deleti | Deletion | 2 | CA | - | 1 | 33529 | 2.98E-03 | 58 | rnr1Y285F | FALSE | CAN | JSY3811 | 1 | rnr1Y285F-R | CCA | nope | >1bp-ins | >1bpdel | none | >1bpdel | ATTGCCCAA | CAAGAAA | GGT | CCA | CA | none | NA | NA_CCA | >1bpdel_CAAGAAA | 8 | FALSE | FALSE |
| 42_v.txt | 33089 | 33089-Deleti | Deletion | 2 | AT | - | 41 | 33529 | 0.1222822 | 50.8292683 | rnr1Y285F | FALSE | CAN | JSY3811 | 1 | rnr1Y285F-R | AAT | nope | >1bp-ins | >1bpdel | none | >1bpdel | AAATAAATA | ATATGAT | TTA | AAT | AT | none | NA | NA_AAT | >1bpdel_ATATGAT | 8 | FALSE | FALSE |
| 42_v.txt | 31663 | 31663-Deleti | Deletion | 1 | A | - | 6 | 11596 | 0.05174198 | 32.5 | rnr1Y285F | FALSE | CAN | JSY3811 | 1 | rnr1Y285F-R | GAA | nope | >1bp-ins | A/T-1 | none | A/T-1 | GCGTGGAAA | AAATGTG | CTT | CTT | T | none | T>none | T>none_CTT | A/T-1_AAATGTG | 8 | FALSE | FALSE |
| 42_v.txt | 31723 | 31723-Deleti | Deletion | 1 | A | - | 5 | 11596 | 0.04311832 | 64 | rnr1Y285F | FALSE | CAN | JSY3811 | 1 | rnr1Y285F-R | CAA | nope | >1bp-ins | A/T-1 | none | A/T-1 | TGTCCCAAA | AAAAAGT | GTT | GTT | T | none | T>none | T>none_GTT | A/T-1_AAAAAGT | 8 | FALSE | FALSE |
| 42_v.txt | 31882 | 31882-Deleti | Deletion | 1 | C | - | 1 | 11596 | 8.62E-03 | 38 | rnr1Y285F | FALSE | CAN | JSY3811 | 1 | rnr1Y285F-R | GCA | nope | >1bp-ins | G/C-1 | none | G/C-1 | AGGCGGCAG | CAGCAAA | CGT | GCA | C | none | nope | nope_GCA | G/C-1_CAGCAAA | 8 | FALSE | FALSE |
| 42_v.txt | 31904 | 31904-Deleti | Deletion | 1 | T | - | 11 | 11596 | 0.0948603 | 39.3636364 | rnr1Y285F | FALSE | CAN | JSY3811 | 1 | rnr1Y285F-R | ATT | nope | >1bp-ins | A/T-1 | none | A/T-1 | ATTGAATTTT | TTTTGGT | TAA | ATT | T | none | nope | nope_ATT | A/T-1_TTTTGGT | 8 | FALSE | FALSE |
| 42_v.txt | 31923 | 31923-Deleti | Deletion | 1 | A | - | 7 | 11597 | 0.06036044 | 37.8571429 | rnr1Y285F | FALSE | CAN | JSY3811 | 1 | rnr1Y285F-R | GAA | nope | >1bp-ins | A/T-1 | none | A/T-1 | GCCGTGAAA | AAACCTT | CTT | CTT | T | none | T>none | T>none_CTT | A/T-1_AAACCTT | 8 | FALSE | FALSE |
| 42_v.txt | 31931 | 31931-Deleti | Deletion | 1 | A | - | 7 | 11597 | 0.06036044 | 37.1428571 | rnr1Y285F | FALSE | CAN | JSY3811 | 1 | rnr1Y285F-R | GAA | nope | >1bp-ins | A/T-1 | none | A/T-1 | ACCTTGAAT | AATAATG | CTT | CTT | T | none | T>none | T>none_CTT | A/T-1_AATAATG | 8 | FALSE | FALSE |
| 42_v.txt | 31979 | 31979-Deleti | Deletion | 1 | G | - | 36 | 23773 | 0.1514323 | 38.9444444 | rnr1Y285F | FALSE | CAN | JSY3811 | 1 | rnr1Y285F-R | CGG | nope | >1bp-ins | G/C-1 | none | G/C-1 | CAAGCCGGG | GGGCATT | GCC | GCC | C | none | C>none | C>none_GCC | G/C-1_GGGCATT | 8 | FALSE | FALSE |
| 42_v.txt | 32022 | 32022-Deleti | Deletion | 1 | C | - | 3 | 23773 | 0.01261936 | 55 | rnr1Y285F | FALSE | CAN | JSY3811 | 1 | rnr1Y285F-R | GCC | nope | >1bp-ins | G/C-1 | none | G/C-1 | GAGATGCCA | CCACGGT | CGG | GCC | C | none | nope | nope_GCC | G/C-1_CCACGGT | 8 | FALSE | FALSE |
| 42_v.txt | 32081 | 32081-Deleti | Deletion | 1 | A | - | 25 | 23771 | 0.10517017 | 63.48 | rnr1Y285F | FALSE | CAN | JSY3811 | 1 | rnr1Y285F-R | CAA | nope | >1bp-ins | A/T-1 | none | A/T-1 | CCATGCAAA | AAAAAAG | GTT | GTT | T | none | T>none | T>none_GTT | A/T-1_AAAAAAG | 8 | FALSE | FALSE |
| 42_v.txt | 32210 | 32210-Deleti | Deletion | 1 | T | - | 8 | 23773 | 0.03365162 | 40.75 | rnr1Y285F | FALSE | CAN | JSY3811 | 1 | rnr1Y285F-R | CTT | nope | >1bp-ins | A/T-1 | none | A/T-1 | ACCACCTTTG | TTTGGTG | GAA | CTT | T | none | nope | nope_CTT | A/T-1_TTTGGTG | 8 | FALSE | FALSE |
| 42_v.txt | 32259 | 32259-Deleti | Deletion | 1 | C | - | 35 | 28271 | 0.12380178 | 37.7714286 | rnr1Y285F | FALSE | CAN | JSY3811 | 1 | rnr1Y285F-R | ACC | nope | >1bp-ins | G/C-1 | none | G/C-1 | GATAGACCA | CCAAATA | TGG | ACC | C | none | nope | nope_ACC | G/C-1_CCAAATA | 8 | FALSE | FALSE |
| 42_v.txt | 32265 | 32265-Deleti | Deletion | 1 | A | - | 76 | 28272 | 0.2688172 | 36.7368421 | rnr1Y285F | FALSE | CAN | JSY3811 | 1 | rnr1Y285F-R | TAA | nope | >1bp-ins | A/T-1 | none | A/T-1 | CCAAATAAA | AAAATAC | ATT | ATT | T | none | T>none | T>none_ATT | A/T-1_AAAATAC | 8 | FALSE | FALSE |
| 42_v.txt | 32343 | 32343-Deleti | Deletion | 1 | G | - | 4 | 28272 | 0.01414827 | 64 | rnr1Y285F | FALSE | CAN | JSY3811 | 1 | rnr1Y285F-R | TGG | nope | >1bp-ins | G/C-1 | none | G/C-1 | ATATGTGGC | GGCAAAA | ACC | ACC | C | none | C>none | C>none_ACC | G/C-1_GGCAAAA | 8 | FALSE | FALSE |
| 42_v.txt | 32346 | 32346-Deleti | Deletion | 1 | A | - | 23 | 28272 | 0.08135258 | 61.6956522 | rnr1Y285F | FALSE | CAN | JSY3811 | 1 | rnr1Y285F-R | CAA | nope | >1bp-ins | A/T-1 | none | A/T-1 | TGTGGCAAA | AAAACCT | GTT | GTT | T | none | T>none | T>none_GTT | A/T-1_AAAACCT | 8 | FALSE | FALSE |
| 42_v.txt | 32416 | 32416-Deleti | Deletion | 1 | T | - | 6 | 28272 | 0.02122241 | 57.1666667 | rnr1Y285F | FALSE | CAN | JSY3811 | 1 | rnr1Y285F-R | GTT | nope | >1bp-ins | A/T-1 | none | A/T-1 | ATTGTGTTAG | TTAGTTT | CAA | GTT | T | none | nope | nope_GTT | A/T-1_TTAGTTT | 8 | FALSE | FALSE |
| 42_v.txt | 32430 | 32430-Deleti | Deletion | 1 | T | - | 22 | 28272 | 0.07781551 | 52.1818182 | rnr1Y285F | FALSE | CAN | JSY3811 | 1 | rnr1Y285F-R | ATT | nope | >1bp-ins | A/T-1 | none | A/T-1 | GGGTCATTG | TTGTATG | TAA | ATT | T | none | nope | nope_ATT | A/T-1_TTGTATG | 8 | FALSE | FALSE |
| 42_v.txt | 32442 | 32442-Deleti | Deletion | 1 | A | - | 10 | 28272 | 0.03537069 | 55.8 | rnr1Y285F | FALSE | CAN | JSY3811 | 1 | rnr1Y285F-R | TAA | nope | >1bp-ins | A/T-1 | none | A/T-1 | GGAACTAAA | AAAAGTC | ATT | ATT | T | none | T>none | T>none_ATT | A/T-1_AAAAGTC | 8 | FALSE | FALSE |
| 42_v.txt | 32490 | 32490-Deleti | Deletion | 1 | A | - | 16 | 28272 | 0.0565931 | 42.6875 | rnr1Y285F | FALSE | CAN | JSY3811 | 1 | rnr1Y285F-R | GAA | nope | >1bp-ins | A/T-1 | none | A/T-1 | ATACGGAAA | AAAACAA | CTT | CTT | T | none | T>none | T>none_CTT | A/T-1_AAAACAA | 8 | FALSE | FALSE |
| 42_v.txt | 32498 | 32498-Deleti | Deletion | 1 | T | - | 18 | 28272 | 0.06366723 | 45.2222222 | rnr1Y285F | FALSE | CAN | JSY3811 | 1 | rnr1Y285F-R | CTT | nope | >1bp-ins | A/T-1 | none | A/T-1 | AACAACTTTT | TTTTTTG | GAA | CTT | T | none | nope | nope_CTT | A/T-1_TTTTTTG | 8 | FALSE | FALSE |
| 42_v.txt | 32514 | 32514-Deleti | Deletion | 1 | G | - | 34 | 28272 | 0.12026033 | 37.0588235 | rnr1Y285F | FALSE | CAN | JSY3811 | 1 | rnr1Y285F-R | TGG | nope | >1bp-ins | G/C-1 | none | G/C-1 | GCTCTTGGA | GGAACGG | ACC | ACC | C | none | C>none | C>none_ACC | G/C-1_GGAACGG | 8 | FALSE | FALSE |
| 42_v.txt | 32522 | 32522-Deleti | Deletion | 1 | T | - | 9 | 28272 | 0.03183362 | 40.8888887 | rnr1Y285F | FALSE | CAN | JSY3811 | 1 | rnr1Y285F-R | ATT | nope | >1bp-ins | A/T-1 | none | A/T-1 | AACGGATTTT | TTTTCTG | TAA | ATT | T | none | nope | nope_ATT | A/T-1_TTTTCTG | 8 | FALSE | FALSE |
| 42_v.txt | 32536 | 32536-Deleti | Deletion | 1 | C | - | 1 | 28272 | 3.54E-03 | 38 | rnr1Y285F | FALSE | CAN | JSY3811 | 1 | rnr1Y285F-R | GCA | nope | >1bp-ins | G/C-1 | none | G/C-1 | GGTTTGCAG | CAGCTTC | CGT | GCA | C | none | nope | nope_GCA | G/C-1_CAGCTTC | 8 | FALSE | FALSE |
| 42_v.txt | 32544 | 32544-Deleti | Deletion | 1 | C | - | 6 | 28272 | 0.02122241 | 37.8333333 | rnr1Y285F | FALSE | CAN | JSY3811 | 1 | rnr1Y285F-R | ACC | nope | >1bp-ins | G/C-1 | none | G/C-1 | GCTTCACCAG | CCAGCAG | TGG | ACC | C | none | nope | nope_ACC | G/C-1_CCAGCAG | 8 | FALSE | FALSE |
| 42_v.txt | 32608 | 32608-Deleti | Deletion | 1 | C | - | 559 | 23846 | 2.34420867 | 44.4096601 | rnr1Y285F | FALSE | CAN | JSY3811 | 1 | rnr1Y285F-R | ACC | nope | >1bp-ins | G/C-1 | none | G/C-1 | AGGAAACCC | CCCAACC | TGG | ACC | C | none | nope | nope_ACC | G/C-1_CCCAACC | 8 | FALSE | FALSE |
| 42_v.txt | 32631 | 32631-Deleti | Deletion | 1 | T | - | 12 | 23846 | 0.05032291 | 48.9166667 | rnr1Y285F | FALSE | CAN | JSY3811 | 1 | rnr1Y285F-R | GTT | nope | >1bp-ins | A/T-1 | none | A/T-1 | CCTTCGTTTT | TTTTTAT | CAA | GTT | T | none | nope | nope_GTT | A/T-1_TTTTTAT | 8 | FALSE | FALSE |
| 42_v.txt | 32658 | 32658-Deleti | Deletion | 1 | C | - | 52 | 23846 | 0.21806592 | 59.1346154 | rnr1Y285F | FALSE | CAN | JSY3811 | 1 | rnr1Y285F-R | ACC | nope | >1bp-ins | G/C-1 | none | G/C-1 | CCTGGACCC | CCCCAGG | TGG | ACC | C | none | nope | nope_ACC | G/C-1_CCCCAGG | 8 | FALSE | FALSE |
| 42_v.txt | 32670 | 32670-Deleti | Deletion | 1 | G | - | 148 | 23846 | 0.62064917 | 58.0405405 | rnr1Y285F | FALSE | CAN | JSY3811 | 1 | rnr1Y285F-R | TGG | nope | >1bp-ins | G/C-1 | none | G/C-1 | GCACCTGGG | GGGTTTC | ACC | ACC | C | none | C>none | C>none_ACC | G/C-1_GGGTTTC | 8 | FALSE | FALSE |
| 42_v.txt | 32707 | 32707-Deleti | Deletion | 1 | C | - | 107 | 23846 | 0.44871257 | 61.3271028 | rnr1Y285F | FALSE | CAN | JSY3811 | 1 | rnr1Y285F-R | ACC | nope | >1bp-ins | G/C-1 | none | G/C-1 | CGGTAACCC | CCCCAGC | TGG | ACC | C | none | nope | nope_ACC | G/C-1_CCCCAGC | 8 | FALSE | FALSE |
| 42_v.txt | 32728 | 32728-Deleti | Deletion | 1 | A | - | 19 | 23846 | 0.07967793 | 60.3157894 | rnr1Y285F | FALSE | CAN | JSY3811 | 1 | rnr1Y285F-R | CAA | nope | >1bp-ins | A/T-1 | none | A/T-1 | CCATACAAA | AAAAACA | GTT | GTT | T | none | T>none | T>none_GTT | A/T-1_AAAAACA | 8 | FALSE | FALSE |
| 42_v.txt | 32760 | 32760-Deleti | Deletion | 1 | A | - | 25 | 23846 | 0.10483939 | 56.6 | rnr1Y285F | FALSE | CAN | JSY3811 | 1 | rnr1Y285F-R | TAA | nope | >1bp-ins | A/T-1 | none | A/T-1 | ATGGCTAAA | AAAACTT | ATT | ATT | T | none | T>none | T>none_ATT | A/T-1_AAAACTT | 8 | FALSE | FALSE |
| 42_v.txt | 32765 | 32765-Deleti | Deletion | 1 | T | - | 2 | 23846 | 8.39E-03 | 64 | rnr1Y285F | FALSE | CAN | JSY3811 | 1 | rnr1Y285F-R | CTT | nope | >1bp-ins | A/T-1 | none | A/T-1 | TAAAACTTTG | TTTGATG | GAA | CTT | T | none | nope | nope_CTT | A/T-1_TTTGATG | 8 | FALSE | FALSE |
| 42_v.txt | 32842 | 32842-Deleti | Deletion | 1 | A | - | 90 | 23846 | 0.37742179 | 42.5777772 | rnr1Y285F | FALSE | CAN | JSY3811 | 1 | rnr1Y285F-R | CAA | nope | >1bp-ins | A/T-1 | none | A/T-1 | TTACCCAAA | AAAAAAT | GTT | GTT | T | none | T>none | T>none_GTT | A/T-1_AAAAAAT | 8 | FALSE | FALSE |
| 42_v.txt | 32884 | 32884-Deleti | Deletion | 1 | A | - | 7 | 33529 | 0.02087745 | 37.7142857 | rnr1Y285F | FALSE | CAN | JSY3811 | 1 | rnr1Y285F-R | CAA | nope | >1bp-ins | A/T-1 | none | A/T-1 | ACGTCCAAA | AAAATTG | GTT | GTT | T | none | T>none | T>none_GTT | A/T-1_AAAATTG | 8 | FALSE | FALSE |
| 42_v.txt | 32917 | 32917-Deleti | Deletion | 1 | C | - | 5 | 33529 | 0.01491246 | 47.4 | rnr1Y285F | FALSE | CAN | JSY3811 | 1 | rnr1Y285F-R | TCC | nope | >1bp-ins | G/C-1 | none | G/C-1 | TAAGTTCCAC | CCAGGGC | AGG | TCC | C | none | nope | nope_TCC | G/C-1_CCAGGGC | 8 | FALSE | FALSE |
| 42_v.txt | 32920 | 32920-Deleti | Deletion | 1 | G | - | 5 | 33529 | 0.01491246 | 58.8 | rnr1Y285F | FALSE | CAN | JSY3811 | 1 | rnr1Y285F-R | AGG | nope | >1bp-ins | G/C-1 | none | G/C-1 | GTTCCAGGG | GGGCAAA | TCC | TCC | C | none | C>none | C>none_TCC | G/C-1_GGGCAAA | 8 | FALSE | FALSE |
| 42_v.txt | 32924 | 32924-Deleti | Deletion | 1 | A | - | 30 | 33529 | 0.08947478 | 51.5666667 | rnr1Y285F | FALSE | CAN | JSY3811 | 1 | rnr1Y285F-R | CAA | nope | >1bp-ins | A/T-1 | none | A/T-1 | CAGGGCAAA | AAAAGTG | GTT | GTT | T | none | T>none | T>none_GTT | A/T-1_AAAAGTG | 8 | FALSE | FALSE |
| 42_v.txt | 32935 | 32935-Deleti | Deletion | 1 | C | - | 5 | 33529 | 0.01491246 | 44.8 | rnr1Y285F | FALSE | CAN | JSY3811 | 1 | rnr1Y285F-R | GCC | nope | >1bp-ins | G/C-1 | none | G/C-1 | TGATTGCCCA | CCCAAGA | CGG | GCC | C | none | nope | nope_GCC | G/C-1_CCCAAGA | 8 | FALSE | FALSE |
| 42_v.txt | 32940 | 32940-Deleti | Deletion | 1 | G | - | 3 | 33529 | 8.95E-03 | 55 | rnr1Y285F | FALSE | CAN | JSY3811 | 1 | rnr1Y285F-R | AGA | nope | >1bp-ins | G/C-1 | none | G/C-1 | GCCCAAGAA | GAAAACC | TCT | TCT | C | none | C>none | C>none_TCT | G/C-1_GAAAACC | 8 | FALSE | FALSE |
| 42_v.txt | 33114 | 33114-Deleti | Deletion | 1 | C | - | 10 | 33529 | 0.02982493 | 45.6000017 | rnr1Y285F | FALSE | CAN | JSY3811 | 1 | rnr1Y285F-R | GCC | nope | >1bp-ins | G/C-1 | none | G/C-1 | ACTGGGCCG | CCGGCGT | CGG | GCC | C | none | nope | nope_GCC | G/C-1_CCGGCGT | 8 | FALSE | FALSE |
| 42_v.txt | 33147 | 33147-Deleti | Deletion | 1 | A | - | 6 | 33529 | 0.01789496 | 36.8333333 | rnr1Y285F | FALSE | CAN | JSY3811 | 1 | rnr1Y285F-R | GAA | nope | >1bp-ins | A/T-1 | none | A/T-1 | CCAATGAAA | AAAAGAC | CTT | CTT | T | none | T>none | T>none_CTT | A/T-1_AAAAGAC | 8 | FALSE | FALSE |
| 42_v.txt | 33175 | 33175-Deleti | Deletion | 1 | G | - | 2 | 33518 | 5.97E-03 | 38 | rnr1Y285F | FALSE | CAN | JSY3811 | 1 | rnr1Y285F-R | AGG | nope | >1bp-ins | G/C-1 | none | G/C-1 | CACCAAGGG | GGGCAAT | TCC | TCC | C | none | C>none | C>none_TCC | G/C-1_GGGCAAT | 8 | FALSE | FALSE |
| 42_v.txt | 33356 | 33356-Deleti | Deletion | 1 | C | - | 28 | 23943 | 0.11694441 | 56.8928571 | rnr1Y285F | FALSE | CAN | JSY3811 | 1 | rnr1Y285F-R | ACC | nope | >1bp-ins | G/C-1 | none | G/C-1 | TATTGACCCA | CCCACGT | TGG | ACC | C | none | nope | nope_ACC | G/C-1_CCCACGT | 8 | FALSE | FALSE |
| 42_v.txt | 33474 | 33474-Deleti | Deletion | 1 | C | - | 66 | 23940 | 0.27568922 | 25.1969697 | rnr1Y285F | FALSE | CAN | JSY3811 | 1 | rnr1Y285F-R | GCC | nope | >1bp-ins | G/C-1 | none | G/C-1 | GCTATGCCTT | CCTTTTT | CGG | GCC | C | none | nope | nope_GCC | G/C-1_CCTTTTT | 8 | FALSE | FALSE |
| 43_v.txt | 31675 | 31675-SNV- | SNV | 1 | A | C | 7 | 10659 | 0.0656722 | 40 | rnr1Y285F | FALSE | CAN | JSY3811 | 2 | rnr1Y285F-R | AAG | A>C | G/C+1 | A/T-1 | TA>GC | TA>GC | GATCAAAGG | AGGTAAT | TTC | TTC | T | G | T>G | T>G_TTC | TA>GC_AGGTAAT | 8 | FALSE | FALSE |
| 43_v.txt | 31723 | 31723-SNV- | SNV | 1 | A | C | 4 | 10659 | 0.03752697 | 55.75 | rnr1Y285F | FALSE | CAN | JSY3811 | 2 | rnr1Y285F-R | CAA | A>C | G/C+1 | A/T-1 | TA>GC | TA>GC | TGTCCCAAA | AAAAAGT | GTT | GTT | T | G | T>G | T>G_GTT | TA>GC_AAAAAGT | 8 | FALSE | FALSE |
| 43_v.txt | 31794 | 31794-SNV-T | SNV | 1 | T | G | 14 | 10659 | 0.1313444 | 53.8571429 | rnr1Y285F | FALSE | CAN | JSY3811 | 2 | rnr1Y285F-R | ATC | T>G | G/C+1 | A/T-1 | TA>GC | TA>GC | TCGACATCTC | TCTCCAA | TAG | ATC | T | C | T>G | T>G_ATC | TA>GC_TCTCCAA | 8 | FALSE | FALSE |
| 43_v.txt | 32109 | 32109-SNV- | SNV | 1 | A | C | 15 | 24741 | 0.06062811 | 60 | rnr1Y285F | FALSE | CAN | JSY3811 | 2 | rnr1Y285F-R | TAA | A>C | G/C+1 | A/T-1 | TA>GC | TA>GC | ATATTTAATA | AATAGCC | ATT | ATT | T | G | T>G | T>G_ATT | TA>GC_AATAGCC | 8 | FALSE | FALSE |
| 43_v.txt | 32337 | 32337-SNV- | SNV | 1 | A | C | 1 | 27533 | 3.63E-03 | 63.999994 | rnr1Y285F | FALSE | CAN | JSY3811 | 2 | rnr1Y285F-R | GAT | A>C | G/C+1 | A/T-1 | TA>GC | TA>GC | TTGAAGATA | ATATGTG | CTA | CTA | T | G | T>G | T>G_CTA | TA>GC_ATATGTG | 8 | FALSE | FALSE |
| 43_v.txt | 32439 | 32439-SNV- | SNV | 1 | A | C | 5 | 27533 | 0.01816003 | 53.4 | rnr1Y285F | FALSE | CAN | JSY3811 | 2 | rnr1Y285F-R | AAC | A>C | G/C+1 | A/T-1 | TA>GC | TA>GC | TATGGAACT | ACTAAAA | TTG | TTG | T | G | T>G | T>G_TTG | TA>GC_ACTAAAA | 8 | FALSE | FALSE |
| 43_v.txt | 32481 | 32481-SNV- | SNV | 1 | A | C | 8 | 27533 | 0.02905604 | 44.125 | rnr1Y285F | FALSE | CAN | JSY3811 | 2 | rnr1Y285F-R | TAA | A>C | G/C+1 | A/T-1 | TA>GC | TA>GC | AAGGTTAAG | AAGATAC | ATT | ATT | T | G | T>G | T>G_ATT | TA>GC_AAGATAC | 8 | FALSE | FALSE |

| | | | | | | | | | | | | | | | | | | | | | | | | | | | | | | | | | | |
|---|---|---|---|---|---|---|---|---|---|---|---|---|---|---|---|---|---|---|---|---|---|---|---|---|---|---|---|---|---|---|---|---|---|---|
| 43_v.txt | 32581 | 32581-SNV-T | SNV | 1 | T | G | 24 | 24364 | 0.09850599 | 34.4166667 | rnr1Y285F | FALSE | CAN | JSY3811 | 2 | rnr1Y285F-R | GTG | T>G | G/C+1 | A/T-1 | TA>GC | TA>GC | GAAATGTGA | TGAAGGC | CAC | GTG | T | C | T>G | T>G_GTG | TA>GC_TGAAGGC | 8 | FALSE | FALSE |
| 43_v.txt | 32785 | 32785-SNV-/ | SNV | 1 | A | C | 11 | 24698 | 0.04453802 | 49.0909091 | rnr1Y285F | FALSE | CAN | JSY3811 | 2 | rnr1Y285F-R | AAC | A>C | G/C+1 | A/T-1 | TA>GC | TA>GC | CCCAGAACT | ACTCGAA | TTG | TTG | T | G | T>G | T>G_TTG | TA>GC_ACTCGAA | 8 | FALSE | FALSE |
| 43_v.txt | 32823 | 32823-SNV-/ | SNV | 1 | A | C | 9 | 24698 | 0.0364402 | 37.1111111 | rnr1Y285F | FALSE | CAN | JSY3811 | 2 | rnr1Y285F-R | CAT | A>C | G/C+1 | A/T-1 | TA>GC | TA>GC | AAGTTCATT | ATTATTG | GTA | GTA | T | G | T>G | T>G_GTA | TA>GC_ATTATTG | 8 | FALSE | FALSE |
| 43_v.txt | 32842 | 32842-SNV-/ | SNV | 1 | A | C | 5 | 24698 | 0.02024455 | 37.9999943 | rnr1Y285F | FALSE | CAN | JSY3811 | 2 | rnr1Y285F-R | CAA | A>C | G/C+1 | A/T-1 | TA>GC | TA>GC | TTACCCAAA | AAAAAAT | GTT | GTT | T | G | T>G | T>G_GTT | TA>GC_AAAAAAT | 8 | FALSE | FALSE |
| 43_v.txt | 32884 | 32884-SNV-/ | SNV | 1 | A | C | 6 | 33567 | 0.0178747 | 30.8333333 | rnr1Y285F | FALSE | CAN | JSY3811 | 2 | rnr1Y285F-R | CAA | A>C | G/C+1 | A/T-1 | TA>GC | TA>GC | ACGTCCAAA | AAAATTG | GTT | GTT | T | G | T>G | T>G_GTT | TA>GC_AAAATTG | 8 | FALSE | FALSE |
| 43_v.txt | 32926 | 32926-SNV-/ | SNV | 1 | A | C | 14 | 33568 | 0.04170639 | 31.5714286 | rnr1Y285F | FALSE | CAN | JSY3811 | 2 | rnr1Y285F-R | AAA | A>C | G/C+1 | A/T-1 | TA>GC | TA>GC | GGGCAAAAC | AAGTGAT | TTT | TTT | T | G | T>G | T>G_TTT | TA>GC_AAGTGAT | 8 | FALSE | FALSE |
| 43_v.txt | 32948 | 32948-SNV-/ | SNV | 1 | A | C | 20 | 33568 | 0.05958055 | 42.3 | rnr1Y285F | FALSE | CAN | JSY3811 | 2 | rnr1Y285F-R | AAT | A>C | G/C+1 | A/T-1 | TA>GC | TA>GC | AAACCAATA | ATACATG | TTA | TTA | T | G | T>G | T>G_TTA | TA>GC_ATACATG | 8 | FALSE | FALSE |
| 43_v.txt | 33095 | 33095-SNV-T | SNV | 1 | T | G | 2 | 33568 | 5.96E-03 | 36.9999776 | rnr1Y285F | FALSE | CAN | JSY3811 | 2 | rnr1Y285F-R | ATA | T>G | G/C+1 | A/T-1 | TA>GC | TA>GC | ATATGATAT | TATAAGA | TAT | ATA | T | C | T>G | T>G_ATA | TA>GC_TATAAGA | 8 | FALSE | FALSE |
| 43_v.txt | 33476 | 33476-SNV-T | SNV | 1 | T | G | 5 | 21680 | 0.02306273 | 36.0002227 | rnr1Y285F | FALSE | CAN | JSY3811 | 2 | rnr1Y285F-R | CTT | T>G | G/C+1 | A/T-1 | TA>GC | TA>GC | TATGCCTTTT | TTTTTTT | GAA | CTT | T | C | T>G | T>G_CTT | TA>GC_TTTTTTT | 8 | FALSE | FALSE |
| 43_v.txt | 32112 | 32112-SNV-/ | SNV | 1 | A | G | 22 | 24741 | 0.08892122 | 55.0909091 | rnr1Y285F | FALSE | CAN | JSY3811 | 2 | rnr1Y285F-R | TAG | A>G | G/C+1 | A/T-1 | TA>CG | TA>CG | TTTAATAGCC | AGCCATT | ATC | ATC | T | C | T>C | T>C_ATC | TA>CG_AGCCATT | 8 | FALSE | FALSE |
| 43_v.txt | 32314 | 32314-SNV-T | SNV | 1 | T | C | 5 | 27533 | 0.01816003 | 38.1999908 | rnr1Y285F | FALSE | CAN | JSY3811 | 2 | rnr1Y285F-R | GTT | T>C | G/C+1 | A/T-1 | TA>CG | TA>CG | TAATGGTTG | TTGTTAA | CAA | GTT | T | G | T>C | T>C_GTT | TA>CG_TTGTTAA | 8 | FALSE | FALSE |
| 43_v.txt | 32346 | 32346-SNV-/ | SNV | 1 | A | G | 8 | 27532 | 0.0290571 | 63.8733236 | rnr1Y285F | FALSE | CAN | JSY3811 | 2 | rnr1Y285F-R | CAA | A>G | G/C+1 | A/T-1 | TA>CG | TA>CG | TGTGGCAAA | AAAACCT | GTT | GTT | T | C | T>C | T>C_GTT | TA>CG_AAAACCT | 8 | FALSE | FALSE |
| 43_v.txt | 32473 | 32473-SNV-/ | SNV | 1 | A | G | 6 | 27533 | 0.02179203 | 37.6674913 | rnr1Y285F | FALSE | CAN | JSY3811 | 2 | rnr1Y285F-R | TAG | A>G | G/C+1 | A/T-1 | TA>CG | TA>CG | CAATGTAGA | AGAAGGT | ATC | ATC | T | C | T>C | T>C_ATC | TA>CG_AGAAGGT | 8 | FALSE | FALSE |
| 43_v.txt | 32938 | 32938-SNV-/ | SNV | 1 | A | G | 116 | 33568 | 0.34556721 | 54.6206897 | rnr1Y285F | FALSE | CAN | JSY3811 | 2 | rnr1Y285F-R | CAA | A>G | G/C+1 | A/T-1 | TA>CG | TA>CG | TTGCCCAAG | AAGAAAA | GTT | GTT | T | C | T>C | T>C_GTT | TA>CG_AAGAAAA | 8 | FALSE | FALSE |
| 43_v.txt | 33476 | 33476-SNV-T | SNV | 1 | T | C | 35 | 21680 | 0.16143911 | 30.3716494 | rnr1Y285F | FALSE | CAN | JSY3811 | 2 | rnr1Y285F-R | CTT | T>C | G/C+1 | A/T-1 | TA>CG | TA>CG | TATGCCTTTT | TTTTTTT | GAA | CTT | T | G | T>C | T>C_CTT | TA>CG_TTTTTTT | 8 | FALSE | FALSE |
| 43_v.txt | 32128 | 32128-SNV-T | SNV | 1 | T | A | 29 | 24741 | 0.11721434 | 58.3448276 | rnr1Y285F | FALSE | CAN | JSY3811 | 2 | rnr1Y285F-R | TTG | T>A | A/T+1 | A/T-1 | TA>AT | TA>AT | AAACTTTGTC | TGTCACC | AAC | TTG | T | T | T>A | T>A_TTG | TA>AT_TGTCACC | 8 | FALSE | FALSE |
| 43_v.txt | 32314 | 32314-SNV-T | SNV | 1 | T | A | 3 | 27533 | 0.01089602 | 31.9999187 | rnr1Y285F | FALSE | CAN | JSY3811 | 2 | rnr1Y285F-R | GTT | T>A | A/T+1 | A/T-1 | TA>AT | TA>AT | TAATGGTTG | TTGTTAA | CAA | GTT | T | T | T>A | T>A_GTT | TA>AT_TTGTTAA | 8 | FALSE | FALSE |
| 43_v.txt | 32338 | 32338-SNV-T | SNV | 1 | T | A | 2 | 27533 | 7.26E-03 | 63.9975357 | rnr1Y285F | FALSE | CAN | JSY3811 | 2 | rnr1Y285F-R | ATA | T>A | A/T+1 | A/T-1 | TA>AT | TA>AT | TGAAGATAT | TATGTGG | TAT | ATA | T | T | T>A | T>A_ATA | TA>AT_TATGTGG | 8 | FALSE | FALSE |
| 43_v.txt | 32346 | 32346-SNV-/ | SNV | 1 | A | T | 4 | 27532 | 0.01452855 | 63.9959122 | rnr1Y285F | FALSE | CAN | JSY3811 | 2 | rnr1Y285F-R | CAA | A>T | A/T+1 | A/T-1 | TA>AT | TA>AT | TGTGGCAAA | AAAACCT | GTT | GTT | T | A | T>A | T>A_GTT | TA>AT_AAAACCT | 8 | FALSE | FALSE |
| 43_v.txt | 32726 | 32726-SNV-/ | SNV | 1 | A | T | 18 | 24698 | 0.0728804 | 60.3333333 | rnr1Y285F | FALSE | CAN | JSY3811 | 2 | rnr1Y285F-R | TAC | A>T | A/T+1 | A/T-1 | TA>AT | TA>AT | AACCATACA | ACAAAAA | ATG | ATG | T | A | T>A | T>A_ATG | TA>AT_ACAAAAA | 8 | FALSE | FALSE |
| 43_v.txt | 32785 | 32785-SNV-/ | SNV | 1 | A | T | 40 | 24698 | 0.16195643 | 47.875 | rnr1Y285F | FALSE | CAN | JSY3811 | 2 | rnr1Y285F-R | AAC | A>T | A/T+1 | A/T-1 | TA>AT | TA>AT | CCCAGAACT | ACTCGAA | TTG | TTG | T | A | T>A | T>A_TTG | TA>AT_ACTCGAA | 8 | FALSE | FALSE |
| 43_v.txt | 33072 | 33072-SNV-/ | SNV | 1 | A | T | 35 | 33567 | 0.10426907 | 47.4842155 | rnr1Y285F | FALSE | CAN | JSY3811 | 2 | rnr1Y285F-R | CAA | A>T | A/T+1 | A/T-1 | TA>AT | TA>AT | TATGCCAAA | AAAGAAC | GTT | GTT | T | A | T>A | T>A_GTT | TA>AT_AAAGAAC | 8 | FALSE | FALSE |
| 43_v.txt | 33095 | 33095-SNV-T | SNV | 1 | T | A | 9 | 33568 | 0.02681125 | 37.6656852 | rnr1Y285F | FALSE | CAN | JSY3811 | 2 | rnr1Y285F-R | ATA | T>A | A/T+1 | A/T-1 | TA>AT | TA>AT | ATATGATAT | TATAAGA | TAT | ATA | T | T | T>A | T>A_ATA | TA>AT_TATAAGA | 8 | FALSE | FALSE |
| 43_v.txt | 33204 | 33204-SNV-/ | SNV | 1 | A | T | 59 | 21681 | 0.27212767 | 39.7966102 | rnr1Y285F | FALSE | CAN | JSY3811 | 2 | rnr1Y285F-R | AAG | A>T | A/T+1 | A/T-1 | TA>AT | TA>AT | TGCTTAAGC | AGCTCTC | TTC | TTC | T | A | T>A | T>A_TTC | TA>AT_AGCTCTC | 8 | FALSE | FALSE |
| 43_v.txt | 31650 | 31650-SNV-( | SNV | 1 | C | T | 26 | 10659 | 0.24392532 | 36.2307692 | rnr1Y285F | FALSE | CAN | JSY3811 | 2 | rnr1Y285F-R | GCG | C>T | A/T+1 | G/C-1 | CG>TA | CG>TA | AGAATGCGA | CGAAATG | CGC | GCG | C | A | C>T | C>T_GCG | CG>TA_CGAAATG | 8 | FALSE | FALSE |
| 43_v.txt | 31658 | 31658-SNV-( | SNV | 1 | C | T | 18 | 10659 | 0.16887138 | 34.4444444 | rnr1Y285F | FALSE | CAN | JSY3811 | 2 | rnr1Y285F-R | GCG | C>T | A/T+1 | G/C-1 | CG>TA | CG>TA | AAATGGCGT | CGTGGAA | CGC | GCG | C | A | C>T | C>T_GCG | CG>TA_CGTGGAA | 8 | FALSE | FALSE |
| 43_v.txt | 31844 | 31844-SNV-( | SNV | 1 | C | T | 26 | 10646 | 0.24422318 | 47.6538462 | rnr1Y285F | FALSE | CAN | JSY3811 | 2 | rnr1Y285F-R | TCC | C>T | A/T+1 | G/C-1 | CG>TA | CG>TA | TAAGATCCA | CCAAACA | AGG | TCC | C | A | C>T | C>T_TCC | CG>TA_CCAAACA | 8 | FALSE | FALSE |
| 43_v.txt | 31884 | 31884-SNV-( | SNV | 1 | G | A | 14 | 10646 | 0.13150479 | 38.5 | rnr1Y285F | FALSE | CAN | JSY3811 | 2 | rnr1Y285F-R | AGC | G>A | A/T+1 | G/C-1 | CG>TA | CG>TA | GCGGCAGCA | GCAAAGC | TCG | TCG | C | T | C>T | C>T_TCG | CG>TA_GCAAAGC | 8 | FALSE | FALSE |
| 43_v.txt | 31911 | 31911-SNV-( | SNV | 1 | G | A | 12 | 10646 | 0.11271839 | 38.1666613 | rnr1Y285F | FALSE | CAN | JSY3811 | 2 | rnr1Y285F-R | TGC | G>A | A/T+1 | G/C-1 | CG>TA | CG>TA | TTTGGTGCA | GCAAAAG | ACG | ACG | C | T | C>T | C>T_ACG | CG>TA_GCAAAAG | 8 | FALSE | FALSE |
| 43_v.txt | 31912 | 31912-SNV-( | SNV | 1 | C | T | 14 | 10646 | 0.13150479 | 39.8569761 | rnr1Y285F | FALSE | CAN | JSY3811 | 2 | rnr1Y285F-R | GCA | C>T | A/T+1 | G/C-1 | CG>TA | CG>TA | TTGGTGCAA | CAAAAGC | CGT | GCA | C | A | C>T | C>T_GCA | CG>TA_CAAAAGC | 8 | FALSE | FALSE |
| 43_v.txt | 31920 | 31920-SNV-( | SNV | 1 | G | A | 12 | 10626 | 0.11293055 | 35.5833333 | rnr1Y285F | FALSE | CAN | JSY3811 | 2 | rnr1Y285F-R | CGT | G>A | A/T+1 | G/C-1 | CG>TA | CG>TA | AAAGCCGTG | GTGAAAC | GCA | GCA | C | T | C>T | C>T_GCA | CG>TA_GTGAAAC | 8 | FALSE | FALSE |
| 43_v.txt | 31962 | 31962-SNV-( | SNV | 1 | G | A | 31 | 24741 | 0.12529809 | 36.8709677 | rnr1Y285F | FALSE | CAN | JSY3811 | 2 | rnr1Y285F-R | CGC | G>A | A/T+1 | G/C-1 | CG>TA | CG>TA | GTGGCCGCA | GCATAAT | GCG | GCG | C | T | C>T | C>T_GCG | CG>TA_GCATAAT | 8 | FALSE | FALSE |
| 43_v.txt | 31972 | 31972-SNV-( | SNV | 1 | C | T | 20 | 24741 | 0.08083748 | 35.35 | rnr1Y285F | FALSE | CAN | JSY3811 | 2 | rnr1Y285F-R | GCC | C>T | A/T+1 | G/C-1 | CG>TA | CG>TA | AATAAGCCA | CCAAGCC | CGG | GCC | C | A | C>T | C>T_GCC | CG>TA_CCAAGCC | 8 | FALSE | FALSE |
| 43_v.txt | 31977 | 31977-SNV-( | SNV | 1 | C | T | 31 | 24741 | 0.12529809 | 46.7096774 | rnr1Y285F | FALSE | CAN | JSY3811 | 2 | rnr1Y285F-R | GCC | C>T | A/T+1 | G/C-1 | CG>TA | CG>TA | GCCAAGCCG | CCGGGCA | CGG | GCC | C | A | C>T | C>T_GCC | CG>TA_CCGGGCA | 8 | FALSE | FALSE |
| 43_v.txt | 31981 | 31981-SNV-( | SNV | 1 | G | A | 174 | 24741 | 0.70328604 | 52.3505747 | rnr1Y285F | FALSE | CAN | JSY3811 | 2 | rnr1Y285F-R | GGC | G>A | A/T+1 | G/C-1 | CG>TA | CG>TA | AGCCGGGCA | GCATTAA | CCG | CCG | C | T | C>T | C>T_CCG | CG>TA_GCATTAA | 8 | FALSE | FALSE |
| 43_v.txt | 32041 | 32041-SNV-( | SNV | 1 | G | A | 54 | 24741 | 0.21826119 | 59.2222222 | rnr1Y285F | FALSE | CAN | JSY3811 | 2 | rnr1Y285F-R | TGC | G>A | A/T+1 | G/C-1 | CG>TA | CG>TA | AAGCTTGCA | GCATAAA | ACG | ACG | C | T | C>T | C>T_ACG | CG>TA_GCATAAA | 8 | FALSE | FALSE |
| 43_v.txt | 32075 | 32075-SNV-( | SNV | 1 | C | T | 25 | 24741 | 0.10104685 | 62.08 | rnr1Y285F | FALSE | CAN | JSY3811 | 2 | rnr1Y285F-R | ACC | C>T | A/T+1 | G/C-1 | CG>TA | CG>TA | AAATAACCA | CCATGCA | TGG | ACC | C | A | C>T | C>T_ACC | CG>TA_CCATGCA | 8 | FALSE | FALSE |
| 43_v.txt | 32088 | 32088-SNV-( | SNV | 1 | C | T | 34 | 24741 | 0.13742371 | 58.6764706 | rnr1Y285F | FALSE | CAN | JSY3811 | 2 | rnr1Y285F-R | GCC | C>T | A/T+1 | G/C-1 | CG>TA | CG>TA | AAAAAGCCT | CCTGCAA | CGG | GCC | C | A | C>T | C>T_GCC | CG>TA_CCTGCAA | 8 | FALSE | FALSE |
| 43_v.txt | 32092 | 32092-SNV-( | SNV | 1 | C | T | 41 | 24741 | 0.16571683 | 60.4634146 | rnr1Y285F | FALSE | CAN | JSY3811 | 2 | rnr1Y285F-R | GCA | C>T | A/T+1 | G/C-1 | CG>TA | CG>TA | AGCCTGCAA | CAACACC | CGT | GCA | C | A | C>T | C>T_GCA | CG>TA_CAACACC | 8 | FALSE | FALSE |
| 43_v.txt | 32114 | 32114-SNV-( | SNV | 1 | C | T | 1839 | 24741 | 7.43300594 | 59.365416 | rnr1Y285F | FALSE | CAN | JSY3811 | 2 | rnr1Y285F-R | GCC | C>T | A/T+1 | G/C-1 | CG>TA | CG>TA | TAATAGCCA | CCATTCG | CGG | GCC | C | A | C>T | C>T_GCC | CG>TA_CCATTCG | 8 | FALSE | FALSE |
| 43_v.txt | 32165 | 32165-SNV-( | SNV | 1 | G | A | 24 | 24741 | 0.09700497 | 42.5841253 | rnr1Y285F | FALSE | CAN | JSY3811 | 2 | rnr1Y285F-R | CGC | G>A | A/T+1 | G/C-1 | CG>TA | CG>TA | CAAAGCGCC | GCCAAAT | GCG | GCG | C | T | C>T | C>T_GCG | CG>TA_GCCAAAT | 8 | FALSE | FALSE |
| 43_v.txt | 32166 | 32166-SNV-( | SNV | 1 | C | T | 31 | 24741 | 0.12529809 | 51.5113211 | rnr1Y285F | FALSE | CAN | JSY3811 | 2 | rnr1Y285F-R | GCC | C>T | A/T+1 | G/C-1 | CG>TA | CG>TA | AAAGCGCCA | CCAAATG | CGG | GCC | C | A | C>T | C>T_GCC | CG>TA_CCAAATG | 8 | FALSE | FALSE |
| 43_v.txt | 32271 | 32271-SNV-( | SNV | 1 | C | T | 38 | 27534 | 0.13801119 | 33.0523256 | rnr1Y285F | FALSE | CAN | JSY3811 | 2 | rnr1Y285F-R | ACG | C>T | A/T+1 | G/C-1 | CG>TA | CG>TA | AAAATACGG | CGGGAAC | TGC | ACG | C | A | C>T | C>T_ACG | CG>TA_CGGGAAC | 8 | FALSE | FALSE |
| 43_v.txt | 32272 | 32272-SNV-( | SNV | 1 | G | A | 10 | 27534 | 0.03631873 | 35.5000268 | rnr1Y285F | FALSE | CAN | JSY3811 | 2 | rnr1Y285F-R | CGG | G>A | A/T+1 | G/C-1 | CG>TA | CG>TA | AAATACGGG | GGGAACC | GCC | GCC | C | T | C>T | C>T_GCC | CG>TA_GGGAACC | 8 | FALSE | FALSE |
| 43_v.txt | 32274 | 32274-SNV-( | SNV | 1 | G | A | 16 | 27534 | 0.05810997 | 35.9375 | rnr1Y285F | FALSE | CAN | JSY3811 | 2 | rnr1Y285F-R | GGA | G>A | A/T+1 | G/C-1 | CG>TA | CG>TA | ATACGGGAA | GAACCAA | CCT | CCT | C | T | C>T | C>T_CCT | CG>TA_GAACCAA | 8 | FALSE | FALSE |
| 43_v.txt | 32299 | 32299-SNV-( | SNV | 1 | C | T | 53 | 27533 | 0.19249628 | 52.9811321 | rnr1Y285F | FALSE | CAN | JSY3811 | 2 | rnr1Y285F-R | GCG | C>T | A/T+1 | G/C-1 | CG>TA | CG>TA | AATTTGCGG | CGGCAGA | CGC | GCG | C | A | C>T | C>T_GCG | CG>TA_CGGCAGA | 8 | FALSE | FALSE |
| 43_v.txt | 32304 | 32304-SNV-( | SNV | 1 | G | A | 26 | 27533 | 0.09443214 | 56.3846154 | rnr1Y285F | FALSE | CAN | JSY3811 | 2 | rnr1Y285F-R | AGA | G>A | A/T+1 | G/C-1 | CG>TA | CG>TA | GCGGCAGAA | GAAATAA | TCT | TCT | C | T | C>T | C>T_TCT | CG>TA_GAAATAA | 8 | FALSE | FALSE |
| 43_v.txt | 32449 | 32449-SNV-( | SNV | 1 | C | T | 28 | 27533 | 0.10169615 | 52.7493702 | rnr1Y285F | FALSE | CAN | JSY3811 | 2 | rnr1Y285F-R | CCA | C>T | A/T+1 | G/C-1 | CG>TA | CG>TA | AAAGTCCAA | CAATGAA | GGT | CCA | C | A | C>T | C>T_CCA | CG>TA_CAATGAA | 8 | FALSE | FALSE |
| 43_v.txt | 32466 | 32466-SNV-( | SNV | 1 | C | T | 37 | 27533 | 0.13438419 | 49.8378378 | rnr1Y285F | FALSE | CAN | JSY3811 | 2 | rnr1Y285F-R | GCC | C>T | A/T+1 | G/C-1 | CG>TA | CG>TA | AGAGAGCCA | CCAATGT | CGG | GCC | C | A | C>T | C>T_GCC | CG>TA_CCAATGT | 8 | FALSE | FALSE |
| 43_v.txt | 32487 | 32487-SNV-( | SNV | 1 | C | T | 27 | 27533 | 0.09806414 | 46.0740741 | rnr1Y285F | FALSE | CAN | JSY3811 | 2 | rnr1Y285F-R | ACG | C>T | A/T+1 | G/C-1 | CG>TA | CG>TA | AAGATACGG | CGGAAAA | TGC | ACG | C | A | C>T | C>T_ACG | CG>TA_CGGAAAA | 8 | FALSE | FALSE |
| 43_v.txt | 32488 | 32488-SNV-( | SNV | 1 | G | A | 11 | 27533 | 0.03995206 | 45.8176136 | rnr1Y285F | FALSE | CAN | JSY3811 | 2 | rnr1Y285F-R | CGG | G>A | A/T+1 | G/C-1 | CG>TA | CG>TA | AGATACGGA | GGAAAAC | GCC | GCC | C | T | C>T | C>T_GCC | CG>TA_GGAAAAC | 8 | FALSE | FALSE |
| 43_v.txt | 32535 | 32535-SNV-( | SNV | 1 | G | A | 45 | 27531 | 0.16345211 | 35.4887762 | rnr1Y285F | FALSE | CAN | JSY3811 | 2 | rnr1Y285F-R | TGC | G>A | A/T+1 | G/C-1 | CG>TA | CG>TA | GGGTTTGCA | GCAGCTT | ACG | ACG | C | T | C>T | C>T_ACG | CG>TA_GCAGCTT | 8 | FALSE | FALSE |
| 43_v.txt | 32536 | 32536-SNV-( | SNV | 1 | C | T | 53 | 27531 | 0.19251026 | 36.2266111 | rnr1Y285F | FALSE | CAN | JSY3811 | 2 | rnr1Y285F-R | GCA | C>T | A/T+1 | G/C-1 | CG>TA | CG>TA | GGTTTGCAG | CAGCTTC | CGT | GCA | C | A | C>T | C>T_GCA | CG>TA_CAGCTTC | 8 | FALSE | FALSE |
| 43_v.txt | 32590 | 32590-SNV-( | SNV | 1 | C | T | 39 | 24364 | 0.16007224 | 36.1025641 | rnr1Y285F | FALSE | CAN | JSY3811 | 2 | rnr1Y285F-R | GCG | C>T | A/T+1 | G/C-1 | CG>TA | CG>TA | AGGCAGCGT | CGTTAAT | CGC | GCG | C | A | C>T | C>T_GCG | CG>TA_CGTTAAT | 8 | FALSE | FALSE |
| 43_v.txt | 32660 | 32660-SNV-( | SNV | 1 | C | T | 30 | 24698 | 0.12146733 | 47.7666667 | rnr1Y285F | FALSE | CAN | JSY3811 | 2 | rnr1Y285F-R | CCC | C>T | A/T+1 | G/C-1 | CG>TA | CG>TA | TGGACCCA | CCAGGCA | GGG | CCC | C | A | C>T | C>T_CCC | CG>TA_CCAGGCA | 8 | FALSE | FALSE |
| 43_v.txt | 32679 | 32679-SNV-( | SNV | 1 | C | T | 35 | 24698 | 0.14171188 | 61.9428571 | rnr1Y285F | FALSE | CAN | JSY3811 | 2 | rnr1Y285F-R | CCA | C>T | A/T+1 | G/C-1 | CG>TA | CG>TA | TTTCTCCAAT | CAATAAC | GGT | CCA | C | A | C>T | C>T_CCA | CG>TA_CAATAAC | 8 | FALSE | FALSE |
| 43_v.txt | 32700 | 32700-SNV-( | SNV | 1 | C | T | 42 | 24698 | 0.17005426 | 59.0952381 | rnr1Y285F | FALSE | CAN | JSY3811 | 2 | rnr1Y285F-R | GCC | C>T | A/T+1 | G/C-1 | CG>TA | CG>TA | ACTGGGCCG | CCGGTAA | CGG | GCC | C | A | C>T | C>T_GCC | CG>TA_CCGGTAA | 8 | FALSE | FALSE |
| 43_v.txt | 32764 | 32764-SNV-( | SNV | 1 | C | T | 216 | 24698 | 0.87456474 | 50.8518519 | rnr1Y285F | FALSE | CAN | JSY3811 | 2 | rnr1Y285F-R | ACT | C>T | A/T+1 | G/C-1 | CG>TA | CG>TA | CTAAAACTTT | CTTTGAT | TGA | ACT | C | A | C>T | C>T_ACT | CG>TA_CTTTGAT | 8 | FALSE | FALSE |
| 43_v.txt | 32776 | 32776-SNV-( | SNV | 1 | C | T | 26 | 24698 | 0.10527168 | 51.6153846 | rnr1Y285F | FALSE | CAN | JSY3811 | 2 | rnr1Y285F-R | GCG | C>T | A/T+1 | G/C-1 | CG>TA | CG>TA | TGGAAGCGA | CGACCCA | CGC | GCG | C | A | C>T | C>T_GCG | CG>TA_CGACCCA | 8 | FALSE | FALSE |
| 43_v.txt | 32781 | 32781-SNV-( | SNV | 1 | C | T | 30 | 24698 | 0.12146733 | 47.9333333 | rnr1Y285F | FALSE | CAN | JSY3811 | 2 | rnr1Y285F-R | CCA | C>T | A/T+1 | G/C-1 | CG>TA | CG>TA | GCGACCCAG | CAGAACT | GGT | CCA | C | A | C>T | C>T_CCA | CG>TA_CAGAACT | 8 | FALSE | FALSE |
| 43_v.txt | 32794 | 32794-SNV-( | SNV | 1 | C | T | 97 | 24698 | 0.39274435 | 45.5154639 | rnr1Y285F | FALSE | CAN | JSY3811 | 2 | rnr1Y285F-R | TCA | C>T | A/T+1 | G/C-1 | CG>TA | CG>TA | CGAATTCACC | CACCGTA | AGT | TCA | C | A | C>T | C>T_TCA | CG>TA_CACCGTA | 8 | FALSE | FALSE |
| 43_v.txt | 32798 | 32798-SNV-( | SNV | 1 | G | A | 34 | 24698 | 0.13766297 | 40.3529412 | rnr1Y285F | FALSE | CAN | JSY3811 | 2 | rnr1Y285F-R | CGT | G>A | A/T+1 | G/C-1 | CG>TA | CG>TA | TTCACCGTAA | GTAATAT | GCA | GCA | C | T | C>T | C>T_GCA | CG>TA_GTAATAT | 8 | FALSE | FALSE |
| 43_v.txt | 32860 | 32860-SNV-( | SNV | 1 | C | T | 19 | 24695 | 0.07693865 | 36.3157895 | rnr1Y285F | FALSE | CAN | JSY3811 | 2 | rnr1Y285F-R | GCC | C>T | A/T+1 | G/C-1 | CG>TA | CG>TA | TCCATGCCGC | CCGCCAG | CGG | GCC | C | A | C>T | C>T_GCC | CG>TA_CCGCCAG | 8 | FALSE | FALSE |
| 43_v.txt | 32900 | 32900-SNV-( | SNV | 1 | G | A | 35 | 33568 | 0.10426597 | 32.6560475 | rnr1Y285F | FALSE | CAN | JSY3811 | 2 | rnr1Y285F-R | GGC | G>A | A/T+1 | G/C-1 | CG>TA | CG>TA | GACTTGGCC | GCCAACT | CCG | CCG | C | T | C>T | C>T_CCG | CG>TA_GCCAACT | 8 | FALSE | FALSE |
| 43_v.txt | 32901 | 32901-SNV-( | SNV | 1 | C | T | 70 | 33568 | 0.20853194 | 31.7573562 | rnr1Y285F | FALSE | CAN | JSY3811 | 2 | rnr1Y285F-R | GCC | C>T | A/T+1 | G/C-1 | CG>TA | CG>TA | ACTTGGCCA | CCAACTA | CGG | GCC | C | A | C>T | C>T_GCC | CG>TA_CCAACTA | 8 | FALSE | FALSE |
| 43_v.txt | 32922 | 32922-SNV-( | SNV | 1 | G | A | 31 | 33568 | 0.09234986 | 42.7096774 | rnr1Y285F | FALSE | CAN | JSY3811 | 2 | rnr1Y285F-R | GGC | G>A | A/T+1 | G/C-1 | CG>TA | CG>TA | TCCAGGGCA | GCAAAAG | CCG | CCG | C | T | C>T | C>T_CCG | CG>TA_GCAAAAG | 8 | FALSE | FALSE |
| 43_v.txt | 32935 | 32935-SNV-( | SNV | 1 | C | T | 55 | 33568 | 0.16384652 | 41.8862218 | rnr1Y285F | FALSE | CAN | JSY3811 | 2 | rnr1Y285F-R | GCC | C>T | A/T+1 | G/C-1 | CG>TA | CG>TA | TGATTGCCCA | CCCAAGA | CGG | GCC | C | A | C>T | C>T_GCC | CG>TA_CCCAAGA | 8 | FALSE | FALSE |
| 43_v.txt | 32936 | 32936-SNV-( | SNV | 1 | C | T | 33 | 33567 | 0.09831084 | 29.3333333 | rnr1Y285F | FALSE | CAN | JSY3811 | 2 | rnr1Y285F-R | CCC | C>T | A/T+1 | G/C-1 | CG>TA | CG>TA | GATTGCCCA | CCAAGAA | GGG | CCC | C | A | C>T | C>T_CCC | CG>TA_CCAAGAA | 8 | FALSE | FALSE |
| 43_v.txt | 32965 | 32965-SNV-( | SNV | 1 | C | T | 35 | 33568 | 0.10426597 | 58.6857143 | rnr1Y285F | FALSE | CAN | JSY3811 | 2 | rnr1Y285F-R | GCC | C>T | A/T+1 | G/C-1 | CG>TA | CG>TA | CATTGGCCG | CCGCACC | CGG | GCC | C | A | C>T | C>T_GCC | CG>TA_CCGCACC | 8 | FALSE | FALSE |
| 43_v.txt | 32967 | 32967-SNV-( | SNV | 1 | G | A | 90 | 33568 | 0.26811249 | 60.7111111 | rnr1Y285F | FALSE | CAN | JSY3811 | 2 | rnr1Y285F-R | CGC | G>A | A/T+1 | G/C-1 | CG>TA | CG>TA | TTGGCCGCA | GCACCAA | GCG | GCG | C | T | C>T | C>T_GCG | CG>TA_GCACCAA | 8 | FALSE | FALSE |
| 43_v.txt | 32977 | 32977-SNV-( | SNV | 1 | C | T | 34 | 33568 | 0.10128694 | 55.2941177 | rnr1Y285F | FALSE | CAN | JSY3811 | 2 | rnr1Y285F-R | GCT | C>T | A/T+1 | G/C-1 | CG>TA | CG>TA | CAAATGCTG | CTGGAGA | CGA | GCT | C | A | C>T | C>T_GCT | CG>TA_CTGGAGA | 8 | FALSE | FALSE |
| 43_v.txt | 33015 | 33015-SNV-( | SNV | 1 | G | A | 639 | 33568 | 1.90359867 | 61.2097027 | rnr1Y285F | FALSE | CAN | JSY3811 | 2 | rnr1Y285F-R | GGA | G>A | A/T+1 | G/C-1 | CG>TA | CG>TA | AAAGAGGAT | GATGTAA | CCT | CCT | C | T | C>T | C>T_CCT | CG>TA_GATGTAA | 8 | FALSE | FALSE |
| 43_v.txt | 33056 | 33056-SNV-( | SNV | 1 | C | T | 56 | 33568 | 0.16682555 | 53.1785714 | rnr1Y285F | FALSE | CAN | JSY3811 | 2 | rnr1Y285F-R | GCG | C>T | A/T+1 | G/C-1 | CG>TA | CG>TA | GGACTGCGT | CGTGACA | CGC | GCG | C | A | C>T | C>T_GCG | CG>TA_CGTGACA | 8 | FALSE | FALSE |
| 43_v.txt | 33069 | 33069-SNV-( | SNV | 1 | G | A | 29 | 33568 | 0.0863918 | 52.06894 | rnr1Y285F | FALSE | CAN | JSY3811 | 2 | rnr1Y285F-R | TGC | G>A | A/T+1 | G/C-1 | CG>TA | CG>TA | GAATATGCC | GCCAAAG | ACG | ACG | C | T | C>T | C>T_ACG | CG>TA_GCCAAAG | 8 | FALSE | FALSE |
| 43_v.txt | 33070 | 33070-SNV-( | SNV | 1 | C | T | 50 | 33568 | 0.14895138 | 55.8990424 | rnr1Y285F | FALSE | CAN | JSY3811 | 2 | rnr1Y285F-R | GCC | C>T | A/T+1 | G/C-1 | CG>TA | CG>TA | AATATGCCA | CCAAAGA | CGG | GCC | C | A | C>T | C>T_GCC | CG>TA_CCAAAGA | 8 | FALSE | FALSE |
| 43_v.txt | 33113 | 33113-SNV-( | SNV | 1 | G | A | 31 | 33568 | 0.09234986 | 43.2588194 | rnr1Y285F | FALSE | CAN | JSY3811 | 2 | rnr1Y285F-R | GGC | G>A | A/T+1 | G/C-1 | CG>TA | CG>TA | CACTGGGCC | GCCGGCG | CCG | CCG | C | T | C>T | C>T_CCG | CG>TA_GCCGGCG | 8 | FALSE | FALSE |
| 43_v.txt | 33114 | 33114-SNV-( | SNV | 1 | C | T | 54 | 33568 | 0.16086749 | 43.1848979 | rnr1Y285F | FALSE | CAN | JSY3811 | 2 | rnr1Y285F-R | GCC | C>T | A/T+1 | G/C-1 | CG>TA | CG>TA | ACTGGGCCG | CCGGCGT | CGG | GCC | C | A | C>T | C>T_GCC | CG>TA_CCGGCGT | 8 | FALSE | FALSE |
| 43_v.txt | 33118 | 33118-SNV-( | SNV | 1 | C | T | 35 | 33567 | 0.10426907 | 42.9709993 | rnr1Y285F | FALSE | CAN | JSY3811 | 2 | rnr1Y285F-R | GCG | C>T | A/T+1 | G/C-1 | CG>TA | CG>TA | GGCCGGCGT | CGTTGGT | CGC | GCG | C | A | C>T | C>T_GCG | CG>TA_CGTTGGT | 8 | FALSE | FALSE |
| 43_v.txt | 33119 | 33119-SNV-( | SNV | 1 | G | A | 33 | 33568 | 0.09830791 | 41.0909091 | rnr1Y285F | FALSE | CAN | JSY3811 | 2 | rnr1Y285F-R | CGT | G>A | A/T+1 | G/C-1 | CG>TA | CG>TA | GCCGGCGTT | GTTGGTC | GCA | GCA | C | T | C>T | C>T_GCA | CG>TA_GTTGGTC | 8 | FALSE | FALSE |
| 43_v.txt | 33153 | 33153-SNV-( | SNV | 1 | C | T | 61 | 33566 | 0.18173151 | 36.868505 | rnr1Y285F | FALSE | CAN | JSY3811 | 2 | rnr1Y285F-R | ACC | C>T | A/T+1 | G/C-1 | CG>TA | CG>TA | AAAAGACCT | CCTGTAC | TGG | ACC | C | A | C>T | C>T_ACC | CG>TA_CCTGTAC | 8 | FALSE | FALSE |
| 43_v.txt | 33258 | 33258-SNV-( | SNV | 1 | C | T | 35 | 21681 | 0.16143167 | 42.6857143 | rnr1Y285F | FALSE | CAN | JSY3811 | 2 | rnr1Y285F-R | GCC | C>T | A/T+1 | G/C-1 | CG>TA | CG>TA | CCTATGCCAT | CCATCCT | CGG | GCC | C | A | C>T | C>T_GCC | CG>TA_CCATCCT | 8 | FALSE | FALSE |
| 43_v.txt | 33276 | 33276-SNV-( | SNV | 1 | G | A | 33 | 21681 | 0.152207 | 39.9090909 | rnr1Y285F | FALSE | CAN | JSY3811 | 2 | rnr1Y285F-R | CGT | G>A | A/T+1 | G/C-1 | CG>TA | CG>TA | GAGAACGTA | GTATCCT | GCA | GCA | C | T | C>T | C>T_GCA | CG>TA_GTATCCT | 8 | FALSE | FALSE |
| 43_v.txt | 33284 | 33284-SNV-( | SNV | 1 | G | A | 25 | 21681 | 0.11530834 | 52 | rnr1Y285F | FALSE | CAN | JSY3811 | 2 | rnr1Y285F-R | CGC | G>A | A/T+1 | G/C-1 | CG>TA | CG>TA | ATCCTCGCCA | GCCATTT | GCG | GCG | C | T | C>T | C>T_GCG | CG>TA_GCCATTT | 8 | FALSE | FALSE |
| 43_v.txt | 33309 | 33309-SNV-( | SNV | 1 | C | T | 31 | 21680 | 0.14298893 | 57.7364783 | rnr1Y285F | FALSE | CAN | JSY3811 | 2 | rnr1Y285F-R | GCG | C>T | A/T+1 | G/C-1 | CG>TA | CG>TA | AAAGAGCGC | CGCAATG | CGC | GCG | C | A | C>T | C>T_GCG | CG>TA_CGCAATG | 8 | FALSE | FALSE |
| 43_v.txt | 33310 | 33310-SNV-( | SNV | 1 | G | A | 25 | 21680 | 0.11531365 | 55.9946018 | rnr1Y285F | FALSE | CAN | JSY3811 | 2 | rnr1Y285F-R | CGC | G>A | A/T+1 | G/C-1 | CG>TA | CG>TA | AAGAGCGCA | GCAATGG | GCG | GCG | C | T | C>T | C>T_GCG | CG>TA_GCAATGG | 8 | FALSE | FALSE |
| 43_v.txt | 33412 | 33412-SNV-( | SNV | 1 | G | A | 30 | 21681 | 0.13837 | 44.3666667 | rnr1Y285F | FALSE | CAN | JSY3811 | 2 | rnr1Y285F-R | GGC | G>A | A/T+1 | G/C-1 | CG>TA | CG>TA | TGACCGGCT | GCTCATT | CCG | CCG | C | T | C>T | C>T_CCG | CG>TA_GCTCATT | 8 | FALSE | FALSE |
| 43_v.txt | 33474 | 33474-SNV-( | SNV | 1 | C | T | 31 | 21681 | 0.14298234 | 34.483871 | rnr1Y285F | FALSE | CAN | JSY3811 | 2 | rnr1Y285F-R | GCC | C>T | A/T+1 | G/C-1 | CG>TA | CG>TA | GCTATGCCTT | CCTTTTT | CGG | GCC | C | A | C>T | C>T_GCC | CG>TA_CCTTTTT | 8 | FALSE | FALSE |
| 43_v.txt | 32027 | 32027-SNV-( | SNV | 1 | G | C | 50 | 24741 | 0.20209369 | 59.28 | rnr1Y285F | FALSE | CAN | JSY3811 | 2 | rnr1Y285F-R | GGT | G>C | G/C+1 | G/C-1 | CG>GC | CG>GC | GCCACGGTA | GTATTTC | CCA | CCA | C | G | C>G | C>G_CCA | CG>GC_GTATTTC | 8 | FALSE | FALSE |
| 43_v.txt | 32033 | 32033-SNV-( | SNV | 1 | C | G | 204 | 24741 | 0.82454226 | 60.9754902 | rnr1Y285F | FALSE | CAN | JSY3811 | 2 | rnr1Y285F-R | TCA | C>G | G/C+1 | G/C-1 | CG>GC | CG>GC | GTATTTCAA | CAAAGCT | AGT | TCA | C | C | C>G | C>G_TCA | CG>GC_CAAAGCT | 8 | FALSE | FALSE |
| 43_v.txt | 32064 | 32064-SNV-( | SNV | 1 | G | C | 39 | 24741 | 0.15763308 | 62.948718 | rnr1Y285F | FALSE | CAN | JSY3811 | 2 | rnr1Y285F-R | TGA | G>C | G/C+1 | G/C-1 | CG>GC | CG>GC | GAGATTGAG | GAGATAA | ACT | ACT | C | G | C>G | C>G_ACT | CG>GC_GAGATAA | 8 | FALSE | FALSE |
| 43_v.txt | 32119 | 32119-SNV-( | SNV | 1 | C | G | 3 | 24741 | 0.01212562 | 54.6666667 | rnr1Y285F | FALSE | CAN | JSY3811 | 2 | rnr1Y285F-R | TCG | C>G | G/C+1 | G/C-1 | CG>GC | CG>GC | GCCATTCGA | CGAAAAC | AGC | TCG | C | C | C>G | C>G_TCG | CG>GC_CGAAAAC | 8 | FALSE | FALSE |
| 43_v.txt | 32165 | 32165-SNV-( | SNV | 1 | G | C | 2 | 24741 | 8.08E-03 | 38.0017602 | rnr1Y285F | FALSE | CAN | JSY3811 | 2 | rnr1Y285F-R | CGC | G>C | G/C+1 | G/C-1 | CG>GC | CG>GC | CAAAGCGCC | GCCAAAT | GCG | GCG | C | G | C>G | C>G_GCG | CG>GC_GCCAAAT | 8 | FALSE | FALSE |
| 43_v.txt | 32271 | 32271-SNV-( | SNV | 1 | C | G | 3 | 27534 | 0.01089562 | 34.3329935 | rnr1Y285F | FALSE | CAN | JSY3811 | 2 | rnr1Y285F-R | ACG | C>G | G/C+1 | G/C-1 | CG>GC | CG>GC | AAAATACGG | CGGGAAC | TGC | ACG | C | C | C>G | C>G_ACG | CG>GC_CGGGAAC | 8 | FALSE | FALSE |
| 43_v.txt | 32471 | 32471-SNV-( | SNV | 1 | G | C | 10 | 27533 | 0.03632005 | 58.8 | rnr1Y285F | FALSE | CAN | JSY3811 | 2 | rnr1Y285F-R | TGT | G>C | G/C+1 | G/C-1 | CG>GC | CG>GC | GCCAATGTA | GTAGAAG | ACA | ACA | C | G | C>G | C>G_ACA | CG>GC_GTAGAAG | 8 | FALSE | FALSE |
| 43_v.txt | 32474 | 32474-SNV-( | SNV | 1 | G | C | 6 | 27533 | 0.02179203 | 59.4921991 | rnr1Y285F | FALSE | CAN | JSY3811 | 2 | rnr1Y285F-R | AGA | G>C | G/C+1 | G/C-1 | CG>GC | CG>GC | AATGTAGAA | GAAGGTT | TCT | TCT | C | G | C>G | C>G_TCT | CG>GC_GAAGGTT | 8 | FALSE | FALSE |
| 43_v.txt | 32488 | 32488-SNV-( | SNV | 1 | G | C | 120 | 27533 | 0.43584063 | 45.6656248 | rnr1Y285F | FALSE | CAN | JSY3811 | 2 | rnr1Y285F-R | CGG | G>C | G/C+1 | G/C-1 | CG>GC | CG>GC | AGATACGGA | GGAAAAC | GCC | GCC | C | G | C>G | C>G_GCC | CG>GC_GGAAAAC | 8 | FALSE | FALSE |
| 43_v.txt | 32936 | 32936-SNV-( | SNV | 1 | C | G | 13 | 33567 | 0.03872851 | 58.5384615 | rnr1Y285F | FALSE | CAN | JSY3811 | 2 | rnr1Y285F-R | CCC | C>G | G/C+1 | G/C-1 | CG>GC | CG>GC | GATTGCCCA | CCAAGAA | GGG | CCC | C | C | C>G | C>G_CCC | CG>GC_CCAAGAA | 8 | FALSE | FALSE |
| 43_v.txt | 32959 | 32959-SNV-( | SNV | 1 | C | G | 22 | 33568 | 0.06553861 | 60.6363636 | rnr1Y285F | FALSE | CAN | JSY3811 | 2 | rnr1Y285F-R | CCA | C>G | G/C+1 | G/C-1 | CG>GC | CG>GC | TGTAACCATT | CATTGGC | GGT | CCA | C | C | C>G | C>G_CCA | CG>GC_CATTGGC | 8 | FALSE | FALSE |
| 43_v.txt | 33024 | 33024-SNV-( | SNV | 1 | G | C | 1047 | 33568 | 3.11904195 | 61.0735435 | rnr1Y285F | FALSE | CAN | JSY3811 | 2 | rnr1Y285F-R | AGG | G>C | G/C+1 | G/C-1 | CG>GC | CG>GC | GTAACAGGG | GGGATGA | TCC | TCC | C | G | C>G | C>G_TCC | CG>GC_GGGATGA | 8 | FALSE | FALSE |
| 43_v.txt | 33093 | 33093-SNV-( | SNV | 1 | G | C | 7 | 33568 | 0.02085319 | 42.998581 | rnr1Y285F | FALSE | CAN | JSY3811 | 2 | rnr1Y285F-R | TGA | G>C | G/C+1 | G/C-1 | CG>GC | CG>GC | AAATATGAT | GATATAA | ACT | ACT | C | G | C>G | C>G_ACT | CG>GC_GATATAA | 8 | FALSE | FALSE |
| 43_v.txt | 33113 | 33113-SNV-( | SNV | 1 | G | C | 3 | 33568 | 8.94E-03 | 43.0009684 | rnr1Y285F | FALSE | CAN | JSY3811 | 2 | rnr1Y285F-R | GGC | G>C | G/C+1 | G/C-1 | CG>GC | CG>GC | CACTGGGCC | GCCGGCG | CCG | CCG | C | G | C>G | C>G_CCG | CG>GC_GCCGGCG | 8 | FALSE | FALSE |
| 43_v.txt | 33114 | 33114-SNV-( | SNV | 1 | C | G | 3 | 33568 | 8.94E-03 | 37.999317 | rnr1Y285F | FALSE | CAN | JSY3811 | 2 | rnr1Y285F-R | GCC | C>G | G/C+1 | G/C-1 | CG>GC | CG>GC | ACTGGGCCG | CCGGCGT | CGG | GCC | C | C | C>G | C>G_GCC | CG>GC_CCGGCGT | 8 | FALSE | FALSE |
| 43_v.txt | 33310 | 33310-SNV-( | SNV | 1 | G | C | 5 | 21680 | 0.02306273 | 18.809853 | rnr1Y285F | FALSE | CAN | JSY3811 | 2 | rnr1Y285F-R | CGC | G>C | G/C+1 | G/C-1 | CG>GC | CG>GC | AAGAGCGCA | GCAATGG | GCG | GCG | C | G | C>G | C>G_GCG | CG>GC_GCAATGG | 8 | FALSE | FALSE |
| 43_v.txt | 33474 | 33474-SNV-( | SNV | 1 | C | G | 2 | 21681 | 9.23E-03 | 18.5 | rnr1Y285F | FALSE | CAN | JSY3811 | 2 | rnr1Y285F-R | GCC | C>G | G/C+1 | G/C-1 | CG>GC | CG>GC | GCTATGCCTT | CCTTTTT | CGG | GCC | C | C | C>G | C>G_GCC | CG>GC_CCTTTTT | 8 | FALSE | FALSE |
| 43_v.txt | 31657 | 31657-SNV-( | SNV | 1 | G | T | 77 | 10659 | 0.72239422 | 18.1354895 | rnr1Y285F | FALSE | CAN | JSY3811 | 2 | rnr1Y285F-R | GGC | G>T | A/T+1 | G/C-1 | CG>AT | CG>AT | GAAATGGCG | GCGTGGA | CCG | CCG | C | A | C>A | C>A_CCG | CG>AT_GCGTGGA | 8 | FALSE | FALSE |
| 43_v.txt | 31658 | 31658-SNV-( | SNV | 1 | C | A | 2 | 10659 | 0.01876349 | 37.5 | rnr1Y285F | FALSE | CAN | JSY3811 | 2 | rnr1Y285F-R | GCG | C>A | A/T+1 | G/C-1 | CG>AT | CG>AT | AAATGGCGT | CGTGGAA | CGC | GCG | C | T | C>A | C>A_GCG | CG>AT_CGTGGAA | 8 | FALSE | FALSE |
| 43_v.txt | 32027 | 32027-SNV-( | SNV | 1 | G | T | 122 | 24741 | 0.49310861 | 59.4918033 | rnr1Y285F | FALSE | CAN | JSY3811 | 2 | rnr1Y285F-R | GGT | G>T | A/T+1 | G/C-1 | CG>AT | CG>AT | GCCACGGTA | GTATTTC | CCA | CCA | C | A | C>A | C>A_CCA | CG>AT_GTATTTC | 8 | FALSE | FALSE |
| 43_v.txt | 32119 | 32119-SNV-( | SNV | 1 | C | A | 174 | 24741 | 0.70328604 | 56.7471264 | rnr1Y285F | FALSE | CAN | JSY3811 | 2 | rnr1Y285F-R | TCG | C>A | A/T+1 | G/C-1 | CG>AT | CG>AT | GCCATTCGA | CGAAAAC | AGC | TCG | C | T | C>A | C>A_TCG | CG>AT_CGAAAAC | 8 | FALSE | FALSE |
| 43_v.txt | 32188 | 32188-SNV-( | SNV | 1 | C | A | 25 | 24741 | 0.10104685 | 42.84 | rnr1Y285F | FALSE | CAN | JSY3811 | 2 | rnr1Y285F-R | ACT | C>A | A/T+1 | G/C-1 | CG>AT | CG>AT | CGAAAACTG | CTGCAAT | TGA | ACT | C | T | C>A | C>A_ACT | CG>AT_CTGCAAT | 8 | FALSE | FALSE |
| 43_v.txt | 32272 | 32272-SNV-( | SNV | 1 | G | T | 24 | 27534 | 0.08716496 | 29.9181432 | rnr1Y285F | FALSE | CAN | JSY3811 | 2 | rnr1Y285F-R | CGG | G>T | A/T+1 | G/C-1 | CG>AT | CG>AT | AAATACGGG | GGGAACC | GCC | GCC | C | A | C>A | C>A_GCC | CG>AT_GGGAACC | 8 | FALSE | FALSE |
| 43_v.txt | 32274 | 32274-SNV-( | SNV | 1 | G | T | 163 | 27534 | 0.59199535 | 36.0306749 | rnr1Y285F | FALSE | CAN | JSY3811 | 2 | rnr1Y285F-R | GGA | G>T | A/T+1 | G/C-1 | CG>AT | CG>AT | ATACGGGAA | GAACCAA | CCT | CCT | C | A | C>A | C>A_CCT | CG>AT_GAACCAA | 8 | FALSE | FALSE |
| 43_v.txt | 32373 | 32373-SNV-( | SNV | 1 | G | T | 55 | 27533 | 0.19976029 | 62.0909091 | rnr1Y285F | FALSE | CAN | JSY3811 | 2 | rnr1Y285F-R | AGC | G>T | A/T+1 | G/C-1 | CG>AT | CG>AT | TCAATAGCA | GCAATAA | TCG | TCG | C | A | C>A | C>A_TCG | CG>AT_GCAATAA | 8 | FALSE | FALSE |
| 43_v.txt | 32432 | 32432-SNV-( | SNV | 1 | G | T | 17 | 27533 | 0.06174409 | 59.1176471 | rnr1Y285F | FALSE | CAN | JSY3811 | 2 | rnr1Y285F-R | TGT | G>T | A/T+1 | G/C-1 | CG>AT | CG>AT | GTCATTGTAT | GTATGGA | ACA | ACA | C | A | C>A | C>A_ACA | CG>AT_GTATGGA | 8 | FALSE | FALSE |
| 43_v.txt | 32487 | 32487-SNV-( | SNV | 1 | C | A | 8 | 27533 | 0.02905604 | 32.75 | rnr1Y285F | FALSE | CAN | JSY3811 | 2 | rnr1Y285F-R | ACG | C>A | A/T+1 | G/C-1 | CG>AT | CG>AT | AAGATACGG | CGGAAAA | TGC | ACG | C | T | C>A | C>A_ACG | CG>AT_CGGAAAA | 8 | FALSE | FALSE |
| 43_v.txt | 32530 | 32530-SNV-( | SNV | 1 | G | T | 21 | 27533 | 0.07627211 | 38.7619048 | rnr1Y285F | FALSE | CAN | JSY3811 | 2 | rnr1Y285F-R | GGG | G>T | A/T+1 | G/C-1 | CG>AT | CG>AT | TTCTGGGGT | GGTTTGC | CCC | CCC | C | A | C>A | C>A_CCC | CG>AT_GGTTTGC | 8 | FALSE | FALSE |
| 43_v.txt | 32545 | 32545-SNV-( | SNV | 1 | C | A | 36 | 27530 | 0.13076644 | 37.3333333 | rnr1Y285F | FALSE | CAN | JSY3811 | 2 | rnr1Y285F-R | CCA | C>A | A/T+1 | G/C-1 | CG>AT | CG>AT | CTTCACCAGC | CAGCAGT | GGT | CCA | C | T | C>A | C>A_CCA | CG>AT_CAGCAGT | 8 | FALSE | FALSE |
| 43_v.txt | 32566 | 32566-SNV-( | SNV | 1 | C | A | 10 | 24356 | 0.04105765 | 37.6 | rnr1Y285F | FALSE | CAN | JSY3811 | 2 | rnr1Y285F-R | TCA | C>A | A/T+1 | G/C-1 | CG>AT | CG>AT | CTAGTTCAGT | CAGTACC | AGT | TCA | C | T | C>A | C>A_TCA | CG>AT_CAGTACC | 8 | FALSE | FALSE |
| 43_v.txt | 32783 | 32783-SNV-( | SNV | 1 | G | T | 28 | 24698 | 0.1133695 | 45.1071429 | rnr1Y285F | FALSE | CAN | JSY3811 | 2 | rnr1Y285F-R | AGA | G>T | A/T+1 | G/C-1 | CG>AT | CG>AT | GACCCAGAA | GAACTCG | TCT | TCT | C | A | C>A | C>A_TCT | CG>AT_GAACTCG | 8 | FALSE | FALSE |
| 43_v.txt | 32917 | 32917-SNV-( | SNV | 1 | C | A | 20 | 33568 | 0.05958055 | 38 | rnr1Y285F | FALSE | CAN | JSY3811 | 2 | rnr1Y285F-R | TCC | C>A | A/T+1 | G/C-1 | CG>AT | CG>AT | TAAGTTCCA | CCAGGGC | AGG | TCC | C | T | C>A | C>A_TCC | CG>AT_CCAGGGC | 8 | FALSE | FALSE |

| | | | | | | | | | | | | | | | | | | | | | | | | | | | | | | | | | | |
|---|---|---|---|---|---|---|---|---|---|---|---|---|---|---|---|---|---|---|---|---|---|---|---|---|---|---|---|---|---|---|---|---|---|---|
| 43_v.txt | 32935 | 32935-SNV-C | SNV | 1 | C | A | 4 | 33568 | 0.01191611 | 24.2492994 | rnr1Y285F | FALSE | CAN | JSY3811 | 2 | rnr1Y285F-R | GCC | C>A | A/T+1 | G/C-1 | CG>AT | CG>AT | TGATTGCCCA | CCCAAGA | CGG | GCC | C | T | C>A | C>A_GCC | CG>AT_CCCAAGA | 8 | FALSE | FALSE |
| 43_v.txt | 32936 | 32936-SNV-C | SNV | 1 | C | A | 26 | 33567 | 0.07745703 | 22.2307692 | rnr1Y285F | FALSE | CAN | JSY3811 | 2 | rnr1Y285F-R | CCC | C>A | A/T+1 | G/C-1 | CG>AT | CG>AT | GATTGCCCA | CCAAGAA | GGG | CCC | C | T | C>A | C>A_CCC | CG>AT_CCAAGAA | 8 | FALSE | FALSE |
| 43_v.txt | 32940 | 32940-SNV-C | SNV | 1 | G | T | 19526 | 33568 | 58.1684938 | 56.2949913 | rnr1Y285F | FALSE | CAN | JSY3811 | 2 | rnr1Y285F-R | AGA | G>T | A/T+1 | G/C-1 | CG>AT | CG>AT | GCCCAAGAA | GAAAACC | TCT | TCT | C | A | C>A | C>A_TCT | CG>AT_GAAAACC | 8 | FALSE | FALSE |
| 43_v.txt | 33012 | 33012-SNV-C | SNV | 1 | G | T | 34 | 33568 | 0.10128694 | 58.7058824 | rnr1Y285F | FALSE | CAN | JSY3811 | 2 | rnr1Y285F-R | AGA | G>T | A/T+1 | G/C-1 | CG>AT | CG>AT | GTGAAAGAG | GAGGATG | TCT | TCT | C | A | C>A | C>A_TCT | CG>AT_GAGGATG | 8 | FALSE | FALSE |
| 43_v.txt | 33024 | 33024-SNV-C | SNV | 1 | G | T | 9 | 33568 | 0.02681125 | 49.2222222 | rnr1Y285F | FALSE | CAN | JSY3811 | 2 | rnr1Y285F-R | AGG | G>T | A/T+1 | G/C-1 | CG>AT | CG>AT | GTAACAGGG | GGGATGA | TCC | TCC | C | A | C>A | C>A_TCC | CG>AT_GGGATGA | 8 | FALSE | FALSE |
| 43_v.txt | 33050 | 33050-SNV-C | SNV | 1 | G | T | 15 | 33568 | 0.04468542 | 44.5333333 | rnr1Y285F | FALSE | CAN | JSY3811 | 2 | rnr1Y285F-R | AGG | G>T | A/T+1 | G/C-1 | CG>AT | CG>AT | ACCCAAGGA | GGACTGC | TCC | TCC | C | A | C>A | C>A_TCC | CG>AT_GGACTGC | 8 | FALSE | FALSE |
| 43_v.txt | 33093 | 33093-SNV-C | SNV | 1 | G | T | 17 | 33568 | 0.05064347 | 44.4112808 | rnr1Y285F | FALSE | CAN | JSY3811 | 2 | rnr1Y285F-R | TGA | G>T | A/T+1 | G/C-1 | CG>AT | CG>AT | AAATATGATA | GATATAA | ACT | ACT | C | A | C>A | C>A_ACT | CG>AT_GATATAA | 8 | FALSE | FALSE |
| 43_v.txt | 33151 | 33151-SNV-C | SNV | 1 | G | T | 29 | 33567 | 0.08639438 | 39.6896552 | rnr1Y285F | FALSE | CAN | JSY3811 | 2 | rnr1Y285F-R | AGA | G>T | A/T+1 | G/C-1 | CG>AT | CG>AT | TGAAAAGAC | GACCTGT | TCT | TCT | C | A | C>A | C>A_TCT | CG>AT_GACCTGT | 8 | FALSE | FALSE |
| 43_v.txt | 33154 | 33154-SNV-C | SNV | 1 | C | A | 14 | 33566 | 0.04170887 | 35.9994491 | rnr1Y285F | FALSE | CAN | JSY3811 | 2 | rnr1Y285F-R | CCT | C>A | A/T+1 | G/C-1 | CG>AT | CG>AT | AAAGACCTG | CTGTACC | GGA | CCT | C | T | C>A | C>A_CCT | CG>AT_CTGTACC | 8 | FALSE | FALSE |
| 43_v.txt | 33175 | 33175-SNV-C | SNV | 1 | G | T | 11 | 33549 | 0.03278786 | 34.2727273 | rnr1Y285F | FALSE | CAN | JSY3811 | 2 | rnr1Y285F-R | AGG | G>T | A/T+1 | G/C-1 | CG>AT | CG>AT | CACCAAGGG | GGGCAAT | TCC | TCC | C | A | C>A | C>A_TCC | CG>AT_GGGCAAT | 8 | FALSE | FALSE |
| 43_v.txt | 33208 | 33208-SNV-C | SNV | 1 | C | A | 30 | 21681 | 0.13837 | 39.4 | rnr1Y285F | FALSE | CAN | JSY3811 | 2 | rnr1Y285F-R | TCT | C>A | A/T+1 | G/C-1 | CG>AT | CG>AT | TAAGCTCTCT | CTCTCTT | AGA | TCT | C | T | C>A | C>A_TCT | CG>AT_CTCTCTT | 8 | FALSE | FALSE |
| 43_v.txt | 33220 | 33220-SNV-C | SNV | 1 | C | A | 39 | 21681 | 0.179881 | 37.5897436 | rnr1Y285F | FALSE | CAN | JSY3811 | 2 | rnr1Y285F-R | TCA | C>A | A/T+1 | G/C-1 | CG>AT | CG>AT | TCACTTCAGC | CAGCGTT | AGT | TCA | C | T | C>A | C>A_TCA | CG>AT_CAGCGTT | 8 | FALSE | FALSE |
| 43_v.txt | 33474 | 33474-SNV-C | SNV | 1 | C | A | 60 | 21681 | 0.27674 | 34.1333333 | rnr1Y285F | FALSE | CAN | JSY3811 | 2 | rnr1Y285F-R | GCC | C>A | A/T+1 | G/C-1 | CG>AT | CG>AT | GCTATGCCTT | CCTTTTT | CGG | GCC | C | T | C>A | C>A_GCC | CG>AT_CCTTTTT | 8 | FALSE | FALSE |
| 43_v.txt | 33475 | 33475-SNV-C | SNV | 1 | C | A | 29 | 21681 | 0.13375767 | 35.3459974 | rnr1Y285F | FALSE | CAN | JSY3811 | 2 | rnr1Y285F-R | CCT | C>A | A/T+1 | G/C-1 | CG>AT | CG>AT | CTATGCCTTT | CTTTTTT | GGA | CCT | C | T | C>A | C>A_CCT | CG>AT_CTTTTTT | 8 | FALSE | FALSE |
| 43_v.txt | 33153 | 33153-MNV- | MNV | 2 | CC | AA | 1 | 33566 | 2.98E-03 | 37.4999988 | rnr1Y285F | FALSE | CAN | JSY3811 | 2 | rnr1Y285F-R | ACC | nope | >1bp-ins | >1bpdel | none | MNV | AAAAGACCT | CCTGTAC | TGG | ACC | CC | none | NA | NA_ACC | MNV_CCTGTAC | 8 | FALSE | FALSE |
| 43_v.txt | 32030 | 32030-Insert | Insertion | 1 | - | T | 204 | 24741 | 0.82454226 | 58.3186275 | rnr1Y285F | FALSE | CAN | JSY3811 | 2 | rnr1Y285F-R | ATT | nope | A/T+1 | >1bpdel | none | A/T+1 | ACGGTATTTC | TTTCAAA | TAA | ATT | - | A | NA | NA_ATT | A/T+1_TTTCAAA | 8 | FALSE | FALSE |
| 43_v.txt | 32081 | 32081-Insert | Insertion | 1 | - | A | 15 | 24741 | 0.06062811 | 63.8666667 | rnr1Y285F | FALSE | CAN | JSY3811 | 2 | rnr1Y285F-R | CAA | nope | A/T+1 | >1bpdel | none | A/T+1 | CCATGCAAA | AAAAAAG | GTT | CAA | - | T | NA | NA_CAA | A/T+1_AAAAAAG | 8 | FALSE | FALSE |
| 43_v.txt | 32346 | 32346-Insert | Insertion | 1 | - | A | 4 | 27533 | 0.01452802 | 64 | rnr1Y285F | FALSE | CAN | JSY3811 | 2 | rnr1Y285F-R | CAA | nope | A/T+1 | >1bpdel | none | A/T+1 | TGTGGCAAA | AAAACCT | GTT | CAA | - | T | NA | NA_CAA | A/T+1_AAAACCT | 8 | FALSE | FALSE |
| 43_v.txt | 32450 | 32450-Insert | Insertion | 1 | - | A | 14 | 27533 | 0.05084807 | 58.4285714 | rnr1Y285F | FALSE | CAN | JSY3811 | 2 | rnr1Y285F-R | CAA | nope | A/T+1 | >1bpdel | none | A/T+1 | AAGTCCAATC | AATGAAT | GTT | CAA | - | T | NA | NA_CAA | A/T+1_AATGAAT | 8 | FALSE | FALSE |
| 43_v.txt | 32736 | 32736-Insert | Insertion | 2 | - | TA | 6 | 24698 | 0.02429347 | 59.3333333 | rnr1Y285F | FALSE | CAN | JSY3811 | 2 | rnr1Y285F-R | GTA | nope | >1bp-ins | >1bpdel | none | >1bp-ins | AAACAGTATA | TATATTA | CAT | GTA | - | none | NA | NA_GTA | >1bp-ins_TATATTA | 8 | FALSE | FALSE |
| 43_v.txt | 32760 | 32760-Insert | Insertion | 1 | - | C | 213 | 24698 | 0.86241801 | 53.0892019 | rnr1Y285F | FALSE | CAN | JSY3811 | 2 | rnr1Y285F-R | TAA | nope | G/C+1 | >1bpdel | none | G/C+1 | ATGGCTAAA | AAAACTT | ATT | TAA | - | G | NA | NA_TAA | G/C+1_AAAACTT | 8 | FALSE | FALSE |
| 43_v.txt | 32842 | 32842-Insert | Insertion | 1 | - | A | 25 | 24698 | 0.10122277 | 43.68 | rnr1Y285F | FALSE | CAN | JSY3811 | 2 | rnr1Y285F-R | CAA | nope | A/T+1 | >1bpdel | none | A/T+1 | TTACCCAAAA | AAAAAAT | GTT | CAA | - | T | NA | NA_CAA | A/T+1_AAAAAAT | 8 | FALSE | FALSE |
| 43_v.txt | 32842 | 32842-Insert | Insertion | 2 | - | AA | 1 | 24698 | 4.05E-03 | 38 | rnr1Y285F | FALSE | CAN | JSY3811 | 2 | rnr1Y285F-R | CAA | nope | >1bp-ins | >1bpdel | none | >1bp-ins | TTACCCAAAA | AAAAAAT | GTT | CAA | - | none | NA | NA_CAA | >1bp-ins_AAAAAAT | 8 | FALSE | FALSE |
| 43_v.txt | 32313 | 32313-Deleti | Deletion | 3 | GTT | - | 8 | 27533 | 0.02905604 | 64 | rnr1Y285F | FALSE | CAN | JSY3811 | 2 | rnr1Y285F-R | GGT | nope | >1bp-ins | >1bpdel | none | >1bpdel | ATAATGGTTC | GTTGTTA | CCA | GGT | GTT | none | NA | NA_GGT | >1bpdel_GTTGTTA | 8 | FALSE | FALSE |
| 43_v.txt | 32337 | 32337-Deleti | Deletion | 2 | AT | - | 27 | 27533 | 0.09806414 | 60.6666667 | rnr1Y285F | FALSE | CAN | JSY3811 | 2 | rnr1Y285F-R | GAT | nope | >1bp-ins | >1bpdel | none | >1bpdel | TTGAAGATAT | ATATGTG | CTA | GAT | AT | none | NA | NA_GAT | >1bpdel_ATATGTG | 8 | FALSE | FALSE |
| 43_v.txt | 32473 | 32473-Deleti | Deletion | 3 | AGA | - | 9 | 27533 | 0.03268805 | 48.7777778 | rnr1Y285F | FALSE | CAN | JSY3811 | 2 | rnr1Y285F-R | TAG | nope | >1bp-ins | >1bpdel | none | >1bpdel | CAATGTAGA | AGAAGGT | ATC | TAG | AGA | none | NA | NA_TAG | >1bpdel_AGAAGGT | 8 | FALSE | FALSE |
| 43_v.txt | 33072 | 33072-Deleti | Deletion | 2 | AA | - | 12 | 33568 | 0.03574833 | 53 | rnr1Y285F | FALSE | CAN | JSY3811 | 2 | rnr1Y285F-R | CAA | nope | >1bp-ins | >1bpdel | none | >1bpdel | TATGCCAAAC | AAAGAAC | GTT | CAA | AA | none | NA | NA_CAA | >1bpdel_AAAGAAC | 8 | FALSE | FALSE |
| 43_v.txt | 33094 | 33094-Deleti | Deletion | 2 | AT | - | 21 | 33568 | 0.06255958 | 54.8571429 | rnr1Y285F | FALSE | CAN | JSY3811 | 2 | rnr1Y285F-R | GAT | nope | >1bp-ins | >1bpdel | none | >1bpdel | AATATGATAT | ATATAAG | CTA | GAT | AT | none | NA | NA_GAT | >1bpdel_ATATAAG | 8 | FALSE | FALSE |
| 43_v.txt | 33206 | 33206-Deleti | Deletion | 2 | CT | - | 6 | 21681 | 0.027674 | 37.8333333 | rnr1Y285F | FALSE | CAN | JSY3811 | 2 | rnr1Y285F-R | GCT | nope | >1bp-ins | >1bpdel | none | >1bpdel | CTTAAGCTCT | CTCTCTC | CGA | GCT | CT | none | NA | NA_GCT | >1bpdel_CTCTCTC | 8 | FALSE | FALSE |
| 43_v.txt | 31663 | 31663-Deleti | Deletion | 1 | A | - | 7 | 10659 | 0.0656722 | 37.5714286 | rnr1Y285F | FALSE | CAN | JSY3811 | 2 | rnr1Y285F-R | GAA | nope | >1bp-ins | A/T-1 | none | A/T-1 | GCGTGGAAA | AAATGTG | CTT | CTT | T | none | T>none | T>none_CTT | A/T-1_AAATGTG | 8 | FALSE | FALSE |
| 43_v.txt | 31723 | 31723-Deleti | Deletion | 1 | A | - | 8 | 10659 | 0.07505395 | 52.875 | rnr1Y285F | FALSE | CAN | JSY3811 | 2 | rnr1Y285F-R | CAA | nope | >1bp-ins | A/T-1 | none | A/T-1 | TGTCCCAAA | AAAAAGT | GTT | GTT | T | none | T>none | T>none_GTT | A/T-1_AAAAAGT | 8 | FALSE | FALSE |
| 43_v.txt | 31863 | 31863-Deleti | Deletion | 1 | A | - | 8 | 10646 | 0.0751456 | 44 | rnr1Y285F | FALSE | CAN | JSY3811 | 2 | rnr1Y285F-R | GAA | nope | >1bp-ins | A/T-1 | none | A/T-1 | AACAGGAAA | AAAATAG | CTT | CTT | T | none | T>none | T>none_CTT | A/T-1_AAAATAG | 8 | FALSE | FALSE |
| 43_v.txt | 31904 | 31904-Deleti | Deletion | 1 | T | - | 6 | 10646 | 0.0563592 | 44 | rnr1Y285F | FALSE | CAN | JSY3811 | 2 | rnr1Y285F-R | ATT | nope | >1bp-ins | A/T-1 | none | A/T-1 | ATTGAATTTT | TTTTGGT | TAA | ATT | T | none | nope | nope_ATT | A/T-1_TTTTGGT | 8 | FALSE | FALSE |
| 43_v.txt | 31913 | 31913-Deleti | Deletion | 1 | A | - | 5 | 10646 | 0.046966 | 37.9999993 | rnr1Y285F | FALSE | CAN | JSY3811 | 2 | rnr1Y285F-R | CAA | nope | >1bp-ins | A/T-1 | none | A/T-1 | TGGTGCAAA | AAAAGCC | GTT | GTT | T | none | T>none | T>none_GTT | A/T-1_AAAAGCC | 8 | FALSE | FALSE |
| 43_v.txt | 31972 | 31972-Deleti | Deletion | 1 | C | - | 7 | 24741 | 0.02829312 | 35 | rnr1Y285F | FALSE | CAN | JSY3811 | 2 | rnr1Y285F-R | GCC | nope | >1bp-ins | G/C-1 | none | G/C-1 | AATAAGCCA | CCAAGCC | CGG | GCC | C | none | nope | nope_GCC | G/C-1_CCAAGCC | 8 | FALSE | FALSE |
| 43_v.txt | 31979 | 31979-Deleti | Deletion | 1 | G | - | 11 | 24741 | 0.04446061 | 43 | rnr1Y285F | FALSE | CAN | JSY3811 | 2 | rnr1Y285F-R | CGG | nope | >1bp-ins | G/C-1 | none | G/C-1 | CAAGCCGGG | GGGCATT | GCC | GCC | C | none | C>none | C>none_GCC | G/C-1_GGGCATT | 8 | FALSE | FALSE |
| 43_v.txt | 32008 | 32008-Deleti | Deletion | 1 | C | - | 10 | 24741 | 0.04041874 | 48.9 | rnr1Y285F | FALSE | CAN | JSY3811 | 2 | rnr1Y285F-R | TCG | nope | >1bp-ins | G/C-1 | none | G/C-1 | GTAACTCGTC | CGTCACG | AGC | TCG | C | none | nope | nope_TCG | G/C-1_CGTCACG | 8 | FALSE | FALSE |
| 43_v.txt | 32045 | 32045-Deleti | Deletion | 1 | A | - | 6 | 24741 | 0.02425124 | 63 | rnr1Y285F | FALSE | CAN | JSY3811 | 2 | rnr1Y285F-R | TAA | nope | >1bp-ins | A/T-1 | none | A/T-1 | TTGCATAAAT | AAATCTG | ATT | ATT | T | none | T>none | T>none_ATT | A/T-1_AAATCTG | 8 | FALSE | FALSE |
| 43_v.txt | 32081 | 32081-Deleti | Deletion | 1 | A | - | 27 | 24741 | 0.10913059 | 63.8148149 | rnr1Y285F | FALSE | CAN | JSY3811 | 2 | rnr1Y285F-R | CAA | nope | >1bp-ins | A/T-1 | none | A/T-1 | CCATGCAAA | AAAAAAG | GTT | GTT | T | none | T>none | T>none_GTT | A/T-1_AAAAAAG | 8 | FALSE | FALSE |
| 43_v.txt | 32346 | 32346-Deleti | Deletion | 1 | A | - | 14 | 27532 | 0.05084992 | 60.4285714 | rnr1Y285F | FALSE | CAN | JSY3811 | 2 | rnr1Y285F-R | CAA | nope | >1bp-ins | A/T-1 | none | A/T-1 | TGTGGCAAA | AAAACCT | GTT | GTT | T | none | T>none | T>none_GTT | A/T-1_AAAACCT | 8 | FALSE | FALSE |
| 43_v.txt | 32356 | 32356-Deleti | Deletion | 1 | T | - | 11 | 27533 | 0.03995206 | 63.5454546 | rnr1Y285F | FALSE | CAN | JSY3811 | 2 | rnr1Y285F-R | GTA | nope | >1bp-ins | A/T-1 | none | A/T-1 | CCTTTGTACC | TACCAGA | CAT | GTA | T | none | nope | nope_GTA | A/T-1_TACCAGA | 8 | FALSE | FALSE |
| 43_v.txt | 32381 | 32381-Deleti | Deletion | 1 | A | - | 8 | 27533 | 0.02905604 | 63.625 | rnr1Y285F | FALSE | CAN | JSY3811 | 2 | rnr1Y285F-R | TAA | nope | >1bp-ins | A/T-1 | none | A/T-1 | AATAATAAA | AAAGGGA | ATT | ATT | T | none | T>none | T>none_ATT | A/T-1_AAAGGGA | 8 | FALSE | FALSE |
| 43_v.txt | 32416 | 32416-Deleti | Deletion | 1 | T | - | 6 | 27533 | 0.02179203 | 64 | rnr1Y285F | FALSE | CAN | JSY3811 | 2 | rnr1Y285F-R | GTT | nope | >1bp-ins | A/T-1 | none | A/T-1 | ATTGTGTTAG | TTAGTTT | CAA | GTT | T | none | nope | nope_GTT | A/T-1_TTAGTTT | 8 | FALSE | FALSE |
| 43_v.txt | 32466 | 32466-Deleti | Deletion | 1 | C | - | 3 | 27533 | 0.01089602 | 46.6666667 | rnr1Y285F | FALSE | CAN | JSY3811 | 2 | rnr1Y285F-R | GCC | nope | >1bp-ins | G/C-1 | none | G/C-1 | AGAGAGCCA | CCAATGT | CGG | GCC | C | none | nope | nope_GCC | G/C-1_CCAATGT | 8 | FALSE | FALSE |
| 43_v.txt | 32490 | 32490-Deleti | Deletion | 1 | A | - | 19 | 27533 | 0.0690081 | 41.5789474 | rnr1Y285F | FALSE | CAN | JSY3811 | 2 | rnr1Y285F-R | GAA | nope | >1bp-ins | A/T-1 | none | A/T-1 | ATACGGAAA | AAAACAA | CTT | CTT | T | none | T>none | T>none_CTT | A/T-1_AAAACAA | 8 | FALSE | FALSE |
| 43_v.txt | 32498 | 32498-Deleti | Deletion | 1 | T | - | 45 | 27533 | 0.16344024 | 43.3777778 | rnr1Y285F | FALSE | CAN | JSY3811 | 2 | rnr1Y285F-R | CTT | nope | >1bp-ins | A/T-1 | none | A/T-1 | AACAACTTTT | TTTTTTG | GAA | CTT | T | none | nope | nope_CTT | A/T-1_TTTTTTG | 8 | FALSE | FALSE |
| 43_v.txt | 32528 | 32528-Deleti | Deletion | 1 | G | - | 5 | 27533 | 0.01816003 | 43.2 | rnr1Y285F | FALSE | CAN | JSY3811 | 2 | rnr1Y285F-R | TGG | nope | >1bp-ins | G/C-1 | none | G/C-1 | TTTTCTGGGG | GGGGTTT | ACC | ACC | C | none | C>none | C>none_ACC | G/C-1_GGGGTTT | 8 | FALSE | FALSE |
| 43_v.txt | 32608 | 32608-Deleti | Deletion | 1 | C | - | 7 | 24364 | 0.02873092 | 37 | rnr1Y285F | FALSE | CAN | JSY3811 | 2 | rnr1Y285F-R | ACC | nope | >1bp-ins | G/C-1 | none | G/C-1 | AGGAAACCC | CCCAACC | TGG | ACC | C | none | nope | nope_ACC | G/C-1_CCCAACC | 8 | FALSE | FALSE |
| 43_v.txt | 32631 | 32631-Deleti | Deletion | 1 | T | - | 13 | 24364 | 0.05335741 | 49.6153846 | rnr1Y285F | FALSE | CAN | JSY3811 | 2 | rnr1Y285F-R | GTT | nope | >1bp-ins | A/T-1 | none | A/T-1 | CCTTCGTTTT | TTTTTAT | CAA | GTT | T | none | nope | nope_GTT | A/T-1_TTTTTAT | 8 | FALSE | FALSE |
| 43_v.txt | 32658 | 32658-Deleti | Deletion | 1 | C | - | 21 | 24698 | 0.08502713 | 61.8095238 | rnr1Y285F | FALSE | CAN | JSY3811 | 2 | rnr1Y285F-R | ACC | nope | >1bp-ins | G/C-1 | none | G/C-1 | CCTGGACCCC | CCCCAGG | TGG | ACC | C | none | nope | nope_ACC | G/C-1_CCCCAGG | 8 | FALSE | FALSE |
| 43_v.txt | 32670 | 32670-Deleti | Deletion | 1 | G | - | 14 | 24698 | 0.05668475 | 57.1428571 | rnr1Y285F | FALSE | CAN | JSY3811 | 2 | rnr1Y285F-R | TGG | nope | >1bp-ins | G/C-1 | none | G/C-1 | GCACCTGGG | GGGTTTC | ACC | ACC | C | none | C>none | C>none_ACC | G/C-1_GGGTTTC | 8 | FALSE | FALSE |
| 43_v.txt | 32697 | 32697-Deleti | Deletion | 1 | G | - | 4 | 24698 | 0.01619564 | 64 | rnr1Y285F | FALSE | CAN | JSY3811 | 2 | rnr1Y285F-R | TGG | nope | >1bp-ins | G/C-1 | none | G/C-1 | CCAACTGGG | GGGCCGG | ACC | ACC | C | none | C>none | C>none_ACC | G/C-1_GGGCCGG | 8 | FALSE | FALSE |
| 43_v.txt | 32707 | 32707-Deleti | Deletion | 1 | C | - | 18 | 24698 | 0.0728804 | 61.7222222 | rnr1Y285F | FALSE | CAN | JSY3811 | 2 | rnr1Y285F-R | ACC | nope | >1bp-ins | G/C-1 | none | G/C-1 | CGGTAACCC | CCCCAGC | TGG | ACC | C | none | nope | nope_ACC | G/C-1_CCCCAGC | 8 | FALSE | FALSE |
| 43_v.txt | 32728 | 32728-Deleti | Deletion | 1 | A | - | 12 | 24698 | 0.04858693 | 61.4166667 | rnr1Y285F | FALSE | CAN | JSY3811 | 2 | rnr1Y285F-R | CAA | nope | >1bp-ins | A/T-1 | none | A/T-1 | CCATACAAA | AAAAACA | GTT | GTT | T | none | T>none | T>none_GTT | A/T-1_AAAAACA | 8 | FALSE | FALSE |
| 43_v.txt | 32779 | 32779-Deleti | Deletion | 1 | C | - | 5 | 24698 | 0.02024455 | 49.4 | rnr1Y285F | FALSE | CAN | JSY3811 | 2 | rnr1Y285F-R | ACC | nope | >1bp-ins | G/C-1 | none | G/C-1 | AAGCGACCC | CCCAGAA | TGG | ACC | C | none | nope | nope_ACC | G/C-1_CCCAGAA | 8 | FALSE | FALSE |
| 43_v.txt | 32804 | 32804-Deleti | Deletion | 1 | T | - | 12 | 24698 | 0.04858693 | 41.1666667 | rnr1Y285F | FALSE | CAN | JSY3811 | 2 | rnr1Y285F-R | ATT | nope | >1bp-ins | A/T-1 | none | A/T-1 | GTAATATTTG | TTTGACA | TAA | ATT | T | none | nope | nope_ATT | A/T-1_TTTGACA | 8 | FALSE | FALSE |
| 43_v.txt | 32839 | 32839-Deleti | Deletion | 1 | C | - | 9 | 24698 | 0.0364402 | 37.6666667 | rnr1Y285F | FALSE | CAN | JSY3811 | 2 | rnr1Y285F-R | ACC | nope | >1bp-ins | G/C-1 | none | G/C-1 | TAATTACCCA | CCCAAAA | TGG | ACC | C | none | nope | nope_ACC | G/C-1_CCCAAAA | 8 | FALSE | FALSE |
| 43_v.txt | 32842 | 32842-Deleti | Deletion | 1 | A | - | 68 | 24698 | 0.27532594 | 41.2205882 | rnr1Y285F | FALSE | CAN | JSY3811 | 2 | rnr1Y285F-R | CAA | nope | >1bp-ins | A/T-1 | none | A/T-1 | TTACCCAAAA | AAAAAAT | GTT | GTT | T | none | T>none | T>none_GTT | A/T-1_AAAAAAT | 8 | FALSE | FALSE |
| 43_v.txt | 32884 | 32884-Deleti | Deletion | 1 | A | - | 33 | 33567 | 0.09831084 | 36.0606061 | rnr1Y285F | FALSE | CAN | JSY3811 | 2 | rnr1Y285F-R | CAA | nope | >1bp-ins | A/T-1 | none | A/T-1 | ACGTCCAAA | AAAATTG | GTT | GTT | T | none | T>none | T>none_GTT | A/T-1_AAAATTG | 8 | FALSE | FALSE |
| 43_v.txt | 32901 | 32901-Deleti | Deletion | 1 | C | - | 1 | 33568 | 2.98E-03 | 38 | rnr1Y285F | FALSE | CAN | JSY3811 | 2 | rnr1Y285F-R | GCC | nope | >1bp-ins | G/C-1 | none | G/C-1 | ACTTGGCCA | CCAACTA | CGG | GCC | C | none | nope | nope_GCC | G/C-1_CCAACTA | 8 | FALSE | FALSE |
| 43_v.txt | 32924 | 32924-Deleti | Deletion | 1 | A | - | 30 | 33568 | 0.08937083 | 53.7666667 | rnr1Y285F | FALSE | CAN | JSY3811 | 2 | rnr1Y285F-R | CAA | nope | >1bp-ins | A/T-1 | none | A/T-1 | CAGGGCAAA | AAAAGTG | GTT | GTT | T | none | T>none | T>none_GTT | A/T-1_AAAAGTG | 8 | FALSE | FALSE |
| 43_v.txt | 32935 | 32935-Deleti | Deletion | 1 | C | - | 5 | 33568 | 0.01489514 | 53.4 | rnr1Y285F | FALSE | CAN | JSY3811 | 2 | rnr1Y285F-R | GCC | nope | >1bp-ins | G/C-1 | none | G/C-1 | TGATTGCCCA | CCCAAGA | CGG | GCC | C | none | nope | nope_GCC | G/C-1_CCCAAGA | 8 | FALSE | FALSE |
| 43_v.txt | 32967 | 32967-Deleti | Deletion | 1 | G | - | 17 | 33568 | 0.05064347 | 62.7058824 | rnr1Y285F | FALSE | CAN | JSY3811 | 2 | rnr1Y285F-R | CGC | nope | >1bp-ins | G/C-1 | none | G/C-1 | TTGGCCGCAC | GCACCAA | GCG | GCG | C | none | C>none | C>none_GCG | G/C-1_GCACCAA | 8 | FALSE | FALSE |
| 43_v.txt | 33000 | 33000-Deleti | Deletion | 1 | A | - | 11 | 33568 | 0.0327693 | 61.5454546 | rnr1Y285F | FALSE | CAN | JSY3811 | 2 | rnr1Y285F-R | GAA | nope | >1bp-ins | A/T-1 | none | A/T-1 | TGTGAGAAA | AAAACTG | CTT | CTT | T | none | T>none | T>none_CTT | A/T-1_AAAACTG | 8 | FALSE | FALSE |
| 43_v.txt | 33070 | 33070-Deleti | Deletion | 1 | C | - | 1 | 33568 | 2.98E-03 | 64 | rnr1Y285F | FALSE | CAN | JSY3811 | 2 | rnr1Y285F-R | GCC | nope | >1bp-ins | G/C-1 | none | G/C-1 | AATATGCCA | CCAAAGA | CGG | GCC | C | none | nope | nope_GCC | G/C-1_CCAAAGA | 8 | FALSE | FALSE |
| 43_v.txt | 33072 | 33072-Deleti | Deletion | 1 | A | - | 2 | 33567 | 5.96E-03 | 63.9999996 | rnr1Y285F | FALSE | CAN | JSY3811 | 2 | rnr1Y285F-R | CAA | nope | >1bp-ins | A/T-1 | none | A/T-1 | TATGCCAAAC | AAAGAAC | GTT | GTT | T | none | T>none | T>none_GTT | A/T-1_AAAGAAC | 8 | FALSE | FALSE |
| 43_v.txt | 33114 | 33114-Deleti | Deletion | 1 | C | - | 1 | 33568 | 2.98E-03 | 64 | rnr1Y285F | FALSE | CAN | JSY3811 | 2 | rnr1Y285F-R | GCC | nope | >1bp-ins | G/C-1 | none | G/C-1 | ACTGGGCCG | CCGGCGT | CGG | GCC | C | none | nope | nope_GCC | G/C-1_CCGGCGT | 8 | FALSE | FALSE |
| 43_v.txt | 33147 | 33147-Deleti | Deletion | 1 | A | - | 8 | 33568 | 0.02383222 | 40.375 | rnr1Y285F | FALSE | CAN | JSY3811 | 2 | rnr1Y285F-R | GAA | nope | >1bp-ins | A/T-1 | none | A/T-1 | CCAATGAAA | AAAAGAC | CTT | CTT | T | none | T>none | T>none_CTT | A/T-1_AAAAGAC | 8 | FALSE | FALSE |
| 43_v.txt | 33175 | 33175-Deleti | Deletion | 1 | G | - | 3 | 33549 | 8.94E-03 | 38 | rnr1Y285F | FALSE | CAN | JSY3811 | 2 | rnr1Y285F-R | AGG | nope | >1bp-ins | G/C-1 | none | G/C-1 | CACCAAGGG | GGGCAAT | TCC | TCC | C | none | C>none | C>none_TCC | G/C-1_GGGCAAT | 8 | FALSE | FALSE |
| 43_v.txt | 33284 | 33284-Deleti | Deletion | 1 | G | - | 2 | 21681 | 9.23E-03 | 51 | rnr1Y285F | FALSE | CAN | JSY3811 | 2 | rnr1Y285F-R | CGC | nope | >1bp-ins | G/C-1 | none | G/C-1 | ATCCTCGCCA | GCCATTT | GCG | GCG | C | none | C>none | C>none_GCG | G/C-1_GCCATTT | 8 | FALSE | FALSE |
| 43_v.txt | 33356 | 33356-Deleti | Deletion | 1 | C | - | 466 | 21681 | 2.14934736 | 55.027897 | rnr1Y285F | FALSE | CAN | JSY3811 | 2 | rnr1Y285F-R | ACC | nope | >1bp-ins | G/C-1 | none | G/C-1 | TATTGACCCA | CCCACGT | TGG | ACC | C | none | nope | nope_ACC | G/C-1_CCCACGT | 8 | FALSE | FALSE |
| 43_v.txt | 33474 | 33474-Deleti | Deletion | 1 | C | - | 65 | 21681 | 0.29980167 | 29.6153846 | rnr1Y285F | FALSE | CAN | JSY3811 | 2 | rnr1Y285F-R | GCC | nope | >1bp-ins | G/C-1 | none | G/C-1 | GCTATGCCTT | CCTTTTT | CGG | GCC | C | none | nope | nope_GCC | G/C-1_CCTTTTT | 8 | FALSE | FALSE |
| 44_v.txt | 31723 | 31723-SNV-A | SNV | 1 | A | C | 7 | 14772 | 0.04738695 | 59.8571429 | rnr1Y285F | FALSE | CAN | JSY3811 | 3 | rnr1Y285F-R | CAA | A>C | G/C+1 | A/T-1 | TA>GC | TA>GC | TGTCCCAAA | AAAAAGT | GTT | GTT | T | G | T>G | T>G_GTT | TA>GC_AAAAAGT | 8 | FALSE | FALSE |
| 44_v.txt | 31839 | 31839-SNV-A | SNV | 1 | A | C | 7 | 14772 | 0.04738695 | 52.8571429 | rnr1Y285F | FALSE | CAN | JSY3811 | 3 | rnr1Y285F-R | TAA | A>C | G/C+1 | A/T-1 | TA>GC | TA>GC | TGAAATAAG | AAGATCC | ATT | ATT | T | G | T>G | T>G_ATT | TA>GC_AAGATCC | 8 | FALSE | FALSE |
| 44_v.txt | 32109 | 32109-SNV-A | SNV | 1 | A | C | 10 | 23899 | 0.04184276 | 63.9 | rnr1Y285F | FALSE | CAN | JSY3811 | 3 | rnr1Y285F-R | TAA | A>C | G/C+1 | A/T-1 | TA>GC | TA>GC | ATATTTAATA | AATAGCC | ATT | ATT | T | G | T>G | T>G_ATT | TA>GC_AATAGCC | 8 | FALSE | FALSE |
| 44_v.txt | 32352 | 32352-SNV-T | SNV | 1 | T | G | 5 | 39917 | 0.01252599 | 55.2 | rnr1Y285F | FALSE | CAN | JSY3811 | 3 | rnr1Y285F-R | CTT | T>G | G/C+1 | A/T-1 | TA>GC | TA>GC | AAAACCTTTG | TTTGTAC | GAA | CTT | T | C | T>G | T>G_CTT | TA>GC_TTTGTAC | 8 | FALSE | FALSE |
| 44_v.txt | 32481 | 32481-SNV-A | SNV | 1 | A | C | 17 | 39917 | 0.04258837 | 45.8235294 | rnr1Y285F | FALSE | CAN | JSY3811 | 3 | rnr1Y285F-R | TAA | A>C | G/C+1 | A/T-1 | TA>GC | TA>GC | AAGGTTAAG | AAGATAC | ATT | ATT | T | G | T>G | T>G_ATT | TA>GC_AAGATAC | 8 | FALSE | FALSE |
| 44_v.txt | 32493 | 32493-SNV-A | SNV | 1 | A | C | 11 | 39917 | 0.02755718 | 25.3650888 | rnr1Y285F | FALSE | CAN | JSY3811 | 3 | rnr1Y285F-R | AAC | A>C | G/C+1 | A/T-1 | TA>GC | TA>GC | CGGAAAACA | ACAACTT | TTG | TTG | T | G | T>G | T>G_TTG | TA>GC_ACAACTT | 8 | FALSE | FALSE |
| 44_v.txt | 32581 | 32581-SNV-T | SNV | 1 | T | G | 32 | 33134 | 0.09657753 | 32.71875 | rnr1Y285F | FALSE | CAN | JSY3811 | 3 | rnr1Y285F-R | GTG | T>G | G/C+1 | A/T-1 | TA>GC | TA>GC | GAAATGTGA | TGAAGGC | CAC | GTG | T | C | T>G | T>G_GTG | TA>GC_TGAAGGC | 8 | FALSE | FALSE |
| 44_v.txt | 32728 | 32728-SNV-A | SNV | 1 | A | C | 3 | 33134 | 9.05E-03 | 37.3334432 | rnr1Y285F | FALSE | CAN | JSY3811 | 3 | rnr1Y285F-R | CAA | A>C | G/C+1 | A/T-1 | TA>GC | TA>GC | CCATACAAA | AAAAACA | GTT | GTT | T | G | T>G | T>G_GTT | TA>GC_AAAAACA | 8 | FALSE | FALSE |
| 44_v.txt | 32739 | 32739-SNV-A | SNV | 1 | A | C | 10 | 33134 | 0.03018048 | 50.8 | rnr1Y285F | FALSE | CAN | JSY3811 | 3 | rnr1Y285F-R | TAT | A>C | G/C+1 | A/T-1 | TA>GC | TA>GC | CAGTATATTA | ATTAGAA | ATA | ATA | T | G | T>G | T>G_ATA | TA>GC_ATTAGAA | 8 | FALSE | FALSE |
| 44_v.txt | 32763 | 32763-SNV-A | SNV | 1 | A | C | 7 | 33134 | 0.02112634 | 49 | rnr1Y285F | FALSE | CAN | JSY3811 | 3 | rnr1Y285F-R | AAC | A>C | G/C+1 | A/T-1 | TA>GC | TA>GC | GCTAAAACTT | ACTTTGA | TTG | TTG | T | G | T>G | T>G_TTG | TA>GC_ACTTTGA | 8 | FALSE | FALSE |
| 44_v.txt | 32842 | 32842-SNV-A | SNV | 1 | A | C | 5 | 33133 | 0.0150907 | 29.2000043 | rnr1Y285F | FALSE | CAN | JSY3811 | 3 | rnr1Y285F-R | CAA | A>C | G/C+1 | A/T-1 | TA>GC | TA>GC | TTACCCAAAA | AAAAAAT | GTT | GTT | T | G | T>G | T>G_GTT | TA>GC_AAAAAAT | 8 | FALSE | FALSE |
| 44_v.txt | 32884 | 32884-SNV-A | SNV | 1 | A | C | 9 | 33681 | 0.0267213 | 31.5555556 | rnr1Y285F | FALSE | CAN | JSY3811 | 3 | rnr1Y285F-R | CAA | A>C | G/C+1 | A/T-1 | TA>GC | TA>GC | ACGTCCAAA | AAAATTG | GTT | GTT | T | G | T>G | T>G_GTT | TA>GC_AAAATTG | 8 | FALSE | FALSE |
| 44_v.txt | 32919 | 32919-SNV-A | SNV | 1 | A | C | 14 | 33681 | 0.04156646 | 36.0714286 | rnr1Y285F | FALSE | CAN | JSY3811 | 3 | rnr1Y285F-R | CAG | A>C | G/C+1 | A/T-1 | TA>GC | TA>GC | AGTTCCAGG | AGGGCAA | GTC | GTC | T | G | T>G | T>G_GTC | TA>GC_AGGGCAA | 8 | FALSE | FALSE |
| 44_v.txt | 32929 | 32929-SNV-T | SNV | 1 | T | G | 14 | 33681 | 0.04156646 | 52.2857143 | rnr1Y285F | FALSE | CAN | JSY3811 | 3 | rnr1Y285F-R | GTG | T>G | G/C+1 | A/T-1 | TA>GC | TA>GC | CAAAAGTGA | TGATTGC | CAC | GTG | T | C | T>G | T>G_GTG | TA>GC_TGATTGC | 8 | FALSE | FALSE |
| 44_v.txt | 33089 | 33089-SNV-A | SNV | 1 | A | C | 16 | 33681 | 0.04750453 | 53.375 | rnr1Y285F | FALSE | CAN | JSY3811 | 3 | rnr1Y285F-R | AAT | A>C | G/C+1 | A/T-1 | TA>GC | TA>GC | AAATAAATAT | ATATGAT | TTA | TTA | T | G | T>G | T>G_TTA | TA>GC_ATATGAT | 8 | FALSE | FALSE |
| 44_v.txt | 33099 | 33099-SNV-A | SNV | 1 | A | C | 9 | 33681 | 0.0267213 | 34.3333333 | rnr1Y285F | FALSE | CAN | JSY3811 | 3 | rnr1Y285F-R | AAG | A>C | G/C+1 | A/T-1 | TA>GC | TA>GC | GATATAAGA | AGAGCGC | TTC | TTC | T | G | T>G | T>G_TTC | TA>GC_AGAGCGC | 8 | FALSE | FALSE |
| 44_v.txt | 33269 | 33269-SNV-A | SNV | 1 | A | C | 203 | 25205 | 0.80539576 | 16.4039409 | rnr1Y285F | FALSE | CAN | JSY3811 | 3 | rnr1Y285F-R | TAG | A>C | G/C+1 | A/T-1 | TA>GC | TA>GC | CTCCATAGAC | AGAGAAC | ATC | ATC | T | G | T>G | T>G_ATC | TA>GC_AGAGAAC | 8 | FALSE | FALSE |
| 44_v.txt | 32081 | 32081-SNV-A | SNV | 1 | A | G | 5 | 23899 | 0.02092138 | 64 | rnr1Y285F | FALSE | CAN | JSY3811 | 3 | rnr1Y285F-R | CAA | A>G | G/C+1 | A/T-1 | TA>CG | TA>CG | CCATGCAAA | AAAAAAG | GTT | GTT | T | C | T>C | T>C_GTT | TA>CG_AAAAAAG | 8 | FALSE | FALSE |
| 44_v.txt | 32202 | 32202-SNV-A | SNV | 1 | A | G | 2 | 23899 | 8.37E-03 | 50.998532 | rnr1Y285F | FALSE | CAN | JSY3811 | 3 | rnr1Y285F-R | AAC | A>G | G/C+1 | A/T-1 | TA>CG | TA>CG | TATGGAACA | ACACCAC | TTG | TTG | T | C | T>C | T>C_TTG | TA>CG_ACACCAC | 8 | FALSE | FALSE |
| 44_v.txt | 32377 | 32377-SNV-T | SNV | 1 | T | C | 6 | 39909 | 0.0150342 | 51.4992689 | rnr1Y285F | FALSE | CAN | JSY3811 | 3 | rnr1Y285F-R | ATA | T>C | G/C+1 | A/T-1 | TA>CG | TA>CG | TAGCAATAAT | TAATAAA | TAT | ATA | T | G | T>C | T>C_ATA | TA>CG_TAATAAA | 8 | FALSE | FALSE |
| 44_v.txt | 32472 | 32472-SNV-T | SNV | 1 | T | C | 25 | 39917 | 0.06262996 | 48.5199999 | rnr1Y285F | FALSE | CAN | JSY3811 | 3 | rnr1Y285F-R | GTA | T>C | G/C+1 | A/T-1 | TA>CG | TA>CG | CCAATGTAGA | TAGAAGG | CAT | GTA | T | G | T>C | T>C_GTA | TA>CG_TAGAAGG | 8 | FALSE | FALSE |
| 44_v.txt | 32919 | 32919-SNV-A | SNV | 1 | A | G | 24 | 33681 | 0.07125679 | 44.6666667 | rnr1Y285F | FALSE | CAN | JSY3811 | 3 | rnr1Y285F-R | CAG | A>G | G/C+1 | A/T-1 | TA>CG | TA>CG | AGTTCCAGG | AGGGCAA | GTC | GTC | T | C | T>C | T>C_GTC | TA>CG_AGGGCAA | 8 | FALSE | FALSE |
| 44_v.txt | 32929 | 32929-SNV-T | SNV | 1 | T | C | 36 | 33681 | 0.10688519 | 43.4166667 | rnr1Y285F | FALSE | CAN | JSY3811 | 3 | rnr1Y285F-R | GTG | T>C | G/C+1 | A/T-1 | TA>CG | TA>CG | CAAAAGTGA | TGATTGC | CAC | GTG | T | G | T>C | T>C_GTG | TA>CG_TGATTGC | 8 | FALSE | FALSE |
| 44_v.txt | 33099 | 33099-SNV-A | SNV | 1 | A | G | 21 | 33681 | 0.06234969 | 50 | rnr1Y285F | FALSE | CAN | JSY3811 | 3 | rnr1Y285F-R | AAG | A>G | G/C+1 | A/T-1 | TA>CG | TA>CG | GATATAAGA | AGAGCGC | TTC | TTC | T | C | T>C | T>C_TTC | TA>CG_AGAGCGC | 8 | FALSE | FALSE |
| 44_v.txt | 32201 | 32201-SNV-A | SNV | 1 | A | T | 2 | 23899 | 8.37E-03 | 35.9999306 | rnr1Y285F | FALSE | CAN | JSY3811 | 3 | rnr1Y285F-R | GAA | A>T | A/T+1 | A/T-1 | TA>AT | TA>AT | GTATGGAAC | AACACCA | CTT | CTT | T | A | T>A | T>A_CTT | TA>AT_AACACCA | 8 | FALSE | FALSE |
| 44_v.txt | 32376 | 32376-SNV-A | SNV | 1 | A | T | 8 | 39917 | 0.02004159 | 63.2490861 | rnr1Y285F | FALSE | CAN | JSY3811 | 3 | rnr1Y285F-R | AAT | A>T | A/T+1 | A/T-1 | TA>AT | TA>AT | ATAGCAATA | ATAATAA | TTA | TTA | T | A | T>A | T>A_TTA | TA>AT_ATAATAA | 8 | FALSE | FALSE |
| 44_v.txt | 32377 | 32377-SNV-T | SNV | 1 | T | A | 5 | 39909 | 0.0125285 | 24.7983567 | rnr1Y285F | FALSE | CAN | JSY3811 | 3 | rnr1Y285F-R | ATA | T>A | A/T+1 | A/T-1 | TA>AT | TA>AT | TAGCAATAAT | TAATAAA | TAT | ATA | T | T | T>A | T>A_ATA | TA>AT_TAATAAA | 8 | FALSE | FALSE |
| 44_v.txt | 32472 | 32472-SNV-T | SNV | 1 | T | A | 8 | 39917 | 0.02004159 | 45.625 | rnr1Y285F | FALSE | CAN | JSY3811 | 3 | rnr1Y285F-R | GTA | T>A | A/T+1 | A/T-1 | TA>AT | TA>AT | CCAATGTAGA | TAGAAGG | CAT | GTA | T | T | T>A | T>A_GTA | TA>AT_TAGAAGG | 8 | FALSE | FALSE |
| 44_v.txt | 32726 | 32726-SNV-A | SNV | 1 | A | T | 19 | 33134 | 0.05734291 | 58.1578947 | rnr1Y285F | FALSE | CAN | JSY3811 | 3 | rnr1Y285F-R | TAC | A>T | A/T+1 | A/T-1 | TA>AT | TA>AT | AACCATACA | ACAAAAA | ATG | ATG | T | A | T>A | T>A_ATG | TA>AT_ACAAAAA | 8 | FALSE | FALSE |
| 44_v.txt | 32763 | 32763-SNV-A | SNV | 1 | A | T | 13 | 33134 | 0.03923462 | 49.6923077 | rnr1Y285F | FALSE | CAN | JSY3811 | 3 | rnr1Y285F-R | AAC | A>T | A/T+1 | A/T-1 | TA>AT | TA>AT | GCTAAAACTT | ACTTTGA | TTG | TTG | T | A | T>A | T>A_TTG | TA>AT_ACTTTGA | 8 | FALSE | FALSE |
| 44_v.txt | 32820 | 32820-SNV-T | SNV | 1 | T | A | 33 | 33134 | 0.09959558 | 40.9951495 | rnr1Y285F | FALSE | CAN | JSY3811 | 3 | rnr1Y285F-R | GTT | T>A | A/T+1 | A/T-1 | TA>AT | TA>AT | AACAAGTTCA | TTCATTA | CAA | GTT | T | T | T>A | T>A_GTT | TA>AT_TTCATTA | 8 | FALSE | FALSE |
| 44_v.txt | 33067 | 33067-SNV-A | SNV | 1 | A | T | 1062 | 33681 | 3.15311303 | 54.4265537 | rnr1Y285F | FALSE | CAN | JSY3811 | 3 | rnr1Y285F-R | TAT | A>T | A/T+1 | A/T-1 | TA>AT | TA>AT | CAGAATATG | ATGCCAA | ATA | ATA | T | A | T>A | T>A_ATA | TA>AT_ATGCCAA | 8 | FALSE | FALSE |
| 44_v.txt | 33138 | 33138-SNV-A | SNV | 1 | A | T | 33 | 33681 | 0.09797809 | 42 | rnr1Y285F | FALSE | CAN | JSY3811 | 3 | rnr1Y285F-R | TAA | A>T | A/T+1 | A/T-1 | TA>AT | TA>AT | GTGGATAAA | AAACCAA | ATT | ATT | T | A | T>A | T>A_ATT | TA>AT_AAACCAA | 8 | FALSE | FALSE |
| 44_v.txt | 31650 | 31650-SNV-C | SNV | 1 | C | T | 30 | 14771 | 0.20310067 | 36.2666667 | rnr1Y285F | FALSE | CAN | JSY3811 | 3 | rnr1Y285F-R | GCG | C>T | A/T+1 | G/C-1 | CG>TA | CG>TA | AGAATGCGA | CGAAATG | CGC | GCG | C | A | C>T | C>T_GCG | CG>TA_CGAAATG | 8 | FALSE | FALSE |
| 44_v.txt | 31658 | 31658-SNV-C | SNV | 1 | C | T | 20 | 14771 | 0.13540045 | 33.1502199 | rnr1Y285F | FALSE | CAN | JSY3811 | 3 | rnr1Y285F-R | GCG | C>T | A/T+1 | G/C-1 | CG>TA | CG>TA | AAATGGCGT | CGTGGAA | CGC | GCG | C | A | C>T | C>T_GCG | CG>TA_CGTGGAA | 8 | FALSE | FALSE |
| 44_v.txt | 31778 | 31778-SNV-C | SNV | 1 | G | A | 15 | 14772 | 0.10154346 | 54.5333333 | rnr1Y285F | FALSE | CAN | JSY3811 | 3 | rnr1Y285F-R | CGG | G>A | A/T+1 | G/C-1 | CG>TA | CG>TA | TCTATCGGA | GGAATCG | GCC | GCC | C | T | C>T | C>T_GCC | CG>TA_GGAATCG | 8 | FALSE | FALSE |
| 44_v.txt | 31829 | 31829-SNV-C | SNV | 1 | G | A | 15 | 14772 | 0.10154346 | 49.2 | rnr1Y285F | FALSE | CAN | JSY3811 | 3 | rnr1Y285F-R | TGC | G>A | A/T+1 | G/C-1 | CG>TA | CG>TA | GAATATGCA | GCATTGA | ACG | ACG | C | T | C>T | C>T_ACG | CG>TA_GCATTGA | 8 | FALSE | FALSE |
| 44_v.txt | 31845 | 31845-SNV-C | SNV | 1 | C | T | 28 | 14772 | 0.18954779 | 51.8214286 | rnr1Y285F | FALSE | CAN | JSY3811 | 3 | rnr1Y285F-R | CCA | C>T | A/T+1 | G/C-1 | CG>TA | CG>TA | AAGATCCAA | CAAACAG | GGT | CCA | C | A | C>T | C>T_CCA | CG>TA_CAAACAG | 8 | FALSE | FALSE |
| 44_v.txt | 31881 | 31881-SNV-C | SNV | 1 | G | A | 14 | 14772 | 0.0947739 | 50.5704789 | rnr1Y285F | FALSE | CAN | JSY3811 | 3 | rnr1Y285F-R | GGC | G>A | A/T+1 | G/C-1 | CG>TA | CG>TA | TAGGCGGCA | GCAGCAA | CCG | CCG | C | T | C>T | C>T_CCG | CG>TA_GCAGCAA | 8 | FALSE | FALSE |
| 44_v.txt | 31882 | 31882-SNV-C | SNV | 1 | C | T | 23 | 14772 | 0.15569997 | 42.9047388 | rnr1Y285F | FALSE | CAN | JSY3811 | 3 | rnr1Y285F-R | GCA | C>T | A/T+1 | G/C-1 | CG>TA | CG>TA | AGGCGGCAG | CAGCAAA | CGT | GCA | C | A | C>T | C>T_GCA | CG>TA_CAGCAAA | 8 | FALSE | FALSE |
| 44_v.txt | 31884 | 31884-SNV-C | SNV | 1 | G | A | 19 | 14772 | 0.12862172 | 34.5263158 | rnr1Y285F | FALSE | CAN | JSY3811 | 3 | rnr1Y285F-R | AGC | G>A | A/T+1 | G/C-1 | CG>TA | CG>TA | GCGGCAGCA | GCAAAGC | TCG | TCG | C | T | C>T | C>T_TCG | CG>TA_GCAAAGC | 8 | FALSE | FALSE |
| 44_v.txt | 31947 | 31947-SNV-C | SNV | 1 | G | A | 14 | 23887 | 0.05860929 | 33.4285714 | rnr1Y285F | FALSE | CAN | JSY3811 | 3 | rnr1Y285F-R | CGT | G>A | A/T+1 | G/C-1 | CG>TA | CG>TA | ATGATCGTCA | GTCATAA | GCA | GCA | C | T | C>T | C>T_GCA | CG>TA_GTCATAA | 8 | FALSE | FALSE |
| 44_v.txt | 31960 | 31960-SNV-C | SNV | 1 | C | T | 36 | 23899 | 0.15063392 | 35.1111111 | rnr1Y285F | FALSE | CAN | JSY3811 | 3 | rnr1Y285F-R | GCC | C>T | A/T+1 | G/C-1 | CG>TA | CG>TA | ATGTGGCCG | CCGCATA | CGG | GCC | C | A | C>T | C>T_GCC | CG>TA_CCGCATA | 8 | FALSE | FALSE |
| 44_v.txt | 31962 | 31962-SNV-C | SNV | 1 | G | A | 42 | 23899 | 0.17573957 | 37.1428571 | rnr1Y285F | FALSE | CAN | JSY3811 | 3 | rnr1Y285F-R | CGC | G>A | A/T+1 | G/C-1 | CG>TA | CG>TA | GTGGCCGCA | GCATAAT | GCG | GCG | C | T | C>T | C>T_GCG | CG>TA_GCATAAT | 8 | FALSE | FALSE |
| 44_v.txt | 31972 | 31972-SNV-C | SNV | 1 | C | T | 16 | 23899 | 0.06694841 | 35.5 | rnr1Y285F | FALSE | CAN | JSY3811 | 3 | rnr1Y285F-R | GCC | C>T | A/T+1 | G/C-1 | CG>TA | CG>TA | AATAAGCCA | CCAAGCC | CGG | GCC | C | A | C>T | C>T_GCC | CG>TA_CCAAGCC | 8 | FALSE | FALSE |
| 44_v.txt | 31981 | 31981-SNV-C | SNV | 1 | G | A | 23 | 23899 | 0.09623834 | 46.7826087 | rnr1Y285F | FALSE | CAN | JSY3811 | 3 | rnr1Y285F-R | GGC | G>A | A/T+1 | G/C-1 | CG>TA | CG>TA | AGCCGGGCA | GCATTAA | CCG | CCG | C | T | C>T | C>T_CCG | CG>TA_GCATTAA | 8 | FALSE | FALSE |
| 44_v.txt | 32022 | 32022-SNV-C | SNV | 1 | C | T | 25 | 23899 | 0.10460689 | 56.8 | rnr1Y285F | FALSE | CAN | JSY3811 | 3 | rnr1Y285F-R | GCC | C>T | A/T+1 | G/C-1 | CG>TA | CG>TA | GAGATGCCA | CCACGGT | CGG | GCC | C | A | C>T | C>T_GCC | CG>TA_CCACGGT | 8 | FALSE | FALSE |

| | | | | | | | | | | | | | | | | | | | | | | | | | | | | | | | | | | |
|---|---|---|---|---|---|---|---|---|---|---|---|---|---|---|---|---|---|---|---|---|---|---|---|---|---|---|---|---|---|---|---|---|---|---|
| 44_v.txt | 32041 | 32041-SNV-( | SNV | 1 | G | A | 21 | 23899 | 0.08786979 | 58.4285714 | rnr1Y285F | FALSE | CAN | JSY3811 | 3 | rnr1Y285F-R | TGC | G>A | A/T+1 | G/C-1 | CG>TA | CG>TA | AAGCTTGCA | GCATAAA | ACG | ACG | C | T | C>T | C>T_ACG | CG>TA_GCATAAA | 8 | FALSE | FALSE |
| 44_v.txt | 32057 | 32057-SNV-( | SNV | 1 | C | T | 23 | 23899 | 0.09623834 | 61.2173913 | rnr1Y285F | FALSE | CAN | JSY3811 | 3 | rnr1Y285F-R | GCG | C>T | A/T+1 | G/C-1 | CG>TA | CG>TA | GATGTGCGA | CGAGATT | CGC | GCG | C | A | C>T | C>T_GCG | CG>TA_CGAGATT | 8 | FALSE | FALSE |
| 44_v.txt | 32088 | 32088-SNV-( | SNV | 1 | C | T | 363 | 23899 | 1.518892 | 60.7245179 | rnr1Y285F | FALSE | CAN | JSY3811 | 3 | rnr1Y285F-R | GCC | C>T | A/T+1 | G/C-1 | CG>TA | CG>TA | AAAAAGCCT | CCTGCAA | CGG | GCC | C | A | C>T | C>T_GCC | CG>TA_CCTGCAA | 8 | FALSE | FALSE |
| 44_v.txt | 32092 | 32092-SNV-( | SNV | 1 | C | T | 40 | 23899 | 0.16737102 | 62.825 | rnr1Y285F | FALSE | CAN | JSY3811 | 3 | rnr1Y285F-R | GCA | C>T | A/T+1 | G/C-1 | CG>TA | CG>TA | AGCCTGCAA | CAACACC | CGT | GCA | C | A | C>T | C>T_GCA | CG>TA_CAACACC | 8 | FALSE | FALSE |
| 44_v.txt | 32114 | 32114-SNV-( | SNV | 1 | C | T | 142 | 23899 | 0.59416712 | 58.6760563 | rnr1Y285F | FALSE | CAN | JSY3811 | 3 | rnr1Y285F-R | GCC | C>T | A/T+1 | G/C-1 | CG>TA | CG>TA | TAATAGCCA | CCATTCG | CGG | GCC | C | A | C>T | C>T_GCC | CG>TA_CCATTCG | 8 | FALSE | FALSE |
| 44_v.txt | 32157 | 32157-SNV-( | SNV | 1 | G | A | 21 | 23899 | 0.08786979 | 47.952381 | rnr1Y285F | FALSE | CAN | JSY3811 | 3 | rnr1Y285F-R | AGC | G>A | A/T+1 | G/C-1 | CG>TA | CG>TA | ATGTAAGCC | GCCAAAG | TCG | TCG | C | T | C>T | C>T_TCG | CG>TA_GCCAAAG | 8 | FALSE | FALSE |
| 44_v.txt | 32165 | 32165-SNV-( | SNV | 1 | G | A | 52 | 23899 | 0.21758233 | 45.4038462 | rnr1Y285F | FALSE | CAN | JSY3811 | 3 | rnr1Y285F-R | CGC | G>A | A/T+1 | G/C-1 | CG>TA | CG>TA | CAAAGCGCC | GCCAAAT | GCG | GCG | C | T | C>T | C>T_GCG | CG>TA_GCCAAAT | 8 | FALSE | FALSE |
| 44_v.txt | 32173 | 32173-SNV-( | SNV | 1 | C | T | 27 | 23899 | 0.11297544 | 45.3333333 | rnr1Y285F | FALSE | CAN | JSY3811 | 3 | rnr1Y285F-R | GCA | C>T | A/T+1 | G/C-1 | CG>TA | CG>TA | CAAATGCAG | CAGCAGT | CGT | GCA | C | A | C>T | C>T_GCA | CG>TA_CAGCAGT | 8 | FALSE | FALSE |
| 44_v.txt | 32191 | 32191-SNV-( | SNV | 1 | C | T | 19 | 23899 | 0.07950123 | 40.5263158 | rnr1Y285F | FALSE | CAN | JSY3811 | 3 | rnr1Y285F-R | GCA | C>T | A/T+1 | G/C-1 | CG>TA | CG>TA | AAACTGCAA | CAATGTA | CGT | GCA | C | A | C>T | C>T_GCA | CG>TA_CAATGTA | 8 | FALSE | FALSE |
| 44_v.txt | 32271 | 32271-SNV-( | SNV | 1 | C | T | 63 | 39917 | 0.15782749 | 29.6666667 | rnr1Y285F | FALSE | CAN | JSY3811 | 3 | rnr1Y285F-R | ACG | C>T | A/T+1 | G/C-1 | CG>TA | CG>TA | AAAATACGG | CGGGAAC | TGC | ACG | C | A | C>T | C>T_ACG | CG>TA_CGGGAAC | 8 | FALSE | FALSE |
| 44_v.txt | 32272 | 32272-SNV-( | SNV | 1 | G | A | 42 | 39917 | 0.10521833 | 36.3810074 | rnr1Y285F | FALSE | CAN | JSY3811 | 3 | rnr1Y285F-R | CGG | G>A | A/T+1 | G/C-1 | CG>TA | CG>TA | AAATACGGG | GGGAACC | GCC | GCC | C | T | C>T | C>T_GCC | CG>TA_GGGAACC | 8 | FALSE | FALSE |
| 44_v.txt | 32299 | 32299-SNV-( | SNV | 1 | C | T | 67 | 39917 | 0.16784829 | 54.641791 | rnr1Y285F | FALSE | CAN | JSY3811 | 3 | rnr1Y285F-R | GCG | C>T | A/T+1 | G/C-1 | CG>TA | CG>TA | AATTTGCGG | CGGCAGA | CGC | GCG | C | A | C>T | C>T_GCG | CG>TA_CGGCAGA | 8 | FALSE | FALSE |
| 44_v.txt | 32344 | 32344-SNV-( | SNV | 1 | G | A | 31 | 39917 | 0.07766115 | 58.0645161 | rnr1Y285F | FALSE | CAN | JSY3811 | 3 | rnr1Y285F-R | GGC | G>A | A/T+1 | G/C-1 | CG>TA | CG>TA | TATGTGGCA | GCAAAAC | CCG | CCG | C | T | C>T | C>T_CCG | CG>TA_GCAAAAC | 8 | FALSE | FALSE |
| 44_v.txt | 32466 | 32466-SNV-( | SNV | 1 | C | T | 62 | 39917 | 0.15532229 | 48.2741936 | rnr1Y285F | FALSE | CAN | JSY3811 | 3 | rnr1Y285F-R | GCC | C>T | A/T+1 | G/C-1 | CG>TA | CG>TA | AGAGAGCCA | CCAATGT | CGG | GCC | C | A | C>T | C>T_GCC | CG>TA_CCAATGT | 8 | FALSE | FALSE |
| 44_v.txt | 32487 | 32487-SNV-( | SNV | 1 | C | T | 26 | 39917 | 0.06513516 | 43.3076923 | rnr1Y285F | FALSE | CAN | JSY3811 | 3 | rnr1Y285F-R | ACG | C>T | A/T+1 | G/C-1 | CG>TA | CG>TA | AAGATACGG | CGGAAAA | TGC | ACG | C | A | C>T | C>T_ACG | CG>TA_CGGAAAA | 8 | FALSE | FALSE |
| 44_v.txt | 32535 | 32535-SNV-( | SNV | 1 | G | A | 55 | 39917 | 0.13778591 | 37.2363636 | rnr1Y285F | FALSE | CAN | JSY3811 | 3 | rnr1Y285F-R | TGC | G>A | A/T+1 | G/C-1 | CG>TA | CG>TA | GGGTTTGCA | GCAGCTT | ACG | ACG | C | T | C>T | C>T_ACG | CG>TA_GCAGCTT | 8 | FALSE | FALSE |
| 44_v.txt | 32536 | 32536-SNV-( | SNV | 1 | C | T | 67 | 39917 | 0.16784829 | 36.2835821 | rnr1Y285F | FALSE | CAN | JSY3811 | 3 | rnr1Y285F-R | GCA | C>T | A/T+1 | G/C-1 | CG>TA | CG>TA | GGTTTGCAG | CAGCTTC | CGT | GCA | C | A | C>T | C>T_GCA | CG>TA_CAGCTTC | 8 | FALSE | FALSE |
| 44_v.txt | 32590 | 32590-SNV-( | SNV | 1 | C | T | 58 | 33134 | 0.17504678 | 35.6551724 | rnr1Y285F | FALSE | CAN | JSY3811 | 3 | rnr1Y285F-R | GCG | C>T | A/T+1 | G/C-1 | CG>TA | CG>TA | AGGCAGCGT | CGTTAAT | CGC | GCG | C | A | C>T | C>T_GCG | CG>TA_CGTTAAT | 8 | FALSE | FALSE |
| 44_v.txt | 32609 | 32609-SNV-( | SNV | 1 | C | T | 73 | 33134 | 0.2203175 | 48.9508285 | rnr1Y285F | FALSE | CAN | JSY3811 | 3 | rnr1Y285F-R | CCC | C>T | A/T+1 | G/C-1 | CG>TA | CG>TA | GGAAACCCA | CCAACCT | GGG | CCC | C | A | C>T | C>T_CCC | CG>TA_CCAACCT | 8 | FALSE | FALSE |
| 44_v.txt | 32700 | 32700-SNV-( | SNV | 1 | C | T | 61 | 33134 | 0.18410092 | 57.9344262 | rnr1Y285F | FALSE | CAN | JSY3811 | 3 | rnr1Y285F-R | GCC | C>T | A/T+1 | G/C-1 | CG>TA | CG>TA | ACTGGGCCG | CCGGTAA | CGG | GCC | C | A | C>T | C>T_GCC | CG>TA_CCGGTAA | 8 | FALSE | FALSE |
| 44_v.txt | 32702 | 32702-SNV-( | SNV | 1 | G | A | 27 | 33134 | 0.08148729 | 54.6666667 | rnr1Y285F | FALSE | CAN | JSY3811 | 3 | rnr1Y285F-R | CGG | G>A | A/T+1 | G/C-1 | CG>TA | CG>TA | TGGGCCGGT | GGTAACC | GCC | GCC | C | T | C>T | C>T_GCC | CG>TA_GGTAACC | 8 | FALSE | FALSE |
| 44_v.txt | 32757 | 32757-SNV-( | SNV | 1 | G | A | 28 | 33134 | 0.08450534 | 51.7142857 | rnr1Y285F | FALSE | CAN | JSY3811 | 3 | rnr1Y285F-R | GGC | G>A | A/T+1 | G/C-1 | CG>TA | CG>TA | ATAATGGCT | GCTAAAA | CCG | CCG | C | T | C>T | C>T_CCG | CG>TA_GCTAAAA | 8 | FALSE | FALSE |
| 44_v.txt | 32776 | 32776-SNV-( | SNV | 1 | C | T | 46 | 33134 | 0.13883021 | 47.2391304 | rnr1Y285F | FALSE | CAN | JSY3811 | 3 | rnr1Y285F-R | GCG | C>T | A/T+1 | G/C-1 | CG>TA | CG>TA | TGGAAGCGA | CGACCCA | CGC | GCG | C | A | C>T | C>T_GCG | CG>TA_CGACCCA | 8 | FALSE | FALSE |
| 44_v.txt | 32796 | 32796-SNV-( | SNV | 1 | C | T | 25 | 33134 | 0.0754512 | 44.48 | rnr1Y285F | FALSE | CAN | JSY3811 | 3 | rnr1Y285F-R | ACC | C>T | A/T+1 | G/C-1 | CG>TA | CG>TA | AATTCACCG | CCGTAAT | TGG | ACC | C | A | C>T | C>T_ACC | CG>TA_CCGTAAT | 8 | FALSE | FALSE |
| 44_v.txt | 32798 | 32798-SNV-( | SNV | 1 | G | A | 49 | 33134 | 0.14788435 | 42.7959184 | rnr1Y285F | FALSE | CAN | JSY3811 | 3 | rnr1Y285F-R | CGT | G>A | A/T+1 | G/C-1 | CG>TA | CG>TA | TTCACCGTA | GTAATAT | GCA | GCA | C | T | C>T | C>T_GCA | CG>TA_GTAATAT | 8 | FALSE | FALSE |
| 44_v.txt | 32855 | 32855-SNV-( | SNV | 1 | C | T | 58 | 33128 | 0.17507848 | 36.5517241 | rnr1Y285F | FALSE | CAN | JSY3811 | 3 | rnr1Y285F-R | TCC | C>T | A/T+1 | G/C-1 | CG>TA | CG>TA | ACTAATCCA | CCATGCC | AGG | TCC | C | A | C>T | C>T_TCC | CG>TA_CCATGCC | 8 | FALSE | FALSE |
| 44_v.txt | 32860 | 32860-SNV-( | SNV | 1 | C | T | 31 | 33128 | 0.09357643 | 35.7419355 | rnr1Y285F | FALSE | CAN | JSY3811 | 3 | rnr1Y285F-R | GCC | C>T | A/T+1 | G/C-1 | CG>TA | CG>TA | TCCATGCCG | CCGCCAG | CGG | GCC | C | A | C>T | C>T_GCC | CG>TA_CCGCCAG | 8 | FALSE | FALSE |
| 44_v.txt | 32862 | 32862-SNV-( | SNV | 1 | G | A | 27 | 33129 | 0.08149959 | 35.5925926 | rnr1Y285F | FALSE | CAN | JSY3811 | 3 | rnr1Y285F-R | CGC | G>A | A/T+1 | G/C-1 | CG>TA | CG>TA | CATGCCGCC | GCCAGTG | GCG | GCG | C | T | C>T | C>T_GCG | CG>TA_GCCAGTG | 8 | FALSE | FALSE |
| 44_v.txt | 32882 | 32882-SNV-( | SNV | 1 | C | T | 103 | 33682 | 0.30580132 | 29.4368932 | rnr1Y285F | FALSE | CAN | JSY3811 | 3 | rnr1Y285F-R | TCC | C>T | A/T+1 | G/C-1 | CG>TA | CG>TA | GTACGTCCA | CCAAAAT | AGG | TCC | C | A | C>T | C>T_TCC | CG>TA_CCAAAAT | 8 | FALSE | FALSE |
| 44_v.txt | 32900 | 32900-SNV-( | SNV | 1 | G | A | 48 | 33681 | 0.14251358 | 35.2065428 | rnr1Y285F | FALSE | CAN | JSY3811 | 3 | rnr1Y285F-R | GGC | G>A | A/T+1 | G/C-1 | CG>TA | CG>TA | GACTTGGCC | GCCAACT | CCG | CCG | C | T | C>T | C>T_CCG | CG>TA_GCCAACT | 8 | FALSE | FALSE |
| 44_v.txt | 32901 | 32901-SNV-( | SNV | 1 | C | T | 74 | 33681 | 0.21970844 | 31.6758869 | rnr1Y285F | FALSE | CAN | JSY3811 | 3 | rnr1Y285F-R | GCC | C>T | A/T+1 | G/C-1 | CG>TA | CG>TA | ACTTGGCCA | CCAACTA | CGG | GCC | C | A | C>T | C>T_GCC | CG>TA_CCAACTA | 8 | FALSE | FALSE |
| 44_v.txt | 32917 | 32917-SNV-( | SNV | 1 | C | T | 107 | 33681 | 0.31768653 | 49.2523365 | rnr1Y285F | FALSE | CAN | JSY3811 | 3 | rnr1Y285F-R | TCC | C>T | A/T+1 | G/C-1 | CG>TA | CG>TA | TAAGTTCCA | CCAGGGC | AGG | TCC | C | A | C>T | C>T_TCC | CG>TA_CCAGGGC | 8 | FALSE | FALSE |
| 44_v.txt | 32922 | 32922-SNV-( | SNV | 1 | G | A | 24 | 33681 | 0.07125679 | 54.6666667 | rnr1Y285F | FALSE | CAN | JSY3811 | 3 | rnr1Y285F-R | GGC | G>A | A/T+1 | G/C-1 | CG>TA | CG>TA | TCCAGGGCA | GCAAAAG | CCG | CCG | C | T | C>T | C>T_CCG | CG>TA_GCAAAAG | 8 | FALSE | FALSE |
| 44_v.txt | 32935 | 32935-SNV-( | SNV | 1 | C | T | 49 | 33681 | 0.14548262 | 37.244898 | rnr1Y285F | FALSE | CAN | JSY3811 | 3 | rnr1Y285F-R | GCC | C>T | A/T+1 | G/C-1 | CG>TA | CG>TA | TGATTGCCC | CCCAAGA | CGG | GCC | C | A | C>T | C>T_GCC | CG>TA_CCCAAGA | 8 | FALSE | FALSE |
| 44_v.txt | 32945 | 32945-SNV-( | SNV | 1 | C | T | 63 | 33681 | 0.18704908 | 58.0634921 | rnr1Y285F | FALSE | CAN | JSY3811 | 3 | rnr1Y285F-R | ACC | C>T | A/T+1 | G/C-1 | CG>TA | CG>TA | AGAAAACCA | CCAATAC | TGG | ACC | C | A | C>T | C>T_ACC | CG>TA_CCAATAC | 8 | FALSE | FALSE |
| 44_v.txt | 32958 | 32958-SNV-( | SNV | 1 | C | T | 236 | 33681 | 0.70069179 | 57.0720339 | rnr1Y285F | FALSE | CAN | JSY3811 | 3 | rnr1Y285F-R | ACC | C>T | A/T+1 | G/C-1 | CG>TA | CG>TA | ATGTAACCA | CCATTGG | TGG | ACC | C | A | C>T | C>T_ACC | CG>TA_CCATTGG | 8 | FALSE | FALSE |
| 44_v.txt | 32965 | 32965-SNV-( | SNV | 1 | C | T | 37 | 33681 | 0.10985422 | 56.6756757 | rnr1Y285F | FALSE | CAN | JSY3811 | 3 | rnr1Y285F-R | GCC | C>T | A/T+1 | G/C-1 | CG>TA | CG>TA | CATTGGCCG | CCGCACC | CGG | GCC | C | A | C>T | C>T_GCC | CG>TA_CCGCACC | 8 | FALSE | FALSE |
| 44_v.txt | 32967 | 32967-SNV-( | SNV | 1 | G | A | 77 | 33681 | 0.22861554 | 60.9220779 | rnr1Y285F | FALSE | CAN | JSY3811 | 3 | rnr1Y285F-R | CGC | G>A | A/T+1 | G/C-1 | CG>TA | CG>TA | TTGGCCGCA | GCACCAA | GCG | GCG | C | T | C>T | C>T_GCG | CG>TA_GCACCAA | 8 | FALSE | FALSE |
| 44_v.txt | 32970 | 32970-SNV-( | SNV | 1 | C | T | 37 | 33681 | 0.10985422 | 57 | rnr1Y285F | FALSE | CAN | JSY3811 | 3 | rnr1Y285F-R | ACC | C>T | A/T+1 | G/C-1 | CG>TA | CG>TA | GCCGCACCA | CCAAATG | TGG | ACC | C | A | C>T | C>T_ACC | CG>TA_CCAAATG | 8 | FALSE | FALSE |
| 44_v.txt | 33037 | 33037-SNV-( | SNV | 1 | C | T | 14 | 33681 | 0.04156646 | 62 | rnr1Y285F | FALSE | CAN | JSY3811 | 3 | rnr1Y285F-R | GCC | C>T | A/T+1 | G/C-1 | CG>TA | CG>TA | ATGTAGCCA | CCATTTC | CGG | GCC | C | A | C>T | C>T_GCC | CG>TA_CCATTTC | 8 | FALSE | FALSE |
| 44_v.txt | 33055 | 33055-SNV-( | SNV | 1 | G | A | 121 | 33681 | 0.35925299 | 57.9465365 | rnr1Y285F | FALSE | CAN | JSY3811 | 3 | rnr1Y285F-R | TGC | G>A | A/T+1 | G/C-1 | CG>TA | CG>TA | AGGACTGCG | GCGTGAC | ACG | ACG | C | T | C>T | C>T_ACG | CG>TA_GCGTGAC | 8 | FALSE | FALSE |
| 44_v.txt | 33056 | 33056-SNV-( | SNV | 1 | C | T | 45 | 33678 | 0.13361839 | 59.196575 | rnr1Y285F | FALSE | CAN | JSY3811 | 3 | rnr1Y285F-R | GCG | C>T | A/T+1 | G/C-1 | CG>TA | CG>TA | GGACTGCGT | CGTGACA | CGC | GCG | C | A | C>T | C>T_GCG | CG>TA_CGTGACA | 8 | FALSE | FALSE |
| 44_v.txt | 33069 | 33069-SNV-( | SNV | 1 | G | A | 26 | 33681 | 0.07719486 | 51.1925071 | rnr1Y285F | FALSE | CAN | JSY3811 | 3 | rnr1Y285F-R | TGC | G>A | A/T+1 | G/C-1 | CG>TA | CG>TA | GAATATGCC | GCCAAAG | ACG | ACG | C | T | C>T | C>T_ACG | CG>TA_GCCAAAG | 8 | FALSE | FALSE |
| 44_v.txt | 33070 | 33070-SNV-( | SNV | 1 | C | T | 47 | 33680 | 0.13954869 | 52.6382979 | rnr1Y285F | FALSE | CAN | JSY3811 | 3 | rnr1Y285F-R | GCC | C>T | A/T+1 | G/C-1 | CG>TA | CG>TA | AATATGCCA | CCAAAGA | CGG | GCC | C | A | C>T | C>T_GCC | CG>TA_CCAAAGA | 8 | FALSE | FALSE |
| 44_v.txt | 33104 | 33104-SNV-( | SNV | 1 | G | A | 32 | 33681 | 0.09500906 | 43.875 | rnr1Y285F | FALSE | CAN | JSY3811 | 3 | rnr1Y285F-R | CGC | G>A | A/T+1 | G/C-1 | CG>TA | CG>TA | AAGAGCGCC | GCCCACT | GCG | GCG | C | T | C>T | C>T_GCG | CG>TA_GCCCACT | 8 | FALSE | FALSE |
| 44_v.txt | 33113 | 33113-SNV-( | SNV | 1 | G | A | 33 | 33681 | 0.09797809 | 42.3501223 | rnr1Y285F | FALSE | CAN | JSY3811 | 3 | rnr1Y285F-R | GGC | G>A | A/T+1 | G/C-1 | CG>TA | CG>TA | CACTGGGCC | GCCGGCG | CCG | CCG | C | T | C>T | C>T_CCG | CG>TA_GCCGGCG | 8 | FALSE | FALSE |
| 44_v.txt | 33114 | 33114-SNV-( | SNV | 1 | C | T | 37 | 33681 | 0.10985422 | 38.918923 | rnr1Y285F | FALSE | CAN | JSY3811 | 3 | rnr1Y285F-R | GCC | C>T | A/T+1 | G/C-1 | CG>TA | CG>TA | ACTGGGCCG | CCGGCGT | CGG | GCC | C | A | C>T | C>T_GCC | CG>TA_CCGGCGT | 8 | FALSE | FALSE |
| 44_v.txt | 33115 | 33115-SNV-( | SNV | 1 | C | T | 42 | 33681 | 0.12469939 | 44.4050615 | rnr1Y285F | FALSE | CAN | JSY3811 | 3 | rnr1Y285F-R | CCG | C>T | A/T+1 | G/C-1 | CG>TA | CG>TA | CTGGGCCGG | CGGCGTT | GGC | CCG | C | A | C>T | C>T_CCG | CG>TA_CGGCGTT | 8 | FALSE | FALSE |
| 44_v.txt | 33118 | 33118-SNV-( | SNV | 1 | C | T | 39 | 33681 | 0.11579229 | 41.3846154 | rnr1Y285F | FALSE | CAN | JSY3811 | 3 | rnr1Y285F-R | GCG | C>T | A/T+1 | G/C-1 | CG>TA | CG>TA | GGCCGGCGT | CGTTGGT | CGC | GCG | C | A | C>T | C>T_GCG | CG>TA_CGTTGGT | 8 | FALSE | FALSE |
| 44_v.txt | 33153 | 33153-SNV-( | SNV | 1 | C | T | 157 | 33681 | 0.46613818 | 36.8407643 | rnr1Y285F | FALSE | CAN | JSY3811 | 3 | rnr1Y285F-R | ACC | C>T | A/T+1 | G/C-1 | CG>TA | CG>TA | AAAAGACCT | CCTGTAC | TGG | ACC | C | A | C>T | C>T_ACC | CG>TA_CCTGTAC | 8 | FALSE | FALSE |
| 44_v.txt | 33156 | 33156-SNV-( | SNV | 1 | G | A | 287 | 33681 | 0.85211247 | 37.2229965 | rnr1Y285F | FALSE | CAN | JSY3811 | 3 | rnr1Y285F-R | TGT | G>A | A/T+1 | G/C-1 | CG>TA | CG>TA | AGACCTGTA | GTACCAA | ACA | ACA | C | T | C>T | C>T_ACA | CG>TA_GTACCAA | 8 | FALSE | FALSE |
| 44_v.txt | 33159 | 33159-SNV-( | SNV | 1 | C | T | 46 | 33680 | 0.13657957 | 37.4340775 | rnr1Y285F | FALSE | CAN | JSY3811 | 3 | rnr1Y285F-R | ACC | C>T | A/T+1 | G/C-1 | CG>TA | CG>TA | CCTGTACCA | CCAATAG | TGG | ACC | C | A | C>T | C>T_ACC | CG>TA_CCAATAG | 8 | FALSE | FALSE |
| 44_v.txt | 33258 | 33258-SNV-( | SNV | 1 | C | T | 37 | 25205 | 0.14679627 | 43.1891892 | rnr1Y285F | FALSE | CAN | JSY3811 | 3 | rnr1Y285F-R | GCC | C>T | A/T+1 | G/C-1 | CG>TA | CG>TA | CCTATGCCAT | CCATCCT | CGG | GCC | C | A | C>T | C>T_GCC | CG>TA_CCATCCT | 8 | FALSE | FALSE |
| 44_v.txt | 33276 | 33276-SNV-( | SNV | 1 | G | A | 50 | 25205 | 0.19837334 | 47.02 | rnr1Y285F | FALSE | CAN | JSY3811 | 3 | rnr1Y285F-R | CGT | G>A | A/T+1 | G/C-1 | CG>TA | CG>TA | GAGAACGTA | GTATCCT | GCA | GCA | C | T | C>T | C>T_GCA | CG>TA_GTATCCT | 8 | FALSE | FALSE |
| 44_v.txt | 33284 | 33284-SNV-( | SNV | 1 | G | A | 31 | 25205 | 0.12299147 | 45.8387097 | rnr1Y285F | FALSE | CAN | JSY3811 | 3 | rnr1Y285F-R | CGC | G>A | A/T+1 | G/C-1 | CG>TA | CG>TA | ATCCTCGCC | GCCATTT | GCG | GCG | C | T | C>T | C>T_GCG | CG>TA_GCCATTT | 8 | FALSE | FALSE |
| 44_v.txt | 33309 | 33309-SNV-( | SNV | 1 | C | T | 38 | 25205 | 0.15076374 | 56.340255 | rnr1Y285F | FALSE | CAN | JSY3811 | 3 | rnr1Y285F-R | GCG | C>T | A/T+1 | G/C-1 | CG>TA | CG>TA | AAAGAGCGC | CGCAATG | CGC | GCG | C | A | C>T | C>T_GCG | CG>TA_CGCAATG | 8 | FALSE | FALSE |
| 44_v.txt | 33310 | 33310-SNV-( | SNV | 1 | G | A | 32 | 25205 | 0.12695894 | 57.0260346 | rnr1Y285F | FALSE | CAN | JSY3811 | 3 | rnr1Y285F-R | CGC | G>A | A/T+1 | G/C-1 | CG>TA | CG>TA | AAGAGCGCA | GCAATGG | GCG | GCG | C | T | C>T | C>T_GCG | CG>TA_GCAATGG | 8 | FALSE | FALSE |
| 44_v.txt | 33392 | 33392-SNV-( | SNV | 1 | G | A | 36 | 25205 | 0.1428288 | 45.3888889 | rnr1Y285F | FALSE | CAN | JSY3811 | 3 | rnr1Y285F-R | CGT | G>A | A/T+1 | G/C-1 | CG>TA | CG>TA | AACGTCGTG | GTGAAAG | GCA | GCA | C | T | C>T | C>T_GCA | CG>TA_GTGAAAG | 8 | FALSE | FALSE |
| 44_v.txt | 33412 | 33412-SNV-( | SNV | 1 | G | A | 29 | 25205 | 0.11505654 | 47.6206897 | rnr1Y285F | FALSE | CAN | JSY3811 | 3 | rnr1Y285F-R | GGC | G>A | A/T+1 | G/C-1 | CG>TA | CG>TA | TGACCGGCT | GCTCATT | CCG | CCG | C | T | C>T | C>T_CCG | CG>TA_GCTCATT | 8 | FALSE | FALSE |
| 44_v.txt | 33474 | 33474-SNV-( | SNV | 1 | C | T | 38 | 25204 | 0.15076972 | 33.5700139 | rnr1Y285F | FALSE | CAN | JSY3811 | 3 | rnr1Y285F-R | GCC | C>T | A/T+1 | G/C-1 | CG>TA | CG>TA | GCTATGCCT | CCTTTTT | CGG | GCC | C | A | C>T | C>T_GCC | CG>TA_CCTTTTT | 8 | FALSE | FALSE |
| 44_v.txt | 31658 | 31658-SNV-( | SNV | 1 | C | G | 2 | 14771 | 0.01354005 | 37.9997272 | rnr1Y285F | FALSE | CAN | JSY3811 | 3 | rnr1Y285F-R | GCG | C>G | G/C+1 | G/C-1 | CG>GC | CG>GC | AAATGGCGT | CGTGGAA | CGC | GCG | C | C | C>G | C>G_GCG | CG>GC_CGTGGAA | 8 | FALSE | FALSE |
| 44_v.txt | 31947 | 31947-SNV-( | SNV | 1 | G | C | 16 | 23887 | 0.06698204 | 37.5 | rnr1Y285F | FALSE | CAN | JSY3811 | 3 | rnr1Y285F-R | CGT | G>C | G/C+1 | G/C-1 | CG>GC | CG>GC | ATGATCGTC | GTCATAA | GCA | GCA | C | G | C>G | C>G_GCA | CG>GC_GTCATAA | 8 | FALSE | FALSE |
| 44_v.txt | 32027 | 32027-SNV-( | SNV | 1 | G | C | 9 | 23899 | 0.03765848 | 63.1111111 | rnr1Y285F | FALSE | CAN | JSY3811 | 3 | rnr1Y285F-R | GGT | G>C | G/C+1 | G/C-1 | CG>GC | CG>GC | GCCACGGTA | GTATTTC | CCA | CCA | C | G | C>G | C>G_CCA | CG>GC_GTATTTC | 8 | FALSE | FALSE |
| 44_v.txt | 32064 | 32064-SNV-( | SNV | 1 | G | C | 9 | 23899 | 0.03765848 | 63.7777778 | rnr1Y285F | FALSE | CAN | JSY3811 | 3 | rnr1Y285F-R | TGA | G>C | G/C+1 | G/C-1 | CG>GC | CG>GC | GAGATTGAG | GAGATAA | ACT | ACT | C | G | C>G | C>G_ACT | CG>GC_GAGATAA | 8 | FALSE | FALSE |
| 44_v.txt | 32292 | 32292-SNV-( | SNV | 1 | G | C | 24 | 39917 | 0.06012476 | 51.0416667 | rnr1Y285F | FALSE | CAN | JSY3811 | 3 | rnr1Y285F-R | TGA | G>C | G/C+1 | G/C-1 | CG>GC | CG>GC | ATATTTGAA | GAATTTG | ACT | ACT | C | G | C>G | C>G_ACT | CG>GC_GAATTTG | 8 | FALSE | FALSE |
| 44_v.txt | 32557 | 32557-SNV-( | SNV | 1 | C | G | 28 | 39904 | 0.0701684 | 37.6068833 | rnr1Y285F | FALSE | CAN | JSY3811 | 3 | rnr1Y285F-R | CCA | C>G | G/C+1 | G/C-1 | CG>GC | CG>GC | TGATACCAA | CAACTAG | GGT | CCA | C | C | C>G | C>G_CCA | CG>GC_CAACTAG | 8 | FALSE | FALSE |
| 44_v.txt | 32566 | 32566-SNV-( | SNV | 1 | C | G | 101 | 33123 | 0.30492407 | 36.5049505 | rnr1Y285F | FALSE | CAN | JSY3811 | 3 | rnr1Y285F-R | TCA | C>G | G/C+1 | G/C-1 | CG>GC | CG>GC | CTAGTTCAG | CAGTACC | AGT | TCA | C | C | C>G | C>G_TCA | CG>GC_CAGTACC | 8 | FALSE | FALSE |
| 44_v.txt | 32940 | 32940-SNV-( | SNV | 1 | G | C | 23 | 33681 | 0.06828776 | 50.6956522 | rnr1Y285F | FALSE | CAN | JSY3811 | 3 | rnr1Y285F-R | AGA | G>C | G/C+1 | G/C-1 | CG>GC | CG>GC | GCCCAAGAA | GAAAACC | TCT | TCT | C | G | C>G | C>G_TCT | CG>GC_GAAAACC | 8 | FALSE | FALSE |
| 44_v.txt | 33018 | 33018-SNV-( | SNV | 1 | G | C | 27 | 33681 | 0.08016389 | 61.6666667 | rnr1Y285F | FALSE | CAN | JSY3811 | 3 | rnr1Y285F-R | TGT | G>C | G/C+1 | G/C-1 | CG>GC | CG>GC | GAGGATGTA | GTAACAG | ACA | ACA | C | G | C>G | C>G_ACA | CG>GC_GTAACAG | 8 | FALSE | FALSE |
| 44_v.txt | 33037 | 33037-SNV-( | SNV | 1 | C | G | 137 | 33681 | 0.40675752 | 59.5036496 | rnr1Y285F | FALSE | CAN | JSY3811 | 3 | rnr1Y285F-R | GCC | C>G | G/C+1 | G/C-1 | CG>GC | CG>GC | ATGTAGCCA | CCATTTC | CGG | GCC | C | C | C>G | C>G_GCC | CG>GC_CCATTTC | 8 | FALSE | FALSE |
| 44_v.txt | 33070 | 33070-SNV-( | SNV | 1 | C | G | 20 | 33680 | 0.05938242 | 51.25 | rnr1Y285F | FALSE | CAN | JSY3811 | 3 | rnr1Y285F-R | GCC | C>G | G/C+1 | G/C-1 | CG>GC | CG>GC | AATATGCCA | CCAAAGA | CGG | GCC | C | C | C>G | C>G_GCC | CG>GC_CCAAAGA | 8 | FALSE | FALSE |
| 44_v.txt | 33114 | 33114-SNV-( | SNV | 1 | C | G | 3 | 33681 | 8.91E-03 | 63.996477 | rnr1Y285F | FALSE | CAN | JSY3811 | 3 | rnr1Y285F-R | GCC | C>G | G/C+1 | G/C-1 | CG>GC | CG>GC | ACTGGGCCG | CCGGCGT | CGG | GCC | C | C | C>G | C>G_GCC | CG>GC_CCGGCGT | 8 | FALSE | FALSE |
| 44_v.txt | 33142 | 33142-SNV-( | SNV | 1 | C | G | 10 | 33681 | 0.02969033 | 41 | rnr1Y285F | FALSE | CAN | JSY3811 | 3 | rnr1Y285F-R | CCA | C>G | G/C+1 | G/C-1 | CG>GC | CG>GC | ATAAACCAA | CAATGAA | GGT | CCA | C | C | C>G | C>G_CCA | CG>GC_CAATGAA | 8 | FALSE | FALSE |
| 44_v.txt | 33153 | 33153-SNV-( | SNV | 1 | C | G | 15 | 33681 | 0.0445355 | 37.9333333 | rnr1Y285F | FALSE | CAN | JSY3811 | 3 | rnr1Y285F-R | ACC | C>G | G/C+1 | G/C-1 | CG>GC | CG>GC | AAAAGACCT | CCTGTAC | TGG | ACC | C | C | C>G | C>G_ACC | CG>GC_CCTGTAC | 8 | FALSE | FALSE |
| 44_v.txt | 33156 | 33156-SNV-( | SNV | 1 | G | C | 6 | 33681 | 0.0178142 | 35.5 | rnr1Y285F | FALSE | CAN | JSY3811 | 3 | rnr1Y285F-R | TGT | G>C | G/C+1 | G/C-1 | CG>GC | CG>GC | AGACCTGTA | GTACCAA | ACA | ACA | C | G | C>G | C>G_ACA | CG>GC_GTACCAA | 8 | FALSE | FALSE |
| 44_v.txt | 33160 | 33160-SNV-( | SNV | 1 | C | G | 11 | 33680 | 0.03266033 | 35.5451112 | rnr1Y285F | FALSE | CAN | JSY3811 | 3 | rnr1Y285F-R | CCA | C>G | G/C+1 | G/C-1 | CG>GC | CG>GC | CTGTACCAA | CAATAGT | GGT | CCA | C | C | C>G | C>G_CCA | CG>GC_CAATAGT | 8 | FALSE | FALSE |
| 44_v.txt | 33310 | 33310-SNV-( | SNV | 1 | G | C | 3 | 25205 | 0.0119024 | 42.9950719 | rnr1Y285F | FALSE | CAN | JSY3811 | 3 | rnr1Y285F-R | CGC | G>C | G/C+1 | G/C-1 | CG>GC | CG>GC | AAGAGCGCA | GCAATGG | GCG | GCG | C | G | C>G | C>G_GCG | CG>GC_GCAATGG | 8 | FALSE | FALSE |
| 44_v.txt | 33456 | 33456-SNV-( | SNV | 1 | G | C | 42 | 25205 | 0.1666336 | 43.2619048 | rnr1Y285F | FALSE | CAN | JSY3811 | 3 | rnr1Y285F-R | TGA | G>C | G/C+1 | G/C-1 | CG>GC | CG>GC | TCTTTTGAAT | GAATTTG | ACT | ACT | C | G | C>G | C>G_ACT | CG>GC_GAATTTG | 8 | FALSE | FALSE |
| 44_v.txt | 31641 | 31641-SNV-( | SNV | 1 | G | T | 45 | 14771 | 0.30465101 | 18.6888889 | rnr1Y285F | FALSE | CAN | JSY3811 | 3 | rnr1Y285F-R | GGT | G>T | A/T+1 | G/C-1 | CG>AT | CG>AT | ATGAGGGTG | GTGAGAA | CCA | CCA | C | A | C>A | C>A_CCA | CG>AT_GTGAGAA | 8 | FALSE | FALSE |
| 44_v.txt | 31657 | 31657-SNV-( | SNV | 1 | G | T | 117 | 14771 | 0.79209261 | 19.1183794 | rnr1Y285F | FALSE | CAN | JSY3811 | 3 | rnr1Y285F-R | GGC | G>T | A/T+1 | G/C-1 | CG>AT | CG>AT | GAAATGGCG | GCGTGGA | CCG | CCG | C | A | C>A | C>A_CCG | CG>AT_GCGTGGA | 8 | FALSE | FALSE |
| 44_v.txt | 32027 | 32027-SNV-( | SNV | 1 | G | T | 22 | 23899 | 0.09205406 | 59.0454546 | rnr1Y285F | FALSE | CAN | JSY3811 | 3 | rnr1Y285F-R | GGT | G>T | A/T+1 | G/C-1 | CG>AT | CG>AT | GCCACGGTA | GTATTTC | CCA | CCA | C | A | C>A | C>A_CCA | CG>AT_GTATTTC | 8 | FALSE | FALSE |
| 44_v.txt | 32064 | 32064-SNV-( | SNV | 1 | G | T | 14 | 23899 | 0.05857986 | 54.5714286 | rnr1Y285F | FALSE | CAN | JSY3811 | 3 | rnr1Y285F-R | TGA | G>T | A/T+1 | G/C-1 | CG>AT | CG>AT | GAGATTGAG | GAGATAA | ACT | ACT | C | A | C>A | C>A_ACT | CG>AT_GAGATAA | 8 | FALSE | FALSE |
| 44_v.txt | 32088 | 32088-SNV-( | SNV | 1 | C | A | 51 | 23899 | 0.21339805 | 61.1568628 | rnr1Y285F | FALSE | CAN | JSY3811 | 3 | rnr1Y285F-R | GCC | C>A | A/T+1 | G/C-1 | CG>AT | CG>AT | AAAAAGCCT | CCTGCAA | CGG | GCC | C | T | C>A | C>A_GCC | CG>AT_CCTGCAA | 8 | FALSE | FALSE |
| 44_v.txt | 32095 | 32095-SNV-( | SNV | 1 | C | A | 617 | 23899 | 2.58169798 | 59.8200972 | rnr1Y285F | FALSE | CAN | JSY3811 | 3 | rnr1Y285F-R | ACA | C>A | A/T+1 | G/C-1 | CG>AT | CG>AT | CTGCAACAC | CACCAGT | TGT | ACA | C | T | C>A | C>A_ACA | CG>AT_CACCAGT | 8 | FALSE | FALSE |
| 44_v.txt | 32149 | 32149-SNV-( | SNV | 1 | C | A | 13 | 23899 | 0.05439558 | 49 | rnr1Y285F | FALSE | CAN | JSY3811 | 3 | rnr1Y285F-R | TCC | C>A | A/T+1 | G/C-1 | CG>AT | CG>AT | ATGTCTCCAT | CCATGTA | AGG | TCC | C | T | C>A | C>A_TCC | CG>AT_CCATGTA | 8 | FALSE | FALSE |
| 44_v.txt | 32195 | 32195-SNV-( | SNV | 1 | G | T | 157 | 23899 | 0.65693125 | 43.2165605 | rnr1Y285F | FALSE | CAN | JSY3811 | 3 | rnr1Y285F-R | TGT | G>T | A/T+1 | G/C-1 | CG>AT | CG>AT | TGCAATGTA | GTATGGA | ACA | ACA | C | A | C>A | C>A_ACA | CG>AT_GTATGGA | 8 | FALSE | FALSE |
| 44_v.txt | 32223 | 32223-SNV-( | SNV | 1 | G | T | 31 | 23899 | 0.12971254 | 37.1290323 | rnr1Y285F | FALSE | CAN | JSY3811 | 3 | rnr1Y285F-R | TGA | G>T | A/T+1 | G/C-1 | CG>AT | CG>AT | GTCCTTGAC | GACAGGA | ACT | ACT | C | A | C>A | C>A_ACT | CG>AT_GACAGGA | 8 | FALSE | FALSE |
| 44_v.txt | 32271 | 32271-SNV-( | SNV | 1 | C | A | 48 | 39917 | 0.12024952 | 34.6875 | rnr1Y285F | FALSE | CAN | JSY3811 | 3 | rnr1Y285F-R | ACG | C>A | A/T+1 | G/C-1 | CG>AT | CG>AT | AAAATACGG | CGGGAAC | TGC | ACG | C | T | C>A | C>A_ACG | CG>AT_CGGGAAC | 8 | FALSE | FALSE |
| 44_v.txt | 32272 | 32272-SNV-( | SNV | 1 | G | T | 27 | 39917 | 0.06764035 | 23.4102211 | rnr1Y285F | FALSE | CAN | JSY3811 | 3 | rnr1Y285F-R | CGG | G>T | A/T+1 | G/C-1 | CG>AT | CG>AT | AAATACGGG | GGGAACC | GCC | GCC | C | A | C>A | C>A_GCC | CG>AT_GGGAACC | 8 | FALSE | FALSE |
| 44_v.txt | 32304 | 32304-SNV-( | SNV | 1 | G | T | 26 | 39917 | 0.06513516 | 49 | rnr1Y285F | FALSE | CAN | JSY3811 | 3 | rnr1Y285F-R | AGA | G>T | A/T+1 | G/C-1 | CG>AT | CG>AT | GCGGCAGAA | GAAATAA | TCT | TCT | C | A | C>A | C>A_TCT | CG>AT_GAAATAA | 8 | FALSE | FALSE |
| 44_v.txt | 32424 | 32424-SNV-( | SNV | 1 | G | T | 8 | 39917 | 0.02004159 | 54.25 | rnr1Y285F | FALSE | CAN | JSY3811 | 3 | rnr1Y285F-R | AGG | G>T | A/T+1 | G/C-1 | CG>AT | CG>AT | AGTTTAGGG | GGGTCAT | TCC | TCC | C | A | C>A | C>A_TCC | CG>AT_GGGTCAT | 8 | FALSE | FALSE |
| 44_v.txt | 32471 | 32471-SNV-( | SNV | 1 | G | T | 23 | 39917 | 0.05761956 | 47.8669079 | rnr1Y285F | FALSE | CAN | JSY3811 | 3 | rnr1Y285F-R | TGT | G>T | A/T+1 | G/C-1 | CG>AT | CG>AT | GCCAATGTA | GTAGAAG | ACA | ACA | C | A | C>A | C>A_ACA | CG>AT_GTAGAAG | 8 | FALSE | FALSE |
| 44_v.txt | 32487 | 32487-SNV-( | SNV | 1 | C | A | 20 | 39917 | 0.05010397 | 45.05 | rnr1Y285F | FALSE | CAN | JSY3811 | 3 | rnr1Y285F-R | ACG | C>A | A/T+1 | G/C-1 | CG>AT | CG>AT | AAGATACGG | CGGAAAA | TGC | ACG | C | T | C>A | C>A_ACG | CG>AT_CGGAAAA | 8 | FALSE | FALSE |
| 44_v.txt | 32536 | 32536-SNV-( | SNV | 1 | C | A | 8 | 39917 | 0.02004159 | 27.375 | rnr1Y285F | FALSE | CAN | JSY3811 | 3 | rnr1Y285F-R | GCA | C>A | A/T+1 | G/C-1 | CG>AT | CG>AT | GGTTTGCAG | CAGCTTC | CGT | GCA | C | T | C>A | C>A_GCA | CG>AT_CAGCTTC | 8 | FALSE | FALSE |
| 44_v.txt | 32556 | 32556-SNV-( | SNV | 1 | C | A | 847 | 39905 | 2.12254104 | 37.3902627 | rnr1Y285F | FALSE | CAN | JSY3811 | 3 | rnr1Y285F-R | ACC | C>A | A/T+1 | G/C-1 | CG>AT | CG>AT | GTGATACCA | CCAACTA | TGG | ACC | C | T | C>A | C>A_ACC | CG>AT_CCAACTA | 8 | FALSE | FALSE |
| 44_v.txt | 32557 | 32557-SNV-( | SNV | 1 | C | A | 5 | 39904 | 0.01253007 | 32.4000897 | rnr1Y285F | FALSE | CAN | JSY3811 | 3 | rnr1Y285F-R | CCA | C>A | A/T+1 | G/C-1 | CG>AT | CG>AT | TGATACCAA | CAACTAG | GGT | CCA | C | T | C>A | C>A_CCA | CG>AT_CAACTAG | 8 | FALSE | FALSE |
| 44_v.txt | 32566 | 32566-SNV-( | SNV | 1 | C | A | 39 | 33123 | 0.11774296 | 36.3076923 | rnr1Y285F | FALSE | CAN | JSY3811 | 3 | rnr1Y285F-R | TCA | C>A | A/T+1 | G/C-1 | CG>AT | CG>AT | CTAGTTCAG | CAGTACC | AGT | TCA | C | T | C>A | C>A_TCA | CG>AT_CAGTACC | 8 | FALSE | FALSE |
| 44_v.txt | 32609 | 32609-SNV-( | SNV | 1 | C | A | 5 | 33134 | 0.01509024 | 40.1976309 | rnr1Y285F | FALSE | CAN | JSY3811 | 3 | rnr1Y285F-R | CCC | C>A | A/T+1 | G/C-1 | CG>AT | CG>AT | GGAAACCCA | CCAACCT | GGG | CCC | C | T | C>A | C>A_CCC | CG>AT_CCAACCT | 8 | FALSE | FALSE |
| 44_v.txt | 32629 | 32629-SNV-( | SNV | 1 | C | A | 25 | 33134 | 0.0754512 | 52.08 | rnr1Y285F | FALSE | CAN | JSY3811 | 3 | rnr1Y285F-R | TCG | C>A | A/T+1 | G/C-1 | CG>AT | CG>AT | TCCCTTCGTT | CGTTTTT | AGC | TCG | C | T | C>A | C>A_TCG | CG>AT_CGTTTTT | 8 | FALSE | FALSE |
| 44_v.txt | 32819 | 32819-SNV-( | SNV | 1 | G | T | 1031 | 33134 | 3.11160741 | 44.0453283 | rnr1Y285F | FALSE | CAN | JSY3811 | 3 | rnr1Y285F-R | AGT | G>T | A/T+1 | G/C-1 | CG>AT | CG>AT | GAACAAGTT | GTTCATT | TCA | TCA | C | A | C>A | C>A_TCA | CG>AT_GTTCATT | 8 | FALSE | FALSE |
| 44_v.txt | 32917 | 32917-SNV-( | SNV | 1 | C | A | 98 | 33681 | 0.29096523 | 50.8571429 | rnr1Y285F | FALSE | CAN | JSY3811 | 3 | rnr1Y285F-R | TCC | C>A | A/T+1 | G/C-1 | CG>AT | CG>AT | TAAGTTCCA | CCAGGGC | AGG | TCC | C | T | C>A | C>A_TCC | CG>AT_CCAGGGC | 8 | FALSE | FALSE |
| 44_v.txt | 32940 | 32940-SNV-( | SNV | 1 | G | T | 21961 | 33681 | 65.2029334 | 56.7795182 | rnr1Y285F | FALSE | CAN | JSY3811 | 3 | rnr1Y285F-R | AGA | G>T | A/T+1 | G/C-1 | CG>AT | CG>AT | GCCCAAGAA | GAAAACC | TCT | TCT | C | A | C>A | C>A_TCT | CG>AT_GAAAACC | 8 | FALSE | FALSE |
| 44_v.txt | 32958 | 32958-SNV-( | SNV | 1 | C | A | 38 | 33681 | 0.11282325 | 54.5263158 | rnr1Y285F | FALSE | CAN | JSY3811 | 3 | rnr1Y285F-R | ACC | C>A | A/T+1 | G/C-1 | CG>AT | CG>AT | ATGTAACCA | CCATTGG | TGG | ACC | C | T | C>A | C>A_ACC | CG>AT_CCATTGG | 8 | FALSE | FALSE |
| 44_v.txt | 33024 | 33024-SNV-( | SNV | 1 | G | T | 14 | 33681 | 0.04156646 | 62.2857143 | rnr1Y285F | FALSE | CAN | JSY3811 | 3 | rnr1Y285F-R | AGG | G>T | A/T+1 | G/C-1 | CG>AT | CG>AT | GTAACAGGG | GGGATGA | TCC | TCC | C | A | C>A | C>A_TCC | CG>AT_GGGATGA | 8 | FALSE | FALSE |
| 44_v.txt | 33070 | 33070-SNV-( | SNV | 1 | C | A | 8 | 33680 | 0.02375297 | 47.5 | rnr1Y285F | FALSE | CAN | JSY3811 | 3 | rnr1Y285F-R | GCC | C>A | A/T+1 | G/C-1 | CG>AT | CG>AT | AATATGCCA | CCAAAGA | CGG | GCC | C | T | C>A | C>A_GCC | CG>AT_CCAAAGA | 8 | FALSE | FALSE |
| 44_v.txt | 33113 | 33113-SNV-( | SNV | 1 | G | T | 1 | 33681 | 2.97E-03 | 37.9971629 | rnr1Y285F | FALSE | CAN | JSY3811 | 3 | rnr1Y285F-R | GGC | G>T | A/T+1 | G/C-1 | CG>AT | CG>AT | CACTGGGCC | GCCGGCG | CCG | CCG | C | A | C>A | C>A_CCG | CG>AT_GCCGGCG | 8 | FALSE | FALSE |
| 44_v.txt | 33151 | 33151-SNV-( | SNV | 1 | G | T | 99 | 33681 | 0.29393427 | 39.8787879 | rnr1Y285F | FALSE | CAN | JSY3811 | 3 | rnr1Y285F-R | AGA | G>T | A/T+1 | G/C-1 | CG>AT | CG>AT | TGAAAAGAC | GACCTGT | TCT | TCT | C | A | C>A | C>A_TCT | CG>AT_GACCTGT | 8 | FALSE | FALSE |
| 44_v.txt | 33186 | 33186-SNV-( | SNV | 1 | C | A | 83 | 25201 | 0.32935201 | 36.7831325 | rnr1Y285F | FALSE | CAN | JSY3811 | 3 | rnr1Y285F-R | ACC | C>A | A/T+1 | G/C-1 | CG>AT | CG>AT | ATCATACCA | CCAATAT | TGG | ACC | C | T | C>A | C>A_ACC | CG>AT_CCAATAT | 8 | FALSE | FALSE |
| 44_v.txt | 33354 | 33354-SNV-( | SNV | 1 | G | T | 125 | 25205 | 0.49593335 | 17.352 | rnr1Y285F | FALSE | CAN | JSY3811 | 3 | rnr1Y285F-R | TGA | G>T | A/T+1 | G/C-1 | CG>AT | CG>AT | GGTATTGAC | GACCCAC | ACT | ACT | C | A | C>A | C>A_ACT | CG>AT_GACCCAC | 8 | FALSE | FALSE |
| 44_v.txt | 33436 | 33436-SNV-( | SNV | 1 | C | A | 17 | 25205 | 0.06744694 | 44.8823529 | rnr1Y285F | FALSE | CAN | JSY3811 | 3 | rnr1Y285F-R | TCT | C>A | A/T+1 | G/C-1 | CG>AT | CG>AT | TCTCCTCTAT | CTATGTC | AGA | TCT | C | T | C>A | C>A_TCT | CG>AT_CTATGTC | 8 | FALSE | FALSE |
| 44_v.txt | 33474 | 33474-SNV-( | SNV | 1 | C | A | 82 | 25204 | 0.32534518 | 33.3601646 | rnr1Y285F | FALSE | CAN | JSY3811 | 3 | rnr1Y285F-R | GCC | C>A | A/T+1 | G/C-1 | CG>AT | CG>AT | GCTATGCCTT | CCTTTTT | CGG | GCC | C | T | C>A | C>A_GCC | CG>AT_CCTTTTT | 8 | FALSE | FALSE |
| 44_v.txt | 33475 | 33475-SNV-( | SNV | 1 | C | A | 27 | 25204 | 0.10712585 | 36.5555556 | rnr1Y285F | FALSE | CAN | JSY3811 | 3 | rnr1Y285F-R | CCT | C>A | A/T+1 | G/C-1 | CG>AT | CG>AT | CTATGCCTTT | CTTTTTT | GGA | CCT | C | T | C>A | C>A_CCT | CG>AT_CTTTTTT | 8 | FALSE | FALSE |
| 44_v.txt | 32556 | 32556-MNV- | MNV | 2 | CC | AA | 1 | 39905 | 2.51E-03 | 37 | rnr1Y285F | FALSE | CAN | JSY3811 | 3 | rnr1Y285F-R | ACC | nope | >1bp-ins | >1bpdel | none | MNV | GTGATACCA | CCAACTA | TGG | ACC | CC | none | NA | NA_ACC | MNV_CCAACTA | 8 | FALSE | FALSE |
| 44_v.txt | 31872 | 31872-Insert | Insertion | 2 | - | AT | 9 | 14772 | 0.06092608 | 39 | rnr1Y285F | FALSE | CAN | JSY3811 | 3 | rnr1Y285F-R | GAT | nope | >1bp-ins | >1bpdel | none | >1bp-ins | ATAGAGATA | ATATAGG | CTA | GAT | - | none | NA | NA_GAT | >1bp-ins_ATATAGG | 8 | FALSE | FALSE |
| 44_v.txt | 32081 | 32081-Insert | Insertion | 1 | - | A | 11 | 23899 | 0.04602703 | 64 | rnr1Y285F | FALSE | CAN | JSY3811 | 3 | rnr1Y285F-R | CAA | nope | A/T+1 | >1bpdel | none | A/T+1 | CCATGCAAA | AAAAAAG | GTT | CAA | - | T | NA | NA_CAA | A/T+1_AAAAAAG | 8 | FALSE | FALSE |
| 44_v.txt | 32631 | 32631-Insert | Insertion | 1 | - | T | 2 | 33134 | 6.04E-03 | 38 | rnr1Y285F | FALSE | CAN | JSY3811 | 3 | rnr1Y285F-R | GTT | nope | A/T+1 | >1bpdel | none | A/T+1 | CCTTCGTTT | TTTTTAT | CAA | GTT | - | A | NA | NA_GTT | A/T+1_TTTTTAT | 8 | FALSE | FALSE |
| 44_v.txt | 32728 | 32728-Insert | Insertion | 1 | - | A | 2 | 33134 | 6.04E-03 | 51 | rnr1Y285F | FALSE | CAN | JSY3811 | 3 | rnr1Y285F-R | CAA | nope | A/T+1 | >1bpdel | none | A/T+1 | CCATACAAA | AAAAACA | GTT | CAA | - | T | NA | NA_CAA | A/T+1_AAAAACA | 8 | FALSE | FALSE |
| 44_v.txt | 32842 | 32842-Insert | Insertion | 1 | - | A | 11 | 33133 | 0.03319953 | 42.3636364 | rnr1Y285F | FALSE | CAN | JSY3811 | 3 | rnr1Y285F-R | CAA | nope | A/T+1 | >1bpdel | none | A/T+1 | TTACCCAAA | AAAAAAT | GTT | CAA | - | T | NA | NA_CAA | A/T+1_AAAAAAT | 8 | FALSE | FALSE |
| 44_v.txt | 33474 | 33474-Insert | Insertion | 1 | - | A | 2 | 25204 | 7.94E-03 | 30 | rnr1Y285F | FALSE | CAN | JSY3811 | 3 | rnr1Y285F-R | GCC | nope | A/T+1 | >1bpdel | none | A/T+1 | GCTATGCCT | CCTTTTT | CGG | GCC | - | T | NA | NA_GCC | A/T+1_CCTTTTT | 8 | FALSE | FALSE |

| | | | | | | | | | | | | | | | | | | | | | | | | | | | | | | | | | | |
|---|---|---|---|---|---|---|---|---|---|---|---|---|---|---|---|---|---|---|---|---|---|---|---|---|---|---|---|---|---|---|---|---|---|---|
| 44_v.txt | 32201 | 32201-Deleti | Deletion | 3 | AAC | - | 6 | 23899 | 0.02510565 | 46.1666667 | rnr1Y285F | FALSE | CAN | JSY3811 | 3 | rnr1Y285F-R | GAA | nope | >1bp-ins | >1bpdel | none | >1bpdel | GTATGGAAC | AACACCA | CTT | GAA | AAC | none | NA | NA_GAA | >1bpdel_AACACCA | 8 | FALSE | FALSE |
| 44_v.txt | 32337 | 32337-Deleti | Deletion | 2 | AT | - | 12 | 39917 | 0.03006238 | 62.5833333 | rnr1Y285F | FALSE | CAN | JSY3811 | 3 | rnr1Y285F-R | GAT | nope | >1bp-ins | >1bpdel | none | >1bpdel | TTGAAGATA | ATATGTG | CTA | GAT | AT | none | NA | NA_GAT | >1bpdel_ATATGTG | 8 | FALSE | FALSE |
| 44_v.txt | 32375 | 32375-Deleti | Deletion | 3 | AAT | - | 22 | 39917 | 0.05511436 | 63.0454546 | rnr1Y285F | FALSE | CAN | JSY3811 | 3 | rnr1Y285F-R | CAA | nope | >1bp-ins | >1bpdel | none | >1bpdel | AATAGCAAT | AATAATA | GTT | CAA | AAT | none | NA | NA_CAA | >1bpdel_AATAATA | 8 | FALSE | FALSE |
| 44_v.txt | 32471 | 32471-Deleti | Deletion | 2 | GT | - | 1 | 39917 | 2.51E-03 | 38 | rnr1Y285F | FALSE | CAN | JSY3811 | 3 | rnr1Y285F-R | TGT | nope | >1bp-ins | >1bpdel | none | >1bpdel | GCCAATGTA | GTAGAAG | ACA | TGT | GT | none | NA | NA_TGT | >1bpdel_GTAGAAG | 8 | FALSE | FALSE |
| 44_v.txt | 32492 | 32492-Deleti | Deletion | 3 | AAC | - | 21 | 39917 | 0.05260916 | 48.9047619 | rnr1Y285F | FALSE | CAN | JSY3811 | 3 | rnr1Y285F-R | AAA | nope | >1bp-ins | >1bpdel | none | >1bpdel | ACGGAAAAC | AACAACT | TTT | AAA | AAC | none | NA | NA_AAA | >1bpdel_AACAACT | 8 | FALSE | FALSE |
| 44_v.txt | 31663 | 31663-Deleti | Deletion | 1 | A | - | 7 | 14772 | 0.04738695 | 36.1428571 | rnr1Y285F | FALSE | CAN | JSY3811 | 3 | rnr1Y285F-R | GAA | nope | >1bp-ins | A/T-1 | none | A/T-1 | GCGTGGAAA | AAATGTG | CTT | CTT | T | none | T>none | T>none_CTT | A/T-1_AAATGTG | 8 | FALSE | FALSE |
| 44_v.txt | 31711 | 31711-Deleti | Deletion | 1 | A | - | 4 | 14772 | 0.02707826 | 64 | rnr1Y285F | FALSE | CAN | JSY3811 | 3 | rnr1Y285F-R | CAA | nope | >1bp-ins | A/T-1 | none | A/T-1 | CATTCCAAA | AAAATTT | GTT | GTT | T | none | T>none | T>none_GTT | A/T-1_AAAATTT | 8 | FALSE | FALSE |
| 44_v.txt | 31723 | 31723-Deleti | Deletion | 1 | A | - | 11 | 14772 | 0.0744652 | 56.8181818 | rnr1Y285F | FALSE | CAN | JSY3811 | 3 | rnr1Y285F-R | CAA | nope | >1bp-ins | A/T-1 | none | A/T-1 | TGTCCCAAA | AAAAAGT | GTT | GTT | T | none | T>none | T>none_GTT | A/T-1_AAAAAGT | 8 | FALSE | FALSE |
| 44_v.txt | 31904 | 31904-Deleti | Deletion | 1 | T | - | 6 | 14772 | 0.04061738 | 46.3333333 | rnr1Y285F | FALSE | CAN | JSY3811 | 3 | rnr1Y285F-R | ATT | nope | >1bp-ins | A/T-1 | none | A/T-1 | ATTGAATTTT | TTTTGGT | TAA | ATT | T | none | nope | nope_ATT | A/T-1_TTTTGGT | 8 | FALSE | FALSE |
| 44_v.txt | 31952 | 31952-Deleti | Deletion | 1 | A | - | 5 | 23899 | 0.02092138 | 37.8 | rnr1Y285F | FALSE | CAN | JSY3811 | 3 | rnr1Y285F-R | TAA | nope | >1bp-ins | A/T-1 | none | A/T-1 | CGTCATAAA | AAATGTG | ATT | ATT | T | none | T>none | T>none_ATT | A/T-1_AAATGTG | 8 | FALSE | FALSE |
| 44_v.txt | 31972 | 31972-Deleti | Deletion | 1 | C | - | 8 | 23899 | 0.0334742 | 35.75 | rnr1Y285F | FALSE | CAN | JSY3811 | 3 | rnr1Y285F-R | GCC | nope | >1bp-ins | G/C-1 | none | G/C-1 | AATAAGCCA | CCAAGCC | CGG | GCC | C | none | nope | nope_GCC | G/C-1_CCAAGCC | 8 | FALSE | FALSE |
| 44_v.txt | 32069 | 32069-Deleti | Deletion | 1 | A | - | 5 | 23899 | 0.02092138 | 58.6 | rnr1Y285F | FALSE | CAN | JSY3811 | 3 | rnr1Y285F-R | TAA | nope | >1bp-ins | A/T-1 | none | A/T-1 | TGAGATAAA | AAATAAC | ATT | ATT | T | none | T>none | T>none_ATT | A/T-1_AAATAAC | 8 | FALSE | FALSE |
| 44_v.txt | 32081 | 32081-Deleti | Deletion | 1 | A | - | 25 | 23899 | 0.10460689 | 63.56 | rnr1Y285F | FALSE | CAN | JSY3811 | 3 | rnr1Y285F-R | CAA | nope | >1bp-ins | A/T-1 | none | A/T-1 | CCATGCAAA | AAAAAAG | GTT | GTT | T | none | T>none | T>none_GTT | A/T-1_AAAAAAG | 8 | FALSE | FALSE |
| 44_v.txt | 32109 | 32109-Deleti | Deletion | 1 | A | - | 28 | 23899 | 0.11715971 | 58.8928571 | rnr1Y285F | FALSE | CAN | JSY3811 | 3 | rnr1Y285F-R | TAA | nope | >1bp-ins | A/T-1 | none | A/T-1 | ATATTTAATA | AATAGCC | ATT | ATT | T | none | T>none | T>none_ATT | A/T-1_AATAGCC | 8 | FALSE | FALSE |
| 44_v.txt | 32149 | 32149-Deleti | Deletion | 1 | C | - | 3 | 23899 | 0.01255283 | 62.3333333 | rnr1Y285F | FALSE | CAN | JSY3811 | 3 | rnr1Y285F-R | TCC | nope | >1bp-ins | G/C-1 | none | G/C-1 | ATGTCTCCAT | CCATGTA | AGG | TCC | C | none | nope | nope_TCC | G/C-1_CCATGTA | 8 | FALSE | FALSE |
| 44_v.txt | 32191 | 32191-Deleti | Deletion | 1 | C | - | 2 | 23899 | 8.37E-03 | 51 | rnr1Y285F | FALSE | CAN | JSY3811 | 3 | rnr1Y285F-R | GCA | nope | >1bp-ins | G/C-1 | none | G/C-1 | AAACTGCAA | CAATGTA | CGT | GCA | C | none | nope | nope_GCA | G/C-1_CAATGTA | 8 | FALSE | FALSE |
| 44_v.txt | 32289 | 32289-Deleti | Deletion | 1 | T | - | 6 | 39917 | 0.01503119 | 54.3333333 | rnr1Y285F | FALSE | CAN | JSY3811 | 3 | rnr1Y285F-R | ATT | nope | >1bp-ins | A/T-1 | none | A/T-1 | TAAATATTTG | TTTGAAT | TAA | ATT | T | none | nope | nope_ATT | A/T-1_TTTGAAT | 8 | FALSE | FALSE |
| 44_v.txt | 32346 | 32346-Deleti | Deletion | 1 | A | - | 47 | 39917 | 0.11774432 | 63.106383 | rnr1Y285F | FALSE | CAN | JSY3811 | 3 | rnr1Y285F-R | CAA | nope | >1bp-ins | A/T-1 | none | A/T-1 | TGTGGCAAA | AAAACCT | GTT | GTT | T | none | T>none | T>none_GTT | A/T-1_AAAACCT | 8 | FALSE | FALSE |
| 44_v.txt | 32352 | 32352-Deleti | Deletion | 1 | T | - | 4 | 39917 | 0.01002079 | 64 | rnr1Y285F | FALSE | CAN | JSY3811 | 3 | rnr1Y285F-R | CTT | nope | >1bp-ins | A/T-1 | none | A/T-1 | AAAACCTTTG | TTTGTAC | GAA | CTT | T | none | nope | nope_CTT | A/T-1_TTTGTAC | 8 | FALSE | FALSE |
| 44_v.txt | 32381 | 32381-Deleti | Deletion | 1 | A | - | 9 | 39917 | 0.02254679 | 59.8888889 | rnr1Y285F | FALSE | CAN | JSY3811 | 3 | rnr1Y285F-R | TAA | nope | >1bp-ins | A/T-1 | none | A/T-1 | AATAATAAA | AAAGGGA | ATT | ATT | T | none | T>none | T>none_ATT | A/T-1_AAAGGGA | 8 | FALSE | FALSE |
| 44_v.txt | 32420 | 32420-Deleti | Deletion | 1 | T | - | 6 | 39917 | 0.01503119 | 59.6666667 | rnr1Y285F | FALSE | CAN | JSY3811 | 3 | rnr1Y285F-R | GTT | nope | >1bp-ins | A/T-1 | none | A/T-1 | TGTTAGTTTA | TTTAGGG | CAA | GTT | T | none | nope | nope_GTT | A/T-1_TTTAGGG | 8 | FALSE | FALSE |
| 44_v.txt | 32424 | 32424-Deleti | Deletion | 1 | G | - | 6 | 39917 | 0.01503119 | 56.6666667 | rnr1Y285F | FALSE | CAN | JSY3811 | 3 | rnr1Y285F-R | AGG | nope | >1bp-ins | G/C-1 | none | G/C-1 | AGTTTAGGG | GGGTCAT | TCC | TCC | C | none | C>none | C>none_TCC | G/C-1_GGGTCAT | 8 | FALSE | FALSE |
| 44_v.txt | 32442 | 32442-Deleti | Deletion | 1 | A | - | 6 | 39917 | 0.01503119 | 50.8333333 | rnr1Y285F | FALSE | CAN | JSY3811 | 3 | rnr1Y285F-R | TAA | nope | >1bp-ins | A/T-1 | none | A/T-1 | GGAACTAAA | AAAAGTC | ATT | ATT | T | none | T>none | T>none_ATT | A/T-1_AAAAGTC | 8 | FALSE | FALSE |
| 44_v.txt | 32475 | 32475-Deleti | Deletion | 1 | A | - | 183 | 39917 | 0.45845129 | 46.8469945 | rnr1Y285F | FALSE | CAN | JSY3811 | 3 | rnr1Y285F-R | GAA | nope | >1bp-ins | A/T-1 | none | A/T-1 | ATGTAGAAG | AAGGTTA | CTT | CTT | T | none | T>none | T>none_CTT | A/T-1_AAGGTTA | 8 | FALSE | FALSE |
| 44_v.txt | 32498 | 32498-Deleti | Deletion | 1 | T | - | 46 | 39917 | 0.11523912 | 44.9130435 | rnr1Y285F | FALSE | CAN | JSY3811 | 3 | rnr1Y285F-R | CTT | nope | >1bp-ins | A/T-1 | none | A/T-1 | AACAACTTTT | TTTTTTG | GAA | CTT | T | none | nope | nope_CTT | A/T-1_TTTTTTG | 8 | FALSE | FALSE |
| 44_v.txt | 32522 | 32522-Deleti | Deletion | 1 | T | - | 30 | 39917 | 0.07515595 | 38.6666667 | rnr1Y285F | FALSE | CAN | JSY3811 | 3 | rnr1Y285F-R | ATT | nope | >1bp-ins | A/T-1 | none | A/T-1 | AACGGATTTT | TTTTCTG | TAA | ATT | T | none | nope | nope_ATT | A/T-1_TTTTCTG | 8 | FALSE | FALSE |
| 44_v.txt | 32528 | 32528-Deleti | Deletion | 1 | G | - | 10 | 39917 | 0.02505198 | 36.3 | rnr1Y285F | FALSE | CAN | JSY3811 | 3 | rnr1Y285F-R | TGG | nope | >1bp-ins | G/C-1 | none | G/C-1 | TTTTCTGGGG | GGGGTTT | ACC | ACC | C | none | C>none | C>none_ACC | G/C-1_GGGGTTT | 8 | FALSE | FALSE |
| 44_v.txt | 32598 | 32598-Deleti | Deletion | 1 | A | - | 5 | 33134 | 0.01509024 | 42.8 | rnr1Y285F | FALSE | CAN | JSY3811 | 3 | rnr1Y285F-R | CAA | nope | >1bp-ins | A/T-1 | none | A/T-1 | TTAATCAAAC | AAAGAGG | GTT | GTT | T | none | T>none | T>none_GTT | A/T-1_AAAGAGG | 8 | FALSE | FALSE |
| 44_v.txt | 32608 | 32608-Deleti | Deletion | 1 | C | - | 48 | 33134 | 0.1448663 | 46.9375022 | rnr1Y285F | FALSE | CAN | JSY3811 | 3 | rnr1Y285F-R | ACC | nope | >1bp-ins | G/C-1 | none | G/C-1 | AGGAAACCC | CCCAACC | TGG | ACC | C | none | nope | nope_ACC | G/C-1_CCCAACC | 8 | FALSE | FALSE |
| 44_v.txt | 32631 | 32631-Deleti | Deletion | 1 | T | - | 10 | 33134 | 0.03018048 | 39.8000057 | rnr1Y285F | FALSE | CAN | JSY3811 | 3 | rnr1Y285F-R | GTT | nope | >1bp-ins | A/T-1 | none | A/T-1 | CCTTCGTTTT | TTTTTAT | CAA | GTT | T | none | nope | nope_GTT | A/T-1_TTTTTAT | 8 | FALSE | FALSE |
| 44_v.txt | 32658 | 32658-Deleti | Deletion | 1 | C | - | 23 | 33134 | 0.0694151 | 55.2173913 | rnr1Y285F | FALSE | CAN | JSY3811 | 3 | rnr1Y285F-R | ACC | nope | >1bp-ins | G/C-1 | none | G/C-1 | CCTGGACCCC | CCCCAGG | TGG | ACC | C | none | nope | nope_ACC | G/C-1_CCCCAGG | 8 | FALSE | FALSE |
| 44_v.txt | 32670 | 32670-Deleti | Deletion | 1 | G | - | 57 | 33134 | 0.17202873 | 58.8245614 | rnr1Y285F | FALSE | CAN | JSY3811 | 3 | rnr1Y285F-R | TGG | nope | >1bp-ins | G/C-1 | none | G/C-1 | GCACCTGGG | GGGTTTC | ACC | ACC | C | none | C>none | C>none_ACC | G/C-1_GGGTTTC | 8 | FALSE | FALSE |
| 44_v.txt | 32707 | 32707-Deleti | Deletion | 1 | C | - | 9 | 33134 | 0.02716243 | 63 | rnr1Y285F | FALSE | CAN | JSY3811 | 3 | rnr1Y285F-R | ACC | nope | >1bp-ins | G/C-1 | none | G/C-1 | CGGTAACCC | CCCCAGC | TGG | ACC | C | none | nope | nope_ACC | G/C-1_CCCCAGC | 8 | FALSE | FALSE |
| 44_v.txt | 32719 | 32719-Deleti | Deletion | 1 | A | - | 13 | 33134 | 0.03923462 | 61.3846154 | rnr1Y285F | FALSE | CAN | JSY3811 | 3 | rnr1Y285F-R | CAA | nope | >1bp-ins | A/T-1 | none | A/T-1 | CACCACAAA | AAACCAT | GTT | GTT | T | none | T>none | T>none_GTT | A/T-1_AAACCAT | 8 | FALSE | FALSE |
| 44_v.txt | 32728 | 32728-Deleti | Deletion | 1 | A | - | 8 | 33134 | 0.02414438 | 60.7500001 | rnr1Y285F | FALSE | CAN | JSY3811 | 3 | rnr1Y285F-R | CAA | nope | >1bp-ins | A/T-1 | none | A/T-1 | CCATACAAA | AAAAACA | GTT | GTT | T | none | T>none | T>none_GTT | A/T-1_AAAAACA | 8 | FALSE | FALSE |
| 44_v.txt | 32779 | 32779-Deleti | Deletion | 1 | C | - | 6 | 33134 | 0.01810829 | 50.5 | rnr1Y285F | FALSE | CAN | JSY3811 | 3 | rnr1Y285F-R | ACC | nope | >1bp-ins | G/C-1 | none | G/C-1 | AAGCGACCC | CCCAGAA | TGG | ACC | C | none | nope | nope_ACC | G/C-1_CCCAGAA | 8 | FALSE | FALSE |
| 44_v.txt | 32842 | 32842-Deleti | Deletion | 1 | A | - | 70 | 33133 | 0.21126973 | 41.342857 | rnr1Y285F | FALSE | CAN | JSY3811 | 3 | rnr1Y285F-R | CAA | nope | >1bp-ins | A/T-1 | none | A/T-1 | TTACCCAAA | AAAAAAT | GTT | GTT | T | none | T>none | T>none_GTT | A/T-1_AAAAAAT | 8 | FALSE | FALSE |
| 44_v.txt | 32882 | 32882-Deleti | Deletion | 1 | C | - | 2 | 33682 | 5.94E-03 | 38 | rnr1Y285F | FALSE | CAN | JSY3811 | 3 | rnr1Y285F-R | TCC | nope | >1bp-ins | G/C-1 | none | G/C-1 | GTACGTCCA | CCAAAAT | AGG | TCC | C | none | nope | nope_TCC | G/C-1_CCAAAAT | 8 | FALSE | FALSE |
| 44_v.txt | 32884 | 32884-Deleti | Deletion | 1 | A | - | 8 | 33681 | 0.02375226 | 37.5 | rnr1Y285F | FALSE | CAN | JSY3811 | 3 | rnr1Y285F-R | CAA | nope | >1bp-ins | A/T-1 | none | A/T-1 | ACGTCCAAA | AAAATTG | GTT | GTT | T | none | T>none | T>none_GTT | A/T-1_AAAATTG | 8 | FALSE | FALSE |
| 44_v.txt | 32901 | 32901-Deleti | Deletion | 1 | C | - | 2 | 33681 | 5.94E-03 | 38 | rnr1Y285F | FALSE | CAN | JSY3811 | 3 | rnr1Y285F-R | GCC | nope | >1bp-ins | G/C-1 | none | G/C-1 | ACTTGGCCA | CCAACTA | CGG | GCC | C | none | nope | nope_GCC | G/C-1_CCAACTA | 8 | FALSE | FALSE |
| 44_v.txt | 32917 | 32917-Deleti | Deletion | 1 | C | - | 2 | 33681 | 5.94E-03 | 38 | rnr1Y285F | FALSE | CAN | JSY3811 | 3 | rnr1Y285F-R | TCC | nope | >1bp-ins | G/C-1 | none | G/C-1 | TAAGTTCCAC | CCAGGGC | AGG | TCC | C | none | nope | nope_TCC | G/C-1_CCAGGGC | 8 | FALSE | FALSE |
| 44_v.txt | 32924 | 32924-Deleti | Deletion | 1 | A | - | 37 | 33681 | 0.10985422 | 52.6486487 | rnr1Y285F | FALSE | CAN | JSY3811 | 3 | rnr1Y285F-R | CAA | nope | >1bp-ins | A/T-1 | none | A/T-1 | CAGGGCAAA | AAAAGTG | GTT | GTT | T | none | T>none | T>none_GTT | A/T-1_AAAAGTG | 8 | FALSE | FALSE |
| 44_v.txt | 32935 | 32935-Deleti | Deletion | 1 | C | - | 6 | 33681 | 0.0178142 | 52.5 | rnr1Y285F | FALSE | CAN | JSY3811 | 3 | rnr1Y285F-R | GCC | nope | >1bp-ins | G/C-1 | none | G/C-1 | TGATTGCCCA | CCCAAGA | CGG | GCC | C | none | nope | nope_GCC | G/C-1_CCCAAGA | 8 | FALSE | FALSE |
| 44_v.txt | 32940 | 32940-Deleti | Deletion | 1 | G | - | 3 | 33681 | 8.91E-03 | 54 | rnr1Y285F | FALSE | CAN | JSY3811 | 3 | rnr1Y285F-R | AGA | nope | >1bp-ins | G/C-1 | none | G/C-1 | GCCCAAGAA | GAAAACC | TCT | TCT | C | none | C>none | C>none_TCT | G/C-1_GAAAACC | 8 | FALSE | FALSE |
| 44_v.txt | 33111 | 33111-Deleti | Deletion | 1 | G | - | 5 | 33681 | 0.01484517 | 34.8 | rnr1Y285F | FALSE | CAN | JSY3811 | 3 | rnr1Y285F-R | TGG | nope | >1bp-ins | G/C-1 | none | G/C-1 | CCCACTGGG | GGGCCGG | ACC | ACC | C | none | C>none | C>none_ACC | G/C-1_GGGCCGG | 8 | FALSE | FALSE |
| 44_v.txt | 33138 | 33138-Deleti | Deletion | 1 | A | - | 7 | 33681 | 0.02078323 | 41.4285714 | rnr1Y285F | FALSE | CAN | JSY3811 | 3 | rnr1Y285F-R | TAA | nope | >1bp-ins | A/T-1 | none | A/T-1 | GTGGATAAA | AAACCAA | ATT | ATT | T | none | T>none | T>none_ATT | A/T-1_AAACCAA | 8 | FALSE | FALSE |
| 44_v.txt | 33147 | 33147-Deleti | Deletion | 1 | A | - | 6 | 33681 | 0.0178142 | 37.8333333 | rnr1Y285F | FALSE | CAN | JSY3811 | 3 | rnr1Y285F-R | GAA | nope | >1bp-ins | A/T-1 | none | A/T-1 | CCAATGAAA | AAAAGAC | CTT | CTT | T | none | T>none | T>none_CTT | A/T-1_AAAAGAC | 8 | FALSE | FALSE |
| 44_v.txt | 33258 | 33258-Deleti | Deletion | 1 | C | - | 2 | 25205 | 7.94E-03 | 38 | rnr1Y285F | FALSE | CAN | JSY3811 | 3 | rnr1Y285F-R | GCC | nope | >1bp-ins | G/C-1 | none | G/C-1 | CCTATGCCAT | CCATCCT | CGG | GCC | C | none | nope | nope_GCC | G/C-1_CCATCCT | 8 | FALSE | FALSE |
| 44_v.txt | 33265 | 33265-Deleti | Deletion | 1 | C | - | 6 | 25205 | 0.0238048 | 42.3333333 | rnr1Y285F | FALSE | CAN | JSY3811 | 3 | rnr1Y285F-R | TCC | nope | >1bp-ins | G/C-1 | none | G/C-1 | CATCCTCCAT | CCATAGA | AGG | TCC | C | none | nope | nope_TCC | G/C-1_CCATAGA | 8 | FALSE | FALSE |
| 44_v.txt | 33356 | 33356-Deleti | Deletion | 1 | C | - | 8 | 25205 | 0.03173973 | 48.75 | rnr1Y285F | FALSE | CAN | JSY3811 | 3 | rnr1Y285F-R | ACC | nope | >1bp-ins | G/C-1 | none | G/C-1 | TATTGACCCA | CCCACGT | TGG | ACC | C | none | nope | nope_ACC | G/C-1_CCCACGT | 8 | FALSE | FALSE |
| 44_v.txt | 33452 | 33452-Deleti | Deletion | 1 | T | - | 6 | 25205 | 0.0238048 | 42 | rnr1Y285F | FALSE | CAN | JSY3811 | 3 | rnr1Y285F-R | CTT | nope | >1bp-ins | A/T-1 | none | A/T-1 | GTCTTCTTTT | TTTTGAA | GAA | CTT | T | none | nope | nope_CTT | A/T-1_TTTTGAA | 8 | FALSE | FALSE |
| 44_v.txt | 33474 | 33474-Deleti | Deletion | 1 | C | - | 45 | 25204 | 0.17854309 | 28.4666667 | rnr1Y285F | FALSE | CAN | JSY3811 | 3 | rnr1Y285F-R | GCC | nope | >1bp-ins | G/C-1 | none | G/C-1 | GCTATGCCTT | CCTTTTT | CGG | GCC | C | none | nope | nope_GCC | G/C-1_CCTTTTT | 8 | FALSE | FALSE |
| 45_v.txt | 31950 | 31950-SNV-A | SNV | 1 | A | C | 11 | 9999 | 0.110011 | 31.6363636 | rnr1Y285F | FALSE | CAN | JSY3811 | 4 | rnr1Y285F-R | CAT | A>C | G/C+1 | A/T-1 | TA>GC | TA>GC | ATCGTCATAA | ATAAATG | GTA | GTA | T | G | T>G | T>G_GTA | TA>GC_ATAAATG | 8 | FALSE | FALSE |
| 45_v.txt | 31964 | 31964-SNV-A | SNV | 1 | A | C | 11 | 9998 | 0.110022 | 36.0909091 | rnr1Y285F | FALSE | CAN | JSY3811 | 4 | rnr1Y285F-R | CAT | A>C | G/C+1 | A/T-1 | TA>GC | TA>GC | GGCCGCATA | ATAATAA | GTA | GTA | T | G | T>G | T>G_GTA | TA>GC_ATAATAA | 8 | FALSE | FALSE |
| 45_v.txt | 31986 | 31986-SNV-A | SNV | 1 | A | C | 4 | 9998 | 0.040008 | 54.75 | rnr1Y285F | FALSE | CAN | JSY3811 | 4 | rnr1Y285F-R | TAA | A>C | G/C+1 | A/T-1 | TA>GC | TA>GC | GGCATTAATT | AATTTAG | ATT | ATT | T | G | T>G | T>G_ATT | TA>GC_AATTTAG | 8 | FALSE | FALSE |
| 45_v.txt | 32109 | 32109-SNV-A | SNV | 1 | A | C | 14 | 9998 | 0.14002801 | 59 | rnr1Y285F | FALSE | CAN | JSY3811 | 4 | rnr1Y285F-R | TAA | A>C | G/C+1 | A/T-1 | TA>GC | TA>GC | ATATTTAATA | AATAGCC | ATT | ATT | T | G | T>G | T>G_ATT | TA>GC_AATAGCC | 8 | FALSE | FALSE |
| 45_v.txt | 32289 | 32289-SNV-T | SNV | 1 | T | G | 10 | 21827 | 0.04581482 | 52.8 | rnr1Y285F | FALSE | CAN | JSY3811 | 4 | rnr1Y285F-R | ATT | T>G | G/C+1 | A/T-1 | TA>GC | TA>GC | TAAATATTTG | TTTGAAT | TAA | ATT | T | C | T>G | T>G_ATT | TA>GC_TTTGAAT | 8 | FALSE | FALSE |
| 45_v.txt | 32445 | 32445-SNV-A | SNV | 1 | A | C | 23 | 21827 | 0.10537408 | 50.6956522 | rnr1Y285F | FALSE | CAN | JSY3811 | 4 | rnr1Y285F-R | AAG | A>C | G/C+1 | A/T-1 | TA>GC | TA>GC | ACTAAAAGT | AGTCCAA | TTC | TTC | T | G | T>G | T>G_TTC | TA>GC_AGTCCAA | 8 | FALSE | FALSE |
| 45_v.txt | 32481 | 32481-SNV-A | SNV | 1 | A | C | 5 | 21827 | 0.02290741 | 52.4 | rnr1Y285F | FALSE | CAN | JSY3811 | 4 | rnr1Y285F-R | TAA | A>C | G/C+1 | A/T-1 | TA>GC | TA>GC | AAGGTTAAG | AAGATAC | ATT | ATT | T | G | T>G | T>G_ATT | TA>GC_AAGATAC | 8 | FALSE | FALSE |
| 45_v.txt | 32800 | 32800-SNV-A | SNV | 1 | A | C | 10 | 13851 | 0.07219695 | 40.4 | rnr1Y285F | FALSE | CAN | JSY3811 | 4 | rnr1Y285F-R | TAA | A>C | G/C+1 | A/T-1 | TA>GC | TA>GC | CACCGTAATA | AATATTT | ATT | ATT | T | G | T>G | T>G_ATT | TA>GC_AATATTT | 8 | FALSE | FALSE |
| 45_v.txt | 32884 | 32884-SNV-A | SNV | 1 | A | C | 7 | 19784 | 0.03538213 | 33.4285714 | rnr1Y285F | FALSE | CAN | JSY3811 | 4 | rnr1Y285F-R | CAA | A>C | G/C+1 | A/T-1 | TA>GC | TA>GC | ACGTCCAAA | AAAATTG | GTT | GTT | T | G | T>G | T>G_GTT | TA>GC_AAAATTG | 8 | FALSE | FALSE |
| 45_v.txt | 32926 | 32926-SNV-A | SNV | 1 | A | C | 13 | 19786 | 0.06570302 | 27.1532131 | rnr1Y285F | FALSE | CAN | JSY3811 | 4 | rnr1Y285F-R | AAA | A>C | G/C+1 | A/T-1 | TA>GC | TA>GC | GGGCAAAAG | AAGTGAT | TTT | TTT | T | G | T>G | T>G_TTT | TA>GC_AAGTGAT | 8 | FALSE | FALSE |
| 45_v.txt | 32927 | 32927-SNV-A | SNV | 1 | A | C | 23 | 19786 | 0.11624381 | 17.5233273 | rnr1Y285F | FALSE | CAN | JSY3811 | 4 | rnr1Y285F-R | AAG | A>C | G/C+1 | A/T-1 | TA>GC | TA>GC | GGCAAAAGT | AGTGATT | TTC | TTC | T | G | T>G | T>G_TTC | TA>GC_AGTGATT | 8 | FALSE | FALSE |
| 45_v.txt | 32929 | 32929-SNV-T | SNV | 1 | T | G | 7 | 19786 | 0.03537855 | 50.4285714 | rnr1Y285F | FALSE | CAN | JSY3811 | 4 | rnr1Y285F-R | GTG | T>G | G/C+1 | A/T-1 | TA>GC | TA>GC | CAAAAGTGA | TGATTGC | CAC | GTG | T | C | T>G | T>G_GTG | TA>GC_TGATTGC | 8 | FALSE | FALSE |
| 45_v.txt | 33189 | 33189-SNV-A | SNV | 1 | A | C | 5 | 11669 | 0.04284857 | 27.4000639 | rnr1Y285F | FALSE | CAN | JSY3811 | 4 | rnr1Y285F-R | AAT | A>C | G/C+1 | A/T-1 | TA>GC | TA>GC | ATACCAATAT | ATATGTC | TTA | TTA | T | G | T>G | T>G_TTA | TA>GC_ATATGTC | 8 | FALSE | FALSE |
| 45_v.txt | 33269 | 33269-SNV-A | SNV | 1 | A | C | 96 | 11670 | 0.82262211 | 17.3854167 | rnr1Y285F | FALSE | CAN | JSY3811 | 4 | rnr1Y285F-R | TAG | A>C | G/C+1 | A/T-1 | TA>GC | TA>GC | CTCCATAGAC | AGAGAAC | ATC | ATC | T | G | T>G | T>G_ATC | TA>GC_AGAGAAC | 8 | FALSE | FALSE |
| 45_v.txt | 32490 | 32490-SNV-A | SNV | 1 | A | G | 18 | 21827 | 0.08246667 | 48.0555556 | rnr1Y285F | FALSE | CAN | JSY3811 | 4 | rnr1Y285F-R | GAA | A>G | G/C+1 | A/T-1 | TA>CG | TA>CG | ATACGGAAA | AAAACAA | CTT | CTT | T | C | T>C | T>C_CTT | TA>CG_AAAACAA | 8 | FALSE | FALSE |
| 45_v.txt | 32842 | 32842-SNV-A | SNV | 1 | A | G | 6 | 13851 | 0.04331817 | 41.8336618 | rnr1Y285F | FALSE | CAN | JSY3811 | 4 | rnr1Y285F-R | CAA | A>G | G/C+1 | A/T-1 | TA>CG | TA>CG | TTACCCAAA | AAAAAAT | GTT | GTT | T | C | T>C | T>C_GTT | TA>CG_AAAAAAT | 8 | FALSE | FALSE |
| 45_v.txt | 32926 | 32926-SNV-A | SNV | 1 | A | G | 3 | 19786 | 0.01516224 | 44.664489 | rnr1Y285F | FALSE | CAN | JSY3811 | 4 | rnr1Y285F-R | AAA | A>G | G/C+1 | A/T-1 | TA>CG | TA>CG | GGGCAAAAG | AAGTGAT | TTT | TTT | T | C | T>C | T>C_TTT | TA>CG_AAGTGAT | 8 | FALSE | FALSE |
| 45_v.txt | 32927 | 32927-SNV-A | SNV | 1 | A | G | 21 | 19786 | 0.10613565 | 49.7138873 | rnr1Y285F | FALSE | CAN | JSY3811 | 4 | rnr1Y285F-R | AAG | A>G | G/C+1 | A/T-1 | TA>CG | TA>CG | GGCAAAAGT | AGTGATT | TTC | TTC | T | C | T>C | T>C_TTC | TA>CG_AGTGATT | 8 | FALSE | FALSE |
| 45_v.txt | 33192 | 33192-SNV-T | SNV | 1 | T | C | 10 | 11670 | 0.0856898 | 30.6 | rnr1Y285F | FALSE | CAN | JSY3811 | 4 | rnr1Y285F-R | ATG | T>C | G/C+1 | A/T-1 | TA>CG | TA>CG | CCAATATGTC | TGTCTTT | TAC | ATG | T | G | T>C | T>C_ATG | TA>CG_TGTCTTT | 8 | FALSE | FALSE |
| 45_v.txt | 33476 | 33476-SNV-T | SNV | 1 | T | C | 19 | 11669 | 0.16282458 | 31.8421053 | rnr1Y285F | FALSE | CAN | JSY3811 | 4 | rnr1Y285F-R | CTT | T>C | G/C+1 | A/T-1 | TA>CG | TA>CG | TATGCCTTTT | TTTTTTT | GAA | CTT | T | G | T>C | T>C_CTT | TA>CG_TTTTTTT | 8 | FALSE | FALSE |
| 45_v.txt | 31964 | 31964-SNV-A | SNV | 1 | A | T | 28 | 9998 | 0.28005601 | 36.5357143 | rnr1Y285F | FALSE | CAN | JSY3811 | 4 | rnr1Y285F-R | CAT | A>T | A/T+1 | A/T-1 | TA>AT | TA>AT | GGCCGCATA | ATAATAA | GTA | GTA | T | A | T>A | T>A_GTA | TA>AT_ATAATAA | 8 | FALSE | FALSE |
| 45_v.txt | 32522 | 32522-SNV-T | SNV | 1 | T | A | 6 | 21827 | 0.02748889 | 36.6669445 | rnr1Y285F | FALSE | CAN | JSY3811 | 4 | rnr1Y285F-R | ATT | T>A | A/T+1 | A/T-1 | TA>AT | TA>AT | AACGGATTTT | TTTTCTG | TAA | ATT | T | T | T>A | T>A_ATT | TA>AT_TTTTCTG | 8 | FALSE | FALSE |
| 45_v.txt | 32929 | 32929-SNV-T | SNV | 1 | T | A | 285 | 19786 | 1.44041241 | 52.3614035 | rnr1Y285F | FALSE | CAN | JSY3811 | 4 | rnr1Y285F-R | GTG | T>A | A/T+1 | A/T-1 | TA>AT | TA>AT | CAAAAGTGA | TGATTGC | CAC | GTG | T | T | T>A | T>A_GTG | TA>AT_TGATTGC | 8 | FALSE | FALSE |
| 45_v.txt | 33067 | 33067-SNV-A | SNV | 1 | A | T | 811 | 19786 | 4.09885778 | 54.6152898 | rnr1Y285F | FALSE | CAN | JSY3811 | 4 | rnr1Y285F-R | TAT | A>T | A/T+1 | A/T-1 | TA>AT | TA>AT | CAGAATATG | ATGCCAA | ATA | ATA | T | A | T>A | T>A_ATA | TA>AT_ATGCCAA | 8 | FALSE | FALSE |
| 45_v.txt | 33192 | 33192-SNV-T | SNV | 1 | T | A | 18 | 11670 | 0.15424165 | 34.8888889 | rnr1Y285F | FALSE | CAN | JSY3811 | 4 | rnr1Y285F-R | ATG | T>A | A/T+1 | A/T-1 | TA>AT | TA>AT | CCAATATGTC | TGTCTTT | TAC | ATG | T | T | T>A | T>A_ATG | TA>AT_TGTCTTT | 8 | FALSE | FALSE |
| 45_v.txt | 33364 | 33364-SNV-T | SNV | 1 | T | A | 30 | 11670 | 0.25706941 | 56.5666667 | rnr1Y285F | FALSE | CAN | JSY3811 | 4 | rnr1Y285F-R | CTG | T>A | A/T+1 | A/T-1 | TA>AT | TA>AT | CACGTCTGTG | TGTGGTG | GAC | CTG | T | T | T>A | T>A_CTG | TA>AT_TGTGGTG | 8 | FALSE | FALSE |
| 45_v.txt | 31650 | 31650-SNV-C | SNV | 1 | C | T | 13 | 6593 | 0.19717883 | 35.8461539 | rnr1Y285F | FALSE | CAN | JSY3811 | 4 | rnr1Y285F-R | GCG | C>T | A/T+1 | G/C-1 | CG>TA | CG>TA | AGAATGCGA | CGAAATG | CGC | GCG | C | A | C>T | C>T_GCG | CG>TA_CGAAATG | 8 | FALSE | FALSE |
| 45_v.txt | 31844 | 31844-SNV-C | SNV | 1 | C | T | 14 | 6593 | 0.21234643 | 47.8571429 | rnr1Y285F | FALSE | CAN | JSY3811 | 4 | rnr1Y285F-R | TCC | C>T | A/T+1 | G/C-1 | CG>TA | CG>TA | TAAGATCCA | CCAAACA | AGG | TCC | C | A | C>T | C>T_TCC | CG>TA_CCAAACA | 8 | FALSE | FALSE |
| 45_v.txt | 31911 | 31911-SNV-G | SNV | 1 | G | A | 10 | 6593 | 0.15167602 | 45.4 | rnr1Y285F | FALSE | CAN | JSY3811 | 4 | rnr1Y285F-R | TGC | G>A | A/T+1 | G/C-1 | CG>TA | CG>TA | TTTGGTGCA | GCAAAAG | ACG | ACG | C | T | C>T | C>T_ACG | CG>TA_GCAAAAG | 8 | FALSE | FALSE |
| 45_v.txt | 31962 | 31962-SNV-G | SNV | 1 | G | A | 20 | 9998 | 0.20004001 | 36.8 | rnr1Y285F | FALSE | CAN | JSY3811 | 4 | rnr1Y285F-R | CGC | G>A | A/T+1 | G/C-1 | CG>TA | CG>TA | GTGGCCGCA | GCATAAT | GCG | GCG | C | T | C>T | C>T_GCG | CG>TA_GCATAAT | 8 | FALSE | FALSE |
| 45_v.txt | 31981 | 31981-SNV-G | SNV | 1 | G | A | 9 | 9998 | 0.090018 | 58.3333333 | rnr1Y285F | FALSE | CAN | JSY3811 | 4 | rnr1Y285F-R | GGC | G>A | A/T+1 | G/C-1 | CG>TA | CG>TA | AGCCGGGCA | GCATTAA | CCG | CCG | C | T | C>T | C>T_CCG | CG>TA_GCATTAA | 8 | FALSE | FALSE |
| 45_v.txt | 32075 | 32075-SNV-C | SNV | 1 | C | T | 22 | 9998 | 0.22004401 | 62 | rnr1Y285F | FALSE | CAN | JSY3811 | 4 | rnr1Y285F-R | ACC | C>T | A/T+1 | G/C-1 | CG>TA | CG>TA | AAATAACCA | CCATGCA | TGG | ACC | C | A | C>T | C>T_ACC | CG>TA_CCATGCA | 8 | FALSE | FALSE |
| 45_v.txt | 32088 | 32088-SNV-C | SNV | 1 | C | T | 52 | 9998 | 0.52010402 | 60.1153846 | rnr1Y285F | FALSE | CAN | JSY3811 | 4 | rnr1Y285F-R | GCC | C>T | A/T+1 | G/C-1 | CG>TA | CG>TA | AAAAAGCCT | CCTGCAA | CGG | GCC | C | A | C>T | C>T_GCC | CG>TA_CCTGCAA | 8 | FALSE | FALSE |
| 45_v.txt | 32092 | 32092-SNV-C | SNV | 1 | C | T | 13 | 9998 | 0.13002601 | 61.3076923 | rnr1Y285F | FALSE | CAN | JSY3811 | 4 | rnr1Y285F-R | GCA | C>T | A/T+1 | G/C-1 | CG>TA | CG>TA | AGCCTGCAA | CAACACC | CGT | GCA | C | A | C>T | C>T_GCA | CG>TA_CAACACC | 8 | FALSE | FALSE |
| 45_v.txt | 32114 | 32114-SNV-C | SNV | 1 | C | T | 74 | 9998 | 0.74014803 | 55.6756757 | rnr1Y285F | FALSE | CAN | JSY3811 | 4 | rnr1Y285F-R | GCC | C>T | A/T+1 | G/C-1 | CG>TA | CG>TA | TAATAGCCAT | CCATTCG | CGG | GCC | C | A | C>T | C>T_GCC | CG>TA_CCATTCG | 8 | FALSE | FALSE |
| 45_v.txt | 32165 | 32165-SNV-G | SNV | 1 | G | A | 16 | 9998 | 0.16003201 | 47.5 | rnr1Y285F | FALSE | CAN | JSY3811 | 4 | rnr1Y285F-R | CGC | G>A | A/T+1 | G/C-1 | CG>TA | CG>TA | CAAAGCGCC | GCCAAAT | GCG | GCG | C | T | C>T | C>T_GCG | CG>TA_GCCAAAT | 8 | FALSE | FALSE |
| 45_v.txt | 32173 | 32173-SNV-C | SNV | 1 | C | T | 13 | 9998 | 0.13002601 | 49.4615385 | rnr1Y285F | FALSE | CAN | JSY3811 | 4 | rnr1Y285F-R | GCA | C>T | A/T+1 | G/C-1 | CG>TA | CG>TA | CAAATGCAG | CAGCAGT | CGT | GCA | C | A | C>T | C>T_GCA | CG>TA_CAGCAGT | 8 | FALSE | FALSE |
| 45_v.txt | 32271 | 32271-SNV-C | SNV | 1 | C | T | 68 | 21827 | 0.31154075 | 32.6617647 | rnr1Y285F | FALSE | CAN | JSY3811 | 4 | rnr1Y285F-R | ACG | C>T | A/T+1 | G/C-1 | CG>TA | CG>TA | AAAATACGG | CGGGAAC | TGC | ACG | C | A | C>T | C>T_ACG | CG>TA_CGGGAAC | 8 | FALSE | FALSE |
| 45_v.txt | 32274 | 32274-SNV-G | SNV | 1 | G | A | 43 | 21827 | 0.19700371 | 36.7441861 | rnr1Y285F | FALSE | CAN | JSY3811 | 4 | rnr1Y285F-R | GGA | G>A | A/T+1 | G/C-1 | CG>TA | CG>TA | ATACGGGAA | GAACCAA | CCT | CCT | C | T | C>T | C>T_CCT | CG>TA_GAACCAA | 8 | FALSE | FALSE |
| 45_v.txt | 32299 | 32299-SNV-C | SNV | 1 | C | T | 37 | 21827 | 0.16951482 | 47.2702703 | rnr1Y285F | FALSE | CAN | JSY3811 | 4 | rnr1Y285F-R | GCG | C>T | A/T+1 | G/C-1 | CG>TA | CG>TA | AATTTGCGG | CGGCAGA | CGC | GCG | C | A | C>T | C>T_GCG | CG>TA_CGGCAGA | 8 | FALSE | FALSE |
| 45_v.txt | 32535 | 32535-SNV-G | SNV | 1 | G | A | 27 | 21826 | 0.12370567 | 36.2957056 | rnr1Y285F | FALSE | CAN | JSY3811 | 4 | rnr1Y285F-R | TGC | G>A | A/T+1 | G/C-1 | CG>TA | CG>TA | GGGTTTGCA | GCAGCTT | ACG | ACG | C | T | C>T | C>T_ACG | CG>TA_GCAGCTT | 8 | FALSE | FALSE |
| 45_v.txt | 32536 | 32536-SNV-C | SNV | 1 | C | T | 43 | 21826 | 0.19701274 | 36.3952951 | rnr1Y285F | FALSE | CAN | JSY3811 | 4 | rnr1Y285F-R | GCA | C>T | A/T+1 | G/C-1 | CG>TA | CG>TA | GGTTTGCAG | CAGCTTC | CGT | GCA | C | A | C>T | C>T_GCA | CG>TA_CAGCTTC | 8 | FALSE | FALSE |
| 45_v.txt | 32571 | 32571-SNV-C | SNV | 1 | C | T | 145 | 13851 | 1.04685582 | 36.3073179 | rnr1Y285F | FALSE | CAN | JSY3811 | 4 | rnr1Y285F-R | ACC | C>T | A/T+1 | G/C-1 | CG>TA | CG>TA | TCAGTACCTT | CCTTGAA | TGG | ACC | C | A | C>T | C>T_ACC | CG>TA_CCTTGAA | 8 | FALSE | FALSE |
| 45_v.txt | 32572 | 32572-SNV-C | SNV | 1 | C | T | 32 | 13851 | 0.23103025 | 34.4671085 | rnr1Y285F | FALSE | CAN | JSY3811 | 4 | rnr1Y285F-R | CCT | C>T | A/T+1 | G/C-1 | CG>TA | CG>TA | CAGTACCTTG | CTTGAAA | GGA | CCT | C | A | C>T | C>T_CCT | CG>TA_CTTGAAA | 8 | FALSE | FALSE |
| 45_v.txt | 32590 | 32590-SNV-C | SNV | 1 | C | T | 28 | 13851 | 0.20215147 | 36.7857143 | rnr1Y285F | FALSE | CAN | JSY3811 | 4 | rnr1Y285F-R | GCG | C>T | A/T+1 | G/C-1 | CG>TA | CG>TA | AGGCAGCGT | CGTTAAT | CGC | GCG | C | A | C>T | C>T_GCG | CG>TA_CGTTAAT | 8 | FALSE | FALSE |
| 45_v.txt | 32678 | 32678-SNV-C | SNV | 1 | C | T | 250 | 13851 | 1.80492383 | 59.368 | rnr1Y285F | FALSE | CAN | JSY3811 | 4 | rnr1Y285F-R | TCC | C>T | A/T+1 | G/C-1 | CG>TA | CG>TA | GTTTCTCCAA | CCAATAA | AGG | TCC | C | A | C>T | C>T_TCC | CG>TA_CCAATAA | 8 | FALSE | FALSE |
| 45_v.txt | 32700 | 32700-SNV-C | SNV | 1 | C | T | 19 | 13851 | 0.13717421 | 55.8947368 | rnr1Y285F | FALSE | CAN | JSY3811 | 4 | rnr1Y285F-R | GCC | C>T | A/T+1 | G/C-1 | CG>TA | CG>TA | ACTGGGCCG | CCGGTAA | CGG | GCC | C | A | C>T | C>T_GCC | CG>TA_CCGGTAA | 8 | FALSE | FALSE |
| 45_v.txt | 32713 | 32713-SNV-C | SNV | 1 | C | T | 18 | 13851 | 0.12995452 | 55.3333333 | rnr1Y285F | FALSE | CAN | JSY3811 | 4 | rnr1Y285F-R | GCA | C>T | A/T+1 | G/C-1 | CG>TA | CG>TA | CCCCAGCAC | CACCACA | CGT | GCA | C | A | C>T | C>T_GCA | CG>TA_CACCACA | 8 | FALSE | FALSE |
| 45_v.txt | 32749 | 32749-SNV-C | SNV | 1 | C | T | 15 | 13851 | 0.10829543 | 47 | rnr1Y285F | FALSE | CAN | JSY3811 | 4 | rnr1Y285F-R | CCG | C>T | A/T+1 | G/C-1 | CG>TA | CG>TA | GAAACCCGA | CGATAAT | GGC | CCG | C | A | C>T | C>T_CCG | CG>TA_CGATAAT | 8 | FALSE | FALSE |
| 45_v.txt | 32788 | 32788-SNV-C | SNV | 1 | C | T | 22 | 13851 | 0.1588333 | 48.6818182 | rnr1Y285F | FALSE | CAN | JSY3811 | 4 | rnr1Y285F-R | TCG | C>T | A/T+1 | G/C-1 | CG>TA | CG>TA | AGAACTCGA | CGAATTC | AGC | TCG | C | A | C>T | C>T_TCG | CG>TA_CGAATTC | 8 | FALSE | FALSE |
| 45_v.txt | 32798 | 32798-SNV-G | SNV | 1 | G | A | 20 | 13851 | 0.14439391 | 49.9 | rnr1Y285F | FALSE | CAN | JSY3811 | 4 | rnr1Y285F-R | CGT | G>A | A/T+1 | G/C-1 | CG>TA | CG>TA | TTCACCGTAA | GTAATAT | GCA | GCA | C | T | C>T | C>T_GCA | CG>TA_GTAATAT | 8 | FALSE | FALSE |
| 45_v.txt | 32840 | 32840-SNV-C | SNV | 1 | C | T | 17 | 13851 | 0.12273482 | 41.2352941 | rnr1Y285F | FALSE | CAN | JSY3811 | 4 | rnr1Y285F-R | CCC | C>T | A/T+1 | G/C-1 | CG>TA | CG>TA | AATTACCCA | CCAAAAA | GGG | CCC | C | A | C>T | C>T_CCC | CG>TA_CCAAAAA | 8 | FALSE | FALSE |
| 45_v.txt | 32860 | 32860-SNV-C | SNV | 1 | C | T | 18 | 13851 | 0.12995452 | 37 | rnr1Y285F | FALSE | CAN | JSY3811 | 4 | rnr1Y285F-R | GCC | C>T | A/T+1 | G/C-1 | CG>TA | CG>TA | TCCATGCCG | CCGCCAG | CGG | GCC | C | A | C>T | C>T_GCC | CG>TA_CCGCCAG | 8 | FALSE | FALSE |
| 45_v.txt | 32882 | 32882-SNV-C | SNV | 1 | C | T | 94 | 19784 | 0.47513142 | 32.4255319 | rnr1Y285F | FALSE | CAN | JSY3811 | 4 | rnr1Y285F-R | TCC | C>T | A/T+1 | G/C-1 | CG>TA | CG>TA | GTACGTCCA | CCAAAAT | AGG | TCC | C | A | C>T | C>T_TCC | CG>TA_CCAAAAT | 8 | FALSE | FALSE |
| 45_v.txt | 32900 | 32900-SNV-G | SNV | 1 | G | A | 22 | 19786 | 0.11118973 | 34.9973375 | rnr1Y285F | FALSE | CAN | JSY3811 | 4 | rnr1Y285F-R | GGC | G>A | A/T+1 | G/C-1 | CG>TA | CG>TA | GACTTGGCC | GCCAACT | CCG | CCG | C | T | C>T | C>T_CCG | CG>TA_GCCAACT | 8 | FALSE | FALSE |
| 45_v.txt | 32901 | 32901-SNV-C | SNV | 1 | C | T | 48 | 19786 | 0.24259578 | 33.229686 | rnr1Y285F | FALSE | CAN | JSY3811 | 4 | rnr1Y285F-R | GCC | C>T | A/T+1 | G/C-1 | CG>TA | CG>TA | ACTTGGCCA | CCAACTA | CGG | GCC | C | A | C>T | C>T_GCC | CG>TA_CCAACTA | 8 | FALSE | FALSE |
| 45_v.txt | 32917 | 32917-SNV-C | SNV | 1 | C | T | 52 | 19786 | 0.26281209 | 46.7115385 | rnr1Y285F | FALSE | CAN | JSY3811 | 4 | rnr1Y285F-R | TCC | C>T | A/T+1 | G/C-1 | CG>TA | CG>TA | TAAGTTCCAC | CCAGGGC | AGG | TCC | C | A | C>T | C>T_TCC | CG>TA_CCAGGGC | 8 | FALSE | FALSE |
| 45_v.txt | 32935 | 32935-SNV-C | SNV | 1 | C | T | 35 | 19786 | 0.17689275 | 42.8830718 | rnr1Y285F | FALSE | CAN | JSY3811 | 4 | rnr1Y285F-R | GCC | C>T | A/T+1 | G/C-1 | CG>TA | CG>TA | TGATTGCCCA | CCCAAGA | CGG | GCC | C | A | C>T | C>T_GCC | CG>TA_CCCAAGA | 8 | FALSE | FALSE |
| 45_v.txt | 32945 | 32945-SNV-C | SNV | 1 | C | T | 60 | 19786 | 0.30324472 | 55.8833333 | rnr1Y285F | FALSE | CAN | JSY3811 | 4 | rnr1Y285F-R | ACC | C>T | A/T+1 | G/C-1 | CG>TA | CG>TA | AGAAAACCA | CCAATAC | TGG | ACC | C | A | C>T | C>T_ACC | CG>TA_CCAATAC | 8 | FALSE | FALSE |
| 45_v.txt | 32958 | 32958-SNV-C | SNV | 1 | C | T | 44 | 19786 | 0.22237946 | 51 | rnr1Y285F | FALSE | CAN | JSY3811 | 4 | rnr1Y285F-R | ACC | C>T | A/T+1 | G/C-1 | CG>TA | CG>TA | ATGTAACCAT | CCATTGG | TGG | ACC | C | A | C>T | C>T_ACC | CG>TA_CCATTGG | 8 | FALSE | FALSE |
| 45_v.txt | 32967 | 32967-SNV-G | SNV | 1 | G | A | 42 | 19786 | 0.2122713 | 59.2619048 | rnr1Y285F | FALSE | CAN | JSY3811 | 4 | rnr1Y285F-R | CGC | G>A | A/T+1 | G/C-1 | CG>TA | CG>TA | TTGGCCGCAC | GCACCAA | GCG | GCG | C | T | C>T | C>T_GCG | CG>TA_GCACCAA | 8 | FALSE | FALSE |
| 45_v.txt | 32977 | 32977-SNV-C | SNV | 1 | C | T | 24 | 19786 | 0.12129789 | 58.7916667 | rnr1Y285F | FALSE | CAN | JSY3811 | 4 | rnr1Y285F-R | GCT | C>T | A/T+1 | G/C-1 | CG>TA | CG>TA | CAAATGCTG | CTGGAGA | CGA | GCT | C | A | C>T | C>T_GCT | CG>TA_CTGGAGA | 8 | FALSE | FALSE |
| 45_v.txt | 33043 | 33043-SNV-C | SNV | 1 | C | T | 23 | 19786 | 0.11624381 | 58.3043478 | rnr1Y285F | FALSE | CAN | JSY3811 | 4 | rnr1Y285F-R | TCA | C>T | A/T+1 | G/C-1 | CG>TA | CG>TA | CCATTTCACC | CACCCAA | AGT | TCA | C | A | C>T | C>T_TCA | CG>TA_CACCCAA | 8 | FALSE | FALSE |
| 45_v.txt | 33056 | 33056-SNV-C | SNV | 1 | C | T | 40 | 19786 | 0.20216315 | 50.2 | rnr1Y285F | FALSE | CAN | JSY3811 | 4 | rnr1Y285F-R | GCG | C>T | A/T+1 | G/C-1 | CG>TA | CG>TA | GGACTGCGT | CGTGACA | CGC | GCG | C | A | C>T | C>T_GCG | CG>TA_CGTGACA | 8 | FALSE | FALSE |
| 45_v.txt | 33069 | 33069-SNV-G | SNV | 1 | G | A | 24 | 19786 | 0.12129789 | 51.4578224 | rnr1Y285F | FALSE | CAN | JSY3811 | 4 | rnr1Y285F-R | TGC | G>A | A/T+1 | G/C-1 | CG>TA | CG>TA | GAATATGCC | GCCAAAG | ACG | ACG | C | T | C>T | C>T_ACG | CG>TA_GCCAAAG | 8 | FALSE | FALSE |
| 45_v.txt | 33070 | 33070-SNV-C | SNV | 1 | C | T | 28 | 19786 | 0.1415142 | 49.8176304 | rnr1Y285F | FALSE | CAN | JSY3811 | 4 | rnr1Y285F-R | GCC | C>T | A/T+1 | G/C-1 | CG>TA | CG>TA | AATATGCCA | CCAAAGA | CGG | GCC | C | A | C>T | C>T_GCC | CG>TA_CCAAAGA | 8 | FALSE | FALSE |
| 45_v.txt | 33104 | 33104-SNV-G | SNV | 1 | G | A | 25 | 19786 | 0.12635197 | 33.84 | rnr1Y285F | FALSE | CAN | JSY3811 | 4 | rnr1Y285F-R | CGC | G>A | A/T+1 | G/C-1 | CG>TA | CG>TA | AAGAGCGCC | GCCCACT | GCG | GCG | C | T | C>T | C>T_GCG | CG>TA_GCCCACT | 8 | FALSE | FALSE |

| | | | | | | | | | | | | | | | | | | | | | | | | | | | | | | | | | | |
|---|---|---|---|---|---|---|---|---|---|---|---|---|---|---|---|---|---|---|---|---|---|---|---|---|---|---|---|---|---|---|---|---|---|---|
| 45_v.txt | 33113 | 33113-SNV-( | SNV | 1 | G | A | 24 | 19786 | 0.12129789 | 42.9166667 | rnr1Y285F | FALSE | CAN | JSY3811 | 4 | rnr1Y285F-R | GGC | G>A | A/T+1 | G/C-1 | CG>TA | CG>TA | CACTGGGCC | GCCGGCG | CCG | CCG | C | T | C>T | C>T_CCG | CG>TA_GCCGGCG | 8 | FALSE | FALSE |
| 45_v.txt | 33153 | 33153-SNV-( | SNV | 1 | C | T | 84 | 19786 | 0.42454261 | 35.8690477 | rnr1Y285F | FALSE | CAN | JSY3811 | 4 | rnr1Y285F-R | ACC | C>T | A/T+1 | G/C-1 | CG>TA | CG>TA | AAAAGACCT | CCTGTAC | TGG | ACC | C | A | C>T | C>T_ACC | CG>TA_CCTGTAC | 8 | FALSE | FALSE |
| 45_v.txt | 33168 | 33168-SNV-( | SNV | 1 | C | T | 19 | 19783 | 0.09604206 | 37.9473684 | rnr1Y285F | FALSE | CAN | JSY3811 | 4 | rnr1Y285F-R | ACC | C>T | A/T+1 | G/C-1 | CG>TA | CG>TA | ATAGTACCA | CCACCAA | TGG | ACC | C | A | C>T | C>T_ACC | CG>TA_CCACCAA | 8 | FALSE | FALSE |
| 45_v.txt | 33258 | 33258-SNV-( | SNV | 1 | C | T | 22 | 11670 | 0.18851757 | 43.6363636 | rnr1Y285F | FALSE | CAN | JSY3811 | 4 | rnr1Y285F-R | GCC | C>T | A/T+1 | G/C-1 | CG>TA | CG>TA | CCTATGCCAT | CCATCCT | CGG | GCC | C | A | C>T | C>T_GCC | CG>TA_CCATCCT | 8 | FALSE | FALSE |
| 45_v.txt | 33310 | 33310-SNV-( | SNV | 1 | G | A | 15 | 11670 | 0.1285347 | 58.2 | rnr1Y285F | FALSE | CAN | JSY3811 | 4 | rnr1Y285F-R | CGC | G>A | A/T+1 | G/C-1 | CG>TA | CG>TA | AAGAGCGCA | GCAATGG | GCG | GCG | C | T | C>T | C>T_GCG | CG>TA_GCAATGG | 8 | FALSE | FALSE |
| 45_v.txt | 33392 | 33392-SNV-( | SNV | 1 | G | A | 17 | 11670 | 0.14567267 | 49.0588235 | rnr1Y285F | FALSE | CAN | JSY3811 | 4 | rnr1Y285F-R | CGT | G>A | A/T+1 | G/C-1 | CG>TA | CG>TA | AACGTCGTG | GTGAAAG | GCA | GCA | C | T | C>T | C>T_GCA | CG>TA_GTGAAAG | 8 | FALSE | FALSE |
| 45_v.txt | 33412 | 33412-SNV-( | SNV | 1 | G | A | 18 | 11670 | 0.15424165 | 46.0555556 | rnr1Y285F | FALSE | CAN | JSY3811 | 4 | rnr1Y285F-R | GGC | G>A | A/T+1 | G/C-1 | CG>TA | CG>TA | TGACCGGCT | GCTCATT | CCG | CCG | C | T | C>T | C>T_CCG | CG>TA_GCTCATT | 8 | FALSE | FALSE |
| 45_v.txt | 33474 | 33474-SNV-( | SNV | 1 | C | T | 18 | 11670 | 0.15424165 | 32.1666667 | rnr1Y285F | FALSE | CAN | JSY3811 | 4 | rnr1Y285F-R | GCC | C>T | A/T+1 | G/C-1 | CG>TA | CG>TA | GCTATGCCTT | CCTTTTT | CGG | GCC | C | A | C>T | C>T_GCC | CG>TA_CCTTTTT | 8 | FALSE | FALSE |
| 45_v.txt | 31956 | 31956-SNV-( | SNV | 1 | G | C | 5 | 9998 | 0.05001 | 34.4 | rnr1Y285F | FALSE | CAN | JSY3811 | 4 | rnr1Y285F-R | TGT | G>C | G/C+1 | G/C-1 | CG>GC | CG>GC | ATAAATGTG | GTGGCCG | ACA | ACA | C | G | C>G | C>G_ACA | CG>GC_GTGGCCG | 8 | FALSE | FALSE |
| 45_v.txt | 31980 | 31980-SNV-( | SNV | 1 | G | C | 13 | 9998 | 0.13002601 | 47.4597392 | rnr1Y285F | FALSE | CAN | JSY3811 | 4 | rnr1Y285F-R | GGG | G>C | G/C+1 | G/C-1 | CG>GC | CG>GC | AAGCCGGGC | GGCATTA | CCC | CCC | C | G | C>G | C>G_CCC | CG>GC_GGCATTA | 8 | FALSE | FALSE |
| 45_v.txt | 32038 | 32038-SNV-( | SNV | 1 | C | G | 11 | 9998 | 0.110022 | 55.0909091 | rnr1Y285F | FALSE | CAN | JSY3811 | 4 | rnr1Y285F-R | GCT | C>G | G/C+1 | G/C-1 | CG>GC | CG>GC | TCAAAGCTTC | CTTGCAT | CGA | GCT | C | C | C>G | C>G_GCT | CG>GC_CTTGCAT | 8 | FALSE | FALSE |
| 45_v.txt | 32292 | 32292-SNV-( | SNV | 1 | G | C | 847 | 21827 | 3.88051496 | 53.2172373 | rnr1Y285F | FALSE | CAN | JSY3811 | 4 | rnr1Y285F-R | TGA | G>C | G/C+1 | G/C-1 | CG>GC | CG>GC | ATATTTGAAT | GAATTTG | ACT | ACT | C | G | C>G | C>G_ACT | CG>GC_GAATTTG | 8 | FALSE | FALSE |
| 45_v.txt | 32432 | 32432-SNV-( | SNV | 1 | G | C | 23 | 21827 | 0.10537408 | 54.0869565 | rnr1Y285F | FALSE | CAN | JSY3811 | 4 | rnr1Y285F-R | TGT | G>C | G/C+1 | G/C-1 | CG>GC | CG>GC | GTCATTGTAT | GTATGGA | ACA | ACA | C | G | C>G | C>G_ACA | CG>GC_GTATGGA | 8 | FALSE | FALSE |
| 45_v.txt | 32571 | 32571-SNV-( | SNV | 1 | C | G | 1 | 13851 | 7.22E-03 | 36.9972092 | rnr1Y285F | FALSE | CAN | JSY3811 | 4 | rnr1Y285F-R | ACC | C>G | G/C+1 | G/C-1 | CG>GC | CG>GC | TCAGTACCTT | CCTTGAA | TGG | ACC | C | C | C>G | C>G_ACC | CG>GC_CCTTGAA | 8 | FALSE | FALSE |
| 45_v.txt | 32749 | 32749-SNV-( | SNV | 1 | C | G | 7 | 13851 | 0.05053787 | 46.7142857 | rnr1Y285F | FALSE | CAN | JSY3811 | 4 | rnr1Y285F-R | CCG | C>G | G/C+1 | G/C-1 | CG>GC | CG>GC | GAAACCCGA | CGATAAT | GGC | CCG | C | C | C>G | C>G_CCG | CG>GC_CGATAAT | 8 | FALSE | FALSE |
| 45_v.txt | 32798 | 32798-SNV-( | SNV | 1 | G | C | 2 | 13851 | 0.01443939 | 51 | rnr1Y285F | FALSE | CAN | JSY3811 | 4 | rnr1Y285F-R | CGT | G>C | G/C+1 | G/C-1 | CG>GC | CG>GC | TTCACCGTAA | GTAATAT | GCA | GCA | C | G | C>G | C>G_GCA | CG>GC_GTAATAT | 8 | FALSE | FALSE |
| 45_v.txt | 32819 | 32819-SNV-( | SNV | 1 | G | C | 4 | 13851 | 0.02887878 | 38 | rnr1Y285F | FALSE | CAN | JSY3811 | 4 | rnr1Y285F-R | AGT | G>C | G/C+1 | G/C-1 | CG>GC | CG>GC | GAACAAGTT | GTTCATT | TCA | TCA | C | G | C>G | C>G_TCA | CG>GC_GTTCATT | 8 | FALSE | FALSE |
| 45_v.txt | 32876 | 32876-SNV-( | SNV | 1 | G | C | 37 | 19779 | 0.18706709 | 37.5405405 | rnr1Y285F | FALSE | CAN | JSY3811 | 4 | rnr1Y285F-R | TGT | G>C | G/C+1 | G/C-1 | CG>GC | CG>GC | AACTTTGTAC | GTACGTC | ACA | ACA | C | G | C>G | C>G_ACA | CG>GC_GTACGTC | 8 | FALSE | FALSE |
| 45_v.txt | 32900 | 32900-SNV-( | SNV | 1 | G | C | 1 | 19786 | 5.05E-03 | 26.997752 | rnr1Y285F | FALSE | CAN | JSY3811 | 4 | rnr1Y285F-R | GGC | G>C | G/C+1 | G/C-1 | CG>GC | CG>GC | GACTTGGCC | GCCAACT | CCG | CCG | C | G | C>G | C>G_CCG | CG>GC_GCCAACT | 8 | FALSE | FALSE |
| 45_v.txt | 32935 | 32935-SNV-( | SNV | 1 | C | G | 13 | 19786 | 0.06570302 | 47.687729 | rnr1Y285F | FALSE | CAN | JSY3811 | 4 | rnr1Y285F-R | GCC | C>G | G/C+1 | G/C-1 | CG>GC | CG>GC | TGATTGCCCA | CCCAAGA | CGG | GCC | C | C | C>G | C>G_GCC | CG>GC_CCCAAGA | 8 | FALSE | FALSE |
| 45_v.txt | 32936 | 32936-SNV-( | SNV | 1 | C | G | 12 | 19786 | 0.06064894 | 48.7608828 | rnr1Y285F | FALSE | CAN | JSY3811 | 4 | rnr1Y285F-R | CCC | C>G | G/C+1 | G/C-1 | CG>GC | CG>GC | GATTGCCCA | CCAAGAA | GGG | CCC | C | C | C>G | C>G_CCC | CG>GC_CCAAGAA | 8 | FALSE | FALSE |
| 45_v.txt | 32971 | 32971-SNV-( | SNV | 1 | C | G | 27 | 19786 | 0.13646012 | 62.1851852 | rnr1Y285F | FALSE | CAN | JSY3811 | 4 | rnr1Y285F-R | CCA | C>G | G/C+1 | G/C-1 | CG>GC | CG>GC | CCGCACCAA | CAAATGC | GGT | CCA | C | C | C>G | C>G_CCA | CG>GC_CAAATGC | 8 | FALSE | FALSE |
| 45_v.txt | 33168 | 33168-SNV-( | SNV | 1 | C | G | 7 | 19783 | 0.03538392 | 37.4285714 | rnr1Y285F | FALSE | CAN | JSY3811 | 4 | rnr1Y285F-R | ACC | C>G | G/C+1 | G/C-1 | CG>GC | CG>GC | ATAGTACCA | CCACCAA | TGG | ACC | C | C | C>G | C>G_ACC | CG>GC_CCACCAA | 8 | FALSE | FALSE |
| 45_v.txt | 33187 | 33187-SNV-( | SNV | 1 | C | G | 16 | 11669 | 0.13711543 | 37.1875 | rnr1Y285F | FALSE | CAN | JSY3811 | 4 | rnr1Y285F-R | CCA | C>G | G/C+1 | G/C-1 | CG>GC | CG>GC | TCATACCAAT | CAATATG | GGT | CCA | C | C | C>G | C>G_CCA | CG>GC_CAATATG | 8 | FALSE | FALSE |
| 45_v.txt | 33205 | 33205-SNV-( | SNV | 1 | G | C | 12 | 11670 | 0.10282776 | 39.5 | rnr1Y285F | FALSE | CAN | JSY3811 | 4 | rnr1Y285F-R | AGC | G>C | G/C+1 | G/C-1 | CG>GC | CG>GC | GCTTAAGCT | GCTCTCT | TCG | TCG | C | G | C>G | C>G_TCG | CG>GC_GCTCTCT | 8 | FALSE | FALSE |
| 45_v.txt | 31657 | 31657-SNV-( | SNV | 1 | G | T | 67 | 6593 | 1.01622933 | 19.4626866 | rnr1Y285F | FALSE | CAN | JSY3811 | 4 | rnr1Y285F-R | GGC | G>T | A/T+1 | G/C-1 | CG>AT | CG>AT | GAAATGGCG | GCGTGGA | CCG | CCG | C | A | C>A | C>A_CCG | CG>AT_GCGTGGA | 8 | FALSE | FALSE |
| 45_v.txt | 31981 | 31981-SNV-( | SNV | 1 | G | T | 24 | 9998 | 0.24004801 | 36.9166667 | rnr1Y285F | FALSE | CAN | JSY3811 | 4 | rnr1Y285F-R | GGC | G>T | A/T+1 | G/C-1 | CG>AT | CG>AT | AGCCGGGCA | GCATTAA | CCG | CCG | C | A | C>A | C>A_CCG | CG>AT_GCATTAA | 8 | FALSE | FALSE |
| 45_v.txt | 32027 | 32027-SNV-( | SNV | 1 | G | T | 22 | 9998 | 0.22004401 | 55.8636364 | rnr1Y285F | FALSE | CAN | JSY3811 | 4 | rnr1Y285F-R | GGT | G>T | A/T+1 | G/C-1 | CG>AT | CG>AT | GCCACGGTA | GTATTTC | CCA | CCA | C | A | C>A | C>A_CCA | CG>AT_GTATTTC | 8 | FALSE | FALSE |
| 45_v.txt | 32188 | 32188-SNV-( | SNV | 1 | C | A | 20 | 9998 | 0.20004001 | 47.7 | rnr1Y285F | FALSE | CAN | JSY3811 | 4 | rnr1Y285F-R | ACT | C>A | A/T+1 | G/C-1 | CG>AT | CG>AT | CGAAAACTG | CTGCAAT | TGA | ACT | C | T | C>A | C>A_ACT | CG>AT_CTGCAAT | 8 | FALSE | FALSE |
| 45_v.txt | 32205 | 32205-SNV-( | SNV | 1 | C | A | 21 | 9998 | 0.21004201 | 44.0476191 | rnr1Y285F | FALSE | CAN | JSY3811 | 4 | rnr1Y285F-R | ACC | C>A | A/T+1 | G/C-1 | CG>AT | CG>AT | GGAACACCA | CCACCTT | TGG | ACC | C | T | C>A | C>A_ACC | CG>AT_CCACCTT | 8 | FALSE | FALSE |
| 45_v.txt | 32271 | 32271-SNV-( | SNV | 1 | C | A | 18 | 21827 | 0.08246667 | 32.7222222 | rnr1Y285F | FALSE | CAN | JSY3811 | 4 | rnr1Y285F-R | ACG | C>A | A/T+1 | G/C-1 | CG>AT | CG>AT | AAAATACGG | CGGGAAC | TGC | ACG | C | T | C>A | C>A_ACG | CG>AT_CGGGAAC | 8 | FALSE | FALSE |
| 45_v.txt | 32274 | 32274-SNV-( | SNV | 1 | G | T | 32 | 21827 | 0.14660741 | 35.125 | rnr1Y285F | FALSE | CAN | JSY3811 | 4 | rnr1Y285F-R | GGA | G>T | A/T+1 | G/C-1 | CG>AT | CG>AT | ATACGGGAA | GAACCAA | CCT | CCT | C | A | C>A | C>A_CCT | CG>AT_GAACCAA | 8 | FALSE | FALSE |
| 45_v.txt | 32292 | 32292-SNV-( | SNV | 1 | G | T | 14 | 21827 | 0.06414074 | 52.1428571 | rnr1Y285F | FALSE | CAN | JSY3811 | 4 | rnr1Y285F-R | TGA | G>T | A/T+1 | G/C-1 | CG>AT | CG>AT | ATATTTGAAT | GAATTTG | ACT | ACT | C | A | C>A | C>A_ACT | CG>AT_GAATTTG | 8 | FALSE | FALSE |
| 45_v.txt | 32373 | 32373-SNV-( | SNV | 1 | G | T | 21 | 21827 | 0.09621112 | 60.2380952 | rnr1Y285F | FALSE | CAN | JSY3811 | 4 | rnr1Y285F-R | AGC | G>T | A/T+1 | G/C-1 | CG>AT | CG>AT | TCAATAGCA | GCAATAA | TCG | TCG | C | A | C>A | C>A_TCG | CG>AT_GCAATAA | 8 | FALSE | FALSE |
| 45_v.txt | 32432 | 32432-SNV-( | SNV | 1 | G | T | 54 | 21827 | 0.24740001 | 56.5555556 | rnr1Y285F | FALSE | CAN | JSY3811 | 4 | rnr1Y285F-R | TGT | G>T | A/T+1 | G/C-1 | CG>AT | CG>AT | GTCATTGTAT | GTATGGA | ACA | ACA | C | A | C>A | C>A_ACA | CG>AT_GTATGGA | 8 | FALSE | FALSE |
| 45_v.txt | 32530 | 32530-SNV-( | SNV | 1 | G | T | 49 | 21827 | 0.2244926 | 37.2857143 | rnr1Y285F | FALSE | CAN | JSY3811 | 4 | rnr1Y285F-R | GGG | G>T | A/T+1 | G/C-1 | CG>AT | CG>AT | TTCTGGGGT | GGTTTGC | CCC | CCC | C | A | C>A | C>A_CCC | CG>AT_GGTTTGC | 8 | FALSE | FALSE |
| 45_v.txt | 32545 | 32545-SNV-( | SNV | 1 | C | A | 21 | 21825 | 0.09621993 | 38 | rnr1Y285F | FALSE | CAN | JSY3811 | 4 | rnr1Y285F-R | CCA | C>A | A/T+1 | G/C-1 | CG>AT | CG>AT | CTTCACCAGC | CAGCAGT | GGT | CCA | C | T | C>A | C>A_CCA | CG>AT_CAGCAGT | 8 | FALSE | FALSE |
| 45_v.txt | 32547 | 32547-SNV-( | SNV | 1 | G | T | 12 | 21823 | 0.05498786 | 37.6666667 | rnr1Y285F | FALSE | CAN | JSY3811 | 4 | rnr1Y285F-R | AGC | G>T | A/T+1 | G/C-1 | CG>AT | CG>AT | TCACCAGCA | GCAGTGA | TCG | TCG | C | A | C>A | C>A_TCG | CG>AT_GCAGTGA | 8 | FALSE | FALSE |
| 45_v.txt | 32556 | 32556-SNV-( | SNV | 1 | C | A | 26 | 21820 | 0.11915674 | 37.6923077 | rnr1Y285F | FALSE | CAN | JSY3811 | 4 | rnr1Y285F-R | ACC | C>A | A/T+1 | G/C-1 | CG>AT | CG>AT | GTGATACCA | CCAACTA | TGG | ACC | C | T | C>A | C>A_ACC | CG>AT_CCAACTA | 8 | FALSE | FALSE |
| 45_v.txt | 32629 | 32629-SNV-( | SNV | 1 | C | A | 21 | 13851 | 0.1516136 | 56.4285714 | rnr1Y285F | FALSE | CAN | JSY3811 | 4 | rnr1Y285F-R | TCG | C>A | A/T+1 | G/C-1 | CG>AT | CG>AT | TCCCTTCGTT | CGTTTTT | AGC | TCG | C | T | C>A | C>A_TCG | CG>AT_CGTTTTT | 8 | FALSE | FALSE |
| 45_v.txt | 32658 | 32658-SNV-( | SNV | 1 | C | A | 36 | 13851 | 0.25990903 | 54.9444444 | rnr1Y285F | FALSE | CAN | JSY3811 | 4 | rnr1Y285F-R | ACC | C>A | A/T+1 | G/C-1 | CG>AT | CG>AT | CCTGGACCC | CCCCAGG | TGG | ACC | C | T | C>A | C>A_ACC | CG>AT_CCCCAGG | 8 | FALSE | FALSE |
| 45_v.txt | 32735 | 32735-SNV-( | SNV | 1 | G | T | 39 | 13851 | 0.28156812 | 56.025641 | rnr1Y285F | FALSE | CAN | JSY3811 | 4 | rnr1Y285F-R | AGT | G>T | A/T+1 | G/C-1 | CG>AT | CG>AT | AAAACAGTA | GTATATT | TCA | TCA | C | A | C>A | C>A_TCA | CG>AT_GTATATT | 8 | FALSE | FALSE |
| 45_v.txt | 32788 | 32788-SNV-( | SNV | 1 | C | A | 6 | 13851 | 0.04331817 | 46 | rnr1Y285F | FALSE | CAN | JSY3811 | 4 | rnr1Y285F-R | TCG | C>A | A/T+1 | G/C-1 | CG>AT | CG>AT | AGAACTCGA | CGAATTC | AGC | TCG | C | T | C>A | C>A_TCG | CG>AT_CGAATTC | 8 | FALSE | FALSE |
| 45_v.txt | 32794 | 32794-SNV-( | SNV | 1 | C | A | 17 | 13851 | 0.12273482 | 41.8235294 | rnr1Y285F | FALSE | CAN | JSY3811 | 4 | rnr1Y285F-R | TCA | C>A | A/T+1 | G/C-1 | CG>AT | CG>AT | CGAATTCACC | CACCGTA | AGT | TCA | C | T | C>A | C>A_TCA | CG>AT_CACCGTA | 8 | FALSE | FALSE |
| 45_v.txt | 32809 | 32809-SNV-( | SNV | 1 | C | A | 14 | 13851 | 0.10107574 | 38.7142857 | rnr1Y285F | FALSE | CAN | JSY3811 | 4 | rnr1Y285F-R | ACA | C>A | A/T+1 | G/C-1 | CG>AT | CG>AT | ATTTGACAG | CAGGGAA | TGT | ACA | C | T | C>A | C>A_ACA | CG>AT_CAGGGAA | 8 | FALSE | FALSE |
| 45_v.txt | 32812 | 32812-SNV-( | SNV | 1 | G | T | 22 | 13851 | 0.1588333 | 46.5909091 | rnr1Y285F | FALSE | CAN | JSY3811 | 4 | rnr1Y285F-R | GGG | G>T | A/T+1 | G/C-1 | CG>AT | CG>AT | TGACAGGGA | GGAACAA | CCC | CCC | C | A | C>A | C>A_CCC | CG>AT_GGAACAA | 8 | FALSE | FALSE |
| 45_v.txt | 32819 | 32819-SNV-( | SNV | 1 | G | T | 53 | 13851 | 0.38264385 | 43.6792453 | rnr1Y285F | FALSE | CAN | JSY3811 | 4 | rnr1Y285F-R | AGT | G>T | A/T+1 | G/C-1 | CG>AT | CG>AT | GAACAAGTT | GTTCATT | TCA | TCA | C | A | C>A | C>A_TCA | CG>AT_GTTCATT | 8 | FALSE | FALSE |
| 45_v.txt | 32917 | 32917-SNV-( | SNV | 1 | C | A | 91 | 19786 | 0.45992116 | 52.021978 | rnr1Y285F | FALSE | CAN | JSY3811 | 4 | rnr1Y285F-R | TCC | C>A | A/T+1 | G/C-1 | CG>AT | CG>AT | TAAGTTCCAC | CCAGGGC | AGG | TCC | C | T | C>A | C>A_TCC | CG>AT_CCAGGGC | 8 | FALSE | FALSE |
| 45_v.txt | 32940 | 32940-SNV-( | SNV | 1 | G | T | 9089 | 19786 | 45.9365208 | 56.2437012 | rnr1Y285F | FALSE | CAN | JSY3811 | 4 | rnr1Y285F-R | AGA | G>T | A/T+1 | G/C-1 | CG>AT | CG>AT | GCCCAAGAA | GAAAACC | TCT | TCT | C | A | C>A | C>A_TCT | CG>AT_GAAAACC | 8 | FALSE | FALSE |
| 45_v.txt | 32958 | 32958-SNV-( | SNV | 1 | C | A | 15 | 19786 | 0.07581118 | 45.0666667 | rnr1Y285F | FALSE | CAN | JSY3811 | 4 | rnr1Y285F-R | ACC | C>A | A/T+1 | G/C-1 | CG>AT | CG>AT | ATGTAACCA | CCATTGG | TGG | ACC | C | T | C>A | C>A_ACC | CG>AT_CCATTGG | 8 | FALSE | FALSE |
| 45_v.txt | 33043 | 33043-SNV-( | SNV | 1 | C | A | 8 | 19786 | 0.04043263 | 60.125 | rnr1Y285F | FALSE | CAN | JSY3811 | 4 | rnr1Y285F-R | TCA | C>A | A/T+1 | G/C-1 | CG>AT | CG>AT | CCATTTCACC | CACCCAA | AGT | TCA | C | T | C>A | C>A_TCA | CG>AT_CACCCAA | 8 | FALSE | FALSE |
| 45_v.txt | 33050 | 33050-SNV-( | SNV | 1 | G | T | 10 | 19786 | 0.05054079 | 52.7 | rnr1Y285F | FALSE | CAN | JSY3811 | 4 | rnr1Y285F-R | AGG | G>T | A/T+1 | G/C-1 | CG>AT | CG>AT | ACCCAAGGA | GGACTGC | TCC | TCC | C | A | C>A | C>A_TCC | CG>AT_GGACTGC | 8 | FALSE | FALSE |
| 45_v.txt | 33151 | 33151-SNV-( | SNV | 1 | G | T | 810 | 19786 | 4.0938037 | 38.7851852 | rnr1Y285F | FALSE | CAN | JSY3811 | 4 | rnr1Y285F-R | AGA | G>T | A/T+1 | G/C-1 | CG>AT | CG>AT | TGAAAAGAC | GACCTGT | TCT | TCT | C | A | C>A | C>A_TCT | CG>AT_GACCTGT | 8 | FALSE | FALSE |
| 45_v.txt | 33153 | 33153-SNV-( | SNV | 1 | C | A | 19 | 19786 | 0.09602749 | 41.8421053 | rnr1Y285F | FALSE | CAN | JSY3811 | 4 | rnr1Y285F-R | ACC | C>A | A/T+1 | G/C-1 | CG>AT | CG>AT | AAAAGACCT | CCTGTAC | TGG | ACC | C | T | C>A | C>A_ACC | CG>AT_CCTGTAC | 8 | FALSE | FALSE |
| 45_v.txt | 33154 | 33154-SNV-( | SNV | 1 | C | A | 20 | 19786 | 0.10108157 | 34.3500921 | rnr1Y285F | FALSE | CAN | JSY3811 | 4 | rnr1Y285F-R | CCT | C>A | A/T+1 | G/C-1 | CG>AT | CG>AT | AAAGACCTG | CTGTACC | GGA | CCT | C | T | C>A | C>A_CCT | CG>AT_CTGTACC | 8 | FALSE | FALSE |
| 45_v.txt | 33160 | 33160-SNV-( | SNV | 1 | C | A | 123 | 19785 | 0.62168309 | 37.5365854 | rnr1Y285F | FALSE | CAN | JSY3811 | 4 | rnr1Y285F-R | CCA | C>A | A/T+1 | G/C-1 | CG>AT | CG>AT | CTGTACCAAT | CAATAGT | GGT | CCA | C | T | C>A | C>A_CCA | CG>AT_CAATAGT | 8 | FALSE | FALSE |
| 45_v.txt | 33195 | 33195-SNV-( | SNV | 1 | C | A | 14 | 11670 | 0.11996572 | 35.9285714 | rnr1Y285F | FALSE | CAN | JSY3811 | 4 | rnr1Y285F-R | TCT | C>A | A/T+1 | G/C-1 | CG>AT | CG>AT | ATATGTCTTT | CTTTGCT | AGA | TCT | C | T | C>A | C>A_TCT | CG>AT_CTTTGCT | 8 | FALSE | FALSE |
| 45_v.txt | 33241 | 33241-SNV-( | SNV | 1 | C | A | 33 | 11670 | 0.28277635 | 42.5454546 | rnr1Y285F | FALSE | CAN | JSY3811 | 4 | rnr1Y285F-R | TCA | C>A | A/T+1 | G/C-1 | CG>AT | CG>AT | CTCCTTCATC | CATCTTC | AGT | TCA | C | T | C>A | C>A_TCA | CG>AT_CATCTTC | 8 | FALSE | FALSE |
| 45_v.txt | 33283 | 33283-SNV-( | SNV | 1 | C | A | 23 | 11670 | 0.19708655 | 52.7826087 | rnr1Y285F | FALSE | CAN | JSY3811 | 4 | rnr1Y285F-R | TCG | C>A | A/T+1 | G/C-1 | CG>AT | CG>AT | TATCCTCGCC | CGCCATT | AGC | TCG | C | T | C>A | C>A_TCG | CG>AT_CGCCATT | 8 | FALSE | FALSE |
| 45_v.txt | 33325 | 33325-SNV-( | SNV | 1 | C | A | 11 | 11670 | 0.09425878 | 44.0909091 | rnr1Y285F | FALSE | CAN | JSY3811 | 4 | rnr1Y285F-R | TCT | C>A | A/T+1 | G/C-1 | CG>AT | CG>AT | ACAATTCTTT | CTTTACT | AGA | TCT | C | T | C>A | C>A_TCT | CG>AT_CTTTACT | 8 | FALSE | FALSE |
| 45_v.txt | 33354 | 33354-SNV-( | SNV | 1 | G | T | 61 | 11670 | 0.5227078 | 17.6065574 | rnr1Y285F | FALSE | CAN | JSY3811 | 4 | rnr1Y285F-R | TGA | G>T | A/T+1 | G/C-1 | CG>AT | CG>AT | GGTATTGAC | GACCCAC | ACT | ACT | C | A | C>A | C>A_ACT | CG>AT_GACCCAC | 8 | FALSE | FALSE |
| 45_v.txt | 33474 | 33474-SNV-( | SNV | 1 | C | A | 40 | 11670 | 0.34275921 | 34.575 | rnr1Y285F | FALSE | CAN | JSY3811 | 4 | rnr1Y285F-R | GCC | C>A | A/T+1 | G/C-1 | CG>AT | CG>AT | GCTATGCCTT | CCTTTTT | CGG | GCC | C | T | C>A | C>A_GCC | CG>AT_CCTTTTT | 8 | FALSE | FALSE |
| 45_v.txt | 33153 | 33153-MNV- | MNV | 2 | CC | AA | 2 | 19786 | 0.01010816 | 35.75 | rnr1Y285F | FALSE | CAN | JSY3811 | 4 | rnr1Y285F-R | ACC | nope | >1bp-ins | >1bpdel | none | MNV | AAAAGACCT | CCTGTAC | TGG | ACC | CC | none | NA | NA_ACC | MNV_CCTGTAC | 8 | FALSE | FALSE |
| 45_v.txt | 31885 | 31885-Insert | Insertion | 1 | - | T | 2 | 6593 | 0.0303352 | 38 | rnr1Y285F | FALSE | CAN | JSY3811 | 4 | rnr1Y285F-R | GCA | nope | A/T+1 | >1bpdel | none | A/T+1 | CGGCAGCAA | CAAAGCT | CGT | GCA | - | A | NA | NA_GCA | A/T+1_CAAAGCT | 8 | FALSE | FALSE |
| 45_v.txt | 32081 | 32081-Insert | Insertion | 1 | - | A | 36 | 9998 | 0.36007201 | 64 | rnr1Y285F | FALSE | CAN | JSY3811 | 4 | rnr1Y285F-R | CAA | nope | A/T+1 | >1bpdel | none | A/T+1 | CCATGCAAA | AAAAAAG | GTT | CAA | - | T | NA | NA_CAA | A/T+1_AAAAAAG | 8 | FALSE | FALSE |
| 45_v.txt | 32468 | 32468-Insert | Insertion | 1 | - | A | 2 | 21827 | 9.16E-03 | 51 | rnr1Y285F | FALSE | CAN | JSY3811 | 4 | rnr1Y285F-R | CAA | nope | A/T+1 | >1bpdel | none | A/T+1 | AGAGCCAAT | AATGTAG | GTT | CAA | - | T | NA | NA_CAA | A/T+1_AATGTAG | 8 | FALSE | FALSE |
| 45_v.txt | 32498 | 32498-Insert | Insertion | 1 | - | T | 2 | 21827 | 9.16E-03 | 38 | rnr1Y285F | FALSE | CAN | JSY3811 | 4 | rnr1Y285F-R | CTT | nope | A/T+1 | >1bpdel | none | A/T+1 | AACAACTTTT | TTTTTTG | GAA | CTT | - | A | NA | NA_CTT | A/T+1_TTTTTTG | 8 | FALSE | FALSE |
| 45_v.txt | 32516 | 32516-Insert | Insertion | 1 | - | A | 2 | 21827 | 9.16E-03 | 38 | rnr1Y285F | FALSE | CAN | JSY3811 | 4 | rnr1Y285F-R | GAA | nope | A/T+1 | >1bpdel | none | A/T+1 | TCTTGGAAC | AACGGAT | CTT | GAA | - | T | NA | NA_GAA | A/T+1_AACGGAT | 8 | FALSE | FALSE |
| 45_v.txt | 32522 | 32522-Insert | Insertion | 1 | - | T | 6 | 21827 | 0.02748889 | 37.1666667 | rnr1Y285F | FALSE | CAN | JSY3811 | 4 | rnr1Y285F-R | ATT | nope | A/T+1 | >1bpdel | none | A/T+1 | AACGGATTT | TTTTCTG | TAA | ATT | - | A | NA | NA_ATT | A/T+1_TTTTCTG | 8 | FALSE | FALSE |
| 45_v.txt | 32728 | 32728-Insert | Insertion | 1 | - | A | 7 | 13851 | 0.05053787 | 64 | rnr1Y285F | FALSE | CAN | JSY3811 | 4 | rnr1Y285F-R | CAA | nope | A/T+1 | >1bpdel | none | A/T+1 | CCATACAAA | AAAAACA | GTT | CAA | - | T | NA | NA_CAA | A/T+1_AAAAACA | 8 | FALSE | FALSE |
| 45_v.txt | 32842 | 32842-Insert | Insertion | 1 | - | A | 26 | 13851 | 0.18771208 | 40 | rnr1Y285F | FALSE | CAN | JSY3811 | 4 | rnr1Y285F-R | CAA | nope | A/T+1 | >1bpdel | none | A/T+1 | TTACCCAAA | AAAAAAT | GTT | CAA | - | T | NA | NA_CAA | A/T+1_AAAAAAT | 8 | FALSE | FALSE |
| 45_v.txt | 33189 | 33189-Deleti | Deletion | 2 | AT | - | 11 | 11669 | 0.09426686 | 37.7272727 | rnr1Y285F | FALSE | CAN | JSY3811 | 4 | rnr1Y285F-R | AAT | nope | >1bp-ins | >1bpdel | none | >1bpdel | ATACCAATA | ATATGTC | TTA | AAT | AT | none | NA | NA_AAT | >1bpdel_ATATGTC | 8 | FALSE | FALSE |
| 45_v.txt | 31723 | 31723-Deleti | Deletion | 1 | A | - | 7 | 6593 | 0.10617321 | 60.2857143 | rnr1Y285F | FALSE | CAN | JSY3811 | 4 | rnr1Y285F-R | CAA | nope | >1bp-ins | A/T-1 | none | A/T-1 | TGTCCCAAA | AAAAAGT | GTT | GTT | T | none | T>none | T>none_GTT | A/T-1_AAAAAGT | 8 | FALSE | FALSE |
| 45_v.txt | 31904 | 31904-Deleti | Deletion | 1 | T | - | 4 | 6593 | 0.06067041 | 38 | rnr1Y285F | FALSE | CAN | JSY3811 | 4 | rnr1Y285F-R | ATT | nope | >1bp-ins | A/T-1 | none | A/T-1 | ATTGAATTT | TTTTGGT | TAA | ATT | T | none | nope | nope_ATT | A/T-1_TTTTGGT | 8 | FALSE | FALSE |
| 45_v.txt | 31913 | 31913-Deleti | Deletion | 1 | A | - | 4 | 6593 | 0.06067041 | 36.5 | rnr1Y285F | FALSE | CAN | JSY3811 | 4 | rnr1Y285F-R | CAA | nope | >1bp-ins | A/T-1 | none | A/T-1 | TGGTGCAAA | AAAAGCC | GTT | GTT | T | none | T>none | T>none_GTT | A/T-1_AAAAGCC | 8 | FALSE | FALSE |
| 45_v.txt | 32081 | 32081-Deleti | Deletion | 1 | A | - | 10 | 9998 | 0.10002 | 64 | rnr1Y285F | FALSE | CAN | JSY3811 | 4 | rnr1Y285F-R | CAA | nope | >1bp-ins | A/T-1 | none | A/T-1 | CCATGCAAA | AAAAAAG | GTT | GTT | T | none | T>none | T>none_GTT | A/T-1_AAAAAAG | 8 | FALSE | FALSE |
| 45_v.txt | 32160 | 32160-Deleti | Deletion | 1 | A | - | 4 | 9998 | 0.040008 | 43.5 | rnr1Y285F | FALSE | CAN | JSY3811 | 4 | rnr1Y285F-R | CAA | nope | >1bp-ins | A/T-1 | none | A/T-1 | TAAGCCAAA | AAAGCGC | GTT | GTT | T | none | T>none | T>none_GTT | A/T-1_AAAGCGC | 8 | FALSE | FALSE |
| 45_v.txt | 32295 | 32295-Deleti | Deletion | 1 | T | - | 30 | 21827 | 0.13744445 | 48.5333333 | rnr1Y285F | FALSE | CAN | JSY3811 | 4 | rnr1Y285F-R | ATT | nope | >1bp-ins | A/T-1 | none | A/T-1 | TTTGAATTTG | TTTGCGG | TAA | ATT | T | none | nope | nope_ATT | A/T-1_TTTGCGG | 8 | FALSE | FALSE |
| 45_v.txt | 32312 | 32312-Deleti | Deletion | 1 | G | - | 5 | 21827 | 0.02290741 | 62.6 | rnr1Y285F | FALSE | CAN | JSY3811 | 4 | rnr1Y285F-R | TGG | nope | >1bp-ins | G/C-1 | none | G/C-1 | AATAATGGT | GGTTGTT | ACC | ACC | C | none | C>none | C>none_ACC | G/C-1_GGTTGTT | 8 | FALSE | FALSE |
| 45_v.txt | 32346 | 32346-Deleti | Deletion | 1 | A | - | 8 | 21827 | 0.03665185 | 59.5 | rnr1Y285F | FALSE | CAN | JSY3811 | 4 | rnr1Y285F-R | CAA | nope | >1bp-ins | A/T-1 | none | A/T-1 | TGTGGCAAA | AAAACCT | GTT | GTT | T | none | T>none | T>none_GTT | A/T-1_AAAACCT | 8 | FALSE | FALSE |
| 45_v.txt | 32481 | 32481-Deleti | Deletion | 1 | A | - | 3 | 21827 | 0.01374445 | 46.6666667 | rnr1Y285F | FALSE | CAN | JSY3811 | 4 | rnr1Y285F-R | TAA | nope | >1bp-ins | A/T-1 | none | A/T-1 | AAGGTTAAG | AAGATAC | ATT | ATT | T | none | T>none | T>none_ATT | A/T-1_AAGATAC | 8 | FALSE | FALSE |
| 45_v.txt | 32490 | 32490-Deleti | Deletion | 1 | A | - | 12 | 21827 | 0.05497778 | 42.3333333 | rnr1Y285F | FALSE | CAN | JSY3811 | 4 | rnr1Y285F-R | GAA | nope | >1bp-ins | A/T-1 | none | A/T-1 | ATACGGAAA | AAAACAA | CTT | CTT | T | none | T>none | T>none_CTT | A/T-1_AAAACAA | 8 | FALSE | FALSE |
| 45_v.txt | 32498 | 32498-Deleti | Deletion | 1 | T | - | 85 | 21827 | 0.38942594 | 44.9882344 | rnr1Y285F | FALSE | CAN | JSY3811 | 4 | rnr1Y285F-R | CTT | nope | >1bp-ins | A/T-1 | none | A/T-1 | AACAACTTTT | TTTTTTG | GAA | CTT | T | none | nope | nope_CTT | A/T-1_TTTTTTG | 8 | FALSE | FALSE |
| 45_v.txt | 32522 | 32522-Deleti | Deletion | 1 | T | - | 16 | 21827 | 0.07330371 | 41.062498 | rnr1Y285F | FALSE | CAN | JSY3811 | 4 | rnr1Y285F-R | ATT | nope | >1bp-ins | A/T-1 | none | A/T-1 | AACGGATTT | TTTTCTG | TAA | ATT | T | none | nope | nope_ATT | A/T-1_TTTTCTG | 8 | FALSE | FALSE |
| 45_v.txt | 32528 | 32528-Deleti | Deletion | 1 | G | - | 7 | 21827 | 0.03207037 | 37.4285714 | rnr1Y285F | FALSE | CAN | JSY3811 | 4 | rnr1Y285F-R | TGG | nope | >1bp-ins | G/C-1 | none | G/C-1 | TTTTCTGGGG | GGGGTTT | ACC | ACC | C | none | C>none | C>none_ACC | G/C-1_GGGGTTT | 8 | FALSE | FALSE |
| 45_v.txt | 32608 | 32608-Deleti | Deletion | 1 | C | - | 15 | 13851 | 0.10829543 | 41.0666667 | rnr1Y285F | FALSE | CAN | JSY3811 | 4 | rnr1Y285F-R | ACC | nope | >1bp-ins | G/C-1 | none | G/C-1 | AGGAAACCC | CCCAACC | TGG | ACC | C | none | nope | nope_ACC | G/C-1_CCCAACC | 8 | FALSE | FALSE |
| 45_v.txt | 32623 | 32623-Deleti | Deletion | 1 | T | - | 14 | 13851 | 0.10107574 | 43 | rnr1Y285F | FALSE | CAN | JSY3811 | 4 | rnr1Y285F-R | CTC | nope | >1bp-ins | A/T-1 | none | A/T-1 | AGAACCTCC | TCCCTTC | GAG | CTC | T | none | nope | nope_CTC | A/T-1_TCCCTTC | 8 | FALSE | FALSE |
| 45_v.txt | 32655 | 32655-Deleti | Deletion | 1 | G | - | 31 | 13851 | 0.22381056 | 58.2258065 | rnr1Y285F | FALSE | CAN | JSY3811 | 4 | rnr1Y285F-R | TGG | nope | >1bp-ins | G/C-1 | none | G/C-1 | ATACCTGGA | GGACCCC | ACC | ACC | C | none | C>none | C>none_ACC | G/C-1_GGACCCC | 8 | FALSE | FALSE |
| 45_v.txt | 32658 | 32658-Deleti | Deletion | 1 | C | - | 58 | 13851 | 0.41874233 | 59.6896552 | rnr1Y285F | FALSE | CAN | JSY3811 | 4 | rnr1Y285F-R | ACC | nope | >1bp-ins | G/C-1 | none | G/C-1 | CCTGGACCC | CCCCAGG | TGG | ACC | C | none | nope | nope_ACC | G/C-1_CCCCAGG | 8 | FALSE | FALSE |
| 45_v.txt | 32670 | 32670-Deleti | Deletion | 1 | G | - | 36 | 13851 | 0.25990903 | 61.4722222 | rnr1Y285F | FALSE | CAN | JSY3811 | 4 | rnr1Y285F-R | TGG | nope | >1bp-ins | G/C-1 | none | G/C-1 | GCACCTGGG | GGGTTTC | ACC | ACC | C | none | C>none | C>none_ACC | G/C-1_GGGTTTC | 8 | FALSE | FALSE |
| 45_v.txt | 32707 | 32707-Deleti | Deletion | 1 | C | - | 24 | 13851 | 0.17327269 | 60.625 | rnr1Y285F | FALSE | CAN | JSY3811 | 4 | rnr1Y285F-R | ACC | nope | >1bp-ins | G/C-1 | none | G/C-1 | CGGTAACCC | CCCCAGC | TGG | ACC | C | none | nope | nope_ACC | G/C-1_CCCCAGC | 8 | FALSE | FALSE |
| 45_v.txt | 32842 | 32842-Deleti | Deletion | 1 | A | - | 50 | 13851 | 0.36098477 | 42.3200011 | rnr1Y285F | FALSE | CAN | JSY3811 | 4 | rnr1Y285F-R | CAA | nope | >1bp-ins | A/T-1 | none | A/T-1 | TTACCCAAA | AAAAAAT | GTT | GTT | T | none | T>none | T>none_GTT | A/T-1_AAAAAAT | 8 | FALSE | FALSE |
| 45_v.txt | 32884 | 32884-Deleti | Deletion | 1 | A | - | 4 | 19784 | 0.02021836 | 38 | rnr1Y285F | FALSE | CAN | JSY3811 | 4 | rnr1Y285F-R | CAA | nope | >1bp-ins | A/T-1 | none | A/T-1 | ACGTCCAAA | AAAATTG | GTT | GTT | T | none | T>none | T>none_GTT | A/T-1_AAAATTG | 8 | FALSE | FALSE |
| 45_v.txt | 32901 | 32901-Deleti | Deletion | 1 | C | - | 1 | 19786 | 5.05E-03 | 38 | rnr1Y285F | FALSE | CAN | JSY3811 | 4 | rnr1Y285F-R | GCC | nope | >1bp-ins | G/C-1 | none | G/C-1 | ACTTGGCCA | CCAACTA | CGG | GCC | C | none | nope | nope_GCC | G/C-1_CCAACTA | 8 | FALSE | FALSE |
| 45_v.txt | 32915 | 32915-Deleti | Deletion | 1 | T | - | 14 | 19786 | 0.0707571 | 49.0714286 | rnr1Y285F | FALSE | CAN | JSY3811 | 4 | rnr1Y285F-R | GTT | nope | >1bp-ins | A/T-1 | none | A/T-1 | ACTAAGTTC | TTCCAGG | CAA | GTT | T | none | nope | nope_GTT | A/T-1_TTCCAGG | 8 | FALSE | FALSE |
| 45_v.txt | 32924 | 32924-Deleti | Deletion | 1 | A | - | 22 | 19786 | 0.11118973 | 56.6363636 | rnr1Y285F | FALSE | CAN | JSY3811 | 4 | rnr1Y285F-R | CAA | nope | >1bp-ins | A/T-1 | none | A/T-1 | CAGGGCAAA | AAAAGTG | GTT | GTT | T | none | T>none | T>none_GTT | A/T-1_AAAAGTG | 8 | FALSE | FALSE |
| 45_v.txt | 32935 | 32935-Deleti | Deletion | 1 | C | - | 48 | 19786 | 0.24259578 | 56.5833324 | rnr1Y285F | FALSE | CAN | JSY3811 | 4 | rnr1Y285F-R | GCC | nope | >1bp-ins | G/C-1 | none | G/C-1 | TGATTGCCCA | CCCAAGA | CGG | GCC | C | none | nope | nope_GCC | G/C-1_CCCAAGA | 8 | FALSE | FALSE |
| 45_v.txt | 32949 | 32949-Deleti | Deletion | 1 | T | - | 5 | 19786 | 0.02527039 | 64 | rnr1Y285F | FALSE | CAN | JSY3811 | 4 | rnr1Y285F-R | ATA | nope | >1bp-ins | A/T-1 | none | A/T-1 | AACCAATAC | TACATGT | TAT | ATA | T | none | nope | nope_ATA | A/T-1_TACATGT | 8 | FALSE | FALSE |
| 45_v.txt | 33147 | 33147-Deleti | Deletion | 1 | A | - | 5 | 19786 | 0.02527039 | 43.2 | rnr1Y285F | FALSE | CAN | JSY3811 | 4 | rnr1Y285F-R | GAA | nope | >1bp-ins | A/T-1 | none | A/T-1 | CCAATGAAA | AAAAGAC | CTT | CTT | T | none | T>none | T>none_CTT | A/T-1_AAAAGAC | 8 | FALSE | FALSE |
| 45_v.txt | 33367 | 33367-Deleti | Deletion | 1 | G | - | 3 | 11670 | 0.02570694 | 64 | rnr1Y285F | FALSE | CAN | JSY3811 | 4 | rnr1Y285F-R | TGG | nope | >1bp-ins | G/C-1 | none | G/C-1 | GTCTGTGGT | GGTGTGT | ACC | ACC | C | none | C>none | C>none_ACC | G/C-1_GGTGTGT | 8 | FALSE | FALSE |
| 45_v.txt | 33474 | 33474-Deleti | Deletion | 1 | C | - | 31 | 11670 | 0.26563839 | 28.1935484 | rnr1Y285F | FALSE | CAN | JSY3811 | 4 | rnr1Y285F-R | GCC | nope | >1bp-ins | G/C-1 | none | G/C-1 | GCTATGCCTT | CCTTTTT | CGG | GCC | C | none | nope | nope_GCC | G/C-1_CCTTTTT | 8 | FALSE | FALSE |
| 161_v.txt | 32256 | 32256-SNV-/ | SNV | 1 | A | C | 14 | 19760 | 0.0708502 | 24.0714286 | rnr1Y285F | FALSE | CAN | JSY3865 | 5 | rnr1Y285F-R | TAG | A>C | G/C+1 | A/T-1 | TA>GC | TA>GC | TTTGATAGA | AGACCAA | ATC | ATC | T | G | T>G | T>G_ATC | TA>GC_AGACCAA | 8 | FALSE | FALSE |
| 161_v.txt | 32283 | 32283-SNV-T | SNV | 1 | T | G | 14 | 19776 | 0.07079288 | 52.1428571 | rnr1Y285F | FALSE | CAN | JSY3865 | 5 | rnr1Y285F-R | GTA | T>G | G/C+1 | A/T-1 | TA>GC | TA>GC | CCAACGTAA | TAAATAT | CAT | GTA | T | C | T>G | T>G_GTA | TA>GC_TAAATAT | 8 | FALSE | FALSE |
| 161_v.txt | 32445 | 32445-SNV-/ | SNV | 1 | A | C | 11 | 19776 | 0.05562298 | 51.5454546 | rnr1Y285F | FALSE | CAN | JSY3865 | 5 | rnr1Y285F-R | AAG | A>C | G/C+1 | A/T-1 | TA>GC | TA>GC | ACTAAAAGT | AGTCCAA | TTC | TTC | T | G | T>G | T>G_TTC | TA>GC_AGTCCAA | 8 | FALSE | FALSE |
| 161_v.txt | 32631 | 32631-SNV-T | SNV | 1 | T | G | 3 | 21855 | 0.01372684 | 55.3333333 | rnr1Y285F | FALSE | CAN | JSY3865 | 5 | rnr1Y285F-R | GTT | T>G | G/C+1 | A/T-1 | TA>GC | TA>GC | CCTTCGTTT | TTTTTAT | CAA | GTT | T | C | T>G | T>G_GTT | TA>GC_TTTTTAT | 8 | FALSE | FALSE |
| 161_v.txt | 32719 | 32719-SNV-/ | SNV | 1 | A | C | 5 | 21854 | 0.02287911 | 21.6476031 | rnr1Y285F | FALSE | CAN | JSY3865 | 5 | rnr1Y285F-R | CAA | A>C | G/C+1 | A/T-1 | TA>GC | TA>GC | CACCACAAA | AAACCAT | GTT | GTT | T | G | T>G | T>G_GTT | TA>GC_AAACCAT | 8 | FALSE | FALSE |
| 161_v.txt | 32721 | 32721-SNV-/ | SNV | 1 | A | C | 5 | 21854 | 0.02287911 | 39.2000603 | rnr1Y285F | FALSE | CAN | JSY3865 | 5 | rnr1Y285F-R | AAC | A>C | G/C+1 | A/T-1 | TA>GC | TA>GC | CCACAAACC | ACCATAC | TTG | TTG | T | G | T>G | T>G_TTG | TA>GC_ACCATAC | 8 | FALSE | FALSE |
| 161_v.txt | 32763 | 32763-SNV-/ | SNV | 1 | A | C | 11 | 21855 | 0.05033173 | 46.7272727 | rnr1Y285F | FALSE | CAN | JSY3865 | 5 | rnr1Y285F-R | AAC | A>C | G/C+1 | A/T-1 | TA>GC | TA>GC | GCTAAAACT | ACTTTGA | TTG | TTG | T | G | T>G | T>G_TTG | TA>GC_ACTTTGA | 8 | FALSE | FALSE |
| 161_v.txt | 32842 | 32842-SNV-/ | SNV | 1 | A | C | 2 | 21855 | 9.15E-03 | 37.9983553 | rnr1Y285F | FALSE | CAN | JSY3865 | 5 | rnr1Y285F-R | CAA | A>C | G/C+1 | A/T-1 | TA>GC | TA>GC | TTACCCAAA | AAAAAAT | GTT | GTT | T | G | T>G | T>G_GTT | TA>GC_AAAAAAT | 8 | FALSE | FALSE |
| 161_v.txt | 33269 | 33269-SNV-/ | SNV | 1 | A | C | 115 | 15799 | 0.72789417 | 15.9478261 | rnr1Y285F | FALSE | CAN | JSY3865 | 5 | rnr1Y285F-R | TAG | A>C | G/C+1 | A/T-1 | TA>GC | TA>GC | CTCCATAGA | AGAGAAC | ATC | ATC | T | G | T>G | T>G_ATC | TA>GC_AGAGAAC | 8 | FALSE | FALSE |
| 161_v.txt | 32055 | 32055-SNV-T | SNV | 1 | T | C | 14 | 18806 | 0.07444433 | 60.5714286 | rnr1Y285F | FALSE | CAN | JSY3865 | 5 | rnr1Y285F-R | GTG | T>C | G/C+1 | A/T-1 | TA>CG | TA>CG | CTGATGTGC | TGCGAGA | CAC | GTG | T | G | T>C | T>C_GTG | TA>CG_TGCGAGA | 8 | FALSE | FALSE |
| 161_v.txt | 32338 | 32338-SNV-T | SNV | 1 | T | C | 2 | 19776 | 0.01011327 | 61.9997496 | rnr1Y285F | FALSE | CAN | JSY3865 | 5 | rnr1Y285F-R | ATA | T>C | G/C+1 | A/T-1 | TA>CG | TA>CG | TGAAGATAT | TATGTGG | TAT | ATA | T | G | T>C | T>C_ATA | TA>CG_TATGTGG | 8 | FALSE | FALSE |
| 161_v.txt | 32719 | 32719-SNV-/ | SNV | 1 | A | G | 9 | 21854 | 0.04118239 | 37.7651106 | rnr1Y285F | FALSE | CAN | JSY3865 | 5 | rnr1Y285F-R | CAA | A>G | G/C+1 | A/T-1 | TA>CG | TA>CG | CACCACAAA | AAACCAT | GTT | GTT | T | C | T>C | T>C_GTT | TA>CG_AAACCAT | 8 | FALSE | FALSE |
| 161_v.txt | 32842 | 32842-SNV-/ | SNV | 1 | A | G | 2 | 21855 | 9.15E-03 | 37.9982323 | rnr1Y285F | FALSE | CAN | JSY3865 | 5 | rnr1Y285F-R | CAA | A>G | G/C+1 | A/T-1 | TA>CG | TA>CG | TTACCCAAA | AAAAAAT | GTT | GTT | T | C | T>C | T>C_GTT | TA>CG_AAAAAAT | 8 | FALSE | FALSE |
| 161_v.txt | 33042 | 33042-SNV-T | SNV | 1 | T | C | 13 | 14700 | 0.08843537 | 55.3846154 | rnr1Y285F | FALSE | CAN | JSY3865 | 5 | rnr1Y285F-R | TTC | T>C | G/C+1 | A/T-1 | TA>CG | TA>CG | GCCATTTCAC | TCACCCA | AAG | TTC | T | G | T>C | T>C_TTC | TA>CG_TCACCCA | 8 | FALSE | FALSE |
| 161_v.txt | 33126 | 33126-SNV-/ | SNV | 1 | A | G | 4 | 14700 | 0.02721088 | 53.4987534 | rnr1Y285F | FALSE | CAN | JSY3865 | 5 | rnr1Y285F-R | CAG | A>G | G/C+1 | A/T-1 | TA>CG | TA>CG | TTGGTCAGA | AGAGGTG | GTC | GTC | T | C | T>C | T>C_GTC | TA>CG_AGAGGTG | 8 | FALSE | FALSE |
| 161_v.txt | 33211 | 33211-SNV-T | SNV | 1 | T | C | 19 | 15799 | 0.12026078 | 36.4210526 | rnr1Y285F | FALSE | CAN | JSY3865 | 5 | rnr1Y285F-R | CTC | T>C | G/C+1 | A/T-1 | TA>CG | TA>CG | GCTCTCTCTT | TCTTCAC | GAG | CTC | T | G | T>C | T>C_CTC | TA>CG_TCTTCAC | 8 | FALSE | FALSE |
| 161_v.txt | 33476 | 33476-SNV-T | SNV | 1 | T | C | 19 | 15797 | 0.120276 | 33.4736842 | rnr1Y285F | FALSE | CAN | JSY3865 | 5 | rnr1Y285F-R | CTT | T>C | G/C+1 | A/T-1 | TA>CG | TA>CG | TATGCCTTTT | TTTTTTT | GAA | CTT | T | G | T>C | T>C_CTT | TA>CG_TTTTTTT | 8 | FALSE | FALSE |
| 161_v.txt | 32265 | 32265-SNV-/ | SNV | 1 | A | T | 15 | 19776 | 0.07584952 | 36.9333333 | rnr1Y285F | FALSE | CAN | JSY3865 | 5 | rnr1Y285F-R | TAA | A>T | A/T+1 | A/T-1 | TA>AT | TA>AT | CCAAATAAA | AAAATAC | ATT | ATT | T | A | T>A | T>A_ATT | TA>AT_AAAATAC | 8 | FALSE | FALSE |

| | | | | | | | | | | | | | | | | | | | | | | | | | | | | | | | | | | |
|---|---|---|---|---|---|---|---|---|---|---|---|---|---|---|---|---|---|---|---|---|---|---|---|---|---|---|---|---|---|---|---|---|---|---|
| 161_v.txt | 32314 | 32314-SNV-T | SNV | 1 | T | A | 2 | 19776 | 0.01011327 | 57.978323 | rnr1Y285F | FALSE | CAN | JSY3865 | 5 | rnr1Y285F-R | GTT | T>A | A/T+1 | A/T-1 | TA>AT | TA>AT | TAATGGTTG | TTGTTAA | CAA | GTT | T | T | T>A | T>A_GTT | TA>AT_TTGTTAA | 8 | FALSE | FALSE |
| 161_v.txt | 32337 | 32337-SNV- | SNV | 1 | A | T | 1 | 19776 | 5.06E-03 | 63.9996364 | rnr1Y285F | FALSE | CAN | JSY3865 | 5 | rnr1Y285F-R | GAT | A>T | A/T+1 | A/T-1 | TA>AT | TA>AT | TTGAAGATA | ATATGTG | CTA | CTA | T | A | T>A | T>A_CTA | TA>AT_ATATGTG | 8 | FALSE | FALSE |
| 161_v.txt | 32338 | 32338-SNV-T | SNV | 1 | T | A | 4 | 19776 | 0.02022654 | 63.9981092 | rnr1Y285F | FALSE | CAN | JSY3865 | 5 | rnr1Y285F-R | ATA | T>A | A/T+1 | A/T-1 | TA>AT | TA>AT | TGAAGATAT | TATGTGG | TAT | ATA | T | T | T>A | T>A_ATA | TA>AT_TATGTGG | 8 | FALSE | FALSE |
| 161_v.txt | 32346 | 32346-SNV- | SNV | 1 | A | T | 394 | 19776 | 1.99231392 | 61.5812183 | rnr1Y285F | FALSE | CAN | JSY3865 | 5 | rnr1Y285F-R | CAA | A>T | A/T+1 | A/T-1 | TA>AT | TA>AT | TGTGGCAAA | AAAACCT | GTT | GTT | T | A | T>A | T>A_GTT | TA>AT_AAAACCT | 8 | FALSE | FALSE |
| 161_v.txt | 32496 | 32496-SNV- | SNV | 1 | A | T | 10 | 19776 | 0.05056634 | 43.1 | rnr1Y285F | FALSE | CAN | JSY3865 | 5 | rnr1Y285F-R | AAC | A>T | A/T+1 | A/T-1 | TA>AT | TA>AT | AAAACAACT | ACTTTTT | TTG | TTG | T | A | T>A | T>A_TTG | TA>AT_ACTTTTT | 8 | FALSE | FALSE |
| 161_v.txt | 32801 | 32801-SNV- | SNV | 1 | A | T | 30 | 21855 | 0.13726836 | 45.1666667 | rnr1Y285F | FALSE | CAN | JSY3865 | 5 | rnr1Y285F-R | AAT | A>T | A/T+1 | A/T-1 | TA>AT | TA>AT | ACCGTAATA | ATATTTG | TTA | TTA | T | A | T>A | T>A_TTA | TA>AT_ATATTTG | 8 | FALSE | FALSE |
| 161_v.txt | 32992 | 32992-SNV-T | SNV | 1 | T | A | 17 | 14700 | 0.11564626 | 56.8823529 | rnr1Y285F | FALSE | CAN | JSY3865 | 5 | rnr1Y285F-R | CTT | T>A | A/T+1 | A/T-1 | TA>AT | TA>AT | GGAATCTTT | TTTGTGA | GAA | CTT | T | T | T>A | T>A_CTT | TA>AT_TTTGTGA | 8 | FALSE | FALSE |
| 161_v.txt | 33060 | 33060-SNV- | SNV | 1 | A | T | 10 | 14700 | 0.06802721 | 61.4 | rnr1Y285F | FALSE | CAN | JSY3865 | 5 | rnr1Y285F-R | GAC | A>T | A/T+1 | A/T-1 | TA>AT | TA>AT | TGCGTGACA | ACAGAAT | CTG | CTG | T | A | T>A | T>A_CTG | TA>AT_ACAGAAT | 8 | FALSE | FALSE |
| 161_v.txt | 33066 | 33066-SNV-T | SNV | 1 | T | A | 1 | 14700 | 6.80E-03 | 63.9970173 | rnr1Y285F | FALSE | CAN | JSY3865 | 5 | rnr1Y285F-R | ATA | T>A | A/T+1 | A/T-1 | TA>AT | TA>AT | ACAGAATAT | TATGCCA | TAT | ATA | T | T | T>A | T>A_ATA | TA>AT_TATGCCA | 8 | FALSE | FALSE |
| 161_v.txt | 33312 | 33312-SNV- | SNV | 1 | A | T | 16 | 15799 | 0.10127223 | 59.1875 | rnr1Y285F | FALSE | CAN | JSY3865 | 5 | rnr1Y285F-R | CAA | A>T | A/T+1 | A/T-1 | TA>AT | TA>AT | GAGCGCAAT | AATGGAT | GTT | GTT | T | A | T>A | T>A_GTT | TA>AT_AATGGAT | 8 | FALSE | FALSE |
| 161_v.txt | 33317 | 33317-SNV- | SNV | 1 | A | T | 25 | 15799 | 0.15823786 | 58.92 | rnr1Y285F | FALSE | CAN | JSY3865 | 5 | rnr1Y285F-R | GAT | A>T | A/T+1 | A/T-1 | TA>AT | TA>AT | CAATGGATA | ATACAAT | CTA | CTA | T | A | T>A | T>A_CTA | TA>AT_ATACAAT | 8 | FALSE | FALSE |
| 161_v.txt | 31844 | 31844-SNV- | SNV | 1 | C | T | 11 | 3276 | 0.33577534 | 47.3636364 | rnr1Y285F | FALSE | CAN | JSY3865 | 5 | rnr1Y285F-R | TCC | C>T | A/T+1 | G/C-1 | CG>TA | CG>TA | TAAGATCCA | CCAAACA | AGG | TCC | C | A | C>T | C>T_TCC | CG>TA_CCAAACA | 8 | FALSE | FALSE |
| 161_v.txt | 31920 | 31920-SNV- | SNV | 1 | G | A | 9 | 3276 | 0.27472528 | 36.4444444 | rnr1Y285F | FALSE | CAN | JSY3865 | 5 | rnr1Y285F-R | CGT | G>A | A/T+1 | G/C-1 | CG>TA | CG>TA | AAAGCCGTG | GTGAAAC | GCA | GCA | C | T | C>T | C>T_GCA | CG>TA_GTGAAAC | 8 | FALSE | FALSE |
| 161_v.txt | 31962 | 31962-SNV- | SNV | 1 | G | A | 24 | 18806 | 0.12761885 | 36.4166667 | rnr1Y285F | FALSE | CAN | JSY3865 | 5 | rnr1Y285F-R | CGC | G>A | A/T+1 | G/C-1 | CG>TA | CG>TA | GTGGCCGCA | GCATAAT | GCG | GCG | C | T | C>T | C>T_GCG | CG>TA_GCATAAT | 8 | FALSE | FALSE |
| 161_v.txt | 31972 | 31972-SNV- | SNV | 1 | C | T | 18 | 18806 | 0.09571413 | 36.6111111 | rnr1Y285F | FALSE | CAN | JSY3865 | 5 | rnr1Y285F-R | GCC | C>T | A/T+1 | G/C-1 | CG>TA | CG>TA | AATAAGCCA | CCAAGCC | CGG | GCC | C | A | C>T | C>T_GCC | CG>TA_CCAAGCC | 8 | FALSE | FALSE |
| 161_v.txt | 31980 | 31980-SNV- | SNV | 1 | G | A | 7 | 18806 | 0.03722216 | 43.4285714 | rnr1Y285F | FALSE | CAN | JSY3865 | 5 | rnr1Y285F-R | GGG | G>A | A/T+1 | G/C-1 | CG>TA | CG>TA | AAGCCGGGC | GGCATTA | CCC | CCC | C | T | C>T | C>T_CCC | CG>TA_GGCATTA | 8 | FALSE | FALSE |
| 161_v.txt | 31981 | 31981-SNV- | SNV | 1 | G | A | 68 | 18806 | 0.36158673 | 50.4411765 | rnr1Y285F | FALSE | CAN | JSY3865 | 5 | rnr1Y285F-R | GGC | G>A | A/T+1 | G/C-1 | CG>TA | CG>TA | AGCCGGGCA | GCATTAA | CCG | CCG | C | T | C>T | C>T_CCG | CG>TA_GCATTAA | 8 | FALSE | FALSE |
| 161_v.txt | 32009 | 32009-SNV- | SNV | 1 | G | A | 8 | 18806 | 0.04253962 | 47.6142049 | rnr1Y285F | FALSE | CAN | JSY3865 | 5 | rnr1Y285F-R | CGT | G>A | A/T+1 | G/C-1 | CG>TA | CG>TA | TAACTCGTC | GTCACGA | GCA | GCA | C | T | C>T | C>T_GCA | CG>TA_GTCACGA | 8 | FALSE | FALSE |
| 161_v.txt | 32057 | 32057-SNV- | SNV | 1 | C | T | 19 | 18806 | 0.10103159 | 59 | rnr1Y285F | FALSE | CAN | JSY3865 | 5 | rnr1Y285F-R | GCG | C>T | A/T+1 | G/C-1 | CG>TA | CG>TA | GATGTGCGA | CGAGATT | CGC | GCG | C | A | C>T | C>T_GCG | CG>TA_CGAGATT | 8 | FALSE | FALSE |
| 161_v.txt | 32092 | 32092-SNV- | SNV | 1 | C | T | 26 | 18806 | 0.13825375 | 57.8461539 | rnr1Y285F | FALSE | CAN | JSY3865 | 5 | rnr1Y285F-R | GCA | C>T | A/T+1 | G/C-1 | CG>TA | CG>TA | AGCCTGCAA | CAACACC | CGT | GCA | C | A | C>T | C>T_GCA | CG>TA_CAACACC | 8 | FALSE | FALSE |
| 161_v.txt | 32114 | 32114-SNV- | SNV | 1 | C | T | 221 | 18806 | 1.17515687 | 59.2895928 | rnr1Y285F | FALSE | CAN | JSY3865 | 5 | rnr1Y285F-R | GCC | C>T | A/T+1 | G/C-1 | CG>TA | CG>TA | TAATAGCCA | CCATTCG | CGG | GCC | C | A | C>T | C>T_GCC | CG>TA_CCATTCG | 8 | FALSE | FALSE |
| 161_v.txt | 32165 | 32165-SNV- | SNV | 1 | G | A | 19 | 18806 | 0.10103159 | 48.631579 | rnr1Y285F | FALSE | CAN | JSY3865 | 5 | rnr1Y285F-R | CGC | G>A | A/T+1 | G/C-1 | CG>TA | CG>TA | CAAAGCGCC | GCCAAAT | GCG | GCG | C | T | C>T | C>T_GCG | CG>TA_GCCAAAT | 8 | FALSE | FALSE |
| 161_v.txt | 32271 | 32271-SNV- | SNV | 1 | C | T | 37 | 19776 | 0.18709547 | 32.1891892 | rnr1Y285F | FALSE | CAN | JSY3865 | 5 | rnr1Y285F-R | ACG | C>T | A/T+1 | G/C-1 | CG>TA | CG>TA | AAAATACGG | CGGGAAC | TGC | ACG | C | A | C>T | C>T_ACG | CG>TA_CGGGAAC | 8 | FALSE | FALSE |
| 161_v.txt | 32274 | 32274-SNV- | SNV | 1 | G | A | 61 | 19776 | 0.30845469 | 35.6885246 | rnr1Y285F | FALSE | CAN | JSY3865 | 5 | rnr1Y285F-R | GGA | G>A | A/T+1 | G/C-1 | CG>TA | CG>TA | ATACGGGAA | GAACCAA | CCT | CCT | C | T | C>T | C>T_CCT | CG>TA_GAACCAA | 8 | FALSE | FALSE |
| 161_v.txt | 32299 | 32299-SNV- | SNV | 1 | C | T | 34 | 19776 | 0.17192557 | 51.2352941 | rnr1Y285F | FALSE | CAN | JSY3865 | 5 | rnr1Y285F-R | GCG | C>T | A/T+1 | G/C-1 | CG>TA | CG>TA | AATTTGCGG | CGGCAGA | CGC | GCG | C | A | C>T | C>T_GCG | CG>TA_CGGCAGA | 8 | FALSE | FALSE |
| 161_v.txt | 32301 | 32301-SNV- | SNV | 1 | G | A | 2277 | 19776 | 11.5139563 | 56.7689943 | rnr1Y285F | FALSE | CAN | JSY3865 | 5 | rnr1Y285F-R | GGC | G>A | A/T+1 | G/C-1 | CG>TA | CG>TA | TTTGCGGCA | GCAGAAA | CCG | CCG | C | T | C>T | C>T_CCG | CG>TA_GCAGAAA | 8 | FALSE | FALSE |
| 161_v.txt | 32449 | 32449-SNV- | SNV | 1 | C | T | 28 | 19776 | 0.14158576 | 54.2857143 | rnr1Y285F | FALSE | CAN | JSY3865 | 5 | rnr1Y285F-R | CCA | C>T | A/T+1 | G/C-1 | CG>TA | CG>TA | AAAGTCCAA | CAATGAA | GGT | CCA | C | A | C>T | C>T_CCA | CG>TA_CAATGAA | 8 | FALSE | FALSE |
| 161_v.txt | 32466 | 32466-SNV- | SNV | 1 | C | T | 27 | 19776 | 0.13652913 | 46.2222222 | rnr1Y285F | FALSE | CAN | JSY3865 | 5 | rnr1Y285F-R | GCC | C>T | A/T+1 | G/C-1 | CG>TA | CG>TA | AGAGAGCCA | CCAATGT | CGG | GCC | C | A | C>T | C>T_GCC | CG>TA_CCAATGT | 8 | FALSE | FALSE |
| 161_v.txt | 32487 | 32487-SNV- | SNV | 1 | C | T | 22 | 19776 | 0.11124596 | 43.2720926 | rnr1Y285F | FALSE | CAN | JSY3865 | 5 | rnr1Y285F-R | ACG | C>T | A/T+1 | G/C-1 | CG>TA | CG>TA | AAGATACGG | CGGAAAA | TGC | ACG | C | A | C>T | C>T_ACG | CG>TA_CGGAAAA | 8 | FALSE | FALSE |
| 161_v.txt | 32508 | 32508-SNV- | SNV | 1 | G | A | 16 | 19776 | 0.08090615 | 36.5625 | rnr1Y285F | FALSE | CAN | JSY3865 | 5 | rnr1Y285F-R | GGC | G>A | A/T+1 | G/C-1 | CG>TA | CG>TA | TTGATGGCT | GCTCTTG | CCG | CCG | C | T | C>T | C>T_CCG | CG>TA_GCTCTTG | 8 | FALSE | FALSE |
| 161_v.txt | 32529 | 32529-SNV- | SNV | 1 | G | A | 32 | 19775 | 0.16182048 | 42.9052617 | rnr1Y285F | FALSE | CAN | JSY3865 | 5 | rnr1Y285F-R | GGG | G>A | A/T+1 | G/C-1 | CG>TA | CG>TA | TTTCTGGGG | GGGTTTG | CCC | CCC | C | T | C>T | C>T_CCC | CG>TA_GGGTTTG | 8 | FALSE | FALSE |
| 161_v.txt | 32530 | 32530-SNV- | SNV | 1 | G | A | 68 | 19776 | 0.34385113 | 39.4264706 | rnr1Y285F | FALSE | CAN | JSY3865 | 5 | rnr1Y285F-R | GGG | G>A | A/T+1 | G/C-1 | CG>TA | CG>TA | TTCTGGGGT | GGTTTGC | CCC | CCC | C | T | C>T | C>T_CCC | CG>TA_GGTTTGC | 8 | FALSE | FALSE |
| 161_v.txt | 32535 | 32535-SNV- | SNV | 1 | G | A | 43 | 19776 | 0.21743528 | 37.9764128 | rnr1Y285F | FALSE | CAN | JSY3865 | 5 | rnr1Y285F-R | TGC | G>A | A/T+1 | G/C-1 | CG>TA | CG>TA | GGGTTTGCA | GCAGCTT | ACG | ACG | C | T | C>T | C>T_ACG | CG>TA_GCAGCTT | 8 | FALSE | FALSE |
| 161_v.txt | 32536 | 32536-SNV- | SNV | 1 | C | T | 40 | 19776 | 0.20226537 | 36.4995066 | rnr1Y285F | FALSE | CAN | JSY3865 | 5 | rnr1Y285F-R | GCA | C>T | A/T+1 | G/C-1 | CG>TA | CG>TA | GGTTTGCAG | CAGCTTC | CGT | GCA | C | A | C>T | C>T_GCA | CG>TA_CAGCTTC | 8 | FALSE | FALSE |
| 161_v.txt | 32545 | 32545-SNV- | SNV | 1 | C | T | 25 | 19776 | 0.12641586 | 36.2 | rnr1Y285F | FALSE | CAN | JSY3865 | 5 | rnr1Y285F-R | CCA | C>T | A/T+1 | G/C-1 | CG>TA | CG>TA | CTTCACCAG | CAGCAGT | GGT | CCA | C | A | C>T | C>T_CCA | CG>TA_CAGCAGT | 8 | FALSE | FALSE |
| 161_v.txt | 32568 | 32568-SNV- | SNV | 1 | G | A | 20 | 21854 | 0.09151643 | 37.25 | rnr1Y285F | FALSE | CAN | JSY3865 | 5 | rnr1Y285F-R | AGT | G>A | A/T+1 | G/C-1 | CG>TA | CG>TA | AGTTCAGTA | GTACCTT | TCA | TCA | C | T | C>T | C>T_TCA | CG>TA_GTACCTT | 8 | FALSE | FALSE |
| 161_v.txt | 32571 | 32571-SNV- | SNV | 1 | C | T | 23 | 21855 | 0.10523908 | 33.6086957 | rnr1Y285F | FALSE | CAN | JSY3865 | 5 | rnr1Y285F-R | ACC | C>T | A/T+1 | G/C-1 | CG>TA | CG>TA | TCAGTACCT | CCTTGAA | TGG | ACC | C | A | C>T | C>T_ACC | CG>TA_CCTTGAA | 8 | FALSE | FALSE |
| 161_v.txt | 32608 | 32608-SNV- | SNV | 1 | C | T | 7 | 21855 | 0.03202928 | 46.0000613 | rnr1Y285F | FALSE | CAN | JSY3865 | 5 | rnr1Y285F-R | ACC | C>T | A/T+1 | G/C-1 | CG>TA | CG>TA | AGGAAACCC | CCCAACC | TGG | ACC | C | A | C>T | C>T_ACC | CG>TA_CCCAACC | 8 | FALSE | FALSE |
| 161_v.txt | 32609 | 32609-SNV- | SNV | 1 | C | T | 100 | 21855 | 0.4575612 | 46.5597082 | rnr1Y285F | FALSE | CAN | JSY3865 | 5 | rnr1Y285F-R | CCC | C>T | A/T+1 | G/C-1 | CG>TA | CG>TA | GGAAACCCA | CCAACCT | GGG | CCC | C | A | C>T | C>T_CCC | CG>TA_CCAACCT | 8 | FALSE | FALSE |
| 161_v.txt | 32749 | 32749-SNV- | SNV | 1 | C | T | 41 | 21855 | 0.18760009 | 54.8048781 | rnr1Y285F | FALSE | CAN | JSY3865 | 5 | rnr1Y285F-R | CCG | C>T | A/T+1 | G/C-1 | CG>TA | CG>TA | GAAACCCGA | CGATAAT | GGC | CCG | C | A | C>T | C>T_CCG | CG>TA_CGATAAT | 8 | FALSE | FALSE |
| 161_v.txt | 32796 | 32796-SNV- | SNV | 1 | C | T | 298 | 21855 | 1.36353237 | 45.9362416 | rnr1Y285F | FALSE | CAN | JSY3865 | 5 | rnr1Y285F-R | ACC | C>T | A/T+1 | G/C-1 | CG>TA | CG>TA | AATTCACCG | CCGTAAT | TGG | ACC | C | A | C>T | C>T_ACC | CG>TA_CCGTAAT | 8 | FALSE | FALSE |
| 161_v.txt | 32798 | 32798-SNV- | SNV | 1 | G | A | 27 | 21855 | 0.12354152 | 46.7407407 | rnr1Y285F | FALSE | CAN | JSY3865 | 5 | rnr1Y285F-R | CGT | G>A | A/T+1 | G/C-1 | CG>TA | CG>TA | TTCACCGTA | GTAATAT | GCA | GCA | C | T | C>T | C>T_GCA | CG>TA_GTAATAT | 8 | FALSE | FALSE |
| 161_v.txt | 32811 | 32811-SNV- | SNV | 1 | G | A | 47 | 21855 | 0.21505376 | 44.5957447 | rnr1Y285F | FALSE | CAN | JSY3865 | 5 | rnr1Y285F-R | AGG | G>A | A/T+1 | G/C-1 | CG>TA | CG>TA | TTGACAGGG | GGGAACA | TCC | TCC | C | T | C>T | C>T_TCC | CG>TA_GGGAACA | 8 | FALSE | FALSE |
| 161_v.txt | 32917 | 32917-SNV- | SNV | 1 | C | T | 33 | 14700 | 0.2244898 | 45.1818182 | rnr1Y285F | FALSE | CAN | JSY3865 | 5 | rnr1Y285F-R | TCC | C>T | A/T+1 | G/C-1 | CG>TA | CG>TA | TAAGTTCCA | CCAGGGC | AGG | TCC | C | A | C>T | C>T_TCC | CG>TA_CCAGGGC | 8 | FALSE | FALSE |
| 161_v.txt | 32958 | 32958-SNV- | SNV | 1 | C | T | 16 | 14700 | 0.10884354 | 39.1875 | rnr1Y285F | FALSE | CAN | JSY3865 | 5 | rnr1Y285F-R | ACC | C>T | A/T+1 | G/C-1 | CG>TA | CG>TA | ATGTAACCA | CCATTGG | TGG | ACC | C | A | C>T | C>T_ACC | CG>TA_CCATTGG | 8 | FALSE | FALSE |
| 161_v.txt | 32965 | 32965-SNV- | SNV | 1 | C | T | 25 | 14700 | 0.17006803 | 60 | rnr1Y285F | FALSE | CAN | JSY3865 | 5 | rnr1Y285F-R | GCC | C>T | A/T+1 | G/C-1 | CG>TA | CG>TA | CATTGGCCG | CCGCACC | CGG | GCC | C | A | C>T | C>T_GCC | CG>TA_CCGCACC | 8 | FALSE | FALSE |
| 161_v.txt | 32967 | 32967-SNV- | SNV | 1 | G | A | 34 | 14700 | 0.23129252 | 61 | rnr1Y285F | FALSE | CAN | JSY3865 | 5 | rnr1Y285F-R | CGC | G>A | A/T+1 | G/C-1 | CG>TA | CG>TA | TTGGCCGCA | GCACCAA | GCG | GCG | C | T | C>T | C>T_GCG | CG>TA_GCACCAA | 8 | FALSE | FALSE |
| 161_v.txt | 32970 | 32970-SNV- | SNV | 1 | C | T | 58 | 14700 | 0.39455782 | 61.7068966 | rnr1Y285F | FALSE | CAN | JSY3865 | 5 | rnr1Y285F-R | ACC | C>T | A/T+1 | G/C-1 | CG>TA | CG>TA | GCCGCACCA | CCAAATG | TGG | ACC | C | A | C>T | C>T_ACC | CG>TA_CCAAATG | 8 | FALSE | FALSE |
| 161_v.txt | 33056 | 33056-SNV- | SNV | 1 | C | T | 25 | 14700 | 0.17006803 | 56.44 | rnr1Y285F | FALSE | CAN | JSY3865 | 5 | rnr1Y285F-R | GCG | C>T | A/T+1 | G/C-1 | CG>TA | CG>TA | GGACTGCGT | CGTGACA | CGC | GCG | C | A | C>T | C>T_GCG | CG>TA_CGTGACA | 8 | FALSE | FALSE |
| 161_v.txt | 33104 | 33104-SNV- | SNV | 1 | G | A | 15 | 14700 | 0.10204082 | 46 | rnr1Y285F | FALSE | CAN | JSY3865 | 5 | rnr1Y285F-R | CGC | G>A | A/T+1 | G/C-1 | CG>TA | CG>TA | AAGAGCGCC | GCCCACT | GCG | GCG | C | T | C>T | C>T_GCG | CG>TA_GCCCACT | 8 | FALSE | FALSE |
| 161_v.txt | 33114 | 33114-SNV- | SNV | 1 | C | T | 769 | 14700 | 5.23129252 | 43.7780869 | rnr1Y285F | FALSE | CAN | JSY3865 | 5 | rnr1Y285F-R | GCC | C>T | A/T+1 | G/C-1 | CG>TA | CG>TA | ACTGGGCCG | CCGGCGT | CGG | GCC | C | A | C>T | C>T_GCC | CG>TA_CCGGCGT | 8 | FALSE | FALSE |
| 161_v.txt | 33115 | 33115-SNV- | SNV | 1 | C | T | 292 | 14700 | 1.98639456 | 44.0302369 | rnr1Y285F | FALSE | CAN | JSY3865 | 5 | rnr1Y285F-R | CCG | C>T | A/T+1 | G/C-1 | CG>TA | CG>TA | CTGGGCCGG | CGGCGTT | GGC | CCG | C | A | C>T | C>T_CCG | CG>TA_CGGCGTT | 8 | FALSE | FALSE |
| 161_v.txt | 33118 | 33118-SNV- | SNV | 1 | C | T | 21 | 14700 | 0.14285714 | 37.8571429 | rnr1Y285F | FALSE | CAN | JSY3865 | 5 | rnr1Y285F-R | GCG | C>T | A/T+1 | G/C-1 | CG>TA | CG>TA | GGCCGGCGT | CGTTGGT | CGC | GCG | C | A | C>T | C>T_GCG | CG>TA_CGTTGGT | 8 | FALSE | FALSE |
| 161_v.txt | 33276 | 33276-SNV- | SNV | 1 | G | A | 30 | 15799 | 0.18988544 | 46.1666667 | rnr1Y285F | FALSE | CAN | JSY3865 | 5 | rnr1Y285F-R | CGT | G>A | A/T+1 | G/C-1 | CG>TA | CG>TA | GAGAACGTA | GTATCCT | GCA | GCA | C | T | C>T | C>T_GCA | CG>TA_GTATCCT | 8 | FALSE | FALSE |
| 161_v.txt | 33284 | 33284-SNV- | SNV | 1 | G | A | 15 | 15799 | 0.09494272 | 44.4666667 | rnr1Y285F | FALSE | CAN | JSY3865 | 5 | rnr1Y285F-R | CGC | G>A | A/T+1 | G/C-1 | CG>TA | CG>TA | ATCCTCGCC | GCCATTT | GCG | GCG | C | T | C>T | C>T_GCG | CG>TA_GCCATTT | 8 | FALSE | FALSE |
| 161_v.txt | 33310 | 33310-SNV- | SNV | 1 | G | A | 22 | 15799 | 0.13924932 | 53.6818182 | rnr1Y285F | FALSE | CAN | JSY3865 | 5 | rnr1Y285F-R | CGC | G>A | A/T+1 | G/C-1 | CG>TA | CG>TA | AAGAGCGCA | GCAATGG | GCG | GCG | C | T | C>T | C>T_GCG | CG>TA_GCAATGG | 8 | FALSE | FALSE |
| 161_v.txt | 33412 | 33412-SNV- | SNV | 1 | G | A | 19 | 15799 | 0.12026078 | 51.5789474 | rnr1Y285F | FALSE | CAN | JSY3865 | 5 | rnr1Y285F-R | GGC | G>A | A/T+1 | G/C-1 | CG>TA | CG>TA | TGACCGGCT | GCTCATT | CCG | CCG | C | T | C>T | C>T_CCG | CG>TA_GCTCATT | 8 | FALSE | FALSE |
| 161_v.txt | 33474 | 33474-SNV- | SNV | 1 | C | T | 24 | 15798 | 0.15191796 | 35 | rnr1Y285F | FALSE | CAN | JSY3865 | 5 | rnr1Y285F-R | GCC | C>T | A/T+1 | G/C-1 | CG>TA | CG>TA | GCTATGCCT | CCTTTTT | CGG | GCC | C | A | C>T | C>T_GCC | CG>TA_CCTTTTT | 8 | FALSE | FALSE |
| 161_v.txt | 31980 | 31980-SNV- | SNV | 1 | G | C | 1 | 18806 | 5.32E-03 | 25 | rnr1Y285F | FALSE | CAN | JSY3865 | 5 | rnr1Y285F-R | GGG | G>C | G/C+1 | G/C-1 | CG>GC | CG>GC | AAGCCGGGC | GGCATTA | CCC | CCC | C | G | C>G | C>G_CCC | CG>GC_GGCATTA | 8 | FALSE | FALSE |
| 161_v.txt | 32038 | 32038-SNV- | SNV | 1 | C | G | 43 | 18806 | 0.22865043 | 56.372093 | rnr1Y285F | FALSE | CAN | JSY3865 | 5 | rnr1Y285F-R | GCT | C>G | G/C+1 | G/C-1 | CG>GC | CG>GC | TCAAAGCTT | CTTGCAT | CGA | GCT | C | C | C>G | C>G_GCT | CG>GC_CTTGCAT | 8 | FALSE | FALSE |
| 161_v.txt | 32199 | 32199-SNV- | SNV | 1 | G | C | 27 | 18806 | 0.1435712 | 42.2942299 | rnr1Y285F | FALSE | CAN | JSY3865 | 5 | rnr1Y285F-R | TGG | G>C | G/C+1 | G/C-1 | CG>GC | CG>GC | ATGTATGGA | GGAACAC | ACC | ACC | C | G | C>G | C>G_ACC | CG>GC_GGAACAC | 8 | FALSE | FALSE |
| 161_v.txt | 32200 | 32200-SNV- | SNV | 1 | G | C | 10 | 18806 | 0.05317452 | 45.3963005 | rnr1Y285F | FALSE | CAN | JSY3865 | 5 | rnr1Y285F-R | GGA | G>C | G/C+1 | G/C-1 | CG>GC | CG>GC | TGTATGGAA | GAACACC | CCT | CCT | C | G | C>G | C>G_CCT | CG>GC_GAACACC | 8 | FALSE | FALSE |
| 161_v.txt | 32223 | 32223-SNV- | SNV | 1 | G | C | 14 | 18806 | 0.07444433 | 36 | rnr1Y285F | FALSE | CAN | JSY3865 | 5 | rnr1Y285F-R | TGA | G>C | G/C+1 | G/C-1 | CG>GC | CG>GC | GTCCTTGAC | GACAGGA | ACT | ACT | C | G | C>G | C>G_ACT | CG>GC_GACAGGA | 8 | FALSE | FALSE |
| 161_v.txt | 32292 | 32292-SNV- | SNV | 1 | G | C | 20 | 19776 | 0.10113269 | 55.25 | rnr1Y285F | FALSE | CAN | JSY3865 | 5 | rnr1Y285F-R | TGA | G>C | G/C+1 | G/C-1 | CG>GC | CG>GC | ATATTTGAA | GAATTTG | ACT | ACT | C | G | C>G | C>G_ACT | CG>GC_GAATTTG | 8 | FALSE | FALSE |
| 161_v.txt | 32471 | 32471-SNV- | SNV | 1 | G | C | 67 | 19776 | 0.3387945 | 48.5074627 | rnr1Y285F | FALSE | CAN | JSY3865 | 5 | rnr1Y285F-R | TGT | G>C | G/C+1 | G/C-1 | CG>GC | CG>GC | GCCAATGTA | GTAGAAG | ACA | ACA | C | G | C>G | C>G_ACA | CG>GC_GTAGAAG | 8 | FALSE | FALSE |
| 161_v.txt | 32535 | 32535-SNV- | SNV | 1 | G | C | 1 | 19776 | 5.06E-03 | 37.9996506 | rnr1Y285F | FALSE | CAN | JSY3865 | 5 | rnr1Y285F-R | TGC | G>C | G/C+1 | G/C-1 | CG>GC | CG>GC | GGGTTTGCA | GCAGCTT | ACG | ACG | C | G | C>G | C>G_ACG | CG>GC_GCAGCTT | 8 | FALSE | FALSE |
| 161_v.txt | 32536 | 32536-SNV- | SNV | 1 | C | G | 3 | 19776 | 0.0151699 | 36.0001506 | rnr1Y285F | FALSE | CAN | JSY3865 | 5 | rnr1Y285F-R | GCA | C>G | G/C+1 | G/C-1 | CG>GC | CG>GC | GGTTTGCAG | CAGCTTC | CGT | GCA | C | C | C>G | C>G_GCA | CG>GC_CAGCTTC | 8 | FALSE | FALSE |
| 161_v.txt | 32571 | 32571-SNV- | SNV | 1 | C | G | 8 | 21855 | 0.0366049 | 37.75 | rnr1Y285F | FALSE | CAN | JSY3865 | 5 | rnr1Y285F-R | ACC | C>G | G/C+1 | G/C-1 | CG>GC | CG>GC | TCAGTACCT | CCTTGAA | TGG | ACC | C | C | C>G | C>G_ACC | CG>GC_CCTTGAA | 8 | FALSE | FALSE |
| 161_v.txt | 32670 | 32670-SNV- | SNV | 1 | G | C | 2 | 21855 | 9.15E-03 | 50 | rnr1Y285F | FALSE | CAN | JSY3865 | 5 | rnr1Y285F-R | TGG | G>C | G/C+1 | G/C-1 | CG>GC | CG>GC | GCACCTGGG | GGGTTTC | ACC | ACC | C | G | C>G | C>G_ACC | CG>GC_GGGTTTC | 8 | FALSE | FALSE |
| 161_v.txt | 32876 | 32876-SNV- | SNV | 1 | G | C | 3 | 14694 | 0.0204165 | 38 | rnr1Y285F | FALSE | CAN | JSY3865 | 5 | rnr1Y285F-R | TGT | G>C | G/C+1 | G/C-1 | CG>GC | CG>GC | AACTTTGTA | GTACGTC | ACA | ACA | C | G | C>G | C>G_ACA | CG>GC_GTACGTC | 8 | FALSE | FALSE |
| 161_v.txt | 32917 | 32917-SNV- | SNV | 1 | C | G | 17 | 14700 | 0.11564626 | 46.9411765 | rnr1Y285F | FALSE | CAN | JSY3865 | 5 | rnr1Y285F-R | TCC | C>G | G/C+1 | G/C-1 | CG>GC | CG>GC | TAAGTTCCA | CCAGGGC | AGG | TCC | C | C | C>G | C>G_TCC | CG>GC_CCAGGGC | 8 | FALSE | FALSE |
| 161_v.txt | 32937 | 32937-SNV- | SNV | 1 | C | G | 9 | 14700 | 0.06122449 | 51.3333333 | rnr1Y285F | FALSE | CAN | JSY3865 | 5 | rnr1Y285F-R | CCA | C>G | G/C+1 | G/C-1 | CG>GC | CG>GC | ATTGCCCAA | CAAGAAA | GGT | CCA | C | C | C>G | C>G_CCA | CG>GC_CAAGAAA | 8 | FALSE | FALSE |
| 161_v.txt | 33024 | 33024-SNV- | SNV | 1 | G | C | 2 | 14700 | 0.01360544 | 64 | rnr1Y285F | FALSE | CAN | JSY3865 | 5 | rnr1Y285F-R | AGG | G>C | G/C+1 | G/C-1 | CG>GC | CG>GC | GTAACAGGG | GGGATGA | TCC | TCC | C | G | C>G | C>G_TCC | CG>GC_GGGATGA | 8 | FALSE | FALSE |
| 161_v.txt | 33114 | 33114-SNV- | SNV | 1 | C | G | 16 | 14700 | 0.10884354 | 44.3092733 | rnr1Y285F | FALSE | CAN | JSY3865 | 5 | rnr1Y285F-R | GCC | C>G | G/C+1 | G/C-1 | CG>GC | CG>GC | ACTGGGCCG | CCGGCGT | CGG | GCC | C | C | C>G | C>G_GCC | CG>GC_CCGGCGT | 8 | FALSE | FALSE |
| 161_v.txt | 33115 | 33115-SNV- | SNV | 1 | C | G | 1 | 14700 | 6.80E-03 | 38 | rnr1Y285F | FALSE | CAN | JSY3865 | 5 | rnr1Y285F-R | CCG | C>G | G/C+1 | G/C-1 | CG>GC | CG>GC | CTGGGCCGG | CGGCGTT | GGC | CCG | C | C | C>G | C>G_CCG | CG>GC_CGGCGTT | 8 | FALSE | FALSE |
| 161_v.txt | 33160 | 33160-SNV- | SNV | 1 | C | G | 20 | 14700 | 0.13605442 | 37.75 | rnr1Y285F | FALSE | CAN | JSY3865 | 5 | rnr1Y285F-R | CCA | C>G | G/C+1 | G/C-1 | CG>GC | CG>GC | CTGTACCAA | CAATAGT | GGT | CCA | C | C | C>G | C>G_CCA | CG>GC_CAATAGT | 8 | FALSE | FALSE |
| 161_v.txt | 33205 | 33205-SNV- | SNV | 1 | G | C | 222 | 15799 | 1.40515223 | 38.9099099 | rnr1Y285F | FALSE | CAN | JSY3865 | 5 | rnr1Y285F-R | AGC | G>C | G/C+1 | G/C-1 | CG>GC | CG>GC | GCTTAAGCT | GCTCTCT | TCG | TCG | C | G | C>G | C>G_TCG | CG>GC_GCTCTCT | 8 | FALSE | FALSE |
| 161_v.txt | 33284 | 33284-SNV- | SNV | 1 | G | C | 3 | 15799 | 0.01898854 | 64 | rnr1Y285F | FALSE | CAN | JSY3865 | 5 | rnr1Y285F-R | CGC | G>C | G/C+1 | G/C-1 | CG>GC | CG>GC | ATCCTCGCC | GCCATTT | GCG | GCG | C | G | C>G | C>G_GCG | CG>GC_GCCATTT | 8 | FALSE | FALSE |
| 161_v.txt | 33456 | 33456-SNV- | SNV | 1 | G | C | 25 | 15799 | 0.15823786 | 42.92 | rnr1Y285F | FALSE | CAN | JSY3865 | 5 | rnr1Y285F-R | TGA | G>C | G/C+1 | G/C-1 | CG>GC | CG>GC | TCTTTTGAA | GAATTTG | ACT | ACT | C | G | C>G | C>G_ACT | CG>GC_GAATTTG | 8 | FALSE | FALSE |
| 161_v.txt | 31817 | 31817-SNV- | SNV | 1 | G | T | 18 | 3276 | 0.54945055 | 63.1666667 | rnr1Y285F | FALSE | CAN | JSY3865 | 5 | rnr1Y285F-R | TGC | G>T | A/T+1 | G/C-1 | CG>AT | CG>AT | AAATCTGCA | GCATCTG | ACG | ACG | C | A | C>A | C>A_ACG | CG>AT_GCATCTG | 8 | FALSE | FALSE |
| 161_v.txt | 31930 | 31930-SNV- | SNV | 1 | G | T | 8 | 3274 | 0.24434942 | 38 | rnr1Y285F | FALSE | CAN | JSY3865 | 5 | rnr1Y285F-R | TGA | G>T | A/T+1 | G/C-1 | CG>AT | CG>AT | AACCTTGAA | GAATAAT | ACT | ACT | C | A | C>A | C>A_ACT | CG>AT_GAATAAT | 8 | FALSE | FALSE |
| 161_v.txt | 31980 | 31980-SNV- | SNV | 1 | G | T | 15 | 18806 | 0.07976178 | 40.4 | rnr1Y285F | FALSE | CAN | JSY3865 | 5 | rnr1Y285F-R | GGG | G>T | A/T+1 | G/C-1 | CG>AT | CG>AT | AAGCCGGGC | GGCATTA | CCC | CCC | C | A | C>A | C>A_CCC | CG>AT_GGCATTA | 8 | FALSE | FALSE |
| 161_v.txt | 31981 | 31981-SNV- | SNV | 1 | G | T | 16 | 18806 | 0.08507923 | 24.3125 | rnr1Y285F | FALSE | CAN | JSY3865 | 5 | rnr1Y285F-R | GGC | G>T | A/T+1 | G/C-1 | CG>AT | CG>AT | AGCCGGGCA | GCATTAA | CCG | CCG | C | A | C>A | C>A_CCG | CG>AT_GCATTAA | 8 | FALSE | FALSE |
| 161_v.txt | 32008 | 32008-SNV- | SNV | 1 | C | A | 29 | 18806 | 0.1542061 | 49.6170279 | rnr1Y285F | FALSE | CAN | JSY3865 | 5 | rnr1Y285F-R | TCG | C>A | A/T+1 | G/C-1 | CG>AT | CG>AT | GTAACTCGT | CGTCACG | AGC | TCG | C | T | C>A | C>A_TCG | CG>AT_CGTCACG | 8 | FALSE | FALSE |
| 161_v.txt | 32027 | 32027-SNV- | SNV | 1 | G | T | 39 | 18806 | 0.20738062 | 59.3846154 | rnr1Y285F | FALSE | CAN | JSY3865 | 5 | rnr1Y285F-R | GGT | G>T | A/T+1 | G/C-1 | CG>AT | CG>AT | GCCACGGTA | GTATTTC | CCA | CCA | C | A | C>A | C>A_CCA | CG>AT_GTATTTC | 8 | FALSE | FALSE |
| 161_v.txt | 32095 | 32095-SNV- | SNV | 1 | C | A | 75 | 18806 | 0.39880889 | 60.04 | rnr1Y285F | FALSE | CAN | JSY3865 | 5 | rnr1Y285F-R | ACA | C>A | A/T+1 | G/C-1 | CG>AT | CG>AT | CTGCAACAC | CACCAGT | TGT | ACA | C | T | C>A | C>A_ACA | CG>AT_CACCAGT | 8 | FALSE | FALSE |
| 161_v.txt | 32223 | 32223-SNV- | SNV | 1 | G | T | 8 | 18806 | 0.04253962 | 38 | rnr1Y285F | FALSE | CAN | JSY3865 | 5 | rnr1Y285F-R | TGA | G>T | A/T+1 | G/C-1 | CG>AT | CG>AT | GTCCTTGAC | GACAGGA | ACT | ACT | C | A | C>A | C>A_ACT | CG>AT_GACAGGA | 8 | FALSE | FALSE |
| 161_v.txt | 32274 | 32274-SNV- | SNV | 1 | G | T | 60 | 19776 | 0.30339806 | 35.7833333 | rnr1Y285F | FALSE | CAN | JSY3865 | 5 | rnr1Y285F-R | GGA | G>T | A/T+1 | G/C-1 | CG>AT | CG>AT | ATACGGGAA | GAACCAA | CCT | CCT | C | A | C>A | C>A_CCT | CG>AT_GAACCAA | 8 | FALSE | FALSE |
| 161_v.txt | 32277 | 32277-SNV- | SNV | 1 | C | A | 31 | 19776 | 0.15675566 | 36.3225807 | rnr1Y285F | FALSE | CAN | JSY3865 | 5 | rnr1Y285F-R | ACC | C>A | A/T+1 | G/C-1 | CG>AT | CG>AT | CGGGAACCA | CCAACGT | TGG | ACC | C | T | C>A | C>A_ACC | CG>AT_CCAACGT | 8 | FALSE | FALSE |
| 161_v.txt | 32304 | 32304-SNV- | SNV | 1 | G | T | 21 | 19776 | 0.10618932 | 48.2857143 | rnr1Y285F | FALSE | CAN | JSY3865 | 5 | rnr1Y285F-R | AGA | G>T | A/T+1 | G/C-1 | CG>AT | CG>AT | GCGGCAGAA | GAAATAA | TCT | TCT | C | A | C>A | C>A_TCT | CG>AT_GAAATAA | 8 | FALSE | FALSE |
| 161_v.txt | 32471 | 32471-SNV- | SNV | 1 | G | T | 50 | 19776 | 0.25283172 | 49.92 | rnr1Y285F | FALSE | CAN | JSY3865 | 5 | rnr1Y285F-R | TGT | G>T | A/T+1 | G/C-1 | CG>AT | CG>AT | GCCAATGTA | GTAGAAG | ACA | ACA | C | A | C>A | C>A_ACA | CG>AT_GTAGAAG | 8 | FALSE | FALSE |
| 161_v.txt | 32488 | 32488-SNV- | SNV | 1 | G | T | 17 | 19776 | 0.08596278 | 45.584592 | rnr1Y285F | FALSE | CAN | JSY3865 | 5 | rnr1Y285F-R | CGG | G>T | A/T+1 | G/C-1 | CG>AT | CG>AT | AGATACGGA | GGAAAAC | GCC | GCC | C | A | C>A | C>A_GCC | CG>AT_GGAAAAC | 8 | FALSE | FALSE |
| 161_v.txt | 32494 | 32494-SNV- | SNV | 1 | C | A | 23 | 19776 | 0.11630259 | 41.2173913 | rnr1Y285F | FALSE | CAN | JSY3865 | 5 | rnr1Y285F-R | ACA | C>A | A/T+1 | G/C-1 | CG>AT | CG>AT | GGAAAACAA | CAACTTT | TGT | ACA | C | T | C>A | C>A_ACA | CG>AT_CAACTTT | 8 | FALSE | FALSE |
| 161_v.txt | 32508 | 32508-SNV- | SNV | 1 | G | T | 12 | 19776 | 0.06067961 | 43.4166667 | rnr1Y285F | FALSE | CAN | JSY3865 | 5 | rnr1Y285F-R | GGC | G>T | A/T+1 | G/C-1 | CG>AT | CG>AT | TTGATGGCT | GCTCTTG | CCG | CCG | C | A | C>A | C>A_CCG | CG>AT_GCTCTTG | 8 | FALSE | FALSE |
| 161_v.txt | 32528 | 32528-SNV- | SNV | 1 | G | T | 1 | 19776 | 5.06E-03 | 33.9998743 | rnr1Y285F | FALSE | CAN | JSY3865 | 5 | rnr1Y285F-R | TGG | G>T | A/T+1 | G/C-1 | CG>AT | CG>AT | TTTTCTGGG | GGGGTTT | ACC | ACC | C | A | C>A | C>A_ACC | CG>AT_GGGGTTT | 8 | FALSE | FALSE |
| 161_v.txt | 32530 | 32530-SNV- | SNV | 1 | G | T | 37 | 19776 | 0.18709547 | 40.3513514 | rnr1Y285F | FALSE | CAN | JSY3865 | 5 | rnr1Y285F-R | GGG | G>T | A/T+1 | G/C-1 | CG>AT | CG>AT | TTCTGGGGT | GGTTTGC | CCC | CCC | C | A | C>A | C>A_CCC | CG>AT_GGTTTGC | 8 | FALSE | FALSE |
| 161_v.txt | 32545 | 32545-SNV- | SNV | 1 | C | A | 9 | 19776 | 0.04550971 | 31.5555556 | rnr1Y285F | FALSE | CAN | JSY3865 | 5 | rnr1Y285F-R | CCA | C>A | A/T+1 | G/C-1 | CG>AT | CG>AT | CTTCACCAG | CAGCAGT | GGT | CCA | C | T | C>A | C>A_CCA | CG>AT_CAGCAGT | 8 | FALSE | FALSE |
| 161_v.txt | 32556 | 32556-SNV- | SNV | 1 | C | A | 38 | 19771 | 0.1922007 | 37.2105263 | rnr1Y285F | FALSE | CAN | JSY3865 | 5 | rnr1Y285F-R | ACC | C>A | A/T+1 | G/C-1 | CG>AT | CG>AT | GTGATACCA | CCAACTA | TGG | ACC | C | T | C>A | C>A_ACC | CG>AT_CCAACTA | 8 | FALSE | FALSE |
| 161_v.txt | 32580 | 32580-SNV- | SNV | 1 | G | T | 16 | 21855 | 0.07320979 | 35.5625 | rnr1Y285F | FALSE | CAN | JSY3865 | 5 | rnr1Y285F-R | TGT | G>T | A/T+1 | G/C-1 | CG>AT | CG>AT | TGAAATGTG | GTGAAGG | ACA | ACA | C | A | C>A | C>A_ACA | CG>AT_GTGAAGG | 8 | FALSE | FALSE |
| 161_v.txt | 32783 | 32783-SNV- | SNV | 1 | G | T | 2931 | 21855 | 13.4111187 | 49.0361651 | rnr1Y285F | FALSE | CAN | JSY3865 | 5 | rnr1Y285F-R | AGA | G>T | A/T+1 | G/C-1 | CG>AT | CG>AT | GACCCAGAA | GAACTCG | TCT | TCT | C | A | C>A | C>A_TCT | CG>AT_GAACTCG | 8 | FALSE | FALSE |
| 161_v.txt | 32876 | 32876-SNV- | SNV | 1 | G | T | 22 | 14694 | 0.14972098 | 33.6363636 | rnr1Y285F | FALSE | CAN | JSY3865 | 5 | rnr1Y285F-R | TGT | G>T | A/T+1 | G/C-1 | CG>AT | CG>AT | AACTTTGTA | GTACGTC | ACA | ACA | C | A | C>A | C>A_ACA | CG>AT_GTACGTC | 8 | FALSE | FALSE |
| 161_v.txt | 32917 | 32917-SNV- | SNV | 1 | C | A | 184 | 14700 | 1.25170068 | 50.7826087 | rnr1Y285F | FALSE | CAN | JSY3865 | 5 | rnr1Y285F-R | TCC | C>A | A/T+1 | G/C-1 | CG>AT | CG>AT | TAAGTTCCA | CCAGGGC | AGG | TCC | C | T | C>A | C>A_TCC | CG>AT_CCAGGGC | 8 | FALSE | FALSE |
| 161_v.txt | 32940 | 32940-SNV- | SNV | 1 | G | T | 36 | 14700 | 0.24489796 | 42.8611111 | rnr1Y285F | FALSE | CAN | JSY3865 | 5 | rnr1Y285F-R | AGA | G>T | A/T+1 | G/C-1 | CG>AT | CG>AT | GCCCAAGAA | GAAAACC | TCT | TCT | C | A | C>A | C>A_TCT | CG>AT_GAAAACC | 8 | FALSE | FALSE |
| 161_v.txt | 32958 | 32958-SNV- | SNV | 1 | C | A | 16 | 14700 | 0.10884354 | 54.25 | rnr1Y285F | FALSE | CAN | JSY3865 | 5 | rnr1Y285F-R | ACC | C>A | A/T+1 | G/C-1 | CG>AT | CG>AT | ATGTAACCA | CCATTGG | TGG | ACC | C | T | C>A | C>A_ACC | CG>AT_CCATTGG | 8 | FALSE | FALSE |
| 161_v.txt | 33024 | 33024-SNV- | SNV | 1 | G | T | 22 | 14700 | 0.14965986 | 61.5909091 | rnr1Y285F | FALSE | CAN | JSY3865 | 5 | rnr1Y285F-R | AGG | G>T | A/T+1 | G/C-1 | CG>AT | CG>AT | GTAACAGGG | GGGATGA | TCC | TCC | C | A | C>A | C>A_TCC | CG>AT_GGGATGA | 8 | FALSE | FALSE |
| 161_v.txt | 33051 | 33051-SNV- | SNV | 1 | G | T | 27 | 14700 | 0.18367347 | 57.9259259 | rnr1Y285F | FALSE | CAN | JSY3865 | 5 | rnr1Y285F-R | GGA | G>T | A/T+1 | G/C-1 | CG>AT | CG>AT | CCCAAGGAC | GACTGCG | CCT | CCT | C | A | C>A | C>A_CCT | CG>AT_GACTGCG | 8 | FALSE | FALSE |
| 161_v.txt | 33127 | 33127-SNV- | SNV | 1 | G | T | 1 | 14700 | 6.80E-03 | 37.9982602 | rnr1Y285F | FALSE | CAN | JSY3865 | 5 | rnr1Y285F-R | AGA | G>T | A/T+1 | G/C-1 | CG>AT | CG>AT | TGGTCAGAG | GAGGTGT | TCT | TCT | C | A | C>A | C>A_TCT | CG>AT_GAGGTGT | 8 | FALSE | FALSE |
| 161_v.txt | 33151 | 33151-SNV- | SNV | 1 | G | T | 26 | 14700 | 0.17687075 | 38.8461539 | rnr1Y285F | FALSE | CAN | JSY3865 | 5 | rnr1Y285F-R | AGA | G>T | A/T+1 | G/C-1 | CG>AT | CG>AT | TGAAAAGAC | GACCTGT | TCT | TCT | C | A | C>A | C>A_TCT | CG>AT_GACCTGT | 8 | FALSE | FALSE |
| 161_v.txt | 33168 | 33168-SNV- | SNV | 1 | C | A | 71 | 14700 | 0.4829932 | 37.6338028 | rnr1Y285F | FALSE | CAN | JSY3865 | 5 | rnr1Y285F-R | ACC | C>A | A/T+1 | G/C-1 | CG>AT | CG>AT | ATAGTACCA | CCACCAA | TGG | ACC | C | T | C>A | C>A_ACC | CG>AT_CCACCAA | 8 | FALSE | FALSE |
| 161_v.txt | 33178 | 33178-SNV- | SNV | 1 | C | A | 13 | 14699 | 0.08844139 | 35.5384615 | rnr1Y285F | FALSE | CAN | JSY3865 | 5 | rnr1Y285F-R | GCA | C>A | A/T+1 | G/C-1 | CG>AT | CG>AT | CAAGGGCAA | CAATCAT | CGT | GCA | C | T | C>A | C>A_GCA | CG>AT_CAATCAT | 8 | FALSE | FALSE |
| 161_v.txt | 33238 | 33238-SNV- | SNV | 1 | C | A | 32 | 15799 | 0.20254447 | 36.21875 | rnr1Y285F | FALSE | CAN | JSY3865 | 5 | rnr1Y285F-R | CCT | C>A | A/T+1 | G/C-1 | CG>AT | CG>AT | CTTCTCCTTC | CTTCATC | GGA | CCT | C | T | C>A | C>A_CCT | CG>AT_CTTCATC | 8 | FALSE | FALSE |
| 161_v.txt | 33325 | 33325-SNV- | SNV | 1 | C | A | 1779 | 15799 | 11.2602063 | 59.7150084 | rnr1Y285F | FALSE | CAN | JSY3865 | 5 | rnr1Y285F-R | TCT | C>A | A/T+1 | G/C-1 | CG>AT | CG>AT | ACAATTCTTT | CTTTACT | AGA | TCT | C | T | C>A | C>A_TCT | CG>AT_CTTTACT | 8 | FALSE | FALSE |
| 161_v.txt | 33474 | 33474-SNV- | SNV | 1 | C | A | 24 | 15798 | 0.15191796 | 35.0416667 | rnr1Y285F | FALSE | CAN | JSY3865 | 5 | rnr1Y285F-R | GCC | C>A | A/T+1 | G/C-1 | CG>AT | CG>AT | GCTATGCCT | CCTTTTT | CGG | GCC | C | T | C>A | C>A_GCC | CG>AT_CCTTTTT | 8 | FALSE | FALSE |
| 161_v.txt | 32008 | 32008-MNV- | MNV | 2 | CG | TT | 10 | 18806 | 0.05317452 | 57.3479623 | rnr1Y285F | FALSE | CAN | JSY3865 | 5 | rnr1Y285F-R | TCG | nope | >1bp-ins | >1bpdel | none | MNV | GTAACTCGT | CGTCACG | AGC | TCG | CG | none | NA | NA_TCG | MNV_CGTCACG | 8 | FALSE | FALSE |
| 161_v.txt | 32007 | 32007-Insert | Insertion | 1 | - | T | 10 | 18806 | 0.05317452 | 58.6 | rnr1Y285F | FALSE | CAN | JSY3865 | 5 | rnr1Y285F-R | CTC | nope | A/T+1 | >1bpdel | none | A/T+1 | GGTAACTCG | TCGTCAC | GAG | CTC | - | A | NA | NA_CTC | A/T+1_TCGTCAC | 8 | FALSE | FALSE |
| 161_v.txt | 32081 | 32081-Insert | Insertion | 1 | - | A | 11 | 18806 | 0.05849197 | 64 | rnr1Y285F | FALSE | CAN | JSY3865 | 5 | rnr1Y285F-R | CAA | nope | A/T+1 | >1bpdel | none | A/T+1 | CCATGCAAA | AAAAAAG | GTT | CAA | - | T | NA | NA_CAA | A/T+1_AAAAAAG | 8 | FALSE | FALSE |
| 161_v.txt | 32490 | 32490-Insert | Insertion | 1 | - | A | 3 | 19776 | 0.0151699 | 45.6666667 | rnr1Y285F | FALSE | CAN | JSY3865 | 5 | rnr1Y285F-R | GAA | nope | A/T+1 | >1bpdel | none | A/T+1 | ATACGGAAA | AAAACAA | CTT | GAA | - | T | NA | NA_GAA | A/T+1_AAAACAA | 8 | FALSE | FALSE |
| 161_v.txt | 32842 | 32842-Insert | Insertion | 1 | - | A | 21 | 21855 | 0.09608785 | 41.5238095 | rnr1Y285F | FALSE | CAN | JSY3865 | 5 | rnr1Y285F-R | CAA | nope | A/T+1 | >1bpdel | none | A/T+1 | TTACCCAAA | AAAAAAT | GTT | CAA | - | T | NA | NA_CAA | A/T+1_AAAAAAT | 8 | FALSE | FALSE |
| 161_v.txt | 33175 | 33175-Insert | Insertion | 1 | - | T | 10 | 14699 | 0.06803184 | 37.8 | rnr1Y285F | FALSE | CAN | JSY3865 | 5 | rnr1Y285F-R | AGG | nope | A/T+1 | >1bpdel | none | A/T+1 | CACCAAGGG | GGGCAAT | TCC | AGG | - | A | NA | NA_AGG | A/T+1_GGGCAAT | 8 | FALSE | FALSE |

| | | | | | | | | | | | | | | | | | | | | | | | | | | | | | | | | | | |
|---|---|---|---|---|---|---|---|---|---|---|---|---|---|---|---|---|---|---|---|---|---|---|---|---|---|---|---|---|---|---|---|---|---|---|
| 161_v.txt | 32313 | 32313-Deleti | Deletion | 3 | GTT | - | 14 | 19776 | 0.07079288 | 53.0714286 | rnr1Y285F | FALSE | CAN | JSY3865 | 5 | rnr1Y285F-R | GGT | nope | >1bp-ins | >1bpdel | none | >1bpdel | ATAATGGTT | GTTGTTA | CCA | GGT | GTT | none | NA | NA_GGT | >1bpdel_GTTGTTA | 8 | FALSE | FALSE |
| 161_v.txt | 32337 | 32337-Deleti | Deletion | 2 | AT | - | 26 | 19776 | 0.13147249 | 61.3076923 | rnr1Y285F | FALSE | CAN | JSY3865 | 5 | rnr1Y285F-R | GAT | nope | >1bp-ins | >1bpdel | none | >1bpdel | TTGAAGATA | ATATGTG | CTA | GAT | AT | none | NA | NA_GAT | >1bpdel_ATATGTG | 8 | FALSE | FALSE |
| 161_v.txt | 32719 | 32719-Deleti | Deletion | 3 | AAA | - | 32 | 21855 | 0.14641958 | 62 | rnr1Y285F | FALSE | CAN | JSY3865 | 5 | rnr1Y285F-R | CAA | nope | >1bp-ins | >1bpdel | none | >1bpdel | CACCACAAA | AAACCAT | GTT | CAA | AAA | none | NA | NA_CAA | >1bpdel_AAACCAT | 8 | FALSE | FALSE |
| 161_v.txt | 33065 | 33065-Deleti | Deletion | 2 | AT | - | 7 | 14700 | 0.04761905 | 64 | rnr1Y285F | FALSE | CAN | JSY3865 | 5 | rnr1Y285F-R | AAT | nope | >1bp-ins | >1bpdel | none | >1bpdel | GACAGAATA | ATATGCC | TTA | AAT | AT | none | NA | NA_AAT | >1bpdel_ATATGCC | 8 | FALSE | FALSE |
| 161_v.txt | 33126 | 33126-Deleti | Deletion | 2 | AG | - | 22 | 14700 | 0.14965986 | 41.2727273 | rnr1Y285F | FALSE | CAN | JSY3865 | 5 | rnr1Y285F-R | CAG | nope | >1bp-ins | >1bpdel | none | >1bpdel | TTGGTCAGA | AGAGGTG | GTC | CAG | AG | none | NA | NA_CAG | >1bpdel_AGAGGTG | 8 | FALSE | FALSE |
| 161_v.txt | 33189 | 33189-Deleti | Deletion | 2 | AT | - | 26 | 15799 | 0.16456738 | 37.8846154 | rnr1Y285F | FALSE | CAN | JSY3865 | 5 | rnr1Y285F-R | AAT | nope | >1bp-ins | >1bpdel | none | >1bpdel | ATACCAATA | ATATGTC | TTA | AAT | AT | none | NA | NA_AAT | >1bpdel_ATATGTC | 8 | FALSE | FALSE |
| 161_v.txt | 31723 | 31723-Deleti | Deletion | 1 | A | - | 4 | 3276 | 0.12210012 | 64 | rnr1Y285F | FALSE | CAN | JSY3865 | 5 | rnr1Y285F-R | CAA | nope | >1bp-ins | A/T-1 | none | A/T-1 | TGTCCCAAA | AAAAAGT | GTT | GTT | T | none | T>none | T>none_GTT | A/T-1_AAAAAGT | 8 | FALSE | FALSE |
| 161_v.txt | 31972 | 31972-Deleti | Deletion | 1 | C | - | 6 | 18806 | 0.03190471 | 35.5 | rnr1Y285F | FALSE | CAN | JSY3865 | 5 | rnr1Y285F-R | GCC | nope | >1bp-ins | G/C-1 | none | G/C-1 | AATAAGCCA | CCAAGCC | CGG | GCC | C | none | nope | nope_GCC | G/C-1_CCAAGCC | 8 | FALSE | FALSE |
| 161_v.txt | 31976 | 31976-Deleti | Deletion | 1 | G | - | 7 | 18806 | 0.03722216 | 39.1428571 | rnr1Y285F | FALSE | CAN | JSY3865 | 5 | rnr1Y285F-R | AGC | nope | >1bp-ins | G/C-1 | none | G/C-1 | AGCCAAGCC | GCCGGGC | TCG | TCG | C | none | C>none | C>none_TCG | G/C-1_GCCGGGC | 8 | FALSE | FALSE |
| 161_v.txt | 32009 | 32009-Deleti | Deletion | 1 | G | - | 2 | 18806 | 0.0106349 | 50.4999855 | rnr1Y285F | FALSE | CAN | JSY3865 | 5 | rnr1Y285F-R | CGT | nope | >1bp-ins | G/C-1 | none | G/C-1 | TAACTCGTCA | GTCACGA | GCA | GCA | C | none | C>none | C>none_GCA | G/C-1_GTCACGA | 8 | FALSE | FALSE |
| 161_v.txt | 32033 | 32033-Deleti | Deletion | 1 | C | - | 6 | 18806 | 0.03190471 | 59.6666667 | rnr1Y285F | FALSE | CAN | JSY3865 | 5 | rnr1Y285F-R | TCA | nope | >1bp-ins | G/C-1 | none | G/C-1 | GTATTTCAAA | CAAAGCT | AGT | TCA | C | none | nope | nope_TCA | G/C-1_CAAAGCT | 8 | FALSE | FALSE |
| 161_v.txt | 32081 | 32081-Deleti | Deletion | 1 | A | - | 16 | 18806 | 0.08507923 | 64 | rnr1Y285F | FALSE | CAN | JSY3865 | 5 | rnr1Y285F-R | CAA | nope | >1bp-ins | A/T-1 | none | A/T-1 | CCATGCAAA | AAAAAAG | GTT | GTT | T | none | T>none | T>none_GTT | A/T-1_AAAAAAG | 8 | FALSE | FALSE |
| 161_v.txt | 32092 | 32092-Deleti | Deletion | 1 | C | - | 3 | 18806 | 0.01595236 | 64 | rnr1Y285F | FALSE | CAN | JSY3865 | 5 | rnr1Y285F-R | GCA | nope | >1bp-ins | G/C-1 | none | G/C-1 | AGCCTGCAA | CAACACC | CGT | GCA | C | none | nope | nope_GCA | G/C-1_CAACACC | 8 | FALSE | FALSE |
| 161_v.txt | 32198 | 32198-Deleti | Deletion | 1 | T | - | 19 | 18806 | 0.10103159 | 43.8947368 | rnr1Y285F | FALSE | CAN | JSY3865 | 5 | rnr1Y285F-R | ATG | nope | >1bp-ins | A/T-1 | none | A/T-1 | AATGTATGG | TGGAACA | TAC | ATG | T | none | nope | nope_ATG | A/T-1_TGGAACA | 8 | FALSE | FALSE |
| 161_v.txt | 32265 | 32265-Deleti | Deletion | 1 | A | - | 3 | 19776 | 0.0151699 | 38 | rnr1Y285F | FALSE | CAN | JSY3865 | 5 | rnr1Y285F-R | TAA | nope | >1bp-ins | A/T-1 | none | A/T-1 | CCAAATAAA | AAAATAC | ATT | ATT | T | none | T>none | T>none_ATT | A/T-1_AAAATAC | 8 | FALSE | FALSE |
| 161_v.txt | 32271 | 32271-Deleti | Deletion | 1 | C | - | 7 | 19776 | 0.03539644 | 37.5714286 | rnr1Y285F | FALSE | CAN | JSY3865 | 5 | rnr1Y285F-R | ACG | nope | >1bp-ins | G/C-1 | none | G/C-1 | AAAATACGG | CGGGAAC | TGC | ACG | C | none | nope | nope_ACG | G/C-1_CGGGAAC | 8 | FALSE | FALSE |
| 161_v.txt | 32346 | 32346-Deleti | Deletion | 1 | A | - | 9 | 19776 | 0.04550971 | 61.5555556 | rnr1Y285F | FALSE | CAN | JSY3865 | 5 | rnr1Y285F-R | CAA | nope | >1bp-ins | A/T-1 | none | A/T-1 | TGTGGCAAA | AAAACCT | GTT | GTT | T | none | T>none | T>none_GTT | A/T-1_AAAACCT | 8 | FALSE | FALSE |
| 161_v.txt | 32381 | 32381-Deleti | Deletion | 1 | A | - | 6 | 19776 | 0.03033981 | 56.8333333 | rnr1Y285F | FALSE | CAN | JSY3865 | 5 | rnr1Y285F-R | TAA | nope | >1bp-ins | A/T-1 | none | A/T-1 | AATAATAAA | AAAGGGA | ATT | ATT | T | none | T>none | T>none_ATT | A/T-1_AAAGGGA | 8 | FALSE | FALSE |
| 161_v.txt | 32440 | 32440-Deleti | Deletion | 1 | C | - | 8 | 19776 | 0.04045307 | 53.125 | rnr1Y285F | FALSE | CAN | JSY3865 | 5 | rnr1Y285F-R | ACT | nope | >1bp-ins | G/C-1 | none | G/C-1 | ATGGAACTA | CTAAAAG | TGA | ACT | C | none | nope | nope_ACT | G/C-1_CTAAAAG | 8 | FALSE | FALSE |
| 161_v.txt | 32466 | 32466-Deleti | Deletion | 1 | C | - | 2 | 19776 | 0.01011327 | 49 | rnr1Y285F | FALSE | CAN | JSY3865 | 5 | rnr1Y285F-R | GCC | nope | >1bp-ins | G/C-1 | none | G/C-1 | AGAGAGCCA | CCAATGT | CGG | GCC | C | none | nope | nope_GCC | G/C-1_CCAATGT | 8 | FALSE | FALSE |
| 161_v.txt | 32498 | 32498-Deleti | Deletion | 1 | T | - | 13 | 19776 | 0.06573625 | 50.4615385 | rnr1Y285F | FALSE | CAN | JSY3865 | 5 | rnr1Y285F-R | CTT | nope | >1bp-ins | A/T-1 | none | A/T-1 | AACAACTTTT | TTTTTTG | GAA | CTT | T | none | nope | nope_CTT | A/T-1_TTTTTTG | 8 | FALSE | FALSE |
| 161_v.txt | 32522 | 32522-Deleti | Deletion | 1 | T | - | 9 | 19776 | 0.04550971 | 40.7777778 | rnr1Y285F | FALSE | CAN | JSY3865 | 5 | rnr1Y285F-R | ATT | nope | >1bp-ins | A/T-1 | none | A/T-1 | AACGGATTTT | TTTTCTG | TAA | ATT | T | none | nope | nope_ATT | A/T-1_TTTTCTG | 8 | FALSE | FALSE |
| 161_v.txt | 32528 | 32528-Deleti | Deletion | 1 | G | - | 4 | 19776 | 0.02022654 | 50.9999974 | rnr1Y285F | FALSE | CAN | JSY3865 | 5 | rnr1Y285F-R | TGG | nope | >1bp-ins | G/C-1 | none | G/C-1 | TTTTCTGGGG | GGGGTTT | ACC | ACC | C | none | C>none | C>none_ACC | G/C-1_GGGGTTT | 8 | FALSE | FALSE |
| 161_v.txt | 32592 | 32592-Deleti | Deletion | 1 | T | - | 13 | 21855 | 0.05948296 | 37.6153846 | rnr1Y285F | FALSE | CAN | JSY3865 | 5 | rnr1Y285F-R | GTT | nope | >1bp-ins | A/T-1 | none | A/T-1 | GCAGCGTTA | TTAATCA | CAA | GTT | T | none | nope | nope_GTT | A/T-1_TTAATCA | 8 | FALSE | FALSE |
| 161_v.txt | 32598 | 32598-Deleti | Deletion | 1 | A | - | 6 | 21855 | 0.02745367 | 45.8333333 | rnr1Y285F | FALSE | CAN | JSY3865 | 5 | rnr1Y285F-R | CAA | nope | >1bp-ins | A/T-1 | none | A/T-1 | TTAATCAAA | AAAGAGG | GTT | GTT | T | none | T>none | T>none_GTT | A/T-1_AAAGAGG | 8 | FALSE | FALSE |
| 161_v.txt | 32608 | 32608-Deleti | Deletion | 1 | C | - | 115 | 21855 | 0.52619538 | 43.8782592 | rnr1Y285F | FALSE | CAN | JSY3865 | 5 | rnr1Y285F-R | ACC | nope | >1bp-ins | G/C-1 | none | G/C-1 | AGGAAACCC | CCCAACC | TGG | ACC | C | none | nope | nope_ACC | G/C-1_CCCAACC | 8 | FALSE | FALSE |
| 161_v.txt | 32631 | 32631-Deleti | Deletion | 1 | T | - | 5 | 21855 | 0.02287806 | 49.8 | rnr1Y285F | FALSE | CAN | JSY3865 | 5 | rnr1Y285F-R | GTT | nope | >1bp-ins | A/T-1 | none | A/T-1 | CCTTCGTTTT | TTTTTAT | CAA | GTT | T | none | nope | nope_GTT | A/T-1_TTTTTAT | 8 | FALSE | FALSE |
| 161_v.txt | 32658 | 32658-Deleti | Deletion | 1 | C | - | 15 | 21855 | 0.06863418 | 61.4 | rnr1Y285F | FALSE | CAN | JSY3865 | 5 | rnr1Y285F-R | ACC | nope | >1bp-ins | G/C-1 | none | G/C-1 | CCTGGACCC | CCCCAGG | TGG | ACC | C | none | nope | nope_ACC | G/C-1_CCCCAGG | 8 | FALSE | FALSE |
| 161_v.txt | 32670 | 32670-Deleti | Deletion | 1 | G | - | 2148 | 21855 | 9.82841455 | 59.7136872 | rnr1Y285F | FALSE | CAN | JSY3865 | 5 | rnr1Y285F-R | TGG | nope | >1bp-ins | G/C-1 | none | G/C-1 | GCACCTGGG | GGGTTTC | ACC | ACC | C | none | C>none | C>none_ACC | G/C-1_GGGTTTC | 8 | FALSE | FALSE |
| 161_v.txt | 32707 | 32707-Deleti | Deletion | 1 | C | - | 10 | 21855 | 0.04575612 | 61.4 | rnr1Y285F | FALSE | CAN | JSY3865 | 5 | rnr1Y285F-R | ACC | nope | >1bp-ins | G/C-1 | none | G/C-1 | CGGTAACCC | CCCCAGC | TGG | ACC | C | none | nope | nope_ACC | G/C-1_CCCCAGC | 8 | FALSE | FALSE |
| 161_v.txt | 32719 | 32719-Deleti | Deletion | 1 | A | - | 2 | 21854 | 9.15E-03 | 63.9999984 | rnr1Y285F | FALSE | CAN | JSY3865 | 5 | rnr1Y285F-R | CAA | nope | >1bp-ins | A/T-1 | none | A/T-1 | CACCACAAA | AAACCAT | GTT | GTT | T | none | T>none | T>none_GTT | A/T-1_AAACCAT | 8 | FALSE | FALSE |
| 161_v.txt | 32728 | 32728-Deleti | Deletion | 1 | A | - | 6 | 21855 | 0.02745367 | 64 | rnr1Y285F | FALSE | CAN | JSY3865 | 5 | rnr1Y285F-R | CAA | nope | >1bp-ins | A/T-1 | none | A/T-1 | CCATACAAA | AAAAACA | GTT | GTT | T | none | T>none | T>none_GTT | A/T-1_AAAAACA | 8 | FALSE | FALSE |
| 161_v.txt | 32744 | 32744-Deleti | Deletion | 1 | A | - | 9 | 21855 | 0.04118051 | 61 | rnr1Y285F | FALSE | CAN | JSY3865 | 5 | rnr1Y285F-R | GAA | nope | >1bp-ins | A/T-1 | none | A/T-1 | TATTAGAAA | AAACCCG | CTT | CTT | T | none | T>none | T>none_CTT | A/T-1_AAACCCG | 8 | FALSE | FALSE |
| 161_v.txt | 32804 | 32804-Deleti | Deletion | 1 | T | - | 6 | 21855 | 0.02745367 | 46 | rnr1Y285F | FALSE | CAN | JSY3865 | 5 | rnr1Y285F-R | ATT | nope | >1bp-ins | A/T-1 | none | A/T-1 | GTAATATTT | TTTGACA | TAA | ATT | T | none | nope | nope_ATT | A/T-1_TTTGACA | 8 | FALSE | FALSE |
| 161_v.txt | 32842 | 32842-Deleti | Deletion | 1 | A | - | 72 | 21855 | 0.32944406 | 43.5277767 | rnr1Y285F | FALSE | CAN | JSY3865 | 5 | rnr1Y285F-R | CAA | nope | >1bp-ins | A/T-1 | none | A/T-1 | TTACCCAAA | AAAAAAT | GTT | GTT | T | none | T>none | T>none_GTT | A/T-1_AAAAAAT | 8 | FALSE | FALSE |
| 161_v.txt | 32924 | 32924-Deleti | Deletion | 1 | A | - | 16 | 14700 | 0.10884354 | 50.375 | rnr1Y285F | FALSE | CAN | JSY3865 | 5 | rnr1Y285F-R | CAA | nope | >1bp-ins | A/T-1 | none | A/T-1 | CAGGGCAAA | AAAAGTG | GTT | GTT | T | none | T>none | T>none_GTT | A/T-1_AAAAGTG | 8 | FALSE | FALSE |
| 161_v.txt | 33168 | 33168-Deleti | Deletion | 1 | C | - | 2 | 14700 | 0.01360544 | 38 | rnr1Y285F | FALSE | CAN | JSY3865 | 5 | rnr1Y285F-R | ACC | nope | >1bp-ins | G/C-1 | none | G/C-1 | ATAGTACCA | CCACCAA | TGG | ACC | C | none | nope | nope_ACC | G/C-1_CCACCAA | 8 | FALSE | FALSE |
| 161_v.txt | 33261 | 33261-Deleti | Deletion | 1 | T | - | 18 | 15799 | 0.11393126 | 37.1111111 | rnr1Y285F | FALSE | CAN | JSY3865 | 5 | rnr1Y285F-R | ATC | nope | >1bp-ins | A/T-1 | none | A/T-1 | ATGCCATCCT | TCCTCCA | TAG | ATC | T | none | nope | nope_ATC | A/T-1_TCCTCCA | 8 | FALSE | FALSE |
| 161_v.txt | 33356 | 33356-Deleti | Deletion | 1 | C | - | 28 | 15799 | 0.17722641 | 57.1428571 | rnr1Y285F | FALSE | CAN | JSY3865 | 5 | rnr1Y285F-R | ACC | nope | >1bp-ins | G/C-1 | none | G/C-1 | TATTGACCCA | CCCACGT | TGG | ACC | C | none | nope | nope_ACC | G/C-1_CCCACGT | 8 | FALSE | FALSE |
| 161_v.txt | 33474 | 33474-Deleti | Deletion | 1 | C | - | 36 | 15798 | 0.22787695 | 29.6944444 | rnr1Y285F | FALSE | CAN | JSY3865 | 5 | rnr1Y285F-R | GCC | nope | >1bp-ins | G/C-1 | none | G/C-1 | GCTATGCCTT | CCTTTTT | CGG | GCC | C | none | nope | nope_GCC | G/C-1_CCTTTTT | 8 | FALSE | FALSE |
| 162_v.txt | 31723 | 31723-SNV-A | SNV | 1 | A | C | 4 | 7952 | 0.05030181 | 64 | rnr1Y285F | FALSE | CAN | JSY3865 | 6 | rnr1Y285F-R | CAA | A>C | G/C+1 | A/T-1 | TA>GC | TA>GC | TGTCCCAAA | AAAAAGT | GTT | GTT | T | G | T>G | T>G_GTT | TA>GC_AAAAAGT | 8 | FALSE | FALSE |
| 162_v.txt | 31950 | 31950-SNV-A | SNV | 1 | A | C | 37 | 21913 | 0.16884954 | 33.8918919 | rnr1Y285F | FALSE | CAN | JSY3865 | 6 | rnr1Y285F-R | CAT | A>C | G/C+1 | A/T-1 | TA>GC | TA>GC | ATCGTCATA | ATAAATG | GTA | GTA | T | G | T>G | T>G_GTA | TA>GC_ATAAATG | 8 | FALSE | FALSE |
| 162_v.txt | 31964 | 31964-SNV-A | SNV | 1 | A | C | 54 | 21913 | 0.24642906 | 36.1111111 | rnr1Y285F | FALSE | CAN | JSY3865 | 6 | rnr1Y285F-R | CAT | A>C | G/C+1 | A/T-1 | TA>GC | TA>GC | GGCCGCATA | ATAATAA | GTA | GTA | T | G | T>G | T>G_GTA | TA>GC_ATAATAA | 8 | FALSE | FALSE |
| 162_v.txt | 32073 | 32073-SNV-A | SNV | 1 | A | C | 20 | 21913 | 0.09127002 | 62.9 | rnr1Y285F | FALSE | CAN | JSY3865 | 6 | rnr1Y285F-R | TAA | A>C | G/C+1 | A/T-1 | TA>GC | TA>GC | ATAAATAAC | AACCATG | ATT | ATT | T | G | T>G | T>G_ATT | TA>GC_AACCATG | 8 | FALSE | FALSE |
| 162_v.txt | 32082 | 32082-SNV-A | SNV | 1 | A | C | 20 | 21912 | 0.09127419 | 62.6501077 | rnr1Y285F | FALSE | CAN | JSY3865 | 6 | rnr1Y285F-R | AAA | A>C | G/C+1 | A/T-1 | TA>GC | TA>GC | CATGCAAAA | AAAAAGC | TTT | TTT | T | G | T>G | T>G_TTT | TA>GC_AAAAAGC | 8 | FALSE | FALSE |
| 162_v.txt | 32094 | 32094-SNV-A | SNV | 1 | A | C | 20 | 21913 | 0.09127002 | 63 | rnr1Y285F | FALSE | CAN | JSY3865 | 6 | rnr1Y285F-R | AAC | A>C | G/C+1 | A/T-1 | TA>GC | TA>GC | CCTGCAACA | ACACCAG | TTG | TTG | T | G | T>G | T>G_TTG | TA>GC_ACACCAG | 8 | FALSE | FALSE |
| 162_v.txt | 32109 | 32109-SNV-A | SNV | 1 | A | C | 20 | 21913 | 0.09127002 | 58.65 | rnr1Y285F | FALSE | CAN | JSY3865 | 6 | rnr1Y285F-R | TAA | A>C | G/C+1 | A/T-1 | TA>GC | TA>GC | ATATTTAATA | AATAGCC | ATT | ATT | T | G | T>G | T>G_ATT | TA>GC_AATAGCC | 8 | FALSE | FALSE |
| 162_v.txt | 32124 | 32124-SNV-A | SNV | 1 | A | C | 8 | 21913 | 0.03650801 | 48 | rnr1Y285F | FALSE | CAN | JSY3865 | 6 | rnr1Y285F-R | AAC | A>C | G/C+1 | A/T-1 | TA>GC | TA>GC | TCGAAAACT | ACTTTGT | TTG | TTG | T | G | T>G | T>G_TTG | TA>GC_ACTTTGT | 8 | FALSE | FALSE |
| 162_v.txt | 32226 | 32226-SNV-A | SNV | 1 | A | C | 23 | 21913 | 0.10496053 | 36.173913 | rnr1Y285F | FALSE | CAN | JSY3865 | 6 | rnr1Y285F-R | CAG | A>C | G/C+1 | A/T-1 | TA>GC | TA>GC | CTTGACAGG | AGGAATT | GTC | GTC | T | G | T>G | T>G_GTC | TA>GC_AGGAATT | 8 | FALSE | FALSE |
| 162_v.txt | 32283 | 32283-SNV-T | SNV | 1 | T | G | 17 | 23484 | 0.07238971 | 57.5294118 | rnr1Y285F | FALSE | CAN | JSY3865 | 6 | rnr1Y285F-R | GTA | T>G | G/C+1 | A/T-1 | TA>GC | TA>GC | CCAACGTAA | TAAATAT | CAT | GTA | T | C | T>G | T>G_GTA | TA>GC_TAAATAT | 8 | FALSE | FALSE |
| 162_v.txt | 32314 | 32314-SNV-T | SNV | 1 | T | G | 5 | 23484 | 0.02129109 | 16.2000709 | rnr1Y285F | FALSE | CAN | JSY3865 | 6 | rnr1Y285F-R | GTT | T>G | G/C+1 | A/T-1 | TA>GC | TA>GC | TAATGGTTG | TTGTTAA | CAA | GTT | T | C | T>G | T>G_GTT | TA>GC_TTGTTAA | 8 | FALSE | FALSE |
| 162_v.txt | 32581 | 32581-SNV-T | SNV | 1 | T | G | 17 | 24455 | 0.06951544 | 34.4706307 | rnr1Y285F | FALSE | CAN | JSY3865 | 6 | rnr1Y285F-R | GTG | T>G | G/C+1 | A/T-1 | TA>GC | TA>GC | GAAATGTGA | TGAAGGC | CAC | GTG | T | C | T>G | T>G_GTG | TA>GC_TGAAGGC | 8 | FALSE | FALSE |
| 162_v.txt | 32631 | 32631-SNV-T | SNV | 1 | T | G | 2 | 24455 | 8.18E-03 | 64 | rnr1Y285F | FALSE | CAN | JSY3865 | 6 | rnr1Y285F-R | GTT | T>G | G/C+1 | A/T-1 | TA>GC | TA>GC | CCTTCGTTTT | TTTTTAT | CAA | GTT | T | C | T>G | T>G_GTT | TA>GC_TTTTTAT | 8 | FALSE | FALSE |
| 162_v.txt | 32681 | 32681-SNV-A | SNV | 1 | A | C | 76 | 24455 | 0.31077489 | 55.8229412 | rnr1Y285F | FALSE | CAN | JSY3865 | 6 | rnr1Y285F-R | AAT | A>C | G/C+1 | A/T-1 | TA>GC | TA>GC | TCTCCAATAA | ATAACGG | TTA | TTA | T | G | T>G | T>G_TTA | TA>GC_ATAACGG | 8 | FALSE | FALSE |
| 162_v.txt | 32802 | 32802-SNV-T | SNV | 1 | T | G | 26 | 24455 | 0.10631773 | 36.7692035 | rnr1Y285F | FALSE | CAN | JSY3865 | 6 | rnr1Y285F-R | ATA | T>G | G/C+1 | A/T-1 | TA>GC | TA>GC | CCGTAATATT | TATTTGA | TAT | ATA | T | C | T>G | T>G_ATA | TA>GC_TATTTGA | 8 | FALSE | FALSE |
| 162_v.txt | 32803 | 32803-SNV-A | SNV | 1 | A | C | 8 | 24455 | 0.03271315 | 43.374501 | rnr1Y285F | FALSE | CAN | JSY3865 | 6 | rnr1Y285F-R | TAT | A>C | G/C+1 | A/T-1 | TA>GC | TA>GC | CGTAATATTT | ATTTGAC | ATA | ATA | T | G | T>G | T>G_ATA | TA>GC_ATTTGAC | 8 | FALSE | FALSE |
| 162_v.txt | 32808 | 32808-SNV-A | SNV | 1 | A | C | 17 | 24455 | 0.06951544 | 39.3538258 | rnr1Y285F | FALSE | CAN | JSY3865 | 6 | rnr1Y285F-R | GAC | A>C | G/C+1 | A/T-1 | TA>GC | TA>GC | TATTTGACA | ACAGGGA | CTG | CTG | T | G | T>G | T>G_CTG | TA>GC_ACAGGGA | 8 | FALSE | FALSE |
| 162_v.txt | 32842 | 32842-SNV-A | SNV | 1 | A | C | 2 | 24455 | 8.18E-03 | 37.9998272 | rnr1Y285F | FALSE | CAN | JSY3865 | 6 | rnr1Y285F-R | CAA | A>C | G/C+1 | A/T-1 | TA>GC | TA>GC | TTACCCAAA | AAAAAAT | GTT | GTT | T | G | T>G | T>G_GTT | TA>GC_AAAAAAT | 8 | FALSE | FALSE |
| 162_v.txt | 32926 | 32926-SNV-A | SNV | 1 | A | C | 10 | 24498 | 0.04081966 | 37.2 | rnr1Y285F | FALSE | CAN | JSY3865 | 6 | rnr1Y285F-R | AAA | A>C | G/C+1 | A/T-1 | TA>GC | TA>GC | GGGCAAAAG | AAGTGAT | TTT | TTT | T | G | T>G | T>G_TTT | TA>GC_AAGTGAT | 8 | FALSE | FALSE |
| 162_v.txt | 32929 | 32929-SNV-T | SNV | 1 | T | G | 17 | 24498 | 0.06939342 | 47.5294118 | rnr1Y285F | FALSE | CAN | JSY3865 | 6 | rnr1Y285F-R | GTG | T>G | G/C+1 | A/T-1 | TA>GC | TA>GC | CAAAAGTGA | TGATTGC | CAC | GTG | T | C | T>G | T>G_GTG | TA>GC_TGATTGC | 8 | FALSE | FALSE |
| 162_v.txt | 33054 | 33054-SNV-T | SNV | 1 | T | G | 250 | 24498 | 1.02049147 | 57.98 | rnr1Y285F | FALSE | CAN | JSY3865 | 6 | rnr1Y285F-R | CTG | T>G | G/C+1 | A/T-1 | TA>GC | TA>GC | AAGGACTGC | TGCGTGA | GAC | CTG | T | C | T>G | T>G_CTG | TA>GC_TGCGTGA | 8 | FALSE | FALSE |
| 162_v.txt | 32077 | 32077-SNV-A | SNV | 1 | A | G | 69 | 21913 | 0.31488158 | 62.3913044 | rnr1Y285F | FALSE | CAN | JSY3865 | 6 | rnr1Y285F-R | CAT | A>G | G/C+1 | A/T-1 | TA>CG | TA>CG | ATAACCATG | ATGCAAA | GTA | GTA | T | C | T>C | T>C_GTA | TA>CG_ATGCAAA | 8 | FALSE | FALSE |
| 162_v.txt | 32081 | 32081-SNV-A | SNV | 1 | A | G | 7 | 21913 | 0.03194451 | 62.7142858 | rnr1Y285F | FALSE | CAN | JSY3865 | 6 | rnr1Y285F-R | CAA | A>G | G/C+1 | A/T-1 | TA>CG | TA>CG | CCATGCAAA | AAAAAAG | GTT | GTT | T | C | T>C | T>C_GTT | TA>CG_AAAAAAG | 8 | FALSE | FALSE |
| 162_v.txt | 32315 | 32315-SNV-T | SNV | 1 | T | C | 8 | 23484 | 0.03406575 | 56.7499993 | rnr1Y285F | FALSE | CAN | JSY3865 | 6 | rnr1Y285F-R | TTG | T>C | G/C+1 | A/T-1 | TA>CG | TA>CG | AATGGTTGT | TGTTAAG | AAC | TTG | T | G | T>C | T>C_TTG | TA>CG_TGTTAAG | 8 | FALSE | FALSE |
| 162_v.txt | 32346 | 32346-SNV-A | SNV | 1 | A | G | 5 | 23484 | 0.02129109 | 54.1994596 | rnr1Y285F | FALSE | CAN | JSY3865 | 6 | rnr1Y285F-R | CAA | A>G | G/C+1 | A/T-1 | TA>CG | TA>CG | TGTGGCAAA | AAAACCT | GTT | GTT | T | C | T>C | T>C_GTT | TA>CG_AAAACCT | 8 | FALSE | FALSE |
| 162_v.txt | 32680 | 32680-SNV-A | SNV | 1 | A | G | 27 | 24454 | 0.11041139 | 57.7409447 | rnr1Y285F | FALSE | CAN | JSY3865 | 6 | rnr1Y285F-R | CAA | A>G | G/C+1 | A/T-1 | TA>CG | TA>CG | TTCTCCAATA | AATAACG | GTT | GTT | T | C | T>C | T>C_GTT | TA>CG_AATAACG | 8 | FALSE | FALSE |
| 162_v.txt | 32808 | 32808-SNV-A | SNV | 1 | A | G | 10 | 24455 | 0.04089143 | 39.1986547 | rnr1Y285F | FALSE | CAN | JSY3865 | 6 | rnr1Y285F-R | GAC | A>G | G/C+1 | A/T-1 | TA>CG | TA>CG | TATTTGACA | ACAGGGA | CTG | CTG | T | C | T>C | T>C_CTG | TA>CG_ACAGGGA | 8 | FALSE | FALSE |
| 162_v.txt | 32929 | 32929-SNV-T | SNV | 1 | T | C | 15 | 24498 | 0.06122949 | 39.2666667 | rnr1Y285F | FALSE | CAN | JSY3865 | 6 | rnr1Y285F-R | GTG | T>C | G/C+1 | A/T-1 | TA>CG | TA>CG | CAAAAGTGA | TGATTGC | CAC | GTG | T | G | T>C | T>C_GTG | TA>CG_TGATTGC | 8 | FALSE | FALSE |
| 162_v.txt | 32314 | 32314-SNV-T | SNV | 1 | T | A | 7 | 23484 | 0.02980753 | 59.8544475 | rnr1Y285F | FALSE | CAN | JSY3865 | 6 | rnr1Y285F-R | GTT | T>A | A/T+1 | A/T-1 | TA>AT | TA>AT | TAATGGTTG | TTGTTAA | CAA | GTT | T | T | T>A | T>A_GTT | TA>AT_TTGTTAA | 8 | FALSE | FALSE |
| 162_v.txt | 32315 | 32315-SNV-T | SNV | 1 | T | A | 3 | 23484 | 0.01277466 | 46.6666666 | rnr1Y285F | FALSE | CAN | JSY3865 | 6 | rnr1Y285F-R | TTG | T>A | A/T+1 | A/T-1 | TA>AT | TA>AT | AATGGTTGT | TGTTAAG | AAC | TTG | T | T | T>A | T>A_TTG | TA>AT_TGTTAAG | 8 | FALSE | FALSE |
| 162_v.txt | 32346 | 32346-SNV-A | SNV | 1 | A | T | 5 | 23484 | 0.02129109 | 60.9970001 | rnr1Y285F | FALSE | CAN | JSY3865 | 6 | rnr1Y285F-R | CAA | A>T | A/T+1 | A/T-1 | TA>AT | TA>AT | TGTGGCAAA | AAAACCT | GTT | GTT | T | A | T>A | T>A_GTT | TA>AT_AAAACCT | 8 | FALSE | FALSE |
| 162_v.txt | 32527 | 32527-SNV-T | SNV | 1 | T | A | 73 | 23484 | 0.31084994 | 42.2286617 | rnr1Y285F | FALSE | CAN | JSY3865 | 6 | rnr1Y285F-R | CTG | T>A | A/T+1 | A/T-1 | TA>AT | TA>AT | ATTTTCTGGG | TGGGGTT | GAC | CTG | T | T | T>A | T>A_CTG | TA>AT_TGGGGTT | 8 | FALSE | FALSE |
| 162_v.txt | 32581 | 32581-SNV-T | SNV | 1 | T | A | 5 | 24455 | 0.02044572 | 35.1998597 | rnr1Y285F | FALSE | CAN | JSY3865 | 6 | rnr1Y285F-R | GTG | T>A | A/T+1 | A/T-1 | TA>AT | TA>AT | GAAATGTGA | TGAAGGC | CAC | GTG | T | T | T>A | T>A_GTG | TA>AT_TGAAGGC | 8 | FALSE | FALSE |
| 162_v.txt | 32916 | 32916-SNV-T | SNV | 1 | T | A | 25 | 24498 | 0.10204915 | 55.1159782 | rnr1Y285F | FALSE | CAN | JSY3865 | 6 | rnr1Y285F-R | TTC | T>A | A/T+1 | A/T-1 | TA>AT | TA>AT | CTAAGTTCCA | TCCAGGG | AAG | TTC | T | T | T>A | T>A_TTC | TA>AT_TCCAGGG | 8 | FALSE | FALSE |
| 162_v.txt | 32929 | 32929-SNV-T | SNV | 1 | T | A | 23 | 24498 | 0.09388522 | 47.4782609 | rnr1Y285F | FALSE | CAN | JSY3865 | 6 | rnr1Y285F-R | GTG | T>A | A/T+1 | A/T-1 | TA>AT | TA>AT | CAAAAGTGA | TGATTGC | CAC | GTG | T | T | T>A | T>A_GTG | TA>AT_TGATTGC | 8 | FALSE | FALSE |
| 162_v.txt | 32983 | 32983-SNV-A | SNV | 1 | A | T | 4 | 24498 | 0.01632786 | 41.4997628 | rnr1Y285F | FALSE | CAN | JSY3865 | 6 | rnr1Y285F-R | GAA | A>T | A/T+1 | A/T-1 | TA>AT | TA>AT | CTGGAGAAA | AAAGGAA | CTT | CTT | T | A | T>A | T>A_CTT | TA>AT_AAAGGAA | 8 | FALSE | FALSE |
| 162_v.txt | 33090 | 33090-SNV-T | SNV | 1 | T | A | 2 | 24498 | 8.16E-03 | 63.9923801 | rnr1Y285F | FALSE | CAN | JSY3865 | 6 | rnr1Y285F-R | ATA | T>A | A/T+1 | A/T-1 | TA>AT | TA>AT | AATAAATAT | TATGATA | TAT | ATA | T | T | T>A | T>A_ATA | TA>AT_TATGATA | 8 | FALSE | FALSE |
| 162_v.txt | 31650 | 31650-SNV-C | SNV | 1 | C | T | 20 | 7952 | 0.25150905 | 36.9 | rnr1Y285F | FALSE | CAN | JSY3865 | 6 | rnr1Y285F-R | GCG | C>T | A/T+1 | G/C-1 | CG>TA | CG>TA | AGAATGCGA | CGAAATG | CGC | GCG | C | A | C>T | C>T_GCG | CG>TA_CGAAATG | 8 | FALSE | FALSE |
| 162_v.txt | 31844 | 31844-SNV-C | SNV | 1 | C | T | 14 | 7952 | 0.17605634 | 42.9285714 | rnr1Y285F | FALSE | CAN | JSY3865 | 6 | rnr1Y285F-R | TCC | C>T | A/T+1 | G/C-1 | CG>TA | CG>TA | TAAGATCCA | CCAAACA | AGG | TCC | C | A | C>T | C>T_TCC | CG>TA_CCAAACA | 8 | FALSE | FALSE |
| 162_v.txt | 31920 | 31920-SNV-G | SNV | 1 | G | A | 9 | 7952 | 0.11317907 | 38 | rnr1Y285F | FALSE | CAN | JSY3865 | 6 | rnr1Y285F-R | CGT | G>A | A/T+1 | G/C-1 | CG>TA | CG>TA | AAAGCCGTG | GTGAAAC | GCA | GCA | C | T | C>T | C>T_GCA | CG>TA_GTGAAAC | 8 | FALSE | FALSE |
| 162_v.txt | 31962 | 31962-SNV-G | SNV | 1 | G | A | 48 | 21913 | 0.21904805 | 35.7083333 | rnr1Y285F | FALSE | CAN | JSY3865 | 6 | rnr1Y285F-R | CGC | G>A | A/T+1 | G/C-1 | CG>TA | CG>TA | GTGGCCGCA | GCATAAT | GCG | GCG | C | T | C>T | C>T_GCG | CG>TA_GCATAAT | 8 | FALSE | FALSE |
| 162_v.txt | 31972 | 31972-SNV-C | SNV | 1 | C | T | 16 | 21913 | 0.07301602 | 36.1875 | rnr1Y285F | FALSE | CAN | JSY3865 | 6 | rnr1Y285F-R | GCC | C>T | A/T+1 | G/C-1 | CG>TA | CG>TA | AATAAGCCA | CCAAGCC | CGG | GCC | C | A | C>T | C>T_GCC | CG>TA_CCAAGCC | 8 | FALSE | FALSE |
| 162_v.txt | 31980 | 31980-SNV-G | SNV | 1 | G | A | 13 | 21913 | 0.05932552 | 49.4615385 | rnr1Y285F | FALSE | CAN | JSY3865 | 6 | rnr1Y285F-R | GGG | G>A | A/T+1 | G/C-1 | CG>TA | CG>TA | AAGCCGGGC | GGCATTA | CCC | CCC | C | T | C>T | C>T_CCC | CG>TA_GGCATTA | 8 | FALSE | FALSE |
| 162_v.txt | 31981 | 31981-SNV-G | SNV | 1 | G | A | 56 | 21913 | 0.25555606 | 50.6788173 | rnr1Y285F | FALSE | CAN | JSY3865 | 6 | rnr1Y285F-R | GGC | G>A | A/T+1 | G/C-1 | CG>TA | CG>TA | AGCCGGGCA | GCATTAA | CCG | CCG | C | T | C>T | C>T_CCG | CG>TA_GCATTAA | 8 | FALSE | FALSE |
| 162_v.txt | 32057 | 32057-SNV-C | SNV | 1 | C | T | 21 | 21913 | 0.09583352 | 63.1418101 | rnr1Y285F | FALSE | CAN | JSY3865 | 6 | rnr1Y285F-R | GCG | C>T | A/T+1 | G/C-1 | CG>TA | CG>TA | GATGTGCGA | CGAGATT | CGC | GCG | C | A | C>T | C>T_GCG | CG>TA_CGAGATT | 8 | FALSE | FALSE |
| 162_v.txt | 32058 | 32058-SNV-G | SNV | 1 | G | A | 10 | 21913 | 0.04563501 | 51.0007109 | rnr1Y285F | FALSE | CAN | JSY3865 | 6 | rnr1Y285F-R | CGA | G>A | A/T+1 | G/C-1 | CG>TA | CG>TA | ATGTGCGAG | GAGATTG | GCT | GCT | C | T | C>T | C>T_GCT | CG>TA_GAGATTG | 8 | FALSE | FALSE |
| 162_v.txt | 32075 | 32075-SNV-C | SNV | 1 | C | T | 101 | 21913 | 0.46091361 | 62.1287129 | rnr1Y285F | FALSE | CAN | JSY3865 | 6 | rnr1Y285F-R | ACC | C>T | A/T+1 | G/C-1 | CG>TA | CG>TA | AAATAACCA | CCATGCA | TGG | ACC | C | A | C>T | C>T_ACC | CG>TA_CCATGCA | 8 | FALSE | FALSE |
| 162_v.txt | 32087 | 32087-SNV-G | SNV | 1 | G | A | 23 | 21912 | 0.10496532 | 58.6957426 | rnr1Y285F | FALSE | CAN | JSY3865 | 6 | rnr1Y285F-R | AGC | G>A | A/T+1 | G/C-1 | CG>TA | CG>TA | AAAAAAGCC | GCCTGCA | TCG | TCG | C | T | C>T | C>T_TCG | CG>TA_GCCTGCA | 8 | FALSE | FALSE |
| 162_v.txt | 32088 | 32088-SNV-C | SNV | 1 | C | T | 129 | 21913 | 0.58869164 | 60.2531831 | rnr1Y285F | FALSE | CAN | JSY3865 | 6 | rnr1Y285F-R | GCC | C>T | A/T+1 | G/C-1 | CG>TA | CG>TA | AAAAAGCCT | CCTGCAA | CGG | GCC | C | A | C>T | C>T_GCC | CG>TA_CCTGCAA | 8 | FALSE | FALSE |
| 162_v.txt | 32092 | 32092-SNV-C | SNV | 1 | C | T | 23 | 21913 | 0.10496053 | 59.7391304 | rnr1Y285F | FALSE | CAN | JSY3865 | 6 | rnr1Y285F-R | GCA | C>T | A/T+1 | G/C-1 | CG>TA | CG>TA | AGCCTGCAA | CAACACC | CGT | GCA | C | A | C>T | C>T_GCA | CG>TA_CAACACC | 8 | FALSE | FALSE |
| 162_v.txt | 32114 | 32114-SNV-C | SNV | 1 | C | T | 271 | 21913 | 1.2367088 | 59.5645757 | rnr1Y285F | FALSE | CAN | JSY3865 | 6 | rnr1Y285F-R | GCC | C>T | A/T+1 | G/C-1 | CG>TA | CG>TA | TAATAGCCA | CCATTCG | CGG | GCC | C | A | C>T | C>T_GCC | CG>TA_CCATTCG | 8 | FALSE | FALSE |
| 162_v.txt | 32165 | 32165-SNV-G | SNV | 1 | G | A | 39 | 21913 | 0.17797654 | 45.129242 | rnr1Y285F | FALSE | CAN | JSY3865 | 6 | rnr1Y285F-R | CGC | G>A | A/T+1 | G/C-1 | CG>TA | CG>TA | CAAAGCGCC | GCCAAAT | GCG | GCG | C | T | C>T | C>T_GCG | CG>TA_GCCAAAT | 8 | FALSE | FALSE |
| 162_v.txt | 32166 | 32166-SNV-C | SNV | 1 | C | T | 35 | 21913 | 0.15972254 | 45.5690743 | rnr1Y285F | FALSE | CAN | JSY3865 | 6 | rnr1Y285F-R | GCC | C>T | A/T+1 | G/C-1 | CG>TA | CG>TA | AAAGCGCCA | CCAAATG | CGG | GCC | C | A | C>T | C>T_GCC | CG>TA_CCAAATG | 8 | FALSE | FALSE |
| 162_v.txt | 32271 | 32271-SNV-C | SNV | 1 | C | T | 35 | 23484 | 0.14903764 | 34.1428571 | rnr1Y285F | FALSE | CAN | JSY3865 | 6 | rnr1Y285F-R | ACG | C>T | A/T+1 | G/C-1 | CG>TA | CG>TA | AAAATACGG | CGGGAAC | TGC | ACG | C | A | C>T | C>T_ACG | CG>TA_CGGGAAC | 8 | FALSE | FALSE |
| 162_v.txt | 32272 | 32272-SNV-G | SNV | 1 | G | A | 16 | 23484 | 0.06813149 | 33.625 | rnr1Y285F | FALSE | CAN | JSY3865 | 6 | rnr1Y285F-R | CGG | G>A | A/T+1 | G/C-1 | CG>TA | CG>TA | AAATACGGG | GGGAACC | GCC | GCC | C | T | C>T | C>T_GCC | CG>TA_GGGAACC | 8 | FALSE | FALSE |
| 162_v.txt | 32299 | 32299-SNV-C | SNV | 1 | C | T | 36 | 23484 | 0.15329586 | 51.9444444 | rnr1Y285F | FALSE | CAN | JSY3865 | 6 | rnr1Y285F-R | GCG | C>T | A/T+1 | G/C-1 | CG>TA | CG>TA | AATTTGCGG | CGGCAGA | CGC | GCG | C | A | C>T | C>T_GCG | CG>TA_CGGCAGA | 8 | FALSE | FALSE |
| 162_v.txt | 32409 | 32409-SNV-G | SNV | 1 | G | A | 525 | 23484 | 2.23556464 | 58.9809524 | rnr1Y285F | FALSE | CAN | JSY3865 | 6 | rnr1Y285F-R | AGA | G>A | A/T+1 | G/C-1 | CG>TA | CG>TA | GAAGTAGAT | GATTGTG | TCT | TCT | C | T | C>T | C>T_TCT | CG>TA_GATTGTG | 8 | FALSE | FALSE |
| 162_v.txt | 32413 | 32413-SNV-G | SNV | 1 | G | A | 74 | 23484 | 0.31510816 | 57.4054054 | rnr1Y285F | FALSE | CAN | JSY3865 | 6 | rnr1Y285F-R | TGT | G>A | A/T+1 | G/C-1 | CG>TA | CG>TA | TAGATTGTG | GTGTTAG | ACA | ACA | C | T | C>T | C>T_ACA | CG>TA_GTGTTAG | 8 | FALSE | FALSE |
| 162_v.txt | 32487 | 32487-SNV-C | SNV | 1 | C | T | 43 | 23484 | 0.18310339 | 44.8139535 | rnr1Y285F | FALSE | CAN | JSY3865 | 6 | rnr1Y285F-R | ACG | C>T | A/T+1 | G/C-1 | CG>TA | CG>TA | AAGATACGG | CGGAAAA | TGC | ACG | C | A | C>T | C>T_ACG | CG>TA_CGGAAAA | 8 | FALSE | FALSE |
| 162_v.txt | 32488 | 32488-SNV-G | SNV | 1 | G | A | 63 | 23484 | 0.26826776 | 45.4285714 | rnr1Y285F | FALSE | CAN | JSY3865 | 6 | rnr1Y285F-R | CGG | G>A | A/T+1 | G/C-1 | CG>TA | CG>TA | AGATACGGA | GGAAAAC | GCC | GCC | C | T | C>T | C>T_GCC | CG>TA_GGAAAAC | 8 | FALSE | FALSE |
| 162_v.txt | 32535 | 32535-SNV-G | SNV | 1 | G | A | 42 | 23484 | 0.17884517 | 36.721295 | rnr1Y285F | FALSE | CAN | JSY3865 | 6 | rnr1Y285F-R | TGC | G>A | A/T+1 | G/C-1 | CG>TA | CG>TA | GGGTTTGCA | GCAGCTT | ACG | ACG | C | T | C>T | C>T_ACG | CG>TA_GCAGCTT | 8 | FALSE | FALSE |
| 162_v.txt | 32536 | 32536-SNV-C | SNV | 1 | C | T | 38 | 23484 | 0.1618123 | 37.6034544 | rnr1Y285F | FALSE | CAN | JSY3865 | 6 | rnr1Y285F-R | GCA | C>T | A/T+1 | G/C-1 | CG>TA | CG>TA | GGTTTGCAG | CAGCTTC | CGT | GCA | C | A | C>T | C>T_GCA | CG>TA_CAGCTTC | 8 | FALSE | FALSE |
| 162_v.txt | 32575 | 32575-SNV-G | SNV | 1 | G | A | 292 | 24455 | 1.19402985 | 37.390411 | rnr1Y285F | FALSE | CAN | JSY3865 | 6 | rnr1Y285F-R | TGA | G>A | A/T+1 | G/C-1 | CG>TA | CG>TA | TACCTTGAA | GAAATGT | ACT | ACT | C | T | C>T | C>T_ACT | CG>TA_GAAATGT | 8 | FALSE | FALSE |
| 162_v.txt | 32590 | 32590-SNV-C | SNV | 1 | C | T | 29 | 24455 | 0.11858516 | 35.5517241 | rnr1Y285F | FALSE | CAN | JSY3865 | 6 | rnr1Y285F-R | GCG | C>T | A/T+1 | G/C-1 | CG>TA | CG>TA | AGGCAGCGT | CGTTAAT | CGC | GCG | C | A | C>T | C>T_GCG | CG>TA_CGTTAAT | 8 | FALSE | FALSE |
| 162_v.txt | 32658 | 32658-SNV-C | SNV | 1 | C | T | 22 | 24455 | 0.08996115 | 50.9545455 | rnr1Y285F | FALSE | CAN | JSY3865 | 6 | rnr1Y285F-R | ACC | C>T | A/T+1 | G/C-1 | CG>TA | CG>TA | CCTGGACCC | CCCCAGG | TGG | ACC | C | A | C>T | C>T_ACC | CG>TA_CCCCAGG | 8 | FALSE | FALSE |
| 162_v.txt | 32700 | 32700-SNV-C | SNV | 1 | C | T | 26 | 24455 | 0.10631773 | 57.1153846 | rnr1Y285F | FALSE | CAN | JSY3865 | 6 | rnr1Y285F-R | GCC | C>T | A/T+1 | G/C-1 | CG>TA | CG>TA | ACTGGGCCG | CCGGTAA | CGG | GCC | C | A | C>T | C>T_GCC | CG>TA_CCGGTAA | 8 | FALSE | FALSE |
| 162_v.txt | 32749 | 32749-SNV-C | SNV | 1 | C | T | 28 | 24455 | 0.11449601 | 50.7142857 | rnr1Y285F | FALSE | CAN | JSY3865 | 6 | rnr1Y285F-R | CCG | C>T | A/T+1 | G/C-1 | CG>TA | CG>TA | GAAACCCGA | CGATAAT | GGC | CCG | C | A | C>T | C>T_CCG | CG>TA_CGATAAT | 8 | FALSE | FALSE |
| 162_v.txt | 32776 | 32776-SNV-C | SNV | 1 | C | T | 24 | 24455 | 0.09813944 | 46.875 | rnr1Y285F | FALSE | CAN | JSY3865 | 6 | rnr1Y285F-R | GCG | C>T | A/T+1 | G/C-1 | CG>TA | CG>TA | TGGAAGCGA | CGACCCA | CGC | GCG | C | A | C>T | C>T_GCG | CG>TA_CGACCCA | 8 | FALSE | FALSE |
| 162_v.txt | 32780 | 32780-SNV-C | SNV | 1 | C | T | 27 | 24455 | 0.11040687 | 45.7260231 | rnr1Y285F | FALSE | CAN | JSY3865 | 6 | rnr1Y285F-R | CCC | C>T | A/T+1 | G/C-1 | CG>TA | CG>TA | AGCGACCCA | CCAGAAC | GGG | CCC | C | A | C>T | C>T_CCC | CG>TA_CCAGAAC | 8 | FALSE | FALSE |
| 162_v.txt | 32796 | 32796-SNV-C | SNV | 1 | C | T | 100 | 24455 | 0.40891433 | 46.1003939 | rnr1Y285F | FALSE | CAN | JSY3865 | 6 | rnr1Y285F-R | ACC | C>T | A/T+1 | G/C-1 | CG>TA | CG>TA | AATTCACCG | CCGTAAT | TGG | ACC | C | A | C>T | C>T_ACC | CG>TA_CCGTAAT | 8 | FALSE | FALSE |
| 162_v.txt | 32797 | 32797-SNV-C | SNV | 1 | C | T | 28 | 24455 | 0.11449601 | 46.6785714 | rnr1Y285F | FALSE | CAN | JSY3865 | 6 | rnr1Y285F-R | CCG | C>T | A/T+1 | G/C-1 | CG>TA | CG>TA | ATTCACCGTA | CGTAATA | GGC | CCG | C | A | C>T | C>T_CCG | CG>TA_CGTAATA | 8 | FALSE | FALSE |
| 162_v.txt | 32798 | 32798-SNV-G | SNV | 1 | G | A | 27 | 24455 | 0.11040687 | 44.037037 | rnr1Y285F | FALSE | CAN | JSY3865 | 6 | rnr1Y285F-R | CGT | G>A | A/T+1 | G/C-1 | CG>TA | CG>TA | TTCACCGTAA | GTAATAT | GCA | GCA | C | T | C>T | C>T_GCA | CG>TA_GTAATAT | 8 | FALSE | FALSE |
| 162_v.txt | 32809 | 32809-SNV-C | SNV | 1 | C | T | 6 | 24455 | 0.02453486 | 46.3314305 | rnr1Y285F | FALSE | CAN | JSY3865 | 6 | rnr1Y285F-R | ACA | C>T | A/T+1 | G/C-1 | CG>TA | CG>TA | ATTTGACAG | CAGGGAA | TGT | ACA | C | A | C>T | C>T_ACA | CG>TA_CAGGGAA | 8 | FALSE | FALSE |
| 162_v.txt | 32840 | 32840-SNV-C | SNV | 1 | C | T | 23 | 24455 | 0.0940503 | 38.65213 | rnr1Y285F | FALSE | CAN | JSY3865 | 6 | rnr1Y285F-R | CCC | C>T | A/T+1 | G/C-1 | CG>TA | CG>TA | AATTACCCA | CCAAAAA | GGG | CCC | C | A | C>T | C>T_CCC | CG>TA_CCAAAAA | 8 | FALSE | FALSE |
| 162_v.txt | 32855 | 32855-SNV-C | SNV | 1 | C | T | 104 | 24455 | 0.42527091 | 36.2596154 | rnr1Y285F | FALSE | CAN | JSY3865 | 6 | rnr1Y285F-R | TCC | C>T | A/T+1 | G/C-1 | CG>TA | CG>TA | ACTAATCCAT | CCATGCC | AGG | TCC | C | A | C>T | C>T_TCC | CG>TA_CCATGCC | 8 | FALSE | FALSE |
| 162_v.txt | 32882 | 32882-SNV-C | SNV | 1 | C | T | 163 | 24498 | 0.66536044 | 32.6809816 | rnr1Y285F | FALSE | CAN | JSY3865 | 6 | rnr1Y285F-R | TCC | C>T | A/T+1 | G/C-1 | CG>TA | CG>TA | GTACGTCCA | CCAAAAT | AGG | TCC | C | A | C>T | C>T_TCC | CG>TA_CCAAAAT | 8 | FALSE | FALSE |
| 162_v.txt | 32901 | 32901-SNV-C | SNV | 1 | C | T | 49 | 24498 | 0.20001633 | 32.5510204 | rnr1Y285F | FALSE | CAN | JSY3865 | 6 | rnr1Y285F-R | GCC | C>T | A/T+1 | G/C-1 | CG>TA | CG>TA | ACTTGGCCA | CCAACTA | CGG | GCC | C | A | C>T | C>T_GCC | CG>TA_CCAACTA | 8 | FALSE | FALSE |
| 162_v.txt | 32917 | 32917-SNV-C | SNV | 1 | C | T | 30 | 24498 | 0.12245898 | 48.5333333 | rnr1Y285F | FALSE | CAN | JSY3865 | 6 | rnr1Y285F-R | TCC | C>T | A/T+1 | G/C-1 | CG>TA | CG>TA | TAAGTTCCAC | CCAGGGC | AGG | TCC | C | A | C>T | C>T_TCC | CG>TA_CCAGGGC | 8 | FALSE | FALSE |
| 162_v.txt | 32935 | 32935-SNV-C | SNV | 1 | C | T | 29 | 24498 | 0.11837701 | 40.6551724 | rnr1Y285F | FALSE | CAN | JSY3865 | 6 | rnr1Y285F-R | GCC | C>T | A/T+1 | G/C-1 | CG>TA | CG>TA | TGATTGCCCA | CCCAAGA | CGG | GCC | C | A | C>T | C>T_GCC | CG>TA_CCCAAGA | 8 | FALSE | FALSE |

| | | | | | | | | | | | | | | | | | | | | | | | | | | | | | | | | | | |
|---|---|---|---|---|---|---|---|---|---|---|---|---|---|---|---|---|---|---|---|---|---|---|---|---|---|---|---|---|---|---|---|---|---|---|
| 162_v.txt | 32945 | 32945-SNV-( | SNV | 1 | C | T | 51 | 24498 | 0.20818026 | 56.0196078 | rnr1Y285F | FALSE | CAN | JSY3865 | 6 | rnr1Y285F-R | ACC | C>T | A/T+1 | G/C-1 | CG>TA | CG>TA | AGAAAACCA | CCAATAC | TGG | ACC | C | A | C>T | C>T_ACC | CG>TA_CCAATAC | 8 | FALSE | FALSE |
| 162_v.txt | 32958 | 32958-SNV-( | SNV | 1 | C | T | 191 | 24498 | 0.77965548 | 58.2146597 | rnr1Y285F | FALSE | CAN | JSY3865 | 6 | rnr1Y285F-R | ACC | C>T | A/T+1 | G/C-1 | CG>TA | CG>TA | ATGTAACCAT | CCATTGG | TGG | ACC | C | A | C>T | C>T_ACC | CG>TA_CCATTGG | 8 | FALSE | FALSE |
| 162_v.txt | 32965 | 32965-SNV-( | SNV | 1 | C | T | 27 | 24498 | 0.11021308 | 59.1851852 | rnr1Y285F | FALSE | CAN | JSY3865 | 6 | rnr1Y285F-R | GCC | C>T | A/T+1 | G/C-1 | CG>TA | CG>TA | CATTGGCCG | CCGCACC | CGG | GCC | C | A | C>T | C>T_GCC | CG>TA_CCGCACC | 8 | FALSE | FALSE |
| 162_v.txt | 32967 | 32967-SNV-( | SNV | 1 | G | A | 58 | 24498 | 0.23675402 | 56.4310345 | rnr1Y285F | FALSE | CAN | JSY3865 | 6 | rnr1Y285F-R | CGC | G>A | A/T+1 | G/C-1 | CG>TA | CG>TA | TTGGCCGCA | GCACCAA | GCG | GCG | C | T | C>T | C>T_GCG | CG>TA_GCACCAA | 8 | FALSE | FALSE |
| 162_v.txt | 33015 | 33015-SNV-( | SNV | 1 | G | A | 19 | 24498 | 0.07755735 | 61.6842105 | rnr1Y285F | FALSE | CAN | JSY3865 | 6 | rnr1Y285F-R | GGA | G>A | A/T+1 | G/C-1 | CG>TA | CG>TA | AAAGAGGAT | GATGTAA | CCT | CCT | C | T | C>T | C>T_CCT | CG>TA_GATGTAA | 8 | FALSE | FALSE |
| 162_v.txt | 33025 | 33025-SNV-( | SNV | 1 | G | A | 25 | 24498 | 0.10204915 | 57.8 | rnr1Y285F | FALSE | CAN | JSY3865 | 6 | rnr1Y285F-R | GGG | G>A | A/T+1 | G/C-1 | CG>TA | CG>TA | TAACAGGGA | GGATGAA | CCC | CCC | C | T | C>T | C>T_CCC | CG>TA_GGATGAA | 8 | FALSE | FALSE |
| 162_v.txt | 33056 | 33056-SNV-( | SNV | 1 | C | T | 33 | 24498 | 0.13470487 | 57.030303 | rnr1Y285F | FALSE | CAN | JSY3865 | 6 | rnr1Y285F-R | GCG | C>T | A/T+1 | G/C-1 | CG>TA | CG>TA | GGACTGCGT | CGTGACA | CGC | GCG | C | A | C>T | C>T_GCG | CG>TA_CGTGACA | 8 | FALSE | FALSE |
| 162_v.txt | 33070 | 33070-SNV-( | SNV | 1 | C | T | 26 | 24498 | 0.10613111 | 53.0769231 | rnr1Y285F | FALSE | CAN | JSY3865 | 6 | rnr1Y285F-R | GCC | C>T | A/T+1 | G/C-1 | CG>TA | CG>TA | AATATGCCA | CCAAAGA | CGG | GCC | C | A | C>T | C>T_GCC | CG>TA_CCAAAGA | 8 | FALSE | FALSE |
| 162_v.txt | 33114 | 33114-SNV-( | SNV | 1 | C | T | 25 | 24498 | 0.10204915 | 43.12 | rnr1Y285F | FALSE | CAN | JSY3865 | 6 | rnr1Y285F-R | GCC | C>T | A/T+1 | G/C-1 | CG>TA | CG>TA | ACTGGGCCG | CCGGCGT | CGG | GCC | C | A | C>T | C>T_GCC | CG>TA_CCGGCGT | 8 | FALSE | FALSE |
| 162_v.txt | 33118 | 33118-SNV-( | SNV | 1 | C | T | 24 | 24498 | 0.09796718 | 37.1666667 | rnr1Y285F | FALSE | CAN | JSY3865 | 6 | rnr1Y285F-R | GCG | C>T | A/T+1 | G/C-1 | CG>TA | CG>TA | GGCCGGCGT | CGTTGGT | CGC | GCG | C | A | C>T | C>T_GCG | CG>TA_CGTTGGT | 8 | FALSE | FALSE |
| 162_v.txt | 33153 | 33153-SNV-( | SNV | 1 | C | T | 26 | 24498 | 0.10613111 | 34.0385757 | rnr1Y285F | FALSE | CAN | JSY3865 | 6 | rnr1Y285F-R | ACC | C>T | A/T+1 | G/C-1 | CG>TA | CG>TA | AAAAGACCT | CCTGTAC | TGG | ACC | C | A | C>T | C>T_ACC | CG>TA_CCTGTAC | 8 | FALSE | FALSE |
| 162_v.txt | 33159 | 33159-SNV-( | SNV | 1 | C | T | 29 | 24496 | 0.11838668 | 33.4827586 | rnr1Y285F | FALSE | CAN | JSY3865 | 6 | rnr1Y285F-R | ACC | C>T | A/T+1 | G/C-1 | CG>TA | CG>TA | CCTGTACCA | CCAATAG | TGG | ACC | C | A | C>T | C>T_ACC | CG>TA_CCAATAG | 8 | FALSE | FALSE |
| 162_v.txt | 33168 | 33168-SNV-( | SNV | 1 | C | T | 131 | 24496 | 0.53478119 | 37.251936 | rnr1Y285F | FALSE | CAN | JSY3865 | 6 | rnr1Y285F-R | ACC | C>T | A/T+1 | G/C-1 | CG>TA | CG>TA | ATAGTACCA | CCACCAA | TGG | ACC | C | A | C>T | C>T_ACC | CG>TA_CCACCAA | 8 | FALSE | FALSE |
| 162_v.txt | 33199 | 33199-SNV-( | SNV | 1 | G | A | 32 | 20807 | 0.1537944 | 37.40625 | rnr1Y285F | FALSE | CAN | JSY3865 | 6 | rnr1Y285F-R | TGC | G>A | A/T+1 | G/C-1 | CG>TA | CG>TA | GTCTTTGCTT | GCTTAAG | ACG | ACG | C | T | C>T | C>T_ACG | CG>TA_GCTTAAG | 8 | FALSE | FALSE |
| 162_v.txt | 33258 | 33258-SNV-( | SNV | 1 | C | T | 41 | 20807 | 0.19704907 | 43.6829268 | rnr1Y285F | FALSE | CAN | JSY3865 | 6 | rnr1Y285F-R | GCC | C>T | A/T+1 | G/C-1 | CG>TA | CG>TA | CCTATGCCAT | CCATCCT | CGG | GCC | C | A | C>T | C>T_GCC | CG>TA_CCATCCT | 8 | FALSE | FALSE |
| 162_v.txt | 33276 | 33276-SNV-( | SNV | 1 | G | A | 32 | 20807 | 0.1537944 | 47.15625 | rnr1Y285F | FALSE | CAN | JSY3865 | 6 | rnr1Y285F-R | CGT | G>A | A/T+1 | G/C-1 | CG>TA | CG>TA | GAGAACGTA | GTATCCT | GCA | GCA | C | T | C>T | C>T_GCA | CG>TA_GTATCCT | 8 | FALSE | FALSE |
| 162_v.txt | 33309 | 33309-SNV-( | SNV | 1 | C | T | 22 | 20806 | 0.10573873 | 59.3603656 | rnr1Y285F | FALSE | CAN | JSY3865 | 6 | rnr1Y285F-R | GCG | C>T | A/T+1 | G/C-1 | CG>TA | CG>TA | AAAGAGCGC | CGCAATG | CGC | GCG | C | A | C>T | C>T_GCG | CG>TA_CGCAATG | 8 | FALSE | FALSE |
| 162_v.txt | 33310 | 33310-SNV-( | SNV | 1 | G | A | 24 | 20805 | 0.11535689 | 60.9957294 | rnr1Y285F | FALSE | CAN | JSY3865 | 6 | rnr1Y285F-R | CGC | G>A | A/T+1 | G/C-1 | CG>TA | CG>TA | AAGAGCGCA | GCAATGG | GCG | GCG | C | T | C>T | C>T_GCG | CG>TA_GCAATGG | 8 | FALSE | FALSE |
| 162_v.txt | 33392 | 33392-SNV-( | SNV | 1 | G | A | 26 | 20807 | 0.12495795 | 52.6153846 | rnr1Y285F | FALSE | CAN | JSY3865 | 6 | rnr1Y285F-R | CGT | G>A | A/T+1 | G/C-1 | CG>TA | CG>TA | AACGTCGTG | GTGAAAG | GCA | GCA | C | T | C>T | C>T_GCA | CG>TA_GTGAAAG | 8 | FALSE | FALSE |
| 162_v.txt | 33412 | 33412-SNV-( | SNV | 1 | G | A | 23 | 20807 | 0.11053972 | 47.5652174 | rnr1Y285F | FALSE | CAN | JSY3865 | 6 | rnr1Y285F-R | GGC | G>A | A/T+1 | G/C-1 | CG>TA | CG>TA | TGACCGGCT | GCTCATT | CCG | CCG | C | T | C>T | C>T_CCG | CG>TA_GCTCATT | 8 | FALSE | FALSE |
| 162_v.txt | 33474 | 33474-SNV-( | SNV | 1 | C | T | 32 | 20804 | 0.15381657 | 35.3125253 | rnr1Y285F | FALSE | CAN | JSY3865 | 6 | rnr1Y285F-R | GCC | C>T | A/T+1 | G/C-1 | CG>TA | CG>TA | GCTATGCCTT | CCTTTTT | CGG | GCC | C | A | C>T | C>T_GCC | CG>TA_CCTTTTT | 8 | FALSE | FALSE |
| 162_v.txt | 32027 | 32027-SNV-( | SNV | 1 | G | C | 10 | 21913 | 0.04563501 | 51.7 | rnr1Y285F | FALSE | CAN | JSY3865 | 6 | rnr1Y285F-R | GGT | G>C | G/C+1 | G/C-1 | CG>GC | CG>GC | GCCACGGTA | GTATTTC | CCA | CCA | C | G | C>G | C>G_CCA | CG>GC_GTATTTC | 8 | FALSE | FALSE |
| 162_v.txt | 32057 | 32057-SNV-( | SNV | 1 | C | G | 1 | 21913 | 4.56E-03 | 47.999462 | rnr1Y285F | FALSE | CAN | JSY3865 | 6 | rnr1Y285F-R | GCG | C>G | G/C+1 | G/C-1 | CG>GC | CG>GC | GATGTGCGA | CGAGATT | CGC | GCG | C | C | C>G | C>G_GCG | CG>GC_CGAGATT | 8 | FALSE | FALSE |
| 162_v.txt | 32064 | 32064-SNV-( | SNV | 1 | G | C | 7 | 21913 | 0.03194451 | 63.7142857 | rnr1Y285F | FALSE | CAN | JSY3865 | 6 | rnr1Y285F-R | TGA | G>C | G/C+1 | G/C-1 | CG>GC | CG>GC | GAGATTGAG | GAGATAA | ACT | ACT | C | G | C>G | C>G_ACT | CG>GC_GAGATAA | 8 | FALSE | FALSE |
| 162_v.txt | 32166 | 32166-SNV-( | SNV | 1 | C | G | 1 | 21913 | 4.56E-03 | 37.9981653 | rnr1Y285F | FALSE | CAN | JSY3865 | 6 | rnr1Y285F-R | GCC | C>G | G/C+1 | G/C-1 | CG>GC | CG>GC | AAAGCGCCA | CCAAATG | CGG | GCC | C | C | C>G | C>G_GCC | CG>GC_CCAAATG | 8 | FALSE | FALSE |
| 162_v.txt | 32199 | 32199-SNV-( | SNV | 1 | G | C | 19 | 21913 | 0.08670652 | 43.5263158 | rnr1Y285F | FALSE | CAN | JSY3865 | 6 | rnr1Y285F-R | TGG | G>C | G/C+1 | G/C-1 | CG>GC | CG>GC | ATGTATGGA | GGAACAC | ACC | ACC | C | G | C>G | C>G_ACC | CG>GC_GGAACAC | 8 | FALSE | FALSE |
| 162_v.txt | 32385 | 32385-SNV-( | SNV | 1 | G | C | 19 | 23484 | 0.08090615 | 53.1052632 | rnr1Y285F | FALSE | CAN | JSY3865 | 6 | rnr1Y285F-R | GGG | G>C | G/C+1 | G/C-1 | CG>GC | CG>GC | ATAAAGGGA | GGAGAAG | CCC | CCC | C | G | C>G | C>G_CCC | CG>GC_GGAGAAG | 8 | FALSE | FALSE |
| 162_v.txt | 32399 | 32399-SNV-( | SNV | 1 | G | C | 11 | 23484 | 0.0468404 | 50.0909091 | rnr1Y285F | FALSE | CAN | JSY3865 | 6 | rnr1Y285F-R | CGT | G>C | G/C+1 | G/C-1 | CG>GC | CG>GC | AGAAACGTA | GTAGGAA | GCA | GCA | C | G | C>G | C>G_GCA | CG>GC_GTAGGAA | 8 | FALSE | FALSE |
| 162_v.txt | 32529 | 32529-SNV-( | SNV | 1 | G | C | 10 | 23484 | 0.04258218 | 40.2961743 | rnr1Y285F | FALSE | CAN | JSY3865 | 6 | rnr1Y285F-R | GGG | G>C | G/C+1 | G/C-1 | CG>GC | CG>GC | TTTCTGGGG | GGGTTTG | CCC | CCC | C | G | C>G | C>G_CCC | CG>GC_GGGTTTG | 8 | FALSE | FALSE |
| 162_v.txt | 32685 | 32685-SNV-( | SNV | 1 | C | G | 7 | 24455 | 0.028624 | 54.4285714 | rnr1Y285F | FALSE | CAN | JSY3865 | 6 | rnr1Y285F-R | ACG | C>G | G/C+1 | G/C-1 | CG>GC | CG>GC | CAATAACGG | CGGAATC | TGC | ACG | C | C | C>G | C>G_ACG | CG>GC_CGGAATC | 8 | FALSE | FALSE |
| 162_v.txt | 32735 | 32735-SNV-( | SNV | 1 | G | C | 4 | 24455 | 0.01635657 | 44.25 | rnr1Y285F | FALSE | CAN | JSY3865 | 6 | rnr1Y285F-R | AGT | G>C | G/C+1 | G/C-1 | CG>GC | CG>GC | AAAACAGTA | GTATATT | TCA | TCA | C | G | C>G | C>G_TCA | CG>GC_GTATATT | 8 | FALSE | FALSE |
| 162_v.txt | 32797 | 32797-SNV-( | SNV | 1 | C | G | 18 | 24455 | 0.07360458 | 49.8888889 | rnr1Y285F | FALSE | CAN | JSY3865 | 6 | rnr1Y285F-R | CCG | C>G | G/C+1 | G/C-1 | CG>GC | CG>GC | ATTCACCGTA | CGTAATA | GGC | CCG | C | C | C>G | C>G_CCG | CG>GC_CGTAATA | 8 | FALSE | FALSE |
| 162_v.txt | 32809 | 32809-SNV-( | SNV | 1 | C | G | 17 | 24455 | 0.06951544 | 39.9976167 | rnr1Y285F | FALSE | CAN | JSY3865 | 6 | rnr1Y285F-R | ACA | C>G | G/C+1 | G/C-1 | CG>GC | CG>GC | ATTTGACAG | CAGGGAA | TGT | ACA | C | C | C>G | C>G_ACA | CG>GC_CAGGGAA | 8 | FALSE | FALSE |
| 162_v.txt | 32829 | 32829-SNV-( | SNV | 1 | G | C | 11 | 24455 | 0.04498058 | 42.7272727 | rnr1Y285F | FALSE | CAN | JSY3865 | 6 | rnr1Y285F-R | TGT | G>C | G/C+1 | G/C-1 | CG>GC | CG>GC | ATTATTGTG | GTGATAA | ACA | ACA | C | G | C>G | C>G_ACA | CG>GC_GTGATAA | 8 | FALSE | FALSE |
| 162_v.txt | 32917 | 32917-SNV-( | SNV | 1 | C | G | 2 | 24498 | 8.16E-03 | 64 | rnr1Y285F | FALSE | CAN | JSY3865 | 6 | rnr1Y285F-R | TCC | C>G | G/C+1 | G/C-1 | CG>GC | CG>GC | TAAGTTCCA | CCAGGGC | AGG | TCC | C | C | C>G | C>G_TCC | CG>GC_CCAGGGC | 8 | FALSE | FALSE |
| 162_v.txt | 32935 | 32935-SNV-( | SNV | 1 | C | G | 12 | 24498 | 0.04898359 | 51.6666667 | rnr1Y285F | FALSE | CAN | JSY3865 | 6 | rnr1Y285F-R | GCC | C>G | G/C+1 | G/C-1 | CG>GC | CG>GC | TGATTGCCCA | CCCAAGA | CGG | GCC | C | C | C>G | C>G_GCC | CG>GC_CCCAAGA | 8 | FALSE | FALSE |
| 162_v.txt | 33114 | 33114-SNV-( | SNV | 1 | C | G | 10 | 24498 | 0.04081966 | 43.2 | rnr1Y285F | FALSE | CAN | JSY3865 | 6 | rnr1Y285F-R | GCC | C>G | G/C+1 | G/C-1 | CG>GC | CG>GC | ACTGGGCCG | CCGGCGT | CGG | GCC | C | C | C>G | C>G_GCC | CG>GC_CCGGCGT | 8 | FALSE | FALSE |
| 162_v.txt | 33154 | 33154-SNV-( | SNV | 1 | C | G | 3 | 24498 | 0.0122459 | 35.9998313 | rnr1Y285F | FALSE | CAN | JSY3865 | 6 | rnr1Y285F-R | CCT | C>G | G/C+1 | G/C-1 | CG>GC | CG>GC | AAAGACCTG | CTGTACC | GGA | CCT | C | C | C>G | C>G_CCT | CG>GC_CTGTACC | 8 | FALSE | FALSE |
| 162_v.txt | 33156 | 33156-SNV-( | SNV | 1 | G | C | 29 | 24496 | 0.11838668 | 35.9655172 | rnr1Y285F | FALSE | CAN | JSY3865 | 6 | rnr1Y285F-R | TGT | G>C | G/C+1 | G/C-1 | CG>GC | CG>GC | AGACCTGTA | GTACCAA | ACA | ACA | C | G | C>G | C>G_ACA | CG>GC_GTACCAA | 8 | FALSE | FALSE |
| 162_v.txt | 33160 | 33160-SNV-( | SNV | 1 | C | G | 3 | 24496 | 0.0122469 | 38 | rnr1Y285F | FALSE | CAN | JSY3865 | 6 | rnr1Y285F-R | CCA | C>G | G/C+1 | G/C-1 | CG>GC | CG>GC | CTGTACCAAT | CAATAGT | GGT | CCA | C | C | C>G | C>G_CCA | CG>GC_CAATAGT | 8 | FALSE | FALSE |
| 162_v.txt | 33169 | 33169-SNV-( | SNV | 1 | C | G | 8 | 24496 | 0.03265839 | 37.9998912 | rnr1Y285F | FALSE | CAN | JSY3865 | 6 | rnr1Y285F-R | CCA | C>G | G/C+1 | G/C-1 | CG>GC | CG>GC | TAGTACCAC | CACCAAG | GGT | CCA | C | C | C>G | C>G_CCA | CG>GC_CACCAAG | 8 | FALSE | FALSE |
| 162_v.txt | 31657 | 31657-SNV-( | SNV | 1 | G | T | 56 | 7952 | 0.70422535 | 20.9642857 | rnr1Y285F | FALSE | CAN | JSY3865 | 6 | rnr1Y285F-R | GGC | G>T | A/T+1 | G/C-1 | CG>AT | CG>AT | GAAATGGCG | GCGTGGA | CCG | CCG | C | A | C>A | C>A_CCG | CG>AT_GCGTGGA | 8 | FALSE | FALSE |
| 162_v.txt | 31869 | 31869-SNV-( | SNV | 1 | G | T | 58 | 7952 | 0.72937626 | 46.2241379 | rnr1Y285F | FALSE | CAN | JSY3865 | 6 | rnr1Y285F-R | AGA | G>T | A/T+1 | G/C-1 | CG>AT | CG>AT | AAAATAGAG | GAGATAT | TCT | TCT | C | A | C>A | C>A_TCT | CG>AT_GAGATAT | 8 | FALSE | FALSE |
| 162_v.txt | 31930 | 31930-SNV-( | SNV | 1 | G | T | 20 | 7952 | 0.25150905 | 37.95 | rnr1Y285F | FALSE | CAN | JSY3865 | 6 | rnr1Y285F-R | TGA | G>T | A/T+1 | G/C-1 | CG>AT | CG>AT | AACCTTGAAT | GAATAAT | ACT | ACT | C | A | C>A | C>A_ACT | CG>AT_GAATAAT | 8 | FALSE | FALSE |
| 162_v.txt | 31980 | 31980-SNV-( | SNV | 1 | G | T | 19 | 21913 | 0.08670652 | 40.2105263 | rnr1Y285F | FALSE | CAN | JSY3865 | 6 | rnr1Y285F-R | GGG | G>T | A/T+1 | G/C-1 | CG>AT | CG>AT | AAGCCGGGC | GGCATTA | CCC | CCC | C | A | C>A | C>A_CCC | CG>AT_GGCATTA | 8 | FALSE | FALSE |
| 162_v.txt | 31981 | 31981-SNV-( | SNV | 1 | G | T | 15 | 21913 | 0.06845252 | 23.7413904 | rnr1Y285F | FALSE | CAN | JSY3865 | 6 | rnr1Y285F-R | GGC | G>T | A/T+1 | G/C-1 | CG>AT | CG>AT | AGCCGGGCA | GCATTAA | CCG | CCG | C | A | C>A | C>A_CCG | CG>AT_GCATTAA | 8 | FALSE | FALSE |
| 162_v.txt | 32027 | 32027-SNV-( | SNV | 1 | G | T | 57 | 21913 | 0.26011956 | 52.2280702 | rnr1Y285F | FALSE | CAN | JSY3865 | 6 | rnr1Y285F-R | GGT | G>T | A/T+1 | G/C-1 | CG>AT | CG>AT | GCCACGGTA | GTATTTC | CCA | CCA | C | A | C>A | C>A_CCA | CG>AT_GTATTTC | 8 | FALSE | FALSE |
| 162_v.txt | 32058 | 32058-SNV-( | SNV | 1 | G | T | 12 | 21913 | 0.05476201 | 60.6677634 | rnr1Y285F | FALSE | CAN | JSY3865 | 6 | rnr1Y285F-R | CGA | G>T | A/T+1 | G/C-1 | CG>AT | CG>AT | ATGTGCGAG | GAGATTG | GCT | GCT | C | A | C>A | C>A_GCT | CG>AT_GAGATTG | 8 | FALSE | FALSE |
| 162_v.txt | 32188 | 32188-SNV-( | SNV | 1 | C | A | 63 | 21913 | 0.28750057 | 44.2380952 | rnr1Y285F | FALSE | CAN | JSY3865 | 6 | rnr1Y285F-R | ACT | C>A | A/T+1 | G/C-1 | CG>AT | CG>AT | CGAAAACTG | CTGCAAT | TGA | ACT | C | T | C>A | C>A_ACT | CG>AT_CTGCAAT | 8 | FALSE | FALSE |
| 162_v.txt | 32271 | 32271-SNV-( | SNV | 1 | C | A | 40 | 23484 | 0.17032874 | 35.625 | rnr1Y285F | FALSE | CAN | JSY3865 | 6 | rnr1Y285F-R | ACG | C>A | A/T+1 | G/C-1 | CG>AT | CG>AT | AAAATACGG | CGGGAAC | TGC | ACG | C | T | C>A | C>A_ACG | CG>AT_CGGGAAC | 8 | FALSE | FALSE |
| 162_v.txt | 32272 | 32272-SNV-( | SNV | 1 | G | T | 27 | 23484 | 0.1149719 | 29.7407407 | rnr1Y285F | FALSE | CAN | JSY3865 | 6 | rnr1Y285F-R | CGG | G>T | A/T+1 | G/C-1 | CG>AT | CG>AT | AAATACGGG | GGGAACC | GCC | GCC | C | A | C>A | C>A_GCC | CG>AT_GGGAACC | 8 | FALSE | FALSE |
| 162_v.txt | 32304 | 32304-SNV-( | SNV | 1 | G | T | 45 | 23484 | 0.19161983 | 58.2222222 | rnr1Y285F | FALSE | CAN | JSY3865 | 6 | rnr1Y285F-R | AGA | G>T | A/T+1 | G/C-1 | CG>AT | CG>AT | GCGGCAGAA | GAAATAA | TCT | TCT | C | A | C>A | C>A_TCT | CG>AT_GAAATAA | 8 | FALSE | FALSE |
| 162_v.txt | 32345 | 32345-SNV-( | SNV | 1 | C | A | 17 | 23484 | 0.07238971 | 48.4716086 | rnr1Y285F | FALSE | CAN | JSY3865 | 6 | rnr1Y285F-R | GCA | C>A | A/T+1 | G/C-1 | CG>AT | CG>AT | ATGTGGCAA | CAAAACC | CGT | GCA | C | T | C>A | C>A_GCA | CG>AT_CAAAACC | 8 | FALSE | FALSE |
| 162_v.txt | 32373 | 32373-SNV-( | SNV | 1 | G | T | 36 | 23484 | 0.15329586 | 57.3888889 | rnr1Y285F | FALSE | CAN | JSY3865 | 6 | rnr1Y285F-R | AGC | G>T | A/T+1 | G/C-1 | CG>AT | CG>AT | TCAATAGCA | GCAATAA | TCG | TCG | C | A | C>A | C>A_TCG | CG>AT_GCAATAA | 8 | FALSE | FALSE |
| 162_v.txt | 32399 | 32399-SNV-( | SNV | 1 | G | T | 9 | 23484 | 0.03832397 | 62.6666667 | rnr1Y285F | FALSE | CAN | JSY3865 | 6 | rnr1Y285F-R | CGT | G>T | A/T+1 | G/C-1 | CG>AT | CG>AT | AGAAACGTA | GTAGGAA | GCA | GCA | C | A | C>A | C>A_GCA | CG>AT_GTAGGAA | 8 | FALSE | FALSE |
| 162_v.txt | 32432 | 32432-SNV-( | SNV | 1 | G | T | 28 | 23484 | 0.11923011 | 55.25 | rnr1Y285F | FALSE | CAN | JSY3865 | 6 | rnr1Y285F-R | TGT | G>T | A/T+1 | G/C-1 | CG>AT | CG>AT | GTCATTGTAT | GTATGGA | ACA | ACA | C | A | C>A | C>A_ACA | CG>AT_GTATGGA | 8 | FALSE | FALSE |
| 162_v.txt | 32487 | 32487-SNV-( | SNV | 1 | C | A | 4 | 23484 | 0.01703287 | 42.75 | rnr1Y285F | FALSE | CAN | JSY3865 | 6 | rnr1Y285F-R | ACG | C>A | A/T+1 | G/C-1 | CG>AT | CG>AT | AAGATACGG | CGGAAAA | TGC | ACG | C | T | C>A | C>A_ACG | CG>AT_CGGAAAA | 8 | FALSE | FALSE |
| 162_v.txt | 32488 | 32488-SNV-( | SNV | 1 | G | T | 10 | 23484 | 0.04258218 | 45.5 | rnr1Y285F | FALSE | CAN | JSY3865 | 6 | rnr1Y285F-R | CGG | G>T | A/T+1 | G/C-1 | CG>AT | CG>AT | AGATACGGA | GGAAAAC | GCC | GCC | C | A | C>A | C>A_GCC | CG>AT_GGAAAAC | 8 | FALSE | FALSE |
| 162_v.txt | 32529 | 32529-SNV-( | SNV | 1 | G | T | 8 | 23484 | 0.03406575 | 36.2484712 | rnr1Y285F | FALSE | CAN | JSY3865 | 6 | rnr1Y285F-R | GGG | G>T | A/T+1 | G/C-1 | CG>AT | CG>AT | TTTCTGGGG | GGGTTTG | CCC | CCC | C | A | C>A | C>A_CCC | CG>AT_GGGTTTG | 8 | FALSE | FALSE |
| 162_v.txt | 32530 | 32530-SNV-( | SNV | 1 | G | T | 68 | 23484 | 0.28955885 | 38.8222437 | rnr1Y285F | FALSE | CAN | JSY3865 | 6 | rnr1Y285F-R | GGG | G>T | A/T+1 | G/C-1 | CG>AT | CG>AT | TTCTGGGGT | GGTTTGC | CCC | CCC | C | A | C>A | C>A_CCC | CG>AT_GGTTTGC | 8 | FALSE | FALSE |
| 162_v.txt | 32531 | 32531-SNV-( | SNV | 1 | G | T | 71 | 23484 | 0.3023335 | 40.3802817 | rnr1Y285F | FALSE | CAN | JSY3865 | 6 | rnr1Y285F-R | GGT | G>T | A/T+1 | G/C-1 | CG>AT | CG>AT | TCTGGGGTT | GTTTGCA | CCA | CCA | C | A | C>A | C>A_CCA | CG>AT_GTTTGCA | 8 | FALSE | FALSE |
| 162_v.txt | 32556 | 32556-SNV-( | SNV | 1 | C | A | 58 | 23477 | 0.24705031 | 36.9827586 | rnr1Y285F | FALSE | CAN | JSY3865 | 6 | rnr1Y285F-R | ACC | C>A | A/T+1 | G/C-1 | CG>AT | CG>AT | GTGATACCA | CCAACTA | TGG | ACC | C | T | C>A | C>A_ACC | CG>AT_CCAACTA | 8 | FALSE | FALSE |
| 162_v.txt | 32572 | 32572-SNV-( | SNV | 1 | C | A | 18 | 24455 | 0.07360458 | 34.4444444 | rnr1Y285F | FALSE | CAN | JSY3865 | 6 | rnr1Y285F-R | CCT | C>A | A/T+1 | G/C-1 | CG>AT | CG>AT | CAGTACCTTG | CTTGAAA | GGA | CCT | C | T | C>A | C>A_CCT | CG>AT_CTTGAAA | 8 | FALSE | FALSE |
| 162_v.txt | 32580 | 32580-SNV-( | SNV | 1 | G | T | 26 | 24455 | 0.10631773 | 36.4590719 | rnr1Y285F | FALSE | CAN | JSY3865 | 6 | rnr1Y285F-R | TGT | G>T | A/T+1 | G/C-1 | CG>AT | CG>AT | TGAAATGTG | GTGAAGG | ACA | ACA | C | A | C>A | C>A_ACA | CG>AT_GTGAAGG | 8 | FALSE | FALSE |
| 162_v.txt | 32621 | 32621-SNV-( | SNV | 1 | C | A | 15 | 24455 | 0.06133715 | 40 | rnr1Y285F | FALSE | CAN | JSY3865 | 6 | rnr1Y285F-R | ACC | C>A | A/T+1 | G/C-1 | CG>AT | CG>AT | TAAGAACCT | CCTCCCT | TGG | ACC | C | T | C>A | C>A_ACC | CG>AT_CCTCCCT | 8 | FALSE | FALSE |
| 162_v.txt | 32629 | 32629-SNV-( | SNV | 1 | C | A | 47 | 24455 | 0.19218974 | 54.0851064 | rnr1Y285F | FALSE | CAN | JSY3865 | 6 | rnr1Y285F-R | TCG | C>A | A/T+1 | G/C-1 | CG>AT | CG>AT | TCCCTTCGTT | CGTTTTT | AGC | TCG | C | T | C>A | C>A_TCG | CG>AT_CGTTTTT | 8 | FALSE | FALSE |
| 162_v.txt | 32735 | 32735-SNV-( | SNV | 1 | G | T | 10 | 24455 | 0.04089143 | 52.8 | rnr1Y285F | FALSE | CAN | JSY3865 | 6 | rnr1Y285F-R | AGT | G>T | A/T+1 | G/C-1 | CG>AT | CG>AT | AAAACAGTA | GTATATT | TCA | TCA | C | A | C>A | C>A_TCA | CG>AT_GTATATT | 8 | FALSE | FALSE |
| 162_v.txt | 32780 | 32780-SNV-( | SNV | 1 | C | A | 8 | 24455 | 0.03271315 | 28.1219446 | rnr1Y285F | FALSE | CAN | JSY3865 | 6 | rnr1Y285F-R | CCC | C>A | A/T+1 | G/C-1 | CG>AT | CG>AT | AGCGACCCA | CCAGAAC | GGG | CCC | C | T | C>A | C>A_CCC | CG>AT_CCAGAAC | 8 | FALSE | FALSE |
| 162_v.txt | 32783 | 32783-SNV-( | SNV | 1 | G | T | 42 | 24455 | 0.17174402 | 44.2380952 | rnr1Y285F | FALSE | CAN | JSY3865 | 6 | rnr1Y285F-R | AGA | G>T | A/T+1 | G/C-1 | CG>AT | CG>AT | GACCCAGAA | GAACTCG | TCT | TCT | C | A | C>A | C>A_TCT | CG>AT_GAACTCG | 8 | FALSE | FALSE |
| 162_v.txt | 32786 | 32786-SNV-( | SNV | 1 | C | A | 38 | 24455 | 0.15538745 | 38.631579 | rnr1Y285F | FALSE | CAN | JSY3865 | 6 | rnr1Y285F-R | ACT | C>A | A/T+1 | G/C-1 | CG>AT | CG>AT | CCAGAACTC | CTCGAAT | TGA | ACT | C | T | C>A | C>A_ACT | CG>AT_CTCGAAT | 8 | FALSE | FALSE |
| 162_v.txt | 32796 | 32796-SNV-( | SNV | 1 | C | A | 4 | 24455 | 0.01635657 | 50.5030917 | rnr1Y285F | FALSE | CAN | JSY3865 | 6 | rnr1Y285F-R | ACC | C>A | A/T+1 | G/C-1 | CG>AT | CG>AT | AATTCACCGT | CCGTAAT | TGG | ACC | C | T | C>A | C>A_ACC | CG>AT_CCGTAAT | 8 | FALSE | FALSE |
| 162_v.txt | 32798 | 32798-SNV-( | SNV | 1 | G | T | 27 | 24455 | 0.11040687 | 50.2592593 | rnr1Y285F | FALSE | CAN | JSY3865 | 6 | rnr1Y285F-R | CGT | G>T | A/T+1 | G/C-1 | CG>AT | CG>AT | TTCACCGTA | GTAATAT | GCA | GCA | C | A | C>A | C>A_GCA | CG>AT_GTAATAT | 8 | FALSE | FALSE |
| 162_v.txt | 32812 | 32812-SNV-( | SNV | 1 | G | T | 9 | 24455 | 0.03680229 | 49.5555556 | rnr1Y285F | FALSE | CAN | JSY3865 | 6 | rnr1Y285F-R | GGG | G>T | A/T+1 | G/C-1 | CG>AT | CG>AT | TGACAGGGA | GGAACAA | CCC | CCC | C | A | C>A | C>A_CCC | CG>AT_GGAACAA | 8 | FALSE | FALSE |
| 162_v.txt | 32840 | 32840-SNV-( | SNV | 1 | C | A | 6 | 24455 | 0.02453486 | 41.1647093 | rnr1Y285F | FALSE | CAN | JSY3865 | 6 | rnr1Y285F-R | CCC | C>A | A/T+1 | G/C-1 | CG>AT | CG>AT | AATTACCCA | CCAAAAA | GGG | CCC | C | T | C>A | C>A_CCC | CG>AT_CCAAAAA | 8 | FALSE | FALSE |
| 162_v.txt | 32917 | 32917-SNV-( | SNV | 1 | C | A | 93 | 24498 | 0.37962283 | 53.7956989 | rnr1Y285F | FALSE | CAN | JSY3865 | 6 | rnr1Y285F-R | TCC | C>A | A/T+1 | G/C-1 | CG>AT | CG>AT | TAAGTTCCA | CCAGGGC | AGG | TCC | C | T | C>A | C>A_TCC | CG>AT_CCAGGGC | 8 | FALSE | FALSE |
| 162_v.txt | 32940 | 32940-SNV-( | SNV | 1 | G | T | 6139 | 24498 | 25.0591885 | 56.3319759 | rnr1Y285F | FALSE | CAN | JSY3865 | 6 | rnr1Y285F-R | AGA | G>T | A/T+1 | G/C-1 | CG>AT | CG>AT | GCCCAAGAA | GAAAACC | TCT | TCT | C | A | C>A | C>A_TCT | CG>AT_GAAAACC | 8 | FALSE | FALSE |
| 162_v.txt | 32954 | 32954-SNV-( | SNV | 1 | G | T | 21 | 24498 | 0.08572128 | 53.9047619 | rnr1Y285F | FALSE | CAN | JSY3865 | 6 | rnr1Y285F-R | TGT | G>T | A/T+1 | G/C-1 | CG>AT | CG>AT | ATACATGTA | GTAACCA | ACA | ACA | C | A | C>A | C>A_ACA | CG>AT_GTAACCA | 8 | FALSE | FALSE |
| 162_v.txt | 32958 | 32958-SNV-( | SNV | 1 | C | A | 16 | 24498 | 0.06531145 | 50.1875 | rnr1Y285F | FALSE | CAN | JSY3865 | 6 | rnr1Y285F-R | ACC | C>A | A/T+1 | G/C-1 | CG>AT | CG>AT | ATGTAACCAT | CCATTGG | TGG | ACC | C | T | C>A | C>A_ACC | CG>AT_CCATTGG | 8 | FALSE | FALSE |
| 162_v.txt | 33015 | 33015-SNV-( | SNV | 1 | G | T | 45 | 24498 | 0.18368846 | 56.0666667 | rnr1Y285F | FALSE | CAN | JSY3865 | 6 | rnr1Y285F-R | GGA | G>T | A/T+1 | G/C-1 | CG>AT | CG>AT | AAAGAGGAT | GATGTAA | CCT | CCT | C | A | C>A | C>A_CCT | CG>AT_GATGTAA | 8 | FALSE | FALSE |
| 162_v.txt | 33024 | 33024-SNV-( | SNV | 1 | G | T | 6 | 24498 | 0.0244918 | 55.1638054 | rnr1Y285F | FALSE | CAN | JSY3865 | 6 | rnr1Y285F-R | AGG | G>T | A/T+1 | G/C-1 | CG>AT | CG>AT | GTAACAGGG | GGGATGA | TCC | TCC | C | A | C>A | C>A_TCC | CG>AT_GGGATGA | 8 | FALSE | FALSE |
| 162_v.txt | 33025 | 33025-SNV-( | SNV | 1 | G | T | 4 | 24498 | 0.01632786 | 64 | rnr1Y285F | FALSE | CAN | JSY3865 | 6 | rnr1Y285F-R | GGG | G>T | A/T+1 | G/C-1 | CG>AT | CG>AT | TAACAGGGA | GGATGAA | CCC | CCC | C | A | C>A | C>A_CCC | CG>AT_GGATGAA | 8 | FALSE | FALSE |
| 162_v.txt | 33093 | 33093-SNV-( | SNV | 1 | G | T | 28 | 24498 | 0.11429505 | 47.0357143 | rnr1Y285F | FALSE | CAN | JSY3865 | 6 | rnr1Y285F-R | TGA | G>T | A/T+1 | G/C-1 | CG>AT | CG>AT | AAATATGAT | GATATAA | ACT | ACT | C | A | C>A | C>A_ACT | CG>AT_GATATAA | 8 | FALSE | FALSE |
| 162_v.txt | 33151 | 33151-SNV-( | SNV | 1 | G | T | 144 | 24498 | 0.58780309 | 38.125 | rnr1Y285F | FALSE | CAN | JSY3865 | 6 | rnr1Y285F-R | AGA | G>T | A/T+1 | G/C-1 | CG>AT | CG>AT | TGAAAAGAC | GACCTGT | TCT | TCT | C | A | C>A | C>A_TCT | CG>AT_GACCTGT | 8 | FALSE | FALSE |
| 162_v.txt | 33153 | 33153-SNV-( | SNV | 1 | C | A | 25 | 24498 | 0.10204915 | 38.4398648 | rnr1Y285F | FALSE | CAN | JSY3865 | 6 | rnr1Y285F-R | ACC | C>A | A/T+1 | G/C-1 | CG>AT | CG>AT | AAAAGACCT | CCTGTAC | TGG | ACC | C | T | C>A | C>A_ACC | CG>AT_CCTGTAC | 8 | FALSE | FALSE |
| 162_v.txt | 33154 | 33154-SNV-( | SNV | 1 | C | A | 63 | 24498 | 0.25716385 | 35.4285043 | rnr1Y285F | FALSE | CAN | JSY3865 | 6 | rnr1Y285F-R | CCT | C>A | A/T+1 | G/C-1 | CG>AT | CG>AT | AAAGACCTG | CTGTACC | GGA | CCT | C | T | C>A | C>A_CCT | CG>AT_CTGTACC | 8 | FALSE | FALSE |
| 162_v.txt | 33159 | 33159-SNV-( | SNV | 1 | C | A | 25 | 24496 | 0.10205748 | 37.96 | rnr1Y285F | FALSE | CAN | JSY3865 | 6 | rnr1Y285F-R | ACC | C>A | A/T+1 | G/C-1 | CG>AT | CG>AT | CCTGTACCA | CCAATAG | TGG | ACC | C | T | C>A | C>A_ACC | CG>AT_CCAATAG | 8 | FALSE | FALSE |
| 162_v.txt | 33160 | 33160-SNV-( | SNV | 1 | C | A | 19 | 24496 | 0.07756368 | 37.4736978 | rnr1Y285F | FALSE | CAN | JSY3865 | 6 | rnr1Y285F-R | CCA | C>A | A/T+1 | G/C-1 | CG>AT | CG>AT | CTGTACCAAT | CAATAGT | GGT | CCA | C | T | C>A | C>A_CCA | CG>AT_CAATAGT | 8 | FALSE | FALSE |
| 162_v.txt | 33168 | 33168-SNV-( | SNV | 1 | C | A | 60 | 24496 | 0.24493795 | 36.9836611 | rnr1Y285F | FALSE | CAN | JSY3865 | 6 | rnr1Y285F-R | ACC | C>A | A/T+1 | G/C-1 | CG>AT | CG>AT | ATAGTACCA | CCACCAA | TGG | ACC | C | T | C>A | C>A_ACC | CG>AT_CCACCAA | 8 | FALSE | FALSE |
| 162_v.txt | 33177 | 33177-SNV-( | SNV | 1 | G | T | 22 | 24490 | 0.08983259 | 34.8636364 | rnr1Y285F | FALSE | CAN | JSY3865 | 6 | rnr1Y285F-R | GGC | G>T | A/T+1 | G/C-1 | CG>AT | CG>AT | CCAAGGGCA | GCAATCA | CCG | CCG | C | A | C>A | C>A_CCG | CG>AT_GCAATCA | 8 | FALSE | FALSE |
| 162_v.txt | 33186 | 33186-SNV-( | SNV | 1 | C | A | 13 | 20803 | 0.06249099 | 34.5384615 | rnr1Y285F | FALSE | CAN | JSY3865 | 6 | rnr1Y285F-R | ACC | C>A | A/T+1 | G/C-1 | CG>AT | CG>AT | ATCATACCA | CCAATAT | TGG | ACC | C | T | C>A | C>A_ACC | CG>AT_CCAATAT | 8 | FALSE | FALSE |
| 162_v.txt | 33325 | 33325-SNV-( | SNV | 1 | C | A | 308 | 20807 | 1.48027106 | 58.2597403 | rnr1Y285F | FALSE | CAN | JSY3865 | 6 | rnr1Y285F-R | TCT | C>A | A/T+1 | G/C-1 | CG>AT | CG>AT | ACAATTCTTT | CTTTACT | AGA | TCT | C | T | C>A | C>A_TCT | CG>AT_CTTTACT | 8 | FALSE | FALSE |
| 162_v.txt | 33354 | 33354-SNV-( | SNV | 1 | G | T | 145 | 20807 | 0.69688086 | 23.1448276 | rnr1Y285F | FALSE | CAN | JSY3865 | 6 | rnr1Y285F-R | TGA | G>T | A/T+1 | G/C-1 | CG>AT | CG>AT | GGTATTGAC | GACCCAC | ACT | ACT | C | A | C>A | C>A_ACT | CG>AT_GACCCAC | 8 | FALSE | FALSE |
| 162_v.txt | 33474 | 33474-SNV-( | SNV | 1 | C | A | 46 | 20804 | 0.22111133 | 34.1309564 | rnr1Y285F | FALSE | CAN | JSY3865 | 6 | rnr1Y285F-R | GCC | C>A | A/T+1 | G/C-1 | CG>AT | CG>AT | GCTATGCCTT | CCTTTTT | CGG | GCC | C | T | C>A | C>A_GCC | CG>AT_CCTTTTT | 8 | FALSE | FALSE |
| 162_v.txt | 33475 | 33475-SNV-( | SNV | 1 | C | A | 17 | 20802 | 0.08172291 | 37.1764706 | rnr1Y285F | FALSE | CAN | JSY3865 | 6 | rnr1Y285F-R | CCT | C>A | A/T+1 | G/C-1 | CG>AT | CG>AT | CTATGCCTTT | CTTTTTT | GGA | CCT | C | T | C>A | C>A_CCT | CG>AT_CTTTTTT | 8 | FALSE | FALSE |
| 162_v.txt | 32839 | 32839-MNV- | MNV | 2 | CC | AA | 1 | 24455 | 4.09E-03 | 26.6668212 | rnr1Y285F | FALSE | CAN | JSY3865 | 6 | rnr1Y285F-R | ACC | nope | >1bp-ins | >1bpdel | none | MNV | TAATTACCCA | CCCAAAA | TGG | ACC | CC | none | NA | NA_ACC | MNV_CCCAAAA | 8 | FALSE | FALSE |
| 162_v.txt | 33168 | 33168-MNV- | MNV | 2 | CC | AA | 1 | 24496 | 4.08E-03 | 32.0000932 | rnr1Y285F | FALSE | CAN | JSY3865 | 6 | rnr1Y285F-R | ACC | nope | >1bp-ins | >1bpdel | none | MNV | ATAGTACCA | CCACCAA | TGG | ACC | CC | none | NA | NA_ACC | MNV_CCACCAA | 8 | FALSE | FALSE |
| 162_v.txt | 33309 | 33309-MNV- | MNV | 2 | CG | TA | 1 | 20807 | 4.81E-03 | 62.9999265 | rnr1Y285F | FALSE | CAN | JSY3865 | 6 | rnr1Y285F-R | GCG | nope | >1bp-ins | >1bpdel | none | MNV | AAAGAGCGC | CGCAATG | CGC | GCG | CG | none | NA | NA_GCG | MNV_CGCAATG | 8 | FALSE | FALSE |
| 162_v.txt | 32081 | 32081-Insert | Insertion | 1 | - | A | 44 | 21913 | 0.20079405 | 62.8636364 | rnr1Y285F | FALSE | CAN | JSY3865 | 6 | rnr1Y285F-R | CAA | nope | A/T+1 | >1bpdel | none | A/T+1 | CCATGCAAA | AAAAAAG | GTT | CAA | - | T | NA | NA_CAA | A/T+1_AAAAAAG | 8 | FALSE | FALSE |
| 162_v.txt | 32183 | 32183-Insert | Insertion | 2 | - | AA | 11 | 21913 | 0.05019851 | 45.2272727 | rnr1Y285F | FALSE | CAN | JSY3865 | 6 | rnr1Y285F-R | CGA | nope | >1bp-ins | >1bpdel | none | >1bp-ins | AGTAACGAA | GAAAACT | GCT | CGA | - | none | NA | NA_CGA | >1bp-ins_GAAAACT | 8 | FALSE | FALSE |
| 162_v.txt | 32344 | 32344-Insert | Insertion | 1 | - | A | 9 | 23484 | 0.03832397 | 60.5555556 | rnr1Y285F | FALSE | CAN | JSY3865 | 6 | rnr1Y285F-R | GGC | nope | A/T+1 | >1bpdel | none | A/T+1 | TATGTGGCA | GCAAAAC | CCG | GGC | - | T | NA | NA_GGC | A/T+1_GCAAAAC | 8 | FALSE | FALSE |
| 162_v.txt | 32407 | 32407-Insert | Insertion | 1 | - | T | 524 | 23484 | 2.23130642 | 60.3912214 | rnr1Y285F | FALSE | CAN | JSY3865 | 6 | rnr1Y285F-R | GTA | nope | A/T+1 | >1bpdel | none | A/T+1 | AGGAAGTAG | TAGATTG | CAT | GTA | - | A | NA | NA_GTA | A/T+1_TAGATTG | 8 | FALSE | FALSE |
| 162_v.txt | 32448 | 32448-Insert | Insertion | 18 | - | CCAATGAAT | 10 | 23484 | 0.04258218 | 52.4555556 | rnr1Y285F | FALSE | CAN | JSY3865 | 6 | rnr1Y285F-R | TCC | nope | >1bp-ins | >1bpdel | none | >1bp-ins | AAAAGTCCA | CCAATGA | AGG | TCC | - | none | NA | NA_TCC | >1bp-ins_CCAATGA | 8 | FALSE | FALSE |
| 162_v.txt | 32448 | 32448-Insert | Insertion | 13 | - | GAATAATAG | 1 | 23484 | 4.26E-03 | 61.3846154 | rnr1Y285F | FALSE | CAN | JSY3865 | 6 | rnr1Y285F-R | TCC | nope | >1bp-ins | >1bpdel | none | >1bp-ins | AAAAGTCCA | CCAATGA | AGG | TCC | - | none | NA | NA_TCC | >1bp-ins_CCAATGA | 8 | FALSE | FALSE |
| 162_v.txt | 32450 | 32450-Insert | Insertion | 1 | - | A | 8 | 23484 | 0.03406575 | 46.375 | rnr1Y285F | FALSE | CAN | JSY3865 | 6 | rnr1Y285F-R | CAA | nope | A/T+1 | >1bpdel | none | A/T+1 | AAGTCCAAT | AATGAAT | GTT | CAA | - | T | NA | NA_CAA | A/T+1_AATGAAT | 8 | FALSE | FALSE |
| 162_v.txt | 32460 | 32460-Insert | Insertion | 2 | - | AG | 14 | 23484 | 0.05961506 | 48.9285714 | rnr1Y285F | FALSE | CAN | JSY3865 | 6 | rnr1Y285F-R | TAG | nope | >1bp-ins | >1bpdel | none | >1bp-ins | AATAATAGA | AGAGAGC | ATC | TAG | - | none | NA | NA_TAG | >1bp-ins_AGAGAGC | 8 | FALSE | FALSE |
| 162_v.txt | 32498 | 32498-Insert | Insertion | 1 | - | T | 18 | 23484 | 0.07664793 | 37.7222222 | rnr1Y285F | FALSE | CAN | JSY3865 | 6 | rnr1Y285F-R | CTT | nope | A/T+1 | >1bpdel | none | A/T+1 | AACAACTTT | TTTTTTG | GAA | CTT | - | A | NA | NA_CTT | A/T+1_TTTTTTG | 8 | FALSE | FALSE |
| 162_v.txt | 32528 | 32528-Insert | Insertion | 1 | - | G | 2 | 23484 | 8.52E-03 | 38 | rnr1Y285F | FALSE | CAN | JSY3865 | 6 | rnr1Y285F-R | TGG | nope | G/C+1 | >1bpdel | none | G/C+1 | TTTTCTGGG | GGGGTTT | ACC | TGG | - | C | NA | NA_TGG | G/C+1_GGGGTTT | 8 | FALSE | FALSE |
| 162_v.txt | 32581 | 32581-Insert | Insertion | 1 | - | T | 1 | 24455 | 4.09E-03 | 11 | rnr1Y285F | FALSE | CAN | JSY3865 | 6 | rnr1Y285F-R | GTG | nope | A/T+1 | >1bpdel | none | A/T+1 | GAAATGTGA | TGAAGGC | CAC | GTG | - | A | NA | NA_GTG | A/T+1_TGAAGGC | 8 | FALSE | FALSE |
| 162_v.txt | 32840 | 32840-Insert | Insertion | 1 | - | T | 1 | 24455 | 4.09E-03 | 37.9999999 | rnr1Y285F | FALSE | CAN | JSY3865 | 6 | rnr1Y285F-R | CCC | nope | A/T+1 | >1bpdel | none | A/T+1 | AATTACCCA | CCAAAAA | GGG | CCC | - | A | NA | NA_CCC | A/T+1_CCAAAAA | 8 | FALSE | FALSE |
| 162_v.txt | 32842 | 32842-Insert | Insertion | 1 | - | A | 73 | 24455 | 0.29850746 | 40.2876712 | rnr1Y285F | FALSE | CAN | JSY3865 | 6 | rnr1Y285F-R | CAA | nope | A/T+1 | >1bpdel | none | A/T+1 | TTACCCAAA | AAAAAAT | GTT | CAA | - | T | NA | NA_CAA | A/T+1_AAAAAAT | 8 | FALSE | FALSE |
| 162_v.txt | 32972 | 32972-Insert | Insertion | 1 | - | A | 10 | 24498 | 0.04081966 | 64 | rnr1Y285F | FALSE | CAN | JSY3865 | 6 | rnr1Y285F-R | CAA | nope | A/T+1 | >1bpdel | none | A/T+1 | CGCACCAAA | AAATGCT | GTT | CAA | - | T | NA | NA_CAA | A/T+1_AAATGCT | 8 | FALSE | FALSE |
| 162_v.txt | 33146 | 33146-Insert | Insertion | 1 | - | C | 6 | 24498 | 0.0244918 | 38 | rnr1Y285F | FALSE | CAN | JSY3865 | 6 | rnr1Y285F-R | TGA | nope | G/C+1 | >1bpdel | none | G/C+1 | ACCAATGAA | GAAAAGA | ACT | TGA | - | G | NA | NA_TGA | G/C+1_GAAAAGA | 8 | FALSE | FALSE |
| 162_v.txt | 33188 | 33188-Insert | Insertion | 1 | - | A | 7 | 20804 | 0.03364738 | 38 | rnr1Y285F | FALSE | CAN | JSY3865 | 6 | rnr1Y285F-R | CAA | nope | A/T+1 | >1bpdel | none | A/T+1 | CATACCAAT | AATATGT | GTT | CAA | - | T | NA | NA_CAA | A/T+1_AATATGT | 8 | FALSE | FALSE |
| 162_v.txt | 33452 | 33452-Insert | Insertion | 1 | - | T | 4 | 20807 | 0.0192243 | 37.5 | rnr1Y285F | FALSE | CAN | JSY3865 | 6 | rnr1Y285F-R | CTT | nope | A/T+1 | >1bpdel | none | A/T+1 | GTCTTCTTTT | TTTTGAA | GAA | CTT | - | A | NA | NA_CTT | A/T+1_TTTTGAA | 8 | FALSE | FALSE |
| 162_v.txt | 32081 | 32081-Deleti | Deletion | 2 | AA | - | 1 | 21913 | 4.56E-03 | 64 | rnr1Y285F | FALSE | CAN | JSY3865 | 6 | rnr1Y285F-R | CAA | nope | >1bp-ins | >1bpdel | none | >1bpdel | CCATGCAAA | AAAAAAG | GTT | CAA | AA | none | NA | NA_CAA | >1bpdel_AAAAAAG | 8 | FALSE | FALSE |
| 162_v.txt | 32314 | 32314-Deleti | Deletion | 2 | TT | - | 5 | 23484 | 0.02129109 | 50.4 | rnr1Y285F | FALSE | CAN | JSY3865 | 6 | rnr1Y285F-R | GTT | nope | >1bp-ins | >1bpdel | none | >1bpdel | TAATGGTTG | TTGTTAA | CAA | GTT | TT | none | NA | NA_GTT | >1bpdel_TTGTTAA | 8 | FALSE | FALSE |
| 162_v.txt | 32460 | 32460-Deleti | Deletion | 2 | AG | - | 10 | 23484 | 0.04258218 | 55.5 | rnr1Y285F | FALSE | CAN | JSY3865 | 6 | rnr1Y285F-R | TAG | nope | >1bp-ins | >1bpdel | none | >1bpdel | AATAATAGA | AGAGAGC | ATC | TAG | AG | none | NA | NA_TAG | >1bpdel_AGAGAGC | 8 | FALSE | FALSE |
| 162_v.txt | 32983 | 32983-Deleti | Deletion | 2 | AA | - | 11 | 24498 | 0.04490163 | 59.6363636 | rnr1Y285F | FALSE | CAN | JSY3865 | 6 | rnr1Y285F-R | GAA | nope | >1bp-ins | >1bpdel | none | >1bpdel | CTGGAGAAA | AAAGGAA | CTT | GAA | AA | none | NA | NA_GAA | >1bpdel_AAAGGAA | 8 | FALSE | FALSE |
| 162_v.txt | 33089 | 33089-Deleti | Deletion | 2 | AT | - | 9 | 24498 | 0.03673769 | 56.6666667 | rnr1Y285F | FALSE | CAN | JSY3865 | 6 | rnr1Y285F-R | AAT | nope | >1bp-ins | >1bpdel | none | >1bpdel | AAATAAATA | ATATGAT | TTA | AAT | AT | none | NA | NA_AAT | >1bpdel_ATATGAT | 8 | FALSE | FALSE |

| | | | | | | | | | | | | | | | | | | | | | | | | | | | | | | | | | | |
|---|---|---|---|---|---|---|---|---|---|---|---|---|---|---|---|---|---|---|---|---|---|---|---|---|---|---|---|---|---|---|---|---|---|---|
| 162_v.txt | 31723 | 31723-Deleti | Deletion | 1 | A | - | 7 | 7952 | 0.08802817 | 56.2857143 | rnr1Y285F | FALSE | CAN | JSY3865 | 6 | rnr1Y285F-R | CAA | nope | >1bp-ins | A/T-1 | none | A/T-1 | TGTCCCAAA | AAAAAGT | GTT | GTT | T | none | T>none | T>none_GTT | A/T-1_AAAAAGT | 8 | FALSE | FALSE |
| 162_v.txt | 31904 | 31904-Deleti | Deletion | 1 | T | - | 5 | 7952 | 0.06287726 | 44.8 | rnr1Y285F | FALSE | CAN | JSY3865 | 6 | rnr1Y285F-R | ATT | nope | >1bp-ins | A/T-1 | none | A/T-1 | ATTGAATTTT | TTTTGGT | TAA | ATT | T | none | nope | nope_ATT | A/T-1_TTTTGGT | 8 | FALSE | FALSE |
| 162_v.txt | 31972 | 31972-Deleti | Deletion | 1 | C | - | 4 | 21913 | 0.01825401 | 38 | rnr1Y285F | FALSE | CAN | JSY3865 | 6 | rnr1Y285F-R | GCC | nope | >1bp-ins | G/C-1 | none | G/C-1 | AATAAGCCA | CCAAGCC | CGG | GCC | C | none | nope | nope_GCC | G/C-1_CCAAGCC | 8 | FALSE | FALSE |
| 162_v.txt | 32057 | 32057-Deleti | Deletion | 1 | C | - | 34 | 21913 | 0.15515904 | 61.8823529 | rnr1Y285F | FALSE | CAN | JSY3865 | 6 | rnr1Y285F-R | GCG | nope | >1bp-ins | G/C-1 | none | G/C-1 | GATGTGCGA | CGAGATT | CGC | GCG | C | none | nope | nope_GCG | G/C-1_CGAGATT | 8 | FALSE | FALSE |
| 162_v.txt | 32075 | 32075-Deleti | Deletion | 1 | C | - | 2 | 21913 | 9.13E-03 | 64 | rnr1Y285F | FALSE | CAN | JSY3865 | 6 | rnr1Y285F-R | ACC | nope | >1bp-ins | G/C-1 | none | G/C-1 | AAATAACCA | CCATGCA | TGG | ACC | C | none | nope | nope_ACC | G/C-1_CCATGCA | 8 | FALSE | FALSE |
| 162_v.txt | 32081 | 32081-Deleti | Deletion | 1 | A | - | 29 | 21913 | 0.13234153 | 63.3448276 | rnr1Y285F | FALSE | CAN | JSY3865 | 6 | rnr1Y285F-R | CAA | nope | >1bp-ins | A/T-1 | none | A/T-1 | CCATGCAAA | AAAAAAG | GTT | GTT | T | none | T>none | T>none_GTT | A/T-1_AAAAAAG | 8 | FALSE | FALSE |
| 162_v.txt | 32154 | 32154-Deleti | Deletion | 1 | T | - | 5 | 21913 | 0.02281751 | 43.2 | rnr1Y285F | FALSE | CAN | JSY3865 | 6 | rnr1Y285F-R | GTA | nope | >1bp-ins | A/T-1 | none | A/T-1 | TCCATGTAAC | TAAGCCA | CAT | GTA | T | none | nope | nope_GTA | A/T-1_TAAGCCA | 8 | FALSE | FALSE |
| 162_v.txt | 32210 | 32210-Deleti | Deletion | 1 | T | - | 5 | 21913 | 0.02281751 | 37.6 | rnr1Y285F | FALSE | CAN | JSY3865 | 6 | rnr1Y285F-R | CTT | nope | >1bp-ins | A/T-1 | none | A/T-1 | ACCACCTTTG | TTTGGTG | GAA | CTT | T | none | nope | nope_CTT | A/T-1_TTTGGTG | 8 | FALSE | FALSE |
| 162_v.txt | 32272 | 32272-Deleti | Deletion | 1 | G | - | 1 | 23484 | 4.26E-03 | 37 | rnr1Y285F | FALSE | CAN | JSY3865 | 6 | rnr1Y285F-R | CGG | nope | >1bp-ins | G/C-1 | none | G/C-1 | AAATACGGG | GGGAACC | GCC | GCC | C | none | C>none | C>none_GCC | G/C-1_GGGAACC | 8 | FALSE | FALSE |
| 162_v.txt | 32346 | 32346-Deleti | Deletion | 1 | A | - | 28 | 23484 | 0.11923011 | 63.2857143 | rnr1Y285F | FALSE | CAN | JSY3865 | 6 | rnr1Y285F-R | CAA | nope | >1bp-ins | A/T-1 | none | A/T-1 | TGTGGCAAA | AAAACCT | GTT | GTT | T | none | T>none | T>none_GTT | A/T-1_AAAACCT | 8 | FALSE | FALSE |
| 162_v.txt | 32381 | 32381-Deleti | Deletion | 1 | A | - | 7 | 23484 | 0.02980753 | 54.7142857 | rnr1Y285F | FALSE | CAN | JSY3865 | 6 | rnr1Y285F-R | TAA | nope | >1bp-ins | A/T-1 | none | A/T-1 | AATAATAAA | AAAGGGA | ATT | ATT | T | none | T>none | T>none_ATT | A/T-1_AAAGGGA | 8 | FALSE | FALSE |
| 162_v.txt | 32424 | 32424-Deleti | Deletion | 1 | G | - | 48 | 23484 | 0.20439448 | 55.5416667 | rnr1Y285F | FALSE | CAN | JSY3865 | 6 | rnr1Y285F-R | AGG | nope | >1bp-ins | G/C-1 | none | G/C-1 | AGTTTAGGG | GGGTCAT | TCC | TCC | C | none | C>none | C>none_TCC | G/C-1_GGGTCAT | 8 | FALSE | FALSE |
| 162_v.txt | 32498 | 32498-Deleti | Deletion | 1 | T | - | 34 | 23484 | 0.14477942 | 46.2647038 | rnr1Y285F | FALSE | CAN | JSY3865 | 6 | rnr1Y285F-R | CTT | nope | >1bp-ins | A/T-1 | none | A/T-1 | AACAACTTTT | TTTTTTG | GAA | CTT | T | none | nope | nope_CTT | A/T-1_TTTTTTG | 8 | FALSE | FALSE |
| 162_v.txt | 32522 | 32522-Deleti | Deletion | 1 | T | - | 16 | 23484 | 0.06813149 | 39.5 | rnr1Y285F | FALSE | CAN | JSY3865 | 6 | rnr1Y285F-R | ATT | nope | >1bp-ins | A/T-1 | none | A/T-1 | AACGGATTTT | TTTTCTG | TAA | ATT | T | none | nope | nope_ATT | A/T-1_TTTTCTG | 8 | FALSE | FALSE |
| 162_v.txt | 32528 | 32528-Deleti | Deletion | 1 | G | - | 7 | 23484 | 0.02980753 | 41.7142808 | rnr1Y285F | FALSE | CAN | JSY3865 | 6 | rnr1Y285F-R | TGG | nope | >1bp-ins | G/C-1 | none | G/C-1 | TTTTCTGGGG | GGGGTTT | ACC | ACC | C | none | C>none | C>none_ACC | G/C-1_GGGGTTT | 8 | FALSE | FALSE |
| 162_v.txt | 32543 | 32543-Deleti | Deletion | 1 | A | - | 690 | 23484 | 2.93817067 | 36.4927536 | rnr1Y285F | FALSE | CAN | JSY3865 | 6 | rnr1Y285F-R | CAC | nope | >1bp-ins | A/T-1 | none | A/T-1 | AGCTTCACCA | ACCAGCA | GTG | GTG | T | none | T>none | T>none_GTG | A/T-1_ACCAGCA | 8 | FALSE | FALSE |
| 162_v.txt | 32608 | 32608-Deleti | Deletion | 1 | C | - | 15 | 24455 | 0.06133715 | 43.5333333 | rnr1Y285F | FALSE | CAN | JSY3865 | 6 | rnr1Y285F-R | ACC | nope | >1bp-ins | G/C-1 | none | G/C-1 | AGGAAACCC | CCCAACC | TGG | ACC | C | none | nope | nope_ACC | G/C-1_CCCAACC | 8 | FALSE | FALSE |
| 162_v.txt | 32618 | 32618-Deleti | Deletion | 1 | G | - | 14 | 24455 | 0.05724801 | 38.3571429 | rnr1Y285F | FALSE | CAN | JSY3865 | 6 | rnr1Y285F-R | AGA | nope | >1bp-ins | G/C-1 | none | G/C-1 | ACCTAAGAA | GAACCTC | TCT | TCT | C | none | C>none | C>none_TCT | G/C-1_GAACCTC | 8 | FALSE | FALSE |
| 162_v.txt | 32631 | 32631-Deleti | Deletion | 1 | T | - | 16 | 24455 | 0.06542629 | 53.25 | rnr1Y285F | FALSE | CAN | JSY3865 | 6 | rnr1Y285F-R | GTT | nope | >1bp-ins | A/T-1 | none | A/T-1 | CCTTCGTTTT | TTTTTAT | CAA | GTT | T | none | nope | nope_GTT | A/T-1_TTTTTAT | 8 | FALSE | FALSE |
| 162_v.txt | 32658 | 32658-Deleti | Deletion | 1 | C | - | 433 | 24455 | 1.77059906 | 60.3418014 | rnr1Y285F | FALSE | CAN | JSY3865 | 6 | rnr1Y285F-R | ACC | nope | >1bp-ins | G/C-1 | none | G/C-1 | CCTGGACCCC | CCCCAGG | TGG | ACC | C | none | nope | nope_ACC | G/C-1_CCCCAGG | 8 | FALSE | FALSE |
| 162_v.txt | 32670 | 32670-Deleti | Deletion | 1 | G | - | 63 | 24455 | 0.25761603 | 58.1111111 | rnr1Y285F | FALSE | CAN | JSY3865 | 6 | rnr1Y285F-R | TGG | nope | >1bp-ins | G/C-1 | none | G/C-1 | GCACCTGGG | GGGTTTC | ACC | ACC | C | none | C>none | C>none_ACC | G/C-1_GGGTTTC | 8 | FALSE | FALSE |
| 162_v.txt | 32707 | 32707-Deleti | Deletion | 1 | C | - | 154 | 24455 | 0.62972807 | 61.987013 | rnr1Y285F | FALSE | CAN | JSY3865 | 6 | rnr1Y285F-R | ACC | nope | >1bp-ins | G/C-1 | none | G/C-1 | CGGTAACCC | CCCCAGC | TGG | ACC | C | none | nope | nope_ACC | G/C-1_CCCCAGC | 8 | FALSE | FALSE |
| 162_v.txt | 32728 | 32728-Deleti | Deletion | 1 | A | - | 6 | 24455 | 0.02453486 | 63.6666667 | rnr1Y285F | FALSE | CAN | JSY3865 | 6 | rnr1Y285F-R | CAA | nope | >1bp-ins | A/T-1 | none | A/T-1 | CCATACAAA | AAAAACA | GTT | GTT | T | none | T>none | T>none_GTT | A/T-1_AAAAACA | 8 | FALSE | FALSE |
| 162_v.txt | 32747 | 32747-Deleti | Deletion | 1 | C | - | 25 | 24455 | 0.10222858 | 55.56 | rnr1Y285F | FALSE | CAN | JSY3865 | 6 | rnr1Y285F-R | ACC | nope | >1bp-ins | G/C-1 | none | G/C-1 | TAGAAACCC | CCCGATA | TGG | ACC | C | none | nope | nope_ACC | G/C-1_CCCGATA | 8 | FALSE | FALSE |
| 162_v.txt | 32760 | 32760-Deleti | Deletion | 1 | A | - | 31 | 24455 | 0.12676344 | 55.9032258 | rnr1Y285F | FALSE | CAN | JSY3865 | 6 | rnr1Y285F-R | TAA | nope | >1bp-ins | A/T-1 | none | A/T-1 | ATGGCTAAA | AAAACTT | ATT | ATT | T | none | T>none | T>none_ATT | A/T-1_AAAACTT | 8 | FALSE | FALSE |
| 162_v.txt | 32779 | 32779-Deleti | Deletion | 1 | C | - | 5 | 24455 | 0.02044572 | 42.999999 | rnr1Y285F | FALSE | CAN | JSY3865 | 6 | rnr1Y285F-R | ACC | nope | >1bp-ins | G/C-1 | none | G/C-1 | AAGCGACCC | CCCAGAA | TGG | ACC | C | none | nope | nope_ACC | G/C-1_CCCAGAA | 8 | FALSE | FALSE |
| 162_v.txt | 32839 | 32839-Deleti | Deletion | 1 | C | - | 8 | 24455 | 0.03271315 | 43.3749983 | rnr1Y285F | FALSE | CAN | JSY3865 | 6 | rnr1Y285F-R | ACC | nope | >1bp-ins | G/C-1 | none | G/C-1 | TAATTACCCA | CCCAAAA | TGG | ACC | C | none | nope | nope_ACC | G/C-1_CCCAAAA | 8 | FALSE | FALSE |
| 162_v.txt | 32842 | 32842-Deleti | Deletion | 1 | A | - | 77 | 24455 | 0.31486404 | 40.9220775 | rnr1Y285F | FALSE | CAN | JSY3865 | 6 | rnr1Y285F-R | CAA | nope | >1bp-ins | A/T-1 | none | A/T-1 | TTACCCAAA | AAAAAAT | GTT | GTT | T | none | T>none | T>none_GTT | A/T-1_AAAAAAT | 8 | FALSE | FALSE |
| 162_v.txt | 32917 | 32917-Deleti | Deletion | 1 | C | - | 12 | 24498 | 0.04898359 | 48.4166667 | rnr1Y285F | FALSE | CAN | JSY3865 | 6 | rnr1Y285F-R | TCC | nope | >1bp-ins | G/C-1 | none | G/C-1 | TAAGTTCCA | CCAGGGC | AGG | TCC | C | none | nope | nope_TCC | G/C-1_CCAGGGC | 8 | FALSE | FALSE |
| 162_v.txt | 32924 | 32924-Deleti | Deletion | 1 | A | - | 19 | 24498 | 0.07755735 | 56.4210526 | rnr1Y285F | FALSE | CAN | JSY3865 | 6 | rnr1Y285F-R | CAA | nope | >1bp-ins | A/T-1 | none | A/T-1 | CAGGGCAAA | AAAAGTG | GTT | GTT | T | none | T>none | T>none_GTT | A/T-1_AAAAGTG | 8 | FALSE | FALSE |
| 162_v.txt | 32932 | 32932-Deleti | Deletion | 1 | T | - | 16 | 24498 | 0.06531145 | 48.75 | rnr1Y285F | FALSE | CAN | JSY3865 | 6 | rnr1Y285F-R | ATT | nope | >1bp-ins | A/T-1 | none | A/T-1 | AAGTGATTG | TTGCCCA | TAA | ATT | T | none | nope | nope_ATT | A/T-1_TTGCCCA | 8 | FALSE | FALSE |
| 162_v.txt | 32945 | 32945-Deleti | Deletion | 1 | C | - | 4 | 24498 | 0.01632786 | 59.75 | rnr1Y285F | FALSE | CAN | JSY3865 | 6 | rnr1Y285F-R | ACC | nope | >1bp-ins | G/C-1 | none | G/C-1 | AGAAAACCA | CCAATAC | TGG | ACC | C | none | nope | nope_ACC | G/C-1_CCAATAC | 8 | FALSE | FALSE |
| 162_v.txt | 32958 | 32958-Deleti | Deletion | 1 | C | - | 2 | 24498 | 8.16E-03 | 64 | rnr1Y285F | FALSE | CAN | JSY3865 | 6 | rnr1Y285F-R | ACC | nope | >1bp-ins | G/C-1 | none | G/C-1 | ATGTAACCAT | CCATTGG | TGG | ACC | C | none | nope | nope_ACC | G/C-1_CCATTGG | 8 | FALSE | FALSE |
| 162_v.txt | 32965 | 32965-Deleti | Deletion | 1 | C | - | 3 | 24498 | 0.0122459 | 55.3333333 | rnr1Y285F | FALSE | CAN | JSY3865 | 6 | rnr1Y285F-R | GCC | nope | >1bp-ins | G/C-1 | none | G/C-1 | CATTGGCCG | CCGCACC | CGG | GCC | C | none | nope | nope_GCC | G/C-1_CCGCACC | 8 | FALSE | FALSE |
| 162_v.txt | 32983 | 32983-Deleti | Deletion | 1 | A | - | 6 | 24498 | 0.0244918 | 63.3333333 | rnr1Y285F | FALSE | CAN | JSY3865 | 6 | rnr1Y285F-R | GAA | nope | >1bp-ins | A/T-1 | none | A/T-1 | CTGGAGAAA | AAAGGAA | CTT | CTT | T | none | T>none | T>none_CTT | A/T-1_AAAGGAA | 8 | FALSE | FALSE |
| 162_v.txt | 32995 | 32995-Deleti | Deletion | 1 | G | - | 64 | 24498 | 0.26124582 | 62.140625 | rnr1Y285F | FALSE | CAN | JSY3865 | 6 | rnr1Y285F-R | TGT | nope | >1bp-ins | G/C-1 | none | G/C-1 | ATCTTTGTGA | GTGAGAA | ACA | ACA | C | none | C>none | C>none_ACA | G/C-1_GTGAGAA | 8 | FALSE | FALSE |
| 162_v.txt | 33024 | 33024-Deleti | Deletion | 1 | G | - | 9 | 24498 | 0.03673769 | 63.9999988 | rnr1Y285F | FALSE | CAN | JSY3865 | 6 | rnr1Y285F-R | AGG | nope | >1bp-ins | G/C-1 | none | G/C-1 | GTAACAGGG | GGGATGA | TCC | TCC | C | none | C>none | C>none_TCC | G/C-1_GGGATGA | 8 | FALSE | FALSE |
| 162_v.txt | 33147 | 33147-Deleti | Deletion | 1 | A | - | 7 | 24498 | 0.02857376 | 41.7142857 | rnr1Y285F | FALSE | CAN | JSY3865 | 6 | rnr1Y285F-R | GAA | nope | >1bp-ins | A/T-1 | none | A/T-1 | CCAATGAAA | AAAAGAC | CTT | CTT | T | none | T>none | T>none_CTT | A/T-1_AAAAGAC | 8 | FALSE | FALSE |
| 162_v.txt | 33168 | 33168-Deleti | Deletion | 1 | C | - | 1 | 24496 | 4.08E-03 | 38 | rnr1Y285F | FALSE | CAN | JSY3865 | 6 | rnr1Y285F-R | ACC | nope | >1bp-ins | G/C-1 | none | G/C-1 | ATAGTACCA | CCACCAA | TGG | ACC | C | none | nope | nope_ACC | G/C-1_CCACCAA | 8 | FALSE | FALSE |
| 162_v.txt | 33356 | 33356-Deleti | Deletion | 1 | C | - | 983 | 20807 | 4.72437161 | 54.4506612 | rnr1Y285F | FALSE | CAN | JSY3865 | 6 | rnr1Y285F-R | ACC | nope | >1bp-ins | G/C-1 | none | G/C-1 | TATTGACCCA | CCCACGT | TGG | ACC | C | none | nope | nope_ACC | G/C-1_CCCACGT | 8 | FALSE | FALSE |
| 162_v.txt | 33474 | 33474-Deleti | Deletion | 1 | C | - | 57 | 20804 | 0.27398577 | 28.5263158 | rnr1Y285F | FALSE | CAN | JSY3865 | 6 | rnr1Y285F-R | GCC | nope | >1bp-ins | G/C-1 | none | G/C-1 | GCTATGCCTT | CCTTTTT | CGG | GCC | C | none | nope | nope_GCC | G/C-1_CCTTTTT | 8 | FALSE | FALSE |
| 163_v.txt | 31839 | 31839-SNV- | SNV | 1 | A | C | 6 | 3349 | 0.17915796 | 51 | rnr1Y285F | FALSE | CAN | JSY3866 | 7 | rnr1Y285F-R | TAA | A>C | G/C+1 | A/T-1 | TA>GC | TA>GC | TGAAATAAG | AAGATCC | ATT | ATT | T | G | T>G | T>G_ATT | TA>GC_AAGATCC | 8 | FALSE | FALSE |
| 163_v.txt | 31986 | 31986-SNV- | SNV | 1 | A | C | 87 | 23392 | 0.37192203 | 51.1494253 | rnr1Y285F | FALSE | CAN | JSY3866 | 7 | rnr1Y285F-R | TAA | A>C | G/C+1 | A/T-1 | TA>GC | TA>GC | GGCATTAATT | AATTTAG | ATT | ATT | T | G | T>G | T>G_ATT | TA>GC_AATTTAG | 8 | FALSE | FALSE |
| 163_v.txt | 32004 | 32004-SNV- | SNV | 1 | A | C | 38 | 23392 | 0.1624487 | 51.7631579 | rnr1Y285F | FALSE | CAN | JSY3866 | 7 | rnr1Y285F-R | TAA | A>C | G/C+1 | A/T-1 | TA>GC | TA>GC | AATGGTAACT | AACTCGT | ATT | ATT | T | G | T>G | T>G_ATT | TA>GC_AACTCGT | 8 | FALSE | FALSE |
| 163_v.txt | 32094 | 32094-SNV- | SNV | 1 | A | C | 18 | 23392 | 0.07694938 | 61.7222222 | rnr1Y285F | FALSE | CAN | JSY3866 | 7 | rnr1Y285F-R | AAC | A>C | G/C+1 | A/T-1 | TA>GC | TA>GC | CCTGCAACA | ACACCAG | TTG | TTG | T | G | T>G | T>G_TTG | TA>GC_ACACCAG | 8 | FALSE | FALSE |
| 163_v.txt | 32284 | 32284-SNV- | SNV | 1 | A | C | 93 | 21737 | 0.42784193 | 37.7634409 | rnr1Y285F | FALSE | CAN | JSY3866 | 7 | rnr1Y285F-R | TAA | A>C | G/C+1 | A/T-1 | TA>GC | TA>GC | CAACGTAAAT | AAATATT | ATT | ATT | T | G | T>G | T>G_ATT | TA>GC_AAATATT | 8 | FALSE | FALSE |
| 163_v.txt | 32310 | 32310-SNV- | SNV | 1 | A | C | 9 | 21737 | 0.04140406 | 50.2222222 | rnr1Y285F | FALSE | CAN | JSY3866 | 7 | rnr1Y285F-R | AAT | A>C | G/C+1 | A/T-1 | TA>GC | TA>GC | GAAATAATG | ATGGTTG | TTA | TTA | T | G | T>G | T>G_TTA | TA>GC_ATGGTTG | 8 | FALSE | FALSE |
| 163_v.txt | 32314 | 32314-SNV-T | SNV | 1 | T | G | 2 | 21737 | 9.20E-03 | 37.5005832 | rnr1Y285F | FALSE | CAN | JSY3866 | 7 | rnr1Y285F-R | GTT | T>G | G/C+1 | A/T-1 | TA>GC | TA>GC | TAATGGTTGT | TTGTTAA | CAA | GTT | T | C | T>G | T>G_GTT | TA>GC_TTGTTAA | 8 | FALSE | FALSE |
| 163_v.txt | 32319 | 32319-SNV- | SNV | 1 | A | C | 25 | 21737 | 0.11501127 | 53.44 | rnr1Y285F | FALSE | CAN | JSY3866 | 7 | rnr1Y285F-R | TAA | A>C | G/C+1 | A/T-1 | TA>GC | TA>GC | GTTGTTAAGA | AAGATAA | ATT | ATT | T | G | T>G | T>G_ATT | TA>GC_AAGATAA | 8 | FALSE | FALSE |
| 163_v.txt | 32475 | 32475-SNV- | SNV | 1 | A | C | 5 | 21737 | 0.02300225 | 53.6 | rnr1Y285F | FALSE | CAN | JSY3866 | 7 | rnr1Y285F-R | GAA | A>C | G/C+1 | A/T-1 | TA>GC | TA>GC | ATGTAGAAG | AAGGTTA | CTT | CTT | T | G | T>G | T>G_CTT | TA>GC_AAGGTTA | 8 | FALSE | FALSE |
| 163_v.txt | 32631 | 32631-SNV-T | SNV | 1 | T | G | 3 | 25230 | 0.01189061 | 50.3333333 | rnr1Y285F | FALSE | CAN | JSY3866 | 7 | rnr1Y285F-R | GTT | T>G | G/C+1 | A/T-1 | TA>GC | TA>GC | CCTTCGTTTT | TTTTTAT | CAA | GTT | T | C | T>G | T>G_GTT | TA>GC_TTTTTAT | 8 | FALSE | FALSE |
| 163_v.txt | 32763 | 32763-SNV- | SNV | 1 | A | C | 26 | 25230 | 0.10305192 | 52.2281481 | rnr1Y285F | FALSE | CAN | JSY3866 | 7 | rnr1Y285F-R | AAC | A>C | G/C+1 | A/T-1 | TA>GC | TA>GC | GCTAAAACTT | ACTTTGA | TTG | TTG | T | G | T>G | T>G_TTG | TA>GC_ACTTTGA | 8 | FALSE | FALSE |
| 163_v.txt | 32842 | 32842-SNV- | SNV | 1 | A | C | 5 | 25230 | 0.01981768 | 34.8 | rnr1Y285F | FALSE | CAN | JSY3866 | 7 | rnr1Y285F-R | CAA | A>C | G/C+1 | A/T-1 | TA>GC | TA>GC | TTACCCAAA | AAAAAAT | GTT | GTT | T | G | T>G | T>G_GTT | TA>GC_AAAAAAT | 8 | FALSE | FALSE |
| 163_v.txt | 33039 | 33039-SNV- | SNV | 1 | A | C | 28 | 16580 | 0.16887817 | 57.1421119 | rnr1Y285F | FALSE | CAN | JSY3866 | 7 | rnr1Y285F-R | CAT | A>C | G/C+1 | A/T-1 | TA>GC | TA>GC | GTAGCCATTT | ATTTCAC | GTA | GTA | T | G | T>G | T>G_GTA | TA>GC_ATTTCAC | 8 | FALSE | FALSE |
| 163_v.txt | 33089 | 33089-SNV- | SNV | 1 | A | C | 28 | 16580 | 0.16887817 | 48.6071429 | rnr1Y285F | FALSE | CAN | JSY3866 | 7 | rnr1Y285F-R | AAT | A>C | G/C+1 | A/T-1 | TA>GC | TA>GC | AAATAAATAT | ATATGAT | TTA | TTA | T | G | T>G | T>G_TTA | TA>GC_ATATGAT | 8 | FALSE | FALSE |
| 163_v.txt | 33189 | 33189-SNV- | SNV | 1 | A | C | 39 | 24849 | 0.15694797 | 36.3333415 | rnr1Y285F | FALSE | CAN | JSY3866 | 7 | rnr1Y285F-R | AAT | A>C | G/C+1 | A/T-1 | TA>GC | TA>GC | ATACCAATAT | ATATGTC | TTA | TTA | T | G | T>G | T>G_TTA | TA>GC_ATATGTC | 8 | FALSE | FALSE |
| 163_v.txt | 33204 | 33204-SNV- | SNV | 1 | A | C | 75 | 24849 | 0.30182301 | 37.1333202 | rnr1Y285F | FALSE | CAN | JSY3866 | 7 | rnr1Y285F-R | AAG | A>C | G/C+1 | A/T-1 | TA>GC | TA>GC | TGCTTAAGCT | AGCTCTC | TTC | TTC | T | G | T>G | T>G_TTC | TA>GC_AGCTCTC | 8 | FALSE | FALSE |
| 163_v.txt | 33269 | 33269-SNV- | SNV | 1 | A | C | 161 | 24849 | 0.6479134 | 15.4782609 | rnr1Y285F | FALSE | CAN | JSY3866 | 7 | rnr1Y285F-R | TAG | A>C | G/C+1 | A/T-1 | TA>GC | TA>GC | CTCCATAGA | AGAGAAC | ATC | ATC | T | G | T>G | T>G_ATC | TA>GC_AGAGAAC | 8 | FALSE | FALSE |
| 163_v.txt | 32081 | 32081-SNV- | SNV | 1 | A | G | 8 | 23392 | 0.03419973 | 52.0000675 | rnr1Y285F | FALSE | CAN | JSY3866 | 7 | rnr1Y285F-R | CAA | A>G | G/C+1 | A/T-1 | TA>CG | TA>CG | CCATGCAAA | AAAAAAG | GTT | GTT | T | C | T>C | T>C_GTT | TA>CG_AAAAAAG | 8 | FALSE | FALSE |
| 163_v.txt | 32337 | 32337-SNV- | SNV | 1 | A | G | 4 | 21737 | 0.0184018 | 50.7499238 | rnr1Y285F | FALSE | CAN | JSY3866 | 7 | rnr1Y285F-R | GAT | A>G | G/C+1 | A/T-1 | TA>CG | TA>CG | TTGAAGATAT | ATATGTG | CTA | CTA | T | C | T>C | T>C_CTA | TA>CG_ATATGTG | 8 | FALSE | FALSE |
| 163_v.txt | 32375 | 32375-SNV- | SNV | 1 | A | G | 3 | 21736 | 0.01380199 | 61.333274 | rnr1Y285F | FALSE | CAN | JSY3866 | 7 | rnr1Y285F-R | CAA | A>G | G/C+1 | A/T-1 | TA>CG | TA>CG | AATAGCAATA | AATAATA | GTT | GTT | T | C | T>C | T>C_GTT | TA>CG_AATAATA | 8 | FALSE | FALSE |
| 163_v.txt | 32377 | 32377-SNV-T | SNV | 1 | T | C | 5 | 21733 | 0.02300649 | 61.5985041 | rnr1Y285F | FALSE | CAN | JSY3866 | 7 | rnr1Y285F-R | ATA | T>C | G/C+1 | A/T-1 | TA>CG | TA>CG | TAGCAATAAT | TAATAAA | TAT | ATA | T | G | T>C | T>C_ATA | TA>CG_TAATAAA | 8 | FALSE | FALSE |
| 163_v.txt | 32454 | 32454-SNV- | SNV | 1 | A | G | 8 | 21737 | 0.03680361 | 41.9981428 | rnr1Y285F | FALSE | CAN | JSY3866 | 7 | rnr1Y285F-R | GAA | A>G | G/C+1 | A/T-1 | TA>CG | TA>CG | CCAATGAATA | AATAATA | CTT | CTT | T | C | T>C | T>C_CTT | TA>CG_AATAATA | 8 | FALSE | FALSE |
| 163_v.txt | 32521 | 32521-SNV- | SNV | 1 | A | G | 7 | 21737 | 0.03220316 | 24.142521 | rnr1Y285F | FALSE | CAN | JSY3866 | 7 | rnr1Y285F-R | GAT | A>G | G/C+1 | A/T-1 | TA>CG | TA>CG | GAACGGATT | ATTTTCT | CTA | CTA | T | C | T>C | T>C_CTA | TA>CG_ATTTTCT | 8 | FALSE | FALSE |
| 163_v.txt | 32522 | 32522-SNV-T | SNV | 1 | T | C | 2 | 21737 | 9.20E-03 | 50.9984438 | rnr1Y285F | FALSE | CAN | JSY3866 | 7 | rnr1Y285F-R | ATT | T>C | G/C+1 | A/T-1 | TA>CG | TA>CG | AACGGATTTT | TTTTCTG | TAA | ATT | T | G | T>C | T>C_ATT | TA>CG_TTTTCTG | 8 | FALSE | FALSE |
| 163_v.txt | 32728 | 32728-SNV- | SNV | 1 | A | G | 15 | 25230 | 0.05945303 | 59.1992157 | rnr1Y285F | FALSE | CAN | JSY3866 | 7 | rnr1Y285F-R | CAA | A>G | G/C+1 | A/T-1 | TA>CG | TA>CG | CCATACAAA | AAAAACA | GTT | GTT | T | C | T>C | T>C_GTT | TA>CG_AAAAACA | 8 | FALSE | FALSE |
| 163_v.txt | 32767 | 32767-SNV-T | SNV | 1 | T | C | 144 | 25230 | 0.57074911 | 53.0625 | rnr1Y285F | FALSE | CAN | JSY3866 | 7 | rnr1Y285F-R | TTG | T>C | G/C+1 | A/T-1 | TA>CG | TA>CG | AAACTTTGAT | TGATGGA | AAC | TTG | T | G | T>C | T>C_TTG | TA>CG_TGATGGA | 8 | FALSE | FALSE |
| 163_v.txt | 32785 | 32785-SNV- | SNV | 1 | A | G | 20 | 25230 | 0.07927071 | 20.549997 | rnr1Y285F | FALSE | CAN | JSY3866 | 7 | rnr1Y285F-R | AAC | A>G | G/C+1 | A/T-1 | TA>CG | TA>CG | CCCAGAACT | ACTCGAA | TTG | TTG | T | C | T>C | T>C_TTG | TA>CG_ACTCGAA | 8 | FALSE | FALSE |
| 163_v.txt | 32842 | 32842-SNV- | SNV | 1 | A | G | 16 | 25230 | 0.06341657 | 40.1875 | rnr1Y285F | FALSE | CAN | JSY3866 | 7 | rnr1Y285F-R | CAA | A>G | G/C+1 | A/T-1 | TA>CG | TA>CG | TTACCCAAA | AAAAAAT | GTT | GTT | T | C | T>C | T>C_GTT | TA>CG_AAAAAAT | 8 | FALSE | FALSE |
| 163_v.txt | 32924 | 32924-SNV- | SNV | 1 | A | G | 7 | 16580 | 0.04221954 | 35.9978755 | rnr1Y285F | FALSE | CAN | JSY3866 | 7 | rnr1Y285F-R | CAA | A>G | G/C+1 | A/T-1 | TA>CG | TA>CG | CAGGGCAAA | AAAAGTG | GTT | GTT | T | C | T>C | T>C_GTT | TA>CG_AAAAGTG | 8 | FALSE | FALSE |
| 163_v.txt | 33040 | 33040-SNV-T | SNV | 1 | T | C | 3 | 16580 | 0.01809409 | 54.9998776 | rnr1Y285F | FALSE | CAN | JSY3866 | 7 | rnr1Y285F-R | ATT | T>C | G/C+1 | A/T-1 | TA>CG | TA>CG | TAGCCATTTC | TTTCACC | TAA | ATT | T | G | T>C | T>C_ATT | TA>CG_TTTCACC | 8 | FALSE | FALSE |
| 163_v.txt | 33094 | 33094-SNV- | SNV | 1 | A | G | 6 | 16580 | 0.03618818 | 45.6675881 | rnr1Y285F | FALSE | CAN | JSY3866 | 7 | rnr1Y285F-R | GAT | A>G | G/C+1 | A/T-1 | TA>CG | TA>CG | AATATGATAT | ATATAAG | CTA | CTA | T | C | T>C | T>C_CTA | TA>CG_ATATAAG | 8 | FALSE | FALSE |
| 163_v.txt | 33095 | 33095-SNV-T | SNV | 1 | T | C | 4 | 16580 | 0.02412545 | 55.999862 | rnr1Y285F | FALSE | CAN | JSY3866 | 7 | rnr1Y285F-R | ATA | T>C | G/C+1 | A/T-1 | TA>CG | TA>CG | ATATGATATA | TATAAGA | TAT | ATA | T | G | T>C | T>C_ATA | TA>CG_TATAAGA | 8 | FALSE | FALSE |
| 163_v.txt | 33476 | 33476-SNV-T | SNV | 1 | T | C | 44 | 24844 | 0.17710514 | 31.9318019 | rnr1Y285F | FALSE | CAN | JSY3866 | 7 | rnr1Y285F-R | CTT | T>C | G/C+1 | A/T-1 | TA>CG | TA>CG | TATGCCTTTT | TTTTTTT | GAA | CTT | T | G | T>C | T>C_CTT | TA>CG_TTTTTTT | 8 | FALSE | FALSE |
| 163_v.txt | 31990 | 31990-SNV-T | SNV | 1 | T | A | 30 | 23392 | 0.12824897 | 53.4333333 | rnr1Y285F | FALSE | CAN | JSY3866 | 7 | rnr1Y285F-R | TTA | T>A | A/T+1 | A/T-1 | TA>AT | TA>AT | TTAATTTAGC | TAGCTTT | AAT | TTA | T | T | T>A | T>A_TTA | TA>AT_TAGCTTT | 8 | FALSE | FALSE |
| 163_v.txt | 32055 | 32055-SNV-T | SNV | 1 | T | A | 18 | 23392 | 0.07694938 | 56.8888889 | rnr1Y285F | FALSE | CAN | JSY3866 | 7 | rnr1Y285F-R | GTG | T>A | A/T+1 | A/T-1 | TA>AT | TA>AT | CTGATGTGC | TGCGAGA | CAC | GTG | T | T | T>A | T>A_GTG | TA>AT_TGCGAGA | 8 | FALSE | FALSE |
| 163_v.txt | 32202 | 32202-SNV- | SNV | 1 | A | T | 194 | 23392 | 0.82934337 | 41.8659794 | rnr1Y285F | FALSE | CAN | JSY3866 | 7 | rnr1Y285F-R | AAC | A>T | A/T+1 | A/T-1 | TA>AT | TA>AT | TATGGAACA | ACACCAC | TTG | TTG | T | A | T>A | T>A_TTG | TA>AT_ACACCAC | 8 | FALSE | FALSE |
| 163_v.txt | 32305 | 32305-SNV- | SNV | 1 | A | T | 3 | 21736 | 0.01380199 | 53.9941423 | rnr1Y285F | FALSE | CAN | JSY3866 | 7 | rnr1Y285F-R | GAA | A>T | A/T+1 | A/T-1 | TA>AT | TA>AT | CGGCAGAAA | AAATAAT | CTT | CTT | T | A | T>A | T>A_CTT | TA>AT_AAATAAT | 8 | FALSE | FALSE |
| 163_v.txt | 32314 | 32314-SNV-T | SNV | 1 | T | A | 3 | 21737 | 0.01380135 | 52.6616399 | rnr1Y285F | FALSE | CAN | JSY3866 | 7 | rnr1Y285F-R | GTT | T>A | A/T+1 | A/T-1 | TA>AT | TA>AT | TAATGGTTGT | TTGTTAA | CAA | GTT | T | T | T>A | T>A_GTT | TA>AT_TTGTTAA | 8 | FALSE | FALSE |
| 163_v.txt | 32315 | 32315-SNV-T | SNV | 1 | T | A | 1 | 21737 | 4.60E-03 | 63.9957366 | rnr1Y285F | FALSE | CAN | JSY3866 | 7 | rnr1Y285F-R | TTG | T>A | A/T+1 | A/T-1 | TA>AT | TA>AT | AATGGTTGTT | TGTTAAG | AAC | TTG | T | T | T>A | T>A_TTG | TA>AT_TGTTAAG | 8 | FALSE | FALSE |
| 163_v.txt | 32337 | 32337-SNV- | SNV | 1 | A | T | 2 | 21737 | 9.20E-03 | 36.9998676 | rnr1Y285F | FALSE | CAN | JSY3866 | 7 | rnr1Y285F-R | GAT | A>T | A/T+1 | A/T-1 | TA>AT | TA>AT | TTGAAGATAT | ATATGTG | CTA | CTA | T | A | T>A | T>A_CTA | TA>AT_ATATGTG | 8 | FALSE | FALSE |
| 163_v.txt | 32375 | 32375-SNV- | SNV | 1 | A | T | 2 | 21736 | 9.20E-03 | 58.4996846 | rnr1Y285F | FALSE | CAN | JSY3866 | 7 | rnr1Y285F-R | CAA | A>T | A/T+1 | A/T-1 | TA>AT | TA>AT | AATAGCAATA | AATAATA | GTT | GTT | T | A | T>A | T>A_GTT | TA>AT_AATAATA | 8 | FALSE | FALSE |
| 163_v.txt | 32377 | 32377-SNV-T | SNV | 1 | T | A | 2 | 21733 | 9.20E-03 | 63.9883491 | rnr1Y285F | FALSE | CAN | JSY3866 | 7 | rnr1Y285F-R | ATA | T>A | A/T+1 | A/T-1 | TA>AT | TA>AT | TAGCAATAAT | TAATAAA | TAT | ATA | T | T | T>A | T>A_ATA | TA>AT_TAATAAA | 8 | FALSE | FALSE |
| 163_v.txt | 32422 | 32422-SNV-T | SNV | 1 | T | A | 23 | 21737 | 0.10581037 | 56.7826087 | rnr1Y285F | FALSE | CAN | JSY3866 | 7 | rnr1Y285F-R | TTA | T>A | A/T+1 | A/T-1 | TA>AT | TA>AT | TTAGTTTAGG | TAGGGTC | AAT | TTA | T | T | T>A | T>A_TTA | TA>AT_TAGGGTC | 8 | FALSE | FALSE |
| 163_v.txt | 32460 | 32460-SNV- | SNV | 1 | A | T | 6 | 21737 | 0.02760271 | 46.6666936 | rnr1Y285F | FALSE | CAN | JSY3866 | 7 | rnr1Y285F-R | TAG | A>T | A/T+1 | A/T-1 | TA>AT | TA>AT | AATAATAGA | AGAGAGC | ATC | ATC | T | A | T>A | T>A_ATC | TA>AT_AGAGAGC | 8 | FALSE | FALSE |
| 163_v.txt | 32728 | 32728-SNV- | SNV | 1 | A | T | 2 | 25230 | 7.93E-03 | 18.999889 | rnr1Y285F | FALSE | CAN | JSY3866 | 7 | rnr1Y285F-R | CAA | A>T | A/T+1 | A/T-1 | TA>AT | TA>AT | CCATACAAA | AAAAACA | GTT | GTT | T | A | T>A | T>A_GTT | TA>AT_AAAAACA | 8 | FALSE | FALSE |
| 163_v.txt | 32763 | 32763-SNV- | SNV | 1 | A | T | 5 | 25230 | 0.01981768 | 52.3782602 | rnr1Y285F | FALSE | CAN | JSY3866 | 7 | rnr1Y285F-R | AAC | A>T | A/T+1 | A/T-1 | TA>AT | TA>AT | GCTAAAACTT | ACTTTGA | TTG | TTG | T | A | T>A | T>A_TTG | TA>AT_ACTTTGA | 8 | FALSE | FALSE |
| 163_v.txt | 32785 | 32785-SNV- | SNV | 1 | A | T | 32 | 25230 | 0.12683314 | 43.8118086 | rnr1Y285F | FALSE | CAN | JSY3866 | 7 | rnr1Y285F-R | AAC | A>T | A/T+1 | A/T-1 | TA>AT | TA>AT | CCCAGAACT | ACTCGAA | TTG | TTG | T | A | T>A | T>A_TTG | TA>AT_ACTCGAA | 8 | FALSE | FALSE |
| 163_v.txt | 33040 | 33040-SNV-T | SNV | 1 | T | A | 3 | 16580 | 0.01809409 | 63.9988431 | rnr1Y285F | FALSE | CAN | JSY3866 | 7 | rnr1Y285F-R | ATT | T>A | A/T+1 | A/T-1 | TA>AT | TA>AT | TAGCCATTTC | TTTCACC | TAA | ATT | T | T | T>A | T>A_ATT | TA>AT_TTTCACC | 8 | FALSE | FALSE |
| 163_v.txt | 33095 | 33095-SNV-T | SNV | 1 | T | A | 3 | 16580 | 0.01809409 | 42.9994037 | rnr1Y285F | FALSE | CAN | JSY3866 | 7 | rnr1Y285F-R | ATA | T>A | A/T+1 | A/T-1 | TA>AT | TA>AT | ATATGATATA | TATAAGA | TAT | ATA | T | T | T>A | T>A_ATA | TA>AT_TATAAGA | 8 | FALSE | FALSE |
| 163_v.txt | 33189 | 33189-SNV- | SNV | 1 | A | T | 6 | 24849 | 0.02414584 | 37.5000013 | rnr1Y285F | FALSE | CAN | JSY3866 | 7 | rnr1Y285F-R | AAT | A>T | A/T+1 | A/T-1 | TA>AT | TA>AT | ATACCAATAT | ATATGTC | TTA | TTA | T | A | T>A | T>A_TTA | TA>AT_ATATGTC | 8 | FALSE | FALSE |
| 163_v.txt | 33204 | 33204-SNV- | SNV | 1 | A | T | 6 | 24849 | 0.02414584 | 36.1652098 | rnr1Y285F | FALSE | CAN | JSY3866 | 7 | rnr1Y285F-R | AAG | A>T | A/T+1 | A/T-1 | TA>AT | TA>AT | TGCTTAAGCT | AGCTCTC | TTC | TTC | T | A | T>A | T>A_TTC | TA>AT_AGCTCTC | 8 | FALSE | FALSE |
| 163_v.txt | 33476 | 33476-SNV-T | SNV | 1 | T | A | 6 | 24844 | 0.0241507 | 33.6656116 | rnr1Y285F | FALSE | CAN | JSY3866 | 7 | rnr1Y285F-R | CTT | T>A | A/T+1 | A/T-1 | TA>AT | TA>AT | TATGCCTTTT | TTTTTTT | GAA | CTT | T | T | T>A | T>A_CTT | TA>AT_TTTTTTT | 8 | FALSE | FALSE |
| 163_v.txt | 31650 | 31650-SNV- | SNV | 1 | C | T | 9 | 3349 | 0.26873694 | 35.2222222 | rnr1Y285F | FALSE | CAN | JSY3866 | 7 | rnr1Y285F-R | GCG | C>T | A/T+1 | G/C-1 | CG>TA | CG>TA | AGAATGCGA | CGAAATG | CGC | GCG | C | A | C>T | C>T_GCG | CG>TA_CGAAATG | 8 | FALSE | FALSE |
| 163_v.txt | 31790 | 31790-SNV- | SNV | 1 | G | A | 4 | 3349 | 0.11943864 | 50.7499995 | rnr1Y285F | FALSE | CAN | JSY3866 | 7 | rnr1Y285F-R | CGA | G>A | A/T+1 | G/C-1 | CG>TA | CG>TA | GATGTCGAC | GACATCT | GCT | GCT | C | T | C>T | C>T_GCT | CG>TA_GACATCT | 8 | FALSE | FALSE |
| 163_v.txt | 31797 | 31797-SNV- | SNV | 1 | C | T | 10 | 3349 | 0.2985966 | 61.2 | rnr1Y285F | FALSE | CAN | JSY3866 | 7 | rnr1Y285F-R | TCC | C>T | A/T+1 | G/C-1 | CG>TA | CG>TA | ACATCTCCAA | CCAATCT | AGG | TCC | C | A | C>T | C>T_TCC | CG>TA_CCAATCT | 8 | FALSE | FALSE |
| 163_v.txt | 31834 | 31834-SNV- | SNV | 1 | G | A | 7 | 3349 | 0.20901762 | 56.4285714 | rnr1Y285F | FALSE | CAN | JSY3866 | 7 | rnr1Y285F-R | TGA | G>A | A/T+1 | G/C-1 | CG>TA | CG>TA | TGCATTGAAA | GAAATAA | ACT | ACT | C | T | C>T | C>T_ACT | CG>TA_GAAATAA | 8 | FALSE | FALSE |
| 163_v.txt | 31960 | 31960-SNV- | SNV | 1 | C | T | 25 | 23392 | 0.10687415 | 36.16 | rnr1Y285F | FALSE | CAN | JSY3866 | 7 | rnr1Y285F-R | GCC | C>T | A/T+1 | G/C-1 | CG>TA | CG>TA | ATGTGGCCG | CCGCATA | CGG | GCC | C | A | C>T | C>T_GCC | CG>TA_CCGCATA | 8 | FALSE | FALSE |
| 163_v.txt | 31962 | 31962-SNV- | SNV | 1 | G | A | 41 | 23392 | 0.1752736 | 36.7073171 | rnr1Y285F | FALSE | CAN | JSY3866 | 7 | rnr1Y285F-R | CGC | G>A | A/T+1 | G/C-1 | CG>TA | CG>TA | GTGGCCGCA | GCATAAT | GCG | GCG | C | T | C>T | C>T_GCG | CG>TA_GCATAAT | 8 | FALSE | FALSE |
| 163_v.txt | 31972 | 31972-SNV- | SNV | 1 | C | T | 14 | 23392 | 0.05984952 | 37.0714286 | rnr1Y285F | FALSE | CAN | JSY3866 | 7 | rnr1Y285F-R | GCC | C>T | A/T+1 | G/C-1 | CG>TA | CG>TA | AATAAGCCA | CCAAGCC | CGG | GCC | C | A | C>T | C>T_GCC | CG>TA_CCAAGCC | 8 | FALSE | FALSE |
| 163_v.txt | 31981 | 31981-SNV- | SNV | 1 | G | A | 38 | 23392 | 0.1624487 | 51.7105263 | rnr1Y285F | FALSE | CAN | JSY3866 | 7 | rnr1Y285F-R | GGC | G>A | A/T+1 | G/C-1 | CG>TA | CG>TA | AGCCGGGCA | GCATTAA | CCG | CCG | C | T | C>T | C>T_CCG | CG>TA_GCATTAA | 8 | FALSE | FALSE |
| 163_v.txt | 32057 | 32057-SNV- | SNV | 1 | C | T | 24 | 23392 | 0.10259918 | 59 | rnr1Y285F | FALSE | CAN | JSY3866 | 7 | rnr1Y285F-R | GCG | C>T | A/T+1 | G/C-1 | CG>TA | CG>TA | GATGTGCGA | CGAGATT | CGC | GCG | C | A | C>T | C>T_GCG | CG>TA_CGAGATT | 8 | FALSE | FALSE |
| 163_v.txt | 32075 | 32075-SNV- | SNV | 1 | C | T | 197 | 23392 | 0.84216826 | 61.1522843 | rnr1Y285F | FALSE | CAN | JSY3866 | 7 | rnr1Y285F-R | ACC | C>T | A/T+1 | G/C-1 | CG>TA | CG>TA | AAATAACCAT | CCATGCA | TGG | ACC | C | A | C>T | C>T_ACC | CG>TA_CCATGCA | 8 | FALSE | FALSE |
| 163_v.txt | 32088 | 32088-SNV- | SNV | 1 | C | T | 138 | 23392 | 0.58994528 | 60.6376812 | rnr1Y285F | FALSE | CAN | JSY3866 | 7 | rnr1Y285F-R | GCC | C>T | A/T+1 | G/C-1 | CG>TA | CG>TA | AAAAAGCCT | CCTGCAA | CGG | GCC | C | A | C>T | C>T_GCC | CG>TA_CCTGCAA | 8 | FALSE | FALSE |
| 163_v.txt | 32092 | 32092-SNV- | SNV | 1 | C | T | 22 | 23392 | 0.09404925 | 62.7272727 | rnr1Y285F | FALSE | CAN | JSY3866 | 7 | rnr1Y285F-R | GCA | C>T | A/T+1 | G/C-1 | CG>TA | CG>TA | AGCCTGCAA | CAACACC | CGT | GCA | C | A | C>T | C>T_GCA | CG>TA_CAACACC | 8 | FALSE | FALSE |
| 163_v.txt | 32097 | 32097-SNV- | SNV | 1 | C | T | 41 | 23392 | 0.1752736 | 60.8780488 | rnr1Y285F | FALSE | CAN | JSY3866 | 7 | rnr1Y285F-R | ACC | C>T | A/T+1 | G/C-1 | CG>TA | CG>TA | GCAACACCA | CCAGTGA | TGG | ACC | C | A | C>T | C>T_ACC | CG>TA_CCAGTGA | 8 | FALSE | FALSE |
| 163_v.txt | 32114 | 32114-SNV- | SNV | 1 | C | T | 446 | 23392 | 1.90663475 | 60.7735426 | rnr1Y285F | FALSE | CAN | JSY3866 | 7 | rnr1Y285F-R | GCC | C>T | A/T+1 | G/C-1 | CG>TA | CG>TA | TAATAGCCAT | CCATTCG | CGG | GCC | C | A | C>T | C>T_GCC | CG>TA_CCATTCG | 8 | FALSE | FALSE |
| 163_v.txt | 32165 | 32165-SNV- | SNV | 1 | G | A | 32 | 23392 | 0.13679891 | 48.3324686 | rnr1Y285F | FALSE | CAN | JSY3866 | 7 | rnr1Y285F-R | CGC | G>A | A/T+1 | G/C-1 | CG>TA | CG>TA | CAAAGCGCC | GCCAAAT | GCG | GCG | C | T | C>T | C>T_GCG | CG>TA_GCCAAAT | 8 | FALSE | FALSE |
| 163_v.txt | 32166 | 32166-SNV- | SNV | 1 | C | T | 121 | 23392 | 0.51727086 | 48.0333927 | rnr1Y285F | FALSE | CAN | JSY3866 | 7 | rnr1Y285F-R | GCC | C>T | A/T+1 | G/C-1 | CG>TA | CG>TA | AAAGCGCCA | CCAAATG | CGG | GCC | C | A | C>T | C>T_GCC | CG>TA_CCAAATG | 8 | FALSE | FALSE |
| 163_v.txt | 32173 | 32173-SNV- | SNV | 1 | C | T | 23 | 23392 | 0.09832421 | 50.1304348 | rnr1Y285F | FALSE | CAN | JSY3866 | 7 | rnr1Y285F-R | GCA | C>T | A/T+1 | G/C-1 | CG>TA | CG>TA | CAAATGCAG | CAGCAGT | CGT | GCA | C | A | C>T | C>T_GCA | CG>TA_CAGCAGT | 8 | FALSE | FALSE |
| 163_v.txt | 32271 | 32271-SNV- | SNV | 1 | C | T | 135 | 21737 | 0.62106086 | 36.2661999 | rnr1Y285F | FALSE | CAN | JSY3866 | 7 | rnr1Y285F-R | ACG | C>T | A/T+1 | G/C-1 | CG>TA | CG>TA | AAAATACGG | CGGGAAC | TGC | ACG | C | A | C>T | C>T_ACG | CG>TA_CGGGAAC | 8 | FALSE | FALSE |
| 163_v.txt | 32272 | 32272-SNV- | SNV | 1 | G | A | 52 | 21737 | 0.23922344 | 35 | rnr1Y285F | FALSE | CAN | JSY3866 | 7 | rnr1Y285F-R | CGG | G>A | A/T+1 | G/C-1 | CG>TA | CG>TA | AAATACGGG | GGGAACC | GCC | GCC | C | T | C>T | C>T_GCC | CG>TA_GGGAACC | 8 | FALSE | FALSE |
| 163_v.txt | 32299 | 32299-SNV- | SNV | 1 | C | T | 40 | 21737 | 0.18401803 | 48.925 | rnr1Y285F | FALSE | CAN | JSY3866 | 7 | rnr1Y285F-R | GCG | C>T | A/T+1 | G/C-1 | CG>TA | CG>TA | AATTTGCGG | CGGCAGA | CGC | GCG | C | A | C>T | C>T_GCG | CG>TA_CGGCAGA | 8 | FALSE | FALSE |
| 163_v.txt | 32304 | 32304-SNV- | SNV | 1 | G | A | 35 | 21737 | 0.16101578 | 50.9137974 | rnr1Y285F | FALSE | CAN | JSY3866 | 7 | rnr1Y285F-R | AGA | G>A | A/T+1 | G/C-1 | CG>TA | CG>TA | GCGGCAGAA | GAAATAA | TCT | TCT | C | T | C>T | C>T_TCT | CG>TA_GAAATAA | 8 | FALSE | FALSE |
| 163_v.txt | 32313 | 32313-SNV- | SNV | 1 | G | A | 17 | 21736 | 0.07821126 | 61.3439899 | rnr1Y285F | FALSE | CAN | JSY3866 | 7 | rnr1Y285F-R | GGT | G>A | A/T+1 | G/C-1 | CG>TA | CG>TA | ATAATGGTT | GTTGTTA | CCA | CCA | C | T | C>T | C>T_CCA | CG>TA_GTTGTTA | 8 | FALSE | FALSE |
| 163_v.txt | 32449 | 32449-SNV- | SNV | 1 | C | T | 118 | 21737 | 0.5428532 | 52.1525424 | rnr1Y285F | FALSE | CAN | JSY3866 | 7 | rnr1Y285F-R | CCA | C>T | A/T+1 | G/C-1 | CG>TA | CG>TA | AAAGTCCAA | CAATGAA | GGT | CCA | C | A | C>T | C>T_CCA | CG>TA_CAATGAA | 8 | FALSE | FALSE |
| 163_v.txt | 32508 | 32508-SNV- | SNV | 1 | G | A | 21 | 21737 | 0.09660947 | 45.5238095 | rnr1Y285F | FALSE | CAN | JSY3866 | 7 | rnr1Y285F-R | GGC | G>A | A/T+1 | G/C-1 | CG>TA | CG>TA | TTGATGGCT | GCTCTTG | CCG | CCG | C | T | C>T | C>T_CCG | CG>TA_GCTCTTG | 8 | FALSE | FALSE |
| 163_v.txt | 32536 | 32536-SNV- | SNV | 1 | C | T | 33 | 21737 | 0.15181488 | 36.5454546 | rnr1Y285F | FALSE | CAN | JSY3866 | 7 | rnr1Y285F-R | GCA | C>T | A/T+1 | G/C-1 | CG>TA | CG>TA | GGTTTGCAG | CAGCTTC | CGT | GCA | C | A | C>T | C>T_GCA | CG>TA_CAGCTTC | 8 | FALSE | FALSE |
| 163_v.txt | 32566 | 32566-SNV- | SNV | 1 | C | T | 50 | 25196 | 0.1984442 | 36.98 | rnr1Y285F | FALSE | CAN | JSY3866 | 7 | rnr1Y285F-R | TCA | C>T | A/T+1 | G/C-1 | CG>TA | CG>TA | CTAGTTCAGT | CAGTACC | AGT | TCA | C | A | C>T | C>T_TCA | CG>TA_CAGTACC | 8 | FALSE | FALSE |

| | | | | | | | | | | | | | | | | | | | | | | | | | | | | | | | | | | |
|---|---|---|---|---|---|---|---|---|---|---|---|---|---|---|---|---|---|---|---|---|---|---|---|---|---|---|---|---|---|---|---|---|---|---|
| 163_v.txt | 32590 | 32590-SNV-( | SNV | 1 | C | T | 53 | 25230 | 0.21006738 | 37.1509434 | rnr1Y285F | FALSE | CAN | JSY3866 | 7 | rnr1Y285F-R | GCG | C>T | A/T+1 | G/C-1 | CG>TA | CG>TA | AGGCAGCGT | CGTTAAT | CGC | GCG | C | A | C>T | C>T_GCG | CG>TA_CGTTAAT | 8 | FALSE | FALSE |
| 163_v.txt | 32658 | 32658-SNV-( | SNV | 1 | C | T | 41 | 25230 | 0.16250495 | 53.9512195 | rnr1Y285F | FALSE | CAN | JSY3866 | 7 | rnr1Y285F-R | ACC | C>T | A/T+1 | G/C-1 | CG>TA | CG>TA | CCTGGACCC | CCCCAGG | TGG | ACC | C | A | C>T | C>T_ACC | CG>TA_CCCCAGG | 8 | FALSE | FALSE |
| 163_v.txt | 32660 | 32660-SNV-( | SNV | 1 | C | T | 34 | 25230 | 0.13476021 | 56.2647059 | rnr1Y285F | FALSE | CAN | JSY3866 | 7 | rnr1Y285F-R | CCC | C>T | A/T+1 | G/C-1 | CG>TA | CG>TA | TGGACCCA | CCAGGCA | GGG | CCC | C | A | C>T | C>T_CCC | CG>TA_CCAGGCA | 8 | FALSE | FALSE |
| 163_v.txt | 32700 | 32700-SNV-( | SNV | 1 | C | T | 25 | 25230 | 0.09908839 | 57.08 | rnr1Y285F | FALSE | CAN | JSY3866 | 7 | rnr1Y285F-R | GCC | C>T | A/T+1 | G/C-1 | CG>TA | CG>TA | ACTGGGCCG | CCGGTAA | CGG | GCC | C | A | C>T | C>T_GCC | CG>TA_CCGGTAA | 8 | FALSE | FALSE |
| 163_v.txt | 32749 | 32749-SNV-( | SNV | 1 | C | T | 67 | 25230 | 0.26555688 | 53.8955224 | rnr1Y285F | FALSE | CAN | JSY3866 | 7 | rnr1Y285F-R | CCG | C>T | A/T+1 | G/C-1 | CG>TA | CG>TA | GAAACCCGA | CGATAAT | GGC | CCG | C | A | C>T | C>T_CCG | CG>TA_CGATAAT | 8 | FALSE | FALSE |
| 163_v.txt | 32757 | 32757-SNV-( | SNV | 1 | G | A | 25 | 25230 | 0.09908839 | 45.64 | rnr1Y285F | FALSE | CAN | JSY3866 | 7 | rnr1Y285F-R | GGC | G>A | A/T+1 | G/C-1 | CG>TA | CG>TA | ATAATGGCT | GCTAAAA | CCG | CCG | C | T | C>T | C>T_CCG | CG>TA_GCTAAAA | 8 | FALSE | FALSE |
| 163_v.txt | 32775 | 32775-SNV-( | SNV | 1 | G | A | 22 | 25230 | 0.08719778 | 50.9553193 | rnr1Y285F | FALSE | CAN | JSY3866 | 7 | rnr1Y285F-R | AGC | G>A | A/T+1 | G/C-1 | CG>TA | CG>TA | ATGGAAGCG | GCGACCC | TCG | TCG | C | T | C>T | C>T_TCG | CG>TA_GCGACCC | 8 | FALSE | FALSE |
| 163_v.txt | 32776 | 32776-SNV-( | SNV | 1 | C | T | 21 | 25229 | 0.08323754 | 47.9028794 | rnr1Y285F | FALSE | CAN | JSY3866 | 7 | rnr1Y285F-R | GCG | C>T | A/T+1 | G/C-1 | CG>TA | CG>TA | TGGAAGCGA | CGACCCA | CGC | GCG | C | A | C>T | C>T_GCG | CG>TA_CGACCCA | 8 | FALSE | FALSE |
| 163_v.txt | 32786 | 32786-SNV-( | SNV | 1 | C | T | 4 | 25230 | 0.01585414 | 46.7493932 | rnr1Y285F | FALSE | CAN | JSY3866 | 7 | rnr1Y285F-R | ACT | C>T | A/T+1 | G/C-1 | CG>TA | CG>TA | CCAGAACTC | CTCGAAT | TGA | ACT | C | A | C>T | C>T_ACT | CG>TA_CTCGAAT | 8 | FALSE | FALSE |
| 163_v.txt | 32794 | 32794-SNV-( | SNV | 1 | C | T | 224 | 25230 | 0.88783195 | 46.0223214 | rnr1Y285F | FALSE | CAN | JSY3866 | 7 | rnr1Y285F-R | TCA | C>T | A/T+1 | G/C-1 | CG>TA | CG>TA | CGAATTCAC | CACCGTA | AGT | TCA | C | A | C>T | C>T_TCA | CG>TA_CACCGTA | 8 | FALSE | FALSE |
| 163_v.txt | 32796 | 32796-SNV-( | SNV | 1 | C | T | 68 | 25230 | 0.26952041 | 45.6176471 | rnr1Y285F | FALSE | CAN | JSY3866 | 7 | rnr1Y285F-R | ACC | C>T | A/T+1 | G/C-1 | CG>TA | CG>TA | AATTCACCGT | CCGTAAT | TGG | ACC | C | A | C>T | C>T_ACC | CG>TA_CCGTAAT | 8 | FALSE | FALSE |
| 163_v.txt | 32798 | 32798-SNV-( | SNV | 1 | G | A | 44 | 25230 | 0.17439556 | 46.5909091 | rnr1Y285F | FALSE | CAN | JSY3866 | 7 | rnr1Y285F-R | CGT | G>A | A/T+1 | G/C-1 | CG>TA | CG>TA | TTCACCGTA | GTAATAT | GCA | GCA | C | T | C>T | C>T_GCA | CG>TA_GTAATAT | 8 | FALSE | FALSE |
| 163_v.txt | 32855 | 32855-SNV-( | SNV | 1 | C | T | 602 | 25229 | 2.38614293 | 36.423588 | rnr1Y285F | FALSE | CAN | JSY3866 | 7 | rnr1Y285F-R | TCC | C>T | A/T+1 | G/C-1 | CG>TA | CG>TA | ACTAATCCAT | CCATGCC | AGG | TCC | C | A | C>T | C>T_TCC | CG>TA_CCATGCC | 8 | FALSE | FALSE |
| 163_v.txt | 32860 | 32860-SNV-( | SNV | 1 | C | T | 30 | 25229 | 0.11891078 | 37.3333333 | rnr1Y285F | FALSE | CAN | JSY3866 | 7 | rnr1Y285F-R | GCC | C>T | A/T+1 | G/C-1 | CG>TA | CG>TA | TCCATGCCG | CCGCCAG | CGG | GCC | C | A | C>T | C>T_GCC | CG>TA_CCGCCAG | 8 | FALSE | FALSE |
| 163_v.txt | 32901 | 32901-SNV-( | SNV | 1 | C | T | 42 | 16580 | 0.25331725 | 32.3809524 | rnr1Y285F | FALSE | CAN | JSY3866 | 7 | rnr1Y285F-R | GCC | C>T | A/T+1 | G/C-1 | CG>TA | CG>TA | ACTTGGCCA | CCAACTA | CGG | GCC | C | A | C>T | C>T_GCC | CG>TA_CCAACTA | 8 | FALSE | FALSE |
| 163_v.txt | 32917 | 32917-SNV-( | SNV | 1 | C | T | 32 | 16580 | 0.19300362 | 40.71875 | rnr1Y285F | FALSE | CAN | JSY3866 | 7 | rnr1Y285F-R | TCC | C>T | A/T+1 | G/C-1 | CG>TA | CG>TA | TAAGTTCCA | CCAGGGC | AGG | TCC | C | A | C>T | C>T_TCC | CG>TA_CCAGGGC | 8 | FALSE | FALSE |
| 163_v.txt | 32936 | 32936-SNV-( | SNV | 1 | C | T | 135 | 16580 | 0.81423402 | 51.6556755 | rnr1Y285F | FALSE | CAN | JSY3866 | 7 | rnr1Y285F-R | CCC | C>T | A/T+1 | G/C-1 | CG>TA | CG>TA | GATTGCCCA | CCAAGAA | GGG | CCC | C | A | C>T | C>T_CCC | CG>TA_CCAAGAA | 8 | FALSE | FALSE |
| 163_v.txt | 32937 | 32937-SNV-( | SNV | 1 | C | T | 52 | 16580 | 0.31363088 | 51.2692308 | rnr1Y285F | FALSE | CAN | JSY3866 | 7 | rnr1Y285F-R | CCA | C>T | A/T+1 | G/C-1 | CG>TA | CG>TA | ATTGCCCAA | CAAGAAA | GGT | CCA | C | A | C>T | C>T_CCA | CG>TA_CAAGAAA | 8 | FALSE | FALSE |
| 163_v.txt | 32945 | 32945-SNV-( | SNV | 1 | C | T | 70 | 16580 | 0.42219542 | 57.3956727 | rnr1Y285F | FALSE | CAN | JSY3866 | 7 | rnr1Y285F-R | ACC | C>T | A/T+1 | G/C-1 | CG>TA | CG>TA | AGAAAACCA | CCAATAC | TGG | ACC | C | A | C>T | C>T_ACC | CG>TA_CCAATAC | 8 | FALSE | FALSE |
| 163_v.txt | 32946 | 32946-SNV-( | SNV | 1 | C | T | 35 | 16580 | 0.21109771 | 57.428758 | rnr1Y285F | FALSE | CAN | JSY3866 | 7 | rnr1Y285F-R | CCA | C>T | A/T+1 | G/C-1 | CG>TA | CG>TA | GAAAACCAA | CAATACA | GGT | CCA | C | A | C>T | C>T_CCA | CG>TA_CAATACA | 8 | FALSE | FALSE |
| 163_v.txt | 32958 | 32958-SNV-( | SNV | 1 | C | T | 69 | 16580 | 0.41616405 | 54.1014493 | rnr1Y285F | FALSE | CAN | JSY3866 | 7 | rnr1Y285F-R | ACC | C>T | A/T+1 | G/C-1 | CG>TA | CG>TA | ATGTAACCAT | CCATTGG | TGG | ACC | C | A | C>T | C>T_ACC | CG>TA_CCATTGG | 8 | FALSE | FALSE |
| 163_v.txt | 32967 | 32967-SNV-( | SNV | 1 | G | A | 43 | 16580 | 0.25934861 | 59.6511628 | rnr1Y285F | FALSE | CAN | JSY3866 | 7 | rnr1Y285F-R | CGC | G>A | A/T+1 | G/C-1 | CG>TA | CG>TA | TTGGCCGCA | GCACCAA | GCG | GCG | C | T | C>T | C>T_GCG | CG>TA_GCACCAA | 8 | FALSE | FALSE |
| 163_v.txt | 32970 | 32970-SNV-( | SNV | 1 | C | T | 124 | 16580 | 0.74788902 | 58.25 | rnr1Y285F | FALSE | CAN | JSY3866 | 7 | rnr1Y285F-R | ACC | C>T | A/T+1 | G/C-1 | CG>TA | CG>TA | GCCGCACCA | CCAAATG | TGG | ACC | C | A | C>T | C>T_ACC | CG>TA_CCAAATG | 8 | FALSE | FALSE |
| 163_v.txt | 33012 | 33012-SNV-( | SNV | 1 | G | A | 43 | 16580 | 0.25934861 | 61.1162791 | rnr1Y285F | FALSE | CAN | JSY3866 | 7 | rnr1Y285F-R | AGA | G>A | A/T+1 | G/C-1 | CG>TA | CG>TA | GTGAAAGAG | GAGGATG | TCT | TCT | C | T | C>T | C>T_TCT | CG>TA_GAGGATG | 8 | FALSE | FALSE |
| 163_v.txt | 33015 | 33015-SNV-( | SNV | 1 | G | A | 50 | 16580 | 0.30156815 | 56.28 | rnr1Y285F | FALSE | CAN | JSY3866 | 7 | rnr1Y285F-R | GGA | G>A | A/T+1 | G/C-1 | CG>TA | CG>TA | AAAGAGGAT | GATGTAA | CCT | CCT | C | T | C>T | C>T_CCT | CG>TA_GATGTAA | 8 | FALSE | FALSE |
| 163_v.txt | 33043 | 33043-SNV-( | SNV | 1 | C | T | 42 | 16580 | 0.25331725 | 61.4285714 | rnr1Y285F | FALSE | CAN | JSY3866 | 7 | rnr1Y285F-R | TCA | C>T | A/T+1 | G/C-1 | CG>TA | CG>TA | CCATTTCACC | CACCCAA | AGT | TCA | C | A | C>T | C>T_TCA | CG>TA_CACCCAA | 8 | FALSE | FALSE |
| 163_v.txt | 33056 | 33056-SNV-( | SNV | 1 | C | T | 21 | 16580 | 0.12665863 | 55.5188587 | rnr1Y285F | FALSE | CAN | JSY3866 | 7 | rnr1Y285F-R | GCG | C>T | A/T+1 | G/C-1 | CG>TA | CG>TA | GGACTGCGT | CGTGACA | CGC | GCG | C | A | C>T | C>T_GCG | CG>TA_CGTGACA | 8 | FALSE | FALSE |
| 163_v.txt | 33057 | 33057-SNV-( | SNV | 1 | G | A | 7 | 16580 | 0.04221954 | 59.1428571 | rnr1Y285F | FALSE | CAN | JSY3866 | 7 | rnr1Y285F-R | CGT | G>A | A/T+1 | G/C-1 | CG>TA | CG>TA | GACTGCGTG | GTGACAG | GCA | GCA | C | T | C>T | C>T_GCA | CG>TA_GTGACAG | 8 | FALSE | FALSE |
| 163_v.txt | 33070 | 33070-SNV-( | SNV | 1 | C | T | 20 | 16580 | 0.12062726 | 54.95 | rnr1Y285F | FALSE | CAN | JSY3866 | 7 | rnr1Y285F-R | GCC | C>T | A/T+1 | G/C-1 | CG>TA | CG>TA | AATATGCCA | CCAAAGA | CGG | GCC | C | A | C>T | C>T_GCC | CG>TA_CCAAAGA | 8 | FALSE | FALSE |
| 163_v.txt | 33103 | 33103-SNV-( | SNV | 1 | C | T | 21 | 16580 | 0.12665863 | 40.6666667 | rnr1Y285F | FALSE | CAN | JSY3866 | 7 | rnr1Y285F-R | GCG | C>T | A/T+1 | G/C-1 | CG>TA | CG>TA | TAAGAGCGC | CGCCCAC | CGC | GCG | C | A | C>T | C>T_GCG | CG>TA_CGCCCAC | 8 | FALSE | FALSE |
| 163_v.txt | 33114 | 33114-SNV-( | SNV | 1 | C | T | 41 | 16580 | 0.24728589 | 42.1463415 | rnr1Y285F | FALSE | CAN | JSY3866 | 7 | rnr1Y285F-R | GCC | C>T | A/T+1 | G/C-1 | CG>TA | CG>TA | ACTGGGCCG | CCGGCGT | CGG | GCC | C | A | C>T | C>T_GCC | CG>TA_CCGGCGT | 8 | FALSE | FALSE |
| 163_v.txt | 33118 | 33118-SNV-( | SNV | 1 | C | T | 18 | 16580 | 0.10856454 | 45.8333333 | rnr1Y285F | FALSE | CAN | JSY3866 | 7 | rnr1Y285F-R | GCG | C>T | A/T+1 | G/C-1 | CG>TA | CG>TA | GGCCGGCGT | CGTTGGT | CGC | GCG | C | A | C>T | C>T_GCG | CG>TA_CGTTGGT | 8 | FALSE | FALSE |
| 163_v.txt | 33153 | 33153-SNV-( | SNV | 1 | C | T | 76 | 16580 | 0.4583836 | 36.6973684 | rnr1Y285F | FALSE | CAN | JSY3866 | 7 | rnr1Y285F-R | ACC | C>T | A/T+1 | G/C-1 | CG>TA | CG>TA | AAAAGACCT | CCTGTAC | TGG | ACC | C | A | C>T | C>T_ACC | CG>TA_CCTGTAC | 8 | FALSE | FALSE |
| 163_v.txt | 33154 | 33154-SNV-( | SNV | 1 | C | T | 13 | 16580 | 0.07840772 | 37.3846154 | rnr1Y285F | FALSE | CAN | JSY3866 | 7 | rnr1Y285F-R | CCT | C>T | A/T+1 | G/C-1 | CG>TA | CG>TA | AAAGACCTG | CTGTACC | GGA | CCT | C | A | C>T | C>T_CCT | CG>TA_CTGTACC | 8 | FALSE | FALSE |
| 163_v.txt | 33171 | 33171-SNV-( | SNV | 1 | C | T | 74 | 16578 | 0.44637471 | 37.0540541 | rnr1Y285F | FALSE | CAN | JSY3866 | 7 | rnr1Y285F-R | ACC | C>T | A/T+1 | G/C-1 | CG>TA | CG>TA | GTACCACCA | CCAAGGG | TGG | ACC | C | A | C>T | C>T_ACC | CG>TA_CCAAGGG | 8 | FALSE | FALSE |
| 163_v.txt | 33258 | 33258-SNV-( | SNV | 1 | C | T | 43 | 24849 | 0.17304519 | 41.0465116 | rnr1Y285F | FALSE | CAN | JSY3866 | 7 | rnr1Y285F-R | GCC | C>T | A/T+1 | G/C-1 | CG>TA | CG>TA | CCTATGCCAT | CCATCCT | CGG | GCC | C | A | C>T | C>T_GCC | CG>TA_CCATCCT | 8 | FALSE | FALSE |
| 163_v.txt | 33276 | 33276-SNV-( | SNV | 1 | G | A | 42 | 24849 | 0.16902089 | 46.5238095 | rnr1Y285F | FALSE | CAN | JSY3866 | 7 | rnr1Y285F-R | CGT | G>A | A/T+1 | G/C-1 | CG>TA | CG>TA | GAGAACGTA | GTATCCT | GCA | GCA | C | T | C>T | C>T_GCA | CG>TA_GTATCCT | 8 | FALSE | FALSE |
| 163_v.txt | 33310 | 33310-SNV-( | SNV | 1 | G | A | 28 | 24849 | 0.11268059 | 54.0357143 | rnr1Y285F | FALSE | CAN | JSY3866 | 7 | rnr1Y285F-R | CGC | G>A | A/T+1 | G/C-1 | CG>TA | CG>TA | AAGAGCGCA | GCAATGG | GCG | GCG | C | T | C>T | C>T_GCG | CG>TA_GCAATGG | 8 | FALSE | FALSE |
| 163_v.txt | 33412 | 33412-SNV-( | SNV | 1 | G | A | 35 | 24849 | 0.14085074 | 47.6 | rnr1Y285F | FALSE | CAN | JSY3866 | 7 | rnr1Y285F-R | GGC | G>A | A/T+1 | G/C-1 | CG>TA | CG>TA | TGACCGGCT | GCTCATT | CCG | CCG | C | T | C>T | C>T_CCG | CG>TA_GCTCATT | 8 | FALSE | FALSE |
| 163_v.txt | 33474 | 33474-SNV-( | SNV | 1 | C | T | 36 | 24845 | 0.14489837 | 33.75 | rnr1Y285F | FALSE | CAN | JSY3866 | 7 | rnr1Y285F-R | GCC | C>T | A/T+1 | G/C-1 | CG>TA | CG>TA | GCTATGCCTT | CCTTTTT | CGG | GCC | C | A | C>T | C>T_GCC | CG>TA_CCTTTTT | 8 | FALSE | FALSE |
| 163_v.txt | 31981 | 31981-SNV-( | SNV | 1 | G | C | 33 | 23392 | 0.14107387 | 44.5757576 | rnr1Y285F | FALSE | CAN | JSY3866 | 7 | rnr1Y285F-R | GGC | G>C | G/C+1 | G/C-1 | CG>GC | CG>GC | AGCCGGGCA | GCATTAA | CCG | CCG | C | G | C>G | C>G_CCG | CG>GC_GCATTAA | 8 | FALSE | FALSE |
| 163_v.txt | 31993 | 31993-SNV-( | SNV | 1 | C | G | 16 | 23392 | 0.06839945 | 52.9310868 | rnr1Y285F | FALSE | CAN | JSY3866 | 7 | rnr1Y285F-R | GCT | C>G | G/C+1 | G/C-1 | CG>GC | CG>GC | ATTTAGCTTT | CTTTAAA | CGA | GCT | C | C | C>G | C>G_GCT | CG>GC_CTTTAAA | 8 | FALSE | FALSE |
| 163_v.txt | 32064 | 32064-SNV-( | SNV | 1 | G | C | 143 | 23392 | 0.61132011 | 61.5314685 | rnr1Y285F | FALSE | CAN | JSY3866 | 7 | rnr1Y285F-R | TGA | G>C | G/C+1 | G/C-1 | CG>GC | CG>GC | GAGATTGAG | GAGATAA | ACT | ACT | C | G | C>G | C>G_ACT | CG>GC_GAGATAA | 8 | FALSE | FALSE |
| 163_v.txt | 32092 | 32092-SNV-( | SNV | 1 | C | G | 22 | 23392 | 0.09404925 | 61.9090909 | rnr1Y285F | FALSE | CAN | JSY3866 | 7 | rnr1Y285F-R | GCA | C>G | G/C+1 | G/C-1 | CG>GC | CG>GC | AGCCTGCAA | CAACACC | CGT | GCA | C | C | C>G | C>G_GCA | CG>GC_CAACACC | 8 | FALSE | FALSE |
| 163_v.txt | 32223 | 32223-SNV-( | SNV | 1 | G | C | 61 | 23392 | 0.26077291 | 36.7704918 | rnr1Y285F | FALSE | CAN | JSY3866 | 7 | rnr1Y285F-R | TGA | G>C | G/C+1 | G/C-1 | CG>GC | CG>GC | GTCCTTGACA | GACAGGA | ACT | ACT | C | G | C>G | C>G_ACT | CG>GC_GACAGGA | 8 | FALSE | FALSE |
| 163_v.txt | 32299 | 32299-SNV-( | SNV | 1 | C | G | 4 | 21737 | 0.0184018 | 57.5 | rnr1Y285F | FALSE | CAN | JSY3866 | 7 | rnr1Y285F-R | GCG | C>G | G/C+1 | G/C-1 | CG>GC | CG>GC | AATTTGCGG | CGGCAGA | CGC | GCG | C | C | C>G | C>G_GCG | CG>GC_CGGCAGA | 8 | FALSE | FALSE |
| 163_v.txt | 32304 | 32304-SNV-( | SNV | 1 | G | C | 5442 | 21737 | 25.0356535 | 56.7185365 | rnr1Y285F | FALSE | CAN | JSY3866 | 7 | rnr1Y285F-R | AGA | G>C | G/C+1 | G/C-1 | CG>GC | CG>GC | GCGGCAGAA | GAAATAA | TCT | TCT | C | G | C>G | C>G_TCT | CG>GC_GAAATAA | 8 | FALSE | FALSE |
| 163_v.txt | 32399 | 32399-SNV-( | SNV | 1 | G | C | 40 | 21737 | 0.18401803 | 59.325 | rnr1Y285F | FALSE | CAN | JSY3866 | 7 | rnr1Y285F-R | CGT | G>C | G/C+1 | G/C-1 | CG>GC | CG>GC | AGAAACGTA | GTAGGAA | GCA | GCA | C | G | C>G | C>G_GCA | CG>GC_GTAGGAA | 8 | FALSE | FALSE |
| 163_v.txt | 32449 | 32449-SNV-( | SNV | 1 | C | G | 35 | 21737 | 0.16101578 | 49.3714286 | rnr1Y285F | FALSE | CAN | JSY3866 | 7 | rnr1Y285F-R | CCA | C>G | G/C+1 | G/C-1 | CG>GC | CG>GC | AAAGTCCAAT | CAATGAA | GGT | CCA | C | C | C>G | C>G_CCA | CG>GC_CAATGAA | 8 | FALSE | FALSE |
| 163_v.txt | 32590 | 32590-SNV-( | SNV | 1 | C | G | 12 | 25230 | 0.04756243 | 34.6666667 | rnr1Y285F | FALSE | CAN | JSY3866 | 7 | rnr1Y285F-R | GCG | C>G | G/C+1 | G/C-1 | CG>GC | CG>GC | AGGCAGCGT | CGTTAAT | CGC | GCG | C | C | C>G | C>G_GCG | CG>GC_CGTTAAT | 8 | FALSE | FALSE |
| 163_v.txt | 32700 | 32700-SNV-( | SNV | 1 | C | G | 3 | 25230 | 0.01189061 | 63.6666667 | rnr1Y285F | FALSE | CAN | JSY3866 | 7 | rnr1Y285F-R | GCC | C>G | G/C+1 | G/C-1 | CG>GC | CG>GC | ACTGGGCCG | CCGGTAA | CGG | GCC | C | C | C>G | C>G_GCC | CG>GC_CCGGTAA | 8 | FALSE | FALSE |
| 163_v.txt | 32776 | 32776-SNV-( | SNV | 1 | C | G | 29 | 25229 | 0.11494709 | 43.6882507 | rnr1Y285F | FALSE | CAN | JSY3866 | 7 | rnr1Y285F-R | GCG | C>G | G/C+1 | G/C-1 | CG>GC | CG>GC | TGGAAGCGA | CGACCCA | CGC | GCG | C | C | C>G | C>G_GCG | CG>GC_CGACCCA | 8 | FALSE | FALSE |
| 163_v.txt | 32786 | 32786-SNV-( | SNV | 1 | C | G | 19 | 25230 | 0.07530717 | 45.6296324 | rnr1Y285F | FALSE | CAN | JSY3866 | 7 | rnr1Y285F-R | ACT | C>G | G/C+1 | G/C-1 | CG>GC | CG>GC | CCAGAACTC | CTCGAAT | TGA | ACT | C | C | C>G | C>G_ACT | CG>GC_CTCGAAT | 8 | FALSE | FALSE |
| 163_v.txt | 32940 | 32940-SNV-( | SNV | 1 | G | C | 3 | 16580 | 0.01809409 | 45.3333333 | rnr1Y285F | FALSE | CAN | JSY3866 | 7 | rnr1Y285F-R | AGA | G>C | G/C+1 | G/C-1 | CG>GC | CG>GC | GCCCAAGAA | GAAAACC | TCT | TCT | C | G | C>G | C>G_TCT | CG>GC_GAAAACC | 8 | FALSE | FALSE |
| 163_v.txt | 33056 | 33056-SNV-( | SNV | 1 | C | G | 1 | 16580 | 6.03E-03 | 26.9993727 | rnr1Y285F | FALSE | CAN | JSY3866 | 7 | rnr1Y285F-R | GCG | C>G | G/C+1 | G/C-1 | CG>GC | CG>GC | GGACTGCGT | CGTGACA | CGC | GCG | C | C | C>G | C>G_GCG | CG>GC_CGTGACA | 8 | FALSE | FALSE |
| 163_v.txt | 33057 | 33057-SNV-( | SNV | 1 | G | C | 8 | 16580 | 0.04825091 | 57.5 | rnr1Y285F | FALSE | CAN | JSY3866 | 7 | rnr1Y285F-R | CGT | G>C | G/C+1 | G/C-1 | CG>GC | CG>GC | GACTGCGTG | GTGACAG | GCA | GCA | C | G | C>G | C>G_GCA | CG>GC_GTGACAG | 8 | FALSE | FALSE |
| 163_v.txt | 33103 | 33103-SNV-( | SNV | 1 | C | G | 6 | 16580 | 0.03618818 | 41.3333333 | rnr1Y285F | FALSE | CAN | JSY3866 | 7 | rnr1Y285F-R | GCG | C>G | G/C+1 | G/C-1 | CG>GC | CG>GC | TAAGAGCGC | CGCCCAC | CGC | GCG | C | C | C>G | C>G_GCG | CG>GC_CGCCCAC | 8 | FALSE | FALSE |
| 163_v.txt | 33154 | 33154-SNV-( | SNV | 1 | C | G | 3 | 16580 | 0.01809409 | 33.3333333 | rnr1Y285F | FALSE | CAN | JSY3866 | 7 | rnr1Y285F-R | CCT | C>G | G/C+1 | G/C-1 | CG>GC | CG>GC | AAAGACCTG | CTGTACC | GGA | CCT | C | C | C>G | C>G_CCT | CG>GC_CTGTACC | 8 | FALSE | FALSE |
| 163_v.txt | 33168 | 33168-SNV-( | SNV | 1 | C | G | 54 | 16578 | 0.3257329 | 37.9814815 | rnr1Y285F | FALSE | CAN | JSY3866 | 7 | rnr1Y285F-R | ACC | C>G | G/C+1 | G/C-1 | CG>GC | CG>GC | ATAGTACCA | CCACCAA | TGG | ACC | C | C | C>G | C>G_ACC | CG>GC_CCACCAA | 8 | FALSE | FALSE |
| 163_v.txt | 33193 | 33193-SNV-( | SNV | 1 | G | C | 16 | 24849 | 0.06438891 | 37.8125 | rnr1Y285F | FALSE | CAN | JSY3866 | 7 | rnr1Y285F-R | TGT | G>C | G/C+1 | G/C-1 | CG>GC | CG>GC | CAATATGTCT | GTCTTTG | ACA | ACA | C | G | C>G | C>G_ACA | CG>GC_GTCTTTG | 8 | FALSE | FALSE |
| 163_v.txt | 31657 | 31657-SNV-( | SNV | 1 | G | T | 23 | 3349 | 0.68677217 | 21.6521739 | rnr1Y285F | FALSE | CAN | JSY3866 | 7 | rnr1Y285F-R | GGC | G>T | A/T+1 | G/C-1 | CG>AT | CG>AT | GAAATGGCG | GCGTGGA | CCG | CCG | C | A | C>A | C>A_CCG | CG>AT_GCGTGGA | 8 | FALSE | FALSE |
| 163_v.txt | 31789 | 31789-SNV-( | SNV | 1 | C | A | 4 | 3349 | 0.11943864 | 28.001681 | rnr1Y285F | FALSE | CAN | JSY3866 | 7 | rnr1Y285F-R | TCG | C>A | A/T+1 | G/C-1 | CG>AT | CG>AT | CGATGTCGA | CGACATC | AGC | TCG | C | T | C>A | C>A_TCG | CG>AT_CGACATC | 8 | FALSE | FALSE |
| 163_v.txt | 31962 | 31962-SNV-( | SNV | 1 | G | T | 4 | 23392 | 0.01709986 | 38 | rnr1Y285F | FALSE | CAN | JSY3866 | 7 | rnr1Y285F-R | CGC | G>T | A/T+1 | G/C-1 | CG>AT | CG>AT | GTGGCCGCA | GCATAAT | GCG | GCG | C | A | C>A | C>A_GCG | CG>AT_GCATAAT | 8 | FALSE | FALSE |
| 163_v.txt | 31963 | 31963-SNV-( | SNV | 1 | C | A | 8 | 23392 | 0.03419973 | 25.2501496 | rnr1Y285F | FALSE | CAN | JSY3866 | 7 | rnr1Y285F-R | GCA | C>A | A/T+1 | G/C-1 | CG>AT | CG>AT | TGGCCGCAT | CATAATA | CGT | GCA | C | T | C>A | C>A_GCA | CG>AT_CATAATA | 8 | FALSE | FALSE |
| 163_v.txt | 31981 | 31981-SNV-( | SNV | 1 | G | T | 27 | 23392 | 0.11542408 | 38.6296296 | rnr1Y285F | FALSE | CAN | JSY3866 | 7 | rnr1Y285F-R | GGC | G>T | A/T+1 | G/C-1 | CG>AT | CG>AT | AGCCGGGCA | GCATTAA | CCG | CCG | C | A | C>A | C>A_CCG | CG>AT_GCATTAA | 8 | FALSE | FALSE |
| 163_v.txt | 32027 | 32027-SNV-( | SNV | 1 | G | T | 11 | 23392 | 0.04702462 | 56.5454546 | rnr1Y285F | FALSE | CAN | JSY3866 | 7 | rnr1Y285F-R | GGT | G>T | A/T+1 | G/C-1 | CG>AT | CG>AT | GCCACGGTA | GTATTTC | CCA | CCA | C | A | C>A | C>A_CCA | CG>AT_GTATTTC | 8 | FALSE | FALSE |
| 163_v.txt | 32064 | 32064-SNV-( | SNV | 1 | G | T | 5 | 23392 | 0.02137483 | 63 | rnr1Y285F | FALSE | CAN | JSY3866 | 7 | rnr1Y285F-R | TGA | G>T | A/T+1 | G/C-1 | CG>AT | CG>AT | GAGATTGAG | GAGATAA | ACT | ACT | C | A | C>A | C>A_ACT | CG>AT_GAGATAA | 8 | FALSE | FALSE |
| 163_v.txt | 32188 | 32188-SNV-( | SNV | 1 | C | A | 24 | 23392 | 0.10259918 | 45.7916667 | rnr1Y285F | FALSE | CAN | JSY3866 | 7 | rnr1Y285F-R | ACT | C>A | A/T+1 | G/C-1 | CG>AT | CG>AT | CGAAAACTG | CTGCAAT | TGA | ACT | C | T | C>A | C>A_ACT | CG>AT_CTGCAAT | 8 | FALSE | FALSE |
| 163_v.txt | 32195 | 32195-SNV-( | SNV | 1 | G | T | 44 | 23392 | 0.1880985 | 47.3863636 | rnr1Y285F | FALSE | CAN | JSY3866 | 7 | rnr1Y285F-R | TGT | G>T | A/T+1 | G/C-1 | CG>AT | CG>AT | TGCAATGTA | GTATGGA | ACA | ACA | C | A | C>A | C>A_ACA | CG>AT_GTATGGA | 8 | FALSE | FALSE |
| 163_v.txt | 32223 | 32223-SNV-( | SNV | 1 | G | T | 28 | 23392 | 0.11969904 | 39.1428571 | rnr1Y285F | FALSE | CAN | JSY3866 | 7 | rnr1Y285F-R | TGA | G>T | A/T+1 | G/C-1 | CG>AT | CG>AT | GTCCTTGACA | GACAGGA | ACT | ACT | C | A | C>A | C>A_ACT | CG>AT_GACAGGA | 8 | FALSE | FALSE |
| 163_v.txt | 32272 | 32272-SNV-( | SNV | 1 | G | T | 7 | 21737 | 0.03220316 | 24.8571429 | rnr1Y285F | FALSE | CAN | JSY3866 | 7 | rnr1Y285F-R | CGG | G>T | A/T+1 | G/C-1 | CG>AT | CG>AT | AAATACGGG | GGGAACC | GCC | GCC | C | A | C>A | C>A_GCC | CG>AT_GGGAACC | 8 | FALSE | FALSE |
| 163_v.txt | 32274 | 32274-SNV-( | SNV | 1 | G | T | 25 | 21737 | 0.11501127 | 34.88 | rnr1Y285F | FALSE | CAN | JSY3866 | 7 | rnr1Y285F-R | GGA | G>T | A/T+1 | G/C-1 | CG>AT | CG>AT | ATACGGGAA | GAACCAA | CCT | CCT | C | A | C>A | C>A_CCT | CG>AT_GAACCAA | 8 | FALSE | FALSE |
| 163_v.txt | 32304 | 32304-SNV-( | SNV | 1 | G | T | 73 | 21737 | 0.33583291 | 55.8073107 | rnr1Y285F | FALSE | CAN | JSY3866 | 7 | rnr1Y285F-R | AGA | G>T | A/T+1 | G/C-1 | CG>AT | CG>AT | GCGGCAGAA | GAAATAA | TCT | TCT | C | A | C>A | C>A_TCT | CG>AT_GAAATAA | 8 | FALSE | FALSE |
| 163_v.txt | 32373 | 32373-SNV-( | SNV | 1 | G | T | 35 | 21737 | 0.16101578 | 58.2857143 | rnr1Y285F | FALSE | CAN | JSY3866 | 7 | rnr1Y285F-R | AGC | G>T | A/T+1 | G/C-1 | CG>AT | CG>AT | TCAATAGCA | GCAATAA | TCG | TCG | C | A | C>A | C>A_TCG | CG>AT_GCAATAA | 8 | FALSE | FALSE |
| 163_v.txt | 32432 | 32432-SNV-( | SNV | 1 | G | T | 15 | 21737 | 0.06900676 | 50.6 | rnr1Y285F | FALSE | CAN | JSY3866 | 7 | rnr1Y285F-R | TGT | G>T | A/T+1 | G/C-1 | CG>AT | CG>AT | GTCATTGTAT | GTATGGA | ACA | ACA | C | A | C>A | C>A_ACA | CG>AT_GTATGGA | 8 | FALSE | FALSE |
| 163_v.txt | 32449 | 32449-SNV-( | SNV | 1 | C | A | 46 | 21737 | 0.21162074 | 54.9130435 | rnr1Y285F | FALSE | CAN | JSY3866 | 7 | rnr1Y285F-R | CCA | C>A | A/T+1 | G/C-1 | CG>AT | CG>AT | AAAGTCCAAT | CAATGAA | GGT | CCA | C | T | C>A | C>A_CCA | CG>AT_CAATGAA | 8 | FALSE | FALSE |
| 163_v.txt | 32515 | 32515-SNV-( | SNV | 1 | G | T | 69 | 21737 | 0.31743111 | 42.8550725 | rnr1Y285F | FALSE | CAN | JSY3866 | 7 | rnr1Y285F-R | GGA | G>T | A/T+1 | G/C-1 | CG>AT | CG>AT | CTCTTGGAA | GAACGGA | CCT | CCT | C | A | C>A | C>A_CCT | CG>AT_GAACGGA | 8 | FALSE | FALSE |
| 163_v.txt | 32530 | 32530-SNV-( | SNV | 1 | G | T | 56 | 21737 | 0.25762525 | 37.875 | rnr1Y285F | FALSE | CAN | JSY3866 | 7 | rnr1Y285F-R | GGG | G>T | A/T+1 | G/C-1 | CG>AT | CG>AT | TTCTGGGGT | GGTTTGC | CCC | CCC | C | A | C>A | C>A_CCC | CG>AT_GGTTTGC | 8 | FALSE | FALSE |
| 163_v.txt | 32538 | 32538-SNV-( | SNV | 1 | G | T | 17 | 21737 | 0.07820766 | 37 | rnr1Y285F | FALSE | CAN | JSY3866 | 7 | rnr1Y285F-R | AGC | G>T | A/T+1 | G/C-1 | CG>AT | CG>AT | TTTGCAGCTT | GCTTCAC | TCG | TCG | C | A | C>A | C>A_TCG | CG>AT_GCTTCAC | 8 | FALSE | FALSE |
| 163_v.txt | 32556 | 32556-SNV-( | SNV | 1 | C | A | 34 | 21733 | 0.15644412 | 36.9411765 | rnr1Y285F | FALSE | CAN | JSY3866 | 7 | rnr1Y285F-R | ACC | C>A | A/T+1 | G/C-1 | CG>AT | CG>AT | GTGATACCA | CCAACTA | TGG | ACC | C | T | C>A | C>A_ACC | CG>AT_CCAACTA | 8 | FALSE | FALSE |
| 163_v.txt | 32629 | 32629-SNV-( | SNV | 1 | C | A | 35 | 25230 | 0.13872374 | 51.4285714 | rnr1Y285F | FALSE | CAN | JSY3866 | 7 | rnr1Y285F-R | TCG | C>A | A/T+1 | G/C-1 | CG>AT | CG>AT | TCCCTTCGTT | CGTTTTT | AGC | TCG | C | T | C>A | C>A_TCG | CG>AT_CGTTTTT | 8 | FALSE | FALSE |
| 163_v.txt | 32735 | 32735-SNV-( | SNV | 1 | G | T | 58 | 25230 | 0.22988506 | 52.5 | rnr1Y285F | FALSE | CAN | JSY3866 | 7 | rnr1Y285F-R | AGT | G>T | A/T+1 | G/C-1 | CG>AT | CG>AT | AAAACAGTA | GTATATT | TCA | TCA | C | A | C>A | C>A_TCA | CG>AT_GTATATT | 8 | FALSE | FALSE |
| 163_v.txt | 32783 | 32783-SNV-( | SNV | 1 | G | T | 33 | 25230 | 0.13079667 | 44.6666667 | rnr1Y285F | FALSE | CAN | JSY3866 | 7 | rnr1Y285F-R | AGA | G>T | A/T+1 | G/C-1 | CG>AT | CG>AT | GACCCAGAA | GAACTCG | TCT | TCT | C | A | C>A | C>A_TCT | CG>AT_GAACTCG | 8 | FALSE | FALSE |
| 163_v.txt | 32794 | 32794-SNV-( | SNV | 1 | C | A | 30 | 25230 | 0.11890606 | 39.6333333 | rnr1Y285F | FALSE | CAN | JSY3866 | 7 | rnr1Y285F-R | TCA | C>A | A/T+1 | G/C-1 | CG>AT | CG>AT | CGAATTCAC | CACCGTA | AGT | TCA | C | T | C>A | C>A_TCA | CG>AT_CACCGTA | 8 | FALSE | FALSE |
| 163_v.txt | 32812 | 32812-SNV-( | SNV | 1 | G | T | 23 | 25230 | 0.09116132 | 42.2173913 | rnr1Y285F | FALSE | CAN | JSY3866 | 7 | rnr1Y285F-R | GGG | G>T | A/T+1 | G/C-1 | CG>AT | CG>AT | TGACAGGGA | GGAACAA | CCC | CCC | C | A | C>A | C>A_CCC | CG>AT_GGAACAA | 8 | FALSE | FALSE |
| 163_v.txt | 32819 | 32819-SNV-( | SNV | 1 | G | T | 14 | 25230 | 0.0554895 | 41.2142857 | rnr1Y285F | FALSE | CAN | JSY3866 | 7 | rnr1Y285F-R | AGT | G>T | A/T+1 | G/C-1 | CG>AT | CG>AT | GAACAAGTT | GTTCATT | TCA | TCA | C | A | C>A | C>A_TCA | CG>AT_GTTCATT | 8 | FALSE | FALSE |
| 163_v.txt | 32876 | 32876-SNV-( | SNV | 1 | G | T | 18 | 16565 | 0.10866284 | 31.2777778 | rnr1Y285F | FALSE | CAN | JSY3866 | 7 | rnr1Y285F-R | TGT | G>T | A/T+1 | G/C-1 | CG>AT | CG>AT | AACTTTGTA | GTACGTC | ACA | ACA | C | A | C>A | C>A_ACA | CG>AT_GTACGTC | 8 | FALSE | FALSE |
| 163_v.txt | 32917 | 32917-SNV-( | SNV | 1 | C | A | 108 | 16580 | 0.65138721 | 52.5740741 | rnr1Y285F | FALSE | CAN | JSY3866 | 7 | rnr1Y285F-R | TCC | C>A | A/T+1 | G/C-1 | CG>AT | CG>AT | TAAGTTCCA | CCAGGGC | AGG | TCC | C | T | C>A | C>A_TCC | CG>AT_CCAGGGC | 8 | FALSE | FALSE |
| 163_v.txt | 32928 | 32928-SNV-( | SNV | 1 | G | T | 111 | 16580 | 0.6694813 | 52.9189189 | rnr1Y285F | FALSE | CAN | JSY3866 | 7 | rnr1Y285F-R | AGT | G>T | A/T+1 | G/C-1 | CG>AT | CG>AT | GCAAAAGTG | GTGATTG | TCA | TCA | C | A | C>A | C>A_TCA | CG>AT_GTGATTG | 8 | FALSE | FALSE |
| 163_v.txt | 32940 | 32940-SNV-( | SNV | 1 | G | T | 208 | 16580 | 1.25452352 | 55.3605769 | rnr1Y285F | FALSE | CAN | JSY3866 | 7 | rnr1Y285F-R | AGA | G>T | A/T+1 | G/C-1 | CG>AT | CG>AT | GCCCAAGAA | GAAAACC | TCT | TCT | C | A | C>A | C>A_TCT | CG>AT_GAAAACC | 8 | FALSE | FALSE |
| 163_v.txt | 32946 | 32946-SNV-( | SNV | 1 | C | A | 4 | 16580 | 0.02412545 | 44.2570357 | rnr1Y285F | FALSE | CAN | JSY3866 | 7 | rnr1Y285F-R | CCA | C>A | A/T+1 | G/C-1 | CG>AT | CG>AT | GAAAACCAA | CAATACA | GGT | CCA | C | T | C>A | C>A_CCA | CG>AT_CAATACA | 8 | FALSE | FALSE |
| 163_v.txt | 32958 | 32958-SNV-( | SNV | 1 | C | A | 24 | 16580 | 0.14475271 | 54.625 | rnr1Y285F | FALSE | CAN | JSY3866 | 7 | rnr1Y285F-R | ACC | C>A | A/T+1 | G/C-1 | CG>AT | CG>AT | ATGTAACCAT | CCATTGG | TGG | ACC | C | T | C>A | C>A_ACC | CG>AT_CCATTGG | 8 | FALSE | FALSE |
| 163_v.txt | 32997 | 32997-SNV-( | SNV | 1 | G | T | 13 | 16580 | 0.07840772 | 58.5384615 | rnr1Y285F | FALSE | CAN | JSY3866 | 7 | rnr1Y285F-R | TGA | G>T | A/T+1 | G/C-1 | CG>AT | CG>AT | CTTTGTGAG | GAGAAAA | ACT | ACT | C | A | C>A | C>A_ACT | CG>AT_GAGAAAA | 8 | FALSE | FALSE |
| 163_v.txt | 33012 | 33012-SNV-( | SNV | 1 | G | T | 14 | 16580 | 0.08443908 | 58.0714286 | rnr1Y285F | FALSE | CAN | JSY3866 | 7 | rnr1Y285F-R | AGA | G>T | A/T+1 | G/C-1 | CG>AT | CG>AT | GTGAAAGAG | GAGGATG | TCT | TCT | C | A | C>A | C>A_TCT | CG>AT_GAGGATG | 8 | FALSE | FALSE |
| 163_v.txt | 33015 | 33015-SNV-( | SNV | 1 | G | T | 36 | 16580 | 0.21712907 | 54.8611111 | rnr1Y285F | FALSE | CAN | JSY3866 | 7 | rnr1Y285F-R | GGA | G>T | A/T+1 | G/C-1 | CG>AT | CG>AT | AAAGAGGAT | GATGTAA | CCT | CCT | C | A | C>A | C>A_CCT | CG>AT_GATGTAA | 8 | FALSE | FALSE |
| 163_v.txt | 33043 | 33043-SNV-( | SNV | 1 | C | A | 9 | 16580 | 0.05428227 | 61.6666667 | rnr1Y285F | FALSE | CAN | JSY3866 | 7 | rnr1Y285F-R | TCA | C>A | A/T+1 | G/C-1 | CG>AT | CG>AT | CCATTTCACC | CACCCAA | AGT | TCA | C | T | C>A | C>A_TCA | CG>AT_CACCCAA | 8 | FALSE | FALSE |
| 163_v.txt | 33093 | 33093-SNV-( | SNV | 1 | G | T | 73 | 16580 | 0.44028951 | 51.2561182 | rnr1Y285F | FALSE | CAN | JSY3866 | 7 | rnr1Y285F-R | TGA | G>T | A/T+1 | G/C-1 | CG>AT | CG>AT | AAATATGAT | GATATAA | ACT | ACT | C | A | C>A | C>A_ACT | CG>AT_GATATAA | 8 | FALSE | FALSE |
| 163_v.txt | 33151 | 33151-SNV-( | SNV | 1 | G | T | 71 | 16580 | 0.42822678 | 38.6901409 | rnr1Y285F | FALSE | CAN | JSY3866 | 7 | rnr1Y285F-R | AGA | G>T | A/T+1 | G/C-1 | CG>AT | CG>AT | TGAAAAGAC | GACCTGT | TCT | TCT | C | A | C>A | C>A_TCT | CG>AT_GACCTGT | 8 | FALSE | FALSE |
| 163_v.txt | 33153 | 33153-SNV-( | SNV | 1 | C | A | 104 | 16580 | 0.62726176 | 42.6923077 | rnr1Y285F | FALSE | CAN | JSY3866 | 7 | rnr1Y285F-R | ACC | C>A | A/T+1 | G/C-1 | CG>AT | CG>AT | AAAAGACCT | CCTGTAC | TGG | ACC | C | T | C>A | C>A_ACC | CG>AT_CCTGTAC | 8 | FALSE | FALSE |
| 163_v.txt | 33154 | 33154-SNV-( | SNV | 1 | C | A | 50 | 16580 | 0.30156815 | 36.64 | rnr1Y285F | FALSE | CAN | JSY3866 | 7 | rnr1Y285F-R | CCT | C>A | A/T+1 | G/C-1 | CG>AT | CG>AT | AAAGACCTG | CTGTACC | GGA | CCT | C | T | C>A | C>A_CCT | CG>AT_CTGTACC | 8 | FALSE | FALSE |
| 163_v.txt | 33168 | 33168-SNV-( | SNV | 1 | C | A | 117 | 16578 | 0.70575462 | 37.2735043 | rnr1Y285F | FALSE | CAN | JSY3866 | 7 | rnr1Y285F-R | ACC | C>A | A/T+1 | G/C-1 | CG>AT | CG>AT | ATAGTACCA | CCACCAA | TGG | ACC | C | T | C>A | C>A_ACC | CG>AT_CCACCAA | 8 | FALSE | FALSE |
| 163_v.txt | 33205 | 33205-SNV-( | SNV | 1 | G | T | 188 | 24848 | 0.75660013 | 39.0557633 | rnr1Y285F | FALSE | CAN | JSY3866 | 7 | rnr1Y285F-R | AGC | G>T | A/T+1 | G/C-1 | CG>AT | CG>AT | GCTTAAGCT | GCTCTCT | TCG | TCG | C | A | C>A | C>A_TCG | CG>AT_GCTCTCT | 8 | FALSE | FALSE |
| 163_v.txt | 33325 | 33325-SNV-( | SNV | 1 | C | A | 43 | 24849 | 0.17304519 | 50.5581395 | rnr1Y285F | FALSE | CAN | JSY3866 | 7 | rnr1Y285F-R | TCT | C>A | A/T+1 | G/C-1 | CG>AT | CG>AT | ACAATTCTTT | CTTTACT | AGA | TCT | C | T | C>A | C>A_TCT | CG>AT_CTTTACT | 8 | FALSE | FALSE |
| 163_v.txt | 33474 | 33474-SNV-( | SNV | 1 | C | A | 65 | 24845 | 0.26162206 | 35.2769231 | rnr1Y285F | FALSE | CAN | JSY3866 | 7 | rnr1Y285F-R | GCC | C>A | A/T+1 | G/C-1 | CG>AT | CG>AT | GCTATGCCTT | CCTTTTT | CGG | GCC | C | T | C>A | C>A_GCC | CG>AT_CCTTTTT | 8 | FALSE | FALSE |
| 163_v.txt | 33475 | 33475-SNV-( | SNV | 1 | C | A | 16 | 24845 | 0.06439928 | 34.9375 | rnr1Y285F | FALSE | CAN | JSY3866 | 7 | rnr1Y285F-R | CCT | C>A | A/T+1 | G/C-1 | CG>AT | CG>AT | CTATGCCTTT | CTTTTTT | GGA | CCT | C | T | C>A | C>A_CCT | CG>AT_CTTTTTT | 8 | FALSE | FALSE |
| 163_v.txt | 32312 | 32312-Repla | Replacement | 2 | GG | A | 5 | 21737 | 0.02300225 | 58.1 | rnr1Y285F | FALSE | CAN | JSY3866 | 7 | rnr1Y285F-R | TGG | nope | A/T+1 | >1bpdel | none | Replac. | AATAATGGT | GGTTGTT | ACC | TGG | GG | T | NA | NA_TGG | Replac._GGTTGTT | 8 | FALSE | FALSE |
| 163_v.txt | 31962 | 31962-MNV- | MNV | 2 | GC | CA | 20 | 23392 | 0.08549932 | 37.65 | rnr1Y285F | FALSE | CAN | JSY3866 | 7 | rnr1Y285F-R | CGC | nope | >1bp-ins | >1bpdel | none | MNV | GTGGCCGCA | GCATAAT | GCG | CGC | GC | none | NA | NA_CGC | MNV_GCATAAT | 8 | FALSE | FALSE |
| 163_v.txt | 33153 | 33153-MNV- | MNV | 2 | CC | AA | 1 | 16580 | 6.03E-03 | 38 | rnr1Y285F | FALSE | CAN | JSY3866 | 7 | rnr1Y285F-R | ACC | nope | >1bp-ins | >1bpdel | none | MNV | AAAAGACCT | CCTGTAC | TGG | ACC | CC | none | NA | NA_ACC | MNV_CCTGTAC | 8 | FALSE | FALSE |
| 163_v.txt | 32081 | 32081-Insert | Insertion | 1 | - | A | 9 | 23392 | 0.03847469 | 61.1111111 | rnr1Y285F | FALSE | CAN | JSY3866 | 7 | rnr1Y285F-R | CAA | nope | A/T+1 | >1bpdel | none | A/T+1 | CCATGCAAA | AAAAAAG | GTT | CAA | - | T | NA | NA_CAA | A/T+1_AAAAAAG | 8 | FALSE | FALSE |
| 163_v.txt | 32312 | 32312-Insert | Insertion | 1 | - | G | 1 | 21737 | 4.60E-03 | 64 | rnr1Y285F | FALSE | CAN | JSY3866 | 7 | rnr1Y285F-R | TGG | nope | G/C+1 | >1bpdel | none | G/C+1 | AATAATGGT | GGTTGTT | ACC | TGG | - | C | NA | NA_TGG | G/C+1_GGTTGTT | 8 | FALSE | FALSE |
| 163_v.txt | 32498 | 32498-Insert | Insertion | 1 | - | T | 3 | 21737 | 0.01380135 | 38 | rnr1Y285F | FALSE | CAN | JSY3866 | 7 | rnr1Y285F-R | CTT | nope | A/T+1 | >1bpdel | none | A/T+1 | AACAACTTTT | TTTTTTG | GAA | CTT | - | A | NA | NA_CTT | A/T+1_TTTTTTG | 8 | FALSE | FALSE |
| 163_v.txt | 32728 | 32728-Insert | Insertion | 1 | - | A | 44 | 25230 | 0.17439556 | 61.9545455 | rnr1Y285F | FALSE | CAN | JSY3866 | 7 | rnr1Y285F-R | CAA | nope | A/T+1 | >1bpdel | none | A/T+1 | CCATACAAA | AAAAACA | GTT | CAA | - | T | NA | NA_CAA | A/T+1_AAAAACA | 8 | FALSE | FALSE |
| 163_v.txt | 32842 | 32842-Insert | Insertion | 1 | - | A | 23 | 25230 | 0.09116132 | 42.6521739 | rnr1Y285F | FALSE | CAN | JSY3866 | 7 | rnr1Y285F-R | CAA | nope | A/T+1 | >1bpdel | none | A/T+1 | TTACCCAAA | AAAAAAT | GTT | CAA | - | T | NA | NA_CAA | A/T+1_AAAAAAT | 8 | FALSE | FALSE |
| 163_v.txt | 32843 | 32843-Insert | Insertion | 1 | - | G | 1 | 25230 | 3.96E-03 | 38 | rnr1Y285F | FALSE | CAN | JSY3866 | 7 | rnr1Y285F-R | AAA | nope | G/C+1 | >1bpdel | none | G/C+1 | TACCCAAAA | AAAAATA | TTT | AAA | - | C | NA | NA_AAA | G/C+1_AAAAATA | 8 | FALSE | FALSE |
| 163_v.txt | 32884 | 32884-Insert | Insertion | 1 | - | A | 117 | 16580 | 0.70566948 | 37.2820513 | rnr1Y285F | FALSE | CAN | JSY3866 | 7 | rnr1Y285F-R | CAA | nope | A/T+1 | >1bpdel | none | A/T+1 | ACGTCCAAA | AAAATTG | GTT | CAA | - | T | NA | NA_CAA | A/T+1_AAAATTG | 8 | FALSE | FALSE |
| 163_v.txt | 32924 | 32924-Insert | Insertion | 1 | - | A | 108 | 16580 | 0.65138721 | 53.0185185 | rnr1Y285F | FALSE | CAN | JSY3866 | 7 | rnr1Y285F-R | CAA | nope | A/T+1 | >1bpdel | none | A/T+1 | CAGGGCAAA | AAAAGTG | GTT | CAA | - | T | NA | NA_CAA | A/T+1_AAAAGTG | 8 | FALSE | FALSE |
| 163_v.txt | 33203 | 33203-Insert | Insertion | 2 | - | AA | 159 | 24849 | 0.63986478 | 38.7578616 | rnr1Y285F | FALSE | CAN | JSY3866 | 7 | rnr1Y285F-R | TAA | nope | >1bp-ins | >1bpdel | none | >1bp-ins | TTGCTTAAG | AAGCTCT | ATT | TAA | - | none | NA | NA_TAA | >1bp-ins_AAGCTCT | 8 | FALSE | FALSE |
| 163_v.txt | 33331 | 33331-Insert | Insertion | 2 | - | TT | 17 | 24849 | 0.06841322 | 60.0294118 | rnr1Y285F | FALSE | CAN | JSY3866 | 7 | rnr1Y285F-R | CTT | nope | >1bp-ins | >1bpdel | none | >1bp-ins | CTTTACTTTT | TTTTCTC | GAA | CTT | - | none | NA | NA_CTT | >1bp-ins_TTTTCTC | 8 | FALSE | FALSE |
| 163_v.txt | 31788 | 31788-Deleti | Deletion | 4 | TCGA | - | 10 | 3349 | 0.2985966 | 58.8 | rnr1Y285F | FALSE | CAN | JSY3866 | 7 | rnr1Y285F-R | GTC | nope | >1bp-ins | >1bpdel | none | >1bpdel | TCGATGTCG | TCGACAT | CAG | GTC | TCGA | none | NA | NA_GTC | >1bpdel_TCGACAT | 8 | FALSE | FALSE |

| | | | | | | | | | | | | | | | | | | | | | | | | | | | | | | | | | | |
|---|---|---|---|---|---|---|---|---|---|---|---|---|---|---|---|---|---|---|---|---|---|---|---|---|---|---|---|---|---|---|---|---|---|---|
| 163_v.txt | 32303 | 32303-Deleti | Deletion | 3 | AGA | - | 16 | 21737 | 0.07360721 | 57.5 | rnr1Y285F | FALSE | CAN | JSY3866 | 7 | rnr1Y285F-R | CAG | nope | >1bp-ins | >1bpdel | none | >1bpdel | TGCGGCAGA | AGAAATA | GTC | CAG | AGA | none | NA | NA_CAG | >1bpdel_AGAAATA | 8 | FALSE | FALSE |
| 163_v.txt | 32313 | 32313-Deleti | Deletion | 3 | GTT | - | 7 | 21737 | 0.03220316 | 62 | rnr1Y285F | FALSE | CAN | JSY3866 | 7 | rnr1Y285F-R | GGT | nope | >1bp-ins | >1bpdel | none | >1bpdel | ATAATGGTT | GTTGTTA | CCA | GGT | GTT | none | NA | NA_GGT | >1bpdel_GTTGTTA | 8 | FALSE | FALSE |
| 163_v.txt | 32337 | 32337-Deleti | Deletion | 2 | AT | - | 31 | 21737 | 0.14261398 | 63.3225807 | rnr1Y285F | FALSE | CAN | JSY3866 | 7 | rnr1Y285F-R | GAT | nope | >1bp-ins | >1bpdel | none | >1bpdel | TTGAAGATA | ATATGTG | CTA | GAT | AT | none | NA | NA_GAT | >1bpdel_ATATGTG | 8 | FALSE | FALSE |
| 163_v.txt | 32375 | 32375-Deleti | Deletion | 3 | AAT | - | 21 | 21737 | 0.09660947 | 62.2380952 | rnr1Y285F | FALSE | CAN | JSY3866 | 7 | rnr1Y285F-R | CAA | nope | >1bp-ins | >1bpdel | none | >1bpdel | AATAGCAAT | AATAATA | GTT | CAA | AAT | none | NA | NA_CAA | >1bpdel_AATAATA | 8 | FALSE | FALSE |
| 163_v.txt | 32454 | 32454-Deleti | Deletion | 3 | AAT | - | 13 | 21737 | 0.05980586 | 56 | rnr1Y285F | FALSE | CAN | JSY3866 | 7 | rnr1Y285F-R | GAA | nope | >1bp-ins | >1bpdel | none | >1bpdel | CCAATGAAT | AATAATA | CTT | GAA | AAT | none | NA | NA_GAA | >1bpdel_AATAATA | 8 | FALSE | FALSE |
| 163_v.txt | 32460 | 32460-Deleti | Deletion | 2 | AG | - | 39 | 21737 | 0.17941758 | 51.7692308 | rnr1Y285F | FALSE | CAN | JSY3866 | 7 | rnr1Y285F-R | TAG | nope | >1bp-ins | >1bpdel | none | >1bpdel | AATAATAGA | AGAGAGC | ATC | TAG | AG | none | NA | NA_TAG | >1bpdel_AGAGAGC | 8 | FALSE | FALSE |
| 163_v.txt | 33094 | 33094-Deleti | Deletion | 2 | AT | - | 12 | 16580 | 0.07237636 | 48.75 | rnr1Y285F | FALSE | CAN | JSY3866 | 7 | rnr1Y285F-R | GAT | nope | >1bp-ins | >1bpdel | none | >1bpdel | AATATGATA | ATATAAG | CTA | GAT | AT | none | NA | NA_GAT | >1bpdel_ATATAAG | 8 | FALSE | FALSE |
| 163_v.txt | 33189 | 33189-Deleti | Deletion | 2 | AT | - | 8 | 24849 | 0.03219446 | 38 | rnr1Y285F | FALSE | CAN | JSY3866 | 7 | rnr1Y285F-R | AAT | nope | >1bp-ins | >1bpdel | none | >1bpdel | ATACCAATA | ATATGTC | TTA | AAT | AT | none | NA | NA_AAT | >1bpdel_ATATGTC | 8 | FALSE | FALSE |
| 163_v.txt | 31952 | 31952-Deleti | Deletion | 1 | A | - | 6 | 23392 | 0.0256498 | 37.8333333 | rnr1Y285F | FALSE | CAN | JSY3866 | 7 | rnr1Y285F-R | TAA | nope | >1bp-ins | A/T-1 | none | A/T-1 | CGTCATAAA | AAATGTG | ATT | ATT | T | none | T>none | T>none_ATT | A/T-1_AAATGTG | 8 | FALSE | FALSE |
| 163_v.txt | 31972 | 31972-Deleti | Deletion | 1 | C | - | 5 | 23392 | 0.02137483 | 37.6 | rnr1Y285F | FALSE | CAN | JSY3866 | 7 | rnr1Y285F-R | GCC | nope | >1bp-ins | G/C-1 | none | G/C-1 | AATAAGCCA | CCAAGCC | CGG | GCC | C | none | nope | nope_GCC | G/C-1_CCAAGCC | 8 | FALSE | FALSE |
| 163_v.txt | 31992 | 31992-Deleti | Deletion | 1 | G | - | 14 | 23392 | 0.05984952 | 57.8571436 | rnr1Y285F | FALSE | CAN | JSY3866 | 7 | rnr1Y285F-R | AGC | nope | >1bp-ins | G/C-1 | none | G/C-1 | AATTTAGCT | GCTTTAA | TCG | TCG | C | none | C>none | C>none_TCG | G/C-1_GCTTTAA | 8 | FALSE | FALSE |
| 163_v.txt | 32081 | 32081-Deleti | Deletion | 1 | A | - | 26 | 23392 | 0.11114911 | 61.8076942 | rnr1Y285F | FALSE | CAN | JSY3866 | 7 | rnr1Y285F-R | CAA | nope | >1bp-ins | A/T-1 | none | A/T-1 | CCATGCAAA | AAAAAAG | GTT | GTT | T | none | T>none | T>none_GTT | A/T-1_AAAAAAG | 8 | FALSE | FALSE |
| 163_v.txt | 32102 | 32102-Deleti | Deletion | 1 | G | - | 35 | 23392 | 0.1496238 | 60.7142857 | rnr1Y285F | FALSE | CAN | JSY3866 | 7 | rnr1Y285F-R | TGA | nope | >1bp-ins | G/C-1 | none | G/C-1 | ACCAGTGAT | GATATTT | ACT | ACT | C | none | C>none | C>none_ACT | G/C-1_GATATTT | 8 | FALSE | FALSE |
| 163_v.txt | 32160 | 32160-Deleti | Deletion | 1 | A | - | 5 | 23392 | 0.02137483 | 58 | rnr1Y285F | FALSE | CAN | JSY3866 | 7 | rnr1Y285F-R | CAA | nope | >1bp-ins | A/T-1 | none | A/T-1 | TAAGCCAAA | AAAGCGC | GTT | GTT | T | none | T>none | T>none_GTT | A/T-1_AAAGCGC | 8 | FALSE | FALSE |
| 163_v.txt | 32210 | 32210-Deleti | Deletion | 1 | T | - | 8 | 23392 | 0.03419973 | 39.875 | rnr1Y285F | FALSE | CAN | JSY3866 | 7 | rnr1Y285F-R | CTT | nope | >1bp-ins | A/T-1 | none | A/T-1 | ACCACCTTTG | TTTGGTG | GAA | CTT | T | none | nope | nope_CTT | A/T-1_TTTGGTG | 8 | FALSE | FALSE |
| 163_v.txt | 32238 | 32238-Deleti | Deletion | 1 | G | - | 14 | 23388 | 0.05985976 | 37.2142857 | rnr1Y285F | FALSE | CAN | JSY3866 | 7 | rnr1Y285F-R | AGC | nope | >1bp-ins | G/C-1 | none | G/C-1 | TTAGGAGCC | GCCAACT | TCG | TCG | C | none | C>none | C>none_TCG | G/C-1_GCCAACT | 8 | FALSE | FALSE |
| 163_v.txt | 32265 | 32265-Deleti | Deletion | 1 | A | - | 6 | 21737 | 0.02760271 | 37.5 | rnr1Y285F | FALSE | CAN | JSY3866 | 7 | rnr1Y285F-R | TAA | nope | >1bp-ins | A/T-1 | none | A/T-1 | CCAAATAAA | AAAATAC | ATT | ATT | T | none | T>none | T>none_ATT | A/T-1_AAAATAC | 8 | FALSE | FALSE |
| 163_v.txt | 32272 | 32272-Deleti | Deletion | 1 | G | - | 1 | 21737 | 4.60E-03 | 37 | rnr1Y285F | FALSE | CAN | JSY3866 | 7 | rnr1Y285F-R | CGG | nope | >1bp-ins | G/C-1 | none | G/C-1 | AAATACGGG | GGGAACC | GCC | GCC | C | none | C>none | C>none_GCC | G/C-1_GGGAACC | 8 | FALSE | FALSE |
| 163_v.txt | 32284 | 32284-Deleti | Deletion | 1 | A | - | 2 | 21737 | 9.20E-03 | 49 | rnr1Y285F | FALSE | CAN | JSY3866 | 7 | rnr1Y285F-R | TAA | nope | >1bp-ins | A/T-1 | none | A/T-1 | CAACGTAAA | AAATATT | ATT | ATT | T | none | T>none | T>none_ATT | A/T-1_AAATATT | 8 | FALSE | FALSE |
| 163_v.txt | 32304 | 32304-Deleti | Deletion | 1 | G | - | 38 | 21737 | 0.17481713 | 52.6592869 | rnr1Y285F | FALSE | CAN | JSY3866 | 7 | rnr1Y285F-R | AGA | nope | >1bp-ins | G/C-1 | none | G/C-1 | GCGGCAGAA | GAAATAA | TCT | TCT | C | none | C>none | C>none_TCT | G/C-1_GAAATAA | 8 | FALSE | FALSE |
| 163_v.txt | 32305 | 32305-Deleti | Deletion | 1 | A | - | 1 | 21736 | 4.60E-03 | 37.9999967 | rnr1Y285F | FALSE | CAN | JSY3866 | 7 | rnr1Y285F-R | GAA | nope | >1bp-ins | A/T-1 | none | A/T-1 | CGGCAGAAA | AAATAAT | CTT | CTT | T | none | T>none | T>none_CTT | A/T-1_AAATAAT | 8 | FALSE | FALSE |
| 163_v.txt | 32346 | 32346-Deleti | Deletion | 1 | A | - | 12 | 21737 | 0.05520541 | 62.5833333 | rnr1Y285F | FALSE | CAN | JSY3866 | 7 | rnr1Y285F-R | CAA | nope | >1bp-ins | A/T-1 | none | A/T-1 | TGTGGCAAA | AAAACCT | GTT | GTT | T | none | T>none | T>none_GTT | A/T-1_AAAACCT | 8 | FALSE | FALSE |
| 163_v.txt | 32381 | 32381-Deleti | Deletion | 1 | A | - | 6 | 21737 | 0.02760271 | 63.1666667 | rnr1Y285F | FALSE | CAN | JSY3866 | 7 | rnr1Y285F-R | TAA | nope | >1bp-ins | A/T-1 | none | A/T-1 | AATAATAAA | AAAGGGA | ATT | ATT | T | none | T>none | T>none_ATT | A/T-1_AAAGGGA | 8 | FALSE | FALSE |
| 163_v.txt | 32429 | 32429-Deleti | Deletion | 1 | A | - | 27 | 21737 | 0.12421217 | 49 | rnr1Y285F | FALSE | CAN | JSY3866 | 7 | rnr1Y285F-R | CAT | nope | >1bp-ins | A/T-1 | none | A/T-1 | AGGGTCATT | ATTGTAT | GTA | GTA | T | none | T>none | T>none_GTA | A/T-1_ATTGTAT | 8 | FALSE | FALSE |
| 163_v.txt | 32442 | 32442-Deleti | Deletion | 1 | A | - | 5 | 21737 | 0.02300225 | 52.6 | rnr1Y285F | FALSE | CAN | JSY3866 | 7 | rnr1Y285F-R | TAA | nope | >1bp-ins | A/T-1 | none | A/T-1 | GGAACTAAA | AAAAGTC | ATT | ATT | T | none | T>none | T>none_ATT | A/T-1_AAAAGTC | 8 | FALSE | FALSE |
| 163_v.txt | 32498 | 32498-Deleti | Deletion | 1 | T | - | 29 | 21737 | 0.13341308 | 46.3448261 | rnr1Y285F | FALSE | CAN | JSY3866 | 7 | rnr1Y285F-R | CTT | nope | >1bp-ins | A/T-1 | none | A/T-1 | AACAACTTTT | TTTTTTG | GAA | CTT | T | none | nope | nope_CTT | A/T-1_TTTTTTG | 8 | FALSE | FALSE |
| 163_v.txt | 32521 | 32521-Deleti | Deletion | 1 | A | - | 74 | 21737 | 0.34043336 | 40.459459 | rnr1Y285F | FALSE | CAN | JSY3866 | 7 | rnr1Y285F-R | GAT | nope | >1bp-ins | A/T-1 | none | A/T-1 | GAACGGATT | ATTTTCT | CTA | CTA | T | none | T>none | T>none_CTA | A/T-1_ATTTTCT | 8 | FALSE | FALSE |
| 163_v.txt | 32522 | 32522-Deleti | Deletion | 1 | T | - | 9 | 21737 | 0.04140406 | 37 | rnr1Y285F | FALSE | CAN | JSY3866 | 7 | rnr1Y285F-R | ATT | nope | >1bp-ins | A/T-1 | none | A/T-1 | AACGGATTT | TTTTCTG | TAA | ATT | T | none | nope | nope_ATT | A/T-1_TTTTCTG | 8 | FALSE | FALSE |
| 163_v.txt | 32556 | 32556-Deleti | Deletion | 1 | C | - | 95 | 21733 | 0.43712327 | 37.7052632 | rnr1Y285F | FALSE | CAN | JSY3866 | 7 | rnr1Y285F-R | ACC | nope | >1bp-ins | G/C-1 | none | G/C-1 | GTGATACCA | CCAACTA | TGG | ACC | C | none | nope | nope_ACC | G/C-1_CCAACTA | 8 | FALSE | FALSE |
| 163_v.txt | 32608 | 32608-Deleti | Deletion | 1 | C | - | 9 | 25230 | 0.03567182 | 43 | rnr1Y285F | FALSE | CAN | JSY3866 | 7 | rnr1Y285F-R | ACC | nope | >1bp-ins | G/C-1 | none | G/C-1 | AGGAAACCC | CCCAACC | TGG | ACC | C | none | nope | nope_ACC | G/C-1_CCCAACC | 8 | FALSE | FALSE |
| 163_v.txt | 32631 | 32631-Deleti | Deletion | 1 | T | - | 10 | 25230 | 0.03963536 | 46.6 | rnr1Y285F | FALSE | CAN | JSY3866 | 7 | rnr1Y285F-R | GTT | nope | >1bp-ins | A/T-1 | none | A/T-1 | CCTTCGTTTT | TTTTTAT | CAA | GTT | T | none | nope | nope_GTT | A/T-1_TTTTTAT | 8 | FALSE | FALSE |
| 163_v.txt | 32658 | 32658-Deleti | Deletion | 1 | C | - | 67 | 25230 | 0.26555688 | 60.1492537 | rnr1Y285F | FALSE | CAN | JSY3866 | 7 | rnr1Y285F-R | ACC | nope | >1bp-ins | G/C-1 | none | G/C-1 | CCTGGACCC | CCCCAGG | TGG | ACC | C | none | nope | nope_ACC | G/C-1_CCCCAGG | 8 | FALSE | FALSE |
| 163_v.txt | 32670 | 32670-Deleti | Deletion | 1 | G | - | 65 | 25230 | 0.25762981 | 61.3846154 | rnr1Y285F | FALSE | CAN | JSY3866 | 7 | rnr1Y285F-R | TGG | nope | >1bp-ins | G/C-1 | none | G/C-1 | GCACCTGGG | GGGTTTC | ACC | ACC | C | none | C>none | C>none_ACC | G/C-1_GGGTTTC | 8 | FALSE | FALSE |
| 163_v.txt | 32707 | 32707-Deleti | Deletion | 1 | C | - | 18 | 25230 | 0.07134364 | 60.1111111 | rnr1Y285F | FALSE | CAN | JSY3866 | 7 | rnr1Y285F-R | ACC | nope | >1bp-ins | G/C-1 | none | G/C-1 | CGGTAACCC | CCCCAGC | TGG | ACC | C | none | nope | nope_ACC | G/C-1_CCCCAGC | 8 | FALSE | FALSE |
| 163_v.txt | 32728 | 32728-Deleti | Deletion | 1 | A | - | 17 | 25230 | 0.0673801 | 60.1764678 | rnr1Y285F | FALSE | CAN | JSY3866 | 7 | rnr1Y285F-R | CAA | nope | >1bp-ins | A/T-1 | none | A/T-1 | CCATACAAA | AAAAACA | GTT | GTT | T | none | T>none | T>none_GTT | A/T-1_AAAAACA | 8 | FALSE | FALSE |
| 163_v.txt | 32760 | 32760-Deleti | Deletion | 1 | A | - | 13 | 25230 | 0.05152596 | 50.1538462 | rnr1Y285F | FALSE | CAN | JSY3866 | 7 | rnr1Y285F-R | TAA | nope | >1bp-ins | A/T-1 | none | A/T-1 | ATGGCTAAA | AAAACTT | ATT | ATT | T | none | T>none | T>none_ATT | A/T-1_AAAACTT | 8 | FALSE | FALSE |
| 163_v.txt | 32764 | 32764-Deleti | Deletion | 1 | C | - | 38 | 25230 | 0.15061435 | 53.5000046 | rnr1Y285F | FALSE | CAN | JSY3866 | 7 | rnr1Y285F-R | ACT | nope | >1bp-ins | G/C-1 | none | G/C-1 | CTAAAACTTT | CTTTGAT | TGA | ACT | C | none | nope | nope_ACT | G/C-1_CTTTGAT | 8 | FALSE | FALSE |
| 163_v.txt | 32779 | 32779-Deleti | Deletion | 1 | C | - | 5 | 25230 | 0.01981768 | 47.6 | rnr1Y285F | FALSE | CAN | JSY3866 | 7 | rnr1Y285F-R | ACC | nope | >1bp-ins | G/C-1 | none | G/C-1 | AAGCGACCC | CCCAGAA | TGG | ACC | C | none | nope | nope_ACC | G/C-1_CCCAGAA | 8 | FALSE | FALSE |
| 163_v.txt | 32842 | 32842-Deleti | Deletion | 1 | A | - | 45 | 25230 | 0.1783591 | 44.6666667 | rnr1Y285F | FALSE | CAN | JSY3866 | 7 | rnr1Y285F-R | CAA | nope | >1bp-ins | A/T-1 | none | A/T-1 | TTACCCAAA | AAAAAAT | GTT | GTT | T | none | T>none | T>none_GTT | A/T-1_AAAAAAT | 8 | FALSE | FALSE |
| 163_v.txt | 32917 | 32917-Deleti | Deletion | 1 | C | - | 2 | 16580 | 0.01206273 | 62.5 | rnr1Y285F | FALSE | CAN | JSY3866 | 7 | rnr1Y285F-R | TCC | nope | >1bp-ins | G/C-1 | none | G/C-1 | TAAGTTCCA | CCAGGGC | AGG | TCC | C | none | nope | nope_TCC | G/C-1_CCAGGGC | 8 | FALSE | FALSE |
| 163_v.txt | 32924 | 32924-Deleti | Deletion | 1 | A | - | 10 | 16580 | 0.06031363 | 49.9999967 | rnr1Y285F | FALSE | CAN | JSY3866 | 7 | rnr1Y285F-R | CAA | nope | >1bp-ins | A/T-1 | none | A/T-1 | CAGGGCAAA | AAAAGTG | GTT | GTT | T | none | T>none | T>none_GTT | A/T-1_AAAAGTG | 8 | FALSE | FALSE |
| 163_v.txt | 33000 | 33000-Deleti | Deletion | 1 | A | - | 5 | 16580 | 0.03015682 | 64 | rnr1Y285F | FALSE | CAN | JSY3866 | 7 | rnr1Y285F-R | GAA | nope | >1bp-ins | A/T-1 | none | A/T-1 | TGTGAGAAA | AAAACTG | CTT | CTT | T | none | T>none | T>none_CTT | A/T-1_AAAACTG | 8 | FALSE | FALSE |
| 163_v.txt | 33040 | 33040-Deleti | Deletion | 1 | T | - | 5 | 16580 | 0.03015682 | 58.6 | rnr1Y285F | FALSE | CAN | JSY3866 | 7 | rnr1Y285F-R | ATT | nope | >1bp-ins | A/T-1 | none | A/T-1 | TAGCCATTTC | TTTCACC | TAA | ATT | T | none | nope | nope_ATT | A/T-1_TTTCACC | 8 | FALSE | FALSE |
| 163_v.txt | 33159 | 33159-Deleti | Deletion | 1 | C | - | 21 | 16579 | 0.12666627 | 38 | rnr1Y285F | FALSE | CAN | JSY3866 | 7 | rnr1Y285F-R | ACC | nope | >1bp-ins | G/C-1 | none | G/C-1 | CCTGTACCA | CCAATAG | TGG | ACC | C | none | nope | nope_ACC | G/C-1_CCAATAG | 8 | FALSE | FALSE |
| 163_v.txt | 33173 | 33173-Deleti | Deletion | 1 | A | - | 28 | 16575 | 0.16892911 | 37.8571429 | rnr1Y285F | FALSE | CAN | JSY3866 | 7 | rnr1Y285F-R | CAA | nope | >1bp-ins | A/T-1 | none | A/T-1 | ACCACCAAG | AAGGGCA | GTT | GTT | T | none | T>none | T>none_GTT | A/T-1_AAGGGCA | 8 | FALSE | FALSE |
| 163_v.txt | 33258 | 33258-Deleti | Deletion | 1 | C | - | 2 | 24849 | 8.05E-03 | 38 | rnr1Y285F | FALSE | CAN | JSY3866 | 7 | rnr1Y285F-R | GCC | nope | >1bp-ins | G/C-1 | none | G/C-1 | CCTATGCCAT | CCATCCT | CGG | GCC | C | none | nope | nope_GCC | G/C-1_CCATCCT | 8 | FALSE | FALSE |
| 163_v.txt | 33288 | 33288-Deleti | Deletion | 1 | T | - | 6 | 24849 | 0.02414584 | 55.3333333 | rnr1Y285F | FALSE | CAN | JSY3866 | 7 | rnr1Y285F-R | ATT | nope | >1bp-ins | A/T-1 | none | A/T-1 | TCGCCATTTA | TTTACTC | TAA | ATT | T | none | nope | nope_ATT | A/T-1_TTTACTC | 8 | FALSE | FALSE |
| 163_v.txt | 33356 | 33356-Deleti | Deletion | 1 | C | - | 24 | 24849 | 0.09658336 | 54.2916667 | rnr1Y285F | FALSE | CAN | JSY3866 | 7 | rnr1Y285F-R | ACC | nope | >1bp-ins | G/C-1 | none | G/C-1 | TATTGACCCA | CCCACGT | TGG | ACC | C | none | nope | nope_ACC | G/C-1_CCCACGT | 8 | FALSE | FALSE |
| 163_v.txt | 33474 | 33474-Deleti | Deletion | 1 | C | - | 72 | 24845 | 0.28979674 | 27.1805556 | rnr1Y285F | FALSE | CAN | JSY3866 | 7 | rnr1Y285F-R | GCC | nope | >1bp-ins | G/C-1 | none | G/C-1 | GCTATGCCTT | CCTTTTT | CGG | GCC | C | none | nope | nope_GCC | G/C-1_CCTTTTT | 8 | FALSE | FALSE |
| 164_v.txt | 32055 | 32055-SNV-T | SNV | 1 | T | G | 27 | 24696 | 0.10932945 | 60.0740741 | rnr1Y285F | FALSE | CAN | JSY3866 | 8 | rnr1Y285F-R | GTG | T>G | G/C+1 | A/T-1 | TA>GC | TA>GC | CTGATGTGC | TGCGAGA | CAC | GTG | T | C | T>G | T>G_GTG | TA>GC_TGCGAGA | 8 | FALSE | FALSE |
| 164_v.txt | 32284 | 32284-SNV-A | SNV | 1 | A | C | 10 | 22423 | 0.04459707 | 32.3 | rnr1Y285F | FALSE | CAN | JSY3866 | 8 | rnr1Y285F-R | TAA | A>C | G/C+1 | A/T-1 | TA>GC | TA>GC | CAACGTAAA | AAATATT | ATT | ATT | T | G | T>G | T>G_ATT | TA>GC_AAATATT | 8 | FALSE | FALSE |
| 164_v.txt | 32288 | 32288-SNV-A | SNV | 1 | A | C | 58 | 22423 | 0.25866298 | 50.8275862 | rnr1Y285F | FALSE | CAN | JSY3866 | 8 | rnr1Y285F-R | TAT | A>C | G/C+1 | A/T-1 | TA>GC | TA>GC | GTAAATATT | ATTTGAA | ATA | ATA | T | G | T>G | T>G_ATA | TA>GC_ATTTGAA | 8 | FALSE | FALSE |
| 164_v.txt | 32439 | 32439-SNV-A | SNV | 1 | A | C | 11 | 22423 | 0.04905677 | 56.6363636 | rnr1Y285F | FALSE | CAN | JSY3866 | 8 | rnr1Y285F-R | AAC | A>C | G/C+1 | A/T-1 | TA>GC | TA>GC | TATGGAACT | ACTAAAA | TTG | TTG | T | G | T>G | T>G_TTG | TA>GC_ACTAAAA | 8 | FALSE | FALSE |
| 164_v.txt | 32445 | 32445-SNV-A | SNV | 1 | A | C | 12 | 22423 | 0.05351648 | 50.1666667 | rnr1Y285F | FALSE | CAN | JSY3866 | 8 | rnr1Y285F-R | AAG | A>C | G/C+1 | A/T-1 | TA>GC | TA>GC | ACTAAAAGT | AGTCCAA | TTC | TTC | T | G | T>G | T>G_TTC | TA>GC_AGTCCAA | 8 | FALSE | FALSE |
| 164_v.txt | 32457 | 32457-SNV-A | SNV | 1 | A | C | 9 | 22423 | 0.04013736 | 51 | rnr1Y285F | FALSE | CAN | JSY3866 | 8 | rnr1Y285F-R | TAA | A>C | G/C+1 | A/T-1 | TA>GC | TA>GC | ATGAATAAT | AATAGAG | ATT | ATT | T | G | T>G | T>G_ATT | TA>GC_AATAGAG | 8 | FALSE | FALSE |
| 164_v.txt | 32565 | 32565-SNV-T | SNV | 1 | T | G | 14 | 24595 | 0.05692214 | 33.0714286 | rnr1Y285F | FALSE | CAN | JSY3866 | 8 | rnr1Y285F-R | TTC | T>G | G/C+1 | A/T-1 | TA>GC | TA>GC | ACTAGTTCA | TCAGTAC | AAG | TTC | T | C | T>G | T>G_TTC | TA>GC_TCAGTAC | 8 | FALSE | FALSE |
| 164_v.txt | 32640 | 32640-SNV-T | SNV | 1 | T | G | 3 | 24610 | 0.01219017 | 64 | rnr1Y285F | FALSE | CAN | JSY3866 | 8 | rnr1Y285F-R | CTT | T>G | G/C+1 | A/T-1 | TA>GC | TA>GC | TTATCCTTAG | TTAGATA | GAA | CTT | T | C | T>G | T>G_CTT | TA>GC_TTAGATA | 8 | FALSE | FALSE |
| 164_v.txt | 32680 | 32680-SNV-A | SNV | 1 | A | C | 6 | 24610 | 0.02438033 | 60.1673132 | rnr1Y285F | FALSE | CAN | JSY3866 | 8 | rnr1Y285F-R | CAA | A>C | G/C+1 | A/T-1 | TA>GC | TA>GC | TTCTCCAATA | AATAACG | GTT | GTT | T | G | T>G | T>G_GTT | TA>GC_AATAACG | 8 | FALSE | FALSE |
| 164_v.txt | 32763 | 32763-SNV-A | SNV | 1 | A | C | 6 | 24610 | 0.02438033 | 50.8333333 | rnr1Y285F | FALSE | CAN | JSY3866 | 8 | rnr1Y285F-R | AAC | A>C | G/C+1 | A/T-1 | TA>GC | TA>GC | GCTAAAACT | ACTTTGA | TTG | TTG | T | G | T>G | T>G_TTG | TA>GC_ACTTTGA | 8 | FALSE | FALSE |
| 164_v.txt | 32803 | 32803-SNV-A | SNV | 1 | A | C | 6 | 24610 | 0.02438033 | 46.6666667 | rnr1Y285F | FALSE | CAN | JSY3866 | 8 | rnr1Y285F-R | TAT | A>C | G/C+1 | A/T-1 | TA>GC | TA>GC | CGTAATATT | ATTTGAC | ATA | ATA | T | G | T>G | T>G_ATA | TA>GC_ATTTGAC | 8 | FALSE | FALSE |
| 164_v.txt | 32919 | 32919-SNV-A | SNV | 1 | A | C | 194 | 19469 | 0.9964559 | 47.8298969 | rnr1Y285F | FALSE | CAN | JSY3866 | 8 | rnr1Y285F-R | CAG | A>C | G/C+1 | A/T-1 | TA>GC | TA>GC | AGTTCCAGG | AGGGCAA | GTC | GTC | T | G | T>G | T>G_GTC | TA>GC_AGGGCAA | 8 | FALSE | FALSE |
| 164_v.txt | 32938 | 32938-SNV-A | SNV | 1 | A | C | 6 | 19469 | 0.03081822 | 49.5 | rnr1Y285F | FALSE | CAN | JSY3866 | 8 | rnr1Y285F-R | CAA | A>C | G/C+1 | A/T-1 | TA>GC | TA>GC | TTGCCCAAG | AAGAAAA | GTT | GTT | T | G | T>G | T>G_GTT | TA>GC_AAGAAAA | 8 | FALSE | FALSE |
| 164_v.txt | 33204 | 33204-SNV-A | SNV | 1 | A | C | 577 | 28651 | 2.01389131 | 37.7244367 | rnr1Y285F | FALSE | CAN | JSY3866 | 8 | rnr1Y285F-R | AAG | A>C | G/C+1 | A/T-1 | TA>GC | TA>GC | TGCTTAAGC | AGCTCTC | TTC | TTC | T | G | T>G | T>G_TTC | TA>GC_AGCTCTC | 8 | FALSE | FALSE |
| 164_v.txt | 32055 | 32055-SNV-T | SNV | 1 | T | C | 21 | 24696 | 0.08503401 | 62.5238095 | rnr1Y285F | FALSE | CAN | JSY3866 | 8 | rnr1Y285F-R | GTG | T>C | G/C+1 | A/T-1 | TA>CG | TA>CG | CTGATGTGC | TGCGAGA | CAC | GTG | T | G | T>C | T>C_GTG | TA>CG_TGCGAGA | 8 | FALSE | FALSE |
| 164_v.txt | 32581 | 32581-SNV-T | SNV | 1 | T | C | 20 | 24611 | 0.08126448 | 33.9 | rnr1Y285F | FALSE | CAN | JSY3866 | 8 | rnr1Y285F-R | GTG | T>C | G/C+1 | A/T-1 | TA>CG | TA>CG | GAAATGTGA | TGAAGGC | CAC | GTG | T | G | T>C | T>C_GTG | TA>CG_TGAAGGC | 8 | FALSE | FALSE |
| 164_v.txt | 32680 | 32680-SNV-A | SNV | 1 | A | G | 8 | 24610 | 0.03250711 | 56.7503442 | rnr1Y285F | FALSE | CAN | JSY3866 | 8 | rnr1Y285F-R | CAA | A>G | G/C+1 | A/T-1 | TA>CG | TA>CG | TTCTCCAATA | AATAACG | GTT | GTT | T | C | T>C | T>C_GTT | TA>CG_AATAACG | 8 | FALSE | FALSE |
| 164_v.txt | 32767 | 32767-SNV-T | SNV | 1 | T | C | 20 | 24610 | 0.08126778 | 54.55 | rnr1Y285F | FALSE | CAN | JSY3866 | 8 | rnr1Y285F-R | TTG | T>C | G/C+1 | A/T-1 | TA>CG | TA>CG | AAACTTTGA | TGATGGA | AAC | TTG | T | G | T>C | T>C_TTG | TA>CG_TGATGGA | 8 | FALSE | FALSE |
| 164_v.txt | 32938 | 32938-SNV-A | SNV | 1 | A | G | 5 | 19469 | 0.02568185 | 51 | rnr1Y285F | FALSE | CAN | JSY3866 | 8 | rnr1Y285F-R | CAA | A>G | G/C+1 | A/T-1 | TA>CG | TA>CG | TTGCCCAAG | AAGAAAA | GTT | GTT | T | C | T>C | T>C_GTT | TA>CG_AAGAAAA | 8 | FALSE | FALSE |
| 164_v.txt | 33192 | 33192-SNV-T | SNV | 1 | T | C | 58 | 28651 | 0.20243622 | 34.1551724 | rnr1Y285F | FALSE | CAN | JSY3866 | 8 | rnr1Y285F-R | ATG | T>C | G/C+1 | A/T-1 | TA>CG | TA>CG | CCAATATGT | TGTCTTT | TAC | ATG | T | G | T>C | T>C_ATG | TA>CG_TGTCTTT | 8 | FALSE | FALSE |
| 164_v.txt | 33326 | 33326-SNV-T | SNV | 1 | T | C | 9 | 28649 | 0.03141471 | 49.3332456 | rnr1Y285F | FALSE | CAN | JSY3866 | 8 | rnr1Y285F-R | CTT | T>C | G/C+1 | A/T-1 | TA>CG | TA>CG | CAATTCTTTA | TTTACTT | GAA | CTT | T | G | T>C | T>C_CTT | TA>CG_TTTACTT | 8 | FALSE | FALSE |
| 164_v.txt | 32073 | 32073-SNV-A | SNV | 1 | A | T | 40 | 24696 | 0.16196955 | 60.225 | rnr1Y285F | FALSE | CAN | JSY3866 | 8 | rnr1Y285F-R | TAA | A>T | A/T+1 | A/T-1 | TA>AT | TA>AT | ATAAATAAC | AACCATG | ATT | ATT | T | A | T>A | T>A_ATT | TA>AT_AACCATG | 8 | FALSE | FALSE |
| 164_v.txt | 32241 | 32241-SNV-A | SNV | 1 | A | T | 30 | 24690 | 0.12150668 | 35.6666667 | rnr1Y285F | FALSE | CAN | JSY3866 | 8 | rnr1Y285F-R | CAA | A>T | A/T+1 | A/T-1 | TA>AT | TA>AT | GGAGCCAAC | AACTTGT | GTT | GTT | T | A | T>A | T>A_GTT | TA>AT_AACTTGT | 8 | FALSE | FALSE |
| 164_v.txt | 32315 | 32315-SNV-T | SNV | 1 | T | A | 5 | 22423 | 0.02229853 | 53.3944047 | rnr1Y285F | FALSE | CAN | JSY3866 | 8 | rnr1Y285F-R | TTG | T>A | A/T+1 | A/T-1 | TA>AT | TA>AT | AATGGTTGT | TGTTAAG | AAC | TTG | T | T | T>A | T>A_TTG | TA>AT_TGTTAAG | 8 | FALSE | FALSE |
| 164_v.txt | 32337 | 32337-SNV-A | SNV | 1 | A | T | 2 | 22423 | 8.92E-03 | 47.4999048 | rnr1Y285F | FALSE | CAN | JSY3866 | 8 | rnr1Y285F-R | GAT | A>T | A/T+1 | A/T-1 | TA>AT | TA>AT | TTGAAGATA | ATATGTG | CTA | CTA | T | A | T>A | T>A_CTA | TA>AT_ATATGTG | 8 | FALSE | FALSE |
| 164_v.txt | 32527 | 32527-SNV-T | SNV | 1 | T | A | 42 | 22423 | 0.18730768 | 42.4047619 | rnr1Y285F | FALSE | CAN | JSY3866 | 8 | rnr1Y285F-R | CTG | T>A | A/T+1 | A/T-1 | TA>AT | TA>AT | ATTTTCTGG | TGGGGTT | GAC | CTG | T | T | T>A | T>A_CTG | TA>AT_TGGGGTT | 8 | FALSE | FALSE |
| 164_v.txt | 32540 | 32540-SNV-T | SNV | 1 | T | A | 19 | 22423 | 0.08473443 | 37.0526316 | rnr1Y285F | FALSE | CAN | JSY3866 | 8 | rnr1Y285F-R | CTT | T>A | A/T+1 | A/T-1 | TA>AT | TA>AT | TGCAGCTTC | TTCACCA | GAA | CTT | T | T | T>A | T>A_CTT | TA>AT_TTCACCA | 8 | FALSE | FALSE |
| 164_v.txt | 33326 | 33326-SNV-T | SNV | 1 | T | A | 4 | 28649 | 0.01396209 | 33.4994988 | rnr1Y285F | FALSE | CAN | JSY3866 | 8 | rnr1Y285F-R | CTT | T>A | A/T+1 | A/T-1 | TA>AT | TA>AT | CAATTCTTTA | TTTACTT | GAA | CTT | T | T | T>A | T>A_CTT | TA>AT_TTTACTT | 8 | FALSE | FALSE |
| 164_v.txt | 31844 | 31844-SNV-C | SNV | 1 | C | T | 9 | 3961 | 0.22721535 | 54.2222222 | rnr1Y285F | FALSE | CAN | JSY3866 | 8 | rnr1Y285F-R | TCC | C>T | A/T+1 | G/C-1 | CG>TA | CG>TA | TAAGATCCA | CCAAACA | AGG | TCC | C | A | C>T | C>T_TCC | CG>TA_CCAAACA | 8 | FALSE | FALSE |
| 164_v.txt | 31960 | 31960-SNV-C | SNV | 1 | C | T | 29 | 24696 | 0.11742792 | 34.8275862 | rnr1Y285F | FALSE | CAN | JSY3866 | 8 | rnr1Y285F-R | GCC | C>T | A/T+1 | G/C-1 | CG>TA | CG>TA | ATGTGGCCG | CCGCATA | CGG | GCC | C | A | C>T | C>T_GCC | CG>TA_CCGCATA | 8 | FALSE | FALSE |
| 164_v.txt | 31962 | 31962-SNV-C | SNV | 1 | G | A | 46 | 24695 | 0.18627253 | 36.5648628 | rnr1Y285F | FALSE | CAN | JSY3866 | 8 | rnr1Y285F-R | CGC | G>A | A/T+1 | G/C-1 | CG>TA | CG>TA | GTGGCCGCA | GCATAAT | GCG | GCG | C | T | C>T | C>T_GCG | CG>TA_GCATAAT | 8 | FALSE | FALSE |
| 164_v.txt | 31963 | 31963-SNV-C | SNV | 1 | C | T | 2 | 24695 | 8.10E-03 | 23.9993716 | rnr1Y285F | FALSE | CAN | JSY3866 | 8 | rnr1Y285F-R | GCA | C>T | A/T+1 | G/C-1 | CG>TA | CG>TA | TGGCCGCAT | CATAATA | CGT | GCA | C | A | C>T | C>T_GCA | CG>TA_CATAATA | 8 | FALSE | FALSE |
| 164_v.txt | 31972 | 31972-SNV-C | SNV | 1 | C | T | 26 | 24695 | 0.10528447 | 37.1923077 | rnr1Y285F | FALSE | CAN | JSY3866 | 8 | rnr1Y285F-R | GCC | C>T | A/T+1 | G/C-1 | CG>TA | CG>TA | AATAAGCCA | CCAAGCC | CGG | GCC | C | A | C>T | C>T_GCC | CG>TA_CCAAGCC | 8 | FALSE | FALSE |
| 164_v.txt | 31981 | 31981-SNV-C | SNV | 1 | G | A | 469 | 24695 | 1.89916987 | 53.4669495 | rnr1Y285F | FALSE | CAN | JSY3866 | 8 | rnr1Y285F-R | GGC | G>A | A/T+1 | G/C-1 | CG>TA | CG>TA | AGCCGGGCA | GCATTAA | CCG | CCG | C | T | C>T | C>T_CCG | CG>TA_GCATTAA | 8 | FALSE | FALSE |
| 164_v.txt | 32057 | 32057-SNV-C | SNV | 1 | C | T | 30 | 24696 | 0.12147716 | 54.1666667 | rnr1Y285F | FALSE | CAN | JSY3866 | 8 | rnr1Y285F-R | GCG | C>T | A/T+1 | G/C-1 | CG>TA | CG>TA | GATGTGCGA | CGAGATT | CGC | GCG | C | A | C>T | C>T_GCG | CG>TA_CGAGATT | 8 | FALSE | FALSE |
| 164_v.txt | 32075 | 32075-SNV-C | SNV | 1 | C | T | 267 | 24696 | 1.08114674 | 61.4868914 | rnr1Y285F | FALSE | CAN | JSY3866 | 8 | rnr1Y285F-R | ACC | C>T | A/T+1 | G/C-1 | CG>TA | CG>TA | AAATAACCA | CCATGCA | TGG | ACC | C | A | C>T | C>T_ACC | CG>TA_CCATGCA | 8 | FALSE | FALSE |
| 164_v.txt | 32088 | 32088-SNV-C | SNV | 1 | C | T | 170 | 24695 | 0.68839846 | 61.2058824 | rnr1Y285F | FALSE | CAN | JSY3866 | 8 | rnr1Y285F-R | GCC | C>T | A/T+1 | G/C-1 | CG>TA | CG>TA | AAAAAGCCT | CCTGCAA | CGG | GCC | C | A | C>T | C>T_GCC | CG>TA_CCTGCAA | 8 | FALSE | FALSE |
| 164_v.txt | 32089 | 32089-SNV-C | SNV | 1 | C | T | 249 | 24694 | 1.00834211 | 61.2207997 | rnr1Y285F | FALSE | CAN | JSY3866 | 8 | rnr1Y285F-R | CCT | C>T | A/T+1 | G/C-1 | CG>TA | CG>TA | AAAAGCCTG | CTGCAAC | GGA | CCT | C | A | C>T | C>T_CCT | CG>TA_CTGCAAC | 8 | FALSE | FALSE |
| 164_v.txt | 32092 | 32092-SNV-C | SNV | 1 | C | T | 36 | 24696 | 0.1457726 | 61.6944444 | rnr1Y285F | FALSE | CAN | JSY3866 | 8 | rnr1Y285F-R | GCA | C>T | A/T+1 | G/C-1 | CG>TA | CG>TA | AGCCTGCAA | CAACACC | CGT | GCA | C | A | C>T | C>T_GCA | CG>TA_CAACACC | 8 | FALSE | FALSE |
| 164_v.txt | 32114 | 32114-SNV-C | SNV | 1 | C | T | 330 | 24696 | 1.33624879 | 59.5545455 | rnr1Y285F | FALSE | CAN | JSY3866 | 8 | rnr1Y285F-R | GCC | C>T | A/T+1 | G/C-1 | CG>TA | CG>TA | TAATAGCCA | CCATTCG | CGG | GCC | C | A | C>T | C>T_GCC | CG>TA_CCATTCG | 8 | FALSE | FALSE |
| 164_v.txt | 32115 | 32115-SNV-C | SNV | 1 | C | T | 48 | 24696 | 0.19436346 | 54.9159017 | rnr1Y285F | FALSE | CAN | JSY3866 | 8 | rnr1Y285F-R | CCA | C>T | A/T+1 | G/C-1 | CG>TA | CG>TA | AATAGCCATT | CATTCGA | GGT | CCA | C | A | C>T | C>T_CCA | CG>TA_CATTCGA | 8 | FALSE | FALSE |
| 164_v.txt | 32166 | 32166-SNV-C | SNV | 1 | C | T | 22 | 24696 | 0.08908325 | 49.9970381 | rnr1Y285F | FALSE | CAN | JSY3866 | 8 | rnr1Y285F-R | GCC | C>T | A/T+1 | G/C-1 | CG>TA | CG>TA | AAAGCGCCA | CCAAATG | CGG | GCC | C | A | C>T | C>T_GCC | CG>TA_CCAAATG | 8 | FALSE | FALSE |
| 164_v.txt | 32259 | 32259-SNV-C | SNV | 1 | C | T | 75 | 22423 | 0.33447799 | 33.1333333 | rnr1Y285F | FALSE | CAN | JSY3866 | 8 | rnr1Y285F-R | ACC | C>T | A/T+1 | G/C-1 | CG>TA | CG>TA | GATAGACCA | CCAAATA | TGG | ACC | C | A | C>T | C>T_ACC | CG>TA_CCAAATA | 8 | FALSE | FALSE |
| 164_v.txt | 32271 | 32271-SNV-C | SNV | 1 | C | T | 81 | 22424 | 0.36122012 | 33.654321 | rnr1Y285F | FALSE | CAN | JSY3866 | 8 | rnr1Y285F-R | ACG | C>T | A/T+1 | G/C-1 | CG>TA | CG>TA | AAAATACGG | CGGGAAC | TGC | ACG | C | A | C>T | C>T_ACG | CG>TA_CGGGAAC | 8 | FALSE | FALSE |
| 164_v.txt | 32272 | 32272-SNV-C | SNV | 1 | G | A | 24 | 22424 | 0.10702818 | 37.25 | rnr1Y285F | FALSE | CAN | JSY3866 | 8 | rnr1Y285F-R | CGG | G>A | A/T+1 | G/C-1 | CG>TA | CG>TA | AAATACGGG | GGGAACC | GCC | GCC | C | T | C>T | C>T_GCC | CG>TA_GGGAACC | 8 | FALSE | FALSE |
| 164_v.txt | 32274 | 32274-SNV-C | SNV | 1 | G | A | 39 | 22423 | 0.17392856 | 36.4358974 | rnr1Y285F | FALSE | CAN | JSY3866 | 8 | rnr1Y285F-R | GGA | G>A | A/T+1 | G/C-1 | CG>TA | CG>TA | ATACGGGAA | GAACCAA | CCT | CCT | C | T | C>T | C>T_CCT | CG>TA_GAACCAA | 8 | FALSE | FALSE |
| 164_v.txt | 32292 | 32292-SNV-C | SNV | 1 | G | A | 24 | 22423 | 0.10703296 | 57.25 | rnr1Y285F | FALSE | CAN | JSY3866 | 8 | rnr1Y285F-R | TGA | G>A | A/T+1 | G/C-1 | CG>TA | CG>TA | ATATTTGAA | GAATTTG | ACT | ACT | C | T | C>T | C>T_ACT | CG>TA_GAATTTG | 8 | FALSE | FALSE |
| 164_v.txt | 32299 | 32299-SNV-C | SNV | 1 | C | T | 38 | 22423 | 0.16946885 | 53.9210526 | rnr1Y285F | FALSE | CAN | JSY3866 | 8 | rnr1Y285F-R | GCG | C>T | A/T+1 | G/C-1 | CG>TA | CG>TA | AATTTGCGG | CGGCAGA | CGC | GCG | C | A | C>T | C>T_GCG | CG>TA_CGGCAGA | 8 | FALSE | FALSE |
| 164_v.txt | 32301 | 32301-SNV-C | SNV | 1 | G | A | 21 | 22423 | 0.09365384 | 53.6190476 | rnr1Y285F | FALSE | CAN | JSY3866 | 8 | rnr1Y285F-R | GGC | G>A | A/T+1 | G/C-1 | CG>TA | CG>TA | TTTGCGGCA | GCAGAAA | CCG | CCG | C | T | C>T | C>T_CCG | CG>TA_GCAGAAA | 8 | FALSE | FALSE |
| 164_v.txt | 32449 | 32449-SNV-C | SNV | 1 | C | T | 5608 | 22423 | 25.0100343 | 53.2960057 | rnr1Y285F | FALSE | CAN | JSY3866 | 8 | rnr1Y285F-R | CCA | C>T | A/T+1 | G/C-1 | CG>TA | CG>TA | AAAGTCCAA | CAATGAA | GGT | CCA | C | A | C>T | C>T_CCA | CG>TA_CAATGAA | 8 | FALSE | FALSE |
| 164_v.txt | 32466 | 32466-SNV-C | SNV | 1 | C | T | 21 | 22423 | 0.09365384 | 52.5238095 | rnr1Y285F | FALSE | CAN | JSY3866 | 8 | rnr1Y285F-R | GCC | C>T | A/T+1 | G/C-1 | CG>TA | CG>TA | AGAGAGCCA | CCAATGT | CGG | GCC | C | A | C>T | C>T_GCC | CG>TA_CCAATGT | 8 | FALSE | FALSE |
| 164_v.txt | 32487 | 32487-SNV-C | SNV | 1 | C | T | 29 | 22423 | 0.12933149 | 44.3450591 | rnr1Y285F | FALSE | CAN | JSY3866 | 8 | rnr1Y285F-R | ACG | C>T | A/T+1 | G/C-1 | CG>TA | CG>TA | AAGATACGG | CGGAAAA | TGC | ACG | C | A | C>T | C>T_ACG | CG>TA_CGGAAAA | 8 | FALSE | FALSE |
| 164_v.txt | 32488 | 32488-SNV-C | SNV | 1 | G | A | 8 | 22423 | 0.03567765 | 47.125 | rnr1Y285F | FALSE | CAN | JSY3866 | 8 | rnr1Y285F-R | CGG | G>A | A/T+1 | G/C-1 | CG>TA | CG>TA | AGATACGGA | GGAAAAC | GCC | GCC | C | T | C>T | C>T_GCC | CG>TA_GGAAAAC | 8 | FALSE | FALSE |
| 164_v.txt | 32535 | 32535-SNV-C | SNV | 1 | G | A | 32 | 22423 | 0.14271061 | 37.2398927 | rnr1Y285F | FALSE | CAN | JSY3866 | 8 | rnr1Y285F-R | TGC | G>A | A/T+1 | G/C-1 | CG>TA | CG>TA | GGGTTTGCA | GCAGCTT | ACG | ACG | C | T | C>T | C>T_ACG | CG>TA_GCAGCTT | 8 | FALSE | FALSE |
| 164_v.txt | 32536 | 32536-SNV-C | SNV | 1 | C | T | 43 | 22423 | 0.19176738 | 36.3482683 | rnr1Y285F | FALSE | CAN | JSY3866 | 8 | rnr1Y285F-R | GCA | C>T | A/T+1 | G/C-1 | CG>TA | CG>TA | GGTTTGCAG | CAGCTTC | CGT | GCA | C | A | C>T | C>T_GCA | CG>TA_CAGCTTC | 8 | FALSE | FALSE |
| 164_v.txt | 32542 | 32542-SNV-C | SNV | 1 | C | T | 27 | 22423 | 0.12041208 | 37.3333333 | rnr1Y285F | FALSE | CAN | JSY3866 | 8 | rnr1Y285F-R | TCA | C>T | A/T+1 | G/C-1 | CG>TA | CG>TA | CAGCTTCACC | CACCAGC | AGT | TCA | C | A | C>T | C>T_TCA | CG>TA_CACCAGC | 8 | FALSE | FALSE |
| 164_v.txt | 32590 | 32590-SNV-C | SNV | 1 | C | T | 46 | 24610 | 0.18691589 | 35.7173913 | rnr1Y285F | FALSE | CAN | JSY3866 | 8 | rnr1Y285F-R | GCG | C>T | A/T+1 | G/C-1 | CG>TA | CG>TA | AGGCAGCGT | CGTTAAT | CGC | GCG | C | A | C>T | C>T_GCG | CG>TA_CGTTAAT | 8 | FALSE | FALSE |
| 164_v.txt | 32609 | 32609-SNV-C | SNV | 1 | C | T | 37 | 24610 | 0.15034539 | 44.7027027 | rnr1Y285F | FALSE | CAN | JSY3866 | 8 | rnr1Y285F-R | CCC | C>T | A/T+1 | G/C-1 | CG>TA | CG>TA | GGAAACCCA | CCAACCT | GGG | CCC | C | A | C>T | C>T_CCC | CG>TA_CCAACCT | 8 | FALSE | FALSE |
| 164_v.txt | 32660 | 32660-SNV-C | SNV | 1 | C | T | 58 | 24610 | 0.23567655 | 57.9137931 | rnr1Y285F | FALSE | CAN | JSY3866 | 8 | rnr1Y285F-R | CCC | C>T | A/T+1 | G/C-1 | CG>TA | CG>TA | TGGACCCCA | CCAGGCA | GGG | CCC | C | A | C>T | C>T_CCC | CG>TA_CCAGGCA | 8 | FALSE | FALSE |
| 164_v.txt | 32679 | 32679-SNV-C | SNV | 1 | C | T | 68 | 24610 | 0.27631044 | 60.9989176 | rnr1Y285F | FALSE | CAN | JSY3866 | 8 | rnr1Y285F-R | CCA | C>T | A/T+1 | G/C-1 | CG>TA | CG>TA | TTTCTCCAAT | CAATAAC | GGT | CCA | C | A | C>T | C>T_CCA | CG>TA_CAATAAC | 8 | FALSE | FALSE |
| 164_v.txt | 32700 | 32700-SNV-C | SNV | 1 | C | T | 25 | 24610 | 0.10158472 | 58.56 | rnr1Y285F | FALSE | CAN | JSY3866 | 8 | rnr1Y285F-R | GCC | C>T | A/T+1 | G/C-1 | CG>TA | CG>TA | ACTGGGCCG | CCGGTAA | CGG | GCC | C | A | C>T | C>T_GCC | CG>TA_CCGGTAA | 8 | FALSE | FALSE |
| 164_v.txt | 32796 | 32796-SNV-C | SNV | 1 | C | T | 32 | 24610 | 0.13002844 | 43.53125 | rnr1Y285F | FALSE | CAN | JSY3866 | 8 | rnr1Y285F-R | ACC | C>T | A/T+1 | G/C-1 | CG>TA | CG>TA | AATTCACCG | CCGTAAT | TGG | ACC | C | A | C>T | C>T_ACC | CG>TA_CCGTAAT | 8 | FALSE | FALSE |
| 164_v.txt | 32798 | 32798-SNV-C | SNV | 1 | G | A | 35 | 24610 | 0.14221861 | 41 | rnr1Y285F | FALSE | CAN | JSY3866 | 8 | rnr1Y285F-R | CGT | G>A | A/T+1 | G/C-1 | CG>TA | CG>TA | TTCACCGTA | GTAATAT | GCA | GCA | C | T | C>T | C>T_GCA | CG>TA_GTAATAT | 8 | FALSE | FALSE |
| 164_v.txt | 32840 | 32840-SNV-C | SNV | 1 | C | T | 33 | 24610 | 0.13409183 | 39.1818182 | rnr1Y285F | FALSE | CAN | JSY3866 | 8 | rnr1Y285F-R | CCC | C>T | A/T+1 | G/C-1 | CG>TA | CG>TA | AATTACCCA | CCAAAAA | GGG | CCC | C | A | C>T | C>T_CCC | CG>TA_CCAAAAA | 8 | FALSE | FALSE |
| 164_v.txt | 32855 | 32855-SNV-C | SNV | 1 | C | T | 440 | 24609 | 1.78796375 | 36.3681818 | rnr1Y285F | FALSE | CAN | JSY3866 | 8 | rnr1Y285F-R | TCC | C>T | A/T+1 | G/C-1 | CG>TA | CG>TA | ACTAATCCAT | CCATGCC | AGG | TCC | C | A | C>T | C>T_TCC | CG>TA_CCATGCC | 8 | FALSE | FALSE |
| 164_v.txt | 32860 | 32860-SNV-C | SNV | 1 | C | T | 22 | 24607 | 0.08940545 | 36.6363636 | rnr1Y285F | FALSE | CAN | JSY3866 | 8 | rnr1Y285F-R | GCC | C>T | A/T+1 | G/C-1 | CG>TA | CG>TA | TCCATGCCG | CCGCCAG | CGG | GCC | C | A | C>T | C>T_GCC | CG>TA_CCGCCAG | 8 | FALSE | FALSE |
| 164_v.txt | 32900 | 32900-SNV-C | SNV | 1 | G | A | 18 | 19468 | 0.09245942 | 35.4408036 | rnr1Y285F | FALSE | CAN | JSY3866 | 8 | rnr1Y285F-R | GGC | G>A | A/T+1 | G/C-1 | CG>TA | CG>TA | GACTTGGCC | GCCAACT | CCG | CCG | C | T | C>T | C>T_CCG | CG>TA_GCCAACT | 8 | FALSE | FALSE |
| 164_v.txt | 32901 | 32901-SNV-C | SNV | 1 | C | T | 42 | 19468 | 0.21573865 | 33.4290289 | rnr1Y285F | FALSE | CAN | JSY3866 | 8 | rnr1Y285F-R | GCC | C>T | A/T+1 | G/C-1 | CG>TA | CG>TA | ACTTGGCCA | CCAACTA | CGG | GCC | C | A | C>T | C>T_GCC | CG>TA_CCAACTA | 8 | FALSE | FALSE |
| 164_v.txt | 32937 | 32937-SNV-C | SNV | 1 | C | T | 24 | 19469 | 0.1232729 | 37.1250608 | rnr1Y285F | FALSE | CAN | JSY3866 | 8 | rnr1Y285F-R | CCA | C>T | A/T+1 | G/C-1 | CG>TA | CG>TA | ATTGCCCAA | CAAGAAA | GGT | CCA | C | A | C>T | C>T_CCA | CG>TA_CAAGAAA | 8 | FALSE | FALSE |
| 164_v.txt | 32940 | 32940-SNV-C | SNV | 1 | G | A | 21 | 19469 | 0.10786378 | 53.3809524 | rnr1Y285F | FALSE | CAN | JSY3866 | 8 | rnr1Y285F-R | AGA | G>A | A/T+1 | G/C-1 | CG>TA | CG>TA | GCCCAAGAA | GAAAACC | TCT | TCT | C | T | C>T | C>T_TCT | CG>TA_GAAAACC | 8 | FALSE | FALSE |

| | | | | | | | | | | | | | | | | | | | | | | | | | | | | | | | | | | |
|---|---|---|---|---|---|---|---|---|---|---|---|---|---|---|---|---|---|---|---|---|---|---|---|---|---|---|---|---|---|---|---|---|---|---|
| 164_v.txt | 32945 | 32945-SNV-( | SNV | 1 | C | T | 108 | 19469 | 0.55472803 | 58.1481482 | rnr1Y285F | FALSE | CAN | JSY3866 | 8 | rnr1Y285F-R | ACC | C>T | A/T+1 | G/C-1 | CG>TA | CG>TA | AGAAAACCA | CCAATAC | TGG | ACC | C | A | C>T | C>T_ACC | CG>TA_CCAATAC | 8 | FALSE | FALSE |
| 164_v.txt | 32958 | 32958-SNV-( | SNV | 1 | C | T | 67 | 19469 | 0.34413683 | 57.8656716 | rnr1Y285F | FALSE | CAN | JSY3866 | 8 | rnr1Y285F-R | ACC | C>T | A/T+1 | G/C-1 | CG>TA | CG>TA | ATGTAACCA | CCATTGG | TGG | ACC | C | A | C>T | C>T_ACC | CG>TA_CCATTGG | 8 | FALSE | FALSE |
| 164_v.txt | 32965 | 32965-SNV-( | SNV | 1 | C | T | 27 | 19469 | 0.13868201 | 58.6666667 | rnr1Y285F | FALSE | CAN | JSY3866 | 8 | rnr1Y285F-R | GCC | C>T | A/T+1 | G/C-1 | CG>TA | CG>TA | CATTGGCCG | CCGCACC | CGG | GCC | C | A | C>T | C>T_GCC | CG>TA_CCGCACC | 8 | FALSE | FALSE |
| 164_v.txt | 32967 | 32967-SNV-( | SNV | 1 | G | A | 49 | 19469 | 0.25168216 | 60.2244898 | rnr1Y285F | FALSE | CAN | JSY3866 | 8 | rnr1Y285F-R | CGC | G>A | A/T+1 | G/C-1 | CG>TA | CG>TA | TTGGCCGCA | GCACCAA | GCG | GCG | C | T | C>T | C>T_GCG | CG>TA_GCACCAA | 8 | FALSE | FALSE |
| 164_v.txt | 33025 | 33025-SNV-( | SNV | 1 | G | A | 20 | 19469 | 0.10272741 | 62.5 | rnr1Y285F | FALSE | CAN | JSY3866 | 8 | rnr1Y285F-R | GGG | G>A | A/T+1 | G/C-1 | CG>TA | CG>TA | TAACAGGGA | GGATGAA | CCC | CCC | C | T | C>T | C>T_CCC | CG>TA_GGATGAA | 8 | FALSE | FALSE |
| 164_v.txt | 33043 | 33043-SNV-( | SNV | 1 | C | T | 21 | 19469 | 0.10786378 | 56.2857143 | rnr1Y285F | FALSE | CAN | JSY3866 | 8 | rnr1Y285F-R | TCA | C>T | A/T+1 | G/C-1 | CG>TA | CG>TA | CCATTTCACC | CACCCAA | AGT | TCA | C | A | C>T | C>T_TCA | CG>TA_CACCCAA | 8 | FALSE | FALSE |
| 164_v.txt | 33056 | 33056-SNV-( | SNV | 1 | C | T | 29 | 19469 | 0.14895475 | 53.8965517 | rnr1Y285F | FALSE | CAN | JSY3866 | 8 | rnr1Y285F-R | GCG | C>T | A/T+1 | G/C-1 | CG>TA | CG>TA | GGACTGCGT | CGTGACA | CGC | GCG | C | A | C>T | C>T_GCG | CG>TA_CGTGACA | 8 | FALSE | FALSE |
| 164_v.txt | 33070 | 33070-SNV-( | SNV | 1 | C | T | 20 | 19469 | 0.10272741 | 60.1 | rnr1Y285F | FALSE | CAN | JSY3866 | 8 | rnr1Y285F-R | GCC | C>T | A/T+1 | G/C-1 | CG>TA | CG>TA | AATATGCCA | CCAAAGA | CGG | GCC | C | A | C>T | C>T_GCC | CG>TA_CCAAAGA | 8 | FALSE | FALSE |
| 164_v.txt | 33114 | 33114-SNV-( | SNV | 1 | C | T | 20 | 19469 | 0.10272741 | 41.5 | rnr1Y285F | FALSE | CAN | JSY3866 | 8 | rnr1Y285F-R | GCC | C>T | A/T+1 | G/C-1 | CG>TA | CG>TA | ACTGGGCCG | CCGGCGT | CGG | GCC | C | A | C>T | C>T_GCC | CG>TA_CCGGCGT | 8 | FALSE | FALSE |
| 164_v.txt | 33118 | 33118-SNV-( | SNV | 1 | C | T | 23 | 19469 | 0.11813653 | 42.6956522 | rnr1Y285F | FALSE | CAN | JSY3866 | 8 | rnr1Y285F-R | GCG | C>T | A/T+1 | G/C-1 | CG>TA | CG>TA | GGCCGGCGT | CGTTGGT | CGC | GCG | C | A | C>T | C>T_GCG | CG>TA_CGTTGGT | 8 | FALSE | FALSE |
| 164_v.txt | 33153 | 33153-SNV-( | SNV | 1 | C | T | 392 | 19469 | 2.01345729 | 37.0331633 | rnr1Y285F | FALSE | CAN | JSY3866 | 8 | rnr1Y285F-R | ACC | C>T | A/T+1 | G/C-1 | CG>TA | CG>TA | AAAAGACCT | CCTGTAC | TGG | ACC | C | A | C>T | C>T_ACC | CG>TA_CCTGTAC | 8 | FALSE | FALSE |
| 164_v.txt | 33159 | 33159-SNV-( | SNV | 1 | C | T | 75 | 19474 | 0.38512889 | 36.48 | rnr1Y285F | FALSE | CAN | JSY3866 | 8 | rnr1Y285F-R | ACC | C>T | A/T+1 | G/C-1 | CG>TA | CG>TA | CCTGTACCA | CCAATAG | TGG | ACC | C | A | C>T | C>T_ACC | CG>TA_CCAATAG | 8 | FALSE | FALSE |
| 164_v.txt | 33168 | 33168-SNV-( | SNV | 1 | C | T | 33 | 19474 | 0.16945671 | 36.0909091 | rnr1Y285F | FALSE | CAN | JSY3866 | 8 | rnr1Y285F-R | ACC | C>T | A/T+1 | G/C-1 | CG>TA | CG>TA | ATAGTACCA | CCACCAA | TGG | ACC | C | A | C>T | C>T_ACC | CG>TA_CCACCAA | 8 | FALSE | FALSE |
| 164_v.txt | 33169 | 33169-SNV-( | SNV | 1 | C | T | 307 | 19474 | 1.57646092 | 37.7785016 | rnr1Y285F | FALSE | CAN | JSY3866 | 8 | rnr1Y285F-R | CCA | C>T | A/T+1 | G/C-1 | CG>TA | CG>TA | TAGTACCAC | CACCAAG | GGT | CCA | C | A | C>T | C>T_CCA | CG>TA_CACCAAG | 8 | FALSE | FALSE |
| 164_v.txt | 33258 | 33258-SNV-( | SNV | 1 | C | T | 42 | 28651 | 0.14659174 | 43.3809524 | rnr1Y285F | FALSE | CAN | JSY3866 | 8 | rnr1Y285F-R | GCC | C>T | A/T+1 | G/C-1 | CG>TA | CG>TA | CCTATGCCAT | CCATCCT | CGG | GCC | C | A | C>T | C>T_GCC | CG>TA_CCATCCT | 8 | FALSE | FALSE |
| 164_v.txt | 33275 | 33275-SNV-( | SNV | 1 | C | T | 32 | 28651 | 0.11168895 | 49.9686065 | rnr1Y285F | FALSE | CAN | JSY3866 | 8 | rnr1Y285F-R | ACG | C>T | A/T+1 | G/C-1 | CG>TA | CG>TA | AGAGAACGT | CGTATCC | TGC | ACG | C | A | C>T | C>T_ACG | CG>TA_CGTATCC | 8 | FALSE | FALSE |
| 164_v.txt | 33276 | 33276-SNV-( | SNV | 1 | G | A | 51 | 28650 | 0.17801047 | 48.5649585 | rnr1Y285F | FALSE | CAN | JSY3866 | 8 | rnr1Y285F-R | CGT | G>A | A/T+1 | G/C-1 | CG>TA | CG>TA | GAGAACGTA | GTATCCT | GCA | GCA | C | T | C>T | C>T_GCA | CG>TA_GTATCCT | 8 | FALSE | FALSE |
| 164_v.txt | 33284 | 33284-SNV-( | SNV | 1 | G | A | 29 | 28651 | 0.10121811 | 48.2758621 | rnr1Y285F | FALSE | CAN | JSY3866 | 8 | rnr1Y285F-R | CGC | G>A | A/T+1 | G/C-1 | CG>TA | CG>TA | ATCCTCGCCA | GCCATTT | GCG | GCG | C | T | C>T | C>T_GCG | CG>TA_GCCATTT | 8 | FALSE | FALSE |
| 164_v.txt | 33309 | 33309-SNV-( | SNV | 1 | C | T | 33 | 28650 | 0.11518325 | 59.121306 | rnr1Y285F | FALSE | CAN | JSY3866 | 8 | rnr1Y285F-R | GCG | C>T | A/T+1 | G/C-1 | CG>TA | CG>TA | AAAGAGCGC | CGCAATG | CGC | GCG | C | A | C>T | C>T_GCG | CG>TA_CGCAATG | 8 | FALSE | FALSE |
| 164_v.txt | 33310 | 33310-SNV-( | SNV | 1 | G | A | 38 | 28648 | 0.13264451 | 51.8917659 | rnr1Y285F | FALSE | CAN | JSY3866 | 8 | rnr1Y285F-R | CGC | G>A | A/T+1 | G/C-1 | CG>TA | CG>TA | AAGAGCGCA | GCAATGG | GCG | GCG | C | T | C>T | C>T_GCG | CG>TA_GCAATGG | 8 | FALSE | FALSE |
| 164_v.txt | 33412 | 33412-SNV-( | SNV | 1 | G | A | 47 | 28651 | 0.16404314 | 39.7446809 | rnr1Y285F | FALSE | CAN | JSY3866 | 8 | rnr1Y285F-R | GGC | G>A | A/T+1 | G/C-1 | CG>TA | CG>TA | TGACCGGCT | GCTCATT | CCG | CCG | C | T | C>T | C>T_CCG | CG>TA_GCTCATT | 8 | FALSE | FALSE |
| 164_v.txt | 33474 | 33474-SNV-( | SNV | 1 | C | T | 31 | 28649 | 0.10820622 | 34.1290323 | rnr1Y285F | FALSE | CAN | JSY3866 | 8 | rnr1Y285F-R | GCC | C>T | A/T+1 | G/C-1 | CG>TA | CG>TA | GCTATGCCT | CCTTTTT | CGG | GCC | C | A | C>T | C>T_GCC | CG>TA_CCTTTTT | 8 | FALSE | FALSE |
| 164_v.txt | 31956 | 31956-SNV-( | SNV | 1 | G | C | 8 | 24696 | 0.03239391 | 37.875 | rnr1Y285F | FALSE | CAN | JSY3866 | 8 | rnr1Y285F-R | TGT | G>C | G/C+1 | G/C-1 | CG>GC | CG>GC | ATAAATGTG | GTGGCCG | ACA | ACA | C | G | C>G | C>G_ACA | CG>GC_GTGGCCG | 8 | FALSE | FALSE |
| 164_v.txt | 31963 | 31963-SNV-( | SNV | 1 | C | G | 9 | 24695 | 0.03644462 | 37.9994122 | rnr1Y285F | FALSE | CAN | JSY3866 | 8 | rnr1Y285F-R | GCA | C>G | G/C+1 | G/C-1 | CG>GC | CG>GC | TGGCCGCAT | CATAATA | CGT | GCA | C | C | C>G | C>G_GCA | CG>GC_CATAATA | 8 | FALSE | FALSE |
| 164_v.txt | 31981 | 31981-SNV-( | SNV | 1 | G | C | 10 | 24695 | 0.04049403 | 41.6 | rnr1Y285F | FALSE | CAN | JSY3866 | 8 | rnr1Y285F-R | GGC | G>C | G/C+1 | G/C-1 | CG>GC | CG>GC | AGCGGGCA | GCATTAA | CCG | CCG | C | G | C>G | C>G_CCG | CG>GC_GCATTAA | 8 | FALSE | FALSE |
| 164_v.txt | 32064 | 32064-SNV-( | SNV | 1 | G | C | 213 | 24696 | 0.86248785 | 62.4600939 | rnr1Y285F | FALSE | CAN | JSY3866 | 8 | rnr1Y285F-R | TGA | G>C | G/C+1 | G/C-1 | CG>GC | CG>GC | GAGATTGAG | GAGATAA | ACT | ACT | C | G | C>G | C>G_ACT | CG>GC_GAGATAA | 8 | FALSE | FALSE |
| 164_v.txt | 32092 | 32092-SNV-( | SNV | 1 | C | G | 8 | 24696 | 0.03239391 | 63.5 | rnr1Y285F | FALSE | CAN | JSY3866 | 8 | rnr1Y285F-R | GCA | C>G | G/C+1 | G/C-1 | CG>GC | CG>GC | AGCCTGCAC | CAACACC | CGT | GCA | C | C | C>G | C>G_GCA | CG>GC_CAACACC | 8 | FALSE | FALSE |
| 164_v.txt | 32115 | 32115-SNV-( | SNV | 1 | C | G | 1 | 24696 | 4.05E-03 | 63.9888147 | rnr1Y285F | FALSE | CAN | JSY3866 | 8 | rnr1Y285F-R | CCA | C>G | G/C+1 | G/C-1 | CG>GC | CG>GC | AATAGCCAT | CATTCGA | GGT | CCA | C | C | C>G | C>G_CCA | CG>GC_CATTCGA | 8 | FALSE | FALSE |
| 164_v.txt | 32167 | 32167-SNV-( | SNV | 1 | C | G | 24 | 24696 | 0.09718173 | 44.6659863 | rnr1Y285F | FALSE | CAN | JSY3866 | 8 | rnr1Y285F-R | CCA | C>G | G/C+1 | G/C-1 | CG>GC | CG>GC | AAGCGCCAA | CAAATGC | GGT | CCA | C | C | C>G | C>G_CCA | CG>GC_CAAATGC | 8 | FALSE | FALSE |
| 164_v.txt | 32304 | 32304-SNV-( | SNV | 1 | G | C | 2217 | 22423 | 9.88716942 | 57.6035183 | rnr1Y285F | FALSE | CAN | JSY3866 | 8 | rnr1Y285F-R | AGA | G>C | G/C+1 | G/C-1 | CG>GC | CG>GC | GCGGCAGAA | GAAATAA | TCT | TCT | C | G | C>G | C>G_TCT | CG>GC_GAAATAA | 8 | FALSE | FALSE |
| 164_v.txt | 32461 | 32461-SNV-( | SNV | 1 | G | C | 1 | 22423 | 4.46E-03 | 31.9998034 | rnr1Y285F | FALSE | CAN | JSY3866 | 8 | rnr1Y285F-R | AGA | G>C | G/C+1 | G/C-1 | CG>GC | CG>GC | ATAATAGAG | GAGAGCC | TCT | TCT | C | G | C>G | C>G_TCT | CG>GC_GAGAGCC | 8 | FALSE | FALSE |
| 164_v.txt | 32488 | 32488-SNV-( | SNV | 1 | G | C | 3 | 22423 | 0.01337912 | 50 | rnr1Y285F | FALSE | CAN | JSY3866 | 8 | rnr1Y285F-R | CGG | G>C | G/C+1 | G/C-1 | CG>GC | CG>GC | AGATACGGA | GGAAAAC | GCC | GCC | C | G | C>G | C>G_GCC | CG>GC_GGAAAAC | 8 | FALSE | FALSE |
| 164_v.txt | 32536 | 32536-SNV-( | SNV | 1 | C | G | 2 | 22423 | 8.92E-03 | 32.4995681 | rnr1Y285F | FALSE | CAN | JSY3866 | 8 | rnr1Y285F-R | GCA | C>G | G/C+1 | G/C-1 | CG>GC | CG>GC | GGTTTGCAG | CAGCTTC | CGT | GCA | C | C | C>G | C>G_GCA | CG>GC_CAGCTTC | 8 | FALSE | FALSE |
| 164_v.txt | 32575 | 32575-SNV-( | SNV | 1 | G | C | 7 | 24611 | 0.02844257 | 38 | rnr1Y285F | FALSE | CAN | JSY3866 | 8 | rnr1Y285F-R | TGA | G>C | G/C+1 | G/C-1 | CG>GC | CG>GC | TACCTTGAA | GAAATGT | ACT | ACT | C | G | C>G | C>G_ACT | CG>GC_GAAATGT | 8 | FALSE | FALSE |
| 164_v.txt | 32629 | 32629-SNV-( | SNV | 1 | C | G | 3 | 24610 | 0.01219017 | 55.3333333 | rnr1Y285F | FALSE | CAN | JSY3866 | 8 | rnr1Y285F-R | TCG | C>G | G/C+1 | G/C-1 | CG>GC | CG>GC | TCCCTTCGTT | CGTTTTT | AGC | TCG | C | C | C>G | C>G_TCG | CG>GC_CGTTTTT | 8 | FALSE | FALSE |
| 164_v.txt | 32679 | 32679-SNV-( | SNV | 1 | C | G | 3 | 24610 | 0.01219017 | 53.3331405 | rnr1Y285F | FALSE | CAN | JSY3866 | 8 | rnr1Y285F-R | CCA | C>G | G/C+1 | G/C-1 | CG>GC | CG>GC | TTTCTCCAAT | CAATAAC | GGT | CCA | C | C | C>G | C>G_CCA | CG>GC_CAATAAC | 8 | FALSE | FALSE |
| 164_v.txt | 32819 | 32819-SNV-( | SNV | 1 | G | C | 6 | 24610 | 0.02438033 | 42.1666667 | rnr1Y285F | FALSE | CAN | JSY3866 | 8 | rnr1Y285F-R | AGT | G>C | G/C+1 | G/C-1 | CG>GC | CG>GC | GAACAAGTT | GTTCATT | TCA | TCA | C | G | C>G | C>G_TCA | CG>GC_GTTCATT | 8 | FALSE | FALSE |
| 164_v.txt | 32876 | 32876-SNV-( | SNV | 1 | G | C | 8 | 19459 | 0.04111208 | 37.5 | rnr1Y285F | FALSE | CAN | JSY3866 | 8 | rnr1Y285F-R | TGT | G>C | G/C+1 | G/C-1 | CG>GC | CG>GC | AACTTTGTA | GTACGTC | ACA | ACA | C | G | C>G | C>G_ACA | CG>GC_GTACGTC | 8 | FALSE | FALSE |
| 164_v.txt | 32937 | 32937-SNV-( | SNV | 1 | C | G | 19 | 19469 | 0.09759104 | 61.9962479 | rnr1Y285F | FALSE | CAN | JSY3866 | 8 | rnr1Y285F-R | CCA | C>G | G/C+1 | G/C-1 | CG>GC | CG>GC | ATTGCCCAA | CAAGAAA | GGT | CCA | C | C | C>G | C>G_CCA | CG>GC_CAAGAAA | 8 | FALSE | FALSE |
| 164_v.txt | 32959 | 32959-SNV-( | SNV | 1 | C | G | 23 | 19469 | 0.11813653 | 55.7789128 | rnr1Y285F | FALSE | CAN | JSY3866 | 8 | rnr1Y285F-R | CCA | C>G | G/C+1 | G/C-1 | CG>GC | CG>GC | TGTAACCATT | CATTGGC | GGT | CCA | C | C | C>G | C>G_CCA | CG>GC_CATTGGC | 8 | FALSE | FALSE |
| 164_v.txt | 32971 | 32971-SNV-( | SNV | 1 | C | G | 109 | 19469 | 0.5598644 | 61.6003861 | rnr1Y285F | FALSE | CAN | JSY3866 | 8 | rnr1Y285F-R | CCA | C>G | G/C+1 | G/C-1 | CG>GC | CG>GC | CCGCACCAA | CAAATGC | GGT | CCA | C | C | C>G | C>G_CCA | CG>GC_CAAATGC | 8 | FALSE | FALSE |
| 164_v.txt | 32997 | 32997-SNV-( | SNV | 1 | G | C | 24 | 19469 | 0.1232729 | 59.7083333 | rnr1Y285F | FALSE | CAN | JSY3866 | 8 | rnr1Y285F-R | TGA | G>C | G/C+1 | G/C-1 | CG>GC | CG>GC | CTTTGTGAG | GAGAAAA | ACT | ACT | C | G | C>G | C>G_ACT | CG>GC_GAGAAAA | 8 | FALSE | FALSE |
| 164_v.txt | 33025 | 33025-SNV-( | SNV | 1 | G | C | 7 | 19469 | 0.0359546 | 60.1428571 | rnr1Y285F | FALSE | CAN | JSY3866 | 8 | rnr1Y285F-R | GGG | G>C | G/C+1 | G/C-1 | CG>GC | CG>GC | TAACAGGGA | GGATGAA | CCC | CCC | C | G | C>G | C>G_CCC | CG>GC_GGATGAA | 8 | FALSE | FALSE |
| 164_v.txt | 33070 | 33070-SNV-( | SNV | 1 | C | G | 4 | 19469 | 0.02054548 | 51 | rnr1Y285F | FALSE | CAN | JSY3866 | 8 | rnr1Y285F-R | GCC | C>G | G/C+1 | G/C-1 | CG>GC | CG>GC | AATATGCCA | CCAAAGA | CGG | GCC | C | C | C>G | C>G_GCC | CG>GC_CCAAAGA | 8 | FALSE | FALSE |
| 164_v.txt | 33106 | 33106-SNV-( | SNV | 1 | C | G | 11 | 19469 | 0.05650008 | 48.5454546 | rnr1Y285F | FALSE | CAN | JSY3866 | 8 | rnr1Y285F-R | CCC | C>G | G/C+1 | G/C-1 | CG>GC | CG>GC | GAGCGCCCA | CCACTGG | GGG | CCC | C | C | C>G | C>G_CCC | CG>GC_CCACTGG | 8 | FALSE | FALSE |
| 164_v.txt | 33378 | 33378-SNV-( | SNV | 1 | G | C | 6 | 28651 | 0.02094168 | 54.6666667 | rnr1Y285F | FALSE | CAN | JSY3866 | 8 | rnr1Y285F-R | TGA | G>C | G/C+1 | G/C-1 | CG>GC | CG>GC | GTTTGTGAA | GAAGCTT | ACT | ACT | C | G | C>G | C>G_ACT | CG>GC_GAAGCTT | 8 | FALSE | FALSE |
| 164_v.txt | 33419 | 33419-SNV-( | SNV | 1 | G | C | 16 | 28651 | 0.05584447 | 49 | rnr1Y285F | FALSE | CAN | JSY3866 | 8 | rnr1Y285F-R | TGT | G>C | G/C+1 | G/C-1 | CG>GC | CG>GC | CTCATTGTAC | GTACATA | ACA | ACA | C | G | C>G | C>G_ACA | CG>GC_GTACATA | 8 | FALSE | FALSE |
| 164_v.txt | 31844 | 31844-SNV-( | SNV | 1 | C | A | 5 | 3961 | 0.12623075 | 35.6 | rnr1Y285F | FALSE | CAN | JSY3866 | 8 | rnr1Y285F-R | TCC | C>A | A/T+1 | G/C-1 | CG>AT | CG>AT | TAAGATCCA | CCAAACA | AGG | TCC | C | T | C>A | C>A_TCC | CG>AT_CCAAACA | 8 | FALSE | FALSE |
| 164_v.txt | 31869 | 31869-SNV-( | SNV | 1 | G | T | 7 | 3961 | 0.17672305 | 54.1428571 | rnr1Y285F | FALSE | CAN | JSY3866 | 8 | rnr1Y285F-R | AGA | G>T | A/T+1 | G/C-1 | CG>AT | CG>AT | AAAATAGAG | GAGATAT | TCT | TCT | C | A | C>A | C>A_TCT | CG>AT_GAGATAT | 8 | FALSE | FALSE |
| 164_v.txt | 31980 | 31980-SNV-( | SNV | 1 | G | T | 22 | 24695 | 0.08908686 | 47.0727656 | rnr1Y285F | FALSE | CAN | JSY3866 | 8 | rnr1Y285F-R | GGG | G>T | A/T+1 | G/C-1 | CG>AT | CG>AT | AAGCCGGGC | GGCATTA | CCC | CCC | C | A | C>A | C>A_CCC | CG>AT_GGCATTA | 8 | FALSE | FALSE |
| 164_v.txt | 31981 | 31981-SNV-( | SNV | 1 | G | T | 29 | 24695 | 0.11743268 | 26.0689656 | rnr1Y285F | FALSE | CAN | JSY3866 | 8 | rnr1Y285F-R | GGC | G>T | A/T+1 | G/C-1 | CG>AT | CG>AT | AGCCGGGCA | GCATTAA | CCG | CCG | C | A | C>A | C>A_CCG | CG>AT_GCATTAA | 8 | FALSE | FALSE |
| 164_v.txt | 32027 | 32027-SNV-( | SNV | 1 | G | T | 61 | 24696 | 0.24700356 | 59.7540984 | rnr1Y285F | FALSE | CAN | JSY3866 | 8 | rnr1Y285F-R | GGT | G>T | A/T+1 | G/C-1 | CG>AT | CG>AT | GCCACGGTA | GTATTTC | CCA | CCA | C | A | C>A | C>A_CCA | CG>AT_GTATTTC | 8 | FALSE | FALSE |
| 164_v.txt | 32088 | 32088-SNV-( | SNV | 1 | C | A | 2 | 24695 | 8.10E-03 | 52 | rnr1Y285F | FALSE | CAN | JSY3866 | 8 | rnr1Y285F-R | GCC | C>A | A/T+1 | G/C-1 | CG>AT | CG>AT | AAAAAGCCT | CCTGCAA | CGG | GCC | C | T | C>A | C>A_GCC | CG>AT_CCTGCAA | 8 | FALSE | FALSE |
| 164_v.txt | 32095 | 32095-SNV-( | SNV | 1 | C | A | 40 | 24696 | 0.16196955 | 59.85 | rnr1Y285F | FALSE | CAN | JSY3866 | 8 | rnr1Y285F-R | ACA | C>A | A/T+1 | G/C-1 | CG>AT | CG>AT | CTGCAACACC | CACCAGT | TGT | ACA | C | T | C>A | C>A_ACA | CG>AT_CACCAGT | 8 | FALSE | FALSE |
| 164_v.txt | 32114 | 32114-SNV-( | SNV | 1 | C | A | 5 | 24696 | 0.02024619 | 43.2 | rnr1Y285F | FALSE | CAN | JSY3866 | 8 | rnr1Y285F-R | GCC | C>A | A/T+1 | G/C-1 | CG>AT | CG>AT | TAATACGCCA | CCATTCG | CGG | GCC | C | T | C>A | C>A_GCC | CG>AT_CCATTCG | 8 | FALSE | FALSE |
| 164_v.txt | 32167 | 32167-SNV-( | SNV | 1 | C | A | 5 | 24696 | 0.02024619 | 31.8006701 | rnr1Y285F | FALSE | CAN | JSY3866 | 8 | rnr1Y285F-R | CCA | C>A | A/T+1 | G/C-1 | CG>AT | CG>AT | AAGCGCCAA | CAAATGC | GGT | CCA | C | T | C>A | C>A_CCA | CG>AT_CAAATGC | 8 | FALSE | FALSE |
| 164_v.txt | 32188 | 32188-SNV-( | SNV | 1 | C | A | 35 | 24696 | 0.14172336 | 46.7428571 | rnr1Y285F | FALSE | CAN | JSY3866 | 8 | rnr1Y285F-R | ACT | C>A | A/T+1 | G/C-1 | CG>AT | CG>AT | CGAAAACTG | CTGCAAT | TGA | ACT | C | T | C>A | C>A_ACT | CG>AT_CTGCAAT | 8 | FALSE | FALSE |
| 164_v.txt | 32272 | 32272-SNV-( | SNV | 1 | G | T | 15 | 22424 | 0.06689262 | 34 | rnr1Y285F | FALSE | CAN | JSY3866 | 8 | rnr1Y285F-R | CGG | G>T | A/T+1 | G/C-1 | CG>AT | CG>AT | AAATACGGG | GGGAACC | GCC | GCC | C | A | C>A | C>A_GCC | CG>AT_GGGAACC | 8 | FALSE | FALSE |
| 164_v.txt | 32274 | 32274-SNV-( | SNV | 1 | G | T | 40 | 22423 | 0.17838826 | 35.125 | rnr1Y285F | FALSE | CAN | JSY3866 | 8 | rnr1Y285F-R | GGA | G>T | A/T+1 | G/C-1 | CG>AT | CG>AT | ATACGGGAA | GAACCAA | CCT | CCT | C | A | C>A | C>A_CCT | CG>AT_GAACCAA | 8 | FALSE | FALSE |
| 164_v.txt | 32368 | 32368-SNV-( | SNV | 1 | C | A | 26 | 22423 | 0.11595237 | 57.9615385 | rnr1Y285F | FALSE | CAN | JSY3866 | 8 | rnr1Y285F-R | TCA | C>A | A/T+1 | G/C-1 | CG>AT | CG>AT | AGTTCTCAAT | CAATAGC | AGT | TCA | C | T | C>A | C>A_TCA | CG>AT_CAATAGC | 8 | FALSE | FALSE |
| 164_v.txt | 32373 | 32373-SNV-( | SNV | 1 | G | T | 26 | 22423 | 0.11595237 | 61.7692308 | rnr1Y285F | FALSE | CAN | JSY3866 | 8 | rnr1Y285F-R | AGC | G>T | A/T+1 | G/C-1 | CG>AT | CG>AT | TCAATAGCA | GCAATAA | TCG | TCG | C | A | C>A | C>A_TCG | CG>AT_GCAATAA | 8 | FALSE | FALSE |
| 164_v.txt | 32424 | 32424-SNV-( | SNV | 1 | G | T | 6 | 22423 | 0.02675824 | 50.3333333 | rnr1Y285F | FALSE | CAN | JSY3866 | 8 | rnr1Y285F-R | AGG | G>T | A/T+1 | G/C-1 | CG>AT | CG>AT | AGTTTAGGG | GGGTCAT | TCC | TCC | C | A | C>A | C>A_TCC | CG>AT_GGGTCAT | 8 | FALSE | FALSE |
| 164_v.txt | 32432 | 32432-SNV-( | SNV | 1 | G | T | 141 | 22423 | 0.62881862 | 57.0851064 | rnr1Y285F | FALSE | CAN | JSY3866 | 8 | rnr1Y285F-R | TGT | G>T | A/T+1 | G/C-1 | CG>AT | CG>AT | GTCATTGTAT | GTATGGA | ACA | ACA | C | A | C>A | C>A_ACA | CG>AT_GTATGGA | 8 | FALSE | FALSE |
| 164_v.txt | 32449 | 32449-SNV-( | SNV | 1 | C | A | 216 | 22423 | 0.96329662 | 52.3657407 | rnr1Y285F | FALSE | CAN | JSY3866 | 8 | rnr1Y285F-R | CCA | C>A | A/T+1 | G/C-1 | CG>AT | CG>AT | AAAGTCCAA | CAATGAA | GGT | CCA | C | T | C>A | C>A_CCA | CG>AT_CAATGAA | 8 | FALSE | FALSE |
| 164_v.txt | 32461 | 32461-SNV-( | SNV | 1 | G | T | 1 | 22423 | 4.46E-03 | 36.9998085 | rnr1Y285F | FALSE | CAN | JSY3866 | 8 | rnr1Y285F-R | AGA | G>T | A/T+1 | G/C-1 | CG>AT | CG>AT | ATAATAGAG | GAGAGCC | TCT | TCT | C | A | C>A | C>A_TCT | CG>AT_GAGAGCC | 8 | FALSE | FALSE |
| 164_v.txt | 32463 | 32463-SNV-( | SNV | 1 | G | T | 86 | 22423 | 0.38353476 | 49.3023256 | rnr1Y285F | FALSE | CAN | JSY3866 | 8 | rnr1Y285F-R | AGA | G>T | A/T+1 | G/C-1 | CG>AT | CG>AT | AATAGAGAG | GAGCCAA | TCT | TCT | C | A | C>A | C>A_TCT | CG>AT_GAGCCAA | 8 | FALSE | FALSE |
| 164_v.txt | 32471 | 32471-SNV-( | SNV | 1 | G | T | 64 | 22423 | 0.28542122 | 48.28125 | rnr1Y285F | FALSE | CAN | JSY3866 | 8 | rnr1Y285F-R | TGT | G>T | A/T+1 | G/C-1 | CG>AT | CG>AT | GCCAATGTA | GTAGAAG | ACA | ACA | C | A | C>A | C>A_ACA | CG>AT_GTAGAAG | 8 | FALSE | FALSE |
| 164_v.txt | 32487 | 32487-SNV-( | SNV | 1 | C | A | 21 | 22423 | 0.09365384 | 43.0997061 | rnr1Y285F | FALSE | CAN | JSY3866 | 8 | rnr1Y285F-R | ACG | C>A | A/T+1 | G/C-1 | CG>AT | CG>AT | AAGATACGG | CGGAAAA | TGC | ACG | C | T | C>A | C>A_ACG | CG>AT_CGGAAAA | 8 | FALSE | FALSE |
| 164_v.txt | 32488 | 32488-SNV-( | SNV | 1 | G | T | 15 | 22423 | 0.0668956 | 44.6666667 | rnr1Y285F | FALSE | CAN | JSY3866 | 8 | rnr1Y285F-R | CGG | G>T | A/T+1 | G/C-1 | CG>AT | CG>AT | AGATACGGA | GGAAAAC | GCC | GCC | C | A | C>A | C>A_GCC | CG>AT_GGAAAAC | 8 | FALSE | FALSE |
| 164_v.txt | 32494 | 32494-SNV-( | SNV | 1 | C | A | 17 | 22423 | 0.07581501 | 45.4705882 | rnr1Y285F | FALSE | CAN | JSY3866 | 8 | rnr1Y285F-R | ACA | C>A | A/T+1 | G/C-1 | CG>AT | CG>AT | GGAAAACAA | CAACTTT | TGT | ACA | C | T | C>A | C>A_ACA | CG>AT_CAACTTT | 8 | FALSE | FALSE |
| 164_v.txt | 32529 | 32529-SNV-( | SNV | 1 | G | T | 65 | 22423 | 0.28988093 | 43.1801599 | rnr1Y285F | FALSE | CAN | JSY3866 | 8 | rnr1Y285F-R | GGG | G>T | A/T+1 | G/C-1 | CG>AT | CG>AT | TTTCTGGGG | GGGTTTG | CCC | CCC | C | A | C>A | C>A_CCC | CG>AT_GGGTTTG | 8 | FALSE | FALSE |
| 164_v.txt | 32530 | 32530-SNV-( | SNV | 1 | G | T | 14 | 22423 | 0.06243589 | 37.5700315 | rnr1Y285F | FALSE | CAN | JSY3866 | 8 | rnr1Y285F-R | GGG | G>T | A/T+1 | G/C-1 | CG>AT | CG>AT | TTCTGGGGT | GGTTTGC | CCC | CCC | C | A | C>A | C>A_CCC | CG>AT_GGTTTGC | 8 | FALSE | FALSE |
| 164_v.txt | 32535 | 32535-SNV-( | SNV | 1 | G | T | 3 | 22423 | 0.01337912 | 37.9818778 | rnr1Y285F | FALSE | CAN | JSY3866 | 8 | rnr1Y285F-R | TGC | G>T | A/T+1 | G/C-1 | CG>AT | CG>AT | GGGTTTGCA | GCAGCTT | ACG | ACG | C | A | C>A | C>A_ACG | CG>AT_GCAGCTT | 8 | FALSE | FALSE |
| 164_v.txt | 32538 | 32538-SNV-( | SNV | 1 | G | T | 33 | 22423 | 0.14717032 | 37.3030303 | rnr1Y285F | FALSE | CAN | JSY3866 | 8 | rnr1Y285F-R | AGC | G>T | A/T+1 | G/C-1 | CG>AT | CG>AT | TTTGCAGCT | GCTTCAC | TCG | TCG | C | A | C>A | C>A_TCG | CG>AT_GCTTCAC | 8 | FALSE | FALSE |
| 164_v.txt | 32544 | 32544-SNV-( | SNV | 1 | C | A | 12 | 22423 | 0.05351648 | 35.6666667 | rnr1Y285F | FALSE | CAN | JSY3866 | 8 | rnr1Y285F-R | ACC | C>A | A/T+1 | G/C-1 | CG>AT | CG>AT | GCTTCACCA | CCAGCAG | TGG | ACC | C | T | C>A | C>A_ACC | CG>AT_CCAGCAG | 8 | FALSE | FALSE |
| 164_v.txt | 32629 | 32629-SNV-( | SNV | 1 | C | A | 14 | 24610 | 0.05688744 | 53.6428571 | rnr1Y285F | FALSE | CAN | JSY3866 | 8 | rnr1Y285F-R | TCG | C>A | A/T+1 | G/C-1 | CG>AT | CG>AT | TCCCTTCGTT | CGTTTTT | AGC | TCG | C | T | C>A | C>A_TCG | CG>AT_CGTTTTT | 8 | FALSE | FALSE |
| 164_v.txt | 32783 | 32783-SNV-( | SNV | 1 | G | T | 14 | 24610 | 0.05688744 | 49.5 | rnr1Y285F | FALSE | CAN | JSY3866 | 8 | rnr1Y285F-R | AGA | G>T | A/T+1 | G/C-1 | CG>AT | CG>AT | GACCCAGAA | GAACTCG | TCT | TCT | C | A | C>A | C>A_TCT | CG>AT_GAACTCG | 8 | FALSE | FALSE |
| 164_v.txt | 32788 | 32788-SNV-( | SNV | 1 | C | A | 30 | 24610 | 0.12190167 | 54.4666667 | rnr1Y285F | FALSE | CAN | JSY3866 | 8 | rnr1Y285F-R | TCG | C>A | A/T+1 | G/C-1 | CG>AT | CG>AT | AGAACTCGA | CGAATTC | AGC | TCG | C | T | C>A | C>A_TCG | CG>AT_CGAATTC | 8 | FALSE | FALSE |
| 164_v.txt | 32798 | 32798-SNV-( | SNV | 1 | G | T | 29 | 24610 | 0.11783828 | 47.3103448 | rnr1Y285F | FALSE | CAN | JSY3866 | 8 | rnr1Y285F-R | CGT | G>T | A/T+1 | G/C-1 | CG>AT | CG>AT | TTCACCGTA | GTAATAT | GCA | GCA | C | A | C>A | C>A_GCA | CG>AT_GTAATAT | 8 | FALSE | FALSE |
| 164_v.txt | 32819 | 32819-SNV-( | SNV | 1 | G | T | 73 | 24610 | 0.29662739 | 41.9315069 | rnr1Y285F | FALSE | CAN | JSY3866 | 8 | rnr1Y285F-R | AGT | G>T | A/T+1 | G/C-1 | CG>AT | CG>AT | GAACAAGTT | GTTCATT | TCA | TCA | C | A | C>A | C>A_TCA | CG>AT_GTTCATT | 8 | FALSE | FALSE |
| 164_v.txt | 32917 | 32917-SNV-( | SNV | 1 | C | A | 68 | 19469 | 0.3492732 | 54.3088235 | rnr1Y285F | FALSE | CAN | JSY3866 | 8 | rnr1Y285F-R | TCC | C>A | A/T+1 | G/C-1 | CG>AT | CG>AT | TAAGTTCCA | CCAGGGC | AGG | TCC | C | T | C>A | C>A_TCC | CG>AT_CCAGGGC | 8 | FALSE | FALSE |
| 164_v.txt | 32940 | 32940-SNV-( | SNV | 1 | G | T | 37 | 19469 | 0.19004571 | 49.2702703 | rnr1Y285F | FALSE | CAN | JSY3866 | 8 | rnr1Y285F-R | AGA | G>T | A/T+1 | G/C-1 | CG>AT | CG>AT | GCCCAAGAA | GAAAACC | TCT | TCT | C | A | C>A | C>A_TCT | CG>AT_GAAAACC | 8 | FALSE | FALSE |
| 164_v.txt | 32958 | 32958-SNV-( | SNV | 1 | C | A | 63 | 19469 | 0.32359135 | 58.1904762 | rnr1Y285F | FALSE | CAN | JSY3866 | 8 | rnr1Y285F-R | ACC | C>A | A/T+1 | G/C-1 | CG>AT | CG>AT | ATGTAACCA | CCATTGG | TGG | ACC | C | T | C>A | C>A_ACC | CG>AT_CCATTGG | 8 | FALSE | FALSE |
| 164_v.txt | 32971 | 32971-SNV-( | SNV | 1 | C | A | 2 | 19469 | 0.01027274 | 63.998394 | rnr1Y285F | FALSE | CAN | JSY3866 | 8 | rnr1Y285F-R | CCA | C>A | A/T+1 | G/C-1 | CG>AT | CG>AT | CCGCACCAA | CAAATGC | GGT | CCA | C | T | C>A | C>A_CCA | CG>AT_CAAATGC | 8 | FALSE | FALSE |
| 164_v.txt | 33033 | 33033-SNV-( | SNV | 1 | G | T | 11 | 19469 | 0.05650008 | 57.7272727 | rnr1Y285F | FALSE | CAN | JSY3866 | 8 | rnr1Y285F-R | TGT | G>T | A/T+1 | G/C-1 | CG>AT | CG>AT | ATGAATGTA | GTAGCCA | ACA | ACA | C | A | C>A | C>A_ACA | CG>AT_GTAGCCA | 8 | FALSE | FALSE |
| 164_v.txt | 33043 | 33043-SNV-( | SNV | 1 | C | A | 15 | 19469 | 0.07704556 | 54.8666667 | rnr1Y285F | FALSE | CAN | JSY3866 | 8 | rnr1Y285F-R | TCA | C>A | A/T+1 | G/C-1 | CG>AT | CG>AT | CCATTTCACC | CACCCAA | AGT | TCA | C | T | C>A | C>A_TCA | CG>AT_CACCCAA | 8 | FALSE | FALSE |
| 164_v.txt | 33050 | 33050-SNV-( | SNV | 1 | G | T | 11 | 19469 | 0.05650008 | 55.4548262 | rnr1Y285F | FALSE | CAN | JSY3866 | 8 | rnr1Y285F-R | AGG | G>T | A/T+1 | G/C-1 | CG>AT | CG>AT | ACCCAAGGA | GGACTGC | TCC | TCC | C | A | C>A | C>A_TCC | CG>AT_GGACTGC | 8 | FALSE | FALSE |
| 164_v.txt | 33051 | 33051-SNV-( | SNV | 1 | G | T | 25 | 19469 | 0.12840927 | 57.7173342 | rnr1Y285F | FALSE | CAN | JSY3866 | 8 | rnr1Y285F-R | GGA | G>T | A/T+1 | G/C-1 | CG>AT | CG>AT | CCCAAGGAC | GACTGCG | CCT | CCT | C | A | C>A | C>A_CCT | CG>AT_GACTGCG | 8 | FALSE | FALSE |
| 164_v.txt | 33102 | 33102-SNV-( | SNV | 1 | G | T | 78 | 19469 | 0.40063691 | 47.5384615 | rnr1Y285F | FALSE | CAN | JSY3866 | 8 | rnr1Y285F-R | AGC | G>T | A/T+1 | G/C-1 | CG>AT | CG>AT | ATAAGAGCG | GCGCCCA | TCG | TCG | C | A | C>A | C>A_TCG | CG>AT_GCGCCCA | 8 | FALSE | FALSE |
| 164_v.txt | 33151 | 33151-SNV-( | SNV | 1 | G | T | 86 | 19469 | 0.44172788 | 38.0232558 | rnr1Y285F | FALSE | CAN | JSY3866 | 8 | rnr1Y285F-R | AGA | G>T | A/T+1 | G/C-1 | CG>AT | CG>AT | TGAAAAGAC | GACCTGT | TCT | TCT | C | A | C>A | C>A_TCT | CG>AT_GACCTGT | 8 | FALSE | FALSE |
| 164_v.txt | 33159 | 33159-SNV-( | SNV | 1 | C | A | 6 | 19474 | 0.03081031 | 33.5 | rnr1Y285F | FALSE | CAN | JSY3866 | 8 | rnr1Y285F-R | ACC | C>A | A/T+1 | G/C-1 | CG>AT | CG>AT | CCTGTACCA | CCAATAG | TGG | ACC | C | T | C>A | C>A_ACC | CG>AT_CCAATAG | 8 | FALSE | FALSE |
| 164_v.txt | 33160 | 33160-SNV-( | SNV | 1 | C | A | 28 | 19474 | 0.14378145 | 37.4642857 | rnr1Y285F | FALSE | CAN | JSY3866 | 8 | rnr1Y285F-R | CCA | C>A | A/T+1 | G/C-1 | CG>AT | CG>AT | CTGTACCAA | CAATAGT | GGT | CCA | C | T | C>A | C>A_CCA | CG>AT_CAATAGT | 8 | FALSE | FALSE |
| 164_v.txt | 33168 | 33168-SNV-( | SNV | 1 | C | A | 18 | 19474 | 0.09243093 | 36.7222222 | rnr1Y285F | FALSE | CAN | JSY3866 | 8 | rnr1Y285F-R | ACC | C>A | A/T+1 | G/C-1 | CG>AT | CG>AT | ATAGTACCA | CCACCAA | TGG | ACC | C | T | C>A | C>A_ACC | CG>AT_CCACCAA | 8 | FALSE | FALSE |
| 164_v.txt | 33186 | 33186-SNV-( | SNV | 1 | C | A | 48 | 28661 | 0.16747497 | 36.0833333 | rnr1Y285F | FALSE | CAN | JSY3866 | 8 | rnr1Y285F-R | ACC | C>A | A/T+1 | G/C-1 | CG>AT | CG>AT | ATCATACCA | CCAATAT | TGG | ACC | C | T | C>A | C>A_ACC | CG>AT_CCAATAT | 8 | FALSE | FALSE |
| 164_v.txt | 33195 | 33195-SNV-( | SNV | 1 | C | A | 17 | 28651 | 0.05933475 | 35.6470588 | rnr1Y285F | FALSE | CAN | JSY3866 | 8 | rnr1Y285F-R | TCT | C>A | A/T+1 | G/C-1 | CG>AT | CG>AT | ATATGTCTT | CTTTGCT | AGA | TCT | C | T | C>A | C>A_TCT | CG>AT_CTTTGCT | 8 | FALSE | FALSE |
| 164_v.txt | 33235 | 33235-SNV-( | SNV | 1 | C | A | 33 | 28651 | 0.11517923 | 38.0606061 | rnr1Y285F | FALSE | CAN | JSY3866 | 8 | rnr1Y285F-R | TCT | C>A | A/T+1 | G/C-1 | CG>AT | CG>AT | GTACTTCTCC | CTCCTTC | AGA | TCT | C | T | C>A | C>A_TCT | CG>AT_CTCCTTC | 8 | FALSE | FALSE |
| 164_v.txt | 33309 | 33309-SNV-( | SNV | 1 | C | A | 9 | 28650 | 0.03141361 | 28.1149535 | rnr1Y285F | FALSE | CAN | JSY3866 | 8 | rnr1Y285F-R | GCG | C>A | A/T+1 | G/C-1 | CG>AT | CG>AT | AAAGAGCGC | CGCAATG | CGC | GCG | C | T | C>A | C>A_GCG | CG>AT_CGCAATG | 8 | FALSE | FALSE |
| 164_v.txt | 33310 | 33310-SNV-( | SNV | 1 | G | T | 13 | 28648 | 0.04537839 | 20.2508682 | rnr1Y285F | FALSE | CAN | JSY3866 | 8 | rnr1Y285F-R | CGC | G>T | A/T+1 | G/C-1 | CG>AT | CG>AT | AAGAGCGCA | GCAATGG | GCG | GCG | C | A | C>A | C>A_GCG | CG>AT_GCAATGG | 8 | FALSE | FALSE |
| 164_v.txt | 33325 | 33325-SNV-( | SNV | 1 | C | A | 96 | 28650 | 0.33507853 | 56.8845804 | rnr1Y285F | FALSE | CAN | JSY3866 | 8 | rnr1Y285F-R | TCT | C>A | A/T+1 | G/C-1 | CG>AT | CG>AT | ACAATTCTTT | CTTTACT | AGA | TCT | C | T | C>A | C>A_TCT | CG>AT_CTTTACT | 8 | FALSE | FALSE |
| 164_v.txt | 33419 | 33419-SNV-( | SNV | 1 | G | T | 348 | 28651 | 1.21461729 | 46.3333333 | rnr1Y285F | FALSE | CAN | JSY3866 | 8 | rnr1Y285F-R | TGT | G>T | A/T+1 | G/C-1 | CG>AT | CG>AT | CTCATTGTAC | GTACATA | ACA | ACA | C | A | C>A | C>A_ACA | CG>AT_GTACATA | 8 | FALSE | FALSE |
| 164_v.txt | 33451 | 33451-SNV-( | SNV | 1 | C | A | 39 | 28651 | 0.1361209 | 41.5337871 | rnr1Y285F | FALSE | CAN | JSY3866 | 8 | rnr1Y285F-R | TCT | C>A | A/T+1 | G/C-1 | CG>AT | CG>AT | CGTCTTCTTT | CTTTTGA | AGA | TCT | C | T | C>A | C>A_TCT | CG>AT_CTTTTGA | 8 | FALSE | FALSE |
| 164_v.txt | 33474 | 33474-SNV-( | SNV | 1 | C | A | 82 | 28649 | 0.28622291 | 34.695122 | rnr1Y285F | FALSE | CAN | JSY3866 | 8 | rnr1Y285F-R | GCC | C>A | A/T+1 | G/C-1 | CG>AT | CG>AT | GCTATGCCTT | CCTTTTT | CGG | GCC | C | T | C>A | C>A_GCC | CG>AT_CCTTTTT | 8 | FALSE | FALSE |
| 164_v.txt | 33475 | 33475-SNV-( | SNV | 1 | C | A | 17 | 28649 | 0.0593389 | 36.8823529 | rnr1Y285F | FALSE | CAN | JSY3866 | 8 | rnr1Y285F-R | CCT | C>A | A/T+1 | G/C-1 | CG>AT | CG>AT | CTATGCCTTT | CTTTTTT | GGA | CCT | C | T | C>A | C>A_CCT | CG>AT_CTTTTTT | 8 | FALSE | FALSE |
| 164_v.txt | 31980 | 31980-MNV- | MNV | 2 | GG | TT | 2 | 24695 | 8.10E-03 | 25.9999998 | rnr1Y285F | FALSE | CAN | JSY3866 | 8 | rnr1Y285F-R | GGG | nope | >1bp-ins | >1bpdel | none | MNV | AAGCCGGGC | GGCATTA | CCC | GGG | GG | none | NA | NA_GGG | MNV_GGCATTA | 8 | FALSE | FALSE |
| 164_v.txt | 31966 | 31966-Insert | Insertion | 1 | - | A | 9 | 24695 | 0.03644462 | 37.1111111 | rnr1Y285F | FALSE | CAN | JSY3866 | 8 | rnr1Y285F-R | TAA | nope | A/T+1 | >1bpdel | none | A/T+1 | CCGCATAAT | AATAAGC | ATT | TAA | - | T | NA | NA_TAA | A/T+1_AATAAGC | 8 | FALSE | FALSE |
| 164_v.txt | 32081 | 32081-Insert | Insertion | 1 | - | A | 19 | 24696 | 0.07693554 | 63.8421053 | rnr1Y285F | FALSE | CAN | JSY3866 | 8 | rnr1Y285F-R | CAA | nope | A/T+1 | >1bpdel | none | A/T+1 | CCATGCAAA | AAAAAAG | GTT | CAA | - | T | NA | NA_CAA | A/T+1_AAAAAAG | 8 | FALSE | FALSE |
| 164_v.txt | 32121 | 32121-Insert | Insertion | 1 | - | A | 6 | 24696 | 0.02429543 | 61.1666667 | rnr1Y285F | FALSE | CAN | JSY3866 | 8 | rnr1Y285F-R | GAA | nope | A/T+1 | >1bpdel | none | A/T+1 | CATTCGAAA | AAAACTT | CTT | GAA | - | T | NA | NA_GAA | A/T+1_AAAACTT | 8 | FALSE | FALSE |
| 164_v.txt | 32378 | 32378-Insert | Insertion | 1 | - | A | 2 | 22423 | 8.92E-03 | 64 | rnr1Y285F | FALSE | CAN | JSY3866 | 8 | rnr1Y285F-R | TAA | nope | A/T+1 | >1bpdel | none | A/T+1 | AGCAATAAT | AATAAAG | ATT | TAA | - | T | NA | NA_TAA | A/T+1_AATAAAG | 8 | FALSE | FALSE |
| 164_v.txt | 32498 | 32498-Insert | Insertion | 1 | - | T | 2 | 22423 | 8.92E-03 | 51 | rnr1Y285F | FALSE | CAN | JSY3866 | 8 | rnr1Y285F-R | CTT | nope | A/T+1 | >1bpdel | none | A/T+1 | AACAACTTT | TTTTTTG | GAA | CTT | - | A | NA | NA_CTT | A/T+1_TTTTTTG | 8 | FALSE | FALSE |
| 164_v.txt | 32728 | 32728-Insert | Insertion | 1 | - | A | 17 | 24610 | 0.06907761 | 62.4705882 | rnr1Y285F | FALSE | CAN | JSY3866 | 8 | rnr1Y285F-R | CAA | nope | A/T+1 | >1bpdel | none | A/T+1 | CCATACAAA | AAAAACA | GTT | CAA | - | T | NA | NA_CAA | A/T+1_AAAAACA | 8 | FALSE | FALSE |
| 164_v.txt | 32842 | 32842-Insert | Insertion | 1 | - | A | 65 | 24610 | 0.26412028 | 44.6 | rnr1Y285F | FALSE | CAN | JSY3866 | 8 | rnr1Y285F-R | CAA | nope | A/T+1 | >1bpdel | none | A/T+1 | TTACCCAAA | AAAAAAT | GTT | CAA | - | T | NA | NA_CAA | A/T+1_AAAAAAT | 8 | FALSE | FALSE |
| 164_v.txt | 32947 | 32947-Insert | Insertion | 1 | - | A | 2 | 19469 | 0.01027274 | 51 | rnr1Y285F | FALSE | CAN | JSY3866 | 8 | rnr1Y285F-R | CAA | nope | A/T+1 | >1bpdel | none | A/T+1 | AAAACCAAT | AATACAT | GTT | CAA | - | T | NA | NA_CAA | A/T+1_AATACAT | 8 | FALSE | FALSE |
| 164_v.txt | 32972 | 32972-Insert | Insertion | 1 | - | A | 2 | 19469 | 0.01027274 | 64 | rnr1Y285F | FALSE | CAN | JSY3866 | 8 | rnr1Y285F-R | CAA | nope | A/T+1 | >1bpdel | none | A/T+1 | CGCACCAAA | AAATGCT | GTT | CAA | - | T | NA | NA_CAA | A/T+1_AAATGCT | 8 | FALSE | FALSE |
| 164_v.txt | 33206 | 33206-Insert | Insertion | 2 | - | CT | 4 | 28651 | 0.01396112 | 37.875 | rnr1Y285F | FALSE | CAN | JSY3866 | 8 | rnr1Y285F-R | GCT | nope | >1bp-ins | >1bpdel | none | >1bp-ins | CTTAAGCTCT | CTCTCTC | CGA | GCT | - | none | NA | NA_GCT | >1bp-ins_CTCTCTC | 8 | FALSE | FALSE |
| 164_v.txt | 33452 | 33452-Insert | Insertion | 1 | - | T | 18 | 28651 | 0.06282503 | 39.2222222 | rnr1Y285F | FALSE | CAN | JSY3866 | 8 | rnr1Y285F-R | CTT | nope | A/T+1 | >1bpdel | none | A/T+1 | GTCTTCTTTT | TTTTGAA | GAA | CTT | - | A | NA | NA_CTT | A/T+1_TTTTGAA | 8 | FALSE | FALSE |
| 164_v.txt | 32313 | 32313-Deleti | Deletion | 3 | GTT | - | 11 | 22423 | 0.04905677 | 58.4545455 | rnr1Y285F | FALSE | CAN | JSY3866 | 8 | rnr1Y285F-R | GGT | nope | >1bp-ins | >1bpdel | none | >1bpdel | ATAATGGTT | GTTGTTA | CCA | GGT | GTT | none | NA | NA_GGT | >1bpdel_GTTGTTA | 8 | FALSE | FALSE |
| 164_v.txt | 32337 | 32337-Deleti | Deletion | 2 | AT | - | 32 | 22423 | 0.14271061 | 58.5625 | rnr1Y285F | FALSE | CAN | JSY3866 | 8 | rnr1Y285F-R | GAT | nope | >1bp-ins | >1bpdel | none | >1bpdel | TTGAAGATA | ATATGTG | CTA | GAT | AT | none | NA | NA_GAT | >1bpdel_ATATGTG | 8 | FALSE | FALSE |
| 164_v.txt | 32460 | 32460-Deleti | Deletion | 2 | AG | - | 134 | 22423 | 0.59760068 | 50.641791 | rnr1Y285F | FALSE | CAN | JSY3866 | 8 | rnr1Y285F-R | TAG | nope | >1bp-ins | >1bpdel | none | >1bpdel | AATAATAGA | AGAGAGC | ATC | TAG | AG | none | NA | NA_TAG | >1bpdel_AGAGAGC | 8 | FALSE | FALSE |
| 164_v.txt | 33168 | 33168-Deleti | Deletion | 2 | CC | - | 1 | 19474 | 5.14E-03 | 37 | rnr1Y285F | FALSE | CAN | JSY3866 | 8 | rnr1Y285F-R | ACC | nope | >1bp-ins | >1bpdel | none | >1bpdel | ATAGTACCA | CCACCAA | TGG | ACC | CC | none | NA | NA_ACC | >1bpdel_CCACCAA | 8 | FALSE | FALSE |

| | | | | | | | | | | | | | | | | | | | | | | | | | | | | | | | | | | |
|---|---|---|---|---|---|---|---|---|---|---|---|---|---|---|---|---|---|---|---|---|---|---|---|---|---|---|---|---|---|---|---|---|---|---|
| 164_v.txt | 33195 | 33195-Deleti | Deletion | 2 | CT | - | 1 | 28651 | 3.49E-03 | 38 | rnr1Y285F | FALSE | CAN | JSY3866 | 8 | rnr1Y285F-R | TCT | nope | >1bp-ins | >1bpdel | none | >1bpdel | ATATGTCTTT | CTTTGCT | AGA | TCT | CT | none | NA | NA_TCT | >1bpdel_CTTTGCT | 8 | FALSE | FALSE |
| 164_v.txt | 31723 | 31723-Deleti | Deletion | 1 | A | - | 4 | 3961 | 0.1009846 | 59.25 | rnr1Y285F | FALSE | CAN | JSY3866 | 8 | rnr1Y285F-R | CAA | nope | >1bp-ins | A/T-1 | none | A/T-1 | TGTCCCAAA | AAAAAGT | GTT | GTT | T | none | T>none | T>none_GTT | A/T-1_AAAAAGT | 8 | FALSE | FALSE |
| 164_v.txt | 31952 | 31952-Deleti | Deletion | 1 | A | - | 8 | 24695 | 0.03239522 | 36.875 | rnr1Y285F | FALSE | CAN | JSY3866 | 8 | rnr1Y285F-R | TAA | nope | >1bp-ins | A/T-1 | none | A/T-1 | CGTCATAAA | AAATGTG | ATT | ATT | T | none | T>none | T>none_ATT | A/T-1_AAATGTG | 8 | FALSE | FALSE |
| 164_v.txt | 31972 | 31972-Deleti | Deletion | 1 | C | - | 6 | 24695 | 0.02429642 | 36.3333333 | rnr1Y285F | FALSE | CAN | JSY3866 | 8 | rnr1Y285F-R | GCC | nope | >1bp-ins | G/C-1 | none | G/C-1 | AATAAGCCA | CCAAGCC | CGG | GCC | C | none | nope | nope_GCC | G/C-1_CCAAGCC | 8 | FALSE | FALSE |
| 164_v.txt | 32069 | 32069-Deleti | Deletion | 1 | A | - | 6 | 24696 | 0.02429543 | 64 | rnr1Y285F | FALSE | CAN | JSY3866 | 8 | rnr1Y285F-R | TAA | nope | >1bp-ins | A/T-1 | none | A/T-1 | TGAGATAAA | AAATAAC | ATT | ATT | T | none | T>none | T>none_ATT | A/T-1_AAATAAC | 8 | FALSE | FALSE |
| 164_v.txt | 32081 | 32081-Deleti | Deletion | 1 | A | - | 41 | 24696 | 0.16601879 | 63.9268293 | rnr1Y285F | FALSE | CAN | JSY3866 | 8 | rnr1Y285F-R | CAA | nope | >1bp-ins | A/T-1 | none | A/T-1 | CCATGCAAA | AAAAAAG | GTT | GTT | T | none | T>none | T>none_GTT | A/T-1_AAAAAAG | 8 | FALSE | FALSE |
| 164_v.txt | 32092 | 32092-Deleti | Deletion | 1 | C | - | 3 | 24696 | 0.01214772 | 64 | rnr1Y285F | FALSE | CAN | JSY3866 | 8 | rnr1Y285F-R | GCA | nope | >1bp-ins | G/C-1 | none | G/C-1 | AGCCTGCAA | CAACACC | CGT | GCA | C | none | nope | nope_GCA | G/C-1_CAACACC | 8 | FALSE | FALSE |
| 164_v.txt | 32114 | 32114-Deleti | Deletion | 1 | C | - | 1 | 24696 | 4.05E-03 | 64 | rnr1Y285F | FALSE | CAN | JSY3866 | 8 | rnr1Y285F-R | GCC | nope | >1bp-ins | G/C-1 | none | G/C-1 | TAATAGCCA | CCATTCG | CGG | GCC | C | none | nope | nope_GCC | G/C-1_CCATTCG | 8 | FALSE | FALSE |
| 164_v.txt | 32184 | 32184-Deleti | Deletion | 1 | A | - | 9 | 24696 | 0.03644315 | 54.4444444 | rnr1Y285F | FALSE | CAN | JSY3866 | 8 | rnr1Y285F-R | GAA | nope | >1bp-ins | A/T-1 | none | A/T-1 | GTAACGAAA | AAAACTG | CTT | CTT | T | none | T>none | T>none_CTT | A/T-1_AAAACTG | 8 | FALSE | FALSE |
| 164_v.txt | 32304 | 32304-Deleti | Deletion | 1 | G | - | 16 | 22423 | 0.07135531 | 62 | rnr1Y285F | FALSE | CAN | JSY3866 | 8 | rnr1Y285F-R | AGA | nope | >1bp-ins | G/C-1 | none | G/C-1 | GCGGCAGAA | GAAATAA | TCT | TCT | C | none | C>none | C>none_TCT | G/C-1_GAAATAA | 8 | FALSE | FALSE |
| 164_v.txt | 32346 | 32346-Deleti | Deletion | 1 | A | - | 15 | 22423 | 0.0668956 | 60.2 | rnr1Y285F | FALSE | CAN | JSY3866 | 8 | rnr1Y285F-R | CAA | nope | >1bp-ins | A/T-1 | none | A/T-1 | TGTGGCAAA | AAAACCT | GTT | GTT | T | none | T>none | T>none_GTT | A/T-1_AAAACCT | 8 | FALSE | FALSE |
| 164_v.txt | 32419 | 32419-Deleti | Deletion | 1 | G | - | 21 | 22423 | 0.09365384 | 58.6666667 | rnr1Y285F | FALSE | CAN | JSY3866 | 8 | rnr1Y285F-R | AGT | nope | >1bp-ins | G/C-1 | none | G/C-1 | GTGTTAGTTT | GTTTAGG | TCA | TCA | C | none | C>none | C>none_TCA | G/C-1_GTTTAGG | 8 | FALSE | FALSE |
| 164_v.txt | 32424 | 32424-Deleti | Deletion | 1 | G | - | 5 | 22423 | 0.02229853 | 48 | rnr1Y285F | FALSE | CAN | JSY3866 | 8 | rnr1Y285F-R | AGG | nope | >1bp-ins | G/C-1 | none | G/C-1 | AGTTTAGGG | GGGTCAT | TCC | TCC | C | none | C>none | C>none_TCC | G/C-1_GGGTCAT | 8 | FALSE | FALSE |
| 164_v.txt | 32457 | 32457-Deleti | Deletion | 1 | A | - | 3 | 22423 | 0.01337912 | 54 | rnr1Y285F | FALSE | CAN | JSY3866 | 8 | rnr1Y285F-R | TAA | nope | >1bp-ins | A/T-1 | none | A/T-1 | ATGAATAAT | AATAGAG | ATT | ATT | T | none | T>none | T>none_ATT | A/T-1_AATAGAG | 8 | FALSE | FALSE |
| 164_v.txt | 32490 | 32490-Deleti | Deletion | 1 | A | - | 22 | 22423 | 0.09811354 | 50.7727273 | rnr1Y285F | FALSE | CAN | JSY3866 | 8 | rnr1Y285F-R | GAA | nope | >1bp-ins | A/T-1 | none | A/T-1 | ATACGGAAA | AAAACAA | CTT | CTT | T | none | T>none | T>none_CTT | A/T-1_AAAACAA | 8 | FALSE | FALSE |
| 164_v.txt | 32498 | 32498-Deleti | Deletion | 1 | T | - | 31 | 22423 | 0.1382509 | 49.7419337 | rnr1Y285F | FALSE | CAN | JSY3866 | 8 | rnr1Y285F-R | CTT | nope | >1bp-ins | A/T-1 | none | A/T-1 | AACAACTTTT | TTTTTTG | GAA | CTT | T | none | nope | nope_CTT | A/T-1_TTTTTTG | 8 | FALSE | FALSE |
| 164_v.txt | 32522 | 32522-Deleti | Deletion | 1 | T | - | 7 | 22423 | 0.03121795 | 40.2857143 | rnr1Y285F | FALSE | CAN | JSY3866 | 8 | rnr1Y285F-R | ATT | nope | >1bp-ins | A/T-1 | none | A/T-1 | AACGGATTTT | TTTTCTG | TAA | ATT | T | none | nope | nope_ATT | A/T-1_TTTTCTG | 8 | FALSE | FALSE |
| 164_v.txt | 32631 | 32631-Deleti | Deletion | 1 | T | - | 10 | 24610 | 0.04063389 | 53 | rnr1Y285F | FALSE | CAN | JSY3866 | 8 | rnr1Y285F-R | GTT | nope | >1bp-ins | A/T-1 | none | A/T-1 | CCTTCGTTTT | TTTTTAT | CAA | GTT | T | none | nope | nope_GTT | A/T-1_TTTTTAT | 8 | FALSE | FALSE |
| 164_v.txt | 32640 | 32640-Deleti | Deletion | 1 | T | - | 5 | 24610 | 0.02031694 | 64 | rnr1Y285F | FALSE | CAN | JSY3866 | 8 | rnr1Y285F-R | CTT | nope | >1bp-ins | A/T-1 | none | A/T-1 | TTATCCTTAG | TTAGATA | GAA | CTT | T | none | nope | nope_CTT | A/T-1_TTAGATA | 8 | FALSE | FALSE |
| 164_v.txt | 32658 | 32658-Deleti | Deletion | 1 | C | - | 15 | 24610 | 0.06095083 | 64 | rnr1Y285F | FALSE | CAN | JSY3866 | 8 | rnr1Y285F-R | ACC | nope | >1bp-ins | G/C-1 | none | G/C-1 | CCTGGACCC | CCCCAGG | TGG | ACC | C | none | nope | nope_ACC | G/C-1_CCCCAGG | 8 | FALSE | FALSE |
| 164_v.txt | 32670 | 32670-Deleti | Deletion | 1 | G | - | 506 | 24610 | 2.05607477 | 59.7509881 | rnr1Y285F | FALSE | CAN | JSY3866 | 8 | rnr1Y285F-R | TGG | nope | >1bp-ins | G/C-1 | none | G/C-1 | GCACCTGGG | GGGTTTC | ACC | ACC | C | none | C>none | C>none_ACC | G/C-1_GGGTTTC | 8 | FALSE | FALSE |
| 164_v.txt | 32680 | 32680-Deleti | Deletion | 1 | A | - | 2 | 24610 | 8.13E-03 | 51.0000054 | rnr1Y285F | FALSE | CAN | JSY3866 | 8 | rnr1Y285F-R | CAA | nope | >1bp-ins | A/T-1 | none | A/T-1 | TTCTCCAATA | AATAACG | GTT | GTT | T | none | T>none | T>none_GTT | A/T-1_AATAACG | 8 | FALSE | FALSE |
| 164_v.txt | 32707 | 32707-Deleti | Deletion | 1 | C | - | 11 | 24610 | 0.04469728 | 61.7272727 | rnr1Y285F | FALSE | CAN | JSY3866 | 8 | rnr1Y285F-R | ACC | nope | >1bp-ins | G/C-1 | none | G/C-1 | CGGTAACCC | CCCCAGC | TGG | ACC | C | none | nope | nope_ACC | G/C-1_CCCCAGC | 8 | FALSE | FALSE |
| 164_v.txt | 32728 | 32728-Deleti | Deletion | 1 | A | - | 46 | 24610 | 0.18691589 | 61.5434784 | rnr1Y285F | FALSE | CAN | JSY3866 | 8 | rnr1Y285F-R | CAA | nope | >1bp-ins | A/T-1 | none | A/T-1 | CCATACAAA | AAAAACA | GTT | GTT | T | none | T>none | T>none_GTT | A/T-1_AAAAACA | 8 | FALSE | FALSE |
| 164_v.txt | 32747 | 32747-Deleti | Deletion | 1 | C | - | 11 | 24610 | 0.04469728 | 61.0909091 | rnr1Y285F | FALSE | CAN | JSY3866 | 8 | rnr1Y285F-R | ACC | nope | >1bp-ins | G/C-1 | none | G/C-1 | TAGAAACCC | CCCGATA | TGG | ACC | C | none | nope | nope_ACC | G/C-1_CCCGATA | 8 | FALSE | FALSE |
| 164_v.txt | 32760 | 32760-Deleti | Deletion | 1 | A | - | 43 | 24610 | 0.17472572 | 53.9302326 | rnr1Y285F | FALSE | CAN | JSY3866 | 8 | rnr1Y285F-R | TAA | nope | >1bp-ins | A/T-1 | none | A/T-1 | ATGGCTAAA | AAAACTT | ATT | ATT | T | none | T>none | T>none_ATT | A/T-1_AAAACTT | 8 | FALSE | FALSE |
| 164_v.txt | 32779 | 32779-Deleti | Deletion | 1 | C | - | 11 | 24610 | 0.04469728 | 49.7272727 | rnr1Y285F | FALSE | CAN | JSY3866 | 8 | rnr1Y285F-R | ACC | nope | >1bp-ins | G/C-1 | none | G/C-1 | AAGCGACCC | CCCAGAA | TGG | ACC | C | none | nope | nope_ACC | G/C-1_CCCAGAA | 8 | FALSE | FALSE |
| 164_v.txt | 32842 | 32842-Deleti | Deletion | 1 | A | - | 178 | 24610 | 0.72328322 | 43.1685393 | rnr1Y285F | FALSE | CAN | JSY3866 | 8 | rnr1Y285F-R | CAA | nope | >1bp-ins | A/T-1 | none | A/T-1 | TTACCCAAA | AAAAAAT | GTT | GTT | T | none | T>none | T>none_GTT | A/T-1_AAAAAAT | 8 | FALSE | FALSE |
| 164_v.txt | 32884 | 32884-Deleti | Deletion | 1 | A | - | 11 | 19468 | 0.05650298 | 36.1818182 | rnr1Y285F | FALSE | CAN | JSY3866 | 8 | rnr1Y285F-R | CAA | nope | >1bp-ins | A/T-1 | none | A/T-1 | ACGTCCAAA | AAAATTG | GTT | GTT | T | none | T>none | T>none_GTT | A/T-1_AAAATTG | 8 | FALSE | FALSE |
| 164_v.txt | 32924 | 32924-Deleti | Deletion | 1 | A | - | 30 | 19469 | 0.15409112 | 55.9 | rnr1Y285F | FALSE | CAN | JSY3866 | 8 | rnr1Y285F-R | CAA | nope | >1bp-ins | A/T-1 | none | A/T-1 | CAGGGCAAA | AAAAGTG | GTT | GTT | T | none | T>none | T>none_GTT | A/T-1_AAAAGTG | 8 | FALSE | FALSE |
| 164_v.txt | 32932 | 32932-Deleti | Deletion | 1 | T | - | 6 | 19469 | 0.03081822 | 59.1666667 | rnr1Y285F | FALSE | CAN | JSY3866 | 8 | rnr1Y285F-R | ATT | nope | >1bp-ins | A/T-1 | none | A/T-1 | AAGTGATTG | TTGCCCA | TAA | ATT | T | none | nope | nope_ATT | A/T-1_TTGCCCA | 8 | FALSE | FALSE |
| 164_v.txt | 33020 | 33020-Deleti | Deletion | 1 | A | - | 5 | 19469 | 0.02568185 | 60.2 | rnr1Y285F | FALSE | CAN | JSY3866 | 8 | rnr1Y285F-R | TAA | nope | >1bp-ins | A/T-1 | none | A/T-1 | GGATGTAAC | AACAGGG | ATT | ATT | T | none | T>none | T>none_ATT | A/T-1_AACAGGG | 8 | FALSE | FALSE |
| 164_v.txt | 33050 | 33050-Deleti | Deletion | 1 | G | - | 1 | 19469 | 5.14E-03 | 38.000001 | rnr1Y285F | FALSE | CAN | JSY3866 | 8 | rnr1Y285F-R | AGG | nope | >1bp-ins | G/C-1 | none | G/C-1 | ACCCAAGGA | GGACTGC | TCC | TCC | C | none | C>none | C>none_TCC | G/C-1_GGACTGC | 8 | FALSE | FALSE |
| 164_v.txt | 33114 | 33114-Deleti | Deletion | 1 | C | - | 7 | 19469 | 0.0359546 | 45.4285714 | rnr1Y285F | FALSE | CAN | JSY3866 | 8 | rnr1Y285F-R | GCC | nope | >1bp-ins | G/C-1 | none | G/C-1 | ACTGGGCCG | CCGGCGT | CGG | GCC | C | none | nope | nope_GCC | G/C-1_CCGGCGT | 8 | FALSE | FALSE |
| 164_v.txt | 33141 | 33141-Deleti | Deletion | 1 | C | - | 6 | 19469 | 0.03081822 | 46.6666667 | rnr1Y285F | FALSE | CAN | JSY3866 | 8 | rnr1Y285F-R | ACC | nope | >1bp-ins | G/C-1 | none | G/C-1 | GATAAACCA | CCAATGA | TGG | ACC | C | none | nope | nope_ACC | G/C-1_CCAATGA | 8 | FALSE | FALSE |
| 164_v.txt | 33196 | 33196-Deleti | Deletion | 1 | T | - | 5 | 28651 | 0.0174514 | 37.9999998 | rnr1Y285F | FALSE | CAN | JSY3866 | 8 | rnr1Y285F-R | CTT | nope | >1bp-ins | A/T-1 | none | A/T-1 | TATGTCTTTG | TTTGCTT | GAA | CTT | T | none | nope | nope_CTT | A/T-1_TTTGCTT | 8 | FALSE | FALSE |
| 164_v.txt | 33326 | 33326-Deleti | Deletion | 1 | T | - | 6 | 28649 | 0.02094314 | 56.6666667 | rnr1Y285F | FALSE | CAN | JSY3866 | 8 | rnr1Y285F-R | CTT | nope | >1bp-ins | A/T-1 | none | A/T-1 | CAATTCTTTA | TTTACTT | GAA | CTT | T | none | nope | nope_CTT | A/T-1_TTTACTT | 8 | FALSE | FALSE |
| 164_v.txt | 33356 | 33356-Deleti | Deletion | 1 | C | - | 8 | 28651 | 0.02792224 | 62.5 | rnr1Y285F | FALSE | CAN | JSY3866 | 8 | rnr1Y285F-R | ACC | nope | >1bp-ins | G/C-1 | none | G/C-1 | TATTGACCCA | CCCACGT | TGG | ACC | C | none | nope | nope_ACC | G/C-1_CCCACGT | 8 | FALSE | FALSE |
| 164_v.txt | 33474 | 33474-Deleti | Deletion | 1 | C | - | 65 | 28649 | 0.22688401 | 31.0615385 | rnr1Y285F | FALSE | CAN | JSY3866 | 8 | rnr1Y285F-R | GCC | nope | >1bp-ins | G/C-1 | none | G/C-1 | GCTATGCCTT | CCTTTTT | CGG | GCC | C | none | nope | nope_GCC | G/C-1_CCTTTTT | 8 | FALSE | FALSE |
| 40_v.txt | 31723 | 31723-SNV- | SNV | 1 | A | C | 12 | 24213 | 0.04956015 | 57.8328534 | rnr1Y285F_pe | FALSE | ARG | JSY3811 | 1 | rnr1Y285F_p | CAA | A>C | G/C+1 | A/T-1 | TA>GC | TA>GC | TGTCCCAAA | AAAAAGT | GTT | GTT | T | G | T>G | T>G_GTT | TA>GC_AAAAAGT | 4 | FALSE | FALSE |
| 40_v.txt | 32493 | 32493-SNV- | SNV | 1 | A | C | 16 | 62778 | 0.02548664 | 33.625 | rnr1Y285F_pe | FALSE | ARG | JSY3811 | 1 | rnr1Y285F_p | AAC | A>C | G/C+1 | A/T-1 | TA>GC | TA>GC | CGGAAAACA | ACAACTT | TTG | TTG | T | G | T>G | T>G_TTG | TA>GC_ACAACTT | 4 | FALSE | FALSE |
| 40_v.txt | 32573 | 32573-SNV- | SNV | 1 | T | G | 6 | 53851 | 0.01114185 | 32.8333333 | rnr1Y285F_pe | FALSE | ARG | JSY3811 | 1 | rnr1Y285F_p | CTT | T>G | G/C+1 | A/T-1 | TA>GC | TA>GC | AGTACCTTG | TTGAAAT | GAA | CTT | T | C | T>G | T>G_CTT | TA>GC_TTGAAAT | 4 | FALSE | FALSE |
| 40_v.txt | 32842 | 32842-SNV- | SNV | 1 | A | C | 8 | 53850 | 0.01485608 | 31.750269 | rnr1Y285F_pe | FALSE | ARG | JSY3811 | 1 | rnr1Y285F_p | CAA | A>C | G/C+1 | A/T-1 | TA>GC | TA>GC | TTACCCAAA | AAAAAAT | GTT | GTT | T | G | T>G | T>G_GTT | TA>GC_AAAAAAT | 4 | FALSE | FALSE |
| 40_v.txt | 32884 | 32884-SNV- | SNV | 1 | A | C | 21 | 87076 | 0.02411686 | 31.1422489 | rnr1Y285F_pe | FALSE | ARG | JSY3811 | 1 | rnr1Y285F_p | CAA | A>C | G/C+1 | A/T-1 | TA>GC | TA>GC | ACGTCCAAA | AAAATTG | GTT | GTT | T | G | T>G | T>G_GTT | TA>GC_AAAATTG | 4 | FALSE | FALSE |
| 40_v.txt | 33269 | 33269-SNV- | SNV | 1 | A | C | 352 | 51277 | 0.68646762 | 16.1931818 | rnr1Y285F_pe | FALSE | ARG | JSY3811 | 1 | rnr1Y285F_p | TAG | A>C | G/C+1 | A/T-1 | TA>GC | TA>GC | CTCCATAGA | AGAGAAC | ATC | ATC | T | G | T>G | T>G_ATC | TA>GC_AGAGAAC | 4 | FALSE | FALSE |
| 40_v.txt | 33476 | 33476-SNV- | SNV | 1 | T | G | 8 | 51275 | 0.01560215 | 29.8759578 | rnr1Y285F_pe | FALSE | ARG | JSY3811 | 1 | rnr1Y285F_p | CTT | T>G | G/C+1 | A/T-1 | TA>GC | TA>GC | TATGCCTTTT | TTTTTTT | GAA | CTT | T | C | T>G | T>G_CTT | TA>GC_TTTTTTT | 4 | FALSE | FALSE |
| 40_v.txt | 32842 | 32842-SNV- | SNV | 1 | A | G | 12 | 53850 | 0.02228412 | 36.5000235 | rnr1Y285F_pe | FALSE | ARG | JSY3811 | 1 | rnr1Y285F_p | CAA | A>G | G/C+1 | A/T-1 | TA>CG | TA>CG | TTACCCAAA | AAAAAAT | GTT | GTT | T | C | T>C | T>C_GTT | TA>CG_AAAAAAT | 4 | FALSE | FALSE |
| 40_v.txt | 32919 | 32919-SNV- | SNV | 1 | A | G | 57 | 87077 | 0.06545931 | 47.2982456 | rnr1Y285F_pe | FALSE | ARG | JSY3811 | 1 | rnr1Y285F_p | CAG | A>G | G/C+1 | A/T-1 | TA>CG | TA>CG | AGTTCCAGG | AGGGCAA | GTC | GTC | T | C | T>C | T>C_GTC | TA>CG_AGGGCAA | 4 | FALSE | FALSE |
| 40_v.txt | 33476 | 33476-SNV- | SNV | 1 | T | C | 68 | 51275 | 0.13261824 | 30.4561676 | rnr1Y285F_pe | FALSE | ARG | JSY3811 | 1 | rnr1Y285F_p | CTT | T>C | G/C+1 | A/T-1 | TA>CG | TA>CG | TATGCCTTTT | TTTTTTT | GAA | CTT | T | G | T>C | T>C_CTT | TA>CG_TTTTTTT | 4 | FALSE | FALSE |
| 40_v.txt | 31650 | 31650-SNV- | SNV | 1 | C | T | 41 | 24213 | 0.16933053 | 36.9512195 | rnr1Y285F_pe | FALSE | ARG | JSY3811 | 1 | rnr1Y285F_p | GCG | C>T | A/T+1 | G/C-1 | CG>TA | CG>TA | AGAATGCGA | CGAAATG | CGC | GCG | C | A | C>T | C>T_GCG | CG>TA_CGAAATG | 4 | FALSE | FALSE |
| 40_v.txt | 31658 | 31658-SNV- | SNV | 1 | C | T | 42 | 24213 | 0.17346054 | 33.6430486 | rnr1Y285F_pe | FALSE | ARG | JSY3811 | 1 | rnr1Y285F_p | GCG | C>T | A/T+1 | G/C-1 | CG>TA | CG>TA | AAATGGCGT | CGTGGAA | CGC | GCG | C | A | C>T | C>T_GCG | CG>TA_CGTGGAA | 4 | FALSE | FALSE |
| 40_v.txt | 31882 | 31882-SNV- | SNV | 1 | C | T | 26 | 24213 | 0.10738033 | 44.2307692 | rnr1Y285F_pe | FALSE | ARG | JSY3811 | 1 | rnr1Y285F_p | GCA | C>T | A/T+1 | G/C-1 | CG>TA | CG>TA | AGGCGGCAG | CAGCAAA | CGT | GCA | C | A | C>T | C>T_GCA | CG>TA_CAGCAAA | 4 | FALSE | FALSE |
| 40_v.txt | 31911 | 31911-SNV- | SNV | 1 | G | A | 30 | 24213 | 0.12390038 | 43.0333333 | rnr1Y285F_pe | FALSE | ARG | JSY3811 | 1 | rnr1Y285F_p | TGC | G>A | A/T+1 | G/C-1 | CG>TA | CG>TA | TTTGGTGCA | GCAAAAG | ACG | ACG | C | T | C>T | C>T_ACG | CG>TA_GCAAAAG | 4 | FALSE | FALSE |
| 40_v.txt | 31920 | 31920-SNV- | SNV | 1 | G | A | 21 | 24211 | 0.08673743 | 37.0476191 | rnr1Y285F_pe | FALSE | ARG | JSY3811 | 1 | rnr1Y285F_p | CGT | G>A | A/T+1 | G/C-1 | CG>TA | CG>TA | AAAGCCGTG | GTGAAAC | GCA | GCA | C | T | C>T | C>T_GCA | CG>TA_GTGAAAC | 4 | FALSE | FALSE |
| 40_v.txt | 31947 | 31947-SNV- | SNV | 1 | G | A | 31 | 47390 | 0.06541464 | 34.7419355 | rnr1Y285F_pe | FALSE | ARG | JSY3811 | 1 | rnr1Y285F_p | CGT | G>A | A/T+1 | G/C-1 | CG>TA | CG>TA | ATGATCGTC | GTCATAA | GCA | GCA | C | T | C>T | C>T_GCA | CG>TA_GTCATAA | 4 | FALSE | FALSE |
| 40_v.txt | 31960 | 31960-SNV- | SNV | 1 | C | T | 47 | 47407 | 0.09914148 | 35.9361702 | rnr1Y285F_pe | FALSE | ARG | JSY3811 | 1 | rnr1Y285F_p | GCC | C>T | A/T+1 | G/C-1 | CG>TA | CG>TA | ATGTGGCCG | CCGCATA | CGG | GCC | C | A | C>T | C>T_GCC | CG>TA_CCGCATA | 4 | FALSE | FALSE |
| 40_v.txt | 31962 | 31962-SNV- | SNV | 1 | G | A | 92 | 47407 | 0.19406417 | 36.7065217 | rnr1Y285F_pe | FALSE | ARG | JSY3811 | 1 | rnr1Y285F_p | CGC | G>A | A/T+1 | G/C-1 | CG>TA | CG>TA | GTGGCCGCA | GCATAAT | GCG | GCG | C | T | C>T | C>T_GCG | CG>TA_GCATAAT | 4 | FALSE | FALSE |
| 40_v.txt | 31972 | 31972-SNV- | SNV | 1 | C | T | 34 | 47407 | 0.07171937 | 37.4117647 | rnr1Y285F_pe | FALSE | ARG | JSY3811 | 1 | rnr1Y285F_p | GCC | C>T | A/T+1 | G/C-1 | CG>TA | CG>TA | AATAAGCCA | CCAAGCC | CGG | GCC | C | A | C>T | C>T_GCC | CG>TA_CCAAGCC | 4 | FALSE | FALSE |
| 40_v.txt | 31981 | 31981-SNV- | SNV | 1 | G | A | 36 | 47407 | 0.07593815 | 50 | rnr1Y285F_pe | FALSE | ARG | JSY3811 | 1 | rnr1Y285F_p | GGC | G>A | A/T+1 | G/C-1 | CG>TA | CG>TA | AGCCGGGCA | GCATTAA | CCG | CCG | C | T | C>T | C>T_CCG | CG>TA_GCATTAA | 4 | FALSE | FALSE |
| 40_v.txt | 32057 | 32057-SNV- | SNV | 1 | C | T | 38 | 47407 | 0.08015694 | 62.2631579 | rnr1Y285F_pe | FALSE | ARG | JSY3811 | 1 | rnr1Y285F_p | GCG | C>T | A/T+1 | G/C-1 | CG>TA | CG>TA | GATGTGCGA | CGAGATT | CGC | GCG | C | A | C>T | C>T_GCG | CG>TA_CGAGATT | 4 | FALSE | FALSE |
| 40_v.txt | 32079 | 32079-SNV- | SNV | 1 | G | A | 27 | 47407 | 0.05695361 | 60.3703704 | rnr1Y285F_pe | FALSE | ARG | JSY3811 | 1 | rnr1Y285F_p | TGC | G>A | A/T+1 | G/C-1 | CG>TA | CG>TA | AACCATGCA | GCAAAAA | ACG | ACG | C | T | C>T | C>T_ACG | CG>TA_GCAAAAA | 4 | FALSE | FALSE |
| 40_v.txt | 32092 | 32092-SNV- | SNV | 1 | C | T | 54 | 47407 | 0.11390723 | 62.0925926 | rnr1Y285F_pe | FALSE | ARG | JSY3811 | 1 | rnr1Y285F_p | GCA | C>T | A/T+1 | G/C-1 | CG>TA | CG>TA | AGCCTGCAA | CAACACC | CGT | GCA | C | A | C>T | C>T_GCA | CG>TA_CAACACC | 4 | FALSE | FALSE |
| 40_v.txt | 32165 | 32165-SNV- | SNV | 1 | G | A | 58 | 47407 | 0.1223448 | 46.4655172 | rnr1Y285F_pe | FALSE | ARG | JSY3811 | 1 | rnr1Y285F_p | CGC | G>A | A/T+1 | G/C-1 | CG>TA | CG>TA | CAAAGCGCC | GCCAAAT | GCG | GCG | C | T | C>T | C>T_GCG | CG>TA_GCCAAAT | 4 | FALSE | FALSE |
| 40_v.txt | 32173 | 32173-SNV- | SNV | 1 | C | T | 37 | 47407 | 0.07804755 | 45.7567568 | rnr1Y285F_pe | FALSE | ARG | JSY3811 | 1 | rnr1Y285F_p | GCA | C>T | A/T+1 | G/C-1 | CG>TA | CG>TA | CAAATGCAG | CAGCAGT | CGT | GCA | C | A | C>T | C>T_GCA | CG>TA_CAGCAGT | 4 | FALSE | FALSE |
| 40_v.txt | 32176 | 32176-SNV- | SNV | 1 | C | T | 35 | 47407 | 0.07382876 | 49.2571429 | rnr1Y285F_pe | FALSE | ARG | JSY3811 | 1 | rnr1Y285F_p | GCA | C>T | A/T+1 | G/C-1 | CG>TA | CG>TA | ATGCAGCAG | CAGTAAC | CGT | GCA | C | A | C>T | C>T_GCA | CG>TA_CAGTAAC | 4 | FALSE | FALSE |
| 40_v.txt | 32271 | 32271-SNV- | SNV | 1 | C | T | 117 | 62778 | 0.18637102 | 32.2131968 | rnr1Y285F_pe | FALSE | ARG | JSY3811 | 1 | rnr1Y285F_p | ACG | C>T | A/T+1 | G/C-1 | CG>TA | CG>TA | AAAATACGG | CGGGAAC | TGC | ACG | C | A | C>T | C>T_ACG | CG>TA_CGGGAAC | 4 | FALSE | FALSE |
| 40_v.txt | 32272 | 32272-SNV- | SNV | 1 | G | A | 15 | 62778 | 0.02389372 | 30.6 | rnr1Y285F_pe | FALSE | ARG | JSY3811 | 1 | rnr1Y285F_p | CGG | G>A | A/T+1 | G/C-1 | CG>TA | CG>TA | AAATACGGG | GGGAACC | GCC | GCC | C | T | C>T | C>T_GCC | CG>TA_GGGAACC | 4 | FALSE | FALSE |
| 40_v.txt | 32299 | 32299-SNV- | SNV | 1 | C | T | 125 | 62778 | 0.19911434 | 51.552 | rnr1Y285F_pe | FALSE | ARG | JSY3811 | 1 | rnr1Y285F_p | GCG | C>T | A/T+1 | G/C-1 | CG>TA | CG>TA | AATTTGCGG | CGGCAGA | CGC | GCG | C | A | C>T | C>T_GCG | CG>TA_CGGCAGA | 4 | FALSE | FALSE |
| 40_v.txt | 32344 | 32344-SNV- | SNV | 1 | G | A | 36 | 62778 | 0.05734493 | 56.8333333 | rnr1Y285F_pe | FALSE | ARG | JSY3811 | 1 | rnr1Y285F_p | GGC | G>A | A/T+1 | G/C-1 | CG>TA | CG>TA | TATGTGGCA | GCAAAAC | CCG | CCG | C | T | C>T | C>T_CCG | CG>TA_GCAAAAC | 4 | FALSE | FALSE |
| 40_v.txt | 32466 | 32466-SNV- | SNV | 1 | C | T | 68 | 62778 | 0.1083182 | 47.6764706 | rnr1Y285F_pe | FALSE | ARG | JSY3811 | 1 | rnr1Y285F_p | GCC | C>T | A/T+1 | G/C-1 | CG>TA | CG>TA | AGAGAGCCA | CCAATGT | CGG | GCC | C | A | C>T | C>T_GCC | CG>TA_CCAATGT | 4 | FALSE | FALSE |
| 40_v.txt | 32487 | 32487-SNV- | SNV | 1 | C | T | 50 | 62778 | 0.07964574 | 46.5 | rnr1Y285F_pe | FALSE | ARG | JSY3811 | 1 | rnr1Y285F_p | ACG | C>T | A/T+1 | G/C-1 | CG>TA | CG>TA | AAGATACGG | CGGAAAA | TGC | ACG | C | A | C>T | C>T_ACG | CG>TA_CGGAAAA | 4 | FALSE | FALSE |
| 40_v.txt | 32535 | 32535-SNV- | SNV | 1 | G | A | 73 | 62776 | 0.11628648 | 37.1370323 | rnr1Y285F_pe | FALSE | ARG | JSY3811 | 1 | rnr1Y285F_p | TGC | G>A | A/T+1 | G/C-1 | CG>TA | CG>TA | GGGTTTGCA | GCAGCTT | ACG | ACG | C | T | C>T | C>T_ACG | CG>TA_GCAGCTT | 4 | FALSE | FALSE |
| 40_v.txt | 32536 | 32536-SNV- | SNV | 1 | C | T | 85 | 62776 | 0.13540207 | 36.5401701 | rnr1Y285F_pe | FALSE | ARG | JSY3811 | 1 | rnr1Y285F_p | GCA | C>T | A/T+1 | G/C-1 | CG>TA | CG>TA | GGTTTGCAG | CAGCTTC | CGT | GCA | C | A | C>T | C>T_GCA | CG>TA_CAGCTTC | 4 | FALSE | FALSE |
| 40_v.txt | 32590 | 32590-SNV- | SNV | 1 | C | T | 90 | 53851 | 0.16712782 | 36.5777778 | rnr1Y285F_pe | FALSE | ARG | JSY3811 | 1 | rnr1Y285F_p | GCG | C>T | A/T+1 | G/C-1 | CG>TA | CG>TA | AGGCAGCGT | CGTTAAT | CGC | GCG | C | A | C>T | C>T_GCG | CG>TA_CGTTAAT | 4 | FALSE | FALSE |
| 40_v.txt | 32700 | 32700-SNV- | SNV | 1 | C | T | 56 | 53851 | 0.10399064 | 58.1607143 | rnr1Y285F_pe | FALSE | ARG | JSY3811 | 1 | rnr1Y285F_p | GCC | C>T | A/T+1 | G/C-1 | CG>TA | CG>TA | ACTGGGCCG | CCGGTAA | CGG | GCC | C | A | C>T | C>T_GCC | CG>TA_CCGGTAA | 4 | FALSE | FALSE |
| 40_v.txt | 32702 | 32702-SNV- | SNV | 1 | G | A | 33 | 53851 | 0.0612802 | 54.3333333 | rnr1Y285F_pe | FALSE | ARG | JSY3811 | 1 | rnr1Y285F_p | CGG | G>A | A/T+1 | G/C-1 | CG>TA | CG>TA | TGGGCCGGT | GGTAACC | GCC | GCC | C | T | C>T | C>T_GCC | CG>TA_GGTAACC | 4 | FALSE | FALSE |
| 40_v.txt | 32757 | 32757-SNV- | SNV | 1 | G | A | 38 | 53851 | 0.07056508 | 50.2631579 | rnr1Y285F_pe | FALSE | ARG | JSY3811 | 1 | rnr1Y285F_p | GGC | G>A | A/T+1 | G/C-1 | CG>TA | CG>TA | ATAATGGCT | GCTAAAA | CCG | CCG | C | T | C>T | C>T_CCG | CG>TA_GCTAAAA | 4 | FALSE | FALSE |
| 40_v.txt | 32776 | 32776-SNV- | SNV | 1 | C | T | 39 | 53851 | 0.07242205 | 49.2051282 | rnr1Y285F_pe | FALSE | ARG | JSY3811 | 1 | rnr1Y285F_p | GCG | C>T | A/T+1 | G/C-1 | CG>TA | CG>TA | TGGAAGCGA | CGACCCA | CGC | GCG | C | A | C>T | C>T_GCG | CG>TA_CGACCCA | 4 | FALSE | FALSE |
| 40_v.txt | 32798 | 32798-SNV- | SNV | 1 | G | A | 75 | 53851 | 0.13927318 | 47.04 | rnr1Y285F_pe | FALSE | ARG | JSY3811 | 1 | rnr1Y285F_p | CGT | G>A | A/T+1 | G/C-1 | CG>TA | CG>TA | TTCACCGTA | GTAATAT | GCA | GCA | C | T | C>T | C>T_GCA | CG>TA_GTAATAT | 4 | FALSE | FALSE |
| 40_v.txt | 32860 | 32860-SNV- | SNV | 1 | C | T | 49 | 53842 | 0.09100702 | 36.8979592 | rnr1Y285F_pe | FALSE | ARG | JSY3811 | 1 | rnr1Y285F_p | GCC | C>T | A/T+1 | G/C-1 | CG>TA | CG>TA | TCCATGCCG | CCGCCAG | CGG | GCC | C | A | C>T | C>T_GCC | CG>TA_CCGCCAG | 4 | FALSE | FALSE |
| 40_v.txt | 32862 | 32862-SNV- | SNV | 1 | G | A | 33 | 53842 | 0.06129044 | 33.3333333 | rnr1Y285F_pe | FALSE | ARG | JSY3811 | 1 | rnr1Y285F_p | CGC | G>A | A/T+1 | G/C-1 | CG>TA | CG>TA | CATGCCGCC | GCCAGTG | GCG | GCG | C | T | C>T | C>T_GCG | CG>TA_GCCAGTG | 4 | FALSE | FALSE |
| 40_v.txt | 32879 | 32879-SNV- | SNV | 1 | C | T | 63 | 87070 | 0.07235558 | 34.9783031 | rnr1Y285F_pe | FALSE | ARG | JSY3811 | 1 | rnr1Y285F_p | ACG | C>T | A/T+1 | G/C-1 | CG>TA | CG>TA | TTTGTACGTC | CGTCCAA | TGC | ACG | C | A | C>T | C>T_ACG | CG>TA_CGTCCAA | 4 | FALSE | FALSE |
| 40_v.txt | 32880 | 32880-SNV- | SNV | 1 | G | A | 49 | 87072 | 0.05627527 | 34.2403471 | rnr1Y285F_pe | FALSE | ARG | JSY3811 | 1 | rnr1Y285F_p | CGT | G>A | A/T+1 | G/C-1 | CG>TA | CG>TA | TTGTACGTC | GTCCAAA | GCA | GCA | C | T | C>T | C>T_GCA | CG>TA_GTCCAAA | 4 | FALSE | FALSE |
| 40_v.txt | 32900 | 32900-SNV- | SNV | 1 | G | A | 82 | 87077 | 0.09416953 | 35.289118 | rnr1Y285F_pe | FALSE | ARG | JSY3811 | 1 | rnr1Y285F_p | GGC | G>A | A/T+1 | G/C-1 | CG>TA | CG>TA | GACTTGGCC | GCCAACT | CCG | CCG | C | T | C>T | C>T_CCG | CG>TA_GCCAACT | 4 | FALSE | FALSE |
| 40_v.txt | 32901 | 32901-SNV- | SNV | 1 | C | T | 166 | 87077 | 0.19063587 | 31.9156627 | rnr1Y285F_pe | FALSE | ARG | JSY3811 | 1 | rnr1Y285F_p | GCC | C>T | A/T+1 | G/C-1 | CG>TA | CG>TA | ACTTGGCCA | CCAACTA | CGG | GCC | C | A | C>T | C>T_GCC | CG>TA_CCAACTA | 4 | FALSE | FALSE |
| 40_v.txt | 32922 | 32922-SNV- | SNV | 1 | G | A | 56 | 87077 | 0.0643109 | 49.7678571 | rnr1Y285F_pe | FALSE | ARG | JSY3811 | 1 | rnr1Y285F_p | GGC | G>A | A/T+1 | G/C-1 | CG>TA | CG>TA | TCCAGGGCA | GCAAAAG | CCG | CCG | C | T | C>T | C>T_CCG | CG>TA_GCAAAAG | 4 | FALSE | FALSE |
| 40_v.txt | 32935 | 32935-SNV- | SNV | 1 | C | T | 114 | 87077 | 0.13091861 | 40.4122807 | rnr1Y285F_pe | FALSE | ARG | JSY3811 | 1 | rnr1Y285F_p | GCC | C>T | A/T+1 | G/C-1 | CG>TA | CG>TA | TGATTGCCC | CCCAAGA | CGG | GCC | C | A | C>T | C>T_GCC | CG>TA_CCCAAGA | 4 | FALSE | FALSE |
| 40_v.txt | 32965 | 32965-SNV- | SNV | 1 | C | T | 82 | 87077 | 0.09416953 | 56.9878049 | rnr1Y285F_pe | FALSE | ARG | JSY3811 | 1 | rnr1Y285F_p | GCC | C>T | A/T+1 | G/C-1 | CG>TA | CG>TA | CATTGGCCG | CCGCACC | CGG | GCC | C | A | C>T | C>T_GCC | CG>TA_CCGCACC | 4 | FALSE | FALSE |
| 40_v.txt | 32967 | 32967-SNV- | SNV | 1 | G | A | 184 | 87077 | 0.21130723 | 59.2554348 | rnr1Y285F_pe | FALSE | ARG | JSY3811 | 1 | rnr1Y285F_p | CGC | G>A | A/T+1 | G/C-1 | CG>TA | CG>TA | TTGGCCGCA | GCACCAA | GCG | GCG | C | T | C>T | C>T_GCG | CG>TA_GCACCAA | 4 | FALSE | FALSE |
| 40_v.txt | 32977 | 32977-SNV- | SNV | 1 | C | T | 57 | 87077 | 0.06545931 | 56.6666667 | rnr1Y285F_pe | FALSE | ARG | JSY3811 | 1 | rnr1Y285F_p | GCT | C>T | A/T+1 | G/C-1 | CG>TA | CG>TA | CAAATGCTG | CTGGAGA | CGA | GCT | C | A | C>T | C>T_GCT | CG>TA_CTGGAGA | 4 | FALSE | FALSE |
| 40_v.txt | 33056 | 33056-SNV- | SNV | 1 | C | T | 122 | 87077 | 0.14010588 | 55.3606557 | rnr1Y285F_pe | FALSE | ARG | JSY3811 | 1 | rnr1Y285F_p | GCG | C>T | A/T+1 | G/C-1 | CG>TA | CG>TA | GGACTGCGT | CGTGACA | CGC | GCG | C | A | C>T | C>T_GCG | CG>TA_CGTGACA | 4 | FALSE | FALSE |
| 40_v.txt | 33069 | 33069-SNV- | SNV | 1 | G | A | 65 | 87077 | 0.07464658 | 54.6608204 | rnr1Y285F_pe | FALSE | ARG | JSY3811 | 1 | rnr1Y285F_p | TGC | G>A | A/T+1 | G/C-1 | CG>TA | CG>TA | GAATATGCC | GCCAAAG | ACG | ACG | C | T | C>T | C>T_ACG | CG>TA_GCCAAAG | 4 | FALSE | FALSE |
| 40_v.txt | 33070 | 33070-SNV- | SNV | 1 | C | T | 95 | 87071 | 0.10910636 | 55.831579 | rnr1Y285F_pe | FALSE | ARG | JSY3811 | 1 | rnr1Y285F_p | GCC | C>T | A/T+1 | G/C-1 | CG>TA | CG>TA | AATATGCCA | CCAAAGA | CGG | GCC | C | A | C>T | C>T_GCC | CG>TA_CCAAAGA | 4 | FALSE | FALSE |
| 40_v.txt | 33104 | 33104-SNV- | SNV | 1 | G | A | 67 | 87077 | 0.0769434 | 44.7014925 | rnr1Y285F_pe | FALSE | ARG | JSY3811 | 1 | rnr1Y285F_p | CGC | G>A | A/T+1 | G/C-1 | CG>TA | CG>TA | AAGAGCGCC | GCCCACT | GCG | GCG | C | T | C>T | C>T_GCG | CG>TA_GCCCACT | 4 | FALSE | FALSE |
| 40_v.txt | 33113 | 33113-SNV- | SNV | 1 | G | A | 44 | 87077 | 0.05052999 | 44.0675833 | rnr1Y285F_pe | FALSE | ARG | JSY3811 | 1 | rnr1Y285F_p | GGC | G>A | A/T+1 | G/C-1 | CG>TA | CG>TA | CACTGGGCC | GCCGGCG | CCG | CCG | C | T | C>T | C>T_CCG | CG>TA_GCCGGCG | 4 | FALSE | FALSE |
| 40_v.txt | 33114 | 33114-SNV- | SNV | 1 | C | T | 88 | 87077 | 0.10105998 | 41.9195044 | rnr1Y285F_pe | FALSE | ARG | JSY3811 | 1 | rnr1Y285F_p | GCC | C>T | A/T+1 | G/C-1 | CG>TA | CG>TA | ACTGGGCCG | CCGGCGT | CGG | GCC | C | A | C>T | C>T_GCC | CG>TA_CCGGCGT | 4 | FALSE | FALSE |
| 40_v.txt | 33118 | 33118-SNV- | SNV | 1 | C | T | 83 | 87077 | 0.09531794 | 39.9512468 | rnr1Y285F_pe | FALSE | ARG | JSY3811 | 1 | rnr1Y285F_p | GCG | C>T | A/T+1 | G/C-1 | CG>TA | CG>TA | GGCCGGCGT | CGTTGGT | CGC | GCG | C | A | C>T | C>T_GCG | CG>TA_CGTTGGT | 4 | FALSE | FALSE |
| 40_v.txt | 33119 | 33119-SNV- | SNV | 1 | G | A | 63 | 87077 | 0.07234976 | 44.284708 | rnr1Y285F_pe | FALSE | ARG | JSY3811 | 1 | rnr1Y285F_p | CGT | G>A | A/T+1 | G/C-1 | CG>TA | CG>TA | GCCGGCGTT | GTTGGTC | GCA | GCA | C | T | C>T | C>T_GCA | CG>TA_GTTGGTC | 4 | FALSE | FALSE |
| 40_v.txt | 33178 | 33178-SNV- | SNV | 1 | C | T | 47 | 87057 | 0.05398762 | 34.7234043 | rnr1Y285F_pe | FALSE | ARG | JSY3811 | 1 | rnr1Y285F_p | GCA | C>T | A/T+1 | G/C-1 | CG>TA | CG>TA | CAAGGGCAA | CAATCAT | CGT | GCA | C | A | C>T | C>T_GCA | CG>TA_CAATCAT | 4 | FALSE | FALSE |
| 40_v.txt | 33258 | 33258-SNV- | SNV | 1 | C | T | 80 | 51277 | 0.15601537 | 43.9375 | rnr1Y285F_pe | FALSE | ARG | JSY3811 | 1 | rnr1Y285F_p | GCC | C>T | A/T+1 | G/C-1 | CG>TA | CG>TA | CCTATGCCAT | CCATCCT | CGG | GCC | C | A | C>T | C>T_GCC | CG>TA_CCATCCT | 4 | FALSE | FALSE |
| 40_v.txt | 33275 | 33275-SNV- | SNV | 1 | C | T | 43 | 51277 | 0.08385826 | 50.6488685 | rnr1Y285F_pe | FALSE | ARG | JSY3811 | 1 | rnr1Y285F_p | ACG | C>T | A/T+1 | G/C-1 | CG>TA | CG>TA | AGAGAACGT | CGTATCC | TGC | ACG | C | A | C>T | C>T_ACG | CG>TA_CGTATCC | 4 | FALSE | FALSE |
| 40_v.txt | 33276 | 33276-SNV- | SNV | 1 | G | A | 61 | 51275 | 0.11896636 | 48.4740806 | rnr1Y285F_pe | FALSE | ARG | JSY3811 | 1 | rnr1Y285F_p | CGT | G>A | A/T+1 | G/C-1 | CG>TA | CG>TA | GAGAACGTA | GTATCCT | GCA | GCA | C | T | C>T | C>T_GCA | CG>TA_GTATCCT | 4 | FALSE | FALSE |
| 40_v.txt | 33284 | 33284-SNV- | SNV | 1 | G | A | 54 | 51277 | 0.10531037 | 45.3703704 | rnr1Y285F_pe | FALSE | ARG | JSY3811 | 1 | rnr1Y285F_p | CGC | G>A | A/T+1 | G/C-1 | CG>TA | CG>TA | ATCCTCGCC | GCCATTT | GCG | GCG | C | T | C>T | C>T_GCG | CG>TA_GCCATTT | 4 | FALSE | FALSE |
| 40_v.txt | 33309 | 33309-SNV- | SNV | 1 | C | T | 52 | 51277 | 0.10140999 | 55.7074295 | rnr1Y285F_pe | FALSE | ARG | JSY3811 | 1 | rnr1Y285F_p | GCG | C>T | A/T+1 | G/C-1 | CG>TA | CG>TA | AAAGAGCGC | CGCAATG | CGC | GCG | C | A | C>T | C>T_GCG | CG>TA_CGCAATG | 4 | FALSE | FALSE |
| 40_v.txt | 33310 | 33310-SNV- | SNV | 1 | G | A | 41 | 51271 | 0.07996723 | 56.3823441 | rnr1Y285F_pe | FALSE | ARG | JSY3811 | 1 | rnr1Y285F_p | CGC | G>A | A/T+1 | G/C-1 | CG>TA | CG>TA | AAGAGCGCA | GCAATGG | GCG | GCG | C | T | C>T | C>T_GCG | CG>TA_GCAATGG | 4 | FALSE | FALSE |
| 40_v.txt | 33392 | 33392-SNV- | SNV | 1 | G | A | 51 | 51277 | 0.0994598 | 52.1960784 | rnr1Y285F_pe | FALSE | ARG | JSY3811 | 1 | rnr1Y285F_p | CGT | G>A | A/T+1 | G/C-1 | CG>TA | CG>TA | AACGTCGTG | GTGAAAG | GCA | GCA | C | T | C>T | C>T_GCA | CG>TA_GTGAAAG | 4 | FALSE | FALSE |
| 40_v.txt | 33412 | 33412-SNV- | SNV | 1 | G | A | 79 | 51277 | 0.15406518 | 47.4303798 | rnr1Y285F_pe | FALSE | ARG | JSY3811 | 1 | rnr1Y285F_p | GGC | G>A | A/T+1 | G/C-1 | CG>TA | CG>TA | TGACCGGCT | GCTCATT | CCG | CCG | C | T | C>T | C>T_CCG | CG>TA_GCTCATT | 4 | FALSE | FALSE |
| 40_v.txt | 33474 | 33474-SNV- | SNV | 1 | C | T | 76 | 51277 | 0.1482146 | 33.916245 | rnr1Y285F_pe | FALSE | ARG | JSY3811 | 1 | rnr1Y285F_p | GCC | C>T | A/T+1 | G/C-1 | CG>TA | CG>TA | GCTATGCCTT | CCTTTTT | CGG | GCC | C | A | C>T | C>T_GCC | CG>TA_CCTTTTT | 4 | FALSE | FALSE |
| 40_v.txt | 31658 | 31658-SNV- | SNV | 1 | C | G | 2 | 24213 | 8.26E-03 | 37.9998926 | rnr1Y285F_pe | FALSE | ARG | JSY3811 | 1 | rnr1Y285F_p | GCG | C>G | G/C+1 | G/C-1 | CG>GC | CG>GC | AAATGGCGT | CGTGGAA | CGC | GCG | C | C | C>G | C>G_GCG | CG>GC_CGTGGAA | 4 | FALSE | FALSE |
| 40_v.txt | 32176 | 32176-SNV- | SNV | 1 | C | G | 6 | 47407 | 0.01265636 | 45.5 | rnr1Y285F_pe | FALSE | ARG | JSY3811 | 1 | rnr1Y285F_p | GCA | C>G | G/C+1 | G/C-1 | CG>GC | CG>GC | ATGCAGCAG | CAGTAAC | CGT | GCA | C | C | C>G | C>G_GCA | CG>GC_CAGTAAC | 4 | FALSE | FALSE |
| 40_v.txt | 32535 | 32535-SNV- | SNV | 1 | G | C | 4 | 62776 | 6.37E-03 | 37.500018 | rnr1Y285F_pe | FALSE | ARG | JSY3811 | 1 | rnr1Y285F_p | TGC | G>C | G/C+1 | G/C-1 | CG>GC | CG>GC | GGGTTTGCA | GCAGCTT | ACG | ACG | C | G | C>G | C>G_ACG | CG>GC_GCAGCTT | 4 | FALSE | FALSE |
| 40_v.txt | 32536 | 32536-SNV- | SNV | 1 | C | G | 4 | 62776 | 6.37E-03 | 37.2489681 | rnr1Y285F_pe | FALSE | ARG | JSY3811 | 1 | rnr1Y285F_p | GCA | C>G | G/C+1 | G/C-1 | CG>GC | CG>GC | GGTTTGCAG | CAGCTTC | CGT | GCA | C | C | C>G | C>G_GCA | CG>GC_CAGCTTC | 4 | FALSE | FALSE |
| 40_v.txt | 32901 | 32901-SNV- | SNV | 1 | C | G | 7 | 87077 | 8.04E-03 | 31.2857143 | rnr1Y285F_pe | FALSE | ARG | JSY3811 | 1 | rnr1Y285F_p | GCC | C>G | G/C+1 | G/C-1 | CG>GC | CG>GC | ACTTGGCCA | CCAACTA | CGG | GCC | C | C | C>G | C>G_GCC | CG>GC_CCAACTA | 4 | FALSE | FALSE |
| 40_v.txt | 33070 | 33070-SNV- | SNV | 1 | C | G | 4 | 87071 | 4.59E-03 | 54.25 | rnr1Y285F_pe | FALSE | ARG | JSY3811 | 1 | rnr1Y285F_p | GCC | C>G | G/C+1 | G/C-1 | CG>GC | CG>GC | AATATGCCA | CCAAAGA | CGG | GCC | C | C | C>G | C>G_GCC | CG>GC_CCAAAGA | 4 | FALSE | FALSE |
| 40_v.txt | 33118 | 33118-SNV- | SNV | 1 | C | G | 6 | 87077 | 6.89E-03 | 33.0002606 | rnr1Y285F_pe | FALSE | ARG | JSY3811 | 1 | rnr1Y285F_p | GCG | C>G | G/C+1 | G/C-1 | CG>GC | CG>GC | GGCCGGCGT | CGTTGGT | CGC | GCG | C | C | C>G | C>G_GCG | CG>GC_CGTTGGT | 4 | FALSE | FALSE |
| 40_v.txt | 33310 | 33310-SNV- | SNV | 1 | G | C | 6 | 51271 | 0.01170252 | 44.6622144 | rnr1Y285F_pe | FALSE | ARG | JSY3811 | 1 | rnr1Y285F_p | CGC | G>C | G/C+1 | G/C-1 | CG>GC | CG>GC | AAGAGCGCA | GCAATGG | GCG | GCG | C | G | C>G | C>G_GCG | CG>GC_GCAATGG | 4 | FALSE | FALSE |
| 40_v.txt | 31657 | 31657-SNV- | SNV | 1 | G | T | 160 | 24213 | 0.66080205 | 18.884815 | rnr1Y285F_pe | FALSE | ARG | JSY3811 | 1 | rnr1Y285F_p | GGC | G>T | A/T+1 | G/C-1 | CG>AT | CG>AT | GAAATGGCG | GCGTGGA | CCG | CCG | C | A | C>A | C>A_CCG | CG>AT_GCGTGGA | 4 | FALSE | FALSE |
| 40_v.txt | 32272 | 32272-SNV- | SNV | 1 | G | T | 18 | 62778 | 0.02867247 | 30.7222222 | rnr1Y285F_pe | FALSE | ARG | JSY3811 | 1 | rnr1Y285F_p | CGG | G>T | A/T+1 | G/C-1 | CG>AT | CG>AT | AAATACGGG | GGGAACC | GCC | GCC | C | A | C>A | C>A_GCC | CG>AT_GGGAACC | 4 | FALSE | FALSE |
| 40_v.txt | 32811 | 32811-SNV- | SNV | 1 | G | T | 17 | 53851 | 0.03156859 | 42.6470588 | rnr1Y285F_pe | FALSE | ARG | JSY3811 | 1 | rnr1Y285F_p | AGG | G>T | A/T+1 | G/C-1 | CG>AT | CG>AT | TTGACAGGG | GGGAACA | TCC | TCC | C | A | C>A | C>A_TCC | CG>AT_GGGAACA | 4 | FALSE | FALSE |

| | | | | | | | | | | | | | | | | | | | | | | | | | | | | | | | | | | |
|---|---|---|---|---|---|---|---|---|---|---|---|---|---|---|---|---|---|---|---|---|---|---|---|---|---|---|---|---|---|---|---|---|---|---|
| 40_v.txt | 32901 | 32901-SNV-( | SNV | 1 | C | A | 6 | 87077 | 6.89E-03 | 38 | rnr1Y285F_pe | FALSE | ARG | JSY3811 | 1 | rnr1Y285F_p | GCC | C>A | A/T+1 | G/C-1 | CG>AT | CG>AT | ACTTGGCCA/ | CCAACTA | CGG | GCC | C | T | C>A | C>A_GCC | CG>AT_CCAACTA | 4 | FALSE | FALSE |
| 40_v.txt | 33050 | 33050-SNV-( | SNV | 1 | G | T | 24 | 87077 | 0.02756181 | 52.375 | rnr1Y285F_pe | FALSE | ARG | JSY3811 | 1 | rnr1Y285F_p | AGG | G>T | A/T+1 | G/C-1 | CG>AT | CG>AT | ACCCAAGGA | GGACTGC | TCC | TCC | C | A | C>A | C>A_TCC | CG>AT_GGACTGC | 4 | FALSE | FALSE |
| 40_v.txt | 33070 | 33070-SNV-( | SNV | 1 | C | A | 17 | 87071 | 0.0195243 | 29.9411765 | rnr1Y285F_pe | FALSE | ARG | JSY3811 | 1 | rnr1Y285F_p | GCC | C>A | A/T+1 | G/C-1 | CG>AT | CG>AT | AATATGCCA/ | CCAAAGA | CGG | GCC | C | T | C>A | C>A_GCC | CG>AT_CCAAAGA | 4 | FALSE | FALSE |
| 40_v.txt | 33175 | 33175-SNV-( | SNV | 1 | G | T | 25 | 87060 | 0.02871583 | 37.8 | rnr1Y285F_pe | FALSE | ARG | JSY3811 | 1 | rnr1Y285F_p | AGG | G>T | A/T+1 | G/C-1 | CG>AT | CG>AT | CACCAAGGG | GGGCAAT | TCC | TCC | C | A | C>A | C>A_TCC | CG>AT_GGGCAAT | 4 | FALSE | FALSE |
| 40_v.txt | 33276 | 33276-SNV-( | SNV | 1 | G | T | 18 | 51275 | 0.03510483 | 14.5547631 | rnr1Y285F_pe | FALSE | ARG | JSY3811 | 1 | rnr1Y285F_p | CGT | G>T | A/T+1 | G/C-1 | CG>AT | CG>AT | GAGAACGTA | GTATCCT | GCA | GCA | C | A | C>A | C>A_GCA | CG>AT_GTATCCT | 4 | FALSE | FALSE |
| 40_v.txt | 33349 | 33349-SNV-( | SNV | 1 | G | T | 230 | 51277 | 0.44854418 | 14.6652174 | rnr1Y285F_pe | FALSE | ARG | JSY3811 | 1 | rnr1Y285F_p | GGT | G>T | A/T+1 | G/C-1 | CG>AT | CG>AT | TCAATGGTA' | GTATTGA | CCA | CCA | C | A | C>A | C>A_CCA | CG>AT_GTATTGA | 4 | FALSE | FALSE |
| 40_v.txt | 33474 | 33474-SNV-( | SNV | 1 | C | A | 109 | 51277 | 0.21257094 | 34.2815949 | rnr1Y285F_pe | FALSE | ARG | JSY3811 | 1 | rnr1Y285F_p | GCC | C>A | A/T+1 | G/C-1 | CG>AT | CG>AT | GCTATGCCTT | CCTTTTT | CGG | GCC | C | T | C>A | C>A_GCC | CG>AT_CCTTTTT | 4 | FALSE | FALSE |
| 40_v.txt | 33475 | 33475-SNV-( | SNV | 1 | C | A | 34 | 51277 | 0.06630653 | 35.7025565 | rnr1Y285F_pe | FALSE | ARG | JSY3811 | 1 | rnr1Y285F_p | CCT | C>A | A/T+1 | G/C-1 | CG>AT | CG>AT | CTATGCCTTT | CTTTTTT | GGA | CCT | C | T | C>A | C>A_CCT | CG>AT_CTTTTTT | 4 | FALSE | FALSE |
| 40_v.txt | 31723 | 31723-Insert | Insertion | 1 | - | A | 2 | 24213 | 8.26E-03 | 56.5 | rnr1Y285F_pe | FALSE | ARG | JSY3811 | 1 | rnr1Y285F_p | CAA | nope | A/T+1 | >1bpdel | none | A/T+1 | TGTCCCAAA/ | AAAAAGT | GTT | CAA | - | T | NA | NA_CAA | A/T+1_AAAAAGT | 4 | FALSE | FALSE |
| 40_v.txt | 32081 | 32081-Insert | Insertion | 1 | - | A | 32 | 47407 | 0.06750058 | 63.875 | rnr1Y285F_pe | FALSE | ARG | JSY3811 | 1 | rnr1Y285F_p | CAA | nope | A/T+1 | >1bpdel | none | A/T+1 | CCATGCAAA/ | AAAAAAG | GTT | CAA | - | T | NA | NA_CAA | A/T+1_AAAAAAG | 4 | FALSE | FALSE |
| 40_v.txt | 32522 | 32522-Insert | Insertion | 1 | - | T | 2 | 62777 | 3.19E-03 | 38 | rnr1Y285F_pe | FALSE | ARG | JSY3811 | 1 | rnr1Y285F_p | ATT | nope | A/T+1 | >1bpdel | none | A/T+1 | AACGGATTT' | TTTTCTG | TAA | ATT | - | A | NA | NA_ATT | A/T+1_TTTTCTG | 4 | FALSE | FALSE |
| 40_v.txt | 32804 | 32804-Insert | Insertion | 1 | - | T | 2 | 53851 | 3.71E-03 | 51 | rnr1Y285F_pe | FALSE | ARG | JSY3811 | 1 | rnr1Y285F_p | ATT | nope | A/T+1 | >1bpdel | none | A/T+1 | GTAATATTTC | TTTGACA | TAA | ATT | - | A | NA | NA_ATT | A/T+1_TTTGACA | 4 | FALSE | FALSE |
| 40_v.txt | 32842 | 32842-Insert | Insertion | 1 | - | A | 19 | 53850 | 0.03528319 | 40.5263158 | rnr1Y285F_pe | FALSE | ARG | JSY3811 | 1 | rnr1Y285F_p | CAA | nope | A/T+1 | >1bpdel | none | A/T+1 | TTACCCAAA/ | AAAAAAT | GTT | CAA | - | T | NA | NA_CAA | A/T+1_AAAAAAT | 4 | FALSE | FALSE |
| 40_v.txt | 32884 | 32884-Insert | Insertion | 1 | - | A | 4 | 87076 | 4.59E-03 | 38 | rnr1Y285F_pe | FALSE | ARG | JSY3811 | 1 | rnr1Y285F_p | CAA | nope | A/T+1 | >1bpdel | none | A/T+1 | ACGTCCAAA/ | AAAATTG | GTT | CAA | - | T | NA | NA_CAA | A/T+1_AAAATTG | 4 | FALSE | FALSE |
| 40_v.txt | 33161 | 33161-Insert | Insertion | 1 | - | A | 2 | 87071 | 2.30E-03 | 38 | rnr1Y285F_pe | FALSE | ARG | JSY3811 | 1 | rnr1Y285F_p | CAA | nope | A/T+1 | >1bpdel | none | A/T+1 | TGTACCAAT/ | AATAGTA | GTT | CAA | - | T | NA | NA_CAA | A/T+1_AATAGTA | 4 | FALSE | FALSE |
| 40_v.txt | 33474 | 33474-Insert | Insertion | 1 | - | A | 1 | 51277 | 1.95E-03 | 34 | rnr1Y285F_pe | FALSE | ARG | JSY3811 | 1 | rnr1Y285F_p | GCC | nope | A/T+1 | >1bpdel | none | A/T+1 | GCTATGCCTT | CCTTTTT | CGG | GCC | - | T | NA | NA_GCC | A/T+1_CCTTTTT | 4 | FALSE | FALSE |
| 40_v.txt | 31679 | 31679-Deleti | Deletion | 1 | A | - | 5 | 24213 | 0.02065006 | 48.4 | rnr1Y285F_pe | FALSE | ARG | JSY3811 | 1 | rnr1Y285F_p | TAA | nope | >1bp-ins | A/T-1 | none | A/T-1 | AAAGGTAAT | AATAAAA | ATT | ATT | T | none | T>none | T>none_ATT | A/T-1_AATAAAA | 4 | FALSE | FALSE |
| 40_v.txt | 31723 | 31723-Deleti | Deletion | 1 | A | - | 24 | 24213 | 0.09912031 | 62.5 | rnr1Y285F_pe | FALSE | ARG | JSY3811 | 1 | rnr1Y285F_p | CAA | nope | >1bp-ins | A/T-1 | none | A/T-1 | TGTCCCAAA/ | AAAAAGT | GTT | GTT | T | none | T>none | T>none_GTT | A/T-1_AAAAAGT | 4 | FALSE | FALSE |
| 40_v.txt | 31904 | 31904-Deleti | Deletion | 1 | T | - | 12 | 24213 | 0.04956015 | 37.4166667 | rnr1Y285F_pe | FALSE | ARG | JSY3811 | 1 | rnr1Y285F_p | ATT | nope | >1bp-ins | A/T-1 | none | A/T-1 | ATTGAATTTT | TTTTGGT | TAA | ATT | T | none | nope | nope_ATT | A/T-1_TTTTGGT | 4 | FALSE | FALSE |
| 40_v.txt | 31913 | 31913-Deleti | Deletion | 1 | A | - | 6 | 24213 | 0.02478008 | 42.3333333 | rnr1Y285F_pe | FALSE | ARG | JSY3811 | 1 | rnr1Y285F_p | CAA | nope | >1bp-ins | A/T-1 | none | A/T-1 | TGGTGCAAA | AAAAGCC | GTT | GTT | T | none | T>none | T>none_GTT | A/T-1_AAAAGCC | 4 | FALSE | FALSE |
| 40_v.txt | 31952 | 31952-Deleti | Deletion | 1 | A | - | 6 | 47407 | 0.01265636 | 37.5 | rnr1Y285F_pe | FALSE | ARG | JSY3811 | 1 | rnr1Y285F_p | TAA | nope | >1bp-ins | A/T-1 | none | A/T-1 | CGTCATAAA' | AAATGTG | ATT | ATT | T | none | T>none | T>none_ATT | A/T-1_AAATGTG | 4 | FALSE | FALSE |
| 40_v.txt | 31972 | 31972-Deleti | Deletion | 1 | C | - | 15 | 47407 | 0.0316409 | 36.1333333 | rnr1Y285F_pe | FALSE | ARG | JSY3811 | 1 | rnr1Y285F_p | GCC | nope | >1bp-ins | G/C-1 | none | G/C-1 | AATAAGCCA. | CCAAGCC | CGG | GCC | C | none | nope | nope_GCC | G/C-1_CCAAGCC | 4 | FALSE | FALSE |
| 40_v.txt | 31979 | 31979-Deleti | Deletion | 1 | G | - | 8 | 47407 | 0.01687515 | 39.5 | rnr1Y285F_pe | FALSE | ARG | JSY3811 | 1 | rnr1Y285F_p | CGG | nope | >1bp-ins | G/C-1 | none | G/C-1 | CAAGCCGGG | GGGCATT | GCC | GCC | C | none | C>none | C>none_GCC | G/C-1_GGGCATT | 4 | FALSE | FALSE |
| 40_v.txt | 32069 | 32069-Deleti | Deletion | 1 | A | - | 10 | 47407 | 0.02109393 | 57.9 | rnr1Y285F_pe | FALSE | ARG | JSY3811 | 1 | rnr1Y285F_p | TAA | nope | >1bp-ins | A/T-1 | none | A/T-1 | TGAGATAAA | AAATAAC | ATT | ATT | T | none | T>none | T>none_ATT | A/T-1_AAATAAC | 4 | FALSE | FALSE |
| 40_v.txt | 32081 | 32081-Deleti | Deletion | 1 | A | - | 37 | 47406 | 0.07804919 | 61.7837841 | rnr1Y285F_pe | FALSE | ARG | JSY3811 | 1 | rnr1Y285F_p | CAA | nope | >1bp-ins | A/T-1 | none | A/T-1 | CCATGCAAA/ | AAAAAAG | GTT | GTT | T | none | T>none | T>none_GTT | A/T-1_AAAAAAG | 4 | FALSE | FALSE |
| 40_v.txt | 32133 | 32133-Deleti | Deletion | 1 | C | - | 6 | 47407 | 0.01265636 | 59.6666667 | rnr1Y285F_pe | FALSE | ARG | JSY3811 | 1 | rnr1Y285F_p | ACC | nope | >1bp-ins | G/C-1 | none | G/C-1 | TTGTCACCAC | CCACCAG | TGG | ACC | C | none | nope | nope_ACC | G/C-1_CCACCAG | 4 | FALSE | FALSE |
| 40_v.txt | 32210 | 32210-Deleti | Deletion | 1 | T | - | 14 | 47407 | 0.0295315 | 44.2857143 | rnr1Y285F_pe | FALSE | ARG | JSY3811 | 1 | rnr1Y285F_p | CTT | nope | >1bp-ins | A/T-1 | none | A/T-1 | ACCACCTTTG | TTTGGTG | GAA | CTT | T | none | nope | nope_CTT | A/T-1_TTTGGTG | 4 | FALSE | FALSE |
| 40_v.txt | 32231 | 32231-Deleti | Deletion | 1 | T | - | 6 | 47407 | 0.01265636 | 37.8333333 | rnr1Y285F_pe | FALSE | ARG | JSY3811 | 1 | rnr1Y285F_p | ATT | nope | >1bp-ins | A/T-1 | none | A/T-1 | CAGGAATTT/ | TTTAGGA | TAA | ATT | T | none | nope | nope_ATT | A/T-1_TTTAGGA | 4 | FALSE | FALSE |
| 40_v.txt | 32272 | 32272-Deleti | Deletion | 1 | G | - | 6 | 62778 | 9.56E-03 | 37.8333333 | rnr1Y285F_pe | FALSE | ARG | JSY3811 | 1 | rnr1Y285F_p | CGG | nope | >1bp-ins | G/C-1 | none | G/C-1 | AAATACGGG | GGGAACC | GCC | GCC | C | none | C>none | C>none_GCC | G/C-1_GGGAACC | 4 | FALSE | FALSE |
| 40_v.txt | 32324 | 32324-Deleti | Deletion | 1 | A | - | 6 | 62778 | 9.56E-03 | 64 | rnr1Y285F_pe | FALSE | ARG | JSY3811 | 1 | rnr1Y285F_p | TAA | nope | >1bp-ins | A/T-1 | none | A/T-1 | TAAGATAAC. | AACAGCG | ATT | ATT | T | none | T>none | T>none_ATT | A/T-1_AACAGCG | 4 | FALSE | FALSE |
| 40_v.txt | 32346 | 32346-Deleti | Deletion | 1 | A | - | 11 | 62778 | 0.01752206 | 55.2727273 | rnr1Y285F_pe | FALSE | ARG | JSY3811 | 1 | rnr1Y285F_p | CAA | nope | >1bp-ins | A/T-1 | none | A/T-1 | TGTGGCAAA | AAAACCT | GTT | GTT | T | none | T>none | T>none_GTT | A/T-1_AAAACCT | 4 | FALSE | FALSE |
| 40_v.txt | 32420 | 32420-Deleti | Deletion | 1 | T | - | 9 | 62778 | 0.01433623 | 61.1111111 | rnr1Y285F_pe | FALSE | ARG | JSY3811 | 1 | rnr1Y285F_p | GTT | nope | >1bp-ins | A/T-1 | none | A/T-1 | TGTTAGTTTA | TTTAGGG | CAA | GTT | T | none | nope | nope_GTT | A/T-1_TTTAGGG | 4 | FALSE | FALSE |
| 40_v.txt | 32442 | 32442-Deleti | Deletion | 1 | A | - | 12 | 62778 | 0.01911498 | 52.4166667 | rnr1Y285F_pe | FALSE | ARG | JSY3811 | 1 | rnr1Y285F_p | TAA | nope | >1bp-ins | A/T-1 | none | A/T-1 | GGAACTAAA | AAAAGTC | ATT | ATT | T | none | T>none | T>none_ATT | A/T-1_AAAAGTC | 4 | FALSE | FALSE |
| 40_v.txt | 32498 | 32498-Deleti | Deletion | 1 | T | - | 70 | 62778 | 0.11150403 | 43.5142857 | rnr1Y285F_pe | FALSE | ARG | JSY3811 | 1 | rnr1Y285F_p | CTT | nope | >1bp-ins | A/T-1 | none | A/T-1 | AACAACTTTT | TTTTTTG | GAA | CTT | T | none | nope | nope_CTT | A/T-1_TTTTTTG | 4 | FALSE | FALSE |
| 40_v.txt | 32528 | 32528-Deleti | Deletion | 1 | G | - | 12 | 62777 | 0.01911528 | 44.5833333 | rnr1Y285F_pe | FALSE | ARG | JSY3811 | 1 | rnr1Y285F_p | TGG | nope | >1bp-ins | G/C-1 | none | G/C-1 | TTTTCTGGGG | GGGGTTT | ACC | ACC | C | none | C>none | C>none_ACC | G/C-1_GGGGTTT | 4 | FALSE | FALSE |
| 40_v.txt | 32573 | 32573-Deleti | Deletion | 1 | T | - | 5 | 53851 | 9.29E-03 | 37.4 | rnr1Y285F_pe | FALSE | ARG | JSY3811 | 1 | rnr1Y285F_p | CTT | nope | >1bp-ins | A/T-1 | none | A/T-1 | AGTACCTTG/ | TTGAAAT | GAA | CTT | T | none | nope | nope_CTT | A/T-1_TTGAAAT | 4 | FALSE | FALSE |
| 40_v.txt | 32598 | 32598-Deleti | Deletion | 1 | A | - | 11 | 53851 | 0.02042673 | 40.0909091 | rnr1Y285F_pe | FALSE | ARG | JSY3811 | 1 | rnr1Y285F_p | CAA | nope | >1bp-ins | A/T-1 | none | A/T-1 | TTAATCAAA( | AAAGAGG | GTT | GTT | T | none | T>none | T>none_GTT | A/T-1_AAAGAGG | 4 | FALSE | FALSE |
| 40_v.txt | 32605 | 32605-Deleti | Deletion | 1 | A | - | 6 | 53851 | 0.01114185 | 37.6666667 | rnr1Y285F_pe | FALSE | ARG | JSY3811 | 1 | rnr1Y285F_p | GAA | nope | >1bp-ins | A/T-1 | none | A/T-1 | AAGAGGAAA | AAACCCA | CTT | CTT | T | none | T>none | T>none_CTT | A/T-1_AAACCCA | 4 | FALSE | FALSE |
| 40_v.txt | 32631 | 32631-Deleti | Deletion | 1 | T | - | 31 | 53851 | 0.05756625 | 45.483871 | rnr1Y285F_pe | FALSE | ARG | JSY3811 | 1 | rnr1Y285F_p | GTT | nope | >1bp-ins | A/T-1 | none | A/T-1 | CCTTCGTTTT | TTTTTAT | CAA | GTT | T | none | nope | nope_GTT | A/T-1_TTTTTAT | 4 | FALSE | FALSE |
| 40_v.txt | 32658 | 32658-Deleti | Deletion | 1 | C | - | 11 | 53851 | 0.02042673 | 60.9090909 | rnr1Y285F_pe | FALSE | ARG | JSY3811 | 1 | rnr1Y285F_p | ACC | nope | >1bp-ins | G/C-1 | none | G/C-1 | CCTGGACCC( | CCCCAGG | TGG | ACC | C | none | nope | nope_ACC | G/C-1_CCCCAGG | 4 | FALSE | FALSE |
| 40_v.txt | 32670 | 32670-Deleti | Deletion | 1 | G | - | 5 | 53851 | 9.29E-03 | 53.6 | rnr1Y285F_pe | FALSE | ARG | JSY3811 | 1 | rnr1Y285F_p | TGG | nope | >1bp-ins | G/C-1 | none | G/C-1 | GCACCTGGG | GGGTTTC | ACC | ACC | C | none | C>none | C>none_ACC | G/C-1_GGGTTTC | 4 | FALSE | FALSE |
| 40_v.txt | 32707 | 32707-Deleti | Deletion | 1 | C | - | 12 | 53851 | 0.02228371 | 62.5833333 | rnr1Y285F_pe | FALSE | ARG | JSY3811 | 1 | rnr1Y285F_p | ACC | nope | >1bp-ins | G/C-1 | none | G/C-1 | CGGTAACCC( | CCCCAGC | TGG | ACC | C | none | nope | nope_ACC | G/C-1_CCCCAGC | 4 | FALSE | FALSE |
| 40_v.txt | 32728 | 32728-Deleti | Deletion | 1 | A | - | 11 | 53851 | 0.02042673 | 56.8181818 | rnr1Y285F_pe | FALSE | ARG | JSY3811 | 1 | rnr1Y285F_p | CAA | nope | >1bp-ins | A/T-1 | none | A/T-1 | CCATACAAA/ | AAAAACA | GTT | GTT | T | none | T>none | T>none_GTT | A/T-1_AAAAACA | 4 | FALSE | FALSE |
| 40_v.txt | 32747 | 32747-Deleti | Deletion | 1 | C | - | 6 | 53851 | 0.01114185 | 54 | rnr1Y285F_pe | FALSE | ARG | JSY3811 | 1 | rnr1Y285F_p | ACC | nope | >1bp-ins | G/C-1 | none | G/C-1 | TAGAAACCC( | CCCGATA | TGG | ACC | C | none | nope | nope_ACC | G/C-1_CCCGATA | 4 | FALSE | FALSE |
| 40_v.txt | 32779 | 32779-Deleti | Deletion | 1 | C | - | 8 | 53851 | 0.01485581 | 55.375 | rnr1Y285F_pe | FALSE | ARG | JSY3811 | 1 | rnr1Y285F_p | ACC | nope | >1bp-ins | G/C-1 | none | G/C-1 | AAGCGACCC. | CCCAGAA | TGG | ACC | C | none | nope | nope_ACC | G/C-1_CCCAGAA | 4 | FALSE | FALSE |
| 40_v.txt | 32842 | 32842-Deleti | Deletion | 1 | A | - | 62 | 53850 | 0.11513463 | 43.8709675 | rnr1Y285F_pe | FALSE | ARG | JSY3811 | 1 | rnr1Y285F_p | CAA | nope | >1bp-ins | A/T-1 | none | A/T-1 | TTACCCAAA/ | AAAAAAT | GTT | GTT | T | none | T>none | T>none_GTT | A/T-1_AAAAAAT | 4 | FALSE | FALSE |
| 40_v.txt | 32884 | 32884-Deleti | Deletion | 1 | A | - | 10 | 87076 | 0.01148422 | 37.5999997 | rnr1Y285F_pe | FALSE | ARG | JSY3811 | 1 | rnr1Y285F_p | CAA | nope | >1bp-ins | A/T-1 | none | A/T-1 | ACGTCCAAA/ | AAAATTG | GTT | GTT | T | none | T>none | T>none_GTT | A/T-1_AAAATTG | 4 | FALSE | FALSE |
| 40_v.txt | 32901 | 32901-Deleti | Deletion | 1 | C | - | 4 | 87077 | 4.59E-03 | 35.75 | rnr1Y285F_pe | FALSE | ARG | JSY3811 | 1 | rnr1Y285F_p | GCC | nope | >1bp-ins | G/C-1 | none | G/C-1 | ACTTGGCCA/ | CCAACTA | CGG | GCC | C | none | nope | nope_GCC | G/C-1_CCAACTA | 4 | FALSE | FALSE |
| 40_v.txt | 32924 | 32924-Deleti | Deletion | 1 | A | - | 79 | 87077 | 0.0907243 | 49.9113924 | rnr1Y285F_pe | FALSE | ARG | JSY3811 | 1 | rnr1Y285F_p | CAA | nope | >1bp-ins | A/T-1 | none | A/T-1 | CAGGGCAAA | AAAAGTG | GTT | GTT | T | none | T>none | T>none_GTT | A/T-1_AAAAGTG | 4 | FALSE | FALSE |
| 40_v.txt | 32935 | 32935-Deleti | Deletion | 1 | C | - | 17 | 87077 | 0.01952295 | 38.8235294 | rnr1Y285F_pe | FALSE | ARG | JSY3811 | 1 | rnr1Y285F_p | GCC | nope | >1bp-ins | G/C-1 | none | G/C-1 | TGATTGCCC/ | CCCAAGA | CGG | GCC | C | none | nope | nope_GCC | G/C-1_CCCAAGA | 4 | FALSE | FALSE |
| 40_v.txt | 33000 | 33000-Deleti | Deletion | 1 | A | - | 9 | 87077 | 0.01033568 | 64 | rnr1Y285F_pe | FALSE | ARG | JSY3811 | 1 | rnr1Y285F_p | GAA | nope | >1bp-ins | A/T-1 | none | A/T-1 | TGTGAGAAA | AAAACTG | CTT | CTT | T | none | T>none | T>none_CTT | A/T-1_AAAACTG | 4 | FALSE | FALSE |
| 40_v.txt | 33083 | 33083-Deleti | Deletion | 1 | A | - | 9 | 87077 | 0.01033568 | 54.5555556 | rnr1Y285F_pe | FALSE | ARG | JSY3811 | 1 | rnr1Y285F_p | TAA | nope | >1bp-ins | A/T-1 | none | A/T-1 | ACCCATAAA' | AAATAAA | ATT | ATT | T | none | T>none | T>none_ATT | A/T-1_AAATAAA | 4 | FALSE | FALSE |
| 40_v.txt | 33147 | 33147-Deleti | Deletion | 1 | A | - | 9 | 87077 | 0.01033568 | 45.8888889 | rnr1Y285F_pe | FALSE | ARG | JSY3811 | 1 | rnr1Y285F_p | GAA | nope | >1bp-ins | A/T-1 | none | A/T-1 | CCAATGAAA. | AAAAGAC | CTT | CTT | T | none | T>none | T>none_CTT | A/T-1_AAAAGAC | 4 | FALSE | FALSE |
| 40_v.txt | 33168 | 33168-Deleti | Deletion | 1 | C | - | 8 | 87069 | 9.19E-03 | 37.75 | rnr1Y285F_pe | FALSE | ARG | JSY3811 | 1 | rnr1Y285F_p | ACC | nope | >1bp-ins | G/C-1 | none | G/C-1 | ATAGTACCA( | CCACCAA | TGG | ACC | C | none | nope | nope_ACC | G/C-1_CCACCAA | 4 | FALSE | FALSE |
| 40_v.txt | 33175 | 33175-Deleti | Deletion | 1 | G | - | 6 | 87060 | 6.89E-03 | 37.8333333 | rnr1Y285F_pe | FALSE | ARG | JSY3811 | 1 | rnr1Y285F_p | AGG | nope | >1bp-ins | G/C-1 | none | G/C-1 | CACCAAGGG | GGGCAAT | TCC | TCC | C | none | C>none | C>none_TCC | G/C-1_GGGCAAT | 4 | FALSE | FALSE |
| 40_v.txt | 33288 | 33288-Deleti | Deletion | 1 | T | - | 7 | 51277 | 0.01365135 | 46.2857143 | rnr1Y285F_pe | FALSE | ARG | JSY3811 | 1 | rnr1Y285F_p | ATT | nope | >1bp-ins | A/T-1 | none | A/T-1 | TCGCCATTTA | TTTACTC | TAA | ATT | T | none | nope | nope_ATT | A/T-1_TTTACTC | 4 | FALSE | FALSE |
| 40_v.txt | 33303 | 33303-Deleti | Deletion | 1 | A | - | 14 | 51277 | 0.02730269 | 61.2857143 | rnr1Y285F_pe | FALSE | ARG | JSY3811 | 1 | rnr1Y285F_p | GAA | nope | >1bp-ins | A/T-1 | none | A/T-1 | GTCGGGAAA | AAAGAGC | CTT | CTT | T | none | T>none | T>none_CTT | A/T-1_AAAGAGC | 4 | FALSE | FALSE |
| 40_v.txt | 33474 | 33474-Deleti | Deletion | 1 | C | - | 138 | 51277 | 0.26912651 | 30.2826083 | rnr1Y285F_pe | FALSE | ARG | JSY3811 | 1 | rnr1Y285F_p | GCC | nope | >1bp-ins | G/C-1 | none | G/C-1 | GCTATGCCTT | CCTTTTT | CGG | GCC | C | none | nope | nope_GCC | G/C-1_CCTTTTT | 4 | FALSE | FALSE |
| 41_v.txt | 31723 | 31723-SNV-/ | SNV | 1 | A | C | 7 | 9117 | 0.07677964 | 60 | rnr1Y285F_pe | FALSE | ARG | JSY3811 | 2 | rnr1Y285F_p | CAA | A>C | G/C+1 | A/T-1 | TA>GC | TA>GC | TGTCCCAAA/ | AAAAAGT | GTT | GTT | T | G | T>G | T>G_GTT | TA>GC_AAAAAGT | 4 | FALSE | FALSE |
| 41_v.txt | 32842 | 32842-SNV-/ | SNV | 1 | A | C | 3 | 25127 | 0.01193935 | 46.666628 | rnr1Y285F_pe | FALSE | ARG | JSY3811 | 2 | rnr1Y285F_p | CAA | A>C | G/C+1 | A/T-1 | TA>GC | TA>GC | TTACCCAAA/ | AAAAAAT | GTT | GTT | T | G | T>G | T>G_GTT | TA>GC_AAAAAAT | 4 | FALSE | FALSE |
| 41_v.txt | 33269 | 33269-SNV-/ | SNV | 1 | A | C | 130 | 17688 | 0.73496156 | 15.4076923 | rnr1Y285F_pe | FALSE | ARG | JSY3811 | 2 | rnr1Y285F_p | TAG | A>C | G/C+1 | A/T-1 | TA>GC | TA>GC | CTCCATAGA( | AGAGAAC | ATC | ATC | T | G | T>G | T>G_ATC | TA>GC_AGAGAAC | 4 | FALSE | FALSE |
| 41_v.txt | 32498 | 32498-SNV-T | SNV | 1 | T | A | 4 | 35122 | 0.01138887 | 44.4999343 | rnr1Y285F_pe | FALSE | ARG | JSY3811 | 2 | rnr1Y285F_p | CTT | T>A | A/T+1 | A/T-1 | TA>AT | TA>AT | AACAACTTTT | TTTTTTG | GAA | CTT | T | T | T>A | T>A_CTT | TA>AT_TTTTTTG | 4 | FALSE | FALSE |
| 41_v.txt | 33099 | 33099-SNV-/ | SNV | 1 | A | T | 11 | 28128 | 0.03910694 | 49.8181818 | rnr1Y285F_pe | FALSE | ARG | JSY3811 | 2 | rnr1Y285F_p | AAG | A>T | A/T+1 | A/T-1 | TA>AT | TA>AT | GATATAAGA | AGAGCGC | TTC | TTC | T | A | T>A | T>A_TTC | TA>AT_AGAGCGC | 4 | FALSE | FALSE |
| 41_v.txt | 31650 | 31650-SNV-( | SNV | 1 | C | T | 21 | 9117 | 0.23033893 | 36.3333333 | rnr1Y285F_pe | FALSE | ARG | JSY3811 | 2 | rnr1Y285F_p | GCG | C>T | A/T+1 | G/C-1 | CG>TA | CG>TA | AGAATGCGA | CGAAATG | CGC | GCG | C | A | C>T | C>T_GCG | CG>TA_CGAAATG | 4 | FALSE | FALSE |
| 41_v.txt | 31658 | 31658-SNV-( | SNV | 1 | C | T | 16 | 9117 | 0.17549633 | 32.9375 | rnr1Y285F_pe | FALSE | ARG | JSY3811 | 2 | rnr1Y285F_p | GCG | C>T | A/T+1 | G/C-1 | CG>TA | CG>TA | AAATGGCGT | CGTGGAA | CGC | GCG | C | A | C>T | C>T_GCG | CG>TA_CGTGGAA | 4 | FALSE | FALSE |
| 41_v.txt | 31882 | 31882-SNV-( | SNV | 1 | C | T | 13 | 9117 | 0.14259077 | 39.4615385 | rnr1Y285F_pe | FALSE | ARG | JSY3811 | 2 | rnr1Y285F_p | GCA | C>T | A/T+1 | G/C-1 | CG>TA | CG>TA | AGGCGGCAG | CAGCAAA | CGT | GCA | C | A | C>T | C>T_GCA | CG>TA_CAGCAAA | 4 | FALSE | FALSE |
| 41_v.txt | 31962 | 31962-SNV-( | SNV | 1 | G | A | 40 | 23281 | 0.17181393 | 36.45 | rnr1Y285F_pe | FALSE | ARG | JSY3811 | 2 | rnr1Y285F_p | CGC | G>A | A/T+1 | G/C-1 | CG>TA | CG>TA | GTGGCCGCA | GCATAAT | GCG | GCG | C | T | C>T | C>T_GCG | CG>TA_GCATAAT | 4 | FALSE | FALSE |
| 41_v.txt | 31972 | 31972-SNV-( | SNV | 1 | C | T | 16 | 23281 | 0.06872557 | 36.6875 | rnr1Y285F_pe | FALSE | ARG | JSY3811 | 2 | rnr1Y285F_p | GCC | C>T | A/T+1 | G/C-1 | CG>TA | CG>TA | AATAAGCCA. | CCAAGCC | CGG | GCC | C | A | C>T | C>T_GCC | CG>TA_CCAAGCC | 4 | FALSE | FALSE |
| 41_v.txt | 31981 | 31981-SNV-( | SNV | 1 | G | A | 19 | 23281 | 0.08161162 | 49.3157895 | rnr1Y285F_pe | FALSE | ARG | JSY3811 | 2 | rnr1Y285F_p | GGC | G>A | A/T+1 | G/C-1 | CG>TA | CG>TA | AGCGGGCA | GCATTAA | CCG | CCG | C | T | C>T | C>T_CCG | CG>TA_GCATTAA | 4 | FALSE | FALSE |
| 41_v.txt | 32092 | 32092-SNV-( | SNV | 1 | C | T | 21 | 23281 | 0.09020231 | 61.3809524 | rnr1Y285F_pe | FALSE | ARG | JSY3811 | 2 | rnr1Y285F_p | GCA | C>T | A/T+1 | G/C-1 | CG>TA | CG>TA | AGCCTGCAA( | CAACACC | CGT | GCA | C | A | C>T | C>T_GCA | CG>TA_CAACACC | 4 | FALSE | FALSE |
| 41_v.txt | 32165 | 32165-SNV-( | SNV | 1 | G | A | 31 | 23281 | 0.13315579 | 44.0967742 | rnr1Y285F_pe | FALSE | ARG | JSY3811 | 2 | rnr1Y285F_p | CGC | G>A | A/T+1 | G/C-1 | CG>TA | CG>TA | CAAAGCGCC | GCCAAAT | GCG | GCG | C | T | C>T | C>T_GCG | CG>TA_GCCAAAT | 4 | FALSE | FALSE |
| 41_v.txt | 32173 | 32173-SNV-( | SNV | 1 | C | T | 22 | 23281 | 0.09449766 | 43.1363636 | rnr1Y285F_pe | FALSE | ARG | JSY3811 | 2 | rnr1Y285F_p | GCA | C>T | A/T+1 | G/C-1 | CG>TA | CG>TA | CAAATGCAG | CAGCAGT | CGT | GCA | C | A | C>T | C>T_GCA | CG>TA_CAGCAGT | 4 | FALSE | FALSE |
| 41_v.txt | 32176 | 32176-SNV-( | SNV | 1 | C | T | 21 | 23281 | 0.09020231 | 49 | rnr1Y285F_pe | FALSE | ARG | JSY3811 | 2 | rnr1Y285F_p | GCA | C>T | A/T+1 | G/C-1 | CG>TA | CG>TA | ATGCAGCAG | CAGTAAC | CGT | GCA | C | A | C>T | C>T_GCA | CG>TA_CAGTAAC | 4 | FALSE | FALSE |
| 41_v.txt | 32271 | 32271-SNV-( | SNV | 1 | C | T | 39 | 35122 | 0.11104151 | 28.6885818 | rnr1Y285F_pe | FALSE | ARG | JSY3811 | 2 | rnr1Y285F_p | ACG | C>T | A/T+1 | G/C-1 | CG>TA | CG>TA | AAAATACGG | CGGGAAC | TGC | ACG | C | A | C>T | C>T_ACG | CG>TA_CGGGAAC | 4 | FALSE | FALSE |
| 41_v.txt | 32299 | 32299-SNV-( | SNV | 1 | C | T | 67 | 35122 | 0.19076362 | 56.5671642 | rnr1Y285F_pe | FALSE | ARG | JSY3811 | 2 | rnr1Y285F_p | GCG | C>T | A/T+1 | G/C-1 | CG>TA | CG>TA | AATTTGCGG( | CGGCAGA | CGC | GCG | C | A | C>T | C>T_GCG | CG>TA_CGGCAGA | 4 | FALSE | FALSE |
| 41_v.txt | 32466 | 32466-SNV-( | SNV | 1 | C | T | 44 | 35122 | 0.1252776 | 48.1818182 | rnr1Y285F_pe | FALSE | ARG | JSY3811 | 2 | rnr1Y285F_p | GCC | C>T | A/T+1 | G/C-1 | CG>TA | CG>TA | AGAGAGCCA | CCAATGT | CGG | GCC | C | A | C>T | C>T_GCC | CG>TA_CCAATGT | 4 | FALSE | FALSE |
| 41_v.txt | 32535 | 32535-SNV-( | SNV | 1 | G | A | 48 | 35122 | 0.13666648 | 37.6448625 | rnr1Y285F_pe | FALSE | ARG | JSY3811 | 2 | rnr1Y285F_p | TGC | G>A | A/T+1 | G/C-1 | CG>TA | CG>TA | GGGTTTGCA( | GCAGCTT | ACG | ACG | C | T | C>T | C>T_ACG | CG>TA_GCAGCTT | 4 | FALSE | FALSE |
| 41_v.txt | 32536 | 32536-SNV-( | SNV | 1 | C | T | 52 | 35122 | 0.14805535 | 37.3451945 | rnr1Y285F_pe | FALSE | ARG | JSY3811 | 2 | rnr1Y285F_p | GCA | C>T | A/T+1 | G/C-1 | CG>TA | CG>TA | GGTTTGCAG( | CAGCTTC | CGT | GCA | C | A | C>T | C>T_GCA | CG>TA_CAGCTTC | 4 | FALSE | FALSE |
| 41_v.txt | 32590 | 32590-SNV-( | SNV | 1 | C | T | 39 | 25127 | 0.15521153 | 36.7179487 | rnr1Y285F_pe | FALSE | ARG | JSY3811 | 2 | rnr1Y285F_p | GCG | C>T | A/T+1 | G/C-1 | CG>TA | CG>TA | AGGCAGCGT | CGTTAAT | CGC | GCG | C | A | C>T | C>T_GCG | CG>TA_CGTTAAT | 4 | FALSE | FALSE |
| 41_v.txt | 32700 | 32700-SNV-( | SNV | 1 | C | T | 30 | 25127 | 0.11939348 | 59.1666667 | rnr1Y285F_pe | FALSE | ARG | JSY3811 | 2 | rnr1Y285F_p | GCC | C>T | A/T+1 | G/C-1 | CG>TA | CG>TA | ACTGGGCCG | CCGGTAA | CGG | GCC | C | A | C>T | C>T_GCC | CG>TA_CCGGTAA | 4 | FALSE | FALSE |
| 41_v.txt | 32702 | 32702-SNV-( | SNV | 1 | G | A | 31 | 25127 | 0.12337326 | 51 | rnr1Y285F_pe | FALSE | ARG | JSY3811 | 2 | rnr1Y285F_p | CGG | G>A | A/T+1 | G/C-1 | CG>TA | CG>TA | TGGGCCGGT. | GGTAACC | GCC | GCC | C | T | C>T | C>T_GCC | CG>TA_GGTAACC | 4 | FALSE | FALSE |
| 41_v.txt | 32798 | 32798-SNV-( | SNV | 1 | G | A | 42 | 25127 | 0.16715087 | 44.5 | rnr1Y285F_pe | FALSE | ARG | JSY3811 | 2 | rnr1Y285F_p | CGT | G>A | A/T+1 | G/C-1 | CG>TA | CG>TA | TTCACCGTA/ | GTAATAT | GCA | GCA | C | T | C>T | C>T_GCA | CG>TA_GTAATAT | 4 | FALSE | FALSE |
| 41_v.txt | 32860 | 32860-SNV-( | SNV | 1 | C | T | 27 | 25126 | 0.10745841 | 37.7777778 | rnr1Y285F_pe | FALSE | ARG | JSY3811 | 2 | rnr1Y285F_p | GCC | C>T | A/T+1 | G/C-1 | CG>TA | CG>TA | TCCATGCCG( | CCGCCAG | CGG | GCC | C | A | C>T | C>T_GCC | CG>TA_CCGCCAG | 4 | FALSE | FALSE |
| 41_v.txt | 32900 | 32900-SNV-( | SNV | 1 | G | A | 27 | 28128 | 0.09598976 | 33.4767442 | rnr1Y285F_pe | FALSE | ARG | JSY3811 | 2 | rnr1Y285F_p | GGC | G>A | A/T+1 | G/C-1 | CG>TA | CG>TA | GACTTGGCC/ | GCCAACT | CCG | CCG | C | T | C>T | C>T_CCG | CG>TA_GCCAACT | 4 | FALSE | FALSE |
| 41_v.txt | 32901 | 32901-SNV-( | SNV | 1 | C | T | 66 | 28128 | 0.23464164 | 31.2730256 | rnr1Y285F_pe | FALSE | ARG | JSY3811 | 2 | rnr1Y285F_p | GCC | C>T | A/T+1 | G/C-1 | CG>TA | CG>TA | ACTTGGCCA/ | CCAACTA | CGG | GCC | C | A | C>T | C>T_GCC | CG>TA_CCAACTA | 4 | FALSE | FALSE |
| 41_v.txt | 32922 | 32922-SNV-( | SNV | 1 | G | A | 24 | 28128 | 0.08532423 | 52.1666667 | rnr1Y285F_pe | FALSE | ARG | JSY3811 | 2 | rnr1Y285F_p | GGC | G>A | A/T+1 | G/C-1 | CG>TA | CG>TA | TCCAGGGCA. | GCAAAAG | CCG | CCG | C | T | C>T | C>T_CCG | CG>TA_GCAAAAG | 4 | FALSE | FALSE |
| 41_v.txt | 32935 | 32935-SNV-( | SNV | 1 | C | T | 43 | 28128 | 0.15287258 | 37.0697674 | rnr1Y285F_pe | FALSE | ARG | JSY3811 | 2 | rnr1Y285F_p | GCC | C>T | A/T+1 | G/C-1 | CG>TA | CG>TA | TGATTGCCC/ | CCCAAGA | CGG | GCC | C | A | C>T | C>T_GCC | CG>TA_CCCAAGA | 4 | FALSE | FALSE |
| 41_v.txt | 32965 | 32965-SNV-( | SNV | 1 | C | T | 27 | 28128 | 0.09598976 | 59.3703704 | rnr1Y285F_pe | FALSE | ARG | JSY3811 | 2 | rnr1Y285F_p | GCC | C>T | A/T+1 | G/C-1 | CG>TA | CG>TA | CATTGGCCG( | CCGCACC | CGG | GCC | C | A | C>T | C>T_GCC | CG>TA_CCGCACC | 4 | FALSE | FALSE |
| 41_v.txt | 32967 | 32967-SNV-( | SNV | 1 | G | A | 55 | 28128 | 0.1955347 | 61.1454546 | rnr1Y285F_pe | FALSE | ARG | JSY3811 | 2 | rnr1Y285F_p | CGC | G>A | A/T+1 | G/C-1 | CG>TA | CG>TA | TTGGCCGCA( | GCACCAA | GCG | GCG | C | T | C>T | C>T_GCG | CG>TA_GCACCAA | 4 | FALSE | FALSE |
| 41_v.txt | 33056 | 33056-SNV-( | SNV | 1 | C | T | 36 | 28128 | 0.12798635 | 53.1944444 | rnr1Y285F_pe | FALSE | ARG | JSY3811 | 2 | rnr1Y285F_p | GCG | C>T | A/T+1 | G/C-1 | CG>TA | CG>TA | GGACTGCGT( | CGTGACA | CGC | GCG | C | A | C>T | C>T_GCG | CG>TA_CGTGACA | 4 | FALSE | FALSE |
| 41_v.txt | 33069 | 33069-SNV-( | SNV | 1 | G | A | 22 | 28128 | 0.07821388 | 58.0907491 | rnr1Y285F_pe | FALSE | ARG | JSY3811 | 2 | rnr1Y285F_p | TGC | G>A | A/T+1 | G/C-1 | CG>TA | CG>TA | GAATATGCC/ | GCCAAAG | ACG | ACG | C | T | C>T | C>T_ACG | CG>TA_GCCAAAG | 4 | FALSE | FALSE |
| 41_v.txt | 33070 | 33070-SNV-( | SNV | 1 | C | T | 30 | 28128 | 0.10665529 | 44.1333333 | rnr1Y285F_pe | FALSE | ARG | JSY3811 | 2 | rnr1Y285F_p | GCC | C>T | A/T+1 | G/C-1 | CG>TA | CG>TA | AATATGCCA/ | CCAAAGA | CGG | GCC | C | A | C>T | C>T_GCC | CG>TA_CCAAAGA | 4 | FALSE | FALSE |
| 41_v.txt | 33113 | 33113-SNV-( | SNV | 1 | G | A | 22 | 28128 | 0.07821388 | 44.8195053 | rnr1Y285F_pe | FALSE | ARG | JSY3811 | 2 | rnr1Y285F_p | GGC | G>A | A/T+1 | G/C-1 | CG>TA | CG>TA | CACTGGGCC | GCCGGCG | CCG | CCG | C | T | C>T | C>T_CCG | CG>TA_GCCGGCG | 4 | FALSE | FALSE |
| 41_v.txt | 33114 | 33114-SNV-( | SNV | 1 | C | T | 23 | 28128 | 0.08176906 | 43.9558486 | rnr1Y285F_pe | FALSE | ARG | JSY3811 | 2 | rnr1Y285F_p | GCC | C>T | A/T+1 | G/C-1 | CG>TA | CG>TA | ACTGGGCCG | CCGGCGT | CGG | GCC | C | A | C>T | C>T_GCC | CG>TA_CCGGCGT | 4 | FALSE | FALSE |
| 41_v.txt | 33118 | 33118-SNV-( | SNV | 1 | C | T | 22 | 28128 | 0.07821388 | 39.453995 | rnr1Y285F_pe | FALSE | ARG | JSY3811 | 2 | rnr1Y285F_p | GCG | C>T | A/T+1 | G/C-1 | CG>TA | CG>TA | GGCCGGCGT | CGTTGGT | CGC | GCG | C | A | C>T | C>T_GCG | CG>TA_CGTTGGT | 4 | FALSE | FALSE |
| 41_v.txt | 33119 | 33119-SNV-( | SNV | 1 | G | A | 22 | 28128 | 0.07821388 | 43.8181818 | rnr1Y285F_pe | FALSE | ARG | JSY3811 | 2 | rnr1Y285F_p | CGT | G>A | A/T+1 | G/C-1 | CG>TA | CG>TA | GCCGGCGTT( | GTTGGTC | GCA | GCA | C | T | C>T | C>T_GCA | CG>TA_GTTGGTC | 4 | FALSE | FALSE |
| 41_v.txt | 33258 | 33258-SNV-( | SNV | 1 | C | T | 26 | 17688 | 0.14699231 | 39.8846154 | rnr1Y285F_pe | FALSE | ARG | JSY3811 | 2 | rnr1Y285F_p | GCC | C>T | A/T+1 | G/C-1 | CG>TA | CG>TA | CCTATGCCAT | CCATCCT | CGG | GCC | C | A | C>T | C>T_GCC | CG>TA_CCATCCT | 4 | FALSE | FALSE |
| 41_v.txt | 33276 | 33276-SNV-( | SNV | 1 | G | A | 27 | 17688 | 0.15264586 | 46.6296296 | rnr1Y285F_pe | FALSE | ARG | JSY3811 | 2 | rnr1Y285F_p | CGT | G>A | A/T+1 | G/C-1 | CG>TA | CG>TA | GAGAACGTA | GTATCCT | GCA | GCA | C | T | C>T | C>T_GCA | CG>TA_GTATCCT | 4 | FALSE | FALSE |
| 41_v.txt | 33284 | 33284-SNV-( | SNV | 1 | G | A | 21 | 17688 | 0.11872456 | 48.6190476 | rnr1Y285F_pe | FALSE | ARG | JSY3811 | 2 | rnr1Y285F_p | CGC | G>A | A/T+1 | G/C-1 | CG>TA | CG>TA | ATCCTCGCCA | GCCATTT | GCG | GCG | C | T | C>T | C>T_GCG | CG>TA_GCCATTT | 4 | FALSE | FALSE |
| 41_v.txt | 33309 | 33309-SNV-( | SNV | 1 | C | T | 23 | 17688 | 0.13003166 | 55.4347826 | rnr1Y285F_pe | FALSE | ARG | JSY3811 | 2 | rnr1Y285F_p | GCG | C>T | A/T+1 | G/C-1 | CG>TA | CG>TA | AAAGAGCGC | CGCAATG | CGC | GCG | C | A | C>T | C>T_GCG | CG>TA_CGCAATG | 4 | FALSE | FALSE |
| 41_v.txt | 33310 | 33310-SNV-( | SNV | 1 | G | A | 19 | 17687 | 0.10742353 | 59.4415719 | rnr1Y285F_pe | FALSE | ARG | JSY3811 | 2 | rnr1Y285F_p | CGC | G>A | A/T+1 | G/C-1 | CG>TA | CG>TA | AAGAGCGCA | GCAATGG | GCG | GCG | C | T | C>T | C>T_GCG | CG>TA_GCAATGG | 4 | FALSE | FALSE |
| 41_v.txt | 33392 | 33392-SNV-( | SNV | 1 | G | A | 22 | 17688 | 0.12437811 | 53.1363636 | rnr1Y285F_pe | FALSE | ARG | JSY3811 | 2 | rnr1Y285F_p | CGT | G>A | A/T+1 | G/C-1 | CG>TA | CG>TA | AACGTCGTG, | GTGAAAG | GCA | GCA | C | T | C>T | C>T_GCA | CG>TA_GTGAAAG | 4 | FALSE | FALSE |
| 41_v.txt | 33412 | 33412-SNV-( | SNV | 1 | G | A | 31 | 17687 | 0.17526997 | 45.1935484 | rnr1Y285F_pe | FALSE | ARG | JSY3811 | 2 | rnr1Y285F_p | GGC | G>A | A/T+1 | G/C-1 | CG>TA | CG>TA | TGACCGGCT( | GCTCATT | CCG | CCG | C | T | C>T | C>T_CCG | CG>TA_GCTCATT | 4 | FALSE | FALSE |
| 41_v.txt | 33474 | 33474-SNV-( | SNV | 1 | C | T | 21 | 17687 | 0.11873127 | 31.8473733 | rnr1Y285F_pe | FALSE | ARG | JSY3811 | 2 | rnr1Y285F_p | GCC | C>T | A/T+1 | G/C-1 | CG>TA | CG>TA | GCTATGCCTT | CCTTTTT | CGG | GCC | C | A | C>T | C>T_GCC | CG>TA_CCTTTTT | 4 | FALSE | FALSE |
| 41_v.txt | 32271 | 32271-SNV-( | SNV | 1 | C | A | 3 | 35122 | 8.54E-03 | 26.9988104 | rnr1Y285F_pe | FALSE | ARG | JSY3811 | 2 | rnr1Y285F_p | ACG | C>A | A/T+1 | G/C-1 | CG>AT | CG>AT | AAAATACGG | CGGGAAC | TGC | ACG | C | T | C>A | C>A_ACG | CG>AT_CGGGAAC | 4 | FALSE | FALSE |
| 41_v.txt | 32272 | 32272-SNV-( | SNV | 1 | G | T | 18 | 35122 | 0.05124993 | 27.5011865 | rnr1Y285F_pe | FALSE | ARG | JSY3811 | 2 | rnr1Y285F_p | CGG | G>T | A/T+1 | G/C-1 | CG>AT | CG>AT | AAATACGGG | GGGAACC | GCC | GCC | C | A | C>A | C>A_GCC | CG>AT_GGGAACC | 4 | FALSE | FALSE |
| 41_v.txt | 33070 | 33070-SNV-( | SNV | 1 | C | A | 5 | 28128 | 0.01777588 | 16.4 | rnr1Y285F_pe | FALSE | ARG | JSY3811 | 2 | rnr1Y285F_p | GCC | C>A | A/T+1 | G/C-1 | CG>AT | CG>AT | AATATGCCA/ | CCAAAGA | CGG | GCC | C | T | C>A | C>A_GCC | CG>AT_CCAAAGA | 4 | FALSE | FALSE |
| 41_v.txt | 33309 | 33309-SNV-( | SNV | 1 | C | A | 9 | 17688 | 0.05088195 | 30 | rnr1Y285F_pe | FALSE | ARG | JSY3811 | 2 | rnr1Y285F_p | GCG | C>A | A/T+1 | G/C-1 | CG>AT | CG>AT | AAAGAGCGC | CGCAATG | CGC | GCG | C | T | C>A | C>A_GCG | CG>AT_CGCAATG | 4 | FALSE | FALSE |
| 41_v.txt | 33474 | 33474-SNV-( | SNV | 1 | C | A | 45 | 17687 | 0.25442415 | 34.3034249 | rnr1Y285F_pe | FALSE | ARG | JSY3811 | 2 | rnr1Y285F_p | GCC | C>A | A/T+1 | G/C-1 | CG>AT | CG>AT | GCTATGCCTT | CCTTTTT | CGG | GCC | C | T | C>A | C>A_GCC | CG>AT_CCTTTTT | 4 | FALSE | FALSE |
| 41_v.txt | 33475 | 33475-SNV-( | SNV | 1 | C | A | 14 | 17687 | 0.07915418 | 33.7857143 | rnr1Y285F_pe | FALSE | ARG | JSY3811 | 2 | rnr1Y285F_p | CCT | C>A | A/T+1 | G/C-1 | CG>AT | CG>AT | CTATGCCTTT | CTTTTTT | GGA | CCT | C | T | C>A | C>A_CCT | CG>AT_CTTTTTT | 4 | FALSE | FALSE |
| 41_v.txt | 32081 | 32081-Insert | Insertion | 1 | - | A | 11 | 23281 | 0.04724883 | 63 | rnr1Y285F_pe | FALSE | ARG | JSY3811 | 2 | rnr1Y285F_p | CAA | nope | A/T+1 | >1bpdel | none | A/T+1 | CCATGCAAA/ | AAAAAAG | GTT | CAA | - | T | NA | NA_CAA | A/T+1_AAAAAAG | 4 | FALSE | FALSE |
| 41_v.txt | 32498 | 32498-Insert | Insertion | 1 | - | T | 2 | 35122 | 5.69E-03 | 37.5 | rnr1Y285F_pe | FALSE | ARG | JSY3811 | 2 | rnr1Y285F_p | CTT | nope | A/T+1 | >1bpdel | none | A/T+1 | AACAACTTTT | TTTTTTG | GAA | CTT | - | A | NA | NA_CTT | A/T+1_TTTTTTG | 4 | FALSE | FALSE |
| 41_v.txt | 32842 | 32842-Insert | Insertion | 1 | - | A | 5 | 25127 | 0.01989891 | 43.2 | rnr1Y285F_pe | FALSE | ARG | JSY3811 | 2 | rnr1Y285F_p | CAA | nope | A/T+1 | >1bpdel | none | A/T+1 | TTACCCAAA/ | AAAAAAT | GTT | CAA | - | T | NA | NA_CAA | A/T+1_AAAAAAT | 4 | FALSE | FALSE |
| 41_v.txt | 33120 | 33120-Insert | Insertion | 1 | - | T | 1 | 28128 | 3.56E-03 | 38 | rnr1Y285F_pe | FALSE | ARG | JSY3811 | 2 | rnr1Y285F_p | GTT | nope | A/T+1 | >1bpdel | none | A/T+1 | CCGGCGTTG( | TTGGTCA | CAA | GTT | - | A | NA | NA_GTT | A/T+1_TTGGTCA | 4 | FALSE | FALSE |
| 41_v.txt | 33474 | 33474-Deleti | Deletion | 2 | CC | - | 1 | 17687 | 5.65E-03 | 25 | rnr1Y285F_pe | FALSE | ARG | JSY3811 | 2 | rnr1Y285F_p | GCC | nope | >1bp-ins | >1bpdel | none | >1bpdel | GCTATGCCTT | CCTTTTT | CGG | GCC | CC | none | NA | NA_GCC | >1bpdel_CCTTTTT | 4 | FALSE | FALSE |
| 41_v.txt | 31663 | 31663-Deleti | Deletion | 1 | A | - | 8 | 9117 | 0.08774816 | 36.875 | rnr1Y285F_pe | FALSE | ARG | JSY3811 | 2 | rnr1Y285F_p | GAA | nope | >1bp-ins | A/T-1 | none | A/T-1 | GCGTGGAAA | AAATGTG | CTT | CTT | T | none | T>none | T>none_CTT | A/T-1_AAATGTG | 4 | FALSE | FALSE |
| 41_v.txt | 31673 | 31673-Deleti | Deletion | 1 | A | - | 4 | 9117 | 0.04387408 | 62.25 | rnr1Y285F_pe | FALSE | ARG | JSY3811 | 2 | rnr1Y285F_p | CAA | nope | >1bp-ins | A/T-1 | none | A/T-1 | GTGATCAAA( | AAAGGTA | GTT | GTT | T | none | T>none | T>none_GTT | A/T-1_AAAGGTA | 4 | FALSE | FALSE |
| 41_v.txt | 31723 | 31723-Deleti | Deletion | 1 | A | - | 14 | 9117 | 0.15355929 | 63.6428571 | rnr1Y285F_pe | FALSE | ARG | JSY3811 | 2 | rnr1Y285F_p | CAA | nope | >1bp-ins | A/T-1 | none | A/T-1 | TGTCCCAAA/ | AAAAAGT | GTT | GTT | T | none | T>none | T>none_GTT | A/T-1_AAAAAGT | 4 | FALSE | FALSE |
| 41_v.txt | 31863 | 31863-Deleti | Deletion | 1 | A | - | 5 | 9117 | 0.0548426 | 52.2 | rnr1Y285F_pe | FALSE | ARG | JSY3811 | 2 | rnr1Y285F_p | GAA | nope | >1bp-ins | A/T-1 | none | A/T-1 | AACAGGAAA | AAAATAG | CTT | CTT | T | none | T>none | T>none_CTT | A/T-1_AAAATAG | 4 | FALSE | FALSE |

| | | | | | | | | | | | | | | | | | | | | | | | | | | | | | | | | | | |
|---|---|---|---|---|---|---|---|---|---|---|---|---|---|---|---|---|---|---|---|---|---|---|---|---|---|---|---|---|---|---|---|---|---|---|
| 41_v.txt | 31972 | 31972-Deleti | Deletion | 1 | C | - | 4 | 23281 | 0.01718139 | 37.25 | rnr1Y285F_pe | FALSE | ARG | JSY3811 | 2 | rnr1Y285F_p | GCC | nope | >1bp-ins | G/C-1 | none | G/C-1 | AATAAGCCA | CCAAGCC | CGG | GCC | C | none | nope | nope_GCC | G/C-1_CCAAGCC | 4 | FALSE | FALSE |
| 41_v.txt | 32081 | 32081-Deleti | Deletion | 1 | A | - | 16 | 23281 | 0.06872557 | 62.7500001 | rnr1Y285F_pe | FALSE | ARG | JSY3811 | 2 | rnr1Y285F_p | CAA | nope | >1bp-ins | A/T-1 | none | A/T-1 | CCATGCAAA | AAAAAAG | GTT | GTT | T | none | T>none | T>none_GTT | A/T-1_AAAAAAG | 4 | FALSE | FALSE |
| 41_v.txt | 32210 | 32210-Deleti | Deletion | 1 | T | - | 7 | 23281 | 0.03006744 | 41.2857143 | rnr1Y285F_pe | FALSE | ARG | JSY3811 | 2 | rnr1Y285F_p | CTT | nope | >1bp-ins | A/T-1 | none | A/T-1 | ACCACCTTTG | TTTGGTG | GAA | CTT | T | none | nope | nope_CTT | A/T-1_TTTGGTG | 4 | FALSE | FALSE |
| 41_v.txt | 32272 | 32272-Deleti | Deletion | 1 | G | - | 2 | 35122 | 5.69E-03 | 38 | rnr1Y285F_pe | FALSE | ARG | JSY3811 | 2 | rnr1Y285F_p | CGG | nope | >1bp-ins | G/C-1 | none | G/C-1 | AAATACGGG | GGGAACC | GCC | GCC | C | none | C>none | C>none_GCC | G/C-1_GGGAACC | 4 | FALSE | FALSE |
| 41_v.txt | 32498 | 32498-Deleti | Deletion | 1 | T | - | 43 | 35122 | 0.12243039 | 47.5348837 | rnr1Y285F_pe | FALSE | ARG | JSY3811 | 2 | rnr1Y285F_p | CTT | nope | >1bp-ins | A/T-1 | none | A/T-1 | AACAACTTTT | TTTTTTG | GAA | CTT | T | none | nope | nope_CTT | A/T-1_TTTTTTG | 4 | FALSE | FALSE |
| 41_v.txt | 32631 | 32631-Deleti | Deletion | 1 | T | - | 9 | 25127 | 0.03581804 | 45.2222222 | rnr1Y285F_pe | FALSE | ARG | JSY3811 | 2 | rnr1Y285F_p | GTT | nope | >1bp-ins | A/T-1 | none | A/T-1 | CCTTCGTTTT | TTTTTAT | CAA | GTT | T | none | nope | nope_GTT | A/T-1_TTTTTAT | 4 | FALSE | FALSE |
| 41_v.txt | 32697 | 32697-Deleti | Deletion | 1 | G | - | 7 | 25127 | 0.02785848 | 64 | rnr1Y285F_pe | FALSE | ARG | JSY3811 | 2 | rnr1Y285F_p | TGG | nope | >1bp-ins | G/C-1 | none | G/C-1 | CCAACTGGG | GGGCCGG | ACC | ACC | C | none | C>none | C>none_ACC | G/C-1_GGGCCGG | 4 | FALSE | FALSE |
| 41_v.txt | 32707 | 32707-Deleti | Deletion | 1 | C | - | 6 | 25127 | 0.0238787 | 58.8333333 | rnr1Y285F_pe | FALSE | ARG | JSY3811 | 2 | rnr1Y285F_p | ACC | nope | >1bp-ins | G/C-1 | none | G/C-1 | CGGTAACCC | CCCCAGC | TGG | ACC | C | none | nope | nope_ACC | G/C-1_CCCCAGC | 4 | FALSE | FALSE |
| 41_v.txt | 32779 | 32779-Deleti | Deletion | 1 | C | - | 5 | 25127 | 0.01989891 | 58.8 | rnr1Y285F_pe | FALSE | ARG | JSY3811 | 2 | rnr1Y285F_p | ACC | nope | >1bp-ins | G/C-1 | none | G/C-1 | AAGCGACCC | CCCAGAA | TGG | ACC | C | none | nope | nope_ACC | G/C-1_CCCAGAA | 4 | FALSE | FALSE |
| 41_v.txt | 32842 | 32842-Deleti | Deletion | 1 | A | - | 28 | 25127 | 0.11143392 | 38.5714286 | rnr1Y285F_pe | FALSE | ARG | JSY3811 | 2 | rnr1Y285F_p | CAA | nope | >1bp-ins | A/T-1 | none | A/T-1 | TTACCCAAA | AAAAAAT | GTT | GTT | T | none | T>none | T>none_GTT | A/T-1_AAAAAAT | 4 | FALSE | FALSE |
| 41_v.txt | 32924 | 32924-Deleti | Deletion | 1 | A | - | 16 | 28128 | 0.05688282 | 52.5625 | rnr1Y285F_pe | FALSE | ARG | JSY3811 | 2 | rnr1Y285F_p | CAA | nope | >1bp-ins | A/T-1 | none | A/T-1 | CAGGGCAAA | AAAAGTG | GTT | GTT | T | none | T>none | T>none_GTT | A/T-1_AAAAGTG | 4 | FALSE | FALSE |
| 41_v.txt | 32935 | 32935-Deleti | Deletion | 1 | C | - | 8 | 28128 | 0.02844141 | 36 | rnr1Y285F_pe | FALSE | ARG | JSY3811 | 2 | rnr1Y285F_p | GCC | nope | >1bp-ins | G/C-1 | none | G/C-1 | TGATTGCCC | CCCAAGA | CGG | GCC | C | none | nope | nope_GCC | G/C-1_CCCAAGA | 4 | FALSE | FALSE |
| 41_v.txt | 32965 | 32965-Deleti | Deletion | 1 | C | - | 3 | 28128 | 0.01066553 | 52.6666667 | rnr1Y285F_pe | FALSE | ARG | JSY3811 | 2 | rnr1Y285F_p | GCC | nope | >1bp-ins | G/C-1 | none | G/C-1 | CATTGGCCG | CCGCACC | CGG | GCC | C | none | nope | nope_GCC | G/C-1_CCGCACC | 4 | FALSE | FALSE |
| 41_v.txt | 33083 | 33083-Deleti | Deletion | 1 | A | - | 7 | 28128 | 0.02488623 | 51.8571429 | rnr1Y285F_pe | FALSE | ARG | JSY3811 | 2 | rnr1Y285F_p | TAA | nope | >1bp-ins | A/T-1 | none | A/T-1 | ACCCATAAA | AAATAAA | ATT | ATT | T | none | T>none | T>none_ATT | A/T-1_AAATAAA | 4 | FALSE | FALSE |
| 41_v.txt | 33474 | 33474-Deleti | Deletion | 1 | C | - | 48 | 17687 | 0.27138576 | 28.1041667 | rnr1Y285F_pe | FALSE | ARG | JSY3811 | 2 | rnr1Y285F_p | GCC | nope | >1bp-ins | G/C-1 | none | G/C-1 | GCTATGCCTT | CCTTTTT | CGG | GCC | C | none | nope | nope_GCC | G/C-1_CCTTTTT | 4 | FALSE | FALSE |
| 165_v.txt | 31723 | 31723-SNV- | SNV | 1 | A | C | 6 | 10074 | 0.05955926 | 50.8352547 | rnr1Y285F_pe | FALSE | ARG | JSY3865 | 3 | rnr1Y285F_p | CAA | A>C | G/C+1 | A/T-1 | TA>GC | TA>GC | TGTCCCAAA | AAAAAGT | GTT | GTT | T | G | T>G | T>G_GTT | TA>GC_AAAAAGT | 4 | FALSE | FALSE |
| 165_v.txt | 32631 | 32631-SNV-T | SNV | 1 | T | G | 5 | 16645 | 0.03003905 | 38 | rnr1Y285F_pe | FALSE | ARG | JSY3865 | 3 | rnr1Y285F_p | GTT | T>G | G/C+1 | A/T-1 | TA>GC | TA>GC | CCTTCGTTTT | TTTTTAT | CAA | GTT | T | C | T>G | T>G_GTT | TA>GC_TTTTTAT | 4 | FALSE | FALSE |
| 165_v.txt | 33269 | 33269-SNV- | SNV | 1 | A | C | 86 | 11988 | 0.71738405 | 15.7906977 | rnr1Y285F_pe | FALSE | ARG | JSY3865 | 3 | rnr1Y285F_p | TAG | A>C | G/C+1 | A/T-1 | TA>GC | TA>GC | CTCCATAGA | AGAGAAC | ATC | ATC | T | G | T>G | T>G_ATC | TA>GC_AGAGAAC | 4 | FALSE | FALSE |
| 165_v.txt | 31650 | 31650-SNV- | SNV | 1 | C | T | 20 | 10074 | 0.19853087 | 36.3 | rnr1Y285F_pe | FALSE | ARG | JSY3865 | 3 | rnr1Y285F_p | GCG | C>T | A/T+1 | G/C-1 | CG>TA | CG>TA | AGAATGCGA | CGAAATG | CGC | GCG | C | A | C>T | C>T_GCG | CG>TA_CGAAATG | 4 | FALSE | FALSE |
| 165_v.txt | 31962 | 31962-SNV- | SNV | 1 | G | A | 18 | 9366 | 0.1921845 | 36 | rnr1Y285F_pe | FALSE | ARG | JSY3865 | 3 | rnr1Y285F_p | CGC | G>A | A/T+1 | G/C-1 | CG>TA | CG>TA | GTGGCCGCA | GCATAAT | GCG | GCG | C | T | C>T | C>T_GCG | CG>TA_GCATAAT | 4 | FALSE | FALSE |
| 165_v.txt | 32092 | 32092-SNV- | SNV | 1 | C | T | 13 | 9366 | 0.13879992 | 61.7692308 | rnr1Y285F_pe | FALSE | ARG | JSY3865 | 3 | rnr1Y285F_p | GCA | C>T | A/T+1 | G/C-1 | CG>TA | CG>TA | AGCCTGCAA | CAACACC | CGT | GCA | C | A | C>T | C>T_GCA | CG>TA_CAACACC | 4 | FALSE | FALSE |
| 165_v.txt | 32165 | 32165-SNV- | SNV | 1 | G | A | 13 | 9366 | 0.13879992 | 54.4615385 | rnr1Y285F_pe | FALSE | ARG | JSY3865 | 3 | rnr1Y285F_p | CGC | G>A | A/T+1 | G/C-1 | CG>TA | CG>TA | CAAAGCGCC | GCCAAAT | GCG | GCG | C | T | C>T | C>T_GCG | CG>TA_GCCAAAT | 4 | FALSE | FALSE |
| 165_v.txt | 32173 | 32173-SNV- | SNV | 1 | C | T | 13 | 9366 | 0.13879992 | 53.9230769 | rnr1Y285F_pe | FALSE | ARG | JSY3865 | 3 | rnr1Y285F_p | GCA | C>T | A/T+1 | G/C-1 | CG>TA | CG>TA | CAAATGCAG | CAGCAGT | CGT | GCA | C | A | C>T | C>T_GCA | CG>TA_CAGCAGT | 4 | FALSE | FALSE |
| 165_v.txt | 32271 | 32271-SNV- | SNV | 1 | C | T | 28 | 11401 | 0.24559249 | 31.75 | rnr1Y285F_pe | FALSE | ARG | JSY3865 | 3 | rnr1Y285F_p | ACG | C>T | A/T+1 | G/C-1 | CG>TA | CG>TA | AAAATACGG | CGGGAAC | TGC | ACG | C | A | C>T | C>T_ACG | CG>TA_CGGGAAC | 4 | FALSE | FALSE |
| 165_v.txt | 32466 | 32466-SNV- | SNV | 1 | C | T | 14 | 11401 | 0.12279625 | 56.1428571 | rnr1Y285F_pe | FALSE | ARG | JSY3865 | 3 | rnr1Y285F_p | GCC | C>T | A/T+1 | G/C-1 | CG>TA | CG>TA | AGAGAGCCA | CCAATGT | CGG | GCC | C | A | C>T | C>T_GCC | CG>TA_CCAATGT | 4 | FALSE | FALSE |
| 165_v.txt | 32487 | 32487-SNV- | SNV | 1 | C | T | 16 | 11401 | 0.14033857 | 45.3125 | rnr1Y285F_pe | FALSE | ARG | JSY3865 | 3 | rnr1Y285F_p | ACG | C>T | A/T+1 | G/C-1 | CG>TA | CG>TA | AAGATACGG | CGGAAAA | TGC | ACG | C | A | C>T | C>T_ACG | CG>TA_CGGAAAA | 4 | FALSE | FALSE |
| 165_v.txt | 32535 | 32535-SNV- | SNV | 1 | G | A | 16 | 11401 | 0.14033857 | 36.4374022 | rnr1Y285F_pe | FALSE | ARG | JSY3865 | 3 | rnr1Y285F_p | TGC | G>A | A/T+1 | G/C-1 | CG>TA | CG>TA | GGGTTTGCA | GCAGCTT | ACG | ACG | C | T | C>T | C>T_ACG | CG>TA_GCAGCTT | 4 | FALSE | FALSE |
| 165_v.txt | 32536 | 32536-SNV- | SNV | 1 | C | T | 17 | 11401 | 0.14910973 | 37.4117647 | rnr1Y285F_pe | FALSE | ARG | JSY3865 | 3 | rnr1Y285F_p | GCA | C>T | A/T+1 | G/C-1 | CG>TA | CG>TA | GGTTTGCAG | CAGCTTC | CGT | GCA | C | A | C>T | C>T_GCA | CG>TA_CAGCTTC | 4 | FALSE | FALSE |
| 165_v.txt | 32590 | 32590-SNV- | SNV | 1 | C | T | 24 | 16645 | 0.14418744 | 34.5 | rnr1Y285F_pe | FALSE | ARG | JSY3865 | 3 | rnr1Y285F_p | GCG | C>T | A/T+1 | G/C-1 | CG>TA | CG>TA | AGGCAGCGT | CGTTAAT | CGC | GCG | C | A | C>T | C>T_GCG | CG>TA_CGTTAAT | 4 | FALSE | FALSE |
| 165_v.txt | 32700 | 32700-SNV- | SNV | 1 | C | T | 20 | 16645 | 0.1201562 | 59.1 | rnr1Y285F_pe | FALSE | ARG | JSY3865 | 3 | rnr1Y285F_p | GCC | C>T | A/T+1 | G/C-1 | CG>TA | CG>TA | ACTGGGCCG | CCGGTAA | CGG | GCC | C | A | C>T | C>T_GCC | CG>TA_CCGGTAA | 4 | FALSE | FALSE |
| 165_v.txt | 32776 | 32776-SNV- | SNV | 1 | C | T | 21 | 16645 | 0.12616401 | 48.7142857 | rnr1Y285F_pe | FALSE | ARG | JSY3865 | 3 | rnr1Y285F_p | GCG | C>T | A/T+1 | G/C-1 | CG>TA | CG>TA | TGGAAGCGA | CGACCCA | CGC | GCG | C | A | C>T | C>T_GCG | CG>TA_CGACCCA | 4 | FALSE | FALSE |
| 165_v.txt | 32798 | 32798-SNV- | SNV | 1 | G | A | 33 | 16645 | 0.19825774 | 43.1212121 | rnr1Y285F_pe | FALSE | ARG | JSY3865 | 3 | rnr1Y285F_p | CGT | G>A | A/T+1 | G/C-1 | CG>TA | CG>TA | TTCACCGTA | GTAATAT | GCA | GCA | C | T | C>T | C>T_GCA | CG>TA_GTAATAT | 4 | FALSE | FALSE |
| 165_v.txt | 32860 | 32860-SNV- | SNV | 1 | C | T | 24 | 16644 | 0.14419611 | 35.125 | rnr1Y285F_pe | FALSE | ARG | JSY3865 | 3 | rnr1Y285F_p | GCC | C>T | A/T+1 | G/C-1 | CG>TA | CG>TA | TCCATGCCG | CCGCCAG | CGG | GCC | C | A | C>T | C>T_GCC | CG>TA_CCGCCAG | 4 | FALSE | FALSE |
| 165_v.txt | 32901 | 32901-SNV- | SNV | 1 | C | T | 28 | 13142 | 0.21305737 | 33.5 | rnr1Y285F_pe | FALSE | ARG | JSY3865 | 3 | rnr1Y285F_p | GCC | C>T | A/T+1 | G/C-1 | CG>TA | CG>TA | ACTTGGCCA | CCAACTA | CGG | GCC | C | A | C>T | C>T_GCC | CG>TA_CCAACTA | 4 | FALSE | FALSE |
| 165_v.txt | 32967 | 32967-SNV- | SNV | 1 | G | A | 37 | 13142 | 0.2815401 | 58.4864865 | rnr1Y285F_pe | FALSE | ARG | JSY3865 | 3 | rnr1Y285F_p | CGC | G>A | A/T+1 | G/C-1 | CG>TA | CG>TA | TTGGCCGCA | GCACCAA | GCG | GCG | C | T | C>T | C>T_GCG | CG>TA_GCACCAA | 4 | FALSE | FALSE |
| 165_v.txt | 33056 | 33056-SNV- | SNV | 1 | C | T | 19 | 13142 | 0.14457465 | 54 | rnr1Y285F_pe | FALSE | ARG | JSY3865 | 3 | rnr1Y285F_p | GCG | C>T | A/T+1 | G/C-1 | CG>TA | CG>TA | GGACTGCGT | CGTGACA | CGC | GCG | C | A | C>T | C>T_GCG | CG>TA_CGTGACA | 4 | FALSE | FALSE |
| 165_v.txt | 33070 | 33070-SNV- | SNV | 1 | C | T | 20 | 13142 | 0.15218384 | 51.8 | rnr1Y285F_pe | FALSE | ARG | JSY3865 | 3 | rnr1Y285F_p | GCC | C>T | A/T+1 | G/C-1 | CG>TA | CG>TA | AATATGCCA | CCAAAGA | CGG | GCC | C | A | C>T | C>T_GCC | CG>TA_CCAAAGA | 4 | FALSE | FALSE |
| 165_v.txt | 33119 | 33119-SNV- | SNV | 1 | G | A | 14 | 13142 | 0.10652869 | 39.7857143 | rnr1Y285F_pe | FALSE | ARG | JSY3865 | 3 | rnr1Y285F_p | CGT | G>A | A/T+1 | G/C-1 | CG>TA | CG>TA | GCCGGCGTT | GTTGGTC | GCA | GCA | C | T | C>T | C>T_GCA | CG>TA_GTTGGTC | 4 | FALSE | FALSE |
| 165_v.txt | 33258 | 33258-SNV- | SNV | 1 | C | T | 22 | 11988 | 0.18351685 | 45.1818182 | rnr1Y285F_pe | FALSE | ARG | JSY3865 | 3 | rnr1Y285F_p | GCC | C>T | A/T+1 | G/C-1 | CG>TA | CG>TA | CCTATGCCAT | CCATCCT | CGG | GCC | C | A | C>T | C>T_GCC | CG>TA_CCATCCT | 4 | FALSE | FALSE |
| 165_v.txt | 33275 | 33275-SNV- | SNV | 1 | C | T | 18 | 11988 | 0.15015015 | 48.8335208 | rnr1Y285F_pe | FALSE | ARG | JSY3865 | 3 | rnr1Y285F_p | ACG | C>T | A/T+1 | G/C-1 | CG>TA | CG>TA | AGAGAACGT | CGTATCC | TGC | ACG | C | A | C>T | C>T_ACG | CG>TA_CGTATCC | 4 | FALSE | FALSE |
| 165_v.txt | 33276 | 33276-SNV- | SNV | 1 | G | A | 31 | 11988 | 0.25859193 | 47.6427574 | rnr1Y285F_pe | FALSE | ARG | JSY3865 | 3 | rnr1Y285F_p | CGT | G>A | A/T+1 | G/C-1 | CG>TA | CG>TA | GAGAACGTA | GTATCCT | GCA | GCA | C | T | C>T | C>T_GCA | CG>TA_GTATCCT | 4 | FALSE | FALSE |
| 165_v.txt | 33309 | 33309-SNV- | SNV | 1 | C | T | 19 | 11988 | 0.15849183 | 55.1052632 | rnr1Y285F_pe | FALSE | ARG | JSY3865 | 3 | rnr1Y285F_p | GCG | C>T | A/T+1 | G/C-1 | CG>TA | CG>TA | AAAGAGCGC | CGCAATG | CGC | GCG | C | A | C>T | C>T_GCG | CG>TA_CGCAATG | 4 | FALSE | FALSE |
| 165_v.txt | 33310 | 33310-SNV- | SNV | 1 | G | A | 16 | 11987 | 0.13347793 | 57.625 | rnr1Y285F_pe | FALSE | ARG | JSY3865 | 3 | rnr1Y285F_p | CGC | G>A | A/T+1 | G/C-1 | CG>TA | CG>TA | AAGAGCGCA | GCAATGG | GCG | GCG | C | T | C>T | C>T_GCG | CG>TA_GCAATGG | 4 | FALSE | FALSE |
| 165_v.txt | 33474 | 33474-SNV- | SNV | 1 | C | T | 19 | 11987 | 0.15850505 | 36.2079323 | rnr1Y285F_pe | FALSE | ARG | JSY3865 | 3 | rnr1Y285F_p | GCC | C>T | A/T+1 | G/C-1 | CG>TA | CG>TA | GCTATGCCTT | CCTTTTT | CGG | GCC | C | A | C>T | C>T_GCC | CG>TA_CCTTTTT | 4 | FALSE | FALSE |
| 165_v.txt | 31650 | 31650-SNV- | SNV | 1 | C | G | 2 | 10074 | 0.01985309 | 37.5 | rnr1Y285F_pe | FALSE | ARG | JSY3865 | 3 | rnr1Y285F_p | GCG | C>G | G/C+1 | G/C-1 | CG>GC | CG>GC | AGAATGCGA | CGAAATG | CGC | GCG | C | C | C>G | C>G_GCG | CG>GC_CGAAATG | 4 | FALSE | FALSE |
| 165_v.txt | 32535 | 32535-SNV- | SNV | 1 | G | C | 1 | 11401 | 8.77E-03 | 37.9974544 | rnr1Y285F_pe | FALSE | ARG | JSY3865 | 3 | rnr1Y285F_p | TGC | G>C | G/C+1 | G/C-1 | CG>GC | CG>GC | GGGTTTGCA | GCAGCTT | ACG | ACG | C | G | C>G | C>G_ACG | CG>GC_GCAGCTT | 4 | FALSE | FALSE |
| 165_v.txt | 33070 | 33070-SNV- | SNV | 1 | C | G | 3 | 13142 | 0.02282758 | 54.6666667 | rnr1Y285F_pe | FALSE | ARG | JSY3865 | 3 | rnr1Y285F_p | GCC | C>G | G/C+1 | G/C-1 | CG>GC | CG>GC | AATATGCCA | CCAAAGA | CGG | GCC | C | C | C>G | C>G_GCC | CG>GC_CCAAAGA | 4 | FALSE | FALSE |
| 165_v.txt | 31657 | 31657-SNV- | SNV | 1 | G | T | 89 | 10074 | 0.88346238 | 19.6067416 | rnr1Y285F_pe | FALSE | ARG | JSY3865 | 3 | rnr1Y285F_p | GGC | G>T | A/T+1 | G/C-1 | CG>AT | CG>AT | GAAATGGCG | GCGTGGA | CCG | CCG | C | A | C>A | C>A_CCG | CG>AT_GCGTGGA | 4 | FALSE | FALSE |
| 165_v.txt | 31710 | 31710-SNV- | SNV | 1 | C | A | 11 | 10074 | 0.10919198 | 44.7272727 | rnr1Y285F_pe | FALSE | ARG | JSY3865 | 3 | rnr1Y285F_p | CCA | C>A | A/T+1 | G/C-1 | CG>AT | CG>AT | ACATTCCAA | CAAAATT | GGT | CCA | C | T | C>A | C>A_CCA | CG>AT_CAAAATT | 4 | FALSE | FALSE |
| 165_v.txt | 32536 | 32536-SNV- | SNV | 1 | C | A | 6 | 11401 | 0.05262696 | 26.6666667 | rnr1Y285F_pe | FALSE | ARG | JSY3865 | 3 | rnr1Y285F_p | GCA | C>A | A/T+1 | G/C-1 | CG>AT | CG>AT | GGTTTGCAG | CAGCTTC | CGT | GCA | C | T | C>A | C>A_GCA | CG>AT_CAGCTTC | 4 | FALSE | FALSE |
| 165_v.txt | 33070 | 33070-SNV- | SNV | 1 | C | A | 6 | 13142 | 0.04565515 | 34.5 | rnr1Y285F_pe | FALSE | ARG | JSY3865 | 3 | rnr1Y285F_p | GCC | C>A | A/T+1 | G/C-1 | CG>AT | CG>AT | AATATGCCA | CCAAAGA | CGG | GCC | C | T | C>A | C>A_GCC | CG>AT_CCAAAGA | 4 | FALSE | FALSE |
| 165_v.txt | 33309 | 33309-SNV- | SNV | 1 | C | A | 2 | 11988 | 0.01668335 | 19 | rnr1Y285F_pe | FALSE | ARG | JSY3865 | 3 | rnr1Y285F_p | GCG | C>A | A/T+1 | G/C-1 | CG>AT | CG>AT | AAAGAGCGC | CGCAATG | CGC | GCG | C | T | C>A | C>A_GCG | CG>AT_CGCAATG | 4 | FALSE | FALSE |
| 165_v.txt | 33310 | 33310-SNV- | SNV | 1 | G | T | 6 | 11987 | 0.05005423 | 19.6666667 | rnr1Y285F_pe | FALSE | ARG | JSY3865 | 3 | rnr1Y285F_p | CGC | G>T | A/T+1 | G/C-1 | CG>AT | CG>AT | AAGAGCGCA | GCAATGG | GCG | GCG | C | A | C>A | C>A_GCG | CG>AT_GCAATGG | 4 | FALSE | FALSE |
| 165_v.txt | 33474 | 33474-SNV- | SNV | 1 | C | A | 43 | 11987 | 0.35872195 | 33.9039679 | rnr1Y285F_pe | FALSE | ARG | JSY3865 | 3 | rnr1Y285F_p | GCC | C>A | A/T+1 | G/C-1 | CG>AT | CG>AT | GCTATGCCTT | CCTTTTT | CGG | GCC | C | T | C>A | C>A_GCC | CG>AT_CCTTTTT | 4 | FALSE | FALSE |
| 165_v.txt | 33475 | 33475-SNV- | SNV | 1 | C | A | 17 | 11987 | 0.14182031 | 35.5294118 | rnr1Y285F_pe | FALSE | ARG | JSY3865 | 3 | rnr1Y285F_p | CCT | C>A | A/T+1 | G/C-1 | CG>AT | CG>AT | CTATGCCTTT | CTTTTTT | GGA | CCT | C | T | C>A | C>A_CCT | CG>AT_CTTTTTT | 4 | FALSE | FALSE |
| 165_v.txt | 31723 | 31723-Insert | Insertion | 1 | - | A | 2 | 10074 | 0.01985309 | 50 | rnr1Y285F_pe | FALSE | ARG | JSY3865 | 3 | rnr1Y285F_p | CAA | nope | A/T+1 | >1bpdel | none | A/T+1 | TGTCCCAAA | AAAAAGT | GTT | CAA | - | T | NA | NA_CAA | A/T+1_AAAAAGT | 4 | FALSE | FALSE |
| 165_v.txt | 32081 | 32081-Insert | Insertion | 1 | - | A | 7 | 9366 | 0.07473842 | 64 | rnr1Y285F_pe | FALSE | ARG | JSY3865 | 3 | rnr1Y285F_p | CAA | nope | A/T+1 | >1bpdel | none | A/T+1 | CCATGCAAA | AAAAAAG | GTT | CAA | - | T | NA | NA_CAA | A/T+1_AAAAAAG | 4 | FALSE | FALSE |
| 165_v.txt | 32842 | 32842-Insert | Insertion | 1 | - | A | 3 | 16645 | 0.01802343 | 55.3333333 | rnr1Y285F_pe | FALSE | ARG | JSY3865 | 3 | rnr1Y285F_p | CAA | nope | A/T+1 | >1bpdel | none | A/T+1 | TTACCCAAA | AAAAAAT | GTT | CAA | - | T | NA | NA_CAA | A/T+1_AAAAAAT | 4 | FALSE | FALSE |
| 165_v.txt | 31723 | 31723-Deleti | Deletion | 1 | A | - | 9 | 10074 | 0.08933889 | 56.1111111 | rnr1Y285F_pe | FALSE | ARG | JSY3865 | 3 | rnr1Y285F_p | CAA | nope | >1bp-ins | A/T-1 | none | A/T-1 | TGTCCCAAA | AAAAAGT | GTT | GTT | T | none | T>none | T>none_GTT | A/T-1_AAAAAGT | 4 | FALSE | FALSE |
| 165_v.txt | 31904 | 31904-Deleti | Deletion | 1 | T | - | 5 | 10074 | 0.04963272 | 48.4 | rnr1Y285F_pe | FALSE | ARG | JSY3865 | 3 | rnr1Y285F_p | ATT | nope | >1bp-ins | A/T-1 | none | A/T-1 | ATTGAATTTT | TTTTGGT | TAA | ATT | T | none | nope | nope_ATT | A/T-1_TTTTGGT | 4 | FALSE | FALSE |
| 165_v.txt | 31913 | 31913-Deleti | Deletion | 1 | A | - | 6 | 10074 | 0.05955926 | 37.8333333 | rnr1Y285F_pe | FALSE | ARG | JSY3865 | 3 | rnr1Y285F_p | CAA | nope | >1bp-ins | A/T-1 | none | A/T-1 | TGGTGCAAA | AAAAGCC | GTT | GTT | T | none | T>none | T>none_GTT | A/T-1_AAAAGCC | 4 | FALSE | FALSE |
| 165_v.txt | 32081 | 32081-Deleti | Deletion | 1 | A | - | 11 | 9365 | 0.11745862 | 63.4545455 | rnr1Y285F_pe | FALSE | ARG | JSY3865 | 3 | rnr1Y285F_p | CAA | nope | >1bp-ins | A/T-1 | none | A/T-1 | CCATGCAAA | AAAAAAG | GTT | GTT | T | none | T>none | T>none_GTT | A/T-1_AAAAAAG | 4 | FALSE | FALSE |
| 165_v.txt | 32442 | 32442-Deleti | Deletion | 1 | A | - | 5 | 11401 | 0.0438558 | 48.2 | rnr1Y285F_pe | FALSE | ARG | JSY3865 | 3 | rnr1Y285F_p | TAA | nope | >1bp-ins | A/T-1 | none | A/T-1 | GGAACTAAA | AAAAGTC | ATT | ATT | T | none | T>none | T>none_ATT | A/T-1_AAAAGTC | 4 | FALSE | FALSE |
| 165_v.txt | 32498 | 32498-Deleti | Deletion | 1 | T | - | 9 | 11401 | 0.07894044 | 55.3333333 | rnr1Y285F_pe | FALSE | ARG | JSY3865 | 3 | rnr1Y285F_p | CTT | nope | >1bp-ins | A/T-1 | none | A/T-1 | AACAACTTTT | TTTTTTG | GAA | CTT | T | none | nope | nope_CTT | A/T-1_TTTTTTG | 4 | FALSE | FALSE |
| 165_v.txt | 32631 | 32631-Deleti | Deletion | 1 | T | - | 6 | 16645 | 0.03604686 | 45.6666667 | rnr1Y285F_pe | FALSE | ARG | JSY3865 | 3 | rnr1Y285F_p | GTT | nope | >1bp-ins | A/T-1 | none | A/T-1 | CCTTCGTTTT | TTTTTAT | CAA | GTT | T | none | nope | nope_GTT | A/T-1_TTTTTAT | 4 | FALSE | FALSE |
| 165_v.txt | 32658 | 32658-Deleti | Deletion | 1 | C | - | 6 | 16645 | 0.03604686 | 53.1666667 | rnr1Y285F_pe | FALSE | ARG | JSY3865 | 3 | rnr1Y285F_p | ACC | nope | >1bp-ins | G/C-1 | none | G/C-1 | CCTGGACCC | CCCCAGG | TGG | ACC | C | none | nope | nope_ACC | G/C-1_CCCCAGG | 4 | FALSE | FALSE |
| 165_v.txt | 32728 | 32728-Deleti | Deletion | 1 | A | - | 6 | 16645 | 0.03604686 | 55.1666667 | rnr1Y285F_pe | FALSE | ARG | JSY3865 | 3 | rnr1Y285F_p | CAA | nope | >1bp-ins | A/T-1 | none | A/T-1 | CCATACAAA | AAAAACA | GTT | GTT | T | none | T>none | T>none_GTT | A/T-1_AAAAACA | 4 | FALSE | FALSE |
| 165_v.txt | 32842 | 32842-Deleti | Deletion | 1 | A | - | 21 | 16645 | 0.12616401 | 41.1428571 | rnr1Y285F_pe | FALSE | ARG | JSY3865 | 3 | rnr1Y285F_p | CAA | nope | >1bp-ins | A/T-1 | none | A/T-1 | TTACCCAAA | AAAAAAT | GTT | GTT | T | none | T>none | T>none_GTT | A/T-1_AAAAAAT | 4 | FALSE | FALSE |
| 165_v.txt | 32924 | 32924-Deleti | Deletion | 1 | A | - | 14 | 13142 | 0.10652869 | 48.2857143 | rnr1Y285F_pe | FALSE | ARG | JSY3865 | 3 | rnr1Y285F_p | CAA | nope | >1bp-ins | A/T-1 | none | A/T-1 | CAGGGCAAA | AAAAGTG | GTT | GTT | T | none | T>none | T>none_GTT | A/T-1_AAAAGTG | 4 | FALSE | FALSE |
| 165_v.txt | 33474 | 33474-Deleti | Deletion | 1 | C | - | 36 | 11987 | 0.30032535 | 30.5277778 | rnr1Y285F_pe | FALSE | ARG | JSY3865 | 3 | rnr1Y285F_p | GCC | nope | >1bp-ins | G/C-1 | none | G/C-1 | GCTATGCCTT | CCTTTTT | CGG | GCC | C | none | nope | nope_GCC | G/C-1_CCTTTTT | 4 | FALSE | FALSE |
| 166_v.txt | 32810 | 32810-SNV- | SNV | 1 | A | C | 5 | 27926 | 0.01790446 | 43.2 | rnr1Y285F_pe | FALSE | ARG | JSY3866 | 4 | rnr1Y285F_p | CAG | A>C | G/C+1 | A/T-1 | TA>GC | TA>GC | TTTGACAGG | AGGGAAC | GTC | GTC | T | G | T>G | T>G_GTC | TA>GC_AGGGAAC | 4 | FALSE | FALSE |
| 166_v.txt | 32842 | 32842-SNV- | SNV | 1 | A | C | 2 | 27926 | 7.16E-03 | 38 | rnr1Y285F_pe | FALSE | ARG | JSY3866 | 4 | rnr1Y285F_p | CAA | A>C | G/C+1 | A/T-1 | TA>GC | TA>GC | TTACCCAAA | AAAAAAT | GTT | GTT | T | G | T>G | T>G_GTT | TA>GC_AAAAAAT | 4 | FALSE | FALSE |
| 166_v.txt | 32884 | 32884-SNV- | SNV | 1 | A | C | 9 | 21126 | 0.04260153 | 27 | rnr1Y285F_pe | FALSE | ARG | JSY3866 | 4 | rnr1Y285F_p | CAA | A>C | G/C+1 | A/T-1 | TA>GC | TA>GC | ACGTCCAAA | AAAATTG | GTT | GTT | T | G | T>G | T>G_GTT | TA>GC_AAAATTG | 4 | FALSE | FALSE |
| 166_v.txt | 33269 | 33269-SNV- | SNV | 1 | A | C | 162 | 18839 | 0.85991826 | 15.845679 | rnr1Y285F_pe | FALSE | ARG | JSY3866 | 4 | rnr1Y285F_p | TAG | A>C | G/C+1 | A/T-1 | TA>GC | TA>GC | CTCCATAGA | AGAGAAC | ATC | ATC | T | G | T>G | T>G_ATC | TA>GC_AGAGAAC | 4 | FALSE | FALSE |
| 166_v.txt | 33476 | 33476-SNV-T | SNV | 1 | T | G | 4 | 18839 | 0.02123255 | 37.500067 | rnr1Y285F_pe | FALSE | ARG | JSY3866 | 4 | rnr1Y285F_p | CTT | T>G | G/C+1 | A/T-1 | TA>GC | TA>GC | TATGCCTTTT | TTTTTTT | GAA | CTT | T | C | T>G | T>G_CTT | TA>GC_TTTTTTT | 4 | FALSE | FALSE |
| 166_v.txt | 32081 | 32081-SNV- | SNV | 1 | A | G | 6 | 16372 | 0.03664794 | 51.1666667 | rnr1Y285F_pe | FALSE | ARG | JSY3866 | 4 | rnr1Y285F_p | CAA | A>G | G/C+1 | A/T-1 | TA>CG | TA>CG | CCATGCAAA | AAAAAAG | GTT | GTT | T | C | T>C | T>C_GTT | TA>CG_AAAAAAG | 4 | FALSE | FALSE |
| 166_v.txt | 32810 | 32810-SNV- | SNV | 1 | A | G | 16 | 27926 | 0.05729428 | 46.8125 | rnr1Y285F_pe | FALSE | ARG | JSY3866 | 4 | rnr1Y285F_p | CAG | A>G | G/C+1 | A/T-1 | TA>CG | TA>CG | TTTGACAGG | AGGGAAC | GTC | GTC | T | C | T>C | T>C_GTC | TA>CG_AGGGAAC | 4 | FALSE | FALSE |
| 166_v.txt | 32842 | 32842-SNV- | SNV | 1 | A | G | 5 | 27926 | 0.01790446 | 48.3998555 | rnr1Y285F_pe | FALSE | ARG | JSY3866 | 4 | rnr1Y285F_p | CAA | A>G | G/C+1 | A/T-1 | TA>CG | TA>CG | TTACCCAAA | AAAAAAT | GTT | GTT | T | C | T>C | T>C_GTT | TA>CG_AAAAAAT | 4 | FALSE | FALSE |
| 166_v.txt | 33476 | 33476-SNV-T | SNV | 1 | T | C | 28 | 18839 | 0.14862785 | 30.4645075 | rnr1Y285F_pe | FALSE | ARG | JSY3866 | 4 | rnr1Y285F_p | CTT | T>C | G/C+1 | A/T-1 | TA>CG | TA>CG | TATGCCTTTT | TTTTTTT | GAA | CTT | T | G | T>C | T>C_CTT | TA>CG_TTTTTTT | 4 | FALSE | FALSE |
| 166_v.txt | 31650 | 31650-SNV- | SNV | 1 | C | T | 22 | 7643 | 0.28784509 | 36.1818182 | rnr1Y285F_pe | FALSE | ARG | JSY3866 | 4 | rnr1Y285F_p | GCG | C>T | A/T+1 | G/C-1 | CG>TA | CG>TA | AGAATGCGA | CGAAATG | CGC | GCG | C | A | C>T | C>T_GCG | CG>TA_CGAAATG | 4 | FALSE | FALSE |
| 166_v.txt | 31830 | 31830-SNV- | SNV | 1 | C | T | 10 | 7643 | 0.13083868 | 60.8 | rnr1Y285F_pe | FALSE | ARG | JSY3866 | 4 | rnr1Y285F_p | GCA | C>T | A/T+1 | G/C-1 | CG>TA | CG>TA | AATATGCAT | CATTGAA | CGT | GCA | C | A | C>T | C>T_GCA | CG>TA_CATTGAA | 4 | FALSE | FALSE |
| 166_v.txt | 31947 | 31947-SNV- | SNV | 1 | G | A | 15 | 16359 | 0.09169265 | 37.9333333 | rnr1Y285F_pe | FALSE | ARG | JSY3866 | 4 | rnr1Y285F_p | CGT | G>A | A/T+1 | G/C-1 | CG>TA | CG>TA | ATGATCGTC | GTCATAA | GCA | GCA | C | T | C>T | C>T_GCA | CG>TA_GTCATAA | 4 | FALSE | FALSE |
| 166_v.txt | 31962 | 31962-SNV- | SNV | 1 | G | A | 22 | 16372 | 0.13437576 | 35.5454546 | rnr1Y285F_pe | FALSE | ARG | JSY3866 | 4 | rnr1Y285F_p | CGC | G>A | A/T+1 | G/C-1 | CG>TA | CG>TA | GTGGCCGCA | GCATAAT | GCG | GCG | C | T | C>T | C>T_GCG | CG>TA_GCATAAT | 4 | FALSE | FALSE |
| 166_v.txt | 32057 | 32057-SNV- | SNV | 1 | C | T | 18 | 16372 | 0.10994381 | 61.8333333 | rnr1Y285F_pe | FALSE | ARG | JSY3866 | 4 | rnr1Y285F_p | GCG | C>T | A/T+1 | G/C-1 | CG>TA | CG>TA | GATGTGCGA | CGAGATT | CGC | GCG | C | A | C>T | C>T_GCG | CG>TA_CGAGATT | 4 | FALSE | FALSE |
| 166_v.txt | 32092 | 32092-SNV- | SNV | 1 | C | T | 17 | 16372 | 0.10383582 | 62.1764706 | rnr1Y285F_pe | FALSE | ARG | JSY3866 | 4 | rnr1Y285F_p | GCA | C>T | A/T+1 | G/C-1 | CG>TA | CG>TA | AGCCTGCAA | CAACACC | CGT | GCA | C | A | C>T | C>T_GCA | CG>TA_CAACACC | 4 | FALSE | FALSE |
| 166_v.txt | 32165 | 32165-SNV- | SNV | 1 | G | A | 30 | 16372 | 0.18323968 | 47.6333333 | rnr1Y285F_pe | FALSE | ARG | JSY3866 | 4 | rnr1Y285F_p | CGC | G>A | A/T+1 | G/C-1 | CG>TA | CG>TA | CAAAGCGCC | GCCAAAT | GCG | GCG | C | T | C>T | C>T_GCG | CG>TA_GCCAAAT | 4 | FALSE | FALSE |
| 166_v.txt | 32173 | 32173-SNV- | SNV | 1 | C | T | 29 | 16372 | 0.17713169 | 47.7586207 | rnr1Y285F_pe | FALSE | ARG | JSY3866 | 4 | rnr1Y285F_p | GCA | C>T | A/T+1 | G/C-1 | CG>TA | CG>TA | CAAATGCAG | CAGCAGT | CGT | GCA | C | A | C>T | C>T_GCA | CG>TA_CAGCAGT | 4 | FALSE | FALSE |
| 166_v.txt | 32271 | 32271-SNV- | SNV | 1 | C | T | 33 | 18011 | 0.18322137 | 32.0606061 | rnr1Y285F_pe | FALSE | ARG | JSY3866 | 4 | rnr1Y285F_p | ACG | C>T | A/T+1 | G/C-1 | CG>TA | CG>TA | AAAATACGG | CGGGAAC | TGC | ACG | C | A | C>T | C>T_ACG | CG>TA_CGGGAAC | 4 | FALSE | FALSE |
| 166_v.txt | 32299 | 32299-SNV- | SNV | 1 | C | T | 33 | 18012 | 0.18321119 | 50.3939394 | rnr1Y285F_pe | FALSE | ARG | JSY3866 | 4 | rnr1Y285F_p | GCG | C>T | A/T+1 | G/C-1 | CG>TA | CG>TA | AATTTGCGG | CGGCAGA | CGC | GCG | C | A | C>T | C>T_GCG | CG>TA_CGGCAGA | 4 | FALSE | FALSE |
| 166_v.txt | 32466 | 32466-SNV- | SNV | 1 | C | T | 34 | 18012 | 0.18876305 | 52.3235294 | rnr1Y285F_pe | FALSE | ARG | JSY3866 | 4 | rnr1Y285F_p | GCC | C>T | A/T+1 | G/C-1 | CG>TA | CG>TA | AGAGAGCCA | CCAATGT | CGG | GCC | C | A | C>T | C>T_GCC | CG>TA_CCAATGT | 4 | FALSE | FALSE |
| 166_v.txt | 32535 | 32535-SNV- | SNV | 1 | G | A | 18 | 18012 | 0.09993338 | 35.9438187 | rnr1Y285F_pe | FALSE | ARG | JSY3866 | 4 | rnr1Y285F_p | TGC | G>A | A/T+1 | G/C-1 | CG>TA | CG>TA | GGGTTTGCA | GCAGCTT | ACG | ACG | C | T | C>T | C>T_ACG | CG>TA_GCAGCTT | 4 | FALSE | FALSE |
| 166_v.txt | 32536 | 32536-SNV- | SNV | 1 | C | T | 26 | 18012 | 0.14434821 | 34.8462685 | rnr1Y285F_pe | FALSE | ARG | JSY3866 | 4 | rnr1Y285F_p | GCA | C>T | A/T+1 | G/C-1 | CG>TA | CG>TA | GGTTTGCAG | CAGCTTC | CGT | GCA | C | A | C>T | C>T_GCA | CG>TA_CAGCTTC | 4 | FALSE | FALSE |
| 166_v.txt | 32590 | 32590-SNV- | SNV | 1 | C | T | 49 | 27926 | 0.17546373 | 36.244898 | rnr1Y285F_pe | FALSE | ARG | JSY3866 | 4 | rnr1Y285F_p | GCG | C>T | A/T+1 | G/C-1 | CG>TA | CG>TA | AGGCAGCGT | CGTTAAT | CGC | GCG | C | A | C>T | C>T_GCG | CG>TA_CGTTAAT | 4 | FALSE | FALSE |
| 166_v.txt | 32700 | 32700-SNV- | SNV | 1 | C | T | 46 | 27926 | 0.16472105 | 59.7826087 | rnr1Y285F_pe | FALSE | ARG | JSY3866 | 4 | rnr1Y285F_p | GCC | C>T | A/T+1 | G/C-1 | CG>TA | CG>TA | ACTGGGCCG | CCGGTAA | CGG | GCC | C | A | C>T | C>T_GCC | CG>TA_CCGGTAA | 4 | FALSE | FALSE |
| 166_v.txt | 32757 | 32757-SNV- | SNV | 1 | G | A | 27 | 27926 | 0.09668409 | 51.5925926 | rnr1Y285F_pe | FALSE | ARG | JSY3866 | 4 | rnr1Y285F_p | GGC | G>A | A/T+1 | G/C-1 | CG>TA | CG>TA | ATAATGGCT | GCTAAAA | CCG | CCG | C | T | C>T | C>T_CCG | CG>TA_GCTAAAA | 4 | FALSE | FALSE |
| 166_v.txt | 32776 | 32776-SNV- | SNV | 1 | C | T | 29 | 27926 | 0.10384588 | 52.4482759 | rnr1Y285F_pe | FALSE | ARG | JSY3866 | 4 | rnr1Y285F_p | GCG | C>T | A/T+1 | G/C-1 | CG>TA | CG>TA | TGGAAGCGA | CGACCCA | CGC | GCG | C | A | C>T | C>T_GCG | CG>TA_CGACCCA | 4 | FALSE | FALSE |
| 166_v.txt | 32798 | 32798-SNV- | SNV | 1 | G | A | 40 | 27926 | 0.14323569 | 41.675 | rnr1Y285F_pe | FALSE | ARG | JSY3866 | 4 | rnr1Y285F_p | CGT | G>A | A/T+1 | G/C-1 | CG>TA | CG>TA | TTCACCGTA | GTAATAT | GCA | GCA | C | T | C>T | C>T_GCA | CG>TA_GTAATAT | 4 | FALSE | FALSE |
| 166_v.txt | 32860 | 32860-SNV- | SNV | 1 | C | T | 39 | 27923 | 0.13966981 | 36.1538462 | rnr1Y285F_pe | FALSE | ARG | JSY3866 | 4 | rnr1Y285F_p | GCC | C>T | A/T+1 | G/C-1 | CG>TA | CG>TA | TCCATGCCG | CCGCCAG | CGG | GCC | C | A | C>T | C>T_GCC | CG>TA_CCGCCAG | 4 | FALSE | FALSE |
| 166_v.txt | 32900 | 32900-SNV- | SNV | 1 | G | A | 24 | 21126 | 0.11360409 | 35.2335103 | rnr1Y285F_pe | FALSE | ARG | JSY3866 | 4 | rnr1Y285F_p | GGC | G>A | A/T+1 | G/C-1 | CG>TA | CG>TA | GACTTGGCC | GCCAACT | CCG | CCG | C | T | C>T | C>T_CCG | CG>TA_GCCAACT | 4 | FALSE | FALSE |
| 166_v.txt | 32901 | 32901-SNV- | SNV | 1 | C | T | 36 | 21126 | 0.17040614 | 30.9725062 | rnr1Y285F_pe | FALSE | ARG | JSY3866 | 4 | rnr1Y285F_p | GCC | C>T | A/T+1 | G/C-1 | CG>TA | CG>TA | ACTTGGCCA | CCAACTA | CGG | GCC | C | A | C>T | C>T_GCC | CG>TA_CCAACTA | 4 | FALSE | FALSE |
| 166_v.txt | 32922 | 32922-SNV- | SNV | 1 | G | A | 24 | 21126 | 0.11360409 | 49.7916667 | rnr1Y285F_pe | FALSE | ARG | JSY3866 | 4 | rnr1Y285F_p | GGC | G>A | A/T+1 | G/C-1 | CG>TA | CG>TA | TCCAGGGCA | GCAAAAG | CCG | CCG | C | T | C>T | C>T_CCG | CG>TA_GCAAAAG | 4 | FALSE | FALSE |
| 166_v.txt | 32935 | 32935-SNV- | SNV | 1 | C | T | 32 | 21126 | 0.15147212 | 42.03125 | rnr1Y285F_pe | FALSE | ARG | JSY3866 | 4 | rnr1Y285F_p | GCC | C>T | A/T+1 | G/C-1 | CG>TA | CG>TA | TGATTGCCC | CCCAAGA | CGG | GCC | C | A | C>T | C>T_GCC | CG>TA_CCCAAGA | 4 | FALSE | FALSE |
| 166_v.txt | 32965 | 32965-SNV- | SNV | 1 | C | T | 24 | 21126 | 0.11360409 | 57.6666667 | rnr1Y285F_pe | FALSE | ARG | JSY3866 | 4 | rnr1Y285F_p | GCC | C>T | A/T+1 | G/C-1 | CG>TA | CG>TA | CATTGGCCG | CCGCACC | CGG | GCC | C | A | C>T | C>T_GCC | CG>TA_CCGCACC | 4 | FALSE | FALSE |
| 166_v.txt | 32967 | 32967-SNV- | SNV | 1 | G | A | 48 | 21126 | 0.22720818 | 60.3541667 | rnr1Y285F_pe | FALSE | ARG | JSY3866 | 4 | rnr1Y285F_p | CGC | G>A | A/T+1 | G/C-1 | CG>TA | CG>TA | TTGGCCGCA | GCACCAA | GCG | GCG | C | T | C>T | C>T_GCG | CG>TA_GCACCAA | 4 | FALSE | FALSE |
| 166_v.txt | 33056 | 33056-SNV- | SNV | 1 | C | T | 21 | 21126 | 0.09940358 | 53.7619048 | rnr1Y285F_pe | FALSE | ARG | JSY3866 | 4 | rnr1Y285F_p | GCG | C>T | A/T+1 | G/C-1 | CG>TA | CG>TA | GGACTGCGT | CGTGACA | CGC | GCG | C | A | C>T | C>T_GCG | CG>TA_CGTGACA | 4 | FALSE | FALSE |
| 166_v.txt | 33070 | 33070-SNV- | SNV | 1 | C | T | 22 | 21126 | 0.10413708 | 52.1818182 | rnr1Y285F_pe | FALSE | ARG | JSY3866 | 4 | rnr1Y285F_p | GCC | C>T | A/T+1 | G/C-1 | CG>TA | CG>TA | AATATGCCA | CCAAAGA | CGG | GCC | C | A | C>T | C>T_GCC | CG>TA_CCAAAGA | 4 | FALSE | FALSE |
| 166_v.txt | 33104 | 33104-SNV- | SNV | 1 | G | A | 27 | 21126 | 0.1278046 | 44.4074074 | rnr1Y285F_pe | FALSE | ARG | JSY3866 | 4 | rnr1Y285F_p | CGC | G>A | A/T+1 | G/C-1 | CG>TA | CG>TA | AAGAGCGCC | GCCCACT | GCG | GCG | C | T | C>T | C>T_GCG | CG>TA_GCCCACT | 4 | FALSE | FALSE |
| 166_v.txt | 33114 | 33114-SNV- | SNV | 1 | C | T | 21 | 21126 | 0.09940358 | 42.952381 | rnr1Y285F_pe | FALSE | ARG | JSY3866 | 4 | rnr1Y285F_p | GCC | C>T | A/T+1 | G/C-1 | CG>TA | CG>TA | ACTGGGCCG | CCGGCGT | CGG | GCC | C | A | C>T | C>T_GCC | CG>TA_CCGGCGT | 4 | FALSE | FALSE |
| 166_v.txt | 33118 | 33118-SNV- | SNV | 1 | C | T | 23 | 21126 | 0.10887059 | 41.6521739 | rnr1Y285F_pe | FALSE | ARG | JSY3866 | 4 | rnr1Y285F_p | GCG | C>T | A/T+1 | G/C-1 | CG>TA | CG>TA | GGCCGGCGT | CGTTGGT | CGC | GCG | C | A | C>T | C>T_GCG | CG>TA_CGTTGGT | 4 | FALSE | FALSE |
| 166_v.txt | 33258 | 33258-SNV- | SNV | 1 | C | T | 24 | 18839 | 0.1273953 | 47.0833333 | rnr1Y285F_pe | FALSE | ARG | JSY3866 | 4 | rnr1Y285F_p | GCC | C>T | A/T+1 | G/C-1 | CG>TA | CG>TA | CCTATGCCAT | CCATCCT | CGG | GCC | C | A | C>T | C>T_GCC | CG>TA_CCATCCT | 4 | FALSE | FALSE |
| 166_v.txt | 33276 | 33276-SNV- | SNV | 1 | G | A | 48 | 18839 | 0.25479059 | 44.5416667 | rnr1Y285F_pe | FALSE | ARG | JSY3866 | 4 | rnr1Y285F_p | CGT | G>A | A/T+1 | G/C-1 | CG>TA | CG>TA | GAGAACGTA | GTATCCT | GCA | GCA | C | T | C>T | C>T_GCA | CG>TA_GTATCCT | 4 | FALSE | FALSE |
| 166_v.txt | 33309 | 33309-SNV- | SNV | 1 | C | T | 21 | 18839 | 0.11147089 | 55.8095238 | rnr1Y285F_pe | FALSE | ARG | JSY3866 | 4 | rnr1Y285F_p | GCG | C>T | A/T+1 | G/C-1 | CG>TA | CG>TA | AAAGAGCGC | CGCAATG | CGC | GCG | C | A | C>T | C>T_GCG | CG>TA_CGCAATG | 4 | FALSE | FALSE |
| 166_v.txt | 33392 | 33392-SNV- | SNV | 1 | G | A | 24 | 18839 | 0.1273953 | 53.8333333 | rnr1Y285F_pe | FALSE | ARG | JSY3866 | 4 | rnr1Y285F_p | CGT | G>A | A/T+1 | G/C-1 | CG>TA | CG>TA | AACGTCGTG | GTGAAAG | GCA | GCA | C | T | C>T | C>T_GCA | CG>TA_GTGAAAG | 4 | FALSE | FALSE |
| 166_v.txt | 33412 | 33412-SNV- | SNV | 1 | G | A | 28 | 18839 | 0.14862785 | 48.9285714 | rnr1Y285F_pe | FALSE | ARG | JSY3866 | 4 | rnr1Y285F_p | GGC | G>A | A/T+1 | G/C-1 | CG>TA | CG>TA | TGACCGGCT | GCTCATT | CCG | CCG | C | T | C>T | C>T_CCG | CG>TA_GCTCATT | 4 | FALSE | FALSE |
| 166_v.txt | 33474 | 33474-SNV- | SNV | 1 | C | T | 25 | 18839 | 0.13270343 | 34.04 | rnr1Y285F_pe | FALSE | ARG | JSY3866 | 4 | rnr1Y285F_p | GCC | C>T | A/T+1 | G/C-1 | CG>TA | CG>TA | GCTATGCCTT | CCTTTTT | CGG | GCC | C | A | C>T | C>T_GCC | CG>TA_CCTTTTT | 4 | FALSE | FALSE |
| 166_v.txt | 32535 | 32535-SNV- | SNV | 1 | G | C | 2 | 18012 | 0.01110371 | 24.9998896 | rnr1Y285F_pe | FALSE | ARG | JSY3866 | 4 | rnr1Y285F_p | TGC | G>C | G/C+1 | G/C-1 | CG>GC | CG>GC | GGGTTTGCA | GCAGCTT | ACG | ACG | C | G | C>G | C>G_ACG | CG>GC_GCAGCTT | 4 | FALSE | FALSE |
| 166_v.txt | 33474 | 33474-SNV- | SNV | 1 | C | A | 63 | 18839 | 0.33441266 | 35.1269841 | rnr1Y285F_pe | FALSE | ARG | JSY3866 | 4 | rnr1Y285F_p | GCC | C>A | A/T+1 | G/C-1 | CG>AT | CG>AT | GCTATGCCTT | CCTTTTT | CGG | GCC | C | T | C>A | C>A_GCC | CG>AT_CCTTTTT | 4 | FALSE | FALSE |
| 166_v.txt | 33475 | 33475-SNV- | SNV | 1 | C | A | 19 | 18839 | 0.10085461 | 35.9473684 | rnr1Y285F_pe | FALSE | ARG | JSY3866 | 4 | rnr1Y285F_p | CCT | C>A | A/T+1 | G/C-1 | CG>AT | CG>AT | CTATGCCTTT | CTTTTTT | GGA | CCT | C | T | C>A | C>A_CCT | CG>AT_CTTTTTT | 4 | FALSE | FALSE |
| 166_v.txt | 32081 | 32081-Insert | Insertion | 1 | - | A | 11 | 16372 | 0.06718788 | 61.6363636 | rnr1Y285F_pe | FALSE | ARG | JSY3866 | 4 | rnr1Y285F_p | CAA | nope | A/T+1 | >1bpdel | none | A/T+1 | CCATGCAAA | AAAAAAG | GTT | CAA | - | T | NA | NA_CAA | A/T+1_AAAAAAG | 4 | FALSE | FALSE |

| | | | | | | | | | | | | | | | | | | | | | | | | | | | | | | | | | | |
|---|---|---|---|---|---|---|---|---|---|---|---|---|---|---|---|---|---|---|---|---|---|---|---|---|---|---|---|---|---|---|---|---|---|---|
| 166_v.txt | 32842 | 32842-Insert | Insertion | 1 | - | A | 10 | 27926 | 0.03580892 | 43.2 | rnr1Y285F_pe | FALSE | ARG | JSY3866 | 4 | rnr1Y285F_p | CAA | nope | A/T+1 | >1bpdel | none | A/T+1 | TTACCCAAA | AAAAAAT | GTT | CAA | - | T | NA | NA_CAA | A/T+1_AAAAAAT | 4 | FALSE | FALSE |
| 166_v.txt | 31723 | 31723-Deleti | Deletion | 1 | A | - | 12 | 7643 | 0.15700641 | 57 | rnr1Y285F_pe | FALSE | ARG | JSY3866 | 4 | rnr1Y285F_p | CAA | nope | >1bp-ins | A/T-1 | none | A/T-1 | TGTCCCAAA | AAAAAGT | GTT | GTT | T | none | T>none | T>none_GTT | A/T-1_AAAAAGT | 4 | FALSE | FALSE |
| 166_v.txt | 31904 | 31904-Deleti | Deletion | 1 | T | - | 6 | 7643 | 0.07850321 | 54.8333333 | rnr1Y285F_pe | FALSE | ARG | JSY3866 | 4 | rnr1Y285F_p | ATT | nope | >1bp-ins | A/T-1 | none | A/T-1 | ATTGAATTTT | TTTTGGT | TAA | ATT | T | none | nope | nope_ATT | A/T-1_TTTTGGT | 4 | FALSE | FALSE |
| 166_v.txt | 32081 | 32081-Deleti | Deletion | 1 | A | - | 13 | 16372 | 0.07940386 | 63.9230769 | rnr1Y285F_pe | FALSE | ARG | JSY3866 | 4 | rnr1Y285F_p | CAA | nope | >1bp-ins | A/T-1 | none | A/T-1 | CCATGCAAA | AAAAAAG | GTT | GTT | T | none | T>none | T>none_GTT | A/T-1_AAAAAAG | 4 | FALSE | FALSE |
| 166_v.txt | 32346 | 32346-Deleti | Deletion | 1 | A | - | 6 | 18012 | 0.03331113 | 63 | rnr1Y285F_pe | FALSE | ARG | JSY3866 | 4 | rnr1Y285F_p | CAA | nope | >1bp-ins | A/T-1 | none | A/T-1 | TGTGGCAAA | AAAACCT | GTT | GTT | T | none | T>none | T>none_GTT | A/T-1_AAAACCT | 4 | FALSE | FALSE |
| 166_v.txt | 32498 | 32498-Deleti | Deletion | 1 | T | - | 26 | 18012 | 0.14434821 | 43.7307692 | rnr1Y285F_pe | FALSE | ARG | JSY3866 | 4 | rnr1Y285F_p | CTT | nope | >1bp-ins | A/T-1 | none | A/T-1 | AACAACTTTT | TTTTTTG | GAA | CTT | T | none | nope | nope_CTT | A/T-1_TTTTTTG | 4 | FALSE | FALSE |
| 166_v.txt | 32631 | 32631-Deleti | Deletion | 1 | T | - | 6 | 27926 | 0.02148535 | 46.3333333 | rnr1Y285F_pe | FALSE | ARG | JSY3866 | 4 | rnr1Y285F_p | GTT | nope | >1bp-ins | A/T-1 | none | A/T-1 | CCTTCGTTTT | TTTTTAT | CAA | GTT | T | none | nope | nope_GTT | A/T-1_TTTTTAT | 4 | FALSE | FALSE |
| 166_v.txt | 32647 | 32647-Deleti | Deletion | 1 | T | - | 6 | 27926 | 0.02148535 | 59 | rnr1Y285F_pe | FALSE | ARG | JSY3866 | 4 | rnr1Y285F_p | ATT | nope | >1bp-ins | A/T-1 | none | A/T-1 | TAGATATTAT | TTATACC | TAA | ATT | T | none | nope | nope_ATT | A/T-1_TTATACC | 4 | FALSE | FALSE |
| 166_v.txt | 32658 | 32658-Deleti | Deletion | 1 | C | - | 12 | 27926 | 0.04297071 | 56.0833333 | rnr1Y285F_pe | FALSE | ARG | JSY3866 | 4 | rnr1Y285F_p | ACC | nope | >1bp-ins | G/C-1 | none | G/C-1 | CCTGGACCC | CCCCAGG | TGG | ACC | C | none | nope | nope_ACC | G/C-1_CCCCAGG | 4 | FALSE | FALSE |
| 166_v.txt | 32728 | 32728-Deleti | Deletion | 1 | A | - | 6 | 27926 | 0.02148535 | 61.5 | rnr1Y285F_pe | FALSE | ARG | JSY3866 | 4 | rnr1Y285F_p | CAA | nope | >1bp-ins | A/T-1 | none | A/T-1 | CCATACAAA | AAAAACA | GTT | GTT | T | none | T>none | T>none_GTT | A/T-1_AAAAACA | 4 | FALSE | FALSE |
| 166_v.txt | 32842 | 32842-Deleti | Deletion | 1 | A | - | 35 | 27926 | 0.12533123 | 40.7428569 | rnr1Y285F_pe | FALSE | ARG | JSY3866 | 4 | rnr1Y285F_p | CAA | nope | >1bp-ins | A/T-1 | none | A/T-1 | TTACCCAAA | AAAAAAT | GTT | GTT | T | none | T>none | T>none_GTT | A/T-1_AAAAAAT | 4 | FALSE | FALSE |
| 166_v.txt | 32884 | 32884-Deleti | Deletion | 1 | A | - | 7 | 21126 | 0.03313453 | 36.5714286 | rnr1Y285F_pe | FALSE | ARG | JSY3866 | 4 | rnr1Y285F_p | CAA | nope | >1bp-ins | A/T-1 | none | A/T-1 | ACGTCCAAA | AAAATTG | GTT | GTT | T | none | T>none | T>none_GTT | A/T-1_AAAATTG | 4 | FALSE | FALSE |
| 166_v.txt | 32924 | 32924-Deleti | Deletion | 1 | A | - | 24 | 21126 | 0.11360409 | 52.1666667 | rnr1Y285F_pe | FALSE | ARG | JSY3866 | 4 | rnr1Y285F_p | CAA | nope | >1bp-ins | A/T-1 | none | A/T-1 | CAGGGCAAA | AAAAGTG | GTT | GTT | T | none | T>none | T>none_GTT | A/T-1_AAAAGTG | 4 | FALSE | FALSE |
| 166_v.txt | 32935 | 32935-Deleti | Deletion | 1 | C | - | 5 | 21126 | 0.02366752 | 50.8 | rnr1Y285F_pe | FALSE | ARG | JSY3866 | 4 | rnr1Y285F_p | GCC | nope | >1bp-ins | G/C-1 | none | G/C-1 | TGATTGCCC | CCCAAGA | CGG | GCC | C | none | nope | nope_GCC | G/C-1_CCCAAGA | 4 | FALSE | FALSE |
| 166_v.txt | 33239 | 33239-Deleti | Deletion | 1 | T | - | 4 | 18839 | 0.02123255 | 43.5 | rnr1Y285F_pe | FALSE | ARG | JSY3866 | 4 | rnr1Y285F_p | CTT | nope | >1bp-ins | A/T-1 | none | A/T-1 | TTCTCCTTCA | TTCATCT | GAA | CTT | T | none | nope | nope_CTT | A/T-1_TTCATCT | 4 | FALSE | FALSE |
| 166_v.txt | 33474 | 33474-Deleti | Deletion | 1 | C | - | 51 | 18839 | 0.27071501 | 28.6470588 | rnr1Y285F_pe | FALSE | ARG | JSY3866 | 4 | rnr1Y285F_p | GCC | nope | >1bp-ins | G/C-1 | none | G/C-1 | GCTATGCCTT | CCTTTTT | CGG | GCC | C | none | nope | nope_GCC | G/C-1_CCTTTTT | 4 | FALSE | FALSE |
| 133_v.txt | 31723 | 31723-SNV-A | SNV | 1 | A | C | 4 | 6373 | 0.06276479 | 64 | rnr1Y285F_pG | TRUE | CAN | JSY2420 | 1 | rnr1Y285F_p | CAA | A>C | G/C+1 | A/T-1 | TA>GC | TA>GC | TGTCCCAAA | AAAAAGT | GTT | GTT | T | G | T>G | T>G_GTT | TA>GC_AAAAAGT | 6 | FALSE | FALSE |
| 133_v.txt | 31854 | 31854-SNV-A | SNV | 1 | A | C | 4 | 6373 | 0.06276479 | 37 | rnr1Y285F_pG | TRUE | CAN | JSY2420 | 1 | rnr1Y285F_p | TAA | A>C | G/C+1 | A/T-1 | TA>GC | TA>GC | ACAGCTAAG | AAGAACA | ATT | ATT | T | G | T>G | T>G_ATT | TA>GC_AAGAACA | 6 | FALSE | FALSE |
| 133_v.txt | 32073 | 32073-SNV-A | SNV | 1 | A | C | 16 | 15057 | 0.10626287 | 63.75 | rnr1Y285F_pG | TRUE | CAN | JSY2420 | 1 | rnr1Y285F_p | TAA | A>C | G/C+1 | A/T-1 | TA>GC | TA>GC | ATAAATAAC | AACCATG | ATT | ATT | T | G | T>G | T>G_ATT | TA>GC_AACCATG | 6 | FALSE | FALSE |
| 133_v.txt | 32294 | 32294-SNV-A | SNV | 1 | A | C | 19 | 21906 | 0.08673423 | 45.8947368 | rnr1Y285F_pG | TRUE | CAN | JSY2420 | 1 | rnr1Y285F_p | AAT | A>C | G/C+1 | A/T-1 | TA>GC | TA>GC | ATTTGAATTT | ATTTGCG | TTA | TTA | T | G | T>G | T>G_TTA | TA>GC_ATTTGCG | 6 | FALSE | FALSE |
| 133_v.txt | 32631 | 32631-SNV-T | SNV | 1 | T | G | 2 | 17261 | 0.01158681 | 53.0000919 | rnr1Y285F_pG | TRUE | CAN | JSY2420 | 1 | rnr1Y285F_p | GTT | T>G | G/C+1 | A/T-1 | TA>GC | TA>GC | CCTTCGTTTT | TTTTTAT | CAA | GTT | T | C | T>G | T>G_GTT | TA>GC_TTTTTAT | 6 | FALSE | FALSE |
| 133_v.txt | 32717 | 32717-SNV-A | SNV | 1 | A | C | 1 | 17260 | 5.79E-03 | 63.9908486 | rnr1Y285F_pG | TRUE | CAN | JSY2420 | 1 | rnr1Y285F_p | CAC | A>C | G/C+1 | A/T-1 | TA>GC | TA>GC | AGCACCACA | ACAAACC | GTG | GTG | T | G | T>G | T>G_GTG | TA>GC_ACAAACC | 6 | FALSE | FALSE |
| 133_v.txt | 32801 | 32801-SNV-A | SNV | 1 | A | C | 6 | 17261 | 0.03476044 | 46.6666667 | rnr1Y285F_pG | TRUE | CAN | JSY2420 | 1 | rnr1Y285F_p | AAT | A>C | G/C+1 | A/T-1 | TA>GC | TA>GC | ACCGTAATAT | ATATTTG | TTA | TTA | T | G | T>G | T>G_TTA | TA>GC_ATATTTG | 6 | FALSE | FALSE |
| 133_v.txt | 32842 | 32842-SNV-A | SNV | 1 | A | C | 5 | 17259 | 0.02897039 | 38 | rnr1Y285F_pG | TRUE | CAN | JSY2420 | 1 | rnr1Y285F_p | CAA | A>C | G/C+1 | A/T-1 | TA>GC | TA>GC | TTACCCAAA | AAAAAAT | GTT | GTT | T | G | T>G | T>G_GTT | TA>GC_AAAAAAT | 6 | FALSE | FALSE |
| 133_v.txt | 32884 | 32884-SNV-A | SNV | 1 | A | C | 2 | 20066 | 9.97E-03 | 37.9960869 | rnr1Y285F_pG | TRUE | CAN | JSY2420 | 1 | rnr1Y285F_p | CAA | A>C | G/C+1 | A/T-1 | TA>GC | TA>GC | ACGTCCAAA | AAAATTG | GTT | GTT | T | G | T>G | T>G_GTT | TA>GC_AAAATTG | 6 | FALSE | FALSE |
| 133_v.txt | 33028 | 33028-SNV-T | SNV | 1 | T | G | 3 | 20066 | 0.01495066 | 33.9993835 | rnr1Y285F_pG | TRUE | CAN | JSY2420 | 1 | rnr1Y285F_p | ATG | T>G | G/C+1 | A/T-1 | TA>GC | TA>GC | CAGGGATGA | TGAATGT | TAC | ATG | T | C | T>G | T>G_ATG | TA>GC_TGAATGT | 6 | FALSE | FALSE |
| 133_v.txt | 33032 | 33032-SNV-T | SNV | 1 | T | G | 2 | 20066 | 9.97E-03 | 63.9974765 | rnr1Y285F_pG | TRUE | CAN | JSY2420 | 1 | rnr1Y285F_p | ATG | T>G | G/C+1 | A/T-1 | TA>GC | TA>GC | GATGAATGT | TGTAGCC | TAC | ATG | T | C | T>G | T>G_ATG | TA>GC_TGTAGCC | 6 | FALSE | FALSE |
| 133_v.txt | 33065 | 33065-SNV-A | SNV | 1 | A | C | 3587 | 20059 | 17.8822474 | 53.574752 | rnr1Y285F_pG | TRUE | CAN | JSY2420 | 1 | rnr1Y285F_p | AAT | A>C | G/C+1 | A/T-1 | TA>GC | TA>GC | GACAGAATA | ATATGCC | TTA | TTA | T | G | T>G | T>G_TTA | TA>GC_ATATGCC | 6 | FALSE | FALSE |
| 133_v.txt | 33089 | 33089-SNV-A | SNV | 1 | A | C | 2 | 20066 | 9.97E-03 | 44.0000549 | rnr1Y285F_pG | TRUE | CAN | JSY2420 | 1 | rnr1Y285F_p | AAT | A>C | G/C+1 | A/T-1 | TA>GC | TA>GC | AAATAAATA | ATATGAT | TTA | TTA | T | G | T>G | T>G_TTA | TA>GC_ATATGAT | 6 | FALSE | FALSE |
| 133_v.txt | 33192 | 33192-SNV-T | SNV | 1 | T | G | 7 | 10944 | 0.06396199 | 37.8571429 | rnr1Y285F_pG | TRUE | CAN | JSY2420 | 1 | rnr1Y285F_p | ATG | T>G | G/C+1 | A/T-1 | TA>GC | TA>GC | CCAATATGT | TGTCTTT | TAC | ATG | T | C | T>G | T>G_ATG | TA>GC_TGTCTTT | 6 | FALSE | FALSE |
| 133_v.txt | 33204 | 33204-SNV-A | SNV | 1 | A | C | 9 | 10943 | 0.08224436 | 29.668696 | rnr1Y285F_pG | TRUE | CAN | JSY2420 | 1 | rnr1Y285F_p | AAG | A>C | G/C+1 | A/T-1 | TA>GC | TA>GC | TGCTTAAGC | AGCTCTC | TTC | TTC | T | G | T>G | T>G_TTC | TA>GC_AGCTCTC | 6 | FALSE | FALSE |
| 133_v.txt | 33269 | 33269-SNV-A | SNV | 1 | A | C | 89 | 10945 | 0.81315669 | 16.1910112 | rnr1Y285F_pG | TRUE | CAN | JSY2420 | 1 | rnr1Y285F_p | TAG | A>C | G/C+1 | A/T-1 | TA>GC | TA>GC | CTCCATAGA | AGAGAAC | ATC | ATC | T | G | T>G | T>G_ATC | TA>GC_AGAGAAC | 6 | FALSE | FALSE |
| 133_v.txt | 33317 | 33317-SNV-A | SNV | 1 | A | C | 10 | 10945 | 0.09136592 | 45.4 | rnr1Y285F_pG | TRUE | CAN | JSY2420 | 1 | rnr1Y285F_p | GAT | A>C | G/C+1 | A/T-1 | TA>GC | TA>GC | CAATGGATA | ATACAAT | CTA | CTA | T | G | T>G | T>G_CTA | TA>GC_ATACAAT | 6 | FALSE | FALSE |
| 133_v.txt | 32346 | 32346-SNV-A | SNV | 1 | A | G | 2 | 21906 | 9.13E-03 | 52.4999228 | rnr1Y285F_pG | TRUE | CAN | JSY2420 | 1 | rnr1Y285F_p | CAA | A>G | G/C+1 | A/T-1 | TA>CG | TA>CG | TGTGGCAAA | AAAACCT | GTT | GTT | T | C | T>C | T>C_GTT | TA>CG_AAAACCT | 6 | FALSE | FALSE |
| 133_v.txt | 32475 | 32475-SNV-A | SNV | 1 | A | G | 5531 | 21906 | 25.2487903 | 46.6050781 | rnr1Y285F_pG | TRUE | CAN | JSY2420 | 1 | rnr1Y285F_p | GAA | A>G | G/C+1 | A/T-1 | TA>CG | TA>CG | ATGTAGAAG | AAGGTTA | CTT | CTT | T | C | T>C | T>C_CTT | TA>CG_AAGGTTA | 6 | FALSE | FALSE |
| 133_v.txt | 32795 | 32795-SNV-A | SNV | 1 | A | G | 6 | 17261 | 0.03476044 | 46.3332627 | rnr1Y285F_pG | TRUE | CAN | JSY2420 | 1 | rnr1Y285F_p | CAC | A>G | G/C+1 | A/T-1 | TA>CG | TA>CG | GAATTCACC | ACCGTAA | GTG | GTG | T | C | T>C | T>C_GTG | TA>CG_ACCGTAA | 6 | FALSE | FALSE |
| 133_v.txt | 32842 | 32842-SNV-A | SNV | 1 | A | G | 7 | 17259 | 0.04055855 | 37.4285714 | rnr1Y285F_pG | TRUE | CAN | JSY2420 | 1 | rnr1Y285F_p | CAA | A>G | G/C+1 | A/T-1 | TA>CG | TA>CG | TTACCCAAA | AAAAAAT | GTT | GTT | T | C | T>C | T>C_GTT | TA>CG_AAAAAAT | 6 | FALSE | FALSE |
| 133_v.txt | 32941 | 32941-SNV-A | SNV | 1 | A | G | 3 | 20066 | 0.01495066 | 49.9965545 | rnr1Y285F_pG | TRUE | CAN | JSY2420 | 1 | rnr1Y285F_p | GAA | A>G | G/C+1 | A/T-1 | TA>CG | TA>CG | CCCAAGAAA | AAAACCA | CTT | CTT | T | C | T>C | T>C_CTT | TA>CG_AAAACCA | 6 | FALSE | FALSE |
| 133_v.txt | 33027 | 33027-SNV-A | SNV | 1 | A | G | 3 | 20066 | 0.01495066 | 63.9997719 | rnr1Y285F_pG | TRUE | CAN | JSY2420 | 1 | rnr1Y285F_p | GAT | A>G | G/C+1 | A/T-1 | TA>CG | TA>CG | ACAGGGATG | ATGAATG | CTA | CTA | T | C | T>C | T>C_CTA | TA>CG_ATGAATG | 6 | FALSE | FALSE |
| 133_v.txt | 33035 | 33035-SNV-A | SNV | 1 | A | G | 9 | 20066 | 0.04485199 | 48.2223783 | rnr1Y285F_pG | TRUE | CAN | JSY2420 | 1 | rnr1Y285F_p | TAG | A>G | G/C+1 | A/T-1 | TA>CG | TA>CG | GAATGTAGC | AGCCATT | ATC | ATC | T | C | T>C | T>C_ATC | TA>CG_AGCCATT | 6 | FALSE | FALSE |
| 133_v.txt | 33090 | 33090-SNV-T | SNV | 1 | T | C | 5 | 20066 | 0.02491777 | 53.3986916 | rnr1Y285F_pG | TRUE | CAN | JSY2420 | 1 | rnr1Y285F_p | ATA | T>C | G/C+1 | A/T-1 | TA>CG | TA>CG | AATAAATAT | TATGATA | TAT | ATA | T | G | T>C | T>C_ATA | TA>CG_TATGATA | 6 | FALSE | FALSE |
| 133_v.txt | 33155 | 33155-SNV-T | SNV | 1 | T | C | 8 | 20065 | 0.03987042 | 37.1248998 | rnr1Y285F_pG | TRUE | CAN | JSY2420 | 1 | rnr1Y285F_p | CTG | T>C | G/C+1 | A/T-1 | TA>CG | TA>CG | AAGACCTGT | TGTACCA | GAC | CTG | T | G | T>C | T>C_CTG | TA>CG_TGTACCA | 6 | FALSE | FALSE |
| 133_v.txt | 31854 | 31854-SNV-A | SNV | 1 | A | T | 18 | 6373 | 0.28244155 | 44.6666667 | rnr1Y285F_pG | TRUE | CAN | JSY2420 | 1 | rnr1Y285F_p | TAA | A>T | A/T+1 | A/T-1 | TA>AT | TA>AT | ACAGCTAAG | AAGAACA | ATT | ATT | T | A | T>A | T>A_ATT | TA>AT_AAGAACA | 6 | FALSE | FALSE |
| 133_v.txt | 32160 | 32160-SNV-A | SNV | 1 | A | T | 18 | 15057 | 0.11954573 | 45.2222222 | rnr1Y285F_pG | TRUE | CAN | JSY2420 | 1 | rnr1Y285F_p | CAA | A>T | A/T+1 | A/T-1 | TA>AT | TA>AT | TAAGCCAAA | AAAGCGC | GTT | GTT | T | A | T>A | T>A_GTT | TA>AT_AAAGCGC | 6 | FALSE | FALSE |
| 133_v.txt | 32346 | 32346-SNV-A | SNV | 1 | A | T | 3 | 21906 | 0.01369488 | 63.9997087 | rnr1Y285F_pG | TRUE | CAN | JSY2420 | 1 | rnr1Y285F_p | CAA | A>T | A/T+1 | A/T-1 | TA>AT | TA>AT | TGTGGCAAA | AAAACCT | GTT | GTT | T | A | T>A | T>A_GTT | TA>AT_AAAACCT | 6 | FALSE | FALSE |
| 133_v.txt | 32456 | 32456-SNV-T | SNV | 1 | T | A | 5 | 21904 | 0.02282688 | 52.0045192 | rnr1Y285F_pG | TRUE | CAN | JSY2420 | 1 | rnr1Y285F_p | ATA | T>A | A/T+1 | A/T-1 | TA>AT | TA>AT | AATGAATAA | TAATAGA | TAT | ATA | T | T | T>A | T>A_ATA | TA>AT_TAATAGA | 6 | FALSE | FALSE |
| 133_v.txt | 32631 | 32631-SNV-T | SNV | 1 | T | A | 5 | 17261 | 0.02896704 | 42.8000437 | rnr1Y285F_pG | TRUE | CAN | JSY2420 | 1 | rnr1Y285F_p | GTT | T>A | A/T+1 | A/T-1 | TA>AT | TA>AT | CCTTCGTTTT | TTTTTAT | CAA | GTT | T | T | T>A | T>A_GTT | TA>AT_TTTTTAT | 6 | FALSE | FALSE |
| 133_v.txt | 32635 | 32635-SNV-T | SNV | 1 | T | A | 22 | 17261 | 0.12745496 | 52.2727273 | rnr1Y285F_pG | TRUE | CAN | JSY2420 | 1 | rnr1Y285F_p | TTA | T>A | A/T+1 | A/T-1 | TA>AT | TA>AT | CGTTTTTATC | TATCCTT | AAT | TTA | T | T | T>A | T>A_TTA | TA>AT_TATCCTT | 6 | FALSE | FALSE |
| 133_v.txt | 32657 | 32657-SNV-A | SNV | 1 | A | T | 3 | 17260 | 0.01738123 | 33.3329256 | rnr1Y285F_pG | TRUE | CAN | JSY2420 | 1 | rnr1Y285F_p | GAC | A>T | A/T+1 | A/T-1 | TA>AT | TA>AT | ACCTGGACC | ACCCCAG | CTG | CTG | T | A | T>A | T>A_CTG | TA>AT_ACCCCAG | 6 | FALSE | FALSE |
| 133_v.txt | 32992 | 32992-SNV-T | SNV | 1 | T | A | 19 | 20066 | 0.09468753 | 56.631579 | rnr1Y285F_pG | TRUE | CAN | JSY2420 | 1 | rnr1Y285F_p | CTT | T>A | A/T+1 | A/T-1 | TA>AT | TA>AT | GGAATCTTT | TTTGTGA | GAA | CTT | T | T | T>A | T>A_CTT | TA>AT_TTTGTGA | 6 | FALSE | FALSE |
| 133_v.txt | 33023 | 33023-SNV-A | SNV | 1 | A | T | 4 | 20066 | 0.01993422 | 63.9999989 | rnr1Y285F_pG | TRUE | CAN | JSY2420 | 1 | rnr1Y285F_p | CAG | A>T | A/T+1 | A/T-1 | TA>AT | TA>AT | TGTAACAGG | AGGGATG | GTC | GTC | T | A | T>A | T>A_GTC | TA>AT_AGGGATG | 6 | FALSE | FALSE |
| 133_v.txt | 33027 | 33027-SNV-A | SNV | 1 | A | T | 2 | 20066 | 9.97E-03 | 47.9997679 | rnr1Y285F_pG | TRUE | CAN | JSY2420 | 1 | rnr1Y285F_p | GAT | A>T | A/T+1 | A/T-1 | TA>AT | TA>AT | ACAGGGATG | ATGAATG | CTA | CTA | T | A | T>A | T>A_CTA | TA>AT_ATGAATG | 6 | FALSE | FALSE |
| 133_v.txt | 33028 | 33028-SNV-T | SNV | 1 | T | A | 8 | 20066 | 0.03986843 | 43.217551 | rnr1Y285F_pG | TRUE | CAN | JSY2420 | 1 | rnr1Y285F_p | ATG | T>A | A/T+1 | A/T-1 | TA>AT | TA>AT | CAGGGATGA | TGAATGT | TAC | ATG | T | T | T>A | T>A_ATG | TA>AT_TGAATGT | 6 | FALSE | FALSE |
| 133_v.txt | 33066 | 33066-SNV-T | SNV | 1 | T | A | 2 | 20066 | 9.97E-03 | 63.9996125 | rnr1Y285F_pG | TRUE | CAN | JSY2420 | 1 | rnr1Y285F_p | ATA | T>A | A/T+1 | A/T-1 | TA>AT | TA>AT | ACAGAATAT | TATGCCA | TAT | ATA | T | T | T>A | T>A_ATA | TA>AT_TATGCCA | 6 | FALSE | FALSE |
| 133_v.txt | 33155 | 33155-SNV-T | SNV | 1 | T | A | 2 | 20065 | 9.97E-03 | 17.9996879 | rnr1Y285F_pG | TRUE | CAN | JSY2420 | 1 | rnr1Y285F_p | CTG | T>A | A/T+1 | A/T-1 | TA>AT | TA>AT | AAGACCTGT | TGTACCA | GAC | CTG | T | T | T>A | T>A_CTG | TA>AT_TGTACCA | 6 | FALSE | FALSE |
| 133_v.txt | 33162 | 33162-SNV-A | SNV | 1 | A | T | 18 | 20064 | 0.08971292 | 34.7777778 | rnr1Y285F_pG | TRUE | CAN | JSY2420 | 1 | rnr1Y285F_p | AAT | A>T | A/T+1 | A/T-1 | TA>AT | TA>AT | GTACCAATA | ATAGTAC | TTA | TTA | T | A | T>A | T>A_TTA | TA>AT_ATAGTAC | 6 | FALSE | FALSE |
| 133_v.txt | 33202 | 33202-SNV-T | SNV | 1 | T | A | 22 | 10943 | 0.20104176 | 37.3636364 | rnr1Y285F_pG | TRUE | CAN | JSY2420 | 1 | rnr1Y285F_p | TTA | T>A | A/T+1 | A/T-1 | TA>AT | TA>AT | TTTGCTTAAG | TAAGCTC | AAT | TTA | T | T | T>A | T>A_TTA | TA>AT_TAAGCTC | 6 | FALSE | FALSE |
| 133_v.txt | 33209 | 33209-SNV-T | SNV | 1 | T | A | 27 | 10945 | 0.24668799 | 38.1111111 | rnr1Y285F_pG | TRUE | CAN | JSY2420 | 1 | rnr1Y285F_p | CTC | T>A | A/T+1 | A/T-1 | TA>AT | TA>AT | AAGCTCTCT | TCTCTTC | GAG | CTC | T | T | T>A | T>A_CTC | TA>AT_TCTCTTC | 6 | FALSE | FALSE |
| 133_v.txt | 33317 | 33317-SNV-A | SNV | 1 | A | T | 58 | 10945 | 0.52992234 | 60.7068966 | rnr1Y285F_pG | TRUE | CAN | JSY2420 | 1 | rnr1Y285F_p | GAT | A>T | A/T+1 | A/T-1 | TA>AT | TA>AT | CAATGGATA | ATACAAT | CTA | CTA | T | A | T>A | T>A_CTA | TA>AT_ATACAAT | 6 | FALSE | FALSE |
| 133_v.txt | 31650 | 31650-SNV-C | SNV | 1 | C | T | 16 | 6373 | 0.25105916 | 32.8125 | rnr1Y285F_pG | TRUE | CAN | JSY2420 | 1 | rnr1Y285F_p | GCG | C>T | A/T+1 | G/C-1 | CG>TA | CG>TA | AGAATGCGA | CGAAATG | CGC | GCG | C | A | C>T | C>T_GCG | CG>TA_CGAAATG | 6 | FALSE | FALSE |
| 133_v.txt | 31844 | 31844-SNV-C | SNV | 1 | C | T | 50 | 6373 | 0.78455986 | 49.8 | rnr1Y285F_pG | TRUE | CAN | JSY2420 | 1 | rnr1Y285F_p | TCC | C>T | A/T+1 | G/C-1 | CG>TA | CG>TA | TAAGATCCA | CCAAACA | AGG | TCC | C | A | C>T | C>T_TCC | CG>TA_CCAAACA | 6 | FALSE | FALSE |
| 133_v.txt | 31882 | 31882-SNV-C | SNV | 1 | C | T | 12 | 6373 | 0.18829437 | 41.5 | rnr1Y285F_pG | TRUE | CAN | JSY2420 | 1 | rnr1Y285F_p | GCA | C>T | A/T+1 | G/C-1 | CG>TA | CG>TA | AGGCGGCAG | CAGCAAA | CGT | GCA | C | A | C>T | C>T_GCA | CG>TA_CAGCAAA | 6 | FALSE | FALSE |
| 133_v.txt | 31962 | 31962-SNV-G | SNV | 1 | G | A | 32 | 15057 | 0.21252574 | 36.84375 | rnr1Y285F_pG | TRUE | CAN | JSY2420 | 1 | rnr1Y285F_p | CGC | G>A | A/T+1 | G/C-1 | CG>TA | CG>TA | GTGGCCGCA | GCATAAT | GCG | GCG | C | T | C>T | C>T_GCG | CG>TA_GCATAAT | 6 | FALSE | FALSE |
| 133_v.txt | 31981 | 31981-SNV-G | SNV | 1 | G | A | 19 | 15057 | 0.12618716 | 48.8421053 | rnr1Y285F_pG | TRUE | CAN | JSY2420 | 1 | rnr1Y285F_p | GGC | G>A | A/T+1 | G/C-1 | CG>TA | CG>TA | AGCCGGGCA | GCATTAA | CCG | CCG | C | T | C>T | C>T_CCG | CG>TA_GCATTAA | 6 | FALSE | FALSE |
| 133_v.txt | 32041 | 32041-SNV-G | SNV | 1 | G | A | 16 | 15057 | 0.10626287 | 60.4375 | rnr1Y285F_pG | TRUE | CAN | JSY2420 | 1 | rnr1Y285F_p | TGC | G>A | A/T+1 | G/C-1 | CG>TA | CG>TA | AAGCTTGCA | GCATAAA | ACG | ACG | C | T | C>T | C>T_ACG | CG>TA_GCATAAA | 6 | FALSE | FALSE |
| 133_v.txt | 32057 | 32057-SNV-C | SNV | 1 | C | T | 19 | 15057 | 0.12618716 | 61.3157895 | rnr1Y285F_pG | TRUE | CAN | JSY2420 | 1 | rnr1Y285F_p | GCG | C>T | A/T+1 | G/C-1 | CG>TA | CG>TA | GATGTGCGA | CGAGATT | CGC | GCG | C | A | C>T | C>T_GCG | CG>TA_CGAGATT | 6 | FALSE | FALSE |
| 133_v.txt | 32075 | 32075-SNV-C | SNV | 1 | C | T | 99 | 15057 | 0.65750149 | 61.8377201 | rnr1Y285F_pG | TRUE | CAN | JSY2420 | 1 | rnr1Y285F_p | ACC | C>T | A/T+1 | G/C-1 | CG>TA | CG>TA | AAATAACCA | CCATGCA | TGG | ACC | C | A | C>T | C>T_ACC | CG>TA_CCATGCA | 6 | FALSE | FALSE |
| 133_v.txt | 32076 | 32076-SNV-C | SNV | 1 | C | T | 22 | 15056 | 0.14612115 | 62.9545455 | rnr1Y285F_pG | TRUE | CAN | JSY2420 | 1 | rnr1Y285F_p | CCA | C>T | A/T+1 | G/C-1 | CG>TA | CG>TA | AATAACCAT | CATGCAA | GGT | CCA | C | A | C>T | C>T_CCA | CG>TA_CATGCAA | 6 | FALSE | FALSE |
| 133_v.txt | 32088 | 32088-SNV-C | SNV | 1 | C | T | 83 | 15057 | 0.55123863 | 59.0481928 | rnr1Y285F_pG | TRUE | CAN | JSY2420 | 1 | rnr1Y285F_p | GCC | C>T | A/T+1 | G/C-1 | CG>TA | CG>TA | AAAAAGCCT | CCTGCAA | CGG | GCC | C | A | C>T | C>T_GCC | CG>TA_CCTGCAA | 6 | FALSE | FALSE |
| 133_v.txt | 32114 | 32114-SNV-C | SNV | 1 | C | T | 59 | 15057 | 0.39184433 | 59.8813559 | rnr1Y285F_pG | TRUE | CAN | JSY2420 | 1 | rnr1Y285F_p | GCC | C>T | A/T+1 | G/C-1 | CG>TA | CG>TA | TAATAGCCA | CCATTCG | CGG | GCC | C | A | C>T | C>T_GCC | CG>TA_CCATTCG | 6 | FALSE | FALSE |
| 133_v.txt | 32165 | 32165-SNV-G | SNV | 1 | G | A | 19 | 15057 | 0.12618716 | 47.6842105 | rnr1Y285F_pG | TRUE | CAN | JSY2420 | 1 | rnr1Y285F_p | CGC | G>A | A/T+1 | G/C-1 | CG>TA | CG>TA | CAAAGCGCC | GCCAAAT | GCG | GCG | C | T | C>T | C>T_GCG | CG>TA_GCCAAAT | 6 | FALSE | FALSE |
| 133_v.txt | 32271 | 32271-SNV-C | SNV | 1 | C | T | 66 | 21906 | 0.30128732 | 32.3476652 | rnr1Y285F_pG | TRUE | CAN | JSY2420 | 1 | rnr1Y285F_p | ACG | C>T | A/T+1 | G/C-1 | CG>TA | CG>TA | AAAATACGG | CGGGAAC | TGC | ACG | C | A | C>T | C>T_ACG | CG>TA_CGGGAAC | 6 | FALSE | FALSE |
| 133_v.txt | 32272 | 32272-SNV-G | SNV | 1 | G | A | 158 | 21906 | 0.72126358 | 36.7151866 | rnr1Y285F_pG | TRUE | CAN | JSY2420 | 1 | rnr1Y285F_p | CGG | G>A | A/T+1 | G/C-1 | CG>TA | CG>TA | AAATACGGG | GGGAACC | GCC | GCC | C | T | C>T | C>T_GCC | CG>TA_GGGAACC | 6 | FALSE | FALSE |
| 133_v.txt | 32299 | 32299-SNV-C | SNV | 1 | C | T | 38 | 21906 | 0.17346846 | 53.8421053 | rnr1Y285F_pG | TRUE | CAN | JSY2420 | 1 | rnr1Y285F_p | GCG | C>T | A/T+1 | G/C-1 | CG>TA | CG>TA | AATTTGCGG | CGGCAGA | CGC | GCG | C | A | C>T | C>T_GCG | CG>TA_CGGCAGA | 6 | FALSE | FALSE |
| 133_v.txt | 32466 | 32466-SNV-C | SNV | 1 | C | T | 39 | 21906 | 0.17803342 | 46.8717949 | rnr1Y285F_pG | TRUE | CAN | JSY2420 | 1 | rnr1Y285F_p | GCC | C>T | A/T+1 | G/C-1 | CG>TA | CG>TA | AGAGAGCCA | CCAATGT | CGG | GCC | C | A | C>T | C>T_GCC | CG>TA_CCAATGT | 6 | FALSE | FALSE |
| 133_v.txt | 32535 | 32535-SNV-G | SNV | 1 | G | A | 33 | 21906 | 0.15064366 | 36.9999448 | rnr1Y285F_pG | TRUE | CAN | JSY2420 | 1 | rnr1Y285F_p | TGC | G>A | A/T+1 | G/C-1 | CG>TA | CG>TA | GGGTTTGCA | GCAGCTT | ACG | ACG | C | T | C>T | C>T_ACG | CG>TA_GCAGCTT | 6 | FALSE | FALSE |
| 133_v.txt | 32536 | 32536-SNV-C | SNV | 1 | C | T | 30 | 21906 | 0.13694878 | 36.8666667 | rnr1Y285F_pG | TRUE | CAN | JSY2420 | 1 | rnr1Y285F_p | GCA | C>T | A/T+1 | G/C-1 | CG>TA | CG>TA | GGTTTGCAG | CAGCTTC | CGT | GCA | C | A | C>T | C>T_GCA | CG>TA_CAGCTTC | 6 | FALSE | FALSE |
| 133_v.txt | 32575 | 32575-SNV-G | SNV | 1 | G | A | 32 | 17262 | 0.18537829 | 35.78125 | rnr1Y285F_pG | TRUE | CAN | JSY2420 | 1 | rnr1Y285F_p | TGA | G>A | A/T+1 | G/C-1 | CG>TA | CG>TA | TACCTTGAA | GAAATGT | ACT | ACT | C | T | C>T | C>T_ACT | CG>TA_GAAATGT | 6 | FALSE | FALSE |
| 133_v.txt | 32590 | 32590-SNV-C | SNV | 1 | C | T | 27 | 17261 | 0.15642199 | 36.3333333 | rnr1Y285F_pG | TRUE | CAN | JSY2420 | 1 | rnr1Y285F_p | GCG | C>T | A/T+1 | G/C-1 | CG>TA | CG>TA | AGGCAGCGT | CGTTAAT | CGC | GCG | C | A | C>T | C>T_GCG | CG>TA_CGTTAAT | 6 | FALSE | FALSE |
| 133_v.txt | 32658 | 32658-SNV-C | SNV | 1 | C | T | 10 | 17261 | 0.05793407 | 51 | rnr1Y285F_pG | TRUE | CAN | JSY2420 | 1 | rnr1Y285F_p | ACC | C>T | A/T+1 | G/C-1 | CG>TA | CG>TA | CCTGGACCC | CCCCAGG | TGG | ACC | C | A | C>T | C>T_ACC | CG>TA_CCCCAGG | 6 | FALSE | FALSE |
| 133_v.txt | 32700 | 32700-SNV-C | SNV | 1 | C | T | 23 | 17261 | 0.13324836 | 54.3913044 | rnr1Y285F_pG | TRUE | CAN | JSY2420 | 1 | rnr1Y285F_p | GCC | C>T | A/T+1 | G/C-1 | CG>TA | CG>TA | ACTGGGCCG | CCGGTAA | CGG | GCC | C | A | C>T | C>T_GCC | CG>TA_CCGGTAA | 6 | FALSE | FALSE |
| 133_v.txt | 32749 | 32749-SNV-C | SNV | 1 | C | T | 19 | 17261 | 0.11007474 | 55.2631579 | rnr1Y285F_pG | TRUE | CAN | JSY2420 | 1 | rnr1Y285F_p | CCG | C>T | A/T+1 | G/C-1 | CG>TA | CG>TA | GAAACCCGA | CGATAAT | GGC | CCG | C | A | C>T | C>T_CCG | CG>TA_CGATAAT | 6 | FALSE | FALSE |
| 133_v.txt | 32776 | 32776-SNV-C | SNV | 1 | C | T | 19 | 17261 | 0.11007474 | 49.7368421 | rnr1Y285F_pG | TRUE | CAN | JSY2420 | 1 | rnr1Y285F_p | GCG | C>T | A/T+1 | G/C-1 | CG>TA | CG>TA | TGGAAGCGA | CGACCCA | CGC | GCG | C | A | C>T | C>T_GCG | CG>TA_CGACCCA | 6 | FALSE | FALSE |
| 133_v.txt | 32796 | 32796-SNV-C | SNV | 1 | C | T | 75 | 17261 | 0.43450553 | 43.8506028 | rnr1Y285F_pG | TRUE | CAN | JSY2420 | 1 | rnr1Y285F_p | ACC | C>T | A/T+1 | G/C-1 | CG>TA | CG>TA | AATTCACCG | CCGTAAT | TGG | ACC | C | A | C>T | C>T_ACC | CG>TA_CCGTAAT | 6 | FALSE | FALSE |
| 133_v.txt | 32797 | 32797-SNV-C | SNV | 1 | C | T | 15 | 17261 | 0.08690111 | 43.8666662 | rnr1Y285F_pG | TRUE | CAN | JSY2420 | 1 | rnr1Y285F_p | CCG | C>T | A/T+1 | G/C-1 | CG>TA | CG>TA | ATTCACCGT | CGTAATA | GGC | CCG | C | A | C>T | C>T_CCG | CG>TA_CGTAATA | 6 | FALSE | FALSE |
| 133_v.txt | 32798 | 32798-SNV-G | SNV | 1 | G | A | 21 | 17261 | 0.12166155 | 46.2857143 | rnr1Y285F_pG | TRUE | CAN | JSY2420 | 1 | rnr1Y285F_p | CGT | G>A | A/T+1 | G/C-1 | CG>TA | CG>TA | TTCACCGTA | GTAATAT | GCA | GCA | C | T | C>T | C>T_GCA | CG>TA_GTAATAT | 6 | FALSE | FALSE |
| 133_v.txt | 32811 | 32811-SNV-G | SNV | 1 | G | A | 21 | 17261 | 0.12166155 | 44.7619048 | rnr1Y285F_pG | TRUE | CAN | JSY2420 | 1 | rnr1Y285F_p | AGG | G>A | A/T+1 | G/C-1 | CG>TA | CG>TA | TTGACAGGG | GGGAACA | TCC | TCC | C | T | C>T | C>T_TCC | CG>TA_GGGAACA | 6 | FALSE | FALSE |
| 133_v.txt | 32855 | 32855-SNV-C | SNV | 1 | C | T | 50 | 17259 | 0.28970392 | 36.6599911 | rnr1Y285F_pG | TRUE | CAN | JSY2420 | 1 | rnr1Y285F_p | TCC | C>T | A/T+1 | G/C-1 | CG>TA | CG>TA | ACTAATCCAT | CCATGCC | AGG | TCC | C | A | C>T | C>T_TCC | CG>TA_CCATGCC | 6 | FALSE | FALSE |
| 133_v.txt | 32856 | 32856-SNV-C | SNV | 1 | C | T | 142 | 17259 | 0.82275914 | 36.6401405 | rnr1Y285F_pG | TRUE | CAN | JSY2420 | 1 | rnr1Y285F_p | CCA | C>T | A/T+1 | G/C-1 | CG>TA | CG>TA | CTAATCCAT | CATGCCG | GGT | CCA | C | A | C>T | C>T_CCA | CG>TA_CATGCCG | 6 | FALSE | FALSE |
| 133_v.txt | 32860 | 32860-SNV-C | SNV | 1 | C | T | 20 | 17259 | 0.11588157 | 36.3 | rnr1Y285F_pG | TRUE | CAN | JSY2420 | 1 | rnr1Y285F_p | GCC | C>T | A/T+1 | G/C-1 | CG>TA | CG>TA | TCCATGCCG | CCGCCAG | CGG | GCC | C | A | C>T | C>T_GCC | CG>TA_CCGCCAG | 6 | FALSE | FALSE |
| 133_v.txt | 32883 | 32883-SNV-C | SNV | 1 | C | T | 42 | 20066 | 0.20930928 | 36.1265825 | rnr1Y285F_pG | TRUE | CAN | JSY2420 | 1 | rnr1Y285F_p | CCA | C>T | A/T+1 | G/C-1 | CG>TA | CG>TA | TACGTCCAA | CAAAATT | GGT | CCA | C | A | C>T | C>T_CCA | CG>TA_CAAAATT | 6 | FALSE | FALSE |
| 133_v.txt | 32901 | 32901-SNV-C | SNV | 1 | C | T | 39 | 20066 | 0.19435862 | 32.6923077 | rnr1Y285F_pG | TRUE | CAN | JSY2420 | 1 | rnr1Y285F_p | GCC | C>T | A/T+1 | G/C-1 | CG>TA | CG>TA | ACTTGGCCA | CCAACTA | CGG | GCC | C | A | C>T | C>T_GCC | CG>TA_CCAACTA | 6 | FALSE | FALSE |
| 133_v.txt | 32945 | 32945-SNV-C | SNV | 1 | C | T | 26 | 20066 | 0.12957241 | 53.3076923 | rnr1Y285F_pG | TRUE | CAN | JSY2420 | 1 | rnr1Y285F_p | ACC | C>T | A/T+1 | G/C-1 | CG>TA | CG>TA | AGAAAACCA | CCAATAC | TGG | ACC | C | A | C>T | C>T_ACC | CG>TA_CCAATAC | 6 | FALSE | FALSE |
| 133_v.txt | 32958 | 32958-SNV-C | SNV | 1 | C | T | 133 | 20066 | 0.66281272 | 54.8421053 | rnr1Y285F_pG | TRUE | CAN | JSY2420 | 1 | rnr1Y285F_p | ACC | C>T | A/T+1 | G/C-1 | CG>TA | CG>TA | ATGTAACCA | CCATTGG | TGG | ACC | C | A | C>T | C>T_ACC | CG>TA_CCATTGG | 6 | FALSE | FALSE |
| 133_v.txt | 32967 | 32967-SNV-G | SNV | 1 | G | A | 35 | 20066 | 0.1744244 | 63 | rnr1Y285F_pG | TRUE | CAN | JSY2420 | 1 | rnr1Y285F_p | CGC | G>A | A/T+1 | G/C-1 | CG>TA | CG>TA | TTGGCCGCA | GCACCAA | GCG | GCG | C | T | C>T | C>T_GCG | CG>TA_GCACCAA | 6 | FALSE | FALSE |
| 133_v.txt | 33037 | 33037-SNV-C | SNV | 1 | C | T | 10 | 20064 | 0.04984051 | 59.6999948 | rnr1Y285F_pG | TRUE | CAN | JSY2420 | 1 | rnr1Y285F_p | GCC | C>T | A/T+1 | G/C-1 | CG>TA | CG>TA | ATGTAGCCA | CCATTTC | CGG | GCC | C | A | C>T | C>T_GCC | CG>TA_CCATTTC | 6 | FALSE | FALSE |
| 133_v.txt | 33069 | 33069-SNV-G | SNV | 1 | G | A | 20 | 20066 | 0.09967109 | 56.7 | rnr1Y285F_pG | TRUE | CAN | JSY2420 | 1 | rnr1Y285F_p | TGC | G>A | A/T+1 | G/C-1 | CG>TA | CG>TA | GAATATGCC | GCCAAAG | ACG | ACG | C | T | C>T | C>T_ACG | CG>TA_GCCAAAG | 6 | FALSE | FALSE |
| 133_v.txt | 33114 | 33114-SNV-C | SNV | 1 | C | T | 17 | 20066 | 0.08472042 | 38.1176421 | rnr1Y285F_pG | TRUE | CAN | JSY2420 | 1 | rnr1Y285F_p | GCC | C>T | A/T+1 | G/C-1 | CG>TA | CG>TA | ACTGGGCCG | CCGGCGT | CGG | GCC | C | A | C>T | C>T_GCC | CG>TA_CCGGCGT | 6 | FALSE | FALSE |
| 133_v.txt | 33115 | 33115-SNV-C | SNV | 1 | C | T | 81 | 20066 | 0.4036679 | 44.1103658 | rnr1Y285F_pG | TRUE | CAN | JSY2420 | 1 | rnr1Y285F_p | CCG | C>T | A/T+1 | G/C-1 | CG>TA | CG>TA | CTGGGCCGG | CGGCGTT | GGC | CCG | C | A | C>T | C>T_CCG | CG>TA_CGGCGTT | 6 | FALSE | FALSE |
| 133_v.txt | 33119 | 33119-SNV-G | SNV | 1 | G | A | 22 | 20066 | 0.10963819 | 40.8636364 | rnr1Y285F_pG | TRUE | CAN | JSY2420 | 1 | rnr1Y285F_p | CGT | G>A | A/T+1 | G/C-1 | CG>TA | CG>TA | GCCGGCGTT | GTTGGTC | GCA | GCA | C | T | C>T | C>T_GCA | CG>TA_GTTGGTC | 6 | FALSE | FALSE |
| 133_v.txt | 33153 | 33153-SNV-C | SNV | 1 | C | T | 40 | 20065 | 0.19935211 | 33.8002908 | rnr1Y285F_pG | TRUE | CAN | JSY2420 | 1 | rnr1Y285F_p | ACC | C>T | A/T+1 | G/C-1 | CG>TA | CG>TA | AAAAGACCT | CCTGTAC | TGG | ACC | C | A | C>T | C>T_ACC | CG>TA_CCTGTAC | 6 | FALSE | FALSE |
| 133_v.txt | 33168 | 33168-SNV-C | SNV | 1 | C | T | 54 | 20063 | 0.26915217 | 37.2222222 | rnr1Y285F_pG | TRUE | CAN | JSY2420 | 1 | rnr1Y285F_p | ACC | C>T | A/T+1 | G/C-1 | CG>TA | CG>TA | ATAGTACCA | CCACCAA | TGG | ACC | C | A | C>T | C>T_ACC | CG>TA_CCACCAA | 6 | FALSE | FALSE |
| 133_v.txt | 33199 | 33199-SNV-G | SNV | 1 | G | A | 30 | 10943 | 0.27414786 | 36.0666667 | rnr1Y285F_pG | TRUE | CAN | JSY2420 | 1 | rnr1Y285F_p | TGC | G>A | A/T+1 | G/C-1 | CG>TA | CG>TA | GTCTTTGCTT | GCTTAAG | ACG | ACG | C | T | C>T | C>T_ACG | CG>TA_GCTTAAG | 6 | FALSE | FALSE |
| 133_v.txt | 33205 | 33205-SNV-G | SNV | 1 | G | A | 3 | 10943 | 0.02741479 | 46.3312938 | rnr1Y285F_pG | TRUE | CAN | JSY2420 | 1 | rnr1Y285F_p | AGC | G>A | A/T+1 | G/C-1 | CG>TA | CG>TA | GCTTAAGCT | GCTCTCT | TCG | TCG | C | T | C>T | C>T_TCG | CG>TA_GCTCTCT | 6 | FALSE | FALSE |
| 133_v.txt | 33258 | 33258-SNV-C | SNV | 1 | C | T | 21 | 10945 | 0.19186843 | 41.8571429 | rnr1Y285F_pG | TRUE | CAN | JSY2420 | 1 | rnr1Y285F_p | GCC | C>T | A/T+1 | G/C-1 | CG>TA | CG>TA | CCTATGCCAT | CCATCCT | CGG | GCC | C | A | C>T | C>T_GCC | CG>TA_CCATCCT | 6 | FALSE | FALSE |
| 133_v.txt | 33474 | 33474-SNV-C | SNV | 1 | C | T | 15 | 10944 | 0.1370614 | 34.8 | rnr1Y285F_pG | TRUE | CAN | JSY2420 | 1 | rnr1Y285F_p | GCC | C>T | A/T+1 | G/C-1 | CG>TA | CG>TA | GCTATGCCTT | CCTTTTT | CGG | GCC | C | A | C>T | C>T_GCC | CG>TA_CCTTTTT | 6 | FALSE | FALSE |
| 133_v.txt | 32064 | 32064-SNV-G | SNV | 1 | G | C | 12 | 15057 | 0.07969715 | 61.3333333 | rnr1Y285F_pG | TRUE | CAN | JSY2420 | 1 | rnr1Y285F_p | TGA | G>C | G/C+1 | G/C-1 | CG>GC | CG>GC | GAGATTGAG | GAGATAA | ACT | ACT | C | G | C>G | C>G_ACT | CG>GC_GAGATAA | 6 | FALSE | FALSE |
| 133_v.txt | 32272 | 32272-SNV-G | SNV | 1 | G | C | 114 | 21906 | 0.52040537 | 35.8421097 | rnr1Y285F_pG | TRUE | CAN | JSY2420 | 1 | rnr1Y285F_p | CGG | G>C | G/C+1 | G/C-1 | CG>GC | CG>GC | AAATACGGG | GGGAACC | GCC | GCC | C | G | C>G | C>G_GCC | CG>GC_GGGAACC | 6 | FALSE | FALSE |
| 133_v.txt | 32304 | 32304-SNV-G | SNV | 1 | G | C | 8 | 21906 | 0.03651968 | 61.125 | rnr1Y285F_pG | TRUE | CAN | JSY2420 | 1 | rnr1Y285F_p | AGA | G>C | G/C+1 | G/C-1 | CG>GC | CG>GC | GCGGCAGAA | GAAATAA | TCT | TCT | C | G | C>G | C>G_TCT | CG>GC_GAAATAA | 6 | FALSE | FALSE |
| 133_v.txt | 32432 | 32432-SNV-G | SNV | 1 | G | C | 28 | 21906 | 0.12781886 | 59.9285714 | rnr1Y285F_pG | TRUE | CAN | JSY2420 | 1 | rnr1Y285F_p | TGT | G>C | G/C+1 | G/C-1 | CG>GC | CG>GC | GTCATTGTAT | GTATGGA | ACA | ACA | C | G | C>G | C>G_ACA | CG>GC_GTATGGA | 6 | FALSE | FALSE |
| 133_v.txt | 32488 | 32488-SNV-G | SNV | 1 | G | C | 16 | 21906 | 0.07303935 | 46 | rnr1Y285F_pG | TRUE | CAN | JSY2420 | 1 | rnr1Y285F_p | CGG | G>C | G/C+1 | G/C-1 | CG>GC | CG>GC | AGATACGGA | GGAAAAC | GCC | GCC | C | G | C>G | C>G_GCC | CG>GC_GGAAAAC | 6 | FALSE | FALSE |
| 133_v.txt | 32514 | 32514-SNV-G | SNV | 1 | G | C | 11 | 21906 | 0.05021455 | 42.1818182 | rnr1Y285F_pG | TRUE | CAN | JSY2420 | 1 | rnr1Y285F_p | TGG | G>C | G/C+1 | G/C-1 | CG>GC | CG>GC | GCTCTTGGA | GGAACGG | ACC | ACC | C | G | C>G | C>G_ACC | CG>GC_GGAACGG | 6 | FALSE | FALSE |
| 133_v.txt | 32529 | 32529-SNV-G | SNV | 1 | G | C | 10 | 21906 | 0.04564959 | 40.7978062 | rnr1Y285F_pG | TRUE | CAN | JSY2420 | 1 | rnr1Y285F_p | GGG | G>C | G/C+1 | G/C-1 | CG>GC | CG>GC | TTTCTGGGG | GGGTTTG | CCC | CCC | C | G | C>G | C>G_CCC | CG>GC_GGGTTTG | 6 | FALSE | FALSE |
| 133_v.txt | 32536 | 32536-SNV-C | SNV | 1 | C | G | 2 | 21906 | 9.13E-03 | 38 | rnr1Y285F_pG | TRUE | CAN | JSY2420 | 1 | rnr1Y285F_p | GCA | C>G | G/C+1 | G/C-1 | CG>GC | CG>GC | GGTTTGCAG | CAGCTTC | CGT | GCA | C | C | C>G | C>G_GCA | CG>GC_CAGCTTC | 6 | FALSE | FALSE |
| 133_v.txt | 32557 | 32557-SNV-C | SNV | 1 | C | G | 8 | 21901 | 0.03652801 | 37.75 | rnr1Y285F_pG | TRUE | CAN | JSY2420 | 1 | rnr1Y285F_p | CCA | C>G | G/C+1 | G/C-1 | CG>GC | CG>GC | TGATACCAA | CAACTAG | GGT | CCA | C | C | C>G | C>G_CCA | CG>GC_CAACTAG | 6 | FALSE | FALSE |
| 133_v.txt | 32580 | 32580-SNV-G | SNV | 1 | G | C | 15 | 17261 | 0.08690111 | 33.2666667 | rnr1Y285F_pG | TRUE | CAN | JSY2420 | 1 | rnr1Y285F_p | TGT | G>C | G/C+1 | G/C-1 | CG>GC | CG>GC | TGAAATGTG | GTGAAGG | ACA | ACA | C | G | C>G | C>G_ACA | CG>GC_GTGAAGG | 6 | FALSE | FALSE |
| 133_v.txt | 32776 | 32776-SNV-C | SNV | 1 | C | G | 6 | 17261 | 0.03476044 | 50.8333333 | rnr1Y285F_pG | TRUE | CAN | JSY2420 | 1 | rnr1Y285F_p | GCG | C>G | G/C+1 | G/C-1 | CG>GC | CG>GC | TGGAAGCGA | CGACCCA | CGC | GCG | C | C | C>G | C>G_GCG | CG>GC_CGACCCA | 6 | FALSE | FALSE |
| 133_v.txt | 32797 | 32797-SNV-C | SNV | 1 | C | G | 1 | 17261 | 5.79E-03 | 37 | rnr1Y285F_pG | TRUE | CAN | JSY2420 | 1 | rnr1Y285F_p | CCG | C>G | G/C+1 | G/C-1 | CG>GC | CG>GC | ATTCACCGT | CGTAATA | GGC | CCG | C | C | C>G | C>G_CCG | CG>GC_CGTAATA | 6 | FALSE | FALSE |
| 133_v.txt | 32809 | 32809-SNV-C | SNV | 1 | C | G | 4 | 17261 | 0.02317363 | 51 | rnr1Y285F_pG | TRUE | CAN | JSY2420 | 1 | rnr1Y285F_p | ACA | C>G | G/C+1 | G/C-1 | CG>GC | CG>GC | ATTTGACAG | CAGGGAA | TGT | ACA | C | C | C>G | C>G_ACA | CG>GC_CAGGGAA | 6 | FALSE | FALSE |

| | | | | | | | | | | | | | | | | | | | | | | | | | | | | | | | | | | |
|---|---|---|---|---|---|---|---|---|---|---|---|---|---|---|---|---|---|---|---|---|---|---|---|---|---|---|---|---|---|---|---|---|---|---|
| 133_v.txt | 32940 | 32940-SNV-( | SNV | 1 | G | C | 3 | 20066 | 0.01495066 | 63.9999999 | rnr1Y285F_pG | TRUE | CAN | JSY2420 | 1 | rnr1Y285F_p | AGA | G>C | G/C+1 | G/C-1 | CG>GC | CG>GC | GCCCAAGAA | GAAAACC | TCT | TCT | C | G | C>G | C>G_TCT | CG>GC_GAAAACC | 6 | FALSE | FALSE |
| 133_v.txt | 32954 | 32954-SNV-( | SNV | 1 | G | C | 14 | 20066 | 0.06976976 | 61.5 | rnr1Y285F_pG | TRUE | CAN | JSY2420 | 1 | rnr1Y285F_p | TGT | G>C | G/C+1 | G/C-1 | CG>GC | CG>GC | ATACATGTA | GTAACCA | ACA | ACA | C | G | C>G | C>G_ACA | CG>GC_GTAACCA | 6 | FALSE | FALSE |
| 133_v.txt | 33024 | 33024-SNV-( | SNV | 1 | G | C | 10 | 20065 | 0.04983803 | 54.0989746 | rnr1Y285F_pG | TRUE | CAN | JSY2420 | 1 | rnr1Y285F_p | AGG | G>C | G/C+1 | G/C-1 | CG>GC | CG>GC | GTAACAGGG | GGGATGA | TCC | TCC | C | G | C>G | C>G_TCC | CG>GC_GGGATGA | 6 | FALSE | FALSE |
| 133_v.txt | 33026 | 33026-SNV-( | SNV | 1 | G | C | 2 | 20066 | 9.97E-03 | 26.4996104 | rnr1Y285F_pG | TRUE | CAN | JSY2420 | 1 | rnr1Y285F_p | GGA | G>C | G/C+1 | G/C-1 | CG>GC | CG>GC | AACAGGGAT | GATGAAT | CCT | CCT | C | G | C>G | C>G_CCT | CG>GC_GATGAAT | 6 | FALSE | FALSE |
| 133_v.txt | 33114 | 33114-SNV-( | SNV | 1 | C | G | 7 | 20066 | 0.03488488 | 41.5707792 | rnr1Y285F_pG | TRUE | CAN | JSY2420 | 1 | rnr1Y285F_p | GCC | C>G | G/C+1 | G/C-1 | CG>GC | CG>GC | ACTGGGCCG | CCGGCGT | CGG | GCC | C | C | C>G | C>G_GCC | CG>GC_CCGGCGT | 6 | FALSE | FALSE |
| 133_v.txt | 33156 | 33156-SNV-( | SNV | 1 | G | C | 71 | 20064 | 0.35386762 | 37.8021948 | rnr1Y285F_pG | TRUE | CAN | JSY2420 | 1 | rnr1Y285F_p | TGT | G>C | G/C+1 | G/C-1 | CG>GC | CG>GC | AGACCTGTA | GTACCAA | ACA | ACA | C | G | C>G | C>G_ACA | CG>GC_GTACCAA | 6 | FALSE | FALSE |
| 133_v.txt | 33168 | 33168-SNV-( | SNV | 1 | C | G | 6 | 20063 | 0.0299058 | 37.6666667 | rnr1Y285F_pG | TRUE | CAN | JSY2420 | 1 | rnr1Y285F_p | ACC | C>G | G/C+1 | G/C-1 | CG>GC | CG>GC | ATAGTACCA | CCACCAA | TGG | ACC | C | C | C>G | C>G_ACC | CG>GC_CCACCAA | 6 | FALSE | FALSE |
| 133_v.txt | 33172 | 33172-SNV-( | SNV | 1 | C | G | 23 | 20063 | 0.11463889 | 38 | rnr1Y285F_pG | TRUE | CAN | JSY2420 | 1 | rnr1Y285F_p | CCA | C>G | G/C+1 | G/C-1 | CG>GC | CG>GC | TACCACCAA | CAAGGGC | GGT | CCA | C | C | C>G | C>G_CCA | CG>GC_CAAGGGC | 6 | FALSE | FALSE |
| 133_v.txt | 33187 | 33187-SNV-( | SNV | 1 | C | G | 26 | 10946 | 0.23752969 | 37.7692308 | rnr1Y285F_pG | TRUE | CAN | JSY2420 | 1 | rnr1Y285F_p | CCA | C>G | G/C+1 | G/C-1 | CG>GC | CG>GC | TCATACCAAT | CAATATG | GGT | CCA | C | C | C>G | C>G_CCA | CG>GC_CAATATG | 6 | FALSE | FALSE |
| 133_v.txt | 33205 | 33205-SNV-( | SNV | 1 | G | C | 8 | 10943 | 0.0731061 | 40.7461232 | rnr1Y285F_pG | TRUE | CAN | JSY2420 | 1 | rnr1Y285F_p | AGC | G>C | G/C+1 | G/C-1 | CG>GC | CG>GC | GCTTAAGCT | GCTCTCT | TCG | TCG | C | G | C>G | C>G_TCG | CG>GC_GCTCTCT | 6 | FALSE | FALSE |
| 133_v.txt | 33419 | 33419-SNV-( | SNV | 1 | G | C | 90 | 10945 | 0.82229329 | 46.4777778 | rnr1Y285F_pG | TRUE | CAN | JSY2420 | 1 | rnr1Y285F_p | TGT | G>C | G/C+1 | G/C-1 | CG>GC | CG>GC | CTCATTGTAC | GTACATA | ACA | ACA | C | G | C>G | C>G_ACA | CG>GC_GTACATA | 6 | FALSE | FALSE |
| 133_v.txt | 33456 | 33456-SNV-( | SNV | 1 | G | C | 14 | 10945 | 0.12791229 | 34.3571429 | rnr1Y285F_pG | TRUE | CAN | JSY2420 | 1 | rnr1Y285F_p | TGA | G>C | G/C+1 | G/C-1 | CG>GC | CG>GC | TCTTTTGAAT | GAATTTG | ACT | ACT | C | G | C>G | C>G_ACT | CG>GC_GAATTTG | 6 | FALSE | FALSE |
| 133_v.txt | 31829 | 31829-SNV-( | SNV | 1 | G | T | 8 | 6373 | 0.12552958 | 53.375 | rnr1Y285F_pG | TRUE | CAN | JSY2420 | 1 | rnr1Y285F_p | TGC | G>T | A/T+1 | G/C-1 | CG>AT | CG>AT | GAATATGCA | GCATTGA | ACG | ACG | C | A | C>A | C>A_ACG | CG>AT_GCATTGA | 6 | FALSE | FALSE |
| 133_v.txt | 32002 | 32002-SNV-( | SNV | 1 | G | T | 7 | 15057 | 0.04649001 | 55.2857143 | rnr1Y285F_pG | TRUE | CAN | JSY2420 | 1 | rnr1Y285F_p | GGT | G>T | A/T+1 | G/C-1 | CG>AT | CG>AT | TAAATGGTA | GTAACTC | CCA | CCA | C | A | C>A | C>A_CCA | CG>AT_GTAACTC | 6 | FALSE | FALSE |
| 133_v.txt | 32027 | 32027-SNV-( | SNV | 1 | G | T | 21 | 15057 | 0.13947001 | 59.5714286 | rnr1Y285F_pG | TRUE | CAN | JSY2420 | 1 | rnr1Y285F_p | GGT | G>T | A/T+1 | G/C-1 | CG>AT | CG>AT | GCCACGGTA | GTATTTC | CCA | CCA | C | A | C>A | C>A_CCA | CG>AT_GTATTTC | 6 | FALSE | FALSE |
| 133_v.txt | 32119 | 32119-SNV-( | SNV | 1 | C | A | 37 | 15057 | 0.24573288 | 55.027027 | rnr1Y285F_pG | TRUE | CAN | JSY2420 | 1 | rnr1Y285F_p | TCG | C>A | A/T+1 | G/C-1 | CG>AT | CG>AT | GCCATTCGA | CGAAAAC | AGC | TCG | C | T | C>A | C>A_TCG | CG>AT_CGAAAAC | 6 | FALSE | FALSE |
| 133_v.txt | 32153 | 32153-SNV-( | SNV | 1 | G | T | 27 | 15057 | 0.17931859 | 53.7777778 | rnr1Y285F_pG | TRUE | CAN | JSY2420 | 1 | rnr1Y285F_p | TGT | G>T | A/T+1 | G/C-1 | CG>AT | CG>AT | CTCCATGTA | GTAAGCC | ACA | ACA | C | A | C>A | C>A_ACA | CG>AT_GTAAGCC | 6 | FALSE | FALSE |
| 133_v.txt | 32199 | 32199-SNV-( | SNV | 1 | G | T | 15 | 15057 | 0.09962144 | 41.2666667 | rnr1Y285F_pG | TRUE | CAN | JSY2420 | 1 | rnr1Y285F_p | TGG | G>T | A/T+1 | G/C-1 | CG>AT | CG>AT | ATGTATGGA | GGAACAC | ACC | ACC | C | A | C>A | C>A_ACC | CG>AT_GGAACAC | 6 | FALSE | FALSE |
| 133_v.txt | 32223 | 32223-SNV-( | SNV | 1 | G | T | 19 | 15057 | 0.12618716 | 40.6842105 | rnr1Y285F_pG | TRUE | CAN | JSY2420 | 1 | rnr1Y285F_p | TGA | G>T | A/T+1 | G/C-1 | CG>AT | CG>AT | GTCCTTGAC | GACAGGA | ACT | ACT | C | A | C>A | C>A_ACT | CG>AT_GACAGGA | 6 | FALSE | FALSE |
| 133_v.txt | 32272 | 32272-SNV-( | SNV | 1 | G | T | 10 | 21906 | 0.04564959 | 24.3000167 | rnr1Y285F_pG | TRUE | CAN | JSY2420 | 1 | rnr1Y285F_p | CGG | G>T | A/T+1 | G/C-1 | CG>AT | CG>AT | AAATACGGG | GGGAACC | GCC | GCC | C | A | C>A | C>A_GCC | CG>AT_GGGAACC | 6 | FALSE | FALSE |
| 133_v.txt | 32277 | 32277-SNV-( | SNV | 1 | C | A | 40 | 21906 | 0.18259838 | 32.025 | rnr1Y285F_pG | TRUE | CAN | JSY2420 | 1 | rnr1Y285F_p | ACC | C>A | A/T+1 | G/C-1 | CG>AT | CG>AT | CGGGAACCA | CCAACGT | TGG | ACC | C | T | C>A | C>A_ACC | CG>AT_CCAACGT | 6 | FALSE | FALSE |
| 133_v.txt | 32282 | 32282-SNV-( | SNV | 1 | G | T | 19 | 21906 | 0.08673423 | 44.631579 | rnr1Y285F_pG | TRUE | CAN | JSY2420 | 1 | rnr1Y285F_p | CGT | G>T | A/T+1 | G/C-1 | CG>AT | CG>AT | ACCAACGTA | GTAAATA | GCA | GCA | C | A | C>A | C>A_GCA | CG>AT_GTAAATA | 6 | FALSE | FALSE |
| 133_v.txt | 32304 | 32304-SNV-( | SNV | 1 | G | T | 10 | 21906 | 0.04564959 | 53.1 | rnr1Y285F_pG | TRUE | CAN | JSY2420 | 1 | rnr1Y285F_p | AGA | G>T | A/T+1 | G/C-1 | CG>AT | CG>AT | GCGGCAGAA | GAAATAA | TCT | TCT | C | A | C>A | C>A_TCT | CG>AT_GAAATAA | 6 | FALSE | FALSE |
| 133_v.txt | 32373 | 32373-SNV-( | SNV | 1 | G | T | 30 | 21906 | 0.13694878 | 58.0666667 | rnr1Y285F_pG | TRUE | CAN | JSY2420 | 1 | rnr1Y285F_p | AGC | G>T | A/T+1 | G/C-1 | CG>AT | CG>AT | TCAATAGCA | GCAATAA | TCG | TCG | C | A | C>A | C>A_TCG | CG>AT_GCAATAA | 6 | FALSE | FALSE |
| 133_v.txt | 32399 | 32399-SNV-( | SNV | 1 | G | T | 43 | 21906 | 0.19629325 | 58.9534884 | rnr1Y285F_pG | TRUE | CAN | JSY2420 | 1 | rnr1Y285F_p | CGT | G>T | A/T+1 | G/C-1 | CG>AT | CG>AT | AGAAACGTA | GTAGGAA | GCA | GCA | C | A | C>A | C>A_GCA | CG>AT_GTAGGAA | 6 | FALSE | FALSE |
| 133_v.txt | 32432 | 32432-SNV-( | SNV | 1 | G | T | 54 | 21906 | 0.24650781 | 58.8703704 | rnr1Y285F_pG | TRUE | CAN | JSY2420 | 1 | rnr1Y285F_p | TGT | G>T | A/T+1 | G/C-1 | CG>AT | CG>AT | GTCATTGTAT | GTATGGA | ACA | ACA | C | A | C>A | C>A_ACA | CG>AT_GTATGGA | 6 | FALSE | FALSE |
| 133_v.txt | 32471 | 32471-SNV-( | SNV | 1 | G | T | 32 | 21906 | 0.1460787 | 42.25 | rnr1Y285F_pG | TRUE | CAN | JSY2420 | 1 | rnr1Y285F_p | TGT | G>T | A/T+1 | G/C-1 | CG>AT | CG>AT | GCCAATGTA | GTAGAAG | ACA | ACA | C | A | C>A | C>A_ACA | CG>AT_GTAGAAG | 6 | FALSE | FALSE |
| 133_v.txt | 32474 | 32474-SNV-( | SNV | 1 | G | T | 12 | 21906 | 0.05477951 | 46.3321907 | rnr1Y285F_pG | TRUE | CAN | JSY2420 | 1 | rnr1Y285F_p | AGA | G>T | A/T+1 | G/C-1 | CG>AT | CG>AT | AATGTAGAA | GAAGGTT | TCT | TCT | C | A | C>A | C>A_TCT | CG>AT_GAAGGTT | 6 | FALSE | FALSE |
| 133_v.txt | 32508 | 32508-SNV-( | SNV | 1 | G | T | 14 | 21906 | 0.06390943 | 39.9285714 | rnr1Y285F_pG | TRUE | CAN | JSY2420 | 1 | rnr1Y285F_p | GGC | G>T | A/T+1 | G/C-1 | CG>AT | CG>AT | TTGATGGCT | GCTCTTG | CCG | CCG | C | A | C>A | C>A_CCG | CG>AT_GCTCTTG | 6 | FALSE | FALSE |
| 133_v.txt | 32529 | 32529-SNV-( | SNV | 1 | G | T | 5 | 21906 | 0.0228248 | 35.7993373 | rnr1Y285F_pG | TRUE | CAN | JSY2420 | 1 | rnr1Y285F_p | GGG | G>T | A/T+1 | G/C-1 | CG>AT | CG>AT | TTTCTGGGG | GGGTTTG | CCC | CCC | C | A | C>A | C>A_CCC | CG>AT_GGGTTTG | 6 | FALSE | FALSE |
| 133_v.txt | 32530 | 32530-SNV-( | SNV | 1 | G | T | 18 | 21906 | 0.08216927 | 39.32238 | rnr1Y285F_pG | TRUE | CAN | JSY2420 | 1 | rnr1Y285F_p | GGG | G>T | A/T+1 | G/C-1 | CG>AT | CG>AT | TTCTGGGGT | GGTTTGC | CCC | CCC | C | A | C>A | C>A_CCC | CG>AT_GGTTTGC | 6 | FALSE | FALSE |
| 133_v.txt | 32575 | 32575-SNV-( | SNV | 1 | G | T | 21 | 17262 | 0.1216545 | 35.4285714 | rnr1Y285F_pG | TRUE | CAN | JSY2420 | 1 | rnr1Y285F_p | TGA | G>T | A/T+1 | G/C-1 | CG>AT | CG>AT | TACCTTGAA | GAAATGT | ACT | ACT | C | A | C>A | C>A_ACT | CG>AT_GAAATGT | 6 | FALSE | FALSE |
| 133_v.txt | 32658 | 32658-SNV-( | SNV | 1 | C | A | 7 | 17261 | 0.04055385 | 25.7142857 | rnr1Y285F_pG | TRUE | CAN | JSY2420 | 1 | rnr1Y285F_p | ACC | C>A | A/T+1 | G/C-1 | CG>AT | CG>AT | CCTGGACCC | CCCCAGG | TGG | ACC | C | T | C>A | C>A_ACC | CG>AT_CCCCAGG | 6 | FALSE | FALSE |
| 133_v.txt | 32676 | 32676-SNV-( | SNV | 1 | C | A | 9 | 17261 | 0.05214066 | 61 | rnr1Y285F_pG | TRUE | CAN | JSY2420 | 1 | rnr1Y285F_p | TCT | C>A | A/T+1 | G/C-1 | CG>AT | CG>AT | GGGTTTCTC | CTCCAAT | AGA | TCT | C | T | C>A | C>A_TCT | CG>AT_CTCCAAT | 6 | FALSE | FALSE |
| 133_v.txt | 32735 | 32735-SNV-( | SNV | 1 | G | T | 12 | 17261 | 0.06952089 | 59.6667406 | rnr1Y285F_pG | TRUE | CAN | JSY2420 | 1 | rnr1Y285F_p | AGT | G>T | A/T+1 | G/C-1 | CG>AT | CG>AT | AAAACAGTA | GTATATT | TCA | TCA | C | A | C>A | C>A_TCA | CG>AT_GTATATT | 6 | FALSE | FALSE |
| 133_v.txt | 32749 | 32749-SNV-( | SNV | 1 | C | A | 6 | 17261 | 0.03476044 | 50.5 | rnr1Y285F_pG | TRUE | CAN | JSY2420 | 1 | rnr1Y285F_p | CCG | C>A | A/T+1 | G/C-1 | CG>AT | CG>AT | GAAACCCGA | CGATAAT | GGC | CCG | C | T | C>A | C>A_CCG | CG>AT_CGATAAT | 6 | FALSE | FALSE |
| 133_v.txt | 32783 | 32783-SNV-( | SNV | 1 | G | T | 19 | 17261 | 0.11007474 | 40.3157895 | rnr1Y285F_pG | TRUE | CAN | JSY2420 | 1 | rnr1Y285F_p | AGA | G>T | A/T+1 | G/C-1 | CG>AT | CG>AT | GACCCAGAA | GAACTCG | TCT | TCT | C | A | C>A | C>A_TCT | CG>AT_GAACTCG | 6 | FALSE | FALSE |
| 133_v.txt | 32797 | 32797-SNV-( | SNV | 1 | C | A | 13 | 17261 | 0.07531429 | 43.3846153 | rnr1Y285F_pG | TRUE | CAN | JSY2420 | 1 | rnr1Y285F_p | CCG | C>A | A/T+1 | G/C-1 | CG>AT | CG>AT | ATTCACCGTA | CGTAATA | GGC | CCG | C | T | C>A | C>A_CCG | CG>AT_CGTAATA | 6 | FALSE | FALSE |
| 133_v.txt | 32809 | 32809-SNV-( | SNV | 1 | C | A | 34 | 17261 | 0.19697584 | 39.7352941 | rnr1Y285F_pG | TRUE | CAN | JSY2420 | 1 | rnr1Y285F_p | ACA | C>A | A/T+1 | G/C-1 | CG>AT | CG>AT | ATTTGACAG | CAGGGAA | TGT | ACA | C | T | C>A | C>A_ACA | CG>AT_CAGGGAA | 6 | FALSE | FALSE |
| 133_v.txt | 32811 | 32811-SNV-( | SNV | 1 | G | T | 8 | 17261 | 0.04634726 | 47.75 | rnr1Y285F_pG | TRUE | CAN | JSY2420 | 1 | rnr1Y285F_p | AGG | G>T | A/T+1 | G/C-1 | CG>AT | CG>AT | TTGACAGGG | GGGAACA | TCC | TCC | C | A | C>A | C>A_TCC | CG>AT_GGGAACA | 6 | FALSE | FALSE |
| 133_v.txt | 32819 | 32819-SNV-( | SNV | 1 | G | T | 32 | 17260 | 0.18539977 | 40.875 | rnr1Y285F_pG | TRUE | CAN | JSY2420 | 1 | rnr1Y285F_p | AGT | G>T | A/T+1 | G/C-1 | CG>AT | CG>AT | GAACAAGTT | GTTCATT | TCA | TCA | C | A | C>A | C>A_TCA | CG>AT_GTTCATT | 6 | FALSE | FALSE |
| 133_v.txt | 32855 | 32855-SNV-( | SNV | 1 | C | A | 7 | 17259 | 0.04055855 | 37.4693157 | rnr1Y285F_pG | TRUE | CAN | JSY2420 | 1 | rnr1Y285F_p | TCC | C>A | A/T+1 | G/C-1 | CG>AT | CG>AT | ACTAATCCAT | CCATGCC | AGG | TCC | C | T | C>A | C>A_TCC | CG>AT_CCATGCC | 6 | FALSE | FALSE |
| 133_v.txt | 32883 | 32883-SNV-( | SNV | 1 | C | A | 3 | 20066 | 0.01495066 | 34.6505704 | rnr1Y285F_pG | TRUE | CAN | JSY2420 | 1 | rnr1Y285F_p | CCA | C>A | A/T+1 | G/C-1 | CG>AT | CG>AT | TACGTCCAA | CAAAATT | GGT | CCA | C | T | C>A | C>A_CCA | CG>AT_CAAAATT | 6 | FALSE | FALSE |
| 133_v.txt | 32917 | 32917-SNV-( | SNV | 1 | C | A | 34 | 20066 | 0.16944085 | 48.6764706 | rnr1Y285F_pG | TRUE | CAN | JSY2420 | 1 | rnr1Y285F_p | TCC | C>A | A/T+1 | G/C-1 | CG>AT | CG>AT | TAAGTTCCA | CCAGGGC | AGG | TCC | C | T | C>A | C>A_TCC | CG>AT_CCAGGGC | 6 | FALSE | FALSE |
| 133_v.txt | 32940 | 32940-SNV-( | SNV | 1 | G | T | 1115 | 20066 | 5.55666301 | 56.7524663 | rnr1Y285F_pG | TRUE | CAN | JSY2420 | 1 | rnr1Y285F_p | AGA | G>T | A/T+1 | G/C-1 | CG>AT | CG>AT | GCCCAAGAA | GAAAACC | TCT | TCT | C | A | C>A | C>A_TCT | CG>AT_GAAAACC | 6 | FALSE | FALSE |
| 133_v.txt | 32954 | 32954-SNV-( | SNV | 1 | G | T | 9 | 20066 | 0.04485199 | 42.3333333 | rnr1Y285F_pG | TRUE | CAN | JSY2420 | 1 | rnr1Y285F_p | TGT | G>T | A/T+1 | G/C-1 | CG>AT | CG>AT | ATACATGTA | GTAACCA | ACA | ACA | C | A | C>A | C>A_ACA | CG>AT_GTAACCA | 6 | FALSE | FALSE |
| 133_v.txt | 33012 | 33012-SNV-( | SNV | 1 | G | T | 4000 | 20066 | 19.9342171 | 60.489 | rnr1Y285F_pG | TRUE | CAN | JSY2420 | 1 | rnr1Y285F_p | AGA | G>T | A/T+1 | G/C-1 | CG>AT | CG>AT | GTGAAAGAG | GAGGATG | TCT | TCT | C | A | C>A | C>A_TCT | CG>AT_GAGGATG | 6 | FALSE | FALSE |
| 133_v.txt | 33015 | 33015-SNV-( | SNV | 1 | G | T | 32 | 20066 | 0.15947374 | 56.96875 | rnr1Y285F_pG | TRUE | CAN | JSY2420 | 1 | rnr1Y285F_p | GGA | G>T | A/T+1 | G/C-1 | CG>AT | CG>AT | AAAGAGGAT | GATGTAA | CCT | CCT | C | A | C>A | C>A_CCT | CG>AT_GATGTAA | 6 | FALSE | FALSE |
| 133_v.txt | 33024 | 33024-SNV-( | SNV | 1 | G | T | 36 | 20065 | 0.1794169 | 63.8564732 | rnr1Y285F_pG | TRUE | CAN | JSY2420 | 1 | rnr1Y285F_p | AGG | G>T | A/T+1 | G/C-1 | CG>AT | CG>AT | GTAACAGGG | GGGATGA | TCC | TCC | C | A | C>A | C>A_TCC | CG>AT_GGGATGA | 6 | FALSE | FALSE |
| 133_v.txt | 33036 | 33036-SNV-( | SNV | 1 | G | T | 2 | 20066 | 9.97E-03 | 63.9650485 | rnr1Y285F_pG | TRUE | CAN | JSY2420 | 1 | rnr1Y285F_p | AGC | G>T | A/T+1 | G/C-1 | CG>AT | CG>AT | AATGTAGCC | GCCATTT | TCG | TCG | C | A | C>A | C>A_TCG | CG>AT_GCCATTT | 6 | FALSE | FALSE |
| 133_v.txt | 33043 | 33043-SNV-( | SNV | 1 | C | A | 16 | 20066 | 0.07973687 | 49.3125 | rnr1Y285F_pG | TRUE | CAN | JSY2420 | 1 | rnr1Y285F_p | TCA | C>A | A/T+1 | G/C-1 | CG>AT | CG>AT | CCATTTCACC | CACCCAA | AGT | TCA | C | T | C>A | C>A_TCA | CG>AT_CACCCAA | 6 | FALSE | FALSE |
| 133_v.txt | 33093 | 33093-SNV-( | SNV | 1 | G | T | 22 | 20066 | 0.10963819 | 52.8181818 | rnr1Y285F_pG | TRUE | CAN | JSY2420 | 1 | rnr1Y285F_p | TGA | G>T | A/T+1 | G/C-1 | CG>AT | CG>AT | AAATATGAT | GATATAA | ACT | ACT | C | A | C>A | C>A_ACT | CG>AT_GATATAA | 6 | FALSE | FALSE |
| 133_v.txt | 33151 | 33151-SNV-( | SNV | 1 | G | T | 37 | 20066 | 0.18439151 | 39.027027 | rnr1Y285F_pG | TRUE | CAN | JSY2420 | 1 | rnr1Y285F_p | AGA | G>T | A/T+1 | G/C-1 | CG>AT | CG>AT | TGAAAAGAC | GACCTGT | TCT | TCT | C | A | C>A | C>A_TCT | CG>AT_GACCTGT | 6 | FALSE | FALSE |
| 133_v.txt | 33154 | 33154-SNV-( | SNV | 1 | C | A | 77 | 20065 | 0.3837528 | 36.8054074 | rnr1Y285F_pG | TRUE | CAN | JSY2420 | 1 | rnr1Y285F_p | CCT | C>A | A/T+1 | G/C-1 | CG>AT | CG>AT | AAAGACCTG | CTGTACC | GGA | CCT | C | T | C>A | C>A_CCT | CG>AT_CTGTACC | 6 | FALSE | FALSE |
| 133_v.txt | 33283 | 33283-SNV-( | SNV | 1 | C | A | 29 | 10945 | 0.26496117 | 48.1724138 | rnr1Y285F_pG | TRUE | CAN | JSY2420 | 1 | rnr1Y285F_p | TCG | C>A | A/T+1 | G/C-1 | CG>AT | CG>AT | TATCCTCGCC | CGCCATT | AGC | TCG | C | T | C>A | C>A_TCG | CG>AT_CGCCATT | 6 | FALSE | FALSE |
| 133_v.txt | 33325 | 33325-SNV-( | SNV | 1 | C | A | 96 | 10945 | 0.87711284 | 59.875 | rnr1Y285F_pG | TRUE | CAN | JSY2420 | 1 | rnr1Y285F_p | TCT | C>A | A/T+1 | G/C-1 | CG>AT | CG>AT | ACAATTCTTT | CTTTACT | AGA | TCT | C | T | C>A | C>A_TCT | CG>AT_CTTTACT | 6 | FALSE | FALSE |
| 133_v.txt | 33354 | 33354-SNV-( | SNV | 1 | G | T | 63 | 10945 | 0.5756053 | 21.8095238 | rnr1Y285F_pG | TRUE | CAN | JSY2420 | 1 | rnr1Y285F_p | TGA | G>T | A/T+1 | G/C-1 | CG>AT | CG>AT | GGTATTGAC | GACCCAC | ACT | ACT | C | A | C>A | C>A_ACT | CG>AT_GACCCAC | 6 | FALSE | FALSE |
| 133_v.txt | 33385 | 33385-SNV-( | SNV | 1 | C | A | 13 | 10945 | 0.1187757 | 47.6923077 | rnr1Y285F_pG | TRUE | CAN | JSY2420 | 1 | rnr1Y285F_p | TCA | C>A | A/T+1 | G/C-1 | CG>AT | CG>AT | AAGCTTCAA | CAACGTC | AGT | TCA | C | T | C>A | C>A_TCA | CG>AT_CAACGTC | 6 | FALSE | FALSE |
| 133_v.txt | 33474 | 33474-SNV-( | SNV | 1 | C | A | 35 | 10944 | 0.31980994 | 34.6285714 | rnr1Y285F_pG | TRUE | CAN | JSY2420 | 1 | rnr1Y285F_p | GCC | C>A | A/T+1 | G/C-1 | CG>AT | CG>AT | GCTATGCCTT | CCTTTTT | CGG | GCC | C | T | C>A | C>A_GCC | CG>AT_CCTTTTT | 6 | FALSE | FALSE |
| 133_v.txt | 33475 | 33475-SNV-( | SNV | 1 | C | A | 11 | 10943 | 0.10052088 | 34.9090909 | rnr1Y285F_pG | TRUE | CAN | JSY2420 | 1 | rnr1Y285F_p | CCT | C>A | A/T+1 | G/C-1 | CG>AT | CG>AT | CTATGCCTTT | CTTTTTT | GGA | CCT | C | T | C>A | C>A_CCT | CG>AT_CTTTTTT | 6 | FALSE | FALSE |
| 133_v.txt | 32271 | 32271-Repla | Replacement | 2 | CG | T | 1 | 21906 | 4.57E-03 | 37.9999999 | rnr1Y285F_pG | TRUE | CAN | JSY2420 | 1 | rnr1Y285F_p | ACG | nope | A/T+1 | >1bpdel | none | Replac. | AAAATACGG | CGGGAAC | TGC | ACG | CG | A | NA | NA_ACG | Replac._CGGGAAC | 6 | FALSE | FALSE |
| 133_v.txt | 32796 | 32796-MNV- | MNV | 2 | CC | AA | 1 | 17261 | 5.79E-03 | 36.9999992 | rnr1Y285F_pG | TRUE | CAN | JSY2420 | 1 | rnr1Y285F_p | ACC | nope | >1bp-ins | >1bpdel | none | MNV | AATTCACCG | CCGTAAT | TGG | ACC | CC | none | NA | NA_ACC | MNV_CCGTAAT | 6 | FALSE | FALSE |
| 133_v.txt | 32940 | 32940-MNV- | MNV | 2 | GA | TG | 1 | 20066 | 4.98E-03 | 63.9999999 | rnr1Y285F_pG | TRUE | CAN | JSY2420 | 1 | rnr1Y285F_p | AGA | nope | >1bp-ins | >1bpdel | none | MNV | GCCCAAGAA | GAAAACC | TCT | AGA | GA | none | NA | NA_AGA | MNV_GAAAACC | 6 | FALSE | FALSE |
| 133_v.txt | 33022 | 33022-MNV- | MNV | 2 | CA | AG | 1 | 20066 | 4.98E-03 | 64 | rnr1Y285F_pG | TRUE | CAN | JSY2420 | 1 | rnr1Y285F_p | ACA | nope | >1bp-ins | >1bpdel | none | MNV | ATGTAACAG | CAGGGAT | TGT | ACA | CA | none | NA | NA_ACA | MNV_CAGGGAT | 6 | FALSE | FALSE |
| 133_v.txt | 33065 | 33065-MNV- | MNV | 2 | AT | CC | 2 | 20066 | 9.97E-03 | 50.9994356 | rnr1Y285F_pG | TRUE | CAN | JSY2420 | 1 | rnr1Y285F_p | AAT | nope | >1bp-ins | >1bpdel | none | MNV | GACAGAATA | ATATGCC | TTA | AAT | AT | none | NA | NA_AAT | MNV_ATATGCC | 6 | FALSE | FALSE |
| 133_v.txt | 32004 | 32004-Insert | Insertion | 1 | - | A | 6 | 15057 | 0.03984858 | 54 | rnr1Y285F_pG | TRUE | CAN | JSY2420 | 1 | rnr1Y285F_p | TAA | nope | A/T+1 | >1bpdel | none | A/T+1 | AATGGTAAC | AACTCGT | ATT | TAA | - | T | NA | NA_TAA | A/T+1_AACTCGT | 6 | FALSE | FALSE |
| 133_v.txt | 32081 | 32081-Insert | Insertion | 1 | - | A | 56 | 15057 | 0.37192004 | 63.5 | rnr1Y285F_pG | TRUE | CAN | JSY2420 | 1 | rnr1Y285F_p | CAA | nope | A/T+1 | >1bpdel | none | A/T+1 | CCATGCAAA | AAAAAAG | GTT | CAA | - | T | NA | NA_CAA | A/T+1_AAAAAAG | 6 | FALSE | FALSE |
| 133_v.txt | 32346 | 32346-Insert | Insertion | 1 | - | A | 13 | 21906 | 0.05934447 | 64 | rnr1Y285F_pG | TRUE | CAN | JSY2420 | 1 | rnr1Y285F_p | CAA | nope | A/T+1 | >1bpdel | none | A/T+1 | TGTGGCAAA | AAAACCT | GTT | CAA | - | T | NA | NA_CAA | A/T+1_AAAACCT | 6 | FALSE | FALSE |
| 133_v.txt | 32381 | 32381-Insert | Insertion | 1 | - | A | 22 | 21906 | 0.10042911 | 62.3181818 | rnr1Y285F_pG | TRUE | CAN | JSY2420 | 1 | rnr1Y285F_p | TAA | nope | A/T+1 | >1bpdel | none | A/T+1 | AATAATAAA | AAAGGGA | ATT | TAA | - | T | NA | NA_TAA | A/T+1_AAAGGGA | 6 | FALSE | FALSE |
| 133_v.txt | 32495 | 32495-Insert | Insertion | 1 | - | A | 2 | 21906 | 9.13E-03 | 38 | rnr1Y285F_pG | TRUE | CAN | JSY2420 | 1 | rnr1Y285F_p | CAA | nope | A/T+1 | >1bpdel | none | A/T+1 | GAAAACAAC | AACTTTT | GTT | CAA | - | T | NA | NA_CAA | A/T+1_AACTTTT | 6 | FALSE | FALSE |
| 133_v.txt | 32528 | 32528-Insert | Insertion | 1 | - | A | 1 | 21906 | 4.57E-03 | 38 | rnr1Y285F_pG | TRUE | CAN | JSY2420 | 1 | rnr1Y285F_p | TGG | nope | A/T+1 | >1bpdel | none | A/T+1 | TTTTCTGGG | GGGGTTT | ACC | TGG | - | T | NA | NA_TGG | A/T+1_GGGGTTT | 6 | FALSE | FALSE |
| 133_v.txt | 32631 | 32631-Insert | Insertion | 1 | - | T | 6 | 17261 | 0.03476044 | 50 | rnr1Y285F_pG | TRUE | CAN | JSY2420 | 1 | rnr1Y285F_p | GTT | nope | A/T+1 | >1bpdel | none | A/T+1 | CCTTCGTTTT | TTTTTAT | CAA | GTT | - | A | NA | NA_GTT | A/T+1_TTTTTAT | 6 | FALSE | FALSE |
| 133_v.txt | 32673 | 32673-Insert | Insertion | 1 | - | T | 8 | 17261 | 0.04634726 | 59.125 | rnr1Y285F_pG | TRUE | CAN | JSY2420 | 1 | rnr1Y285F_p | GTT | nope | A/T+1 | >1bpdel | none | A/T+1 | CCTGGGTTT | TTTCTCC | CAA | GTT | - | A | NA | NA_GTT | A/T+1_TTTCTCC | 6 | FALSE | FALSE |
| 133_v.txt | 32728 | 32728-Insert | Insertion | 1 | - | A | 3 | 17261 | 0.01738022 | 48.3333333 | rnr1Y285F_pG | TRUE | CAN | JSY2420 | 1 | rnr1Y285F_p | CAA | nope | A/T+1 | >1bpdel | none | A/T+1 | CCATACAAA | AAAAACA | GTT | CAA | - | T | NA | NA_CAA | A/T+1_AAAAACA | 6 | FALSE | FALSE |
| 133_v.txt | 32760 | 32760-Insert | Insertion | 1 | - | A | 7 | 17261 | 0.04055385 | 52.8571429 | rnr1Y285F_pG | TRUE | CAN | JSY2420 | 1 | rnr1Y285F_p | TAA | nope | A/T+1 | >1bpdel | none | A/T+1 | ATGGCTAAA | AAAACTT | ATT | TAA | - | T | NA | NA_TAA | A/T+1_AAAACTT | 6 | FALSE | FALSE |
| 133_v.txt | 32842 | 32842-Insert | Insertion | 1 | - | A | 32 | 17259 | 0.18541051 | 41.5 | rnr1Y285F_pG | TRUE | CAN | JSY2420 | 1 | rnr1Y285F_p | CAA | nope | A/T+1 | >1bpdel | none | A/T+1 | TTACCCAAA | AAAAAAT | GTT | CAA | - | T | NA | NA_CAA | A/T+1_AAAAAAT | 6 | FALSE | FALSE |
| 133_v.txt | 33326 | 33326-Insert | Insertion | 1 | - | T | 6 | 10945 | 0.05481955 | 64 | rnr1Y285F_pG | TRUE | CAN | JSY2420 | 1 | rnr1Y285F_p | CTT | nope | A/T+1 | >1bpdel | none | A/T+1 | CAATTCTTTA | TTTACTT | GAA | CTT | - | A | NA | NA_CTT | A/T+1_TTTACTT | 6 | FALSE | FALSE |
| 133_v.txt | 32454 | 32454-Deleti | Deletion | 3 | AAT | - | 11 | 21906 | 0.05021455 | 51.4545455 | rnr1Y285F_pG | TRUE | CAN | JSY2420 | 1 | rnr1Y285F_p | GAA | nope | >1bp-ins | >1bpdel | none | >1bpdel | CCAATGAAT | AATAATA | CTT | GAA | AAT | none | NA | NA_GAA | >1bpdel_AATAATA | 6 | FALSE | FALSE |
| 133_v.txt | 32716 | 32716-Deleti | Deletion | 2 | CA | - | 13 | 17261 | 0.07531429 | 62 | rnr1Y285F_pG | TRUE | CAN | JSY2420 | 1 | rnr1Y285F_p | CCA | nope | >1bp-ins | >1bpdel | none | >1bpdel | CAGCACCAC | CACAAAC | GGT | CCA | CA | none | NA | NA_CCA | >1bpdel_CACAAAC | 6 | FALSE | FALSE |
| 133_v.txt | 32736 | 32736-Deleti | Deletion | 2 | TA | - | 28 | 17261 | 0.1622154 | 59.6785714 | rnr1Y285F_pG | TRUE | CAN | JSY2420 | 1 | rnr1Y285F_p | GTA | nope | >1bp-ins | >1bpdel | none | >1bpdel | AAACAGTAT | TATATTA | CAT | GTA | TA | none | NA | NA_GTA | >1bpdel_TATATTA | 6 | FALSE | FALSE |
| 133_v.txt | 32855 | 32855-Deleti | Deletion | 2 | CC | - | 1 | 17259 | 5.79E-03 | 38 | rnr1Y285F_pG | TRUE | CAN | JSY2420 | 1 | rnr1Y285F_p | TCC | nope | >1bp-ins | >1bpdel | none | >1bpdel | ACTAATCCAT | CCATGCC | AGG | TCC | CC | none | NA | NA_TCC | >1bpdel_CCATGCC | 6 | FALSE | FALSE |
| 133_v.txt | 33022 | 33022-Deleti | Deletion | 2 | CA | - | 1 | 20066 | 4.98E-03 | 64 | rnr1Y285F_pG | TRUE | CAN | JSY2420 | 1 | rnr1Y285F_p | ACA | nope | >1bp-ins | >1bpdel | none | >1bpdel | ATGTAACAG | CAGGGAT | TGT | ACA | CA | none | NA | NA_ACA | >1bpdel_CAGGGAT | 6 | FALSE | FALSE |
| 133_v.txt | 33022 | 33022-Deleti | Deletion | 16 | CAGGGATGA | - | 103 | 20066 | 0.51330609 | 61.1359223 | rnr1Y285F_pG | TRUE | CAN | JSY2420 | 1 | rnr1Y285F_p | ACA | nope | >1bp-ins | >1bpdel | none | >1bpdel | ATGTAACAG | CAGGGAT | TGT | ACA | CAGGGATGA | none | NA | NA_ACA | >1bpdel_CAGGGAT | 6 | FALSE | FALSE |
| 133_v.txt | 33025 | 33025-Deleti | Deletion | 3 | GGA | - | 1 | 20066 | 4.98E-03 | 64 | rnr1Y285F_pG | TRUE | CAN | JSY2420 | 1 | rnr1Y285F_p | GGG | nope | >1bp-ins | >1bpdel | none | >1bpdel | TAACAGGGA | GGATGAA | CCC | GGG | GGA | none | NA | NA_GGG | >1bpdel_GGATGAA | 6 | FALSE | FALSE |
| 133_v.txt | 33031 | 33031-Deleti | Deletion | 2 | AT | - | 1 | 20066 | 4.98E-03 | 64 | rnr1Y285F_pG | TRUE | CAN | JSY2420 | 1 | rnr1Y285F_p | AAT | nope | >1bp-ins | >1bpdel | none | >1bpdel | GGATGAATG | ATGTAGC | TTA | AAT | AT | none | NA | NA_AAT | >1bpdel_ATGTAGC | 6 | FALSE | FALSE |
| 133_v.txt | 33032 | 33032-Deleti | Deletion | 6 | TGTAGC | - | 1 | 20066 | 4.98E-03 | 64 | rnr1Y285F_pG | TRUE | CAN | JSY2420 | 1 | rnr1Y285F_p | ATG | nope | >1bp-ins | >1bpdel | none | >1bpdel | GATGAATGT | TGTAGCC | TAC | ATG | TGTAGC | none | NA | NA_ATG | >1bpdel_TGTAGCC | 6 | FALSE | FALSE |
| 133_v.txt | 33065 | 33065-Deleti | Deletion | 2 | AT | - | 10 | 20066 | 0.04983554 | 52.9 | rnr1Y285F_pG | TRUE | CAN | JSY2420 | 1 | rnr1Y285F_p | AAT | nope | >1bp-ins | >1bpdel | none | >1bpdel | GACAGAATA | ATATGCC | TTA | AAT | AT | none | NA | NA_AAT | >1bpdel_ATATGCC | 6 | FALSE | FALSE |
| 133_v.txt | 33089 | 33089-Deleti | Deletion | 2 | AT | - | 39 | 20066 | 0.19435862 | 51.2564103 | rnr1Y285F_pG | TRUE | CAN | JSY2420 | 1 | rnr1Y285F_p | AAT | nope | >1bp-ins | >1bpdel | none | >1bpdel | AAATAAATA | ATATGAT | TTA | AAT | AT | none | NA | NA_AAT | >1bpdel_ATATGAT | 6 | FALSE | FALSE |
| 133_v.txt | 33155 | 33155-Deleti | Deletion | 2 | TG | - | 21 | 20065 | 0.10465986 | 37.3333333 | rnr1Y285F_pG | TRUE | CAN | JSY2420 | 1 | rnr1Y285F_p | CTG | nope | >1bp-ins | >1bpdel | none | >1bpdel | AAGACCTGT | TGTACCA | GAC | CTG | TG | none | NA | NA_CTG | >1bpdel_TGTACCA | 6 | FALSE | FALSE |
| 133_v.txt | 31723 | 31723-Deleti | Deletion | 1 | A | - | 7 | 6373 | 0.10983838 | 64 | rnr1Y285F_pG | TRUE | CAN | JSY2420 | 1 | rnr1Y285F_p | CAA | nope | >1bp-ins | A/T-1 | none | A/T-1 | TGTCCCAAA | AAAAAGT | GTT | GTT | T | none | T>none | T>none_GTT | A/T-1_AAAAAGT | 6 | FALSE | FALSE |
| 133_v.txt | 31863 | 31863-Deleti | Deletion | 1 | A | - | 9 | 6373 | 0.14122078 | 47.7777778 | rnr1Y285F_pG | TRUE | CAN | JSY2420 | 1 | rnr1Y285F_p | GAA | nope | >1bp-ins | A/T-1 | none | A/T-1 | AACAGGAAA | AAAATAG | CTT | CTT | T | none | T>none | T>none_CTT | A/T-1_AAAATAG | 6 | FALSE | FALSE |
| 133_v.txt | 31904 | 31904-Deleti | Deletion | 1 | T | - | 9 | 6371 | 0.14126511 | 41.4444444 | rnr1Y285F_pG | TRUE | CAN | JSY2420 | 1 | rnr1Y285F_p | ATT | nope | >1bp-ins | A/T-1 | none | A/T-1 | ATTGAATTTT | TTTTGGT | TAA | ATT | T | none | nope | nope_ATT | A/T-1_TTTTGGT | 6 | FALSE | FALSE |
| 133_v.txt | 32073 | 32073-Deleti | Deletion | 1 | A | - | 3 | 15057 | 0.01992429 | 56.6666667 | rnr1Y285F_pG | TRUE | CAN | JSY2420 | 1 | rnr1Y285F_p | TAA | nope | >1bp-ins | A/T-1 | none | A/T-1 | ATAAATAAC | AACCATG | ATT | ATT | T | none | T>none | T>none_ATT | A/T-1_AACCATG | 6 | FALSE | FALSE |
| 133_v.txt | 32075 | 32075-Deleti | Deletion | 1 | C | - | 1 | 15057 | 6.64E-03 | 64 | rnr1Y285F_pG | TRUE | CAN | JSY2420 | 1 | rnr1Y285F_p | ACC | nope | >1bp-ins | G/C-1 | none | G/C-1 | AAATAACCA | CCATGCA | TGG | ACC | C | none | nope | nope_ACC | G/C-1_CCATGCA | 6 | FALSE | FALSE |
| 133_v.txt | 32081 | 32081-Deleti | Deletion | 1 | A | - | 13 | 15057 | 0.08633858 | 63.1538462 | rnr1Y285F_pG | TRUE | CAN | JSY2420 | 1 | rnr1Y285F_p | CAA | nope | >1bp-ins | A/T-1 | none | A/T-1 | CCATGCAAA | AAAAAAG | GTT | GTT | T | none | T>none | T>none_GTT | A/T-1_AAAAAAG | 6 | FALSE | FALSE |
| 133_v.txt | 32216 | 32216-Deleti | Deletion | 1 | G | - | 7 | 15057 | 0.04649001 | 38 | rnr1Y285F_pG | TRUE | CAN | JSY2420 | 1 | rnr1Y285F_p | TGG | nope | >1bp-ins | G/C-1 | none | G/C-1 | TTTGGTGGT | GGTCCTT | ACC | ACC | C | none | C>none | C>none_ACC | G/C-1_GGTCCTT | 6 | FALSE | FALSE |
| 133_v.txt | 32346 | 32346-Deleti | Deletion | 1 | A | - | 54 | 21906 | 0.24650781 | 61.3703704 | rnr1Y285F_pG | TRUE | CAN | JSY2420 | 1 | rnr1Y285F_p | CAA | nope | >1bp-ins | A/T-1 | none | A/T-1 | TGTGGCAAA | AAAACCT | GTT | GTT | T | none | T>none | T>none_GTT | A/T-1_AAAACCT | 6 | FALSE | FALSE |
| 133_v.txt | 32419 | 32419-Deleti | Deletion | 1 | G | - | 9 | 21906 | 0.04108463 | 60.7777778 | rnr1Y285F_pG | TRUE | CAN | JSY2420 | 1 | rnr1Y285F_p | AGT | nope | >1bp-ins | G/C-1 | none | G/C-1 | GTGTTAGTTT | GTTTAGG | TCA | TCA | C | none | C>none | C>none_TCA | G/C-1_GTTTAGG | 6 | FALSE | FALSE |
| 133_v.txt | 32424 | 32424-Deleti | Deletion | 1 | G | - | 9 | 21906 | 0.04108463 | 64 | rnr1Y285F_pG | TRUE | CAN | JSY2420 | 1 | rnr1Y285F_p | AGG | nope | >1bp-ins | G/C-1 | none | G/C-1 | AGTTTAGGG | GGGTCAT | TCC | TCC | C | none | C>none | C>none_TCC | G/C-1_GGGTCAT | 6 | FALSE | FALSE |
| 133_v.txt | 32490 | 32490-Deleti | Deletion | 1 | A | - | 10 | 21906 | 0.04564959 | 45.7 | rnr1Y285F_pG | TRUE | CAN | JSY2420 | 1 | rnr1Y285F_p | GAA | nope | >1bp-ins | A/T-1 | none | A/T-1 | ATACGGAAA | AAAACAA | CTT | CTT | T | none | T>none | T>none_CTT | A/T-1_AAAACAA | 6 | FALSE | FALSE |
| 133_v.txt | 32498 | 32498-Deleti | Deletion | 1 | T | - | 58 | 21906 | 0.26476764 | 43.637931 | rnr1Y285F_pG | TRUE | CAN | JSY2420 | 1 | rnr1Y285F_p | CTT | nope | >1bp-ins | A/T-1 | none | A/T-1 | AACAACTTTT | TTTTTTG | GAA | CTT | T | none | nope | nope_CTT | A/T-1_TTTTTTG | 6 | FALSE | FALSE |
| 133_v.txt | 32528 | 32528-Deleti | Deletion | 1 | G | - | 7 | 21906 | 0.03195472 | 38.7142857 | rnr1Y285F_pG | TRUE | CAN | JSY2420 | 1 | rnr1Y285F_p | TGG | nope | >1bp-ins | G/C-1 | none | G/C-1 | TTTTCTGGG | GGGGTTT | ACC | ACC | C | none | C>none | C>none_ACC | G/C-1_GGGGTTT | 6 | FALSE | FALSE |
| 133_v.txt | 32571 | 32571-Deleti | Deletion | 1 | C | - | 16 | 17262 | 0.09268914 | 36.8125 | rnr1Y285F_pG | TRUE | CAN | JSY2420 | 1 | rnr1Y285F_p | ACC | nope | >1bp-ins | G/C-1 | none | G/C-1 | TCAGTACCTT | CCTTGAA | TGG | ACC | C | none | nope | nope_ACC | G/C-1_CCTTGAA | 6 | FALSE | FALSE |
| 133_v.txt | 32631 | 32631-Deleti | Deletion | 1 | T | - | 12 | 17261 | 0.06952089 | 52.5833338 | rnr1Y285F_pG | TRUE | CAN | JSY2420 | 1 | rnr1Y285F_p | GTT | nope | >1bp-ins | A/T-1 | none | A/T-1 | CCTTCGTTTT | TTTTTAT | CAA | GTT | T | none | nope | nope_GTT | A/T-1_TTTTTAT | 6 | FALSE | FALSE |
| 133_v.txt | 32657 | 32657-Deleti | Deletion | 1 | A | - | 15 | 17260 | 0.08690614 | 58.9333333 | rnr1Y285F_pG | TRUE | CAN | JSY2420 | 1 | rnr1Y285F_p | GAC | nope | >1bp-ins | A/T-1 | none | A/T-1 | ACCTGGACC | ACCCCAG | CTG | CTG | T | none | T>none | T>none_CTG | A/T-1_ACCCCAG | 6 | FALSE | FALSE |
| 133_v.txt | 32658 | 32658-Deleti | Deletion | 1 | C | - | 7 | 17261 | 0.04055385 | 52.1428571 | rnr1Y285F_pG | TRUE | CAN | JSY2420 | 1 | rnr1Y285F_p | ACC | nope | >1bp-ins | G/C-1 | none | G/C-1 | CCTGGACCC | CCCCAGG | TGG | ACC | C | none | nope | nope_ACC | G/C-1_CCCCAGG | 6 | FALSE | FALSE |
| 133_v.txt | 32705 | 32705-Deleti | Deletion | 1 | A | - | 9 | 17261 | 0.05214066 | 59.4444444 | rnr1Y285F_pG | TRUE | CAN | JSY2420 | 1 | rnr1Y285F_p | TAA | nope | >1bp-ins | A/T-1 | none | A/T-1 | GCCGGTAAC | AACCCCA | ATT | ATT | T | none | T>none | T>none_ATT | A/T-1_AACCCCA | 6 | FALSE | FALSE |
| 133_v.txt | 32707 | 32707-Deleti | Deletion | 1 | C | - | 17 | 17261 | 0.09848792 | 59.7647059 | rnr1Y285F_pG | TRUE | CAN | JSY2420 | 1 | rnr1Y285F_p | ACC | nope | >1bp-ins | G/C-1 | none | G/C-1 | CGGTAACCC | CCCCAGC | TGG | ACC | C | none | nope | nope_ACC | G/C-1_CCCCAGC | 6 | FALSE | FALSE |
| 133_v.txt | 32728 | 32728-Deleti | Deletion | 1 | A | - | 20 | 17261 | 0.11586814 | 61.35 | rnr1Y285F_pG | TRUE | CAN | JSY2420 | 1 | rnr1Y285F_p | CAA | nope | >1bp-ins | A/T-1 | none | A/T-1 | CCATACAAA | AAAAACA | GTT | GTT | T | none | T>none | T>none_GTT | A/T-1_AAAAACA | 6 | FALSE | FALSE |
| 133_v.txt | 32795 | 32795-Deleti | Deletion | 1 | A | - | 31 | 17261 | 0.17959562 | 47.0645161 | rnr1Y285F_pG | TRUE | CAN | JSY2420 | 1 | rnr1Y285F_p | CAC | nope | >1bp-ins | A/T-1 | none | A/T-1 | GAATTCACC | ACCGTAA | GTG | GTG | T | none | T>none | T>none_GTG | A/T-1_ACCGTAA | 6 | FALSE | FALSE |
| 133_v.txt | 32842 | 32842-Deleti | Deletion | 1 | A | - | 114 | 17259 | 0.66052494 | 42.2807018 | rnr1Y285F_pG | TRUE | CAN | JSY2420 | 1 | rnr1Y285F_p | CAA | nope | >1bp-ins | A/T-1 | none | A/T-1 | TTACCCAAA | AAAAAAT | GTT | GTT | T | none | T>none | T>none_GTT | A/T-1_AAAAAAT | 6 | FALSE | FALSE |
| 133_v.txt | 32855 | 32855-Deleti | Deletion | 1 | C | - | 1 | 17259 | 5.79E-03 | 38 | rnr1Y285F_pG | TRUE | CAN | JSY2420 | 1 | rnr1Y285F_p | TCC | nope | >1bp-ins | G/C-1 | none | G/C-1 | ACTAATCCAT | CCATGCC | AGG | TCC | C | none | nope | nope_TCC | G/C-1_CCATGCC | 6 | FALSE | FALSE |
| 133_v.txt | 32884 | 32884-Deleti | Deletion | 1 | A | - | 10 | 20066 | 0.04983554 | 36.0999989 | rnr1Y285F_pG | TRUE | CAN | JSY2420 | 1 | rnr1Y285F_p | CAA | nope | >1bp-ins | A/T-1 | none | A/T-1 | ACGTCCAAA | AAAATTG | GTT | GTT | T | none | T>none | T>none_GTT | A/T-1_AAAATTG | 6 | FALSE | FALSE |
| 133_v.txt | 32924 | 32924-Deleti | Deletion | 1 | A | - | 79 | 20066 | 0.39370079 | 55.1772152 | rnr1Y285F_pG | TRUE | CAN | JSY2420 | 1 | rnr1Y285F_p | CAA | nope | >1bp-ins | A/T-1 | none | A/T-1 | CAGGGCAAA | AAAAGTG | GTT | GTT | T | none | T>none | T>none_GTT | A/T-1_AAAAGTG | 6 | FALSE | FALSE |
| 133_v.txt | 32941 | 32941-Deleti | Deletion | 1 | A | - | 22 | 20066 | 0.10963819 | 57.727269 | rnr1Y285F_pG | TRUE | CAN | JSY2420 | 1 | rnr1Y285F_p | GAA | nope | >1bp-ins | A/T-1 | none | A/T-1 | CCCAAGAAA | AAAACCA | CTT | CTT | T | none | T>none | T>none_CTT | A/T-1_AAAACCA | 6 | FALSE | FALSE |
| 133_v.txt | 33000 | 33000-Deleti | Deletion | 1 | A | - | 6 | 20066 | 0.02990133 | 63.6666667 | rnr1Y285F_pG | TRUE | CAN | JSY2420 | 1 | rnr1Y285F_p | GAA | nope | >1bp-ins | A/T-1 | none | A/T-1 | TGTGAGAAA | AAAACTG | CTT | CTT | T | none | T>none | T>none_CTT | A/T-1_AAAACTG | 6 | FALSE | FALSE |
| 133_v.txt | 33008 | 33008-Deleti | Deletion | 1 | G | - | 12 | 20066 | 0.05980265 | 60.5833333 | rnr1Y285F_pG | TRUE | CAN | JSY2420 | 1 | rnr1Y285F_p | TGA | nope | >1bp-ins | G/C-1 | none | G/C-1 | AACTGTGAA | GAAAGAG | ACT | ACT | C | none | C>none | C>none_ACT | G/C-1_GAAAGAG | 6 | FALSE | FALSE |
| 133_v.txt | 33029 | 33029-Deleti | Deletion | 1 | G | - | 1 | 20066 | 4.98E-03 | 64 | rnr1Y285F_pG | TRUE | CAN | JSY2420 | 1 | rnr1Y285F_p | TGA | nope | >1bp-ins | G/C-1 | none | G/C-1 | AGGGATGAA | GAATGTA | ACT | ACT | C | none | C>none | C>none_ACT | G/C-1_GAATGTA | 6 | FALSE | FALSE |
| 133_v.txt | 33037 | 33037-Deleti | Deletion | 1 | C | - | 1 | 20064 | 4.98E-03 | 64 | rnr1Y285F_pG | TRUE | CAN | JSY2420 | 1 | rnr1Y285F_p | GCC | nope | >1bp-ins | G/C-1 | none | G/C-1 | ATGTAGCCA | CCATTTC | CGG | GCC | C | none | nope | nope_GCC | G/C-1_CCATTTC | 6 | FALSE | FALSE |

| | | | | | | | | | | | | | | | | | | | | | | | | | | | | | | | | | |
|---|---|---|---|---|---|---|---|---|---|---|---|---|---|---|---|---|---|---|---|---|---|---|---|---|---|---|---|---|---|---|---|---|---|
| 133_v.txt | 33147 | 33147-Deleti | Deletion | 1 | A | - | 6 | 20066 | 0.02990133 | 37.6666667 | rnr1Y285F_pG | TRUE | CAN | JSY2420 | 1 | rnr1Y285F_p GAA | nope | >1bp-ins | A/T-1 | none | A/T-1 | CCAATGAAA | AAAAGAC | CTT | CTT | T | none | T>none | T>none_CTT | A/T-1_AAAAGAC | 6 | FALSE | FALSE |
| 133_v.txt | 33168 | 33168-Deleti | Deletion | 1 | C | - | 2 | 20063 | 9.97E-03 | 38 | rnr1Y285F_pG | TRUE | CAN | JSY2420 | 1 | rnr1Y285F_p ACC | nope | >1bp-ins | G/C-1 | none | G/C-1 | ATAGTACCA | CCACCAA | TGG | ACC | C | none | nope | nope_ACC | G/C-1_CCACCAA | 6 | FALSE | FALSE |
| 133_v.txt | 33474 | 33474-Deleti | Deletion | 1 | C | - | 38 | 10944 | 0.34722222 | 29.1578947 | rnr1Y285F_pG | TRUE | CAN | JSY2420 | 1 | rnr1Y285F_p GCC | nope | >1bp-ins | G/C-1 | none | G/C-1 | GCTATGCCT | CCTTTTT | CGG | GCC | C | none | nope | nope_GCC | G/C-1_CCTTTTT | 6 | FALSE | FALSE |
| 134_v.txt | 31723 | 31723-SNV-/ | SNV | 1 | A | C | 6 | 8528 | 0.07035647 | 61.1666953 | rnr1Y285F_pG | TRUE | CAN | JSY2420 | 2 | rnr1Y285F_p CAA | A>C | G/C+1 | A/T-1 | TA>GC | TA>GC | TGTCCCAAA | AAAAAGT | GTT | GTT | T | G | T>G | T>G_GTT | TA>GC_AAAAAGT | 6 | FALSE | FALSE |
| 134_v.txt | 31907 | 31907-SNV-T | SNV | 1 | T | G | 4 | 8528 | 0.04690432 | 37.4998982 | rnr1Y285F_pG | TRUE | CAN | JSY2420 | 2 | rnr1Y285F_p TTG | T>G | G/C+1 | A/T-1 | TA>GC | TA>GC | GAATTTTGG | TGGTGCA | AAC | TTG | T | C | T>G | T>G_TTG | TA>GC_TGGTGCA | 6 | FALSE | FALSE |
| 134_v.txt | 31967 | 31967-SNV-/ | SNV | 1 | A | C | 16 | 19468 | 0.08218615 | 25.1875 | rnr1Y285F_pG | TRUE | CAN | JSY2420 | 2 | rnr1Y285F_p AAT | A>C | G/C+1 | A/T-1 | TA>GC | TA>GC | CGCATAATA | ATAAGCC | TTA | TTA | T | G | T>G | T>G_TTA | TA>GC_ATAAGCC | 6 | FALSE | FALSE |
| 134_v.txt | 32265 | 32265-SNV-/ | SNV | 1 | A | C | 17 | 25529 | 0.06659094 | 33.4117647 | rnr1Y285F_pG | TRUE | CAN | JSY2420 | 2 | rnr1Y285F_p TAA | A>C | G/C+1 | A/T-1 | TA>GC | TA>GC | CCAAATAAA | AAAATAC | ATT | ATT | T | G | T>G | T>G_ATT | TA>GC_AAAATAC | 6 | FALSE | FALSE |
| 134_v.txt | 32283 | 32283-SNV-T | SNV | 1 | T | G | 5 | 25529 | 0.01958557 | 60.4 | rnr1Y285F_pG | TRUE | CAN | JSY2420 | 2 | rnr1Y285F_p GTA | T>G | G/C+1 | A/T-1 | TA>GC | TA>GC | CCAACGTAA | TAAATAT | CAT | GTA | T | C | T>G | T>G_GTA | TA>GC_TAAATAT | 6 | FALSE | FALSE |
| 134_v.txt | 32309 | 32309-SNV-/ | SNV | 1 | A | C | 1 | 25529 | 3.92E-03 | 64 | rnr1Y285F_pG | TRUE | CAN | JSY2420 | 2 | rnr1Y285F_p TAA | A>C | G/C+1 | A/T-1 | TA>GC | TA>GC | AGAAATAAT | AATGGTT | ATT | ATT | T | G | T>G | T>G_ATT | TA>GC_AATGGTT | 6 | FALSE | FALSE |
| 134_v.txt | 32310 | 32310-SNV-/ | SNV | 1 | A | C | 17 | 25529 | 0.06659094 | 50.9999309 | rnr1Y285F_pG | TRUE | CAN | JSY2420 | 2 | rnr1Y285F_p AAT | A>C | G/C+1 | A/T-1 | TA>GC | TA>GC | GAAATAATG | ATGGTTG | TTA | TTA | T | G | T>G | T>G_TTA | TA>GC_ATGGTTG | 6 | FALSE | FALSE |
| 134_v.txt | 32352 | 32352-SNV-T | SNV | 1 | T | G | 5 | 25529 | 0.01958557 | 62.6 | rnr1Y285F_pG | TRUE | CAN | JSY2420 | 2 | rnr1Y285F_p CTT | T>G | G/C+1 | A/T-1 | TA>GC | TA>GC | AAAACCTTT | TTTGTAC | GAA | CTT | T | C | T>G | T>G_CTT | TA>GC_TTTGTAC | 6 | FALSE | FALSE |
| 134_v.txt | 32493 | 32493-SNV-/ | SNV | 1 | A | C | 9 | 25529 | 0.03525403 | 39.8887821 | rnr1Y285F_pG | TRUE | CAN | JSY2420 | 2 | rnr1Y285F_p AAC | A>C | G/C+1 | A/T-1 | TA>GC | TA>GC | CGGAAAACA | ACAACTT | TTG | TTG | T | G | T>G | T>G_TTG | TA>GC_ACAACTT | 6 | FALSE | FALSE |
| 134_v.txt | 32631 | 32631-SNV-T | SNV | 1 | T | G | 4 | 23621 | 0.01693408 | 43.0006551 | rnr1Y285F_pG | TRUE | CAN | JSY2420 | 2 | rnr1Y285F_p GTT | T>G | G/C+1 | A/T-1 | TA>GC | TA>GC | CCTTCGTTTT | TTTTTAT | CAA | GTT | T | C | T>G | T>G_GTT | TA>GC_TTTTTAT | 6 | FALSE | FALSE |
| 134_v.txt | 32763 | 32763-SNV-/ | SNV | 1 | A | C | 19 | 23621 | 0.0804369 | 51.7368421 | rnr1Y285F_pG | TRUE | CAN | JSY2420 | 2 | rnr1Y285F_p AAC | A>C | G/C+1 | A/T-1 | TA>GC | TA>GC | GCTAAAACT | ACTTTGA | TTG | TTG | T | G | T>G | T>G_TTG | TA>GC_ACTTTGA | 6 | FALSE | FALSE |
| 134_v.txt | 32767 | 32767-SNV-T | SNV | 1 | T | G | 18 | 23621 | 0.07620338 | 30 | rnr1Y285F_pG | TRUE | CAN | JSY2420 | 2 | rnr1Y285F_p TTG | T>G | G/C+1 | A/T-1 | TA>GC | TA>GC | AAACTTTGA | TGATGGA | AAC | TTG | T | C | T>G | T>G_TTG | TA>GC_TGATGGA | 6 | FALSE | FALSE |
| 134_v.txt | 32785 | 32785-SNV-/ | SNV | 1 | A | C | 8 | 23621 | 0.03386817 | 36.375 | rnr1Y285F_pG | TRUE | CAN | JSY2420 | 2 | rnr1Y285F_p AAC | A>C | G/C+1 | A/T-1 | TA>GC | TA>GC | CCCAGAACT | ACTCGAA | TTG | TTG | T | G | T>G | T>G_TTG | TA>GC_ACTCGAA | 6 | FALSE | FALSE |
| 134_v.txt | 32842 | 32842-SNV-/ | SNV | 1 | A | C | 5 | 23621 | 0.02116761 | 37.8000159 | rnr1Y285F_pG | TRUE | CAN | JSY2420 | 2 | rnr1Y285F_p CAA | A>C | G/C+1 | A/T-1 | TA>GC | TA>GC | TTACCCAAA | AAAAAAT | GTT | GTT | T | G | T>G | T>G_GTT | TA>GC_AAAAAAT | 6 | FALSE | FALSE |
| 134_v.txt | 32903 | 32903-SNV-/ | SNV | 1 | A | C | 52 | 26566 | 0.19573891 | 44.4038462 | rnr1Y285F_pG | TRUE | CAN | JSY2420 | 2 | rnr1Y285F_p CAA | A>C | G/C+1 | A/T-1 | TA>GC | TA>GC | TTGGCCAAC | AACTACA | GTT | GTT | T | G | T>G | T>G_GTT | TA>GC_AACTACA | 6 | FALSE | FALSE |
| 134_v.txt | 32913 | 32913-SNV-/ | SNV | 1 | A | C | 54 | 26566 | 0.20326733 | 34.1851852 | rnr1Y285F_pG | TRUE | CAN | JSY2420 | 2 | rnr1Y285F_p AAG | A>C | G/C+1 | A/T-1 | TA>GC | TA>GC | ACACTAAGT | AGTTCCA | TTC | TTC | T | G | T>G | T>G_TTC | TA>GC_AGTTCCA | 6 | FALSE | FALSE |
| 134_v.txt | 32938 | 32938-SNV-/ | SNV | 1 | A | C | 41 | 26566 | 0.15433261 | 53.7272366 | rnr1Y285F_pG | TRUE | CAN | JSY2420 | 2 | rnr1Y285F_p CAA | A>C | G/C+1 | A/T-1 | TA>GC | TA>GC | TTGCCCAAG | AAGAAAA | GTT | GTT | T | G | T>G | T>G_GTT | TA>GC_AAGAAAA | 6 | FALSE | FALSE |
| 134_v.txt | 33126 | 33126-SNV-/ | SNV | 1 | A | C | 25 | 26566 | 0.09410525 | 41.6 | rnr1Y285F_pG | TRUE | CAN | JSY2420 | 2 | rnr1Y285F_p CAG | A>C | G/C+1 | A/T-1 | TA>GC | TA>GC | TTGGTCAGA | AGAGGTG | GTC | GTC | T | G | T>G | T>G_GTC | TA>GC_AGAGGTG | 6 | FALSE | FALSE |
| 134_v.txt | 33204 | 33204-SNV-/ | SNV | 1 | A | C | 147 | 19947 | 0.73695293 | 36.4557823 | rnr1Y285F_pG | TRUE | CAN | JSY2420 | 2 | rnr1Y285F_p AAG | A>C | G/C+1 | A/T-1 | TA>GC | TA>GC | TGCTTAAGC | AGCTCTC | TTC | TTC | T | G | T>G | T>G_TTC | TA>GC_AGCTCTC | 6 | FALSE | FALSE |
| 134_v.txt | 33269 | 33269-SNV-/ | SNV | 1 | A | C | 156 | 19947 | 0.78207249 | 15.3461539 | rnr1Y285F_pG | TRUE | CAN | JSY2420 | 2 | rnr1Y285F_p TAG | A>C | G/C+1 | A/T-1 | TA>GC | TA>GC | CTCCATAGA | AGAGAAC | ATC | ATC | T | G | T>G | T>G_ATC | TA>GC_AGAGAAC | 6 | FALSE | FALSE |
| 134_v.txt | 32309 | 32309-SNV-/ | SNV | 1 | A | G | 24 | 25529 | 0.09401073 | 58.9169485 | rnr1Y285F_pG | TRUE | CAN | JSY2420 | 2 | rnr1Y285F_p TAA | A>G | G/C+1 | A/T-1 | TA>CG | TA>CG | AGAAATAAT | AATGGTT | ATT | ATT | T | C | T>C | T>C_ATT | TA>CG_AATGGTT | 6 | FALSE | FALSE |
| 134_v.txt | 32337 | 32337-SNV-/ | SNV | 1 | A | G | 2 | 25529 | 7.83E-03 | 47.9999809 | rnr1Y285F_pG | TRUE | CAN | JSY2420 | 2 | rnr1Y285F_p GAT | A>G | G/C+1 | A/T-1 | TA>CG | TA>CG | TTGAAGATA | ATATGTG | CTA | CTA | T | C | T>C | T>C_CTA | TA>CG_ATATGTG | 6 | FALSE | FALSE |
| 134_v.txt | 32375 | 32375-SNV-/ | SNV | 1 | A | G | 4 | 25529 | 0.01566846 | 54.9998948 | rnr1Y285F_pG | TRUE | CAN | JSY2420 | 2 | rnr1Y285F_p CAA | A>G | G/C+1 | A/T-1 | TA>CG | TA>CG | AATAGCAAT | AATAATA | GTT | GTT | T | C | T>C | T>C_GTT | TA>CG_AATAATA | 6 | FALSE | FALSE |
| 134_v.txt | 32377 | 32377-SNV-T | SNV | 1 | T | C | 8 | 25526 | 0.03134059 | 56.3747154 | rnr1Y285F_pG | TRUE | CAN | JSY2420 | 2 | rnr1Y285F_p ATA | T>C | G/C+1 | A/T-1 | TA>CG | TA>CG | TAGCAATAA | TAATAAA | TAT | ATA | T | G | T>C | T>C_ATA | TA>CG_TAATAAA | 6 | FALSE | FALSE |
| 134_v.txt | 32475 | 32475-SNV-/ | SNV | 1 | A | G | 2238 | 25529 | 8.76650084 | 46.8771767 | rnr1Y285F_pG | TRUE | CAN | JSY2420 | 2 | rnr1Y285F_p GAA | A>G | G/C+1 | A/T-1 | TA>CG | TA>CG | ATGTAGAAG | AAGGTTA | CTT | CTT | T | C | T>C | T>C_CTT | TA>CG_AAGGTTA | 6 | FALSE | FALSE |
| 134_v.txt | 32766 | 32766-SNV-T | SNV | 1 | T | C | 5 | 23621 | 0.02116761 | 52.5998429 | rnr1Y285F_pG | TRUE | CAN | JSY2420 | 2 | rnr1Y285F_p TTT | T>C | G/C+1 | A/T-1 | TA>CG | TA>CG | AAAACTTTG | TTGATGG | AAA | TTT | T | G | T>C | T>C_TTT | TA>CG_TTGATGG | 6 | FALSE | FALSE |
| 134_v.txt | 32767 | 32767-SNV-T | SNV | 1 | T | C | 12 | 23621 | 0.05080225 | 60.25 | rnr1Y285F_pG | TRUE | CAN | JSY2420 | 2 | rnr1Y285F_p TTG | T>C | G/C+1 | A/T-1 | TA>CG | TA>CG | AAACTTTGA | TGATGGA | AAC | TTG | T | G | T>C | T>C_TTG | TA>CG_TGATGGA | 6 | FALSE | FALSE |
| 134_v.txt | 32820 | 32820-SNV-T | SNV | 1 | T | C | 5 | 23621 | 0.02116761 | 34.5999605 | rnr1Y285F_pG | TRUE | CAN | JSY2420 | 2 | rnr1Y285F_p GTT | T>C | G/C+1 | A/T-1 | TA>CG | TA>CG | AACAAGTTC | TTCATTA | CAA | GTT | T | G | T>C | T>C_GTT | TA>CG_TTCATTA | 6 | FALSE | FALSE |
| 134_v.txt | 32927 | 32927-SNV-/ | SNV | 1 | A | G | 26 | 26566 | 0.09786946 | 48.9209916 | rnr1Y285F_pG | TRUE | CAN | JSY2420 | 2 | rnr1Y285F_p AAG | A>G | G/C+1 | A/T-1 | TA>CG | TA>CG | GGCAAAAGT | AGTGATT | TTC | TTC | T | C | T>C | T>C_TTC | TA>CG_AGTGATT | 6 | FALSE | FALSE |
| 134_v.txt | 32929 | 32929-SNV-T | SNV | 1 | T | C | 14 | 26566 | 0.05269894 | 42.071297 | rnr1Y285F_pG | TRUE | CAN | JSY2420 | 2 | rnr1Y285F_p GTG | T>C | G/C+1 | A/T-1 | TA>CG | TA>CG | CAAAAGTGA | TGATTGC | CAC | GTG | T | G | T>C | T>C_GTG | TA>CG_TGATTGC | 6 | FALSE | FALSE |
| 134_v.txt | 32938 | 32938-SNV-/ | SNV | 1 | A | G | 164 | 26566 | 0.61733042 | 54.5820195 | rnr1Y285F_pG | TRUE | CAN | JSY2420 | 2 | rnr1Y285F_p CAA | A>G | G/C+1 | A/T-1 | TA>CG | TA>CG | TTGCCCAAG | AAGAAAA | GTT | GTT | T | C | T>C | T>C_GTT | TA>CG_AAGAAAA | 6 | FALSE | FALSE |
| 134_v.txt | 33040 | 33040-SNV-T | SNV | 1 | T | C | 50 | 26566 | 0.1882105 | 58.3204534 | rnr1Y285F_pG | TRUE | CAN | JSY2420 | 2 | rnr1Y285F_p ATT | T>C | G/C+1 | A/T-1 | TA>CG | TA>CG | TAGCCATTTC | TTTCACC | TAA | ATT | T | G | T>C | T>C_ATT | TA>CG_TTTCACC | 6 | FALSE | FALSE |
| 134_v.txt | 33204 | 33204-SNV-/ | SNV | 1 | A | G | 32 | 19947 | 0.16042513 | 35.4375 | rnr1Y285F_pG | TRUE | CAN | JSY2420 | 2 | rnr1Y285F_p AAG | A>G | G/C+1 | A/T-1 | TA>CG | TA>CG | TGCTTAAGC | AGCTCTC | TTC | TTC | T | C | T>C | T>C_TTC | TA>CG_AGCTCTC | 6 | FALSE | FALSE |
| 134_v.txt | 31906 | 31906-SNV-T | SNV | 1 | T | A | 41 | 8528 | 0.48076923 | 41.9493866 | rnr1Y285F_pG | TRUE | CAN | JSY2420 | 2 | rnr1Y285F_p TTT | T>A | A/T+1 | A/T-1 | TA>AT | TA>AT | TGAATTTTG | TTGGTGC | AAA | TTT | T | T | T>A | T>A_TTT | TA>AT_TTGGTGC | 6 | FALSE | FALSE |
| 134_v.txt | 32004 | 32004-SNV-/ | SNV | 1 | A | T | 24 | 19468 | 0.12327923 | 52.875 | rnr1Y285F_pG | TRUE | CAN | JSY2420 | 2 | rnr1Y285F_p TAA | A>T | A/T+1 | A/T-1 | TA>AT | TA>AT | AATGGTAAC | AACTCGT | ATT | ATT | T | A | T>A | T>A_ATT | TA>AT_AACTCGT | 6 | FALSE | FALSE |
| 134_v.txt | 32128 | 32128-SNV-T | SNV | 1 | T | A | 143 | 19468 | 0.73453873 | 56.986014 | rnr1Y285F_pG | TRUE | CAN | JSY2420 | 2 | rnr1Y285F_p TTG | T>A | A/T+1 | A/T-1 | TA>AT | TA>AT | AAACTTTGT | TGTCACC | AAC | TTG | T | T | T>A | T>A_TTG | TA>AT_TGTCACC | 6 | FALSE | FALSE |
| 134_v.txt | 32310 | 32310-SNV-/ | SNV | 1 | A | T | 2 | 25529 | 7.83E-03 | 39.4997138 | rnr1Y285F_pG | TRUE | CAN | JSY2420 | 2 | rnr1Y285F_p AAT | A>T | A/T+1 | A/T-1 | TA>AT | TA>AT | GAAATAATG | ATGGTTG | TTA | TTA | T | A | T>A | T>A_TTA | TA>AT_ATGGTTG | 6 | FALSE | FALSE |
| 134_v.txt | 32337 | 32337-SNV-/ | SNV | 1 | A | T | 3 | 25529 | 0.01175134 | 27.6664313 | rnr1Y285F_pG | TRUE | CAN | JSY2420 | 2 | rnr1Y285F_p GAT | A>T | A/T+1 | A/T-1 | TA>AT | TA>AT | TTGAAGATA | ATATGTG | CTA | CTA | T | A | T>A | T>A_CTA | TA>AT_ATATGTG | 6 | FALSE | FALSE |
| 134_v.txt | 32338 | 32338-SNV-T | SNV | 1 | T | A | 3 | 25529 | 0.01175134 | 63.996659 | rnr1Y285F_pG | TRUE | CAN | JSY2420 | 2 | rnr1Y285F_p ATA | T>A | A/T+1 | A/T-1 | TA>AT | TA>AT | TGAAGATAT | TATGTGG | TAT | ATA | T | T | T>A | T>A_ATA | TA>AT_TATGTGG | 6 | FALSE | FALSE |
| 134_v.txt | 32375 | 32375-SNV-/ | SNV | 1 | A | T | 2 | 25529 | 7.83E-03 | 47.4993676 | rnr1Y285F_pG | TRUE | CAN | JSY2420 | 2 | rnr1Y285F_p CAA | A>T | A/T+1 | A/T-1 | TA>AT | TA>AT | AATAGCAAT | AATAATA | GTT | GTT | T | A | T>A | T>A_GTT | TA>AT_AATAATA | 6 | FALSE | FALSE |
| 134_v.txt | 32377 | 32377-SNV-T | SNV | 1 | T | A | 4 | 25526 | 0.0156703 | 37.4977734 | rnr1Y285F_pG | TRUE | CAN | JSY2420 | 2 | rnr1Y285F_p ATA | T>A | A/T+1 | A/T-1 | TA>AT | TA>AT | TAGCAATAA | TAATAAA | TAT | ATA | T | T | T>A | T>A_ATA | TA>AT_TAATAAA | 6 | FALSE | FALSE |
| 134_v.txt | 32442 | 32442-SNV-/ | SNV | 1 | A | T | 15 | 25529 | 0.05875671 | 51.2 | rnr1Y285F_pG | TRUE | CAN | JSY2420 | 2 | rnr1Y285F_p TAA | A>T | A/T+1 | A/T-1 | TA>AT | TA>AT | GGAACTAAA | AAAAGTC | ATT | ATT | T | A | T>A | T>A_ATT | TA>AT_AAAAGTC | 6 | FALSE | FALSE |
| 134_v.txt | 32457 | 32457-SNV-/ | SNV | 1 | A | T | 29 | 25529 | 0.1135963 | 53.6896552 | rnr1Y285F_pG | TRUE | CAN | JSY2420 | 2 | rnr1Y285F_p TAA | A>T | A/T+1 | A/T-1 | TA>AT | TA>AT | ATGAATAAT | AATAGAG | ATT | ATT | T | A | T>A | T>A_ATT | TA>AT_AATAGAG | 6 | FALSE | FALSE |
| 134_v.txt | 32498 | 32498-SNV-T | SNV | 1 | T | A | 4 | 25529 | 0.01566846 | 44.4986323 | rnr1Y285F_pG | TRUE | CAN | JSY2420 | 2 | rnr1Y285F_p CTT | T>A | A/T+1 | A/T-1 | TA>AT | TA>AT | AACAACTTT | TTTTTTG | GAA | CTT | T | T | T>A | T>A_CTT | TA>AT_TTTTTTG | 6 | FALSE | FALSE |
| 134_v.txt | 32766 | 32766-SNV-T | SNV | 1 | T | A | 17 | 23621 | 0.07196986 | 46.70452 | rnr1Y285F_pG | TRUE | CAN | JSY2420 | 2 | rnr1Y285F_p TTT | T>A | A/T+1 | A/T-1 | TA>AT | TA>AT | AAAACTTTG | TTGATGG | AAA | TTT | T | T | T>A | T>A_TTT | TA>AT_TTGATGG | 6 | FALSE | FALSE |
| 134_v.txt | 32767 | 32767-SNV-T | SNV | 1 | T | A | 5 | 23621 | 0.02116761 | 48.2 | rnr1Y285F_pG | TRUE | CAN | JSY2420 | 2 | rnr1Y285F_p TTG | T>A | A/T+1 | A/T-1 | TA>AT | TA>AT | AAACTTTGA | TGATGGA | AAC | TTG | T | T | T>A | T>A_TTG | TA>AT_TGATGGA | 6 | FALSE | FALSE |
| 134_v.txt | 32820 | 32820-SNV-T | SNV | 1 | T | A | 18 | 23621 | 0.07620338 | 41.1660747 | rnr1Y285F_pG | TRUE | CAN | JSY2420 | 2 | rnr1Y285F_p GTT | T>A | A/T+1 | A/T-1 | TA>AT | TA>AT | AACAAGTTC | TTCATTA | CAA | GTT | T | T | T>A | T>A_GTT | TA>AT_TTCATTA | 6 | FALSE | FALSE |
| 134_v.txt | 32927 | 32927-SNV-/ | SNV | 1 | A | T | 5 | 26566 | 0.01882105 | 42.5918355 | rnr1Y285F_pG | TRUE | CAN | JSY2420 | 2 | rnr1Y285F_p AAG | A>T | A/T+1 | A/T-1 | TA>AT | TA>AT | GGCAAAAGT | AGTGATT | TTC | TTC | T | A | T>A | T>A_TTC | TA>AT_AGTGATT | 6 | FALSE | FALSE |
| 134_v.txt | 32929 | 32929-SNV-T | SNV | 1 | T | A | 26 | 26566 | 0.09786946 | 44.2295647 | rnr1Y285F_pG | TRUE | CAN | JSY2420 | 2 | rnr1Y285F_p GTG | T>A | A/T+1 | A/T-1 | TA>AT | TA>AT | CAAAAGTGA | TGATTGC | CAC | GTG | T | T | T>A | T>A_GTG | TA>AT_TGATTGC | 6 | FALSE | FALSE |
| 134_v.txt | 33039 | 33039-SNV-/ | SNV | 1 | A | T | 33 | 26565 | 0.1242236 | 61.3924349 | rnr1Y285F_pG | TRUE | CAN | JSY2420 | 2 | rnr1Y285F_p CAT | A>T | A/T+1 | A/T-1 | TA>AT | TA>AT | GTAGCCATT | ATTTCAC | GTA | GTA | T | A | T>A | T>A_GTA | TA>AT_ATTTCAC | 6 | FALSE | FALSE |
| 134_v.txt | 33317 | 33317-SNV-/ | SNV | 1 | A | T | 26 | 19947 | 0.13034542 | 60.2307692 | rnr1Y285F_pG | TRUE | CAN | JSY2420 | 2 | rnr1Y285F_p GAT | A>T | A/T+1 | A/T-1 | TA>AT | TA>AT | CAATGGATA | ATACAAT | CTA | CTA | T | A | T>A | T>A_CTA | TA>AT_ATACAAT | 6 | FALSE | FALSE |
| 134_v.txt | 33321 | 33321-SNV-/ | SNV | 1 | A | T | 11 | 19947 | 0.05514614 | 61.9090909 | rnr1Y285F_pG | TRUE | CAN | JSY2420 | 2 | rnr1Y285F_p CAA | A>T | A/T+1 | A/T-1 | TA>AT | TA>AT | GGATACAAT | AATTCTT | GTT | GTT | T | A | T>A | T>A_GTT | TA>AT_AATTCTT | 6 | FALSE | FALSE |
| 134_v.txt | 31784 | 31784-SNV-( | SNV | 1 | G | A | 12 | 8528 | 0.14071295 | 54.9166667 | rnr1Y285F_pG | TRUE | CAN | JSY2420 | 2 | rnr1Y285F_p CGA | G>A | A/T+1 | G/C-1 | CG>TA | CG>TA | GGAATCGAT | GATGTCG | GCT | GCT | C | T | C>T | C>T_GCT | CG>TA_GATGTCG | 6 | FALSE | FALSE |
| 134_v.txt | 31829 | 31829-SNV-( | SNV | 1 | G | A | 4 | 8528 | 0.04690432 | 57.25 | rnr1Y285F_pG | TRUE | CAN | JSY2420 | 2 | rnr1Y285F_p TGC | G>A | A/T+1 | G/C-1 | CG>TA | CG>TA | GAATATGCA | GCATTGA | ACG | ACG | C | T | C>T | C>T_ACG | CG>TA_GCATTGA | 6 | FALSE | FALSE |
| 134_v.txt | 31830 | 31830-SNV-( | SNV | 1 | C | T | 11 | 8528 | 0.12898687 | 56.9063984 | rnr1Y285F_pG | TRUE | CAN | JSY2420 | 2 | rnr1Y285F_p GCA | C>T | A/T+1 | G/C-1 | CG>TA | CG>TA | AATATGCAT | CATTGAA | CGT | GCA | C | A | C>T | C>T_GCA | CG>TA_CATTGAA | 6 | FALSE | FALSE |
| 134_v.txt | 31879 | 31879-SNV-( | SNV | 1 | C | T | 13 | 8528 | 0.15243902 | 41.0769231 | rnr1Y285F_pG | TRUE | CAN | JSY2420 | 2 | rnr1Y285F_p GCG | C>T | A/T+1 | G/C-1 | CG>TA | CG>TA | TATAGGCGG | CGGCAGC | CGC | GCG | C | A | C>T | C>T_GCG | CG>TA_CGGCAGC | 6 | FALSE | FALSE |
| 134_v.txt | 31882 | 31882-SNV-( | SNV | 1 | C | T | 16 | 8528 | 0.18761726 | 45.75 | rnr1Y285F_pG | TRUE | CAN | JSY2420 | 2 | rnr1Y285F_p GCA | C>T | A/T+1 | G/C-1 | CG>TA | CG>TA | AGGCGGCAG | CAGCAAA | CGT | GCA | C | A | C>T | C>T_GCA | CG>TA_CAGCAAA | 6 | FALSE | FALSE |
| 134_v.txt | 31962 | 31962-SNV-( | SNV | 1 | G | A | 42 | 19468 | 0.21573865 | 37.9285714 | rnr1Y285F_pG | TRUE | CAN | JSY2420 | 2 | rnr1Y285F_p CGC | G>A | A/T+1 | G/C-1 | CG>TA | CG>TA | GTGGCCGCA | GCATAAT | GCG | GCG | C | T | C>T | C>T_GCG | CG>TA_GCATAAT | 6 | FALSE | FALSE |
| 134_v.txt | 31972 | 31972-SNV-( | SNV | 1 | C | T | 15 | 19468 | 0.07704952 | 37.5333333 | rnr1Y285F_pG | TRUE | CAN | JSY2420 | 2 | rnr1Y285F_p GCC | C>T | A/T+1 | G/C-1 | CG>TA | CG>TA | AATAAGCCA | CCAAGCC | CGG | GCC | C | A | C>T | C>T_GCC | CG>TA_CCAAGCC | 6 | FALSE | FALSE |
| 134_v.txt | 31981 | 31981-SNV-( | SNV | 1 | G | A | 45 | 19468 | 0.23114855 | 52.0444444 | rnr1Y285F_pG | TRUE | CAN | JSY2420 | 2 | rnr1Y285F_p GGC | G>A | A/T+1 | G/C-1 | CG>TA | CG>TA | AGCCGGGCA | GCATTAA | CCG | CCG | C | T | C>T | C>T_CCG | CG>TA_GCATTAA | 6 | FALSE | FALSE |
| 134_v.txt | 32041 | 32041-SNV-( | SNV | 1 | G | A | 27 | 19468 | 0.13868913 | 62.9259259 | rnr1Y285F_pG | TRUE | CAN | JSY2420 | 2 | rnr1Y285F_p TGC | G>A | A/T+1 | G/C-1 | CG>TA | CG>TA | AAGCTTGCA | GCATAAA | ACG | ACG | C | T | C>T | C>T_ACG | CG>TA_GCATAAA | 6 | FALSE | FALSE |
| 134_v.txt | 32057 | 32057-SNV-( | SNV | 1 | C | T | 23 | 19468 | 0.11814259 | 62.4347826 | rnr1Y285F_pG | TRUE | CAN | JSY2420 | 2 | rnr1Y285F_p GCG | C>T | A/T+1 | G/C-1 | CG>TA | CG>TA | GATGTGCGA | CGAGATT | CGC | GCG | C | A | C>T | C>T_GCG | CG>TA_CGAGATT | 6 | FALSE | FALSE |
| 134_v.txt | 32075 | 32075-SNV-( | SNV | 1 | C | T | 94 | 19467 | 0.48286844 | 62.3723492 | rnr1Y285F_pG | TRUE | CAN | JSY2420 | 2 | rnr1Y285F_p ACC | C>T | A/T+1 | G/C-1 | CG>TA | CG>TA | AAATAACCA | CCATGCA | TGG | ACC | C | A | C>T | C>T_ACC | CG>TA_CCATGCA | 6 | FALSE | FALSE |
| 134_v.txt | 32076 | 32076-SNV-( | SNV | 1 | C | T | 20 | 19465 | 0.10274852 | 56.0988687 | rnr1Y285F_pG | TRUE | CAN | JSY2420 | 2 | rnr1Y285F_p CCA | C>T | A/T+1 | G/C-1 | CG>TA | CG>TA | AATAACCAT | CATGCAA | GGT | CCA | C | A | C>T | C>T_CCA | CG>TA_CATGCAA | 6 | FALSE | FALSE |
| 134_v.txt | 32088 | 32088-SNV-( | SNV | 1 | C | T | 102 | 19468 | 0.52393672 | 60.5 | rnr1Y285F_pG | TRUE | CAN | JSY2420 | 2 | rnr1Y285F_p GCC | C>T | A/T+1 | G/C-1 | CG>TA | CG>TA | AAAAAGCCT | CCTGCAA | CGG | GCC | C | A | C>T | C>T_GCC | CG>TA_CCTGCAA | 6 | FALSE | FALSE |
| 134_v.txt | 32092 | 32092-SNV-( | SNV | 1 | C | T | 25 | 19468 | 0.12841586 | 62.08 | rnr1Y285F_pG | TRUE | CAN | JSY2420 | 2 | rnr1Y285F_p GCA | C>T | A/T+1 | G/C-1 | CG>TA | CG>TA | AGCCTGCAA | CAACACC | CGT | GCA | C | A | C>T | C>T_GCA | CG>TA_CAACACC | 6 | FALSE | FALSE |
| 134_v.txt | 32114 | 32114-SNV-( | SNV | 1 | C | T | 330 | 19467 | 1.69517645 | 58.9999994 | rnr1Y285F_pG | TRUE | CAN | JSY2420 | 2 | rnr1Y285F_p GCC | C>T | A/T+1 | G/C-1 | CG>TA | CG>TA | TAATAGCCA | CCATTCG | CGG | GCC | C | A | C>T | C>T_GCC | CG>TA_CCATTCG | 6 | FALSE | FALSE |
| 134_v.txt | 32115 | 32115-SNV-( | SNV | 1 | C | T | 31 | 19467 | 0.15924385 | 59.0275029 | rnr1Y285F_pG | TRUE | CAN | JSY2420 | 2 | rnr1Y285F_p CCA | C>T | A/T+1 | G/C-1 | CG>TA | CG>TA | AATAGCCAT | CATTCGA | GGT | CCA | C | A | C>T | C>T_CCA | CG>TA_CATTCGA | 6 | FALSE | FALSE |
| 134_v.txt | 32165 | 32165-SNV-( | SNV | 1 | G | A | 24 | 19468 | 0.12327923 | 53.7493038 | rnr1Y285F_pG | TRUE | CAN | JSY2420 | 2 | rnr1Y285F_p CGC | G>A | A/T+1 | G/C-1 | CG>TA | CG>TA | CAAAGCGCC | GCCAAAT | GCG | GCG | C | T | C>T | C>T_GCG | CG>TA_GCCAAAT | 6 | FALSE | FALSE |
| 134_v.txt | 32166 | 32166-SNV-( | SNV | 1 | C | T | 98 | 19468 | 0.50339018 | 47.2812393 | rnr1Y285F_pG | TRUE | CAN | JSY2420 | 2 | rnr1Y285F_p GCC | C>T | A/T+1 | G/C-1 | CG>TA | CG>TA | AAAGCGCCA | CCAAATG | CGG | GCC | C | A | C>T | C>T_GCC | CG>TA_CCAAATG | 6 | FALSE | FALSE |
| 134_v.txt | 32173 | 32173-SNV-( | SNV | 1 | C | T | 22 | 19468 | 0.11300596 | 48.5454546 | rnr1Y285F_pG | TRUE | CAN | JSY2420 | 2 | rnr1Y285F_p GCA | C>T | A/T+1 | G/C-1 | CG>TA | CG>TA | CAAATGCAG | CAGCAGT | CGT | GCA | C | A | C>T | C>T_GCA | CG>TA_CAGCAGT | 6 | FALSE | FALSE |
| 134_v.txt | 32271 | 32271-SNV-( | SNV | 1 | C | T | 80 | 25529 | 0.31336911 | 35.7875 | rnr1Y285F_pG | TRUE | CAN | JSY2420 | 2 | rnr1Y285F_p ACG | C>T | A/T+1 | G/C-1 | CG>TA | CG>TA | AAAATACGG | CGGGAAC | TGC | ACG | C | A | C>T | C>T_ACG | CG>TA_CGGGAAC | 6 | FALSE | FALSE |
| 134_v.txt | 32274 | 32274-SNV-( | SNV | 1 | G | A | 207 | 25529 | 0.81084257 | 36.7971015 | rnr1Y285F_pG | TRUE | CAN | JSY2420 | 2 | rnr1Y285F_p GGA | G>A | A/T+1 | G/C-1 | CG>TA | CG>TA | ATACGGGAA | GAACCAA | CCT | CCT | C | T | C>T | C>T_CCT | CG>TA_GAACCAA | 6 | FALSE | FALSE |
| 134_v.txt | 32299 | 32299-SNV-( | SNV | 1 | C | T | 49 | 25529 | 0.19193858 | 51.6326531 | rnr1Y285F_pG | TRUE | CAN | JSY2420 | 2 | rnr1Y285F_p GCG | C>T | A/T+1 | G/C-1 | CG>TA | CG>TA | AATTTGCGG | CGGCAGA | CGC | GCG | C | A | C>T | C>T_GCG | CG>TA_CGGCAGA | 6 | FALSE | FALSE |
| 134_v.txt | 32304 | 32304-SNV-( | SNV | 1 | G | A | 37 | 25529 | 0.14493321 | 59.1081081 | rnr1Y285F_pG | TRUE | CAN | JSY2420 | 2 | rnr1Y285F_p AGA | G>A | A/T+1 | G/C-1 | CG>TA | CG>TA | GCGGCAGAA | GAAATAA | TCT | TCT | C | T | C>T | C>T_TCT | CG>TA_GAAATAA | 6 | FALSE | FALSE |
| 134_v.txt | 32413 | 32413-SNV-( | SNV | 1 | G | A | 32 | 25529 | 0.12534764 | 54.125 | rnr1Y285F_pG | TRUE | CAN | JSY2420 | 2 | rnr1Y285F_p TGT | G>A | A/T+1 | G/C-1 | CG>TA | CG>TA | TAGATTGTG | GTGTTAG | ACA | ACA | C | T | C>T | C>T_ACA | CG>TA_GTGTTAG | 6 | FALSE | FALSE |
| 134_v.txt | 32449 | 32449-SNV-( | SNV | 1 | C | T | 35 | 25529 | 0.13709899 | 53.8571429 | rnr1Y285F_pG | TRUE | CAN | JSY2420 | 2 | rnr1Y285F_p CCA | C>T | A/T+1 | G/C-1 | CG>TA | CG>TA | AAAGTCCAA | CAATGAA | GGT | CCA | C | A | C>T | C>T_CCA | CG>TA_CAATGAA | 6 | FALSE | FALSE |
| 134_v.txt | 32466 | 32466-SNV-( | SNV | 1 | C | T | 35 | 25529 | 0.13709899 | 48.9294654 | rnr1Y285F_pG | TRUE | CAN | JSY2420 | 2 | rnr1Y285F_p GCC | C>T | A/T+1 | G/C-1 | CG>TA | CG>TA | AGAGAGCCA | CCAATGT | CGG | GCC | C | A | C>T | C>T_GCC | CG>TA_CCAATGT | 6 | FALSE | FALSE |
| 134_v.txt | 32488 | 32488-SNV-( | SNV | 1 | G | A | 32 | 25529 | 0.12534764 | 47.03125 | rnr1Y285F_pG | TRUE | CAN | JSY2420 | 2 | rnr1Y285F_p CGG | G>A | A/T+1 | G/C-1 | CG>TA | CG>TA | AGATACGGA | GGAAAAC | GCC | GCC | C | T | C>T | C>T_GCC | CG>TA_GGAAAAC | 6 | FALSE | FALSE |
| 134_v.txt | 32508 | 32508-SNV-( | SNV | 1 | G | A | 28 | 25529 | 0.10967919 | 39.2857143 | rnr1Y285F_pG | TRUE | CAN | JSY2420 | 2 | rnr1Y285F_p GGC | G>A | A/T+1 | G/C-1 | CG>TA | CG>TA | TTGATGGCT | GCTCTTG | CCG | CCG | C | T | C>T | C>T_CCG | CG>TA_GCTCTTG | 6 | FALSE | FALSE |
| 134_v.txt | 32535 | 32535-SNV-( | SNV | 1 | G | A | 27 | 25529 | 0.10576208 | 37.1850728 | rnr1Y285F_pG | TRUE | CAN | JSY2420 | 2 | rnr1Y285F_p TGC | G>A | A/T+1 | G/C-1 | CG>TA | CG>TA | GGGTTTGCA | GCAGCTT | ACG | ACG | C | T | C>T | C>T_ACG | CG>TA_GCAGCTT | 6 | FALSE | FALSE |
| 134_v.txt | 32536 | 32536-SNV-( | SNV | 1 | C | T | 43 | 25529 | 0.1684359 | 37.2085239 | rnr1Y285F_pG | TRUE | CAN | JSY2420 | 2 | rnr1Y285F_p GCA | C>T | A/T+1 | G/C-1 | CG>TA | CG>TA | GGTTTGCAG | CAGCTTC | CGT | GCA | C | A | C>T | C>T_GCA | CG>TA_CAGCTTC | 6 | FALSE | FALSE |
| 134_v.txt | 32542 | 32542-SNV-( | SNV | 1 | C | T | 105 | 25529 | 0.41129696 | 37.1809524 | rnr1Y285F_pG | TRUE | CAN | JSY2420 | 2 | rnr1Y285F_p TCA | C>T | A/T+1 | G/C-1 | CG>TA | CG>TA | CAGCTTCAC | CACCAGC | AGT | TCA | C | A | C>T | C>T_TCA | CG>TA_CACCAGC | 6 | FALSE | FALSE |
| 134_v.txt | 32545 | 32545-SNV-( | SNV | 1 | C | T | 37 | 25529 | 0.14493321 | 36.6756757 | rnr1Y285F_pG | TRUE | CAN | JSY2420 | 2 | rnr1Y285F_p CCA | C>T | A/T+1 | G/C-1 | CG>TA | CG>TA | CTTCACCAG | CAGCAGT | GGT | CCA | C | A | C>T | C>T_CCA | CG>TA_CAGCAGT | 6 | FALSE | FALSE |
| 134_v.txt | 32568 | 32568-SNV-( | SNV | 1 | G | A | 21 | 23619 | 0.08891147 | 37.5238095 | rnr1Y285F_pG | TRUE | CAN | JSY2420 | 2 | rnr1Y285F_p AGT | G>A | A/T+1 | G/C-1 | CG>TA | CG>TA | AGTTCAGTA | GTACCTT | TCA | TCA | C | T | C>T | C>T_TCA | CG>TA_GTACCTT | 6 | FALSE | FALSE |
| 134_v.txt | 32575 | 32575-SNV-( | SNV | 1 | G | A | 69 | 23621 | 0.29211295 | 36.4782609 | rnr1Y285F_pG | TRUE | CAN | JSY2420 | 2 | rnr1Y285F_p TGA | G>A | A/T+1 | G/C-1 | CG>TA | CG>TA | TACCTTGAA | GAAATGT | ACT | ACT | C | T | C>T | C>T_ACT | CG>TA_GAAATGT | 6 | FALSE | FALSE |
| 134_v.txt | 32590 | 32590-SNV-( | SNV | 1 | C | T | 44 | 23621 | 0.18627493 | 36.5454546 | rnr1Y285F_pG | TRUE | CAN | JSY2420 | 2 | rnr1Y285F_p GCG | C>T | A/T+1 | G/C-1 | CG>TA | CG>TA | AGGCAGCGT | CGTTAAT | CGC | GCG | C | A | C>T | C>T_GCG | CG>TA_CGTTAAT | 6 | FALSE | FALSE |
| 134_v.txt | 32629 | 32629-SNV-( | SNV | 1 | C | T | 7 | 23621 | 0.02963465 | 56.0002449 | rnr1Y285F_pG | TRUE | CAN | JSY2420 | 2 | rnr1Y285F_p TCG | C>T | A/T+1 | G/C-1 | CG>TA | CG>TA | TCCCTTCGTT | CGTTTTT | AGC | TCG | C | A | C>T | C>T_TCG | CG>TA_CGTTTTT | 6 | FALSE | FALSE |
| 134_v.txt | 32630 | 32630-SNV-( | SNV | 1 | G | A | 4 | 23621 | 0.01693408 | 36.9998573 | rnr1Y285F_pG | TRUE | CAN | JSY2420 | 2 | rnr1Y285F_p CGT | G>A | A/T+1 | G/C-1 | CG>TA | CG>TA | CCCTTCGTTT | GTTTTTA | GCA | GCA | C | T | C>T | C>T_GCA | CG>TA_GTTTTTA | 6 | FALSE | FALSE |
| 134_v.txt | 32658 | 32658-SNV-( | SNV | 1 | C | T | 17 | 23621 | 0.07196986 | 56.6470588 | rnr1Y285F_pG | TRUE | CAN | JSY2420 | 2 | rnr1Y285F_p ACC | C>T | A/T+1 | G/C-1 | CG>TA | CG>TA | CCTGGACCC | CCCCAGG | TGG | ACC | C | A | C>T | C>T_ACC | CG>TA_CCCCAGG | 6 | FALSE | FALSE |
| 134_v.txt | 32661 | 32661-SNV-( | SNV | 1 | C | T | 96 | 23621 | 0.40641802 | 59.5520833 | rnr1Y285F_pG | TRUE | CAN | JSY2420 | 2 | rnr1Y285F_p CCA | C>T | A/T+1 | G/C-1 | CG>TA | CG>TA | GGACCCCAG | CAGGCAC | GGT | CCA | C | A | C>T | C>T_CCA | CG>TA_CAGGCAC | 6 | FALSE | FALSE |
| 134_v.txt | 32700 | 32700-SNV-( | SNV | 1 | C | T | 23 | 23621 | 0.09737098 | 61.6956522 | rnr1Y285F_pG | TRUE | CAN | JSY2420 | 2 | rnr1Y285F_p GCC | C>T | A/T+1 | G/C-1 | CG>TA | CG>TA | ACTGGGCCG | CCGGTAA | CGG | GCC | C | A | C>T | C>T_GCC | CG>TA_CCGGTAA | 6 | FALSE | FALSE |
| 134_v.txt | 32796 | 32796-SNV-( | SNV | 1 | C | T | 54 | 23621 | 0.22861014 | 44.8146286 | rnr1Y285F_pG | TRUE | CAN | JSY2420 | 2 | rnr1Y285F_p ACC | C>T | A/T+1 | G/C-1 | CG>TA | CG>TA | AATTCACCG | CCGTAAT | TGG | ACC | C | A | C>T | C>T_ACC | CG>TA_CCGTAAT | 6 | FALSE | FALSE |
| 134_v.txt | 32797 | 32797-SNV-( | SNV | 1 | C | T | 28 | 23621 | 0.11853859 | 47.8222178 | rnr1Y285F_pG | TRUE | CAN | JSY2420 | 2 | rnr1Y285F_p CCG | C>T | A/T+1 | G/C-1 | CG>TA | CG>TA | ATTCACCGT | CGTAATA | GGC | CCG | C | A | C>T | C>T_CCG | CG>TA_CGTAATA | 6 | FALSE | FALSE |
| 134_v.txt | 32798 | 32798-SNV-( | SNV | 1 | G | A | 45 | 23621 | 0.19050845 | 41.2437697 | rnr1Y285F_pG | TRUE | CAN | JSY2420 | 2 | rnr1Y285F_p CGT | G>A | A/T+1 | G/C-1 | CG>TA | CG>TA | TTCACCGTA | GTAATAT | GCA | GCA | C | T | C>T | C>T_GCA | CG>TA_GTAATAT | 6 | FALSE | FALSE |
| 134_v.txt | 32840 | 32840-SNV-( | SNV | 1 | C | T | 28 | 23621 | 0.11853859 | 39.4285714 | rnr1Y285F_pG | TRUE | CAN | JSY2420 | 2 | rnr1Y285F_p CCC | C>T | A/T+1 | G/C-1 | CG>TA | CG>TA | AATTACCCA | CCAAAAA | GGG | CCC | C | A | C>T | C>T_CCC | CG>TA_CCAAAAA | 6 | FALSE | FALSE |
| 134_v.txt | 32855 | 32855-SNV-( | SNV | 1 | C | T | 211 | 23621 | 0.89327294 | 36.6872038 | rnr1Y285F_pG | TRUE | CAN | JSY2420 | 2 | rnr1Y285F_p TCC | C>T | A/T+1 | G/C-1 | CG>TA | CG>TA | ACTAATCCAT | CCATGCC | AGG | TCC | C | A | C>T | C>T_TCC | CG>TA_CCATGCC | 6 | FALSE | FALSE |
| 134_v.txt | 32860 | 32860-SNV-( | SNV | 1 | C | T | 22 | 23619 | 0.09314535 | 37.4090909 | rnr1Y285F_pG | TRUE | CAN | JSY2420 | 2 | rnr1Y285F_p GCC | C>T | A/T+1 | G/C-1 | CG>TA | CG>TA | TCCATGCCG | CCGCCAG | CGG | GCC | C | A | C>T | C>T_GCC | CG>TA_CCGCCAG | 6 | FALSE | FALSE |
| 134_v.txt | 32862 | 32862-SNV-( | SNV | 1 | G | A | 25 | 23618 | 0.10585147 | 34 | rnr1Y285F_pG | TRUE | CAN | JSY2420 | 2 | rnr1Y285F_p CGC | G>A | A/T+1 | G/C-1 | CG>TA | CG>TA | CATGCCGCC | GCCAGTG | GCG | GCG | C | T | C>T | C>T_GCG | CG>TA_GCCAGTG | 6 | FALSE | FALSE |
| 134_v.txt | 32883 | 32883-SNV-( | SNV | 1 | C | T | 83 | 26567 | 0.31241766 | 36.1204819 | rnr1Y285F_pG | TRUE | CAN | JSY2420 | 2 | rnr1Y285F_p CCA | C>T | A/T+1 | G/C-1 | CG>TA | CG>TA | TACGTCCAA | CAAAATT | GGT | CCA | C | A | C>T | C>T_CCA | CG>TA_CAAAATT | 6 | FALSE | FALSE |
| 134_v.txt | 32901 | 32901-SNV-( | SNV | 1 | C | T | 66 | 26566 | 0.24843785 | 30.5 | rnr1Y285F_pG | TRUE | CAN | JSY2420 | 2 | rnr1Y285F_p GCC | C>T | A/T+1 | G/C-1 | CG>TA | CG>TA | ACTTGGCCA | CCAACTA | CGG | GCC | C | A | C>T | C>T_GCC | CG>TA_CCAACTA | 6 | FALSE | FALSE |
| 134_v.txt | 32917 | 32917-SNV-( | SNV | 1 | C | T | 484 | 26566 | 1.82187759 | 48.7066116 | rnr1Y285F_pG | TRUE | CAN | JSY2420 | 2 | rnr1Y285F_p TCC | C>T | A/T+1 | G/C-1 | CG>TA | CG>TA | TAAGTTCCA | CCAGGGC | AGG | TCC | C | A | C>T | C>T_TCC | CG>TA_CCAGGGC | 6 | FALSE | FALSE |
| 134_v.txt | 32922 | 32922-SNV-( | SNV | 1 | G | A | 20 | 26566 | 0.0752842 | 50.5995732 | rnr1Y285F_pG | TRUE | CAN | JSY2420 | 2 | rnr1Y285F_p GGC | G>A | A/T+1 | G/C-1 | CG>TA | CG>TA | TCCAGGGCA | GCAAAAG | CCG | CCG | C | T | C>T | C>T_CCG | CG>TA_GCAAAAG | 6 | FALSE | FALSE |
| 134_v.txt | 32965 | 32965-SNV-( | SNV | 1 | C | T | 31 | 26566 | 0.11669051 | 58.1935484 | rnr1Y285F_pG | TRUE | CAN | JSY2420 | 2 | rnr1Y285F_p GCC | C>T | A/T+1 | G/C-1 | CG>TA | CG>TA | CATTGGCCG | CCGCACC | CGG | GCC | C | A | C>T | C>T_GCC | CG>TA_CCGCACC | 6 | FALSE | FALSE |
| 134_v.txt | 32967 | 32967-SNV-( | SNV | 1 | G | A | 49 | 26566 | 0.18444629 | 59.3469388 | rnr1Y285F_pG | TRUE | CAN | JSY2420 | 2 | rnr1Y285F_p CGC | G>A | A/T+1 | G/C-1 | CG>TA | CG>TA | TTGGCCGCA | GCACCAA | GCG | GCG | C | T | C>T | C>T_GCG | CG>TA_GCACCAA | 6 | FALSE | FALSE |
| 134_v.txt | 32995 | 32995-SNV-( | SNV | 1 | G | A | 22 | 26566 | 0.08281262 | 59.3181818 | rnr1Y285F_pG | TRUE | CAN | JSY2420 | 2 | rnr1Y285F_p TGT | G>A | A/T+1 | G/C-1 | CG>TA | CG>TA | ATCTTTGTGA | GTGAGAA | ACA | ACA | C | T | C>T | C>T_ACA | CG>TA_GTGAGAA | 6 | FALSE | FALSE |
| 134_v.txt | 33015 | 33015-SNV-( | SNV | 1 | G | A | 28 | 26566 | 0.10539788 | 54.75 | rnr1Y285F_pG | TRUE | CAN | JSY2420 | 2 | rnr1Y285F_p GGA | G>A | A/T+1 | G/C-1 | CG>TA | CG>TA | AAAGAGGAT | GATGTAA | CCT | CCT | C | T | C>T | C>T_CCT | CG>TA_GATGTAA | 6 | FALSE | FALSE |
| 134_v.txt | 33047 | 33047-SNV-( | SNV | 1 | C | T | 6 | 26565 | 0.02258611 | 59.325457 | rnr1Y285F_pG | TRUE | CAN | JSY2420 | 2 | rnr1Y285F_p CCA | C>T | A/T+1 | G/C-1 | CG>TA | CG>TA | TTCACCCAA | CAAGGAC | GGT | CCA | C | A | C>T | C>T_CCA | CG>TA_CAAGGAC | 6 | FALSE | FALSE |
| 134_v.txt | 33056 | 33056-SNV-( | SNV | 1 | C | T | 38 | 26566 | 0.14303998 | 52.0263158 | rnr1Y285F_pG | TRUE | CAN | JSY2420 | 2 | rnr1Y285F_p GCG | C>T | A/T+1 | G/C-1 | CG>TA | CG>TA | GGACTGCGT | CGTGACA | CGC | GCG | C | A | C>T | C>T_GCG | CG>TA_CGTGACA | 6 | FALSE | FALSE |
| 134_v.txt | 33069 | 33069-SNV-( | SNV | 1 | G | A | 23 | 26566 | 0.08657683 | 55.2616314 | rnr1Y285F_pG | TRUE | CAN | JSY2420 | 2 | rnr1Y285F_p TGC | G>A | A/T+1 | G/C-1 | CG>TA | CG>TA | GAATATGCC | GCCAAAG | ACG | ACG | C | T | C>T | C>T_ACG | CG>TA_GCCAAAG | 6 | FALSE | FALSE |
| 134_v.txt | 33070 | 33070-SNV-( | SNV | 1 | C | T | 35 | 26565 | 0.13175231 | 52.7714286 | rnr1Y285F_pG | TRUE | CAN | JSY2420 | 2 | rnr1Y285F_p GCC | C>T | A/T+1 | G/C-1 | CG>TA | CG>TA | AATATGCCA | CCAAAGA | CGG | GCC | C | A | C>T | C>T_GCC | CG>TA_CCAAAGA | 6 | FALSE | FALSE |
| 134_v.txt | 33113 | 33113-SNV-( | SNV | 1 | G | A | 25 | 26566 | 0.09410525 | 38.2799999 | rnr1Y285F_pG | TRUE | CAN | JSY2420 | 2 | rnr1Y285F_p GGC | G>A | A/T+1 | G/C-1 | CG>TA | CG>TA | CACTGGGCC | GCCGGCG | CCG | CCG | C | T | C>T | C>T_CCG | CG>TA_GCCGGCG | 6 | FALSE | FALSE |
| 134_v.txt | 33114 | 33114-SNV-( | SNV | 1 | C | T | 26 | 26566 | 0.09786946 | 41.8839985 | rnr1Y285F_pG | TRUE | CAN | JSY2420 | 2 | rnr1Y285F_p GCC | C>T | A/T+1 | G/C-1 | CG>TA | CG>TA | ACTGGGCCG | CCGGCGT | CGG | GCC | C | A | C>T | C>T_GCC | CG>TA_CCGGCGT | 6 | FALSE | FALSE |
| 134_v.txt | 33115 | 33115-SNV-( | SNV | 1 | C | T | 11 | 26566 | 0.04140631 | 34.5452186 | rnr1Y285F_pG | TRUE | CAN | JSY2420 | 2 | rnr1Y285F_p CCG | C>T | A/T+1 | G/C-1 | CG>TA | CG>TA | CTGGGCCGG | CGGCGTT | GGC | CCG | C | A | C>T | C>T_CCG | CG>TA_CGGCGTT | 6 | FALSE | FALSE |
| 134_v.txt | 33118 | 33118-SNV-( | SNV | 1 | C | T | 23 | 26566 | 0.08657683 | 40.5652174 | rnr1Y285F_pG | TRUE | CAN | JSY2420 | 2 | rnr1Y285F_p GCG | C>T | A/T+1 | G/C-1 | CG>TA | CG>TA | GGCCGGCGT | CGTTGGT | CGC | GCG | C | A | C>T | C>T_GCG | CG>TA_CGTTGGT | 6 | FALSE | FALSE |
| 134_v.txt | 33153 | 33153-SNV-( | SNV | 1 | C | T | 95 | 26566 | 0.35759994 | 36.6210526 | rnr1Y285F_pG | TRUE | CAN | JSY2420 | 2 | rnr1Y285F_p ACC | C>T | A/T+1 | G/C-1 | CG>TA | CG>TA | AAAAGACCT | CCTGTAC | TGG | ACC | C | A | C>T | C>T_ACC | CG>TA_CCTGTAC | 6 | FALSE | FALSE |

| | | | | | | | | | | | | | | | | | | | | | | | | | | | | | | | | | | |
|---|---|---|---|---|---|---|---|---|---|---|---|---|---|---|---|---|---|---|---|---|---|---|---|---|---|---|---|---|---|---|---|---|---|---|
| 134_v.txt | 33154 | 33154-SNV- | SNV | 1 | C | T | 57 | 26566 | 0.21455996 | 36.631579 | rnr1Y285F_pG | TRUE | CAN | JSY2420 | 2 | rnr1Y285F_p | CCT | C>T | A/T+1 | G/C-1 | CG>TA | CG>TA | AAAGACCTG | CTGTACC | GGA | CCT | C | A | C>T | C>T_CCT | CG>TA_CTGTACC | 6 | FALSE | FALSE |
| 134_v.txt | 33159 | 33159-SNV- | SNV | 1 | C | T | 44 | 26565 | 0.16563147 | 36.5681401 | rnr1Y285F_pG | TRUE | CAN | JSY2420 | 2 | rnr1Y285F_p | ACC | C>T | A/T+1 | G/C-1 | CG>TA | CG>TA | CCTGTACCA | CCAATAG | TGG | ACC | C | A | C>T | C>T_ACC | CG>TA_CCAATAG | 6 | FALSE | FALSE |
| 134_v.txt | 33168 | 33168-SNV- | SNV | 1 | C | T | 176 | 26561 | 0.66262565 | 37.4715909 | rnr1Y285F_pG | TRUE | CAN | JSY2420 | 2 | rnr1Y285F_p | ACC | C>T | A/T+1 | G/C-1 | CG>TA | CG>TA | ATAGTACCA | CCACCAA | TGG | ACC | C | A | C>T | C>T_ACC | CG>TA_CCACCAA | 6 | FALSE | FALSE |
| 134_v.txt | 33258 | 33258-SNV- | SNV | 1 | C | T | 23 | 19947 | 0.11530556 | 45.0869565 | rnr1Y285F_pG | TRUE | CAN | JSY2420 | 2 | rnr1Y285F_p | GCC | C>T | A/T+1 | G/C-1 | CG>TA | CG>TA | CCTATGCCAT | CCATCCT | CGG | GCC | C | A | C>T | C>T_GCC | CG>TA_CCATCCT | 6 | FALSE | FALSE |
| 134_v.txt | 33275 | 33275-SNV- | SNV | 1 | C | T | 24 | 19947 | 0.12031885 | 44.9547851 | rnr1Y285F_pG | TRUE | CAN | JSY2420 | 2 | rnr1Y285F_p | ACG | C>T | A/T+1 | G/C-1 | CG>TA | CG>TA | AGAGAACGT | CGTATCC | TGC | ACG | C | A | C>T | C>T_ACG | CG>TA_CGTATCC | 6 | FALSE | FALSE |
| 134_v.txt | 33276 | 33276-SNV- | SNV | 1 | G | A | 32 | 19946 | 0.16043317 | 44.9618912 | rnr1Y285F_pG | TRUE | CAN | JSY2420 | 2 | rnr1Y285F_p | CGT | G>A | A/T+1 | G/C-1 | CG>TA | CG>TA | GAGAACGTA | GTATCCT | GCA | GCA | C | T | C>T | C>T_GCA | CG>TA_GTATCCT | 6 | FALSE | FALSE |
| 134_v.txt | 33284 | 33284-SNV- | SNV | 1 | G | A | 24 | 19947 | 0.12031885 | 46.2916667 | rnr1Y285F_pG | TRUE | CAN | JSY2420 | 2 | rnr1Y285F_p | CGC | G>A | A/T+1 | G/C-1 | CG>TA | CG>TA | ATCCTCGCC | GCCATTT | GCG | GCG | C | T | C>T | C>T_GCG | CG>TA_GCCATTT | 6 | FALSE | FALSE |
| 134_v.txt | 33309 | 33309-SNV- | SNV | 1 | C | T | 29 | 19946 | 0.14539256 | 53.8275862 | rnr1Y285F_pG | TRUE | CAN | JSY2420 | 2 | rnr1Y285F_p | GCG | C>T | A/T+1 | G/C-1 | CG>TA | CG>TA | AAAGAGCGC | CGCAATG | CGC | GCG | C | A | C>T | C>T_GCG | CG>TA_CGCAATG | 6 | FALSE | FALSE |
| 134_v.txt | 33310 | 33310-SNV- | SNV | 1 | G | A | 28 | 19940 | 0.14042126 | 61.0149404 | rnr1Y285F_pG | TRUE | CAN | JSY2420 | 2 | rnr1Y285F_p | CGC | G>A | A/T+1 | G/C-1 | CG>TA | CG>TA | AAGAGCGCA | GCAATGG | GCG | GCG | C | T | C>T | C>T_GCG | CG>TA_GCAATGG | 6 | FALSE | FALSE |
| 134_v.txt | 33412 | 33412-SNV- | SNV | 1 | G | A | 26 | 19948 | 0.13033888 | 51.3846154 | rnr1Y285F_pG | TRUE | CAN | JSY2420 | 2 | rnr1Y285F_p | GGC | G>A | A/T+1 | G/C-1 | CG>TA | CG>TA | TGACCGGCT | GCTCATT | CCG | CCG | C | T | C>T | C>T_CCG | CG>TA_GCTCATT | 6 | FALSE | FALSE |
| 134_v.txt | 33474 | 33474-SNV- | SNV | 1 | C | T | 32 | 19946 | 0.16043317 | 33.5750596 | rnr1Y285F_pG | TRUE | CAN | JSY2420 | 2 | rnr1Y285F_p | GCC | C>T | A/T+1 | G/C-1 | CG>TA | CG>TA | GCTATGCCTT | CCTTTTT | CGG | GCC | C | A | C>T | C>T_GCC | CG>TA_CCTTTTT | 6 | FALSE | FALSE |
| 134_v.txt | 31981 | 31981-SNV- | SNV | 1 | G | C | 18 | 19468 | 0.09245942 | 45.8888889 | rnr1Y285F_pG | TRUE | CAN | JSY2420 | 2 | rnr1Y285F_p | GGC | G>C | G/C+1 | G/C-1 | CG>GC | CG>GC | AGCCGGGCA | GCATTAA | CCG | CCG | C | G | C>G | C>G_CCG | CG>GC_GCATTAA | 6 | FALSE | FALSE |
| 134_v.txt | 32064 | 32064-SNV- | SNV | 1 | G | C | 41 | 19468 | 0.21060201 | 62.7560976 | rnr1Y285F_pG | TRUE | CAN | JSY2420 | 2 | rnr1Y285F_p | TGA | G>C | G/C+1 | G/C-1 | CG>GC | CG>GC | GAGATTGAG | GAGATAA | ACT | ACT | C | G | C>G | C>G_ACT | CG>GC_GAGATAA | 6 | FALSE | FALSE |
| 134_v.txt | 32115 | 32115-SNV- | SNV | 1 | C | G | 4 | 19467 | 0.02054759 | 63.9974213 | rnr1Y285F_pG | TRUE | CAN | JSY2420 | 2 | rnr1Y285F_p | CCA | C>G | G/C+1 | G/C-1 | CG>GC | CG>GC | AATAGCCAT | CATTCGA | GGT | CCA | C | C | C>G | C>G_CCA | CG>GC_CATTCGA | 6 | FALSE | FALSE |
| 134_v.txt | 32200 | 32200-SNV- | SNV | 1 | G | C | 12 | 19468 | 0.06163961 | 44.25 | rnr1Y285F_pG | TRUE | CAN | JSY2420 | 2 | rnr1Y285F_p | GGA | G>C | G/C+1 | G/C-1 | CG>GC | CG>GC | TGTATGGAA | GAACACC | CCT | CCT | C | G | C>G | C>G_CCT | CG>GC_GAACACC | 6 | FALSE | FALSE |
| 134_v.txt | 32278 | 32278-SNV- | SNV | 1 | C | G | 56 | 25529 | 0.21935838 | 36.875 | rnr1Y285F_pG | TRUE | CAN | JSY2420 | 2 | rnr1Y285F_p | CCA | C>G | G/C+1 | G/C-1 | CG>GC | CG>GC | GGGAACCAA | CAACGTA | GGT | CCA | C | C | C>G | C>G_CCA | CG>GC_CAACGTA | 6 | FALSE | FALSE |
| 134_v.txt | 32292 | 32292-SNV- | SNV | 1 | G | C | 39 | 25529 | 0.15276744 | 51.025641 | rnr1Y285F_pG | TRUE | CAN | JSY2420 | 2 | rnr1Y285F_p | TGA | G>C | G/C+1 | G/C-1 | CG>GC | CG>GC | ATATTTGAAT | GAATTTG | ACT | ACT | C | G | C>G | C>G_ACT | CG>GC_GAATTTG | 6 | FALSE | FALSE |
| 134_v.txt | 32467 | 32467-SNV- | SNV | 1 | C | G | 129 | 25528 | 0.50532748 | 47.2219927 | rnr1Y285F_pG | TRUE | CAN | JSY2420 | 2 | rnr1Y285F_p | CCA | C>G | G/C+1 | G/C-1 | CG>GC | CG>GC | GAGAGCCAA | CAATGTA | GGT | CCA | C | C | C>G | C>G_CCA | CG>GC_CAATGTA | 6 | FALSE | FALSE |
| 134_v.txt | 32536 | 32536-SNV- | SNV | 1 | C | G | 2 | 25529 | 7.83E-03 | 35.9993125 | rnr1Y285F_pG | TRUE | CAN | JSY2420 | 2 | rnr1Y285F_p | GCA | C>G | G/C+1 | G/C-1 | CG>GC | CG>GC | GGTTTGCAG | CAGCTTC | CGT | GCA | C | C | C>G | C>G_GCA | CG>GC_CAGCTTC | 6 | FALSE | FALSE |
| 134_v.txt | 32829 | 32829-SNV- | SNV | 1 | G | C | 153 | 23621 | 0.64772872 | 44.2352941 | rnr1Y285F_pG | TRUE | CAN | JSY2420 | 2 | rnr1Y285F_p | TGT | G>C | G/C+1 | G/C-1 | CG>GC | CG>GC | ATTATTGTGA | GTGATAA | ACA | ACA | C | G | C>G | C>G_ACA | CG>GC_GTGATAA | 6 | FALSE | FALSE |
| 134_v.txt | 32876 | 32876-SNV- | SNV | 1 | G | C | 28 | 26557 | 0.1054336 | 36.5357143 | rnr1Y285F_pG | TRUE | CAN | JSY2420 | 2 | rnr1Y285F_p | TGT | G>C | G/C+1 | G/C-1 | CG>GC | CG>GC | AACTTTGTAC | GTACGTC | ACA | ACA | C | G | C>G | C>G_ACA | CG>GC_GTACGTC | 6 | FALSE | FALSE |
| 134_v.txt | 32917 | 32917-SNV- | SNV | 1 | C | G | 39 | 26566 | 0.14680419 | 54.974359 | rnr1Y285F_pG | TRUE | CAN | JSY2420 | 2 | rnr1Y285F_p | TCC | C>G | G/C+1 | G/C-1 | CG>GC | CG>GC | TAAGTTCCA | CCAGGGC | AGG | TCC | C | C | C>G | C>G_TCC | CG>GC_CCAGGGC | 6 | FALSE | FALSE |
| 134_v.txt | 32922 | 32922-SNV- | SNV | 1 | G | C | 2 | 26566 | 7.53E-03 | 35.9999573 | rnr1Y285F_pG | TRUE | CAN | JSY2420 | 2 | rnr1Y285F_p | GGC | G>C | G/C+1 | G/C-1 | CG>GC | CG>GC | TCCAGGGCA | GCAAAAG | CCG | CCG | C | G | C>G | C>G_CCG | CG>GC_GCAAAAG | 6 | FALSE | FALSE |
| 134_v.txt | 32937 | 32937-SNV- | SNV | 1 | C | G | 93 | 26566 | 0.35007152 | 56.449658 | rnr1Y285F_pG | TRUE | CAN | JSY2420 | 2 | rnr1Y285F_p | CCA | C>G | G/C+1 | G/C-1 | CG>GC | CG>GC | ATTGCCCAA | CAAGAAA | GGT | CCA | C | C | C>G | C>G_CCA | CG>GC_CAAGAAA | 6 | FALSE | FALSE |
| 134_v.txt | 33024 | 33024-SNV- | SNV | 1 | G | C | 42 | 26566 | 0.15809682 | 60.6904762 | rnr1Y285F_pG | TRUE | CAN | JSY2420 | 2 | rnr1Y285F_p | AGG | G>C | G/C+1 | G/C-1 | CG>GC | CG>GC | GTAACAGGG | GGGATGA | TCC | TCC | C | G | C>G | C>G_TCC | CG>GC_GGGATGA | 6 | FALSE | FALSE |
| 134_v.txt | 33070 | 33070-SNV- | SNV | 1 | C | G | 72 | 26565 | 0.27103332 | 53.4027778 | rnr1Y285F_pG | TRUE | CAN | JSY2420 | 2 | rnr1Y285F_p | GCC | C>G | G/C+1 | G/C-1 | CG>GC | CG>GC | AATATGCCA | CCAAAGA | CGG | GCC | C | C | C>G | C>G_GCC | CG>GC_CCAAAGA | 6 | FALSE | FALSE |
| 134_v.txt | 33079 | 33079-SNV- | SNV | 1 | C | G | 36 | 26566 | 0.13551156 | 54 | rnr1Y285F_pG | TRUE | CAN | JSY2420 | 2 | rnr1Y285F_p | CCC | C>G | G/C+1 | G/C-1 | CG>GC | CG>GC | AAGAACCCA | CCATAAA | GGG | CCC | C | C | C>G | C>G_CCC | CG>GC_CCATAAA | 6 | FALSE | FALSE |
| 134_v.txt | 33114 | 33114-SNV- | SNV | 1 | C | G | 55 | 26566 | 0.20703154 | 44.9785601 | rnr1Y285F_pG | TRUE | CAN | JSY2420 | 2 | rnr1Y285F_p | GCC | C>G | G/C+1 | G/C-1 | CG>GC | CG>GC | ACTGGGCCG | CCGGCGT | CGG | GCC | C | C | C>G | C>G_GCC | CG>GC_CCGGCGT | 6 | FALSE | FALSE |
| 134_v.txt | 33115 | 33115-SNV- | SNV | 1 | C | G | 24 | 26566 | 0.09034104 | 46.0813357 | rnr1Y285F_pG | TRUE | CAN | JSY2420 | 2 | rnr1Y285F_p | CCG | C>G | G/C+1 | G/C-1 | CG>GC | CG>GC | CTGGGCCGG | CGGCGTT | GGC | CCG | C | C | C>G | C>G_CCG | CG>GC_CGGCGTT | 6 | FALSE | FALSE |
| 134_v.txt | 33118 | 33118-SNV- | SNV | 1 | C | G | 163 | 26566 | 0.61356621 | 43.2392638 | rnr1Y285F_pG | TRUE | CAN | JSY2420 | 2 | rnr1Y285F_p | GCG | C>G | G/C+1 | G/C-1 | CG>GC | CG>GC | GGCCGGCGT | CGTTGGT | CGC | GCG | C | C | C>G | C>G_GCG | CG>GC_CGTTGGT | 6 | FALSE | FALSE |
| 134_v.txt | 33160 | 33160-SNV- | SNV | 1 | C | G | 64 | 26565 | 0.2409185 | 37.8433102 | rnr1Y285F_pG | TRUE | CAN | JSY2420 | 2 | rnr1Y285F_p | CCA | C>G | G/C+1 | G/C-1 | CG>GC | CG>GC | CTGTACCAAT | CAATAGT | GGT | CCA | C | C | C>G | C>G_CCA | CG>GC_CAATAGT | 6 | FALSE | FALSE |
| 134_v.txt | 33168 | 33168-SNV- | SNV | 1 | C | G | 23 | 26561 | 0.08659313 | 36.9565217 | rnr1Y285F_pG | TRUE | CAN | JSY2420 | 2 | rnr1Y285F_p | ACC | C>G | G/C+1 | G/C-1 | CG>GC | CG>GC | ATAGTACCA | CCACCAA | TGG | ACC | C | C | C>G | C>G_ACC | CG>GC_CCACCAA | 6 | FALSE | FALSE |
| 134_v.txt | 33419 | 33419-SNV- | SNV | 1 | G | C | 23 | 19948 | 0.11529978 | 43.0869565 | rnr1Y285F_pG | TRUE | CAN | JSY2420 | 2 | rnr1Y285F_p | TGT | G>C | G/C+1 | G/C-1 | CG>GC | CG>GC | CTCATTGTAC | GTACATA | ACA | ACA | C | G | C>G | C>G_ACA | CG>GC_GTACATA | 6 | FALSE | FALSE |
| 134_v.txt | 31657 | 31657-SNV- | SNV | 1 | G | T | 91 | 8528 | 1.06707317 | 17.6263736 | rnr1Y285F_pG | TRUE | CAN | JSY2420 | 2 | rnr1Y285F_p | GGC | G>T | A/T+1 | G/C-1 | CG>AT | CG>AT | GAAATGGCG | GCGTGGA | CCG | CCG | C | A | C>A | C>A_CCG | CG>AT_GCGTGGA | 6 | FALSE | FALSE |
| 134_v.txt | 31829 | 31829-SNV- | SNV | 1 | G | T | 7 | 8528 | 0.08208255 | 56 | rnr1Y285F_pG | TRUE | CAN | JSY2420 | 2 | rnr1Y285F_p | TGC | G>T | A/T+1 | G/C-1 | CG>AT | CG>AT | GAATATGCA | GCATTGA | ACG | ACG | C | A | C>A | C>A_ACG | CG>AT_GCATTGA | 6 | FALSE | FALSE |
| 134_v.txt | 31869 | 31869-SNV- | SNV | 1 | G | T | 16 | 8528 | 0.18761726 | 46.5 | rnr1Y285F_pG | TRUE | CAN | JSY2420 | 2 | rnr1Y285F_p | AGA | G>T | A/T+1 | G/C-1 | CG>AT | CG>AT | AAAATAGAG | GAGATAT | TCT | TCT | C | A | C>A | C>A_TCT | CG>AT_GAGATAT | 6 | FALSE | FALSE |
| 134_v.txt | 31980 | 31980-SNV- | SNV | 1 | G | T | 22 | 19468 | 0.11300596 | 45.8950658 | rnr1Y285F_pG | TRUE | CAN | JSY2420 | 2 | rnr1Y285F_p | GGG | G>T | A/T+1 | G/C-1 | CG>AT | CG>AT | AAGCCGGGC | GGCATTA | CCC | CCC | C | A | C>A | C>A_CCC | CG>AT_GGCATTA | 6 | FALSE | FALSE |
| 134_v.txt | 31981 | 31981-SNV- | SNV | 1 | G | T | 14 | 19468 | 0.07191288 | 32.0714286 | rnr1Y285F_pG | TRUE | CAN | JSY2420 | 2 | rnr1Y285F_p | GGC | G>T | A/T+1 | G/C-1 | CG>AT | CG>AT | AGCCGGGCA | GCATTAA | CCG | CCG | C | A | C>A | C>A_CCG | CG>AT_GCATTAA | 6 | FALSE | FALSE |
| 134_v.txt | 32008 | 32008-SNV- | SNV | 1 | C | A | 4770 | 19468 | 24.5017465 | 57.587631 | rnr1Y285F_pG | TRUE | CAN | JSY2420 | 2 | rnr1Y285F_p | TCG | C>A | A/T+1 | G/C-1 | CG>AT | CG>AT | GTAACTCGT | CGTCACG | AGC | TCG | C | T | C>A | C>A_TCG | CG>AT_CGTCACG | 6 | FALSE | FALSE |
| 134_v.txt | 32027 | 32027-SNV- | SNV | 1 | G | T | 18 | 19468 | 0.09245942 | 54.1666667 | rnr1Y285F_pG | TRUE | CAN | JSY2420 | 2 | rnr1Y285F_p | GGT | G>T | A/T+1 | G/C-1 | CG>AT | CG>AT | GCCACGGTA | GTATTTC | CCA | CCA | C | A | C>A | C>A_CCA | CG>AT_GTATTTC | 6 | FALSE | FALSE |
| 134_v.txt | 32088 | 32088-SNV- | SNV | 1 | C | A | 197 | 19468 | 1.01191699 | 61.35533 | rnr1Y285F_pG | TRUE | CAN | JSY2420 | 2 | rnr1Y285F_p | GCC | C>A | A/T+1 | G/C-1 | CG>AT | CG>AT | AAAAAGCCT | CCTGCAA | CGG | GCC | C | T | C>A | C>A_GCC | CG>AT_CCTGCAA | 6 | FALSE | FALSE |
| 134_v.txt | 32095 | 32095-SNV- | SNV | 1 | C | A | 217 | 19468 | 1.11464968 | 60.7327189 | rnr1Y285F_pG | TRUE | CAN | JSY2420 | 2 | rnr1Y285F_p | ACA | C>A | A/T+1 | G/C-1 | CG>AT | CG>AT | CTGCAACAC | CACCAGT | TGT | ACA | C | T | C>A | C>A_ACA | CG>AT_CACCAGT | 6 | FALSE | FALSE |
| 134_v.txt | 32114 | 32114-SNV- | SNV | 1 | C | A | 5 | 19467 | 0.02568449 | 29.8 | rnr1Y285F_pG | TRUE | CAN | JSY2420 | 2 | rnr1Y285F_p | GCC | C>A | A/T+1 | G/C-1 | CG>AT | CG>AT | TAATAGCCA | CCATTCG | CGG | GCC | C | T | C>A | C>A_GCC | CG>AT_CCATTCG | 6 | FALSE | FALSE |
| 134_v.txt | 32153 | 32153-SNV- | SNV | 1 | G | T | 52 | 19468 | 0.26710499 | 53.2884615 | rnr1Y285F_pG | TRUE | CAN | JSY2420 | 2 | rnr1Y285F_p | TGT | G>T | A/T+1 | G/C-1 | CG>AT | CG>AT | CTCCATGTAA | GTAAGCC | ACA | ACA | C | A | C>A | C>A_ACA | CG>AT_GTAAGCC | 6 | FALSE | FALSE |
| 134_v.txt | 32188 | 32188-SNV- | SNV | 1 | C | A | 42 | 19468 | 0.21573865 | 45.3095238 | rnr1Y285F_pG | TRUE | CAN | JSY2420 | 2 | rnr1Y285F_p | ACT | C>A | A/T+1 | G/C-1 | CG>AT | CG>AT | CGAAAACTG | CTGCAAT | TGA | ACT | C | T | C>A | C>A_ACT | CG>AT_CTGCAAT | 6 | FALSE | FALSE |
| 134_v.txt | 32238 | 32238-SNV- | SNV | 1 | G | T | 23 | 19463 | 0.11817294 | 37.3478261 | rnr1Y285F_pG | TRUE | CAN | JSY2420 | 2 | rnr1Y285F_p | AGC | G>T | A/T+1 | G/C-1 | CG>AT | CG>AT | TTAGGAGCC | GCCAACT | TCG | TCG | C | A | C>A | C>A_TCG | CG>AT_GCCAACT | 6 | FALSE | FALSE |
| 134_v.txt | 32274 | 32274-SNV- | SNV | 1 | G | T | 574 | 25529 | 2.24842336 | 36.5400697 | rnr1Y285F_pG | TRUE | CAN | JSY2420 | 2 | rnr1Y285F_p | GGA | G>T | A/T+1 | G/C-1 | CG>AT | CG>AT | ATACGGGAA | GAACCAA | CCT | CCT | C | A | C>A | C>A_CCT | CG>AT_GAACCAA | 6 | FALSE | FALSE |
| 134_v.txt | 32304 | 32304-SNV- | SNV | 1 | G | T | 15 | 25529 | 0.05875671 | 51.6 | rnr1Y285F_pG | TRUE | CAN | JSY2420 | 2 | rnr1Y285F_p | AGA | G>T | A/T+1 | G/C-1 | CG>AT | CG>AT | GCGGCAGAA | GAAATAA | TCT | TCT | C | A | C>A | C>A_TCT | CG>AT_GAAATAA | 6 | FALSE | FALSE |
| 134_v.txt | 32373 | 32373-SNV- | SNV | 1 | G | T | 31 | 25529 | 0.12143053 | 56 | rnr1Y285F_pG | TRUE | CAN | JSY2420 | 2 | rnr1Y285F_p | AGC | G>T | A/T+1 | G/C-1 | CG>AT | CG>AT | TCAATAGCA | GCAATAA | TCG | TCG | C | A | C>A | C>A_TCG | CG>AT_GCAATAA | 6 | FALSE | FALSE |
| 134_v.txt | 32432 | 32432-SNV- | SNV | 1 | G | T | 16 | 25529 | 0.06267382 | 60.25 | rnr1Y285F_pG | TRUE | CAN | JSY2420 | 2 | rnr1Y285F_p | TGT | G>T | A/T+1 | G/C-1 | CG>AT | CG>AT | GTCATTGTAT | GTATGGA | ACA | ACA | C | A | C>A | C>A_ACA | CG>AT_GTATGGA | 6 | FALSE | FALSE |
| 134_v.txt | 32463 | 32463-SNV- | SNV | 1 | G | T | 49 | 25529 | 0.19193858 | 48.2040816 | rnr1Y285F_pG | TRUE | CAN | JSY2420 | 2 | rnr1Y285F_p | AGA | G>T | A/T+1 | G/C-1 | CG>AT | CG>AT | AATAGAGAG | GAGCCAA | TCT | TCT | C | A | C>A | C>A_TCT | CG>AT_GAGCCAA | 6 | FALSE | FALSE |
| 134_v.txt | 32466 | 32466-SNV- | SNV | 1 | C | A | 4 | 25529 | 0.01566846 | 52.9909815 | rnr1Y285F_pG | TRUE | CAN | JSY2420 | 2 | rnr1Y285F_p | GCC | C>A | A/T+1 | G/C-1 | CG>AT | CG>AT | AGAGAGCCA | CCAATGT | CGG | GCC | C | T | C>A | C>A_GCC | CG>AT_CCAATGT | 6 | FALSE | FALSE |
| 134_v.txt | 32467 | 32467-SNV- | SNV | 1 | C | A | 8 | 25528 | 0.03133814 | 32.7511902 | rnr1Y285F_pG | TRUE | CAN | JSY2420 | 2 | rnr1Y285F_p | CCA | C>A | A/T+1 | G/C-1 | CG>AT | CG>AT | GAGAGCCAA | CAATGTA | GGT | CCA | C | T | C>A | C>A_CCA | CG>AT_CAATGTA | 6 | FALSE | FALSE |
| 134_v.txt | 32474 | 32474-SNV- | SNV | 1 | G | T | 17 | 25529 | 0.06659094 | 50.107324 | rnr1Y285F_pG | TRUE | CAN | JSY2420 | 2 | rnr1Y285F_p | AGA | G>T | A/T+1 | G/C-1 | CG>AT | CG>AT | AATGTAGAA | GAAGGTT | TCT | TCT | C | A | C>A | C>A_TCT | CG>AT_GAAGGTT | 6 | FALSE | FALSE |
| 134_v.txt | 32494 | 32494-SNV- | SNV | 1 | C | A | 28 | 25529 | 0.10967919 | 46.071055 | rnr1Y285F_pG | TRUE | CAN | JSY2420 | 2 | rnr1Y285F_p | ACA | C>A | A/T+1 | G/C-1 | CG>AT | CG>AT | GGAAAACAA | CAACTTT | TGT | ACA | C | T | C>A | C>A_ACA | CG>AT_CAACTTT | 6 | FALSE | FALSE |
| 134_v.txt | 32530 | 32530-SNV- | SNV | 1 | G | T | 32 | 25529 | 0.12534764 | 36.6875 | rnr1Y285F_pG | TRUE | CAN | JSY2420 | 2 | rnr1Y285F_p | GGG | G>T | A/T+1 | G/C-1 | CG>AT | CG>AT | TTCTGGGGT | GGTTTGC | CCC | CCC | C | A | C>A | C>A_CCC | CG>AT_GGTTTGC | 6 | FALSE | FALSE |
| 134_v.txt | 32547 | 32547-SNV- | SNV | 1 | G | T | 25 | 25527 | 0.09793552 | 37 | rnr1Y285F_pG | TRUE | CAN | JSY2420 | 2 | rnr1Y285F_p | AGC | G>T | A/T+1 | G/C-1 | CG>AT | CG>AT | TCACCAGCA | GCAGTGA | TCG | TCG | C | A | C>A | C>A_TCG | CG>AT_GCAGTGA | 6 | FALSE | FALSE |
| 134_v.txt | 32556 | 32556-SNV- | SNV | 1 | C | A | 21 | 25522 | 0.08228195 | 37.8095238 | rnr1Y285F_pG | TRUE | CAN | JSY2420 | 2 | rnr1Y285F_p | ACC | C>A | A/T+1 | G/C-1 | CG>AT | CG>AT | GTGATACCA | CCAACTA | TGG | ACC | C | T | C>A | C>A_ACC | CG>AT_CCAACTA | 6 | FALSE | FALSE |
| 134_v.txt | 32566 | 32566-SNV- | SNV | 1 | C | A | 38 | 23613 | 0.1609283 | 37.5526316 | rnr1Y285F_pG | TRUE | CAN | JSY2420 | 2 | rnr1Y285F_p | TCA | C>A | A/T+1 | G/C-1 | CG>AT | CG>AT | CTAGTTCAGT | CAGTACC | AGT | TCA | C | T | C>A | C>A_TCA | CG>AT_CAGTACC | 6 | FALSE | FALSE |
| 134_v.txt | 32572 | 32572-SNV- | SNV | 1 | C | A | 69 | 23621 | 0.29211295 | 36.7101449 | rnr1Y285F_pG | TRUE | CAN | JSY2420 | 2 | rnr1Y285F_p | CCT | C>A | A/T+1 | G/C-1 | CG>AT | CG>AT | CAGTACCTTG | CTTGAAA | GGA | CCT | C | T | C>A | C>A_CCT | CG>AT_CTTGAAA | 6 | FALSE | FALSE |
| 134_v.txt | 32580 | 32580-SNV- | SNV | 1 | G | T | 64 | 23621 | 0.27094535 | 35.65625 | rnr1Y285F_pG | TRUE | CAN | JSY2420 | 2 | rnr1Y285F_p | TGT | G>T | A/T+1 | G/C-1 | CG>AT | CG>AT | TGAAATGTG | GTGAAGG | ACA | ACA | C | A | C>A | C>A_ACA | CG>AT_GTGAAGG | 6 | FALSE | FALSE |
| 134_v.txt | 32629 | 32629-SNV- | SNV | 1 | C | A | 18 | 23621 | 0.07620338 | 49.1176248 | rnr1Y285F_pG | TRUE | CAN | JSY2420 | 2 | rnr1Y285F_p | TCG | C>A | A/T+1 | G/C-1 | CG>AT | CG>AT | TCCCTTCGTT | CGTTTTT | AGC | TCG | C | T | C>A | C>A_TCG | CG>AT_CGTTTTT | 6 | FALSE | FALSE |
| 134_v.txt | 32692 | 32692-SNV- | SNV | 1 | C | A | 11 | 23621 | 0.04656873 | 55.8181818 | rnr1Y285F_pG | TRUE | CAN | JSY2420 | 2 | rnr1Y285F_p | CCA | C>A | A/T+1 | G/C-1 | CG>AT | CG>AT | GGAATCCAA | CAACTGG | GGT | CCA | C | T | C>A | C>A_CCA | CG>AT_CAACTGG | 6 | FALSE | FALSE |
| 134_v.txt | 32819 | 32819-SNV- | SNV | 1 | G | T | 27 | 23621 | 0.11430507 | 39.8147926 | rnr1Y285F_pG | TRUE | CAN | JSY2420 | 2 | rnr1Y285F_p | AGT | G>T | A/T+1 | G/C-1 | CG>AT | CG>AT | GAACAAGTT | GTTCATT | TCA | TCA | C | A | C>A | C>A_TCA | CG>AT_GTTCATT | 6 | FALSE | FALSE |
| 134_v.txt | 32917 | 32917-SNV- | SNV | 1 | C | A | 134 | 26566 | 0.50440413 | 50.9477612 | rnr1Y285F_pG | TRUE | CAN | JSY2420 | 2 | rnr1Y285F_p | TCC | C>A | A/T+1 | G/C-1 | CG>AT | CG>AT | TAAGTTCCA | CCAGGGC | AGG | TCC | C | T | C>A | C>A_TCC | CG>AT_CCAGGGC | 6 | FALSE | FALSE |
| 134_v.txt | 32928 | 32928-SNV- | SNV | 1 | G | T | 207 | 26566 | 0.77919145 | 53.8199535 | rnr1Y285F_pG | TRUE | CAN | JSY2420 | 2 | rnr1Y285F_p | AGT | G>T | A/T+1 | G/C-1 | CG>AT | CG>AT | GCAAAAGTG | GTGATTG | TCA | TCA | C | A | C>A | C>A_TCA | CG>AT_GTGATTG | 6 | FALSE | FALSE |
| 134_v.txt | 32937 | 32937-SNV- | SNV | 1 | C | A | 9 | 26566 | 0.03387789 | 41.2127675 | rnr1Y285F_pG | TRUE | CAN | JSY2420 | 2 | rnr1Y285F_p | CCA | C>A | A/T+1 | G/C-1 | CG>AT | CG>AT | ATTGCCCAA | CAAGAAA | GGT | CCA | C | T | C>A | C>A_CCA | CG>AT_CAAGAAA | 6 | FALSE | FALSE |
| 134_v.txt | 32940 | 32940-SNV- | SNV | 1 | G | T | 372 | 26566 | 1.40028608 | 56.5322581 | rnr1Y285F_pG | TRUE | CAN | JSY2420 | 2 | rnr1Y285F_p | AGA | G>T | A/T+1 | G/C-1 | CG>AT | CG>AT | GCCCAAGAA | GAAAACC | TCT | TCT | C | A | C>A | C>A_TCT | CG>AT_GAAAACC | 6 | FALSE | FALSE |
| 134_v.txt | 33006 | 33006-SNV- | SNV | 1 | G | T | 60 | 26566 | 0.22585259 | 61.5666667 | rnr1Y285F_pG | TRUE | CAN | JSY2420 | 2 | rnr1Y285F_p | TGT | G>T | A/T+1 | G/C-1 | CG>AT | CG>AT | AAAACTGTG | GTGAAAG | ACA | ACA | C | A | C>A | C>A_ACA | CG>AT_GTGAAAG | 6 | FALSE | FALSE |
| 134_v.txt | 33024 | 33024-SNV- | SNV | 1 | G | T | 41 | 26566 | 0.15433261 | 63.195122 | rnr1Y285F_pG | TRUE | CAN | JSY2420 | 2 | rnr1Y285F_p | AGG | G>T | A/T+1 | G/C-1 | CG>AT | CG>AT | GTAACAGGG | GGGATGA | TCC | TCC | C | A | C>A | C>A_TCC | CG>AT_GGGATGA | 6 | FALSE | FALSE |
| 134_v.txt | 33043 | 33043-SNV- | SNV | 1 | C | A | 23 | 26566 | 0.08657683 | 56.3043478 | rnr1Y285F_pG | TRUE | CAN | JSY2420 | 2 | rnr1Y285F_p | TCA | C>A | A/T+1 | G/C-1 | CG>AT | CG>AT | CCATTTCACC | CACCCAA | AGT | TCA | C | T | C>A | C>A_TCA | CG>AT_CACCCAA | 6 | FALSE | FALSE |
| 134_v.txt | 33070 | 33070-SNV- | SNV | 1 | C | A | 3 | 26565 | 0.01129306 | 24.3333333 | rnr1Y285F_pG | TRUE | CAN | JSY2420 | 2 | rnr1Y285F_p | GCC | C>A | A/T+1 | G/C-1 | CG>AT | CG>AT | AATATGCCA | CCAAAGA | CGG | GCC | C | T | C>A | C>A_GCC | CG>AT_CCAAAGA | 6 | FALSE | FALSE |
| 134_v.txt | 33079 | 33079-SNV- | SNV | 1 | C | A | 13 | 26566 | 0.04893473 | 41 | rnr1Y285F_pG | TRUE | CAN | JSY2420 | 2 | rnr1Y285F_p | CCC | C>A | A/T+1 | G/C-1 | CG>AT | CG>AT | AAGAACCCA | CCATAAA | GGG | CCC | C | T | C>A | C>A_CCC | CG>AT_CCATAAA | 6 | FALSE | FALSE |
| 134_v.txt | 33112 | 33112-SNV- | SNV | 1 | G | T | 17 | 26566 | 0.06399157 | 40.9264308 | rnr1Y285F_pG | TRUE | CAN | JSY2420 | 2 | rnr1Y285F_p | GGG | G>T | A/T+1 | G/C-1 | CG>AT | CG>AT | CCACTGGGC | GGCCGGC | CCC | CCC | C | A | C>A | C>A_CCC | CG>AT_GGCCGGC | 6 | FALSE | FALSE |
| 134_v.txt | 33151 | 33151-SNV- | SNV | 1 | G | T | 26 | 26566 | 0.09786946 | 37.5384615 | rnr1Y285F_pG | TRUE | CAN | JSY2420 | 2 | rnr1Y285F_p | AGA | G>T | A/T+1 | G/C-1 | CG>AT | CG>AT | TGAAAAGAC | GACCTGT | TCT | TCT | C | A | C>A | C>A_TCT | CG>AT_GACCTGT | 6 | FALSE | FALSE |
| 134_v.txt | 33154 | 33154-SNV- | SNV | 1 | C | A | 14 | 26566 | 0.05269894 | 33.7142857 | rnr1Y285F_pG | TRUE | CAN | JSY2420 | 2 | rnr1Y285F_p | CCT | C>A | A/T+1 | G/C-1 | CG>AT | CG>AT | AAAGACCTG | CTGTACC | GGA | CCT | C | T | C>A | C>A_CCT | CG>AT_CTGTACC | 6 | FALSE | FALSE |
| 134_v.txt | 33160 | 33160-SNV- | SNV | 1 | C | A | 6 | 26565 | 0.02258611 | 29.0000462 | rnr1Y285F_pG | TRUE | CAN | JSY2420 | 2 | rnr1Y285F_p | CCA | C>A | A/T+1 | G/C-1 | CG>AT | CG>AT | CTGTACCAAT | CAATAGT | GGT | CCA | C | T | C>A | C>A_CCA | CG>AT_CAATAGT | 6 | FALSE | FALSE |
| 134_v.txt | 33186 | 33186-SNV- | SNV | 1 | C | A | 22 | 19945 | 0.11030333 | 35.7727273 | rnr1Y285F_pG | TRUE | CAN | JSY2420 | 2 | rnr1Y285F_p | ACC | C>A | A/T+1 | G/C-1 | CG>AT | CG>AT | ATCATACCA | CCAATAT | TGG | ACC | C | T | C>A | C>A_ACC | CG>AT_CCAATAT | 6 | FALSE | FALSE |
| 134_v.txt | 33195 | 33195-SNV- | SNV | 1 | C | A | 19 | 19947 | 0.09525242 | 37.3157895 | rnr1Y285F_pG | TRUE | CAN | JSY2420 | 2 | rnr1Y285F_p | TCT | C>A | A/T+1 | G/C-1 | CG>AT | CG>AT | ATATGTCTTT | CTTTGCT | AGA | TCT | C | T | C>A | C>A_TCT | CG>AT_CTTTGCT | 6 | FALSE | FALSE |
| 134_v.txt | 33241 | 33241-SNV- | SNV | 1 | C | A | 33 | 19947 | 0.16543841 | 45.4545455 | rnr1Y285F_pG | TRUE | CAN | JSY2420 | 2 | rnr1Y285F_p | TCA | C>A | A/T+1 | G/C-1 | CG>AT | CG>AT | CTCCTTCATC | CATCTTC | AGT | TCA | C | T | C>A | C>A_TCA | CG>AT_CATCTTC | 6 | FALSE | FALSE |
| 134_v.txt | 33247 | 33247-SNV- | SNV | 1 | C | A | 63 | 19947 | 0.31583697 | 39.4761905 | rnr1Y285F_pG | TRUE | CAN | JSY2420 | 2 | rnr1Y285F_p | TCA | C>A | A/T+1 | G/C-1 | CG>AT | CG>AT | CATCTTCATC | CATCACC | AGT | TCA | C | T | C>A | C>A_TCA | CG>AT_CATCACC | 6 | FALSE | FALSE |
| 134_v.txt | 33309 | 33309-SNV- | SNV | 1 | C | A | 8 | 19946 | 0.04010829 | 26.75 | rnr1Y285F_pG | TRUE | CAN | JSY2420 | 2 | rnr1Y285F_p | GCG | C>A | A/T+1 | G/C-1 | CG>AT | CG>AT | AAAGAGCGC | CGCAATG | CGC | GCG | C | T | C>A | C>A_GCG | CG>AT_CGCAATG | 6 | FALSE | FALSE |
| 134_v.txt | 33325 | 33325-SNV- | SNV | 1 | C | A | 40 | 19947 | 0.20053141 | 55.75 | rnr1Y285F_pG | TRUE | CAN | JSY2420 | 2 | rnr1Y285F_p | TCT | C>A | A/T+1 | G/C-1 | CG>AT | CG>AT | ACAATTCTTT | CTTTACT | AGA | TCT | C | T | C>A | C>A_TCT | CG>AT_CTTTACT | 6 | FALSE | FALSE |
| 134_v.txt | 33419 | 33419-SNV- | SNV | 1 | G | T | 656 | 19948 | 3.28855023 | 47.160061 | rnr1Y285F_pG | TRUE | CAN | JSY2420 | 2 | rnr1Y285F_p | TGT | G>T | A/T+1 | G/C-1 | CG>AT | CG>AT | CTCATTGTAC | GTACATA | ACA | ACA | C | A | C>A | C>A_ACA | CG>AT_GTACATA | 6 | FALSE | FALSE |
| 134_v.txt | 33433 | 33433-SNV- | SNV | 1 | C | A | 22 | 19948 | 0.11028675 | 42.0454546 | rnr1Y285F_pG | TRUE | CAN | JSY2420 | 2 | rnr1Y285F_p | TCC | C>A | A/T+1 | G/C-1 | CG>AT | CG>AT | GCTTCTCCTC | CCTCTAT | AGG | TCC | C | T | C>A | C>A_TCC | CG>AT_CCTCTAT | 6 | FALSE | FALSE |
| 134_v.txt | 33474 | 33474-SNV- | SNV | 1 | C | A | 57 | 19946 | 0.28577158 | 35.0092444 | rnr1Y285F_pG | TRUE | CAN | JSY2420 | 2 | rnr1Y285F_p | GCC | C>A | A/T+1 | G/C-1 | CG>AT | CG>AT | GCTATGCCTT | CCTTTTT | CGG | GCC | C | T | C>A | C>A_GCC | CG>AT_CCTTTTT | 6 | FALSE | FALSE |
| 134_v.txt | 33475 | 33475-SNV- | SNV | 1 | C | A | 10 | 19946 | 0.05013537 | 37.9000374 | rnr1Y285F_pG | TRUE | CAN | JSY2420 | 2 | rnr1Y285F_p | CCT | C>A | A/T+1 | G/C-1 | CG>AT | CG>AT | CTATGCCTTT | CTTTTTT | GGA | CCT | C | T | C>A | C>A_CCT | CG>AT_CTTTTTT | 6 | FALSE | FALSE |
| 134_v.txt | 32630 | 32630-Repla | Replacement | 2 | G | AT | 13 | 23621 | 0.05503577 | 50.5769193 | rnr1Y285F_pG | TRUE | CAN | JSY2420 | 2 | rnr1Y285F_p | CGT | nope | >1bp-ins | G/C-1 | none | Replac. | CCCTTCGTTT | GTTTTTA | GCA | GCA | C | none | C>none | C>none_GCA | Replac._GTTTTTA | 6 | FALSE | FALSE |
| 134_v.txt | 32922 | 32922-Repla | Replacement | 2 | G | AA | 7 | 26566 | 0.02634947 | 63.7142856 | rnr1Y285F_pG | TRUE | CAN | JSY2420 | 2 | rnr1Y285F_p | GGC | nope | >1bp-ins | G/C-1 | none | Replac. | TCCAGGGCA | GCAAAAG | CCG | CCG | C | none | C>none | C>none_CCG | Replac._GCAAAAG | 6 | FALSE | FALSE |
| 134_v.txt | 32114 | 32114-MNV- | MNV | 2 | CC | AA | 1 | 19468 | 5.14E-03 | 32.9999996 | rnr1Y285F_pG | TRUE | CAN | JSY2420 | 2 | rnr1Y285F_p | GCC | nope | >1bp-ins | >1bpdel | none | MNV | TAATAGCCA | CCATTCG | CGG | GCC | CC | none | NA | NA_GCC | MNV_CCATTCG | 6 | FALSE | FALSE |
| 134_v.txt | 32796 | 32796-MNV- | MNV | 2 | CC | TT | 1 | 23621 | 4.23E-03 | 31.0000002 | rnr1Y285F_pG | TRUE | CAN | JSY2420 | 2 | rnr1Y285F_p | ACC | nope | >1bp-ins | >1bpdel | none | MNV | AATTCACCGT | CCGTAAT | TGG | ACC | CC | none | NA | NA_ACC | MNV_CCGTAAT | 6 | FALSE | FALSE |
| 134_v.txt | 33113 | 33113-MNV- | MNV | 2 | GC | AG | 1 | 26566 | 3.76E-03 | 37.9999998 | rnr1Y285F_pG | TRUE | CAN | JSY2420 | 2 | rnr1Y285F_p | GGC | nope | >1bp-ins | >1bpdel | none | MNV | CACTGGGCC | GCCGGCG | CCG | GGC | GC | none | NA | NA_GGC | MNV_GCCGGCG | 6 | FALSE | FALSE |
| 134_v.txt | 33114 | 33114-MNV- | MNV | 2 | CC | AA | 1 | 26566 | 3.76E-03 | 36.5000428 | rnr1Y285F_pG | TRUE | CAN | JSY2420 | 2 | rnr1Y285F_p | GCC | nope | >1bp-ins | >1bpdel | none | MNV | ACTGGGCCG | CCGGCGT | CGG | GCC | CC | none | NA | NA_GCC | MNV_CCGGCGT | 6 | FALSE | FALSE |
| 134_v.txt | 33114 | 33114-MNV- | MNV | 2 | CC | TG | 1 | 26566 | 3.76E-03 | 37.4999998 | rnr1Y285F_pG | TRUE | CAN | JSY2420 | 2 | rnr1Y285F_p | GCC | nope | >1bp-ins | >1bpdel | none | MNV | ACTGGGCCG | CCGGCGT | CGG | GCC | CC | none | NA | NA_GCC | MNV_CCGGCGT | 6 | FALSE | FALSE |
| 134_v.txt | 33153 | 33153-MNV- | MNV | 2 | CC | AA | 1 | 26566 | 3.76E-03 | 37 | rnr1Y285F_pG | TRUE | CAN | JSY2420 | 2 | rnr1Y285F_p | ACC | nope | >1bp-ins | >1bpdel | none | MNV | AAAAGACCT | CCTGTAC | TGG | ACC | CC | none | NA | NA_ACC | MNV_CCTGTAC | 6 | FALSE | FALSE |
| 134_v.txt | 33159 | 33159-MNV- | MNV | 2 | CC | AA | 2 | 26565 | 7.53E-03 | 32.7501142 | rnr1Y285F_pG | TRUE | CAN | JSY2420 | 2 | rnr1Y285F_p | ACC | nope | >1bp-ins | >1bpdel | none | MNV | CCTGTACCA | CCAATAG | TGG | ACC | CC | none | NA | NA_ACC | MNV_CCAATAG | 6 | FALSE | FALSE |
| 134_v.txt | 31723 | 31723-Insert | Insertion | 1 | - | A | 2 | 8528 | 0.02345216 | 64 | rnr1Y285F_pG | TRUE | CAN | JSY2420 | 2 | rnr1Y285F_p | CAA | nope | A/T+1 | >1bpdel | none | A/T+1 | TGTCCCAAA | AAAAAGT | GTT | CAA | - | T | NA | NA_CAA | A/T+1_AAAAAGT | 6 | FALSE | FALSE |
| 134_v.txt | 31904 | 31904-Insert | Insertion | 1 | - | T | 17 | 8528 | 0.19934334 | 42.2941177 | rnr1Y285F_pG | TRUE | CAN | JSY2420 | 2 | rnr1Y285F_p | ATT | nope | A/T+1 | >1bpdel | none | A/T+1 | ATTGAATTTT | TTTTGGT | TAA | ATT | - | A | NA | NA_ATT | A/T+1_TTTTGGT | 6 | FALSE | FALSE |
| 134_v.txt | 31912 | 31912-Insert | Insertion | 1 | - | T | 2 | 8528 | 0.02345216 | 38 | rnr1Y285F_pG | TRUE | CAN | JSY2420 | 2 | rnr1Y285F_p | GCA | nope | A/T+1 | >1bpdel | none | A/T+1 | TTGGTGCAA | CAAAAGC | CGT | GCA | - | A | NA | NA_GCA | A/T+1_CAAAAGC | 6 | FALSE | FALSE |
| 134_v.txt | 32081 | 32081-Insert | Insertion | 1 | - | A | 2060 | 19468 | 10.581467 | 63.4135922 | rnr1Y285F_pG | TRUE | CAN | JSY2420 | 2 | rnr1Y285F_p | CAA | nope | A/T+1 | >1bpdel | none | A/T+1 | CCATGCAAA | AAAAAAG | GTT | CAA | - | T | NA | NA_CAA | A/T+1_AAAAAAG | 6 | FALSE | FALSE |
| 134_v.txt | 32081 | 32081-Insert | Insertion | 2 | - | AA | 2 | 19468 | 0.01027327 | 62.5 | rnr1Y285F_pG | TRUE | CAN | JSY2420 | 2 | rnr1Y285F_p | CAA | nope | >1bp-ins | >1bpdel | none | >1bp-ins | CCATGCAAA | AAAAAAG | GTT | CAA | - | none | NA | NA_CAA | >1bp-ins_AAAAAAG | 6 | FALSE | FALSE |
| 134_v.txt | 32087 | 32087-Insert | Insertion | 1 | - | G | 4 | 19468 | 0.02054654 | 57.25 | rnr1Y285F_pG | TRUE | CAN | JSY2420 | 2 | rnr1Y285F_p | AGC | nope | G/C+1 | >1bpdel | none | G/C+1 | AAAAAAGCC | GCCTGCA | TCG | AGC | - | C | NA | NA_AGC | G/C+1_GCCTGCA | 6 | FALSE | FALSE |
| 134_v.txt | 32498 | 32498-Insert | Insertion | 1 | - | T | 3 | 25529 | 0.01175134 | 38 | rnr1Y285F_pG | TRUE | CAN | JSY2420 | 2 | rnr1Y285F_p | CTT | nope | A/T+1 | >1bpdel | none | A/T+1 | AACAACTTTT | TTTTTTG | GAA | CTT | - | A | NA | NA_CTT | A/T+1_TTTTTTG | 6 | FALSE | FALSE |
| 134_v.txt | 32728 | 32728-Insert | Insertion | 1 | - | A | 59 | 23621 | 0.24977774 | 62 | rnr1Y285F_pG | TRUE | CAN | JSY2420 | 2 | rnr1Y285F_p | CAA | nope | A/T+1 | >1bpdel | none | A/T+1 | CCATACAAA | AAAAACA | GTT | CAA | - | T | NA | NA_CAA | A/T+1_AAAAACA | 6 | FALSE | FALSE |
| 134_v.txt | 32842 | 32842-Insert | Insertion | 1 | - | A | 226 | 23621 | 0.95677575 | 41.9823009 | rnr1Y285F_pG | TRUE | CAN | JSY2420 | 2 | rnr1Y285F_p | CAA | nope | A/T+1 | >1bpdel | none | A/T+1 | TTACCCAAA | AAAAAAT | GTT | CAA | - | T | NA | NA_CAA | A/T+1_AAAAAAT | 6 | FALSE | FALSE |
| 134_v.txt | 33036 | 33036-Insert | Insertion | 33 | - | GCCATTTCAC | 37 | 26566 | 0.13927577 | 59.6134316 | rnr1Y285F_pG | TRUE | CAN | JSY2420 | 2 | rnr1Y285F_p | AGC | nope | >1bp-ins | >1bpdel | none | >1bp-ins | AATGTAGCC | GCCATTT | TCG | AGC | - | none | NA | NA_AGC | >1bp-ins_GCCATTT | 6 | FALSE | FALSE |
| 134_v.txt | 33170 | 33170-Insert | Insertion | 1 | - | A | 40 | 26561 | 0.15059674 | 37.975 | rnr1Y285F_pG | TRUE | CAN | JSY2420 | 2 | rnr1Y285F_p | CAC | nope | A/T+1 | >1bpdel | none | A/T+1 | AGTACCACC | ACCAAGG | GTG | CAC | - | T | NA | NA_CAC | A/T+1_ACCAAGG | 6 | FALSE | FALSE |
| 134_v.txt | 33206 | 33206-Insert | Insertion | 2 | - | CT | 14 | 19947 | 0.07018599 | 37.75 | rnr1Y285F_pG | TRUE | CAN | JSY2420 | 2 | rnr1Y285F_p | GCT | nope | >1bp-ins | >1bpdel | none | >1bp-ins | CTTAAGCTCT | CTCTCTC | CGA | GCT | - | none | NA | NA_GCT | >1bp-ins_CTCTCTC | 6 | FALSE | FALSE |
| 134_v.txt | 31913 | 31913-Deleti | Deletion | 2 | AA | - | 11 | 8528 | 0.12898687 | 37.8181818 | rnr1Y285F_pG | TRUE | CAN | JSY2420 | 2 | rnr1Y285F_p | CAA | nope | >1bp-ins | >1bpdel | none | >1bpdel | TGGTGCAAA | AAAAGCC | GTT | CAA | AA | none | NA | NA_CAA | >1bpdel_AAAAGCC | 6 | FALSE | FALSE |
| 134_v.txt | 32337 | 32337-Deleti | Deletion | 2 | AT | - | 123 | 25529 | 0.48180501 | 61.7642276 | rnr1Y285F_pG | TRUE | CAN | JSY2420 | 2 | rnr1Y285F_p | GAT | nope | >1bp-ins | >1bpdel | none | >1bpdel | TTGAAGATA | ATATGTG | CTA | GAT | AT | none | NA | NA_GAT | >1bpdel_ATATGTG | 6 | FALSE | FALSE |
| 134_v.txt | 32375 | 32375-Deleti | Deletion | 3 | AAT | - | 15 | 25529 | 0.05875671 | 60.0666667 | rnr1Y285F_pG | TRUE | CAN | JSY2420 | 2 | rnr1Y285F_p | CAA | nope | >1bp-ins | >1bpdel | none | >1bpdel | AATAGCAAT | AATAATA | GTT | CAA | AAT | none | NA | NA_CAA | >1bpdel_AATAATA | 6 | FALSE | FALSE |
| 134_v.txt | 32804 | 32804-Deleti | Deletion | 2 | TT | - | 26 | 23621 | 0.11007155 | 40.7307692 | rnr1Y285F_pG | TRUE | CAN | JSY2420 | 2 | rnr1Y285F_p | ATT | nope | >1bp-ins | >1bpdel | none | >1bpdel | GTAATATTT | TTTGACA | TAA | ATT | TT | none | NA | NA_ATT | >1bpdel_TTTGACA | 6 | FALSE | FALSE |
| 134_v.txt | 33045 | 33045-Deleti | Deletion | 3 | CCC | - | 47 | 26566 | 0.17691787 | 57.7659575 | rnr1Y285F_pG | TRUE | CAN | JSY2420 | 2 | rnr1Y285F_p | ACC | nope | >1bp-ins | >1bpdel | none | >1bpdel | ATTTCACCCA | CCCAAGG | TGG | ACC | CCC | none | NA | NA_ACC | >1bpdel_CCCAAGG | 6 | FALSE | FALSE |
| 134_v.txt | 31723 | 31723-Deleti | Deletion | 1 | A | - | 10 | 8528 | 0.11726079 | 63.9000001 | rnr1Y285F_pG | TRUE | CAN | JSY2420 | 2 | rnr1Y285F_p | CAA | nope | >1bp-ins | A/T-1 | none | A/T-1 | TGTCCCAAA | AAAAAGT | GTT | GTT | T | none | T>none | T>none_GTT | A/T-1_AAAAAGT | 6 | FALSE | FALSE |
| 134_v.txt | 31904 | 31904-Deleti | Deletion | 1 | T | - | 11 | 8528 | 0.12898687 | 41.8181809 | rnr1Y285F_pG | TRUE | CAN | JSY2420 | 2 | rnr1Y285F_p | ATT | nope | >1bp-ins | A/T-1 | none | A/T-1 | ATTGAATTTT | TTTTGGT | TAA | ATT | T | none | nope | nope_ATT | A/T-1_TTTTGGT | 6 | FALSE | FALSE |
| 134_v.txt | 31913 | 31913-Deleti | Deletion | 1 | A | - | 4 | 8528 | 0.04690432 | 44.4999964 | rnr1Y285F_pG | TRUE | CAN | JSY2420 | 2 | rnr1Y285F_p | CAA | nope | >1bp-ins | A/T-1 | none | A/T-1 | TGGTGCAAA | AAAAGCC | GTT | GTT | T | none | T>none | T>none_GTT | A/T-1_AAAAGCC | 6 | FALSE | FALSE |
| 134_v.txt | 31972 | 31972-Deleti | Deletion | 1 | C | - | 6 | 19468 | 0.03081981 | 35.1666667 | rnr1Y285F_pG | TRUE | CAN | JSY2420 | 2 | rnr1Y285F_p | GCC | nope | >1bp-ins | G/C-1 | none | G/C-1 | AATAAGCCA | CCAAGCC | CGG | GCC | C | none | nope | nope_GCC | G/C-1_CCAAGCC | 6 | FALSE | FALSE |
| 134_v.txt | 31997 | 31997-Deleti | Deletion | 1 | A | - | 37 | 19468 | 0.19005548 | 52.3783784 | rnr1Y285F_pG | TRUE | CAN | JSY2420 | 2 | rnr1Y285F_p | TAA | nope | >1bp-ins | A/T-1 | none | A/T-1 | AGCTTTAAA | AAATGGT | ATT | ATT | T | none | T>none | T>none_ATT | A/T-1_AAATGGT | 6 | FALSE | FALSE |
| 134_v.txt | 32004 | 32004-Deleti | Deletion | 1 | A | - | 2 | 19468 | 0.01027327 | 64 | rnr1Y285F_pG | TRUE | CAN | JSY2420 | 2 | rnr1Y285F_p | TAA | nope | >1bp-ins | A/T-1 | none | A/T-1 | AATGGTAAC | AACTCGT | ATT | ATT | T | none | T>none | T>none_ATT | A/T-1_AACTCGT | 6 | FALSE | FALSE |

| | | | | | | | | | | | | | | | | | | | | | | | | | | | | | | | | | | |
|---|---|---|---|---|---|---|---|---|---|---|---|---|---|---|---|---|---|---|---|---|---|---|---|---|---|---|---|---|---|---|---|---|---|---|
| 134_v.txt | 32081 | 32081-Deleti | Deletion | 1 | A | - | 69 | 19468 | 0.35442778 | 63.7101451 | rnr1Y285F_pG | TRUE | CAN | JSY2420 | 2 | rnr1Y285F_p | CAA | nope | >1bp-ins | A/T-1 | none | A/T-1 | CCATGCAAA | AAAAAAG | GTT | GTT | T | none | T>none | T>none_GTT | A/T-1_AAAAAAG | 6 | FALSE | FALSE |
| 134_v.txt | 32114 | 32114-Deleti | Deletion | 1 | C | - | 2 | 19467 | 0.0102738 | 62 | rnr1Y285F_pG | TRUE | CAN | JSY2420 | 2 | rnr1Y285F_p | GCC | nope | >1bp-ins | G/C-1 | none | G/C-1 | TAATAGCCA | CCATTCG | CGG | GCC | C | none | nope | nope_GCC | G/C-1_CCATTCG | 6 | FALSE | FALSE |
| 134_v.txt | 32166 | 32166-Deleti | Deletion | 1 | C | - | 1 | 19468 | 5.14E-03 | 63.9999941 | rnr1Y285F_pG | TRUE | CAN | JSY2420 | 2 | rnr1Y285F_p | GCC | nope | >1bp-ins | G/C-1 | none | G/C-1 | AAAGCGCCA | CCAAATG | CGG | GCC | C | none | nope | nope_GCC | G/C-1_CCAAATG | 6 | FALSE | FALSE |
| 134_v.txt | 32210 | 32210-Deleti | Deletion | 1 | T | - | 7 | 19467 | 0.03595829 | 41.2857143 | rnr1Y285F_pG | TRUE | CAN | JSY2420 | 2 | rnr1Y285F_p | CTT | nope | >1bp-ins | A/T-1 | none | A/T-1 | ACCACCTTTG | TTTGGTG | GAA | CTT | T | none | nope | nope_CTT | A/T-1_TTTGGTG | 6 | FALSE | FALSE |
| 134_v.txt | 32265 | 32265-Deleti | Deletion | 1 | A | - | 2 | 25529 | 7.83E-03 | 38 | rnr1Y285F_pG | TRUE | CAN | JSY2420 | 2 | rnr1Y285F_p | TAA | nope | >1bp-ins | A/T-1 | none | A/T-1 | CCAAATAAA | AAAATAC | ATT | ATT | T | none | T>none | T>none_ATT | A/T-1_AAAATAC | 6 | FALSE | FALSE |
| 134_v.txt | 32304 | 32304-Deleti | Deletion | 1 | G | - | 38 | 25529 | 0.14885033 | 60.4736842 | rnr1Y285F_pG | TRUE | CAN | JSY2420 | 2 | rnr1Y285F_p | AGA | nope | >1bp-ins | G/C-1 | none | G/C-1 | GCGGCAGAA | GAAATAA | TCT | TCT | C | none | C>none | C>none_TCT | G/C-1_GAAATAA | 6 | FALSE | FALSE |
| 134_v.txt | 32352 | 32352-Deleti | Deletion | 1 | T | - | 2 | 25529 | 7.83E-03 | 63.5 | rnr1Y285F_pG | TRUE | CAN | JSY2420 | 2 | rnr1Y285F_p | CTT | nope | >1bp-ins | A/T-1 | none | A/T-1 | AAAACCTTT | TTTGTAC | GAA | CTT | T | none | nope | nope_CTT | A/T-1_TTTGTAC | 6 | FALSE | FALSE |
| 134_v.txt | 32357 | 32357-Deleti | Deletion | 1 | A | - | 194 | 25529 | 0.75992009 | 61.1082474 | rnr1Y285F_pG | TRUE | CAN | JSY2420 | 2 | rnr1Y285F_p | TAC | nope | >1bp-ins | A/T-1 | none | A/T-1 | CTTTGTACCA | ACCAGAG | ATG | ATG | T | none | T>none | T>none_ATG | A/T-1_ACCAGAG | 6 | FALSE | FALSE |
| 134_v.txt | 32381 | 32381-Deleti | Deletion | 1 | A | - | 7 | 25529 | 0.0274198 | 64 | rnr1Y285F_pG | TRUE | CAN | JSY2420 | 2 | rnr1Y285F_p | TAA | nope | >1bp-ins | A/T-1 | none | A/T-1 | AATAATAAA | AAAGGGA | ATT | ATT | T | none | T>none | T>none_ATT | A/T-1_AAAGGGA | 6 | FALSE | FALSE |
| 134_v.txt | 32424 | 32424-Deleti | Deletion | 1 | G | - | 162 | 25529 | 0.63457245 | 57.3703704 | rnr1Y285F_pG | TRUE | CAN | JSY2420 | 2 | rnr1Y285F_p | AGG | nope | >1bp-ins | G/C-1 | none | G/C-1 | AGTTTAGGG | GGGTCAT | TCC | TCC | C | none | C>none | C>none_TCC | G/C-1_GGGTCAT | 6 | FALSE | FALSE |
| 134_v.txt | 32427 | 32427-Deleti | Deletion | 1 | T | - | 25 | 25529 | 0.09792785 | 60.44 | rnr1Y285F_pG | TRUE | CAN | JSY2420 | 2 | rnr1Y285F_p | GTC | nope | >1bp-ins | A/T-1 | none | A/T-1 | TTAGGGTCA | TCATTGT | CAG | GTC | T | none | nope | nope_GTC | A/T-1_TCATTGT | 6 | FALSE | FALSE |
| 134_v.txt | 32442 | 32442-Deleti | Deletion | 1 | A | - | 4 | 25529 | 0.01566846 | 64 | rnr1Y285F_pG | TRUE | CAN | JSY2420 | 2 | rnr1Y285F_p | TAA | nope | >1bp-ins | A/T-1 | none | A/T-1 | GGAACTAAA | AAAAGTC | ATT | ATT | T | none | T>none | T>none_ATT | A/T-1_AAAAGTC | 6 | FALSE | FALSE |
| 134_v.txt | 32498 | 32498-Deleti | Deletion | 1 | T | - | 31 | 25529 | 0.12143053 | 44.4193483 | rnr1Y285F_pG | TRUE | CAN | JSY2420 | 2 | rnr1Y285F_p | CTT | nope | >1bp-ins | A/T-1 | none | A/T-1 | AACAACTTTT | TTTTTTG | GAA | CTT | T | none | nope | nope_CTT | A/T-1_TTTTTTG | 6 | FALSE | FALSE |
| 134_v.txt | 32522 | 32522-Deleti | Deletion | 1 | T | - | 25 | 25529 | 0.09792785 | 39.92 | rnr1Y285F_pG | TRUE | CAN | JSY2420 | 2 | rnr1Y285F_p | ATT | nope | >1bp-ins | A/T-1 | none | A/T-1 | AACGGATTT | TTTTCTG | TAA | ATT | T | none | nope | nope_ATT | A/T-1_TTTTCTG | 6 | FALSE | FALSE |
| 134_v.txt | 32528 | 32528-Deleti | Deletion | 1 | G | - | 5 | 25529 | 0.01958557 | 37.2 | rnr1Y285F_pG | TRUE | CAN | JSY2420 | 2 | rnr1Y285F_p | TGG | nope | >1bp-ins | G/C-1 | none | G/C-1 | TTTTCTGGG | GGGGTTT | ACC | ACC | C | none | C>none | C>none_ACC | G/C-1_GGGGTTT | 6 | FALSE | FALSE |
| 134_v.txt | 32535 | 32535-Deleti | Deletion | 1 | G | - | 22 | 25529 | 0.08617651 | 37.8636364 | rnr1Y285F_pG | TRUE | CAN | JSY2420 | 2 | rnr1Y285F_p | TGC | nope | >1bp-ins | G/C-1 | none | G/C-1 | GGGTTTGCA | GCAGCTT | ACG | ACG | C | none | C>none | C>none_ACG | G/C-1_GCAGCTT | 6 | FALSE | FALSE |
| 134_v.txt | 32608 | 32608-Deleti | Deletion | 1 | C | - | 306 | 23621 | 1.29545743 | 45.3954248 | rnr1Y285F_pG | TRUE | CAN | JSY2420 | 2 | rnr1Y285F_p | ACC | nope | >1bp-ins | G/C-1 | none | G/C-1 | AGGAAACCC | CCCAACC | TGG | ACC | C | none | nope | nope_ACC | G/C-1_CCCAACC | 6 | FALSE | FALSE |
| 134_v.txt | 32613 | 32613-Deleti | Deletion | 1 | C | - | 78 | 23621 | 0.33021464 | 45.6282051 | rnr1Y285F_pG | TRUE | CAN | JSY2420 | 2 | rnr1Y285F_p | ACC | nope | >1bp-ins | G/C-1 | none | G/C-1 | ACCCAACCT | CCTAAGA | TGG | ACC | C | none | nope | nope_ACC | G/C-1_CCTAAGA | 6 | FALSE | FALSE |
| 134_v.txt | 32615 | 32615-Deleti | Deletion | 1 | T | - | 140 | 23621 | 0.59269294 | 45.8142857 | rnr1Y285F_pG | TRUE | CAN | JSY2420 | 2 | rnr1Y285F_p | CTA | nope | >1bp-ins | A/T-1 | none | A/T-1 | CCAACCTAA | TAAGAAC | GAT | CTA | T | none | nope | nope_CTA | A/T-1_TAAGAAC | 6 | FALSE | FALSE |
| 134_v.txt | 32619 | 32619-Deleti | Deletion | 1 | A | - | 32 | 23621 | 0.13547267 | 45.8125 | rnr1Y285F_pG | TRUE | CAN | JSY2420 | 2 | rnr1Y285F_p | GAA | nope | >1bp-ins | A/T-1 | none | A/T-1 | CCTAAGAAC | AACCTCC | CTT | CTT | T | none | T>none | T>none_CTT | A/T-1_AACCTCC | 6 | FALSE | FALSE |
| 134_v.txt | 32631 | 32631-Deleti | Deletion | 1 | T | - | 192 | 23621 | 0.81283604 | 51.4166713 | rnr1Y285F_pG | TRUE | CAN | JSY2420 | 2 | rnr1Y285F_p | GTT | nope | >1bp-ins | A/T-1 | none | A/T-1 | CCTTCGTTTT | TTTTTAT | CAA | GTT | T | none | nope | nope_GTT | A/T-1_TTTTTAT | 6 | FALSE | FALSE |
| 134_v.txt | 32658 | 32658-Deleti | Deletion | 1 | C | - | 8 | 23621 | 0.03386817 | 56.5 | rnr1Y285F_pG | TRUE | CAN | JSY2420 | 2 | rnr1Y285F_p | ACC | nope | >1bp-ins | G/C-1 | none | G/C-1 | CCTGGACCC | CCCCAGG | TGG | ACC | C | none | nope | nope_ACC | G/C-1_CCCCAGG | 6 | FALSE | FALSE |
| 134_v.txt | 32707 | 32707-Deleti | Deletion | 1 | C | - | 154 | 23621 | 0.65196224 | 62.8311688 | rnr1Y285F_pG | TRUE | CAN | JSY2420 | 2 | rnr1Y285F_p | ACC | nope | >1bp-ins | G/C-1 | none | G/C-1 | CGGTAACCC | CCCCAGC | TGG | ACC | C | none | nope | nope_ACC | G/C-1_CCCCAGC | 6 | FALSE | FALSE |
| 134_v.txt | 32714 | 32714-Deleti | Deletion | 1 | A | - | 24 | 23621 | 0.10160451 | 63.375 | rnr1Y285F_pG | TRUE | CAN | JSY2420 | 2 | rnr1Y285F_p | CAC | nope | >1bp-ins | A/T-1 | none | A/T-1 | CCCAGCACC | ACCACAA | GTG | GTG | T | none | T>none | T>none_GTG | A/T-1_ACCACAA | 6 | FALSE | FALSE |
| 134_v.txt | 32728 | 32728-Deleti | Deletion | 1 | A | - | 37 | 23621 | 0.15664028 | 61.9999897 | rnr1Y285F_pG | TRUE | CAN | JSY2420 | 2 | rnr1Y285F_p | CAA | nope | >1bp-ins | A/T-1 | none | A/T-1 | CCATACAAA | AAAAACA | GTT | GTT | T | none | T>none | T>none_GTT | A/T-1_AAAAACA | 6 | FALSE | FALSE |
| 134_v.txt | 32740 | 32740-Deleti | Deletion | 1 | T | - | 5 | 23621 | 0.02116761 | 48.4 | rnr1Y285F_pG | TRUE | CAN | JSY2420 | 2 | rnr1Y285F_p | ATT | nope | >1bp-ins | A/T-1 | none | A/T-1 | AGTATATTA | TTAGAAA | TAA | ATT | T | none | nope | nope_ATT | A/T-1_TTAGAAA | 6 | FALSE | FALSE |
| 134_v.txt | 32744 | 32744-Deleti | Deletion | 1 | A | - | 802 | 23621 | 3.39528386 | 57.0822943 | rnr1Y285F_pG | TRUE | CAN | JSY2420 | 2 | rnr1Y285F_p | GAA | nope | >1bp-ins | A/T-1 | none | A/T-1 | TATTAGAAA | AAACCCG | CTT | CTT | T | none | T>none | T>none_CTT | A/T-1_AAACCCG | 6 | FALSE | FALSE |
| 134_v.txt | 32804 | 32804-Deleti | Deletion | 1 | T | - | 2 | 23621 | 8.47E-03 | 38.0000226 | rnr1Y285F_pG | TRUE | CAN | JSY2420 | 2 | rnr1Y285F_p | ATT | nope | >1bp-ins | A/T-1 | none | A/T-1 | GTAATATTT | TTTGACA | TAA | ATT | T | none | nope | nope_ATT | A/T-1_TTTGACA | 6 | FALSE | FALSE |
| 134_v.txt | 32824 | 32824-Deleti | Deletion | 1 | T | - | 5 | 23621 | 0.02116761 | 37.8 | rnr1Y285F_pG | TRUE | CAN | JSY2420 | 2 | rnr1Y285F_p | ATT | nope | >1bp-ins | A/T-1 | none | A/T-1 | AGTTCATTAT | TTATTGT | TAA | ATT | T | none | nope | nope_ATT | A/T-1_TTATTGT | 6 | FALSE | FALSE |
| 134_v.txt | 32842 | 32842-Deleti | Deletion | 1 | A | - | 135 | 23621 | 0.57152534 | 42.6666624 | rnr1Y285F_pG | TRUE | CAN | JSY2420 | 2 | rnr1Y285F_p | CAA | nope | >1bp-ins | A/T-1 | none | A/T-1 | TTACCCAAA | AAAAAAT | GTT | GTT | T | none | T>none | T>none_GTT | A/T-1_AAAAAAT | 6 | FALSE | FALSE |
| 134_v.txt | 32899 | 32899-Deleti | Deletion | 1 | G | - | 43 | 26566 | 0.16186103 | 35.8139535 | rnr1Y285F_pG | TRUE | CAN | JSY2420 | 2 | rnr1Y285F_p | TGG | nope | >1bp-ins | G/C-1 | none | G/C-1 | TGACTTGGC | GGCCAAC | ACC | ACC | C | none | C>none | C>none_ACC | G/C-1_GGCCAAC | 6 | FALSE | FALSE |
| 134_v.txt | 32924 | 32924-Deleti | Deletion | 1 | A | - | 24 | 26566 | 0.09034104 | 57.4166667 | rnr1Y285F_pG | TRUE | CAN | JSY2420 | 2 | rnr1Y285F_p | CAA | nope | >1bp-ins | A/T-1 | none | A/T-1 | CAGGGCAAA | AAAAGTG | GTT | GTT | T | none | T>none | T>none_GTT | A/T-1_AAAAGTG | 6 | FALSE | FALSE |
| 134_v.txt | 32995 | 32995-Deleti | Deletion | 1 | G | - | 21 | 26566 | 0.07904841 | 60.6666667 | rnr1Y285F_pG | TRUE | CAN | JSY2420 | 2 | rnr1Y285F_p | TGT | nope | >1bp-ins | G/C-1 | none | G/C-1 | ATCTTTGTGA | GTGAGAA | ACA | ACA | C | none | C>none | C>none_ACA | G/C-1_GTGAGAA | 6 | FALSE | FALSE |
| 134_v.txt | 33000 | 33000-Deleti | Deletion | 1 | A | - | 29 | 26566 | 0.10916209 | 59.9655172 | rnr1Y285F_pG | TRUE | CAN | JSY2420 | 2 | rnr1Y285F_p | GAA | nope | >1bp-ins | A/T-1 | none | A/T-1 | TGTGAGAAA | AAAACTG | CTT | CTT | T | none | T>none | T>none_CTT | A/T-1_AAAACTG | 6 | FALSE | FALSE |
| 134_v.txt | 33040 | 33040-Deleti | Deletion | 1 | T | - | 2 | 26566 | 7.53E-03 | 64 | rnr1Y285F_pG | TRUE | CAN | JSY2420 | 2 | rnr1Y285F_p | ATT | nope | >1bp-ins | A/T-1 | none | A/T-1 | TAGCCATTTC | TTTCACC | TAA | ATT | T | none | nope | nope_ATT | A/T-1_TTTCACC | 6 | FALSE | FALSE |
| 134_v.txt | 33111 | 33111-Deleti | Deletion | 1 | G | - | 14 | 26566 | 0.05269894 | 37.0000057 | rnr1Y285F_pG | TRUE | CAN | JSY2420 | 2 | rnr1Y285F_p | TGG | nope | >1bp-ins | G/C-1 | none | G/C-1 | CCCACTGGG | GGGCCGG | ACC | ACC | C | none | C>none | C>none_ACC | G/C-1_GGGCCGG | 6 | FALSE | FALSE |
| 134_v.txt | 33147 | 33147-Deleti | Deletion | 1 | A | - | 6 | 26566 | 0.02258526 | 37.8333333 | rnr1Y285F_pG | TRUE | CAN | JSY2420 | 2 | rnr1Y285F_p | GAA | nope | >1bp-ins | A/T-1 | none | A/T-1 | CCAATGAAA | AAAAGAC | CTT | CTT | T | none | T>none | T>none_CTT | A/T-1_AAAAGAC | 6 | FALSE | FALSE |
| 134_v.txt | 33153 | 33153-Deleti | Deletion | 1 | C | - | 1 | 26566 | 3.76E-03 | 37 | rnr1Y285F_pG | TRUE | CAN | JSY2420 | 2 | rnr1Y285F_p | ACC | nope | >1bp-ins | G/C-1 | none | G/C-1 | AAAAGACCT | CCTGTAC | TGG | ACC | C | none | nope | nope_ACC | G/C-1_CCTGTAC | 6 | FALSE | FALSE |
| 134_v.txt | 33350 | 33350-Deleti | Deletion | 1 | T | - | 37 | 19948 | 0.18548225 | 50.7027027 | rnr1Y285F_pG | TRUE | CAN | JSY2420 | 2 | rnr1Y285F_p | GTA | nope | >1bp-ins | A/T-1 | none | A/T-1 | CAATGGTAT | TATTGAC | CAT | GTA | T | none | nope | nope_GTA | A/T-1_TATTGAC | 6 | FALSE | FALSE |
| 134_v.txt | 33368 | 33368-Deleti | Deletion | 1 | G | - | 4 | 19948 | 0.02005214 | 57.5 | rnr1Y285F_pG | TRUE | CAN | JSY2420 | 2 | rnr1Y285F_p | GGT | nope | >1bp-ins | G/C-1 | none | G/C-1 | TCTGTGGTG | GTGTGTT | CCA | CCA | C | none | C>none | C>none_CCA | G/C-1_GTGTGTT | 6 | FALSE | FALSE |
| 134_v.txt | 33474 | 33474-Deleti | Deletion | 1 | C | - | 35 | 19946 | 0.17547378 | 30.0856776 | rnr1Y285F_pG | TRUE | CAN | JSY2420 | 2 | rnr1Y285F_p | GCC | nope | >1bp-ins | G/C-1 | none | G/C-1 | GCTATGCCT | CCTTTTT | CGG | GCC | C | none | nope | nope_GCC | G/C-1_CCTTTTT | 6 | FALSE | FALSE |
| 135_v.txt | 31723 | 31723-SNV-A | SNV | 1 | A | C | 4 | 10907 | 0.0366737 | 63.9994538 | rnr1Y285F_pG | TRUE | CAN | JSY2421 | 3 | rnr1Y285F_p | CAA | A>C | G/C+1 | A/T-1 | TA>GC | TA>GC | TGTCCCAAA | AAAAAGT | GTT | GTT | T | G | T>G | T>G_GTT | TA>GC_AAAAAGT | 6 | FALSE | FALSE |
| 135_v.txt | 31986 | 31986-SNV-A | SNV | 1 | A | C | 46 | 23199 | 0.19828441 | 51.173913 | rnr1Y285F_pG | TRUE | CAN | JSY2421 | 3 | rnr1Y285F_p | TAA | A>C | G/C+1 | A/T-1 | TA>GC | TA>GC | GGCATTAAT | AATTTAG | ATT | ATT | T | G | T>G | T>G_ATT | TA>GC_AATTTAG | 6 | FALSE | FALSE |
| 135_v.txt | 32073 | 32073-SNV-A | SNV | 1 | A | C | 20 | 23199 | 0.08621061 | 60.85 | rnr1Y285F_pG | TRUE | CAN | JSY2421 | 3 | rnr1Y285F_p | TAA | A>C | G/C+1 | A/T-1 | TA>GC | TA>GC | ATAAATAAC | AACCATG | ATT | ATT | T | G | T>G | T>G_ATT | TA>GC_AACCATG | 6 | FALSE | FALSE |
| 135_v.txt | 32081 | 32081-SNV-A | SNV | 1 | A | C | 1 | 23199 | 4.31E-03 | 15.0004277 | rnr1Y285F_pG | TRUE | CAN | JSY2421 | 3 | rnr1Y285F_p | CAA | A>C | G/C+1 | A/T-1 | TA>GC | TA>GC | CCATGCAAA | AAAAAAG | GTT | GTT | T | G | T>G | T>G_GTT | TA>GC_AAAAAAG | 6 | FALSE | FALSE |
| 135_v.txt | 32226 | 32226-SNV-A | SNV | 1 | A | C | 42 | 23199 | 0.18104229 | 35.7857728 | rnr1Y285F_pG | TRUE | CAN | JSY2421 | 3 | rnr1Y285F_p | CAG | A>C | G/C+1 | A/T-1 | TA>GC | TA>GC | CTTGACAGG | AGGAATT | GTC | GTC | T | G | T>G | T>G_GTC | TA>GC_AGGAATT | 6 | FALSE | FALSE |
| 135_v.txt | 32445 | 32445-SNV-A | SNV | 1 | A | C | 67 | 33161 | 0.20204457 | 52.3880597 | rnr1Y285F_pG | TRUE | CAN | JSY2421 | 3 | rnr1Y285F_p | AAG | A>C | G/C+1 | A/T-1 | TA>GC | TA>GC | ACTAAAAGT | AGTCCAA | TTC | TTC | T | G | T>G | T>G_TTC | TA>GC_AGTCCAA | 6 | FALSE | FALSE |
| 135_v.txt | 32481 | 32481-SNV-A | SNV | 1 | A | C | 35 | 33161 | 0.10554567 | 45.8857143 | rnr1Y285F_pG | TRUE | CAN | JSY2421 | 3 | rnr1Y285F_p | TAA | A>C | G/C+1 | A/T-1 | TA>GC | TA>GC | AAGGTTAAG | AAGATAC | ATT | ATT | T | G | T>G | T>G_ATT | TA>GC_AAGATAC | 6 | FALSE | FALSE |
| 135_v.txt | 32493 | 32493-SNV-A | SNV | 1 | A | C | 7 | 33161 | 0.02110913 | 37.4253015 | rnr1Y285F_pG | TRUE | CAN | JSY2421 | 3 | rnr1Y285F_p | AAC | A>C | G/C+1 | A/T-1 | TA>GC | TA>GC | CGGAAAACA | ACAACTT | TTG | TTG | T | G | T>G | T>G_TTG | TA>GC_ACAACTT | 6 | FALSE | FALSE |
| 135_v.txt | 32581 | 32581-SNV-T | SNV | 1 | T | G | 46 | 27973 | 0.16444429 | 35.4565217 | rnr1Y285F_pG | TRUE | CAN | JSY2421 | 3 | rnr1Y285F_p | GTG | T>G | G/C+1 | A/T-1 | TA>GC | TA>GC | GAAATGTGA | TGAAGGC | CAC | GTG | T | C | T>G | T>G_GTG | TA>GC_TGAAGGC | 6 | FALSE | FALSE |
| 135_v.txt | 32642 | 32642-SNV-A | SNV | 1 | A | C | 4 | 27973 | 0.0142995 | 31.5005609 | rnr1Y285F_pG | TRUE | CAN | JSY2421 | 3 | rnr1Y285F_p | TAG | A>C | G/C+1 | A/T-1 | TA>GC | TA>GC | ATCCTTAGAT | AGATATT | ATC | ATC | T | G | T>G | T>G_ATC | TA>GC_AGATATT | 6 | FALSE | FALSE |
| 135_v.txt | 32680 | 32680-SNV-A | SNV | 1 | A | C | 9 | 27973 | 0.03217388 | 58.5523079 | rnr1Y285F_pG | TRUE | CAN | JSY2421 | 3 | rnr1Y285F_p | CAA | A>C | G/C+1 | A/T-1 | TA>GC | TA>GC | TTCTCCAATA | AATAACG | GTT | GTT | T | G | T>G | T>G_GTT | TA>GC_AATAACG | 6 | FALSE | FALSE |
| 135_v.txt | 32739 | 32739-SNV-A | SNV | 1 | A | C | 56 | 27973 | 0.20019304 | 60.4107143 | rnr1Y285F_pG | TRUE | CAN | JSY2421 | 3 | rnr1Y285F_p | TAT | A>C | G/C+1 | A/T-1 | TA>GC | TA>GC | CAGTATATTA | ATTAGAA | ATA | ATA | T | G | T>G | T>G_ATA | TA>GC_ATTAGAA | 6 | FALSE | FALSE |
| 135_v.txt | 32763 | 32763-SNV-A | SNV | 1 | A | C | 22 | 27973 | 0.07864727 | 50.3636364 | rnr1Y285F_pG | TRUE | CAN | JSY2421 | 3 | rnr1Y285F_p | AAC | A>C | G/C+1 | A/T-1 | TA>GC | TA>GC | GCTAAAACT | ACTTTGA | TTG | TTG | T | G | T>G | T>G_TTG | TA>GC_ACTTTGA | 6 | FALSE | FALSE |
| 135_v.txt | 32799 | 32799-SNV-T | SNV | 1 | T | G | 229 | 27973 | 0.81864655 | 45.0408867 | rnr1Y285F_pG | TRUE | CAN | JSY2421 | 3 | rnr1Y285F_p | GTA | T>G | G/C+1 | A/T-1 | TA>GC | TA>GC | TCACCGTAAT | TAATATT | CAT | GTA | T | C | T>G | T>G_GTA | TA>GC_TAATATT | 6 | FALSE | FALSE |
| 135_v.txt | 32800 | 32800-SNV-A | SNV | 1 | A | C | 4 | 27973 | 0.0142995 | 38 | rnr1Y285F_pG | TRUE | CAN | JSY2421 | 3 | rnr1Y285F_p | TAA | A>C | G/C+1 | A/T-1 | TA>GC | TA>GC | CACCGTAAT | AATATTT | ATT | ATT | T | G | T>G | T>G_ATT | TA>GC_AATATTT | 6 | FALSE | FALSE |
| 135_v.txt | 32801 | 32801-SNV-A | SNV | 1 | A | C | 27 | 27973 | 0.09652165 | 46.0736893 | rnr1Y285F_pG | TRUE | CAN | JSY2421 | 3 | rnr1Y285F_p | AAT | A>C | G/C+1 | A/T-1 | TA>GC | TA>GC | ACCGTAATA | ATATTTG | TTA | TTA | T | G | T>G | T>G_TTA | TA>GC_ATATTTG | 6 | FALSE | FALSE |
| 135_v.txt | 32808 | 32808-SNV-A | SNV | 1 | A | C | 9 | 27973 | 0.03217388 | 49.4392071 | rnr1Y285F_pG | TRUE | CAN | JSY2421 | 3 | rnr1Y285F_p | GAC | A>C | G/C+1 | A/T-1 | TA>GC | TA>GC | TATTTGACAC | ACAGGGA | CTG | CTG | T | G | T>G | T>G_CTG | TA>GC_ACAGGGA | 6 | FALSE | FALSE |
| 135_v.txt | 32842 | 32842-SNV-A | SNV | 1 | A | C | 4 | 27973 | 0.0142995 | 34.2510898 | rnr1Y285F_pG | TRUE | CAN | JSY2421 | 3 | rnr1Y285F_p | CAA | A>C | G/C+1 | A/T-1 | TA>GC | TA>GC | TTACCCAAA | AAAAAAT | GTT | GTT | T | G | T>G | T>G_GTT | TA>GC_AAAAAAT | 6 | FALSE | FALSE |
| 135_v.txt | 32884 | 32884-SNV-A | SNV | 1 | A | C | 9 | 29054 | 0.0309768 | 31.5555556 | rnr1Y285F_pG | TRUE | CAN | JSY2421 | 3 | rnr1Y285F_p | CAA | A>C | G/C+1 | A/T-1 | TA>GC | TA>GC | ACGTCCAAA | AAAATTG | GTT | GTT | T | G | T>G | T>G_GTT | TA>GC_AAAATTG | 6 | FALSE | FALSE |
| 135_v.txt | 32915 | 32915-SNV-T | SNV | 1 | T | G | 25 | 29052 | 0.0860526 | 44.16 | rnr1Y285F_pG | TRUE | CAN | JSY2421 | 3 | rnr1Y285F_p | GTT | T>G | G/C+1 | A/T-1 | TA>GC | TA>GC | ACTAAGTTC | TTCCAGG | CAA | GTT | T | C | T>G | T>G_GTT | TA>GC_TTCCAGG | 6 | FALSE | FALSE |
| 135_v.txt | 32929 | 32929-SNV-T | SNV | 1 | T | G | 28 | 29052 | 0.09637891 | 51.2142524 | rnr1Y285F_pG | TRUE | CAN | JSY2421 | 3 | rnr1Y285F_p | GTG | T>G | G/C+1 | A/T-1 | TA>GC | TA>GC | CAAAAGTGA | TGATTGC | CAC | GTG | T | C | T>G | T>G_GTG | TA>GC_TGATTGC | 6 | FALSE | FALSE |
| 135_v.txt | 33000 | 33000-SNV-A | SNV | 1 | A | C | 3 | 29054 | 0.0103256 | 22.000719 | rnr1Y285F_pG | TRUE | CAN | JSY2421 | 3 | rnr1Y285F_p | GAA | A>C | G/C+1 | A/T-1 | TA>GC | TA>GC | TGTGAGAAA | AAAACTG | CTT | CTT | T | G | T>G | T>G_CTT | TA>GC_AAAACTG | 6 | FALSE | FALSE |
| 135_v.txt | 33019 | 33019-SNV-T | SNV | 1 | T | G | 6 | 29054 | 0.0206512 | 59 | rnr1Y285F_pG | TRUE | CAN | JSY2421 | 3 | rnr1Y285F_p | GTA | T>G | G/C+1 | A/T-1 | TA>GC | TA>GC | AGGATGTAA | TAACAGG | CAT | GTA | T | C | T>G | T>G_GTA | TA>GC_TAACAGG | 6 | FALSE | FALSE |
| 135_v.txt | 33065 | 33065-SNV-A | SNV | 1 | A | C | 15 | 29051 | 0.05163334 | 47.2 | rnr1Y285F_pG | TRUE | CAN | JSY2421 | 3 | rnr1Y285F_p | AAT | A>C | G/C+1 | A/T-1 | TA>GC | TA>GC | GACAGAATA | ATATGCC | TTA | TTA | T | G | T>G | T>G_TTA | TA>GC_ATATGCC | 6 | FALSE | FALSE |
| 135_v.txt | 33089 | 33089-SNV-A | SNV | 1 | A | C | 29 | 29051 | 0.09982445 | 48.1724138 | rnr1Y285F_pG | TRUE | CAN | JSY2421 | 3 | rnr1Y285F_p | AAT | A>C | G/C+1 | A/T-1 | TA>GC | TA>GC | AAATAAATA | ATATGAT | TTA | TTA | T | G | T>G | T>G_TTA | TA>GC_ATATGAT | 6 | FALSE | FALSE |
| 135_v.txt | 33138 | 33138-SNV-A | SNV | 1 | A | C | 77 | 29051 | 0.26505112 | 39.6363636 | rnr1Y285F_pG | TRUE | CAN | JSY2421 | 3 | rnr1Y285F_p | TAA | A>C | G/C+1 | A/T-1 | TA>GC | TA>GC | GTGGATAAA | AAACCAA | ATT | ATT | T | G | T>G | T>G_ATT | TA>GC_AAACCAA | 6 | FALSE | FALSE |
| 135_v.txt | 33269 | 33269-SNV-A | SNV | 1 | A | C | 155 | 19836 | 0.78140754 | 16.3225807 | rnr1Y285F_pG | TRUE | CAN | JSY2421 | 3 | rnr1Y285F_p | TAG | A>C | G/C+1 | A/T-1 | TA>GC | TA>GC | CTCCATAGAC | AGAGAAC | ATC | ATC | T | G | T>G | T>G_ATC | TA>GC_AGAGAAC | 6 | FALSE | FALSE |
| 135_v.txt | 32048 | 32048-SNV-T | SNV | 1 | T | C | 2 | 23199 | 8.62E-03 | 63.9957497 | rnr1Y285F_pG | TRUE | CAN | JSY2421 | 3 | rnr1Y285F_p | ATC | T>C | G/C+1 | A/T-1 | TA>CG | TA>CG | CATAAATCT | TCTGATG | TAG | ATC | T | G | T>C | T>C_ATC | TA>CG_TCTGATG | 6 | FALSE | FALSE |
| 135_v.txt | 32055 | 32055-SNV-T | SNV | 1 | T | C | 50 | 23199 | 0.21552653 | 59.52 | rnr1Y285F_pG | TRUE | CAN | JSY2421 | 3 | rnr1Y285F_p | GTG | T>C | G/C+1 | A/T-1 | TA>CG | TA>CG | CTGATGTGC | TGCGAGA | CAC | GTG | T | G | T>C | T>C_GTG | TA>CG_TGCGAGA | 6 | FALSE | FALSE |
| 135_v.txt | 32126 | 32126-SNV-T | SNV | 1 | T | C | 4 | 23199 | 0.01724212 | 56.7495817 | rnr1Y285F_pG | TRUE | CAN | JSY2421 | 3 | rnr1Y285F_p | CTT | T>C | G/C+1 | A/T-1 | TA>CG | TA>CG | GAAAACTTT | TTTGTCA | GAA | CTT | T | G | T>C | T>C_CTT | TA>CG_TTTGTCA | 6 | FALSE | FALSE |
| 135_v.txt | 32256 | 32256-SNV-A | SNV | 1 | A | G | 40 | 33111 | 0.12080578 | 36.45 | rnr1Y285F_pG | TRUE | CAN | JSY2421 | 3 | rnr1Y285F_p | TAG | A>G | G/C+1 | A/T-1 | TA>CG | TA>CG | TTTGATAGA | AGACCAA | ATC | ATC | T | C | T>C | T>C_ATC | TA>CG_AGACCAA | 6 | FALSE | FALSE |
| 135_v.txt | 32303 | 32303-SNV-A | SNV | 1 | A | G | 9 | 33160 | 0.02714113 | 57.7765838 | rnr1Y285F_pG | TRUE | CAN | JSY2421 | 3 | rnr1Y285F_p | CAG | A>G | G/C+1 | A/T-1 | TA>CG | TA>CG | TGCGGCAGA | AGAAATA | GTC | GTC | T | C | T>C | T>C_GTC | TA>CG_AGAAATA | 6 | FALSE | FALSE |
| 135_v.txt | 32346 | 32346-SNV-A | SNV | 1 | A | G | 8 | 33160 | 0.02412545 | 53.1247768 | rnr1Y285F_pG | TRUE | CAN | JSY2421 | 3 | rnr1Y285F_p | CAA | A>G | G/C+1 | A/T-1 | TA>CG | TA>CG | TGTGGCAAA | AAAACCT | GTT | GTT | T | C | T>C | T>C_GTT | TA>CG_AAAACCT | 6 | FALSE | FALSE |
| 135_v.txt | 32445 | 32445-SNV-A | SNV | 1 | A | G | 79 | 33161 | 0.23823166 | 51.6708861 | rnr1Y285F_pG | TRUE | CAN | JSY2421 | 3 | rnr1Y285F_p | AAG | A>G | G/C+1 | A/T-1 | TA>CG | TA>CG | ACTAAAAGT | AGTCCAA | TTC | TTC | T | C | T>C | T>C_TTC | TA>CG_AGTCCAA | 6 | FALSE | FALSE |
| 135_v.txt | 32472 | 32472-SNV-T | SNV | 1 | T | C | 31 | 33161 | 0.09348331 | 50.5134403 | rnr1Y285F_pG | TRUE | CAN | JSY2421 | 3 | rnr1Y285F_p | GTA | T>C | G/C+1 | A/T-1 | TA>CG | TA>CG | CCAATGTAG | TAGAAGG | CAT | GTA | T | G | T>C | T>C_GTA | TA>CG_TAGAAGG | 6 | FALSE | FALSE |
| 135_v.txt | 32473 | 32473-SNV-A | SNV | 1 | A | G | 10 | 33161 | 0.03015591 | 43.1 | rnr1Y285F_pG | TRUE | CAN | JSY2421 | 3 | rnr1Y285F_p | TAG | A>G | G/C+1 | A/T-1 | TA>CG | TA>CG | CAATGTAGA | AGAAGGT | ATC | ATC | T | C | T>C | T>C_ATC | TA>CG_AGAAGGT | 6 | FALSE | FALSE |
| 135_v.txt | 32490 | 32490-SNV-A | SNV | 1 | A | G | 31 | 33161 | 0.09348331 | 44.6774194 | rnr1Y285F_pG | TRUE | CAN | JSY2421 | 3 | rnr1Y285F_p | GAA | A>G | G/C+1 | A/T-1 | TA>CG | TA>CG | ATACGGAAA | AAAACAA | CTT | CTT | T | C | T>C | T>C_CTT | TA>CG_AAAACAA | 6 | FALSE | FALSE |
| 135_v.txt | 32734 | 32734-SNV-A | SNV | 1 | A | G | 27 | 27972 | 0.0965251 | 60.1481933 | rnr1Y285F_pG | TRUE | CAN | JSY2421 | 3 | rnr1Y285F_p | CAG | A>G | G/C+1 | A/T-1 | TA>CG | TA>CG | AAAAACAGT | AGTATAT | GTC | GTC | T | C | T>C | T>C_GTC | TA>CG_AGTATAT | 6 | FALSE | FALSE |
| 135_v.txt | 32752 | 32752-SNV-T | SNV | 1 | T | C | 4 | 27973 | 0.0142995 | 54.2499149 | rnr1Y285F_pG | TRUE | CAN | JSY2421 | 3 | rnr1Y285F_p | ATA | T>C | G/C+1 | A/T-1 | TA>CG | TA>CG | ACCCGATAA | TAATGGC | TAT | ATA | T | G | T>C | T>C_ATA | TA>CG_TAATGGC | 6 | FALSE | FALSE |
| 135_v.txt | 32753 | 32753-SNV-A | SNV | 1 | A | G | 8 | 27973 | 0.02859901 | 21.0000501 | rnr1Y285F_pG | TRUE | CAN | JSY2421 | 3 | rnr1Y285F_p | TAA | A>G | G/C+1 | A/T-1 | TA>CG | TA>CG | CCCGATAAT | AATGGCT | ATT | ATT | T | C | T>C | T>C_ATT | TA>CG_AATGGCT | 6 | FALSE | FALSE |
| 135_v.txt | 32767 | 32767-SNV-T | SNV | 1 | T | C | 24 | 27973 | 0.08579702 | 50 | rnr1Y285F_pG | TRUE | CAN | JSY2421 | 3 | rnr1Y285F_p | TTG | T>C | G/C+1 | A/T-1 | TA>CG | TA>CG | AAACTTTGA | TGATGGA | AAC | TTG | T | G | T>C | T>C_TTG | TA>CG_TGATGGA | 6 | FALSE | FALSE |
| 135_v.txt | 32800 | 32800-SNV-A | SNV | 1 | A | G | 9 | 27973 | 0.03217388 | 38.7777778 | rnr1Y285F_pG | TRUE | CAN | JSY2421 | 3 | rnr1Y285F_p | TAA | A>G | G/C+1 | A/T-1 | TA>CG | TA>CG | CACCGTAAT | AATATTT | ATT | ATT | T | C | T>C | T>C_ATT | TA>CG_AATATTT | 6 | FALSE | FALSE |
| 135_v.txt | 32929 | 32929-SNV-T | SNV | 1 | T | C | 94 | 29052 | 0.32355776 | 51.0510767 | rnr1Y285F_pG | TRUE | CAN | JSY2421 | 3 | rnr1Y285F_p | GTG | T>C | G/C+1 | A/T-1 | TA>CG | TA>CG | CAAAAGTGA | TGATTGC | CAC | GTG | T | G | T>C | T>C_GTG | TA>CG_TGATTGC | 6 | FALSE | FALSE |
| 135_v.txt | 33189 | 33189-SNV-A | SNV | 1 | A | G | 3 | 19828 | 0.01513012 | 37.999855 | rnr1Y285F_pG | TRUE | CAN | JSY2421 | 3 | rnr1Y285F_p | AAT | A>G | G/C+1 | A/T-1 | TA>CG | TA>CG | ATACCAATA | ATATGTC | TTA | TTA | T | C | T>C | T>C_TTA | TA>CG_ATATGTC | 6 | FALSE | FALSE |
| 135_v.txt | 33192 | 33192-SNV-T | SNV | 1 | T | C | 20 | 19828 | 0.10086746 | 34.05 | rnr1Y285F_pG | TRUE | CAN | JSY2421 | 3 | rnr1Y285F_p | ATG | T>C | G/C+1 | A/T-1 | TA>CG | TA>CG | CCAATATGT | TGTCTTT | TAC | ATG | T | G | T>C | T>C_ATG | TA>CG_TGTCTTT | 6 | FALSE | FALSE |
| 135_v.txt | 33207 | 33207-SNV-T | SNV | 1 | T | C | 14 | 19825 | 0.07061791 | 18.8590713 | rnr1Y285F_pG | TRUE | CAN | JSY2421 | 3 | rnr1Y285F_p | CTC | T>C | G/C+1 | A/T-1 | TA>CG | TA>CG | TTAAGCTCT | TCTCTCT | GAG | CTC | T | G | T>C | T>C_CTC | TA>CG_TCTCTCT | 6 | FALSE | FALSE |
| 135_v.txt | 31974 | 31974-SNV-A | SNV | 1 | A | T | 29 | 23199 | 0.12500539 | 39.7199936 | rnr1Y285F_pG | TRUE | CAN | JSY2421 | 3 | rnr1Y285F_p | CAA | A>T | A/T+1 | A/T-1 | TA>AT | TA>AT | TAAGCCAAG | AAGCCGG | GTT | GTT | T | A | T>A | T>A_GTT | TA>AT_AAGCCGG | 6 | FALSE | FALSE |
| 135_v.txt | 32073 | 32073-SNV-A | SNV | 1 | A | T | 28 | 23199 | 0.12069486 | 61.8214286 | rnr1Y285F_pG | TRUE | CAN | JSY2421 | 3 | rnr1Y285F_p | TAA | A>T | A/T+1 | A/T-1 | TA>AT | TA>AT | ATAAATAAC | AACCATG | ATT | ATT | T | A | T>A | T>A_ATT | TA>AT_AACCATG | 6 | FALSE | FALSE |
| 135_v.txt | 32105 | 32105-SNV-A | SNV | 1 | A | T | 19 | 23199 | 0.08190008 | 61.8947368 | rnr1Y285F_pG | TRUE | CAN | JSY2421 | 3 | rnr1Y285F_p | TAT | A>T | A/T+1 | A/T-1 | TA>AT | TA>AT | AGTGATATT | ATTTAAT | ATA | ATA | T | A | T>A | T>A_ATA | TA>AT_ATTTAAT | 6 | FALSE | FALSE |
| 135_v.txt | 32110 | 32110-SNV-A | SNV | 1 | A | T | 20 | 23199 | 0.08621061 | 60.55 | rnr1Y285F_pG | TRUE | CAN | JSY2421 | 3 | rnr1Y285F_p | AAT | A>T | A/T+1 | A/T-1 | TA>AT | TA>AT | TATTTAATAG | ATAGCCA | TTA | TTA | T | A | T>A | T>A_TTA | TA>AT_ATAGCCA | 6 | FALSE | FALSE |
| 135_v.txt | 32126 | 32126-SNV-T | SNV | 1 | T | A | 8 | 23199 | 0.03448425 | 20.9997889 | rnr1Y285F_pG | TRUE | CAN | JSY2421 | 3 | rnr1Y285F_p | CTT | T>A | A/T+1 | A/T-1 | TA>AT | TA>AT | GAAAACTTT | TTTGTCA | GAA | CTT | T | T | T>A | T>A_CTT | TA>AT_TTTGTCA | 6 | FALSE | FALSE |
| 135_v.txt | 32127 | 32127-SNV-T | SNV | 1 | T | A | 2 | 23199 | 8.62E-03 | 50.9985028 | rnr1Y285F_pG | TRUE | CAN | JSY2421 | 3 | rnr1Y285F_p | TTT | T>A | A/T+1 | A/T-1 | TA>AT | TA>AT | AAAACTTTG | TTGTCAC | AAA | TTT | T | T | T>A | T>A_TTT | TA>AT_TTGTCAC | 6 | FALSE | FALSE |
| 135_v.txt | 32212 | 32212-SNV-T | SNV | 1 | T | A | 17 | 23199 | 0.07327902 | 45.8823529 | rnr1Y285F_pG | TRUE | CAN | JSY2421 | 3 | rnr1Y285F_p | TTG | T>A | A/T+1 | A/T-1 | TA>AT | TA>AT | CACCTTTGGT | TGGTGGT | AAC | TTG | T | T | T>A | T>A_TTG | TA>AT_TGGTGGT | 6 | FALSE | FALSE |
| 135_v.txt | 32303 | 32303-SNV-A | SNV | 1 | A | T | 3 | 33160 | 9.05E-03 | 42.9957521 | rnr1Y285F_pG | TRUE | CAN | JSY2421 | 3 | rnr1Y285F_p | CAG | A>T | A/T+1 | A/T-1 | TA>AT | TA>AT | TGCGGCAGA | AGAAATA | GTC | GTC | T | A | T>A | T>A_GTC | TA>AT_AGAAATA | 6 | FALSE | FALSE |
| 135_v.txt | 32319 | 32319-SNV-A | SNV | 1 | A | T | 124 | 33161 | 0.37393324 | 56.983871 | rnr1Y285F_pG | TRUE | CAN | JSY2421 | 3 | rnr1Y285F_p | TAA | A>T | A/T+1 | A/T-1 | TA>AT | TA>AT | GTTGTTAAG | AAGATAA | ATT | ATT | T | A | T>A | T>A_ATT | TA>AT_AAGATAA | 6 | FALSE | FALSE |
| 135_v.txt | 32346 | 32346-SNV-A | SNV | 1 | A | T | 4 | 33160 | 0.01206273 | 13.7499423 | rnr1Y285F_pG | TRUE | CAN | JSY2421 | 3 | rnr1Y285F_p | CAA | A>T | A/T+1 | A/T-1 | TA>AT | TA>AT | TGTGGCAAA | AAAACCT | GTT | GTT | T | A | T>A | T>A_GTT | TA>AT_AAAACCT | 6 | FALSE | FALSE |
| 135_v.txt | 32408 | 32408-SNV-A | SNV | 1 | A | T | 98 | 33161 | 0.29552788 | 56.9591837 | rnr1Y285F_pG | TRUE | CAN | JSY2421 | 3 | rnr1Y285F_p | TAG | A>T | A/T+1 | A/T-1 | TA>AT | TA>AT | GGAAGTAGA | AGATTGT | ATC | ATC | T | A | T>A | T>A_ATC | TA>AT_AGATTGT | 6 | FALSE | FALSE |
| 135_v.txt | 32473 | 32473-SNV-A | SNV | 1 | A | T | 22 | 33161 | 0.06634299 | 47.6363636 | rnr1Y285F_pG | TRUE | CAN | JSY2421 | 3 | rnr1Y285F_p | TAG | A>T | A/T+1 | A/T-1 | TA>AT | TA>AT | CAATGTAGA | AGAAGGT | ATC | ATC | T | A | T>A | T>A_ATC | TA>AT_AGAAGGT | 6 | FALSE | FALSE |
| 135_v.txt | 32498 | 32498-SNV-T | SNV | 1 | T | A | 4 | 33161 | 0.01206236 | 50.7477794 | rnr1Y285F_pG | TRUE | CAN | JSY2421 | 3 | rnr1Y285F_p | CTT | T>A | A/T+1 | A/T-1 | TA>AT | TA>AT | AACAACTTTT | TTTTTTG | GAA | CTT | T | T | T>A | T>A_CTT | TA>AT_TTTTTTG | 6 | FALSE | FALSE |
| 135_v.txt | 32517 | 32517-SNV-A | SNV | 1 | A | T | 75 | 33160 | 0.22617612 | 41.96 | rnr1Y285F_pG | TRUE | CAN | JSY2421 | 3 | rnr1Y285F_p | AAC | A>T | A/T+1 | A/T-1 | TA>AT | TA>AT | CTTGGAACG | ACGGATT | TTG | TTG | T | A | T>A | T>A_TTG | TA>AT_ACGGATT | 6 | FALSE | FALSE |
| 135_v.txt | 32524 | 32524-SNV-T | SNV | 1 | T | A | 45 | 33160 | 0.13570567 | 42.1555556 | rnr1Y285F_pG | TRUE | CAN | JSY2421 | 3 | rnr1Y285F_p | TTT | T>A | A/T+1 | A/T-1 | TA>AT | TA>AT | CGGATTTTCT | TTCTGGG | AAA | TTT | T | T | T>A | T>A_TTT | TA>AT_TTCTGGG | 6 | FALSE | FALSE |
| 135_v.txt | 32751 | 32751-SNV-A | SNV | 1 | A | T | 4 | 27973 | 0.0142995 | 50.7493135 | rnr1Y285F_pG | TRUE | CAN | JSY2421 | 3 | rnr1Y285F_p | GAT | A>T | A/T+1 | A/T-1 | TA>AT | TA>AT | AACCCGATA | ATAATGG | CTA | CTA | T | A | T>A | T>A_CTA | TA>AT_ATAATGG | 6 | FALSE | FALSE |
| 135_v.txt | 32752 | 32752-SNV-T | SNV | 1 | T | A | 2 | 27973 | 7.15E-03 | 62.9992128 | rnr1Y285F_pG | TRUE | CAN | JSY2421 | 3 | rnr1Y285F_p | ATA | T>A | A/T+1 | A/T-1 | TA>AT | TA>AT | ACCCGATAA | TAATGGC | TAT | ATA | T | T | T>A | T>A_ATA | TA>AT_TAATGGC | 6 | FALSE | FALSE |
| 135_v.txt | 32753 | 32753-SNV-A | SNV | 1 | A | T | 2 | 27973 | 7.15E-03 | 21.0000069 | rnr1Y285F_pG | TRUE | CAN | JSY2421 | 3 | rnr1Y285F_p | TAA | A>T | A/T+1 | A/T-1 | TA>AT | TA>AT | CCCGATAAT | AATGGCT | ATT | ATT | T | A | T>A | T>A_ATT | TA>AT_AATGGCT | 6 | FALSE | FALSE |
| 135_v.txt | 32763 | 32763-SNV-A | SNV | 1 | A | T | 59 | 27973 | 0.21091767 | 45.8474576 | rnr1Y285F_pG | TRUE | CAN | JSY2421 | 3 | rnr1Y285F_p | AAC | A>T | A/T+1 | A/T-1 | TA>AT | TA>AT | GCTAAAACT | ACTTTGA | TTG | TTG | T | A | T>A | T>A_TTG | TA>AT_ACTTTGA | 6 | FALSE | FALSE |
| 135_v.txt | 32787 | 32787-SNV-T | SNV | 1 | T | A | 52 | 27973 | 0.18589354 | 49.4037486 | rnr1Y285F_pG | TRUE | CAN | JSY2421 | 3 | rnr1Y285F_p | CTC | T>A | A/T+1 | A/T-1 | TA>AT | TA>AT | CAGAACTCG | TCGAATT | GAG | CTC | T | T | T>A | T>A_CTC | TA>AT_TCGAATT | 6 | FALSE | FALSE |
| 135_v.txt | 32947 | 32947-SNV-A | SNV | 1 | A | T | 58 | 29052 | 0.19964202 | 58.7758621 | rnr1Y285F_pG | TRUE | CAN | JSY2421 | 3 | rnr1Y285F_p | CAA | A>T | A/T+1 | A/T-1 | TA>AT | TA>AT | AAAACCAAT | AATACAT | GTT | GTT | T | A | T>A | T>A_GTT | TA>AT_AATACAT | 6 | FALSE | FALSE |
| 135_v.txt | 33019 | 33019-SNV-T | SNV | 1 | T | A | 10 | 29054 | 0.03441867 | 55.9 | rnr1Y285F_pG | TRUE | CAN | JSY2421 | 3 | rnr1Y285F_p | GTA | T>A | A/T+1 | A/T-1 | TA>AT | TA>AT | AGGATGTAA | TAACAGG | CAT | GTA | T | T | T>A | T>A_GTA | TA>AT_TAACAGG | 6 | FALSE | FALSE |
| 135_v.txt | 33072 | 33072-SNV-A | SNV | 1 | A | T | 20 | 29051 | 0.06884445 | 49.1 | rnr1Y285F_pG | TRUE | CAN | JSY2421 | 3 | rnr1Y285F_p | CAA | A>T | A/T+1 | A/T-1 | TA>AT | TA>AT | TATGCCAAA | AAAGAAC | GTT | GTT | T | A | T>A | T>A_GTT | TA>AT_AAAGAAC | 6 | FALSE | FALSE |
| 135_v.txt | 33089 | 33089-SNV-A | SNV | 1 | A | T | 53 | 29051 | 0.18243778 | 47.509434 | rnr1Y285F_pG | TRUE | CAN | JSY2421 | 3 | rnr1Y285F_p | AAT | A>T | A/T+1 | A/T-1 | TA>AT | TA>AT | AAATAAATA | ATATGAT | TTA | TTA | T | A | T>A | T>A_TTA | TA>AT_ATATGAT | 6 | FALSE | FALSE |
| 135_v.txt | 33189 | 33189-SNV-A | SNV | 1 | A | T | 3 | 19828 | 0.01513012 | 24.3332311 | rnr1Y285F_pG | TRUE | CAN | JSY2421 | 3 | rnr1Y285F_p | AAT | A>T | A/T+1 | A/T-1 | TA>AT | TA>AT | ATACCAATA | ATATGTC | TTA | TTA | T | A | T>A | T>A_TTA | TA>AT_ATATGTC | 6 | FALSE | FALSE |
| 135_v.txt | 33196 | 33196-SNV-T | SNV | 1 | T | A | 70 | 19827 | 0.35305392 | 36.9 | rnr1Y285F_pG | TRUE | CAN | JSY2421 | 3 | rnr1Y285F_p | CTT | T>A | A/T+1 | A/T-1 | TA>AT | TA>AT | TATGTCTTTG | TTTGCTT | GAA | CTT | T | T | T>A | T>A_CTT | TA>AT_TTTGCTT | 6 | FALSE | FALSE |
| 135_v.txt | 31650 | 31650-SNV-C | SNV | 1 | C | T | 19 | 10907 | 0.17420006 | 35.2105263 | rnr1Y285F_pG | TRUE | CAN | JSY2421 | 3 | rnr1Y285F_p | GCG | C>T | A/T+1 | G/C-1 | CG>TA | CG>TA | AGAATGCGA | CGAAATG | CGC | GCG | C | A | C>T | C>T_GCG | CG>TA_CGAAATG | 6 | FALSE | FALSE |
| 135_v.txt | 31658 | 31658-SNV-C | SNV | 1 | C | T | 19 | 10907 | 0.17420006 | 37.1578947 | rnr1Y285F_pG | TRUE | CAN | JSY2421 | 3 | rnr1Y285F_p | GCG | C>T | A/T+1 | G/C-1 | CG>TA | CG>TA | AAATGGCGT | CGTGGAA | CGC | GCG | C | A | C>T | C>T_GCG | CG>TA_CGTGGAA | 6 | FALSE | FALSE |
| 135_v.txt | 31834 | 31834-SNV-C | SNV | 1 | G | A | 18 | 10907 | 0.16503163 | 50 | rnr1Y285F_pG | TRUE | CAN | JSY2421 | 3 | rnr1Y285F_p | TGA | G>A | A/T+1 | G/C-1 | CG>TA | CG>TA | TGCATTGAA | GAAATAA | ACT | ACT | C | T | C>T | C>T_ACT | CG>TA_GAAATAA | 6 | FALSE | FALSE |
| 135_v.txt | 31844 | 31844-SNV-C | SNV | 1 | C | T | 111 | 10907 | 1.01769506 | 49.1801802 | rnr1Y285F_pG | TRUE | CAN | JSY2421 | 3 | rnr1Y285F_p | TCC | C>T | A/T+1 | G/C-1 | CG>TA | CG>TA | TAAGATCCA | CCAAACA | AGG | TCC | C | A | C>T | C>T_TCC | CG>TA_CCAAACA | 6 | FALSE | FALSE |
| 135_v.txt | 31911 | 31911-SNV-C | SNV | 1 | G | A | 11 | 10906 | 0.10086191 | 33.9090909 | rnr1Y285F_pG | TRUE | CAN | JSY2421 | 3 | rnr1Y285F_p | TGC | G>A | A/T+1 | G/C-1 | CG>TA | CG>TA | TTTGGTGCA | GCAAAAG | ACG | ACG | C | T | C>T | C>T_ACG | CG>TA_GCAAAAG | 6 | FALSE | FALSE |
| 135_v.txt | 31912 | 31912-SNV-C | SNV | 1 | C | T | 5 | 10906 | 0.04584632 | 36.9992338 | rnr1Y285F_pG | TRUE | CAN | JSY2421 | 3 | rnr1Y285F_p | GCA | C>T | A/T+1 | G/C-1 | CG>TA | CG>TA | TTGGTGCAA | CAAAAGC | CGT | GCA | C | A | C>T | C>T_GCA | CG>TA_CAAAAGC | 6 | FALSE | FALSE |

| | | | | | | | | | | | | | | | | | | | | | | | | | | | | | | | | | | |
|---|---|---|---|---|---|---|---|---|---|---|---|---|---|---|---|---|---|---|---|---|---|---|---|---|---|---|---|---|---|---|---|---|---|---|
| 135_v.txt | 31920 | 31920-SNV-( | SNV | 1 | G | A | 7 | 10906 | 0.06418485 | 38 | rnr1Y285F_pG | TRUE | CAN | JSY2421 | 3 | rnr1Y285F_p | CGT | G>A | A/T+1 | G/C-1 | CG>TA | CG>TA | AAAGCCGTG | GTGAAAC | GCA | GCA | C | T | C>T | C>T_GCA | CG>TA_GTGAAAC | 6 | FALSE | FALSE |
| 135_v.txt | 31922 | 31922-SNV-( | SNV | 1 | G | A | 34 | 10906 | 0.311755 | 36.9698074 | rnr1Y285F_pG | TRUE | CAN | JSY2421 | 3 | rnr1Y285F_p | TGA | G>A | A/T+1 | G/C-1 | CG>TA | CG>TA | AGCCGTGAA | GAAACCT | ACT | ACT | C | T | C>T | C>T_ACT | CG>TA_GAAACCT | 6 | FALSE | FALSE |
| 135_v.txt | 31962 | 31962-SNV-( | SNV | 1 | G | A | 46 | 23199 | 0.19828441 | 36.4347826 | rnr1Y285F_pG | TRUE | CAN | JSY2421 | 3 | rnr1Y285F_p | CGC | G>A | A/T+1 | G/C-1 | CG>TA | CG>TA | GTGGCCGCA | GCATAAT | GCG | GCG | C | T | C>T | C>T_GCG | CG>TA_GCATAAT | 6 | FALSE | FALSE |
| 135_v.txt | 31977 | 31977-SNV-( | SNV | 1 | C | T | 21 | 23199 | 0.09052114 | 38.3333333 | rnr1Y285F_pG | TRUE | CAN | JSY2421 | 3 | rnr1Y285F_p | GCC | C>T | A/T+1 | G/C-1 | CG>TA | CG>TA | GCCAAGCCG | CCGGGCA | CGG | GCC | C | A | C>T | C>T_GCC | CG>TA_CCGGGCA | 6 | FALSE | FALSE |
| 135_v.txt | 31980 | 31980-SNV-( | SNV | 1 | G | A | 26 | 23199 | 0.1120738 | 50.2884778 | rnr1Y285F_pG | TRUE | CAN | JSY2421 | 3 | rnr1Y285F_p | GGG | G>A | A/T+1 | G/C-1 | CG>TA | CG>TA | AAGCCGGGC | GGCATTA | CCC | CCC | C | T | C>T | C>T_CCC | CG>TA_GGCATTA | 6 | FALSE | FALSE |
| 135_v.txt | 31981 | 31981-SNV-( | SNV | 1 | G | A | 35 | 23199 | 0.15086857 | 51.5428571 | rnr1Y285F_pG | TRUE | CAN | JSY2421 | 3 | rnr1Y285F_p | GGC | G>A | A/T+1 | G/C-1 | CG>TA | CG>TA | AGCCGGGCA | GCATTAA | CCG | CCG | C | T | C>T | C>T_CCG | CG>TA_GCATTAA | 6 | FALSE | FALSE |
| 135_v.txt | 32041 | 32041-SNV-( | SNV | 1 | G | A | 21 | 23198 | 0.09052505 | 55.952381 | rnr1Y285F_pG | TRUE | CAN | JSY2421 | 3 | rnr1Y285F_p | TGC | G>A | A/T+1 | G/C-1 | CG>TA | CG>TA | AAGCTTGCA | GCATAAA | ACG | ACG | C | T | C>T | C>T_ACG | CG>TA_GCATAAA | 6 | FALSE | FALSE |
| 135_v.txt | 32042 | 32042-SNV-( | SNV | 1 | C | T | 12 | 23196 | 0.05173306 | 52.8318025 | rnr1Y285F_pG | TRUE | CAN | JSY2421 | 3 | rnr1Y285F_p | GCA | C>T | A/T+1 | G/C-1 | CG>TA | CG>TA | AGCTTGCAT | CATAAAT | CGT | GCA | C | A | C>T | C>T_GCA | CG>TA_CATAAAT | 6 | FALSE | FALSE |
| 135_v.txt | 32075 | 32075-SNV-( | SNV | 1 | C | T | 296 | 23199 | 1.27591707 | 61.9797297 | rnr1Y285F_pG | TRUE | CAN | JSY2421 | 3 | rnr1Y285F_p | ACC | C>T | A/T+1 | G/C-1 | CG>TA | CG>TA | AAATAACCA | CCATGCA | TGG | ACC | C | A | C>T | C>T_ACC | CG>TA_CCATGCA | 6 | FALSE | FALSE |
| 135_v.txt | 32088 | 32088-SNV-( | SNV | 1 | C | T | 323 | 23198 | 1.39236141 | 61.407111 | rnr1Y285F_pG | TRUE | CAN | JSY2421 | 3 | rnr1Y285F_p | GCC | C>T | A/T+1 | G/C-1 | CG>TA | CG>TA | AAAAAGCCT | CCTGCAA | CGG | GCC | C | A | C>T | C>T_GCC | CG>TA_CCTGCAA | 6 | FALSE | FALSE |
| 135_v.txt | 32092 | 32092-SNV-( | SNV | 1 | C | T | 32 | 23199 | 0.13793698 | 60.375 | rnr1Y285F_pG | TRUE | CAN | JSY2421 | 3 | rnr1Y285F_p | GCA | C>T | A/T+1 | G/C-1 | CG>TA | CG>TA | AGCCTGCAA | CAACACC | CGT | GCA | C | A | C>T | C>T_GCA | CG>TA_CAACACC | 6 | FALSE | FALSE |
| 135_v.txt | 32114 | 32114-SNV-( | SNV | 1 | C | T | 303 | 23197 | 1.30620339 | 59.7917506 | rnr1Y285F_pG | TRUE | CAN | JSY2421 | 3 | rnr1Y285F_p | GCC | C>T | A/T+1 | G/C-1 | CG>TA | CG>TA | TAATAGCCA | CCATTCG | CGG | GCC | C | A | C>T | C>T_GCC | CG>TA_CCATTCG | 6 | FALSE | FALSE |
| 135_v.txt | 32165 | 32165-SNV-( | SNV | 1 | G | A | 28 | 23199 | 0.12069486 | 48.8928571 | rnr1Y285F_pG | TRUE | CAN | JSY2421 | 3 | rnr1Y285F_p | CGC | G>A | A/T+1 | G/C-1 | CG>TA | CG>TA | CAAAGCGCC | GCCAAAT | GCG | GCG | C | T | C>T | C>T_GCG | CG>TA_GCCAAAT | 6 | FALSE | FALSE |
| 135_v.txt | 32271 | 32271-SNV-( | SNV | 1 | C | T | 86 | 33139 | 0.25951296 | 33.1277046 | rnr1Y285F_pG | TRUE | CAN | JSY2421 | 3 | rnr1Y285F_p | ACG | C>T | A/T+1 | G/C-1 | CG>TA | CG>TA | AAAATACGG | CGGGAAC | TGC | ACG | C | A | C>T | C>T_ACG | CG>TA_CGGGAAC | 6 | FALSE | FALSE |
| 135_v.txt | 32272 | 32272-SNV-( | SNV | 1 | G | A | 68 | 33139 | 0.20519629 | 35.9264706 | rnr1Y285F_pG | TRUE | CAN | JSY2421 | 3 | rnr1Y285F_p | CGG | G>A | A/T+1 | G/C-1 | CG>TA | CG>TA | AAATACGGG | GGGAACC | GCC | GCC | C | T | C>T | C>T_GCC | CG>TA_GGGAACC | 6 | FALSE | FALSE |
| 135_v.txt | 32299 | 32299-SNV-( | SNV | 1 | C | T | 48 | 33161 | 0.14474835 | 52.9583333 | rnr1Y285F_pG | TRUE | CAN | JSY2421 | 3 | rnr1Y285F_p | GCG | C>T | A/T+1 | G/C-1 | CG>TA | CG>TA | AATTTGCGG | CGGCAGA | CGC | GCG | C | A | C>T | C>T_GCG | CG>TA_CGGCAGA | 6 | FALSE | FALSE |
| 135_v.txt | 32301 | 32301-SNV-( | SNV | 1 | G | A | 16 | 33161 | 0.04824945 | 46.6873694 | rnr1Y285F_pG | TRUE | CAN | JSY2421 | 3 | rnr1Y285F_p | GGC | G>A | A/T+1 | G/C-1 | CG>TA | CG>TA | TTTGCGGCA | GCAGAAA | CCG | CCG | C | T | C>T | C>T_CCG | CG>TA_GCAGAAA | 6 | FALSE | FALSE |
| 135_v.txt | 32302 | 32302-SNV-( | SNV | 1 | C | T | 16 | 33161 | 0.04824945 | 54.4992281 | rnr1Y285F_pG | TRUE | CAN | JSY2421 | 3 | rnr1Y285F_p | GCA | C>T | A/T+1 | G/C-1 | CG>TA | CG>TA | TTGCGGCAG | CAGAAAT | CGT | GCA | C | A | C>T | C>T_GCA | CG>TA_CAGAAAT | 6 | FALSE | FALSE |
| 135_v.txt | 32304 | 32304-SNV-( | SNV | 1 | G | A | 10 | 33161 | 0.03015591 | 54.7999015 | rnr1Y285F_pG | TRUE | CAN | JSY2421 | 3 | rnr1Y285F_p | AGA | G>A | A/T+1 | G/C-1 | CG>TA | CG>TA | GCGGCAGAA | GAAATAA | TCT | TCT | C | T | C>T | C>T_TCT | CG>TA_GAAATAA | 6 | FALSE | FALSE |
| 135_v.txt | 32313 | 32313-SNV-( | SNV | 1 | G | A | 65 | 33161 | 0.19601339 | 58.0307692 | rnr1Y285F_pG | TRUE | CAN | JSY2421 | 3 | rnr1Y285F_p | GGT | G>A | A/T+1 | G/C-1 | CG>TA | CG>TA | ATAATGGTT | GTTGTTA | CCA | CCA | C | T | C>T | C>T_CCA | CG>TA_GTTGTTA | 6 | FALSE | FALSE |
| 135_v.txt | 32345 | 32345-SNV-( | SNV | 1 | C | T | 11 | 33161 | 0.0331715 | 58.8163839 | rnr1Y285F_pG | TRUE | CAN | JSY2421 | 3 | rnr1Y285F_p | GCA | C>T | A/T+1 | G/C-1 | CG>TA | CG>TA | ATGTGGCAA | CAAAACC | CGT | GCA | C | A | C>T | C>T_GCA | CG>TA_CAAAACC | 6 | FALSE | FALSE |
| 135_v.txt | 32413 | 32413-SNV-( | SNV | 1 | G | A | 1133 | 33161 | 3.41666415 | 58.7714034 | rnr1Y285F_pG | TRUE | CAN | JSY2421 | 3 | rnr1Y285F_p | TGT | G>A | A/T+1 | G/C-1 | CG>TA | CG>TA | TAGATTGTG | GTGTTAG | ACA | ACA | C | T | C>T | C>T_ACA | CG>TA_GTGTTAG | 6 | FALSE | FALSE |
| 135_v.txt | 32436 | 32436-SNV-( | SNV | 1 | G | A | 45 | 33161 | 0.13570158 | 47.3111111 | rnr1Y285F_pG | TRUE | CAN | JSY2421 | 3 | rnr1Y285F_p | TGG | G>A | A/T+1 | G/C-1 | CG>TA | CG>TA | TTGTATGGA | GGAACTA | ACC | ACC | C | T | C>T | C>T_ACC | CG>TA_GGAACTA | 6 | FALSE | FALSE |
| 135_v.txt | 32448 | 32448-SNV-( | SNV | 1 | C | T | 94 | 33161 | 0.28346552 | 52.4660314 | rnr1Y285F_pG | TRUE | CAN | JSY2421 | 3 | rnr1Y285F_p | TCC | C>T | A/T+1 | G/C-1 | CG>TA | CG>TA | AAAAGTCCA | CCAATGA | AGG | TCC | C | A | C>T | C>T_TCC | CG>TA_CCAATGA | 6 | FALSE | FALSE |
| 135_v.txt | 32449 | 32449-SNV-( | SNV | 1 | C | T | 87 | 33161 | 0.26235638 | 49.0804598 | rnr1Y285F_pG | TRUE | CAN | JSY2421 | 3 | rnr1Y285F_p | CCA | C>T | A/T+1 | G/C-1 | CG>TA | CG>TA | AAAGTCCAA | CAATGAA | GGT | CCA | C | A | C>T | C>T_CCA | CG>TA_CAATGAA | 6 | FALSE | FALSE |
| 135_v.txt | 32466 | 32466-SNV-( | SNV | 1 | C | T | 42 | 33161 | 0.12665481 | 48.1666667 | rnr1Y285F_pG | TRUE | CAN | JSY2421 | 3 | rnr1Y285F_p | GCC | C>T | A/T+1 | G/C-1 | CG>TA | CG>TA | AGAGAGCCA | CCAATGT | CGG | GCC | C | A | C>T | C>T_GCC | CG>TA_CCAATGT | 6 | FALSE | FALSE |
| 135_v.txt | 32487 | 32487-SNV-( | SNV | 1 | C | T | 51 | 33161 | 0.15379512 | 46.1176471 | rnr1Y285F_pG | TRUE | CAN | JSY2421 | 3 | rnr1Y285F_p | ACG | C>T | A/T+1 | G/C-1 | CG>TA | CG>TA | AAGATACGG | CGGAAAA | TGC | ACG | C | A | C>T | C>T_ACG | CG>TA_CGGAAAA | 6 | FALSE | FALSE |
| 135_v.txt | 32529 | 32529-SNV-( | SNV | 1 | G | A | 49 | 33160 | 0.1477684 | 38.6335814 | rnr1Y285F_pG | TRUE | CAN | JSY2421 | 3 | rnr1Y285F_p | GGG | G>A | A/T+1 | G/C-1 | CG>TA | CG>TA | TTTCTGGGG | GGGTTTG | CCC | CCC | C | T | C>T | C>T_CCC | CG>TA_GGGTTTG | 6 | FALSE | FALSE |
| 135_v.txt | 32530 | 32530-SNV-( | SNV | 1 | G | A | 30 | 33160 | 0.09047045 | 37.9666667 | rnr1Y285F_pG | TRUE | CAN | JSY2421 | 3 | rnr1Y285F_p | GGG | G>A | A/T+1 | G/C-1 | CG>TA | CG>TA | TTCTGGGGT | GGTTTGC | CCC | CCC | C | T | C>T | C>T_CCC | CG>TA_GGTTTGC | 6 | FALSE | FALSE |
| 135_v.txt | 32535 | 32535-SNV-( | SNV | 1 | G | A | 38 | 33160 | 0.1145959 | 37.7890554 | rnr1Y285F_pG | TRUE | CAN | JSY2421 | 3 | rnr1Y285F_p | TGC | G>A | A/T+1 | G/C-1 | CG>TA | CG>TA | GGGTTTGCA | GCAGCTT | ACG | ACG | C | T | C>T | C>T_ACG | CG>TA_GCAGCTT | 6 | FALSE | FALSE |
| 135_v.txt | 32536 | 32536-SNV-( | SNV | 1 | C | T | 60 | 33160 | 0.18094089 | 36.8158274 | rnr1Y285F_pG | TRUE | CAN | JSY2421 | 3 | rnr1Y285F_p | GCA | C>T | A/T+1 | G/C-1 | CG>TA | CG>TA | GGTTTGCAG | CAGCTTC | CGT | GCA | C | A | C>T | C>T_GCA | CG>TA_CAGCTTC | 6 | FALSE | FALSE |
| 135_v.txt | 32545 | 32545-SNV-( | SNV | 1 | C | T | 82 | 33159 | 0.24729334 | 36.8658537 | rnr1Y285F_pG | TRUE | CAN | JSY2421 | 3 | rnr1Y285F_p | CCA | C>T | A/T+1 | G/C-1 | CG>TA | CG>TA | CTTCACCAG | CAGCAGT | GGT | CCA | C | A | C>T | C>T_CCA | CG>TA_CAGCAGT | 6 | FALSE | FALSE |
| 135_v.txt | 32566 | 32566-SNV-( | SNV | 1 | C | T | 20 | 27962 | 0.07152564 | 37.15 | rnr1Y285F_pG | TRUE | CAN | JSY2421 | 3 | rnr1Y285F_p | TCA | C>T | A/T+1 | G/C-1 | CG>TA | CG>TA | CTAGTTCAG | CAGTACC | AGT | TCA | C | A | C>T | C>T_TCA | CG>TA_CAGTACC | 6 | FALSE | FALSE |
| 135_v.txt | 32571 | 32571-SNV-( | SNV | 1 | C | T | 61 | 27974 | 0.21805963 | 36.6126505 | rnr1Y285F_pG | TRUE | CAN | JSY2421 | 3 | rnr1Y285F_p | ACC | C>T | A/T+1 | G/C-1 | CG>TA | CG>TA | TCAGTACCT | CCTTGAA | TGG | ACC | C | A | C>T | C>T_ACC | CG>TA_CCTTGAA | 6 | FALSE | FALSE |
| 135_v.txt | 32572 | 32572-SNV-( | SNV | 1 | C | T | 65 | 27974 | 0.23235862 | 34.5516246 | rnr1Y285F_pG | TRUE | CAN | JSY2421 | 3 | rnr1Y285F_p | CCT | C>T | A/T+1 | G/C-1 | CG>TA | CG>TA | CAGTACCTT | CTTGAAA | GGA | CCT | C | A | C>T | C>T_CCT | CG>TA_CTTGAAA | 6 | FALSE | FALSE |
| 135_v.txt | 32590 | 32590-SNV-( | SNV | 1 | C | T | 46 | 27973 | 0.16444429 | 37.4565217 | rnr1Y285F_pG | TRUE | CAN | JSY2421 | 3 | rnr1Y285F_p | GCG | C>T | A/T+1 | G/C-1 | CG>TA | CG>TA | AGGCAGCGT | CGTTAAT | CGC | GCG | C | A | C>T | C>T_GCG | CG>TA_CGTTAAT | 6 | FALSE | FALSE |
| 135_v.txt | 32610 | 32610-SNV-( | SNV | 1 | C | T | 26 | 27973 | 0.09294677 | 51.6538462 | rnr1Y285F_pG | TRUE | CAN | JSY2421 | 3 | rnr1Y285F_p | CCA | C>T | A/T+1 | G/C-1 | CG>TA | CG>TA | GAAACCCAA | CAACCTA | GGT | CCA | C | A | C>T | C>T_CCA | CG>TA_CAACCTA | 6 | FALSE | FALSE |
| 135_v.txt | 32643 | 32643-SNV-( | SNV | 1 | G | A | 5 | 27973 | 0.01787438 | 36.9998338 | rnr1Y285F_pG | TRUE | CAN | JSY2421 | 3 | rnr1Y285F_p | AGA | G>A | A/T+1 | G/C-1 | CG>TA | CG>TA | TCCTTAGAT | GATATTA | TCT | TCT | C | T | C>T | C>T_TCT | CG>TA_GATATTA | 6 | FALSE | FALSE |
| 135_v.txt | 32658 | 32658-SNV-( | SNV | 1 | C | T | 67 | 27973 | 0.23951668 | 55.5522388 | rnr1Y285F_pG | TRUE | CAN | JSY2421 | 3 | rnr1Y285F_p | ACC | C>T | A/T+1 | G/C-1 | CG>TA | CG>TA | CCTGGACCC | CCCCAGG | TGG | ACC | C | A | C>T | C>T_ACC | CG>TA_CCCCAGG | 6 | FALSE | FALSE |
| 135_v.txt | 32678 | 32678-SNV-( | SNV | 1 | C | T | 485 | 27972 | 1.73387673 | 60.0942329 | rnr1Y285F_pG | TRUE | CAN | JSY2421 | 3 | rnr1Y285F_p | TCC | C>T | A/T+1 | G/C-1 | CG>TA | CG>TA | GTTTCTCCAA | CCAATAA | AGG | TCC | C | A | C>T | C>T_TCC | CG>TA_CCAATAA | 6 | FALSE | FALSE |
| 135_v.txt | 32679 | 32679-SNV-( | SNV | 1 | C | T | 145 | 27971 | 0.51839405 | 59.8166661 | rnr1Y285F_pG | TRUE | CAN | JSY2421 | 3 | rnr1Y285F_p | CCA | C>T | A/T+1 | G/C-1 | CG>TA | CG>TA | TTTCTCCAAT | CAATAAC | GGT | CCA | C | A | C>T | C>T_CCA | CG>TA_CAATAAC | 6 | FALSE | FALSE |
| 135_v.txt | 32685 | 32685-SNV-( | SNV | 1 | C | T | 8 | 27973 | 0.02859901 | 54.9991578 | rnr1Y285F_pG | TRUE | CAN | JSY2421 | 3 | rnr1Y285F_p | ACG | C>T | A/T+1 | G/C-1 | CG>TA | CG>TA | CAATAACGG | CGGAATC | TGC | ACG | C | A | C>T | C>T_ACG | CG>TA_CGGAATC | 6 | FALSE | FALSE |
| 135_v.txt | 32686 | 32686-SNV-( | SNV | 1 | G | A | 11 | 27973 | 0.03932363 | 45.0770326 | rnr1Y285F_pG | TRUE | CAN | JSY2421 | 3 | rnr1Y285F_p | CGG | G>A | A/T+1 | G/C-1 | CG>TA | CG>TA | AATAACGGA | GGAATCC | GCC | GCC | C | T | C>T | C>T_GCC | CG>TA_GGAATCC | 6 | FALSE | FALSE |
| 135_v.txt | 32700 | 32700-SNV-( | SNV | 1 | C | T | 29 | 27973 | 0.1036714 | 58.137931 | rnr1Y285F_pG | TRUE | CAN | JSY2421 | 3 | rnr1Y285F_p | GCC | C>T | A/T+1 | G/C-1 | CG>TA | CG>TA | ACTGGGCCG | CCGGTAA | CGG | GCC | C | A | C>T | C>T_GCC | CG>TA_CCGGTAA | 6 | FALSE | FALSE |
| 135_v.txt | 32757 | 32757-SNV-( | SNV | 1 | G | A | 23 | 27973 | 0.08222214 | 55.5217391 | rnr1Y285F_pG | TRUE | CAN | JSY2421 | 3 | rnr1Y285F_p | GGC | G>A | A/T+1 | G/C-1 | CG>TA | CG>TA | ATAATGGCT | GCTAAAA | CCG | CCG | C | T | C>T | C>T_CCG | CG>TA_GCTAAAA | 6 | FALSE | FALSE |
| 135_v.txt | 32788 | 32788-SNV-( | SNV | 1 | C | T | 54 | 27973 | 0.19304329 | 43.4214695 | rnr1Y285F_pG | TRUE | CAN | JSY2421 | 3 | rnr1Y285F_p | TCG | C>T | A/T+1 | G/C-1 | CG>TA | CG>TA | AGAACTCGA | CGAATTC | AGC | TCG | C | A | C>T | C>T_TCG | CG>TA_CGAATTC | 6 | FALSE | FALSE |
| 135_v.txt | 32798 | 32798-SNV-( | SNV | 1 | G | A | 50 | 27973 | 0.17874379 | 40.919976 | rnr1Y285F_pG | TRUE | CAN | JSY2421 | 3 | rnr1Y285F_p | CGT | G>A | A/T+1 | G/C-1 | CG>TA | CG>TA | TTCACCGTA | GTAATAT | GCA | GCA | C | T | C>T | C>T_GCA | CG>TA_GTAATAT | 6 | FALSE | FALSE |
| 135_v.txt | 32809 | 32809-SNV-( | SNV | 1 | C | T | 17 | 27973 | 0.06077289 | 48.2941177 | rnr1Y285F_pG | TRUE | CAN | JSY2421 | 3 | rnr1Y285F_p | ACA | C>T | A/T+1 | G/C-1 | CG>TA | CG>TA | ATTTGACAG | CAGGGAA | TGT | ACA | C | A | C>T | C>T_ACA | CG>TA_CAGGGAA | 6 | FALSE | FALSE |
| 135_v.txt | 32840 | 32840-SNV-( | SNV | 1 | C | T | 37 | 27973 | 0.1322704 | 38.7153917 | rnr1Y285F_pG | TRUE | CAN | JSY2421 | 3 | rnr1Y285F_p | CCC | C>T | A/T+1 | G/C-1 | CG>TA | CG>TA | AATTACCCA | CCAAAAA | GGG | CCC | C | A | C>T | C>T_CCC | CG>TA_CCAAAAA | 6 | FALSE | FALSE |
| 135_v.txt | 32841 | 32841-SNV-( | SNV | 1 | C | T | 445 | 27973 | 1.59081972 | 42.1469613 | rnr1Y285F_pG | TRUE | CAN | JSY2421 | 3 | rnr1Y285F_p | CCA | C>T | A/T+1 | G/C-1 | CG>TA | CG>TA | ATTACCCAA | CAAAAAA | GGT | CCA | C | A | C>T | C>T_CCA | CG>TA_CAAAAAA | 6 | FALSE | FALSE |
| 135_v.txt | 32855 | 32855-SNV-( | SNV | 1 | C | T | 173 | 27972 | 0.61847562 | 36.6473988 | rnr1Y285F_pG | TRUE | CAN | JSY2421 | 3 | rnr1Y285F_p | TCC | C>T | A/T+1 | G/C-1 | CG>TA | CG>TA | ACTAATCCAT | CCATGCC | AGG | TCC | C | A | C>T | C>T_TCC | CG>TA_CCATGCC | 6 | FALSE | FALSE |
| 135_v.txt | 32860 | 32860-SNV-( | SNV | 1 | C | T | 30 | 27972 | 0.10725011 | 37.1666667 | rnr1Y285F_pG | TRUE | CAN | JSY2421 | 3 | rnr1Y285F_p | GCC | C>T | A/T+1 | G/C-1 | CG>TA | CG>TA | TCCATGCCG | CCGCCAG | CGG | GCC | C | A | C>T | C>T_GCC | CG>TA_CCGCCAG | 6 | FALSE | FALSE |
| 135_v.txt | 32899 | 32899-SNV-( | SNV | 1 | G | A | 40 | 29052 | 0.13768415 | 37.2378106 | rnr1Y285F_pG | TRUE | CAN | JSY2421 | 3 | rnr1Y285F_p | TGG | G>A | A/T+1 | G/C-1 | CG>TA | CG>TA | TGACTTGGC | GGCCAAC | ACC | ACC | C | T | C>T | C>T_ACC | CG>TA_GGCCAAC | 6 | FALSE | FALSE |
| 135_v.txt | 32900 | 32900-SNV-( | SNV | 1 | G | A | 26 | 29052 | 0.0894947 | 34.9596766 | rnr1Y285F_pG | TRUE | CAN | JSY2421 | 3 | rnr1Y285F_p | GGC | G>A | A/T+1 | G/C-1 | CG>TA | CG>TA | GACTTGGCC | GCCAACT | CCG | CCG | C | T | C>T | C>T_CCG | CG>TA_GCCAACT | 6 | FALSE | FALSE |
| 135_v.txt | 32901 | 32901-SNV-( | SNV | 1 | C | T | 61 | 29052 | 0.20996833 | 32.9833977 | rnr1Y285F_pG | TRUE | CAN | JSY2421 | 3 | rnr1Y285F_p | GCC | C>T | A/T+1 | G/C-1 | CG>TA | CG>TA | ACTTGGCCA | CCAACTA | CGG | GCC | C | A | C>T | C>T_GCC | CG>TA_CCAACTA | 6 | FALSE | FALSE |
| 135_v.txt | 32917 | 32917-SNV-( | SNV | 1 | C | T | 74 | 29052 | 0.25471568 | 46.0810811 | rnr1Y285F_pG | TRUE | CAN | JSY2421 | 3 | rnr1Y285F_p | TCC | C>T | A/T+1 | G/C-1 | CG>TA | CG>TA | TAAGTTCCA | CCAGGGC | AGG | TCC | C | A | C>T | C>T_TCC | CG>TA_CCAGGGC | 6 | FALSE | FALSE |
| 135_v.txt | 32922 | 32922-SNV-( | SNV | 1 | G | A | 29 | 29052 | 0.09982101 | 50.6206897 | rnr1Y285F_pG | TRUE | CAN | JSY2421 | 3 | rnr1Y285F_p | GGC | G>A | A/T+1 | G/C-1 | CG>TA | CG>TA | TCCAGGGCA | GCAAAAG | CCG | CCG | C | T | C>T | C>T_CCG | CG>TA_GCAAAAG | 6 | FALSE | FALSE |
| 135_v.txt | 32935 | 32935-SNV-( | SNV | 1 | C | T | 32 | 29052 | 0.11014732 | 33.7509765 | rnr1Y285F_pG | TRUE | CAN | JSY2421 | 3 | rnr1Y285F_p | GCC | C>T | A/T+1 | G/C-1 | CG>TA | CG>TA | TGATTGCCC | CCCAAGA | CGG | GCC | C | A | C>T | C>T_GCC | CG>TA_CCCAAGA | 6 | FALSE | FALSE |
| 135_v.txt | 32936 | 32936-SNV-( | SNV | 1 | C | T | 33 | 29052 | 0.11358943 | 33.538794 | rnr1Y285F_pG | TRUE | CAN | JSY2421 | 3 | rnr1Y285F_p | CCC | C>T | A/T+1 | G/C-1 | CG>TA | CG>TA | GATTGCCCA | CCAAGAA | GGG | CCC | C | A | C>T | C>T_CCC | CG>TA_CCAAGAA | 6 | FALSE | FALSE |
| 135_v.txt | 32940 | 32940-SNV-( | SNV | 1 | G | A | 38 | 29052 | 0.13079995 | 50.4210526 | rnr1Y285F_pG | TRUE | CAN | JSY2421 | 3 | rnr1Y285F_p | AGA | G>A | A/T+1 | G/C-1 | CG>TA | CG>TA | GCCCAAGAA | GAAAACC | TCT | TCT | C | T | C>T | C>T_TCT | CG>TA_GAAAACC | 6 | FALSE | FALSE |
| 135_v.txt | 32945 | 32945-SNV-( | SNV | 1 | C | T | 539 | 29052 | 1.85529396 | 57.4972171 | rnr1Y285F_pG | TRUE | CAN | JSY2421 | 3 | rnr1Y285F_p | ACC | C>T | A/T+1 | G/C-1 | CG>TA | CG>TA | AGAAAACCA | CCAATAC | TGG | ACC | C | A | C>T | C>T_ACC | CG>TA_CCAATAC | 6 | FALSE | FALSE |
| 135_v.txt | 32958 | 32958-SNV-( | SNV | 1 | C | T | 91 | 29052 | 0.31323145 | 52.3736264 | rnr1Y285F_pG | TRUE | CAN | JSY2421 | 3 | rnr1Y285F_p | ACC | C>T | A/T+1 | G/C-1 | CG>TA | CG>TA | ATGTAACCA | CCATTGG | TGG | ACC | C | A | C>T | C>T_ACC | CG>TA_CCATTGG | 6 | FALSE | FALSE |
| 135_v.txt | 32965 | 32965-SNV-( | SNV | 1 | C | T | 26 | 29052 | 0.0894947 | 58.0769231 | rnr1Y285F_pG | TRUE | CAN | JSY2421 | 3 | rnr1Y285F_p | GCC | C>T | A/T+1 | G/C-1 | CG>TA | CG>TA | CATTGGCCG | CCGCACC | CGG | GCC | C | A | C>T | C>T_GCC | CG>TA_CCGCACC | 6 | FALSE | FALSE |
| 135_v.txt | 32967 | 32967-SNV-( | SNV | 1 | G | A | 64 | 29052 | 0.22029464 | 58.859375 | rnr1Y285F_pG | TRUE | CAN | JSY2421 | 3 | rnr1Y285F_p | CGC | G>A | A/T+1 | G/C-1 | CG>TA | CG>TA | TTGGCCGCA | GCACCAA | GCG | GCG | C | T | C>T | C>T_GCG | CG>TA_GCACCAA | 6 | FALSE | FALSE |
| 135_v.txt | 33015 | 33015-SNV-( | SNV | 1 | G | A | 46 | 29054 | 0.15832588 | 55.1086957 | rnr1Y285F_pG | TRUE | CAN | JSY2421 | 3 | rnr1Y285F_p | GGA | G>A | A/T+1 | G/C-1 | CG>TA | CG>TA | AAAGAGGAT | GATGTAA | CCT | CCT | C | T | C>T | C>T_CCT | CG>TA_GATGTAA | 6 | FALSE | FALSE |
| 135_v.txt | 33043 | 33043-SNV-( | SNV | 1 | C | T | 29 | 29054 | 0.09981414 | 59.9310345 | rnr1Y285F_pG | TRUE | CAN | JSY2421 | 3 | rnr1Y285F_p | TCA | C>T | A/T+1 | G/C-1 | CG>TA | CG>TA | CCATTTCACC | CACCCAA | AGT | TCA | C | A | C>T | C>T_TCA | CG>TA_CACCCAA | 6 | FALSE | FALSE |
| 135_v.txt | 33056 | 33056-SNV-( | SNV | 1 | C | T | 38 | 29054 | 0.13079094 | 53.2368421 | rnr1Y285F_pG | TRUE | CAN | JSY2421 | 3 | rnr1Y285F_p | GCG | C>T | A/T+1 | G/C-1 | CG>TA | CG>TA | GGACTGCGT | CGTGACA | CGC | GCG | C | A | C>T | C>T_GCG | CG>TA_CGTGACA | 6 | FALSE | FALSE |
| 135_v.txt | 33069 | 33069-SNV-( | SNV | 1 | G | A | 32 | 29051 | 0.11015111 | 55.8125 | rnr1Y285F_pG | TRUE | CAN | JSY2421 | 3 | rnr1Y285F_p | TGC | G>A | A/T+1 | G/C-1 | CG>TA | CG>TA | GAATATGCC | GCCAAAG | ACG | ACG | C | T | C>T | C>T_ACG | CG>TA_GCCAAAG | 6 | FALSE | FALSE |
| 135_v.txt | 33070 | 33070-SNV-( | SNV | 1 | C | T | 23 | 29051 | 0.07917111 | 54.1301459 | rnr1Y285F_pG | TRUE | CAN | JSY2421 | 3 | rnr1Y285F_p | GCC | C>T | A/T+1 | G/C-1 | CG>TA | CG>TA | AATATGCCA | CCAAAGA | CGG | GCC | C | A | C>T | C>T_GCC | CG>TA_CCAAAGA | 6 | FALSE | FALSE |
| 135_v.txt | 33114 | 33114-SNV-( | SNV | 1 | C | T | 45 | 29051 | 0.1549 | 42.5060658 | rnr1Y285F_pG | TRUE | CAN | JSY2421 | 3 | rnr1Y285F_p | GCC | C>T | A/T+1 | G/C-1 | CG>TA | CG>TA | ACTGGGCCG | CCGGCGT | CGG | GCC | C | A | C>T | C>T_GCC | CG>TA_CCGGCGT | 6 | FALSE | FALSE |
| 135_v.txt | 33115 | 33115-SNV-( | SNV | 1 | C | T | 93 | 29051 | 0.32012667 | 45.0000808 | rnr1Y285F_pG | TRUE | CAN | JSY2421 | 3 | rnr1Y285F_p | CCG | C>T | A/T+1 | G/C-1 | CG>TA | CG>TA | CTGGGCCGG | CGGCGTT | GGC | CCG | C | A | C>T | C>T_CCG | CG>TA_CGGCGTT | 6 | FALSE | FALSE |
| 135_v.txt | 33153 | 33153-SNV-( | SNV | 1 | C | T | 221 | 29051 | 0.76073113 | 36.7149321 | rnr1Y285F_pG | TRUE | CAN | JSY2421 | 3 | rnr1Y285F_p | ACC | C>T | A/T+1 | G/C-1 | CG>TA | CG>TA | AAAAGACCT | CCTGTAC | TGG | ACC | C | A | C>T | C>T_ACC | CG>TA_CCTGTAC | 6 | FALSE | FALSE |
| 135_v.txt | 33159 | 33159-SNV-( | SNV | 1 | C | T | 152 | 29050 | 0.5232358 | 36.5197368 | rnr1Y285F_pG | TRUE | CAN | JSY2421 | 3 | rnr1Y285F_p | ACC | C>T | A/T+1 | G/C-1 | CG>TA | CG>TA | CCTGTACCA | CCAATAG | TGG | ACC | C | A | C>T | C>T_ACC | CG>TA_CCAATAG | 6 | FALSE | FALSE |
| 135_v.txt | 33169 | 33169-SNV-( | SNV | 1 | C | T | 21 | 29049 | 0.07229165 | 36.2857143 | rnr1Y285F_pG | TRUE | CAN | JSY2421 | 3 | rnr1Y285F_p | CCA | C>T | A/T+1 | G/C-1 | CG>TA | CG>TA | TAGTACCAC | CACCAAG | GGT | CCA | C | A | C>T | C>T_CCA | CG>TA_CACCAAG | 6 | FALSE | FALSE |
| 135_v.txt | 33171 | 33171-SNV-( | SNV | 1 | C | T | 116 | 29049 | 0.39932528 | 36.7844828 | rnr1Y285F_pG | TRUE | CAN | JSY2421 | 3 | rnr1Y285F_p | ACC | C>T | A/T+1 | G/C-1 | CG>TA | CG>TA | GTACCACCA | CCAAGGG | TGG | ACC | C | A | C>T | C>T_ACC | CG>TA_CCAAGGG | 6 | FALSE | FALSE |
| 135_v.txt | 33177 | 33177-SNV-( | SNV | 1 | G | A | 18 | 29044 | 0.06197494 | 36.4998835 | rnr1Y285F_pG | TRUE | CAN | JSY2421 | 3 | rnr1Y285F_p | GGC | G>A | A/T+1 | G/C-1 | CG>TA | CG>TA | CCAAGGGCA | GCAATCA | CCG | CCG | C | T | C>T | C>T_CCG | CG>TA_GCAATCA | 6 | FALSE | FALSE |
| 135_v.txt | 33178 | 33178-SNV-( | SNV | 1 | C | T | 10 | 29043 | 0.03443171 | 34.6998465 | rnr1Y285F_pG | TRUE | CAN | JSY2421 | 3 | rnr1Y285F_p | GCA | C>T | A/T+1 | G/C-1 | CG>TA | CG>TA | CAAGGGCAA | CAATCAT | CGT | GCA | C | A | C>T | C>T_GCA | CG>TA_CAATCAT | 6 | FALSE | FALSE |
| 135_v.txt | 33193 | 33193-SNV-( | SNV | 1 | G | A | 92 | 19828 | 0.46399032 | 35.8817648 | rnr1Y285F_pG | TRUE | CAN | JSY2421 | 3 | rnr1Y285F_p | TGT | G>A | A/T+1 | G/C-1 | CG>TA | CG>TA | CAATATGTC | GTCTTTG | ACA | ACA | C | T | C>T | C>T_ACA | CG>TA_GTCTTTG | 6 | FALSE | FALSE |
| 135_v.txt | 33258 | 33258-SNV-( | SNV | 1 | C | T | 39 | 19836 | 0.19661222 | 42.8461539 | rnr1Y285F_pG | TRUE | CAN | JSY2421 | 3 | rnr1Y285F_p | GCC | C>T | A/T+1 | G/C-1 | CG>TA | CG>TA | CCTATGCCAT | CCATCCT | CGG | GCC | C | A | C>T | C>T_GCC | CG>TA_CCATCCT | 6 | FALSE | FALSE |
| 135_v.txt | 33276 | 33276-SNV-( | SNV | 1 | G | A | 36 | 19836 | 0.1814882 | 49.7222222 | rnr1Y285F_pG | TRUE | CAN | JSY2421 | 3 | rnr1Y285F_p | CGT | G>A | A/T+1 | G/C-1 | CG>TA | CG>TA | GAGAACGTA | GTATCCT | GCA | GCA | C | T | C>T | C>T_GCA | CG>TA_GTATCCT | 6 | FALSE | FALSE |
| 135_v.txt | 33310 | 33310-SNV-( | SNV | 1 | G | A | 28 | 19836 | 0.14115749 | 54.5714286 | rnr1Y285F_pG | TRUE | CAN | JSY2421 | 3 | rnr1Y285F_p | CGC | G>A | A/T+1 | G/C-1 | CG>TA | CG>TA | AAGAGCGCA | GCAATGG | GCG | GCG | C | T | C>T | C>T_GCG | CG>TA_GCAATGG | 6 | FALSE | FALSE |
| 135_v.txt | 33376 | 33376-SNV-( | SNV | 1 | G | A | 25 | 19836 | 0.12603348 | 52.2 | rnr1Y285F_pG | TRUE | CAN | JSY2421 | 3 | rnr1Y285F_p | TGT | G>A | A/T+1 | G/C-1 | CG>TA | CG>TA | GTGTTTGTG | GTGAAGC | ACA | ACA | C | T | C>T | C>T_ACA | CG>TA_GTGAAGC | 6 | FALSE | FALSE |
| 135_v.txt | 33392 | 33392-SNV-( | SNV | 1 | G | A | 35 | 19836 | 0.17644686 | 53.1142857 | rnr1Y285F_pG | TRUE | CAN | JSY2421 | 3 | rnr1Y285F_p | CGT | G>A | A/T+1 | G/C-1 | CG>TA | CG>TA | AACGTCGTG | GTGAAAG | GCA | GCA | C | T | C>T | C>T_GCA | CG>TA_GTGAAAG | 6 | FALSE | FALSE |
| 135_v.txt | 33412 | 33412-SNV-( | SNV | 1 | G | A | 29 | 19836 | 0.14619883 | 48.1724138 | rnr1Y285F_pG | TRUE | CAN | JSY2421 | 3 | rnr1Y285F_p | GGC | G>A | A/T+1 | G/C-1 | CG>TA | CG>TA | TGACCGGCT | GCTCATT | CCG | CCG | C | T | C>T | C>T_CCG | CG>TA_GCTCATT | 6 | FALSE | FALSE |
| 135_v.txt | 33474 | 33474-SNV-( | SNV | 1 | C | T | 27 | 19833 | 0.13613674 | 31.4444444 | rnr1Y285F_pG | TRUE | CAN | JSY2421 | 3 | rnr1Y285F_p | GCC | C>T | A/T+1 | G/C-1 | CG>TA | CG>TA | GCTATGCCT | CCTTTTT | CGG | GCC | C | A | C>T | C>T_GCC | CG>TA_CCTTTTT | 6 | FALSE | FALSE |
| 135_v.txt | 31912 | 31912-SNV-( | SNV | 1 | C | G | 1 | 10906 | 9.17E-03 | 37.9987913 | rnr1Y285F_pG | TRUE | CAN | JSY2421 | 3 | rnr1Y285F_p | GCA | C>G | G/C+1 | G/C-1 | CG>GC | CG>GC | TTGGTGCAA | CAAAAGC | CGT | GCA | C | C | C>G | C>G_GCA | CG>GC_CAAAAGC | 6 | FALSE | FALSE |
| 135_v.txt | 31917 | 31917-SNV-( | SNV | 1 | G | C | 38 | 10906 | 0.34843206 | 37.5263158 | rnr1Y285F_pG | TRUE | CAN | JSY2421 | 3 | rnr1Y285F_p | AGC | G>C | G/C+1 | G/C-1 | CG>GC | CG>GC | GCAAAAGCC | GCCGTGA | TCG | TCG | C | G | C>G | C>G_TCG | CG>GC_GCCGTGA | 6 | FALSE | FALSE |
| 135_v.txt | 31981 | 31981-SNV-( | SNV | 1 | G | C | 15 | 23199 | 0.06465796 | 48.9333333 | rnr1Y285F_pG | TRUE | CAN | JSY2421 | 3 | rnr1Y285F_p | GGC | G>C | G/C+1 | G/C-1 | CG>GC | CG>GC | AGCCGGGCA | GCATTAA | CCG | CCG | C | G | C>G | C>G_CCG | CG>GC_GCATTAA | 6 | FALSE | FALSE |
| 135_v.txt | 32027 | 32027-SNV-( | SNV | 1 | G | C | 24 | 23199 | 0.10345274 | 57.375 | rnr1Y285F_pG | TRUE | CAN | JSY2421 | 3 | rnr1Y285F_p | GGT | G>C | G/C+1 | G/C-1 | CG>GC | CG>GC | GCCACGGTA | GTATTTC | CCA | CCA | C | G | C>G | C>G_CCA | CG>GC_GTATTTC | 6 | FALSE | FALSE |
| 135_v.txt | 32042 | 32042-SNV-( | SNV | 1 | C | G | 3 | 23196 | 0.01293326 | 37.9980572 | rnr1Y285F_pG | TRUE | CAN | JSY2421 | 3 | rnr1Y285F_p | GCA | C>G | G/C+1 | G/C-1 | CG>GC | CG>GC | AGCTTGCAT | CATAAAT | CGT | GCA | C | C | C>G | C>G_GCA | CG>GC_CATAAAT | 6 | FALSE | FALSE |
| 135_v.txt | 32064 | 32064-SNV-( | SNV | 1 | G | C | 42 | 23199 | 0.18104229 | 62.7380952 | rnr1Y285F_pG | TRUE | CAN | JSY2421 | 3 | rnr1Y285F_p | TGA | G>C | G/C+1 | G/C-1 | CG>GC | CG>GC | GAGATTGAG | GAGATAA | ACT | ACT | C | G | C>G | C>G_ACT | CG>GC_GAGATAA | 6 | FALSE | FALSE |
| 135_v.txt | 32182 | 32182-SNV-( | SNV | 1 | C | G | 15 | 23199 | 0.06465796 | 49.8666667 | rnr1Y285F_pG | TRUE | CAN | JSY2421 | 3 | rnr1Y285F_p | ACG | C>G | G/C+1 | G/C-1 | CG>GC | CG>GC | CAGTAACGA | CGAAAAC | TGC | ACG | C | C | C>G | C>G_ACG | CG>GC_CGAAAAC | 6 | FALSE | FALSE |
| 135_v.txt | 32272 | 32272-SNV-( | SNV | 1 | G | C | 118 | 33139 | 0.35607592 | 36 | rnr1Y285F_pG | TRUE | CAN | JSY2421 | 3 | rnr1Y285F_p | CGG | G>C | G/C+1 | G/C-1 | CG>GC | CG>GC | AAATACGGG | GGGAACC | GCC | GCC | C | G | C>G | C>G_GCC | CG>GC_GGGAACC | 6 | FALSE | FALSE |
| 135_v.txt | 32292 | 32292-SNV-( | SNV | 1 | G | C | 112 | 33161 | 0.33774615 | 52.2053571 | rnr1Y285F_pG | TRUE | CAN | JSY2421 | 3 | rnr1Y285F_p | TGA | G>C | G/C+1 | G/C-1 | CG>GC | CG>GC | ATATTTGAA | GAATTTG | ACT | ACT | C | G | C>G | C>G_ACT | CG>GC_GAATTTG | 6 | FALSE | FALSE |
| 135_v.txt | 32304 | 32304-SNV-( | SNV | 1 | G | C | 14 | 33161 | 0.04221827 | 53.8571348 | rnr1Y285F_pG | TRUE | CAN | JSY2421 | 3 | rnr1Y285F_p | AGA | G>C | G/C+1 | G/C-1 | CG>GC | CG>GC | GCGGCAGAA | GAAATAA | TCT | TCT | C | G | C>G | C>G_TCT | CG>GC_GAAATAA | 6 | FALSE | FALSE |
| 135_v.txt | 32316 | 32316-SNV-( | SNV | 1 | G | C | 24 | 33161 | 0.07237418 | 61.1666667 | rnr1Y285F_pG | TRUE | CAN | JSY2421 | 3 | rnr1Y285F_p | TGT | G>C | G/C+1 | G/C-1 | CG>GC | CG>GC | ATGGTTGTT | GTTAAGA | ACA | ACA | C | G | C>G | C>G_ACA | CG>GC_GTTAAGA | 6 | FALSE | FALSE |
| 135_v.txt | 32345 | 32345-SNV-( | SNV | 1 | C | G | 2 | 33161 | 6.03E-03 | 44.9988695 | rnr1Y285F_pG | TRUE | CAN | JSY2421 | 3 | rnr1Y285F_p | GCA | C>G | G/C+1 | G/C-1 | CG>GC | CG>GC | ATGTGGCAA | CAAAACC | CGT | GCA | C | C | C>G | C>G_GCA | CG>GC_CAAAACC | 6 | FALSE | FALSE |
| 135_v.txt | 32385 | 32385-SNV-( | SNV | 1 | G | C | 12 | 33161 | 0.03618709 | 62.6666667 | rnr1Y285F_pG | TRUE | CAN | JSY2421 | 3 | rnr1Y285F_p | GGG | G>C | G/C+1 | G/C-1 | CG>GC | CG>GC | ATAAAGGGA | GGAGAAG | CCC | CCC | C | G | C>G | C>G_CCC | CG>GC_GGAGAAG | 6 | FALSE | FALSE |
| 135_v.txt | 32474 | 32474-SNV-( | SNV | 1 | G | C | 1 | 33161 | 3.02E-03 | 63.9980145 | rnr1Y285F_pG | TRUE | CAN | JSY2421 | 3 | rnr1Y285F_p | AGA | G>C | G/C+1 | G/C-1 | CG>GC | CG>GC | AATGTAGAA | GAAGGTT | TCT | TCT | C | G | C>G | C>G_TCT | CG>GC_GAAGGTT | 6 | FALSE | FALSE |
| 135_v.txt | 32487 | 32487-SNV-( | SNV | 1 | C | G | 53 | 33161 | 0.1598263 | 46.1509434 | rnr1Y285F_pG | TRUE | CAN | JSY2421 | 3 | rnr1Y285F_p | ACG | C>G | G/C+1 | G/C-1 | CG>GC | CG>GC | AAGATACGG | CGGAAAA | TGC | ACG | C | C | C>G | C>G_ACG | CG>GC_CGGAAAA | 6 | FALSE | FALSE |
| 135_v.txt | 32529 | 32529-SNV-( | SNV | 1 | G | C | 18 | 33160 | 0.05428227 | 43.1678432 | rnr1Y285F_pG | TRUE | CAN | JSY2421 | 3 | rnr1Y285F_p | GGG | G>C | G/C+1 | G/C-1 | CG>GC | CG>GC | TTTCTGGGG | GGGTTTG | CCC | CCC | C | G | C>G | C>G_CCC | CG>GC_GGGTTTG | 6 | FALSE | FALSE |
| 135_v.txt | 32539 | 32539-SNV-( | SNV | 1 | C | G | 28 | 33160 | 0.08443908 | 36.75 | rnr1Y285F_pG | TRUE | CAN | JSY2421 | 3 | rnr1Y285F_p | GCT | C>G | G/C+1 | G/C-1 | CG>GC | CG>GC | TTGCAGCTTC | CTTCACC | CGA | GCT | C | C | C>G | C>G_GCT | CG>GC_CTTCACC | 6 | FALSE | FALSE |
| 135_v.txt | 32566 | 32566-SNV-( | SNV | 1 | C | G | 19 | 27962 | 0.06794936 | 33.9473684 | rnr1Y285F_pG | TRUE | CAN | JSY2421 | 3 | rnr1Y285F_p | TCA | C>G | G/C+1 | G/C-1 | CG>GC | CG>GC | CTAGTTCAG | CAGTACC | AGT | TCA | C | C | C>G | C>G_TCA | CG>GC_CAGTACC | 6 | FALSE | FALSE |
| 135_v.txt | 32685 | 32685-SNV-( | SNV | 1 | C | G | 37 | 27973 | 0.1322704 | 61.8361655 | rnr1Y285F_pG | TRUE | CAN | JSY2421 | 3 | rnr1Y285F_p | ACG | C>G | G/C+1 | G/C-1 | CG>GC | CG>GC | CAATAACGG | CGGAATC | TGC | ACG | C | C | C>G | C>G_ACG | CG>GC_CGGAATC | 6 | FALSE | FALSE |
| 135_v.txt | 32749 | 32749-SNV-( | SNV | 1 | C | G | 47 | 27973 | 0.16801916 | 56.212766 | rnr1Y285F_pG | TRUE | CAN | JSY2421 | 3 | rnr1Y285F_p | CCG | C>G | G/C+1 | G/C-1 | CG>GC | CG>GC | GAAACCCGA | CGATAAT | GGC | CCG | C | C | C>G | C>G_CCG | CG>GC_CGATAAT | 6 | FALSE | FALSE |
| 135_v.txt | 32788 | 32788-SNV-( | SNV | 1 | C | G | 2 | 27973 | 7.15E-03 | 46.9952745 | rnr1Y285F_pG | TRUE | CAN | JSY2421 | 3 | rnr1Y285F_p | TCG | C>G | G/C+1 | G/C-1 | CG>GC | CG>GC | AGAACTCGA | CGAATTC | AGC | TCG | C | C | C>G | C>G_TCG | CG>GC_CGAATTC | 6 | FALSE | FALSE |
| 135_v.txt | 32797 | 32797-SNV-( | SNV | 1 | C | G | 23 | 27973 | 0.08222214 | 47.6479934 | rnr1Y285F_pG | TRUE | CAN | JSY2421 | 3 | rnr1Y285F_p | CCG | C>G | G/C+1 | G/C-1 | CG>GC | CG>GC | ATTCACCGT | CGTAATA | GGC | CCG | C | C | C>G | C>G_CCG | CG>GC_CGTAATA | 6 | FALSE | FALSE |
| 135_v.txt | 32809 | 32809-SNV-( | SNV | 1 | C | G | 14 | 27973 | 0.05004826 | 39.9285714 | rnr1Y285F_pG | TRUE | CAN | JSY2421 | 3 | rnr1Y285F_p | ACA | C>G | G/C+1 | G/C-1 | CG>GC | CG>GC | ATTTGACAG | CAGGGAA | TGT | ACA | C | C | C>G | C>G_ACA | CG>GC_CAGGGAA | 6 | FALSE | FALSE |
| 135_v.txt | 32937 | 32937-SNV-( | SNV | 1 | C | G | 23 | 29052 | 0.07916839 | 57.54691 | rnr1Y285F_pG | TRUE | CAN | JSY2421 | 3 | rnr1Y285F_p | CCA | C>G | G/C+1 | G/C-1 | CG>GC | CG>GC | ATTGCCCAA | CAAGAAA | GGT | CCA | C | C | C>G | C>G_CCA | CG>GC_CAAGAAA | 6 | FALSE | FALSE |
| 135_v.txt | 32997 | 32997-SNV-( | SNV | 1 | G | C | 20 | 29054 | 0.06883734 | 61.6 | rnr1Y285F_pG | TRUE | CAN | JSY2421 | 3 | rnr1Y285F_p | TGA | G>C | G/C+1 | G/C-1 | CG>GC | CG>GC | CTTTGTGAG | GAGAAAA | ACT | ACT | C | G | C>G | C>G_ACT | CG>GC_GAGAAAA | 6 | FALSE | FALSE |
| 135_v.txt | 33070 | 33070-SNV-( | SNV | 1 | C | G | 18 | 29051 | 0.06196 | 57.7777575 | rnr1Y285F_pG | TRUE | CAN | JSY2421 | 3 | rnr1Y285F_p | GCC | C>G | G/C+1 | G/C-1 | CG>GC | CG>GC | AATATGCCA | CCAAAGA | CGG | GCC | C | C | C>G | C>G_GCC | CG>GC_CCAAAGA | 6 | FALSE | FALSE |
| 135_v.txt | 33103 | 33103-SNV-( | SNV | 1 | C | G | 9 | 29051 | 0.03098 | 39.6666667 | rnr1Y285F_pG | TRUE | CAN | JSY2421 | 3 | rnr1Y285F_p | GCG | C>G | G/C+1 | G/C-1 | CG>GC | CG>GC | TAAGAGCGC | CGCCCAC | CGC | GCG | C | C | C>G | C>G_GCG | CG>GC_CGCCCAC | 6 | FALSE | FALSE |
| 135_v.txt | 33153 | 33153-SNV-( | SNV | 1 | C | G | 23 | 29051 | 0.07917111 | 37.173913 | rnr1Y285F_pG | TRUE | CAN | JSY2421 | 3 | rnr1Y285F_p | ACC | C>G | G/C+1 | G/C-1 | CG>GC | CG>GC | AAAAGACCT | CCTGTAC | TGG | ACC | C | C | C>G | C>G_ACC | CG>GC_CCTGTAC | 6 | FALSE | FALSE |
| 135_v.txt | 33154 | 33154-SNV-( | SNV | 1 | C | G | 3 | 29051 | 0.01032667 | 38 | rnr1Y285F_pG | TRUE | CAN | JSY2421 | 3 | rnr1Y285F_p | CCT | C>G | G/C+1 | G/C-1 | CG>GC | CG>GC | AAAGACCTG | CTGTACC | GGA | CCT | C | C | C>G | C>G_CCT | CG>GC_CTGTACC | 6 | FALSE | FALSE |
| 135_v.txt | 33178 | 33178-SNV-( | SNV | 1 | C | G | 7 | 29043 | 0.02410219 | 37.1421496 | rnr1Y285F_pG | TRUE | CAN | JSY2421 | 3 | rnr1Y285F_p | GCA | C>G | G/C+1 | G/C-1 | CG>GC | CG>GC | CAAGGGCAA | CAATCAT | CGT | GCA | C | C | C>G | C>G_GCA | CG>GC_CAATCAT | 6 | FALSE | FALSE |
| 135_v.txt | 33187 | 33187-SNV-( | SNV | 1 | C | G | 6 | 19828 | 0.03026024 | 38 | rnr1Y285F_pG | TRUE | CAN | JSY2421 | 3 | rnr1Y285F_p | CCA | C>G | G/C+1 | G/C-1 | CG>GC | CG>GC | TCATACCAAT | CAATATG | GGT | CCA | C | C | C>G | C>G_CCA | CG>GC_CAATATG | 6 | FALSE | FALSE |
| 135_v.txt | 33210 | 33210-SNV-( | SNV | 1 | C | G | 113 | 19827 | 0.56992989 | 39.0088496 | rnr1Y285F_pG | TRUE | CAN | JSY2421 | 3 | rnr1Y285F_p | TCT | C>G | G/C+1 | G/C-1 | CG>GC | CG>GC | AGCTCTCTCT | CTCTTCA | AGA | TCT | C | C | C>G | C>G_TCT | CG>GC_CTCTTCA | 6 | FALSE | FALSE |
| 135_v.txt | 33456 | 33456-SNV-( | SNV | 1 | G | C | 24 | 19836 | 0.12099214 | 39.25 | rnr1Y285F_pG | TRUE | CAN | JSY2421 | 3 | rnr1Y285F_p | TGA | G>C | G/C+1 | G/C-1 | CG>GC | CG>GC | TCTTTTGAAT | GAATTTG | ACT | ACT | C | G | C>G | C>G_ACT | CG>GC_GAATTTG | 6 | FALSE | FALSE |
| 135_v.txt | 31817 | 31817-SNV-( | SNV | 1 | G | T | 18 | 10907 | 0.16503163 | 58.9444444 | rnr1Y285F_pG | TRUE | CAN | JSY2421 | 3 | rnr1Y285F_p | TGC | G>T | A/T+1 | G/C-1 | CG>AT | CG>AT | AAATCTGCA | GCATCTG | ACG | ACG | C | A | C>A | C>A_ACG | CG>AT_GCATCTG | 6 | FALSE | FALSE |

| | | | | | | | | | | | | | | | | | | | | | | | | | | | | | | | | | | |
|---|---|---|---|---|---|---|---|---|---|---|---|---|---|---|---|---|---|---|---|---|---|---|---|---|---|---|---|---|---|---|---|---|---|---|
| 135_v.txt | 31829 | 31829-SNV- | SNV | 1 | G | T | 9 | 10907 | 0.08251582 | 56.4444444 | rnr1Y285F_pG | TRUE | CAN | JSY2421 | 3 | rnr1Y285F_p | TGC | G>T | A/T+1 | G/C-1 | CG>AT | CG>AT | GAATATGCA | GCATTGA | ACG | ACG | C | A | C>A | C>A_ACG | CG>AT_GCATTGA | 6 | FALSE | FALSE |
| 135_v.txt | 31869 | 31869-SNV- | SNV | 1 | G | T | 9 | 10907 | 0.08251582 | 46 | rnr1Y285F_pG | TRUE | CAN | JSY2421 | 3 | rnr1Y285F_p | AGA | G>T | A/T+1 | G/C-1 | CG>AT | CG>AT | AAAATAGAG | GAGATAT | TCT | TCT | C | A | C>A | C>A_TCT | CG>AT_GAGATAT | 6 | FALSE | FALSE |
| 135_v.txt | 31920 | 31920-SNV- | SNV | 1 | G | T | 33 | 10906 | 0.30258573 | 36.7575758 | rnr1Y285F_pG | TRUE | CAN | JSY2421 | 3 | rnr1Y285F_p | CGT | G>T | A/T+1 | G/C-1 | CG>AT | CG>AT | AAAGCCGTG | GTGAAAC | GCA | GCA | C | A | C>A | C>A_GCA | CG>AT_GTGAAAC | 6 | FALSE | FALSE |
| 135_v.txt | 31973 | 31973-SNV- | SNV | 1 | C | A | 37 | 23199 | 0.15948963 | 31.445638 | rnr1Y285F_pG | TRUE | CAN | JSY2421 | 3 | rnr1Y285F_p | CCA | C>A | A/T+1 | G/C-1 | CG>AT | CG>AT | ATAAGCCAA | CAAGCCG | GGT | CCA | C | T | C>A | C>A_CCA | CG>AT_CAAGCCG | 6 | FALSE | FALSE |
| 135_v.txt | 31980 | 31980-SNV- | SNV | 1 | G | T | 17 | 23199 | 0.07327902 | 37.403875 | rnr1Y285F_pG | TRUE | CAN | JSY2421 | 3 | rnr1Y285F_p | GGG | G>T | A/T+1 | G/C-1 | CG>AT | CG>AT | AAGCCGGGC | GGCATTA | CCC | CCC | C | A | C>A | C>A_CCC | CG>AT_GGCATTA | 6 | FALSE | FALSE |
| 135_v.txt | 31981 | 31981-SNV- | SNV | 1 | G | T | 27 | 23199 | 0.11638433 | 30.6296296 | rnr1Y285F_pG | TRUE | CAN | JSY2421 | 3 | rnr1Y285F_p | GGC | G>T | A/T+1 | G/C-1 | CG>AT | CG>AT | AGCCGGGCA | GCATTAA | CCG | CCG | C | A | C>A | C>A_CCG | CG>AT_GCATTAA | 6 | FALSE | FALSE |
| 135_v.txt | 32027 | 32027-SNV- | SNV | 1 | G | T | 112 | 23199 | 0.48277943 | 59.2678571 | rnr1Y285F_pG | TRUE | CAN | JSY2421 | 3 | rnr1Y285F_p | GGT | G>T | A/T+1 | G/C-1 | CG>AT | CG>AT | GCCACGGTA | GTATTTC | CCA | CCA | C | A | C>A | C>A_CCA | CG>AT_GTATTTC | 6 | FALSE | FALSE |
| 135_v.txt | 32089 | 32089-SNV- | SNV | 1 | C | A | 30 | 23198 | 0.12932149 | 57.1664825 | rnr1Y285F_pG | TRUE | CAN | JSY2421 | 3 | rnr1Y285F_p | CCT | C>A | A/T+1 | G/C-1 | CG>AT | CG>AT | AAAAGCCTG | CTGCAAC | GGA | CCT | C | T | C>A | C>A_CCT | CG>AT_CTGCAAC | 6 | FALSE | FALSE |
| 135_v.txt | 32119 | 32119-SNV- | SNV | 1 | C | A | 33 | 23199 | 0.14224751 | 55.969697 | rnr1Y285F_pG | TRUE | CAN | JSY2421 | 3 | rnr1Y285F_p | TCG | C>A | A/T+1 | G/C-1 | CG>AT | CG>AT | GCCATTCGA | CGAAAAC | AGC | TCG | C | T | C>A | C>A_TCG | CG>AT_CGAAAAC | 6 | FALSE | FALSE |
| 135_v.txt | 32149 | 32149-SNV- | SNV | 1 | C | A | 21 | 23199 | 0.09052114 | 49.4761905 | rnr1Y285F_pG | TRUE | CAN | JSY2421 | 3 | rnr1Y285F_p | TCC | C>A | A/T+1 | G/C-1 | CG>AT | CG>AT | ATGTCTCCAT | CCATGTA | AGG | TCC | C | T | C>A | C>A_TCC | CG>AT_CCATGTA | 6 | FALSE | FALSE |
| 135_v.txt | 32153 | 32153-SNV- | SNV | 1 | G | T | 78 | 23199 | 0.33622139 | 50.1282051 | rnr1Y285F_pG | TRUE | CAN | JSY2421 | 3 | rnr1Y285F_p | TGT | G>T | A/T+1 | G/C-1 | CG>AT | CG>AT | CTCCATGTA | GTAAGCC | ACA | ACA | C | A | C>A | C>A_ACA | CG>AT_GTAAGCC | 6 | FALSE | FALSE |
| 135_v.txt | 32188 | 32188-SNV- | SNV | 1 | C | A | 14 | 23199 | 0.06034743 | 46.0714286 | rnr1Y285F_pG | TRUE | CAN | JSY2421 | 3 | rnr1Y285F_p | ACT | C>A | A/T+1 | G/C-1 | CG>AT | CG>AT | CGAAAACTG | CTGCAAT | TGA | ACT | C | T | C>A | C>A_ACT | CG>AT_CTGCAAT | 6 | FALSE | FALSE |
| 135_v.txt | 32195 | 32195-SNV- | SNV | 1 | G | T | 99 | 23199 | 0.42674253 | 45.010101 | rnr1Y285F_pG | TRUE | CAN | JSY2421 | 3 | rnr1Y285F_p | TGT | G>T | A/T+1 | G/C-1 | CG>AT | CG>AT | TGCAATGTA | GTATGGA | ACA | ACA | C | A | C>A | C>A_ACA | CG>AT_GTATGGA | 6 | FALSE | FALSE |
| 135_v.txt | 32199 | 32199-SNV- | SNV | 1 | G | T | 102 | 23199 | 0.43967412 | 41.2156863 | rnr1Y285F_pG | TRUE | CAN | JSY2421 | 3 | rnr1Y285F_p | TGG | G>T | A/T+1 | G/C-1 | CG>AT | CG>AT | ATGTATGGA | GGAACAC | ACC | ACC | C | A | C>A | C>A_ACC | CG>AT_GGAACAC | 6 | FALSE | FALSE |
| 135_v.txt | 32223 | 32223-SNV- | SNV | 1 | G | T | 30 | 23199 | 0.12931592 | 40.8 | rnr1Y285F_pG | TRUE | CAN | JSY2421 | 3 | rnr1Y285F_p | TGA | G>T | A/T+1 | G/C-1 | CG>AT | CG>AT | GTCCTTGAC | GACAGGA | ACT | ACT | C | A | C>A | C>A_ACT | CG>AT_GACAGGA | 6 | FALSE | FALSE |
| 135_v.txt | 32272 | 32272-SNV- | SNV | 1 | G | T | 76 | 33139 | 0.22933704 | 35.6184211 | rnr1Y285F_pG | TRUE | CAN | JSY2421 | 3 | rnr1Y285F_p | CGG | G>T | A/T+1 | G/C-1 | CG>AT | CG>AT | AAATACGGG | GGGAACC | GCC | GCC | C | A | C>A | C>A_GCC | CG>AT_GGGAACC | 6 | FALSE | FALSE |
| 135_v.txt | 32274 | 32274-SNV- | SNV | 1 | G | T | 49 | 33139 | 0.14786204 | 35.3469388 | rnr1Y285F_pG | TRUE | CAN | JSY2421 | 3 | rnr1Y285F_p | GGA | G>T | A/T+1 | G/C-1 | CG>AT | CG>AT | ATACGGGAA | GAACCAA | CCT | CCT | C | A | C>A | C>A_CCT | CG>AT_GAACCAA | 6 | FALSE | FALSE |
| 135_v.txt | 32282 | 32282-SNV- | SNV | 1 | G | T | 55 | 33139 | 0.16596759 | 41.7272727 | rnr1Y285F_pG | TRUE | CAN | JSY2421 | 3 | rnr1Y285F_p | CGT | G>T | A/T+1 | G/C-1 | CG>AT | CG>AT | ACCAACGTA | GTAAATA | GCA | GCA | C | A | C>A | C>A_GCA | CG>AT_GTAAATA | 6 | FALSE | FALSE |
| 135_v.txt | 32292 | 32292-SNV- | SNV | 1 | G | T | 20 | 33161 | 0.06031181 | 44 | rnr1Y285F_pG | TRUE | CAN | JSY2421 | 3 | rnr1Y285F_p | TGA | G>T | A/T+1 | G/C-1 | CG>AT | CG>AT | ATATTTGAAT | GAATTTG | ACT | ACT | C | A | C>A | C>A_ACT | CG>AT_GAATTTG | 6 | FALSE | FALSE |
| 135_v.txt | 32301 | 32301-SNV- | SNV | 1 | G | T | 34 | 33161 | 0.10253008 | 49.9114148 | rnr1Y285F_pG | TRUE | CAN | JSY2421 | 3 | rnr1Y285F_p | GGC | G>T | A/T+1 | G/C-1 | CG>AT | CG>AT | TTTGCGGCA | GCAGAAA | CCG | CCG | C | A | C>A | C>A_CCG | CG>AT_GCAGAAA | 6 | FALSE | FALSE |
| 135_v.txt | 32304 | 32304-SNV- | SNV | 1 | G | T | 133 | 33161 | 0.40107355 | 59.7712231 | rnr1Y285F_pG | TRUE | CAN | JSY2421 | 3 | rnr1Y285F_p | AGA | G>T | A/T+1 | G/C-1 | CG>AT | CG>AT | GCGGCAGAA | GAAATAA | TCT | TCT | C | A | C>A | C>A_TCT | CG>AT_GAAATAA | 6 | FALSE | FALSE |
| 135_v.txt | 32373 | 32373-SNV- | SNV | 1 | G | T | 46 | 33161 | 0.13871717 | 59.7391304 | rnr1Y285F_pG | TRUE | CAN | JSY2421 | 3 | rnr1Y285F_p | AGC | G>T | A/T+1 | G/C-1 | CG>AT | CG>AT | TCAATAGCA | GCAATAA | TCG | TCG | C | A | C>A | C>A_TCG | CG>AT_GCAATAA | 6 | FALSE | FALSE |
| 135_v.txt | 32399 | 32399-SNV- | SNV | 1 | G | T | 243 | 33161 | 0.73278852 | 58.7736626 | rnr1Y285F_pG | TRUE | CAN | JSY2421 | 3 | rnr1Y285F_p | CGT | G>T | A/T+1 | G/C-1 | CG>AT | CG>AT | AGAAACGTA | GTAGGAA | GCA | GCA | C | A | C>A | C>A_GCA | CG>AT_GTAGGAA | 6 | FALSE | FALSE |
| 135_v.txt | 32432 | 32432-SNV- | SNV | 1 | G | T | 48 | 33161 | 0.14474835 | 58.3125 | rnr1Y285F_pG | TRUE | CAN | JSY2421 | 3 | rnr1Y285F_p | TGT | G>T | A/T+1 | G/C-1 | CG>AT | CG>AT | GTCATTGTAT | GTATGGA | ACA | ACA | C | A | C>A | C>A_ACA | CG>AT_GTATGGA | 6 | FALSE | FALSE |
| 135_v.txt | 32449 | 32449-SNV- | SNV | 1 | C | A | 4 | 33161 | 0.01206236 | 39.25 | rnr1Y285F_pG | TRUE | CAN | JSY2421 | 3 | rnr1Y285F_p | CCA | C>A | A/T+1 | G/C-1 | CG>AT | CG>AT | AAAGTCCAA | CAATGAA | GGT | CCA | C | T | C>A | C>A_CCA | CG>AT_CAATGAA | 6 | FALSE | FALSE |
| 135_v.txt | 32474 | 32474-SNV- | SNV | 1 | G | T | 27 | 33161 | 0.08142095 | 53.5914515 | rnr1Y285F_pG | TRUE | CAN | JSY2421 | 3 | rnr1Y285F_p | AGA | G>T | A/T+1 | G/C-1 | CG>AT | CG>AT | AATGTAGAA | GAAGGTT | TCT | TCT | C | A | C>A | C>A_TCT | CG>AT_GAAGGTT | 6 | FALSE | FALSE |
| 135_v.txt | 32487 | 32487-SNV- | SNV | 1 | C | A | 40 | 33161 | 0.12062362 | 44.475 | rnr1Y285F_pG | TRUE | CAN | JSY2421 | 3 | rnr1Y285F_p | ACG | C>A | A/T+1 | G/C-1 | CG>AT | CG>AT | AAGATACGG | CGGAAAA | TGC | ACG | C | T | C>A | C>A_ACG | CG>AT_CGGAAAA | 6 | FALSE | FALSE |
| 135_v.txt | 32530 | 32530-SNV- | SNV | 1 | G | T | 36 | 33160 | 0.10856454 | 38.0833333 | rnr1Y285F_pG | TRUE | CAN | JSY2421 | 3 | rnr1Y285F_p | GGG | G>T | A/T+1 | G/C-1 | CG>AT | CG>AT | TTCTGGGGT | GGTTTGC | CCC | CCC | C | A | C>A | C>A_CCC | CG>AT_GGTTTGC | 6 | FALSE | FALSE |
| 135_v.txt | 32556 | 32556-SNV- | SNV | 1 | C | A | 21 | 33156 | 0.06333695 | 33.9047619 | rnr1Y285F_pG | TRUE | CAN | JSY2421 | 3 | rnr1Y285F_p | ACC | C>A | A/T+1 | G/C-1 | CG>AT | CG>AT | GTGATACCA | CCAACTA | TGG | ACC | C | T | C>A | C>A_ACC | CG>AT_CCAACTA | 6 | FALSE | FALSE |
| 135_v.txt | 32566 | 32566-SNV- | SNV | 1 | C | A | 40 | 27962 | 0.14305128 | 35.45 | rnr1Y285F_pG | TRUE | CAN | JSY2421 | 3 | rnr1Y285F_p | TCA | C>A | A/T+1 | G/C-1 | CG>AT | CG>AT | CTAGTTCAGT | CAGTACC | AGT | TCA | C | T | C>A | C>A_TCA | CG>AT_CAGTACC | 6 | FALSE | FALSE |
| 135_v.txt | 32572 | 32572-SNV- | SNV | 1 | C | A | 36 | 27974 | 0.12869093 | 34.7209335 | rnr1Y285F_pG | TRUE | CAN | JSY2421 | 3 | rnr1Y285F_p | CCT | C>A | A/T+1 | G/C-1 | CG>AT | CG>AT | CAGTACCTTG | CTTGAAA | GGA | CCT | C | T | C>A | C>A_CCT | CG>AT_CTTGAAA | 6 | FALSE | FALSE |
| 135_v.txt | 32575 | 32575-SNV- | SNV | 1 | G | T | 80 | 27973 | 0.28599006 | 36.275 | rnr1Y285F_pG | TRUE | CAN | JSY2421 | 3 | rnr1Y285F_p | TGA | G>T | A/T+1 | G/C-1 | CG>AT | CG>AT | TACCTTGAA | GAAATGT | ACT | ACT | C | A | C>A | C>A_ACT | CG>AT_GAAATGT | 6 | FALSE | FALSE |
| 135_v.txt | 32629 | 32629-SNV- | SNV | 1 | C | A | 190 | 27973 | 0.6792264 | 54.1105263 | rnr1Y285F_pG | TRUE | CAN | JSY2421 | 3 | rnr1Y285F_p | TCG | C>A | A/T+1 | G/C-1 | CG>AT | CG>AT | TCCCTTCGTT | CGTTTTT | AGC | TCG | C | T | C>A | C>A_TCG | CG>AT_CGTTTTT | 6 | FALSE | FALSE |
| 135_v.txt | 32686 | 32686-SNV- | SNV | 1 | G | T | 5 | 27973 | 0.01787438 | 21.1981151 | rnr1Y285F_pG | TRUE | CAN | JSY2421 | 3 | rnr1Y285F_p | CGG | G>T | A/T+1 | G/C-1 | CG>AT | CG>AT | AATAACGGA | GGAATCC | GCC | GCC | C | A | C>A | C>A_GCC | CG>AT_GGAATCC | 6 | FALSE | FALSE |
| 135_v.txt | 32735 | 32735-SNV- | SNV | 1 | G | T | 48 | 27973 | 0.17159404 | 58.2077945 | rnr1Y285F_pG | TRUE | CAN | JSY2421 | 3 | rnr1Y285F_p | AGT | G>T | A/T+1 | G/C-1 | CG>AT | CG>AT | AAAACAGTA | GTATATT | TCA | TCA | C | A | C>A | C>A_TCA | CG>AT_GTATATT | 6 | FALSE | FALSE |
| 135_v.txt | 32783 | 32783-SNV- | SNV | 1 | G | T | 25 | 27973 | 0.08937189 | 43.56 | rnr1Y285F_pG | TRUE | CAN | JSY2421 | 3 | rnr1Y285F_p | AGA | G>T | A/T+1 | G/C-1 | CG>AT | CG>AT | GACCCAGAA | GAACTCG | TCT | TCT | C | A | C>A | C>A_TCT | CG>AT_GAACTCG | 6 | FALSE | FALSE |
| 135_v.txt | 32809 | 32809-SNV- | SNV | 1 | C | A | 49 | 27973 | 0.17516891 | 42.2653061 | rnr1Y285F_pG | TRUE | CAN | JSY2421 | 3 | rnr1Y285F_p | ACA | C>A | A/T+1 | G/C-1 | CG>AT | CG>AT | ATTTGACAG | CAGGGAA | TGT | ACA | C | T | C>A | C>A_ACA | CG>AT_CAGGGAA | 6 | FALSE | FALSE |
| 135_v.txt | 32819 | 32819-SNV- | SNV | 1 | G | T | 52 | 27973 | 0.18589354 | 43.4038462 | rnr1Y285F_pG | TRUE | CAN | JSY2421 | 3 | rnr1Y285F_p | AGT | G>T | A/T+1 | G/C-1 | CG>AT | CG>AT | GAACAAGTT | GTTCATT | TCA | TCA | C | A | C>A | C>A_TCA | CG>AT_GTTCATT | 6 | FALSE | FALSE |
| 135_v.txt | 32840 | 32840-SNV- | SNV | 1 | C | A | 2 | 27973 | 7.15E-03 | 30.497997 | rnr1Y285F_pG | TRUE | CAN | JSY2421 | 3 | rnr1Y285F_p | CCC | C>A | A/T+1 | G/C-1 | CG>AT | CG>AT | AATTACCCA | CCAAAAA | GGG | CCC | C | T | C>A | C>A_CCC | CG>AT_CCAAAAA | 6 | FALSE | FALSE |
| 135_v.txt | 32876 | 32876-SNV- | SNV | 1 | G | T | 85 | 29043 | 0.29266949 | 35.6588235 | rnr1Y285F_pG | TRUE | CAN | JSY2421 | 3 | rnr1Y285F_p | TGT | G>T | A/T+1 | G/C-1 | CG>AT | CG>AT | AACTTTGTAC | GTACGTC | ACA | ACA | C | A | C>A | C>A_ACA | CG>AT_GTACGTC | 6 | FALSE | FALSE |
| 135_v.txt | 32899 | 32899-SNV- | SNV | 1 | G | T | 71 | 29052 | 0.24438937 | 14.8730259 | rnr1Y285F_pG | TRUE | CAN | JSY2421 | 3 | rnr1Y285F_p | TGG | G>T | A/T+1 | G/C-1 | CG>AT | CG>AT | TGACTTGGC | GGCCAAC | ACC | ACC | C | A | C>A | C>A_ACC | CG>AT_GGCCAAC | 6 | FALSE | FALSE |
| 135_v.txt | 32917 | 32917-SNV- | SNV | 1 | C | A | 322 | 29052 | 1.10835743 | 51.4192547 | rnr1Y285F_pG | TRUE | CAN | JSY2421 | 3 | rnr1Y285F_p | TCC | C>A | A/T+1 | G/C-1 | CG>AT | CG>AT | TAAGTTCCA | CCAGGGC | AGG | TCC | C | T | C>A | C>A_TCC | CG>AT_CCAGGGC | 6 | FALSE | FALSE |
| 135_v.txt | 32928 | 32928-SNV- | SNV | 1 | G | T | 15 | 29052 | 0.05163156 | 43.5288304 | rnr1Y285F_pG | TRUE | CAN | JSY2421 | 3 | rnr1Y285F_p | AGT | G>T | A/T+1 | G/C-1 | CG>AT | CG>AT | GCAAAAGTG | GTGATTG | TCA | TCA | C | A | C>A | C>A_TCA | CG>AT_GTGATTG | 6 | FALSE | FALSE |
| 135_v.txt | 32936 | 32936-SNV- | SNV | 1 | C | A | 53 | 29052 | 0.1824315 | 36.2089862 | rnr1Y285F_pG | TRUE | CAN | JSY2421 | 3 | rnr1Y285F_p | CCC | C>A | A/T+1 | G/C-1 | CG>AT | CG>AT | GATTGCCCA | CCAAGAA | GGG | CCC | C | T | C>A | C>A_CCC | CG>AT_CCAAGAA | 6 | FALSE | FALSE |
| 135_v.txt | 32940 | 32940-SNV- | SNV | 1 | G | T | 259 | 29052 | 0.89150489 | 54.1389961 | rnr1Y285F_pG | TRUE | CAN | JSY2421 | 3 | rnr1Y285F_p | AGA | G>T | A/T+1 | G/C-1 | CG>AT | CG>AT | GCCCAAGAA | GAAAACC | TCT | TCT | C | A | C>A | C>A_TCT | CG>AT_GAAAACC | 6 | FALSE | FALSE |
| 135_v.txt | 32954 | 32954-SNV- | SNV | 1 | G | T | 145 | 29052 | 0.49910505 | 55.937931 | rnr1Y285F_pG | TRUE | CAN | JSY2421 | 3 | rnr1Y285F_p | TGT | G>T | A/T+1 | G/C-1 | CG>AT | CG>AT | ATACATGTA | GTAACCA | ACA | ACA | C | A | C>A | C>A_ACA | CG>AT_GTAACCA | 6 | FALSE | FALSE |
| 135_v.txt | 33006 | 33006-SNV- | SNV | 1 | G | T | 17 | 29054 | 0.05851174 | 58.6470588 | rnr1Y285F_pG | TRUE | CAN | JSY2421 | 3 | rnr1Y285F_p | TGT | G>T | A/T+1 | G/C-1 | CG>AT | CG>AT | AAAACTGTG | GTGAAAG | ACA | ACA | C | A | C>A | C>A_ACA | CG>AT_GTGAAAG | 6 | FALSE | FALSE |
| 135_v.txt | 33015 | 33015-SNV- | SNV | 1 | G | T | 33 | 29054 | 0.11358161 | 58 | rnr1Y285F_pG | TRUE | CAN | JSY2421 | 3 | rnr1Y285F_p | GGA | G>T | A/T+1 | G/C-1 | CG>AT | CG>AT | AAAGAGGAT | GATGTAA | CCT | CCT | C | A | C>A | C>A_CCT | CG>AT_GATGTAA | 6 | FALSE | FALSE |
| 135_v.txt | 33024 | 33024-SNV- | SNV | 1 | G | T | 71 | 29054 | 0.24437255 | 60.1267606 | rnr1Y285F_pG | TRUE | CAN | JSY2421 | 3 | rnr1Y285F_p | AGG | G>T | A/T+1 | G/C-1 | CG>AT | CG>AT | GTAACAGGG | GGGATGA | TCC | TCC | C | A | C>A | C>A_TCC | CG>AT_GGGATGA | 6 | FALSE | FALSE |
| 135_v.txt | 33043 | 33043-SNV- | SNV | 1 | C | A | 77 | 29054 | 0.26502375 | 59.5844156 | rnr1Y285F_pG | TRUE | CAN | JSY2421 | 3 | rnr1Y285F_p | TCA | C>A | A/T+1 | G/C-1 | CG>AT | CG>AT | CCATTTCACC | CACCCAA | AGT | TCA | C | T | C>A | C>A_TCA | CG>AT_CACCCAA | 6 | FALSE | FALSE |
| 135_v.txt | 33093 | 33093-SNV- | SNV | 1 | G | T | 104 | 29051 | 0.35799112 | 50.1730769 | rnr1Y285F_pG | TRUE | CAN | JSY2421 | 3 | rnr1Y285F_p | TGA | G>T | A/T+1 | G/C-1 | CG>AT | CG>AT | AAATATGAT | GATATAA | ACT | ACT | C | A | C>A | C>A_ACT | CG>AT_GATATAA | 6 | FALSE | FALSE |
| 135_v.txt | 33114 | 33114-SNV- | SNV | 1 | C | A | 56 | 29051 | 0.19276445 | 43.7844231 | rnr1Y285F_pG | TRUE | CAN | JSY2421 | 3 | rnr1Y285F_p | GCC | C>A | A/T+1 | G/C-1 | CG>AT | CG>AT | ACTGGGCCG | CCGGCGT | CGG | GCC | C | T | C>A | C>A_GCC | CG>AT_CCGGCGT | 6 | FALSE | FALSE |
| 135_v.txt | 33151 | 33151-SNV- | SNV | 1 | G | T | 213 | 29051 | 0.73319335 | 40.1079812 | rnr1Y285F_pG | TRUE | CAN | JSY2421 | 3 | rnr1Y285F_p | AGA | G>T | A/T+1 | G/C-1 | CG>AT | CG>AT | TGAAAAGAC | GACCTGT | TCT | TCT | C | A | C>A | C>A_TCT | CG>AT_GACCTGT | 6 | FALSE | FALSE |
| 135_v.txt | 33153 | 33153-SNV- | SNV | 1 | C | A | 90 | 29051 | 0.30980001 | 38.9 | rnr1Y285F_pG | TRUE | CAN | JSY2421 | 3 | rnr1Y285F_p | ACC | C>A | A/T+1 | G/C-1 | CG>AT | CG>AT | AAAAGACCT | CCTGTAC | TGG | ACC | C | T | C>A | C>A_ACC | CG>AT_CCTGTAC | 6 | FALSE | FALSE |
| 135_v.txt | 33154 | 33154-SNV- | SNV | 1 | C | A | 140 | 29051 | 0.48191112 | 36.6072456 | rnr1Y285F_pG | TRUE | CAN | JSY2421 | 3 | rnr1Y285F_p | CCT | C>A | A/T+1 | G/C-1 | CG>AT | CG>AT | AAAGACCTG | CTGTACC | GGA | CCT | C | T | C>A | C>A_CCT | CG>AT_CTGTACC | 6 | FALSE | FALSE |
| 135_v.txt | 33169 | 33169-SNV- | SNV | 1 | C | A | 48 | 29049 | 0.16523805 | 37.5833333 | rnr1Y285F_pG | TRUE | CAN | JSY2421 | 3 | rnr1Y285F_p | CCA | C>A | A/T+1 | G/C-1 | CG>AT | CG>AT | TAGTACCAC | CACCAAG | GGT | CCA | C | T | C>A | C>A_CCA | CG>AT_CACCAAG | 6 | FALSE | FALSE |
| 135_v.txt | 33177 | 33177-SNV- | SNV | 1 | G | T | 21 | 29044 | 0.07230409 | 37.7133921 | rnr1Y285F_pG | TRUE | CAN | JSY2421 | 3 | rnr1Y285F_p | GGC | G>T | A/T+1 | G/C-1 | CG>AT | CG>AT | CCAAGGGCA | GCAATCA | CCG | CCG | C | A | C>A | C>A_CCG | CG>AT_GCAATCA | 6 | FALSE | FALSE |
| 135_v.txt | 33208 | 33208-SNV- | SNV | 1 | C | A | 20 | 19827 | 0.10087255 | 41.9975714 | rnr1Y285F_pG | TRUE | CAN | JSY2421 | 3 | rnr1Y285F_p | TCT | C>A | A/T+1 | G/C-1 | CG>AT | CG>AT | TAAGCTCTCT | CTCTCTT | AGA | TCT | C | T | C>A | C>A_TCT | CG>AT_CTCTCTT | 6 | FALSE | FALSE |
| 135_v.txt | 33210 | 33210-SNV- | SNV | 1 | C | A | 10 | 19827 | 0.05043627 | 37.4 | rnr1Y285F_pG | TRUE | CAN | JSY2421 | 3 | rnr1Y285F_p | TCT | C>A | A/T+1 | G/C-1 | CG>AT | CG>AT | AGCTCTCTCT | CTCTTCA | AGA | TCT | C | T | C>A | C>A_TCT | CG>AT_CTCTTCA | 6 | FALSE | FALSE |
| 135_v.txt | 33220 | 33220-SNV- | SNV | 1 | C | A | 63 | 19836 | 0.31760436 | 39.031746 | rnr1Y285F_pG | TRUE | CAN | JSY2421 | 3 | rnr1Y285F_p | TCA | C>A | A/T+1 | G/C-1 | CG>AT | CG>AT | TCACTTCAGC | CAGCGTT | AGT | TCA | C | T | C>A | C>A_TCA | CG>AT_CAGCGTT | 6 | FALSE | FALSE |
| 135_v.txt | 33238 | 33238-SNV- | SNV | 1 | C | A | 21 | 19836 | 0.10586812 | 37.3809524 | rnr1Y285F_pG | TRUE | CAN | JSY2421 | 3 | rnr1Y285F_p | CCT | C>A | A/T+1 | G/C-1 | CG>AT | CG>AT | CTTCTCCTTC | CTTCATC | GGA | CCT | C | T | C>A | C>A_CCT | CG>AT_CTTCATC | 6 | FALSE | FALSE |
| 135_v.txt | 33241 | 33241-SNV- | SNV | 1 | C | A | 13 | 19836 | 0.06553741 | 44 | rnr1Y285F_pG | TRUE | CAN | JSY2421 | 3 | rnr1Y285F_p | TCA | C>A | A/T+1 | G/C-1 | CG>AT | CG>AT | CTCCTTCATC | CATCTTC | AGT | TCA | C | T | C>A | C>A_TCA | CG>AT_CATCTTC | 6 | FALSE | FALSE |
| 135_v.txt | 33325 | 33325-SNV- | SNV | 1 | C | A | 94 | 19836 | 0.47388586 | 57.0638298 | rnr1Y285F_pG | TRUE | CAN | JSY2421 | 3 | rnr1Y285F_p | TCT | C>A | A/T+1 | G/C-1 | CG>AT | CG>AT | ACAATTCTTT | CTTTACT | AGA | TCT | C | T | C>A | C>A_TCT | CG>AT_CTTTACT | 6 | FALSE | FALSE |
| 135_v.txt | 33354 | 33354-SNV- | SNV | 1 | G | T | 206 | 19836 | 1.03851583 | 37.5194175 | rnr1Y285F_pG | TRUE | CAN | JSY2421 | 3 | rnr1Y285F_p | TGA | G>T | A/T+1 | G/C-1 | CG>AT | CG>AT | GGTATTGAC | GACCCAC | ACT | ACT | C | A | C>A | C>A_ACT | CG>AT_GACCCAC | 6 | FALSE | FALSE |
| 135_v.txt | 33451 | 33451-SNV- | SNV | 1 | C | A | 65 | 19836 | 0.32768703 | 41.8153846 | rnr1Y285F_pG | TRUE | CAN | JSY2421 | 3 | rnr1Y285F_p | TCT | C>A | A/T+1 | G/C-1 | CG>AT | CG>AT | CGTCTTCTTT | CTTTTGA | AGA | TCT | C | T | C>A | C>A_TCT | CG>AT_CTTTTGA | 6 | FALSE | FALSE |
| 135_v.txt | 33474 | 33474-SNV- | SNV | 1 | C | A | 51 | 19833 | 0.25714718 | 32.9215686 | rnr1Y285F_pG | TRUE | CAN | JSY2421 | 3 | rnr1Y285F_p | GCC | C>A | A/T+1 | G/C-1 | CG>AT | CG>AT | GCTATGCCTT | CCTTTTT | CGG | GCC | C | T | C>A | C>A_GCC | CG>AT_CCTTTTT | 6 | FALSE | FALSE |
| 135_v.txt | 31911 | 31911-Repla | Replacement | 2 | GC | A | 6 | 10906 | 0.05501559 | 45.4444444 | rnr1Y285F_pG | TRUE | CAN | JSY2421 | 3 | rnr1Y285F_p | TGC | nope | A/T+1 | >1bpdel | none | Replac. | TTTGGTGCA | GCAAAAG | ACG | TGC | GC | T | NA | NA_TGC | Replac._GCAAAAG | 6 | FALSE | FALSE |
| 135_v.txt | 31912 | 31912-Repla | Replacement | 2 | C | AA | 32 | 10906 | 0.29341647 | 41.5468179 | rnr1Y285F_pG | TRUE | CAN | JSY2421 | 3 | rnr1Y285F_p | GCA | nope | >1bp-ins | G/C-1 | none | Replac. | TTGGTGCAA | CAAAAGC | CGT | GCA | C | none | nope | nope_GCA | Replac._CAAAAGC | 6 | FALSE | FALSE |
| 135_v.txt | 32497 | 32497-Repla | Replacement | 2 | C | TT | 43 | 33161 | 0.1296704 | 46.2674335 | rnr1Y285F_pG | TRUE | CAN | JSY2421 | 3 | rnr1Y285F_p | ACT | nope | >1bp-ins | G/C-1 | none | Replac. | AAACAACTTT | CTTTTTT | TGA | ACT | C | none | nope | nope_ACT | Replac._CTTTTTT | 6 | FALSE | FALSE |
| 135_v.txt | 32899 | 32899-Repla | Replacement | 2 | GG | A | 1 | 29052 | 3.44E-03 | 34 | rnr1Y285F_pG | TRUE | CAN | JSY2421 | 3 | rnr1Y285F_p | TGG | nope | A/T+1 | >1bpdel | none | Replac. | TGACTTGGC | GGCCAAC | ACC | TGG | GG | T | NA | NA_TGG | Replac._GGCCAAC | 6 | FALSE | FALSE |
| 135_v.txt | 32048 | 32048-MNV- | MNV | 2 | TC | AA | 52 | 23199 | 0.22414759 | 62.7496962 | rnr1Y285F_pG | TRUE | CAN | JSY2421 | 3 | rnr1Y285F_p | ATC | nope | >1bp-ins | >1bpdel | none | MNV | CATAAATCTC | TCTGATG | TAG | ATC | TC | none | NA | NA_ATC | MNV_TCTGATG | 6 | FALSE | FALSE |
| 135_v.txt | 32303 | 32303-MNV- | MNV | 2 | AG | GT | 71 | 33161 | 0.21410693 | 57.2484474 | rnr1Y285F_pG | TRUE | CAN | JSY2421 | 3 | rnr1Y285F_p | CAG | nope | >1bp-ins | >1bpdel | none | MNV | TGCGGCAGA | AGAAATA | GTC | CAG | AG | none | NA | NA_CAG | MNV_AGAAATA | 6 | FALSE | FALSE |
| 135_v.txt | 32571 | 32571-MNV- | MNV | 2 | CC | GT | 1 | 27974 | 3.58E-03 | 37.9999578 | rnr1Y285F_pG | TRUE | CAN | JSY2421 | 3 | rnr1Y285F_p | ACC | nope | >1bp-ins | >1bpdel | none | MNV | TCAGTACCTT | CCTTGAA | TGG | ACC | CC | none | NA | NA_ACC | MNV_CCTTGAA | 6 | FALSE | FALSE |
| 135_v.txt | 32798 | 32798-MNV- | MNV | 2 | GT | AG | 1 | 27973 | 3.58E-03 | 37.9999954 | rnr1Y285F_pG | TRUE | CAN | JSY2421 | 3 | rnr1Y285F_p | CGT | nope | >1bp-ins | >1bpdel | none | MNV | TTCACCGTAA | GTAATAT | GCA | CGT | GT | none | NA | NA_CGT | MNV_GTAATAT | 6 | FALSE | FALSE |
| 135_v.txt | 32800 | 32800-MNV- | MNV | 2 | AA | CT | 1 | 27973 | 3.58E-03 | 37.5000084 | rnr1Y285F_pG | TRUE | CAN | JSY2421 | 3 | rnr1Y285F_p | TAA | nope | >1bp-ins | >1bpdel | none | MNV | CACCGTAATA | AATATTT | ATT | TAA | AA | none | NA | NA_TAA | MNV_AATATTT | 6 | FALSE | FALSE |
| 135_v.txt | 32840 | 32840-MNV- | MNV | 2 | CC | AA | 1 | 27973 | 3.58E-03 | 31.000468 | rnr1Y285F_pG | TRUE | CAN | JSY2421 | 3 | rnr1Y285F_p | CCC | nope | >1bp-ins | >1bpdel | none | MNV | AATTACCCA | CCAAAAA | GGG | CCC | CC | none | NA | NA_CCC | MNV_CCAAAAA | 6 | FALSE | FALSE |
| 135_v.txt | 33153 | 33153-MNV- | MNV | 2 | CC | TA | 1 | 29052 | 3.44E-03 | 38 | rnr1Y285F_pG | TRUE | CAN | JSY2421 | 3 | rnr1Y285F_p | ACC | nope | >1bp-ins | >1bpdel | none | MNV | AAAAGACCT | CCTGTAC | TGG | ACC | CC | none | NA | NA_ACC | MNV_CCTGTAC | 6 | FALSE | FALSE |
| 135_v.txt | 31723 | 31723-Insert | Insertion | 1 | - | A | 2 | 10907 | 0.01833685 | 64 | rnr1Y285F_pG | TRUE | CAN | JSY2421 | 3 | rnr1Y285F_p | CAA | nope | A/T+1 | >1bpdel | none | A/T+1 | TGTCCCAAA | AAAAAGT | GTT | CAA | - | T | NA | NA_CAA | A/T+1_AAAAAGT | 6 | FALSE | FALSE |
| 135_v.txt | 31892 | 31892-Insert | Insertion | 4 | - | AACA | 24 | 10907 | 0.22004218 | 40.7395833 | rnr1Y285F_pG | TRUE | CAN | JSY2421 | 3 | rnr1Y285F_p | TAA | nope | >1bp-ins | >1bpdel | none | >1bp-ins | AAAGCTAAC | AACACCA | ATT | TAA | - | none | NA | NA_TAA | >1bp-ins_AACACCA | 6 | FALSE | FALSE |
| 135_v.txt | 31964 | 31964-Insert | Insertion | 1 | - | A | 32 | 23199 | 0.13793698 | 37.46875 | rnr1Y285F_pG | TRUE | CAN | JSY2421 | 3 | rnr1Y285F_p | CAT | nope | A/T+1 | >1bpdel | none | A/T+1 | GGCCGCATA | ATAATAA | GTA | CAT | - | T | NA | NA_CAT | A/T+1_ATAATAA | 6 | FALSE | FALSE |
| 135_v.txt | 31966 | 31966-Insert | Insertion | 1 | - | A | 6 | 23199 | 0.02586318 | 37.3333333 | rnr1Y285F_pG | TRUE | CAN | JSY2421 | 3 | rnr1Y285F_p | TAA | nope | A/T+1 | >1bpdel | none | A/T+1 | CCGCATAATA | AATAAGC | ATT | TAA | - | T | NA | NA_TAA | A/T+1_AATAAGC | 6 | FALSE | FALSE |
| 135_v.txt | 31967 | 31967-Insert | Insertion | 2 | - | AT | 19 | 23199 | 0.08190008 | 37.2894737 | rnr1Y285F_pG | TRUE | CAN | JSY2421 | 3 | rnr1Y285F_p | AAT | nope | >1bp-ins | >1bpdel | none | >1bp-ins | CGCATAATA | ATAAGCC | TTA | AAT | - | none | NA | NA_AAT | >1bp-ins_ATAAGCC | 6 | FALSE | FALSE |
| 135_v.txt | 32081 | 32081-Insert | Insertion | 1 | - | A | 2491 | 23199 | 10.7375318 | 63.593336 | rnr1Y285F_pG | TRUE | CAN | JSY2421 | 3 | rnr1Y285F_p | CAA | nope | A/T+1 | >1bpdel | none | A/T+1 | CCATGCAAA | AAAAAAG | GTT | CAA | - | T | NA | NA_CAA | A/T+1_AAAAAAG | 6 | FALSE | FALSE |
| 135_v.txt | 32081 | 32081-Insert | Insertion | 2 | - | AA | 2 | 23199 | 8.62E-03 | 64 | rnr1Y285F_pG | TRUE | CAN | JSY2421 | 3 | rnr1Y285F_p | CAA | nope | >1bp-ins | >1bpdel | none | >1bp-ins | CCATGCAAA | AAAAAAG | GTT | CAA | - | none | NA | NA_CAA | >1bp-ins_AAAAAAG | 6 | FALSE | FALSE |
| 135_v.txt | 32082 | 32082-Insert | Insertion | 1 | - | C | 1 | 23199 | 4.31E-03 | 2 | rnr1Y285F_pG | TRUE | CAN | JSY2421 | 3 | rnr1Y285F_p | AAA | nope | G/C+1 | >1bpdel | none | G/C+1 | CATGCAAAA | AAAAAGC | TTT | AAA | - | G | NA | NA_AAA | G/C+1_AAAAAGC | 6 | FALSE | FALSE |
| 135_v.txt | 32087 | 32087-Insert | Insertion | 1 | - | G | 2 | 23199 | 8.62E-03 | 64 | rnr1Y285F_pG | TRUE | CAN | JSY2421 | 3 | rnr1Y285F_p | AGC | nope | G/C+1 | >1bpdel | none | G/C+1 | AAAAAAGCC | GCCTGCA | TCG | AGC | - | C | NA | NA_AGC | G/C+1_GCCTGCA | 6 | FALSE | FALSE |
| 135_v.txt | 32475 | 32475-Insert | Insertion | 1 | - | A | 34 | 33161 | 0.10253008 | 46.9117647 | rnr1Y285F_pG | TRUE | CAN | JSY2421 | 3 | rnr1Y285F_p | GAA | nope | A/T+1 | >1bpdel | none | A/T+1 | ATGTAGAAG | AAGGTTA | CTT | GAA | - | T | NA | NA_GAA | A/T+1_AAGGTTA | 6 | FALSE | FALSE |
| 135_v.txt | 32498 | 32498-Insert | Insertion | 1 | - | T | 4 | 33161 | 0.01206236 | 37 | rnr1Y285F_pG | TRUE | CAN | JSY2421 | 3 | rnr1Y285F_p | CTT | nope | A/T+1 | >1bpdel | none | A/T+1 | AACAACTTTT | TTTTTTG | GAA | CTT | - | A | NA | NA_CTT | A/T+1_TTTTTTG | 6 | FALSE | FALSE |
| 135_v.txt | 32528 | 32528-Insert | Insertion | 1 | - | G | 2 | 33160 | 6.03E-03 | 51 | rnr1Y285F_pG | TRUE | CAN | JSY2421 | 3 | rnr1Y285F_p | TGG | nope | G/C+1 | >1bpdel | none | G/C+1 | TTTTCTGGGG | GGGGTTT | ACC | TGG | - | C | NA | NA_TGG | G/C+1_GGGGTTT | 6 | FALSE | FALSE |
| 135_v.txt | 32532 | 32532-Insert | Insertion | 1 | - | T | 40 | 33160 | 0.12062726 | 41.225 | rnr1Y285F_pG | TRUE | CAN | JSY2421 | 3 | rnr1Y285F_p | GTT | nope | A/T+1 | >1bpdel | none | A/T+1 | CTGGGGTTTC | TTTGCAG | CAA | GTT | - | A | NA | NA_GTT | A/T+1_TTTGCAG | 6 | FALSE | FALSE |
| 135_v.txt | 32665 | 32665-Insert | Insertion | 27 | - | CACCTGGGT | 4 | 27973 | 0.0142995 | 56.7962963 | rnr1Y285F_pG | TRUE | CAN | JSY2421 | 3 | rnr1Y285F_p | GCA | nope | >1bp-ins | >1bpdel | none | >1bp-ins | CCCAGGCAC | CACCTGG | CGT | GCA | - | none | NA | NA_GCA | >1bp-ins_CACCTGG | 6 | FALSE | FALSE |
| 135_v.txt | 32677 | 32677-Insert | Insertion | 1 | - | T | 21 | 27973 | 0.07507239 | 59.8571429 | rnr1Y285F_pG | TRUE | CAN | JSY2421 | 3 | rnr1Y285F_p | CTC | nope | A/T+1 | >1bpdel | none | A/T+1 | GGTTTCTCCA | TCCAATA | GAG | CTC | - | A | NA | NA_CTC | A/T+1_TCCAATA | 6 | FALSE | FALSE |
| 135_v.txt | 32728 | 32728-Insert | Insertion | 1 | - | A | 6 | 27973 | 0.02144926 | 63.5 | rnr1Y285F_pG | TRUE | CAN | JSY2421 | 3 | rnr1Y285F_p | CAA | nope | A/T+1 | >1bpdel | none | A/T+1 | CCATACAAA | AAAAACA | GTT | CAA | - | T | NA | NA_CAA | A/T+1_AAAAACA | 6 | FALSE | FALSE |
| 135_v.txt | 32842 | 32842-Insert | Insertion | 1 | - | A | 210 | 27973 | 0.75072391 | 43.6857143 | rnr1Y285F_pG | TRUE | CAN | JSY2421 | 3 | rnr1Y285F_p | CAA | nope | A/T+1 | >1bpdel | none | A/T+1 | TTACCCAAA | AAAAAAT | GTT | CAA | - | T | NA | NA_CAA | A/T+1_AAAAAAT | 6 | FALSE | FALSE |
| 135_v.txt | 33013 | 33013-Insert | Insertion | 10 | - | AGGATGTAA | 25 | 29054 | 0.08604667 | 62.624 | rnr1Y285F_pG | TRUE | CAN | JSY2421 | 3 | rnr1Y285F_p | GAG | nope | >1bp-ins | >1bpdel | none | >1bp-ins | TGAAAGAGG | AGGATGT | CTC | GAG | - | none | NA | NA_GAG | >1bp-ins_AGGATGT | 6 | FALSE | FALSE |
| 135_v.txt | 33036 | 33036-Insert | Insertion | 33 | - | GCCATTTCAC | 14 | 29054 | 0.04818614 | 59.465368 | rnr1Y285F_pG | TRUE | CAN | JSY2421 | 3 | rnr1Y285F_p | AGC | nope | >1bp-ins | >1bpdel | none | >1bp-ins | AATGTAGCC | GCCATTT | TCG | AGC | - | none | NA | NA_AGC | >1bp-ins_GCCATTT | 6 | FALSE | FALSE |
| 135_v.txt | 33096 | 33096-Insert | Insertion | 3 | - | ATA | 18 | 29051 | 0.06196 | 53.462963 | rnr1Y285F_pG | TRUE | CAN | JSY2421 | 3 | rnr1Y285F_p | TAT | nope | >1bp-ins | >1bpdel | none | >1bp-ins | TATGATATA | ATAAGAG | ATA | TAT | - | none | NA | NA_TAT | >1bp-ins_ATAAGAG | 6 | FALSE | FALSE |
| 135_v.txt | 33206 | 33206-Insert | Insertion | 2 | - | AT | 3 | 19827 | 0.01513088 | 37.8333333 | rnr1Y285F_pG | TRUE | CAN | JSY2421 | 3 | rnr1Y285F_p | GCT | nope | >1bp-ins | >1bpdel | none | >1bp-ins | CTTAAGCTCT | CTCTCTC | CGA | GCT | - | none | NA | NA_GCT | >1bp-ins_CTCTCTC | 6 | FALSE | FALSE |
| 135_v.txt | 33206 | 33206-Insert | Insertion | 2 | - | CT | 24 | 19827 | 0.12104706 | 37.5625 | rnr1Y285F_pG | TRUE | CAN | JSY2421 | 3 | rnr1Y285F_p | GCT | nope | >1bp-ins | >1bpdel | none | >1bp-ins | CTTAAGCTCT | CTCTCTC | CGA | GCT | - | none | NA | NA_GCT | >1bp-ins_CTCTCTC | 6 | FALSE | FALSE |
| 135_v.txt | 33384 | 33384-Insert | Insertion | 2 | - | TC | 9 | 19836 | 0.04537205 | 52.0555556 | rnr1Y285F_pG | TRUE | CAN | JSY2421 | 3 | rnr1Y285F_p | TTC | nope | >1bp-ins | >1bpdel | none | >1bp-ins | GAAGCTTCA | TCAACGT | AAG | TTC | - | none | NA | NA_TTC | >1bp-ins_TCAACGT | 6 | FALSE | FALSE |
| 135_v.txt | 32126 | 32126-Deleti | Deletion | 2 | TT | - | 30 | 23199 | 0.12931592 | 57.8666667 | rnr1Y285F_pG | TRUE | CAN | JSY2421 | 3 | rnr1Y285F_p | CTT | nope | >1bp-ins | >1bpdel | none | >1bpdel | GAAAACTTT | TTTGTCA | GAA | CTT | TT | none | NA | NA_CTT | >1bpdel_TTTGTCA | 6 | FALSE | FALSE |
| 135_v.txt | 32335 | 32335-Deleti | Deletion | 2 | AG | - | 28 | 33161 | 0.08443654 | 63.3214286 | rnr1Y285F_pG | TRUE | CAN | JSY2421 | 3 | rnr1Y285F_p | AAG | nope | >1bp-ins | >1bpdel | none | >1bpdel | CGTTGAAGA | AGATATG | TTC | AAG | AG | none | NA | NA_AAG | >1bpdel_AGATATG | 6 | FALSE | FALSE |
| 135_v.txt | 32454 | 32454-Deleti | Deletion | 3 | AAT | - | 37 | 33161 | 0.11157685 | 52.1621622 | rnr1Y285F_pG | TRUE | CAN | JSY2421 | 3 | rnr1Y285F_p | GAA | nope | >1bp-ins | >1bpdel | none | >1bpdel | CCAATGAAT | AATAATA | CTT | GAA | AAT | none | NA | NA_GAA | >1bpdel_AATAATA | 6 | FALSE | FALSE |
| 135_v.txt | 32642 | 32642-Deleti | Deletion | 2 | AG | - | 184 | 27973 | 0.65777714 | 51.7065217 | rnr1Y285F_pG | TRUE | CAN | JSY2421 | 3 | rnr1Y285F_p | TAG | nope | >1bp-ins | >1bpdel | none | >1bpdel | ATCCTTAGAT | AGATATT | ATC | TAG | AG | none | NA | NA_TAG | >1bpdel_AGATATT | 6 | FALSE | FALSE |
| 135_v.txt | 32751 | 32751-Deleti | Deletion | 3 | ATA | - | 20 | 27973 | 0.07149752 | 59.7 | rnr1Y285F_pG | TRUE | CAN | JSY2421 | 3 | rnr1Y285F_p | GAT | nope | >1bp-ins | >1bpdel | none | >1bpdel | AACCCGATA | ATAATGG | CTA | GAT | ATA | none | NA | NA_GAT | >1bpdel_ATAATGG | 6 | FALSE | FALSE |
| 135_v.txt | 32994 | 32994-Deleti | Deletion | 2 | TG | - | 11 | 29054 | 0.03786054 | 60.5454546 | rnr1Y285F_pG | TRUE | CAN | JSY2421 | 3 | rnr1Y285F_p | TTG | nope | >1bp-ins | >1bpdel | none | >1bpdel | AATCTTTGTG | TGTGAGA | AAC | TTG | TG | none | NA | NA_TTG | >1bpdel_TGTGAGA | 6 | FALSE | FALSE |
| 135_v.txt | 33000 | 33000-Deleti | Deletion | 2 | AA | - | 18 | 29054 | 0.0619536 | 62.3888889 | rnr1Y285F_pG | TRUE | CAN | JSY2421 | 3 | rnr1Y285F_p | GAA | nope | >1bp-ins | >1bpdel | none | >1bpdel | TGTGAGAAA | AAAACTG | CTT | GAA | AA | none | NA | NA_GAA | >1bpdel_AAAACTG | 6 | FALSE | FALSE |
| 135_v.txt | 33189 | 33189-Deleti | Deletion | 2 | AT | - | 14 | 19828 | 0.07060722 | 35.5714286 | rnr1Y285F_pG | TRUE | CAN | JSY2421 | 3 | rnr1Y285F_p | AAT | nope | >1bp-ins | >1bpdel | none | >1bpdel | ATACCAATAT | ATATGTC | TTA | AAT | AT | none | NA | NA_AAT | >1bpdel_ATATGTC | 6 | FALSE | FALSE |
| 135_v.txt | 33206 | 33206-Deleti | Deletion | 2 | CT | - | 26 | 19827 | 0.13113431 | 37.6153846 | rnr1Y285F_pG | TRUE | CAN | JSY2421 | 3 | rnr1Y285F_p | GCT | nope | >1bp-ins | >1bpdel | none | >1bpdel | CTTAAGCTCT | CTCTCTC | CGA | GCT | CT | none | NA | NA_GCT | >1bpdel_CTCTCTC | 6 | FALSE | FALSE |
| 135_v.txt | 31723 | 31723-Deleti | Deletion | 1 | A | - | 9 | 10907 | 0.08251582 | 63.2222226 | rnr1Y285F_pG | TRUE | CAN | JSY2421 | 3 | rnr1Y285F_p | CAA | nope | >1bp-ins | A/T-1 | none | A/T-1 | TGTCCCAAA | AAAAAGT | GTT | GTT | T | none | T>none | T>none_GTT | A/T-1_AAAAAGT | 6 | FALSE | FALSE |
| 135_v.txt | 31890 | 31890-Deleti | Deletion | 1 | C | - | 16 | 10907 | 0.14669478 | 44.0625 | rnr1Y285F_pG | TRUE | CAN | JSY2421 | 3 | rnr1Y285F_p | GCT | nope | >1bp-ins | G/C-1 | none | G/C-1 | GCAAAGCTA | CTAACAC | CGA | GCT | C | none | nope | nope_GCT | G/C-1_CTAACAC | 6 | FALSE | FALSE |
| 135_v.txt | 31904 | 31904-Deleti | Deletion | 1 | T | - | 8 | 10906 | 0.07335412 | 42.375 | rnr1Y285F_pG | TRUE | CAN | JSY2421 | 3 | rnr1Y285F_p | ATT | nope | >1bp-ins | A/T-1 | none | A/T-1 | ATTGAATTTT | TTTTGGT | TAA | ATT | T | none | nope | nope_ATT | A/T-1_TTTTGGT | 6 | FALSE | FALSE |
| 135_v.txt | 31923 | 31923-Deleti | Deletion | 1 | A | - | 22 | 10906 | 0.20172382 | 37.8181673 | rnr1Y285F_pG | TRUE | CAN | JSY2421 | 3 | rnr1Y285F_p | GAA | nope | >1bp-ins | A/T-1 | none | A/T-1 | GCCGTGAAA | AAACCTT | CTT | CTT | T | none | T>none | T>none_CTT | A/T-1_AAACCTT | 6 | FALSE | FALSE |
| 135_v.txt | 31952 | 31952-Deleti | Deletion | 1 | A | - | 18 | 23199 | 0.07758955 | 36.9444444 | rnr1Y285F_pG | TRUE | CAN | JSY2421 | 3 | rnr1Y285F_p | TAA | nope | >1bp-ins | A/T-1 | none | A/T-1 | CGTCATAAAT | AAATGTG | ATT | ATT | T | none | T>none | T>none_ATT | A/T-1_AAATGTG | 6 | FALSE | FALSE |
| 135_v.txt | 31977 | 31977-Deleti | Deletion | 1 | C | - | 21 | 23199 | 0.09052114 | 44.3333333 | rnr1Y285F_pG | TRUE | CAN | JSY2421 | 3 | rnr1Y285F_p | GCC | nope | >1bp-ins | G/C-1 | none | G/C-1 | GCCAAGCCG | CCGGGCA | CGG | GCC | C | none | nope | nope_GCC | G/C-1_CCGGGCA | 6 | FALSE | FALSE |
| 135_v.txt | 31979 | 31979-Deleti | Deletion | 1 | G | - | 23 | 23199 | 0.0991422 | 41.6087071 | rnr1Y285F_pG | TRUE | CAN | JSY2421 | 3 | rnr1Y285F_p | CGG | nope | >1bp-ins | G/C-1 | none | G/C-1 | CAAGCCGGG | GGGCATT | GCC | GCC | C | none | C>none | C>none_GCC | G/C-1_GGGCATT | 6 | FALSE | FALSE |
| 135_v.txt | 31988 | 31988-Deleti | Deletion | 1 | T | - | 5 | 23199 | 0.02155265 | 51.4 | rnr1Y285F_pG | TRUE | CAN | JSY2421 | 3 | rnr1Y285F_p | ATT | nope | >1bp-ins | A/T-1 | none | A/T-1 | CATTAATTTA | TTTAGCT | TAA | ATT | T | none | nope | nope_ATT | A/T-1_TTTAGCT | 6 | FALSE | FALSE |
| 135_v.txt | 32024 | 32024-Deleti | Deletion | 1 | A | - | 7 | 23199 | 0.03017371 | 41.2857143 | rnr1Y285F_pG | TRUE | CAN | JSY2421 | 3 | rnr1Y285F_p | CAC | nope | >1bp-ins | A/T-1 | none | A/T-1 | GATGCCACG | ACGGTAT | GTG | GTG | T | none | T>none | T>none_GTG | A/T-1_ACGGTAT | 6 | FALSE | FALSE |
| 135_v.txt | 32034 | 32034-Deleti | Deletion | 1 | A | - | 46 | 23199 | 0.19828441 | 58.8478261 | rnr1Y285F_pG | TRUE | CAN | JSY2421 | 3 | rnr1Y285F_p | CAA | nope | >1bp-ins | A/T-1 | none | A/T-1 | TATTTCAAAG | AAAGCTT | GTT | GTT | T | none | T>none | T>none_GTT | A/T-1_AAAGCTT | 6 | FALSE | FALSE |

| | | | | | | | | | | | | | | | | | | | | | | | | | | | | | | | | | | |
|---|---|---|---|---|---|---|---|---|---|---|---|---|---|---|---|---|---|---|---|---|---|---|---|---|---|---|---|---|---|---|---|---|---|---|
| 135_v.txt | 32042 | 32042-Deleti | Deletion | 1 | C | - | 52 | 23196 | 0.22417658 | 61.2499992 | rnr1Y285F_pG | TRUE | CAN | JSY2421 | 3 | rnr1Y285F_p | GCA | nope | >1bp-ins | G/C-1 | none | G/C-1 | AGCTTGCAT | CATAAAT | CGT | GCA | C | none | nope | nope_GCA | G/C-1_CATAAAT | 6 | FALSE | FALSE |
| 135_v.txt | 32068 | 32068-Deleti | Deletion | 1 | T | - | 24 | 23199 | 0.10345274 | 61.7916667 | rnr1Y285F_pG | TRUE | CAN | JSY2421 | 3 | rnr1Y285F_p | ATA | nope | >1bp-ins | A/T-1 | none | A/T-1 | TTGAGATAA | TAAATAA | TAT | ATA | T | none | nope | nope_ATA | A/T-1_TAAATAA | 6 | FALSE | FALSE |
| 135_v.txt | 32073 | 32073-Deleti | Deletion | 1 | A | - | 2 | 23199 | 8.62E-03 | 64 | rnr1Y285F_pG | TRUE | CAN | JSY2421 | 3 | rnr1Y285F_p | TAA | nope | >1bp-ins | A/T-1 | none | A/T-1 | ATAAATAAC | AACCATG | ATT | ATT | T | none | T>none | T>none_ATT | A/T-1_AACCATG | 6 | FALSE | FALSE |
| 135_v.txt | 32075 | 32075-Deleti | Deletion | 1 | C | - | 2 | 23199 | 8.62E-03 | 64 | rnr1Y285F_pG | TRUE | CAN | JSY2421 | 3 | rnr1Y285F_p | ACC | nope | >1bp-ins | G/C-1 | none | G/C-1 | AAATAACCA | CCATGCA | TGG | ACC | C | none | nope | nope_ACC | G/C-1_CCATGCA | 6 | FALSE | FALSE |
| 135_v.txt | 32081 | 32081-Deleti | Deletion | 1 | A | - | 16 | 23199 | 0.06896849 | 63.0625 | rnr1Y285F_pG | TRUE | CAN | JSY2421 | 3 | rnr1Y285F_p | CAA | nope | >1bp-ins | A/T-1 | none | A/T-1 | CCATGCAAA | AAAAAAG | GTT | GTT | T | none | T>none | T>none_GTT | A/T-1_AAAAAAG | 6 | FALSE | FALSE |
| 135_v.txt | 32113 | 32113-Deleti | Deletion | 1 | G | - | 18 | 23199 | 0.07758955 | 60.6111131 | rnr1Y285F_pG | TRUE | CAN | JSY2421 | 3 | rnr1Y285F_p | AGC | nope | >1bp-ins | G/C-1 | none | G/C-1 | TTAATAGCC | GCCATTC | TCG | TCG | C | none | C>none | C>none_TCG | G/C-1_GCCATTC | 6 | FALSE | FALSE |
| 135_v.txt | 32168 | 32168-Deleti | Deletion | 1 | A | - | 32 | 23199 | 0.13793698 | 48.6875 | rnr1Y285F_pG | TRUE | CAN | JSY2421 | 3 | rnr1Y285F_p | CAA | nope | >1bp-ins | A/T-1 | none | A/T-1 | AGCGCCAAA | AAATGCA | GTT | GTT | T | none | T>none | T>none_GTT | A/T-1_AAATGCA | 6 | FALSE | FALSE |
| 135_v.txt | 32184 | 32184-Deleti | Deletion | 1 | A | - | 139 | 23199 | 0.59916376 | 48.7769784 | rnr1Y285F_pG | TRUE | CAN | JSY2421 | 3 | rnr1Y285F_p | GAA | nope | >1bp-ins | A/T-1 | none | A/T-1 | GTAACGAAA | AAAACTG | CTT | CTT | T | none | T>none | T>none_CTT | A/T-1_AAAACTG | 6 | FALSE | FALSE |
| 135_v.txt | 32188 | 32188-Deleti | Deletion | 1 | C | - | 15 | 23199 | 0.06465796 | 49.2666667 | rnr1Y285F_pG | TRUE | CAN | JSY2421 | 3 | rnr1Y285F_p | ACT | nope | >1bp-ins | G/C-1 | none | G/C-1 | CGAAAACTG | CTGCAAT | TGA | ACT | C | none | nope | nope_ACT | G/C-1_CTGCAAT | 6 | FALSE | FALSE |
| 135_v.txt | 32205 | 32205-Deleti | Deletion | 1 | C | - | 6 | 23199 | 0.02586318 | 49.1666667 | rnr1Y285F_pG | TRUE | CAN | JSY2421 | 3 | rnr1Y285F_p | ACC | nope | >1bp-ins | G/C-1 | none | G/C-1 | GGAACACCA | CCACCTT | TGG | ACC | C | none | nope | nope_ACC | G/C-1_CCACCTT | 6 | FALSE | FALSE |
| 135_v.txt | 32210 | 32210-Deleti | Deletion | 1 | T | - | 6 | 23199 | 0.02586318 | 50.6666667 | rnr1Y285F_pG | TRUE | CAN | JSY2421 | 3 | rnr1Y285F_p | CTT | nope | >1bp-ins | A/T-1 | none | A/T-1 | ACCACCTTTG | TTTGGTG | GAA | CTT | T | none | nope | nope_CTT | A/T-1_TTTGGTG | 6 | FALSE | FALSE |
| 135_v.txt | 32225 | 32225-Deleti | Deletion | 1 | C | - | 9 | 23199 | 0.03879478 | 37.5555553 | rnr1Y285F_pG | TRUE | CAN | JSY2421 | 3 | rnr1Y285F_p | ACA | nope | >1bp-ins | G/C-1 | none | G/C-1 | CCTTGACAG | CAGGAAT | TGT | ACA | C | none | nope | nope_ACA | G/C-1_CAGGAAT | 6 | FALSE | FALSE |
| 135_v.txt | 32265 | 32265-Deleti | Deletion | 1 | A | - | 6 | 33139 | 0.01810556 | 38 | rnr1Y285F_pG | TRUE | CAN | JSY2421 | 3 | rnr1Y285F_p | TAA | nope | >1bp-ins | A/T-1 | none | A/T-1 | CCAAATAAA | AAAATAC | ATT | ATT | T | none | T>none | T>none_ATT | A/T-1_AAAATAC | 6 | FALSE | FALSE |
| 135_v.txt | 32277 | 32277-Deleti | Deletion | 1 | C | - | 12 | 33139 | 0.03621111 | 35.25 | rnr1Y285F_pG | TRUE | CAN | JSY2421 | 3 | rnr1Y285F_p | ACC | nope | >1bp-ins | G/C-1 | none | G/C-1 | CGGGAACCA | CCAACGT | TGG | ACC | C | none | nope | nope_ACC | G/C-1_CCAACGT | 6 | FALSE | FALSE |
| 135_v.txt | 32302 | 32302-Deleti | Deletion | 1 | C | - | 42 | 33161 | 0.12665481 | 56.2619063 | rnr1Y285F_pG | TRUE | CAN | JSY2421 | 3 | rnr1Y285F_p | GCA | nope | >1bp-ins | G/C-1 | none | G/C-1 | TTGCGGCAG | CAGAAAT | CGT | GCA | C | none | nope | nope_GCA | G/C-1_CAGAAAT | 6 | FALSE | FALSE |
| 135_v.txt | 32345 | 32345-Deleti | Deletion | 1 | C | - | 14 | 33161 | 0.04221827 | 59.3571413 | rnr1Y285F_pG | TRUE | CAN | JSY2421 | 3 | rnr1Y285F_p | GCA | nope | >1bp-ins | G/C-1 | none | G/C-1 | ATGTGGCAA | CAAAACC | CGT | GCA | C | none | nope | nope_GCA | G/C-1_CAAAACC | 6 | FALSE | FALSE |
| 135_v.txt | 32346 | 32346-Deleti | Deletion | 1 | A | - | 224 | 33160 | 0.67551267 | 61.4241056 | rnr1Y285F_pG | TRUE | CAN | JSY2421 | 3 | rnr1Y285F_p | CAA | nope | >1bp-ins | A/T-1 | none | A/T-1 | TGTGGCAAA | AAAACCT | GTT | GTT | T | none | T>none | T>none_GTT | A/T-1_AAAACCT | 6 | FALSE | FALSE |
| 135_v.txt | 32381 | 32381-Deleti | Deletion | 1 | A | - | 11 | 33161 | 0.0331715 | 63.1818182 | rnr1Y285F_pG | TRUE | CAN | JSY2421 | 3 | rnr1Y285F_p | TAA | nope | >1bp-ins | A/T-1 | none | A/T-1 | AATAATAAA | AAAGGGA | ATT | ATT | T | none | T>none | T>none_ATT | A/T-1_AAAGGGA | 6 | FALSE | FALSE |
| 135_v.txt | 32419 | 32419-Deleti | Deletion | 1 | G | - | 25 | 33161 | 0.07538977 | 56.84 | rnr1Y285F_pG | TRUE | CAN | JSY2421 | 3 | rnr1Y285F_p | AGT | nope | >1bp-ins | G/C-1 | none | G/C-1 | GTGTTAGTTT | GTTTAGG | TCA | TCA | C | none | C>none | C>none_TCA | G/C-1_GTTTAGG | 6 | FALSE | FALSE |
| 135_v.txt | 32436 | 32436-Deleti | Deletion | 1 | G | - | 2 | 33161 | 6.03E-03 | 50.5 | rnr1Y285F_pG | TRUE | CAN | JSY2421 | 3 | rnr1Y285F_p | TGG | nope | >1bp-ins | G/C-1 | none | G/C-1 | TTGTATGGA | GGAACTA | ACC | ACC | C | none | C>none | C>none_ACC | G/C-1_GGAACTA | 6 | FALSE | FALSE |
| 135_v.txt | 32442 | 32442-Deleti | Deletion | 1 | A | - | 17 | 33161 | 0.05126504 | 56.3529412 | rnr1Y285F_pG | TRUE | CAN | JSY2421 | 3 | rnr1Y285F_p | TAA | nope | >1bp-ins | A/T-1 | none | A/T-1 | GGAACTAAA | AAAAGTC | ATT | ATT | T | none | T>none | T>none_ATT | A/T-1_AAAAGTC | 6 | FALSE | FALSE |
| 135_v.txt | 32448 | 32448-Deleti | Deletion | 1 | C | - | 2 | 33161 | 6.03E-03 | 50.0000001 | rnr1Y285F_pG | TRUE | CAN | JSY2421 | 3 | rnr1Y285F_p | TCC | nope | >1bp-ins | G/C-1 | none | G/C-1 | AAAAGTCCA | CCAATGA | AGG | TCC | C | none | nope | nope_TCC | G/C-1_CCAATGA | 6 | FALSE | FALSE |
| 135_v.txt | 32490 | 32490-Deleti | Deletion | 1 | A | - | 28 | 33161 | 0.08443654 | 50.1428571 | rnr1Y285F_pG | TRUE | CAN | JSY2421 | 3 | rnr1Y285F_p | GAA | nope | >1bp-ins | A/T-1 | none | A/T-1 | ATACGGAAA | AAAACAA | CTT | CTT | T | none | T>none | T>none_CTT | A/T-1_AAAACAA | 6 | FALSE | FALSE |
| 135_v.txt | 32494 | 32494-Deleti | Deletion | 1 | C | - | 115 | 33160 | 0.34680338 | 46.6260871 | rnr1Y285F_pG | TRUE | CAN | JSY2421 | 3 | rnr1Y285F_p | ACA | nope | >1bp-ins | G/C-1 | none | G/C-1 | GGAAAACAA | CAACTTT | TGT | ACA | C | none | nope | nope_ACA | G/C-1_CAACTTT | 6 | FALSE | FALSE |
| 135_v.txt | 32498 | 32498-Deleti | Deletion | 1 | T | - | 311 | 33161 | 0.93784868 | 44.344052 | rnr1Y285F_pG | TRUE | CAN | JSY2421 | 3 | rnr1Y285F_p | CTT | nope | >1bp-ins | A/T-1 | none | A/T-1 | AACAACTTTT | TTTTTTG | GAA | CTT | T | none | nope | nope_CTT | A/T-1_TTTTTTG | 6 | FALSE | FALSE |
| 135_v.txt | 32522 | 32522-Deleti | Deletion | 1 | T | - | 66 | 33160 | 0.19903498 | 40.1212121 | rnr1Y285F_pG | TRUE | CAN | JSY2421 | 3 | rnr1Y285F_p | ATT | nope | >1bp-ins | A/T-1 | none | A/T-1 | AACGGATTTT | TTTTCTG | TAA | ATT | T | none | nope | nope_ATT | A/T-1_TTTTCTG | 6 | FALSE | FALSE |
| 135_v.txt | 32528 | 32528-Deleti | Deletion | 1 | G | - | 9 | 33160 | 0.02714113 | 38.8888875 | rnr1Y285F_pG | TRUE | CAN | JSY2421 | 3 | rnr1Y285F_p | TGG | nope | >1bp-ins | G/C-1 | none | G/C-1 | TTTTCTGGGG | GGGGTTT | ACC | ACC | C | none | C>none | C>none_ACC | G/C-1_GGGGTTT | 6 | FALSE | FALSE |
| 135_v.txt | 32598 | 32598-Deleti | Deletion | 1 | A | - | 6 | 27973 | 0.02144926 | 38 | rnr1Y285F_pG | TRUE | CAN | JSY2421 | 3 | rnr1Y285F_p | CAA | nope | >1bp-ins | A/T-1 | none | A/T-1 | TTAATCAAA | AAAGAGG | GTT | GTT | T | none | T>none | T>none_GTT | A/T-1_AAAGAGG | 6 | FALSE | FALSE |
| 135_v.txt | 32608 | 32608-Deleti | Deletion | 1 | C | - | 28 | 27973 | 0.10009652 | 44.4642857 | rnr1Y285F_pG | TRUE | CAN | JSY2421 | 3 | rnr1Y285F_p | ACC | nope | >1bp-ins | G/C-1 | none | G/C-1 | AGGAAACCC | CCCAACC | TGG | ACC | C | none | nope | nope_ACC | G/C-1_CCCAACC | 6 | FALSE | FALSE |
| 135_v.txt | 32631 | 32631-Deleti | Deletion | 1 | T | - | 231 | 27973 | 0.8257963 | 49.2727273 | rnr1Y285F_pG | TRUE | CAN | JSY2421 | 3 | rnr1Y285F_p | GTT | nope | >1bp-ins | A/T-1 | none | A/T-1 | CCTTCGTTTT | TTTTTAT | CAA | GTT | T | none | nope | nope_GTT | A/T-1_TTTTTAT | 6 | FALSE | FALSE |
| 135_v.txt | 32658 | 32658-Deleti | Deletion | 1 | C | - | 119 | 27973 | 0.42541022 | 58.7563025 | rnr1Y285F_pG | TRUE | CAN | JSY2421 | 3 | rnr1Y285F_p | ACC | nope | >1bp-ins | G/C-1 | none | G/C-1 | CCTGGACCC | CCCCAGG | TGG | ACC | C | none | nope | nope_ACC | G/C-1_CCCCAGG | 6 | FALSE | FALSE |
| 135_v.txt | 32670 | 32670-Deleti | Deletion | 1 | G | - | 109 | 27973 | 0.38966146 | 59.5412844 | rnr1Y285F_pG | TRUE | CAN | JSY2421 | 3 | rnr1Y285F_p | TGG | nope | >1bp-ins | G/C-1 | none | G/C-1 | GCACCTGGG | GGGTTTC | ACC | ACC | C | none | C>none | C>none_ACC | G/C-1_GGGTTTC | 6 | FALSE | FALSE |
| 135_v.txt | 32680 | 32680-Deleti | Deletion | 1 | A | - | 2 | 27973 | 7.15E-03 | 38.0000089 | rnr1Y285F_pG | TRUE | CAN | JSY2421 | 3 | rnr1Y285F_p | CAA | nope | >1bp-ins | A/T-1 | none | A/T-1 | TTCTCCAATA | AATAACG | GTT | GTT | T | none | T>none | T>none_GTT | A/T-1_AATAACG | 6 | FALSE | FALSE |
| 135_v.txt | 32686 | 32686-Deleti | Deletion | 1 | G | - | 28 | 27973 | 0.10009652 | 63.5 | rnr1Y285F_pG | TRUE | CAN | JSY2421 | 3 | rnr1Y285F_p | CGG | nope | >1bp-ins | G/C-1 | none | G/C-1 | AATAACGGA | GGAATCC | GCC | GCC | C | none | C>none | C>none_GCC | G/C-1_GGAATCC | 6 | FALSE | FALSE |
| 135_v.txt | 32707 | 32707-Deleti | Deletion | 1 | C | - | 80 | 27973 | 0.28599006 | 62.6375 | rnr1Y285F_pG | TRUE | CAN | JSY2421 | 3 | rnr1Y285F_p | ACC | nope | >1bp-ins | G/C-1 | none | G/C-1 | CGGTAACCC | CCCCAGC | TGG | ACC | C | none | nope | nope_ACC | G/C-1_CCCCAGC | 6 | FALSE | FALSE |
| 135_v.txt | 32728 | 32728-Deleti | Deletion | 1 | A | - | 74 | 27973 | 0.26454081 | 60.0405308 | rnr1Y285F_pG | TRUE | CAN | JSY2421 | 3 | rnr1Y285F_p | CAA | nope | >1bp-ins | A/T-1 | none | A/T-1 | CCATACAAA | AAAAACA | GTT | GTT | T | none | T>none | T>none_GTT | A/T-1_AAAAACA | 6 | FALSE | FALSE |
| 135_v.txt | 32752 | 32752-Deleti | Deletion | 1 | T | - | 3 | 27973 | 0.01072463 | 55.3333318 | rnr1Y285F_pG | TRUE | CAN | JSY2421 | 3 | rnr1Y285F_p | ATA | nope | >1bp-ins | A/T-1 | none | A/T-1 | ACCCGATAA | TAATGGC | TAT | ATA | T | none | nope | nope_ATA | A/T-1_TAATGGC | 6 | FALSE | FALSE |
| 135_v.txt | 32804 | 32804-Deleti | Deletion | 1 | T | - | 22 | 27973 | 0.07864727 | 46.1363636 | rnr1Y285F_pG | TRUE | CAN | JSY2421 | 3 | rnr1Y285F_p | ATT | nope | >1bp-ins | A/T-1 | none | A/T-1 | GTAATATTTG | TTTGACA | TAA | ATT | T | none | nope | nope_ATT | A/T-1_TTTGACA | 6 | FALSE | FALSE |
| 135_v.txt | 32842 | 32842-Deleti | Deletion | 1 | A | - | 142 | 27973 | 0.50763236 | 42.2887354 | rnr1Y285F_pG | TRUE | CAN | JSY2421 | 3 | rnr1Y285F_p | CAA | nope | >1bp-ins | A/T-1 | none | A/T-1 | TTACCCAAA | AAAAAAT | GTT | GTT | T | none | T>none | T>none_GTT | A/T-1_AAAAAAT | 6 | FALSE | FALSE |
| 135_v.txt | 32884 | 32884-Deleti | Deletion | 1 | A | - | 29 | 29054 | 0.09981414 | 36.5517241 | rnr1Y285F_pG | TRUE | CAN | JSY2421 | 3 | rnr1Y285F_p | CAA | nope | >1bp-ins | A/T-1 | none | A/T-1 | ACGTCCAAA | AAAATTG | GTT | GTT | T | none | T>none | T>none_GTT | A/T-1_AAAATTG | 6 | FALSE | FALSE |
| 135_v.txt | 32901 | 32901-Deleti | Deletion | 1 | C | - | 1 | 29052 | 3.44E-03 | 37.9999997 | rnr1Y285F_pG | TRUE | CAN | JSY2421 | 3 | rnr1Y285F_p | GCC | nope | >1bp-ins | G/C-1 | none | G/C-1 | ACTTGGCCA | CCAACTA | CGG | GCC | C | none | nope | nope_GCC | G/C-1_CCAACTA | 6 | FALSE | FALSE |
| 135_v.txt | 32917 | 32917-Deleti | Deletion | 1 | C | - | 101 | 29052 | 0.34765249 | 46.980198 | rnr1Y285F_pG | TRUE | CAN | JSY2421 | 3 | rnr1Y285F_p | TCC | nope | >1bp-ins | G/C-1 | none | G/C-1 | TAAGTTCCA | CCAGGGC | AGG | TCC | C | none | nope | nope_TCC | G/C-1_CCAGGGC | 6 | FALSE | FALSE |
| 135_v.txt | 32924 | 32924-Deleti | Deletion | 1 | A | - | 22 | 29052 | 0.07572628 | 52.0909091 | rnr1Y285F_pG | TRUE | CAN | JSY2421 | 3 | rnr1Y285F_p | CAA | nope | >1bp-ins | A/T-1 | none | A/T-1 | CAGGGCAAA | AAAAGTG | GTT | GTT | T | none | T>none | T>none_GTT | A/T-1_AAAAGTG | 6 | FALSE | FALSE |
| 135_v.txt | 32935 | 32935-Deleti | Deletion | 1 | C | - | 6 | 29052 | 0.02065262 | 54.1666678 | rnr1Y285F_pG | TRUE | CAN | JSY2421 | 3 | rnr1Y285F_p | GCC | nope | >1bp-ins | G/C-1 | none | G/C-1 | TGATTGCCC | CCCAAGA | CGG | GCC | C | none | nope | nope_GCC | G/C-1_CCCAAGA | 6 | FALSE | FALSE |
| 135_v.txt | 33000 | 33000-Deleti | Deletion | 1 | A | - | 66 | 29054 | 0.22716321 | 63.3181819 | rnr1Y285F_pG | TRUE | CAN | JSY2421 | 3 | rnr1Y285F_p | GAA | nope | >1bp-ins | A/T-1 | none | A/T-1 | TGTGAGAAA | AAAACTG | CTT | CTT | T | none | T>none | T>none_CTT | A/T-1_AAAACTG | 6 | FALSE | FALSE |
| 135_v.txt | 33019 | 33019-Deleti | Deletion | 1 | T | - | 2 | 29054 | 6.88E-03 | 64 | rnr1Y285F_pG | TRUE | CAN | JSY2421 | 3 | rnr1Y285F_p | GTA | nope | >1bp-ins | A/T-1 | none | A/T-1 | AGGATGTAA | TAACAGG | CAT | GTA | T | none | nope | nope_GTA | A/T-1_TAACAGG | 6 | FALSE | FALSE |
| 135_v.txt | 33045 | 33045-Deleti | Deletion | 1 | C | - | 7 | 29054 | 0.02409307 | 64 | rnr1Y285F_pG | TRUE | CAN | JSY2421 | 3 | rnr1Y285F_p | ACC | nope | >1bp-ins | G/C-1 | none | G/C-1 | ATTTCACCCA | CCCAAGG | TGG | ACC | C | none | nope | nope_ACC | G/C-1_CCCAAGG | 6 | FALSE | FALSE |
| 135_v.txt | 33105 | 33105-Deleti | Deletion | 1 | C | - | 71 | 29051 | 0.24439778 | 48.2253521 | rnr1Y285F_pG | TRUE | CAN | JSY2421 | 3 | rnr1Y285F_p | GCC | nope | >1bp-ins | G/C-1 | none | G/C-1 | AGAGCGCCC | CCCACTG | CGG | GCC | C | none | nope | nope_GCC | G/C-1_CCCACTG | 6 | FALSE | FALSE |
| 135_v.txt | 33114 | 33114-Deleti | Deletion | 1 | C | - | 1 | 29051 | 3.44E-03 | 63.9999302 | rnr1Y285F_pG | TRUE | CAN | JSY2421 | 3 | rnr1Y285F_p | GCC | nope | >1bp-ins | G/C-1 | none | G/C-1 | ACTGGGCCG | CCGGCGT | CGG | GCC | C | none | nope | nope_GCC | G/C-1_CCGGCGT | 6 | FALSE | FALSE |
| 135_v.txt | 33138 | 33138-Deleti | Deletion | 1 | A | - | 3 | 29051 | 0.01032667 | 46.6666667 | rnr1Y285F_pG | TRUE | CAN | JSY2421 | 3 | rnr1Y285F_p | TAA | nope | >1bp-ins | A/T-1 | none | A/T-1 | GTGGATAAA | AAACCAA | ATT | ATT | T | none | T>none | T>none_ATT | A/T-1_AAACCAA | 6 | FALSE | FALSE |
| 135_v.txt | 33157 | 33157-Deleti | Deletion | 1 | T | - | 7 | 29051 | 0.02409556 | 37.8571429 | rnr1Y285F_pG | TRUE | CAN | JSY2421 | 3 | rnr1Y285F_p | GTA | nope | >1bp-ins | A/T-1 | none | A/T-1 | GACCTGTAC | TACCAAT | CAT | GTA | T | none | nope | nope_GTA | A/T-1_TACCAAT | 6 | FALSE | FALSE |
| 135_v.txt | 33171 | 33171-Deleti | Deletion | 1 | C | - | 3 | 29049 | 0.01032738 | 38 | rnr1Y285F_pG | TRUE | CAN | JSY2421 | 3 | rnr1Y285F_p | ACC | nope | >1bp-ins | G/C-1 | none | G/C-1 | GTACCACCA | CCAAGGG | TGG | ACC | C | none | nope | nope_ACC | G/C-1_CCAAGGG | 6 | FALSE | FALSE |
| 135_v.txt | 33178 | 33178-Deleti | Deletion | 1 | C | - | 12 | 29043 | 0.04131805 | 37 | rnr1Y285F_pG | TRUE | CAN | JSY2421 | 3 | rnr1Y285F_p | GCA | nope | >1bp-ins | G/C-1 | none | G/C-1 | CAAGGGCAA | CAATCAT | CGT | GCA | C | none | nope | nope_GCA | G/C-1_CAATCAT | 6 | FALSE | FALSE |
| 135_v.txt | 33230 | 33230-Deleti | Deletion | 1 | T | - | 59 | 19836 | 0.297439 | 37.1186441 | rnr1Y285F_pG | TRUE | CAN | JSY2421 | 3 | rnr1Y285F_p | GTA | nope | >1bp-ins | A/T-1 | none | A/T-1 | GTTCTGTACT | TACTTCT | CAT | GTA | T | none | nope | nope_GTA | A/T-1_TACTTCT | 6 | FALSE | FALSE |
| 135_v.txt | 33356 | 33356-Deleti | Deletion | 1 | C | - | 7 | 19836 | 0.03528937 | 56.2857143 | rnr1Y285F_pG | TRUE | CAN | JSY2421 | 3 | rnr1Y285F_p | ACC | nope | >1bp-ins | G/C-1 | none | G/C-1 | TATTGACCCA | CCCACGT | TGG | ACC | C | none | nope | nope_ACC | G/C-1_CCCACGT | 6 | FALSE | FALSE |
| 135_v.txt | 33474 | 33474-Deleti | Deletion | 1 | C | - | 46 | 19833 | 0.23193667 | 29.1086957 | rnr1Y285F_pG | TRUE | CAN | JSY2421 | 3 | rnr1Y285F_p | GCC | nope | >1bp-ins | G/C-1 | none | G/C-1 | GCTATGCCTT | CCTTTTT | CGG | GCC | C | none | nope | nope_GCC | G/C-1_CCTTTTT | 6 | FALSE | FALSE |
| 136_v.txt | 31723 | 31723-SNV- | SNV | 1 | A | C | 2 | 11836 | 0.0168976 | 64 | rnr1Y285F_pG | TRUE | CAN | JSY2421 | 4 | rnr1Y285F_p | CAA | A>C | G/C+1 | A/T-1 | TA>GC | TA>GC | TGTCCCAAA | AAAAAGT | GTT | GTT | T | G | T>G | T>G_GTT | TA>GC_AAAAAGT | 6 | FALSE | FALSE |
| 136_v.txt | 31854 | 31854-SNV- | SNV | 1 | A | C | 13 | 11836 | 0.1098344 | 47.5384615 | rnr1Y285F_pG | TRUE | CAN | JSY2421 | 4 | rnr1Y285F_p | TAA | A>C | G/C+1 | A/T-1 | TA>GC | TA>GC | ACAGCTAAG | AAGAACA | ATT | ATT | T | G | T>G | T>G_ATT | TA>GC_AAGAACA | 6 | FALSE | FALSE |
| 136_v.txt | 31986 | 31986-SNV- | SNV | 1 | A | C | 4222 | 24887 | 16.9646804 | 51.4909995 | rnr1Y285F_pG | TRUE | CAN | JSY2421 | 4 | rnr1Y285F_p | TAA | A>C | G/C+1 | A/T-1 | TA>GC | TA>GC | GGCATTAAT | AATTTAG | ATT | ATT | T | G | T>G | T>G_ATT | TA>GC_AATTTAG | 6 | FALSE | FALSE |
| 136_v.txt | 32073 | 32073-SNV- | SNV | 1 | A | C | 25 | 24887 | 0.10045405 | 58.92 | rnr1Y285F_pG | TRUE | CAN | JSY2421 | 4 | rnr1Y285F_p | TAA | A>C | G/C+1 | A/T-1 | TA>GC | TA>GC | ATAAATAAC | AACCATG | ATT | ATT | T | G | T>G | T>G_ATT | TA>GC_AACCATG | 6 | FALSE | FALSE |
| 136_v.txt | 32124 | 32124-SNV- | SNV | 1 | A | C | 7 | 24887 | 0.02812714 | 52.1428571 | rnr1Y285F_pG | TRUE | CAN | JSY2421 | 4 | rnr1Y285F_p | AAC | A>C | G/C+1 | A/T-1 | TA>GC | TA>GC | TCGAAAACT | ACTTTGT | TTG | TTG | T | G | T>G | T>G_TTG | TA>GC_ACTTTGT | 6 | FALSE | FALSE |
| 136_v.txt | 32288 | 32288-SNV- | SNV | 1 | A | C | 41 | 27455 | 0.14933528 | 38.5121951 | rnr1Y285F_pG | TRUE | CAN | JSY2421 | 4 | rnr1Y285F_p | TAT | A>C | G/C+1 | A/T-1 | TA>GC | TA>GC | GTAAATATT | ATTTGAA | ATA | ATA | T | G | T>G | T>G_ATA | TA>GC_ATTTGAA | 6 | FALSE | FALSE |
| 136_v.txt | 32441 | 32441-SNV-T | SNV | 1 | T | G | 7 | 27455 | 0.02549627 | 60.2857143 | rnr1Y285F_pG | TRUE | CAN | JSY2421 | 4 | rnr1Y285F_p | CTA | T>G | G/C+1 | A/T-1 | TA>GC | TA>GC | TGGAACTAA | TAAAAGT | GAT | CTA | T | C | T>G | T>G_CTA | TA>GC_TAAAAGT | 6 | FALSE | FALSE |
| 136_v.txt | 32445 | 32445-SNV- | SNV | 1 | A | C | 21 | 27455 | 0.0764888 | 57.0952381 | rnr1Y285F_pG | TRUE | CAN | JSY2421 | 4 | rnr1Y285F_p | AAG | A>C | G/C+1 | A/T-1 | TA>GC | TA>GC | ACTAAAAGT | AGTCCAA | TTC | TTC | T | G | T>G | T>G_TTC | TA>GC_AGTCCAA | 6 | FALSE | FALSE |
| 136_v.txt | 32450 | 32450-SNV- | SNV | 1 | A | C | 8 | 27455 | 0.02913859 | 54.1239371 | rnr1Y285F_pG | TRUE | CAN | JSY2421 | 4 | rnr1Y285F_p | CAA | A>C | G/C+1 | A/T-1 | TA>GC | TA>GC | AAGTCCAAT | AATGAAT | GTT | GTT | T | G | T>G | T>G_GTT | TA>GC_AATGAAT | 6 | FALSE | FALSE |
| 136_v.txt | 32493 | 32493-SNV- | SNV | 1 | A | C | 6 | 27455 | 0.02185394 | 42.3333333 | rnr1Y285F_pG | TRUE | CAN | JSY2421 | 4 | rnr1Y285F_p | AAC | A>C | G/C+1 | A/T-1 | TA>GC | TA>GC | CGGAAAACA | ACAACTT | TTG | TTG | T | G | T>G | T>G_TTG | TA>GC_ACAACTT | 6 | FALSE | FALSE |
| 136_v.txt | 32581 | 32581-SNV-T | SNV | 1 | T | G | 35 | 29096 | 0.12029145 | 36.7428571 | rnr1Y285F_pG | TRUE | CAN | JSY2421 | 4 | rnr1Y285F_p | GTG | T>G | G/C+1 | A/T-1 | TA>GC | TA>GC | GAAATGTGA | TGAAGGC | CAC | GTG | T | C | T>G | T>G_GTG | TA>GC_TGAAGGC | 6 | FALSE | FALSE |
| 136_v.txt | 32631 | 32631-SNV-T | SNV | 1 | T | G | 6 | 29096 | 0.02062139 | 46.6666667 | rnr1Y285F_pG | TRUE | CAN | JSY2421 | 4 | rnr1Y285F_p | GTT | T>G | G/C+1 | A/T-1 | TA>GC | TA>GC | CCTTCGTTTT | TTTTTAT | CAA | GTT | T | C | T>G | T>G_GTT | TA>GC_TTTTTAT | 6 | FALSE | FALSE |
| 136_v.txt | 32766 | 32766-SNV-T | SNV | 1 | T | G | 13 | 29096 | 0.04467968 | 37.6923077 | rnr1Y285F_pG | TRUE | CAN | JSY2421 | 4 | rnr1Y285F_p | TTT | T>G | G/C+1 | A/T-1 | TA>GC | TA>GC | AAAACTTTG | TTGATGG | AAA | TTT | T | C | T>G | T>G_TTT | TA>GC_TTGATGG | 6 | FALSE | FALSE |
| 136_v.txt | 32801 | 32801-SNV- | SNV | 1 | A | C | 12 | 29096 | 0.04124278 | 45.3333333 | rnr1Y285F_pG | TRUE | CAN | JSY2421 | 4 | rnr1Y285F_p | AAT | A>C | G/C+1 | A/T-1 | TA>GC | TA>GC | ACCGTAATA | ATATTTG | TTA | TTA | T | G | T>G | T>G_TTA | TA>GC_ATATTTG | 6 | FALSE | FALSE |
| 136_v.txt | 32842 | 32842-SNV- | SNV | 1 | A | C | 2 | 29096 | 6.87E-03 | 35.9999426 | rnr1Y285F_pG | TRUE | CAN | JSY2421 | 4 | rnr1Y285F_p | CAA | A>C | G/C+1 | A/T-1 | TA>GC | TA>GC | TTACCCAAA | AAAAAAT | GTT | GTT | T | G | T>G | T>G_GTT | TA>GC_AAAAAAT | 6 | FALSE | FALSE |
| 136_v.txt | 32884 | 32884-SNV- | SNV | 1 | A | C | 9 | 34459 | 0.026118 | 32 | rnr1Y285F_pG | TRUE | CAN | JSY2421 | 4 | rnr1Y285F_p | CAA | A>C | G/C+1 | A/T-1 | TA>GC | TA>GC | ACGTCCAAA | AAAATTG | GTT | GTT | T | G | T>G | T>G_GTT | TA>GC_AAAATTG | 6 | FALSE | FALSE |
| 136_v.txt | 32919 | 32919-SNV- | SNV | 1 | A | C | 10 | 34459 | 0.02902 | 31.7 | rnr1Y285F_pG | TRUE | CAN | JSY2421 | 4 | rnr1Y285F_p | CAG | A>C | G/C+1 | A/T-1 | TA>GC | TA>GC | AGTTCCAGG | AGGGCAA | GTC | GTC | T | G | T>G | T>G_GTC | TA>GC_AGGGCAA | 6 | FALSE | FALSE |
| 136_v.txt | 32952 | 32952-SNV- | SNV | 1 | A | C | 20 | 34459 | 0.05803999 | 36.7 | rnr1Y285F_pG | TRUE | CAN | JSY2421 | 4 | rnr1Y285F_p | CAT | A>C | G/C+1 | A/T-1 | TA>GC | TA>GC | CAATACATG | ATGTAAC | GTA | GTA | T | G | T>G | T>G_GTA | TA>GC_ATGTAAC | 6 | FALSE | FALSE |
| 136_v.txt | 33054 | 33054-SNV-T | SNV | 1 | T | G | 39 | 34459 | 0.11317798 | 53.8974359 | rnr1Y285F_pG | TRUE | CAN | JSY2421 | 4 | rnr1Y285F_p | CTG | T>G | G/C+1 | A/T-1 | TA>GC | TA>GC | AAGGACTGC | TGCGTGA | GAC | CTG | T | C | T>G | T>G_CTG | TA>GC_TGCGTGA | 6 | FALSE | FALSE |
| 136_v.txt | 33099 | 33099-SNV- | SNV | 1 | A | C | 8 | 34459 | 0.023216 | 40.6249994 | rnr1Y285F_pG | TRUE | CAN | JSY2421 | 4 | rnr1Y285F_p | AAG | A>C | G/C+1 | A/T-1 | TA>GC | TA>GC | GATATAAGA | AGAGCGC | TTC | TTC | T | G | T>G | T>G_TTC | TA>GC_AGAGCGC | 6 | FALSE | FALSE |
| 136_v.txt | 33166 | 33166-SNV-T | SNV | 1 | T | G | 54 | 34459 | 0.15670797 | 37.7221077 | rnr1Y285F_pG | TRUE | CAN | JSY2421 | 4 | rnr1Y285F_p | GTA | T>G | G/C+1 | A/T-1 | TA>GC | TA>GC | CAATAGTAC | TACCACC | CAT | GTA | T | C | T>G | T>G_GTA | TA>GC_TACCACC | 6 | FALSE | FALSE |
| 136_v.txt | 33192 | 33192-SNV-T | SNV | 1 | T | G | 4 | 15059 | 0.02656219 | 38 | rnr1Y285F_pG | TRUE | CAN | JSY2421 | 4 | rnr1Y285F_p | ATG | T>G | G/C+1 | A/T-1 | TA>GC | TA>GC | CCAATATGT | TGTCTTT | TAC | ATG | T | C | T>G | T>G_ATG | TA>GC_TGTCTTT | 6 | FALSE | FALSE |
| 136_v.txt | 33214 | 33214-SNV-T | SNV | 1 | T | G | 26 | 17622 | 0.14754284 | 38.303464 | rnr1Y285F_pG | TRUE | CAN | JSY2421 | 4 | rnr1Y285F_p | TTC | T>G | G/C+1 | A/T-1 | TA>GC | TA>GC | CTCTCTTCAC | TCACTTC | AAG | TTC | T | C | T>G | T>G_TTC | TA>GC_TCACTTC | 6 | FALSE | FALSE |
| 136_v.txt | 32160 | 32160-SNV- | SNV | 1 | A | G | 5 | 24887 | 0.02009081 | 52.7964981 | rnr1Y285F_pG | TRUE | CAN | JSY2421 | 4 | rnr1Y285F_p | CAA | A>G | G/C+1 | A/T-1 | TA>CG | TA>CG | TAAGCCAAA | AAAGCGC | GTT | GTT | T | C | T>C | T>C_GTT | TA>CG_AAAGCGC | 6 | FALSE | FALSE |
| 136_v.txt | 32256 | 32256-SNV- | SNV | 1 | A | G | 52 | 27447 | 0.18945604 | 36.2115385 | rnr1Y285F_pG | TRUE | CAN | JSY2421 | 4 | rnr1Y285F_p | TAG | A>G | G/C+1 | A/T-1 | TA>CG | TA>CG | TTTGATAGA | AGACCAA | ATC | ATC | T | C | T>C | T>C_ATC | TA>CG_AGACCAA | 6 | FALSE | FALSE |
| 136_v.txt | 32919 | 32919-SNV- | SNV | 1 | A | G | 25 | 34459 | 0.07254999 | 45.68 | rnr1Y285F_pG | TRUE | CAN | JSY2421 | 4 | rnr1Y285F_p | CAG | A>G | G/C+1 | A/T-1 | TA>CG | TA>CG | AGTTCCAGG | AGGGCAA | GTC | GTC | T | C | T>C | T>C_GTC | TA>CG_AGGGCAA | 6 | FALSE | FALSE |
| 136_v.txt | 32941 | 32941-SNV- | SNV | 1 | A | G | 6 | 34459 | 0.017412 | 52.9836925 | rnr1Y285F_pG | TRUE | CAN | JSY2421 | 4 | rnr1Y285F_p | GAA | A>G | G/C+1 | A/T-1 | TA>CG | TA>CG | CCCAAGAAA | AAAACCA | CTT | CTT | T | C | T>C | T>C_CTT | TA>CG_AAAACCA | 6 | FALSE | FALSE |
| 136_v.txt | 33052 | 33052-SNV- | SNV | 1 | A | G | 31 | 34459 | 0.08996198 | 53.1938781 | rnr1Y285F_pG | TRUE | CAN | JSY2421 | 4 | rnr1Y285F_p | GAC | A>G | G/C+1 | A/T-1 | TA>CG | TA>CG | CCAAGGACT | ACTGCGT | CTG | CTG | T | C | T>C | T>C_CTG | TA>CG_ACTGCGT | 6 | FALSE | FALSE |
| 136_v.txt | 33054 | 33054-SNV-T | SNV | 1 | T | C | 14 | 34459 | 0.04062799 | 52.8571429 | rnr1Y285F_pG | TRUE | CAN | JSY2421 | 4 | rnr1Y285F_p | CTG | T>C | G/C+1 | A/T-1 | TA>CG | TA>CG | AAGGACTGC | TGCGTGA | GAC | CTG | T | G | T>C | T>C_CTG | TA>CG_TGCGTGA | 6 | FALSE | FALSE |
| 136_v.txt | 33095 | 33095-SNV-T | SNV | 1 | T | C | 4 | 34459 | 0.011608 | 32.4999946 | rnr1Y285F_pG | TRUE | CAN | JSY2421 | 4 | rnr1Y285F_p | ATA | T>C | G/C+1 | A/T-1 | TA>CG | TA>CG | ATATGATAT | TATAAGA | TAT | ATA | T | G | T>C | T>C_ATA | TA>CG_TATAAGA | 6 | FALSE | FALSE |
| 136_v.txt | 33099 | 33099-SNV- | SNV | 1 | A | G | 21 | 34459 | 0.06094199 | 48.9047608 | rnr1Y285F_pG | TRUE | CAN | JSY2421 | 4 | rnr1Y285F_p | AAG | A>G | G/C+1 | A/T-1 | TA>CG | TA>CG | GATATAAGA | AGAGCGC | TTC | TTC | T | C | T>C | T>C_TTC | TA>CG_AGAGCGC | 6 | FALSE | FALSE |
| 136_v.txt | 33167 | 33167-SNV- | SNV | 1 | A | G | 7 | 34459 | 0.020314 | 37.5708441 | rnr1Y285F_pG | TRUE | CAN | JSY2421 | 4 | rnr1Y285F_p | TAC | A>G | G/C+1 | A/T-1 | TA>CG | TA>CG | AATAGTACC | ACCACCA | ATG | ATG | T | C | T>C | T>C_ATG | TA>CG_ACCACCA | 6 | FALSE | FALSE |
| 136_v.txt | 33192 | 33192-SNV-T | SNV | 1 | T | C | 17 | 15059 | 0.1128893 | 29.2941177 | rnr1Y285F_pG | TRUE | CAN | JSY2421 | 4 | rnr1Y285F_p | ATG | T>C | G/C+1 | A/T-1 | TA>CG | TA>CG | CCAATATGT | TGTCTTT | TAC | ATG | T | G | T>C | T>C_ATG | TA>CG_TGTCTTT | 6 | FALSE | FALSE |
| 136_v.txt | 33207 | 33207-SNV-T | SNV | 1 | T | C | 12 | 17621 | 0.06810056 | 21.8333112 | rnr1Y285F_pG | TRUE | CAN | JSY2421 | 4 | rnr1Y285F_p | CTC | T>C | G/C+1 | A/T-1 | TA>CG | TA>CG | TTAAGCTCTC | TCTCTCT | GAG | CTC | T | G | T>C | T>C_CTC | TA>CG_TCTCTCT | 6 | FALSE | FALSE |
| 136_v.txt | 31839 | 31839-SNV- | SNV | 1 | A | T | 41 | 11836 | 0.34640081 | 49.2926829 | rnr1Y285F_pG | TRUE | CAN | JSY2421 | 4 | rnr1Y285F_p | TAA | A>T | A/T+1 | A/T-1 | TA>AT | TA>AT | TGAAATAAG | AAGATCC | ATT | ATT | T | A | T>A | T>A_ATT | TA>AT_AAGATCC | 6 | FALSE | FALSE |
| 136_v.txt | 32109 | 32109-SNV- | SNV | 1 | A | T | 27 | 24887 | 0.10849038 | 55.5185185 | rnr1Y285F_pG | TRUE | CAN | JSY2421 | 4 | rnr1Y285F_p | TAA | A>T | A/T+1 | A/T-1 | TA>AT | TA>AT | ATATTTAATA | AATAGCC | ATT | ATT | T | A | T>A | T>A_ATT | TA>AT_AATAGCC | 6 | FALSE | FALSE |
| 136_v.txt | 32288 | 32288-SNV- | SNV | 1 | A | T | 385 | 27455 | 1.40229466 | 49.9454546 | rnr1Y285F_pG | TRUE | CAN | JSY2421 | 4 | rnr1Y285F_p | TAT | A>T | A/T+1 | A/T-1 | TA>AT | TA>AT | GTAAATATTT | ATTTGAA | ATA | ATA | T | A | T>A | T>A_ATA | TA>AT_ATTTGAA | 6 | FALSE | FALSE |
| 136_v.txt | 32338 | 32338-SNV-T | SNV | 1 | T | A | 4 | 27455 | 0.0145693 | 59.4986012 | rnr1Y285F_pG | TRUE | CAN | JSY2421 | 4 | rnr1Y285F_p | ATA | T>A | A/T+1 | A/T-1 | TA>AT | TA>AT | TGAAGATAT | TATGTGG | TAT | ATA | T | T | T>A | T>A_ATA | TA>AT_TATGTGG | 6 | FALSE | FALSE |
| 136_v.txt | 32473 | 32473-SNV- | SNV | 1 | A | T | 13 | 27455 | 0.04735021 | 39.8461539 | rnr1Y285F_pG | TRUE | CAN | JSY2421 | 4 | rnr1Y285F_p | TAG | A>T | A/T+1 | A/T-1 | TA>AT | TA>AT | CAATGTAGA | AGAAGGT | ATC | ATC | T | A | T>A | T>A_ATC | TA>AT_AGAAGGT | 6 | FALSE | FALSE |
| 136_v.txt | 32728 | 32728-SNV- | SNV | 1 | A | T | 3 | 29096 | 0.0103107 | 49.9947637 | rnr1Y285F_pG | TRUE | CAN | JSY2421 | 4 | rnr1Y285F_p | CAA | A>T | A/T+1 | A/T-1 | TA>AT | TA>AT | CCATACAAA | AAAAACA | GTT | GTT | T | A | T>A | T>A_GTT | TA>AT_AAAAACA | 6 | FALSE | FALSE |
| 136_v.txt | 32948 | 32948-SNV- | SNV | 1 | A | T | 112 | 34459 | 0.32502394 | 57.8303571 | rnr1Y285F_pG | TRUE | CAN | JSY2421 | 4 | rnr1Y285F_p | AAT | A>T | A/T+1 | A/T-1 | TA>AT | TA>AT | AAACCAATA | ATACATG | TTA | TTA | T | A | T>A | T>A_TTA | TA>AT_ATACATG | 6 | FALSE | FALSE |
| 136_v.txt | 33089 | 33089-SNV- | SNV | 1 | A | T | 2 | 34459 | 5.80E-03 | 50.9995886 | rnr1Y285F_pG | TRUE | CAN | JSY2421 | 4 | rnr1Y285F_p | AAT | A>T | A/T+1 | A/T-1 | TA>AT | TA>AT | AAATAAATA | ATATGAT | TTA | TTA | T | A | T>A | T>A_TTA | TA>AT_ATATGAT | 6 | FALSE | FALSE |
| 136_v.txt | 33090 | 33090-SNV-T | SNV | 1 | T | A | 5 | 34459 | 0.01451 | 42.796669 | rnr1Y285F_pG | TRUE | CAN | JSY2421 | 4 | rnr1Y285F_p | ATA | T>A | A/T+1 | A/T-1 | TA>AT | TA>AT | AATAAATAT | TATGATA | TAT | ATA | T | T | T>A | T>A_ATA | TA>AT_TATGATA | 6 | FALSE | FALSE |
| 136_v.txt | 33095 | 33095-SNV-T | SNV | 1 | T | A | 13 | 34459 | 0.03772599 | 42.5381081 | rnr1Y285F_pG | TRUE | CAN | JSY2421 | 4 | rnr1Y285F_p | ATA | T>A | A/T+1 | A/T-1 | TA>AT | TA>AT | ATATGATAT | TATAAGA | TAT | ATA | T | T | T>A | T>A_ATA | TA>AT_TATAAGA | 6 | FALSE | FALSE |
| 136_v.txt | 33096 | 33096-SNV- | SNV | 1 | A | T | 6 | 34459 | 0.017412 | 40.9998862 | rnr1Y285F_pG | TRUE | CAN | JSY2421 | 4 | rnr1Y285F_p | TAT | A>T | A/T+1 | A/T-1 | TA>AT | TA>AT | TATGATATA | ATAAGAG | ATA | ATA | T | A | T>A | T>A_ATA | TA>AT_ATAAGAG | 6 | FALSE | FALSE |
| 136_v.txt | 33097 | 33097-SNV-T | SNV | 1 | T | A | 8 | 34459 | 0.023216 | 37.748788 | rnr1Y285F_pG | TRUE | CAN | JSY2421 | 4 | rnr1Y285F_p | ATA | T>A | A/T+1 | A/T-1 | TA>AT | TA>AT | ATGATATAA | TAAGAGC | TAT | ATA | T | T | T>A | T>A_ATA | TA>AT_TAAGAGC | 6 | FALSE | FALSE |
| 136_v.txt | 33099 | 33099-SNV- | SNV | 1 | A | T | 9 | 34459 | 0.026118 | 55.1111106 | rnr1Y285F_pG | TRUE | CAN | JSY2421 | 4 | rnr1Y285F_p | AAG | A>T | A/T+1 | A/T-1 | TA>AT | TA>AT | GATATAAGA | AGAGCGC | TTC | TTC | T | A | T>A | T>A_TTC | TA>AT_AGAGCGC | 6 | FALSE | FALSE |
| 136_v.txt | 33194 | 33194-SNV-T | SNV | 1 | T | A | 15 | 17620 | 0.08513053 | 35.5312154 | rnr1Y285F_pG | TRUE | CAN | JSY2421 | 4 | rnr1Y285F_p | GTC | T>A | A/T+1 | A/T-1 | TA>AT | TA>AT | AATATGTCTT | TCTTTGC | CAG | GTC | T | T | T>A | T>A_GTC | TA>AT_TCTTTGC | 6 | FALSE | FALSE |
| 136_v.txt | 33207 | 33207-SNV-T | SNV | 1 | T | A | 1 | 17621 | 5.68E-03 | 36.9993269 | rnr1Y285F_pG | TRUE | CAN | JSY2421 | 4 | rnr1Y285F_p | CTC | T>A | A/T+1 | A/T-1 | TA>AT | TA>AT | TTAAGCTCTC | TCTCTCT | GAG | CTC | T | T | T>A | T>A_CTC | TA>AT_TCTCTCT | 6 | FALSE | FALSE |
| 136_v.txt | 31650 | 31650-SNV- | SNV | 1 | C | T | 33 | 11836 | 0.27881041 | 36.5151515 | rnr1Y285F_pG | TRUE | CAN | JSY2421 | 4 | rnr1Y285F_p | GCG | C>T | A/T+1 | G/C-1 | CG>TA | CG>TA | AGAATGCGA | CGAAATG | CGC | GCG | C | A | C>T | C>T_GCG | CG>TA_CGAAATG | 6 | FALSE | FALSE |
| 136_v.txt | 31830 | 31830-SNV- | SNV | 1 | C | T | 13 | 11836 | 0.1098344 | 57.5384615 | rnr1Y285F_pG | TRUE | CAN | JSY2421 | 4 | rnr1Y285F_p | GCA | C>T | A/T+1 | G/C-1 | CG>TA | CG>TA | AATATGCATT | CATTGAA | CGT | GCA | C | A | C>T | C>T_GCA | CG>TA_CATTGAA | 6 | FALSE | FALSE |
| 136_v.txt | 31844 | 31844-SNV- | SNV | 1 | C | T | 66 | 11836 | 0.55762082 | 50.9393939 | rnr1Y285F_pG | TRUE | CAN | JSY2421 | 4 | rnr1Y285F_p | TCC | C>T | A/T+1 | G/C-1 | CG>TA | CG>TA | TAAGATCCA | CCAAACA | AGG | TCC | C | A | C>T | C>T_TCC | CG>TA_CCAAACA | 6 | FALSE | FALSE |
| 136_v.txt | 31881 | 31881-SNV- | SNV | 1 | G | A | 12 | 11836 | 0.1013856 | 44.9166667 | rnr1Y285F_pG | TRUE | CAN | JSY2421 | 4 | rnr1Y285F_p | GGC | G>A | A/T+1 | G/C-1 | CG>TA | CG>TA | TAGGCGGCA | GCAGCAA | CCG | CCG | C | T | C>T | C>T_CCG | CG>TA_GCAGCAA | 6 | FALSE | FALSE |
| 136_v.txt | 31882 | 31882-SNV- | SNV | 1 | C | T | 13 | 11836 | 0.1098344 | 49.1448449 | rnr1Y285F_pG | TRUE | CAN | JSY2421 | 4 | rnr1Y285F_p | GCA | C>T | A/T+1 | G/C-1 | CG>TA | CG>TA | AGGCGGCAG | CAGCAAA | CGT | GCA | C | A | C>T | C>T_GCA | CG>TA_CAGCAAA | 6 | FALSE | FALSE |
| 136_v.txt | 31920 | 31920-SNV- | SNV | 1 | G | A | 12 | 11834 | 0.10140274 | 37.9166667 | rnr1Y285F_pG | TRUE | CAN | JSY2421 | 4 | rnr1Y285F_p | CGT | G>A | A/T+1 | G/C-1 | CG>TA | CG>TA | AAAGCCGTG | GTGAAAC | GCA | GCA | C | T | C>T | C>T_GCA | CG>TA_GTGAAAC | 6 | FALSE | FALSE |
| 136_v.txt | 31962 | 31962-SNV- | SNV | 1 | G | A | 50 | 24887 | 0.20090811 | 35.62 | rnr1Y285F_pG | TRUE | CAN | JSY2421 | 4 | rnr1Y285F_p | CGC | G>A | A/T+1 | G/C-1 | CG>TA | CG>TA | GTGGCCGCA | GCATAAT | GCG | GCG | C | T | C>T | C>T_GCG | CG>TA_GCATAAT | 6 | FALSE | FALSE |
| 136_v.txt | 31981 | 31981-SNV- | SNV | 1 | G | A | 39 | 24887 | 0.15670832 | 47.4871795 | rnr1Y285F_pG | TRUE | CAN | JSY2421 | 4 | rnr1Y285F_p | GGC | G>A | A/T+1 | G/C-1 | CG>TA | CG>TA | AGCCGGGCA | GCATTAA | CCG | CCG | C | T | C>T | C>T_CCG | CG>TA_GCATTAA | 6 | FALSE | FALSE |
| 136_v.txt | 32041 | 32041-SNV- | SNV | 1 | G | A | 25 | 24887 | 0.10045405 | 62.88 | rnr1Y285F_pG | TRUE | CAN | JSY2421 | 4 | rnr1Y285F_p | TGC | G>A | A/T+1 | G/C-1 | CG>TA | CG>TA | AAGCTTGCA | GCATAAA | ACG | ACG | C | T | C>T | C>T_ACG | CG>TA_GCATAAA | 6 | FALSE | FALSE |
| 136_v.txt | 32075 | 32075-SNV- | SNV | 1 | C | T | 492 | 24882 | 1.97733301 | 62.0061965 | rnr1Y285F_pG | TRUE | CAN | JSY2421 | 4 | rnr1Y285F_p | ACC | C>T | A/T+1 | G/C-1 | CG>TA | CG>TA | AAATAACCA | CCATGCA | TGG | ACC | C | A | C>T | C>T_ACC | CG>TA_CCATGCA | 6 | FALSE | FALSE |
| 136_v.txt | 32076 | 32076-SNV- | SNV | 1 | C | T | 59 | 24886 | 0.23708109 | 61.966295 | rnr1Y285F_pG | TRUE | CAN | JSY2421 | 4 | rnr1Y285F_p | CCA | C>T | A/T+1 | G/C-1 | CG>TA | CG>TA | AATAACCAT | CATGCAA | GGT | CCA | C | A | C>T | C>T_CCA | CG>TA_CATGCAA | 6 | FALSE | FALSE |
| 136_v.txt | 32079 | 32079-SNV- | SNV | 1 | G | A | 13 | 24887 | 0.05223611 | 63.3846154 | rnr1Y285F_pG | TRUE | CAN | JSY2421 | 4 | rnr1Y285F_p | TGC | G>A | A/T+1 | G/C-1 | CG>TA | CG>TA | AACCATGCA | GCAAAAA | ACG | ACG | C | T | C>T | C>T_ACG | CG>TA_GCAAAAA | 6 | FALSE | FALSE |
| 136_v.txt | 32080 | 32080-SNV- | SNV | 1 | C | T | 12 | 24887 | 0.04821795 | 55.8334109 | rnr1Y285F_pG | TRUE | CAN | JSY2421 | 4 | rnr1Y285F_p | GCA | C>T | A/T+1 | G/C-1 | CG>TA | CG>TA | ACCATGCAA | CAAAAAA | CGT | GCA | C | A | C>T | C>T_GCA | CG>TA_CAAAAAA | 6 | FALSE | FALSE |
| 136_v.txt | 32088 | 32088-SNV- | SNV | 1 | C | T | 194 | 24887 | 0.77952345 | 61.314433 | rnr1Y285F_pG | TRUE | CAN | JSY2421 | 4 | rnr1Y285F_p | GCC | C>T | A/T+1 | G/C-1 | CG>TA | CG>TA | AAAAAGCCT | CCTGCAA | CGG | GCC | C | A | C>T | C>T_GCC | CG>TA_CCTGCAA | 6 | FALSE | FALSE |

| | | | | | | | | | | | | | | | | | | | | | | | | | | | | | | | | | | |
|---|---|---|---|---|---|---|---|---|---|---|---|---|---|---|---|---|---|---|---|---|---|---|---|---|---|---|---|---|---|---|---|---|---|---|
| 136_v.txt | 32092 | 32092-SNV-( | SNV | 1 | C | T | 28 | 24887 | 0.11250854 | 61.8571429 | rnr1Y285F_pG | TRUE | CAN | JSY2421 | 4 | rnr1Y285F_p | GCA | C>T | A/T+1 | G/C-1 | CG>TA | CG>TA | AGCCTGCAA( | CAACACC | CGT | GCA | C | A | C>T | C>T_GCA | CG>TA_CAACACC | 6 | FALSE | FALSE |
| 136_v.txt | 32114 | 32114-SNV-( | SNV | 1 | C | T | 123 | 24887 | 0.49423394 | 60.504065 | rnr1Y285F_pG | TRUE | CAN | JSY2421 | 4 | rnr1Y285F_p | GCC | C>T | A/T+1 | G/C-1 | CG>TA | CG>TA | TAATAGCCAT | CCATTCG | CGG | GCC | C | A | C>T | C>T_GCC | CG>TA_CCATTCG | 6 | FALSE | FALSE |
| 136_v.txt | 32157 | 32157-SNV-( | SNV | 1 | G | A | 19 | 24887 | 0.07634508 | 53.5254791 | rnr1Y285F_pG | TRUE | CAN | JSY2421 | 4 | rnr1Y285F_p | AGC | G>A | A/T+1 | G/C-1 | CG>TA | CG>TA | ATGTAAGCC/ | GCCAAAG | TCG | TCG | C | T | C>T | C>T_TCG | CG>TA_GCCAAAG | 6 | FALSE | FALSE |
| 136_v.txt | 32158 | 32158-SNV-( | SNV | 1 | C | T | 7 | 24887 | 0.02812714 | 56.9999901 | rnr1Y285F_pG | TRUE | CAN | JSY2421 | 4 | rnr1Y285F_p | GCC | C>T | A/T+1 | G/C-1 | CG>TA | CG>TA | TGTAAGCCA/ | CCAAAGC | CGG | GCC | C | A | C>T | C>T_GCC | CG>TA_CCAAAGC | 6 | FALSE | FALSE |
| 136_v.txt | 32159 | 32159-SNV-( | SNV | 1 | C | T | 4 | 24887 | 0.01607265 | 50.7486739 | rnr1Y285F_pG | TRUE | CAN | JSY2421 | 4 | rnr1Y285F_p | CCA | C>T | A/T+1 | G/C-1 | CG>TA | CG>TA | GTAAGCCAA. | CAAAGCG | GGT | CCA | C | A | C>T | C>T_CCA | CG>TA_CAAAGCG | 6 | FALSE | FALSE |
| 136_v.txt | 32164 | 32164-SNV-( | SNV | 1 | C | T | 10 | 24887 | 0.04018162 | 49.799388 | rnr1Y285F_pG | TRUE | CAN | JSY2421 | 4 | rnr1Y285F_p | GCG | C>T | A/T+1 | G/C-1 | CG>TA | CG>TA | CCAAAGCGC( | CGCCAAA | CGC | GCG | C | A | C>T | C>T_GCG | CG>TA_CGCCAAA | 6 | FALSE | FALSE |
| 136_v.txt | 32165 | 32165-SNV-( | SNV | 1 | G | A | 35 | 24887 | 0.14063567 | 44.6856491 | rnr1Y285F_pG | TRUE | CAN | JSY2421 | 4 | rnr1Y285F_p | CGC | G>A | A/T+1 | G/C-1 | CG>TA | CG>TA | CAAAGCGCC. | GCCAAAT | GCG | GCG | C | T | C>T | C>T_GCG | CG>TA_GCCAAAT | 6 | FALSE | FALSE |
| 136_v.txt | 32176 | 32176-SNV-( | SNV | 1 | C | T | 26 | 24887 | 0.10447221 | 44.3461539 | rnr1Y285F_pG | TRUE | CAN | JSY2421 | 4 | rnr1Y285F_p | GCA | C>T | A/T+1 | G/C-1 | CG>TA | CG>TA | ATGCAGCAG | CAGTAAC | CGT | GCA | C | A | C>T | C>T_GCA | CG>TA_CAGTAAC | 6 | FALSE | FALSE |
| 136_v.txt | 32271 | 32271-SNV-( | SNV | 1 | C | T | 50 | 27455 | 0.18211619 | 33.16 | rnr1Y285F_pG | TRUE | CAN | JSY2421 | 4 | rnr1Y285F_p | ACG | C>T | A/T+1 | G/C-1 | CG>TA | CG>TA | AAAATACGG | CGGGAAC | TGC | ACG | C | A | C>T | C>T_ACG | CG>TA_CGGGAAC | 6 | FALSE | FALSE |
| 136_v.txt | 32274 | 32274-SNV-( | SNV | 1 | G | A | 19 | 27455 | 0.06920415 | 34.8947368 | rnr1Y285F_pG | TRUE | CAN | JSY2421 | 4 | rnr1Y285F_p | GGA | G>A | A/T+1 | G/C-1 | CG>TA | CG>TA | ATACGGGAA | GAACCAA | CCT | CCT | C | T | C>T | C>T_CCT | CG>TA_GAACCAA | 6 | FALSE | FALSE |
| 136_v.txt | 32299 | 32299-SNV-( | SNV | 1 | C | T | 58 | 27455 | 0.21125478 | 52.0862069 | rnr1Y285F_pG | TRUE | CAN | JSY2421 | 4 | rnr1Y285F_p | GCG | C>T | A/T+1 | G/C-1 | CG>TA | CG>TA | AATTTGCGG( | CGGCAGA | CGC | GCG | C | A | C>T | C>T_GCG | CG>TA_CGGCAGA | 6 | FALSE | FALSE |
| 136_v.txt | 32463 | 32463-SNV-( | SNV | 1 | G | A | 28 | 27455 | 0.10198507 | 42.7142857 | rnr1Y285F_pG | TRUE | CAN | JSY2421 | 4 | rnr1Y285F_p | AGA | G>A | A/T+1 | G/C-1 | CG>TA | CG>TA | AATAGAGAG | GAGCCAA | TCT | TCT | C | T | C>T | C>T_TCT | CG>TA_GAGCCAA | 6 | FALSE | FALSE |
| 136_v.txt | 32466 | 32466-SNV-( | SNV | 1 | C | T | 29 | 27455 | 0.10562739 | 51.4827586 | rnr1Y285F_pG | TRUE | CAN | JSY2421 | 4 | rnr1Y285F_p | GCC | C>T | A/T+1 | G/C-1 | CG>TA | CG>TA | AGAGAGCCA | CCAATGT | CGG | GCC | C | A | C>T | C>T_GCC | CG>TA_CCAATGT | 6 | FALSE | FALSE |
| 136_v.txt | 32487 | 32487-SNV-( | SNV | 1 | C | T | 31 | 27455 | 0.11291204 | 43.7419355 | rnr1Y285F_pG | TRUE | CAN | JSY2421 | 4 | rnr1Y285F_p | ACG | C>T | A/T+1 | G/C-1 | CG>TA | CG>TA | AAGATACGG | CGGAAAA | TGC | ACG | C | A | C>T | C>T_ACG | CG>TA_CGGAAAA | 6 | FALSE | FALSE |
| 136_v.txt | 32508 | 32508-SNV-( | SNV | 1 | G | A | 24 | 27455 | 0.08741577 | 38.4583333 | rnr1Y285F_pG | TRUE | CAN | JSY2421 | 4 | rnr1Y285F_p | GGC | G>A | A/T+1 | G/C-1 | CG>TA | CG>TA | TTGATGGCT( | GCTCTTG | CCG | CCG | C | T | C>T | C>T_CCG | CG>TA_GCTCTTG | 6 | FALSE | FALSE |
| 136_v.txt | 32529 | 32529-SNV-( | SNV | 1 | G | A | 29 | 27455 | 0.10562739 | 39.1724138 | rnr1Y285F_pG | TRUE | CAN | JSY2421 | 4 | rnr1Y285F_p | GGG | G>A | A/T+1 | G/C-1 | CG>TA | CG>TA | TTTCTGGGGT | GGGTTTG | CCC | CCC | C | T | C>T | C>T_CCC | CG>TA_GGGTTTG | 6 | FALSE | FALSE |
| 136_v.txt | 32535 | 32535-SNV-( | SNV | 1 | G | A | 26 | 27455 | 0.09470042 | 37.4230769 | rnr1Y285F_pG | TRUE | CAN | JSY2421 | 4 | rnr1Y285F_p | TGC | G>A | A/T+1 | G/C-1 | CG>TA | CG>TA | GGGTTTGCA( | GCAGCTT | ACG | ACG | C | T | C>T | C>T_ACG | CG>TA_GCAGCTT | 6 | FALSE | FALSE |
| 136_v.txt | 32536 | 32536-SNV-( | SNV | 1 | C | T | 56 | 27455 | 0.20397013 | 36.3925841 | rnr1Y285F_pG | TRUE | CAN | JSY2421 | 4 | rnr1Y285F_p | GCA | C>T | A/T+1 | G/C-1 | CG>TA | CG>TA | GGTTTGCAG( | CAGCTTC | CGT | GCA | C | A | C>T | C>T_GCA | CG>TA_CAGCTTC | 6 | FALSE | FALSE |
| 136_v.txt | 32571 | 32571-SNV-( | SNV | 1 | C | T | 53 | 29096 | 0.18215562 | 35.1698113 | rnr1Y285F_pG | TRUE | CAN | JSY2421 | 4 | rnr1Y285F_p | ACC | C>T | A/T+1 | G/C-1 | CG>TA | CG>TA | TCAGTACCTT | CCTTGAA | TGG | ACC | C | A | C>T | C>T_ACC | CG>TA_CCTTGAA | 6 | FALSE | FALSE |
| 136_v.txt | 32590 | 32590-SNV-( | SNV | 1 | C | T | 51 | 29096 | 0.17528183 | 37.0980392 | rnr1Y285F_pG | TRUE | CAN | JSY2421 | 4 | rnr1Y285F_p | GCG | C>T | A/T+1 | G/C-1 | CG>TA | CG>TA | AGGCAGCGT | CGTTAAT | CGC | GCG | C | A | C>T | C>T_GCG | CG>TA_CGTTAAT | 6 | FALSE | FALSE |
| 136_v.txt | 32660 | 32660-SNV-( | SNV | 1 | C | T | 51 | 29096 | 0.17528183 | 55.1960784 | rnr1Y285F_pG | TRUE | CAN | JSY2421 | 4 | rnr1Y285F_p | CCC | C>T | A/T+1 | G/C-1 | CG>TA | CG>TA | TGGACCCCA( | CCAGGCA | GGG | CCC | C | A | C>T | C>T_CCC | CG>TA_CCAGGCA | 6 | FALSE | FALSE |
| 136_v.txt | 32678 | 32678-SNV-( | SNV | 1 | C | T | 68 | 29096 | 0.2337091 | 58.9558824 | rnr1Y285F_pG | TRUE | CAN | JSY2421 | 4 | rnr1Y285F_p | TCC | C>T | A/T+1 | G/C-1 | CG>TA | CG>TA | GTTTCTCCAA | CCAATAA | AGG | TCC | C | A | C>T | C>T_TCC | CG>TA_CCAATAA | 6 | FALSE | FALSE |
| 136_v.txt | 32700 | 32700-SNV-( | SNV | 1 | C | T | 30 | 29096 | 0.10310696 | 56.0666667 | rnr1Y285F_pG | TRUE | CAN | JSY2421 | 4 | rnr1Y285F_p | GCC | C>T | A/T+1 | G/C-1 | CG>TA | CG>TA | ACTGGGCCG | CCGGTAA | CGG | GCC | C | A | C>T | C>T_GCC | CG>TA_CCGGTAA | 6 | FALSE | FALSE |
| 136_v.txt | 32713 | 32713-SNV-( | SNV | 1 | C | T | 23 | 29096 | 0.07904867 | 60.0434783 | rnr1Y285F_pG | TRUE | CAN | JSY2421 | 4 | rnr1Y285F_p | GCA | C>T | A/T+1 | G/C-1 | CG>TA | CG>TA | CCCCAGCAC( | CACCACA | CGT | GCA | C | A | C>T | C>T_GCA | CG>TA_CACCACA | 6 | FALSE | FALSE |
| 136_v.txt | 32750 | 32750-SNV-( | SNV | 1 | G | A | 11 | 29096 | 0.03780588 | 48.5453383 | rnr1Y285F_pG | TRUE | CAN | JSY2421 | 4 | rnr1Y285F_p | CGA | G>A | A/T+1 | G/C-1 | CG>TA | CG>TA | AAACCCGAT/ | GATAATG | GCT | GCT | C | T | C>T | C>T_GCT | CG>TA_GATAATG | 6 | FALSE | FALSE |
| 136_v.txt | 32776 | 32776-SNV-( | SNV | 1 | C | T | 26 | 29096 | 0.08935936 | 49.9230769 | rnr1Y285F_pG | TRUE | CAN | JSY2421 | 4 | rnr1Y285F_p | GCG | C>T | A/T+1 | G/C-1 | CG>TA | CG>TA | TGGAAGCGA | CGACCCA | CGC | GCG | C | A | C>T | C>T_GCG | CG>TA_CGACCCA | 6 | FALSE | FALSE |
| 136_v.txt | 32788 | 32788-SNV-( | SNV | 1 | C | T | 228 | 29096 | 0.78361287 | 47.2675439 | rnr1Y285F_pG | TRUE | CAN | JSY2421 | 4 | rnr1Y285F_p | TCG | C>T | A/T+1 | G/C-1 | CG>TA | CG>TA | AGAACTCGA. | CGAATTC | AGC | TCG | C | A | C>T | C>T_TCG | CG>TA_CGAATTC | 6 | FALSE | FALSE |
| 136_v.txt | 32797 | 32797-SNV-( | SNV | 1 | C | T | 260 | 29096 | 0.89359362 | 47.7076923 | rnr1Y285F_pG | TRUE | CAN | JSY2421 | 4 | rnr1Y285F_p | CCG | C>T | A/T+1 | G/C-1 | CG>TA | CG>TA | ATTCACCGTA | CGTAATA | GGC | CCG | C | A | C>T | C>T_CCG | CG>TA_CGTAATA | 6 | FALSE | FALSE |
| 136_v.txt | 32798 | 32798-SNV-( | SNV | 1 | G | A | 46 | 29096 | 0.15809733 | 42.8693925 | rnr1Y285F_pG | TRUE | CAN | JSY2421 | 4 | rnr1Y285F_p | CGT | G>A | A/T+1 | G/C-1 | CG>TA | CG>TA | TTCACCGTAA | GTAATAT | GCA | GCA | C | T | C>T | C>T_GCA | CG>TA_GTAATAT | 6 | FALSE | FALSE |
| 136_v.txt | 32855 | 32855-SNV-( | SNV | 1 | C | T | 218 | 29094 | 0.74929539 | 36.4357798 | rnr1Y285F_pG | TRUE | CAN | JSY2421 | 4 | rnr1Y285F_p | TCC | C>T | A/T+1 | G/C-1 | CG>TA | CG>TA | ACTAATCCAT | CCATGCC | AGG | TCC | C | A | C>T | C>T_TCC | CG>TA_CCATGCC | 6 | FALSE | FALSE |
| 136_v.txt | 32860 | 32860-SNV-( | SNV | 1 | C | T | 39 | 29094 | 0.13404826 | 37.1282051 | rnr1Y285F_pG | TRUE | CAN | JSY2421 | 4 | rnr1Y285F_p | GCC | C>T | A/T+1 | G/C-1 | CG>TA | CG>TA | TCCATGCCG( | CCGCCAG | CGG | GCC | C | A | C>T | C>T_GCC | CG>TA_CCGCCAG | 6 | FALSE | FALSE |
| 136_v.txt | 32862 | 32862-SNV-( | SNV | 1 | G | A | 15 | 29094 | 0.05155702 | 36.0666667 | rnr1Y285F_pG | TRUE | CAN | JSY2421 | 4 | rnr1Y285F_p | CGC | G>A | A/T+1 | G/C-1 | CG>TA | CG>TA | CATGCCGCC/ | GCCAGTG | GCG | GCG | C | T | C>T | C>T_GCG | CG>TA_GCCAGTG | 6 | FALSE | FALSE |
| 136_v.txt | 32900 | 32900-SNV-( | SNV | 1 | G | A | 42 | 34459 | 0.12188398 | 34.7594761 | rnr1Y285F_pG | TRUE | CAN | JSY2421 | 4 | rnr1Y285F_p | GGC | G>A | A/T+1 | G/C-1 | CG>TA | CG>TA | GACTTGGCC/ | GCCAACT | CCG | CCG | C | T | C>T | C>T_CCG | CG>TA_GCCAACT | 6 | FALSE | FALSE |
| 136_v.txt | 32901 | 32901-SNV-( | SNV | 1 | C | T | 62 | 34459 | 0.17992397 | 33.4193375 | rnr1Y285F_pG | TRUE | CAN | JSY2421 | 4 | rnr1Y285F_p | GCC | C>T | A/T+1 | G/C-1 | CG>TA | CG>TA | ACTTGGCCA/ | CCAACTA | CGG | GCC | C | A | C>T | C>T_GCC | CG>TA_CCAACTA | 6 | FALSE | FALSE |
| 136_v.txt | 32917 | 32917-SNV-( | SNV | 1 | C | T | 103 | 34459 | 0.29890595 | 45.2621359 | rnr1Y285F_pG | TRUE | CAN | JSY2421 | 4 | rnr1Y285F_p | TCC | C>T | A/T+1 | G/C-1 | CG>TA | CG>TA | TAAGTTCCA( | CCAGGGC | AGG | TCC | C | A | C>T | C>T_TCC | CG>TA_CCAGGGC | 6 | FALSE | FALSE |
| 136_v.txt | 32922 | 32922-SNV-( | SNV | 1 | G | A | 27 | 34459 | 0.07835399 | 49.1481482 | rnr1Y285F_pG | TRUE | CAN | JSY2421 | 4 | rnr1Y285F_p | GGC | G>A | A/T+1 | G/C-1 | CG>TA | CG>TA | TCCAGGGCA. | GCAAAAG | CCG | CCG | C | T | C>T | C>T_CCG | CG>TA_GCAAAAG | 6 | FALSE | FALSE |
| 136_v.txt | 32945 | 32945-SNV-( | SNV | 1 | C | T | 39 | 34459 | 0.11317798 | 54.331774 | rnr1Y285F_pG | TRUE | CAN | JSY2421 | 4 | rnr1Y285F_p | ACC | C>T | A/T+1 | G/C-1 | CG>TA | CG>TA | AGAAAACCA | CCAATAC | TGG | ACC | C | A | C>T | C>T_ACC | CG>TA_CCAATAC | 6 | FALSE | FALSE |
| 136_v.txt | 32946 | 32946-SNV-( | SNV | 1 | C | T | 97 | 34459 | 0.28149395 | 57.5766653 | rnr1Y285F_pG | TRUE | CAN | JSY2421 | 4 | rnr1Y285F_p | CCA | C>T | A/T+1 | G/C-1 | CG>TA | CG>TA | GAAAACCAA | CAATACA | GGT | CCA | C | A | C>T | C>T_CCA | CG>TA_CAATACA | 6 | FALSE | FALSE |
| 136_v.txt | 32958 | 32958-SNV-( | SNV | 1 | C | T | 167 | 34459 | 0.48463391 | 56.4610778 | rnr1Y285F_pG | TRUE | CAN | JSY2421 | 4 | rnr1Y285F_p | ACC | C>T | A/T+1 | G/C-1 | CG>TA | CG>TA | ATGTAACCAT | CCATTGG | TGG | ACC | C | A | C>T | C>T_ACC | CG>TA_CCATTGG | 6 | FALSE | FALSE |
| 136_v.txt | 32965 | 32965-SNV-( | SNV | 1 | C | T | 36 | 34459 | 0.10447198 | 55.6111111 | rnr1Y285F_pG | TRUE | CAN | JSY2421 | 4 | rnr1Y285F_p | GCC | C>T | A/T+1 | G/C-1 | CG>TA | CG>TA | CATTGGCCG( | CCGCACC | CGG | GCC | C | A | C>T | C>T_GCC | CG>TA_CCGCACC | 6 | FALSE | FALSE |
| 136_v.txt | 32967 | 32967-SNV-( | SNV | 1 | G | A | 89 | 34459 | 0.25827795 | 58.2696629 | rnr1Y285F_pG | TRUE | CAN | JSY2421 | 4 | rnr1Y285F_p | CGC | G>A | A/T+1 | G/C-1 | CG>TA | CG>TA | TTGGCCGCA( | GCACCAA | GCG | GCG | C | T | C>T | C>T_GCG | CG>TA_GCACCAA | 6 | FALSE | FALSE |
| 136_v.txt | 33015 | 33015-SNV-( | SNV | 1 | G | A | 21 | 34459 | 0.06094199 | 46 | rnr1Y285F_pG | TRUE | CAN | JSY2421 | 4 | rnr1Y285F_p | GGA | G>A | A/T+1 | G/C-1 | CG>TA | CG>TA | AAAGAGGAT | GATGTAA | CCT | CCT | C | T | C>T | C>T_CCT | CG>TA_GATGTAA | 6 | FALSE | FALSE |
| 136_v.txt | 33051 | 33051-SNV-( | SNV | 1 | G | A | 24 | 34459 | 0.06964799 | 46.8751775 | rnr1Y285F_pG | TRUE | CAN | JSY2421 | 4 | rnr1Y285F_p | GGA | G>A | A/T+1 | G/C-1 | CG>TA | CG>TA | CCCAAGGACT | GACTGCG | CCT | CCT | C | T | C>T | C>T_CCT | CG>TA_GACTGCG | 6 | FALSE | FALSE |
| 136_v.txt | 33056 | 33056-SNV-( | SNV | 1 | C | T | 39 | 34459 | 0.11317798 | 58.3333333 | rnr1Y285F_pG | TRUE | CAN | JSY2421 | 4 | rnr1Y285F_p | GCG | C>T | A/T+1 | G/C-1 | CG>TA | CG>TA | GGACTGCGT( | CGTGACA | CGC | GCG | C | A | C>T | C>T_GCG | CG>TA_CGTGACA | 6 | FALSE | FALSE |
| 136_v.txt | 33070 | 33070-SNV-( | SNV | 1 | C | T | 38 | 34459 | 0.11027598 | 48.7894737 | rnr1Y285F_pG | TRUE | CAN | JSY2421 | 4 | rnr1Y285F_p | GCC | C>T | A/T+1 | G/C-1 | CG>TA | CG>TA | AATATGCCA/ | CCAAAGA | CGG | GCC | C | A | C>T | C>T_GCC | CG>TA_CCAAAGA | 6 | FALSE | FALSE |
| 136_v.txt | 33103 | 33103-SNV-( | SNV | 1 | C | T | 14 | 34459 | 0.04062799 | 44.9978647 | rnr1Y285F_pG | TRUE | CAN | JSY2421 | 4 | rnr1Y285F_p | GCG | C>T | A/T+1 | G/C-1 | CG>TA | CG>TA | TAAGAGCGC | CGCCCAC | CGC | GCG | C | A | C>T | C>T_GCG | CG>TA_CGCCCAC | 6 | FALSE | FALSE |
| 136_v.txt | 33104 | 33104-SNV-( | SNV | 1 | G | A | 30 | 34459 | 0.08705998 | 36.7661752 | rnr1Y285F_pG | TRUE | CAN | JSY2421 | 4 | rnr1Y285F_p | CGC | G>A | A/T+1 | G/C-1 | CG>TA | CG>TA | AAGAGCGCC | GCCCACT | GCG | GCG | C | T | C>T | C>T_GCG | CG>TA_GCCCACT | 6 | FALSE | FALSE |
| 136_v.txt | 33113 | 33113-SNV-( | SNV | 1 | G | A | 22 | 34459 | 0.06384399 | 45.4090909 | rnr1Y285F_pG | TRUE | CAN | JSY2421 | 4 | rnr1Y285F_p | GGC | G>A | A/T+1 | G/C-1 | CG>TA | CG>TA | CACTGGGCC( | GCCGGCG | CCG | CCG | C | T | C>T | C>T_CCG | CG>TA_GCCGGCG | 6 | FALSE | FALSE |
| 136_v.txt | 33114 | 33114-SNV-( | SNV | 1 | C | T | 56 | 34459 | 0.16251197 | 39.9084526 | rnr1Y285F_pG | TRUE | CAN | JSY2421 | 4 | rnr1Y285F_p | GCC | C>T | A/T+1 | G/C-1 | CG>TA | CG>TA | ACTGGGCCG | CCGGCGT | CGG | GCC | C | A | C>T | C>T_GCC | CG>TA_CCGGCGT | 6 | FALSE | FALSE |
| 136_v.txt | 33115 | 33115-SNV-( | SNV | 1 | C | T | 64 | 34459 | 0.18572797 | 44.1848646 | rnr1Y285F_pG | TRUE | CAN | JSY2421 | 4 | rnr1Y285F_p | CCG | C>T | A/T+1 | G/C-1 | CG>TA | CG>TA | CTGGGCCGG | CGGCGTT | GGC | CCG | C | A | C>T | C>T_CCG | CG>TA_CGGCGTT | 6 | FALSE | FALSE |
| 136_v.txt | 33118 | 33118-SNV-( | SNV | 1 | C | T | 30 | 34459 | 0.08705998 | 43.4663433 | rnr1Y285F_pG | TRUE | CAN | JSY2421 | 4 | rnr1Y285F_p | GCG | C>T | A/T+1 | G/C-1 | CG>TA | CG>TA | GGCCGGCGT | CGTTGGT | CGC | GCG | C | A | C>T | C>T_GCG | CG>TA_CGTTGGT | 6 | FALSE | FALSE |
| 136_v.txt | 33119 | 33119-SNV-( | SNV | 1 | G | A | 26 | 34459 | 0.07545199 | 44.9612846 | rnr1Y285F_pG | TRUE | CAN | JSY2421 | 4 | rnr1Y285F_p | CGT | G>A | A/T+1 | G/C-1 | CG>TA | CG>TA | GCCGGCGTT( | GTTGGTC | GCA | GCA | C | T | C>T | C>T_GCA | CG>TA_GTTGGTC | 6 | FALSE | FALSE |
| 136_v.txt | 33153 | 33153-SNV-( | SNV | 1 | C | T | 15 | 34459 | 0.04352999 | 33.3333334 | rnr1Y285F_pG | TRUE | CAN | JSY2421 | 4 | rnr1Y285F_p | ACC | C>T | A/T+1 | G/C-1 | CG>TA | CG>TA | AAAAGACCT( | CCTGTAC | TGG | ACC | C | A | C>T | C>T_ACC | CG>TA_CCTGTAC | 6 | FALSE | FALSE |
| 136_v.txt | 33159 | 33159-SNV-( | SNV | 1 | C | T | 140 | 34462 | 0.40624456 | 37.0428873 | rnr1Y285F_pG | TRUE | CAN | JSY2421 | 4 | rnr1Y285F_p | ACC | C>T | A/T+1 | G/C-1 | CG>TA | CG>TA | CCTGTACCA/ | CCAATAG | TGG | ACC | C | A | C>T | C>T_ACC | CG>TA_CCAATAG | 6 | FALSE | FALSE |
| 136_v.txt | 33169 | 33169-SNV-( | SNV | 1 | C | T | 9 | 34458 | 0.02611875 | 34.8889694 | rnr1Y285F_pG | TRUE | CAN | JSY2421 | 4 | rnr1Y285F_p | CCA | C>T | A/T+1 | G/C-1 | CG>TA | CG>TA | TAGTACCAC( | CACCAAG | GGT | CCA | C | A | C>T | C>T_CCA | CG>TA_CACCAAG | 6 | FALSE | FALSE |
| 136_v.txt | 33171 | 33171-SNV-( | SNV | 1 | C | T | 4346 | 34457 | 12.6128218 | 37.3693051 | rnr1Y285F_pG | TRUE | CAN | JSY2421 | 4 | rnr1Y285F_p | ACC | C>T | A/T+1 | G/C-1 | CG>TA | CG>TA | GTACCACCA/ | CCAAGGG | TGG | ACC | C | A | C>T | C>T_ACC | CG>TA_CCAAGGG | 6 | FALSE | FALSE |
| 136_v.txt | 33206 | 33206-SNV-( | SNV | 1 | C | T | 1 | 17621 | 5.68E-03 | 37.9997287 | rnr1Y285F_pG | TRUE | CAN | JSY2421 | 4 | rnr1Y285F_p | GCT | C>T | A/T+1 | G/C-1 | CG>TA | CG>TA | CTTAAGCTCT | CTCTCTC | CGA | GCT | C | A | C>T | C>T_GCT | CG>TA_CTCTCTC | 6 | FALSE | FALSE |
| 136_v.txt | 33258 | 33258-SNV-( | SNV | 1 | C | T | 35 | 17622 | 0.19861537 | 42.0571429 | rnr1Y285F_pG | TRUE | CAN | JSY2421 | 4 | rnr1Y285F_p | GCC | C>T | A/T+1 | G/C-1 | CG>TA | CG>TA | CCTATGCCAT | CCATCCT | CGG | GCC | C | A | C>T | C>T_GCC | CG>TA_CCATCCT | 6 | FALSE | FALSE |
| 136_v.txt | 33276 | 33276-SNV-( | SNV | 1 | G | A | 35 | 17622 | 0.19861537 | 44.5142857 | rnr1Y285F_pG | TRUE | CAN | JSY2421 | 4 | rnr1Y285F_p | CGT | G>A | A/T+1 | G/C-1 | CG>TA | CG>TA | GAGAACGTA | GTATCCT | GCA | GCA | C | T | C>T | C>T_GCA | CG>TA_GTATCCT | 6 | FALSE | FALSE |
| 136_v.txt | 33310 | 33310-SNV-( | SNV | 1 | G | A | 22 | 17622 | 0.12484395 | 58.1818182 | rnr1Y285F_pG | TRUE | CAN | JSY2421 | 4 | rnr1Y285F_p | CGC | G>A | A/T+1 | G/C-1 | CG>TA | CG>TA | AAGAGCGCA | GCAATGG | GCG | GCG | C | T | C>T | C>T_GCG | CG>TA_GCAATGG | 6 | FALSE | FALSE |
| 136_v.txt | 33412 | 33412-SNV-( | SNV | 1 | G | A | 38 | 17622 | 0.21563954 | 47.6842105 | rnr1Y285F_pG | TRUE | CAN | JSY2421 | 4 | rnr1Y285F_p | GGC | G>A | A/T+1 | G/C-1 | CG>TA | CG>TA | TGACCGGCT( | GCTCATT | CCG | CCG | C | T | C>T | C>T_CCG | CG>TA_GCTCATT | 6 | FALSE | FALSE |
| 136_v.txt | 33474 | 33474-SNV-( | SNV | 1 | C | T | 24 | 17622 | 0.1361934 | 35.4166667 | rnr1Y285F_pG | TRUE | CAN | JSY2421 | 4 | rnr1Y285F_p | GCC | C>T | A/T+1 | G/C-1 | CG>TA | CG>TA | GCTATGCCTT | CCTTTTT | CGG | GCC | C | A | C>T | C>T_GCC | CG>TA_CCTTTTT | 6 | FALSE | FALSE |
| 136_v.txt | 31657 | 31657-SNV-( | SNV | 1 | G | C | 7 | 11836 | 0.0591416 | 20.1428571 | rnr1Y285F_pG | TRUE | CAN | JSY2421 | 4 | rnr1Y285F_p | GGC | G>C | G/C+1 | G/C-1 | CG>GC | CG>GC | GAAATGGCG | GCGTGGA | CCG | CCG | C | G | C>G | C>G_CCG | CG>GC_GCGTGGA | 6 | FALSE | FALSE |
| 136_v.txt | 31956 | 31956-SNV-( | SNV | 1 | G | C | 13 | 24887 | 0.05223611 | 37.7692308 | rnr1Y285F_pG | TRUE | CAN | JSY2421 | 4 | rnr1Y285F_p | TGT | G>C | G/C+1 | G/C-1 | CG>GC | CG>GC | ATAAATGTG( | GTGGCCG | ACA | ACA | C | G | C>G | C>G_ACA | CG>GC_GTGGCCG | 6 | FALSE | FALSE |
| 136_v.txt | 32027 | 32027-SNV-( | SNV | 1 | G | C | 35 | 24886 | 0.14064132 | 60.053933 | rnr1Y285F_pG | TRUE | CAN | JSY2421 | 4 | rnr1Y285F_p | GGT | G>C | G/C+1 | G/C-1 | CG>GC | CG>GC | GCCACGGTA' | GTATTTC | CCA | CCA | C | G | C>G | C>G_CCA | CG>GC_GTATTTC | 6 | FALSE | FALSE |
| 136_v.txt | 32064 | 32064-SNV-( | SNV | 1 | G | C | 42 | 24887 | 0.16876281 | 61.7380952 | rnr1Y285F_pG | TRUE | CAN | JSY2421 | 4 | rnr1Y285F_p | TGA | G>C | G/C+1 | G/C-1 | CG>GC | CG>GC | GAGATTGAG | GAGATAA | ACT | ACT | C | G | C>G | C>G_ACT | CG>GC_GAGATAA | 6 | FALSE | FALSE |
| 136_v.txt | 32076 | 32076-SNV-( | SNV | 1 | C | G | 2 | 24886 | 8.04E-03 | 62.5004069 | rnr1Y285F_pG | TRUE | CAN | JSY2421 | 4 | rnr1Y285F_p | CCA | C>G | G/C+1 | G/C-1 | CG>GC | CG>GC | AATAACCAT( | CATGCAA | GGT | CCA | C | C | C>G | C>G_CCA | CG>GC_CATGCAA | 6 | FALSE | FALSE |
| 136_v.txt | 32159 | 32159-SNV-( | SNV | 1 | C | G | 1 | 24887 | 4.02E-03 | 61.9966447 | rnr1Y285F_pG | TRUE | CAN | JSY2421 | 4 | rnr1Y285F_p | CCA | C>G | G/C+1 | G/C-1 | CG>GC | CG>GC | GTAAGCCAA. | CAAAGCG | GGT | CCA | C | C | C>G | C>G_CCA | CG>GC_CAAAGCG | 6 | FALSE | FALSE |
| 136_v.txt | 32164 | 32164-SNV-( | SNV | 1 | C | G | 2 | 24887 | 8.04E-03 | 42.9995767 | rnr1Y285F_pG | TRUE | CAN | JSY2421 | 4 | rnr1Y285F_p | GCG | C>G | G/C+1 | G/C-1 | CG>GC | CG>GC | CCAAAGCGC( | CGCCAAA | CGC | GCG | C | C | C>G | C>G_GCG | CG>GC_CGCCAAA | 6 | FALSE | FALSE |
| 136_v.txt | 32195 | 32195-SNV-( | SNV | 1 | G | C | 9 | 24887 | 0.03616346 | 40.6666667 | rnr1Y285F_pG | TRUE | CAN | JSY2421 | 4 | rnr1Y285F_p | TGT | G>C | G/C+1 | G/C-1 | CG>GC | CG>GC | TGCAATGTAT | GTATGGA | ACA | ACA | C | G | C>G | C>G_ACA | CG>GC_GTATGGA | 6 | FALSE | FALSE |
| 136_v.txt | 32223 | 32223-SNV-( | SNV | 1 | G | C | 32 | 24887 | 0.12858119 | 36.09375 | rnr1Y285F_pG | TRUE | CAN | JSY2421 | 4 | rnr1Y285F_p | TGA | G>C | G/C+1 | G/C-1 | CG>GC | CG>GC | GTCCTTGAC/ | GACAGGA | ACT | ACT | C | G | C>G | C>G_ACT | CG>GC_GACAGGA | 6 | FALSE | FALSE |
| 136_v.txt | 32292 | 32292-SNV-( | SNV | 1 | G | C | 138 | 27455 | 0.50264069 | 51.8768116 | rnr1Y285F_pG | TRUE | CAN | JSY2421 | 4 | rnr1Y285F_p | TGA | G>C | G/C+1 | G/C-1 | CG>GC | CG>GC | ATATTTGAAT | GAATTTG | ACT | ACT | C | G | C>G | C>G_ACT | CG>GC_GAATTTG | 6 | FALSE | FALSE |
| 136_v.txt | 32536 | 32536-SNV-( | SNV | 1 | C | G | 2 | 27455 | 7.29E-03 | 37.9990972 | rnr1Y285F_pG | TRUE | CAN | JSY2421 | 4 | rnr1Y285F_p | GCA | C>G | G/C+1 | G/C-1 | CG>GC | CG>GC | GGTTTGCAG( | CAGCTTC | CGT | GCA | C | C | C>G | C>G_GCA | CG>GC_CAGCTTC | 6 | FALSE | FALSE |
| 136_v.txt | 32587 | 32587-SNV-( | SNV | 1 | C | G | 10 | 29096 | 0.03436899 | 34.7 | rnr1Y285F_pG | TRUE | CAN | JSY2421 | 4 | rnr1Y285F_p | GCA | C>G | G/C+1 | G/C-1 | CG>GC | CG>GC | TGAAGGCAG | CAGCGTT | CGT | GCA | C | C | C>G | C>G_GCA | CG>GC_CAGCGTT | 6 | FALSE | FALSE |
| 136_v.txt | 32735 | 32735-SNV-( | SNV | 1 | G | C | 7 | 29096 | 0.02405829 | 56 | rnr1Y285F_pG | TRUE | CAN | JSY2421 | 4 | rnr1Y285F_p | AGT | G>C | G/C+1 | G/C-1 | CG>GC | CG>GC | AAAACAGTA | GTATATT | TCA | TCA | C | G | C>G | C>G_TCA | CG>GC_GTATATT | 6 | FALSE | FALSE |
| 136_v.txt | 32749 | 32749-SNV-( | SNV | 1 | C | G | 39 | 29096 | 0.13403904 | 55.3838137 | rnr1Y285F_pG | TRUE | CAN | JSY2421 | 4 | rnr1Y285F_p | CCG | C>G | G/C+1 | G/C-1 | CG>GC | CG>GC | GAAACCCGA | CGATAAT | GGC | CCG | C | C | C>G | C>G_CCG | CG>GC_CGATAAT | 6 | FALSE | FALSE |
| 136_v.txt | 32797 | 32797-SNV-( | SNV | 1 | C | G | 16 | 29096 | 0.05499038 | 46.0625 | rnr1Y285F_pG | TRUE | CAN | JSY2421 | 4 | rnr1Y285F_p | CCG | C>G | G/C+1 | G/C-1 | CG>GC | CG>GC | ATTCACCGTA | CGTAATA | GGC | CCG | C | C | C>G | C>G_CCG | CG>GC_CGTAATA | 6 | FALSE | FALSE |
| 136_v.txt | 32862 | 32862-SNV-( | SNV | 1 | G | C | 9 | 29094 | 0.03093421 | 37.6666667 | rnr1Y285F_pG | TRUE | CAN | JSY2421 | 4 | rnr1Y285F_p | CGC | G>C | G/C+1 | G/C-1 | CG>GC | CG>GC | CATGCCGCC/ | GCCAGTG | GCG | GCG | C | G | C>G | C>G_GCG | CG>GC_GCCAGTG | 6 | FALSE | FALSE |
| 136_v.txt | 32936 | 32936-SNV-( | SNV | 1 | C | G | 11 | 34459 | 0.03192199 | 56.5051531 | rnr1Y285F_pG | TRUE | CAN | JSY2421 | 4 | rnr1Y285F_p | CCC | C>G | G/C+1 | G/C-1 | CG>GC | CG>GC | GATTGCCCA/ | CCAAGAA | GGG | CCC | C | C | C>G | C>G_CCC | CG>GC_CCAAGAA | 6 | FALSE | FALSE |
| 136_v.txt | 32937 | 32937-SNV-( | SNV | 1 | C | G | 23 | 34459 | 0.06674599 | 50.6904076 | rnr1Y285F_pG | TRUE | CAN | JSY2421 | 4 | rnr1Y285F_p | CCA | C>G | G/C+1 | G/C-1 | CG>GC | CG>GC | ATTGCCCAA( | CAAGAAA | GGT | CCA | C | C | C>G | C>G_CCA | CG>GC_CAAGAAA | 6 | FALSE | FALSE |
| 136_v.txt | 33018 | 33018-SNV-( | SNV | 1 | G | C | 20 | 34459 | 0.05803999 | 59.1 | rnr1Y285F_pG | TRUE | CAN | JSY2421 | 4 | rnr1Y285F_p | TGT | G>C | G/C+1 | G/C-1 | CG>GC | CG>GC | GAGGATGTA | GTAACAG | ACA | ACA | C | G | C>G | C>G_ACA | CG>GC_GTAACAG | 6 | FALSE | FALSE |
| 136_v.txt | 33024 | 33024-SNV-( | SNV | 1 | G | C | 34 | 34459 | 0.09866798 | 60.1470588 | rnr1Y285F_pG | TRUE | CAN | JSY2421 | 4 | rnr1Y285F_p | AGG | G>C | G/C+1 | G/C-1 | CG>GC | CG>GC | GTAACAGGG | GGGATGA | TCC | TCC | C | G | C>G | C>G_TCC | CG>GC_GGGATGA | 6 | FALSE | FALSE |
| 136_v.txt | 33070 | 33070-SNV-( | SNV | 1 | C | G | 127 | 34459 | 0.36855393 | 55.6377953 | rnr1Y285F_pG | TRUE | CAN | JSY2421 | 4 | rnr1Y285F_p | GCC | C>G | G/C+1 | G/C-1 | CG>GC | CG>GC | AATATGCCA/ | CCAAAGA | CGG | GCC | C | C | C>G | C>G_GCC | CG>GC_CCAAAGA | 6 | FALSE | FALSE |
| 136_v.txt | 33093 | 33093-SNV-( | SNV | 1 | G | C | 24 | 34459 | 0.06964799 | 46.2915793 | rnr1Y285F_pG | TRUE | CAN | JSY2421 | 4 | rnr1Y285F_p | TGA | G>C | G/C+1 | G/C-1 | CG>GC | CG>GC | AAATATGAT/ | GATATAA | ACT | ACT | C | G | C>G | C>G_ACT | CG>GC_GATATAA | 6 | FALSE | FALSE |
| 136_v.txt | 33103 | 33103-SNV-( | SNV | 1 | C | G | 34 | 34459 | 0.09866798 | 40.2034018 | rnr1Y285F_pG | TRUE | CAN | JSY2421 | 4 | rnr1Y285F_p | GCG | C>G | G/C+1 | G/C-1 | CG>GC | CG>GC | TAAGAGCGC | CGCCCAC | CGC | GCG | C | C | C>G | C>G_GCG | CG>GC_CGCCCAC | 6 | FALSE | FALSE |
| 136_v.txt | 33115 | 33115-SNV-( | SNV | 1 | C | G | 7 | 34459 | 0.020314 | 41.7110776 | rnr1Y285F_pG | TRUE | CAN | JSY2421 | 4 | rnr1Y285F_p | CCG | C>G | G/C+1 | G/C-1 | CG>GC | CG>GC | CTGGGCCGG | CGGCGTT | GGC | CCG | C | C | C>G | C>G_CCG | CG>GC_CGGCGTT | 6 | FALSE | FALSE |
| 136_v.txt | 33118 | 33118-SNV-( | SNV | 1 | C | G | 2 | 34459 | 5.80E-03 | 50.9988752 | rnr1Y285F_pG | TRUE | CAN | JSY2421 | 4 | rnr1Y285F_p | GCG | C>G | G/C+1 | G/C-1 | CG>GC | CG>GC | GGCCGGCGT | CGTTGGT | CGC | GCG | C | C | C>G | C>G_GCG | CG>GC_CGTTGGT | 6 | FALSE | FALSE |
| 136_v.txt | 33154 | 33154-SNV-( | SNV | 1 | C | G | 2 | 34458 | 5.80E-03 | 37.5000002 | rnr1Y285F_pG | TRUE | CAN | JSY2421 | 4 | rnr1Y285F_p | CCT | C>G | G/C+1 | G/C-1 | CG>GC | CG>GC | AAAGACCTG' | CTGTACC | GGA | CCT | C | C | C>G | C>G_CCT | CG>GC_CTGTACC | 6 | FALSE | FALSE |
| 136_v.txt | 33160 | 33160-SNV-( | SNV | 1 | C | G | 21 | 34462 | 0.06093668 | 36.5231614 | rnr1Y285F_pG | TRUE | CAN | JSY2421 | 4 | rnr1Y285F_p | CCA | C>G | G/C+1 | G/C-1 | CG>GC | CG>GC | CTGTACCAAT | CAATAGT | GGT | CCA | C | C | C>G | C>G_CCA | CG>GC_CAATAGT | 6 | FALSE | FALSE |
| 136_v.txt | 33195 | 33195-SNV-( | SNV | 1 | C | G | 1 | 17622 | 5.68E-03 | 37.9975516 | rnr1Y285F_pG | TRUE | CAN | JSY2421 | 4 | rnr1Y285F_p | TCT | C>G | G/C+1 | G/C-1 | CG>GC | CG>GC | ATATGTCTTT | CTTTGCT | AGA | TCT | C | C | C>G | C>G_TCT | CG>GC_CTTTGCT | 6 | FALSE | FALSE |
| 136_v.txt | 33276 | 33276-SNV-( | SNV | 1 | G | C | 5 | 17622 | 0.02837362 | 18.8 | rnr1Y285F_pG | TRUE | CAN | JSY2421 | 4 | rnr1Y285F_p | CGT | G>C | G/C+1 | G/C-1 | CG>GC | CG>GC | GAGAACGTA | GTATCCT | GCA | GCA | C | G | C>G | C>G_GCA | CG>GC_GTATCCT | 6 | FALSE | FALSE |
| 136_v.txt | 33378 | 33378-SNV-( | SNV | 1 | G | C | 143 | 17622 | 0.81148564 | 55.6923077 | rnr1Y285F_pG | TRUE | CAN | JSY2421 | 4 | rnr1Y285F_p | TGA | G>C | G/C+1 | G/C-1 | CG>GC | CG>GC | GTTTGTGAA( | GAAGCTT | ACT | ACT | C | G | C>G | C>G_ACT | CG>GC_GAAGCTT | 6 | FALSE | FALSE |
| 136_v.txt | 31657 | 31657-SNV-( | SNV | 1 | G | T | 86 | 11836 | 0.72659682 | 20.0348837 | rnr1Y285F_pG | TRUE | CAN | JSY2421 | 4 | rnr1Y285F_p | GGC | G>T | A/T+1 | G/C-1 | CG>AT | CG>AT | GAAATGGCG | GCGTGGA | CCG | CCG | C | A | C>A | C>A_CCG | CG>AT_GCGTGGA | 6 | FALSE | FALSE |
| 136_v.txt | 31930 | 31930-SNV-( | SNV | 1 | G | T | 11 | 11834 | 0.09295251 | 37.6363636 | rnr1Y285F_pG | TRUE | CAN | JSY2421 | 4 | rnr1Y285F_p | TGA | G>T | A/T+1 | G/C-1 | CG>AT | CG>AT | AACCTTGAAT | GAATAAT | ACT | ACT | C | A | C>A | C>A_ACT | CG>AT_GAATAAT | 6 | FALSE | FALSE |
| 136_v.txt | 31981 | 31981-SNV-( | SNV | 1 | G | T | 37 | 24887 | 0.148672 | 31.1891892 | rnr1Y285F_pG | TRUE | CAN | JSY2421 | 4 | rnr1Y285F_p | GGC | G>T | A/T+1 | G/C-1 | CG>AT | CG>AT | AGCCGGGCA | GCATTAA | CCG | CCG | C | A | C>A | C>A_CCG | CG>AT_GCATTAA | 6 | FALSE | FALSE |
| 136_v.txt | 32014 | 32014-SNV-( | SNV | 1 | G | T | 9 | 24887 | 0.03616346 | 27.5574456 | rnr1Y285F_pG | TRUE | CAN | JSY2421 | 4 | rnr1Y285F_p | CGA | G>T | A/T+1 | G/C-1 | CG>AT | CG>AT | CGTCACGAG. | GAGAGAT | GCT | GCT | C | A | C>A | C>A_GCT | CG>AT_GAGAGAT | 6 | FALSE | FALSE |
| 136_v.txt | 32026 | 32026-SNV-( | SNV | 1 | G | T | 5 | 24887 | 0.02009081 | 42.7974538 | rnr1Y285F_pG | TRUE | CAN | JSY2421 | 4 | rnr1Y285F_p | CGG | G>T | A/T+1 | G/C-1 | CG>AT | CG>AT | TGCCACGGT/ | GGTATTT | GCC | GCC | C | A | C>A | C>A_GCC | CG>AT_GGTATTT | 6 | FALSE | FALSE |
| 136_v.txt | 32027 | 32027-SNV-( | SNV | 1 | G | T | 448 | 24886 | 1.80020895 | 60.4694593 | rnr1Y285F_pG | TRUE | CAN | JSY2421 | 4 | rnr1Y285F_p | GGT | G>T | A/T+1 | G/C-1 | CG>AT | CG>AT | GCCACGGTA' | GTATTTC | CCA | CCA | C | A | C>A | C>A_CCA | CG>AT_GTATTTC | 6 | FALSE | FALSE |
| 136_v.txt | 32058 | 32058-SNV-( | SNV | 1 | G | T | 84 | 24887 | 0.33752562 | 59.9166667 | rnr1Y285F_pG | TRUE | CAN | JSY2421 | 4 | rnr1Y285F_p | CGA | G>T | A/T+1 | G/C-1 | CG>AT | CG>AT | ATGTGCGAG | GAGATTG | GCT | GCT | C | A | C>A | C>A_GCT | CG>AT_GAGATTG | 6 | FALSE | FALSE |
| 136_v.txt | 32274 | 32274-SNV-( | SNV | 1 | G | T | 31 | 27455 | 0.11291204 | 34.2258065 | rnr1Y285F_pG | TRUE | CAN | JSY2421 | 4 | rnr1Y285F_p | GGA | G>T | A/T+1 | G/C-1 | CG>AT | CG>AT | ATACGGGAA | GAACCAA | CCT | CCT | C | A | C>A | C>A_CCT | CG>AT_GAACCAA | 6 | FALSE | FALSE |
| 136_v.txt | 32304 | 32304-SNV-( | SNV | 1 | G | T | 206 | 27455 | 0.7503187 | 56.6636918 | rnr1Y285F_pG | TRUE | CAN | JSY2421 | 4 | rnr1Y285F_p | AGA | G>T | A/T+1 | G/C-1 | CG>AT | CG>AT | GCGGCAGAA | GAAATAA | TCT | TCT | C | A | C>A | C>A_TCT | CG>AT_GAAATAA | 6 | FALSE | FALSE |
| 136_v.txt | 32368 | 32368-SNV-( | SNV | 1 | C | A | 30 | 27455 | 0.10926971 | 57.7 | rnr1Y285F_pG | TRUE | CAN | JSY2421 | 4 | rnr1Y285F_p | TCA | C>A | A/T+1 | G/C-1 | CG>AT | CG>AT | AGTTCTCAAT | CAATAGC | AGT | TCA | C | T | C>A | C>A_TCA | CG>AT_CAATAGC | 6 | FALSE | FALSE |
| 136_v.txt | 32373 | 32373-SNV-( | SNV | 1 | G | T | 32 | 27455 | 0.11655436 | 59.9375 | rnr1Y285F_pG | TRUE | CAN | JSY2421 | 4 | rnr1Y285F_p | AGC | G>T | A/T+1 | G/C-1 | CG>AT | CG>AT | TCAATAGCA/ | GCAATAA | TCG | TCG | C | A | C>A | C>A_TCG | CG>AT_GCAATAA | 6 | FALSE | FALSE |
| 136_v.txt | 32432 | 32432-SNV-( | SNV | 1 | G | T | 71 | 27455 | 0.25860499 | 57.4225352 | rnr1Y285F_pG | TRUE | CAN | JSY2421 | 4 | rnr1Y285F_p | TGT | G>T | A/T+1 | G/C-1 | CG>AT | CG>AT | GTCATTGTAT | GTATGGA | ACA | ACA | C | A | C>A | C>A_ACA | CG>AT_GTATGGA | 6 | FALSE | FALSE |
| 136_v.txt | 32461 | 32461-SNV-( | SNV | 1 | G | T | 3 | 27455 | 0.01092697 | 38 | rnr1Y285F_pG | TRUE | CAN | JSY2421 | 4 | rnr1Y285F_p | AGA | G>T | A/T+1 | G/C-1 | CG>AT | CG>AT | ATAATAGAG | GAGAGCC | TCT | TCT | C | A | C>A | C>A_TCT | CG>AT_GAGAGCC | 6 | FALSE | FALSE |
| 136_v.txt | 32471 | 32471-SNV-( | SNV | 1 | G | T | 31 | 27455 | 0.11291204 | 48.1935484 | rnr1Y285F_pG | TRUE | CAN | JSY2421 | 4 | rnr1Y285F_p | TGT | G>T | A/T+1 | G/C-1 | CG>AT | CG>AT | GCCAATGTA( | GTAGAAG | ACA | ACA | C | A | C>A | C>A_ACA | CG>AT_GTAGAAG | 6 | FALSE | FALSE |
| 136_v.txt | 32529 | 32529-SNV-( | SNV | 1 | G | T | 7 | 27455 | 0.02549627 | 34.2857143 | rnr1Y285F_pG | TRUE | CAN | JSY2421 | 4 | rnr1Y285F_p | GGG | G>T | A/T+1 | G/C-1 | CG>AT | CG>AT | TTTCTGGGGT | GGGTTTG | CCC | CCC | C | A | C>A | C>A_CCC | CG>AT_GGGTTTG | 6 | FALSE | FALSE |
| 136_v.txt | 32530 | 32530-SNV-( | SNV | 1 | G | T | 30 | 27455 | 0.10926971 | 35.1000345 | rnr1Y285F_pG | TRUE | CAN | JSY2421 | 4 | rnr1Y285F_p | GGG | G>T | A/T+1 | G/C-1 | CG>AT | CG>AT | TTCTGGGGTT | GGTTTGC | CCC | CCC | C | A | C>A | C>A_CCC | CG>AT_GGTTTGC | 6 | FALSE | FALSE |
| 136_v.txt | 32538 | 32538-SNV-( | SNV | 1 | G | T | 23 | 27455 | 0.08377345 | 37.3913044 | rnr1Y285F_pG | TRUE | CAN | JSY2421 | 4 | rnr1Y285F_p | AGC | G>T | A/T+1 | G/C-1 | CG>AT | CG>AT | TTTGCAGCTT | GCTTCAC | TCG | TCG | C | A | C>A | C>A_TCG | CG>AT_GCTTCAC | 6 | FALSE | FALSE |
| 136_v.txt | 32542 | 32542-SNV-( | SNV | 1 | C | A | 39 | 27454 | 0.1420558 | 36.974359 | rnr1Y285F_pG | TRUE | CAN | JSY2421 | 4 | rnr1Y285F_p | TCA | C>A | A/T+1 | G/C-1 | CG>AT | CG>AT | CAGCTTCAC( | CACCAGC | AGT | TCA | C | T | C>A | C>A_TCA | CG>AT_CACCAGC | 6 | FALSE | FALSE |
| 136_v.txt | 32585 | 32585-SNV-( | SNV | 1 | G | T | 8 | 29096 | 0.02749519 | 32.875 | rnr1Y285F_pG | TRUE | CAN | JSY2421 | 4 | rnr1Y285F_p | AGG | G>T | A/T+1 | G/C-1 | CG>AT | CG>AT | TGTGAAGGC | GGCAGCG | TCC | TCC | C | A | C>A | C>A_TCC | CG>AT_GGCAGCG | 6 | FALSE | FALSE |
| 136_v.txt | 32692 | 32692-SNV-( | SNV | 1 | C | A | 33 | 29096 | 0.11341765 | 63.1515152 | rnr1Y285F_pG | TRUE | CAN | JSY2421 | 4 | rnr1Y285F_p | CCA | C>A | A/T+1 | G/C-1 | CG>AT | CG>AT | GGAATCCAA' | CAACTGG | GGT | CCA | C | T | C>A | C>A_CCA | CG>AT_CAACTGG | 6 | FALSE | FALSE |
| 136_v.txt | 32749 | 32749-SNV-( | SNV | 1 | C | A | 5 | 29096 | 0.01718449 | 58.7976259 | rnr1Y285F_pG | TRUE | CAN | JSY2421 | 4 | rnr1Y285F_p | CCG | C>A | A/T+1 | G/C-1 | CG>AT | CG>AT | GAAACCCGA | CGATAAT | GGC | CCG | C | T | C>A | C>A_CCG | CG>AT_CGATAAT | 6 | FALSE | FALSE |
| 136_v.txt | 32783 | 32783-SNV-( | SNV | 1 | G | T | 46 | 29096 | 0.15809733 | 44.9130435 | rnr1Y285F_pG | TRUE | CAN | JSY2421 | 4 | rnr1Y285F_p | AGA | G>T | A/T+1 | G/C-1 | CG>AT | CG>AT | GACCCAGAA | GAACTCG | TCT | TCT | C | A | C>A | C>A_TCT | CG>AT_GAACTCG | 6 | FALSE | FALSE |
| 136_v.txt | 32788 | 32788-SNV-( | SNV | 1 | C | A | 55 | 29096 | 0.18902942 | 46.8727273 | rnr1Y285F_pG | TRUE | CAN | JSY2421 | 4 | rnr1Y285F_p | TCG | C>A | A/T+1 | G/C-1 | CG>AT | CG>AT | AGAACTCGA. | CGAATTC | AGC | TCG | C | T | C>A | C>A_TCG | CG>AT_CGAATTC | 6 | FALSE | FALSE |
| 136_v.txt | 32797 | 32797-SNV-( | SNV | 1 | C | A | 3 | 29096 | 0.0103107 | 38 | rnr1Y285F_pG | TRUE | CAN | JSY2421 | 4 | rnr1Y285F_p | CCG | C>A | A/T+1 | G/C-1 | CG>AT | CG>AT | ATTCACCGTA | CGTAATA | GGC | CCG | C | T | C>A | C>A_CCG | CG>AT_CGTAATA | 6 | FALSE | FALSE |
| 136_v.txt | 32809 | 32809-SNV-( | SNV | 1 | C | A | 13 | 29096 | 0.04467968 | 41.8461539 | rnr1Y285F_pG | TRUE | CAN | JSY2421 | 4 | rnr1Y285F_p | ACA | C>A | A/T+1 | G/C-1 | CG>AT | CG>AT | ATTTGACAG( | CAGGGAA | TGT | ACA | C | T | C>A | C>A_ACA | CG>AT_CAGGGAA | 6 | FALSE | FALSE |
| 136_v.txt | 32819 | 32819-SNV-( | SNV | 1 | G | T | 65 | 29096 | 0.22339841 | 45.4 | rnr1Y285F_pG | TRUE | CAN | JSY2421 | 4 | rnr1Y285F_p | AGT | G>T | A/T+1 | G/C-1 | CG>AT | CG>AT | GAACAAGTT( | GTTCATT | TCA | TCA | C | A | C>A | C>A_TCA | CG>AT_GTTCATT | 6 | FALSE | FALSE |
| 136_v.txt | 32917 | 32917-SNV-( | SNV | 1 | C | A | 237 | 34459 | 0.68777388 | 51.535865 | rnr1Y285F_pG | TRUE | CAN | JSY2421 | 4 | rnr1Y285F_p | TCC | C>A | A/T+1 | G/C-1 | CG>AT | CG>AT | TAAGTTCCA( | CCAGGGC | AGG | TCC | C | T | C>A | C>A_TCC | CG>AT_CCAGGGC | 6 | FALSE | FALSE |

| | | | | | | | | | | | | | | | | | | | | | | | | | | | | | | | | | | |
|---|---|---|---|---|---|---|---|---|---|---|---|---|---|---|---|---|---|---|---|---|---|---|---|---|---|---|---|---|---|---|---|---|---|---|
| 136_v.txt | 32928 | 32928-SNV-C | SNV | 1 | G | T | 13 | 34459 | 0.03772599 | 47.1538462 | rnr1Y285F_pG | TRUE | CAN | JSY2421 | 4 | rnr1Y285F_p | AGT | G>T | A/T+1 | G/C-1 | CG>AT | CG>AT | GCAAAAGTG | GTGATTG | TCA | TCA | C | A | C>A | C>A_TCA | CG>AT_GTGATTG | 6 | FALSE | FALSE |
| 136_v.txt | 32936 | 32936-SNV-C | SNV | 1 | C | A | 34 | 34459 | 0.09866798 | 21.5293256 | rnr1Y285F_pG | TRUE | CAN | JSY2421 | 4 | rnr1Y285F_p | CCC | C>A | A/T+1 | G/C-1 | CG>AT | CG>AT | GATTGCCCA | CCAAGAA | GGG | CCC | C | T | C>A | C>A_CCC | CG>AT_CCAAGAA | 6 | FALSE | FALSE |
| 136_v.txt | 32937 | 32937-SNV-C | SNV | 1 | C | A | 21 | 34459 | 0.06094199 | 43.1304044 | rnr1Y285F_pG | TRUE | CAN | JSY2421 | 4 | rnr1Y285F_p | CCA | C>A | A/T+1 | G/C-1 | CG>AT | CG>AT | ATTGCCCAA | CAAGAAA | GGT | CCA | C | T | C>A | C>A_CCA | CG>AT_CAAGAAA | 6 | FALSE | FALSE |
| 136_v.txt | 32940 | 32940-SNV-C | SNV | 1 | G | T | 38 | 34459 | 0.11027598 | 45.5264455 | rnr1Y285F_pG | TRUE | CAN | JSY2421 | 4 | rnr1Y285F_p | AGA | G>T | A/T+1 | G/C-1 | CG>AT | CG>AT | GCCCAAGAA | GAAAACC | TCT | TCT | C | A | C>A | C>A_TCT | CG>AT_GAAAACC | 6 | FALSE | FALSE |
| 136_v.txt | 33012 | 33012-SNV-C | SNV | 1 | G | T | 237 | 34459 | 0.68777388 | 59.9535865 | rnr1Y285F_pG | TRUE | CAN | JSY2421 | 4 | rnr1Y285F_p | AGA | G>T | A/T+1 | G/C-1 | CG>AT | CG>AT | GTGAAAGAG | GAGGATG | TCT | TCT | C | A | C>A | C>A_TCT | CG>AT_GAGGATG | 6 | FALSE | FALSE |
| 136_v.txt | 33015 | 33015-SNV-C | SNV | 1 | G | T | 41 | 34459 | 0.11898198 | 57.6097561 | rnr1Y285F_pG | TRUE | CAN | JSY2421 | 4 | rnr1Y285F_p | GGA | G>T | A/T+1 | G/C-1 | CG>AT | CG>AT | AAAGAGGAT | GATGTAA | CCT | CCT | C | A | C>A | C>A_CCT | CG>AT_GATGTAA | 6 | FALSE | FALSE |
| 136_v.txt | 33043 | 33043-SNV-C | SNV | 1 | C | A | 52 | 34459 | 0.15090397 | 57.7884615 | rnr1Y285F_pG | TRUE | CAN | JSY2421 | 4 | rnr1Y285F_p | TCA | C>A | A/T+1 | G/C-1 | CG>AT | CG>AT | CCATTTCACC | CACCCAA | AGT | TCA | C | T | C>A | C>A_TCA | CG>AT_CACCCAA | 6 | FALSE | FALSE |
| 136_v.txt | 33050 | 33050-SNV-C | SNV | 1 | G | T | 17 | 34459 | 0.04933399 | 51.2934975 | rnr1Y285F_pG | TRUE | CAN | JSY2421 | 4 | rnr1Y285F_p | AGG | G>T | A/T+1 | G/C-1 | CG>AT | CG>AT | ACCCAAGGA | GGACTGC | TCC | TCC | C | A | C>A | C>A_TCC | CG>AT_GGACTGC | 6 | FALSE | FALSE |
| 136_v.txt | 33051 | 33051-SNV-C | SNV | 1 | G | T | 33 | 34459 | 0.09576598 | 53.1844484 | rnr1Y285F_pG | TRUE | CAN | JSY2421 | 4 | rnr1Y285F_p | GGA | G>T | A/T+1 | G/C-1 | CG>AT | CG>AT | CCCAAGGAC | GACTGCG | CCT | CCT | C | A | C>A | C>A_CCT | CG>AT_GACTGCG | 6 | FALSE | FALSE |
| 136_v.txt | 33093 | 33093-SNV-C | SNV | 1 | G | T | 99 | 34459 | 0.28729795 | 50.2726249 | rnr1Y285F_pG | TRUE | CAN | JSY2421 | 4 | rnr1Y285F_p | TGA | G>T | A/T+1 | G/C-1 | CG>AT | CG>AT | AAATATGAT | GATATAA | ACT | ACT | C | A | C>A | C>A_ACT | CG>AT_GATATAA | 6 | FALSE | FALSE |
| 136_v.txt | 33151 | 33151-SNV-C | SNV | 1 | G | T | 248 | 34459 | 0.71969587 | 39.7258065 | rnr1Y285F_pG | TRUE | CAN | JSY2421 | 4 | rnr1Y285F_p | AGA | G>T | A/T+1 | G/C-1 | CG>AT | CG>AT | TGAAAAGAC | GACCTGT | TCT | TCT | C | A | C>A | C>A_TCT | CG>AT_GACCTGT | 6 | FALSE | FALSE |
| 136_v.txt | 33153 | 33153-SNV-C | SNV | 1 | C | A | 23 | 34459 | 0.06674599 | 39.0434783 | rnr1Y285F_pG | TRUE | CAN | JSY2421 | 4 | rnr1Y285F_p | ACC | C>A | A/T+1 | G/C-1 | CG>AT | CG>AT | AAAAGACCT | CCTGTAC | TGG | ACC | C | T | C>A | C>A_ACC | CG>AT_CCTGTAC | 6 | FALSE | FALSE |
| 136_v.txt | 33154 | 33154-SNV-C | SNV | 1 | C | A | 133 | 34458 | 0.38597713 | 37.2782063 | rnr1Y285F_pG | TRUE | CAN | JSY2421 | 4 | rnr1Y285F_p | CCT | C>A | A/T+1 | G/C-1 | CG>AT | CG>AT | AAAGACCTG | CTGTACC | GGA | CCT | C | T | C>A | C>A_CCT | CG>AT_CTGTACC | 6 | FALSE | FALSE |
| 136_v.txt | 33159 | 33159-SNV-C | SNV | 1 | C | A | 20 | 34462 | 0.05803494 | 35.6500526 | rnr1Y285F_pG | TRUE | CAN | JSY2421 | 4 | rnr1Y285F_p | ACC | C>A | A/T+1 | G/C-1 | CG>AT | CG>AT | CCTGTACCA | CCAATAG | TGG | ACC | C | T | C>A | C>A_ACC | CG>AT_CCAATAG | 6 | FALSE | FALSE |
| 136_v.txt | 33168 | 33168-SNV-C | SNV | 1 | C | A | 185 | 34458 | 0.53688548 | 37.108163 | rnr1Y285F_pG | TRUE | CAN | JSY2421 | 4 | rnr1Y285F_p | ACC | C>A | A/T+1 | G/C-1 | CG>AT | CG>AT | ATAGTACCA | CCACCAA | TGG | ACC | C | T | C>A | C>A_ACC | CG>AT_CCACCAA | 6 | FALSE | FALSE |
| 136_v.txt | 33195 | 33195-SNV-C | SNV | 1 | C | A | 33 | 17622 | 0.18726592 | 34.8476643 | rnr1Y285F_pG | TRUE | CAN | JSY2421 | 4 | rnr1Y285F_p | TCT | C>A | A/T+1 | G/C-1 | CG>AT | CG>AT | ATATGTCTTT | CTTTGCT | AGA | TCT | C | T | C>A | C>A_TCT | CG>AT_CTTTGCT | 6 | FALSE | FALSE |
| 136_v.txt | 33210 | 33210-SNV-C | SNV | 1 | C | A | 38 | 17622 | 0.21563954 | 40.4210526 | rnr1Y285F_pG | TRUE | CAN | JSY2421 | 4 | rnr1Y285F_p | TCT | C>A | A/T+1 | G/C-1 | CG>AT | CG>AT | AGCTCTCTCT | CTCTTCA | AGA | TCT | C | T | C>A | C>A_TCT | CG>AT_CTCTTCA | 6 | FALSE | FALSE |
| 136_v.txt | 33235 | 33235-SNV-C | SNV | 1 | C | A | 69 | 17622 | 0.39155601 | 37 | rnr1Y285F_pG | TRUE | CAN | JSY2421 | 4 | rnr1Y285F_p | TCT | C>A | A/T+1 | G/C-1 | CG>AT | CG>AT | GTACTTCTCC | CTCCTTC | AGA | TCT | C | T | C>A | C>A_TCT | CG>AT_CTCCTTC | 6 | FALSE | FALSE |
| 136_v.txt | 33238 | 33238-SNV-C | SNV | 1 | C | A | 37 | 17622 | 0.20996482 | 36.8378378 | rnr1Y285F_pG | TRUE | CAN | JSY2421 | 4 | rnr1Y285F_p | CCT | C>A | A/T+1 | G/C-1 | CG>AT | CG>AT | CTTCTCCTTC | CTTCATC | GGA | CCT | C | T | C>A | C>A_CCT | CG>AT_CTTCATC | 6 | FALSE | FALSE |
| 136_v.txt | 33283 | 33283-SNV-C | SNV | 1 | C | A | 34 | 17622 | 0.19294064 | 45.5 | rnr1Y285F_pG | TRUE | CAN | JSY2421 | 4 | rnr1Y285F_p | TCG | C>A | A/T+1 | G/C-1 | CG>AT | CG>AT | TATCCTCGCC | CGCCATT | AGC | TCG | C | T | C>A | C>A_TCG | CG>AT_CGCCATT | 6 | FALSE | FALSE |
| 136_v.txt | 33385 | 33385-SNV-C | SNV | 1 | C | A | 56 | 17622 | 0.31778459 | 50.3571429 | rnr1Y285F_pG | TRUE | CAN | JSY2421 | 4 | rnr1Y285F_p | TCA | C>A | A/T+1 | G/C-1 | CG>AT | CG>AT | AAGCTTCAA | CAACGTC | AGT | TCA | C | T | C>A | C>A_TCA | CG>AT_CAACGTC | 6 | FALSE | FALSE |
| 136_v.txt | 33433 | 33433-SNV-C | SNV | 1 | C | A | 52 | 17622 | 0.29508569 | 42.5384615 | rnr1Y285F_pG | TRUE | CAN | JSY2421 | 4 | rnr1Y285F_p | TCC | C>A | A/T+1 | G/C-1 | CG>AT | CG>AT | GCTTCTCCTC | CCTCTAT | AGG | TCC | C | T | C>A | C>A_TCC | CG>AT_CCTCTAT | 6 | FALSE | FALSE |
| 136_v.txt | 33456 | 33456-SNV-C | SNV | 1 | G | T | 145 | 17622 | 0.82283509 | 42.3655172 | rnr1Y285F_pG | TRUE | CAN | JSY2421 | 4 | rnr1Y285F_p | TGA | G>T | A/T+1 | G/C-1 | CG>AT | CG>AT | TCTTTTGAAT | GAATTTG | ACT | ACT | C | A | C>A | C>A_ACT | CG>AT_GAATTTG | 6 | FALSE | FALSE |
| 136_v.txt | 33474 | 33474-SNV-C | SNV | 1 | C | A | 57 | 17622 | 0.32345931 | 33.3157895 | rnr1Y285F_pG | TRUE | CAN | JSY2421 | 4 | rnr1Y285F_p | GCC | C>A | A/T+1 | G/C-1 | CG>AT | CG>AT | GCTATGCCTT | CCTTTTT | CGG | GCC | C | T | C>A | C>A_GCC | CG>AT_CCTTTTT | 6 | FALSE | FALSE |
| 136_v.txt | 32304 | 32304-Repla | Replacement | 2 | G | CA | 6 | 27455 | 0.02185394 | 63.0832259 | rnr1Y285F_pG | TRUE | CAN | JSY2421 | 4 | rnr1Y285F_p | AGA | nope | >1bp-ins | G/C-1 | none | Replac. | GCGGCAGAA | GAAATAA | TCT | TCT | C | none | C>none | C>none_TCT | Replac._GAAATAA | 6 | FALSE | FALSE |
| 136_v.txt | 32079 | 32079-MNV- | MNV | 2 | GC | AA | 18 | 24887 | 0.07232692 | 63.8889088 | rnr1Y285F_pG | TRUE | CAN | JSY2421 | 4 | rnr1Y285F_p | TGC | nope | >1bp-ins | >1bpdel | none | MNV | AACCATGCA | GCAAAAA | ACG | TGC | GC | none | NA | NA_TGC | MNV_GCAAAAA | 6 | FALSE | FALSE |
| 136_v.txt | 32748 | 32748-MNV- | MNV | 2 | CC | AA | 1 | 29096 | 3.44E-03 | 36.0001943 | rnr1Y285F_pG | TRUE | CAN | JSY2421 | 4 | rnr1Y285F_p | CCC | nope | >1bp-ins | >1bpdel | none | MNV | AGAAACCCG | CCGATAA | GGG | CCC | CC | none | NA | NA_CCC | MNV_CCGATAA | 6 | FALSE | FALSE |
| 136_v.txt | 33153 | 33153-MNV- | MNV | 2 | CC | AA | 1 | 34459 | 2.90E-03 | 32.5 | rnr1Y285F_pG | TRUE | CAN | JSY2421 | 4 | rnr1Y285F_p | ACC | nope | >1bp-ins | >1bpdel | none | MNV | AAAAGACCT | CCTGTAC | TGG | ACC | CC | none | NA | NA_ACC | MNV_CCTGTAC | 6 | FALSE | FALSE |
| 136_v.txt | 33153 | 33153-MNV- | MNV | 2 | CC | GA | 1 | 34459 | 2.90E-03 | 38 | rnr1Y285F_pG | TRUE | CAN | JSY2421 | 4 | rnr1Y285F_p | ACC | nope | >1bp-ins | >1bpdel | none | MNV | AAAAGACCT | CCTGTAC | TGG | ACC | CC | none | NA | NA_ACC | MNV_CCTGTAC | 6 | FALSE | FALSE |
| 136_v.txt | 33159 | 33159-MNV- | MNV | 2 | CC | AA | 1 | 34462 | 2.90E-03 | 27.0000753 | rnr1Y285F_pG | TRUE | CAN | JSY2421 | 4 | rnr1Y285F_p | ACC | nope | >1bp-ins | >1bpdel | none | MNV | CCTGTACCA | CCAATAG | TGG | ACC | CC | none | NA | NA_ACC | MNV_CCAATAG | 6 | FALSE | FALSE |
| 136_v.txt | 33168 | 33168-MNV- | MNV | 2 | CC | AA | 1 | 34458 | 2.90E-03 | 36.5000086 | rnr1Y285F_pG | TRUE | CAN | JSY2421 | 4 | rnr1Y285F_p | ACC | nope | >1bp-ins | >1bpdel | none | MNV | ATAGTACCA | CCACCAA | TGG | ACC | CC | none | NA | NA_ACC | MNV_CCACCAA | 6 | FALSE | FALSE |
| 136_v.txt | 32081 | 32081-Insert | Insertion | 1 | - | A | 179 | 24887 | 0.71925102 | 63.8156425 | rnr1Y285F_pG | TRUE | CAN | JSY2421 | 4 | rnr1Y285F_p | CAA | nope | A/T+1 | >1bpdel | none | A/T+1 | CCATGCAAA | AAAAAAG | GTT | CAA | - | T | NA | NA_CAA | A/T+1_AAAAAAG | 6 | FALSE | FALSE |
| 136_v.txt | 32231 | 32231-Insert | Insertion | 1 | - | T | 56 | 24887 | 0.22501708 | 37.5 | rnr1Y285F_pG | TRUE | CAN | JSY2421 | 4 | rnr1Y285F_p | ATT | nope | A/T+1 | >1bpdel | none | A/T+1 | CAGGAATTT | TTTAGGA | TAA | ATT | - | A | NA | NA_ATT | A/T+1_TTTAGGA | 6 | FALSE | FALSE |
| 136_v.txt | 32265 | 32265-Insert | Insertion | 1 | - | A | 3 | 27455 | 0.01092697 | 38 | rnr1Y285F_pG | TRUE | CAN | JSY2421 | 4 | rnr1Y285F_p | TAA | nope | A/T+1 | >1bpdel | none | A/T+1 | CCAAATAAA | AAAATAC | ATT | TAA | - | T | NA | NA_TAA | A/T+1_AAAATAC | 6 | FALSE | FALSE |
| 136_v.txt | 32305 | 32305-Insert | Insertion | 1 | - | A | 1 | 27455 | 3.64E-03 | 61 | rnr1Y285F_pG | TRUE | CAN | JSY2421 | 4 | rnr1Y285F_p | GAA | nope | A/T+1 | >1bpdel | none | A/T+1 | CGGCAGAAA | AAATAAT | CTT | GAA | - | T | NA | NA_GAA | A/T+1_AAATAAT | 6 | FALSE | FALSE |
| 136_v.txt | 32450 | 32450-Insert | Insertion | 1 | - | A | 10 | 27455 | 0.03642324 | 54.7 | rnr1Y285F_pG | TRUE | CAN | JSY2421 | 4 | rnr1Y285F_p | CAA | nope | A/T+1 | >1bpdel | none | A/T+1 | AAGTCCAAT | AATGAAT | GTT | CAA | - | T | NA | NA_CAA | A/T+1_AATGAAT | 6 | FALSE | FALSE |
| 136_v.txt | 32498 | 32498-Insert | Insertion | 1 | - | T | 3 | 27455 | 0.01092697 | 44 | rnr1Y285F_pG | TRUE | CAN | JSY2421 | 4 | rnr1Y285F_p | CTT | nope | A/T+1 | >1bpdel | none | A/T+1 | AACAACTTTT | TTTTTTG | GAA | CTT | - | A | NA | NA_CTT | A/T+1_TTTTTTG | 6 | FALSE | FALSE |
| 136_v.txt | 32728 | 32728-Insert | Insertion | 1 | - | A | 28 | 29096 | 0.09623316 | 61.8928571 | rnr1Y285F_pG | TRUE | CAN | JSY2421 | 4 | rnr1Y285F_p | CAA | nope | A/T+1 | >1bpdel | none | A/T+1 | CCATACAAA | AAAAACA | GTT | CAA | - | T | NA | NA_CAA | A/T+1_AAAAACA | 6 | FALSE | FALSE |
| 136_v.txt | 32842 | 32842-Insert | Insertion | 1 | - | A | 25 | 29096 | 0.08592246 | 44.88 | rnr1Y285F_pG | TRUE | CAN | JSY2421 | 4 | rnr1Y285F_p | CAA | nope | A/T+1 | >1bpdel | none | A/T+1 | TTACCCAAA | AAAAAAT | GTT | CAA | - | T | NA | NA_CAA | A/T+1_AAAAAAT | 6 | FALSE | FALSE |
| 136_v.txt | 33205 | 33205-Insert | Insertion | 5 | - | GCTCT | 5 | 17622 | 0.02837362 | 41.64 | rnr1Y285F_pG | TRUE | CAN | JSY2421 | 4 | rnr1Y285F_p | AGC | nope | >1bp-ins | >1bpdel | none | >1bp-ins | GCTTAAGCT | GCTCTCT | TCG | AGC | - | none | NA | NA_AGC | >1bp-ins_GCTCTCT | 6 | FALSE | FALSE |
| 136_v.txt | 33206 | 33206-Insert | Insertion | 2 | - | CT | 658 | 17622 | 3.7339689 | 38.4490882 | rnr1Y285F_pG | TRUE | CAN | JSY2421 | 4 | rnr1Y285F_p | GCT | nope | >1bp-ins | >1bpdel | none | >1bp-ins | CTTAAGCTCT | CTCTCTC | CGA | GCT | - | none | NA | NA_GCT | >1bp-ins_CTCTCTC | 6 | FALSE | FALSE |
| 136_v.txt | 33214 | 33214-Insert | Insertion | 1 | - | C | 2 | 17622 | 0.01134945 | 38 | rnr1Y285F_pG | TRUE | CAN | JSY2421 | 4 | rnr1Y285F_p | TTC | nope | G/C+1 | >1bpdel | none | G/C+1 | CTCTCTTCAC | TCACTTC | AAG | TTC | - | G | NA | NA_TTC | G/C+1_TCACTTC | 6 | FALSE | FALSE |
| 136_v.txt | 33214 | 33214-Insert | Insertion | 2 | - | CC | 1 | 17622 | 5.68E-03 | 24.5 | rnr1Y285F_pG | TRUE | CAN | JSY2421 | 4 | rnr1Y285F_p | TTC | nope | >1bp-ins | >1bpdel | none | >1bp-ins | CTCTCTTCAC | TCACTTC | AAG | TTC | - | none | NA | NA_TTC | >1bp-ins_TCACTTC | 6 | FALSE | FALSE |
| 136_v.txt | 33214 | 33214-Insert | Insertion | 2 | - | CG | 1 | 17622 | 5.68E-03 | 24.5 | rnr1Y285F_pG | TRUE | CAN | JSY2421 | 4 | rnr1Y285F_p | TTC | nope | >1bp-ins | >1bpdel | none | >1bp-ins | CTCTCTTCAC | TCACTTC | AAG | TTC | - | none | NA | NA_TTC | >1bp-ins_TCACTTC | 6 | FALSE | FALSE |
| 136_v.txt | 33215 | 33215-Insert | Insertion | 1 | - | C | 2 | 17622 | 0.01134945 | 24.5 | rnr1Y285F_pG | TRUE | CAN | JSY2421 | 4 | rnr1Y285F_p | TCA | nope | G/C+1 | >1bpdel | none | G/C+1 | TCTCTTCACT | CACTTCA | AGT | TCA | - | G | NA | NA_TCA | G/C+1_CACTTCA | 6 | FALSE | FALSE |
| 136_v.txt | 33341 | 33341-Insert | Insertion | 1 | - | T | 4 | 17622 | 0.0226989 | 63.75 | rnr1Y285F_pG | TRUE | CAN | JSY2421 | 4 | rnr1Y285F_p | CTT | nope | A/T+1 | >1bpdel | none | A/T+1 | CTCATCTTTC | TTTCAAT | GAA | CTT | - | A | NA | NA_CTT | A/T+1_TTTCAAT | 6 | FALSE | FALSE |
| 136_v.txt | 32014 | 32014-Deleti | Deletion | 2 | GA | - | 42 | 24887 | 0.16876281 | 55.7380952 | rnr1Y285F_pG | TRUE | CAN | JSY2421 | 4 | rnr1Y285F_p | CGA | nope | >1bp-ins | >1bpdel | none | >1bpdel | CGTCACGAG | GAGAGAT | GCT | CGA | GA | none | NA | NA_CGA | >1bpdel_GAGAGAT | 6 | FALSE | FALSE |
| 136_v.txt | 32157 | 32157-Deleti | Deletion | 8 | GCCAAAGC | - | 33 | 24887 | 0.13259935 | 48.9393939 | rnr1Y285F_pG | TRUE | CAN | JSY2421 | 4 | rnr1Y285F_p | AGC | nope | >1bp-ins | >1bpdel | none | >1bpdel | ATGTAAGCC | GCCAAAG | TCG | AGC | GCCAAAGC | none | NA | NA_AGC | >1bpdel_GCCAAAG | 6 | FALSE | FALSE |
| 136_v.txt | 32337 | 32337-Deleti | Deletion | 2 | AT | - | 25 | 27455 | 0.0910581 | 61.2 | rnr1Y285F_pG | TRUE | CAN | JSY2421 | 4 | rnr1Y285F_p | GAT | nope | >1bp-ins | >1bpdel | none | >1bpdel | TTGAAGATA | ATATGTG | CTA | GAT | AT | none | NA | NA_GAT | >1bpdel_ATATGTG | 6 | FALSE | FALSE |
| 136_v.txt | 32415 | 32415-Deleti | Deletion | 4 | GTTA | - | 216 | 27455 | 0.78674194 | 57.3796296 | rnr1Y285F_pG | TRUE | CAN | JSY2421 | 4 | rnr1Y285F_p | TGT | nope | >1bp-ins | >1bpdel | none | >1bpdel | GATTGTGTT | GTTAGTT | ACA | TGT | GTTA | none | NA | NA_TGT | >1bpdel_GTTAGTT | 6 | FALSE | FALSE |
| 136_v.txt | 32453 | 32453-Deleti | Deletion | 2 | GA | - | 8 | 27455 | 0.02913859 | 54.25 | rnr1Y285F_pG | TRUE | CAN | JSY2421 | 4 | rnr1Y285F_p | TGA | nope | >1bp-ins | >1bpdel | none | >1bpdel | TCCAATGAA | GAATAAT | ACT | TGA | GA | none | NA | NA_TGA | >1bpdel_GAATAAT | 6 | FALSE | FALSE |
| 136_v.txt | 32460 | 32460-Deleti | Deletion | 2 | AG | - | 24 | 27455 | 0.08741577 | 50.375 | rnr1Y285F_pG | TRUE | CAN | JSY2421 | 4 | rnr1Y285F_p | TAG | nope | >1bp-ins | >1bpdel | none | >1bpdel | AATAATAGA | AGAGAGC | ATC | TAG | AG | none | NA | NA_TAG | >1bpdel_AGAGAGC | 6 | FALSE | FALSE |
| 136_v.txt | 32748 | 32748-Deleti | Deletion | 3 | CCG | - | 27 | 29096 | 0.09279626 | 59.2222222 | rnr1Y285F_pG | TRUE | CAN | JSY2421 | 4 | rnr1Y285F_p | CCC | nope | >1bp-ins | >1bpdel | none | >1bpdel | AGAAACCCG | CCGATAA | GGG | CCC | CCG | none | NA | NA_CCC | >1bpdel_CCGATAA | 6 | FALSE | FALSE |
| 136_v.txt | 33089 | 33089-Deleti | Deletion | 2 | AT | - | 211 | 34459 | 0.61232189 | 52.0663507 | rnr1Y285F_pG | TRUE | CAN | JSY2421 | 4 | rnr1Y285F_p | AAT | nope | >1bp-ins | >1bpdel | none | >1bpdel | AAATAAATA | ATATGAT | TTA | AAT | AT | none | NA | NA_AAT | >1bpdel_ATATGAT | 6 | FALSE | FALSE |
| 136_v.txt | 33093 | 33093-Deleti | Deletion | 7 | GATATAA | - | 71 | 34459 | 0.20604196 | 54.1267606 | rnr1Y285F_pG | TRUE | CAN | JSY2421 | 4 | rnr1Y285F_p | TGA | nope | >1bp-ins | >1bpdel | none | >1bpdel | AAATATGAT | GATATAA | ACT | TGA | GATATAA | none | NA | NA_TGA | >1bpdel_GATATAA | 6 | FALSE | FALSE |
| 136_v.txt | 33167 | 33167-Deleti | Deletion | 3 | ACC | - | 10 | 34459 | 0.02902 | 37.6 | rnr1Y285F_pG | TRUE | CAN | JSY2421 | 4 | rnr1Y285F_p | TAC | nope | >1bp-ins | >1bpdel | none | >1bpdel | AATAGTACC | ACCACCA | ATG | TAC | ACC | none | NA | NA_TAC | >1bpdel_ACCACCA | 6 | FALSE | FALSE |
| 136_v.txt | 33189 | 33189-Deleti | Deletion | 2 | AT | - | 200 | 15080 | 1.32625995 | 37.51 | rnr1Y285F_pG | TRUE | CAN | JSY2421 | 4 | rnr1Y285F_p | AAT | nope | >1bp-ins | >1bpdel | none | >1bpdel | ATACCAATA | ATATGTC | TTA | AAT | AT | none | NA | NA_AAT | >1bpdel_ATATGTC | 6 | FALSE | FALSE |
| 136_v.txt | 33206 | 33206-Deleti | Deletion | 2 | CT | - | 26 | 17622 | 0.14754284 | 37.3461539 | rnr1Y285F_pG | TRUE | CAN | JSY2421 | 4 | rnr1Y285F_p | GCT | nope | >1bp-ins | >1bpdel | none | >1bpdel | CTTAAGCTCT | CTCTCTC | CGA | GCT | CT | none | NA | NA_GCT | >1bpdel_CTCTCTC | 6 | FALSE | FALSE |
| 136_v.txt | 31663 | 31663-Deleti | Deletion | 1 | A | - | 5 | 11836 | 0.042244 | 37.2 | rnr1Y285F_pG | TRUE | CAN | JSY2421 | 4 | rnr1Y285F_p | GAA | nope | >1bp-ins | A/T-1 | none | A/T-1 | GCGTGGAAA | AAATGTG | CTT | CTT | T | none | T>none | T>none_CTT | A/T-1_AAATGTG | 6 | FALSE | FALSE |
| 136_v.txt | 31723 | 31723-Deleti | Deletion | 1 | A | - | 13 | 11836 | 0.1098344 | 62.8461539 | rnr1Y285F_pG | TRUE | CAN | JSY2421 | 4 | rnr1Y285F_p | CAA | nope | >1bp-ins | A/T-1 | none | A/T-1 | TGTCCCAAA | AAAAAGT | GTT | GTT | T | none | T>none | T>none_GTT | A/T-1_AAAAAGT | 6 | FALSE | FALSE |
| 136_v.txt | 31904 | 31904-Deleti | Deletion | 1 | T | - | 12 | 11836 | 0.1013856 | 45.6666667 | rnr1Y285F_pG | TRUE | CAN | JSY2421 | 4 | rnr1Y285F_p | ATT | nope | >1bp-ins | A/T-1 | none | A/T-1 | ATTGAATTTT | TTTTGGT | TAA | ATT | T | none | nope | nope_ATT | A/T-1_TTTTGGT | 6 | FALSE | FALSE |
| 136_v.txt | 31913 | 31913-Deleti | Deletion | 1 | A | - | 5 | 11836 | 0.042244 | 36.6 | rnr1Y285F_pG | TRUE | CAN | JSY2421 | 4 | rnr1Y285F_p | CAA | nope | >1bp-ins | A/T-1 | none | A/T-1 | TGGTGCAAA | AAAAGCC | GTT | GTT | T | none | T>none | T>none_GTT | A/T-1_AAAAGCC | 6 | FALSE | FALSE |
| 136_v.txt | 31923 | 31923-Deleti | Deletion | 1 | A | - | 39 | 11835 | 0.32953105 | 37.7179487 | rnr1Y285F_pG | TRUE | CAN | JSY2421 | 4 | rnr1Y285F_p | GAA | nope | >1bp-ins | A/T-1 | none | A/T-1 | GCCGTGAAA | AAACCTT | CTT | CTT | T | none | T>none | T>none_CTT | A/T-1_AAACCTT | 6 | FALSE | FALSE |
| 136_v.txt | 31966 | 31966-Deleti | Deletion | 1 | A | - | 6 | 24887 | 0.02410897 | 38 | rnr1Y285F_pG | TRUE | CAN | JSY2421 | 4 | rnr1Y285F_p | TAA | nope | >1bp-ins | A/T-1 | none | A/T-1 | CCGCATAAT | AATAAGC | ATT | ATT | T | none | T>none | T>none_ATT | A/T-1_AATAAGC | 6 | FALSE | FALSE |
| 136_v.txt | 31984 | 31984-Deleti | Deletion | 1 | T | - | 167 | 24887 | 0.67103307 | 51.9820359 | rnr1Y285F_pG | TRUE | CAN | JSY2421 | 4 | rnr1Y285F_p | ATT | nope | >1bp-ins | A/T-1 | none | A/T-1 | CGGGCATTA | TTAATTT | TAA | ATT | T | none | nope | nope_ATT | A/T-1_TTAATTT | 6 | FALSE | FALSE |
| 136_v.txt | 32026 | 32026-Deleti | Deletion | 1 | G | - | 21 | 24887 | 0.0843814 | 58.0476202 | rnr1Y285F_pG | TRUE | CAN | JSY2421 | 4 | rnr1Y285F_p | CGG | nope | >1bp-ins | G/C-1 | none | G/C-1 | TGCCACGGT | GGTATTT | GCC | GCC | C | none | C>none | C>none_GCC | G/C-1_GGTATTT | 6 | FALSE | FALSE |
| 136_v.txt | 32044 | 32044-Deleti | Deletion | 1 | T | - | 28 | 24887 | 0.11250854 | 62.7857143 | rnr1Y285F_pG | TRUE | CAN | JSY2421 | 4 | rnr1Y285F_p | ATA | nope | >1bp-ins | A/T-1 | none | A/T-1 | CTTGCATAA | TAAATCT | TAT | ATA | T | none | nope | nope_ATA | A/T-1_TAAATCT | 6 | FALSE | FALSE |
| 136_v.txt | 32081 | 32081-Deleti | Deletion | 1 | A | - | 24 | 24885 | 0.09644364 | 63.6666667 | rnr1Y285F_pG | TRUE | CAN | JSY2421 | 4 | rnr1Y285F_p | CAA | nope | >1bp-ins | A/T-1 | none | A/T-1 | CCATGCAAA | AAAAAAG | GTT | GTT | T | none | T>none | T>none_GTT | A/T-1_AAAAAAG | 6 | FALSE | FALSE |
| 136_v.txt | 32160 | 32160-Deleti | Deletion | 1 | A | - | 3 | 24887 | 0.01205449 | 38 | rnr1Y285F_pG | TRUE | CAN | JSY2421 | 4 | rnr1Y285F_p | CAA | nope | >1bp-ins | A/T-1 | none | A/T-1 | TAAGCCAAA | AAAGCGC | GTT | GTT | T | none | T>none | T>none_GTT | A/T-1_AAAGCGC | 6 | FALSE | FALSE |
| 136_v.txt | 32184 | 32184-Deleti | Deletion | 1 | A | - | 11 | 24887 | 0.04419978 | 39.9090909 | rnr1Y285F_pG | TRUE | CAN | JSY2421 | 4 | rnr1Y285F_p | GAA | nope | >1bp-ins | A/T-1 | none | A/T-1 | GTAACGAAA | AAAACTG | CTT | CTT | T | none | T>none | T>none_CTT | A/T-1_AAAACTG | 6 | FALSE | FALSE |
| 136_v.txt | 32265 | 32265-Deleti | Deletion | 1 | A | - | 6 | 27455 | 0.02185394 | 37.6666651 | rnr1Y285F_pG | TRUE | CAN | JSY2421 | 4 | rnr1Y285F_p | TAA | nope | >1bp-ins | A/T-1 | none | A/T-1 | CCAAATAAA | AAAATAC | ATT | ATT | T | none | T>none | T>none_ATT | A/T-1_AAAATAC | 6 | FALSE | FALSE |
| 136_v.txt | 32304 | 32304-Deleti | Deletion | 1 | G | - | 3 | 27455 | 0.01092697 | 54.9999991 | rnr1Y285F_pG | TRUE | CAN | JSY2421 | 4 | rnr1Y285F_p | AGA | nope | >1bp-ins | G/C-1 | none | G/C-1 | GCGGCAGAA | GAAATAA | TCT | TCT | C | none | C>none | C>none_TCT | G/C-1_GAAATAA | 6 | FALSE | FALSE |
| 136_v.txt | 32346 | 32346-Deleti | Deletion | 1 | A | - | 38 | 27455 | 0.13840831 | 61.9210526 | rnr1Y285F_pG | TRUE | CAN | JSY2421 | 4 | rnr1Y285F_p | CAA | nope | >1bp-ins | A/T-1 | none | A/T-1 | TGTGGCAAA | AAAACCT | GTT | GTT | T | none | T>none | T>none_GTT | A/T-1_AAAACCT | 6 | FALSE | FALSE |
| 136_v.txt | 32381 | 32381-Deleti | Deletion | 1 | A | - | 10 | 27455 | 0.03642324 | 63.3 | rnr1Y285F_pG | TRUE | CAN | JSY2421 | 4 | rnr1Y285F_p | TAA | nope | >1bp-ins | A/T-1 | none | A/T-1 | AATAATAAA | AAAGGGA | ATT | ATT | T | none | T>none | T>none_ATT | A/T-1_AAAGGGA | 6 | FALSE | FALSE |
| 136_v.txt | 32416 | 32416-Deleti | Deletion | 1 | T | - | 2 | 27455 | 7.29E-03 | 64 | rnr1Y285F_pG | TRUE | CAN | JSY2421 | 4 | rnr1Y285F_p | GTT | nope | >1bp-ins | A/T-1 | none | A/T-1 | ATTGTGTTAC | TTAGTTT | CAA | GTT | T | none | nope | nope_GTT | A/T-1_TTAGTTT | 6 | FALSE | FALSE |
| 136_v.txt | 32424 | 32424-Deleti | Deletion | 1 | G | - | 22 | 27455 | 0.08013112 | 56.7272727 | rnr1Y285F_pG | TRUE | CAN | JSY2421 | 4 | rnr1Y285F_p | AGG | nope | >1bp-ins | G/C-1 | none | G/C-1 | AGTTTAGGG | GGGTCAT | TCC | TCC | C | none | C>none | C>none_TCC | G/C-1_GGGTCAT | 6 | FALSE | FALSE |
| 136_v.txt | 32460 | 32460-Deleti | Deletion | 1 | A | - | 8 | 27455 | 0.02913859 | 49 | rnr1Y285F_pG | TRUE | CAN | JSY2421 | 4 | rnr1Y285F_p | TAG | nope | >1bp-ins | A/T-1 | none | A/T-1 | AATAATAGA | AGAGAGC | ATC | ATC | T | none | T>none | T>none_ATC | A/T-1_AGAGAGC | 6 | FALSE | FALSE |
| 136_v.txt | 32490 | 32490-Deleti | Deletion | 1 | A | - | 8 | 27455 | 0.02913859 | 51 | rnr1Y285F_pG | TRUE | CAN | JSY2421 | 4 | rnr1Y285F_p | GAA | nope | >1bp-ins | A/T-1 | none | A/T-1 | ATACGGAAA | AAAACAA | CTT | CTT | T | none | T>none | T>none_CTT | A/T-1_AAAACAA | 6 | FALSE | FALSE |
| 136_v.txt | 32498 | 32498-Deleti | Deletion | 1 | T | - | 65 | 27455 | 0.23675105 | 45.9538456 | rnr1Y285F_pG | TRUE | CAN | JSY2421 | 4 | rnr1Y285F_p | CTT | nope | >1bp-ins | A/T-1 | none | A/T-1 | AACAACTTTT | TTTTTTG | GAA | CTT | T | none | nope | nope_CTT | A/T-1_TTTTTTG | 6 | FALSE | FALSE |
| 136_v.txt | 32522 | 32522-Deleti | Deletion | 1 | T | - | 10 | 27455 | 0.03642324 | 36 | rnr1Y285F_pG | TRUE | CAN | JSY2421 | 4 | rnr1Y285F_p | ATT | nope | >1bp-ins | A/T-1 | none | A/T-1 | AACGGATTTT | TTTTCTG | TAA | ATT | T | none | nope | nope_ATT | A/T-1_TTTTCTG | 6 | FALSE | FALSE |
| 136_v.txt | 32576 | 32576-Deleti | Deletion | 1 | A | - | 12 | 29096 | 0.04124278 | 37.9166667 | rnr1Y285F_pG | TRUE | CAN | JSY2421 | 4 | rnr1Y285F_p | GAA | nope | >1bp-ins | A/T-1 | none | A/T-1 | ACCTTGAAA | AAATGTG | CTT | CTT | T | none | T>none | T>none_CTT | A/T-1_AAATGTG | 6 | FALSE | FALSE |
| 136_v.txt | 32585 | 32585-Deleti | Deletion | 1 | G | - | 33 | 29096 | 0.11341765 | 37.7272727 | rnr1Y285F_pG | TRUE | CAN | JSY2421 | 4 | rnr1Y285F_p | AGG | nope | >1bp-ins | G/C-1 | none | G/C-1 | TGTGAAGGC | GGCAGCG | TCC | TCC | C | none | C>none | C>none_TCC | G/C-1_GGCAGCG | 6 | FALSE | FALSE |
| 136_v.txt | 32631 | 32631-Deleti | Deletion | 1 | T | - | 9 | 29096 | 0.03093209 | 52.2222222 | rnr1Y285F_pG | TRUE | CAN | JSY2421 | 4 | rnr1Y285F_p | GTT | nope | >1bp-ins | A/T-1 | none | A/T-1 | CCTTCGTTTT | TTTTTAT | CAA | GTT | T | none | nope | nope_GTT | A/T-1_TTTTTAT | 6 | FALSE | FALSE |
| 136_v.txt | 32658 | 32658-Deleti | Deletion | 1 | C | - | 251 | 29096 | 0.86266153 | 60.12749 | rnr1Y285F_pG | TRUE | CAN | JSY2421 | 4 | rnr1Y285F_p | ACC | nope | >1bp-ins | G/C-1 | none | G/C-1 | CCTGGACCC | CCCCAGG | TGG | ACC | C | none | nope | nope_ACC | G/C-1_CCCCAGG | 6 | FALSE | FALSE |
| 136_v.txt | 32670 | 32670-Deleti | Deletion | 1 | G | - | 71 | 29096 | 0.2440198 | 59.2112676 | rnr1Y285F_pG | TRUE | CAN | JSY2421 | 4 | rnr1Y285F_p | TGG | nope | >1bp-ins | G/C-1 | none | G/C-1 | GCACCTGGG | GGGTTTC | ACC | ACC | C | none | C>none | C>none_ACC | G/C-1_GGGTTTC | 6 | FALSE | FALSE |
| 136_v.txt | 32707 | 32707-Deleti | Deletion | 1 | C | - | 18 | 29096 | 0.06186417 | 64 | rnr1Y285F_pG | TRUE | CAN | JSY2421 | 4 | rnr1Y285F_p | ACC | nope | >1bp-ins | G/C-1 | none | G/C-1 | CGGTAACCC | CCCCAGC | TGG | ACC | C | none | nope | nope_ACC | G/C-1_CCCCAGC | 6 | FALSE | FALSE |
| 136_v.txt | 32719 | 32719-Deleti | Deletion | 1 | A | - | 12 | 29096 | 0.04124278 | 63.5833333 | rnr1Y285F_pG | TRUE | CAN | JSY2421 | 4 | rnr1Y285F_p | CAA | nope | >1bp-ins | A/T-1 | none | A/T-1 | CACCACAAA | AAACCAT | GTT | GTT | T | none | T>none | T>none_GTT | A/T-1_AAACCAT | 6 | FALSE | FALSE |
| 136_v.txt | 32728 | 32728-Deleti | Deletion | 1 | A | - | 19 | 29096 | 0.06530107 | 61.9473661 | rnr1Y285F_pG | TRUE | CAN | JSY2421 | 4 | rnr1Y285F_p | CAA | nope | >1bp-ins | A/T-1 | none | A/T-1 | CCATACAAA | AAAAACA | GTT | GTT | T | none | T>none | T>none_GTT | A/T-1_AAAAACA | 6 | FALSE | FALSE |
| 136_v.txt | 32760 | 32760-Deleti | Deletion | 1 | A | - | 12 | 29096 | 0.04124278 | 53.3333333 | rnr1Y285F_pG | TRUE | CAN | JSY2421 | 4 | rnr1Y285F_p | TAA | nope | >1bp-ins | A/T-1 | none | A/T-1 | ATGGCTAAA | AAAACTT | ATT | ATT | T | none | T>none | T>none_ATT | A/T-1_AAAACTT | 6 | FALSE | FALSE |
| 136_v.txt | 32842 | 32842-Deleti | Deletion | 1 | A | - | 185 | 29096 | 0.63582623 | 41.1297282 | rnr1Y285F_pG | TRUE | CAN | JSY2421 | 4 | rnr1Y285F_p | CAA | nope | >1bp-ins | A/T-1 | none | A/T-1 | TTACCCAAA | AAAAAAT | GTT | GTT | T | none | T>none | T>none_GTT | A/T-1_AAAAAAT | 6 | FALSE | FALSE |
| 136_v.txt | 32868 | 32868-Deleti | Deletion | 1 | G | - | 86 | 29086 | 0.2956749 | 37.7790698 | rnr1Y285F_pG | TRUE | CAN | JSY2421 | 4 | rnr1Y285F_p | TGG | nope | >1bp-ins | G/C-1 | none | G/C-1 | GCCAGTGGA | GGAACTT | ACC | ACC | C | none | C>none | C>none_ACC | G/C-1_GGAACTT | 6 | FALSE | FALSE |
| 136_v.txt | 32884 | 32884-Deleti | Deletion | 1 | A | - | 11 | 34459 | 0.03192199 | 36.3636364 | rnr1Y285F_pG | TRUE | CAN | JSY2421 | 4 | rnr1Y285F_p | CAA | nope | >1bp-ins | A/T-1 | none | A/T-1 | ACGTCCAAA | AAAATTG | GTT | GTT | T | none | T>none | T>none_GTT | A/T-1_AAAATTG | 6 | FALSE | FALSE |
| 136_v.txt | 32924 | 32924-Deleti | Deletion | 1 | A | - | 36 | 34459 | 0.10447198 | 51.4166667 | rnr1Y285F_pG | TRUE | CAN | JSY2421 | 4 | rnr1Y285F_p | CAA | nope | >1bp-ins | A/T-1 | none | A/T-1 | CAGGGCAAA | AAAAGTG | GTT | GTT | T | none | T>none | T>none_GTT | A/T-1_AAAAGTG | 6 | FALSE | FALSE |
| 136_v.txt | 32941 | 32941-Deleti | Deletion | 1 | A | - | 23 | 34459 | 0.06674599 | 58.1739123 | rnr1Y285F_pG | TRUE | CAN | JSY2421 | 4 | rnr1Y285F_p | GAA | nope | >1bp-ins | A/T-1 | none | A/T-1 | CCCAAGAAA | AAAACCA | CTT | CTT | T | none | T>none | T>none_CTT | A/T-1_AAAACCA | 6 | FALSE | FALSE |
| 136_v.txt | 33000 | 33000-Deleti | Deletion | 1 | A | - | 27 | 34459 | 0.07835399 | 63.8518519 | rnr1Y285F_pG | TRUE | CAN | JSY2421 | 4 | rnr1Y285F_p | GAA | nope | >1bp-ins | A/T-1 | none | A/T-1 | TGTGAGAAA | AAAACTG | CTT | CTT | T | none | T>none | T>none_CTT | A/T-1_AAAACTG | 6 | FALSE | FALSE |
| 136_v.txt | 33078 | 33078-Deleti | Deletion | 1 | C | - | 106 | 34459 | 0.30761195 | 51.9433962 | rnr1Y285F_pG | TRUE | CAN | JSY2421 | 4 | rnr1Y285F_p | ACC | nope | >1bp-ins | G/C-1 | none | G/C-1 | AAAGAACCC | CCCATAA | TGG | ACC | C | none | nope | nope_ACC | G/C-1_CCCATAA | 6 | FALSE | FALSE |
| 136_v.txt | 33083 | 33083-Deleti | Deletion | 1 | A | - | 83 | 34459 | 0.24086596 | 52.9879518 | rnr1Y285F_pG | TRUE | CAN | JSY2421 | 4 | rnr1Y285F_p | TAA | nope | >1bp-ins | A/T-1 | none | A/T-1 | ACCCATAAA | AAATAAA | ATT | ATT | T | none | T>none | T>none_ATT | A/T-1_AAATAAA | 6 | FALSE | FALSE |
| 136_v.txt | 33093 | 33093-Deleti | Deletion | 1 | G | - | 1 | 34459 | 2.90E-03 | 64 | rnr1Y285F_pG | TRUE | CAN | JSY2421 | 4 | rnr1Y285F_p | TGA | nope | >1bp-ins | G/C-1 | none | G/C-1 | AAATATGAT | GATATAA | ACT | ACT | C | none | C>none | C>none_ACT | G/C-1_GATATAA | 6 | FALSE | FALSE |
| 136_v.txt | 33095 | 33095-Deleti | Deletion | 1 | T | - | 1 | 34459 | 2.90E-03 | 64 | rnr1Y285F_pG | TRUE | CAN | JSY2421 | 4 | rnr1Y285F_p | ATA | nope | >1bp-ins | A/T-1 | none | A/T-1 | ATATGATAT | TATAAGA | TAT | ATA | T | none | nope | nope_ATA | A/T-1_TATAAGA | 6 | FALSE | FALSE |
| 136_v.txt | 33098 | 33098-Deleti | Deletion | 1 | A | - | 1 | 34459 | 2.90E-03 | 56.9999961 | rnr1Y285F_pG | TRUE | CAN | JSY2421 | 4 | rnr1Y285F_p | TAA | nope | >1bp-ins | A/T-1 | none | A/T-1 | TGATATAAG | AAGAGCG | ATT | ATT | T | none | T>none | T>none_ATT | A/T-1_AAGAGCG | 6 | FALSE | FALSE |
| 136_v.txt | 33153 | 33153-Deleti | Deletion | 1 | C | - | 2 | 34459 | 5.80E-03 | 37.5 | rnr1Y285F_pG | TRUE | CAN | JSY2421 | 4 | rnr1Y285F_p | ACC | nope | >1bp-ins | G/C-1 | none | G/C-1 | AAAAGACCT | CCTGTAC | TGG | ACC | C | none | nope | nope_ACC | G/C-1_CCTGTAC | 6 | FALSE | FALSE |
| 136_v.txt | 33279 | 33279-Deleti | Deletion | 1 | T | - | 19 | 17622 | 0.10781977 | 48.1578947 | rnr1Y285F_pG | TRUE | CAN | JSY2421 | 4 | rnr1Y285F_p | ATC | nope | >1bp-ins | A/T-1 | none | A/T-1 | AACGTATCC | TCCTCGC | TAG | ATC | T | none | nope | nope_ATC | A/T-1_TCCTCGC | 6 | FALSE | FALSE |
| 136_v.txt | 33330 | 33330-Deleti | Deletion | 1 | C | - | 7 | 17622 | 0.03972307 | 59.7142857 | rnr1Y285F_pG | TRUE | CAN | JSY2421 | 4 | rnr1Y285F_p | ACT | nope | >1bp-ins | G/C-1 | none | G/C-1 | TCTTTACTTT | CTTTTCT | TGA | ACT | C | none | nope | nope_ACT | G/C-1_CTTTTCT | 6 | FALSE | FALSE |
| 136_v.txt | 33356 | 33356-Deleti | Deletion | 1 | C | - | 30 | 17622 | 0.17024174 | 56.8666667 | rnr1Y285F_pG | TRUE | CAN | JSY2421 | 4 | rnr1Y285F_p | ACC | nope | >1bp-ins | G/C-1 | none | G/C-1 | TATTGACCCA | CCCACGT | TGG | ACC | C | none | nope | nope_ACC | G/C-1_CCCACGT | 6 | FALSE | FALSE |
| 136_v.txt | 33474 | 33474-Deleti | Deletion | 1 | C | - | 51 | 17622 | 0.28941096 | 28.4117647 | rnr1Y285F_pG | TRUE | CAN | JSY2421 | 4 | rnr1Y285F_p | GCC | nope | >1bp-ins | G/C-1 | none | G/C-1 | GCTATGCCTT | CCTTTTT | CGG | GCC | C | none | nope | nope_GCC | G/C-1_CCTTTTT | 6 | FALSE | FALSE |
| 174_v.txt | 31854 | 31854-SNV-A | SNV | 1 | A | C | 6 | 7006 | 0.08564088 | 45.6666667 | rnr1Y285F_pG | TRUE | CAN | JSY2420 | 5 | rnr1Y285F_p | TAA | A>C | G/C+1 | A/T-1 | TA>GC | TA>GC | ACAGCTAAG | AAGAACA | ATT | ATT | T | G | T>G | T>G_ATT | TA>GC_AAGAACA | 6 | FALSE | FALSE |
| 174_v.txt | 31950 | 31950-SNV-A | SNV | 1 | A | C | 14 | 11591 | 0.12078337 | 35.7142857 | rnr1Y285F_pG | TRUE | CAN | JSY2420 | 5 | rnr1Y285F_p | CAT | A>C | G/C+1 | A/T-1 | TA>GC | TA>GC | ATCGTCATA | ATAAATG | GTA | GTA | T | G | T>G | T>G_GTA | TA>GC_ATAAATG | 6 | FALSE | FALSE |
| 174_v.txt | 31986 | 31986-SNV-A | SNV | 1 | A | C | 21 | 11593 | 0.18114379 | 51.0952381 | rnr1Y285F_pG | TRUE | CAN | JSY2420 | 5 | rnr1Y285F_p | TAA | A>C | G/C+1 | A/T-1 | TA>GC | TA>GC | GGCATTAAT | AATTTAG | ATT | ATT | T | G | T>G | T>G_ATT | TA>GC_AATTTAG | 6 | FALSE | FALSE |
| 174_v.txt | 32226 | 32226-SNV-A | SNV | 1 | A | C | 15 | 11593 | 0.12938842 | 37.2666667 | rnr1Y285F_pG | TRUE | CAN | JSY2420 | 5 | rnr1Y285F_p | CAG | A>C | G/C+1 | A/T-1 | TA>GC | TA>GC | CTTGACAGG | AGGAATT | GTC | GTC | T | G | T>G | T>G_GTC | TA>GC_AGGAATT | 6 | FALSE | FALSE |
| 174_v.txt | 32283 | 32283-SNV-T | SNV | 1 | T | G | 9 | 21310 | 0.04223369 | 35.5553852 | rnr1Y285F_pG | TRUE | CAN | JSY2420 | 5 | rnr1Y285F_p | GTA | T>G | G/C+1 | A/T-1 | TA>GC | TA>GC | CCAACGTAA | TAAATAT | CAT | GTA | T | C | T>G | T>G_GTA | TA>GC_TAAATAT | 6 | FALSE | FALSE |
| 174_v.txt | 32288 | 32288-SNV-A | SNV | 1 | A | C | 46 | 21310 | 0.2158611 | 46.1525732 | rnr1Y285F_pG | TRUE | CAN | JSY2420 | 5 | rnr1Y285F_p | TAT | A>C | G/C+1 | A/T-1 | TA>GC | TA>GC | GTAAATATT | ATTTGAA | ATA | ATA | T | G | T>G | T>G_ATA | TA>GC_ATTTGAA | 6 | FALSE | FALSE |
| 174_v.txt | 32416 | 32416-SNV-T | SNV | 1 | T | G | 2 | 21310 | 9.39E-03 | 51 | rnr1Y285F_pG | TRUE | CAN | JSY2420 | 5 | rnr1Y285F_p | GTT | T>G | G/C+1 | A/T-1 | TA>GC | TA>GC | ATTGTGTTAC | TTAGTTT | CAA | GTT | T | C | T>G | T>G_GTT | TA>GC_TTAGTTT | 6 | FALSE | FALSE |
| 174_v.txt | 32439 | 32439-SNV-A | SNV | 1 | A | C | 24 | 21310 | 0.11262318 | 57.5833333 | rnr1Y285F_pG | TRUE | CAN | JSY2420 | 5 | rnr1Y285F_p | AAC | A>C | G/C+1 | A/T-1 | TA>GC | TA>GC | TATGGAACT | ACTAAAA | TTG | TTG | T | G | T>G | T>G_TTG | TA>GC_ACTAAAA | 6 | FALSE | FALSE |
| 174_v.txt | 32445 | 32445-SNV-A | SNV | 1 | A | C | 115 | 21310 | 0.53965275 | 53.5565217 | rnr1Y285F_pG | TRUE | CAN | JSY2420 | 5 | rnr1Y285F_p | AAG | A>C | G/C+1 | A/T-1 | TA>GC | TA>GC | ACTAAAAGT | AGTCCAA | TTC | TTC | T | G | T>G | T>G_TTC | TA>GC_AGTCCAA | 6 | FALSE | FALSE |
| 174_v.txt | 32457 | 32457-SNV-A | SNV | 1 | A | C | 11 | 21310 | 0.05161896 | 53.9090909 | rnr1Y285F_pG | TRUE | CAN | JSY2420 | 5 | rnr1Y285F_p | TAA | A>C | G/C+1 | A/T-1 | TA>GC | TA>GC | ATGAATAAT | AATAGAG | ATT | ATT | T | G | T>G | T>G_ATT | TA>GC_AATAGAG | 6 | FALSE | FALSE |
| 174_v.txt | 32573 | 32573-SNV-T | SNV | 1 | T | G | 6 | 22591 | 0.02655925 | 24.5 | rnr1Y285F_pG | TRUE | CAN | JSY2420 | 5 | rnr1Y285F_p | CTT | T>G | G/C+1 | A/T-1 | TA>GC | TA>GC | AGTACCTTG | TTGAAAT | GAA | CTT | T | C | T>G | T>G_CTT | TA>GC_TTGAAAT | 6 | FALSE | FALSE |
| 174_v.txt | 32760 | 32760-SNV-A | SNV | 1 | A | C | 11 | 22590 | 0.04869411 | 49.3636364 | rnr1Y285F_pG | TRUE | CAN | JSY2420 | 5 | rnr1Y285F_p | TAA | A>C | G/C+1 | A/T-1 | TA>GC | TA>GC | ATGGCTAAA | AAAACTT | ATT | ATT | T | G | T>G | T>G_ATT | TA>GC_AAAACTT | 6 | FALSE | FALSE |
| 174_v.txt | 32792 | 32792-SNV-T | SNV | 1 | T | G | 9 | 22590 | 0.03984064 | 54.2222222 | rnr1Y285F_pG | TRUE | CAN | JSY2420 | 5 | rnr1Y285F_p | ATT | T>G | G/C+1 | A/T-1 | TA>GC | TA>GC | CTCGAATTCA | TTCACCG | TAA | ATT | T | C | T>G | T>G_ATT | TA>GC_TTCACCG | 6 | FALSE | FALSE |
| 174_v.txt | 32803 | 32803-SNV-A | SNV | 1 | A | C | 16 | 22590 | 0.0708278 | 46.125 | rnr1Y285F_pG | TRUE | CAN | JSY2420 | 5 | rnr1Y285F_p | TAT | A>C | G/C+1 | A/T-1 | TA>GC | TA>GC | CGTAATATTT | ATTTGAC | ATA | ATA | T | G | T>G | T>G_ATA | TA>GC_ATTTGAC | 6 | FALSE | FALSE |

| | | | | | | | | | | | | | | | | | | | | | | | | | | | | | | | | | | |
|---|---|---|---|---|---|---|---|---|---|---|---|---|---|---|---|---|---|---|---|---|---|---|---|---|---|---|---|---|---|---|---|---|---|---|
| 174_v.txt | 32808 | 32808-SNV-A | SNV | 1 | A | C | 504 | 22590 | 2.2310757 | 44.9928975 | rnr1Y285F_pG | TRUE | CAN | JSY2420 | 5 | rnr1Y285F_p | GAC | A>C | G/C+1 | A/T-1 | TA>GC | TA>GC | TATTTGACA | ACAGGGA | CTG | CTG | T | G | T>G | T>G_CTG | TA>GC_ACAGGGA | 6 | FALSE | FALSE |
| 174_v.txt | 32823 | 32823-SNV-A | SNV | 1 | A | C | 7 | 22590 | 0.03098716 | 48.1428571 | rnr1Y285F_pG | TRUE | CAN | JSY2420 | 5 | rnr1Y285F_p | CAT | A>C | G/C+1 | A/T-1 | TA>GC | TA>GC | AAGTTCATT | ATTATTG | GTA | GTA | T | G | T>G | T>G_GTA | TA>GC_ATTATTG | 6 | FALSE | FALSE |
| 174_v.txt | 32842 | 32842-SNV-A | SNV | 1 | A | C | 6 | 22590 | 0.02656043 | 37.8333414 | rnr1Y285F_pG | TRUE | CAN | JSY2420 | 5 | rnr1Y285F_p | CAA | A>C | G/C+1 | A/T-1 | TA>GC | TA>GC | TTACCCAAA | AAAAAAT | GTT | GTT | T | G | T>G | T>G_GTT | TA>GC_AAAAAAT | 6 | FALSE | FALSE |
| 174_v.txt | 32844 | 32844-SNV-A | SNV | 1 | A | C | 14 | 22590 | 0.06197433 | 37.9285714 | rnr1Y285F_pG | TRUE | CAN | JSY2420 | 5 | rnr1Y285F_p | AAA | A>C | G/C+1 | A/T-1 | TA>GC | TA>GC | ACCCAAAAA | AAAATAC | TTT | TTT | T | G | T>G | T>G_TTT | TA>GC_AAAATAC | 6 | FALSE | FALSE |
| 174_v.txt | 32884 | 32884-SNV-A | SNV | 1 | A | C | 3 | 23328 | 0.01286008 | 37.999058 | rnr1Y285F_pG | TRUE | CAN | JSY2420 | 5 | rnr1Y285F_p | CAA | A>C | G/C+1 | A/T-1 | TA>GC | TA>GC | ACGTCCAAA | AAAATTG | GTT | GTT | T | G | T>G | T>G_GTT | TA>GC_AAAATTG | 6 | FALSE | FALSE |
| 174_v.txt | 32915 | 32915-SNV-T | SNV | 1 | T | G | 9 | 23328 | 0.03858025 | 39.9988077 | rnr1Y285F_pG | TRUE | CAN | JSY2420 | 5 | rnr1Y285F_p | GTT | T>G | G/C+1 | A/T-1 | TA>GC | TA>GC | ACTAAGTTC | TTCCAGG | CAA | GTT | T | C | T>G | T>G_GTT | TA>GC_TTCCAGG | 6 | FALSE | FALSE |
| 174_v.txt | 32919 | 32919-SNV-A | SNV | 1 | A | C | 7 | 23328 | 0.03000686 | 32.8571429 | rnr1Y285F_pG | TRUE | CAN | JSY2420 | 5 | rnr1Y285F_p | CAG | A>C | G/C+1 | A/T-1 | TA>GC | TA>GC | AGTTCCAGG | AGGGCAA | GTC | GTC | T | G | T>G | T>G_GTC | TA>GC_AGGGCAA | 6 | FALSE | FALSE |
| 174_v.txt | 32929 | 32929-SNV-T | SNV | 1 | T | G | 14 | 23328 | 0.06001372 | 57.7142857 | rnr1Y285F_pG | TRUE | CAN | JSY2420 | 5 | rnr1Y285F_p | GTG | T>G | G/C+1 | A/T-1 | TA>GC | TA>GC | CAAAAGTGA | TGATTGC | CAC | GTG | T | C | T>G | T>G_GTG | TA>GC_TGATTGC | 6 | FALSE | FALSE |
| 174_v.txt | 33039 | 33039-SNV-A | SNV | 1 | A | C | 27 | 23328 | 0.11574074 | 60.5925926 | rnr1Y285F_pG | TRUE | CAN | JSY2420 | 5 | rnr1Y285F_p | CAT | A>C | G/C+1 | A/T-1 | TA>GC | TA>GC | GTAGCCATT | ATTTCAC | GTA | GTA | T | G | T>G | T>G_GTA | TA>GC_ATTTCAC | 6 | FALSE | FALSE |
| 174_v.txt | 33138 | 33138-SNV-A | SNV | 1 | A | C | 7 | 23328 | 0.03000686 | 42.5714286 | rnr1Y285F_pG | TRUE | CAN | JSY2420 | 5 | rnr1Y285F_p | TAA | A>C | G/C+1 | A/T-1 | TA>GC | TA>GC | GTGGATAAA | AAACCAA | ATT | ATT | T | G | T>G | T>G_ATT | TA>GC_AAACCAA | 6 | FALSE | FALSE |
| 174_v.txt | 33162 | 33162-SNV-A | SNV | 1 | A | C | 10 | 23325 | 0.04287245 | 36.6 | rnr1Y285F_pG | TRUE | CAN | JSY2420 | 5 | rnr1Y285F_p | AAT | A>C | G/C+1 | A/T-1 | TA>GC | TA>GC | GTACCAATA | ATAGTAC | TTA | TTA | T | G | T>G | T>G_TTA | TA>GC_ATAGTAC | 6 | FALSE | FALSE |
| 174_v.txt | 33269 | 33269-SNV-A | SNV | 1 | A | C | 85 | 12706 | 0.66897529 | 16.3647059 | rnr1Y285F_pG | TRUE | CAN | JSY2420 | 5 | rnr1Y285F_p | TAG | A>C | G/C+1 | A/T-1 | TA>GC | TA>GC | CTCCATAGA | AGAGAAC | ATC | ATC | T | G | T>G | T>G_ATC | TA>GC_AGAGAAC | 6 | FALSE | FALSE |
| 174_v.txt | 32174 | 32174-SNV-A | SNV | 1 | A | G | 8 | 11593 | 0.06900716 | 51.1249991 | rnr1Y285F_pG | TRUE | CAN | JSY2420 | 5 | rnr1Y285F_p | CAG | A>G | G/C+1 | A/T-1 | TA>CG | TA>CG | AAATGCAGC | AGCAGTA | GTC | GTC | T | C | T>C | T>C_GTC | TA>CG_AGCAGTA | 6 | FALSE | FALSE |
| 174_v.txt | 32226 | 32226-SNV-A | SNV | 1 | A | G | 7 | 11593 | 0.06038127 | 38 | rnr1Y285F_pG | TRUE | CAN | JSY2420 | 5 | rnr1Y285F_p | CAG | A>G | G/C+1 | A/T-1 | TA>CG | TA>CG | CTTGACAGG | AGGAATT | GTC | GTC | T | C | T>C | T>C_GTC | TA>CG_AGGAATT | 6 | FALSE | FALSE |
| 174_v.txt | 32284 | 32284-SNV-A | SNV | 1 | A | G | 5 | 21310 | 0.02346316 | 39.1995347 | rnr1Y285F_pG | TRUE | CAN | JSY2420 | 5 | rnr1Y285F_p | TAA | A>G | G/C+1 | A/T-1 | TA>CG | TA>CG | CAACGTAAA | AAATATT | ATT | ATT | T | C | T>C | T>C_ATT | TA>CG_AAATATT | 6 | FALSE | FALSE |
| 174_v.txt | 32337 | 32337-SNV-A | SNV | 1 | A | G | 4 | 21310 | 0.01877053 | 58.4998696 | rnr1Y285F_pG | TRUE | CAN | JSY2420 | 5 | rnr1Y285F_p | GAT | A>G | G/C+1 | A/T-1 | TA>CG | TA>CG | TTGAAGATA | ATATGTG | CTA | CTA | T | C | T>C | T>C_CTA | TA>CG_ATATGTG | 6 | FALSE | FALSE |
| 174_v.txt | 32445 | 32445-SNV-A | SNV | 1 | A | G | 23 | 21310 | 0.10793055 | 52.3913044 | rnr1Y285F_pG | TRUE | CAN | JSY2420 | 5 | rnr1Y285F_p | AAG | A>G | G/C+1 | A/T-1 | TA>CG | TA>CG | ACTAAAAGT | AGTCCAA | TTC | TTC | T | C | T>C | T>C_TTC | TA>CG_AGTCCAA | 6 | FALSE | FALSE |
| 174_v.txt | 32472 | 32472-SNV-T | SNV | 1 | T | C | 35 | 21310 | 0.16424214 | 47.6857143 | rnr1Y285F_pG | TRUE | CAN | JSY2420 | 5 | rnr1Y285F_p | GTA | T>C | G/C+1 | A/T-1 | TA>CG | TA>CG | CCAATGTAG | TAGAAGG | CAT | GTA | T | G | T>C | T>C_GTA | TA>CG_TAGAAGG | 6 | FALSE | FALSE |
| 174_v.txt | 32490 | 32490-SNV-A | SNV | 1 | A | G | 7 | 21310 | 0.03284843 | 44.4263664 | rnr1Y285F_pG | TRUE | CAN | JSY2420 | 5 | rnr1Y285F_p | GAA | A>G | G/C+1 | A/T-1 | TA>CG | TA>CG | ATACGGAAA | AAAACAA | CTT | CTT | T | C | T>C | T>C_CTT | TA>CG_AAAACAA | 6 | FALSE | FALSE |
| 174_v.txt | 32492 | 32492-SNV-A | SNV | 1 | A | G | 5 | 21310 | 0.02346316 | 27.1992913 | rnr1Y285F_pG | TRUE | CAN | JSY2420 | 5 | rnr1Y285F_p | AAA | A>G | G/C+1 | A/T-1 | TA>CG | TA>CG | ACGGAAAAC | AACAACT | TTT | TTT | T | C | T>C | T>C_TTT | TA>CG_AACAACT | 6 | FALSE | FALSE |
| 174_v.txt | 32573 | 32573-SNV-T | SNV | 1 | T | C | 6 | 22591 | 0.02655925 | 33.3333333 | rnr1Y285F_pG | TRUE | CAN | JSY2420 | 5 | rnr1Y285F_p | CTT | T>C | G/C+1 | A/T-1 | TA>CG | TA>CG | AGTACCTTG | TTGAAAT | GAA | CTT | T | G | T>C | T>C_CTT | TA>CG_TTGAAAT | 6 | FALSE | FALSE |
| 174_v.txt | 32648 | 32648-SNV-T | SNV | 1 | T | C | 5 | 22590 | 0.02213369 | 62.5998723 | rnr1Y285F_pG | TRUE | CAN | JSY2420 | 5 | rnr1Y285F_p | TTA | T>C | G/C+1 | A/T-1 | TA>CG | TA>CG | AGATATTAT | TATACCT | AAT | TTA | T | G | T>C | T>C_TTA | TA>CG_TATACCT | 6 | FALSE | FALSE |
| 174_v.txt | 32649 | 32649-SNV-A | SNV | 1 | A | G | 3 | 22590 | 0.01328021 | 50.6665193 | rnr1Y285F_pG | TRUE | CAN | JSY2420 | 5 | rnr1Y285F_p | TAT | A>G | G/C+1 | A/T-1 | TA>CG | TA>CG | GATATTATA | ATACCTG | ATA | ATA | T | C | T>C | T>C_ATA | TA>CG_ATACCTG | 6 | FALSE | FALSE |
| 174_v.txt | 32680 | 32680-SNV-A | SNV | 1 | A | G | 21 | 22590 | 0.09296149 | 57.9047619 | rnr1Y285F_pG | TRUE | CAN | JSY2420 | 5 | rnr1Y285F_p | CAA | A>G | G/C+1 | A/T-1 | TA>CG | TA>CG | TTCTCCAATA | AATAACG | GTT | GTT | T | C | T>C | T>C_GTT | TA>CG_AATAACG | 6 | FALSE | FALSE |
| 174_v.txt | 32696 | 32696-SNV-T | SNV | 1 | T | C | 4 | 22590 | 0.01770695 | 58.9996793 | rnr1Y285F_pG | TRUE | CAN | JSY2420 | 5 | rnr1Y285F_p | CTG | T>C | G/C+1 | A/T-1 | TA>CG | TA>CG | TCCAACTGG | TGGGCCG | GAC | CTG | T | G | T>C | T>C_CTG | TA>CG_TGGGCCG | 6 | FALSE | FALSE |
| 174_v.txt | 32740 | 32740-SNV-T | SNV | 1 | T | C | 6 | 22589 | 0.0265616 | 53.9981402 | rnr1Y285F_pG | TRUE | CAN | JSY2420 | 5 | rnr1Y285F_p | ATT | T>C | G/C+1 | A/T-1 | TA>CG | TA>CG | AGTATATTA | TTAGAAA | TAA | ATT | T | G | T>C | T>C_ATT | TA>CG_TTAGAAA | 6 | FALSE | FALSE |
| 174_v.txt | 32808 | 32808-SNV-A | SNV | 1 | A | G | 8 | 22590 | 0.0354139 | 50.8749505 | rnr1Y285F_pG | TRUE | CAN | JSY2420 | 5 | rnr1Y285F_p | GAC | A>G | G/C+1 | A/T-1 | TA>CG | TA>CG | TATTTGACA | ACAGGGA | CTG | CTG | T | C | T>C | T>C_CTG | TA>CG_ACAGGGA | 6 | FALSE | FALSE |
| 174_v.txt | 32842 | 32842-SNV-A | SNV | 1 | A | G | 5 | 22590 | 0.02213369 | 43.1998826 | rnr1Y285F_pG | TRUE | CAN | JSY2420 | 5 | rnr1Y285F_p | CAA | A>G | G/C+1 | A/T-1 | TA>CG | TA>CG | TTACCCAAA | AAAAAAT | GTT | GTT | T | C | T>C | T>C_GTT | TA>CG_AAAAAAT | 6 | FALSE | FALSE |
| 174_v.txt | 32919 | 32919-SNV-A | SNV | 1 | A | G | 20 | 23328 | 0.08573388 | 50.8 | rnr1Y285F_pG | TRUE | CAN | JSY2420 | 5 | rnr1Y285F_p | CAG | A>G | G/C+1 | A/T-1 | TA>CG | TA>CG | AGTTCCAGG | AGGGCAA | GTC | GTC | T | C | T>C | T>C_GTC | TA>CG_AGGGCAA | 6 | FALSE | FALSE |
| 174_v.txt | 32927 | 32927-SNV-A | SNV | 1 | A | G | 24 | 23328 | 0.10288066 | 47.7916667 | rnr1Y285F_pG | TRUE | CAN | JSY2420 | 5 | rnr1Y285F_p | AAG | A>G | G/C+1 | A/T-1 | TA>CG | TA>CG | GGCAAAAGT | AGTGATT | TTC | TTC | T | C | T>C | T>C_TTC | TA>CG_AGTGATT | 6 | FALSE | FALSE |
| 174_v.txt | 32283 | 32283-SNV-T | SNV | 1 | T | A | 10 | 21310 | 0.04692633 | 24.0982967 | rnr1Y285F_pG | TRUE | CAN | JSY2420 | 5 | rnr1Y285F_p | GTA | T>A | A/T+1 | A/T-1 | TA>AT | TA>AT | CCAACGTAA | TAAATAT | CAT | GTA | T | T | T>A | T>A_GTA | TA>AT_TAAATAT | 6 | FALSE | FALSE |
| 174_v.txt | 32284 | 32284-SNV-A | SNV | 1 | A | T | 129 | 21310 | 0.6053496 | 45.5078042 | rnr1Y285F_pG | TRUE | CAN | JSY2420 | 5 | rnr1Y285F_p | TAA | A>T | A/T+1 | A/T-1 | TA>AT | TA>AT | CAACGTAAA | AAATATT | ATT | ATT | T | A | T>A | T>A_ATT | TA>AT_AAATATT | 6 | FALSE | FALSE |
| 174_v.txt | 32289 | 32289-SNV-T | SNV | 1 | T | A | 21 | 21310 | 0.09854528 | 51.9487454 | rnr1Y285F_pG | TRUE | CAN | JSY2420 | 5 | rnr1Y285F_p | ATT | T>A | A/T+1 | A/T-1 | TA>AT | TA>AT | TAAATATTT | TTTGAAT | TAA | ATT | T | T | T>A | T>A_ATT | TA>AT_TTTGAAT | 6 | FALSE | FALSE |
| 174_v.txt | 32290 | 32290-SNV-T | SNV | 1 | T | A | 28 | 21310 | 0.13139371 | 53.309984 | rnr1Y285F_pG | TRUE | CAN | JSY2420 | 5 | rnr1Y285F_p | TTT | T>A | A/T+1 | A/T-1 | TA>AT | TA>AT | AAATATTTG | TTGAATT | AAA | TTT | T | T | T>A | T>A_TTT | TA>AT_TTGAATT | 6 | FALSE | FALSE |
| 174_v.txt | 32337 | 32337-SNV-A | SNV | 1 | A | T | 2 | 21310 | 9.39E-03 | 48.9997849 | rnr1Y285F_pG | TRUE | CAN | JSY2420 | 5 | rnr1Y285F_p | GAT | A>T | A/T+1 | A/T-1 | TA>AT | TA>AT | TTGAAGATA | ATATGTG | CTA | CTA | T | A | T>A | T>A_CTA | TA>AT_ATATGTG | 6 | FALSE | FALSE |
| 174_v.txt | 32349 | 32349-SNV-A | SNV | 1 | A | T | 10 | 21310 | 0.04692633 | 60.9 | rnr1Y285F_pG | TRUE | CAN | JSY2420 | 5 | rnr1Y285F_p | AAC | A>T | A/T+1 | A/T-1 | TA>AT | TA>AT | GGCAAAACC | ACCTTTG | TTG | TTG | T | A | T>A | T>A_TTG | TA>AT_ACCTTTG | 6 | FALSE | FALSE |
| 174_v.txt | 32356 | 32356-SNV-T | SNV | 1 | T | A | 23 | 21310 | 0.10793055 | 62.2608696 | rnr1Y285F_pG | TRUE | CAN | JSY2420 | 5 | rnr1Y285F_p | GTA | T>A | A/T+1 | A/T-1 | TA>AT | TA>AT | CCTTTGTACC | TACCAGA | CAT | GTA | T | T | T>A | T>A_GTA | TA>AT_TACCAGA | 6 | FALSE | FALSE |
| 174_v.txt | 32422 | 32422-SNV-T | SNV | 1 | T | A | 33 | 21310 | 0.15485688 | 50.6969697 | rnr1Y285F_pG | TRUE | CAN | JSY2420 | 5 | rnr1Y285F_p | TTA | T>A | A/T+1 | A/T-1 | TA>AT | TA>AT | TTAGTTTAG | TAGGGTC | AAT | TTA | T | T | T>A | T>A_TTA | TA>AT_TAGGGTC | 6 | FALSE | FALSE |
| 174_v.txt | 32472 | 32472-SNV-T | SNV | 1 | T | A | 6 | 21310 | 0.0281558 | 48 | rnr1Y285F_pG | TRUE | CAN | JSY2420 | 5 | rnr1Y285F_p | GTA | T>A | A/T+1 | A/T-1 | TA>AT | TA>AT | CCAATGTAG | TAGAAGG | CAT | GTA | T | T | T>A | T>A_GTA | TA>AT_TAGAAGG | 6 | FALSE | FALSE |
| 174_v.txt | 32481 | 32481-SNV-A | SNV | 1 | A | T | 24 | 21310 | 0.11262318 | 48.0416667 | rnr1Y285F_pG | TRUE | CAN | JSY2420 | 5 | rnr1Y285F_p | TAA | A>T | A/T+1 | A/T-1 | TA>AT | TA>AT | AAGGTTAAG | AAGATAC | ATT | ATT | T | A | T>A | T>A_ATT | TA>AT_AAGATAC | 6 | FALSE | FALSE |
| 174_v.txt | 32573 | 32573-SNV-T | SNV | 1 | T | A | 17 | 22591 | 0.07525121 | 37.4705882 | rnr1Y285F_pG | TRUE | CAN | JSY2420 | 5 | rnr1Y285F_p | CTT | T>A | A/T+1 | A/T-1 | TA>AT | TA>AT | AGTACCTTG | TTGAAAT | GAA | CTT | T | T | T>A | T>A_CTT | TA>AT_TTGAAAT | 6 | FALSE | FALSE |
| 174_v.txt | 32616 | 32616-SNV-A | SNV | 1 | A | T | 27 | 22590 | 0.11952191 | 43.4074074 | rnr1Y285F_pG | TRUE | CAN | JSY2420 | 5 | rnr1Y285F_p | TAA | A>T | A/T+1 | A/T-1 | TA>AT | TA>AT | CAACCTAAG | AAGAACC | ATT | ATT | T | A | T>A | T>A_ATT | TA>AT_AAGAACC | 6 | FALSE | FALSE |
| 174_v.txt | 32649 | 32649-SNV-A | SNV | 1 | A | T | 1 | 22590 | 4.43E-03 | 63.9995283 | rnr1Y285F_pG | TRUE | CAN | JSY2420 | 5 | rnr1Y285F_p | TAT | A>T | A/T+1 | A/T-1 | TA>AT | TA>AT | GATATTATA | ATACCTG | ATA | ATA | T | A | T>A | T>A_ATA | TA>AT_ATACCTG | 6 | FALSE | FALSE |
| 174_v.txt | 32696 | 32696-SNV-T | SNV | 1 | T | A | 2 | 22590 | 8.85E-03 | 63.9987612 | rnr1Y285F_pG | TRUE | CAN | JSY2420 | 5 | rnr1Y285F_p | CTG | T>A | A/T+1 | A/T-1 | TA>AT | TA>AT | TCCAACTGG | TGGGCCG | GAC | CTG | T | T | T>A | T>A_CTG | TA>AT_TGGGCCG | 6 | FALSE | FALSE |
| 174_v.txt | 32717 | 32717-SNV-A | SNV | 1 | A | T | 31 | 22590 | 0.13722886 | 59 | rnr1Y285F_pG | TRUE | CAN | JSY2420 | 5 | rnr1Y285F_p | CAC | A>T | A/T+1 | A/T-1 | TA>AT | TA>AT | AGCACCACA | ACAAACC | GTG | GTG | T | A | T>A | T>A_GTG | TA>AT_ACAAACC | 6 | FALSE | FALSE |
| 174_v.txt | 32736 | 32736-SNV-T | SNV | 1 | T | A | 2 | 22590 | 8.85E-03 | 50.9970748 | rnr1Y285F_pG | TRUE | CAN | JSY2420 | 5 | rnr1Y285F_p | GTA | T>A | A/T+1 | A/T-1 | TA>AT | TA>AT | AAACAGTAT | TATATTA | CAT | GTA | T | T | T>A | T>A_GTA | TA>AT_TATATTA | 6 | FALSE | FALSE |
| 174_v.txt | 32740 | 32740-SNV-T | SNV | 1 | T | A | 5 | 22589 | 0.02213467 | 52.9865429 | rnr1Y285F_pG | TRUE | CAN | JSY2420 | 5 | rnr1Y285F_p | ATT | T>A | A/T+1 | A/T-1 | TA>AT | TA>AT | AGTATATTA | TTAGAAA | TAA | ATT | T | T | T>A | T>A_ATT | TA>AT_TTAGAAA | 6 | FALSE | FALSE |
| 174_v.txt | 32760 | 32760-SNV-A | SNV | 1 | A | T | 24 | 22590 | 0.1062417 | 52.5416667 | rnr1Y285F_pG | TRUE | CAN | JSY2420 | 5 | rnr1Y285F_p | TAA | A>T | A/T+1 | A/T-1 | TA>AT | TA>AT | ATGGCTAAA | AAAACTT | ATT | ATT | T | A | T>A | T>A_ATT | TA>AT_AAAACTT | 6 | FALSE | FALSE |
| 174_v.txt | 32915 | 32915-SNV-T | SNV | 1 | T | A | 14 | 23328 | 0.06001372 | 19.0708073 | rnr1Y285F_pG | TRUE | CAN | JSY2420 | 5 | rnr1Y285F_p | GTT | T>A | A/T+1 | A/T-1 | TA>AT | TA>AT | ACTAAGTTC | TTCCAGG | CAA | GTT | T | T | T>A | T>A_GTT | TA>AT_TTCCAGG | 6 | FALSE | FALSE |
| 174_v.txt | 32916 | 32916-SNV-T | SNV | 1 | T | A | 41 | 23327 | 0.17576199 | 50.046109 | rnr1Y285F_pG | TRUE | CAN | JSY2420 | 5 | rnr1Y285F_p | TTC | T>A | A/T+1 | A/T-1 | TA>AT | TA>AT | CTAAGTTCC | TCCAGGG | AAG | TTC | T | T | T>A | T>A_TTC | TA>AT_TCCAGGG | 6 | FALSE | FALSE |
| 174_v.txt | 33039 | 33039-SNV-A | SNV | 1 | A | T | 19 | 23328 | 0.08144719 | 62.0526316 | rnr1Y285F_pG | TRUE | CAN | JSY2420 | 5 | rnr1Y285F_p | CAT | A>T | A/T+1 | A/T-1 | TA>AT | TA>AT | GTAGCCATT | ATTTCAC | GTA | GTA | T | A | T>A | T>A_GTA | TA>AT_ATTTCAC | 6 | FALSE | FALSE |
| 174_v.txt | 33065 | 33065-SNV-A | SNV | 1 | A | T | 14 | 23328 | 0.06001372 | 54.3571429 | rnr1Y285F_pG | TRUE | CAN | JSY2420 | 5 | rnr1Y285F_p | AAT | A>T | A/T+1 | A/T-1 | TA>AT | TA>AT | GACAGAATA | ATATGCC | TTA | TTA | T | A | T>A | T>A_TTA | TA>AT_ATATGCC | 6 | FALSE | FALSE |
| 174_v.txt | 33089 | 33089-SNV-A | SNV | 1 | A | T | 28 | 23328 | 0.12002744 | 49.9999497 | rnr1Y285F_pG | TRUE | CAN | JSY2420 | 5 | rnr1Y285F_p | AAT | A>T | A/T+1 | A/T-1 | TA>AT | TA>AT | AAATAAATA | ATATGAT | TTA | TTA | T | A | T>A | T>A_TTA | TA>AT_ATATGAT | 6 | FALSE | FALSE |
| 174_v.txt | 33090 | 33090-SNV-T | SNV | 1 | T | A | 4 | 23328 | 0.01714678 | 50.49578 | rnr1Y285F_pG | TRUE | CAN | JSY2420 | 5 | rnr1Y285F_p | ATA | T>A | A/T+1 | A/T-1 | TA>AT | TA>AT | AATAAATAT | TATGATA | TAT | ATA | T | T | T>A | T>A_ATA | TA>AT_TATGATA | 6 | FALSE | FALSE |
| 174_v.txt | 33192 | 33192-SNV-T | SNV | 1 | T | A | 17 | 12689 | 0.13397431 | 35.7058824 | rnr1Y285F_pG | TRUE | CAN | JSY2420 | 5 | rnr1Y285F_p | ATG | T>A | A/T+1 | A/T-1 | TA>AT | TA>AT | CCAATATGT | TGTCTTT | TAC | ATG | T | T | T>A | T>A_ATG | TA>AT_TGTCTTT | 6 | FALSE | FALSE |
| 174_v.txt | 33202 | 33202-SNV-T | SNV | 1 | T | A | 19 | 12697 | 0.14964165 | 36.7894737 | rnr1Y285F_pG | TRUE | CAN | JSY2420 | 5 | rnr1Y285F_p | TTA | T>A | A/T+1 | A/T-1 | TA>AT | TA>AT | TTTGCTTAAG | TAAGCTC | AAT | TTA | T | T | T>A | T>A_TTA | TA>AT_TAAGCTC | 6 | FALSE | FALSE |
| 174_v.txt | 33295 | 33295-SNV-T | SNV | 1 | T | A | 30 | 12706 | 0.23610893 | 54.7 | rnr1Y285F_pG | TRUE | CAN | JSY2420 | 5 | rnr1Y285F_p | CTC | T>A | A/T+1 | A/T-1 | TA>AT | TA>AT | TTACTCTCGT | TCGTCGG | GAG | CTC | T | T | T>A | T>A_CTC | TA>AT_TCGTCGG | 6 | FALSE | FALSE |
| 174_v.txt | 33317 | 33317-SNV-A | SNV | 1 | A | T | 13 | 12706 | 0.10231387 | 63.7692308 | rnr1Y285F_pG | TRUE | CAN | JSY2420 | 5 | rnr1Y285F_p | GAT | A>T | A/T+1 | A/T-1 | TA>AT | TA>AT | CAATGGATA | ATACAAT | CTA | CTA | T | A | T>A | T>A_CTA | TA>AT_ATACAAT | 6 | FALSE | FALSE |
| 174_v.txt | 31658 | 31658-SNV-C | SNV | 1 | C | T | 15 | 7006 | 0.2141022 | 33.2666667 | rnr1Y285F_pG | TRUE | CAN | JSY2420 | 5 | rnr1Y285F_p | GCG | C>T | A/T+1 | G/C-1 | CG>TA | CG>TA | AAATGGCGT | CGTGGAA | CGC | GCG | C | A | C>T | C>T_GCG | CG>TA_CGTGGAA | 6 | FALSE | FALSE |
| 174_v.txt | 31844 | 31844-SNV-C | SNV | 1 | C | T | 16 | 7006 | 0.22837568 | 45.6875 | rnr1Y285F_pG | TRUE | CAN | JSY2420 | 5 | rnr1Y285F_p | TCC | C>T | A/T+1 | G/C-1 | CG>TA | CG>TA | TAAGATCCA | CCAAACA | AGG | TCC | C | A | C>T | C>T_TCC | CG>TA_CCAAACA | 6 | FALSE | FALSE |
| 174_v.txt | 31930 | 31930-SNV-G | SNV | 1 | G | A | 19 | 7006 | 0.27119612 | 36.0526316 | rnr1Y285F_pG | TRUE | CAN | JSY2420 | 5 | rnr1Y285F_p | TGA | G>A | A/T+1 | G/C-1 | CG>TA | CG>TA | AACCTTGAA | GAATAAT | ACT | ACT | C | T | C>T | C>T_ACT | CG>TA_GAATAAT | 6 | FALSE | FALSE |
| 174_v.txt | 31947 | 31947-SNV-G | SNV | 1 | G | A | 10 | 11583 | 0.08633342 | 36 | rnr1Y285F_pG | TRUE | CAN | JSY2420 | 5 | rnr1Y285F_p | CGT | G>A | A/T+1 | G/C-1 | CG>TA | CG>TA | ATGATCGTC | GTCATAA | GCA | GCA | C | T | C>T | C>T_GCA | CG>TA_GTCATAA | 6 | FALSE | FALSE |
| 174_v.txt | 31962 | 31962-SNV-G | SNV | 1 | G | A | 26 | 11593 | 0.22427327 | 35.7692308 | rnr1Y285F_pG | TRUE | CAN | JSY2420 | 5 | rnr1Y285F_p | CGC | G>A | A/T+1 | G/C-1 | CG>TA | CG>TA | GTGGCCGCA | GCATAAT | GCG | GCG | C | T | C>T | C>T_GCG | CG>TA_GCATAAT | 6 | FALSE | FALSE |
| 174_v.txt | 31972 | 31972-SNV-C | SNV | 1 | C | T | 11 | 11593 | 0.09488484 | 36.7272727 | rnr1Y285F_pG | TRUE | CAN | JSY2420 | 5 | rnr1Y285F_p | GCC | C>T | A/T+1 | G/C-1 | CG>TA | CG>TA | AATAAGCCA | CCAAGCC | CGG | GCC | C | A | C>T | C>T_GCC | CG>TA_CCAAGCC | 6 | FALSE | FALSE |
| 174_v.txt | 31981 | 31981-SNV-G | SNV | 1 | G | A | 23 | 11593 | 0.19839558 | 44.6086957 | rnr1Y285F_pG | TRUE | CAN | JSY2420 | 5 | rnr1Y285F_p | GGC | G>A | A/T+1 | G/C-1 | CG>TA | CG>TA | AGCCGGGCA | GCATTAA | CCG | CCG | C | T | C>T | C>T_CCG | CG>TA_GCATTAA | 6 | FALSE | FALSE |
| 174_v.txt | 32075 | 32075-SNV-C | SNV | 1 | C | T | 165 | 11593 | 1.42327267 | 61.3878788 | rnr1Y285F_pG | TRUE | CAN | JSY2420 | 5 | rnr1Y285F_p | ACC | C>T | A/T+1 | G/C-1 | CG>TA | CG>TA | AAATAACCA | CCATGCA | TGG | ACC | C | A | C>T | C>T_ACC | CG>TA_CCATGCA | 6 | FALSE | FALSE |
| 174_v.txt | 32088 | 32088-SNV-C | SNV | 1 | C | T | 49 | 11593 | 0.42266885 | 61.3265306 | rnr1Y285F_pG | TRUE | CAN | JSY2420 | 5 | rnr1Y285F_p | GCC | C>T | A/T+1 | G/C-1 | CG>TA | CG>TA | AAAAAGCCT | CCTGCAA | CGG | GCC | C | A | C>T | C>T_GCC | CG>TA_CCTGCAA | 6 | FALSE | FALSE |
| 174_v.txt | 32092 | 32092-SNV-C | SNV | 1 | C | T | 17 | 11592 | 0.14665286 | 51.8212546 | rnr1Y285F_pG | TRUE | CAN | JSY2420 | 5 | rnr1Y285F_p | GCA | C>T | A/T+1 | G/C-1 | CG>TA | CG>TA | AGCCTGCAA | CAACACC | CGT | GCA | C | A | C>T | C>T_GCA | CG>TA_CAACACC | 6 | FALSE | FALSE |
| 174_v.txt | 32114 | 32114-SNV-C | SNV | 1 | C | T | 86 | 11593 | 0.74182697 | 59.4186047 | rnr1Y285F_pG | TRUE | CAN | JSY2420 | 5 | rnr1Y285F_p | GCC | C>T | A/T+1 | G/C-1 | CG>TA | CG>TA | TAATAGCCA | CCATTCG | CGG | GCC | C | A | C>T | C>T_GCC | CG>TA_CCATTCG | 6 | FALSE | FALSE |
| 174_v.txt | 32165 | 32165-SNV-G | SNV | 1 | G | A | 16 | 11593 | 0.13801432 | 51.9375 | rnr1Y285F_pG | TRUE | CAN | JSY2420 | 5 | rnr1Y285F_p | CGC | G>A | A/T+1 | G/C-1 | CG>TA | CG>TA | CAAAGCGCC | GCCAAAT | GCG | GCG | C | T | C>T | C>T_GCG | CG>TA_GCCAAAT | 6 | FALSE | FALSE |
| 174_v.txt | 32166 | 32166-SNV-C | SNV | 1 | C | T | 34 | 11593 | 0.29328043 | 48.2336068 | rnr1Y285F_pG | TRUE | CAN | JSY2420 | 5 | rnr1Y285F_p | GCC | C>T | A/T+1 | G/C-1 | CG>TA | CG>TA | AAAGCGCCA | CCAAATG | CGG | GCC | C | A | C>T | C>T_GCC | CG>TA_CCAAATG | 6 | FALSE | FALSE |
| 174_v.txt | 32173 | 32173-SNV-C | SNV | 1 | C | T | 14 | 11593 | 0.12076253 | 41.4989097 | rnr1Y285F_pG | TRUE | CAN | JSY2420 | 5 | rnr1Y285F_p | GCA | C>T | A/T+1 | G/C-1 | CG>TA | CG>TA | CAAATGCAG | CAGCAGT | CGT | GCA | C | A | C>T | C>T_GCA | CG>TA_CAGCAGT | 6 | FALSE | FALSE |
| 174_v.txt | 32176 | 32176-SNV-C | SNV | 1 | C | T | 9 | 11593 | 0.07763305 | 46.000824 | rnr1Y285F_pG | TRUE | CAN | JSY2420 | 5 | rnr1Y285F_p | GCA | C>T | A/T+1 | G/C-1 | CG>TA | CG>TA | ATGCAGCAG | CAGTAAC | CGT | GCA | C | A | C>T | C>T_GCA | CG>TA_CAGTAAC | 6 | FALSE | FALSE |
| 174_v.txt | 32205 | 32205-SNV-C | SNV | 1 | C | T | 14 | 11593 | 0.12076253 | 44.5714286 | rnr1Y285F_pG | TRUE | CAN | JSY2420 | 5 | rnr1Y285F_p | ACC | C>T | A/T+1 | G/C-1 | CG>TA | CG>TA | GGAACACCA | CCACCTT | TGG | ACC | C | A | C>T | C>T_ACC | CG>TA_CCACCTT | 6 | FALSE | FALSE |
| 174_v.txt | 32271 | 32271-SNV-C | SNV | 1 | C | T | 34 | 21310 | 0.15954951 | 33.7352941 | rnr1Y285F_pG | TRUE | CAN | JSY2420 | 5 | rnr1Y285F_p | ACG | C>T | A/T+1 | G/C-1 | CG>TA | CG>TA | AAAATACGG | CGGGAAC | TGC | ACG | C | A | C>T | C>T_ACG | CG>TA_CGGGAAC | 6 | FALSE | FALSE |
| 174_v.txt | 32272 | 32272-SNV-G | SNV | 1 | G | A | 25 | 21310 | 0.11731581 | 34.3574649 | rnr1Y285F_pG | TRUE | CAN | JSY2420 | 5 | rnr1Y285F_p | CGG | G>A | A/T+1 | G/C-1 | CG>TA | CG>TA | AAATACGGG | GGGAACC | GCC | GCC | C | T | C>T | C>T_GCC | CG>TA_GGGAACC | 6 | FALSE | FALSE |
| 174_v.txt | 32274 | 32274-SNV-G | SNV | 1 | G | A | 78 | 21310 | 0.36602534 | 36.6794872 | rnr1Y285F_pG | TRUE | CAN | JSY2420 | 5 | rnr1Y285F_p | GGA | G>A | A/T+1 | G/C-1 | CG>TA | CG>TA | ATACGGGAA | GAACCAA | CCT | CCT | C | T | C>T | C>T_CCT | CG>TA_GAACCAA | 6 | FALSE | FALSE |
| 174_v.txt | 32277 | 32277-SNV-C | SNV | 1 | C | T | 31 | 21310 | 0.14547161 | 15.8706665 | rnr1Y285F_pG | TRUE | CAN | JSY2420 | 5 | rnr1Y285F_p | ACC | C>T | A/T+1 | G/C-1 | CG>TA | CG>TA | CGGGAACCA | CCAACGT | TGG | ACC | C | A | C>T | C>T_ACC | CG>TA_CCAACGT | 6 | FALSE | FALSE |
| 174_v.txt | 32299 | 32299-SNV-C | SNV | 1 | C | T | 51 | 21310 | 0.23932426 | 53.4509804 | rnr1Y285F_pG | TRUE | CAN | JSY2420 | 5 | rnr1Y285F_p | GCG | C>T | A/T+1 | G/C-1 | CG>TA | CG>TA | AATTTGCGG | CGGCAGA | CGC | GCG | C | A | C>T | C>T_GCG | CG>TA_CGGCAGA | 6 | FALSE | FALSE |
| 174_v.txt | 32301 | 32301-SNV-G | SNV | 1 | G | A | 16 | 21310 | 0.07508212 | 55.5625 | rnr1Y285F_pG | TRUE | CAN | JSY2420 | 5 | rnr1Y285F_p | GGC | G>A | A/T+1 | G/C-1 | CG>TA | CG>TA | TTTGCGGCA | GCAGAAA | CCG | CCG | C | T | C>T | C>T_CCG | CG>TA_GCAGAAA | 6 | FALSE | FALSE |
| 174_v.txt | 32413 | 32413-SNV-G | SNV | 1 | G | A | 40 | 21310 | 0.1877053 | 57.125 | rnr1Y285F_pG | TRUE | CAN | JSY2420 | 5 | rnr1Y285F_p | TGT | G>A | A/T+1 | G/C-1 | CG>TA | CG>TA | TAGATTGTG | GTGTTAG | ACA | ACA | C | T | C>T | C>T_ACA | CG>TA_GTGTTAG | 6 | FALSE | FALSE |
| 174_v.txt | 32436 | 32436-SNV-G | SNV | 1 | G | A | 26 | 21310 | 0.12200845 | 53 | rnr1Y285F_pG | TRUE | CAN | JSY2420 | 5 | rnr1Y285F_p | TGG | G>A | A/T+1 | G/C-1 | CG>TA | CG>TA | TTGTATGGA | GGAACTA | ACC | ACC | C | T | C>T | C>T_ACC | CG>TA_GGAACTA | 6 | FALSE | FALSE |
| 174_v.txt | 32449 | 32449-SNV-C | SNV | 1 | C | T | 37 | 21310 | 0.17362741 | 51.3243243 | rnr1Y285F_pG | TRUE | CAN | JSY2420 | 5 | rnr1Y285F_p | CCA | C>T | A/T+1 | G/C-1 | CG>TA | CG>TA | AAAGTCCAA | CAATGAA | GGT | CCA | C | A | C>T | C>T_CCA | CG>TA_CAATGAA | 6 | FALSE | FALSE |
| 174_v.txt | 32466 | 32466-SNV-C | SNV | 1 | C | T | 44 | 21310 | 0.20647583 | 46.9545455 | rnr1Y285F_pG | TRUE | CAN | JSY2420 | 5 | rnr1Y285F_p | GCC | C>T | A/T+1 | G/C-1 | CG>TA | CG>TA | AGAGAGCCA | CCAATGT | CGG | GCC | C | A | C>T | C>T_GCC | CG>TA_CCAATGT | 6 | FALSE | FALSE |
| 174_v.txt | 32487 | 32487-SNV-C | SNV | 1 | C | T | 21 | 21310 | 0.09854528 | 42.5238095 | rnr1Y285F_pG | TRUE | CAN | JSY2420 | 5 | rnr1Y285F_p | ACG | C>T | A/T+1 | G/C-1 | CG>TA | CG>TA | AAGATACGG | CGGAAAA | TGC | ACG | C | A | C>T | C>T_ACG | CG>TA_CGGAAAA | 6 | FALSE | FALSE |
| 174_v.txt | 32535 | 32535-SNV-G | SNV | 1 | G | A | 27 | 21309 | 0.12670703 | 36.5550685 | rnr1Y285F_pG | TRUE | CAN | JSY2420 | 5 | rnr1Y285F_p | TGC | G>A | A/T+1 | G/C-1 | CG>TA | CG>TA | GGGTTTGCA | GCAGCTT | ACG | ACG | C | T | C>T | C>T_ACG | CG>TA_GCAGCTT | 6 | FALSE | FALSE |
| 174_v.txt | 32536 | 32536-SNV-C | SNV | 1 | C | T | 41 | 21309 | 0.19240696 | 36.7067664 | rnr1Y285F_pG | TRUE | CAN | JSY2420 | 5 | rnr1Y285F_p | GCA | C>T | A/T+1 | G/C-1 | CG>TA | CG>TA | GGTTTGCAG | CAGCTTC | CGT | GCA | C | A | C>T | C>T_GCA | CG>TA_CAGCTTC | 6 | FALSE | FALSE |
| 174_v.txt | 32539 | 32539-SNV-C | SNV | 1 | C | T | 6 | 21309 | 0.02815712 | 37.9989375 | rnr1Y285F_pG | TRUE | CAN | JSY2420 | 5 | rnr1Y285F_p | GCT | C>T | A/T+1 | G/C-1 | CG>TA | CG>TA | TTGCAGCTT | CTTCACC | CGA | GCT | C | A | C>T | C>T_GCT | CG>TA_CTTCACC | 6 | FALSE | FALSE |
| 174_v.txt | 32566 | 32566-SNV-C | SNV | 1 | C | T | 38 | 22585 | 0.16825327 | 36 | rnr1Y285F_pG | TRUE | CAN | JSY2420 | 5 | rnr1Y285F_p | TCA | C>T | A/T+1 | G/C-1 | CG>TA | CG>TA | CTAGTTCAG | CAGTACC | AGT | TCA | C | A | C>T | C>T_TCA | CG>TA_CAGTACC | 6 | FALSE | FALSE |
| 174_v.txt | 32568 | 32568-SNV-G | SNV | 1 | G | A | 22 | 22592 | 0.0973796 | 37.4545455 | rnr1Y285F_pG | TRUE | CAN | JSY2420 | 5 | rnr1Y285F_p | AGT | G>A | A/T+1 | G/C-1 | CG>TA | CG>TA | AGTTCAGTA | GTACCTT | TCA | TCA | C | T | C>T | C>T_TCA | CG>TA_GTACCTT | 6 | FALSE | FALSE |
| 174_v.txt | 32571 | 32571-SNV-C | SNV | 1 | C | T | 111 | 22592 | 0.49132436 | 35.5844645 | rnr1Y285F_pG | TRUE | CAN | JSY2420 | 5 | rnr1Y285F_p | ACC | C>T | A/T+1 | G/C-1 | CG>TA | CG>TA | TCAGTACCT | CCTTGAA | TGG | ACC | C | A | C>T | C>T_ACC | CG>TA_CCTTGAA | 6 | FALSE | FALSE |
| 174_v.txt | 32572 | 32572-SNV-C | SNV | 1 | C | T | 12 | 22591 | 0.0531185 | 24.5000139 | rnr1Y285F_pG | TRUE | CAN | JSY2420 | 5 | rnr1Y285F_p | CCT | C>T | A/T+1 | G/C-1 | CG>TA | CG>TA | CAGTACCTT | CTTGAAA | GGA | CCT | C | A | C>T | C>T_CCT | CG>TA_CTTGAAA | 6 | FALSE | FALSE |
| 174_v.txt | 32590 | 32590-SNV-C | SNV | 1 | C | T | 57 | 22590 | 0.25232404 | 35.3157895 | rnr1Y285F_pG | TRUE | CAN | JSY2420 | 5 | rnr1Y285F_p | GCG | C>T | A/T+1 | G/C-1 | CG>TA | CG>TA | AGGCAGCGT | CGTTAAT | CGC | GCG | C | A | C>T | C>T_GCG | CG>TA_CGTTAAT | 6 | FALSE | FALSE |
| 174_v.txt | 32658 | 32658-SNV-C | SNV | 1 | C | T | 25 | 22590 | 0.11066844 | 51.56 | rnr1Y285F_pG | TRUE | CAN | JSY2420 | 5 | rnr1Y285F_p | ACC | C>T | A/T+1 | G/C-1 | CG>TA | CG>TA | CCTGGACCC | CCCCAGG | TGG | ACC | C | A | C>T | C>T_ACC | CG>TA_CCCCAGG | 6 | FALSE | FALSE |
| 174_v.txt | 32660 | 32660-SNV-C | SNV | 1 | C | T | 29 | 22590 | 0.12837539 | 56.7931035 | rnr1Y285F_pG | TRUE | CAN | JSY2420 | 5 | rnr1Y285F_p | CCC | C>T | A/T+1 | G/C-1 | CG>TA | CG>TA | TGGACCCCA | CCAGGCA | GGG | CCC | C | A | C>T | C>T_CCC | CG>TA_CCAGGCA | 6 | FALSE | FALSE |
| 174_v.txt | 32697 | 32697-SNV-G | SNV | 1 | G | A | 2 | 22590 | 8.85E-03 | 17.4999982 | rnr1Y285F_pG | TRUE | CAN | JSY2420 | 5 | rnr1Y285F_p | TGG | G>A | A/T+1 | G/C-1 | CG>TA | CG>TA | CCAACTGGG | GGGCCGG | ACC | ACC | C | T | C>T | C>T_ACC | CG>TA_GGGCCGG | 6 | FALSE | FALSE |
| 174_v.txt | 32698 | 32698-SNV-G | SNV | 1 | G | A | 16 | 22590 | 0.0708278 | 20.934512 | rnr1Y285F_pG | TRUE | CAN | JSY2420 | 5 | rnr1Y285F_p | GGG | G>A | A/T+1 | G/C-1 | CG>TA | CG>TA | CAACTGGGC | GGCCGGT | CCC | CCC | C | T | C>T | C>T_CCC | CG>TA_GGCCGGT | 6 | FALSE | FALSE |
| 174_v.txt | 32699 | 32699-SNV-G | SNV | 1 | G | A | 11 | 22590 | 0.04869411 | 51.7276255 | rnr1Y285F_pG | TRUE | CAN | JSY2420 | 5 | rnr1Y285F_p | GGC | G>A | A/T+1 | G/C-1 | CG>TA | CG>TA | AACTGGGCC | GCCGGTA | CCG | CCG | C | T | C>T | C>T_CCG | CG>TA_GCCGGTA | 6 | FALSE | FALSE |
| 174_v.txt | 32700 | 32700-SNV-C | SNV | 1 | C | T | 23 | 22590 | 0.10181496 | 56.4109612 | rnr1Y285F_pG | TRUE | CAN | JSY2420 | 5 | rnr1Y285F_p | GCC | C>T | A/T+1 | G/C-1 | CG>TA | CG>TA | ACTGGGCCG | CCGGTAA | CGG | GCC | C | A | C>T | C>T_GCC | CG>TA_CCGGTAA | 6 | FALSE | FALSE |
| 174_v.txt | 32701 | 32701-SNV-C | SNV | 1 | C | T | 8 | 22589 | 0.03541547 | 54.7503081 | rnr1Y285F_pG | TRUE | CAN | JSY2420 | 5 | rnr1Y285F_p | CCG | C>T | A/T+1 | G/C-1 | CG>TA | CG>TA | CTGGGCCGG | CGGTAAC | GGC | CCG | C | A | C>T | C>T_CCG | CG>TA_CGGTAAC | 6 | FALSE | FALSE |
| 174_v.txt | 32702 | 32702-SNV-G | SNV | 1 | G | A | 47 | 22590 | 0.20805666 | 60.2977522 | rnr1Y285F_pG | TRUE | CAN | JSY2420 | 5 | rnr1Y285F_p | CGG | G>A | A/T+1 | G/C-1 | CG>TA | CG>TA | TGGGCCGGT | GGTAACC | GCC | GCC | C | T | C>T | C>T_GCC | CG>TA_GGTAACC | 6 | FALSE | FALSE |
| 174_v.txt | 32703 | 32703-SNV-G | SNV | 1 | G | A | 10 | 22590 | 0.04426738 | 46.7909563 | rnr1Y285F_pG | TRUE | CAN | JSY2420 | 5 | rnr1Y285F_p | GGT | G>A | A/T+1 | G/C-1 | CG>TA | CG>TA | GGGCCGGTA | GTAACCC | CCA | CCA | C | T | C>T | C>T_CCA | CG>TA_GTAACCC | 6 | FALSE | FALSE |
| 174_v.txt | 32749 | 32749-SNV-C | SNV | 1 | C | T | 29 | 22590 | 0.12837539 | 54.2413793 | rnr1Y285F_pG | TRUE | CAN | JSY2420 | 5 | rnr1Y285F_p | CCG | C>T | A/T+1 | G/C-1 | CG>TA | CG>TA | GAAACCCGA | CGATAAT | GGC | CCG | C | A | C>T | C>T_CCG | CG>TA_CGATAAT | 6 | FALSE | FALSE |
| 174_v.txt | 32757 | 32757-SNV-G | SNV | 1 | G | A | 26 | 22590 | 0.11509518 | 59.0384615 | rnr1Y285F_pG | TRUE | CAN | JSY2420 | 5 | rnr1Y285F_p | GGC | G>A | A/T+1 | G/C-1 | CG>TA | CG>TA | ATAATGGCT | GCTAAAA | CCG | CCG | C | T | C>T | C>T_CCG | CG>TA_GCTAAAA | 6 | FALSE | FALSE |
| 174_v.txt | 32776 | 32776-SNV-C | SNV | 1 | C | T | 26 | 22590 | 0.11509518 | 52.3461539 | rnr1Y285F_pG | TRUE | CAN | JSY2420 | 5 | rnr1Y285F_p | GCG | C>T | A/T+1 | G/C-1 | CG>TA | CG>TA | TGGAAGCGA | CGACCCA | CGC | GCG | C | A | C>T | C>T_GCG | CG>TA_CGACCCA | 6 | FALSE | FALSE |
| 174_v.txt | 32779 | 32779-SNV-C | SNV | 1 | C | T | 5 | 22590 | 0.02213369 | 45.9993779 | rnr1Y285F_pG | TRUE | CAN | JSY2420 | 5 | rnr1Y285F_p | ACC | C>T | A/T+1 | G/C-1 | CG>TA | CG>TA | AAGCGACCC | CCCAGAA | TGG | ACC | C | A | C>T | C>T_ACC | CG>TA_CCCAGAA | 6 | FALSE | FALSE |
| 174_v.txt | 32780 | 32780-SNV-C | SNV | 1 | C | T | 53 | 22590 | 0.23461709 | 46.5096937 | rnr1Y285F_pG | TRUE | CAN | JSY2420 | 5 | rnr1Y285F_p | CCC | C>T | A/T+1 | G/C-1 | CG>TA | CG>TA | AGCGACCCA | CCAGAAC | GGG | CCC | C | A | C>T | C>T_CCC | CG>TA_CCAGAAC | 6 | FALSE | FALSE |
| 174_v.txt | 32788 | 32788-SNV-C | SNV | 1 | C | T | 41 | 22590 | 0.18149624 | 48.3658537 | rnr1Y285F_pG | TRUE | CAN | JSY2420 | 5 | rnr1Y285F_p | TCG | C>T | A/T+1 | G/C-1 | CG>TA | CG>TA | AGAACTCGA | CGAATTC | AGC | TCG | C | A | C>T | C>T_TCG | CG>TA_CGAATTC | 6 | FALSE | FALSE |
| 174_v.txt | 32796 | 32796-SNV-C | SNV | 1 | C | T | 31 | 22590 | 0.13722886 | 40.516129 | rnr1Y285F_pG | TRUE | CAN | JSY2420 | 5 | rnr1Y285F_p | ACC | C>T | A/T+1 | G/C-1 | CG>TA | CG>TA | AATTCACCG | CCGTAAT | TGG | ACC | C | A | C>T | C>T_ACC | CG>TA_CCGTAAT | 6 | FALSE | FALSE |
| 174_v.txt | 32798 | 32798-SNV-G | SNV | 1 | G | A | 46 | 22590 | 0.20362993 | 41.9130435 | rnr1Y285F_pG | TRUE | CAN | JSY2420 | 5 | rnr1Y285F_p | CGT | G>A | A/T+1 | G/C-1 | CG>TA | CG>TA | TTCACCGTA | GTAATAT | GCA | GCA | C | T | C>T | C>T_GCA | CG>TA_GTAATAT | 6 | FALSE | FALSE |
| 174_v.txt | 32809 | 32809-SNV-C | SNV | 1 | C | T | 18 | 22590 | 0.07968128 | 45.5 | rnr1Y285F_pG | TRUE | CAN | JSY2420 | 5 | rnr1Y285F_p | ACA | C>T | A/T+1 | G/C-1 | CG>TA | CG>TA | ATTTGACAG | CAGGGAA | TGT | ACA | C | A | C>T | C>T_ACA | CG>TA_CAGGGAA | 6 | FALSE | FALSE |
| 174_v.txt | 32855 | 32855-SNV-C | SNV | 1 | C | T | 3439 | 22589 | 15.2242242 | 36.2305903 | rnr1Y285F_pG | TRUE | CAN | JSY2420 | 5 | rnr1Y285F_p | TCC | C>T | A/T+1 | G/C-1 | CG>TA | CG>TA | ACTAATCCA | CCATGCC | AGG | TCC | C | A | C>T | C>T_TCC | CG>TA_CCATGCC | 6 | FALSE | FALSE |
| 174_v.txt | 32862 | 32862-SNV-G | SNV | 1 | G | A | 20 | 22588 | 0.08854259 | 36.35 | rnr1Y285F_pG | TRUE | CAN | JSY2420 | 5 | rnr1Y285F_p | CGC | G>A | A/T+1 | G/C-1 | CG>TA | CG>TA | CATGCCGCC | GCCAGTG | GCG | GCG | C | T | C>T | C>T_GCG | CG>TA_GCCAGTG | 6 | FALSE | FALSE |
| 174_v.txt | 32882 | 32882-SNV-C | SNV | 1 | C | T | 53 | 23328 | 0.22719479 | 29.6603774 | rnr1Y285F_pG | TRUE | CAN | JSY2420 | 5 | rnr1Y285F_p | TCC | C>T | A/T+1 | G/C-1 | CG>TA | CG>TA | GTACGTCCA | CCAAAAT | AGG | TCC | C | A | C>T | C>T_TCC | CG>TA_CCAAAAT | 6 | FALSE | FALSE |
| 174_v.txt | 32900 | 32900-SNV-G | SNV | 1 | G | A | 25 | 23328 | 0.10716735 | 37.3980127 | rnr1Y285F_pG | TRUE | CAN | JSY2420 | 5 | rnr1Y285F_p | GGC | G>A | A/T+1 | G/C-1 | CG>TA | CG>TA | GACTTGGCC | GCCAACT | CCG | CCG | C | T | C>T | C>T_CCG | CG>TA_GCCAACT | 6 | FALSE | FALSE |
| 174_v.txt | 32901 | 32901-SNV-C | SNV | 1 | C | T | 54 | 23328 | 0.23148148 | 34.6855943 | rnr1Y285F_pG | TRUE | CAN | JSY2420 | 5 | rnr1Y285F_p | GCC | C>T | A/T+1 | G/C-1 | CG>TA | CG>TA | ACTTGGCCA | CCAACTA | CGG | GCC | C | A | C>T | C>T_GCC | CG>TA_CCAACTA | 6 | FALSE | FALSE |
| 174_v.txt | 32917 | 32917-SNV-C | SNV | 1 | C | T | 113 | 23328 | 0.48439643 | 47.539823 | rnr1Y285F_pG | TRUE | CAN | JSY2420 | 5 | rnr1Y285F_p | TCC | C>T | A/T+1 | G/C-1 | CG>TA | CG>TA | TAAGTTCCA | CCAGGGC | AGG | TCC | C | A | C>T | C>T_TCC | CG>TA_CCAGGGC | 6 | FALSE | FALSE |
| 174_v.txt | 32922 | 32922-SNV-G | SNV | 1 | G | A | 23 | 23328 | 0.09859396 | 51.4347826 | rnr1Y285F_pG | TRUE | CAN | JSY2420 | 5 | rnr1Y285F_p | GGC | G>A | A/T+1 | G/C-1 | CG>TA | CG>TA | TCCAGGGCA | GCAAAAG | CCG | CCG | C | T | C>T | C>T_CCG | CG>TA_GCAAAAG | 6 | FALSE | FALSE |
| 174_v.txt | 32937 | 32937-SNV-C | SNV | 1 | C | T | 104 | 23328 | 0.44581619 | 47.6634615 | rnr1Y285F_pG | TRUE | CAN | JSY2420 | 5 | rnr1Y285F_p | CCA | C>T | A/T+1 | G/C-1 | CG>TA | CG>TA | ATTGCCCAA | CAAGAAA | GGT | CCA | C | A | C>T | C>T_CCA | CG>TA_CAAGAAA | 6 | FALSE | FALSE |

| | | | | | | | | | | | | | | | | | | | | | | | | | | | | | | | | | | |
|---|---|---|---|---|---|---|---|---|---|---|---|---|---|---|---|---|---|---|---|---|---|---|---|---|---|---|---|---|---|---|---|---|---|---|
| 174_v.txt | 32945 | 32945-SNV- | SNV | 1 | C | T | 107 | 23328 | 0.45867627 | 56.5327103 | rnr1Y285F_pG | TRUE | CAN | JSY2420 | 5 | rnr1Y285F_p | ACC | C>T | A/T+1 | G/C-1 | CG>TA | CG>TA | AGAAAACCA | CCAATAC | TGG | ACC | C | A | C>T | C>T_ACC | CG>TA_CCAATAC | 6 | FALSE | FALSE |
| 174_v.txt | 32958 | 32958-SNV- | SNV | 1 | C | T | 168 | 23328 | 0.72016461 | 56.0654762 | rnr1Y285F_pG | TRUE | CAN | JSY2420 | 5 | rnr1Y285F_p | ACC | C>T | A/T+1 | G/C-1 | CG>TA | CG>TA | ATGTAACCA | CCATTGG | TGG | ACC | C | A | C>T | C>T_ACC | CG>TA_CCATTGG | 6 | FALSE | FALSE |
| 174_v.txt | 32967 | 32967-SNV- | SNV | 1 | G | A | 60 | 23328 | 0.25720165 | 57.8333333 | rnr1Y285F_pG | TRUE | CAN | JSY2420 | 5 | rnr1Y285F_p | CGC | G>A | A/T+1 | G/C-1 | CG>TA | CG>TA | TTGGCCGCA | GCACCAA | GCG | GCG | C | T | C>T | C>T_GCG | CG>TA_GCACCAA | 6 | FALSE | FALSE |
| 174_v.txt | 32970 | 32970-SNV- | SNV | 1 | C | T | 30 | 23328 | 0.12860082 | 63.2333333 | rnr1Y285F_pG | TRUE | CAN | JSY2420 | 5 | rnr1Y285F_p | ACC | C>T | A/T+1 | G/C-1 | CG>TA | CG>TA | GCCGCACCA | CCAAATG | TGG | ACC | C | A | C>T | C>T_ACC | CG>TA_CCAAATG | 6 | FALSE | FALSE |
| 174_v.txt | 33015 | 33015-SNV- | SNV | 1 | G | A | 22 | 23328 | 0.09430727 | 59.8636364 | rnr1Y285F_pG | TRUE | CAN | JSY2420 | 5 | rnr1Y285F_p | GGA | G>A | A/T+1 | G/C-1 | CG>TA | CG>TA | AAAGAGGAT | GATGTAA | CCT | CCT | C | T | C>T | C>T_CCT | CG>TA_GATGTAA | 6 | FALSE | FALSE |
| 174_v.txt | 33056 | 33056-SNV- | SNV | 1 | C | T | 26 | 23328 | 0.11145405 | 57.8042599 | rnr1Y285F_pG | TRUE | CAN | JSY2420 | 5 | rnr1Y285F_p | GCG | C>T | A/T+1 | G/C-1 | CG>TA | CG>TA | GGACTGCGT | CGTGACA | CGC | GCG | C | A | C>T | C>T_GCG | CG>TA_CGTGACA | 6 | FALSE | FALSE |
| 174_v.txt | 33057 | 33057-SNV- | SNV | 1 | G | A | 12 | 23328 | 0.05144033 | 52.0827415 | rnr1Y285F_pG | TRUE | CAN | JSY2420 | 5 | rnr1Y285F_p | CGT | G>A | A/T+1 | G/C-1 | CG>TA | CG>TA | GACTGCGTG | GTGACAG | GCA | GCA | C | T | C>T | C>T_GCA | CG>TA_GTGACAG | 6 | FALSE | FALSE |
| 174_v.txt | 33070 | 33070-SNV- | SNV | 1 | C | T | 28 | 23328 | 0.12002744 | 58.7142857 | rnr1Y285F_pG | TRUE | CAN | JSY2420 | 5 | rnr1Y285F_p | GCC | C>T | A/T+1 | G/C-1 | CG>TA | CG>TA | AATATGCCA | CCAAAGA | CGG | GCC | C | A | C>T | C>T_GCC | CG>TA_CCAAAGA | 6 | FALSE | FALSE |
| 174_v.txt | 33075 | 33075-SNV- | SNV | 1 | G | A | 26 | 23328 | 0.11145405 | 47.4230769 | rnr1Y285F_pG | TRUE | CAN | JSY2420 | 5 | rnr1Y285F_p | AGA | G>A | A/T+1 | G/C-1 | CG>TA | CG>TA | GCCAAAGAA | GAACCCA | TCT | TCT | C | T | C>T | C>T_TCT | CG>TA_GAACCCA | 6 | FALSE | FALSE |
| 174_v.txt | 33078 | 33078-SNV- | SNV | 1 | C | T | 5 | 23328 | 0.02143347 | 52.7993867 | rnr1Y285F_pG | TRUE | CAN | JSY2420 | 5 | rnr1Y285F_p | ACC | C>T | A/T+1 | G/C-1 | CG>TA | CG>TA | AAAGAACCC | CCCATAA | TGG | ACC | C | A | C>T | C>T_ACC | CG>TA_CCCATAA | 6 | FALSE | FALSE |
| 174_v.txt | 33111 | 33111-SNV- | SNV | 1 | G | A | 35 | 23328 | 0.15003429 | 38.4857143 | rnr1Y285F_pG | TRUE | CAN | JSY2420 | 5 | rnr1Y285F_p | TGG | G>A | A/T+1 | G/C-1 | CG>TA | CG>TA | CCCACTGGG | GGGCCGG | ACC | ACC | C | T | C>T | C>T_ACC | CG>TA_GGGCCGG | 6 | FALSE | FALSE |
| 174_v.txt | 33114 | 33114-SNV- | SNV | 1 | C | T | 19 | 23328 | 0.08144719 | 42.3684211 | rnr1Y285F_pG | TRUE | CAN | JSY2420 | 5 | rnr1Y285F_p | GCC | C>T | A/T+1 | G/C-1 | CG>TA | CG>TA | ACTGGGCCG | CCGGCGT | CGG | GCC | C | A | C>T | C>T_GCC | CG>TA_CCGGCGT | 6 | FALSE | FALSE |
| 174_v.txt | 33119 | 33119-SNV- | SNV | 1 | G | A | 32 | 23328 | 0.13717421 | 38.9375 | rnr1Y285F_pG | TRUE | CAN | JSY2420 | 5 | rnr1Y285F_p | CGT | G>A | A/T+1 | G/C-1 | CG>TA | CG>TA | GCCGGCGTT | GTTGGTC | GCA | GCA | C | T | C>T | C>T_GCA | CG>TA_GTTGGTC | 6 | FALSE | FALSE |
| 174_v.txt | 33153 | 33153-SNV- | SNV | 1 | C | T | 264 | 23327 | 1.13173576 | 36.7804088 | rnr1Y285F_pG | TRUE | CAN | JSY2420 | 5 | rnr1Y285F_p | ACC | C>T | A/T+1 | G/C-1 | CG>TA | CG>TA | AAAAGACCT | CCTGTAC | TGG | ACC | C | A | C>T | C>T_ACC | CG>TA_CCTGTAC | 6 | FALSE | FALSE |
| 174_v.txt | 33154 | 33154-SNV- | SNV | 1 | C | T | 17 | 23327 | 0.07287692 | 36.3529412 | rnr1Y285F_pG | TRUE | CAN | JSY2420 | 5 | rnr1Y285F_p | CCT | C>T | A/T+1 | G/C-1 | CG>TA | CG>TA | AAAGACCTG | CTGTACC | GGA | CCT | C | A | C>T | C>T_CCT | CG>TA_CTGTACC | 6 | FALSE | FALSE |
| 174_v.txt | 33156 | 33156-SNV- | SNV | 1 | G | A | 21 | 23327 | 0.09002444 | 35.952381 | rnr1Y285F_pG | TRUE | CAN | JSY2420 | 5 | rnr1Y285F_p | TGT | G>A | A/T+1 | G/C-1 | CG>TA | CG>TA | AGACCTGTA | GTACCAA | ACA | ACA | C | T | C>T | C>T_ACA | CG>TA_GTACCAA | 6 | FALSE | FALSE |
| 174_v.txt | 33159 | 33159-SNV- | SNV | 1 | C | T | 48 | 23326 | 0.20577896 | 37 | rnr1Y285F_pG | TRUE | CAN | JSY2420 | 5 | rnr1Y285F_p | ACC | C>T | A/T+1 | G/C-1 | CG>TA | CG>TA | CCTGTACCA | CCAATAG | TGG | ACC | C | A | C>T | C>T_ACC | CG>TA_CCAATAG | 6 | FALSE | FALSE |
| 174_v.txt | 33168 | 33168-SNV- | SNV | 1 | C | T | 168 | 23325 | 0.72025724 | 37.2321786 | rnr1Y285F_pG | TRUE | CAN | JSY2420 | 5 | rnr1Y285F_p | ACC | C>T | A/T+1 | G/C-1 | CG>TA | CG>TA | ATAGTACCA | CCACCAA | TGG | ACC | C | A | C>T | C>T_ACC | CG>TA_CCACCAA | 6 | FALSE | FALSE |
| 174_v.txt | 33171 | 33171-SNV- | SNV | 1 | C | T | 203 | 23325 | 0.87031083 | 37.408867 | rnr1Y285F_pG | TRUE | CAN | JSY2420 | 5 | rnr1Y285F_p | ACC | C>T | A/T+1 | G/C-1 | CG>TA | CG>TA | GTACCACCA | CCAAGGG | TGG | ACC | C | A | C>T | C>T_ACC | CG>TA_CCAAGGG | 6 | FALSE | FALSE |
| 174_v.txt | 33177 | 33177-SNV- | SNV | 1 | G | A | 14 | 23323 | 0.06002658 | 37.8571429 | rnr1Y285F_pG | TRUE | CAN | JSY2420 | 5 | rnr1Y285F_p | GGC | G>A | A/T+1 | G/C-1 | CG>TA | CG>TA | CCAAGGGCA | GCAATCA | CCG | CCG | C | T | C>T | C>T_CCG | CG>TA_GCAATCA | 6 | FALSE | FALSE |
| 174_v.txt | 33199 | 33199-SNV- | SNV | 1 | G | A | 22 | 12689 | 0.17337852 | 36.6818182 | rnr1Y285F_pG | TRUE | CAN | JSY2420 | 5 | rnr1Y285F_p | TGC | G>A | A/T+1 | G/C-1 | CG>TA | CG>TA | GTCTTTGCTT | GCTTAAG | ACG | ACG | C | T | C>T | C>T_ACG | CG>TA_GCTTAAG | 6 | FALSE | FALSE |
| 174_v.txt | 33205 | 33205-SNV- | SNV | 1 | G | A | 8 | 12697 | 0.06300701 | 38.6239045 | rnr1Y285F_pG | TRUE | CAN | JSY2420 | 5 | rnr1Y285F_p | AGC | G>A | A/T+1 | G/C-1 | CG>TA | CG>TA | GCTTAAGCT | GCTCTCT | TCG | TCG | C | T | C>T | C>T_TCG | CG>TA_GCTCTCT | 6 | FALSE | FALSE |
| 174_v.txt | 33258 | 33258-SNV- | SNV | 1 | C | T | 24 | 12706 | 0.18888714 | 43.7083333 | rnr1Y285F_pG | TRUE | CAN | JSY2420 | 5 | rnr1Y285F_p | GCC | C>T | A/T+1 | G/C-1 | CG>TA | CG>TA | CCTATGCCAT | CCATCCT | CGG | GCC | C | A | C>T | C>T_GCC | CG>TA_CCATCCT | 6 | FALSE | FALSE |
| 174_v.txt | 33276 | 33276-SNV- | SNV | 1 | G | A | 14 | 12706 | 0.11018417 | 50.7857143 | rnr1Y285F_pG | TRUE | CAN | JSY2420 | 5 | rnr1Y285F_p | CGT | G>A | A/T+1 | G/C-1 | CG>TA | CG>TA | GAGAACGTA | GTATCCT | GCA | GCA | C | T | C>T | C>T_GCA | CG>TA_GTATCCT | 6 | FALSE | FALSE |
| 174_v.txt | 33310 | 33310-SNV- | SNV | 1 | G | A | 20 | 12706 | 0.15740595 | 60.1 | rnr1Y285F_pG | TRUE | CAN | JSY2420 | 5 | rnr1Y285F_p | CGC | G>A | A/T+1 | G/C-1 | CG>TA | CG>TA | AAGAGCGCA | GCAATGG | GCG | GCG | C | T | C>T | C>T_GCG | CG>TA_GCAATGG | 6 | FALSE | FALSE |
| 174_v.txt | 33412 | 33412-SNV- | SNV | 1 | G | A | 19 | 12706 | 0.14953565 | 44.4210526 | rnr1Y285F_pG | TRUE | CAN | JSY2420 | 5 | rnr1Y285F_p | GGC | G>A | A/T+1 | G/C-1 | CG>TA | CG>TA | TGACCGGCT | GCTCATT | CCG | CCG | C | T | C>T | C>T_CCG | CG>TA_GCTCATT | 6 | FALSE | FALSE |
| 174_v.txt | 33474 | 33474-SNV- | SNV | 1 | C | T | 18 | 12705 | 0.14167651 | 33.8888889 | rnr1Y285F_pG | TRUE | CAN | JSY2420 | 5 | rnr1Y285F_p | GCC | C>T | A/T+1 | G/C-1 | CG>TA | CG>TA | GCTATGCCTT | CCTTTTT | CGG | GCC | C | A | C>T | C>T_GCC | CG>TA_CCTTTTT | 6 | FALSE | FALSE |
| 174_v.txt | 31947 | 31947-SNV- | SNV | 1 | G | C | 5 | 11583 | 0.04316671 | 38 | rnr1Y285F_pG | TRUE | CAN | JSY2420 | 5 | rnr1Y285F_p | CGT | G>C | G/C+1 | G/C-1 | CG>GC | CG>GC | ATGATCGTC | GTCATAA | GCA | GCA | C | G | C>G | C>G_GCA | CG>GC_GTCATAA | 6 | FALSE | FALSE |
| 174_v.txt | 31981 | 31981-SNV- | SNV | 1 | G | C | 9 | 11593 | 0.07763305 | 40.8888889 | rnr1Y285F_pG | TRUE | CAN | JSY2420 | 5 | rnr1Y285F_p | GGC | G>C | G/C+1 | G/C-1 | CG>GC | CG>GC | AGCCGGGCA | GCATTAA | CCG | CCG | C | G | C>G | C>G_CCG | CG>GC_GCATTAA | 6 | FALSE | FALSE |
| 174_v.txt | 32027 | 32027-SNV- | SNV | 1 | G | C | 2 | 11591 | 0.01725477 | 45.5003745 | rnr1Y285F_pG | TRUE | CAN | JSY2420 | 5 | rnr1Y285F_p | GGT | G>C | G/C+1 | G/C-1 | CG>GC | CG>GC | GCCACGGTA | GTATTTC | CCA | CCA | C | G | C>G | C>G_CCA | CG>GC_GTATTTC | 6 | FALSE | FALSE |
| 174_v.txt | 32038 | 32038-SNV- | SNV | 1 | C | G | 31 | 11593 | 0.26740274 | 57.0967742 | rnr1Y285F_pG | TRUE | CAN | JSY2420 | 5 | rnr1Y285F_p | GCT | C>G | G/C+1 | G/C-1 | CG>GC | CG>GC | TCAAAGCTT | CTTGCAT | CGA | GCT | C | C | C>G | C>G_GCT | CG>GC_CTTGCAT | 6 | FALSE | FALSE |
| 174_v.txt | 32064 | 32064-SNV- | SNV | 1 | G | C | 7 | 11593 | 0.06038127 | 54.5714286 | rnr1Y285F_pG | TRUE | CAN | JSY2420 | 5 | rnr1Y285F_p | TGA | G>C | G/C+1 | G/C-1 | CG>GC | CG>GC | GAGATTGAG | GAGATAA | ACT | ACT | C | G | C>G | C>G_ACT | CG>GC_GAGATAA | 6 | FALSE | FALSE |
| 174_v.txt | 32092 | 32092-SNV- | SNV | 1 | C | G | 5 | 11592 | 0.0431332 | 63.992479 | rnr1Y285F_pG | TRUE | CAN | JSY2420 | 5 | rnr1Y285F_p | GCA | C>G | G/C+1 | G/C-1 | CG>GC | CG>GC | AGCCTGCAA | CAACACC | CGT | GCA | C | C | C>G | C>G_GCA | CG>GC_CAACACC | 6 | FALSE | FALSE |
| 174_v.txt | 32176 | 32176-SNV- | SNV | 1 | C | G | 1 | 11593 | 8.63E-03 | 38.0051272 | rnr1Y285F_pG | TRUE | CAN | JSY2420 | 5 | rnr1Y285F_p | GCA | C>G | G/C+1 | G/C-1 | CG>GC | CG>GC | ATGCAGCAG | CAGTAAC | CGT | GCA | C | C | C>G | C>G_GCA | CG>GC_CAGTAAC | 6 | FALSE | FALSE |
| 174_v.txt | 32271 | 32271-SNV- | SNV | 1 | C | G | 145 | 21310 | 0.68043172 | 35.937931 | rnr1Y285F_pG | TRUE | CAN | JSY2420 | 5 | rnr1Y285F_p | ACG | C>G | G/C+1 | G/C-1 | CG>GC | CG>GC | AAAATACGG | CGGGAAC | TGC | ACG | C | C | C>G | C>G_ACG | CG>GC_CGGGAAC | 6 | FALSE | FALSE |
| 174_v.txt | 32272 | 32272-SNV- | SNV | 1 | G | C | 12 | 21310 | 0.05631159 | 37.4111702 | rnr1Y285F_pG | TRUE | CAN | JSY2420 | 5 | rnr1Y285F_p | CGG | G>C | G/C+1 | G/C-1 | CG>GC | CG>GC | AAATACGGG | GGGAACC | GCC | GCC | C | G | C>G | C>G_GCC | CG>GC_GGGAACC | 6 | FALSE | FALSE |
| 174_v.txt | 32292 | 32292-SNV- | SNV | 1 | G | C | 37 | 21310 | 0.17362741 | 52.8378378 | rnr1Y285F_pG | TRUE | CAN | JSY2420 | 5 | rnr1Y285F_p | TGA | G>C | G/C+1 | G/C-1 | CG>GC | CG>GC | ATATTTGAAT | GAATTTG | ACT | ACT | C | G | C>G | C>G_ACT | CG>GC_GAATTTG | 6 | FALSE | FALSE |
| 174_v.txt | 32301 | 32301-SNV- | SNV | 1 | G | C | 6 | 21310 | 0.0281558 | 50 | rnr1Y285F_pG | TRUE | CAN | JSY2420 | 5 | rnr1Y285F_p | GGC | G>C | G/C+1 | G/C-1 | CG>GC | CG>GC | TTTGCGGCA | GCAGAAA | CCG | CCG | C | G | C>G | C>G_CCG | CG>GC_GCAGAAA | 6 | FALSE | FALSE |
| 174_v.txt | 32316 | 32316-SNV- | SNV | 1 | G | C | 14 | 21310 | 0.06569686 | 61.1428571 | rnr1Y285F_pG | TRUE | CAN | JSY2420 | 5 | rnr1Y285F_p | TGT | G>C | G/C+1 | G/C-1 | CG>GC | CG>GC | ATGGTTGTT | GTTAAGA | ACA | ACA | C | G | C>G | C>G_ACA | CG>GC_GTTAAGA | 6 | FALSE | FALSE |
| 174_v.txt | 32432 | 32432-SNV- | SNV | 1 | G | C | 3 | 21310 | 0.0140779 | 58.6666667 | rnr1Y285F_pG | TRUE | CAN | JSY2420 | 5 | rnr1Y285F_p | TGT | G>C | G/C+1 | G/C-1 | CG>GC | CG>GC | GTCATTGTAT | GTATGGA | ACA | ACA | C | G | C>G | C>G_ACA | CG>GC_GTATGGA | 6 | FALSE | FALSE |
| 174_v.txt | 32474 | 32474-SNV- | SNV | 1 | G | C | 64 | 21310 | 0.30032848 | 47.75 | rnr1Y285F_pG | TRUE | CAN | JSY2420 | 5 | rnr1Y285F_p | AGA | G>C | G/C+1 | G/C-1 | CG>GC | CG>GC | AATGTAGAA | GAAGGTT | TCT | TCT | C | G | C>G | C>G_TCT | CG>GC_GAAGGTT | 6 | FALSE | FALSE |
| 174_v.txt | 32487 | 32487-SNV- | SNV | 1 | C | G | 9 | 21310 | 0.04223369 | 49.4444444 | rnr1Y285F_pG | TRUE | CAN | JSY2420 | 5 | rnr1Y285F_p | ACG | C>G | G/C+1 | G/C-1 | CG>GC | CG>GC | AAGATACGG | CGGAAAA | TGC | ACG | C | C | C>G | C>G_ACG | CG>GC_CGGAAAA | 6 | FALSE | FALSE |
| 174_v.txt | 32538 | 32538-SNV- | SNV | 1 | G | C | 6 | 21309 | 0.02815712 | 37.996809 | rnr1Y285F_pG | TRUE | CAN | JSY2420 | 5 | rnr1Y285F_p | AGC | G>C | G/C+1 | G/C-1 | CG>GC | CG>GC | TTTGCAGCTT | GCTTCAC | TCG | TCG | C | G | C>G | C>G_TCG | CG>GC_GCTTCAC | 6 | FALSE | FALSE |
| 174_v.txt | 32539 | 32539-SNV- | SNV | 1 | C | G | 32 | 21309 | 0.15017129 | 36.342454 | rnr1Y285F_pG | TRUE | CAN | JSY2420 | 5 | rnr1Y285F_p | GCT | C>G | G/C+1 | G/C-1 | CG>GC | CG>GC | TTGCAGCTTC | CTTCACC | CGA | GCT | C | C | C>G | C>G_GCT | CG>GC_CTTCACC | 6 | FALSE | FALSE |
| 174_v.txt | 32557 | 32557-SNV- | SNV | 1 | C | G | 17 | 21302 | 0.07980471 | 37.6463366 | rnr1Y285F_pG | TRUE | CAN | JSY2420 | 5 | rnr1Y285F_p | CCA | C>G | G/C+1 | G/C-1 | CG>GC | CG>GC | TGATACCAA | CAACTAG | GGT | CCA | C | C | C>G | C>G_CCA | CG>GC_CAACTAG | 6 | FALSE | FALSE |
| 174_v.txt | 32566 | 32566-SNV- | SNV | 1 | C | G | 54 | 22585 | 0.23909675 | 37.0185185 | rnr1Y285F_pG | TRUE | CAN | JSY2420 | 5 | rnr1Y285F_p | TCA | C>G | G/C+1 | G/C-1 | CG>GC | CG>GC | CTAGTTCAG | CAGTACC | AGT | TCA | C | C | C>G | C>G_TCA | CG>GC_CAGTACC | 6 | FALSE | FALSE |
| 174_v.txt | 32572 | 32572-SNV- | SNV | 1 | C | G | 28 | 22591 | 0.12394316 | 37.3233104 | rnr1Y285F_pG | TRUE | CAN | JSY2420 | 5 | rnr1Y285F_p | CCT | C>G | G/C+1 | G/C-1 | CG>GC | CG>GC | CAGTACCTT | CTTGAAA | GGA | CCT | C | C | C>G | C>G_CCT | CG>GC_CTTGAAA | 6 | FALSE | FALSE |
| 174_v.txt | 32580 | 32580-SNV- | SNV | 1 | G | C | 57 | 22591 | 0.25231287 | 37.6666667 | rnr1Y285F_pG | TRUE | CAN | JSY2420 | 5 | rnr1Y285F_p | TGT | G>C | G/C+1 | G/C-1 | CG>GC | CG>GC | TGAAATGTG | GTGAAGG | ACA | ACA | C | G | C>G | C>G_ACA | CG>GC_GTGAAGG | 6 | FALSE | FALSE |
| 174_v.txt | 32710 | 32710-SNV- | SNV | 1 | C | G | 4 | 22590 | 0.01770695 | 63.9979898 | rnr1Y285F_pG | TRUE | CAN | JSY2420 | 5 | rnr1Y285F_p | CCA | C>G | G/C+1 | G/C-1 | CG>GC | CG>GC | TAACCCCAG | CAGCACC | GGT | CCA | C | C | C>G | C>G_CCA | CG>GC_CAGCACC | 6 | FALSE | FALSE |
| 174_v.txt | 32735 | 32735-SNV- | SNV | 1 | G | C | 22 | 22590 | 0.09738823 | 59.8956224 | rnr1Y285F_pG | TRUE | CAN | JSY2420 | 5 | rnr1Y285F_p | AGT | G>C | G/C+1 | G/C-1 | CG>GC | CG>GC | AAAACAGTA | GTATATT | TCA | TCA | C | G | C>G | C>G_TCA | CG>GC_GTATATT | 6 | FALSE | FALSE |
| 174_v.txt | 32776 | 32776-SNV- | SNV | 1 | C | G | 99 | 22590 | 0.43824701 | 50.5151515 | rnr1Y285F_pG | TRUE | CAN | JSY2420 | 5 | rnr1Y285F_p | GCG | C>G | G/C+1 | G/C-1 | CG>GC | CG>GC | TGGAAGCGA | CGACCCA | CGC | GCG | C | C | C>G | C>G_GCG | CG>GC_CGACCCA | 6 | FALSE | FALSE |
| 174_v.txt | 32786 | 32786-SNV- | SNV | 1 | C | G | 5 | 22590 | 0.02213369 | 32.6 | rnr1Y285F_pG | TRUE | CAN | JSY2420 | 5 | rnr1Y285F_p | ACT | C>G | G/C+1 | G/C-1 | CG>GC | CG>GC | CCAGAACTC | CTCGAAT | TGA | ACT | C | C | C>G | C>G_ACT | CG>GC_CTCGAAT | 6 | FALSE | FALSE |
| 174_v.txt | 32797 | 32797-SNV- | SNV | 1 | C | G | 23 | 22590 | 0.10181496 | 46.3919138 | rnr1Y285F_pG | TRUE | CAN | JSY2420 | 5 | rnr1Y285F_p | CCG | C>G | G/C+1 | G/C-1 | CG>GC | CG>GC | ATTCACCGT | CGTAATA | GGC | CCG | C | C | C>G | C>G_CCG | CG>GC_CGTAATA | 6 | FALSE | FALSE |
| 174_v.txt | 32798 | 32798-SNV- | SNV | 1 | G | C | 6 | 22590 | 0.02656043 | 51 | rnr1Y285F_pG | TRUE | CAN | JSY2420 | 5 | rnr1Y285F_p | CGT | G>C | G/C+1 | G/C-1 | CG>GC | CG>GC | TTCACCGTA | GTAATAT | GCA | GCA | C | G | C>G | C>G_GCA | CG>GC_GTAATAT | 6 | FALSE | FALSE |
| 174_v.txt | 32819 | 32819-SNV- | SNV | 1 | G | C | 76 | 22590 | 0.33643205 | 43.1842105 | rnr1Y285F_pG | TRUE | CAN | JSY2420 | 5 | rnr1Y285F_p | AGT | G>C | G/C+1 | G/C-1 | CG>GC | CG>GC | GAACAAGTT | GTTCATT | TCA | TCA | C | G | C>G | C>G_TCA | CG>GC_GTTCATT | 6 | FALSE | FALSE |
| 174_v.txt | 32876 | 32876-SNV- | SNV | 1 | G | C | 33 | 23316 | 0.14153371 | 36.7878788 | rnr1Y285F_pG | TRUE | CAN | JSY2420 | 5 | rnr1Y285F_p | TGT | G>C | G/C+1 | G/C-1 | CG>GC | CG>GC | AACTTTGTA | GTACGTC | ACA | ACA | C | G | C>G | C>G_ACA | CG>GC_GTACGTC | 6 | FALSE | FALSE |
| 174_v.txt | 32917 | 32917-SNV- | SNV | 1 | C | G | 35 | 23328 | 0.15003429 | 48.4 | rnr1Y285F_pG | TRUE | CAN | JSY2420 | 5 | rnr1Y285F_p | TCC | C>G | G/C+1 | G/C-1 | CG>GC | CG>GC | TAAGTTCCA | CCAGGGC | AGG | TCC | C | C | C>G | C>G_TCC | CG>GC_CCAGGGC | 6 | FALSE | FALSE |
| 174_v.txt | 32940 | 32940-SNV- | SNV | 1 | G | C | 9 | 23328 | 0.03858025 | 54.1111111 | rnr1Y285F_pG | TRUE | CAN | JSY2420 | 5 | rnr1Y285F_p | AGA | G>C | G/C+1 | G/C-1 | CG>GC | CG>GC | GCCCAAGAA | GAAAACC | TCT | TCT | C | G | C>G | C>G_TCT | CG>GC_GAAAACC | 6 | FALSE | FALSE |
| 174_v.txt | 32967 | 32967-SNV- | SNV | 1 | G | C | 5 | 23328 | 0.02143347 | 41 | rnr1Y285F_pG | TRUE | CAN | JSY2420 | 5 | rnr1Y285F_p | CGC | G>C | G/C+1 | G/C-1 | CG>GC | CG>GC | TTGGCCGCA | GCACCAA | GCG | GCG | C | G | C>G | C>G_GCG | CG>GC_GCACCAA | 6 | FALSE | FALSE |
| 174_v.txt | 32997 | 32997-SNV- | SNV | 1 | G | C | 20 | 23328 | 0.08573388 | 60.45 | rnr1Y285F_pG | TRUE | CAN | JSY2420 | 5 | rnr1Y285F_p | TGA | G>C | G/C+1 | G/C-1 | CG>GC | CG>GC | CTTTGTGAG | GAGAAAA | ACT | ACT | C | G | C>G | C>G_ACT | CG>GC_GAGAAAA | 6 | FALSE | FALSE |
| 174_v.txt | 33037 | 33037-SNV- | SNV | 1 | C | G | 75 | 23328 | 0.32150206 | 59.88 | rnr1Y285F_pG | TRUE | CAN | JSY2420 | 5 | rnr1Y285F_p | GCC | C>G | G/C+1 | G/C-1 | CG>GC | CG>GC | ATGTAGCCA | CCATTTC | CGG | GCC | C | C | C>G | C>G_GCC | CG>GC_CCATTTC | 6 | FALSE | FALSE |
| 174_v.txt | 33057 | 33057-SNV- | SNV | 1 | G | C | 35 | 23328 | 0.15003429 | 55.5129104 | rnr1Y285F_pG | TRUE | CAN | JSY2420 | 5 | rnr1Y285F_p | CGT | G>C | G/C+1 | G/C-1 | CG>GC | CG>GC | GACTGCGTG | GTGACAG | GCA | GCA | C | G | C>G | C>G_GCA | CG>GC_GTGACAG | 6 | FALSE | FALSE |
| 174_v.txt | 33070 | 33070-SNV- | SNV | 1 | C | G | 30 | 23328 | 0.12860082 | 55.3333333 | rnr1Y285F_pG | TRUE | CAN | JSY2420 | 5 | rnr1Y285F_p | GCC | C>G | G/C+1 | G/C-1 | CG>GC | CG>GC | AATATGCCA | CCAAAGA | CGG | GCC | C | C | C>G | C>G_GCC | CG>GC_CCAAAGA | 6 | FALSE | FALSE |
| 174_v.txt | 33079 | 33079-SNV- | SNV | 1 | C | G | 17 | 23328 | 0.0728738 | 53.3537256 | rnr1Y285F_pG | TRUE | CAN | JSY2420 | 5 | rnr1Y285F_p | CCC | C>G | G/C+1 | G/C-1 | CG>GC | CG>GC | AAGAACCCA | CCATAAA | GGG | CCC | C | C | C>G | C>G_CCC | CG>GC_CCATAAA | 6 | FALSE | FALSE |
| 174_v.txt | 33093 | 33093-SNV- | SNV | 1 | G | C | 6 | 23328 | 0.02572017 | 45.5 | rnr1Y285F_pG | TRUE | CAN | JSY2420 | 5 | rnr1Y285F_p | TGA | G>C | G/C+1 | G/C-1 | CG>GC | CG>GC | AAATATGAT | GATATAA | ACT | ACT | C | G | C>G | C>G_ACT | CG>GC_GATATAA | 6 | FALSE | FALSE |
| 174_v.txt | 33111 | 33111-SNV- | SNV | 1 | G | C | 106 | 23328 | 0.45438958 | 42.4811321 | rnr1Y285F_pG | TRUE | CAN | JSY2420 | 5 | rnr1Y285F_p | TGG | G>C | G/C+1 | G/C-1 | CG>GC | CG>GC | CCCACTGGG | GGGCCGG | ACC | ACC | C | G | C>G | C>G_ACC | CG>GC_GGGCCGG | 6 | FALSE | FALSE |
| 174_v.txt | 33115 | 33115-SNV- | SNV | 1 | C | G | 14 | 23328 | 0.06001372 | 43.2854241 | rnr1Y285F_pG | TRUE | CAN | JSY2420 | 5 | rnr1Y285F_p | CCG | C>G | G/C+1 | G/C-1 | CG>GC | CG>GC | CTGGGCCGG | CGGCGTT | GGC | CCG | C | C | C>G | C>G_CCG | CG>GC_CGGCGTT | 6 | FALSE | FALSE |
| 174_v.txt | 33151 | 33151-SNV- | SNV | 1 | G | C | 13 | 23328 | 0.05572702 | 38.7692308 | rnr1Y285F_pG | TRUE | CAN | JSY2420 | 5 | rnr1Y285F_p | AGA | G>C | G/C+1 | G/C-1 | CG>GC | CG>GC | TGAAAAGAC | GACCTGT | TCT | TCT | C | G | C>G | C>G_TCT | CG>GC_GACCTGT | 6 | FALSE | FALSE |
| 174_v.txt | 33154 | 33154-SNV- | SNV | 1 | C | G | 10 | 23327 | 0.04286878 | 37.9 | rnr1Y285F_pG | TRUE | CAN | JSY2420 | 5 | rnr1Y285F_p | CCT | C>G | G/C+1 | G/C-1 | CG>GC | CG>GC | AAAGACCTG | CTGTACC | GGA | CCT | C | C | C>G | C>G_CCT | CG>GC_CTGTACC | 6 | FALSE | FALSE |
| 174_v.txt | 33156 | 33156-SNV- | SNV | 1 | G | C | 47 | 23327 | 0.20148326 | 37.0638298 | rnr1Y285F_pG | TRUE | CAN | JSY2420 | 5 | rnr1Y285F_p | TGT | G>C | G/C+1 | G/C-1 | CG>GC | CG>GC | AGACCTGTA | GTACCAA | ACA | ACA | C | G | C>G | C>G_ACA | CG>GC_GTACCAA | 6 | FALSE | FALSE |
| 174_v.txt | 33205 | 33205-SNV- | SNV | 1 | G | C | 66 | 12697 | 0.51980783 | 38.7249202 | rnr1Y285F_pG | TRUE | CAN | JSY2420 | 5 | rnr1Y285F_p | AGC | G>C | G/C+1 | G/C-1 | CG>GC | CG>GC | GCTTAAGCT | GCTCTCT | TCG | TCG | C | G | C>G | C>G_TCG | CG>GC_GCTCTCT | 6 | FALSE | FALSE |
| 174_v.txt | 33210 | 33210-SNV- | SNV | 1 | C | G | 4 | 12697 | 0.03150351 | 37.75 | rnr1Y285F_pG | TRUE | CAN | JSY2420 | 5 | rnr1Y285F_p | TCT | C>G | G/C+1 | G/C-1 | CG>GC | CG>GC | AGCTCTCTCT | CTCTTCA | AGA | TCT | C | C | C>G | C>G_TCT | CG>GC_CTCTTCA | 6 | FALSE | FALSE |
| 174_v.txt | 33354 | 33354-SNV- | SNV | 1 | G | C | 2 | 12706 | 0.0157406 | 64 | rnr1Y285F_pG | TRUE | CAN | JSY2420 | 5 | rnr1Y285F_p | TGA | G>C | G/C+1 | G/C-1 | CG>GC | CG>GC | GGTATTGAC | GACCCAC | ACT | ACT | C | G | C>G | C>G_ACT | CG>GC_GACCCAC | 6 | FALSE | FALSE |
| 174_v.txt | 33419 | 33419-SNV- | SNV | 1 | G | C | 40 | 12706 | 0.3148119 | 50.9 | rnr1Y285F_pG | TRUE | CAN | JSY2420 | 5 | rnr1Y285F_p | TGT | G>C | G/C+1 | G/C-1 | CG>GC | CG>GC | CTCATTGTAC | GTACATA | ACA | ACA | C | G | C>G | C>G_ACA | CG>GC_GTACATA | 6 | FALSE | FALSE |
| 174_v.txt | 31911 | 31911-SNV- | SNV | 1 | G | T | 6 | 7006 | 0.08564088 | 38 | rnr1Y285F_pG | TRUE | CAN | JSY2420 | 5 | rnr1Y285F_p | TGC | G>T | A/T+1 | G/C-1 | CG>AT | CG>AT | TTTGGTGCA | GCAAAAG | ACG | ACG | C | A | C>A | C>A_ACG | CG>AT_GCAAAAG | 6 | FALSE | FALSE |
| 174_v.txt | 31930 | 31930-SNV- | SNV | 1 | G | T | 9 | 7006 | 0.12846132 | 37.7777778 | rnr1Y285F_pG | TRUE | CAN | JSY2420 | 5 | rnr1Y285F_p | TGA | G>T | A/T+1 | G/C-1 | CG>AT | CG>AT | AACCTTGAA | GAATAAT | ACT | ACT | C | A | C>A | C>A_ACT | CG>AT_GAATAAT | 6 | FALSE | FALSE |
| 174_v.txt | 32026 | 32026-SNV- | SNV | 1 | G | T | 2 | 11593 | 0.01725179 | 33.4993136 | rnr1Y285F_pG | TRUE | CAN | JSY2420 | 5 | rnr1Y285F_p | CGG | G>T | A/T+1 | G/C-1 | CG>AT | CG>AT | TGCCACGGT | GGTATTT | GCC | GCC | C | A | C>A | C>A_GCC | CG>AT_GGTATTT | 6 | FALSE | FALSE |
| 174_v.txt | 32027 | 32027-SNV- | SNV | 1 | G | T | 58 | 11591 | 0.50038823 | 59.4651178 | rnr1Y285F_pG | TRUE | CAN | JSY2420 | 5 | rnr1Y285F_p | GGT | G>T | A/T+1 | G/C-1 | CG>AT | CG>AT | GCCACGGTA | GTATTTC | CCA | CCA | C | A | C>A | C>A_CCA | CG>AT_GTATTTC | 6 | FALSE | FALSE |
| 174_v.txt | 32064 | 32064-SNV- | SNV | 1 | G | T | 5 | 11593 | 0.04312948 | 64 | rnr1Y285F_pG | TRUE | CAN | JSY2420 | 5 | rnr1Y285F_p | TGA | G>T | A/T+1 | G/C-1 | CG>AT | CG>AT | GAGATTGAG | GAGATAA | ACT | ACT | C | A | C>A | C>A_ACT | CG>AT_GAGATAA | 6 | FALSE | FALSE |
| 174_v.txt | 32088 | 32088-SNV- | SNV | 1 | C | A | 2 | 11593 | 0.01725179 | 20.0061038 | rnr1Y285F_pG | TRUE | CAN | JSY2420 | 5 | rnr1Y285F_p | GCC | C>A | A/T+1 | G/C-1 | CG>AT | CG>AT | AAAAAGCCT | CCTGCAA | CGG | GCC | C | T | C>A | C>A_GCC | CG>AT_CCTGCAA | 6 | FALSE | FALSE |
| 174_v.txt | 32089 | 32089-SNV- | SNV | 1 | C | A | 12 | 11593 | 0.10351074 | 45.25 | rnr1Y285F_pG | TRUE | CAN | JSY2420 | 5 | rnr1Y285F_p | CCT | C>A | A/T+1 | G/C-1 | CG>AT | CG>AT | AAAAGCCTG | CTGCAAC | GGA | CCT | C | T | C>A | C>A_CCT | CG>AT_CTGCAAC | 6 | FALSE | FALSE |
| 174_v.txt | 32188 | 32188-SNV- | SNV | 1 | C | A | 39 | 11593 | 0.3364099 | 39.4358974 | rnr1Y285F_pG | TRUE | CAN | JSY2420 | 5 | rnr1Y285F_p | ACT | C>A | A/T+1 | G/C-1 | CG>AT | CG>AT | CGAAAACTG | CTGCAAT | TGA | ACT | C | T | C>A | C>A_ACT | CG>AT_CTGCAAT | 6 | FALSE | FALSE |
| 174_v.txt | 32195 | 32195-SNV- | SNV | 1 | G | T | 25 | 11593 | 0.21564737 | 39 | rnr1Y285F_pG | TRUE | CAN | JSY2420 | 5 | rnr1Y285F_p | TGT | G>T | A/T+1 | G/C-1 | CG>AT | CG>AT | TGCAATGTA | GTATGGA | ACA | ACA | C | A | C>A | C>A_ACA | CG>AT_GTATGGA | 6 | FALSE | FALSE |
| 174_v.txt | 32205 | 32205-SNV- | SNV | 1 | C | A | 11 | 11593 | 0.09488484 | 34.7272727 | rnr1Y285F_pG | TRUE | CAN | JSY2420 | 5 | rnr1Y285F_p | ACC | C>A | A/T+1 | G/C-1 | CG>AT | CG>AT | GGAACACCA | CCACCTT | TGG | ACC | C | T | C>A | C>A_ACC | CG>AT_CCACCTT | 6 | FALSE | FALSE |
| 174_v.txt | 32223 | 32223-SNV- | SNV | 1 | G | T | 18 | 11593 | 0.15526611 | 39.2222222 | rnr1Y285F_pG | TRUE | CAN | JSY2420 | 5 | rnr1Y285F_p | TGA | G>T | A/T+1 | G/C-1 | CG>AT | CG>AT | GTCCTTGAC | GACAGGA | ACT | ACT | C | A | C>A | C>A_ACT | CG>AT_GACAGGA | 6 | FALSE | FALSE |
| 174_v.txt | 32271 | 32271-SNV- | SNV | 1 | C | A | 7 | 21310 | 0.03284843 | 34.1428571 | rnr1Y285F_pG | TRUE | CAN | JSY2420 | 5 | rnr1Y285F_p | ACG | C>A | A/T+1 | G/C-1 | CG>AT | CG>AT | AAAATACGG | CGGGAAC | TGC | ACG | C | T | C>A | C>A_ACG | CG>AT_CGGGAAC | 6 | FALSE | FALSE |
| 174_v.txt | 32274 | 32274-SNV- | SNV | 1 | G | T | 120 | 21310 | 0.56311591 | 36.65 | rnr1Y285F_pG | TRUE | CAN | JSY2420 | 5 | rnr1Y285F_p | GGA | G>T | A/T+1 | G/C-1 | CG>AT | CG>AT | ATACGGGAA | GAACCAA | CCT | CCT | C | A | C>A | C>A_CCT | CG>AT_GAACCAA | 6 | FALSE | FALSE |
| 174_v.txt | 32277 | 32277-SNV- | SNV | 1 | C | A | 21 | 21310 | 0.09854528 | 31.7096383 | rnr1Y285F_pG | TRUE | CAN | JSY2420 | 5 | rnr1Y285F_p | ACC | C>A | A/T+1 | G/C-1 | CG>AT | CG>AT | CGGGAACCA | CCAACGT | TGG | ACC | C | T | C>A | C>A_ACC | CG>AT_CCAACGT | 6 | FALSE | FALSE |
| 174_v.txt | 32292 | 32292-SNV- | SNV | 1 | G | T | 50 | 21310 | 0.23463163 | 52.32 | rnr1Y285F_pG | TRUE | CAN | JSY2420 | 5 | rnr1Y285F_p | TGA | G>T | A/T+1 | G/C-1 | CG>AT | CG>AT | ATATTTGAA | GAATTTG | ACT | ACT | C | A | C>A | C>A_ACT | CG>AT_GAATTTG | 6 | FALSE | FALSE |
| 174_v.txt | 32304 | 32304-SNV- | SNV | 1 | G | T | 14 | 21310 | 0.06569686 | 48.7142857 | rnr1Y285F_pG | TRUE | CAN | JSY2420 | 5 | rnr1Y285F_p | AGA | G>T | A/T+1 | G/C-1 | CG>AT | CG>AT | GCGGCAGAA | GAAATAA | TCT | TCT | C | A | C>A | C>A_TCT | CG>AT_GAAATAA | 6 | FALSE | FALSE |
| 174_v.txt | 32373 | 32373-SNV- | SNV | 1 | G | T | 21 | 21310 | 0.09854528 | 60.9047619 | rnr1Y285F_pG | TRUE | CAN | JSY2420 | 5 | rnr1Y285F_p | AGC | G>T | A/T+1 | G/C-1 | CG>AT | CG>AT | TCAATAGCA | GCAATAA | TCG | TCG | C | A | C>A | C>A_TCG | CG>AT_GCAATAA | 6 | FALSE | FALSE |
| 174_v.txt | 32399 | 32399-SNV- | SNV | 1 | G | T | 736 | 21310 | 3.45377757 | 59.2649457 | rnr1Y285F_pG | TRUE | CAN | JSY2420 | 5 | rnr1Y285F_p | CGT | G>T | A/T+1 | G/C-1 | CG>AT | CG>AT | AGAAACGTA | GTAGGAA | GCA | GCA | C | A | C>A | C>A_GCA | CG>AT_GTAGGAA | 6 | FALSE | FALSE |
| 174_v.txt | 32432 | 32432-SNV- | SNV | 1 | G | T | 45 | 21310 | 0.21116847 | 58.5111111 | rnr1Y285F_pG | TRUE | CAN | JSY2420 | 5 | rnr1Y285F_p | TGT | G>T | A/T+1 | G/C-1 | CG>AT | CG>AT | GTCATTGTAT | GTATGGA | ACA | ACA | C | A | C>A | C>A_ACA | CG>AT_GTATGGA | 6 | FALSE | FALSE |
| 174_v.txt | 32449 | 32449-SNV- | SNV | 1 | C | A | 23 | 21310 | 0.10793055 | 50.173913 | rnr1Y285F_pG | TRUE | CAN | JSY2420 | 5 | rnr1Y285F_p | CCA | C>A | A/T+1 | G/C-1 | CG>AT | CG>AT | AAAGTCCAA | CAATGAA | GGT | CCA | C | T | C>A | C>A_CCA | CG>AT_CAATGAA | 6 | FALSE | FALSE |
| 174_v.txt | 32463 | 32463-SNV- | SNV | 1 | G | T | 20 | 21310 | 0.09385265 | 53.55 | rnr1Y285F_pG | TRUE | CAN | JSY2420 | 5 | rnr1Y285F_p | AGA | G>T | A/T+1 | G/C-1 | CG>AT | CG>AT | AATAGAGAG | GAGCCAA | TCT | TCT | C | A | C>A | C>A_TCT | CG>AT_GAGCCAA | 6 | FALSE | FALSE |
| 174_v.txt | 32471 | 32471-SNV- | SNV | 1 | G | T | 28 | 21310 | 0.13139371 | 47.3914748 | rnr1Y285F_pG | TRUE | CAN | JSY2420 | 5 | rnr1Y285F_p | TGT | G>T | A/T+1 | G/C-1 | CG>AT | CG>AT | GCCAATGTA | GTAGAAG | ACA | ACA | C | A | C>A | C>A_ACA | CG>AT_GTAGAAG | 6 | FALSE | FALSE |
| 174_v.txt | 32474 | 32474-SNV- | SNV | 1 | G | T | 25 | 21310 | 0.11731581 | 49.48 | rnr1Y285F_pG | TRUE | CAN | JSY2420 | 5 | rnr1Y285F_p | AGA | G>T | A/T+1 | G/C-1 | CG>AT | CG>AT | AATGTAGAA | GAAGGTT | TCT | TCT | C | A | C>A | C>A_TCT | CG>AT_GAAGGTT | 6 | FALSE | FALSE |
| 174_v.txt | 32487 | 32487-SNV- | SNV | 1 | C | A | 13 | 21310 | 0.06100422 | 51.0769231 | rnr1Y285F_pG | TRUE | CAN | JSY2420 | 5 | rnr1Y285F_p | ACG | C>A | A/T+1 | G/C-1 | CG>AT | CG>AT | AAGATACGG | CGGAAAA | TGC | ACG | C | T | C>A | C>A_ACG | CG>AT_CGGAAAA | 6 | FALSE | FALSE |
| 174_v.txt | 32494 | 32494-SNV- | SNV | 1 | C | A | 4 | 21310 | 0.01877053 | 44.500282 | rnr1Y285F_pG | TRUE | CAN | JSY2420 | 5 | rnr1Y285F_p | ACA | C>A | A/T+1 | G/C-1 | CG>AT | CG>AT | GGAAAACAA | CAACTTT | TGT | ACA | C | T | C>A | C>A_ACA | CG>AT_CAACTTT | 6 | FALSE | FALSE |
| 174_v.txt | 32508 | 32508-SNV- | SNV | 1 | G | T | 60 | 21310 | 0.28155795 | 39.5333333 | rnr1Y285F_pG | TRUE | CAN | JSY2420 | 5 | rnr1Y285F_p | GGC | G>T | A/T+1 | G/C-1 | CG>AT | CG>AT | TTGATGGCT | GCTCTTG | CCG | CCG | C | A | C>A | C>A_CCG | CG>AT_GCTCTTG | 6 | FALSE | FALSE |
| 174_v.txt | 32530 | 32530-SNV- | SNV | 1 | G | T | 12 | 21310 | 0.05631159 | 35.6666667 | rnr1Y285F_pG | TRUE | CAN | JSY2420 | 5 | rnr1Y285F_p | GGG | G>T | A/T+1 | G/C-1 | CG>AT | CG>AT | TTCTGGGGT | GGTTTGC | CCC | CCC | C | A | C>A | C>A_CCC | CG>AT_GGTTTGC | 6 | FALSE | FALSE |
| 174_v.txt | 32542 | 32542-SNV- | SNV | 1 | C | A | 46 | 21309 | 0.21587123 | 36.7391304 | rnr1Y285F_pG | TRUE | CAN | JSY2420 | 5 | rnr1Y285F_p | TCA | C>A | A/T+1 | G/C-1 | CG>AT | CG>AT | CAGCTTCAC | CACCAGC | AGT | TCA | C | T | C>A | C>A_TCA | CG>AT_CACCAGC | 6 | FALSE | FALSE |
| 174_v.txt | 32556 | 32556-SNV- | SNV | 1 | C | A | 32 | 21303 | 0.15021359 | 36.5619292 | rnr1Y285F_pG | TRUE | CAN | JSY2420 | 5 | rnr1Y285F_p | ACC | C>A | A/T+1 | G/C-1 | CG>AT | CG>AT | GTGATACCA | CCAACTA | TGG | ACC | C | T | C>A | C>A_ACC | CG>AT_CCAACTA | 6 | FALSE | FALSE |
| 174_v.txt | 32566 | 32566-SNV- | SNV | 1 | C | A | 10 | 22585 | 0.04427718 | 36 | rnr1Y285F_pG | TRUE | CAN | JSY2420 | 5 | rnr1Y285F_p | TCA | C>A | A/T+1 | G/C-1 | CG>AT | CG>AT | CTAGTTCAG | CAGTACC | AGT | TCA | C | T | C>A | C>A_TCA | CG>AT_CAGTACC | 6 | FALSE | FALSE |
| 174_v.txt | 32571 | 32571-SNV- | SNV | 1 | C | A | 27 | 22592 | 0.11951133 | 36.4814712 | rnr1Y285F_pG | TRUE | CAN | JSY2420 | 5 | rnr1Y285F_p | ACC | C>A | A/T+1 | G/C-1 | CG>AT | CG>AT | TCAGTACCTT | CCTTGAA | TGG | ACC | C | T | C>A | C>A_ACC | CG>AT_CCTTGAA | 6 | FALSE | FALSE |
| 174_v.txt | 32572 | 32572-SNV- | SNV | 1 | C | A | 5 | 22591 | 0.02213271 | 32.2000155 | rnr1Y285F_pG | TRUE | CAN | JSY2420 | 5 | rnr1Y285F_p | CCT | C>A | A/T+1 | G/C-1 | CG>AT | CG>AT | CAGTACCTT | CTTGAAA | GGA | CCT | C | T | C>A | C>A_CCT | CG>AT_CTTGAAA | 6 | FALSE | FALSE |
| 174_v.txt | 32575 | 32575-SNV- | SNV | 1 | G | T | 17 | 22591 | 0.07525121 | 36.1764706 | rnr1Y285F_pG | TRUE | CAN | JSY2420 | 5 | rnr1Y285F_p | TGA | G>T | A/T+1 | G/C-1 | CG>AT | CG>AT | TACCTTGAA | GAAATGT | ACT | ACT | C | A | C>A | C>A_ACT | CG>AT_GAAATGT | 6 | FALSE | FALSE |
| 174_v.txt | 32629 | 32629-SNV- | SNV | 1 | C | A | 23 | 22590 | 0.10181496 | 54.1304348 | rnr1Y285F_pG | TRUE | CAN | JSY2420 | 5 | rnr1Y285F_p | TCG | C>A | A/T+1 | G/C-1 | CG>AT | CG>AT | TCCCTTCGTT | CGTTTTT | AGC | TCG | C | T | C>A | C>A_TCG | CG>AT_CGTTTTT | 6 | FALSE | FALSE |
| 174_v.txt | 32698 | 32698-SNV- | SNV | 1 | G | T | 29 | 22590 | 0.12837539 | 14.7929235 | rnr1Y285F_pG | TRUE | CAN | JSY2420 | 5 | rnr1Y285F_p | GGG | G>T | A/T+1 | G/C-1 | CG>AT | CG>AT | CAACTGGGC | GGCCGGT | CCC | CCC | C | A | C>A | C>A_CCC | CG>AT_GGCCGGT | 6 | FALSE | FALSE |
| 174_v.txt | 32702 | 32702-SNV- | SNV | 1 | G | T | 4 | 22590 | 0.01770695 | 54.2499282 | rnr1Y285F_pG | TRUE | CAN | JSY2420 | 5 | rnr1Y285F_p | CGG | G>T | A/T+1 | G/C-1 | CG>AT | CG>AT | TGGGCCGGT | GGTAACC | GCC | GCC | C | A | C>A | C>A_GCC | CG>AT_GGTAACC | 6 | FALSE | FALSE |
| 174_v.txt | 32709 | 32709-SNV- | SNV | 1 | C | A | 7 | 22588 | 0.03098991 | 43.575205 | rnr1Y285F_pG | TRUE | CAN | JSY2420 | 5 | rnr1Y285F_p | CCC | C>A | A/T+1 | G/C-1 | CG>AT | CG>AT | GTAACCCCA | CCAGCAC | GGG | CCC | C | T | C>A | C>A_CCC | CG>AT_CCAGCAC | 6 | FALSE | FALSE |
| 174_v.txt | 32735 | 32735-SNV- | SNV | 1 | G | T | 96 | 22590 | 0.4249668 | 56.4534424 | rnr1Y285F_pG | TRUE | CAN | JSY2420 | 5 | rnr1Y285F_p | AGT | G>T | A/T+1 | G/C-1 | CG>AT | CG>AT | AAAACAGTA | GTATATT | TCA | TCA | C | A | C>A | C>A_TCA | CG>AT_GTATATT | 6 | FALSE | FALSE |
| 174_v.txt | 32783 | 32783-SNV- | SNV | 1 | G | T | 30 | 22590 | 0.13280213 | 49.3 | rnr1Y285F_pG | TRUE | CAN | JSY2420 | 5 | rnr1Y285F_p | AGA | G>T | A/T+1 | G/C-1 | CG>AT | CG>AT | GACCCAGAA | GAACTCG | TCT | TCT | C | A | C>A | C>A_TCT | CG>AT_GAACTCG | 6 | FALSE | FALSE |
| 174_v.txt | 32786 | 32786-SNV- | SNV | 1 | C | A | 30 | 22590 | 0.13280213 | 54.0333333 | rnr1Y285F_pG | TRUE | CAN | JSY2420 | 5 | rnr1Y285F_p | ACT | C>A | A/T+1 | G/C-1 | CG>AT | CG>AT | CCAGAACTC | CTCGAAT | TGA | ACT | C | T | C>A | C>A_ACT | CG>AT_CTCGAAT | 6 | FALSE | FALSE |
| 174_v.txt | 32794 | 32794-SNV- | SNV | 1 | C | A | 169 | 22590 | 0.74811864 | 44.4023669 | rnr1Y285F_pG | TRUE | CAN | JSY2420 | 5 | rnr1Y285F_p | TCA | C>A | A/T+1 | G/C-1 | CG>AT | CG>AT | CGAATTCAC | CACCGTA | AGT | TCA | C | T | C>A | C>A_TCA | CG>AT_CACCGTA | 6 | FALSE | FALSE |
| 174_v.txt | 32796 | 32796-SNV- | SNV | 1 | C | A | 29 | 22590 | 0.12837539 | 47.7931035 | rnr1Y285F_pG | TRUE | CAN | JSY2420 | 5 | rnr1Y285F_p | ACC | C>A | A/T+1 | G/C-1 | CG>AT | CG>AT | AATTCACCG | CCGTAAT | TGG | ACC | C | T | C>A | C>A_ACC | CG>AT_CCGTAAT | 6 | FALSE | FALSE |
| 174_v.txt | 32797 | 32797-SNV- | SNV | 1 | C | A | 10 | 22590 | 0.04426738 | 51.400644 | rnr1Y285F_pG | TRUE | CAN | JSY2420 | 5 | rnr1Y285F_p | CCG | C>A | A/T+1 | G/C-1 | CG>AT | CG>AT | ATTCACCGT | CGTAATA | GGC | CCG | C | T | C>A | C>A_CCG | CG>AT_CGTAATA | 6 | FALSE | FALSE |
| 174_v.txt | 32798 | 32798-SNV- | SNV | 1 | G | T | 33 | 22590 | 0.14608234 | 42.4545455 | rnr1Y285F_pG | TRUE | CAN | JSY2420 | 5 | rnr1Y285F_p | CGT | G>T | A/T+1 | G/C-1 | CG>AT | CG>AT | TTCACCGTA | GTAATAT | GCA | GCA | C | A | C>A | C>A_GCA | CG>AT_GTAATAT | 6 | FALSE | FALSE |
| 174_v.txt | 32809 | 32809-SNV- | SNV | 1 | C | A | 36 | 22590 | 0.15936255 | 45.0833333 | rnr1Y285F_pG | TRUE | CAN | JSY2420 | 5 | rnr1Y285F_p | ACA | C>A | A/T+1 | G/C-1 | CG>AT | CG>AT | ATTTGACAG | CAGGGAA | TGT | ACA | C | T | C>A | C>A_ACA | CG>AT_CAGGGAA | 6 | FALSE | FALSE |

| | | | | | | | | | | | | | | | | | | | | | | | | | | | | | | | | | | |
|---|---|---|---|---|---|---|---|---|---|---|---|---|---|---|---|---|---|---|---|---|---|---|---|---|---|---|---|---|---|---|---|---|---|---|
| 174_v.txt | 32811 | 32811-SNV-( | SNV | 1 | G | T | 12 | 22590 | 0.05312085 | 50.75 | rnr1Y285F_pG | TRUE | CAN | JSY2420 | 5 | rnr1Y285F_p | AGG | G>T | A/T+1 | G/C-1 | CG>AT | CG>AT | TTGACAGGG | GGGAACA | TCC | TCC | C | A | C>A | C>A_TCC | CG>AT_GGGAACA | 6 | FALSE | FALSE |
| 174_v.txt | 32819 | 32819-SNV-( | SNV | 1 | G | T | 111 | 22590 | 0.49136786 | 44.8018018 | rnr1Y285F_pG | TRUE | CAN | JSY2420 | 5 | rnr1Y285F_p | AGT | G>T | A/T+1 | G/C-1 | CG>AT | CG>AT | GAACAAGTT( | GTTCATT | TCA | TCA | C | A | C>A | C>A_TCA | CG>AT_GTTCATT | 6 | FALSE | FALSE |
| 174_v.txt | 32876 | 32876-SNV-( | SNV | 1 | G | T | 27 | 23316 | 0.11580031 | 37.5925926 | rnr1Y285F_pG | TRUE | CAN | JSY2420 | 5 | rnr1Y285F_p | TGT | G>T | A/T+1 | G/C-1 | CG>AT | CG>AT | AACTTTGTA( | GTACGTC | ACA | ACA | C | A | C>A | C>A_ACA | CG>AT_GTACGTC | 6 | FALSE | FALSE |
| 174_v.txt | 32917 | 32917-SNV-( | SNV | 1 | C | A | 213 | 23328 | 0.91306584 | 53.3661972 | rnr1Y285F_pG | TRUE | CAN | JSY2420 | 5 | rnr1Y285F_p | TCC | C>A | A/T+1 | G/C-1 | CG>AT | CG>AT | TAAGTTCCA( | CCAGGGC | AGG | TCC | C | T | C>A | C>A_TCC | CG>AT_CCAGGGC | 6 | FALSE | FALSE |
| 174_v.txt | 32940 | 32940-SNV-( | SNV | 1 | G | T | 114 | 23328 | 0.48868313 | 54.8684211 | rnr1Y285F_pG | TRUE | CAN | JSY2420 | 5 | rnr1Y285F_p | AGA | G>T | A/T+1 | G/C-1 | CG>AT | CG>AT | GCCCAAGAA | GAAAACC | TCT | TCT | C | A | C>A | C>A_TCT | CG>AT_GAAAACC | 6 | FALSE | FALSE |
| 174_v.txt | 33012 | 33012-SNV-( | SNV | 1 | G | T | 41 | 23328 | 0.17575446 | 62.3658537 | rnr1Y285F_pG | TRUE | CAN | JSY2420 | 5 | rnr1Y285F_p | AGA | G>T | A/T+1 | G/C-1 | CG>AT | CG>AT | GTGAAAGAG | GAGGATG | TCT | TCT | C | A | C>A | C>A_TCT | CG>AT_GAGGATG | 6 | FALSE | FALSE |
| 174_v.txt | 33024 | 33024-SNV-( | SNV | 1 | G | T | 14 | 23328 | 0.06001372 | 61.5 | rnr1Y285F_pG | TRUE | CAN | JSY2420 | 5 | rnr1Y285F_p | AGG | G>T | A/T+1 | G/C-1 | CG>AT | CG>AT | GTAACAGGG | GGGATGA | TCC | TCC | C | A | C>A | C>A_TCC | CG>AT_GGGATGA | 6 | FALSE | FALSE |
| 174_v.txt | 33033 | 33033-SNV-( | SNV | 1 | G | T | 34 | 23328 | 0.1457476 | 59.1470588 | rnr1Y285F_pG | TRUE | CAN | JSY2420 | 5 | rnr1Y285F_p | TGT | G>T | A/T+1 | G/C-1 | CG>AT | CG>AT | ATGAATGTA( | GTAGCCA | ACA | ACA | C | A | C>A | C>A_ACA | CG>AT_GTAGCCA | 6 | FALSE | FALSE |
| 174_v.txt | 33043 | 33043-SNV-( | SNV | 1 | C | A | 96 | 23328 | 0.41152263 | 61.46875 | rnr1Y285F_pG | TRUE | CAN | JSY2420 | 5 | rnr1Y285F_p | TCA | C>A | A/T+1 | G/C-1 | CG>AT | CG>AT | CCATTTCACC | CACCCAA | AGT | TCA | C | T | C>A | C>A_TCA | CG>AT_CACCCAA | 6 | FALSE | FALSE |
| 174_v.txt | 33051 | 33051-SNV-( | SNV | 1 | G | T | 14 | 23328 | 0.06001372 | 57.2857143 | rnr1Y285F_pG | TRUE | CAN | JSY2420 | 5 | rnr1Y285F_p | GGA | G>T | A/T+1 | G/C-1 | CG>AT | CG>AT | CCCAAGGAC | GACTGCG | CCT | CCT | C | A | C>A | C>A_CCT | CG>AT_GACTGCG | 6 | FALSE | FALSE |
| 174_v.txt | 33093 | 33093-SNV-( | SNV | 1 | G | T | 66 | 23328 | 0.28292181 | 48.530303 | rnr1Y285F_pG | TRUE | CAN | JSY2420 | 5 | rnr1Y285F_p | TGA | G>T | A/T+1 | G/C-1 | CG>AT | CG>AT | AAATATGAT/ | GATATAA | ACT | ACT | C | A | C>A | C>A_ACT | CG>AT_GATATAA | 6 | FALSE | FALSE |
| 174_v.txt | 33114 | 33114-SNV-( | SNV | 1 | C | A | 98 | 23328 | 0.42009602 | 44.7346939 | rnr1Y285F_pG | TRUE | CAN | JSY2420 | 5 | rnr1Y285F_p | GCC | C>A | A/T+1 | G/C-1 | CG>AT | CG>AT | ACTGGGCCG | CCGGCGT | CGG | GCC | C | T | C>A | C>A_GCC | CG>AT_CCGGCGT | 6 | FALSE | FALSE |
| 174_v.txt | 33115 | 33115-SNV-( | SNV | 1 | C | A | 13 | 23328 | 0.05572702 | 35.3855371 | rnr1Y285F_pG | TRUE | CAN | JSY2420 | 5 | rnr1Y285F_p | CCG | C>A | A/T+1 | G/C-1 | CG>AT | CG>AT | CTGGGCCGG | CGGCGTT | GGC | CCG | C | T | C>A | C>A_CCG | CG>AT_CGGCGTT | 6 | FALSE | FALSE |
| 174_v.txt | 33151 | 33151-SNV-( | SNV | 1 | G | T | 27 | 23328 | 0.11574074 | 39.4814815 | rnr1Y285F_pG | TRUE | CAN | JSY2420 | 5 | rnr1Y285F_p | AGA | G>T | A/T+1 | G/C-1 | CG>AT | CG>AT | TGAAAAGAC | GACCTGT | TCT | TCT | C | A | C>A | C>A_TCT | CG>AT_GACCTGT | 6 | FALSE | FALSE |
| 174_v.txt | 33153 | 33153-SNV-( | SNV | 1 | C | A | 54 | 23327 | 0.23149141 | 40.5544604 | rnr1Y285F_pG | TRUE | CAN | JSY2420 | 5 | rnr1Y285F_p | ACC | C>A | A/T+1 | G/C-1 | CG>AT | CG>AT | AAAAGACCT( | CCTGTAC | TGG | ACC | C | T | C>A | C>A_ACC | CG>AT_CCTGTAC | 6 | FALSE | FALSE |
| 174_v.txt | 33154 | 33154-SNV-( | SNV | 1 | C | A | 40 | 23327 | 0.17147512 | 36.525 | rnr1Y285F_pG | TRUE | CAN | JSY2420 | 5 | rnr1Y285F_p | CCT | C>A | A/T+1 | G/C-1 | CG>AT | CG>AT | AAAGACCTG | CTGTACC | GGA | CCT | C | T | C>A | C>A_CCT | CG>AT_CTGTACC | 6 | FALSE | FALSE |
| 174_v.txt | 33156 | 33156-SNV-( | SNV | 1 | G | T | 9 | 23327 | 0.0385819 | 37.3333333 | rnr1Y285F_pG | TRUE | CAN | JSY2420 | 5 | rnr1Y285F_p | TGT | G>T | A/T+1 | G/C-1 | CG>AT | CG>AT | AGACCTGTA( | GTACCAA | ACA | ACA | C | A | C>A | C>A_ACA | CG>AT_GTACCAA | 6 | FALSE | FALSE |
| 174_v.txt | 33159 | 33159-SNV-( | SNV | 1 | C | A | 23 | 23326 | 0.09860242 | 37.9565217 | rnr1Y285F_pG | TRUE | CAN | JSY2420 | 5 | rnr1Y285F_p | ACC | C>A | A/T+1 | G/C-1 | CG>AT | CG>AT | CCTGTACCA( | CCAATAG | TGG | ACC | C | T | C>A | C>A_ACC | CG>AT_CCAATAG | 6 | FALSE | FALSE |
| 174_v.txt | 33168 | 33168-SNV-( | SNV | 1 | C | A | 29 | 23325 | 0.12433012 | 35.9941286 | rnr1Y285F_pG | TRUE | CAN | JSY2420 | 5 | rnr1Y285F_p | ACC | C>A | A/T+1 | G/C-1 | CG>AT | CG>AT | ATAGTACCA( | CCACCAA | TGG | ACC | C | T | C>A | C>A_ACC | CG>AT_CCACCAA | 6 | FALSE | FALSE |
| 174_v.txt | 33177 | 33177-SNV-( | SNV | 1 | G | T | 14 | 23323 | 0.06002658 | 38 | rnr1Y285F_pG | TRUE | CAN | JSY2420 | 5 | rnr1Y285F_p | GGC | G>T | A/T+1 | G/C-1 | CG>AT | CG>AT | CCAAGGGCA | GCAATCA | CCG | CCG | C | A | C>A | C>A_CCG | CG>AT_GCAATCA | 6 | FALSE | FALSE |
| 174_v.txt | 33208 | 33208-SNV-( | SNV | 1 | C | A | 316 | 12697 | 2.48877688 | 39.4841772 | rnr1Y285F_pG | TRUE | CAN | JSY2420 | 5 | rnr1Y285F_p | TCT | C>A | A/T+1 | G/C-1 | CG>AT | CG>AT | TAAGCTCTCT | CTCTCTT | AGA | TCT | C | T | C>A | C>A_TCT | CG>AT_CTCTCTT | 6 | FALSE | FALSE |
| 174_v.txt | 33210 | 33210-SNV-( | SNV | 1 | C | A | 15 | 12697 | 0.11813814 | 35.6 | rnr1Y285F_pG | TRUE | CAN | JSY2420 | 5 | rnr1Y285F_p | TCT | C>A | A/T+1 | G/C-1 | CG>AT | CG>AT | AGCTCTCTCT | CTCTTCA | AGA | TCT | C | T | C>A | C>A_TCT | CG>AT_CTCTTCA | 6 | FALSE | FALSE |
| 174_v.txt | 33238 | 33238-SNV-( | SNV | 1 | C | A | 15 | 12706 | 0.11805446 | 38.3333333 | rnr1Y285F_pG | TRUE | CAN | JSY2420 | 5 | rnr1Y285F_p | CCT | C>A | A/T+1 | G/C-1 | CG>AT | CG>AT | CTTCTCCTTC/ | CTTCATC | GGA | CCT | C | T | C>A | C>A_CCT | CG>AT_CTTCATC | 6 | FALSE | FALSE |
| 174_v.txt | 33241 | 33241-SNV-( | SNV | 1 | C | A | 11 | 12706 | 0.08657327 | 41.6363636 | rnr1Y285F_pG | TRUE | CAN | JSY2420 | 5 | rnr1Y285F_p | TCA | C>A | A/T+1 | G/C-1 | CG>AT | CG>AT | CTCCTTCATC | CATCTTC | AGT | TCA | C | T | C>A | C>A_TCA | CG>AT_CATCTTC | 6 | FALSE | FALSE |
| 174_v.txt | 33265 | 33265-SNV-( | SNV | 1 | C | A | 36 | 12706 | 0.28333071 | 43.8333333 | rnr1Y285F_pG | TRUE | CAN | JSY2420 | 5 | rnr1Y285F_p | TCC | C>A | A/T+1 | G/C-1 | CG>AT | CG>AT | CATCCTCCAT | CCATAGA | AGG | TCC | C | T | C>A | C>A_TCC | CG>AT_CCATAGA | 6 | FALSE | FALSE |
| 174_v.txt | 33325 | 33325-SNV-( | SNV | 1 | C | A | 82 | 12706 | 0.6453644 | 56.3536585 | rnr1Y285F_pG | TRUE | CAN | JSY2420 | 5 | rnr1Y285F_p | TCT | C>A | A/T+1 | G/C-1 | CG>AT | CG>AT | ACAATTCTTT | CTTTACT | AGA | TCT | C | T | C>A | C>A_TCT | CG>AT_CTTTACT | 6 | FALSE | FALSE |
| 174_v.txt | 33354 | 33354-SNV-( | SNV | 1 | G | T | 89 | 12706 | 0.70045648 | 19.8202247 | rnr1Y285F_pG | TRUE | CAN | JSY2420 | 5 | rnr1Y285F_p | TGA | G>T | A/T+1 | G/C-1 | CG>AT | CG>AT | GGTATTGAC( | GACCCAC | ACT | ACT | C | A | C>A | C>A_ACT | CG>AT_GACCCAC | 6 | FALSE | FALSE |
| 174_v.txt | 33474 | 33474-SNV-( | SNV | 1 | C | A | 37 | 12705 | 0.29122393 | 34.6216216 | rnr1Y285F_pG | TRUE | CAN | JSY2420 | 5 | rnr1Y285F_p | GCC | C>A | A/T+1 | G/C-1 | CG>AT | CG>AT | GCTATGCCTT | CCTTTTT | CGG | GCC | C | T | C>A | C>A_GCC | CG>AT_CCTTTTT | 6 | FALSE | FALSE |
| 174_v.txt | 33475 | 33475-SNV-( | SNV | 1 | C | A | 12 | 12705 | 0.094451 | 35.25 | rnr1Y285F_pG | TRUE | CAN | JSY2420 | 5 | rnr1Y285F_p | CCT | C>A | A/T+1 | G/C-1 | CG>AT | CG>AT | CTATGCCTTT | CTTTTTT | GGA | CCT | C | T | C>A | C>A_CCT | CG>AT_CTTTTTT | 6 | FALSE | FALSE |
| 174_v.txt | 32091 | 32091-MNV- | MNV | 2 | GC | TT | 12 | 11593 | 0.10351074 | 62.5416623 | rnr1Y285F_pG | TRUE | CAN | JSY2420 | 5 | rnr1Y285F_p | TGC | nope | >1bp-ins | >1bpdel | none | MNV | AAGCCTGCA, | GCAACAC | ACG | TGC | GC | none | NA | NA_TGC | MNV_GCAACAC | 6 | FALSE | FALSE |
| 174_v.txt | 32556 | 32556-MNV- | MNV | 2 | CC | AA | 2 | 21303 | 9.39E-03 | 34.5000386 | rnr1Y285F_pG | TRUE | CAN | JSY2420 | 5 | rnr1Y285F_p | ACC | nope | >1bp-ins | >1bpdel | none | MNV | GTGATACCA/ | CCAACTA | TGG | ACC | CC | none | NA | NA_ACC | MNV_CCAACTA | 6 | FALSE | FALSE |
| 174_v.txt | 32709 | 32709-MNV- | MNV | 2 | CC | AT | 52 | 22590 | 0.23019035 | 62.7980342 | rnr1Y285F_pG | TRUE | CAN | JSY2420 | 5 | rnr1Y285F_p | CCC | nope | >1bp-ins | >1bpdel | none | MNV | GTAACCCCA( | CCAGCAC | GGG | CCC | CC | none | NA | NA_CCC | MNV_CCAGCAC | 6 | FALSE | FALSE |
| 174_v.txt | 31819 | 31819-Insert | Insertion | 23 | - | ATCTGAATAT | 3 | 7006 | 0.04282044 | 54.4637681 | rnr1Y285F_pG | TRUE | CAN | JSY2420 | 5 | rnr1Y285F_p | CAT | nope | >1bp-ins | >1bpdel | none | >1bp-ins | ATCTGCATCT | ATCTGAA | GTA | CAT | - | none | NA | NA_CAT | >1bp-ins_ATCTGAA | 6 | FALSE | FALSE |
| 174_v.txt | 31863 | 31863-Insert | Insertion | 1 | - | A | 5 | 7006 | 0.0713674 | 43.2 | rnr1Y285F_pG | TRUE | CAN | JSY2420 | 5 | rnr1Y285F_p | GAA | nope | A/T+1 | >1bpdel | none | A/T+1 | AACAGGAAA | AAAATAG | CTT | GAA | - | T | NA | NA_GAA | A/T+1_AAAATAG | 6 | FALSE | FALSE |
| 174_v.txt | 31909 | 31909-Insert | Insertion | 1 | - | C | 1 | 7006 | 0.01427348 | 11 | rnr1Y285F_pG | TRUE | CAN | JSY2420 | 5 | rnr1Y285F_p | GGT | nope | G/C+1 | >1bpdel | none | G/C+1 | ATTTTGGTG( | GTGCAAA | CCA | GGT | - | G | NA | NA_GGT | G/C+1_GTGCAAA | 6 | FALSE | FALSE |
| 174_v.txt | 32081 | 32081-Insert | Insertion | 1 | - | A | 25 | 11593 | 0.21564737 | 62.4 | rnr1Y285F_pG | TRUE | CAN | JSY2420 | 5 | rnr1Y285F_p | CAA | nope | A/T+1 | >1bpdel | none | A/T+1 | CCATGCAAA/ | AAAAAAG | GTT | CAA | - | T | NA | NA_CAA | A/T+1_AAAAAAG | 6 | FALSE | FALSE |
| 174_v.txt | 32081 | 32081-Insert | Insertion | 2 | - | AA | 4 | 11593 | 0.03450358 | 59.875 | rnr1Y285F_pG | TRUE | CAN | JSY2420 | 5 | rnr1Y285F_p | CAA | nope | >1bp-ins | >1bpdel | none | >1bp-ins | CCATGCAAA/ | AAAAAAG | GTT | CAA | - | none | NA | NA_CAA | >1bp-ins_AAAAAAG | 6 | FALSE | FALSE |
| 174_v.txt | 32351 | 32351-Insert | Insertion | 1 | - | A | 3 | 21310 | 0.0140779 | 62.3333333 | rnr1Y285F_pG | TRUE | CAN | JSY2420 | 5 | rnr1Y285F_p | CCT | nope | A/T+1 | >1bpdel | none | A/T+1 | CAAAACCTTT | CTTTGTA | GGA | CCT | - | T | NA | NA_CCT | A/T+1_CTTTGTA | 6 | FALSE | FALSE |
| 174_v.txt | 32352 | 32352-Insert | Insertion | 1 | - | T | 18 | 21310 | 0.08446739 | 61 | rnr1Y285F_pG | TRUE | CAN | JSY2420 | 5 | rnr1Y285F_p | CTT | nope | A/T+1 | >1bpdel | none | A/T+1 | AAAACCTTTC | TTTGTAC | GAA | CTT | - | A | NA | NA_CTT | A/T+1_TTTGTAC | 6 | FALSE | FALSE |
| 174_v.txt | 32375 | 32375-Insert | Insertion | 1 | - | A | 11 | 21310 | 0.05161896 | 64 | rnr1Y285F_pG | TRUE | CAN | JSY2420 | 5 | rnr1Y285F_p | CAA | nope | A/T+1 | >1bpdel | none | A/T+1 | AATAGCAAT/ | AATAATA | GTT | CAA | - | T | NA | NA_CAA | A/T+1_AATAATA | 6 | FALSE | FALSE |
| 174_v.txt | 32402 | 32402-Insert | Insertion | 34 | - | GGAAGTAGA | 3 | 21310 | 0.0140779 | 63.3137255 | rnr1Y285F_pG | TRUE | CAN | JSY2420 | 5 | rnr1Y285F_p | AGG | nope | >1bp-ins | >1bpdel | none | >1bp-ins | AACGTAGGA | GGAAGTA | TCC | AGG | - | none | NA | NA_AGG | >1bp-ins_GGAAGTA | 6 | FALSE | FALSE |
| 174_v.txt | 32490 | 32490-Insert | Insertion | 1 | - | A | 3 | 21310 | 0.0140779 | 46.6666667 | rnr1Y285F_pG | TRUE | CAN | JSY2420 | 5 | rnr1Y285F_p | GAA | nope | A/T+1 | >1bpdel | none | A/T+1 | ATACGGAAA | AAAACAA | CTT | GAA | - | T | NA | NA_GAA | A/T+1_AAAACAA | 6 | FALSE | FALSE |
| 174_v.txt | 32498 | 32498-Insert | Insertion | 1 | - | T | 6 | 21310 | 0.0281558 | 44.3333333 | rnr1Y285F_pG | TRUE | CAN | JSY2420 | 5 | rnr1Y285F_p | CTT | nope | A/T+1 | >1bpdel | none | A/T+1 | AACAACTTTT | TTTTTTG | GAA | CTT | - | A | NA | NA_CTT | A/T+1_TTTTTTG | 6 | FALSE | FALSE |
| 174_v.txt | 32608 | 32608-Insert | Insertion | 1 | - | C | 7 | 22590 | 0.03098716 | 49.1428571 | rnr1Y285F_pG | TRUE | CAN | JSY2420 | 5 | rnr1Y285F_p | ACC | nope | G/C+1 | >1bpdel | none | G/C+1 | AGGAAACCC | CCCAACC | TGG | ACC | - | G | NA | NA_ACC | G/C+1_CCCAACC | 6 | FALSE | FALSE |
| 174_v.txt | 32842 | 32842-Insert | Insertion | 1 | - | A | 225 | 22590 | 0.99601594 | 44.0977778 | rnr1Y285F_pG | TRUE | CAN | JSY2420 | 5 | rnr1Y285F_p | CAA | nope | A/T+1 | >1bpdel | none | A/T+1 | TTACCCAAA/ | AAAAAAT | GTT | CAA | - | T | NA | NA_CAA | A/T+1_AAAAAAT | 6 | FALSE | FALSE |
| 174_v.txt | 32842 | 32842-Insert | Insertion | 2 | - | AA | 1 | 22590 | 4.43E-03 | 38 | rnr1Y285F_pG | TRUE | CAN | JSY2420 | 5 | rnr1Y285F_p | CAA | nope | >1bp-ins | >1bpdel | none | >1bp-ins | TTACCCAAA/ | AAAAAAT | GTT | CAA | - | none | NA | NA_CAA | >1bp-ins_AAAAAAT | 6 | FALSE | FALSE |
| 174_v.txt | 32881 | 32881-Insert | Insertion | 1 | - | T | 2 | 23328 | 8.57E-03 | 38 | rnr1Y285F_pG | TRUE | CAN | JSY2420 | 5 | rnr1Y285F_p | GTC | nope | A/T+1 | >1bpdel | none | A/T+1 | TGTACGTCC/ | TCCAAAA | CAG | GTC | - | A | NA | NA_GTC | A/T+1_TCCAAAA | 6 | FALSE | FALSE |
| 174_v.txt | 32884 | 32884-Insert | Insertion | 1 | - | A | 17 | 23328 | 0.0728738 | 36.2352941 | rnr1Y285F_pG | TRUE | CAN | JSY2420 | 5 | rnr1Y285F_p | CAA | nope | A/T+1 | >1bpdel | none | A/T+1 | ACGTCCAAA/ | AAAATTG | GTT | CAA | - | T | NA | NA_CAA | A/T+1_AAAATTG | 6 | FALSE | FALSE |
| 174_v.txt | 33072 | 33072-Insert | Insertion | 2 | - | AA | 20 | 23328 | 0.08573388 | 58.125 | rnr1Y285F_pG | TRUE | CAN | JSY2420 | 5 | rnr1Y285F_p | CAA | nope | >1bp-ins | >1bpdel | none | >1bp-ins | TATGCCAAA( | AAAGAAC | GTT | CAA | - | none | NA | NA_CAA | >1bp-ins_AAAGAAC | 6 | FALSE | FALSE |
| 174_v.txt | 33147 | 33147-Insert | Insertion | 1 | - | A | 23 | 23328 | 0.09859396 | 39.0434783 | rnr1Y285F_pG | TRUE | CAN | JSY2420 | 5 | rnr1Y285F_p | GAA | nope | A/T+1 | >1bpdel | none | A/T+1 | CCAATGAAA. | AAAAGAC | CTT | GAA | - | T | NA | NA_GAA | A/T+1_AAAAGAC | 6 | FALSE | FALSE |
| 174_v.txt | 33168 | 33168-Insert | Insertion | 1 | - | C | 20 | 23325 | 0.08574491 | 37.4 | rnr1Y285F_pG | TRUE | CAN | JSY2420 | 5 | rnr1Y285F_p | ACC | nope | G/C+1 | >1bpdel | none | G/C+1 | ATAGTACCA( | CCACCAA | TGG | ACC | - | G | NA | NA_ACC | G/C+1_CCACCAA | 6 | FALSE | FALSE |
| 174_v.txt | 33206 | 33206-Insert | Insertion | 2 | - | CC | 1 | 12697 | 7.88E-03 | 32.5 | rnr1Y285F_pG | TRUE | CAN | JSY2420 | 5 | rnr1Y285F_p | GCT | nope | >1bp-ins | >1bpdel | none | >1bp-ins | CTTAAGCTCT | CTCTCTC | CGA | GCT | - | none | NA | NA_GCT | >1bp-ins_CTCTCTC | 6 | FALSE | FALSE |
| 174_v.txt | 33206 | 33206-Insert | Insertion | 2 | - | CT | 10 | 12697 | 0.07875876 | 40.4 | rnr1Y285F_pG | TRUE | CAN | JSY2420 | 5 | rnr1Y285F_p | GCT | nope | >1bp-ins | >1bpdel | none | >1bp-ins | CTTAAGCTCT | CTCTCTC | CGA | GCT | - | none | NA | NA_GCT | >1bp-ins_CTCTCTC | 6 | FALSE | FALSE |
| 174_v.txt | 33376 | 33376-Insert | Insertion | 2 | - | TC | 22 | 12706 | 0.17314655 | 56.6818182 | rnr1Y285F_pG | TRUE | CAN | JSY2420 | 5 | rnr1Y285F_p | TGT | nope | >1bp-ins | >1bpdel | none | >1bp-ins | GTGTTTGTG/ | GTGAAGC | ACA | TGT | - | none | NA | NA_TGT | >1bp-ins_GTGAAGC | 6 | FALSE | FALSE |
| 174_v.txt | 33417 | 33417-Insert | Insertion | 1 | - | T | 6 | 12706 | 0.04722179 | 43.1666667 | rnr1Y285F_pG | TRUE | CAN | JSY2420 | 5 | rnr1Y285F_p | ATT | nope | A/T+1 | >1bpdel | none | A/T+1 | GGCTCATTG1 | TTGTACA | TAA | ATT | - | A | NA | NA_ATT | A/T+1_TTGTACA | 6 | FALSE | FALSE |
| 174_v.txt | 31893 | 31893-Deleti | Deletion | 2 | AC | - | 12 | 7006 | 0.17128176 | 40.8333333 | rnr1Y285F_pG | TRUE | CAN | JSY2420 | 5 | rnr1Y285F_p | AAC | nope | >1bp-ins | >1bpdel | none | >1bpdel | AAGCTAACA( | ACACCAT | TTG | AAC | AC | none | NA | NA_AAC | >1bpdel_ACACCAT | 6 | FALSE | FALSE |
| 174_v.txt | 31908 | 31908-Deleti | Deletion | 2 | GG | - | 6 | 7006 | 0.08564088 | 42.3333333 | rnr1Y285F_pG | TRUE | CAN | JSY2420 | 5 | rnr1Y285F_p | TGG | nope | >1bp-ins | >1bpdel | none | >1bpdel | AATTTTGGTG | GGTGCAA | ACC | TGG | GG | none | NA | NA_TGG | >1bpdel_GGTGCAA | 6 | FALSE | FALSE |
| 174_v.txt | 32173 | 32173-Deleti | Deletion | 2 | CA | - | 3 | 11593 | 0.02587769 | 38 | rnr1Y285F_pG | TRUE | CAN | JSY2420 | 5 | rnr1Y285F_p | GCA | nope | >1bp-ins | >1bpdel | none | >1bpdel | CAAATGCAG | CAGCAGT | CGT | GCA | CA | none | NA | NA_GCA | >1bpdel_CAGCAGT | 6 | FALSE | FALSE |
| 174_v.txt | 32173 | 32173-Deleti | Deletion | 4 | CAGC | - | 13 | 11593 | 0.11213663 | 50.9230769 | rnr1Y285F_pG | TRUE | CAN | JSY2420 | 5 | rnr1Y285F_p | GCA | nope | >1bp-ins | >1bpdel | none | >1bpdel | CAAATGCAG | CAGCAGT | CGT | GCA | CAGC | none | NA | NA_GCA | >1bpdel_CAGCAGT | 6 | FALSE | FALSE |
| 174_v.txt | 32277 | 32277-Deleti | Deletion | 2 | CC | - | 18 | 21310 | 0.08446739 | 35.3333333 | rnr1Y285F_pG | TRUE | CAN | JSY2420 | 5 | rnr1Y285F_p | ACC | nope | >1bp-ins | >1bpdel | none | >1bpdel | CGGGAACCA | CCAACGT | TGG | ACC | CC | none | NA | NA_ACC | >1bpdel_CCAACGT | 6 | FALSE | FALSE |
| 174_v.txt | 32337 | 32337-Deleti | Deletion | 2 | AT | - | 6 | 21310 | 0.0281558 | 64 | rnr1Y285F_pG | TRUE | CAN | JSY2420 | 5 | rnr1Y285F_p | GAT | nope | >1bp-ins | >1bpdel | none | >1bpdel | TTGAAGATA | ATATGTG | CTA | GAT | AT | none | NA | NA_GAT | >1bpdel_ATATGTG | 6 | FALSE | FALSE |
| 174_v.txt | 32492 | 32492-Deleti | Deletion | 3 | AAC | - | 37 | 21310 | 0.17362741 | 44.4594595 | rnr1Y285F_pG | TRUE | CAN | JSY2420 | 5 | rnr1Y285F_p | AAA | nope | >1bp-ins | >1bpdel | none | >1bpdel | ACGGAAAAC | AACAACT | TTT | AAA | AAC | none | NA | NA_AAA | >1bpdel_AACAACT | 6 | FALSE | FALSE |
| 174_v.txt | 32571 | 32571-Deleti | Deletion | 2 | CC | - | 1 | 22592 | 4.43E-03 | 38 | rnr1Y285F_pG | TRUE | CAN | JSY2420 | 5 | rnr1Y285F_p | ACC | nope | >1bp-ins | >1bpdel | none | >1bpdel | TCAGTACCTT | CCTTGAA | TGG | ACC | CC | none | NA | NA_ACC | >1bpdel_CCTTGAA | 6 | FALSE | FALSE |
| 174_v.txt | 32648 | 32648-Deleti | Deletion | 2 | TA | - | 11 | 22590 | 0.04869411 | 57.7272727 | rnr1Y285F_pG | TRUE | CAN | JSY2420 | 5 | rnr1Y285F_p | TTA | nope | >1bp-ins | >1bpdel | none | >1bpdel | AGATATTAT/ | TATACCT | AAT | TTA | TA | none | NA | NA_TTA | >1bpdel_TATACCT | 6 | FALSE | FALSE |
| 174_v.txt | 32696 | 32696-Deleti | Deletion | 8 | TGGGCCGG | - | 52 | 22590 | 0.23019035 | 61.7115385 | rnr1Y285F_pG | TRUE | CAN | JSY2420 | 5 | rnr1Y285F_p | CTG | nope | >1bp-ins | >1bpdel | none | >1bpdel | TCCAACTGG( | TGGGCCG | GAC | CTG | TGGGCCGG | none | NA | NA_CTG | >1bpdel_TGGGCCG | 6 | FALSE | FALSE |
| 174_v.txt | 32736 | 32736-Deleti | Deletion | 2 | TA | - | 17 | 22590 | 0.07525454 | 58.5882353 | rnr1Y285F_pG | TRUE | CAN | JSY2420 | 5 | rnr1Y285F_p | GTA | nope | >1bp-ins | >1bpdel | none | >1bpdel | AAACAGTAT. | TATATTA | CAT | GTA | TA | none | NA | NA_GTA | >1bpdel_TATATTA | 6 | FALSE | FALSE |
| 174_v.txt | 32739 | 32739-Deleti | Deletion | 2 | AT | - | 5 | 22590 | 0.02213369 | 64 | rnr1Y285F_pG | TRUE | CAN | JSY2420 | 5 | rnr1Y285F_p | TAT | nope | >1bp-ins | >1bpdel | none | >1bpdel | CAGTATATTA | ATTAGAA | ATA | TAT | AT | none | NA | NA_TAT | >1bpdel_ATTAGAA | 6 | FALSE | FALSE |
| 174_v.txt | 32779 | 32779-Deleti | Deletion | 2 | CC | - | 1 | 22590 | 4.43E-03 | 64 | rnr1Y285F_pG | TRUE | CAN | JSY2420 | 5 | rnr1Y285F_p | ACC | nope | >1bp-ins | >1bpdel | none | >1bpdel | AAGCGACCC. | CCCAGAA | TGG | ACC | CC | none | NA | NA_ACC | >1bpdel_CCCAGAA | 6 | FALSE | FALSE |
| 174_v.txt | 32994 | 32994-Deleti | Deletion | 2 | TG | - | 8 | 23328 | 0.03429355 | 60.75 | rnr1Y285F_pG | TRUE | CAN | JSY2420 | 5 | rnr1Y285F_p | TTG | nope | >1bp-ins | >1bpdel | none | >1bpdel | AATCTTTGTG | TGTGAGA | AAC | TTG | TG | none | NA | NA_TTG | >1bpdel_TGTGAGA | 6 | FALSE | FALSE |
| 174_v.txt | 33057 | 33057-Deleti | Deletion | 2 | GT | - | 28 | 23328 | 0.12002744 | 61.5 | rnr1Y285F_pG | TRUE | CAN | JSY2420 | 5 | rnr1Y285F_p | CGT | nope | >1bp-ins | >1bpdel | none | >1bpdel | GACTGCGTG. | GTGACAG | GCA | CGT | GT | none | NA | NA_CGT | >1bpdel_GTGACAG | 6 | FALSE | FALSE |
| 174_v.txt | 33089 | 33089-Deleti | Deletion | 2 | AT | - | 23 | 23328 | 0.09859396 | 51.2608696 | rnr1Y285F_pG | TRUE | CAN | JSY2420 | 5 | rnr1Y285F_p | AAT | nope | >1bp-ins | >1bpdel | none | >1bpdel | AAATAAATA | ATATGAT | TTA | AAT | AT | none | NA | NA_AAT | >1bpdel_ATATGAT | 6 | FALSE | FALSE |
| 174_v.txt | 31682 | 31682-Deleti | Deletion | 1 | A | - | 4 | 7006 | 0.05709392 | 51 | rnr1Y285F_pG | TRUE | CAN | JSY2420 | 5 | rnr1Y285F_p | TAA | nope | >1bp-ins | A/T-1 | none | A/T-1 | GGTAATAAA | AAAACGT | ATT | ATT | T | none | T>none | T>none_ATT | A/T-1_AAAACGT | 6 | FALSE | FALSE |
| 174_v.txt | 31723 | 31723-Deleti | Deletion | 1 | A | - | 12 | 7006 | 0.17128176 | 58.1666667 | rnr1Y285F_pG | TRUE | CAN | JSY2420 | 5 | rnr1Y285F_p | CAA | nope | >1bp-ins | A/T-1 | none | A/T-1 | TGTCCCAAA/ | AAAAAGT | GTT | GTT | T | none | T>none | T>none_GTT | A/T-1_AAAAAGT | 6 | FALSE | FALSE |
| 174_v.txt | 31904 | 31904-Deleti | Deletion | 1 | T | - | 18 | 7006 | 0.25692264 | 46.5555556 | rnr1Y285F_pG | TRUE | CAN | JSY2420 | 5 | rnr1Y285F_p | ATT | nope | >1bp-ins | A/T-1 | none | A/T-1 | ATTGAATTTT | TTTTGGT | TAA | ATT | T | none | nope | nope_ATT | A/T-1_TTTTGGT | 6 | FALSE | FALSE |
| 174_v.txt | 31972 | 31972-Deleti | Deletion | 1 | C | - | 8 | 11593 | 0.06900716 | 36.5 | rnr1Y285F_pG | TRUE | CAN | JSY2420 | 5 | rnr1Y285F_p | GCC | nope | >1bp-ins | G/C-1 | none | G/C-1 | AATAAGCCA. | CCAAGCC | CGG | GCC | C | none | nope | nope_GCC | G/C-1_CCAAGCC | 6 | FALSE | FALSE |
| 174_v.txt | 31979 | 31979-Deleti | Deletion | 1 | G | - | 77 | 11593 | 0.66419391 | 43.2467533 | rnr1Y285F_pG | TRUE | CAN | JSY2420 | 5 | rnr1Y285F_p | CGG | nope | >1bp-ins | G/C-1 | none | G/C-1 | CAAGCCGGG | GGGCATT | GCC | GCC | C | none | C>none | C>none_GCC | G/C-1_GGGCATT | 6 | FALSE | FALSE |
| 174_v.txt | 32026 | 32026-Deleti | Deletion | 1 | G | - | 16 | 11593 | 0.13801432 | 53.4999988 | rnr1Y285F_pG | TRUE | CAN | JSY2420 | 5 | rnr1Y285F_p | CGG | nope | >1bp-ins | G/C-1 | none | G/C-1 | TGCCACGGT/ | GGTATTT | GCC | GCC | C | none | C>none | C>none_GCC | G/C-1_GGTATTT | 6 | FALSE | FALSE |
| 174_v.txt | 32075 | 32075-Deleti | Deletion | 1 | C | - | 16 | 11593 | 0.13801432 | 60.75 | rnr1Y285F_pG | TRUE | CAN | JSY2420 | 5 | rnr1Y285F_p | ACC | nope | >1bp-ins | G/C-1 | none | G/C-1 | AAATAACCA | CCATGCA | TGG | ACC | C | none | nope | nope_ACC | G/C-1_CCATGCA | 6 | FALSE | FALSE |
| 174_v.txt | 32081 | 32081-Deleti | Deletion | 1 | A | - | 21 | 11593 | 0.18114379 | 62.7619046 | rnr1Y285F_pG | TRUE | CAN | JSY2420 | 5 | rnr1Y285F_p | CAA | nope | >1bp-ins | A/T-1 | none | A/T-1 | CCATGCAAA/ | AAAAAAG | GTT | GTT | T | none | T>none | T>none_GTT | A/T-1_AAAAAAG | 6 | FALSE | FALSE |
| 174_v.txt | 32087 | 32087-Deleti | Deletion | 1 | G | - | 17 | 11592 | 0.14665286 | 63.0588231 | rnr1Y285F_pG | TRUE | CAN | JSY2420 | 5 | rnr1Y285F_p | AGC | nope | >1bp-ins | G/C-1 | none | G/C-1 | AAAAAAGCC | GCCTGCA | TCG | TCG | C | none | C>none | C>none_TCG | G/C-1_GCCTGCA | 6 | FALSE | FALSE |
| 174_v.txt | 32125 | 32125-Deleti | Deletion | 1 | C | - | 17 | 11593 | 0.14664021 | 58.9411765 | rnr1Y285F_pG | TRUE | CAN | JSY2420 | 5 | rnr1Y285F_p | ACT | nope | >1bp-ins | G/C-1 | none | G/C-1 | CGAAAACTT | CTTTGTC | TGA | ACT | C | none | nope | nope_ACT | G/C-1_CTTTGTC | 6 | FALSE | FALSE |
| 174_v.txt | 32216 | 32216-Deleti | Deletion | 1 | G | - | 5 | 11593 | 0.04312948 | 38 | rnr1Y285F_pG | TRUE | CAN | JSY2420 | 5 | rnr1Y285F_p | TGG | nope | >1bp-ins | G/C-1 | none | G/C-1 | TTTGGTGGT( | GGTCCTT | ACC | ACC | C | none | C>none | C>none_ACC | G/C-1_GGTCCTT | 6 | FALSE | FALSE |
| 174_v.txt | 32272 | 32272-Deleti | Deletion | 1 | G | - | 2 | 21310 | 9.39E-03 | 37.4999971 | rnr1Y285F_pG | TRUE | CAN | JSY2420 | 5 | rnr1Y285F_p | CGG | nope | >1bp-ins | G/C-1 | none | G/C-1 | AAATACGGG | GGGAACC | GCC | GCC | C | none | C>none | C>none_GCC | G/C-1_GGGAACC | 6 | FALSE | FALSE |
| 174_v.txt | 32277 | 32277-Deleti | Deletion | 1 | C | - | 3 | 21310 | 0.0140779 | 36.9999888 | rnr1Y285F_pG | TRUE | CAN | JSY2420 | 5 | rnr1Y285F_p | ACC | nope | >1bp-ins | G/C-1 | none | G/C-1 | CGGGAACCA | CCAACGT | TGG | ACC | C | none | nope | nope_ACC | G/C-1_CCAACGT | 6 | FALSE | FALSE |
| 174_v.txt | 32284 | 32284-Deleti | Deletion | 1 | A | - | 30 | 21310 | 0.14077898 | 48.7333306 | rnr1Y285F_pG | TRUE | CAN | JSY2420 | 5 | rnr1Y285F_p | TAA | nope | >1bp-ins | A/T-1 | none | A/T-1 | CAACGTAAA | AAATATT | ATT | ATT | T | none | T>none | T>none_ATT | A/T-1_AAATATT | 6 | FALSE | FALSE |
| 174_v.txt | 32288 | 32288-Deleti | Deletion | 1 | A | - | 20 | 21310 | 0.09385265 | 47.1 | rnr1Y285F_pG | TRUE | CAN | JSY2420 | 5 | rnr1Y285F_p | TAT | nope | >1bp-ins | A/T-1 | none | A/T-1 | GTAAATATTT | ATTTGAA | ATA | ATA | T | none | T>none | T>none_ATA | A/T-1_ATTTGAA | 6 | FALSE | FALSE |
| 174_v.txt | 32308 | 32308-Deleti | Deletion | 1 | T | - | 18 | 21310 | 0.08446739 | 61.8333333 | rnr1Y285F_pG | TRUE | CAN | JSY2420 | 5 | rnr1Y285F_p | ATA | nope | >1bp-ins | A/T-1 | none | A/T-1 | CAGAAATAA | TAATGGT | TAT | ATA | T | none | nope | nope_ATA | A/T-1_TAATGGT | 6 | FALSE | FALSE |
| 174_v.txt | 32346 | 32346-Deleti | Deletion | 1 | A | - | 16 | 21310 | 0.07508212 | 62.375 | rnr1Y285F_pG | TRUE | CAN | JSY2420 | 5 | rnr1Y285F_p | CAA | nope | >1bp-ins | A/T-1 | none | A/T-1 | TGTGGCAAA | AAAACCT | GTT | GTT | T | none | T>none | T>none_GTT | A/T-1_AAAACCT | 6 | FALSE | FALSE |
| 174_v.txt | 32358 | 32358-Deleti | Deletion | 1 | C | - | 23 | 21310 | 0.10793055 | 61.4347826 | rnr1Y285F_pG | TRUE | CAN | JSY2420 | 5 | rnr1Y285F_p | ACC | nope | >1bp-ins | G/C-1 | none | G/C-1 | TTTGTACCAC | CCAGAGT | TGG | ACC | C | none | nope | nope_ACC | G/C-1_CCAGAGT | 6 | FALSE | FALSE |
| 174_v.txt | 32416 | 32416-Deleti | Deletion | 1 | T | - | 6 | 21310 | 0.0281558 | 50 | rnr1Y285F_pG | TRUE | CAN | JSY2420 | 5 | rnr1Y285F_p | GTT | nope | >1bp-ins | A/T-1 | none | A/T-1 | ATTGTGTTAC | TTAGTTT | CAA | GTT | T | none | nope | nope_GTT | A/T-1_TTAGTTT | 6 | FALSE | FALSE |
| 174_v.txt | 32419 | 32419-Deleti | Deletion | 1 | G | - | 12 | 21310 | 0.05631159 | 59.4166667 | rnr1Y285F_pG | TRUE | CAN | JSY2420 | 5 | rnr1Y285F_p | AGT | nope | >1bp-ins | G/C-1 | none | G/C-1 | GTGTTAGTTT | GTTTAGG | TCA | TCA | C | none | C>none | C>none_TCA | G/C-1_GTTTAGG | 6 | FALSE | FALSE |
| 174_v.txt | 32481 | 32481-Deleti | Deletion | 1 | A | - | 6 | 21310 | 0.0281558 | 46 | rnr1Y285F_pG | TRUE | CAN | JSY2420 | 5 | rnr1Y285F_p | TAA | nope | >1bp-ins | A/T-1 | none | A/T-1 | AAGGTTAAG | AAGATAC | ATT | ATT | T | none | T>none | T>none_ATT | A/T-1_AAGATAC | 6 | FALSE | FALSE |
| 174_v.txt | 32487 | 32487-Deleti | Deletion | 1 | C | - | 3 | 21310 | 0.0140779 | 46.6666667 | rnr1Y285F_pG | TRUE | CAN | JSY2420 | 5 | rnr1Y285F_p | ACG | nope | >1bp-ins | G/C-1 | none | G/C-1 | AAGATACGG | CGGAAAA | TGC | ACG | C | none | nope | nope_ACG | G/C-1_CGGAAAA | 6 | FALSE | FALSE |
| 174_v.txt | 32490 | 32490-Deleti | Deletion | 1 | A | - | 27 | 21310 | 0.12670108 | 45.2962963 | rnr1Y285F_pG | TRUE | CAN | JSY2420 | 5 | rnr1Y285F_p | GAA | nope | >1bp-ins | A/T-1 | none | A/T-1 | ATACGGAAA | AAAACAA | CTT | CTT | T | none | T>none | T>none_CTT | A/T-1_AAAACAA | 6 | FALSE | FALSE |
| 174_v.txt | 32498 | 32498-Deleti | Deletion | 1 | T | - | 60 | 21310 | 0.28155795 | 44.2833328 | rnr1Y285F_pG | TRUE | CAN | JSY2420 | 5 | rnr1Y285F_p | CTT | nope | >1bp-ins | A/T-1 | none | A/T-1 | AACAACTTTT | TTTTTTG | GAA | CTT | T | none | nope | nope_CTT | A/T-1_TTTTTTG | 6 | FALSE | FALSE |
| 174_v.txt | 32522 | 32522-Deleti | Deletion | 1 | T | - | 7 | 21310 | 0.03284843 | 37.7142857 | rnr1Y285F_pG | TRUE | CAN | JSY2420 | 5 | rnr1Y285F_p | ATT | nope | >1bp-ins | A/T-1 | none | A/T-1 | AACGGATTT | TTTTCTG | TAA | ATT | T | none | nope | nope_ATT | A/T-1_TTTTCTG | 6 | FALSE | FALSE |
| 174_v.txt | 32528 | 32528-Deleti | Deletion | 1 | G | - | 7 | 21310 | 0.03284843 | 35.8571429 | rnr1Y285F_pG | TRUE | CAN | JSY2420 | 5 | rnr1Y285F_p | TGG | nope | >1bp-ins | G/C-1 | none | G/C-1 | TTTTCTGGGC | GGGGTTT | ACC | ACC | C | none | C>none | C>none_ACC | G/C-1_GGGGTTT | 6 | FALSE | FALSE |
| 174_v.txt | 32608 | 32608-Deleti | Deletion | 1 | C | - | 199 | 22590 | 0.88092076 | 44.3819091 | rnr1Y285F_pG | TRUE | CAN | JSY2420 | 5 | rnr1Y285F_p | ACC | nope | >1bp-ins | G/C-1 | none | G/C-1 | AGGAAACCC | CCCAACC | TGG | ACC | C | none | nope | nope_ACC | G/C-1_CCCAACC | 6 | FALSE | FALSE |
| 174_v.txt | 32611 | 32611-Deleti | Deletion | 1 | A | - | 28 | 22590 | 0.12394865 | 39.6785714 | rnr1Y285F_pG | TRUE | CAN | JSY2420 | 5 | rnr1Y285F_p | CAA | nope | >1bp-ins | A/T-1 | none | A/T-1 | AAACCCAAC( | AACCTAA | GTT | GTT | T | none | T>none | T>none_GTT | A/T-1_AACCTAA | 6 | FALSE | FALSE |
| 174_v.txt | 32631 | 32631-Deleti | Deletion | 1 | T | - | 12 | 22590 | 0.05312085 | 47.6666667 | rnr1Y285F_pG | TRUE | CAN | JSY2420 | 5 | rnr1Y285F_p | GTT | nope | >1bp-ins | A/T-1 | none | A/T-1 | CCTTCGTTTT | TTTTTAT | CAA | GTT | T | none | nope | nope_GTT | A/T-1_TTTTTAT | 6 | FALSE | FALSE |
| 174_v.txt | 32647 | 32647-Deleti | Deletion | 1 | T | - | 6 | 22590 | 0.02656043 | 59.6666651 | rnr1Y285F_pG | TRUE | CAN | JSY2420 | 5 | rnr1Y285F_p | ATT | nope | >1bp-ins | A/T-1 | none | A/T-1 | TAGATATTAT | TTATACC | TAA | ATT | T | none | nope | nope_ATT | A/T-1_TTATACC | 6 | FALSE | FALSE |
| 174_v.txt | 32658 | 32658-Deleti | Deletion | 1 | C | - | 93 | 22590 | 0.41168659 | 60.9892473 | rnr1Y285F_pG | TRUE | CAN | JSY2420 | 5 | rnr1Y285F_p | ACC | nope | >1bp-ins | G/C-1 | none | G/C-1 | CCTGGACCCC | CCCCAGG | TGG | ACC | C | none | nope | nope_ACC | G/C-1_CCCCAGG | 6 | FALSE | FALSE |
| 174_v.txt | 32670 | 32670-Deleti | Deletion | 1 | G | - | 279 | 22590 | 1.23505976 | 60.3835125 | rnr1Y285F_pG | TRUE | CAN | JSY2420 | 5 | rnr1Y285F_p | TGG | nope | >1bp-ins | G/C-1 | none | G/C-1 | GCACCTGGG | GGGTTTC | ACC | ACC | C | none | C>none | C>none_ACC | G/C-1_GGGTTTC | 6 | FALSE | FALSE |
| 174_v.txt | 32697 | 32697-Deleti | Deletion | 1 | G | - | 1 | 22590 | 4.43E-03 | 18.9999999 | rnr1Y285F_pG | TRUE | CAN | JSY2420 | 5 | rnr1Y285F_p | TGG | nope | >1bp-ins | G/C-1 | none | G/C-1 | CCAACTGGG | GGGCCGG | ACC | ACC | C | none | C>none | C>none_ACC | G/C-1_GGGCCGG | 6 | FALSE | FALSE |
| 174_v.txt | 32705 | 32705-Deleti | Deletion | 1 | A | - | 29 | 22590 | 0.12837539 | 61.0689655 | rnr1Y285F_pG | TRUE | CAN | JSY2420 | 5 | rnr1Y285F_p | TAA | nope | >1bp-ins | A/T-1 | none | A/T-1 | GCCGGTAAC | AACCCCA | ATT | ATT | T | none | T>none | T>none_ATT | A/T-1_AACCCCA | 6 | FALSE | FALSE |
| 174_v.txt | 32707 | 32707-Deleti | Deletion | 1 | C | - | 24 | 22590 | 0.1062417 | 60.6666667 | rnr1Y285F_pG | TRUE | CAN | JSY2420 | 5 | rnr1Y285F_p | ACC | nope | >1bp-ins | G/C-1 | none | G/C-1 | CGGTAACCC( | CCCCAGC | TGG | ACC | C | none | nope | nope_ACC | G/C-1_CCCCAGC | 6 | FALSE | FALSE |
| 174_v.txt | 32719 | 32719-Deleti | Deletion | 1 | A | - | 18 | 22590 | 0.07968128 | 63.6666667 | rnr1Y285F_pG | TRUE | CAN | JSY2420 | 5 | rnr1Y285F_p | CAA | nope | >1bp-ins | A/T-1 | none | A/T-1 | CACCACAAA( | AAACCAT | GTT | GTT | T | none | T>none | T>none_GTT | A/T-1_AAACCAT | 6 | FALSE | FALSE |
| 174_v.txt | 32728 | 32728-Deleti | Deletion | 1 | A | - | 18 | 22590 | 0.07968128 | 58.0555556 | rnr1Y285F_pG | TRUE | CAN | JSY2420 | 5 | rnr1Y285F_p | CAA | nope | >1bp-ins | A/T-1 | none | A/T-1 | CCATACAAA/ | AAAAACA | GTT | GTT | T | none | T>none | T>none_GTT | A/T-1_AAAAACA | 6 | FALSE | FALSE |
| 174_v.txt | 32740 | 32740-Deleti | Deletion | 1 | T | - | 3 | 22589 | 0.0132808 | 63.9999897 | rnr1Y285F_pG | TRUE | CAN | JSY2420 | 5 | rnr1Y285F_p | ATT | nope | >1bp-ins | A/T-1 | none | A/T-1 | AGTATATTA( | TTAGAAA | TAA | ATT | T | none | nope | nope_ATT | A/T-1_TTAGAAA | 6 | FALSE | FALSE |
| 174_v.txt | 32747 | 32747-Deleti | Deletion | 1 | C | - | 20 | 22590 | 0.08853475 | 58.2 | rnr1Y285F_pG | TRUE | CAN | JSY2420 | 5 | rnr1Y285F_p | ACC | nope | >1bp-ins | G/C-1 | none | G/C-1 | TAGAAACCC( | CCCGATA | TGG | ACC | C | none | nope | nope_ACC | G/C-1_CCCGATA | 6 | FALSE | FALSE |
| 174_v.txt | 32760 | 32760-Deleti | Deletion | 1 | A | - | 11 | 22590 | 0.04869411 | 61.1818182 | rnr1Y285F_pG | TRUE | CAN | JSY2420 | 5 | rnr1Y285F_p | TAA | nope | >1bp-ins | A/T-1 | none | A/T-1 | ATGGCTAAA. | AAAACTT | ATT | ATT | T | none | T>none | T>none_ATT | A/T-1_AAAACTT | 6 | FALSE | FALSE |
| 174_v.txt | 32779 | 32779-Deleti | Deletion | 1 | C | - | 37 | 22590 | 0.16378929 | 52.4594587 | rnr1Y285F_pG | TRUE | CAN | JSY2420 | 5 | rnr1Y285F_p | ACC | nope | >1bp-ins | G/C-1 | none | G/C-1 | AAGCGACCC. | CCCAGAA | TGG | ACC | C | none | nope | nope_ACC | G/C-1_CCCAGAA | 6 | FALSE | FALSE |
| 174_v.txt | 32814 | 32814-Deleti | Deletion | 1 | A | - | 33 | 22590 | 0.14608234 | 41.3333333 | rnr1Y285F_pG | TRUE | CAN | JSY2420 | 5 | rnr1Y285F_p | GAA | nope | >1bp-ins | A/T-1 | none | A/T-1 | ACAGGGAAC | AACAAGT | CTT | CTT | T | none | T>none | T>none_CTT | A/T-1_AACAAGT | 6 | FALSE | FALSE |
| 174_v.txt | 32819 | 32819-Deleti | Deletion | 1 | G | - | 3 | 22590 | 0.01328021 | 37.6666667 | rnr1Y285F_pG | TRUE | CAN | JSY2420 | 5 | rnr1Y285F_p | AGT | nope | >1bp-ins | G/C-1 | none | G/C-1 | GAACAAGTT( | GTTCATT | TCA | TCA | C | none | C>none | C>none_TCA | G/C-1_GTTCATT | 6 | FALSE | FALSE |
| 174_v.txt | 32842 | 32842-Deleti | Deletion | 1 | A | - | 304 | 22590 | 1.3457282 | 43.0986837 | rnr1Y285F_pG | TRUE | CAN | JSY2420 | 5 | rnr1Y285F_p | CAA | nope | >1bp-ins | A/T-1 | none | A/T-1 | TTACCCAAA/ | AAAAAAT | GTT | GTT | T | none | T>none | T>none_GTT | A/T-1_AAAAAAT | 6 | FALSE | FALSE |
| 174_v.txt | 32848 | 32848-Deleti | Deletion | 1 | T | - | 32 | 22590 | 0.1416556 | 37.71875 | rnr1Y285F_pG | TRUE | CAN | JSY2420 | 5 | rnr1Y285F_p | ATA | nope | >1bp-ins | A/T-1 | none | A/T-1 | AAAAAATAC | TACTAAT | TAT | ATA | T | none | nope | nope_ATA | A/T-1_TACTAAT | 6 | FALSE | FALSE |
| 174_v.txt | 32876 | 32876-Deleti | Deletion | 1 | G | - | 2 | 23316 | 8.58E-03 | 38 | rnr1Y285F_pG | TRUE | CAN | JSY2420 | 5 | rnr1Y285F_p | TGT | nope | >1bp-ins | G/C-1 | none | G/C-1 | AACTTTGTA( | GTACGTC | ACA | ACA | C | none | C>none | C>none_ACA | G/C-1_GTACGTC | 6 | FALSE | FALSE |
| 174_v.txt | 32884 | 32884-Deleti | Deletion | 1 | A | - | 5 | 23328 | 0.02143347 | 36.6 | rnr1Y285F_pG | TRUE | CAN | JSY2420 | 5 | rnr1Y285F_p | CAA | nope | >1bp-ins | A/T-1 | none | A/T-1 | ACGTCCAAA/ | AAAATTG | GTT | GTT | T | none | T>none | T>none_GTT | A/T-1_AAAATTG | 6 | FALSE | FALSE |
| 174_v.txt | 32917 | 32917-Deleti | Deletion | 1 | C | - | 82 | 23328 | 0.35150892 | 49.902439 | rnr1Y285F_pG | TRUE | CAN | JSY2420 | 5 | rnr1Y285F_p | TCC | nope | >1bp-ins | G/C-1 | none | G/C-1 | TAAGTTCCA( | CCAGGGC | AGG | TCC | C | none | nope | nope_TCC | G/C-1_CCAGGGC | 6 | FALSE | FALSE |

| | | | | | | | | | | | | | | | | | | | | | | | | | | | | | | | | | | |
|---|---|---|---|---|---|---|---|---|---|---|---|---|---|---|---|---|---|---|---|---|---|---|---|---|---|---|---|---|---|---|---|---|---|---|
| 174_v.txt | 32919 | 32919-Deleti | Deletion | 1 | A | - | 7 | 23328 | 0.03000686 | 58.8571429 | rnr1Y285F_pG | TRUE | CAN | JSY2420 | 5 | rnr1Y285F_p | CAG | nope | >1bp-ins | A/T-1 | none | A/T-1 | AGTTCCAGG | AGGGCAA | GTC | GTC | T | none | T>none | T>none_GTC | A/T-1_AGGGCAA | 6 | FALSE | FALSE |
| 174_v.txt | 32924 | 32924-Deleti | Deletion | 1 | A | - | 21 | 23328 | 0.09002058 | 49.5238095 | rnr1Y285F_pG | TRUE | CAN | JSY2420 | 5 | rnr1Y285F_p | CAA | nope | >1bp-ins | A/T-1 | none | A/T-1 | CAGGGCAAA | AAAAGTG | GTT | GTT | T | none | T>none | T>none_GTT | A/T-1_AAAAGTG | 6 | FALSE | FALSE |
| 174_v.txt | 32945 | 32945-Deleti | Deletion | 1 | C | - | 16 | 23328 | 0.06858711 | 61.6875 | rnr1Y285F_pG | TRUE | CAN | JSY2420 | 5 | rnr1Y285F_p | ACC | nope | >1bp-ins | G/C-1 | none | G/C-1 | AGAAAACCA | CCAATAC | TGG | ACC | C | none | nope | nope_ACC | G/C-1_CCAATAC | 6 | FALSE | FALSE |
| 174_v.txt | 33000 | 33000-Deleti | Deletion | 1 | A | - | 23 | 23328 | 0.09859396 | 63.3478261 | rnr1Y285F_pG | TRUE | CAN | JSY2420 | 5 | rnr1Y285F_p | GAA | nope | >1bp-ins | A/T-1 | none | A/T-1 | TGTGAGAAA | AAAACTG | CTT | CTT | T | none | T>none | T>none_CTT | A/T-1_AAAACTG | 6 | FALSE | FALSE |
| 174_v.txt | 33078 | 33078-Deleti | Deletion | 1 | C | - | 26 | 23328 | 0.11145405 | 57.4230765 | rnr1Y285F_pG | TRUE | CAN | JSY2420 | 5 | rnr1Y285F_p | ACC | nope | >1bp-ins | G/C-1 | none | G/C-1 | AAAGAACCC | CCCATAA | TGG | ACC | C | none | nope | nope_ACC | G/C-1_CCCATAA | 6 | FALSE | FALSE |
| 174_v.txt | 33111 | 33111-Deleti | Deletion | 1 | G | - | 3 | 23328 | 0.01286008 | 54 | rnr1Y285F_pG | TRUE | CAN | JSY2420 | 5 | rnr1Y285F_p | TGG | nope | >1bp-ins | G/C-1 | none | G/C-1 | CCCACTGGG | GGGCCGG | ACC | ACC | C | none | C>none | C>none_ACC | G/C-1_GGGCCGG | 6 | FALSE | FALSE |
| 174_v.txt | 33138 | 33138-Deleti | Deletion | 1 | A | - | 3 | 23328 | 0.01286008 | 36.3333333 | rnr1Y285F_pG | TRUE | CAN | JSY2420 | 5 | rnr1Y285F_p | TAA | nope | >1bp-ins | A/T-1 | none | A/T-1 | GTGGATAAA | AAACCAA | ATT | ATT | T | none | T>none | T>none_ATT | A/T-1_AAACCAA | 6 | FALSE | FALSE |
| 174_v.txt | 33215 | 33215-Deleti | Deletion | 1 | C | - | 5 | 12706 | 0.03935149 | 38 | rnr1Y285F_pG | TRUE | CAN | JSY2420 | 5 | rnr1Y285F_p | TCA | nope | >1bp-ins | G/C-1 | none | G/C-1 | TCTCTTCACT | CACTTCA | AGT | TCA | C | none | nope | nope_TCA | G/C-1_CACTTCA | 6 | FALSE | FALSE |
| 174_v.txt | 33356 | 33356-Deleti | Deletion | 1 | C | - | 22 | 12706 | 0.17314655 | 52.2727273 | rnr1Y285F_pG | TRUE | CAN | JSY2420 | 5 | rnr1Y285F_p | ACC | nope | >1bp-ins | G/C-1 | none | G/C-1 | TATTGACCC | CCCACGT | TGG | ACC | C | none | nope | nope_ACC | G/C-1_CCCACGT | 6 | FALSE | FALSE |
| 174_v.txt | 33474 | 33474-Deleti | Deletion | 1 | C | - | 32 | 12705 | 0.25186934 | 27.71875 | rnr1Y285F_pG | TRUE | CAN | JSY2420 | 5 | rnr1Y285F_p | GCC | nope | >1bp-ins | G/C-1 | none | G/C-1 | GCTATGCCTT | CCTTTTT | CGG | GCC | C | none | nope | nope_GCC | G/C-1_CCTTTTT | 6 | FALSE | FALSE |
| 175_v.txt | 31723 | 31723-SNV-A | SNV | 1 | A | C | 7 | 9519 | 0.07353714 | 56.1428571 | rnr1Y285F_pG | TRUE | CAN | JSY2421 | 6 | rnr1Y285F_p | CAA | A>C | G/C+1 | A/T-1 | TA>GC | TA>GC | TGTCCCAAA | AAAAAGT | GTT | GTT | T | G | T>G | T>G_GTT | TA>GC_AAAAAGT | 6 | FALSE | FALSE |
| 175_v.txt | 31839 | 31839-SNV-A | SNV | 1 | A | C | 18 | 9519 | 0.18909549 | 47.3888889 | rnr1Y285F_pG | TRUE | CAN | JSY2421 | 6 | rnr1Y285F_p | TAA | A>C | G/C+1 | A/T-1 | TA>GC | TA>GC | TGAAATAAG | AAGATCC | ATT | ATT | T | G | T>G | T>G_ATT | TA>GC_AAGATCC | 6 | FALSE | FALSE |
| 175_v.txt | 32109 | 32109-SNV-A | SNV | 1 | A | C | 8 | 13710 | 0.05835157 | 58.5 | rnr1Y285F_pG | TRUE | CAN | JSY2421 | 6 | rnr1Y285F_p | TAA | A>C | G/C+1 | A/T-1 | TA>GC | TA>GC | ATATTTAATA | AATAGCC | ATT | ATT | T | G | T>G | T>G_ATT | TA>GC_AATAGCC | 6 | FALSE | FALSE |
| 175_v.txt | 32256 | 32256-SNV-A | SNV | 1 | A | C | 8 | 18151 | 0.04407471 | 17.75 | rnr1Y285F_pG | TRUE | CAN | JSY2421 | 6 | rnr1Y285F_p | TAG | A>C | G/C+1 | A/T-1 | TA>GC | TA>GC | TTTGATAGA | AGACCAA | ATC | ATC | T | G | T>G | T>G_ATC | TA>GC_AGACCAA | 6 | FALSE | FALSE |
| 175_v.txt | 32294 | 32294-SNV-A | SNV | 1 | A | C | 13 | 18153 | 0.07161351 | 46.2307692 | rnr1Y285F_pG | TRUE | CAN | JSY2421 | 6 | rnr1Y285F_p | AAT | A>C | G/C+1 | A/T-1 | TA>GC | TA>GC | ATTTGAATTT | ATTTGCG | TTA | TTA | T | G | T>G | T>G_TTA | TA>GC_ATTTGCG | 6 | FALSE | FALSE |
| 175_v.txt | 32314 | 32314-SNV-T | SNV | 1 | T | G | 11 | 18153 | 0.06059605 | 58 | rnr1Y285F_pG | TRUE | CAN | JSY2421 | 6 | rnr1Y285F_p | GTT | T>G | G/C+1 | A/T-1 | TA>GC | TA>GC | TAATGGTTG | TTGTTAA | CAA | GTT | T | C | T>G | T>G_GTT | TA>GC_TTGTTAA | 6 | FALSE | FALSE |
| 175_v.txt | 32439 | 32439-SNV-A | SNV | 1 | A | C | 19 | 18153 | 0.1046659 | 49.4736842 | rnr1Y285F_pG | TRUE | CAN | JSY2421 | 6 | rnr1Y285F_p | AAC | A>C | G/C+1 | A/T-1 | TA>GC | TA>GC | TATGGAACT | ACTAAAA | TTG | TTG | T | G | T>G | T>G_TTG | TA>GC_ACTAAAA | 6 | FALSE | FALSE |
| 175_v.txt | 32541 | 32541-SNV-T | SNV | 1 | T | G | 41 | 18153 | 0.22585799 | 37.1463415 | rnr1Y285F_pG | TRUE | CAN | JSY2421 | 6 | rnr1Y285F_p | TTC | T>G | G/C+1 | A/T-1 | TA>GC | TA>GC | GCAGCTTCA | TCACCAG | AAG | TTC | T | C | T>G | T>G_TTC | TA>GC_TCACCAG | 6 | FALSE | FALSE |
| 175_v.txt | 32680 | 32680-SNV-A | SNV | 1 | A | C | 10 | 25011 | 0.03998241 | 58.9007475 | rnr1Y285F_pG | TRUE | CAN | JSY2421 | 6 | rnr1Y285F_p | CAA | A>C | G/C+1 | A/T-1 | TA>GC | TA>GC | TTCTCCAATA | AATAACG | GTT | GTT | T | G | T>G | T>G_GTT | TA>GC_AATAACG | 6 | FALSE | FALSE |
| 175_v.txt | 32681 | 32681-SNV-A | SNV | 1 | A | C | 22 | 25011 | 0.0879613 | 43.0924226 | rnr1Y285F_pG | TRUE | CAN | JSY2421 | 6 | rnr1Y285F_p | AAT | A>C | G/C+1 | A/T-1 | TA>GC | TA>GC | TCTCCAATAA | ATAACGG | TTA | TTA | T | G | T>G | T>G_TTA | TA>GC_ATAACGG | 6 | FALSE | FALSE |
| 175_v.txt | 32717 | 32717-SNV-A | SNV | 1 | A | C | 1 | 25011 | 4.00E-03 | 60.0009152 | rnr1Y285F_pG | TRUE | CAN | JSY2421 | 6 | rnr1Y285F_p | CAC | A>C | G/C+1 | A/T-1 | TA>GC | TA>GC | AGCACCACA | ACAAACC | GTG | GTG | T | G | T>G | T>G_GTG | TA>GC_ACAAACC | 6 | FALSE | FALSE |
| 175_v.txt | 32728 | 32728-SNV-A | SNV | 1 | A | C | 6 | 25011 | 0.02398945 | 14.166687 | rnr1Y285F_pG | TRUE | CAN | JSY2421 | 6 | rnr1Y285F_p | CAA | A>C | G/C+1 | A/T-1 | TA>GC | TA>GC | CCATACAAA | AAAAACA | GTT | GTT | T | G | T>G | T>G_GTT | TA>GC_AAAAACA | 6 | FALSE | FALSE |
| 175_v.txt | 32763 | 32763-SNV-A | SNV | 1 | A | C | 8 | 25011 | 0.03198593 | 40.25 | rnr1Y285F_pG | TRUE | CAN | JSY2421 | 6 | rnr1Y285F_p | AAC | A>C | G/C+1 | A/T-1 | TA>GC | TA>GC | GCTAAAACT | ACTTTGA | TTG | TTG | T | G | T>G | T>G_TTG | TA>GC_ACTTTGA | 6 | FALSE | FALSE |
| 175_v.txt | 32801 | 32801-SNV-A | SNV | 1 | A | C | 45 | 25011 | 0.17992084 | 42.5333333 | rnr1Y285F_pG | TRUE | CAN | JSY2421 | 6 | rnr1Y285F_p | AAT | A>C | G/C+1 | A/T-1 | TA>GC | TA>GC | ACCGTAATA | ATATTTG | TTA | TTA | T | G | T>G | T>G_TTA | TA>GC_ATATTTG | 6 | FALSE | FALSE |
| 175_v.txt | 32842 | 32842-SNV-A | SNV | 1 | A | C | 4 | 25011 | 0.01599296 | 38.0018763 | rnr1Y285F_pG | TRUE | CAN | JSY2421 | 6 | rnr1Y285F_p | CAA | A>C | G/C+1 | A/T-1 | TA>GC | TA>GC | TTACCCAAA | AAAAAAT | GTT | GTT | T | G | T>G | T>G_GTT | TA>GC_AAAAAAT | 6 | FALSE | FALSE |
| 175_v.txt | 32884 | 32884-SNV-A | SNV | 1 | A | C | 2 | 24207 | 8.26E-03 | 37.9971988 | rnr1Y285F_pG | TRUE | CAN | JSY2421 | 6 | rnr1Y285F_p | CAA | A>C | G/C+1 | A/T-1 | TA>GC | TA>GC | ACGTCCAAA | AAAATTG | GTT | GTT | T | G | T>G | T>G_GTT | TA>GC_AAAATTG | 6 | FALSE | FALSE |
| 175_v.txt | 32929 | 32929-SNV-T | SNV | 1 | T | G | 17 | 24207 | 0.07022762 | 46.8823529 | rnr1Y285F_pG | TRUE | CAN | JSY2421 | 6 | rnr1Y285F_p | GTG | T>G | G/C+1 | A/T-1 | TA>GC | TA>GC | CAAAAGTGA | TGATTGC | CAC | GTG | T | C | T>G | T>G_GTG | TA>GC_TGATTGC | 6 | FALSE | FALSE |
| 175_v.txt | 32952 | 32952-SNV-A | SNV | 1 | A | C | 12 | 24207 | 0.04957244 | 56.0833333 | rnr1Y285F_pG | TRUE | CAN | JSY2421 | 6 | rnr1Y285F_p | CAT | A>C | G/C+1 | A/T-1 | TA>GC | TA>GC | CAATACATG | ATGTAAC | GTA | GTA | T | G | T>G | T>G_GTA | TA>GC_ATGTAAC | 6 | FALSE | FALSE |
| 175_v.txt | 33089 | 33089-SNV-A | SNV | 1 | A | C | 7 | 24205 | 0.02891965 | 44.7142928 | rnr1Y285F_pG | TRUE | CAN | JSY2421 | 6 | rnr1Y285F_p | AAT | A>C | G/C+1 | A/T-1 | TA>GC | TA>GC | AAATAAATA | ATATGAT | TTA | TTA | T | G | T>G | T>G_TTA | TA>GC_ATATGAT | 6 | FALSE | FALSE |
| 175_v.txt | 33099 | 33099-SNV-A | SNV | 1 | A | C | 17 | 24207 | 0.07022762 | 29.2941177 | rnr1Y285F_pG | TRUE | CAN | JSY2421 | 6 | rnr1Y285F_p | AAG | A>C | G/C+1 | A/T-1 | TA>GC | TA>GC | GATATAAGA | AGAGCGC | TTC | TTC | T | G | T>G | T>G_TTC | TA>GC_AGAGCGC | 6 | FALSE | FALSE |
| 175_v.txt | 33166 | 33166-SNV-T | SNV | 1 | T | G | 95 | 24206 | 0.39246468 | 37.7263158 | rnr1Y285F_pG | TRUE | CAN | JSY2421 | 6 | rnr1Y285F_p | GTA | T>G | G/C+1 | A/T-1 | TA>GC | TA>GC | CAATAGTAC | TACCACC | CAT | GTA | T | C | T>G | T>G_GTA | TA>GC_TACCACC | 6 | FALSE | FALSE |
| 175_v.txt | 33269 | 33269-SNV-A | SNV | 1 | A | C | 91 | 13039 | 0.69790628 | 16.5494506 | rnr1Y285F_pG | TRUE | CAN | JSY2421 | 6 | rnr1Y285F_p | TAG | A>C | G/C+1 | A/T-1 | TA>GC | TA>GC | CTCCATAGA | AGAGAAC | ATC | ATC | T | G | T>G | T>G_ATC | TA>GC_AGAGAAC | 6 | FALSE | FALSE |
| 175_v.txt | 32256 | 32256-SNV-A | SNV | 1 | A | G | 21 | 18151 | 0.11569611 | 36.3333333 | rnr1Y285F_pG | TRUE | CAN | JSY2421 | 6 | rnr1Y285F_p | TAG | A>G | G/C+1 | A/T-1 | TA>CG | TA>CG | TTTGATAGA | AGACCAA | ATC | ATC | T | C | T>C | T>C_ATC | TA>CG_AGACCAA | 6 | FALSE | FALSE |
| 175_v.txt | 32320 | 32320-SNV-A | SNV | 1 | A | G | 2 | 18153 | 0.01101746 | 63.9997021 | rnr1Y285F_pG | TRUE | CAN | JSY2421 | 6 | rnr1Y285F_p | AAG | A>G | G/C+1 | A/T-1 | TA>CG | TA>CG | TTGTTAAGAT | AGATAAC | TTC | TTC | T | C | T>C | T>C_TTC | TA>CG_AGATAAC | 6 | FALSE | FALSE |
| 175_v.txt | 32322 | 32322-SNV-A | SNV | 1 | A | G | 5 | 18153 | 0.02754366 | 33.1999532 | rnr1Y285F_pG | TRUE | CAN | JSY2421 | 6 | rnr1Y285F_p | GAT | A>G | G/C+1 | A/T-1 | TA>CG | TA>CG | GTTAAGATA | ATAACAG | CTA | CTA | T | C | T>C | T>C_CTA | TA>CG_ATAACAG | 6 | FALSE | FALSE |
| 175_v.txt | 32324 | 32324-SNV-A | SNV | 1 | A | G | 7 | 18153 | 0.03856112 | 56.4285701 | rnr1Y285F_pG | TRUE | CAN | JSY2421 | 6 | rnr1Y285F_p | TAA | A>G | G/C+1 | A/T-1 | TA>CG | TA>CG | TAAGATAAC | AACAGCG | ATT | ATT | T | C | T>C | T>C_ATT | TA>CG_AACAGCG | 6 | FALSE | FALSE |
| 175_v.txt | 32325 | 32325-SNV-A | SNV | 1 | A | G | 3 | 18153 | 0.01652619 | 63.9997989 | rnr1Y285F_pG | TRUE | CAN | JSY2421 | 6 | rnr1Y285F_p | AAC | A>G | G/C+1 | A/T-1 | TA>CG | TA>CG | AAGATAACA | ACAGCGT | TTG | TTG | T | C | T>C | T>C_TTG | TA>CG_ACAGCGT | 6 | FALSE | FALSE |
| 175_v.txt | 32327 | 32327-SNV-A | SNV | 1 | A | G | 6 | 18153 | 0.03305239 | 54.8328433 | rnr1Y285F_pG | TRUE | CAN | JSY2421 | 6 | rnr1Y285F_p | CAG | A>G | G/C+1 | A/T-1 | TA>CG | TA>CG | GATAACAGC | AGCGTTG | GTC | GTC | T | C | T>C | T>C_GTC | TA>CG_AGCGTTG | 6 | FALSE | FALSE |
| 175_v.txt | 32387 | 32387-SNV-A | SNV | 1 | A | G | 9 | 18153 | 0.04957858 | 63.3333333 | rnr1Y285F_pG | TRUE | CAN | JSY2421 | 6 | rnr1Y285F_p | GAG | A>G | G/C+1 | A/T-1 | TA>CG | TA>CG | AAAGGGAGA | AGAAGTA | CTC | CTC | T | C | T>C | T>C_CTC | TA>CG_AGAAGTA | 6 | FALSE | FALSE |
| 175_v.txt | 32490 | 32490-SNV-A | SNV | 1 | A | G | 120 | 18153 | 0.66104776 | 44.775 | rnr1Y285F_pG | TRUE | CAN | JSY2421 | 6 | rnr1Y285F_p | GAA | A>G | G/C+1 | A/T-1 | TA>CG | TA>CG | ATACGGAAA | AAAACAA | CTT | CTT | T | C | T>C | T>C_CTT | TA>CG_AAAACAA | 6 | FALSE | FALSE |
| 175_v.txt | 32541 | 32541-SNV-T | SNV | 1 | T | C | 23 | 18153 | 0.12670082 | 34.5217391 | rnr1Y285F_pG | TRUE | CAN | JSY2421 | 6 | rnr1Y285F_p | TTC | T>C | G/C+1 | A/T-1 | TA>CG | TA>CG | GCAGCTTCA | TCACCAG | AAG | TTC | T | G | T>C | T>C_TTC | TA>CG_TCACCAG | 6 | FALSE | FALSE |
| 175_v.txt | 32574 | 32574-SNV-T | SNV | 1 | T | C | 107 | 25012 | 0.42779466 | 37.1868158 | rnr1Y285F_pG | TRUE | CAN | JSY2421 | 6 | rnr1Y285F_p | TTG | T>C | G/C+1 | A/T-1 | TA>CG | TA>CG | GTACCTTGA | TGAAATG | AAC | TTG | T | G | T>C | T>C_TTG | TA>CG_TGAAATG | 6 | FALSE | FALSE |
| 175_v.txt | 32642 | 32642-SNV-A | SNV | 1 | A | G | 3 | 25011 | 0.01199472 | 55.305768 | rnr1Y285F_pG | TRUE | CAN | JSY2421 | 6 | rnr1Y285F_p | TAG | A>G | G/C+1 | A/T-1 | TA>CG | TA>CG | ATCCTTAGAT | AGATATT | ATC | ATC | T | C | T>C | T>C_ATC | TA>CG_AGATATT | 6 | FALSE | FALSE |
| 175_v.txt | 32680 | 32680-SNV-A | SNV | 1 | A | G | 7 | 25011 | 0.02798769 | 56.286033 | rnr1Y285F_pG | TRUE | CAN | JSY2421 | 6 | rnr1Y285F_p | CAA | A>G | G/C+1 | A/T-1 | TA>CG | TA>CG | TTCTCCAATA | AATAACG | GTT | GTT | T | C | T>C | T>C_GTT | TA>CG_AATAACG | 6 | FALSE | FALSE |
| 175_v.txt | 32717 | 32717-SNV-A | SNV | 1 | A | G | 6 | 25011 | 0.02398945 | 59.5002722 | rnr1Y285F_pG | TRUE | CAN | JSY2421 | 6 | rnr1Y285F_p | CAC | A>G | G/C+1 | A/T-1 | TA>CG | TA>CG | AGCACCACA | ACAAACC | GTG | GTG | T | C | T>C | T>C_GTG | TA>CG_ACAAACC | 6 | FALSE | FALSE |
| 175_v.txt | 32787 | 32787-SNV-T | SNV | 1 | T | C | 11 | 25011 | 0.04398065 | 40.9090909 | rnr1Y285F_pG | TRUE | CAN | JSY2421 | 6 | rnr1Y285F_p | CTC | T>C | G/C+1 | A/T-1 | TA>CG | TA>CG | CAGAACTCG | TCGAATT | GAG | CTC | T | G | T>C | T>C_CTC | TA>CG_TCGAATT | 6 | FALSE | FALSE |
| 175_v.txt | 32884 | 32884-SNV-A | SNV | 1 | A | G | 4 | 24207 | 0.01652415 | 37.9961007 | rnr1Y285F_pG | TRUE | CAN | JSY2421 | 6 | rnr1Y285F_p | CAA | A>G | G/C+1 | A/T-1 | TA>CG | TA>CG | ACGTCCAAA | AAAATTG | GTT | GTT | T | C | T>C | T>C_GTT | TA>CG_AAAATTG | 6 | FALSE | FALSE |
| 175_v.txt | 32924 | 32924-SNV-A | SNV | 1 | A | G | 5 | 24207 | 0.02065518 | 39.7999482 | rnr1Y285F_pG | TRUE | CAN | JSY2421 | 6 | rnr1Y285F_p | CAA | A>G | G/C+1 | A/T-1 | TA>CG | TA>CG | CAGGGCAAA | AAAAGTG | GTT | GTT | T | C | T>C | T>C_GTT | TA>CG_AAAAGTG | 6 | FALSE | FALSE |
| 175_v.txt | 32941 | 32941-SNV-A | SNV | 1 | A | G | 7 | 24207 | 0.02891726 | 40.4281334 | rnr1Y285F_pG | TRUE | CAN | JSY2421 | 6 | rnr1Y285F_p | GAA | A>G | G/C+1 | A/T-1 | TA>CG | TA>CG | CCCAAGAAA | AAAACCA | CTT | CTT | T | C | T>C | T>C_CTT | TA>CG_AAAACCA | 6 | FALSE | FALSE |
| 175_v.txt | 31867 | 31867-SNV-T | SNV | 1 | T | A | 11 | 9519 | 0.11555836 | 40.2722925 | rnr1Y285F_pG | TRUE | CAN | JSY2421 | 6 | rnr1Y285F_p | ATA | T>A | A/T+1 | A/T-1 | TA>AT | TA>AT | GGAAAATAG | TAGAGAT | TAT | ATA | T | T | T>A | T>A_ATA | TA>AT_TAGAGAT | 6 | FALSE | FALSE |
| 175_v.txt | 31986 | 31986-SNV-A | SNV | 1 | A | T | 33 | 13700 | 0.24087591 | 50.9393939 | rnr1Y285F_pG | TRUE | CAN | JSY2421 | 6 | rnr1Y285F_p | TAA | A>T | A/T+1 | A/T-1 | TA>AT | TA>AT | GGCATTAATT | AATTTAG | ATT | ATT | T | A | T>A | T>A_ATT | TA>AT_AATTTAG | 6 | FALSE | FALSE |
| 175_v.txt | 32055 | 32055-SNV-T | SNV | 1 | T | A | 29 | 13710 | 0.21152444 | 58.6551724 | rnr1Y285F_pG | TRUE | CAN | JSY2421 | 6 | rnr1Y285F_p | GTG | T>A | A/T+1 | A/T-1 | TA>AT | TA>AT | CTGATGTGC | TGCGAGA | CAC | GTG | T | T | T>A | T>A_GTG | TA>AT_TGCGAGA | 6 | FALSE | FALSE |
| 175_v.txt | 32109 | 32109-SNV-A | SNV | 1 | A | T | 20 | 13710 | 0.14587892 | 58.75 | rnr1Y285F_pG | TRUE | CAN | JSY2421 | 6 | rnr1Y285F_p | TAA | A>T | A/T+1 | A/T-1 | TA>AT | TA>AT | ATATTTAATA | AATAGCC | ATT | ATT | T | A | T>A | T>A_ATT | TA>AT_AATAGCC | 6 | FALSE | FALSE |
| 175_v.txt | 32169 | 32169-SNV-A | SNV | 1 | A | T | 14 | 13710 | 0.10211524 | 46.7142857 | rnr1Y285F_pG | TRUE | CAN | JSY2421 | 6 | rnr1Y285F_p | AAA | A>T | A/T+1 | A/T-1 | TA>AT | TA>AT | GCGCCAAAT | AATGCAG | TTT | TTT | T | A | T>A | T>A_TTT | TA>AT_AATGCAG | 6 | FALSE | FALSE |
| 175_v.txt | 32319 | 32319-SNV-A | SNV | 1 | A | T | 14 | 18153 | 0.07712224 | 58.6378798 | rnr1Y285F_pG | TRUE | CAN | JSY2421 | 6 | rnr1Y285F_p | TAA | A>T | A/T+1 | A/T-1 | TA>AT | TA>AT | GTTGTTAAG | AAGATAA | ATT | ATT | T | A | T>A | T>A_ATT | TA>AT_AAGATAA | 6 | FALSE | FALSE |
| 175_v.txt | 32320 | 32320-SNV-A | SNV | 1 | A | T | 4 | 18153 | 0.02203493 | 36.7495625 | rnr1Y285F_pG | TRUE | CAN | JSY2421 | 6 | rnr1Y285F_p | AAG | A>T | A/T+1 | A/T-1 | TA>AT | TA>AT | TTGTTAAGAT | AGATAAC | TTC | TTC | T | A | T>A | T>A_TTC | TA>AT_AGATAAC | 6 | FALSE | FALSE |
| 175_v.txt | 32325 | 32325-SNV-A | SNV | 1 | A | T | 2 | 18153 | 0.01101746 | 50.4997074 | rnr1Y285F_pG | TRUE | CAN | JSY2421 | 6 | rnr1Y285F_p | AAC | A>T | A/T+1 | A/T-1 | TA>AT | TA>AT | AAGATAACA | ACAGCGT | TTG | TTG | T | A | T>A | T>A_TTG | TA>AT_ACAGCGT | 6 | FALSE | FALSE |
| 175_v.txt | 32442 | 32442-SNV-A | SNV | 1 | A | T | 9 | 18153 | 0.04957858 | 55 | rnr1Y285F_pG | TRUE | CAN | JSY2421 | 6 | rnr1Y285F_p | TAA | A>T | A/T+1 | A/T-1 | TA>AT | TA>AT | GGAACTAAA | AAAAGTC | ATT | ATT | T | A | T>A | T>A_ATT | TA>AT_AAAAGTC | 6 | FALSE | FALSE |
| 175_v.txt | 32574 | 32574-SNV-T | SNV | 1 | T | A | 5 | 25012 | 0.01999041 | 31.3997214 | rnr1Y285F_pG | TRUE | CAN | JSY2421 | 6 | rnr1Y285F_p | TTG | T>A | A/T+1 | A/T-1 | TA>AT | TA>AT | GTACCTTGA | TGAAATG | AAC | TTG | T | T | T>A | T>A_TTG | TA>AT_TGAAATG | 6 | FALSE | FALSE |
| 175_v.txt | 32654 | 32654-SNV-T | SNV | 1 | T | A | 17 | 25011 | 0.06797009 | 58.9411765 | rnr1Y285F_pG | TRUE | CAN | JSY2421 | 6 | rnr1Y285F_p | CTG | T>A | A/T+1 | A/T-1 | TA>AT | TA>AT | TATACCTGG | TGGACCC | GAC | CTG | T | T | T>A | T>A_CTG | TA>AT_TGGACCC | 6 | FALSE | FALSE |
| 175_v.txt | 32732 | 32732-SNV-A | SNV | 1 | A | T | 32 | 25011 | 0.12794371 | 57.375 | rnr1Y285F_pG | TRUE | CAN | JSY2421 | 6 | rnr1Y285F_p | AAC | A>T | A/T+1 | A/T-1 | TA>AT | TA>AT | ACAAAAACA | ACAGTAT | TTG | TTG | T | A | T>A | T>A_TTG | TA>AT_ACAGTAT | 6 | FALSE | FALSE |
| 175_v.txt | 32787 | 32787-SNV-T | SNV | 1 | T | A | 25 | 25011 | 0.09995602 | 48.36 | rnr1Y285F_pG | TRUE | CAN | JSY2421 | 6 | rnr1Y285F_p | CTC | T>A | A/T+1 | A/T-1 | TA>AT | TA>AT | CAGAACTCG | TCGAATT | GAG | CTC | T | T | T>A | T>A_CTC | TA>AT_TCGAATT | 6 | FALSE | FALSE |
| 175_v.txt | 32924 | 32924-SNV-A | SNV | 1 | A | T | 3 | 24207 | 0.01239311 | 37.3330949 | rnr1Y285F_pG | TRUE | CAN | JSY2421 | 6 | rnr1Y285F_p | CAA | A>T | A/T+1 | A/T-1 | TA>AT | TA>AT | CAGGGCAAA | AAAAGTG | GTT | GTT | T | A | T>A | T>A_GTT | TA>AT_AAAAGTG | 6 | FALSE | FALSE |
| 175_v.txt | 32941 | 32941-SNV-A | SNV | 1 | A | T | 3 | 24207 | 0.01239311 | 28.6663583 | rnr1Y285F_pG | TRUE | CAN | JSY2421 | 6 | rnr1Y285F_p | GAA | A>T | A/T+1 | A/T-1 | TA>AT | TA>AT | CCCAAGAAA | AAAACCA | CTT | CTT | T | A | T>A | T>A_CTT | TA>AT_AAAACCA | 6 | FALSE | FALSE |
| 175_v.txt | 33087 | 33087-SNV-A | SNV | 1 | A | T | 30 | 24207 | 0.12393109 | 57.2 | rnr1Y285F_pG | TRUE | CAN | JSY2421 | 6 | rnr1Y285F_p | TAA | A>T | A/T+1 | A/T-1 | TA>AT | TA>AT | ATAAATAAAT | AAATATG | ATT | ATT | T | A | T>A | T>A_ATT | TA>AT_AAATATG | 6 | FALSE | FALSE |
| 175_v.txt | 33089 | 33089-SNV-A | SNV | 1 | A | T | 420 | 24205 | 1.73517868 | 50.1878144 | rnr1Y285F_pG | TRUE | CAN | JSY2421 | 6 | rnr1Y285F_p | AAT | A>T | A/T+1 | A/T-1 | TA>AT | TA>AT | AAATAAATA | ATATGAT | TTA | TTA | T | A | T>A | T>A_TTA | TA>AT_ATATGAT | 6 | FALSE | FALSE |
| 175_v.txt | 33317 | 33317-SNV-A | SNV | 1 | A | T | 11 | 13039 | 0.0843623 | 54.4545455 | rnr1Y285F_pG | TRUE | CAN | JSY2421 | 6 | rnr1Y285F_p | GAT | A>T | A/T+1 | A/T-1 | TA>AT | TA>AT | CAATGGATA | ATACAAT | CTA | CTA | T | A | T>A | T>A_CTA | TA>AT_ATACAAT | 6 | FALSE | FALSE |
| 175_v.txt | 31650 | 31650-SNV-C | SNV | 1 | C | T | 27 | 9519 | 0.28364324 | 35.2592593 | rnr1Y285F_pG | TRUE | CAN | JSY2421 | 6 | rnr1Y285F_p | GCG | C>T | A/T+1 | G/C-1 | CG>TA | CG>TA | AGAATGCGA | CGAAATG | CGC | GCG | C | A | C>T | C>T_GCG | CG>TA_CGAAATG | 6 | FALSE | FALSE |
| 175_v.txt | 31830 | 31830-SNV-C | SNV | 1 | C | T | 13 | 9519 | 0.13656897 | 57.1538462 | rnr1Y285F_pG | TRUE | CAN | JSY2421 | 6 | rnr1Y285F_p | GCA | C>T | A/T+1 | G/C-1 | CG>TA | CG>TA | AATATGCATT | CATTGAA | CGT | GCA | C | A | C>T | C>T_GCA | CG>TA_CATTGAA | 6 | FALSE | FALSE |
| 175_v.txt | 31834 | 31834-SNV-C | SNV | 1 | G | A | 16 | 9519 | 0.16808488 | 54.9375 | rnr1Y285F_pG | TRUE | CAN | JSY2421 | 6 | rnr1Y285F_p | TGA | G>A | A/T+1 | G/C-1 | CG>TA | CG>TA | TGCATTGAA | GAAATAA | ACT | ACT | C | T | C>T | C>T_ACT | CG>TA_GAAATAA | 6 | FALSE | FALSE |
| 175_v.txt | 31844 | 31844-SNV-C | SNV | 1 | C | T | 16 | 9519 | 0.16808488 | 51.375 | rnr1Y285F_pG | TRUE | CAN | JSY2421 | 6 | rnr1Y285F_p | TCC | C>T | A/T+1 | G/C-1 | CG>TA | CG>TA | TAAGATCCA | CCAAACA | AGG | TCC | C | A | C>T | C>T_TCC | CG>TA_CCAAACA | 6 | FALSE | FALSE |
| 175_v.txt | 31869 | 31869-SNV-C | SNV | 1 | G | A | 12 | 9519 | 0.12606366 | 39.5 | rnr1Y285F_pG | TRUE | CAN | JSY2421 | 6 | rnr1Y285F_p | AGA | G>A | A/T+1 | G/C-1 | CG>TA | CG>TA | AAAATAGAG | GAGATAT | TCT | TCT | C | T | C>T | C>T_TCT | CG>TA_GAGATAT | 6 | FALSE | FALSE |
| 175_v.txt | 31962 | 31962-SNV-C | SNV | 1 | G | A | 30 | 13700 | 0.2189781 | 35.5333333 | rnr1Y285F_pG | TRUE | CAN | JSY2421 | 6 | rnr1Y285F_p | CGC | G>A | A/T+1 | G/C-1 | CG>TA | CG>TA | GTGGCCGCA | GCATAAT | GCG | GCG | C | T | C>T | C>T_GCG | CG>TA_GCATAAT | 6 | FALSE | FALSE |
| 175_v.txt | 31979 | 31979-SNV-C | SNV | 1 | G | A | 8 | 13700 | 0.05839416 | 52.9953731 | rnr1Y285F_pG | TRUE | CAN | JSY2421 | 6 | rnr1Y285F_p | CGG | G>A | A/T+1 | G/C-1 | CG>TA | CG>TA | CAAGCCGGG | GGGCATT | GCC | GCC | C | T | C>T | C>T_GCC | CG>TA_GGGCATT | 6 | FALSE | FALSE |
| 175_v.txt | 32057 | 32057-SNV-C | SNV | 1 | C | T | 16 | 13710 | 0.11670314 | 59.0625 | rnr1Y285F_pG | TRUE | CAN | JSY2421 | 6 | rnr1Y285F_p | GCG | C>T | A/T+1 | G/C-1 | CG>TA | CG>TA | GATGTGCGA | CGAGATT | CGC | GCG | C | A | C>T | C>T_GCG | CG>TA_CGAGATT | 6 | FALSE | FALSE |
| 175_v.txt | 32075 | 32075-SNV-C | SNV | 1 | C | T | 232 | 13710 | 1.69219548 | 60.8793103 | rnr1Y285F_pG | TRUE | CAN | JSY2421 | 6 | rnr1Y285F_p | ACC | C>T | A/T+1 | G/C-1 | CG>TA | CG>TA | AAATAACCA | CCATGCA | TGG | ACC | C | A | C>T | C>T_ACC | CG>TA_CCATGCA | 6 | FALSE | FALSE |
| 175_v.txt | 32088 | 32088-SNV-C | SNV | 1 | C | T | 244 | 13710 | 1.77972283 | 61.1803279 | rnr1Y285F_pG | TRUE | CAN | JSY2421 | 6 | rnr1Y285F_p | GCC | C>T | A/T+1 | G/C-1 | CG>TA | CG>TA | AAAAAGCCT | CCTGCAA | CGG | GCC | C | A | C>T | C>T_GCC | CG>TA_CCTGCAA | 6 | FALSE | FALSE |
| 175_v.txt | 32092 | 32092-SNV-C | SNV | 1 | C | T | 18 | 13710 | 0.13129103 | 60.1666667 | rnr1Y285F_pG | TRUE | CAN | JSY2421 | 6 | rnr1Y285F_p | GCA | C>T | A/T+1 | G/C-1 | CG>TA | CG>TA | AGCCTGCAA | CAACACC | CGT | GCA | C | A | C>T | C>T_GCA | CG>TA_CAACACC | 6 | FALSE | FALSE |
| 175_v.txt | 32114 | 32114-SNV-C | SNV | 1 | C | T | 70 | 13710 | 0.51057622 | 60.2 | rnr1Y285F_pG | TRUE | CAN | JSY2421 | 6 | rnr1Y285F_p | GCC | C>T | A/T+1 | G/C-1 | CG>TA | CG>TA | TAATAGCCA | CCATTCG | CGG | GCC | C | A | C>T | C>T_GCC | CG>TA_CCATTCG | 6 | FALSE | FALSE |
| 175_v.txt | 32165 | 32165-SNV-C | SNV | 1 | G | A | 23 | 13710 | 0.16776076 | 48.6086957 | rnr1Y285F_pG | TRUE | CAN | JSY2421 | 6 | rnr1Y285F_p | CGC | G>A | A/T+1 | G/C-1 | CG>TA | CG>TA | CAAAGCGCC | GCCAAAT | GCG | GCG | C | T | C>T | C>T_GCG | CG>TA_GCCAAAT | 6 | FALSE | FALSE |
| 175_v.txt | 32190 | 32190-SNV-C | SNV | 1 | G | A | 16 | 13710 | 0.11670314 | 42 | rnr1Y285F_pG | TRUE | CAN | JSY2421 | 6 | rnr1Y285F_p | TGC | G>A | A/T+1 | G/C-1 | CG>TA | CG>TA | AAAACTGCA | GCAATGT | ACG | ACG | C | T | C>T | C>T_ACG | CG>TA_GCAATGT | 6 | FALSE | FALSE |
| 175_v.txt | 32271 | 32271-SNV-C | SNV | 1 | C | T | 66 | 18153 | 0.36357627 | 35.5909091 | rnr1Y285F_pG | TRUE | CAN | JSY2421 | 6 | rnr1Y285F_p | ACG | C>T | A/T+1 | G/C-1 | CG>TA | CG>TA | AAAATACGG | CGGGAAC | TGC | ACG | C | A | C>T | C>T_ACG | CG>TA_CGGGAAC | 6 | FALSE | FALSE |
| 175_v.txt | 32274 | 32274-SNV-C | SNV | 1 | G | A | 179 | 18153 | 0.98606291 | 36.8379888 | rnr1Y285F_pG | TRUE | CAN | JSY2421 | 6 | rnr1Y285F_p | GGA | G>A | A/T+1 | G/C-1 | CG>TA | CG>TA | ATACGGGAA | GAACCAA | CCT | CCT | C | T | C>T | C>T_CCT | CG>TA_GAACCAA | 6 | FALSE | FALSE |
| 175_v.txt | 32299 | 32299-SNV-C | SNV | 1 | C | T | 42 | 18153 | 0.23136672 | 51.5714286 | rnr1Y285F_pG | TRUE | CAN | JSY2421 | 6 | rnr1Y285F_p | GCG | C>T | A/T+1 | G/C-1 | CG>TA | CG>TA | AATTTGCGG | CGGCAGA | CGC | GCG | C | A | C>T | C>T_GCG | CG>TA_CGGCAGA | 6 | FALSE | FALSE |
| 175_v.txt | 32304 | 32304-SNV-C | SNV | 1 | G | A | 15 | 18153 | 0.08263097 | 55.2 | rnr1Y285F_pG | TRUE | CAN | JSY2421 | 6 | rnr1Y285F_p | AGA | G>A | A/T+1 | G/C-1 | CG>TA | CG>TA | GCGGCAGAA | GAAATAA | TCT | TCT | C | T | C>T | C>T_TCT | CG>TA_GAAATAA | 6 | FALSE | FALSE |
| 175_v.txt | 32326 | 32326-SNV-C | SNV | 1 | C | T | 6 | 18153 | 0.03305239 | 61.9990493 | rnr1Y285F_pG | TRUE | CAN | JSY2421 | 6 | rnr1Y285F_p | ACA | C>T | A/T+1 | G/C-1 | CG>TA | CG>TA | AGATAACAG | CAGCGTT | TGT | ACA | C | A | C>T | C>T_ACA | CG>TA_CAGCGTT | 6 | FALSE | FALSE |
| 175_v.txt | 32449 | 32449-SNV-C | SNV | 1 | C | T | 64 | 18153 | 0.35255881 | 53.125 | rnr1Y285F_pG | TRUE | CAN | JSY2421 | 6 | rnr1Y285F_p | CCA | C>T | A/T+1 | G/C-1 | CG>TA | CG>TA | AAAGTCCAA | CAATGAA | GGT | CCA | C | A | C>T | C>T_CCA | CG>TA_CAATGAA | 6 | FALSE | FALSE |
| 175_v.txt | 32466 | 32466-SNV-C | SNV | 1 | C | T | 28 | 18153 | 0.15424448 | 49.7142857 | rnr1Y285F_pG | TRUE | CAN | JSY2421 | 6 | rnr1Y285F_p | GCC | C>T | A/T+1 | G/C-1 | CG>TA | CG>TA | AGAGAGCCA | CCAATGT | CGG | GCC | C | A | C>T | C>T_GCC | CG>TA_CCAATGT | 6 | FALSE | FALSE |
| 175_v.txt | 32487 | 32487-SNV-C | SNV | 1 | C | T | 32 | 18153 | 0.1762794 | 46.7191546 | rnr1Y285F_pG | TRUE | CAN | JSY2421 | 6 | rnr1Y285F_p | ACG | C>T | A/T+1 | G/C-1 | CG>TA | CG>TA | AAGATACGG | CGGAAAA | TGC | ACG | C | A | C>T | C>T_ACG | CG>TA_CGGAAAA | 6 | FALSE | FALSE |
| 175_v.txt | 32488 | 32488-SNV-C | SNV | 1 | G | A | 32 | 18153 | 0.1762794 | 45.8403671 | rnr1Y285F_pG | TRUE | CAN | JSY2421 | 6 | rnr1Y285F_p | CGG | G>A | A/T+1 | G/C-1 | CG>TA | CG>TA | AGATACGGA | GGAAAAC | GCC | GCC | C | T | C>T | C>T_GCC | CG>TA_GGAAAAC | 6 | FALSE | FALSE |
| 175_v.txt | 32529 | 32529-SNV-C | SNV | 1 | G | A | 25 | 18153 | 0.13771828 | 36.8803282 | rnr1Y285F_pG | TRUE | CAN | JSY2421 | 6 | rnr1Y285F_p | GGG | G>A | A/T+1 | G/C-1 | CG>TA | CG>TA | TTTCTGGGG | GGGTTTG | CCC | CCC | C | T | C>T | C>T_CCC | CG>TA_GGGTTTG | 6 | FALSE | FALSE |
| 175_v.txt | 32535 | 32535-SNV-C | SNV | 1 | G | A | 21 | 18153 | 0.11568336 | 37.7989395 | rnr1Y285F_pG | TRUE | CAN | JSY2421 | 6 | rnr1Y285F_p | TGC | G>A | A/T+1 | G/C-1 | CG>TA | CG>TA | GGGTTTGCA | GCAGCTT | ACG | ACG | C | T | C>T | C>T_ACG | CG>TA_GCAGCTT | 6 | FALSE | FALSE |
| 175_v.txt | 32536 | 32536-SNV-C | SNV | 1 | C | T | 22 | 18153 | 0.12119209 | 36.5455782 | rnr1Y285F_pG | TRUE | CAN | JSY2421 | 6 | rnr1Y285F_p | GCA | C>T | A/T+1 | G/C-1 | CG>TA | CG>TA | GGTTTGCAG | CAGCTTC | CGT | GCA | C | A | C>T | C>T_GCA | CG>TA_CAGCTTC | 6 | FALSE | FALSE |
| 175_v.txt | 32566 | 32566-SNV-C | SNV | 1 | C | T | 38 | 25009 | 0.1519453 | 36.5263158 | rnr1Y285F_pG | TRUE | CAN | JSY2421 | 6 | rnr1Y285F_p | TCA | C>T | A/T+1 | G/C-1 | CG>TA | CG>TA | CTAGTTCAG | CAGTACC | AGT | TCA | C | A | C>T | C>T_TCA | CG>TA_CAGTACC | 6 | FALSE | FALSE |
| 175_v.txt | 32571 | 32571-SNV-C | SNV | 1 | C | T | 108 | 25012 | 0.43179274 | 35.4814815 | rnr1Y285F_pG | TRUE | CAN | JSY2421 | 6 | rnr1Y285F_p | ACC | C>T | A/T+1 | G/C-1 | CG>TA | CG>TA | TCAGTACCT | CCTTGAA | TGG | ACC | C | A | C>T | C>T_ACC | CG>TA_CCTTGAA | 6 | FALSE | FALSE |
| 175_v.txt | 32587 | 32587-SNV-C | SNV | 1 | C | T | 27 | 25011 | 0.1079525 | 35 | rnr1Y285F_pG | TRUE | CAN | JSY2421 | 6 | rnr1Y285F_p | GCA | C>T | A/T+1 | G/C-1 | CG>TA | CG>TA | TGAAGGCAG | CAGCGTT | CGT | GCA | C | A | C>T | C>T_GCA | CG>TA_CAGCGTT | 6 | FALSE | FALSE |
| 175_v.txt | 32590 | 32590-SNV-C | SNV | 1 | C | T | 47 | 25011 | 0.18791732 | 36.4680851 | rnr1Y285F_pG | TRUE | CAN | JSY2421 | 6 | rnr1Y285F_p | GCG | C>T | A/T+1 | G/C-1 | CG>TA | CG>TA | AGGCAGCGT | CGTTAAT | CGC | GCG | C | A | C>T | C>T_GCG | CG>TA_CGTTAAT | 6 | FALSE | FALSE |
| 175_v.txt | 32609 | 32609-SNV-C | SNV | 1 | C | T | 47 | 25011 | 0.18791732 | 44.8291564 | rnr1Y285F_pG | TRUE | CAN | JSY2421 | 6 | rnr1Y285F_p | CCC | C>T | A/T+1 | G/C-1 | CG>TA | CG>TA | GGAAACCCA | CCAACCT | GGG | CCC | C | A | C>T | C>T_CCC | CG>TA_CCAACCT | 6 | FALSE | FALSE |
| 175_v.txt | 32658 | 32658-SNV-C | SNV | 1 | C | T | 26 | 25011 | 0.10395426 | 51.3846154 | rnr1Y285F_pG | TRUE | CAN | JSY2421 | 6 | rnr1Y285F_p | ACC | C>T | A/T+1 | G/C-1 | CG>TA | CG>TA | CCTGGACCC | CCCCAGG | TGG | ACC | C | A | C>T | C>T_ACC | CG>TA_CCCCAGG | 6 | FALSE | FALSE |
| 175_v.txt | 32660 | 32660-SNV-C | SNV | 1 | C | T | 30 | 25011 | 0.11994722 | 54.0322311 | rnr1Y285F_pG | TRUE | CAN | JSY2421 | 6 | rnr1Y285F_p | CCC | C>T | A/T+1 | G/C-1 | CG>TA | CG>TA | TGGACCCCA | CCAGGCA | GGG | CCC | C | A | C>T | C>T_CCC | CG>TA_CCAGGCA | 6 | FALSE | FALSE |
| 175_v.txt | 32661 | 32661-SNV-C | SNV | 1 | C | T | 34 | 25011 | 0.13594019 | 57.9397111 | rnr1Y285F_pG | TRUE | CAN | JSY2421 | 6 | rnr1Y285F_p | CCA | C>T | A/T+1 | G/C-1 | CG>TA | CG>TA | GGACCCCAG | CAGGCAC | GGT | CCA | C | A | C>T | C>T_CCA | CG>TA_CAGGCAC | 6 | FALSE | FALSE |
| 175_v.txt | 32678 | 32678-SNV-C | SNV | 1 | C | T | 27 | 25011 | 0.1079525 | 54.9259259 | rnr1Y285F_pG | TRUE | CAN | JSY2421 | 6 | rnr1Y285F_p | TCC | C>T | A/T+1 | G/C-1 | CG>TA | CG>TA | GTTTCTCCAA | CCAATAA | AGG | TCC | C | A | C>T | C>T_TCC | CG>TA_CCAATAA | 6 | FALSE | FALSE |
| 175_v.txt | 32700 | 32700-SNV-C | SNV | 1 | C | T | 35 | 25011 | 0.13993843 | 60.2857143 | rnr1Y285F_pG | TRUE | CAN | JSY2421 | 6 | rnr1Y285F_p | GCC | C>T | A/T+1 | G/C-1 | CG>TA | CG>TA | ACTGGGCCG | CCGGTAA | CGG | GCC | C | A | C>T | C>T_GCC | CG>TA_CCGGTAA | 6 | FALSE | FALSE |
| 175_v.txt | 32749 | 32749-SNV-C | SNV | 1 | C | T | 39 | 25011 | 0.15593139 | 57.2820513 | rnr1Y285F_pG | TRUE | CAN | JSY2421 | 6 | rnr1Y285F_p | CCG | C>T | A/T+1 | G/C-1 | CG>TA | CG>TA | GAAACCCGA | CGATAAT | GGC | CCG | C | A | C>T | C>T_CCG | CG>TA_CGATAAT | 6 | FALSE | FALSE |
| 175_v.txt | 32757 | 32757-SNV-C | SNV | 1 | G | A | 28 | 25011 | 0.11195074 | 53.6071429 | rnr1Y285F_pG | TRUE | CAN | JSY2421 | 6 | rnr1Y285F_p | GGC | G>A | A/T+1 | G/C-1 | CG>TA | CG>TA | ATAATGGCT | GCTAAAA | CCG | CCG | C | T | C>T | C>T_CCG | CG>TA_GCTAAAA | 6 | FALSE | FALSE |
| 175_v.txt | 32776 | 32776-SNV-C | SNV | 1 | C | T | 32 | 25011 | 0.12794371 | 51.625 | rnr1Y285F_pG | TRUE | CAN | JSY2421 | 6 | rnr1Y285F_p | GCG | C>T | A/T+1 | G/C-1 | CG>TA | CG>TA | TGGAAGCGA | CGACCCA | CGC | GCG | C | A | C>T | C>T_GCG | CG>TA_CGACCCA | 6 | FALSE | FALSE |
| 175_v.txt | 32780 | 32780-SNV-C | SNV | 1 | C | T | 88 | 25011 | 0.35184519 | 47.125 | rnr1Y285F_pG | TRUE | CAN | JSY2421 | 6 | rnr1Y285F_p | CCC | C>T | A/T+1 | G/C-1 | CG>TA | CG>TA | AGCGACCCA | CCAGAAC | GGG | CCC | C | A | C>T | C>T_CCC | CG>TA_CCAGAAC | 6 | FALSE | FALSE |
| 175_v.txt | 32788 | 32788-SNV-C | SNV | 1 | C | T | 410 | 25011 | 1.63927872 | 47.1442677 | rnr1Y285F_pG | TRUE | CAN | JSY2421 | 6 | rnr1Y285F_p | TCG | C>T | A/T+1 | G/C-1 | CG>TA | CG>TA | AGAACTCGA | CGAATTC | AGC | TCG | C | A | C>T | C>T_TCG | CG>TA_CGAATTC | 6 | FALSE | FALSE |
| 175_v.txt | 32796 | 32796-SNV-C | SNV | 1 | C | T | 109 | 25011 | 0.43580824 | 47.3944954 | rnr1Y285F_pG | TRUE | CAN | JSY2421 | 6 | rnr1Y285F_p | ACC | C>T | A/T+1 | G/C-1 | CG>TA | CG>TA | AATTCACCG | CCGTAAT | TGG | ACC | C | A | C>T | C>T_ACC | CG>TA_CCGTAAT | 6 | FALSE | FALSE |
| 175_v.txt | 32797 | 32797-SNV-C | SNV | 1 | C | T | 26 | 25011 | 0.10395426 | 49.3845249 | rnr1Y285F_pG | TRUE | CAN | JSY2421 | 6 | rnr1Y285F_p | CCG | C>T | A/T+1 | G/C-1 | CG>TA | CG>TA | ATTCACCGTA | CGTAATA | GGC | CCG | C | A | C>T | C>T_CCG | CG>TA_CGTAATA | 6 | FALSE | FALSE |
| 175_v.txt | 32798 | 32798-SNV-C | SNV | 1 | G | A | 46 | 25011 | 0.18391908 | 43.0216503 | rnr1Y285F_pG | TRUE | CAN | JSY2421 | 6 | rnr1Y285F_p | CGT | G>A | A/T+1 | G/C-1 | CG>TA | CG>TA | TTCACCGTA | GTAATAT | GCA | GCA | C | T | C>T | C>T_GCA | CG>TA_GTAATAT | 6 | FALSE | FALSE |
| 175_v.txt | 32855 | 32855-SNV-C | SNV | 1 | C | T | 245 | 25010 | 0.97960816 | 35.6897959 | rnr1Y285F_pG | TRUE | CAN | JSY2421 | 6 | rnr1Y285F_p | TCC | C>T | A/T+1 | G/C-1 | CG>TA | CG>TA | ACTAATCCA | CCATGCC | AGG | TCC | C | A | C>T | C>T_TCC | CG>TA_CCATGCC | 6 | FALSE | FALSE |
| 175_v.txt | 32859 | 32859-SNV-C | SNV | 1 | G | A | 28 | 25007 | 0.11196865 | 35.570685 | rnr1Y285F_pG | TRUE | CAN | JSY2421 | 6 | rnr1Y285F_p | TGC | G>A | A/T+1 | G/C-1 | CG>TA | CG>TA | ATCCATGCC | GCCGCCA | ACG | ACG | C | T | C>T | C>T_ACG | CG>TA_GCCGCCA | 6 | FALSE | FALSE |
| 175_v.txt | 32860 | 32860-SNV-C | SNV | 1 | C | T | 31 | 25007 | 0.12396529 | 33.7096438 | rnr1Y285F_pG | TRUE | CAN | JSY2421 | 6 | rnr1Y285F_p | GCC | C>T | A/T+1 | G/C-1 | CG>TA | CG>TA | TCCATGCCG | CCGCCAG | CGG | GCC | C | A | C>T | C>T_GCC | CG>TA_CCGCCAG | 6 | FALSE | FALSE |
| 175_v.txt | 32890 | 32890-SNV-C | SNV | 1 | G | A | 33 | 24207 | 0.1363242 | 35.5757576 | rnr1Y285F_pG | TRUE | CAN | JSY2421 | 6 | rnr1Y285F_p | TGA | G>A | A/T+1 | G/C-1 | CG>TA | CG>TA | AAAATTGAA | GAATGAC | ACT | ACT | C | T | C>T | C>T_ACT | CG>TA_GAATGAC | 6 | FALSE | FALSE |
| 175_v.txt | 32900 | 32900-SNV-C | SNV | 1 | G | A | 44 | 24207 | 0.18176561 | 34.8159444 | rnr1Y285F_pG | TRUE | CAN | JSY2421 | 6 | rnr1Y285F_p | GGC | G>A | A/T+1 | G/C-1 | CG>TA | CG>TA | GACTTGGCC | GCCAACT | CCG | CCG | C | T | C>T | C>T_CCG | CG>TA_GCCAACT | 6 | FALSE | FALSE |
| 175_v.txt | 32901 | 32901-SNV-C | SNV | 1 | C | T | 41 | 24207 | 0.1693725 | 34.4636434 | rnr1Y285F_pG | TRUE | CAN | JSY2421 | 6 | rnr1Y285F_p | GCC | C>T | A/T+1 | G/C-1 | CG>TA | CG>TA | ACTTGGCCA | CCAACTA | CGG | GCC | C | A | C>T | C>T_GCC | CG>TA_CCAACTA | 6 | FALSE | FALSE |
| 175_v.txt | 32917 | 32917-SNV-C | SNV | 1 | C | T | 86 | 24207 | 0.35526914 | 47.1627907 | rnr1Y285F_pG | TRUE | CAN | JSY2421 | 6 | rnr1Y285F_p | TCC | C>T | A/T+1 | G/C-1 | CG>TA | CG>TA | TAAGTTCCA | CCAGGGC | AGG | TCC | C | A | C>T | C>T_TCC | CG>TA_CCAGGGC | 6 | FALSE | FALSE |
| 175_v.txt | 32945 | 32945-SNV-C | SNV | 1 | C | T | 56 | 24207 | 0.23133804 | 54.7851439 | rnr1Y285F_pG | TRUE | CAN | JSY2421 | 6 | rnr1Y285F_p | ACC | C>T | A/T+1 | G/C-1 | CG>TA | CG>TA | AGAAAACCA | CCAATAC | TGG | ACC | C | A | C>T | C>T_ACC | CG>TA_CCAATAC | 6 | FALSE | FALSE |

| | | | | | | | | | | | | | | | | | | | | | | | | | | | | | | | | | | |
|---|---|---|---|---|---|---|---|---|---|---|---|---|---|---|---|---|---|---|---|---|---|---|---|---|---|---|---|---|---|---|---|---|---|---|
| 175_v.txt | 32946 | 32946-SNV-( | SNV | 1 | C | T | 89 | 24207 | 0.36766225 | 55.8604954 | rnr1Y285F_pG | TRUE | CAN | JSY2421 | 6 | rnr1Y285F_p | CCA | C>T | A/T+1 | G/C-1 | CG>TA | CG>TA | GAAAACCAA | CAATACA | GGT | CCA | C | A | C>T | C>T_CCA | CG>TA_CAATACA | 6 | FALSE | FALSE |
| 175_v.txt | 32958 | 32958-SNV-( | SNV | 1 | C | T | 134 | 24207 | 0.55355889 | 54.984403 | rnr1Y285F_pG | TRUE | CAN | JSY2421 | 6 | rnr1Y285F_p | ACC | C>T | A/T+1 | G/C-1 | CG>TA | CG>TA | ATGTAACCAT | CCATTGG | TGG | ACC | C | A | C>T | C>T_ACC | CG>TA_CCATTGG | 6 | FALSE | FALSE |
| 175_v.txt | 32965 | 32965-SNV-( | SNV | 1 | C | T | 26 | 24207 | 0.10740695 | 57.1153846 | rnr1Y285F_pG | TRUE | CAN | JSY2421 | 6 | rnr1Y285F_p | GCC | C>T | A/T+1 | G/C-1 | CG>TA | CG>TA | CATTGGCCG | CCGCACC | CGG | GCC | C | A | C>T | C>T_GCC | CG>TA_CCGCACC | 6 | FALSE | FALSE |
| 175_v.txt | 32967 | 32967-SNV-( | SNV | 1 | G | A | 60 | 24207 | 0.24786219 | 58.7666667 | rnr1Y285F_pG | TRUE | CAN | JSY2421 | 6 | rnr1Y285F_p | CGC | G>A | A/T+1 | G/C-1 | CG>TA | CG>TA | TTGGCCGCA | GCACCAA | GCG | GCG | C | T | C>T | C>T_GCG | CG>TA_GCACCAA | 6 | FALSE | FALSE |
| 175_v.txt | 32970 | 32970-SNV-( | SNV | 1 | C | T | 46 | 24207 | 0.19002768 | 58.3695652 | rnr1Y285F_pG | TRUE | CAN | JSY2421 | 6 | rnr1Y285F_p | ACC | C>T | A/T+1 | G/C-1 | CG>TA | CG>TA | GCCGCACCA | CCAAATG | TGG | ACC | C | A | C>T | C>T_ACC | CG>TA_CCAAATG | 6 | FALSE | FALSE |
| 175_v.txt | 33006 | 33006-SNV-( | SNV | 1 | G | A | 13 | 24207 | 0.05370347 | 59.7692308 | rnr1Y285F_pG | TRUE | CAN | JSY2421 | 6 | rnr1Y285F_p | TGT | G>A | A/T+1 | G/C-1 | CG>TA | CG>TA | AAAACTGTG | GTGAAAG | ACA | ACA | C | T | C>T | C>T_ACA | CG>TA_GTGAAAG | 6 | FALSE | FALSE |
| 175_v.txt | 33037 | 33037-SNV-( | SNV | 1 | C | T | 15 | 24207 | 0.06196555 | 63.4 | rnr1Y285F_pG | TRUE | CAN | JSY2421 | 6 | rnr1Y285F_p | GCC | C>T | A/T+1 | G/C-1 | CG>TA | CG>TA | ATGTAGCCAT | CCATTTC | CGG | GCC | C | A | C>T | C>T_GCC | CG>TA_CCATTTC | 6 | FALSE | FALSE |
| 175_v.txt | 33056 | 33056-SNV-( | SNV | 1 | C | T | 35 | 24207 | 0.14458628 | 59.1369574 | rnr1Y285F_pG | TRUE | CAN | JSY2421 | 6 | rnr1Y285F_p | GCG | C>T | A/T+1 | G/C-1 | CG>TA | CG>TA | GGACTGCGT | CGTGACA | CGC | GCG | C | A | C>T | C>T_GCG | CG>TA_CGTGACA | 6 | FALSE | FALSE |
| 175_v.txt | 33070 | 33070-SNV-( | SNV | 1 | C | T | 37 | 24207 | 0.15284835 | 49.1621622 | rnr1Y285F_pG | TRUE | CAN | JSY2421 | 6 | rnr1Y285F_p | GCC | C>T | A/T+1 | G/C-1 | CG>TA | CG>TA | AATATGCCA | CCAAAGA | CGG | GCC | C | A | C>T | C>T_GCC | CG>TA_CCAAAGA | 6 | FALSE | FALSE |
| 175_v.txt | 33104 | 33104-SNV-( | SNV | 1 | G | A | 38 | 24207 | 0.15697939 | 48.1842105 | rnr1Y285F_pG | TRUE | CAN | JSY2421 | 6 | rnr1Y285F_p | CGC | G>A | A/T+1 | G/C-1 | CG>TA | CG>TA | AAGAGCGCC | GCCCACT | GCG | GCG | C | T | C>T | C>T_GCG | CG>TA_GCCCACT | 6 | FALSE | FALSE |
| 175_v.txt | 33114 | 33114-SNV-( | SNV | 1 | C | T | 43 | 24207 | 0.17763457 | 44.1162791 | rnr1Y285F_pG | TRUE | CAN | JSY2421 | 6 | rnr1Y285F_p | GCC | C>T | A/T+1 | G/C-1 | CG>TA | CG>TA | ACTGGGCCG | CCGGCGT | CGG | GCC | C | A | C>T | C>T_GCC | CG>TA_CCGGCGT | 6 | FALSE | FALSE |
| 175_v.txt | 33115 | 33115-SNV-( | SNV | 1 | C | T | 50 | 24207 | 0.20655182 | 41.24 | rnr1Y285F_pG | TRUE | CAN | JSY2421 | 6 | rnr1Y285F_p | CCG | C>T | A/T+1 | G/C-1 | CG>TA | CG>TA | CTGGGCCGG | CGGCGTT | GGC | CCG | C | A | C>T | C>T_CCG | CG>TA_CGGCGTT | 6 | FALSE | FALSE |
| 175_v.txt | 33118 | 33118-SNV-( | SNV | 1 | C | T | 29 | 24207 | 0.11980006 | 43.8275862 | rnr1Y285F_pG | TRUE | CAN | JSY2421 | 6 | rnr1Y285F_p | GCG | C>T | A/T+1 | G/C-1 | CG>TA | CG>TA | GGCCGGCGT | CGTTGGT | CGC | GCG | C | A | C>T | C>T_GCG | CG>TA_CGTTGGT | 6 | FALSE | FALSE |
| 175_v.txt | 33153 | 33153-SNV-( | SNV | 1 | C | T | 220 | 24207 | 0.90882803 | 36.4727273 | rnr1Y285F_pG | TRUE | CAN | JSY2421 | 6 | rnr1Y285F_p | ACC | C>T | A/T+1 | G/C-1 | CG>TA | CG>TA | AAAAGACCT | CCTGTAC | TGG | ACC | C | A | C>T | C>T_ACC | CG>TA_CCTGTAC | 6 | FALSE | FALSE |
| 175_v.txt | 33154 | 33154-SNV-( | SNV | 1 | C | T | 7 | 24207 | 0.02891726 | 37.1428571 | rnr1Y285F_pG | TRUE | CAN | JSY2421 | 6 | rnr1Y285F_p | CCT | C>T | A/T+1 | G/C-1 | CG>TA | CG>TA | AAAGACCTG | CTGTACC | GGA | CCT | C | A | C>T | C>T_CCT | CG>TA_CTGTACC | 6 | FALSE | FALSE |
| 175_v.txt | 33159 | 33159-SNV-( | SNV | 1 | C | T | 60 | 24206 | 0.24787243 | 37.3833333 | rnr1Y285F_pG | TRUE | CAN | JSY2421 | 6 | rnr1Y285F_p | ACC | C>T | A/T+1 | G/C-1 | CG>TA | CG>TA | CCTGTACCA | CCAATAG | TGG | ACC | C | A | C>T | C>T_ACC | CG>TA_CCAATAG | 6 | FALSE | FALSE |
| 175_v.txt | 33168 | 33168-SNV-( | SNV | 1 | C | T | 60 | 24206 | 0.24787243 | 37.3666667 | rnr1Y285F_pG | TRUE | CAN | JSY2421 | 6 | rnr1Y285F_p | ACC | C>T | A/T+1 | G/C-1 | CG>TA | CG>TA | ATAGTACCA | CCACCAA | TGG | ACC | C | A | C>T | C>T_ACC | CG>TA_CCACCAA | 6 | FALSE | FALSE |
| 175_v.txt | 33177 | 33177-SNV-( | SNV | 1 | G | A | 17 | 24200 | 0.07024793 | 35.3529412 | rnr1Y285F_pG | TRUE | CAN | JSY2421 | 6 | rnr1Y285F_p | GGC | G>A | A/T+1 | G/C-1 | CG>TA | CG>TA | CCAAGGGCA | GCAATCA | CCG | CCG | C | T | C>T | C>T_CCG | CG>TA_GCAATCA | 6 | FALSE | FALSE |
| 175_v.txt | 33199 | 33199-SNV-( | SNV | 1 | G | A | 71 | 13006 | 0.54590189 | 36.5492958 | rnr1Y285F_pG | TRUE | CAN | JSY2421 | 6 | rnr1Y285F_p | TGC | G>A | A/T+1 | G/C-1 | CG>TA | CG>TA | GTCTTTGCTT | GCTTAAG | ACG | ACG | C | T | C>T | C>T_ACG | CG>TA_GCTTAAG | 6 | FALSE | FALSE |
| 175_v.txt | 33205 | 33205-SNV-( | SNV | 1 | G | A | 8 | 13005 | 0.0615148 | 37.1244585 | rnr1Y285F_pG | TRUE | CAN | JSY2421 | 6 | rnr1Y285F_p | AGC | G>A | A/T+1 | G/C-1 | CG>TA | CG>TA | GCTTAAGCT | GCTCTCT | TCG | TCG | C | T | C>T | C>T_TCG | CG>TA_GCTCTCT | 6 | FALSE | FALSE |
| 175_v.txt | 33276 | 33276-SNV-( | SNV | 1 | G | A | 23 | 13039 | 0.1763939 | 46.4782609 | rnr1Y285F_pG | TRUE | CAN | JSY2421 | 6 | rnr1Y285F_p | CGT | G>A | A/T+1 | G/C-1 | CG>TA | CG>TA | GAGAACGTA | GTATCCT | GCA | GCA | C | T | C>T | C>T_GCA | CG>TA_GTATCCT | 6 | FALSE | FALSE |
| 175_v.txt | 33309 | 33309-SNV-( | SNV | 1 | C | T | 18 | 13039 | 0.1380474 | 60.2741763 | rnr1Y285F_pG | TRUE | CAN | JSY2421 | 6 | rnr1Y285F_p | GCG | C>T | A/T+1 | G/C-1 | CG>TA | CG>TA | AAAGAGCGC | CGCAATG | CGC | GCG | C | A | C>T | C>T_GCG | CG>TA_CGCAATG | 6 | FALSE | FALSE |
| 175_v.txt | 33310 | 33310-SNV-( | SNV | 1 | G | A | 19 | 13038 | 0.14572787 | 56.924377 | rnr1Y285F_pG | TRUE | CAN | JSY2421 | 6 | rnr1Y285F_p | CGC | G>A | A/T+1 | G/C-1 | CG>TA | CG>TA | AAGAGCGCA | GCAATGG | GCG | GCG | C | T | C>T | C>T_GCG | CG>TA_GCAATGG | 6 | FALSE | FALSE |
| 175_v.txt | 33412 | 33412-SNV-( | SNV | 1 | G | A | 23 | 13039 | 0.1763939 | 48.4347826 | rnr1Y285F_pG | TRUE | CAN | JSY2421 | 6 | rnr1Y285F_p | GGC | G>A | A/T+1 | G/C-1 | CG>TA | CG>TA | TGACCGGCT | GCTCATT | CCG | CCG | C | T | C>T | C>T_CCG | CG>TA_GCTCATT | 6 | FALSE | FALSE |
| 175_v.txt | 33474 | 33474-SNV-( | SNV | 1 | C | T | 23 | 13038 | 0.17640743 | 34.3478261 | rnr1Y285F_pG | TRUE | CAN | JSY2421 | 6 | rnr1Y285F_p | GCC | C>T | A/T+1 | G/C-1 | CG>TA | CG>TA | GCTATGCCTT | CCTTTTT | CGG | GCC | C | A | C>T | C>T_GCC | CG>TA_CCTTTTT | 6 | FALSE | FALSE |
| 175_v.txt | 31979 | 31979-SNV-( | SNV | 1 | G | C | 1 | 13700 | 7.30E-03 | 34.9980893 | rnr1Y285F_pG | TRUE | CAN | JSY2421 | 6 | rnr1Y285F_p | CGG | G>C | G/C+1 | G/C-1 | CG>GC | CG>GC | CAAGCCGGG | GGGCATT | GCC | GCC | C | G | C>G | C>G_GCC | CG>GC_GGGCATT | 6 | FALSE | FALSE |
| 175_v.txt | 31980 | 31980-SNV-( | SNV | 1 | G | C | 5 | 13700 | 0.03649635 | 42.7860041 | rnr1Y285F_pG | TRUE | CAN | JSY2421 | 6 | rnr1Y285F_p | GGG | G>C | G/C+1 | G/C-1 | CG>GC | CG>GC | AAGCCGGGC | GGCATTA | CCC | CCC | C | G | C>G | C>G_CCC | CG>GC_GGCATTA | 6 | FALSE | FALSE |
| 175_v.txt | 32027 | 32027-SNV-( | SNV | 1 | G | C | 28 | 13710 | 0.20423049 | 58.6428571 | rnr1Y285F_pG | TRUE | CAN | JSY2421 | 6 | rnr1Y285F_p | GGT | G>C | G/C+1 | G/C-1 | CG>GC | CG>GC | GCCACGGTA | GTATTTC | CCA | CCA | C | G | C>G | C>G_CCA | CG>GC_GTATTTC | 6 | FALSE | FALSE |
| 175_v.txt | 32038 | 32038-SNV-( | SNV | 1 | C | G | 19 | 13710 | 0.13858498 | 59.9473684 | rnr1Y285F_pG | TRUE | CAN | JSY2421 | 6 | rnr1Y285F_p | GCT | C>G | G/C+1 | G/C-1 | CG>GC | CG>GC | TCAAAGCTT | CTTGCAT | CGA | GCT | C | C | C>G | C>G_GCT | CG>GC_CTTGCAT | 6 | FALSE | FALSE |
| 175_v.txt | 32190 | 32190-SNV-( | SNV | 1 | G | C | 2 | 13710 | 0.01458789 | 38 | rnr1Y285F_pG | TRUE | CAN | JSY2421 | 6 | rnr1Y285F_p | TGC | G>C | G/C+1 | G/C-1 | CG>GC | CG>GC | AAAACTGCA | GCAATGT | ACG | ACG | C | G | C>G | C>G_ACG | CG>GC_GCAATGT | 6 | FALSE | FALSE |
| 175_v.txt | 32292 | 32292-SNV-( | SNV | 1 | G | C | 41 | 18153 | 0.22585799 | 52.902439 | rnr1Y285F_pG | TRUE | CAN | JSY2421 | 6 | rnr1Y285F_p | TGA | G>C | G/C+1 | G/C-1 | CG>GC | CG>GC | ATATTTGAAT | GAATTTG | ACT | ACT | C | G | C>G | C>G_ACT | CG>GC_GAATTTG | 6 | FALSE | FALSE |
| 175_v.txt | 32432 | 32432-SNV-( | SNV | 1 | G | C | 9 | 18153 | 0.04957858 | 57.4444444 | rnr1Y285F_pG | TRUE | CAN | JSY2421 | 6 | rnr1Y285F_p | TGT | G>C | G/C+1 | G/C-1 | CG>GC | CG>GC | GTCATTGTAT | GTATGGA | ACA | ACA | C | G | C>G | C>G_ACA | CG>GC_GTATGGA | 6 | FALSE | FALSE |
| 175_v.txt | 32539 | 32539-SNV-( | SNV | 1 | C | G | 41 | 18153 | 0.22585799 | 37.8292683 | rnr1Y285F_pG | TRUE | CAN | JSY2421 | 6 | rnr1Y285F_p | GCT | C>G | G/C+1 | G/C-1 | CG>GC | CG>GC | TTGCAGCTTC | CTTCACC | CGA | GCT | C | C | C>G | C>G_GCT | CG>GC_CTTCACC | 6 | FALSE | FALSE |
| 175_v.txt | 32557 | 32557-SNV-( | SNV | 1 | C | G | 15 | 18147 | 0.08265829 | 33.2652818 | rnr1Y285F_pG | TRUE | CAN | JSY2421 | 6 | rnr1Y285F_p | CCA | C>G | G/C+1 | G/C-1 | CG>GC | CG>GC | TGATACCAA | CAACTAG | GGT | CCA | C | C | C>G | C>G_CCA | CG>GC_CAACTAG | 6 | FALSE | FALSE |
| 175_v.txt | 32566 | 32566-SNV-( | SNV | 1 | C | G | 3 | 25009 | 0.01199568 | 37 | rnr1Y285F_pG | TRUE | CAN | JSY2421 | 6 | rnr1Y285F_p | TCA | C>G | G/C+1 | G/C-1 | CG>GC | CG>GC | CTAGTTCAGT | CAGTACC | AGT | TCA | C | C | C>G | C>G_TCA | CG>GC_CAGTACC | 6 | FALSE | FALSE |
| 175_v.txt | 32575 | 32575-SNV-( | SNV | 1 | G | C | 3 | 25012 | 0.01199424 | 37.9968542 | rnr1Y285F_pG | TRUE | CAN | JSY2421 | 6 | rnr1Y285F_p | TGA | G>C | G/C+1 | G/C-1 | CG>GC | CG>GC | TACCTTGAA | GAAATGT | ACT | ACT | C | G | C>G | C>G_ACT | CG>GC_GAAATGT | 6 | FALSE | FALSE |
| 175_v.txt | 32776 | 32776-SNV-( | SNV | 1 | C | G | 18 | 25011 | 0.07196833 | 43.2777778 | rnr1Y285F_pG | TRUE | CAN | JSY2421 | 6 | rnr1Y285F_p | GCG | C>G | G/C+1 | G/C-1 | CG>GC | CG>GC | TGGAAGCGA | CGACCCA | CGC | GCG | C | C | C>G | C>G_GCG | CG>GC_CGACCCA | 6 | FALSE | FALSE |
| 175_v.txt | 32786 | 32786-SNV-( | SNV | 1 | C | G | 22 | 25011 | 0.0879613 | 49.7264393 | rnr1Y285F_pG | TRUE | CAN | JSY2421 | 6 | rnr1Y285F_p | ACT | C>G | G/C+1 | G/C-1 | CG>GC | CG>GC | CCAGAACTC | CTCGAAT | TGA | ACT | C | C | C>G | C>G_ACT | CG>GC_CTCGAAT | 6 | FALSE | FALSE |
| 175_v.txt | 32797 | 32797-SNV-( | SNV | 1 | C | G | 2 | 25011 | 8.00E-03 | 36.5030985 | rnr1Y285F_pG | TRUE | CAN | JSY2421 | 6 | rnr1Y285F_p | CCG | C>G | G/C+1 | G/C-1 | CG>GC | CG>GC | ATTCACCGTA | CGTAATA | GGC | CCG | C | C | C>G | C>G_CCG | CG>GC_CGTAATA | 6 | FALSE | FALSE |
| 175_v.txt | 32819 | 32819-SNV-( | SNV | 1 | G | C | 15 | 25011 | 0.05997361 | 43.1333333 | rnr1Y285F_pG | TRUE | CAN | JSY2421 | 6 | rnr1Y285F_p | AGT | G>C | G/C+1 | G/C-1 | CG>GC | CG>GC | GAACAAGTT | GTTCATT | TCA | TCA | C | G | C>G | C>G_TCA | CG>GC_GTTCATT | 6 | FALSE | FALSE |
| 175_v.txt | 32829 | 32829-SNV-( | SNV | 1 | G | C | 8 | 25011 | 0.03198593 | 50.75 | rnr1Y285F_pG | TRUE | CAN | JSY2421 | 6 | rnr1Y285F_p | TGT | G>C | G/C+1 | G/C-1 | CG>GC | CG>GC | ATTATTGTGA | GTGATAA | ACA | ACA | C | G | C>G | C>G_ACA | CG>GC_GTGATAA | 6 | FALSE | FALSE |
| 175_v.txt | 32876 | 32876-SNV-( | SNV | 1 | G | C | 60 | 24198 | 0.24795438 | 37.0833333 | rnr1Y285F_pG | TRUE | CAN | JSY2421 | 6 | rnr1Y285F_p | TGT | G>C | G/C+1 | G/C-1 | CG>GC | CG>GC | AACTTTGTAC | GTACGTC | ACA | ACA | C | G | C>G | C>G_ACA | CG>GC_GTACGTC | 6 | FALSE | FALSE |
| 175_v.txt | 32917 | 32917-SNV-( | SNV | 1 | C | G | 91 | 24207 | 0.37592432 | 44.4175824 | rnr1Y285F_pG | TRUE | CAN | JSY2421 | 6 | rnr1Y285F_p | TCC | C>G | G/C+1 | G/C-1 | CG>GC | CG>GC | TAAGTTCCAC | CCAGGGC | AGG | TCC | C | C | C>G | C>G_TCC | CG>GC_CCAGGGC | 6 | FALSE | FALSE |
| 175_v.txt | 32936 | 32936-SNV-( | SNV | 1 | C | G | 21 | 24207 | 0.08675177 | 47.3333333 | rnr1Y285F_pG | TRUE | CAN | JSY2421 | 6 | rnr1Y285F_p | CCC | C>G | G/C+1 | G/C-1 | CG>GC | CG>GC | GATTGCCCA | CCAAGAA | GGG | CCC | C | C | C>G | C>G_CCC | CG>GC_CCAAGAA | 6 | FALSE | FALSE |
| 175_v.txt | 32959 | 32959-SNV-( | SNV | 1 | C | G | 20 | 24207 | 0.08262073 | 54.8946737 | rnr1Y285F_pG | TRUE | CAN | JSY2421 | 6 | rnr1Y285F_p | CCA | C>G | G/C+1 | G/C-1 | CG>GC | CG>GC | TGTAACCATT | CATTGGC | GGT | CCA | C | C | C>G | C>G_CCA | CG>GC_CATTGGC | 6 | FALSE | FALSE |
| 175_v.txt | 32997 | 32997-SNV-( | SNV | 1 | G | C | 14 | 24207 | 0.05783451 | 62.4285714 | rnr1Y285F_pG | TRUE | CAN | JSY2421 | 6 | rnr1Y285F_p | TGA | G>C | G/C+1 | G/C-1 | CG>GC | CG>GC | CTTTGTGAGA | GAGAAAA | ACT | ACT | C | G | C>G | C>G_ACT | CG>GC_GAGAAAA | 6 | FALSE | FALSE |
| 175_v.txt | 33037 | 33037-SNV-( | SNV | 1 | C | G | 47 | 24207 | 0.19415871 | 60.7446809 | rnr1Y285F_pG | TRUE | CAN | JSY2421 | 6 | rnr1Y285F_p | GCC | C>G | G/C+1 | G/C-1 | CG>GC | CG>GC | ATGTAGCCAT | CCATTTC | CGG | GCC | C | C | C>G | C>G_GCC | CG>GC_CCATTTC | 6 | FALSE | FALSE |
| 175_v.txt | 33057 | 33057-SNV-( | SNV | 1 | G | C | 12 | 24207 | 0.04957244 | 51.832694 | rnr1Y285F_pG | TRUE | CAN | JSY2421 | 6 | rnr1Y285F_p | CGT | G>C | G/C+1 | G/C-1 | CG>GC | CG>GC | GACTGCGTG | GTGACAG | GCA | GCA | C | G | C>G | C>G_GCA | CG>GC_GTGACAG | 6 | FALSE | FALSE |
| 175_v.txt | 33070 | 33070-SNV-( | SNV | 1 | C | G | 68 | 24207 | 0.28091048 | 56.3529412 | rnr1Y285F_pG | TRUE | CAN | JSY2421 | 6 | rnr1Y285F_p | GCC | C>G | G/C+1 | G/C-1 | CG>GC | CG>GC | AATATGCCA | CCAAAGA | CGG | GCC | C | C | C>G | C>G_GCC | CG>GC_CCAAAGA | 6 | FALSE | FALSE |
| 175_v.txt | 33156 | 33156-SNV-( | SNV | 1 | G | C | 223 | 24206 | 0.92125919 | 37.1748879 | rnr1Y285F_pG | TRUE | CAN | JSY2421 | 6 | rnr1Y285F_p | TGT | G>C | G/C+1 | G/C-1 | CG>GC | CG>GC | AGACCTGTA | GTACCAA | ACA | ACA | C | G | C>G | C>G_ACA | CG>GC_GTACCAA | 6 | FALSE | FALSE |
| 175_v.txt | 33168 | 33168-SNV-( | SNV | 1 | C | G | 2 | 24206 | 8.26E-03 | 38 | rnr1Y285F_pG | TRUE | CAN | JSY2421 | 6 | rnr1Y285F_p | ACC | C>G | G/C+1 | G/C-1 | CG>GC | CG>GC | ATAGTACCA | CCACCAA | TGG | ACC | C | C | C>G | C>G_ACC | CG>GC_CCACCAA | 6 | FALSE | FALSE |
| 175_v.txt | 33169 | 33169-SNV-( | SNV | 1 | C | G | 26 | 24206 | 0.10741139 | 37.5741991 | rnr1Y285F_pG | TRUE | CAN | JSY2421 | 6 | rnr1Y285F_p | CCA | C>G | G/C+1 | G/C-1 | CG>GC | CG>GC | TAGTACCAC | CACCAAG | GGT | CCA | C | C | C>G | C>G_CCA | CG>GC_CACCAAG | 6 | FALSE | FALSE |
| 175_v.txt | 33193 | 33193-SNV-( | SNV | 1 | G | C | 23 | 13005 | 0.17685506 | 37.2608696 | rnr1Y285F_pG | TRUE | CAN | JSY2421 | 6 | rnr1Y285F_p | TGT | G>C | G/C+1 | G/C-1 | CG>GC | CG>GC | CAATATGTCT | GTCTTTG | ACA | ACA | C | G | C>G | C>G_ACA | CG>GC_GTCTTTG | 6 | FALSE | FALSE |
| 175_v.txt | 33205 | 33205-SNV-( | SNV | 1 | G | C | 58 | 13005 | 0.44598232 | 39.0161857 | rnr1Y285F_pG | TRUE | CAN | JSY2421 | 6 | rnr1Y285F_p | AGC | G>C | G/C+1 | G/C-1 | CG>GC | CG>GC | GCTTAAGCT | GCTCTCT | TCG | TCG | C | G | C>G | C>G_TCG | CG>GC_GCTCTCT | 6 | FALSE | FALSE |
| 175_v.txt | 33210 | 33210-SNV-( | SNV | 1 | C | G | 12 | 13039 | 0.0920316 | 37.5 | rnr1Y285F_pG | TRUE | CAN | JSY2421 | 6 | rnr1Y285F_p | TCT | C>G | G/C+1 | G/C-1 | CG>GC | CG>GC | AGCTCTCTCT | CTCTTCA | AGA | TCT | C | C | C>G | C>G_TCT | CG>GC_CTCTTCA | 6 | FALSE | FALSE |
| 175_v.txt | 33276 | 33276-SNV-( | SNV | 1 | G | C | 3 | 13039 | 0.0230079 | 55.3333333 | rnr1Y285F_pG | TRUE | CAN | JSY2421 | 6 | rnr1Y285F_p | CGT | G>C | G/C+1 | G/C-1 | CG>GC | CG>GC | GAGAACGTA | GTATCCT | GCA | GCA | C | G | C>G | C>G_GCA | CG>GC_GTATCCT | 6 | FALSE | FALSE |
| 175_v.txt | 33354 | 33354-SNV-( | SNV | 1 | G | C | 5 | 13039 | 0.0383465 | 51.4 | rnr1Y285F_pG | TRUE | CAN | JSY2421 | 6 | rnr1Y285F_p | TGA | G>C | G/C+1 | G/C-1 | CG>GC | CG>GC | GGTATTGAC | GACCCAC | ACT | ACT | C | G | C>G | C>G_ACT | CG>GC_GACCCAC | 6 | FALSE | FALSE |
| 175_v.txt | 33378 | 33378-SNV-( | SNV | 1 | G | C | 10 | 13039 | 0.076693 | 52 | rnr1Y285F_pG | TRUE | CAN | JSY2421 | 6 | rnr1Y285F_p | TGA | G>C | G/C+1 | G/C-1 | CG>GC | CG>GC | GTTTGTGAA | GAAGCTT | ACT | ACT | C | G | C>G | C>G_ACT | CG>GC_GAAGCTT | 6 | FALSE | FALSE |
| 175_v.txt | 33412 | 33412-SNV-( | SNV | 1 | G | C | 17 | 13039 | 0.1303781 | 50.5882353 | rnr1Y285F_pG | TRUE | CAN | JSY2421 | 6 | rnr1Y285F_p | GGC | G>C | G/C+1 | G/C-1 | CG>GC | CG>GC | TGACCGGCT | GCTCATT | CCG | CCG | C | G | C>G | C>G_CCG | CG>GC_GCTCATT | 6 | FALSE | FALSE |
| 175_v.txt | 33419 | 33419-SNV-( | SNV | 1 | G | C | 74 | 13039 | 0.56752819 | 48.5405405 | rnr1Y285F_pG | TRUE | CAN | JSY2421 | 6 | rnr1Y285F_p | TGT | G>C | G/C+1 | G/C-1 | CG>GC | CG>GC | CTCATTGTAC | GTACATA | ACA | ACA | C | G | C>G | C>G_ACA | CG>GC_GTACATA | 6 | FALSE | FALSE |
| 175_v.txt | 31869 | 31869-SNV-( | SNV | 1 | G | T | 4 | 9519 | 0.04202122 | 50.75 | rnr1Y285F_pG | TRUE | CAN | JSY2421 | 6 | rnr1Y285F_p | AGA | G>T | A/T+1 | G/C-1 | CG>AT | CG>AT | AAAATAGAG | GAGATAT | TCT | TCT | C | A | C>A | C>A_TCT | CG>AT_GAGATAT | 6 | FALSE | FALSE |
| 175_v.txt | 31980 | 31980-SNV-( | SNV | 1 | G | T | 10 | 13700 | 0.0729927 | 40.4869396 | rnr1Y285F_pG | TRUE | CAN | JSY2421 | 6 | rnr1Y285F_p | GGG | G>T | A/T+1 | G/C-1 | CG>AT | CG>AT | AAGCCGGGC | GGCATTA | CCC | CCC | C | A | C>A | C>A_CCC | CG>AT_GGCATTA | 6 | FALSE | FALSE |
| 175_v.txt | 32008 | 32008-SNV-( | SNV | 1 | C | A | 19 | 13700 | 0.13868613 | 38.6842105 | rnr1Y285F_pG | TRUE | CAN | JSY2421 | 6 | rnr1Y285F_p | TCG | C>A | A/T+1 | G/C-1 | CG>AT | CG>AT | GTAACTCGT | CGTCACG | AGC | TCG | C | T | C>A | C>A_TCG | CG>AT_CGTCACG | 6 | FALSE | FALSE |
| 175_v.txt | 32027 | 32027-SNV-( | SNV | 1 | G | T | 37 | 13710 | 0.269876 | 57.4594595 | rnr1Y285F_pG | TRUE | CAN | JSY2421 | 6 | rnr1Y285F_p | GGT | G>T | A/T+1 | G/C-1 | CG>AT | CG>AT | GCCACGGTA | GTATTTC | CCA | CCA | C | A | C>A | C>A_CCA | CG>AT_GTATTTC | 6 | FALSE | FALSE |
| 175_v.txt | 32153 | 32153-SNV-( | SNV | 1 | G | T | 58 | 13710 | 0.42304887 | 55.5172414 | rnr1Y285F_pG | TRUE | CAN | JSY2421 | 6 | rnr1Y285F_p | TGT | G>T | A/T+1 | G/C-1 | CG>AT | CG>AT | CTCCATGTAA | GTAAGCC | ACA | ACA | C | A | C>A | C>A_ACA | CG>AT_GTAAGCC | 6 | FALSE | FALSE |
| 175_v.txt | 32188 | 32188-SNV-( | SNV | 1 | C | A | 79 | 13710 | 0.57622174 | 44.9873418 | rnr1Y285F_pG | TRUE | CAN | JSY2421 | 6 | rnr1Y285F_p | ACT | C>A | A/T+1 | G/C-1 | CG>AT | CG>AT | CGAAAACTG | CTGCAAT | TGA | ACT | C | T | C>A | C>A_ACT | CG>AT_CTGCAAT | 6 | FALSE | FALSE |
| 175_v.txt | 32190 | 32190-SNV-( | SNV | 1 | G | T | 4 | 13710 | 0.02917578 | 51 | rnr1Y285F_pG | TRUE | CAN | JSY2421 | 6 | rnr1Y285F_p | TGC | G>T | A/T+1 | G/C-1 | CG>AT | CG>AT | AAAACTGCA | GCAATGT | ACG | ACG | C | A | C>A | C>A_ACG | CG>AT_GCAATGT | 6 | FALSE | FALSE |
| 175_v.txt | 32195 | 32195-SNV-( | SNV | 1 | G | T | 34 | 13710 | 0.24799417 | 42.9117647 | rnr1Y285F_pG | TRUE | CAN | JSY2421 | 6 | rnr1Y285F_p | TGT | G>T | A/T+1 | G/C-1 | CG>AT | CG>AT | TGCAATGTAT | GTATGGA | ACA | ACA | C | A | C>A | C>A_ACA | CG>AT_GTATGGA | 6 | FALSE | FALSE |
| 175_v.txt | 32274 | 32274-SNV-( | SNV | 1 | G | T | 141 | 18153 | 0.77673112 | 36.4539007 | rnr1Y285F_pG | TRUE | CAN | JSY2421 | 6 | rnr1Y285F_p | GGA | G>T | A/T+1 | G/C-1 | CG>AT | CG>AT | ATACGGGAA | GAACCAA | CCT | CCT | C | A | C>A | C>A_CCT | CG>AT_GAACCAA | 6 | FALSE | FALSE |
| 175_v.txt | 32282 | 32282-SNV-( | SNV | 1 | G | T | 63 | 18153 | 0.34705007 | 41.7301587 | rnr1Y285F_pG | TRUE | CAN | JSY2421 | 6 | rnr1Y285F_p | CGT | G>T | A/T+1 | G/C-1 | CG>AT | CG>AT | ACCAACGTA | GTAAATA | GCA | GCA | C | A | C>A | C>A_GCA | CG>AT_GTAAATA | 6 | FALSE | FALSE |
| 175_v.txt | 32292 | 32292-SNV-( | SNV | 1 | G | T | 8 | 18153 | 0.04406985 | 35.625 | rnr1Y285F_pG | TRUE | CAN | JSY2421 | 6 | rnr1Y285F_p | TGA | G>T | A/T+1 | G/C-1 | CG>AT | CG>AT | ATATTTGAAT | GAATTTG | ACT | ACT | C | A | C>A | C>A_ACT | CG>AT_GAATTTG | 6 | FALSE | FALSE |
| 175_v.txt | 32304 | 32304-SNV-( | SNV | 1 | G | T | 48 | 18153 | 0.2644191 | 55.0833333 | rnr1Y285F_pG | TRUE | CAN | JSY2421 | 6 | rnr1Y285F_p | AGA | G>T | A/T+1 | G/C-1 | CG>AT | CG>AT | GCGGCAGAA | GAAATAA | TCT | TCT | C | A | C>A | C>A_TCT | CG>AT_GAAATAA | 6 | FALSE | FALSE |
| 175_v.txt | 32373 | 32373-SNV-( | SNV | 1 | G | T | 240 | 18153 | 1.32209552 | 60.7 | rnr1Y285F_pG | TRUE | CAN | JSY2421 | 6 | rnr1Y285F_p | AGC | G>T | A/T+1 | G/C-1 | CG>AT | CG>AT | TCAATAGCA | GCAATAA | TCG | TCG | C | A | C>A | C>A_TCG | CG>AT_GCAATAA | 6 | FALSE | FALSE |
| 175_v.txt | 32399 | 32399-SNV-( | SNV | 1 | G | T | 83 | 18153 | 0.4572247 | 56.8795181 | rnr1Y285F_pG | TRUE | CAN | JSY2421 | 6 | rnr1Y285F_p | CGT | G>T | A/T+1 | G/C-1 | CG>AT | CG>AT | AGAAACGTA | GTAGGAA | GCA | GCA | C | A | C>A | C>A_GCA | CG>AT_GTAGGAA | 6 | FALSE | FALSE |
| 175_v.txt | 32432 | 32432-SNV-( | SNV | 1 | G | T | 78 | 18153 | 0.42968105 | 56.5769231 | rnr1Y285F_pG | TRUE | CAN | JSY2421 | 6 | rnr1Y285F_p | TGT | G>T | A/T+1 | G/C-1 | CG>AT | CG>AT | GTCATTGTAT | GTATGGA | ACA | ACA | C | A | C>A | C>A_ACA | CG>AT_GTATGGA | 6 | FALSE | FALSE |
| 175_v.txt | 32449 | 32449-SNV-( | SNV | 1 | C | A | 12 | 18153 | 0.06610478 | 57.1666667 | rnr1Y285F_pG | TRUE | CAN | JSY2421 | 6 | rnr1Y285F_p | CCA | C>A | A/T+1 | G/C-1 | CG>AT | CG>AT | AAAGTCCAA | CAATGAA | GGT | CCA | C | T | C>A | C>A_CCA | CG>AT_CAATGAA | 6 | FALSE | FALSE |
| 175_v.txt | 32463 | 32463-SNV-( | SNV | 1 | G | T | 127 | 18153 | 0.69960888 | 50.5173779 | rnr1Y285F_pG | TRUE | CAN | JSY2421 | 6 | rnr1Y285F_p | AGA | G>T | A/T+1 | G/C-1 | CG>AT | CG>AT | AATAGAGAG | GAGCCAA | TCT | TCT | C | A | C>A | C>A_TCT | CG>AT_GAGCCAA | 6 | FALSE | FALSE |
| 175_v.txt | 32471 | 32471-SNV-( | SNV | 1 | G | T | 45 | 18153 | 0.24789291 | 47.2444444 | rnr1Y285F_pG | TRUE | CAN | JSY2421 | 6 | rnr1Y285F_p | TGT | G>T | A/T+1 | G/C-1 | CG>AT | CG>AT | GCCAATGTA | GTAGAAG | ACA | ACA | C | A | C>A | C>A_ACA | CG>AT_GTAGAAG | 6 | FALSE | FALSE |
| 175_v.txt | 32487 | 32487-SNV-( | SNV | 1 | C | A | 8 | 18153 | 0.04406985 | 50.7539092 | rnr1Y285F_pG | TRUE | CAN | JSY2421 | 6 | rnr1Y285F_p | ACG | C>A | A/T+1 | G/C-1 | CG>AT | CG>AT | AAGATACGG | CGGAAAA | TGC | ACG | C | T | C>A | C>A_ACG | CG>AT_CGGAAAA | 6 | FALSE | FALSE |
| 175_v.txt | 32494 | 32494-SNV-( | SNV | 1 | C | A | 16 | 18153 | 0.0881397 | 49.5625 | rnr1Y285F_pG | TRUE | CAN | JSY2421 | 6 | rnr1Y285F_p | ACA | C>A | A/T+1 | G/C-1 | CG>AT | CG>AT | GGAAAACAA | CAACTTT | TGT | ACA | C | T | C>A | C>A_ACA | CG>AT_CAACTTT | 6 | FALSE | FALSE |
| 175_v.txt | 32508 | 32508-SNV-( | SNV | 1 | G | T | 29 | 18153 | 0.15975321 | 42.3793103 | rnr1Y285F_pG | TRUE | CAN | JSY2421 | 6 | rnr1Y285F_p | GGC | G>T | A/T+1 | G/C-1 | CG>AT | CG>AT | TTGATGGCT | GCTCTTG | CCG | CCG | C | A | C>A | C>A_CCG | CG>AT_GCTCTTG | 6 | FALSE | FALSE |
| 175_v.txt | 32530 | 32530-SNV-( | SNV | 1 | G | T | 22 | 18153 | 0.12119209 | 37.4986013 | rnr1Y285F_pG | TRUE | CAN | JSY2421 | 6 | rnr1Y285F_p | GGG | G>T | A/T+1 | G/C-1 | CG>AT | CG>AT | TTCTGGGGT | GGTTTGC | CCC | CCC | C | A | C>A | C>A_CCC | CG>AT_GGTTTGC | 6 | FALSE | FALSE |
| 175_v.txt | 32547 | 32547-SNV-( | SNV | 1 | G | T | 87 | 18152 | 0.47928603 | 36.4712644 | rnr1Y285F_pG | TRUE | CAN | JSY2421 | 6 | rnr1Y285F_p | AGC | G>T | A/T+1 | G/C-1 | CG>AT | CG>AT | TCACCAGCA | GCAGTGA | TCG | TCG | C | A | C>A | C>A_TCG | CG>AT_GCAGTGA | 6 | FALSE | FALSE |
| 175_v.txt | 32556 | 32556-SNV-( | SNV | 1 | C | A | 32 | 18148 | 0.17632797 | 37.1875 | rnr1Y285F_pG | TRUE | CAN | JSY2421 | 6 | rnr1Y285F_p | ACC | C>A | A/T+1 | G/C-1 | CG>AT | CG>AT | GTGATACCA | CCAACTA | TGG | ACC | C | T | C>A | C>A_ACC | CG>AT_CCAACTA | 6 | FALSE | FALSE |
| 175_v.txt | 32566 | 32566-SNV-( | SNV | 1 | C | A | 8 | 25009 | 0.03198848 | 38 | rnr1Y285F_pG | TRUE | CAN | JSY2421 | 6 | rnr1Y285F_p | TCA | C>A | A/T+1 | G/C-1 | CG>AT | CG>AT | CTAGTTCAGT | CAGTACC | AGT | TCA | C | T | C>A | C>A_TCA | CG>AT_CAGTACC | 6 | FALSE | FALSE |
| 175_v.txt | 32575 | 32575-SNV-( | SNV | 1 | G | T | 15 | 25012 | 0.05997121 | 35.3985236 | rnr1Y285F_pG | TRUE | CAN | JSY2421 | 6 | rnr1Y285F_p | TGA | G>T | A/T+1 | G/C-1 | CG>AT | CG>AT | TACCTTGAA | GAAATGT | ACT | ACT | C | A | C>A | C>A_ACT | CG>AT_GAAATGT | 6 | FALSE | FALSE |
| 175_v.txt | 32629 | 32629-SNV-( | SNV | 1 | C | A | 40 | 25011 | 0.15992963 | 48.5 | rnr1Y285F_pG | TRUE | CAN | JSY2421 | 6 | rnr1Y285F_p | TCG | C>A | A/T+1 | G/C-1 | CG>AT | CG>AT | TCCCTTCGTT | CGTTTTT | AGC | TCG | C | T | C>A | C>A_TCG | CG>AT_CGTTTTT | 6 | FALSE | FALSE |
| 175_v.txt | 32692 | 32692-SNV-( | SNV | 1 | C | A | 45 | 25011 | 0.17992084 | 57.6444444 | rnr1Y285F_pG | TRUE | CAN | JSY2421 | 6 | rnr1Y285F_p | CCA | C>A | A/T+1 | G/C-1 | CG>AT | CG>AT | GGAATCCAA | CAACTGG | GGT | CCA | C | T | C>A | C>A_CCA | CG>AT_CAACTGG | 6 | FALSE | FALSE |
| 175_v.txt | 32735 | 32735-SNV-( | SNV | 1 | G | T | 14 | 25011 | 0.05597537 | 59.2142857 | rnr1Y285F_pG | TRUE | CAN | JSY2421 | 6 | rnr1Y285F_p | AGT | G>T | A/T+1 | G/C-1 | CG>AT | CG>AT | AAAACAGTA | GTATATT | TCA | TCA | C | A | C>A | C>A_TCA | CG>AT_GTATATT | 6 | FALSE | FALSE |
| 175_v.txt | 32783 | 32783-SNV-( | SNV | 1 | G | T | 14 | 25011 | 0.05597537 | 54.2142857 | rnr1Y285F_pG | TRUE | CAN | JSY2421 | 6 | rnr1Y285F_p | AGA | G>T | A/T+1 | G/C-1 | CG>AT | CG>AT | GACCCAGAA | GAACTCG | TCT | TCT | C | A | C>A | C>A_TCT | CG>AT_GAACTCG | 6 | FALSE | FALSE |
| 175_v.txt | 32794 | 32794-SNV-( | SNV | 1 | C | A | 200 | 25011 | 0.79964816 | 43.11 | rnr1Y285F_pG | TRUE | CAN | JSY2421 | 6 | rnr1Y285F_p | TCA | C>A | A/T+1 | G/C-1 | CG>AT | CG>AT | CGAATTCAC | CACCGTA | AGT | TCA | C | T | C>A | C>A_TCA | CG>AT_CACCGTA | 6 | FALSE | FALSE |
| 175_v.txt | 32796 | 32796-SNV-( | SNV | 1 | C | A | 59 | 25011 | 0.23589621 | 44.779661 | rnr1Y285F_pG | TRUE | CAN | JSY2421 | 6 | rnr1Y285F_p | ACC | C>A | A/T+1 | G/C-1 | CG>AT | CG>AT | AATTCACCGT | CCGTAAT | TGG | ACC | C | T | C>A | C>A_ACC | CG>AT_CCGTAAT | 6 | FALSE | FALSE |
| 175_v.txt | 32819 | 32819-SNV-( | SNV | 1 | G | T | 25 | 25011 | 0.09995602 | 41.28 | rnr1Y285F_pG | TRUE | CAN | JSY2421 | 6 | rnr1Y285F_p | AGT | G>T | A/T+1 | G/C-1 | CG>AT | CG>AT | GAACAAGTT | GTTCATT | TCA | TCA | C | A | C>A | C>A_TCA | CG>AT_GTTCATT | 6 | FALSE | FALSE |
| 175_v.txt | 32876 | 32876-SNV-( | SNV | 1 | G | T | 31 | 24198 | 0.12810976 | 37.483871 | rnr1Y285F_pG | TRUE | CAN | JSY2421 | 6 | rnr1Y285F_p | TGT | G>T | A/T+1 | G/C-1 | CG>AT | CG>AT | AACTTTGTAC | GTACGTC | ACA | ACA | C | A | C>A | C>A_ACA | CG>AT_GTACGTC | 6 | FALSE | FALSE |
| 175_v.txt | 32917 | 32917-SNV-( | SNV | 1 | C | A | 840 | 24207 | 3.47007064 | 51.202381 | rnr1Y285F_pG | TRUE | CAN | JSY2421 | 6 | rnr1Y285F_p | TCC | C>A | A/T+1 | G/C-1 | CG>AT | CG>AT | TAAGTTCCAC | CCAGGGC | AGG | TCC | C | T | C>A | C>A_TCC | CG>AT_CCAGGGC | 6 | FALSE | FALSE |
| 175_v.txt | 32940 | 32940-SNV-( | SNV | 1 | G | T | 69 | 24206 | 0.28505329 | 49.5921772 | rnr1Y285F_pG | TRUE | CAN | JSY2421 | 6 | rnr1Y285F_p | AGA | G>T | A/T+1 | G/C-1 | CG>AT | CG>AT | GCCCAAGAA | GAAAACC | TCT | TCT | C | A | C>A | C>A_TCT | CG>AT_GAAAACC | 6 | FALSE | FALSE |
| 175_v.txt | 33006 | 33006-SNV-( | SNV | 1 | G | T | 52 | 24207 | 0.2148139 | 60.3076923 | rnr1Y285F_pG | TRUE | CAN | JSY2421 | 6 | rnr1Y285F_p | TGT | G>T | A/T+1 | G/C-1 | CG>AT | CG>AT | AAAACTGTG | GTGAAAG | ACA | ACA | C | A | C>A | C>A_ACA | CG>AT_GTGAAAG | 6 | FALSE | FALSE |
| 175_v.txt | 33024 | 33024-SNV-( | SNV | 1 | G | T | 35 | 24207 | 0.14458628 | 60.6285714 | rnr1Y285F_pG | TRUE | CAN | JSY2421 | 6 | rnr1Y285F_p | AGG | G>T | A/T+1 | G/C-1 | CG>AT | CG>AT | GTAACAGGG | GGGATGA | TCC | TCC | C | A | C>A | C>A_TCC | CG>AT_GGGATGA | 6 | FALSE | FALSE |
| 175_v.txt | 33093 | 33093-SNV-( | SNV | 1 | G | T | 57 | 24207 | 0.23546908 | 48.3859649 | rnr1Y285F_pG | TRUE | CAN | JSY2421 | 6 | rnr1Y285F_p | TGA | G>T | A/T+1 | G/C-1 | CG>AT | CG>AT | AAATATGAT | GATATAA | ACT | ACT | C | A | C>A | C>A_ACT | CG>AT_GATATAA | 6 | FALSE | FALSE |
| 175_v.txt | 33112 | 33112-SNV-( | SNV | 1 | G | T | 37 | 24207 | 0.15284835 | 43.6486487 | rnr1Y285F_pG | TRUE | CAN | JSY2421 | 6 | rnr1Y285F_p | GGG | G>T | A/T+1 | G/C-1 | CG>AT | CG>AT | CCACTGGGC | GGCCGGC | CCC | CCC | C | A | C>A | C>A_CCC | CG>AT_GGCCGGC | 6 | FALSE | FALSE |
| 175_v.txt | 33114 | 33114-SNV-( | SNV | 1 | C | A | 13 | 24207 | 0.05370347 | 42.3076923 | rnr1Y285F_pG | TRUE | CAN | JSY2421 | 6 | rnr1Y285F_p | GCC | C>A | A/T+1 | G/C-1 | CG>AT | CG>AT | ACTGGGCCG | CCGGCGT | CGG | GCC | C | T | C>A | C>A_GCC | CG>AT_CCGGCGT | 6 | FALSE | FALSE |
| 175_v.txt | 33115 | 33115-SNV-( | SNV | 1 | C | A | 29 | 24207 | 0.11980006 | 45.9310345 | rnr1Y285F_pG | TRUE | CAN | JSY2421 | 6 | rnr1Y285F_p | CCG | C>A | A/T+1 | G/C-1 | CG>AT | CG>AT | CTGGGCCGG | CGGCGTT | GGC | CCG | C | T | C>A | C>A_CCG | CG>AT_CGGCGTT | 6 | FALSE | FALSE |
| 175_v.txt | 33151 | 33151-SNV-( | SNV | 1 | G | T | 50 | 24207 | 0.20655182 | 38.08 | rnr1Y285F_pG | TRUE | CAN | JSY2421 | 6 | rnr1Y285F_p | AGA | G>T | A/T+1 | G/C-1 | CG>AT | CG>AT | TGAAAAGAC | GACCTGT | TCT | TCT | C | A | C>A | C>A_TCT | CG>AT_GACCTGT | 6 | FALSE | FALSE |
| 175_v.txt | 33153 | 33153-SNV-( | SNV | 1 | C | A | 11 | 24207 | 0.0454414 | 37.0909091 | rnr1Y285F_pG | TRUE | CAN | JSY2421 | 6 | rnr1Y285F_p | ACC | C>A | A/T+1 | G/C-1 | CG>AT | CG>AT | AAAAGACCT | CCTGTAC | TGG | ACC | C | T | C>A | C>A_ACC | CG>AT_CCTGTAC | 6 | FALSE | FALSE |
| 175_v.txt | 33154 | 33154-SNV-( | SNV | 1 | C | A | 68 | 24207 | 0.28091048 | 36.8676471 | rnr1Y285F_pG | TRUE | CAN | JSY2421 | 6 | rnr1Y285F_p | CCT | C>A | A/T+1 | G/C-1 | CG>AT | CG>AT | AAAGACCTG | CTGTACC | GGA | CCT | C | T | C>A | C>A_CCT | CG>AT_CTGTACC | 6 | FALSE | FALSE |
| 175_v.txt | 33156 | 33156-SNV-( | SNV | 1 | G | T | 14 | 24206 | 0.0578369 | 37.8571429 | rnr1Y285F_pG | TRUE | CAN | JSY2421 | 6 | rnr1Y285F_p | TGT | G>T | A/T+1 | G/C-1 | CG>AT | CG>AT | AGACCTGTA | GTACCAA | ACA | ACA | C | A | C>A | C>A_ACA | CG>AT_GTACCAA | 6 | FALSE | FALSE |
| 175_v.txt | 33159 | 33159-SNV-( | SNV | 1 | C | A | 45 | 24206 | 0.18590432 | 37.8222222 | rnr1Y285F_pG | TRUE | CAN | JSY2421 | 6 | rnr1Y285F_p | ACC | C>A | A/T+1 | G/C-1 | CG>AT | CG>AT | CCTGTACCA | CCAATAG | TGG | ACC | C | T | C>A | C>A_ACC | CG>AT_CCAATAG | 6 | FALSE | FALSE |
| 175_v.txt | 33168 | 33168-SNV-( | SNV | 1 | C | A | 126 | 24206 | 0.5205321 | 36.9047619 | rnr1Y285F_pG | TRUE | CAN | JSY2421 | 6 | rnr1Y285F_p | ACC | C>A | A/T+1 | G/C-1 | CG>AT | CG>AT | ATAGTACCA | CCACCAA | TGG | ACC | C | T | C>A | C>A_ACC | CG>AT_CCACCAA | 6 | FALSE | FALSE |
| 175_v.txt | 33177 | 33177-SNV-( | SNV | 1 | G | T | 26 | 24200 | 0.10743802 | 37.6153846 | rnr1Y285F_pG | TRUE | CAN | JSY2421 | 6 | rnr1Y285F_p | GGC | G>T | A/T+1 | G/C-1 | CG>AT | CG>AT | CCAAGGGCA | GCAATCA | CCG | CCG | C | A | C>A | C>A_CCG | CG>AT_GCAATCA | 6 | FALSE | FALSE |
| 175_v.txt | 33186 | 33186-SNV-( | SNV | 1 | C | A | 12 | 13000 | 0.09230769 | 35.5 | rnr1Y285F_pG | TRUE | CAN | JSY2421 | 6 | rnr1Y285F_p | ACC | C>A | A/T+1 | G/C-1 | CG>AT | CG>AT | ATCATACCA | CCAATAT | TGG | ACC | C | T | C>A | C>A_ACC | CG>AT_CCAATAT | 6 | FALSE | FALSE |
| 175_v.txt | 33210 | 33210-SNV-( | SNV | 1 | C | A | 84 | 13039 | 0.64422118 | 38.0714286 | rnr1Y285F_pG | TRUE | CAN | JSY2421 | 6 | rnr1Y285F_p | TCT | C>A | A/T+1 | G/C-1 | CG>AT | CG>AT | AGCTCTCTCT | CTCTTCA | AGA | TCT | C | T | C>A | C>A_TCT | CG>AT_CTCTTCA | 6 | FALSE | FALSE |
| 175_v.txt | 33241 | 33241-SNV-( | SNV | 1 | C | A | 42 | 13039 | 0.32211059 | 40.3333333 | rnr1Y285F_pG | TRUE | CAN | JSY2421 | 6 | rnr1Y285F_p | TCA | C>A | A/T+1 | G/C-1 | CG>AT | CG>AT | CTCCTTCATC | CATCTTC | AGT | TCA | C | T | C>A | C>A_TCA | CG>AT_CATCTTC | 6 | FALSE | FALSE |
| 175_v.txt | 33325 | 33325-SNV-( | SNV | 1 | C | A | 144 | 13039 | 1.10437917 | 56.625 | rnr1Y285F_pG | TRUE | CAN | JSY2421 | 6 | rnr1Y285F_p | TCT | C>A | A/T+1 | G/C-1 | CG>AT | CG>AT | ACAATTCTTT | CTTTACT | AGA | TCT | C | T | C>A | C>A_TCT | CG>AT_CTTTACT | 6 | FALSE | FALSE |
| 175_v.txt | 33354 | 33354-SNV-( | SNV | 1 | G | T | 68 | 13039 | 0.52151239 | 18.8970588 | rnr1Y285F_pG | TRUE | CAN | JSY2421 | 6 | rnr1Y285F_p | TGA | G>T | A/T+1 | G/C-1 | CG>AT | CG>AT | GGTATTGAC | GACCCAC | ACT | ACT | C | A | C>A | C>A_ACT | CG>AT_GACCCAC | 6 | FALSE | FALSE |
| 175_v.txt | 33474 | 33474-SNV-( | SNV | 1 | C | A | 50 | 13038 | 0.3834944 | 34.92 | rnr1Y285F_pG | TRUE | CAN | JSY2421 | 6 | rnr1Y285F_p | GCC | C>A | A/T+1 | G/C-1 | CG>AT | CG>AT | GCTATGCCTT | CCTTTTT | CGG | GCC | C | T | C>A | C>A_GCC | CG>AT_CCTTTTT | 6 | FALSE | FALSE |
| 175_v.txt | 31872 | 31872-MNV- | MNV | 2 | AT | GA | 10 | 9519 | 0.10505305 | 40.5532106 | rnr1Y285F_pG | TRUE | CAN | JSY2421 | 6 | rnr1Y285F_p | GAT | nope | >1bp-ins | >1bpdel | none | MNV | ATAGAGATA | ATATAGG | CTA | GAT | AT | none | NA | NA_GAT | MNV_ATATAGG | 6 | FALSE | FALSE |
| 175_v.txt | 32940 | 32940-MNV- | MNV | 2 | GA | AG | 523 | 24207 | 2.16053208 | 54.6657249 | rnr1Y285F_pG | TRUE | CAN | JSY2421 | 6 | rnr1Y285F_p | AGA | nope | >1bp-ins | >1bpdel | none | MNV | GCCCAAGAA | GAAAACC | TCT | AGA | GA | none | NA | NA_AGA | MNV_GAAAACC | 6 | FALSE | FALSE |
| 175_v.txt | 33171 | 33171-MNV- | MNV | 2 | CC | TA | 42 | 24206 | 0.1735107 | 37.4163819 | rnr1Y285F_pG | TRUE | CAN | JSY2421 | 6 | rnr1Y285F_p | ACC | nope | >1bp-ins | >1bpdel | none | MNV | GTACCACCA | CCAAGGG | TGG | ACC | CC | none | NA | NA_ACC | MNV_CCAAGGG | 6 | FALSE | FALSE |
| 175_v.txt | 32081 | 32081-Insert | Insertion | 1 | - | A | 32 | 13710 | 0.23340627 | 61.625 | rnr1Y285F_pG | TRUE | CAN | JSY2421 | 6 | rnr1Y285F_p | CAA | nope | A/T+1 | >1bpdel | none | A/T+1 | CCATGCAAA | AAAAAAG | GTT | CAA | - | T | NA | NA_CAA | A/T+1_AAAAAAG | 6 | FALSE | FALSE |

| | | | | | | | | | | | | | | | | | | | | | | | | | | | | | | | | | | |
|---|---|---|---|---|---|---|---|---|---|---|---|---|---|---|---|---|---|---|---|---|---|---|---|---|---|---|---|---|---|---|---|---|---|---|
| 175_v.txt | 32180 | 32180-Insert | Insertion | 1 | - | A | 13 | 13710 | 0.0948213 | 49 | rnr1Y285F_pG | TRUE | CAN | JSY2421 | 6 | rnr1Y285F_p | TAA | nope | A/T+1 | >1bpdel | none | A/T+1 | AGCAGTAAC | AACGAAA | ATT | TAA | - | T | NA | NA_TAA | A/T+1_AACGAAA | 6 | FALSE | FALSE |
| 175_v.txt | 32498 | 32498-Insert | Insertion | 1 | - | T | 3 | 18153 | 0.01652619 | 54 | rnr1Y285F_pG | TRUE | CAN | JSY2421 | 6 | rnr1Y285F_p | CTT | nope | A/T+1 | >1bpdel | none | A/T+1 | AACAACTTTT | TTTTTTG | GAA | CTT | - | A | NA | NA_CTT | A/T+1_TTTTTTG | 6 | FALSE | FALSE |
| 175_v.txt | 32528 | 32528-Insert | Insertion | 1 | - | G | 2 | 18153 | 0.01101746 | 64 | rnr1Y285F_pG | TRUE | CAN | JSY2421 | 6 | rnr1Y285F_p | TGG | nope | G/C+1 | >1bpdel | none | G/C+1 | TTTTCTGGG | GGGGTTT | ACC | TGG | - | C | NA | NA_TGG | G/C+1_GGGGTTT | 6 | FALSE | FALSE |
| 175_v.txt | 32656 | 32656-Insert | Insertion | 1 | - | A | 11 | 25011 | 0.04398065 | 59.1818182 | rnr1Y285F_pG | TRUE | CAN | JSY2421 | 6 | rnr1Y285F_p | GGA | nope | A/T+1 | >1bpdel | none | A/T+1 | TACCTGGAC | GACCCCA | CCT | GGA | - | T | NA | NA_GGA | A/T+1_GACCCCA | 6 | FALSE | FALSE |
| 175_v.txt | 32673 | 32673-Insert | Insertion | 1 | - | T | 26 | 25011 | 0.10395426 | 58.2692308 | rnr1Y285F_pG | TRUE | CAN | JSY2421 | 6 | rnr1Y285F_p | GTT | nope | A/T+1 | >1bpdel | none | A/T+1 | CCTGGGTTTC | TTTCTCC | CAA | GTT | - | A | NA | NA_GTT | A/T+1_TTTCTCC | 6 | FALSE | FALSE |
| 175_v.txt | 32728 | 32728-Insert | Insertion | 1 | - | A | 168 | 25011 | 0.67170445 | 62.0178571 | rnr1Y285F_pG | TRUE | CAN | JSY2421 | 6 | rnr1Y285F_p | CAA | nope | A/T+1 | >1bpdel | none | A/T+1 | CCATACAAA | AAAAACA | GTT | CAA | - | T | NA | NA_CAA | A/T+1_AAAAACA | 6 | FALSE | FALSE |
| 175_v.txt | 32728 | 32728-Insert | Insertion | 1 | - | G | 4 | 25011 | 0.01599296 | 59.5 | rnr1Y285F_pG | TRUE | CAN | JSY2421 | 6 | rnr1Y285F_p | CAA | nope | G/C+1 | >1bpdel | none | G/C+1 | CCATACAAA | AAAAACA | GTT | CAA | - | C | NA | NA_CAA | G/C+1_AAAAACA | 6 | FALSE | FALSE |
| 175_v.txt | 32760 | 32760-Insert | Insertion | 1 | - | A | 2 | 25011 | 8.00E-03 | 51 | rnr1Y285F_pG | TRUE | CAN | JSY2421 | 6 | rnr1Y285F_p | TAA | nope | A/T+1 | >1bpdel | none | A/T+1 | ATGGCTAAA | AAAACTT | ATT | TAA | - | T | NA | NA_TAA | A/T+1_AAAACTT | 6 | FALSE | FALSE |
| 175_v.txt | 32840 | 32840-Insert | Insertion | 1 | - | A | 4 | 25011 | 0.01599296 | 37.75 | rnr1Y285F_pG | TRUE | CAN | JSY2421 | 6 | rnr1Y285F_p | CCC | nope | A/T+1 | >1bpdel | none | A/T+1 | AATTACCCA | CCAAAAA | GGG | CCC | - | T | NA | NA_CCC | A/T+1_CCAAAAA | 6 | FALSE | FALSE |
| 175_v.txt | 32842 | 32842-Insert | Insertion | 1 | - | A | 123 | 25011 | 0.49178362 | 43.4227642 | rnr1Y285F_pG | TRUE | CAN | JSY2421 | 6 | rnr1Y285F_p | CAA | nope | A/T+1 | >1bpdel | none | A/T+1 | TTACCCAAA | AAAAAAT | GTT | CAA | - | T | NA | NA_CAA | A/T+1_AAAAAAT | 6 | FALSE | FALSE |
| 175_v.txt | 32842 | 32842-Insert | Insertion | 1 | - | G | 20 | 25011 | 0.07996482 | 38.85 | rnr1Y285F_pG | TRUE | CAN | JSY2421 | 6 | rnr1Y285F_p | CAA | nope | G/C+1 | >1bpdel | none | G/C+1 | TTACCCAAA | AAAAAAT | GTT | CAA | - | C | NA | NA_CAA | G/C+1_AAAAAAT | 6 | FALSE | FALSE |
| 175_v.txt | 32884 | 32884-Insert | Insertion | 1 | - | A | 27 | 24207 | 0.11153799 | 37.2592593 | rnr1Y285F_pG | TRUE | CAN | JSY2421 | 6 | rnr1Y285F_p | CAA | nope | A/T+1 | >1bpdel | none | A/T+1 | ACGTCCAAA | AAAATTG | GTT | CAA | - | T | NA | NA_CAA | A/T+1_AAAATTG | 6 | FALSE | FALSE |
| 175_v.txt | 32912 | 32912-Insert | Insertion | 1 | - | A | 127 | 24207 | 0.52464163 | 46.9212598 | rnr1Y285F_pG | TRUE | CAN | JSY2421 | 6 | rnr1Y285F_p | TAA | nope | A/T+1 | >1bpdel | none | A/T+1 | TACACTAAG | AAGTTCC | ATT | TAA | - | T | NA | NA_TAA | A/T+1_AAGTTCC | 6 | FALSE | FALSE |
| 175_v.txt | 32924 | 32924-Insert | Insertion | 1 | - | A | 6 | 24207 | 0.02478622 | 55.3333333 | rnr1Y285F_pG | TRUE | CAN | JSY2421 | 6 | rnr1Y285F_p | CAA | nope | A/T+1 | >1bpdel | none | A/T+1 | CAGGGCAAA | AAAAGTG | GTT | CAA | - | T | NA | NA_CAA | A/T+1_AAAAGTG | 6 | FALSE | FALSE |
| 175_v.txt | 32975 | 32975-Insert | Insertion | 1 | - | T | 7 | 24207 | 0.02891726 | 60.1428571 | rnr1Y285F_pG | TRUE | CAN | JSY2421 | 6 | rnr1Y285F_p | ATG | nope | A/T+1 | >1bpdel | none | A/T+1 | ACCAAATGC | TGCTGGA | TAC | ATG | - | A | NA | NA_ATG | A/T+1_TGCTGGA | 6 | FALSE | FALSE |
| 175_v.txt | 33126 | 33126-Insert | Insertion | 1 | - | A | 10 | 24207 | 0.04131037 | 48.2 | rnr1Y285F_pG | TRUE | CAN | JSY2421 | 6 | rnr1Y285F_p | CAG | nope | A/T+1 | >1bpdel | none | A/T+1 | TTGGTCAGA | AGAGGTG | GTC | CAG | - | T | NA | NA_CAG | A/T+1_AGAGGTG | 6 | FALSE | FALSE |
| 175_v.txt | 33206 | 33206-Insert | Insertion | 2 | - | CT | 17 | 13005 | 0.13071895 | 37.6764706 | rnr1Y285F_pG | TRUE | CAN | JSY2421 | 6 | rnr1Y285F_p | GCT | nope | >1bp-ins | >1bpdel | none | >1bp-ins | CTTAAGCTCT | CTCTCTC | CGA | GCT | - | none | NA | NA_GCT | >1bp-ins_CTCTCTC | 6 | FALSE | FALSE |
| 175_v.txt | 33303 | 33303-Insert | Insertion | 1 | - | A | 3 | 13039 | 0.0230079 | 64 | rnr1Y285F_pG | TRUE | CAN | JSY2421 | 6 | rnr1Y285F_p | GAA | nope | A/T+1 | >1bpdel | none | A/T+1 | GTCGGGAAA | AAAGAGC | CTT | GAA | - | T | NA | NA_GAA | A/T+1_AAAGAGC | 6 | FALSE | FALSE |
| 175_v.txt | 33424 | 33424-Insert | Insertion | 14 | - | TATGCTTCTC | 5 | 13039 | 0.0383465 | 51.1142857 | rnr1Y285F_pG | TRUE | CAN | JSY2421 | 6 | rnr1Y285F_p | ATA | nope | >1bp-ins | >1bpdel | none | >1bp-ins | TGTACATATG | TATGCTT | TAT | ATA | - | none | NA | NA_ATA | >1bp-ins_TATGCTT | 6 | FALSE | FALSE |
| 175_v.txt | 31868 | 31868-Deleti | Deletion | 2 | AG | - | 5 | 9519 | 0.05252653 | 48.4 | rnr1Y285F_pG | TRUE | CAN | JSY2421 | 6 | rnr1Y285F_p | TAG | nope | >1bp-ins | >1bpdel | none | >1bpdel | GAAAATAGA | AGAGATA | ATC | TAG | AG | none | NA | NA_TAG | >1bpdel_AGAGATA | 6 | FALSE | FALSE |
| 175_v.txt | 32318 | 32318-Deleti | Deletion | 3 | TAA | - | 17 | 18153 | 0.09364843 | 51.0588235 | rnr1Y285F_pG | TRUE | CAN | JSY2421 | 6 | rnr1Y285F_p | TTA | nope | >1bp-ins | >1bpdel | none | >1bpdel | GGTTGTTAA | TAAGATA | AAT | TTA | TAA | none | NA | NA_TTA | >1bpdel_TAAGATA | 6 | FALSE | FALSE |
| 175_v.txt | 32322 | 32322-Deleti | Deletion | 6 | ATAACA | - | 17 | 18153 | 0.09364843 | 49.8235294 | rnr1Y285F_pG | TRUE | CAN | JSY2421 | 6 | rnr1Y285F_p | GAT | nope | >1bp-ins | >1bpdel | none | >1bpdel | GTTAAGATA | ATAACAG | CTA | GAT | ATAACA | none | NA | NA_GAT | >1bpdel_ATAACAG | 6 | FALSE | FALSE |
| 175_v.txt | 32346 | 32346-Deleti | Deletion | 2 | AA | - | 20 | 18153 | 0.11017463 | 60.55 | rnr1Y285F_pG | TRUE | CAN | JSY2421 | 6 | rnr1Y285F_p | CAA | nope | >1bp-ins | >1bpdel | none | >1bpdel | TGTGGCAAA | AAAACCT | GTT | CAA | AA | none | NA | NA_CAA | >1bpdel_AAAACCT | 6 | FALSE | FALSE |
| 175_v.txt | 32642 | 32642-Deleti | Deletion | 2 | AG | - | 12 | 25011 | 0.04797889 | 48.249993 | rnr1Y285F_pG | TRUE | CAN | JSY2421 | 6 | rnr1Y285F_p | TAG | nope | >1bp-ins | >1bpdel | none | >1bpdel | ATCCTTAGAT | AGATATT | ATC | TAG | AG | none | NA | NA_TAG | >1bpdel_AGATATT | 6 | FALSE | FALSE |
| 175_v.txt | 32716 | 32716-Deleti | Deletion | 2 | CA | - | 15 | 25011 | 0.05997361 | 64 | rnr1Y285F_pG | TRUE | CAN | JSY2421 | 6 | rnr1Y285F_p | CCA | nope | >1bp-ins | >1bpdel | none | >1bpdel | CAGCACCAC | CACAAAC | GGT | CCA | CA | none | NA | NA_CCA | >1bpdel_CACAAAC | 6 | FALSE | FALSE |
| 175_v.txt | 33089 | 33089-Deleti | Deletion | 2 | AT | - | 9 | 24207 | 0.03717933 | 53.8888889 | rnr1Y285F_pG | TRUE | CAN | JSY2421 | 6 | rnr1Y285F_p | AAT | nope | >1bp-ins | >1bpdel | none | >1bpdel | AAATAAATA | ATATGAT | TTA | AAT | AT | none | NA | NA_AAT | >1bpdel_ATATGAT | 6 | FALSE | FALSE |
| 175_v.txt | 33189 | 33189-Deleti | Deletion | 2 | AT | - | 22 | 13004 | 0.16917871 | 37.7272727 | rnr1Y285F_pG | TRUE | CAN | JSY2421 | 6 | rnr1Y285F_p | AAT | nope | >1bp-ins | >1bpdel | none | >1bpdel | ATACCAATA | ATATGTC | TTA | AAT | AT | none | NA | NA_AAT | >1bpdel_ATATGTC | 6 | FALSE | FALSE |
| 175_v.txt | 31723 | 31723-Deleti | Deletion | 1 | A | - | 17 | 9519 | 0.17859019 | 63.7058824 | rnr1Y285F_pG | TRUE | CAN | JSY2421 | 6 | rnr1Y285F_p | CAA | nope | >1bp-ins | A/T-1 | none | A/T-1 | TGTCCCAAA | AAAAAGT | GTT | GTT | T | none | T>none | T>none_GTT | A/T-1_AAAAAGT | 6 | FALSE | FALSE |
| 175_v.txt | 31834 | 31834-Deleti | Deletion | 1 | G | - | 3 | 9519 | 0.03151592 | 64 | rnr1Y285F_pG | TRUE | CAN | JSY2421 | 6 | rnr1Y285F_p | TGA | nope | >1bp-ins | G/C-1 | none | G/C-1 | TGCATTGAA | GAAATAA | ACT | ACT | C | none | C>none | C>none_ACT | G/C-1_GAAATAA | 6 | FALSE | FALSE |
| 175_v.txt | 31861 | 31861-Deleti | Deletion | 1 | G | - | 10 | 9519 | 0.10505305 | 40.2 | rnr1Y285F_pG | TRUE | CAN | JSY2421 | 6 | rnr1Y285F_p | AGG | nope | >1bp-ins | G/C-1 | none | G/C-1 | AGAACAGGA | GGAAAAT | TCC | TCC | C | none | C>none | C>none_TCC | G/C-1_GGAAAAT | 6 | FALSE | FALSE |
| 175_v.txt | 31904 | 31904-Deleti | Deletion | 1 | T | - | 8 | 9519 | 0.08404244 | 51 | rnr1Y285F_pG | TRUE | CAN | JSY2421 | 6 | rnr1Y285F_p | ATT | nope | >1bp-ins | A/T-1 | none | A/T-1 | ATTGAATTTT | TTTTGGT | TAA | ATT | T | none | nope | nope_ATT | A/T-1_TTTTGGT | 6 | FALSE | FALSE |
| 175_v.txt | 31913 | 31913-Deleti | Deletion | 1 | A | - | 6 | 9519 | 0.06303183 | 46.6666667 | rnr1Y285F_pG | TRUE | CAN | JSY2421 | 6 | rnr1Y285F_p | CAA | nope | >1bp-ins | A/T-1 | none | A/T-1 | TGGTGCAAA | AAAAGCC | GTT | GTT | T | none | T>none | T>none_GTT | A/T-1_AAAAGCC | 6 | FALSE | FALSE |
| 175_v.txt | 31979 | 31979-Deleti | Deletion | 1 | G | - | 34 | 13700 | 0.24817518 | 38.2647038 | rnr1Y285F_pG | TRUE | CAN | JSY2421 | 6 | rnr1Y285F_p | CGG | nope | >1bp-ins | G/C-1 | none | G/C-1 | CAAGCCGGG | GGGCATT | GCC | GCC | C | none | C>none | C>none_GCC | G/C-1_GGGCATT | 6 | FALSE | FALSE |
| 175_v.txt | 32075 | 32075-Deleti | Deletion | 1 | C | - | 7 | 13710 | 0.05105762 | 63.5714286 | rnr1Y285F_pG | TRUE | CAN | JSY2421 | 6 | rnr1Y285F_p | ACC | nope | >1bp-ins | G/C-1 | none | G/C-1 | AAATAACCA | CCATGCA | TGG | ACC | C | none | nope | nope_ACC | G/C-1_CCATGCA | 6 | FALSE | FALSE |
| 175_v.txt | 32081 | 32081-Deleti | Deletion | 1 | A | - | 23 | 13710 | 0.16776076 | 62.1739134 | rnr1Y285F_pG | TRUE | CAN | JSY2421 | 6 | rnr1Y285F_p | CAA | nope | >1bp-ins | A/T-1 | none | A/T-1 | CCATGCAAA | AAAAAAG | GTT | GTT | T | none | T>none | T>none_GTT | A/T-1_AAAAAAG | 6 | FALSE | FALSE |
| 175_v.txt | 32092 | 32092-Deleti | Deletion | 1 | C | - | 2 | 13710 | 0.01458789 | 63 | rnr1Y285F_pG | TRUE | CAN | JSY2421 | 6 | rnr1Y285F_p | GCA | nope | >1bp-ins | G/C-1 | none | G/C-1 | AGCCTGCAA | CAACACC | CGT | GCA | C | none | nope | nope_GCA | G/C-1_CAACACC | 6 | FALSE | FALSE |
| 175_v.txt | 32121 | 32121-Deleti | Deletion | 1 | A | - | 7 | 13710 | 0.05105762 | 59.2857143 | rnr1Y285F_pG | TRUE | CAN | JSY2421 | 6 | rnr1Y285F_p | GAA | nope | >1bp-ins | A/T-1 | none | A/T-1 | CATTCGAAA | AAAACTT | CTT | CTT | T | none | T>none | T>none_CTT | A/T-1_AAAACTT | 6 | FALSE | FALSE |
| 175_v.txt | 32184 | 32184-Deleti | Deletion | 1 | A | - | 16 | 13710 | 0.11670314 | 45.6875 | rnr1Y285F_pG | TRUE | CAN | JSY2421 | 6 | rnr1Y285F_p | GAA | nope | >1bp-ins | A/T-1 | none | A/T-1 | GTAACGAAA | AAAACTG | CTT | CTT | T | none | T>none | T>none_CTT | A/T-1_AAAACTG | 6 | FALSE | FALSE |
| 175_v.txt | 32314 | 32314-Deleti | Deletion | 1 | T | - | 2 | 18153 | 0.01101746 | 61.5 | rnr1Y285F_pG | TRUE | CAN | JSY2421 | 6 | rnr1Y285F_p | GTT | nope | >1bp-ins | A/T-1 | none | A/T-1 | TAATGGTTG | TTGTTAA | CAA | GTT | T | none | nope | nope_GTT | A/T-1_TTGTTAA | 6 | FALSE | FALSE |
| 175_v.txt | 32346 | 32346-Deleti | Deletion | 1 | A | - | 46 | 18153 | 0.25340164 | 59.173913 | rnr1Y285F_pG | TRUE | CAN | JSY2421 | 6 | rnr1Y285F_p | CAA | nope | >1bp-ins | A/T-1 | none | A/T-1 | TGTGGCAAA | AAAACCT | GTT | GTT | T | none | T>none | T>none_GTT | A/T-1_AAAACCT | 6 | FALSE | FALSE |
| 175_v.txt | 32377 | 32377-Deleti | Deletion | 1 | T | - | 6 | 18153 | 0.03305239 | 64 | rnr1Y285F_pG | TRUE | CAN | JSY2421 | 6 | rnr1Y285F_p | ATA | nope | >1bp-ins | A/T-1 | none | A/T-1 | TAGCAATAA | TAATAAA | TAT | ATA | T | none | nope | nope_ATA | A/T-1_TAATAAA | 6 | FALSE | FALSE |
| 175_v.txt | 32387 | 32387-Deleti | Deletion | 1 | A | - | 36 | 18153 | 0.19831433 | 61.6666667 | rnr1Y285F_pG | TRUE | CAN | JSY2421 | 6 | rnr1Y285F_p | GAG | nope | >1bp-ins | A/T-1 | none | A/T-1 | AAAGGGAGA | AGAAGTA | CTC | CTC | T | none | T>none | T>none_CTC | A/T-1_AGAAGTA | 6 | FALSE | FALSE |
| 175_v.txt | 32420 | 32420-Deleti | Deletion | 1 | T | - | 12 | 18153 | 0.06610478 | 59.5833333 | rnr1Y285F_pG | TRUE | CAN | JSY2421 | 6 | rnr1Y285F_p | GTT | nope | >1bp-ins | A/T-1 | none | A/T-1 | TGTTAGTTTA | TTTAGGG | CAA | GTT | T | none | nope | nope_GTT | A/T-1_TTTAGGG | 6 | FALSE | FALSE |
| 175_v.txt | 32424 | 32424-Deleti | Deletion | 1 | G | - | 5 | 18153 | 0.02754366 | 63 | rnr1Y285F_pG | TRUE | CAN | JSY2421 | 6 | rnr1Y285F_p | AGG | nope | >1bp-ins | G/C-1 | none | G/C-1 | AGTTTAGGG | GGGTCAT | TCC | TCC | C | none | C>none | C>none_TCC | G/C-1_GGGTCAT | 6 | FALSE | FALSE |
| 175_v.txt | 32442 | 32442-Deleti | Deletion | 1 | A | - | 3 | 18153 | 0.01652619 | 63 | rnr1Y285F_pG | TRUE | CAN | JSY2421 | 6 | rnr1Y285F_p | TAA | nope | >1bp-ins | A/T-1 | none | A/T-1 | GGAACTAAA | AAAAGTC | ATT | ATT | T | none | T>none | T>none_ATT | A/T-1_AAAAGTC | 6 | FALSE | FALSE |
| 175_v.txt | 32464 | 32464-Deleti | Deletion | 1 | A | - | 9 | 18153 | 0.04957858 | 51.1111016 | rnr1Y285F_pG | TRUE | CAN | JSY2421 | 6 | rnr1Y285F_p | GAG | nope | >1bp-ins | A/T-1 | none | A/T-1 | ATAGAGAGC | AGCCAAT | CTC | CTC | T | none | T>none | T>none_CTC | A/T-1_AGCCAAT | 6 | FALSE | FALSE |
| 175_v.txt | 32494 | 32494-Deleti | Deletion | 1 | C | - | 4 | 18153 | 0.02203493 | 44.5 | rnr1Y285F_pG | TRUE | CAN | JSY2421 | 6 | rnr1Y285F_p | ACA | nope | >1bp-ins | G/C-1 | none | G/C-1 | GGAAAACAA | CAACTTT | TGT | ACA | C | none | nope | nope_ACA | G/C-1_CAACTTT | 6 | FALSE | FALSE |
| 175_v.txt | 32498 | 32498-Deleti | Deletion | 1 | T | - | 38 | 18153 | 0.20933179 | 45.5789445 | rnr1Y285F_pG | TRUE | CAN | JSY2421 | 6 | rnr1Y285F_p | CTT | nope | >1bp-ins | A/T-1 | none | A/T-1 | AACAACTTTT | TTTTTTG | GAA | CTT | T | none | nope | nope_CTT | A/T-1_TTTTTTG | 6 | FALSE | FALSE |
| 175_v.txt | 32514 | 32514-Deleti | Deletion | 1 | G | - | 5 | 18153 | 0.02754366 | 38 | rnr1Y285F_pG | TRUE | CAN | JSY2421 | 6 | rnr1Y285F_p | TGG | nope | >1bp-ins | G/C-1 | none | G/C-1 | GCTCTTGGA | GGAACGG | ACC | ACC | C | none | C>none | C>none_ACC | G/C-1_GGAACGG | 6 | FALSE | FALSE |
| 175_v.txt | 32608 | 32608-Deleti | Deletion | 1 | C | - | 11 | 25010 | 0.04398241 | 38.7272724 | rnr1Y285F_pG | TRUE | CAN | JSY2421 | 6 | rnr1Y285F_p | ACC | nope | >1bp-ins | G/C-1 | none | G/C-1 | AGGAAACCC | CCCAACC | TGG | ACC | C | none | nope | nope_ACC | G/C-1_CCCAACC | 6 | FALSE | FALSE |
| 175_v.txt | 32631 | 32631-Deleti | Deletion | 1 | T | - | 7 | 25011 | 0.02798769 | 41.5714286 | rnr1Y285F_pG | TRUE | CAN | JSY2421 | 6 | rnr1Y285F_p | GTT | nope | >1bp-ins | A/T-1 | none | A/T-1 | CCTTCGTTTT | TTTTTAT | CAA | GTT | T | none | nope | nope_GTT | A/T-1_TTTTTAT | 6 | FALSE | FALSE |
| 175_v.txt | 32658 | 32658-Deleti | Deletion | 1 | C | - | 33 | 25011 | 0.13194195 | 56.3636364 | rnr1Y285F_pG | TRUE | CAN | JSY2421 | 6 | rnr1Y285F_p | ACC | nope | >1bp-ins | G/C-1 | none | G/C-1 | CCTGGACCC | CCCCAGG | TGG | ACC | C | none | nope | nope_ACC | G/C-1_CCCCAGG | 6 | FALSE | FALSE |
| 175_v.txt | 32670 | 32670-Deleti | Deletion | 1 | G | - | 628 | 25011 | 2.51089521 | 59.0031847 | rnr1Y285F_pG | TRUE | CAN | JSY2421 | 6 | rnr1Y285F_p | TGG | nope | >1bp-ins | G/C-1 | none | G/C-1 | GCACCTGGG | GGGTTTC | ACC | ACC | C | none | C>none | C>none_ACC | G/C-1_GGGTTTC | 6 | FALSE | FALSE |
| 175_v.txt | 32673 | 32673-Deleti | Deletion | 1 | T | - | 10 | 25011 | 0.03998241 | 56.1000002 | rnr1Y285F_pG | TRUE | CAN | JSY2421 | 6 | rnr1Y285F_p | GTT | nope | >1bp-ins | A/T-1 | none | A/T-1 | CCTGGGTTTC | TTTCTCC | CAA | GTT | T | none | nope | nope_GTT | A/T-1_TTTCTCC | 6 | FALSE | FALSE |
| 175_v.txt | 32678 | 32678-Deleti | Deletion | 1 | C | - | 3 | 25011 | 0.01199472 | 62.3333333 | rnr1Y285F_pG | TRUE | CAN | JSY2421 | 6 | rnr1Y285F_p | TCC | nope | >1bp-ins | G/C-1 | none | G/C-1 | GTTTCTCCAA | CCAATAA | AGG | TCC | C | none | nope | nope_TCC | G/C-1_CCAATAA | 6 | FALSE | FALSE |
| 175_v.txt | 32707 | 32707-Deleti | Deletion | 1 | C | - | 49 | 25011 | 0.1959138 | 62 | rnr1Y285F_pG | TRUE | CAN | JSY2421 | 6 | rnr1Y285F_p | ACC | nope | >1bp-ins | G/C-1 | none | G/C-1 | CGGTAACCC | CCCCAGC | TGG | ACC | C | none | nope | nope_ACC | G/C-1_CCCCAGC | 6 | FALSE | FALSE |
| 175_v.txt | 32719 | 32719-Deleti | Deletion | 1 | A | - | 19 | 25011 | 0.07596658 | 60.631579 | rnr1Y285F_pG | TRUE | CAN | JSY2421 | 6 | rnr1Y285F_p | CAA | nope | >1bp-ins | A/T-1 | none | A/T-1 | CACCACAAA | AAACCAT | GTT | GTT | T | none | T>none | T>none_GTT | A/T-1_AAACCAT | 6 | FALSE | FALSE |
| 175_v.txt | 32728 | 32728-Deleti | Deletion | 1 | A | - | 11 | 25011 | 0.04398065 | 60.7272696 | rnr1Y285F_pG | TRUE | CAN | JSY2421 | 6 | rnr1Y285F_p | CAA | nope | >1bp-ins | A/T-1 | none | A/T-1 | CCATACAAA | AAAAACA | GTT | GTT | T | none | T>none | T>none_GTT | A/T-1_AAAAACA | 6 | FALSE | FALSE |
| 175_v.txt | 32760 | 32760-Deleti | Deletion | 1 | A | - | 12 | 25011 | 0.04797889 | 57.999997 | rnr1Y285F_pG | TRUE | CAN | JSY2421 | 6 | rnr1Y285F_p | TAA | nope | >1bp-ins | A/T-1 | none | A/T-1 | ATGGCTAAA | AAAACTT | ATT | ATT | T | none | T>none | T>none_ATT | A/T-1_AAAACTT | 6 | FALSE | FALSE |
| 175_v.txt | 32804 | 32804-Deleti | Deletion | 1 | T | - | 6 | 25011 | 0.02398945 | 55.3333333 | rnr1Y285F_pG | TRUE | CAN | JSY2421 | 6 | rnr1Y285F_p | ATT | nope | >1bp-ins | A/T-1 | none | A/T-1 | GTAATATTTG | TTTGACA | TAA | ATT | T | none | nope | nope_ATT | A/T-1_TTTGACA | 6 | FALSE | FALSE |
| 175_v.txt | 32842 | 32842-Deleti | Deletion | 1 | A | - | 114 | 25011 | 0.45579945 | 44.6842096 | rnr1Y285F_pG | TRUE | CAN | JSY2421 | 6 | rnr1Y285F_p | CAA | nope | >1bp-ins | A/T-1 | none | A/T-1 | TTACCCAAA | AAAAAAT | GTT | GTT | T | none | T>none | T>none_GTT | A/T-1_AAAAAAT | 6 | FALSE | FALSE |
| 175_v.txt | 32860 | 32860-Deleti | Deletion | 1 | C | - | 124 | 25007 | 0.49586116 | 37.1854835 | rnr1Y285F_pG | TRUE | CAN | JSY2421 | 6 | rnr1Y285F_p | GCC | nope | >1bp-ins | G/C-1 | none | G/C-1 | TCCATGCCGC | CCGCCAG | CGG | GCC | C | none | nope | nope_GCC | G/C-1_CCGCCAG | 6 | FALSE | FALSE |
| 175_v.txt | 32884 | 32884-Deleti | Deletion | 1 | A | - | 10 | 24207 | 0.04131037 | 36.7999971 | rnr1Y285F_pG | TRUE | CAN | JSY2421 | 6 | rnr1Y285F_p | CAA | nope | >1bp-ins | A/T-1 | none | A/T-1 | ACGTCCAAA | AAAATTG | GTT | GTT | T | none | T>none | T>none_GTT | A/T-1_AAAATTG | 6 | FALSE | FALSE |
| 175_v.txt | 32924 | 32924-Deleti | Deletion | 1 | A | - | 36 | 24207 | 0.14871731 | 55.7222216 | rnr1Y285F_pG | TRUE | CAN | JSY2421 | 6 | rnr1Y285F_p | CAA | nope | >1bp-ins | A/T-1 | none | A/T-1 | CAGGGCAAA | AAAAGTG | GTT | GTT | T | none | T>none | T>none_GTT | A/T-1_AAAAGTG | 6 | FALSE | FALSE |
| 175_v.txt | 32932 | 32932-Deleti | Deletion | 1 | T | - | 13 | 24207 | 0.05370347 | 54.3846154 | rnr1Y285F_pG | TRUE | CAN | JSY2421 | 6 | rnr1Y285F_p | ATT | nope | >1bp-ins | A/T-1 | none | A/T-1 | AAGTGATTG | TTGCCCA | TAA | ATT | T | none | nope | nope_ATT | A/T-1_TTGCCCA | 6 | FALSE | FALSE |
| 175_v.txt | 32941 | 32941-Deleti | Deletion | 1 | A | - | 4 | 24207 | 0.01652415 | 55.7499959 | rnr1Y285F_pG | TRUE | CAN | JSY2421 | 6 | rnr1Y285F_p | GAA | nope | >1bp-ins | A/T-1 | none | A/T-1 | CCCAAGAAA | AAAACCA | CTT | CTT | T | none | T>none | T>none_CTT | A/T-1_AAAACCA | 6 | FALSE | FALSE |
| 175_v.txt | 33000 | 33000-Deleti | Deletion | 1 | A | - | 33 | 24207 | 0.1363242 | 62.7575758 | rnr1Y285F_pG | TRUE | CAN | JSY2421 | 6 | rnr1Y285F_p | GAA | nope | >1bp-ins | A/T-1 | none | A/T-1 | TGTGAGAAA | AAAACTG | CTT | CTT | T | none | T>none | T>none_CTT | A/T-1_AAAACTG | 6 | FALSE | FALSE |
| 175_v.txt | 33072 | 33072-Deleti | Deletion | 1 | A | - | 40 | 24207 | 0.16524146 | 50.775 | rnr1Y285F_pG | TRUE | CAN | JSY2421 | 6 | rnr1Y285F_p | CAA | nope | >1bp-ins | A/T-1 | none | A/T-1 | TATGCCAAA | AAAGAAC | GTT | GTT | T | none | T>none | T>none_GTT | A/T-1_AAAGAAC | 6 | FALSE | FALSE |
| 175_v.txt | 33087 | 33087-Deleti | Deletion | 1 | A | - | 3 | 24207 | 0.01239311 | 63 | rnr1Y285F_pG | TRUE | CAN | JSY2421 | 6 | rnr1Y285F_p | TAA | nope | >1bp-ins | A/T-1 | none | A/T-1 | ATAAATAAA | AAATATG | ATT | ATT | T | none | T>none | T>none_ATT | A/T-1_AAATATG | 6 | FALSE | FALSE |
| 175_v.txt | 33104 | 33104-Deleti | Deletion | 1 | G | - | 13 | 24207 | 0.05370347 | 46.3846154 | rnr1Y285F_pG | TRUE | CAN | JSY2421 | 6 | rnr1Y285F_p | CGC | nope | >1bp-ins | G/C-1 | none | G/C-1 | AAGAGCGCC | GCCCACT | GCG | GCG | C | none | C>none | C>none_GCG | G/C-1_GCCCACT | 6 | FALSE | FALSE |
| 175_v.txt | 33114 | 33114-Deleti | Deletion | 1 | C | - | 1 | 24207 | 4.13E-03 | 38 | rnr1Y285F_pG | TRUE | CAN | JSY2421 | 6 | rnr1Y285F_p | GCC | nope | >1bp-ins | G/C-1 | none | G/C-1 | ACTGGGCCG | CCGGCGT | CGG | GCC | C | none | nope | nope_GCC | G/C-1_CCGGCGT | 6 | FALSE | FALSE |
| 175_v.txt | 33141 | 33141-Deleti | Deletion | 1 | C | - | 7 | 24207 | 0.02891726 | 41.1428571 | rnr1Y285F_pG | TRUE | CAN | JSY2421 | 6 | rnr1Y285F_p | ACC | nope | >1bp-ins | G/C-1 | none | G/C-1 | GATAAACCA | CCAATGA | TGG | ACC | C | none | nope | nope_ACC | G/C-1_CCAATGA | 6 | FALSE | FALSE |
| 175_v.txt | 33147 | 33147-Deleti | Deletion | 1 | A | - | 7 | 24207 | 0.02891726 | 37 | rnr1Y285F_pG | TRUE | CAN | JSY2421 | 6 | rnr1Y285F_p | GAA | nope | >1bp-ins | A/T-1 | none | A/T-1 | CCAATGAAA | AAAAGAC | CTT | CTT | T | none | T>none | T>none_CTT | A/T-1_AAAAGAC | 6 | FALSE | FALSE |
| 175_v.txt | 33159 | 33159-Deleti | Deletion | 1 | C | - | 2 | 24206 | 8.26E-03 | 38 | rnr1Y285F_pG | TRUE | CAN | JSY2421 | 6 | rnr1Y285F_p | ACC | nope | >1bp-ins | G/C-1 | none | G/C-1 | CCTGTACCA | CCAATAG | TGG | ACC | C | none | nope | nope_ACC | G/C-1_CCAATAG | 6 | FALSE | FALSE |
| 175_v.txt | 33168 | 33168-Deleti | Deletion | 1 | C | - | 4 | 24206 | 0.01652483 | 38 | rnr1Y285F_pG | TRUE | CAN | JSY2421 | 6 | rnr1Y285F_p | ACC | nope | >1bp-ins | G/C-1 | none | G/C-1 | ATAGTACCA | CCACCAA | TGG | ACC | C | none | nope | nope_ACC | G/C-1_CCACCAA | 6 | FALSE | FALSE |
| 175_v.txt | 33171 | 33171-Deleti | Deletion | 1 | C | - | 1 | 24206 | 4.13E-03 | 37.9999999 | rnr1Y285F_pG | TRUE | CAN | JSY2421 | 6 | rnr1Y285F_p | ACC | nope | >1bp-ins | G/C-1 | none | G/C-1 | GTACCACCA | CCAAGGG | TGG | ACC | C | none | nope | nope_ACC | G/C-1_CCAAGGG | 6 | FALSE | FALSE |
| 175_v.txt | 33210 | 33210-Deleti | Deletion | 1 | C | - | 4 | 13039 | 0.0306772 | 37.75 | rnr1Y285F_pG | TRUE | CAN | JSY2421 | 6 | rnr1Y285F_p | TCT | nope | >1bp-ins | G/C-1 | none | G/C-1 | AGCTCTCTCT | CTCTTCA | AGA | TCT | C | none | nope | nope_TCT | G/C-1_CTCTTCA | 6 | FALSE | FALSE |
| 175_v.txt | 33356 | 33356-Deleti | Deletion | 1 | C | - | 37 | 13039 | 0.28376409 | 52.7567568 | rnr1Y285F_pG | TRUE | CAN | JSY2421 | 6 | rnr1Y285F_p | ACC | nope | >1bp-ins | G/C-1 | none | G/C-1 | TATTGACCC | CCCACGT | TGG | ACC | C | none | nope | nope_ACC | G/C-1_CCCACGT | 6 | FALSE | FALSE |
| 175_v.txt | 33474 | 33474-Deleti | Deletion | 1 | C | - | 25 | 13038 | 0.1917472 | 27.96 | rnr1Y285F_pG | TRUE | CAN | JSY2421 | 6 | rnr1Y285F_p | GCC | nope | >1bp-ins | G/C-1 | none | G/C-1 | GCTATGCCTT | CCTTTTT | CGG | GCC | C | none | nope | nope_GCC | G/C-1_CCTTTTT | 6 | FALSE | FALSE |
| 5_v.txt | 31723 | 31723-SNV-A | SNV | 1 | A | C | 4 | 9499 | 0.0421097 | 64 | WT | FALSE | CAN | JSY13 | 1 | WT-Rep1 | CAA | A>C | G/C+1 | A/T-1 | TA>GC | TA>GC | TGTCCCAAA | AAAAAGT | GTT | GTT | T | G | T>G | T>G_GTT | TA>GC_AAAAAGT | 9 | FALSE | FALSE |
| 5_v.txt | 31839 | 31839-SNV-A | SNV | 1 | A | C | 6 | 9499 | 0.06316454 | 48.3333333 | WT | FALSE | CAN | JSY13 | 1 | WT-Rep1 | TAA | A>C | G/C+1 | A/T-1 | TA>GC | TA>GC | TGAAATAAG | AAGATCC | ATT | ATT | T | G | T>G | T>G_ATT | TA>GC_AAGATCC | 9 | FALSE | FALSE |
| 5_v.txt | 31854 | 31854-SNV-A | SNV | 1 | A | C | 12 | 9499 | 0.12632909 | 43.1666667 | WT | FALSE | CAN | JSY13 | 1 | WT-Rep1 | TAA | A>C | G/C+1 | A/T-1 | TA>GC | TA>GC | ACAGCTAAG | AAGAACA | ATT | ATT | T | G | T>G | T>G_ATT | TA>GC_AAGAACA | 9 | FALSE | FALSE |
| 5_v.txt | 31860 | 31860-SNV-A | SNV | 1 | A | C | 17 | 9499 | 0.17896621 | 47.5294118 | WT | FALSE | CAN | JSY13 | 1 | WT-Rep1 | CAG | A>C | G/C+1 | A/T-1 | TA>GC | TA>GC | AAGAACAGG | AGGAAAA | GTC | GTC | T | G | T>G | T>G_GTC | TA>GC_AGGAAAA | 9 | FALSE | FALSE |
| 5_v.txt | 31874 | 31874-SNV-A | SNV | 1 | A | C | 18 | 9499 | 0.18949363 | 43.2222222 | WT | FALSE | CAN | JSY13 | 1 | WT-Rep1 | TAT | A>C | G/C+1 | A/T-1 | TA>GC | TA>GC | AGAGATATA | ATAGGCG | ATA | ATA | T | G | T>G | T>G_ATA | TA>GC_ATAGGCG | 9 | FALSE | FALSE |
| 5_v.txt | 31950 | 31950-SNV-A | SNV | 1 | A | C | 36 | 18263 | 0.19711986 | 34.2777778 | WT | FALSE | CAN | JSY13 | 1 | WT-Rep1 | CAT | A>C | G/C+1 | A/T-1 | TA>GC | TA>GC | ATCGTCATA | ATAAATG | GTA | GTA | T | G | T>G | T>G_GTA | TA>GC_ATAAATG | 9 | FALSE | FALSE |
| 5_v.txt | 31964 | 31964-SNV-A | SNV | 1 | A | C | 6 | 18279 | 0.03282455 | 28.6666667 | WT | FALSE | CAN | JSY13 | 1 | WT-Rep1 | CAT | A>C | G/C+1 | A/T-1 | TA>GC | TA>GC | GGCCGCATA | ATAATAA | GTA | GTA | T | G | T>G | T>G_GTA | TA>GC_ATAATAA | 9 | FALSE | FALSE |
| 5_v.txt | 31986 | 31986-SNV-A | SNV | 1 | A | C | 157 | 18279 | 0.85890913 | 51.7324841 | WT | FALSE | CAN | JSY13 | 1 | WT-Rep1 | TAA | A>C | G/C+1 | A/T-1 | TA>GC | TA>GC | GGCATTAAT | AATTTAG | ATT | ATT | T | G | T>G | T>G_ATT | TA>GC_AATTTAG | 9 | FALSE | FALSE |
| 5_v.txt | 32015 | 32015-SNV-A | SNV | 1 | A | C | 2 | 18279 | 0.01094152 | 27.0014518 | WT | FALSE | CAN | JSY13 | 1 | WT-Rep1 | GAG | A>C | G/C+1 | A/T-1 | TA>GC | TA>GC | GTCACGAGA | AGAGATG | CTC | CTC | T | G | T>G | T>G_CTC | TA>GC_AGAGATG | 9 | FALSE | FALSE |
| 5_v.txt | 32101 | 32101-SNV-T | SNV | 1 | T | G | 10 | 18279 | 0.05470759 | 55.6 | WT | FALSE | CAN | JSY13 | 1 | WT-Rep1 | GTG | T>G | G/C+1 | A/T-1 | TA>GC | TA>GC | CACCAGTGA | TGATATT | CAC | GTG | T | C | T>G | T>G_GTG | TA>GC_TGATATT | 9 | FALSE | FALSE |
| 5_v.txt | 32109 | 32109-SNV-A | SNV | 1 | A | C | 17 | 18279 | 0.0930029 | 61.5882353 | WT | FALSE | CAN | JSY13 | 1 | WT-Rep1 | TAA | A>C | G/C+1 | A/T-1 | TA>GC | TA>GC | ATATTTAATA | AATAGCC | ATT | ATT | T | G | T>G | T>G_ATT | TA>GC_AATAGCC | 9 | FALSE | FALSE |
| 5_v.txt | 32256 | 32256-SNV-A | SNV | 1 | A | C | 11 | 21411 | 0.05137546 | 26.5454546 | WT | FALSE | CAN | JSY13 | 1 | WT-Rep1 | TAG | A>C | G/C+1 | A/T-1 | TA>GC | TA>GC | TTTGATAGA | AGACCAA | ATC | ATC | T | G | T>G | T>G_ATC | TA>GC_AGACCAA | 9 | FALSE | FALSE |
| 5_v.txt | 32265 | 32265-SNV-A | SNV | 1 | A | C | 27 | 21420 | 0.12605042 | 32.5925926 | WT | FALSE | CAN | JSY13 | 1 | WT-Rep1 | TAA | A>C | G/C+1 | A/T-1 | TA>GC | TA>GC | CCAAATAAA | AAAATAC | ATT | ATT | T | G | T>G | T>G_ATT | TA>GC_AAAATAC | 9 | FALSE | FALSE |
| 5_v.txt | 32283 | 32283-SNV-T | SNV | 1 | T | G | 2810 | 21420 | 13.1185808 | 52.9231794 | WT | FALSE | CAN | JSY13 | 1 | WT-Rep1 | GTA | T>G | G/C+1 | A/T-1 | TA>GC | TA>GC | CCAACGTAA | TAAATAT | CAT | GTA | T | C | T>G | T>G_GTA | TA>GC_TAAATAT | 9 | FALSE | FALSE |
| 5_v.txt | 32284 | 32284-SNV-A | SNV | 1 | A | C | 12 | 21421 | 0.05601979 | 39.083132 | WT | FALSE | CAN | JSY13 | 1 | WT-Rep1 | TAA | A>C | G/C+1 | A/T-1 | TA>GC | TA>GC | CAACGTAAA | AAATATT | ATT | ATT | T | G | T>G | T>G_ATT | TA>GC_AAATATT | 9 | FALSE | FALSE |
| 5_v.txt | 32288 | 32288-SNV-A | SNV | 1 | A | C | 58 | 21421 | 0.27076234 | 49.4137931 | WT | FALSE | CAN | JSY13 | 1 | WT-Rep1 | TAT | A>C | G/C+1 | A/T-1 | TA>GC | TA>GC | GTAAATATTT | ATTTGAA | ATA | ATA | T | G | T>G | T>G_ATA | TA>GC_ATTTGAA | 9 | FALSE | FALSE |
| 5_v.txt | 32293 | 32293-SNV-A | SNV | 1 | A | C | 21 | 21421 | 0.09803464 | 17.1424858 | WT | FALSE | CAN | JSY13 | 1 | WT-Rep1 | GAA | A>C | G/C+1 | A/T-1 | TA>GC | TA>GC | TATTTGAATT | AATTTGC | CTT | CTT | T | G | T>G | T>G_CTT | TA>GC_AATTTGC | 9 | FALSE | FALSE |
| 5_v.txt | 32294 | 32294-SNV-A | SNV | 1 | A | C | 21 | 21421 | 0.09803464 | 46.5714286 | WT | FALSE | CAN | JSY13 | 1 | WT-Rep1 | AAT | A>C | G/C+1 | A/T-1 | TA>GC | TA>GC | ATTTGAATTT | ATTTGCG | TTA | TTA | T | G | T>G | T>G_TTA | TA>GC_ATTTGCG | 9 | FALSE | FALSE |
| 5_v.txt | 32311 | 32311-SNV-T | SNV | 1 | T | G | 4 | 21421 | 0.01867327 | 44.25 | WT | FALSE | CAN | JSY13 | 1 | WT-Rep1 | ATG | T>G | G/C+1 | A/T-1 | TA>GC | TA>GC | AAATAATGG | TGGTTGT | TAC | ATG | T | C | T>G | T>G_ATG | TA>GC_TGGTTGT | 9 | FALSE | FALSE |
| 5_v.txt | 32319 | 32319-SNV-A | SNV | 1 | A | C | 49 | 21421 | 0.22874749 | 57.7069005 | WT | FALSE | CAN | JSY13 | 1 | WT-Rep1 | TAA | A>C | G/C+1 | A/T-1 | TA>GC | TA>GC | GTTGTTAAG | AAGATAA | ATT | ATT | T | G | T>G | T>G_ATT | TA>GC_AAGATAA | 9 | FALSE | FALSE |
| 5_v.txt | 32414 | 32414-SNV-T | SNV | 1 | T | G | 3 | 21420 | 0.0140056 | 28.6667912 | WT | FALSE | CAN | JSY13 | 1 | WT-Rep1 | GTG | T>G | G/C+1 | A/T-1 | TA>GC | TA>GC | AGATTGTGT | TGTTAGT | CAC | GTG | T | C | T>G | T>G_GTG | TA>GC_TGTTAGT | 9 | FALSE | FALSE |
| 5_v.txt | 32439 | 32439-SNV-A | SNV | 1 | A | C | 52 | 21421 | 0.24275244 | 51.3461539 | WT | FALSE | CAN | JSY13 | 1 | WT-Rep1 | AAC | A>C | G/C+1 | A/T-1 | TA>GC | TA>GC | TATGGAACT | ACTAAAA | TTG | TTG | T | G | T>G | T>G_TTG | TA>GC_ACTAAAA | 9 | FALSE | FALSE |
| 5_v.txt | 32445 | 32445-SNV-A | SNV | 1 | A | C | 22 | 21421 | 0.10270296 | 50.3181818 | WT | FALSE | CAN | JSY13 | 1 | WT-Rep1 | AAG | A>C | G/C+1 | A/T-1 | TA>GC | TA>GC | ACTAAAAGT | AGTCCAA | TTC | TTC | T | G | T>G | T>G_TTC | TA>GC_AGTCCAA | 9 | FALSE | FALSE |
| 5_v.txt | 32455 | 32455-SNV-A | SNV | 1 | A | C | 1 | 21421 | 4.67E-03 | 38.0013 | WT | FALSE | CAN | JSY13 | 1 | WT-Rep1 | AAT | A>C | G/C+1 | A/T-1 | TA>GC | TA>GC | CAATGAATA | ATAATAG | TTA | TTA | T | G | T>G | T>G_TTA | TA>GC_ATAATAG | 9 | FALSE | FALSE |
| 5_v.txt | 32472 | 32472-SNV-T | SNV | 1 | T | G | 57 | 21421 | 0.26609402 | 47.2955327 | WT | FALSE | CAN | JSY13 | 1 | WT-Rep1 | GTA | T>G | G/C+1 | A/T-1 | TA>GC | TA>GC | CCAATGTAG | TAGAAGG | CAT | GTA | T | C | T>G | T>G_GTA | TA>GC_TAGAAGG | 9 | FALSE | FALSE |
| 5_v.txt | 32473 | 32473-SNV-A | SNV | 1 | A | C | 10 | 21421 | 0.04668316 | 50.0963849 | WT | FALSE | CAN | JSY13 | 1 | WT-Rep1 | TAG | A>C | G/C+1 | A/T-1 | TA>GC | TA>GC | CAATGTAGA | AGAAGGT | ATC | ATC | T | G | T>G | T>G_ATC | TA>GC_AGAAGGT | 9 | FALSE | FALSE |
| 5_v.txt | 32481 | 32481-SNV-A | SNV | 1 | A | C | 20 | 21421 | 0.09336632 | 49.65 | WT | FALSE | CAN | JSY13 | 1 | WT-Rep1 | TAA | A>C | G/C+1 | A/T-1 | TA>GC | TA>GC | AAGGTTAAG | AAGATAC | ATT | ATT | T | G | T>G | T>G_ATT | TA>GC_AAGATAC | 9 | FALSE | FALSE |
| 5_v.txt | 32493 | 32493-SNV-A | SNV | 1 | A | C | 2 | 21421 | 9.34E-03 | 37.4983208 | WT | FALSE | CAN | JSY13 | 1 | WT-Rep1 | AAC | A>C | G/C+1 | A/T-1 | TA>GC | TA>GC | CGGAAAACA | ACAACTT | TTG | TTG | T | G | T>G | T>G_TTG | TA>GC_ACAACTT | 9 | FALSE | FALSE |
| 5_v.txt | 32541 | 32541-SNV-T | SNV | 1 | T | G | 64 | 21421 | 0.29877223 | 36.4375149 | WT | FALSE | CAN | JSY13 | 1 | WT-Rep1 | TTC | T>G | G/C+1 | A/T-1 | TA>GC | TA>GC | GCAGCTTCA | TCACCAG | AAG | TTC | T | C | T>G | T>G_TTC | TA>GC_TCACCAG | 9 | FALSE | FALSE |
| 5_v.txt | 32573 | 32573-SNV-T | SNV | 1 | T | G | 9 | 22153 | 0.04062655 | 34.6666667 | WT | FALSE | CAN | JSY13 | 1 | WT-Rep1 | CTT | T>G | G/C+1 | A/T-1 | TA>GC | TA>GC | AGTACCTTG | TTGAAAT | GAA | CTT | T | C | T>G | T>G_CTT | TA>GC_TTGAAAT | 9 | FALSE | FALSE |
| 5_v.txt | 32581 | 32581-SNV-T | SNV | 1 | T | G | 16 | 22153 | 0.07222498 | 36.8125 | WT | FALSE | CAN | JSY13 | 1 | WT-Rep1 | GTG | T>G | G/C+1 | A/T-1 | TA>GC | TA>GC | GAAATGTGA | TGAAGGC | CAC | GTG | T | C | T>G | T>G_GTG | TA>GC_TGAAGGC | 9 | FALSE | FALSE |
| 5_v.txt | 32616 | 32616-SNV-A | SNV | 1 | A | C | 22 | 22156 | 0.0992959 | 48.5 | WT | FALSE | CAN | JSY13 | 1 | WT-Rep1 | TAA | A>C | G/C+1 | A/T-1 | TA>GC | TA>GC | CAACCTAAG | AAGAACC | ATT | ATT | T | G | T>G | T>G_ATT | TA>GC_AAGAACC | 9 | FALSE | FALSE |
| 5_v.txt | 32657 | 32657-SNV-A | SNV | 1 | A | C | 2 | 22156 | 9.03E-03 | 59.9990318 | WT | FALSE | CAN | JSY13 | 1 | WT-Rep1 | GAC | A>C | G/C+1 | A/T-1 | TA>GC | TA>GC | ACCTGGACC | ACCCCAG | CTG | CTG | T | G | T>G | T>G_CTG | TA>GC_ACCCCAG | 9 | FALSE | FALSE |
| 5_v.txt | 32680 | 32680-SNV-A | SNV | 1 | A | C | 38 | 22156 | 0.1715111 | 58.5762946 | WT | FALSE | CAN | JSY13 | 1 | WT-Rep1 | CAA | A>C | G/C+1 | A/T-1 | TA>GC | TA>GC | TTCTCCAATA | AATAACG | GTT | GTT | T | G | T>G | T>G_GTT | TA>GC_AATAACG | 9 | FALSE | FALSE |
| 5_v.txt | 32742 | 32742-SNV-A | SNV | 1 | A | C | 10 | 22177 | 0.04509176 | 54 | WT | FALSE | CAN | JSY13 | 1 | WT-Rep1 | TAG | A>C | G/C+1 | A/T-1 | TA>GC | TA>GC | TATATTAGA | AGAAACC | ATC | ATC | T | G | T>G | T>G_ATC | TA>GC_AGAAACC | 9 | FALSE | FALSE |
| 5_v.txt | 32760 | 32760-SNV-A | SNV | 1 | A | C | 1 | 22176 | 4.51E-03 | 63.9993335 | WT | FALSE | CAN | JSY13 | 1 | WT-Rep1 | TAA | A>C | G/C+1 | A/T-1 | TA>GC | TA>GC | ATGGCTAAA | AAAACTT | ATT | ATT | T | G | T>G | T>G_ATT | TA>GC_AAAACTT | 9 | FALSE | FALSE |
| 5_v.txt | 32763 | 32763-SNV-A | SNV | 1 | A | C | 10 | 22177 | 0.04509176 | 46.7 | WT | FALSE | CAN | JSY13 | 1 | WT-Rep1 | AAC | A>C | G/C+1 | A/T-1 | TA>GC | TA>GC | GCTAAAACT | ACTTTGA | TTG | TTG | T | G | T>G | T>G_TTG | TA>GC_ACTTTGA | 9 | FALSE | FALSE |
| 5_v.txt | 32769 | 32769-SNV-A | SNV | 1 | A | C | 9 | 22177 | 0.04058259 | 56.6666667 | WT | FALSE | CAN | JSY13 | 1 | WT-Rep1 | GAT | A>C | G/C+1 | A/T-1 | TA>GC | TA>GC | ACTTTGATG | ATGGAAG | CTA | CTA | T | G | T>G | T>G_CTA | TA>GC_ATGGAAG | 9 | FALSE | FALSE |
| 5_v.txt | 32787 | 32787-SNV-T | SNV | 1 | T | G | 16 | 22177 | 0.07214682 | 46.0034884 | WT | FALSE | CAN | JSY13 | 1 | WT-Rep1 | CTC | T>G | G/C+1 | A/T-1 | TA>GC | TA>GC | CAGAACTCG | TCGAATT | GAG | CTC | T | C | T>G | T>G_CTC | TA>GC_TCGAATT | 9 | FALSE | FALSE |
| 5_v.txt | 32792 | 32792-SNV-T | SNV | 1 | T | G | 14 | 22177 | 0.06312847 | 50.2142857 | WT | FALSE | CAN | JSY13 | 1 | WT-Rep1 | ATT | T>G | G/C+1 | A/T-1 | TA>GC | TA>GC | CTCGAATTC | TTCACCG | TAA | ATT | T | C | T>G | T>G_ATT | TA>GC_TTCACCG | 9 | FALSE | FALSE |

| | | | | | | | | | | | | | | | | | | | | | | | | | | | | | | | | | | |
|---|---|---|---|---|---|---|---|---|---|---|---|---|---|---|---|---|---|---|---|---|---|---|---|---|---|---|---|---|---|---|---|---|---|---|
| 5_v.txt | 32800 | 32800-SNV-A | SNV | 1 | A | C | 17 | 22177 | 0.076656 | 48.5878638 | WT | FALSE | CAN | JSY13 | 1 | WT-Rep1 | TAA | A>C | G/C+1 | A/T-1 | TA>GC | TA>GC | CACCGTAAT | AATATTT | ATT | ATT | T | G | T>G | T>G_ATT | TA>GC_AATATTT | 9 | FALSE | FALSE |
| 5_v.txt | 32801 | 32801-SNV-A | SNV | 1 | A | C | 29 | 22177 | 0.13076611 | 43.5874225 | WT | FALSE | CAN | JSY13 | 1 | WT-Rep1 | AAT | A>C | G/C+1 | A/T-1 | TA>GC | TA>GC | ACCGTAATA | ATATTTG | TTA | TTA | T | G | T>G | T>G_TTA | TA>GC_ATATTTG | 9 | FALSE | FALSE |
| 5_v.txt | 32803 | 32803-SNV-A | SNV | 1 | A | C | 8 | 22177 | 0.03607341 | 37.2502567 | WT | FALSE | CAN | JSY13 | 1 | WT-Rep1 | TAT | A>C | G/C+1 | A/T-1 | TA>GC | TA>GC | CGTAATATTT | ATTTGAC | ATA | ATA | T | G | T>G | T>G_ATA | TA>GC_ATTTGAC | 9 | FALSE | FALSE |
| 5_v.txt | 32823 | 32823-SNV-A | SNV | 1 | A | C | 34 | 22177 | 0.15331199 | 42.6176471 | WT | FALSE | CAN | JSY13 | 1 | WT-Rep1 | CAT | A>C | G/C+1 | A/T-1 | TA>GC | TA>GC | AAGTTCATT | ATTATTG | GTA | GTA | T | G | T>G | T>G_GTA | TA>GC_ATTATTG | 9 | FALSE | FALSE |
| 5_v.txt | 32842 | 32842-SNV-A | SNV | 1 | A | C | 5 | 22177 | 0.02254588 | 37.199776 | WT | FALSE | CAN | JSY13 | 1 | WT-Rep1 | CAA | A>C | G/C+1 | A/T-1 | TA>GC | TA>GC | TTACCCAAA | AAAAAAT | GTT | GTT | T | G | T>G | T>G_GTT | TA>GC_AAAAAAT | 9 | FALSE | FALSE |
| 5_v.txt | 32884 | 32884-SNV-A | SNV | 1 | A | C | 5 | 24305 | 0.0205719 | 38 | WT | FALSE | CAN | JSY13 | 1 | WT-Rep1 | CAA | A>C | G/C+1 | A/T-1 | TA>GC | TA>GC | ACGTCCAAA | AAAATTG | GTT | GTT | T | G | T>G | T>G_GTT | TA>GC_AAAATTG | 9 | FALSE | FALSE |
| 5_v.txt | 32915 | 32915-SNV-T | SNV | 1 | T | G | 10 | 24305 | 0.0411438 | 48.5 | WT | FALSE | CAN | JSY13 | 1 | WT-Rep1 | GTT | T>G | G/C+1 | A/T-1 | TA>GC | TA>GC | ACTAAGTTC | TTCCAGG | CAA | GTT | T | C | T>G | T>G_GTT | TA>GC_TTCCAGG | 9 | FALSE | FALSE |
| 5_v.txt | 32919 | 32919-SNV-A | SNV | 1 | A | C | 13 | 24305 | 0.05348694 | 27.1554331 | WT | FALSE | CAN | JSY13 | 1 | WT-Rep1 | CAG | A>C | G/C+1 | A/T-1 | TA>GC | TA>GC | AGTTCCAGG | AGGGCAA | GTC | GTC | T | G | T>G | T>G_GTC | TA>GC_AGGGCAA | 9 | FALSE | FALSE |
| 5_v.txt | 32929 | 32929-SNV-T | SNV | 1 | T | G | 10 | 24343 | 0.04107957 | 51.9989009 | WT | FALSE | CAN | JSY13 | 1 | WT-Rep1 | GTG | T>G | G/C+1 | A/T-1 | TA>GC | TA>GC | CAAAAGTGA | TGATTGC | CAC | GTG | T | C | T>G | T>G_GTG | TA>GC_TGATTGC | 9 | FALSE | FALSE |
| 5_v.txt | 32938 | 32938-SNV-A | SNV | 1 | A | C | 75 | 24343 | 0.30809678 | 50.4930066 | WT | FALSE | CAN | JSY13 | 1 | WT-Rep1 | CAA | A>C | G/C+1 | A/T-1 | TA>GC | TA>GC | TTGCCCAAG | AAGAAAA | GTT | GTT | T | G | T>G | T>G_GTT | TA>GC_AAGAAAA | 9 | FALSE | FALSE |
| 5_v.txt | 32941 | 32941-SNV-A | SNV | 1 | A | C | 9 | 24343 | 0.03697161 | 16.8905127 | WT | FALSE | CAN | JSY13 | 1 | WT-Rep1 | GAA | A>C | G/C+1 | A/T-1 | TA>GC | TA>GC | CCCAAGAAA | AAAACCA | CTT | CTT | T | G | T>G | T>G_CTT | TA>GC_AAAACCA | 9 | FALSE | FALSE |
| 5_v.txt | 33054 | 33054-SNV-T | SNV | 1 | T | G | 64 | 24341 | 0.26293086 | 52.9373832 | WT | FALSE | CAN | JSY13 | 1 | WT-Rep1 | CTG | T>G | G/C+1 | A/T-1 | TA>GC | TA>GC | AAGGACTGC | TGCGTGA | GAC | CTG | T | C | T>G | T>G_CTG | TA>GC_TGCGTGA | 9 | FALSE | FALSE |
| 5_v.txt | 33084 | 33084-SNV-A | SNV | 1 | A | C | 1 | 24343 | 4.11E-03 | 63.9995854 | WT | FALSE | CAN | JSY13 | 1 | WT-Rep1 | AAA | A>C | G/C+1 | A/T-1 | TA>GC | TA>GC | CCCATAAAT | AATAAAT | TTT | TTT | T | G | T>G | T>G_TTT | TA>GC_AATAAAT | 9 | FALSE | FALSE |
| 5_v.txt | 33150 | 33150-SNV-A | SNV | 1 | A | C | 40 | 24343 | 0.16431829 | 38.475 | WT | FALSE | CAN | JSY13 | 1 | WT-Rep1 | AAG | A>C | G/C+1 | A/T-1 | TA>GC | TA>GC | ATGAAAAGA | AGACCTG | TTC | TTC | T | G | T>G | T>G_TTC | TA>GC_AGACCTG | 9 | FALSE | FALSE |
| 5_v.txt | 33157 | 33157-SNV-T | SNV | 1 | T | G | 7 | 24342 | 0.02875688 | 29.430795 | WT | FALSE | CAN | JSY13 | 1 | WT-Rep1 | GTA | T>G | G/C+1 | A/T-1 | TA>GC | TA>GC | GACCTGTAC | TACCAAT | CAT | GTA | T | C | T>G | T>G_GTA | TA>GC_TACCAAT | 9 | FALSE | FALSE |
| 5_v.txt | 33166 | 33166-SNV-T | SNV | 1 | T | G | 22 | 24300 | 0.09053498 | 37.1818182 | WT | FALSE | CAN | JSY13 | 1 | WT-Rep1 | GTA | T>G | G/C+1 | A/T-1 | TA>GC | TA>GC | CAATAGTAC | TACCACC | CAT | GTA | T | C | T>G | T>G_GTA | TA>GC_TACCACC | 9 | FALSE | FALSE |
| 5_v.txt | 33192 | 33192-SNV-T | SNV | 1 | T | G | 4 | 7277 | 0.05496771 | 36.9994487 | WT | FALSE | CAN | JSY13 | 1 | WT-Rep1 | ATG | T>G | G/C+1 | A/T-1 | TA>GC | TA>GC | CCAATATGT | TGTCTTT | TAC | ATG | T | C | T>G | T>G_ATG | TA>GC_TGTCTTT | 9 | FALSE | FALSE |
| 5_v.txt | 33204 | 33204-SNV-A | SNV | 1 | A | C | 8 | 7277 | 0.10993541 | 34.0038098 | WT | FALSE | CAN | JSY13 | 1 | WT-Rep1 | AAG | A>C | G/C+1 | A/T-1 | TA>GC | TA>GC | TGCTTAAGC | AGCTCTC | TTC | TTC | T | G | T>G | T>G_TTC | TA>GC_AGCTCTC | 9 | FALSE | FALSE |
| 5_v.txt | 33287 | 33287-SNV-A | SNV | 1 | A | C | 14 | 7277 | 0.19238697 | 11.9997734 | WT | FALSE | CAN | JSY13 | 1 | WT-Rep1 | CAT | A>C | G/C+1 | A/T-1 | TA>GC | TA>GC | CTCGCCATTT | ATTTACT | GTA | GTA | T | G | T>G | T>G_GTA | TA>GC_ATTTACT | 9 | FALSE | FALSE |
| 5_v.txt | 33291 | 33291-SNV-A | SNV | 1 | A | C | 3 | 7277 | 0.04122578 | 37.6665848 | WT | FALSE | CAN | JSY13 | 1 | WT-Rep1 | TAC | A>C | G/C+1 | A/T-1 | TA>GC | TA>GC | CCATTTACTC | ACTCTCG | ATG | ATG | T | G | T>G | T>G_ATG | TA>GC_ACTCTCG | 9 | FALSE | FALSE |
| 5_v.txt | 32070 | 32070-SNV-A | SNV | 1 | A | G | 2 | 18278 | 0.01094212 | 54.9999744 | WT | FALSE | CAN | JSY13 | 1 | WT-Rep1 | AAA | A>G | G/C+1 | A/T-1 | TA>CG | TA>CG | GAGATAAAT | AATAACC | TTT | TTT | T | C | T>C | T>C_TTT | TA>CG_AATAACC | 9 | FALSE | FALSE |
| 5_v.txt | 32073 | 32073-SNV-A | SNV | 1 | A | G | 5 | 18279 | 0.02735379 | 63.999318 | WT | FALSE | CAN | JSY13 | 1 | WT-Rep1 | TAA | A>G | G/C+1 | A/T-1 | TA>CG | TA>CG | ATAAATAAC | AACCATG | ATT | ATT | T | C | T>C | T>C_ATT | TA>CG_AACCATG | 9 | FALSE | FALSE |
| 5_v.txt | 32077 | 32077-SNV-A | SNV | 1 | A | G | 21 | 18278 | 0.11489222 | 63.2381494 | WT | FALSE | CAN | JSY13 | 1 | WT-Rep1 | CAT | A>G | G/C+1 | A/T-1 | TA>CG | TA>CG | ATAACCATG | ATGCAAA | GTA | GTA | T | C | T>C | T>C_GTA | TA>CG_ATGCAAA | 9 | FALSE | FALSE |
| 5_v.txt | 32081 | 32081-SNV-A | SNV | 1 | A | G | 6 | 18279 | 0.03282455 | 48.9999618 | WT | FALSE | CAN | JSY13 | 1 | WT-Rep1 | CAA | A>G | G/C+1 | A/T-1 | TA>CG | TA>CG | CCATGCAAA | AAAAAAG | GTT | GTT | T | C | T>C | T>C_GTT | TA>CG_AAAAAAG | 9 | FALSE | FALSE |
| 5_v.txt | 32112 | 32112-SNV-A | SNV | 1 | A | G | 26 | 18279 | 0.14223973 | 57.5 | WT | FALSE | CAN | JSY13 | 1 | WT-Rep1 | TAG | A>G | G/C+1 | A/T-1 | TA>CG | TA>CG | TTTAATAGC | AGCCATT | ATC | ATC | T | C | T>C | T>C_ATC | TA>CG_AGCCATT | 9 | FALSE | FALSE |
| 5_v.txt | 32256 | 32256-SNV-A | SNV | 1 | A | G | 57 | 21411 | 0.2662183 | 35.5964912 | WT | FALSE | CAN | JSY13 | 1 | WT-Rep1 | TAG | A>G | G/C+1 | A/T-1 | TA>CG | TA>CG | TTTGATAGA | AGACCAA | ATC | ATC | T | C | T>C | T>C_ATC | TA>CG_AGACCAA | 9 | FALSE | FALSE |
| 5_v.txt | 32275 | 32275-SNV-A | SNV | 1 | A | G | 18 | 21420 | 0.08403361 | 32.9368442 | WT | FALSE | CAN | JSY13 | 1 | WT-Rep1 | GAA | A>G | G/C+1 | A/T-1 | TA>CG | TA>CG | TACGGGAAC | AACCAAC | CTT | CTT | T | C | T>C | T>C_CTT | TA>CG_AACCAAC | 9 | FALSE | FALSE |
| 5_v.txt | 32293 | 32293-SNV-A | SNV | 1 | A | G | 90 | 21421 | 0.42014845 | 54.1168936 | WT | FALSE | CAN | JSY13 | 1 | WT-Rep1 | GAA | A>G | G/C+1 | A/T-1 | TA>CG | TA>CG | TATTTGAATT | AATTTGC | CTT | CTT | T | C | T>C | T>C_CTT | TA>CG_AATTTGC | 9 | FALSE | FALSE |
| 5_v.txt | 32294 | 32294-SNV-A | SNV | 1 | A | G | 5 | 21421 | 0.02334158 | 47.8 | WT | FALSE | CAN | JSY13 | 1 | WT-Rep1 | AAT | A>G | G/C+1 | A/T-1 | TA>CG | TA>CG | ATTTGAATTT | ATTTGCG | TTA | TTA | T | C | T>C | T>C_TTA | TA>CG_ATTTGCG | 9 | FALSE | FALSE |
| 5_v.txt | 32320 | 32320-SNV-A | SNV | 1 | A | G | 5 | 21421 | 0.02334158 | 38.799684 | WT | FALSE | CAN | JSY13 | 1 | WT-Rep1 | AAG | A>G | G/C+1 | A/T-1 | TA>CG | TA>CG | TTGTTAAGA | AGATAAC | TTC | TTC | T | C | T>C | T>C_TTC | TA>CG_AGATAAC | 9 | FALSE | FALSE |
| 5_v.txt | 32338 | 32338-SNV-T | SNV | 1 | T | C | 1 | 21421 | 4.67E-03 | 54.9999904 | WT | FALSE | CAN | JSY13 | 1 | WT-Rep1 | ATA | T>C | G/C+1 | A/T-1 | TA>CG | TA>CG | TGAAGATAT | TATGTGG | TAT | ATA | T | G | T>C | T>C_ATA | TA>CG_TATGTGG | 9 | FALSE | FALSE |
| 5_v.txt | 32445 | 32445-SNV-A | SNV | 1 | A | G | 132 | 21421 | 0.61621773 | 51.6666667 | WT | FALSE | CAN | JSY13 | 1 | WT-Rep1 | AAG | A>G | G/C+1 | A/T-1 | TA>CG | TA>CG | ACTAAAAGT | AGTCCAA | TTC | TTC | T | C | T>C | T>C_TTC | TA>CG_AGTCCAA | 9 | FALSE | FALSE |
| 5_v.txt | 32454 | 32454-SNV-A | SNV | 1 | A | G | 2 | 21421 | 9.34E-03 | 50.9999808 | WT | FALSE | CAN | JSY13 | 1 | WT-Rep1 | GAA | A>G | G/C+1 | A/T-1 | TA>CG | TA>CG | CCAATGAAT | AATAATA | CTT | CTT | T | C | T>C | T>C_CTT | TA>CG_AATAATA | 9 | FALSE | FALSE |
| 5_v.txt | 32459 | 32459-SNV-T | SNV | 1 | T | C | 2 | 21421 | 9.34E-03 | 63.9999482 | WT | FALSE | CAN | JSY13 | 1 | WT-Rep1 | ATA | T>C | G/C+1 | A/T-1 | TA>CG | TA>CG | GAATAATAG | TAGAGAG | TAT | ATA | T | G | T>C | T>C_ATA | TA>CG_TAGAGAG | 9 | FALSE | FALSE |
| 5_v.txt | 32472 | 32472-SNV-T | SNV | 1 | T | C | 38 | 21421 | 0.17739601 | 47.815039 | WT | FALSE | CAN | JSY13 | 1 | WT-Rep1 | GTA | T>C | G/C+1 | A/T-1 | TA>CG | TA>CG | CCAATGTAG | TAGAAGG | CAT | GTA | T | G | T>C | T>C_GTA | TA>CG_TAGAAGG | 9 | FALSE | FALSE |
| 5_v.txt | 32490 | 32490-SNV-A | SNV | 1 | A | G | 56 | 21421 | 0.2614257 | 41.9285714 | WT | FALSE | CAN | JSY13 | 1 | WT-Rep1 | GAA | A>G | G/C+1 | A/T-1 | TA>CG | TA>CG | ATACGGAAA | AAAACAA | CTT | CTT | T | C | T>C | T>C_CTT | TA>CG_AAAACAA | 9 | FALSE | FALSE |
| 5_v.txt | 32493 | 32493-SNV-A | SNV | 1 | A | G | 4 | 21421 | 0.01867327 | 17.4997265 | WT | FALSE | CAN | JSY13 | 1 | WT-Rep1 | AAC | A>G | G/C+1 | A/T-1 | TA>CG | TA>CG | CGGAAAACA | ACAACTT | TTG | TTG | T | C | T>C | T>C_TTG | TA>CG_ACAACTT | 9 | FALSE | FALSE |
| 5_v.txt | 32527 | 32527-SNV-T | SNV | 1 | T | C | 6 | 21421 | 0.0280099 | 36.3331607 | WT | FALSE | CAN | JSY13 | 1 | WT-Rep1 | CTG | T>C | G/C+1 | A/T-1 | TA>CG | TA>CG | ATTTTCTGG | TGGGGTT | GAC | CTG | T | G | T>C | T>C_CTG | TA>CG_TGGGGTT | 9 | FALSE | FALSE |
| 5_v.txt | 32573 | 32573-SNV-T | SNV | 1 | T | C | 6 | 22153 | 0.02708437 | 37.6666667 | WT | FALSE | CAN | JSY13 | 1 | WT-Rep1 | CTT | T>C | G/C+1 | A/T-1 | TA>CG | TA>CG | AGTACCTTG | TTGAAAT | GAA | CTT | T | G | T>C | T>C_CTT | TA>CG_TTGAAAT | 9 | FALSE | FALSE |
| 5_v.txt | 32583 | 32583-SNV-A | SNV | 1 | A | G | 74 | 22153 | 0.33404054 | 36.5135135 | WT | FALSE | CAN | JSY13 | 1 | WT-Rep1 | GAA | A>G | G/C+1 | A/T-1 | TA>CG | TA>CG | AATGTGAAG | AAGGCAG | CTT | CTT | T | C | T>C | T>C_CTT | TA>CG_AAGGCAG | 9 | FALSE | FALSE |
| 5_v.txt | 32662 | 32662-SNV-A | SNV | 1 | A | G | 22 | 22156 | 0.0992959 | 55.1817678 | WT | FALSE | CAN | JSY13 | 1 | WT-Rep1 | CAG | A>G | G/C+1 | A/T-1 | TA>CG | TA>CG | GACCCCAGG | AGGCACC | GTC | GTC | T | C | T>C | T>C_GTC | TA>CG_AGGCACC | 9 | FALSE | FALSE |
| 5_v.txt | 32680 | 32680-SNV-A | SNV | 1 | A | G | 61 | 22156 | 0.27532046 | 59.6670016 | WT | FALSE | CAN | JSY13 | 1 | WT-Rep1 | CAA | A>G | G/C+1 | A/T-1 | TA>CG | TA>CG | TTCTCCAATA | AATAACG | GTT | GTT | T | C | T>C | T>C_GTT | TA>CG_AATAACG | 9 | FALSE | FALSE |
| 5_v.txt | 32767 | 32767-SNV-T | SNV | 1 | T | C | 35 | 22177 | 0.15782117 | 45.7714286 | WT | FALSE | CAN | JSY13 | 1 | WT-Rep1 | TTG | T>C | G/C+1 | A/T-1 | TA>CG | TA>CG | AAACTTTGA | TGATGGA | AAC | TTG | T | G | T>C | T>C_TTG | TA>CG_TGATGGA | 9 | FALSE | FALSE |
| 5_v.txt | 32787 | 32787-SNV-T | SNV | 1 | T | C | 10 | 22177 | 0.04509176 | 40.5018527 | WT | FALSE | CAN | JSY13 | 1 | WT-Rep1 | CTC | T>C | G/C+1 | A/T-1 | TA>CG | TA>CG | CAGAACTCG | TCGAATT | GAG | CTC | T | G | T>C | T>C_CTC | TA>CG_TCGAATT | 9 | FALSE | FALSE |
| 5_v.txt | 32808 | 32808-SNV-A | SNV | 1 | A | G | 222 | 22177 | 1.00103711 | 44.0225225 | WT | FALSE | CAN | JSY13 | 1 | WT-Rep1 | GAC | A>G | G/C+1 | A/T-1 | TA>CG | TA>CG | TATTTGACA | ACAGGGA | CTG | CTG | T | C | T>C | T>C_CTG | TA>CG_ACAGGGA | 9 | FALSE | FALSE |
| 5_v.txt | 32919 | 32919-SNV-A | SNV | 1 | A | G | 22 | 24305 | 0.09051636 | 46.1371545 | WT | FALSE | CAN | JSY13 | 1 | WT-Rep1 | CAG | A>G | G/C+1 | A/T-1 | TA>CG | TA>CG | AGTTCCAGG | AGGGCAA | GTC | GTC | T | C | T>C | T>C_GTC | TA>CG_AGGGCAA | 9 | FALSE | FALSE |
| 5_v.txt | 32927 | 32927-SNV-A | SNV | 1 | A | G | 26 | 24343 | 0.10680689 | 46.6128345 | WT | FALSE | CAN | JSY13 | 1 | WT-Rep1 | AAG | A>G | G/C+1 | A/T-1 | TA>CG | TA>CG | GGCAAAAGT | AGTGATT | TTC | TTC | T | C | T>C | T>C_TTC | TA>CG_AGTGATT | 9 | FALSE | FALSE |
| 5_v.txt | 32941 | 32941-SNV-A | SNV | 1 | A | G | 6 | 24343 | 0.02464774 | 53.166837 | WT | FALSE | CAN | JSY13 | 1 | WT-Rep1 | GAA | A>G | G/C+1 | A/T-1 | TA>CG | TA>CG | CCCAAGAAA | AAAACCA | CTT | CTT | T | C | T>C | T>C_CTT | TA>CG_AAAACCA | 9 | FALSE | FALSE |
| 5_v.txt | 33052 | 33052-SNV-A | SNV | 1 | A | G | 25 | 24343 | 0.10269893 | 57.1960022 | WT | FALSE | CAN | JSY13 | 1 | WT-Rep1 | GAC | A>G | G/C+1 | A/T-1 | TA>CG | TA>CG | CCAAGGACT | ACTGCGT | CTG | CTG | T | C | T>C | T>C_CTG | TA>CG_ACTGCGT | 9 | FALSE | FALSE |
| 5_v.txt | 33065 | 33065-SNV-A | SNV | 1 | A | G | 5 | 24343 | 0.02053979 | 58.7997073 | WT | FALSE | CAN | JSY13 | 1 | WT-Rep1 | AAT | A>G | G/C+1 | A/T-1 | TA>CG | TA>CG | GACAGAATA | ATATGCC | TTA | TTA | T | C | T>C | T>C_TTA | TA>CG_ATATGCC | 9 | FALSE | FALSE |
| 5_v.txt | 33126 | 33126-SNV-A | SNV | 1 | A | G | 19 | 24343 | 0.07805119 | 38.2105263 | WT | FALSE | CAN | JSY13 | 1 | WT-Rep1 | CAG | A>G | G/C+1 | A/T-1 | TA>CG | TA>CG | TTGGTCAGA | AGAGGTG | GTC | GTC | T | C | T>C | T>C_GTC | TA>CG_AGAGGTG | 9 | FALSE | FALSE |
| 5_v.txt | 33150 | 33150-SNV-A | SNV | 1 | A | G | 17 | 24343 | 0.06983527 | 34.6470588 | WT | FALSE | CAN | JSY13 | 1 | WT-Rep1 | AAG | A>G | G/C+1 | A/T-1 | TA>CG | TA>CG | ATGAAAAGA | AGACCTG | TTC | TTC | T | C | T>C | T>C_TTC | TA>CG_AGACCTG | 9 | FALSE | FALSE |
| 5_v.txt | 33157 | 33157-SNV-T | SNV | 1 | T | C | 25 | 24342 | 0.10270315 | 35.8000862 | WT | FALSE | CAN | JSY13 | 1 | WT-Rep1 | GTA | T>C | G/C+1 | A/T-1 | TA>CG | TA>CG | GACCTGTAC | TACCAAT | CAT | GTA | T | G | T>C | T>C_GTA | TA>CG_TACCAAT | 9 | FALSE | FALSE |
| 5_v.txt | 33293 | 33293-SNV-T | SNV | 1 | T | C | 2 | 7277 | 0.02748385 | 29.4866419 | WT | FALSE | CAN | JSY13 | 1 | WT-Rep1 | CTC | T>C | G/C+1 | A/T-1 | TA>CG | TA>CG | ATTTACTCTC | TCTCGTC | GAG | CTC | T | G | T>C | T>C_CTC | TA>CG_TCTCGTC | 9 | FALSE | FALSE |
| 5_v.txt | 33476 | 33476-SNV-T | SNV | 1 | T | C | 17 | 7275 | 0.23367698 | 30.9411765 | WT | FALSE | CAN | JSY13 | 1 | WT-Rep1 | CTT | T>C | G/C+1 | A/T-1 | TA>CG | TA>CG | TATGCCTTTT | TTTTTTT | GAA | CTT | T | G | T>C | T>C_CTT | TA>CG_TTTTTTT | 9 | FALSE | FALSE |
| 5_v.txt | 31804 | 31804-SNV-T | SNV | 1 | T | A | 12 | 9497 | 0.12635569 | 51.577002 | WT | FALSE | CAN | JSY13 | 1 | WT-Rep1 | TTC | T>A | A/T+1 | A/T-1 | TA>AT | TA>AT | CAATCTTCCA | TCCAAAT | AAG | TTC | T | T | T>A | T>A_TTC | TA>AT_TCCAAAT | 9 | FALSE | FALSE |
| 5_v.txt | 31816 | 31816-SNV-T | SNV | 1 | T | A | 17 | 9499 | 0.17896621 | 52.2922597 | WT | FALSE | CAN | JSY13 | 1 | WT-Rep1 | CTG | T>A | A/T+1 | A/T-1 | TA>AT | TA>AT | TAAATCTGC | TGCATCT | GAC | CTG | T | T | T>A | T>A_CTG | TA>AT_TGCATCT | 9 | FALSE | FALSE |
| 5_v.txt | 31822 | 31822-SNV-T | SNV | 1 | T | A | 36 | 9499 | 0.37898726 | 53.7777778 | WT | FALSE | CAN | JSY13 | 1 | WT-Rep1 | CTG | T>A | A/T+1 | A/T-1 | TA>AT | TA>AT | TGCATCTGA | TGAATAT | GAC | CTG | T | T | T>A | T>A_CTG | TA>AT_TGAATAT | 9 | FALSE | FALSE |
| 5_v.txt | 31854 | 31854-SNV-A | SNV | 1 | A | T | 16 | 9499 | 0.16843878 | 53.4375 | WT | FALSE | CAN | JSY13 | 1 | WT-Rep1 | TAA | A>T | A/T+1 | A/T-1 | TA>AT | TA>AT | ACAGCTAAG | AAGAACA | ATT | ATT | T | A | T>A | T>A_ATT | TA>AT_AAGAACA | 9 | FALSE | FALSE |
| 5_v.txt | 31906 | 31906-SNV-T | SNV | 1 | T | A | 23 | 9484 | 0.24251371 | 39.2173913 | WT | FALSE | CAN | JSY13 | 1 | WT-Rep1 | TTT | T>A | A/T+1 | A/T-1 | TA>AT | TA>AT | TGAATTTTG | TTGGTGC | AAA | TTT | T | T | T>A | T>A_TTT | TA>AT_TTGGTGC | 9 | FALSE | FALSE |
| 5_v.txt | 31964 | 31964-SNV-A | SNV | 1 | A | T | 19 | 18279 | 0.10394442 | 35.0526316 | WT | FALSE | CAN | JSY13 | 1 | WT-Rep1 | CAT | A>T | A/T+1 | A/T-1 | TA>AT | TA>AT | GGCCGCATA | ATAATAA | GTA | GTA | T | A | T>A | T>A_GTA | TA>AT_ATAATAA | 9 | FALSE | FALSE |
| 5_v.txt | 31974 | 31974-SNV-A | SNV | 1 | A | T | 12 | 18279 | 0.06564911 | 41.4967915 | WT | FALSE | CAN | JSY13 | 1 | WT-Rep1 | CAA | A>T | A/T+1 | A/T-1 | TA>AT | TA>AT | TAAGCCAAG | AAGCCGG | GTT | GTT | T | A | T>A | T>A_GTT | TA>AT_AAGCCGG | 9 | FALSE | FALSE |
| 5_v.txt | 31986 | 31986-SNV-A | SNV | 1 | A | T | 10 | 18279 | 0.05470759 | 47.7 | WT | FALSE | CAN | JSY13 | 1 | WT-Rep1 | TAA | A>T | A/T+1 | A/T-1 | TA>AT | TA>AT | GGCATTAAT | AATTTAG | ATT | ATT | T | A | T>A | T>A_ATT | TA>AT_AATTTAG | 9 | FALSE | FALSE |
| 5_v.txt | 31990 | 31990-SNV-T | SNV | 1 | T | A | 25 | 18279 | 0.13676897 | 56.32 | WT | FALSE | CAN | JSY13 | 1 | WT-Rep1 | TTA | T>A | A/T+1 | A/T-1 | TA>AT | TA>AT | TTAATTTAG | TAGCTTT | AAT | TTA | T | T | T>A | T>A_TTA | TA>AT_TAGCTTT | 9 | FALSE | FALSE |
| 5_v.txt | 31996 | 31996-SNV-T | SNV | 1 | T | A | 69 | 18279 | 0.37748236 | 55.4927536 | WT | FALSE | CAN | JSY13 | 1 | WT-Rep1 | TTA | T>A | A/T+1 | A/T-1 | TA>AT | TA>AT | TAGCTTTAA | TAAATGG | AAT | TTA | T | T | T>A | T>A_TTA | TA>AT_TAAATGG | 9 | FALSE | FALSE |
| 5_v.txt | 32050 | 32050-SNV-T | SNV | 1 | T | A | 28 | 18279 | 0.15318125 | 62.2857143 | WT | FALSE | CAN | JSY13 | 1 | WT-Rep1 | CTG | T>A | A/T+1 | A/T-1 | TA>AT | TA>AT | TAAATCTGA | TGATGTG | GAC | CTG | T | T | T>A | T>A_CTG | TA>AT_TGATGTG | 9 | FALSE | FALSE |
| 5_v.txt | 32070 | 32070-SNV-A | SNV | 1 | A | T | 2 | 18278 | 0.01094212 | 32.4998101 | WT | FALSE | CAN | JSY13 | 1 | WT-Rep1 | AAA | A>T | A/T+1 | A/T-1 | TA>AT | TA>AT | GAGATAAAT | AATAACC | TTT | TTT | T | A | T>A | T>A_TTT | TA>AT_AATAACC | 9 | FALSE | FALSE |
| 5_v.txt | 32072 | 32072-SNV-T | SNV | 1 | T | A | 1 | 18279 | 5.47E-03 | 63.9990076 | WT | FALSE | CAN | JSY13 | 1 | WT-Rep1 | ATA | T>A | A/T+1 | A/T-1 | TA>AT | TA>AT | GATAAATAA | TAACCAT | TAT | ATA | T | T | T>A | T>A_ATA | TA>AT_TAACCAT | 9 | FALSE | FALSE |
| 5_v.txt | 32073 | 32073-SNV-A | SNV | 1 | A | T | 16 | 18279 | 0.08753214 | 58.9344662 | WT | FALSE | CAN | JSY13 | 1 | WT-Rep1 | TAA | A>T | A/T+1 | A/T-1 | TA>AT | TA>AT | ATAAATAAC | AACCATG | ATT | ATT | T | A | T>A | T>A_ATT | TA>AT_AACCATG | 9 | FALSE | FALSE |
| 5_v.txt | 32081 | 32081-SNV-A | SNV | 1 | A | T | 1 | 18279 | 5.47E-03 | 63.9993624 | WT | FALSE | CAN | JSY13 | 1 | WT-Rep1 | CAA | A>T | A/T+1 | A/T-1 | TA>AT | TA>AT | CCATGCAAA | AAAAAAG | GTT | GTT | T | A | T>A | T>A_GTT | TA>AT_AAAAAAG | 9 | FALSE | FALSE |
| 5_v.txt | 32294 | 32294-SNV-A | SNV | 1 | A | T | 5 | 21421 | 0.02334158 | 46.8 | WT | FALSE | CAN | JSY13 | 1 | WT-Rep1 | AAT | A>T | A/T+1 | A/T-1 | TA>AT | TA>AT | ATTTGAATTT | ATTTGCG | TTA | TTA | T | A | T>A | T>A_TTA | TA>AT_ATTTGCG | 9 | FALSE | FALSE |
| 5_v.txt | 32314 | 32314-SNV-T | SNV | 1 | T | A | 3 | 21421 | 0.01400495 | 49.6621564 | WT | FALSE | CAN | JSY13 | 1 | WT-Rep1 | GTT | T>A | A/T+1 | A/T-1 | TA>AT | TA>AT | TAATGGTTG | TTGTTAA | CAA | GTT | T | T | T>A | T>A_GTT | TA>AT_TTGTTAA | 9 | FALSE | FALSE |
| 5_v.txt | 32315 | 32315-SNV-T | SNV | 1 | T | A | 1 | 21421 | 4.67E-03 | 63.9998588 | WT | FALSE | CAN | JSY13 | 1 | WT-Rep1 | TTG | T>A | A/T+1 | A/T-1 | TA>AT | TA>AT | AATGGTTGT | TGTTAAG | AAC | TTG | T | T | T>A | T>A_TTG | TA>AT_TGTTAAG | 9 | FALSE | FALSE |
| 5_v.txt | 32317 | 32317-SNV-T | SNV | 1 | T | A | 4 | 21421 | 0.01867327 | 62.7041293 | WT | FALSE | CAN | JSY13 | 1 | WT-Rep1 | GTT | T>A | A/T+1 | A/T-1 | TA>AT | TA>AT | TGGTTGTTA | TTAAGAT | CAA | GTT | T | T | T>A | T>A_GTT | TA>AT_TTAAGAT | 9 | FALSE | FALSE |
| 5_v.txt | 32319 | 32319-SNV-A | SNV | 1 | A | T | 8 | 21421 | 0.03734653 | 56.8740145 | WT | FALSE | CAN | JSY13 | 1 | WT-Rep1 | TAA | A>T | A/T+1 | A/T-1 | TA>AT | TA>AT | GTTGTTAAG | AAGATAA | ATT | ATT | T | A | T>A | T>A_ATT | TA>AT_AAGATAA | 9 | FALSE | FALSE |
| 5_v.txt | 32320 | 32320-SNV-A | SNV | 1 | A | T | 2 | 21421 | 9.34E-03 | 32.9987096 | WT | FALSE | CAN | JSY13 | 1 | WT-Rep1 | AAG | A>T | A/T+1 | A/T-1 | TA>AT | TA>AT | TTGTTAAGA | AGATAAC | TTC | TTC | T | A | T>A | T>A_TTC | TA>AT_AGATAAC | 9 | FALSE | FALSE |
| 5_v.txt | 32338 | 32338-SNV-T | SNV | 1 | T | A | 3 | 21421 | 0.01400495 | 53.3315947 | WT | FALSE | CAN | JSY13 | 1 | WT-Rep1 | ATA | T>A | A/T+1 | A/T-1 | TA>AT | TA>AT | TGAAGATAT | TATGTGG | TAT | ATA | T | T | T>A | T>A_ATA | TA>AT_TATGTGG | 9 | FALSE | FALSE |
| 5_v.txt | 32414 | 32414-SNV-T | SNV | 1 | T | A | 6 | 21420 | 0.02801121 | 42.4979134 | WT | FALSE | CAN | JSY13 | 1 | WT-Rep1 | GTG | T>A | A/T+1 | A/T-1 | TA>AT | TA>AT | AGATTGTGT | TGTTAGT | CAC | GTG | T | T | T>A | T>A_GTG | TA>AT_TGTTAGT | 9 | FALSE | FALSE |
| 5_v.txt | 32445 | 32445-SNV-A | SNV | 1 | A | T | 13 | 21421 | 0.06068811 | 50 | WT | FALSE | CAN | JSY13 | 1 | WT-Rep1 | AAG | A>T | A/T+1 | A/T-1 | TA>AT | TA>AT | ACTAAAAGT | AGTCCAA | TTC | TTC | T | A | T>A | T>A_TTC | TA>AT_AGTCCAA | 9 | FALSE | FALSE |
| 5_v.txt | 32454 | 32454-SNV-A | SNV | 1 | A | T | 3 | 21421 | 0.01400495 | 53.3328283 | WT | FALSE | CAN | JSY13 | 1 | WT-Rep1 | GAA | A>T | A/T+1 | A/T-1 | TA>AT | TA>AT | CCAATGAAT | AATAATA | CTT | CTT | T | A | T>A | T>A_CTT | TA>AT_AATAATA | 9 | FALSE | FALSE |
| 5_v.txt | 32456 | 32456-SNV-T | SNV | 1 | T | A | 5 | 21421 | 0.02334158 | 37.7995404 | WT | FALSE | CAN | JSY13 | 1 | WT-Rep1 | ATA | T>A | A/T+1 | A/T-1 | TA>AT | TA>AT | AATGAATAA | TAATAGA | TAT | ATA | T | T | T>A | T>A_ATA | TA>AT_TAATAGA | 9 | FALSE | FALSE |
| 5_v.txt | 32459 | 32459-SNV-T | SNV | 1 | T | A | 3 | 21421 | 0.01400495 | 62.3324283 | WT | FALSE | CAN | JSY13 | 1 | WT-Rep1 | ATA | T>A | A/T+1 | A/T-1 | TA>AT | TA>AT | GAATAATAG | TAGAGAG | TAT | ATA | T | T | T>A | T>A_ATA | TA>AT_TAGAGAG | 9 | FALSE | FALSE |
| 5_v.txt | 32481 | 32481-SNV-A | SNV | 1 | A | T | 8 | 21421 | 0.03734653 | 41.5 | WT | FALSE | CAN | JSY13 | 1 | WT-Rep1 | TAA | A>T | A/T+1 | A/T-1 | TA>AT | TA>AT | AAGGTTAAG | AAGATAC | ATT | ATT | T | A | T>A | T>A_ATT | TA>AT_AAGATAC | 9 | FALSE | FALSE |
| 5_v.txt | 32498 | 32498-SNV-T | SNV | 1 | T | A | 5 | 21421 | 0.02334158 | 37.5996978 | WT | FALSE | CAN | JSY13 | 1 | WT-Rep1 | CTT | T>A | A/T+1 | A/T-1 | TA>AT | TA>AT | AACAACTTTT | TTTTTTG | GAA | CTT | T | T | T>A | T>A_CTT | TA>AT_TTTTTTG | 9 | FALSE | FALSE |
| 5_v.txt | 32527 | 32527-SNV-T | SNV | 1 | T | A | 1 | 21421 | 4.67E-03 | 37.9980322 | WT | FALSE | CAN | JSY13 | 1 | WT-Rep1 | CTG | T>A | A/T+1 | A/T-1 | TA>AT | TA>AT | ATTTTCTGG | TGGGGTT | GAC | CTG | T | T | T>A | T>A_CTG | TA>AT_TGGGGTT | 9 | FALSE | FALSE |
| 5_v.txt | 32541 | 32541-SNV-T | SNV | 1 | T | A | 4 | 21421 | 0.01867327 | 38 | WT | FALSE | CAN | JSY13 | 1 | WT-Rep1 | TTC | T>A | A/T+1 | A/T-1 | TA>AT | TA>AT | GCAGCTTCA | TCACCAG | AAG | TTC | T | T | T>A | T>A_TTC | TA>AT_TCACCAG | 9 | FALSE | FALSE |
| 5_v.txt | 32573 | 32573-SNV-T | SNV | 1 | T | A | 7 | 22153 | 0.03159843 | 37.8571429 | WT | FALSE | CAN | JSY13 | 1 | WT-Rep1 | CTT | T>A | A/T+1 | A/T-1 | TA>AT | TA>AT | AGTACCTTG | TTGAAAT | GAA | CTT | T | T | T>A | T>A_CTT | TA>AT_TTGAAAT | 9 | FALSE | FALSE |
| 5_v.txt | 32598 | 32598-SNV-A | SNV | 1 | A | T | 13 | 22156 | 0.05867485 | 52.3076923 | WT | FALSE | CAN | JSY13 | 1 | WT-Rep1 | CAA | A>T | A/T+1 | A/T-1 | TA>AT | TA>AT | TTAATCAAA | AAAGAGG | GTT | GTT | T | A | T>A | T>A_GTT | TA>AT_AAAGAGG | 9 | FALSE | FALSE |
| 5_v.txt | 32631 | 32631-SNV-T | SNV | 1 | T | A | 6 | 22156 | 0.0270807 | 43.1582607 | WT | FALSE | CAN | JSY13 | 1 | WT-Rep1 | GTT | T>A | A/T+1 | A/T-1 | TA>AT | TA>AT | CCTTCGTTTT | TTTTTAT | CAA | GTT | T | T | T>A | T>A_GTT | TA>AT_TTTTTAT | 9 | FALSE | FALSE |
| 5_v.txt | 32635 | 32635-SNV-T | SNV | 1 | T | A | 41 | 22156 | 0.18505145 | 57.0243902 | WT | FALSE | CAN | JSY13 | 1 | WT-Rep1 | TTA | T>A | A/T+1 | A/T-1 | TA>AT | TA>AT | CGTTTTATC | TATCCTT | AAT | TTA | T | T | T>A | T>A_TTA | TA>AT_TATCCTT | 9 | FALSE | FALSE |
| 5_v.txt | 32662 | 32662-SNV-A | SNV | 1 | A | T | 5 | 22156 | 0.02256725 | 31.1996736 | WT | FALSE | CAN | JSY13 | 1 | WT-Rep1 | CAG | A>T | A/T+1 | A/T-1 | TA>AT | TA>AT | GACCCCAGG | AGGCACC | GTC | GTC | T | A | T>A | T>A_GTC | TA>AT_AGGCACC | 9 | FALSE | FALSE |
| 5_v.txt | 32726 | 32726-SNV-A | SNV | 1 | A | T | 118 | 22177 | 0.53208279 | 56.7457627 | WT | FALSE | CAN | JSY13 | 1 | WT-Rep1 | TAC | A>T | A/T+1 | A/T-1 | TA>AT | TA>AT | AACCATACA | ACAAAAA | ATG | ATG | T | A | T>A | T>A_ATG | TA>AT_ACAAAAA | 9 | FALSE | FALSE |
| 5_v.txt | 32734 | 32734-SNV-A | SNV | 1 | A | T | 33 | 22177 | 0.14880281 | 55.0909091 | WT | FALSE | CAN | JSY13 | 1 | WT-Rep1 | CAG | A>T | A/T+1 | A/T-1 | TA>AT | TA>AT | AAAAACAGT | AGTATAT | GTC | GTC | T | A | T>A | T>A_GTC | TA>AT_AGTATAT | 9 | FALSE | FALSE |
| 5_v.txt | 32760 | 32760-SNV-A | SNV | 1 | A | T | 38 | 22176 | 0.17135642 | 52.4444134 | WT | FALSE | CAN | JSY13 | 1 | WT-Rep1 | TAA | A>T | A/T+1 | A/T-1 | TA>AT | TA>AT | ATGGCTAAA | AAAACTT | ATT | ATT | T | A | T>A | T>A_ATT | TA>AT_AAAACTT | 9 | FALSE | FALSE |
| 5_v.txt | 32804 | 32804-SNV-T | SNV | 1 | T | A | 223 | 22177 | 1.00554629 | 45.4616827 | WT | FALSE | CAN | JSY13 | 1 | WT-Rep1 | ATT | T>A | A/T+1 | A/T-1 | TA>AT | TA>AT | GTAATATTTG | TTTGACA | TAA | ATT | T | T | T>A | T>A_ATT | TA>AT_TTTGACA | 9 | FALSE | FALSE |
| 5_v.txt | 32805 | 32805-SNV-T | SNV | 1 | T | A | 1 | 22177 | 4.51E-03 | 37.99821 | WT | FALSE | CAN | JSY13 | 1 | WT-Rep1 | TTT | T>A | A/T+1 | A/T-1 | TA>AT | TA>AT | TAATATTTGA | TTGACAG | AAA | TTT | T | T | T>A | T>A_TTT | TA>AT_TTGACAG | 9 | FALSE | FALSE |
| 5_v.txt | 32875 | 32875-SNV-T | SNV | 1 | T | A | 23 | 24288 | 0.09469697 | 37.6064704 | WT | FALSE | CAN | JSY13 | 1 | WT-Rep1 | TTG | T>A | A/T+1 | A/T-1 | TA>AT | TA>AT | GAACTTTGT | TGTACGT | AAC | TTG | T | T | T>A | T>A_TTG | TA>AT_TGTACGT | 9 | FALSE | FALSE |
| 5_v.txt | 32929 | 32929-SNV-T | SNV | 1 | T | A | 7 | 24343 | 0.0287557 | 42.8536669 | WT | FALSE | CAN | JSY13 | 1 | WT-Rep1 | GTG | T>A | A/T+1 | A/T-1 | TA>AT | TA>AT | CAAAAGTGA | TGATTGC | CAC | GTG | T | T | T>A | T>A_GTG | TA>AT_TGATTGC | 9 | FALSE | FALSE |
| 5_v.txt | 32932 | 32932-SNV-T | SNV | 1 | T | A | 30 | 24343 | 0.12323871 | 51.4333333 | WT | FALSE | CAN | JSY13 | 1 | WT-Rep1 | ATT | T>A | A/T+1 | A/T-1 | TA>AT | TA>AT | AAGTGATTG | TTGCCCA | TAA | ATT | T | T | T>A | T>A_ATT | TA>AT_TTGCCCA | 9 | FALSE | FALSE |
| 5_v.txt | 32938 | 32938-SNV-A | SNV | 1 | A | T | 22 | 24343 | 0.09037506 | 53.7252921 | WT | FALSE | CAN | JSY13 | 1 | WT-Rep1 | CAA | A>T | A/T+1 | A/T-1 | TA>AT | TA>AT | TTGCCCAAG | AAGAAAA | GTT | GTT | T | A | T>A | T>A_GTT | TA>AT_AAGAAAA | 9 | FALSE | FALSE |
| 5_v.txt | 32948 | 32948-SNV-A | SNV | 1 | A | T | 50 | 24343 | 0.20539786 | 56.62 | WT | FALSE | CAN | JSY13 | 1 | WT-Rep1 | AAT | A>T | A/T+1 | A/T-1 | TA>AT | TA>AT | AAACCAATA | ATACATG | TTA | TTA | T | A | T>A | T>A_TTA | TA>AT_ATACATG | 9 | FALSE | FALSE |
| 5_v.txt | 32992 | 32992-SNV-T | SNV | 1 | T | A | 19 | 24343 | 0.07805119 | 57.3092996 | WT | FALSE | CAN | JSY13 | 1 | WT-Rep1 | CTT | T>A | A/T+1 | A/T-1 | TA>AT | TA>AT | GGAATCTTT | TTTGTGA | GAA | CTT | T | T | T>A | T>A_CTT | TA>AT_TTTGTGA | 9 | FALSE | FALSE |
| 5_v.txt | 33048 | 33048-SNV-A | SNV | 1 | A | T | 17 | 24343 | 0.06983527 | 60.3529412 | WT | FALSE | CAN | JSY13 | 1 | WT-Rep1 | CAA | A>T | A/T+1 | A/T-1 | TA>AT | TA>AT | TCACCCAAG | AAGGACT | GTT | GTT | T | A | T>A | T>A_GTT | TA>AT_AAGGACT | 9 | FALSE | FALSE |
| 5_v.txt | 33066 | 33066-SNV-T | SNV | 1 | T | A | 1 | 24343 | 4.11E-03 | 63.9974811 | WT | FALSE | CAN | JSY13 | 1 | WT-Rep1 | ATA | T>A | A/T+1 | A/T-1 | TA>AT | TA>AT | ACAGAATAT | TATGCCA | TAT | ATA | T | T | T>A | T>A_ATA | TA>AT_TATGCCA | 9 | FALSE | FALSE |
| 5_v.txt | 33082 | 33082-SNV-T | SNV | 1 | T | A | 6 | 24343 | 0.02464774 | 24.3323908 | WT | FALSE | CAN | JSY13 | 1 | WT-Rep1 | ATA | T>A | A/T+1 | A/T-1 | TA>AT | TA>AT | AACCCATAA | TAAATAA | TAT | ATA | T | T | T>A | T>A_ATA | TA>AT_TAAATAA | 9 | FALSE | FALSE |
| 5_v.txt | 33089 | 33089-SNV-A | SNV | 1 | A | T | 31 | 24343 | 0.12734667 | 42.9677419 | WT | FALSE | CAN | JSY13 | 1 | WT-Rep1 | AAT | A>T | A/T+1 | A/T-1 | TA>AT | TA>AT | AAATAAATA | ATATGAT | TTA | TTA | T | A | T>A | T>A_TTA | TA>AT_ATATGAT | 9 | FALSE | FALSE |
| 5_v.txt | 33150 | 33150-SNV-A | SNV | 1 | A | T | 17 | 24343 | 0.06983527 | 37.6470588 | WT | FALSE | CAN | JSY13 | 1 | WT-Rep1 | AAG | A>T | A/T+1 | A/T-1 | TA>AT | TA>AT | ATGAAAAGA | AGACCTG | TTC | TTC | T | A | T>A | T>A_TTC | TA>AT_AGACCTG | 9 | FALSE | FALSE |
| 5_v.txt | 33162 | 33162-SNV-A | SNV | 1 | A | T | 34 | 24311 | 0.13985439 | 36.4117647 | WT | FALSE | CAN | JSY13 | 1 | WT-Rep1 | AAT | A>T | A/T+1 | A/T-1 | TA>AT | TA>AT | GTACCAATA | ATAGTAC | TTA | TTA | T | A | T>A | T>A_TTA | TA>AT_ATAGTAC | 9 | FALSE | FALSE |
| 5_v.txt | 33192 | 33192-SNV-T | SNV | 1 | T | A | 8 | 7277 | 0.10993541 | 25.4998739 | WT | FALSE | CAN | JSY13 | 1 | WT-Rep1 | ATG | T>A | A/T+1 | A/T-1 | TA>AT | TA>AT | CCAATATGT | TGTCTTT | TAC | ATG | T | T | T>A | T>A_ATG | TA>AT_TGTCTTT | 9 | FALSE | FALSE |
| 5_v.txt | 33287 | 33287-SNV-A | SNV | 1 | A | T | 2 | 7277 | 0.02748385 | 63.9959814 | WT | FALSE | CAN | JSY13 | 1 | WT-Rep1 | CAT | A>T | A/T+1 | A/T-1 | TA>AT | TA>AT | CTCGCCATTT | ATTTACT | GTA | GTA | T | A | T>A | T>A_GTA | TA>AT_ATTTACT | 9 | FALSE | FALSE |
| 5_v.txt | 33293 | 33293-SNV-T | SNV | 1 | T | A | 3 | 7277 | 0.04122578 | 28.8307038 | WT | FALSE | CAN | JSY13 | 1 | WT-Rep1 | CTC | T>A | A/T+1 | A/T-1 | TA>AT | TA>AT | ATTTACTCTC | TCTCGTC | GAG | CTC | T | T | T>A | T>A_CTC | TA>AT_TCTCGTC | 9 | FALSE | FALSE |
| 5_v.txt | 33364 | 33364-SNV-T | SNV | 1 | T | A | 22 | 7277 | 0.30232239 | 51.5454546 | WT | FALSE | CAN | JSY13 | 1 | WT-Rep1 | CTG | T>A | A/T+1 | A/T-1 | TA>AT | TA>AT | CACGTCTGT | TGTGGTG | GAC | CTG | T | T | T>A | T>A_CTG | TA>AT_TGTGGTG | 9 | FALSE | FALSE |
| 5_v.txt | 31805 | 31805-SNV-C | SNV | 1 | C | T | 15 | 9499 | 0.15791136 | 58.2666667 | WT | FALSE | CAN | JSY13 | 1 | WT-Rep1 | TCC | C>T | A/T+1 | G/C-1 | CG>TA | CG>TA | AATCTTCCA | CCAAATA | AGG | TCC | C | A | C>T | C>T_TCC | CG>TA_CCAAATA | 9 | FALSE | FALSE |
| 5_v.txt | 31834 | 31834-SNV-G | SNV | 1 | G | A | 20 | 9499 | 0.21054848 | 49.7 | WT | FALSE | CAN | JSY13 | 1 | WT-Rep1 | TGA | G>A | A/T+1 | G/C-1 | CG>TA | CG>TA | TGCATTGAA | GAAATAA | ACT | ACT | C | T | C>T | C>T_ACT | CG>TA_GAAATAA | 9 | FALSE | FALSE |
| 5_v.txt | 31845 | 31845-SNV-C | SNV | 1 | C | T | 46 | 9499 | 0.4842615 | 52.2391304 | WT | FALSE | CAN | JSY13 | 1 | WT-Rep1 | CCA | C>T | A/T+1 | G/C-1 | CG>TA | CG>TA | AAGATCCAA | CAAACAG | GGT | CCA | C | A | C>T | C>T_CCA | CG>TA_CAAACAG | 9 | FALSE | FALSE |
| 5_v.txt | 31882 | 31882-SNV-C | SNV | 1 | C | T | 15 | 9499 | 0.15791136 | 41.8 | WT | FALSE | CAN | JSY13 | 1 | WT-Rep1 | GCA | C>T | A/T+1 | G/C-1 | CG>TA | CG>TA | AGGCGGCAG | CAGCAAA | CGT | GCA | C | A | C>T | C>T_GCA | CG>TA_CAGCAAA | 9 | FALSE | FALSE |
| 5_v.txt | 31885 | 31885-SNV-C | SNV | 1 | C | T | 5 | 9499 | 0.05263712 | 42.998418 | WT | FALSE | CAN | JSY13 | 1 | WT-Rep1 | GCA | C>T | A/T+1 | G/C-1 | CG>TA | CG>TA | CGGCAGCAA | CAAAGCT | CGT | GCA | C | A | C>T | C>T_GCA | CG>TA_CAAAGCT | 9 | FALSE | FALSE |
| 5_v.txt | 31930 | 31930-SNV-G | SNV | 1 | G | A | 11 | 9485 | 0.11597259 | 37.7272727 | WT | FALSE | CAN | JSY13 | 1 | WT-Rep1 | TGA | G>A | A/T+1 | G/C-1 | CG>TA | CG>TA | AACCTTGAA | GAATAAT | ACT | ACT | C | T | C>T | C>T_ACT | CG>TA_GAATAAT | 9 | FALSE | FALSE |
| 5_v.txt | 31962 | 31962-SNV-G | SNV | 1 | G | A | 24 | 18279 | 0.13129821 | 34.9583333 | WT | FALSE | CAN | JSY13 | 1 | WT-Rep1 | CGC | G>A | A/T+1 | G/C-1 | CG>TA | CG>TA | GTGGCCGCA | GCATAAT | GCG | GCG | C | T | C>T | C>T_GCG | CG>TA_GCATAAT | 9 | FALSE | FALSE |

| | | | | | | | | | | | | | | | | | | | | | | | | | | | | | | | | | | |
|---|---|---|---|---|---|---|---|---|---|---|---|---|---|---|---|---|---|---|---|---|---|---|---|---|---|---|---|---|---|---|---|---|---|---|
| 5_v.txt | 31972 | 31972-SNV- | SNV | 1 | C | T | 18 | 18279 | 0.09847366 | 36.9437097 | WT | FALSE | CAN | JSY13 | 1 | WT-Rep1 | GCC | C>T | A/T+1 | G/C-1 | CG>TA | CG>TA | AATAAGCCA | CCAAGCC | CGG | GCC | C | A | C>T | C>T_GCC | CG>TA_CCAAGCC | 9 | FALSE | FALSE |
| 5_v.txt | 31980 | 31980-SNV- | SNV | 1 | G | A | 64 | 18279 | 0.35012856 | 46.3640664 | WT | FALSE | CAN | JSY13 | 1 | WT-Rep1 | GGG | G>A | A/T+1 | G/C-1 | CG>TA | CG>TA | AAGCCGGGC | GGCATTA | CCC | CCC | C | T | C>T | C>T_CCC | CG>TA_GGCATTA | 9 | FALSE | FALSE |
| 5_v.txt | 31981 | 31981-SNV- | SNV | 1 | G | A | 24 | 18279 | 0.13129821 | 46.5313038 | WT | FALSE | CAN | JSY13 | 1 | WT-Rep1 | GGC | G>A | A/T+1 | G/C-1 | CG>TA | CG>TA | AGCCGGGCA | GCATTAA | CCG | CCG | C | T | C>T | C>T_CCG | CG>TA_GCATTAA | 9 | FALSE | FALSE |
| 5_v.txt | 32041 | 32041-SNV- | SNV | 1 | G | A | 51 | 18279 | 0.2790087 | 59.9215686 | WT | FALSE | CAN | JSY13 | 1 | WT-Rep1 | TGC | G>A | A/T+1 | G/C-1 | CG>TA | CG>TA | AAGCTTGCA | GCATAAA | ACG | ACG | C | T | C>T | C>T_ACG | CG>TA_GCATAAA | 9 | FALSE | FALSE |
| 5_v.txt | 32075 | 32075-SNV- | SNV | 1 | C | T | 27 | 18279 | 0.14771049 | 61.2951451 | WT | FALSE | CAN | JSY13 | 1 | WT-Rep1 | ACC | C>T | A/T+1 | G/C-1 | CG>TA | CG>TA | AAATAACCA | CCATGCA | TGG | ACC | C | A | C>T | C>T_ACC | CG>TA_CCATGCA | 9 | FALSE | FALSE |
| 5_v.txt | 32076 | 32076-SNV- | SNV | 1 | C | T | 48 | 18278 | 0.26261079 | 60.7905667 | WT | FALSE | CAN | JSY13 | 1 | WT-Rep1 | CCA | C>T | A/T+1 | G/C-1 | CG>TA | CG>TA | AATAACCAT | CATGCAA | GGT | CCA | C | A | C>T | C>T_CCA | CG>TA_CATGCAA | 9 | FALSE | FALSE |
| 5_v.txt | 32080 | 32080-SNV- | SNV | 1 | C | T | 8 | 18279 | 0.04376607 | 57.25 | WT | FALSE | CAN | JSY13 | 1 | WT-Rep1 | GCA | C>T | A/T+1 | G/C-1 | CG>TA | CG>TA | ACCATGCAA | CAAAAAA | CGT | GCA | C | A | C>T | C>T_GCA | CG>TA_CAAAAAA | 9 | FALSE | FALSE |
| 5_v.txt | 32088 | 32088-SNV- | SNV | 1 | C | T | 46 | 18278 | 0.25166867 | 62.7391304 | WT | FALSE | CAN | JSY13 | 1 | WT-Rep1 | GCC | C>T | A/T+1 | G/C-1 | CG>TA | CG>TA | AAAAAGCCT | CCTGCAA | CGG | GCC | C | A | C>T | C>T_GCC | CG>TA_CCTGCAA | 9 | FALSE | FALSE |
| 5_v.txt | 32092 | 32092-SNV- | SNV | 1 | C | T | 21 | 18279 | 0.11488594 | 62.952381 | WT | FALSE | CAN | JSY13 | 1 | WT-Rep1 | GCA | C>T | A/T+1 | G/C-1 | CG>TA | CG>TA | AGCCTGCAA | CAACACC | CGT | GCA | C | A | C>T | C>T_GCA | CG>TA_CAACACC | 9 | FALSE | FALSE |
| 5_v.txt | 32165 | 32165-SNV- | SNV | 1 | G | A | 23 | 18279 | 0.12582745 | 51.9130435 | WT | FALSE | CAN | JSY13 | 1 | WT-Rep1 | CGC | G>A | A/T+1 | G/C-1 | CG>TA | CG>TA | CAAAGCGCC | GCCAAAT | GCG | GCG | C | T | C>T | C>T_GCG | CG>TA_GCCAAAT | 9 | FALSE | FALSE |
| 5_v.txt | 32271 | 32271-SNV- | SNV | 1 | C | T | 59 | 21420 | 0.27544351 | 32.0847458 | WT | FALSE | CAN | JSY13 | 1 | WT-Rep1 | ACG | C>T | A/T+1 | G/C-1 | CG>TA | CG>TA | AAAATACGG | CGGGAAC | TGC | ACG | C | A | C>T | C>T_ACG | CG>TA_CGGGAAC | 9 | FALSE | FALSE |
| 5_v.txt | 32272 | 32272-SNV- | SNV | 1 | G | A | 45 | 21420 | 0.21008403 | 37.2444444 | WT | FALSE | CAN | JSY13 | 1 | WT-Rep1 | CGG | G>A | A/T+1 | G/C-1 | CG>TA | CG>TA | AAATACGGG | GGGAACC | GCC | GCC | C | T | C>T | C>T_GCC | CG>TA_GGGAACC | 9 | FALSE | FALSE |
| 5_v.txt | 32274 | 32274-SNV- | SNV | 1 | G | A | 13 | 21420 | 0.06069094 | 37.3846121 | WT | FALSE | CAN | JSY13 | 1 | WT-Rep1 | GGA | G>A | A/T+1 | G/C-1 | CG>TA | CG>TA | ATACGGGAA | GAACCAA | CCT | CCT | C | T | C>T | C>T_CCT | CG>TA_GAACCAA | 9 | FALSE | FALSE |
| 5_v.txt | 32292 | 32292-SNV- | SNV | 1 | G | A | 34 | 21421 | 0.15872275 | 51.3529412 | WT | FALSE | CAN | JSY13 | 1 | WT-Rep1 | TGA | G>A | A/T+1 | G/C-1 | CG>TA | CG>TA | ATATTTGAA | GAATTTG | ACT | ACT | C | T | C>T | C>T_ACT | CG>TA_GAATTTG | 9 | FALSE | FALSE |
| 5_v.txt | 32299 | 32299-SNV- | SNV | 1 | C | T | 42 | 21421 | 0.19606928 | 51.9285714 | WT | FALSE | CAN | JSY13 | 1 | WT-Rep1 | GCG | C>T | A/T+1 | G/C-1 | CG>TA | CG>TA | AATTTGCGG | CGGCAGA | CGC | GCG | C | A | C>T | C>T_GCG | CG>TA_CGGCAGA | 9 | FALSE | FALSE |
| 5_v.txt | 32344 | 32344-SNV- | SNV | 1 | G | A | 27 | 21421 | 0.12604454 | 60.3333333 | WT | FALSE | CAN | JSY13 | 1 | WT-Rep1 | GGC | G>A | A/T+1 | G/C-1 | CG>TA | CG>TA | TATGTGGCA | GCAAAAC | CCG | CCG | C | T | C>T | C>T_CCG | CG>TA_GCAAAAC | 9 | FALSE | FALSE |
| 5_v.txt | 32413 | 32413-SNV- | SNV | 1 | G | A | 17 | 21421 | 0.07936137 | 57.2347025 | WT | FALSE | CAN | JSY13 | 1 | WT-Rep1 | TGT | G>A | A/T+1 | G/C-1 | CG>TA | CG>TA | TAGATTGTG | GTGTTAG | ACA | ACA | C | T | C>T | C>T_ACA | CG>TA_GTGTTAG | 9 | FALSE | FALSE |
| 5_v.txt | 32449 | 32449-SNV- | SNV | 1 | C | T | 172 | 21421 | 0.80295038 | 52.25 | WT | FALSE | CAN | JSY13 | 1 | WT-Rep1 | CCA | C>T | A/T+1 | G/C-1 | CG>TA | CG>TA | AAAGTCCAA | CAATGAA | GGT | CCA | C | A | C>T | C>T_CCA | CG>TA_CAATGAA | 9 | FALSE | FALSE |
| 5_v.txt | 32463 | 32463-SNV- | SNV | 1 | G | A | 38 | 21421 | 0.17739601 | 50.4210526 | WT | FALSE | CAN | JSY13 | 1 | WT-Rep1 | AGA | G>A | A/T+1 | G/C-1 | CG>TA | CG>TA | AATAGAGAG | GAGCCAA | TCT | TCT | C | T | C>T | C>T_TCT | CG>TA_GAGCCAA | 9 | FALSE | FALSE |
| 5_v.txt | 32487 | 32487-SNV- | SNV | 1 | C | T | 54 | 21421 | 0.25208907 | 44.8518519 | WT | FALSE | CAN | JSY13 | 1 | WT-Rep1 | ACG | C>T | A/T+1 | G/C-1 | CG>TA | CG>TA | AAGATACGG | CGGAAAA | TGC | ACG | C | A | C>T | C>T_ACG | CG>TA_CGGAAAA | 9 | FALSE | FALSE |
| 5_v.txt | 32488 | 32488-SNV- | SNV | 1 | G | A | 103 | 21421 | 0.48083656 | 50.0580131 | WT | FALSE | CAN | JSY13 | 1 | WT-Rep1 | CGG | G>A | A/T+1 | G/C-1 | CG>TA | CG>TA | AGATACGGA | GGAAAAC | GCC | GCC | C | T | C>T | C>T_GCC | CG>TA_GGAAAAC | 9 | FALSE | FALSE |
| 5_v.txt | 32508 | 32508-SNV- | SNV | 1 | G | A | 20 | 21421 | 0.09336632 | 38.75 | WT | FALSE | CAN | JSY13 | 1 | WT-Rep1 | GGC | G>A | A/T+1 | G/C-1 | CG>TA | CG>TA | TTGATGGCT | GCTCTTG | CCG | CCG | C | T | C>T | C>T_CCG | CG>TA_GCTCTTG | 9 | FALSE | FALSE |
| 5_v.txt | 32528 | 32528-SNV- | SNV | 1 | G | A | 2 | 21421 | 9.34E-03 | 32.4998694 | WT | FALSE | CAN | JSY13 | 1 | WT-Rep1 | TGG | G>A | A/T+1 | G/C-1 | CG>TA | CG>TA | TTTTCTGGG | GGGGTTT | ACC | ACC | C | T | C>T | C>T_ACC | CG>TA_GGGGTTT | 9 | FALSE | FALSE |
| 5_v.txt | 32529 | 32529-SNV- | SNV | 1 | G | A | 12 | 21421 | 0.05601979 | 37.9166667 | WT | FALSE | CAN | JSY13 | 1 | WT-Rep1 | GGG | G>A | A/T+1 | G/C-1 | CG>TA | CG>TA | TTTCTGGGG | GGGTTTG | CCC | CCC | C | T | C>T | C>T_CCC | CG>TA_GGGTTTG | 9 | FALSE | FALSE |
| 5_v.txt | 32530 | 32530-SNV- | SNV | 1 | G | A | 54 | 21421 | 0.25208907 | 38.2592593 | WT | FALSE | CAN | JSY13 | 1 | WT-Rep1 | GGG | G>A | A/T+1 | G/C-1 | CG>TA | CG>TA | TTCTGGGGT | GGTTTGC | CCC | CCC | C | T | C>T | C>T_CCC | CG>TA_GGTTTGC | 9 | FALSE | FALSE |
| 5_v.txt | 32535 | 32535-SNV- | SNV | 1 | G | A | 33 | 21421 | 0.15405443 | 36.0000969 | WT | FALSE | CAN | JSY13 | 1 | WT-Rep1 | TGC | G>A | A/T+1 | G/C-1 | CG>TA | CG>TA | GGGTTTGCA | GCAGCTT | ACG | ACG | C | T | C>T | C>T_ACG | CG>TA_GCAGCTT | 9 | FALSE | FALSE |
| 5_v.txt | 32536 | 32536-SNV- | SNV | 1 | C | T | 32 | 21421 | 0.14938612 | 36.5174184 | WT | FALSE | CAN | JSY13 | 1 | WT-Rep1 | GCA | C>T | A/T+1 | G/C-1 | CG>TA | CG>TA | GGTTTGCAG | CAGCTTC | CGT | GCA | C | A | C>T | C>T_GCA | CG>TA_CAGCTTC | 9 | FALSE | FALSE |
| 5_v.txt | 32545 | 32545-SNV- | SNV | 1 | C | T | 63 | 21421 | 0.29410392 | 37.3333333 | WT | FALSE | CAN | JSY13 | 1 | WT-Rep1 | CCA | C>T | A/T+1 | G/C-1 | CG>TA | CG>TA | CTTCACCAG | CAGCAGT | GGT | CCA | C | A | C>T | C>T_CCA | CG>TA_CAGCAGT | 9 | FALSE | FALSE |
| 5_v.txt | 32566 | 32566-SNV- | SNV | 1 | C | T | 21 | 22145 | 0.09482953 | 35.4285714 | WT | FALSE | CAN | JSY13 | 1 | WT-Rep1 | TCA | C>T | A/T+1 | G/C-1 | CG>TA | CG>TA | CTAGTTCAG | CAGTACC | AGT | TCA | C | A | C>T | C>T_TCA | CG>TA_CAGTACC | 9 | FALSE | FALSE |
| 5_v.txt | 32572 | 32572-SNV- | SNV | 1 | C | T | 20 | 22153 | 0.09028123 | 30.9001398 | WT | FALSE | CAN | JSY13 | 1 | WT-Rep1 | CCT | C>T | A/T+1 | G/C-1 | CG>TA | CG>TA | CAGTACCTT | CTTGAAA | GGA | CCT | C | A | C>T | C>T_CCT | CG>TA_CTTGAAA | 9 | FALSE | FALSE |
| 5_v.txt | 32575 | 32575-SNV- | SNV | 1 | G | A | 36 | 22153 | 0.16250621 | 36.3333333 | WT | FALSE | CAN | JSY13 | 1 | WT-Rep1 | TGA | G>A | A/T+1 | G/C-1 | CG>TA | CG>TA | TACCTTGAA | GAAATGT | ACT | ACT | C | T | C>T | C>T_ACT | CG>TA_GAAATGT | 9 | FALSE | FALSE |
| 5_v.txt | 32590 | 32590-SNV- | SNV | 1 | C | T | 39 | 22156 | 0.17602455 | 35.7435897 | WT | FALSE | CAN | JSY13 | 1 | WT-Rep1 | GCG | C>T | A/T+1 | G/C-1 | CG>TA | CG>TA | AGGCAGCGT | CGTTAAT | CGC | GCG | C | A | C>T | C>T_GCG | CG>TA_CGTTAAT | 9 | FALSE | FALSE |
| 5_v.txt | 32629 | 32629-SNV- | SNV | 1 | C | T | 37 | 22156 | 0.16699765 | 54.6466807 | WT | FALSE | CAN | JSY13 | 1 | WT-Rep1 | TCG | C>T | A/T+1 | G/C-1 | CG>TA | CG>TA | TCCCTTCGT | CGTTTTT | AGC | TCG | C | A | C>T | C>T_TCG | CG>TA_CGTTTTT | 9 | FALSE | FALSE |
| 5_v.txt | 32658 | 32658-SNV- | SNV | 1 | C | T | 34 | 22156 | 0.1534573 | 56.3654373 | WT | FALSE | CAN | JSY13 | 1 | WT-Rep1 | ACC | C>T | A/T+1 | G/C-1 | CG>TA | CG>TA | CCTGGACCC | CCCCAGG | TGG | ACC | C | A | C>T | C>T_ACC | CG>TA_CCCCAGG | 9 | FALSE | FALSE |
| 5_v.txt | 32661 | 32661-SNV- | SNV | 1 | C | T | 33 | 22156 | 0.14894385 | 58.3588636 | WT | FALSE | CAN | JSY13 | 1 | WT-Rep1 | CCA | C>T | A/T+1 | G/C-1 | CG>TA | CG>TA | GGACCCCAG | CAGGCAC | GGT | CCA | C | A | C>T | C>T_CCA | CG>TA_CAGGCAC | 9 | FALSE | FALSE |
| 5_v.txt | 32679 | 32679-SNV- | SNV | 1 | C | T | 39 | 22155 | 0.1760325 | 57.5087737 | WT | FALSE | CAN | JSY13 | 1 | WT-Rep1 | CCA | C>T | A/T+1 | G/C-1 | CG>TA | CG>TA | TTTCTCCAA | CAATAAC | GGT | CCA | C | A | C>T | C>T_CCA | CG>TA_CAATAAC | 9 | FALSE | FALSE |
| 5_v.txt | 32700 | 32700-SNV- | SNV | 1 | C | T | 30 | 22156 | 0.1354035 | 57.7 | WT | FALSE | CAN | JSY13 | 1 | WT-Rep1 | GCC | C>T | A/T+1 | G/C-1 | CG>TA | CG>TA | ACTGGGCCG | CCGGTAA | CGG | GCC | C | A | C>T | C>T_GCC | CG>TA_CCGGTAA | 9 | FALSE | FALSE |
| 5_v.txt | 32749 | 32749-SNV- | SNV | 1 | C | T | 31 | 22177 | 0.13978446 | 54.7088886 | WT | FALSE | CAN | JSY13 | 1 | WT-Rep1 | CCG | C>T | A/T+1 | G/C-1 | CG>TA | CG>TA | GAAACCCGA | CGATAAT | GGC | CCG | C | A | C>T | C>T_CCG | CG>TA_CGATAAT | 9 | FALSE | FALSE |
| 5_v.txt | 32757 | 32757-SNV- | SNV | 1 | G | A | 16 | 22177 | 0.07214682 | 50.3114928 | WT | FALSE | CAN | JSY13 | 1 | WT-Rep1 | GGC | G>A | A/T+1 | G/C-1 | CG>TA | CG>TA | ATAATGGCT | GCTAAAA | CCG | CCG | C | T | C>T | C>T_CCG | CG>TA_GCTAAAA | 9 | FALSE | FALSE |
| 5_v.txt | 32776 | 32776-SNV- | SNV | 1 | C | T | 25 | 22177 | 0.1127294 | 48.08 | WT | FALSE | CAN | JSY13 | 1 | WT-Rep1 | GCG | C>T | A/T+1 | G/C-1 | CG>TA | CG>TA | TGGAAGCGA | CGACCCA | CGC | GCG | C | A | C>T | C>T_GCG | CG>TA_CGACCCA | 9 | FALSE | FALSE |
| 5_v.txt | 32780 | 32780-SNV- | SNV | 1 | C | T | 28 | 22177 | 0.12625693 | 43.3571429 | WT | FALSE | CAN | JSY13 | 1 | WT-Rep1 | CCC | C>T | A/T+1 | G/C-1 | CG>TA | CG>TA | AGCGACCCA | CCAGAAC | GGG | CCC | C | A | C>T | C>T_CCC | CG>TA_CCAGAAC | 9 | FALSE | FALSE |
| 5_v.txt | 32788 | 32788-SNV- | SNV | 1 | C | T | 55 | 22176 | 0.24801587 | 45.6373976 | WT | FALSE | CAN | JSY13 | 1 | WT-Rep1 | TCG | C>T | A/T+1 | G/C-1 | CG>TA | CG>TA | AGAACTCGA | CGAATTC | AGC | TCG | C | A | C>T | C>T_TCG | CG>TA_CGAATTC | 9 | FALSE | FALSE |
| 5_v.txt | 32797 | 32797-SNV- | SNV | 1 | C | T | 111 | 22177 | 0.50051856 | 47.2522523 | WT | FALSE | CAN | JSY13 | 1 | WT-Rep1 | CCG | C>T | A/T+1 | G/C-1 | CG>TA | CG>TA | ATTCACCGT | CGTAATA | GGC | CCG | C | A | C>T | C>T_CCG | CG>TA_CGTAATA | 9 | FALSE | FALSE |
| 5_v.txt | 32798 | 32798-SNV- | SNV | 1 | G | A | 26 | 22177 | 0.11723858 | 41.7307692 | WT | FALSE | CAN | JSY13 | 1 | WT-Rep1 | CGT | G>A | A/T+1 | G/C-1 | CG>TA | CG>TA | TTCACCGTA | GTAATAT | GCA | GCA | C | T | C>T | C>T_GCA | CG>TA_GTAATAT | 9 | FALSE | FALSE |
| 5_v.txt | 32811 | 32811-SNV- | SNV | 1 | G | A | 60 | 22177 | 0.27055057 | 42.5166667 | WT | FALSE | CAN | JSY13 | 1 | WT-Rep1 | AGG | G>A | A/T+1 | G/C-1 | CG>TA | CG>TA | TTGACAGGG | GGGAACA | TCC | TCC | C | T | C>T | C>T_TCC | CG>TA_GGGAACA | 9 | FALSE | FALSE |
| 5_v.txt | 32840 | 32840-SNV- | SNV | 1 | C | T | 33 | 22177 | 0.14880281 | 41.666499 | WT | FALSE | CAN | JSY13 | 1 | WT-Rep1 | CCC | C>T | A/T+1 | G/C-1 | CG>TA | CG>TA | AATTACCCA | CCAAAAA | GGG | CCC | C | A | C>T | C>T_CCC | CG>TA_CCAAAAA | 9 | FALSE | FALSE |
| 5_v.txt | 32855 | 32855-SNV- | SNV | 1 | C | T | 46 | 22175 | 0.20744081 | 35.1089228 | WT | FALSE | CAN | JSY13 | 1 | WT-Rep1 | TCC | C>T | A/T+1 | G/C-1 | CG>TA | CG>TA | ACTAATCCA | CCATGCC | AGG | TCC | C | A | C>T | C>T_TCC | CG>TA_CCATGCC | 9 | FALSE | FALSE |
| 5_v.txt | 32856 | 32856-SNV- | SNV | 1 | C | T | 85 | 22175 | 0.38331454 | 37.0471512 | WT | FALSE | CAN | JSY13 | 1 | WT-Rep1 | CCA | C>T | A/T+1 | G/C-1 | CG>TA | CG>TA | CTAATCCAT | CATGCCG | GGT | CCA | C | A | C>T | C>T_CCA | CG>TA_CATGCCG | 9 | FALSE | FALSE |
| 5_v.txt | 32860 | 32860-SNV- | SNV | 1 | C | T | 24 | 22175 | 0.10822999 | 35.4166667 | WT | FALSE | CAN | JSY13 | 1 | WT-Rep1 | GCC | C>T | A/T+1 | G/C-1 | CG>TA | CG>TA | TCCATGCCG | CCGCCAG | CGG | GCC | C | A | C>T | C>T_GCC | CG>TA_CCGCCAG | 9 | FALSE | FALSE |
| 5_v.txt | 32882 | 32882-SNV- | SNV | 1 | C | T | 60 | 24305 | 0.24686279 | 27 | WT | FALSE | CAN | JSY13 | 1 | WT-Rep1 | TCC | C>T | A/T+1 | G/C-1 | CG>TA | CG>TA | GTACGTCCA | CCAAAAT | AGG | TCC | C | A | C>T | C>T_TCC | CG>TA_CCAAAAT | 9 | FALSE | FALSE |
| 5_v.txt | 32899 | 32899-SNV- | SNV | 1 | G | A | 53 | 24305 | 0.21806213 | 33.7033039 | WT | FALSE | CAN | JSY13 | 1 | WT-Rep1 | TGG | G>A | A/T+1 | G/C-1 | CG>TA | CG>TA | TGACTTGGC | GGCCAAC | ACC | ACC | C | T | C>T | C>T_ACC | CG>TA_GGCCAAC | 9 | FALSE | FALSE |
| 5_v.txt | 32900 | 32900-SNV- | SNV | 1 | G | A | 34 | 24305 | 0.13988891 | 36.3189414 | WT | FALSE | CAN | JSY13 | 1 | WT-Rep1 | GGC | G>A | A/T+1 | G/C-1 | CG>TA | CG>TA | GACTTGGCC | GCCAACT | CCG | CCG | C | T | C>T | C>T_CCG | CG>TA_GCCAACT | 9 | FALSE | FALSE |
| 5_v.txt | 32901 | 32901-SNV- | SNV | 1 | C | T | 62 | 24305 | 0.25509155 | 29.593554 | WT | FALSE | CAN | JSY13 | 1 | WT-Rep1 | GCC | C>T | A/T+1 | G/C-1 | CG>TA | CG>TA | ACTTGGCCA | CCAACTA | CGG | GCC | C | A | C>T | C>T_GCC | CG>TA_CCAACTA | 9 | FALSE | FALSE |
| 5_v.txt | 32917 | 32917-SNV- | SNV | 1 | C | T | 31 | 24305 | 0.12754577 | 41.483871 | WT | FALSE | CAN | JSY13 | 1 | WT-Rep1 | TCC | C>T | A/T+1 | G/C-1 | CG>TA | CG>TA | TAAGTTCCA | CCAGGGC | AGG | TCC | C | A | C>T | C>T_TCC | CG>TA_CCAGGGC | 9 | FALSE | FALSE |
| 5_v.txt | 32935 | 32935-SNV- | SNV | 1 | C | T | 36 | 24343 | 0.14788646 | 33.0285344 | WT | FALSE | CAN | JSY13 | 1 | WT-Rep1 | GCC | C>T | A/T+1 | G/C-1 | CG>TA | CG>TA | TGATTGCCC | CCCAAGA | CGG | GCC | C | A | C>T | C>T_GCC | CG>TA_CCCAAGA | 9 | FALSE | FALSE |
| 5_v.txt | 32937 | 32937-SNV- | SNV | 1 | C | T | 33 | 24343 | 0.13556259 | 25.2121219 | WT | FALSE | CAN | JSY13 | 1 | WT-Rep1 | CCA | C>T | A/T+1 | G/C-1 | CG>TA | CG>TA | ATTGCCCAA | CAAGAAA | GGT | CCA | C | A | C>T | C>T_CCA | CG>TA_CAAGAAA | 9 | FALSE | FALSE |
| 5_v.txt | 32940 | 32940-SNV- | SNV | 1 | G | A | 83 | 24343 | 0.34096044 | 52.7831325 | WT | FALSE | CAN | JSY13 | 1 | WT-Rep1 | AGA | G>A | A/T+1 | G/C-1 | CG>TA | CG>TA | GCCCAAGAA | GAAAACC | TCT | TCT | C | T | C>T | C>T_TCT | CG>TA_GAAAACC | 9 | FALSE | FALSE |
| 5_v.txt | 32945 | 32945-SNV- | SNV | 1 | C | T | 113 | 24343 | 0.46419915 | 57.9675013 | WT | FALSE | CAN | JSY13 | 1 | WT-Rep1 | ACC | C>T | A/T+1 | G/C-1 | CG>TA | CG>TA | AGAAAACCA | CCAATAC | TGG | ACC | C | A | C>T | C>T_ACC | CG>TA_CCAATAC | 9 | FALSE | FALSE |
| 5_v.txt | 32946 | 32946-SNV- | SNV | 1 | C | T | 49 | 24343 | 0.2012899 | 55.9999606 | WT | FALSE | CAN | JSY13 | 1 | WT-Rep1 | CCA | C>T | A/T+1 | G/C-1 | CG>TA | CG>TA | GAAAACCAA | CAATACA | GGT | CCA | C | A | C>T | C>T_CCA | CG>TA_CAATACA | 9 | FALSE | FALSE |
| 5_v.txt | 32958 | 32958-SNV- | SNV | 1 | C | T | 108 | 24343 | 0.44365937 | 53.3240741 | WT | FALSE | CAN | JSY13 | 1 | WT-Rep1 | ACC | C>T | A/T+1 | G/C-1 | CG>TA | CG>TA | ATGTAACCA | CCATTGG | TGG | ACC | C | A | C>T | C>T_ACC | CG>TA_CCATTGG | 9 | FALSE | FALSE |
| 5_v.txt | 32965 | 32965-SNV- | SNV | 1 | C | T | 29 | 24343 | 0.11913076 | 58.0689655 | WT | FALSE | CAN | JSY13 | 1 | WT-Rep1 | GCC | C>T | A/T+1 | G/C-1 | CG>TA | CG>TA | CATTGGCCG | CCGCACC | CGG | GCC | C | A | C>T | C>T_GCC | CG>TA_CCGCACC | 9 | FALSE | FALSE |
| 5_v.txt | 32967 | 32967-SNV- | SNV | 1 | G | A | 64 | 24343 | 0.26290926 | 58.3125 | WT | FALSE | CAN | JSY13 | 1 | WT-Rep1 | CGC | G>A | A/T+1 | G/C-1 | CG>TA | CG>TA | TTGGCCGCA | GCACCAA | GCG | GCG | C | T | C>T | C>T_GCG | CG>TA_GCACCAA | 9 | FALSE | FALSE |
| 5_v.txt | 32982 | 32982-SNV- | SNV | 1 | G | A | 178 | 24343 | 0.73121637 | 61.1460674 | WT | FALSE | CAN | JSY13 | 1 | WT-Rep1 | AGA | G>A | A/T+1 | G/C-1 | CG>TA | CG>TA | GCTGGAGAA | GAAAGGA | TCT | TCT | C | T | C>T | C>T_TCT | CG>TA_GAAAGGA | 9 | FALSE | FALSE |
| 5_v.txt | 32995 | 32995-SNV- | SNV | 1 | G | A | 59 | 24343 | 0.24236947 | 61.4576271 | WT | FALSE | CAN | JSY13 | 1 | WT-Rep1 | TGT | G>A | A/T+1 | G/C-1 | CG>TA | CG>TA | ATCTTTGTG | GTGAGAA | ACA | ACA | C | T | C>T | C>T_ACA | CG>TA_GTGAGAA | 9 | FALSE | FALSE |
| 5_v.txt | 33012 | 33012-SNV- | SNV | 1 | G | A | 27 | 24343 | 0.11091484 | 61 | WT | FALSE | CAN | JSY13 | 1 | WT-Rep1 | AGA | G>A | A/T+1 | G/C-1 | CG>TA | CG>TA | GTGAAAGAG | GAGGATG | TCT | TCT | C | T | C>T | C>T_TCT | CG>TA_GAGGATG | 9 | FALSE | FALSE |
| 5_v.txt | 33015 | 33015-SNV- | SNV | 1 | G | A | 39 | 24343 | 0.16021033 | 53.6153846 | WT | FALSE | CAN | JSY13 | 1 | WT-Rep1 | GGA | G>A | A/T+1 | G/C-1 | CG>TA | CG>TA | AAAGAGGAT | GATGTAA | CCT | CCT | C | T | C>T | C>T_CCT | CG>TA_GATGTAA | 9 | FALSE | FALSE |
| 5_v.txt | 33024 | 33024-SNV- | SNV | 1 | G | A | 14 | 24343 | 0.0575114 | 61.8571429 | WT | FALSE | CAN | JSY13 | 1 | WT-Rep1 | AGG | G>A | A/T+1 | G/C-1 | CG>TA | CG>TA | GTAACAGGG | GGGATGA | TCC | TCC | C | T | C>T | C>T_TCC | CG>TA_GGGATGA | 9 | FALSE | FALSE |
| 5_v.txt | 33043 | 33043-SNV- | SNV | 1 | C | T | 21 | 24343 | 0.0862671 | 55.952381 | WT | FALSE | CAN | JSY13 | 1 | WT-Rep1 | TCA | C>T | A/T+1 | G/C-1 | CG>TA | CG>TA | CCATTTCACC | CACCCAA | AGT | TCA | C | A | C>T | C>T_TCA | CG>TA_CACCCAA | 9 | FALSE | FALSE |
| 5_v.txt | 33051 | 33051-SNV- | SNV | 1 | G | A | 41 | 24343 | 0.16842624 | 55.8292683 | WT | FALSE | CAN | JSY13 | 1 | WT-Rep1 | GGA | G>A | A/T+1 | G/C-1 | CG>TA | CG>TA | CCCAAGGAC | GACTGCG | CCT | CCT | C | T | C>T | C>T_CCT | CG>TA_GACTGCG | 9 | FALSE | FALSE |
| 5_v.txt | 33055 | 33055-SNV- | SNV | 1 | G | A | 74 | 24343 | 0.30398883 | 58.2290967 | WT | FALSE | CAN | JSY13 | 1 | WT-Rep1 | TGC | G>A | A/T+1 | G/C-1 | CG>TA | CG>TA | AGGACTGCG | GCGTGAC | ACG | ACG | C | T | C>T | C>T_ACG | CG>TA_GCGTGAC | 9 | FALSE | FALSE |
| 5_v.txt | 33056 | 33056-SNV- | SNV | 1 | C | T | 24 | 24343 | 0.09859097 | 51.8150134 | WT | FALSE | CAN | JSY13 | 1 | WT-Rep1 | GCG | C>T | A/T+1 | G/C-1 | CG>TA | CG>TA | GGACTGCGT | CGTGACA | CGC | GCG | C | A | C>T | C>T_GCG | CG>TA_CGTGACA | 9 | FALSE | FALSE |
| 5_v.txt | 33057 | 33057-SNV- | SNV | 1 | G | A | 5 | 24343 | 0.02053979 | 42.8004149 | WT | FALSE | CAN | JSY13 | 1 | WT-Rep1 | CGT | G>A | A/T+1 | G/C-1 | CG>TA | CG>TA | GACTGCGTG | GTGACAG | GCA | GCA | C | T | C>T | C>T_GCA | CG>TA_GTGACAG | 9 | FALSE | FALSE |
| 5_v.txt | 33069 | 33069-SNV- | SNV | 1 | G | A | 23 | 24343 | 0.09448301 | 51.0869565 | WT | FALSE | CAN | JSY13 | 1 | WT-Rep1 | TGC | G>A | A/T+1 | G/C-1 | CG>TA | CG>TA | GAATATGCC | GCCAAAG | ACG | ACG | C | T | C>T | C>T_ACG | CG>TA_GCCAAAG | 9 | FALSE | FALSE |
| 5_v.txt | 33078 | 33078-SNV- | SNV | 1 | C | T | 18 | 24343 | 0.07394323 | 52.2771873 | WT | FALSE | CAN | JSY13 | 1 | WT-Rep1 | ACC | C>T | A/T+1 | G/C-1 | CG>TA | CG>TA | AAAGAACCC | CCCATAA | TGG | ACC | C | A | C>T | C>T_ACC | CG>TA_CCCATAA | 9 | FALSE | FALSE |
| 5_v.txt | 33111 | 33111-SNV- | SNV | 1 | G | A | 62 | 24343 | 0.25469334 | 43.5967742 | WT | FALSE | CAN | JSY13 | 1 | WT-Rep1 | TGG | G>A | A/T+1 | G/C-1 | CG>TA | CG>TA | CCCACTGGG | GGGCCGG | ACC | ACC | C | T | C>T | C>T_ACC | CG>TA_GGGCCGG | 9 | FALSE | FALSE |
| 5_v.txt | 33114 | 33114-SNV- | SNV | 1 | C | T | 50 | 24343 | 0.20539786 | 42.3198175 | WT | FALSE | CAN | JSY13 | 1 | WT-Rep1 | GCC | C>T | A/T+1 | G/C-1 | CG>TA | CG>TA | ACTGGGCCG | CCGGCGT | CGG | GCC | C | A | C>T | C>T_GCC | CG>TA_CCGGCGT | 9 | FALSE | FALSE |
| 5_v.txt | 33115 | 33115-SNV- | SNV | 1 | C | T | 46 | 24343 | 0.18896603 | 44.1304348 | WT | FALSE | CAN | JSY13 | 1 | WT-Rep1 | CCG | C>T | A/T+1 | G/C-1 | CG>TA | CG>TA | CTGGGCCGG | CGGCGTT | GGC | CCG | C | A | C>T | C>T_CCG | CG>TA_CGGCGTT | 9 | FALSE | FALSE |
| 5_v.txt | 33118 | 33118-SNV- | SNV | 1 | C | T | 29 | 24343 | 0.11913076 | 40.2413793 | WT | FALSE | CAN | JSY13 | 1 | WT-Rep1 | GCG | C>T | A/T+1 | G/C-1 | CG>TA | CG>TA | GGCCGGCGT | CGTTGGT | CGC | GCG | C | A | C>T | C>T_GCG | CG>TA_CGTTGGT | 9 | FALSE | FALSE |
| 5_v.txt | 33153 | 33153-SNV- | SNV | 1 | C | T | 26 | 24343 | 0.10680689 | 37.1154052 | WT | FALSE | CAN | JSY13 | 1 | WT-Rep1 | ACC | C>T | A/T+1 | G/C-1 | CG>TA | CG>TA | AAAAGACCT | CCTGTAC | TGG | ACC | C | A | C>T | C>T_ACC | CG>TA_CCTGTAC | 9 | FALSE | FALSE |
| 5_v.txt | 33168 | 33168-SNV- | SNV | 1 | C | T | 26 | 24300 | 0.10699589 | 36.4230738 | WT | FALSE | CAN | JSY13 | 1 | WT-Rep1 | ACC | C>T | A/T+1 | G/C-1 | CG>TA | CG>TA | ATAGTACCA | CCACCAA | TGG | ACC | C | A | C>T | C>T_ACC | CG>TA_CCACCAA | 9 | FALSE | FALSE |
| 5_v.txt | 33171 | 33171-SNV- | SNV | 1 | C | T | 111 | 24300 | 0.45679012 | 37.2791175 | WT | FALSE | CAN | JSY13 | 1 | WT-Rep1 | ACC | C>T | A/T+1 | G/C-1 | CG>TA | CG>TA | GTACCACCA | CCAAGGG | TGG | ACC | C | A | C>T | C>T_ACC | CG>TA_CCAAGGG | 9 | FALSE | FALSE |
| 5_v.txt | 33172 | 33172-SNV- | SNV | 1 | C | T | 21 | 24300 | 0.08641975 | 36.5238095 | WT | FALSE | CAN | JSY13 | 1 | WT-Rep1 | CCA | C>T | A/T+1 | G/C-1 | CG>TA | CG>TA | TACCACCAA | CAAGGGC | GGT | CCA | C | A | C>T | C>T_CCA | CG>TA_CAAGGGC | 9 | FALSE | FALSE |
| 5_v.txt | 33177 | 33177-SNV- | SNV | 1 | G | A | 19 | 24298 | 0.07819574 | 37.8421053 | WT | FALSE | CAN | JSY13 | 1 | WT-Rep1 | GGC | G>A | A/T+1 | G/C-1 | CG>TA | CG>TA | CCAAGGGCA | GCAATCA | CCG | CCG | C | T | C>T | C>T_CCG | CG>TA_GCAATCA | 9 | FALSE | FALSE |
| 5_v.txt | 33178 | 33178-SNV- | SNV | 1 | C | T | 14 | 24298 | 0.05761791 | 32.2858977 | WT | FALSE | CAN | JSY13 | 1 | WT-Rep1 | GCA | C>T | A/T+1 | G/C-1 | CG>TA | CG>TA | CAAGGGCAA | CAATCAT | CGT | GCA | C | A | C>T | C>T_GCA | CG>TA_CAATCAT | 9 | FALSE | FALSE |
| 5_v.txt | 33284 | 33284-SNV- | SNV | 1 | G | A | 7 | 7276 | 0.09620671 | 43.7013153 | WT | FALSE | CAN | JSY13 | 1 | WT-Rep1 | CGC | G>A | A/T+1 | G/C-1 | CG>TA | CG>TA | ATCCTCGCC | GCCATTT | GCG | GCG | C | T | C>T | C>T_GCG | CG>TA_GCCATTT | 9 | FALSE | FALSE |
| 5_v.txt | 33376 | 33376-SNV- | SNV | 1 | G | A | 18 | 7277 | 0.24735468 | 56.3888889 | WT | FALSE | CAN | JSY13 | 1 | WT-Rep1 | TGT | G>A | A/T+1 | G/C-1 | CG>TA | CG>TA | GTGTTTGTG | GTGAAGC | ACA | ACA | C | T | C>T | C>T_ACA | CG>TA_GTGAAGC | 9 | FALSE | FALSE |
| 5_v.txt | 33412 | 33412-SNV- | SNV | 1 | G | A | 15 | 7277 | 0.2061289 | 39.6 | WT | FALSE | CAN | JSY13 | 1 | WT-Rep1 | GGC | G>A | A/T+1 | G/C-1 | CG>TA | CG>TA | TGACCGGCT | GCTCATT | CCG | CCG | C | T | C>T | C>T_CCG | CG>TA_GCTCATT | 9 | FALSE | FALSE |
| 5_v.txt | 31805 | 31805-SNV- | SNV | 1 | C | G | 1 | 9499 | 0.01052742 | 64 | WT | FALSE | CAN | JSY13 | 1 | WT-Rep1 | TCC | C>G | G/C+1 | G/C-1 | CG>GC | CG>GC | AATCTTCCA | CCAAATA | AGG | TCC | C | C | C>G | C>G_TCC | CG>GC_CCAAATA | 9 | FALSE | FALSE |
| 5_v.txt | 31947 | 31947-SNV- | SNV | 1 | G | C | 11 | 18256 | 0.06025416 | 34.7272727 | WT | FALSE | CAN | JSY13 | 1 | WT-Rep1 | CGT | G>C | G/C+1 | G/C-1 | CG>GC | CG>GC | ATGATCGTC | GTCATAA | GCA | GCA | C | G | C>G | C>G_GCA | CG>GC_GTCATAA | 9 | FALSE | FALSE |
| 5_v.txt | 31956 | 31956-SNV- | SNV | 1 | G | C | 13 | 18263 | 0.07118217 | 35.3076923 | WT | FALSE | CAN | JSY13 | 1 | WT-Rep1 | TGT | G>C | G/C+1 | G/C-1 | CG>GC | CG>GC | ATAAATGTG | GTGGCCG | ACA | ACA | C | G | C>G | C>G_ACA | CG>GC_GTGGCCG | 9 | FALSE | FALSE |
| 5_v.txt | 31981 | 31981-SNV- | SNV | 1 | G | C | 3 | 18279 | 0.01641228 | 53.9775915 | WT | FALSE | CAN | JSY13 | 1 | WT-Rep1 | GGC | G>C | G/C+1 | G/C-1 | CG>GC | CG>GC | AGCCGGGCA | GCATTAA | CCG | CCG | C | G | C>G | C>G_CCG | CG>GC_GCATTAA | 9 | FALSE | FALSE |
| 5_v.txt | 32027 | 32027-SNV- | SNV | 1 | G | C | 51 | 18279 | 0.2790087 | 57.1372549 | WT | FALSE | CAN | JSY13 | 1 | WT-Rep1 | GGT | G>C | G/C+1 | G/C-1 | CG>GC | CG>GC | GCCACGGTA | GTATTTC | CCA | CCA | C | G | C>G | C>G_CCA | CG>GC_GTATTTC | 9 | FALSE | FALSE |
| 5_v.txt | 32058 | 32058-SNV- | SNV | 1 | G | C | 6 | 18279 | 0.03282455 | 64 | WT | FALSE | CAN | JSY13 | 1 | WT-Rep1 | CGA | G>C | G/C+1 | G/C-1 | CG>GC | CG>GC | ATGTGCGAG | GAGATTG | GCT | GCT | C | G | C>G | C>G_GCT | CG>GC_GAGATTG | 9 | FALSE | FALSE |
| 5_v.txt | 32064 | 32064-SNV- | SNV | 1 | G | C | 31 | 18279 | 0.16959352 | 60.1612903 | WT | FALSE | CAN | JSY13 | 1 | WT-Rep1 | TGA | G>C | G/C+1 | G/C-1 | CG>GC | CG>GC | GAGATTGAG | GAGATAA | ACT | ACT | C | G | C>G | C>G_ACT | CG>GC_GAGATAA | 9 | FALSE | FALSE |
| 5_v.txt | 32075 | 32075-SNV- | SNV | 1 | C | G | 1 | 18279 | 5.47E-03 | 63.9971482 | WT | FALSE | CAN | JSY13 | 1 | WT-Rep1 | ACC | C>G | G/C+1 | G/C-1 | CG>GC | CG>GC | AAATAACCA | CCATGCA | TGG | ACC | C | C | C>G | C>G_ACC | CG>GC_CCATGCA | 9 | FALSE | FALSE |
| 5_v.txt | 32088 | 32088-SNV- | SNV | 1 | C | G | 2 | 18278 | 0.01094212 | 43.5 | WT | FALSE | CAN | JSY13 | 1 | WT-Rep1 | GCC | C>G | G/C+1 | G/C-1 | CG>GC | CG>GC | AAAAAGCCT | CCTGCAA | CGG | GCC | C | C | C>G | C>G_GCC | CG>GC_CCTGCAA | 9 | FALSE | FALSE |
| 5_v.txt | 32089 | 32089-SNV- | SNV | 1 | C | G | 33 | 18278 | 0.18054492 | 59.2428619 | WT | FALSE | CAN | JSY13 | 1 | WT-Rep1 | CCT | C>G | G/C+1 | G/C-1 | CG>GC | CG>GC | AAAAGCCTG | CTGCAAC | GGA | CCT | C | C | C>G | C>G_CCT | CG>GC_CTGCAAC | 9 | FALSE | FALSE |
| 5_v.txt | 32092 | 32092-SNV- | SNV | 1 | C | G | 4 | 18279 | 0.02188304 | 62.75 | WT | FALSE | CAN | JSY13 | 1 | WT-Rep1 | GCA | C>G | G/C+1 | G/C-1 | CG>GC | CG>GC | AGCCTGCAA | CAACACC | CGT | GCA | C | C | C>G | C>G_GCA | CG>GC_CAACACC | 9 | FALSE | FALSE |
| 5_v.txt | 32153 | 32153-SNV- | SNV | 1 | G | C | 23 | 18279 | 0.12582745 | 49.4347826 | WT | FALSE | CAN | JSY13 | 1 | WT-Rep1 | TGT | G>C | G/C+1 | G/C-1 | CG>GC | CG>GC | CTCCATGTA | GTAAGCC | ACA | ACA | C | G | C>G | C>G_ACA | CG>GC_GTAAGCC | 9 | FALSE | FALSE |
| 5_v.txt | 32199 | 32199-SNV- | SNV | 1 | G | C | 33 | 18270 | 0.18062397 | 45.3333333 | WT | FALSE | CAN | JSY13 | 1 | WT-Rep1 | TGG | G>C | G/C+1 | G/C-1 | CG>GC | CG>GC | ATGTATGGA | GGAACAC | ACC | ACC | C | G | C>G | C>G_ACC | CG>GC_GGAACAC | 9 | FALSE | FALSE |
| 5_v.txt | 32223 | 32223-SNV- | SNV | 1 | G | C | 31 | 18270 | 0.16967707 | 37.0967742 | WT | FALSE | CAN | JSY13 | 1 | WT-Rep1 | TGA | G>C | G/C+1 | G/C-1 | CG>GC | CG>GC | GTCCTTGAC | GACAGGA | ACT | ACT | C | G | C>G | C>G_ACT | CG>GC_GACAGGA | 9 | FALSE | FALSE |
| 5_v.txt | 32271 | 32271-SNV- | SNV | 1 | C | G | 9 | 21420 | 0.04201681 | 36.4444444 | WT | FALSE | CAN | JSY13 | 1 | WT-Rep1 | ACG | C>G | G/C+1 | G/C-1 | CG>GC | CG>GC | AAAATACGG | CGGGAAC | TGC | ACG | C | C | C>G | C>G_ACG | CG>GC_CGGGAAC | 9 | FALSE | FALSE |
| 5_v.txt | 32272 | 32272-SNV- | SNV | 1 | G | C | 9 | 21420 | 0.04201681 | 35.8888889 | WT | FALSE | CAN | JSY13 | 1 | WT-Rep1 | CGG | G>C | G/C+1 | G/C-1 | CG>GC | CG>GC | AAATACGGG | GGGAACC | GCC | GCC | C | G | C>G | C>G_GCC | CG>GC_GGGAACC | 9 | FALSE | FALSE |
| 5_v.txt | 32278 | 32278-SNV- | SNV | 1 | C | G | 22 | 21420 | 0.10270775 | 34.3181818 | WT | FALSE | CAN | JSY13 | 1 | WT-Rep1 | CCA | C>G | G/C+1 | G/C-1 | CG>GC | CG>GC | GGGAACCAA | CAACGTA | GGT | CCA | C | C | C>G | C>G_CCA | CG>GC_CAACGTA | 9 | FALSE | FALSE |
| 5_v.txt | 32282 | 32282-SNV- | SNV | 1 | G | C | 16 | 21420 | 0.07469655 | 36.6241485 | WT | FALSE | CAN | JSY13 | 1 | WT-Rep1 | CGT | G>C | G/C+1 | G/C-1 | CG>GC | CG>GC | ACCAACGTA | GTAAATA | GCA | GCA | C | G | C>G | C>G_GCA | CG>GC_GTAAATA | 9 | FALSE | FALSE |
| 5_v.txt | 32292 | 32292-SNV- | SNV | 1 | G | C | 34 | 21421 | 0.15872275 | 52.5588235 | WT | FALSE | CAN | JSY13 | 1 | WT-Rep1 | TGA | G>C | G/C+1 | G/C-1 | CG>GC | CG>GC | ATATTTGAA | GAATTTG | ACT | ACT | C | G | C>G | C>G_ACT | CG>GC_GAATTTG | 9 | FALSE | FALSE |
| 5_v.txt | 32304 | 32304-SNV- | SNV | 1 | G | C | 4 | 21421 | 0.01867327 | 62.25 | WT | FALSE | CAN | JSY13 | 1 | WT-Rep1 | AGA | G>C | G/C+1 | G/C-1 | CG>GC | CG>GC | GCGGCAGAA | GAAATAA | TCT | TCT | C | G | C>G | C>G_TCT | CG>GC_GAAATAA | 9 | FALSE | FALSE |
| 5_v.txt | 32316 | 32316-SNV- | SNV | 1 | G | C | 4 | 21421 | 0.01867327 | 28.2538116 | WT | FALSE | CAN | JSY13 | 1 | WT-Rep1 | TGT | G>C | G/C+1 | G/C-1 | CG>GC | CG>GC | ATGGTTGTT | GTTAAGA | ACA | ACA | C | G | C>G | C>G_ACA | CG>GC_GTTAAGA | 9 | FALSE | FALSE |
| 5_v.txt | 32385 | 32385-SNV- | SNV | 1 | G | C | 13 | 21419 | 0.06069378 | 59.4531251 | WT | FALSE | CAN | JSY13 | 1 | WT-Rep1 | GGG | G>C | G/C+1 | G/C-1 | CG>GC | CG>GC | ATAAAGGGA | GGAGAAG | CCC | CCC | C | G | C>G | C>G_CCC | CG>GC_GGAGAAG | 9 | FALSE | FALSE |
| 5_v.txt | 32432 | 32432-SNV- | SNV | 1 | G | C | 21 | 21421 | 0.09803464 | 56.2380952 | WT | FALSE | CAN | JSY13 | 1 | WT-Rep1 | TGT | G>C | G/C+1 | G/C-1 | CG>GC | CG>GC | GTCATTGTA | GTATGGA | ACA | ACA | C | G | C>G | C>G_ACA | CG>GC_GTATGGA | 9 | FALSE | FALSE |
| 5_v.txt | 32449 | 32449-SNV- | SNV | 1 | C | G | 4 | 21421 | 0.01867327 | 56 | WT | FALSE | CAN | JSY13 | 1 | WT-Rep1 | CCA | C>G | G/C+1 | G/C-1 | CG>GC | CG>GC | AAAGTCCAA | CAATGAA | GGT | CCA | C | C | C>G | C>G_CCA | CG>GC_CAATGAA | 9 | FALSE | FALSE |
| 5_v.txt | 32471 | 32471-SNV- | SNV | 1 | G | C | 3 | 21421 | 0.01400495 | 46.6669318 | WT | FALSE | CAN | JSY13 | 1 | WT-Rep1 | TGT | G>C | G/C+1 | G/C-1 | CG>GC | CG>GC | GCCAATGTA | GTAGAAG | ACA | ACA | C | G | C>G | C>G_ACA | CG>GC_GTAGAAG | 9 | FALSE | FALSE |
| 5_v.txt | 32487 | 32487-SNV- | SNV | 1 | C | G | 25 | 21421 | 0.1167079 | 43.28 | WT | FALSE | CAN | JSY13 | 1 | WT-Rep1 | ACG | C>G | G/C+1 | G/C-1 | CG>GC | CG>GC | AAGATACGG | CGGAAAA | TGC | ACG | C | C | C>G | C>G_ACG | CG>GC_CGGAAAA | 9 | FALSE | FALSE |
| 5_v.txt | 32488 | 32488-SNV- | SNV | 1 | G | C | 40 | 21421 | 0.18673265 | 44.9747342 | WT | FALSE | CAN | JSY13 | 1 | WT-Rep1 | CGG | G>C | G/C+1 | G/C-1 | CG>GC | CG>GC | AGATACGGA | GGAAAAC | GCC | GCC | C | G | C>G | C>G_GCC | CG>GC_GGAAAAC | 9 | FALSE | FALSE |
| 5_v.txt | 32520 | 32520-SNV- | SNV | 1 | G | C | 38 | 21421 | 0.17739601 | 37.3421053 | WT | FALSE | CAN | JSY13 | 1 | WT-Rep1 | GGA | G>C | G/C+1 | G/C-1 | CG>GC | CG>GC | GGAACGGAT | GATTTTC | CCT | CCT | C | G | C>G | C>G_CCT | CG>GC_GATTTTC | 9 | FALSE | FALSE |
| 5_v.txt | 32528 | 32528-SNV- | SNV | 1 | G | C | 1 | 21421 | 4.67E-03 | 37.9995872 | WT | FALSE | CAN | JSY13 | 1 | WT-Rep1 | TGG | G>C | G/C+1 | G/C-1 | CG>GC | CG>GC | TTTTCTGGG | GGGGTTT | ACC | ACC | C | G | C>G | C>G_ACC | CG>GC_GGGGTTT | 9 | FALSE | FALSE |
| 5_v.txt | 32529 | 32529-SNV- | SNV | 1 | G | C | 12 | 21421 | 0.05601979 | 38.75 | WT | FALSE | CAN | JSY13 | 1 | WT-Rep1 | GGG | G>C | G/C+1 | G/C-1 | CG>GC | CG>GC | TTTCTGGGG | GGGTTTG | CCC | CCC | C | G | C>G | C>G_CCC | CG>GC_GGGTTTG | 9 | FALSE | FALSE |
| 5_v.txt | 32539 | 32539-SNV- | SNV | 1 | C | G | 22 | 21421 | 0.10270296 | 37.2727273 | WT | FALSE | CAN | JSY13 | 1 | WT-Rep1 | GCT | C>G | G/C+1 | G/C-1 | CG>GC | CG>GC | TTGCAGCTT | CTTCACC | CGA | GCT | C | C | C>G | C>G_GCT | CG>GC_CTTCACC | 9 | FALSE | FALSE |
| 5_v.txt | 32566 | 32566-SNV- | SNV | 1 | C | G | 32 | 22145 | 0.14450215 | 37.5625 | WT | FALSE | CAN | JSY13 | 1 | WT-Rep1 | TCA | C>G | G/C+1 | G/C-1 | CG>GC | CG>GC | CTAGTTCAG | CAGTACC | AGT | TCA | C | C | C>G | C>G_TCA | CG>GC_CAGTACC | 9 | FALSE | FALSE |

| | | | | | | | | | | | | | | | | | | | | | | | | | | | | | | | | | | |
|---|---|---|---|---|---|---|---|---|---|---|---|---|---|---|---|---|---|---|---|---|---|---|---|---|---|---|---|---|---|---|---|---|---|---|
| 5_v.txt | 32679 | 32679-SNV-( | SNV | 1 | C | G | 4 | 22155 | 0.01805462 | 63.9977614 | WT | FALSE | CAN | JSY13 | 1 | WT-Rep1 | CCA | C>G | G/C+1 | G/C-1 | CG>GC | CG>GC | TTTCTCCAAT | CAATAAC | GGT | CCA | C | C | C>G | C>G_CCA | CG>GC_CAATAAC | 9 | FALSE | FALSE |
| 5_v.txt | 32685 | 32685-SNV-( | SNV | 1 | C | G | 39 | 22156 | 0.17602455 | 60.5128205 | WT | FALSE | CAN | JSY13 | 1 | WT-Rep1 | ACG | C>G | G/C+1 | G/C-1 | CG>GC | CG>GC | CAATAACGG | CGGAATC | TGC | ACG | C | C | C>G | C>G_ACG | CG>GC_CGGAATC | 9 | FALSE | FALSE |
| 5_v.txt | 32749 | 32749-SNV-( | SNV | 1 | C | G | 26 | 22177 | 0.11723858 | 56.767728 | WT | FALSE | CAN | JSY13 | 1 | WT-Rep1 | CCG | C>G | G/C+1 | G/C-1 | CG>GC | CG>GC | GAAACCCGA | CGATAAT | GGC | CCG | C | C | C>G | C>G_CCG | CG>GC_CGATAAT | 9 | FALSE | FALSE |
| 5_v.txt | 32757 | 32757-SNV-( | SNV | 1 | G | C | 7 | 22177 | 0.03156423 | 45.7144545 | WT | FALSE | CAN | JSY13 | 1 | WT-Rep1 | GGC | G>C | G/C+1 | G/C-1 | CG>GC | CG>GC | ATAATGGCT | GCTAAAA | CCG | CCG | C | G | C>G | C>G_CCG | CG>GC_GCTAAAA | 9 | FALSE | FALSE |
| 5_v.txt | 32758 | 32758-SNV-( | SNV | 1 | C | G | 2 | 22177 | 9.02E-03 | 37.4970113 | WT | FALSE | CAN | JSY13 | 1 | WT-Rep1 | GCT | C>G | G/C+1 | G/C-1 | CG>GC | CG>GC | TAATGGCTA | CTAAAAC | CGA | GCT | C | C | C>G | C>G_GCT | CG>GC_CTAAAAC | 9 | FALSE | FALSE |
| 5_v.txt | 32776 | 32776-SNV-( | SNV | 1 | C | G | 10 | 22177 | 0.04509176 | 43.4 | WT | FALSE | CAN | JSY13 | 1 | WT-Rep1 | GCG | C>G | G/C+1 | G/C-1 | CG>GC | CG>GC | TGGAAGCGA | CGACCCA | CGC | GCG | C | C | C>G | C>G_GCG | CG>GC_CGACCCA | 9 | FALSE | FALSE |
| 5_v.txt | 32786 | 32786-SNV-( | SNV | 1 | C | G | 49 | 22177 | 0.22094963 | 48.9596618 | WT | FALSE | CAN | JSY13 | 1 | WT-Rep1 | ACT | C>G | G/C+1 | G/C-1 | CG>GC | CG>GC | CCAGAACTC | CTCGAAT | TGA | ACT | C | C | C>G | C>G_ACT | CG>GC_CTCGAAT | 9 | FALSE | FALSE |
| 5_v.txt | 32797 | 32797-SNV-( | SNV | 1 | C | G | 31 | 22177 | 0.13978446 | 47.2258065 | WT | FALSE | CAN | JSY13 | 1 | WT-Rep1 | CCG | C>G | G/C+1 | G/C-1 | CG>GC | CG>GC | ATTCACCGTA | CGTAATA | GGC | CCG | C | C | C>G | C>G_CCG | CG>GC_CGTAATA | 9 | FALSE | FALSE |
| 5_v.txt | 32798 | 32798-SNV-( | SNV | 1 | G | C | 42 | 22177 | 0.1893854 | 45.2380952 | WT | FALSE | CAN | JSY13 | 1 | WT-Rep1 | CGT | G>C | G/C+1 | G/C-1 | CG>GC | CG>GC | TTCACCGTAA | GTAATAT | GCA | GCA | C | G | C>G | C>G_GCA | CG>GC_GTAATAT | 9 | FALSE | FALSE |
| 5_v.txt | 32819 | 32819-SNV-( | SNV | 1 | G | C | 10 | 22177 | 0.04509176 | 42.5 | WT | FALSE | CAN | JSY13 | 1 | WT-Rep1 | AGT | G>C | G/C+1 | G/C-1 | CG>GC | CG>GC | GAACAAGTT | GTTCATT | TCA | TCA | C | G | C>G | C>G_TCA | CG>GC_GTTCATT | 9 | FALSE | FALSE |
| 5_v.txt | 32876 | 32876-SNV-( | SNV | 1 | G | C | 86 | 24300 | 0.35390947 | 37.3720086 | WT | FALSE | CAN | JSY13 | 1 | WT-Rep1 | TGT | G>C | G/C+1 | G/C-1 | CG>GC | CG>GC | AACTTTGTAC | GTACGTC | ACA | ACA | C | G | C>G | C>G_ACA | CG>GC_GTACGTC | 9 | FALSE | FALSE |
| 5_v.txt | 32900 | 32900-SNV-( | SNV | 1 | G | C | 4 | 24305 | 0.01645752 | 29.2467426 | WT | FALSE | CAN | JSY13 | 1 | WT-Rep1 | GGC | G>C | G/C+1 | G/C-1 | CG>GC | CG>GC | GACTTGGCC | GCCAACT | CCG | CCG | C | G | C>G | C>G_CCG | CG>GC_GCCAACT | 9 | FALSE | FALSE |
| 5_v.txt | 32917 | 32917-SNV-( | SNV | 1 | C | G | 8 | 24305 | 0.03291504 | 51.75 | WT | FALSE | CAN | JSY13 | 1 | WT-Rep1 | TCC | C>G | G/C+1 | G/C-1 | CG>GC | CG>GC | TAAGTTCCAC | CCAGGGC | AGG | TCC | C | C | C>G | C>G_TCC | CG>GC_CCAGGGC | 9 | FALSE | FALSE |
| 5_v.txt | 32920 | 32920-SNV-( | SNV | 1 | G | C | 1 | 24305 | 4.11E-03 | 43.9973062 | WT | FALSE | CAN | JSY13 | 1 | WT-Rep1 | AGG | G>C | G/C+1 | G/C-1 | CG>GC | CG>GC | GTTCCAGGG | GGGCAAA | TCC | TCC | C | G | C>G | C>G_TCC | CG>GC_GGGCAAA | 9 | FALSE | FALSE |
| 5_v.txt | 32935 | 32935-SNV-( | SNV | 1 | C | G | 3 | 24343 | 0.01232387 | 50.3344459 | WT | FALSE | CAN | JSY13 | 1 | WT-Rep1 | GCC | C>G | G/C+1 | G/C-1 | CG>GC | CG>GC | TGATTGCCCA | CCCAAGA | CGG | GCC | C | C | C>G | C>G_GCC | CG>GC_CCCAAGA | 9 | FALSE | FALSE |
| 5_v.txt | 32936 | 32936-SNV-( | SNV | 1 | C | G | 17 | 24341 | 0.06984101 | 48.1545043 | WT | FALSE | CAN | JSY13 | 1 | WT-Rep1 | CCC | C>G | G/C+1 | G/C-1 | CG>GC | CG>GC | GATTGCCCA | CCAAGAA | GGG | CCC | C | C | C>G | C>G_CCC | CG>GC_CCAAGAA | 9 | FALSE | FALSE |
| 5_v.txt | 32945 | 32945-SNV-( | SNV | 1 | C | G | 7 | 24343 | 0.0287557 | 60.2774053 | WT | FALSE | CAN | JSY13 | 1 | WT-Rep1 | ACC | C>G | G/C+1 | G/C-1 | CG>GC | CG>GC | AGAAAACCA | CCAATAC | TGG | ACC | C | C | C>G | C>G_ACC | CG>GC_CCAATAC | 9 | FALSE | FALSE |
| 5_v.txt | 32971 | 32971-SNV-( | SNV | 1 | C | G | 21 | 24343 | 0.0862671 | 63.9975002 | WT | FALSE | CAN | JSY13 | 1 | WT-Rep1 | CCA | C>G | G/C+1 | G/C-1 | CG>GC | CG>GC | CCGCACCAA | CAAATGC | GGT | CCA | C | C | C>G | C>G_CCA | CG>GC_CAAATGC | 9 | FALSE | FALSE |
| 5_v.txt | 32997 | 32997-SNV-( | SNV | 1 | G | C | 107 | 24343 | 0.43955141 | 60.9906542 | WT | FALSE | CAN | JSY13 | 1 | WT-Rep1 | TGA | G>C | G/C+1 | G/C-1 | CG>GC | CG>GC | CTTTGTGAG | GAGAAAA | ACT | ACT | C | G | C>G | C>G_ACT | CG>GC_GAGAAAA | 9 | FALSE | FALSE |
| 5_v.txt | 33024 | 33024-SNV-( | SNV | 1 | G | C | 6 | 24343 | 0.02464774 | 58.6666667 | WT | FALSE | CAN | JSY13 | 1 | WT-Rep1 | AGG | G>C | G/C+1 | G/C-1 | CG>GC | CG>GC | GTAACAGGG | GGGATGA | TCC | TCC | C | G | C>G | C>G_TCC | CG>GC_GGGATGA | 9 | FALSE | FALSE |
| 5_v.txt | 33037 | 33037-SNV-( | SNV | 1 | C | G | 19 | 24343 | 0.07805119 | 56.3157895 | WT | FALSE | CAN | JSY13 | 1 | WT-Rep1 | GCC | C>G | G/C+1 | G/C-1 | CG>GC | CG>GC | ATGTAGCCA | CCATTTC | CGG | GCC | C | C | C>G | C>G_GCC | CG>GC_CCATTTC | 9 | FALSE | FALSE |
| 5_v.txt | 33057 | 33057-SNV-( | SNV | 1 | G | C | 9 | 24343 | 0.03697161 | 57.6662277 | WT | FALSE | CAN | JSY13 | 1 | WT-Rep1 | CGT | G>C | G/C+1 | G/C-1 | CG>GC | CG>GC | GACTGCGTG | GTGACAG | GCA | GCA | C | G | C>G | C>G_GCA | CG>GC_GTGACAG | 9 | FALSE | FALSE |
| 5_v.txt | 33078 | 33078-SNV-( | SNV | 1 | C | G | 1 | 24343 | 4.11E-03 | 63.9986878 | WT | FALSE | CAN | JSY13 | 1 | WT-Rep1 | ACC | C>G | G/C+1 | G/C-1 | CG>GC | CG>GC | AAAGAACCC | CCCATAA | TGG | ACC | C | C | C>G | C>G_ACC | CG>GC_CCCATAA | 9 | FALSE | FALSE |
| 5_v.txt | 33079 | 33079-SNV-( | SNV | 1 | C | G | 14 | 24343 | 0.0575114 | 51.2124771 | WT | FALSE | CAN | JSY13 | 1 | WT-Rep1 | CCC | C>G | G/C+1 | G/C-1 | CG>GC | CG>GC | AAGAACCCA | CCATAAA | GGG | CCC | C | C | C>G | C>G_CCC | CG>GC_CCATAAA | 9 | FALSE | FALSE |
| 5_v.txt | 33093 | 33093-SNV-( | SNV | 1 | G | C | 23 | 24343 | 0.09448301 | 51.4348237 | WT | FALSE | CAN | JSY13 | 1 | WT-Rep1 | TGA | G>C | G/C+1 | G/C-1 | CG>GC | CG>GC | AAATATGAT | GATATAA | ACT | ACT | C | G | C>G | C>G_ACT | CG>GC_GATATAA | 9 | FALSE | FALSE |
| 5_v.txt | 33114 | 33114-SNV-( | SNV | 1 | C | G | 6 | 24343 | 0.02464774 | 46.3321944 | WT | FALSE | CAN | JSY13 | 1 | WT-Rep1 | GCC | C>G | G/C+1 | G/C-1 | CG>GC | CG>GC | ACTGGGCCG | CCGGCGT | CGG | GCC | C | C | C>G | C>G_GCC | CG>GC_CCGGCGT | 9 | FALSE | FALSE |
| 5_v.txt | 33115 | 33115-SNV-( | SNV | 1 | C | G | 4 | 24343 | 0.01643183 | 31.75 | WT | FALSE | CAN | JSY13 | 1 | WT-Rep1 | CCG | C>G | G/C+1 | G/C-1 | CG>GC | CG>GC | CTGGGCCGG | CGGCGTT | GGC | CCG | C | C | C>G | C>G_CCG | CG>GC_CGGCGTT | 9 | FALSE | FALSE |
| 5_v.txt | 33118 | 33118-SNV-( | SNV | 1 | C | G | 24 | 24343 | 0.09859097 | 42.2916667 | WT | FALSE | CAN | JSY13 | 1 | WT-Rep1 | GCG | C>G | G/C+1 | G/C-1 | CG>GC | CG>GC | GGCCGGCGT | CGTTGGT | CGC | GCG | C | C | C>G | C>G_GCG | CG>GC_CGTTGGT | 9 | FALSE | FALSE |
| 5_v.txt | 33154 | 33154-SNV-( | SNV | 1 | C | G | 25 | 24343 | 0.10269893 | 36.9600774 | WT | FALSE | CAN | JSY13 | 1 | WT-Rep1 | CCT | C>G | G/C+1 | G/C-1 | CG>GC | CG>GC | AAAGACCTG | CTGTACC | GGA | CCT | C | C | C>G | C>G_CCT | CG>GC_CTGTACC | 9 | FALSE | FALSE |
| 5_v.txt | 33156 | 33156-SNV-( | SNV | 1 | G | C | 18 | 24342 | 0.07394627 | 37.4998824 | WT | FALSE | CAN | JSY13 | 1 | WT-Rep1 | TGT | G>C | G/C+1 | G/C-1 | CG>GC | CG>GC | AGACCTGTA | GTACCAA | ACA | ACA | C | G | C>G | C>G_ACA | CG>GC_GTACCAA | 9 | FALSE | FALSE |
| 5_v.txt | 33160 | 33160-SNV-( | SNV | 1 | C | G | 9 | 24341 | 0.03697465 | 35.7777778 | WT | FALSE | CAN | JSY13 | 1 | WT-Rep1 | CCA | C>G | G/C+1 | G/C-1 | CG>GC | CG>GC | CTGTACCAAT | CAATAGT | GGT | CCA | C | C | C>G | C>G_CCA | CG>GC_CAATAGT | 9 | FALSE | FALSE |
| 5_v.txt | 33168 | 33168-SNV-( | SNV | 1 | C | G | 32 | 24300 | 0.13168724 | 37.8437489 | WT | FALSE | CAN | JSY13 | 1 | WT-Rep1 | ACC | C>G | G/C+1 | G/C-1 | CG>GC | CG>GC | ATAGTACCA | CCACCAA | TGG | ACC | C | C | C>G | C>G_ACC | CG>GC_CCACCAA | 9 | FALSE | FALSE |
| 5_v.txt | 33169 | 33169-SNV-( | SNV | 1 | C | G | 3 | 24300 | 0.01234568 | 33.3337496 | WT | FALSE | CAN | JSY13 | 1 | WT-Rep1 | CCA | C>G | G/C+1 | G/C-1 | CG>GC | CG>GC | TAGTACCAC | CACCAAG | GGT | CCA | C | C | C>G | C>G_CCA | CG>GC_CACCAAG | 9 | FALSE | FALSE |
| 5_v.txt | 33172 | 33172-SNV-( | SNV | 1 | C | G | 34 | 24300 | 0.1399177 | 36.8235294 | WT | FALSE | CAN | JSY13 | 1 | WT-Rep1 | CCA | C>G | G/C+1 | G/C-1 | CG>GC | CG>GC | TACCACCAA | CAAGGGC | GGT | CCA | C | C | C>G | C>G_CCA | CG>GC_CAAGGGC | 9 | FALSE | FALSE |
| 5_v.txt | 33178 | 33178-SNV-( | SNV | 1 | C | G | 51 | 24298 | 0.20989382 | 37.3920541 | WT | FALSE | CAN | JSY13 | 1 | WT-Rep1 | GCA | C>G | G/C+1 | G/C-1 | CG>GC | CG>GC | CAAGGGCAA | CAATCAT | CGT | GCA | C | C | C>G | C>G_GCA | CG>GC_CAATCAT | 9 | FALSE | FALSE |
| 5_v.txt | 33193 | 33193-SNV-( | SNV | 1 | G | C | 9 | 7277 | 0.12367734 | 34.8865409 | WT | FALSE | CAN | JSY13 | 1 | WT-Rep1 | TGT | G>C | G/C+1 | G/C-1 | CG>GC | CG>GC | CAATATGTCT | GTCTTTG | ACA | ACA | C | G | C>G | C>G_ACA | CG>GC_GTCTTTG | 9 | FALSE | FALSE |
| 5_v.txt | 33205 | 33205-SNV-( | SNV | 1 | G | C | 9 | 7277 | 0.12367734 | 37.2209243 | WT | FALSE | CAN | JSY13 | 1 | WT-Rep1 | AGC | G>C | G/C+1 | G/C-1 | CG>GC | CG>GC | GCTTAAGCT | GCTCTCT | TCG | TCG | C | G | C>G | C>G_TCG | CG>GC_GCTCTCT | 9 | FALSE | FALSE |
| 5_v.txt | 33210 | 33210-SNV-( | SNV | 1 | C | G | 25 | 7277 | 0.34354817 | 39.68 | WT | FALSE | CAN | JSY13 | 1 | WT-Rep1 | TCT | C>G | G/C+1 | G/C-1 | CG>GC | CG>GC | AGCTCTCTCT | CTCTTCA | AGA | TCT | C | C | C>G | C>G_TCT | CG>GC_CTCTTCA | 9 | FALSE | FALSE |
| 5_v.txt | 33283 | 33283-SNV-( | SNV | 1 | C | G | 1 | 7277 | 0.01374193 | 63.9999632 | WT | FALSE | CAN | JSY13 | 1 | WT-Rep1 | TCG | C>G | G/C+1 | G/C-1 | CG>GC | CG>GC | TATCCTCGCC | CGCCATT | AGC | TCG | C | C | C>G | C>G_TCG | CG>GC_CGCCATT | 9 | FALSE | FALSE |
| 5_v.txt | 33285 | 33285-SNV-( | SNV | 1 | C | G | 1 | 7277 | 0.01374193 | 37.9882864 | WT | FALSE | CAN | JSY13 | 1 | WT-Rep1 | GCC | C>G | G/C+1 | G/C-1 | CG>GC | CG>GC | TCCTCGCCAT | CCATTTA | CGG | GCC | C | C | C>G | C>G_GCC | CG>GC_CCATTTA | 9 | FALSE | FALSE |
| 5_v.txt | 33354 | 33354-SNV-( | SNV | 1 | G | C | 4 | 7277 | 0.05496771 | 49.25 | WT | FALSE | CAN | JSY13 | 1 | WT-Rep1 | TGA | G>C | G/C+1 | G/C-1 | CG>GC | CG>GC | GGTATTGAC | GACCCAC | ACT | ACT | C | G | C>G | C>G_ACT | CG>GC_GACCCAC | 9 | FALSE | FALSE |
| 5_v.txt | 33419 | 33419-SNV-( | SNV | 1 | G | C | 15 | 7277 | 0.2061289 | 41.5333333 | WT | FALSE | CAN | JSY13 | 1 | WT-Rep1 | TGT | G>C | G/C+1 | G/C-1 | CG>GC | CG>GC | CTCATTGTAC | GTACATA | ACA | ACA | C | G | C>G | C>G_ACA | CG>GC_GTACATA | 9 | FALSE | FALSE |
| 5_v.txt | 33456 | 33456-SNV-( | SNV | 1 | G | C | 9 | 7276 | 0.12369434 | 42.5555556 | WT | FALSE | CAN | JSY13 | 1 | WT-Rep1 | TGA | G>C | G/C+1 | G/C-1 | CG>GC | CG>GC | TCTTTTGAAT | GAATTTG | ACT | ACT | C | G | C>G | C>G_ACT | CG>GC_GAATTTG | 9 | FALSE | FALSE |
| 5_v.txt | 31657 | 31657-SNV-( | SNV | 1 | G | T | 94 | 9499 | 0.98957785 | 18.9468085 | WT | FALSE | CAN | JSY13 | 1 | WT-Rep1 | GGC | G>T | A/T+1 | G/C-1 | CG>AT | CG>AT | GAAATGGCG | GCGTGGA | CCG | CCG | C | A | C>A | C>A_CCG | CG>AT_GCGTGGA | 9 | FALSE | FALSE |
| 5_v.txt | 31798 | 31798-SNV-( | SNV | 1 | C | A | 36 | 9499 | 0.37898726 | 54.3333333 | WT | FALSE | CAN | JSY13 | 1 | WT-Rep1 | CCA | C>A | A/T+1 | G/C-1 | CG>AT | CG>AT | CATCTCCAAT | CAATCTT | GGT | CCA | C | T | C>A | C>A_CCA | CG>AT_CAATCTT | 9 | FALSE | FALSE |
| 5_v.txt | 31817 | 31817-SNV-( | SNV | 1 | G | T | 12 | 9499 | 0.12632909 | 52.416976 | WT | FALSE | CAN | JSY13 | 1 | WT-Rep1 | TGC | G>T | A/T+1 | G/C-1 | CG>AT | CG>AT | AAATCTGCAT | GCATCTG | ACG | ACG | C | A | C>A | C>A_ACG | CG>AT_GCATCTG | 9 | FALSE | FALSE |
| 5_v.txt | 31845 | 31845-SNV-( | SNV | 1 | C | A | 8 | 9499 | 0.08421939 | 25.75 | WT | FALSE | CAN | JSY13 | 1 | WT-Rep1 | CCA | C>A | A/T+1 | G/C-1 | CG>AT | CG>AT | AAGATCCAA | CAAACAG | GGT | CCA | C | T | C>A | C>A_CCA | CG>AT_CAAACAG | 9 | FALSE | FALSE |
| 5_v.txt | 31869 | 31869-SNV-( | SNV | 1 | G | T | 128 | 9499 | 1.34751026 | 47.890625 | WT | FALSE | CAN | JSY13 | 1 | WT-Rep1 | AGA | G>T | A/T+1 | G/C-1 | CG>AT | CG>AT | AAAATAGAG | GAGATAT | TCT | TCT | C | A | C>A | C>A_TCT | CG>AT_GAGATAT | 9 | FALSE | FALSE |
| 5_v.txt | 31930 | 31930-SNV-( | SNV | 1 | G | T | 14 | 9485 | 0.14760148 | 37.2857143 | WT | FALSE | CAN | JSY13 | 1 | WT-Rep1 | TGA | G>T | A/T+1 | G/C-1 | CG>AT | CG>AT | AACCTTGAAT | GAATAAT | ACT | ACT | C | A | C>A | C>A_ACT | CG>AT_GAATAAT | 9 | FALSE | FALSE |
| 5_v.txt | 31973 | 31973-SNV-( | SNV | 1 | C | A | 16 | 18279 | 0.08753214 | 20.8096257 | WT | FALSE | CAN | JSY13 | 1 | WT-Rep1 | CCA | C>A | A/T+1 | G/C-1 | CG>AT | CG>AT | ATAAGCCAA | CAAGCCG | GGT | CCA | C | T | C>A | C>A_CCA | CG>AT_CAAGCCG | 9 | FALSE | FALSE |
| 5_v.txt | 31980 | 31980-SNV-( | SNV | 1 | G | T | 13 | 18279 | 0.07111986 | 35.7544055 | WT | FALSE | CAN | JSY13 | 1 | WT-Rep1 | GGG | G>T | A/T+1 | G/C-1 | CG>AT | CG>AT | AAGCCGGGC | GGCATTA | CCC | CCC | C | A | C>A | C>A_CCC | CG>AT_GGCATTA | 9 | FALSE | FALSE |
| 5_v.txt | 32008 | 32008-SNV-( | SNV | 1 | C | A | 116 | 18279 | 0.63460802 | 53.3706897 | WT | FALSE | CAN | JSY13 | 1 | WT-Rep1 | TCG | C>A | A/T+1 | G/C-1 | CG>AT | CG>AT | GTAACTCGTC | CGTCACG | AGC | TCG | C | T | C>A | C>A_TCG | CG>AT_CGTCACG | 9 | FALSE | FALSE |
| 5_v.txt | 32027 | 32027-SNV-( | SNV | 1 | G | T | 15 | 18279 | 0.08206138 | 49.9333333 | WT | FALSE | CAN | JSY13 | 1 | WT-Rep1 | GGT | G>T | A/T+1 | G/C-1 | CG>AT | CG>AT | GCCACGGTA | GTATTTC | CCA | CCA | C | A | C>A | C>A_CCA | CG>AT_GTATTTC | 9 | FALSE | FALSE |
| 5_v.txt | 32058 | 32058-SNV-( | SNV | 1 | G | T | 13 | 18279 | 0.07111986 | 58.7692308 | WT | FALSE | CAN | JSY13 | 1 | WT-Rep1 | CGA | G>T | A/T+1 | G/C-1 | CG>AT | CG>AT | ATGTGCGAG | GAGATTG | GCT | GCT | C | A | C>A | C>A_GCT | CG>AT_GAGATTG | 9 | FALSE | FALSE |
| 5_v.txt | 32064 | 32064-SNV-( | SNV | 1 | G | T | 17 | 18279 | 0.0930029 | 62.1764706 | WT | FALSE | CAN | JSY13 | 1 | WT-Rep1 | TGA | G>T | A/T+1 | G/C-1 | CG>AT | CG>AT | GAGATTGAG | GAGATAA | ACT | ACT | C | A | C>A | C>A_ACT | CG>AT_GAGATAA | 9 | FALSE | FALSE |
| 5_v.txt | 32080 | 32080-SNV-( | SNV | 1 | C | A | 3 | 18279 | 0.01641228 | 64 | WT | FALSE | CAN | JSY13 | 1 | WT-Rep1 | GCA | C>A | A/T+1 | G/C-1 | CG>AT | CG>AT | ACCATGCAA | CAAAAAA | CGT | GCA | C | T | C>A | C>A_GCA | CG>AT_CAAAAAA | 9 | FALSE | FALSE |
| 5_v.txt | 32088 | 32088-SNV-( | SNV | 1 | C | A | 41 | 18278 | 0.22431338 | 59.6829268 | WT | FALSE | CAN | JSY13 | 1 | WT-Rep1 | GCC | C>A | A/T+1 | G/C-1 | CG>AT | CG>AT | AAAAAGCCT | CCTGCAA | CGG | GCC | C | T | C>A | C>A_GCC | CG>AT_CCTGCAA | 9 | FALSE | FALSE |
| 5_v.txt | 32089 | 32089-SNV-( | SNV | 1 | C | A | 10 | 18278 | 0.05471058 | 48.3025863 | WT | FALSE | CAN | JSY13 | 1 | WT-Rep1 | CCT | C>A | A/T+1 | G/C-1 | CG>AT | CG>AT | AAAAGCCTG | CTGCAAC | GGA | CCT | C | T | C>A | C>A_CCT | CG>AT_CTGCAAC | 9 | FALSE | FALSE |
| 5_v.txt | 32095 | 32095-SNV-( | SNV | 1 | C | A | 112 | 18279 | 0.61272499 | 60.0982143 | WT | FALSE | CAN | JSY13 | 1 | WT-Rep1 | ACA | C>A | A/T+1 | G/C-1 | CG>AT | CG>AT | CTGCAACAC | CACCAGT | TGT | ACA | C | T | C>A | C>A_ACA | CG>AT_CACCAGT | 9 | FALSE | FALSE |
| 5_v.txt | 32119 | 32119-SNV-( | SNV | 1 | C | A | 30 | 18279 | 0.16412276 | 58.1666667 | WT | FALSE | CAN | JSY13 | 1 | WT-Rep1 | TCG | C>A | A/T+1 | G/C-1 | CG>AT | CG>AT | GCCATTCGA | CGAAAAC | AGC | TCG | C | T | C>A | C>A_TCG | CG>AT_CGAAAAC | 9 | FALSE | FALSE |
| 5_v.txt | 32149 | 32149-SNV-( | SNV | 1 | C | A | 62 | 18279 | 0.33918705 | 51.9193548 | WT | FALSE | CAN | JSY13 | 1 | WT-Rep1 | TCC | C>A | A/T+1 | G/C-1 | CG>AT | CG>AT | ATGTCTCCAT | CCATGTA | AGG | TCC | C | T | C>A | C>A_TCC | CG>AT_CCATGTA | 9 | FALSE | FALSE |
| 5_v.txt | 32188 | 32188-SNV-( | SNV | 1 | C | A | 30 | 18279 | 0.16412276 | 43.7333333 | WT | FALSE | CAN | JSY13 | 1 | WT-Rep1 | ACT | C>A | A/T+1 | G/C-1 | CG>AT | CG>AT | CGAAAACTG | CTGCAAT | TGA | ACT | C | T | C>A | C>A_ACT | CG>AT_CTGCAAT | 9 | FALSE | FALSE |
| 5_v.txt | 32199 | 32199-SNV-( | SNV | 1 | G | T | 12 | 18270 | 0.06568145 | 44.1666667 | WT | FALSE | CAN | JSY13 | 1 | WT-Rep1 | TGG | G>T | A/T+1 | G/C-1 | CG>AT | CG>AT | ATGTATGGA | GGAACAC | ACC | ACC | C | A | C>A | C>A_ACC | CG>AT_GGAACAC | 9 | FALSE | FALSE |
| 5_v.txt | 32203 | 32203-SNV-( | SNV | 1 | C | A | 34 | 18270 | 0.18609743 | 45.3529412 | WT | FALSE | CAN | JSY13 | 1 | WT-Rep1 | ACA | C>A | A/T+1 | G/C-1 | CG>AT | CG>AT | ATGGAACAC | CACCACC | TGT | ACA | C | T | C>A | C>A_ACA | CG>AT_CACCACC | 9 | FALSE | FALSE |
| 5_v.txt | 32223 | 32223-SNV-( | SNV | 1 | G | T | 19 | 18270 | 0.10399562 | 40.7894737 | WT | FALSE | CAN | JSY13 | 1 | WT-Rep1 | TGA | G>T | A/T+1 | G/C-1 | CG>AT | CG>AT | GTCCTTGACA | GACAGGA | ACT | ACT | C | A | C>A | C>A_ACT | CG>AT_GACAGGA | 9 | FALSE | FALSE |
| 5_v.txt | 32246 | 32246-SNV-( | SNV | 1 | G | T | 20 | 18266 | 0.10949305 | 36.45 | WT | FALSE | CAN | JSY13 | 1 | WT-Rep1 | TGT | G>T | A/T+1 | G/C-1 | CG>AT | CG>AT | CAACTTGTTC | GTTCTTT | ACA | ACA | C | A | C>A | C>A_ACA | CG>AT_GTTCTTT | 9 | FALSE | FALSE |
| 5_v.txt | 32271 | 32271-SNV-( | SNV | 1 | C | A | 14 | 21420 | 0.06535948 | 37.5714286 | WT | FALSE | CAN | JSY13 | 1 | WT-Rep1 | ACG | C>A | A/T+1 | G/C-1 | CG>AT | CG>AT | AAAATACGG | CGGGAAC | TGC | ACG | C | T | C>A | C>A_ACG | CG>AT_CGGGAAC | 9 | FALSE | FALSE |
| 5_v.txt | 32272 | 32272-SNV-( | SNV | 1 | G | T | 32 | 21420 | 0.14939309 | 34.875 | WT | FALSE | CAN | JSY13 | 1 | WT-Rep1 | CGG | G>T | A/T+1 | G/C-1 | CG>AT | CG>AT | AAATACGGG | GGGAACC | GCC | GCC | C | A | C>A | C>A_GCC | CG>AT_GGGAACC | 9 | FALSE | FALSE |
| 5_v.txt | 32274 | 32274-SNV-( | SNV | 1 | G | T | 31 | 21420 | 0.14472456 | 35.5810971 | WT | FALSE | CAN | JSY13 | 1 | WT-Rep1 | GGA | G>T | A/T+1 | G/C-1 | CG>AT | CG>AT | ATACGGGAA | GAACCAA | CCT | CCT | C | A | C>A | C>A_CCT | CG>AT_GAACCAA | 9 | FALSE | FALSE |
| 5_v.txt | 32282 | 32282-SNV-( | SNV | 1 | G | T | 159 | 21420 | 0.74229692 | 45.3936161 | WT | FALSE | CAN | JSY13 | 1 | WT-Rep1 | CGT | G>T | A/T+1 | G/C-1 | CG>AT | CG>AT | ACCAACGTA | GTAAATA | GCA | GCA | C | A | C>A | C>A_GCA | CG>AT_GTAAATA | 9 | FALSE | FALSE |
| 5_v.txt | 32292 | 32292-SNV-( | SNV | 1 | G | T | 31 | 21421 | 0.1447178 | 48.8709677 | WT | FALSE | CAN | JSY13 | 1 | WT-Rep1 | TGA | G>T | A/T+1 | G/C-1 | CG>AT | CG>AT | ATATTTGAAT | GAATTTG | ACT | ACT | C | A | C>A | C>A_ACT | CG>AT_GAATTTG | 9 | FALSE | FALSE |
| 5_v.txt | 32301 | 32301-SNV-( | SNV | 1 | G | T | 21 | 21421 | 0.09803464 | 50.7619048 | WT | FALSE | CAN | JSY13 | 1 | WT-Rep1 | GGC | G>T | A/T+1 | G/C-1 | CG>AT | CG>AT | TTTGCGGCA | GCAGAAA | CCG | CCG | C | A | C>A | C>A_CCG | CG>AT_GCAGAAA | 9 | FALSE | FALSE |
| 5_v.txt | 32304 | 32304-SNV-( | SNV | 1 | G | T | 17 | 21421 | 0.07936137 | 46.7058824 | WT | FALSE | CAN | JSY13 | 1 | WT-Rep1 | AGA | G>T | A/T+1 | G/C-1 | CG>AT | CG>AT | GCGGCAGAA | GAAATAA | TCT | TCT | C | A | C>A | C>A_TCT | CG>AT_GAAATAA | 9 | FALSE | FALSE |
| 5_v.txt | 32313 | 32313-SNV-( | SNV | 1 | G | T | 1 | 21421 | 4.67E-03 | 63.9998094 | WT | FALSE | CAN | JSY13 | 1 | WT-Rep1 | GGT | G>T | A/T+1 | G/C-1 | CG>AT | CG>AT | ATAATGGTT | GTTGTTA | CCA | CCA | C | A | C>A | C>A_CCA | CG>AT_GTTGTTA | 9 | FALSE | FALSE |
| 5_v.txt | 32368 | 32368-SNV-( | SNV | 1 | C | A | 31 | 21421 | 0.1447178 | 57.7096774 | WT | FALSE | CAN | JSY13 | 1 | WT-Rep1 | TCA | C>A | A/T+1 | G/C-1 | CG>AT | CG>AT | AGTTCTCAAT | CAATAGC | AGT | TCA | C | T | C>A | C>A_TCA | CG>AT_CAATAGC | 9 | FALSE | FALSE |
| 5_v.txt | 32373 | 32373-SNV-( | SNV | 1 | G | T | 23 | 21421 | 0.10737127 | 61.9130435 | WT | FALSE | CAN | JSY13 | 1 | WT-Rep1 | AGC | G>T | A/T+1 | G/C-1 | CG>AT | CG>AT | TCAATAGCA | GCAATAA | TCG | TCG | C | A | C>A | C>A_TCG | CG>AT_GCAATAA | 9 | FALSE | FALSE |
| 5_v.txt | 32399 | 32399-SNV-( | SNV | 1 | G | T | 197 | 21421 | 0.91965828 | 58.7411168 | WT | FALSE | CAN | JSY13 | 1 | WT-Rep1 | CGT | G>T | A/T+1 | G/C-1 | CG>AT | CG>AT | AGAAACGTA | GTAGGAA | GCA | GCA | C | A | C>A | C>A_GCA | CG>AT_GTAGGAA | 9 | FALSE | FALSE |
| 5_v.txt | 32432 | 32432-SNV-( | SNV | 1 | G | T | 76 | 21421 | 0.35479203 | 56.5921053 | WT | FALSE | CAN | JSY13 | 1 | WT-Rep1 | TGT | G>T | A/T+1 | G/C-1 | CG>AT | CG>AT | GTCATTGTAT | GTATGGA | ACA | ACA | C | A | C>A | C>A_ACA | CG>AT_GTATGGA | 9 | FALSE | FALSE |
| 5_v.txt | 32449 | 32449-SNV-( | SNV | 1 | C | A | 10 | 21421 | 0.04668316 | 42.5 | WT | FALSE | CAN | JSY13 | 1 | WT-Rep1 | CCA | C>A | A/T+1 | G/C-1 | CG>AT | CG>AT | AAAGTCCAAT | CAATGAA | GGT | CCA | C | T | C>A | C>A_CCA | CG>AT_CAATGAA | 9 | FALSE | FALSE |
| 5_v.txt | 32463 | 32463-SNV-( | SNV | 1 | G | T | 19 | 21421 | 0.08869801 | 50.1052632 | WT | FALSE | CAN | JSY13 | 1 | WT-Rep1 | AGA | G>T | A/T+1 | G/C-1 | CG>AT | CG>AT | AATAGAGAG | GAGCCAA | TCT | TCT | C | A | C>A | C>A_TCT | CG>AT_GAGCCAA | 9 | FALSE | FALSE |
| 5_v.txt | 32471 | 32471-SNV-( | SNV | 1 | G | T | 24 | 21421 | 0.11203959 | 46.8751866 | WT | FALSE | CAN | JSY13 | 1 | WT-Rep1 | TGT | G>T | A/T+1 | G/C-1 | CG>AT | CG>AT | GCCAATGTA | GTAGAAG | ACA | ACA | C | A | C>A | C>A_ACA | CG>AT_GTAGAAG | 9 | FALSE | FALSE |
| 5_v.txt | 32474 | 32474-SNV-( | SNV | 1 | G | T | 19 | 21421 | 0.08869801 | 51.5818194 | WT | FALSE | CAN | JSY13 | 1 | WT-Rep1 | AGA | G>T | A/T+1 | G/C-1 | CG>AT | CG>AT | AATGTAGAA | GAAGGTT | TCT | TCT | C | A | C>A | C>A_TCT | CG>AT_GAAGGTT | 9 | FALSE | FALSE |
| 5_v.txt | 32487 | 32487-SNV-( | SNV | 1 | C | A | 45 | 21421 | 0.21007423 | 46.1555556 | WT | FALSE | CAN | JSY13 | 1 | WT-Rep1 | ACG | C>A | A/T+1 | G/C-1 | CG>AT | CG>AT | AAGATACGG | CGGAAAA | TGC | ACG | C | T | C>A | C>A_ACG | CG>AT_CGGAAAA | 9 | FALSE | FALSE |
| 5_v.txt | 32488 | 32488-SNV-( | SNV | 1 | G | T | 47 | 21421 | 0.21941086 | 44.531707 | WT | FALSE | CAN | JSY13 | 1 | WT-Rep1 | CGG | G>T | A/T+1 | G/C-1 | CG>AT | CG>AT | AGATACGGA | GGAAAAC | GCC | GCC | C | A | C>A | C>A_GCC | CG>AT_GGAAAAC | 9 | FALSE | FALSE |
| 5_v.txt | 32494 | 32494-SNV-( | SNV | 1 | C | A | 70 | 21421 | 0.32678213 | 45.2556714 | WT | FALSE | CAN | JSY13 | 1 | WT-Rep1 | ACA | C>A | A/T+1 | G/C-1 | CG>AT | CG>AT | GGAAAACAA | CAACTTT | TGT | ACA | C | T | C>A | C>A_ACA | CG>AT_CAACTTT | 9 | FALSE | FALSE |
| 5_v.txt | 32508 | 32508-SNV-( | SNV | 1 | G | T | 12 | 21421 | 0.05601979 | 41.4166667 | WT | FALSE | CAN | JSY13 | 1 | WT-Rep1 | GGC | G>T | A/T+1 | G/C-1 | CG>AT | CG>AT | TTGATGGCT | GCTCTTG | CCG | CCG | C | A | C>A | C>A_CCG | CG>AT_GCTCTTG | 9 | FALSE | FALSE |
| 5_v.txt | 32529 | 32529-SNV-( | SNV | 1 | G | T | 7 | 21421 | 0.03267821 | 49 | WT | FALSE | CAN | JSY13 | 1 | WT-Rep1 | GGG | G>T | A/T+1 | G/C-1 | CG>AT | CG>AT | TTTCTGGGG | GGGTTTG | CCC | CCC | C | A | C>A | C>A_CCC | CG>AT_GGGTTTG | 9 | FALSE | FALSE |
| 5_v.txt | 32530 | 32530-SNV-( | SNV | 1 | G | T | 54 | 21421 | 0.25208907 | 36.7592593 | WT | FALSE | CAN | JSY13 | 1 | WT-Rep1 | GGG | G>T | A/T+1 | G/C-1 | CG>AT | CG>AT | TTCTGGGGT | GGTTTGC | CCC | CCC | C | A | C>A | C>A_CCC | CG>AT_GGTTTGC | 9 | FALSE | FALSE |
| 5_v.txt | 32542 | 32542-SNV-( | SNV | 1 | C | A | 76 | 21421 | 0.35479203 | 36.0653762 | WT | FALSE | CAN | JSY13 | 1 | WT-Rep1 | TCA | C>A | A/T+1 | G/C-1 | CG>AT | CG>AT | CAGCTTCACC | CACCAGC | AGT | TCA | C | T | C>A | C>A_TCA | CG>AT_CACCAGC | 9 | FALSE | FALSE |
| 5_v.txt | 32566 | 32566-SNV-( | SNV | 1 | C | A | 35 | 22145 | 0.15804922 | 37.4285714 | WT | FALSE | CAN | JSY13 | 1 | WT-Rep1 | TCA | C>A | A/T+1 | G/C-1 | CG>AT | CG>AT | CTAGTTCAGT | CAGTACC | AGT | TCA | C | T | C>A | C>A_TCA | CG>AT_CAGTACC | 9 | FALSE | FALSE |
| 5_v.txt | 32572 | 32572-SNV-( | SNV | 1 | C | A | 29 | 22153 | 0.13090778 | 36.9313184 | WT | FALSE | CAN | JSY13 | 1 | WT-Rep1 | CCT | C>A | A/T+1 | G/C-1 | CG>AT | CG>AT | CAGTACCTTG | CTTGAAA | GGA | CCT | C | T | C>A | C>A_CCT | CG>AT_CTTGAAA | 9 | FALSE | FALSE |
| 5_v.txt | 32575 | 32575-SNV-( | SNV | 1 | G | T | 87 | 22153 | 0.39272333 | 36.7816092 | WT | FALSE | CAN | JSY13 | 1 | WT-Rep1 | TGA | G>T | A/T+1 | G/C-1 | CG>AT | CG>AT | TACCTTGAA | GAAATGT | ACT | ACT | C | A | C>A | C>A_ACT | CG>AT_GAAATGT | 9 | FALSE | FALSE |
| 5_v.txt | 32629 | 32629-SNV-( | SNV | 1 | C | A | 151 | 22156 | 0.68153096 | 52.9521823 | WT | FALSE | CAN | JSY13 | 1 | WT-Rep1 | TCG | C>A | A/T+1 | G/C-1 | CG>AT | CG>AT | TCCCTTCGTT | CGTTTTT | AGC | TCG | C | T | C>A | C>A_TCG | CG>AT_CGTTTTT | 9 | FALSE | FALSE |
| 5_v.txt | 32630 | 32630-SNV-( | SNV | 1 | G | T | 72 | 22156 | 0.32496841 | 49.9444495 | WT | FALSE | CAN | JSY13 | 1 | WT-Rep1 | CGT | G>T | A/T+1 | G/C-1 | CG>AT | CG>AT | CCCTTCGTTT | GTTTTTA | GCA | GCA | C | A | C>A | C>A_GCA | CG>AT_GTTTTTA | 9 | FALSE | FALSE |
| 5_v.txt | 32692 | 32692-SNV-( | SNV | 1 | C | A | 84 | 22156 | 0.37912981 | 58.7857143 | WT | FALSE | CAN | JSY13 | 1 | WT-Rep1 | CCA | C>A | A/T+1 | G/C-1 | CG>AT | CG>AT | GGAATCCAA | CAACTGG | GGT | CCA | C | T | C>A | C>A_CCA | CG>AT_CAACTGG | 9 | FALSE | FALSE |
| 5_v.txt | 32788 | 32788-SNV-( | SNV | 1 | C | A | 66 | 22176 | 0.29761905 | 46.3981145 | WT | FALSE | CAN | JSY13 | 1 | WT-Rep1 | TCG | C>A | A/T+1 | G/C-1 | CG>AT | CG>AT | AGAACTCGA | CGAATTC | AGC | TCG | C | T | C>A | C>A_TCG | CG>AT_CGAATTC | 9 | FALSE | FALSE |
| 5_v.txt | 32794 | 32794-SNV-( | SNV | 1 | C | A | 32 | 22177 | 0.14429364 | 38.34375 | WT | FALSE | CAN | JSY13 | 1 | WT-Rep1 | TCA | C>A | A/T+1 | G/C-1 | CG>AT | CG>AT | CGAATTCACC | CACCGTA | AGT | TCA | C | T | C>A | C>A_TCA | CG>AT_CACCGTA | 9 | FALSE | FALSE |
| 5_v.txt | 32797 | 32797-SNV-( | SNV | 1 | C | A | 19 | 22177 | 0.08567435 | 39.9473684 | WT | FALSE | CAN | JSY13 | 1 | WT-Rep1 | CCG | C>A | A/T+1 | G/C-1 | CG>AT | CG>AT | ATTCACCGTA | CGTAATA | GGC | CCG | C | T | C>A | C>A_CCG | CG>AT_CGTAATA | 9 | FALSE | FALSE |
| 5_v.txt | 32798 | 32798-SNV-( | SNV | 1 | G | T | 31 | 22177 | 0.13978446 | 42.3548387 | WT | FALSE | CAN | JSY13 | 1 | WT-Rep1 | CGT | G>T | A/T+1 | G/C-1 | CG>AT | CG>AT | TTCACCGTAA | GTAATAT | GCA | GCA | C | A | C>A | C>A_GCA | CG>AT_GTAATAT | 9 | FALSE | FALSE |
| 5_v.txt | 32819 | 32819-SNV-( | SNV | 1 | G | T | 44 | 22177 | 0.19840375 | 41.5 | WT | FALSE | CAN | JSY13 | 1 | WT-Rep1 | AGT | G>T | A/T+1 | G/C-1 | CG>AT | CG>AT | GAACAAGTT | GTTCATT | TCA | TCA | C | A | C>A | C>A_TCA | CG>AT_GTTCATT | 9 | FALSE | FALSE |
| 5_v.txt | 32855 | 32855-SNV-( | SNV | 1 | C | A | 7 | 22175 | 0.03156708 | 36.4286384 | WT | FALSE | CAN | JSY13 | 1 | WT-Rep1 | TCC | C>A | A/T+1 | G/C-1 | CG>AT | CG>AT | ACTAATCCAT | CCATGCC | AGG | TCC | C | T | C>A | C>A_TCC | CG>AT_CCATGCC | 9 | FALSE | FALSE |
| 5_v.txt | 32876 | 32876-SNV-( | SNV | 1 | G | T | 81 | 24300 | 0.33333333 | 35.9260967 | WT | FALSE | CAN | JSY13 | 1 | WT-Rep1 | TGT | G>T | A/T+1 | G/C-1 | CG>AT | CG>AT | AACTTTGTAC | GTACGTC | ACA | ACA | C | A | C>A | C>A_ACA | CG>AT_GTACGTC | 9 | FALSE | FALSE |
| 5_v.txt | 32917 | 32917-SNV-( | SNV | 1 | C | A | 81 | 24305 | 0.33326476 | 53.8271605 | WT | FALSE | CAN | JSY13 | 1 | WT-Rep1 | TCC | C>A | A/T+1 | G/C-1 | CG>AT | CG>AT | TAAGTTCCAC | CCAGGGC | AGG | TCC | C | T | C>A | C>A_TCC | CG>AT_CCAGGGC | 9 | FALSE | FALSE |
| 5_v.txt | 32928 | 32928-SNV-( | SNV | 1 | G | T | 23 | 24343 | 0.09448301 | 44.2519507 | WT | FALSE | CAN | JSY13 | 1 | WT-Rep1 | AGT | G>T | A/T+1 | G/C-1 | CG>AT | CG>AT | GCAAAAGTG | GTGATTG | TCA | TCA | C | A | C>A | C>A_TCA | CG>AT_GTGATTG | 9 | FALSE | FALSE |
| 5_v.txt | 32936 | 32936-SNV-( | SNV | 1 | C | A | 42 | 24341 | 0.17254838 | 36.4239099 | WT | FALSE | CAN | JSY13 | 1 | WT-Rep1 | CCC | C>A | A/T+1 | G/C-1 | CG>AT | CG>AT | GATTGCCCA | CCAAGAA | GGG | CCC | C | T | C>A | C>A_CCC | CG>AT_CCAAGAA | 9 | FALSE | FALSE |
| 5_v.txt | 32937 | 32937-SNV-( | SNV | 1 | C | A | 29 | 24343 | 0.11913076 | 52.4137931 | WT | FALSE | CAN | JSY13 | 1 | WT-Rep1 | CCA | C>A | A/T+1 | G/C-1 | CG>AT | CG>AT | ATTGCCCAA | CAAGAAA | GGT | CCA | C | T | C>A | C>A_CCA | CG>AT_CAAGAAA | 9 | FALSE | FALSE |
| 5_v.txt | 32940 | 32940-SNV-( | SNV | 1 | G | T | 42 | 24343 | 0.1725342 | 45.3095238 | WT | FALSE | CAN | JSY13 | 1 | WT-Rep1 | AGA | G>T | A/T+1 | G/C-1 | CG>AT | CG>AT | GCCCAAGAA | GAAAACC | TCT | TCT | C | A | C>A | C>A_TCT | CG>AT_GAAAACC | 9 | FALSE | FALSE |
| 5_v.txt | 32946 | 32946-SNV-( | SNV | 1 | C | A | 14 | 24343 | 0.0575114 | 52.2142444 | WT | FALSE | CAN | JSY13 | 1 | WT-Rep1 | CCA | C>A | A/T+1 | G/C-1 | CG>AT | CG>AT | GAAAACCAA | CAATACA | GGT | CCA | C | T | C>A | C>A_CCA | CG>AT_CAATACA | 9 | FALSE | FALSE |
| 5_v.txt | 32997 | 32997-SNV-( | SNV | 1 | G | T | 59 | 24343 | 0.24236947 | 58.220339 | WT | FALSE | CAN | JSY13 | 1 | WT-Rep1 | TGA | G>T | A/T+1 | G/C-1 | CG>AT | CG>AT | CTTTGTGAG | GAGAAAA | ACT | ACT | C | A | C>A | C>A_ACT | CG>AT_GAGAAAA | 9 | FALSE | FALSE |
| 5_v.txt | 33006 | 33006-SNV-( | SNV | 1 | G | T | 36 | 24343 | 0.14788646 | 60.0555556 | WT | FALSE | CAN | JSY13 | 1 | WT-Rep1 | TGT | G>T | A/T+1 | G/C-1 | CG>AT | CG>AT | AAAACTGTG | GTGAAAG | ACA | ACA | C | A | C>A | C>A_ACA | CG>AT_GTGAAAG | 9 | FALSE | FALSE |
| 5_v.txt | 33015 | 33015-SNV-( | SNV | 1 | G | T | 17 | 24343 | 0.06983527 | 52.2352941 | WT | FALSE | CAN | JSY13 | 1 | WT-Rep1 | GGA | G>T | A/T+1 | G/C-1 | CG>AT | CG>AT | AAAGAGGAT | GATGTAA | CCT | CCT | C | A | C>A | C>A_CCT | CG>AT_GATGTAA | 9 | FALSE | FALSE |
| 5_v.txt | 33024 | 33024-SNV-( | SNV | 1 | G | T | 80 | 24343 | 0.32863657 | 60.125 | WT | FALSE | CAN | JSY13 | 1 | WT-Rep1 | AGG | G>T | A/T+1 | G/C-1 | CG>AT | CG>AT | GTAACAGGG | GGGATGA | TCC | TCC | C | A | C>A | C>A_TCC | CG>AT_GGGATGA | 9 | FALSE | FALSE |
| 5_v.txt | 33043 | 33043-SNV-( | SNV | 1 | C | A | 73 | 24343 | 0.29988087 | 60.3150685 | WT | FALSE | CAN | JSY13 | 1 | WT-Rep1 | TCA | C>A | A/T+1 | G/C-1 | CG>AT | CG>AT | CCATTTCACC | CACCCAA | AGT | TCA | C | T | C>A | C>A_TCA | CG>AT_CACCCAA | 9 | FALSE | FALSE |
| 5_v.txt | 33051 | 33051-SNV-( | SNV | 1 | G | T | 207 | 24343 | 0.85034712 | 57.2318841 | WT | FALSE | CAN | JSY13 | 1 | WT-Rep1 | GGA | G>T | A/T+1 | G/C-1 | CG>AT | CG>AT | CCCAAGGAC | GACTGCG | CCT | CCT | C | A | C>A | C>A_CCT | CG>AT_GACTGCG | 9 | FALSE | FALSE |
| 5_v.txt | 33056 | 33056-SNV-( | SNV | 1 | C | A | 6 | 24343 | 0.02464774 | 41.3269966 | WT | FALSE | CAN | JSY13 | 1 | WT-Rep1 | GCG | C>A | A/T+1 | G/C-1 | CG>AT | CG>AT | GGACTGCGT | CGTGACA | CGC | GCG | C | T | C>A | C>A_GCG | CG>AT_CGTGACA | 9 | FALSE | FALSE |
| 5_v.txt | 33079 | 33079-SNV-( | SNV | 1 | C | A | 8 | 24343 | 0.03286366 | 39.1247653 | WT | FALSE | CAN | JSY13 | 1 | WT-Rep1 | CCC | C>A | A/T+1 | G/C-1 | CG>AT | CG>AT | AAGAACCCA | CCATAAA | GGG | CCC | C | T | C>A | C>A_CCC | CG>AT_CCATAAA | 9 | FALSE | FALSE |
| 5_v.txt | 33093 | 33093-SNV-( | SNV | 1 | G | T | 52 | 24343 | 0.21361377 | 49.5577131 | WT | FALSE | CAN | JSY13 | 1 | WT-Rep1 | TGA | G>T | A/T+1 | G/C-1 | CG>AT | CG>AT | AAATATGAT | GATATAA | ACT | ACT | C | A | C>A | C>A_ACT | CG>AT_GATATAA | 9 | FALSE | FALSE |
| 5_v.txt | 33111 | 33111-SNV-( | SNV | 1 | G | T | 6 | 24343 | 0.02464774 | 46 | WT | FALSE | CAN | JSY13 | 1 | WT-Rep1 | TGG | G>T | A/T+1 | G/C-1 | CG>AT | CG>AT | CCCACTGGG | GGGCCGG | ACC | ACC | C | A | C>A | C>A_ACC | CG>AT_GGGCCGG | 9 | FALSE | FALSE |
| 5_v.txt | 33112 | 33112-SNV-( | SNV | 1 | G | T | 19 | 24343 | 0.07805119 | 45.8327382 | WT | FALSE | CAN | JSY13 | 1 | WT-Rep1 | GGG | G>T | A/T+1 | G/C-1 | CG>AT | CG>AT | CCACTGGGC | GGCCGGC | CCC | CCC | C | A | C>A | C>A_CCC | CG>AT_GGCCGGC | 9 | FALSE | FALSE |

| | | | | | | | | | | | | | | | | | | | | | | | | | | | | | | | | | | |
|---|---|---|---|---|---|---|---|---|---|---|---|---|---|---|---|---|---|---|---|---|---|---|---|---|---|---|---|---|---|---|---|---|---|---|
| 5_v.txt | 33115 | 33115-SNV-( | SNV | 1 | C | A | 48 | 24343 | 0.19718194 | 42.6875 | WT | FALSE | CAN | JSY13 | 1 | WT-Rep1 | CCG | C>A | A/T+1 | G/C-1 | CG>AT | CG>AT | CTGGGCCGG | CGGCGTT | GGC | CCG | C | T | C>A | C>A_CCG | CG>AT_CGGCGTT | 9 | FALSE | FALSE |
| 5_v.txt | 33151 | 33151-SNV-( | SNV | 1 | G | T | 82 | 24343 | 0.33685248 | 40.1828994 | WT | FALSE | CAN | JSY13 | 1 | WT-Rep1 | AGA | G>T | A/T+1 | G/C-1 | CG>AT | CG>AT | TGAAAAGAC | GACCTGT | TCT | TCT | C | A | C>A | C>A_TCT | CG>AT_GACCTGT | 9 | FALSE | FALSE |
| 5_v.txt | 33154 | 33154-SNV-( | SNV | 1 | C | A | 32 | 24343 | 0.13145463 | 36.9063624 | WT | FALSE | CAN | JSY13 | 1 | WT-Rep1 | CCT | C>A | A/T+1 | G/C-1 | CG>AT | CG>AT | AAAGACCTG | CTGTACC | GGA | CCT | C | T | C>A | C>A_CCT | CG>AT_CTGTACC | 9 | FALSE | FALSE |
| 5_v.txt | 33160 | 33160-SNV-( | SNV | 1 | C | A | 32 | 24341 | 0.13146543 | 36.1875 | WT | FALSE | CAN | JSY13 | 1 | WT-Rep1 | CCA | C>A | A/T+1 | G/C-1 | CG>AT | CG>AT | CTGTACCAAT | CAATAGT | GGT | CCA | C | T | C>A | C>A_CCA | CG>AT_CAATAGT | 9 | FALSE | FALSE |
| 5_v.txt | 33168 | 33168-SNV-( | SNV | 1 | C | A | 6 | 24300 | 0.02469136 | 29.4999995 | WT | FALSE | CAN | JSY13 | 1 | WT-Rep1 | ACC | C>A | A/T+1 | G/C-1 | CG>AT | CG>AT | ATAGTACCA( | CCACCAA | TGG | ACC | C | T | C>A | C>A_ACC | CG>AT_CCACCAA | 9 | FALSE | FALSE |
| 5_v.txt | 33169 | 33169-SNV-( | SNV | 1 | C | A | 16 | 24300 | 0.06584362 | 37.7499319 | WT | FALSE | CAN | JSY13 | 1 | WT-Rep1 | CCA | C>A | A/T+1 | G/C-1 | CG>AT | CG>AT | TAGTACCAC( | CACCAAG | GGT | CCA | C | T | C>A | C>A_CCA | CG>AT_CACCAAG | 9 | FALSE | FALSE |
| 5_v.txt | 33172 | 33172-SNV-( | SNV | 1 | C | A | 6 | 24300 | 0.02469136 | 33.5 | WT | FALSE | CAN | JSY13 | 1 | WT-Rep1 | CCA | C>A | A/T+1 | G/C-1 | CG>AT | CG>AT | TACCACCAA( | CAAGGGC | GGT | CCA | C | T | C>A | C>A_CCA | CG>AT_CAAGGGC | 9 | FALSE | FALSE |
| 5_v.txt | 33175 | 33175-SNV-( | SNV | 1 | G | T | 7 | 24298 | 0.02880896 | 37.7142857 | WT | FALSE | CAN | JSY13 | 1 | WT-Rep1 | AGG | G>T | A/T+1 | G/C-1 | CG>AT | CG>AT | CACCAAGGG | GGGCAAT | TCC | TCC | C | A | C>A | C>A_TCC | CG>AT_GGGCAAT | 9 | FALSE | FALSE |
| 5_v.txt | 33177 | 33177-SNV-( | SNV | 1 | G | T | 43 | 24298 | 0.1769693 | 36.9534884 | WT | FALSE | CAN | JSY13 | 1 | WT-Rep1 | GGC | G>T | A/T+1 | G/C-1 | CG>AT | CG>AT | CCAAGGGCA | GCAATCA | CCG | CCG | C | A | C>A | C>A_CCG | CG>AT_GCAATCA | 9 | FALSE | FALSE |
| 5_v.txt | 33186 | 33186-SNV-( | SNV | 1 | C | A | 20 | 7277 | 0.27483853 | 36.75 | WT | FALSE | CAN | JSY13 | 1 | WT-Rep1 | ACC | C>A | A/T+1 | G/C-1 | CG>AT | CG>AT | ATCATACCA/ | CCAATAT | TGG | ACC | C | T | C>A | C>A_ACC | CG>AT_CCAATAT | 9 | FALSE | FALSE |
| 5_v.txt | 33235 | 33235-SNV-( | SNV | 1 | C | A | 23 | 7277 | 0.31606431 | 33.0434783 | WT | FALSE | CAN | JSY13 | 1 | WT-Rep1 | TCT | C>A | A/T+1 | G/C-1 | CG>AT | CG>AT | GTACTTCTCC | CTCCTTC | AGA | TCT | C | T | C>A | C>A_TCT | CG>AT_CTCCTTC | 9 | FALSE | FALSE |
| 5_v.txt | 33241 | 33241-SNV-( | SNV | 1 | C | A | 15 | 7277 | 0.2061289 | 42.8666667 | WT | FALSE | CAN | JSY13 | 1 | WT-Rep1 | TCA | C>A | A/T+1 | G/C-1 | CG>AT | CG>AT | CTCCTTCATC | CATCTTC | AGT | TCA | C | T | C>A | C>A_TCA | CG>AT_CATCTTC | 9 | FALSE | FALSE |
| 5_v.txt | 33247 | 33247-SNV-( | SNV | 1 | C | A | 14 | 7277 | 0.19238697 | 33.5 | WT | FALSE | CAN | JSY13 | 1 | WT-Rep1 | TCA | C>A | A/T+1 | G/C-1 | CG>AT | CG>AT | CATCTTCATC | CATCACC | AGT | TCA | C | T | C>A | C>A_TCA | CG>AT_CATCACC | 9 | FALSE | FALSE |
| 5_v.txt | 33283 | 33283-SNV-( | SNV | 1 | C | A | 13 | 7277 | 0.17864505 | 37.6941 | WT | FALSE | CAN | JSY13 | 1 | WT-Rep1 | TCG | C>A | A/T+1 | G/C-1 | CG>AT | CG>AT | TATCCTCGCC | CGCCATT | AGC | TCG | C | T | C>A | C>A_TCG | CG>AT_CGCCATT | 9 | FALSE | FALSE |
| 5_v.txt | 33284 | 33284-SNV-( | SNV | 1 | G | T | 1 | 7276 | 0.01374382 | 62.999584 | WT | FALSE | CAN | JSY13 | 1 | WT-Rep1 | CGC | G>T | A/T+1 | G/C-1 | CG>AT | CG>AT | ATCCTCGCCA | GCCATTT | GCG | GCG | C | A | C>A | C>A_GCG | CG>AT_GCCATTT | 9 | FALSE | FALSE |
| 5_v.txt | 33285 | 33285-SNV-( | SNV | 1 | C | A | 5 | 7277 | 0.06870963 | 40.9977289 | WT | FALSE | CAN | JSY13 | 1 | WT-Rep1 | GCC | C>A | A/T+1 | G/C-1 | CG>AT | CG>AT | TCCTCGCCAT | CCATTTA | CGG | GCC | C | T | C>A | C>A_GCC | CG>AT_CCATTTA | 9 | FALSE | FALSE |
| 5_v.txt | 33286 | 33286-SNV-( | SNV | 1 | C | A | 1 | 7276 | 0.01374382 | 63.9996307 | WT | FALSE | CAN | JSY13 | 1 | WT-Rep1 | CCA | C>A | A/T+1 | G/C-1 | CG>AT | CG>AT | CCTCGCCATT | CATTTAC | GGT | CCA | C | T | C>A | C>A_CCA | CG>AT_CATTTAC | 9 | FALSE | FALSE |
| 5_v.txt | 33325 | 33325-SNV-( | SNV | 1 | C | A | 18 | 7277 | 0.24735468 | 54.0555556 | WT | FALSE | CAN | JSY13 | 1 | WT-Rep1 | TCT | C>A | A/T+1 | G/C-1 | CG>AT | CG>AT | ACAATTCTTT | CTTTACT | AGA | TCT | C | T | C>A | C>A_TCT | CG>AT_CTTTACT | 9 | FALSE | FALSE |
| 5_v.txt | 33337 | 33337-SNV-( | SNV | 1 | C | A | 9 | 7277 | 0.12367734 | 52.2222222 | WT | FALSE | CAN | JSY13 | 1 | WT-Rep1 | TCA | C>A | A/T+1 | G/C-1 | CG>AT | CG>AT | TTTTCTCATC | CATCTTT | AGT | TCA | C | T | C>A | C>A_TCA | CG>AT_CATCTTT | 9 | FALSE | FALSE |
| 5_v.txt | 33354 | 33354-SNV-( | SNV | 1 | G | T | 39 | 7277 | 0.53593514 | 28.7179487 | WT | FALSE | CAN | JSY13 | 1 | WT-Rep1 | TGA | G>T | A/T+1 | G/C-1 | CG>AT | CG>AT | GGTATTGAC( | GACCCAC | ACT | ACT | C | A | C>A | C>A_ACT | CG>AT_GACCCAC | 9 | FALSE | FALSE |
| 5_v.txt | 33385 | 33385-SNV-( | SNV | 1 | C | A | 10 | 7277 | 0.13741927 | 47.7 | WT | FALSE | CAN | JSY13 | 1 | WT-Rep1 | TCA | C>A | A/T+1 | G/C-1 | CG>AT | CG>AT | AAGCTTCAA( | CAACGTC | AGT | TCA | C | T | C>A | C>A_TCA | CG>AT_CAACGTC | 9 | FALSE | FALSE |
| 5_v.txt | 33419 | 33419-SNV-( | SNV | 1 | G | T | 104 | 7277 | 1.42916037 | 47.6730769 | WT | FALSE | CAN | JSY13 | 1 | WT-Rep1 | TGT | G>T | A/T+1 | G/C-1 | CG>AT | CG>AT | CTCATTGTAC | GTACATA | ACA | ACA | C | A | C>A | C>A_ACA | CG>AT_GTACATA | 9 | FALSE | FALSE |
| 5_v.txt | 33474 | 33474-SNV-( | SNV | 1 | C | A | 14 | 7275 | 0.19243986 | 33.7857143 | WT | FALSE | CAN | JSY13 | 1 | WT-Rep1 | GCC | C>A | A/T+1 | G/C-1 | CG>AT | CG>AT | GCTATGCCTT | CCTTTTT | CGG | GCC | C | T | C>A | C>A_GCC | CG>AT_CCTTTTT | 9 | FALSE | FALSE |
| 5_v.txt | 31885 | 31885-Repla | Replacement | 2 | C | AA | 19 | 9499 | 0.20002106 | 44.7629363 | WT | FALSE | CAN | JSY13 | 1 | WT-Rep1 | GCA | nope | >1bp-ins | G/C-1 | none | Replac. | CGGCAGCAA | CAAAGCT | CGT | GCA | C | none | nope | nope_GCA | Replac._CAAAGCT | 9 | FALSE | FALSE |
| 5_v.txt | 31972 | 31972-Repla | Replacement | 2 | CC | T | 24 | 18279 | 0.13129821 | 39.3333256 | WT | FALSE | CAN | JSY13 | 1 | WT-Rep1 | GCC | nope | A/T+1 | >1bpdel | none | Replac. | AATAAGCCA | CCAAGCC | CGG | GCC | CC | A | NA | NA_GCC | Replac._CCAAGCC | 9 | FALSE | FALSE |
| 5_v.txt | 32080 | 32080-Repla | Replacement | 2 | C | AA | 9 | 18279 | 0.04923683 | 64 | WT | FALSE | CAN | JSY13 | 1 | WT-Rep1 | GCA | nope | >1bp-ins | G/C-1 | none | Replac. | ACCATGCAA/ | CAAAAAA | CGT | GCA | C | none | nope | nope_GCA | Replac._CAAAAAA | 9 | FALSE | FALSE |
| 5_v.txt | 32498 | 32498-Repla | Replacement | 3 | T | AAC | 1 | 21421 | 4.67E-03 | 38 | WT | FALSE | CAN | JSY13 | 1 | WT-Rep1 | CTT | nope | >1bp-ins | A/T-1 | none | Replac. | AACAACTTTT | TTTTTTG | GAA | CTT | T | none | nope | nope_CTT | Replac._TTTTTTG | 9 | FALSE | FALSE |
| 5_v.txt | 32526 | 32526-Repla | Replacement | 2 | CT | A | 20 | 21421 | 0.09336632 | 38.9496539 | WT | FALSE | CAN | JSY13 | 1 | WT-Rep1 | TCT | nope | A/T+1 | >1bpdel | none | Replac. | GATTTTCTGG | CTGGGGT | AGA | TCT | CT | T | NA | NA_TCT | Replac._CTGGGGT | 9 | FALSE | FALSE |
| 5_v.txt | 32934 | 32934-Repla | Replacement | 2 | GC | T | 2 | 24343 | 8.22E-03 | 49.25 | WT | FALSE | CAN | JSY13 | 1 | WT-Rep1 | TGC | nope | A/T+1 | >1bpdel | none | Replac. | GTGATTGCC( | GCCCAAG | ACG | TGC | GC | A | NA | NA_TGC | Replac._GCCCAAG | 9 | FALSE | FALSE |
| 5_v.txt | 33168 | 33168-Repla | Replacement | 3 | CCA | TT | 20 | 24300 | 0.08230453 | 37.65 | WT | FALSE | CAN | JSY13 | 1 | WT-Rep1 | ACC | nope | >1bp-ins | >1bpdel | none | Replac. | ATAGTACCA( | CCACCAA | TGG | ACC | CCA | none | NA | NA_ACC | Replac._CCACCAA | 9 | FALSE | FALSE |
| 5_v.txt | 32274 | 32274-MNV- | MNV | 2 | GA | TG | 1 | 21420 | 4.67E-03 | 36.4999899 | WT | FALSE | CAN | JSY13 | 1 | WT-Rep1 | GGA | nope | >1bp-ins | >1bpdel | none | MNV | ATACGGGAA | GAACCAA | CCT | GGA | GA | none | NA | NA_GGA | MNV_GAACCAA | 9 | FALSE | FALSE |
| 5_v.txt | 32413 | 32413-MNV- | MNV | 2 | GT | AG | 69 | 21421 | 0.32211381 | 58.1159055 | WT | FALSE | CAN | JSY13 | 1 | WT-Rep1 | TGT | nope | >1bp-ins | >1bpdel | none | MNV | TAGATTGTG | GTGTTAG | ACA | TGT | GT | none | NA | NA_TGT | MNV_GTGTTAG | 9 | FALSE | FALSE |
| 5_v.txt | 32541 | 32541-MNV- | MNV | 2 | TC | GA | 1 | 21421 | 4.67E-03 | 36.4999998 | WT | FALSE | CAN | JSY13 | 1 | WT-Rep1 | TTC | nope | >1bp-ins | >1bpdel | none | MNV | GCAGCTTCA( | TCACCAG | AAG | TTC | TC | none | NA | NA_TTC | MNV_TCACCAG | 9 | FALSE | FALSE |
| 5_v.txt | 32937 | 32937-MNV- | MNV | 2 | CA | AG | 25 | 24343 | 0.10269893 | 53.659938 | WT | FALSE | CAN | JSY13 | 1 | WT-Rep1 | CCA | nope | >1bp-ins | >1bpdel | none | MNV | ATTGCCCAA( | CAAGAAA | GGT | CCA | CA | none | NA | NA_CCA | MNV_CAAGAAA | 9 | FALSE | FALSE |
| 5_v.txt | 32945 | 32945-MNV- | MNV | 2 | CC | AA | 1 | 24343 | 4.11E-03 | 63.9987657 | WT | FALSE | CAN | JSY13 | 1 | WT-Rep1 | ACC | nope | >1bp-ins | >1bpdel | none | MNV | AGAAAACCA | CCAATAC | TGG | ACC | CC | none | NA | NA_ACC | MNV_CCAATAC | 9 | FALSE | FALSE |
| 5_v.txt | 33153 | 33153-MNV- | MNV | 2 | CC | AA | 1 | 24343 | 4.11E-03 | 32.5 | WT | FALSE | CAN | JSY13 | 1 | WT-Rep1 | ACC | nope | >1bp-ins | >1bpdel | none | MNV | AAAAGACCT | CCTGTAC | TGG | ACC | CC | none | NA | NA_ACC | MNV_CCTGTAC | 9 | FALSE | FALSE |
| 5_v.txt | 33168 | 33168-MNV- | MNV | 2 | CC | AT | 14 | 24300 | 0.05761317 | 37.7500021 | WT | FALSE | CAN | JSY13 | 1 | WT-Rep1 | ACC | nope | >1bp-ins | >1bpdel | none | MNV | ATAGTACCA( | CCACCAA | TGG | ACC | CC | none | NA | NA_ACC | MNV_CCACCAA | 9 | FALSE | FALSE |
| 5_v.txt | 31769 | 31769-Insert | Insertion | 32 | - | TCTTCTATCG | 2 | 9499 | 0.02105485 | 48.84375 | WT | FALSE | CAN | JSY13 | 1 | WT-Rep1 | CTC | nope | >1bp-ins | >1bpdel | none | >1bp-ins | AATGTCTCTT | TCTTCTA | GAG | CTC | - | none | NA | NA_CTC | >1bp-ins_TCTTCTA | 9 | FALSE | FALSE |
| 5_v.txt | 31892 | 31892-Insert | Insertion | 1 | - | A | 3 | 9498 | 0.0315856 | 46.6666667 | WT | FALSE | CAN | JSY13 | 1 | WT-Rep1 | TAA | nope | A/T+1 | >1bpdel | none | A/T+1 | AAAGCTAAC | AACACCA | ATT | TAA | - | T | NA | NA_TAA | A/T+1_AACACCA | 9 | FALSE | FALSE |
| 5_v.txt | 31903 | 31903-Insert | Insertion | 1 | - | C | 2 | 9484 | 0.02108815 | 38 | WT | FALSE | CAN | JSY13 | 1 | WT-Rep1 | AAT | nope | G/C+1 | >1bpdel | none | G/C+1 | CATTGAATTT | ATTTTGG | TTA | AAT | - | G | NA | NA_AAT | G/C+1_ATTTTGG | 9 | FALSE | FALSE |
| 5_v.txt | 31984 | 31984-Insert | Insertion | 5 | - | TTAAT | 18 | 18279 | 0.09847366 | 51.4888889 | WT | FALSE | CAN | JSY13 | 1 | WT-Rep1 | ATT | nope | >1bp-ins | >1bpdel | none | >1bp-ins | CGGGCATTA | TTAATTT | TAA | ATT | - | none | NA | NA_ATT | >1bp-ins_TTAATTT | 9 | FALSE | FALSE |
| 5_v.txt | 32005 | 32005-Insert | Insertion | 3 | - | ACT | 7 | 18279 | 0.03829531 | 53.0952381 | WT | FALSE | CAN | JSY13 | 1 | WT-Rep1 | AAC | nope | >1bp-ins | >1bpdel | none | >1bp-ins | ATGGTAACT( | ACTCGTC | TTG | AAC | - | none | NA | NA_AAC | >1bp-ins_ACTCGTC | 9 | FALSE | FALSE |
| 5_v.txt | 32034 | 32034-Insert | Insertion | 1 | - | A | 8 | 18279 | 0.04376607 | 64 | WT | FALSE | CAN | JSY13 | 1 | WT-Rep1 | CAA | nope | A/T+1 | >1bpdel | none | A/T+1 | TATTTCAAA | AAAGCTT | GTT | CAA | - | T | NA | NA_CAA | A/T+1_AAAGCTT | 9 | FALSE | FALSE |
| 5_v.txt | 32081 | 32081-Insert | Insertion | 1 | - | A | 54 | 18279 | 0.29542098 | 63.3333333 | WT | FALSE | CAN | JSY13 | 1 | WT-Rep1 | CAA | nope | A/T+1 | >1bpdel | none | A/T+1 | CCATGCAAA/ | AAAAAAG | GTT | CAA | - | T | NA | NA_CAA | A/T+1_AAAAAAG | 9 | FALSE | FALSE |
| 5_v.txt | 32081 | 32081-Insert | Insertion | 2 | - | AA | 1 | 18279 | 5.47E-03 | 64 | WT | FALSE | CAN | JSY13 | 1 | WT-Rep1 | CAA | nope | >1bp-ins | >1bpdel | none | >1bp-ins | CCATGCAAA/ | AAAAAAG | GTT | CAA | - | none | NA | NA_CAA | >1bp-ins_AAAAAAG | 9 | FALSE | FALSE |
| 5_v.txt | 32081 | 32081-Insert | Insertion | 3 | - | AAA | 12 | 18279 | 0.06564911 | 60.3611111 | WT | FALSE | CAN | JSY13 | 1 | WT-Rep1 | CAA | nope | >1bp-ins | >1bpdel | none | >1bp-ins | CCATGCAAA/ | AAAAAAG | GTT | CAA | - | none | NA | NA_CAA | >1bp-ins_AAAAAAG | 9 | FALSE | FALSE |
| 5_v.txt | 32081 | 32081-Insert | Insertion | 1 | - | G | 1 | 18279 | 5.47E-03 | 62 | WT | FALSE | CAN | JSY13 | 1 | WT-Rep1 | CAA | nope | G/C+1 | >1bpdel | none | G/C+1 | CCATGCAAA/ | AAAAAAG | GTT | CAA | - | C | NA | NA_CAA | G/C+1_AAAAAAG | 9 | FALSE | FALSE |
| 5_v.txt | 32168 | 32168-Insert | Insertion | 1 | - | A | 6 | 18279 | 0.03282455 | 48 | WT | FALSE | CAN | JSY13 | 1 | WT-Rep1 | CAA | nope | A/T+1 | >1bpdel | none | A/T+1 | AGCGCCAAA | AAATGCA | GTT | CAA | - | T | NA | NA_CAA | A/T+1_AAATGCA | 9 | FALSE | FALSE |
| 5_v.txt | 32229 | 32229-Insert | Insertion | 1 | - | T | 5 | 18270 | 0.02736727 | 37.8 | WT | FALSE | CAN | JSY13 | 1 | WT-Rep1 | GAA | nope | A/T+1 | >1bpdel | none | A/T+1 | GACAGGAAT | AATTTAG | CTT | GAA | - | A | NA | NA_GAA | A/T+1_AATTTAG | 9 | FALSE | FALSE |
| 5_v.txt | 32261 | 32261-Insert | Insertion | 1 | - | A | 3 | 21420 | 0.0140056 | 38 | WT | FALSE | CAN | JSY13 | 1 | WT-Rep1 | CAA | nope | A/T+1 | >1bpdel | none | A/T+1 | TAGACCAAA | AAATAAA | GTT | CAA | - | T | NA | NA_CAA | A/T+1_AAATAAA | 9 | FALSE | FALSE |
| 5_v.txt | 32346 | 32346-Insert | Insertion | 1 | - | A | 19 | 21421 | 0.08869801 | 60.1578947 | WT | FALSE | CAN | JSY13 | 1 | WT-Rep1 | CAA | nope | A/T+1 | >1bpdel | none | A/T+1 | TGTGGCAAA | AAAACCT | GTT | CAA | - | T | NA | NA_CAA | A/T+1_AAAACCT | 9 | FALSE | FALSE |
| 5_v.txt | 32402 | 32402-Insert | Insertion | 11 | - | GGAAGTAGA | 15 | 21421 | 0.07002474 | 55.5090909 | WT | FALSE | CAN | JSY13 | 1 | WT-Rep1 | AGG | nope | >1bp-ins | >1bpdel | none | >1bp-ins | AACGTAGGA | GGAAGTA | TCC | AGG | - | none | NA | NA_AGG | >1bp-ins_GGAAGTA | 9 | FALSE | FALSE |
| 5_v.txt | 32476 | 32476-Insert | Insertion | 14 | - | AGGTTAAGA | 35 | 21421 | 0.16339107 | 43.4367347 | WT | FALSE | CAN | JSY13 | 1 | WT-Rep1 | AAG | nope | >1bp-ins | >1bpdel | none | >1bp-ins | TGTAGAAGG | AGGTTAA | TTC | AAG | - | none | NA | NA_AAG | >1bp-ins_AGGTTAA | 9 | FALSE | FALSE |
| 5_v.txt | 32488 | 32488-Insert | Insertion | 14 | - | GGAGGTTAA | 1 | 21421 | 4.67E-03 | 36.9285714 | WT | FALSE | CAN | JSY13 | 1 | WT-Rep1 | CGG | nope | >1bp-ins | >1bpdel | none | >1bp-ins | AGATACGGA | GGAAAAC | GCC | CGG | - | none | NA | NA_CGG | >1bp-ins_GGAAAAC | 9 | FALSE | FALSE |
| 5_v.txt | 32493 | 32493-Insert | Insertion | 3 | - | GAT | 1 | 21421 | 4.67E-03 | 38 | WT | FALSE | CAN | JSY13 | 1 | WT-Rep1 | AAC | nope | >1bp-ins | >1bpdel | none | >1bp-ins | CGGAAAACA | ACAACTT | TTG | AAC | - | none | NA | NA_AAC | >1bp-ins_ACAACTT | 9 | FALSE | FALSE |
| 5_v.txt | 32495 | 32495-Insert | Insertion | 4 | - | GGAA | 1 | 21421 | 4.67E-03 | 38 | WT | FALSE | CAN | JSY13 | 1 | WT-Rep1 | CAA | nope | >1bp-ins | >1bpdel | none | >1bp-ins | GAAAACAAC | AACTTTT | GTT | CAA | - | none | NA | NA_CAA | >1bp-ins_AACTTTT | 9 | FALSE | FALSE |
| 5_v.txt | 32498 | 32498-Insert | Insertion | 1 | - | T | 9 | 21421 | 0.04201485 | 49.4444441 | WT | FALSE | CAN | JSY13 | 1 | WT-Rep1 | CTT | nope | A/T+1 | >1bpdel | none | A/T+1 | AACAACTTTT | TTTTTTG | GAA | CTT | - | A | NA | NA_CTT | A/T+1_TTTTTTG | 9 | FALSE | FALSE |
| 5_v.txt | 32528 | 32528-Insert | Insertion | 1 | - | G | 25 | 21421 | 0.1167079 | 39.56 | WT | FALSE | CAN | JSY13 | 1 | WT-Rep1 | TGG | nope | G/C+1 | >1bpdel | none | G/C+1 | TTTTCTGGG( | GGGGTTT | ACC | TGG | - | C | NA | NA_TGG | G/C+1_GGGGTTT | 9 | FALSE | FALSE |
| 5_v.txt | 32629 | 32629-Insert | Insertion | 1 | - | C | 70 | 22156 | 0.31594151 | 51.1285714 | WT | FALSE | CAN | JSY13 | 1 | WT-Rep1 | TCG | nope | G/C+1 | >1bpdel | none | G/C+1 | TCCCTTCGTT | CGTTTTT | AGC | TCG | - | G | NA | NA_TCG | G/C+1_CGTTTTT | 9 | FALSE | FALSE |
| 5_v.txt | 32646 | 32646-Insert | Insertion | 16 | - | ATTATACCTG | 3 | 22156 | 0.01354035 | 51.625 | WT | FALSE | CAN | JSY13 | 1 | WT-Rep1 | TAT | nope | >1bp-ins | >1bpdel | none | >1bp-ins | TTAGATATTA | ATTATAC | ATA | TAT | - | none | NA | NA_TAT | >1bp-ins_ATTATAC | 9 | FALSE | FALSE |
| 5_v.txt | 32647 | 32647-Insert | Insertion | 1 | - | T | 8 | 22156 | 0.0361076 | 64 | WT | FALSE | CAN | JSY13 | 1 | WT-Rep1 | ATT | nope | A/T+1 | >1bpdel | none | A/T+1 | TAGATATTAT | TTATACC | TAA | ATT | - | A | NA | NA_ATT | A/T+1_TTATACC | 9 | FALSE | FALSE |
| 5_v.txt | 32728 | 32728-Insert | Insertion | 1 | - | A | 58 | 22177 | 0.26153222 | 60.3448276 | WT | FALSE | CAN | JSY13 | 1 | WT-Rep1 | CAA | nope | A/T+1 | >1bpdel | none | A/T+1 | CCATACAAA/ | AAAAACA | GTT | CAA | - | T | NA | NA_CAA | A/T+1_AAAAACA | 9 | FALSE | FALSE |
| 5_v.txt | 32750 | 32750-Insert | Insertion | 4 | - | GATA | 12 | 22177 | 0.05411011 | 58.8958333 | WT | FALSE | CAN | JSY13 | 1 | WT-Rep1 | CGA | nope | >1bp-ins | >1bpdel | none | >1bp-ins | AAACCCGAT/ | GATAATG | GCT | CGA | - | none | NA | NA_CGA | >1bp-ins_GATAATG | 9 | FALSE | FALSE |
| 5_v.txt | 32760 | 32760-Insert | Insertion | 1 | - | A | 11 | 22177 | 0.04960094 | 51.8181818 | WT | FALSE | CAN | JSY13 | 1 | WT-Rep1 | TAA | nope | A/T+1 | >1bpdel | none | A/T+1 | ATGGCTAAA | AAAACTT | ATT | TAA | - | T | NA | NA_TAA | A/T+1_AAAACTT | 9 | FALSE | FALSE |
| 5_v.txt | 32841 | 32841-Insert | Insertion | 1 | - | G | 10 | 22177 | 0.04509176 | 36.2816549 | WT | FALSE | CAN | JSY13 | 1 | WT-Rep1 | CCA | nope | G/C+1 | >1bpdel | none | G/C+1 | ATTACCCAA/ | CAAAAAA | GGT | CCA | - | C | NA | NA_CCA | G/C+1_CAAAAAA | 9 | FALSE | FALSE |
| 5_v.txt | 32842 | 32842-Insert | Insertion | 1 | - | A | 414 | 22177 | 1.86679894 | 42.3333333 | WT | FALSE | CAN | JSY13 | 1 | WT-Rep1 | CAA | nope | A/T+1 | >1bpdel | none | A/T+1 | TTACCCAAA/ | AAAAAAT | GTT | CAA | - | T | NA | NA_CAA | A/T+1_AAAAAAT | 9 | FALSE | FALSE |
| 5_v.txt | 32842 | 32842-Insert | Insertion | 2 | - | AA | 1 | 22177 | 4.51E-03 | 38 | WT | FALSE | CAN | JSY13 | 1 | WT-Rep1 | CAA | nope | >1bp-ins | >1bpdel | none | >1bp-ins | TTACCCAAA/ | AAAAAAT | GTT | CAA | - | none | NA | NA_CAA | >1bp-ins_AAAAAAT | 9 | FALSE | FALSE |
| 5_v.txt | 32924 | 32924-Insert | Insertion | 1 | - | A | 26 | 24305 | 0.10697387 | 53.6923077 | WT | FALSE | CAN | JSY13 | 1 | WT-Rep1 | CAA | nope | A/T+1 | >1bpdel | none | A/T+1 | CAGGGCAAA | AAAAGTG | GTT | CAA | - | T | NA | NA_CAA | A/T+1_AAAAGTG | 9 | FALSE | FALSE |
| 5_v.txt | 32953 | 32953-Insert | Insertion | 3 | - | TGT | 8 | 24343 | 0.03286366 | 51.4166667 | WT | FALSE | CAN | JSY13 | 1 | WT-Rep1 | ATG | nope | >1bp-ins | >1bpdel | none | >1bp-ins | AATACATGT/ | TGTAACC | TAC | ATG | - | none | NA | NA_ATG | >1bp-ins_TGTAACC | 9 | FALSE | FALSE |
| 5_v.txt | 32972 | 32972-Insert | Insertion | 1 | - | A | 36 | 24343 | 0.14788646 | 61.2222222 | WT | FALSE | CAN | JSY13 | 1 | WT-Rep1 | CAA | nope | A/T+1 | >1bpdel | none | A/T+1 | CGCACCAAA | AAATGCT | GTT | CAA | - | T | NA | NA_CAA | A/T+1_AAATGCT | 9 | FALSE | FALSE |
| 5_v.txt | 32992 | 32992-Insert | Insertion | 1 | - | T | 7 | 24343 | 0.0287557 | 63.1428571 | WT | FALSE | CAN | JSY13 | 1 | WT-Rep1 | CTT | nope | A/T+1 | >1bpdel | none | A/T+1 | GGAATCTTT( | TTTGTGA | GAA | CTT | - | A | NA | NA_CTT | A/T+1_TTTGTGA | 9 | FALSE | FALSE |
| 5_v.txt | 33009 | 33009-Insert | Insertion | 1 | - | A | 10 | 24343 | 0.04107957 | 58.8 | WT | FALSE | CAN | JSY13 | 1 | WT-Rep1 | GAA | nope | A/T+1 | >1bpdel | none | A/T+1 | ACTGTGAAA | AAAGAGG | CTT | GAA | - | T | NA | NA_GAA | A/T+1_AAAGAGG | 9 | FALSE | FALSE |
| 5_v.txt | 33036 | 33036-Insert | Insertion | 33 | - | GCCATTTCAC | 8 | 24343 | 0.03286366 | 56.2916667 | WT | FALSE | CAN | JSY13 | 1 | WT-Rep1 | AGC | nope | >1bp-ins | >1bpdel | none | >1bp-ins | AATGTAGCC/ | GCCATTT | TCG | AGC | - | none | NA | NA_AGC | >1bp-ins_GCCATTT | 9 | FALSE | FALSE |
| 5_v.txt | 33045 | 33045-Insert | Insertion | 1 | - | C | 12 | 24343 | 0.04929549 | 59.3333333 | WT | FALSE | CAN | JSY13 | 1 | WT-Rep1 | ACC | nope | G/C+1 | >1bpdel | none | G/C+1 | ATTTCACCCA | CCCAAGG | TGG | ACC | - | G | NA | NA_ACC | G/C+1_CCCAAGG | 9 | FALSE | FALSE |
| 5_v.txt | 33046 | 33046-Insert | Insertion | 24 | - | CCAAGGACT( | 3 | 24343 | 0.01232387 | 54.375 | WT | FALSE | CAN | JSY13 | 1 | WT-Rep1 | CCC | nope | >1bp-ins | >1bpdel | none | >1bp-ins | TTTCACCCAA | CCAAGGA | GGG | CCC | - | none | NA | NA_CCC | >1bp-ins_CCAAGGA | 9 | FALSE | FALSE |
| 5_v.txt | 33078 | 33078-Insert | Insertion | 1 | - | C | 1 | 24343 | 4.11E-03 | 38 | WT | FALSE | CAN | JSY13 | 1 | WT-Rep1 | ACC | nope | G/C+1 | >1bpdel | none | G/C+1 | AAAGAACCC | CCCATAA | TGG | ACC | - | G | NA | NA_ACC | G/C+1_CCCATAA | 9 | FALSE | FALSE |
| 5_v.txt | 33147 | 33147-Insert | Insertion | 2 | - | AA | 3 | 24343 | 0.01232387 | 36.8333333 | WT | FALSE | CAN | JSY13 | 1 | WT-Rep1 | GAA | nope | >1bp-ins | >1bpdel | none | >1bp-ins | CCAATGAAA | AAAAGAC | CTT | GAA | - | none | NA | NA_GAA | >1bp-ins_AAAAGAC | 9 | FALSE | FALSE |
| 5_v.txt | 33148 | 33148-Insert | Insertion | 2 | - | AG | 1 | 24343 | 4.11E-03 | 64 | WT | FALSE | CAN | JSY13 | 1 | WT-Rep1 | AAA | nope | >1bp-ins | >1bpdel | none | >1bp-ins | CAATGAAAA | AAAGACC | TTT | AAA | - | none | NA | NA_AAA | >1bp-ins_AAAGACC | 9 | FALSE | FALSE |
| 5_v.txt | 33206 | 33206-Insert | Insertion | 2 | - | CT | 7 | 7277 | 0.09619349 | 36.7142857 | WT | FALSE | CAN | JSY13 | 1 | WT-Rep1 | GCT | nope | >1bp-ins | >1bpdel | none | >1bp-ins | CTTAAGCTCT | CTCTCTC | CGA | GCT | - | none | NA | NA_GCT | >1bp-ins_CTCTCTC | 9 | FALSE | FALSE |
| 5_v.txt | 33254 | 33254-Insert | Insertion | 4 | - | TATG | 6 | 7277 | 0.08245156 | 36.6666667 | WT | FALSE | CAN | JSY13 | 1 | WT-Rep1 | CTA | nope | >1bp-ins | >1bpdel | none | >1bp-ins | ATCACCTATG | TATGCCA | GAT | CTA | - | none | NA | NA_CTA | >1bp-ins_TATGCCA | 9 | FALSE | FALSE |
| 5_v.txt | 33437 | 33437-Insert | Insertion | 1 | - | T | 8 | 7277 | 0.10993541 | 41 | WT | FALSE | CAN | JSY13 | 1 | WT-Rep1 | CTA | nope | A/T+1 | >1bpdel | none | A/T+1 | CTCCTCTATG | TATGTCG | GAT | CTA | - | A | NA | NA_CTA | A/T+1_TATGTCG | 9 | FALSE | FALSE |
| 5_v.txt | 32014 | 32014-Deleti | Deletion | 2 | GA | - | 28 | 18279 | 0.15318125 | 57.2142857 | WT | FALSE | CAN | JSY13 | 1 | WT-Rep1 | CGA | nope | >1bp-ins | >1bpdel | none | >1bpdel | CGTCACGAG | GAGAGAT | GCT | CGA | GA | none | NA | NA_CGA | >1bpdel_GAGAGAT | 9 | FALSE | FALSE |
| 5_v.txt | 32070 | 32070-Deleti | Deletion | 3 | AAT | - | 42 | 18279 | 0.22977187 | 63.0238095 | WT | FALSE | CAN | JSY13 | 1 | WT-Rep1 | AAA | nope | >1bp-ins | >1bpdel | none | >1bpdel | GAGATAAAT | AATAACC | TTT | AAA | AAT | none | NA | NA_AAA | >1bpdel_AATAACC | 9 | FALSE | FALSE |
| 5_v.txt | 32313 | 32313-Deleti | Deletion | 3 | GTT | - | 23 | 21421 | 0.10737127 | 61.0869565 | WT | FALSE | CAN | JSY13 | 1 | WT-Rep1 | GGT | nope | >1bp-ins | >1bpdel | none | >1bpdel | ATAATGGTT( | GTTGTTA | CCA | GGT | GTT | none | NA | NA_GGT | >1bpdel_GTTGTTA | 9 | FALSE | FALSE |
| 5_v.txt | 32316 | 32316-Deleti | Deletion | 5 | GTTAA | - | 11 | 21421 | 0.05135148 | 61.5454546 | WT | FALSE | CAN | JSY13 | 1 | WT-Rep1 | TGT | nope | >1bp-ins | >1bpdel | none | >1bpdel | ATGGTTGTT/ | GTTAAGA | ACA | TGT | GTTAA | none | NA | NA_TGT | >1bpdel_GTTAAGA | 9 | FALSE | FALSE |
| 5_v.txt | 32337 | 32337-Deleti | Deletion | 2 | AT | - | 148 | 21421 | 0.69091079 | 62.3378378 | WT | FALSE | CAN | JSY13 | 1 | WT-Rep1 | GAT | nope | >1bp-ins | >1bpdel | none | >1bpdel | TTGAAGATA | ATATGTG | CTA | GAT | AT | none | NA | NA_GAT | >1bpdel_ATATGTG | 9 | FALSE | FALSE |
| 5_v.txt | 32454 | 32454-Deleti | Deletion | 3 | AAT | - | 17 | 21421 | 0.07936137 | 46.8823529 | WT | FALSE | CAN | JSY13 | 1 | WT-Rep1 | GAA | nope | >1bp-ins | >1bpdel | none | >1bpdel | CCAATGAAT/ | AATAATA | CTT | GAA | AAT | none | NA | NA_GAA | >1bpdel_AATAATA | 9 | FALSE | FALSE |
| 5_v.txt | 32458 | 32458-Deleti | Deletion | 2 | AT | - | 20 | 21421 | 0.09336632 | 46 | WT | FALSE | CAN | JSY13 | 1 | WT-Rep1 | AAT | nope | >1bp-ins | >1bpdel | none | >1bpdel | TGAATAATA( | ATAGAGA | TTA | AAT | AT | none | NA | NA_AAT | >1bpdel_ATAGAGA | 9 | FALSE | FALSE |
| 5_v.txt | 32473 | 32473-Deleti | Deletion | 3 | AGA | - | 11 | 21421 | 0.05135148 | 55.1818182 | WT | FALSE | CAN | JSY13 | 1 | WT-Rep1 | TAG | nope | >1bp-ins | >1bpdel | none | >1bpdel | CAATGTAGA | AGAAGGT | ATC | TAG | AGA | none | NA | NA_TAG | >1bpdel_AGAAGGT | 9 | FALSE | FALSE |
| 5_v.txt | 32492 | 32492-Deleti | Deletion | 3 | AAC | - | 647 | 21421 | 3.02040054 | 44.3802164 | WT | FALSE | CAN | JSY13 | 1 | WT-Rep1 | AAA | nope | >1bp-ins | >1bpdel | none | >1bpdel | ACGGAAAAC | AACAACT | TTT | AAA | AAC | none | NA | NA_AAA | >1bpdel_AACAACT | 9 | FALSE | FALSE |
| 5_v.txt | 32493 | 32493-Deleti | Deletion | 2 | AC | - | 1 | 21421 | 4.67E-03 | 38 | WT | FALSE | CAN | JSY13 | 1 | WT-Rep1 | AAC | nope | >1bp-ins | >1bpdel | none | >1bpdel | CGGAAAACA | ACAACTT | TTG | AAC | AC | none | NA | NA_AAC | >1bpdel_ACAACTT | 9 | FALSE | FALSE |
| 5_v.txt | 32648 | 32648-Deleti | Deletion | 2 | TA | - | 8 | 22156 | 0.0361076 | 50.5000923 | WT | FALSE | CAN | JSY13 | 1 | WT-Rep1 | TTA | nope | >1bp-ins | >1bpdel | none | >1bpdel | AGATATTAT/ | TATACCT | AAT | TTA | TA | none | NA | NA_TTA | >1bpdel_TATACCT | 9 | FALSE | FALSE |
| 5_v.txt | 32804 | 32804-Deleti | Deletion | 2 | TT | - | 6 | 22177 | 0.02705506 | 37.166669 | WT | FALSE | CAN | JSY13 | 1 | WT-Rep1 | ATT | nope | >1bp-ins | >1bpdel | none | >1bpdel | GTAATATTTG | TTTGACA | TAA | ATT | TT | none | NA | NA_ATT | >1bpdel_TTTGACA | 9 | FALSE | FALSE |
| 5_v.txt | 32919 | 32919-Deleti | Deletion | 2 | AG | - | 1 | 24305 | 4.11E-03 | 11 | WT | FALSE | CAN | JSY13 | 1 | WT-Rep1 | CAG | nope | >1bp-ins | >1bpdel | none | >1bpdel | AGTTCCAGG | AGGGCAA | GTC | CAG | AG | none | NA | NA_CAG | >1bpdel_AGGGCAA | 9 | FALSE | FALSE |
| 5_v.txt | 32927 | 32927-Deleti | Deletion | 3 | AGT | - | 1 | 24343 | 4.11E-03 | 11 | WT | FALSE | CAN | JSY13 | 1 | WT-Rep1 | AAG | nope | >1bp-ins | >1bpdel | none | >1bpdel | GGCAAAAGT | AGTGATT | TTC | AAG | AGT | none | NA | NA_AAG | >1bpdel_AGTGATT | 9 | FALSE | FALSE |
| 5_v.txt | 33055 | 33055-Deleti | Deletion | 2 | GC | - | 1 | 24343 | 4.11E-03 | 64 | WT | FALSE | CAN | JSY13 | 1 | WT-Rep1 | TGC | nope | >1bp-ins | >1bpdel | none | >1bpdel | AGGACTGCG | GCGTGAC | ACG | TGC | GC | none | NA | NA_TGC | >1bpdel_GCGTGAC | 9 | FALSE | FALSE |
| 5_v.txt | 33065 | 33065-Deleti | Deletion | 2 | AT | - | 11 | 24343 | 0.04518753 | 61.5454599 | WT | FALSE | CAN | JSY13 | 1 | WT-Rep1 | AAT | nope | >1bp-ins | >1bpdel | none | >1bpdel | GACAGAATA | ATATGCC | TTA | AAT | AT | none | NA | NA_AAT | >1bpdel_ATATGCC | 9 | FALSE | FALSE |
| 5_v.txt | 33081 | 33081-Deleti | Deletion | 4 | ATAA | - | 12 | 24343 | 0.04929549 | 47.4166667 | WT | FALSE | CAN | JSY13 | 1 | WT-Rep1 | CAT | nope | >1bp-ins | >1bpdel | none | >1bpdel | GAACCCATA/ | ATAAATA | GTA | CAT | ATAA | none | NA | NA_CAT | >1bpdel_ATAAATA | 9 | FALSE | FALSE |
| 5_v.txt | 33094 | 33094-Deleti | Deletion | 2 | AT | - | 119 | 24343 | 0.4888469 | 48.6302521 | WT | FALSE | CAN | JSY13 | 1 | WT-Rep1 | GAT | nope | >1bp-ins | >1bpdel | none | >1bpdel | AATATGATA | ATATAAG | CTA | GAT | AT | none | NA | NA_GAT | >1bpdel_ATATAAG | 9 | FALSE | FALSE |
| 5_v.txt | 33281 | 33281-Deleti | Deletion | 13 | CTCGCCATTT | - | 19 | 7277 | 0.26109661 | 56.8947368 | WT | FALSE | CAN | JSY13 | 1 | WT-Rep1 | CCT | nope | >1bp-ins | >1bpdel | none | >1bpdel | CGTATCCTCG | CTCGCCA | GGA | CCT | CTCGCCATTT | none | NA | NA_CCT | >1bpdel_CTCGCCA | 9 | FALSE | FALSE |
| 5_v.txt | 31723 | 31723-Deleti | Deletion | 1 | A | - | 14 | 9499 | 0.14738394 | 59.1428571 | WT | FALSE | CAN | JSY13 | 1 | WT-Rep1 | CAA | nope | >1bp-ins | A/T-1 | none | A/T-1 | TGTCCCAAA/ | AAAAAGT | GTT | GTT | T | none | T>none | T>none_GTT | A/T-1_AAAAAGT | 9 | FALSE | FALSE |
| 5_v.txt | 31805 | 31805-Deleti | Deletion | 1 | C | - | 3 | 9499 | 0.03158227 | 53 | WT | FALSE | CAN | JSY13 | 1 | WT-Rep1 | TCC | nope | >1bp-ins | G/C-1 | none | G/C-1 | AATCTTCCAA | CCAAATA | AGG | TCC | C | none | nope | nope_TCC | G/C-1_CCAAATA | 9 | FALSE | FALSE |
| 5_v.txt | 31846 | 31846-Deleti | Deletion | 1 | A | - | 6 | 9499 | 0.06316454 | 54.833355 | WT | FALSE | CAN | JSY13 | 1 | WT-Rep1 | CAA | nope | >1bp-ins | A/T-1 | none | A/T-1 | AGATCCAAA | AAACAGC | GTT | GTT | T | none | T>none | T>none_GTT | A/T-1_AAACAGC | 9 | FALSE | FALSE |
| 5_v.txt | 31854 | 31854-Deleti | Deletion | 1 | A | - | 3 | 9499 | 0.03158227 | 64 | WT | FALSE | CAN | JSY13 | 1 | WT-Rep1 | TAA | nope | >1bp-ins | A/T-1 | none | A/T-1 | ACAGCTAAG | AAGAACA | ATT | ATT | T | none | T>none | T>none_ATT | A/T-1_AAGAACA | 9 | FALSE | FALSE |
| 5_v.txt | 31863 | 31863-Deleti | Deletion | 1 | A | - | 13 | 9499 | 0.13685651 | 44.3076923 | WT | FALSE | CAN | JSY13 | 1 | WT-Rep1 | GAA | nope | >1bp-ins | A/T-1 | none | A/T-1 | AACAGGAAA | AAAATAG | CTT | CTT | T | none | T>none | T>none_CTT | A/T-1_AAAATAG | 9 | FALSE | FALSE |
| 5_v.txt | 31904 | 31904-Deleti | Deletion | 1 | T | - | 56 | 9484 | 0.59046816 | 43.3035714 | WT | FALSE | CAN | JSY13 | 1 | WT-Rep1 | ATT | nope | >1bp-ins | A/T-1 | none | A/T-1 | ATTGAATTTT | TTTTGGT | TAA | ATT | T | none | nope | nope_ATT | A/T-1_TTTTGGT | 9 | FALSE | FALSE |
| 5_v.txt | 31913 | 31913-Deleti | Deletion | 1 | A | - | 5 | 9484 | 0.05272037 | 38 | WT | FALSE | CAN | JSY13 | 1 | WT-Rep1 | CAA | nope | >1bp-ins | A/T-1 | none | A/T-1 | TGGTGCAAA | AAAAGCC | GTT | GTT | T | none | T>none | T>none_GTT | A/T-1_AAAAGCC | 9 | FALSE | FALSE |
| 5_v.txt | 31926 | 31926-Deleti | Deletion | 1 | C | - | 5 | 9485 | 0.05271481 | 37.8 | WT | FALSE | CAN | JSY13 | 1 | WT-Rep1 | ACC | nope | >1bp-ins | G/C-1 | none | G/C-1 | GTGAAACCT | CCTTGAA | TGG | ACC | C | none | nope | nope_ACC | G/C-1_CCTTGAA | 9 | FALSE | FALSE |
| 5_v.txt | 31952 | 31952-Deleti | Deletion | 1 | A | - | 13 | 18263 | 0.07118217 | 37.4615385 | WT | FALSE | CAN | JSY13 | 1 | WT-Rep1 | TAA | nope | >1bp-ins | A/T-1 | none | A/T-1 | CGTCATAAAT | AAATGTG | ATT | ATT | T | none | T>none | T>none_ATT | A/T-1_AAATGTG | 9 | FALSE | FALSE |
| 5_v.txt | 31972 | 31972-Deleti | Deletion | 1 | C | - | 9 | 18279 | 0.04923683 | 34.8888897 | WT | FALSE | CAN | JSY13 | 1 | WT-Rep1 | GCC | nope | >1bp-ins | G/C-1 | none | G/C-1 | AATAAGCCA | CCAAGCC | CGG | GCC | C | none | nope | nope_GCC | G/C-1_CCAAGCC | 9 | FALSE | FALSE |
| 5_v.txt | 32015 | 32015-Deleti | Deletion | 1 | A | - | 3 | 18279 | 0.01641228 | 63.333316 | WT | FALSE | CAN | JSY13 | 1 | WT-Rep1 | GAG | nope | >1bp-ins | A/T-1 | none | A/T-1 | GTCACGAGA | AGAGATG | CTC | CTC | T | none | T>none | T>none_CTC | A/T-1_AGAGATG | 9 | FALSE | FALSE |
| 5_v.txt | 32068 | 32068-Deleti | Deletion | 1 | T | - | 6 | 18279 | 0.03282455 | 64 | WT | FALSE | CAN | JSY13 | 1 | WT-Rep1 | ATA | nope | >1bp-ins | A/T-1 | none | A/T-1 | TTGAGATAA | TAAATAA | TAT | ATA | T | none | nope | nope_ATA | A/T-1_TAAATAA | 9 | FALSE | FALSE |
| 5_v.txt | 32070 | 32070-Deleti | Deletion | 1 | A | - | 6 | 18278 | 0.03282635 | 63.9999973 | WT | FALSE | CAN | JSY13 | 1 | WT-Rep1 | AAA | nope | >1bp-ins | A/T-1 | none | A/T-1 | GAGATAAAT | AATAACC | TTT | TTT | T | none | T>none | T>none_TTT | A/T-1_AATAACC | 9 | FALSE | FALSE |
| 5_v.txt | 32073 | 32073-Deleti | Deletion | 1 | A | - | 6 | 18279 | 0.03282455 | 64 | WT | FALSE | CAN | JSY13 | 1 | WT-Rep1 | TAA | nope | >1bp-ins | A/T-1 | none | A/T-1 | ATAAATAAC( | AACCATG | ATT | ATT | T | none | T>none | T>none_ATT | A/T-1_AACCATG | 9 | FALSE | FALSE |
| 5_v.txt | 32075 | 32075-Deleti | Deletion | 1 | C | - | 1 | 18279 | 5.47E-03 | 63.9999415 | WT | FALSE | CAN | JSY13 | 1 | WT-Rep1 | ACC | nope | >1bp-ins | G/C-1 | none | G/C-1 | AAATAACCA | CCATGCA | TGG | ACC | C | none | nope | nope_ACC | G/C-1_CCATGCA | 9 | FALSE | FALSE |
| 5_v.txt | 32080 | 32080-Deleti | Deletion | 1 | C | - | 2 | 18279 | 0.01094152 | 64 | WT | FALSE | CAN | JSY13 | 1 | WT-Rep1 | GCA | nope | >1bp-ins | G/C-1 | none | G/C-1 | ACCATGCAA/ | CAAAAAA | CGT | GCA | C | none | nope | nope_GCA | G/C-1_CAAAAAA | 9 | FALSE | FALSE |
| 5_v.txt | 32081 | 32081-Deleti | Deletion | 1 | A | - | 93 | 18279 | 0.50878057 | 63.1397822 | WT | FALSE | CAN | JSY13 | 1 | WT-Rep1 | CAA | nope | >1bp-ins | A/T-1 | none | A/T-1 | CCATGCAAA/ | AAAAAAG | GTT | GTT | T | none | T>none | T>none_GTT | A/T-1_AAAAAAG | 9 | FALSE | FALSE |
| 5_v.txt | 32092 | 32092-Deleti | Deletion | 1 | C | - | 3 | 18279 | 0.01641228 | 62.3333333 | WT | FALSE | CAN | JSY13 | 1 | WT-Rep1 | GCA | nope | >1bp-ins | G/C-1 | none | G/C-1 | AGCCTGCAA( | CAACACC | CGT | GCA | C | none | nope | nope_GCA | G/C-1_CAACACC | 9 | FALSE | FALSE |
| 5_v.txt | 32145 | 32145-Deleti | Deletion | 1 | G | - | 8 | 18279 | 0.04376607 | 52.125 | WT | FALSE | CAN | JSY13 | 1 | WT-Rep1 | TGT | nope | >1bp-ins | G/C-1 | none | G/C-1 | GTAGATGTC | GTCTCCA | ACA | ACA | C | none | C>none | C>none_ACA | G/C-1_GTCTCCA | 9 | FALSE | FALSE |

| | | | | | | | | | | | | | | | | | | | | | | | | | | | | | | | | | | |
|---|---|---|---|---|---|---|---|---|---|---|---|---|---|---|---|---|---|---|---|---|---|---|---|---|---|---|---|---|---|---|---|---|---|---|
| 5_v.txt | 32147 | 32147-Deleti | Deletion | 1 | C | - | 8 | 18279 | 0.04376607 | 54.125 | WT | FALSE | CAN | JSY13 | 1 | WT-Rep1 | TCT | nope | >1bp-ins | G/C-1 | none | G/C-1 | AGATGTCTC | CTCCATG | AGA | TCT | C | none | nope | nope_TCT | G/C-1_CTCCATG | 9 | FALSE | FALSE |
| 5_v.txt | 32149 | 32149-Deleti | Deletion | 1 | C | - | 10 | 18279 | 0.05470759 | 56.1 | WT | FALSE | CAN | JSY13 | 1 | WT-Rep1 | TCC | nope | >1bp-ins | G/C-1 | none | G/C-1 | ATGTCTCCAT | CCATGTA | AGG | TCC | C | none | nope | nope_TCC | G/C-1_CCATGTA | 9 | FALSE | FALSE |
| 5_v.txt | 32160 | 32160-Deleti | Deletion | 1 | A | - | 11 | 18279 | 0.06017835 | 54.5454546 | WT | FALSE | CAN | JSY13 | 1 | WT-Rep1 | CAA | nope | >1bp-ins | A/T-1 | none | A/T-1 | TAAGCCAAA | AAAGCGC | GTT | GTT | T | none | T>none | T>none_GTT | A/T-1_AAAGCGC | 9 | FALSE | FALSE |
| 5_v.txt | 32191 | 32191-Deleti | Deletion | 1 | C | - | 9 | 18279 | 0.04923683 | 49.3333333 | WT | FALSE | CAN | JSY13 | 1 | WT-Rep1 | GCA | nope | >1bp-ins | G/C-1 | none | G/C-1 | AAACTGCAA | CAATGTA | CGT | GCA | C | none | nope | nope_GCA | G/C-1_CAATGTA | 9 | FALSE | FALSE |
| 5_v.txt | 32227 | 32227-Deleti | Deletion | 1 | G | - | 19 | 18270 | 0.10399562 | 38 | WT | FALSE | CAN | JSY13 | 1 | WT-Rep1 | AGG | nope | >1bp-ins | G/C-1 | none | G/C-1 | TTGACAGGA | GGAATTT | TCC | TCC | C | none | C>none | C>none_TCC | G/C-1_GGAATTT | 9 | FALSE | FALSE |
| 5_v.txt | 32235 | 32235-Deleti | Deletion | 1 | G | - | 15 | 18270 | 0.08210181 | 36.9333333 | WT | FALSE | CAN | JSY13 | 1 | WT-Rep1 | AGG | nope | >1bp-ins | G/C-1 | none | G/C-1 | AATTTAGGA | GGAGCCA | TCC | TCC | C | none | C>none | C>none_TCC | G/C-1_GGAGCCA | 9 | FALSE | FALSE |
| 5_v.txt | 32239 | 32239-Deleti | Deletion | 1 | C | - | 14 | 18268 | 0.07663674 | 38 | WT | FALSE | CAN | JSY13 | 1 | WT-Rep1 | GCC | nope | >1bp-ins | G/C-1 | none | G/C-1 | TAGGAGCCA | CCAACTT | CGG | GCC | C | none | nope | nope_GCC | G/C-1_CCAACTT | 9 | FALSE | FALSE |
| 5_v.txt | 32265 | 32265-Deleti | Deletion | 1 | A | - | 5 | 21420 | 0.02334267 | 37.8 | WT | FALSE | CAN | JSY13 | 1 | WT-Rep1 | TAA | nope | >1bp-ins | A/T-1 | none | A/T-1 | CCAAATAAA | AAAATAC | ATT | ATT | T | none | T>none | T>none_ATT | A/T-1_AAAATAC | 9 | FALSE | FALSE |
| 5_v.txt | 32275 | 32275-Deleti | Deletion | 1 | A | - | 24 | 21420 | 0.11204482 | 35.7083318 | WT | FALSE | CAN | JSY13 | 1 | WT-Rep1 | GAA | nope | >1bp-ins | A/T-1 | none | A/T-1 | TACGGGAAC | AACCAAC | CTT | CTT | T | none | T>none | T>none_CTT | A/T-1_AACCAAC | 9 | FALSE | FALSE |
| 5_v.txt | 32283 | 32283-Deleti | Deletion | 1 | T | - | 9 | 21420 | 0.04201681 | 56.9999358 | WT | FALSE | CAN | JSY13 | 1 | WT-Rep1 | GTA | nope | >1bp-ins | A/T-1 | none | A/T-1 | CCAACGTAA | TAAATAT | CAT | GTA | T | none | nope | nope_GTA | A/T-1_TAAATAT | 9 | FALSE | FALSE |
| 5_v.txt | 32304 | 32304-Deleti | Deletion | 1 | G | - | 24 | 21421 | 0.11203959 | 57.8333333 | WT | FALSE | CAN | JSY13 | 1 | WT-Rep1 | AGA | nope | >1bp-ins | G/C-1 | none | G/C-1 | GCGGCAGAA | GAAATAA | TCT | TCT | C | none | C>none | C>none_TCT | G/C-1_GAAATAA | 9 | FALSE | FALSE |
| 5_v.txt | 32311 | 32311-Deleti | Deletion | 1 | T | - | 50 | 21421 | 0.23341581 | 56.1 | WT | FALSE | CAN | JSY13 | 1 | WT-Rep1 | ATG | nope | >1bp-ins | A/T-1 | none | A/T-1 | AAATAATGG | TGGTTGT | TAC | ATG | T | none | nope | nope_ATG | A/T-1_TGGTTGT | 9 | FALSE | FALSE |
| 5_v.txt | 32377 | 32377-Deleti | Deletion | 1 | T | - | 7 | 21421 | 0.03267821 | 64 | WT | FALSE | CAN | JSY13 | 1 | WT-Rep1 | ATA | nope | >1bp-ins | A/T-1 | none | A/T-1 | TAGCAATAA | TAATAAA | TAT | ATA | T | none | nope | nope_ATA | A/T-1_TAATAAA | 9 | FALSE | FALSE |
| 5_v.txt | 32384 | 32384-Deleti | Deletion | 1 | G | - | 7 | 21421 | 0.03267821 | 61.8571377 | WT | FALSE | CAN | JSY13 | 1 | WT-Rep1 | AGG | nope | >1bp-ins | G/C-1 | none | G/C-1 | AATAAAGGG | GGGAGAA | TCC | TCC | C | none | C>none | C>none_TCC | G/C-1_GGGAGAA | 9 | FALSE | FALSE |
| 5_v.txt | 32442 | 32442-Deleti | Deletion | 1 | A | - | 8 | 21421 | 0.03734653 | 56.625 | WT | FALSE | CAN | JSY13 | 1 | WT-Rep1 | TAA | nope | >1bp-ins | A/T-1 | none | A/T-1 | GGAACTAAA | AAAAGTC | ATT | ATT | T | none | T>none | T>none_ATT | A/T-1_AAAAGTC | 9 | FALSE | FALSE |
| 5_v.txt | 32449 | 32449-Deleti | Deletion | 1 | C | - | 3 | 21421 | 0.01400495 | 38 | WT | FALSE | CAN | JSY13 | 1 | WT-Rep1 | CCA | nope | >1bp-ins | G/C-1 | none | G/C-1 | AAAGTCCAA | CAATGAA | GGT | CCA | C | none | nope | nope_CCA | G/C-1_CAATGAA | 9 | FALSE | FALSE |
| 5_v.txt | 32454 | 32454-Deleti | Deletion | 1 | A | - | 202 | 21421 | 0.94299986 | 51.8316813 | WT | FALSE | CAN | JSY13 | 1 | WT-Rep1 | GAA | nope | >1bp-ins | A/T-1 | none | A/T-1 | CCAATGAAT | AATAATA | CTT | CTT | T | none | T>none | T>none_CTT | A/T-1_AATAATA | 9 | FALSE | FALSE |
| 5_v.txt | 32461 | 32461-Deleti | Deletion | 1 | G | - | 5 | 21421 | 0.02334158 | 43.2 | WT | FALSE | CAN | JSY13 | 1 | WT-Rep1 | AGA | nope | >1bp-ins | G/C-1 | none | G/C-1 | ATAATAGAG | GAGAGCC | TCT | TCT | C | none | C>none | C>none_TCT | G/C-1_GAGAGCC | 9 | FALSE | FALSE |
| 5_v.txt | 32463 | 32463-Deleti | Deletion | 1 | G | - | 3 | 21421 | 0.01400495 | 38 | WT | FALSE | CAN | JSY13 | 1 | WT-Rep1 | AGA | nope | >1bp-ins | G/C-1 | none | G/C-1 | AATAGAGAG | GAGCCAA | TCT | TCT | C | none | C>none | C>none_TCT | G/C-1_GAGCCAA | 9 | FALSE | FALSE |
| 5_v.txt | 32498 | 32498-Deleti | Deletion | 1 | T | - | 44 | 21421 | 0.20540591 | 45.3409063 | WT | FALSE | CAN | JSY13 | 1 | WT-Rep1 | CTT | nope | >1bp-ins | A/T-1 | none | A/T-1 | AACAACTTTT | TTTTTTG | GAA | CTT | T | none | nope | nope_CTT | A/T-1_TTTTTTG | 9 | FALSE | FALSE |
| 5_v.txt | 32528 | 32528-Deleti | Deletion | 1 | G | - | 49 | 21421 | 0.22874749 | 42.9591833 | WT | FALSE | CAN | JSY13 | 1 | WT-Rep1 | TGG | nope | >1bp-ins | G/C-1 | none | G/C-1 | TTTTCTGGG | GGGGTTT | ACC | ACC | C | none | C>none | C>none_ACC | G/C-1_GGGGTTT | 9 | FALSE | FALSE |
| 5_v.txt | 32545 | 32545-Deleti | Deletion | 1 | C | - | 4 | 21421 | 0.01867327 | 38 | WT | FALSE | CAN | JSY13 | 1 | WT-Rep1 | CCA | nope | >1bp-ins | G/C-1 | none | G/C-1 | CTTCACCAG | CAGCAGT | GGT | CCA | C | none | nope | nope_CCA | G/C-1_CAGCAGT | 9 | FALSE | FALSE |
| 5_v.txt | 32594 | 32594-Deleti | Deletion | 1 | A | - | 14 | 22156 | 0.0631883 | 40.7857143 | WT | FALSE | CAN | JSY13 | 1 | WT-Rep1 | TAA | nope | >1bp-ins | A/T-1 | none | A/T-1 | AGCGTTAAT | AATCAAA | ATT | ATT | T | none | T>none | T>none_ATT | A/T-1_AATCAAA | 9 | FALSE | FALSE |
| 5_v.txt | 32624 | 32624-Deleti | Deletion | 1 | C | - | 22 | 22156 | 0.0992959 | 47.4545455 | WT | FALSE | CAN | JSY13 | 1 | WT-Rep1 | TCC | nope | >1bp-ins | G/C-1 | none | G/C-1 | GAACCTCCC | CCCTTCG | AGG | TCC | C | none | nope | nope_TCC | G/C-1_CCCTTCG | 9 | FALSE | FALSE |
| 5_v.txt | 32631 | 32631-Deleti | Deletion | 1 | T | - | 19 | 22156 | 0.08575555 | 48.3157891 | WT | FALSE | CAN | JSY13 | 1 | WT-Rep1 | GTT | nope | >1bp-ins | A/T-1 | none | A/T-1 | CCTTCGTTT | TTTTTAT | CAA | GTT | T | none | nope | nope_GTT | A/T-1_TTTTTAT | 9 | FALSE | FALSE |
| 5_v.txt | 32657 | 32657-Deleti | Deletion | 1 | A | - | 19 | 22156 | 0.08575555 | 58.4736615 | WT | FALSE | CAN | JSY13 | 1 | WT-Rep1 | GAC | nope | >1bp-ins | A/T-1 | none | A/T-1 | ACCTGGACC | ACCCCAG | CTG | CTG | T | none | T>none | T>none_CTG | A/T-1_ACCCCAG | 9 | FALSE | FALSE |
| 5_v.txt | 32658 | 32658-Deleti | Deletion | 1 | C | - | 37 | 22156 | 0.16699765 | 57.9999937 | WT | FALSE | CAN | JSY13 | 1 | WT-Rep1 | ACC | nope | >1bp-ins | G/C-1 | none | G/C-1 | CCTGGACCC | CCCCAGG | TGG | ACC | C | none | nope | nope_ACC | G/C-1_CCCCAGG | 9 | FALSE | FALSE |
| 5_v.txt | 32667 | 32667-Deleti | Deletion | 1 | C | - | 10 | 22156 | 0.0451345 | 58.2 | WT | FALSE | CAN | JSY13 | 1 | WT-Rep1 | ACC | nope | >1bp-ins | G/C-1 | none | G/C-1 | CAGGCACCT | CCTGGGT | TGG | ACC | C | none | nope | nope_ACC | G/C-1_CCTGGGT | 9 | FALSE | FALSE |
| 5_v.txt | 32677 | 32677-Deleti | Deletion | 1 | T | - | 17 | 22156 | 0.07672865 | 56.2941177 | WT | FALSE | CAN | JSY13 | 1 | WT-Rep1 | CTC | nope | >1bp-ins | A/T-1 | none | A/T-1 | GGTTTCTCCA | TCCAATA | GAG | CTC | T | none | nope | nope_CTC | A/T-1_TCCAATA | 9 | FALSE | FALSE |
| 5_v.txt | 32707 | 32707-Deleti | Deletion | 1 | C | - | 41 | 22177 | 0.18487622 | 62.9512195 | WT | FALSE | CAN | JSY13 | 1 | WT-Rep1 | ACC | nope | >1bp-ins | G/C-1 | none | G/C-1 | CGGTAACCC | CCCCAGC | TGG | ACC | C | none | nope | nope_ACC | G/C-1_CCCCAGC | 9 | FALSE | FALSE |
| 5_v.txt | 32715 | 32715-Deleti | Deletion | 1 | C | - | 28 | 22177 | 0.12625693 | 59.6785714 | WT | FALSE | CAN | JSY13 | 1 | WT-Rep1 | ACC | nope | >1bp-ins | G/C-1 | none | G/C-1 | CCAGCACCA | CCACAAA | TGG | ACC | C | none | nope | nope_ACC | G/C-1_CCACAAA | 9 | FALSE | FALSE |
| 5_v.txt | 32719 | 32719-Deleti | Deletion | 1 | A | - | 30 | 22177 | 0.13527529 | 62.6 | WT | FALSE | CAN | JSY13 | 1 | WT-Rep1 | CAA | nope | >1bp-ins | A/T-1 | none | A/T-1 | CACCACAAA | AAACCAT | GTT | GTT | T | none | T>none | T>none_GTT | A/T-1_AAACCAT | 9 | FALSE | FALSE |
| 5_v.txt | 32728 | 32728-Deleti | Deletion | 1 | A | - | 76 | 22177 | 0.34269739 | 60.9078963 | WT | FALSE | CAN | JSY13 | 1 | WT-Rep1 | CAA | nope | >1bp-ins | A/T-1 | none | A/T-1 | CCATACAAA | AAAAACA | GTT | GTT | T | none | T>none | T>none_GTT | A/T-1_AAAAACA | 9 | FALSE | FALSE |
| 5_v.txt | 32758 | 32758-Deleti | Deletion | 1 | C | - | 8 | 22177 | 0.03607341 | 57.2499967 | WT | FALSE | CAN | JSY13 | 1 | WT-Rep1 | GCT | nope | >1bp-ins | G/C-1 | none | G/C-1 | TAATGGCTA | CTAAAAC | CGA | GCT | C | none | nope | nope_GCT | G/C-1_CTAAAAC | 9 | FALSE | FALSE |
| 5_v.txt | 32760 | 32760-Deleti | Deletion | 1 | A | - | 16 | 22176 | 0.07215007 | 47.9999872 | WT | FALSE | CAN | JSY13 | 1 | WT-Rep1 | TAA | nope | >1bp-ins | A/T-1 | none | A/T-1 | ATGGCTAAA | AAAACTT | ATT | ATT | T | none | T>none | T>none_ATT | A/T-1_AAAACTT | 9 | FALSE | FALSE |
| 5_v.txt | 32788 | 32788-Deleti | Deletion | 1 | C | - | 2 | 22176 | 9.02E-03 | 50.9999955 | WT | FALSE | CAN | JSY13 | 1 | WT-Rep1 | TCG | nope | >1bp-ins | G/C-1 | none | G/C-1 | AGAACTCGA | CGAATTC | AGC | TCG | C | none | nope | nope_TCG | G/C-1_CGAATTC | 9 | FALSE | FALSE |
| 5_v.txt | 32811 | 32811-Deleti | Deletion | 1 | G | - | 22 | 22177 | 0.09920188 | 45.7272727 | WT | FALSE | CAN | JSY13 | 1 | WT-Rep1 | AGG | nope | >1bp-ins | G/C-1 | none | G/C-1 | TTGACAGGG | GGGAACA | TCC | TCC | C | none | C>none | C>none_TCC | G/C-1_GGGAACA | 9 | FALSE | FALSE |
| 5_v.txt | 32816 | 32816-Deleti | Deletion | 1 | C | - | 11 | 22177 | 0.04960094 | 40.9090909 | WT | FALSE | CAN | JSY13 | 1 | WT-Rep1 | ACA | nope | >1bp-ins | G/C-1 | none | G/C-1 | AGGGAACAA | CAAGTTC | TGT | ACA | C | none | nope | nope_ACA | G/C-1_CAAGTTC | 9 | FALSE | FALSE |
| 5_v.txt | 32839 | 32839-Deleti | Deletion | 1 | C | - | 7 | 22177 | 0.03156423 | 40.2857261 | WT | FALSE | CAN | JSY13 | 1 | WT-Rep1 | ACC | nope | >1bp-ins | G/C-1 | none | G/C-1 | TAATTACCCA | CCCAAAA | TGG | ACC | C | none | nope | nope_ACC | G/C-1_CCCAAAA | 9 | FALSE | FALSE |
| 5_v.txt | 32842 | 32842-Deleti | Deletion | 1 | A | - | 203 | 22177 | 0.91536276 | 43.1527036 | WT | FALSE | CAN | JSY13 | 1 | WT-Rep1 | CAA | nope | >1bp-ins | A/T-1 | none | A/T-1 | TTACCCAAA | AAAAAAT | GTT | GTT | T | none | T>none | T>none_GTT | A/T-1_AAAAAAT | 9 | FALSE | FALSE |
| 5_v.txt | 32884 | 32884-Deleti | Deletion | 1 | A | - | 25 | 24305 | 0.10285949 | 36.88 | WT | FALSE | CAN | JSY13 | 1 | WT-Rep1 | CAA | nope | >1bp-ins | A/T-1 | none | A/T-1 | ACGTCCAAA | AAAATTG | GTT | GTT | T | none | T>none | T>none_GTT | A/T-1_AAAATTG | 9 | FALSE | FALSE |
| 5_v.txt | 32920 | 32920-Deleti | Deletion | 1 | G | - | 12 | 24305 | 0.04937256 | 51.9166831 | WT | FALSE | CAN | JSY13 | 1 | WT-Rep1 | AGG | nope | >1bp-ins | G/C-1 | none | G/C-1 | GTTCCAGGG | GGGCAAA | TCC | TCC | C | none | C>none | C>none_TCC | G/C-1_GGGCAAA | 9 | FALSE | FALSE |
| 5_v.txt | 32924 | 32924-Deleti | Deletion | 1 | A | - | 24 | 24305 | 0.09874511 | 48.5833391 | WT | FALSE | CAN | JSY13 | 1 | WT-Rep1 | CAA | nope | >1bp-ins | A/T-1 | none | A/T-1 | CAGGGCAAA | AAAAGTG | GTT | GTT | T | none | T>none | T>none_GTT | A/T-1_AAAAGTG | 9 | FALSE | FALSE |
| 5_v.txt | 32934 | 32934-Deleti | Deletion | 1 | G | - | 33 | 24343 | 0.13556259 | 52.6968872 | WT | FALSE | CAN | JSY13 | 1 | WT-Rep1 | TGC | nope | >1bp-ins | G/C-1 | none | G/C-1 | GTGATTGCC | GCCCAAG | ACG | ACG | C | none | C>none | C>none_ACG | G/C-1_GCCCAAG | 9 | FALSE | FALSE |
| 5_v.txt | 32935 | 32935-Deleti | Deletion | 1 | C | - | 27 | 24343 | 0.11091484 | 52.7037216 | WT | FALSE | CAN | JSY13 | 1 | WT-Rep1 | GCC | nope | >1bp-ins | G/C-1 | none | G/C-1 | TGATTGCCC | CCCAAGA | CGG | GCC | C | none | nope | nope_GCC | G/C-1_CCCAAGA | 9 | FALSE | FALSE |
| 5_v.txt | 32941 | 32941-Deleti | Deletion | 1 | A | - | 11 | 24343 | 0.04518753 | 54.3636458 | WT | FALSE | CAN | JSY13 | 1 | WT-Rep1 | GAA | nope | >1bp-ins | A/T-1 | none | A/T-1 | CCCAAGAAA | AAAACCA | CTT | CTT | T | none | T>none | T>none_CTT | A/T-1_AAAACCA | 9 | FALSE | FALSE |
| 5_v.txt | 32986 | 32986-Deleti | Deletion | 1 | G | - | 18 | 24343 | 0.07394323 | 59.0555556 | WT | FALSE | CAN | JSY13 | 1 | WT-Rep1 | AGG | nope | >1bp-ins | G/C-1 | none | G/C-1 | GAGAAAGGA | GGAATCT | TCC | TCC | C | none | C>none | C>none_TCC | G/C-1_GGAATCT | 9 | FALSE | FALSE |
| 5_v.txt | 33000 | 33000-Deleti | Deletion | 1 | A | - | 63 | 24343 | 0.2588013 | 62.5555556 | WT | FALSE | CAN | JSY13 | 1 | WT-Rep1 | GAA | nope | >1bp-ins | A/T-1 | none | A/T-1 | TGTGAGAAA | AAAACTG | CTT | CTT | T | none | T>none | T>none_CTT | A/T-1_AAAACTG | 9 | FALSE | FALSE |
| 5_v.txt | 33072 | 33072-Deleti | Deletion | 1 | A | - | 18 | 24343 | 0.07394323 | 49 | WT | FALSE | CAN | JSY13 | 1 | WT-Rep1 | CAA | nope | >1bp-ins | A/T-1 | none | A/T-1 | TATGCCAAA | AAAGAAC | GTT | GTT | T | none | T>none | T>none_GTT | A/T-1_AAAGAAC | 9 | FALSE | FALSE |
| 5_v.txt | 33075 | 33075-Deleti | Deletion | 1 | G | - | 8 | 24343 | 0.03286366 | 53.6250032 | WT | FALSE | CAN | JSY13 | 1 | WT-Rep1 | AGA | nope | >1bp-ins | G/C-1 | none | G/C-1 | GCCAAAGAA | GAACCCA | TCT | TCT | C | none | C>none | C>none_TCT | G/C-1_GAACCCA | 9 | FALSE | FALSE |
| 5_v.txt | 33076 | 33076-Deleti | Deletion | 1 | A | - | 8 | 24343 | 0.03286366 | 59.9998672 | WT | FALSE | CAN | JSY13 | 1 | WT-Rep1 | GAA | nope | >1bp-ins | A/T-1 | none | A/T-1 | CCAAAGAAC | AACCCAT | CTT | CTT | T | none | T>none | T>none_CTT | A/T-1_AACCCAT | 9 | FALSE | FALSE |
| 5_v.txt | 33078 | 33078-Deleti | Deletion | 1 | C | - | 12 | 24343 | 0.04929549 | 52.7499999 | WT | FALSE | CAN | JSY13 | 1 | WT-Rep1 | ACC | nope | >1bp-ins | G/C-1 | none | G/C-1 | AAAGAACCC | CCCATAA | TGG | ACC | C | none | nope | nope_ACC | G/C-1_CCCATAA | 9 | FALSE | FALSE |
| 5_v.txt | 33083 | 33083-Deleti | Deletion | 1 | A | - | 20 | 24343 | 0.08215914 | 49.2500348 | WT | FALSE | CAN | JSY13 | 1 | WT-Rep1 | TAA | nope | >1bp-ins | A/T-1 | none | A/T-1 | ACCCATAAA | AAATAAA | ATT | ATT | T | none | T>none | T>none_ATT | A/T-1_AAATAAA | 9 | FALSE | FALSE |
| 5_v.txt | 33111 | 33111-Deleti | Deletion | 1 | G | - | 82 | 24343 | 0.33685248 | 41.1097561 | WT | FALSE | CAN | JSY13 | 1 | WT-Rep1 | TGG | nope | >1bp-ins | G/C-1 | none | G/C-1 | CCCACTGGG | GGGCCGG | ACC | ACC | C | none | C>none | C>none_ACC | G/C-1_GGGCCGG | 9 | FALSE | FALSE |
| 5_v.txt | 33114 | 33114-Deleti | Deletion | 1 | C | - | 1 | 24343 | 4.11E-03 | 63.9999887 | WT | FALSE | CAN | JSY13 | 1 | WT-Rep1 | GCC | nope | >1bp-ins | G/C-1 | none | G/C-1 | ACTGGGCCG | CCGGCGT | CGG | GCC | C | none | nope | nope_GCC | G/C-1_CCGGCGT | 9 | FALSE | FALSE |
| 5_v.txt | 33147 | 33147-Deleti | Deletion | 1 | A | - | 47 | 24343 | 0.19307398 | 39.31916 | WT | FALSE | CAN | JSY13 | 1 | WT-Rep1 | GAA | nope | >1bp-ins | A/T-1 | none | A/T-1 | CCAATGAAA | AAAAGAC | CTT | CTT | T | none | T>none | T>none_CTT | A/T-1_AAAAGAC | 9 | FALSE | FALSE |
| 5_v.txt | 33153 | 33153-Deleti | Deletion | 1 | C | - | 5 | 24343 | 0.02053979 | 37.6000001 | WT | FALSE | CAN | JSY13 | 1 | WT-Rep1 | ACC | nope | >1bp-ins | G/C-1 | none | G/C-1 | AAAAGACCT | CCTGTAC | TGG | ACC | C | none | nope | nope_ACC | G/C-1_CCTGTAC | 9 | FALSE | FALSE |
| 5_v.txt | 33168 | 33168-Deleti | Deletion | 1 | C | - | 4 | 24300 | 0.01646091 | 36.5 | WT | FALSE | CAN | JSY13 | 1 | WT-Rep1 | ACC | nope | >1bp-ins | G/C-1 | none | G/C-1 | ATAGTACCA | CCACCAA | TGG | ACC | C | none | nope | nope_ACC | G/C-1_CCACCAA | 9 | FALSE | FALSE |
| 5_v.txt | 33171 | 33171-Deleti | Deletion | 1 | C | - | 2 | 24300 | 8.23E-03 | 36.5000006 | WT | FALSE | CAN | JSY13 | 1 | WT-Rep1 | ACC | nope | >1bp-ins | G/C-1 | none | G/C-1 | GTACCACCA | CCAAGGG | TGG | ACC | C | none | nope | nope_ACC | G/C-1_CCAAGGG | 9 | FALSE | FALSE |
| 5_v.txt | 33175 | 33175-Deleti | Deletion | 1 | G | - | 7 | 24298 | 0.02880896 | 37.7142857 | WT | FALSE | CAN | JSY13 | 1 | WT-Rep1 | AGG | nope | >1bp-ins | G/C-1 | none | G/C-1 | CACCAAGGG | GGGCAAT | TCC | TCC | C | none | C>none | C>none_TCC | G/C-1_GGGCAAT | 9 | FALSE | FALSE |
| 5_v.txt | 33178 | 33178-Deleti | Deletion | 1 | C | - | 12 | 24298 | 0.04938678 | 37.9166654 | WT | FALSE | CAN | JSY13 | 1 | WT-Rep1 | GCA | nope | >1bp-ins | G/C-1 | none | G/C-1 | CAAGGGCAA | CAATCAT | CGT | GCA | C | none | nope | nope_GCA | G/C-1_CAATCAT | 9 | FALSE | FALSE |
| 5_v.txt | 33207 | 33207-Deleti | Deletion | 1 | T | - | 4 | 7277 | 0.05496771 | 36.5 | WT | FALSE | CAN | JSY13 | 1 | WT-Rep1 | CTC | nope | >1bp-ins | A/T-1 | none | A/T-1 | TTAAGCTCTC | TCTCTCT | GAG | CTC | T | none | nope | nope_CTC | A/T-1_TCTCTCT | 9 | FALSE | FALSE |
| 5_v.txt | 33373 | 33373-Deleti | Deletion | 1 | T | - | 8 | 7277 | 0.10993541 | 49.75 | WT | FALSE | CAN | JSY13 | 1 | WT-Rep1 | GTT | nope | >1bp-ins | A/T-1 | none | A/T-1 | GGTGTGTTT | TTTGTGA | CAA | GTT | T | none | nope | nope_GTT | A/T-1_TTTGTGA | 9 | FALSE | FALSE |
| 5_v.txt | 33474 | 33474-Deleti | Deletion | 1 | C | - | 19 | 7275 | 0.26116839 | 29 | WT | FALSE | CAN | JSY13 | 1 | WT-Rep1 | GCC | nope | >1bp-ins | G/C-1 | none | G/C-1 | GCTATGCCTT | CCTTTTT | CGG | GCC | C | none | nope | nope_GCC | G/C-1_CCTTTTT | 9 | FALSE | FALSE |
| 6_v.txt | 31986 | 31986-SNV-A | SNV | 1 | A | C | 16 | 21739 | 0.07360044 | 52.1875 | WT | FALSE | CAN | JSY13 | 2 | WT-Rep2 | TAA | A>C | G/C+1 | A/T-1 | TA>GC | TA>GC | GGCATTAAT | AATTTAG | ATT | ATT | T | G | T>G | T>G_ATT | TA>GC_AATTTAG | 9 | FALSE | FALSE |
| 6_v.txt | 32162 | 32162-SNV-A | SNV | 1 | A | C | 4 | 21737 | 0.0184018 | 16.9999328 | WT | FALSE | CAN | JSY13 | 2 | WT-Rep2 | AAG | A>C | G/C+1 | A/T-1 | TA>GC | TA>GC | AGCCAAAGC | AGCGCCA | TTC | TTC | T | G | T>G | T>G_TTC | TA>GC_AGCGCCA | 9 | FALSE | FALSE |
| 6_v.txt | 32319 | 32319-SNV-A | SNV | 1 | A | C | 6 | 32056 | 0.01871725 | 54 | WT | FALSE | CAN | JSY13 | 2 | WT-Rep2 | TAA | A>C | G/C+1 | A/T-1 | TA>GC | TA>GC | GTTGTTAAG | AAGATAA | ATT | ATT | T | G | T>G | T>G_ATT | TA>GC_AAGATAA | 9 | FALSE | FALSE |
| 6_v.txt | 32441 | 32441-SNV-T | SNV | 1 | T | G | 11 | 32055 | 0.03431602 | 51.5442688 | WT | FALSE | CAN | JSY13 | 2 | WT-Rep2 | CTA | T>G | G/C+1 | A/T-1 | TA>GC | TA>GC | TGGAACTAA | TAAAAGT | GAT | CTA | T | C | T>G | T>G_CTA | TA>GC_TAAAAGT | 9 | FALSE | FALSE |
| 6_v.txt | 32445 | 32445-SNV-A | SNV | 1 | A | C | 11 | 32056 | 0.03431495 | 45.3636364 | WT | FALSE | CAN | JSY13 | 2 | WT-Rep2 | AAG | A>C | G/C+1 | A/T-1 | TA>GC | TA>GC | ACTAAAAGT | AGTCCAA | TTC | TTC | T | G | T>G | T>G_TTC | TA>GC_AGTCCAA | 9 | FALSE | FALSE |
| 6_v.txt | 32472 | 32472-SNV-T | SNV | 1 | T | G | 38 | 32056 | 0.11854255 | 42.6328391 | WT | FALSE | CAN | JSY13 | 2 | WT-Rep2 | GTA | T>G | G/C+1 | A/T-1 | TA>GC | TA>GC | CCAATGTAG | TAGAAGG | CAT | GTA | T | C | T>G | T>G_GTA | TA>GC_TAGAAGG | 9 | FALSE | FALSE |
| 6_v.txt | 32473 | 32473-SNV-A | SNV | 1 | A | C | 6 | 32056 | 0.01871725 | 40.167766 | WT | FALSE | CAN | JSY13 | 2 | WT-Rep2 | TAG | A>C | G/C+1 | A/T-1 | TA>GC | TA>GC | CAATGTAGA | AGAAGGT | ATC | ATC | T | G | T>G | T>G_ATC | TA>GC_AGAAGGT | 9 | FALSE | FALSE |
| 6_v.txt | 32573 | 32573-SNV-T | SNV | 1 | T | G | 62 | 25283 | 0.24522406 | 36.0483871 | WT | FALSE | CAN | JSY13 | 2 | WT-Rep2 | CTT | T>G | G/C+1 | A/T-1 | TA>GC | TA>GC | AGTACCTTG | TTGAAAT | GAA | CTT | T | C | T>G | T>G_CTT | TA>GC_TTGAAAT | 9 | FALSE | FALSE |
| 6_v.txt | 32574 | 32574-SNV-T | SNV | 1 | T | G | 15 | 25283 | 0.0593284 | 36.6656349 | WT | FALSE | CAN | JSY13 | 2 | WT-Rep2 | TTG | T>G | G/C+1 | A/T-1 | TA>GC | TA>GC | GTACCTTGA | TGAAATG | AAC | TTG | T | C | T>G | T>G_TTG | TA>GC_TGAAATG | 9 | FALSE | FALSE |
| 6_v.txt | 32631 | 32631-SNV-T | SNV | 1 | T | G | 2 | 25283 | 7.91E-03 | 63.9992793 | WT | FALSE | CAN | JSY13 | 2 | WT-Rep2 | GTT | T>G | G/C+1 | A/T-1 | TA>GC | TA>GC | CCTTCGTTT | TTTTTAT | CAA | GTT | T | C | T>G | T>G_GTT | TA>GC_TTTTTAT | 9 | FALSE | FALSE |
| 6_v.txt | 32705 | 32705-SNV-A | SNV | 1 | A | C | 9 | 25283 | 0.03559704 | 50.2222222 | WT | FALSE | CAN | JSY13 | 2 | WT-Rep2 | TAA | A>C | G/C+1 | A/T-1 | TA>GC | TA>GC | GCCGGTAAC | AACCCCA | ATT | ATT | T | G | T>G | T>G_ATT | TA>GC_AACCCCA | 9 | FALSE | FALSE |
| 6_v.txt | 32842 | 32842-SNV-A | SNV | 1 | A | C | 5 | 25283 | 0.01977613 | 31.6003198 | WT | FALSE | CAN | JSY13 | 2 | WT-Rep2 | CAA | A>C | G/C+1 | A/T-1 | TA>GC | TA>GC | TTACCCAAA | AAAAAAT | GTT | GTT | T | G | T>G | T>G_GTT | TA>GC_AAAAAAT | 9 | FALSE | FALSE |
| 6_v.txt | 32884 | 32884-SNV-A | SNV | 1 | A | C | 7 | 20701 | 0.03381479 | 32.8574681 | WT | FALSE | CAN | JSY13 | 2 | WT-Rep2 | CAA | A>C | G/C+1 | A/T-1 | TA>GC | TA>GC | ACGTCCAAA | AAAATTG | GTT | GTT | T | G | T>G | T>G_GTT | TA>GC_AAAATTG | 9 | FALSE | FALSE |
| 6_v.txt | 32915 | 32915-SNV-T | SNV | 1 | T | G | 11 | 20702 | 0.05313496 | 33 | WT | FALSE | CAN | JSY13 | 2 | WT-Rep2 | GTT | T>G | G/C+1 | A/T-1 | TA>GC | TA>GC | ACTAAGTTC | TTCCAGG | CAA | GTT | T | C | T>G | T>G_GTT | TA>GC_TTCCAGG | 9 | FALSE | FALSE |
| 6_v.txt | 32919 | 32919-SNV-A | SNV | 1 | A | C | 27 | 20702 | 0.13042218 | 41.4814815 | WT | FALSE | CAN | JSY13 | 2 | WT-Rep2 | CAG | A>C | G/C+1 | A/T-1 | TA>GC | TA>GC | AGTTCCAGG | AGGGCAA | GTC | GTC | T | G | T>G | T>G_GTC | TA>GC_AGGGCAA | 9 | FALSE | FALSE |
| 6_v.txt | 33099 | 33099-SNV-A | SNV | 1 | A | C | 7 | 20702 | 0.03381316 | 48.1428571 | WT | FALSE | CAN | JSY13 | 2 | WT-Rep2 | AAG | A>C | G/C+1 | A/T-1 | TA>GC | TA>GC | GATATAAGA | AGAGCGC | TTC | TTC | T | G | T>G | T>G_TTC | TA>GC_AGAGCGC | 9 | FALSE | FALSE |
| 6_v.txt | 33204 | 33204-SNV-A | SNV | 1 | A | C | 19 | 13775 | 0.13793104 | 32.631579 | WT | FALSE | CAN | JSY13 | 2 | WT-Rep2 | AAG | A>C | G/C+1 | A/T-1 | TA>GC | TA>GC | TGCTTAAGC | AGCTCTC | TTC | TTC | T | G | T>G | T>G_TTC | TA>GC_AGCTCTC | 9 | FALSE | FALSE |
| 6_v.txt | 33207 | 33207-SNV-T | SNV | 1 | T | G | 1 | 13775 | 7.26E-03 | 38.0000001 | WT | FALSE | CAN | JSY13 | 2 | WT-Rep2 | CTC | T>G | G/C+1 | A/T-1 | TA>GC | TA>GC | TTAAGCTCTC | TCTCTCT | GAG | CTC | T | C | T>G | T>G_CTC | TA>GC_TCTCTCT | 9 | FALSE | FALSE |
| 6_v.txt | 31965 | 31965-SNV-T | SNV | 1 | T | C | 6 | 21739 | 0.02760017 | 37.3333455 | WT | FALSE | CAN | JSY13 | 2 | WT-Rep2 | ATA | T>C | G/C+1 | A/T-1 | TA>CG | TA>CG | GCCGCATAA | TAATAAG | TAT | ATA | T | G | T>C | T>C_ATA | TA>CG_TAATAAG | 9 | FALSE | FALSE |
| 6_v.txt | 32081 | 32081-SNV-A | SNV | 1 | A | G | 3 | 21739 | 0.01380008 | 56.6666252 | WT | FALSE | CAN | JSY13 | 2 | WT-Rep2 | CAA | A>G | G/C+1 | A/T-1 | TA>CG | TA>CG | CCATGCAAA | AAAAAAG | GTT | GTT | T | C | T>C | T>C_GTT | TA>CG_AAAAAAG | 9 | FALSE | FALSE |
| 6_v.txt | 32162 | 32162-SNV-A | SNV | 1 | A | G | 2 | 21737 | 9.20E-03 | 50.9984205 | WT | FALSE | CAN | JSY13 | 2 | WT-Rep2 | AAG | A>G | G/C+1 | A/T-1 | TA>CG | TA>CG | AGCCAAAGC | AGCGCCA | TTC | TTC | T | C | T>C | T>C_TTC | TA>CG_AGCGCCA | 9 | FALSE | FALSE |
| 6_v.txt | 32337 | 32337-SNV-A | SNV | 1 | A | G | 4 | 32056 | 0.01247816 | 39.7499874 | WT | FALSE | CAN | JSY13 | 2 | WT-Rep2 | GAT | A>G | G/C+1 | A/T-1 | TA>CG | TA>CG | TTGAAGATA | ATATGTG | CTA | CTA | T | C | T>C | T>C_CTA | TA>CG_ATATGTG | 9 | FALSE | FALSE |
| 6_v.txt | 32441 | 32441-SNV-T | SNV | 1 | T | C | 10 | 32055 | 0.03119638 | 46.9999309 | WT | FALSE | CAN | JSY13 | 2 | WT-Rep2 | CTA | T>C | G/C+1 | A/T-1 | TA>CG | TA>CG | TGGAACTAA | TAAAAGT | GAT | CTA | T | G | T>C | T>C_CTA | TA>CG_TAAAAGT | 9 | FALSE | FALSE |
| 6_v.txt | 32445 | 32445-SNV-A | SNV | 1 | A | G | 49 | 32056 | 0.1528575 | 46 | WT | FALSE | CAN | JSY13 | 2 | WT-Rep2 | AAG | A>G | G/C+1 | A/T-1 | TA>CG | TA>CG | ACTAAAAGT | AGTCCAA | TTC | TTC | T | C | T>C | T>C_TTC | TA>CG_AGTCCAA | 9 | FALSE | FALSE |
| 6_v.txt | 32473 | 32473-SNV-A | SNV | 1 | A | G | 12 | 32056 | 0.03743449 | 42.6670311 | WT | FALSE | CAN | JSY13 | 2 | WT-Rep2 | TAG | A>G | G/C+1 | A/T-1 | TA>CG | TA>CG | CAATGTAGA | AGAAGGT | ATC | ATC | T | C | T>C | T>C_ATC | TA>CG_AGAAGGT | 9 | FALSE | FALSE |
| 6_v.txt | 32476 | 32476-SNV-A | SNV | 1 | A | G | 26 | 32056 | 0.08110806 | 42.4999061 | WT | FALSE | CAN | JSY13 | 2 | WT-Rep2 | AAG | A>G | G/C+1 | A/T-1 | TA>CG | TA>CG | TGTAGAAGG | AGGTTAA | TTC | TTC | T | C | T>C | T>C_TTC | TA>CG_AGGTTAA | 9 | FALSE | FALSE |
| 6_v.txt | 32479 | 32479-SNV-T | SNV | 1 | T | C | 5 | 32056 | 0.0155977 | 36.8000373 | WT | FALSE | CAN | JSY13 | 2 | WT-Rep2 | GTT | T>C | G/C+1 | A/T-1 | TA>CG | TA>CG | AGAAGGTTA | TTAAGAT | CAA | GTT | T | G | T>C | T>C_GTT | TA>CG_TTAAGAT | 9 | FALSE | FALSE |
| 6_v.txt | 32573 | 32573-SNV-T | SNV | 1 | T | C | 11 | 25283 | 0.0435075 | 35.4545455 | WT | FALSE | CAN | JSY13 | 2 | WT-Rep2 | CTT | T>C | G/C+1 | A/T-1 | TA>CG | TA>CG | AGTACCTTG | TTGAAAT | GAA | CTT | T | G | T>C | T>C_CTT | TA>CG_TTGAAAT | 9 | FALSE | FALSE |
| 6_v.txt | 32583 | 32583-SNV-A | SNV | 1 | A | G | 22 | 25283 | 0.08701499 | 34.5454546 | WT | FALSE | CAN | JSY13 | 2 | WT-Rep2 | GAA | A>G | G/C+1 | A/T-1 | TA>CG | TA>CG | AATGTGAAG | AAGGCAG | CTT | CTT | T | C | T>C | T>C_CTT | TA>CG_AAGGCAG | 9 | FALSE | FALSE |
| 6_v.txt | 32605 | 32605-SNV-A | SNV | 1 | A | G | 7 | 25281 | 0.02768878 | 30.9999078 | WT | FALSE | CAN | JSY13 | 2 | WT-Rep2 | GAA | A>G | G/C+1 | A/T-1 | TA>CG | TA>CG | AAGAGGAAA | AAACCCA | CTT | CTT | T | C | T>C | T>C_CTT | TA>CG_AAACCCA | 9 | FALSE | FALSE |
| 6_v.txt | 32719 | 32719-SNV-A | SNV | 1 | A | G | 7 | 25282 | 0.02768768 | 27.8571345 | WT | FALSE | CAN | JSY13 | 2 | WT-Rep2 | CAA | A>G | G/C+1 | A/T-1 | TA>CG | TA>CG | CACCACAAA | AAACCAT | GTT | GTT | T | C | T>C | T>C_GTT | TA>CG_AAACCAT | 9 | FALSE | FALSE |
| 6_v.txt | 32767 | 32767-SNV-T | SNV | 1 | T | C | 13 | 25283 | 0.05141795 | 53.3841207 | WT | FALSE | CAN | JSY13 | 2 | WT-Rep2 | TTG | T>C | G/C+1 | A/T-1 | TA>CG | TA>CG | AAACTTTGA | TGATGGA | AAC | TTG | T | G | T>C | T>C_TTG | TA>CG_TGATGGA | 9 | FALSE | FALSE |
| 6_v.txt | 32769 | 32769-SNV-A | SNV | 1 | A | G | 8 | 25283 | 0.03164182 | 44.372856 | WT | FALSE | CAN | JSY13 | 2 | WT-Rep2 | GAT | A>G | G/C+1 | A/T-1 | TA>CG | TA>CG | ACTTTGATG | ATGGAAG | CTA | CTA | T | C | T>C | T>C_CTA | TA>CG_ATGGAAG | 9 | FALSE | FALSE |
| 6_v.txt | 32884 | 32884-SNV-A | SNV | 1 | A | G | 9 | 20701 | 0.04347616 | 30.5557672 | WT | FALSE | CAN | JSY13 | 2 | WT-Rep2 | CAA | A>G | G/C+1 | A/T-1 | TA>CG | TA>CG | ACGTCCAAA | AAAATTG | GTT | GTT | T | C | T>C | T>C_GTT | TA>CG_AAAATTG | 9 | FALSE | FALSE |
| 6_v.txt | 32919 | 32919-SNV-A | SNV | 1 | A | G | 14 | 20702 | 0.06762632 | 45.2857143 | WT | FALSE | CAN | JSY13 | 2 | WT-Rep2 | CAG | A>G | G/C+1 | A/T-1 | TA>CG | TA>CG | AGTTCCAGG | AGGGCAA | GTC | GTC | T | C | T>C | T>C_GTC | TA>CG_AGGGCAA | 9 | FALSE | FALSE |
| 6_v.txt | 32947 | 32947-SNV-A | SNV | 1 | A | G | 26 | 20702 | 0.12559173 | 56.1923077 | WT | FALSE | CAN | JSY13 | 2 | WT-Rep2 | CAA | A>G | G/C+1 | A/T-1 | TA>CG | TA>CG | AAAACCAAT | AATACAT | GTT | GTT | T | C | T>C | T>C_GTT | TA>CG_AATACAT | 9 | FALSE | FALSE |
| 6_v.txt | 33099 | 33099-SNV-A | SNV | 1 | A | G | 13 | 20702 | 0.06279587 | 47 | WT | FALSE | CAN | JSY13 | 2 | WT-Rep2 | AAG | A>G | G/C+1 | A/T-1 | TA>CG | TA>CG | GATATAAGA | AGAGCGC | TTC | TTC | T | C | T>C | T>C_TTC | TA>CG_AGAGCGC | 9 | FALSE | FALSE |
| 6_v.txt | 33126 | 33126-SNV-A | SNV | 1 | A | G | 21 | 20702 | 0.10143947 | 41.9045379 | WT | FALSE | CAN | JSY13 | 2 | WT-Rep2 | CAG | A>G | G/C+1 | A/T-1 | TA>CG | TA>CG | TTGGTCAGA | AGAGGTG | GTC | GTC | T | C | T>C | T>C_GTC | TA>CG_AGAGGTG | 9 | FALSE | FALSE |
| 6_v.txt | 33207 | 33207-SNV-T | SNV | 1 | T | C | 10 | 13775 | 0.07259528 | 25.7000004 | WT | FALSE | CAN | JSY13 | 2 | WT-Rep2 | CTC | T>C | G/C+1 | A/T-1 | TA>CG | TA>CG | TTAAGCTCTC | TCTCTCT | GAG | CTC | T | G | T>C | T>C_CTC | TA>CG_TCTCTCT | 9 | FALSE | FALSE |
| 6_v.txt | 33209 | 33209-SNV-T | SNV | 1 | T | C | 6 | 13775 | 0.04355717 | 38.3332984 | WT | FALSE | CAN | JSY13 | 2 | WT-Rep2 | CTC | T>C | G/C+1 | A/T-1 | TA>CG | TA>CG | AAGCTCTCTC | TCTCTTC | GAG | CTC | T | G | T>C | T>C_CTC | TA>CG_TCTCTTC | 9 | FALSE | FALSE |
| 6_v.txt | 33211 | 33211-SNV-T | SNV | 1 | T | C | 10 | 13776 | 0.07259001 | 34.6001251 | WT | FALSE | CAN | JSY13 | 2 | WT-Rep2 | CTC | T>C | G/C+1 | A/T-1 | TA>CG | TA>CG | GCTCTCTCTT | TCTTCAC | GAG | CTC | T | G | T>C | T>C_CTC | TA>CG_TCTTCAC | 9 | FALSE | FALSE |
| 6_v.txt | 33476 | 33476-SNV-T | SNV | 1 | T | C | 25 | 13774 | 0.18150138 | 29.52 | WT | FALSE | CAN | JSY13 | 2 | WT-Rep2 | CTT | T>C | G/C+1 | A/T-1 | TA>CG | TA>CG | TATGCCTTTT | TTTTTTT | GAA | CTT | T | G | T>C | T>C_CTT | TA>CG_TTTTTTT | 9 | FALSE | FALSE |
| 6_v.txt | 31965 | 31965-SNV-T | SNV | 1 | T | A | 4 | 21739 | 0.01840011 | 31.0001671 | WT | FALSE | CAN | JSY13 | 2 | WT-Rep2 | ATA | T>A | A/T+1 | A/T-1 | TA>AT | TA>AT | GCCGCATAA | TAATAAG | TAT | ATA | T | T | T>A | T>A_ATA | TA>AT_TAATAAG | 9 | FALSE | FALSE |
| 6_v.txt | 32081 | 32081-SNV-A | SNV | 1 | A | T | 1 | 21739 | 4.60E-03 | 61.9997491 | WT | FALSE | CAN | JSY13 | 2 | WT-Rep2 | CAA | A>T | A/T+1 | A/T-1 | TA>AT | TA>AT | CCATGCAAA | AAAAAAG | GTT | GTT | T | A | T>A | T>A_GTT | TA>AT_AAAAAAG | 9 | FALSE | FALSE |
| 6_v.txt | 32265 | 32265-SNV-A | SNV | 1 | A | T | 25 | 32056 | 0.07798852 | 37.4 | WT | FALSE | CAN | JSY13 | 2 | WT-Rep2 | TAA | A>T | A/T+1 | A/T-1 | TA>AT | TA>AT | CCAAATAAA | AAAATAC | ATT | ATT | T | A | T>A | T>A_ATT | TA>AT_AAAATAC | 9 | FALSE | FALSE |
| 6_v.txt | 32319 | 32319-SNV-A | SNV | 1 | A | T | 239 | 32056 | 0.74557025 | 58.6736402 | WT | FALSE | CAN | JSY13 | 2 | WT-Rep2 | TAA | A>T | A/T+1 | A/T-1 | TA>AT | TA>AT | GTTGTTAAG | AAGATAA | ATT | ATT | T | A | T>A | T>A_ATT | TA>AT_AAGATAA | 9 | FALSE | FALSE |
| 6_v.txt | 32337 | 32337-SNV-A | SNV | 1 | A | T | 5 | 32056 | 0.0155977 | 58.5995433 | WT | FALSE | CAN | JSY13 | 2 | WT-Rep2 | GAT | A>T | A/T+1 | A/T-1 | TA>AT | TA>AT | TTGAAGATA | ATATGTG | CTA | CTA | T | A | T>A | T>A_CTA | TA>AT_ATATGTG | 9 | FALSE | FALSE |
| 6_v.txt | 32422 | 32422-SNV-T | SNV | 1 | T | A | 30 | 32056 | 0.09358622 | 53.9666667 | WT | FALSE | CAN | JSY13 | 2 | WT-Rep2 | TTA | T>A | A/T+1 | A/T-1 | TA>AT | TA>AT | TTAGTTTAG | TAGGGTC | AAT | TTA | T | T | T>A | T>A_TTA | TA>AT_TAGGGTC | 9 | FALSE | FALSE |
| 6_v.txt | 32479 | 32479-SNV-T | SNV | 1 | T | A | 3 | 32056 | 9.36E-03 | 29.0000266 | WT | FALSE | CAN | JSY13 | 2 | WT-Rep2 | GTT | T>A | A/T+1 | A/T-1 | TA>AT | TA>AT | AGAAGGTTA | TTAAGAT | CAA | GTT | T | T | T>A | T>A_GTT | TA>AT_TTAAGAT | 9 | FALSE | FALSE |
| 6_v.txt | 32498 | 32498-SNV-T | SNV | 1 | T | A | 6 | 32056 | 0.01871725 | 37.9973218 | WT | FALSE | CAN | JSY13 | 2 | WT-Rep2 | CTT | T>A | A/T+1 | A/T-1 | TA>AT | TA>AT | AACAACTTTT | TTTTTTG | GAA | CTT | T | T | T>A | T>A_CTT | TA>AT_TTTTTTG | 9 | FALSE | FALSE |
| 6_v.txt | 32541 | 32541-SNV-T | SNV | 1 | T | A | 20 | 32056 | 0.06239082 | 37.7 | WT | FALSE | CAN | JSY13 | 2 | WT-Rep2 | TTC | T>A | A/T+1 | A/T-1 | TA>AT | TA>AT | GCAGCTTCA | TCACCAG | AAG | TTC | T | T | T>A | T>A_TTC | TA>AT_TCACCAG | 9 | FALSE | FALSE |
| 6_v.txt | 32573 | 32573-SNV-T | SNV | 1 | T | A | 11 | 25283 | 0.0435075 | 36.3636364 | WT | FALSE | CAN | JSY13 | 2 | WT-Rep2 | CTT | T>A | A/T+1 | A/T-1 | TA>AT | TA>AT | AGTACCTTG | TTGAAAT | GAA | CTT | T | T | T>A | T>A_CTT | TA>AT_TTGAAAT | 9 | FALSE | FALSE |
| 6_v.txt | 32605 | 32605-SNV-A | SNV | 1 | A | T | 11 | 25281 | 0.04351094 | 15.4544035 | WT | FALSE | CAN | JSY13 | 2 | WT-Rep2 | GAA | A>T | A/T+1 | A/T-1 | TA>AT | TA>AT | AAGAGGAAA | AAACCCA | CTT | CTT | T | A | T>A | T>A_CTT | TA>AT_AAACCCA | 9 | FALSE | FALSE |
| 6_v.txt | 32719 | 32719-SNV-A | SNV | 1 | A | T | 1 | 25282 | 3.96E-03 | 63.9996537 | WT | FALSE | CAN | JSY13 | 2 | WT-Rep2 | CAA | A>T | A/T+1 | A/T-1 | TA>AT | TA>AT | CACCACAAA | AAACCAT | GTT | GTT | T | A | T>A | T>A_GTT | TA>AT_AAACCAT | 9 | FALSE | FALSE |
| 6_v.txt | 32767 | 32767-SNV-T | SNV | 1 | T | A | 3 | 25283 | 0.01186568 | 28.6660485 | WT | FALSE | CAN | JSY13 | 2 | WT-Rep2 | TTG | T>A | A/T+1 | A/T-1 | TA>AT | TA>AT | AAACTTTGA | TGATGGA | AAC | TTG | T | T | T>A | T>A_TTG | TA>AT_TGATGGA | 9 | FALSE | FALSE |

| | | | | | | | | | | | | | | | | | | | | | | | | | | | | | | | | | | |
|---|---|---|---|---|---|---|---|---|---|---|---|---|---|---|---|---|---|---|---|---|---|---|---|---|---|---|---|---|---|---|---|---|---|---|
| 6_v.txt | 32929 | 32929-SNV-T | SNV | 1 | T | A | 16 | 20702 | 0.07728722 | 53 | WT | FALSE | CAN | JSY13 | 2 | WT-Rep2 | GTG | T>A | A/T+1 | A/T-1 | TA>AT | TA>AT | CAAAAGTGA | TGATTGC | CAC | GTG | T | T | T>A | T>A_GTG | TA>AT_TGATTGC | 9 | FALSE | FALSE |
| 6_v.txt | 33030 | 33030-SNV- | SNV | 1 | A | T | 2 | 20702 | 9.66E-03 | 30.9999692 | WT | FALSE | CAN | JSY13 | 2 | WT-Rep2 | GAA | A>T | A/T+1 | A/T-1 | TA>AT | TA>AT | GGGATGAAT | AATGTAG | CTT | CTT | T | A | T>A | T>A_CTT | TA>AT_AATGTAG | 9 | FALSE | FALSE |
| 6_v.txt | 33072 | 33072-SNV- | SNV | 1 | A | T | 10 | 20702 | 0.04830451 | 50.9 | WT | FALSE | CAN | JSY13 | 2 | WT-Rep2 | CAA | A>T | A/T+1 | A/T-1 | TA>AT | TA>AT | TATGCCAAA | AAAGAAC | GTT | GTT | T | A | T>A | T>A_GTT | TA>AT_AAAGAAC | 9 | FALSE | FALSE |
| 6_v.txt | 33204 | 33204-SNV- | SNV | 1 | A | T | 10 | 13775 | 0.07259528 | 37.5 | WT | FALSE | CAN | JSY13 | 2 | WT-Rep2 | AAG | A>T | A/T+1 | A/T-1 | TA>AT | TA>AT | TGCTTAAGC | AGCTCTC | TTC | TTC | T | A | T>A | T>A_TTC | TA>AT_AGCTCTC | 9 | FALSE | FALSE |
| 6_v.txt | 31650 | 31650-SNV- | SNV | 1 | C | T | 32 | 11614 | 0.27552953 | 36.5625 | WT | FALSE | CAN | JSY13 | 2 | WT-Rep2 | GCG | C>T | A/T+1 | G/C-1 | CG>TA | CG>TA | AGAATGCGA | CGAAATG | CGC | GCG | C | A | C>T | C>T_GCG | CG>TA_CGAAATG | 9 | FALSE | FALSE |
| 6_v.txt | 31806 | 31806-SNV- | SNV | 1 | C | T | 545 | 11614 | 4.69261236 | 58.3761468 | WT | FALSE | CAN | JSY13 | 2 | WT-Rep2 | CCA | C>T | A/T+1 | G/C-1 | CG>TA | CG>TA | ATCTTCCAAA | CAAATAA | GGT | CCA | C | A | C>T | C>T_CCA | CG>TA_CAAATAA | 9 | FALSE | FALSE |
| 6_v.txt | 31844 | 31844-SNV- | SNV | 1 | C | T | 19 | 11614 | 0.16359566 | 49.1578947 | WT | FALSE | CAN | JSY13 | 2 | WT-Rep2 | TCC | C>T | A/T+1 | G/C-1 | CG>TA | CG>TA | TAAGATCCA | CCAAACA | AGG | TCC | C | A | C>T | C>T_TCC | CG>TA_CCAAACA | 9 | FALSE | FALSE |
| 6_v.txt | 31962 | 31962-SNV- | SNV | 1 | G | A | 41 | 21739 | 0.18860113 | 35.902439 | WT | FALSE | CAN | JSY13 | 2 | WT-Rep2 | CGC | G>A | A/T+1 | G/C-1 | CG>TA | CG>TA | GTGGCCGCA | GCATAAT | GCG | GCG | C | T | C>T | C>T_GCG | CG>TA_GCATAAT | 9 | FALSE | FALSE |
| 6_v.txt | 31981 | 31981-SNV- | SNV | 1 | G | A | 22 | 21739 | 0.10120061 | 47.2727273 | WT | FALSE | CAN | JSY13 | 2 | WT-Rep2 | GGC | G>A | A/T+1 | G/C-1 | CG>TA | CG>TA | AGCCGGGCA | GCATTAA | CCG | CCG | C | T | C>T | C>T_CCG | CG>TA_GCATTAA | 9 | FALSE | FALSE |
| 6_v.txt | 32041 | 32041-SNV- | SNV | 1 | G | A | 22 | 21739 | 0.10120061 | 56 | WT | FALSE | CAN | JSY13 | 2 | WT-Rep2 | TGC | G>A | A/T+1 | G/C-1 | CG>TA | CG>TA | AAGCTTGCA | GCATAAA | ACG | ACG | C | T | C>T | C>T_ACG | CG>TA_GCATAAA | 9 | FALSE | FALSE |
| 6_v.txt | 32057 | 32057-SNV- | SNV | 1 | C | T | 12 | 21739 | 0.05520033 | 57 | WT | FALSE | CAN | JSY13 | 2 | WT-Rep2 | GCG | C>T | A/T+1 | G/C-1 | CG>TA | CG>TA | GATGTGCGA | CGAGATT | CGC | GCG | C | A | C>T | C>T_GCG | CG>TA_CGAGATT | 9 | FALSE | FALSE |
| 6_v.txt | 32076 | 32076-SNV- | SNV | 1 | C | T | 22 | 21739 | 0.10120061 | 60.4090909 | WT | FALSE | CAN | JSY13 | 2 | WT-Rep2 | CCA | C>T | A/T+1 | G/C-1 | CG>TA | CG>TA | AATAACCAT | CATGCAA | GGT | CCA | C | A | C>T | C>T_CCA | CG>TA_CATGCAA | 9 | FALSE | FALSE |
| 6_v.txt | 32089 | 32089-SNV- | SNV | 1 | C | T | 269 | 21739 | 1.23740742 | 62.1226766 | WT | FALSE | CAN | JSY13 | 2 | WT-Rep2 | CCT | C>T | A/T+1 | G/C-1 | CG>TA | CG>TA | AAAAGCCTG | CTGCAAC | GGA | CCT | C | A | C>T | C>T_CCT | CG>TA_CTGCAAC | 9 | FALSE | FALSE |
| 6_v.txt | 32092 | 32092-SNV- | SNV | 1 | C | T | 15 | 21739 | 0.06900041 | 59.7333333 | WT | FALSE | CAN | JSY13 | 2 | WT-Rep2 | GCA | C>T | A/T+1 | G/C-1 | CG>TA | CG>TA | AGCCTGCAA | CAACACC | CGT | GCA | C | A | C>T | C>T_GCA | CG>TA_CAACACC | 9 | FALSE | FALSE |
| 6_v.txt | 32114 | 32114-SNV- | SNV | 1 | C | T | 43 | 21739 | 0.19780119 | 61.4651163 | WT | FALSE | CAN | JSY13 | 2 | WT-Rep2 | GCC | C>T | A/T+1 | G/C-1 | CG>TA | CG>TA | TAATAGCCA | CCATTCG | CGG | GCC | C | A | C>T | C>T_GCC | CG>TA_CCATTCG | 9 | FALSE | FALSE |
| 6_v.txt | 32157 | 32157-SNV- | SNV | 1 | G | A | 18 | 21737 | 0.08280812 | 46.5558381 | WT | FALSE | CAN | JSY13 | 2 | WT-Rep2 | AGC | G>A | A/T+1 | G/C-1 | CG>TA | CG>TA | ATGTAAGCC | GCCAAAG | TCG | TCG | C | T | C>T | C>T_TCG | CG>TA_GCCAAAG | 9 | FALSE | FALSE |
| 6_v.txt | 32158 | 32158-SNV- | SNV | 1 | C | T | 8 | 21737 | 0.03680361 | 43.6235783 | WT | FALSE | CAN | JSY13 | 2 | WT-Rep2 | GCC | C>T | A/T+1 | G/C-1 | CG>TA | CG>TA | TGTAAGCCA | CCAAAGC | CGG | GCC | C | A | C>T | C>T_GCC | CG>TA_CCAAAGC | 9 | FALSE | FALSE |
| 6_v.txt | 32164 | 32164-SNV- | SNV | 1 | C | T | 1 | 21737 | 4.60E-03 | 24.0057609 | WT | FALSE | CAN | JSY13 | 2 | WT-Rep2 | GCG | C>T | A/T+1 | G/C-1 | CG>TA | CG>TA | CCAAAGCGC | CGCCAAA | CGC | GCG | C | A | C>T | C>T_GCG | CG>TA_CGCCAAA | 9 | FALSE | FALSE |
| 6_v.txt | 32165 | 32165-SNV- | SNV | 1 | G | A | 29 | 21737 | 0.13341308 | 49.9665422 | WT | FALSE | CAN | JSY13 | 2 | WT-Rep2 | CGC | G>A | A/T+1 | G/C-1 | CG>TA | CG>TA | CAAAGCGCC | GCCAAAT | GCG | GCG | C | T | C>T | C>T_GCG | CG>TA_GCCAAAT | 9 | FALSE | FALSE |
| 6_v.txt | 32176 | 32176-SNV- | SNV | 1 | C | T | 20 | 21737 | 0.09200902 | 50.85 | WT | FALSE | CAN | JSY13 | 2 | WT-Rep2 | GCA | C>T | A/T+1 | G/C-1 | CG>TA | CG>TA | ATGCAGCAG | CAGTAAC | CGT | GCA | C | A | C>T | C>T_GCA | CG>TA_CAGTAAC | 9 | FALSE | FALSE |
| 6_v.txt | 32271 | 32271-SNV- | SNV | 1 | C | T | 51 | 32056 | 0.15909658 | 31.5882353 | WT | FALSE | CAN | JSY13 | 2 | WT-Rep2 | ACG | C>T | A/T+1 | G/C-1 | CG>TA | CG>TA | AAAATACGG | CGGGAAC | TGC | ACG | C | A | C>T | C>T_ACG | CG>TA_CGGGAAC | 9 | FALSE | FALSE |
| 6_v.txt | 32299 | 32299-SNV- | SNV | 1 | C | T | 33 | 32056 | 0.10294485 | 48.6893485 | WT | FALSE | CAN | JSY13 | 2 | WT-Rep2 | GCG | C>T | A/T+1 | G/C-1 | CG>TA | CG>TA | AATTTGCGG | CGGCAGA | CGC | GCG | C | A | C>T | C>T_GCG | CG>TA_CGGCAGA | 9 | FALSE | FALSE |
| 6_v.txt | 32449 | 32449-SNV- | SNV | 1 | C | T | 25 | 32056 | 0.07798852 | 55.52 | WT | FALSE | CAN | JSY13 | 2 | WT-Rep2 | CCA | C>T | A/T+1 | G/C-1 | CG>TA | CG>TA | AAAGTCCAA | CAATGAA | GGT | CCA | C | A | C>T | C>T_CCA | CG>TA_CAATGAA | 9 | FALSE | FALSE |
| 6_v.txt | 32466 | 32466-SNV- | SNV | 1 | C | T | 34 | 32056 | 0.10606439 | 48.6764706 | WT | FALSE | CAN | JSY13 | 2 | WT-Rep2 | GCC | C>T | A/T+1 | G/C-1 | CG>TA | CG>TA | AGAGAGCCA | CCAATGT | CGG | GCC | C | A | C>T | C>T_GCC | CG>TA_CCAATGT | 9 | FALSE | FALSE |
| 6_v.txt | 32487 | 32487-SNV- | SNV | 1 | C | T | 16 | 32056 | 0.04991265 | 45.6243938 | WT | FALSE | CAN | JSY13 | 2 | WT-Rep2 | ACG | C>T | A/T+1 | G/C-1 | CG>TA | CG>TA | AAGATACGG | CGGAAAA | TGC | ACG | C | A | C>T | C>T_ACG | CG>TA_CGGAAAA | 9 | FALSE | FALSE |
| 6_v.txt | 32488 | 32488-SNV- | SNV | 1 | G | A | 66 | 32056 | 0.20588969 | 46.5010408 | WT | FALSE | CAN | JSY13 | 2 | WT-Rep2 | CGG | G>A | A/T+1 | G/C-1 | CG>TA | CG>TA | AGATACGGA | GGAAAAC | GCC | GCC | C | T | C>T | C>T_GCC | CG>TA_GGAAAAC | 9 | FALSE | FALSE |
| 6_v.txt | 32535 | 32535-SNV- | SNV | 1 | G | A | 37 | 32056 | 0.11542301 | 37.0000497 | WT | FALSE | CAN | JSY13 | 2 | WT-Rep2 | TGC | G>A | A/T+1 | G/C-1 | CG>TA | CG>TA | GGGTTTGCA | GCAGCTT | ACG | ACG | C | T | C>T | C>T_ACG | CG>TA_GCAGCTT | 9 | FALSE | FALSE |
| 6_v.txt | 32536 | 32536-SNV- | SNV | 1 | C | T | 64 | 32056 | 0.19965061 | 37.0946433 | WT | FALSE | CAN | JSY13 | 2 | WT-Rep2 | GCA | C>T | A/T+1 | G/C-1 | CG>TA | CG>TA | GGTTTGCAG | CAGCTTC | CGT | GCA | C | A | C>T | C>T_GCA | CG>TA_CAGCTTC | 9 | FALSE | FALSE |
| 6_v.txt | 32590 | 32590-SNV- | SNV | 1 | C | T | 31 | 25283 | 0.12261203 | 37.1612903 | WT | FALSE | CAN | JSY13 | 2 | WT-Rep2 | GCG | C>T | A/T+1 | G/C-1 | CG>TA | CG>TA | AGGCAGCGT | CGTTAAT | CGC | GCG | C | A | C>T | C>T_GCG | CG>TA_CGTTAAT | 9 | FALSE | FALSE |
| 6_v.txt | 32604 | 32604-SNV- | SNV | 1 | G | A | 2 | 25283 | 7.91E-03 | 37.999736 | WT | FALSE | CAN | JSY13 | 2 | WT-Rep2 | GGA | G>A | A/T+1 | G/C-1 | CG>TA | CG>TA | AAAGAGGA | GAAACCC | CCT | CCT | C | T | C>T | C>T_CCT | CG>TA_GAAACCC | 9 | FALSE | FALSE |
| 6_v.txt | 32679 | 32679-SNV- | SNV | 1 | C | T | 41 | 25283 | 0.1621643 | 59.5121951 | WT | FALSE | CAN | JSY13 | 2 | WT-Rep2 | CCA | C>T | A/T+1 | G/C-1 | CG>TA | CG>TA | TTTCTCCAAT | CAATAAC | GGT | CCA | C | A | C>T | C>T_CCA | CG>TA_CAATAAC | 9 | FALSE | FALSE |
| 6_v.txt | 32700 | 32700-SNV- | SNV | 1 | C | T | 35 | 25283 | 0.13843294 | 61.7688697 | WT | FALSE | CAN | JSY13 | 2 | WT-Rep2 | GCC | C>T | A/T+1 | G/C-1 | CG>TA | CG>TA | ACTGGGCCG | CCGGTAA | CGG | GCC | C | A | C>T | C>T_GCC | CG>TA_CCGGTAA | 9 | FALSE | FALSE |
| 6_v.txt | 32701 | 32701-SNV- | SNV | 1 | C | T | 6 | 25283 | 0.02373136 | 63.8323653 | WT | FALSE | CAN | JSY13 | 2 | WT-Rep2 | CCG | C>T | A/T+1 | G/C-1 | CG>TA | CG>TA | CTGGGCCGG | CGGTAAC | GGC | CCG | C | A | C>T | C>T_CCG | CG>TA_CGGTAAC | 9 | FALSE | FALSE |
| 6_v.txt | 32757 | 32757-SNV- | SNV | 1 | G | A | 27 | 25283 | 0.10679113 | 56.8148148 | WT | FALSE | CAN | JSY13 | 2 | WT-Rep2 | GGC | G>A | A/T+1 | G/C-1 | CG>TA | CG>TA | ATAATGGCT | GCTAAAA | CCG | CCG | C | T | C>T | C>T_CCG | CG>TA_GCTAAAA | 9 | FALSE | FALSE |
| 6_v.txt | 32776 | 32776-SNV- | SNV | 1 | C | T | 29 | 25283 | 0.11470158 | 49.7586207 | WT | FALSE | CAN | JSY13 | 2 | WT-Rep2 | GCG | C>T | A/T+1 | G/C-1 | CG>TA | CG>TA | TGGAAGCGA | CGACCCA | CGC | GCG | C | A | C>T | C>T_GCG | CG>TA_CGACCCA | 9 | FALSE | FALSE |
| 6_v.txt | 32798 | 32798-SNV- | SNV | 1 | G | A | 41 | 25283 | 0.1621643 | 44.5609756 | WT | FALSE | CAN | JSY13 | 2 | WT-Rep2 | CGT | G>A | A/T+1 | G/C-1 | CG>TA | CG>TA | TTCACCGTA | GTAATAT | GCA | GCA | C | T | C>T | C>T_GCA | CG>TA_GTAATAT | 9 | FALSE | FALSE |
| 6_v.txt | 32860 | 32860-SNV- | SNV | 1 | C | T | 31 | 25282 | 0.12261688 | 37.2580645 | WT | FALSE | CAN | JSY13 | 2 | WT-Rep2 | GCC | C>T | A/T+1 | G/C-1 | CG>TA | CG>TA | TCCATGCCG | CCGCCAG | CGG | GCC | C | A | C>T | C>T_GCC | CG>TA_CCGCCAG | 9 | FALSE | FALSE |
| 6_v.txt | 32883 | 32883-SNV- | SNV | 1 | C | T | 42 | 20701 | 0.20288875 | 36.5 | WT | FALSE | CAN | JSY13 | 2 | WT-Rep2 | CCA | C>T | A/T+1 | G/C-1 | CG>TA | CG>TA | TACGTCCAA | CAAAATT | GGT | CCA | C | A | C>T | C>T_CCA | CG>TA_CAAAATT | 9 | FALSE | FALSE |
| 6_v.txt | 32899 | 32899-SNV- | SNV | 1 | G | A | 94 | 20702 | 0.45406241 | 36.2270658 | WT | FALSE | CAN | JSY13 | 2 | WT-Rep2 | TGG | G>A | A/T+1 | G/C-1 | CG>TA | CG>TA | TGACTTGGC | GGCCAAC | ACC | ACC | C | T | C>T | C>T_ACC | CG>TA_GGCCAAC | 9 | FALSE | FALSE |
| 6_v.txt | 32900 | 32900-SNV- | SNV | 1 | G | A | 26 | 20702 | 0.12559173 | 33.9206952 | WT | FALSE | CAN | JSY13 | 2 | WT-Rep2 | GGC | G>A | A/T+1 | G/C-1 | CG>TA | CG>TA | GACTTGGCC | GCCAACT | CCG | CCG | C | T | C>T | C>T_CCG | CG>TA_GCCAACT | 9 | FALSE | FALSE |
| 6_v.txt | 32901 | 32901-SNV- | SNV | 1 | C | T | 30 | 20702 | 0.14491354 | 32.5994947 | WT | FALSE | CAN | JSY13 | 2 | WT-Rep2 | GCC | C>T | A/T+1 | G/C-1 | CG>TA | CG>TA | ACTTGGCCA | CCAACTA | CGG | GCC | C | A | C>T | C>T_GCC | CG>TA_CCAACTA | 9 | FALSE | FALSE |
| 6_v.txt | 32935 | 32935-SNV- | SNV | 1 | C | T | 27 | 20702 | 0.13042218 | 30.962963 | WT | FALSE | CAN | JSY13 | 2 | WT-Rep2 | GCC | C>T | A/T+1 | G/C-1 | CG>TA | CG>TA | TGATTGCCCA | CCCAAGA | CGG | GCC | C | A | C>T | C>T_GCC | CG>TA_CCCAAGA | 9 | FALSE | FALSE |
| 6_v.txt | 32958 | 32958-SNV- | SNV | 1 | C | T | 31 | 20702 | 0.14974399 | 43.7096774 | WT | FALSE | CAN | JSY13 | 2 | WT-Rep2 | ACC | C>T | A/T+1 | G/C-1 | CG>TA | CG>TA | ATGTAACCA | CCATTGG | TGG | ACC | C | A | C>T | C>T_ACC | CG>TA_CCATTGG | 9 | FALSE | FALSE |
| 6_v.txt | 32965 | 32965-SNV- | SNV | 1 | C | T | 20 | 20702 | 0.09660902 | 59.35 | WT | FALSE | CAN | JSY13 | 2 | WT-Rep2 | GCC | C>T | A/T+1 | G/C-1 | CG>TA | CG>TA | CATTGGCCG | CCGCACC | CGG | GCC | C | A | C>T | C>T_GCC | CG>TA_CCGCACC | 9 | FALSE | FALSE |
| 6_v.txt | 32967 | 32967-SNV- | SNV | 1 | G | A | 44 | 20702 | 0.21253985 | 62.4318182 | WT | FALSE | CAN | JSY13 | 2 | WT-Rep2 | CGC | G>A | A/T+1 | G/C-1 | CG>TA | CG>TA | TTGGCCGCA | GCACCAA | GCG | GCG | C | T | C>T | C>T_GCG | CG>TA_GCACCAA | 9 | FALSE | FALSE |
| 6_v.txt | 32970 | 32970-SNV- | SNV | 1 | C | T | 36 | 20702 | 0.17389624 | 57.9444444 | WT | FALSE | CAN | JSY13 | 2 | WT-Rep2 | ACC | C>T | A/T+1 | G/C-1 | CG>TA | CG>TA | GCCGCACCA | CCAAATG | TGG | ACC | C | A | C>T | C>T_ACC | CG>TA_CCAAATG | 9 | FALSE | FALSE |
| 6_v.txt | 33012 | 33012-SNV- | SNV | 1 | G | A | 90 | 20702 | 0.43474061 | 61.7444444 | WT | FALSE | CAN | JSY13 | 2 | WT-Rep2 | AGA | G>A | A/T+1 | G/C-1 | CG>TA | CG>TA | GTGAAAGAG | GAGGATG | TCT | TCT | C | T | C>T | C>T_TCT | CG>TA_GAGGATG | 9 | FALSE | FALSE |
| 6_v.txt | 33055 | 33055-SNV- | SNV | 1 | G | A | 132 | 20702 | 0.63761955 | 58.3181818 | WT | FALSE | CAN | JSY13 | 2 | WT-Rep2 | TGC | G>A | A/T+1 | G/C-1 | CG>TA | CG>TA | AGGACTGCG | GCGTGAC | ACG | ACG | C | T | C>T | C>T_ACG | CG>TA_GCGTGAC | 9 | FALSE | FALSE |
| 6_v.txt | 33056 | 33056-SNV- | SNV | 1 | C | T | 31 | 20702 | 0.14974399 | 57.9032258 | WT | FALSE | CAN | JSY13 | 2 | WT-Rep2 | GCG | C>T | A/T+1 | G/C-1 | CG>TA | CG>TA | GGACTGCGT | CGTGACA | CGC | GCG | C | A | C>T | C>T_GCG | CG>TA_CGTGACA | 9 | FALSE | FALSE |
| 6_v.txt | 33070 | 33070-SNV- | SNV | 1 | C | T | 20 | 20702 | 0.09660902 | 53.1 | WT | FALSE | CAN | JSY13 | 2 | WT-Rep2 | GCC | C>T | A/T+1 | G/C-1 | CG>TA | CG>TA | AATATGCCA | CCAAAGA | CGG | GCC | C | A | C>T | C>T_GCC | CG>TA_CCAAAGA | 9 | FALSE | FALSE |
| 6_v.txt | 33113 | 33113-SNV- | SNV | 1 | G | A | 21 | 20702 | 0.10143947 | 40.8095238 | WT | FALSE | CAN | JSY13 | 2 | WT-Rep2 | GGC | G>A | A/T+1 | G/C-1 | CG>TA | CG>TA | CACTGGGCC | GCCGGCG | CCG | CCG | C | T | C>T | C>T_CCG | CG>TA_GCCGGCG | 9 | FALSE | FALSE |
| 6_v.txt | 33114 | 33114-SNV- | SNV | 1 | C | T | 29 | 20702 | 0.14008308 | 43.3759549 | WT | FALSE | CAN | JSY13 | 2 | WT-Rep2 | GCC | C>T | A/T+1 | G/C-1 | CG>TA | CG>TA | ACTGGGCCG | CCGGCGT | CGG | GCC | C | A | C>T | C>T_GCC | CG>TA_CCGGCGT | 9 | FALSE | FALSE |
| 6_v.txt | 33115 | 33115-SNV- | SNV | 1 | C | T | 9 | 20702 | 0.04347406 | 40.8888889 | WT | FALSE | CAN | JSY13 | 2 | WT-Rep2 | CCG | C>T | A/T+1 | G/C-1 | CG>TA | CG>TA | CTGGGCCGG | CGGCGTT | GGC | CCG | C | A | C>T | C>T_CCG | CG>TA_CGGCGTT | 9 | FALSE | FALSE |
| 6_v.txt | 33118 | 33118-SNV- | SNV | 1 | C | T | 25 | 20702 | 0.12076128 | 39.44 | WT | FALSE | CAN | JSY13 | 2 | WT-Rep2 | GCG | C>T | A/T+1 | G/C-1 | CG>TA | CG>TA | GGCCGGCGT | CGTTGGT | CGC | GCG | C | A | C>T | C>T_GCG | CG>TA_CGTTGGT | 9 | FALSE | FALSE |
| 6_v.txt | 33169 | 33169-SNV- | SNV | 1 | C | T | 9 | 20695 | 0.04348877 | 37.6666667 | WT | FALSE | CAN | JSY13 | 2 | WT-Rep2 | CCA | C>T | A/T+1 | G/C-1 | CG>TA | CG>TA | TAGTACCAC | CACCAAG | GGT | CCA | C | A | C>T | C>T_CCA | CG>TA_CACCAAG | 9 | FALSE | FALSE |
| 6_v.txt | 33199 | 33199-SNV- | SNV | 1 | G | A | 11 | 13775 | 0.07985481 | 36.6360027 | WT | FALSE | CAN | JSY13 | 2 | WT-Rep2 | TGC | G>A | A/T+1 | G/C-1 | CG>TA | CG>TA | GTCTTTGCTT | GCTTAAG | ACG | ACG | C | T | C>T | C>T_ACG | CG>TA_GCTTAAG | 9 | FALSE | FALSE |
| 6_v.txt | 33258 | 33258-SNV- | SNV | 1 | C | T | 19 | 13776 | 0.13792102 | 45.8947368 | WT | FALSE | CAN | JSY13 | 2 | WT-Rep2 | GCC | C>T | A/T+1 | G/C-1 | CG>TA | CG>TA | CCTATGCCAT | CCATCCT | CGG | GCC | C | A | C>T | C>T_GCC | CG>TA_CCATCCT | 9 | FALSE | FALSE |
| 6_v.txt | 33474 | 33474-SNV- | SNV | 1 | C | T | 21 | 13775 | 0.15245009 | 31.5714286 | WT | FALSE | CAN | JSY13 | 2 | WT-Rep2 | GCC | C>T | A/T+1 | G/C-1 | CG>TA | CG>TA | GCTATGCCTT | CCTTTTT | CGG | GCC | C | A | C>T | C>T_GCC | CG>TA_CCTTTTT | 9 | FALSE | FALSE |
| 6_v.txt | 31981 | 31981-SNV- | SNV | 1 | G | C | 111 | 21739 | 0.51060306 | 44.5585586 | WT | FALSE | CAN | JSY13 | 2 | WT-Rep2 | GGC | G>C | G/C+1 | G/C-1 | CG>GC | CG>GC | AGCCGGGCA | GCATTAA | CCG | CCG | C | G | C>G | C>G_CCG | CG>GC_GCATTAA | 9 | FALSE | FALSE |
| 6_v.txt | 32057 | 32057-SNV- | SNV | 1 | C | G | 2 | 21739 | 9.20E-03 | 59.5 | WT | FALSE | CAN | JSY13 | 2 | WT-Rep2 | GCG | C>G | G/C+1 | G/C-1 | CG>GC | CG>GC | GATGTGCGA | CGAGATT | CGC | GCG | C | C | C>G | C>G_GCG | CG>GC_CGAGATT | 9 | FALSE | FALSE |
| 6_v.txt | 32092 | 32092-SNV- | SNV | 1 | C | G | 7 | 21739 | 0.03220019 | 64 | WT | FALSE | CAN | JSY13 | 2 | WT-Rep2 | GCA | C>G | G/C+1 | G/C-1 | CG>GC | CG>GC | AGCCTGCAA | CAACACC | CGT | GCA | C | C | C>G | C>G_GCA | CG>GC_CAACACC | 9 | FALSE | FALSE |
| 6_v.txt | 32164 | 32164-SNV- | SNV | 1 | C | G | 2 | 21737 | 9.20E-03 | 50.5009279 | WT | FALSE | CAN | JSY13 | 2 | WT-Rep2 | GCG | C>G | G/C+1 | G/C-1 | CG>GC | CG>GC | CCAAAGCGC | CGCCAAA | CGC | GCG | C | C | C>G | C>G_GCG | CG>GC_CGCCAAA | 9 | FALSE | FALSE |
| 6_v.txt | 32282 | 32282-SNV- | SNV | 1 | G | C | 6 | 32056 | 0.01871725 | 37.3333333 | WT | FALSE | CAN | JSY13 | 2 | WT-Rep2 | CGT | G>C | G/C+1 | G/C-1 | CG>GC | CG>GC | ACCAACGTA | GTAAATA | GCA | GCA | C | G | C>G | C>G_GCA | CG>GC_GTAAATA | 9 | FALSE | FALSE |
| 6_v.txt | 32298 | 32298-SNV- | SNV | 1 | G | C | 21 | 32056 | 0.06551036 | 28.9061135 | WT | FALSE | CAN | JSY13 | 2 | WT-Rep2 | TGC | G>C | G/C+1 | G/C-1 | CG>GC | CG>GC | GAATTTGCG | GCGGCAG | ACG | ACG | C | G | C>G | C>G_ACG | CG>GC_GCGGCAG | 9 | FALSE | FALSE |
| 6_v.txt | 32471 | 32471-SNV- | SNV | 1 | G | C | 17 | 32056 | 0.05303219 | 47.0569488 | WT | FALSE | CAN | JSY13 | 2 | WT-Rep2 | TGT | G>C | G/C+1 | G/C-1 | CG>GC | CG>GC | GCCAATGTA | GTAGAAG | ACA | ACA | C | G | C>G | C>G_ACA | CG>GC_GTAGAAG | 9 | FALSE | FALSE |
| 6_v.txt | 32478 | 32478-SNV- | SNV | 1 | G | C | 1 | 32056 | 3.12E-03 | 26.999628 | WT | FALSE | CAN | JSY13 | 2 | WT-Rep2 | GGT | G>C | G/C+1 | G/C-1 | CG>GC | CG>GC | TAGAAGGTT | GTTAAGA | CCA | CCA | C | G | C>G | C>G_CCA | CG>GC_GTTAAGA | 9 | FALSE | FALSE |
| 6_v.txt | 32487 | 32487-SNV- | SNV | 1 | C | G | 9 | 32056 | 0.02807587 | 48.5540726 | WT | FALSE | CAN | JSY13 | 2 | WT-Rep2 | ACG | C>G | G/C+1 | G/C-1 | CG>GC | CG>GC | AAGATACGG | CGGAAAA | TGC | ACG | C | C | C>G | C>G_ACG | CG>GC_CGGAAAA | 9 | FALSE | FALSE |
| 6_v.txt | 32488 | 32488-SNV- | SNV | 1 | G | C | 13 | 32056 | 0.04055403 | 47.7697995 | WT | FALSE | CAN | JSY13 | 2 | WT-Rep2 | CGG | G>C | G/C+1 | G/C-1 | CG>GC | CG>GC | AGATACGGA | GGAAAAC | GCC | GCC | C | G | C>G | C>G_GCC | CG>GC_GGAAAAC | 9 | FALSE | FALSE |
| 6_v.txt | 32557 | 32557-SNV- | SNV | 1 | C | G | 140 | 32042 | 0.43692653 | 37.0285714 | WT | FALSE | CAN | JSY13 | 2 | WT-Rep2 | CCA | C>G | G/C+1 | G/C-1 | CG>GC | CG>GC | TGATACCAA | CAACTAG | GGT | CCA | C | C | C>G | C>G_CCA | CG>GC_CAACTAG | 9 | FALSE | FALSE |
| 6_v.txt | 32566 | 32566-SNV- | SNV | 1 | C | G | 34 | 25271 | 0.13454157 | 37.6176471 | WT | FALSE | CAN | JSY13 | 2 | WT-Rep2 | TCA | C>G | G/C+1 | G/C-1 | CG>GC | CG>GC | CTAGTTCAG | CAGTACC | AGT | TCA | C | C | C>G | C>G_TCA | CG>GC_CAGTACC | 9 | FALSE | FALSE |
| 6_v.txt | 32587 | 32587-SNV- | SNV | 1 | C | G | 12 | 25283 | 0.04746272 | 36.3333333 | WT | FALSE | CAN | JSY13 | 2 | WT-Rep2 | GCA | C>G | G/C+1 | G/C-1 | CG>GC | CG>GC | TGAAGGCAG | CAGCGTT | CGT | GCA | C | C | C>G | C>G_GCA | CG>GC_CAGCGTT | 9 | FALSE | FALSE |
| 6_v.txt | 32604 | 32604-SNV- | SNV | 1 | G | C | 1 | 25283 | 3.96E-03 | 63.998545 | WT | FALSE | CAN | JSY13 | 2 | WT-Rep2 | GGA | G>C | G/C+1 | G/C-1 | CG>GC | CG>GC | AAAGAGGA | GAAACCC | CCT | CCT | C | G | C>G | C>G_CCT | CG>GC_GAAACCC | 9 | FALSE | FALSE |
| 6_v.txt | 32811 | 32811-SNV- | SNV | 1 | G | C | 21 | 25283 | 0.08305976 | 46.4761905 | WT | FALSE | CAN | JSY13 | 2 | WT-Rep2 | AGG | G>C | G/C+1 | G/C-1 | CG>GC | CG>GC | TTGACAGGG | GGGAACA | TCC | TCC | C | G | C>G | C>G_TCC | CG>GC_GGGAACA | 9 | FALSE | FALSE |
| 6_v.txt | 32876 | 32876-SNV- | SNV | 1 | G | C | 7 | 20695 | 0.0338246 | 37.7142857 | WT | FALSE | CAN | JSY13 | 2 | WT-Rep2 | TGT | G>C | G/C+1 | G/C-1 | CG>GC | CG>GC | AACTTTGTA | GTACGTC | ACA | ACA | C | G | C>G | C>G_ACA | CG>GC_GTACGTC | 9 | FALSE | FALSE |
| 6_v.txt | 32883 | 32883-SNV- | SNV | 1 | C | G | 3 | 20701 | 0.01449205 | 28.3333333 | WT | FALSE | CAN | JSY13 | 2 | WT-Rep2 | CCA | C>G | G/C+1 | G/C-1 | CG>GC | CG>GC | TACGTCCAA | CAAAATT | GGT | CCA | C | C | C>G | C>G_CCA | CG>GC_CAAAATT | 9 | FALSE | FALSE |
| 6_v.txt | 32901 | 32901-SNV- | SNV | 1 | C | G | 2 | 20702 | 9.66E-03 | 37.9993163 | WT | FALSE | CAN | JSY13 | 2 | WT-Rep2 | GCC | C>G | G/C+1 | G/C-1 | CG>GC | CG>GC | ACTTGGCCA | CCAACTA | CGG | GCC | C | C | C>G | C>G_GCC | CG>GC_CCAACTA | 9 | FALSE | FALSE |
| 6_v.txt | 32937 | 32937-SNV- | SNV | 1 | C | G | 9 | 20702 | 0.04347406 | 54.2222222 | WT | FALSE | CAN | JSY13 | 2 | WT-Rep2 | CCA | C>G | G/C+1 | G/C-1 | CG>GC | CG>GC | ATTGCCCAA | CAAGAAA | GGT | CCA | C | C | C>G | C>G_CCA | CG>GC_CAAGAAA | 9 | FALSE | FALSE |
| 6_v.txt | 33111 | 33111-SNV- | SNV | 1 | G | C | 18 | 20702 | 0.08694812 | 41.6111111 | WT | FALSE | CAN | JSY13 | 2 | WT-Rep2 | TGG | G>C | G/C+1 | G/C-1 | CG>GC | CG>GC | CCCACTGGG | GGGCCGG | ACC | ACC | C | G | C>G | C>G_ACC | CG>GC_GGGCCGG | 9 | FALSE | FALSE |
| 6_v.txt | 33114 | 33114-SNV- | SNV | 1 | C | G | 3 | 20702 | 0.01449135 | 55.3303389 | WT | FALSE | CAN | JSY13 | 2 | WT-Rep2 | GCC | C>G | G/C+1 | G/C-1 | CG>GC | CG>GC | ACTGGGCCG | CCGGCGT | CGG | GCC | C | C | C>G | C>G_GCC | CG>GC_CCGGCGT | 9 | FALSE | FALSE |
| 6_v.txt | 33168 | 33168-SNV- | SNV | 1 | C | G | 23 | 20695 | 0.11113796 | 35.9977 | WT | FALSE | CAN | JSY13 | 2 | WT-Rep2 | ACC | C>G | G/C+1 | G/C-1 | CG>GC | CG>GC | ATAGTACCA | CCACCAA | TGG | ACC | C | C | C>G | C>G_ACC | CG>GC_CCACCAA | 9 | FALSE | FALSE |
| 6_v.txt | 33169 | 33169-SNV- | SNV | 1 | C | G | 2 | 20695 | 9.66E-03 | 38 | WT | FALSE | CAN | JSY13 | 2 | WT-Rep2 | CCA | C>G | G/C+1 | G/C-1 | CG>GC | CG>GC | TAGTACCAC | CACCAAG | GGT | CCA | C | C | C>G | C>G_CCA | CG>GC_CACCAAG | 9 | FALSE | FALSE |
| 6_v.txt | 31657 | 31657-SNV- | SNV | 1 | G | T | 93 | 11614 | 0.80075771 | 19.483871 | WT | FALSE | CAN | JSY13 | 2 | WT-Rep2 | GGC | G>T | A/T+1 | G/C-1 | CG>AT | CG>AT | GAAATGGCG | GCGTGGA | CCG | CCG | C | A | C>A | C>A_CCG | CG>AT_GCGTGGA | 9 | FALSE | FALSE |
| 6_v.txt | 32057 | 32057-SNV- | SNV | 1 | C | A | 5 | 21739 | 0.02300014 | 45.4 | WT | FALSE | CAN | JSY13 | 2 | WT-Rep2 | GCG | C>A | A/T+1 | G/C-1 | CG>AT | CG>AT | GATGTGCGA | CGAGATT | CGC | GCG | C | T | C>A | C>A_GCG | CG>AT_CGAGATT | 9 | FALSE | FALSE |
| 6_v.txt | 32058 | 32058-SNV- | SNV | 1 | G | T | 341 | 21739 | 1.56860941 | 61.0842456 | WT | FALSE | CAN | JSY13 | 2 | WT-Rep2 | CGA | G>T | A/T+1 | G/C-1 | CG>AT | CG>AT | ATGTGCGAG | GAGATTG | GCT | GCT | C | A | C>A | C>A_GCT | CG>AT_GAGATTG | 9 | FALSE | FALSE |
| 6_v.txt | 32149 | 32149-SNV- | SNV | 1 | C | A | 13 | 21739 | 0.05980036 | 48.6153846 | WT | FALSE | CAN | JSY13 | 2 | WT-Rep2 | TCC | C>A | A/T+1 | G/C-1 | CG>AT | CG>AT | ATGTCTCCAT | CCATGTA | AGG | TCC | C | T | C>A | C>A_TCC | CG>AT_CCATGTA | 9 | FALSE | FALSE |
| 6_v.txt | 32223 | 32223-SNV- | SNV | 1 | G | T | 11 | 21737 | 0.05060496 | 35.5454546 | WT | FALSE | CAN | JSY13 | 2 | WT-Rep2 | TGA | G>T | A/T+1 | G/C-1 | CG>AT | CG>AT | GTCCTTGAC | GACAGGA | ACT | ACT | C | A | C>A | C>A_ACT | CG>AT_GACAGGA | 9 | FALSE | FALSE |
| 6_v.txt | 32282 | 32282-SNV- | SNV | 1 | G | T | 48 | 32056 | 0.14973796 | 42.4791667 | WT | FALSE | CAN | JSY13 | 2 | WT-Rep2 | CGT | G>T | A/T+1 | G/C-1 | CG>AT | CG>AT | ACCAACGTA | GTAAATA | GCA | GCA | C | A | C>A | C>A_GCA | CG>AT_GTAAATA | 9 | FALSE | FALSE |
| 6_v.txt | 32399 | 32399-SNV- | SNV | 1 | G | T | 92 | 32056 | 0.28699775 | 57.5217391 | WT | FALSE | CAN | JSY13 | 2 | WT-Rep2 | CGT | G>T | A/T+1 | G/C-1 | CG>AT | CG>AT | AGAAACGTA | GTAGGAA | GCA | GCA | C | A | C>A | C>A_GCA | CG>AT_GTAGGAA | 9 | FALSE | FALSE |
| 6_v.txt | 32432 | 32432-SNV- | SNV | 1 | G | T | 127 | 32056 | 0.39618168 | 57.3228347 | WT | FALSE | CAN | JSY13 | 2 | WT-Rep2 | TGT | G>T | A/T+1 | G/C-1 | CG>AT | CG>AT | GTCATTGTAT | GTATGGA | ACA | ACA | C | A | C>A | C>A_ACA | CG>AT_GTATGGA | 9 | FALSE | FALSE |
| 6_v.txt | 32474 | 32474-SNV- | SNV | 1 | G | T | 113 | 32056 | 0.35250811 | 48.5046273 | WT | FALSE | CAN | JSY13 | 2 | WT-Rep2 | AGA | G>T | A/T+1 | G/C-1 | CG>AT | CG>AT | AATGTAGAA | GAAGGTT | TCT | TCT | C | A | C>A | C>A_TCT | CG>AT_GAAGGTT | 9 | FALSE | FALSE |
| 6_v.txt | 32530 | 32530-SNV- | SNV | 1 | G | T | 153 | 32056 | 0.47728974 | 38.379085 | WT | FALSE | CAN | JSY13 | 2 | WT-Rep2 | GGG | G>T | A/T+1 | G/C-1 | CG>AT | CG>AT | TTCTGGGGT | GGTTTGC | CCC | CCC | C | A | C>A | C>A_CCC | CG>AT_GGTTTGC | 9 | FALSE | FALSE |
| 6_v.txt | 32538 | 32538-SNV- | SNV | 1 | G | T | 20 | 32056 | 0.06239082 | 37.95 | WT | FALSE | CAN | JSY13 | 2 | WT-Rep2 | AGC | G>T | A/T+1 | G/C-1 | CG>AT | CG>AT | TTTGCAGCTT | GCTTCAC | TCG | TCG | C | A | C>A | C>A_TCG | CG>AT_GCTTCAC | 9 | FALSE | FALSE |
| 6_v.txt | 32794 | 32794-SNV- | SNV | 1 | C | A | 33 | 25283 | 0.13052249 | 44.1212121 | WT | FALSE | CAN | JSY13 | 2 | WT-Rep2 | TCA | C>A | A/T+1 | G/C-1 | CG>AT | CG>AT | CGAATTCAC | CACCGTA | AGT | TCA | C | T | C>A | C>A_TCA | CG>AT_CACCGTA | 9 | FALSE | FALSE |
| 6_v.txt | 32876 | 32876-SNV- | SNV | 1 | G | T | 13 | 20695 | 0.06281711 | 31.6923077 | WT | FALSE | CAN | JSY13 | 2 | WT-Rep2 | TGT | G>T | A/T+1 | G/C-1 | CG>AT | CG>AT | AACTTTGTA | GTACGTC | ACA | ACA | C | A | C>A | C>A_ACA | CG>AT_GTACGTC | 9 | FALSE | FALSE |
| 6_v.txt | 32883 | 32883-SNV- | SNV | 1 | C | A | 4 | 20701 | 0.01932274 | 31 | WT | FALSE | CAN | JSY13 | 2 | WT-Rep2 | CCA | C>A | A/T+1 | G/C-1 | CG>AT | CG>AT | TACGTCCAA | CAAAATT | GGT | CCA | C | T | C>A | C>A_CCA | CG>AT_CAAAATT | 9 | FALSE | FALSE |
| 6_v.txt | 32899 | 32899-SNV- | SNV | 1 | G | T | 55 | 20702 | 0.26567481 | 14.9634986 | WT | FALSE | CAN | JSY13 | 2 | WT-Rep2 | TGG | G>T | A/T+1 | G/C-1 | CG>AT | CG>AT | TGACTTGGC | GGCCAAC | ACC | ACC | C | A | C>A | C>A_ACC | CG>AT_GGCCAAC | 9 | FALSE | FALSE |
| 6_v.txt | 32954 | 32954-SNV- | SNV | 1 | G | T | 13 | 20702 | 0.06279587 | 48.2307692 | WT | FALSE | CAN | JSY13 | 2 | WT-Rep2 | TGT | G>T | A/T+1 | G/C-1 | CG>AT | CG>AT | ATACATGTA | GTAACCA | ACA | ACA | C | A | C>A | C>A_ACA | CG>AT_GTAACCA | 9 | FALSE | FALSE |
| 6_v.txt | 33006 | 33006-SNV- | SNV | 1 | G | T | 133 | 20702 | 0.64245001 | 60.6240602 | WT | FALSE | CAN | JSY13 | 2 | WT-Rep2 | TGT | G>T | A/T+1 | G/C-1 | CG>AT | CG>AT | AAAACTGTG | GTGAAAG | ACA | ACA | C | A | C>A | C>A_ACA | CG>AT_GTGAAAG | 9 | FALSE | FALSE |
| 6_v.txt | 33018 | 33018-SNV- | SNV | 1 | G | T | 117 | 20700 | 0.56521739 | 60.2225227 | WT | FALSE | CAN | JSY13 | 2 | WT-Rep2 | TGT | G>T | A/T+1 | G/C-1 | CG>AT | CG>AT | GAGGATGTA | GTAACAG | ACA | ACA | C | A | C>A | C>A_ACA | CG>AT_GTAACAG | 9 | FALSE | FALSE |
| 6_v.txt | 33033 | 33033-SNV- | SNV | 1 | G | T | 1581 | 20702 | 7.63694329 | 60.1391524 | WT | FALSE | CAN | JSY13 | 2 | WT-Rep2 | TGT | G>T | A/T+1 | G/C-1 | CG>AT | CG>AT | ATGAATGTA | GTAGCCA | ACA | ACA | C | A | C>A | C>A_ACA | CG>AT_GTAGCCA | 9 | FALSE | FALSE |
| 6_v.txt | 33043 | 33043-SNV- | SNV | 1 | C | A | 72 | 20702 | 0.34779248 | 61.4722222 | WT | FALSE | CAN | JSY13 | 2 | WT-Rep2 | TCA | C>A | A/T+1 | G/C-1 | CG>AT | CG>AT | CCATTTCACC | CACCCAA | AGT | TCA | C | T | C>A | C>A_TCA | CG>AT_CACCCAA | 9 | FALSE | FALSE |
| 6_v.txt | 33056 | 33056-SNV- | SNV | 1 | C | A | 7 | 20702 | 0.03381316 | 27.7142857 | WT | FALSE | CAN | JSY13 | 2 | WT-Rep2 | GCG | C>A | A/T+1 | G/C-1 | CG>AT | CG>AT | GGACTGCGT | CGTGACA | CGC | GCG | C | T | C>A | C>A_GCG | CG>AT_CGTGACA | 9 | FALSE | FALSE |
| 6_v.txt | 33115 | 33115-SNV- | SNV | 1 | C | A | 142 | 20702 | 0.68592407 | 42.1901409 | WT | FALSE | CAN | JSY13 | 2 | WT-Rep2 | CCG | C>A | A/T+1 | G/C-1 | CG>AT | CG>AT | CTGGGCCGG | CGGCGTT | GGC | CCG | C | T | C>A | C>A_CCG | CG>AT_CGGCGTT | 9 | FALSE | FALSE |
| 6_v.txt | 33151 | 33151-SNV- | SNV | 1 | G | T | 12 | 20702 | 0.05796541 | 34.5 | WT | FALSE | CAN | JSY13 | 2 | WT-Rep2 | AGA | G>T | A/T+1 | G/C-1 | CG>AT | CG>AT | TGAAAAGAC | GACCTGT | TCT | TCT | C | A | C>A | C>A_TCT | CG>AT_GACCTGT | 9 | FALSE | FALSE |
| 6_v.txt | 33160 | 33160-SNV- | SNV | 1 | C | A | 16 | 20709 | 0.07726109 | 37.625 | WT | FALSE | CAN | JSY13 | 2 | WT-Rep2 | CCA | C>A | A/T+1 | G/C-1 | CG>AT | CG>AT | CTGTACCAAT | CAATAGT | GGT | CCA | C | T | C>A | C>A_CCA | CG>AT_CAATAGT | 9 | FALSE | FALSE |
| 6_v.txt | 33168 | 33168-SNV- | SNV | 1 | C | A | 597 | 20695 | 2.88475477 | 37.1516139 | WT | FALSE | CAN | JSY13 | 2 | WT-Rep2 | ACC | C>A | A/T+1 | G/C-1 | CG>AT | CG>AT | ATAGTACCA | CCACCAA | TGG | ACC | C | T | C>A | C>A_ACC | CG>AT_CCACCAA | 9 | FALSE | FALSE |
| 6_v.txt | 33169 | 33169-SNV- | SNV | 1 | C | A | 17 | 20695 | 0.08214545 | 37.2941177 | WT | FALSE | CAN | JSY13 | 2 | WT-Rep2 | CCA | C>A | A/T+1 | G/C-1 | CG>AT | CG>AT | TAGTACCAC | CACCAAG | GGT | CCA | C | T | C>A | C>A_CCA | CG>AT_CACCAAG | 9 | FALSE | FALSE |
| 6_v.txt | 33186 | 33186-SNV- | SNV | 1 | C | A | 9 | 13763 | 0.06539272 | 38 | WT | FALSE | CAN | JSY13 | 2 | WT-Rep2 | ACC | C>A | A/T+1 | G/C-1 | CG>AT | CG>AT | ATCATACCA | CCAATAT | TGG | ACC | C | T | C>A | C>A_ACC | CG>AT_CCAATAT | 9 | FALSE | FALSE |
| 6_v.txt | 33208 | 33208-SNV- | SNV | 1 | C | A | 2 | 13775 | 0.01451906 | 38 | WT | FALSE | CAN | JSY13 | 2 | WT-Rep2 | TCT | C>A | A/T+1 | G/C-1 | CG>AT | CG>AT | TAAGCTCTCT | CTCTCTT | AGA | TCT | C | T | C>A | C>A_TCT | CG>AT_CTCTCTT | 9 | FALSE | FALSE |
| 6_v.txt | 33210 | 33210-SNV- | SNV | 1 | C | A | 4 | 13775 | 0.02903811 | 33.4985316 | WT | FALSE | CAN | JSY13 | 2 | WT-Rep2 | TCT | C>A | A/T+1 | G/C-1 | CG>AT | CG>AT | AGCTCTCTCT | CTCTTCA | AGA | TCT | C | T | C>A | C>A_TCT | CG>AT_CTCTTCA | 9 | FALSE | FALSE |
| 6_v.txt | 33354 | 33354-SNV- | SNV | 1 | G | T | 79 | 13776 | 0.57346109 | 18.278481 | WT | FALSE | CAN | JSY13 | 2 | WT-Rep2 | TGA | G>T | A/T+1 | G/C-1 | CG>AT | CG>AT | GGTATTGAC | GACCCAC | ACT | ACT | C | A | C>A | C>A_ACT | CG>AT_GACCCAC | 9 | FALSE | FALSE |
| 6_v.txt | 33415 | 33415-SNV- | SNV | 1 | C | A | 26 | 13776 | 0.18873403 | 38.7692308 | WT | FALSE | CAN | JSY13 | 2 | WT-Rep2 | TCA | C>A | A/T+1 | G/C-1 | CG>AT | CG>AT | CCGGCTCATT | CATTGTA | AGT | TCA | C | T | C>A | C>A_TCA | CG>AT_CATTGTA | 9 | FALSE | FALSE |
| 6_v.txt | 33474 | 33474-SNV- | SNV | 1 | C | A | 38 | 13775 | 0.27586207 | 33.9473684 | WT | FALSE | CAN | JSY13 | 2 | WT-Rep2 | GCC | C>A | A/T+1 | G/C-1 | CG>AT | CG>AT | GCTATGCCTT | CCTTTTT | CGG | GCC | C | T | C>A | C>A_GCC | CG>AT_CCTTTTT | 9 | FALSE | FALSE |
| 6_v.txt | 32488 | 32488-Repla | Replacement | 2 | G | AC | 1 | 32056 | 3.12E-03 | 38 | WT | FALSE | CAN | JSY13 | 2 | WT-Rep2 | CGG | nope | >1bp-ins | G/C-1 | none | Replac. | AGATACGGA | GGAAAAC | GCC | GCC | C | none | C>none | C>none_GCC | Replac._GGAAAAC | 9 | FALSE | FALSE |
| 6_v.txt | 32604 | 32604-Repla | Replacement | 2 | G | CA | 577 | 25283 | 2.28216588 | 42.5987866 | WT | FALSE | CAN | JSY13 | 2 | WT-Rep2 | GGA | nope | >1bp-ins | G/C-1 | none | Replac. | AAAGAGGA | GAAACCC | CCT | CCT | C | none | C>none | C>none_CCT | Replac._GAAACCC | 9 | FALSE | FALSE |
| 6_v.txt | 32700 | 32700-Repla | Replacement | 9 | CC | TTGTGTTGT | 1 | 25283 | 3.96E-03 | 64 | WT | FALSE | CAN | JSY13 | 2 | WT-Rep2 | GCC | nope | >1bp-ins | >1bpdel | none | Replac. | ACTGGGCCG | CCGGTAA | CGG | GCC | CC | none | NA | NA_GCC | Replac._CCGGTAA | 9 | FALSE | FALSE |
| 6_v.txt | 33030 | 33030-Repla | Replacement | 2 | AA | C | 10 | 20702 | 0.04830451 | 63.5 | WT | FALSE | CAN | JSY13 | 2 | WT-Rep2 | GAA | nope | G/C+1 | >1bpdel | none | Replac. | GGGATGAAT | AATGTAG | CTT | GAA | AA | G | NA | NA_GAA | Replac._AATGTAG | 9 | FALSE | FALSE |
| 6_v.txt | 31964 | 31964-MNV- | MNV | 2 | AT | TA | 22 | 21739 | 0.10120061 | 37.613488 | WT | FALSE | CAN | JSY13 | 2 | WT-Rep2 | CAT | nope | >1bp-ins | >1bpdel | none | MNV | GGCCGCATA | ATAATAA | GTA | CAT | AT | none | NA | NA_CAT | MNV_ATAATAA | 9 | FALSE | FALSE |
| 6_v.txt | 32298 | 32298-MNV- | MNV | 2 | GC | CA | 5278 | 32056 | 16.4649364 | 55.8531804 | WT | FALSE | CAN | JSY13 | 2 | WT-Rep2 | TGC | nope | >1bp-ins | >1bpdel | none | MNV | GAATTTGCG | GCGGCAG | ACG | TGC | GC | none | NA | NA_TGC | MNV_GCGGCAG | 9 | FALSE | FALSE |

| | | | | | | | | | | | | | | | | | | | | | | | | | | | | | | | | | | |
|---|---|---|---|---|---|---|---|---|---|---|---|---|---|---|---|---|---|---|---|---|---|---|---|---|---|---|---|---|---|---|---|---|---|---|
| 6_v.txt | 32298 | 32298-MNV- | MNV | 2 | GC | CG | 6 | 32056 | 0.01871725 | 55.7509243 | WT | FALSE | CAN | JSY13 | 2 | WT-Rep2 | TGC | nope | >1bp-ins | >1bpdel | none | MNV | GAATTTGCG | GCGGCAG | ACG | TGC | GC | none | NA | NA_TGC | MNV_GCGGCAG | 9 | FALSE | FALSE |
| 6_v.txt | 33168 | 33168-MNV- | MNV | 2 | CC | AA | 1 | 20695 | 4.83E-03 | 36 | WT | FALSE | CAN | JSY13 | 2 | WT-Rep2 | ACC | nope | >1bp-ins | >1bpdel | none | MNV | ATAGTACCA | CCACCAA | TGG | ACC | CC | none | NA | NA_ACC | MNV_CCACCAA | 9 | FALSE | FALSE |
| 6_v.txt | 33198 | 33198-MNV- | MNV | 2 | TG | CA | 12 | 13775 | 0.08711434 | 37.0626475 | WT | FALSE | CAN | JSY13 | 2 | WT-Rep2 | TTG | nope | >1bp-ins | >1bpdel | none | MNV | TGTCTTTGCT | TGCTTAA | AAC | TTG | TG | none | NA | NA_TTG | MNV_TGCTTAA | 9 | FALSE | FALSE |
| 6_v.txt | 31964 | 31964-Insert | Insertion | 3 | - | ATA | 3 | 21739 | 0.01380008 | 37.7777778 | WT | FALSE | CAN | JSY13 | 2 | WT-Rep2 | CAT | nope | >1bp-ins | >1bpdel | none | >1bp-ins | GGCCGCATA | ATAATAA | GTA | CAT | - | none | NA | NA_CAT | >1bp-ins_ATAATAA | 9 | FALSE | FALSE |
| 6_v.txt | 31969 | 31969-Insert | Insertion | 2 | - | TA | 22 | 21739 | 0.10120061 | 37.3181818 | WT | FALSE | CAN | JSY13 | 2 | WT-Rep2 | TAA | nope | >1bp-ins | >1bpdel | none | >1bp-ins | CATAATAAG | AAGCCAA | ATT | TAA | - | none | NA | NA_TAA | >1bp-ins_AAGCCAA | 9 | FALSE | FALSE |
| 6_v.txt | 31996 | 31996-Insert | Insertion | 4 | - | TAAA | 25 | 21739 | 0.11500069 | 48.14 | WT | FALSE | CAN | JSY13 | 2 | WT-Rep2 | TTA | nope | >1bp-ins | >1bpdel | none | >1bp-ins | TAGCTTTAA | TAAATGG | AAT | TTA | - | none | NA | NA_TTA | >1bp-ins_TAAATGG | 9 | FALSE | FALSE |
| 6_v.txt | 32033 | 32033-Insert | Insertion | 1 | - | A | 8 | 21739 | 0.03680022 | 64 | WT | FALSE | CAN | JSY13 | 2 | WT-Rep2 | TCA | nope | A/T+1 | >1bpdel | none | A/T+1 | GTATTTCAA | CAAAGCT | AGT | TCA | - | T | NA | NA_TCA | A/T+1_CAAAGCT | 9 | FALSE | FALSE |
| 6_v.txt | 32068 | 32068-Insert | Insertion | 1 | - | T | 4 | 21739 | 0.01840011 | 56.5 | WT | FALSE | CAN | JSY13 | 2 | WT-Rep2 | ATA | nope | A/T+1 | >1bpdel | none | A/T+1 | TTGAGATAA | TAAATAA | TAT | ATA | - | A | NA | NA_ATA | A/T+1_TAAATAA | 9 | FALSE | FALSE |
| 6_v.txt | 32081 | 32081-Insert | Insertion | 1 | - | A | 38 | 21739 | 0.17480105 | 63.8684211 | WT | FALSE | CAN | JSY13 | 2 | WT-Rep2 | CAA | nope | A/T+1 | >1bpdel | none | A/T+1 | CCATGCAAA | AAAAAAG | GTT | CAA | - | T | NA | NA_CAA | A/T+1_AAAAAAG | 9 | FALSE | FALSE |
| 6_v.txt | 32164 | 32164-Insert | Insertion | 18 | - | CGCCAAATG | 1 | 21737 | 4.60E-03 | 38 | WT | FALSE | CAN | JSY13 | 2 | WT-Rep2 | GCG | nope | >1bp-ins | >1bpdel | none | >1bp-ins | CCAAAGCGC | CGCCAAA | CGC | GCG | - | none | NA | NA_GCG | >1bp-ins_CGCCAAA | 9 | FALSE | FALSE |
| 6_v.txt | 32277 | 32277-Insert | Insertion | 1 | - | C | 22 | 32056 | 0.0686299 | 37 | WT | FALSE | CAN | JSY13 | 2 | WT-Rep2 | ACC | nope | G/C+1 | >1bpdel | none | G/C+1 | CGGGAACCA | CCAACGT | TGG | ACC | - | G | NA | NA_ACC | G/C+1_CCAACGT | 9 | FALSE | FALSE |
| 6_v.txt | 32300 | 32300-Insert | Insertion | 1 | - | A | 1 | 32056 | 3.12E-03 | 34 | WT | FALSE | CAN | JSY13 | 2 | WT-Rep2 | CGG | nope | A/T+1 | >1bpdel | none | A/T+1 | ATTTGCGGC | GGCAGAA | GCC | CGG | - | T | NA | NA_CGG | A/T+1_GGCAGAA | 9 | FALSE | FALSE |
| 6_v.txt | 32375 | 32375-Insert | Insertion | 1 | - | A | 9 | 32056 | 0.02807587 | 61 | WT | FALSE | CAN | JSY13 | 2 | WT-Rep2 | CAA | nope | A/T+1 | >1bpdel | none | A/T+1 | AATAGCAAT | AATAATA | GTT | CAA | - | T | NA | NA_CAA | A/T+1_AATAATA | 9 | FALSE | FALSE |
| 6_v.txt | 32468 | 32468-Insert | Insertion | 1 | - | A | 4 | 32056 | 0.01247816 | 57.5 | WT | FALSE | CAN | JSY13 | 2 | WT-Rep2 | CAA | nope | A/T+1 | >1bpdel | none | A/T+1 | AGAGCCAAT | AATGTAG | GTT | CAA | - | T | NA | NA_CAA | A/T+1_AATGTAG | 9 | FALSE | FALSE |
| 6_v.txt | 32489 | 32489-Insert | Insertion | 1 | - | C | 12 | 32056 | 0.03743449 | 46.9166667 | WT | FALSE | CAN | JSY13 | 2 | WT-Rep2 | GGA | nope | G/C+1 | >1bpdel | none | G/C+1 | GATACGGAA | GAAAACA | CCT | GGA | - | G | NA | NA_GGA | G/C+1_GAAAACA | 9 | FALSE | FALSE |
| 6_v.txt | 32498 | 32498-Insert | Insertion | 1 | - | T | 11 | 32056 | 0.03431495 | 42.3636364 | WT | FALSE | CAN | JSY13 | 2 | WT-Rep2 | CTT | nope | A/T+1 | >1bpdel | none | A/T+1 | AACAACTTTT | TTTTTTG | GAA | CTT | - | A | NA | NA_CTT | A/T+1_TTTTTTG | 9 | FALSE | FALSE |
| 6_v.txt | 32605 | 32605-Insert | Insertion | 1 | - | A | 1 | 25283 | 3.96E-03 | 33 | WT | FALSE | CAN | JSY13 | 2 | WT-Rep2 | GAA | nope | A/T+1 | >1bpdel | none | A/T+1 | AAGAGGAAA | AAACCCA | CTT | GAA | - | T | NA | NA_GAA | A/T+1_AAACCCA | 9 | FALSE | FALSE |
| 6_v.txt | 32631 | 32631-Insert | Insertion | 1 | - | T | 25 | 25283 | 0.09888067 | 47.56 | WT | FALSE | CAN | JSY13 | 2 | WT-Rep2 | GTT | nope | A/T+1 | >1bpdel | none | A/T+1 | CCTTCGTTTT | TTTTTAT | CAA | GTT | - | A | NA | NA_GTT | A/T+1_TTTTTAT | 9 | FALSE | FALSE |
| 6_v.txt | 32648 | 32648-Insert | Insertion | 2 | - | TA | 1 | 25283 | 3.96E-03 | 38 | WT | FALSE | CAN | JSY13 | 2 | WT-Rep2 | TTA | nope | >1bp-ins | >1bpdel | none | >1bp-ins | AGATATTATA | TATACCT | AAT | TTA | - | none | NA | NA_TTA | >1bp-ins_TATACCT | 9 | FALSE | FALSE |
| 6_v.txt | 32648 | 32648-Insert | Insertion | 4 | - | TATA | 5 | 25283 | 0.01977613 | 58.65 | WT | FALSE | CAN | JSY13 | 2 | WT-Rep2 | TTA | nope | >1bp-ins | >1bpdel | none | >1bp-ins | AGATATTATA | TATACCT | AAT | TTA | - | none | NA | NA_TTA | >1bp-ins_TATACCT | 9 | FALSE | FALSE |
| 6_v.txt | 32719 | 32719-Insert | Insertion | 1 | - | A | 6 | 25283 | 0.02373136 | 59.6666667 | WT | FALSE | CAN | JSY13 | 2 | WT-Rep2 | CAA | nope | A/T+1 | >1bpdel | none | A/T+1 | CACCACAAA | AAACCAT | GTT | CAA | - | T | NA | NA_CAA | A/T+1_AAACCAT | 9 | FALSE | FALSE |
| 6_v.txt | 32841 | 32841-Insert | Insertion | 1 | - | A | 6 | 25283 | 0.02373136 | 50.1666667 | WT | FALSE | CAN | JSY13 | 2 | WT-Rep2 | CCA | nope | A/T+1 | >1bpdel | none | A/T+1 | ATTACCCAA | CAAAAAA | GGT | CCA | - | T | NA | NA_CCA | A/T+1_CAAAAAA | 9 | FALSE | FALSE |
| 6_v.txt | 32842 | 32842-Insert | Insertion | 1 | - | A | 72 | 25283 | 0.28477633 | 43.3194444 | WT | FALSE | CAN | JSY13 | 2 | WT-Rep2 | CAA | nope | A/T+1 | >1bpdel | none | A/T+1 | TTACCCAAA | AAAAAAT | GTT | CAA | - | T | NA | NA_CAA | A/T+1_AAAAAAT | 9 | FALSE | FALSE |
| 6_v.txt | 32901 | 32901-Insert | Insertion | 1 | - | T | 1 | 20702 | 4.83E-03 | 38 | WT | FALSE | CAN | JSY13 | 2 | WT-Rep2 | GCC | nope | A/T+1 | >1bpdel | none | A/T+1 | ACTTGGCCA | CCAACTA | CGG | GCC | - | A | NA | NA_GCC | A/T+1_CCAACTA | 9 | FALSE | FALSE |
| 6_v.txt | 32903 | 32903-Insert | Insertion | 1 | - | A | 21 | 20702 | 0.10143947 | 42.6666667 | WT | FALSE | CAN | JSY13 | 2 | WT-Rep2 | CAA | nope | A/T+1 | >1bpdel | none | A/T+1 | TTGGCCAAC | AACTACA | GTT | CAA | - | T | NA | NA_CAA | A/T+1_AACTACA | 9 | FALSE | FALSE |
| 6_v.txt | 33018 | 33018-Insert | Insertion | 6 | - | GTAACA | 2 | 20702 | 9.66E-03 | 51 | WT | FALSE | CAN | JSY13 | 2 | WT-Rep2 | TGT | nope | >1bp-ins | >1bpdel | none | >1bp-ins | GAGGATGTA | GTAACAG | ACA | TGT | - | none | NA | NA_TGT | >1bp-ins_GTAACAG | 9 | FALSE | FALSE |
| 6_v.txt | 33428 | 33428-Insert | Insertion | 1 | - | A | 3 | 13776 | 0.021777 | 50.6666667 | WT | FALSE | CAN | JSY13 | 2 | WT-Rep2 | GCT | nope | A/T+1 | >1bpdel | none | A/T+1 | CATATGCTTC | CTTCTCC | CGA | GCT | - | T | NA | NA_GCT | A/T+1_CTTCTCC | 9 | FALSE | FALSE |
| 6_v.txt | 32157 | 32157-Deleti | Deletion | 8 | GCCAAAGC | - | 11 | 21737 | 0.05060496 | 56.3636364 | WT | FALSE | CAN | JSY13 | 2 | WT-Rep2 | AGC | nope | >1bp-ins | >1bpdel | none | >1bpdel | ATGTAAGCC | GCCAAAG | TCG | AGC | GCCAAAGC | none | NA | NA_AGC | >1bpdel_GCCAAAG | 9 | FALSE | FALSE |
| 6_v.txt | 32160 | 32160-Deleti | Deletion | 2 | AA | - | 1 | 21737 | 4.60E-03 | 38 | WT | FALSE | CAN | JSY13 | 2 | WT-Rep2 | CAA | nope | >1bp-ins | >1bpdel | none | >1bpdel | TAAGCCAAA | AAAGCGC | GTT | CAA | AA | none | NA | NA_CAA | >1bpdel_AAAGCGC | 9 | FALSE | FALSE |
| 6_v.txt | 32337 | 32337-Deleti | Deletion | 2 | AT | - | 149 | 32056 | 0.46481158 | 62.3758389 | WT | FALSE | CAN | JSY13 | 2 | WT-Rep2 | GAT | nope | >1bp-ins | >1bpdel | none | >1bpdel | TTGAAGATA | ATATGTG | CTA | GAT | AT | none | NA | NA_GAT | >1bpdel_ATATGTG | 9 | FALSE | FALSE |
| 6_v.txt | 32472 | 32472-Deleti | Deletion | 2 | TA | - | 18 | 32056 | 0.05615174 | 48.8888889 | WT | FALSE | CAN | JSY13 | 2 | WT-Rep2 | GTA | nope | >1bp-ins | >1bpdel | none | >1bpdel | CCAATGTAG | TAGAAGG | CAT | GTA | TA | none | NA | NA_GTA | >1bpdel_TAGAAGG | 9 | FALSE | FALSE |
| 6_v.txt | 32476 | 32476-Deleti | Deletion | 4 | AGGT | - | 18 | 32056 | 0.05615174 | 48.6666667 | WT | FALSE | CAN | JSY13 | 2 | WT-Rep2 | AAG | nope | >1bp-ins | >1bpdel | none | >1bpdel | TGTAGAAGG | AGGTTAA | TTC | AAG | AGGT | none | NA | NA_AAG | >1bpdel_AGGTTAA | 9 | FALSE | FALSE |
| 6_v.txt | 32700 | 32700-Deleti | Deletion | 2 | CC | - | 7 | 25283 | 0.02768659 | 64 | WT | FALSE | CAN | JSY13 | 2 | WT-Rep2 | GCC | nope | >1bp-ins | >1bpdel | none | >1bpdel | ACTGGGCCG | CCGGTAA | CGG | GCC | CC | none | NA | NA_GCC | >1bpdel_CCGGTAA | 9 | FALSE | FALSE |
| 6_v.txt | 32767 | 32767-Deleti | Deletion | 3 | TGA | - | 8 | 25283 | 0.03164182 | 47.25 | WT | FALSE | CAN | JSY13 | 2 | WT-Rep2 | TTG | nope | >1bp-ins | >1bpdel | none | >1bpdel | AAACTTTGAT | TGATGGA | AAC | TTG | TGA | none | NA | NA_TTG | >1bpdel_TGATGGA | 9 | FALSE | FALSE |
| 6_v.txt | 33126 | 33126-Deleti | Deletion | 2 | AG | - | 10 | 20702 | 0.04830451 | 44.3 | WT | FALSE | CAN | JSY13 | 2 | WT-Rep2 | CAG | nope | >1bp-ins | >1bpdel | none | >1bpdel | TTGGTCAGA | AGAGGTG | GTC | CAG | AG | none | NA | NA_CAG | >1bpdel_AGAGGTG | 9 | FALSE | FALSE |
| 6_v.txt | 33206 | 33206-Deleti | Deletion | 2 | CT | - | 1 | 13775 | 7.26E-03 | 38 | WT | FALSE | CAN | JSY13 | 2 | WT-Rep2 | GCT | nope | >1bp-ins | >1bpdel | none | >1bpdel | CTTAAGCTCT | CTCTCTC | CGA | GCT | CT | none | NA | NA_GCT | >1bpdel_CTCTCTC | 9 | FALSE | FALSE |
| 6_v.txt | 33206 | 33206-Deleti | Deletion | 6 | CTCTCT | - | 8 | 13776 | 0.05807201 | 37.5 | WT | FALSE | CAN | JSY13 | 2 | WT-Rep2 | GCT | nope | >1bp-ins | >1bpdel | none | >1bpdel | CTTAAGCTCT | CTCTCTC | CGA | GCT | CTCTCT | none | NA | NA_GCT | >1bpdel_CTCTCTC | 9 | FALSE | FALSE |
| 6_v.txt | 31723 | 31723-Deleti | Deletion | 1 | A | - | 10 | 11614 | 0.08610298 | 62.6 | WT | FALSE | CAN | JSY13 | 2 | WT-Rep2 | CAA | nope | >1bp-ins | A/T-1 | none | A/T-1 | TGTCCCAAA | AAAAAGT | GTT | GTT | T | none | T>none | T>none_GTT | A/T-1_AAAAAGT | 9 | FALSE | FALSE |
| 6_v.txt | 31863 | 31863-Deleti | Deletion | 1 | A | - | 7 | 11607 | 0.06030844 | 56.1428571 | WT | FALSE | CAN | JSY13 | 2 | WT-Rep2 | GAA | nope | >1bp-ins | A/T-1 | none | A/T-1 | AACAGGAAA | AAAATAG | CTT | CTT | T | none | T>none | T>none_CTT | A/T-1_AAAATAG | 9 | FALSE | FALSE |
| 6_v.txt | 31899 | 31899-Deleti | Deletion | 1 | T | - | 7 | 11607 | 0.06030844 | 42.5714286 | WT | FALSE | CAN | JSY13 | 2 | WT-Rep2 | ATT | nope | >1bp-ins | A/T-1 | none | A/T-1 | ACACCATTG | TTGAATT | TAA | ATT | T | none | nope | nope_ATT | A/T-1_TTGAATT | 9 | FALSE | FALSE |
| 6_v.txt | 31904 | 31904-Deleti | Deletion | 1 | T | - | 6 | 11607 | 0.05169294 | 49.1666667 | WT | FALSE | CAN | JSY13 | 2 | WT-Rep2 | ATT | nope | >1bp-ins | A/T-1 | none | A/T-1 | ATTGAATTTT | TTTTGGT | TAA | ATT | T | none | nope | nope_ATT | A/T-1_TTTTGGT | 9 | FALSE | FALSE |
| 6_v.txt | 31913 | 31913-Deleti | Deletion | 1 | A | - | 15 | 11607 | 0.12923236 | 37.6 | WT | FALSE | CAN | JSY13 | 2 | WT-Rep2 | CAA | nope | >1bp-ins | A/T-1 | none | A/T-1 | TGGTGCAAA | AAAAGCC | GTT | GTT | T | none | T>none | T>none_GTT | A/T-1_AAAAGCC | 9 | FALSE | FALSE |
| 6_v.txt | 31918 | 31918-Deleti | Deletion | 1 | C | - | 9 | 11607 | 0.07753942 | 37.2222222 | WT | FALSE | CAN | JSY13 | 2 | WT-Rep2 | GCC | nope | >1bp-ins | G/C-1 | none | G/C-1 | CAAAAGCCG | CCGTGAA | CGG | GCC | C | none | nope | nope_GCC | G/C-1_CCGTGAA | 9 | FALSE | FALSE |
| 6_v.txt | 31923 | 31923-Deleti | Deletion | 1 | A | - | 10 | 11607 | 0.08615491 | 38 | WT | FALSE | CAN | JSY13 | 2 | WT-Rep2 | GAA | nope | >1bp-ins | A/T-1 | none | A/T-1 | GCCGTGAAA | AAACCTT | CTT | CTT | T | none | T>none | T>none_CTT | A/T-1_AAACCTT | 9 | FALSE | FALSE |
| 6_v.txt | 32057 | 32057-Deleti | Deletion | 1 | C | - | 16 | 21739 | 0.07360044 | 59.75 | WT | FALSE | CAN | JSY13 | 2 | WT-Rep2 | GCG | nope | >1bp-ins | G/C-1 | none | G/C-1 | GATGTGCGA | CGAGATT | CGC | GCG | C | none | nope | nope_GCG | G/C-1_CGAGATT | 9 | FALSE | FALSE |
| 6_v.txt | 32081 | 32081-Deleti | Deletion | 1 | A | - | 30 | 21739 | 0.13800083 | 62.4999982 | WT | FALSE | CAN | JSY13 | 2 | WT-Rep2 | CAA | nope | >1bp-ins | A/T-1 | none | A/T-1 | CCATGCAAA | AAAAAAG | GTT | GTT | T | none | T>none | T>none_GTT | A/T-1_AAAAAAG | 9 | FALSE | FALSE |
| 6_v.txt | 32160 | 32160-Deleti | Deletion | 1 | A | - | 4 | 21737 | 0.0184018 | 57.4999958 | WT | FALSE | CAN | JSY13 | 2 | WT-Rep2 | CAA | nope | >1bp-ins | A/T-1 | none | A/T-1 | TAAGCCAAA | AAAGCGC | GTT | GTT | T | none | T>none | T>none_GTT | A/T-1_AAAGCGC | 9 | FALSE | FALSE |
| 6_v.txt | 32180 | 32180-Deleti | Deletion | 1 | A | - | 38 | 21737 | 0.17481713 | 47.0526316 | WT | FALSE | CAN | JSY13 | 2 | WT-Rep2 | TAA | nope | >1bp-ins | A/T-1 | none | A/T-1 | AGCAGTAAC | AACGAAA | ATT | ATT | T | none | T>none | T>none_ATT | A/T-1_AACGAAA | 9 | FALSE | FALSE |
| 6_v.txt | 32184 | 32184-Deleti | Deletion | 1 | A | - | 35 | 21737 | 0.16101578 | 45.6285714 | WT | FALSE | CAN | JSY13 | 2 | WT-Rep2 | GAA | nope | >1bp-ins | A/T-1 | none | A/T-1 | GTAACGAAA | AAAACTG | CTT | CTT | T | none | T>none | T>none_CTT | A/T-1_AAAACTG | 9 | FALSE | FALSE |
| 6_v.txt | 32238 | 32238-Deleti | Deletion | 1 | G | - | 31 | 21732 | 0.14264679 | 37.8064516 | WT | FALSE | CAN | JSY13 | 2 | WT-Rep2 | AGC | nope | >1bp-ins | G/C-1 | none | G/C-1 | TTAGGAGCC | GCCAACT | TCG | TCG | C | none | C>none | C>none_TCG | G/C-1_GCCAACT | 9 | FALSE | FALSE |
| 6_v.txt | 32249 | 32249-Deleti | Deletion | 1 | C | - | 5 | 21717 | 0.02302344 | 38 | WT | FALSE | CAN | JSY13 | 2 | WT-Rep2 | TCT | nope | >1bp-ins | G/C-1 | none | G/C-1 | CTTGTTCTTT | CTTTGAT | AGA | TCT | C | none | nope | nope_TCT | G/C-1_CTTTGAT | 9 | FALSE | FALSE |
| 6_v.txt | 32298 | 32298-Deleti | Deletion | 1 | G | - | 4 | 32056 | 0.01247816 | 43.2500039 | WT | FALSE | CAN | JSY13 | 2 | WT-Rep2 | TGC | nope | >1bp-ins | G/C-1 | none | G/C-1 | GAATTTGCG | GCGGCAG | ACG | ACG | C | none | C>none | C>none_ACG | G/C-1_GCGGCAG | 9 | FALSE | FALSE |
| 6_v.txt | 32299 | 32299-Deleti | Deletion | 1 | C | - | 1 | 32056 | 3.12E-03 | 54.0000071 | WT | FALSE | CAN | JSY13 | 2 | WT-Rep2 | GCG | nope | >1bp-ins | G/C-1 | none | G/C-1 | AATTTGCGG | CGGCAGA | CGC | GCG | C | none | nope | nope_GCG | G/C-1_CGGCAGA | 9 | FALSE | FALSE |
| 6_v.txt | 32328 | 32328-Deleti | Deletion | 1 | G | - | 8 | 32056 | 0.02495633 | 57.125 | WT | FALSE | CAN | JSY13 | 2 | WT-Rep2 | AGC | nope | >1bp-ins | G/C-1 | none | G/C-1 | ATAACAGCG | GCGTTGA | TCG | TCG | C | none | C>none | C>none_TCG | G/C-1_GCGTTGA | 9 | FALSE | FALSE |
| 6_v.txt | 32387 | 32387-Deleti | Deletion | 1 | A | - | 21 | 32056 | 0.06551036 | 59.0952381 | WT | FALSE | CAN | JSY13 | 2 | WT-Rep2 | GAG | nope | >1bp-ins | A/T-1 | none | A/T-1 | AAAGGGAGA | AGAAGTA | CTC | CTC | T | none | T>none | T>none_CTC | A/T-1_AGAAGTA | 9 | FALSE | FALSE |
| 6_v.txt | 32436 | 32436-Deleti | Deletion | 1 | G | - | 59 | 32056 | 0.18405291 | 56.8983051 | WT | FALSE | CAN | JSY13 | 2 | WT-Rep2 | TGG | nope | >1bp-ins | G/C-1 | none | G/C-1 | TTGTATGGA | GGAACTA | ACC | ACC | C | none | C>none | C>none_ACC | G/C-1_GGAACTA | 9 | FALSE | FALSE |
| 6_v.txt | 32442 | 32442-Deleti | Deletion | 1 | A | - | 59 | 32056 | 0.18405291 | 52.9322038 | WT | FALSE | CAN | JSY13 | 2 | WT-Rep2 | TAA | nope | >1bp-ins | A/T-1 | none | A/T-1 | GGAACTAAA | AAAAGTC | ATT | ATT | T | none | T>none | T>none_ATT | A/T-1_AAAAGTC | 9 | FALSE | FALSE |
| 6_v.txt | 32470 | 32470-Deleti | Deletion | 1 | T | - | 18 | 32056 | 0.05615174 | 49.3333113 | WT | FALSE | CAN | JSY13 | 2 | WT-Rep2 | ATG | nope | >1bp-ins | A/T-1 | none | A/T-1 | AGCCAATGT | TGTAGAA | TAC | ATG | T | none | nope | nope_ATG | A/T-1_TGTAGAA | 9 | FALSE | FALSE |
| 6_v.txt | 32498 | 32498-Deleti | Deletion | 1 | T | - | 68 | 32056 | 0.21212878 | 43.7794104 | WT | FALSE | CAN | JSY13 | 2 | WT-Rep2 | CTT | nope | >1bp-ins | A/T-1 | none | A/T-1 | AACAACTTTT | TTTTTTG | GAA | CTT | T | none | nope | nope_CTT | A/T-1_TTTTTTG | 9 | FALSE | FALSE |
| 6_v.txt | 32532 | 32532-Deleti | Deletion | 1 | T | - | 22 | 32056 | 0.0686299 | 39.0909091 | WT | FALSE | CAN | JSY13 | 2 | WT-Rep2 | GTT | nope | >1bp-ins | A/T-1 | none | A/T-1 | CTGGGGTTT | TTTGCAG | CAA | GTT | T | none | nope | nope_GTT | A/T-1_TTTGCAG | 9 | FALSE | FALSE |
| 6_v.txt | 32573 | 32573-Deleti | Deletion | 1 | T | - | 3 | 25283 | 0.01186568 | 36.6666667 | WT | FALSE | CAN | JSY13 | 2 | WT-Rep2 | CTT | nope | >1bp-ins | A/T-1 | none | A/T-1 | AGTACCTTG | TTGAAAT | GAA | CTT | T | none | nope | nope_CTT | A/T-1_TTGAAAT | 9 | FALSE | FALSE |
| 6_v.txt | 32583 | 32583-Deleti | Deletion | 1 | A | - | 4 | 25283 | 0.01582091 | 30.75 | WT | FALSE | CAN | JSY13 | 2 | WT-Rep2 | GAA | nope | >1bp-ins | A/T-1 | none | A/T-1 | AATGTGAAG | AAGGCAG | CTT | CTT | T | none | T>none | T>none_CTT | A/T-1_AAGGCAG | 9 | FALSE | FALSE |
| 6_v.txt | 32598 | 32598-Deleti | Deletion | 1 | A | - | 4681 | 25283 | 18.5144168 | 41.4142277 | WT | FALSE | CAN | JSY13 | 2 | WT-Rep2 | CAA | nope | >1bp-ins | A/T-1 | none | A/T-1 | TTAATCAAA | AAAGAGG | GTT | GTT | T | none | T>none | T>none_GTT | A/T-1_AAAGAGG | 9 | FALSE | FALSE |
| 6_v.txt | 32605 | 32605-Deleti | Deletion | 1 | A | - | 7 | 25281 | 0.02768878 | 36.9999994 | WT | FALSE | CAN | JSY13 | 2 | WT-Rep2 | GAA | nope | >1bp-ins | A/T-1 | none | A/T-1 | AAGAGGAAA | AAACCCA | CTT | CTT | T | none | T>none | T>none_CTT | A/T-1_AAACCCA | 9 | FALSE | FALSE |
| 6_v.txt | 32631 | 32631-Deleti | Deletion | 1 | T | - | 8 | 25283 | 0.03164182 | 43.5000113 | WT | FALSE | CAN | JSY13 | 2 | WT-Rep2 | GTT | nope | >1bp-ins | A/T-1 | none | A/T-1 | CCTTCGTTTT | TTTTTAT | CAA | GTT | T | none | nope | nope_GTT | A/T-1_TTTTTAT | 9 | FALSE | FALSE |
| 6_v.txt | 32640 | 32640-Deleti | Deletion | 1 | T | - | 7 | 25283 | 0.02768659 | 52.8571429 | WT | FALSE | CAN | JSY13 | 2 | WT-Rep2 | CTT | nope | >1bp-ins | A/T-1 | none | A/T-1 | TTATCCTTAG | TTAGATA | GAA | CTT | T | none | nope | nope_CTT | A/T-1_TTAGATA | 9 | FALSE | FALSE |
| 6_v.txt | 32658 | 32658-Deleti | Deletion | 1 | C | - | 9 | 25283 | 0.03559704 | 51.6666667 | WT | FALSE | CAN | JSY13 | 2 | WT-Rep2 | ACC | nope | >1bp-ins | G/C-1 | none | G/C-1 | CCTGGACCC | CCCCAGG | TGG | ACC | C | none | nope | nope_ACC | G/C-1_CCCCAGG | 9 | FALSE | FALSE |
| 6_v.txt | 32662 | 32662-Deleti | Deletion | 1 | A | - | 10 | 25283 | 0.03955227 | 63.3 | WT | FALSE | CAN | JSY13 | 2 | WT-Rep2 | CAG | nope | >1bp-ins | A/T-1 | none | A/T-1 | GACCCCAGG | AGGCACC | GTC | GTC | T | none | T>none | T>none_GTC | A/T-1_AGGCACC | 9 | FALSE | FALSE |
| 6_v.txt | 32683 | 32683-Deleti | Deletion | 1 | A | - | 53 | 25283 | 0.20962702 | 60.9811321 | WT | FALSE | CAN | JSY13 | 2 | WT-Rep2 | TAA | nope | >1bp-ins | A/T-1 | none | A/T-1 | TCCAATAAC | AACGGAA | ATT | ATT | T | none | T>none | T>none_ATT | A/T-1_AACGGAA | 9 | FALSE | FALSE |
| 6_v.txt | 32719 | 32719-Deleti | Deletion | 1 | A | - | 46 | 25282 | 0.18194763 | 59.7608681 | WT | FALSE | CAN | JSY13 | 2 | WT-Rep2 | CAA | nope | >1bp-ins | A/T-1 | none | A/T-1 | CACCACAAA | AAACCAT | GTT | GTT | T | none | T>none | T>none_GTT | A/T-1_AAACCAT | 9 | FALSE | FALSE |
| 6_v.txt | 32728 | 32728-Deleti | Deletion | 1 | A | - | 13 | 25283 | 0.05141795 | 60.9230769 | WT | FALSE | CAN | JSY13 | 2 | WT-Rep2 | CAA | nope | >1bp-ins | A/T-1 | none | A/T-1 | CCATACAAA | AAAAACA | GTT | GTT | T | none | T>none | T>none_GTT | A/T-1_AAAAACA | 9 | FALSE | FALSE |
| 6_v.txt | 32779 | 32779-Deleti | Deletion | 1 | C | - | 8 | 25283 | 0.03164182 | 57.25 | WT | FALSE | CAN | JSY13 | 2 | WT-Rep2 | ACC | nope | >1bp-ins | G/C-1 | none | G/C-1 | AAGCGACCC | CCCAGAA | TGG | ACC | C | none | nope | nope_ACC | G/C-1_CCCAGAA | 9 | FALSE | FALSE |
| 6_v.txt | 32842 | 32842-Deleti | Deletion | 1 | A | - | 858 | 25283 | 3.39358462 | 43.5594353 | WT | FALSE | CAN | JSY13 | 2 | WT-Rep2 | CAA | nope | >1bp-ins | A/T-1 | none | A/T-1 | TTACCCAAA | AAAAAAT | GTT | GTT | T | none | T>none | T>none_GTT | A/T-1_AAAAAAT | 9 | FALSE | FALSE |
| 6_v.txt | 32884 | 32884-Deleti | Deletion | 1 | A | - | 13 | 20701 | 0.0627989 | 37.1538469 | WT | FALSE | CAN | JSY13 | 2 | WT-Rep2 | CAA | nope | >1bp-ins | A/T-1 | none | A/T-1 | ACGTCCAAA | AAAATTG | GTT | GTT | T | none | T>none | T>none_GTT | A/T-1_AAAATTG | 9 | FALSE | FALSE |
| 6_v.txt | 32901 | 32901-Deleti | Deletion | 1 | C | - | 1 | 20702 | 4.83E-03 | 38 | WT | FALSE | CAN | JSY13 | 2 | WT-Rep2 | GCC | nope | >1bp-ins | G/C-1 | none | G/C-1 | ACTTGGCCA | CCAACTA | CGG | GCC | C | none | nope | nope_GCC | G/C-1_CCAACTA | 9 | FALSE | FALSE |
| 6_v.txt | 32924 | 32924-Deleti | Deletion | 1 | A | - | 23 | 20702 | 0.11110038 | 51.4782609 | WT | FALSE | CAN | JSY13 | 2 | WT-Rep2 | CAA | nope | >1bp-ins | A/T-1 | none | A/T-1 | CAGGGCAAA | AAAAGTG | GTT | GTT | T | none | T>none | T>none_GTT | A/T-1_AAAAGTG | 9 | FALSE | FALSE |
| 6_v.txt | 32935 | 32935-Deleti | Deletion | 1 | C | - | 7 | 20702 | 0.03381316 | 48.4285714 | WT | FALSE | CAN | JSY13 | 2 | WT-Rep2 | GCC | nope | >1bp-ins | G/C-1 | none | G/C-1 | TGATTGCCC | CCCAAGA | CGG | GCC | C | none | nope | nope_GCC | G/C-1_CCCAAGA | 9 | FALSE | FALSE |
| 6_v.txt | 33111 | 33111-Deleti | Deletion | 1 | G | - | 3 | 20702 | 0.01449135 | 45.6666667 | WT | FALSE | CAN | JSY13 | 2 | WT-Rep2 | TGG | nope | >1bp-ins | G/C-1 | none | G/C-1 | CCCACTGGG | GGGCCGG | ACC | ACC | C | none | C>none | C>none_ACC | G/C-1_GGGCCGG | 9 | FALSE | FALSE |
| 6_v.txt | 33168 | 33168-Deleti | Deletion | 1 | C | - | 1 | 20695 | 4.83E-03 | 37.9999991 | WT | FALSE | CAN | JSY13 | 2 | WT-Rep2 | ACC | nope | >1bp-ins | G/C-1 | none | G/C-1 | ATAGTACCA | CCACCAA | TGG | ACC | C | none | nope | nope_ACC | G/C-1_CCACCAA | 9 | FALSE | FALSE |
| 6_v.txt | 33317 | 33317-Deleti | Deletion | 1 | A | - | 7 | 13776 | 0.05081301 | 56.8571429 | WT | FALSE | CAN | JSY13 | 2 | WT-Rep2 | GAT | nope | >1bp-ins | A/T-1 | none | A/T-1 | CAATGGATA | ATACAAT | CTA | CTA | T | none | T>none | T>none_CTA | A/T-1_ATACAAT | 9 | FALSE | FALSE |
| 6_v.txt | 33474 | 33474-Deleti | Deletion | 1 | C | - | 29 | 13775 | 0.21052632 | 27.3448276 | WT | FALSE | CAN | JSY13 | 2 | WT-Rep2 | GCC | nope | >1bp-ins | G/C-1 | none | G/C-1 | GCTATGCCTT | CCTTTTT | CGG | GCC | C | none | nope | nope_GCC | G/C-1_CCTTTTT | 9 | FALSE | FALSE |
| 7_v.txt | 31986 | 31986-SNV-A | SNV | 1 | A | C | 10164 | 22492 | 45.1894007 | 50.4585793 | WT | FALSE | CAN | JSY13 | 3 | WT-Rep3 | TAA | A>C | G/C+1 | A/T-1 | TA>GC | TA>GC | GGCATTAAT | AATTTAG | ATT | ATT | T | G | T>G | T>G_ATT | TA>GC_AATTTAG | 9 | FALSE | FALSE |
| 7_v.txt | 32162 | 32162-SNV-A | SNV | 1 | A | C | 3 | 22492 | 0.01333808 | 19.3339905 | WT | FALSE | CAN | JSY13 | 3 | WT-Rep3 | AAG | A>C | G/C+1 | A/T-1 | TA>GC | TA>GC | AGCCAAAGC | AGCGCCA | TTC | TTC | T | G | T>G | T>G_TTC | TA>GC_AGCGCCA | 9 | FALSE | FALSE |
| 7_v.txt | 32265 | 32265-SNV-A | SNV | 1 | A | C | 13 | 53882 | 0.0241268 | 18.2306866 | WT | FALSE | CAN | JSY13 | 3 | WT-Rep3 | TAA | A>C | G/C+1 | A/T-1 | TA>GC | TA>GC | CCAAATAAA | AAAATAC | ATT | ATT | T | G | T>G | T>G_ATT | TA>GC_AAAATAC | 9 | FALSE | FALSE |
| 7_v.txt | 32439 | 32439-SNV-A | SNV | 1 | A | C | 12 | 53882 | 0.02227089 | 49.0833333 | WT | FALSE | CAN | JSY13 | 3 | WT-Rep3 | AAC | A>C | G/C+1 | A/T-1 | TA>GC | TA>GC | TATGGAACT | ACTAAAA | TTG | TTG | T | G | T>G | T>G_TTG | TA>GC_ACTAAAA | 9 | FALSE | FALSE |
| 7_v.txt | 32445 | 32445-SNV-A | SNV | 1 | A | C | 27 | 53882 | 0.0501095 | 46.1481482 | WT | FALSE | CAN | JSY13 | 3 | WT-Rep3 | AAG | A>C | G/C+1 | A/T-1 | TA>GC | TA>GC | ACTAAAAGT | AGTCCAA | TTC | TTC | T | G | T>G | T>G_TTC | TA>GC_AGTCCAA | 9 | FALSE | FALSE |
| 7_v.txt | 32457 | 32457-SNV-A | SNV | 1 | A | C | 8 | 53882 | 0.01484726 | 44.75 | WT | FALSE | CAN | JSY13 | 3 | WT-Rep3 | TAA | A>C | G/C+1 | A/T-1 | TA>GC | TA>GC | ATGAATAAT | AATAGAG | ATT | ATT | T | G | T>G | T>G_ATT | TA>GC_AATAGAG | 9 | FALSE | FALSE |
| 7_v.txt | 32493 | 32493-SNV-A | SNV | 1 | A | C | 9 | 53882 | 0.01670317 | 47.4427857 | WT | FALSE | CAN | JSY13 | 3 | WT-Rep3 | AAC | A>C | G/C+1 | A/T-1 | TA>GC | TA>GC | CGGAAAACA | ACAACTT | TTG | TTG | T | G | T>G | T>G_TTG | TA>GC_ACAACTT | 9 | FALSE | FALSE |
| 7_v.txt | 32616 | 32616-SNV-A | SNV | 1 | A | C | 10 | 14183 | 0.07050695 | 42.5 | WT | FALSE | CAN | JSY13 | 3 | WT-Rep3 | TAA | A>C | G/C+1 | A/T-1 | TA>GC | TA>GC | CAACCTAAG | AAGAACC | ATT | ATT | T | G | T>G | T>G_ATT | TA>GC_AAGAACC | 9 | FALSE | FALSE |
| 7_v.txt | 32760 | 32760-SNV-A | SNV | 1 | A | C | 1 | 14183 | 7.05E-03 | 63.9985588 | WT | FALSE | CAN | JSY13 | 3 | WT-Rep3 | TAA | A>C | G/C+1 | A/T-1 | TA>GC | TA>GC | ATGGCTAAA | AAAACTT | ATT | ATT | T | G | T>G | T>G_ATT | TA>GC_AAAACTT | 9 | FALSE | FALSE |
| 7_v.txt | 32801 | 32801-SNV-A | SNV | 1 | A | C | 12 | 14183 | 0.08460833 | 46.1666667 | WT | FALSE | CAN | JSY13 | 3 | WT-Rep3 | AAT | A>C | G/C+1 | A/T-1 | TA>GC | TA>GC | ACCGTAATAT | ATATTTG | TTA | TTA | T | G | T>G | T>G_TTA | TA>GC_ATATTTG | 9 | FALSE | FALSE |
| 7_v.txt | 32823 | 32823-SNV-A | SNV | 1 | A | C | 30 | 14183 | 0.21152084 | 44.3333333 | WT | FALSE | CAN | JSY13 | 3 | WT-Rep3 | CAT | A>C | G/C+1 | A/T-1 | TA>GC | TA>GC | AAGTTCATTA | ATTATTG | GTA | GTA | T | G | T>G | T>G_GTA | TA>GC_ATTATTG | 9 | FALSE | FALSE |
| 7_v.txt | 32884 | 32884-SNV-A | SNV | 1 | A | C | 6 | 34274 | 0.01750598 | 34.831601 | WT | FALSE | CAN | JSY13 | 3 | WT-Rep3 | CAA | A>C | G/C+1 | A/T-1 | TA>GC | TA>GC | ACGTCCAAA | AAAATTG | GTT | GTT | T | G | T>G | T>G_GTT | TA>GC_AAAATTG | 9 | FALSE | FALSE |
| 7_v.txt | 33269 | 33269-SNV-A | SNV | 1 | A | C | 154 | 21999 | 0.70003182 | 15.987013 | WT | FALSE | CAN | JSY13 | 3 | WT-Rep3 | TAG | A>C | G/C+1 | A/T-1 | TA>GC | TA>GC | CTCCATAGA | AGAGAAC | ATC | ATC | T | G | T>G | T>G_ATC | TA>GC_AGAGAAC | 9 | FALSE | FALSE |
| 7_v.txt | 33476 | 33476-SNV-T | SNV | 1 | T | G | 3 | 21996 | 0.01363884 | 36.3330251 | WT | FALSE | CAN | JSY13 | 3 | WT-Rep3 | CTT | T>G | G/C+1 | A/T-1 | TA>GC | TA>GC | TATGCCTTTT | TTTTTTT | GAA | CTT | T | C | T>G | T>G_CTT | TA>GC_TTTTTTT | 9 | FALSE | FALSE |
| 7_v.txt | 32161 | 32161-SNV-A | SNV | 1 | A | G | 3 | 22492 | 0.01333808 | 27.658572 | WT | FALSE | CAN | JSY13 | 3 | WT-Rep3 | AAA | A>G | G/C+1 | A/T-1 | TA>CG | TA>CG | AAGCCAAAG | AAGCGCC | TTT | TTT | T | C | T>C | T>C_TTT | TA>CG_AAGCGCC | 9 | FALSE | FALSE |
| 7_v.txt | 32162 | 32162-SNV-A | SNV | 1 | A | G | 6 | 22492 | 0.02667615 | 44.329883 | WT | FALSE | CAN | JSY13 | 3 | WT-Rep3 | AAG | A>G | G/C+1 | A/T-1 | TA>CG | TA>CG | AGCCAAAGC | AGCGCCA | TTC | TTC | T | C | T>C | T>C_TTC | TA>CG_AGCGCCA | 9 | FALSE | FALSE |
| 7_v.txt | 32265 | 32265-SNV-A | SNV | 1 | A | G | 8 | 53882 | 0.01484726 | 29.7495215 | WT | FALSE | CAN | JSY13 | 3 | WT-Rep3 | TAA | A>G | G/C+1 | A/T-1 | TA>CG | TA>CG | CCAAATAAA | AAAATAC | ATT | ATT | T | C | T>C | T>C_ATT | TA>CG_AAAATAC | 9 | FALSE | FALSE |
| 7_v.txt | 32275 | 32275-SNV-A | SNV | 1 | A | G | 42 | 53882 | 0.07794811 | 33.0713264 | WT | FALSE | CAN | JSY13 | 3 | WT-Rep3 | GAA | A>G | G/C+1 | A/T-1 | TA>CG | TA>CG | TACGGGAAC | AACCAAC | CTT | CTT | T | C | T>C | T>C_CTT | TA>CG_AACCAAC | 9 | FALSE | FALSE |
| 7_v.txt | 32337 | 32337-SNV-A | SNV | 1 | A | G | 5 | 53882 | 9.28E-03 | 45.9999524 | WT | FALSE | CAN | JSY13 | 3 | WT-Rep3 | GAT | A>G | G/C+1 | A/T-1 | TA>CG | TA>CG | TTGAAGATA | ATATGTG | CTA | CTA | T | C | T>C | T>C_CTA | TA>CG_ATATGTG | 9 | FALSE | FALSE |
| 7_v.txt | 32346 | 32346-SNV-A | SNV | 1 | A | G | 13 | 53882 | 0.0241268 | 51.1537925 | WT | FALSE | CAN | JSY13 | 3 | WT-Rep3 | CAA | A>G | G/C+1 | A/T-1 | TA>CG | TA>CG | TGTGGCAAA | AAAACCT | GTT | GTT | T | C | T>C | T>C_GTT | TA>CG_AAAACCT | 9 | FALSE | FALSE |
| 7_v.txt | 32445 | 32445-SNV-A | SNV | 1 | A | G | 31 | 53882 | 0.05753313 | 49.1935484 | WT | FALSE | CAN | JSY13 | 3 | WT-Rep3 | AAG | A>G | G/C+1 | A/T-1 | TA>CG | TA>CG | ACTAAAAGT | AGTCCAA | TTC | TTC | T | C | T>C | T>C_TTC | TA>CG_AGTCCAA | 9 | FALSE | FALSE |
| 7_v.txt | 32492 | 32492-SNV-A | SNV | 1 | A | G | 15 | 53882 | 0.02783861 | 21.1333279 | WT | FALSE | CAN | JSY13 | 3 | WT-Rep3 | AAA | A>G | G/C+1 | A/T-1 | TA>CG | TA>CG | ACGGAAAAC | AACAACT | TTT | TTT | T | C | T>C | T>C_TTT | TA>CG_AACAACT | 9 | FALSE | FALSE |
| 7_v.txt | 32734 | 32734-SNV-A | SNV | 1 | A | G | 53 | 14183 | 0.37368681 | 56.8867925 | WT | FALSE | CAN | JSY13 | 3 | WT-Rep3 | CAG | A>G | G/C+1 | A/T-1 | TA>CG | TA>CG | AAAAACAGT | AGTATAT | GTC | GTC | T | C | T>C | T>C_GTC | TA>CG_AGTATAT | 9 | FALSE | FALSE |
| 7_v.txt | 32754 | 32754-SNV-A | SNV | 1 | A | G | 2 | 14183 | 0.01410139 | 50.9996322 | WT | FALSE | CAN | JSY13 | 3 | WT-Rep3 | AAT | A>G | G/C+1 | A/T-1 | TA>CG | TA>CG | CCGATAATG | ATGGCTA | TTA | TTA | T | C | T>C | T>C_TTA | TA>CG_ATGGCTA | 9 | FALSE | FALSE |
| 7_v.txt | 32759 | 32759-SNV-T | SNV | 1 | T | C | 7 | 14183 | 0.04935486 | 43.2861827 | WT | FALSE | CAN | JSY13 | 3 | WT-Rep3 | CTA | T>C | G/C+1 | A/T-1 | TA>CG | TA>CG | AATGGCTAA | TAAAACT | GAT | CTA | T | G | T>C | T>C_CTA | TA>CG_TAAAACT | 9 | FALSE | FALSE |
| 7_v.txt | 32763 | 32763-SNV-A | SNV | 1 | A | G | 4 | 14182 | 0.02820477 | 37.711316 | WT | FALSE | CAN | JSY13 | 3 | WT-Rep3 | AAC | A>G | G/C+1 | A/T-1 | TA>CG | TA>CG | GCTAAAACT | ACTTTGA | TTG | TTG | T | C | T>C | T>C_TTG | TA>CG_ACTTTGA | 9 | FALSE | FALSE |
| 7_v.txt | 32938 | 32938-SNV-A | SNV | 1 | A | G | 61 | 34274 | 0.17797748 | 55.3934426 | WT | FALSE | CAN | JSY13 | 3 | WT-Rep3 | CAA | A>G | G/C+1 | A/T-1 | TA>CG | TA>CG | TTGCCCAAG | AAGAAAA | GTT | GTT | T | C | T>C | T>C_GTT | TA>CG_AAGAAAA | 9 | FALSE | FALSE |
| 7_v.txt | 33150 | 33150-SNV-A | SNV | 1 | A | G | 25 | 34303 | 0.07287992 | 34.52 | WT | FALSE | CAN | JSY13 | 3 | WT-Rep3 | AAG | A>G | G/C+1 | A/T-1 | TA>CG | TA>CG | ATGAAAAGA | AGACCTG | TTC | TTC | T | C | T>C | T>C_TTC | TA>CG_AGACCTG | 9 | FALSE | FALSE |
| 7_v.txt | 33476 | 33476-SNV-T | SNV | 1 | T | C | 31 | 21996 | 0.14093472 | 31.7416986 | WT | FALSE | CAN | JSY13 | 3 | WT-Rep3 | CTT | T>C | G/C+1 | A/T-1 | TA>CG | TA>CG | TATGCCTTTT | TTTTTTT | GAA | CTT | T | G | T>C | T>C_CTT | TA>CG_TTTTTTT | 9 | FALSE | FALSE |
| 7_v.txt | 31983 | 31983-SNV-A | SNV | 1 | A | T | 28 | 22492 | 0.12448871 | 44 | WT | FALSE | CAN | JSY13 | 3 | WT-Rep3 | CAT | A>T | A/T+1 | A/T-1 | TA>AT | TA>AT | CCGGGCATT | ATTAATT | GTA | GTA | T | A | T>A | T>A_GTA | TA>AT_ATTAATT | 9 | FALSE | FALSE |
| 7_v.txt | 32160 | 32160-SNV-A | SNV | 1 | A | T | 1 | 22492 | 4.45E-03 | 63.9962305 | WT | FALSE | CAN | JSY13 | 3 | WT-Rep3 | CAA | A>T | A/T+1 | A/T-1 | TA>AT | TA>AT | TAAGCCAAA | AAAGCGC | GTT | GTT | T | A | T>A | T>A_GTT | TA>AT_AAAGCGC | 9 | FALSE | FALSE |
| 7_v.txt | 32161 | 32161-SNV-A | SNV | 1 | A | T | 2 | 22492 | 8.89E-03 | 49.4973466 | WT | FALSE | CAN | JSY13 | 3 | WT-Rep3 | AAA | A>T | A/T+1 | A/T-1 | TA>AT | TA>AT | AAGCCAAAG | AAGCGCC | TTT | TTT | T | A | T>A | T>A_TTT | TA>AT_AAGCGCC | 9 | FALSE | FALSE |
| 7_v.txt | 32266 | 32266-SNV-A | SNV | 1 | A | T | 5 | 53882 | 9.28E-03 | 24.1999344 | WT | FALSE | CAN | JSY13 | 3 | WT-Rep3 | AAA | A>T | A/T+1 | A/T-1 | TA>AT | TA>AT | CAAATAAAA | AAATACG | TTT | TTT | T | A | T>A | T>A_TTT | TA>AT_AAATACG | 9 | FALSE | FALSE |
| 7_v.txt | 32275 | 32275-SNV-A | SNV | 1 | A | T | 12 | 53882 | 0.02227089 | 26.165778 | WT | FALSE | CAN | JSY13 | 3 | WT-Rep3 | GAA | A>T | A/T+1 | A/T-1 | TA>AT | TA>AT | TACGGGAAC | AACCAAC | CTT | CTT | T | A | T>A | T>A_CTT | TA>AT_AACCAAC | 9 | FALSE | FALSE |
| 7_v.txt | 32337 | 32337-SNV-A | SNV | 1 | A | T | 8 | 53882 | 0.01484726 | 44.9996717 | WT | FALSE | CAN | JSY13 | 3 | WT-Rep3 | GAT | A>T | A/T+1 | A/T-1 | TA>AT | TA>AT | TTGAAGATA | ATATGTG | CTA | CTA | T | A | T>A | T>A_CTA | TA>AT_ATATGTG | 9 | FALSE | FALSE |
| 7_v.txt | 32346 | 32346-SNV-A | SNV | 1 | A | T | 235 | 53882 | 0.43613823 | 61.2828323 | WT | FALSE | CAN | JSY13 | 3 | WT-Rep3 | CAA | A>T | A/T+1 | A/T-1 | TA>AT | TA>AT | TGTGGCAAA | AAAACCT | GTT | GTT | T | A | T>A | T>A_GTT | TA>AT_AAAACCT | 9 | FALSE | FALSE |
| 7_v.txt | 32492 | 32492-SNV-A | SNV | 1 | A | T | 3 | 53882 | 5.57E-03 | 19.9999941 | WT | FALSE | CAN | JSY13 | 3 | WT-Rep3 | AAA | A>T | A/T+1 | A/T-1 | TA>AT | TA>AT | ACGGAAAAC | AACAACT | TTT | TTT | T | A | T>A | T>A_TTT | TA>AT_AACAACT | 9 | FALSE | FALSE |
| 7_v.txt | 32496 | 32496-SNV-A | SNV | 1 | A | T | 91 | 53882 | 0.16888757 | 43.2417582 | WT | FALSE | CAN | JSY13 | 3 | WT-Rep3 | AAC | A>T | A/T+1 | A/T-1 | TA>AT | TA>AT | AAAACAACT | ACTTTTT | TTG | TTG | T | A | T>A | T>A_TTG | TA>AT_ACTTTTT | 9 | FALSE | FALSE |

| | | | | | | | | | | | | | | | | | | | | | | | | | | | | | | | | | | |
|---|---|---|---|---|---|---|---|---|---|---|---|---|---|---|---|---|---|---|---|---|---|---|---|---|---|---|---|---|---|---|---|---|---|---|
| 7_v.txt | 32499 | 32499-SNV-T | SNV | 1 | T | A | 4 | 53882 | 7.42E-03 | 44.2488316 | WT | FALSE | CAN | JSY13 | 3 | WT-Rep3 | TTT | T>A | A/T+1 | A/T-1 | TA>AT | TA>AT | ACAACTTTTT | TTTTTGA | AAA | TTT | T | T | T>A | T>A_TTT | TA>AT_TTTTTGA | 9 | FALSE | FALSE |
| 7_v.txt | 32524 | 32524-SNV-T | SNV | 1 | T | A | 27 | 53882 | 0.0501095 | 41.4814815 | WT | FALSE | CAN | JSY13 | 3 | WT-Rep3 | TTT | T>A | A/T+1 | A/T-1 | TA>AT | TA>AT | CGGATTTTCT | TTCTGGG | AAA | TTT | T | T | T>A | T>A_TTT | TA>AT_TTCTGGG | 9 | FALSE | FALSE |
| 7_v.txt | 32598 | 32598-SNV-/ | SNV | 1 | A | T | 5 | 14183 | 0.03525347 | 55 | WT | FALSE | CAN | JSY13 | 3 | WT-Rep3 | CAA | A>T | A/T+1 | A/T-1 | TA>AT | TA>AT | TTAATCAAA( | AAAGAGG | GTT | GTT | T | A | T>A | T>A_GTT | TA>AT_AAAGAGG | 9 | FALSE | FALSE |
| 7_v.txt | 32755 | 32755-SNV-T | SNV | 1 | T | A | 1 | 14181 | 7.05E-03 | 37.9990727 | WT | FALSE | CAN | JSY13 | 3 | WT-Rep3 | ATG | T>A | A/T+1 | A/T-1 | TA>AT | TA>AT | CGATAATGG | TGGCTAA | TAC | ATG | T | T | T>A | T>A_ATG | TA>AT_TGGCTAA | 9 | FALSE | FALSE |
| 7_v.txt | 32767 | 32767-SNV-T | SNV | 1 | T | A | 7 | 14183 | 0.04935486 | 44.8519549 | WT | FALSE | CAN | JSY13 | 3 | WT-Rep3 | TTG | T>A | A/T+1 | A/T-1 | TA>AT | TA>AT | AAACTTTGAT | TGATGGA | AAC | TTG | T | T | T>A | T>A_TTG | TA>AT_TGATGGA | 9 | FALSE | FALSE |
| 7_v.txt | 32821 | 32821-SNV-T | SNV | 1 | T | A | 20 | 14183 | 0.14101389 | 41.25 | WT | FALSE | CAN | JSY13 | 3 | WT-Rep3 | TTC | T>A | A/T+1 | A/T-1 | TA>AT | TA>AT | ACAAGTTCAT | TCATTAT | AAG | TTC | T | T | T>A | T>A_TTC | TA>AT_TCATTAT | 9 | FALSE | FALSE |
| 7_v.txt | 33067 | 33067-SNV-/ | SNV | 1 | A | T | 207 | 34303 | 0.60344576 | 55.1111111 | WT | FALSE | CAN | JSY13 | 3 | WT-Rep3 | TAT | A>T | A/T+1 | A/T-1 | TA>AT | TA>AT | CAGAATATG( | ATGCCAA | ATA | ATA | T | A | T>A | T>A_ATA | TA>AT_ATGCCAA | 9 | FALSE | FALSE |
| 7_v.txt | 33192 | 33192-SNV-T | SNV | 1 | T | A | 22 | 21998 | 0.10000909 | 34.5909091 | WT | FALSE | CAN | JSY13 | 3 | WT-Rep3 | ATG | T>A | A/T+1 | A/T-1 | TA>AT | TA>AT | CCAATATGT( | TGTCTTT | TAC | ATG | T | T | T>A | T>A_ATG | TA>AT_TGTCTTT | 9 | FALSE | FALSE |
| 7_v.txt | 33321 | 33321-SNV-/ | SNV | 1 | A | T | 593 | 21999 | 2.69557707 | 59.3490725 | WT | FALSE | CAN | JSY13 | 3 | WT-Rep3 | CAA | A>T | A/T+1 | A/T-1 | TA>AT | TA>AT | GGATACAAT | AATTCTT | GTT | GTT | T | A | T>A | T>A_GTT | TA>AT_AATTCTT | 9 | FALSE | FALSE |
| 7_v.txt | 33345 | 33345-SNV-/ | SNV | 1 | A | T | 38 | 21999 | 0.17273512 | 41.5526316 | WT | FALSE | CAN | JSY13 | 3 | WT-Rep3 | CAA | A>T | A/T+1 | A/T-1 | TA>AT | TA>AT | TCTTTCAATG | AATGGTA | GTT | GTT | T | A | T>A | T>A_GTT | TA>AT_AATGGTA | 9 | FALSE | FALSE |
| 7_v.txt | 31650 | 31650-SNV-( | SNV | 1 | C | T | 11 | 6282 | 0.17510347 | 37.0909091 | WT | FALSE | CAN | JSY13 | 3 | WT-Rep3 | GCG | C>T | A/T+1 | G/C-1 | CG>TA | CG>TA | AGAATGCGA | CGAAATG | CGC | GCG | C | A | C>T | C>T_GCG | CG>TA_CGAAATG | 9 | FALSE | FALSE |
| 7_v.txt | 31806 | 31806-SNV-( | SNV | 1 | C | T | 24 | 6282 | 0.38204394 | 54.7083333 | WT | FALSE | CAN | JSY13 | 3 | WT-Rep3 | CCA | C>T | A/T+1 | G/C-1 | CG>TA | CG>TA | ATCTTCCAAA | CAAATAA | GGT | CCA | C | A | C>T | C>T_CCA | CG>TA_CAAATAA | 9 | FALSE | FALSE |
| 7_v.txt | 31882 | 31882-SNV-( | SNV | 1 | C | T | 10 | 6282 | 0.15918497 | 39.8 | WT | FALSE | CAN | JSY13 | 3 | WT-Rep3 | GCA | C>T | A/T+1 | G/C-1 | CG>TA | CG>TA | AGGCGGCAG | CAGCAAA | CGT | GCA | C | A | C>T | C>T_GCA | CG>TA_CAGCAAA | 9 | FALSE | FALSE |
| 7_v.txt | 31960 | 31960-SNV-( | SNV | 1 | C | T | 22 | 22492 | 0.09781256 | 37.1363636 | WT | FALSE | CAN | JSY13 | 3 | WT-Rep3 | GCC | C>T | A/T+1 | G/C-1 | CG>TA | CG>TA | ATGTGGCCG( | CCGCATA | CGG | GCC | C | A | C>T | C>T_GCC | CG>TA_CCGCATA | 9 | FALSE | FALSE |
| 7_v.txt | 31962 | 31962-SNV-( | SNV | 1 | G | A | 34 | 22492 | 0.15116486 | 36.5 | WT | FALSE | CAN | JSY13 | 3 | WT-Rep3 | CGC | G>A | A/T+1 | G/C-1 | CG>TA | CG>TA | GTGGCCGCA | GCATAAT | GCG | GCG | C | T | C>T | C>T_GCG | CG>TA_GCATAAT | 9 | FALSE | FALSE |
| 7_v.txt | 31972 | 31972-SNV-( | SNV | 1 | C | T | 18 | 22492 | 0.08002846 | 36.2222222 | WT | FALSE | CAN | JSY13 | 3 | WT-Rep3 | GCC | C>T | A/T+1 | G/C-1 | CG>TA | CG>TA | AATAAGCCA. | CCAAGCC | CGG | GCC | C | A | C>T | C>T_GCC | CG>TA_CCAAGCC | 9 | FALSE | FALSE |
| 7_v.txt | 31981 | 31981-SNV-( | SNV | 1 | G | A | 20 | 22492 | 0.08892051 | 45.9 | WT | FALSE | CAN | JSY13 | 3 | WT-Rep3 | GGC | G>A | A/T+1 | G/C-1 | CG>TA | CG>TA | AGCCGGGCA | GCATTAA | CCG | CCG | C | T | C>T | C>T_CCG | CG>TA_GCATTAA | 9 | FALSE | FALSE |
| 7_v.txt | 32057 | 32057-SNV-( | SNV | 1 | C | T | 20 | 22489 | 0.08893237 | 56.7501527 | WT | FALSE | CAN | JSY13 | 3 | WT-Rep3 | GCG | C>T | A/T+1 | G/C-1 | CG>TA | CG>TA | GATGTGCGA | CGAGATT | CGC | GCG | C | A | C>T | C>T_GCG | CG>TA_CGAGATT | 9 | FALSE | FALSE |
| 7_v.txt | 32058 | 32058-SNV-( | SNV | 1 | G | A | 4 | 22492 | 0.0177841 | 53.7496465 | WT | FALSE | CAN | JSY13 | 3 | WT-Rep3 | CGA | G>A | A/T+1 | G/C-1 | CG>TA | CG>TA | ATGTGCGAG | GAGATTG | GCT | GCT | C | T | C>T | C>T_GCT | CG>TA_GAGATTG | 9 | FALSE | FALSE |
| 7_v.txt | 32092 | 32092-SNV-( | SNV | 1 | C | T | 28 | 22492 | 0.12448871 | 58.8571429 | WT | FALSE | CAN | JSY13 | 3 | WT-Rep3 | GCA | C>T | A/T+1 | G/C-1 | CG>TA | CG>TA | AGCCTGCAA( | CAACACC | CGT | GCA | C | A | C>T | C>T_GCA | CG>TA_CAACACC | 9 | FALSE | FALSE |
| 7_v.txt | 32157 | 32157-SNV-( | SNV | 1 | G | A | 14 | 22491 | 0.06224712 | 37.0713925 | WT | FALSE | CAN | JSY13 | 3 | WT-Rep3 | AGC | G>A | A/T+1 | G/C-1 | CG>TA | CG>TA | ATGTAAGCC/ | GCCAAAG | TCG | TCG | C | T | C>T | C>T_TCG | CG>TA_GCCAAAG | 9 | FALSE | FALSE |
| 7_v.txt | 32158 | 32158-SNV-( | SNV | 1 | C | T | 6 | 22492 | 0.02667615 | 52.8327204 | WT | FALSE | CAN | JSY13 | 3 | WT-Rep3 | GCC | C>T | A/T+1 | G/C-1 | CG>TA | CG>TA | TGTAAGCCA/ | CCAAAGC | CGG | GCC | C | A | C>T | C>T_GCC | CG>TA_CCAAAGC | 9 | FALSE | FALSE |
| 7_v.txt | 32159 | 32159-SNV-( | SNV | 1 | C | T | 5 | 22492 | 0.02223013 | 46.5989739 | WT | FALSE | CAN | JSY13 | 3 | WT-Rep3 | CCA | C>T | A/T+1 | G/C-1 | CG>TA | CG>TA | GTAAGCCAA. | CAAAGCG | GGT | CCA | C | A | C>T | C>T_CCA | CG>TA_CAAAGCG | 9 | FALSE | FALSE |
| 7_v.txt | 32165 | 32165-SNV-( | SNV | 1 | G | A | 22 | 22492 | 0.09781256 | 46.5922906 | WT | FALSE | CAN | JSY13 | 3 | WT-Rep3 | CGC | G>A | A/T+1 | G/C-1 | CG>TA | CG>TA | CAAAGCGCC. | GCCAAAT | GCG | GCG | C | T | C>T | C>T_GCG | CG>TA_GCCAAAT | 9 | FALSE | FALSE |
| 7_v.txt | 32271 | 32271-SNV-( | SNV | 1 | C | T | 48 | 53882 | 0.08908355 | 32.661816 | WT | FALSE | CAN | JSY13 | 3 | WT-Rep3 | ACG | C>T | A/T+1 | G/C-1 | CG>TA | CG>TA | AAAATACGG | CGGGAAC | TGC | ACG | C | A | C>T | C>T_ACG | CG>TA_CGGGAAC | 9 | FALSE | FALSE |
| 7_v.txt | 32272 | 32272-SNV-( | SNV | 1 | G | A | 39 | 53882 | 0.07238039 | 34.2820741 | WT | FALSE | CAN | JSY13 | 3 | WT-Rep3 | CGG | G>A | A/T+1 | G/C-1 | CG>TA | CG>TA | AAATACGGG | GGGAACC | GCC | GCC | C | T | C>T | C>T_GCC | CG>TA_GGGAACC | 9 | FALSE | FALSE |
| 7_v.txt | 32274 | 32274-SNV-( | SNV | 1 | G | A | 492 | 53882 | 0.91310642 | 37.1869739 | WT | FALSE | CAN | JSY13 | 3 | WT-Rep3 | GGA | G>A | A/T+1 | G/C-1 | CG>TA | CG>TA | ATACGGGAA | GAACCAA | CCT | CCT | C | T | C>T | C>T_CCT | CG>TA_GAACCAA | 9 | FALSE | FALSE |
| 7_v.txt | 32299 | 32299-SNV-( | SNV | 1 | C | T | 95 | 53882 | 0.1763112 | 51.1894737 | WT | FALSE | CAN | JSY13 | 3 | WT-Rep3 | GCG | C>T | A/T+1 | G/C-1 | CG>TA | CG>TA | AATTTGCGG( | CGGCAGA | CGC | GCG | C | A | C>T | C>T_GCG | CG>TA_CGGCAGA | 9 | FALSE | FALSE |
| 7_v.txt | 32463 | 32463-SNV-( | SNV | 1 | G | A | 71 | 53882 | 0.13176942 | 45.6619718 | WT | FALSE | CAN | JSY13 | 3 | WT-Rep3 | AGA | G>A | A/T+1 | G/C-1 | CG>TA | CG>TA | AATAGAGAG | GAGCCAA | TCT | TCT | C | T | C>T | C>T_TCT | CG>TA_GAGCCAA | 9 | FALSE | FALSE |
| 7_v.txt | 32466 | 32466-SNV-( | SNV | 1 | C | T | 49 | 53882 | 0.09093946 | 47.1511094 | WT | FALSE | CAN | JSY13 | 3 | WT-Rep3 | GCC | C>T | A/T+1 | G/C-1 | CG>TA | CG>TA | AGAGAGCCA | CCAATGT | CGG | GCC | C | A | C>T | C>T_GCC | CG>TA_CCAATGT | 9 | FALSE | FALSE |
| 7_v.txt | 32467 | 32467-SNV-( | SNV | 1 | C | T | 13 | 53881 | 0.02412724 | 43.6118099 | WT | FALSE | CAN | JSY13 | 3 | WT-Rep3 | CCA | C>T | A/T+1 | G/C-1 | CG>TA | CG>TA | GAGAGCCAA | CAATGTA | GGT | CCA | C | A | C>T | C>T_CCA | CG>TA_CAATGTA | 9 | FALSE | FALSE |
| 7_v.txt | 32487 | 32487-SNV-( | SNV | 1 | C | T | 44 | 53881 | 0.08166144 | 40.5681337 | WT | FALSE | CAN | JSY13 | 3 | WT-Rep3 | ACG | C>T | A/T+1 | G/C-1 | CG>TA | CG>TA | AAGATACGG | CGGAAAA | TGC | ACG | C | A | C>T | C>T_ACG | CG>TA_CGGAAAA | 9 | FALSE | FALSE |
| 7_v.txt | 32488 | 32488-SNV-( | SNV | 1 | G | A | 53 | 53882 | 0.09836309 | 42.3205202 | WT | FALSE | CAN | JSY13 | 3 | WT-Rep3 | CGG | G>A | A/T+1 | G/C-1 | CG>TA | CG>TA | AGATACGGA | GGAAAAC | GCC | GCC | C | T | C>T | C>T_GCC | CG>TA_GGAAAAC | 9 | FALSE | FALSE |
| 7_v.txt | 32535 | 32535-SNV-( | SNV | 1 | G | A | 59 | 53882 | 0.10949853 | 37.0332503 | WT | FALSE | CAN | JSY13 | 3 | WT-Rep3 | TGC | G>A | A/T+1 | G/C-1 | CG>TA | CG>TA | GGGTTTGCA( | GCAGCTT | ACG | ACG | C | T | C>T | C>T_ACG | CG>TA_GCAGCTT | 9 | FALSE | FALSE |
| 7_v.txt | 32536 | 32536-SNV-( | SNV | 1 | C | T | 80 | 53882 | 0.14847259 | 37.299712 | WT | FALSE | CAN | JSY13 | 3 | WT-Rep3 | GCA | C>T | A/T+1 | G/C-1 | CG>TA | CG>TA | GGTTTGCAG( | CAGCTTC | CGT | GCA | C | A | C>T | C>T_GCA | CG>TA_CAGCTTC | 9 | FALSE | FALSE |
| 7_v.txt | 32545 | 32545-SNV-( | SNV | 1 | C | T | 177 | 53882 | 0.3284956 | 37.180791 | WT | FALSE | CAN | JSY13 | 3 | WT-Rep3 | CCA | C>T | A/T+1 | G/C-1 | CG>TA | CG>TA | CTTCACCAGC | CAGCAGT | GGT | CCA | C | A | C>T | C>T_CCA | CG>TA_CAGCAGT | 9 | FALSE | FALSE |
| 7_v.txt | 32749 | 32749-SNV-( | SNV | 1 | C | T | 18 | 14183 | 0.1269125 | 59 | WT | FALSE | CAN | JSY13 | 3 | WT-Rep3 | CCG | C>T | A/T+1 | G/C-1 | CG>TA | CG>TA | GAAACCCGA | CGATAAT | GGC | CCG | C | A | C>T | C>T_CCG | CG>TA_CGATAAT | 9 | FALSE | FALSE |
| 7_v.txt | 32756 | 32756-SNV-( | SNV | 1 | G | A | 6 | 14183 | 0.04230417 | 36.3333341 | WT | FALSE | CAN | JSY13 | 3 | WT-Rep3 | TGG | G>A | A/T+1 | G/C-1 | CG>TA | CG>TA | GATAATGGC | GGCTAAA | ACC | ACC | C | T | C>T | C>T_ACC | CG>TA_GGCTAAA | 9 | FALSE | FALSE |
| 7_v.txt | 32757 | 32757-SNV-( | SNV | 1 | G | A | 5 | 14183 | 0.03525347 | 52.600867 | WT | FALSE | CAN | JSY13 | 3 | WT-Rep3 | GGC | G>A | A/T+1 | G/C-1 | CG>TA | CG>TA | ATAATGGCT/ | GCTAAAA | CCG | CCG | C | T | C>T | C>T_CCG | CG>TA_GCTAAAA | 9 | FALSE | FALSE |
| 7_v.txt | 32776 | 32776-SNV-( | SNV | 1 | C | T | 11 | 14183 | 0.07755764 | 48.9090909 | WT | FALSE | CAN | JSY13 | 3 | WT-Rep3 | GCG | C>T | A/T+1 | G/C-1 | CG>TA | CG>TA | TGGAAGCGA | CGACCCA | CGC | GCG | C | A | C>T | C>T_GCG | CG>TA_CGACCCA | 9 | FALSE | FALSE |
| 7_v.txt | 32797 | 32797-SNV-( | SNV | 1 | C | T | 13 | 14182 | 0.09166549 | 49.6923077 | WT | FALSE | CAN | JSY13 | 3 | WT-Rep3 | CCG | C>T | A/T+1 | G/C-1 | CG>TA | CG>TA | ATTCACCGTA | CGTAATA | GGC | CCG | C | A | C>T | C>T_CCG | CG>TA_CGTAATA | 9 | FALSE | FALSE |
| 7_v.txt | 32798 | 32798-SNV-( | SNV | 1 | G | A | 14 | 14183 | 0.09870972 | 41.4996563 | WT | FALSE | CAN | JSY13 | 3 | WT-Rep3 | CGT | G>A | A/T+1 | G/C-1 | CG>TA | CG>TA | TTCACCGTA/ | GTAATAT | GCA | GCA | C | T | C>T | C>T_GCA | CG>TA_GTAATAT | 9 | FALSE | FALSE |
| 7_v.txt | 32855 | 32855-SNV-( | SNV | 1 | C | T | 381 | 14182 | 2.68650402 | 36.0656168 | WT | FALSE | CAN | JSY13 | 3 | WT-Rep3 | TCC | C>T | A/T+1 | G/C-1 | CG>TA | CG>TA | ACTAATCCAT | CCATGCC | AGG | TCC | C | A | C>T | C>T_TCC | CG>TA_CCATGCC | 9 | FALSE | FALSE |
| 7_v.txt | 32882 | 32882-SNV-( | SNV | 1 | C | T | 72 | 34274 | 0.21007178 | 26.0532273 | WT | FALSE | CAN | JSY13 | 3 | WT-Rep3 | TCC | C>T | A/T+1 | G/C-1 | CG>TA | CG>TA | GTACGTCCA/ | CCAAAAT | AGG | TCC | C | A | C>T | C>T_TCC | CG>TA_CCAAAAT | 9 | FALSE | FALSE |
| 7_v.txt | 32883 | 32883-SNV-( | SNV | 1 | C | T | 54 | 34274 | 0.15755383 | 35.962963 | WT | FALSE | CAN | JSY13 | 3 | WT-Rep3 | CCA | C>T | A/T+1 | G/C-1 | CG>TA | CG>TA | TACGTCCAA/ | CAAAATT | GGT | CCA | C | A | C>T | C>T_CCA | CG>TA_CAAAATT | 9 | FALSE | FALSE |
| 7_v.txt | 32901 | 32901-SNV-( | SNV | 1 | C | T | 66 | 34274 | 0.19256579 | 31.1363636 | WT | FALSE | CAN | JSY13 | 3 | WT-Rep3 | GCC | C>T | A/T+1 | G/C-1 | CG>TA | CG>TA | ACTTGGCCA/ | CCAACTA | CGG | GCC | C | A | C>T | C>T_GCC | CG>TA_CCAACTA | 9 | FALSE | FALSE |
| 7_v.txt | 32922 | 32922-SNV-( | SNV | 1 | G | A | 25 | 34274 | 0.07294159 | 46.32 | WT | FALSE | CAN | JSY13 | 3 | WT-Rep3 | GGC | G>A | A/T+1 | G/C-1 | CG>TA | CG>TA | TCCAGGGCA. | GCAAAAG | CCG | CCG | C | T | C>T | C>T_CCG | CG>TA_GCAAAAG | 9 | FALSE | FALSE |
| 7_v.txt | 32934 | 32934-SNV-( | SNV | 1 | G | A | 12 | 34274 | 0.03501196 | 41.1682927 | WT | FALSE | CAN | JSY13 | 3 | WT-Rep3 | TGC | G>A | A/T+1 | G/C-1 | CG>TA | CG>TA | GTGATTGCC( | GCCCAAG | ACG | ACG | C | T | C>T | C>T_ACG | CG>TA_GCCCAAG | 9 | FALSE | FALSE |
| 7_v.txt | 32935 | 32935-SNV-( | SNV | 1 | C | T | 35 | 34274 | 0.10211822 | 36.2008538 | WT | FALSE | CAN | JSY13 | 3 | WT-Rep3 | GCC | C>T | A/T+1 | G/C-1 | CG>TA | CG>TA | TGATTGCCC/ | CCCAAGA | CGG | GCC | C | A | C>T | C>T_GCC | CG>TA_CCCAAGA | 9 | FALSE | FALSE |
| 7_v.txt | 32958 | 32958-SNV-( | SNV | 1 | C | T | 101 | 34303 | 0.29443489 | 49.6732673 | WT | FALSE | CAN | JSY13 | 3 | WT-Rep3 | ACC | C>T | A/T+1 | G/C-1 | CG>TA | CG>TA | ATGTAACCAT | CCATTGG | TGG | ACC | C | A | C>T | C>T_ACC | CG>TA_CCATTGG | 9 | FALSE | FALSE |
| 7_v.txt | 32965 | 32965-SNV-( | SNV | 1 | C | T | 32 | 34303 | 0.0932863 | 57.84375 | WT | FALSE | CAN | JSY13 | 3 | WT-Rep3 | GCC | C>T | A/T+1 | G/C-1 | CG>TA | CG>TA | CATTGGCCG( | CCGCACC | CGG | GCC | C | A | C>T | C>T_GCC | CG>TA_CCGCACC | 9 | FALSE | FALSE |
| 7_v.txt | 32967 | 32967-SNV-( | SNV | 1 | G | A | 64 | 34303 | 0.1865726 | 57.203125 | WT | FALSE | CAN | JSY13 | 3 | WT-Rep3 | CGC | G>A | A/T+1 | G/C-1 | CG>TA | CG>TA | TTGGCCGCA( | GCACCAA | GCG | GCG | C | T | C>T | C>T_GCG | CG>TA_GCACCAA | 9 | FALSE | FALSE |
| 7_v.txt | 32995 | 32995-SNV-( | SNV | 1 | G | A | 25 | 34303 | 0.07287992 | 58.48 | WT | FALSE | CAN | JSY13 | 3 | WT-Rep3 | TGT | G>A | A/T+1 | G/C-1 | CG>TA | CG>TA | ATCTTTGTGA | GTGAGAA | ACA | ACA | C | T | C>T | C>T_ACA | CG>TA_GTGAGAA | 9 | FALSE | FALSE |
| 7_v.txt | 33043 | 33043-SNV-( | SNV | 1 | C | T | 47 | 34303 | 0.13701426 | 58.5957447 | WT | FALSE | CAN | JSY13 | 3 | WT-Rep3 | TCA | C>T | A/T+1 | G/C-1 | CG>TA | CG>TA | CCATTTCACC | CACCCAA | AGT | TCA | C | A | C>T | C>T_TCA | CG>TA_CACCCAA | 9 | FALSE | FALSE |
| 7_v.txt | 33056 | 33056-SNV-( | SNV | 1 | C | T | 46 | 34303 | 0.13409906 | 55.4782609 | WT | FALSE | CAN | JSY13 | 3 | WT-Rep3 | GCG | C>T | A/T+1 | G/C-1 | CG>TA | CG>TA | GGACTGCGT( | CGTGACA | CGC | GCG | C | A | C>T | C>T_GCG | CG>TA_CGTGACA | 9 | FALSE | FALSE |
| 7_v.txt | 33070 | 33070-SNV-( | SNV | 1 | C | T | 37 | 34303 | 0.10786229 | 53.8648649 | WT | FALSE | CAN | JSY13 | 3 | WT-Rep3 | GCC | C>T | A/T+1 | G/C-1 | CG>TA | CG>TA | AATATGCCA/ | CCAAAGA | CGG | GCC | C | A | C>T | C>T_GCC | CG>TA_CCAAAGA | 9 | FALSE | FALSE |
| 7_v.txt | 33078 | 33078-SNV-( | SNV | 1 | C | T | 61 | 34303 | 0.17782701 | 52.704918 | WT | FALSE | CAN | JSY13 | 3 | WT-Rep3 | ACC | C>T | A/T+1 | G/C-1 | CG>TA | CG>TA | AAAGAACCC. | CCCATAA | TGG | ACC | C | A | C>T | C>T_ACC | CG>TA_CCCATAA | 9 | FALSE | FALSE |
| 7_v.txt | 33118 | 33118-SNV-( | SNV | 1 | C | T | 28 | 34303 | 0.08162551 | 39.8928571 | WT | FALSE | CAN | JSY13 | 3 | WT-Rep3 | GCG | C>T | A/T+1 | G/C-1 | CG>TA | CG>TA | GGCCGGCGT | CGTTGGT | CGC | GCG | C | A | C>T | C>T_GCG | CG>TA_CGTTGGT | 9 | FALSE | FALSE |
| 7_v.txt | 33168 | 33168-SNV-( | SNV | 1 | C | T | 16 | 34299 | 0.04664859 | 36.8749567 | WT | FALSE | CAN | JSY13 | 3 | WT-Rep3 | ACC | C>T | A/T+1 | G/C-1 | CG>TA | CG>TA | ATAGTACCA( | CCACCAA | TGG | ACC | C | A | C>T | C>T_ACC | CG>TA_CCACCAA | 9 | FALSE | FALSE |
| 7_v.txt | 33258 | 33258-SNV-( | SNV | 1 | C | T | 30 | 21999 | 0.13636984 | 40.9333333 | WT | FALSE | CAN | JSY13 | 3 | WT-Rep3 | GCC | C>T | A/T+1 | G/C-1 | CG>TA | CG>TA | CCTATGCCAT | CCATCCT | CGG | GCC | C | A | C>T | C>T_GCC | CG>TA_CCATCCT | 9 | FALSE | FALSE |
| 7_v.txt | 33276 | 33276-SNV-( | SNV | 1 | G | A | 31 | 21999 | 0.1409155 | 45.0322581 | WT | FALSE | CAN | JSY13 | 3 | WT-Rep3 | CGT | G>A | A/T+1 | G/C-1 | CG>TA | CG>TA | GAGAACGTA | GTATCCT | GCA | GCA | C | T | C>T | C>T_GCA | CG>TA_GTATCCT | 9 | FALSE | FALSE |
| 7_v.txt | 33310 | 33310-SNV-( | SNV | 1 | G | A | 20 | 21999 | 0.09091322 | 57.45 | WT | FALSE | CAN | JSY13 | 3 | WT-Rep3 | CGC | G>A | A/T+1 | G/C-1 | CG>TA | CG>TA | AAGAGCGCA | GCAATGG | GCG | GCG | C | T | C>T | C>T_GCG | CG>TA_GCAATGG | 9 | FALSE | FALSE |
| 7_v.txt | 33412 | 33412-SNV-( | SNV | 1 | G | A | 24 | 21999 | 0.10909587 | 46.5833333 | WT | FALSE | CAN | JSY13 | 3 | WT-Rep3 | GGC | G>A | A/T+1 | G/C-1 | CG>TA | CG>TA | TGACCGGCT( | GCTCATT | CCG | CCG | C | T | C>T | C>T_CCG | CG>TA_GCTCATT | 9 | FALSE | FALSE |
| 7_v.txt | 33474 | 33474-SNV-( | SNV | 1 | C | T | 40 | 21997 | 0.18184298 | 35.375 | WT | FALSE | CAN | JSY13 | 3 | WT-Rep3 | GCC | C>T | A/T+1 | G/C-1 | CG>TA | CG>TA | GCTATGCCTT | CCTTTTT | CGG | GCC | C | A | C>T | C>T_GCC | CG>TA_CCTTTTT | 9 | FALSE | FALSE |
| 7_v.txt | 32058 | 32058-SNV-( | SNV | 1 | G | C | 10 | 22492 | 0.04446025 | 51.9990393 | WT | FALSE | CAN | JSY13 | 3 | WT-Rep3 | CGA | G>C | G/C+1 | G/C-1 | CG>GC | CG>GC | ATGTGCGAG | GAGATTG | GCT | GCT | C | G | C>G | C>G_GCT | CG>GC_GAGATTG | 9 | FALSE | FALSE |
| 7_v.txt | 32064 | 32064-SNV-( | SNV | 1 | G | C | 34 | 22492 | 0.15116486 | 62.8823529 | WT | FALSE | CAN | JSY13 | 3 | WT-Rep3 | TGA | G>C | G/C+1 | G/C-1 | CG>GC | CG>GC | GAGATTGAG | GAGATAA | ACT | ACT | C | G | C>G | C>G_ACT | CG>GC_GAGATAA | 9 | FALSE | FALSE |
| 7_v.txt | 32153 | 32153-SNV-( | SNV | 1 | G | C | 8 | 22492 | 0.0355682 | 42 | WT | FALSE | CAN | JSY13 | 3 | WT-Rep3 | TGT | G>C | G/C+1 | G/C-1 | CG>GC | CG>GC | CTCCATGTA/ | GTAAGCC | ACA | ACA | C | G | C>G | C>G_ACA | CG>GC_GTAAGCC | 9 | FALSE | FALSE |
| 7_v.txt | 32165 | 32165-SNV-( | SNV | 1 | G | C | 1 | 22492 | 4.45E-03 | 29.0034363 | WT | FALSE | CAN | JSY13 | 3 | WT-Rep3 | CGC | G>C | G/C+1 | G/C-1 | CG>GC | CG>GC | CAAAGCGCC. | GCCAAAT | GCG | GCG | C | G | C>G | C>G_GCG | CG>GC_GCCAAAT | 9 | FALSE | FALSE |
| 7_v.txt | 32200 | 32200-SNV-( | SNV | 1 | G | C | 76 | 22492 | 0.33789792 | 45.131579 | WT | FALSE | CAN | JSY13 | 3 | WT-Rep3 | GGA | G>C | G/C+1 | G/C-1 | CG>GC | CG>GC | TGTATGGAA( | GAACACC | CCT | CCT | C | G | C>G | C>G_CCT | CG>GC_GAACACC | 9 | FALSE | FALSE |
| 7_v.txt | 32282 | 32282-SNV-( | SNV | 1 | G | C | 14 | 53882 | 0.0259827 | 39.8571429 | WT | FALSE | CAN | JSY13 | 3 | WT-Rep3 | CGT | G>C | G/C+1 | G/C-1 | CG>GC | CG>GC | ACCAACGTA/ | GTAAATA | GCA | GCA | C | G | C>G | C>G_GCA | CG>GC_GTAAATA | 9 | FALSE | FALSE |
| 7_v.txt | 32292 | 32292-SNV-( | SNV | 1 | G | C | 32 | 53882 | 0.05938904 | 56.90625 | WT | FALSE | CAN | JSY13 | 3 | WT-Rep3 | TGA | G>C | G/C+1 | G/C-1 | CG>GC | CG>GC | ATATTTGAAT | GAATTTG | ACT | ACT | C | G | C>G | C>G_ACT | CG>GC_GAATTTG | 9 | FALSE | FALSE |
| 7_v.txt | 32302 | 32302-SNV-( | SNV | 1 | C | G | 31 | 53882 | 0.05753313 | 54.516129 | WT | FALSE | CAN | JSY13 | 3 | WT-Rep3 | GCA | C>G | G/C+1 | G/C-1 | CG>GC | CG>GC | TTGCGGCAG. | CAGAAAT | CGT | GCA | C | C | C>G | C>G_GCA | CG>GC_CAGAAAT | 9 | FALSE | FALSE |
| 7_v.txt | 32432 | 32432-SNV-( | SNV | 1 | G | C | 28 | 53882 | 0.05196541 | 56.8571429 | WT | FALSE | CAN | JSY13 | 3 | WT-Rep3 | TGT | G>C | G/C+1 | G/C-1 | CG>GC | CG>GC | GTCATTGTAT | GTATGGA | ACA | ACA | C | G | C>G | C>G_ACA | CG>GC_GTATGGA | 9 | FALSE | FALSE |
| 7_v.txt | 32467 | 32467-SNV-( | SNV | 1 | C | G | 32 | 53881 | 0.05939014 | 50.2726741 | WT | FALSE | CAN | JSY13 | 3 | WT-Rep3 | CCA | C>G | G/C+1 | G/C-1 | CG>GC | CG>GC | GAGAGCCAA | CAATGTA | GGT | CCA | C | C | C>G | C>G_CCA | CG>GC_CAATGTA | 9 | FALSE | FALSE |
| 7_v.txt | 32471 | 32471-SNV-( | SNV | 1 | G | C | 10 | 53882 | 0.01855907 | 42.3 | WT | FALSE | CAN | JSY13 | 3 | WT-Rep3 | TGT | G>C | G/C+1 | G/C-1 | CG>GC | CG>GC | GCCAATGTA( | GTAGAAG | ACA | ACA | C | G | C>G | C>G_ACA | CG>GC_GTAGAAG | 9 | FALSE | FALSE |
| 7_v.txt | 32474 | 32474-SNV-( | SNV | 1 | G | C | 14 | 53882 | 0.0259827 | 42.7142857 | WT | FALSE | CAN | JSY13 | 3 | WT-Rep3 | AGA | G>C | G/C+1 | G/C-1 | CG>GC | CG>GC | AATGTAGAA | GAAGGTT | TCT | TCT | C | G | C>G | C>G_TCT | CG>GC_GAAGGTT | 9 | FALSE | FALSE |
| 7_v.txt | 32487 | 32487-SNV-( | SNV | 1 | C | G | 13 | 53881 | 0.02412724 | 42.4614733 | WT | FALSE | CAN | JSY13 | 3 | WT-Rep3 | ACG | C>G | G/C+1 | G/C-1 | CG>GC | CG>GC | AAGATACGG | CGGAAAA | TGC | ACG | C | C | C>G | C>G_ACG | CG>GC_CGGAAAA | 9 | FALSE | FALSE |
| 7_v.txt | 32536 | 32536-SNV-( | SNV | 1 | C | G | 4 | 53882 | 7.42E-03 | 36.9999086 | WT | FALSE | CAN | JSY13 | 3 | WT-Rep3 | GCA | C>G | G/C+1 | G/C-1 | CG>GC | CG>GC | GGTTTGCAG( | CAGCTTC | CGT | GCA | C | C | C>G | C>G_GCA | CG>GC_CAGCTTC | 9 | FALSE | FALSE |
| 7_v.txt | 32758 | 32758-SNV-( | SNV | 1 | C | G | 1 | 14183 | 7.05E-03 | 63.9986641 | WT | FALSE | CAN | JSY13 | 3 | WT-Rep3 | GCT | C>G | G/C+1 | G/C-1 | CG>GC | CG>GC | TAATGGCTA/ | CTAAAAC | CGA | GCT | C | C | C>G | C>G_GCT | CG>GC_CTAAAAC | 9 | FALSE | FALSE |
| 7_v.txt | 32776 | 32776-SNV-( | SNV | 1 | C | G | 5 | 14183 | 0.03525347 | 48.4 | WT | FALSE | CAN | JSY13 | 3 | WT-Rep3 | GCG | C>G | G/C+1 | G/C-1 | CG>GC | CG>GC | TGGAAGCGA | CGACCCA | CGC | GCG | C | C | C>G | C>G_GCG | CG>GC_CGACCCA | 9 | FALSE | FALSE |
| 7_v.txt | 32788 | 32788-SNV-( | SNV | 1 | C | G | 8 | 14183 | 0.05640556 | 47.5 | WT | FALSE | CAN | JSY13 | 3 | WT-Rep3 | TCG | C>G | G/C+1 | G/C-1 | CG>GC | CG>GC | AGAACTCGA. | CGAATTC | AGC | TCG | C | C | C>G | C>G_TCG | CG>GC_CGAATTC | 9 | FALSE | FALSE |
| 7_v.txt | 32797 | 32797-SNV-( | SNV | 1 | C | G | 5 | 14182 | 0.03525596 | 40.2 | WT | FALSE | CAN | JSY13 | 3 | WT-Rep3 | CCG | C>G | G/C+1 | G/C-1 | CG>GC | CG>GC | ATTCACCGTA | CGTAATA | GGC | CCG | C | C | C>G | C>G_CCG | CG>GC_CGTAATA | 9 | FALSE | FALSE |
| 7_v.txt | 32934 | 32934-SNV-( | SNV | 1 | G | C | 3 | 34274 | 8.75E-03 | 40.3332456 | WT | FALSE | CAN | JSY13 | 3 | WT-Rep3 | TGC | G>C | G/C+1 | G/C-1 | CG>GC | CG>GC | GTGATTGCC( | GCCCAAG | ACG | ACG | C | G | C>G | C>G_ACG | CG>GC_GCCCAAG | 9 | FALSE | FALSE |
| 7_v.txt | 33037 | 33037-SNV-( | SNV | 1 | C | G | 12 | 34303 | 0.03498236 | 64 | WT | FALSE | CAN | JSY13 | 3 | WT-Rep3 | GCC | C>G | G/C+1 | G/C-1 | CG>GC | CG>GC | ATGTAGCCAT | CCATTTC | CGG | GCC | C | C | C>G | C>G_GCC | CG>GC_CCATTTC | 9 | FALSE | FALSE |
| 7_v.txt | 33070 | 33070-SNV-( | SNV | 1 | C | G | 112 | 34303 | 0.32650206 | 55.9285714 | WT | FALSE | CAN | JSY13 | 3 | WT-Rep3 | GCC | C>G | G/C+1 | G/C-1 | CG>GC | CG>GC | AATATGCCA/ | CCAAAGA | CGG | GCC | C | C | C>G | C>G_GCC | CG>GC_CCAAAGA | 9 | FALSE | FALSE |
| 7_v.txt | 33118 | 33118-SNV-( | SNV | 1 | C | G | 5 | 34303 | 0.01457599 | 42.6 | WT | FALSE | CAN | JSY13 | 3 | WT-Rep3 | GCG | C>G | G/C+1 | G/C-1 | CG>GC | CG>GC | GGCCGGCGT | CGTTGGT | CGC | GCG | C | C | C>G | C>G_GCG | CG>GC_CGTTGGT | 9 | FALSE | FALSE |
| 7_v.txt | 33172 | 33172-SNV-( | SNV | 1 | C | G | 18 | 34299 | 0.05247966 | 37.8333333 | WT | FALSE | CAN | JSY13 | 3 | WT-Rep3 | CCA | C>G | G/C+1 | G/C-1 | CG>GC | CG>GC | TACCACCAA( | CAAGGGC | GGT | CCA | C | C | C>G | C>G_CCA | CG>GC_CAAGGGC | 9 | FALSE | FALSE |
| 7_v.txt | 33354 | 33354-SNV-( | SNV | 1 | G | C | 19 | 21988 | 0.08641077 | 50.2601657 | WT | FALSE | CAN | JSY13 | 3 | WT-Rep3 | TGA | G>C | G/C+1 | G/C-1 | CG>GC | CG>GC | GGTATTGAC( | GACCCAC | ACT | ACT | C | G | C>G | C>G_ACT | CG>GC_GACCCAC | 9 | FALSE | FALSE |
| 7_v.txt | 33378 | 33378-SNV-( | SNV | 1 | G | C | 24 | 21999 | 0.10909587 | 53.9166667 | WT | FALSE | CAN | JSY13 | 3 | WT-Rep3 | TGA | G>C | G/C+1 | G/C-1 | CG>GC | CG>GC | GTTTGTGAA( | GAAGCTT | ACT | ACT | C | G | C>G | C>G_ACT | CG>GC_GAAGCTT | 9 | FALSE | FALSE |
| 7_v.txt | 33419 | 33419-SNV-( | SNV | 1 | G | C | 8 | 21999 | 0.03636529 | 46.5 | WT | FALSE | CAN | JSY13 | 3 | WT-Rep3 | TGT | G>C | G/C+1 | G/C-1 | CG>GC | CG>GC | CTCATTGTAC | GTACATA | ACA | ACA | C | G | C>G | C>G_ACA | CG>GC_GTACATA | 9 | FALSE | FALSE |
| 7_v.txt | 31657 | 31657-SNV-( | SNV | 1 | G | T | 57 | 6282 | 0.90735435 | 20.122807 | WT | FALSE | CAN | JSY13 | 3 | WT-Rep3 | GGC | G>T | A/T+1 | G/C-1 | CG>AT | CG>AT | GAAATGGCG | GCGTGGA | CCG | CCG | C | A | C>A | C>A_CCG | CG>AT_GCGTGGA | 9 | FALSE | FALSE |
| 7_v.txt | 32158 | 32158-SNV-( | SNV | 1 | C | A | 5 | 22492 | 0.02223013 | 26.9955322 | WT | FALSE | CAN | JSY13 | 3 | WT-Rep3 | GCC | C>A | A/T+1 | G/C-1 | CG>AT | CG>AT | TGTAAGCCA/ | CCAAAGC | CGG | GCC | C | T | C>A | C>A_GCC | CG>AT_CCAAAGC | 9 | FALSE | FALSE |
| 7_v.txt | 32272 | 32272-SNV-( | SNV | 1 | G | T | 26 | 53882 | 0.04825359 | 28.0010995 | WT | FALSE | CAN | JSY13 | 3 | WT-Rep3 | CGG | G>T | A/T+1 | G/C-1 | CG>AT | CG>AT | AAATACGGG | GGGAACC | GCC | GCC | C | A | C>A | C>A_GCC | CG>AT_GGGAACC | 9 | FALSE | FALSE |
| 7_v.txt | 32274 | 32274-SNV-( | SNV | 1 | G | T | 41 | 53882 | 0.0760922 | 34.4147617 | WT | FALSE | CAN | JSY13 | 3 | WT-Rep3 | GGA | G>T | A/T+1 | G/C-1 | CG>AT | CG>AT | ATACGGGAA | GAACCAA | CCT | CCT | C | A | C>A | C>A_CCT | CG>AT_GAACCAA | 9 | FALSE | FALSE |
| 7_v.txt | 32282 | 32282-SNV-( | SNV | 1 | G | T | 14 | 53882 | 0.0259827 | 34.3571429 | WT | FALSE | CAN | JSY13 | 3 | WT-Rep3 | CGT | G>T | A/T+1 | G/C-1 | CG>AT | CG>AT | ACCAACGTA/ | GTAAATA | GCA | GCA | C | A | C>A | C>A_GCA | CG>AT_GTAAATA | 9 | FALSE | FALSE |
| 7_v.txt | 32368 | 32368-SNV-( | SNV | 1 | C | A | 31 | 53882 | 0.05753313 | 49.6774194 | WT | FALSE | CAN | JSY13 | 3 | WT-Rep3 | TCA | C>A | A/T+1 | G/C-1 | CG>AT | CG>AT | AGTTCTCAAT | CAATAGC | AGT | TCA | C | T | C>A | C>A_TCA | CG>AT_CAATAGC | 9 | FALSE | FALSE |
| 7_v.txt | 32373 | 32373-SNV-( | SNV | 1 | G | T | 26 | 53882 | 0.04825359 | 55.9615385 | WT | FALSE | CAN | JSY13 | 3 | WT-Rep3 | AGC | G>T | A/T+1 | G/C-1 | CG>AT | CG>AT | TCAATAGCA/ | GCAATAA | TCG | TCG | C | A | C>A | C>A_TCG | CG>AT_GCAATAA | 9 | FALSE | FALSE |
| 7_v.txt | 32399 | 32399-SNV-( | SNV | 1 | G | T | 27 | 53882 | 0.0501095 | 59.7037037 | WT | FALSE | CAN | JSY13 | 3 | WT-Rep3 | CGT | G>T | A/T+1 | G/C-1 | CG>AT | CG>AT | AGAAACGTA | GTAGGAA | GCA | GCA | C | A | C>A | C>A_GCA | CG>AT_GTAGGAA | 9 | FALSE | FALSE |
| 7_v.txt | 32432 | 32432-SNV-( | SNV | 1 | G | T | 97 | 53882 | 0.18002301 | 56.9587629 | WT | FALSE | CAN | JSY13 | 3 | WT-Rep3 | TGT | G>T | A/T+1 | G/C-1 | CG>AT | CG>AT | GTCATTGTAT | GTATGGA | ACA | ACA | C | A | C>A | C>A_ACA | CG>AT_GTATGGA | 9 | FALSE | FALSE |
| 7_v.txt | 32487 | 32487-SNV-( | SNV | 1 | C | A | 13 | 53881 | 0.02412724 | 42.6152726 | WT | FALSE | CAN | JSY13 | 3 | WT-Rep3 | ACG | C>A | A/T+1 | G/C-1 | CG>AT | CG>AT | AAGATACGG | CGGAAAA | TGC | ACG | C | T | C>A | C>A_ACG | CG>AT_CGGAAAA | 9 | FALSE | FALSE |
| 7_v.txt | 32530 | 32530-SNV-( | SNV | 1 | G | T | 51 | 53882 | 0.09465128 | 37.7254902 | WT | FALSE | CAN | JSY13 | 3 | WT-Rep3 | GGG | G>T | A/T+1 | G/C-1 | CG>AT | CG>AT | TTCTGGGGTT | GGTTTGC | CCC | CCC | C | A | C>A | C>A_CCC | CG>AT_GGTTTGC | 9 | FALSE | FALSE |
| 7_v.txt | 32536 | 32536-SNV-( | SNV | 1 | C | A | 11 | 53882 | 0.02041498 | 28.1820715 | WT | FALSE | CAN | JSY13 | 3 | WT-Rep3 | GCA | C>A | A/T+1 | G/C-1 | CG>AT | CG>AT | GGTTTGCAG( | CAGCTTC | CGT | GCA | C | T | C>A | C>A_GCA | CG>AT_CAGCTTC | 9 | FALSE | FALSE |
| 7_v.txt | 32629 | 32629-SNV-( | SNV | 1 | C | A | 205 | 14183 | 1.44539237 | 52.7317073 | WT | FALSE | CAN | JSY13 | 3 | WT-Rep3 | TCG | C>A | A/T+1 | G/C-1 | CG>AT | CG>AT | TCCCTTCGTT | CGTTTTT | AGC | TCG | C | T | C>A | C>A_TCG | CG>AT_CGTTTTT | 9 | FALSE | FALSE |
| 7_v.txt | 32788 | 32788-SNV-( | SNV | 1 | C | A | 6 | 14183 | 0.04230417 | 55.1666667 | WT | FALSE | CAN | JSY13 | 3 | WT-Rep3 | TCG | C>A | A/T+1 | G/C-1 | CG>AT | CG>AT | AGAACTCGA. | CGAATTC | AGC | TCG | C | T | C>A | C>A_TCG | CG>AT_CGAATTC | 9 | FALSE | FALSE |
| 7_v.txt | 32797 | 32797-SNV-( | SNV | 1 | C | A | 4 | 14182 | 0.02820477 | 41 | WT | FALSE | CAN | JSY13 | 3 | WT-Rep3 | CCG | C>A | A/T+1 | G/C-1 | CG>AT | CG>AT | ATTCACCGTA | CGTAATA | GGC | CCG | C | T | C>A | C>A_CCG | CG>AT_CGTAATA | 9 | FALSE | FALSE |
| 7_v.txt | 32809 | 32809-SNV-( | SNV | 1 | C | A | 19 | 14183 | 0.1339632 | 43.2105263 | WT | FALSE | CAN | JSY13 | 3 | WT-Rep3 | ACA | C>A | A/T+1 | G/C-1 | CG>AT | CG>AT | ATTTGACAG( | CAGGGAA | TGT | ACA | C | T | C>A | C>A_ACA | CG>AT_CAGGGAA | 9 | FALSE | FALSE |
| 7_v.txt | 32840 | 32840-SNV-( | SNV | 1 | C | A | 3 | 14183 | 0.02115208 | 29 | WT | FALSE | CAN | JSY13 | 3 | WT-Rep3 | CCC | C>A | A/T+1 | G/C-1 | CG>AT | CG>AT | AATTACCCA/ | CCAAAAA | GGG | CCC | C | T | C>A | C>A_CCC | CG>AT_CCAAAAA | 9 | FALSE | FALSE |
| 7_v.txt | 32876 | 32876-SNV-( | SNV | 1 | G | T | 17 | 34260 | 0.04962055 | 32.7647059 | WT | FALSE | CAN | JSY13 | 3 | WT-Rep3 | TGT | G>T | A/T+1 | G/C-1 | CG>AT | CG>AT | AACTTTGTA( | GTACGTC | ACA | ACA | C | A | C>A | C>A_ACA | CG>AT_GTACGTC | 9 | FALSE | FALSE |
| 7_v.txt | 32883 | 32883-SNV-( | SNV | 1 | C | A | 5 | 34274 | 0.01458832 | 27.2 | WT | FALSE | CAN | JSY13 | 3 | WT-Rep3 | CCA | C>A | A/T+1 | G/C-1 | CG>AT | CG>AT | TACGTCCAA/ | CAAAATT | GGT | CCA | C | T | C>A | C>A_CCA | CG>AT_CAAAATT | 9 | FALSE | FALSE |
| 7_v.txt | 32917 | 32917-SNV-( | SNV | 1 | C | A | 20 | 34274 | 0.05835327 | 44.25 | WT | FALSE | CAN | JSY13 | 3 | WT-Rep3 | TCC | C>A | A/T+1 | G/C-1 | CG>AT | CG>AT | TAAGTTCCA( | CCAGGGC | AGG | TCC | C | T | C>A | C>A_TCC | CG>AT_CCAGGGC | 9 | FALSE | FALSE |
| 7_v.txt | 32997 | 32997-SNV-( | SNV | 1 | G | T | 15 | 34303 | 0.04372795 | 62.2666667 | WT | FALSE | CAN | JSY13 | 3 | WT-Rep3 | TGA | G>T | A/T+1 | G/C-1 | CG>AT | CG>AT | CTTTGTGAG/ | GAGAAAA | ACT | ACT | C | A | C>A | C>A_ACT | CG>AT_GAGAAAA | 9 | FALSE | FALSE |
| 7_v.txt | 33033 | 33033-SNV-( | SNV | 1 | G | T | 2900 | 34303 | 8.45407107 | 60.8872414 | WT | FALSE | CAN | JSY13 | 3 | WT-Rep3 | TGT | G>T | A/T+1 | G/C-1 | CG>AT | CG>AT | ATGAATGTA( | GTAGCCA | ACA | ACA | C | A | C>A | C>A_ACA | CG>AT_GTAGCCA | 9 | FALSE | FALSE |
| 7_v.txt | 33159 | 33159-SNV-( | SNV | 1 | C | A | 34 | 34302 | 0.09911959 | 37.8823529 | WT | FALSE | CAN | JSY13 | 3 | WT-Rep3 | ACC | C>A | A/T+1 | G/C-1 | CG>AT | CG>AT | CCTGTACCA/ | CCAATAG | TGG | ACC | C | T | C>A | C>A_ACC | CG>AT_CCAATAG | 9 | FALSE | FALSE |
| 7_v.txt | 33169 | 33169-SNV-( | SNV | 1 | C | A | 3 | 34299 | 8.75E-03 | 37.9997521 | WT | FALSE | CAN | JSY13 | 3 | WT-Rep3 | CCA | C>A | A/T+1 | G/C-1 | CG>AT | CG>AT | TAGTACCAC( | CACCAAG | GGT | CCA | C | T | C>A | C>A_CCA | CG>AT_CACCAAG | 9 | FALSE | FALSE |
| 7_v.txt | 33283 | 33283-SNV-( | SNV | 1 | C | A | 14 | 21999 | 0.06363926 | 46.4285714 | WT | FALSE | CAN | JSY13 | 3 | WT-Rep3 | TCG | C>A | A/T+1 | G/C-1 | CG>AT | CG>AT | TATCCTCGCC | CGCCATT | AGC | TCG | C | T | C>A | C>A_TCG | CG>AT_CGCCATT | 9 | FALSE | FALSE |
| 7_v.txt | 33378 | 33378-SNV-( | SNV | 1 | G | T | 14 | 21999 | 0.06363926 | 45.8571429 | WT | FALSE | CAN | JSY13 | 3 | WT-Rep3 | TGA | G>T | A/T+1 | G/C-1 | CG>AT | CG>AT | GTTTGTGAA( | GAAGCTT | ACT | ACT | C | A | C>A | C>A_ACT | CG>AT_GAAGCTT | 9 | FALSE | FALSE |
| 7_v.txt | 33474 | 33474-SNV-( | SNV | 1 | C | A | 49 | 21997 | 0.22275765 | 31.4081633 | WT | FALSE | CAN | JSY13 | 3 | WT-Rep3 | GCC | C>A | A/T+1 | G/C-1 | CG>AT | CG>AT | GCTATGCCTT | CCTTTTT | CGG | GCC | C | T | C>A | C>A_GCC | CG>AT_CCTTTTT | 9 | FALSE | FALSE |
| 7_v.txt | 33475 | 33475-SNV-( | SNV | 1 | C | A | 16 | 21997 | 0.07273719 | 34.9375 | WT | FALSE | CAN | JSY13 | 3 | WT-Rep3 | CCT | C>A | A/T+1 | G/C-1 | CG>AT | CG>AT | CTATGCCTTT | CTTTTTT | GGA | CCT | C | T | C>A | C>A_CCT | CG>AT_CTTTTTT | 9 | FALSE | FALSE |
| 7_v.txt | 32765 | 32765-MNV- | MNV | 2 | TT | AG | 6 | 14183 | 0.04230417 | 60.4166559 | WT | FALSE | CAN | JSY13 | 3 | WT-Rep3 | CTT | nope | >1bp-ins | >1bpdel | none | MNV | TAAAACTTT( | TTTGATG | GAA | CTT | TT | none | NA | NA_CTT | MNV_TTTGATG | 9 | FALSE | FALSE |

| | | | | | | | | | | | | | | | | | | | | | | | | | | | | | | | | | | |
|---|---|---|---|---|---|---|---|---|---|---|---|---|---|---|---|---|---|---|---|---|---|---|---|---|---|---|---|---|---|---|---|---|---|---|
| 7_v.txt | 33168 | 33168-MNV- | MNV | 2 | CC | TT | 1 | 34299 | 2.92E-03 | 37.9999535 | WT | FALSE | CAN | JSY13 | 3 | WT-Rep3 | ACC | nope | >1bp-ins | >1bpdel | none | MNV | ATAGTACCA( | CCACCAA | TGG | ACC | CC | none | NA | NA_ACC | MNV_CCACCAA | 9 | FALSE | FALSE |
| 7_v.txt | 32080 | 32080-Insert | Insertion | 1 | - | C | 3 | 22492 | 0.01333808 | 62.3333333 | WT | FALSE | CAN | JSY13 | 3 | WT-Rep3 | GCA | nope | G/C+1 | >1bpdel | none | G/C+1 | ACCATGCAA/ | CAAAAAA | CGT | GCA | - | G | NA | NA_GCA | G/C+1_CAAAAAA | 9 | FALSE | FALSE |
| 7_v.txt | 32081 | 32081-Insert | Insertion | 1 | - | A | 18 | 22492 | 0.08002846 | 63.8888889 | WT | FALSE | CAN | JSY13 | 3 | WT-Rep3 | CAA | nope | A/T+1 | >1bpdel | none | A/T+1 | CCATGCAAA/ | AAAAAAG | GTT | CAA | - | T | NA | NA_CAA | A/T+1_AAAAAAG | 9 | FALSE | FALSE |
| 7_v.txt | 32346 | 32346-Insert | Insertion | 1 | - | A | 6 | 53882 | 0.01113544 | 59.3333333 | WT | FALSE | CAN | JSY13 | 3 | WT-Rep3 | CAA | nope | A/T+1 | >1bpdel | none | A/T+1 | TGTGGCAAA | AAAACCT | GTT | CAA | - | T | NA | NA_CAA | A/T+1_AAAACCT | 9 | FALSE | FALSE |
| 7_v.txt | 32498 | 32498-Insert | Insertion | 1 | - | T | 11 | 53882 | 0.02041498 | 44.7272727 | WT | FALSE | CAN | JSY13 | 3 | WT-Rep3 | CTT | nope | A/T+1 | >1bpdel | none | A/T+1 | AACAACTTTT | TTTTTTG | GAA | CTT | - | A | NA | NA_CTT | A/T+1_TTTTTTG | 9 | FALSE | FALSE |
| 7_v.txt | 32658 | 32658-Insert | Insertion | 1 | - | C | 4 | 14183 | 0.02820278 | 57.25 | WT | FALSE | CAN | JSY13 | 3 | WT-Rep3 | ACC | nope | G/C+1 | >1bpdel | none | G/C+1 | CCTGGACCC( | CCCCAGG | TGG | ACC | - | G | NA | NA_ACC | G/C+1_CCCCAGG | 9 | FALSE | FALSE |
| 7_v.txt | 32728 | 32728-Insert | Insertion | 1 | - | A | 5 | 14183 | 0.03525347 | 58.4 | WT | FALSE | CAN | JSY13 | 3 | WT-Rep3 | CAA | nope | A/T+1 | >1bpdel | none | A/T+1 | CCATACAAA/ | AAAAACA | GTT | CAA | - | T | NA | NA_CAA | A/T+1_AAAAACA | 9 | FALSE | FALSE |
| 7_v.txt | 32760 | 32760-Insert | Insertion | 1 | - | A | 1 | 14183 | 7.05E-03 | 63.9999996 | WT | FALSE | CAN | JSY13 | 3 | WT-Rep3 | TAA | nope | A/T+1 | >1bpdel | none | A/T+1 | ATGGCTAAA. | AAAACTT | ATT | TAA | - | T | NA | NA_TAA | A/T+1_AAAACTT | 9 | FALSE | FALSE |
| 7_v.txt | 32842 | 32842-Insert | Insertion | 1 | - | A | 25 | 14183 | 0.17626736 | 45.28 | WT | FALSE | CAN | JSY13 | 3 | WT-Rep3 | CAA | nope | A/T+1 | >1bpdel | none | A/T+1 | TTACCCAAAA | AAAAAAT | GTT | CAA | - | T | NA | NA_CAA | A/T+1_AAAAAAT | 9 | FALSE | FALSE |
| 7_v.txt | 32884 | 32884-Insert | Insertion | 1 | - | A | 10 | 34274 | 0.02917664 | 37.6 | WT | FALSE | CAN | JSY13 | 3 | WT-Rep3 | CAA | nope | A/T+1 | >1bpdel | none | A/T+1 | ACGTCCAAA/ | AAAATTG | GTT | CAA | - | T | NA | NA_CAA | A/T+1_AAAATTG | 9 | FALSE | FALSE |
| 7_v.txt | 32970 | 32970-Insert | Insertion | 1 | - | C | 4 | 34303 | 0.01166079 | 57.5 | WT | FALSE | CAN | JSY13 | 3 | WT-Rep3 | ACC | nope | G/C+1 | >1bpdel | none | G/C+1 | GCCGCACCA( | CCAAATG | TGG | ACC | - | G | NA | NA_ACC | G/C+1_CCAAATG | 9 | FALSE | FALSE |
| 7_v.txt | 33009 | 33009-Insert | Insertion | 1 | - | A | 3 | 34303 | 8.75E-03 | 64 | WT | FALSE | CAN | JSY13 | 3 | WT-Rep3 | GAA | nope | A/T+1 | >1bpdel | none | A/T+1 | ACTGTGAAA( | AAAGAGG | CTT | GAA | - | T | NA | NA_GAA | A/T+1_AAAGAGG | 9 | FALSE | FALSE |
| 7_v.txt | 33019 | 33019-Insert | Insertion | 1 | - | T | 16 | 34303 | 0.04664315 | 60.3125 | WT | FALSE | CAN | JSY13 | 3 | WT-Rep3 | GTA | nope | A/T+1 | >1bpdel | none | A/T+1 | AGGATGTAA | TAACAGG | CAT | GTA | - | A | NA | NA_GTA | A/T+1_TAACAGG | 9 | FALSE | FALSE |
| 7_v.txt | 33281 | 33281-Insert | Insertion | 13 | - | CTCGCCATTT | 2 | 21999 | 9.09E-03 | 43.3076923 | WT | FALSE | CAN | JSY13 | 3 | WT-Rep3 | CCT | nope | >1bp-ins | >1bpdel | none | >1bp-ins | CGTATCCTCC | CTCGCCA | GGA | CCT | - | none | NA | NA_CCT | >1bp-ins_CTCGCCA | 9 | FALSE | FALSE |
| 7_v.txt | 33355 | 33355-Insert | Insertion | 4 | - | ACCC | 2 | 21999 | 9.09E-03 | 63 | WT | FALSE | CAN | JSY13 | 3 | WT-Rep3 | GAC | nope | >1bp-ins | >1bpdel | none | >1bp-ins | GTATTGACC( | ACCCACG | CTG | GAC | - | none | NA | NA_GAC | >1bp-ins_ACCCACG | 9 | FALSE | FALSE |
| 7_v.txt | 33356 | 33356-Insert | Insertion | 1 | - | C | 15 | 21999 | 0.06818492 | 61.5333333 | WT | FALSE | CAN | JSY13 | 3 | WT-Rep3 | ACC | nope | G/C+1 | >1bpdel | none | G/C+1 | TATTGACCCA | CCCACGT | TGG | ACC | - | G | NA | NA_ACC | G/C+1_CCCACGT | 9 | FALSE | FALSE |
| 7_v.txt | 32157 | 32157-Deleti | Deletion | 8 | GCCAAAGC | - | 18 | 22492 | 0.08002846 | 47.6666667 | WT | FALSE | CAN | JSY13 | 3 | WT-Rep3 | AGC | nope | >1bp-ins | >1bpdel | none | >1bpdel | ATGTAAGCC/ | GCCAAAG | TCG | AGC | GCCAAAGC | none | NA | NA_AGC | >1bpdel_GCCAAAG | 9 | FALSE | FALSE |
| 7_v.txt | 32265 | 32265-Deleti | Deletion | 2 | AA | - | 12 | 53882 | 0.02227089 | 36.8333333 | WT | FALSE | CAN | JSY13 | 3 | WT-Rep3 | TAA | nope | >1bp-ins | >1bpdel | none | >1bpdel | CCAAATAAA/ | AAAATAC | ATT | TAA | AA | none | NA | NA_TAA | >1bpdel_AAAATAC | 9 | FALSE | FALSE |
| 7_v.txt | 32337 | 32337-Deleti | Deletion | 2 | AT | - | 31 | 53882 | 0.05753313 | 61.483871 | WT | FALSE | CAN | JSY13 | 3 | WT-Rep3 | GAT | nope | >1bp-ins | >1bpdel | none | >1bpdel | TTGAAGATA' | ATATGTG | CTA | GAT | AT | none | NA | NA_GAT | >1bpdel_ATATGTG | 9 | FALSE | FALSE |
| 7_v.txt | 32492 | 32492-Deleti | Deletion | 3 | AAC | - | 87 | 53882 | 0.16146394 | 43.7816092 | WT | FALSE | CAN | JSY13 | 3 | WT-Rep3 | AAA | nope | >1bp-ins | >1bpdel | none | >1bpdel | ACGGAAAAC | AACAACT | TTT | AAA | AAC | none | NA | NA_AAA | >1bpdel_AACAACT | 9 | FALSE | FALSE |
| 7_v.txt | 32498 | 32498-Deleti | Deletion | 2 | TT | - | 178 | 53882 | 0.33035151 | 44.6460674 | WT | FALSE | CAN | JSY13 | 3 | WT-Rep3 | CTT | nope | >1bp-ins | >1bpdel | none | >1bpdel | AACAACTTTT | TTTTTTG | GAA | CTT | TT | none | NA | NA_CTT | >1bpdel_TTTTTTG | 9 | FALSE | FALSE |
| 7_v.txt | 32754 | 32754-Deleti | Deletion | 15 | ATGGCTAAA. | - | 19 | 14183 | 0.1339632 | 54 | WT | FALSE | CAN | JSY13 | 3 | WT-Rep3 | AAT | nope | >1bp-ins | >1bpdel | none | >1bpdel | CCGATAATG( | ATGGCTA | TTA | AAT | ATGGCTAAA. | none | NA | NA_AAT | >1bpdel_ATGGCTA | 9 | FALSE | FALSE |
| 7_v.txt | 32804 | 32804-Deleti | Deletion | 2 | TT | - | 5 | 14183 | 0.03525347 | 47.2 | WT | FALSE | CAN | JSY13 | 3 | WT-Rep3 | ATT | nope | >1bp-ins | >1bpdel | none | >1bpdel | GTAATATTT( | TTTGACA | TAA | ATT | TT | none | NA | NA_ATT | >1bpdel_TTTGACA | 9 | FALSE | FALSE |
| 7_v.txt | 32839 | 32839-Deleti | Deletion | 2 | CC | - | 5 | 14183 | 0.03525347 | 48.2 | WT | FALSE | CAN | JSY13 | 3 | WT-Rep3 | ACC | nope | >1bp-ins | >1bpdel | none | >1bpdel | TAATTACCCA | CCCAAAA | TGG | ACC | CC | none | NA | NA_ACC | >1bpdel_CCCAAAA | 9 | FALSE | FALSE |
| 7_v.txt | 33168 | 33168-Deleti | Deletion | 2 | CC | - | 9 | 34299 | 0.02623983 | 37.7777778 | WT | FALSE | CAN | JSY13 | 3 | WT-Rep3 | ACC | nope | >1bp-ins | >1bpdel | none | >1bpdel | ATAGTACCA( | CCACCAA | TGG | ACC | CC | none | NA | NA_ACC | >1bpdel_CCACCAA | 9 | FALSE | FALSE |
| 7_v.txt | 31723 | 31723-Deleti | Deletion | 1 | A | - | 6 | 6282 | 0.09551098 | 64 | WT | FALSE | CAN | JSY13 | 3 | WT-Rep3 | CAA | nope | >1bp-ins | A/T-1 | none | A/T-1 | TGTCCCAAA/ | AAAAAGT | GTT | GTT | T | none | T>none | T>none_GTT | A/T-1_AAAAAGT | 9 | FALSE | FALSE |
| 7_v.txt | 31960 | 31960-Deleti | Deletion | 1 | C | - | 2 | 22492 | 8.89E-03 | 38 | WT | FALSE | CAN | JSY13 | 3 | WT-Rep3 | GCC | nope | >1bp-ins | G/C-1 | none | G/C-1 | ATGTGGCCG( | CCGCATA | CGG | GCC | C | none | nope | nope_GCC | G/C-1_CCGCATA | 9 | FALSE | FALSE |
| 7_v.txt | 31972 | 31972-Deleti | Deletion | 1 | C | - | 2 | 22492 | 8.89E-03 | 38 | WT | FALSE | CAN | JSY13 | 3 | WT-Rep3 | GCC | nope | >1bp-ins | G/C-1 | none | G/C-1 | AATAAGCCA. | CCAAGCC | CGG | GCC | C | none | nope | nope_GCC | G/C-1_CCAAGCC | 9 | FALSE | FALSE |
| 7_v.txt | 32081 | 32081-Deleti | Deletion | 1 | A | - | 20 | 22491 | 0.08892446 | 64 | WT | FALSE | CAN | JSY13 | 3 | WT-Rep3 | CAA | nope | >1bp-ins | A/T-1 | none | A/T-1 | CCATGCAAA/ | AAAAAAG | GTT | GTT | T | none | T>none | T>none_GTT | A/T-1_AAAAAAG | 9 | FALSE | FALSE |
| 7_v.txt | 32106 | 32106-Deleti | Deletion | 1 | T | - | 8 | 22492 | 0.0355682 | 60.375 | WT | FALSE | CAN | JSY13 | 3 | WT-Rep3 | ATT | nope | >1bp-ins | A/T-1 | none | A/T-1 | GTGATATTTA | TTTAATA | TAA | ATT | T | none | nope | nope_ATT | A/T-1_TTTAATA | 9 | FALSE | FALSE |
| 7_v.txt | 32265 | 32265-Deleti | Deletion | 1 | A | - | 7 | 53882 | 0.01299135 | 38 | WT | FALSE | CAN | JSY13 | 3 | WT-Rep3 | TAA | nope | >1bp-ins | A/T-1 | none | A/T-1 | CCAAATAAA/ | AAAATAC | ATT | ATT | T | none | T>none | T>none_ATT | A/T-1_AAAATAC | 9 | FALSE | FALSE |
| 7_v.txt | 32346 | 32346-Deleti | Deletion | 1 | A | - | 10 | 53882 | 0.01855907 | 63.3 | WT | FALSE | CAN | JSY13 | 3 | WT-Rep3 | CAA | nope | >1bp-ins | A/T-1 | none | A/T-1 | TGTGGCAAA | AAAACCT | GTT | GTT | T | none | T>none | T>none_GTT | A/T-1_AAAACCT | 9 | FALSE | FALSE |
| 7_v.txt | 32350 | 32350-Deleti | Deletion | 1 | C | - | 12 | 53882 | 0.02227089 | 60.0833333 | WT | FALSE | CAN | JSY13 | 3 | WT-Rep3 | ACC | nope | >1bp-ins | G/C-1 | none | G/C-1 | GCAAAACCT' | CCTTTGT | TGG | ACC | C | none | nope | nope_ACC | G/C-1_CCTTTGT | 9 | FALSE | FALSE |
| 7_v.txt | 32352 | 32352-Deleti | Deletion | 1 | T | - | 10 | 53882 | 0.01855907 | 56.9 | WT | FALSE | CAN | JSY13 | 3 | WT-Rep3 | CTT | nope | >1bp-ins | A/T-1 | none | A/T-1 | AAAACCTTT( | TTTGTAC | GAA | CTT | T | none | nope | nope_CTT | A/T-1_TTTGTAC | 9 | FALSE | FALSE |
| 7_v.txt | 32381 | 32381-Deleti | Deletion | 1 | A | - | 9 | 53882 | 0.01670317 | 62.2222222 | WT | FALSE | CAN | JSY13 | 3 | WT-Rep3 | TAA | nope | >1bp-ins | A/T-1 | none | A/T-1 | AATAATAAA( | AAAGGGA | ATT | ATT | T | none | T>none | T>none_ATT | A/T-1_AAAGGGA | 9 | FALSE | FALSE |
| 7_v.txt | 32442 | 32442-Deleti | Deletion | 1 | A | - | 13 | 53882 | 0.0241268 | 56.4615385 | WT | FALSE | CAN | JSY13 | 3 | WT-Rep3 | TAA | nope | >1bp-ins | A/T-1 | none | A/T-1 | GGAACTAAA | AAAAGTC | ATT | ATT | T | none | T>none | T>none_ATT | A/T-1_AAAAGTC | 9 | FALSE | FALSE |
| 7_v.txt | 32457 | 32457-Deleti | Deletion | 1 | A | - | 4 | 53882 | 7.42E-03 | 44.5 | WT | FALSE | CAN | JSY13 | 3 | WT-Rep3 | TAA | nope | >1bp-ins | A/T-1 | none | A/T-1 | ATGAATAAT/ | AATAGAG | ATT | ATT | T | none | T>none | T>none_ATT | A/T-1_AATAGAG | 9 | FALSE | FALSE |
| 7_v.txt | 32466 | 32466-Deleti | Deletion | 1 | C | - | 5 | 53882 | 9.28E-03 | 41.7999996 | WT | FALSE | CAN | JSY13 | 3 | WT-Rep3 | GCC | nope | >1bp-ins | G/C-1 | none | G/C-1 | AGAGAGCCA | CCAATGT | CGG | GCC | C | none | nope | nope_GCC | G/C-1_CCAATGT | 9 | FALSE | FALSE |
| 7_v.txt | 32498 | 32498-Deleti | Deletion | 1 | T | - | 108 | 53882 | 0.20043799 | 48.7777775 | WT | FALSE | CAN | JSY13 | 3 | WT-Rep3 | CTT | nope | >1bp-ins | A/T-1 | none | A/T-1 | AACAACTTTT | TTTTTTG | GAA | CTT | T | none | nope | nope_CTT | A/T-1_TTTTTTG | 9 | FALSE | FALSE |
| 7_v.txt | 32528 | 32528-Deleti | Deletion | 1 | G | - | 7 | 53882 | 0.01299135 | 48.7142857 | WT | FALSE | CAN | JSY13 | 3 | WT-Rep3 | TGG | nope | >1bp-ins | G/C-1 | none | G/C-1 | TTTTCTGGG( | GGGGTTT | ACC | ACC | C | none | C>none | C>none_ACC | G/C-1_GGGGTTT | 9 | FALSE | FALSE |
| 7_v.txt | 32598 | 32598-Deleti | Deletion | 1 | A | - | 1164 | 14183 | 8.20700839 | 41.7817869 | WT | FALSE | CAN | JSY13 | 3 | WT-Rep3 | CAA | nope | >1bp-ins | A/T-1 | none | A/T-1 | TTAATCAAA( | AAAGAGG | GTT | GTT | T | none | T>none | T>none_GTT | A/T-1_AAAGAGG | 9 | FALSE | FALSE |
| 7_v.txt | 32631 | 32631-Deleti | Deletion | 1 | T | - | 6 | 14183 | 0.04230417 | 50.8333333 | WT | FALSE | CAN | JSY13 | 3 | WT-Rep3 | GTT | nope | >1bp-ins | A/T-1 | none | A/T-1 | CCTTCGTTTT | TTTTTAT | CAA | GTT | T | none | nope | nope_GTT | A/T-1_TTTTTAT | 9 | FALSE | FALSE |
| 7_v.txt | 32658 | 32658-Deleti | Deletion | 1 | C | - | 15 | 14183 | 0.10576042 | 54.5333371 | WT | FALSE | CAN | JSY13 | 3 | WT-Rep3 | ACC | nope | >1bp-ins | G/C-1 | none | G/C-1 | CCTGGACCC( | CCCCAGG | TGG | ACC | C | none | nope | nope_ACC | G/C-1_CCCCAGG | 9 | FALSE | FALSE |
| 7_v.txt | 32707 | 32707-Deleti | Deletion | 1 | C | - | 5 | 14183 | 0.03525347 | 55.4 | WT | FALSE | CAN | JSY13 | 3 | WT-Rep3 | ACC | nope | >1bp-ins | G/C-1 | none | G/C-1 | CGGTAACCC( | CCCCAGC | TGG | ACC | C | none | nope | nope_ACC | G/C-1_CCCCAGC | 9 | FALSE | FALSE |
| 7_v.txt | 32728 | 32728-Deleti | Deletion | 1 | A | - | 11 | 14182 | 0.07756311 | 61.5454546 | WT | FALSE | CAN | JSY13 | 3 | WT-Rep3 | CAA | nope | >1bp-ins | A/T-1 | none | A/T-1 | CCATACAAA/ | AAAAACA | GTT | GTT | T | none | T>none | T>none_GTT | A/T-1_AAAAACA | 9 | FALSE | FALSE |
| 7_v.txt | 32758 | 32758-Deleti | Deletion | 1 | C | - | 6 | 14183 | 0.04230417 | 55.1666627 | WT | FALSE | CAN | JSY13 | 3 | WT-Rep3 | GCT | nope | >1bp-ins | G/C-1 | none | G/C-1 | TAATGGCTA/ | CTAAAAC | CGA | GCT | C | none | nope | nope_GCT | G/C-1_CTAAAAC | 9 | FALSE | FALSE |
| 7_v.txt | 32760 | 32760-Deleti | Deletion | 1 | A | - | 2 | 14183 | 0.01410139 | 36.5 | WT | FALSE | CAN | JSY13 | 3 | WT-Rep3 | TAA | nope | >1bp-ins | A/T-1 | none | A/T-1 | ATGGCTAAA. | AAAACTT | ATT | ATT | T | none | T>none | T>none_ATT | A/T-1_AAAACTT | 9 | FALSE | FALSE |
| 7_v.txt | 32779 | 32779-Deleti | Deletion | 1 | C | - | 4 | 14183 | 0.02820278 | 49.5 | WT | FALSE | CAN | JSY13 | 3 | WT-Rep3 | ACC | nope | >1bp-ins | G/C-1 | none | G/C-1 | AAGCGACCC. | CCCAGAA | TGG | ACC | C | none | nope | nope_ACC | G/C-1_CCCAGAA | 9 | FALSE | FALSE |
| 7_v.txt | 32839 | 32839-Deleti | Deletion | 1 | C | - | 1 | 14183 | 7.05E-03 | 38 | WT | FALSE | CAN | JSY13 | 3 | WT-Rep3 | ACC | nope | >1bp-ins | G/C-1 | none | G/C-1 | TAATTACCCA | CCCAAAA | TGG | ACC | C | none | nope | nope_ACC | G/C-1_CCCAAAA | 9 | FALSE | FALSE |
| 7_v.txt | 32842 | 32842-Deleti | Deletion | 1 | A | - | 45 | 14183 | 0.31728125 | 40.0222222 | WT | FALSE | CAN | JSY13 | 3 | WT-Rep3 | CAA | nope | >1bp-ins | A/T-1 | none | A/T-1 | TTACCCAAA/ | AAAAAAT | GTT | GTT | T | none | T>none | T>none_GTT | A/T-1_AAAAAAT | 9 | FALSE | FALSE |
| 7_v.txt | 32882 | 32882-Deleti | Deletion | 1 | C | - | 2 | 34274 | 5.84E-03 | 35.9999994 | WT | FALSE | CAN | JSY13 | 3 | WT-Rep3 | TCC | nope | >1bp-ins | G/C-1 | none | G/C-1 | GTACGTCCA/ | CCAAAAT | AGG | TCC | C | none | nope | nope_TCC | G/C-1_CCAAAAT | 9 | FALSE | FALSE |
| 7_v.txt | 32884 | 32884-Deleti | Deletion | 1 | A | - | 9 | 34274 | 0.02625897 | 37.5555556 | WT | FALSE | CAN | JSY13 | 3 | WT-Rep3 | CAA | nope | >1bp-ins | A/T-1 | none | A/T-1 | ACGTCCAAA/ | AAAATTG | GTT | GTT | T | none | T>none | T>none_GTT | A/T-1_AAAATTG | 9 | FALSE | FALSE |
| 7_v.txt | 32924 | 32924-Deleti | Deletion | 1 | A | - | 31 | 34274 | 0.09044757 | 51.8709677 | WT | FALSE | CAN | JSY13 | 3 | WT-Rep3 | CAA | nope | >1bp-ins | A/T-1 | none | A/T-1 | CAGGGCAAA | AAAAGTG | GTT | GTT | T | none | T>none | T>none_GTT | A/T-1_AAAAGTG | 9 | FALSE | FALSE |
| 7_v.txt | 32934 | 32934-Deleti | Deletion | 1 | G | - | 10 | 34274 | 0.02917664 | 55.5 | WT | FALSE | CAN | JSY13 | 3 | WT-Rep3 | TGC | nope | >1bp-ins | G/C-1 | none | G/C-1 | GTGATTGCC( | GCCCAAG | ACG | ACG | C | none | C>none | C>none_ACG | G/C-1_GCCCAAG | 9 | FALSE | FALSE |
| 7_v.txt | 32935 | 32935-Deleti | Deletion | 1 | C | - | 9 | 34274 | 0.02625897 | 46.333335 | WT | FALSE | CAN | JSY13 | 3 | WT-Rep3 | GCC | nope | >1bp-ins | G/C-1 | none | G/C-1 | TGATTGCCCA | CCCAAGA | CGG | GCC | C | none | nope | nope_GCC | G/C-1_CCCAAGA | 9 | FALSE | FALSE |
| 7_v.txt | 32965 | 32965-Deleti | Deletion | 1 | C | - | 7 | 34303 | 0.02040638 | 56.8571429 | WT | FALSE | CAN | JSY13 | 3 | WT-Rep3 | GCC | nope | >1bp-ins | G/C-1 | none | G/C-1 | CATTGGCCG( | CCGCACC | CGG | GCC | C | none | nope | nope_GCC | G/C-1_CCGCACC | 9 | FALSE | FALSE |
| 7_v.txt | 33000 | 33000-Deleti | Deletion | 1 | A | - | 7 | 34303 | 0.02040638 | 60.4285714 | WT | FALSE | CAN | JSY13 | 3 | WT-Rep3 | GAA | nope | >1bp-ins | A/T-1 | none | A/T-1 | TGTGAGAAA | AAAACTG | CTT | CTT | T | none | T>none | T>none_CTT | A/T-1_AAAACTG | 9 | FALSE | FALSE |
| 7_v.txt | 33014 | 33014-Deleti | Deletion | 1 | G | - | 11 | 34303 | 0.03206717 | 63.0909091 | WT | FALSE | CAN | JSY13 | 3 | WT-Rep3 | AGG | nope | >1bp-ins | G/C-1 | none | G/C-1 | GAAAGAGG/ | GGATGTA | TCC | TCC | C | none | C>none | C>none_TCC | G/C-1_GGATGTA | 9 | FALSE | FALSE |
| 7_v.txt | 33018 | 33018-Deleti | Deletion | 1 | G | - | 6 | 34302 | 0.01749169 | 61.3333342 | WT | FALSE | CAN | JSY13 | 3 | WT-Rep3 | TGT | nope | >1bp-ins | G/C-1 | none | G/C-1 | GAGGATGTA | GTAACAG | ACA | ACA | C | none | C>none | C>none_ACA | G/C-1_GTAACAG | 9 | FALSE | FALSE |
| 7_v.txt | 33138 | 33138-Deleti | Deletion | 1 | A | - | 8 | 34303 | 0.02332158 | 44.25 | WT | FALSE | CAN | JSY13 | 3 | WT-Rep3 | TAA | nope | >1bp-ins | A/T-1 | none | A/T-1 | GTGGATAAA | AAACCAA | ATT | ATT | T | none | T>none | T>none_ATT | A/T-1_AAACCAA | 9 | FALSE | FALSE |
| 7_v.txt | 33147 | 33147-Deleti | Deletion | 1 | A | - | 10 | 34303 | 0.02915197 | 39.5 | WT | FALSE | CAN | JSY13 | 3 | WT-Rep3 | GAA | nope | >1bp-ins | A/T-1 | none | A/T-1 | CCAATGAAA. | AAAAGAC | CTT | CTT | T | none | T>none | T>none_CTT | A/T-1_AAAAGAC | 9 | FALSE | FALSE |
| 7_v.txt | 33168 | 33168-Deleti | Deletion | 1 | C | - | 6 | 34299 | 0.01749322 | 37.8333333 | WT | FALSE | CAN | JSY13 | 3 | WT-Rep3 | ACC | nope | >1bp-ins | G/C-1 | none | G/C-1 | ATAGTACCA( | CCACCAA | TGG | ACC | C | none | nope | nope_ACC | G/C-1_CCACCAA | 9 | FALSE | FALSE |
| 7_v.txt | 33474 | 33474-Deleti | Deletion | 1 | C | - | 61 | 21997 | 0.27731054 | 26.8032787 | WT | FALSE | CAN | JSY13 | 3 | WT-Rep3 | GCC | nope | >1bp-ins | G/C-1 | none | G/C-1 | GCTATGCCTT | CCTTTTT | CGG | GCC | C | none | nope | nope_GCC | G/C-1_CCTTTTT | 9 | FALSE | FALSE |
| 169_v.txt | 31964 | 31964-SNV-/ | SNV | 1 | A | C | 19 | 19698 | 0.09645649 | 35.3157895 | WT | FALSE | CAN | JSY13 | 4 | WT-Rep4 | CAT | A>C | G/C+1 | A/T-1 | TA>GC | TA>GC | GGCCGCATA. | ATAATAA | GTA | GTA | T | G | T>G | T>G_GTA | TA>GC_ATAATAA | 9 | FALSE | FALSE |
| 169_v.txt | 32109 | 32109-SNV-/ | SNV | 1 | A | C | 10 | 19698 | 0.05076658 | 51.5 | WT | FALSE | CAN | JSY13 | 4 | WT-Rep4 | TAA | A>C | G/C+1 | A/T-1 | TA>GC | TA>GC | ATATTTAATA | AATAGCC | ATT | ATT | T | G | T>G | T>G_ATT | TA>GC_AATAGCC | 9 | FALSE | FALSE |
| 169_v.txt | 32226 | 32226-SNV-/ | SNV | 1 | A | C | 10 | 19698 | 0.05076658 | 37.9 | WT | FALSE | CAN | JSY13 | 4 | WT-Rep4 | CAG | A>C | G/C+1 | A/T-1 | TA>GC | TA>GC | CTTGACAGG. | AGGAATT | GTC | GTC | T | G | T>G | T>G_GTC | TA>GC_AGGAATT | 9 | FALSE | FALSE |
| 169_v.txt | 32445 | 32445-SNV-/ | SNV | 1 | A | C | 24 | 51504 | 0.04659832 | 51.75 | WT | FALSE | CAN | JSY13 | 4 | WT-Rep4 | AAG | A>C | G/C+1 | A/T-1 | TA>GC | TA>GC | ACTAAAAGT( | AGTCCAA | TTC | TTC | T | G | T>G | T>G_TTC | TA>GC_AGTCCAA | 9 | FALSE | FALSE |
| 169_v.txt | 32473 | 32473-SNV-/ | SNV | 1 | A | C | 10 | 51504 | 0.01941597 | 50.9967781 | WT | FALSE | CAN | JSY13 | 4 | WT-Rep4 | TAG | A>C | G/C+1 | A/T-1 | TA>GC | TA>GC | CAATGTAGA. | AGAAGGT | ATC | ATC | T | G | T>G | T>G_ATC | TA>GC_AGAAGGT | 9 | FALSE | FALSE |
| 169_v.txt | 32574 | 32574-SNV-T | SNV | 1 | T | G | 7 | 12118 | 0.05776531 | 37.8569431 | WT | FALSE | CAN | JSY13 | 4 | WT-Rep4 | TTG | T>G | G/C+1 | A/T-1 | TA>GC | TA>GC | GTACCTTGA/ | TGAAATG | AAC | TTG | T | C | T>G | T>G_TTG | TA>GC_TGAAATG | 9 | FALSE | FALSE |
| 169_v.txt | 32742 | 32742-SNV-/ | SNV | 1 | A | C | 5 | 12117 | 0.04126434 | 53.2 | WT | FALSE | CAN | JSY13 | 4 | WT-Rep4 | TAG | A>C | G/C+1 | A/T-1 | TA>GC | TA>GC | TATATTAGA/ | AGAAACC | ATC | ATC | T | G | T>G | T>G_ATC | TA>GC_AGAAACC | 9 | FALSE | FALSE |
| 169_v.txt | 32800 | 32800-SNV-/ | SNV | 1 | A | C | 11 | 12117 | 0.09078155 | 44.3638157 | WT | FALSE | CAN | JSY13 | 4 | WT-Rep4 | TAA | A>C | G/C+1 | A/T-1 | TA>GC | TA>GC | CACCGTAAT/ | AATATTT | ATT | ATT | T | G | T>G | T>G_ATT | TA>GC_AATATTT | 9 | FALSE | FALSE |
| 169_v.txt | 32801 | 32801-SNV-/ | SNV | 1 | A | C | 7 | 12116 | 0.05777484 | 45.142777 | WT | FALSE | CAN | JSY13 | 4 | WT-Rep4 | AAT | A>C | G/C+1 | A/T-1 | TA>GC | TA>GC | ACCGTAATA' | ATATTTG | TTA | TTA | T | G | T>G | T>G_TTA | TA>GC_ATATTTG | 9 | FALSE | FALSE |
| 169_v.txt | 32842 | 32842-SNV-/ | SNV | 1 | A | C | 3 | 12117 | 0.0247586 | 32.3333333 | WT | FALSE | CAN | JSY13 | 4 | WT-Rep4 | CAA | A>C | G/C+1 | A/T-1 | TA>GC | TA>GC | TTACCCAAA/ | AAAAAAT | GTT | GTT | T | G | T>G | T>G_GTT | TA>GC_AAAAAAT | 9 | FALSE | FALSE |
| 169_v.txt | 32919 | 32919-SNV-/ | SNV | 1 | A | C | 21 | 26544 | 0.07911392 | 38.1428571 | WT | FALSE | CAN | JSY13 | 4 | WT-Rep4 | CAG | A>C | G/C+1 | A/T-1 | TA>GC | TA>GC | AGTTCCAGG( | AGGGCAA | GTC | GTC | T | G | T>G | T>G_GTC | TA>GC_AGGGCAA | 9 | FALSE | FALSE |
| 169_v.txt | 32985 | 32985-SNV-/ | SNV | 1 | A | C | 10 | 26544 | 0.0376733 | 63.8 | WT | FALSE | CAN | JSY13 | 4 | WT-Rep4 | AAG | A>C | G/C+1 | A/T-1 | TA>GC | TA>GC | GGAGAAAG( | AGGAATC | TTC | TTC | T | G | T>G | T>G_TTC | TA>GC_AGGAATC | 9 | FALSE | FALSE |
| 169_v.txt | 33039 | 33039-SNV-/ | SNV | 1 | A | C | 17 | 26544 | 0.06404461 | 61.8823529 | WT | FALSE | CAN | JSY13 | 4 | WT-Rep4 | CAT | A>C | G/C+1 | A/T-1 | TA>GC | TA>GC | GTAGCCATTT | ATTTCAC | GTA | GTA | T | G | T>G | T>G_GTA | TA>GC_ATTTCAC | 9 | FALSE | FALSE |
| 169_v.txt | 33166 | 33166-SNV-T | SNV | 1 | T | G | 8 | 26546 | 0.03013637 | 36.5 | WT | FALSE | CAN | JSY13 | 4 | WT-Rep4 | GTA | T>G | G/C+1 | A/T-1 | TA>GC | TA>GC | CAATAGTAC( | TACCACC | CAT | GTA | T | C | T>G | T>G_GTA | TA>GC_TACCACC | 9 | FALSE | FALSE |
| 169_v.txt | 33204 | 33204-SNV-/ | SNV | 1 | A | C | 273 | 16226 | 1.6824849 | 37.3956044 | WT | FALSE | CAN | JSY13 | 4 | WT-Rep4 | AAG | A>C | G/C+1 | A/T-1 | TA>GC | TA>GC | TGCTTAAGCT | AGCTCTC | TTC | TTC | T | G | T>G | T>G_TTC | TA>GC_AGCTCTC | 9 | FALSE | FALSE |
| 169_v.txt | 33207 | 33207-SNV-T | SNV | 1 | T | G | 4 | 16226 | 0.02465179 | 33.9995464 | WT | FALSE | CAN | JSY13 | 4 | WT-Rep4 | CTC | T>G | G/C+1 | A/T-1 | TA>GC | TA>GC | TTAAGCTCTC | TCTCTCT | GAG | CTC | T | C | T>G | T>G_CTC | TA>GC_TCTCTCT | 9 | FALSE | FALSE |
| 169_v.txt | 33465 | 33465-SNV-/ | SNV | 1 | A | C | 64 | 16227 | 0.39440439 | 15.3125 | WT | FALSE | CAN | JSY13 | 4 | WT-Rep4 | CAT | A>C | G/C+1 | A/T-1 | TA>GC | TA>GC | TTTGTCATTG | ATTGCTA | GTA | GTA | T | G | T>G | T>G_GTA | TA>GC_ATTGCTA | 9 | FALSE | FALSE |
| 169_v.txt | 33476 | 33476-SNV-T | SNV | 1 | T | G | 3 | 16225 | 0.01848999 | 32 | WT | FALSE | CAN | JSY13 | 4 | WT-Rep4 | CTT | T>G | G/C+1 | A/T-1 | TA>GC | TA>GC | TATGCCTTTT | TTTTTTT | GAA | CTT | T | C | T>G | T>G_CTT | TA>GC_TTTTTTT | 9 | FALSE | FALSE |
| 169_v.txt | 32256 | 32256-SNV-/ | SNV | 1 | A | G | 52 | 51480 | 0.1010101 | 35.7884615 | WT | FALSE | CAN | JSY13 | 4 | WT-Rep4 | TAG | A>G | G/C+1 | A/T-1 | TA>CG | TA>CG | TTTGATAGA( | AGACCAA | ATC | ATC | T | C | T>C | T>C_ATC | TA>CG_AGACCAA | 9 | FALSE | FALSE |
| 169_v.txt | 32262 | 32262-SNV-/ | SNV | 1 | A | G | 37 | 51504 | 0.07183908 | 36.027027 | WT | FALSE | CAN | JSY13 | 4 | WT-Rep4 | AAA | A>G | G/C+1 | A/T-1 | TA>CG | TA>CG | AGACCAAAT. | AATAAAA | TTT | TTT | T | C | T>C | T>C_TTT | TA>CG_AATAAAA | 9 | FALSE | FALSE |
| 169_v.txt | 32293 | 32293-SNV-/ | SNV | 1 | A | G | 93 | 51504 | 0.1805685 | 53.3010753 | WT | FALSE | CAN | JSY13 | 4 | WT-Rep4 | GAA | A>G | G/C+1 | A/T-1 | TA>CG | TA>CG | TATTTGAATT | AATTTGC | CTT | CTT | T | C | T>C | T>C_CTT | TA>CG_AATTTGC | 9 | FALSE | FALSE |
| 169_v.txt | 32337 | 32337-SNV-/ | SNV | 1 | A | G | 5 | 51504 | 9.71E-03 | 55.1999764 | WT | FALSE | CAN | JSY13 | 4 | WT-Rep4 | GAT | A>G | G/C+1 | A/T-1 | TA>CG | TA>CG | TTGAAGATA' | ATATGTG | CTA | CTA | T | C | T>C | T>C_CTA | TA>CG_ATATGTG | 9 | FALSE | FALSE |
| 169_v.txt | 32338 | 32338-SNV-T | SNV | 1 | T | C | 7 | 51504 | 0.01359118 | 46.4285232 | WT | FALSE | CAN | JSY13 | 4 | WT-Rep4 | ATA | T>C | G/C+1 | A/T-1 | TA>CG | TA>CG | TGAAGATAT( | TATGTGG | TAT | ATA | T | G | T>C | T>C_ATA | TA>CG_TATGTGG | 9 | FALSE | FALSE |
| 169_v.txt | 32445 | 32445-SNV-/ | SNV | 1 | A | G | 49 | 51504 | 0.09513824 | 53.1020408 | WT | FALSE | CAN | JSY13 | 4 | WT-Rep4 | AAG | A>G | G/C+1 | A/T-1 | TA>CG | TA>CG | ACTAAAAGT( | AGTCCAA | TTC | TTC | T | C | T>C | T>C_TTC | TA>CG_AGTCCAA | 9 | FALSE | FALSE |
| 169_v.txt | 32472 | 32472-SNV-T | SNV | 1 | T | C | 47 | 51504 | 0.09125505 | 49.3617021 | WT | FALSE | CAN | JSY13 | 4 | WT-Rep4 | GTA | T>C | G/C+1 | A/T-1 | TA>CG | TA>CG | CCAATGTAG/ | TAGAAGG | CAT | GTA | T | G | T>C | T>C_GTA | TA>CG_TAGAAGG | 9 | FALSE | FALSE |
| 169_v.txt | 32680 | 32680-SNV-/ | SNV | 1 | A | G | 17 | 12117 | 0.14029875 | 63.2941177 | WT | FALSE | CAN | JSY13 | 4 | WT-Rep4 | CAA | A>G | G/C+1 | A/T-1 | TA>CG | TA>CG | TTCTCCAATA | AATAACG | GTT | GTT | T | C | T>C | T>C_GTT | TA>CG_AATAACG | 9 | FALSE | FALSE |
| 169_v.txt | 32919 | 32919-SNV-/ | SNV | 1 | A | G | 19 | 26544 | 0.07157927 | 46.9473684 | WT | FALSE | CAN | JSY13 | 4 | WT-Rep4 | CAG | A>G | G/C+1 | A/T-1 | TA>CG | TA>CG | AGTTCCAGG( | AGGGCAA | GTC | GTC | T | C | T>C | T>C_GTC | TA>CG_AGGGCAA | 9 | FALSE | FALSE |
| 169_v.txt | 32990 | 32990-SNV-T | SNV | 1 | T | C | 26 | 26544 | 0.09795057 | 61.9615385 | WT | FALSE | CAN | JSY13 | 4 | WT-Rep4 | ATC | T>C | G/C+1 | A/T-1 | TA>CG | TA>CG | AAGGAATCT | TCTTTGT | TAG | ATC | T | G | T>C | T>C_ATC | TA>CG_TCTTTGT | 9 | FALSE | FALSE |
| 169_v.txt | 33476 | 33476-SNV-T | SNV | 1 | T | C | 21 | 16225 | 0.12942989 | 33.952381 | WT | FALSE | CAN | JSY13 | 4 | WT-Rep4 | CTT | T>C | G/C+1 | A/T-1 | TA>CG | TA>CG | TATGCCTTTT | TTTTTTT | GAA | CTT | T | G | T>C | T>C_CTT | TA>CG_TTTTTTT | 9 | FALSE | FALSE |
| 169_v.txt | 32050 | 32050-SNV-T | SNV | 1 | T | A | 16 | 19698 | 0.08122652 | 63.5 | WT | FALSE | CAN | JSY13 | 4 | WT-Rep4 | CTG | T>A | A/T+1 | A/T-1 | TA>AT | TA>AT | TAAATCTGA1 | TGATGTG | GAC | CTG | T | T | T>A | T>A_CTG | TA>AT_TGATGTG | 9 | FALSE | FALSE |
| 169_v.txt | 32289 | 32289-SNV-T | SNV | 1 | T | A | 69 | 51504 | 0.13397018 | 52.9710145 | WT | FALSE | CAN | JSY13 | 4 | WT-Rep4 | ATT | T>A | A/T+1 | A/T-1 | TA>AT | TA>AT | TAAATATTTG | TTTGAAT | TAA | ATT | T | T | T>A | T>A_ATT | TA>AT_TTTGAAT | 9 | FALSE | FALSE |
| 169_v.txt | 32337 | 32337-SNV-/ | SNV | 1 | A | T | 6 | 51504 | 0.01164958 | 44.6665727 | WT | FALSE | CAN | JSY13 | 4 | WT-Rep4 | GAT | A>T | A/T+1 | A/T-1 | TA>AT | TA>AT | TTGAAGATA' | ATATGTG | CTA | CTA | T | A | T>A | T>A_CTA | TA>AT_ATATGTG | 9 | FALSE | FALSE |
| 169_v.txt | 32338 | 32338-SNV-T | SNV | 1 | T | A | 5 | 51504 | 9.71E-03 | 56.599077 | WT | FALSE | CAN | JSY13 | 4 | WT-Rep4 | ATA | T>A | A/T+1 | A/T-1 | TA>AT | TA>AT | TGAAGATAT( | TATGTGG | TAT | ATA | T | T | T>A | T>A_ATA | TA>AT_TATGTGG | 9 | FALSE | FALSE |
| 169_v.txt | 32354 | 32354-SNV-T | SNV | 1 | T | A | 63 | 51504 | 0.1223206 | 63.5714286 | WT | FALSE | CAN | JSY13 | 4 | WT-Rep4 | TTG | T>A | A/T+1 | A/T-1 | TA>AT | TA>AT | AACCTTTGTA | TGTACCA | AAC | TTG | T | T | T>A | T>A_TTG | TA>AT_TGTACCA | 9 | FALSE | FALSE |
| 169_v.txt | 32472 | 32472-SNV-T | SNV | 1 | T | A | 10 | 51504 | 0.01941597 | 41.9 | WT | FALSE | CAN | JSY13 | 4 | WT-Rep4 | GTA | T>A | A/T+1 | A/T-1 | TA>AT | TA>AT | CCAATGTAG/ | TAGAAGG | CAT | GTA | T | T | T>A | T>A_GTA | TA>AT_TAGAAGG | 9 | FALSE | FALSE |
| 169_v.txt | 33065 | 33065-SNV-/ | SNV | 1 | A | T | 16 | 26544 | 0.06027728 | 50.0625 | WT | FALSE | CAN | JSY13 | 4 | WT-Rep4 | AAT | A>T | A/T+1 | A/T-1 | TA>AT | TA>AT | GACAGAATA | ATATGCC | TTA | TTA | T | A | T>A | T>A_TTA | TA>AT_ATATGCC | 9 | FALSE | FALSE |
| 169_v.txt | 33067 | 33067-SNV-/ | SNV | 1 | A | T | 682 | 26544 | 2.56931887 | 55.5645161 | WT | FALSE | CAN | JSY13 | 4 | WT-Rep4 | TAT | A>T | A/T+1 | A/T-1 | TA>AT | TA>AT | CAGAATATG( | ATGCCAA | ATA | ATA | T | A | T>A | T>A_ATA | TA>AT_ATGCCAA | 9 | FALSE | FALSE |
| 169_v.txt | 33089 | 33089-SNV-/ | SNV | 1 | A | T | 41 | 26544 | 0.15446052 | 50.0722485 | WT | FALSE | CAN | JSY13 | 4 | WT-Rep4 | AAT | A>T | A/T+1 | A/T-1 | TA>AT | TA>AT | AAATAAATA' | ATATGAT | TTA | TTA | T | A | T>A | T>A_TTA | TA>AT_ATATGAT | 9 | FALSE | FALSE |
| 169_v.txt | 33090 | 33090-SNV-T | SNV | 1 | T | A | 3 | 26544 | 0.01130199 | 37.6654469 | WT | FALSE | CAN | JSY13 | 4 | WT-Rep4 | ATA | T>A | A/T+1 | A/T-1 | TA>AT | TA>AT | AATAAATAT( | TATGATA | TAT | ATA | T | T | T>A | T>A_ATA | TA>AT_TATGATA | 9 | FALSE | FALSE |
| 169_v.txt | 31806 | 31806-SNV-( | SNV | 1 | C | T | 351 | 3301 | 10.6331415 | 59.2905983 | WT | FALSE | CAN | JSY13 | 4 | WT-Rep4 | CCA | C>T | A/T+1 | G/C-1 | CG>TA | CG>TA | ATCTTCCAAA | CAAATAA | GGT | CCA | C | A | C>T | C>T_CCA | CG>TA_CAAATAA | 9 | FALSE | FALSE |
| 169_v.txt | 31962 | 31962-SNV-( | SNV | 1 | G | A | 25 | 19698 | 0.12691644 | 37.48 | WT | FALSE | CAN | JSY13 | 4 | WT-Rep4 | CGC | G>A | A/T+1 | G/C-1 | CG>TA | CG>TA | GTGGCCGCA | GCATAAT | GCG | GCG | C | T | C>T | C>T_GCG | CG>TA_GCATAAT | 9 | FALSE | FALSE |
| 169_v.txt | 31972 | 31972-SNV-( | SNV | 1 | C | T | 14 | 19698 | 0.07107321 | 37.7142857 | WT | FALSE | CAN | JSY13 | 4 | WT-Rep4 | GCC | C>T | A/T+1 | G/C-1 | CG>TA | CG>TA | AATAAGCCA. | CCAAGCC | CGG | GCC | C | A | C>T | C>T_GCC | CG>TA_CCAAGCC | 9 | FALSE | FALSE |
| 169_v.txt | 32075 | 32075-SNV-( | SNV | 1 | C | T | 79 | 19698 | 0.40105595 | 61.443038 | WT | FALSE | CAN | JSY13 | 4 | WT-Rep4 | ACC | C>T | A/T+1 | G/C-1 | CG>TA | CG>TA | AAATAACCA' | CCATGCA | TGG | ACC | C | A | C>T | C>T_ACC | CG>TA_CCATGCA | 9 | FALSE | FALSE |
| 169_v.txt | 32092 | 32092-SNV-( | SNV | 1 | C | T | 29 | 19698 | 0.14722307 | 57.7241379 | WT | FALSE | CAN | JSY13 | 4 | WT-Rep4 | GCA | C>T | A/T+1 | G/C-1 | CG>TA | CG>TA | AGCCTGCAA( | CAACACC | CGT | GCA | C | A | C>T | C>T_GCA | CG>TA_CAACACC | 9 | FALSE | FALSE |
| 169_v.txt | 32165 | 32165-SNV-( | SNV | 1 | G | A | 18 | 19698 | 0.09137984 | 45.5555556 | WT | FALSE | CAN | JSY13 | 4 | WT-Rep4 | CGC | G>A | A/T+1 | G/C-1 | CG>TA | CG>TA | CAAAGCGCC. | GCCAAAT | GCG | GCG | C | T | C>T | C>T_GCG | CG>TA_GCCAAAT | 9 | FALSE | FALSE |
| 169_v.txt | 32205 | 32205-SNV-( | SNV | 1 | C | T | 47 | 19698 | 0.2386029 | 40.3617021 | WT | FALSE | CAN | JSY13 | 4 | WT-Rep4 | ACC | C>T | A/T+1 | G/C-1 | CG>TA | CG>TA | GGAACACCA | CCACCTT | TGG | ACC | C | A | C>T | C>T_ACC | CG>TA_CCACCTT | 9 | FALSE | FALSE |
| 169_v.txt | 32259 | 32259-SNV-( | SNV | 1 | C | T | 64 | 51502 | 0.12426702 | 31.109375 | WT | FALSE | CAN | JSY13 | 4 | WT-Rep4 | ACC | C>T | A/T+1 | G/C-1 | CG>TA | CG>TA | GATAGACCA. | CCAAATA | TGG | ACC | C | A | C>T | C>T_ACC | CG>TA_CCAAATA | 9 | FALSE | FALSE |
| 169_v.txt | 32271 | 32271-SNV-( | SNV | 1 | C | T | 95 | 51504 | 0.18445169 | 34.0524853 | WT | FALSE | CAN | JSY13 | 4 | WT-Rep4 | ACG | C>T | A/T+1 | G/C-1 | CG>TA | CG>TA | AAAATACGG | CGGGAAC | TGC | ACG | C | A | C>T | C>T_ACG | CG>TA_CGGGAAC | 9 | FALSE | FALSE |
| 169_v.txt | 32272 | 32272-SNV-( | SNV | 1 | G | A | 87 | 51504 | 0.16891892 | 36.5517241 | WT | FALSE | CAN | JSY13 | 4 | WT-Rep4 | CGG | G>A | A/T+1 | G/C-1 | CG>TA | CG>TA | AAATACGGG | GGGAACC | GCC | GCC | C | T | C>T | C>T_GCC | CG>TA_GGGAACC | 9 | FALSE | FALSE |
| 169_v.txt | 32299 | 32299-SNV-( | SNV | 1 | C | T | 84 | 51504 | 0.16309413 | 51.0476191 | WT | FALSE | CAN | JSY13 | 4 | WT-Rep4 | GCG | C>T | A/T+1 | G/C-1 | CG>TA | CG>TA | AATTTGCGG( | CGGCAGA | CGC | GCG | C | A | C>T | C>T_GCG | CG>TA_CGGCAGA | 9 | FALSE | FALSE |
| 169_v.txt | 32301 | 32301-SNV-( | SNV | 1 | G | A | 74 | 51504 | 0.14367816 | 52.0945946 | WT | FALSE | CAN | JSY13 | 4 | WT-Rep4 | GGC | G>A | A/T+1 | G/C-1 | CG>TA | CG>TA | TTTGCGGCA( | GCAGAAA | CCG | CCG | C | T | C>T | C>T_CCG | CG>TA_GCAGAAA | 9 | FALSE | FALSE |
| 169_v.txt | 32466 | 32466-SNV-( | SNV | 1 | C | T | 73 | 51504 | 0.14173656 | 48.3420949 | WT | FALSE | CAN | JSY13 | 4 | WT-Rep4 | GCC | C>T | A/T+1 | G/C-1 | CG>TA | CG>TA | AGAGAGCCA | CCAATGT | CGG | GCC | C | A | C>T | C>T_GCC | CG>TA_CCAATGT | 9 | FALSE | FALSE |
| 169_v.txt | 32467 | 32467-SNV-( | SNV | 1 | C | T | 13 | 51503 | 0.02524125 | 35.8450036 | WT | FALSE | CAN | JSY13 | 4 | WT-Rep4 | CCA | C>T | A/T+1 | G/C-1 | CG>TA | CG>TA | GAGAGCCAA | CAATGTA | GGT | CCA | C | A | C>T | C>T_CCA | CG>TA_CAATGTA | 9 | FALSE | FALSE |
| 169_v.txt | 32487 | 32487-SNV-( | SNV | 1 | C | T | 31 | 51504 | 0.0601895 | 47.6451613 | WT | FALSE | CAN | JSY13 | 4 | WT-Rep4 | ACG | C>T | A/T+1 | G/C-1 | CG>TA | CG>TA | AAGATACGG | CGGAAAA | TGC | ACG | C | A | C>T | C>T_ACG | CG>TA_CGGAAAA | 9 | FALSE | FALSE |
| 169_v.txt | 32529 | 32529-SNV-( | SNV | 1 | G | A | 125 | 51504 | 0.2426996 | 41.568 | WT | FALSE | CAN | JSY13 | 4 | WT-Rep4 | GGG | G>A | A/T+1 | G/C-1 | CG>TA | CG>TA | TTTCTGGGG1 | GGGTTTG | CCC | CCC | C | T | C>T | C>T_CCC | CG>TA_GGGTTTG | 9 | FALSE | FALSE |
| 169_v.txt | 32530 | 32530-SNV-( | SNV | 1 | G | A | 14 | 51504 | 0.02718236 | 38.5690795 | WT | FALSE | CAN | JSY13 | 4 | WT-Rep4 | GGG | G>A | A/T+1 | G/C-1 | CG>TA | CG>TA | TTCTGGGGT | GGTTTGC | CCC | CCC | C | T | C>T | C>T_CCC | CG>TA_GGTTTGC | 9 | FALSE | FALSE |

| | | | | | | | | | | | | | | | | | | | | | | | | | | | | | | | | | | |
|---|---|---|---|---|---|---|---|---|---|---|---|---|---|---|---|---|---|---|---|---|---|---|---|---|---|---|---|---|---|---|---|---|---|---|
| 169_v.txt | 32535 | 32535-SNV-( | SNV | 1 | G | A | 59 | 51504 | 0.11455421 | 36.7790816 | WT | FALSE | CAN | JSY13 | 4 | WT-Rep4 | TGC | G>A | A/T+1 | G/C-1 | CG>TA | CG>TA | GGGTTTGCA | GCAGCTT | ACG | ACG | C | T | C>T | C>T_ACG | CG>TA_GCAGCTT | 9 | FALSE | FALSE |
| 169_v.txt | 32536 | 32536-SNV-( | SNV | 1 | C | T | 72 | 51504 | 0.13979497 | 35.3194248 | WT | FALSE | CAN | JSY13 | 4 | WT-Rep4 | GCA | C>T | A/T+1 | G/C-1 | CG>TA | CG>TA | GGTTTGCAG | CAGCTTC | CGT | GCA | C | A | C>T | C>T_GCA | CG>TA_CAGCTTC | 9 | FALSE | FALSE |
| 169_v.txt | 32545 | 32545-SNV-( | SNV | 1 | C | T | 555 | 51503 | 1.07760713 | 37.4576577 | WT | FALSE | CAN | JSY13 | 4 | WT-Rep4 | CCA | C>T | A/T+1 | G/C-1 | CG>TA | CG>TA | CTTCACCAG | CAGCAGT | GGT | CCA | C | A | C>T | C>T_CCA | CG>TA_CAGCAGT | 9 | FALSE | FALSE |
| 169_v.txt | 32575 | 32575-SNV-( | SNV | 1 | G | A | 17 | 12118 | 0.14028718 | 36.2941177 | WT | FALSE | CAN | JSY13 | 4 | WT-Rep4 | TGA | G>A | A/T+1 | G/C-1 | CG>TA | CG>TA | TACCTTGAA | GAAATGT | ACT | ACT | C | T | C>T | C>T_ACT | CG>TA_GAAATGT | 9 | FALSE | FALSE |
| 169_v.txt | 32590 | 32590-SNV-( | SNV | 1 | C | T | 15 | 12117 | 0.12379302 | 36.6666667 | WT | FALSE | CAN | JSY13 | 4 | WT-Rep4 | GCG | C>T | A/T+1 | G/C-1 | CG>TA | CG>TA | AGGCAGCGT | CGTTAAT | CGC | GCG | C | A | C>T | C>T_GCG | CG>TA_CGTTAAT | 9 | FALSE | FALSE |
| 169_v.txt | 32757 | 32757-SNV-( | SNV | 1 | G | A | 15 | 12117 | 0.12379302 | 56.2666667 | WT | FALSE | CAN | JSY13 | 4 | WT-Rep4 | GGC | G>A | A/T+1 | G/C-1 | CG>TA | CG>TA | ATAATGGCT | GCTAAAA | CCG | CCG | C | T | C>T | C>T_CCG | CG>TA_GCTAAAA | 9 | FALSE | FALSE |
| 169_v.txt | 32776 | 32776-SNV-( | SNV | 1 | C | T | 16 | 12117 | 0.13204589 | 46.3125 | WT | FALSE | CAN | JSY13 | 4 | WT-Rep4 | GCG | C>T | A/T+1 | G/C-1 | CG>TA | CG>TA | TGGAAGCGA | CGACCCA | CGC | GCG | C | A | C>T | C>T_GCG | CG>TA_CGACCCA | 9 | FALSE | FALSE |
| 169_v.txt | 32797 | 32797-SNV-( | SNV | 1 | C | T | 6 | 12117 | 0.04951721 | 49.1666667 | WT | FALSE | CAN | JSY13 | 4 | WT-Rep4 | CCG | C>T | A/T+1 | G/C-1 | CG>TA | CG>TA | ATTCACCGT | CGTAATA | GGC | CCG | C | A | C>T | C>T_CCG | CG>TA_CGTAATA | 9 | FALSE | FALSE |
| 169_v.txt | 32798 | 32798-SNV-( | SNV | 1 | G | A | 12 | 12117 | 0.09903442 | 48.1666667 | WT | FALSE | CAN | JSY13 | 4 | WT-Rep4 | CGT | G>A | A/T+1 | G/C-1 | CG>TA | CG>TA | TTCACCGTA | GTAATAT | GCA | GCA | C | T | C>T | C>T_GCA | CG>TA_GTAATAT | 9 | FALSE | FALSE |
| 169_v.txt | 32840 | 32840-SNV-( | SNV | 1 | C | T | 22 | 12117 | 0.18156309 | 37.5454546 | WT | FALSE | CAN | JSY13 | 4 | WT-Rep4 | CCC | C>T | A/T+1 | G/C-1 | CG>TA | CG>TA | AATTACCCA | CCAAAAA | GGG | CCC | C | A | C>T | C>T_CCC | CG>TA_CCAAAAA | 9 | FALSE | FALSE |
| 169_v.txt | 32855 | 32855-SNV-( | SNV | 1 | C | T | 15 | 12117 | 0.12379302 | 36.5333333 | WT | FALSE | CAN | JSY13 | 4 | WT-Rep4 | TCC | C>T | A/T+1 | G/C-1 | CG>TA | CG>TA | ACTAATCCAT | CCATGCC | AGG | TCC | C | A | C>T | C>T_TCC | CG>TA_CCATGCC | 9 | FALSE | FALSE |
| 169_v.txt | 32860 | 32860-SNV-( | SNV | 1 | C | T | 16 | 12117 | 0.13204589 | 37.375 | WT | FALSE | CAN | JSY13 | 4 | WT-Rep4 | GCC | C>T | A/T+1 | G/C-1 | CG>TA | CG>TA | TCCATGCCG | CCGCCAG | CGG | GCC | C | A | C>T | C>T_GCC | CG>TA_CCGCCAG | 9 | FALSE | FALSE |
| 169_v.txt | 32900 | 32900-SNV-( | SNV | 1 | G | A | 27 | 26544 | 0.1017179 | 35.6220616 | WT | FALSE | CAN | JSY13 | 4 | WT-Rep4 | GGC | G>A | A/T+1 | G/C-1 | CG>TA | CG>TA | GACTTGGCC | GCCAACT | CCG | CCG | C | T | C>T | C>T_CCG | CG>TA_GCCAACT | 9 | FALSE | FALSE |
| 169_v.txt | 32901 | 32901-SNV-( | SNV | 1 | C | T | 46 | 26544 | 0.17329717 | 31.8042305 | WT | FALSE | CAN | JSY13 | 4 | WT-Rep4 | GCC | C>T | A/T+1 | G/C-1 | CG>TA | CG>TA | ACTTGGCCA | CCAACTA | CGG | GCC | C | A | C>T | C>T_GCC | CG>TA_CCAACTA | 9 | FALSE | FALSE |
| 169_v.txt | 32936 | 32936-SNV-( | SNV | 1 | C | T | 63 | 26544 | 0.23734177 | 45.031746 | WT | FALSE | CAN | JSY13 | 4 | WT-Rep4 | CCC | C>T | A/T+1 | G/C-1 | CG>TA | CG>TA | GATTGCCCA | CCAAGAA | GGG | CCC | C | A | C>T | C>T_CCC | CG>TA_CCAAGAA | 9 | FALSE | FALSE |
| 169_v.txt | 32965 | 32965-SNV-( | SNV | 1 | C | T | 26 | 26544 | 0.09795057 | 56.4615385 | WT | FALSE | CAN | JSY13 | 4 | WT-Rep4 | GCC | C>T | A/T+1 | G/C-1 | CG>TA | CG>TA | CATTGGCCG | CCGCACC | CGG | GCC | C | A | C>T | C>T_GCC | CG>TA_CCGCACC | 9 | FALSE | FALSE |
| 169_v.txt | 32967 | 32967-SNV-( | SNV | 1 | G | A | 50 | 26544 | 0.18836649 | 60.18 | WT | FALSE | CAN | JSY13 | 4 | WT-Rep4 | CGC | G>A | A/T+1 | G/C-1 | CG>TA | CG>TA | TTGGCCGCA | GCACCAA | GCG | GCG | C | T | C>T | C>T_GCG | CG>TA_GCACCAA | 9 | FALSE | FALSE |
| 169_v.txt | 32970 | 32970-SNV-( | SNV | 1 | C | T | 24 | 26544 | 0.09041591 | 63.0416667 | WT | FALSE | CAN | JSY13 | 4 | WT-Rep4 | ACC | C>T | A/T+1 | G/C-1 | CG>TA | CG>TA | GCCGCACCA | CCAAATG | TGG | ACC | C | A | C>T | C>T_ACC | CG>TA_CCAAATG | 9 | FALSE | FALSE |
| 169_v.txt | 32977 | 32977-SNV-( | SNV | 1 | C | T | 24 | 26544 | 0.09041591 | 61.2083333 | WT | FALSE | CAN | JSY13 | 4 | WT-Rep4 | GCT | C>T | A/T+1 | G/C-1 | CG>TA | CG>TA | CAAATGCTG | CTGGAGA | CGA | GCT | C | A | C>T | C>T_GCT | CG>TA_CTGGAGA | 9 | FALSE | FALSE |
| 169_v.txt | 32995 | 32995-SNV-( | SNV | 1 | G | A | 26 | 26544 | 0.09795057 | 60.6538462 | WT | FALSE | CAN | JSY13 | 4 | WT-Rep4 | TGT | G>A | A/T+1 | G/C-1 | CG>TA | CG>TA | ATCTTTGTGA | GTGAGAA | ACA | ACA | C | T | C>T | C>T_ACA | CG>TA_GTGAGAA | 9 | FALSE | FALSE |
| 169_v.txt | 33015 | 33015-SNV-( | SNV | 1 | G | A | 16 | 26544 | 0.06027728 | 51.625 | WT | FALSE | CAN | JSY13 | 4 | WT-Rep4 | GGA | G>A | A/T+1 | G/C-1 | CG>TA | CG>TA | AAAGAGGAT | GATGTAA | CCT | CCT | C | T | C>T | C>T_CCT | CG>TA_GATGTAA | 9 | FALSE | FALSE |
| 169_v.txt | 33043 | 33043-SNV-( | SNV | 1 | C | T | 272 | 26544 | 1.02471368 | 59.7536765 | WT | FALSE | CAN | JSY13 | 4 | WT-Rep4 | TCA | C>T | A/T+1 | G/C-1 | CG>TA | CG>TA | CCATTTCACC | CACCCAA | AGT | TCA | C | A | C>T | C>T_TCA | CG>TA_CACCCAA | 9 | FALSE | FALSE |
| 169_v.txt | 33055 | 33055-SNV-( | SNV | 1 | G | A | 604 | 26543 | 2.27555288 | 57.9648201 | WT | FALSE | CAN | JSY13 | 4 | WT-Rep4 | TGC | G>A | A/T+1 | G/C-1 | CG>TA | CG>TA | AGGACTGCG | GCGTGAC | ACG | ACG | C | T | C>T | C>T_ACG | CG>TA_GCGTGAC | 9 | FALSE | FALSE |
| 169_v.txt | 33056 | 33056-SNV-( | SNV | 1 | C | T | 31 | 26544 | 0.11678722 | 55.8683028 | WT | FALSE | CAN | JSY13 | 4 | WT-Rep4 | GCG | C>T | A/T+1 | G/C-1 | CG>TA | CG>TA | GGACTGCGT | CGTGACA | CGC | GCG | C | A | C>T | C>T_GCG | CG>TA_CGTGACA | 9 | FALSE | FALSE |
| 169_v.txt | 33114 | 33114-SNV-( | SNV | 1 | C | T | 36 | 26544 | 0.13562387 | 43.7481118 | WT | FALSE | CAN | JSY13 | 4 | WT-Rep4 | GCC | C>T | A/T+1 | G/C-1 | CG>TA | CG>TA | ACTGGGCCG | CCGGCGT | CGG | GCC | C | A | C>T | C>T_GCC | CG>TA_CCGGCGT | 9 | FALSE | FALSE |
| 169_v.txt | 33115 | 33115-SNV-( | SNV | 1 | C | T | 44 | 26544 | 0.16576251 | 42.3635183 | WT | FALSE | CAN | JSY13 | 4 | WT-Rep4 | CCG | C>T | A/T+1 | G/C-1 | CG>TA | CG>TA | CTGGGCCGG | CGGCGTT | GGC | CCG | C | A | C>T | C>T_CCG | CG>TA_CGGCGTT | 9 | FALSE | FALSE |
| 169_v.txt | 33206 | 33206-SNV-( | SNV | 1 | C | T | 1 | 16226 | 6.16E-03 | 37.9996973 | WT | FALSE | CAN | JSY13 | 4 | WT-Rep4 | GCT | C>T | A/T+1 | G/C-1 | CG>TA | CG>TA | CTTAAGCTCT | CTCTCTC | CGA | GCT | C | A | C>T | C>T_GCT | CG>TA_CTCTCTC | 9 | FALSE | FALSE |
| 169_v.txt | 33229 | 33229-SNV-( | SNV | 1 | G | A | 65 | 16226 | 0.40059164 | 38.7076923 | WT | FALSE | CAN | JSY13 | 4 | WT-Rep4 | TGT | G>A | A/T+1 | G/C-1 | CG>TA | CG>TA | CGTTCTGTAC | GTACTTC | ACA | ACA | C | T | C>T | C>T_ACA | CG>TA_GTACTTC | 9 | FALSE | FALSE |
| 169_v.txt | 33258 | 33258-SNV-( | SNV | 1 | C | T | 18 | 16227 | 0.11092623 | 43.4444444 | WT | FALSE | CAN | JSY13 | 4 | WT-Rep4 | GCC | C>T | A/T+1 | G/C-1 | CG>TA | CG>TA | CCTATGCCAT | CCATCCT | CGG | GCC | C | A | C>T | C>T_GCC | CG>TA_CCATCCT | 9 | FALSE | FALSE |
| 169_v.txt | 33276 | 33276-SNV-( | SNV | 1 | G | A | 21 | 16227 | 0.12941394 | 46.0952381 | WT | FALSE | CAN | JSY13 | 4 | WT-Rep4 | CGT | G>A | A/T+1 | G/C-1 | CG>TA | CG>TA | GAGAACGTA | GTATCCT | GCA | GCA | C | T | C>T | C>T_GCA | CG>TA_GTATCCT | 9 | FALSE | FALSE |
| 169_v.txt | 33310 | 33310-SNV-( | SNV | 1 | G | A | 22 | 16227 | 0.13557651 | 59.0454546 | WT | FALSE | CAN | JSY13 | 4 | WT-Rep4 | CGC | G>A | A/T+1 | G/C-1 | CG>TA | CG>TA | AAGAGCGCA | GCAATGG | GCG | GCG | C | T | C>T | C>T_GCG | CG>TA_GCAATGG | 9 | FALSE | FALSE |
| 169_v.txt | 33412 | 33412-SNV-( | SNV | 1 | G | A | 25 | 16227 | 0.15406421 | 47.44 | WT | FALSE | CAN | JSY13 | 4 | WT-Rep4 | GGC | G>A | A/T+1 | G/C-1 | CG>TA | CG>TA | TGACCGGCT | GCTCATT | CCG | CCG | C | T | C>T | C>T_CCG | CG>TA_GCTCATT | 9 | FALSE | FALSE |
| 169_v.txt | 33474 | 33474-SNV-( | SNV | 1 | C | T | 22 | 16227 | 0.13557651 | 35.8636452 | WT | FALSE | CAN | JSY13 | 4 | WT-Rep4 | GCC | C>T | A/T+1 | G/C-1 | CG>TA | CG>TA | GCTATGCCTT | CCTTTTT | CGG | GCC | C | A | C>T | C>T_GCC | CG>TA_CCTTTTT | 9 | FALSE | FALSE |
| 169_v.txt | 31956 | 31956-SNV-( | SNV | 1 | G | C | 24 | 19698 | 0.12183978 | 34.9583333 | WT | FALSE | CAN | JSY13 | 4 | WT-Rep4 | TGT | G>C | G/C+1 | G/C-1 | CG>GC | CG>GC | ATAAATGTG | GTGGCCG | ACA | ACA | C | G | C>G | C>G_ACA | CG>GC_GTGGCCG | 9 | FALSE | FALSE |
| 169_v.txt | 32092 | 32092-SNV-( | SNV | 1 | C | G | 5 | 19698 | 0.02538329 | 64 | WT | FALSE | CAN | JSY13 | 4 | WT-Rep4 | GCA | C>G | G/C+1 | G/C-1 | CG>GC | CG>GC | AGCCTGCAA | CAACACC | CGT | GCA | C | C | C>G | C>G_GCA | CG>GC_CAACACC | 9 | FALSE | FALSE |
| 169_v.txt | 32188 | 32188-SNV-( | SNV | 1 | C | G | 7 | 19698 | 0.0355366 | 41.7142857 | WT | FALSE | CAN | JSY13 | 4 | WT-Rep4 | ACT | C>G | G/C+1 | G/C-1 | CG>GC | CG>GC | CGAAAACTG | CTGCAAT | TGA | ACT | C | C | C>G | C>G_ACT | CG>GC_CTGCAAT | 9 | FALSE | FALSE |
| 169_v.txt | 32223 | 32223-SNV-( | SNV | 1 | G | C | 21 | 19698 | 0.10660981 | 34.6672334 | WT | FALSE | CAN | JSY13 | 4 | WT-Rep4 | TGA | G>C | G/C+1 | G/C-1 | CG>GC | CG>GC | GTCCTTGAC | GACAGGA | ACT | ACT | C | G | C>G | C>G_ACT | CG>GC_GACAGGA | 9 | FALSE | FALSE |
| 169_v.txt | 32271 | 32271-SNV-( | SNV | 1 | C | G | 1372 | 51504 | 2.66387077 | 36.2043152 | WT | FALSE | CAN | JSY13 | 4 | WT-Rep4 | ACG | C>G | G/C+1 | G/C-1 | CG>GC | CG>GC | AAAATACGG | CGGGAAC | TGC | ACG | C | C | C>G | C>G_ACG | CG>GC_CGGGAAC | 9 | FALSE | FALSE |
| 169_v.txt | 32316 | 32316-SNV-( | SNV | 1 | G | C | 22 | 51504 | 0.04271513 | 55.6818182 | WT | FALSE | CAN | JSY13 | 4 | WT-Rep4 | TGT | G>C | G/C+1 | G/C-1 | CG>GC | CG>GC | ATGGTTGTT | GTTAAGA | ACA | ACA | C | G | C>G | C>G_ACA | CG>GC_GTTAAGA | 9 | FALSE | FALSE |
| 169_v.txt | 32358 | 32358-SNV-( | SNV | 1 | C | G | 3 | 51503 | 5.83E-03 | 52.6520588 | WT | FALSE | CAN | JSY13 | 4 | WT-Rep4 | ACC | C>G | G/C+1 | G/C-1 | CG>GC | CG>GC | TTTGTACCAG | CCAGAGT | TGG | ACC | C | C | C>G | C>G_ACC | CG>GC_CCAGAGT | 9 | FALSE | FALSE |
| 169_v.txt | 32432 | 32432-SNV-( | SNV | 1 | G | C | 12 | 51504 | 0.02329916 | 45.8333333 | WT | FALSE | CAN | JSY13 | 4 | WT-Rep4 | TGT | G>C | G/C+1 | G/C-1 | CG>GC | CG>GC | GTCATTGTAT | GTATGGA | ACA | ACA | C | G | C>G | C>G_ACA | CG>GC_GTATGGA | 9 | FALSE | FALSE |
| 169_v.txt | 32467 | 32467-SNV-( | SNV | 1 | C | G | 19 | 51503 | 0.03689106 | 47.7336305 | WT | FALSE | CAN | JSY13 | 4 | WT-Rep4 | CCA | C>G | G/C+1 | G/C-1 | CG>GC | CG>GC | GAGAGCCAA | CAATGTA | GGT | CCA | C | C | C>G | C>G_CCA | CG>GC_CAATGTA | 9 | FALSE | FALSE |
| 169_v.txt | 32474 | 32474-SNV-( | SNV | 1 | G | C | 59 | 51504 | 0.11455421 | 47.9304545 | WT | FALSE | CAN | JSY13 | 4 | WT-Rep4 | AGA | G>C | G/C+1 | G/C-1 | CG>GC | CG>GC | AATGTAGAA | GAAGGTT | TCT | TCT | C | G | C>G | C>G_TCT | CG>GC_GAAGGTT | 9 | FALSE | FALSE |
| 169_v.txt | 32487 | 32487-SNV-( | SNV | 1 | C | G | 1513 | 51504 | 2.93763591 | 46.4600132 | WT | FALSE | CAN | JSY13 | 4 | WT-Rep4 | ACG | C>G | G/C+1 | G/C-1 | CG>GC | CG>GC | AAGATACGG | CGGAAAA | TGC | ACG | C | C | C>G | C>G_ACG | CG>GC_CGGAAAA | 9 | FALSE | FALSE |
| 169_v.txt | 32529 | 32529-SNV-( | SNV | 1 | G | C | 4 | 51504 | 7.77E-03 | 37.25 | WT | FALSE | CAN | JSY13 | 4 | WT-Rep4 | GGG | G>C | G/C+1 | G/C-1 | CG>GC | CG>GC | TTTCTGGGG | GGGTTTG | CCC | CCC | C | G | C>G | C>G_CCC | CG>GC_GGGTTTG | 9 | FALSE | FALSE |
| 169_v.txt | 32539 | 32539-SNV-( | SNV | 1 | C | G | 18 | 51504 | 0.03494874 | 37.3888889 | WT | FALSE | CAN | JSY13 | 4 | WT-Rep4 | GCT | C>G | G/C+1 | G/C-1 | CG>GC | CG>GC | TTGCAGCTT | CTTCACC | CGA | GCT | C | C | C>G | C>G_GCT | CG>GC_CTTCACC | 9 | FALSE | FALSE |
| 169_v.txt | 32776 | 32776-SNV-( | SNV | 1 | C | G | 16 | 12117 | 0.13204589 | 42.5625 | WT | FALSE | CAN | JSY13 | 4 | WT-Rep4 | GCG | C>G | G/C+1 | G/C-1 | CG>GC | CG>GC | TGGAAGCGA | CGACCCA | CGC | GCG | C | C | C>G | C>G_GCG | CG>GC_CGACCCA | 9 | FALSE | FALSE |
| 169_v.txt | 32797 | 32797-SNV-( | SNV | 1 | C | G | 123 | 12117 | 1.01510275 | 45.4227642 | WT | FALSE | CAN | JSY13 | 4 | WT-Rep4 | CCG | C>G | G/C+1 | G/C-1 | CG>GC | CG>GC | ATTCACCGT | CGTAATA | GGC | CCG | C | C | C>G | C>G_CCG | CG>GC_CGTAATA | 9 | FALSE | FALSE |
| 169_v.txt | 32798 | 32798-SNV-( | SNV | 1 | G | C | 2 | 12117 | 0.01650574 | 51 | WT | FALSE | CAN | JSY13 | 4 | WT-Rep4 | CGT | G>C | G/C+1 | G/C-1 | CG>GC | CG>GC | TTCACCGTA | GTAATAT | GCA | GCA | C | G | C>G | C>G_GCA | CG>GC_GTAATAT | 9 | FALSE | FALSE |
| 169_v.txt | 32809 | 32809-SNV-( | SNV | 1 | C | G | 21 | 12117 | 0.17331023 | 47.5714286 | WT | FALSE | CAN | JSY13 | 4 | WT-Rep4 | ACA | C>G | G/C+1 | G/C-1 | CG>GC | CG>GC | ATTTGACAG | CAGGGAA | TGT | ACA | C | C | C>G | C>G_ACA | CG>GC_CAGGGAA | 9 | FALSE | FALSE |
| 169_v.txt | 32917 | 32917-SNV-( | SNV | 1 | C | G | 9 | 26544 | 0.03390597 | 50.1111111 | WT | FALSE | CAN | JSY13 | 4 | WT-Rep4 | TCC | C>G | G/C+1 | G/C-1 | CG>GC | CG>GC | TAAGTTCCA | CCAGGGC | AGG | TCC | C | C | C>G | C>G_TCC | CG>GC_CCAGGGC | 9 | FALSE | FALSE |
| 169_v.txt | 32936 | 32936-SNV-( | SNV | 1 | C | G | 121 | 26544 | 0.4558469 | 49.1157025 | WT | FALSE | CAN | JSY13 | 4 | WT-Rep4 | CCC | C>G | G/C+1 | G/C-1 | CG>GC | CG>GC | GATTGCCCA | CCAAGAA | GGG | CCC | C | C | C>G | C>G_CCC | CG>GC_CCAAGAA | 9 | FALSE | FALSE |
| 169_v.txt | 32959 | 32959-SNV-( | SNV | 1 | C | G | 14 | 26544 | 0.05274262 | 61.7857143 | WT | FALSE | CAN | JSY13 | 4 | WT-Rep4 | CCA | C>G | G/C+1 | G/C-1 | CG>GC | CG>GC | TGTAACCATT | CATTGGC | GGT | CCA | C | C | C>G | C>G_CCA | CG>GC_CATTGGC | 9 | FALSE | FALSE |
| 169_v.txt | 32997 | 32997-SNV-( | SNV | 1 | G | C | 14 | 26544 | 0.05274262 | 59.9285714 | WT | FALSE | CAN | JSY13 | 4 | WT-Rep4 | TGA | G>C | G/C+1 | G/C-1 | CG>GC | CG>GC | CTTTGTGAG | GAGAAAA | ACT | ACT | C | G | C>G | C>G_ACT | CG>GC_GAGAAAA | 9 | FALSE | FALSE |
| 169_v.txt | 33056 | 33056-SNV-( | SNV | 1 | C | G | 2 | 26544 | 7.54E-03 | 63.9922377 | WT | FALSE | CAN | JSY13 | 4 | WT-Rep4 | GCG | C>G | G/C+1 | G/C-1 | CG>GC | CG>GC | GGACTGCGT | CGTGACA | CGC | GCG | C | C | C>G | C>G_GCG | CG>GC_CGTGACA | 9 | FALSE | FALSE |
| 169_v.txt | 33156 | 33156-SNV-( | SNV | 1 | G | C | 26 | 26542 | 0.09795795 | 36.3461539 | WT | FALSE | CAN | JSY13 | 4 | WT-Rep4 | TGT | G>C | G/C+1 | G/C-1 | CG>GC | CG>GC | AGACCTGTA | GTACCAA | ACA | ACA | C | G | C>G | C>G_ACA | CG>GC_GTACCAA | 9 | FALSE | FALSE |
| 169_v.txt | 33160 | 33160-SNV-( | SNV | 1 | C | G | 8 | 26546 | 0.03013637 | 37.9953255 | WT | FALSE | CAN | JSY13 | 4 | WT-Rep4 | CCA | C>G | G/C+1 | G/C-1 | CG>GC | CG>GC | CTGTACCAAT | CAATAGT | GGT | CCA | C | C | C>G | C>G_CCA | CG>GC_CAATAGT | 9 | FALSE | FALSE |
| 169_v.txt | 33193 | 33193-SNV-( | SNV | 1 | G | C | 9 | 16223 | 0.05547679 | 37.7777778 | WT | FALSE | CAN | JSY13 | 4 | WT-Rep4 | TGT | G>C | G/C+1 | G/C-1 | CG>GC | CG>GC | CAATATGTCT | GTCTTTG | ACA | ACA | C | G | C>G | C>G_ACA | CG>GC_GTCTTTG | 9 | FALSE | FALSE |
| 169_v.txt | 32088 | 32088-SNV-( | SNV | 1 | C | A | 19 | 19698 | 0.09645649 | 56.6842105 | WT | FALSE | CAN | JSY13 | 4 | WT-Rep4 | GCC | C>A | A/T+1 | G/C-1 | CG>AT | CG>AT | AAAAAGCCT | CCTGCAA | CGG | GCC | C | T | C>A | C>A_GCC | CG>AT_CCTGCAA | 9 | FALSE | FALSE |
| 169_v.txt | 32119 | 32119-SNV-( | SNV | 1 | C | A | 15 | 19698 | 0.07614986 | 52.8666667 | WT | FALSE | CAN | JSY13 | 4 | WT-Rep4 | TCG | C>A | A/T+1 | G/C-1 | CG>AT | CG>AT | GCCATTCGA | CGAAAAC | AGC | TCG | C | T | C>A | C>A_TCG | CG>AT_CGAAAAC | 9 | FALSE | FALSE |
| 169_v.txt | 32188 | 32188-SNV-( | SNV | 1 | C | A | 131 | 19698 | 0.66504214 | 45.7175573 | WT | FALSE | CAN | JSY13 | 4 | WT-Rep4 | ACT | C>A | A/T+1 | G/C-1 | CG>AT | CG>AT | CGAAAACTG | CTGCAAT | TGA | ACT | C | T | C>A | C>A_ACT | CG>AT_CTGCAAT | 9 | FALSE | FALSE |
| 169_v.txt | 32272 | 32272-SNV-( | SNV | 1 | G | T | 16 | 51504 | 0.03106555 | 25.5 | WT | FALSE | CAN | JSY13 | 4 | WT-Rep4 | CGG | G>T | A/T+1 | G/C-1 | CG>AT | CG>AT | AAATACGGG | GGGAACC | GCC | GCC | C | A | C>A | C>A_GCC | CG>AT_GGGAACC | 9 | FALSE | FALSE |
| 169_v.txt | 32274 | 32274-SNV-( | SNV | 1 | G | T | 24 | 51504 | 0.04659832 | 33.7916667 | WT | FALSE | CAN | JSY13 | 4 | WT-Rep4 | GGA | G>T | A/T+1 | G/C-1 | CG>AT | CG>AT | ATACGGGAA | GAACCAA | CCT | CCT | C | A | C>A | C>A_CCT | CG>AT_GAACCAA | 9 | FALSE | FALSE |
| 169_v.txt | 32282 | 32282-SNV-( | SNV | 1 | G | T | 55 | 51504 | 0.10678782 | 38.9454546 | WT | FALSE | CAN | JSY13 | 4 | WT-Rep4 | CGT | G>T | A/T+1 | G/C-1 | CG>AT | CG>AT | ACCAACGTA | GTAAATA | GCA | GCA | C | A | C>A | C>A_GCA | CG>AT_GTAAATA | 9 | FALSE | FALSE |
| 169_v.txt | 32304 | 32304-SNV-( | SNV | 1 | G | T | 29 | 51504 | 0.05630631 | 52.137931 | WT | FALSE | CAN | JSY13 | 4 | WT-Rep4 | AGA | G>T | A/T+1 | G/C-1 | CG>AT | CG>AT | GCGGCAGAA | GAAATAA | TCT | TCT | C | A | C>A | C>A_TCT | CG>AT_GAAATAA | 9 | FALSE | FALSE |
| 169_v.txt | 32359 | 32359-SNV-( | SNV | 1 | C | A | 11 | 51504 | 0.02135757 | 32.5469036 | WT | FALSE | CAN | JSY13 | 4 | WT-Rep4 | CCA | C>A | A/T+1 | G/C-1 | CG>AT | CG>AT | TTGTACCAG | CAGAGTT | GGT | CCA | C | T | C>A | C>A_CCA | CG>AT_CAGAGTT | 9 | FALSE | FALSE |
| 169_v.txt | 32399 | 32399-SNV-( | SNV | 1 | G | T | 132 | 51504 | 0.25629077 | 59.4924242 | WT | FALSE | CAN | JSY13 | 4 | WT-Rep4 | CGT | G>T | A/T+1 | G/C-1 | CG>AT | CG>AT | AGAAACGTA | GTAGGAA | GCA | GCA | C | A | C>A | C>A_GCA | CG>AT_GTAGGAA | 9 | FALSE | FALSE |
| 169_v.txt | 32463 | 32463-SNV-( | SNV | 1 | G | T | 25 | 51504 | 0.04853992 | 46.28 | WT | FALSE | CAN | JSY13 | 4 | WT-Rep4 | AGA | G>T | A/T+1 | G/C-1 | CG>AT | CG>AT | AATAGAGAG | GAGCCAA | TCT | TCT | C | A | C>A | C>A_TCT | CG>AT_GAGCCAA | 9 | FALSE | FALSE |
| 169_v.txt | 32529 | 32529-SNV-( | SNV | 1 | G | T | 11 | 51504 | 0.02135757 | 33.4545455 | WT | FALSE | CAN | JSY13 | 4 | WT-Rep4 | GGG | G>T | A/T+1 | G/C-1 | CG>AT | CG>AT | TTTCTGGGG | GGGTTTG | CCC | CCC | C | A | C>A | C>A_CCC | CG>AT_GGGTTTG | 9 | FALSE | FALSE |
| 169_v.txt | 32530 | 32530-SNV-( | SNV | 1 | G | T | 16 | 51504 | 0.03106555 | 35.3732228 | WT | FALSE | CAN | JSY13 | 4 | WT-Rep4 | GGG | G>T | A/T+1 | G/C-1 | CG>AT | CG>AT | TTCTGGGGT | GGTTTGC | CCC | CCC | C | A | C>A | C>A_CCC | CG>AT_GGTTTGC | 9 | FALSE | FALSE |
| 169_v.txt | 32566 | 32566-SNV-( | SNV | 1 | C | A | 14 | 12118 | 0.11553062 | 37.4285714 | WT | FALSE | CAN | JSY13 | 4 | WT-Rep4 | TCA | C>A | A/T+1 | G/C-1 | CG>AT | CG>AT | CTAGTTCAG | CAGTACC | AGT | TCA | C | T | C>A | C>A_TCA | CG>AT_CAGTACC | 9 | FALSE | FALSE |
| 169_v.txt | 32580 | 32580-SNV-( | SNV | 1 | G | T | 8 | 12117 | 0.06602294 | 35.75 | WT | FALSE | CAN | JSY13 | 4 | WT-Rep4 | TGT | G>T | A/T+1 | G/C-1 | CG>AT | CG>AT | TGAAATGTG | GTGAAGG | ACA | ACA | C | A | C>A | C>A_ACA | CG>AT_GTGAAGG | 9 | FALSE | FALSE |
| 169_v.txt | 32786 | 32786-SNV-( | SNV | 1 | C | A | 11 | 12117 | 0.09078155 | 54.5454546 | WT | FALSE | CAN | JSY13 | 4 | WT-Rep4 | ACT | C>A | A/T+1 | G/C-1 | CG>AT | CG>AT | CCAGAACTC | CTCGAAT | TGA | ACT | C | T | C>A | C>A_ACT | CG>AT_CTCGAAT | 9 | FALSE | FALSE |
| 169_v.txt | 32794 | 32794-SNV-( | SNV | 1 | C | A | 9 | 12117 | 0.07427581 | 49.3333333 | WT | FALSE | CAN | JSY13 | 4 | WT-Rep4 | TCA | C>A | A/T+1 | G/C-1 | CG>AT | CG>AT | CGAATTCAC | CACCGTA | AGT | TCA | C | T | C>A | C>A_TCA | CG>AT_CACCGTA | 9 | FALSE | FALSE |
| 169_v.txt | 32797 | 32797-SNV-( | SNV | 1 | C | A | 18 | 12117 | 0.14855162 | 46.6666667 | WT | FALSE | CAN | JSY13 | 4 | WT-Rep4 | CCG | C>A | A/T+1 | G/C-1 | CG>AT | CG>AT | ATTCACCGT | CGTAATA | GGC | CCG | C | T | C>A | C>A_CCG | CG>AT_CGTAATA | 9 | FALSE | FALSE |
| 169_v.txt | 32798 | 32798-SNV-( | SNV | 1 | G | T | 5 | 12117 | 0.04126434 | 42.8 | WT | FALSE | CAN | JSY13 | 4 | WT-Rep4 | CGT | G>T | A/T+1 | G/C-1 | CG>AT | CG>AT | TTCACCGTA | GTAATAT | GCA | GCA | C | A | C>A | C>A_GCA | CG>AT_GTAATAT | 9 | FALSE | FALSE |
| 169_v.txt | 32940 | 32940-SNV-( | SNV | 1 | G | T | 36 | 26544 | 0.13562387 | 33.8611111 | WT | FALSE | CAN | JSY13 | 4 | WT-Rep4 | AGA | G>T | A/T+1 | G/C-1 | CG>AT | CG>AT | GCCCAAGAA | GAAAACC | TCT | TCT | C | A | C>A | C>A_TCT | CG>AT_GAAAACC | 9 | FALSE | FALSE |
| 169_v.txt | 32954 | 32954-SNV-( | SNV | 1 | G | T | 8 | 26544 | 0.03013864 | 47.123908 | WT | FALSE | CAN | JSY13 | 4 | WT-Rep4 | TGT | G>T | A/T+1 | G/C-1 | CG>AT | CG>AT | ATACATGTA | GTAACCA | ACA | ACA | C | A | C>A | C>A_ACA | CG>AT_GTAACCA | 9 | FALSE | FALSE |
| 169_v.txt | 33015 | 33015-SNV-( | SNV | 1 | G | T | 139 | 26544 | 0.52365883 | 57.6834532 | WT | FALSE | CAN | JSY13 | 4 | WT-Rep4 | GGA | G>T | A/T+1 | G/C-1 | CG>AT | CG>AT | AAAGAGGAT | GATGTAA | CCT | CCT | C | A | C>A | C>A_CCT | CG>AT_GATGTAA | 9 | FALSE | FALSE |
| 169_v.txt | 33033 | 33033-SNV-( | SNV | 1 | G | T | 4994 | 26544 | 18.8140446 | 60.4279135 | WT | FALSE | CAN | JSY13 | 4 | WT-Rep4 | TGT | G>T | A/T+1 | G/C-1 | CG>AT | CG>AT | ATGAATGTA | GTAGCCA | ACA | ACA | C | A | C>A | C>A_ACA | CG>AT_GTAGCCA | 9 | FALSE | FALSE |
| 169_v.txt | 33051 | 33051-SNV-( | SNV | 1 | G | T | 166 | 26544 | 0.62537673 | 57.6325301 | WT | FALSE | CAN | JSY13 | 4 | WT-Rep4 | GGA | G>T | A/T+1 | G/C-1 | CG>AT | CG>AT | CCCAAGGAC | GACTGCG | CCT | CCT | C | A | C>A | C>A_CCT | CG>AT_GACTGCG | 9 | FALSE | FALSE |
| 169_v.txt | 33055 | 33055-SNV-( | SNV | 1 | G | T | 2 | 26543 | 7.54E-03 | 50.9947784 | WT | FALSE | CAN | JSY13 | 4 | WT-Rep4 | TGC | G>T | A/T+1 | G/C-1 | CG>AT | CG>AT | AGGACTGCG | GCGTGAC | ACG | ACG | C | A | C>A | C>A_ACG | CG>AT_GCGTGAC | 9 | FALSE | FALSE |
| 169_v.txt | 33093 | 33093-SNV-( | SNV | 1 | G | T | 13 | 26544 | 0.04897529 | 51.2307692 | WT | FALSE | CAN | JSY13 | 4 | WT-Rep4 | TGA | G>T | A/T+1 | G/C-1 | CG>AT | CG>AT | AAATATGAT | GATATAA | ACT | ACT | C | A | C>A | C>A_ACT | CG>AT_GATATAA | 9 | FALSE | FALSE |
| 169_v.txt | 33112 | 33112-SNV-( | SNV | 1 | G | T | 15 | 26544 | 0.05650995 | 47.1333333 | WT | FALSE | CAN | JSY13 | 4 | WT-Rep4 | GGG | G>T | A/T+1 | G/C-1 | CG>AT | CG>AT | CCACTGGGC | GGCCGGC | CCC | CCC | C | A | C>A | C>A_CCC | CG>AT_GGCCGGC | 9 | FALSE | FALSE |
| 169_v.txt | 33115 | 33115-SNV-( | SNV | 1 | C | A | 13 | 26544 | 0.04897529 | 39.9226889 | WT | FALSE | CAN | JSY13 | 4 | WT-Rep4 | CCG | C>A | A/T+1 | G/C-1 | CG>AT | CG>AT | CTGGGCCGG | CGGCGTT | GGC | CCG | C | T | C>A | C>A_CCG | CG>AT_CGGCGTT | 9 | FALSE | FALSE |
| 169_v.txt | 33151 | 33151-SNV-( | SNV | 1 | G | T | 17 | 26544 | 0.06404461 | 37.2352941 | WT | FALSE | CAN | JSY13 | 4 | WT-Rep4 | AGA | G>T | A/T+1 | G/C-1 | CG>AT | CG>AT | TGAAAAGAC | GACCTGT | TCT | TCT | C | A | C>A | C>A_TCT | CG>AT_GACCTGT | 9 | FALSE | FALSE |
| 169_v.txt | 33153 | 33153-SNV-( | SNV | 1 | C | A | 14 | 26543 | 0.0527446 | 35.7142857 | WT | FALSE | CAN | JSY13 | 4 | WT-Rep4 | ACC | C>A | A/T+1 | G/C-1 | CG>AT | CG>AT | AAAAGACCT | CCTGTAC | TGG | ACC | C | T | C>A | C>A_ACC | CG>AT_CCTGTAC | 9 | FALSE | FALSE |
| 169_v.txt | 33159 | 33159-SNV-( | SNV | 1 | C | A | 10 | 26546 | 0.03767046 | 37.8991716 | WT | FALSE | CAN | JSY13 | 4 | WT-Rep4 | ACC | C>A | A/T+1 | G/C-1 | CG>AT | CG>AT | CCTGTACCA | CCAATAG | TGG | ACC | C | T | C>A | C>A_ACC | CG>AT_CCAATAG | 9 | FALSE | FALSE |
| 169_v.txt | 33160 | 33160-SNV-( | SNV | 1 | C | A | 3 | 26546 | 0.01130114 | 37.3323284 | WT | FALSE | CAN | JSY13 | 4 | WT-Rep4 | CCA | C>A | A/T+1 | G/C-1 | CG>AT | CG>AT | CTGTACCAAT | CAATAGT | GGT | CCA | C | T | C>A | C>A_CCA | CG>AT_CAATAGT | 9 | FALSE | FALSE |
| 169_v.txt | 33169 | 33169-SNV-( | SNV | 1 | C | A | 12 | 26545 | 0.04520625 | 37.75 | WT | FALSE | CAN | JSY13 | 4 | WT-Rep4 | CCA | C>A | A/T+1 | G/C-1 | CG>AT | CG>AT | TAGTACCAC | CACCAAG | GGT | CCA | C | T | C>A | C>A_CCA | CG>AT_CACCAAG | 9 | FALSE | FALSE |
| 169_v.txt | 33419 | 33419-SNV-( | SNV | 1 | G | T | 12 | 16227 | 0.07395082 | 38.6666667 | WT | FALSE | CAN | JSY13 | 4 | WT-Rep4 | TGT | G>T | A/T+1 | G/C-1 | CG>AT | CG>AT | CTCATTGTAC | GTACATA | ACA | ACA | C | A | C>A | C>A_ACA | CG>AT_GTACATA | 9 | FALSE | FALSE |
| 169_v.txt | 33474 | 33474-SNV-( | SNV | 1 | C | A | 48 | 16227 | 0.29580329 | 34.5838269 | WT | FALSE | CAN | JSY13 | 4 | WT-Rep4 | GCC | C>A | A/T+1 | G/C-1 | CG>AT | CG>AT | GCTATGCCTT | CCTTTTT | CGG | GCC | C | T | C>A | C>A_GCC | CG>AT_CCTTTTT | 9 | FALSE | FALSE |
| 169_v.txt | 33475 | 33475-SNV-( | SNV | 1 | C | A | 17 | 16227 | 0.10476367 | 36.116238 | WT | FALSE | CAN | JSY13 | 4 | WT-Rep4 | CCT | C>A | A/T+1 | G/C-1 | CG>AT | CG>AT | CTATGCCTTT | CTTTTTT | GGA | CCT | C | T | C>A | C>A_CCT | CG>AT_CTTTTTT | 9 | FALSE | FALSE |
| 169_v.txt | 33206 | 33206-Repla | Replacement | 2 | CT | A | 1 | 16226 | 6.16E-03 | 38 | WT | FALSE | CAN | JSY13 | 4 | WT-Rep4 | GCT | nope | A/T+1 | >1bpdel | none | Replac. | CTTAAGCTCT | CTCTCTC | CGA | GCT | CT | T | NA | NA_GCT | Replac._CTCTCTC | 9 | FALSE | FALSE |
| 169_v.txt | 33114 | 33114-MNV- | MNV | 2 | CC | AA | 1 | 26544 | 3.77E-03 | 31.4999999 | WT | FALSE | CAN | JSY13 | 4 | WT-Rep4 | GCC | nope | >1bp-ins | >1bpdel | none | MNV | ACTGGGCCG | CCGGCGT | CGG | GCC | CC | none | NA | NA_GCC | MNV_CCGGCGT | 9 | FALSE | FALSE |
| 169_v.txt | 33159 | 33159-MNV- | MNV | 2 | CC | AA | 2 | 26546 | 7.53E-03 | 30.9996881 | WT | FALSE | CAN | JSY13 | 4 | WT-Rep4 | ACC | nope | >1bp-ins | >1bpdel | none | MNV | CCTGTACCA | CCAATAG | TGG | ACC | CC | none | NA | NA_ACC | MNV_CCAATAG | 9 | FALSE | FALSE |
| 169_v.txt | 31886 | 31886-Insert | Insertion | 1 | - | A | 4 | 3301 | 0.1211754 | 44.5 | WT | FALSE | CAN | JSY13 | 4 | WT-Rep4 | CAA | nope | A/T+1 | >1bpdel | none | A/T+1 | GGCAGCAAA | AAAGCTA | GTT | CAA | - | T | NA | NA_CAA | A/T+1_AAAGCTA | 9 | FALSE | FALSE |
| 169_v.txt | 31974 | 31974-Insert | Insertion | 2 | - | AA | 3 | 19698 | 0.01522997 | 56.5 | WT | FALSE | CAN | JSY13 | 4 | WT-Rep4 | CAA | nope | >1bp-ins | >1bpdel | none | >1bp-ins | TAAGCCAAG | AAGCCGG | GTT | CAA | - | none | NA | NA_CAA | >1bp-ins_AAGCCGG | 9 | FALSE | FALSE |
| 169_v.txt | 32080 | 32080-Insert | Insertion | 1 | - | A | 6 | 19698 | 0.03045995 | 64 | WT | FALSE | CAN | JSY13 | 4 | WT-Rep4 | GCA | nope | A/T+1 | >1bpdel | none | A/T+1 | ACCATGCAA | CAAAAAA | CGT | GCA | - | T | NA | NA_GCA | A/T+1_CAAAAAA | 9 | FALSE | FALSE |
| 169_v.txt | 32081 | 32081-Insert | Insertion | 1 | - | A | 55 | 19698 | 0.27921616 | 63.7454546 | WT | FALSE | CAN | JSY13 | 4 | WT-Rep4 | CAA | nope | A/T+1 | >1bpdel | none | A/T+1 | CCATGCAAA | AAAAAAG | GTT | CAA | - | T | NA | NA_CAA | A/T+1_AAAAAAG | 9 | FALSE | FALSE |
| 169_v.txt | 32126 | 32126-Insert | Insertion | 2 | - | TT | 132 | 19698 | 0.67011879 | 57.7110266 | WT | FALSE | CAN | JSY13 | 4 | WT-Rep4 | CTT | nope | >1bp-ins | >1bpdel | none | >1bp-ins | GAAAACTTT | TTTGTCA | GAA | CTT | - | none | NA | NA_CTT | >1bp-ins_TTTGTCA | 9 | FALSE | FALSE |
| 169_v.txt | 32160 | 32160-Insert | Insertion | 1 | - | A | 6 | 19698 | 0.03045995 | 48.6666667 | WT | FALSE | CAN | JSY13 | 4 | WT-Rep4 | CAA | nope | A/T+1 | >1bpdel | none | A/T+1 | TAAGCCAAA | AAAGCGC | GTT | CAA | - | T | NA | NA_CAA | A/T+1_AAAGCGC | 9 | FALSE | FALSE |
| 169_v.txt | 32160 | 32160-Insert | Insertion | 2 | - | AA | 44 | 19698 | 0.22337293 | 57.1818182 | WT | FALSE | CAN | JSY13 | 4 | WT-Rep4 | CAA | nope | >1bp-ins | >1bpdel | none | >1bp-ins | TAAGCCAAA | AAAGCGC | GTT | CAA | - | none | NA | NA_CAA | >1bp-ins_AAAGCGC | 9 | FALSE | FALSE |
| 169_v.txt | 32184 | 32184-Insert | Insertion | 1 | - | A | 6 | 19698 | 0.03045995 | 42.3333333 | WT | FALSE | CAN | JSY13 | 4 | WT-Rep4 | GAA | nope | A/T+1 | >1bpdel | none | A/T+1 | GTAACGAAA | AAAACTG | CTT | GAA | - | T | NA | NA_GAA | A/T+1_AAAACTG | 9 | FALSE | FALSE |
| 169_v.txt | 32498 | 32498-Insert | Insertion | 1 | - | T | 28 | 51504 | 0.05436471 | 45 | WT | FALSE | CAN | JSY13 | 4 | WT-Rep4 | CTT | nope | A/T+1 | >1bpdel | none | A/T+1 | AACAACTTT | TTTTTTG | GAA | CTT | - | A | NA | NA_CTT | A/T+1_TTTTTTG | 9 | FALSE | FALSE |
| 169_v.txt | 32676 | 32676-Insert | Insertion | 31 | - | CTCCAATAA | 3 | 12117 | 0.0247586 | 63.6451613 | WT | FALSE | CAN | JSY13 | 4 | WT-Rep4 | TCT | nope | >1bp-ins | >1bpdel | none | >1bp-ins | GGGTTTCTC | CTCCAAT | AGA | TCT | - | none | NA | NA_TCT | >1bp-ins_CTCCAAT | 9 | FALSE | FALSE |
| 169_v.txt | 32842 | 32842-Insert | Insertion | 1 | - | A | 15 | 12117 | 0.12379302 | 39.5333333 | WT | FALSE | CAN | JSY13 | 4 | WT-Rep4 | CAA | nope | A/T+1 | >1bpdel | none | A/T+1 | TTACCCAAA | AAAAAAT | GTT | CAA | - | T | NA | NA_CAA | A/T+1_AAAAAAT | 9 | FALSE | FALSE |
| 169_v.txt | 33046 | 33046-Insert | Insertion | 24 | - | CCAAGGACT | 9 | 26544 | 0.03390597 | 62.9166667 | WT | FALSE | CAN | JSY13 | 4 | WT-Rep4 | CCC | nope | >1bp-ins | >1bpdel | none | >1bp-ins | TTTCACCCAA | CCAAGGA | GGG | CCC | - | none | NA | NA_CCC | >1bp-ins_CCAAGGA | 9 | FALSE | FALSE |
| 169_v.txt | 33138 | 33138-Insert | Insertion | 1 | - | A | 15 | 26544 | 0.05650995 | 41.3333333 | WT | FALSE | CAN | JSY13 | 4 | WT-Rep4 | TAA | nope | A/T+1 | >1bpdel | none | A/T+1 | GTGGATAAA | AAACCAA | ATT | TAA | - | T | NA | NA_TAA | A/T+1_AAACCAA | 9 | FALSE | FALSE |
| 169_v.txt | 33206 | 33206-Insert | Insertion | 2 | - | CT | 15 | 16226 | 0.09244423 | 37.4 | WT | FALSE | CAN | JSY13 | 4 | WT-Rep4 | GCT | nope | >1bp-ins | >1bpdel | none | >1bp-ins | CTTAAGCTCT | CTCTCTC | CGA | GCT | - | none | NA | NA_GCT | >1bp-ins_CTCTCTC | 9 | FALSE | FALSE |
| 169_v.txt | 32337 | 32337-Deleti | Deletion | 2 | AT | - | 21 | 51504 | 0.04077353 | 62.3333333 | WT | FALSE | CAN | JSY13 | 4 | WT-Rep4 | GAT | nope | >1bp-ins | >1bpdel | none | >1bpdel | TTGAAGATA | ATATGTG | CTA | GAT | AT | none | NA | NA_GAT | >1bpdel_ATATGTG | 9 | FALSE | FALSE |
| 169_v.txt | 32358 | 32358-Deleti | Deletion | 2 | CC | - | 58 | 51504 | 0.11261261 | 62.7068966 | WT | FALSE | CAN | JSY13 | 4 | WT-Rep4 | ACC | nope | >1bp-ins | >1bpdel | none | >1bpdel | TTTGTACCAG | CCAGAGT | TGG | ACC | CC | none | NA | NA_ACC | >1bpdel_CCAGAGT | 9 | FALSE | FALSE |
| 169_v.txt | 32498 | 32498-Deleti | Deletion | 2 | TT | - | 60 | 51504 | 0.11649581 | 45.3833333 | WT | FALSE | CAN | JSY13 | 4 | WT-Rep4 | CTT | nope | >1bp-ins | >1bpdel | none | >1bpdel | AACAACTTTT | TTTTTTG | GAA | CTT | TT | none | NA | NA_CTT | >1bpdel_TTTTTTG | 9 | FALSE | FALSE |
| 169_v.txt | 32953 | 32953-Deleti | Deletion | 2 | TG | - | 25 | 26544 | 0.09418324 | 55.36 | WT | FALSE | CAN | JSY13 | 4 | WT-Rep4 | ATG | nope | >1bp-ins | >1bpdel | none | >1bpdel | AATACATGT | TGTAACC | TAC | ATG | TG | none | NA | NA_ATG | >1bpdel_TGTAACC | 9 | FALSE | FALSE |
| 169_v.txt | 33089 | 33089-Deleti | Deletion | 2 | AT | - | 10 | 26544 | 0.0376733 | 47.7000363 | WT | FALSE | CAN | JSY13 | 4 | WT-Rep4 | AAT | nope | >1bp-ins | >1bpdel | none | >1bpdel | AAATAAATA | ATATGAT | TTA | AAT | AT | none | NA | NA_AAT | >1bpdel_ATATGAT | 9 | FALSE | FALSE |
| 169_v.txt | 33206 | 33206-Deleti | Deletion | 2 | CT | - | 14 | 16226 | 0.08628128 | 37.6428571 | WT | FALSE | CAN | JSY13 | 4 | WT-Rep4 | GCT | nope | >1bp-ins | >1bpdel | none | >1bpdel | CTTAAGCTCT | CTCTCTC | CGA | GCT | CT | none | NA | NA_GCT | >1bpdel_CTCTCTC | 9 | FALSE | FALSE |
| 169_v.txt | 31806 | 31806-Deleti | Deletion | 1 | C | - | 2 | 3301 | 0.0605877 | 51 | WT | FALSE | CAN | JSY13 | 4 | WT-Rep4 | CCA | nope | >1bp-ins | G/C-1 | none | G/C-1 | ATCTTCCAAA | CAAATAA | GGT | CCA | C | none | nope | nope_CCA | G/C-1_CAAATAA | 9 | FALSE | FALSE |
| 169_v.txt | 31863 | 31863-Deleti | Deletion | 1 | A | - | 5 | 3301 | 0.15146925 | 42.2 | WT | FALSE | CAN | JSY13 | 4 | WT-Rep4 | GAA | nope | >1bp-ins | A/T-1 | none | A/T-1 | AACAGGAAA | AAAATAG | CTT | CTT | T | none | T>none | T>none_CTT | A/T-1_AAAATAG | 9 | FALSE | FALSE |
| 169_v.txt | 31972 | 31972-Deleti | Deletion | 1 | C | - | 6 | 19698 | 0.03045995 | 35.8333333 | WT | FALSE | CAN | JSY13 | 4 | WT-Rep4 | GCC | nope | >1bp-ins | G/C-1 | none | G/C-1 | AATAAGCCA | CCAAGCC | CGG | GCC | C | none | nope | nope_GCC | G/C-1_CCAAGCC | 9 | FALSE | FALSE |

| | | | | | | | | | | | | | | | | | | | | | | | | | | | | | | | | | | |
|---|---|---|---|---|---|---|---|---|---|---|---|---|---|---|---|---|---|---|---|---|---|---|---|---|---|---|---|---|---|---|---|---|---|---|
| 169_v.txt | 32081 | 32081-Deleti | Deletion | 1 | A | - | 19 | 19698 | 0.09645649 | 63.8947369 | WT | FALSE | CAN | JSY13 | 4 | WT-Rep4 | CAA | nope | >1bp-ins | A/T-1 | none | A/T-1 | CCATGCAAA | AAAAAAG | GTT | GTT | T | none | T>none | T>none_GTT | A/T-1_AAAAAAG | 9 | FALSE | FALSE |
| 169_v.txt | 32224 | 32224-Deleti | Deletion | 1 | A | - | 6 | 19698 | 0.03045995 | 37.8333336 | WT | FALSE | CAN | JSY13 | 4 | WT-Rep4 | GAC | nope | >1bp-ins | A/T-1 | none | A/T-1 | TCCTTGACAG | ACAGGAA | CTG | CTG | T | none | T>none | T>none_CTG | A/T-1_ACAGGAA | 9 | FALSE | FALSE |
| 169_v.txt | 32289 | 32289-Deleti | Deletion | 1 | T | - | 7 | 51504 | 0.01359118 | 48.7142857 | WT | FALSE | CAN | JSY13 | 4 | WT-Rep4 | ATT | nope | >1bp-ins | A/T-1 | none | A/T-1 | TAAATATTTG | TTTGAAT | TAA | ATT | T | none | nope | nope_ATT | A/T-1_TTTGAAT | 9 | FALSE | FALSE |
| 169_v.txt | 32313 | 32313-Deleti | Deletion | 1 | G | - | 8 | 51504 | 0.01553277 | 62.25 | WT | FALSE | CAN | JSY13 | 4 | WT-Rep4 | GGT | nope | >1bp-ins | G/C-1 | none | G/C-1 | ATAATGGTT | GTTGTTA | CCA | CCA | C | none | C>none | C>none_CCA | G/C-1_GTTGTTA | 9 | FALSE | FALSE |
| 169_v.txt | 32346 | 32346-Deleti | Deletion | 1 | A | - | 8 | 51504 | 0.01553277 | 62.625 | WT | FALSE | CAN | JSY13 | 4 | WT-Rep4 | CAA | nope | >1bp-ins | A/T-1 | none | A/T-1 | TGTGGCAAA | AAAACCT | GTT | GTT | T | none | T>none | T>none_GTT | A/T-1_AAAACCT | 9 | FALSE | FALSE |
| 169_v.txt | 32358 | 32358-Deleti | Deletion | 1 | C | - | 5 | 51503 | 9.71E-03 | 58.8000053 | WT | FALSE | CAN | JSY13 | 4 | WT-Rep4 | ACC | nope | >1bp-ins | G/C-1 | none | G/C-1 | TTTGTACCAG | CCAGAGT | TGG | ACC | C | none | nope | nope_ACC | G/C-1_CCAGAGT | 9 | FALSE | FALSE |
| 169_v.txt | 32369 | 32369-Deleti | Deletion | 1 | A | - | 26 | 51504 | 0.05048152 | 62.9230769 | WT | FALSE | CAN | JSY13 | 4 | WT-Rep4 | CAA | nope | >1bp-ins | A/T-1 | none | A/T-1 | GTTCTCAATA | AATAGCA | GTT | GTT | T | none | T>none | T>none_GTT | A/T-1_AATAGCA | 9 | FALSE | FALSE |
| 169_v.txt | 32442 | 32442-Deleti | Deletion | 1 | A | - | 11 | 51504 | 0.02135757 | 56.8181818 | WT | FALSE | CAN | JSY13 | 4 | WT-Rep4 | TAA | nope | >1bp-ins | A/T-1 | none | A/T-1 | GGAACTAAA | AAAAGTC | ATT | ATT | T | none | T>none | T>none_ATT | A/T-1_AAAAGTC | 9 | FALSE | FALSE |
| 169_v.txt | 32473 | 32473-Deleti | Deletion | 1 | A | - | 3 | 51504 | 5.83E-03 | 38.0000038 | WT | FALSE | CAN | JSY13 | 4 | WT-Rep4 | TAG | nope | >1bp-ins | A/T-1 | none | A/T-1 | CAATGTAGA | AGAAGGT | ATC | ATC | T | none | T>none | T>none_ATC | A/T-1_AGAAGGT | 9 | FALSE | FALSE |
| 169_v.txt | 32490 | 32490-Deleti | Deletion | 1 | A | - | 10 | 51504 | 0.01941597 | 46.2 | WT | FALSE | CAN | JSY13 | 4 | WT-Rep4 | GAA | nope | >1bp-ins | A/T-1 | none | A/T-1 | ATACGGAAA | AAAACAA | CTT | CTT | T | none | T>none | T>none_CTT | A/T-1_AAAACAA | 9 | FALSE | FALSE |
| 169_v.txt | 32498 | 32498-Deleti | Deletion | 1 | T | - | 89 | 51504 | 0.17280211 | 43.3370733 | WT | FALSE | CAN | JSY13 | 4 | WT-Rep4 | CTT | nope | >1bp-ins | A/T-1 | none | A/T-1 | AACAACTTTT | TTTTTTG | GAA | CTT | T | none | nope | nope_CTT | A/T-1_TTTTTTG | 9 | FALSE | FALSE |
| 169_v.txt | 32528 | 32528-Deleti | Deletion | 1 | G | - | 9 | 51504 | 0.01747437 | 42.4444435 | WT | FALSE | CAN | JSY13 | 4 | WT-Rep4 | TGG | nope | >1bp-ins | G/C-1 | none | G/C-1 | TTTTCTGGGG | GGGGTTT | ACC | ACC | C | none | C>none | C>none_ACC | G/C-1_GGGGTTT | 9 | FALSE | FALSE |
| 169_v.txt | 32598 | 32598-Deleti | Deletion | 1 | A | - | 1790 | 12117 | 14.7726335 | 39.9162011 | WT | FALSE | CAN | JSY13 | 4 | WT-Rep4 | CAA | nope | >1bp-ins | A/T-1 | none | A/T-1 | TTAATCAAA | AAAGAGG | GTT | GTT | T | none | T>none | T>none_GTT | A/T-1_AAAGAGG | 9 | FALSE | FALSE |
| 169_v.txt | 32631 | 32631-Deleti | Deletion | 1 | T | - | 4 | 12117 | 0.03301147 | 44.25 | WT | FALSE | CAN | JSY13 | 4 | WT-Rep4 | GTT | nope | >1bp-ins | A/T-1 | none | A/T-1 | CCTTCGTTTT | TTTTTAT | CAA | GTT | T | none | nope | nope_GTT | A/T-1_TTTTTAT | 9 | FALSE | FALSE |
| 169_v.txt | 32658 | 32658-Deleti | Deletion | 1 | C | - | 5 | 12117 | 0.04126434 | 58.8 | WT | FALSE | CAN | JSY13 | 4 | WT-Rep4 | ACC | nope | >1bp-ins | G/C-1 | none | G/C-1 | CCTGGACCC | CCCCAGG | TGG | ACC | C | none | nope | nope_ACC | G/C-1_CCCCAGG | 9 | FALSE | FALSE |
| 169_v.txt | 32677 | 32677-Deleti | Deletion | 1 | T | - | 10 | 12117 | 0.08252868 | 60.2 | WT | FALSE | CAN | JSY13 | 4 | WT-Rep4 | CTC | nope | >1bp-ins | A/T-1 | none | A/T-1 | GGTTTCTCCA | TCCAATA | GAG | CTC | T | none | nope | nope_CTC | A/T-1_TCCAATA | 9 | FALSE | FALSE |
| 169_v.txt | 32728 | 32728-Deleti | Deletion | 1 | A | - | 5 | 12117 | 0.04126434 | 61 | WT | FALSE | CAN | JSY13 | 4 | WT-Rep4 | CAA | nope | >1bp-ins | A/T-1 | none | A/T-1 | CCATACAAA | AAAAACA | GTT | GTT | T | none | T>none | T>none_GTT | A/T-1_AAAAACA | 9 | FALSE | FALSE |
| 169_v.txt | 32811 | 32811-Deleti | Deletion | 1 | G | - | 12 | 12117 | 0.09903442 | 50.8333333 | WT | FALSE | CAN | JSY13 | 4 | WT-Rep4 | AGG | nope | >1bp-ins | G/C-1 | none | G/C-1 | TTGACAGGG | GGGAACA | TCC | TCC | C | none | C>none | C>none_TCC | G/C-1_GGGAACA | 9 | FALSE | FALSE |
| 169_v.txt | 32842 | 32842-Deleti | Deletion | 1 | A | - | 166 | 12117 | 1.36997607 | 42.1325301 | WT | FALSE | CAN | JSY13 | 4 | WT-Rep4 | CAA | nope | >1bp-ins | A/T-1 | none | A/T-1 | TTACCCAAA | AAAAAAT | GTT | GTT | T | none | T>none | T>none_GTT | A/T-1_AAAAAAT | 9 | FALSE | FALSE |
| 169_v.txt | 32884 | 32884-Deleti | Deletion | 1 | A | - | 29 | 26544 | 0.10925256 | 36.2758621 | WT | FALSE | CAN | JSY13 | 4 | WT-Rep4 | CAA | nope | >1bp-ins | A/T-1 | none | A/T-1 | ACGTCCAAA | AAAATTG | GTT | GTT | T | none | T>none | T>none_GTT | A/T-1_AAAATTG | 9 | FALSE | FALSE |
| 169_v.txt | 32901 | 32901-Deleti | Deletion | 1 | C | - | 3 | 26544 | 0.01130199 | 38 | WT | FALSE | CAN | JSY13 | 4 | WT-Rep4 | GCC | nope | >1bp-ins | G/C-1 | none | G/C-1 | ACTTGGCCA | CCAACTA | CGG | GCC | C | none | nope | nope_GCC | G/C-1_CCAACTA | 9 | FALSE | FALSE |
| 169_v.txt | 32924 | 32924-Deleti | Deletion | 1 | A | - | 31 | 26544 | 0.11678722 | 51.2580645 | WT | FALSE | CAN | JSY13 | 4 | WT-Rep4 | CAA | nope | >1bp-ins | A/T-1 | none | A/T-1 | CAGGGCAAA | AAAAGTG | GTT | GTT | T | none | T>none | T>none_GTT | A/T-1_AAAAGTG | 9 | FALSE | FALSE |
| 169_v.txt | 32967 | 32967-Deleti | Deletion | 1 | G | - | 3 | 26544 | 0.01130199 | 61.6666667 | WT | FALSE | CAN | JSY13 | 4 | WT-Rep4 | CGC | nope | >1bp-ins | G/C-1 | none | G/C-1 | TTGGCCGCA | GCACCAA | GCG | GCG | C | none | C>none | C>none_GCG | G/C-1_GCACCAA | 9 | FALSE | FALSE |
| 169_v.txt | 32995 | 32995-Deleti | Deletion | 1 | G | - | 26 | 26544 | 0.09795057 | 62.3076923 | WT | FALSE | CAN | JSY13 | 4 | WT-Rep4 | TGT | nope | >1bp-ins | G/C-1 | none | G/C-1 | ATCTTTGTGA | GTGAGAA | ACA | ACA | C | none | C>none | C>none_ACA | G/C-1_GTGAGAA | 9 | FALSE | FALSE |
| 169_v.txt | 33116 | 33116-Deleti | Deletion | 1 | G | - | 20 | 26544 | 0.07534659 | 37.1000042 | WT | FALSE | CAN | JSY13 | 4 | WT-Rep4 | CGG | nope | >1bp-ins | G/C-1 | none | G/C-1 | TGGGCCGGC | GGCGTTG | GCC | GCC | C | none | C>none | C>none_GCC | G/C-1_GGCGTTG | 9 | FALSE | FALSE |
| 169_v.txt | 33474 | 33474-Deleti | Deletion | 1 | C | - | 37 | 16227 | 0.22801504 | 30.5675676 | WT | FALSE | CAN | JSY13 | 4 | WT-Rep4 | GCC | nope | >1bp-ins | G/C-1 | none | G/C-1 | GCTATGCCTT | CCTTTTT | CGG | GCC | C | none | nope | nope_GCC | G/C-1_CCTTTTT | 9 | FALSE | FALSE |
| 188_v.txt | 31723 | 31723-SNV-A | SNV | 1 | A | C | 2 | 4236 | 0.04721435 | 64 | WT | FALSE | CAN | JSY13 | 5 | WT-Rep5 | CAA | A>C | G/C+1 | A/T-1 | TA>GC | TA>GC | TGTCCCAAA | AAAAAGT | GTT | GTT | T | G | T>G | T>G_GTT | TA>GC_AAAAAGT | 9 | FALSE | FALSE |
| 188_v.txt | 31967 | 31967-SNV-A | SNV | 1 | A | C | 17 | 12088 | 0.14063534 | 29.1176471 | WT | FALSE | CAN | JSY13 | 5 | WT-Rep5 | AAT | A>C | G/C+1 | A/T-1 | TA>GC | TA>GC | CGCATAATA | ATAAGCC | TTA | TTA | T | G | T>G | T>G_TTA | TA>GC_ATAAGCC | 9 | FALSE | FALSE |
| 188_v.txt | 32288 | 32288-SNV-A | SNV | 1 | A | C | 38 | 31738 | 0.11973029 | 45.1052632 | WT | FALSE | CAN | JSY13 | 5 | WT-Rep5 | TAT | A>C | G/C+1 | A/T-1 | TA>GC | TA>GC | GTAAATATTT | ATTTGAA | ATA | ATA | T | G | T>G | T>G_ATA | TA>GC_ATTTGAA | 9 | FALSE | FALSE |
| 188_v.txt | 32498 | 32498-SNV-T | SNV | 1 | T | G | 2 | 31738 | 6.30E-03 | 37.5014345 | WT | FALSE | CAN | JSY13 | 5 | WT-Rep5 | CTT | T>G | G/C+1 | A/T-1 | TA>GC | TA>GC | AACAACTTTT | TTTTTTG | GAA | CTT | T | C | T>G | T>G_CTT | TA>GC_TTTTTTG | 9 | FALSE | FALSE |
| 188_v.txt | 32631 | 32631-SNV-T | SNV | 1 | T | G | 1 | 30744 | 3.25E-03 | 35.999566 | WT | FALSE | CAN | JSY13 | 5 | WT-Rep5 | GTT | T>G | G/C+1 | A/T-1 | TA>GC | TA>GC | CCTTCGTTTT | TTTTTAT | CAA | GTT | T | C | T>G | T>G_GTT | TA>GC_TTTTTAT | 9 | FALSE | FALSE |
| 188_v.txt | 32640 | 32640-SNV-T | SNV | 1 | T | G | 2 | 30744 | 6.51E-03 | 51 | WT | FALSE | CAN | JSY13 | 5 | WT-Rep5 | CTT | T>G | G/C+1 | A/T-1 | TA>GC | TA>GC | TTATCCTTAG | TTAGATA | GAA | CTT | T | C | T>G | T>G_CTT | TA>GC_TTAGATA | 9 | FALSE | FALSE |
| 188_v.txt | 32662 | 32662-SNV-A | SNV | 1 | A | C | 3 | 30744 | 9.76E-03 | 40.3349832 | WT | FALSE | CAN | JSY13 | 5 | WT-Rep5 | CAG | A>C | G/C+1 | A/T-1 | TA>GC | TA>GC | GACCCCAGG | AGGCACC | GTC | GTC | T | G | T>G | T>G_GTC | TA>GC_AGGCACC | 9 | FALSE | FALSE |
| 188_v.txt | 32884 | 32884-SNV-A | SNV | 1 | A | C | 5 | 25521 | 0.01959171 | 36.9996477 | WT | FALSE | CAN | JSY13 | 5 | WT-Rep5 | CAA | A>C | G/C+1 | A/T-1 | TA>GC | TA>GC | ACGTCCAAA | AAAATTG | GTT | GTT | T | G | T>G | T>G_GTT | TA>GC_AAAATTG | 9 | FALSE | FALSE |
| 188_v.txt | 32913 | 32913-SNV-A | SNV | 1 | A | C | 24 | 25520 | 0.09404389 | 26.375 | WT | FALSE | CAN | JSY13 | 5 | WT-Rep5 | AAG | A>C | G/C+1 | A/T-1 | TA>GC | TA>GC | ACACTAAGT | AGTTCCA | TTC | TTC | T | G | T>G | T>G_TTC | TA>GC_AGTTCCA | 9 | FALSE | FALSE |
| 188_v.txt | 33003 | 33003-SNV-A | SNV | 1 | A | C | 4 | 25520 | 0.01567398 | 64 | WT | FALSE | CAN | JSY13 | 5 | WT-Rep5 | AAC | A>C | G/C+1 | A/T-1 | TA>GC | TA>GC | GAGAAAACT | ACTGTGA | TTG | TTG | T | G | T>G | T>G_TTG | TA>GC_ACTGTGA | 9 | FALSE | FALSE |
| 188_v.txt | 33099 | 33099-SNV-A | SNV | 1 | A | C | 16 | 25520 | 0.06269593 | 43.3125 | WT | FALSE | CAN | JSY13 | 5 | WT-Rep5 | AAG | A>C | G/C+1 | A/T-1 | TA>GC | TA>GC | GATATAAGA | AGAGCGC | TTC | TTC | T | G | T>G | T>G_TTC | TA>GC_AGAGCGC | 9 | FALSE | FALSE |
| 188_v.txt | 33204 | 33204-SNV-A | SNV | 1 | A | C | 147 | 17251 | 0.85212452 | 36.462585 | WT | FALSE | CAN | JSY13 | 5 | WT-Rep5 | AAG | A>C | G/C+1 | A/T-1 | TA>GC | TA>GC | TGCTTAAGCT | AGCTCTC | TTC | TTC | T | G | T>G | T>G_TTC | TA>GC_AGCTCTC | 9 | FALSE | FALSE |
| 188_v.txt | 33269 | 33269-SNV-A | SNV | 1 | A | C | 138 | 17251 | 0.79995363 | 15.7608696 | WT | FALSE | CAN | JSY13 | 5 | WT-Rep5 | TAG | A>C | G/C+1 | A/T-1 | TA>GC | TA>GC | CTCCATAGA | AGAGAAC | ATC | ATC | T | G | T>G | T>G_ATC | TA>GC_AGAGAAC | 9 | FALSE | FALSE |
| 188_v.txt | 33477 | 33477-SNV-T | SNV | 1 | T | G | 5 | 17247 | 0.02899055 | 36.6 | WT | FALSE | CAN | JSY13 | 5 | WT-Rep5 | TTT | T>G | G/C+1 | A/T-1 | TA>GC | TA>GC | ATGCCTTTTT | TTTTTTT | AAA | TTT | T | C | T>G | T>G_TTT | TA>GC_TTTTTTT | 9 | FALSE | FALSE |
| 188_v.txt | 32081 | 32081-SNV-A | SNV | 1 | A | G | 6 | 12088 | 0.049636 | 63.1663697 | WT | FALSE | CAN | JSY13 | 5 | WT-Rep5 | CAA | A>G | G/C+1 | A/T-1 | TA>CG | TA>CG | CCATGCAAA | AAAAAAG | GTT | GTT | T | C | T>C | T>C_GTT | TA>CG_AAAAAAG | 9 | FALSE | FALSE |
| 188_v.txt | 32293 | 32293-SNV-A | SNV | 1 | A | G | 22 | 31738 | 0.06931754 | 55.7727273 | WT | FALSE | CAN | JSY13 | 5 | WT-Rep5 | GAA | A>G | G/C+1 | A/T-1 | TA>CG | TA>CG | TATTTGAATT | AATTTGC | CTT | CTT | T | C | T>C | T>C_CTT | TA>CG_AATTTGC | 9 | FALSE | FALSE |
| 188_v.txt | 32381 | 32381-SNV-A | SNV | 1 | A | G | 9 | 31736 | 0.02835896 | 53.7773338 | WT | FALSE | CAN | JSY13 | 5 | WT-Rep5 | TAA | A>G | G/C+1 | A/T-1 | TA>CG | TA>CG | AATAATAAA | AAAGGGA | ATT | ATT | T | C | T>C | T>C_ATT | TA>CG_AAAGGGA | 9 | FALSE | FALSE |
| 188_v.txt | 32382 | 32382-SNV-A | SNV | 1 | A | G | 3 | 31738 | 9.45E-03 | 46.3332754 | WT | FALSE | CAN | JSY13 | 5 | WT-Rep5 | AAA | A>G | G/C+1 | A/T-1 | TA>CG | TA>CG | ATAATAAAG | AAGGGAG | TTT | TTT | T | C | T>C | T>C_TTT | TA>CG_AAGGGAG | 9 | FALSE | FALSE |
| 188_v.txt | 32383 | 32383-SNV-A | SNV | 1 | A | G | 5 | 31738 | 0.01575399 | 63.9990916 | WT | FALSE | CAN | JSY13 | 5 | WT-Rep5 | AAG | A>G | G/C+1 | A/T-1 | TA>CG | TA>CG | TAATAAAGG | AGGGAGA | TTC | TTC | T | C | T>C | T>C_TTC | TA>CG_AGGGAGA | 9 | FALSE | FALSE |
| 188_v.txt | 32387 | 32387-SNV-A | SNV | 1 | A | G | 18 | 31737 | 0.05671614 | 51.6066409 | WT | FALSE | CAN | JSY13 | 5 | WT-Rep5 | GAG | A>G | G/C+1 | A/T-1 | TA>CG | TA>CG | AAAGGGAGA | AGAAGTA | CTC | CTC | T | C | T>C | T>C_CTC | TA>CG_AGAAGTA | 9 | FALSE | FALSE |
| 188_v.txt | 32389 | 32389-SNV-A | SNV | 1 | A | G | 13 | 31738 | 0.04096036 | 30.6888679 | WT | FALSE | CAN | JSY13 | 5 | WT-Rep5 | GAA | A>G | G/C+1 | A/T-1 | TA>CG | TA>CG | AGGGAGAA | AAGTAGA | CTT | CTT | T | C | T>C | T>C_CTT | TA>CG_AAGTAGA | 9 | FALSE | FALSE |
| 188_v.txt | 32390 | 32390-SNV-A | SNV | 1 | A | G | 5 | 31738 | 0.01575399 | 58.8000409 | WT | FALSE | CAN | JSY13 | 5 | WT-Rep5 | AAG | A>G | G/C+1 | A/T-1 | TA>CG | TA>CG | GGGAGAAGT | AGTAGAA | TTC | TTC | T | C | T>C | T>C_TTC | TA>CG_AGTAGAA | 9 | FALSE | FALSE |
| 188_v.txt | 32392 | 32392-SNV-T | SNV | 1 | T | C | 3 | 31738 | 9.45E-03 | 43.9999619 | WT | FALSE | CAN | JSY13 | 5 | WT-Rep5 | GTA | T>C | G/C+1 | A/T-1 | TA>CG | TA>CG | GAGAAGTAG | TAGAAAC | CAT | GTA | T | G | T>C | T>C_GTA | TA>CG_TAGAAAC | 9 | FALSE | FALSE |
| 188_v.txt | 32393 | 32393-SNV-A | SNV | 1 | A | G | 9 | 31738 | 0.02835717 | 55.4441858 | WT | FALSE | CAN | JSY13 | 5 | WT-Rep5 | TAG | A>G | G/C+1 | A/T-1 | TA>CG | TA>CG | AGAAGTAGA | AGAAACG | ATC | ATC | T | C | T>C | T>C_ATC | TA>CG_AGAAACG | 9 | FALSE | FALSE |
| 188_v.txt | 32445 | 32445-SNV-A | SNV | 1 | A | G | 40 | 31738 | 0.12603189 | 52.4 | WT | FALSE | CAN | JSY13 | 5 | WT-Rep5 | AAG | A>G | G/C+1 | A/T-1 | TA>CG | TA>CG | ACTAAAAGT | AGTCCAA | TTC | TTC | T | C | T>C | T>C_TTC | TA>CG_AGTCCAA | 9 | FALSE | FALSE |
| 188_v.txt | 32631 | 32631-SNV-T | SNV | 1 | T | C | 6 | 30744 | 0.019516 | 19.999939 | WT | FALSE | CAN | JSY13 | 5 | WT-Rep5 | GTT | T>C | G/C+1 | A/T-1 | TA>CG | TA>CG | CCTTCGTTTT | TTTTTAT | CAA | GTT | T | G | T>C | T>C_GTT | TA>CG_TTTTTAT | 9 | FALSE | FALSE |
| 188_v.txt | 32662 | 32662-SNV-A | SNV | 1 | A | G | 10 | 30744 | 0.03252667 | 51.1978678 | WT | FALSE | CAN | JSY13 | 5 | WT-Rep5 | CAG | A>G | G/C+1 | A/T-1 | TA>CG | TA>CG | GACCCCAGG | AGGCACC | GTC | GTC | T | C | T>C | T>C_GTC | TA>CG_AGGCACC | 9 | FALSE | FALSE |
| 188_v.txt | 32680 | 32680-SNV-A | SNV | 1 | A | G | 75 | 30744 | 0.24395004 | 60.466159 | WT | FALSE | CAN | JSY13 | 5 | WT-Rep5 | CAA | A>G | G/C+1 | A/T-1 | TA>CG | TA>CG | TTCTCCAATA | AATAACG | GTT | GTT | T | C | T>C | T>C_GTT | TA>CG_AATAACG | 9 | FALSE | FALSE |
| 188_v.txt | 32728 | 32728-SNV-A | SNV | 1 | A | G | 6 | 30742 | 0.01951727 | 56.9991407 | WT | FALSE | CAN | JSY13 | 5 | WT-Rep5 | CAA | A>G | G/C+1 | A/T-1 | TA>CG | TA>CG | CCATACAAA | AAAAACA | GTT | GTT | T | C | T>C | T>C_GTT | TA>CG_AAAAACA | 9 | FALSE | FALSE |
| 188_v.txt | 32799 | 32799-SNV-T | SNV | 1 | T | C | 45 | 30744 | 0.14637002 | 41.5343478 | WT | FALSE | CAN | JSY13 | 5 | WT-Rep5 | GTA | T>C | G/C+1 | A/T-1 | TA>CG | TA>CG | TCACCGTAAT | TAATATT | CAT | GTA | T | G | T>C | T>C_GTA | TA>CG_TAATATT | 9 | FALSE | FALSE |
| 188_v.txt | 33204 | 33204-SNV-A | SNV | 1 | A | G | 13 | 17251 | 0.07535795 | 35.9230769 | WT | FALSE | CAN | JSY13 | 5 | WT-Rep5 | AAG | A>G | G/C+1 | A/T-1 | TA>CG | TA>CG | TGCTTAAGCT | AGCTCTC | TTC | TTC | T | C | T>C | T>C_TTC | TA>CG_AGCTCTC | 9 | FALSE | FALSE |
| 188_v.txt | 32381 | 32381-SNV-A | SNV | 1 | A | T | 3 | 31736 | 9.45E-03 | 63.9967262 | WT | FALSE | CAN | JSY13 | 5 | WT-Rep5 | TAA | A>T | A/T+1 | A/T-1 | TA>AT | TA>AT | AATAATAAA | AAAGGGA | ATT | ATT | T | A | T>A | T>A_ATT | TA>AT_AAAGGGA | 9 | FALSE | FALSE |
| 188_v.txt | 32382 | 32382-SNV-A | SNV | 1 | A | T | 3 | 31738 | 9.45E-03 | 53.9995293 | WT | FALSE | CAN | JSY13 | 5 | WT-Rep5 | AAA | A>T | A/T+1 | A/T-1 | TA>AT | TA>AT | ATAATAAAG | AAGGGAG | TTT | TTT | T | A | T>A | T>A_TTT | TA>AT_AAGGGAG | 9 | FALSE | FALSE |
| 188_v.txt | 32389 | 32389-SNV-A | SNV | 1 | A | T | 2 | 31738 | 6.30E-03 | 63.9994279 | WT | FALSE | CAN | JSY13 | 5 | WT-Rep5 | GAA | A>T | A/T+1 | A/T-1 | TA>AT | TA>AT | AGGGAGAA | AAGTAGA | CTT | CTT | T | A | T>A | T>A_CTT | TA>AT_AAGTAGA | 9 | FALSE | FALSE |
| 188_v.txt | 32392 | 32392-SNV-T | SNV | 1 | T | A | 5 | 31738 | 0.01575399 | 49.3983424 | WT | FALSE | CAN | JSY13 | 5 | WT-Rep5 | GTA | T>A | A/T+1 | A/T-1 | TA>AT | TA>AT | GAGAAGTAG | TAGAAAC | CAT | GTA | T | T | T>A | T>A_GTA | TA>AT_TAGAAAC | 9 | FALSE | FALSE |
| 188_v.txt | 32680 | 32680-SNV-A | SNV | 1 | A | T | 1 | 30744 | 3.25E-03 | 63.9995157 | WT | FALSE | CAN | JSY13 | 5 | WT-Rep5 | CAA | A>T | A/T+1 | A/T-1 | TA>AT | TA>AT | TTCTCCAATA | AATAACG | GTT | GTT | T | A | T>A | T>A_GTT | TA>AT_AATAACG | 9 | FALSE | FALSE |
| 188_v.txt | 33065 | 33065-SNV-A | SNV | 1 | A | T | 14 | 25520 | 0.05485893 | 53.7857143 | WT | FALSE | CAN | JSY13 | 5 | WT-Rep5 | AAT | A>T | A/T+1 | A/T-1 | TA>AT | TA>AT | GACAGAATA | ATATGCC | TTA | TTA | T | A | T>A | T>A_TTA | TA>AT_ATATGCC | 9 | FALSE | FALSE |
| 188_v.txt | 33430 | 33430-SNV-T | SNV | 1 | T | A | 36 | 17251 | 0.20868356 | 40.8888889 | WT | FALSE | CAN | JSY13 | 5 | WT-Rep5 | TTC | T>A | A/T+1 | A/T-1 | TA>AT | TA>AT | TATGCTTCTC | TCTCCTC | AAG | TTC | T | T | T>A | T>A_TTC | TA>AT_TCTCCTC | 9 | FALSE | FALSE |
| 188_v.txt | 31844 | 31844-SNV-C | SNV | 1 | C | T | 28 | 4236 | 0.66100094 | 50.9285714 | WT | FALSE | CAN | JSY13 | 5 | WT-Rep5 | TCC | C>T | A/T+1 | G/C-1 | CG>TA | CG>TA | TAAGATCCA | CCAAACA | AGG | TCC | C | A | C>T | C>T_TCC | CG>TA_CCAAACA | 9 | FALSE | FALSE |
| 188_v.txt | 31962 | 31962-SNV-G | SNV | 1 | G | A | 15 | 12088 | 0.12409001 | 37.8 | WT | FALSE | CAN | JSY13 | 5 | WT-Rep5 | CGC | G>A | A/T+1 | G/C-1 | CG>TA | CG>TA | GTGGCCGCA | GCATAAT | GCG | GCG | C | T | C>T | C>T_GCG | CG>TA_GCATAAT | 9 | FALSE | FALSE |
| 188_v.txt | 31981 | 31981-SNV-G | SNV | 1 | G | A | 17 | 12088 | 0.14063534 | 47.2352941 | WT | FALSE | CAN | JSY13 | 5 | WT-Rep5 | GGC | G>A | A/T+1 | G/C-1 | CG>TA | CG>TA | AGCCGGGCA | GCATTAA | CCG | CCG | C | T | C>T | C>T_CCG | CG>TA_GCATTAA | 9 | FALSE | FALSE |
| 188_v.txt | 32041 | 32041-SNV-G | SNV | 1 | G | A | 16 | 12088 | 0.13236267 | 59.25 | WT | FALSE | CAN | JSY13 | 5 | WT-Rep5 | TGC | G>A | A/T+1 | G/C-1 | CG>TA | CG>TA | AAGCTTGCA | GCATAAA | ACG | ACG | C | T | C>T | C>T_ACG | CG>TA_GCATAAA | 9 | FALSE | FALSE |
| 188_v.txt | 32075 | 32075-SNV-C | SNV | 1 | C | T | 120 | 12088 | 0.99272005 | 61.9416667 | WT | FALSE | CAN | JSY13 | 5 | WT-Rep5 | ACC | C>T | A/T+1 | G/C-1 | CG>TA | CG>TA | AAATAACCA | CCATGCA | TGG | ACC | C | A | C>T | C>T_ACC | CG>TA_CCATGCA | 9 | FALSE | FALSE |
| 188_v.txt | 32088 | 32088-SNV-C | SNV | 1 | C | T | 45 | 12088 | 0.37227002 | 62.6444444 | WT | FALSE | CAN | JSY13 | 5 | WT-Rep5 | GCC | C>T | A/T+1 | G/C-1 | CG>TA | CG>TA | AAAAAGCCT | CCTGCAA | CGG | GCC | C | A | C>T | C>T_GCC | CG>TA_CCTGCAA | 9 | FALSE | FALSE |
| 188_v.txt | 32092 | 32092-SNV-C | SNV | 1 | C | T | 15 | 12088 | 0.12409001 | 59.8 | WT | FALSE | CAN | JSY13 | 5 | WT-Rep5 | GCA | C>T | A/T+1 | G/C-1 | CG>TA | CG>TA | AGCCTGCAA | CAACACC | CGT | GCA | C | A | C>T | C>T_GCA | CG>TA_CAACACC | 9 | FALSE | FALSE |
| 188_v.txt | 32114 | 32114-SNV-C | SNV | 1 | C | T | 72 | 12088 | 0.59563203 | 59.7361111 | WT | FALSE | CAN | JSY13 | 5 | WT-Rep5 | GCC | C>T | A/T+1 | G/C-1 | CG>TA | CG>TA | TAATAGCCAT | CCATTCG | CGG | GCC | C | A | C>T | C>T_GCC | CG>TA_CCATTCG | 9 | FALSE | FALSE |
| 188_v.txt | 32259 | 32259-SNV-C | SNV | 1 | C | T | 66 | 31737 | 0.20795916 | 31 | WT | FALSE | CAN | JSY13 | 5 | WT-Rep5 | ACC | C>T | A/T+1 | G/C-1 | CG>TA | CG>TA | GATAGACCA | CCAAATA | TGG | ACC | C | A | C>T | C>T_ACC | CG>TA_CCAAATA | 9 | FALSE | FALSE |
| 188_v.txt | 32271 | 32271-SNV-C | SNV | 1 | C | T | 131 | 31738 | 0.41275443 | 36.0458015 | WT | FALSE | CAN | JSY13 | 5 | WT-Rep5 | ACG | C>T | A/T+1 | G/C-1 | CG>TA | CG>TA | AAAATACGG | CGGGAAC | TGC | ACG | C | A | C>T | C>T_ACG | CG>TA_CGGGAAC | 9 | FALSE | FALSE |
| 188_v.txt | 32299 | 32299-SNV-C | SNV | 1 | C | T | 59 | 31738 | 0.18589703 | 52.661017 | WT | FALSE | CAN | JSY13 | 5 | WT-Rep5 | GCG | C>T | A/T+1 | G/C-1 | CG>TA | CG>TA | AATTTGCGG | CGGCAGA | CGC | GCG | C | A | C>T | C>T_GCG | CG>TA_CGGCAGA | 9 | FALSE | FALSE |
| 188_v.txt | 32385 | 32385-SNV-G | SNV | 1 | G | A | 6 | 31738 | 0.01890478 | 41.9999585 | WT | FALSE | CAN | JSY13 | 5 | WT-Rep5 | GGG | G>A | A/T+1 | G/C-1 | CG>TA | CG>TA | ATAAAGGGA | GGAGAAG | CCC | CCC | C | T | C>T | C>T_CCC | CG>TA_GGAGAAG | 9 | FALSE | FALSE |
| 188_v.txt | 32388 | 32388-SNV-G | SNV | 1 | G | A | 6 | 31738 | 0.01890478 | 37.4948587 | WT | FALSE | CAN | JSY13 | 5 | WT-Rep5 | AGA | G>A | A/T+1 | G/C-1 | CG>TA | CG>TA | AAGGGAGA | GAAGTAG | TCT | TCT | C | T | C>T | C>T_TCT | CG>TA_GAAGTAG | 9 | FALSE | FALSE |
| 188_v.txt | 32394 | 32394-SNV-G | SNV | 1 | G | A | 2 | 31738 | 6.30E-03 | 32.4999721 | WT | FALSE | CAN | JSY13 | 5 | WT-Rep5 | AGA | G>A | A/T+1 | G/C-1 | CG>TA | CG>TA | GAAGTAGAA | GAAACGT | TCT | TCT | C | T | C>T | C>T_TCT | CG>TA_GAAACGT | 9 | FALSE | FALSE |
| 188_v.txt | 32449 | 32449-SNV-C | SNV | 1 | C | T | 44 | 31738 | 0.13863508 | 49.8863636 | WT | FALSE | CAN | JSY13 | 5 | WT-Rep5 | CCA | C>T | A/T+1 | G/C-1 | CG>TA | CG>TA | AAAGTCCAA | CAATGAA | GGT | CCA | C | A | C>T | C>T_CCA | CG>TA_CAATGAA | 9 | FALSE | FALSE |
| 188_v.txt | 32466 | 32466-SNV-C | SNV | 1 | C | T | 43 | 31738 | 0.13548428 | 49.6744186 | WT | FALSE | CAN | JSY13 | 5 | WT-Rep5 | GCC | C>T | A/T+1 | G/C-1 | CG>TA | CG>TA | AGAGAGCCA | CCAATGT | CGG | GCC | C | A | C>T | C>T_GCC | CG>TA_CCAATGT | 9 | FALSE | FALSE |
| 188_v.txt | 32487 | 32487-SNV-C | SNV | 1 | C | T | 128 | 31738 | 0.40330204 | 46.0231783 | WT | FALSE | CAN | JSY13 | 5 | WT-Rep5 | ACG | C>T | A/T+1 | G/C-1 | CG>TA | CG>TA | AAGATACGG | CGGAAAA | TGC | ACG | C | A | C>T | C>T_ACG | CG>TA_CGGAAAA | 9 | FALSE | FALSE |
| 188_v.txt | 32488 | 32488-SNV-G | SNV | 1 | G | A | 19 | 31738 | 0.05986515 | 40.4736842 | WT | FALSE | CAN | JSY13 | 5 | WT-Rep5 | CGG | G>A | A/T+1 | G/C-1 | CG>TA | CG>TA | AGATACGGA | GGAAAAC | GCC | GCC | C | T | C>T | C>T_GCC | CG>TA_GGAAAAC | 9 | FALSE | FALSE |
| 188_v.txt | 32535 | 32535-SNV-G | SNV | 1 | G | A | 32 | 31738 | 0.10082551 | 37.3372585 | WT | FALSE | CAN | JSY13 | 5 | WT-Rep5 | TGC | G>A | A/T+1 | G/C-1 | CG>TA | CG>TA | GGGTTTGCA | GCAGCTT | ACG | ACG | C | T | C>T | C>T_ACG | CG>TA_GCAGCTT | 9 | FALSE | FALSE |
| 188_v.txt | 32536 | 32536-SNV-C | SNV | 1 | C | T | 43 | 31738 | 0.13548428 | 37.4879243 | WT | FALSE | CAN | JSY13 | 5 | WT-Rep5 | GCA | C>T | A/T+1 | G/C-1 | CG>TA | CG>TA | GGTTTGCAG | CAGCTTC | CGT | GCA | C | A | C>T | C>T_GCA | CG>TA_CAGCTTC | 9 | FALSE | FALSE |
| 188_v.txt | 32566 | 32566-SNV-C | SNV | 1 | C | T | 33 | 30736 | 0.10736596 | 36.030303 | WT | FALSE | CAN | JSY13 | 5 | WT-Rep5 | TCA | C>T | A/T+1 | G/C-1 | CG>TA | CG>TA | CTAGTTCAGT | CAGTACC | AGT | TCA | C | A | C>T | C>T_TCA | CG>TA_CAGTACC | 9 | FALSE | FALSE |
| 188_v.txt | 32590 | 32590-SNV-C | SNV | 1 | C | T | 44 | 30744 | 0.14311736 | 36.8181818 | WT | FALSE | CAN | JSY13 | 5 | WT-Rep5 | GCG | C>T | A/T+1 | G/C-1 | CG>TA | CG>TA | AGGCAGCGT | CGTTAAT | CGC | GCG | C | A | C>T | C>T_GCG | CG>TA_CGTTAAT | 9 | FALSE | FALSE |
| 188_v.txt | 32609 | 32609-SNV-C | SNV | 1 | C | T | 111 | 30744 | 0.36104606 | 46.8647311 | WT | FALSE | CAN | JSY13 | 5 | WT-Rep5 | CCC | C>T | A/T+1 | G/C-1 | CG>TA | CG>TA | GGAAACCCA | CCAACCT | GGG | CCC | C | A | C>T | C>T_CCC | CG>TA_CCAACCT | 9 | FALSE | FALSE |
| 188_v.txt | 32610 | 32610-SNV-C | SNV | 1 | C | T | 47 | 30744 | 0.15287536 | 46.314434 | WT | FALSE | CAN | JSY13 | 5 | WT-Rep5 | CCA | C>T | A/T+1 | G/C-1 | CG>TA | CG>TA | GAAACCCAA | CAACCTA | GGT | CCA | C | A | C>T | C>T_CCA | CG>TA_CAACCTA | 9 | FALSE | FALSE |
| 188_v.txt | 32679 | 32679-SNV-C | SNV | 1 | C | T | 119 | 30741 | 0.38710517 | 59.9521346 | WT | FALSE | CAN | JSY13 | 5 | WT-Rep5 | CCA | C>T | A/T+1 | G/C-1 | CG>TA | CG>TA | TTTCTCCAAT | CAATAAC | GGT | CCA | C | A | C>T | C>T_CCA | CG>TA_CAATAAC | 9 | FALSE | FALSE |
| 188_v.txt | 32700 | 32700-SNV-C | SNV | 1 | C | T | 34 | 30744 | 0.11059068 | 59.9411765 | WT | FALSE | CAN | JSY13 | 5 | WT-Rep5 | GCC | C>T | A/T+1 | G/C-1 | CG>TA | CG>TA | ACTGGGCCG | CCGGTAA | CGG | GCC | C | A | C>T | C>T_GCC | CG>TA_CCGGTAA | 9 | FALSE | FALSE |
| 188_v.txt | 32749 | 32749-SNV-C | SNV | 1 | C | T | 148 | 30744 | 0.48139474 | 54.8918919 | WT | FALSE | CAN | JSY13 | 5 | WT-Rep5 | CCG | C>T | A/T+1 | G/C-1 | CG>TA | CG>TA | GAAACCCGA | CGATAAT | GGC | CCG | C | A | C>T | C>T_CCG | CG>TA_CGATAAT | 9 | FALSE | FALSE |
| 188_v.txt | 32757 | 32757-SNV-G | SNV | 1 | G | A | 25 | 30744 | 0.08131668 | 52.84 | WT | FALSE | CAN | JSY13 | 5 | WT-Rep5 | GGC | G>A | A/T+1 | G/C-1 | CG>TA | CG>TA | ATAATGGCT | GCTAAAA | CCG | CCG | C | T | C>T | C>T_CCG | CG>TA_GCTAAAA | 9 | FALSE | FALSE |
| 188_v.txt | 32776 | 32776-SNV-C | SNV | 1 | C | T | 35 | 30744 | 0.11384335 | 45.7714286 | WT | FALSE | CAN | JSY13 | 5 | WT-Rep5 | GCG | C>T | A/T+1 | G/C-1 | CG>TA | CG>TA | TGGAAGCGA | CGACCCA | CGC | GCG | C | A | C>T | C>T_GCG | CG>TA_CGACCCA | 9 | FALSE | FALSE |
| 188_v.txt | 32780 | 32780-SNV-C | SNV | 1 | C | T | 50 | 30744 | 0.16263336 | 47.0179858 | WT | FALSE | CAN | JSY13 | 5 | WT-Rep5 | CCC | C>T | A/T+1 | G/C-1 | CG>TA | CG>TA | AGCGACCCA | CCAGAAC | GGG | CCC | C | A | C>T | C>T_CCC | CG>TA_CCAGAAC | 9 | FALSE | FALSE |
| 188_v.txt | 32781 | 32781-SNV-C | SNV | 1 | C | T | 350 | 30744 | 1.13843352 | 49.6364793 | WT | FALSE | CAN | JSY13 | 5 | WT-Rep5 | CCA | C>T | A/T+1 | G/C-1 | CG>TA | CG>TA | GCGACCCAG | CAGAACT | GGT | CCA | C | A | C>T | C>T_CCA | CG>TA_CAGAACT | 9 | FALSE | FALSE |
| 188_v.txt | 32796 | 32796-SNV-C | SNV | 1 | C | T | 262 | 30744 | 0.8521988 | 47.129771 | WT | FALSE | CAN | JSY13 | 5 | WT-Rep5 | ACC | C>T | A/T+1 | G/C-1 | CG>TA | CG>TA | AATTCACCGT | CCGTAAT | TGG | ACC | C | A | C>T | C>T_ACC | CG>TA_CCGTAAT | 9 | FALSE | FALSE |
| 188_v.txt | 32797 | 32797-SNV-C | SNV | 1 | C | T | 140 | 30744 | 0.45537341 | 48.5637517 | WT | FALSE | CAN | JSY13 | 5 | WT-Rep5 | CCG | C>T | A/T+1 | G/C-1 | CG>TA | CG>TA | ATTCACCGTA | CGTAATA | GGC | CCG | C | A | C>T | C>T_CCG | CG>TA_CGTAATA | 9 | FALSE | FALSE |
| 188_v.txt | 32798 | 32798-SNV-G | SNV | 1 | G | A | 53 | 30744 | 0.17239136 | 40.1434543 | WT | FALSE | CAN | JSY13 | 5 | WT-Rep5 | CGT | G>A | A/T+1 | G/C-1 | CG>TA | CG>TA | TTCACCGTAA | GTAATAT | GCA | GCA | C | T | C>T | C>T_GCA | CG>TA_GTAATAT | 9 | FALSE | FALSE |
| 188_v.txt | 32840 | 32840-SNV-C | SNV | 1 | C | T | 37 | 30744 | 0.12034869 | 37.5675676 | WT | FALSE | CAN | JSY13 | 5 | WT-Rep5 | CCC | C>T | A/T+1 | G/C-1 | CG>TA | CG>TA | AATTACCCA | CCAAAAA | GGG | CCC | C | A | C>T | C>T_CCC | CG>TA_CCAAAAA | 9 | FALSE | FALSE |
| 188_v.txt | 32855 | 32855-SNV-C | SNV | 1 | C | T | 109 | 30742 | 0.35456379 | 35.8899083 | WT | FALSE | CAN | JSY13 | 5 | WT-Rep5 | TCC | C>T | A/T+1 | G/C-1 | CG>TA | CG>TA | ACTAATCCAT | CCATGCC | AGG | TCC | C | A | C>T | C>T_TCC | CG>TA_CCATGCC | 9 | FALSE | FALSE |
| 188_v.txt | 32860 | 32860-SNV-C | SNV | 1 | C | T | 34 | 30739 | 0.11060867 | 37.7647059 | WT | FALSE | CAN | JSY13 | 5 | WT-Rep5 | GCC | C>T | A/T+1 | G/C-1 | CG>TA | CG>TA | TCCATGCCG | CCGCCAG | CGG | GCC | C | A | C>T | C>T_GCC | CG>TA_CCGCCAG | 9 | FALSE | FALSE |
| 188_v.txt | 32862 | 32862-SNV-G | SNV | 1 | G | A | 25 | 30739 | 0.08132991 | 36.08 | WT | FALSE | CAN | JSY13 | 5 | WT-Rep5 | CGC | G>A | A/T+1 | G/C-1 | CG>TA | CG>TA | CATGCCGCC | GCCAGTG | GCG | GCG | C | T | C>T | C>T_GCG | CG>TA_GCCAGTG | 9 | FALSE | FALSE |
| 188_v.txt | 32900 | 32900-SNV-G | SNV | 1 | G | A | 26 | 25520 | 0.10188088 | 35.8770834 | WT | FALSE | CAN | JSY13 | 5 | WT-Rep5 | GGC | G>A | A/T+1 | G/C-1 | CG>TA | CG>TA | GACTTGGCC | GCCAACT | CCG | CCG | C | T | C>T | C>T_CCG | CG>TA_GCCAACT | 9 | FALSE | FALSE |
| 188_v.txt | 32901 | 32901-SNV-C | SNV | 1 | C | T | 51 | 25520 | 0.19984326 | 32.0199512 | WT | FALSE | CAN | JSY13 | 5 | WT-Rep5 | GCC | C>T | A/T+1 | G/C-1 | CG>TA | CG>TA | ACTTGGCCA | CCAACTA | CGG | GCC | C | A | C>T | C>T_GCC | CG>TA_CCAACTA | 9 | FALSE | FALSE |
| 188_v.txt | 32917 | 32917-SNV-C | SNV | 1 | C | T | 552 | 25520 | 2.1630094 | 49.1358696 | WT | FALSE | CAN | JSY13 | 5 | WT-Rep5 | TCC | C>T | A/T+1 | G/C-1 | CG>TA | CG>TA | TAAGTTCCA | CCAGGGC | AGG | TCC | C | A | C>T | C>T_TCC | CG>TA_CCAGGGC | 9 | FALSE | FALSE |
| 188_v.txt | 32922 | 32922-SNV-G | SNV | 1 | G | A | 19 | 25520 | 0.07445141 | 44.3157895 | WT | FALSE | CAN | JSY13 | 5 | WT-Rep5 | GGC | G>A | A/T+1 | G/C-1 | CG>TA | CG>TA | TCCAGGGCA | GCAAAAG | CCG | CCG | C | T | C>T | C>T_CCG | CG>TA_GCAAAAG | 9 | FALSE | FALSE |
| 188_v.txt | 32940 | 32940-SNV-G | SNV | 1 | G | A | 42 | 25520 | 0.1645768 | 50.4890563 | WT | FALSE | CAN | JSY13 | 5 | WT-Rep5 | AGA | G>A | A/T+1 | G/C-1 | CG>TA | CG>TA | GCCCAAGAA | GAAAACC | TCT | TCT | C | T | C>T | C>T_TCT | CG>TA_GAAAACC | 9 | FALSE | FALSE |
| 188_v.txt | 32945 | 32945-SNV-C | SNV | 1 | C | T | 71 | 25520 | 0.27821317 | 57.5915493 | WT | FALSE | CAN | JSY13 | 5 | WT-Rep5 | ACC | C>T | A/T+1 | G/C-1 | CG>TA | CG>TA | AGAAAACCA | CCAATAC | TGG | ACC | C | A | C>T | C>T_ACC | CG>TA_CCAATAC | 9 | FALSE | FALSE |
| 188_v.txt | 32958 | 32958-SNV-C | SNV | 1 | C | T | 123 | 25520 | 0.48197492 | 56.2195122 | WT | FALSE | CAN | JSY13 | 5 | WT-Rep5 | ACC | C>T | A/T+1 | G/C-1 | CG>TA | CG>TA | ATGTAACCAT | CCATTGG | TGG | ACC | C | A | C>T | C>T_ACC | CG>TA_CCATTGG | 9 | FALSE | FALSE |
| 188_v.txt | 32967 | 32967-SNV-G | SNV | 1 | G | A | 60 | 25520 | 0.23510972 | 59.8666667 | WT | FALSE | CAN | JSY13 | 5 | WT-Rep5 | CGC | G>A | A/T+1 | G/C-1 | CG>TA | CG>TA | TTGGCCGCA | GCACCAA | GCG | GCG | C | T | C>T | C>T_GCG | CG>TA_GCACCAA | 9 | FALSE | FALSE |
| 188_v.txt | 32970 | 32970-SNV-C | SNV | 1 | C | T | 191 | 25520 | 0.7484326 | 58.513089 | WT | FALSE | CAN | JSY13 | 5 | WT-Rep5 | ACC | C>T | A/T+1 | G/C-1 | CG>TA | CG>TA | GCCGCACCA | CCAAATG | TGG | ACC | C | A | C>T | C>T_ACC | CG>TA_CCAAATG | 9 | FALSE | FALSE |
| 188_v.txt | 33055 | 33055-SNV-G | SNV | 1 | G | A | 35 | 25520 | 0.13714734 | 55.4282997 | WT | FALSE | CAN | JSY13 | 5 | WT-Rep5 | TGC | G>A | A/T+1 | G/C-1 | CG>TA | CG>TA | AGGACTGCG | GCGTGAC | ACG | ACG | C | T | C>T | C>T_ACG | CG>TA_GCGTGAC | 9 | FALSE | FALSE |
| 188_v.txt | 33056 | 33056-SNV-C | SNV | 1 | C | T | 30 | 25520 | 0.11755486 | 55.1613919 | WT | FALSE | CAN | JSY13 | 5 | WT-Rep5 | GCG | C>T | A/T+1 | G/C-1 | CG>TA | CG>TA | GGACTGCGT | CGTGACA | CGC | GCG | C | A | C>T | C>T_GCG | CG>TA_CGTGACA | 9 | FALSE | FALSE |
| 188_v.txt | 33069 | 33069-SNV-G | SNV | 1 | G | A | 25 | 25520 | 0.09796238 | 50.7006508 | WT | FALSE | CAN | JSY13 | 5 | WT-Rep5 | TGC | G>A | A/T+1 | G/C-1 | CG>TA | CG>TA | GAATATGCC | GCCAAAG | ACG | ACG | C | T | C>T | C>T_ACG | CG>TA_GCCAAAG | 9 | FALSE | FALSE |
| 188_v.txt | 33070 | 33070-SNV-C | SNV | 1 | C | T | 36 | 25520 | 0.14106583 | 53.2469773 | WT | FALSE | CAN | JSY13 | 5 | WT-Rep5 | GCC | C>T | A/T+1 | G/C-1 | CG>TA | CG>TA | AATATGCCA | CCAAAGA | CGG | GCC | C | A | C>T | C>T_GCC | CG>TA_CCAAAGA | 9 | FALSE | FALSE |
| 188_v.txt | 33104 | 33104-SNV-G | SNV | 1 | G | A | 27 | 25520 | 0.10579937 | 40.037037 | WT | FALSE | CAN | JSY13 | 5 | WT-Rep5 | CGC | G>A | A/T+1 | G/C-1 | CG>TA | CG>TA | AAGAGCGCC | GCCCACT | GCG | GCG | C | T | C>T | C>T_GCG | CG>TA_GCCCACT | 9 | FALSE | FALSE |
| 188_v.txt | 33111 | 33111-SNV-G | SNV | 1 | G | A | 32 | 25520 | 0.12539185 | 39.3420834 | WT | FALSE | CAN | JSY13 | 5 | WT-Rep5 | TGG | G>A | A/T+1 | G/C-1 | CG>TA | CG>TA | CCCACTGGG | GGGCCGG | ACC | ACC | C | T | C>T | C>T_ACC | CG>TA_GGGCCGG | 9 | FALSE | FALSE |
| 188_v.txt | 33114 | 33114-SNV-C | SNV | 1 | C | T | 59 | 25520 | 0.23119122 | 41.5432139 | WT | FALSE | CAN | JSY13 | 5 | WT-Rep5 | GCC | C>T | A/T+1 | G/C-1 | CG>TA | CG>TA | ACTGGGCCG | CCGGCGT | CGG | GCC | C | A | C>T | C>T_GCC | CG>TA_CCGGCGT | 9 | FALSE | FALSE |
| 188_v.txt | 33115 | 33115-SNV-C | SNV | 1 | C | T | 32 | 25520 | 0.12539185 | 46.4366482 | WT | FALSE | CAN | JSY13 | 5 | WT-Rep5 | CCG | C>T | A/T+1 | G/C-1 | CG>TA | CG>TA | CTGGGCCGG | CGGCGTT | GGC | CCG | C | A | C>T | C>T_CCG | CG>TA_CGGCGTT | 9 | FALSE | FALSE |

| | | | | | | | | | | | | | | | | | | | | | | | | | | | | | | | | | | |
|---|---|---|---|---|---|---|---|---|---|---|---|---|---|---|---|---|---|---|---|---|---|---|---|---|---|---|---|---|---|---|---|---|---|---|
| 188_v.txt | 33119 | 33119-SNV- | SNV | 1 | G | A | 21 | 25520 | 0.0822884 | 43.5714286 | WT | FALSE | CAN | JSY13 | 5 | WT-Rep5 | CGT | G>A | A/T+1 | G/C-1 | CG>TA | CG>TA | GCCGGCGTT | GTTGGTC | GCA | GCA | C | T | C>T | C>T_GCA | CG>TA_GTTGGTC | 9 | FALSE | FALSE |
| 188_v.txt | 33153 | 33153-SNV- | SNV | 1 | C | T | 56 | 25519 | 0.21944434 | 35.8559475 | WT | FALSE | CAN | JSY13 | 5 | WT-Rep5 | ACC | C>T | A/T+1 | G/C-1 | CG>TA | CG>TA | AAAAGACCT | CCTGTAC | TGG | ACC | C | A | C>T | C>T_ACC | CG>TA_CCTGTAC | 9 | FALSE | FALSE |
| 188_v.txt | 33154 | 33154-SNV- | SNV | 1 | C | T | 11 | 25519 | 0.04310514 | 35.0000522 | WT | FALSE | CAN | JSY13 | 5 | WT-Rep5 | CCT | C>T | A/T+1 | G/C-1 | CG>TA | CG>TA | AAAGACCTG | CTGTACC | GGA | CCT | C | A | C>T | C>T_CCT | CG>TA_CTGTACC | 9 | FALSE | FALSE |
| 188_v.txt | 33156 | 33156-SNV- | SNV | 1 | G | A | 36 | 25518 | 0.14107689 | 37.6944444 | WT | FALSE | CAN | JSY13 | 5 | WT-Rep5 | TGT | G>A | A/T+1 | G/C-1 | CG>TA | CG>TA | AGACCTGTA | GTACCAA | ACA | ACA | C | T | C>T | C>T_ACA | CG>TA_GTACCAA | 9 | FALSE | FALSE |
| 188_v.txt | 33159 | 33159-SNV- | SNV | 1 | C | T | 38 | 25518 | 0.14891449 | 36.5263158 | WT | FALSE | CAN | JSY13 | 5 | WT-Rep5 | ACC | C>T | A/T+1 | G/C-1 | CG>TA | CG>TA | CCTGTACCA | CCAATAG | TGG | ACC | C | A | C>T | C>T_ACC | CG>TA_CCAATAG | 9 | FALSE | FALSE |
| 188_v.txt | 33168 | 33168-SNV- | SNV | 1 | C | T | 44 | 25518 | 0.17242731 | 37.1591722 | WT | FALSE | CAN | JSY13 | 5 | WT-Rep5 | ACC | C>T | A/T+1 | G/C-1 | CG>TA | CG>TA | ATAGTACCA | CCACCAA | TGG | ACC | C | A | C>T | C>T_ACC | CG>TA_CCACCAA | 9 | FALSE | FALSE |
| 188_v.txt | 33169 | 33169-SNV- | SNV | 1 | C | T | 77 | 25518 | 0.30174779 | 37.5847731 | WT | FALSE | CAN | JSY13 | 5 | WT-Rep5 | CCA | C>T | A/T+1 | G/C-1 | CG>TA | CG>TA | TAGTACCAC | CACCAAG | GGT | CCA | C | A | C>T | C>T_CCA | CG>TA_CACCAAG | 9 | FALSE | FALSE |
| 188_v.txt | 33171 | 33171-SNV- | SNV | 1 | C | T | 43 | 25518 | 0.1685085 | 36.1162791 | WT | FALSE | CAN | JSY13 | 5 | WT-Rep5 | ACC | C>T | A/T+1 | G/C-1 | CG>TA | CG>TA | GTACCACCA | CCAAGGG | TGG | ACC | C | A | C>T | C>T_ACC | CG>TA_CCAAGGG | 9 | FALSE | FALSE |
| 188_v.txt | 33229 | 33229-SNV- | SNV | 1 | G | A | 28 | 17251 | 0.16230943 | 33.8928571 | WT | FALSE | CAN | JSY13 | 5 | WT-Rep5 | TGT | G>A | A/T+1 | G/C-1 | CG>TA | CG>TA | CGTTCTGTAC | GTACTTC | ACA | ACA | C | T | C>T | C>T_ACA | CG>TA_GTACTTC | 9 | FALSE | FALSE |
| 188_v.txt | 33258 | 33258-SNV- | SNV | 1 | C | T | 34 | 17251 | 0.19709002 | 45.2058824 | WT | FALSE | CAN | JSY13 | 5 | WT-Rep5 | GCC | C>T | A/T+1 | G/C-1 | CG>TA | CG>TA | CCTATGCCAT | CCATCCT | CGG | GCC | C | A | C>T | C>T_GCC | CG>TA_CCATCCT | 9 | FALSE | FALSE |
| 188_v.txt | 33275 | 33275-SNV- | SNV | 1 | C | T | 20 | 17251 | 0.11593531 | 43.6974655 | WT | FALSE | CAN | JSY13 | 5 | WT-Rep5 | ACG | C>T | A/T+1 | G/C-1 | CG>TA | CG>TA | AGAGAACGT | CGTATCC | TGC | ACG | C | A | C>T | C>T_ACG | CG>TA_CGTATCC | 9 | FALSE | FALSE |
| 188_v.txt | 33276 | 33276-SNV- | SNV | 1 | G | A | 24 | 17250 | 0.13913044 | 49.3401058 | WT | FALSE | CAN | JSY13 | 5 | WT-Rep5 | CGT | G>A | A/T+1 | G/C-1 | CG>TA | CG>TA | GAGAACGTA | GTATCCT | GCA | GCA | C | T | C>T | C>T_GCA | CG>TA_GTATCCT | 9 | FALSE | FALSE |
| 188_v.txt | 33284 | 33284-SNV- | SNV | 1 | G | A | 23 | 17251 | 0.1333256 | 47.6086957 | WT | FALSE | CAN | JSY13 | 5 | WT-Rep5 | CGC | G>A | A/T+1 | G/C-1 | CG>TA | CG>TA | ATCCTCGCCA | GCCATTT | GCG | GCG | C | T | C>T | C>T_GCG | CG>TA_GCCATTT | 9 | FALSE | FALSE |
| 188_v.txt | 33310 | 33310-SNV- | SNV | 1 | G | A | 32 | 17251 | 0.18549649 | 57.4375 | WT | FALSE | CAN | JSY13 | 5 | WT-Rep5 | CGC | G>A | A/T+1 | G/C-1 | CG>TA | CG>TA | AAGAGCGCA | GCAATGG | GCG | GCG | C | T | C>T | C>T_GCG | CG>TA_GCAATGG | 9 | FALSE | FALSE |
| 188_v.txt | 33392 | 33392-SNV- | SNV | 1 | G | A | 22 | 17251 | 0.12752884 | 51.8181818 | WT | FALSE | CAN | JSY13 | 5 | WT-Rep5 | CGT | G>A | A/T+1 | G/C-1 | CG>TA | CG>TA | AACGTCGTG | GTGAAAG | GCA | GCA | C | T | C>T | C>T_GCA | CG>TA_GTGAAAG | 9 | FALSE | FALSE |
| 188_v.txt | 33412 | 33412-SNV- | SNV | 1 | G | A | 29 | 17251 | 0.1681062 | 51.3793103 | WT | FALSE | CAN | JSY13 | 5 | WT-Rep5 | GGC | G>A | A/T+1 | G/C-1 | CG>TA | CG>TA | TGACCGGCT | GCTCATT | CCG | CCG | C | T | C>T | C>T_CCG | CG>TA_GCTCATT | 9 | FALSE | FALSE |
| 188_v.txt | 33474 | 33474-SNV- | SNV | 1 | C | T | 21 | 17249 | 0.12174619 | 34.1904762 | WT | FALSE | CAN | JSY13 | 5 | WT-Rep5 | GCC | C>T | A/T+1 | G/C-1 | CG>TA | CG>TA | GCTATGCCTT | CCTTTTT | CGG | GCC | C | A | C>T | C>T_GCC | CG>TA_CCTTTTT | 9 | FALSE | FALSE |
| 188_v.txt | 32304 | 32304-SNV- | SNV | 1 | G | C | 5 | 31738 | 0.01575399 | 44.8 | WT | FALSE | CAN | JSY13 | 5 | WT-Rep5 | AGA | G>C | G/C+1 | G/C-1 | CG>GC | CG>GC | GCGGCAGAA | GAAATAA | TCT | TCT | C | G | C>G | C>G_TCT | CG>GC_GAAATAA | 9 | FALSE | FALSE |
| 188_v.txt | 32316 | 32316-SNV- | SNV | 1 | G | C | 57 | 31738 | 0.17959544 | 59.9122807 | WT | FALSE | CAN | JSY13 | 5 | WT-Rep5 | TGT | G>C | G/C+1 | G/C-1 | CG>GC | CG>GC | ATGGTTGTT | GTTAAGA | ACA | ACA | C | G | C>G | C>G_ACA | CG>GC_GTTAAGA | 9 | FALSE | FALSE |
| 188_v.txt | 32488 | 32488-SNV- | SNV | 1 | G | C | 3 | 31738 | 9.45E-03 | 41.3333333 | WT | FALSE | CAN | JSY13 | 5 | WT-Rep5 | CGG | G>C | G/C+1 | G/C-1 | CG>GC | CG>GC | AGATACGGA | GGAAAAC | GCC | GCC | C | G | C>G | C>G_GCC | CG>GC_GGAAAAC | 9 | FALSE | FALSE |
| 188_v.txt | 32536 | 32536-SNV- | SNV | 1 | C | G | 1 | 31738 | 3.15E-03 | 30.9997347 | WT | FALSE | CAN | JSY13 | 5 | WT-Rep5 | GCA | C>G | G/C+1 | G/C-1 | CG>GC | CG>GC | GGTTTGCAG | CAGCTTC | CGT | GCA | C | C | C>G | C>G_GCA | CG>GC_CAGCTTC | 9 | FALSE | FALSE |
| 188_v.txt | 32557 | 32557-SNV- | SNV | 1 | C | G | 19 | 31728 | 0.05988401 | 37.6842105 | WT | FALSE | CAN | JSY13 | 5 | WT-Rep5 | CCA | C>G | G/C+1 | G/C-1 | CG>GC | CG>GC | TGATACCAA | CAACTAG | GGT | CCA | C | C | C>G | C>G_CCA | CG>GC_CAACTAG | 9 | FALSE | FALSE |
| 188_v.txt | 32691 | 32691-SNV- | SNV | 1 | C | G | 1 | 30743 | 3.25E-03 | 63.9938979 | WT | FALSE | CAN | JSY13 | 5 | WT-Rep5 | TCC | C>G | G/C+1 | G/C-1 | CG>GC | CG>GC | CGGAATCCA | CCAACTG | AGG | TCC | C | C | C>G | C>G_TCC | CG>GC_CCAACTG | 9 | FALSE | FALSE |
| 188_v.txt | 32735 | 32735-SNV- | SNV | 1 | G | C | 13 | 30744 | 0.04228467 | 60 | WT | FALSE | CAN | JSY13 | 5 | WT-Rep5 | AGT | G>C | G/C+1 | G/C-1 | CG>GC | CG>GC | AAAACAGTA | GTATATT | TCA | TCA | C | G | C>G | C>G_TCA | CG>GC_GTATATT | 9 | FALSE | FALSE |
| 188_v.txt | 32796 | 32796-SNV- | SNV | 1 | C | G | 2 | 30744 | 6.51E-03 | 27 | WT | FALSE | CAN | JSY13 | 5 | WT-Rep5 | ACC | C>G | G/C+1 | G/C-1 | CG>GC | CG>GC | AATTCACCG | CCGTAAT | TGG | ACC | C | C | C>G | C>G_ACC | CG>GC_CCGTAAT | 9 | FALSE | FALSE |
| 188_v.txt | 32811 | 32811-SNV- | SNV | 1 | G | C | 2 | 30744 | 6.51E-03 | 31.4995388 | WT | FALSE | CAN | JSY13 | 5 | WT-Rep5 | AGG | G>C | G/C+1 | G/C-1 | CG>GC | CG>GC | TTGACAGGG | GGGAACA | TCC | TCC | C | G | C>G | C>G_TCC | CG>GC_GGGAACA | 9 | FALSE | FALSE |
| 188_v.txt | 32917 | 32917-SNV- | SNV | 1 | C | G | 3 | 25520 | 0.01175549 | 37 | WT | FALSE | CAN | JSY13 | 5 | WT-Rep5 | TCC | C>G | G/C+1 | G/C-1 | CG>GC | CG>GC | TAAGTTCCA | CCAGGGC | AGG | TCC | C | C | C>G | C>G_TCC | CG>GC_CCAGGGC | 9 | FALSE | FALSE |
| 188_v.txt | 33070 | 33070-SNV- | SNV | 1 | C | G | 3 | 25520 | 0.01175549 | 63.9840987 | WT | FALSE | CAN | JSY13 | 5 | WT-Rep5 | GCC | C>G | G/C+1 | G/C-1 | CG>GC | CG>GC | AATATGCCA | CCAAAGA | CGG | GCC | C | C | C>G | C>G_GCC | CG>GC_CCAAAGA | 9 | FALSE | FALSE |
| 188_v.txt | 33079 | 33079-SNV- | SNV | 1 | C | G | 4515 | 25520 | 17.6920063 | 53.2555925 | WT | FALSE | CAN | JSY13 | 5 | WT-Rep5 | CCC | C>G | G/C+1 | G/C-1 | CG>GC | CG>GC | AAGAACCCA | CCATAAA | GGG | CCC | C | C | C>G | C>G_CCC | CG>GC_CCATAAA | 9 | FALSE | FALSE |
| 188_v.txt | 33119 | 33119-SNV- | SNV | 1 | G | C | 2 | 25520 | 7.84E-03 | 38 | WT | FALSE | CAN | JSY13 | 5 | WT-Rep5 | CGT | G>C | G/C+1 | G/C-1 | CG>GC | CG>GC | GCCGGCGTT | GTTGGTC | GCA | GCA | C | G | C>G | C>G_GCA | CG>GC_GTTGGTC | 9 | FALSE | FALSE |
| 188_v.txt | 33153 | 33153-SNV- | SNV | 1 | C | G | 1 | 25519 | 3.92E-03 | 37.9982825 | WT | FALSE | CAN | JSY13 | 5 | WT-Rep5 | ACC | C>G | G/C+1 | G/C-1 | CG>GC | CG>GC | AAAAGACCT | CCTGTAC | TGG | ACC | C | C | C>G | C>G_ACC | CG>GC_CCTGTAC | 9 | FALSE | FALSE |
| 188_v.txt | 32058 | 32058-SNV- | SNV | 1 | G | T | 56 | 12088 | 0.46326936 | 62.0178571 | WT | FALSE | CAN | JSY13 | 5 | WT-Rep5 | CGA | G>T | A/T+1 | G/C-1 | CG>AT | CG>AT | ATGTGCGAG | GAGATTG | GCT | GCT | C | A | C>A | C>A_GCT | CG>AT_GAGATTG | 9 | FALSE | FALSE |
| 188_v.txt | 32088 | 32088-SNV- | SNV | 1 | C | A | 48 | 12088 | 0.39708802 | 59.875 | WT | FALSE | CAN | JSY13 | 5 | WT-Rep5 | GCC | C>A | A/T+1 | G/C-1 | CG>AT | CG>AT | AAAAAGCCT | CCTGCAA | CGG | GCC | C | T | C>A | C>A_GCC | CG>AT_CCTGCAA | 9 | FALSE | FALSE |
| 188_v.txt | 32089 | 32089-SNV- | SNV | 1 | C | A | 59 | 12088 | 0.48808736 | 59.9157267 | WT | FALSE | CAN | JSY13 | 5 | WT-Rep5 | CCT | C>A | A/T+1 | G/C-1 | CG>AT | CG>AT | AAAAGCCTG | CTGCAAC | GGA | CCT | C | T | C>A | C>A_CCT | CG>AT_CTGCAAC | 9 | FALSE | FALSE |
| 188_v.txt | 32095 | 32095-SNV- | SNV | 1 | C | A | 12 | 12088 | 0.09927201 | 54.5 | WT | FALSE | CAN | JSY13 | 5 | WT-Rep5 | ACA | C>A | A/T+1 | G/C-1 | CG>AT | CG>AT | CTGCAACAC | CACCAGT | TGT | ACA | C | T | C>A | C>A_ACA | CG>AT_CACCAGT | 9 | FALSE | FALSE |
| 188_v.txt | 32188 | 32188-SNV- | SNV | 1 | C | A | 28 | 12088 | 0.23163468 | 44.2142857 | WT | FALSE | CAN | JSY13 | 5 | WT-Rep5 | ACT | C>A | A/T+1 | G/C-1 | CG>AT | CG>AT | CGAAAACTG | CTGCAAT | TGA | ACT | C | T | C>A | C>A_ACT | CG>AT_CTGCAAT | 9 | FALSE | FALSE |
| 188_v.txt | 32195 | 32195-SNV- | SNV | 1 | G | T | 17 | 12088 | 0.14063534 | 42.5882353 | WT | FALSE | CAN | JSY13 | 5 | WT-Rep5 | TGT | G>T | A/T+1 | G/C-1 | CG>AT | CG>AT | TGCAATGTA | GTATGGA | ACA | ACA | C | A | C>A | C>A_ACA | CG>AT_GTATGGA | 9 | FALSE | FALSE |
| 188_v.txt | 32238 | 32238-SNV- | SNV | 1 | G | T | 10 | 12087 | 0.08273352 | 35.6 | WT | FALSE | CAN | JSY13 | 5 | WT-Rep5 | AGC | G>T | A/T+1 | G/C-1 | CG>AT | CG>AT | TTAGGAGCC | GCCAACT | TCG | TCG | C | A | C>A | C>A_TCG | CG>AT_GCCAACT | 9 | FALSE | FALSE |
| 188_v.txt | 32271 | 32271-SNV- | SNV | 1 | C | A | 100 | 31738 | 0.31507972 | 35.36 | WT | FALSE | CAN | JSY13 | 5 | WT-Rep5 | ACG | C>A | A/T+1 | G/C-1 | CG>AT | CG>AT | AAAATACGG | CGGGAAC | TGC | ACG | C | T | C>A | C>A_ACG | CG>AT_CGGGAAC | 9 | FALSE | FALSE |
| 188_v.txt | 32272 | 32272-SNV- | SNV | 1 | G | T | 65 | 31738 | 0.20480182 | 35.2615612 | WT | FALSE | CAN | JSY13 | 5 | WT-Rep5 | CGG | G>T | A/T+1 | G/C-1 | CG>AT | CG>AT | AAATACGGG | GGGAACC | GCC | GCC | C | A | C>A | C>A_GCC | CG>AT_GGGAACC | 9 | FALSE | FALSE |
| 188_v.txt | 32304 | 32304-SNV- | SNV | 1 | G | T | 22 | 31738 | 0.06931754 | 53.6363636 | WT | FALSE | CAN | JSY13 | 5 | WT-Rep5 | AGA | G>T | A/T+1 | G/C-1 | CG>AT | CG>AT | GCGGCAGAA | GAAATAA | TCT | TCT | C | A | C>A | C>A_TCT | CG>AT_GAAATAA | 9 | FALSE | FALSE |
| 188_v.txt | 32373 | 32373-SNV- | SNV | 1 | G | T | 17 | 31738 | 0.05356355 | 49.2352941 | WT | FALSE | CAN | JSY13 | 5 | WT-Rep5 | AGC | G>T | A/T+1 | G/C-1 | CG>AT | CG>AT | TCAATAGCA | GCAATAA | TCG | TCG | C | A | C>A | C>A_TCG | CG>AT_GCAATAA | 9 | FALSE | FALSE |
| 188_v.txt | 32384 | 32384-SNV- | SNV | 1 | G | T | 7 | 31737 | 0.02205628 | 57.7122149 | WT | FALSE | CAN | JSY13 | 5 | WT-Rep5 | AGG | G>T | A/T+1 | G/C-1 | CG>AT | CG>AT | AATAAAGGG | GGGAGAA | TCC | TCC | C | A | C>A | C>A_TCC | CG>AT_GGGAGAA | 9 | FALSE | FALSE |
| 188_v.txt | 32385 | 32385-SNV- | SNV | 1 | G | T | 6 | 31738 | 0.01890478 | 25.9999744 | WT | FALSE | CAN | JSY13 | 5 | WT-Rep5 | GGG | G>T | A/T+1 | G/C-1 | CG>AT | CG>AT | ATAAAGGGA | GGAGAAG | CCC | CCC | C | A | C>A | C>A_CCC | CG>AT_GGAGAAG | 9 | FALSE | FALSE |
| 188_v.txt | 32449 | 32449-SNV- | SNV | 1 | C | A | 63 | 31738 | 0.19850022 | 52.8571429 | WT | FALSE | CAN | JSY13 | 5 | WT-Rep5 | CCA | C>A | A/T+1 | G/C-1 | CG>AT | CG>AT | AAAGTCCAA | CAATGAA | GGT | CCA | C | T | C>A | C>A_CCA | CG>AT_CAATGAA | 9 | FALSE | FALSE |
| 188_v.txt | 32474 | 32474-SNV- | SNV | 1 | G | T | 13 | 31738 | 0.04096036 | 45.6153846 | WT | FALSE | CAN | JSY13 | 5 | WT-Rep5 | AGA | G>T | A/T+1 | G/C-1 | CG>AT | CG>AT | AATGTAGAA | GAAGGTT | TCT | TCT | C | A | C>A | C>A_TCT | CG>AT_GAAGGTT | 9 | FALSE | FALSE |
| 188_v.txt | 32487 | 32487-SNV- | SNV | 1 | C | A | 24 | 31738 | 0.07561913 | 45.1643375 | WT | FALSE | CAN | JSY13 | 5 | WT-Rep5 | ACG | C>A | A/T+1 | G/C-1 | CG>AT | CG>AT | AAGATACGG | CGGAAAA | TGC | ACG | C | T | C>A | C>A_ACG | CG>AT_CGGAAAA | 9 | FALSE | FALSE |
| 188_v.txt | 32488 | 32488-SNV- | SNV | 1 | G | T | 24 | 31738 | 0.07561913 | 41.7916667 | WT | FALSE | CAN | JSY13 | 5 | WT-Rep5 | CGG | G>T | A/T+1 | G/C-1 | CG>AT | CG>AT | AGATACGGA | GGAAAAC | GCC | GCC | C | A | C>A | C>A_GCC | CG>AT_GGAAAAC | 9 | FALSE | FALSE |
| 188_v.txt | 32508 | 32508-SNV- | SNV | 1 | G | T | 52 | 31738 | 0.16384145 | 44.6153846 | WT | FALSE | CAN | JSY13 | 5 | WT-Rep5 | GGC | G>T | A/T+1 | G/C-1 | CG>AT | CG>AT | TTGATGGCT | GCTCTTG | CCG | CCG | C | A | C>A | C>A_CCG | CG>AT_GCTCTTG | 9 | FALSE | FALSE |
| 188_v.txt | 32530 | 32530-SNV- | SNV | 1 | G | T | 33 | 31738 | 0.10397631 | 36.4242424 | WT | FALSE | CAN | JSY13 | 5 | WT-Rep5 | GGG | G>T | A/T+1 | G/C-1 | CG>AT | CG>AT | TTCTGGGGT | GGTTTGC | CCC | CCC | C | A | C>A | C>A_CCC | CG>AT_GGTTTGC | 9 | FALSE | FALSE |
| 188_v.txt | 32538 | 32538-SNV- | SNV | 1 | G | T | 19 | 31738 | 0.05986515 | 34.2631579 | WT | FALSE | CAN | JSY13 | 5 | WT-Rep5 | AGC | G>T | A/T+1 | G/C-1 | CG>AT | CG>AT | TTTGCAGCTT | GCTTCAC | TCG | TCG | C | A | C>A | C>A_TCG | CG>AT_GCTTCAC | 9 | FALSE | FALSE |
| 188_v.txt | 32547 | 32547-SNV- | SNV | 1 | G | T | 47 | 31733 | 0.1481108 | 37.2765957 | WT | FALSE | CAN | JSY13 | 5 | WT-Rep5 | AGC | G>T | A/T+1 | G/C-1 | CG>AT | CG>AT | TCACCAGCA | GCAGTGA | TCG | TCG | C | A | C>A | C>A_TCG | CG>AT_GCAGTGA | 9 | FALSE | FALSE |
| 188_v.txt | 32663 | 32663-SNV- | SNV | 1 | G | T | 8 | 30744 | 0.02602134 | 52.623954 | WT | FALSE | CAN | JSY13 | 5 | WT-Rep5 | AGG | G>T | A/T+1 | G/C-1 | CG>AT | CG>AT | ACCCCAGGC | GGCACCT | TCC | TCC | C | A | C>A | C>A_TCC | CG>AT_GGCACCT | 9 | FALSE | FALSE |
| 188_v.txt | 32691 | 32691-SNV- | SNV | 1 | C | A | 4 | 30743 | 0.01301109 | 39.2495728 | WT | FALSE | CAN | JSY13 | 5 | WT-Rep5 | TCC | C>A | A/T+1 | G/C-1 | CG>AT | CG>AT | CGGAATCCA | CCAACTG | AGG | TCC | C | T | C>A | C>A_TCC | CG>AT_CCAACTG | 9 | FALSE | FALSE |
| 188_v.txt | 32692 | 32692-SNV- | SNV | 1 | C | A | 29 | 30743 | 0.09433042 | 38.6806588 | WT | FALSE | CAN | JSY13 | 5 | WT-Rep5 | CCA | C>A | A/T+1 | G/C-1 | CG>AT | CG>AT | GGAATCCAA | CAACTGG | GGT | CCA | C | T | C>A | C>A_CCA | CG>AT_CAACTGG | 9 | FALSE | FALSE |
| 188_v.txt | 32735 | 32735-SNV- | SNV | 1 | G | T | 46 | 30744 | 0.14962269 | 58.6956522 | WT | FALSE | CAN | JSY13 | 5 | WT-Rep5 | AGT | G>T | A/T+1 | G/C-1 | CG>AT | CG>AT | AAAACAGTA | GTATATT | TCA | TCA | C | A | C>A | C>A_TCA | CG>AT_GTATATT | 9 | FALSE | FALSE |
| 188_v.txt | 32781 | 32781-SNV- | SNV | 1 | C | A | 9 | 30744 | 0.02927401 | 23.5593051 | WT | FALSE | CAN | JSY13 | 5 | WT-Rep5 | CCA | C>A | A/T+1 | G/C-1 | CG>AT | CG>AT | GCGACCCAG | CAGAACT | GGT | CCA | C | T | C>A | C>A_CCA | CG>AT_CAGAACT | 9 | FALSE | FALSE |
| 188_v.txt | 32796 | 32796-SNV- | SNV | 1 | C | A | 8 | 30744 | 0.02602134 | 22.75 | WT | FALSE | CAN | JSY13 | 5 | WT-Rep5 | ACC | C>A | A/T+1 | G/C-1 | CG>AT | CG>AT | AATTCACCG | CCGTAAT | TGG | ACC | C | T | C>A | C>A_ACC | CG>AT_CCGTAAT | 9 | FALSE | FALSE |
| 188_v.txt | 32797 | 32797-SNV- | SNV | 1 | C | A | 6 | 30744 | 0.019516 | 37.4959462 | WT | FALSE | CAN | JSY13 | 5 | WT-Rep5 | CCG | C>A | A/T+1 | G/C-1 | CG>AT | CG>AT | ATTCACCGTA | CGTAATA | GGC | CCG | C | T | C>A | C>A_CCG | CG>AT_CGTAATA | 9 | FALSE | FALSE |
| 188_v.txt | 32809 | 32809-SNV- | SNV | 1 | C | A | 92 | 30744 | 0.29924538 | 45.1956522 | WT | FALSE | CAN | JSY13 | 5 | WT-Rep5 | ACA | C>A | A/T+1 | G/C-1 | CG>AT | CG>AT | ATTTGACAG | CAGGGAA | TGT | ACA | C | T | C>A | C>A_ACA | CG>AT_CAGGGAA | 9 | FALSE | FALSE |
| 188_v.txt | 32811 | 32811-SNV- | SNV | 1 | G | T | 16 | 30744 | 0.05204268 | 44.1240845 | WT | FALSE | CAN | JSY13 | 5 | WT-Rep5 | AGG | G>T | A/T+1 | G/C-1 | CG>AT | CG>AT | TTGACAGGG | GGGAACA | TCC | TCC | C | A | C>A | C>A_TCC | CG>AT_GGGAACA | 9 | FALSE | FALSE |
| 188_v.txt | 32812 | 32812-SNV- | SNV | 1 | G | T | 21 | 30744 | 0.06830601 | 46.4751714 | WT | FALSE | CAN | JSY13 | 5 | WT-Rep5 | GGG | G>T | A/T+1 | G/C-1 | CG>AT | CG>AT | TGACAGGGA | GGAACAA | CCC | CCC | C | A | C>A | C>A_CCC | CG>AT_GGAACAA | 9 | FALSE | FALSE |
| 188_v.txt | 32819 | 32819-SNV- | SNV | 1 | G | T | 15 | 30744 | 0.04879001 | 42.3333333 | WT | FALSE | CAN | JSY13 | 5 | WT-Rep5 | AGT | G>T | A/T+1 | G/C-1 | CG>AT | CG>AT | GAACAAGTT | GTTCATT | TCA | TCA | C | A | C>A | C>A_TCA | CG>AT_GTTCATT | 9 | FALSE | FALSE |
| 188_v.txt | 32840 | 32840-SNV- | SNV | 1 | C | A | 5 | 30744 | 0.01626334 | 32.4 | WT | FALSE | CAN | JSY13 | 5 | WT-Rep5 | CCC | C>A | A/T+1 | G/C-1 | CG>AT | CG>AT | AATTACCCA | CCAAAAA | GGG | CCC | C | T | C>A | C>A_CCC | CG>AT_CCAAAAA | 9 | FALSE | FALSE |
| 188_v.txt | 32917 | 32917-SNV- | SNV | 1 | C | A | 39 | 25520 | 0.15282132 | 52.2307692 | WT | FALSE | CAN | JSY13 | 5 | WT-Rep5 | TCC | C>A | A/T+1 | G/C-1 | CG>AT | CG>AT | TAAGTTCCA | CCAGGGC | AGG | TCC | C | T | C>A | C>A_TCC | CG>AT_CCAGGGC | 9 | FALSE | FALSE |
| 188_v.txt | 32936 | 32936-SNV- | SNV | 1 | C | A | 35 | 25520 | 0.13714734 | 34.7142857 | WT | FALSE | CAN | JSY13 | 5 | WT-Rep5 | CCC | C>A | A/T+1 | G/C-1 | CG>AT | CG>AT | GATTGCCCA | CCAAGAA | GGG | CCC | C | T | C>A | C>A_CCC | CG>AT_CCAAGAA | 9 | FALSE | FALSE |
| 188_v.txt | 32940 | 32940-SNV- | SNV | 1 | G | T | 49 | 25520 | 0.19200627 | 49.0779764 | WT | FALSE | CAN | JSY13 | 5 | WT-Rep5 | AGA | G>T | A/T+1 | G/C-1 | CG>AT | CG>AT | GCCCAAGAA | GAAAACC | TCT | TCT | C | A | C>A | C>A_TCT | CG>AT_GAAAACC | 9 | FALSE | FALSE |
| 188_v.txt | 33012 | 33012-SNV- | SNV | 1 | G | T | 44 | 25520 | 0.17241379 | 58.9545455 | WT | FALSE | CAN | JSY13 | 5 | WT-Rep5 | AGA | G>T | A/T+1 | G/C-1 | CG>AT | CG>AT | GTGAAAGAG | GAGGATG | TCT | TCT | C | A | C>A | C>A_TCT | CG>AT_GAGGATG | 9 | FALSE | FALSE |
| 188_v.txt | 33024 | 33024-SNV- | SNV | 1 | G | T | 32 | 25520 | 0.12539185 | 57.5 | WT | FALSE | CAN | JSY13 | 5 | WT-Rep5 | AGG | G>T | A/T+1 | G/C-1 | CG>AT | CG>AT | GTAACAGGG | GGGATGA | TCC | TCC | C | A | C>A | C>A_TCC | CG>AT_GGGATGA | 9 | FALSE | FALSE |
| 188_v.txt | 33033 | 33033-SNV- | SNV | 1 | G | T | 15 | 25520 | 0.05877743 | 55.1333333 | WT | FALSE | CAN | JSY13 | 5 | WT-Rep5 | TGT | G>T | A/T+1 | G/C-1 | CG>AT | CG>AT | ATGAATGTA | GTAGCCA | ACA | ACA | C | A | C>A | C>A_ACA | CG>AT_GTAGCCA | 9 | FALSE | FALSE |
| 188_v.txt | 33069 | 33069-SNV- | SNV | 1 | G | T | 1 | 25520 | 3.92E-03 | 63.993211 | WT | FALSE | CAN | JSY13 | 5 | WT-Rep5 | TGC | G>T | A/T+1 | G/C-1 | CG>AT | CG>AT | GAATATGCC | GCCAAAG | ACG | ACG | C | A | C>A | C>A_ACG | CG>AT_GCCAAAG | 9 | FALSE | FALSE |
| 188_v.txt | 33079 | 33079-SNV- | SNV | 1 | C | A | 9 | 25520 | 0.03526646 | 50.3333333 | WT | FALSE | CAN | JSY13 | 5 | WT-Rep5 | CCC | C>A | A/T+1 | G/C-1 | CG>AT | CG>AT | AAGAACCCA | CCATAAA | GGG | CCC | C | T | C>A | C>A_CCC | CG>AT_CCATAAA | 9 | FALSE | FALSE |
| 188_v.txt | 33093 | 33093-SNV- | SNV | 1 | G | T | 11 | 25520 | 0.04310345 | 48.7272727 | WT | FALSE | CAN | JSY13 | 5 | WT-Rep5 | TGA | G>T | A/T+1 | G/C-1 | CG>AT | CG>AT | AAATATGAT | GATATAA | ACT | ACT | C | A | C>A | C>A_ACT | CG>AT_GATATAA | 9 | FALSE | FALSE |
| 188_v.txt | 33111 | 33111-SNV- | SNV | 1 | G | T | 5 | 25520 | 0.01959248 | 41.9960607 | WT | FALSE | CAN | JSY13 | 5 | WT-Rep5 | TGG | G>T | A/T+1 | G/C-1 | CG>AT | CG>AT | CCCACTGGG | GGGCCGG | ACC | ACC | C | A | C>A | C>A_ACC | CG>AT_GGGCCGG | 9 | FALSE | FALSE |
| 188_v.txt | 33112 | 33112-SNV- | SNV | 1 | G | T | 46 | 25520 | 0.18025078 | 42.6799137 | WT | FALSE | CAN | JSY13 | 5 | WT-Rep5 | GGG | G>T | A/T+1 | G/C-1 | CG>AT | CG>AT | CCACTGGGC | GGCCGGC | CCC | CCC | C | A | C>A | C>A_CCC | CG>AT_GGCCGGC | 9 | FALSE | FALSE |
| 188_v.txt | 33154 | 33154-SNV- | SNV | 1 | C | A | 1819 | 25519 | 7.12802226 | 37.4498367 | WT | FALSE | CAN | JSY13 | 5 | WT-Rep5 | CCT | C>A | A/T+1 | G/C-1 | CG>AT | CG>AT | AAAGACCTG | CTGTACC | GGA | CCT | C | T | C>A | C>A_CCT | CG>AT_CTGTACC | 9 | FALSE | FALSE |
| 188_v.txt | 33169 | 33169-SNV- | SNV | 1 | C | A | 16 | 25518 | 0.06270084 | 37.9927127 | WT | FALSE | CAN | JSY13 | 5 | WT-Rep5 | CCA | C>A | A/T+1 | G/C-1 | CG>AT | CG>AT | TAGTACCAC | CACCAAG | GGT | CCA | C | T | C>A | C>A_CCA | CG>AT_CACCAAG | 9 | FALSE | FALSE |
| 188_v.txt | 33175 | 33175-SNV- | SNV | 1 | G | T | 9 | 25512 | 0.03527752 | 35 | WT | FALSE | CAN | JSY13 | 5 | WT-Rep5 | AGG | G>T | A/T+1 | G/C-1 | CG>AT | CG>AT | CACCAAGGG | GGGCAAT | TCC | TCC | C | A | C>A | C>A_TCC | CG>AT_GGGCAAT | 9 | FALSE | FALSE |
| 188_v.txt | 33238 | 33238-SNV- | SNV | 1 | C | A | 55 | 17251 | 0.3188221 | 37.0727273 | WT | FALSE | CAN | JSY13 | 5 | WT-Rep5 | CCT | C>A | A/T+1 | G/C-1 | CG>AT | CG>AT | CTTCTCCTTC | CTTCATC | GGA | CCT | C | T | C>A | C>A_CCT | CG>AT_CTTCATC | 9 | FALSE | FALSE |
| 188_v.txt | 33276 | 33276-SNV- | SNV | 1 | G | T | 7 | 17250 | 0.04057971 | 24.1388998 | WT | FALSE | CAN | JSY13 | 5 | WT-Rep5 | CGT | G>T | A/T+1 | G/C-1 | CG>AT | CG>AT | GAGAACGTA | GTATCCT | GCA | GCA | C | A | C>A | C>A_GCA | CG>AT_GTATCCT | 9 | FALSE | FALSE |
| 188_v.txt | 33325 | 33325-SNV- | SNV | 1 | C | A | 88 | 17251 | 0.51011536 | 57.0454546 | WT | FALSE | CAN | JSY13 | 5 | WT-Rep5 | TCT | C>A | A/T+1 | G/C-1 | CG>AT | CG>AT | ACAATTCTTT | CTTTACT | AGA | TCT | C | T | C>A | C>A_TCT | CG>AT_CTTTACT | 9 | FALSE | FALSE |
| 188_v.txt | 33436 | 33436-SNV- | SNV | 1 | C | A | 27 | 17251 | 0.15651267 | 46.7777778 | WT | FALSE | CAN | JSY13 | 5 | WT-Rep5 | TCT | C>A | A/T+1 | G/C-1 | CG>AT | CG>AT | TCTCCTCTAT | CTATGTC | AGA | TCT | C | T | C>A | C>A_TCT | CG>AT_CTATGTC | 9 | FALSE | FALSE |
| 188_v.txt | 33474 | 33474-SNV- | SNV | 1 | C | A | 52 | 17249 | 0.30146675 | 32.6153846 | WT | FALSE | CAN | JSY13 | 5 | WT-Rep5 | GCC | C>A | A/T+1 | G/C-1 | CG>AT | CG>AT | GCTATGCCTT | CCTTTTT | CGG | GCC | C | T | C>A | C>A_GCC | CG>AT_CCTTTTT | 9 | FALSE | FALSE |
| 188_v.txt | 33475 | 33475-SNV- | SNV | 1 | C | A | 11 | 17249 | 0.06377181 | 35.3636364 | WT | FALSE | CAN | JSY13 | 5 | WT-Rep5 | CCT | C>A | A/T+1 | G/C-1 | CG>AT | CG>AT | CTATGCCTTT | CTTTTTT | GGA | CCT | C | T | C>A | C>A_CCT | CG>AT_CTTTTTT | 9 | FALSE | FALSE |
| 188_v.txt | 32798 | 32798-MNV- | MNV | 2 | GT | AC | 1 | 30744 | 3.25E-03 | 37.9999732 | WT | FALSE | CAN | JSY13 | 5 | WT-Rep5 | CGT | nope | >1bp-ins | >1bpdel | none | MNV | TTCACCGTAA | GTAATAT | GCA | CGT | GT | none | NA | NA_CGT | MNV_GTAATAT | 9 | FALSE | FALSE |
| 188_v.txt | 33114 | 33114-MNV- | MNV | 2 | CC | TA | 1 | 25520 | 3.92E-03 | 36.0002879 | WT | FALSE | CAN | JSY13 | 5 | WT-Rep5 | GCC | nope | >1bp-ins | >1bpdel | none | MNV | ACTGGGCCG | CCGGCGT | CGG | GCC | CC | none | NA | NA_GCC | MNV_CCGGCGT | 9 | FALSE | FALSE |
| 188_v.txt | 33168 | 33168-MNV- | MNV | 2 | CC | AA | 3 | 25518 | 0.01175641 | 31.3333783 | WT | FALSE | CAN | JSY13 | 5 | WT-Rep5 | ACC | nope | >1bp-ins | >1bpdel | none | MNV | ATAGTACCA | CCACCAA | TGG | ACC | CC | none | NA | NA_ACC | MNV_CCACCAA | 9 | FALSE | FALSE |
| 188_v.txt | 31904 | 31904-Insert | Insertion | 1 | - | T | 3 | 4236 | 0.07082153 | 46.6666667 | WT | FALSE | CAN | JSY13 | 5 | WT-Rep5 | ATT | nope | A/T+1 | >1bpdel | none | A/T+1 | ATTGAATTTT | TTTTGGT | TAA | ATT | - | A | NA | NA_ATT | A/T+1_TTTTGGT | 9 | FALSE | FALSE |
| 188_v.txt | 32081 | 32081-Insert | Insertion | 1 | - | A | 239 | 12088 | 1.97716744 | 63.6485356 | WT | FALSE | CAN | JSY13 | 5 | WT-Rep5 | CAA | nope | A/T+1 | >1bpdel | none | A/T+1 | CCATGCAAA | AAAAAAG | GTT | CAA | - | T | NA | NA_CAA | A/T+1_AAAAAAG | 9 | FALSE | FALSE |
| 188_v.txt | 32121 | 32121-Insert | Insertion | 1 | - | A | 8 | 12088 | 0.06618134 | 60.75 | WT | FALSE | CAN | JSY13 | 5 | WT-Rep5 | GAA | nope | A/T+1 | >1bpdel | none | A/T+1 | CATTCGAAA | AAAACTT | CTT | GAA | - | T | NA | NA_GAA | A/T+1_AAAACTT | 9 | FALSE | FALSE |
| 188_v.txt | 32265 | 32265-Insert | Insertion | 1 | - | A | 2 | 31738 | 6.30E-03 | 37.5 | WT | FALSE | CAN | JSY13 | 5 | WT-Rep5 | TAA | nope | A/T+1 | >1bpdel | none | A/T+1 | CCAAATAAA | AAAATAC | ATT | TAA | - | T | NA | NA_TAA | A/T+1_AAAATAC | 9 | FALSE | FALSE |
| 188_v.txt | 32498 | 32498-Insert | Insertion | 1 | - | T | 149 | 31738 | 0.46946878 | 44.4966443 | WT | FALSE | CAN | JSY13 | 5 | WT-Rep5 | CTT | nope | A/T+1 | >1bpdel | none | A/T+1 | AACAACTTTT | TTTTTTG | GAA | CTT | - | A | NA | NA_CTT | A/T+1_TTTTTTG | 9 | FALSE | FALSE |
| 188_v.txt | 32528 | 32528-Insert | Insertion | 1 | - | G | 5 | 31738 | 0.01575399 | 38 | WT | FALSE | CAN | JSY13 | 5 | WT-Rep5 | TGG | nope | G/C+1 | >1bpdel | none | G/C+1 | TTTTCTGGG | GGGGTTT | ACC | TGG | - | C | NA | NA_TGG | G/C+1_GGGGTTT | 9 | FALSE | FALSE |
| 188_v.txt | 32631 | 32631-Insert | Insertion | 1 | - | T | 50 | 30744 | 0.16263336 | 49.82 | WT | FALSE | CAN | JSY13 | 5 | WT-Rep5 | GTT | nope | A/T+1 | >1bpdel | none | A/T+1 | CCTTCGTTTT | TTTTTAT | CAA | GTT | - | A | NA | NA_GTT | A/T+1_TTTTTAT | 9 | FALSE | FALSE |
| 188_v.txt | 32680 | 32680-Insert | Insertion | 1 | - | A | 11 | 30744 | 0.03577934 | 64 | WT | FALSE | CAN | JSY13 | 5 | WT-Rep5 | CAA | nope | A/T+1 | >1bpdel | none | A/T+1 | TTCTCCAATA | AATAACG | GTT | CAA | - | T | NA | NA_CAA | A/T+1_AATAACG | 9 | FALSE | FALSE |
| 188_v.txt | 32728 | 32728-Insert | Insertion | 1 | - | A | 132 | 30744 | 0.42935207 | 62.2045455 | WT | FALSE | CAN | JSY13 | 5 | WT-Rep5 | CAA | nope | A/T+1 | >1bpdel | none | A/T+1 | CCATACAAA | AAAAACA | GTT | CAA | - | T | NA | NA_CAA | A/T+1_AAAAACA | 9 | FALSE | FALSE |
| 188_v.txt | 32760 | 32760-Insert | Insertion | 1 | - | A | 4 | 30744 | 0.01301067 | 64 | WT | FALSE | CAN | JSY13 | 5 | WT-Rep5 | TAA | nope | A/T+1 | >1bpdel | none | A/T+1 | ATGGCTAAA | AAAACTT | ATT | TAA | - | T | NA | NA_TAA | A/T+1_AAAACTT | 9 | FALSE | FALSE |
| 188_v.txt | 32839 | 32839-Insert | Insertion | 1 | - | T | 1 | 30744 | 3.25E-03 | 3 | WT | FALSE | CAN | JSY13 | 5 | WT-Rep5 | ACC | nope | A/T+1 | >1bpdel | none | A/T+1 | TAATTACCCA | CCCAAAA | TGG | ACC | - | A | NA | NA_ACC | A/T+1_CCCAAAA | 9 | FALSE | FALSE |
| 188_v.txt | 32841 | 32841-Insert | Insertion | 1 | - | A | 1 | 30744 | 3.25E-03 | 38 | WT | FALSE | CAN | JSY13 | 5 | WT-Rep5 | CCA | nope | A/T+1 | >1bpdel | none | A/T+1 | ATTACCCAA | CAAAAAA | GGT | CCA | - | T | NA | NA_CCA | A/T+1_CAAAAAA | 9 | FALSE | FALSE |
| 188_v.txt | 32842 | 32842-Insert | Insertion | 1 | - | A | 1500 | 30744 | 4.87900078 | 42.0733333 | WT | FALSE | CAN | JSY13 | 5 | WT-Rep5 | CAA | nope | A/T+1 | >1bpdel | none | A/T+1 | TTACCCAAA | AAAAAAT | GTT | CAA | - | T | NA | NA_CAA | A/T+1_AAAAAAT | 9 | FALSE | FALSE |
| 188_v.txt | 32843 | 32843-Insert | Insertion | 1 | - | G | 1 | 30744 | 3.25E-03 | 38 | WT | FALSE | CAN | JSY13 | 5 | WT-Rep5 | AAA | nope | G/C+1 | >1bpdel | none | G/C+1 | TACCCAAAA | AAAAATA | TTT | AAA | - | C | NA | NA_AAA | G/C+1_AAAAATA | 9 | FALSE | FALSE |
| 188_v.txt | 32884 | 32884-Insert | Insertion | 1 | - | A | 3 | 25521 | 0.01175503 | 35 | WT | FALSE | CAN | JSY13 | 5 | WT-Rep5 | CAA | nope | A/T+1 | >1bpdel | none | A/T+1 | ACGTCCAAA | AAAATTG | GTT | CAA | - | T | NA | NA_CAA | A/T+1_AAAATTG | 9 | FALSE | FALSE |
| 188_v.txt | 32983 | 32983-Insert | Insertion | 1 | - | A | 3 | 25520 | 0.01175549 | 60.6666667 | WT | FALSE | CAN | JSY13 | 5 | WT-Rep5 | GAA | nope | A/T+1 | >1bpdel | none | A/T+1 | CTGGAGAAA | AAAGGAA | CTT | GAA | - | T | NA | NA_GAA | A/T+1_AAAGGAA | 9 | FALSE | FALSE |
| 188_v.txt | 33206 | 33206-Insert | Insertion | 2 | - | CT | 28 | 17251 | 0.16230943 | 37.2142857 | WT | FALSE | CAN | JSY13 | 5 | WT-Rep5 | GCT | nope | >1bp-ins | >1bpdel | none | >1bp-ins | CTTAAGCTCT | CTCTCTC | CGA | GCT | - | none | NA | NA_GCT | >1bp-ins_CTCTCTC | 9 | FALSE | FALSE |
| 188_v.txt | 33331 | 33331-Insert | Insertion | 1 | - | T | 3 | 17251 | 0.0173903 | 62.3333333 | WT | FALSE | CAN | JSY13 | 5 | WT-Rep5 | CTT | nope | A/T+1 | >1bpdel | none | A/T+1 | CTTTACTTTT | TTTTCTC | GAA | CTT | - | A | NA | NA_CTT | A/T+1_TTTTCTC | 9 | FALSE | FALSE |
| 188_v.txt | 33452 | 33452-Insert | Insertion | 1 | - | T | 3 | 17251 | 0.0173903 | 46.6666667 | WT | FALSE | CAN | JSY13 | 5 | WT-Rep5 | CTT | nope | A/T+1 | >1bpdel | none | A/T+1 | GTCTTCTTTT | TTTTGAA | GAA | CTT | - | A | NA | NA_CTT | A/T+1_TTTTGAA | 9 | FALSE | FALSE |
| 188_v.txt | 32381 | 32381-Deleti | Deletion | 14 | AAAGGGAG | - | 31 | 31738 | 0.09767471 | 63.3548387 | WT | FALSE | CAN | JSY13 | 5 | WT-Rep5 | TAA | nope | >1bp-ins | >1bpdel | none | >1bpdel | AATAATAAA | AAAGGGA | ATT | TAA | AAAGGGAG | none | NA | NA_TAA | >1bpdel_AAAGGGA | 9 | FALSE | FALSE |
| 188_v.txt | 33213 | 33213-Deleti | Deletion | 2 | TT | - | 14 | 17251 | 0.08115472 | 36.9285714 | WT | FALSE | CAN | JSY13 | 5 | WT-Rep5 | CTT | nope | >1bp-ins | >1bpdel | none | >1bpdel | TCTCTCTTCA | TTCACTT | GAA | CTT | TT | none | NA | NA_CTT | >1bpdel_TTCACTT | 9 | FALSE | FALSE |
| 188_v.txt | 31723 | 31723-Deleti | Deletion | 1 | A | - | 7 | 4236 | 0.16525024 | 64 | WT | FALSE | CAN | JSY13 | 5 | WT-Rep5 | CAA | nope | >1bp-ins | A/T-1 | none | A/T-1 | TGTCCCAAA | AAAAAGT | GTT | GTT | T | none | T>none | T>none_GTT | A/T-1_AAAAAGT | 9 | FALSE | FALSE |
| 188_v.txt | 31846 | 31846-Deleti | Deletion | 1 | A | - | 32 | 4236 | 0.75542965 | 51.9375 | WT | FALSE | CAN | JSY13 | 5 | WT-Rep5 | CAA | nope | >1bp-ins | A/T-1 | none | A/T-1 | AGATCCAAA | AAACAGC | GTT | GTT | T | none | T>none | T>none_GTT | A/T-1_AAACAGC | 9 | FALSE | FALSE |
| 188_v.txt | 31857 | 31857-Deleti | Deletion | 1 | A | - | 9 | 4236 | 0.21246459 | 52.1111111 | WT | FALSE | CAN | JSY13 | 5 | WT-Rep5 | GAA | nope | >1bp-ins | A/T-1 | none | A/T-1 | GCTAAGAAC | AACAGGA | CTT | CTT | T | none | T>none | T>none_CTT | A/T-1_AACAGGA | 9 | FALSE | FALSE |
| 188_v.txt | 31863 | 31863-Deleti | Deletion | 1 | A | - | 18 | 4236 | 0.42492918 | 44.4444444 | WT | FALSE | CAN | JSY13 | 5 | WT-Rep5 | GAA | nope | >1bp-ins | A/T-1 | none | A/T-1 | AACAGGAAA | AAAATAG | CTT | CTT | T | none | T>none | T>none_CTT | A/T-1_AAAATAG | 9 | FALSE | FALSE |
| 188_v.txt | 31904 | 31904-Deleti | Deletion | 1 | T | - | 13 | 4236 | 0.3068933 | 38.9999981 | WT | FALSE | CAN | JSY13 | 5 | WT-Rep5 | ATT | nope | >1bp-ins | A/T-1 | none | A/T-1 | ATTGAATTTT | TTTTGGT | TAA | ATT | T | none | nope | nope_ATT | A/T-1_TTTTGGT | 9 | FALSE | FALSE |
| 188_v.txt | 31913 | 31913-Deleti | Deletion | 1 | A | - | 5 | 4236 | 0.11803588 | 43.2 | WT | FALSE | CAN | JSY13 | 5 | WT-Rep5 | CAA | nope | >1bp-ins | A/T-1 | none | A/T-1 | TGGTGCAAA | AAAAGCC | GTT | GTT | T | none | T>none | T>none_GTT | A/T-1_AAAAGCC | 9 | FALSE | FALSE |
| 188_v.txt | 31979 | 31979-Deleti | Deletion | 1 | G | - | 24 | 12088 | 0.19854401 | 43.125 | WT | FALSE | CAN | JSY13 | 5 | WT-Rep5 | CGG | nope | >1bp-ins | G/C-1 | none | G/C-1 | CAAGCCGGG | GGGCATT | GCC | GCC | C | none | C>none | C>none_GCC | G/C-1_GGGCATT | 9 | FALSE | FALSE |
| 188_v.txt | 32030 | 32030-Deleti | Deletion | 1 | T | - | 13 | 12088 | 0.10754467 | 62 | WT | FALSE | CAN | JSY13 | 5 | WT-Rep5 | ATT | nope | >1bp-ins | A/T-1 | none | A/T-1 | ACGGTATTT | TTTCAAA | TAA | ATT | T | none | nope | nope_ATT | A/T-1_TTTCAAA | 9 | FALSE | FALSE |
| 188_v.txt | 32075 | 32075-Deleti | Deletion | 1 | C | - | 3 | 12088 | 0.024818 | 64 | WT | FALSE | CAN | JSY13 | 5 | WT-Rep5 | ACC | nope | >1bp-ins | G/C-1 | none | G/C-1 | AAATAACCA | CCATGCA | TGG | ACC | C | none | nope | nope_ACC | G/C-1_CCATGCA | 9 | FALSE | FALSE |
| 188_v.txt | 32081 | 32081-Deleti | Deletion | 1 | A | - | 389 | 12088 | 3.21806751 | 62.1773779 | WT | FALSE | CAN | JSY13 | 5 | WT-Rep5 | CAA | nope | >1bp-ins | A/T-1 | none | A/T-1 | CCATGCAAA | AAAAAAG | GTT | GTT | T | none | T>none | T>none_GTT | A/T-1_AAAAAAG | 9 | FALSE | FALSE |
| 188_v.txt | 32145 | 32145-Deleti | Deletion | 1 | G | - | 10 | 12088 | 0.08272667 | 51.1 | WT | FALSE | CAN | JSY13 | 5 | WT-Rep5 | TGT | nope | >1bp-ins | G/C-1 | none | G/C-1 | GTAGATGTC | GTCTCCA | ACA | ACA | C | none | C>none | C>none_ACA | G/C-1_GTCTCCA | 9 | FALSE | FALSE |

| | | | | | | | | | | | | | | | | | | | | | | | | | | | | | | | | | | |
|---|---|---|---|---|---|---|---|---|---|---|---|---|---|---|---|---|---|---|---|---|---|---|---|---|---|---|---|---|---|---|---|---|---|---|
| 188_v.txt | 32184 | 32184-Deleti | Deletion | 1 | A | - | 5 | 12088 | 0.04136334 | 45.4 | WT | FALSE | CAN | JSY13 | 5 | WT-Rep5 | GAA | nope | >1bp-ins | A/T-1 | none | A/T-1 | GTAACGAAA | AAAACTG | CTT | CTT | T | none | T>none | T>none_CTT | A/T-1_AAAACTG | 9 | FALSE | FALSE |
| 188_v.txt | 32199 | 32199-Deleti | Deletion | 1 | G | - | 5 | 12088 | 0.04136334 | 43 | WT | FALSE | CAN | JSY13 | 5 | WT-Rep5 | TGG | nope | >1bp-ins | G/C-1 | none | G/C-1 | ATGTATGGA | GGAACAC | ACC | ACC | C | none | C>none | C>none_ACC | G/C-1_GGAACAC | 9 | FALSE | FALSE |
| 188_v.txt | 32231 | 32231-Deleti | Deletion | 1 | T | - | 8 | 12088 | 0.06618134 | 37.875 | WT | FALSE | CAN | JSY13 | 5 | WT-Rep5 | ATT | nope | >1bp-ins | A/T-1 | none | A/T-1 | CAGGAATTT | TTTAGGA | TAA | ATT | T | none | nope | nope_ATT | A/T-1_TTTAGGA | 9 | FALSE | FALSE |
| 188_v.txt | 32272 | 32272-Deleti | Deletion | 1 | G | - | 3 | 31738 | 9.45E-03 | 37.0000001 | WT | FALSE | CAN | JSY13 | 5 | WT-Rep5 | CGG | nope | >1bp-ins | G/C-1 | none | G/C-1 | AAATACGGG | GGGAACC | GCC | GCC | C | none | C>none | C>none_GCC | G/C-1_GGGAACC | 9 | FALSE | FALSE |
| 188_v.txt | 32275 | 32275-Deleti | Deletion | 1 | A | - | 11 | 31738 | 0.03465877 | 37.1818182 | WT | FALSE | CAN | JSY13 | 5 | WT-Rep5 | GAA | nope | >1bp-ins | A/T-1 | none | A/T-1 | TACGGGAAC | AACCAAC | CTT | CTT | T | none | T>none | T>none_CTT | A/T-1_AACCAAC | 9 | FALSE | FALSE |
| 188_v.txt | 32334 | 32334-Deleti | Deletion | 1 | A | - | 66 | 31738 | 0.20795261 | 59.9545455 | WT | FALSE | CAN | JSY13 | 5 | WT-Rep5 | GAA | nope | >1bp-ins | A/T-1 | none | A/T-1 | GCGTTGAAG | AAGATAT | CTT | CTT | T | none | T>none | T>none_CTT | A/T-1_AAGATAT | 9 | FALSE | FALSE |
| 188_v.txt | 32346 | 32346-Deleti | Deletion | 1 | A | - | 81 | 31738 | 0.25521457 | 61.4197531 | WT | FALSE | CAN | JSY13 | 5 | WT-Rep5 | CAA | nope | >1bp-ins | A/T-1 | none | A/T-1 | TGTGGCAAA | AAAACCT | GTT | GTT | T | none | T>none | T>none_GTT | A/T-1_AAAACCT | 9 | FALSE | FALSE |
| 188_v.txt | 32381 | 32381-Deleti | Deletion | 1 | A | - | 5 | 31736 | 0.01575498 | 61.3999995 | WT | FALSE | CAN | JSY13 | 5 | WT-Rep5 | TAA | nope | >1bp-ins | A/T-1 | none | A/T-1 | AATAATAAA | AAAGGGA | ATT | ATT | T | none | T>none | T>none_ATT | A/T-1_AAAGGGA | 9 | FALSE | FALSE |
| 188_v.txt | 32424 | 32424-Deleti | Deletion | 1 | G | - | 11 | 31738 | 0.03465877 | 55.2727273 | WT | FALSE | CAN | JSY13 | 5 | WT-Rep5 | AGG | nope | >1bp-ins | G/C-1 | none | G/C-1 | AGTTTAGGG | GGGTCAT | TCC | TCC | C | none | C>none | C>none_TCC | G/C-1_GGGTCAT | 9 | FALSE | FALSE |
| 188_v.txt | 32442 | 32442-Deleti | Deletion | 1 | A | - | 9 | 31738 | 0.02835717 | 55.2222222 | WT | FALSE | CAN | JSY13 | 5 | WT-Rep5 | TAA | nope | >1bp-ins | A/T-1 | none | A/T-1 | GGAACTAAA | AAAAGTC | ATT | ATT | T | none | T>none | T>none_ATT | A/T-1_AAAAGTC | 9 | FALSE | FALSE |
| 188_v.txt | 32490 | 32490-Deleti | Deletion | 1 | A | - | 557 | 31738 | 1.75499401 | 45.3393178 | WT | FALSE | CAN | JSY13 | 5 | WT-Rep5 | GAA | nope | >1bp-ins | A/T-1 | none | A/T-1 | ATACGGAAA | AAAACAA | CTT | CTT | T | none | T>none | T>none_CTT | A/T-1_AAAACAA | 9 | FALSE | FALSE |
| 188_v.txt | 32498 | 32498-Deleti | Deletion | 1 | T | - | 947 | 31738 | 2.9838049 | 45.2460385 | WT | FALSE | CAN | JSY13 | 5 | WT-Rep5 | CTT | nope | >1bp-ins | A/T-1 | none | A/T-1 | AACAACTTTT | TTTTTTG | GAA | CTT | T | none | nope | nope_CTT | A/T-1_TTTTTTG | 9 | FALSE | FALSE |
| 188_v.txt | 32514 | 32514-Deleti | Deletion | 1 | G | - | 25 | 31738 | 0.07876993 | 40 | WT | FALSE | CAN | JSY13 | 5 | WT-Rep5 | TGG | nope | >1bp-ins | G/C-1 | none | G/C-1 | GCTCTTGGA | GGAACGG | ACC | ACC | C | none | C>none | C>none_ACC | G/C-1_GGAACGG | 9 | FALSE | FALSE |
| 188_v.txt | 32522 | 32522-Deleti | Deletion | 1 | T | - | 13 | 31738 | 0.04096036 | 38 | WT | FALSE | CAN | JSY13 | 5 | WT-Rep5 | ATT | nope | >1bp-ins | A/T-1 | none | A/T-1 | AACGGATTT | TTTTCTG | TAA | ATT | T | none | nope | nope_ATT | A/T-1_TTTTCTG | 9 | FALSE | FALSE |
| 188_v.txt | 32528 | 32528-Deleti | Deletion | 1 | G | - | 5 | 31738 | 0.01575399 | 41.7999982 | WT | FALSE | CAN | JSY13 | 5 | WT-Rep5 | TGG | nope | >1bp-ins | G/C-1 | none | G/C-1 | TTTTCTGGG | GGGGTTT | ACC | ACC | C | none | C>none | C>none_ACC | G/C-1_GGGGTTT | 9 | FALSE | FALSE |
| 188_v.txt | 32605 | 32605-Deleti | Deletion | 1 | A | - | 9 | 30744 | 0.02927401 | 40.6666667 | WT | FALSE | CAN | JSY13 | 5 | WT-Rep5 | GAA | nope | >1bp-ins | A/T-1 | none | A/T-1 | AAGAGGAAA | AAACCCA | CTT | CTT | T | none | T>none | T>none_CTT | A/T-1_AAACCCA | 9 | FALSE | FALSE |
| 188_v.txt | 32608 | 32608-Deleti | Deletion | 1 | C | - | 42 | 30744 | 0.13661202 | 43.7619027 | WT | FALSE | CAN | JSY13 | 5 | WT-Rep5 | ACC | nope | >1bp-ins | G/C-1 | none | G/C-1 | AGGAAACCC | CCCAACC | TGG | ACC | C | none | nope | nope_ACC | G/C-1_CCCAACC | 9 | FALSE | FALSE |
| 188_v.txt | 32624 | 32624-Deleti | Deletion | 1 | C | - | 25 | 30744 | 0.08131668 | 47.28 | WT | FALSE | CAN | JSY13 | 5 | WT-Rep5 | TCC | nope | >1bp-ins | G/C-1 | none | G/C-1 | GAACCTCCCT | CCCTTCG | AGG | TCC | C | none | nope | nope_TCC | G/C-1_CCCTTCG | 9 | FALSE | FALSE |
| 188_v.txt | 32631 | 32631-Deleti | Deletion | 1 | T | - | 89 | 30744 | 0.28948738 | 49.5730334 | WT | FALSE | CAN | JSY13 | 5 | WT-Rep5 | GTT | nope | >1bp-ins | A/T-1 | none | A/T-1 | CCTTCGTTTT | TTTTTAT | CAA | GTT | T | none | nope | nope_GTT | A/T-1_TTTTTAT | 9 | FALSE | FALSE |
| 188_v.txt | 32640 | 32640-Deleti | Deletion | 1 | T | - | 8 | 30744 | 0.02602134 | 56.625 | WT | FALSE | CAN | JSY13 | 5 | WT-Rep5 | CTT | nope | >1bp-ins | A/T-1 | none | A/T-1 | TTATCCTTAG | TTAGATA | GAA | CTT | T | none | nope | nope_CTT | A/T-1_TTAGATA | 9 | FALSE | FALSE |
| 188_v.txt | 32658 | 32658-Deleti | Deletion | 1 | C | - | 33 | 30744 | 0.10733802 | 58.9090909 | WT | FALSE | CAN | JSY13 | 5 | WT-Rep5 | ACC | nope | >1bp-ins | G/C-1 | none | G/C-1 | CCTGGACCC | CCCCAGG | TGG | ACC | C | none | nope | nope_ACC | G/C-1_CCCCAGG | 9 | FALSE | FALSE |
| 188_v.txt | 32662 | 32662-Deleti | Deletion | 1 | A | - | 12 | 30744 | 0.03903201 | 59.5833285 | WT | FALSE | CAN | JSY13 | 5 | WT-Rep5 | CAG | nope | >1bp-ins | A/T-1 | none | A/T-1 | GACCCCAGG | AGGCACC | GTC | GTC | T | none | T>none | T>none_GTC | A/T-1_AGGCACC | 9 | FALSE | FALSE |
| 188_v.txt | 32663 | 32663-Deleti | Deletion | 1 | G | - | 10 | 30744 | 0.03252667 | 62.5000001 | WT | FALSE | CAN | JSY13 | 5 | WT-Rep5 | AGG | nope | >1bp-ins | G/C-1 | none | G/C-1 | ACCCCAGGC | GGCACCT | TCC | TCC | C | none | C>none | C>none_TCC | G/C-1_GGCACCT | 9 | FALSE | FALSE |
| 188_v.txt | 32670 | 32670-Deleti | Deletion | 1 | G | - | 93 | 30744 | 0.30249805 | 58.0322581 | WT | FALSE | CAN | JSY13 | 5 | WT-Rep5 | TGG | nope | >1bp-ins | G/C-1 | none | G/C-1 | GCACCTGGG | GGGTTTC | ACC | ACC | C | none | C>none | C>none_ACC | G/C-1_GGGTTTC | 9 | FALSE | FALSE |
| 188_v.txt | 32680 | 32680-Deleti | Deletion | 1 | A | - | 1 | 30744 | 3.25E-03 | 26 | WT | FALSE | CAN | JSY13 | 5 | WT-Rep5 | CAA | nope | >1bp-ins | A/T-1 | none | A/T-1 | TTCTCCAATA | AATAACG | GTT | GTT | T | none | T>none | T>none_GTT | A/T-1_AATAACG | 9 | FALSE | FALSE |
| 188_v.txt | 32691 | 32691-Deleti | Deletion | 1 | C | - | 27 | 30743 | 0.08782487 | 60.3333333 | WT | FALSE | CAN | JSY13 | 5 | WT-Rep5 | TCC | nope | >1bp-ins | G/C-1 | none | G/C-1 | CGGAATCCA | CCAACTG | AGG | TCC | C | none | nope | nope_TCC | G/C-1_CCAACTG | 9 | FALSE | FALSE |
| 188_v.txt | 32697 | 32697-Deleti | Deletion | 1 | G | - | 6 | 30744 | 0.019516 | 64 | WT | FALSE | CAN | JSY13 | 5 | WT-Rep5 | TGG | nope | >1bp-ins | G/C-1 | none | G/C-1 | CCAACTGGG | GGGCCGG | ACC | ACC | C | none | C>none | C>none_ACC | G/C-1_GGGCCGG | 9 | FALSE | FALSE |
| 188_v.txt | 32707 | 32707-Deleti | Deletion | 1 | C | - | 58 | 30744 | 0.1886547 | 62.7413793 | WT | FALSE | CAN | JSY13 | 5 | WT-Rep5 | ACC | nope | >1bp-ins | G/C-1 | none | G/C-1 | CGGTAACCC | CCCCAGC | TGG | ACC | C | none | nope | nope_ACC | G/C-1_CCCCAGC | 9 | FALSE | FALSE |
| 188_v.txt | 32719 | 32719-Deleti | Deletion | 1 | A | - | 34 | 30744 | 0.11059068 | 60.9411765 | WT | FALSE | CAN | JSY13 | 5 | WT-Rep5 | CAA | nope | >1bp-ins | A/T-1 | none | A/T-1 | CACCACAAA | AAACCAT | GTT | GTT | T | none | T>none | T>none_GTT | A/T-1_AAACCAT | 9 | FALSE | FALSE |
| 188_v.txt | 32728 | 32728-Deleti | Deletion | 1 | A | - | 4412 | 30742 | 14.3517013 | 60.9111492 | WT | FALSE | CAN | JSY13 | 5 | WT-Rep5 | CAA | nope | >1bp-ins | A/T-1 | none | A/T-1 | CCATACAAA | AAAAACA | GTT | GTT | T | none | T>none | T>none_GTT | A/T-1_AAAAACA | 9 | FALSE | FALSE |
| 188_v.txt | 32740 | 32740-Deleti | Deletion | 1 | T | - | 5 | 30744 | 0.01626334 | 47.6 | WT | FALSE | CAN | JSY13 | 5 | WT-Rep5 | ATT | nope | >1bp-ins | A/T-1 | none | A/T-1 | AGTATATTA | TTAGAAA | TAA | ATT | T | none | nope | nope_ATT | A/T-1_TTAGAAA | 9 | FALSE | FALSE |
| 188_v.txt | 32744 | 32744-Deleti | Deletion | 1 | A | - | 22 | 30744 | 0.07155868 | 59.9545455 | WT | FALSE | CAN | JSY13 | 5 | WT-Rep5 | GAA | nope | >1bp-ins | A/T-1 | none | A/T-1 | TATTAGAAA | AAACCCG | CTT | CTT | T | none | T>none | T>none_CTT | A/T-1_AAACCCG | 9 | FALSE | FALSE |
| 188_v.txt | 32760 | 32760-Deleti | Deletion | 1 | A | - | 300 | 30744 | 0.97580016 | 52.909999 | WT | FALSE | CAN | JSY13 | 5 | WT-Rep5 | TAA | nope | >1bp-ins | A/T-1 | none | A/T-1 | ATGGCTAAA | AAAACTT | ATT | ATT | T | none | T>none | T>none_ATT | A/T-1_AAAACTT | 9 | FALSE | FALSE |
| 188_v.txt | 32779 | 32779-Deleti | Deletion | 1 | C | - | 7 | 30743 | 0.02276941 | 52.8571356 | WT | FALSE | CAN | JSY13 | 5 | WT-Rep5 | ACC | nope | >1bp-ins | G/C-1 | none | G/C-1 | AAGCGACCC | CCCAGAA | TGG | ACC | C | none | nope | nope_ACC | G/C-1_CCCAGAA | 9 | FALSE | FALSE |
| 188_v.txt | 32839 | 32839-Deleti | Deletion | 1 | C | - | 23 | 30744 | 0.07481135 | 40.086952 | WT | FALSE | CAN | JSY13 | 5 | WT-Rep5 | ACC | nope | >1bp-ins | G/C-1 | none | G/C-1 | TAATTACCCA | CCCAAAA | TGG | ACC | C | none | nope | nope_ACC | G/C-1_CCCAAAA | 9 | FALSE | FALSE |
| 188_v.txt | 32842 | 32842-Deleti | Deletion | 1 | A | - | 2457 | 30744 | 7.99180328 | 41.963775 | WT | FALSE | CAN | JSY13 | 5 | WT-Rep5 | CAA | nope | >1bp-ins | A/T-1 | none | A/T-1 | TTACCCAAA | AAAAAAT | GTT | GTT | T | none | T>none | T>none_GTT | A/T-1_AAAAAAT | 9 | FALSE | FALSE |
| 188_v.txt | 32884 | 32884-Deleti | Deletion | 1 | A | - | 8 | 25521 | 0.03134673 | 37.1250001 | WT | FALSE | CAN | JSY13 | 5 | WT-Rep5 | CAA | nope | >1bp-ins | A/T-1 | none | A/T-1 | ACGTCCAAA | AAAATTG | GTT | GTT | T | none | T>none | T>none_GTT | A/T-1_AAAATTG | 9 | FALSE | FALSE |
| 188_v.txt | 32901 | 32901-Deleti | Deletion | 1 | C | - | 1 | 25520 | 3.92E-03 | 38 | WT | FALSE | CAN | JSY13 | 5 | WT-Rep5 | GCC | nope | >1bp-ins | G/C-1 | none | G/C-1 | ACTTGGCCA | CCAACTA | CGG | GCC | C | none | nope | nope_GCC | G/C-1_CCAACTA | 9 | FALSE | FALSE |
| 188_v.txt | 32924 | 32924-Deleti | Deletion | 1 | A | - | 526 | 25520 | 2.06112853 | 53.6806084 | WT | FALSE | CAN | JSY13 | 5 | WT-Rep5 | CAA | nope | >1bp-ins | A/T-1 | none | A/T-1 | CAGGGCAAA | AAAAGTG | GTT | GTT | T | none | T>none | T>none_GTT | A/T-1_AAAAGTG | 9 | FALSE | FALSE |
| 188_v.txt | 32941 | 32941-Deleti | Deletion | 1 | A | - | 122 | 25520 | 0.47805643 | 56.8196654 | WT | FALSE | CAN | JSY13 | 5 | WT-Rep5 | GAA | nope | >1bp-ins | A/T-1 | none | A/T-1 | CCCAAGAAA | AAAACCA | CTT | CTT | T | none | T>none | T>none_CTT | A/T-1_AAAACCA | 9 | FALSE | FALSE |
| 188_v.txt | 32972 | 32972-Deleti | Deletion | 1 | A | - | 19 | 25520 | 0.07445141 | 63.0526316 | WT | FALSE | CAN | JSY13 | 5 | WT-Rep5 | CAA | nope | >1bp-ins | A/T-1 | none | A/T-1 | CGCACCAAA | AAATGCT | GTT | GTT | T | none | T>none | T>none_GTT | A/T-1_AAATGCT | 9 | FALSE | FALSE |
| 188_v.txt | 33000 | 33000-Deleti | Deletion | 1 | A | - | 284 | 25520 | 1.11285267 | 62.4401409 | WT | FALSE | CAN | JSY13 | 5 | WT-Rep5 | GAA | nope | >1bp-ins | A/T-1 | none | A/T-1 | TGTGAGAAA | AAAACTG | CTT | CTT | T | none | T>none | T>none_CTT | A/T-1_AAAACTG | 9 | FALSE | FALSE |
| 188_v.txt | 33040 | 33040-Deleti | Deletion | 1 | T | - | 18 | 25520 | 0.07053292 | 56.8888889 | WT | FALSE | CAN | JSY13 | 5 | WT-Rep5 | ATT | nope | >1bp-ins | A/T-1 | none | A/T-1 | TAGCCATTTC | TTTCACC | TAA | ATT | T | none | nope | nope_ATT | A/T-1_TTTCACC | 9 | FALSE | FALSE |
| 188_v.txt | 33147 | 33147-Deleti | Deletion | 1 | A | - | 19 | 25520 | 0.07445141 | 40.1578947 | WT | FALSE | CAN | JSY13 | 5 | WT-Rep5 | GAA | nope | >1bp-ins | A/T-1 | none | A/T-1 | CCAATGAAA | AAAAGAC | CTT | CTT | T | none | T>none | T>none_CTT | A/T-1_AAAAGAC | 9 | FALSE | FALSE |
| 188_v.txt | 33154 | 33154-Deleti | Deletion | 1 | C | - | 1 | 25519 | 3.92E-03 | 38 | WT | FALSE | CAN | JSY13 | 5 | WT-Rep5 | CCT | nope | >1bp-ins | G/C-1 | none | G/C-1 | AAAGACCTG | CTGTACC | GGA | CCT | C | none | nope | nope_CCT | G/C-1_CTGTACC | 9 | FALSE | FALSE |
| 188_v.txt | 33168 | 33168-Deleti | Deletion | 1 | C | - | 2 | 25518 | 7.84E-03 | 38 | WT | FALSE | CAN | JSY13 | 5 | WT-Rep5 | ACC | nope | >1bp-ins | G/C-1 | none | G/C-1 | ATAGTACCA | CCACCAA | TGG | ACC | C | none | nope | nope_ACC | G/C-1_CCACCAA | 9 | FALSE | FALSE |
| 188_v.txt | 33175 | 33175-Deleti | Deletion | 1 | G | - | 2 | 25512 | 7.84E-03 | 38 | WT | FALSE | CAN | JSY13 | 5 | WT-Rep5 | AGG | nope | >1bp-ins | G/C-1 | none | G/C-1 | CACCAAGGG | GGGCAAT | TCC | TCC | C | none | C>none | C>none_TCC | G/C-1_GGGCAAT | 9 | FALSE | FALSE |
| 188_v.txt | 33196 | 33196-Deleti | Deletion | 1 | T | - | 8 | 17251 | 0.04637412 | 35.75 | WT | FALSE | CAN | JSY13 | 5 | WT-Rep5 | CTT | nope | >1bp-ins | A/T-1 | none | A/T-1 | TATGTCTTTG | TTTGCTT | GAA | CTT | T | none | nope | nope_CTT | A/T-1_TTTGCTT | 9 | FALSE | FALSE |
| 188_v.txt | 33474 | 33474-Deleti | Deletion | 1 | C | - | 40 | 17249 | 0.2318975 | 30.6 | WT | FALSE | CAN | JSY13 | 5 | WT-Rep5 | GCC | nope | >1bp-ins | G/C-1 | none | G/C-1 | GCTATGCCTT | CCTTTTT | CGG | GCC | C | none | nope | nope_GCC | G/C-1_CCTTTTT | 9 | FALSE | FALSE |
| 129_v.txt | 32112 | 32112-SNV- | SNV | 1 | A | C | 8 | 18543 | 0.04314297 | 57.25 | WT | FALSE | CAN | JSY2424 | 6 | WT-Rep6 | TAG | A>C | G/C+1 | A/T-1 | TA>GC | TA>GC | TTTAATAGC | AGCCATT | ATC | ATC | T | G | T>G | T>G_ATC | TA>GC_AGCCATT | 9 | FALSE | FALSE |
| 129_v.txt | 32283 | 32283-SNV-T | SNV | 1 | T | G | 233 | 34188 | 0.68152568 | 50.8525316 | WT | FALSE | CAN | JSY2424 | 6 | WT-Rep6 | GTA | T>G | G/C+1 | A/T-1 | TA>GC | TA>GC | CCAACGTAA | TAAATAT | CAT | GTA | T | C | T>G | T>G_GTA | TA>GC_TAAATAT | 9 | FALSE | FALSE |
| 129_v.txt | 32369 | 32369-SNV- | SNV | 1 | A | C | 11 | 34188 | 0.03217503 | 54.0002365 | WT | FALSE | CAN | JSY2424 | 6 | WT-Rep6 | CAA | A>C | G/C+1 | A/T-1 | TA>GC | TA>GC | GTTCTCAATA | AATAGCA | GTT | GTT | T | G | T>G | T>G_GTT | TA>GC_AATAGCA | 9 | FALSE | FALSE |
| 129_v.txt | 32493 | 32493-SNV- | SNV | 1 | A | C | 21 | 34188 | 0.06142506 | 22.7143474 | WT | FALSE | CAN | JSY2424 | 6 | WT-Rep6 | AAC | A>C | G/C+1 | A/T-1 | TA>GC | TA>GC | CGGAAAACA | ACAACTT | TTG | TTG | T | G | T>G | T>G_TTG | TA>GC_ACAACTT | 9 | FALSE | FALSE |
| 129_v.txt | 32573 | 32573-SNV-T | SNV | 1 | T | G | 6 | 30732 | 0.01952362 | 37.6666667 | WT | FALSE | CAN | JSY2424 | 6 | WT-Rep6 | CTT | T>G | G/C+1 | A/T-1 | TA>GC | TA>GC | AGTACCTTG | TTGAAAT | GAA | CTT | T | C | T>G | T>G_CTT | TA>GC_TTGAAAT | 9 | FALSE | FALSE |
| 129_v.txt | 32792 | 32792-SNV-T | SNV | 1 | T | G | 24 | 30731 | 0.07809704 | 43.375 | WT | FALSE | CAN | JSY2424 | 6 | WT-Rep6 | ATT | T>G | G/C+1 | A/T-1 | TA>GC | TA>GC | CTCGAATTCA | TTCACCG | TAA | ATT | T | C | T>G | T>G_ATT | TA>GC_TTCACCG | 9 | FALSE | FALSE |
| 129_v.txt | 32799 | 32799-SNV-T | SNV | 1 | T | G | 82 | 30731 | 0.26683154 | 44.4036514 | WT | FALSE | CAN | JSY2424 | 6 | WT-Rep6 | GTA | T>G | G/C+1 | A/T-1 | TA>GC | TA>GC | TCACCGTAAT | TAATATT | CAT | GTA | T | C | T>G | T>G_GTA | TA>GC_TAATATT | 9 | FALSE | FALSE |
| 129_v.txt | 32842 | 32842-SNV- | SNV | 1 | A | C | 4 | 30731 | 0.01301617 | 37.0002254 | WT | FALSE | CAN | JSY2424 | 6 | WT-Rep6 | CAA | A>C | G/C+1 | A/T-1 | TA>GC | TA>GC | TTACCCAAA | AAAAAAT | GTT | GTT | T | G | T>G | T>G_GTT | TA>GC_AAAAAAT | 9 | FALSE | FALSE |
| 129_v.txt | 32884 | 32884-SNV- | SNV | 1 | A | C | 4 | 27506 | 0.01454228 | 30.5003814 | WT | FALSE | CAN | JSY2424 | 6 | WT-Rep6 | CAA | A>C | G/C+1 | A/T-1 | TA>GC | TA>GC | ACGTCCAAA | AAAATTG | GTT | GTT | T | G | T>G | T>G_GTT | TA>GC_AAAATTG | 9 | FALSE | FALSE |
| 129_v.txt | 32938 | 32938-SNV- | SNV | 1 | A | C | 35 | 27506 | 0.12724497 | 49.6285714 | WT | FALSE | CAN | JSY2424 | 6 | WT-Rep6 | CAA | A>C | G/C+1 | A/T-1 | TA>GC | TA>GC | TTGCCCAAG | AAGAAAA | GTT | GTT | T | G | T>G | T>G_GTT | TA>GC_AAGAAAA | 9 | FALSE | FALSE |
| 129_v.txt | 33065 | 33065-SNV- | SNV | 1 | A | C | 90 | 27506 | 0.32720134 | 54.6222222 | WT | FALSE | CAN | JSY2424 | 6 | WT-Rep6 | AAT | A>C | G/C+1 | A/T-1 | TA>GC | TA>GC | GACAGAATA | ATATGCC | TTA | TTA | T | G | T>G | T>G_TTA | TA>GC_ATATGCC | 9 | FALSE | FALSE |
| 129_v.txt | 33081 | 33081-SNV- | SNV | 1 | A | C | 20 | 27506 | 0.07271141 | 56.65 | WT | FALSE | CAN | JSY2424 | 6 | WT-Rep6 | CAT | A>C | G/C+1 | A/T-1 | TA>GC | TA>GC | GAACCCATA | ATAAATA | GTA | GTA | T | G | T>G | T>G_GTA | TA>GC_ATAAATA | 9 | FALSE | FALSE |
| 129_v.txt | 33099 | 33099-SNV- | SNV | 1 | A | C | 18 | 27506 | 0.06544027 | 39.6111111 | WT | FALSE | CAN | JSY2424 | 6 | WT-Rep6 | AAG | A>C | G/C+1 | A/T-1 | TA>GC | TA>GC | GATATAAGA | AGAGCGC | TTC | TTC | T | G | T>G | T>G_TTC | TA>GC_AGAGCGC | 9 | FALSE | FALSE |
| 129_v.txt | 33126 | 33126-SNV- | SNV | 1 | A | C | 11 | 27506 | 0.03999128 | 37.7272727 | WT | FALSE | CAN | JSY2424 | 6 | WT-Rep6 | CAG | A>C | G/C+1 | A/T-1 | TA>GC | TA>GC | TTGGTCAGA | AGAGGTG | GTC | GTC | T | G | T>G | T>G_GTC | TA>GC_AGAGGTG | 9 | FALSE | FALSE |
| 129_v.txt | 33204 | 33204-SNV- | SNV | 1 | A | C | 32 | 27156 | 0.11783768 | 36.75 | WT | FALSE | CAN | JSY2424 | 6 | WT-Rep6 | AAG | A>C | G/C+1 | A/T-1 | TA>GC | TA>GC | TGCTTAAGC | AGCTCTC | TTC | TTC | T | G | T>G | T>G_TTC | TA>GC_AGCTCTC | 9 | FALSE | FALSE |
| 129_v.txt | 33269 | 33269-SNV- | SNV | 1 | A | C | 192 | 27157 | 0.70700004 | 17.6614583 | WT | FALSE | CAN | JSY2424 | 6 | WT-Rep6 | TAG | A>C | G/C+1 | A/T-1 | TA>GC | TA>GC | CTCCATAGA | AGAGAAC | ATC | ATC | T | G | T>G | T>G_ATC | TA>GC_AGAGAAC | 9 | FALSE | FALSE |
| 129_v.txt | 32275 | 32275-SNV- | SNV | 1 | A | G | 36 | 34188 | 0.10530011 | 33.2777078 | WT | FALSE | CAN | JSY2424 | 6 | WT-Rep6 | GAA | A>G | G/C+1 | A/T-1 | TA>CG | TA>CG | TACGGGAAC | AACCAAC | CTT | CTT | T | C | T>C | T>C_CTT | TA>CG_AACCAAC | 9 | FALSE | FALSE |
| 129_v.txt | 32338 | 32338-SNV-T | SNV | 1 | T | C | 4 | 34188 | 0.01170001 | 63.9997488 | WT | FALSE | CAN | JSY2424 | 6 | WT-Rep6 | ATA | T>C | G/C+1 | A/T-1 | TA>CG | TA>CG | TGAAGATAT | TATGTGG | TAT | ATA | T | G | T>C | T>C_ATA | TA>CG_TATGTGG | 9 | FALSE | FALSE |
| 129_v.txt | 32573 | 32573-SNV-T | SNV | 1 | T | C | 15 | 30732 | 0.04880906 | 35.2 | WT | FALSE | CAN | JSY2424 | 6 | WT-Rep6 | CTT | T>C | G/C+1 | A/T-1 | TA>CG | TA>CG | AGTACCTTG | TTGAAAT | GAA | CTT | T | G | T>C | T>C_CTT | TA>CG_TTGAAAT | 9 | FALSE | FALSE |
| 129_v.txt | 32799 | 32799-SNV-T | SNV | 1 | T | C | 18 | 30731 | 0.05857278 | 46.4823238 | WT | FALSE | CAN | JSY2424 | 6 | WT-Rep6 | GTA | T>C | G/C+1 | A/T-1 | TA>CG | TA>CG | TCACCGTAAT | TAATATT | CAT | GTA | T | G | T>C | T>C_GTA | TA>CG_TAATATT | 9 | FALSE | FALSE |
| 129_v.txt | 33139 | 33139-SNV- | SNV | 1 | A | G | 30 | 27506 | 0.10906711 | 39.3666667 | WT | FALSE | CAN | JSY2424 | 6 | WT-Rep6 | AAA | A>G | G/C+1 | A/T-1 | TA>CG | TA>CG | TGGATAAAC | AACCAAT | TTT | TTT | T | C | T>C | T>C_TTT | TA>CG_AACCAAT | 9 | FALSE | FALSE |
| 129_v.txt | 33476 | 33476-SNV-T | SNV | 1 | T | C | 28 | 27153 | 0.10311936 | 33 | WT | FALSE | CAN | JSY2424 | 6 | WT-Rep6 | CTT | T>C | G/C+1 | A/T-1 | TA>CG | TA>CG | TATGCCTTTT | TTTTTTT | GAA | CTT | T | G | T>C | T>C_CTT | TA>CG_TTTTTTT | 9 | FALSE | FALSE |
| 129_v.txt | 31874 | 31874-SNV- | SNV | 1 | A | T | 115 | 10450 | 1.10047847 | 42.3652174 | WT | FALSE | CAN | JSY2424 | 6 | WT-Rep6 | TAT | A>T | A/T+1 | A/T-1 | TA>AT | TA>AT | AGAGATATA | ATAGGCG | ATA | ATA | T | A | T>A | T>A_ATA | TA>AT_ATAGGCG | 9 | FALSE | FALSE |
| 129_v.txt | 31990 | 31990-SNV-T | SNV | 1 | T | A | 28 | 18543 | 0.15100038 | 52.8214286 | WT | FALSE | CAN | JSY2424 | 6 | WT-Rep6 | TTA | T>A | A/T+1 | A/T-1 | TA>AT | TA>AT | TTAATTTAGC | TAGCTTT | AAT | TTA | T | T | T>A | T>A_TTA | TA>AT_TAGCTTT | 9 | FALSE | FALSE |
| 129_v.txt | 32275 | 32275-SNV- | SNV | 1 | A | T | 5 | 34188 | 0.01462502 | 14.200003 | WT | FALSE | CAN | JSY2424 | 6 | WT-Rep6 | GAA | A>T | A/T+1 | A/T-1 | TA>AT | TA>AT | TACGGGAAC | AACCAAC | CTT | CTT | T | A | T>A | T>A_CTT | TA>AT_AACCAAC | 9 | FALSE | FALSE |
| 129_v.txt | 32284 | 32284-SNV- | SNV | 1 | A | T | 45 | 34188 | 0.13162513 | 42.6139636 | WT | FALSE | CAN | JSY2424 | 6 | WT-Rep6 | TAA | A>T | A/T+1 | A/T-1 | TA>AT | TA>AT | CAACGTAAA | AAATATT | ATT | ATT | T | A | T>A | T>A_ATT | TA>AT_AAATATT | 9 | FALSE | FALSE |
| 129_v.txt | 32337 | 32337-SNV- | SNV | 1 | A | T | 12 | 34188 | 0.03510004 | 36.8331755 | WT | FALSE | CAN | JSY2424 | 6 | WT-Rep6 | GAT | A>T | A/T+1 | A/T-1 | TA>AT | TA>AT | TTGAAGATA | ATATGTG | CTA | CTA | T | A | T>A | T>A_CTA | TA>AT_ATATGTG | 9 | FALSE | FALSE |
| 129_v.txt | 32338 | 32338-SNV-T | SNV | 1 | T | A | 4 | 34188 | 0.01170001 | 63.9978421 | WT | FALSE | CAN | JSY2424 | 6 | WT-Rep6 | ATA | T>A | A/T+1 | A/T-1 | TA>AT | TA>AT | TGAAGATAT | TATGTGG | TAT | ATA | T | T | T>A | T>A_ATA | TA>AT_TATGTGG | 9 | FALSE | FALSE |
| 129_v.txt | 32573 | 32573-SNV-T | SNV | 1 | T | A | 8 | 30732 | 0.0260315 | 35.875 | WT | FALSE | CAN | JSY2424 | 6 | WT-Rep6 | CTT | T>A | A/T+1 | A/T-1 | TA>AT | TA>AT | AGTACCTTG | TTGAAAT | GAA | CTT | T | T | T>A | T>A_CTT | TA>AT_TTGAAAT | 9 | FALSE | FALSE |
| 129_v.txt | 33089 | 33089-SNV- | SNV | 1 | A | T | 14 | 27506 | 0.05089799 | 52.0714286 | WT | FALSE | CAN | JSY2424 | 6 | WT-Rep6 | AAT | A>T | A/T+1 | A/T-1 | TA>AT | TA>AT | AAATAAATA | ATATGAT | TTA | TTA | T | A | T>A | T>A_TTA | TA>AT_ATATGAT | 9 | FALSE | FALSE |
| 129_v.txt | 33162 | 33162-SNV- | SNV | 1 | A | T | 26 | 27475 | 0.09463148 | 36.1923077 | WT | FALSE | CAN | JSY2424 | 6 | WT-Rep6 | AAT | A>T | A/T+1 | A/T-1 | TA>AT | TA>AT | GTACCAATA | ATAGTAC | TTA | TTA | T | A | T>A | T>A_TTA | TA>AT_ATAGTAC | 9 | FALSE | FALSE |
| 129_v.txt | 33303 | 33303-SNV- | SNV | 1 | A | T | 11 | 27157 | 0.04050521 | 49.7272727 | WT | FALSE | CAN | JSY2424 | 6 | WT-Rep6 | GAA | A>T | A/T+1 | A/T-1 | TA>AT | TA>AT | GTCGGGAAA | AAAGAGC | CTT | CTT | T | A | T>A | T>A_CTT | TA>AT_AAAGAGC | 9 | FALSE | FALSE |
| 129_v.txt | 31650 | 31650-SNV- | SNV | 1 | C | T | 19 | 10450 | 0.18181818 | 36.6842105 | WT | FALSE | CAN | JSY2424 | 6 | WT-Rep6 | GCG | C>T | A/T+1 | G/C-1 | CG>TA | CG>TA | AGAATGCGA | CGAAATG | CGC | GCG | C | A | C>T | C>T_GCG | CG>TA_CGAAATG | 9 | FALSE | FALSE |
| 129_v.txt | 31806 | 31806-SNV- | SNV | 1 | C | T | 11 | 10450 | 0.10526316 | 56.4545455 | WT | FALSE | CAN | JSY2424 | 6 | WT-Rep6 | CCA | C>T | A/T+1 | G/C-1 | CG>TA | CG>TA | ATCTTCCAAA | CAAATAA | GGT | CCA | C | A | C>T | C>T_CCA | CG>TA_CAAATAA | 9 | FALSE | FALSE |
| 129_v.txt | 31817 | 31817-SNV- | SNV | 1 | G | A | 10 | 10450 | 0.09569378 | 56.1 | WT | FALSE | CAN | JSY2424 | 6 | WT-Rep6 | TGC | G>A | A/T+1 | G/C-1 | CG>TA | CG>TA | AAATCTGCA | GCATCTG | ACG | ACG | C | T | C>T | C>T_ACG | CG>TA_GCATCTG | 9 | FALSE | FALSE |
| 129_v.txt | 31830 | 31830-SNV- | SNV | 1 | C | T | 14 | 10450 | 0.13397129 | 49 | WT | FALSE | CAN | JSY2424 | 6 | WT-Rep6 | GCA | C>T | A/T+1 | G/C-1 | CG>TA | CG>TA | AATATGCAT | CATTGAA | CGT | GCA | C | A | C>T | C>T_GCA | CG>TA_CATTGAA | 9 | FALSE | FALSE |
| 129_v.txt | 31882 | 31882-SNV- | SNV | 1 | C | T | 14 | 10450 | 0.13397129 | 40.9285714 | WT | FALSE | CAN | JSY2424 | 6 | WT-Rep6 | GCA | C>T | A/T+1 | G/C-1 | CG>TA | CG>TA | AGGCGGCAG | CAGCAAA | CGT | GCA | C | A | C>T | C>T_GCA | CG>TA_CAGCAAA | 9 | FALSE | FALSE |
| 129_v.txt | 31912 | 31912-SNV- | SNV | 1 | C | T | 12 | 10450 | 0.11483254 | 39.0833333 | WT | FALSE | CAN | JSY2424 | 6 | WT-Rep6 | GCA | C>T | A/T+1 | G/C-1 | CG>TA | CG>TA | TTGGTGCAA | CAAAAGC | CGT | GCA | C | A | C>T | C>T_GCA | CG>TA_CAAAAGC | 9 | FALSE | FALSE |
| 129_v.txt | 31930 | 31930-SNV- | SNV | 1 | G | A | 15 | 10448 | 0.14356815 | 36.2 | WT | FALSE | CAN | JSY2424 | 6 | WT-Rep6 | TGA | G>A | A/T+1 | G/C-1 | CG>TA | CG>TA | AACCTTGAA | GAATAAT | ACT | ACT | C | T | C>T | C>T_ACT | CG>TA_GAATAAT | 9 | FALSE | FALSE |
| 129_v.txt | 31962 | 31962-SNV- | SNV | 1 | G | A | 33 | 18543 | 0.17796473 | 36.3333333 | WT | FALSE | CAN | JSY2424 | 6 | WT-Rep6 | CGC | G>A | A/T+1 | G/C-1 | CG>TA | CG>TA | GTGGCCGCA | GCATAAT | GCG | GCG | C | T | C>T | C>T_GCG | CG>TA_GCATAAT | 9 | FALSE | FALSE |
| 129_v.txt | 31972 | 31972-SNV- | SNV | 1 | C | T | 18 | 18543 | 0.09707167 | 37.7777778 | WT | FALSE | CAN | JSY2424 | 6 | WT-Rep6 | GCC | C>T | A/T+1 | G/C-1 | CG>TA | CG>TA | AATAAGCCA | CCAAGCC | CGG | GCC | C | A | C>T | C>T_GCC | CG>TA_CCAAGCC | 9 | FALSE | FALSE |
| 129_v.txt | 31980 | 31980-SNV- | SNV | 1 | G | A | 13 | 18543 | 0.07010732 | 45.6153846 | WT | FALSE | CAN | JSY2424 | 6 | WT-Rep6 | GGG | G>A | A/T+1 | G/C-1 | CG>TA | CG>TA | AAGCCGGGC | GGCATTA | CCC | CCC | C | T | C>T | C>T_CCC | CG>TA_GGCATTA | 9 | FALSE | FALSE |
| 129_v.txt | 31981 | 31981-SNV- | SNV | 1 | G | A | 23 | 18543 | 0.12403602 | 40.3190489 | WT | FALSE | CAN | JSY2424 | 6 | WT-Rep6 | GGC | G>A | A/T+1 | G/C-1 | CG>TA | CG>TA | AGCCGGGCA | GCATTAA | CCG | CCG | C | T | C>T | C>T_CCG | CG>TA_GCATTAA | 9 | FALSE | FALSE |
| 129_v.txt | 32092 | 32092-SNV- | SNV | 1 | C | T | 19 | 18543 | 0.10246454 | 63.4210526 | WT | FALSE | CAN | JSY2424 | 6 | WT-Rep6 | GCA | C>T | A/T+1 | G/C-1 | CG>TA | CG>TA | AGCCTGCAA | CAACACC | CGT | GCA | C | A | C>T | C>T_GCA | CG>TA_CAACACC | 9 | FALSE | FALSE |
| 129_v.txt | 32114 | 32114-SNV- | SNV | 1 | C | T | 1345 | 18543 | 7.25341099 | 59.3167286 | WT | FALSE | CAN | JSY2424 | 6 | WT-Rep6 | GCC | C>T | A/T+1 | G/C-1 | CG>TA | CG>TA | TAATAGCCA | CCATTCG | CGG | GCC | C | A | C>T | C>T_GCC | CG>TA_CCATTCG | 9 | FALSE | FALSE |
| 129_v.txt | 32165 | 32165-SNV- | SNV | 1 | G | A | 29 | 18543 | 0.15639325 | 47.1724138 | WT | FALSE | CAN | JSY2424 | 6 | WT-Rep6 | CGC | G>A | A/T+1 | G/C-1 | CG>TA | CG>TA | CAAAGCGCC | GCCAAAT | GCG | GCG | C | T | C>T | C>T_GCG | CG>TA_GCCAAAT | 9 | FALSE | FALSE |
| 129_v.txt | 32271 | 32271-SNV- | SNV | 1 | C | T | 52 | 34188 | 0.15210015 | 30.5191905 | WT | FALSE | CAN | JSY2424 | 6 | WT-Rep6 | ACG | C>T | A/T+1 | G/C-1 | CG>TA | CG>TA | AAAATACGG | CGGGAAC | TGC | ACG | C | A | C>T | C>T_ACG | CG>TA_CGGGAAC | 9 | FALSE | FALSE |
| 129_v.txt | 32272 | 32272-SNV- | SNV | 1 | G | A | 103 | 34188 | 0.3012753 | 35.930724 | WT | FALSE | CAN | JSY2424 | 6 | WT-Rep6 | CGG | G>A | A/T+1 | G/C-1 | CG>TA | CG>TA | AAATACGGG | GGGAACC | GCC | GCC | C | T | C>T | C>T_GCC | CG>TA_GGGAACC | 9 | FALSE | FALSE |
| 129_v.txt | 32274 | 32274-SNV- | SNV | 1 | G | A | 73 | 34188 | 0.21352521 | 36.6027959 | WT | FALSE | CAN | JSY2424 | 6 | WT-Rep6 | GGA | G>A | A/T+1 | G/C-1 | CG>TA | CG>TA | ATACGGGAA | GAACCAA | CCT | CCT | C | T | C>T | C>T_CCT | CG>TA_GAACCAA | 9 | FALSE | FALSE |
| 129_v.txt | 32292 | 32292-SNV- | SNV | 1 | G | A | 232 | 34188 | 0.67860068 | 53.3448276 | WT | FALSE | CAN | JSY2424 | 6 | WT-Rep6 | TGA | G>A | A/T+1 | G/C-1 | CG>TA | CG>TA | ATATTTGAAT | GAATTTG | ACT | ACT | C | T | C>T | C>T_ACT | CG>TA_GAATTTG | 9 | FALSE | FALSE |
| 129_v.txt | 32299 | 32299-SNV- | SNV | 1 | C | T | 53 | 34188 | 0.15502516 | 50.6037736 | WT | FALSE | CAN | JSY2424 | 6 | WT-Rep6 | GCG | C>T | A/T+1 | G/C-1 | CG>TA | CG>TA | AATTTGCGG | CGGCAGA | CGC | GCG | C | A | C>T | C>T_GCG | CG>TA_CGGCAGA | 9 | FALSE | FALSE |
| 129_v.txt | 32344 | 32344-SNV- | SNV | 1 | G | A | 22 | 34188 | 0.06435006 | 62.6363636 | WT | FALSE | CAN | JSY2424 | 6 | WT-Rep6 | GGC | G>A | A/T+1 | G/C-1 | CG>TA | CG>TA | TATGTGGCA | GCAAAAC | CCG | CCG | C | T | C>T | C>T_CCG | CG>TA_GCAAAAC | 9 | FALSE | FALSE |
| 129_v.txt | 32413 | 32413-SNV- | SNV | 1 | G | A | 66 | 34188 | 0.19305019 | 57.5757576 | WT | FALSE | CAN | JSY2424 | 6 | WT-Rep6 | TGT | G>A | A/T+1 | G/C-1 | CG>TA | CG>TA | TAGATTGTG | GTGTTAG | ACA | ACA | C | T | C>T | C>T_ACA | CG>TA_GTGTTAG | 9 | FALSE | FALSE |
| 129_v.txt | 32449 | 32449-SNV- | SNV | 1 | C | T | 50 | 34188 | 0.14625015 | 52.6 | WT | FALSE | CAN | JSY2424 | 6 | WT-Rep6 | CCA | C>T | A/T+1 | G/C-1 | CG>TA | CG>TA | AAAGTCCAA | CAATGAA | GGT | CCA | C | A | C>T | C>T_CCA | CG>TA_CAATGAA | 9 | FALSE | FALSE |
| 129_v.txt | 32466 | 32466-SNV- | SNV | 1 | C | T | 51 | 34188 | 0.14917515 | 44 | WT | FALSE | CAN | JSY2424 | 6 | WT-Rep6 | GCC | C>T | A/T+1 | G/C-1 | CG>TA | CG>TA | AGAGAGCCA | CCAATGT | CGG | GCC | C | A | C>T | C>T_GCC | CG>TA_CCAATGT | 9 | FALSE | FALSE |
| 129_v.txt | 32488 | 32488-SNV- | SNV | 1 | G | A | 81 | 34188 | 0.23692524 | 40.691358 | WT | FALSE | CAN | JSY2424 | 6 | WT-Rep6 | CGG | G>A | A/T+1 | G/C-1 | CG>TA | CG>TA | AGATACGGA | GGAAAAC | GCC | GCC | C | T | C>T | C>T_GCC | CG>TA_GGAAAAC | 9 | FALSE | FALSE |
| 129_v.txt | 32520 | 32520-SNV- | SNV | 1 | G | A | 13493 | 34187 | 39.4682189 | 42.5529534 | WT | FALSE | CAN | JSY2424 | 6 | WT-Rep6 | GGA | G>A | A/T+1 | G/C-1 | CG>TA | CG>TA | GGAACGGAT | GATTTTC | CCT | CCT | C | T | C>T | C>T_CCT | CG>TA_GATTTTC | 9 | FALSE | FALSE |
| 129_v.txt | 32535 | 32535-SNV- | SNV | 1 | G | A | 28 | 34185 | 0.08190727 | 36.1423637 | WT | FALSE | CAN | JSY2424 | 6 | WT-Rep6 | TGC | G>A | A/T+1 | G/C-1 | CG>TA | CG>TA | GGGTTTGCA | GCAGCTT | ACG | ACG | C | T | C>T | C>T_ACG | CG>TA_GCAGCTT | 9 | FALSE | FALSE |
| 129_v.txt | 32536 | 32536-SNV- | SNV | 1 | C | T | 53 | 34185 | 0.15503876 | 36.754775 | WT | FALSE | CAN | JSY2424 | 6 | WT-Rep6 | GCA | C>T | A/T+1 | G/C-1 | CG>TA | CG>TA | GGTTTGCAG | CAGCTTC | CGT | GCA | C | A | C>T | C>T_GCA | CG>TA_CAGCTTC | 9 | FALSE | FALSE |
| 129_v.txt | 32566 | 32566-SNV- | SNV | 1 | C | T | 70 | 30720 | 0.22786458 | 35.3428571 | WT | FALSE | CAN | JSY2424 | 6 | WT-Rep6 | TCA | C>T | A/T+1 | G/C-1 | CG>TA | CG>TA | CTAGTTCAG | CAGTACC | AGT | TCA | C | A | C>T | C>T_TCA | CG>TA_CAGTACC | 9 | FALSE | FALSE |
| 129_v.txt | 32572 | 32572-SNV- | SNV | 1 | C | T | 39 | 30732 | 0.12690355 | 30.8461539 | WT | FALSE | CAN | JSY2424 | 6 | WT-Rep6 | CCT | C>T | A/T+1 | G/C-1 | CG>TA | CG>TA | CAGTACCTT | CTTGAAA | GGA | CCT | C | A | C>T | C>T_CCT | CG>TA_CTTGAAA | 9 | FALSE | FALSE |
| 129_v.txt | 32575 | 32575-SNV- | SNV | 1 | G | A | 45 | 30731 | 0.14643194 | 35.5555556 | WT | FALSE | CAN | JSY2424 | 6 | WT-Rep6 | TGA | G>A | A/T+1 | G/C-1 | CG>TA | CG>TA | TACCTTGAA | GAAATGT | ACT | ACT | C | T | C>T | C>T_ACT | CG>TA_GAAATGT | 9 | FALSE | FALSE |
| 129_v.txt | 32590 | 32590-SNV- | SNV | 1 | C | T | 43 | 30731 | 0.13992386 | 36.3953488 | WT | FALSE | CAN | JSY2424 | 6 | WT-Rep6 | GCG | C>T | A/T+1 | G/C-1 | CG>TA | CG>TA | AGGCAGCGT | CGTTAAT | CGC | GCG | C | A | C>T | C>T_GCG | CG>TA_CGTTAAT | 9 | FALSE | FALSE |
| 129_v.txt | 32609 | 32609-SNV- | SNV | 1 | C | T | 33 | 30731 | 0.10738342 | 47.4242424 | WT | FALSE | CAN | JSY2424 | 6 | WT-Rep6 | CCC | C>T | A/T+1 | G/C-1 | CG>TA | CG>TA | GGAAACCCA | CCAACCT | GGG | CCC | C | A | C>T | C>T_CCC | CG>TA_CCAACCT | 9 | FALSE | FALSE |
| 129_v.txt | 32700 | 32700-SNV- | SNV | 1 | C | T | 31 | 30731 | 0.10087534 | 54.3225807 | WT | FALSE | CAN | JSY2424 | 6 | WT-Rep6 | GCC | C>T | A/T+1 | G/C-1 | CG>TA | CG>TA | ACTGGGCCG | CCGGTAA | CGG | GCC | C | A | C>T | C>T_GCC | CG>TA_CCGGTAA | 9 | FALSE | FALSE |
| 129_v.txt | 32733 | 32733-SNV- | SNV | 1 | C | T | 39 | 30731 | 0.12690768 | 55.6410256 | WT | FALSE | CAN | JSY2424 | 6 | WT-Rep6 | ACA | C>T | A/T+1 | G/C-1 | CG>TA | CG>TA | CAAAAACAG | CAGTATA | TGT | ACA | C | A | C>T | C>T_ACA | CG>TA_CAGTATA | 9 | FALSE | FALSE |
| 129_v.txt | 32749 | 32749-SNV- | SNV | 1 | C | T | 29 | 30731 | 0.09436725 | 52.7241379 | WT | FALSE | CAN | JSY2424 | 6 | WT-Rep6 | CCG | C>T | A/T+1 | G/C-1 | CG>TA | CG>TA | GAAACCCGA | CGATAAT | GGC | CCG | C | A | C>T | C>T_CCG | CG>TA_CGATAAT | 9 | FALSE | FALSE |
| 129_v.txt | 32757 | 32757-SNV- | SNV | 1 | G | A | 31 | 30731 | 0.10087534 | 53.6451613 | WT | FALSE | CAN | JSY2424 | 6 | WT-Rep6 | GGC | G>A | A/T+1 | G/C-1 | CG>TA | CG>TA | ATAATGGCT | GCTAAAA | CCG | CCG | C | T | C>T | C>T_CCG | CG>TA_GCTAAAA | 9 | FALSE | FALSE |
| 129_v.txt | 32776 | 32776-SNV- | SNV | 1 | C | T | 26 | 30731 | 0.08460512 | 51.1538462 | WT | FALSE | CAN | JSY2424 | 6 | WT-Rep6 | GCG | C>T | A/T+1 | G/C-1 | CG>TA | CG>TA | TGGAAGCGA | CGACCCA | CGC | GCG | C | A | C>T | C>T_GCG | CG>TA_CGACCCA | 9 | FALSE | FALSE |
| 129_v.txt | 32788 | 32788-SNV- | SNV | 1 | C | T | 450 | 30731 | 1.46431942 | 45.4644444 | WT | FALSE | CAN | JSY2424 | 6 | WT-Rep6 | TCG | C>T | A/T+1 | G/C-1 | CG>TA | CG>TA | AGAACTCGA | CGAATTC | AGC | TCG | C | A | C>T | C>T_TCG | CG>TA_CGAATTC | 9 | FALSE | FALSE |

| | | | | | | | | | | | | | | | | | | | | | | | | | | | | | | | | | | |
|---|---|---|---|---|---|---|---|---|---|---|---|---|---|---|---|---|---|---|---|---|---|---|---|---|---|---|---|---|---|---|---|---|---|---|
| 129_v.txt | 32796 | 32796-SNV-( | SNV | 1 | C | T | 24 | 30731 | 0.07809704 | 44.1953023 | WT | FALSE | CAN | JSY2424 | 6 | WT-Rep6 | ACC | C>T | A/T+1 | G/C-1 | CG>TA | CG>TA | AATTCACCG1 | CCGTAAT | TGG | ACC | C | A | C>T | C>T_ACC | CG>TA_CCGTAAT | 9 | FALSE | FALSE |
| 129_v.txt | 32798 | 32798-SNV-( | SNV | 1 | G | A | 35 | 30731 | 0.11389151 | 46.6 | WT | FALSE | CAN | JSY2424 | 6 | WT-Rep6 | CGT | G>A | A/T+1 | G/C-1 | CG>TA | CG>TA | TTCACCGTA/ | GTAATAT | GCA | GCA | C | T | C>T | C>T_GCA | CG>TA_GTAATAT | 9 | FALSE | FALSE |
| 129_v.txt | 32855 | 32855-SNV-( | SNV | 1 | C | T | 114 | 30731 | 0.37096092 | 35.3859649 | WT | FALSE | CAN | JSY2424 | 6 | WT-Rep6 | TCC | C>T | A/T+1 | G/C-1 | CG>TA | CG>TA | ACTAATCCA1 | CCATGCC | AGG | TCC | C | A | C>T | C>T_TCC | CG>TA_CCATGCC | 9 | FALSE | FALSE |
| 129_v.txt | 32860 | 32860-SNV-( | SNV | 1 | C | T | 28 | 30728 | 0.0911221 | 36.3214286 | WT | FALSE | CAN | JSY2424 | 6 | WT-Rep6 | GCC | C>T | A/T+1 | G/C-1 | CG>TA | CG>TA | TCCATGCCG( | CCGCCAG | CGG | GCC | C | A | C>T | C>T_GCC | CG>TA_CCGCCAG | 9 | FALSE | FALSE |
| 129_v.txt | 32862 | 32862-SNV-( | SNV | 1 | G | A | 24 | 30728 | 0.07810466 | 35.5416667 | WT | FALSE | CAN | JSY2424 | 6 | WT-Rep6 | CGC | G>A | A/T+1 | G/C-1 | CG>TA | CG>TA | CATGCCGCC/ | GCCAGTG | GCG | GCG | C | T | C>T | C>T_GCG | CG>TA_GCCAGTG | 9 | FALSE | FALSE |
| 129_v.txt | 32879 | 32879-SNV-( | SNV | 1 | C | T | 22 | 27505 | 0.07998546 | 35.6363636 | WT | FALSE | CAN | JSY2424 | 6 | WT-Rep6 | ACG | C>T | A/T+1 | G/C-1 | CG>TA | CG>TA | TTTGTACGTC | CGTCCAA | TGC | ACG | C | A | C>T | C>T_ACG | CG>TA_CGTCCAA | 9 | FALSE | FALSE |
| 129_v.txt | 32882 | 32882-SNV-( | SNV | 1 | C | T | 235 | 27506 | 0.85435905 | 34.4893617 | WT | FALSE | CAN | JSY2424 | 6 | WT-Rep6 | TCC | C>T | A/T+1 | G/C-1 | CG>TA | CG>TA | GTACGTCCA/ | CCAAAAT | AGG | TCC | C | A | C>T | C>T_TCC | CG>TA_CCAAAAT | 9 | FALSE | FALSE |
| 129_v.txt | 32890 | 32890-SNV-( | SNV | 1 | G | A | 29 | 27506 | 0.10543154 | 36.9655172 | WT | FALSE | CAN | JSY2424 | 6 | WT-Rep6 | TGA | G>A | A/T+1 | G/C-1 | CG>TA | CG>TA | AAAATTGAA | GAATGAC | ACT | ACT | C | T | C>T | C>T_ACT | CG>TA_GAATGAC | 9 | FALSE | FALSE |
| 129_v.txt | 32901 | 32901-SNV-( | SNV | 1 | C | T | 60 | 27506 | 0.21813423 | 32.6166667 | WT | FALSE | CAN | JSY2424 | 6 | WT-Rep6 | GCC | C>T | A/T+1 | G/C-1 | CG>TA | CG>TA | ACTTGGCCA/ | CCAACTA | CGG | GCC | C | A | C>T | C>T_GCC | CG>TA_CCAACTA | 9 | FALSE | FALSE |
| 129_v.txt | 32922 | 32922-SNV-( | SNV | 1 | G | A | 26 | 27506 | 0.09452483 | 55.7692308 | WT | FALSE | CAN | JSY2424 | 6 | WT-Rep6 | GGC | G>A | A/T+1 | G/C-1 | CG>TA | CG>TA | TCCAGGGCA. | GCAAAAG | CCG | CCG | C | T | C>T | C>T_CCG | CG>TA_GCAAAAG | 9 | FALSE | FALSE |
| 129_v.txt | 32935 | 32935-SNV-( | SNV | 1 | C | T | 26 | 27506 | 0.09452483 | 38.6538462 | WT | FALSE | CAN | JSY2424 | 6 | WT-Rep6 | GCC | C>T | A/T+1 | G/C-1 | CG>TA | CG>TA | TGATTGCCCA | CCCAAGA | CGG | GCC | C | A | C>T | C>T_GCC | CG>TA_CCCAAGA | 9 | FALSE | FALSE |
| 129_v.txt | 32965 | 32965-SNV-( | SNV | 1 | C | T | 28 | 27506 | 0.10179597 | 58.8928571 | WT | FALSE | CAN | JSY2424 | 6 | WT-Rep6 | GCC | C>T | A/T+1 | G/C-1 | CG>TA | CG>TA | CATTGGCCG( | CCGCACC | CGG | GCC | C | A | C>T | C>T_GCC | CG>TA_CCGCACC | 9 | FALSE | FALSE |
| 129_v.txt | 32967 | 32967-SNV-( | SNV | 1 | G | A | 67 | 27506 | 0.24358322 | 57.4179105 | WT | FALSE | CAN | JSY2424 | 6 | WT-Rep6 | CGC | G>A | A/T+1 | G/C-1 | CG>TA | CG>TA | TTGGCCGCA( | GCACCAA | GCG | GCG | C | T | C>T | C>T_GCG | CG>TA_GCACCAA | 9 | FALSE | FALSE |
| 129_v.txt | 32977 | 32977-SNV-( | SNV | 1 | C | T | 28 | 27506 | 0.10179597 | 52.4642857 | WT | FALSE | CAN | JSY2424 | 6 | WT-Rep6 | GCT | C>T | A/T+1 | G/C-1 | CG>TA | CG>TA | CAAATGCTG( | CTGGAGA | CGA | GCT | C | A | C>T | C>T_GCT | CG>TA_CTGGAGA | 9 | FALSE | FALSE |
| 129_v.txt | 33037 | 33037-SNV-( | SNV | 1 | C | T | 15 | 27506 | 0.05453356 | 58.4 | WT | FALSE | CAN | JSY2424 | 6 | WT-Rep6 | GCC | C>T | A/T+1 | G/C-1 | CG>TA | CG>TA | ATGTAGCCA' | CCATTTC | CGG | GCC | C | A | C>T | C>T_GCC | CG>TA_CCATTTC | 9 | FALSE | FALSE |
| 129_v.txt | 33043 | 33043-SNV-( | SNV | 1 | C | T | 25 | 27506 | 0.09088926 | 57.4 | WT | FALSE | CAN | JSY2424 | 6 | WT-Rep6 | TCA | C>T | A/T+1 | G/C-1 | CG>TA | CG>TA | CCATTTCACC | CACCCAA | AGT | TCA | C | A | C>T | C>T_TCA | CG>TA_CACCCAA | 9 | FALSE | FALSE |
| 129_v.txt | 33056 | 33056-SNV-( | SNV | 1 | C | T | 37 | 27506 | 0.13451611 | 58.1348495 | WT | FALSE | CAN | JSY2424 | 6 | WT-Rep6 | GCG | C>T | A/T+1 | G/C-1 | CG>TA | CG>TA | GGACTGCGT( | CGTGACA | CGC | GCG | C | A | C>T | C>T_GCG | CG>TA_CGTGACA | 9 | FALSE | FALSE |
| 129_v.txt | 33057 | 33057-SNV-( | SNV | 1 | G | A | 7 | 27506 | 0.02544899 | 60.4263946 | WT | FALSE | CAN | JSY2424 | 6 | WT-Rep6 | CGT | G>A | A/T+1 | G/C-1 | CG>TA | CG>TA | GACTGCGTG. | GTGACAG | GCA | GCA | C | T | C>T | C>T_GCA | CG>TA_GTGACAG | 9 | FALSE | FALSE |
| 129_v.txt | 33069 | 33069-SNV-( | SNV | 1 | G | A | 23 | 27506 | 0.08361812 | 56.173913 | WT | FALSE | CAN | JSY2424 | 6 | WT-Rep6 | TGC | G>A | A/T+1 | G/C-1 | CG>TA | CG>TA | GAATATGCC/ | GCCAAAG | ACG | ACG | C | T | C>T | C>T_ACG | CG>TA_GCCAAAG | 9 | FALSE | FALSE |
| 129_v.txt | 33070 | 33070-SNV-( | SNV | 1 | C | T | 29 | 27506 | 0.10543154 | 50.7234738 | WT | FALSE | CAN | JSY2424 | 6 | WT-Rep6 | GCC | C>T | A/T+1 | G/C-1 | CG>TA | CG>TA | AATATGCCA/ | CCAAAGA | CGG | GCC | C | A | C>T | C>T_GCC | CG>TA_CCAAAGA | 9 | FALSE | FALSE |
| 129_v.txt | 33078 | 33078-SNV-( | SNV | 1 | C | T | 34 | 27506 | 0.12360939 | 50.9117647 | WT | FALSE | CAN | JSY2424 | 6 | WT-Rep6 | ACC | C>T | A/T+1 | G/C-1 | CG>TA | CG>TA | AAAGAACCC | CCCATAA | TGG | ACC | C | A | C>T | C>T_ACC | CG>TA_CCCATAA | 9 | FALSE | FALSE |
| 129_v.txt | 33114 | 33114-SNV-( | SNV | 1 | C | T | 24 | 27506 | 0.08725369 | 37.0833333 | WT | FALSE | CAN | JSY2424 | 6 | WT-Rep6 | GCC | C>T | A/T+1 | G/C-1 | CG>TA | CG>TA | ACTGGGCCG | CCGGCGT | CGG | GCC | C | A | C>T | C>T_GCC | CG>TA_CCGGCGT | 9 | FALSE | FALSE |
| 129_v.txt | 33118 | 33118-SNV-( | SNV | 1 | C | T | 27 | 27506 | 0.0981604 | 41.4426264 | WT | FALSE | CAN | JSY2424 | 6 | WT-Rep6 | GCG | C>T | A/T+1 | G/C-1 | CG>TA | CG>TA | GGCCGGCGT | CGTTGGT | CGC | GCG | C | A | C>T | C>T_GCG | CG>TA_CGTTGGT | 9 | FALSE | FALSE |
| 129_v.txt | 33119 | 33119-SNV-( | SNV | 1 | G | A | 24 | 27506 | 0.08725369 | 41.7083333 | WT | FALSE | CAN | JSY2424 | 6 | WT-Rep6 | CGT | G>A | A/T+1 | G/C-1 | CG>TA | CG>TA | GCCGGCGTT( | GTTGGTC | GCA | GCA | C | T | C>T | C>T_GCA | CG>TA_GTTGGTC | 9 | FALSE | FALSE |
| 129_v.txt | 33177 | 33177-SNV-( | SNV | 1 | G | A | 14 | 27468 | 0.0509684 | 37.2857143 | WT | FALSE | CAN | JSY2424 | 6 | WT-Rep6 | GGC | G>A | A/T+1 | G/C-1 | CG>TA | CG>TA | CCAAGGGCA | GCAATCA | CCG | CCG | C | T | C>T | C>T_CCG | CG>TA_GCAATCA | 9 | FALSE | FALSE |
| 129_v.txt | 33258 | 33258-SNV-( | SNV | 1 | C | T | 31 | 27157 | 0.11415105 | 40.5806452 | WT | FALSE | CAN | JSY2424 | 6 | WT-Rep6 | GCC | C>T | A/T+1 | G/C-1 | CG>TA | CG>TA | CCTATGCCAT | CCATCCT | CGG | GCC | C | A | C>T | C>T_GCC | CG>TA_CCATCCT | 9 | FALSE | FALSE |
| 129_v.txt | 33275 | 33275-SNV-( | SNV | 1 | C | T | 26 | 27157 | 0.09573959 | 52.6523307 | WT | FALSE | CAN | JSY2424 | 6 | WT-Rep6 | ACG | C>T | A/T+1 | G/C-1 | CG>TA | CG>TA | AGAGAACGT | CGTATCC | TGC | ACG | C | A | C>T | C>T_ACG | CG>TA_CGTATCC | 9 | FALSE | FALSE |
| 129_v.txt | 33276 | 33276-SNV-( | SNV | 1 | G | A | 44 | 27156 | 0.16202681 | 46.8825997 | WT | FALSE | CAN | JSY2424 | 6 | WT-Rep6 | CGT | G>A | A/T+1 | G/C-1 | CG>TA | CG>TA | GAGAACGTA | GTATCCT | GCA | GCA | C | T | C>T | C>T_GCA | CG>TA_GTATCCT | 9 | FALSE | FALSE |
| 129_v.txt | 33284 | 33284-SNV-( | SNV | 1 | G | A | 33 | 27157 | 0.12151563 | 44 | WT | FALSE | CAN | JSY2424 | 6 | WT-Rep6 | CGC | G>A | A/T+1 | G/C-1 | CG>TA | CG>TA | ATCCTCGCCA | GCCATTT | GCG | GCG | C | T | C>T | C>T_GCG | CG>TA_GCCATTT | 9 | FALSE | FALSE |
| 129_v.txt | 33310 | 33310-SNV-( | SNV | 1 | G | A | 39 | 27157 | 0.14360938 | 53.5384615 | WT | FALSE | CAN | JSY2424 | 6 | WT-Rep6 | CGC | G>A | A/T+1 | G/C-1 | CG>TA | CG>TA | AAGAGCGCA | GCAATGG | GCG | GCG | C | T | C>T | C>T_GCG | CG>TA_GCAATGG | 9 | FALSE | FALSE |
| 129_v.txt | 33392 | 33392-SNV-( | SNV | 1 | G | A | 27 | 27157 | 0.09942188 | 54.7037037 | WT | FALSE | CAN | JSY2424 | 6 | WT-Rep6 | CGT | G>A | A/T+1 | G/C-1 | CG>TA | CG>TA | AACGTCGTG. | GTGAAAG | GCA | GCA | C | T | C>T | C>T_GCA | CG>TA_GTGAAAG | 9 | FALSE | FALSE |
| 129_v.txt | 33412 | 33412-SNV-( | SNV | 1 | G | A | 32 | 27157 | 0.11783334 | 49.5 | WT | FALSE | CAN | JSY2424 | 6 | WT-Rep6 | GGC | G>A | A/T+1 | G/C-1 | CG>TA | CG>TA | TGACCGGCT( | GCTCATT | CCG | CCG | C | T | C>T | C>T_CCG | CG>TA_GCTCATT | 9 | FALSE | FALSE |
| 129_v.txt | 33474 | 33474-SNV-( | SNV | 1 | C | T | 32 | 27155 | 0.11784202 | 31.375 | WT | FALSE | CAN | JSY2424 | 6 | WT-Rep6 | GCC | C>T | A/T+1 | G/C-1 | CG>TA | CG>TA | GCTATGCCTT | CCTTTTT | CGG | GCC | C | A | C>T | C>T_GCC | CG>TA_CCTTTTT | 9 | FALSE | FALSE |
| 129_v.txt | 31980 | 31980-SNV-( | SNV | 1 | G | C | 2 | 18543 | 0.01078574 | 57.5 | WT | FALSE | CAN | JSY2424 | 6 | WT-Rep6 | GGG | G>C | G/C+1 | G/C-1 | CG>GC | CG>GC | AAGCCGGGC | GGCATTA | CCC | CCC | C | G | C>G | C>G_CCC | CG>GC_GGCATTA | 9 | FALSE | FALSE |
| 129_v.txt | 32058 | 32058-SNV-( | SNV | 1 | G | C | 9 | 18543 | 0.04853584 | 61.4444444 | WT | FALSE | CAN | JSY2424 | 6 | WT-Rep6 | CGA | G>C | G/C+1 | G/C-1 | CG>GC | CG>GC | ATGTGCGAG | GAGATTG | GCT | GCT | C | G | C>G | C>G_GCT | CG>GC_GAGATTG | 9 | FALSE | FALSE |
| 129_v.txt | 32282 | 32282-SNV-( | SNV | 1 | G | C | 33 | 34188 | 0.0965251 | 41.4515645 | WT | FALSE | CAN | JSY2424 | 6 | WT-Rep6 | CGT | G>C | G/C+1 | G/C-1 | CG>GC | CG>GC | ACCAACGTA/ | GTAAATA | GCA | GCA | C | G | C>G | C>G_GCA | CG>GC_GTAAATA | 9 | FALSE | FALSE |
| 129_v.txt | 32292 | 32292-SNV-( | SNV | 1 | G | C | 6 | 34188 | 0.01755002 | 55.1666667 | WT | FALSE | CAN | JSY2424 | 6 | WT-Rep6 | TGA | G>C | G/C+1 | G/C-1 | CG>GC | CG>GC | ATATTTGAA1 | GAATTTG | ACT | ACT | C | G | C>G | C>G_ACT | CG>GC_GAATTTG | 9 | FALSE | FALSE |
| 129_v.txt | 32316 | 32316-SNV-( | SNV | 1 | G | C | 24 | 34188 | 0.07020007 | 58.2916667 | WT | FALSE | CAN | JSY2424 | 6 | WT-Rep6 | TGT | G>C | G/C+1 | G/C-1 | CG>GC | CG>GC | ATGGTTGTT/ | GTTAAGA | ACA | ACA | C | G | C>G | C>G_ACA | CG>GC_GTTAAGA | 9 | FALSE | FALSE |
| 129_v.txt | 32528 | 32528-SNV-( | SNV | 1 | G | C | 4 | 34187 | 0.01170035 | 34 | WT | FALSE | CAN | JSY2424 | 6 | WT-Rep6 | TGG | G>C | G/C+1 | G/C-1 | CG>GC | CG>GC | TTTTCTGGGG | GGGGTTT | ACC | ACC | C | G | C>G | C>G_ACC | CG>GC_GGGGTTT | 9 | FALSE | FALSE |
| 129_v.txt | 32538 | 32538-SNV-( | SNV | 1 | G | C | 17 | 34185 | 0.04972941 | 35.4705882 | WT | FALSE | CAN | JSY2424 | 6 | WT-Rep6 | AGC | G>C | G/C+1 | G/C-1 | CG>GC | CG>GC | TTTGCAGCTT | GCTTCAC | TCG | TCG | C | G | C>G | C>G_TCG | CG>GC_GCTTCAC | 9 | FALSE | FALSE |
| 129_v.txt | 32557 | 32557-SNV-( | SNV | 1 | C | G | 101 | 34173 | 0.29555497 | 37.6732673 | WT | FALSE | CAN | JSY2424 | 6 | WT-Rep6 | CCA | C>G | G/C+1 | G/C-1 | CG>GC | CG>GC | TGATACCAA( | CAACTAG | GGT | CCA | C | C | C>G | C>G_CCA | CG>GC_CAACTAG | 9 | FALSE | FALSE |
| 129_v.txt | 32566 | 32566-SNV-( | SNV | 1 | C | G | 109 | 30720 | 0.35481771 | 36.4311927 | WT | FALSE | CAN | JSY2424 | 6 | WT-Rep6 | TCA | C>G | G/C+1 | G/C-1 | CG>GC | CG>GC | CTAGTTCAGT | CAGTACC | AGT | TCA | C | C | C>G | C>G_TCA | CG>GC_CAGTACC | 9 | FALSE | FALSE |
| 129_v.txt | 32572 | 32572-SNV-( | SNV | 1 | C | G | 25 | 30732 | 0.08134843 | 36.68 | WT | FALSE | CAN | JSY2424 | 6 | WT-Rep6 | CCT | C>G | G/C+1 | G/C-1 | CG>GC | CG>GC | CAGTACCTTC | CTTGAAA | GGA | CCT | C | C | C>G | C>G_CCT | CG>GC_CTTGAAA | 9 | FALSE | FALSE |
| 129_v.txt | 32685 | 32685-SNV-( | SNV | 1 | C | G | 28 | 30731 | 0.09111321 | 62.0357143 | WT | FALSE | CAN | JSY2424 | 6 | WT-Rep6 | ACG | C>G | G/C+1 | G/C-1 | CG>GC | CG>GC | CAATAACGG | CGGAATC | TGC | ACG | C | C | C>G | C>G_ACG | CG>GC_CGGAATC | 9 | FALSE | FALSE |
| 129_v.txt | 32735 | 32735-SNV-( | SNV | 1 | G | C | 26 | 30731 | 0.08460512 | 57.2692308 | WT | FALSE | CAN | JSY2424 | 6 | WT-Rep6 | AGT | G>C | G/C+1 | G/C-1 | CG>GC | CG>GC | AAAACAGTA | GTATATT | TCA | TCA | C | G | C>G | C>G_TCA | CG>GC_GTATATT | 9 | FALSE | FALSE |
| 129_v.txt | 32794 | 32794-SNV-( | SNV | 1 | C | G | 12 | 30731 | 0.03904852 | 44.9166667 | WT | FALSE | CAN | JSY2424 | 6 | WT-Rep6 | TCA | C>G | G/C+1 | G/C-1 | CG>GC | CG>GC | CGAATTCAC( | CACCGTA | AGT | TCA | C | C | C>G | C>G_TCA | CG>GC_CACCGTA | 9 | FALSE | FALSE |
| 129_v.txt | 32811 | 32811-SNV-( | SNV | 1 | G | C | 25 | 30731 | 0.08135108 | 44.88 | WT | FALSE | CAN | JSY2424 | 6 | WT-Rep6 | AGG | G>C | G/C+1 | G/C-1 | CG>GC | CG>GC | TTGACAGGG | GGGAACA | TCC | TCC | C | G | C>G | C>G_TCC | CG>GC_GGGAACA | 9 | FALSE | FALSE |
| 129_v.txt | 32971 | 32971-SNV-( | SNV | 1 | C | G | 16 | 27506 | 0.05816913 | 56.6875 | WT | FALSE | CAN | JSY2424 | 6 | WT-Rep6 | CCA | C>G | G/C+1 | G/C-1 | CG>GC | CG>GC | CCGCACCAA/ | CAAATGC | GGT | CCA | C | C | C>G | C>G_CCA | CG>GC_CAAATGC | 9 | FALSE | FALSE |
| 129_v.txt | 32977 | 32977-SNV-( | SNV | 1 | C | G | 4 | 27506 | 0.01454228 | 62.75 | WT | FALSE | CAN | JSY2424 | 6 | WT-Rep6 | GCT | C>G | G/C+1 | G/C-1 | CG>GC | CG>GC | CAAATGCTG( | CTGGAGA | CGA | GCT | C | C | C>G | C>G_GCT | CG>GC_CTGGAGA | 9 | FALSE | FALSE |
| 129_v.txt | 33037 | 33037-SNV-( | SNV | 1 | C | G | 5 | 27506 | 0.01817785 | 64 | WT | FALSE | CAN | JSY2424 | 6 | WT-Rep6 | GCC | C>G | G/C+1 | G/C-1 | CG>GC | CG>GC | ATGTAGCCA' | CCATTTC | CGG | GCC | C | C | C>G | C>G_GCC | CG>GC_CCATTTC | 9 | FALSE | FALSE |
| 129_v.txt | 33057 | 33057-SNV-( | SNV | 1 | G | C | 111 | 27506 | 0.40354832 | 57.8053738 | WT | FALSE | CAN | JSY2424 | 6 | WT-Rep6 | CGT | G>C | G/C+1 | G/C-1 | CG>GC | CG>GC | GACTGCGTG. | GTGACAG | GCA | GCA | C | G | C>G | C>G_GCA | CG>GC_GTGACAG | 9 | FALSE | FALSE |
| 129_v.txt | 33070 | 33070-SNV-( | SNV | 1 | C | G | 2 | 27506 | 7.27E-03 | 24.4997955 | WT | FALSE | CAN | JSY2424 | 6 | WT-Rep6 | GCC | C>G | G/C+1 | G/C-1 | CG>GC | CG>GC | AATATGCCA/ | CCAAAGA | CGG | GCC | C | C | C>G | C>G_GCC | CG>GC_CCAAAGA | 9 | FALSE | FALSE |
| 129_v.txt | 33093 | 33093-SNV-( | SNV | 1 | G | C | 21 | 27506 | 0.07634698 | 55.5714286 | WT | FALSE | CAN | JSY2424 | 6 | WT-Rep6 | TGA | G>C | G/C+1 | G/C-1 | CG>GC | CG>GC | AAATATGAT/ | GATATAA | ACT | ACT | C | G | C>G | C>G_ACT | CG>GC_GATATAA | 9 | FALSE | FALSE |
| 129_v.txt | 33118 | 33118-SNV-( | SNV | 1 | C | G | 2 | 27506 | 7.27E-03 | 37.9987654 | WT | FALSE | CAN | JSY2424 | 6 | WT-Rep6 | GCG | C>G | G/C+1 | G/C-1 | CG>GC | CG>GC | GGCCGGCGT | CGTTGGT | CGC | GCG | C | C | C>G | C>G_GCG | CG>GC_CGTTGGT | 9 | FALSE | FALSE |
| 129_v.txt | 33156 | 33156-SNV-( | SNV | 1 | G | C | 27 | 27476 | 0.09826758 | 37.7407407 | WT | FALSE | CAN | JSY2424 | 6 | WT-Rep6 | TGT | G>C | G/C+1 | G/C-1 | CG>GC | CG>GC | AGACCTGTA( | GTACCAA | ACA | ACA | C | G | C>G | C>G_ACA | CG>GC_GTACCAA | 9 | FALSE | FALSE |
| 129_v.txt | 33169 | 33169-SNV-( | SNV | 1 | C | G | 121 | 27472 | 0.44044846 | 37.6198347 | WT | FALSE | CAN | JSY2424 | 6 | WT-Rep6 | CCA | C>G | G/C+1 | G/C-1 | CG>GC | CG>GC | TAGTACCAC( | CACCAAG | GGT | CCA | C | C | C>G | C>G_CCA | CG>GC_CACCAAG | 9 | FALSE | FALSE |
| 129_v.txt | 33172 | 33172-SNV-( | SNV | 1 | C | G | 23 | 27472 | 0.08372161 | 35.5217391 | WT | FALSE | CAN | JSY2424 | 6 | WT-Rep6 | CCA | C>G | G/C+1 | G/C-1 | CG>GC | CG>GC | TACCACCAA( | CAAGGGC | GGT | CCA | C | C | C>G | C>G_CCA | CG>GC_CAAGGGC | 9 | FALSE | FALSE |
| 129_v.txt | 33275 | 33275-SNV-( | SNV | 1 | C | G | 1 | 27157 | 3.68E-03 | 44.9977584 | WT | FALSE | CAN | JSY2424 | 6 | WT-Rep6 | ACG | C>G | G/C+1 | G/C-1 | CG>GC | CG>GC | AGAGAACGT | CGTATCC | TGC | ACG | C | C | C>G | C>G_ACG | CG>GC_CGTATCC | 9 | FALSE | FALSE |
| 129_v.txt | 33276 | 33276-SNV-( | SNV | 1 | G | C | 3 | 27156 | 0.01104728 | 44.6635088 | WT | FALSE | CAN | JSY2424 | 6 | WT-Rep6 | CGT | G>C | G/C+1 | G/C-1 | CG>GC | CG>GC | GAGAACGTA | GTATCCT | GCA | GCA | C | G | C>G | C>G_GCA | CG>GC_GTATCCT | 9 | FALSE | FALSE |
| 129_v.txt | 33419 | 33419-SNV-( | SNV | 1 | G | C | 9 | 27157 | 0.03314063 | 51.6666667 | WT | FALSE | CAN | JSY2424 | 6 | WT-Rep6 | TGT | G>C | G/C+1 | G/C-1 | CG>GC | CG>GC | CTCATTGTAC | GTACATA | ACA | ACA | C | G | C>G | C>G_ACA | CG>GC_GTACATA | 9 | FALSE | FALSE |
| 129_v.txt | 31806 | 31806-SNV-( | SNV | 1 | C | A | 5 | 10450 | 0.04784689 | 57.8 | WT | FALSE | CAN | JSY2424 | 6 | WT-Rep6 | CCA | C>A | A/T+1 | G/C-1 | CG>AT | CG>AT | ATCTTCCAAA | CAAATAA | GGT | CCA | C | T | C>A | C>A_CCA | CG>AT_CAAATAA | 9 | FALSE | FALSE |
| 129_v.txt | 31817 | 31817-SNV-( | SNV | 1 | G | T | 8 | 10450 | 0.07655502 | 45.375 | WT | FALSE | CAN | JSY2424 | 6 | WT-Rep6 | TGC | G>T | A/T+1 | G/C-1 | CG>AT | CG>AT | AAATCTGCA' | GCATCTG | ACG | ACG | C | A | C>A | C>A_ACG | CG>AT_GCATCTG | 9 | FALSE | FALSE |
| 129_v.txt | 31980 | 31980-SNV-( | SNV | 1 | G | T | 14 | 18543 | 0.07550019 | 40.1428571 | WT | FALSE | CAN | JSY2424 | 6 | WT-Rep6 | GGG | G>T | A/T+1 | G/C-1 | CG>AT | CG>AT | AAGCCGGGC | GGCATTA | CCC | CCC | C | A | C>A | C>A_CCC | CG>AT_GGCATTA | 9 | FALSE | FALSE |
| 129_v.txt | 32095 | 32095-SNV-( | SNV | 1 | C | A | 29 | 18543 | 0.15639325 | 57.2413793 | WT | FALSE | CAN | JSY2424 | 6 | WT-Rep6 | ACA | C>A | A/T+1 | G/C-1 | CG>AT | CG>AT | CTGCAACAC( | CACCAGT | TGT | ACA | C | T | C>A | C>A_ACA | CG>AT_CACCAGT | 9 | FALSE | FALSE |
| 129_v.txt | 32282 | 32282-SNV-( | SNV | 1 | G | T | 38 | 34188 | 0.11115011 | 45.603328 | WT | FALSE | CAN | JSY2424 | 6 | WT-Rep6 | CGT | G>T | A/T+1 | G/C-1 | CG>AT | CG>AT | ACCAACGTA/ | GTAAATA | GCA | GCA | C | A | C>A | C>A_GCA | CG>AT_GTAAATA | 9 | FALSE | FALSE |
| 129_v.txt | 32292 | 32292-SNV-( | SNV | 1 | G | T | 57 | 34188 | 0.16672517 | 53.5614035 | WT | FALSE | CAN | JSY2424 | 6 | WT-Rep6 | TGA | G>T | A/T+1 | G/C-1 | CG>AT | CG>AT | ATATTTGAA1 | GAATTTG | ACT | ACT | C | A | C>A | C>A_ACT | CG>AT_GAATTTG | 9 | FALSE | FALSE |
| 129_v.txt | 32368 | 32368-SNV-( | SNV | 1 | C | A | 69 | 34186 | 0.20183701 | 58.1596135 | WT | FALSE | CAN | JSY2424 | 6 | WT-Rep6 | TCA | C>A | A/T+1 | G/C-1 | CG>AT | CG>AT | AGTTCTCAA1 | CAATAGC | AGT | TCA | C | T | C>A | C>A_TCA | CG>AT_CAATAGC | 9 | FALSE | FALSE |
| 129_v.txt | 32384 | 32384-SNV-( | SNV | 1 | G | T | 13 | 34188 | 0.03802504 | 58.6923077 | WT | FALSE | CAN | JSY2424 | 6 | WT-Rep6 | AGG | G>T | A/T+1 | G/C-1 | CG>AT | CG>AT | AATAAAGGG | GGGAGAA | TCC | TCC | C | A | C>A | C>A_TCC | CG>AT_GGGAGAA | 9 | FALSE | FALSE |
| 129_v.txt | 32399 | 32399-SNV-( | SNV | 1 | G | T | 226 | 34188 | 0.66105066 | 58.6460177 | WT | FALSE | CAN | JSY2424 | 6 | WT-Rep6 | CGT | G>T | A/T+1 | G/C-1 | CG>AT | CG>AT | AGAAACGTA | GTAGGAA | GCA | GCA | C | A | C>A | C>A_GCA | CG>AT_GTAGGAA | 9 | FALSE | FALSE |
| 129_v.txt | 32432 | 32432-SNV-( | SNV | 1 | G | T | 39 | 34188 | 0.11407511 | 51.6153846 | WT | FALSE | CAN | JSY2424 | 6 | WT-Rep6 | TGT | G>T | A/T+1 | G/C-1 | CG>AT | CG>AT | GTCATTGTAT | GTATGGA | ACA | ACA | C | A | C>A | C>A_ACA | CG>AT_GTATGGA | 9 | FALSE | FALSE |
| 129_v.txt | 32463 | 32463-SNV-( | SNV | 1 | G | T | 29 | 34188 | 0.08482509 | 49.7241379 | WT | FALSE | CAN | JSY2424 | 6 | WT-Rep6 | AGA | G>T | A/T+1 | G/C-1 | CG>AT | CG>AT | AATAGAGAG | GAGCCAA | TCT | TCT | C | A | C>A | C>A_TCT | CG>AT_GAGCCAA | 9 | FALSE | FALSE |
| 129_v.txt | 32471 | 32471-SNV-( | SNV | 1 | G | T | 79 | 34188 | 0.23107523 | 49.0253165 | WT | FALSE | CAN | JSY2424 | 6 | WT-Rep6 | TGT | G>T | A/T+1 | G/C-1 | CG>AT | CG>AT | GCCAATGTA( | GTAGAAG | ACA | ACA | C | A | C>A | C>A_ACA | CG>AT_GTAGAAG | 9 | FALSE | FALSE |
| 129_v.txt | 32488 | 32488-SNV-( | SNV | 1 | G | T | 27 | 34188 | 0.07897508 | 44.4074074 | WT | FALSE | CAN | JSY2424 | 6 | WT-Rep6 | CGG | G>T | A/T+1 | G/C-1 | CG>AT | CG>AT | AGATACGGA | GGAAAAC | GCC | GCC | C | A | C>A | C>A_GCC | CG>AT_GGAAAAC | 9 | FALSE | FALSE |
| 129_v.txt | 32494 | 32494-SNV-( | SNV | 1 | C | A | 22 | 34188 | 0.06435006 | 47.5454699 | WT | FALSE | CAN | JSY2424 | 6 | WT-Rep6 | ACA | C>A | A/T+1 | G/C-1 | CG>AT | CG>AT | GGAAAACAA | CAACTTT | TGT | ACA | C | T | C>A | C>A_ACA | CG>AT_CAACTTT | 9 | FALSE | FALSE |
| 129_v.txt | 32508 | 32508-SNV-( | SNV | 1 | G | T | 22 | 34188 | 0.06435006 | 46.4545455 | WT | FALSE | CAN | JSY2424 | 6 | WT-Rep6 | GGC | G>T | A/T+1 | G/C-1 | CG>AT | CG>AT | TTGATGGCT( | GCTCTTG | CCG | CCG | C | A | C>A | C>A_CCG | CG>AT_GCTCTTG | 9 | FALSE | FALSE |
| 129_v.txt | 32572 | 32572-SNV-( | SNV | 1 | C | A | 20 | 30732 | 0.06507875 | 35 | WT | FALSE | CAN | JSY2424 | 6 | WT-Rep6 | CCT | C>A | A/T+1 | G/C-1 | CG>AT | CG>AT | CAGTACCTTC | CTTGAAA | GGA | CCT | C | T | C>A | C>A_CCT | CG>AT_CTTGAAA | 9 | FALSE | FALSE |
| 129_v.txt | 32658 | 32658-SNV-( | SNV | 1 | C | A | 19 | 30730 | 0.06182883 | 20.2628332 | WT | FALSE | CAN | JSY2424 | 6 | WT-Rep6 | ACC | C>A | A/T+1 | G/C-1 | CG>AT | CG>AT | CCTGGACCCC | CCCCAGG | TGG | ACC | C | T | C>A | C>A_ACC | CG>AT_CCCCAGG | 9 | FALSE | FALSE |
| 129_v.txt | 32735 | 32735-SNV-( | SNV | 1 | G | T | 9 | 30731 | 0.02928639 | 55 | WT | FALSE | CAN | JSY2424 | 6 | WT-Rep6 | AGT | G>T | A/T+1 | G/C-1 | CG>AT | CG>AT | AAAACAGTA | GTATATT | TCA | TCA | C | A | C>A | C>A_TCA | CG>AT_GTATATT | 9 | FALSE | FALSE |
| 129_v.txt | 32794 | 32794-SNV-( | SNV | 1 | C | A | 26 | 30731 | 0.08460512 | 46.1538462 | WT | FALSE | CAN | JSY2424 | 6 | WT-Rep6 | TCA | C>A | A/T+1 | G/C-1 | CG>AT | CG>AT | CGAATTCAC( | CACCGTA | AGT | TCA | C | T | C>A | C>A_TCA | CG>AT_CACCGTA | 9 | FALSE | FALSE |
| 129_v.txt | 32796 | 32796-SNV-( | SNV | 1 | C | A | 28 | 30731 | 0.09111321 | 46.5966814 | WT | FALSE | CAN | JSY2424 | 6 | WT-Rep6 | ACC | C>A | A/T+1 | G/C-1 | CG>AT | CG>AT | AATTCACCG1 | CCGTAAT | TGG | ACC | C | T | C>A | C>A_ACC | CG>AT_CCGTAAT | 9 | FALSE | FALSE |
| 129_v.txt | 32797 | 32797-SNV-( | SNV | 1 | C | A | 44 | 30731 | 0.1431779 | 42.2953657 | WT | FALSE | CAN | JSY2424 | 6 | WT-Rep6 | CCG | C>A | A/T+1 | G/C-1 | CG>AT | CG>AT | ATTCACCGT/ | CGTAATA | GGC | CCG | C | T | C>A | C>A_CCG | CG>AT_CGTAATA | 9 | FALSE | FALSE |
| 129_v.txt | 32798 | 32798-SNV-( | SNV | 1 | G | T | 46 | 30731 | 0.14968599 | 43.6304348 | WT | FALSE | CAN | JSY2424 | 6 | WT-Rep6 | CGT | G>T | A/T+1 | G/C-1 | CG>AT | CG>AT | TTCACCGTA/ | GTAATAT | GCA | GCA | C | A | C>A | C>A_GCA | CG>AT_GTAATAT | 9 | FALSE | FALSE |
| 129_v.txt | 32811 | 32811-SNV-( | SNV | 1 | G | T | 7 | 30731 | 0.0227783 | 37.5714286 | WT | FALSE | CAN | JSY2424 | 6 | WT-Rep6 | AGG | G>T | A/T+1 | G/C-1 | CG>AT | CG>AT | TTGACAGGG | GGGAACA | TCC | TCC | C | A | C>A | C>A_TCC | CG>AT_GGGAACA | 9 | FALSE | FALSE |
| 129_v.txt | 32928 | 32928-SNV-( | SNV | 1 | G | T | 12 | 27506 | 0.04362685 | 52.9166667 | WT | FALSE | CAN | JSY2424 | 6 | WT-Rep6 | AGT | G>T | A/T+1 | G/C-1 | CG>AT | CG>AT | GCAAAAGTG | GTGATTG | TCA | TCA | C | A | C>A | C>A_TCA | CG>AT_GTGATTG | 9 | FALSE | FALSE |
| 129_v.txt | 33006 | 33006-SNV-( | SNV | 1 | G | T | 9 | 27506 | 0.03272013 | 64 | WT | FALSE | CAN | JSY2424 | 6 | WT-Rep6 | TGT | G>T | A/T+1 | G/C-1 | CG>AT | CG>AT | AAAACTGTG. | GTGAAAG | ACA | ACA | C | A | C>A | C>A_ACA | CG>AT_GTGAAAG | 9 | FALSE | FALSE |
| 129_v.txt | 33012 | 33012-SNV-( | SNV | 1 | G | T | 33 | 27506 | 0.11997382 | 56.2121212 | WT | FALSE | CAN | JSY2424 | 6 | WT-Rep6 | AGA | G>T | A/T+1 | G/C-1 | CG>AT | CG>AT | GTGAAAGAG | GAGGATG | TCT | TCT | C | A | C>A | C>A_TCT | CG>AT_GAGGATG | 9 | FALSE | FALSE |
| 129_v.txt | 33018 | 33018-SNV-( | SNV | 1 | G | T | 27 | 27506 | 0.0981604 | 61 | WT | FALSE | CAN | JSY2424 | 6 | WT-Rep6 | TGT | G>T | A/T+1 | G/C-1 | CG>AT | CG>AT | GAGGATGTA | GTAACAG | ACA | ACA | C | A | C>A | C>A_ACA | CG>AT_GTAACAG | 9 | FALSE | FALSE |
| 129_v.txt | 33151 | 33151-SNV-( | SNV | 1 | G | T | 31 | 27477 | 0.11282163 | 37.1290323 | WT | FALSE | CAN | JSY2424 | 6 | WT-Rep6 | AGA | G>T | A/T+1 | G/C-1 | CG>AT | CG>AT | TGAAAAGAC | GACCTGT | TCT | TCT | C | A | C>A | C>A_TCT | CG>AT_GACCTGT | 9 | FALSE | FALSE |
| 129_v.txt | 33175 | 33175-SNV-( | SNV | 1 | G | T | 5 | 27469 | 0.01820234 | 38 | WT | FALSE | CAN | JSY2424 | 6 | WT-Rep6 | AGG | G>T | A/T+1 | G/C-1 | CG>AT | CG>AT | CACCAAGGG | GGGCAAT | TCC | TCC | C | A | C>A | C>A_TCC | CG>AT_GGGCAAT | 9 | FALSE | FALSE |
| 129_v.txt | 33177 | 33177-SNV-( | SNV | 1 | G | T | 12 | 27468 | 0.0436872 | 37.25 | WT | FALSE | CAN | JSY2424 | 6 | WT-Rep6 | GGC | G>T | A/T+1 | G/C-1 | CG>AT | CG>AT | CCAAGGGCA | GCAATCA | CCG | CCG | C | A | C>A | C>A_CCG | CG>AT_GCAATCA | 9 | FALSE | FALSE |
| 129_v.txt | 33210 | 33210-SNV-( | SNV | 1 | C | A | 18 | 27157 | 0.06628125 | 34.4444444 | WT | FALSE | CAN | JSY2424 | 6 | WT-Rep6 | TCT | C>A | A/T+1 | G/C-1 | CG>AT | CG>AT | AGCTCTCTCT | CTCTTCA | AGA | TCT | C | T | C>A | C>A_TCT | CG>AT_CTCTTCA | 9 | FALSE | FALSE |
| 129_v.txt | 33378 | 33378-SNV-( | SNV | 1 | G | T | 184 | 27157 | 0.6775417 | 56.25 | WT | FALSE | CAN | JSY2424 | 6 | WT-Rep6 | TGA | G>T | A/T+1 | G/C-1 | CG>AT | CG>AT | GTTTGTGAA( | GAAGCTT | ACT | ACT | C | A | C>A | C>A_ACT | CG>AT_GAAGCTT | 9 | FALSE | FALSE |
| 129_v.txt | 33474 | 33474-SNV-( | SNV | 1 | C | A | 58 | 27155 | 0.21358866 | 35.1724138 | WT | FALSE | CAN | JSY2424 | 6 | WT-Rep6 | GCC | C>A | A/T+1 | G/C-1 | CG>AT | CG>AT | GCTATGCCTT | CCTTTTT | CGG | GCC | C | T | C>A | C>A_GCC | CG>AT_CCTTTTT | 9 | FALSE | FALSE |
| 129_v.txt | 32743 | 32743-Repla | Replacement | 2 | G | CA | 22 | 30731 | 0.07158895 | 62.6136358 | WT | FALSE | CAN | JSY2424 | 6 | WT-Rep6 | AGA | nope | >1bp-ins | G/C-1 | none | Replac. | ATATTAGAA/ | GAAACCC | TCT | TCT | C | none | C>none | C>none_TCT | Replac._GAAACCC | 9 | FALSE | FALSE |
| 129_v.txt | 32274 | 32274-MNV- | MNV | 2 | GA | AT | 1 | 34188 | 2.93E-03 | 37.9999963 | WT | FALSE | CAN | JSY2424 | 6 | WT-Rep6 | GGA | nope | >1bp-ins | >1bpdel | none | MNV | ATACGGGAA | GAACCAA | CCT | GGA | GA | none | NA | NA_GGA | MNV_GAACCAA | 9 | FALSE | FALSE |
| 129_v.txt | 32081 | 32081-Insert | Insertion | 1 | - | A | 48 | 18543 | 0.25885779 | 63.4375 | WT | FALSE | CAN | JSY2424 | 6 | WT-Rep6 | CAA | nope | A/T+1 | >1bpdel | none | A/T+1 | CCATGCAAA/ | AAAAAAG | GTT | CAA | - | T | NA | NA_CAA | A/T+1_AAAAAAG | 9 | FALSE | FALSE |
| 129_v.txt | 32442 | 32442-Insert | Insertion | 1 | - | A | 4 | 34188 | 0.01170001 | 50.25 | WT | FALSE | CAN | JSY2424 | 6 | WT-Rep6 | TAA | nope | A/T+1 | >1bpdel | none | A/T+1 | GGAACTAAA | AAAAGTC | ATT | TAA | - | T | NA | NA_TAA | A/T+1_AAAAGTC | 9 | FALSE | FALSE |
| 129_v.txt | 32468 | 32468-Insert | Insertion | 1 | - | A | 32 | 34188 | 0.09360009 | 45.25 | WT | FALSE | CAN | JSY2424 | 6 | WT-Rep6 | CAA | nope | A/T+1 | >1bpdel | none | A/T+1 | AGAGCCAAT | AATGTAG | GTT | CAA | - | T | NA | NA_CAA | A/T+1_AATGTAG | 9 | FALSE | FALSE |
| 129_v.txt | 32494 | 32494-Insert | Insertion | 1 | - | C | 1 | 34188 | 2.93E-03 | 11 | WT | FALSE | CAN | JSY2424 | 6 | WT-Rep6 | ACA | nope | G/C+1 | >1bpdel | none | G/C+1 | GGAAAACAA | CAACTTT | TGT | ACA | - | G | NA | NA_ACA | G/C+1_CAACTTT | 9 | FALSE | FALSE |
| 129_v.txt | 32658 | 32658-Insert | Insertion | 1 | - | C | 24 | 30731 | 0.07809704 | 62.3333333 | WT | FALSE | CAN | JSY2424 | 6 | WT-Rep6 | ACC | nope | G/C+1 | >1bpdel | none | G/C+1 | CCTGGACCCC | CCCCAGG | TGG | ACC | - | G | NA | NA_ACC | G/C+1_CCCCAGG | 9 | FALSE | FALSE |
| 129_v.txt | 32728 | 32728-Insert | Insertion | 1 | - | A | 22 | 30731 | 0.07158895 | 59.6818182 | WT | FALSE | CAN | JSY2424 | 6 | WT-Rep6 | CAA | nope | A/T+1 | >1bpdel | none | A/T+1 | CCATACAAA/ | AAAAACA | GTT | CAA | - | T | NA | NA_CAA | A/T+1_AAAAACA | 9 | FALSE | FALSE |
| 129_v.txt | 32842 | 32842-Insert | Insertion | 1 | - | A | 131 | 30731 | 0.42627965 | 42.5267176 | WT | FALSE | CAN | JSY2424 | 6 | WT-Rep6 | CAA | nope | A/T+1 | >1bpdel | none | A/T+1 | TTACCCAAA/ | AAAAAAT | GTT | CAA | - | T | NA | NA_CAA | A/T+1_AAAAAAT | 9 | FALSE | FALSE |
| 129_v.txt | 32884 | 32884-Insert | Insertion | 1 | - | A | 2 | 27506 | 7.27E-03 | 38 | WT | FALSE | CAN | JSY2424 | 6 | WT-Rep6 | CAA | nope | A/T+1 | >1bpdel | none | A/T+1 | ACGTCCAAA/ | AAAATTG | GTT | CAA | - | T | NA | NA_CAA | A/T+1_AAAATTG | 9 | FALSE | FALSE |
| 129_v.txt | 33014 | 33014-Insert | Insertion | 11 | - | GGATGTAAC | 3 | 27506 | 0.01090671 | 63.9393939 | WT | FALSE | CAN | JSY2424 | 6 | WT-Rep6 | AGG | nope | >1bp-ins | >1bpdel | none | >1bp-ins | GAAAGAGG/ | GGATGTA | TCC | AGG | - | none | NA | NA_AGG | >1bp-ins_GGATGTA | 9 | FALSE | FALSE |
| 129_v.txt | 33072 | 33072-Insert | Insertion | 1 | - | A | 27 | 27506 | 0.0981604 | 57.7037037 | WT | FALSE | CAN | JSY2424 | 6 | WT-Rep6 | CAA | nope | A/T+1 | >1bpdel | none | A/T+1 | TATGCCAAA( | AAAGAAC | GTT | CAA | - | T | NA | NA_CAA | A/T+1_AAAGAAC | 9 | FALSE | FALSE |
| 129_v.txt | 33088 | 33088-Insert | Insertion | 10 | - | AATATGATA' | 18 | 27506 | 0.06544027 | 53.7833333 | WT | FALSE | CAN | JSY2424 | 6 | WT-Rep6 | AAA | nope | >1bp-ins | >1bpdel | none | >1bp-ins | TAAATAAAT/ | AATATGA | TTT | AAA | - | none | NA | NA_AAA | >1bp-ins_AATATGA | 9 | FALSE | FALSE |
| 129_v.txt | 33230 | 33230-Insert | Insertion | 1 | - | A | 3 | 27157 | 0.01104688 | 36.3333333 | WT | FALSE | CAN | JSY2424 | 6 | WT-Rep6 | GTA | nope | A/T+1 | >1bpdel | none | A/T+1 | GTTCTGTACT | TACTTCT | CAT | GTA | - | T | NA | NA_GTA | A/T+1_TACTTCT | 9 | FALSE | FALSE |
| 129_v.txt | 32014 | 32014-Deleti | Deletion | 2 | GA | - | 17 | 18543 | 0.0916788 | 58.9411765 | WT | FALSE | CAN | JSY2424 | 6 | WT-Rep6 | CGA | nope | >1bp-ins | >1bpdel | none | >1bpdel | CGTCACGAG. | GAGAGAT | GCT | CGA | GA | none | NA | NA_CGA | >1bpdel_GAGAGAT | 9 | FALSE | FALSE |
| 129_v.txt | 32337 | 32337-Deleti | Deletion | 2 | AT | - | 43 | 34188 | 0.12577513 | 60.9069767 | WT | FALSE | CAN | JSY2424 | 6 | WT-Rep6 | GAT | nope | >1bp-ins | >1bpdel | none | >1bpdel | TTGAAGATA' | ATATGTG | CTA | GAT | AT | none | NA | NA_GAT | >1bpdel_ATATGTG | 9 | FALSE | FALSE |
| 129_v.txt | 33189 | 33189-Deleti | Deletion | 2 | AT | - | 23 | 27153 | 0.08470519 | 37.7826087 | WT | FALSE | CAN | JSY2424 | 6 | WT-Rep6 | AAT | nope | >1bp-ins | >1bpdel | none | >1bpdel | ATACCAATA1 | ATATGTC | TTA | AAT | AT | none | NA | NA_AAT | >1bpdel_ATATGTC | 9 | FALSE | FALSE |
| 129_v.txt | 31723 | 31723-Deleti | Deletion | 1 | A | - | 6 | 10450 | 0.05741627 | 61.3333333 | WT | FALSE | CAN | JSY2424 | 6 | WT-Rep6 | CAA | nope | >1bp-ins | A/T-1 | none | A/T-1 | TGTCCCAAA/ | AAAAAGT | GTT | GTT | T | none | T>none | T>none_GTT | A/T-1_AAAAAGT | 9 | FALSE | FALSE |
| 129_v.txt | 31874 | 31874-Deleti | Deletion | 1 | A | - | 2 | 10450 | 0.01913876 | 38 | WT | FALSE | CAN | JSY2424 | 6 | WT-Rep6 | TAT | nope | >1bp-ins | A/T-1 | none | A/T-1 | AGAGATATA | ATAGGCG | ATA | ATA | T | none | T>none | T>none_ATA | A/T-1_ATAGGCG | 9 | FALSE | FALSE |
| 129_v.txt | 31958 | 31958-Deleti | Deletion | 1 | G | - | 12 | 18543 | 0.06471445 | 37.5 | WT | FALSE | CAN | JSY2424 | 6 | WT-Rep6 | TGG | nope | >1bp-ins | G/C-1 | none | G/C-1 | AAATGTGGC | GGCCGCA | ACC | ACC | C | none | C>none | C>none_ACC | G/C-1_GGCCGCA | 9 | FALSE | FALSE |
| 129_v.txt | 32081 | 32081-Deleti | Deletion | 1 | A | - | 32 | 18543 | 0.17257186 | 62.3750004 | WT | FALSE | CAN | JSY2424 | 6 | WT-Rep6 | CAA | nope | >1bp-ins | A/T-1 | none | A/T-1 | CCATGCAAA/ | AAAAAAG | GTT | GTT | T | none | T>none | T>none_GTT | A/T-1_AAAAAAG | 9 | FALSE | FALSE |
| 129_v.txt | 32149 | 32149-Deleti | Deletion | 1 | C | - | 1984 | 18543 | 10.6994553 | 51.9637097 | WT | FALSE | CAN | JSY2424 | 6 | WT-Rep6 | TCC | nope | >1bp-ins | G/C-1 | none | G/C-1 | ATGTCTCCAT | CCATGTA | AGG | TCC | C | none | nope | nope_TCC | G/C-1_CCATGTA | 9 | FALSE | FALSE |
| 129_v.txt | 32168 | 32168-Deleti | Deletion | 1 | A | - | 8 | 18543 | 0.04314297 | 53.125 | WT | FALSE | CAN | JSY2424 | 6 | WT-Rep6 | CAA | nope | >1bp-ins | A/T-1 | none | A/T-1 | AGCGCCAAA | AAATGCA | GTT | GTT | T | none | T>none | T>none_GTT | A/T-1_AAATGCA | 9 | FALSE | FALSE |
| 129_v.txt | 32210 | 32210-Deleti | Deletion | 1 | T | - | 5 | 18543 | 0.02696435 | 34.8 | WT | FALSE | CAN | JSY2424 | 6 | WT-Rep6 | CTT | nope | >1bp-ins | A/T-1 | none | A/T-1 | ACCACCTTTG | TTTGGTG | GAA | CTT | T | none | nope | nope_CTT | A/T-1_TTTGGTG | 9 | FALSE | FALSE |
| 129_v.txt | 32282 | 32282-Deleti | Deletion | 1 | G | - | 47 | 34188 | 0.13747514 | 36.5957446 | WT | FALSE | CAN | JSY2424 | 6 | WT-Rep6 | CGT | nope | >1bp-ins | G/C-1 | none | G/C-1 | ACCAACGTA/ | GTAAATA | GCA | GCA | C | none | C>none | C>none_GCA | G/C-1_GTAAATA | 9 | FALSE | FALSE |
| 129_v.txt | 32284 | 32284-Deleti | Deletion | 1 | A | - | 8 | 34188 | 0.02340002 | 40.375 | WT | FALSE | CAN | JSY2424 | 6 | WT-Rep6 | TAA | nope | >1bp-ins | A/T-1 | none | A/T-1 | CAACGTAAA' | AAATATT | ATT | ATT | T | none | T>none | T>none_ATT | A/T-1_AAATATT | 9 | FALSE | FALSE |

| | | | | | | | | | | | | | | | | | | | | | | | | | | | | | | | | | | |
|---|---|---|---|---|---|---|---|---|---|---|---|---|---|---|---|---|---|---|---|---|---|---|---|---|---|---|---|---|---|---|---|---|---|---|
| 129_v.txt | 32341 | 32341-Deleti | Deletion | 1 | G | - | 8 | 34188 | 0.02340002 | 61 | WT | FALSE | CAN | JSY2424 | 6 | WT-Rep6 | TGT | nope | >1bp-ins | G/C-1 | none | G/C-1 | AGATATGTG | GTGGCAA | ACA | ACA | C | none | C>none | C>none_ACA | G/C-1_GTGGCAA | 9 | FALSE | FALSE |
| 129_v.txt | 32346 | 32346-Deleti | Deletion | 1 | A | - | 22 | 34188 | 0.06435006 | 61.1363636 | WT | FALSE | CAN | JSY2424 | 6 | WT-Rep6 | CAA | nope | >1bp-ins | A/T-1 | none | A/T-1 | TGTGGCAAA | AAAACCT | GTT | GTT | T | none | T>none | T>none_GTT | A/T-1_AAAACCT | 9 | FALSE | FALSE |
| 129_v.txt | 32352 | 32352-Deleti | Deletion | 1 | T | - | 7 | 34188 | 0.02047502 | 60.8571429 | WT | FALSE | CAN | JSY2424 | 6 | WT-Rep6 | CTT | nope | >1bp-ins | A/T-1 | none | A/T-1 | AAAACCTTT | TTTGTAC | GAA | CTT | T | none | nope | nope_CTT | A/T-1_TTTGTAC | 9 | FALSE | FALSE |
| 129_v.txt | 32369 | 32369-Deleti | Deletion | 1 | A | - | 2 | 34188 | 5.85E-03 | 64 | WT | FALSE | CAN | JSY2424 | 6 | WT-Rep6 | CAA | nope | >1bp-ins | A/T-1 | none | A/T-1 | GTTCTCAATA | AATAGCA | GTT | GTT | T | none | T>none | T>none_GTT | A/T-1_AATAGCA | 9 | FALSE | FALSE |
| 129_v.txt | 32381 | 32381-Deleti | Deletion | 1 | A | - | 7 | 34188 | 0.02047502 | 61.2857143 | WT | FALSE | CAN | JSY2424 | 6 | WT-Rep6 | TAA | nope | >1bp-ins | A/T-1 | none | A/T-1 | AATAATAAA | AAAGGGA | ATT | ATT | T | none | T>none | T>none_ATT | A/T-1_AAAGGGA | 9 | FALSE | FALSE |
| 129_v.txt | 32387 | 32387-Deleti | Deletion | 1 | A | - | 9 | 34188 | 0.02632503 | 64 | WT | FALSE | CAN | JSY2424 | 6 | WT-Rep6 | GAG | nope | >1bp-ins | A/T-1 | none | A/T-1 | AAAGGGAGA | AGAAGTA | CTC | CTC | T | none | T>none | T>none_CTC | A/T-1_AGAAGTA | 9 | FALSE | FALSE |
| 129_v.txt | 32498 | 32498-Deleti | Deletion | 1 | T | - | 153 | 34188 | 0.44752545 | 44.2222222 | WT | FALSE | CAN | JSY2424 | 6 | WT-Rep6 | CTT | nope | >1bp-ins | A/T-1 | none | A/T-1 | AACAACTTT | TTTTTTG | GAA | CTT | T | none | nope | nope_CTT | A/T-1_TTTTTTG | 9 | FALSE | FALSE |
| 129_v.txt | 32522 | 32522-Deleti | Deletion | 1 | T | - | 12 | 34187 | 0.03510106 | 37.75 | WT | FALSE | CAN | JSY2424 | 6 | WT-Rep6 | ATT | nope | >1bp-ins | A/T-1 | none | A/T-1 | AACGGATTT | TTTTCTG | TAA | ATT | T | none | nope | nope_ATT | A/T-1_TTTTCTG | 9 | FALSE | FALSE |
| 129_v.txt | 32528 | 32528-Deleti | Deletion | 1 | G | - | 13488 | 34187 | 39.4535935 | 41.8104982 | WT | FALSE | CAN | JSY2424 | 6 | WT-Rep6 | TGG | nope | >1bp-ins | G/C-1 | none | G/C-1 | TTTTCTGGG | GGGGTTT | ACC | ACC | C | none | C>none | C>none_ACC | G/C-1_GGGGTTT | 9 | FALSE | FALSE |
| 129_v.txt | 32573 | 32573-Deleti | Deletion | 1 | T | - | 3 | 30732 | 9.76E-03 | 38 | WT | FALSE | CAN | JSY2424 | 6 | WT-Rep6 | CTT | nope | >1bp-ins | A/T-1 | none | A/T-1 | AGTACCTTG | TTGAAAT | GAA | CTT | T | none | nope | nope_CTT | A/T-1_TTGAAAT | 9 | FALSE | FALSE |
| 129_v.txt | 32631 | 32631-Deleti | Deletion | 1 | T | - | 23 | 30731 | 0.07484299 | 47.173913 | WT | FALSE | CAN | JSY2424 | 6 | WT-Rep6 | GTT | nope | >1bp-ins | A/T-1 | none | A/T-1 | CCTTCGTTTT | TTTTTAT | CAA | GTT | T | none | nope | nope_GTT | A/T-1_TTTTTAT | 9 | FALSE | FALSE |
| 129_v.txt | 32658 | 32658-Deleti | Deletion | 1 | C | - | 7 | 30730 | 0.02277904 | 61.2857103 | WT | FALSE | CAN | JSY2424 | 6 | WT-Rep6 | ACC | nope | >1bp-ins | G/C-1 | none | G/C-1 | CCTGGACCC | CCCCAGG | TGG | ACC | C | none | nope | nope_ACC | G/C-1_CCCCAGG | 9 | FALSE | FALSE |
| 129_v.txt | 32707 | 32707-Deleti | Deletion | 1 | C | - | 9 | 30731 | 0.02928639 | 57.5555556 | WT | FALSE | CAN | JSY2424 | 6 | WT-Rep6 | ACC | nope | >1bp-ins | G/C-1 | none | G/C-1 | CGGTAACCC | CCCCAGC | TGG | ACC | C | none | nope | nope_ACC | G/C-1_CCCCAGC | 9 | FALSE | FALSE |
| 129_v.txt | 32740 | 32740-Deleti | Deletion | 1 | T | - | 5 | 30731 | 0.01627022 | 57.4 | WT | FALSE | CAN | JSY2424 | 6 | WT-Rep6 | ATT | nope | >1bp-ins | A/T-1 | none | A/T-1 | AGTATATTA | TTAGAAA | TAA | ATT | T | none | nope | nope_ATT | A/T-1_TTAGAAA | 9 | FALSE | FALSE |
| 129_v.txt | 32842 | 32842-Deleti | Deletion | 1 | A | - | 50 | 30731 | 0.16270216 | 39.74 | WT | FALSE | CAN | JSY2424 | 6 | WT-Rep6 | CAA | nope | >1bp-ins | A/T-1 | none | A/T-1 | TTACCCAAA | AAAAAAT | GTT | GTT | T | none | T>none | T>none_GTT | A/T-1_AAAAAAT | 9 | FALSE | FALSE |
| 129_v.txt | 32884 | 32884-Deleti | Deletion | 1 | A | - | 8 | 27506 | 0.02908456 | 37.3749998 | WT | FALSE | CAN | JSY2424 | 6 | WT-Rep6 | CAA | nope | >1bp-ins | A/T-1 | none | A/T-1 | ACGTCCAAA | AAAATTG | GTT | GTT | T | none | T>none | T>none_GTT | A/T-1_AAAATTG | 9 | FALSE | FALSE |
| 129_v.txt | 32920 | 32920-Deleti | Deletion | 1 | G | - | 9 | 27506 | 0.03272013 | 52.7777778 | WT | FALSE | CAN | JSY2424 | 6 | WT-Rep6 | AGG | nope | >1bp-ins | G/C-1 | none | G/C-1 | GTTCCAGGG | GGGCAAA | TCC | TCC | C | none | C>none | C>none_TCC | G/C-1_GGGCAAA | 9 | FALSE | FALSE |
| 129_v.txt | 32924 | 32924-Deleti | Deletion | 1 | A | - | 17 | 27506 | 0.0618047 | 51.6470588 | WT | FALSE | CAN | JSY2424 | 6 | WT-Rep6 | CAA | nope | >1bp-ins | A/T-1 | none | A/T-1 | CAGGGCAAA | AAAAGTG | GTT | GTT | T | none | T>none | T>none_GTT | A/T-1_AAAAGTG | 9 | FALSE | FALSE |
| 129_v.txt | 32935 | 32935-Deleti | Deletion | 1 | C | - | 7 | 27506 | 0.02544899 | 46.7142857 | WT | FALSE | CAN | JSY2424 | 6 | WT-Rep6 | GCC | nope | >1bp-ins | G/C-1 | none | G/C-1 | TGATTGCCCA | CCCAAGA | CGG | GCC | C | none | nope | nope_GCC | G/C-1_CCCAAGA | 9 | FALSE | FALSE |
| 129_v.txt | 32941 | 32941-Deleti | Deletion | 1 | A | - | 8 | 27506 | 0.02908456 | 63.25 | WT | FALSE | CAN | JSY2424 | 6 | WT-Rep6 | GAA | nope | >1bp-ins | A/T-1 | none | A/T-1 | CCCAAGAAA | AAAACCA | CTT | CTT | T | none | T>none | T>none_CTT | A/T-1_AAAACCA | 9 | FALSE | FALSE |
| 129_v.txt | 32965 | 32965-Deleti | Deletion | 1 | C | - | 12 | 27506 | 0.04362685 | 59.6666667 | WT | FALSE | CAN | JSY2424 | 6 | WT-Rep6 | GCC | nope | >1bp-ins | G/C-1 | none | G/C-1 | CATTGGCCG | CCGCACC | CGG | GCC | C | none | nope | nope_GCC | G/C-1_CCGCACC | 9 | FALSE | FALSE |
| 129_v.txt | 33000 | 33000-Deleti | Deletion | 1 | A | - | 27 | 27506 | 0.0981604 | 61.1851852 | WT | FALSE | CAN | JSY2424 | 6 | WT-Rep6 | GAA | nope | >1bp-ins | A/T-1 | none | A/T-1 | TGTGAGAAA | AAAACTG | CTT | CTT | T | none | T>none | T>none_CTT | A/T-1_AAAACTG | 9 | FALSE | FALSE |
| 129_v.txt | 33037 | 33037-Deleti | Deletion | 1 | C | - | 4 | 27506 | 0.01454228 | 57.25 | WT | FALSE | CAN | JSY2424 | 6 | WT-Rep6 | GCC | nope | >1bp-ins | G/C-1 | none | G/C-1 | ATGTAGCCA | CCATTTC | CGG | GCC | C | none | nope | nope_GCC | G/C-1_CCATTTC | 9 | FALSE | FALSE |
| 129_v.txt | 33040 | 33040-Deleti | Deletion | 1 | T | - | 5 | 27506 | 0.01817785 | 58.8 | WT | FALSE | CAN | JSY2424 | 6 | WT-Rep6 | ATT | nope | >1bp-ins | A/T-1 | none | A/T-1 | TAGCCATTTC | TTTCACC | TAA | ATT | T | none | nope | nope_ATT | A/T-1_TTTCACC | 9 | FALSE | FALSE |
| 129_v.txt | 33122 | 33122-Deleti | Deletion | 1 | G | - | 9 | 27506 | 0.03272013 | 40.6666667 | WT | FALSE | CAN | JSY2424 | 6 | WT-Rep6 | TGG | nope | >1bp-ins | G/C-1 | none | G/C-1 | GGCGTTGGT | GGTCAGA | ACC | ACC | C | none | C>none | C>none_ACC | G/C-1_GGTCAGA | 9 | FALSE | FALSE |
| 129_v.txt | 33169 | 33169-Deleti | Deletion | 1 | C | - | 3 | 27472 | 0.01092021 | 37.6666667 | WT | FALSE | CAN | JSY2424 | 6 | WT-Rep6 | CCA | nope | >1bp-ins | G/C-1 | none | G/C-1 | TAGTACCAC | CACCAAG | GGT | CCA | C | none | nope | nope_CCA | G/C-1_CACCAAG | 9 | FALSE | FALSE |
| 129_v.txt | 33175 | 33175-Deleti | Deletion | 1 | G | - | 7 | 27469 | 0.02548327 | 37.4285714 | WT | FALSE | CAN | JSY2424 | 6 | WT-Rep6 | AGG | nope | >1bp-ins | G/C-1 | none | G/C-1 | CACCAAGGG | GGGCAAT | TCC | TCC | C | none | C>none | C>none_TCC | G/C-1_GGGCAAT | 9 | FALSE | FALSE |
| 129_v.txt | 33194 | 33194-Deleti | Deletion | 1 | T | - | 10 | 27156 | 0.03682428 | 37.8 | WT | FALSE | CAN | JSY2424 | 6 | WT-Rep6 | GTC | nope | >1bp-ins | A/T-1 | none | A/T-1 | AATATGTCTT | TCTTTGC | CAG | GTC | T | none | nope | nope_GTC | A/T-1_TCTTTGC | 9 | FALSE | FALSE |
| 129_v.txt | 33303 | 33303-Deleti | Deletion | 1 | A | - | 6 | 27157 | 0.02209375 | 54.3333333 | WT | FALSE | CAN | JSY2424 | 6 | WT-Rep6 | GAA | nope | >1bp-ins | A/T-1 | none | A/T-1 | GTCGGGAAA | AAAGAGC | CTT | CTT | T | none | T>none | T>none_CTT | A/T-1_AAAGAGC | 9 | FALSE | FALSE |
| 129_v.txt | 33400 | 33400-Deleti | Deletion | 1 | G | - | 9 | 27157 | 0.03314063 | 58.1111111 | WT | FALSE | CAN | JSY2424 | 6 | WT-Rep6 | AGG | nope | >1bp-ins | G/C-1 | none | G/C-1 | GAAAGAGGG | GGGTTGT | TCC | TCC | C | none | C>none | C>none_TCC | G/C-1_GGGTTGT | 9 | FALSE | FALSE |
| 129_v.txt | 33474 | 33474-Deleti | Deletion | 1 | C | - | 70 | 27155 | 0.25777942 | 30.0285714 | WT | FALSE | CAN | JSY2424 | 6 | WT-Rep6 | GCC | nope | >1bp-ins | G/C-1 | none | G/C-1 | GCTATGCCTT | CCTTTTT | CGG | GCC | C | none | nope | nope_GCC | G/C-1_CCTTTTT | 9 | FALSE | FALSE |
| 130_v.txt | 31663 | 31663-SNV-A | SNV | 1 | A | C | 8 | 11604 | 0.06894174 | 17 | WT | FALSE | CAN | JSY2424 | 7 | WT-Rep7 | GAA | A>C | G/C+1 | A/T-1 | TA>GC | TA>GC | GCGTGGAAA | AAATGTG | CTT | CTT | T | G | T>G | T>G_CTT | TA>GC_AAATGTG | 9 | FALSE | FALSE |
| 130_v.txt | 31723 | 31723-SNV-A | SNV | 1 | A | C | 3 | 11604 | 0.02585315 | 33.3333333 | WT | FALSE | CAN | JSY2424 | 7 | WT-Rep7 | CAA | A>C | G/C+1 | A/T-1 | TA>GC | TA>GC | TGTCCCAAA | AAAAAGT | GTT | GTT | T | G | T>G | T>G_GTT | TA>GC_AAAAAGT | 9 | FALSE | FALSE |
| 130_v.txt | 31950 | 31950-SNV-A | SNV | 1 | A | C | 28 | 31094 | 0.09004953 | 32.0714286 | WT | FALSE | CAN | JSY2424 | 7 | WT-Rep7 | CAT | A>C | G/C+1 | A/T-1 | TA>GC | TA>GC | ATCGTCATA | ATAAATG | GTA | GTA | T | G | T>G | T>G_GTA | TA>GC_ATAAATG | 9 | FALSE | FALSE |
| 130_v.txt | 31964 | 31964-SNV-A | SNV | 1 | A | C | 37 | 31094 | 0.11899402 | 36.2971344 | WT | FALSE | CAN | JSY2424 | 7 | WT-Rep7 | CAT | A>C | G/C+1 | A/T-1 | TA>GC | TA>GC | GGCCGCATA | ATAATAA | GTA | GTA | T | G | T>G | T>G_GTA | TA>GC_ATAATAA | 9 | FALSE | FALSE |
| 130_v.txt | 31986 | 31986-SNV-A | SNV | 1 | A | C | 40 | 31094 | 0.12864218 | 48.85 | WT | FALSE | CAN | JSY2424 | 7 | WT-Rep7 | TAA | A>C | G/C+1 | A/T-1 | TA>GC | TA>GC | GGCATTAAT | AATTTAG | ATT | ATT | T | G | T>G | T>G_ATT | TA>GC_AATTTAG | 9 | FALSE | FALSE |
| 130_v.txt | 32067 | 32067-SNV-A | SNV | 1 | A | C | 53 | 31094 | 0.17045089 | 56.9811321 | WT | FALSE | CAN | JSY2424 | 7 | WT-Rep7 | GAT | A>C | G/C+1 | A/T-1 | TA>GC | TA>GC | ATTGAGATA | ATAAATA | CTA | CTA | T | G | T>G | T>G_CTA | TA>GC_ATAAATA | 9 | FALSE | FALSE |
| 130_v.txt | 32109 | 32109-SNV-A | SNV | 1 | A | C | 10 | 31094 | 0.03216055 | 61.3 | WT | FALSE | CAN | JSY2424 | 7 | WT-Rep7 | TAA | A>C | G/C+1 | A/T-1 | TA>GC | TA>GC | ATATTTAATA | AATAGCC | ATT | ATT | T | G | T>G | T>G_ATT | TA>GC_AATAGCC | 9 | FALSE | FALSE |
| 130_v.txt | 32226 | 32226-SNV-A | SNV | 1 | A | C | 8 | 31093 | 0.02572926 | 37.125 | WT | FALSE | CAN | JSY2424 | 7 | WT-Rep7 | CAG | A>C | G/C+1 | A/T-1 | TA>GC | TA>GC | CTTGACAGG | AGGAATT | GTC | GTC | T | G | T>G | T>G_GTC | TA>GC_AGGAATT | 9 | FALSE | FALSE |
| 130_v.txt | 32265 | 32265-SNV-A | SNV | 1 | A | C | 14 | 43326 | 0.03231316 | 23.3572177 | WT | FALSE | CAN | JSY2424 | 7 | WT-Rep7 | TAA | A>C | G/C+1 | A/T-1 | TA>GC | TA>GC | CCAAATAAA | AAAATAC | ATT | ATT | T | G | T>G | T>G_ATT | TA>GC_AAAATAC | 9 | FALSE | FALSE |
| 130_v.txt | 32283 | 32283-SNV-T | SNV | 1 | T | G | 28 | 43326 | 0.06462632 | 57.101901 | WT | FALSE | CAN | JSY2424 | 7 | WT-Rep7 | GTA | T>G | G/C+1 | A/T-1 | TA>GC | TA>GC | CCAACGTAA | TAAATAT | CAT | GTA | T | C | T>G | T>G_GTA | TA>GC_TAAATAT | 9 | FALSE | FALSE |
| 130_v.txt | 32317 | 32317-SNV-T | SNV | 1 | T | G | 4 | 43325 | 9.23E-03 | 22.5005149 | WT | FALSE | CAN | JSY2424 | 7 | WT-Rep7 | GTT | T>G | G/C+1 | A/T-1 | TA>GC | TA>GC | TGGTTGTTA | TTAAGAT | CAA | GTT | T | C | T>G | T>G_GTT | TA>GC_TTAAGAT | 9 | FALSE | FALSE |
| 130_v.txt | 32319 | 32319-SNV-A | SNV | 1 | A | C | 22 | 43326 | 0.05077782 | 58.5868473 | WT | FALSE | CAN | JSY2424 | 7 | WT-Rep7 | TAA | A>C | G/C+1 | A/T-1 | TA>GC | TA>GC | GTTGTTAAG | AAGATAA | ATT | ATT | T | G | T>G | T>G_ATT | TA>GC_AAGATAA | 9 | FALSE | FALSE |
| 130_v.txt | 32327 | 32327-SNV-A | SNV | 1 | A | C | 1 | 43326 | 2.31E-03 | 63.9999256 | WT | FALSE | CAN | JSY2424 | 7 | WT-Rep7 | CAG | A>C | G/C+1 | A/T-1 | TA>GC | TA>GC | GATAACAGC | AGCGTTG | GTC | GTC | T | G | T>G | T>G_GTC | TA>GC_AGCGTTG | 9 | FALSE | FALSE |
| 130_v.txt | 32439 | 32439-SNV-A | SNV | 1 | A | C | 19 | 43326 | 0.04385358 | 60.4736842 | WT | FALSE | CAN | JSY2424 | 7 | WT-Rep7 | AAC | A>C | G/C+1 | A/T-1 | TA>GC | TA>GC | TATGGAACT | ACTAAAA | TTG | TTG | T | G | T>G | T>G_TTG | TA>GC_ACTAAAA | 9 | FALSE | FALSE |
| 130_v.txt | 32457 | 32457-SNV-A | SNV | 1 | A | C | 61 | 43326 | 0.14079306 | 53.8852459 | WT | FALSE | CAN | JSY2424 | 7 | WT-Rep7 | TAA | A>C | G/C+1 | A/T-1 | TA>GC | TA>GC | ATGAATAAT | AATAGAG | ATT | ATT | T | G | T>G | T>G_ATT | TA>GC_AATAGAG | 9 | FALSE | FALSE |
| 130_v.txt | 32475 | 32475-SNV-A | SNV | 1 | A | C | 64 | 43326 | 0.14771731 | 48.265625 | WT | FALSE | CAN | JSY2424 | 7 | WT-Rep7 | GAA | A>C | G/C+1 | A/T-1 | TA>GC | TA>GC | ATGTAGAAG | AAGGTTA | CTT | CTT | T | G | T>G | T>G_CTT | TA>GC_AAGGTTA | 9 | FALSE | FALSE |
| 130_v.txt | 32541 | 32541-SNV-T | SNV | 1 | T | G | 90 | 43325 | 0.20773226 | 37.6660083 | WT | FALSE | CAN | JSY2424 | 7 | WT-Rep7 | TTC | T>G | G/C+1 | A/T-1 | TA>GC | TA>GC | GCAGCTTCA | TCACCAG | AAG | TTC | T | C | T>G | T>G_TTC | TA>GC_TCACCAG | 9 | FALSE | FALSE |
| 130_v.txt | 32616 | 32616-SNV-A | SNV | 1 | A | C | 22 | 31688 | 0.06942691 | 41.6817224 | WT | FALSE | CAN | JSY2424 | 7 | WT-Rep7 | TAA | A>C | G/C+1 | A/T-1 | TA>GC | TA>GC | CAACCTAAG | AAGAACC | ATT | ATT | T | G | T>G | T>G_ATT | TA>GC_AAGAACC | 9 | FALSE | FALSE |
| 130_v.txt | 32680 | 32680-SNV-A | SNV | 1 | A | C | 20 | 31687 | 0.06311737 | 54.750398 | WT | FALSE | CAN | JSY2424 | 7 | WT-Rep7 | CAA | A>C | G/C+1 | A/T-1 | TA>GC | TA>GC | TTCTCCAATA | AATAACG | GTT | GTT | T | G | T>G | T>G_GTT | TA>GC_AATAACG | 9 | FALSE | FALSE |
| 130_v.txt | 32681 | 32681-SNV-A | SNV | 1 | A | C | 15 | 31688 | 0.04733653 | 41.7326755 | WT | FALSE | CAN | JSY2424 | 7 | WT-Rep7 | AAT | A>C | G/C+1 | A/T-1 | TA>GC | TA>GC | TCTCCAATA | ATAACGG | TTA | TTA | T | G | T>G | T>G_TTA | TA>GC_ATAACGG | 9 | FALSE | FALSE |
| 130_v.txt | 32760 | 32760-SNV-A | SNV | 1 | A | C | 3 | 31688 | 9.47E-03 | 46.6664078 | WT | FALSE | CAN | JSY2424 | 7 | WT-Rep7 | TAA | A>C | G/C+1 | A/T-1 | TA>GC | TA>GC | ATGGCTAAA | AAAACTT | ATT | ATT | T | G | T>G | T>G_ATT | TA>GC_AAAACTT | 9 | FALSE | FALSE |
| 130_v.txt | 32763 | 32763-SNV-A | SNV | 1 | A | C | 7 | 31688 | 0.02209038 | 54.4285714 | WT | FALSE | CAN | JSY2424 | 7 | WT-Rep7 | AAC | A>C | G/C+1 | A/T-1 | TA>GC | TA>GC | GCTAAAACT | ACTTTGA | TTG | TTG | T | G | T>G | T>G_TTG | TA>GC_ACTTTGA | 9 | FALSE | FALSE |
| 130_v.txt | 32799 | 32799-SNV-T | SNV | 1 | T | G | 8 | 31688 | 0.02524615 | 37.8747052 | WT | FALSE | CAN | JSY2424 | 7 | WT-Rep7 | GTA | T>G | G/C+1 | A/T-1 | TA>GC | TA>GC | TCACCGTAA | TAATATT | CAT | GTA | T | C | T>G | T>G_GTA | TA>GC_TAATATT | 9 | FALSE | FALSE |
| 130_v.txt | 32801 | 32801-SNV-A | SNV | 1 | A | C | 14 | 31688 | 0.04418076 | 44.2857143 | WT | FALSE | CAN | JSY2424 | 7 | WT-Rep7 | AAT | A>C | G/C+1 | A/T-1 | TA>GC | TA>GC | ACCGTAATA | ATATTTG | TTA | TTA | T | G | T>G | T>G_TTA | TA>GC_ATATTTG | 9 | FALSE | FALSE |
| 130_v.txt | 32823 | 32823-SNV-A | SNV | 1 | A | C | 9 | 31687 | 0.02840282 | 46.5555556 | WT | FALSE | CAN | JSY2424 | 7 | WT-Rep7 | CAT | A>C | G/C+1 | A/T-1 | TA>GC | TA>GC | AAGTTCATT | ATTATTG | GTA | GTA | T | G | T>G | T>G_GTA | TA>GC_ATTATTG | 9 | FALSE | FALSE |
| 130_v.txt | 32842 | 32842-SNV-A | SNV | 1 | A | C | 7 | 31687 | 0.02209108 | 39.5712585 | WT | FALSE | CAN | JSY2424 | 7 | WT-Rep7 | CAA | A>C | G/C+1 | A/T-1 | TA>GC | TA>GC | TTACCCAAA | AAAAAAT | GTT | GTT | T | G | T>G | T>G_GTT | TA>GC_AAAAAAT | 9 | FALSE | FALSE |
| 130_v.txt | 32857 | 32857-SNV-A | SNV | 1 | A | C | 2 | 31686 | 6.31E-03 | 23.5003597 | WT | FALSE | CAN | JSY2424 | 7 | WT-Rep7 | CAT | A>C | G/C+1 | A/T-1 | TA>GC | TA>GC | TAATCCATG | ATGCCGC | GTA | GTA | T | G | T>G | T>G_GTA | TA>GC_ATGCCGC | 9 | FALSE | FALSE |
| 130_v.txt | 32858 | 32858-SNV-T | SNV | 1 | T | G | 1 | 31686 | 3.16E-03 | 22.9999998 | WT | FALSE | CAN | JSY2424 | 7 | WT-Rep7 | ATG | T>G | G/C+1 | A/T-1 | TA>GC | TA>GC | AATCCATGC | TGCCGCC | TAC | ATG | T | C | T>G | T>G_ATG | TA>GC_TGCCGCC | 9 | FALSE | FALSE |
| 130_v.txt | 32913 | 32913-SNV-A | SNV | 1 | A | C | 28 | 33860 | 0.08269344 | 30.1071429 | WT | FALSE | CAN | JSY2424 | 7 | WT-Rep7 | AAG | A>C | G/C+1 | A/T-1 | TA>GC | TA>GC | ACACTAAGT | AGTTCCA | TTC | TTC | T | G | T>G | T>G_TTC | TA>GC_AGTTCCA | 9 | FALSE | FALSE |
| 130_v.txt | 32916 | 32916-SNV-T | SNV | 1 | T | G | 49 | 33860 | 0.14471353 | 44.795305 | WT | FALSE | CAN | JSY2424 | 7 | WT-Rep7 | TTC | T>G | G/C+1 | A/T-1 | TA>GC | TA>GC | CTAAGTTCCA | TCCAGGG | AAG | TTC | T | C | T>G | T>G_TTC | TA>GC_TCCAGGG | 9 | FALSE | FALSE |
| 130_v.txt | 32929 | 32929-SNV-T | SNV | 1 | T | G | 34 | 33860 | 0.10041347 | 51.7941177 | WT | FALSE | CAN | JSY2424 | 7 | WT-Rep7 | GTG | T>G | G/C+1 | A/T-1 | TA>GC | TA>GC | CAAAAGTGA | TGATTGC | CAC | GTG | T | C | T>G | T>G_GTG | TA>GC_TGATTGC | 9 | FALSE | FALSE |
| 130_v.txt | 32938 | 32938-SNV-A | SNV | 1 | A | C | 14 | 33860 | 0.04134672 | 46.2142857 | WT | FALSE | CAN | JSY2424 | 7 | WT-Rep7 | CAA | A>C | G/C+1 | A/T-1 | TA>GC | TA>GC | TTGCCCAAG | AAGAAAA | GTT | GTT | T | G | T>G | T>G_GTT | TA>GC_AAGAAAA | 9 | FALSE | FALSE |
| 130_v.txt | 32948 | 32948-SNV-A | SNV | 1 | A | C | 23 | 33860 | 0.06792676 | 42 | WT | FALSE | CAN | JSY2424 | 7 | WT-Rep7 | AAT | A>C | G/C+1 | A/T-1 | TA>GC | TA>GC | AAACCAATA | ATACATG | TTA | TTA | T | G | T>G | T>G_TTA | TA>GC_ATACATG | 9 | FALSE | FALSE |
| 130_v.txt | 33039 | 33039-SNV-A | SNV | 1 | A | C | 10 | 33860 | 0.02953337 | 59.3990081 | WT | FALSE | CAN | JSY2424 | 7 | WT-Rep7 | CAT | A>C | G/C+1 | A/T-1 | TA>GC | TA>GC | GTAGCCATT | ATTTCAC | GTA | GTA | T | G | T>G | T>G_GTA | TA>GC_ATTTCAC | 9 | FALSE | FALSE |
| 130_v.txt | 33054 | 33054-SNV-T | SNV | 1 | T | G | 9 | 33860 | 0.02658004 | 55 | WT | FALSE | CAN | JSY2424 | 7 | WT-Rep7 | CTG | T>G | G/C+1 | A/T-1 | TA>GC | TA>GC | AAGGACTGC | TGCGTGA | GAC | CTG | T | C | T>G | T>G_CTG | TA>GC_TGCGTGA | 9 | FALSE | FALSE |
| 130_v.txt | 33058 | 33058-SNV-T | SNV | 1 | T | G | 5 | 33860 | 0.01476669 | 45.2 | WT | FALSE | CAN | JSY2424 | 7 | WT-Rep7 | GTG | T>G | G/C+1 | A/T-1 | TA>GC | TA>GC | ACTGCGTGA | TGACAGA | CAC | GTG | T | C | T>G | T>G_GTG | TA>GC_TGACAGA | 9 | FALSE | FALSE |
| 130_v.txt | 33065 | 33065-SNV-A | SNV | 1 | A | C | 2 | 33860 | 5.91E-03 | 56.4958697 | WT | FALSE | CAN | JSY2424 | 7 | WT-Rep7 | AAT | A>C | G/C+1 | A/T-1 | TA>GC | TA>GC | GACAGAATA | ATATGCC | TTA | TTA | T | G | T>G | T>G_TTA | TA>GC_ATATGCC | 9 | FALSE | FALSE |
| 130_v.txt | 33157 | 33157-SNV-T | SNV | 1 | T | G | 8 | 33843 | 0.02363857 | 36.125 | WT | FALSE | CAN | JSY2424 | 7 | WT-Rep7 | GTA | T>G | G/C+1 | A/T-1 | TA>GC | TA>GC | GACCTGTAC | TACCAAT | CAT | GTA | T | C | T>G | T>G_GTA | TA>GC_TACCAAT | 9 | FALSE | FALSE |
| 130_v.txt | 33189 | 33189-SNV-A | SNV | 1 | A | C | 5 | 32447 | 0.01540975 | 22.4006635 | WT | FALSE | CAN | JSY2424 | 7 | WT-Rep7 | AAT | A>C | G/C+1 | A/T-1 | TA>GC | TA>GC | ATACCAATA | ATATGTC | TTA | TTA | T | G | T>G | T>G_TTA | TA>GC_ATATGTC | 9 | FALSE | FALSE |
| 130_v.txt | 33207 | 33207-SNV-T | SNV | 1 | T | G | 4 | 32447 | 0.0123278 | 24.0009626 | WT | FALSE | CAN | JSY2424 | 7 | WT-Rep7 | CTC | T>G | G/C+1 | A/T-1 | TA>GC | TA>GC | TTAAGCTCTC | TCTCTCT | GAG | CTC | T | C | T>G | T>G_CTC | TA>GC_TCTCTCT | 9 | FALSE | FALSE |
| 130_v.txt | 33269 | 33269-SNV-A | SNV | 1 | A | C | 230 | 32448 | 0.70882643 | 15.5217391 | WT | FALSE | CAN | JSY2424 | 7 | WT-Rep7 | TAG | A>C | G/C+1 | A/T-1 | TA>GC | TA>GC | CTCCATAGA | AGAGAAC | ATC | ATC | T | G | T>G | T>G_ATC | TA>GC_AGAGAAC | 9 | FALSE | FALSE |
| 130_v.txt | 33465 | 33465-SNV-A | SNV | 1 | A | C | 197 | 32448 | 0.60712525 | 14.4213198 | WT | FALSE | CAN | JSY2424 | 7 | WT-Rep7 | CAT | A>C | G/C+1 | A/T-1 | TA>GC | TA>GC | TTTGTCATTG | ATTGCTA | GTA | GTA | T | G | T>G | T>G_GTA | TA>GC_ATTGCTA | 9 | FALSE | FALSE |
| 130_v.txt | 33476 | 33476-SNV-T | SNV | 1 | T | G | 4 | 32447 | 0.0123278 | 33.75 | WT | FALSE | CAN | JSY2424 | 7 | WT-Rep7 | CTT | T>G | G/C+1 | A/T-1 | TA>GC | TA>GC | TATGCCTTTT | TTTTTTT | GAA | CTT | T | C | T>G | T>G_CTT | TA>GC_TTTTTTT | 9 | FALSE | FALSE |
| 130_v.txt | 32015 | 32015-SNV-A | SNV | 1 | A | G | 3 | 31094 | 9.65E-03 | 52.33322 | WT | FALSE | CAN | JSY2424 | 7 | WT-Rep7 | GAG | A>G | G/C+1 | A/T-1 | TA>CG | TA>CG | GTCACGAGA | AGAGATG | CTC | CTC | T | C | T>C | T>C_CTC | TA>CG_AGAGATG | 9 | FALSE | FALSE |
| 130_v.txt | 32055 | 32055-SNV-T | SNV | 1 | T | C | 20 | 31094 | 0.06432109 | 58.1496476 | WT | FALSE | CAN | JSY2424 | 7 | WT-Rep7 | GTG | T>C | G/C+1 | A/T-1 | TA>CG | TA>CG | CTGATGTGC | TGCGAGA | CAC | GTG | T | G | T>C | T>C_GTG | TA>CG_TGCGAGA | 9 | FALSE | FALSE |
| 130_v.txt | 32069 | 32069-SNV-A | SNV | 1 | A | G | 4 | 31094 | 0.01286422 | 63.9994135 | WT | FALSE | CAN | JSY2424 | 7 | WT-Rep7 | TAA | A>G | G/C+1 | A/T-1 | TA>CG | TA>CG | TGAGATAAA | AAATAAC | ATT | ATT | T | C | T>C | T>C_ATT | TA>CG_AAATAAC | 9 | FALSE | FALSE |
| 130_v.txt | 32081 | 32081-SNV-A | SNV | 1 | A | G | 11 | 31094 | 0.0353766 | 63.7271748 | WT | FALSE | CAN | JSY2424 | 7 | WT-Rep7 | CAA | A>G | G/C+1 | A/T-1 | TA>CG | TA>CG | CCATGCAAA | AAAAAAG | GTT | GTT | T | C | T>C | T>C_GTT | TA>CG_AAAAAAG | 9 | FALSE | FALSE |
| 130_v.txt | 32319 | 32319-SNV-A | SNV | 1 | A | G | 9 | 43326 | 0.02077275 | 55.3310479 | WT | FALSE | CAN | JSY2424 | 7 | WT-Rep7 | TAA | A>G | G/C+1 | A/T-1 | TA>CG | TA>CG | GTTGTTAAG | AAGATAA | ATT | ATT | T | C | T>C | T>C_ATT | TA>CG_AAGATAA | 9 | FALSE | FALSE |
| 130_v.txt | 32320 | 32320-SNV-A | SNV | 1 | A | G | 12 | 43326 | 0.027697 | 50.8327411 | WT | FALSE | CAN | JSY2424 | 7 | WT-Rep7 | AAG | A>G | G/C+1 | A/T-1 | TA>CG | TA>CG | TTGTTAAGA | AGATAAC | TTC | TTC | T | C | T>C | T>C_TTC | TA>CG_AGATAAC | 9 | FALSE | FALSE |
| 130_v.txt | 32322 | 32322-SNV-A | SNV | 1 | A | G | 3 | 43326 | 6.92E-03 | 63.9999169 | WT | FALSE | CAN | JSY2424 | 7 | WT-Rep7 | GAT | A>G | G/C+1 | A/T-1 | TA>CG | TA>CG | GTTAAGATA | ATAACAG | CTA | CTA | T | C | T>C | T>C_CTA | TA>CG_ATAACAG | 9 | FALSE | FALSE |
| 130_v.txt | 32324 | 32324-SNV-A | SNV | 1 | A | G | 3 | 43326 | 6.92E-03 | 58.999989 | WT | FALSE | CAN | JSY2424 | 7 | WT-Rep7 | TAA | A>G | G/C+1 | A/T-1 | TA>CG | TA>CG | TAAGATAAC | AACAGCG | ATT | ATT | T | C | T>C | T>C_ATT | TA>CG_AACAGCG | 9 | FALSE | FALSE |
| 130_v.txt | 32327 | 32327-SNV-A | SNV | 1 | A | G | 9 | 43326 | 0.02077275 | 53.5556286 | WT | FALSE | CAN | JSY2424 | 7 | WT-Rep7 | CAG | A>G | G/C+1 | A/T-1 | TA>CG | TA>CG | GATAACAGC | AGCGTTG | GTC | GTC | T | C | T>C | T>C_GTC | TA>CG_AGCGTTG | 9 | FALSE | FALSE |
| 130_v.txt | 32331 | 32331-SNV-T | SNV | 1 | T | C | 4 | 43325 | 9.23E-03 | 36.9999679 | WT | FALSE | CAN | JSY2424 | 7 | WT-Rep7 | GTT | T>C | G/C+1 | A/T-1 | TA>CG | TA>CG | ACAGCGTTG | TTGAAGA | CAA | GTT | T | G | T>C | T>C_GTT | TA>CG_TTGAAGA | 9 | FALSE | FALSE |
| 130_v.txt | 32335 | 32335-SNV-A | SNV | 1 | A | G | 6 | 43326 | 0.0138485 | 47.993295 | WT | FALSE | CAN | JSY2424 | 7 | WT-Rep7 | AAG | A>G | G/C+1 | A/T-1 | TA>CG | TA>CG | CGTTGAAGA | AGATATG | TTC | TTC | T | C | T>C | T>C_TTC | TA>CG_AGATATG | 9 | FALSE | FALSE |
| 130_v.txt | 32338 | 32338-SNV-T | SNV | 1 | T | C | 8 | 43326 | 0.01846466 | 42.9999115 | WT | FALSE | CAN | JSY2424 | 7 | WT-Rep7 | ATA | T>C | G/C+1 | A/T-1 | TA>CG | TA>CG | TGAAGATAT | TATGTGG | TAT | ATA | T | G | T>C | T>C_ATA | TA>CG_TATGTGG | 9 | FALSE | FALSE |
| 130_v.txt | 32381 | 32381-SNV-A | SNV | 1 | A | G | 7 | 43325 | 0.01615695 | 50.2834328 | WT | FALSE | CAN | JSY2424 | 7 | WT-Rep7 | TAA | A>G | G/C+1 | A/T-1 | TA>CG | TA>CG | AATAATAAA | AAAGGGA | ATT | ATT | T | C | T>C | T>C_ATT | TA>CG_AAAGGGA | 9 | FALSE | FALSE |
| 130_v.txt | 32445 | 32445-SNV-A | SNV | 1 | A | G | 31 | 43326 | 0.07155057 | 47.1935484 | WT | FALSE | CAN | JSY2424 | 7 | WT-Rep7 | AAG | A>G | G/C+1 | A/T-1 | TA>CG | TA>CG | ACTAAAAGT | AGTCCAA | TTC | TTC | T | C | T>C | T>C_TTC | TA>CG_AGTCCAA | 9 | FALSE | FALSE |
| 130_v.txt | 32490 | 32490-SNV-A | SNV | 1 | A | G | 171 | 43326 | 0.39468218 | 43.9182405 | WT | FALSE | CAN | JSY2424 | 7 | WT-Rep7 | GAA | A>G | G/C+1 | A/T-1 | TA>CG | TA>CG | ATACGGAAA | AAAACAA | CTT | CTT | T | C | T>C | T>C_CTT | TA>CG_AAAACAA | 9 | FALSE | FALSE |
| 130_v.txt | 32541 | 32541-SNV-T | SNV | 1 | T | C | 52 | 43325 | 0.12002308 | 36.3068245 | WT | FALSE | CAN | JSY2424 | 7 | WT-Rep7 | TTC | T>C | G/C+1 | A/T-1 | TA>CG | TA>CG | GCAGCTTCA | TCACCAG | AAG | TTC | T | G | T>C | T>C_TTC | TA>CG_TCACCAG | 9 | FALSE | FALSE |
| 130_v.txt | 32680 | 32680-SNV-A | SNV | 1 | A | G | 94 | 31687 | 0.29665162 | 61.4253687 | WT | FALSE | CAN | JSY2424 | 7 | WT-Rep7 | CAA | A>G | G/C+1 | A/T-1 | TA>CG | TA>CG | TTCTCCAATA | AATAACG | GTT | GTT | T | C | T>C | T>C_GTT | TA>CG_AATAACG | 9 | FALSE | FALSE |
| 130_v.txt | 32728 | 32728-SNV-A | SNV | 1 | A | G | 9 | 31688 | 0.02840192 | 52.110856 | WT | FALSE | CAN | JSY2424 | 7 | WT-Rep7 | CAA | A>G | G/C+1 | A/T-1 | TA>CG | TA>CG | CCATACAAA | AAAAACA | GTT | GTT | T | C | T>C | T>C_GTT | TA>CG_AAAAACA | 9 | FALSE | FALSE |
| 130_v.txt | 32787 | 32787-SNV-T | SNV | 1 | T | C | 11 | 31688 | 0.03471346 | 43.0908277 | WT | FALSE | CAN | JSY2424 | 7 | WT-Rep7 | CTC | T>C | G/C+1 | A/T-1 | TA>CG | TA>CG | CAGAACTCG | TCGAATT | GAG | CTC | T | G | T>C | T>C_CTC | TA>CG_TCGAATT | 9 | FALSE | FALSE |
| 130_v.txt | 32858 | 32858-SNV-T | SNV | 1 | T | C | 7 | 31686 | 0.02209178 | 25.999985 | WT | FALSE | CAN | JSY2424 | 7 | WT-Rep7 | ATG | T>C | G/C+1 | A/T-1 | TA>CG | TA>CG | AATCCATGC | TGCCGCC | TAC | ATG | T | G | T>C | T>C_ATG | TA>CG_TGCCGCC | 9 | FALSE | FALSE |
| 130_v.txt | 32919 | 32919-SNV-A | SNV | 1 | A | G | 28 | 33860 | 0.08269344 | 45.5357143 | WT | FALSE | CAN | JSY2424 | 7 | WT-Rep7 | CAG | A>G | G/C+1 | A/T-1 | TA>CG | TA>CG | AGTTCCAGG | AGGGCAA | GTC | GTC | T | C | T>C | T>C_GTC | TA>CG_AGGGCAA | 9 | FALSE | FALSE |
| 130_v.txt | 32938 | 32938-SNV-A | SNV | 1 | A | G | 7 | 33860 | 0.02067336 | 52.1428571 | WT | FALSE | CAN | JSY2424 | 7 | WT-Rep7 | CAA | A>G | G/C+1 | A/T-1 | TA>CG | TA>CG | TTGCCCAAG | AAGAAAA | GTT | GTT | T | C | T>C | T>C_GTT | TA>CG_AAGAAAA | 9 | FALSE | FALSE |
| 130_v.txt | 32941 | 32941-SNV-A | SNV | 1 | A | G | 11 | 33860 | 0.03248671 | 53.818084 | WT | FALSE | CAN | JSY2424 | 7 | WT-Rep7 | GAA | A>G | G/C+1 | A/T-1 | TA>CG | TA>CG | CCCAAGAAA | AAAACCA | CTT | CTT | T | C | T>C | T>C_CTT | TA>CG_AAAACCA | 9 | FALSE | FALSE |
| 130_v.txt | 32994 | 32994-SNV-T | SNV | 1 | T | C | 8 | 33860 | 0.0236267 | 36.9999931 | WT | FALSE | CAN | JSY2424 | 7 | WT-Rep7 | TTG | T>C | G/C+1 | A/T-1 | TA>CG | TA>CG | AATCTTTGTG | TGTGAGA | AAC | TTG | T | G | T>C | T>C_TTG | TA>CG_TGTGAGA | 9 | FALSE | FALSE |
| 130_v.txt | 33000 | 33000-SNV-A | SNV | 1 | A | G | 4 | 33860 | 0.01181335 | 40.4986976 | WT | FALSE | CAN | JSY2424 | 7 | WT-Rep7 | GAA | A>G | G/C+1 | A/T-1 | TA>CG | TA>CG | TGTGAGAAA | AAAACTG | CTT | CTT | T | C | T>C | T>C_CTT | TA>CG_AAAACTG | 9 | FALSE | FALSE |
| 130_v.txt | 33039 | 33039-SNV-A | SNV | 1 | A | G | 6 | 33860 | 0.01772002 | 62.995093 | WT | FALSE | CAN | JSY2424 | 7 | WT-Rep7 | CAT | A>G | G/C+1 | A/T-1 | TA>CG | TA>CG | GTAGCCATT | ATTTCAC | GTA | GTA | T | C | T>C | T>C_GTA | TA>CG_ATTTCAC | 9 | FALSE | FALSE |
| 130_v.txt | 33065 | 33065-SNV-A | SNV | 1 | A | G | 12 | 33860 | 0.03544005 | 52.2464832 | WT | FALSE | CAN | JSY2424 | 7 | WT-Rep7 | AAT | A>G | G/C+1 | A/T-1 | TA>CG | TA>CG | GACAGAATA | ATATGCC | TTA | TTA | T | C | T>C | T>C_TTA | TA>CG_ATATGCC | 9 | FALSE | FALSE |
| 130_v.txt | 33092 | 33092-SNV-T | SNV | 1 | T | C | 5 | 33860 | 0.01476669 | 58.5998018 | WT | FALSE | CAN | JSY2424 | 7 | WT-Rep7 | ATG | T>C | G/C+1 | A/T-1 | TA>CG | TA>CG | TAAATATGA | TGATATA | TAC | ATG | T | G | T>C | T>C_ATG | TA>CG_TGATATA | 9 | FALSE | FALSE |
| 130_v.txt | 33157 | 33157-SNV-T | SNV | 1 | T | C | 29 | 33843 | 0.0856898 | 37.5172414 | WT | FALSE | CAN | JSY2424 | 7 | WT-Rep7 | GTA | T>C | G/C+1 | A/T-1 | TA>CG | TA>CG | GACCTGTAC | TACCAAT | CAT | GTA | T | G | T>C | T>C_GTA | TA>CG_TACCAAT | 9 | FALSE | FALSE |
| 130_v.txt | 33207 | 33207-SNV-T | SNV | 1 | T | C | 20 | 32447 | 0.06163898 | 22.3493909 | WT | FALSE | CAN | JSY2424 | 7 | WT-Rep7 | CTC | T>C | G/C+1 | A/T-1 | TA>CG | TA>CG | TTAAGCTCTC | TCTCTCT | GAG | CTC | T | G | T>C | T>C_CTC | TA>CG_TCTCTCT | 9 | FALSE | FALSE |
| 130_v.txt | 33476 | 33476-SNV-T | SNV | 1 | T | C | 52 | 32447 | 0.16026135 | 31.75 | WT | FALSE | CAN | JSY2424 | 7 | WT-Rep7 | CTT | T>C | G/C+1 | A/T-1 | TA>CG | TA>CG | TATGCCTTTT | TTTTTTT | GAA | CTT | T | G | T>C | T>C_CTT | TA>CG_TTTTTTT | 9 | FALSE | FALSE |
| 130_v.txt | 31904 | 31904-SNV-T | SNV | 1 | T | A | 1 | 11585 | 8.63E-03 | 37.9983285 | WT | FALSE | CAN | JSY2424 | 7 | WT-Rep7 | ATT | T>A | A/T+1 | A/T-1 | TA>AT | TA>AT | ATTGAATTTT | TTTTGGT | TAA | ATT | T | T | T>A | T>A_ATT | TA>AT_TTTTGGT | 9 | FALSE | FALSE |
| 130_v.txt | 31967 | 31967-SNV-A | SNV | 1 | A | T | 31 | 31094 | 0.09969769 | 31.6129032 | WT | FALSE | CAN | JSY2424 | 7 | WT-Rep7 | AAT | A>T | A/T+1 | A/T-1 | TA>AT | TA>AT | CGCATAATA | ATAAGCC | TTA | TTA | T | A | T>A | T>A_TTA | TA>AT_ATAAGCC | 9 | FALSE | FALSE |
| 130_v.txt | 32015 | 32015-SNV-A | SNV | 1 | A | T | 3 | 31094 | 9.65E-03 | 28.6666397 | WT | FALSE | CAN | JSY2424 | 7 | WT-Rep7 | GAG | A>T | A/T+1 | A/T-1 | TA>AT | TA>AT | GTCACGAGA | AGAGATG | CTC | CTC | T | A | T>A | T>A_CTC | TA>AT_AGAGATG | 9 | FALSE | FALSE |
| 130_v.txt | 32055 | 32055-SNV-T | SNV | 1 | T | A | 6 | 31094 | 0.01929633 | 57.9981227 | WT | FALSE | CAN | JSY2424 | 7 | WT-Rep7 | GTG | T>A | A/T+1 | A/T-1 | TA>AT | TA>AT | CTGATGTGC | TGCGAGA | CAC | GTG | T | T | T>A | T>A_GTG | TA>AT_TGCGAGA | 9 | FALSE | FALSE |
| 130_v.txt | 32069 | 32069-SNV-A | SNV | 1 | A | T | 1 | 31094 | 3.22E-03 | 63.9994087 | WT | FALSE | CAN | JSY2424 | 7 | WT-Rep7 | TAA | A>T | A/T+1 | A/T-1 | TA>AT | TA>AT | TGAGATAAA | AAATAAC | ATT | ATT | T | A | T>A | T>A_ATT | TA>AT_AAATAAC | 9 | FALSE | FALSE |
| 130_v.txt | 32245 | 32245-SNV-T | SNV | 1 | T | A | 24 | 31092 | 0.07719027 | 38 | WT | FALSE | CAN | JSY2424 | 7 | WT-Rep7 | TTG | T>A | A/T+1 | A/T-1 | TA>AT | TA>AT | CCAACTTGTT | TGTTCTT | AAC | TTG | T | T | T>A | T>A_TTG | TA>AT_TGTTCTT | 9 | FALSE | FALSE |
| 130_v.txt | 32265 | 32265-SNV-A | SNV | 1 | A | T | 55 | 43326 | 0.12694456 | 35.763649 | WT | FALSE | CAN | JSY2424 | 7 | WT-Rep7 | TAA | A>T | A/T+1 | A/T-1 | TA>AT | TA>AT | CCAAATAAA | AAAATAC | ATT | ATT | T | A | T>A | T>A_ATT | TA>AT_AAAATAC | 9 | FALSE | FALSE |
| 130_v.txt | 32284 | 32284-SNV-A | SNV | 1 | A | T | 12 | 43326 | 0.027697 | 39.4958535 | WT | FALSE | CAN | JSY2424 | 7 | WT-Rep7 | TAA | A>T | A/T+1 | A/T-1 | TA>AT | TA>AT | CAACGTAAA | AAATATT | ATT | ATT | T | A | T>A | T>A_ATT | TA>AT_AAATATT | 9 | FALSE | FALSE |
| 130_v.txt | 32287 | 32287-SNV-T | SNV | 1 | T | A | 29 | 43326 | 0.0669344 | 46.1724138 | WT | FALSE | CAN | JSY2424 | 7 | WT-Rep7 | ATA | T>A | A/T+1 | A/T-1 | TA>AT | TA>AT | CGTAAATATT | TATTTGA | TAT | ATA | T | T | T>A | T>A_ATA | TA>AT_TATTTGA | 9 | FALSE | FALSE |
| 130_v.txt | 32317 | 32317-SNV-T | SNV | 1 | T | A | 9 | 43325 | 0.02077323 | 50.4341392 | WT | FALSE | CAN | JSY2424 | 7 | WT-Rep7 | GTT | T>A | A/T+1 | A/T-1 | TA>AT | TA>AT | TGGTTGTTA | TTAAGAT | CAA | GTT | T | T | T>A | T>A_GTT | TA>AT_TTAAGAT | 9 | FALSE | FALSE |
| 130_v.txt | 32320 | 32320-SNV-A | SNV | 1 | A | T | 5 | 43326 | 0.01154042 | 56.3971178 | WT | FALSE | CAN | JSY2424 | 7 | WT-Rep7 | AAG | A>T | A/T+1 | A/T-1 | TA>AT | TA>AT | TTGTTAAGA | AGATAAC | TTC | TTC | T | A | T>A | T>A_TTC | TA>AT_AGATAAC | 9 | FALSE | FALSE |
| 130_v.txt | 32322 | 32322-SNV-A | SNV | 1 | A | T | 6 | 43326 | 0.0138485 | 43.6665594 | WT | FALSE | CAN | JSY2424 | 7 | WT-Rep7 | GAT | A>T | A/T+1 | A/T-1 | TA>AT | TA>AT | GTTAAGATA | ATAACAG | CTA | CTA | T | A | T>A | T>A_CTA | TA>AT_ATAACAG | 9 | FALSE | FALSE |
| 130_v.txt | 32323 | 32323-SNV-T | SNV | 1 | T | A | 4 | 43326 | 9.23E-03 | 63.99867 | WT | FALSE | CAN | JSY2424 | 7 | WT-Rep7 | ATA | T>A | A/T+1 | A/T-1 | TA>AT | TA>AT | TTAAGATAA | TAACAGC | TAT | ATA | T | T | T>A | T>A_ATA | TA>AT_TAACAGC | 9 | FALSE | FALSE |
| 130_v.txt | 32325 | 32325-SNV-A | SNV | 1 | A | T | 8 | 43326 | 0.01846466 | 48.4993705 | WT | FALSE | CAN | JSY2424 | 7 | WT-Rep7 | AAC | A>T | A/T+1 | A/T-1 | TA>AT | TA>AT | AAGATAACA | ACAGCGT | TTG | TTG | T | A | T>A | T>A_TTG | TA>AT_ACAGCGT | 9 | FALSE | FALSE |

| | | | | | | | | | | | | | | | | | | | | | | | | | | | | | | | | | | |
|---|---|---|---|---|---|---|---|---|---|---|---|---|---|---|---|---|---|---|---|---|---|---|---|---|---|---|---|---|---|---|---|---|---|---|
| 130_v.txt | 32331 | 32331-SNV-1 | SNV | 1 | T | A | 4 | 43325 | 9.23E-03 | 63.9971596 | WT | FALSE | CAN | JSY2424 | 7 | WT-Rep7 | GTT | T>A | A/T+1 | A/T-1 | TA>AT | TA>AT | ACAGCGTTG | TTGAAGA | CAA | GTT | T | T | T>A | T>A_GTT | TA>AT_TTGAAGA | 9 | FALSE | FALSE |
| 130_v.txt | 32332 | 32332-SNV-1 | SNV | 1 | T | A | 6 | 43326 | 0.0138485 | 51.9950336 | WT | FALSE | CAN | JSY2424 | 7 | WT-Rep7 | TTG | T>A | A/T+1 | A/T-1 | TA>AT | TA>AT | CAGCGTTGA | TGAAGAT | AAC | TTG | T | T | T>A | T>A_TTG | TA>AT_TGAAGAT | 9 | FALSE | FALSE |
| 130_v.txt | 32338 | 32338-SNV-1 | SNV | 1 | T | A | 3 | 43326 | 6.92E-03 | 58.6651607 | WT | FALSE | CAN | JSY2424 | 7 | WT-Rep7 | ATA | T>A | A/T+1 | A/T-1 | TA>AT | TA>AT | TGAAGATAT | TATGTGG | TAT | ATA | T | T | T>A | T>A_ATA | TA>AT_TATGTGG | 9 | FALSE | FALSE |
| 130_v.txt | 32380 | 32380-SNV-1 | SNV | 1 | T | A | 29 | 43326 | 0.0669344 | 60.1011179 | WT | FALSE | CAN | JSY2424 | 7 | WT-Rep7 | ATA | T>A | A/T+1 | A/T-1 | TA>AT | TA>AT | CAATAATAA | TAAAGGG | TAT | ATA | T | T | T>A | T>A_ATA | TA>AT_TAAAGGG | 9 | FALSE | FALSE |
| 130_v.txt | 32407 | 32407-SNV-1 | SNV | 1 | T | A | 5 | 43324 | 0.01154095 | 28.9999818 | WT | FALSE | CAN | JSY2424 | 7 | WT-Rep7 | GTA | T>A | A/T+1 | A/T-1 | TA>AT | TA>AT | AGGAAGTAC | TAGATTG | CAT | GTA | T | T | T>A | T>A_GTA | TA>AT_TAGATTG | 9 | FALSE | FALSE |
| 130_v.txt | 32408 | 32408-SNV-/ | SNV | 1 | A | T | 17 | 43326 | 0.03923741 | 27.2344738 | WT | FALSE | CAN | JSY2424 | 7 | WT-Rep7 | TAG | A>T | A/T+1 | A/T-1 | TA>AT | TA>AT | GGAAGTAGA | AGATTGT | ATC | ATC | T | A | T>A | T>A_ATC | TA>AT_AGATTGT | 9 | FALSE | FALSE |
| 130_v.txt | 32410 | 32410-SNV-/ | SNV | 1 | A | T | 16 | 43326 | 0.03692933 | 58.4375 | WT | FALSE | CAN | JSY2424 | 7 | WT-Rep7 | GAT | A>T | A/T+1 | A/T-1 | TA>AT | TA>AT | AAGTAGATT | ATTGTGT | CTA | CTA | T | A | T>A | T>A_CTA | TA>AT_ATTGTGT | 9 | FALSE | FALSE |
| 130_v.txt | 32498 | 32498-SNV-1 | SNV | 1 | T | A | 8 | 43326 | 0.01846466 | 43.9999489 | WT | FALSE | CAN | JSY2424 | 7 | WT-Rep7 | CTT | T>A | A/T+1 | A/T-1 | TA>AT | TA>AT | AACAACTTT | TTTTTTG | GAA | CTT | T | T | T>A | T>A_CTT | TA>AT_TTTTTTG | 9 | FALSE | FALSE |
| 130_v.txt | 32503 | 32503-SNV-1 | SNV | 1 | T | A | 45 | 43326 | 0.10386373 | 47.6 | WT | FALSE | CAN | JSY2424 | 7 | WT-Rep7 | TTG | T>A | A/T+1 | A/T-1 | TA>AT | TA>AT | CTTTTTTGAT | TGATGGC | AAC | TTG | T | T | T>A | T>A_TTG | TA>AT_TGATGGC | 9 | FALSE | FALSE |
| 130_v.txt | 32760 | 32760-SNV-/ | SNV | 1 | A | T | 15 | 31688 | 0.04733653 | 48.9950198 | WT | FALSE | CAN | JSY2424 | 7 | WT-Rep7 | TAA | A>T | A/T+1 | A/T-1 | TA>AT | TA>AT | ATGGCTAAA | AAAACTT | ATT | ATT | T | A | T>A | T>A_ATT | TA>AT_AAAACTT | 9 | FALSE | FALSE |
| 130_v.txt | 32767 | 32767-SNV-1 | SNV | 1 | T | A | 39 | 31688 | 0.12307498 | 50.7948718 | WT | FALSE | CAN | JSY2424 | 7 | WT-Rep7 | TTG | T>A | A/T+1 | A/T-1 | TA>AT | TA>AT | AAACTTTGA | TGATGGA | AAC | TTG | T | T | T>A | T>A_TTG | TA>AT_TGATGGA | 9 | FALSE | FALSE |
| 130_v.txt | 32787 | 32787-SNV-1 | SNV | 1 | T | A | 1 | 31688 | 3.16E-03 | 63.9999548 | WT | FALSE | CAN | JSY2424 | 7 | WT-Rep7 | CTC | T>A | A/T+1 | A/T-1 | TA>AT | TA>AT | CAGAACTCG | TCGAATT | GAG | CTC | T | T | T>A | T>A_CTC | TA>AT_TCGAATT | 9 | FALSE | FALSE |
| 130_v.txt | 32799 | 32799-SNV-1 | SNV | 1 | T | A | 5 | 31688 | 0.01577884 | 43.1969413 | WT | FALSE | CAN | JSY2424 | 7 | WT-Rep7 | GTA | T>A | A/T+1 | A/T-1 | TA>AT | TA>AT | TCACCGTAA | TAATATT | CAT | GTA | T | T | T>A | T>A_GTA | TA>AT_TAATATT | 9 | FALSE | FALSE |
| 130_v.txt | 32858 | 32858-SNV-1 | SNV | 1 | T | A | 8 | 31686 | 0.02524774 | 35.1241977 | WT | FALSE | CAN | JSY2424 | 7 | WT-Rep7 | ATG | T>A | A/T+1 | A/T-1 | TA>AT | TA>AT | AATCCATGC | TGCCGCC | TAC | ATG | T | T | T>A | T>A_ATG | TA>AT_TGCCGCC | 9 | FALSE | FALSE |
| 130_v.txt | 32916 | 32916-SNV-1 | SNV | 1 | T | A | 8 | 33860 | 0.0236267 | 38.6240404 | WT | FALSE | CAN | JSY2424 | 7 | WT-Rep7 | TTC | T>A | A/T+1 | A/T-1 | TA>AT | TA>AT | CTAAGTTCC | TCCAGGG | AAG | TTC | T | T | T>A | T>A_TTC | TA>AT_TCCAGGG | 9 | FALSE | FALSE |
| 130_v.txt | 32938 | 32938-SNV-/ | SNV | 1 | A | T | 14 | 33860 | 0.04134672 | 51.6428571 | WT | FALSE | CAN | JSY2424 | 7 | WT-Rep7 | CAA | A>T | A/T+1 | A/T-1 | TA>AT | TA>AT | TTGCCCAAG | AAGAAAA | GTT | GTT | T | A | T>A | T>A_GTT | TA>AT_AAGAAAA | 9 | FALSE | FALSE |
| 130_v.txt | 32941 | 32941-SNV-/ | SNV | 1 | A | T | 6 | 33860 | 0.01772002 | 24.0000237 | WT | FALSE | CAN | JSY2424 | 7 | WT-Rep7 | GAA | A>T | A/T+1 | A/T-1 | TA>AT | TA>AT | CCCAAGAAA | AAAACCA | CTT | CTT | T | A | T>A | T>A_CTT | TA>AT_AAAACCA | 9 | FALSE | FALSE |
| 130_v.txt | 32994 | 32994-SNV-1 | SNV | 1 | T | A | 2 | 33860 | 5.91E-03 | 63.9996002 | WT | FALSE | CAN | JSY2424 | 7 | WT-Rep7 | TTG | T>A | A/T+1 | A/T-1 | TA>AT | TA>AT | AATCTTGTG | TGTGAGA | AAC | TTG | T | T | T>A | T>A_TTG | TA>AT_TGTGAGA | 9 | FALSE | FALSE |
| 130_v.txt | 33040 | 33040-SNV-1 | SNV | 1 | T | A | 5 | 33859 | 0.01476712 | 53.3981607 | WT | FALSE | CAN | JSY2424 | 7 | WT-Rep7 | ATT | T>A | A/T+1 | A/T-1 | TA>AT | TA>AT | TAGCCATTTC | TTTCACC | TAA | ATT | T | T | T>A | T>A_ATT | TA>AT_TTTCACC | 9 | FALSE | FALSE |
| 130_v.txt | 33066 | 33066-SNV-1 | SNV | 1 | T | A | 2 | 33860 | 5.91E-03 | 63.9958636 | WT | FALSE | CAN | JSY2424 | 7 | WT-Rep7 | ATA | T>A | A/T+1 | A/T-1 | TA>AT | TA>AT | ACAGAATAT | TATGCCA | TAT | ATA | T | T | T>A | T>A_ATA | TA>AT_TATGCCA | 9 | FALSE | FALSE |
| 130_v.txt | 33092 | 33092-SNV-1 | SNV | 1 | T | A | 7 | 33860 | 0.02067336 | 45.1416442 | WT | FALSE | CAN | JSY2424 | 7 | WT-Rep7 | ATG | T>A | A/T+1 | A/T-1 | TA>AT | TA>AT | TAAATATGT | TGATATA | TAC | ATG | T | T | T>A | T>A_ATG | TA>AT_TGATATA | 9 | FALSE | FALSE |
| 130_v.txt | 33095 | 33095-SNV-1 | SNV | 1 | T | A | 4 | 33860 | 0.01181335 | 44.503611 | WT | FALSE | CAN | JSY2424 | 7 | WT-Rep7 | ATA | T>A | A/T+1 | A/T-1 | TA>AT | TA>AT | ATATGATAT | TATAAGA | TAT | ATA | T | T | T>A | T>A_ATA | TA>AT_TATAAGA | 9 | FALSE | FALSE |
| 130_v.txt | 33157 | 33157-SNV-1 | SNV | 1 | T | A | 10 | 33843 | 0.02954821 | 37.3 | WT | FALSE | CAN | JSY2424 | 7 | WT-Rep7 | GTA | T>A | A/T+1 | A/T-1 | TA>AT | TA>AT | GACCTGTAC | TACCAAT | CAT | GTA | T | T | T>A | T>A_GTA | TA>AT_TACCAAT | 9 | FALSE | FALSE |
| 130_v.txt | 31650 | 31650-SNV-( | SNV | 1 | C | T | 27 | 11604 | 0.23267839 | 36.6296296 | WT | FALSE | CAN | JSY2424 | 7 | WT-Rep7 | GCG | C>T | A/T+1 | G/C-1 | CG>TA | CG>TA | AGAATGCGA | CGAAATG | CGC | GCG | C | A | C>T | C>T_GCG | CG>TA_CGAAATG | 9 | FALSE | FALSE |
| 130_v.txt | 31806 | 31806-SNV-( | SNV | 1 | C | T | 28 | 11604 | 0.24129611 | 56.0363389 | WT | FALSE | CAN | JSY2424 | 7 | WT-Rep7 | CCA | C>T | A/T+1 | G/C-1 | CG>TA | CG>TA | ATCTTCCAA | CAAATAA | GGT | CCA | C | A | C>T | C>T_CCA | CG>TA_CAAATAA | 9 | FALSE | FALSE |
| 130_v.txt | 31834 | 31834-SNV-( | SNV | 1 | G | A | 55 | 11604 | 0.47397449 | 50.7636364 | WT | FALSE | CAN | JSY2424 | 7 | WT-Rep7 | TGA | G>A | A/T+1 | G/C-1 | CG>TA | CG>TA | TGCATTGAA | GAAATAA | ACT | ACT | C | T | C>T | C>T_ACT | CG>TA_GAAATAA | 9 | FALSE | FALSE |
| 130_v.txt | 31884 | 31884-SNV-( | SNV | 1 | G | A | 20 | 11604 | 0.17235436 | 32.45 | WT | FALSE | CAN | JSY2424 | 7 | WT-Rep7 | AGC | G>A | A/T+1 | G/C-1 | CG>TA | CG>TA | GCGGCAGCA | GCAAAGC | TCG | TCG | C | T | C>T | C>T_TCG | CG>TA_GCAAAGC | 9 | FALSE | FALSE |
| 130_v.txt | 31960 | 31960-SNV-( | SNV | 1 | C | T | 27 | 31094 | 0.08683347 | 35.2962963 | WT | FALSE | CAN | JSY2424 | 7 | WT-Rep7 | GCC | C>T | A/T+1 | G/C-1 | CG>TA | CG>TA | ATGTGGCG | CCGCATA | CGG | GCC | C | A | C>T | C>T_GCC | CG>TA_CCGCATA | 9 | FALSE | FALSE |
| 130_v.txt | 31962 | 31962-SNV-( | SNV | 1 | G | A | 48 | 31094 | 0.15437062 | 37.5416667 | WT | FALSE | CAN | JSY2424 | 7 | WT-Rep7 | CGC | G>A | A/T+1 | G/C-1 | CG>TA | CG>TA | GTGGCCGCA | GCATAAT | GCG | GCG | C | T | C>T | C>T_GCG | CG>TA_GCATAAT | 9 | FALSE | FALSE |
| 130_v.txt | 31972 | 31972-SNV-( | SNV | 1 | C | T | 18 | 31094 | 0.05788898 | 37.1111111 | WT | FALSE | CAN | JSY2424 | 7 | WT-Rep7 | GCC | C>T | A/T+1 | G/C-1 | CG>TA | CG>TA | AATAAGCCA | CCAAGCC | CGG | GCC | C | A | C>T | C>T_GCC | CG>TA_CCAAGCC | 9 | FALSE | FALSE |
| 130_v.txt | 31981 | 31981-SNV-( | SNV | 1 | G | A | 60 | 31094 | 0.19296327 | 51.8166667 | WT | FALSE | CAN | JSY2424 | 7 | WT-Rep7 | GGC | G>A | A/T+1 | G/C-1 | CG>TA | CG>TA | AGCCGGGCA | GCATTAA | CCG | CCG | C | T | C>T | C>T_CCG | CG>TA_GCATTAA | 9 | FALSE | FALSE |
| 130_v.txt | 32014 | 32014-SNV-( | SNV | 1 | G | A | 4 | 31094 | 0.01286422 | 39.9998751 | WT | FALSE | CAN | JSY2424 | 7 | WT-Rep7 | CGA | G>A | A/T+1 | G/C-1 | CG>TA | CG>TA | CGTCACGAG | GAGAGAT | GCT | GCT | C | T | C>T | C>T_GCT | CG>TA_GAGAGAT | 9 | FALSE | FALSE |
| 130_v.txt | 32041 | 32041-SNV-( | SNV | 1 | G | A | 52 | 31094 | 0.16723484 | 59.8846154 | WT | FALSE | CAN | JSY2424 | 7 | WT-Rep7 | TGC | G>A | A/T+1 | G/C-1 | CG>TA | CG>TA | AAGCTTGCA | GCATAAA | ACG | ACG | C | T | C>T | C>T_ACG | CG>TA_GCATAAA | 9 | FALSE | FALSE |
| 130_v.txt | 32049 | 32049-SNV-( | SNV | 1 | C | T | 2 | 31094 | 6.43E-03 | 63.999135 | WT | FALSE | CAN | JSY2424 | 7 | WT-Rep7 | TCT | C>T | A/T+1 | G/C-1 | CG>TA | CG>TA | ATAAATCTG | CTGATGT | AGA | TCT | C | A | C>T | C>T_TCT | CG>TA_CTGATGT | 9 | FALSE | FALSE |
| 130_v.txt | 32054 | 32054-SNV-( | SNV | 1 | G | A | 5 | 31094 | 0.01608027 | 62.3992237 | WT | FALSE | CAN | JSY2424 | 7 | WT-Rep7 | TGT | G>A | A/T+1 | G/C-1 | CG>TA | CG>TA | TCTGATGTG | GTGCGAG | ACA | ACA | C | T | C>T | C>T_ACA | CG>TA_GTGCGAG | 9 | FALSE | FALSE |
| 130_v.txt | 32057 | 32057-SNV-( | SNV | 1 | C | T | 32 | 31093 | 0.10291706 | 61.0927408 | WT | FALSE | CAN | JSY2424 | 7 | WT-Rep7 | GCG | C>T | A/T+1 | G/C-1 | CG>TA | CG>TA | GATGTGCGA | CGAGATT | CGC | GCG | C | A | C>T | C>T_GCG | CG>TA_CGAGATT | 9 | FALSE | FALSE |
| 130_v.txt | 32076 | 32076-SNV-( | SNV | 1 | C | T | 26 | 31094 | 0.08361742 | 61.8846154 | WT | FALSE | CAN | JSY2424 | 7 | WT-Rep7 | CCA | C>T | A/T+1 | G/C-1 | CG>TA | CG>TA | AATAACCAT | CATGCAA | GGT | CCA | C | A | C>T | C>T_CCA | CG>TA_CATGCAA | 9 | FALSE | FALSE |
| 130_v.txt | 32088 | 32088-SNV-( | SNV | 1 | C | T | 14 | 31094 | 0.04502476 | 62.6428571 | WT | FALSE | CAN | JSY2424 | 7 | WT-Rep7 | GCC | C>T | A/T+1 | G/C-1 | CG>TA | CG>TA | AAAAAGCCT | CCTGCAA | CGG | GCC | C | A | C>T | C>T_GCC | CG>TA_CCTGCAA | 9 | FALSE | FALSE |
| 130_v.txt | 32092 | 32092-SNV-( | SNV | 1 | C | T | 28 | 31094 | 0.09004953 | 56.6785714 | WT | FALSE | CAN | JSY2424 | 7 | WT-Rep7 | GCA | C>T | A/T+1 | G/C-1 | CG>TA | CG>TA | AGCCTGCAA | CAACACC | CGT | GCA | C | A | C>T | C>T_GCA | CG>TA_CAACACC | 9 | FALSE | FALSE |
| 130_v.txt | 32115 | 32115-SNV-( | SNV | 1 | C | T | 45 | 31094 | 0.14472246 | 55.3111111 | WT | FALSE | CAN | JSY2424 | 7 | WT-Rep7 | CCA | C>T | A/T+1 | G/C-1 | CG>TA | CG>TA | AATAGCCAT | CATTCGA | GGT | CCA | C | A | C>T | C>T_CCA | CG>TA_CATTCGA | 9 | FALSE | FALSE |
| 130_v.txt | 32129 | 32129-SNV-( | SNV | 1 | G | A | 8 | 31093 | 0.02572926 | 41.999785 | WT | FALSE | CAN | JSY2424 | 7 | WT-Rep7 | TGT | G>A | A/T+1 | G/C-1 | CG>TA | CG>TA | AACTTGTCA | GTCACCA | ACA | ACA | C | T | C>T | C>T_ACA | CG>TA_GTCACCA | 9 | FALSE | FALSE |
| 130_v.txt | 32165 | 32165-SNV-( | SNV | 1 | G | A | 25 | 31094 | 0.08040136 | 50.32 | WT | FALSE | CAN | JSY2424 | 7 | WT-Rep7 | CGC | G>A | A/T+1 | G/C-1 | CG>TA | CG>TA | CAAAGCGCC | GCCAAAT | GCG | GCG | C | T | C>T | C>T_GCG | CG>TA_GCCAAAT | 9 | FALSE | FALSE |
| 130_v.txt | 32190 | 32190-SNV-( | SNV | 1 | G | A | 11 | 31094 | 0.0353376 | 42.3634697 | WT | FALSE | CAN | JSY2424 | 7 | WT-Rep7 | TGC | G>A | A/T+1 | G/C-1 | CG>TA | CG>TA | AAAACTGCA | GCAATGT | ACG | ACG | C | T | C>T | C>T_ACG | CG>TA_GCAATGT | 9 | FALSE | FALSE |
| 130_v.txt | 32191 | 32191-SNV-( | SNV | 1 | C | T | 9 | 31094 | 0.02894449 | 52.1101701 | WT | FALSE | CAN | JSY2424 | 7 | WT-Rep7 | GCA | C>T | A/T+1 | G/C-1 | CG>TA | CG>TA | AAACTGCAA | CAATGTA | CGT | GCA | C | A | C>T | C>T_GCA | CG>TA_CAATGTA | 9 | FALSE | FALSE |
| 130_v.txt | 32271 | 32271-SNV-( | SNV | 1 | C | T | 95 | 43326 | 0.21926788 | 33.6947368 | WT | FALSE | CAN | JSY2424 | 7 | WT-Rep7 | ACG | C>T | A/T+1 | G/C-1 | CG>TA | CG>TA | AAAATACGG | CGGGAAC | TGC | ACG | C | A | C>T | C>T_ACG | CG>TA_CGGGAAC | 9 | FALSE | FALSE |
| 130_v.txt | 32274 | 32274-SNV-( | SNV | 1 | G | A | 24 | 43326 | 0.05539399 | 36.1666667 | WT | FALSE | CAN | JSY2424 | 7 | WT-Rep7 | GGA | G>A | A/T+1 | G/C-1 | CG>TA | CG>TA | ATACGGGAA | GAACCAA | CCT | CCT | C | T | C>T | C>T_CCT | CG>TA_GAACCAA | 9 | FALSE | FALSE |
| 130_v.txt | 32282 | 32282-SNV-( | SNV | 1 | G | A | 11 | 43326 | 0.02538891 | 43.9084755 | WT | FALSE | CAN | JSY2424 | 7 | WT-Rep7 | CGT | G>A | A/T+1 | G/C-1 | CG>TA | CG>TA | ACCAACGTA | GTAAATA | GCA | GCA | C | T | C>T | C>T_GCA | CG>TA_GTAAATA | 9 | FALSE | FALSE |
| 130_v.txt | 32292 | 32292-SNV-( | SNV | 1 | G | A | 44 | 43326 | 0.10155565 | 58.0681818 | WT | FALSE | CAN | JSY2424 | 7 | WT-Rep7 | TGA | G>A | A/T+1 | G/C-1 | CG>TA | CG>TA | ATATTTGAA | GAATTTG | ACT | ACT | C | T | C>T | C>T_ACT | CG>TA_GAATTTG | 9 | FALSE | FALSE |
| 130_v.txt | 32299 | 32299-SNV-( | SNV | 1 | C | T | 64 | 43326 | 0.14771731 | 56.625 | WT | FALSE | CAN | JSY2424 | 7 | WT-Rep7 | GCG | C>T | A/T+1 | G/C-1 | CG>TA | CG>TA | AATTTGCGG | CGGCAGA | CGC | GCG | C | A | C>T | C>T_GCG | CG>TA_CGGCAGA | 9 | FALSE | FALSE |
| 130_v.txt | 32301 | 32301-SNV-( | SNV | 1 | G | A | 44 | 43326 | 0.10155565 | 57.8409091 | WT | FALSE | CAN | JSY2424 | 7 | WT-Rep7 | GGC | G>A | A/T+1 | G/C-1 | CG>TA | CG>TA | TTTGCGGCA | GCAGAAA | CCG | CCG | C | T | C>T | C>T_CCG | CG>TA_GCAGAAA | 9 | FALSE | FALSE |
| 130_v.txt | 32321 | 32321-SNV-( | SNV | 1 | G | A | 5 | 43326 | 0.01154042 | 45.3999172 | WT | FALSE | CAN | JSY2424 | 7 | WT-Rep7 | AGA | G>A | A/T+1 | G/C-1 | CG>TA | CG>TA | TGTTAAGAT | GATAACA | TCT | TCT | C | T | C>T | C>T_TCT | CG>TA_GATAACA | 9 | FALSE | FALSE |
| 130_v.txt | 32328 | 32328-SNV-( | SNV | 1 | G | A | 3 | 43326 | 6.92E-03 | 63.9960184 | WT | FALSE | CAN | JSY2424 | 7 | WT-Rep7 | AGC | G>A | A/T+1 | G/C-1 | CG>TA | CG>TA | ATAACAGCG | GCGTTGA | TCG | TCG | C | T | C>T | C>T_TCG | CG>TA_GCGTTGA | 9 | FALSE | FALSE |
| 130_v.txt | 32329 | 32329-SNV-( | SNV | 1 | C | T | 22 | 43325 | 0.050779 | 43.1278544 | WT | FALSE | CAN | JSY2424 | 7 | WT-Rep7 | GCG | C>T | A/T+1 | G/C-1 | CG>TA | CG>TA | TAACAGCGT | CGTTGAA | CGC | GCG | C | A | C>T | C>T_GCG | CG>TA_CGTTGAA | 9 | FALSE | FALSE |
| 130_v.txt | 32330 | 32330-SNV-( | SNV | 1 | G | A | 12 | 43326 | 0.027697 | 59.582486 | WT | FALSE | CAN | JSY2424 | 7 | WT-Rep7 | CGT | G>A | A/T+1 | G/C-1 | CG>TA | CG>TA | AACAGCGTT | GTTGAAG | GCA | GCA | C | T | C>T | C>T_GCA | CG>TA_GTTGAAG | 9 | FALSE | FALSE |
| 130_v.txt | 32336 | 32336-SNV-( | SNV | 1 | G | A | 5 | 43326 | 0.01154042 | 54.9947261 | WT | FALSE | CAN | JSY2424 | 7 | WT-Rep7 | AGA | G>A | A/T+1 | G/C-1 | CG>TA | CG>TA | GTTGAAGAT | GATATGT | TCT | TCT | C | T | C>T | C>T_TCT | CG>TA_GATATGT | 9 | FALSE | FALSE |
| 130_v.txt | 32341 | 32341-SNV-( | SNV | 1 | G | A | 34 | 43326 | 0.07847482 | 59.4117647 | WT | FALSE | CAN | JSY2424 | 7 | WT-Rep7 | TGT | G>A | A/T+1 | G/C-1 | CG>TA | CG>TA | AGATATGTG | GTGGCAA | ACA | ACA | C | T | C>T | C>T_ACA | CG>TA_GTGGCAA | 9 | FALSE | FALSE |
| 130_v.txt | 32449 | 32449-SNV-( | SNV | 1 | C | T | 28 | 43326 | 0.06462632 | 44.9642857 | WT | FALSE | CAN | JSY2424 | 7 | WT-Rep7 | CCA | C>T | A/T+1 | G/C-1 | CG>TA | CG>TA | AAAGTCCAA | CAATGAA | GGT | CCA | C | A | C>T | C>T_CCA | CG>TA_CAATGAA | 9 | FALSE | FALSE |
| 130_v.txt | 32463 | 32463-SNV-( | SNV | 1 | G | A | 37 | 43326 | 0.08539907 | 44.1081081 | WT | FALSE | CAN | JSY2424 | 7 | WT-Rep7 | AGA | G>A | A/T+1 | G/C-1 | CG>TA | CG>TA | AATAGAGAG | GAGCCAA | TCT | TCT | C | T | C>T | C>T_TCT | CG>TA_GAGCCAA | 9 | FALSE | FALSE |
| 130_v.txt | 32466 | 32466-SNV-( | SNV | 1 | C | T | 50 | 43326 | 0.11540415 | 47.6767958 | WT | FALSE | CAN | JSY2424 | 7 | WT-Rep7 | GCC | C>T | A/T+1 | G/C-1 | CG>TA | CG>TA | AGAGAGCCA | CCAATGT | CGG | GCC | C | A | C>T | C>T_GCC | CG>TA_CCAATGT | 9 | FALSE | FALSE |
| 130_v.txt | 32467 | 32467-SNV-( | SNV | 1 | C | T | 9 | 43326 | 0.02077275 | 50.8861154 | WT | FALSE | CAN | JSY2424 | 7 | WT-Rep7 | CCA | C>T | A/T+1 | G/C-1 | CG>TA | CG>TA | GAGAGCCAA | CAATGTA | GGT | CCA | C | A | C>T | C>T_CCA | CG>TA_CAATGTA | 9 | FALSE | FALSE |
| 130_v.txt | 32487 | 32487-SNV-( | SNV | 1 | C | T | 110 | 43325 | 0.25389498 | 45.4818182 | WT | FALSE | CAN | JSY2424 | 7 | WT-Rep7 | ACG | C>T | A/T+1 | G/C-1 | CG>TA | CG>TA | AAGATACGG | CGGAAAA | TGC | ACG | C | A | C>T | C>T_ACG | CG>TA_CGGAAAA | 9 | FALSE | FALSE |
| 130_v.txt | 32488 | 32488-SNV-( | SNV | 1 | G | A | 22 | 43326 | 0.05077782 | 44.3181027 | WT | FALSE | CAN | JSY2424 | 7 | WT-Rep7 | CGG | G>A | A/T+1 | G/C-1 | CG>TA | CG>TA | AGATACGGA | GGAAAAC | GCC | GCC | C | T | C>T | C>T_GCC | CG>TA_GGAAAAC | 9 | FALSE | FALSE |
| 130_v.txt | 32508 | 32508-SNV-( | SNV | 1 | G | A | 45 | 43326 | 0.10386373 | 40.9333333 | WT | FALSE | CAN | JSY2424 | 7 | WT-Rep7 | GGC | G>A | A/T+1 | G/C-1 | CG>TA | CG>TA | TTGATGGCT | GCTCTTG | CCG | CCG | C | T | C>T | C>T_CCG | CG>TA_GCTCTTG | 9 | FALSE | FALSE |
| 130_v.txt | 32529 | 32529-SNV-( | SNV | 1 | G | A | 98 | 43326 | 0.22619213 | 43.3978852 | WT | FALSE | CAN | JSY2424 | 7 | WT-Rep7 | GGG | G>A | A/T+1 | G/C-1 | CG>TA | CG>TA | TTTCTGGGG | GGGTTTG | CCC | CCC | C | T | C>T | C>T_CCC | CG>TA_GGGTTTG | 9 | FALSE | FALSE |
| 130_v.txt | 32535 | 32535-SNV-( | SNV | 1 | G | A | 70 | 43325 | 0.16156953 | 37.0822986 | WT | FALSE | CAN | JSY2424 | 7 | WT-Rep7 | TGC | G>A | A/T+1 | G/C-1 | CG>TA | CG>TA | GGGTTTGCA | GCAGCTT | ACG | ACG | C | T | C>T | C>T_ACG | CG>TA_GCAGCTT | 9 | FALSE | FALSE |
| 130_v.txt | 32536 | 32536-SNV-( | SNV | 1 | C | T | 44 | 43325 | 0.10155799 | 37.3181597 | WT | FALSE | CAN | JSY2424 | 7 | WT-Rep7 | GCA | C>T | A/T+1 | G/C-1 | CG>TA | CG>TA | GGTTTGCAG | CAGCTTC | CGT | GCA | C | A | C>T | C>T_GCA | CG>TA_CAGCTTC | 9 | FALSE | FALSE |
| 130_v.txt | 32542 | 32542-SNV-( | SNV | 1 | C | T | 43 | 43325 | 0.09924986 | 36.0689518 | WT | FALSE | CAN | JSY2424 | 7 | WT-Rep7 | TCA | C>T | A/T+1 | G/C-1 | CG>TA | CG>TA | CAGCTTCAC | CACCAGC | AGT | TCA | C | A | C>T | C>T_TCA | CG>TA_CACCAGC | 9 | FALSE | FALSE |
| 130_v.txt | 32566 | 32566-SNV-( | SNV | 1 | C | T | 41 | 31633 | 0.12961148 | 35.0243902 | WT | FALSE | CAN | JSY2424 | 7 | WT-Rep7 | TCA | C>T | A/T+1 | G/C-1 | CG>TA | CG>TA | CTAGTTCAG | CAGTACC | AGT | TCA | C | A | C>T | C>T_TCA | CG>TA_CAGTACC | 9 | FALSE | FALSE |
| 130_v.txt | 32571 | 32571-SNV-( | SNV | 1 | C | T | 65 | 31647 | 0.20539072 | 35.1230769 | WT | FALSE | CAN | JSY2424 | 7 | WT-Rep7 | ACC | C>T | A/T+1 | G/C-1 | CG>TA | CG>TA | TCAGTACCT | CCTTGAA | TGG | ACC | C | A | C>T | C>T_ACC | CG>TA_CCTTGAA | 9 | FALSE | FALSE |
| 130_v.txt | 32590 | 32590-SNV-( | SNV | 1 | C | T | 31 | 31688 | 0.09782883 | 36.0967742 | WT | FALSE | CAN | JSY2424 | 7 | WT-Rep7 | GCG | C>T | A/T+1 | G/C-1 | CG>TA | CG>TA | AGGCAGCGT | CGTTAAT | CGC | GCG | C | A | C>T | C>T_GCG | CG>TA_CGTTAAT | 9 | FALSE | FALSE |
| 130_v.txt | 32658 | 32658-SNV-( | SNV | 1 | C | T | 80 | 31688 | 0.2524615 | 55.6875 | WT | FALSE | CAN | JSY2424 | 7 | WT-Rep7 | ACC | C>T | A/T+1 | G/C-1 | CG>TA | CG>TA | CCTGGACCC | CCCCAGG | TGG | ACC | C | A | C>T | C>T_ACC | CG>TA_CCCCAGG | 9 | FALSE | FALSE |
| 130_v.txt | 32679 | 32679-SNV-( | SNV | 1 | C | T | 39 | 31687 | 0.12307887 | 59.2051282 | WT | FALSE | CAN | JSY2424 | 7 | WT-Rep7 | CCA | C>T | A/T+1 | G/C-1 | CG>TA | CG>TA | TTTCTCCAA | CAATAAC | GGT | CCA | C | A | C>T | C>T_CCA | CG>TA_CAATAAC | 9 | FALSE | FALSE |
| 130_v.txt | 32700 | 32700-SNV-( | SNV | 1 | C | T | 35 | 31688 | 0.11045191 | 56.5142857 | WT | FALSE | CAN | JSY2424 | 7 | WT-Rep7 | GCC | C>T | A/T+1 | G/C-1 | CG>TA | CG>TA | ACTGGGCCG | CCGGTAA | CGG | GCC | C | A | C>T | C>T_GCC | CG>TA_CCGGTAA | 9 | FALSE | FALSE |
| 130_v.txt | 32749 | 32749-SNV-( | SNV | 1 | C | T | 203 | 31688 | 0.64062106 | 52.6600985 | WT | FALSE | CAN | JSY2424 | 7 | WT-Rep7 | CCG | C>T | A/T+1 | G/C-1 | CG>TA | CG>TA | GAAACCCGA | CGATAAT | GGC | CCG | C | A | C>T | C>T_CCG | CG>TA_CGATAAT | 9 | FALSE | FALSE |
| 130_v.txt | 32788 | 32788-SNV-( | SNV | 1 | C | T | 1482 | 31688 | 4.67684928 | 47.5623321 | WT | FALSE | CAN | JSY2424 | 7 | WT-Rep7 | TCG | C>T | A/T+1 | G/C-1 | CG>TA | CG>TA | AGAACTCGA | CGAATTC | AGC | TCG | C | A | C>T | C>T_TCG | CG>TA_CGAATTC | 9 | FALSE | FALSE |
| 130_v.txt | 32794 | 32794-SNV-( | SNV | 1 | C | T | 67 | 31688 | 0.21143651 | 43.9701493 | WT | FALSE | CAN | JSY2424 | 7 | WT-Rep7 | TCA | C>T | A/T+1 | G/C-1 | CG>TA | CG>TA | CGAATTCAC | CACCGTA | AGT | TCA | C | A | C>T | C>T_TCA | CG>TA_CACCGTA | 9 | FALSE | FALSE |
| 130_v.txt | 32797 | 32797-SNV-( | SNV | 1 | C | T | 53 | 31688 | 0.16725574 | 48.0377359 | WT | FALSE | CAN | JSY2424 | 7 | WT-Rep7 | CCG | C>T | A/T+1 | G/C-1 | CG>TA | CG>TA | ATTCACCGT | CGTAATA | GGC | CCG | C | A | C>T | C>T_CCG | CG>TA_CGTAATA | 9 | FALSE | FALSE |
| 130_v.txt | 32798 | 32798-SNV-( | SNV | 1 | G | A | 39 | 31688 | 0.12307498 | 45.8461539 | WT | FALSE | CAN | JSY2424 | 7 | WT-Rep7 | CGT | G>A | A/T+1 | G/C-1 | CG>TA | CG>TA | TTCACCGTA | GTAATAT | GCA | GCA | C | T | C>T | C>T_GCA | CG>TA_GTAATAT | 9 | FALSE | FALSE |
| 130_v.txt | 32809 | 32809-SNV-( | SNV | 1 | C | T | 27 | 31688 | 0.08520576 | 43.4444444 | WT | FALSE | CAN | JSY2424 | 7 | WT-Rep7 | ACA | C>T | A/T+1 | G/C-1 | CG>TA | CG>TA | ATTTGACAG | CAGGGAA | TGT | ACA | C | A | C>T | C>T_ACA | CG>TA_CAGGGAA | 9 | FALSE | FALSE |
| 130_v.txt | 32811 | 32811-SNV-( | SNV | 1 | G | A | 22 | 31688 | 0.06942691 | 44.5 | WT | FALSE | CAN | JSY2424 | 7 | WT-Rep7 | AGG | G>A | A/T+1 | G/C-1 | CG>TA | CG>TA | TTGACAGGG | GGGAACA | TCC | TCC | C | T | C>T | C>T_TCC | CG>TA_GGGAACA | 9 | FALSE | FALSE |
| 130_v.txt | 32841 | 32841-SNV-( | SNV | 1 | C | T | 41 | 31687 | 0.1293906 | 42.5843361 | WT | FALSE | CAN | JSY2424 | 7 | WT-Rep7 | CCA | C>T | A/T+1 | G/C-1 | CG>TA | CG>TA | ATTACCCAA | CAAAAAA | GGT | CCA | C | A | C>T | C>T_CCA | CG>TA_CAAAAAA | 9 | FALSE | FALSE |
| 130_v.txt | 32859 | 32859-SNV-( | SNV | 1 | G | A | 9 | 31686 | 0.02840371 | 34.105872 | WT | FALSE | CAN | JSY2424 | 7 | WT-Rep7 | TGC | G>A | A/T+1 | G/C-1 | CG>TA | CG>TA | ATCCATGCC | GCCGCCA | ACG | ACG | C | T | C>T | C>T_ACG | CG>TA_GCCGCCA | 9 | FALSE | FALSE |
| 130_v.txt | 32860 | 32860-SNV-( | SNV | 1 | C | T | 30 | 31686 | 0.09467904 | 37.0666667 | WT | FALSE | CAN | JSY2424 | 7 | WT-Rep7 | GCC | C>T | A/T+1 | G/C-1 | CG>TA | CG>TA | TCCATGCCG | CCGCCAG | CGG | GCC | C | A | C>T | C>T_GCC | CG>TA_CCGCCAG | 9 | FALSE | FALSE |
| 130_v.txt | 32862 | 32862-SNV-( | SNV | 1 | G | A | 26 | 31685 | 0.08205776 | 35.8076923 | WT | FALSE | CAN | JSY2424 | 7 | WT-Rep7 | CGC | G>A | A/T+1 | G/C-1 | CG>TA | CG>TA | CATGCCGCC | GCCAGTG | GCG | GCG | C | T | C>T | C>T_GCG | CG>TA_GCCAGTG | 9 | FALSE | FALSE |
| 130_v.txt | 32890 | 32890-SNV-( | SNV | 1 | G | A | 324 | 33859 | 0.95690954 | 37.2006173 | WT | FALSE | CAN | JSY2424 | 7 | WT-Rep7 | TGA | G>A | A/T+1 | G/C-1 | CG>TA | CG>TA | AAAATTGAA | GAATGAC | ACT | ACT | C | T | C>T | C>T_ACT | CG>TA_GAATGAC | 9 | FALSE | FALSE |
| 130_v.txt | 32901 | 32901-SNV-( | SNV | 1 | C | T | 62 | 33859 | 0.18311232 | 34.2419355 | WT | FALSE | CAN | JSY2424 | 7 | WT-Rep7 | GCC | C>T | A/T+1 | G/C-1 | CG>TA | CG>TA | ACTTGGCCA | CCAACTA | CGG | GCC | C | A | C>T | C>T_GCC | CG>TA_CCAACTA | 9 | FALSE | FALSE |
| 130_v.txt | 32917 | 32917-SNV-( | SNV | 1 | C | T | 43 | 33860 | 0.1269935 | 46.9483554 | WT | FALSE | CAN | JSY2424 | 7 | WT-Rep7 | TCC | C>T | A/T+1 | G/C-1 | CG>TA | CG>TA | TAAGTTCCA | CCAGGGC | AGG | TCC | C | A | C>T | C>T_TCC | CG>TA_CCAGGGC | 9 | FALSE | FALSE |
| 130_v.txt | 32940 | 32940-SNV-( | SNV | 1 | G | A | 21 | 33860 | 0.06202008 | 46.8512909 | WT | FALSE | CAN | JSY2424 | 7 | WT-Rep7 | AGA | G>A | A/T+1 | G/C-1 | CG>TA | CG>TA | GCCCAAGAA | GAAAACC | TCT | TCT | C | T | C>T | C>T_TCT | CG>TA_GAAAACC | 9 | FALSE | FALSE |
| 130_v.txt | 32946 | 32946-SNV-( | SNV | 1 | C | T | 47 | 33860 | 0.13880685 | 56.9361702 | WT | FALSE | CAN | JSY2424 | 7 | WT-Rep7 | CCA | C>T | A/T+1 | G/C-1 | CG>TA | CG>TA | GAAAACCAA | CAATACA | GGT | CCA | C | A | C>T | C>T_CCA | CG>TA_CAATACA | 9 | FALSE | FALSE |
| 130_v.txt | 32965 | 32965-SNV-( | SNV | 1 | C | T | 35 | 33860 | 0.10336681 | 59.8 | WT | FALSE | CAN | JSY2424 | 7 | WT-Rep7 | GCC | C>T | A/T+1 | G/C-1 | CG>TA | CG>TA | CATTGGCCG | CCGCACC | CGG | GCC | C | A | C>T | C>T_GCC | CG>TA_CCGCACC | 9 | FALSE | FALSE |
| 130_v.txt | 32967 | 32967-SNV-( | SNV | 1 | G | A | 60 | 33860 | 0.17720024 | 61.65 | WT | FALSE | CAN | JSY2424 | 7 | WT-Rep7 | CGC | G>A | A/T+1 | G/C-1 | CG>TA | CG>TA | TTGGCCGCA | GCACCAA | GCG | GCG | C | T | C>T | C>T_GCG | CG>TA_GCACCAA | 9 | FALSE | FALSE |
| 130_v.txt | 32995 | 32995-SNV-( | SNV | 1 | G | A | 172 | 33860 | 0.50797401 | 62.4360464 | WT | FALSE | CAN | JSY2424 | 7 | WT-Rep7 | TGT | G>A | A/T+1 | G/C-1 | CG>TA | CG>TA | ATCTTTGTGA | GTGAGAA | ACA | ACA | C | T | C>T | C>T_ACA | CG>TA_GTGAGAA | 9 | FALSE | FALSE |
| 130_v.txt | 33015 | 33015-SNV-( | SNV | 1 | G | A | 63 | 33860 | 0.18606025 | 58.4603175 | WT | FALSE | CAN | JSY2424 | 7 | WT-Rep7 | GGA | G>A | A/T+1 | G/C-1 | CG>TA | CG>TA | AAAGAGGAT | GATGTAA | CCT | CCT | C | T | C>T | C>T_CCT | CG>TA_GATGTAA | 9 | FALSE | FALSE |
| 130_v.txt | 33056 | 33056-SNV-( | SNV | 1 | C | T | 33 | 33860 | 0.09746013 | 50.7878788 | WT | FALSE | CAN | JSY2424 | 7 | WT-Rep7 | GCG | C>T | A/T+1 | G/C-1 | CG>TA | CG>TA | GGACTGCGT | CGTGACA | CGC | GCG | C | A | C>T | C>T_GCG | CG>TA_CGTGACA | 9 | FALSE | FALSE |
| 130_v.txt | 33070 | 33070-SNV-( | SNV | 1 | C | T | 23 | 33860 | 0.06792676 | 51.5217391 | WT | FALSE | CAN | JSY2424 | 7 | WT-Rep7 | GCC | C>T | A/T+1 | G/C-1 | CG>TA | CG>TA | AATATGCCA | CCAAAGA | CGG | GCC | C | A | C>T | C>T_GCC | CG>TA_CCAAAGA | 9 | FALSE | FALSE |
| 130_v.txt | 33114 | 33114-SNV-( | SNV | 1 | C | T | 64 | 33860 | 0.18901359 | 41.9999175 | WT | FALSE | CAN | JSY2424 | 7 | WT-Rep7 | GCC | C>T | A/T+1 | G/C-1 | CG>TA | CG>TA | ACTGGGCCG | CCGGCGT | CGG | GCC | C | A | C>T | C>T_GCC | CG>TA_CCGGCGT | 9 | FALSE | FALSE |
| 130_v.txt | 33115 | 33115-SNV-( | SNV | 1 | C | T | 144 | 33860 | 0.42528057 | 42.589159 | WT | FALSE | CAN | JSY2424 | 7 | WT-Rep7 | CCG | C>T | A/T+1 | G/C-1 | CG>TA | CG>TA | CTGGGCCGG | CGGCGTT | GGC | CCG | C | A | C>T | C>T_CCG | CG>TA_CGGCGTT | 9 | FALSE | FALSE |
| 130_v.txt | 33118 | 33118-SNV-( | SNV | 1 | C | T | 27 | 33860 | 0.07974011 | 42 | WT | FALSE | CAN | JSY2424 | 7 | WT-Rep7 | GCG | C>T | A/T+1 | G/C-1 | CG>TA | CG>TA | GGCCGGCGT | CGTTGGT | CGC | GCG | C | A | C>T | C>T_GCG | CG>TA_CGTTGGT | 9 | FALSE | FALSE |
| 130_v.txt | 33153 | 33153-SNV-( | SNV | 1 | C | T | 36 | 33859 | 0.10632328 | 36.1389515 | WT | FALSE | CAN | JSY2424 | 7 | WT-Rep7 | ACC | C>T | A/T+1 | G/C-1 | CG>TA | CG>TA | AAAAGACCT | CCTGTAC | TGG | ACC | C | A | C>T | C>T_ACC | CG>TA_CCTGTAC | 9 | FALSE | FALSE |
| 130_v.txt | 33156 | 33156-SNV-( | SNV | 1 | G | A | 9 | 33843 | 0.02659339 | 32.0000596 | WT | FALSE | CAN | JSY2424 | 7 | WT-Rep7 | TGT | G>A | A/T+1 | G/C-1 | CG>TA | CG>TA | AGACCTGTA | GTACCAA | ACA | ACA | C | T | C>T | C>T_ACA | CG>TA_GTACCAA | 9 | FALSE | FALSE |
| 130_v.txt | 33168 | 33168-SNV-( | SNV | 1 | C | T | 13 | 33840 | 0.03841608 | 34.3845846 | WT | FALSE | CAN | JSY2424 | 7 | WT-Rep7 | ACC | C>T | A/T+1 | G/C-1 | CG>TA | CG>TA | ATAGTACCA | CCACCAA | TGG | ACC | C | A | C>T | C>T_ACC | CG>TA_CCACCAA | 9 | FALSE | FALSE |
| 130_v.txt | 33169 | 33169-SNV-( | SNV | 1 | C | T | 10 | 33840 | 0.02955083 | 37.8000054 | WT | FALSE | CAN | JSY2424 | 7 | WT-Rep7 | CCA | C>T | A/T+1 | G/C-1 | CG>TA | CG>TA | TAGTACCAC | CACCAAG | GGT | CCA | C | A | C>T | C>T_CCA | CG>TA_CACCAAG | 9 | FALSE | FALSE |
| 130_v.txt | 33206 | 33206-SNV-( | SNV | 1 | C | T | 2 | 32448 | 6.16E-03 | 50.9989621 | WT | FALSE | CAN | JSY2424 | 7 | WT-Rep7 | GCT | C>T | A/T+1 | G/C-1 | CG>TA | CG>TA | CTTAAGCTC | CTCTCTC | CGA | GCT | C | A | C>T | C>T_GCT | CG>TA_CTCTCTC | 9 | FALSE | FALSE |
| 130_v.txt | 33229 | 33229-SNV-( | SNV | 1 | G | A | 31 | 32448 | 0.09553748 | 36.8709677 | WT | FALSE | CAN | JSY2424 | 7 | WT-Rep7 | TGT | G>A | A/T+1 | G/C-1 | CG>TA | CG>TA | CGTTCTGTA | GTACTTC | ACA | ACA | C | T | C>T | C>T_ACA | CG>TA_GTACTTC | 9 | FALSE | FALSE |
| 130_v.txt | 33258 | 33258-SNV-( | SNV | 1 | C | T | 34 | 32448 | 0.10478304 | 45.4117647 | WT | FALSE | CAN | JSY2424 | 7 | WT-Rep7 | GCC | C>T | A/T+1 | G/C-1 | CG>TA | CG>TA | CCTATGCCA | CCATCCT | CGG | GCC | C | A | C>T | C>T_GCC | CG>TA_CCATCCT | 9 | FALSE | FALSE |
| 130_v.txt | 33275 | 33275-SNV-( | SNV | 1 | C | T | 25 | 32448 | 0.07704635 | 49.9192559 | WT | FALSE | CAN | JSY2424 | 7 | WT-Rep7 | ACG | C>T | A/T+1 | G/C-1 | CG>TA | CG>TA | AGAGAACGT | CGTATCC | TGC | ACG | C | A | C>T | C>T_ACG | CG>TA_CGTATCC | 9 | FALSE | FALSE |
| 130_v.txt | 33276 | 33276-SNV-( | SNV | 1 | G | A | 30 | 32448 | 0.09245562 | 48.7650885 | WT | FALSE | CAN | JSY2424 | 7 | WT-Rep7 | CGT | G>A | A/T+1 | G/C-1 | CG>TA | CG>TA | GAGAACGTA | GTATCCT | GCA | GCA | C | T | C>T | C>T_GCA | CG>TA_GTATCCT | 9 | FALSE | FALSE |
| 130_v.txt | 33284 | 33284-SNV-( | SNV | 1 | G | A | 29 | 32448 | 0.08937377 | 43.2068966 | WT | FALSE | CAN | JSY2424 | 7 | WT-Rep7 | CGC | G>A | A/T+1 | G/C-1 | CG>TA | CG>TA | ATCCTCGCC | GCCATTT | GCG | GCG | C | T | C>T | C>T_GCG | CG>TA_GCCATTT | 9 | FALSE | FALSE |
| 130_v.txt | 33309 | 33309-SNV-( | SNV | 1 | C | T | 25 | 32448 | 0.07704635 | 58.2632411 | WT | FALSE | CAN | JSY2424 | 7 | WT-Rep7 | GCG | C>T | A/T+1 | G/C-1 | CG>TA | CG>TA | AAAGAGCGC | CGCAATG | CGC | GCG | C | A | C>T | C>T_GCG | CG>TA_CGCAATG | 9 | FALSE | FALSE |
| 130_v.txt | 33310 | 33310-SNV-( | SNV | 1 | G | A | 25 | 32442 | 0.0770606 | 58.0095194 | WT | FALSE | CAN | JSY2424 | 7 | WT-Rep7 | CGC | G>A | A/T+1 | G/C-1 | CG>TA | CG>TA | AAGAGCGCA | GCAATGG | GCG | GCG | C | T | C>T | C>T_GCG | CG>TA_GCAATGG | 9 | FALSE | FALSE |
| 130_v.txt | 33376 | 33376-SNV-( | SNV | 1 | G | A | 49 | 32448 | 0.15101085 | 53.5102041 | WT | FALSE | CAN | JSY2424 | 7 | WT-Rep7 | TGT | G>A | A/T+1 | G/C-1 | CG>TA | CG>TA | GTGTTTGTG | GTGAAGC | ACA | ACA | C | T | C>T | C>T_ACA | CG>TA_GTGAAGC | 9 | FALSE | FALSE |
| 130_v.txt | 33392 | 33392-SNV-( | SNV | 1 | G | A | 28 | 32448 | 0.08629191 | 51.5357143 | WT | FALSE | CAN | JSY2424 | 7 | WT-Rep7 | CGT | G>A | A/T+1 | G/C-1 | CG>TA | CG>TA | AACGTCGTG | GTGAAAG | GCA | GCA | C | T | C>T | C>T_GCA | CG>TA_GTGAAAG | 9 | FALSE | FALSE |
| 130_v.txt | 33412 | 33412-SNV-( | SNV | 1 | G | A | 46 | 32448 | 0.14176529 | 52.2826087 | WT | FALSE | CAN | JSY2424 | 7 | WT-Rep7 | GGC | G>A | A/T+1 | G/C-1 | CG>TA | CG>TA | TGACCGGCT | GCTCATT | CCG | CCG | C | T | C>T | C>T_CCG | CG>TA_GCTCATT | 9 | FALSE | FALSE |
| 130_v.txt | 33474 | 33474-SNV-( | SNV | 1 | C | T | 47 | 32448 | 0.14484714 | 36.2245019 | WT | FALSE | CAN | JSY2424 | 7 | WT-Rep7 | GCC | C>T | A/T+1 | G/C-1 | CG>TA | CG>TA | GCTATGCCT | CCTTTTT | CGG | GCC | C | A | C>T | C>T_GCC | CG>TA_CCTTTTT | 9 | FALSE | FALSE |
| 130_v.txt | 31947 | 31947-SNV-( | SNV | 1 | G | C | 8 | 31052 | 0.02576324 | 34.625 | WT | FALSE | CAN | JSY2424 | 7 | WT-Rep7 | CGT | G>C | G/C+1 | G/C-1 | CG>GC | CG>GC | ATGATCGTC | GTCATAA | GCA | GCA | C | G | C>G | C>G_GCA | CG>GC_GTCATAA | 9 | FALSE | FALSE |
| 130_v.txt | 32027 | 32027-SNV-( | SNV | 1 | G | C | 14 | 31094 | 0.04502476 | 57.0714286 | WT | FALSE | CAN | JSY2424 | 7 | WT-Rep7 | GGT | G>C | G/C+1 | G/C-1 | CG>GC | CG>GC | GCCACGGTA | GTATTTC | CCA | CCA | C | G | C>G | C>G_CCA | CG>GC_GTATTTC | 9 | FALSE | FALSE |
| 130_v.txt | 32033 | 32033-SNV-( | SNV | 1 | C | G | 2 | 31094 | 6.43E-03 | 63.9940302 | WT | FALSE | CAN | JSY2424 | 7 | WT-Rep7 | TCA | C>G | G/C+1 | G/C-1 | CG>GC | CG>GC | GTATTTCAA | CAAAGCT | AGT | TCA | C | C | C>G | C>G_TCA | CG>GC_CAAAGCT | 9 | FALSE | FALSE |
| 130_v.txt | 32038 | 32038-SNV-( | SNV | 1 | C | G | 24 | 31094 | 0.07718531 | 61.625 | WT | FALSE | CAN | JSY2424 | 7 | WT-Rep7 | GCT | C>G | G/C+1 | G/C-1 | CG>GC | CG>GC | TCAAAGCTC | CTTGCAT | CGA | GCT | C | C | C>G | C>G_GCT | CG>GC_CTTGCAT | 9 | FALSE | FALSE |
| 130_v.txt | 32049 | 32049-SNV-( | SNV | 1 | C | G | 1 | 31094 | 3.22E-03 | 63.9979604 | WT | FALSE | CAN | JSY2424 | 7 | WT-Rep7 | TCT | C>G | G/C+1 | G/C-1 | CG>GC | CG>GC | ATAAATCTG | CTGATGT | AGA | TCT | C | C | C>G | C>G_TCT | CG>GC_CTGATGT | 9 | FALSE | FALSE |

| | | | | | | | | | | | | | | | | | | | | | | | | | | | | | | | | | | |
|---|---|---|---|---|---|---|---|---|---|---|---|---|---|---|---|---|---|---|---|---|---|---|---|---|---|---|---|---|---|---|---|---|---|---|
| 130_v.txt | 32058 | 32058-SNV-( | SNV | 1 | G | C | 6 | 31094 | 0.01929633 | 63.9739828 | WT | FALSE | CAN | JSY2424 | 7 | WT-Rep7 | CGA | G>C | G/C+1 | G/C-1 | CG>GC | CG>GC | ATGTGCGAG | GAGATTG | GCT | GCT | C | G | C>G | C>G_GCT | CG>GC_GAGATTG | 9 | FALSE | FALSE |
| 130_v.txt | 32064 | 32064-SNV-( | SNV | 1 | G | C | 22 | 31094 | 0.0707532 | 60.9545455 | WT | FALSE | CAN | JSY2424 | 7 | WT-Rep7 | TGA | G>C | G/C+1 | G/C-1 | CG>GC | CG>GC | GAGATTGAG | GAGATAA | ACT | ACT | C | G | C>G | C>G_ACT | CG>GC_GAGATAA | 9 | FALSE | FALSE |
| 130_v.txt | 32095 | 32095-SNV-( | SNV | 1 | C | G | 11 | 31094 | 0.0353766 | 61.6363636 | WT | FALSE | CAN | JSY2424 | 7 | WT-Rep7 | ACA | C>G | G/C+1 | G/C-1 | CG>GC | CG>GC | CTGCAACAC | CACCAGT | TGT | ACA | C | C | C>G | C>G_ACA | CG>GC_CACCAGT | 9 | FALSE | FALSE |
| 130_v.txt | 32098 | 32098-SNV-( | SNV | 1 | C | G | 63 | 31094 | 0.20261144 | 61.0952381 | WT | FALSE | CAN | JSY2424 | 7 | WT-Rep7 | CCA | C>G | G/C+1 | G/C-1 | CG>GC | CG>GC | CAACACCAG | CAGTGAT | GGT | CCA | C | C | C>G | C>G_CCA | CG>GC_CAGTGAT | 9 | FALSE | FALSE |
| 130_v.txt | 32129 | 32129-SNV-( | SNV | 1 | G | C | 2 | 31093 | 6.43E-03 | 35.9994681 | WT | FALSE | CAN | JSY2424 | 7 | WT-Rep7 | TGT | G>C | G/C+1 | G/C-1 | CG>GC | CG>GC | AACTTTGTCA | GTCACCA | ACA | ACA | C | G | C>G | C>G_ACA | CG>GC_GTCACCA | 9 | FALSE | FALSE |
| 130_v.txt | 32200 | 32200-SNV-( | SNV | 1 | G | C | 10 | 31094 | 0.03216055 | 43.3 | WT | FALSE | CAN | JSY2424 | 7 | WT-Rep7 | GGA | G>C | G/C+1 | G/C-1 | CG>GC | CG>GC | TGTATGGAA | GAACACC | CCT | CCT | C | G | C>G | C>G_CCT | CG>GC_GAACACC | 9 | FALSE | FALSE |
| 130_v.txt | 32223 | 32223-SNV-( | SNV | 1 | G | C | 12 | 31094 | 0.03859266 | 37.5 | WT | FALSE | CAN | JSY2424 | 7 | WT-Rep7 | TGA | G>C | G/C+1 | G/C-1 | CG>GC | CG>GC | GTCCTTGACA | GACAGGA | ACT | ACT | C | G | C>G | C>G_ACT | CG>GC_GACAGGA | 9 | FALSE | FALSE |
| 130_v.txt | 32271 | 32271-SNV-( | SNV | 1 | C | G | 12 | 43326 | 0.027697 | 35.1666667 | WT | FALSE | CAN | JSY2424 | 7 | WT-Rep7 | ACG | C>G | G/C+1 | G/C-1 | CG>GC | CG>GC | AAAATACGG | CGGGAAC | TGC | ACG | C | C | C>G | C>G_ACG | CG>GC_CGGGAAC | 9 | FALSE | FALSE |
| 130_v.txt | 32278 | 32278-SNV-( | SNV | 1 | C | G | 71 | 43326 | 0.16387389 | 36.1549296 | WT | FALSE | CAN | JSY2424 | 7 | WT-Rep7 | CCA | C>G | G/C+1 | G/C-1 | CG>GC | CG>GC | GGGAACCAA | CAACGTA | GGT | CCA | C | C | C>G | C>G_CCA | CG>GC_CAACGTA | 9 | FALSE | FALSE |
| 130_v.txt | 32282 | 32282-SNV-( | SNV | 1 | G | C | 27 | 43326 | 0.06231824 | 39.5545189 | WT | FALSE | CAN | JSY2424 | 7 | WT-Rep7 | CGT | G>C | G/C+1 | G/C-1 | CG>GC | CG>GC | ACCAACGTA | GTAAATA | GCA | GCA | C | G | C>G | C>G_GCA | CG>GC_GTAAATA | 9 | FALSE | FALSE |
| 130_v.txt | 32292 | 32292-SNV-( | SNV | 1 | G | C | 3 | 43326 | 6.92E-03 | 55.3333333 | WT | FALSE | CAN | JSY2424 | 7 | WT-Rep7 | TGA | G>C | G/C+1 | G/C-1 | CG>GC | CG>GC | ATATTTGAAT | GAATTTG | ACT | ACT | C | G | C>G | C>G_ACT | CG>GC_GAATTTG | 9 | FALSE | FALSE |
| 130_v.txt | 32304 | 32304-SNV-( | SNV | 1 | G | C | 54 | 43326 | 0.12463648 | 54.9074074 | WT | FALSE | CAN | JSY2424 | 7 | WT-Rep7 | AGA | G>C | G/C+1 | G/C-1 | CG>GC | CG>GC | GCGGCAGAA | GAAATAA | TCT | TCT | C | G | C>G | C>G_TCT | CG>GC_GAAATAA | 9 | FALSE | FALSE |
| 130_v.txt | 32316 | 32316-SNV-( | SNV | 1 | G | C | 107 | 43326 | 0.24696487 | 59.6760675 | WT | FALSE | CAN | JSY2424 | 7 | WT-Rep7 | TGT | G>C | G/C+1 | G/C-1 | CG>GC | CG>GC | ATGGTTGTTA | GTTAAGA | ACA | ACA | C | G | C>G | C>G_ACA | CG>GC_GTTAAGA | 9 | FALSE | FALSE |
| 130_v.txt | 32326 | 32326-SNV-( | SNV | 1 | C | G | 1 | 43326 | 2.31E-03 | 56.9929268 | WT | FALSE | CAN | JSY2424 | 7 | WT-Rep7 | ACA | C>G | G/C+1 | G/C-1 | CG>GC | CG>GC | AGATAACAG | CAGCGTT | TGT | ACA | C | C | C>G | C>G_ACA | CG>GC_CAGCGTT | 9 | FALSE | FALSE |
| 130_v.txt | 32328 | 32328-SNV-( | SNV | 1 | G | C | 3 | 43326 | 6.92E-03 | 63.9999349 | WT | FALSE | CAN | JSY2424 | 7 | WT-Rep7 | AGC | G>C | G/C+1 | G/C-1 | CG>GC | CG>GC | ATAACAGCG | GCGTTGA | TCG | TCG | C | G | C>G | C>G_TCG | CG>GC_GCGTTGA | 9 | FALSE | FALSE |
| 130_v.txt | 32333 | 32333-SNV-( | SNV | 1 | G | C | 6 | 43326 | 0.0138485 | 46.330542 | WT | FALSE | CAN | JSY2424 | 7 | WT-Rep7 | TGA | G>C | G/C+1 | G/C-1 | CG>GC | CG>GC | AGCGTTGAA | GAAGATA | ACT | ACT | C | G | C>G | C>G_ACT | CG>GC_GAAGATA | 9 | FALSE | FALSE |
| 130_v.txt | 32399 | 32399-SNV-( | SNV | 1 | G | C | 23 | 43326 | 0.05308591 | 57.173913 | WT | FALSE | CAN | JSY2424 | 7 | WT-Rep7 | CGT | G>C | G/C+1 | G/C-1 | CG>GC | CG>GC | AGAAACGTA | GTAGGAA | GCA | GCA | C | G | C>G | C>G_GCA | CG>GC_GTAGGAA | 9 | FALSE | FALSE |
| 130_v.txt | 32432 | 32432-SNV-( | SNV | 1 | G | C | 14 | 43326 | 0.03231316 | 57.2857143 | WT | FALSE | CAN | JSY2424 | 7 | WT-Rep7 | TGT | G>C | G/C+1 | G/C-1 | CG>GC | CG>GC | GTCATTGTAT | GTATGGA | ACA | ACA | C | G | C>G | C>G_ACA | CG>GC_GTATGGA | 9 | FALSE | FALSE |
| 130_v.txt | 32449 | 32449-SNV-( | SNV | 1 | C | G | 8 | 43326 | 0.01846466 | 46.625 | WT | FALSE | CAN | JSY2424 | 7 | WT-Rep7 | CCA | C>G | G/C+1 | G/C-1 | CG>GC | CG>GC | AAAGTCCAA | CAATGAA | GGT | CCA | C | C | C>G | C>G_CCA | CG>GC_CAATGAA | 9 | FALSE | FALSE |
| 130_v.txt | 32467 | 32467-SNV-( | SNV | 1 | C | G | 9 | 43326 | 0.02077275 | 49.2168001 | WT | FALSE | CAN | JSY2424 | 7 | WT-Rep7 | CCA | C>G | G/C+1 | G/C-1 | CG>GC | CG>GC | GAGAGCCAA | CAATGTA | GGT | CCA | C | C | C>G | C>G_CCA | CG>GC_CAATGTA | 9 | FALSE | FALSE |
| 130_v.txt | 32487 | 32487-SNV-( | SNV | 1 | C | G | 3 | 43325 | 6.92E-03 | 50.6666667 | WT | FALSE | CAN | JSY2424 | 7 | WT-Rep7 | ACG | C>G | G/C+1 | G/C-1 | CG>GC | CG>GC | AAGATACGG | CGGAAAA | TGC | ACG | C | C | C>G | C>G_ACG | CG>GC_CGGAAAA | 9 | FALSE | FALSE |
| 130_v.txt | 32488 | 32488-SNV-( | SNV | 1 | G | C | 4 | 43326 | 9.23E-03 | 30.7500866 | WT | FALSE | CAN | JSY2424 | 7 | WT-Rep7 | CGG | G>C | G/C+1 | G/C-1 | CG>GC | CG>GC | AGATACGGA | GGAAAAC | GCC | GCC | C | G | C>G | C>G_GCC | CG>GC_GGAAAAC | 9 | FALSE | FALSE |
| 130_v.txt | 32535 | 32535-SNV-( | SNV | 1 | G | C | 5 | 43325 | 0.01154068 | 22.200287 | WT | FALSE | CAN | JSY2424 | 7 | WT-Rep7 | TGC | G>C | G/C+1 | G/C-1 | CG>GC | CG>GC | GGGTTTGCA | GCAGCTT | ACG | ACG | C | G | C>G | C>G_ACG | CG>GC_GCAGCTT | 9 | FALSE | FALSE |
| 130_v.txt | 32539 | 32539-SNV-( | SNV | 1 | C | G | 61 | 43325 | 0.14079631 | 37.0983607 | WT | FALSE | CAN | JSY2424 | 7 | WT-Rep7 | GCT | C>G | G/C+1 | G/C-1 | CG>GC | CG>GC | TTGCAGCTTC | CTTCACC | CGA | GCT | C | C | C>G | C>G_GCT | CG>GC_CTTCACC | 9 | FALSE | FALSE |
| 130_v.txt | 32542 | 32542-SNV-( | SNV | 1 | C | G | 2 | 43325 | 4.62E-03 | 37.9990237 | WT | FALSE | CAN | JSY2424 | 7 | WT-Rep7 | TCA | C>G | G/C+1 | G/C-1 | CG>GC | CG>GC | CAGCTTCACC | CACCAGC | AGT | TCA | C | C | C>G | C>G_TCA | CG>GC_CACCAGC | 9 | FALSE | FALSE |
| 130_v.txt | 32557 | 32557-SNV-( | SNV | 1 | C | G | 13 | 43311 | 0.03001547 | 37.5384615 | WT | FALSE | CAN | JSY2424 | 7 | WT-Rep7 | CCA | C>G | G/C+1 | G/C-1 | CG>GC | CG>GC | TGATACCAA | CAACTAG | GGT | CCA | C | C | C>G | C>G_CCA | CG>GC_CAACTAG | 9 | FALSE | FALSE |
| 130_v.txt | 32566 | 32566-SNV-( | SNV | 1 | C | G | 34 | 31633 | 0.10748269 | 35.7647059 | WT | FALSE | CAN | JSY2424 | 7 | WT-Rep7 | TCA | C>G | G/C+1 | G/C-1 | CG>GC | CG>GC | CTAGTTCAGT | CAGTACC | AGT | TCA | C | C | C>G | C>G_TCA | CG>GC_CAGTACC | 9 | FALSE | FALSE |
| 130_v.txt | 32679 | 32679-SNV-( | SNV | 1 | C | G | 1 | 31687 | 3.16E-03 | 57 | WT | FALSE | CAN | JSY2424 | 7 | WT-Rep7 | CCA | C>G | G/C+1 | G/C-1 | CG>GC | CG>GC | TTTCTCCAAT | CAATAAC | GGT | CCA | C | C | C>G | C>G_CCA | CG>GC_CAATAAC | 9 | FALSE | FALSE |
| 130_v.txt | 32685 | 32685-SNV-( | SNV | 1 | C | G | 13 | 31688 | 0.04102499 | 63.3076923 | WT | FALSE | CAN | JSY2424 | 7 | WT-Rep7 | ACG | C>G | G/C+1 | G/C-1 | CG>GC | CG>GC | CAATAACGG | CGGAATC | TGC | ACG | C | C | C>G | C>G_ACG | CG>GC_CGGAATC | 9 | FALSE | FALSE |
| 130_v.txt | 32749 | 32749-SNV-( | SNV | 1 | C | G | 11 | 31688 | 0.03471346 | 56.2727273 | WT | FALSE | CAN | JSY2424 | 7 | WT-Rep7 | CCG | C>G | G/C+1 | G/C-1 | CG>GC | CG>GC | GAAACCCGA | CGATAAT | GGC | CCG | C | C | C>G | C>G_CCG | CG>GC_CGATAAT | 9 | FALSE | FALSE |
| 130_v.txt | 32786 | 32786-SNV-( | SNV | 1 | C | G | 15 | 31688 | 0.04733653 | 48.1317477 | WT | FALSE | CAN | JSY2424 | 7 | WT-Rep7 | ACT | C>G | G/C+1 | G/C-1 | CG>GC | CG>GC | CCAGAACTC | CTCGAAT | TGA | ACT | C | C | C>G | C>G_ACT | CG>GC_CTCGAAT | 9 | FALSE | FALSE |
| 130_v.txt | 32809 | 32809-SNV-( | SNV | 1 | C | G | 4 | 31688 | 0.01262308 | 42.5 | WT | FALSE | CAN | JSY2424 | 7 | WT-Rep7 | ACA | C>G | G/C+1 | G/C-1 | CG>GC | CG>GC | ATTTGACAG | CAGGGAA | TGT | ACA | C | C | C>G | C>G_ACA | CG>GC_CAGGGAA | 9 | FALSE | FALSE |
| 130_v.txt | 32819 | 32819-SNV-( | SNV | 1 | G | C | 93 | 31687 | 0.29349576 | 45.0967742 | WT | FALSE | CAN | JSY2424 | 7 | WT-Rep7 | AGT | G>C | G/C+1 | G/C-1 | CG>GC | CG>GC | GAACAAGTT | GTTCATT | TCA | TCA | C | G | C>G | C>G_TCA | CG>GC_GTTCATT | 9 | FALSE | FALSE |
| 130_v.txt | 32856 | 32856-SNV-( | SNV | 1 | C | G | 1 | 31688 | 3.16E-03 | 24.0000004 | WT | FALSE | CAN | JSY2424 | 7 | WT-Rep7 | CCA | C>G | G/C+1 | G/C-1 | CG>GC | CG>GC | CTAATCCATG | CATGCCG | GGT | CCA | C | C | C>G | C>G_CCA | CG>GC_CATGCCG | 9 | FALSE | FALSE |
| 130_v.txt | 32860 | 32860-SNV-( | SNV | 1 | C | G | 1 | 31686 | 3.16E-03 | 38 | WT | FALSE | CAN | JSY2424 | 7 | WT-Rep7 | GCC | C>G | G/C+1 | G/C-1 | CG>GC | CG>GC | TCCATGCCG | CCGCCAG | CGG | GCC | C | C | C>G | C>G_GCC | CG>GC_CCGCCAG | 9 | FALSE | FALSE |
| 130_v.txt | 32876 | 32876-SNV-( | SNV | 1 | G | C | 170 | 33843 | 0.50231953 | 37.1882353 | WT | FALSE | CAN | JSY2424 | 7 | WT-Rep7 | TGT | G>C | G/C+1 | G/C-1 | CG>GC | CG>GC | AACTTTGTAC | GTACGTC | ACA | ACA | C | G | C>G | C>G_ACA | CG>GC_GTACGTC | 9 | FALSE | FALSE |
| 130_v.txt | 32937 | 32937-SNV-( | SNV | 1 | C | G | 11 | 33860 | 0.03248671 | 55.9532134 | WT | FALSE | CAN | JSY2424 | 7 | WT-Rep7 | CCA | C>G | G/C+1 | G/C-1 | CG>GC | CG>GC | ATTGCCCAA | CAAGAAA | GGT | CCA | C | C | C>G | C>G_CCA | CG>GC_CAAGAAA | 9 | FALSE | FALSE |
| 130_v.txt | 32959 | 32959-SNV-( | SNV | 1 | C | G | 19 | 33860 | 0.05611341 | 56.2631579 | WT | FALSE | CAN | JSY2424 | 7 | WT-Rep7 | CCA | C>G | G/C+1 | G/C-1 | CG>GC | CG>GC | TGTAACCATT | CATTGGC | GGT | CCA | C | C | C>G | C>G_CCA | CG>GC_CATTGGC | 9 | FALSE | FALSE |
| 130_v.txt | 32997 | 32997-SNV-( | SNV | 1 | G | C | 16 | 33860 | 0.0472534 | 60.9375 | WT | FALSE | CAN | JSY2424 | 7 | WT-Rep7 | TGA | G>C | G/C+1 | G/C-1 | CG>GC | CG>GC | CTTTGTGAG | GAGAAAA | ACT | ACT | C | G | C>G | C>G_ACT | CG>GC_GAGAAAA | 9 | FALSE | FALSE |
| 130_v.txt | 33070 | 33070-SNV-( | SNV | 1 | C | G | 66 | 33860 | 0.19492026 | 56.5909091 | WT | FALSE | CAN | JSY2424 | 7 | WT-Rep7 | GCC | C>G | G/C+1 | G/C-1 | CG>GC | CG>GC | AATATGCCA | CCAAAGA | CGG | GCC | C | C | C>G | C>G_GCC | CG>GC_CCAAAGA | 9 | FALSE | FALSE |
| 130_v.txt | 33079 | 33079-SNV-( | SNV | 1 | C | G | 74 | 33860 | 0.21854696 | 53.4324324 | WT | FALSE | CAN | JSY2424 | 7 | WT-Rep7 | CCC | C>G | G/C+1 | G/C-1 | CG>GC | CG>GC | AAGAACCCA | CCATAAA | GGG | CCC | C | C | C>G | C>G_CCC | CG>GC_CCATAAA | 9 | FALSE | FALSE |
| 130_v.txt | 33111 | 33111-SNV-( | SNV | 1 | G | C | 17 | 33860 | 0.05020673 | 44.4117647 | WT | FALSE | CAN | JSY2424 | 7 | WT-Rep7 | TGG | G>C | G/C+1 | G/C-1 | CG>GC | CG>GC | CCCACTGGG | GGGCCGG | ACC | ACC | C | G | C>G | C>G_ACC | CG>GC_GGGCCGG | 9 | FALSE | FALSE |
| 130_v.txt | 33115 | 33115-SNV-( | SNV | 1 | C | G | 5 | 33860 | 0.01476669 | 48.1995927 | WT | FALSE | CAN | JSY2424 | 7 | WT-Rep7 | CCG | C>G | G/C+1 | G/C-1 | CG>GC | CG>GC | CTGGGCCGG | CGGCGTT | GGC | CCG | C | C | C>G | C>G_CCG | CG>GC_CGGCGTT | 9 | FALSE | FALSE |
| 130_v.txt | 33154 | 33154-SNV-( | SNV | 1 | C | G | 25 | 33859 | 0.07383561 | 37.519914 | WT | FALSE | CAN | JSY2424 | 7 | WT-Rep7 | CCT | C>G | G/C+1 | G/C-1 | CG>GC | CG>GC | AAAGACCTG | CTGTACC | GGA | CCT | C | C | C>G | C>G_CCT | CG>GC_CTGTACC | 9 | FALSE | FALSE |
| 130_v.txt | 33156 | 33156-SNV-( | SNV | 1 | G | C | 84 | 33843 | 0.24820495 | 37.1785698 | WT | FALSE | CAN | JSY2424 | 7 | WT-Rep7 | TGT | G>C | G/C+1 | G/C-1 | CG>GC | CG>GC | AGACCTGTA | GTACCAA | ACA | ACA | C | G | C>G | C>G_ACA | CG>GC_GTACCAA | 9 | FALSE | FALSE |
| 130_v.txt | 33160 | 33160-SNV-( | SNV | 1 | C | G | 11 | 33843 | 0.03250303 | 36.8181818 | WT | FALSE | CAN | JSY2424 | 7 | WT-Rep7 | CCA | C>G | G/C+1 | G/C-1 | CG>GC | CG>GC | CTGTACCAAT | CAATAGT | GGT | CCA | C | C | C>G | C>G_CCA | CG>GC_CAATAGT | 9 | FALSE | FALSE |
| 130_v.txt | 33168 | 33168-SNV-( | SNV | 1 | C | G | 16 | 33840 | 0.04728132 | 37.9374926 | WT | FALSE | CAN | JSY2424 | 7 | WT-Rep7 | ACC | C>G | G/C+1 | G/C-1 | CG>GC | CG>GC | ATAGTACCA | CCACCAA | TGG | ACC | C | C | C>G | C>G_ACC | CG>GC_CCACCAA | 9 | FALSE | FALSE |
| 130_v.txt | 33169 | 33169-SNV-( | SNV | 1 | C | G | 15 | 33840 | 0.04432624 | 36.0670521 | WT | FALSE | CAN | JSY2424 | 7 | WT-Rep7 | CCA | C>G | G/C+1 | G/C-1 | CG>GC | CG>GC | TAGTACCAC | CACCAAG | GGT | CCA | C | C | C>G | C>G_CCA | CG>GC_CACCAAG | 9 | FALSE | FALSE |
| 130_v.txt | 33172 | 33172-SNV-( | SNV | 1 | C | G | 20 | 33839 | 0.0591034 | 36.65 | WT | FALSE | CAN | JSY2424 | 7 | WT-Rep7 | CCA | C>G | G/C+1 | G/C-1 | CG>GC | CG>GC | TACCACCAA | CAAGGGC | GGT | CCA | C | C | C>G | C>G_CCA | CG>GC_CAAGGGC | 9 | FALSE | FALSE |
| 130_v.txt | 33193 | 33193-SNV-( | SNV | 1 | G | C | 240 | 32448 | 0.73964497 | 36.9083333 | WT | FALSE | CAN | JSY2424 | 7 | WT-Rep7 | TGT | G>C | G/C+1 | G/C-1 | CG>GC | CG>GC | CAATATGTCT | GTCTTTG | ACA | ACA | C | G | C>G | C>G_ACA | CG>GC_GTCTTTG | 9 | FALSE | FALSE |
| 130_v.txt | 33276 | 33276-SNV-( | SNV | 1 | G | C | 3 | 32448 | 9.25E-03 | 24.6681391 | WT | FALSE | CAN | JSY2424 | 7 | WT-Rep7 | CGT | G>C | G/C+1 | G/C-1 | CG>GC | CG>GC | GAGAACGTA | GTATCCT | GCA | GCA | C | G | C>G | C>G_GCA | CG>GC_GTATCCT | 9 | FALSE | FALSE |
| 130_v.txt | 33419 | 33419-SNV-( | SNV | 1 | G | C | 13 | 32448 | 0.0400641 | 47.7692308 | WT | FALSE | CAN | JSY2424 | 7 | WT-Rep7 | TGT | G>C | G/C+1 | G/C-1 | CG>GC | CG>GC | CTCATTGTAC | GTACATA | ACA | ACA | C | G | C>G | C>G_ACA | CG>GC_GTACATA | 9 | FALSE | FALSE |
| 130_v.txt | 33456 | 33456-SNV-( | SNV | 1 | G | C | 133 | 32448 | 0.40988659 | 42.3157895 | WT | FALSE | CAN | JSY2424 | 7 | WT-Rep7 | TGA | G>C | G/C+1 | G/C-1 | CG>GC | CG>GC | TCTTTTGAAT | GAATTTG | ACT | ACT | C | G | C>G | C>G_ACT | CG>GC_GAATTTG | 9 | FALSE | FALSE |
| 130_v.txt | 31869 | 31869-SNV-( | SNV | 1 | G | T | 20 | 11604 | 0.17235436 | 45.05 | WT | FALSE | CAN | JSY2424 | 7 | WT-Rep7 | AGA | G>T | A/T+1 | G/C-1 | CG>AT | CG>AT | AAAATAGAG | GAGATAT | TCT | TCT | C | A | C>A | C>A_TCT | CG>AT_GAGATAT | 9 | FALSE | FALSE |
| 130_v.txt | 31871 | 31871-SNV-( | SNV | 1 | G | T | 15 | 11604 | 0.12926577 | 45.1333333 | WT | FALSE | CAN | JSY2424 | 7 | WT-Rep7 | AGA | G>T | A/T+1 | G/C-1 | CG>AT | CG>AT | AATAGAGAT | GATATAG | TCT | TCT | C | A | C>A | C>A_TCT | CG>AT_GATATAG | 9 | FALSE | FALSE |
| 130_v.txt | 32058 | 32058-SNV-( | SNV | 1 | G | T | 21 | 31094 | 0.06753715 | 56.8912191 | WT | FALSE | CAN | JSY2424 | 7 | WT-Rep7 | CGA | G>T | A/T+1 | G/C-1 | CG>AT | CG>AT | ATGTGCGAG | GAGATTG | GCT | GCT | C | A | C>A | C>A_GCT | CG>AT_GAGATTG | 9 | FALSE | FALSE |
| 130_v.txt | 32088 | 32088-SNV-( | SNV | 1 | C | A | 21 | 31094 | 0.06753715 | 59.8095238 | WT | FALSE | CAN | JSY2424 | 7 | WT-Rep7 | GCC | C>A | A/T+1 | G/C-1 | CG>AT | CG>AT | AAAAAGCCT | CCTGCAA | CGG | GCC | C | T | C>A | C>A_GCC | CG>AT_CCTGCAA | 9 | FALSE | FALSE |
| 130_v.txt | 32095 | 32095-SNV-( | SNV | 1 | C | A | 24 | 31094 | 0.07718531 | 54.6666667 | WT | FALSE | CAN | JSY2424 | 7 | WT-Rep7 | ACA | C>A | A/T+1 | G/C-1 | CG>AT | CG>AT | CTGCAACAC | CACCAGT | TGT | ACA | C | T | C>A | C>A_ACA | CG>AT_CACCAGT | 9 | FALSE | FALSE |
| 130_v.txt | 32119 | 32119-SNV-( | SNV | 1 | C | A | 40 | 31094 | 0.12864218 | 59.075 | WT | FALSE | CAN | JSY2424 | 7 | WT-Rep7 | TCG | C>A | A/T+1 | G/C-1 | CG>AT | CG>AT | GCCATTCGA | CGAAAAC | AGC | TCG | C | T | C>A | C>A_TCG | CG>AT_CGAAAAC | 9 | FALSE | FALSE |
| 130_v.txt | 32153 | 32153-SNV-( | SNV | 1 | G | T | 12 | 31094 | 0.03859266 | 52.5833333 | WT | FALSE | CAN | JSY2424 | 7 | WT-Rep7 | TGT | G>T | A/T+1 | G/C-1 | CG>AT | CG>AT | CTCCATGTA | GTAAGCC | ACA | ACA | C | A | C>A | C>A_ACA | CG>AT_GTAAGCC | 9 | FALSE | FALSE |
| 130_v.txt | 32188 | 32188-SNV-( | SNV | 1 | C | A | 57 | 31094 | 0.18331511 | 41.4561404 | WT | FALSE | CAN | JSY2424 | 7 | WT-Rep7 | ACT | C>A | A/T+1 | G/C-1 | CG>AT | CG>AT | CGAAAACTG | CTGCAAT | TGA | ACT | C | T | C>A | C>A_ACT | CG>AT_CTGCAAT | 9 | FALSE | FALSE |
| 130_v.txt | 32203 | 32203-SNV-( | SNV | 1 | C | A | 16 | 31094 | 0.05145687 | 49.625 | WT | FALSE | CAN | JSY2424 | 7 | WT-Rep7 | ACA | C>A | A/T+1 | G/C-1 | CG>AT | CG>AT | ATGGAACAC | CACCACC | TGT | ACA | C | T | C>A | C>A_ACA | CG>AT_CACCACC | 9 | FALSE | FALSE |
| 130_v.txt | 32260 | 32260-SNV-( | SNV | 1 | C | A | 13 | 43327 | 0.03000439 | 20.8468543 | WT | FALSE | CAN | JSY2424 | 7 | WT-Rep7 | CCA | C>A | A/T+1 | G/C-1 | CG>AT | CG>AT | ATAGACCAA | CAAATAA | GGT | CCA | C | T | C>A | C>A_CCA | CG>AT_CAAATAA | 9 | FALSE | FALSE |
| 130_v.txt | 32271 | 32271-SNV-( | SNV | 1 | C | A | 57 | 43326 | 0.13156073 | 36.3684211 | WT | FALSE | CAN | JSY2424 | 7 | WT-Rep7 | ACG | C>A | A/T+1 | G/C-1 | CG>AT | CG>AT | AAAATACGG | CGGGAAC | TGC | ACG | C | T | C>A | C>A_ACG | CG>AT_CGGGAAC | 9 | FALSE | FALSE |
| 130_v.txt | 32274 | 32274-SNV-( | SNV | 1 | G | T | 18 | 43326 | 0.04154549 | 33.2777778 | WT | FALSE | CAN | JSY2424 | 7 | WT-Rep7 | GGA | G>T | A/T+1 | G/C-1 | CG>AT | CG>AT | ATACGGGAA | GAACCAA | CCT | CCT | C | A | C>A | C>A_CCT | CG>AT_GAACCAA | 9 | FALSE | FALSE |
| 130_v.txt | 32292 | 32292-SNV-( | SNV | 1 | G | T | 19 | 43326 | 0.04385358 | 33.7894737 | WT | FALSE | CAN | JSY2424 | 7 | WT-Rep7 | TGA | G>T | A/T+1 | G/C-1 | CG>AT | CG>AT | ATATTTGAAT | GAATTTG | ACT | ACT | C | A | C>A | C>A_ACT | CG>AT_GAATTTG | 9 | FALSE | FALSE |
| 130_v.txt | 32313 | 32313-SNV-( | SNV | 1 | G | T | 28 | 43326 | 0.06462632 | 47.3928571 | WT | FALSE | CAN | JSY2424 | 7 | WT-Rep7 | GGT | G>T | A/T+1 | G/C-1 | CG>AT | CG>AT | ATAATGGTT | GTTGTTA | CCA | CCA | C | A | C>A | C>A_CCA | CG>AT_GTTGTTA | 9 | FALSE | FALSE |
| 130_v.txt | 32329 | 32329-SNV-( | SNV | 1 | C | A | 12 | 43325 | 0.02769763 | 28.8333461 | WT | FALSE | CAN | JSY2424 | 7 | WT-Rep7 | GCG | C>A | A/T+1 | G/C-1 | CG>AT | CG>AT | TAACAGCGT | CGTTGAA | CGC | GCG | C | T | C>A | C>A_GCG | CG>AT_CGTTGAA | 9 | FALSE | FALSE |
| 130_v.txt | 32336 | 32336-SNV-( | SNV | 1 | G | T | 2 | 43326 | 4.62E-03 | 19.9993937 | WT | FALSE | CAN | JSY2424 | 7 | WT-Rep7 | AGA | G>T | A/T+1 | G/C-1 | CG>AT | CG>AT | GTTGAAGAT | GATATGT | TCT | TCT | C | A | C>A | C>A_TCT | CG>AT_GATATGT | 9 | FALSE | FALSE |
| 130_v.txt | 32368 | 32368-SNV-( | SNV | 1 | C | A | 50 | 43326 | 0.11540415 | 55.54 | WT | FALSE | CAN | JSY2424 | 7 | WT-Rep7 | TCA | C>A | A/T+1 | G/C-1 | CG>AT | CG>AT | AGTTCTCAAT | CAATAGC | AGT | TCA | C | T | C>A | C>A_TCA | CG>AT_CAATAGC | 9 | FALSE | FALSE |
| 130_v.txt | 32373 | 32373-SNV-( | SNV | 1 | G | T | 53 | 43326 | 0.12232839 | 61.4150943 | WT | FALSE | CAN | JSY2424 | 7 | WT-Rep7 | AGC | G>T | A/T+1 | G/C-1 | CG>AT | CG>AT | TCAATAGCA | GCAATAA | TCG | TCG | C | A | C>A | C>A_TCG | CG>AT_GCAATAA | 9 | FALSE | FALSE |
| 130_v.txt | 32399 | 32399-SNV-( | SNV | 1 | G | T | 255 | 43326 | 0.58856114 | 59.6588235 | WT | FALSE | CAN | JSY2424 | 7 | WT-Rep7 | CGT | G>T | A/T+1 | G/C-1 | CG>AT | CG>AT | AGAAACGTA | GTAGGAA | GCA | GCA | C | A | C>A | C>A_GCA | CG>AT_GTAGGAA | 9 | FALSE | FALSE |
| 130_v.txt | 32432 | 32432-SNV-( | SNV | 1 | G | T | 996 | 43326 | 2.29885058 | 56.5582329 | WT | FALSE | CAN | JSY2424 | 7 | WT-Rep7 | TGT | G>T | A/T+1 | G/C-1 | CG>AT | CG>AT | GTCATTGTAT | GTATGGA | ACA | ACA | C | A | C>A | C>A_ACA | CG>AT_GTATGGA | 9 | FALSE | FALSE |
| 130_v.txt | 32449 | 32449-SNV-( | SNV | 1 | C | A | 17 | 43326 | 0.03923741 | 46 | WT | FALSE | CAN | JSY2424 | 7 | WT-Rep7 | CCA | C>A | A/T+1 | G/C-1 | CG>AT | CG>AT | AAAGTCCAA | CAATGAA | GGT | CCA | C | T | C>A | C>A_CCA | CG>AT_CAATGAA | 9 | FALSE | FALSE |
| 130_v.txt | 32471 | 32471-SNV-( | SNV | 1 | G | T | 30 | 43326 | 0.06924249 | 44.5666667 | WT | FALSE | CAN | JSY2424 | 7 | WT-Rep7 | TGT | G>T | A/T+1 | G/C-1 | CG>AT | CG>AT | GCCAATGTA | GTAGAAG | ACA | ACA | C | A | C>A | C>A_ACA | CG>AT_GTAGAAG | 9 | FALSE | FALSE |
| 130_v.txt | 32487 | 32487-SNV-( | SNV | 1 | C | A | 18 | 43325 | 0.04154645 | 35.4444444 | WT | FALSE | CAN | JSY2424 | 7 | WT-Rep7 | ACG | C>A | A/T+1 | G/C-1 | CG>AT | CG>AT | AAGATACGG | CGGAAAA | TGC | ACG | C | T | C>A | C>A_ACG | CG>AT_CGGAAAA | 9 | FALSE | FALSE |
| 130_v.txt | 32488 | 32488-SNV-( | SNV | 1 | G | T | 36 | 43326 | 0.08309099 | 48.555374 | WT | FALSE | CAN | JSY2424 | 7 | WT-Rep7 | CGG | G>T | A/T+1 | G/C-1 | CG>AT | CG>AT | AGATACGGA | GGAAAAC | GCC | GCC | C | A | C>A | C>A_GCC | CG>AT_GGAAAAC | 9 | FALSE | FALSE |
| 130_v.txt | 32494 | 32494-SNV-( | SNV | 1 | C | A | 154 | 43326 | 0.35544477 | 45.0909091 | WT | FALSE | CAN | JSY2424 | 7 | WT-Rep7 | ACA | C>A | A/T+1 | G/C-1 | CG>AT | CG>AT | GGAAAACAA | CAACTTT | TGT | ACA | C | T | C>A | C>A_ACA | CG>AT_CAACTTT | 9 | FALSE | FALSE |
| 130_v.txt | 32508 | 32508-SNV-( | SNV | 1 | G | T | 9 | 43326 | 0.02077275 | 38.8888889 | WT | FALSE | CAN | JSY2424 | 7 | WT-Rep7 | GGC | G>T | A/T+1 | G/C-1 | CG>AT | CG>AT | TTGATGGCT | GCTCTTG | CCG | CCG | C | A | C>A | C>A_CCG | CG>AT_GCTCTTG | 9 | FALSE | FALSE |
| 130_v.txt | 32529 | 32529-SNV-( | SNV | 1 | G | T | 5 | 43326 | 0.01154042 | 47.7987494 | WT | FALSE | CAN | JSY2424 | 7 | WT-Rep7 | GGG | G>T | A/T+1 | G/C-1 | CG>AT | CG>AT | TTTCTGGGG | GGGTTTG | CCC | CCC | C | A | C>A | C>A_CCC | CG>AT_GGGTTTG | 9 | FALSE | FALSE |
| 130_v.txt | 32530 | 32530-SNV-( | SNV | 1 | G | T | 30 | 43326 | 0.06924249 | 31.4671618 | WT | FALSE | CAN | JSY2424 | 7 | WT-Rep7 | GGG | G>T | A/T+1 | G/C-1 | CG>AT | CG>AT | TTCTGGGGT | GGTTTGC | CCC | CCC | C | A | C>A | C>A_CCC | CG>AT_GGTTTGC | 9 | FALSE | FALSE |
| 130_v.txt | 32536 | 32536-SNV-( | SNV | 1 | C | A | 6 | 43325 | 0.01384882 | 31.6671613 | WT | FALSE | CAN | JSY2424 | 7 | WT-Rep7 | GCA | C>A | A/T+1 | G/C-1 | CG>AT | CG>AT | GGTTTGCAG | CAGCTTC | CGT | GCA | C | T | C>A | C>A_GCA | CG>AT_CAGCTTC | 9 | FALSE | FALSE |
| 130_v.txt | 32544 | 32544-SNV-( | SNV | 1 | C | A | 27 | 43325 | 0.06231968 | 36.8148148 | WT | FALSE | CAN | JSY2424 | 7 | WT-Rep7 | ACC | C>A | A/T+1 | G/C-1 | CG>AT | CG>AT | GCTTCACCAG | CCAGCAG | TGG | ACC | C | T | C>A | C>A_ACC | CG>AT_CCAGCAG | 9 | FALSE | FALSE |
| 130_v.txt | 32575 | 32575-SNV-( | SNV | 1 | G | T | 13 | 31689 | 0.0410237 | 36.1538462 | WT | FALSE | CAN | JSY2424 | 7 | WT-Rep7 | TGA | G>T | A/T+1 | G/C-1 | CG>AT | CG>AT | TACCTTGAA | GAAATGT | ACT | ACT | C | A | C>A | C>A_ACT | CG>AT_GAAATGT | 9 | FALSE | FALSE |
| 130_v.txt | 32629 | 32629-SNV-( | SNV | 1 | C | A | 55 | 31688 | 0.17356728 | 55.5454546 | WT | FALSE | CAN | JSY2424 | 7 | WT-Rep7 | TCG | C>A | A/T+1 | G/C-1 | CG>AT | CG>AT | TCCCTTCGTT | CGTTTTT | AGC | TCG | C | T | C>A | C>A_TCG | CG>AT_CGTTTTT | 9 | FALSE | FALSE |
| 130_v.txt | 32679 | 32679-SNV-( | SNV | 1 | C | A | 9 | 31687 | 0.02840282 | 32.2222222 | WT | FALSE | CAN | JSY2424 | 7 | WT-Rep7 | CCA | C>A | A/T+1 | G/C-1 | CG>AT | CG>AT | TTTCTCCAAT | CAATAAC | GGT | CCA | C | T | C>A | C>A_CCA | CG>AT_CAATAAC | 9 | FALSE | FALSE |
| 130_v.txt | 32692 | 32692-SNV-( | SNV | 1 | C | A | 132 | 31688 | 0.41656147 | 58.8257576 | WT | FALSE | CAN | JSY2424 | 7 | WT-Rep7 | CCA | C>A | A/T+1 | G/C-1 | CG>AT | CG>AT | GGAATCCAA | CAACTGG | GGT | CCA | C | T | C>A | C>A_CCA | CG>AT_CAACTGG | 9 | FALSE | FALSE |
| 130_v.txt | 32735 | 32735-SNV-( | SNV | 1 | G | T | 56 | 31688 | 0.17672305 | 56.6430854 | WT | FALSE | CAN | JSY2424 | 7 | WT-Rep7 | AGT | G>T | A/T+1 | G/C-1 | CG>AT | CG>AT | AAAACAGTA | GTATATT | TCA | TCA | C | A | C>A | C>A_TCA | CG>AT_GTATATT | 9 | FALSE | FALSE |
| 130_v.txt | 32794 | 32794-SNV-( | SNV | 1 | C | A | 57 | 31688 | 0.17987882 | 40.6140351 | WT | FALSE | CAN | JSY2424 | 7 | WT-Rep7 | TCA | C>A | A/T+1 | G/C-1 | CG>AT | CG>AT | CGAATTCAC | CACCGTA | AGT | TCA | C | T | C>A | C>A_TCA | CG>AT_CACCGTA | 9 | FALSE | FALSE |
| 130_v.txt | 32797 | 32797-SNV-( | SNV | 1 | C | A | 130 | 31688 | 0.41024994 | 46.4769231 | WT | FALSE | CAN | JSY2424 | 7 | WT-Rep7 | CCG | C>A | A/T+1 | G/C-1 | CG>AT | CG>AT | ATTCACCGTA | CGTAATA | GGC | CCG | C | T | C>A | C>A_CCG | CG>AT_CGTAATA | 9 | FALSE | FALSE |
| 130_v.txt | 32798 | 32798-SNV-( | SNV | 1 | G | T | 5 | 31688 | 0.01577884 | 37.4 | WT | FALSE | CAN | JSY2424 | 7 | WT-Rep7 | CGT | G>T | A/T+1 | G/C-1 | CG>AT | CG>AT | TTCACCGTA | GTAATAT | GCA | GCA | C | A | C>A | C>A_GCA | CG>AT_GTAATAT | 9 | FALSE | FALSE |
| 130_v.txt | 32811 | 32811-SNV-( | SNV | 1 | G | T | 32 | 31688 | 0.1009846 | 41.78125 | WT | FALSE | CAN | JSY2424 | 7 | WT-Rep7 | AGG | G>T | A/T+1 | G/C-1 | CG>AT | CG>AT | TTGACAGGG | GGGAACA | TCC | TCC | C | A | C>A | C>A_TCC | CG>AT_GGGAACA | 9 | FALSE | FALSE |
| 130_v.txt | 32856 | 32856-SNV-( | SNV | 1 | C | A | 5 | 31688 | 0.01577884 | 35.7999331 | WT | FALSE | CAN | JSY2424 | 7 | WT-Rep7 | CCA | C>A | A/T+1 | G/C-1 | CG>AT | CG>AT | CTAATCCATG | CATGCCG | GGT | CCA | C | T | C>A | C>A_CCA | CG>AT_CATGCCG | 9 | FALSE | FALSE |
| 130_v.txt | 32859 | 32859-SNV-( | SNV | 1 | G | T | 2 | 31686 | 6.31E-03 | 22.0004767 | WT | FALSE | CAN | JSY2424 | 7 | WT-Rep7 | TGC | G>T | A/T+1 | G/C-1 | CG>AT | CG>AT | ATCCATGCC | GCCGCCA | ACG | ACG | C | A | C>A | C>A_ACG | CG>AT_GCCGCCA | 9 | FALSE | FALSE |
| 130_v.txt | 32860 | 32860-SNV-( | SNV | 1 | C | A | 4 | 31686 | 0.01262387 | 20 | WT | FALSE | CAN | JSY2424 | 7 | WT-Rep7 | GCC | C>A | A/T+1 | G/C-1 | CG>AT | CG>AT | TCCATGCCG | CCGCCAG | CGG | GCC | C | T | C>A | C>A_GCC | CG>AT_CCGCCAG | 9 | FALSE | FALSE |
| 130_v.txt | 32876 | 32876-SNV-( | SNV | 1 | G | T | 16 | 33843 | 0.04727713 | 28.5625 | WT | FALSE | CAN | JSY2424 | 7 | WT-Rep7 | TGT | G>T | A/T+1 | G/C-1 | CG>AT | CG>AT | AACTTTGTAC | GTACGTC | ACA | ACA | C | A | C>A | C>A_ACA | CG>AT_GTACGTC | 9 | FALSE | FALSE |
| 130_v.txt | 32917 | 32917-SNV-( | SNV | 1 | C | A | 27 | 33860 | 0.07974011 | 54.2039039 | WT | FALSE | CAN | JSY2424 | 7 | WT-Rep7 | TCC | C>A | A/T+1 | G/C-1 | CG>AT | CG>AT | TAAGTTCCA | CCAGGGC | AGG | TCC | C | T | C>A | C>A_TCC | CG>AT_CCAGGGC | 9 | FALSE | FALSE |
| 130_v.txt | 32937 | 32937-SNV-( | SNV | 1 | C | A | 24 | 33860 | 0.0708801 | 48.239044 | WT | FALSE | CAN | JSY2424 | 7 | WT-Rep7 | CCA | C>A | A/T+1 | G/C-1 | CG>AT | CG>AT | ATTGCCCAA | CAAGAAA | GGT | CCA | C | T | C>A | C>A_CCA | CG>AT_CAAGAAA | 9 | FALSE | FALSE |
| 130_v.txt | 32940 | 32940-SNV-( | SNV | 1 | G | T | 57 | 33860 | 0.16834023 | 45.5573321 | WT | FALSE | CAN | JSY2424 | 7 | WT-Rep7 | AGA | G>T | A/T+1 | G/C-1 | CG>AT | CG>AT | GCCCAAGAA | GAAAACC | TCT | TCT | C | A | C>A | C>A_TCT | CG>AT_GAAAACC | 9 | FALSE | FALSE |
| 130_v.txt | 32970 | 32970-SNV-( | SNV | 1 | C | A | 106 | 33860 | 0.31305375 | 61.1226415 | WT | FALSE | CAN | JSY2424 | 7 | WT-Rep7 | ACC | C>A | A/T+1 | G/C-1 | CG>AT | CG>AT | GCCGCACCA | CCAAATG | TGG | ACC | C | T | C>A | C>A_ACC | CG>AT_CCAAATG | 9 | FALSE | FALSE |
| 130_v.txt | 33006 | 33006-SNV-( | SNV | 1 | G | T | 47 | 33860 | 0.13880685 | 59.5106383 | WT | FALSE | CAN | JSY2424 | 7 | WT-Rep7 | TGT | G>T | A/T+1 | G/C-1 | CG>AT | CG>AT | AAAACTGTG | GTGAAAG | ACA | ACA | C | A | C>A | C>A_ACA | CG>AT_GTGAAAG | 9 | FALSE | FALSE |
| 130_v.txt | 33015 | 33015-SNV-( | SNV | 1 | G | T | 73 | 33860 | 0.21559362 | 59.2191781 | WT | FALSE | CAN | JSY2424 | 7 | WT-Rep7 | GGA | G>T | A/T+1 | G/C-1 | CG>AT | CG>AT | AAAGAGGAT | GATGTAA | CCT | CCT | C | A | C>A | C>A_CCT | CG>AT_GATGTAA | 9 | FALSE | FALSE |
| 130_v.txt | 33018 | 33018-SNV-( | SNV | 1 | G | T | 28 | 33860 | 0.08269344 | 59.6785714 | WT | FALSE | CAN | JSY2424 | 7 | WT-Rep7 | TGT | G>T | A/T+1 | G/C-1 | CG>AT | CG>AT | GAGGATGTA | GTAACAG | ACA | ACA | C | A | C>A | C>A_ACA | CG>AT_GTAACAG | 9 | FALSE | FALSE |
| 130_v.txt | 33024 | 33024-SNV-( | SNV | 1 | G | T | 26 | 33860 | 0.07678677 | 60.4615385 | WT | FALSE | CAN | JSY2424 | 7 | WT-Rep7 | AGG | G>T | A/T+1 | G/C-1 | CG>AT | CG>AT | GTAACAGGG | GGGATGA | TCC | TCC | C | A | C>A | C>A_TCC | CG>AT_GGGATGA | 9 | FALSE | FALSE |
| 130_v.txt | 33051 | 33051-SNV-( | SNV | 1 | G | T | 17 | 33860 | 0.05020673 | 56 | WT | FALSE | CAN | JSY2424 | 7 | WT-Rep7 | GGA | G>T | A/T+1 | G/C-1 | CG>AT | CG>AT | CCCAAGGAC | GACTGCG | CCT | CCT | C | A | C>A | C>A_CCT | CG>AT_GACTGCG | 9 | FALSE | FALSE |
| 130_v.txt | 33114 | 33114-SNV-( | SNV | 1 | C | A | 106 | 33860 | 0.31305375 | 45.0362364 | WT | FALSE | CAN | JSY2424 | 7 | WT-Rep7 | GCC | C>A | A/T+1 | G/C-1 | CG>AT | CG>AT | ACTGGGCCG | CCGGCGT | CGG | GCC | C | T | C>A | C>A_GCC | CG>AT_CCGGCGT | 9 | FALSE | FALSE |
| 130_v.txt | 33151 | 33151-SNV-( | SNV | 1 | G | T | 63 | 33859 | 0.18606574 | 39.2857143 | WT | FALSE | CAN | JSY2424 | 7 | WT-Rep7 | AGA | G>T | A/T+1 | G/C-1 | CG>AT | CG>AT | TGAAAAGAC | GACCTGT | TCT | TCT | C | A | C>A | C>A_TCT | CG>AT_GACCTGT | 9 | FALSE | FALSE |
| 130_v.txt | 33154 | 33154-SNV-( | SNV | 1 | C | A | 7 | 33859 | 0.02067397 | 24.8575724 | WT | FALSE | CAN | JSY2424 | 7 | WT-Rep7 | CCT | C>A | A/T+1 | G/C-1 | CG>AT | CG>AT | AAAGACCTG | CTGTACC | GGA | CCT | C | T | C>A | C>A_CCT | CG>AT_CTGTACC | 9 | FALSE | FALSE |
| 130_v.txt | 33156 | 33156-SNV-( | SNV | 1 | G | T | 14 | 33843 | 0.04136749 | 37.4999935 | WT | FALSE | CAN | JSY2424 | 7 | WT-Rep7 | TGT | G>T | A/T+1 | G/C-1 | CG>AT | CG>AT | AGACCTGTA | GTACCAA | ACA | ACA | C | A | C>A | C>A_ACA | CG>AT_GTACCAA | 9 | FALSE | FALSE |
| 130_v.txt | 33168 | 33168-SNV-( | SNV | 1 | C | A | 9 | 33840 | 0.02659575 | 36.1110572 | WT | FALSE | CAN | JSY2424 | 7 | WT-Rep7 | ACC | C>A | A/T+1 | G/C-1 | CG>AT | CG>AT | ATAGTACCA | CCACCAA | TGG | ACC | C | T | C>A | C>A_ACC | CG>AT_CCACCAA | 9 | FALSE | FALSE |
| 130_v.txt | 33193 | 33193-SNV-( | SNV | 1 | G | T | 31 | 32448 | 0.09553748 | 33.1612903 | WT | FALSE | CAN | JSY2424 | 7 | WT-Rep7 | TGT | G>T | A/T+1 | G/C-1 | CG>AT | CG>AT | CAATATGTCT | GTCTTTG | ACA | ACA | C | A | C>A | C>A_ACA | CG>AT_GTCTTTG | 9 | FALSE | FALSE |
| 130_v.txt | 33208 | 33208-SNV-( | SNV | 1 | C | A | 33 | 32448 | 0.10170118 | 36.9676092 | WT | FALSE | CAN | JSY2424 | 7 | WT-Rep7 | TCT | C>A | A/T+1 | G/C-1 | CG>AT | CG>AT | TAAGCTCTCT | CTCTCTT | AGA | TCT | C | T | C>A | C>A_TCT | CG>AT_CTCTCTT | 9 | FALSE | FALSE |
| 130_v.txt | 33238 | 33238-SNV-( | SNV | 1 | C | A | 92 | 32448 | 0.28353057 | 38.076087 | WT | FALSE | CAN | JSY2424 | 7 | WT-Rep7 | CCT | C>A | A/T+1 | G/C-1 | CG>AT | CG>AT | CTTCTCCTTC | CTTCATC | GGA | CCT | C | T | C>A | C>A_CCT | CG>AT_CTTCATC | 9 | FALSE | FALSE |
| 130_v.txt | 33309 | 33309-SNV-( | SNV | 1 | C | A | 8 | 32448 | 0.02465483 | 19.6247199 | WT | FALSE | CAN | JSY2424 | 7 | WT-Rep7 | GCG | C>A | A/T+1 | G/C-1 | CG>AT | CG>AT | AAAGAGCGC | CGCAATG | CGC | GCG | C | T | C>A | C>A_GCG | CG>AT_CGCAATG | 9 | FALSE | FALSE |
| 130_v.txt | 33310 | 33310-SNV-( | SNV | 1 | G | T | 28 | 32442 | 0.08630787 | 11.7886755 | WT | FALSE | CAN | JSY2424 | 7 | WT-Rep7 | CGC | G>T | A/T+1 | G/C-1 | CG>AT | CG>AT | AAGAGCGCA | GCAATGG | GCG | GCG | C | A | C>A | C>A_GCG | CG>AT_GCAATGG | 9 | FALSE | FALSE |
| 130_v.txt | 33325 | 33325-SNV-( | SNV | 1 | C | A | 34 | 32448 | 0.10478304 | 57.0882353 | WT | FALSE | CAN | JSY2424 | 7 | WT-Rep7 | TCT | C>A | A/T+1 | G/C-1 | CG>AT | CG>AT | ACAATTCTTT | CTTTACT | AGA | TCT | C | T | C>A | C>A_TCT | CG>AT_CTTTACT | 9 | FALSE | FALSE |
| 130_v.txt | 33337 | 33337-SNV-( | SNV | 1 | C | A | 23 | 32448 | 0.07088264 | 44.8695652 | WT | FALSE | CAN | JSY2424 | 7 | WT-Rep7 | TCA | C>A | A/T+1 | G/C-1 | CG>AT | CG>AT | TTTTCTCATC | CATCTTT | AGT | TCA | C | T | C>A | C>A_TCA | CG>AT_CATCTTT | 9 | FALSE | FALSE |
| 130_v.txt | 33354 | 33354-SNV-( | SNV | 1 | G | T | 150 | 32448 | 0.46227811 | 17.3533333 | WT | FALSE | CAN | JSY2424 | 7 | WT-Rep7 | TGA | G>T | A/T+1 | G/C-1 | CG>AT | CG>AT | GGTATTGAC | GACCCAC | ACT | ACT | C | A | C>A | C>A_ACT | CG>AT_GACCCAC | 9 | FALSE | FALSE |
| 130_v.txt | 33415 | 33415-SNV-( | SNV | 1 | C | A | 48 | 32448 | 0.14792899 | 41.6458333 | WT | FALSE | CAN | JSY2424 | 7 | WT-Rep7 | TCA | C>A | A/T+1 | G/C-1 | CG>AT | CG>AT | CCGGCTCATT | CATTGTA | AGT | TCA | C | T | C>A | C>A_TCA | CG>AT_CATTGTA | 9 | FALSE | FALSE |

| | | | | | | | | | | | | | | | | | | | | | | | | | | | | | | | | | | | |
|---|---|---|---|---|---|---|---|---|---|---|---|---|---|---|---|---|---|---|---|---|---|---|---|---|---|---|---|---|---|---|---|---|---|---|---|
| 130_v.txt | 33419 | 33419-SNV-( | SNV | 1 | G | T | 18 | 32448 | 0.05547337 | 40.8888889 | WT | | FALSE | CAN | JSY2424 | 7 | WT-Rep7 | TGT | G>T | A/T+1 | G/C-1 | CG>AT | CG>AT | CTCATTGTAC | GTACATA | ACA | ACA | C | A | C>A | C>A_ACA | CG>AT_GTACATA | 9 | FALSE | FALSE |
| 130_v.txt | 33436 | 33436-SNV-( | SNV | 1 | C | A | 19 | 32448 | 0.05855523 | 40.5263158 | WT | | FALSE | CAN | JSY2424 | 7 | WT-Rep7 | TCT | C>A | A/T+1 | G/C-1 | CG>AT | CG>AT | TCTCCTCTAT | CTATGTC | AGA | TCT | C | T | C>A | C>A_TCT | CG>AT_CTATGTC | 9 | FALSE | FALSE |
| 130_v.txt | 33474 | 33474-SNV-( | SNV | 1 | C | A | 67 | 32448 | 0.20648422 | 34.9054172 | WT | | FALSE | CAN | JSY2424 | 7 | WT-Rep7 | GCC | C>A | A/T+1 | G/C-1 | CG>AT | CG>AT | GCTATGCCTT | CCTTTTT | CGG | GCC | C | T | C>A | C>A_GCC | CG>AT_CCTTTTT | 9 | FALSE | FALSE |
| 130_v.txt | 33475 | 33475-SNV-( | SNV | 1 | C | A | 17 | 32448 | 0.05239152 | 34.7647059 | WT | | FALSE | CAN | JSY2424 | 7 | WT-Rep7 | CCT | C>A | A/T+1 | G/C-1 | CG>AT | CG>AT | CTATGCCTTT | CTTTTTT | GGA | CCT | C | T | C>A | C>A_CCT | CG>AT_CTTTTTT | 9 | FALSE | FALSE |
| 130_v.txt | 32033 | 32033-Repla | Replacement | 2 | C | AA | 9 | 31094 | 0.02894449 | 61.1110697 | WT | | FALSE | CAN | JSY2424 | 7 | WT-Rep7 | TCA | nope | >1bp-ins | G/C-1 | none | Replac. | GTATTTCAAA | CAAAGCT | AGT | TCA | C | none | nope | nope_TCA | Replac._CAAAGCT | 9 | FALSE | FALSE |
| 130_v.txt | 32489 | 32489-Repla | Replacement | 2 | G | AA | 5 | 43326 | 0.01154042 | 38 | WT | | FALSE | CAN | JSY2424 | 7 | WT-Rep7 | GGA | nope | >1bp-ins | G/C-1 | none | Replac. | GATACGGAA | GAAAACA | CCT | CCT | C | none | C>none | C>none_CCT | Replac._GAAAACA | 9 | FALSE | FALSE |
| 130_v.txt | 32489 | 32489-Repla | Replacement | 2 | G | CA | 72 | 43326 | 0.16618197 | 46.2361008 | WT | | FALSE | CAN | JSY2424 | 7 | WT-Rep7 | GGA | nope | >1bp-ins | G/C-1 | none | Replac. | GATACGGAA | GAAAACA | CCT | CCT | C | none | C>none | C>none_CCT | Replac._GAAAACA | 9 | FALSE | FALSE |
| 130_v.txt | 32940 | 32940-Repla | Replacement | 2 | GA | C | 6 | 33860 | 0.01772002 | 62.8333333 | WT | | FALSE | CAN | JSY2424 | 7 | WT-Rep7 | AGA | nope | G/C+1 | >1bpdel | none | Replac. | GCCCAAGAA | GAAAACC | TCT | AGA | GA | G | NA | NA_AGA | Replac._GAAAACC | 9 | FALSE | FALSE |
| 130_v.txt | 33114 | 33114-Repla | Replacement | 2 | CC | T | 1 | 33860 | 2.95E-03 | 38 | WT | | FALSE | CAN | JSY2424 | 7 | WT-Rep7 | GCC | nope | A/T+1 | >1bpdel | none | Replac. | ACTGGGCCG | CCGGCGT | CGG | GCC | CC | A | NA | NA_GCC | Replac._CCGGCGT | 9 | FALSE | FALSE |
| 130_v.txt | 32190 | 32190-MNV- | MNV | 2 | GC | CT | 23 | 31094 | 0.07396926 | 43.2173912 | WT | | FALSE | CAN | JSY2424 | 7 | WT-Rep7 | TGC | nope | >1bp-ins | >1bpdel | none | MNV | AAAACTGCA | GCAATGT | ACG | TGC | GC | none | NA | NA_TGC | MNV_GCAATGT | 9 | FALSE | FALSE |
| 130_v.txt | 32259 | 32259-MNV- | MNV | 2 | CC | AA | 15 | 43327 | 0.03462045 | 37.0999821 | WT | | FALSE | CAN | JSY2424 | 7 | WT-Rep7 | ACC | nope | >1bp-ins | >1bpdel | none | MNV | GATAGACCA | CCAAATA | TGG | ACC | CC | none | NA | NA_ACC | MNV_CCAAATA | 9 | FALSE | FALSE |
| 130_v.txt | 32316 | 32316-MNV- | MNV | 2 | GT | TA | 20 | 43326 | 0.04616166 | 60.5001633 | WT | | FALSE | CAN | JSY2424 | 7 | WT-Rep7 | TGT | nope | >1bp-ins | >1bpdel | none | MNV | ATGGTTGTTA | GTTAAGA | ACA | TGT | GT | none | NA | NA_TGT | MNV_GTTAAGA | 9 | FALSE | FALSE |
| 130_v.txt | 32541 | 32541-MNV- | MNV | 2 | TC | GA | 3 | 43325 | 6.92E-03 | 30.0000568 | WT | | FALSE | CAN | JSY2424 | 7 | WT-Rep7 | TTC | nope | >1bp-ins | >1bpdel | none | MNV | GCAGCTTCA | TCACCAG | AAG | TTC | TC | none | NA | NA_TTC | MNV_TCACCAG | 9 | FALSE | FALSE |
| 130_v.txt | 32787 | 32787-MNV- | MNV | 2 | TC | AA | 37 | 31688 | 0.11676344 | 46.3361058 | WT | | FALSE | CAN | JSY2424 | 7 | WT-Rep7 | CTC | nope | >1bp-ins | >1bpdel | none | MNV | CAGAACTCGA | TCGAATT | GAG | CTC | TC | none | NA | NA_CTC | MNV_TCGAATT | 9 | FALSE | FALSE |
| 130_v.txt | 33153 | 33153-MNV- | MNV | 2 | CC | TT | 1 | 33859 | 2.95E-03 | 37.9999494 | WT | | FALSE | CAN | JSY2424 | 7 | WT-Rep7 | ACC | nope | >1bp-ins | >1bpdel | none | MNV | AAAAGACCT | CCTGTAC | TGG | ACC | CC | none | NA | NA_ACC | MNV_CCTGTAC | 9 | FALSE | FALSE |
| 130_v.txt | 33168 | 33168-MNV- | MNV | 2 | CC | AA | 1 | 33840 | 2.96E-03 | 35.0000504 | WT | | FALSE | CAN | JSY2424 | 7 | WT-Rep7 | ACC | nope | >1bp-ins | >1bpdel | none | MNV | ATAGTACCA | CCACCAA | TGG | ACC | CC | none | NA | NA_ACC | MNV_CCACCAA | 9 | FALSE | FALSE |
| 130_v.txt | 31904 | 31904-Insert | Insertion | 2 | - | TT | 6 | 11585 | 0.05179111 | 37.3333333 | WT | | FALSE | CAN | JSY2424 | 7 | WT-Rep7 | ATT | nope | >1bp-ins | >1bpdel | none | >1bp-ins | ATTGAATTTT | TTTTGGT | TAA | ATT | - | none | NA | NA_ATT | >1bp-ins_TTTTGGT | 9 | FALSE | FALSE |
| 130_v.txt | 31920 | 31920-Insert | Insertion | 2 | - | GT | 4 | 11584 | 0.03453039 | 37.25 | WT | | FALSE | CAN | JSY2424 | 7 | WT-Rep7 | CGT | nope | >1bp-ins | >1bpdel | none | >1bp-ins | AAAGCCGTG | GTGAAAC | GCA | CGT | - | none | NA | NA_CGT | >1bp-ins_GTGAAAC | 9 | FALSE | FALSE |
| 130_v.txt | 31964 | 31964-Insert | Insertion | 3 | - | ATA | 19 | 31094 | 0.06110504 | 36.9649123 | WT | | FALSE | CAN | JSY2424 | 7 | WT-Rep7 | CAT | nope | >1bp-ins | >1bpdel | none | >1bp-ins | GGCCGCATA | ATAATAA | GTA | CAT | - | none | NA | NA_CAT | >1bp-ins_ATAATAA | 9 | FALSE | FALSE |
| 130_v.txt | 32069 | 32069-Insert | Insertion | 1 | - | A | 48 | 31094 | 0.15437062 | 63.0416667 | WT | | FALSE | CAN | JSY2424 | 7 | WT-Rep7 | TAA | nope | A/T+1 | >1bpdel | none | A/T+1 | TGAGATAAA | AAATAAC | ATT | TAA | - | T | NA | NA_TAA | A/T+1_AAATAAC | 9 | FALSE | FALSE |
| 130_v.txt | 32081 | 32081-Insert | Insertion | 1 | - | A | 22 | 31094 | 0.0707532 | 64 | WT | | FALSE | CAN | JSY2424 | 7 | WT-Rep7 | CAA | nope | A/T+1 | >1bpdel | none | A/T+1 | CCATGCAAA | AAAAAAG | GTT | CAA | - | T | NA | NA_CAA | A/T+1_AAAAAAG | 9 | FALSE | FALSE |
| 130_v.txt | 32265 | 32265-Insert | Insertion | 1 | - | A | 3 | 43326 | 6.92E-03 | 36.3333333 | WT | | FALSE | CAN | JSY2424 | 7 | WT-Rep7 | TAA | nope | A/T+1 | >1bpdel | none | A/T+1 | CCAAATAAA | AAAATAC | ATT | TAA | - | T | NA | NA_TAA | A/T+1_AAAATAC | 9 | FALSE | FALSE |
| 130_v.txt | 32284 | 32284-Insert | Insertion | 1 | - | A | 1 | 43326 | 2.31E-03 | 64 | WT | | FALSE | CAN | JSY2424 | 7 | WT-Rep7 | TAA | nope | A/T+1 | >1bpdel | none | A/T+1 | CAACGTAAA | AAATATT | ATT | TAA | - | T | NA | NA_TAA | A/T+1_AAATATT | 9 | FALSE | FALSE |
| 130_v.txt | 32309 | 32309-Insert | Insertion | 1 | - | A | 20 | 43326 | 0.04616166 | 59.75 | WT | | FALSE | CAN | JSY2424 | 7 | WT-Rep7 | TAA | nope | A/T+1 | >1bpdel | none | A/T+1 | AGAAATAAT | AATGGTT | ATT | TAA | - | T | NA | NA_TAA | A/T+1_AATGGTT | 9 | FALSE | FALSE |
| 130_v.txt | 32324 | 32324-Insert | Insertion | 1 | - | A | 1 | 43326 | 2.31E-03 | 63.9999998 | WT | | FALSE | CAN | JSY2424 | 7 | WT-Rep7 | TAA | nope | A/T+1 | >1bpdel | none | A/T+1 | TAAGATAAC | AACAGCG | ATT | TAA | - | T | NA | NA_TAA | A/T+1_AACAGCG | 9 | FALSE | FALSE |
| 130_v.txt | 32448 | 32448-Insert | Insertion | 18 | - | CCAATGAAT | 20 | 43326 | 0.04616166 | 48.8583333 | WT | | FALSE | CAN | JSY2424 | 7 | WT-Rep7 | TCC | nope | >1bp-ins | >1bpdel | none | >1bp-ins | AAAAGTCCA | CCAATGA | AGG | TCC | - | none | NA | NA_TCC | >1bp-ins_CCAATGA | 9 | FALSE | FALSE |
| 130_v.txt | 32468 | 32468-Insert | Insertion | 1 | - | A | 1 | 43326 | 2.31E-03 | 38 | WT | | FALSE | CAN | JSY2424 | 7 | WT-Rep7 | CAA | nope | A/T+1 | >1bpdel | none | A/T+1 | AGAGCCAAT | AATGTAG | GTT | CAA | - | T | NA | NA_CAA | A/T+1_AATGTAG | 9 | FALSE | FALSE |
| 130_v.txt | 32490 | 32490-Insert | Insertion | 1 | - | A | 39 | 43326 | 0.09001523 | 45.1025641 | WT | | FALSE | CAN | JSY2424 | 7 | WT-Rep7 | GAA | nope | A/T+1 | >1bpdel | none | A/T+1 | ATACGGAAA | AAAACAA | CTT | GAA | - | T | NA | NA_GAA | A/T+1_AAAACAA | 9 | FALSE | FALSE |
| 130_v.txt | 32498 | 32498-Insert | Insertion | 1 | - | T | 18 | 43326 | 0.04154549 | 47.1666667 | WT | | FALSE | CAN | JSY2424 | 7 | WT-Rep7 | CTT | nope | A/T+1 | >1bpdel | none | A/T+1 | AACAACTTTT | TTTTTTG | GAA | CTT | - | A | NA | NA_CTT | A/T+1_TTTTTTG | 9 | FALSE | FALSE |
| 130_v.txt | 32619 | 32619-Insert | Insertion | 1 | - | A | 4 | 31688 | 0.01262308 | 44.5 | WT | | FALSE | CAN | JSY2424 | 7 | WT-Rep7 | GAA | nope | A/T+1 | >1bpdel | none | A/T+1 | CCTAAGAAC | AACCTCC | CTT | GAA | - | T | NA | NA_GAA | A/T+1_AACCTCC | 9 | FALSE | FALSE |
| 130_v.txt | 32631 | 32631-Insert | Insertion | 1 | - | T | 47 | 31688 | 0.14832113 | 50.7021277 | WT | | FALSE | CAN | JSY2424 | 7 | WT-Rep7 | GTT | nope | A/T+1 | >1bpdel | none | A/T+1 | CCTTCGTTTT | TTTTTAT | CAA | GTT | - | A | NA | NA_GTT | A/T+1_TTTTTAT | 9 | FALSE | FALSE |
| 130_v.txt | 32638 | 32638-Insert | Insertion | 1 | - | A | 4 | 31688 | 0.01262308 | 44.25 | WT | | FALSE | CAN | JSY2424 | 7 | WT-Rep7 | TCC | nope | A/T+1 | >1bpdel | none | A/T+1 | TTTTATCCTT | CCTTAGA | AGG | TCC | - | T | NA | NA_TCC | A/T+1_CCTTAGA | 9 | FALSE | FALSE |
| 130_v.txt | 32728 | 32728-Insert | Insertion | 1 | - | A | 9 | 31688 | 0.02840192 | 57.8888889 | WT | | FALSE | CAN | JSY2424 | 7 | WT-Rep7 | CAA | nope | A/T+1 | >1bpdel | none | A/T+1 | CCATACAAA | AAAAACA | GTT | CAA | - | T | NA | NA_CAA | A/T+1_AAAAACA | 9 | FALSE | FALSE |
| 130_v.txt | 32736 | 32736-Insert | Insertion | 2 | - | CA | 1 | 31688 | 3.16E-03 | 60.5 | WT | | FALSE | CAN | JSY2424 | 7 | WT-Rep7 | GTA | nope | >1bp-ins | >1bpdel | none | >1bp-ins | AAACAGTAT | TATATTA | CAT | GTA | - | none | NA | NA_GTA | >1bp-ins_TATATTA | 9 | FALSE | FALSE |
| 130_v.txt | 32736 | 32736-Insert | Insertion | 2 | - | TA | 21 | 31688 | 0.06627114 | 58.6428571 | WT | | FALSE | CAN | JSY2424 | 7 | WT-Rep7 | GTA | nope | >1bp-ins | >1bpdel | none | >1bp-ins | AAACAGTAT | TATATTA | CAT | GTA | - | none | NA | NA_GTA | >1bp-ins_TATATTA | 9 | FALSE | FALSE |
| 130_v.txt | 32760 | 32760-Insert | Insertion | 1 | - | A | 25 | 31688 | 0.07889422 | 54.92 | WT | | FALSE | CAN | JSY2424 | 7 | WT-Rep7 | TAA | nope | A/T+1 | >1bpdel | none | A/T+1 | ATGGCTAAA | AAAACTT | ATT | TAA | - | T | NA | NA_TAA | A/T+1_AAAACTT | 9 | FALSE | FALSE |
| 130_v.txt | 32842 | 32842-Insert | Insertion | 1 | - | A | 74 | 31687 | 0.23353426 | 44.3918919 | WT | | FALSE | CAN | JSY2424 | 7 | WT-Rep7 | CAA | nope | A/T+1 | >1bpdel | none | A/T+1 | TTACCCAAA | AAAAAAT | GTT | CAA | - | T | NA | NA_CAA | A/T+1_AAAAAAT | 9 | FALSE | FALSE |
| 130_v.txt | 33166 | 33166-Insert | Insertion | 4 | - | TACC | 13 | 33840 | 0.03841608 | 37.7884615 | WT | | FALSE | CAN | JSY2424 | 7 | WT-Rep7 | GTA | nope | >1bp-ins | >1bpdel | none | >1bp-ins | CAATAGTAC | TACCACC | CAT | GTA | - | none | NA | NA_GTA | >1bp-ins_TACCACC | 9 | FALSE | FALSE |
| 130_v.txt | 33236 | 33236-Insert | Insertion | 13 | - | TCCTTCATCT | 6 | 32448 | 0.01849112 | 38.9230769 | WT | | FALSE | CAN | JSY2424 | 7 | WT-Rep7 | CTC | nope | >1bp-ins | >1bpdel | none | >1bp-ins | TACTTCTCCT | TCCTTCA | GAG | CTC | - | none | NA | NA_CTC | >1bp-ins_TCCTTCA | 9 | FALSE | FALSE |
| 130_v.txt | 33475 | 33475-Insert | Insertion | 1 | - | T | 1 | 32448 | 3.08E-03 | 31 | WT | | FALSE | CAN | JSY2424 | 7 | WT-Rep7 | CCT | nope | A/T+1 | >1bpdel | none | A/T+1 | CTATGCCTTT | CTTTTTT | GGA | CCT | - | A | NA | NA_CCT | A/T+1_CTTTTTT | 9 | FALSE | FALSE |
| 130_v.txt | 32014 | 32014-Deleti | Deletion | 2 | GA | - | 9 | 31094 | 0.02894449 | 58 | WT | | FALSE | CAN | JSY2424 | 7 | WT-Rep7 | CGA | nope | >1bp-ins | >1bpdel | none | >1bpdel | CGTCACGAG | GAGAGAT | GCT | CGA | GA | none | NA | NA_CGA | >1bpdel_GAGAGAT | 9 | FALSE | FALSE |
| 130_v.txt | 32048 | 32048-Deleti | Deletion | 2 | TC | - | 31 | 31094 | 0.09969769 | 61.3548387 | WT | | FALSE | CAN | JSY2424 | 7 | WT-Rep7 | ATC | nope | >1bp-ins | >1bpdel | none | >1bpdel | CATAAATCT | TCTGATG | TAG | ATC | TC | none | NA | NA_ATC | >1bpdel_TCTGATG | 9 | FALSE | FALSE |
| 130_v.txt | 32053 | 32053-Deleti | Deletion | 2 | TG | - | 31 | 31094 | 0.09969769 | 61.8709677 | WT | | FALSE | CAN | JSY2424 | 7 | WT-Rep7 | ATG | nope | >1bp-ins | >1bpdel | none | >1bpdel | ATCTGATGT | TGTGCGA | TAC | ATG | TG | none | NA | NA_ATG | >1bpdel_TGTGCGA | 9 | FALSE | FALSE |
| 130_v.txt | 32128 | 32128-Deleti | Deletion | 2 | TG | - | 10 | 31094 | 0.03216055 | 54.1 | WT | | FALSE | CAN | JSY2424 | 7 | WT-Rep7 | TTG | nope | >1bp-ins | >1bpdel | none | >1bpdel | AAACTTTGT | TGTCACC | AAC | TTG | TG | none | NA | NA_TTG | >1bpdel_TGTCACC | 9 | FALSE | FALSE |
| 130_v.txt | 32316 | 32316-Deleti | Deletion | 2 | GT | - | 1 | 43326 | 2.31E-03 | 63.9999999 | WT | | FALSE | CAN | JSY2424 | 7 | WT-Rep7 | TGT | nope | >1bp-ins | >1bpdel | none | >1bpdel | ATGGTTGTTA | GTTAAGA | ACA | TGT | GT | none | NA | NA_TGT | >1bpdel_GTTAAGA | 9 | FALSE | FALSE |
| 130_v.txt | 32319 | 32319-Deleti | Deletion | 3 | AAG | - | 1 | 43326 | 2.31E-03 | 64 | WT | | FALSE | CAN | JSY2424 | 7 | WT-Rep7 | TAA | nope | >1bp-ins | >1bpdel | none | >1bpdel | GTTGTTAAGA | AAGATAA | ATT | TAA | AAG | none | NA | NA_TAA | >1bpdel_AAGATAA | 9 | FALSE | FALSE |
| 130_v.txt | 32319 | 32319-Deleti | Deletion | 15 | AAGATAACA | - | 110 | 43326 | 0.25388912 | 62.7909091 | WT | | FALSE | CAN | JSY2424 | 7 | WT-Rep7 | TAA | nope | >1bp-ins | >1bpdel | none | >1bpdel | GTTGTTAAGA | AAGATAA | ATT | TAA | AAGATAACA | none | NA | NA_TAA | >1bpdel_AAGATAA | 9 | FALSE | FALSE |
| 130_v.txt | 32324 | 32324-Deleti | Deletion | 7 | AACAGCG | - | 1 | 43326 | 2.31E-03 | 58 | WT | | FALSE | CAN | JSY2424 | 7 | WT-Rep7 | TAA | nope | >1bp-ins | >1bpdel | none | >1bpdel | TAAGATAAC | AACAGCG | ATT | TAA | AACAGCG | none | NA | NA_TAA | >1bpdel_AACAGCG | 9 | FALSE | FALSE |
| 130_v.txt | 32332 | 32332-Deleti | Deletion | 2 | TG | - | 1 | 43326 | 2.31E-03 | 64 | WT | | FALSE | CAN | JSY2424 | 7 | WT-Rep7 | TTG | nope | >1bp-ins | >1bpdel | none | >1bpdel | CAGCGTTGA | TGAAGAT | AAC | TTG | TG | none | NA | NA_TTG | >1bpdel_TGAAGAT | 9 | FALSE | FALSE |
| 130_v.txt | 32335 | 32335-Deleti | Deletion | 2 | AG | - | 25 | 43326 | 0.05770207 | 64 | WT | | FALSE | CAN | JSY2424 | 7 | WT-Rep7 | AAG | nope | >1bp-ins | >1bpdel | none | >1bpdel | CGTTGAAGA | AGATATG | TTC | AAG | AG | none | NA | NA_AAG | >1bpdel_AGATATG | 9 | FALSE | FALSE |
| 130_v.txt | 32337 | 32337-Deleti | Deletion | 2 | AT | - | 174 | 43326 | 0.40160643 | 61.9195402 | WT | | FALSE | CAN | JSY2424 | 7 | WT-Rep7 | GAT | nope | >1bp-ins | >1bpdel | none | >1bpdel | TTGAAGATA | ATATGTG | CTA | GAT | AT | none | NA | NA_GAT | >1bpdel_ATATGTG | 9 | FALSE | FALSE |
| 130_v.txt | 32407 | 32407-Deleti | Deletion | 2 | TA | - | 13 | 43326 | 0.03000508 | 61.6923077 | WT | | FALSE | CAN | JSY2424 | 7 | WT-Rep7 | GTA | nope | >1bp-ins | >1bpdel | none | >1bpdel | AGGAAGTAG | TAGATTG | CAT | GTA | TA | none | NA | NA_GTA | >1bpdel_TAGATTG | 9 | FALSE | FALSE |
| 130_v.txt | 32856 | 32856-Deleti | Deletion | 4 | CATG | - | 15 | 31688 | 0.04733653 | 37.6 | WT | | FALSE | CAN | JSY2424 | 7 | WT-Rep7 | CCA | nope | >1bp-ins | >1bpdel | none | >1bpdel | CTAATCCATG | CATGCCG | GGT | CCA | CATG | none | NA | NA_CCA | >1bpdel_CATGCCG | 9 | FALSE | FALSE |
| 130_v.txt | 32994 | 32994-Deleti | Deletion | 2 | TG | - | 35 | 33860 | 0.10336681 | 62.5714286 | WT | | FALSE | CAN | JSY2424 | 7 | WT-Rep7 | TTG | nope | >1bp-ins | >1bpdel | none | >1bpdel | AATCTTTGTG | TGTGAGA | AAC | TTG | TG | none | NA | NA_TTG | >1bpdel_TGTGAGA | 9 | FALSE | FALSE |
| 130_v.txt | 33065 | 33065-Deleti | Deletion | 2 | AT | - | 26 | 33860 | 0.07678677 | 56.8461548 | WT | | FALSE | CAN | JSY2424 | 7 | WT-Rep7 | AAT | nope | >1bp-ins | >1bpdel | none | >1bpdel | GACAGAATA | ATATGCC | TTA | AAT | AT | none | NA | NA_AAT | >1bpdel_ATATGCC | 9 | FALSE | FALSE |
| 130_v.txt | 33089 | 33089-Deleti | Deletion | 5 | ATATG | - | 58 | 33860 | 0.17129356 | 54.5689655 | WT | | FALSE | CAN | JSY2424 | 7 | WT-Rep7 | AAT | nope | >1bp-ins | >1bpdel | none | >1bpdel | AAATAAATA | ATATGAT | TTA | AAT | ATATG | none | NA | NA_AAT | >1bpdel_ATATGAT | 9 | FALSE | FALSE |
| 130_v.txt | 33092 | 33092-Deleti | Deletion | 4 | TGAT | - | 1 | 33860 | 2.95E-03 | 64 | WT | | FALSE | CAN | JSY2424 | 7 | WT-Rep7 | ATG | nope | >1bp-ins | >1bpdel | none | >1bpdel | TAAATATGA | TGATATA | TAC | ATG | TGAT | none | NA | NA_ATG | >1bpdel_TGATATA | 9 | FALSE | FALSE |
| 130_v.txt | 33094 | 33094-Deleti | Deletion | 2 | AT | - | 32 | 33860 | 0.09450679 | 52.5 | WT | | FALSE | CAN | JSY2424 | 7 | WT-Rep7 | GAT | nope | >1bp-ins | >1bpdel | none | >1bpdel | AATATGATA | ATATAAG | CTA | GAT | AT | none | NA | NA_GAT | >1bpdel_ATATAAG | 9 | FALSE | FALSE |
| 130_v.txt | 33167 | 33167-Deleti | Deletion | 3 | ACC | - | 15 | 33840 | 0.04432624 | 38 | WT | | FALSE | CAN | JSY2424 | 7 | WT-Rep7 | TAC | nope | >1bp-ins | >1bpdel | none | >1bpdel | AATAGTACC | ACCACCA | ATG | TAC | ACC | none | NA | NA_TAC | >1bpdel_ACCACCA | 9 | FALSE | FALSE |
| 130_v.txt | 33189 | 33189-Deleti | Deletion | 2 | AT | - | 19 | 32449 | 0.05855342 | 37.5789476 | WT | | FALSE | CAN | JSY2424 | 7 | WT-Rep7 | AAT | nope | >1bp-ins | >1bpdel | none | >1bpdel | ATACCAATA | ATATGTC | TTA | AAT | AT | none | NA | NA_AAT | >1bpdel_ATATGTC | 9 | FALSE | FALSE |
| 130_v.txt | 33206 | 33206-Deleti | Deletion | 2 | CT | - | 35 | 32448 | 0.10786489 | 37.6857143 | WT | | FALSE | CAN | JSY2424 | 7 | WT-Rep7 | GCT | nope | >1bp-ins | >1bpdel | none | >1bpdel | CTTAAGCTCT | CTCTCTC | CGA | GCT | CT | none | NA | NA_GCT | >1bpdel_CTCTCTC | 9 | FALSE | FALSE |
| 130_v.txt | 33474 | 33474-Deleti | Deletion | 2 | CC | - | 1 | 32448 | 3.08E-03 | 31 | WT | | FALSE | CAN | JSY2424 | 7 | WT-Rep7 | GCC | nope | >1bp-ins | >1bpdel | none | >1bpdel | GCTATGCCTT | CCTTTTT | CGG | GCC | CC | none | NA | NA_GCC | >1bpdel_CCTTTTT | 9 | FALSE | FALSE |
| 130_v.txt | 31663 | 31663-Deleti | Deletion | 1 | A | - | 5 | 11604 | 0.04308859 | 34.2 | WT | | FALSE | CAN | JSY2424 | 7 | WT-Rep7 | GAA | nope | >1bp-ins | A/T-1 | none | A/T-1 | GCGTGGAAA | AAATGTG | CTT | CTT | T | none | T>none | T>none_CTT | A/T-1_AAATGTG | 9 | FALSE | FALSE |
| 130_v.txt | 31723 | 31723-Deleti | Deletion | 1 | A | - | 12 | 11604 | 0.10341262 | 63.1666667 | WT | | FALSE | CAN | JSY2424 | 7 | WT-Rep7 | CAA | nope | >1bp-ins | A/T-1 | none | A/T-1 | TGTCCCAAA | AAAAAGT | GTT | GTT | T | none | T>none | T>none_GTT | A/T-1_AAAAAGT | 9 | FALSE | FALSE |
| 130_v.txt | 31747 | 31747-Deleti | Deletion | 1 | C | - | 5 | 11604 | 0.04308859 | 58 | WT | | FALSE | CAN | JSY2424 | 7 | WT-Rep7 | TCC | nope | >1bp-ins | G/C-1 | none | G/C-1 | GATCTTCCCA | CCCATAC | AGG | TCC | C | none | nope | nope_TCC | G/C-1_CCCATAC | 9 | FALSE | FALSE |
| 130_v.txt | 31807 | 31807-Deleti | Deletion | 1 | A | - | 1808 | 11604 | 15.5808342 | 58.9773233 | WT | | FALSE | CAN | JSY2424 | 7 | WT-Rep7 | CAA | nope | >1bp-ins | A/T-1 | none | A/T-1 | TCTTCCAAAT | AAATAAA | GTT | GTT | T | none | T>none | T>none_GTT | A/T-1_AAATAAA | 9 | FALSE | FALSE |
| 130_v.txt | 31824 | 31824-Deleti | Deletion | 1 | A | - | 11 | 11604 | 0.0947949 | 56 | WT | | FALSE | CAN | JSY2424 | 7 | WT-Rep7 | GAA | nope | >1bp-ins | A/T-1 | none | A/T-1 | CATCTGAAT | AATATGC | CTT | CTT | T | none | T>none | T>none_CTT | A/T-1_AATATGC | 9 | FALSE | FALSE |
| 130_v.txt | 31863 | 31863-Deleti | Deletion | 1 | A | - | 13 | 11604 | 0.11203033 | 46.5384615 | WT | | FALSE | CAN | JSY2424 | 7 | WT-Rep7 | GAA | nope | >1bp-ins | A/T-1 | none | A/T-1 | AACAGGAAA | AAAATAG | CTT | CTT | T | none | T>none | T>none_CTT | A/T-1_AAAATAG | 9 | FALSE | FALSE |
| 130_v.txt | 31904 | 31904-Deleti | Deletion | 1 | T | - | 6 | 11585 | 0.05179111 | 44.833333 | WT | | FALSE | CAN | JSY2424 | 7 | WT-Rep7 | ATT | nope | >1bp-ins | A/T-1 | none | A/T-1 | ATTGAATTTT | TTTTGGT | TAA | ATT | T | none | nope | nope_ATT | A/T-1_TTTTGGT | 9 | FALSE | FALSE |
| 130_v.txt | 31972 | 31972-Deleti | Deletion | 1 | C | - | 9 | 31094 | 0.02894449 | 34.5555556 | WT | | FALSE | CAN | JSY2424 | 7 | WT-Rep7 | GCC | nope | >1bp-ins | G/C-1 | none | G/C-1 | AATAAGCCA | CCAAGCC | CGG | GCC | C | none | nope | nope_GCC | G/C-1_CCAAGCC | 9 | FALSE | FALSE |
| 130_v.txt | 31988 | 31988-Deleti | Deletion | 1 | T | - | 7 | 31094 | 0.02251238 | 48.8571429 | WT | | FALSE | CAN | JSY2424 | 7 | WT-Rep7 | ATT | nope | >1bp-ins | A/T-1 | none | A/T-1 | CATTAATTTA | TTTAGCT | TAA | ATT | T | none | nope | nope_ATT | A/T-1_TTTAGCT | 9 | FALSE | FALSE |
| 130_v.txt | 32057 | 32057-Deleti | Deletion | 1 | C | - | 32 | 31093 | 0.10291706 | 62.5937503 | WT | | FALSE | CAN | JSY2424 | 7 | WT-Rep7 | GCG | nope | >1bp-ins | G/C-1 | none | G/C-1 | GATGTGCGA | CGAGATT | CGC | GCG | C | none | nope | nope_GCG | G/C-1_CGAGATT | 9 | FALSE | FALSE |
| 130_v.txt | 32064 | 32064-Deleti | Deletion | 1 | G | - | 17 | 31094 | 0.05467293 | 60.2352941 | WT | | FALSE | CAN | JSY2424 | 7 | WT-Rep7 | TGA | nope | >1bp-ins | G/C-1 | none | G/C-1 | GAGATTGAG | GAGATAA | ACT | ACT | C | none | C>none | C>none_ACT | G/C-1_GAGATAA | 9 | FALSE | FALSE |
| 130_v.txt | 32069 | 32069-Deleti | Deletion | 1 | A | - | 6 | 31094 | 0.01929633 | 63.6666667 | WT | | FALSE | CAN | JSY2424 | 7 | WT-Rep7 | TAA | nope | >1bp-ins | A/T-1 | none | A/T-1 | TGAGATAAA | AAATAAC | ATT | ATT | T | none | T>none | T>none_ATT | A/T-1_AAATAAC | 9 | FALSE | FALSE |
| 130_v.txt | 32081 | 32081-Deleti | Deletion | 1 | A | - | 66 | 31094 | 0.2122596 | 63.2424242 | WT | | FALSE | CAN | JSY2424 | 7 | WT-Rep7 | CAA | nope | >1bp-ins | A/T-1 | none | A/T-1 | CCATGCAAA | AAAAAAG | GTT | GTT | T | none | T>none | T>none_GTT | A/T-1_AAAAAAG | 9 | FALSE | FALSE |
| 130_v.txt | 32113 | 32113-Deleti | Deletion | 1 | G | - | 8 | 31094 | 0.02572844 | 55.5 | WT | | FALSE | CAN | JSY2424 | 7 | WT-Rep7 | AGC | nope | >1bp-ins | G/C-1 | none | G/C-1 | TTAATAGCC | GCCATTC | TCG | TCG | C | none | C>none | C>none_TCG | G/C-1_GCCATTC | 9 | FALSE | FALSE |
| 130_v.txt | 32210 | 32210-Deleti | Deletion | 1 | T | - | 8 | 31094 | 0.02572844 | 42.75 | WT | | FALSE | CAN | JSY2424 | 7 | WT-Rep7 | CTT | nope | >1bp-ins | A/T-1 | none | A/T-1 | ACCACCTTTG | TTTGGTG | GAA | CTT | T | none | nope | nope_CTT | A/T-1_TTTGGTG | 9 | FALSE | FALSE |
| 130_v.txt | 32259 | 32259-Deleti | Deletion | 1 | C | - | 9 | 43327 | 0.02077227 | 36.6666669 | WT | | FALSE | CAN | JSY2424 | 7 | WT-Rep7 | ACC | nope | >1bp-ins | G/C-1 | none | G/C-1 | GATAGACCA | CCAAATA | TGG | ACC | C | none | nope | nope_ACC | G/C-1_CCAAATA | 9 | FALSE | FALSE |
| 130_v.txt | 32265 | 32265-Deleti | Deletion | 1 | A | - | 16 | 43326 | 0.03692933 | 37.5625 | WT | | FALSE | CAN | JSY2424 | 7 | WT-Rep7 | TAA | nope | >1bp-ins | A/T-1 | none | A/T-1 | CCAAATAAA | AAAATAC | ATT | ATT | T | none | T>none | T>none_ATT | A/T-1_AAAATAC | 9 | FALSE | FALSE |
| 130_v.txt | 32282 | 32282-Deleti | Deletion | 1 | G | - | 12 | 43326 | 0.027697 | 36.8333335 | WT | | FALSE | CAN | JSY2424 | 7 | WT-Rep7 | CGT | nope | >1bp-ins | G/C-1 | none | G/C-1 | ACCAACGTA | GTAAATA | GCA | GCA | C | none | C>none | C>none_GCA | G/C-1_GTAAATA | 9 | FALSE | FALSE |
| 130_v.txt | 32284 | 32284-Deleti | Deletion | 1 | A | - | 5 | 43326 | 0.01154042 | 47.9999913 | WT | | FALSE | CAN | JSY2424 | 7 | WT-Rep7 | TAA | nope | >1bp-ins | A/T-1 | none | A/T-1 | CAACGTAAA | AAATATT | ATT | ATT | T | none | T>none | T>none_ATT | A/T-1_AAATATT | 9 | FALSE | FALSE |
| 130_v.txt | 32304 | 32304-Deleti | Deletion | 1 | G | - | 17 | 43326 | 0.03923741 | 59.6470588 | WT | | FALSE | CAN | JSY2424 | 7 | WT-Rep7 | AGA | nope | >1bp-ins | G/C-1 | none | G/C-1 | GCGGCAGAA | GAAATAA | TCT | TCT | C | none | C>none | C>none_TCT | G/C-1_GAAATAA | 9 | FALSE | FALSE |
| 130_v.txt | 32323 | 32323-Deleti | Deletion | 1 | T | - | 4 | 43326 | 9.23E-03 | 57.25 | WT | | FALSE | CAN | JSY2424 | 7 | WT-Rep7 | ATA | nope | >1bp-ins | A/T-1 | none | A/T-1 | TTAAGATAA | TAACAGC | TAT | ATA | T | none | nope | nope_ATA | A/T-1_TAACAGC | 9 | FALSE | FALSE |
| 130_v.txt | 32358 | 32358-Deleti | Deletion | 1 | C | - | 8 | 43326 | 0.01846466 | 63.625 | WT | | FALSE | CAN | JSY2424 | 7 | WT-Rep7 | ACC | nope | >1bp-ins | G/C-1 | none | G/C-1 | TTTGTACCAG | CCAGAGT | TGG | ACC | C | none | nope | nope_ACC | G/C-1_CCAGAGT | 9 | FALSE | FALSE |
| 130_v.txt | 32373 | 32373-Deleti | Deletion | 1 | G | - | 62 | 43326 | 0.14310114 | 62.9193548 | WT | | FALSE | CAN | JSY2424 | 7 | WT-Rep7 | AGC | nope | >1bp-ins | G/C-1 | none | G/C-1 | TCAATAGCA | GCAATAA | TCG | TCG | C | none | C>none | C>none_TCG | G/C-1_GCAATAA | 9 | FALSE | FALSE |
| 130_v.txt | 32381 | 32381-Deleti | Deletion | 1 | A | - | 9 | 43325 | 0.02077323 | 62.111109 | WT | | FALSE | CAN | JSY2424 | 7 | WT-Rep7 | TAA | nope | >1bp-ins | A/T-1 | none | A/T-1 | AATAATAAA | AAAGGGA | ATT | ATT | T | none | T>none | T>none_ATT | A/T-1_AAAGGGA | 9 | FALSE | FALSE |
| 130_v.txt | 32407 | 32407-Deleti | Deletion | 1 | T | - | 2 | 43324 | 4.62E-03 | 50.9999998 | WT | | FALSE | CAN | JSY2424 | 7 | WT-Rep7 | GTA | nope | >1bp-ins | A/T-1 | none | A/T-1 | AGGAAGTAG | TAGATTG | CAT | GTA | T | none | nope | nope_GTA | A/T-1_TAGATTG | 9 | FALSE | FALSE |
| 130_v.txt | 32420 | 32420-Deleti | Deletion | 1 | T | - | 10 | 43326 | 0.02308083 | 54.5 | WT | | FALSE | CAN | JSY2424 | 7 | WT-Rep7 | GTT | nope | >1bp-ins | A/T-1 | none | A/T-1 | TGTTAGTTTA | TTTAGGG | CAA | GTT | T | none | nope | nope_GTT | A/T-1_TTTAGGG | 9 | FALSE | FALSE |
| 130_v.txt | 32490 | 32490-Deleti | Deletion | 1 | A | - | 67 | 43326 | 0.15464156 | 45.2388062 | WT | | FALSE | CAN | JSY2424 | 7 | WT-Rep7 | GAA | nope | >1bp-ins | A/T-1 | none | A/T-1 | ATACGGAAA | AAAACAA | CTT | CTT | T | none | T>none | T>none_CTT | A/T-1_AAAACAA | 9 | FALSE | FALSE |
| 130_v.txt | 32498 | 32498-Deleti | Deletion | 1 | T | - | 137 | 43326 | 0.31620736 | 45.7372262 | WT | | FALSE | CAN | JSY2424 | 7 | WT-Rep7 | CTT | nope | >1bp-ins | A/T-1 | none | A/T-1 | AACAACTTTT | TTTTTTG | GAA | CTT | T | none | nope | nope_CTT | A/T-1_TTTTTTG | 9 | FALSE | FALSE |
| 130_v.txt | 32615 | 32615-Deleti | Deletion | 1 | T | - | 25 | 31688 | 0.07889422 | 44.4799988 | WT | | FALSE | CAN | JSY2424 | 7 | WT-Rep7 | CTA | nope | >1bp-ins | A/T-1 | none | A/T-1 | CCAACCTAA | TAAGAAC | GAT | CTA | T | none | nope | nope_CTA | A/T-1_TAAGAAC | 9 | FALSE | FALSE |
| 130_v.txt | 32616 | 32616-Deleti | Deletion | 1 | A | - | 2 | 31688 | 6.31E-03 | 38.0000004 | WT | | FALSE | CAN | JSY2424 | 7 | WT-Rep7 | TAA | nope | >1bp-ins | A/T-1 | none | A/T-1 | CAACCTAAG | AAGAACC | ATT | ATT | T | none | T>none | T>none_ATT | A/T-1_AAGAACC | 9 | FALSE | FALSE |
| 130_v.txt | 32631 | 32631-Deleti | Deletion | 1 | T | - | 14 | 31688 | 0.04418076 | 52.2142863 | WT | | FALSE | CAN | JSY2424 | 7 | WT-Rep7 | GTT | nope | >1bp-ins | A/T-1 | none | A/T-1 | CCTTCGTTTT | TTTTTAT | CAA | GTT | T | none | nope | nope_GTT | A/T-1_TTTTTAT | 9 | FALSE | FALSE |
| 130_v.txt | 32658 | 32658-Deleti | Deletion | 1 | C | - | 16 | 31688 | 0.0504923 | 57.875 | WT | | FALSE | CAN | JSY2424 | 7 | WT-Rep7 | ACC | nope | >1bp-ins | G/C-1 | none | G/C-1 | CCTGGACCC | CCCCAGG | TGG | ACC | C | none | nope | nope_ACC | G/C-1_CCCCAGG | 9 | FALSE | FALSE |
| 130_v.txt | 32670 | 32670-Deleti | Deletion | 1 | G | - | 9 | 31688 | 0.02840192 | 56.5555556 | WT | | FALSE | CAN | JSY2424 | 7 | WT-Rep7 | TGG | nope | >1bp-ins | G/C-1 | none | G/C-1 | GCACCTGGG | GGGTTTC | ACC | ACC | C | none | C>none | C>none_ACC | G/C-1_GGGTTTC | 9 | FALSE | FALSE |
| 130_v.txt | 32688 | 32688-Deleti | Deletion | 1 | A | - | 8 | 31688 | 0.02524615 | 58.75 | WT | | FALSE | CAN | JSY2424 | 7 | WT-Rep7 | GAA | nope | >1bp-ins | A/T-1 | none | A/T-1 | TAACGGAAT | AATCCAA | CTT | CTT | T | none | T>none | T>none_CTT | A/T-1_AATCCAA | 9 | FALSE | FALSE |
| 130_v.txt | 32697 | 32697-Deleti | Deletion | 1 | G | - | 8 | 31688 | 0.02524615 | 63.5 | WT | | FALSE | CAN | JSY2424 | 7 | WT-Rep7 | TGG | nope | >1bp-ins | G/C-1 | none | G/C-1 | CCAACTGGG | GGGCCGG | ACC | ACC | C | none | C>none | C>none_ACC | G/C-1_GGGCCGG | 9 | FALSE | FALSE |
| 130_v.txt | 32707 | 32707-Deleti | Deletion | 1 | C | - | 10 | 31688 | 0.03155769 | 61.3 | WT | | FALSE | CAN | JSY2424 | 7 | WT-Rep7 | ACC | nope | >1bp-ins | G/C-1 | none | G/C-1 | CGGTAACCC | CCCCAGC | TGG | ACC | C | none | nope | nope_ACC | G/C-1_CCCCAGC | 9 | FALSE | FALSE |
| 130_v.txt | 32728 | 32728-Deleti | Deletion | 1 | A | - | 20 | 31688 | 0.06311538 | 61.1000002 | WT | | FALSE | CAN | JSY2424 | 7 | WT-Rep7 | CAA | nope | >1bp-ins | A/T-1 | none | A/T-1 | CCATACAAA | AAAAACA | GTT | GTT | T | none | T>none | T>none_GTT | A/T-1_AAAAACA | 9 | FALSE | FALSE |
| 130_v.txt | 32760 | 32760-Deleti | Deletion | 1 | A | - | 3 | 31688 | 9.47E-03 | 53.9999993 | WT | | FALSE | CAN | JSY2424 | 7 | WT-Rep7 | TAA | nope | >1bp-ins | A/T-1 | none | A/T-1 | ATGGCTAAA | AAAACTT | ATT | ATT | T | none | T>none | T>none_ATT | A/T-1_AAAACTT | 9 | FALSE | FALSE |
| 130_v.txt | 32811 | 32811-Deleti | Deletion | 1 | G | - | 10 | 31688 | 0.03155769 | 46.5 | WT | | FALSE | CAN | JSY2424 | 7 | WT-Rep7 | AGG | nope | >1bp-ins | G/C-1 | none | G/C-1 | TTGACAGGG | GGGAACA | TCC | TCC | C | none | C>none | C>none_TCC | G/C-1_GGGAACA | 9 | FALSE | FALSE |
| 130_v.txt | 32839 | 32839-Deleti | Deletion | 1 | C | - | 23 | 31687 | 0.07258497 | 39.8695652 | WT | | FALSE | CAN | JSY2424 | 7 | WT-Rep7 | ACC | nope | >1bp-ins | G/C-1 | none | G/C-1 | TAATTACCCA | CCCAAAA | TGG | ACC | C | none | nope | nope_ACC | G/C-1_CCCAAAA | 9 | FALSE | FALSE |
| 130_v.txt | 32842 | 32842-Deleti | Deletion | 1 | A | - | 110 | 31687 | 0.34714552 | 42.8090897 | WT | | FALSE | CAN | JSY2424 | 7 | WT-Rep7 | CAA | nope | >1bp-ins | A/T-1 | none | A/T-1 | TTACCCAAA | AAAAAAT | GTT | GTT | T | none | T>none | T>none_GTT | A/T-1_AAAAAAT | 9 | FALSE | FALSE |
| 130_v.txt | 32860 | 32860-Deleti | Deletion | 1 | C | - | 2 | 31686 | 6.31E-03 | 38 | WT | | FALSE | CAN | JSY2424 | 7 | WT-Rep7 | GCC | nope | >1bp-ins | G/C-1 | none | G/C-1 | TCCATGCCG | CCGCCAG | CGG | GCC | C | none | nope | nope_GCC | G/C-1_CCGCCAG | 9 | FALSE | FALSE |
| 130_v.txt | 32882 | 32882-Deleti | Deletion | 1 | C | - | 36 | 33858 | 0.10632642 | 36.3611111 | WT | | FALSE | CAN | JSY2424 | 7 | WT-Rep7 | TCC | nope | >1bp-ins | G/C-1 | none | G/C-1 | GTACGTCCA | CCAAAAT | AGG | TCC | C | none | nope | nope_TCC | G/C-1_CCAAAAT | 9 | FALSE | FALSE |
| 130_v.txt | 32884 | 32884-Deleti | Deletion | 1 | A | - | 20 | 33859 | 0.05906849 | 37.05 | WT | | FALSE | CAN | JSY2424 | 7 | WT-Rep7 | CAA | nope | >1bp-ins | A/T-1 | none | A/T-1 | ACGTCCAAA | AAAATTG | GTT | GTT | T | none | T>none | T>none_GTT | A/T-1_AAAATTG | 9 | FALSE | FALSE |
| 130_v.txt | 32917 | 32917-Deleti | Deletion | 1 | C | - | 1 | 33860 | 2.95E-03 | 61 | WT | | FALSE | CAN | JSY2424 | 7 | WT-Rep7 | TCC | nope | >1bp-ins | G/C-1 | none | G/C-1 | TAAGTTCCA | CCAGGGC | AGG | TCC | C | none | nope | nope_TCC | G/C-1_CCAGGGC | 9 | FALSE | FALSE |
| 130_v.txt | 32924 | 32924-Deleti | Deletion | 1 | A | - | 16 | 33860 | 0.0472534 | 57.1875 | WT | | FALSE | CAN | JSY2424 | 7 | WT-Rep7 | CAA | nope | >1bp-ins | A/T-1 | none | A/T-1 | CAGGGCAAA | AAAAGTG | GTT | GTT | T | none | T>none | T>none_GTT | A/T-1_AAAAGTG | 9 | FALSE | FALSE |
| 130_v.txt | 32934 | 32934-Deleti | Deletion | 1 | G | - | 25 | 33860 | 0.07383343 | 51.08 | WT | | FALSE | CAN | JSY2424 | 7 | WT-Rep7 | TGC | nope | >1bp-ins | G/C-1 | none | G/C-1 | GTGATTGCC | GCCCAAG | ACG | ACG | C | none | C>none | C>none_ACG | G/C-1_GCCCAAG | 9 | FALSE | FALSE |
| 130_v.txt | 32972 | 32972-Deleti | Deletion | 1 | A | - | 8 | 33860 | 0.0236267 | 64 | WT | | FALSE | CAN | JSY2424 | 7 | WT-Rep7 | CAA | nope | >1bp-ins | A/T-1 | none | A/T-1 | CGCACCAAA | AAATGCT | GTT | GTT | T | none | T>none | T>none_GTT | A/T-1_AAATGCT | 9 | FALSE | FALSE |
| 130_v.txt | 32999 | 32999-Deleti | Deletion | 1 | G | - | 24 | 33859 | 0.07088219 | 63.9999993 | WT | | FALSE | CAN | JSY2424 | 7 | WT-Rep7 | AGA | nope | >1bp-ins | G/C-1 | none | G/C-1 | TTGTGAGAA | GAAAACT | TCT | TCT | C | none | C>none | C>none_TCT | G/C-1_GAAAACT | 9 | FALSE | FALSE |
| 130_v.txt | 33000 | 33000-Deleti | Deletion | 1 | A | - | 30 | 33860 | 0.08860012 | 59.5 | WT | | FALSE | CAN | JSY2424 | 7 | WT-Rep7 | GAA | nope | >1bp-ins | A/T-1 | none | A/T-1 | TGTGAGAAA | AAAACTG | CTT | CTT | T | none | T>none | T>none_CTT | A/T-1_AAAACTG | 9 | FALSE | FALSE |
| 130_v.txt | 33040 | 33040-Deleti | Deletion | 1 | T | - | 22 | 33859 | 0.06497534 | 60.5909088 | WT | | FALSE | CAN | JSY2424 | 7 | WT-Rep7 | ATT | nope | >1bp-ins | A/T-1 | none | A/T-1 | TAGCCATTTC | TTTCACC | TAA | ATT | T | none | nope | nope_ATT | A/T-1_TTTCACC | 9 | FALSE | FALSE |
| 130_v.txt | 33058 | 33058-Deleti | Deletion | 1 | T | - | 6 | 33860 | 0.01772002 | 59.6666667 | WT | | FALSE | CAN | JSY2424 | 7 | WT-Rep7 | GTG | nope | >1bp-ins | A/T-1 | none | A/T-1 | ACTGCGTGA | TGACAGA | CAC | GTG | T | none | nope | nope_GTG | A/T-1_TGACAGA | 9 | FALSE | FALSE |
| 130_v.txt | 33090 | 33090-Deleti | Deletion | 1 | T | - | 1 | 33860 | 2.95E-03 | 64 | WT | | FALSE | CAN | JSY2424 | 7 | WT-Rep7 | ATA | nope | >1bp-ins | A/T-1 | none | A/T-1 | AATAAATAT | TATGATA | TAT | ATA | T | none | nope | nope_ATA | A/T-1_TATGATA | 9 | FALSE | FALSE |
| 130_v.txt | 33111 | 33111-Deleti | Deletion | 1 | G | - | 4 | 33860 | 0.01181335 | 51 | WT | | FALSE | CAN | JSY2424 | 7 | WT-Rep7 | TGG | nope | >1bp-ins | G/C-1 | none | G/C-1 | CCCACTGGG | GGGCCGG | ACC | ACC | C | none | C>none | C>none_ACC | G/C-1_GGGCCGG | 9 | FALSE | FALSE |
| 130_v.txt | 33114 | 33114-Deleti | Deletion | 1 | C | - | 14 | 33860 | 0.04134672 | 50.5 | WT | | FALSE | CAN | JSY2424 | 7 | WT-Rep7 | GCC | nope | >1bp-ins | G/C-1 | none | G/C-1 | ACTGGGCCG | CCGGCGT | CGG | GCC | C | none | nope | nope_GCC | G/C-1_CCGGCGT | 9 | FALSE | FALSE |
| 130_v.txt | 33147 | 33147-Deleti | Deletion | 1 | A | - | 50 | 33860 | 0.14766686 | 39.9 | WT | | FALSE | CAN | JSY2424 | 7 | WT-Rep7 | GAA | nope | >1bp-ins | A/T-1 | none | A/T-1 | CCAATGAAA | AAAAGAC | CTT | CTT | T | none | T>none | T>none_CTT | A/T-1_AAAAGAC | 9 | FALSE | FALSE |
| 130_v.txt | 33157 | 33157-Deleti | Deletion | 1 | T | - | 11 | 33843 | 0.03250303 | 38 | WT | | FALSE | CAN | JSY2424 | 7 | WT-Rep7 | GTA | nope | >1bp-ins | A/T-1 | none | A/T-1 | GACCTGTAC | TACCAAT | CAT | GTA | T | none | nope | nope_GTA | A/T-1_TACCAAT | 9 | FALSE | FALSE |

| | | | | | | | | | | | | | | | | | | | | | | | | | | | | | | | | | | |
|---|---|---|---|---|---|---|---|---|---|---|---|---|---|---|---|---|---|---|---|---|---|---|---|---|---|---|---|---|---|---|---|---|---|---|
| 130_v.txt | 33167 | 33167-Deleti | Deletion | 1 | A | - | 8 | 33840 | 0.02364066 | 35.7500001 | WT | FALSE | CAN | JSY2424 | 7 | WT-Rep7 | TAC | nope | >1bp-ins | A/T-1 | none | A/T-1 | AATAGTACC | ACCACCA | ATG | ATG | T | none | T>none | T>none_ATG | A/T-1_ACCACCA | 9 | FALSE | FALSE |
| 130_v.txt | 33168 | 33168-Deleti | Deletion | 1 | C | - | 3 | 33840 | 8.87E-03 | 37.6666667 | WT | FALSE | CAN | JSY2424 | 7 | WT-Rep7 | ACC | nope | >1bp-ins | G/C-1 | none | G/C-1 | ATAGTACCA | CCACCAA | TGG | ACC | C | none | nope | nope_ACC | G/C-1_CCACCAA | 9 | FALSE | FALSE |
| 130_v.txt | 33199 | 33199-Deleti | Deletion | 1 | G | - | 112 | 32448 | 0.34516765 | 37.0625 | WT | FALSE | CAN | JSY2424 | 7 | WT-Rep7 | TGC | nope | >1bp-ins | G/C-1 | none | G/C-1 | GTCTTTGCTT | GCTTAAG | ACG | ACG | C | none | C>none | C>none_ACG | G/C-1_GCTTAAG | 9 | FALSE | FALSE |
| 130_v.txt | 33417 | 33417-Deleti | Deletion | 1 | T | - | 22 | 32448 | 0.06780079 | 47.2272727 | WT | FALSE | CAN | JSY2424 | 7 | WT-Rep7 | ATT | nope | >1bp-ins | A/T-1 | none | A/T-1 | GGCTCATTG | TTGTACA | TAA | ATT | T | none | nope | nope_ATT | A/T-1_TTGTACA | 9 | FALSE | FALSE |
| 130_v.txt | 33459 | 33459-Deleti | Deletion | 1 | T | - | 33 | 32448 | 0.10170118 | 37.8181818 | WT | FALSE | CAN | JSY2424 | 7 | WT-Rep7 | ATT | nope | >1bp-ins | A/T-1 | none | A/T-1 | TTTGAATTTG | TTTGTCA | TAA | ATT | T | none | nope | nope_ATT | A/T-1_TTTGTCA | 9 | FALSE | FALSE |
| 130_v.txt | 33474 | 33474-Deleti | Deletion | 1 | C | - | 94 | 32448 | 0.28969428 | 29.0638288 | WT | FALSE | CAN | JSY2424 | 7 | WT-Rep7 | GCC | nope | >1bp-ins | G/C-1 | none | G/C-1 | GCTATGCCTT | CCTTTTT | CGG | GCC | C | none | nope | nope_GCC | G/C-1_CCTTTTT | 9 | FALSE | FALSE |
| 131_v.txt | 31723 | 31723-SNV- | SNV | 1 | A | C | 8 | 16915 | 0.0472953 | 53 | WT | FALSE | CAN | JSY2424 | 8 | WT-Rep8 | CAA | A>C | G/C+1 | A/T-1 | TA>GC | TA>GC | TGTCCCAAA | AAAAAGT | GTT | GTT | T | G | T>G | T>G_GTT | TA>GC_AAAAAGT | 9 | FALSE | FALSE |
| 131_v.txt | 31950 | 31950-SNV- | SNV | 1 | A | C | 6 | 27413 | 0.02188743 | 37.5 | WT | FALSE | CAN | JSY2424 | 8 | WT-Rep8 | CAT | A>C | G/C+1 | A/T-1 | TA>GC | TA>GC | ATCGTCATA | ATAAATG | GTA | GTA | T | G | T>G | T>G_GTA | TA>GC_ATAAATG | 9 | FALSE | FALSE |
| 131_v.txt | 31986 | 31986-SNV- | SNV | 1 | A | C | 8 | 27414 | 0.02918217 | 45.75 | WT | FALSE | CAN | JSY2424 | 8 | WT-Rep8 | TAA | A>C | G/C+1 | A/T-1 | TA>GC | TA>GC | GGCATTAAT | AATTTAG | ATT | ATT | T | G | T>G | T>G_ATT | TA>GC_AATTTAG | 9 | FALSE | FALSE |
| 131_v.txt | 32126 | 32126-SNV-T | SNV | 1 | T | G | 7 | 27414 | 0.0255344 | 17.5713563 | WT | FALSE | CAN | JSY2424 | 8 | WT-Rep8 | CTT | T>G | G/C+1 | A/T-1 | TA>GC | TA>GC | GAAAACTTT | TTTGTCA | GAA | CTT | T | C | T>G | T>G_CTT | TA>GC_TTTGTCA | 9 | FALSE | FALSE |
| 131_v.txt | 32187 | 32187-SNV- | SNV | 1 | A | C | 8 | 27410 | 0.02918643 | 36.75 | WT | FALSE | CAN | JSY2424 | 8 | WT-Rep8 | AAC | A>C | G/C+1 | A/T-1 | TA>GC | TA>GC | ACGAAAACT | ACTGCAA | TTG | TTG | T | G | T>G | T>G_TTG | TA>GC_ACTGCAA | 9 | FALSE | FALSE |
| 131_v.txt | 32294 | 32294-SNV- | SNV | 1 | A | C | 72 | 35671 | 0.20184464 | 49.0416068 | WT | FALSE | CAN | JSY2424 | 8 | WT-Rep8 | AAT | A>C | G/C+1 | A/T-1 | TA>GC | TA>GC | ATTTGAATTT | ATTTGCG | TTA | TTA | T | G | T>G | T>G_TTA | TA>GC_ATTTGCG | 9 | FALSE | FALSE |
| 131_v.txt | 32473 | 32473-SNV- | SNV | 1 | A | C | 21 | 35672 | 0.0588697 | 43.381584 | WT | FALSE | CAN | JSY2424 | 8 | WT-Rep8 | TAG | A>C | G/C+1 | A/T-1 | TA>GC | TA>GC | CAATGTAGA | AGAAGGT | ATC | ATC | T | G | T>G | T>G_ATC | TA>GC_AGAAGGT | 9 | FALSE | FALSE |
| 131_v.txt | 32481 | 32481-SNV- | SNV | 1 | A | C | 13 | 35672 | 0.03644315 | 42.8461539 | WT | FALSE | CAN | JSY2424 | 8 | WT-Rep8 | TAA | A>C | G/C+1 | A/T-1 | TA>GC | TA>GC | AAGGTTAAG | AAGATAC | ATT | ATT | T | G | T>G | T>G_ATT | TA>GC_AAGATAC | 9 | FALSE | FALSE |
| 131_v.txt | 32681 | 32681-SNV- | SNV | 1 | A | C | 9 | 20964 | 0.04293074 | 52.2222222 | WT | FALSE | CAN | JSY2424 | 8 | WT-Rep8 | AAT | A>C | G/C+1 | A/T-1 | TA>GC | TA>GC | TCTCCAATAA | ATAACGG | TTA | TTA | T | G | T>G | T>G_TTA | TA>GC_ATAACGG | 9 | FALSE | FALSE |
| 131_v.txt | 32800 | 32800-SNV- | SNV | 1 | A | C | 6 | 20964 | 0.02862049 | 46.6666667 | WT | FALSE | CAN | JSY2424 | 8 | WT-Rep8 | TAA | A>C | G/C+1 | A/T-1 | TA>GC | TA>GC | CACCGTAAT | AATATTT | ATT | ATT | T | G | T>G | T>G_ATT | TA>GC_AATATTT | 9 | FALSE | FALSE |
| 131_v.txt | 32842 | 32842-SNV- | SNV | 1 | A | C | 3 | 20964 | 0.01431025 | 46.3331356 | WT | FALSE | CAN | JSY2424 | 8 | WT-Rep8 | CAA | A>C | G/C+1 | A/T-1 | TA>GC | TA>GC | TTACCCAAA | AAAAAAT | GTT | GTT | T | G | T>G | T>G_GTT | TA>GC_AAAAAAT | 9 | FALSE | FALSE |
| 131_v.txt | 32884 | 32884-SNV- | SNV | 1 | A | C | 16 | 31752 | 0.05039053 | 34.625 | WT | FALSE | CAN | JSY2424 | 8 | WT-Rep8 | CAA | A>C | G/C+1 | A/T-1 | TA>GC | TA>GC | ACGTCCAAA | AAAATTG | GTT | GTT | T | G | T>G | T>G_GTT | TA>GC_AAAATTG | 9 | FALSE | FALSE |
| 131_v.txt | 32919 | 32919-SNV- | SNV | 1 | A | C | 6 | 31752 | 0.01889645 | 24.1680762 | WT | FALSE | CAN | JSY2424 | 8 | WT-Rep8 | CAG | A>C | G/C+1 | A/T-1 | TA>GC | TA>GC | AGTTCCAGG | AGGGCAA | GTC | GTC | T | G | T>G | T>G_GTC | TA>GC_AGGGCAA | 9 | FALSE | FALSE |
| 131_v.txt | 33189 | 33189-SNV- | SNV | 1 | A | C | 11 | 31292 | 0.03515276 | 35.4545455 | WT | FALSE | CAN | JSY2424 | 8 | WT-Rep8 | AAT | A>C | G/C+1 | A/T-1 | TA>GC | TA>GC | ATACCAATA | ATATGTC | TTA | TTA | T | G | T>G | T>G_TTA | TA>GC_ATATGTC | 9 | FALSE | FALSE |
| 131_v.txt | 33269 | 33269-SNV- | SNV | 1 | A | C | 223 | 31295 | 0.71257389 | 15.1748879 | WT | FALSE | CAN | JSY2424 | 8 | WT-Rep8 | TAG | A>C | G/C+1 | A/T-1 | TA>GC | TA>GC | CTCCATAGA | AGAGAAC | ATC | ATC | T | G | T>G | T>G_ATC | TA>GC_AGAGAAC | 9 | FALSE | FALSE |
| 131_v.txt | 33476 | 33476-SNV-T | SNV | 1 | T | G | 9 | 31291 | 0.02876226 | 32.2222222 | WT | FALSE | CAN | JSY2424 | 8 | WT-Rep8 | CTT | T>G | G/C+1 | A/T-1 | TA>GC | TA>GC | TATGCCTTTT | TTTTTTT | GAA | CTT | T | C | T>G | T>G_CTT | TA>GC_TTTTTTT | 9 | FALSE | FALSE |
| 131_v.txt | 31819 | 31819-SNV- | SNV | 1 | A | G | 7 | 16915 | 0.04138339 | 35.1428016 | WT | FALSE | CAN | JSY2424 | 8 | WT-Rep8 | CAT | A>G | G/C+1 | A/T-1 | TA>CG | TA>CG | ATCTGCATCT | ATCTGAA | GTA | GTA | T | C | T>C | T>C_GTA | TA>CG_ATCTGAA | 9 | FALSE | FALSE |
| 131_v.txt | 31820 | 31820-SNV-T | SNV | 1 | T | C | 3 | 16915 | 0.01773574 | 55.3332203 | WT | FALSE | CAN | JSY2424 | 8 | WT-Rep8 | ATC | T>C | G/C+1 | A/T-1 | TA>CG | TA>CG | TCTGCATCTG | TCTGAAT | TAG | ATC | T | G | T>C | T>C_ATC | TA>CG_TCTGAAT | 9 | FALSE | FALSE |
| 131_v.txt | 31822 | 31822-SNV-T | SNV | 1 | T | C | 6 | 16915 | 0.03547148 | 49.4994858 | WT | FALSE | CAN | JSY2424 | 8 | WT-Rep8 | CTG | T>C | G/C+1 | A/T-1 | TA>CG | TA>CG | TGCATCTGA | TGAATAT | GAC | CTG | T | G | T>C | T>C_CTG | TA>CG_TGAATAT | 9 | FALSE | FALSE |
| 131_v.txt | 32055 | 32055-SNV-T | SNV | 1 | T | C | 25 | 27414 | 0.09119428 | 62.6 | WT | FALSE | CAN | JSY2424 | 8 | WT-Rep8 | GTG | T>C | G/C+1 | A/T-1 | TA>CG | TA>CG | CTGATGTGC | TGCGAGA | CAC | GTG | T | G | T>C | T>C_GTG | TA>CG_TGCGAGA | 9 | FALSE | FALSE |
| 131_v.txt | 32081 | 32081-SNV- | SNV | 1 | A | G | 10 | 27414 | 0.03647771 | 49.5001476 | WT | FALSE | CAN | JSY2424 | 8 | WT-Rep8 | CAA | A>G | G/C+1 | A/T-1 | TA>CG | TA>CG | CCATGCAAA | AAAAAAG | GTT | GTT | T | C | T>C | T>C_GTT | TA>CG_AAAAAAG | 9 | FALSE | FALSE |
| 131_v.txt | 32126 | 32126-SNV-T | SNV | 1 | T | C | 6 | 27414 | 0.02188663 | 51.4984144 | WT | FALSE | CAN | JSY2424 | 8 | WT-Rep8 | CTT | T>C | G/C+1 | A/T-1 | TA>CG | TA>CG | GAAAACTTT | TTTGTCA | GAA | CTT | T | G | T>C | T>C_CTT | TA>CG_TTTGTCA | 9 | FALSE | FALSE |
| 131_v.txt | 32293 | 32293-SNV- | SNV | 1 | A | G | 58 | 35671 | 0.16259707 | 50.9598239 | WT | FALSE | CAN | JSY2424 | 8 | WT-Rep8 | GAA | A>G | G/C+1 | A/T-1 | TA>CG | TA>CG | TATTTGAATT | AATTTGC | CTT | CTT | T | C | T>C | T>C_CTT | TA>CG_AATTTGC | 9 | FALSE | FALSE |
| 131_v.txt | 32314 | 32314-SNV-T | SNV | 1 | T | C | 11 | 35671 | 0.03083738 | 33.4466661 | WT | FALSE | CAN | JSY2424 | 8 | WT-Rep8 | GTT | T>C | G/C+1 | A/T-1 | TA>CG | TA>CG | TAATGGTTG | TTGTTAA | CAA | GTT | T | G | T>C | T>C_GTT | TA>CG_TTGTTAA | 9 | FALSE | FALSE |
| 131_v.txt | 32315 | 32315-SNV-T | SNV | 1 | T | C | 6 | 35671 | 0.01682039 | 53.8333037 | WT | FALSE | CAN | JSY2424 | 8 | WT-Rep8 | TTG | T>C | G/C+1 | A/T-1 | TA>CG | TA>CG | AATGGTTGT | TGTTAAG | AAC | TTG | T | G | T>C | T>C_TTG | TA>CG_TGTTAAG | 9 | FALSE | FALSE |
| 131_v.txt | 32473 | 32473-SNV- | SNV | 1 | A | G | 11 | 35672 | 0.03083651 | 47.3636258 | WT | FALSE | CAN | JSY2424 | 8 | WT-Rep8 | TAG | A>G | G/C+1 | A/T-1 | TA>CG | TA>CG | CAATGTAGA | AGAAGGT | ATC | ATC | T | C | T>C | T>C_ATC | TA>CG_AGAAGGT | 9 | FALSE | FALSE |
| 131_v.txt | 32527 | 32527-SNV-T | SNV | 1 | T | C | 12 | 35672 | 0.03363983 | 36.4165793 | WT | FALSE | CAN | JSY2424 | 8 | WT-Rep8 | CTG | T>C | G/C+1 | A/T-1 | TA>CG | TA>CG | ATTTTCTGG | TGGGGTT | GAC | CTG | T | G | T>C | T>C_CTG | TA>CG_TGGGGTT | 9 | FALSE | FALSE |
| 131_v.txt | 32734 | 32734-SNV- | SNV | 1 | A | G | 27 | 20964 | 0.12879222 | 53.962963 | WT | FALSE | CAN | JSY2424 | 8 | WT-Rep8 | CAG | A>G | G/C+1 | A/T-1 | TA>CG | TA>CG | AAAAACAGT | AGTATAT | GTC | GTC | T | C | T>C | T>C_GTC | TA>CG_AGTATAT | 9 | FALSE | FALSE |
| 131_v.txt | 32919 | 32919-SNV- | SNV | 1 | A | G | 25 | 31752 | 0.0787352 | 45.2803231 | WT | FALSE | CAN | JSY2424 | 8 | WT-Rep8 | CAG | A>G | G/C+1 | A/T-1 | TA>CG | TA>CG | AGTTCCAGG | AGGGCAA | GTC | GTC | T | C | T>C | T>C_GTC | TA>CG_AGGGCAA | 9 | FALSE | FALSE |
| 131_v.txt | 33214 | 33214-SNV-T | SNV | 1 | T | C | 9 | 31295 | 0.02875859 | 28.9999885 | WT | FALSE | CAN | JSY2424 | 8 | WT-Rep8 | TTC | T>C | G/C+1 | A/T-1 | TA>CG | TA>CG | CTCTCTTCAC | TCACTTC | AAG | TTC | T | G | T>C | T>C_TTC | TA>CG_TCACTTC | 9 | FALSE | FALSE |
| 131_v.txt | 33476 | 33476-SNV-T | SNV | 1 | T | C | 25 | 31291 | 0.07989518 | 28.48 | WT | FALSE | CAN | JSY2424 | 8 | WT-Rep8 | CTT | T>C | G/C+1 | A/T-1 | TA>CG | TA>CG | TATGCCTTTT | TTTTTTT | GAA | CTT | T | G | T>C | T>C_CTT | TA>CG_TTTTTTT | 9 | FALSE | FALSE |
| 131_v.txt | 31814 | 31814-SNV-T | SNV | 1 | T | A | 5 | 16915 | 0.02955956 | 33.9993665 | WT | FALSE | CAN | JSY2424 | 8 | WT-Rep8 | ATC | T>A | A/T+1 | A/T-1 | TA>AT | TA>AT | AATAAATCT | TCTGCAT | TAG | ATC | T | T | T>A | T>A_ATC | TA>AT_TCTGCAT | 9 | FALSE | FALSE |
| 131_v.txt | 31819 | 31819-SNV- | SNV | 1 | A | T | 3 | 16915 | 0.01773574 | 63.9989698 | WT | FALSE | CAN | JSY2424 | 8 | WT-Rep8 | CAT | A>T | A/T+1 | A/T-1 | TA>AT | TA>AT | ATCTGCATCT | ATCTGAA | GTA | GTA | T | A | T>A | T>A_GTA | TA>AT_ATCTGAA | 9 | FALSE | FALSE |
| 131_v.txt | 31820 | 31820-SNV-T | SNV | 1 | T | A | 5 | 16915 | 0.02955956 | 49.1991974 | WT | FALSE | CAN | JSY2424 | 8 | WT-Rep8 | ATC | T>A | A/T+1 | A/T-1 | TA>AT | TA>AT | TCTGCATCTG | TCTGAAT | TAG | ATC | T | T | T>A | T>A_ATC | TA>AT_TCTGAAT | 9 | FALSE | FALSE |
| 131_v.txt | 31822 | 31822-SNV-T | SNV | 1 | T | A | 6 | 16915 | 0.03547148 | 37.4986292 | WT | FALSE | CAN | JSY2424 | 8 | WT-Rep8 | CTG | T>A | A/T+1 | A/T-1 | TA>AT | TA>AT | TGCATCTGA | TGAATAT | GAC | CTG | T | T | T>A | T>A_CTG | TA>AT_TGAATAT | 9 | FALSE | FALSE |
| 131_v.txt | 32073 | 32073-SNV- | SNV | 1 | A | T | 22 | 27414 | 0.08025097 | 59.9545455 | WT | FALSE | CAN | JSY2424 | 8 | WT-Rep8 | TAA | A>T | A/T+1 | A/T-1 | TA>AT | TA>AT | ATAAATAAC | AACCATG | ATT | ATT | T | A | T>A | T>A_ATT | TA>AT_AACCATG | 9 | FALSE | FALSE |
| 131_v.txt | 32127 | 32127-SNV-T | SNV | 1 | T | A | 2 | 27414 | 7.30E-03 | 33.9998466 | WT | FALSE | CAN | JSY2424 | 8 | WT-Rep8 | TTT | T>A | A/T+1 | A/T-1 | TA>AT | TA>AT | AAAACTTTG | TTGTCAC | AAA | TTT | T | T | T>A | T>A_TTT | TA>AT_TTGTCAC | 9 | FALSE | FALSE |
| 131_v.txt | 32314 | 32314-SNV-T | SNV | 1 | T | A | 6 | 35671 | 0.01682039 | 45.5019272 | WT | FALSE | CAN | JSY2424 | 8 | WT-Rep8 | GTT | T>A | A/T+1 | A/T-1 | TA>AT | TA>AT | TAATGGTTG | TTGTTAA | CAA | GTT | T | T | T>A | T>A_GTT | TA>AT_TTGTTAA | 9 | FALSE | FALSE |
| 131_v.txt | 32315 | 32315-SNV-T | SNV | 1 | T | A | 4 | 35671 | 0.01121359 | 49.2497048 | WT | FALSE | CAN | JSY2424 | 8 | WT-Rep8 | TTG | T>A | A/T+1 | A/T-1 | TA>AT | TA>AT | AATGGTTGT | TGTTAAG | AAC | TTG | T | T | T>A | T>A_TTG | TA>AT_TGTTAAG | 9 | FALSE | FALSE |
| 131_v.txt | 32338 | 32338-SNV-T | SNV | 1 | T | A | 5 | 35671 | 0.01401699 | 56.5996091 | WT | FALSE | CAN | JSY2424 | 8 | WT-Rep8 | ATA | T>A | A/T+1 | A/T-1 | TA>AT | TA>AT | TGAAGATAT | TATGTGG | TAT | ATA | T | T | T>A | T>A_ATA | TA>AT_TATGTGG | 9 | FALSE | FALSE |
| 131_v.txt | 32429 | 32429-SNV- | SNV | 1 | A | T | 33 | 35672 | 0.09250953 | 49.9090909 | WT | FALSE | CAN | JSY2424 | 8 | WT-Rep8 | CAT | A>T | A/T+1 | A/T-1 | TA>AT | TA>AT | AGGGTCATT | ATTGTAT | GTA | GTA | T | A | T>A | T>A_GTA | TA>AT_ATTGTAT | 9 | FALSE | FALSE |
| 131_v.txt | 32527 | 32527-SNV-T | SNV | 1 | T | A | 30 | 35672 | 0.08409957 | 39.2971873 | WT | FALSE | CAN | JSY2424 | 8 | WT-Rep8 | CTG | T>A | A/T+1 | A/T-1 | TA>AT | TA>AT | ATTTTCTGG | TGGGGTT | GAC | CTG | T | T | T>A | T>A_CTG | TA>AT_TGGGGTT | 9 | FALSE | FALSE |
| 131_v.txt | 32820 | 32820-SNV-T | SNV | 1 | T | A | 28 | 20964 | 0.1335623 | 45 | WT | FALSE | CAN | JSY2424 | 8 | WT-Rep8 | GTT | T>A | A/T+1 | A/T-1 | TA>AT | TA>AT | AACAAGTTC | TTCATTA | CAA | GTT | T | T | T>A | T>A_GTT | TA>AT_TTCATTA | 9 | FALSE | FALSE |
| 131_v.txt | 33213 | 33213-SNV-T | SNV | 1 | T | A | 4 | 31295 | 0.0127816 | 36.9959619 | WT | FALSE | CAN | JSY2424 | 8 | WT-Rep8 | CTT | T>A | A/T+1 | A/T-1 | TA>AT | TA>AT | TCTCTCTTCA | TTCACTT | GAA | CTT | T | T | T>A | T>A_CTT | TA>AT_TTCACTT | 9 | FALSE | FALSE |
| 131_v.txt | 33214 | 33214-SNV-T | SNV | 1 | T | A | 1 | 31295 | 3.20E-03 | 37.9998963 | WT | FALSE | CAN | JSY2424 | 8 | WT-Rep8 | TTC | T>A | A/T+1 | A/T-1 | TA>AT | TA>AT | CTCTCTTCAC | TCACTTC | AAG | TTC | T | T | T>A | T>A_TTC | TA>AT_TCACTTC | 9 | FALSE | FALSE |
| 131_v.txt | 33345 | 33345-SNV- | SNV | 1 | A | T | 45 | 31295 | 0.14379294 | 35.9333333 | WT | FALSE | CAN | JSY2424 | 8 | WT-Rep8 | CAA | A>T | A/T+1 | A/T-1 | TA>AT | TA>AT | TCTTTCAATG | AATGGTA | GTT | GTT | T | A | T>A | T>A_GTT | TA>AT_AATGGTA | 9 | FALSE | FALSE |
| 131_v.txt | 33476 | 33476-SNV-T | SNV | 1 | T | A | 5 | 31291 | 0.01597904 | 36.2 | WT | FALSE | CAN | JSY2424 | 8 | WT-Rep8 | CTT | T>A | A/T+1 | A/T-1 | TA>AT | TA>AT | TATGCCTTTT | TTTTTTT | GAA | CTT | T | T | T>A | T>A_CTT | TA>AT_TTTTTTT | 9 | FALSE | FALSE |
| 131_v.txt | 31650 | 31650-SNV- | SNV | 1 | C | T | 40 | 16915 | 0.2364765 | 35.875 | WT | FALSE | CAN | JSY2424 | 8 | WT-Rep8 | GCG | C>T | A/T+1 | G/C-1 | CG>TA | CG>TA | AGAATGCGA | CGAAATG | CGC | GCG | C | A | C>T | C>T_GCG | CG>TA_CGAAATG | 9 | FALSE | FALSE |
| 131_v.txt | 31658 | 31658-SNV- | SNV | 1 | C | T | 22 | 16915 | 0.13006208 | 37.6818458 | WT | FALSE | CAN | JSY2424 | 8 | WT-Rep8 | GCG | C>T | A/T+1 | G/C-1 | CG>TA | CG>TA | AAATGGCGT | CGTGGAA | CGC | GCG | C | A | C>T | C>T_GCG | CG>TA_CGTGGAA | 9 | FALSE | FALSE |
| 131_v.txt | 31817 | 31817-SNV- | SNV | 1 | G | A | 5 | 16915 | 0.02955956 | 63.3999951 | WT | FALSE | CAN | JSY2424 | 8 | WT-Rep8 | TGC | G>A | A/T+1 | G/C-1 | CG>TA | CG>TA | AAATCTGCA | GCATCTG | ACG | ACG | C | T | C>T | C>T_ACG | CG>TA_GCATCTG | 9 | FALSE | FALSE |
| 131_v.txt | 31821 | 31821-SNV- | SNV | 1 | C | T | 1 | 16915 | 5.91E-03 | 33.9995686 | WT | FALSE | CAN | JSY2424 | 8 | WT-Rep8 | TCT | C>T | A/T+1 | G/C-1 | CG>TA | CG>TA | CTGCATCTG | CTGAATA | AGA | TCT | C | A | C>T | C>T_TCT | CG>TA_CTGAATA | 9 | FALSE | FALSE |
| 131_v.txt | 31830 | 31830-SNV- | SNV | 1 | C | T | 21 | 16915 | 0.12415016 | 47.6190476 | WT | FALSE | CAN | JSY2424 | 8 | WT-Rep8 | GCA | C>T | A/T+1 | G/C-1 | CG>TA | CG>TA | AATATGCAT | CATTGAA | CGT | GCA | C | A | C>T | C>T_GCA | CG>TA_CATTGAA | 9 | FALSE | FALSE |
| 131_v.txt | 31881 | 31881-SNV- | SNV | 1 | G | A | 15 | 16915 | 0.08867869 | 43.532821 | WT | FALSE | CAN | JSY2424 | 8 | WT-Rep8 | GGC | G>A | A/T+1 | G/C-1 | CG>TA | CG>TA | TAGGCGGCA | GCAGCAA | CCG | CCG | C | T | C>T | C>T_CCG | CG>TA_GCAGCAA | 9 | FALSE | FALSE |
| 131_v.txt | 31882 | 31882-SNV- | SNV | 1 | C | T | 17 | 16915 | 0.10050251 | 41.2263001 | WT | FALSE | CAN | JSY2424 | 8 | WT-Rep8 | GCA | C>T | A/T+1 | G/C-1 | CG>TA | CG>TA | AGGCGGCAG | CAGCAAA | CGT | GCA | C | A | C>T | C>T_GCA | CG>TA_CAGCAAA | 9 | FALSE | FALSE |
| 131_v.txt | 31920 | 31920-SNV- | SNV | 1 | G | A | 16 | 16911 | 0.09461297 | 36.6875 | WT | FALSE | CAN | JSY2424 | 8 | WT-Rep8 | CGT | G>A | A/T+1 | G/C-1 | CG>TA | CG>TA | AAAGCCGTG | GTGAAAC | GCA | GCA | C | T | C>T | C>T_GCA | CG>TA_GTGAAAC | 9 | FALSE | FALSE |
| 131_v.txt | 31960 | 31960-SNV- | SNV | 1 | C | T | 27 | 27414 | 0.09848982 | 35.2962963 | WT | FALSE | CAN | JSY2424 | 8 | WT-Rep8 | GCC | C>T | A/T+1 | G/C-1 | CG>TA | CG>TA | ATGTGGCCG | CCGCATA | CGG | GCC | C | A | C>T | C>T_GCC | CG>TA_CCGCATA | 9 | FALSE | FALSE |
| 131_v.txt | 31962 | 31962-SNV- | SNV | 1 | G | A | 58 | 27414 | 0.21157073 | 37.5862069 | WT | FALSE | CAN | JSY2424 | 8 | WT-Rep8 | CGC | G>A | A/T+1 | G/C-1 | CG>TA | CG>TA | GTGGCCGCA | GCATAAT | GCG | GCG | C | T | C>T | C>T_GCG | CG>TA_GCATAAT | 9 | FALSE | FALSE |
| 131_v.txt | 31972 | 31972-SNV- | SNV | 1 | C | T | 20 | 27414 | 0.07295542 | 35.65 | WT | FALSE | CAN | JSY2424 | 8 | WT-Rep8 | GCC | C>T | A/T+1 | G/C-1 | CG>TA | CG>TA | AATAAGCCA | CCAAGCC | CGG | GCC | C | A | C>T | C>T_GCC | CG>TA_CCAAGCC | 9 | FALSE | FALSE |
| 131_v.txt | 31981 | 31981-SNV- | SNV | 1 | G | A | 31 | 27414 | 0.11308091 | 49.7419355 | WT | FALSE | CAN | JSY2424 | 8 | WT-Rep8 | GGC | G>A | A/T+1 | G/C-1 | CG>TA | CG>TA | AGCCGGGCA | GCATTAA | CCG | CCG | C | T | C>T | C>T_CCG | CG>TA_GCATTAA | 9 | FALSE | FALSE |
| 131_v.txt | 32057 | 32057-SNV- | SNV | 1 | C | T | 22 | 27414 | 0.08025097 | 60.4090909 | WT | FALSE | CAN | JSY2424 | 8 | WT-Rep8 | GCG | C>T | A/T+1 | G/C-1 | CG>TA | CG>TA | GATGTGCGA | CGAGATT | CGC | GCG | C | A | C>T | C>T_GCG | CG>TA_CGAGATT | 9 | FALSE | FALSE |
| 131_v.txt | 32114 | 32114-SNV- | SNV | 1 | C | T | 36 | 27414 | 0.13131976 | 52.5555556 | WT | FALSE | CAN | JSY2424 | 8 | WT-Rep8 | GCC | C>T | A/T+1 | G/C-1 | CG>TA | CG>TA | TAATAGCCA | CCATTCG | CGG | GCC | C | A | C>T | C>T_GCC | CG>TA_CCATTCG | 9 | FALSE | FALSE |
| 131_v.txt | 32165 | 32165-SNV- | SNV | 1 | G | A | 43 | 27410 | 0.15687705 | 44.0232558 | WT | FALSE | CAN | JSY2424 | 8 | WT-Rep8 | CGC | G>A | A/T+1 | G/C-1 | CG>TA | CG>TA | CAAAGCGCC | GCCAAAT | GCG | GCG | C | T | C>T | C>T_GCG | CG>TA_GCCAAAT | 9 | FALSE | FALSE |
| 131_v.txt | 32166 | 32166-SNV- | SNV | 1 | C | T | 27 | 27410 | 0.0985042 | 40.7034475 | WT | FALSE | CAN | JSY2424 | 8 | WT-Rep8 | GCC | C>T | A/T+1 | G/C-1 | CG>TA | CG>TA | AAAGCGCCA | CCAAATG | CGG | GCC | C | A | C>T | C>T_GCC | CG>TA_CCAAATG | 9 | FALSE | FALSE |
| 131_v.txt | 32173 | 32173-SNV- | SNV | 1 | C | T | 35 | 27410 | 0.12769062 | 48.7428571 | WT | FALSE | CAN | JSY2424 | 8 | WT-Rep8 | GCA | C>T | A/T+1 | G/C-1 | CG>TA | CG>TA | CAAATGCAG | CAGCAGT | CGT | GCA | C | A | C>T | C>T_GCA | CG>TA_CAGCAGT | 9 | FALSE | FALSE |
| 131_v.txt | 32176 | 32176-SNV- | SNV | 1 | C | T | 21 | 27410 | 0.07661437 | 43.3333333 | WT | FALSE | CAN | JSY2424 | 8 | WT-Rep8 | GCA | C>T | A/T+1 | G/C-1 | CG>TA | CG>TA | ATGCAGCAG | CAGTAAC | CGT | GCA | C | A | C>T | C>T_GCA | CG>TA_CAGTAAC | 9 | FALSE | FALSE |
| 131_v.txt | 32271 | 32271-SNV- | SNV | 1 | C | T | 47 | 35671 | 0.13175969 | 30.212766 | WT | FALSE | CAN | JSY2424 | 8 | WT-Rep8 | ACG | C>T | A/T+1 | G/C-1 | CG>TA | CG>TA | AAAATACGG | CGGGAAC | TGC | ACG | C | A | C>T | C>T_ACG | CG>TA_CGGGAAC | 9 | FALSE | FALSE |
| 131_v.txt | 32299 | 32299-SNV- | SNV | 1 | C | T | 59 | 35671 | 0.16540047 | 53.220339 | WT | FALSE | CAN | JSY2424 | 8 | WT-Rep8 | GCG | C>T | A/T+1 | G/C-1 | CG>TA | CG>TA | AATTTGCGG | CGGCAGA | CGC | GCG | C | A | C>T | C>T_GCG | CG>TA_CGGCAGA | 9 | FALSE | FALSE |
| 131_v.txt | 32436 | 32436-SNV- | SNV | 1 | G | A | 968 | 35672 | 2.71361292 | 55.2479339 | WT | FALSE | CAN | JSY2424 | 8 | WT-Rep8 | TGG | G>A | A/T+1 | G/C-1 | CG>TA | CG>TA | TTGTATGGA | GGAACTA | ACC | ACC | C | T | C>T | C>T_ACC | CG>TA_GGAACTA | 9 | FALSE | FALSE |
| 131_v.txt | 32449 | 32449-SNV- | SNV | 1 | C | T | 27 | 35672 | 0.07568962 | 42.9259259 | WT | FALSE | CAN | JSY2424 | 8 | WT-Rep8 | CCA | C>T | A/T+1 | G/C-1 | CG>TA | CG>TA | AAAGTCCAA | CAATGAA | GGT | CCA | C | A | C>T | C>T_CCA | CG>TA_CAATGAA | 9 | FALSE | FALSE |
| 131_v.txt | 32463 | 32463-SNV- | SNV | 1 | G | A | 61 | 35672 | 0.17100247 | 47.2295082 | WT | FALSE | CAN | JSY2424 | 8 | WT-Rep8 | AGA | G>A | A/T+1 | G/C-1 | CG>TA | CG>TA | AATAGAGAG | GAGCCAA | TCT | TCT | C | T | C>T | C>T_TCT | CG>TA_GAGCCAA | 9 | FALSE | FALSE |
| 131_v.txt | 32466 | 32466-SNV- | SNV | 1 | C | T | 55 | 35672 | 0.15418255 | 42.0363636 | WT | FALSE | CAN | JSY2424 | 8 | WT-Rep8 | GCC | C>T | A/T+1 | G/C-1 | CG>TA | CG>TA | AGAGAGCCA | CCAATGT | CGG | GCC | C | A | C>T | C>T_GCC | CG>TA_CCAATGT | 9 | FALSE | FALSE |
| 131_v.txt | 32487 | 32487-SNV- | SNV | 1 | C | T | 34 | 35672 | 0.09531285 | 40.8823529 | WT | FALSE | CAN | JSY2424 | 8 | WT-Rep8 | ACG | C>T | A/T+1 | G/C-1 | CG>TA | CG>TA | AAGATACGG | CGGAAAA | TGC | ACG | C | A | C>T | C>T_ACG | CG>TA_CGGAAAA | 9 | FALSE | FALSE |
| 131_v.txt | 32528 | 32528-SNV- | SNV | 1 | G | A | 7 | 35672 | 0.01962323 | 38.4283634 | WT | FALSE | CAN | JSY2424 | 8 | WT-Rep8 | TGG | G>A | A/T+1 | G/C-1 | CG>TA | CG>TA | TTTTCTGGG | GGGGTTT | ACC | ACC | C | T | C>T | C>T_ACC | CG>TA_GGGGTTT | 9 | FALSE | FALSE |
| 131_v.txt | 32530 | 32530-SNV- | SNV | 1 | G | A | 17 | 35672 | 0.04765643 | 36.6470588 | WT | FALSE | CAN | JSY2424 | 8 | WT-Rep8 | GGG | G>A | A/T+1 | G/C-1 | CG>TA | CG>TA | TTCTGGGGT | GGTTTGC | CCC | CCC | C | T | C>T | C>T_CCC | CG>TA_GGTTTGC | 9 | FALSE | FALSE |
| 131_v.txt | 32535 | 32535-SNV- | SNV | 1 | G | A | 47 | 35672 | 0.131756 | 36.7233686 | WT | FALSE | CAN | JSY2424 | 8 | WT-Rep8 | TGC | G>A | A/T+1 | G/C-1 | CG>TA | CG>TA | GGGTTTGCA | GCAGCTT | ACG | ACG | C | T | C>T | C>T_ACG | CG>TA_GCAGCTT | 9 | FALSE | FALSE |
| 131_v.txt | 32536 | 32536-SNV- | SNV | 1 | C | T | 49 | 35672 | 0.13736264 | 36.8360604 | WT | FALSE | CAN | JSY2424 | 8 | WT-Rep8 | GCA | C>T | A/T+1 | G/C-1 | CG>TA | CG>TA | GGTTTGCAG | CAGCTTC | CGT | GCA | C | A | C>T | C>T_GCA | CG>TA_CAGCTTC | 9 | FALSE | FALSE |
| 131_v.txt | 32575 | 32575-SNV- | SNV | 1 | G | A | 23 | 20965 | 0.10970665 | 35.1304348 | WT | FALSE | CAN | JSY2424 | 8 | WT-Rep8 | TGA | G>A | A/T+1 | G/C-1 | CG>TA | CG>TA | TACCTTGAA | GAAATGT | ACT | ACT | C | T | C>T | C>T_ACT | CG>TA_GAAATGT | 9 | FALSE | FALSE |
| 131_v.txt | 32590 | 32590-SNV- | SNV | 1 | C | T | 36 | 20964 | 0.17172295 | 34.8055556 | WT | FALSE | CAN | JSY2424 | 8 | WT-Rep8 | GCG | C>T | A/T+1 | G/C-1 | CG>TA | CG>TA | AGGCAGCGT | CGTTAAT | CGC | GCG | C | A | C>T | C>T_GCG | CG>TA_CGTTAAT | 9 | FALSE | FALSE |
| 131_v.txt | 32700 | 32700-SNV- | SNV | 1 | C | T | 33 | 20964 | 0.15741271 | 57.9090909 | WT | FALSE | CAN | JSY2424 | 8 | WT-Rep8 | GCC | C>T | A/T+1 | G/C-1 | CG>TA | CG>TA | ACTGGGCCG | CCGGTAA | CGG | GCC | C | A | C>T | C>T_GCC | CG>TA_CCGGTAA | 9 | FALSE | FALSE |
| 131_v.txt | 32702 | 32702-SNV- | SNV | 1 | G | A | 25 | 20964 | 0.11925205 | 43.76 | WT | FALSE | CAN | JSY2424 | 8 | WT-Rep8 | CGG | G>A | A/T+1 | G/C-1 | CG>TA | CG>TA | TGGGCCGGT | GGTAACC | GCC | GCC | C | T | C>T | C>T_GCC | CG>TA_GGTAACC | 9 | FALSE | FALSE |
| 131_v.txt | 32713 | 32713-SNV- | SNV | 1 | C | T | 20 | 20964 | 0.09540164 | 58.7 | WT | FALSE | CAN | JSY2424 | 8 | WT-Rep8 | GCA | C>T | A/T+1 | G/C-1 | CG>TA | CG>TA | CCCCAGCAC | CACCACA | CGT | GCA | C | A | C>T | C>T_GCA | CG>TA_CACCACA | 9 | FALSE | FALSE |
| 131_v.txt | 32798 | 32798-SNV- | SNV | 1 | G | A | 25 | 20964 | 0.11925205 | 40.48 | WT | FALSE | CAN | JSY2424 | 8 | WT-Rep8 | CGT | G>A | A/T+1 | G/C-1 | CG>TA | CG>TA | TTCACCGTA | GTAATAT | GCA | GCA | C | T | C>T | C>T_GCA | CG>TA_GTAATAT | 9 | FALSE | FALSE |
| 131_v.txt | 32860 | 32860-SNV- | SNV | 1 | C | T | 20 | 20961 | 0.0954153 | 34.3 | WT | FALSE | CAN | JSY2424 | 8 | WT-Rep8 | GCC | C>T | A/T+1 | G/C-1 | CG>TA | CG>TA | TCCATGCCG | CCGCCAG | CGG | GCC | C | A | C>T | C>T_GCC | CG>TA_CCGCCAG | 9 | FALSE | FALSE |
| 131_v.txt | 32900 | 32900-SNV- | SNV | 1 | G | A | 27 | 31752 | 0.08503401 | 36.4797809 | WT | FALSE | CAN | JSY2424 | 8 | WT-Rep8 | GGC | G>A | A/T+1 | G/C-1 | CG>TA | CG>TA | GACTTGGCC | GCCAACT | CCG | CCG | C | T | C>T | C>T_CCG | CG>TA_GCCAACT | 9 | FALSE | FALSE |
| 131_v.txt | 32901 | 32901-SNV- | SNV | 1 | C | T | 65 | 31752 | 0.20471151 | 32.7230152 | WT | FALSE | CAN | JSY2424 | 8 | WT-Rep8 | GCC | C>T | A/T+1 | G/C-1 | CG>TA | CG>TA | ACTTGGCCA | CCAACTA | CGG | GCC | C | A | C>T | C>T_GCC | CG>TA_CCAACTA | 9 | FALSE | FALSE |
| 131_v.txt | 32922 | 32922-SNV- | SNV | 1 | G | A | 25 | 31752 | 0.0787352 | 48.44 | WT | FALSE | CAN | JSY2424 | 8 | WT-Rep8 | GGC | G>A | A/T+1 | G/C-1 | CG>TA | CG>TA | TCCAGGGCA | GCAAAAG | CCG | CCG | C | T | C>T | C>T_CCG | CG>TA_GCAAAAG | 9 | FALSE | FALSE |
| 131_v.txt | 32935 | 32935-SNV- | SNV | 1 | C | T | 39 | 31752 | 0.12282691 | 39.7948718 | WT | FALSE | CAN | JSY2424 | 8 | WT-Rep8 | GCC | C>T | A/T+1 | G/C-1 | CG>TA | CG>TA | TGATTGCCC | CCCAAGA | CGG | GCC | C | A | C>T | C>T_GCC | CG>TA_CCCAAGA | 9 | FALSE | FALSE |
| 131_v.txt | 32937 | 32937-SNV- | SNV | 1 | C | T | 90 | 31752 | 0.28344671 | 42.2888889 | WT | FALSE | CAN | JSY2424 | 8 | WT-Rep8 | CCA | C>T | A/T+1 | G/C-1 | CG>TA | CG>TA | ATTGCCCAA | CAAGAAA | GGT | CCA | C | A | C>T | C>T_CCA | CG>TA_CAAGAAA | 9 | FALSE | FALSE |
| 131_v.txt | 32946 | 32946-SNV- | SNV | 1 | C | T | 32 | 31752 | 0.10078105 | 54.0625 | WT | FALSE | CAN | JSY2424 | 8 | WT-Rep8 | CCA | C>T | A/T+1 | G/C-1 | CG>TA | CG>TA | GAAAACCAA | CAATACA | GGT | CCA | C | A | C>T | C>T_CCA | CG>TA_CAATACA | 9 | FALSE | FALSE |
| 131_v.txt | 32965 | 32965-SNV- | SNV | 1 | C | T | 42 | 31752 | 0.13227513 | 57.7857143 | WT | FALSE | CAN | JSY2424 | 8 | WT-Rep8 | GCC | C>T | A/T+1 | G/C-1 | CG>TA | CG>TA | CATTGGCCG | CCGCACC | CGG | GCC | C | A | C>T | C>T_GCC | CG>TA_CCGCACC | 9 | FALSE | FALSE |
| 131_v.txt | 32967 | 32967-SNV- | SNV | 1 | G | A | 74 | 31752 | 0.23305619 | 57.2432432 | WT | FALSE | CAN | JSY2424 | 8 | WT-Rep8 | CGC | G>A | A/T+1 | G/C-1 | CG>TA | CG>TA | TTGGCCGCA | GCACCAA | GCG | GCG | C | T | C>T | C>T_GCG | CG>TA_GCACCAA | 9 | FALSE | FALSE |
| 131_v.txt | 32977 | 32977-SNV- | SNV | 1 | C | T | 35 | 31752 | 0.11022928 | 58.8857143 | WT | FALSE | CAN | JSY2424 | 8 | WT-Rep8 | GCT | C>T | A/T+1 | G/C-1 | CG>TA | CG>TA | CAAATGCTG | CTGGAGA | CGA | GCT | C | A | C>T | C>T_GCT | CG>TA_CTGGAGA | 9 | FALSE | FALSE |
| 131_v.txt | 33056 | 33056-SNV- | SNV | 1 | C | T | 43 | 31752 | 0.13542454 | 56.2093023 | WT | FALSE | CAN | JSY2424 | 8 | WT-Rep8 | GCG | C>T | A/T+1 | G/C-1 | CG>TA | CG>TA | GGACTGCGT | CGTGACA | CGC | GCG | C | A | C>T | C>T_GCG | CG>TA_CGTGACA | 9 | FALSE | FALSE |
| 131_v.txt | 33070 | 33070-SNV- | SNV | 1 | C | T | 38 | 31752 | 0.1196775 | 58.1842105 | WT | FALSE | CAN | JSY2424 | 8 | WT-Rep8 | GCC | C>T | A/T+1 | G/C-1 | CG>TA | CG>TA | AATATGCCA | CCAAAGA | CGG | GCC | C | A | C>T | C>T_GCC | CG>TA_CCAAAGA | 9 | FALSE | FALSE |
| 131_v.txt | 33104 | 33104-SNV- | SNV | 1 | G | A | 48 | 31752 | 0.15117158 | 36.7083333 | WT | FALSE | CAN | JSY2424 | 8 | WT-Rep8 | CGC | G>A | A/T+1 | G/C-1 | CG>TA | CG>TA | AAGAGCGCC | GCCCACT | GCG | GCG | C | T | C>T | C>T_GCG | CG>TA_GCCCACT | 9 | FALSE | FALSE |
| 131_v.txt | 33113 | 33113-SNV- | SNV | 1 | G | A | 24 | 31752 | 0.07558579 | 39.4179311 | WT | FALSE | CAN | JSY2424 | 8 | WT-Rep8 | GGC | G>A | A/T+1 | G/C-1 | CG>TA | CG>TA | CACTGGGCC | GCCGGCG | CCG | CCG | C | T | C>T | C>T_CCG | CG>TA_GCCGGCG | 9 | FALSE | FALSE |
| 131_v.txt | 33114 | 33114-SNV- | SNV | 1 | C | T | 24 | 31752 | 0.07558579 | 42.9587533 | WT | FALSE | CAN | JSY2424 | 8 | WT-Rep8 | GCC | C>T | A/T+1 | G/C-1 | CG>TA | CG>TA | ACTGGGCCG | CCGGCGT | CGG | GCC | C | A | C>T | C>T_GCC | CG>TA_CCGGCGT | 9 | FALSE | FALSE |
| 131_v.txt | 33118 | 33118-SNV- | SNV | 1 | C | T | 33 | 31752 | 0.10393046 | 39.9086262 | WT | FALSE | CAN | JSY2424 | 8 | WT-Rep8 | GCG | C>T | A/T+1 | G/C-1 | CG>TA | CG>TA | GGCCGGCGT | CGTTGGT | CGC | GCG | C | A | C>T | C>T_GCG | CG>TA_CGTTGGT | 9 | FALSE | FALSE |
| 131_v.txt | 33119 | 33119-SNV- | SNV | 1 | G | A | 25 | 31752 | 0.0787352 | 44.9197142 | WT | FALSE | CAN | JSY2424 | 8 | WT-Rep8 | CGT | G>A | A/T+1 | G/C-1 | CG>TA | CG>TA | GCCGGCGTT | GTTGGTC | GCA | GCA | C | T | C>T | C>T_GCA | CG>TA_GTTGGTC | 9 | FALSE | FALSE |
| 131_v.txt | 33168 | 33168-SNV- | SNV | 1 | C | T | 32 | 31746 | 0.1008001 | 36.7185517 | WT | FALSE | CAN | JSY2424 | 8 | WT-Rep8 | ACC | C>T | A/T+1 | G/C-1 | CG>TA | CG>TA | ATAGTACCA | CCACCAA | TGG | ACC | C | A | C>T | C>T_ACC | CG>TA_CCACCAA | 9 | FALSE | FALSE |
| 131_v.txt | 33171 | 33171-SNV- | SNV | 1 | C | T | 49 | 31746 | 0.15435015 | 36.3877551 | WT | FALSE | CAN | JSY2424 | 8 | WT-Rep8 | ACC | C>T | A/T+1 | G/C-1 | CG>TA | CG>TA | GTACCACCA | CCAAGGG | TGG | ACC | C | A | C>T | C>T_ACC | CG>TA_CCAAGGG | 9 | FALSE | FALSE |
| 131_v.txt | 33205 | 33205-SNV- | SNV | 1 | G | A | 10 | 31295 | 0.03195399 | 37.5002328 | WT | FALSE | CAN | JSY2424 | 8 | WT-Rep8 | AGC | G>A | A/T+1 | G/C-1 | CG>TA | CG>TA | GCTTAAGCT | GCTCTCT | TCG | TCG | C | T | C>T | C>T_TCG | CG>TA_GCTCTCT | 9 | FALSE | FALSE |
| 131_v.txt | 33215 | 33215-SNV- | SNV | 1 | C | T | 6 | 31294 | 0.019173 | 31.8310314 | WT | FALSE | CAN | JSY2424 | 8 | WT-Rep8 | TCA | C>T | A/T+1 | G/C-1 | CG>TA | CG>TA | TCTCTTCACT | CACTTCA | AGT | TCA | C | A | C>T | C>T_TCA | CG>TA_CACTTCA | 9 | FALSE | FALSE |
| 131_v.txt | 33258 | 33258-SNV- | SNV | 1 | C | T | 33 | 31295 | 0.10544816 | 37.8484849 | WT | FALSE | CAN | JSY2424 | 8 | WT-Rep8 | GCC | C>T | A/T+1 | G/C-1 | CG>TA | CG>TA | CCTATGCCAT | CCATCCT | CGG | GCC | C | A | C>T | C>T_GCC | CG>TA_CCATCCT | 9 | FALSE | FALSE |
| 131_v.txt | 33275 | 33275-SNV- | SNV | 1 | C | T | 27 | 31295 | 0.08627576 | 42.4813392 | WT | FALSE | CAN | JSY2424 | 8 | WT-Rep8 | ACG | C>T | A/T+1 | G/C-1 | CG>TA | CG>TA | AGAGAACGT | CGTATCC | TGC | ACG | C | A | C>T | C>T_ACG | CG>TA_CGTATCC | 9 | FALSE | FALSE |
| 131_v.txt | 33276 | 33276-SNV- | SNV | 1 | G | A | 43 | 31295 | 0.13740214 | 46.5697574 | WT | FALSE | CAN | JSY2424 | 8 | WT-Rep8 | CGT | G>A | A/T+1 | G/C-1 | CG>TA | CG>TA | GAGAACGTA | GTATCCT | GCA | GCA | C | T | C>T | C>T_GCA | CG>TA_GTATCCT | 9 | FALSE | FALSE |
| 131_v.txt | 33309 | 33309-SNV- | SNV | 1 | C | T | 31 | 31295 | 0.09905736 | 55.6097725 | WT | FALSE | CAN | JSY2424 | 8 | WT-Rep8 | GCG | C>T | A/T+1 | G/C-1 | CG>TA | CG>TA | AAAGAGCGC | CGCAATG | CGC | GCG | C | A | C>T | C>T_GCG | CG>TA_CGCAATG | 9 | FALSE | FALSE |
| 131_v.txt | 33310 | 33310-SNV- | SNV | 1 | G | A | 37 | 31292 | 0.11824108 | 54.5815313 | WT | FALSE | CAN | JSY2424 | 8 | WT-Rep8 | CGC | G>A | A/T+1 | G/C-1 | CG>TA | CG>TA | AAGAGCGCA | GCAATGG | GCG | GCG | C | T | C>T | C>T_GCG | CG>TA_GCAATGG | 9 | FALSE | FALSE |
| 131_v.txt | 33392 | 33392-SNV- | SNV | 1 | G | A | 37 | 31295 | 0.11822975 | 49.6756757 | WT | FALSE | CAN | JSY2424 | 8 | WT-Rep8 | CGT | G>A | A/T+1 | G/C-1 | CG>TA | CG>TA | AACGTCGTG | GTGAAAG | GCA | GCA | C | T | C>T | C>T_GCA | CG>TA_GTGAAAG | 9 | FALSE | FALSE |
| 131_v.txt | 33412 | 33412-SNV- | SNV | 1 | G | A | 55 | 31295 | 0.17574692 | 45.2181818 | WT | FALSE | CAN | JSY2424 | 8 | WT-Rep8 | GGC | G>A | A/T+1 | G/C-1 | CG>TA | CG>TA | TGACCGGCT | GCTCATT | CCG | CCG | C | T | C>T | C>T_CCG | CG>TA_GCTCATT | 9 | FALSE | FALSE |
| 131_v.txt | 33474 | 33474-SNV- | SNV | 1 | C | T | 32 | 31292 | 0.10226256 | 34.0937658 | WT | FALSE | CAN | JSY2424 | 8 | WT-Rep8 | GCC | C>T | A/T+1 | G/C-1 | CG>TA | CG>TA | GCTATGCCTT | CCTTTTT | CGG | GCC | C | A | C>T | C>T_GCC | CG>TA_CCTTTTT | 9 | FALSE | FALSE |
| 131_v.txt | 31657 | 31657-SNV- | SNV | 1 | G | C | 8 | 16915 | 0.0472953 | 13.8748603 | WT | FALSE | CAN | JSY2424 | 8 | WT-Rep8 | GGC | G>C | G/C+1 | G/C-1 | CG>GC | CG>GC | GAAATGGCG | GCGTGGA | CCG | CCG | C | G | C>G | C>G_CCG | CG>GC_GCGTGGA | 9 | FALSE | FALSE |
| 131_v.txt | 31823 | 31823-SNV- | SNV | 1 | G | C | 1 | 16915 | 5.91E-03 | 58.9821734 | WT | FALSE | CAN | JSY2424 | 8 | WT-Rep8 | TGA | G>C | G/C+1 | G/C-1 | CG>GC | CG>GC | GCATCTGAA | GAATATG | ACT | ACT | C | G | C>G | C>G_ACT | CG>GC_GAATATG | 9 | FALSE | FALSE |

| | | | | | | | | | | | | | | | | | | | | | | | | | | | | | | | | | | |
|---|---|---|---|---|---|---|---|---|---|---|---|---|---|---|---|---|---|---|---|---|---|---|---|---|---|---|---|---|---|---|---|---|---|---|
| 131_v.txt | 32271 | 32271-SNV-C | SNV | 1 | C | G | 33 | 35671 | 0.09251213 | 35.6666667 | WT | FALSE | CAN | JSY2424 | 8 | WT-Rep8 | ACG | C>G | G/C+1 | G/C-1 | CG>GC | CG>GC | AAAATACGG | CGGGAAC | TGC | ACG | C | C | C>G | C>G_ACG | CG>GC_CGGGAAC | 9 | FALSE | FALSE |
| 131_v.txt | 32302 | 32302-SNV-C | SNV | 1 | C | G | 11 | 35671 | 0.03083738 | 58 | WT | FALSE | CAN | JSY2424 | 8 | WT-Rep8 | GCA | C>G | G/C+1 | G/C-1 | CG>GC | CG>GC | TTGCGGCAG | CAGAAAT | CGT | GCA | C | C | C>G | C>G_GCA | CG>GC_CAGAAAT | 9 | FALSE | FALSE |
| 131_v.txt | 32399 | 32399-SNV-C | SNV | 1 | G | C | 125 | 35672 | 0.35041489 | 58.264 | WT | FALSE | CAN | JSY2424 | 8 | WT-Rep8 | CGT | G>C | G/C+1 | G/C-1 | CG>GC | CG>GC | AGAAACGTA | GTAGGAA | GCA | GCA | C | G | C>G | C>G_GCA | CG>GC_GTAGGAA | 9 | FALSE | FALSE |
| 131_v.txt | 32449 | 32449-SNV-C | SNV | 1 | C | G | 9 | 35672 | 0.02522987 | 40 | WT | FALSE | CAN | JSY2424 | 8 | WT-Rep8 | CCA | C>G | G/C+1 | G/C-1 | CG>GC | CG>GC | AAAGTCCAA | CAATGAA | GGT | CCA | C | C | C>G | C>G_CCA | CG>GC_CAATGAA | 9 | FALSE | FALSE |
| 131_v.txt | 32528 | 32528-SNV-C | SNV | 1 | G | C | 1 | 35672 | 2.80E-03 | 37.9995297 | WT | FALSE | CAN | JSY2424 | 8 | WT-Rep8 | TGG | G>C | G/C+1 | G/C-1 | CG>GC | CG>GC | TTTTCTGGG | GGGGTTT | ACC | ACC | C | G | C>G | C>G_ACC | CG>GC_GGGGTTT | 9 | FALSE | FALSE |
| 131_v.txt | 32536 | 32536-SNV-C | SNV | 1 | C | G | 3 | 35672 | 8.41E-03 | 37.665734 | WT | FALSE | CAN | JSY2424 | 8 | WT-Rep8 | GCA | C>G | G/C+1 | G/C-1 | CG>GC | CG>GC | GGTTTGCAG | CAGCTTC | CGT | GCA | C | C | C>G | C>G_GCA | CG>GC_CAGCTTC | 9 | FALSE | FALSE |
| 131_v.txt | 32786 | 32786-SNV-C | SNV | 1 | C | G | 9 | 20964 | 0.04293074 | 46.3333333 | WT | FALSE | CAN | JSY2424 | 8 | WT-Rep8 | ACT | C>G | G/C+1 | G/C-1 | CG>GC | CG>GC | CCAGAACTC | CTCGAAT | TGA | ACT | C | C | C>G | C>G_ACT | CG>GC_CTCGAAT | 9 | FALSE | FALSE |
| 131_v.txt | 32788 | 32788-SNV-C | SNV | 1 | C | G | 97 | 20964 | 0.46269796 | 45.6391753 | WT | FALSE | CAN | JSY2424 | 8 | WT-Rep8 | TCG | C>G | G/C+1 | G/C-1 | CG>GC | CG>GC | AGAACTCGA | CGAATTC | AGC | TCG | C | C | C>G | C>G_TCG | CG>GC_CGAATTC | 9 | FALSE | FALSE |
| 131_v.txt | 32794 | 32794-SNV-C | SNV | 1 | C | G | 9 | 20964 | 0.04293074 | 41.7777778 | WT | FALSE | CAN | JSY2424 | 8 | WT-Rep8 | TCA | C>G | G/C+1 | G/C-1 | CG>GC | CG>GC | CGAATTCAC | CACCGTA | AGT | TCA | C | C | C>G | C>G_TCA | CG>GC_CACCGTA | 9 | FALSE | FALSE |
| 131_v.txt | 32876 | 32876-SNV-C | SNV | 1 | G | C | 26 | 31735 | 0.08192847 | 35.8076923 | WT | FALSE | CAN | JSY2424 | 8 | WT-Rep8 | TGT | G>C | G/C+1 | G/C-1 | CG>GC | CG>GC | AACTTTGTA | GTACGTC | ACA | ACA | C | G | C>G | C>G_ACA | CG>GC_GTACGTC | 9 | FALSE | FALSE |
| 131_v.txt | 32946 | 32946-SNV-C | SNV | 1 | C | G | 3 | 31752 | 9.45E-03 | 61.6666667 | WT | FALSE | CAN | JSY2424 | 8 | WT-Rep8 | CCA | C>G | G/C+1 | G/C-1 | CG>GC | CG>GC | GAAAACCAA | CAATACA | GGT | CCA | C | C | C>G | C>G_CCA | CG>GC_CAATACA | 9 | FALSE | FALSE |
| 131_v.txt | 33118 | 33118-SNV-C | SNV | 1 | C | G | 2 | 31752 | 6.30E-03 | 50.998015 | WT | FALSE | CAN | JSY2424 | 8 | WT-Rep8 | GCG | C>G | G/C+1 | G/C-1 | CG>GC | CG>GC | GGCCGGCGT | CGTTGGT | CGC | GCG | C | C | C>G | C>G_GCG | CG>GC_CGTTGGT | 9 | FALSE | FALSE |
| 131_v.txt | 33169 | 33169-SNV-C | SNV | 1 | C | G | 53 | 31746 | 0.16695017 | 37.4322891 | WT | FALSE | CAN | JSY2424 | 8 | WT-Rep8 | CCA | C>G | G/C+1 | G/C-1 | CG>GC | CG>GC | TAGTACCAC | CACCAAG | GGT | CCA | C | C | C>G | C>G_CCA | CG>GC_CACCAAG | 9 | FALSE | FALSE |
| 131_v.txt | 33187 | 33187-SNV-C | SNV | 1 | C | G | 39 | 31283 | 0.12466835 | 36.5641026 | WT | FALSE | CAN | JSY2424 | 8 | WT-Rep8 | CCA | C>G | G/C+1 | G/C-1 | CG>GC | CG>GC | TCATACCAA | CAATATG | GGT | CCA | C | C | C>G | C>G_CCA | CG>GC_CAATATG | 9 | FALSE | FALSE |
| 131_v.txt | 33276 | 33276-SNV-C | SNV | 1 | G | C | 1 | 31295 | 3.20E-03 | 43.9908643 | WT | FALSE | CAN | JSY2424 | 8 | WT-Rep8 | CGT | G>C | G/C+1 | G/C-1 | CG>GC | CG>GC | GAGAACGTA | GTATCCT | GCA | GCA | C | G | C>G | C>G_GCA | CG>GC_GTATCCT | 9 | FALSE | FALSE |
| 131_v.txt | 33309 | 33309-SNV-C | SNV | 1 | C | G | 1 | 31295 | 3.20E-03 | 59.9967028 | WT | FALSE | CAN | JSY2424 | 8 | WT-Rep8 | GCG | C>G | G/C+1 | G/C-1 | CG>GC | CG>GC | AAAGAGCGC | CGCAATG | CGC | GCG | C | C | C>G | C>G_GCG | CG>GC_CGCAATG | 9 | FALSE | FALSE |
| 131_v.txt | 33354 | 33354-SNV-C | SNV | 1 | G | C | 194 | 31295 | 0.61990733 | 54.6340206 | WT | FALSE | CAN | JSY2424 | 8 | WT-Rep8 | TGA | G>C | G/C+1 | G/C-1 | CG>GC | CG>GC | GGTATTGAC | GACCCAC | ACT | ACT | C | G | C>G | C>G_ACT | CG>GC_GACCCAC | 9 | FALSE | FALSE |
| 131_v.txt | 33378 | 33378-SNV-C | SNV | 1 | G | C | 32 | 31295 | 0.10225276 | 54.875 | WT | FALSE | CAN | JSY2424 | 8 | WT-Rep8 | TGA | G>C | G/C+1 | G/C-1 | CG>GC | CG>GC | GTTTGTGAA | GAAGCTT | ACT | ACT | C | G | C>G | C>G_ACT | CG>GC_GAAGCTT | 9 | FALSE | FALSE |
| 131_v.txt | 31657 | 31657-SNV-C | SNV | 1 | G | T | 122 | 16915 | 0.72125333 | 20.4994669 | WT | FALSE | CAN | JSY2424 | 8 | WT-Rep8 | GGC | G>T | A/T+1 | G/C-1 | CG>AT | CG>AT | GAAATGGCG | GCGTGGA | CCG | CCG | C | A | C>A | C>A_CCG | CG>AT_GCGTGGA | 9 | FALSE | FALSE |
| 131_v.txt | 31821 | 31821-SNV-C | SNV | 1 | C | A | 2 | 16915 | 0.01182383 | 17.4999978 | WT | FALSE | CAN | JSY2424 | 8 | WT-Rep8 | TCT | C>A | A/T+1 | G/C-1 | CG>AT | CG>AT | CTGCATCTG | CTGAATA | AGA | TCT | C | T | C>A | C>A_TCT | CG>AT_CTGAATA | 9 | FALSE | FALSE |
| 131_v.txt | 31877 | 31877-SNV-C | SNV | 1 | G | T | 10 | 16915 | 0.05911913 | 46.9 | WT | FALSE | CAN | JSY2424 | 8 | WT-Rep8 | AGG | G>T | A/T+1 | G/C-1 | CG>AT | CG>AT | GATATAGGC | GGCGGCA | TCC | TCC | C | A | C>A | C>A_TCC | CG>AT_GGCGGCA | 9 | FALSE | FALSE |
| 131_v.txt | 31981 | 31981-SNV-C | SNV | 1 | G | T | 37 | 27414 | 0.13496754 | 21.027027 | WT | FALSE | CAN | JSY2424 | 8 | WT-Rep8 | GGC | G>T | A/T+1 | G/C-1 | CG>AT | CG>AT | AGCCGGGCA | GCATTAA | CCG | CCG | C | A | C>A | C>A_CCG | CG>AT_GCATTAA | 9 | FALSE | FALSE |
| 131_v.txt | 32238 | 32238-SNV-C | SNV | 1 | G | T | 15 | 27408 | 0.05472855 | 36.0666667 | WT | FALSE | CAN | JSY2424 | 8 | WT-Rep8 | AGC | G>T | A/T+1 | G/C-1 | CG>AT | CG>AT | TTAGGAGCC | GCCAACT | TCG | TCG | C | A | C>A | C>A_TCG | CG>AT_GCCAACT | 9 | FALSE | FALSE |
| 131_v.txt | 32368 | 32368-SNV-C | SNV | 1 | C | A | 179 | 35671 | 0.50180819 | 60.6256983 | WT | FALSE | CAN | JSY2424 | 8 | WT-Rep8 | TCA | C>A | A/T+1 | G/C-1 | CG>AT | CG>AT | AGTTCTCAA | CAATAGC | AGT | TCA | C | T | C>A | C>A_TCA | CG>AT_CAATAGC | 9 | FALSE | FALSE |
| 131_v.txt | 32373 | 32373-SNV-C | SNV | 1 | G | T | 31 | 35671 | 0.08690533 | 55.0967742 | WT | FALSE | CAN | JSY2424 | 8 | WT-Rep8 | AGC | G>T | A/T+1 | G/C-1 | CG>AT | CG>AT | TCAATAGCA | GCAATAA | TCG | TCG | C | A | C>A | C>A_TCG | CG>AT_GCAATAA | 9 | FALSE | FALSE |
| 131_v.txt | 32399 | 32399-SNV-C | SNV | 1 | G | T | 17 | 35672 | 0.04765643 | 55.7058824 | WT | FALSE | CAN | JSY2424 | 8 | WT-Rep8 | CGT | G>T | A/T+1 | G/C-1 | CG>AT | CG>AT | AGAAACGTA | GTAGGAA | GCA | GCA | C | A | C>A | C>A_GCA | CG>AT_GTAGGAA | 9 | FALSE | FALSE |
| 131_v.txt | 32432 | 32432-SNV-C | SNV | 1 | G | T | 31 | 35672 | 0.08690289 | 51.4516129 | WT | FALSE | CAN | JSY2424 | 8 | WT-Rep8 | TGT | G>T | A/T+1 | G/C-1 | CG>AT | CG>AT | GTCATTGTA | GTATGGA | ACA | ACA | C | A | C>A | C>A_ACA | CG>AT_GTATGGA | 9 | FALSE | FALSE |
| 131_v.txt | 32471 | 32471-SNV-C | SNV | 1 | G | T | 39 | 35672 | 0.10932945 | 49.5128205 | WT | FALSE | CAN | JSY2424 | 8 | WT-Rep8 | TGT | G>T | A/T+1 | G/C-1 | CG>AT | CG>AT | GCCAATGTA | GTAGAAG | ACA | ACA | C | A | C>A | C>A_ACA | CG>AT_GTAGAAG | 9 | FALSE | FALSE |
| 131_v.txt | 32530 | 32530-SNV-C | SNV | 1 | G | T | 21079 | 35672 | 59.0911639 | 39.3918118 | WT | FALSE | CAN | JSY2424 | 8 | WT-Rep8 | GGG | G>T | A/T+1 | G/C-1 | CG>AT | CG>AT | TTCTGGGGT | GGTTTGC | CCC | CCC | C | A | C>A | C>A_CCC | CG>AT_GGTTTGC | 9 | FALSE | FALSE |
| 131_v.txt | 32575 | 32575-SNV-C | SNV | 1 | G | T | 422 | 20965 | 2.01287861 | 36.8815166 | WT | FALSE | CAN | JSY2424 | 8 | WT-Rep8 | TGA | G>T | A/T+1 | G/C-1 | CG>AT | CG>AT | TACCTTGAA | GAAATGT | ACT | ACT | C | A | C>A | C>A_ACT | CG>AT_GAAATGT | 9 | FALSE | FALSE |
| 131_v.txt | 32794 | 32794-SNV-C | SNV | 1 | C | A | 36 | 20964 | 0.17172295 | 39.5277778 | WT | FALSE | CAN | JSY2424 | 8 | WT-Rep8 | TCA | C>A | A/T+1 | G/C-1 | CG>AT | CG>AT | CGAATTCAC | CACCGTA | AGT | TCA | C | T | C>A | C>A_TCA | CG>AT_CACCGTA | 9 | FALSE | FALSE |
| 131_v.txt | 32811 | 32811-SNV-C | SNV | 1 | G | T | 10 | 20964 | 0.04770082 | 36.7 | WT | FALSE | CAN | JSY2424 | 8 | WT-Rep8 | AGG | G>T | A/T+1 | G/C-1 | CG>AT | CG>AT | TTGACAGGG | GGGAACA | TCC | TCC | C | A | C>A | C>A_TCC | CG>AT_GGGAACA | 9 | FALSE | FALSE |
| 131_v.txt | 33033 | 33033-SNV-C | SNV | 1 | G | T | 37 | 31752 | 0.11652809 | 55.8378378 | WT | FALSE | CAN | JSY2424 | 8 | WT-Rep8 | TGT | G>T | A/T+1 | G/C-1 | CG>AT | CG>AT | ATGAATGTA | GTAGCCA | ACA | ACA | C | A | C>A | C>A_ACA | CG>AT_GTAGCCA | 9 | FALSE | FALSE |
| 131_v.txt | 33114 | 33114-SNV-C | SNV | 1 | C | A | 54 | 31752 | 0.17006803 | 41.3384032 | WT | FALSE | CAN | JSY2424 | 8 | WT-Rep8 | GCC | C>A | A/T+1 | G/C-1 | CG>AT | CG>AT | ACTGGGCCG | CCGGCGT | CGG | GCC | C | T | C>A | C>A_GCC | CG>AT_CCGGCGT | 9 | FALSE | FALSE |
| 131_v.txt | 33151 | 33151-SNV-C | SNV | 1 | G | T | 52 | 31752 | 0.16376921 | 37.9807692 | WT | FALSE | CAN | JSY2424 | 8 | WT-Rep8 | AGA | G>T | A/T+1 | G/C-1 | CG>AT | CG>AT | TGAAAAGAC | GACCTGT | TCT | TCT | C | A | C>A | C>A_TCT | CG>AT_GACCTGT | 9 | FALSE | FALSE |
| 131_v.txt | 33169 | 33169-SNV-C | SNV | 1 | C | A | 4 | 31746 | 0.01260001 | 36.2497341 | WT | FALSE | CAN | JSY2424 | 8 | WT-Rep8 | CCA | C>A | A/T+1 | G/C-1 | CG>AT | CG>AT | TAGTACCAC | CACCAAG | GGT | CCA | C | T | C>A | C>A_CCA | CG>AT_CACCAAG | 9 | FALSE | FALSE |
| 131_v.txt | 33385 | 33385-SNV-C | SNV | 1 | C | A | 24 | 31295 | 0.07668957 | 49.875 | WT | FALSE | CAN | JSY2424 | 8 | WT-Rep8 | TCA | C>A | A/T+1 | G/C-1 | CG>AT | CG>AT | AAGCTTCAA | CAACGTC | AGT | TCA | C | T | C>A | C>A_TCA | CG>AT_CAACGTC | 9 | FALSE | FALSE |
| 131_v.txt | 33474 | 33474-SNV-C | SNV | 1 | C | A | 76 | 31292 | 0.24287358 | 34.5264371 | WT | FALSE | CAN | JSY2424 | 8 | WT-Rep8 | GCC | C>A | A/T+1 | G/C-1 | CG>AT | CG>AT | GCTATGCCTT | CCTTTTT | CGG | GCC | C | T | C>A | C>A_GCC | CG>AT_CCTTTTT | 9 | FALSE | FALSE |
| 131_v.txt | 33475 | 33475-SNV-C | SNV | 1 | C | A | 26 | 31292 | 0.08308833 | 37.3076923 | WT | FALSE | CAN | JSY2424 | 8 | WT-Rep8 | CCT | C>A | A/T+1 | G/C-1 | CG>AT | CG>AT | CTATGCCTTT | CTTTTTT | GGA | CCT | C | T | C>A | C>A_CCT | CG>AT_CTTTTTT | 9 | FALSE | FALSE |
| 131_v.txt | 32337 | 32337-Repla | Replacement | 2 | AT | G | 1 | 35671 | 2.80E-03 | 42 | WT | FALSE | CAN | JSY2424 | 8 | WT-Rep8 | GAT | nope | G/C+1 | >1bpdel | none | Replac. | TTGAAGATA | ATATGTG | CTA | GAT | AT | C | NA | NA_GAT | Replac._ATATGTG | 9 | FALSE | FALSE |
| 131_v.txt | 33205 | 33205-Repla | Replacement | 2 | G | CC | 6 | 31295 | 0.01917239 | 37.5 | WT | FALSE | CAN | JSY2424 | 8 | WT-Rep8 | AGC | nope | >1bp-ins | G/C-1 | none | Replac. | GCTTAAGCT | GCTCTCT | TCG | TCG | C | none | C>none | C>none_TCG | Replac._GCTCTCT | 9 | FALSE | FALSE |
| 131_v.txt | 33168 | 33168-MNV- | MNV | 2 | CC | TG | 1 | 31746 | 3.15E-03 | 37.9999996 | WT | FALSE | CAN | JSY2424 | 8 | WT-Rep8 | ACC | nope | >1bp-ins | >1bpdel | none | MNV | ATAGTACCA | CCACCAA | TGG | ACC | CC | none | NA | NA_ACC | MNV_CCACCAA | 9 | FALSE | FALSE |
| 131_v.txt | 32081 | 32081-Insert | Insertion | 1 | - | A | 207 | 27414 | 0.75508864 | 63.826087 | WT | FALSE | CAN | JSY2424 | 8 | WT-Rep8 | CAA | nope | A/T+1 | >1bpdel | none | A/T+1 | CCATGCAAA | AAAAAAG | GTT | CAA | - | T | NA | NA_CAA | A/T+1_AAAAAAG | 9 | FALSE | FALSE |
| 131_v.txt | 32081 | 32081-Insert | Insertion | 2 | - | AA | 1 | 27414 | 3.65E-03 | 64 | WT | FALSE | CAN | JSY2424 | 8 | WT-Rep8 | CAA | nope | >1bp-ins | >1bpdel | none | >1bp-ins | CCATGCAAA | AAAAAAG | GTT | CAA | - | none | NA | NA_CAA | >1bp-ins_AAAAAAG | 9 | FALSE | FALSE |
| 131_v.txt | 32295 | 32295-Insert | Insertion | 1 | - | T | 1 | 35671 | 2.80E-03 | 64 | WT | FALSE | CAN | JSY2424 | 8 | WT-Rep8 | ATT | nope | A/T+1 | >1bpdel | none | A/T+1 | TTTGAATTTG | TTTGCGG | TAA | ATT | - | A | NA | NA_ATT | A/T+1_TTTGCGG | 9 | FALSE | FALSE |
| 131_v.txt | 32473 | 32473-Insert | Insertion | 2 | - | AG | 40 | 35672 | 0.11213277 | 46.5125 | WT | FALSE | CAN | JSY2424 | 8 | WT-Rep8 | TAG | nope | >1bp-ins | >1bpdel | none | >1bp-ins | CAATGTAGA | AGAAGGT | ATC | TAG | - | none | NA | NA_TAG | >1bp-ins_AGAAGGT | 9 | FALSE | FALSE |
| 131_v.txt | 32705 | 32705-Insert | Insertion | 1 | - | A | 1 | 20964 | 4.77E-03 | 10 | WT | FALSE | CAN | JSY2424 | 8 | WT-Rep8 | TAA | nope | A/T+1 | >1bpdel | none | A/T+1 | GCCGGTAAC | AACCCCA | ATT | TAA | - | T | NA | NA_TAA | A/T+1_AACCCCA | 9 | FALSE | FALSE |
| 131_v.txt | 32705 | 32705-Insert | Insertion | 40 | - | AACCCCAGC | 3 | 20964 | 0.01431025 | 58.325 | WT | FALSE | CAN | JSY2424 | 8 | WT-Rep8 | TAA | nope | >1bp-ins | >1bpdel | none | >1bp-ins | GCCGGTAAC | AACCCCA | ATT | TAA | - | none | NA | NA_TAA | >1bp-ins_AACCCCA | 9 | FALSE | FALSE |
| 131_v.txt | 32842 | 32842-Insert | Insertion | 1 | - | A | 7 | 20964 | 0.03339057 | 41.7142857 | WT | FALSE | CAN | JSY2424 | 8 | WT-Rep8 | CAA | nope | A/T+1 | >1bpdel | none | A/T+1 | TTACCCAAA | AAAAAAT | GTT | CAA | - | T | NA | NA_CAA | A/T+1_AAAAAAT | 9 | FALSE | FALSE |
| 131_v.txt | 32942 | 32942-Insert | Insertion | 5 | - | AAACC | 4 | 31752 | 0.01259763 | 50.35 | WT | FALSE | CAN | JSY2424 | 8 | WT-Rep8 | AAA | nope | >1bp-ins | >1bpdel | none | >1bp-ins | CCAAGAAAA | AAACCAA | TTT | AAA | - | none | NA | NA_AAA | >1bp-ins_AAACCAA | 9 | FALSE | FALSE |
| 131_v.txt | 33048 | 33048-Insert | Insertion | 1 | - | A | 3 | 31752 | 9.45E-03 | 64 | WT | FALSE | CAN | JSY2424 | 8 | WT-Rep8 | CAA | nope | A/T+1 | >1bpdel | none | A/T+1 | TCACCCAAG | AAGGACT | GTT | CAA | - | T | NA | NA_CAA | A/T+1_AAGGACT | 9 | FALSE | FALSE |
| 131_v.txt | 33081 | 33081-Insert | Insertion | 1 | - | A | 25 | 31752 | 0.0787352 | 55.68 | WT | FALSE | CAN | JSY2424 | 8 | WT-Rep8 | CAT | nope | A/T+1 | >1bpdel | none | A/T+1 | GAACCCATA | ATAAATA | GTA | CAT | - | T | NA | NA_CAT | A/T+1_ATAAATA | 9 | FALSE | FALSE |
| 131_v.txt | 33475 | 33475-Insert | Insertion | 1 | - | A | 1 | 31292 | 3.20E-03 | 38 | WT | FALSE | CAN | JSY2424 | 8 | WT-Rep8 | CCT | nope | A/T+1 | >1bpdel | none | A/T+1 | CTATGCCTTT | CTTTTTT | GGA | CCT | - | T | NA | NA_CCT | A/T+1_CTTTTTT | 9 | FALSE | FALSE |
| 131_v.txt | 31812 | 31812-Deleti | Deletion | 12 | AATCTGCAT | - | 16 | 16915 | 0.0945906 | 53.875 | WT | FALSE | CAN | JSY2424 | 8 | WT-Rep8 | AAA | nope | >1bp-ins | >1bpdel | none | >1bpdel | CAAATAAAT | AATCTGC | TTT | AAA | AATCTGCAT | none | NA | NA_AAA | >1bpdel_AATCTGC | 9 | FALSE | FALSE |
| 131_v.txt | 32126 | 32126-Deleti | Deletion | 2 | TT | - | 27 | 27414 | 0.09848982 | 58.4444444 | WT | FALSE | CAN | JSY2424 | 8 | WT-Rep8 | CTT | nope | >1bp-ins | >1bpdel | none | >1bpdel | GAAAACTTT | TTTGTCA | GAA | CTT | TT | none | NA | NA_CTT | >1bpdel_TTTGTCA | 9 | FALSE | FALSE |
| 131_v.txt | 32313 | 32313-Deleti | Deletion | 3 | GTT | - | 10 | 35671 | 0.02803398 | 52.6 | WT | FALSE | CAN | JSY2424 | 8 | WT-Rep8 | GGT | nope | >1bp-ins | >1bpdel | none | >1bpdel | ATAATGGTT | GTTGTTA | CCA | GGT | GTT | none | NA | NA_GGT | >1bpdel_GTTGTTA | 9 | FALSE | FALSE |
| 131_v.txt | 32337 | 32337-Deleti | Deletion | 2 | AT | - | 68 | 35671 | 0.19063105 | 62.8970588 | WT | FALSE | CAN | JSY2424 | 8 | WT-Rep8 | GAT | nope | >1bp-ins | >1bpdel | none | >1bpdel | TTGAAGATA | ATATGTG | CTA | GAT | AT | none | NA | NA_GAT | >1bpdel_ATATGTG | 9 | FALSE | FALSE |
| 131_v.txt | 33212 | 33212-Deleti | Deletion | 5 | CTTCA | - | 8 | 31295 | 0.02556319 | 37.875 | WT | FALSE | CAN | JSY2424 | 8 | WT-Rep8 | TCT | nope | >1bp-ins | >1bpdel | none | >1bpdel | CTCTCTCTTC | CTTCACT | AGA | TCT | CTTCA | none | NA | NA_TCT | >1bpdel_CTTCACT | 9 | FALSE | FALSE |
| 131_v.txt | 33474 | 33474-Deleti | Deletion | 2 | CC | - | 1 | 31292 | 3.20E-03 | 22 | WT | FALSE | CAN | JSY2424 | 8 | WT-Rep8 | GCC | nope | >1bp-ins | >1bpdel | none | >1bpdel | GCTATGCCTT | CCTTTTT | CGG | GCC | CC | none | NA | NA_GCC | >1bpdel_CCTTTTT | 9 | FALSE | FALSE |
| 131_v.txt | 31663 | 31663-Deleti | Deletion | 1 | A | - | 10 | 16915 | 0.05911913 | 35.9 | WT | FALSE | CAN | JSY2424 | 8 | WT-Rep8 | GAA | nope | >1bp-ins | A/T-1 | none | A/T-1 | GCGTGGAAA | AAATGTG | CTT | CTT | T | none | T>none | T>none_CTT | A/T-1_AAATGTG | 9 | FALSE | FALSE |
| 131_v.txt | 31723 | 31723-Deleti | Deletion | 1 | A | - | 21 | 16915 | 0.12415016 | 63.2857143 | WT | FALSE | CAN | JSY2424 | 8 | WT-Rep8 | CAA | nope | >1bp-ins | A/T-1 | none | A/T-1 | TGTCCCAAA | AAAAAGT | GTT | GTT | T | none | T>none | T>none_GTT | A/T-1_AAAAAGT | 9 | FALSE | FALSE |
| 131_v.txt | 31904 | 31904-Deleti | Deletion | 1 | T | - | 6 | 16912 | 0.03547777 | 37.8333333 | WT | FALSE | CAN | JSY2424 | 8 | WT-Rep8 | ATT | nope | >1bp-ins | A/T-1 | none | A/T-1 | ATTGAATTTT | TTTTGGT | TAA | ATT | T | none | nope | nope_ATT | A/T-1_TTTTGGT | 9 | FALSE | FALSE |
| 131_v.txt | 31920 | 31920-Deleti | Deletion | 1 | G | - | 2 | 16911 | 0.01182662 | 38 | WT | FALSE | CAN | JSY2424 | 8 | WT-Rep8 | CGT | nope | >1bp-ins | G/C-1 | none | G/C-1 | AAAGCCGTG | GTGAAAC | GCA | GCA | C | none | C>none | C>none_GCA | G/C-1_GTGAAAC | 9 | FALSE | FALSE |
| 131_v.txt | 31972 | 31972-Deleti | Deletion | 1 | C | - | 7 | 27414 | 0.0255344 | 35.8571429 | WT | FALSE | CAN | JSY2424 | 8 | WT-Rep8 | GCC | nope | >1bp-ins | G/C-1 | none | G/C-1 | AATAAGCCA | CCAAGCC | CGG | GCC | C | none | nope | nope_GCC | G/C-1_CCAAGCC | 9 | FALSE | FALSE |
| 131_v.txt | 32057 | 32057-Deleti | Deletion | 1 | C | - | 4 | 27414 | 0.01459109 | 62.75 | WT | FALSE | CAN | JSY2424 | 8 | WT-Rep8 | GCG | nope | >1bp-ins | G/C-1 | none | G/C-1 | GATGTGCGA | CGAGATT | CGC | GCG | C | none | nope | nope_GCG | G/C-1_CGAGATT | 9 | FALSE | FALSE |
| 131_v.txt | 32073 | 32073-Deleti | Deletion | 1 | A | - | 3 | 27414 | 0.01094331 | 64 | WT | FALSE | CAN | JSY2424 | 8 | WT-Rep8 | TAA | nope | >1bp-ins | A/T-1 | none | A/T-1 | ATAAATAAC | AACCATG | ATT | ATT | T | none | T>none | T>none_ATT | A/T-1_AACCATG | 9 | FALSE | FALSE |
| 131_v.txt | 32081 | 32081-Deleti | Deletion | 1 | A | - | 21 | 27414 | 0.0766032 | 63.7142858 | WT | FALSE | CAN | JSY2424 | 8 | WT-Rep8 | CAA | nope | >1bp-ins | A/T-1 | none | A/T-1 | CCATGCAAA | AAAAAAG | GTT | GTT | T | none | T>none | T>none_GTT | A/T-1_AAAAAAG | 9 | FALSE | FALSE |
| 131_v.txt | 32126 | 32126-Deleti | Deletion | 1 | T | - | 2 | 27414 | 7.30E-03 | 50.9999995 | WT | FALSE | CAN | JSY2424 | 8 | WT-Rep8 | CTT | nope | >1bp-ins | A/T-1 | none | A/T-1 | GAAAACTTT | TTTGTCA | GAA | CTT | T | none | nope | nope_CTT | A/T-1_TTTGTCA | 9 | FALSE | FALSE |
| 131_v.txt | 32420 | 32420-Deleti | Deletion | 1 | T | - | 8 | 35672 | 0.02242655 | 56.375 | WT | FALSE | CAN | JSY2424 | 8 | WT-Rep8 | GTT | nope | >1bp-ins | A/T-1 | none | A/T-1 | TGTTAGTTTA | TTTAGGG | CAA | GTT | T | none | nope | nope_GTT | A/T-1_TTTAGGG | 9 | FALSE | FALSE |
| 131_v.txt | 32436 | 32436-Deleti | Deletion | 1 | G | - | 2 | 35672 | 5.61E-03 | 51 | WT | FALSE | CAN | JSY2424 | 8 | WT-Rep8 | TGG | nope | >1bp-ins | G/C-1 | none | G/C-1 | TTGTATGGA | GGAACTA | ACC | ACC | C | none | C>none | C>none_ACC | G/C-1_GGAACTA | 9 | FALSE | FALSE |
| 131_v.txt | 32442 | 32442-Deleti | Deletion | 1 | A | - | 9 | 35672 | 0.02522987 | 57.7777778 | WT | FALSE | CAN | JSY2424 | 8 | WT-Rep8 | TAA | nope | >1bp-ins | A/T-1 | none | A/T-1 | GGAACTAAA | AAAAGTC | ATT | ATT | T | none | T>none | T>none_ATT | A/T-1_AAAAGTC | 9 | FALSE | FALSE |
| 131_v.txt | 32498 | 32498-Deleti | Deletion | 1 | T | - | 87 | 35672 | 0.24388876 | 43.8735632 | WT | FALSE | CAN | JSY2424 | 8 | WT-Rep8 | CTT | nope | >1bp-ins | A/T-1 | none | A/T-1 | AACAACTTTT | TTTTTTG | GAA | CTT | T | none | nope | nope_CTT | A/T-1_TTTTTTG | 9 | FALSE | FALSE |
| 131_v.txt | 32522 | 32522-Deleti | Deletion | 1 | T | - | 9 | 35672 | 0.02522987 | 40.8888889 | WT | FALSE | CAN | JSY2424 | 8 | WT-Rep8 | ATT | nope | >1bp-ins | A/T-1 | none | A/T-1 | AACGGATTTT | TTTTCTG | TAA | ATT | T | none | nope | nope_ATT | A/T-1_TTTTCTG | 9 | FALSE | FALSE |
| 131_v.txt | 32528 | 32528-Deleti | Deletion | 1 | G | - | 6 | 35672 | 0.01681992 | 41.4999998 | WT | FALSE | CAN | JSY2424 | 8 | WT-Rep8 | TGG | nope | >1bp-ins | G/C-1 | none | G/C-1 | TTTTCTGGG | GGGGTTT | ACC | ACC | C | none | C>none | C>none_ACC | G/C-1_GGGGTTT | 9 | FALSE | FALSE |
| 131_v.txt | 32631 | 32631-Deleti | Deletion | 1 | T | - | 12 | 20964 | 0.05724099 | 45.5833333 | WT | FALSE | CAN | JSY2424 | 8 | WT-Rep8 | GTT | nope | >1bp-ins | A/T-1 | none | A/T-1 | CCTTCGTTTT | TTTTTAT | CAA | GTT | T | none | nope | nope_GTT | A/T-1_TTTTTAT | 9 | FALSE | FALSE |
| 131_v.txt | 32658 | 32658-Deleti | Deletion | 1 | C | - | 9 | 20964 | 0.04293074 | 55.8888889 | WT | FALSE | CAN | JSY2424 | 8 | WT-Rep8 | ACC | nope | >1bp-ins | G/C-1 | none | G/C-1 | CCTGGACCC | CCCCAGG | TGG | ACC | C | none | nope | nope_ACC | G/C-1_CCCCAGG | 9 | FALSE | FALSE |
| 131_v.txt | 32697 | 32697-Deleti | Deletion | 1 | G | - | 4 | 20964 | 0.01908033 | 64 | WT | FALSE | CAN | JSY2424 | 8 | WT-Rep8 | TGG | nope | >1bp-ins | G/C-1 | none | G/C-1 | CCAACTGGG | GGGCCGG | ACC | ACC | C | none | C>none | C>none_ACC | G/C-1_GGGCCGG | 9 | FALSE | FALSE |
| 131_v.txt | 32707 | 32707-Deleti | Deletion | 1 | C | - | 5 | 20964 | 0.02385041 | 58.8 | WT | FALSE | CAN | JSY2424 | 8 | WT-Rep8 | ACC | nope | >1bp-ins | G/C-1 | none | G/C-1 | CGGTAACCC | CCCCAGC | TGG | ACC | C | none | nope | nope_ACC | G/C-1_CCCCAGC | 9 | FALSE | FALSE |
| 131_v.txt | 32744 | 32744-Deleti | Deletion | 1 | A | - | 14 | 20964 | 0.06678115 | 59.6428571 | WT | FALSE | CAN | JSY2424 | 8 | WT-Rep8 | GAA | nope | >1bp-ins | A/T-1 | none | A/T-1 | TATTAGAAA | AAACCCG | CTT | CTT | T | none | T>none | T>none_CTT | A/T-1_AAACCCG | 9 | FALSE | FALSE |
| 131_v.txt | 32779 | 32779-Deleti | Deletion | 1 | C | - | 8 | 20964 | 0.03816066 | 43.5 | WT | FALSE | CAN | JSY2424 | 8 | WT-Rep8 | ACC | nope | >1bp-ins | G/C-1 | none | G/C-1 | AAGCGACCC | CCCAGAA | TGG | ACC | C | none | nope | nope_ACC | G/C-1_CCCAGAA | 9 | FALSE | FALSE |
| 131_v.txt | 32800 | 32800-Deleti | Deletion | 1 | A | - | 3 | 20964 | 0.01431025 | 46 | WT | FALSE | CAN | JSY2424 | 8 | WT-Rep8 | TAA | nope | >1bp-ins | A/T-1 | none | A/T-1 | CACCGTAAT | AATATTT | ATT | ATT | T | none | T>none | T>none_ATT | A/T-1_AATATTT | 9 | FALSE | FALSE |
| 131_v.txt | 32842 | 32842-Deleti | Deletion | 1 | A | - | 36 | 20964 | 0.17172295 | 43.3888874 | WT | FALSE | CAN | JSY2424 | 8 | WT-Rep8 | CAA | nope | >1bp-ins | A/T-1 | none | A/T-1 | TTACCCAAA | AAAAAAT | GTT | GTT | T | none | T>none | T>none_GTT | A/T-1_AAAAAAT | 9 | FALSE | FALSE |
| 131_v.txt | 32884 | 32884-Deleti | Deletion | 1 | A | - | 6 | 31752 | 0.01889645 | 36.8333333 | WT | FALSE | CAN | JSY2424 | 8 | WT-Rep8 | CAA | nope | >1bp-ins | A/T-1 | none | A/T-1 | ACGTCCAAA | AAAATTG | GTT | GTT | T | none | T>none | T>none_GTT | A/T-1_AAAATTG | 9 | FALSE | FALSE |
| 131_v.txt | 32901 | 32901-Deleti | Deletion | 1 | C | - | 1 | 31752 | 3.15E-03 | 37.9999999 | WT | FALSE | CAN | JSY2424 | 8 | WT-Rep8 | GCC | nope | >1bp-ins | G/C-1 | none | G/C-1 | ACTTGGCCA | CCAACTA | CGG | GCC | C | none | nope | nope_GCC | G/C-1_CCAACTA | 9 | FALSE | FALSE |
| 131_v.txt | 32919 | 32919-Deleti | Deletion | 1 | A | - | 2065 | 31752 | 6.50352734 | 50.4305107 | WT | FALSE | CAN | JSY2424 | 8 | WT-Rep8 | CAG | nope | >1bp-ins | A/T-1 | none | A/T-1 | AGTTCCAGG | AGGGCAA | GTC | GTC | T | none | T>none | T>none_GTC | A/T-1_AGGGCAA | 9 | FALSE | FALSE |
| 131_v.txt | 32920 | 32920-Deleti | Deletion | 1 | G | - | 5 | 31752 | 0.01574704 | 53.6000109 | WT | FALSE | CAN | JSY2424 | 8 | WT-Rep8 | AGG | nope | >1bp-ins | G/C-1 | none | G/C-1 | GTTCCAGGG | GGGCAAA | TCC | TCC | C | none | C>none | C>none_TCC | G/C-1_GGGCAAA | 9 | FALSE | FALSE |
| 131_v.txt | 32924 | 32924-Deleti | Deletion | 1 | A | - | 31 | 31752 | 0.09763165 | 55.5483871 | WT | FALSE | CAN | JSY2424 | 8 | WT-Rep8 | CAA | nope | >1bp-ins | A/T-1 | none | A/T-1 | CAGGGCAAA | AAAAGTG | GTT | GTT | T | none | T>none | T>none_GTT | A/T-1_AAAAGTG | 9 | FALSE | FALSE |
| 131_v.txt | 32935 | 32935-Deleti | Deletion | 1 | C | - | 2 | 31752 | 6.30E-03 | 64 | WT | FALSE | CAN | JSY2424 | 8 | WT-Rep8 | GCC | nope | >1bp-ins | G/C-1 | none | G/C-1 | TGATTGCCC | CCCAAGA | CGG | GCC | C | none | nope | nope_GCC | G/C-1_CCCAAGA | 9 | FALSE | FALSE |
| 131_v.txt | 32965 | 32965-Deleti | Deletion | 1 | C | - | 3 | 31752 | 9.45E-03 | 53.6666667 | WT | FALSE | CAN | JSY2424 | 8 | WT-Rep8 | GCC | nope | >1bp-ins | G/C-1 | none | G/C-1 | CATTGGCCG | CCGCACC | CGG | GCC | C | none | nope | nope_GCC | G/C-1_CCGCACC | 9 | FALSE | FALSE |
| 131_v.txt | 33000 | 33000-Deleti | Deletion | 1 | A | - | 6 | 31752 | 0.01889645 | 64 | WT | FALSE | CAN | JSY2424 | 8 | WT-Rep8 | GAA | nope | >1bp-ins | A/T-1 | none | A/T-1 | TGTGAGAAA | AAAACTG | CTT | CTT | T | none | T>none | T>none_CTT | A/T-1_AAAACTG | 9 | FALSE | FALSE |
| 131_v.txt | 33147 | 33147-Deleti | Deletion | 1 | A | - | 9 | 31752 | 0.02834467 | 37.8888889 | WT | FALSE | CAN | JSY2424 | 8 | WT-Rep8 | GAA | nope | >1bp-ins | A/T-1 | none | A/T-1 | CCAATGAAA | AAAAGAC | CTT | CTT | T | none | T>none | T>none_CTT | A/T-1_AAAAGAC | 9 | FALSE | FALSE |
| 131_v.txt | 33168 | 33168-Deleti | Deletion | 1 | C | - | 1 | 31746 | 3.15E-03 | 37.9999999 | WT | FALSE | CAN | JSY2424 | 8 | WT-Rep8 | ACC | nope | >1bp-ins | G/C-1 | none | G/C-1 | ATAGTACCA | CCACCAA | TGG | ACC | C | none | nope | nope_ACC | G/C-1_CCACCAA | 9 | FALSE | FALSE |
| 131_v.txt | 33239 | 33239-Deleti | Deletion | 1 | T | - | 6 | 31295 | 0.01917239 | 38 | WT | FALSE | CAN | JSY2424 | 8 | WT-Rep8 | CTT | nope | >1bp-ins | A/T-1 | none | A/T-1 | TTCTCCTTCA | TTCATCT | GAA | CTT | T | none | nope | nope_CTT | A/T-1_TTCATCT | 9 | FALSE | FALSE |
| 131_v.txt | 33288 | 33288-Deleti | Deletion | 1 | T | - | 7 | 31295 | 0.02236779 | 43.8571429 | WT | FALSE | CAN | JSY2424 | 8 | WT-Rep8 | ATT | nope | >1bp-ins | A/T-1 | none | A/T-1 | TCGCCATTTA | TTTACTC | TAA | ATT | T | none | nope | nope_ATT | A/T-1_TTTACTC | 9 | FALSE | FALSE |
| 131_v.txt | 33408 | 33408-Deleti | Deletion | 1 | A | - | 12 | 31295 | 0.03834478 | 45.0833333 | WT | FALSE | CAN | JSY2424 | 8 | WT-Rep8 | GAC | nope | >1bp-ins | A/T-1 | none | A/T-1 | GTTGTGACC | ACCGGCT | CTG | CTG | T | none | T>none | T>none_CTG | A/T-1_ACCGGCT | 9 | FALSE | FALSE |
| 131_v.txt | 33474 | 33474-Deleti | Deletion | 1 | C | - | 88 | 31292 | 0.28122204 | 30.4545455 | WT | FALSE | CAN | JSY2424 | 8 | WT-Rep8 | GCC | nope | >1bp-ins | G/C-1 | none | G/C-1 | GCTATGCCTT | CCTTTTT | CGG | GCC | C | none | nope | nope_GCC | G/C-1_CCTTTTT | 9 | FALSE | FALSE |
| 132_v.txt | 31723 | 31723-SNV-A | SNV | 1 | A | C | 5 | 10432 | 0.04792945 | 56.4 | WT | FALSE | CAN | JSY2424 | 9 | WT-Rep9 | CAA | A>C | G/C+1 | A/T-1 | TA>GC | TA>GC | TGTCCCAAA | AAAAAGT | GTT | GTT | T | G | T>G | T>G_GTT | TA>GC_AAAAAGT | 9 | FALSE | FALSE |
| 132_v.txt | 31839 | 31839-SNV-A | SNV | 1 | A | C | 5 | 10432 | 0.04792945 | 53.6 | WT | FALSE | CAN | JSY2424 | 9 | WT-Rep9 | TAA | A>C | G/C+1 | A/T-1 | TA>GC | TA>GC | TGAAATAAG | AAGATCC | ATT | ATT | T | G | T>G | T>G_ATT | TA>GC_AAGATCC | 9 | FALSE | FALSE |
| 132_v.txt | 32081 | 32081-SNV-A | SNV | 1 | A | C | 1 | 24990 | 4.00E-03 | 63.9996748 | WT | FALSE | CAN | JSY2424 | 9 | WT-Rep9 | CAA | A>C | G/C+1 | A/T-1 | TA>GC | TA>GC | CCATGCAAA | AAAAAAG | GTT | GTT | T | G | T>G | T>G_GTT | TA>GC_AAAAAAG | 9 | FALSE | FALSE |
| 132_v.txt | 32284 | 32284-SNV-A | SNV | 1 | A | C | 63 | 33196 | 0.1897819 | 37.6666667 | WT | FALSE | CAN | JSY2424 | 9 | WT-Rep9 | TAA | A>C | G/C+1 | A/T-1 | TA>GC | TA>GC | CAACGTAAA | AAATATT | ATT | ATT | T | G | T>G | T>G_ATT | TA>GC_AAATATT | 9 | FALSE | FALSE |
| 132_v.txt | 32481 | 32481-SNV-A | SNV | 1 | A | C | 10 | 33196 | 0.03012411 | 47.4 | WT | FALSE | CAN | JSY2424 | 9 | WT-Rep9 | TAA | A>C | G/C+1 | A/T-1 | TA>GC | TA>GC | AAGGTTAAG | AAGATAC | ATT | ATT | T | G | T>G | T>G_ATT | TA>GC_AAGATAC | 9 | FALSE | FALSE |
| 132_v.txt | 32662 | 32662-SNV-A | SNV | 1 | A | C | 2 | 30600 | 6.54E-03 | 64 | WT | FALSE | CAN | JSY2424 | 9 | WT-Rep9 | CAG | A>C | G/C+1 | A/T-1 | TA>GC | TA>GC | GACCCCAGG | AGGCACC | GTC | GTC | T | G | T>G | T>G_GTC | TA>GC_AGGCACC | 9 | FALSE | FALSE |
| 132_v.txt | 32823 | 32823-SNV-A | SNV | 1 | A | C | 27 | 30600 | 0.08823529 | 42.0740741 | WT | FALSE | CAN | JSY2424 | 9 | WT-Rep9 | CAT | A>C | G/C+1 | A/T-1 | TA>GC | TA>GC | AAGTTCATTA | ATTATTG | GTA | GTA | T | G | T>G | T>G_GTA | TA>GC_ATTATTG | 9 | FALSE | FALSE |
| 132_v.txt | 32842 | 32842-SNV-A | SNV | 1 | A | C | 3 | 30600 | 9.80E-03 | 37.3333358 | WT | FALSE | CAN | JSY2424 | 9 | WT-Rep9 | CAA | A>C | G/C+1 | A/T-1 | TA>GC | TA>GC | TTACCCAAA | AAAAAAT | GTT | GTT | T | G | T>G | T>G_GTT | TA>GC_AAAAAAT | 9 | FALSE | FALSE |
| 132_v.txt | 32884 | 32884-SNV-A | SNV | 1 | A | C | 11 | 30857 | 0.03564831 | 35 | WT | FALSE | CAN | JSY2424 | 9 | WT-Rep9 | CAA | A>C | G/C+1 | A/T-1 | TA>GC | TA>GC | ACGTCCAAA | AAAATTG | GTT | GTT | T | G | T>G | T>G_GTT | TA>GC_AAAATTG | 9 | FALSE | FALSE |
| 132_v.txt | 32913 | 32913-SNV-A | SNV | 1 | A | C | 33 | 30857 | 0.10694494 | 29.1515152 | WT | FALSE | CAN | JSY2424 | 9 | WT-Rep9 | AAG | A>C | G/C+1 | A/T-1 | TA>GC | TA>GC | ACACTAAGT | AGTTCCA | TTC | TTC | T | G | T>G | T>G_TTC | TA>GC_AGTTCCA | 9 | FALSE | FALSE |
| 132_v.txt | 33039 | 33039-SNV-A | SNV | 1 | A | C | 18 | 30857 | 0.0583336 | 55.7777778 | WT | FALSE | CAN | JSY2424 | 9 | WT-Rep9 | CAT | A>C | G/C+1 | A/T-1 | TA>GC | TA>GC | GTAGCCATT | ATTTCAC | GTA | GTA | T | G | T>G | T>G_GTA | TA>GC_ATTTCAC | 9 | FALSE | FALSE |
| 132_v.txt | 33166 | 33166-SNV-T | SNV | 1 | T | G | 8 | 30853 | 0.02592941 | 38 | WT | FALSE | CAN | JSY2424 | 9 | WT-Rep9 | GTA | T>G | G/C+1 | A/T-1 | TA>GC | TA>GC | CAATAGTAC | TACCACC | CAT | GTA | T | C | T>G | T>G_GTA | TA>GC_TACCACC | 9 | FALSE | FALSE |
| 132_v.txt | 33269 | 33269-SNV-A | SNV | 1 | A | C | 207 | 33366 | 0.62039202 | 16.468599 | WT | FALSE | CAN | JSY2424 | 9 | WT-Rep9 | TAG | A>C | G/C+1 | A/T-1 | TA>GC | TA>GC | CTCCATAGA | AGAGAAC | ATC | ATC | T | G | T>G | T>G_ATC | TA>GC_AGAGAAC | 9 | FALSE | FALSE |
| 132_v.txt | 32081 | 32081-SNV-A | SNV | 1 | A | G | 7 | 24990 | 0.02801121 | 52.2859665 | WT | FALSE | CAN | JSY2424 | 9 | WT-Rep9 | CAA | A>G | G/C+1 | A/T-1 | TA>CG | TA>CG | CCATGCAAA | AAAAAAG | GTT | GTT | T | C | T>C | T>C_GTT | TA>CG_AAAAAAG | 9 | FALSE | FALSE |
| 132_v.txt | 32256 | 32256-SNV-A | SNV | 1 | A | G | 34 | 33178 | 0.10247755 | 36.7352941 | WT | FALSE | CAN | JSY2424 | 9 | WT-Rep9 | TAG | A>G | G/C+1 | A/T-1 | TA>CG | TA>CG | TTTGATAGA | AGACCAA | ATC | ATC | T | C | T>C | T>C_ATC | TA>CG_AGACCAA | 9 | FALSE | FALSE |
| 132_v.txt | 32473 | 32473-SNV-A | SNV | 1 | A | G | 27 | 33196 | 0.0813351 | 44.8888877 | WT | FALSE | CAN | JSY2424 | 9 | WT-Rep9 | TAG | A>G | G/C+1 | A/T-1 | TA>CG | TA>CG | CAATGTAGA | AGAAGGT | ATC | ATC | T | C | T>C | T>C_ATC | TA>CG_AGAAGGT | 9 | FALSE | FALSE |
| 132_v.txt | 32490 | 32490-SNV-A | SNV | 1 | A | G | 173 | 33196 | 0.52114713 | 45.1269599 | WT | FALSE | CAN | JSY2424 | 9 | WT-Rep9 | GAA | A>G | G/C+1 | A/T-1 | TA>CG | TA>CG | ATACGGAAA | AAAACAA | CTT | CTT | T | C | T>C | T>C_CTT | TA>CG_AAAACAA | 9 | FALSE | FALSE |
| 132_v.txt | 32662 | 32662-SNV-A | SNV | 1 | A | G | 19 | 30600 | 0.0620915 | 52.0526316 | WT | FALSE | CAN | JSY2424 | 9 | WT-Rep9 | CAG | A>G | G/C+1 | A/T-1 | TA>CG | TA>CG | GACCCCAGG | AGGCACC | GTC | GTC | T | C | T>C | T>C_GTC | TA>CG_AGGCACC | 9 | FALSE | FALSE |
| 132_v.txt | 33150 | 33150-SNV-A | SNV | 1 | A | G | 27 | 30856 | 0.08750324 | 32.925935 | WT | FALSE | CAN | JSY2424 | 9 | WT-Rep9 | AAG | A>G | G/C+1 | A/T-1 | TA>CG | TA>CG | ATGAAAAGA | AGACCTG | TTC | TTC | T | C | T>C | T>C_TTC | TA>CG_AGACCTG | 9 | FALSE | FALSE |
| 132_v.txt | 33192 | 33192-SNV-T | SNV | 1 | T | C | 87 | 33364 | 0.2607601 | 34.4598656 | WT | FALSE | CAN | JSY2424 | 9 | WT-Rep9 | ATG | T>C | G/C+1 | A/T-1 | TA>CG | TA>CG | CCAATATGT | TGTCTTT | TAC | ATG | T | G | T>C | T>C_ATG | TA>CG_TGTCTTT | 9 | FALSE | FALSE |
| 132_v.txt | 33331 | 33331-SNV-T | SNV | 1 | T | C | 5 | 33365 | 0.01498576 | 58.399922 | WT | FALSE | CAN | JSY2424 | 9 | WT-Rep9 | CTT | T>C | G/C+1 | A/T-1 | TA>CG | TA>CG | CTTTACTTT | TTTTCTC | GAA | CTT | T | G | T>C | T>C_CTT | TA>CG_TTTTCTC | 9 | FALSE | FALSE |
| 132_v.txt | 33476 | 33476-SNV-T | SNV | 1 | T | C | 49 | 33362 | 0.14687369 | 30.4089041 | WT | FALSE | CAN | JSY2424 | 9 | WT-Rep9 | CTT | T>C | G/C+1 | A/T-1 | TA>CG | TA>CG | TATGCCTTTT | TTTTTTT | GAA | CTT | T | G | T>C | T>C_CTT | TA>CG_TTTTTTT | 9 | FALSE | FALSE |
| 132_v.txt | 32294 | 32294-SNV-A | SNV | 1 | A | T | 19 | 33196 | 0.05723581 | 50.0526316 | WT | FALSE | CAN | JSY2424 | 9 | WT-Rep9 | AAT | A>T | A/T+1 | A/T-1 | TA>AT | TA>AT | ATTTGAATTT | ATTTGCG | TTA | TTA | T | A | T>A | T>A_TTA | TA>AT_ATTTGCG | 9 | FALSE | FALSE |

| | | | | | | | | | | | | | | | | | | | | | | | | | | | | | | | | | | |
|---|---|---|---|---|---|---|---|---|---|---|---|---|---|---|---|---|---|---|---|---|---|---|---|---|---|---|---|---|---|---|---|---|---|---|
| 132_v.txt | 32314 | 32314-SNV-T | SNV | 1 | T | A | 8 | 33196 | 0.02409929 | 49.2409652 | WT | FALSE | CAN | JSY2424 | 9 | WT-Rep9 | GTT | T>A | A/T+1 | A/T-1 | TA>AT | TA>AT | TAATGGTTG | TTGTTAA | CAA | GTT | T | T | T>A | T>A_GTT | TA>AT_TTGTTAA | 9 | FALSE | FALSE |
| 132_v.txt | 32315 | 32315-SNV-T | SNV | 1 | T | A | 7 | 33196 | 0.02108688 | 51.9979933 | WT | FALSE | CAN | JSY2424 | 9 | WT-Rep9 | TTG | T>A | A/T+1 | A/T-1 | TA>AT | TA>AT | AATGGTTGT | TGTTAAG | AAC | TTG | T | T | T>A | T>A_TTG | TA>AT_TGTTAAG | 9 | FALSE | FALSE |
| 132_v.txt | 32947 | 32947-SNV-A | SNV | 1 | A | T | 37 | 30857 | 0.11990796 | 59.5945946 | WT | FALSE | CAN | JSY2424 | 9 | WT-Rep9 | CAA | A>T | A/T+1 | A/T-1 | TA>AT | TA>AT | AAAACCAAT | AATACAT | GTT | GTT | T | A | T>A | T>A_GTT | TA>AT_AATACAT | 9 | FALSE | FALSE |
| 132_v.txt | 33192 | 33192-SNV-T | SNV | 1 | T | A | 66 | 33364 | 0.19781801 | 34.0921233 | WT | FALSE | CAN | JSY2424 | 9 | WT-Rep9 | ATG | T>A | A/T+1 | A/T-1 | TA>AT | TA>AT | CCAATATGT | TGTCTTT | TAC | ATG | T | T | T>A | T>A_ATG | TA>AT_TGTCTTT | 9 | FALSE | FALSE |
| 132_v.txt | 33321 | 33321-SNV-A | SNV | 1 | A | T | 42 | 33366 | 0.12587664 | 60.9047619 | WT | FALSE | CAN | JSY2424 | 9 | WT-Rep9 | CAA | A>T | A/T+1 | A/T-1 | TA>AT | TA>AT | GGATACAAT | AATTCTT | GTT | GTT | T | A | T>A | T>A_GTT | TA>AT_AATTCTT | 9 | FALSE | FALSE |
| 132_v.txt | 33328 | 33328-SNV-T | SNV | 1 | T | A | 46 | 33366 | 0.13786489 | 59 | WT | FALSE | CAN | JSY2424 | 9 | WT-Rep9 | TTA | T>A | A/T+1 | A/T-1 | TA>AT | TA>AT | ATTCTTTACT | TACTTTT | AAT | TTA | T | T | T>A | T>A_TTA | TA>AT_TACTTTT | 9 | FALSE | FALSE |
| 132_v.txt | 33331 | 33331-SNV-T | SNV | 1 | T | A | 3 | 33365 | 8.99E-03 | 51.6661354 | WT | FALSE | CAN | JSY2424 | 9 | WT-Rep9 | CTT | T>A | A/T+1 | A/T-1 | TA>AT | TA>AT | CTTTACTTTT | TTTTCTC | GAA | CTT | T | T | T>A | T>A_CTT | TA>AT_TTTTCTC | 9 | FALSE | FALSE |
| 132_v.txt | 31650 | 31650-SNV-C | SNV | 1 | C | T | 22 | 10432 | 0.21088957 | 35.2727273 | WT | FALSE | CAN | JSY2424 | 9 | WT-Rep9 | GCG | C>T | A/T+1 | G/C-1 | CG>TA | CG>TA | AGAATGCGA | CGAAATG | CGC | GCG | C | A | C>T | C>T_GCG | CG>TA_CGAAATG | 9 | FALSE | FALSE |
| 132_v.txt | 31947 | 31947-SNV-G | SNV | 1 | G | A | 26 | 24974 | 0.10410827 | 36.2307692 | WT | FALSE | CAN | JSY2424 | 9 | WT-Rep9 | CGT | G>A | A/T+1 | G/C-1 | CG>TA | CG>TA | ATGATCGTC | GTCATAA | GCA | GCA | C | T | C>T | C>T_GCA | CG>TA_GTCATAA | 9 | FALSE | FALSE |
| 132_v.txt | 31962 | 31962-SNV-G | SNV | 1 | G | A | 42 | 24990 | 0.16806723 | 36.9761905 | WT | FALSE | CAN | JSY2424 | 9 | WT-Rep9 | CGC | G>A | A/T+1 | G/C-1 | CG>TA | CG>TA | GTGGCCGCA | GCATAAT | GCG | GCG | C | T | C>T | C>T_GCG | CG>TA_GCATAAT | 9 | FALSE | FALSE |
| 132_v.txt | 31972 | 31972-SNV-C | SNV | 1 | C | T | 19 | 24990 | 0.07603041 | 37.7368421 | WT | FALSE | CAN | JSY2424 | 9 | WT-Rep9 | GCC | C>T | A/T+1 | G/C-1 | CG>TA | CG>TA | AATAAGCCA | CCAAGCC | CGG | GCC | C | A | C>T | C>T_GCC | CG>TA_CCAAGCC | 9 | FALSE | FALSE |
| 132_v.txt | 31977 | 31977-SNV-C | SNV | 1 | C | T | 26 | 24990 | 0.10404162 | 41.8461539 | WT | FALSE | CAN | JSY2424 | 9 | WT-Rep9 | GCC | C>T | A/T+1 | G/C-1 | CG>TA | CG>TA | GCCAAGCCG | CCGGGCA | CGG | GCC | C | A | C>T | C>T_GCC | CG>TA_CCGGGCA | 9 | FALSE | FALSE |
| 132_v.txt | 31981 | 31981-SNV-G | SNV | 1 | G | A | 26 | 24990 | 0.10404162 | 47.8076923 | WT | FALSE | CAN | JSY2424 | 9 | WT-Rep9 | GGC | G>A | A/T+1 | G/C-1 | CG>TA | CG>TA | AGCCGGGCA | GCATTAA | CCG | CCG | C | T | C>T | C>T_CCG | CG>TA_GCATTAA | 9 | FALSE | FALSE |
| 132_v.txt | 32092 | 32092-SNV-C | SNV | 1 | C | T | 26 | 24990 | 0.10404162 | 60.9230769 | WT | FALSE | CAN | JSY2424 | 9 | WT-Rep9 | GCA | C>T | A/T+1 | G/C-1 | CG>TA | CG>TA | AGCCTGCAA | CAACACC | CGT | GCA | C | A | C>T | C>T_GCA | CG>TA_CAACACC | 9 | FALSE | FALSE |
| 132_v.txt | 32114 | 32114-SNV-C | SNV | 1 | C | T | 12414 | 24990 | 49.6758704 | 59.6063316 | WT | FALSE | CAN | JSY2424 | 9 | WT-Rep9 | GCC | C>T | A/T+1 | G/C-1 | CG>TA | CG>TA | TAATAGCCA | CCATTCG | CGG | GCC | C | A | C>T | C>T_GCC | CG>TA_CCATTCG | 9 | FALSE | FALSE |
| 132_v.txt | 32165 | 32165-SNV-G | SNV | 1 | G | A | 30 | 24990 | 0.12004802 | 50.3333333 | WT | FALSE | CAN | JSY2424 | 9 | WT-Rep9 | CGC | G>A | A/T+1 | G/C-1 | CG>TA | CG>TA | CAAAGCGCC | GCCAAAT | GCG | GCG | C | T | C>T | C>T_GCG | CG>TA_GCCAAAT | 9 | FALSE | FALSE |
| 132_v.txt | 32271 | 32271-SNV-C | SNV | 1 | C | T | 54 | 33196 | 0.1626702 | 32.0740741 | WT | FALSE | CAN | JSY2424 | 9 | WT-Rep9 | ACG | C>T | A/T+1 | G/C-1 | CG>TA | CG>TA | AAAATACGG | CGGGAAC | TGC | ACG | C | A | C>T | C>T_ACG | CG>TA_CGGGAAC | 9 | FALSE | FALSE |
| 132_v.txt | 32292 | 32292-SNV-G | SNV | 1 | G | A | 322 | 33196 | 0.96999639 | 54.3881988 | WT | FALSE | CAN | JSY2424 | 9 | WT-Rep9 | TGA | G>A | A/T+1 | G/C-1 | CG>TA | CG>TA | ATATTTGAA | GAATTTG | ACT | ACT | C | T | C>T | C>T_ACT | CG>TA_GAATTTG | 9 | FALSE | FALSE |
| 132_v.txt | 32299 | 32299-SNV-C | SNV | 1 | C | T | 64 | 33196 | 0.19279431 | 54.46875 | WT | FALSE | CAN | JSY2424 | 9 | WT-Rep9 | GCG | C>T | A/T+1 | G/C-1 | CG>TA | CG>TA | AATTTGCGG | CGGCAGA | CGC | GCG | C | A | C>T | C>T_GCG | CG>TA_CGGCAGA | 9 | FALSE | FALSE |
| 132_v.txt | 32304 | 32304-SNV-G | SNV | 1 | G | A | 23 | 33196 | 0.06928546 | 60.7391304 | WT | FALSE | CAN | JSY2424 | 9 | WT-Rep9 | AGA | G>A | A/T+1 | G/C-1 | CG>TA | CG>TA | GCGGCAGAA | GAAATAA | TCT | TCT | C | T | C>T | C>T_TCT | CG>TA_GAAATAA | 9 | FALSE | FALSE |
| 132_v.txt | 32344 | 32344-SNV-G | SNV | 1 | G | A | 26 | 33196 | 0.07832269 | 58.5 | WT | FALSE | CAN | JSY2424 | 9 | WT-Rep9 | GGC | G>A | A/T+1 | G/C-1 | CG>TA | CG>TA | TATGTGGCA | GCAAAAC | CCG | CCG | C | T | C>T | C>T_CCG | CG>TA_GCAAAAC | 9 | FALSE | FALSE |
| 132_v.txt | 32466 | 32466-SNV-C | SNV | 1 | C | T | 50 | 33196 | 0.15062056 | 42.78 | WT | FALSE | CAN | JSY2424 | 9 | WT-Rep9 | GCC | C>T | A/T+1 | G/C-1 | CG>TA | CG>TA | AGAGAGCCA | CCAATGT | CGG | GCC | C | A | C>T | C>T_GCC | CG>TA_CCAATGT | 9 | FALSE | FALSE |
| 132_v.txt | 32487 | 32487-SNV-C | SNV | 1 | C | T | 26 | 33196 | 0.07832269 | 39.7307692 | WT | FALSE | CAN | JSY2424 | 9 | WT-Rep9 | ACG | C>T | A/T+1 | G/C-1 | CG>TA | CG>TA | AAGATACGG | CGGAAAA | TGC | ACG | C | A | C>T | C>T_ACG | CG>TA_CGGAAAA | 9 | FALSE | FALSE |
| 132_v.txt | 32535 | 32535-SNV-G | SNV | 1 | G | A | 39 | 33196 | 0.11748403 | 36.9735627 | WT | FALSE | CAN | JSY2424 | 9 | WT-Rep9 | TGC | G>A | A/T+1 | G/C-1 | CG>TA | CG>TA | GGGTTTGCA | GCAGCTT | ACG | ACG | C | T | C>T | C>T_ACG | CG>TA_GCAGCTT | 9 | FALSE | FALSE |
| 132_v.txt | 32536 | 32536-SNV-C | SNV | 1 | C | T | 53 | 33196 | 0.15965779 | 37.2906867 | WT | FALSE | CAN | JSY2424 | 9 | WT-Rep9 | GCA | C>T | A/T+1 | G/C-1 | CG>TA | CG>TA | GGTTTGCAG | CAGCTTC | CGT | GCA | C | A | C>T | C>T_GCA | CG>TA_CAGCTTC | 9 | FALSE | FALSE |
| 132_v.txt | 32572 | 32572-SNV-C | SNV | 1 | C | T | 51 | 30600 | 0.16666667 | 32.9803922 | WT | FALSE | CAN | JSY2424 | 9 | WT-Rep9 | CCT | C>T | A/T+1 | G/C-1 | CG>TA | CG>TA | CAGTACCTT | CTTGAAA | GGA | CCT | C | A | C>T | C>T_CCT | CG>TA_CTTGAAA | 9 | FALSE | FALSE |
| 132_v.txt | 32590 | 32590-SNV-C | SNV | 1 | C | T | 62 | 30600 | 0.20261438 | 36.7903226 | WT | FALSE | CAN | JSY2424 | 9 | WT-Rep9 | GCG | C>T | A/T+1 | G/C-1 | CG>TA | CG>TA | AGGCAGCGT | CGTTAAT | CGC | GCG | C | A | C>T | C>T_GCG | CG>TA_CGTTAAT | 9 | FALSE | FALSE |
| 132_v.txt | 32658 | 32658-SNV-C | SNV | 1 | C | T | 50 | 30600 | 0.16339869 | 55.54 | WT | FALSE | CAN | JSY2424 | 9 | WT-Rep9 | ACC | C>T | A/T+1 | G/C-1 | CG>TA | CG>TA | CCTGGACCC | CCCCAGG | TGG | ACC | C | A | C>T | C>T_ACC | CG>TA_CCCCAGG | 9 | FALSE | FALSE |
| 132_v.txt | 32700 | 32700-SNV-C | SNV | 1 | C | T | 41 | 30600 | 0.13398693 | 61.097561 | WT | FALSE | CAN | JSY2424 | 9 | WT-Rep9 | GCC | C>T | A/T+1 | G/C-1 | CG>TA | CG>TA | ACTGGGCCG | CCGGTAA | CGG | GCC | C | A | C>T | C>T_GCC | CG>TA_CCGGTAA | 9 | FALSE | FALSE |
| 132_v.txt | 32713 | 32713-SNV-C | SNV | 1 | C | T | 22 | 30600 | 0.07189543 | 63.1818182 | WT | FALSE | CAN | JSY2424 | 9 | WT-Rep9 | GCA | C>T | A/T+1 | G/C-1 | CG>TA | CG>TA | CCCCAGCAC | CACCACA | CGT | GCA | C | A | C>T | C>T_GCA | CG>TA_CACCACA | 9 | FALSE | FALSE |
| 132_v.txt | 32757 | 32757-SNV-G | SNV | 1 | G | A | 24 | 30600 | 0.07843137 | 54.375 | WT | FALSE | CAN | JSY2424 | 9 | WT-Rep9 | GGC | G>A | A/T+1 | G/C-1 | CG>TA | CG>TA | ATAATGGCT | GCTAAAA | CCG | CCG | C | T | C>T | C>T_CCG | CG>TA_GCTAAAA | 9 | FALSE | FALSE |
| 132_v.txt | 32775 | 32775-SNV-G | SNV | 1 | G | A | 30 | 30600 | 0.09803922 | 48.3661716 | WT | FALSE | CAN | JSY2424 | 9 | WT-Rep9 | AGC | G>A | A/T+1 | G/C-1 | CG>TA | CG>TA | ATGGAAGCG | GCGACCC | TCG | TCG | C | T | C>T | C>T_TCG | CG>TA_GCGACCC | 9 | FALSE | FALSE |
| 132_v.txt | 32776 | 32776-SNV-C | SNV | 1 | C | T | 33 | 30599 | 0.10784666 | 47.939206 | WT | FALSE | CAN | JSY2424 | 9 | WT-Rep9 | GCG | C>T | A/T+1 | G/C-1 | CG>TA | CG>TA | TGGAAGCGA | CGACCCA | CGC | GCG | C | A | C>T | C>T_GCG | CG>TA_CGACCCA | 9 | FALSE | FALSE |
| 132_v.txt | 32797 | 32797-SNV-C | SNV | 1 | C | T | 145 | 30600 | 0.47385621 | 44.1862069 | WT | FALSE | CAN | JSY2424 | 9 | WT-Rep9 | CCG | C>T | A/T+1 | G/C-1 | CG>TA | CG>TA | ATTCACCGT | CGTAATA | GGC | CCG | C | A | C>T | C>T_CCG | CG>TA_CGTAATA | 9 | FALSE | FALSE |
| 132_v.txt | 32798 | 32798-SNV-G | SNV | 1 | G | A | 53 | 30600 | 0.17320261 | 42.7929047 | WT | FALSE | CAN | JSY2424 | 9 | WT-Rep9 | CGT | G>A | A/T+1 | G/C-1 | CG>TA | CG>TA | TTCACCGTA | GTAATAT | GCA | GCA | C | T | C>T | C>T_GCA | CG>TA_GTAATAT | 9 | FALSE | FALSE |
| 132_v.txt | 32856 | 32856-SNV-C | SNV | 1 | C | T | 27 | 30599 | 0.08823818 | 34.7407407 | WT | FALSE | CAN | JSY2424 | 9 | WT-Rep9 | CCA | C>T | A/T+1 | G/C-1 | CG>TA | CG>TA | CTAATCCAT | CATGCCG | GGT | CCA | C | A | C>T | C>T_CCA | CG>TA_CATGCCG | 9 | FALSE | FALSE |
| 132_v.txt | 32860 | 32860-SNV-C | SNV | 1 | C | T | 35 | 30596 | 0.11439404 | 37.1714286 | WT | FALSE | CAN | JSY2424 | 9 | WT-Rep9 | GCC | C>T | A/T+1 | G/C-1 | CG>TA | CG>TA | TCCATGCCG | CCGCCAG | CGG | GCC | C | A | C>T | C>T_GCC | CG>TA_CCGCCAG | 9 | FALSE | FALSE |
| 132_v.txt | 32879 | 32879-SNV-C | SNV | 1 | C | T | 24 | 30856 | 0.07778066 | 35.4166667 | WT | FALSE | CAN | JSY2424 | 9 | WT-Rep9 | ACG | C>T | A/T+1 | G/C-1 | CG>TA | CG>TA | TTTGTACGT | CGTCCAA | TGC | ACG | C | A | C>T | C>T_ACG | CG>TA_CGTCCAA | 9 | FALSE | FALSE |
| 132_v.txt | 32890 | 32890-SNV-G | SNV | 1 | G | A | 132 | 30857 | 0.42777976 | 36.9848485 | WT | FALSE | CAN | JSY2424 | 9 | WT-Rep9 | TGA | G>A | A/T+1 | G/C-1 | CG>TA | CG>TA | AAAATTGAA | GAATGAC | ACT | ACT | C | T | C>T | C>T_ACT | CG>TA_GAATGAC | 9 | FALSE | FALSE |
| 132_v.txt | 32900 | 32900-SNV-G | SNV | 1 | G | A | 29 | 30857 | 0.09398192 | 36.9964792 | WT | FALSE | CAN | JSY2424 | 9 | WT-Rep9 | GGC | G>A | A/T+1 | G/C-1 | CG>TA | CG>TA | GACTTGGCC | GCCAACT | CCG | CCG | C | T | C>T | C>T_CCG | CG>TA_GCCAACT | 9 | FALSE | FALSE |
| 132_v.txt | 32901 | 32901-SNV-C | SNV | 1 | C | T | 66 | 30857 | 0.21388988 | 32.8791985 | WT | FALSE | CAN | JSY2424 | 9 | WT-Rep9 | GCC | C>T | A/T+1 | G/C-1 | CG>TA | CG>TA | ACTTGGCCA | CCAACTA | CGG | GCC | C | A | C>T | C>T_GCC | CG>TA_CCAACTA | 9 | FALSE | FALSE |
| 132_v.txt | 32935 | 32935-SNV-C | SNV | 1 | C | T | 41 | 30857 | 0.13287099 | 36.2927441 | WT | FALSE | CAN | JSY2424 | 9 | WT-Rep9 | GCC | C>T | A/T+1 | G/C-1 | CG>TA | CG>TA | TGATTGCCC | CCCAAGA | CGG | GCC | C | A | C>T | C>T_GCC | CG>TA_CCCAAGA | 9 | FALSE | FALSE |
| 132_v.txt | 32965 | 32965-SNV-C | SNV | 1 | C | T | 35 | 30857 | 0.11342645 | 56.8285714 | WT | FALSE | CAN | JSY2424 | 9 | WT-Rep9 | GCC | C>T | A/T+1 | G/C-1 | CG>TA | CG>TA | CATTGGCCG | CCGCACC | CGG | GCC | C | A | C>T | C>T_GCC | CG>TA_CCGCACC | 9 | FALSE | FALSE |
| 132_v.txt | 32967 | 32967-SNV-G | SNV | 1 | G | A | 80 | 30857 | 0.25926046 | 58.75 | WT | FALSE | CAN | JSY2424 | 9 | WT-Rep9 | CGC | G>A | A/T+1 | G/C-1 | CG>TA | CG>TA | TTGGCCGCA | GCACCAA | GCG | GCG | C | T | C>T | C>T_GCG | CG>TA_GCACCAA | 9 | FALSE | FALSE |
| 132_v.txt | 32977 | 32977-SNV-C | SNV | 1 | C | T | 26 | 30857 | 0.08425965 | 57.9615385 | WT | FALSE | CAN | JSY2424 | 9 | WT-Rep9 | GCT | C>T | A/T+1 | G/C-1 | CG>TA | CG>TA | CAAATGCTG | CTGGAGA | CGA | GCT | C | A | C>T | C>T_GCT | CG>TA_CTGGAGA | 9 | FALSE | FALSE |
| 132_v.txt | 32995 | 32995-SNV-G | SNV | 1 | G | A | 77 | 30857 | 0.24953819 | 61.8441558 | WT | FALSE | CAN | JSY2424 | 9 | WT-Rep9 | TGT | G>A | A/T+1 | G/C-1 | CG>TA | CG>TA | ATCTTTGTG | GTGAGAA | ACA | ACA | C | T | C>T | C>T_ACA | CG>TA_GTGAGAA | 9 | FALSE | FALSE |
| 132_v.txt | 33012 | 33012-SNV-G | SNV | 1 | G | A | 74 | 30857 | 0.23981593 | 60.2432432 | WT | FALSE | CAN | JSY2424 | 9 | WT-Rep9 | AGA | G>A | A/T+1 | G/C-1 | CG>TA | CG>TA | GTGAAAGAG | GAGGATG | TCT | TCT | C | T | C>T | C>T_TCT | CG>TA_GAGGATG | 9 | FALSE | FALSE |
| 132_v.txt | 33056 | 33056-SNV-C | SNV | 1 | C | T | 44 | 30857 | 0.14259325 | 56.5 | WT | FALSE | CAN | JSY2424 | 9 | WT-Rep9 | GCG | C>T | A/T+1 | G/C-1 | CG>TA | CG>TA | GGACTGCGT | CGTGACA | CGC | GCG | C | A | C>T | C>T_GCG | CG>TA_CGTGACA | 9 | FALSE | FALSE |
| 132_v.txt | 33070 | 33070-SNV-C | SNV | 1 | C | T | 30 | 30857 | 0.09722267 | 50.7333333 | WT | FALSE | CAN | JSY2424 | 9 | WT-Rep9 | GCC | C>T | A/T+1 | G/C-1 | CG>TA | CG>TA | AATATGCCA | CCAAAGA | CGG | GCC | C | A | C>T | C>T_GCC | CG>TA_CCAAAGA | 9 | FALSE | FALSE |
| 132_v.txt | 33104 | 33104-SNV-G | SNV | 1 | G | A | 37 | 30857 | 0.11990796 | 43.1891892 | WT | FALSE | CAN | JSY2424 | 9 | WT-Rep9 | CGC | G>A | A/T+1 | G/C-1 | CG>TA | CG>TA | AAGAGCGCC | GCCCACT | GCG | GCG | C | T | C>T | C>T_GCG | CG>TA_GCCCACT | 9 | FALSE | FALSE |
| 132_v.txt | 33111 | 33111-SNV-G | SNV | 1 | G | A | 31 | 30857 | 0.10046343 | 45.483871 | WT | FALSE | CAN | JSY2424 | 9 | WT-Rep9 | TGG | G>A | A/T+1 | G/C-1 | CG>TA | CG>TA | CCCACTGGG | GGGCCGG | ACC | ACC | C | T | C>T | C>T_ACC | CG>TA_GGGCCGG | 9 | FALSE | FALSE |
| 132_v.txt | 33114 | 33114-SNV-C | SNV | 1 | C | T | 31 | 30857 | 0.10046343 | 39.0322581 | WT | FALSE | CAN | JSY2424 | 9 | WT-Rep9 | GCC | C>T | A/T+1 | G/C-1 | CG>TA | CG>TA | ACTGGGCCG | CCGGCGT | CGG | GCC | C | A | C>T | C>T_GCC | CG>TA_CCGGCGT | 9 | FALSE | FALSE |
| 132_v.txt | 33118 | 33118-SNV-C | SNV | 1 | C | T | 27 | 30857 | 0.08750041 | 38.8514716 | WT | FALSE | CAN | JSY2424 | 9 | WT-Rep9 | GCG | C>T | A/T+1 | G/C-1 | CG>TA | CG>TA | GGCCGGCGT | CGTTGGT | CGC | GCG | C | A | C>T | C>T_GCG | CG>TA_CGTTGGT | 9 | FALSE | FALSE |
| 132_v.txt | 33119 | 33119-SNV-G | SNV | 1 | G | A | 27 | 30857 | 0.08750041 | 45.1477543 | WT | FALSE | CAN | JSY2424 | 9 | WT-Rep9 | CGT | G>A | A/T+1 | G/C-1 | CG>TA | CG>TA | GCCGGCGTT | GTTGGTC | GCA | GCA | C | T | C>T | C>T_GCA | CG>TA_GTTGGTC | 9 | FALSE | FALSE |
| 132_v.txt | 33159 | 33159-SNV-C | SNV | 1 | C | T | 33 | 30855 | 0.10695187 | 35.6060606 | WT | FALSE | CAN | JSY2424 | 9 | WT-Rep9 | ACC | C>T | A/T+1 | G/C-1 | CG>TA | CG>TA | CCTGTACCA | CCAATAG | TGG | ACC | C | A | C>T | C>T_ACC | CG>TA_CCAATAG | 9 | FALSE | FALSE |
| 132_v.txt | 33193 | 33193-SNV-G | SNV | 1 | G | A | 150 | 33365 | 0.44957291 | 36.5056913 | WT | FALSE | CAN | JSY2424 | 9 | WT-Rep9 | TGT | G>A | A/T+1 | G/C-1 | CG>TA | CG>TA | CAATATGTC | GTCTTTG | ACA | ACA | C | T | C>T | C>T_ACA | CG>TA_GTCTTTG | 9 | FALSE | FALSE |
| 132_v.txt | 33258 | 33258-SNV-C | SNV | 1 | C | T | 50 | 33366 | 0.14985314 | 42.9 | WT | FALSE | CAN | JSY2424 | 9 | WT-Rep9 | GCC | C>T | A/T+1 | G/C-1 | CG>TA | CG>TA | CCTATGCCAT | CCATCCT | CGG | GCC | C | A | C>T | C>T_GCC | CG>TA_CCATCCT | 9 | FALSE | FALSE |
| 132_v.txt | 33276 | 33276-SNV-G | SNV | 1 | G | A | 57 | 33366 | 0.17083258 | 46.5614035 | WT | FALSE | CAN | JSY2424 | 9 | WT-Rep9 | CGT | G>A | A/T+1 | G/C-1 | CG>TA | CG>TA | GAGAACGTA | GTATCCT | GCA | GCA | C | T | C>T | C>T_GCA | CG>TA_GTATCCT | 9 | FALSE | FALSE |
| 132_v.txt | 33284 | 33284-SNV-G | SNV | 1 | G | A | 32 | 33366 | 0.09590601 | 52 | WT | FALSE | CAN | JSY2424 | 9 | WT-Rep9 | CGC | G>A | A/T+1 | G/C-1 | CG>TA | CG>TA | ATCCTCGCC | GCCATTT | GCG | GCG | C | T | C>T | C>T_GCG | CG>TA_GCCATTT | 9 | FALSE | FALSE |
| 132_v.txt | 33309 | 33309-SNV-C | SNV | 1 | C | T | 40 | 33366 | 0.11988252 | 55.1726908 | WT | FALSE | CAN | JSY2424 | 9 | WT-Rep9 | GCG | C>T | A/T+1 | G/C-1 | CG>TA | CG>TA | AAAGAGCGC | CGCAATG | CGC | GCG | C | A | C>T | C>T_GCG | CG>TA_CGCAATG | 9 | FALSE | FALSE |
| 132_v.txt | 33310 | 33310-SNV-G | SNV | 1 | G | A | 38 | 33361 | 0.11390546 | 55.5263158 | WT | FALSE | CAN | JSY2424 | 9 | WT-Rep9 | CGC | G>A | A/T+1 | G/C-1 | CG>TA | CG>TA | AAGAGCGCA | GCAATGG | GCG | GCG | C | T | C>T | C>T_GCG | CG>TA_GCAATGG | 9 | FALSE | FALSE |
| 132_v.txt | 33376 | 33376-SNV-G | SNV | 1 | G | A | 85 | 33366 | 0.25475035 | 54.7764706 | WT | FALSE | CAN | JSY2424 | 9 | WT-Rep9 | TGT | G>A | A/T+1 | G/C-1 | CG>TA | CG>TA | GTGTTTGTG | GTGAAGC | ACA | ACA | C | T | C>T | C>T_ACA | CG>TA_GTGAAGC | 9 | FALSE | FALSE |
| 132_v.txt | 33392 | 33392-SNV-G | SNV | 1 | G | A | 39 | 33366 | 0.11688545 | 47.7435897 | WT | FALSE | CAN | JSY2424 | 9 | WT-Rep9 | CGT | G>A | A/T+1 | G/C-1 | CG>TA | CG>TA | AACGTCGTG | GTGAAAG | GCA | GCA | C | T | C>T | C>T_GCA | CG>TA_GTGAAAG | 9 | FALSE | FALSE |
| 132_v.txt | 33412 | 33412-SNV-G | SNV | 1 | G | A | 49 | 33366 | 0.14685608 | 43.4081633 | WT | FALSE | CAN | JSY2424 | 9 | WT-Rep9 | GGC | G>A | A/T+1 | G/C-1 | CG>TA | CG>TA | TGACCGGCT | GCTCATT | CCG | CCG | C | T | C>T | C>T_CCG | CG>TA_GCTCATT | 9 | FALSE | FALSE |
| 132_v.txt | 33474 | 33474-SNV-C | SNV | 1 | C | T | 49 | 33362 | 0.14687369 | 34.6452232 | WT | FALSE | CAN | JSY2424 | 9 | WT-Rep9 | GCC | C>T | A/T+1 | G/C-1 | CG>TA | CG>TA | GCTATGCCT | CCTTTTT | CGG | GCC | C | A | C>T | C>T_GCC | CG>TA_CCTTTTT | 9 | FALSE | FALSE |
| 132_v.txt | 32033 | 32033-SNV-C | SNV | 1 | C | G | 1 | 24990 | 4.00E-03 | 63.9960154 | WT | FALSE | CAN | JSY2424 | 9 | WT-Rep9 | TCA | C>G | G/C+1 | G/C-1 | CG>GC | CG>GC | GTATTTCAA | CAAAGCT | AGT | TCA | C | C | C>G | C>G_TCA | CG>GC_CAAAGCT | 9 | FALSE | FALSE |
| 132_v.txt | 32199 | 32199-SNV-G | SNV | 1 | G | C | 44 | 24990 | 0.17607043 | 38.8863636 | WT | FALSE | CAN | JSY2424 | 9 | WT-Rep9 | TGG | G>C | G/C+1 | G/C-1 | CG>GC | CG>GC | ATGTATGGA | GGAACAC | ACC | ACC | C | G | C>G | C>G_ACC | CG>GC_GGAACAC | 9 | FALSE | FALSE |
| 132_v.txt | 32278 | 32278-SNV-C | SNV | 1 | C | G | 21 | 33196 | 0.06326063 | 35.8571429 | WT | FALSE | CAN | JSY2424 | 9 | WT-Rep9 | CCA | C>G | G/C+1 | G/C-1 | CG>GC | CG>GC | GGGAACCAA | CAACGTA | GGT | CCA | C | C | C>G | C>G_CCA | CG>GC_CAACGTA | 9 | FALSE | FALSE |
| 132_v.txt | 32282 | 32282-SNV-G | SNV | 1 | G | C | 15 | 33196 | 0.04518617 | 41.3333333 | WT | FALSE | CAN | JSY2424 | 9 | WT-Rep9 | CGT | G>C | G/C+1 | G/C-1 | CG>GC | CG>GC | ACCAACGTA | GTAAATA | GCA | GCA | C | G | C>G | C>G_GCA | CG>GC_GTAAATA | 9 | FALSE | FALSE |
| 132_v.txt | 32474 | 32474-SNV-G | SNV | 1 | G | C | 3 | 33196 | 9.04E-03 | 44.9985948 | WT | FALSE | CAN | JSY2424 | 9 | WT-Rep9 | AGA | G>C | G/C+1 | G/C-1 | CG>GC | CG>GC | AATGTAGAA | GAAGGTT | TCT | TCT | C | G | C>G | C>G_TCT | CG>GC_GAAGGTT | 9 | FALSE | FALSE |
| 132_v.txt | 32529 | 32529-SNV-G | SNV | 1 | G | C | 2530 | 33196 | 7.62140017 | 41.5810277 | WT | FALSE | CAN | JSY2424 | 9 | WT-Rep9 | GGG | G>C | G/C+1 | G/C-1 | CG>GC | CG>GC | TTTCTGGGG | GGGTTTG | CCC | CCC | C | G | C>G | C>G_CCC | CG>GC_GGGTTTG | 9 | FALSE | FALSE |
| 132_v.txt | 32535 | 32535-SNV-G | SNV | 1 | G | C | 1 | 33196 | 3.01E-03 | 37.9993729 | WT | FALSE | CAN | JSY2424 | 9 | WT-Rep9 | TGC | G>C | G/C+1 | G/C-1 | CG>GC | CG>GC | GGGTTTGCA | GCAGCTT | ACG | ACG | C | G | C>G | C>G_ACG | CG>GC_GCAGCTT | 9 | FALSE | FALSE |
| 132_v.txt | 32757 | 32757-SNV-G | SNV | 1 | G | C | 4 | 30600 | 0.0130719 | 53.75 | WT | FALSE | CAN | JSY2424 | 9 | WT-Rep9 | GGC | G>C | G/C+1 | G/C-1 | CG>GC | CG>GC | ATAATGGCT | GCTAAAA | CCG | CCG | C | G | C>G | C>G_CCG | CG>GC_GCTAAAA | 9 | FALSE | FALSE |
| 132_v.txt | 32794 | 32794-SNV-C | SNV | 1 | C | G | 29 | 30600 | 0.09477124 | 42.2758621 | WT | FALSE | CAN | JSY2424 | 9 | WT-Rep9 | TCA | C>G | G/C+1 | G/C-1 | CG>GC | CG>GC | CGAATTCAC | CACCGTA | AGT | TCA | C | C | C>G | C>G_TCA | CG>GC_CACCGTA | 9 | FALSE | FALSE |
| 132_v.txt | 32901 | 32901-SNV-C | SNV | 1 | C | G | 2 | 30857 | 6.48E-03 | 37.5000965 | WT | FALSE | CAN | JSY2424 | 9 | WT-Rep9 | GCC | C>G | G/C+1 | G/C-1 | CG>GC | CG>GC | ACTTGGCCA | CCAACTA | CGG | GCC | C | C | C>G | C>G_GCC | CG>GC_CCAACTA | 9 | FALSE | FALSE |
| 132_v.txt | 32928 | 32928-SNV-G | SNV | 1 | G | C | 3 | 30857 | 9.72E-03 | 62.6666667 | WT | FALSE | CAN | JSY2424 | 9 | WT-Rep9 | AGT | G>C | G/C+1 | G/C-1 | CG>GC | CG>GC | GCAAAAGTG | GTGATTG | TCA | TCA | C | G | C>G | C>G_TCA | CG>GC_GTGATTG | 9 | FALSE | FALSE |
| 132_v.txt | 32935 | 32935-SNV-C | SNV | 1 | C | G | 1 | 30857 | 3.24E-03 | 42.0009558 | WT | FALSE | CAN | JSY2424 | 9 | WT-Rep9 | GCC | C>G | G/C+1 | G/C-1 | CG>GC | CG>GC | TGATTGCCC | CCCAAGA | CGG | GCC | C | C | C>G | C>G_GCC | CG>GC_CCCAAGA | 9 | FALSE | FALSE |
| 132_v.txt | 32936 | 32936-SNV-C | SNV | 1 | C | G | 31 | 30857 | 0.10046343 | 53.7488997 | WT | FALSE | CAN | JSY2424 | 9 | WT-Rep9 | CCC | C>G | G/C+1 | G/C-1 | CG>GC | CG>GC | GATTGCCCA | CCAAGAA | GGG | CCC | C | C | C>G | C>G_CCC | CG>GC_CCAAGAA | 9 | FALSE | FALSE |
| 132_v.txt | 32997 | 32997-SNV-G | SNV | 1 | G | C | 16 | 30857 | 0.05185209 | 63.1875 | WT | FALSE | CAN | JSY2424 | 9 | WT-Rep9 | TGA | G>C | G/C+1 | G/C-1 | CG>GC | CG>GC | CTTTGTGAG | GAGAAAA | ACT | ACT | C | G | C>G | C>G_ACT | CG>GC_GAGAAAA | 9 | FALSE | FALSE |
| 132_v.txt | 33070 | 33070-SNV-C | SNV | 1 | C | G | 4 | 30857 | 0.01296302 | 55.5 | WT | FALSE | CAN | JSY2424 | 9 | WT-Rep9 | GCC | C>G | G/C+1 | G/C-1 | CG>GC | CG>GC | AATATGCCA | CCAAAGA | CGG | GCC | C | C | C>G | C>G_GCC | CG>GC_CCAAAGA | 9 | FALSE | FALSE |
| 132_v.txt | 33118 | 33118-SNV-C | SNV | 1 | C | G | 4 | 30857 | 0.01296302 | 37.9995925 | WT | FALSE | CAN | JSY2424 | 9 | WT-Rep9 | GCG | C>G | G/C+1 | G/C-1 | CG>GC | CG>GC | GGCCGGCGT | CGTTGGT | CGC | GCG | C | C | C>G | C>G_GCG | CG>GC_CGTTGGT | 9 | FALSE | FALSE |
| 132_v.txt | 33310 | 33310-SNV-G | SNV | 1 | G | C | 7 | 33361 | 0.02098259 | 25 | WT | FALSE | CAN | JSY2424 | 9 | WT-Rep9 | CGC | G>C | G/C+1 | G/C-1 | CG>GC | CG>GC | AAGAGCGCA | GCAATGG | GCG | GCG | C | G | C>G | C>G_GCG | CG>GC_GCAATGG | 9 | FALSE | FALSE |
| 132_v.txt | 31657 | 31657-SNV-G | SNV | 1 | G | T | 94 | 10432 | 0.90107362 | 18.9148936 | WT | FALSE | CAN | JSY2424 | 9 | WT-Rep9 | GGC | G>T | A/T+1 | G/C-1 | CG>AT | CG>AT | GAAATGGCG | GCGTGGA | CCG | CCG | C | A | C>A | C>A_CCG | CG>AT_GCGTGGA | 9 | FALSE | FALSE |
| 132_v.txt | 31829 | 31829-SNV-G | SNV | 1 | G | T | 15 | 10432 | 0.14378834 | 50.1333333 | WT | FALSE | CAN | JSY2424 | 9 | WT-Rep9 | TGC | G>T | A/T+1 | G/C-1 | CG>AT | CG>AT | GAATATGCA | GCATTGA | ACG | ACG | C | A | C>A | C>A_ACG | CG>AT_GCATTGA | 9 | FALSE | FALSE |
| 132_v.txt | 32064 | 32064-SNV-G | SNV | 1 | G | T | 11 | 24990 | 0.04401761 | 54.2727273 | WT | FALSE | CAN | JSY2424 | 9 | WT-Rep9 | TGA | G>T | A/T+1 | G/C-1 | CG>AT | CG>AT | GAGATTGAG | GAGATAA | ACT | ACT | C | A | C>A | C>A_ACT | CG>AT_GAGATAA | 9 | FALSE | FALSE |
| 132_v.txt | 32095 | 32095-SNV-C | SNV | 1 | C | A | 20 | 24990 | 0.08003201 | 58.7 | WT | FALSE | CAN | JSY2424 | 9 | WT-Rep9 | ACA | C>A | A/T+1 | G/C-1 | CG>AT | CG>AT | CTGCAACAC | CACCAGT | TGT | ACA | C | T | C>A | C>A_ACA | CG>AT_CACCAGT | 9 | FALSE | FALSE |
| 132_v.txt | 32220 | 32220-SNV-C | SNV | 1 | C | A | 13 | 24990 | 0.05202081 | 42 | WT | FALSE | CAN | JSY2424 | 9 | WT-Rep9 | CCT | C>A | A/T+1 | G/C-1 | CG>AT | CG>AT | GTGGTCCTT | CTTGACA | GGA | CCT | C | T | C>A | C>A_CCT | CG>AT_CTTGACA | 9 | FALSE | FALSE |
| 132_v.txt | 32282 | 32282-SNV-G | SNV | 1 | G | T | 33 | 33196 | 0.09940957 | 40.2424242 | WT | FALSE | CAN | JSY2424 | 9 | WT-Rep9 | CGT | G>T | A/T+1 | G/C-1 | CG>AT | CG>AT | ACCAACGTA | GTAAATA | GCA | GCA | C | A | C>A | C>A_GCA | CG>AT_GTAAATA | 9 | FALSE | FALSE |
| 132_v.txt | 32432 | 32432-SNV-G | SNV | 1 | G | T | 865 | 33196 | 2.60573563 | 56.1757225 | WT | FALSE | CAN | JSY2424 | 9 | WT-Rep9 | TGT | G>T | A/T+1 | G/C-1 | CG>AT | CG>AT | GTCATTGTA | GTATGGA | ACA | ACA | C | A | C>A | C>A_ACA | CG>AT_GTATGGA | 9 | FALSE | FALSE |
| 132_v.txt | 32474 | 32474-SNV-G | SNV | 1 | G | T | 16 | 33196 | 0.04819858 | 43.749217 | WT | FALSE | CAN | JSY2424 | 9 | WT-Rep9 | AGA | G>T | A/T+1 | G/C-1 | CG>AT | CG>AT | AATGTAGAA | GAAGGTT | TCT | TCT | C | A | C>A | C>A_TCT | CG>AT_GAAGGTT | 9 | FALSE | FALSE |
| 132_v.txt | 32529 | 32529-SNV-G | SNV | 1 | G | T | 33 | 33196 | 0.09940957 | 43.9090909 | WT | FALSE | CAN | JSY2424 | 9 | WT-Rep9 | GGG | G>T | A/T+1 | G/C-1 | CG>AT | CG>AT | TTTCTGGGG | GGGTTTG | CCC | CCC | C | A | C>A | C>A_CCC | CG>AT_GGGTTTG | 9 | FALSE | FALSE |
| 132_v.txt | 32536 | 32536-SNV-C | SNV | 1 | C | A | 8 | 33196 | 0.02409929 | 23.4954457 | WT | FALSE | CAN | JSY2424 | 9 | WT-Rep9 | GCA | C>A | A/T+1 | G/C-1 | CG>AT | CG>AT | GGTTTGCAG | CAGCTTC | CGT | GCA | C | T | C>A | C>A_GCA | CG>AT_CAGCTTC | 9 | FALSE | FALSE |
| 132_v.txt | 32776 | 32776-SNV-C | SNV | 1 | C | A | 6 | 30599 | 0.01960848 | 30.1679424 | WT | FALSE | CAN | JSY2424 | 9 | WT-Rep9 | GCG | C>A | A/T+1 | G/C-1 | CG>AT | CG>AT | TGGAAGCGA | CGACCCA | CGC | GCG | C | T | C>A | C>A_GCG | CG>AT_CGACCCA | 9 | FALSE | FALSE |
| 132_v.txt | 32797 | 32797-SNV-C | SNV | 1 | C | A | 37 | 30600 | 0.12091503 | 45.8378378 | WT | FALSE | CAN | JSY2424 | 9 | WT-Rep9 | CCG | C>A | A/T+1 | G/C-1 | CG>AT | CG>AT | ATTCACCGT | CGTAATA | GGC | CCG | C | T | C>A | C>A_CCG | CG>AT_CGTAATA | 9 | FALSE | FALSE |
| 132_v.txt | 32811 | 32811-SNV-G | SNV | 1 | G | T | 12 | 30600 | 0.03921569 | 48.7493651 | WT | FALSE | CAN | JSY2424 | 9 | WT-Rep9 | AGG | G>T | A/T+1 | G/C-1 | CG>AT | CG>AT | TTGACAGGG | GGGAACA | TCC | TCC | C | A | C>A | C>A_TCC | CG>AT_GGGAACA | 9 | FALSE | FALSE |
| 132_v.txt | 32928 | 32928-SNV-G | SNV | 1 | G | T | 51 | 30857 | 0.16527854 | 48.3921569 | WT | FALSE | CAN | JSY2424 | 9 | WT-Rep9 | AGT | G>T | A/T+1 | G/C-1 | CG>AT | CG>AT | GCAAAAGTG | GTGATTG | TCA | TCA | C | A | C>A | C>A_TCA | CG>AT_GTGATTG | 9 | FALSE | FALSE |
| 132_v.txt | 32958 | 32958-SNV-C | SNV | 1 | C | A | 18 | 30857 | 0.0583336 | 53.8888889 | WT | FALSE | CAN | JSY2424 | 9 | WT-Rep9 | ACC | C>A | A/T+1 | G/C-1 | CG>AT | CG>AT | ATGTAACCA | CCATTGG | TGG | ACC | C | T | C>A | C>A_ACC | CG>AT_CCATTGG | 9 | FALSE | FALSE |
| 132_v.txt | 33024 | 33024-SNV-G | SNV | 1 | G | T | 31 | 30857 | 0.10046343 | 61.5483871 | WT | FALSE | CAN | JSY2424 | 9 | WT-Rep9 | AGG | G>T | A/T+1 | G/C-1 | CG>AT | CG>AT | GTAACAGGG | GGGATGA | TCC | TCC | C | A | C>A | C>A_TCC | CG>AT_GGGATGA | 9 | FALSE | FALSE |
| 132_v.txt | 33051 | 33051-SNV-G | SNV | 1 | G | T | 56 | 30857 | 0.18148232 | 56.5535714 | WT | FALSE | CAN | JSY2424 | 9 | WT-Rep9 | GGA | G>T | A/T+1 | G/C-1 | CG>AT | CG>AT | CCCAAGGAC | GACTGCG | CCT | CCT | C | A | C>A | C>A_CCT | CG>AT_GACTGCG | 9 | FALSE | FALSE |
| 132_v.txt | 33151 | 33151-SNV-G | SNV | 1 | G | T | 169 | 30856 | 0.54770547 | 39.2958419 | WT | FALSE | CAN | JSY2424 | 9 | WT-Rep9 | AGA | G>T | A/T+1 | G/C-1 | CG>AT | CG>AT | TGAAAAGAC | GACCTGT | TCT | TCT | C | A | C>A | C>A_TCT | CG>AT_GACCTGT | 9 | FALSE | FALSE |
| 132_v.txt | 33169 | 33169-SNV-C | SNV | 1 | C | A | 81 | 30853 | 0.26253525 | 37.9506173 | WT | FALSE | CAN | JSY2424 | 9 | WT-Rep9 | CCA | C>A | A/T+1 | G/C-1 | CG>AT | CG>AT | TAGTACCAC | CACCAAG | GGT | CCA | C | T | C>A | C>A_CCA | CG>AT_CACCAAG | 9 | FALSE | FALSE |
| 132_v.txt | 33310 | 33310-SNV-G | SNV | 1 | G | T | 17 | 33361 | 0.05095771 | 21.5882353 | WT | FALSE | CAN | JSY2424 | 9 | WT-Rep9 | CGC | G>T | A/T+1 | G/C-1 | CG>AT | CG>AT | AAGAGCGCA | GCAATGG | GCG | GCG | C | A | C>A | C>A_GCG | CG>AT_GCAATGG | 9 | FALSE | FALSE |
| 132_v.txt | 33330 | 33330-SNV-C | SNV | 1 | C | A | 41 | 33366 | 0.12287958 | 59.7563922 | WT | FALSE | CAN | JSY2424 | 9 | WT-Rep9 | ACT | C>A | A/T+1 | G/C-1 | CG>AT | CG>AT | TCTTTACTTT | CTTTTCT | TGA | ACT | C | T | C>A | C>A_ACT | CG>AT_CTTTTCT | 9 | FALSE | FALSE |
| 132_v.txt | 33419 | 33419-SNV-G | SNV | 1 | G | T | 45 | 33366 | 0.13486783 | 39.5111111 | WT | FALSE | CAN | JSY2424 | 9 | WT-Rep9 | TGT | G>T | A/T+1 | G/C-1 | CG>AT | CG>AT | CTCATTGTAC | GTACATA | ACA | ACA | C | A | C>A | C>A_ACA | CG>AT_GTACATA | 9 | FALSE | FALSE |
| 132_v.txt | 33433 | 33433-SNV-C | SNV | 1 | C | A | 26 | 33366 | 0.07792364 | 44.6923077 | WT | FALSE | CAN | JSY2424 | 9 | WT-Rep9 | TCC | C>A | A/T+1 | G/C-1 | CG>AT | CG>AT | GCTTCTCCTC | CCTCTAT | AGG | TCC | C | T | C>A | C>A_TCC | CG>AT_CCTCTAT | 9 | FALSE | FALSE |
| 132_v.txt | 33456 | 33456-SNV-G | SNV | 1 | G | T | 51 | 33365 | 0.15285479 | 28.3529412 | WT | FALSE | CAN | JSY2424 | 9 | WT-Rep9 | TGA | G>T | A/T+1 | G/C-1 | CG>AT | CG>AT | TCTTTTGAAT | GAATTTG | ACT | ACT | C | A | C>A | C>A_ACT | CG>AT_GAATTTG | 9 | FALSE | FALSE |
| 132_v.txt | 33474 | 33474-SNV-C | SNV | 1 | C | A | 96 | 33362 | 0.28775253 | 34.1410894 | WT | FALSE | CAN | JSY2424 | 9 | WT-Rep9 | GCC | C>A | A/T+1 | G/C-1 | CG>AT | CG>AT | GCTATGCCTT | CCTTTTT | CGG | GCC | C | T | C>A | C>A_GCC | CG>AT_CCTTTTT | 9 | FALSE | FALSE |
| 132_v.txt | 33475 | 33475-SNV-C | SNV | 1 | C | A | 22 | 33362 | 0.06594329 | 34.2269903 | WT | FALSE | CAN | JSY2424 | 9 | WT-Rep9 | CCT | C>A | A/T+1 | G/C-1 | CG>AT | CG>AT | CTATGCCTTT | CTTTTTT | GGA | CCT | C | T | C>A | C>A_CCT | CG>AT_CTTTTTT | 9 | FALSE | FALSE |
| 132_v.txt | 32811 | 32811-MNV- | MNV | 2 | GG | AA | 112 | 30600 | 0.36601307 | 44.1744332 | WT | FALSE | CAN | JSY2424 | 9 | WT-Rep9 | AGG | nope | >1bp-ins | >1bpdel | none | MNV | TTGACAGGG | GGGAACA | TCC | AGG | GG | none | NA | NA_AGG | MNV_GGGAACA | 9 | FALSE | FALSE |
| 132_v.txt | 33330 | 33330-MNV- | MNV | 2 | CT | AC | 1 | 33366 | 3.00E-03 | 63.999973 | WT | FALSE | CAN | JSY2424 | 9 | WT-Rep9 | ACT | nope | >1bp-ins | >1bpdel | none | MNV | TCTTTACTTT | CTTTTCT | TGA | ACT | CT | none | NA | NA_ACT | MNV_CTTTTCT | 9 | FALSE | FALSE |
| 132_v.txt | 32081 | 32081-Insert | Insertion | 1 | - | A | 15 | 24990 | 0.06002401 | 62.7333333 | WT | FALSE | CAN | JSY2424 | 9 | WT-Rep9 | CAA | nope | A/T+1 | >1bpdel | none | A/T+1 | CCATGCAAA | AAAAAAG | GTT | CAA | - | T | NA | NA_CAA | A/T+1_AAAAAAG | 9 | FALSE | FALSE |
| 132_v.txt | 32490 | 32490-Insert | Insertion | 1 | - | A | 19 | 33196 | 0.05723581 | 42.1052632 | WT | FALSE | CAN | JSY2424 | 9 | WT-Rep9 | GAA | nope | A/T+1 | >1bpdel | none | A/T+1 | ATACGGAAA | AAAACAA | CTT | GAA | - | T | NA | NA_GAA | A/T+1_AAAACAA | 9 | FALSE | FALSE |
| 132_v.txt | 32705 | 32705-Insert | Insertion | 1 | - | A | 4 | 30600 | 0.0130719 | 63.25 | WT | FALSE | CAN | JSY2424 | 9 | WT-Rep9 | TAA | nope | A/T+1 | >1bpdel | none | A/T+1 | GCCGGTAAC | AACCCCA | ATT | TAA | - | T | NA | NA_TAA | A/T+1_AACCCCA | 9 | FALSE | FALSE |
| 132_v.txt | 32734 | 32734-Insert | Insertion | 3 | - | AGT | 98 | 30600 | 0.32026144 | 60.4965986 | WT | FALSE | CAN | JSY2424 | 9 | WT-Rep9 | CAG | nope | >1bp-ins | >1bpdel | none | >1bp-ins | AAAAACAGT | AGTATAT | GTC | CAG | - | none | NA | NA_CAG | >1bp-ins_AGTATAT | 9 | FALSE | FALSE |
| 132_v.txt | 32775 | 32775-Insert | Insertion | 1 | - | G | 1 | 30600 | 3.27E-03 | 11 | WT | FALSE | CAN | JSY2424 | 9 | WT-Rep9 | AGC | nope | G/C+1 | >1bpdel | none | G/C+1 | ATGGAAGCG | GCGACCC | TCG | AGC | - | C | NA | NA_AGC | G/C+1_GCGACCC | 9 | FALSE | FALSE |
| 132_v.txt | 32842 | 32842-Insert | Insertion | 1 | - | A | 458 | 30600 | 1.49673203 | 41.709607 | WT | FALSE | CAN | JSY2424 | 9 | WT-Rep9 | CAA | nope | A/T+1 | >1bpdel | none | A/T+1 | TTACCCAAA | AAAAAAT | GTT | CAA | - | T | NA | NA_CAA | A/T+1_AAAAAAT | 9 | FALSE | FALSE |
| 132_v.txt | 33015 | 33015-Insert | Insertion | 9 | - | GATGTAACA | 47 | 30857 | 0.15231552 | 62.7257683 | WT | FALSE | CAN | JSY2424 | 9 | WT-Rep9 | GGA | nope | >1bp-ins | >1bpdel | none | >1bp-ins | AAAGAGGAT | GATGTAA | CCT | GGA | - | none | NA | NA_GGA | >1bp-ins_GATGTAA | 9 | FALSE | FALSE |
| 132_v.txt | 33223 | 33223-Insert | Insertion | 11 | - | CGTTCTGTAC | 9 | 33366 | 0.02697357 | 38.2626263 | WT | FALSE | CAN | JSY2424 | 9 | WT-Rep9 | GCG | nope | >1bp-ins | >1bpdel | none | >1bp-ins | CTTCAGCGTT | CGTTCTG | CGC | GCG | - | none | NA | NA_GCG | >1bp-ins_CGTTCTG | 9 | FALSE | FALSE |
| 132_v.txt | 32032 | 32032-Deleti | Deletion | 2 | TC | - | 10 | 24990 | 0.04001601 | 64 | WT | FALSE | CAN | JSY2424 | 9 | WT-Rep9 | TTC | nope | >1bp-ins | >1bpdel | none | >1bpdel | GGTATTTCA | TCAAAGC | AAG | TTC | TC | none | NA | NA_TTC | >1bpdel_TCAAAGC | 9 | FALSE | FALSE |

| | | | | | | | | | | | | | | | | | | | | | | | | | | | | | | | | | | |
|---|---|---|---|---|---|---|---|---|---|---|---|---|---|---|---|---|---|---|---|---|---|---|---|---|---|---|---|---|---|---|---|---|---|---|
| 132_v.txt | 32313 | 32313-Deleti | Deletion | 3 | GTT | - | 31 | 33196 | 0.09338475 | 60.1935484 | WT | FALSE | CAN | JSY2424 | 9 | WT-Rep9 | GGT | nope | >1bp-ins | >1bpdel | none | >1bpdel | ATAATGGTT | GTTGTTA | CCA | GGT | GTT | none | NA | NA_GGT | >1bpdel_GTTGTTA | 9 | FALSE | FALSE |
| 132_v.txt | 32337 | 32337-Deleti | Deletion | 2 | AT | - | 77 | 33196 | 0.23195566 | 62.6883117 | WT | FALSE | CAN | JSY2424 | 9 | WT-Rep9 | GAT | nope | >1bp-ins | >1bpdel | none | >1bpdel | TTGAAGATA | ATATGTG | CTA | GAT | AT | none | NA | NA_GAT | >1bpdel_ATATGTG | 9 | FALSE | FALSE |
| 132_v.txt | 32473 | 32473-Deleti | Deletion | 2 | AG | - | 1 | 33196 | 3.01E-03 | 32 | WT | FALSE | CAN | JSY2424 | 9 | WT-Rep9 | TAG | nope | >1bp-ins | >1bpdel | none | >1bpdel | CAATGTAGA | AGAAGGT | ATC | TAG | AG | none | NA | NA_TAG | >1bpdel_AGAAGGT | 9 | FALSE | FALSE |
| 132_v.txt | 33089 | 33089-Deleti | Deletion | 2 | AT | - | 83 | 30857 | 0.26898273 | 53.5903615 | WT | FALSE | CAN | JSY2424 | 9 | WT-Rep9 | AAT | nope | >1bp-ins | >1bpdel | none | >1bpdel | AAATAAATA | ATATGAT | TTA | AAT | AT | none | NA | NA_AAT | >1bpdel_ATATGAT | 9 | FALSE | FALSE |
| 132_v.txt | 33474 | 33474-Deleti | Deletion | 2 | CC | - | 1 | 33362 | 3.00E-03 | 10 | WT | FALSE | CAN | JSY2424 | 9 | WT-Rep9 | GCC | nope | >1bp-ins | >1bpdel | none | >1bpdel | GCTATGCCT | CCTTTTT | CGG | GCC | CC | none | NA | NA_GCC | >1bpdel_CCTTTTT | 9 | FALSE | FALSE |
| 132_v.txt | 31663 | 31663-Deleti | Deletion | 1 | A | - | 6 | 10432 | 0.05751534 | 37.3333333 | WT | FALSE | CAN | JSY2424 | 9 | WT-Rep9 | GAA | nope | >1bp-ins | A/T-1 | none | A/T-1 | GCGTGGAAA | AAATGTG | CTT | CTT | T | none | T>none | T>none_CTT | A/T-1_AAATGTG | 9 | FALSE | FALSE |
| 132_v.txt | 31723 | 31723-Deleti | Deletion | 1 | A | - | 17 | 10432 | 0.16296012 | 59.1764706 | WT | FALSE | CAN | JSY2424 | 9 | WT-Rep9 | CAA | nope | >1bp-ins | A/T-1 | none | A/T-1 | TGTCCCAAA | AAAAAGT | GTT | GTT | T | none | T>none | T>none_GTT | A/T-1_AAAAAGT | 9 | FALSE | FALSE |
| 132_v.txt | 31904 | 31904-Deleti | Deletion | 1 | T | - | 8 | 10423 | 0.07675333 | 43.875 | WT | FALSE | CAN | JSY2424 | 9 | WT-Rep9 | ATT | nope | >1bp-ins | A/T-1 | none | A/T-1 | ATTGAATTT | TTTTGGT | TAA | ATT | T | none | nope | nope_ATT | A/T-1_TTTTGGT | 9 | FALSE | FALSE |
| 132_v.txt | 31972 | 31972-Deleti | Deletion | 1 | C | - | 6 | 24990 | 0.0240096 | 37.1666667 | WT | FALSE | CAN | JSY2424 | 9 | WT-Rep9 | GCC | nope | >1bp-ins | G/C-1 | none | G/C-1 | AATAAGCCA | CCAAGCC | CGG | GCC | C | none | nope | nope_GCC | G/C-1_CCAAGCC | 9 | FALSE | FALSE |
| 132_v.txt | 32033 | 32033-Deleti | Deletion | 1 | C | - | 1 | 24990 | 4.00E-03 | 64 | WT | FALSE | CAN | JSY2424 | 9 | WT-Rep9 | TCA | nope | >1bp-ins | G/C-1 | none | G/C-1 | GTATTTCAA | CAAAGCT | AGT | TCA | C | none | nope | nope_TCA | G/C-1_CAAAGCT | 9 | FALSE | FALSE |
| 132_v.txt | 32069 | 32069-Deleti | Deletion | 1 | A | - | 5 | 24990 | 0.020008 | 64 | WT | FALSE | CAN | JSY2424 | 9 | WT-Rep9 | TAA | nope | >1bp-ins | A/T-1 | none | A/T-1 | TGAGATAAA | AAATAAC | ATT | ATT | T | none | T>none | T>none_ATT | A/T-1_AAATAAC | 9 | FALSE | FALSE |
| 132_v.txt | 32081 | 32081-Deleti | Deletion | 1 | A | - | 77 | 24990 | 0.30812325 | 62.7402597 | WT | FALSE | CAN | JSY2424 | 9 | WT-Rep9 | CAA | nope | >1bp-ins | A/T-1 | none | A/T-1 | CCATGCAAA | AAAAAAG | GTT | GTT | T | none | T>none | T>none_GTT | A/T-1_AAAAAAG | 9 | FALSE | FALSE |
| 132_v.txt | 32210 | 32210-Deleti | Deletion | 1 | T | - | 9 | 24990 | 0.03601441 | 40.8888889 | WT | FALSE | CAN | JSY2424 | 9 | WT-Rep9 | CTT | nope | >1bp-ins | A/T-1 | none | A/T-1 | ACCACCTTTG | TTTGGTG | GAA | CTT | T | none | nope | nope_CTT | A/T-1_TTTGGTG | 9 | FALSE | FALSE |
| 132_v.txt | 32218 | 32218-Deleti | Deletion | 1 | T | - | 51 | 24990 | 0.20408163 | 39.8235294 | WT | FALSE | CAN | JSY2424 | 9 | WT-Rep9 | GTC | nope | >1bp-ins | A/T-1 | none | A/T-1 | TGGTGGTCC | TCCTTGA | CAG | GTC | T | none | nope | nope_GTC | A/T-1_TCCTTGA | 9 | FALSE | FALSE |
| 132_v.txt | 32231 | 32231-Deleti | Deletion | 1 | T | - | 5 | 24990 | 0.020008 | 37.8 | WT | FALSE | CAN | JSY2424 | 9 | WT-Rep9 | ATT | nope | >1bp-ins | A/T-1 | none | A/T-1 | CAGGAATTT | TTTAGGA | TAA | ATT | T | none | nope | nope_ATT | A/T-1_TTTAGGA | 9 | FALSE | FALSE |
| 132_v.txt | 32261 | 32261-Deleti | Deletion | 1 | A | - | 99 | 33195 | 0.29823769 | 37.4646465 | WT | FALSE | CAN | JSY2424 | 9 | WT-Rep9 | CAA | nope | >1bp-ins | A/T-1 | none | A/T-1 | TAGACCAAA | AAATAAA | GTT | GTT | T | none | T>none | T>none_GTT | A/T-1_AAATAAA | 9 | FALSE | FALSE |
| 132_v.txt | 32284 | 32284-Deleti | Deletion | 1 | A | - | 6 | 33196 | 0.01807447 | 51 | WT | FALSE | CAN | JSY2424 | 9 | WT-Rep9 | TAA | nope | >1bp-ins | A/T-1 | none | A/T-1 | CAACGTAAA | AAATATT | ATT | ATT | T | none | T>none | T>none_ATT | A/T-1_AAATATT | 9 | FALSE | FALSE |
| 132_v.txt | 32346 | 32346-Deleti | Deletion | 1 | A | - | 6 | 33196 | 0.01807447 | 63.1666667 | WT | FALSE | CAN | JSY2424 | 9 | WT-Rep9 | CAA | nope | >1bp-ins | A/T-1 | none | A/T-1 | TGTGGCAAA | AAAACCT | GTT | GTT | T | none | T>none | T>none_GTT | A/T-1_AAAACCT | 9 | FALSE | FALSE |
| 132_v.txt | 32381 | 32381-Deleti | Deletion | 1 | A | - | 6 | 33196 | 0.01807447 | 63.3333333 | WT | FALSE | CAN | JSY2424 | 9 | WT-Rep9 | TAA | nope | >1bp-ins | A/T-1 | none | A/T-1 | AATAATAAA | AAAGGGA | ATT | ATT | T | none | T>none | T>none_ATT | A/T-1_AAAGGGA | 9 | FALSE | FALSE |
| 132_v.txt | 32416 | 32416-Deleti | Deletion | 1 | T | - | 6 | 33196 | 0.01807447 | 50.8333333 | WT | FALSE | CAN | JSY2424 | 9 | WT-Rep9 | GTT | nope | >1bp-ins | A/T-1 | none | A/T-1 | ATTGTGTTAG | TTAGTTT | CAA | GTT | T | none | nope | nope_GTT | A/T-1_TTAGTTT | 9 | FALSE | FALSE |
| 132_v.txt | 32420 | 32420-Deleti | Deletion | 1 | T | - | 8 | 33196 | 0.02409929 | 63.75 | WT | FALSE | CAN | JSY2424 | 9 | WT-Rep9 | GTT | nope | >1bp-ins | A/T-1 | none | A/T-1 | TGTTAGTTTA | TTTAGGG | CAA | GTT | T | none | nope | nope_GTT | A/T-1_TTTAGGG | 9 | FALSE | FALSE |
| 132_v.txt | 32442 | 32442-Deleti | Deletion | 1 | A | - | 9 | 33196 | 0.0271117 | 49 | WT | FALSE | CAN | JSY2424 | 9 | WT-Rep9 | TAA | nope | >1bp-ins | A/T-1 | none | A/T-1 | GGAACTAAA | AAAAGTC | ATT | ATT | T | none | T>none | T>none_ATT | A/T-1_AAAAGTC | 9 | FALSE | FALSE |
| 132_v.txt | 32490 | 32490-Deleti | Deletion | 1 | A | - | 2 | 33196 | 6.03E-03 | 38 | WT | FALSE | CAN | JSY2424 | 9 | WT-Rep9 | GAA | nope | >1bp-ins | A/T-1 | none | A/T-1 | ATACGGAAA | AAAACAA | CTT | CTT | T | none | T>none | T>none_CTT | A/T-1_AAAACAA | 9 | FALSE | FALSE |
| 132_v.txt | 32498 | 32498-Deleti | Deletion | 1 | T | - | 34 | 33196 | 0.10242198 | 46.6176471 | WT | FALSE | CAN | JSY2424 | 9 | WT-Rep9 | CTT | nope | >1bp-ins | A/T-1 | none | A/T-1 | AACAACTTTT | TTTTTTG | GAA | CTT | T | none | nope | nope_CTT | A/T-1_TTTTTTG | 9 | FALSE | FALSE |
| 132_v.txt | 32514 | 32514-Deleti | Deletion | 1 | G | - | 18 | 33196 | 0.0542234 | 39.3333333 | WT | FALSE | CAN | JSY2424 | 9 | WT-Rep9 | TGG | nope | >1bp-ins | G/C-1 | none | G/C-1 | GCTCTTGGA | GGAACGG | ACC | ACC | C | none | C>none | C>none_ACC | G/C-1_GGAACGG | 9 | FALSE | FALSE |
| 132_v.txt | 32630 | 32630-Deleti | Deletion | 1 | G | - | 20 | 30600 | 0.06535948 | 50.2999936 | WT | FALSE | CAN | JSY2424 | 9 | WT-Rep9 | CGT | nope | >1bp-ins | G/C-1 | none | G/C-1 | CCCTTCGTTT | GTTTTTA | GCA | GCA | C | none | C>none | C>none_GCA | G/C-1_GTTTTTA | 9 | FALSE | FALSE |
| 132_v.txt | 32631 | 32631-Deleti | Deletion | 1 | T | - | 11 | 30600 | 0.03594771 | 49.3636037 | WT | FALSE | CAN | JSY2424 | 9 | WT-Rep9 | GTT | nope | >1bp-ins | A/T-1 | none | A/T-1 | CCTTCGTTTT | TTTTTAT | CAA | GTT | T | none | nope | nope_GTT | A/T-1_TTTTTAT | 9 | FALSE | FALSE |
| 132_v.txt | 32637 | 32637-Deleti | Deletion | 1 | T | - | 16 | 30600 | 0.05228758 | 56.5 | WT | FALSE | CAN | JSY2424 | 9 | WT-Rep9 | ATC | nope | >1bp-ins | A/T-1 | none | A/T-1 | TTTTTATCCT | TCCTTAG | TAG | ATC | T | none | nope | nope_ATC | A/T-1_TCCTTAG | 9 | FALSE | FALSE |
| 132_v.txt | 32640 | 32640-Deleti | Deletion | 1 | T | - | 8 | 30600 | 0.02614379 | 58.5 | WT | FALSE | CAN | JSY2424 | 9 | WT-Rep9 | CTT | nope | >1bp-ins | A/T-1 | none | A/T-1 | TTATCCTTAG | TTAGATA | GAA | CTT | T | none | nope | nope_CTT | A/T-1_TTAGATA | 9 | FALSE | FALSE |
| 132_v.txt | 32658 | 32658-Deleti | Deletion | 1 | C | - | 8 | 30600 | 0.02614379 | 57.375 | WT | FALSE | CAN | JSY2424 | 9 | WT-Rep9 | ACC | nope | >1bp-ins | G/C-1 | none | G/C-1 | CCTGGACCC | CCCCAGG | TGG | ACC | C | none | nope | nope_ACC | G/C-1_CCCCAGG | 9 | FALSE | FALSE |
| 132_v.txt | 32662 | 32662-Deleti | Deletion | 1 | A | - | 34 | 30600 | 0.11111111 | 62.1176471 | WT | FALSE | CAN | JSY2424 | 9 | WT-Rep9 | CAG | nope | >1bp-ins | A/T-1 | none | A/T-1 | GACCCCAGG | AGGCACC | GTC | GTC | T | none | T>none | T>none_GTC | A/T-1_AGGCACC | 9 | FALSE | FALSE |
| 132_v.txt | 32728 | 32728-Deleti | Deletion | 1 | A | - | 10 | 30600 | 0.03267974 | 62.6 | WT | FALSE | CAN | JSY2424 | 9 | WT-Rep9 | CAA | nope | >1bp-ins | A/T-1 | none | A/T-1 | CCATACAAA | AAAAACA | GTT | GTT | T | none | T>none | T>none_GTT | A/T-1_AAAAACA | 9 | FALSE | FALSE |
| 132_v.txt | 32779 | 32779-Deleti | Deletion | 1 | C | - | 8 | 30600 | 0.02614379 | 45.625 | WT | FALSE | CAN | JSY2424 | 9 | WT-Rep9 | ACC | nope | >1bp-ins | G/C-1 | none | G/C-1 | AAGCGACCC | CCCAGAA | TGG | ACC | C | none | nope | nope_ACC | G/C-1_CCCAGAA | 9 | FALSE | FALSE |
| 132_v.txt | 32811 | 32811-Deleti | Deletion | 1 | G | - | 2 | 30600 | 6.54E-03 | 63.9999977 | WT | FALSE | CAN | JSY2424 | 9 | WT-Rep9 | AGG | nope | >1bp-ins | G/C-1 | none | G/C-1 | TTGACAGGG | GGGAACA | TCC | TCC | C | none | C>none | C>none_TCC | G/C-1_GGGAACA | 9 | FALSE | FALSE |
| 132_v.txt | 32836 | 32836-Deleti | Deletion | 1 | T | - | 18 | 30600 | 0.05882353 | 43.0555556 | WT | FALSE | CAN | JSY2424 | 9 | WT-Rep9 | ATT | nope | >1bp-ins | A/T-1 | none | A/T-1 | TGATAATTA | TTACCCA | TAA | ATT | T | none | nope | nope_ATT | A/T-1_TTACCCA | 9 | FALSE | FALSE |
| 132_v.txt | 32842 | 32842-Deleti | Deletion | 1 | A | - | 77 | 30600 | 0.25163399 | 41.7012985 | WT | FALSE | CAN | JSY2424 | 9 | WT-Rep9 | CAA | nope | >1bp-ins | A/T-1 | none | A/T-1 | TTACCCAAA | AAAAAAT | GTT | GTT | T | none | T>none | T>none_GTT | A/T-1_AAAAAAT | 9 | FALSE | FALSE |
| 132_v.txt | 32884 | 32884-Deleti | Deletion | 1 | A | - | 12 | 30857 | 0.03888907 | 37.0833333 | WT | FALSE | CAN | JSY2424 | 9 | WT-Rep9 | CAA | nope | >1bp-ins | A/T-1 | none | A/T-1 | ACGTCCAAA | AAAATTG | GTT | GTT | T | none | T>none | T>none_GTT | A/T-1_AAAATTG | 9 | FALSE | FALSE |
| 132_v.txt | 32924 | 32924-Deleti | Deletion | 1 | A | - | 38 | 30857 | 0.12314872 | 54.9210526 | WT | FALSE | CAN | JSY2424 | 9 | WT-Rep9 | CAA | nope | >1bp-ins | A/T-1 | none | A/T-1 | CAGGGCAAA | AAAAGTG | GTT | GTT | T | none | T>none | T>none_GTT | A/T-1_AAAAGTG | 9 | FALSE | FALSE |
| 132_v.txt | 32935 | 32935-Deleti | Deletion | 1 | C | - | 7 | 30857 | 0.02268529 | 41.2857164 | WT | FALSE | CAN | JSY2424 | 9 | WT-Rep9 | GCC | nope | >1bp-ins | G/C-1 | none | G/C-1 | TGATTGCCC | CCCAAGA | CGG | GCC | C | none | nope | nope_GCC | G/C-1_CCCAAGA | 9 | FALSE | FALSE |
| 132_v.txt | 32992 | 32992-Deleti | Deletion | 1 | T | - | 7 | 30857 | 0.02268529 | 61.1428571 | WT | FALSE | CAN | JSY2424 | 9 | WT-Rep9 | CTT | nope | >1bp-ins | A/T-1 | none | A/T-1 | GGAATCTTT | TTTGTGA | GAA | CTT | T | none | nope | nope_CTT | A/T-1_TTTGTGA | 9 | FALSE | FALSE |
| 132_v.txt | 33027 | 33027-Deleti | Deletion | 1 | A | - | 34 | 30857 | 0.1101857 | 60.5294118 | WT | FALSE | CAN | JSY2424 | 9 | WT-Rep9 | GAT | nope | >1bp-ins | A/T-1 | none | A/T-1 | ACAGGGATG | ATGAATG | CTA | CTA | T | none | T>none | T>none_CTA | A/T-1_ATGAATG | 9 | FALSE | FALSE |
| 132_v.txt | 33138 | 33138-Deleti | Deletion | 1 | A | - | 7 | 30857 | 0.02268529 | 40.1428571 | WT | FALSE | CAN | JSY2424 | 9 | WT-Rep9 | TAA | nope | >1bp-ins | A/T-1 | none | A/T-1 | GTGGATAAA | AAACCAA | ATT | ATT | T | none | T>none | T>none_ATT | A/T-1_AAACCAA | 9 | FALSE | FALSE |
| 132_v.txt | 33147 | 33147-Deleti | Deletion | 1 | A | - | 6 | 30857 | 0.01944454 | 38 | WT | FALSE | CAN | JSY2424 | 9 | WT-Rep9 | GAA | nope | >1bp-ins | A/T-1 | none | A/T-1 | CCAATGAAA | AAAAGAC | CTT | CTT | T | none | T>none | T>none_CTT | A/T-1_AAAAGAC | 9 | FALSE | FALSE |
| 132_v.txt | 33166 | 33166-Deleti | Deletion | 1 | T | - | 30 | 30853 | 0.09723528 | 37.9 | WT | FALSE | CAN | JSY2424 | 9 | WT-Rep9 | GTA | nope | >1bp-ins | A/T-1 | none | A/T-1 | CAATAGTAC | TACCACC | CAT | GTA | T | none | nope | nope_GTA | A/T-1_TACCACC | 9 | FALSE | FALSE |
| 132_v.txt | 33239 | 33239-Deleti | Deletion | 1 | T | - | 5 | 33366 | 0.01498531 | 43.2 | WT | FALSE | CAN | JSY2424 | 9 | WT-Rep9 | CTT | nope | >1bp-ins | A/T-1 | none | A/T-1 | TTCTCCTTCA | TTCATCT | GAA | CTT | T | none | nope | nope_CTT | A/T-1_TTCATCT | 9 | FALSE | FALSE |
| 132_v.txt | 33326 | 33326-Deleti | Deletion | 1 | T | - | 99 | 33366 | 0.29670923 | 61.3636364 | WT | FALSE | CAN | JSY2424 | 9 | WT-Rep9 | CTT | nope | >1bp-ins | A/T-1 | none | A/T-1 | CAATTCTTTA | TTTACTT | GAA | CTT | T | none | nope | nope_CTT | A/T-1_TTTACTT | 9 | FALSE | FALSE |
| 132_v.txt | 33331 | 33331-Deleti | Deletion | 1 | T | - | 12 | 33365 | 0.03596583 | 62.8333333 | WT | FALSE | CAN | JSY2424 | 9 | WT-Rep9 | CTT | nope | >1bp-ins | A/T-1 | none | A/T-1 | CTTTACTTTT | TTTTCTC | GAA | CTT | T | none | nope | nope_CTT | A/T-1_TTTTCTC | 9 | FALSE | FALSE |
| 132_v.txt | 33474 | 33474-Deleti | Deletion | 1 | C | - | 76 | 33362 | 0.22780409 | 29.6184211 | WT | FALSE | CAN | JSY2424 | 9 | WT-Rep9 | GCC | nope | >1bp-ins | G/C-1 | none | G/C-1 | GCTATGCCT | CCTTTTT | CGG | GCC | C | none | nope | nope_GCC | G/C-1_CCTTTTT | 9 | FALSE | FALSE |
| 168_v.txt | 31723 | 31723-SNV-A | SNV | 1 | A | C | 2 | 3138 | 0.06373486 | 51 | WT | FALSE | CAN | JSY13 | 1A | WT-Rep1A | CAA | A>C | G/C+1 | A/T-1 | TA>GC | TA>GC | TGTCCCAAA | AAAAAGT | GTT | GTT | T | G | T>G | T>G_GTT | TA>GC_AAAAAGT | 9 | TRUE | FALSE |
| 168_v.txt | 31950 | 31950-SNV-A | SNV | 1 | A | C | 29 | 14348 | 0.20211876 | 35.8275862 | WT | FALSE | CAN | JSY13 | 1A | WT-Rep1A | CAT | A>C | G/C+1 | A/T-1 | TA>GC | TA>GC | ATCGTCATA | ATAAATG | GTA | GTA | T | G | T>G | T>G_GTA | TA>GC_ATAAATG | 9 | TRUE | FALSE |
| 168_v.txt | 31966 | 31966-SNV-A | SNV | 1 | A | C | 9 | 14355 | 0.06269593 | 32.7777778 | WT | FALSE | CAN | JSY13 | 1A | WT-Rep1A | TAA | A>C | G/C+1 | A/T-1 | TA>GC | TA>GC | CCGCATAAT | AATAAGC | ATT | ATT | T | G | T>G | T>G_ATT | TA>GC_AATAAGC | 9 | TRUE | FALSE |
| 168_v.txt | 31986 | 31986-SNV-A | SNV | 1 | A | C | 127 | 14355 | 0.88470916 | 51.2913386 | WT | FALSE | CAN | JSY13 | 1A | WT-Rep1A | TAA | A>C | G/C+1 | A/T-1 | TA>GC | TA>GC | GGCATTAAT | AATTTAG | ATT | ATT | T | G | T>G | T>G_ATT | TA>GC_AATTTAG | 9 | TRUE | FALSE |
| 168_v.txt | 32055 | 32055-SNV-T | SNV | 1 | T | G | 3 | 14355 | 0.02089864 | 63.3333333 | WT | FALSE | CAN | JSY13 | 1A | WT-Rep1A | GTG | T>G | G/C+1 | A/T-1 | TA>GC | TA>GC | CTGATGTGC | TGCGAGA | CAC | GTG | T | C | T>G | T>G_GTG | TA>GC_TGCGAGA | 9 | TRUE | FALSE |
| 168_v.txt | 32101 | 32101-SNV-T | SNV | 1 | T | G | 6 | 14355 | 0.04179728 | 59.5 | WT | FALSE | CAN | JSY13 | 1A | WT-Rep1A | GTG | T>G | G/C+1 | A/T-1 | TA>GC | TA>GC | CACCAGTGA | TGATATT | CAC | GTG | T | C | T>G | T>G_GTG | TA>GC_TGATATT | 9 | TRUE | FALSE |
| 168_v.txt | 32109 | 32109-SNV-A | SNV | 1 | A | C | 13 | 14355 | 0.09056078 | 61.8461539 | WT | FALSE | CAN | JSY13 | 1A | WT-Rep1A | TAA | A>C | G/C+1 | A/T-1 | TA>GC | TA>GC | ATATTTAATA | AATAGCC | ATT | ATT | T | G | T>G | T>G_ATT | TA>GC_AATAGCC | 9 | TRUE | FALSE |
| 168_v.txt | 32226 | 32226-SNV-A | SNV | 1 | A | C | 13 | 14341 | 0.09064919 | 36.7693539 | WT | FALSE | CAN | JSY13 | 1A | WT-Rep1A | CAG | A>C | G/C+1 | A/T-1 | TA>GC | TA>GC | CTTGACAGG | AGGAATT | GTC | GTC | T | G | T>G | T>G_GTC | TA>GC_AGGAATT | 9 | TRUE | FALSE |
| 168_v.txt | 32256 | 32256-SNV-A | SNV | 1 | A | C | 23 | 46462 | 0.04950282 | 18.4782609 | WT | FALSE | CAN | JSY13 | 1A | WT-Rep1A | TAG | A>C | G/C+1 | A/T-1 | TA>GC | TA>GC | TTTGATAGA | AGACCAA | ATC | ATC | T | G | T>G | T>G_ATC | TA>GC_AGACCAA | 9 | TRUE | FALSE |
| 168_v.txt | 32265 | 32265-SNV-A | SNV | 1 | A | C | 45 | 46483 | 0.09680959 | 30.6888889 | WT | FALSE | CAN | JSY13 | 1A | WT-Rep1A | TAA | A>C | G/C+1 | A/T-1 | TA>GC | TA>GC | CCAAATAAA | AAAATAC | ATT | ATT | T | G | T>G | T>G_ATT | TA>GC_AAAATAC | 9 | TRUE | FALSE |
| 168_v.txt | 32283 | 32283-SNV-T | SNV | 1 | T | G | 6029 | 46483 | 12.9703332 | 52.7436166 | WT | FALSE | CAN | JSY13 | 1A | WT-Rep1A | GTA | T>G | G/C+1 | A/T-1 | TA>GC | TA>GC | CCAACGTAA | TAAATAT | CAT | GTA | T | C | T>G | T>G_GTA | TA>GC_TAAATAT | 9 | TRUE | FALSE |
| 168_v.txt | 32284 | 32284-SNV-A | SNV | 1 | A | C | 23 | 46483 | 0.04948046 | 28.2612947 | WT | FALSE | CAN | JSY13 | 1A | WT-Rep1A | TAA | A>C | G/C+1 | A/T-1 | TA>GC | TA>GC | CAACGTAAA | AAATATT | ATT | ATT | T | G | T>G | T>G_ATT | TA>GC_AAATATT | 9 | TRUE | FALSE |
| 168_v.txt | 32288 | 32288-SNV-A | SNV | 1 | A | C | 106 | 46483 | 0.22804036 | 52.3657481 | WT | FALSE | CAN | JSY13 | 1A | WT-Rep1A | TAT | A>C | G/C+1 | A/T-1 | TA>GC | TA>GC | GTAAATATTT | ATTTGAA | ATA | ATA | T | G | T>G | T>G_ATA | TA>GC_ATTTGAA | 9 | TRUE | FALSE |
| 168_v.txt | 32294 | 32294-SNV-A | SNV | 1 | A | C | 53 | 46483 | 0.11402018 | 49.4909499 | WT | FALSE | CAN | JSY13 | 1A | WT-Rep1A | AAT | A>C | G/C+1 | A/T-1 | TA>GC | TA>GC | ATTTGAATTT | ATTTGCG | TTA | TTA | T | G | T>G | T>G_TTA | TA>GC_ATTTGCG | 9 | TRUE | FALSE |
| 168_v.txt | 32310 | 32310-SNV-A | SNV | 1 | A | C | 15 | 46483 | 0.03226986 | 46.7326453 | WT | FALSE | CAN | JSY13 | 1A | WT-Rep1A | AAT | A>C | G/C+1 | A/T-1 | TA>GC | TA>GC | GAAATAATG | ATGGTTG | TTA | TTA | T | G | T>G | T>G_TTA | TA>GC_ATGGTTG | 9 | TRUE | FALSE |
| 168_v.txt | 32319 | 32319-SNV-A | SNV | 1 | A | C | 93 | 46483 | 0.20007315 | 56.6200455 | WT | FALSE | CAN | JSY13 | 1A | WT-Rep1A | TAA | A>C | G/C+1 | A/T-1 | TA>GC | TA>GC | GTTGTTAAG | AAGATAA | ATT | ATT | T | G | T>G | T>G_ATT | TA>GC_AAGATAA | 9 | TRUE | FALSE |
| 168_v.txt | 32439 | 32439-SNV-A | SNV | 1 | A | C | 112 | 46483 | 0.2409483 | 53.6339286 | WT | FALSE | CAN | JSY13 | 1A | WT-Rep1A | AAC | A>C | G/C+1 | A/T-1 | TA>GC | TA>GC | TATGGAACT | ACTAAAA | TTG | TTG | T | G | T>G | T>G_TTG | TA>GC_ACTAAAA | 9 | TRUE | FALSE |
| 168_v.txt | 32445 | 32445-SNV-A | SNV | 1 | A | C | 49 | 46483 | 0.10541488 | 52.3469388 | WT | FALSE | CAN | JSY13 | 1A | WT-Rep1A | AAG | A>C | G/C+1 | A/T-1 | TA>GC | TA>GC | ACTAAAAGT | AGTCCAA | TTC | TTC | T | G | T>G | T>G_TTC | TA>GC_AGTCCAA | 9 | TRUE | FALSE |
| 168_v.txt | 32472 | 32472-SNV-T | SNV | 1 | T | G | 111 | 46483 | 0.23879698 | 47.1891831 | WT | FALSE | CAN | JSY13 | 1A | WT-Rep1A | GTA | T>G | G/C+1 | A/T-1 | TA>GC | TA>GC | CCAATGTAG | TAGAAGG | CAT | GTA | T | C | T>G | T>G_GTA | TA>GC_TAGAAGG | 9 | TRUE | FALSE |
| 168_v.txt | 32473 | 32473-SNV-A | SNV | 1 | A | C | 29 | 46483 | 0.0623884 | 45.7222951 | WT | FALSE | CAN | JSY13 | 1A | WT-Rep1A | TAG | A>C | G/C+1 | A/T-1 | TA>GC | TA>GC | CAATGTAGA | AGAAGGT | ATC | ATC | T | G | T>G | T>G_ATC | TA>GC_AGAAGGT | 9 | TRUE | FALSE |
| 168_v.txt | 32481 | 32481-SNV-A | SNV | 1 | A | C | 60 | 46483 | 0.12907945 | 45.3333333 | WT | FALSE | CAN | JSY13 | 1A | WT-Rep1A | TAA | A>C | G/C+1 | A/T-1 | TA>GC | TA>GC | AAGGTTAAG | AAGATAC | ATT | ATT | T | G | T>G | T>G_ATT | TA>GC_AAGATAC | 9 | TRUE | FALSE |
| 168_v.txt | 32493 | 32493-SNV-A | SNV | 1 | A | C | 8 | 46483 | 0.01721059 | 30.2521875 | WT | FALSE | CAN | JSY13 | 1A | WT-Rep1A | AAC | A>C | G/C+1 | A/T-1 | TA>GC | TA>GC | CGGAAAACA | ACAACTT | TTG | TTG | T | G | T>G | T>G_TTG | TA>GC_ACAACTT | 9 | TRUE | FALSE |
| 168_v.txt | 32527 | 32527-SNV-T | SNV | 1 | T | G | 7 | 46483 | 0.01505927 | 19.4643235 | WT | FALSE | CAN | JSY13 | 1A | WT-Rep1A | CTG | T>G | G/C+1 | A/T-1 | TA>GC | TA>GC | ATTTTCTGG | TGGGGTT | GAC | CTG | T | C | T>G | T>G_CTG | TA>GC_TGGGGTT | 9 | TRUE | FALSE |
| 168_v.txt | 32541 | 32541-SNV-T | SNV | 1 | T | G | 139 | 46481 | 0.29904692 | 37.2942535 | WT | FALSE | CAN | JSY13 | 1A | WT-Rep1A | TTC | T>G | G/C+1 | A/T-1 | TA>GC | TA>GC | GCAGCTTCA | TCACCAG | AAG | TTC | T | C | T>G | T>G_TTC | TA>GC_TCACCAG | 9 | TRUE | FALSE |
| 168_v.txt | 32581 | 32581-SNV-T | SNV | 1 | T | G | 6 | 7613 | 0.07881256 | 32.6666667 | WT | FALSE | CAN | JSY13 | 1A | WT-Rep1A | GTG | T>G | G/C+1 | A/T-1 | TA>GC | TA>GC | GAAATGTGA | TGAAGGC | CAC | GTG | T | C | T>G | T>G_GTG | TA>GC_TGAAGGC | 9 | TRUE | FALSE |
| 168_v.txt | 32616 | 32616-SNV-A | SNV | 1 | A | C | 4 | 7614 | 0.0525348 | 50.25 | WT | FALSE | CAN | JSY13 | 1A | WT-Rep1A | TAA | A>C | G/C+1 | A/T-1 | TA>GC | TA>GC | CAACCTAAG | AAGAACC | ATT | ATT | T | G | T>G | T>G_ATT | TA>GC_AAGAACC | 9 | TRUE | FALSE |
| 168_v.txt | 32680 | 32680-SNV-A | SNV | 1 | A | C | 9 | 7614 | 0.11820331 | 55 | WT | FALSE | CAN | JSY13 | 1A | WT-Rep1A | CAA | A>C | G/C+1 | A/T-1 | TA>GC | TA>GC | TTCTCCAATA | AATAACG | GTT | GTT | T | G | T>G | T>G_GTT | TA>GC_AATAACG | 9 | TRUE | FALSE |
| 168_v.txt | 32728 | 32728-SNV-A | SNV | 1 | A | C | 2 | 7622 | 0.02623983 | 20.5006044 | WT | FALSE | CAN | JSY13 | 1A | WT-Rep1A | CAA | A>C | G/C+1 | A/T-1 | TA>GC | TA>GC | CCATACAAA | AAAAACA | GTT | GTT | T | G | T>G | T>G_GTT | TA>GC_AAAAACA | 9 | TRUE | FALSE |
| 168_v.txt | 32760 | 32760-SNV-A | SNV | 1 | A | C | 2 | 7622 | 0.02623983 | 24.5001226 | WT | FALSE | CAN | JSY13 | 1A | WT-Rep1A | TAA | A>C | G/C+1 | A/T-1 | TA>GC | TA>GC | ATGGCTAAA | AAAACTT | ATT | ATT | T | G | T>G | T>G_ATT | TA>GC_AAAACTT | 9 | TRUE | FALSE |
| 168_v.txt | 32787 | 32787-SNV-T | SNV | 1 | T | G | 11 | 7622 | 0.14431908 | 41.9090909 | WT | FALSE | CAN | JSY13 | 1A | WT-Rep1A | CTC | T>G | G/C+1 | A/T-1 | TA>GC | TA>GC | CAGAACTCG | TCGAATT | GAG | CTC | T | C | T>G | T>G_CTC | TA>GC_TCGAATT | 9 | TRUE | FALSE |
| 168_v.txt | 32792 | 32792-SNV-T | SNV | 1 | T | G | 6 | 7622 | 0.0787195 | 47.1666667 | WT | FALSE | CAN | JSY13 | 1A | WT-Rep1A | ATT | T>G | G/C+1 | A/T-1 | TA>GC | TA>GC | CTCGAATTCA | TTCACCG | TAA | ATT | T | C | T>G | T>G_ATT | TA>GC_TTCACCG | 9 | TRUE | FALSE |
| 168_v.txt | 32800 | 32800-SNV-A | SNV | 1 | A | C | 6 | 7622 | 0.0787195 | 43.8333333 | WT | FALSE | CAN | JSY13 | 1A | WT-Rep1A | TAA | A>C | G/C+1 | A/T-1 | TA>GC | TA>GC | CACCGTAAT | AATATTT | ATT | ATT | T | G | T>G | T>G_ATT | TA>GC_AATATTT | 9 | TRUE | FALSE |
| 168_v.txt | 32801 | 32801-SNV-A | SNV | 1 | A | C | 7 | 7622 | 0.09183941 | 49.4309924 | WT | FALSE | CAN | JSY13 | 1A | WT-Rep1A | AAT | A>C | G/C+1 | A/T-1 | TA>GC | TA>GC | ACCGTAATA | ATATTTG | TTA | TTA | T | G | T>G | T>G_TTA | TA>GC_ATATTTG | 9 | TRUE | FALSE |
| 168_v.txt | 32803 | 32803-SNV-A | SNV | 1 | A | C | 4 | 7622 | 0.05247966 | 57.5 | WT | FALSE | CAN | JSY13 | 1A | WT-Rep1A | TAT | A>C | G/C+1 | A/T-1 | TA>GC | TA>GC | CGTAATATTT | ATTTGAC | ATA | ATA | T | G | T>G | T>G_ATA | TA>GC_ATTTGAC | 9 | TRUE | FALSE |
| 168_v.txt | 32823 | 32823-SNV-A | SNV | 1 | A | C | 5 | 7622 | 0.06559958 | 40.4 | WT | FALSE | CAN | JSY13 | 1A | WT-Rep1A | CAT | A>C | G/C+1 | A/T-1 | TA>GC | TA>GC | AAGTTCATTA | ATTATTG | GTA | GTA | T | G | T>G | T>G_GTA | TA>GC_ATTATTG | 9 | TRUE | FALSE |
| 168_v.txt | 32884 | 32884-SNV-A | SNV | 1 | A | C | 4 | 22850 | 0.01750547 | 37.5 | WT | FALSE | CAN | JSY13 | 1A | WT-Rep1A | CAA | A>C | G/C+1 | A/T-1 | TA>GC | TA>GC | ACGTCCAAA | AAAATTG | GTT | GTT | T | G | T>G | T>G_GTT | TA>GC_AAAATTG | 9 | TRUE | FALSE |
| 168_v.txt | 32915 | 32915-SNV-T | SNV | 1 | T | G | 6 | 22850 | 0.02625821 | 58.3333333 | WT | FALSE | CAN | JSY13 | 1A | WT-Rep1A | GTT | T>G | G/C+1 | A/T-1 | TA>GC | TA>GC | ACTAAGTTC | TTCCAGG | CAA | GTT | T | C | T>G | T>G_GTT | TA>GC_TTCCAGG | 9 | TRUE | FALSE |
| 168_v.txt | 32919 | 32919-SNV-A | SNV | 1 | A | C | 9 | 22850 | 0.03938731 | 39.4450033 | WT | FALSE | CAN | JSY13 | 1A | WT-Rep1A | CAG | A>C | G/C+1 | A/T-1 | TA>GC | TA>GC | AGTTCCAGG | AGGGCAA | GTC | GTC | T | G | T>G | T>G_GTC | TA>GC_AGGGCAA | 9 | TRUE | FALSE |
| 168_v.txt | 32929 | 32929-SNV-T | SNV | 1 | T | G | 14 | 22885 | 0.06117544 | 53.8581076 | WT | FALSE | CAN | JSY13 | 1A | WT-Rep1A | GTG | T>G | G/C+1 | A/T-1 | TA>GC | TA>GC | CAAAAGTGA | TGATTGC | CAC | GTG | T | C | T>G | T>G_GTG | TA>GC_TGATTGC | 9 | TRUE | FALSE |
| 168_v.txt | 32938 | 32938-SNV-A | SNV | 1 | A | C | 60 | 22886 | 0.26216901 | 53.3166667 | WT | FALSE | CAN | JSY13 | 1A | WT-Rep1A | CAA | A>C | G/C+1 | A/T-1 | TA>GC | TA>GC | TTGCCCAAG | AAGAAAA | GTT | GTT | T | G | T>G | T>G_GTT | TA>GC_AAGAAAA | 9 | TRUE | FALSE |
| 168_v.txt | 33054 | 33054-SNV-T | SNV | 1 | T | G | 55 | 22886 | 0.24032159 | 58.5613067 | WT | FALSE | CAN | JSY13 | 1A | WT-Rep1A | CTG | T>G | G/C+1 | A/T-1 | TA>GC | TA>GC | AAGGACTGC | TGCGTGA | GAC | CTG | T | C | T>G | T>G_CTG | TA>GC_TGCGTGA | 9 | TRUE | FALSE |
| 168_v.txt | 33150 | 33150-SNV-A | SNV | 1 | A | C | 42 | 22886 | 0.18351831 | 38.9285714 | WT | FALSE | CAN | JSY13 | 1A | WT-Rep1A | AAG | A>C | G/C+1 | A/T-1 | TA>GC | TA>GC | ATGAAAAGA | AGACCTG | TTC | TTC | T | G | T>G | T>G_TTC | TA>GC_AGACCTG | 9 | TRUE | FALSE |
| 168_v.txt | 33157 | 33157-SNV-T | SNV | 1 | T | G | 3 | 22885 | 0.01310902 | 38 | WT | FALSE | CAN | JSY13 | 1A | WT-Rep1A | GTA | T>G | G/C+1 | A/T-1 | TA>GC | TA>GC | GACCTGTAC | TACCAAT | CAT | GTA | T | C | T>G | T>G_GTA | TA>GC_TACCAAT | 9 | TRUE | FALSE |
| 168_v.txt | 33166 | 33166-SNV-T | SNV | 1 | T | G | 25 | 22848 | 0.10941877 | 35.76 | WT | FALSE | CAN | JSY13 | 1A | WT-Rep1A | GTA | T>G | G/C+1 | A/T-1 | TA>GC | TA>GC | CAATAGTAC | TACCACC | CAT | GTA | T | C | T>G | T>G_GTA | TA>GC_TACCACC | 9 | TRUE | FALSE |
| 168_v.txt | 33192 | 33192-SNV-T | SNV | 1 | T | G | 5 | 23357 | 0.02140686 | 37.2 | WT | FALSE | CAN | JSY13 | 1A | WT-Rep1A | ATG | T>G | G/C+1 | A/T-1 | TA>GC | TA>GC | CCAATATGT | TGTCTTT | TAC | ATG | T | C | T>G | T>G_ATG | TA>GC_TGTCTTT | 9 | TRUE | FALSE |
| 168_v.txt | 33204 | 33204-SNV-A | SNV | 1 | A | C | 29 | 23355 | 0.12417041 | 34.9996639 | WT | FALSE | CAN | JSY13 | 1A | WT-Rep1A | AAG | A>C | G/C+1 | A/T-1 | TA>GC | TA>GC | TGCTTAAGC | AGCTCTC | TTC | TTC | T | G | T>G | T>G_TTC | TA>GC_AGCTCTC | 9 | TRUE | FALSE |
| 168_v.txt | 33269 | 33269-SNV-A | SNV | 1 | A | C | 176 | 23355 | 0.75358596 | 16.2727273 | WT | FALSE | CAN | JSY13 | 1A | WT-Rep1A | TAG | A>C | G/C+1 | A/T-1 | TA>GC | TA>GC | CTCCATAGA | AGAGAAC | ATC | ATC | T | G | T>G | T>G_ATC | TA>GC_AGAGAAC | 9 | TRUE | FALSE |
| 168_v.txt | 33287 | 33287-SNV-A | SNV | 1 | A | C | 45 | 23355 | 0.19267823 | 14.6667338 | WT | FALSE | CAN | JSY13 | 1A | WT-Rep1A | CAT | A>C | G/C+1 | A/T-1 | TA>GC | TA>GC | CTCGCCATTT | ATTTACT | GTA | GTA | T | G | T>G | T>G_GTA | TA>GC_ATTTACT | 9 | TRUE | FALSE |
| 168_v.txt | 33291 | 33291-SNV-A | SNV | 1 | A | C | 28 | 23355 | 0.11988868 | 42.3606716 | WT | FALSE | CAN | JSY13 | 1A | WT-Rep1A | TAC | A>C | G/C+1 | A/T-1 | TA>GC | TA>GC | CCATTTACTC | ACTCTCG | ATG | ATG | T | G | T>G | T>G_ATG | TA>GC_ACTCTCG | 9 | TRUE | FALSE |
| 168_v.txt | 32055 | 32055-SNV-T | SNV | 1 | T | C | 10 | 14355 | 0.06966214 | 46.6 | WT | FALSE | CAN | JSY13 | 1A | WT-Rep1A | GTG | T>C | G/C+1 | A/T-1 | TA>CG | TA>CG | CTGATGTGC | TGCGAGA | CAC | GTG | T | G | T>C | T>C_GTG | TA>CG_TGCGAGA | 9 | TRUE | FALSE |
| 168_v.txt | 32072 | 32072-SNV-T | SNV | 1 | T | C | 4 | 14355 | 0.02786486 | 40.7498438 | WT | FALSE | CAN | JSY13 | 1A | WT-Rep1A | ATA | T>C | G/C+1 | A/T-1 | TA>CG | TA>CG | GATAAATAA | TAACCAT | TAT | ATA | T | G | T>C | T>C_ATA | TA>CG_TAACCAT | 9 | TRUE | FALSE |
| 168_v.txt | 32081 | 32081-SNV-A | SNV | 1 | A | G | 6 | 14355 | 0.04179728 | 63.8333333 | WT | FALSE | CAN | JSY13 | 1A | WT-Rep1A | CAA | A>G | G/C+1 | A/T-1 | TA>CG | TA>CG | CCATGCAAA | AAAAAAG | GTT | GTT | T | C | T>C | T>C_GTT | TA>CG_AAAAAAG | 9 | TRUE | FALSE |
| 168_v.txt | 32226 | 32226-SNV-A | SNV | 1 | A | G | 4 | 14341 | 0.02789206 | 38 | WT | FALSE | CAN | JSY13 | 1A | WT-Rep1A | CAG | A>G | G/C+1 | A/T-1 | TA>CG | TA>CG | CTTGACAGG | AGGAATT | GTC | GTC | T | C | T>C | T>C_GTC | TA>CG_AGGAATT | 9 | TRUE | FALSE |
| 168_v.txt | 32256 | 32256-SNV-A | SNV | 1 | A | G | 94 | 46462 | 0.20231587 | 36.2978723 | WT | FALSE | CAN | JSY13 | 1A | WT-Rep1A | TAG | A>G | G/C+1 | A/T-1 | TA>CG | TA>CG | TTTGATAGA | AGACCAA | ATC | ATC | T | C | T>C | T>C_ATC | TA>CG_AGACCAA | 9 | TRUE | FALSE |
| 168_v.txt | 32275 | 32275-SNV-A | SNV | 1 | A | G | 33 | 46483 | 0.0709937 | 31.574812 | WT | FALSE | CAN | JSY13 | 1A | WT-Rep1A | GAA | A>G | G/C+1 | A/T-1 | TA>CG | TA>CG | TACGGGAAC | AACCAAC | CTT | CTT | T | C | T>C | T>C_CTT | TA>CG_AACCAAC | 9 | TRUE | FALSE |
| 168_v.txt | 32293 | 32293-SNV-A | SNV | 1 | A | G | 213 | 46483 | 0.45823204 | 54.4841994 | WT | FALSE | CAN | JSY13 | 1A | WT-Rep1A | GAA | A>G | G/C+1 | A/T-1 | TA>CG | TA>CG | TATTTGAATT | AATTTGC | CTT | CTT | T | C | T>C | T>C_CTT | TA>CG_AATTTGC | 9 | TRUE | FALSE |
| 168_v.txt | 32317 | 32317-SNV-T | SNV | 1 | T | C | 11 | 46483 | 0.02366457 | 22.3638598 | WT | FALSE | CAN | JSY13 | 1A | WT-Rep1A | GTT | T>C | G/C+1 | A/T-1 | TA>CG | TA>CG | TGGTTGTTA | TTAAGAT | CAA | GTT | T | G | T>C | T>C_GTT | TA>CG_TTAAGAT | 9 | TRUE | FALSE |
| 168_v.txt | 32319 | 32319-SNV-A | SNV | 1 | A | G | 13 | 46483 | 0.02796721 | 54.4579246 | WT | FALSE | CAN | JSY13 | 1A | WT-Rep1A | TAA | A>G | G/C+1 | A/T-1 | TA>CG | TA>CG | GTTGTTAAG | AAGATAA | ATT | ATT | T | C | T>C | T>C_ATT | TA>CG_AAGATAA | 9 | TRUE | FALSE |
| 168_v.txt | 32337 | 32337-SNV-A | SNV | 1 | A | G | 8 | 46483 | 0.01721059 | 56.3749263 | WT | FALSE | CAN | JSY13 | 1A | WT-Rep1A | GAT | A>G | G/C+1 | A/T-1 | TA>CG | TA>CG | TTGAAGATA | ATATGTG | CTA | CTA | T | C | T>C | T>C_CTA | TA>CG_ATATGTG | 9 | TRUE | FALSE |
| 168_v.txt | 32375 | 32375-SNV-A | SNV | 1 | A | G | 8 | 46481 | 0.01721133 | 54.7500859 | WT | FALSE | CAN | JSY13 | 1A | WT-Rep1A | CAA | A>G | G/C+1 | A/T-1 | TA>CG | TA>CG | AATAGCAAT | AATAATA | GTT | GTT | T | C | T>C | T>C_GTT | TA>CG_AATAATA | 9 | TRUE | FALSE |
| 168_v.txt | 32376 | 32376-SNV-A | SNV | 1 | A | G | 10 | 46482 | 0.0215137 | 37.199934 | WT | FALSE | CAN | JSY13 | 1A | WT-Rep1A | AAT | A>G | G/C+1 | A/T-1 | TA>CG | TA>CG | ATAGCAATA | ATAATAA | TTA | TTA | T | C | T>C | T>C_TTA | TA>CG_ATAATAA | 9 | TRUE | FALSE |
| 168_v.txt | 32445 | 32445-SNV-A | SNV | 1 | A | G | 301 | 46483 | 0.64754857 | 53.6478405 | WT | FALSE | CAN | JSY13 | 1A | WT-Rep1A | AAG | A>G | G/C+1 | A/T-1 | TA>CG | TA>CG | ACTAAAAGT | AGTCCAA | TTC | TTC | T | C | T>C | T>C_TTC | TA>CG_AGTCCAA | 9 | TRUE | FALSE |
| 168_v.txt | 32454 | 32454-SNV-A | SNV | 1 | A | G | 10 | 46482 | 0.0215137 | 40.1996046 | WT | FALSE | CAN | JSY13 | 1A | WT-Rep1A | GAA | A>G | G/C+1 | A/T-1 | TA>CG | TA>CG | CCAATGAAT | AATAATA | CTT | CTT | T | C | T>C | T>C_CTT | TA>CG_AATAATA | 9 | TRUE | FALSE |
| 168_v.txt | 32456 | 32456-SNV-T | SNV | 1 | T | C | 4 | 46482 | 8.61E-03 | 49.7500226 | WT | FALSE | CAN | JSY13 | 1A | WT-Rep1A | ATA | T>C | G/C+1 | A/T-1 | TA>CG | TA>CG | AATGAATAA | TAATAGA | TAT | ATA | T | G | T>C | T>C_ATA | TA>CG_TAATAGA | 9 | TRUE | FALSE |
| 168_v.txt | 32460 | 32460-SNV-A | SNV | 1 | A | G | 14 | 46483 | 0.03011854 | 50.4267645 | WT | FALSE | CAN | JSY13 | 1A | WT-Rep1A | TAG | A>G | G/C+1 | A/T-1 | TA>CG | TA>CG | AATAATAGA | AGAGAGC | ATC | ATC | T | C | T>C | T>C_ATC | TA>CG_AGAGAGC | 9 | TRUE | FALSE |
| 168_v.txt | 32472 | 32472-SNV-T | SNV | 1 | T | C | 76 | 46483 | 0.16350064 | 47.9736785 | WT | FALSE | CAN | JSY13 | 1A | WT-Rep1A | GTA | T>C | G/C+1 | A/T-1 | TA>CG | TA>CG | CCAATGTAG | TAGAAGG | CAT | GTA | T | G | T>C | T>C_GTA | TA>CG_TAGAAGG | 9 | TRUE | FALSE |
| 168_v.txt | 32473 | 32473-SNV-A | SNV | 1 | A | G | 22 | 46483 | 0.04732913 | 48.6148489 | WT | FALSE | CAN | JSY13 | 1A | WT-Rep1A | TAG | A>G | G/C+1 | A/T-1 | TA>CG | TA>CG | CAATGTAGA | AGAAGGT | ATC | ATC | T | C | T>C | T>C_ATC | TA>CG_AGAAGGT | 9 | TRUE | FALSE |
| 168_v.txt | 32490 | 32490-SNV-A | SNV | 1 | A | G | 92 | 46483 | 0.19792182 | 47.7282609 | WT | FALSE | CAN | JSY13 | 1A | WT-Rep1A | GAA | A>G | G/C+1 | A/T-1 | TA>CG | TA>CG | ATACGGAAA | AAAACAA | CTT | CTT | T | C | T>C | T>C_CTT | TA>CG_AAAACAA | 9 | TRUE | FALSE |
| 168_v.txt | 32527 | 32527-SNV-T | SNV | 1 | T | C | 14 | 46483 | 0.03011854 | 35.4711746 | WT | FALSE | CAN | JSY13 | 1A | WT-Rep1A | CTG | T>C | G/C+1 | A/T-1 | TA>CG | TA>CG | ATTTTCTGG | TGGGGTT | GAC | CTG | T | G | T>C | T>C_CTG | TA>CG_TGGGGTT | 9 | TRUE | FALSE |
| 168_v.txt | 32583 | 32583-SNV-A | SNV | 1 | A | G | 15 | 7613 | 0.19703139 | 32.9333333 | WT | FALSE | CAN | JSY13 | 1A | WT-Rep1A | GAA | A>G | G/C+1 | A/T-1 | TA>CG | TA>CG | AATGTGAAG | AAGGCAG | CTT | CTT | T | C | T>C | T>C_CTT | TA>CG_AAGGCAG | 9 | TRUE | FALSE |
| 168_v.txt | 32680 | 32680-SNV-A | SNV | 1 | A | G | 22 | 7614 | 0.28894142 | 61.5909091 | WT | FALSE | CAN | JSY13 | 1A | WT-Rep1A | CAA | A>G | G/C+1 | A/T-1 | TA>CG | TA>CG | TTCTCCAATA | AATAACG | GTT | GTT | T | C | T>C | T>C_GTT | TA>CG_AATAACG | 9 | TRUE | FALSE |

| | | | | | | | | | | | | | | | | | | | | | | | | | | | | | | | | | | |
|---|---|---|---|---|---|---|---|---|---|---|---|---|---|---|---|---|---|---|---|---|---|---|---|---|---|---|---|---|---|---|---|---|---|---|
| 168_v.txt | 32767 | 32767-SNV-T | SNV | 1 | T | C | 15 | 7622 | 0.19679874 | 56 | WT | FALSE | CAN | JSY13 | 1A | WT-Rep1A | TTG | T>C | G/C+1 | A/T-1 | TA>CG | TA>CG | AAACTTTGA | TGATGGA | AAC | TTG | T | G | T>C | T>C_TTG | TA>CG_TGATGGA | 9 | TRUE | FALSE |
| 168_v.txt | 32787 | 32787-SNV-T | SNV | 1 | T | C | 6 | 7622 | 0.0787195 | 38.8333333 | WT | FALSE | CAN | JSY13 | 1A | WT-Rep1A | CTC | T>C | G/C+1 | A/T-1 | TA>CG | TA>CG | CAGAACTCG | TCGAATT | GAG | CTC | T | G | T>C | T>C_CTC | TA>CG_TCGAATT | 9 | TRUE | FALSE |
| 168_v.txt | 32800 | 32800-SNV-A | SNV | 1 | A | G | 3 | 7622 | 0.03935975 | 55.3333333 | WT | FALSE | CAN | JSY13 | 1A | WT-Rep1A | TAA | A>G | G/C+1 | A/T-1 | TA>CG | TA>CG | CACCGTAAT | AATATTT | ATT | ATT | T | C | T>C | T>C_ATT | TA>CG_AATATTT | 9 | TRUE | FALSE |
| 168_v.txt | 32808 | 32808-SNV-A | SNV | 1 | A | G | 87 | 7622 | 1.1414327 | 44.1494253 | WT | FALSE | CAN | JSY13 | 1A | WT-Rep1A | GAC | A>G | G/C+1 | A/T-1 | TA>CG | TA>CG | TATTTGACA | ACAGGGA | CTG | CTG | T | C | T>C | T>C_CTG | TA>CG_ACAGGGA | 9 | TRUE | FALSE |
| 168_v.txt | 32919 | 32919-SNV-A | SNV | 1 | A | G | 22 | 22850 | 0.09628009 | 47.0915179 | WT | FALSE | CAN | JSY13 | 1A | WT-Rep1A | CAG | A>G | G/C+1 | A/T-1 | TA>CG | TA>CG | AGTTCCAGG | AGGGCAA | GTC | GTC | T | C | T>C | T>C_GTC | TA>CG_AGGGCAA | 9 | TRUE | FALSE |
| 168_v.txt | 32929 | 32929-SNV-T | SNV | 1 | T | C | 8 | 22885 | 0.0349574 | 43.1248763 | WT | FALSE | CAN | JSY13 | 1A | WT-Rep1A | GTG | T>C | G/C+1 | A/T-1 | TA>CG | TA>CG | CAAAAGTGA | TGATTGC | CAC | GTG | T | G | T>C | T>C_GTG | TA>CG_TGATTGC | 9 | TRUE | FALSE |
| 168_v.txt | 32938 | 32938-SNV-A | SNV | 1 | A | G | 6 | 22886 | 0.0262169 | 53.1666667 | WT | FALSE | CAN | JSY13 | 1A | WT-Rep1A | CAA | A>G | G/C+1 | A/T-1 | TA>CG | TA>CG | TTGCCCAAG | AAGAAAA | GTT | GTT | T | C | T>C | T>C_GTT | TA>CG_AAGAAAA | 9 | TRUE | FALSE |
| 168_v.txt | 32941 | 32941-SNV-A | SNV | 1 | A | G | 8 | 22886 | 0.03495587 | 55.4874086 | WT | FALSE | CAN | JSY13 | 1A | WT-Rep1A | GAA | A>G | G/C+1 | A/T-1 | TA>CG | TA>CG | CCCAAGAAA | AAAACCA | CTT | CTT | T | C | T>C | T>C_CTT | TA>CG_AAAACCA | 9 | TRUE | FALSE |
| 168_v.txt | 33065 | 33065-SNV-A | SNV | 1 | A | G | 3 | 22886 | 0.01310845 | 63.9998938 | WT | FALSE | CAN | JSY13 | 1A | WT-Rep1A | AAT | A>G | G/C+1 | A/T-1 | TA>CG | TA>CG | GACAGAATA | ATATGCC | TTA | TTA | T | C | T>C | T>C_TTA | TA>CG_ATATGCC | 9 | TRUE | FALSE |
| 168_v.txt | 33066 | 33066-SNV-T | SNV | 1 | T | C | 4 | 22886 | 0.01747793 | 56.4998711 | WT | FALSE | CAN | JSY13 | 1A | WT-Rep1A | ATA | T>C | G/C+1 | A/T-1 | TA>CG | TA>CG | ACAGAATAT | TATGCCA | TAT | ATA | T | G | T>C | T>C_ATA | TA>CG_TATGCCA | 9 | TRUE | FALSE |
| 168_v.txt | 33076 | 33076-SNV-A | SNV | 1 | A | G | 7 | 22886 | 0.03058639 | 50.2856771 | WT | FALSE | CAN | JSY13 | 1A | WT-Rep1A | GAA | A>G | G/C+1 | A/T-1 | TA>CG | TA>CG | CCAAAGAAC | AACCCAT | CTT | CTT | T | C | T>C | T>C_CTT | TA>CG_AACCCAT | 9 | TRUE | FALSE |
| 168_v.txt | 33077 | 33077-SNV-A | SNV | 1 | A | G | 6 | 22886 | 0.0262169 | 24.4999818 | WT | FALSE | CAN | JSY13 | 1A | WT-Rep1A | AAC | A>G | G/C+1 | A/T-1 | TA>CG | TA>CG | CAAAGAACC | ACCCATA | TTG | TTG | T | C | T>C | T>C_TTG | TA>CG_ACCCATA | 9 | TRUE | FALSE |
| 168_v.txt | 33150 | 33150-SNV-A | SNV | 1 | A | G | 29 | 22886 | 0.12671502 | 37.2413793 | WT | FALSE | CAN | JSY13 | 1A | WT-Rep1A | AAG | A>G | G/C+1 | A/T-1 | TA>CG | TA>CG | ATGAAAAGA | AGACCTG | TTC | TTC | T | C | T>C | T>C_TTC | TA>CG_AGACCTG | 9 | TRUE | FALSE |
| 168_v.txt | 33157 | 33157-SNV-T | SNV | 1 | T | C | 22 | 22885 | 0.09613284 | 36 | WT | FALSE | CAN | JSY13 | 1A | WT-Rep1A | GTA | T>C | G/C+1 | A/T-1 | TA>CG | TA>CG | GACCTGTAC | TACCAAT | CAT | GTA | T | G | T>C | T>C_GTA | TA>CG_TACCAAT | 9 | TRUE | FALSE |
| 168_v.txt | 33170 | 33170-SNV-A | SNV | 1 | A | G | 8 | 22847 | 0.03501554 | 35.75 | WT | FALSE | CAN | JSY13 | 1A | WT-Rep1A | CAC | A>G | G/C+1 | A/T-1 | TA>CG | TA>CG | AGTACCACC | ACCAAGG | GTG | GTG | T | C | T>C | T>C_GTG | TA>CG_ACCAAGG | 9 | TRUE | FALSE |
| 168_v.txt | 33192 | 33192-SNV-T | SNV | 1 | T | C | 15 | 23357 | 0.06422058 | 33.5333333 | WT | FALSE | CAN | JSY13 | 1A | WT-Rep1A | ATG | T>C | G/C+1 | A/T-1 | TA>CG | TA>CG | CCAATATGT | TGTCTTT | TAC | ATG | T | G | T>C | T>C_ATG | TA>CG_TGTCTTT | 9 | TRUE | FALSE |
| 168_v.txt | 33207 | 33207-SNV-T | SNV | 1 | T | C | 14 | 23354 | 0.0599469 | 21.4284596 | WT | FALSE | CAN | JSY13 | 1A | WT-Rep1A | CTC | T>C | G/C+1 | A/T-1 | TA>CG | TA>CG | TTAAGCTCTC | TCTCTCT | GAG | CTC | T | G | T>C | T>C_CTC | TA>CG_TCTCTCT | 9 | TRUE | FALSE |
| 168_v.txt | 33291 | 33291-SNV-A | SNV | 1 | A | G | 6 | 23355 | 0.02569043 | 52.8251813 | WT | FALSE | CAN | JSY13 | 1A | WT-Rep1A | TAC | A>G | G/C+1 | A/T-1 | TA>CG | TA>CG | CCATTTACTC | ACTCTCG | ATG | ATG | T | C | T>C | T>C_ATG | TA>CG_ACTCTCG | 9 | TRUE | FALSE |
| 168_v.txt | 33293 | 33293-SNV-T | SNV | 1 | T | C | 12 | 23355 | 0.05138086 | 34.2499677 | WT | FALSE | CAN | JSY13 | 1A | WT-Rep1A | CTC | T>C | G/C+1 | A/T-1 | TA>CG | TA>CG | ATTTACTCTC | TCTCGTC | GAG | CTC | T | G | T>C | T>C_CTC | TA>CG_TCTCGTC | 9 | TRUE | FALSE |
| 168_v.txt | 33476 | 33476-SNV-T | SNV | 1 | T | C | 34 | 23348 | 0.14562275 | 31.1176471 | WT | FALSE | CAN | JSY13 | 1A | WT-Rep1A | CTT | T>C | G/C+1 | A/T-1 | TA>CG | TA>CG | TATGCCTTTT | TTTTTTT | GAA | CTT | T | G | T>C | T>C_CTT | TA>CG_TTTTTTT | 9 | TRUE | FALSE |
| 168_v.txt | 31816 | 31816-SNV-T | SNV | 1 | T | A | 11 | 3138 | 0.35054175 | 62.6363636 | WT | FALSE | CAN | JSY13 | 1A | WT-Rep1A | CTG | T>A | A/T+1 | A/T-1 | TA>AT | TA>AT | TAAATCTGC | TGCATCT | GAC | CTG | T | T | T>A | T>A_CTG | TA>AT_TGCATCT | 9 | TRUE | FALSE |
| 168_v.txt | 31822 | 31822-SNV-T | SNV | 1 | T | A | 18 | 3138 | 0.57361377 | 51.8888889 | WT | FALSE | CAN | JSY13 | 1A | WT-Rep1A | CTG | T>A | A/T+1 | A/T-1 | TA>AT | TA>AT | TGCATCTGA | TGAATAT | GAC | CTG | T | T | T>A | T>A_CTG | TA>AT_TGAATAT | 9 | TRUE | FALSE |
| 168_v.txt | 31906 | 31906-SNV-T | SNV | 1 | T | A | 10 | 3127 | 0.31979533 | 37.3 | WT | FALSE | CAN | JSY13 | 1A | WT-Rep1A | TTT | T>A | A/T+1 | A/T-1 | TA>AT | TA>AT | TGAATTTTG | TTGGTGC | AAA | TTT | T | T | T>A | T>A_TTT | TA>AT_TTGGTGC | 9 | TRUE | FALSE |
| 168_v.txt | 31964 | 31964-SNV-A | SNV | 1 | A | T | 23 | 14355 | 0.16022292 | 34.8695652 | WT | FALSE | CAN | JSY13 | 1A | WT-Rep1A | CAT | A>T | A/T+1 | A/T-1 | TA>AT | TA>AT | GGCCGCATA | ATAATAA | GTA | GTA | T | A | T>A | T>A_GTA | TA>AT_ATAATAA | 9 | TRUE | FALSE |
| 168_v.txt | 31986 | 31986-SNV-A | SNV | 1 | A | T | 7 | 14355 | 0.0487635 | 44.7142857 | WT | FALSE | CAN | JSY13 | 1A | WT-Rep1A | TAA | A>T | A/T+1 | A/T-1 | TA>AT | TA>AT | GGCATTAAT | AATTTAG | ATT | ATT | T | A | T>A | T>A_ATT | TA>AT_AATTTAG | 9 | TRUE | FALSE |
| 168_v.txt | 31996 | 31996-SNV-T | SNV | 1 | T | A | 75 | 14355 | 0.52246604 | 53.4533333 | WT | FALSE | CAN | JSY13 | 1A | WT-Rep1A | TTA | T>A | A/T+1 | A/T-1 | TA>AT | TA>AT | TAGCTTTAA | TAAATGG | AAT | TTA | T | T | T>A | T>A_TTA | TA>AT_TAAATGG | 9 | TRUE | FALSE |
| 168_v.txt | 32072 | 32072-SNV-T | SNV | 1 | T | A | 1 | 14355 | 6.97E-03 | 63.9966641 | WT | FALSE | CAN | JSY13 | 1A | WT-Rep1A | ATA | T>A | A/T+1 | A/T-1 | TA>AT | TA>AT | GATAAATAA | TAACCAT | TAT | ATA | T | T | T>A | T>A_ATA | TA>AT_TAACCAT | 9 | TRUE | FALSE |
| 168_v.txt | 32073 | 32073-SNV-A | SNV | 1 | A | T | 19 | 14355 | 0.13235806 | 61.1049862 | WT | FALSE | CAN | JSY13 | 1A | WT-Rep1A | TAA | A>T | A/T+1 | A/T-1 | TA>AT | TA>AT | ATAAATAAC | AACCATG | ATT | ATT | T | A | T>A | T>A_ATT | TA>AT_AACCATG | 9 | TRUE | FALSE |
| 168_v.txt | 32230 | 32230-SNV-A | SNV | 1 | A | T | 10 | 14341 | 0.06973014 | 36.6 | WT | FALSE | CAN | JSY13 | 1A | WT-Rep1A | AAT | A>T | A/T+1 | A/T-1 | TA>AT | TA>AT | ACAGGAATT | ATTTAGG | TTA | TTA | T | A | T>A | T>A_TTA | TA>AT_ATTTAGG | 9 | TRUE | FALSE |
| 168_v.txt | 32247 | 32247-SNV-T | SNV | 1 | T | A | 3 | 14341 | 0.02091904 | 32.9998993 | WT | FALSE | CAN | JSY13 | 1A | WT-Rep1A | GTT | T>A | A/T+1 | A/T-1 | TA>AT | TA>AT | AACTTGTTCT | TTCTTTG | CAA | GTT | T | T | T>A | T>A_GTT | TA>AT_TTCTTTG | 9 | TRUE | FALSE |
| 168_v.txt | 32314 | 32314-SNV-T | SNV | 1 | T | A | 8 | 46483 | 0.01721059 | 55.6150058 | WT | FALSE | CAN | JSY13 | 1A | WT-Rep1A | GTT | T>A | A/T+1 | A/T-1 | TA>AT | TA>AT | TAATGGTTG | TTGTTAA | CAA | GTT | T | T | T>A | T>A_GTT | TA>AT_TTGTTAA | 9 | TRUE | FALSE |
| 168_v.txt | 32317 | 32317-SNV-T | SNV | 1 | T | A | 7 | 46483 | 0.01505927 | 48.7134724 | WT | FALSE | CAN | JSY13 | 1A | WT-Rep1A | GTT | T>A | A/T+1 | A/T-1 | TA>AT | TA>AT | TGGTTGTTA | TTAAGAT | CAA | GTT | T | T | T>A | T>A_GTT | TA>AT_TTAAGAT | 9 | TRUE | FALSE |
| 168_v.txt | 32318 | 32318-SNV-T | SNV | 1 | T | A | 6 | 46483 | 0.01290795 | 36.1651614 | WT | FALSE | CAN | JSY13 | 1A | WT-Rep1A | TTA | T>A | A/T+1 | A/T-1 | TA>AT | TA>AT | GGTTGTTAA | TAAGATA | AAT | TTA | T | T | T>A | T>A_TTA | TA>AT_TAAGATA | 9 | TRUE | FALSE |
| 168_v.txt | 32337 | 32337-SNV-A | SNV | 1 | A | T | 6 | 46483 | 0.01290795 | 28.8333044 | WT | FALSE | CAN | JSY13 | 1A | WT-Rep1A | GAT | A>T | A/T+1 | A/T-1 | TA>AT | TA>AT | TTGAAGATA | ATATGTG | CTA | CTA | T | A | T>A | T>A_CTA | TA>AT_ATATGTG | 9 | TRUE | FALSE |
| 168_v.txt | 32376 | 32376-SNV-A | SNV | 1 | A | T | 8 | 46482 | 0.01721096 | 43.1246307 | WT | FALSE | CAN | JSY13 | 1A | WT-Rep1A | AAT | A>T | A/T+1 | A/T-1 | TA>AT | TA>AT | ATAGCAATA | ATAATAA | TTA | TTA | T | A | T>A | T>A_TTA | TA>AT_ATAATAA | 9 | TRUE | FALSE |
| 168_v.txt | 32377 | 32377-SNV-T | SNV | 1 | T | A | 3 | 46475 | 6.46E-03 | 55.3223715 | WT | FALSE | CAN | JSY13 | 1A | WT-Rep1A | ATA | T>A | A/T+1 | A/T-1 | TA>AT | TA>AT | TAGCAATAA | TAATAAA | TAT | ATA | T | T | T>A | T>A_ATA | TA>AT_TAATAAA | 9 | TRUE | FALSE |
| 168_v.txt | 32414 | 32414-SNV-T | SNV | 1 | T | A | 12 | 46481 | 0.025817 | 51.4994004 | WT | FALSE | CAN | JSY13 | 1A | WT-Rep1A | GTG | T>A | A/T+1 | A/T-1 | TA>AT | TA>AT | AGATTGTGT | TGTTAGT | CAC | GTG | T | T | T>A | T>A_GTG | TA>AT_TGTTAGT | 9 | TRUE | FALSE |
| 168_v.txt | 32445 | 32445-SNV-A | SNV | 1 | A | T | 25 | 46483 | 0.0537831 | 55.16 | WT | FALSE | CAN | JSY13 | 1A | WT-Rep1A | AAG | A>T | A/T+1 | A/T-1 | TA>AT | TA>AT | ACTAAAAGT | AGTCCAA | TTC | TTC | T | A | T>A | T>A_TTC | TA>AT_AGTCCAA | 9 | TRUE | FALSE |
| 168_v.txt | 32455 | 32455-SNV-A | SNV | 1 | A | T | 4 | 46483 | 8.61E-03 | 38.4993796 | WT | FALSE | CAN | JSY13 | 1A | WT-Rep1A | AAT | A>T | A/T+1 | A/T-1 | TA>AT | TA>AT | CAATGAATA | ATAATAG | TTA | TTA | T | A | T>A | T>A_TTA | TA>AT_ATAATAG | 9 | TRUE | FALSE |
| 168_v.txt | 32456 | 32456-SNV-T | SNV | 1 | T | A | 8 | 46482 | 0.01721096 | 37.6250715 | WT | FALSE | CAN | JSY13 | 1A | WT-Rep1A | ATA | T>A | A/T+1 | A/T-1 | TA>AT | TA>AT | AATGAATAA | TAATAGA | TAT | ATA | T | T | T>A | T>A_ATA | TA>AT_TAATAGA | 9 | TRUE | FALSE |
| 168_v.txt | 32459 | 32459-SNV-T | SNV | 1 | T | A | 7 | 46483 | 0.01505927 | 45.2839956 | WT | FALSE | CAN | JSY13 | 1A | WT-Rep1A | ATA | T>A | A/T+1 | A/T-1 | TA>AT | TA>AT | GAATAATAG | TAGAGAG | TAT | ATA | T | T | T>A | T>A_ATA | TA>AT_TAGAGAG | 9 | TRUE | FALSE |
| 168_v.txt | 32472 | 32472-SNV-T | SNV | 1 | T | A | 8 | 46483 | 0.01721059 | 47.3749984 | WT | FALSE | CAN | JSY13 | 1A | WT-Rep1A | GTA | T>A | A/T+1 | A/T-1 | TA>AT | TA>AT | CCAATGTAG | TAGAAGG | CAT | GTA | T | T | T>A | T>A_GTA | TA>AT_TAGAAGG | 9 | TRUE | FALSE |
| 168_v.txt | 32481 | 32481-SNV-A | SNV | 1 | A | T | 25 | 46483 | 0.0537831 | 44.4 | WT | FALSE | CAN | JSY13 | 1A | WT-Rep1A | TAA | A>T | A/T+1 | A/T-1 | TA>AT | TA>AT | AAGGTTAAG | AAGATAC | ATT | ATT | T | A | T>A | T>A_ATT | TA>AT_AAGATAC | 9 | TRUE | FALSE |
| 168_v.txt | 32512 | 32512-SNV-T | SNV | 1 | T | A | 22 | 46483 | 0.04732913 | 41.2727273 | WT | FALSE | CAN | JSY13 | 1A | WT-Rep1A | CTT | T>A | A/T+1 | A/T-1 | TA>AT | TA>AT | TGGCTCTTG | TTGGAAC | GAA | CTT | T | T | T>A | T>A_CTT | TA>AT_TTGGAAC | 9 | TRUE | FALSE |
| 168_v.txt | 32635 | 32635-SNV-T | SNV | 1 | T | A | 11 | 7614 | 0.14447071 | 55.0909091 | WT | FALSE | CAN | JSY13 | 1A | WT-Rep1A | TTA | T>A | A/T+1 | A/T-1 | TA>AT | TA>AT | CGTTTTTATC | TATCCTT | AAT | TTA | T | T | T>A | T>A_TTA | TA>AT_TATCCTT | 9 | TRUE | FALSE |
| 168_v.txt | 32726 | 32726-SNV-A | SNV | 1 | A | T | 39 | 7622 | 0.51167673 | 54.1282051 | WT | FALSE | CAN | JSY13 | 1A | WT-Rep1A | TAC | A>T | A/T+1 | A/T-1 | TA>AT | TA>AT | AACCATACA | ACAAAAA | ATG | ATG | T | A | T>A | T>A_ATG | TA>AT_ACAAAAA | 9 | TRUE | FALSE |
| 168_v.txt | 32734 | 32734-SNV-A | SNV | 1 | A | T | 7 | 7622 | 0.09183941 | 64 | WT | FALSE | CAN | JSY13 | 1A | WT-Rep1A | CAG | A>T | A/T+1 | A/T-1 | TA>AT | TA>AT | AAAAACAGT | AGTATAT | GTC | GTC | T | A | T>A | T>A_GTC | TA>AT_AGTATAT | 9 | TRUE | FALSE |
| 168_v.txt | 32760 | 32760-SNV-A | SNV | 1 | A | T | 10 | 7622 | 0.13119916 | 56.1997367 | WT | FALSE | CAN | JSY13 | 1A | WT-Rep1A | TAA | A>T | A/T+1 | A/T-1 | TA>AT | TA>AT | ATGGCTAAA | AAAACTT | ATT | ATT | T | A | T>A | T>A_ATT | TA>AT_AAAACTT | 9 | TRUE | FALSE |
| 168_v.txt | 32804 | 32804-SNV-T | SNV | 1 | T | A | 87 | 7622 | 1.1414327 | 47.1500462 | WT | FALSE | CAN | JSY13 | 1A | WT-Rep1A | ATT | T>A | A/T+1 | A/T-1 | TA>AT | TA>AT | GTAATATTT | TTTGACA | TAA | ATT | T | T | T>A | T>A_ATT | TA>AT_TTTGACA | 9 | TRUE | FALSE |
| 168_v.txt | 32915 | 32915-SNV-T | SNV | 1 | T | A | 20 | 22850 | 0.08752735 | 29.15 | WT | FALSE | CAN | JSY13 | 1A | WT-Rep1A | GTT | T>A | A/T+1 | A/T-1 | TA>AT | TA>AT | ACTAAGTTC | TTCCAGG | CAA | GTT | T | T | T>A | T>A_GTT | TA>AT_TTCCAGG | 9 | TRUE | FALSE |
| 168_v.txt | 32924 | 32924-SNV-A | SNV | 1 | A | T | 2 | 22850 | 8.75E-03 | 50.9984129 | WT | FALSE | CAN | JSY13 | 1A | WT-Rep1A | CAA | A>T | A/T+1 | A/T-1 | TA>AT | TA>AT | CAGGGCAAA | AAAAGTG | GTT | GTT | T | A | T>A | T>A_GTT | TA>AT_AAAAGTG | 9 | TRUE | FALSE |
| 168_v.txt | 32932 | 32932-SNV-T | SNV | 1 | T | A | 24 | 22886 | 0.10486761 | 57.4583333 | WT | FALSE | CAN | JSY13 | 1A | WT-Rep1A | ATT | T>A | A/T+1 | A/T-1 | TA>AT | TA>AT | AAGTGATTG | TTGCCCA | TAA | ATT | T | T | T>A | T>A_ATT | TA>AT_TTGCCCA | 9 | TRUE | FALSE |
| 168_v.txt | 32938 | 32938-SNV-A | SNV | 1 | A | T | 22 | 22886 | 0.09612864 | 45.2272727 | WT | FALSE | CAN | JSY13 | 1A | WT-Rep1A | CAA | A>T | A/T+1 | A/T-1 | TA>AT | TA>AT | TTGCCCAAG | AAGAAAA | GTT | GTT | T | A | T>A | T>A_GTT | TA>AT_AAGAAAA | 9 | TRUE | FALSE |
| 168_v.txt | 32948 | 32948-SNV-A | SNV | 1 | A | T | 42 | 22886 | 0.18351831 | 60.5476191 | WT | FALSE | CAN | JSY13 | 1A | WT-Rep1A | AAT | A>T | A/T+1 | A/T-1 | TA>AT | TA>AT | AAACCAATA | ATACATG | TTA | TTA | T | A | T>A | T>A_TTA | TA>AT_ATACATG | 9 | TRUE | FALSE |
| 168_v.txt | 32992 | 32992-SNV-T | SNV | 1 | T | A | 24 | 22886 | 0.10486761 | 54.4533402 | WT | FALSE | CAN | JSY13 | 1A | WT-Rep1A | CTT | T>A | A/T+1 | A/T-1 | TA>AT | TA>AT | GGAATCTTT | TTTGTGA | GAA | CTT | T | T | T>A | T>A_CTT | TA>AT_TTTGTGA | 9 | TRUE | FALSE |
| 168_v.txt | 33039 | 33039-SNV-A | SNV | 1 | A | T | 13 | 22886 | 0.05680329 | 58.3846533 | WT | FALSE | CAN | JSY13 | 1A | WT-Rep1A | CAT | A>T | A/T+1 | A/T-1 | TA>AT | TA>AT | GTAGCCATT | ATTTCAC | GTA | GTA | T | A | T>A | T>A_GTA | TA>AT_ATTTCAC | 9 | TRUE | FALSE |
| 168_v.txt | 33040 | 33040-SNV-T | SNV | 1 | T | A | 3 | 22885 | 0.01310902 | 49.9994029 | WT | FALSE | CAN | JSY13 | 1A | WT-Rep1A | ATT | T>A | A/T+1 | A/T-1 | TA>AT | TA>AT | TAGCCATTTC | TTTCACC | TAA | ATT | T | T | T>A | T>A_ATT | TA>AT_TTTCACC | 9 | TRUE | FALSE |
| 168_v.txt | 33065 | 33065-SNV-A | SNV | 1 | A | T | 3 | 22886 | 0.01310845 | 63.9994754 | WT | FALSE | CAN | JSY13 | 1A | WT-Rep1A | AAT | A>T | A/T+1 | A/T-1 | TA>AT | TA>AT | GACAGAATA | ATATGCC | TTA | TTA | T | A | T>A | T>A_TTA | TA>AT_ATATGCC | 9 | TRUE | FALSE |
| 168_v.txt | 33066 | 33066-SNV-T | SNV | 1 | T | A | 3 | 22886 | 0.01310845 | 53.3319218 | WT | FALSE | CAN | JSY13 | 1A | WT-Rep1A | ATA | T>A | A/T+1 | A/T-1 | TA>AT | TA>AT | ACAGAATAT | TATGCCA | TAT | ATA | T | T | T>A | T>A_ATA | TA>AT_TATGCCA | 9 | TRUE | FALSE |
| 168_v.txt | 33077 | 33077-SNV-A | SNV | 1 | A | T | 5 | 22886 | 0.02184742 | 31.7996878 | WT | FALSE | CAN | JSY13 | 1A | WT-Rep1A | AAC | A>T | A/T+1 | A/T-1 | TA>AT | TA>AT | CAAAGAACC | ACCCATA | TTG | TTG | T | A | T>A | T>A_TTG | TA>AT_ACCCATA | 9 | TRUE | FALSE |
| 168_v.txt | 33089 | 33089-SNV-A | SNV | 1 | A | T | 27 | 22886 | 0.11797606 | 48.962963 | WT | FALSE | CAN | JSY13 | 1A | WT-Rep1A | AAT | A>T | A/T+1 | A/T-1 | TA>AT | TA>AT | AAATAAATA | ATATGAT | TTA | TTA | T | A | T>A | T>A_TTA | TA>AT_ATATGAT | 9 | TRUE | FALSE |
| 168_v.txt | 33094 | 33094-SNV-A | SNV | 1 | A | T | 5 | 22886 | 0.02184742 | 53.0000026 | WT | FALSE | CAN | JSY13 | 1A | WT-Rep1A | GAT | A>T | A/T+1 | A/T-1 | TA>AT | TA>AT | AATATGATA | ATATAAG | CTA | CTA | T | A | T>A | T>A_CTA | TA>AT_ATATAAG | 9 | TRUE | FALSE |
| 168_v.txt | 33095 | 33095-SNV-T | SNV | 1 | T | A | 5 | 22886 | 0.02184742 | 47.7999741 | WT | FALSE | CAN | JSY13 | 1A | WT-Rep1A | ATA | T>A | A/T+1 | A/T-1 | TA>AT | TA>AT | ATATGATAT | TATAAGA | TAT | ATA | T | T | T>A | T>A_ATA | TA>AT_TATAAGA | 9 | TRUE | FALSE |
| 168_v.txt | 33150 | 33150-SNV-A | SNV | 1 | A | T | 19 | 22886 | 0.08302019 | 37.9473684 | WT | FALSE | CAN | JSY13 | 1A | WT-Rep1A | AAG | A>T | A/T+1 | A/T-1 | TA>AT | TA>AT | ATGAAAAGA | AGACCTG | TTC | TTC | T | A | T>A | T>A_TTC | TA>AT_AGACCTG | 9 | TRUE | FALSE |
| 168_v.txt | 33162 | 33162-SNV-A | SNV | 1 | A | T | 36 | 22858 | 0.15749409 | 37.0277778 | WT | FALSE | CAN | JSY13 | 1A | WT-Rep1A | AAT | A>T | A/T+1 | A/T-1 | TA>AT | TA>AT | GTACCAATA | ATAGTAC | TTA | TTA | T | A | T>A | T>A_TTA | TA>AT_ATAGTAC | 9 | TRUE | FALSE |
| 168_v.txt | 33170 | 33170-SNV-A | SNV | 1 | A | T | 5 | 22847 | 0.02188471 | 37.1998412 | WT | FALSE | CAN | JSY13 | 1A | WT-Rep1A | CAC | A>T | A/T+1 | A/T-1 | TA>AT | TA>AT | AGTACCACC | ACCAAGG | GTG | GTG | T | A | T>A | T>A_GTG | TA>AT_ACCAAGG | 9 | TRUE | FALSE |
| 168_v.txt | 33192 | 33192-SNV-T | SNV | 1 | T | A | 12 | 23357 | 0.05137646 | 29.0833333 | WT | FALSE | CAN | JSY13 | 1A | WT-Rep1A | ATG | T>A | A/T+1 | A/T-1 | TA>AT | TA>AT | CCAATATGT | TGTCTTT | TAC | ATG | T | T | T>A | T>A_ATG | TA>AT_TGTCTTT | 9 | TRUE | FALSE |
| 168_v.txt | 33209 | 33209-SNV-T | SNV | 1 | T | A | 15 | 23355 | 0.06422608 | 36.4592237 | WT | FALSE | CAN | JSY13 | 1A | WT-Rep1A | CTC | T>A | A/T+1 | A/T-1 | TA>AT | TA>AT | AAGCTCTCTC | TCTCTTC | GAG | CTC | T | T | T>A | T>A_CTC | TA>AT_TCTCTTC | 9 | TRUE | FALSE |
| 168_v.txt | 33211 | 33211-SNV-T | SNV | 1 | T | A | 38 | 23355 | 0.16270606 | 38.0511992 | WT | FALSE | CAN | JSY13 | 1A | WT-Rep1A | CTC | T>A | A/T+1 | A/T-1 | TA>AT | TA>AT | GCTCTCTCTT | TCTTCAC | GAG | CTC | T | T | T>A | T>A_CTC | TA>AT_TCTTCAC | 9 | TRUE | FALSE |
| 168_v.txt | 33282 | 33282-SNV-T | SNV | 1 | T | A | 2 | 23355 | 8.56E-03 | 37.9968763 | WT | FALSE | CAN | JSY13 | 1A | WT-Rep1A | CTC | T>A | A/T+1 | A/T-1 | TA>AT | TA>AT | GTATCCTCG | TCGCCAT | GAG | CTC | T | T | T>A | T>A_CTC | TA>AT_TCGCCAT | 9 | TRUE | FALSE |
| 168_v.txt | 33290 | 33290-SNV-T | SNV | 1 | T | A | 3 | 23355 | 0.01284522 | 53.332766 | WT | FALSE | CAN | JSY13 | 1A | WT-Rep1A | TTA | T>A | A/T+1 | A/T-1 | TA>AT | TA>AT | GCCATTTACT | TACTCTC | AAT | TTA | T | T | T>A | T>A_TTA | TA>AT_TACTCTC | 9 | TRUE | FALSE |
| 168_v.txt | 33293 | 33293-SNV-T | SNV | 1 | T | A | 1 | 23355 | 4.28E-03 | 63.9996229 | WT | FALSE | CAN | JSY13 | 1A | WT-Rep1A | CTC | T>A | A/T+1 | A/T-1 | TA>AT | TA>AT | ATTTACTCTC | TCTCGTC | GAG | CTC | T | T | T>A | T>A_CTC | TA>AT_TCTCGTC | 9 | TRUE | FALSE |
| 168_v.txt | 33317 | 33317-SNV-A | SNV | 1 | A | T | 17 | 23355 | 0.07278955 | 54.8235294 | WT | FALSE | CAN | JSY13 | 1A | WT-Rep1A | GAT | A>T | A/T+1 | A/T-1 | TA>AT | TA>AT | CAATGGATA | ATACAAT | CTA | CTA | T | A | T>A | T>A_CTA | TA>AT_ATACAAT | 9 | TRUE | FALSE |
| 168_v.txt | 33364 | 33364-SNV-T | SNV | 1 | T | A | 69 | 23355 | 0.29543995 | 53.7246377 | WT | FALSE | CAN | JSY13 | 1A | WT-Rep1A | CTG | T>A | A/T+1 | A/T-1 | TA>AT | TA>AT | CACGTCTGT | TGTGGTG | GAC | CTG | T | T | T>A | T>A_CTG | TA>AT_TGTGGTG | 9 | TRUE | FALSE |
| 168_v.txt | 31834 | 31834-SNV-G | SNV | 1 | G | A | 8 | 3138 | 0.25493945 | 54.125 | WT | FALSE | CAN | JSY13 | 1A | WT-Rep1A | TGA | G>A | A/T+1 | G/C-1 | CG>TA | CG>TA | TGCATTGAA | GAAATAA | ACT | ACT | C | T | C>T | C>T_ACT | CG>TA_GAAATAA | 9 | TRUE | FALSE |
| 168_v.txt | 31845 | 31845-SNV-C | SNV | 1 | C | T | 9 | 3138 | 0.28680688 | 53.8888889 | WT | FALSE | CAN | JSY13 | 1A | WT-Rep1A | CCA | C>T | A/T+1 | G/C-1 | CG>TA | CG>TA | AAGATCCAA | CAAACAG | GGT | CCA | C | A | C>T | C>T_CCA | CG>TA_CAAACAG | 9 | TRUE | FALSE |
| 168_v.txt | 31962 | 31962-SNV-G | SNV | 1 | G | A | 20 | 14355 | 0.13932428 | 33.95 | WT | FALSE | CAN | JSY13 | 1A | WT-Rep1A | CGC | G>A | A/T+1 | G/C-1 | CG>TA | CG>TA | GTGGCCGCA | GCATAAT | GCG | GCG | C | T | C>T | C>T_GCG | CG>TA_GCATAAT | 9 | TRUE | FALSE |
| 168_v.txt | 31972 | 31972-SNV-C | SNV | 1 | C | T | 16 | 14355 | 0.11145942 | 37.1866386 | WT | FALSE | CAN | JSY13 | 1A | WT-Rep1A | GCC | C>T | A/T+1 | G/C-1 | CG>TA | CG>TA | AATAAGCCA | CCAAGCC | CGG | GCC | C | A | C>T | C>T_GCC | CG>TA_CCAAGCC | 9 | TRUE | FALSE |
| 168_v.txt | 31980 | 31980-SNV-G | SNV | 1 | G | A | 29 | 14354 | 0.20203428 | 49.5172414 | WT | FALSE | CAN | JSY13 | 1A | WT-Rep1A | GGG | G>A | A/T+1 | G/C-1 | CG>TA | CG>TA | AAGCCGGGC | GGCATTA | CCC | CCC | C | T | C>T | C>T_CCC | CG>TA_GGCATTA | 9 | TRUE | FALSE |
| 168_v.txt | 31981 | 31981-SNV-G | SNV | 1 | G | A | 25 | 14355 | 0.17415535 | 47.9923363 | WT | FALSE | CAN | JSY13 | 1A | WT-Rep1A | GGC | G>A | A/T+1 | G/C-1 | CG>TA | CG>TA | AGCCGGGCA | GCATTAA | CCG | CCG | C | T | C>T | C>T_CCG | CG>TA_GCATTAA | 9 | TRUE | FALSE |
| 168_v.txt | 32014 | 32014-SNV-G | SNV | 1 | G | A | 4 | 14355 | 0.02786486 | 57.4993646 | WT | FALSE | CAN | JSY13 | 1A | WT-Rep1A | CGA | G>A | A/T+1 | G/C-1 | CG>TA | CG>TA | CGTCACGAG | GAGAGAT | GCT | GCT | C | T | C>T | C>T_GCT | CG>TA_GAGAGAT | 9 | TRUE | FALSE |
| 168_v.txt | 32041 | 32041-SNV-G | SNV | 1 | G | A | 31 | 14355 | 0.21595263 | 63.8064516 | WT | FALSE | CAN | JSY13 | 1A | WT-Rep1A | TGC | G>A | A/T+1 | G/C-1 | CG>TA | CG>TA | AAGCTTGCA | GCATAAA | ACG | ACG | C | T | C>T | C>T_ACG | CG>TA_GCATAAA | 9 | TRUE | FALSE |
| 168_v.txt | 32075 | 32075-SNV-C | SNV | 1 | C | T | 28 | 14354 | 0.19506758 | 63.7819306 | WT | FALSE | CAN | JSY13 | 1A | WT-Rep1A | ACC | C>T | A/T+1 | G/C-1 | CG>TA | CG>TA | AAATAACCA | CCATGCA | TGG | ACC | C | A | C>T | C>T_ACC | CG>TA_CCATGCA | 9 | TRUE | FALSE |
| 168_v.txt | 32076 | 32076-SNV-C | SNV | 1 | C | T | 28 | 14355 | 0.19505399 | 62.3571429 | WT | FALSE | CAN | JSY13 | 1A | WT-Rep1A | CCA | C>T | A/T+1 | G/C-1 | CG>TA | CG>TA | AATAACCAT | CATGCAA | GGT | CCA | C | A | C>T | C>T_CCA | CG>TA_CATGCAA | 9 | TRUE | FALSE |
| 168_v.txt | 32080 | 32080-SNV-C | SNV | 1 | C | T | 5 | 14355 | 0.03483107 | 63.9976501 | WT | FALSE | CAN | JSY13 | 1A | WT-Rep1A | GCA | C>T | A/T+1 | G/C-1 | CG>TA | CG>TA | ACCATGCAA | CAAAAAA | CGT | GCA | C | A | C>T | C>T_GCA | CG>TA_CAAAAAA | 9 | TRUE | FALSE |
| 168_v.txt | 32088 | 32088-SNV-C | SNV | 1 | C | T | 37 | 14355 | 0.25774991 | 60.945946 | WT | FALSE | CAN | JSY13 | 1A | WT-Rep1A | GCC | C>T | A/T+1 | G/C-1 | CG>TA | CG>TA | AAAAAGCCT | CCTGCAA | CGG | GCC | C | A | C>T | C>T_GCC | CG>TA_CCTGCAA | 9 | TRUE | FALSE |
| 168_v.txt | 32092 | 32092-SNV-C | SNV | 1 | C | T | 17 | 14355 | 0.11842564 | 54.2352941 | WT | FALSE | CAN | JSY13 | 1A | WT-Rep1A | GCA | C>T | A/T+1 | G/C-1 | CG>TA | CG>TA | AGCCTGCAA | CAACACC | CGT | GCA | C | A | C>T | C>T_GCA | CG>TA_CAACACC | 9 | TRUE | FALSE |
| 168_v.txt | 32165 | 32165-SNV-G | SNV | 1 | G | A | 19 | 14355 | 0.13235806 | 39.8947368 | WT | FALSE | CAN | JSY13 | 1A | WT-Rep1A | CGC | G>A | A/T+1 | G/C-1 | CG>TA | CG>TA | CAAAGCGCC | GCCAAAT | GCG | GCG | C | T | C>T | C>T_GCG | CG>TA_GCCAAAT | 9 | TRUE | FALSE |
| 168_v.txt | 32271 | 32271-SNV-C | SNV | 1 | C | T | 95 | 46483 | 0.20437579 | 33.6210526 | WT | FALSE | CAN | JSY13 | 1A | WT-Rep1A | ACG | C>T | A/T+1 | G/C-1 | CG>TA | CG>TA | AAAATACGG | CGGGAAC | TGC | ACG | C | A | C>T | C>T_ACG | CG>TA_CGGGAAC | 9 | TRUE | FALSE |
| 168_v.txt | 32272 | 32272-SNV-G | SNV | 1 | G | A | 119 | 46483 | 0.25600757 | 36.9495943 | WT | FALSE | CAN | JSY13 | 1A | WT-Rep1A | CGG | G>A | A/T+1 | G/C-1 | CG>TA | CG>TA | AAATACGGG | GGGAACC | GCC | GCC | C | T | C>T | C>T_GCC | CG>TA_GGGAACC | 9 | TRUE | FALSE |
| 168_v.txt | 32274 | 32274-SNV-G | SNV | 1 | G | A | 45 | 46483 | 0.09680959 | 35.5777778 | WT | FALSE | CAN | JSY13 | 1A | WT-Rep1A | GGA | G>A | A/T+1 | G/C-1 | CG>TA | CG>TA | ATACGGGAA | GAACCAA | CCT | CCT | C | T | C>T | C>T_CCT | CG>TA_GAACCAA | 9 | TRUE | FALSE |
| 168_v.txt | 32292 | 32292-SNV-G | SNV | 1 | G | A | 106 | 46483 | 0.22804036 | 52.7358491 | WT | FALSE | CAN | JSY13 | 1A | WT-Rep1A | TGA | G>A | A/T+1 | G/C-1 | CG>TA | CG>TA | ATATTTGAA | GAATTTG | ACT | ACT | C | T | C>T | C>T_ACT | CG>TA_GAATTTG | 9 | TRUE | FALSE |
| 168_v.txt | 32299 | 32299-SNV-C | SNV | 1 | C | T | 79 | 46483 | 0.16995461 | 51.8987342 | WT | FALSE | CAN | JSY13 | 1A | WT-Rep1A | GCG | C>T | A/T+1 | G/C-1 | CG>TA | CG>TA | AATTTGCGG | CGGCAGA | CGC | GCG | C | A | C>T | C>T_GCG | CG>TA_CGGCAGA | 9 | TRUE | FALSE |
| 168_v.txt | 32316 | 32316-SNV-G | SNV | 1 | G | A | 5 | 46483 | 0.01075662 | 39.3995911 | WT | FALSE | CAN | JSY13 | 1A | WT-Rep1A | TGT | G>A | A/T+1 | G/C-1 | CG>TA | CG>TA | ATGGTTGTT | GTTAAGA | ACA | ACA | C | T | C>T | C>T_ACA | CG>TA_GTTAAGA | 9 | TRUE | FALSE |
| 168_v.txt | 32344 | 32344-SNV-G | SNV | 1 | G | A | 70 | 46483 | 0.15059269 | 60 | WT | FALSE | CAN | JSY13 | 1A | WT-Rep1A | GGC | G>A | A/T+1 | G/C-1 | CG>TA | CG>TA | TATGTGGCA | GCAAAAC | CCG | CCG | C | T | C>T | C>T_CCG | CG>TA_GCAAAAC | 9 | TRUE | FALSE |
| 168_v.txt | 32398 | 32398-SNV-C | SNV | 1 | C | T | 10 | 46483 | 0.02151324 | 48.4993371 | WT | FALSE | CAN | JSY13 | 1A | WT-Rep1A | ACG | C>T | A/T+1 | G/C-1 | CG>TA | CG>TA | TAGAAACGT | CGTAGGA | TGC | ACG | C | A | C>T | C>T_ACG | CG>TA_CGTAGGA | 9 | TRUE | FALSE |
| 168_v.txt | 32413 | 32413-SNV-G | SNV | 1 | G | A | 48 | 46482 | 0.10326578 | 53.2705127 | WT | FALSE | CAN | JSY13 | 1A | WT-Rep1A | TGT | G>A | A/T+1 | G/C-1 | CG>TA | CG>TA | TAGATTGTG | GTGTTAG | ACA | ACA | C | T | C>T | C>T_ACA | CG>TA_GTGTTAG | 9 | TRUE | FALSE |
| 168_v.txt | 32436 | 32436-SNV-G | SNV | 1 | G | A | 61 | 46483 | 0.13123077 | 52.5409836 | WT | FALSE | CAN | JSY13 | 1A | WT-Rep1A | TGG | G>A | A/T+1 | G/C-1 | CG>TA | CG>TA | TTGTATGGA | GGAACTA | ACC | ACC | C | T | C>T | C>T_ACC | CG>TA_GGAACTA | 9 | TRUE | FALSE |
| 168_v.txt | 32449 | 32449-SNV-C | SNV | 1 | C | T | 321 | 46483 | 0.69057505 | 52.271028 | WT | FALSE | CAN | JSY13 | 1A | WT-Rep1A | CCA | C>T | A/T+1 | G/C-1 | CG>TA | CG>TA | AAAGTCCAA | CAATGAA | GGT | CCA | C | A | C>T | C>T_CCA | CG>TA_CAATGAA | 9 | TRUE | FALSE |
| 168_v.txt | 32463 | 32463-SNV-G | SNV | 1 | G | A | 53 | 46483 | 0.11402018 | 46.2075472 | WT | FALSE | CAN | JSY13 | 1A | WT-Rep1A | AGA | G>A | A/T+1 | G/C-1 | CG>TA | CG>TA | AATAGAGAG | GAGCCAA | TCT | TCT | C | T | C>T | C>T_TCT | CG>TA_GAGCCAA | 9 | TRUE | FALSE |
| 168_v.txt | 32466 | 32466-SNV-C | SNV | 1 | C | T | 61 | 46483 | 0.13123077 | 46.5901639 | WT | FALSE | CAN | JSY13 | 1A | WT-Rep1A | GCC | C>T | A/T+1 | G/C-1 | CG>TA | CG>TA | AGAGAGCCA | CCAATGT | CGG | GCC | C | A | C>T | C>T_GCC | CG>TA_CCAATGT | 9 | TRUE | FALSE |
| 168_v.txt | 32487 | 32487-SNV-C | SNV | 1 | C | T | 105 | 46483 | 0.22588904 | 46.9434109 | WT | FALSE | CAN | JSY13 | 1A | WT-Rep1A | ACG | C>T | A/T+1 | G/C-1 | CG>TA | CG>TA | AAGATACGG | CGGAAAA | TGC | ACG | C | A | C>T | C>T_ACG | CG>TA_CGGAAAA | 9 | TRUE | FALSE |
| 168_v.txt | 32488 | 32488-SNV-G | SNV | 1 | G | A | 216 | 46483 | 0.46468601 | 46.6712963 | WT | FALSE | CAN | JSY13 | 1A | WT-Rep1A | CGG | G>A | A/T+1 | G/C-1 | CG>TA | CG>TA | AGATACGGA | GGAAAAC | GCC | GCC | C | T | C>T | C>T_GCC | CG>TA_GGAAAAC | 9 | TRUE | FALSE |
| 168_v.txt | 32508 | 32508-SNV-G | SNV | 1 | G | A | 25 | 46483 | 0.0537831 | 31.4000001 | WT | FALSE | CAN | JSY13 | 1A | WT-Rep1A | GGC | G>A | A/T+1 | G/C-1 | CG>TA | CG>TA | TTGATGGCT | GCTCTTG | CCG | CCG | C | T | C>T | C>T_CCG | CG>TA_GCTCTTG | 9 | TRUE | FALSE |
| 168_v.txt | 32509 | 32509-SNV-C | SNV | 1 | C | T | 11 | 46483 | 0.02366457 | 49.7266234 | WT | FALSE | CAN | JSY13 | 1A | WT-Rep1A | GCT | C>T | A/T+1 | G/C-1 | CG>TA | CG>TA | TGATGGCTC | CTCTTGG | CGA | GCT | C | A | C>T | C>T_GCT | CG>TA_CTCTTGG | 9 | TRUE | FALSE |
| 168_v.txt | 32529 | 32529-SNV-G | SNV | 1 | G | A | 20 | 46483 | 0.04302648 | 34.2 | WT | FALSE | CAN | JSY13 | 1A | WT-Rep1A | GGG | G>A | A/T+1 | G/C-1 | CG>TA | CG>TA | TTTCTGGGG | GGGTTTG | CCC | CCC | C | T | C>T | C>T_CCC | CG>TA_GGGTTTG | 9 | TRUE | FALSE |
| 168_v.txt | 32530 | 32530-SNV-G | SNV | 1 | G | A | 95 | 46483 | 0.20437579 | 38.7578947 | WT | FALSE | CAN | JSY13 | 1A | WT-Rep1A | GGG | G>A | A/T+1 | G/C-1 | CG>TA | CG>TA | TTCTGGGGT | GGTTTGC | CCC | CCC | C | T | C>T | C>T_CCC | CG>TA_GGTTTGC | 9 | TRUE | FALSE |
| 168_v.txt | 32535 | 32535-SNV-G | SNV | 1 | G | A | 47 | 46481 | 0.10111659 | 36.360648 | WT | FALSE | CAN | JSY13 | 1A | WT-Rep1A | TGC | G>A | A/T+1 | G/C-1 | CG>TA | CG>TA | GGGTTTGCA | GCAGCTT | ACG | ACG | C | T | C>T | C>T_ACG | CG>TA_GCAGCTT | 9 | TRUE | FALSE |
| 168_v.txt | 32536 | 32536-SNV-C | SNV | 1 | C | T | 69 | 46481 | 0.14844775 | 37.4049131 | WT | FALSE | CAN | JSY13 | 1A | WT-Rep1A | GCA | C>T | A/T+1 | G/C-1 | CG>TA | CG>TA | GGTTTGCAG | CAGCTTC | CGT | GCA | C | A | C>T | C>T_GCA | CG>TA_CAGCTTC | 9 | TRUE | FALSE |
| 168_v.txt | 32545 | 32545-SNV-C | SNV | 1 | C | T | 112 | 46481 | 0.24095867 | 36.8655702 | WT | FALSE | CAN | JSY13 | 1A | WT-Rep1A | CCA | C>T | A/T+1 | G/C-1 | CG>TA | CG>TA | CTTCACCAG | CAGCAGT | GGT | CCA | C | A | C>T | C>T_CCA | CG>TA_CAGCAGT | 9 | TRUE | FALSE |
| 168_v.txt | 32547 | 32547-SNV-G | SNV | 1 | G | A | 20 | 46480 | 0.04302926 | 37.2992781 | WT | FALSE | CAN | JSY13 | 1A | WT-Rep1A | AGC | G>A | A/T+1 | G/C-1 | CG>TA | CG>TA | TCACCAGCA | GCAGTGA | TCG | TCG | C | T | C>T | C>T_TCG | CG>TA_GCAGTGA | 9 | TRUE | FALSE |
| 168_v.txt | 32566 | 32566-SNV-C | SNV | 1 | C | T | 8 | 7614 | 0.10506961 | 34.625 | WT | FALSE | CAN | JSY13 | 1A | WT-Rep1A | TCA | C>T | A/T+1 | G/C-1 | CG>TA | CG>TA | CTAGTTCAG | CAGTACC | AGT | TCA | C | A | C>T | C>T_TCA | CG>TA_CAGTACC | 9 | TRUE | FALSE |
| 168_v.txt | 32575 | 32575-SNV-G | SNV | 1 | G | A | 12 | 7614 | 0.15760441 | 38 | WT | FALSE | CAN | JSY13 | 1A | WT-Rep1A | TGA | G>A | A/T+1 | G/C-1 | CG>TA | CG>TA | TACCTTGAA | GAAATGT | ACT | ACT | C | T | C>T | C>T_ACT | CG>TA_GAAATGT | 9 | TRUE | FALSE |
| 168_v.txt | 32629 | 32629-SNV-C | SNV | 1 | C | T | 11 | 7614 | 0.14447071 | 53.2727273 | WT | FALSE | CAN | JSY13 | 1A | WT-Rep1A | TCG | C>T | A/T+1 | G/C-1 | CG>TA | CG>TA | TCCCTTCGT | CGTTTTT | AGC | TCG | C | A | C>T | C>T_TCG | CG>TA_CGTTTTT | 9 | TRUE | FALSE |
| 168_v.txt | 32630 | 32630-SNV-G | SNV | 1 | G | A | 3 | 7614 | 0.0394011 | 47 | WT | FALSE | CAN | JSY13 | 1A | WT-Rep1A | CGT | G>A | A/T+1 | G/C-1 | CG>TA | CG>TA | CCCTTCGTT | GTTTTTA | GCA | GCA | C | T | C>T | C>T_GCA | CG>TA_GTTTTTA | 9 | TRUE | FALSE |
| 168_v.txt | 32658 | 32658-SNV-C | SNV | 1 | C | T | 9 | 7614 | 0.11820331 | 62.6666667 | WT | FALSE | CAN | JSY13 | 1A | WT-Rep1A | ACC | C>T | A/T+1 | G/C-1 | CG>TA | CG>TA | CCTGGACCC | CCCCAGG | TGG | ACC | C | A | C>T | C>T_ACC | CG>TA_CCCCAGG | 9 | TRUE | FALSE |
| 168_v.txt | 32661 | 32661-SNV-C | SNV | 1 | C | T | 17 | 7614 | 0.22327292 | 54.8235294 | WT | FALSE | CAN | JSY13 | 1A | WT-Rep1A | CCA | C>T | A/T+1 | G/C-1 | CG>TA | CG>TA | GGACCCCAG | CAGGCAC | GGT | CCA | C | A | C>T | C>T_CCA | CG>TA_CAGGCAC | 9 | TRUE | FALSE |
| 168_v.txt | 32749 | 32749-SNV-C | SNV | 1 | C | T | 7 | 7622 | 0.09183941 | 43.7125154 | WT | FALSE | CAN | JSY13 | 1A | WT-Rep1A | CCG | C>T | A/T+1 | G/C-1 | CG>TA | CG>TA | GAAACCCGA | CGATAAT | GGC | CCG | C | A | C>T | C>T_CCG | CG>TA_CGATAAT | 9 | TRUE | FALSE |
| 168_v.txt | 32776 | 32776-SNV-C | SNV | 1 | C | T | 14 | 7622 | 0.18367883 | 44.5714286 | WT | FALSE | CAN | JSY13 | 1A | WT-Rep1A | GCG | C>T | A/T+1 | G/C-1 | CG>TA | CG>TA | TGGAAGCGA | CGACCCA | CGC | GCG | C | A | C>T | C>T_GCG | CG>TA_CGACCCA | 9 | TRUE | FALSE |

| | | | | | | | | | | | | | | | | | | | | | | | | | | | | | | | | | | |
|---|---|---|---|---|---|---|---|---|---|---|---|---|---|---|---|---|---|---|---|---|---|---|---|---|---|---|---|---|---|---|---|---|---|---|
| 168_v.txt | 32780 | 32780-SNV-( | SNV | 1 | C | T | 18 | 7622 | 0.23615849 | 46.7222222 | WT | FALSE | CAN | JSY13 | 1A | WT-Rep1A | CCC | C>T | A/T+1 | G/C-1 | CG>TA | CG>TA | AGCGACCCA | CCAGAAC | GGG | CCC | C | A | C>T | C>T_CCC | CG>TA_CCAGAAC | 9 | TRUE | FALSE |
| 168_v.txt | 32788 | 32788-SNV-( | SNV | 1 | C | T | 24 | 7622 | 0.31487799 | 45.375 | WT | FALSE | CAN | JSY13 | 1A | WT-Rep1A | TCG | C>T | A/T+1 | G/C-1 | CG>TA | CG>TA | AGAACTCGA | CGAATTC | AGC | TCG | C | A | C>T | C>T_TCG | CG>TA_CGAATTC | 9 | TRUE | FALSE |
| 168_v.txt | 32794 | 32794-SNV-( | SNV | 1 | C | T | 7 | 7622 | 0.09183941 | 38.2857143 | WT | FALSE | CAN | JSY13 | 1A | WT-Rep1A | TCA | C>T | A/T+1 | G/C-1 | CG>TA | CG>TA | CGAATTCAC | CACCGTA | AGT | TCA | C | A | C>T | C>T_TCA | CG>TA_CACCGTA | 9 | TRUE | FALSE |
| 168_v.txt | 32797 | 32797-SNV-( | SNV | 1 | C | T | 47 | 7622 | 0.61663605 | 44.2582056 | WT | FALSE | CAN | JSY13 | 1A | WT-Rep1A | CCG | C>T | A/T+1 | G/C-1 | CG>TA | CG>TA | ATTCACCGTA | CGTAATA | GGC | CCG | C | A | C>T | C>T_CCG | CG>TA_CGTAATA | 9 | TRUE | FALSE |
| 168_v.txt | 32798 | 32798-SNV-( | SNV | 1 | G | A | 9 | 7622 | 0.11807924 | 43.5555556 | WT | FALSE | CAN | JSY13 | 1A | WT-Rep1A | CGT | G>A | A/T+1 | G/C-1 | CG>TA | CG>TA | TTCACCGTA | GTAATAT | GCA | GCA | C | T | C>T | C>T_GCA | CG>TA_GTAATAT | 9 | TRUE | FALSE |
| 168_v.txt | 32811 | 32811-SNV-( | SNV | 1 | G | A | 22 | 7622 | 0.28863815 | 40.1818182 | WT | FALSE | CAN | JSY13 | 1A | WT-Rep1A | AGG | G>A | A/T+1 | G/C-1 | CG>TA | CG>TA | TTGACAGGG | GGGAACA | TCC | TCC | C | T | C>T | C>T_TCC | CG>TA_GGGAACA | 9 | TRUE | FALSE |
| 168_v.txt | 32855 | 32855-SNV-( | SNV | 1 | C | T | 14 | 7622 | 0.18367883 | 37.5714286 | WT | FALSE | CAN | JSY13 | 1A | WT-Rep1A | TCC | C>T | A/T+1 | G/C-1 | CG>TA | CG>TA | ACTAATCCAT | CCATGCC | AGG | TCC | C | A | C>T | C>T_TCC | CG>TA_CCATGCC | 9 | TRUE | FALSE |
| 168_v.txt | 32856 | 32856-SNV-( | SNV | 1 | C | T | 39 | 7622 | 0.51167673 | 37.7692308 | WT | FALSE | CAN | JSY13 | 1A | WT-Rep1A | CCA | C>T | A/T+1 | G/C-1 | CG>TA | CG>TA | CTAATCCATC | CATGCCG | GGT | CCA | C | A | C>T | C>T_CCA | CG>TA_CATGCCG | 9 | TRUE | FALSE |
| 168_v.txt | 32882 | 32882-SNV-( | SNV | 1 | C | T | 40 | 22850 | 0.17505471 | 27.575 | WT | FALSE | CAN | JSY13 | 1A | WT-Rep1A | TCC | C>T | A/T+1 | G/C-1 | CG>TA | CG>TA | GTACGTCCA | CCAAAAT | AGG | TCC | C | A | C>T | C>T_TCC | CG>TA_CCAAAAT | 9 | TRUE | FALSE |
| 168_v.txt | 32899 | 32899-SNV-( | SNV | 1 | G | A | 49 | 22850 | 0.21444201 | 35.1033221 | WT | FALSE | CAN | JSY13 | 1A | WT-Rep1A | TGG | G>A | A/T+1 | G/C-1 | CG>TA | CG>TA | TGACTTGGC | GGCCAAC | ACC | ACC | C | T | C>T | C>T_ACC | CG>TA_GGCCAAC | 9 | TRUE | FALSE |
| 168_v.txt | 32900 | 32900-SNV-( | SNV | 1 | G | A | 23 | 22850 | 0.10065646 | 37.2142043 | WT | FALSE | CAN | JSY13 | 1A | WT-Rep1A | GGC | G>A | A/T+1 | G/C-1 | CG>TA | CG>TA | GACTTGGCC | GCCAACT | CCG | CCG | C | T | C>T | C>T_CCG | CG>TA_GCCAACT | 9 | TRUE | FALSE |
| 168_v.txt | 32901 | 32901-SNV-( | SNV | 1 | C | T | 44 | 22850 | 0.19256018 | 30.7275885 | WT | FALSE | CAN | JSY13 | 1A | WT-Rep1A | GCC | C>T | A/T+1 | G/C-1 | CG>TA | CG>TA | ACTTGGCCA | CCAACTA | CGG | GCC | C | A | C>T | C>T_GCC | CG>TA_CCAACTA | 9 | TRUE | FALSE |
| 168_v.txt | 32922 | 32922-SNV-( | SNV | 1 | G | A | 25 | 22850 | 0.10940919 | 50.72 | WT | FALSE | CAN | JSY13 | 1A | WT-Rep1A | GGC | G>A | A/T+1 | G/C-1 | CG>TA | CG>TA | TCCAGGGCA | GCAAAAG | CCG | CCG | C | T | C>T | C>T_CCG | CG>TA_GCAAAAG | 9 | TRUE | FALSE |
| 168_v.txt | 32934 | 32934-SNV-( | SNV | 1 | G | A | 10 | 22886 | 0.04369484 | 44.9994527 | WT | FALSE | CAN | JSY13 | 1A | WT-Rep1A | TGC | G>A | A/T+1 | G/C-1 | CG>TA | CG>TA | GTGATTGCC | GCCCAAG | ACG | ACG | C | T | C>T | C>T_ACG | CG>TA_GCCCAAG | 9 | TRUE | FALSE |
| 168_v.txt | 32935 | 32935-SNV-( | SNV | 1 | C | T | 22 | 22886 | 0.09612864 | 32.7726603 | WT | FALSE | CAN | JSY13 | 1A | WT-Rep1A | GCC | C>T | A/T+1 | G/C-1 | CG>TA | CG>TA | TGATTGCCCA | CCCAAGA | CGG | GCC | C | A | C>T | C>T_GCC | CG>TA_CCCAAGA | 9 | TRUE | FALSE |
| 168_v.txt | 32937 | 32937-SNV-( | SNV | 1 | C | T | 27 | 22886 | 0.11797606 | 29.8148148 | WT | FALSE | CAN | JSY13 | 1A | WT-Rep1A | CCA | C>T | A/T+1 | G/C-1 | CG>TA | CG>TA | ATTGCCCAAC | CAAGAAA | GGT | CCA | C | A | C>T | C>T_CCA | CG>TA_CAAGAAA | 9 | TRUE | FALSE |
| 168_v.txt | 32940 | 32940-SNV-( | SNV | 1 | G | A | 78 | 22886 | 0.34081972 | 50.7564103 | WT | FALSE | CAN | JSY13 | 1A | WT-Rep1A | AGA | G>A | A/T+1 | G/C-1 | CG>TA | CG>TA | GCCCAAGAA | GAAAACC | TCT | TCT | C | T | C>T | C>T_TCT | CG>TA_GAAAACC | 9 | TRUE | FALSE |
| 168_v.txt | 32945 | 32945-SNV-( | SNV | 1 | C | T | 112 | 22886 | 0.48938216 | 57.6531767 | WT | FALSE | CAN | JSY13 | 1A | WT-Rep1A | ACC | C>T | A/T+1 | G/C-1 | CG>TA | CG>TA | AGAAAACCA | CCAATAC | TGG | ACC | C | A | C>T | C>T_ACC | CG>TA_CCAATAC | 9 | TRUE | FALSE |
| 168_v.txt | 32946 | 32946-SNV-( | SNV | 1 | C | T | 42 | 22886 | 0.18351831 | 58.6666667 | WT | FALSE | CAN | JSY13 | 1A | WT-Rep1A | CCA | C>T | A/T+1 | G/C-1 | CG>TA | CG>TA | GAAAACCAA | CAATACA | GGT | CCA | C | A | C>T | C>T_CCA | CG>TA_CAATACA | 9 | TRUE | FALSE |
| 168_v.txt | 32958 | 32958-SNV-( | SNV | 1 | C | T | 84 | 22886 | 0.36703662 | 56.1547619 | WT | FALSE | CAN | JSY13 | 1A | WT-Rep1A | ACC | C>T | A/T+1 | G/C-1 | CG>TA | CG>TA | ATGTAACCA | CCATTGG | TGG | ACC | C | A | C>T | C>T_ACC | CG>TA_CCATTGG | 9 | TRUE | FALSE |
| 168_v.txt | 32965 | 32965-SNV-( | SNV | 1 | C | T | 29 | 22886 | 0.12671502 | 57.4827586 | WT | FALSE | CAN | JSY13 | 1A | WT-Rep1A | GCC | C>T | A/T+1 | G/C-1 | CG>TA | CG>TA | CATTGGCCG | CCGCACC | CGG | GCC | C | A | C>T | C>T_GCC | CG>TA_CCGCACC | 9 | TRUE | FALSE |
| 168_v.txt | 32967 | 32967-SNV-( | SNV | 1 | G | A | 48 | 22886 | 0.20973521 | 56.25 | WT | FALSE | CAN | JSY13 | 1A | WT-Rep1A | CGC | G>A | A/T+1 | G/C-1 | CG>TA | CG>TA | TTGGCCGCA | GCACCAA | GCG | GCG | C | T | C>T | C>T_GCG | CG>TA_GCACCAA | 9 | TRUE | FALSE |
| 168_v.txt | 32977 | 32977-SNV-( | SNV | 1 | C | T | 24 | 22886 | 0.10486761 | 58.2083333 | WT | FALSE | CAN | JSY13 | 1A | WT-Rep1A | GCT | C>T | A/T+1 | G/C-1 | CG>TA | CG>TA | CAAATGCTG | CTGGAGA | CGA | GCT | C | A | C>T | C>T_GCT | CG>TA_CTGGAGA | 9 | TRUE | FALSE |
| 168_v.txt | 32982 | 32982-SNV-( | SNV | 1 | G | A | 139 | 22886 | 0.60735821 | 60.057554 | WT | FALSE | CAN | JSY13 | 1A | WT-Rep1A | AGA | G>A | A/T+1 | G/C-1 | CG>TA | CG>TA | GCTGGAGAA | GAAAGGA | TCT | TCT | C | T | C>T | C>T_TCT | CG>TA_GAAAGGA | 9 | TRUE | FALSE |
| 168_v.txt | 32995 | 32995-SNV-( | SNV | 1 | G | A | 57 | 22886 | 0.24906056 | 63.3333333 | WT | FALSE | CAN | JSY13 | 1A | WT-Rep1A | TGT | G>A | A/T+1 | G/C-1 | CG>TA | CG>TA | ATCTTTGTGA | GTGAGAA | ACA | ACA | C | T | C>T | C>T_ACA | CG>TA_GTGAGAA | 9 | TRUE | FALSE |
| 168_v.txt | 33012 | 33012-SNV-( | SNV | 1 | G | A | 19 | 22886 | 0.08302019 | 63.2105263 | WT | FALSE | CAN | JSY13 | 1A | WT-Rep1A | AGA | G>A | A/T+1 | G/C-1 | CG>TA | CG>TA | GTGAAAGAG | GAGGATG | TCT | TCT | C | T | C>T | C>T_TCT | CG>TA_GAGGATG | 9 | TRUE | FALSE |
| 168_v.txt | 33015 | 33015-SNV-( | SNV | 1 | G | A | 32 | 22886 | 0.13982347 | 59.59375 | WT | FALSE | CAN | JSY13 | 1A | WT-Rep1A | GGA | G>A | A/T+1 | G/C-1 | CG>TA | CG>TA | AAAGAGGAT | GATGTAA | CCT | CCT | C | T | C>T | C>T_CCT | CG>TA_GATGTAA | 9 | TRUE | FALSE |
| 168_v.txt | 33024 | 33024-SNV-( | SNV | 1 | G | A | 10 | 22886 | 0.04369484 | 64 | WT | FALSE | CAN | JSY13 | 1A | WT-Rep1A | AGG | G>A | A/T+1 | G/C-1 | CG>TA | CG>TA | GTAACAGGG | GGGATGA | TCC | TCC | C | T | C>T | C>T_TCC | CG>TA_GGGATGA | 9 | TRUE | FALSE |
| 168_v.txt | 33037 | 33037-SNV-( | SNV | 1 | C | T | 16 | 22886 | 0.06991174 | 54.875 | WT | FALSE | CAN | JSY13 | 1A | WT-Rep1A | GCC | C>T | A/T+1 | G/C-1 | CG>TA | CG>TA | ATGTAGCCA | CCATTTC | CGG | GCC | C | A | C>T | C>T_GCC | CG>TA_CCATTTC | 9 | TRUE | FALSE |
| 168_v.txt | 33051 | 33051-SNV-( | SNV | 1 | G | A | 48 | 22886 | 0.20973521 | 57.8541667 | WT | FALSE | CAN | JSY13 | 1A | WT-Rep1A | GGA | G>A | A/T+1 | G/C-1 | CG>TA | CG>TA | CCCAAGGAC | GACTGCG | CCT | CCT | C | T | C>T | C>T_CCT | CG>TA_GACTGCG | 9 | TRUE | FALSE |
| 168_v.txt | 33055 | 33055-SNV-( | SNV | 1 | G | A | 51 | 22886 | 0.22284366 | 52.9605784 | WT | FALSE | CAN | JSY13 | 1A | WT-Rep1A | TGC | G>A | A/T+1 | G/C-1 | CG>TA | CG>TA | AGGACTGCG | GCGTGAC | ACG | ACG | C | T | C>T | C>T_ACG | CG>TA_GCGTGAC | 9 | TRUE | FALSE |
| 168_v.txt | 33056 | 33056-SNV-( | SNV | 1 | C | T | 27 | 22886 | 0.11797606 | 56.1481482 | WT | FALSE | CAN | JSY13 | 1A | WT-Rep1A | GCG | C>T | A/T+1 | G/C-1 | CG>TA | CG>TA | GGACTGCGT | CGTGACA | CGC | GCG | C | A | C>T | C>T_GCG | CG>TA_CGTGACA | 9 | TRUE | FALSE |
| 168_v.txt | 33069 | 33069-SNV-( | SNV | 1 | G | A | 20 | 22886 | 0.08738967 | 52.8002766 | WT | FALSE | CAN | JSY13 | 1A | WT-Rep1A | TGC | G>A | A/T+1 | G/C-1 | CG>TA | CG>TA | GAATATGCC | GCCAAAG | ACG | ACG | C | T | C>T | C>T_ACG | CG>TA_GCCAAAG | 9 | TRUE | FALSE |
| 168_v.txt | 33070 | 33070-SNV-( | SNV | 1 | C | T | 32 | 22886 | 0.13982347 | 54.15625 | WT | FALSE | CAN | JSY13 | 1A | WT-Rep1A | GCC | C>T | A/T+1 | G/C-1 | CG>TA | CG>TA | AATATGCCA | CCAAAGA | CGG | GCC | C | A | C>T | C>T_GCC | CG>TA_CCAAAGA | 9 | TRUE | FALSE |
| 168_v.txt | 33111 | 33111-SNV-( | SNV | 1 | G | A | 63 | 22886 | 0.27527746 | 41.4401253 | WT | FALSE | CAN | JSY13 | 1A | WT-Rep1A | TGG | G>A | A/T+1 | G/C-1 | CG>TA | CG>TA | CCCACTGGG | GGGCCGG | ACC | ACC | C | T | C>T | C>T_ACC | CG>TA_GGGCCGG | 9 | TRUE | FALSE |
| 168_v.txt | 33114 | 33114-SNV-( | SNV | 1 | C | T | 46 | 22886 | 0.20099624 | 41.1955102 | WT | FALSE | CAN | JSY13 | 1A | WT-Rep1A | GCC | C>T | A/T+1 | G/C-1 | CG>TA | CG>TA | ACTGGGCCG | CCGGCGT | CGG | GCC | C | A | C>T | C>T_GCC | CG>TA_CCGGCGT | 9 | TRUE | FALSE |
| 168_v.txt | 33115 | 33115-SNV-( | SNV | 1 | C | T | 50 | 22886 | 0.21847418 | 46.78 | WT | FALSE | CAN | JSY13 | 1A | WT-Rep1A | CCG | C>T | A/T+1 | G/C-1 | CG>TA | CG>TA | CTGGGCCGG | CGGCGTT | GGC | CCG | C | A | C>T | C>T_CCG | CG>TA_CGGCGTT | 9 | TRUE | FALSE |
| 168_v.txt | 33118 | 33118-SNV-( | SNV | 1 | C | T | 21 | 22886 | 0.09175915 | 43.7142857 | WT | FALSE | CAN | JSY13 | 1A | WT-Rep1A | GCG | C>T | A/T+1 | G/C-1 | CG>TA | CG>TA | GGCCGGCGT | CGTTGGT | CGC | GCG | C | A | C>T | C>T_GCG | CG>TA_CGTTGGT | 9 | TRUE | FALSE |
| 168_v.txt | 33153 | 33153-SNV-( | SNV | 1 | C | T | 27 | 22886 | 0.11797606 | 36.6294331 | WT | FALSE | CAN | JSY13 | 1A | WT-Rep1A | ACC | C>T | A/T+1 | G/C-1 | CG>TA | CG>TA | AAAAGACCT | CCTGTAC | TGG | ACC | C | A | C>T | C>T_ACC | CG>TA_CCTGTAC | 9 | TRUE | FALSE |
| 168_v.txt | 33168 | 33168-SNV-( | SNV | 1 | C | T | 22 | 22847 | 0.09629273 | 37.8172784 | WT | FALSE | CAN | JSY13 | 1A | WT-Rep1A | ACC | C>T | A/T+1 | G/C-1 | CG>TA | CG>TA | ATAGTACCA | CCACCAA | TGG | ACC | C | A | C>T | C>T_ACC | CG>TA_CCACCAA | 9 | TRUE | FALSE |
| 168_v.txt | 33169 | 33169-SNV-( | SNV | 1 | C | T | 12 | 22847 | 0.05252331 | 37.9999976 | WT | FALSE | CAN | JSY13 | 1A | WT-Rep1A | CCA | C>T | A/T+1 | G/C-1 | CG>TA | CG>TA | TAGTACCAC | CACCAAG | GGT | CCA | C | A | C>T | C>T_CCA | CG>TA_CACCAAG | 9 | TRUE | FALSE |
| 168_v.txt | 33171 | 33171-SNV-( | SNV | 1 | C | T | 101 | 22847 | 0.44207117 | 37.3662261 | WT | FALSE | CAN | JSY13 | 1A | WT-Rep1A | ACC | C>T | A/T+1 | G/C-1 | CG>TA | CG>TA | GTACCACCA | CCAAGGG | TGG | ACC | C | A | C>T | C>T_ACC | CG>TA_CCAAGGG | 9 | TRUE | FALSE |
| 168_v.txt | 33172 | 33172-SNV-( | SNV | 1 | C | T | 23 | 22847 | 0.10066967 | 36.8255322 | WT | FALSE | CAN | JSY13 | 1A | WT-Rep1A | CCA | C>T | A/T+1 | G/C-1 | CG>TA | CG>TA | TACCACCAA | CAAGGGC | GGT | CCA | C | A | C>T | C>T_CCA | CG>TA_CAAGGGC | 9 | TRUE | FALSE |
| 168_v.txt | 33177 | 33177-SNV-( | SNV | 1 | G | A | 11 | 22843 | 0.0481548 | 37.8181818 | WT | FALSE | CAN | JSY13 | 1A | WT-Rep1A | GGC | G>A | A/T+1 | G/C-1 | CG>TA | CG>TA | CCAAGGGCA | GCAATCA | CCG | CCG | C | T | C>T | C>T_CCG | CG>TA_GCAATCA | 9 | TRUE | FALSE |
| 168_v.txt | 33205 | 33205-SNV-( | SNV | 1 | G | A | 12 | 23355 | 0.05138086 | 34.1656589 | WT | FALSE | CAN | JSY13 | 1A | WT-Rep1A | AGC | G>A | A/T+1 | G/C-1 | CG>TA | CG>TA | GCTTAAGCT | GCTCTCT | TCG | TCG | C | T | C>T | C>T_TCG | CG>TA_GCTCTCT | 9 | TRUE | FALSE |
| 168_v.txt | 33258 | 33258-SNV-( | SNV | 1 | C | T | 32 | 23355 | 0.13701563 | 41.0625 | WT | FALSE | CAN | JSY13 | 1A | WT-Rep1A | GCC | C>T | A/T+1 | G/C-1 | CG>TA | CG>TA | CCTATGCCAT | CCATCCT | CGG | GCC | C | A | C>T | C>T_GCC | CG>TA_CCATCCT | 9 | TRUE | FALSE |
| 168_v.txt | 33284 | 33284-SNV-( | SNV | 1 | G | A | 20 | 23355 | 0.08563477 | 49.8396159 | WT | FALSE | CAN | JSY13 | 1A | WT-Rep1A | CGC | G>A | A/T+1 | G/C-1 | CG>TA | CG>TA | ATCCTCGCCA | GCCATTT | GCG | GCG | C | T | C>T | C>T_GCG | CG>TA_GCCATTT | 9 | TRUE | FALSE |
| 168_v.txt | 33286 | 33286-SNV-( | SNV | 1 | C | T | 1 | 23354 | 4.28E-03 | 63.9997558 | WT | FALSE | CAN | JSY13 | 1A | WT-Rep1A | CCA | C>T | A/T+1 | G/C-1 | CG>TA | CG>TA | CCTCGCCATT | CATTTAC | GGT | CCA | C | A | C>T | C>T_CCA | CG>TA_CATTTAC | 9 | TRUE | FALSE |
| 168_v.txt | 33292 | 33292-SNV-( | SNV | 1 | C | T | 5 | 23355 | 0.02140869 | 42.3997985 | WT | FALSE | CAN | JSY13 | 1A | WT-Rep1A | ACT | C>T | A/T+1 | G/C-1 | CG>TA | CG>TA | CATTTACTCT | CTCTCGT | TGA | ACT | C | A | C>T | C>T_ACT | CG>TA_CTCTCGT | 9 | TRUE | FALSE |
| 168_v.txt | 33309 | 33309-SNV-( | SNV | 1 | C | T | 27 | 23353 | 0.11561684 | 58.5149004 | WT | FALSE | CAN | JSY13 | 1A | WT-Rep1A | GCG | C>T | A/T+1 | G/C-1 | CG>TA | CG>TA | AAAGAGCGC | CGCAATG | CGC | GCG | C | A | C>T | C>T_GCG | CG>TA_CGCAATG | 9 | TRUE | FALSE |
| 168_v.txt | 33310 | 33310-SNV-( | SNV | 1 | G | A | 33 | 23352 | 0.14131552 | 58.7161682 | WT | FALSE | CAN | JSY13 | 1A | WT-Rep1A | CGC | G>A | A/T+1 | G/C-1 | CG>TA | CG>TA | AAGAGCGCA | GCAATGG | GCG | GCG | C | T | C>T | C>T_GCG | CG>TA_GCAATGG | 9 | TRUE | FALSE |
| 168_v.txt | 33376 | 33376-SNV-( | SNV | 1 | G | A | 37 | 23355 | 0.15842432 | 51.4324324 | WT | FALSE | CAN | JSY13 | 1A | WT-Rep1A | TGT | G>A | A/T+1 | G/C-1 | CG>TA | CG>TA | GTGTTTGTG | GTGAAGC | ACA | ACA | C | T | C>T | C>T_ACA | CG>TA_GTGAAGC | 9 | TRUE | FALSE |
| 168_v.txt | 33412 | 33412-SNV-( | SNV | 1 | G | A | 24 | 23355 | 0.10276172 | 49.4583333 | WT | FALSE | CAN | JSY13 | 1A | WT-Rep1A | GGC | G>A | A/T+1 | G/C-1 | CG>TA | CG>TA | TGACCGGCT | GCTCATT | CCG | CCG | C | T | C>T | C>T_CCG | CG>TA_GCTCATT | 9 | TRUE | FALSE |
| 168_v.txt | 33436 | 33436-SNV-( | SNV | 1 | C | T | 8 | 23355 | 0.03425391 | 31.3751338 | WT | FALSE | CAN | JSY13 | 1A | WT-Rep1A | TCT | C>T | A/T+1 | G/C-1 | CG>TA | CG>TA | TCTCCTCTAT | CTATGTC | AGA | TCT | C | A | C>T | C>T_TCT | CG>TA_CTATGTC | 9 | TRUE | FALSE |
| 168_v.txt | 33474 | 33474-SNV-( | SNV | 1 | C | T | 37 | 23349 | 0.15846503 | 34.8918919 | WT | FALSE | CAN | JSY13 | 1A | WT-Rep1A | GCC | C>T | A/T+1 | G/C-1 | CG>TA | CG>TA | GCTATGCCTT | CCTTTTT | CGG | GCC | C | A | C>T | C>T_GCC | CG>TA_CCTTTTT | 9 | TRUE | FALSE |
| 168_v.txt | 31956 | 31956-SNV-( | SNV | 1 | G | C | 14 | 14348 | 0.09757458 | 37.6428571 | WT | FALSE | CAN | JSY13 | 1A | WT-Rep1A | TGT | G>C | G/C+1 | G/C-1 | CG>GC | CG>GC | ATAAATGTG | GTGGCCG | ACA | ACA | C | G | C>G | C>G_ACA | CG>GC_GTGGCCG | 9 | TRUE | FALSE |
| 168_v.txt | 31981 | 31981-SNV-( | SNV | 1 | G | C | 3 | 14355 | 0.02089864 | 58.3243794 | WT | FALSE | CAN | JSY13 | 1A | WT-Rep1A | GGC | G>C | G/C+1 | G/C-1 | CG>GC | CG>GC | AGCCGGGCA | GCATTAA | CCG | CCG | C | G | C>G | C>G_CCG | CG>GC_GCATTAA | 9 | TRUE | FALSE |
| 168_v.txt | 32027 | 32027-SNV-( | SNV | 1 | G | C | 60 | 14355 | 0.41797283 | 59.2166667 | WT | FALSE | CAN | JSY13 | 1A | WT-Rep1A | GGT | G>C | G/C+1 | G/C-1 | CG>GC | CG>GC | GCCACGGTA | GTATTTC | CCA | CCA | C | G | C>G | C>G_CCA | CG>GC_GTATTTC | 9 | TRUE | FALSE |
| 168_v.txt | 32038 | 32038-SNV-( | SNV | 1 | C | G | 5 | 14355 | 0.03483107 | 64 | WT | FALSE | CAN | JSY13 | 1A | WT-Rep1A | GCT | C>G | G/C+1 | G/C-1 | CG>GC | CG>GC | TCAAAGCTTC | CTTGCAT | CGA | GCT | C | C | C>G | C>G_GCT | CG>GC_CTTGCAT | 9 | TRUE | FALSE |
| 168_v.txt | 32058 | 32058-SNV-( | SNV | 1 | G | C | 4 | 14355 | 0.02786486 | 58.25 | WT | FALSE | CAN | JSY13 | 1A | WT-Rep1A | CGA | G>C | G/C+1 | G/C-1 | CG>GC | CG>GC | ATGTGCGAG | GAGATTG | GCT | GCT | C | G | C>G | C>G_GCT | CG>GC_GAGATTG | 9 | TRUE | FALSE |
| 168_v.txt | 32064 | 32064-SNV-( | SNV | 1 | G | C | 17 | 14355 | 0.11842564 | 61.5882353 | WT | FALSE | CAN | JSY13 | 1A | WT-Rep1A | TGA | G>C | G/C+1 | G/C-1 | CG>GC | CG>GC | GAGATTGAG | GAGATAA | ACT | ACT | C | G | C>G | C>G_ACT | CG>GC_GAGATAA | 9 | TRUE | FALSE |
| 168_v.txt | 32080 | 32080-SNV-( | SNV | 1 | C | G | 2 | 14355 | 0.01393243 | 39.4986716 | WT | FALSE | CAN | JSY13 | 1A | WT-Rep1A | GCA | C>G | G/C+1 | G/C-1 | CG>GC | CG>GC | ACCATGCAA | CAAAAAA | CGT | GCA | C | C | C>G | C>G_GCA | CG>GC_CAAAAAA | 9 | TRUE | FALSE |
| 168_v.txt | 32088 | 32088-SNV-( | SNV | 1 | C | G | 2 | 14355 | 0.01393243 | 64 | WT | FALSE | CAN | JSY13 | 1A | WT-Rep1A | GCC | C>G | G/C+1 | G/C-1 | CG>GC | CG>GC | AAAAAGCCT | CCTGCAA | CGG | GCC | C | C | C>G | C>G_GCC | CG>GC_CCTGCAA | 9 | TRUE | FALSE |
| 168_v.txt | 32089 | 32089-SNV-( | SNV | 1 | C | G | 18 | 14355 | 0.12539185 | 59.2781592 | WT | FALSE | CAN | JSY13 | 1A | WT-Rep1A | CCT | C>G | G/C+1 | G/C-1 | CG>GC | CG>GC | AAAAGCCTG | CTGCAAC | GGA | CCT | C | C | C>G | C>G_CCT | CG>GC_CTGCAAC | 9 | TRUE | FALSE |
| 168_v.txt | 32092 | 32092-SNV-( | SNV | 1 | C | G | 10 | 14355 | 0.06966214 | 58.8 | WT | FALSE | CAN | JSY13 | 1A | WT-Rep1A | GCA | C>G | G/C+1 | G/C-1 | CG>GC | CG>GC | AGCCTGCAA | CAACACC | CGT | GCA | C | C | C>G | C>G_GCA | CG>GC_CAACACC | 9 | TRUE | FALSE |
| 168_v.txt | 32153 | 32153-SNV-( | SNV | 1 | G | C | 10 | 14355 | 0.06966214 | 47.9 | WT | FALSE | CAN | JSY13 | 1A | WT-Rep1A | TGT | G>C | G/C+1 | G/C-1 | CG>GC | CG>GC | CTCCATGTA | GTAAGCC | ACA | ACA | C | G | C>G | C>G_ACA | CG>GC_GTAAGCC | 9 | TRUE | FALSE |
| 168_v.txt | 32199 | 32199-SNV-( | SNV | 1 | G | C | 35 | 14341 | 0.24405551 | 43.3714286 | WT | FALSE | CAN | JSY13 | 1A | WT-Rep1A | TGG | G>C | G/C+1 | G/C-1 | CG>GC | CG>GC | ATGTATGGA | GGAACAC | ACC | ACC | C | G | C>G | C>G_ACC | CG>GC_GGAACAC | 9 | TRUE | FALSE |
| 168_v.txt | 32223 | 32223-SNV-( | SNV | 1 | G | C | 20 | 14341 | 0.13946029 | 37.8 | WT | FALSE | CAN | JSY13 | 1A | WT-Rep1A | TGA | G>C | G/C+1 | G/C-1 | CG>GC | CG>GC | GTCCTTGAC | GACAGGA | ACT | ACT | C | G | C>G | C>G_ACT | CG>GC_GACAGGA | 9 | TRUE | FALSE |
| 168_v.txt | 32271 | 32271-SNV-( | SNV | 1 | C | G | 24 | 46483 | 0.05163178 | 36.875 | WT | FALSE | CAN | JSY13 | 1A | WT-Rep1A | ACG | C>G | G/C+1 | G/C-1 | CG>GC | CG>GC | AAAATACGG | CGGGAAC | TGC | ACG | C | C | C>G | C>G_ACG | CG>GC_CGGGAAC | 9 | TRUE | FALSE |
| 168_v.txt | 32272 | 32272-SNV-( | SNV | 1 | G | C | 26 | 46483 | 0.05593443 | 36.65386 | WT | FALSE | CAN | JSY13 | 1A | WT-Rep1A | CGG | G>C | G/C+1 | G/C-1 | CG>GC | CG>GC | AAATACGGG | GGGAACC | GCC | GCC | C | G | C>G | C>G_GCC | CG>GC_GGGAACC | 9 | TRUE | FALSE |
| 168_v.txt | 32278 | 32278-SNV-( | SNV | 1 | C | G | 34 | 46483 | 0.07314502 | 36.0882353 | WT | FALSE | CAN | JSY13 | 1A | WT-Rep1A | CCA | C>G | G/C+1 | G/C-1 | CG>GC | CG>GC | GGGAACCAA | CAACGTA | GGT | CCA | C | C | C>G | C>G_CCA | CG>GC_CAACGTA | 9 | TRUE | FALSE |
| 168_v.txt | 32282 | 32282-SNV-( | SNV | 1 | G | C | 30 | 46483 | 0.06453972 | 40.1327964 | WT | FALSE | CAN | JSY13 | 1A | WT-Rep1A | CGT | G>C | G/C+1 | G/C-1 | CG>GC | CG>GC | ACCAACGTA | GTAAATA | GCA | GCA | C | G | C>G | C>G_GCA | CG>GC_GTAAATA | 9 | TRUE | FALSE |
| 168_v.txt | 32292 | 32292-SNV-( | SNV | 1 | G | C | 68 | 46483 | 0.14629004 | 56.7058824 | WT | FALSE | CAN | JSY13 | 1A | WT-Rep1A | TGA | G>C | G/C+1 | G/C-1 | CG>GC | CG>GC | ATATTTGAAT | GAATTTG | ACT | ACT | C | G | C>G | C>G_ACT | CG>GC_GAATTTG | 9 | TRUE | FALSE |
| 168_v.txt | 32304 | 32304-SNV-( | SNV | 1 | G | C | 9 | 46483 | 0.01936192 | 55.2222222 | WT | FALSE | CAN | JSY13 | 1A | WT-Rep1A | AGA | G>C | G/C+1 | G/C-1 | CG>GC | CG>GC | GCGGCAGAA | GAAATAA | TCT | TCT | C | G | C>G | C>G_TCT | CG>GC_GAAATAA | 9 | TRUE | FALSE |
| 168_v.txt | 32316 | 32316-SNV-( | SNV | 1 | G | C | 17 | 46483 | 0.03657251 | 52.5275414 | WT | FALSE | CAN | JSY13 | 1A | WT-Rep1A | TGT | G>C | G/C+1 | G/C-1 | CG>GC | CG>GC | ATGGTTGTT | GTTAAGA | ACA | ACA | C | G | C>G | C>G_ACA | CG>GC_GTTAAGA | 9 | TRUE | FALSE |
| 168_v.txt | 32385 | 32385-SNV-( | SNV | 1 | G | C | 41 | 46482 | 0.08820619 | 59.649047 | WT | FALSE | CAN | JSY13 | 1A | WT-Rep1A | GGG | G>C | G/C+1 | G/C-1 | CG>GC | CG>GC | ATAAAGGGA | GGAGAAG | CCC | CCC | C | G | C>G | C>G_CCC | CG>GC_GGAGAAG | 9 | TRUE | FALSE |
| 168_v.txt | 32432 | 32432-SNV-( | SNV | 1 | G | C | 42 | 46483 | 0.09035561 | 53.0476191 | WT | FALSE | CAN | JSY13 | 1A | WT-Rep1A | TGT | G>C | G/C+1 | G/C-1 | CG>GC | CG>GC | GTCATTGTAT | GTATGGA | ACA | ACA | C | G | C>G | C>G_ACA | CG>GC_GTATGGA | 9 | TRUE | FALSE |
| 168_v.txt | 32471 | 32471-SNV-( | SNV | 1 | G | C | 6 | 46483 | 0.01290795 | 59.6531572 | WT | FALSE | CAN | JSY13 | 1A | WT-Rep1A | TGT | G>C | G/C+1 | G/C-1 | CG>GC | CG>GC | GCCAATGTA | GTAGAAG | ACA | ACA | C | G | C>G | C>G_ACA | CG>GC_GTAGAAG | 9 | TRUE | FALSE |
| 168_v.txt | 32474 | 32474-SNV-( | SNV | 1 | G | C | 1 | 46483 | 2.15E-03 | 63.9934355 | WT | FALSE | CAN | JSY13 | 1A | WT-Rep1A | AGA | G>C | G/C+1 | G/C-1 | CG>GC | CG>GC | AATGTAGAA | GAAGGTT | TCT | TCT | C | G | C>G | C>G_TCT | CG>GC_GAAGGTT | 9 | TRUE | FALSE |
| 168_v.txt | 32487 | 32487-SNV-( | SNV | 1 | C | G | 42 | 46483 | 0.09035561 | 48.0719816 | WT | FALSE | CAN | JSY13 | 1A | WT-Rep1A | ACG | C>G | G/C+1 | G/C-1 | CG>GC | CG>GC | AAGATACGG | CGGAAAA | TGC | ACG | C | C | C>G | C>G_ACG | CG>GC_CGGAAAA | 9 | TRUE | FALSE |
| 168_v.txt | 32488 | 32488-SNV-( | SNV | 1 | G | C | 82 | 46483 | 0.17640858 | 46.8292683 | WT | FALSE | CAN | JSY13 | 1A | WT-Rep1A | CGG | G>C | G/C+1 | G/C-1 | CG>GC | CG>GC | AGATACGGA | GGAAAAC | GCC | GCC | C | G | C>G | C>G_GCC | CG>GC_GGAAAAC | 9 | TRUE | FALSE |
| 168_v.txt | 32494 | 32494-SNV-( | SNV | 1 | C | G | 1 | 46483 | 2.15E-03 | 37.9998778 | WT | FALSE | CAN | JSY13 | 1A | WT-Rep1A | ACA | C>G | G/C+1 | G/C-1 | CG>GC | CG>GC | GGAAAACAA | CAACTTT | TGT | ACA | C | C | C>G | C>G_ACA | CG>GC_CAACTTT | 9 | TRUE | FALSE |
| 168_v.txt | 32509 | 32509-SNV-( | SNV | 1 | C | G | 4 | 46483 | 8.61E-03 | 44.999487 | WT | FALSE | CAN | JSY13 | 1A | WT-Rep1A | GCT | C>G | G/C+1 | G/C-1 | CG>GC | CG>GC | TGATGGCTC | CTCTTGG | CGA | GCT | C | C | C>G | C>G_GCT | CG>GC_CTCTTGG | 9 | TRUE | FALSE |
| 168_v.txt | 32520 | 32520-SNV-( | SNV | 1 | G | C | 90 | 46483 | 0.19361917 | 37.5111111 | WT | FALSE | CAN | JSY13 | 1A | WT-Rep1A | GGA | G>C | G/C+1 | G/C-1 | CG>GC | CG>GC | GGAACGGAT | GATTTTC | CCT | CCT | C | G | C>G | C>G_CCT | CG>GC_GATTTTC | 9 | TRUE | FALSE |
| 168_v.txt | 32529 | 32529-SNV-( | SNV | 1 | G | C | 21 | 46483 | 0.04517781 | 43.6190476 | WT | FALSE | CAN | JSY13 | 1A | WT-Rep1A | GGG | G>C | G/C+1 | G/C-1 | CG>GC | CG>GC | TTTCTGGGG | GGGTTTG | CCC | CCC | C | G | C>G | C>G_CCC | CG>GC_GGGTTTG | 9 | TRUE | FALSE |
| 168_v.txt | 32536 | 32536-SNV-( | SNV | 1 | C | G | 3 | 46481 | 6.45E-03 | 37.6658416 | WT | FALSE | CAN | JSY13 | 1A | WT-Rep1A | GCA | C>G | G/C+1 | G/C-1 | CG>GC | CG>GC | GGTTTGCAG | CAGCTTC | CGT | GCA | C | C | C>G | C>G_GCA | CG>GC_CAGCTTC | 9 | TRUE | FALSE |
| 168_v.txt | 32539 | 32539-SNV-( | SNV | 1 | C | G | 29 | 46481 | 0.06239109 | 37.6896552 | WT | FALSE | CAN | JSY13 | 1A | WT-Rep1A | GCT | C>G | G/C+1 | G/C-1 | CG>GC | CG>GC | TTGCAGCTTC | CTTCACC | CGA | GCT | C | C | C>G | C>G_GCT | CG>GC_CTTCACC | 9 | TRUE | FALSE |
| 168_v.txt | 32566 | 32566-SNV-( | SNV | 1 | C | G | 8 | 7614 | 0.10506961 | 37.875 | WT | FALSE | CAN | JSY13 | 1A | WT-Rep1A | TCA | C>G | G/C+1 | G/C-1 | CG>GC | CG>GC | CTAGTTCAG | CAGTACC | AGT | TCA | C | C | C>G | C>G_TCA | CG>GC_CAGTACC | 9 | TRUE | FALSE |
| 168_v.txt | 32685 | 32685-SNV-( | SNV | 1 | C | G | 9 | 7614 | 0.11820331 | 61.8888889 | WT | FALSE | CAN | JSY13 | 1A | WT-Rep1A | ACG | C>G | G/C+1 | G/C-1 | CG>GC | CG>GC | CAATAACGG | CGGAATC | TGC | ACG | C | C | C>G | C>G_ACG | CG>GC_CGGAATC | 9 | TRUE | FALSE |
| 168_v.txt | 32749 | 32749-SNV-( | SNV | 1 | C | G | 11 | 7622 | 0.14431908 | 51.3609238 | WT | FALSE | CAN | JSY13 | 1A | WT-Rep1A | CCG | C>G | G/C+1 | G/C-1 | CG>GC | CG>GC | GAAACCCGA | CGATAAT | GGC | CCG | C | C | C>G | C>G_CCG | CG>GC_CGATAAT | 9 | TRUE | FALSE |
| 168_v.txt | 32776 | 32776-SNV-( | SNV | 1 | C | G | 4 | 7622 | 0.05247966 | 36.5 | WT | FALSE | CAN | JSY13 | 1A | WT-Rep1A | GCG | C>G | G/C+1 | G/C-1 | CG>GC | CG>GC | TGGAAGCGA | CGACCCA | CGC | GCG | C | C | C>G | C>G_GCG | CG>GC_CGACCCA | 9 | TRUE | FALSE |
| 168_v.txt | 32786 | 32786-SNV-( | SNV | 1 | C | G | 22 | 7622 | 0.28863815 | 47.3157078 | WT | FALSE | CAN | JSY13 | 1A | WT-Rep1A | ACT | C>G | G/C+1 | G/C-1 | CG>GC | CG>GC | CCAGAACTC | CTCGAAT | TGA | ACT | C | C | C>G | C>G_ACT | CG>GC_CTCGAAT | 9 | TRUE | FALSE |
| 168_v.txt | 32797 | 32797-SNV-( | SNV | 1 | C | G | 13 | 7622 | 0.17055891 | 45.7773219 | WT | FALSE | CAN | JSY13 | 1A | WT-Rep1A | CCG | C>G | G/C+1 | G/C-1 | CG>GC | CG>GC | ATTCACCGTA | CGTAATA | GGC | CCG | C | C | C>G | C>G_CCG | CG>GC_CGTAATA | 9 | TRUE | FALSE |
| 168_v.txt | 32798 | 32798-SNV-( | SNV | 1 | G | C | 12 | 7622 | 0.15743899 | 46.3333333 | WT | FALSE | CAN | JSY13 | 1A | WT-Rep1A | CGT | G>C | G/C+1 | G/C-1 | CG>GC | CG>GC | TTCACCGTA | GTAATAT | GCA | GCA | C | G | C>G | C>G_GCA | CG>GC_GTAATAT | 9 | TRUE | FALSE |
| 168_v.txt | 32819 | 32819-SNV-( | SNV | 1 | G | C | 5 | 7622 | 0.06559958 | 48.4 | WT | FALSE | CAN | JSY13 | 1A | WT-Rep1A | AGT | G>C | G/C+1 | G/C-1 | CG>GC | CG>GC | GAACAAGTT | GTTCATT | TCA | TCA | C | G | C>G | C>G_TCA | CG>GC_GTTCATT | 9 | TRUE | FALSE |
| 168_v.txt | 32876 | 32876-SNV-( | SNV | 1 | G | C | 67 | 22839 | 0.29335785 | 37.6567164 | WT | FALSE | CAN | JSY13 | 1A | WT-Rep1A | TGT | G>C | G/C+1 | G/C-1 | CG>GC | CG>GC | AACTTTGTAC | GTACGTC | ACA | ACA | C | G | C>G | C>G_ACA | CG>GC_GTACGTC | 9 | TRUE | FALSE |
| 168_v.txt | 32900 | 32900-SNV-( | SNV | 1 | G | C | 2 | 22850 | 8.75E-03 | 37.9953795 | WT | FALSE | CAN | JSY13 | 1A | WT-Rep1A | GGC | G>C | G/C+1 | G/C-1 | CG>GC | CG>GC | GACTTGGCC | GCCAACT | CCG | CCG | C | G | C>G | C>G_CCG | CG>GC_GCCAACT | 9 | TRUE | FALSE |
| 168_v.txt | 32917 | 32917-SNV-( | SNV | 1 | C | G | 8 | 22850 | 0.03501094 | 55.25 | WT | FALSE | CAN | JSY13 | 1A | WT-Rep1A | TCC | C>G | G/C+1 | G/C-1 | CG>GC | CG>GC | TAAGTTCCAC | CCAGGGC | AGG | TCC | C | C | C>G | C>G_TCC | CG>GC_CCAGGGC | 9 | TRUE | FALSE |
| 168_v.txt | 32935 | 32935-SNV-( | SNV | 1 | C | G | 3 | 22886 | 0.01310845 | 40.9997512 | WT | FALSE | CAN | JSY13 | 1A | WT-Rep1A | GCC | C>G | G/C+1 | G/C-1 | CG>GC | CG>GC | TGATTGCCCA | CCCAAGA | CGG | GCC | C | C | C>G | C>G_GCC | CG>GC_CCCAAGA | 9 | TRUE | FALSE |
| 168_v.txt | 32936 | 32936-SNV-( | SNV | 1 | C | G | 15 | 22886 | 0.06554225 | 45.5099213 | WT | FALSE | CAN | JSY13 | 1A | WT-Rep1A | CCC | C>G | G/C+1 | G/C-1 | CG>GC | CG>GC | GATTGCCCA | CCAAGAA | GGG | CCC | C | C | C>G | C>G_CCC | CG>GC_CCAAGAA | 9 | TRUE | FALSE |
| 168_v.txt | 32945 | 32945-SNV-( | SNV | 1 | C | G | 5 | 22886 | 0.02184742 | 63.9919823 | WT | FALSE | CAN | JSY13 | 1A | WT-Rep1A | ACC | C>G | G/C+1 | G/C-1 | CG>GC | CG>GC | AGAAAACCA | CCAATAC | TGG | ACC | C | C | C>G | C>G_ACC | CG>GC_CCAATAC | 9 | TRUE | FALSE |
| 168_v.txt | 32971 | 32971-SNV-( | SNV | 1 | C | G | 25 | 22886 | 0.10923709 | 63.7947587 | WT | FALSE | CAN | JSY13 | 1A | WT-Rep1A | CCA | C>G | G/C+1 | G/C-1 | CG>GC | CG>GC | CCGCACCAA | CAAATGC | GGT | CCA | C | C | C>G | C>G_CCA | CG>GC_CAAATGC | 9 | TRUE | FALSE |
| 168_v.txt | 32997 | 32997-SNV-( | SNV | 1 | G | C | 90 | 22886 | 0.39325352 | 61.4333333 | WT | FALSE | CAN | JSY13 | 1A | WT-Rep1A | TGA | G>C | G/C+1 | G/C-1 | CG>GC | CG>GC | CTTTGTGAG | GAGAAAA | ACT | ACT | C | G | C>G | C>G_ACT | CG>GC_GAGAAAA | 9 | TRUE | FALSE |
| 168_v.txt | 33024 | 33024-SNV-( | SNV | 1 | G | C | 3 | 22886 | 0.01310845 | 64 | WT | FALSE | CAN | JSY13 | 1A | WT-Rep1A | AGG | G>C | G/C+1 | G/C-1 | CG>GC | CG>GC | GTAACAGGG | GGGATGA | TCC | TCC | C | G | C>G | C>G_TCC | CG>GC_GGGATGA | 9 | TRUE | FALSE |
| 168_v.txt | 33037 | 33037-SNV-( | SNV | 1 | C | G | 17 | 22886 | 0.07428122 | 59.3529412 | WT | FALSE | CAN | JSY13 | 1A | WT-Rep1A | GCC | C>G | G/C+1 | G/C-1 | CG>GC | CG>GC | ATGTAGCCA | CCATTTC | CGG | GCC | C | C | C>G | C>G_GCC | CG>GC_CCATTTC | 9 | TRUE | FALSE |
| 168_v.txt | 33056 | 33056-SNV-( | SNV | 1 | C | G | 2 | 22886 | 8.74E-03 | 40 | WT | FALSE | CAN | JSY13 | 1A | WT-Rep1A | GCG | C>G | G/C+1 | G/C-1 | CG>GC | CG>GC | GGACTGCGT | CGTGACA | CGC | GCG | C | C | C>G | C>G_GCG | CG>GC_CGTGACA | 9 | TRUE | FALSE |
| 168_v.txt | 33057 | 33057-SNV-( | SNV | 1 | G | C | 9 | 22886 | 0.03932535 | 61.1106744 | WT | FALSE | CAN | JSY13 | 1A | WT-Rep1A | CGT | G>C | G/C+1 | G/C-1 | CG>GC | CG>GC | GACTGCGTG | GTGACAG | GCA | GCA | C | G | C>G | C>G_GCA | CG>GC_GTGACAG | 9 | TRUE | FALSE |
| 168_v.txt | 33070 | 33070-SNV-( | SNV | 1 | C | G | 10 | 22886 | 0.04369484 | 55.8 | WT | FALSE | CAN | JSY13 | 1A | WT-Rep1A | GCC | C>G | G/C+1 | G/C-1 | CG>GC | CG>GC | AATATGCCA | CCAAAGA | CGG | GCC | C | C | C>G | C>G_GCC | CG>GC_CCAAAGA | 9 | TRUE | FALSE |
| 168_v.txt | 33079 | 33079-SNV-( | SNV | 1 | C | G | 17 | 22886 | 0.07428122 | 50.1764706 | WT | FALSE | CAN | JSY13 | 1A | WT-Rep1A | CCC | C>G | G/C+1 | G/C-1 | CG>GC | CG>GC | AAGAACCCA | CCATAAA | GGG | CCC | C | C | C>G | C>G_CCC | CG>GC_CCATAAA | 9 | TRUE | FALSE |
| 168_v.txt | 33093 | 33093-SNV-( | SNV | 1 | G | C | 11 | 22886 | 0.04806432 | 41.271338 | WT | FALSE | CAN | JSY13 | 1A | WT-Rep1A | TGA | G>C | G/C+1 | G/C-1 | CG>GC | CG>GC | AAATATGAT | GATATAA | ACT | ACT | C | G | C>G | C>G_ACT | CG>GC_GATATAA | 9 | TRUE | FALSE |
| 168_v.txt | 33114 | 33114-SNV-( | SNV | 1 | C | G | 6 | 22886 | 0.0262169 | 46.6654545 | WT | FALSE | CAN | JSY13 | 1A | WT-Rep1A | GCC | C>G | G/C+1 | G/C-1 | CG>GC | CG>GC | ACTGGGCCG | CCGGCGT | CGG | GCC | C | C | C>G | C>G_GCC | CG>GC_CCGGCGT | 9 | TRUE | FALSE |
| 168_v.txt | 33115 | 33115-SNV-( | SNV | 1 | C | G | 3 | 22886 | 0.01310845 | 37.3333333 | WT | FALSE | CAN | JSY13 | 1A | WT-Rep1A | CCG | C>G | G/C+1 | G/C-1 | CG>GC | CG>GC | CTGGGCCGG | CGGCGTT | GGC | CCG | C | C | C>G | C>G_CCG | CG>GC_CGGCGTT | 9 | TRUE | FALSE |
| 168_v.txt | 33118 | 33118-SNV-( | SNV | 1 | C | G | 31 | 22886 | 0.13545399 | 41.8064516 | WT | FALSE | CAN | JSY13 | 1A | WT-Rep1A | GCG | C>G | G/C+1 | G/C-1 | CG>GC | CG>GC | GGCCGGCGT | CGTTGGT | CGC | GCG | C | C | C>G | C>G_GCG | CG>GC_CGTTGGT | 9 | TRUE | FALSE |
| 168_v.txt | 33154 | 33154-SNV-( | SNV | 1 | C | G | 22 | 22886 | 0.09612864 | 37.9978816 | WT | FALSE | CAN | JSY13 | 1A | WT-Rep1A | CCT | C>G | G/C+1 | G/C-1 | CG>GC | CG>GC | AAAGACCTG | CTGTACC | GGA | CCT | C | C | C>G | C>G_CCT | CG>GC_CTGTACC | 9 | TRUE | FALSE |
| 168_v.txt | 33156 | 33156-SNV-( | SNV | 1 | G | C | 12 | 22885 | 0.05243609 | 38 | WT | FALSE | CAN | JSY13 | 1A | WT-Rep1A | TGT | G>C | G/C+1 | G/C-1 | CG>GC | CG>GC | AGACCTGTA | GTACCAA | ACA | ACA | C | G | C>G | C>G_ACA | CG>GC_GTACCAA | 9 | TRUE | FALSE |
| 168_v.txt | 33160 | 33160-SNV-( | SNV | 1 | C | G | 16 | 22885 | 0.06991479 | 37.6875 | WT | FALSE | CAN | JSY13 | 1A | WT-Rep1A | CCA | C>G | G/C+1 | G/C-1 | CG>GC | CG>GC | CTGTACCAAT | CAATAGT | GGT | CCA | C | C | C>G | C>G_CCA | CG>GC_CAATAGT | 9 | TRUE | FALSE |
| 168_v.txt | 33168 | 33168-SNV-( | SNV | 1 | C | G | 35 | 22847 | 0.15319298 | 36.9130124 | WT | FALSE | CAN | JSY13 | 1A | WT-Rep1A | ACC | C>G | G/C+1 | G/C-1 | CG>GC | CG>GC | ATAGTACCA | CCACCAA | TGG | ACC | C | C | C>G | C>G_ACC | CG>GC_CCACCAA | 9 | TRUE | FALSE |
| 168_v.txt | 33169 | 33169-SNV-( | SNV | 1 | C | G | 3 | 22847 | 0.01313083 | 37.6666663 | WT | FALSE | CAN | JSY13 | 1A | WT-Rep1A | CCA | C>G | G/C+1 | G/C-1 | CG>GC | CG>GC | TAGTACCAC | CACCAAG | GGT | CCA | C | C | C>G | C>G_CCA | CG>GC_CACCAAG | 9 | TRUE | FALSE |
| 168_v.txt | 33172 | 33172-SNV-( | SNV | 1 | C | G | 32 | 22847 | 0.14006215 | 37.4993992 | WT | FALSE | CAN | JSY13 | 1A | WT-Rep1A | CCA | C>G | G/C+1 | G/C-1 | CG>GC | CG>GC | TACCACCAA | CAAGGGC | GGT | CCA | C | C | C>G | C>G_CCA | CG>GC_CAAGGGC | 9 | TRUE | FALSE |

| | | | | | | | | | | | | | | | | | | | | | | | | | | | | | | | | | | |
|---|---|---|---|---|---|---|---|---|---|---|---|---|---|---|---|---|---|---|---|---|---|---|---|---|---|---|---|---|---|---|---|---|---|---|
| 168_v.txt | 33178 | 33178-SNV-( | SNV | 1 | C | G | 36 | 22843 | 0.15759751 | 36.541014 | WT | FALSE | CAN | JSY13 | 1A | WT-Rep1A | GCA | C>G | G/C+1 | G/C-1 | CG>GC | CG>GC | CAAGGGCAA | CAATCAT | CGT | GCA | C | C | C>G | C>G_GCA | CG>GC_CAATCAT | 9 | TRUE | FALSE |
| 168_v.txt | 33193 | 33193-SNV-( | SNV | 1 | G | C | 21 | 23357 | 0.08990881 | 36.9988741 | WT | FALSE | CAN | JSY13 | 1A | WT-Rep1A | TGT | G>C | G/C+1 | G/C-1 | CG>GC | CG>GC | CAATATGTC | GTCTTTG | ACA | ACA | C | G | C>G | C>G_ACA | CG>GC_GTCTTTG | 9 | TRUE | FALSE |
| 168_v.txt | 33205 | 33205-SNV-( | SNV | 1 | G | C | 26 | 23355 | 0.1113252 | 39.035616 | WT | FALSE | CAN | JSY13 | 1A | WT-Rep1A | AGC | G>C | G/C+1 | G/C-1 | CG>GC | CG>GC | GCTTAAGCT | GCTCTCT | TCG | TCG | C | G | C>G | C>G_TCG | CG>GC_GCTCTCT | 9 | TRUE | FALSE |
| 168_v.txt | 33210 | 33210-SNV-( | SNV | 1 | C | G | 49 | 23354 | 0.20981417 | 40.6506523 | WT | FALSE | CAN | JSY13 | 1A | WT-Rep1A | TCT | C>G | G/C+1 | G/C-1 | CG>GC | CG>GC | AGCTCTCTCT | CTCTTCA | AGA | TCT | C | C | C>G | C>G_TCT | CG>GC_CTCTTCA | 9 | TRUE | FALSE |
| 168_v.txt | 33284 | 33284-SNV-( | SNV | 1 | G | C | 5 | 23355 | 0.02140869 | 48.3813238 | WT | FALSE | CAN | JSY13 | 1A | WT-Rep1A | CGC | G>C | G/C+1 | G/C-1 | CG>GC | CG>GC | ATCCTCGCC | GCCATTT | GCG | GCG | C | G | C>G | C>G_GCG | CG>GC_GCCATTT | 9 | TRUE | FALSE |
| 168_v.txt | 33286 | 33286-SNV-( | SNV | 1 | C | G | 1 | 23354 | 4.28E-03 | 63.9997212 | WT | FALSE | CAN | JSY13 | 1A | WT-Rep1A | CCA | C>G | G/C+1 | G/C-1 | CG>GC | CG>GC | CCTCGCCATT | CATTTAC | GGT | CCA | C | C | C>G | C>G_CCA | CG>GC_CATTTAC | 9 | TRUE | FALSE |
| 168_v.txt | 33309 | 33309-SNV-( | SNV | 1 | C | G | 2 | 23353 | 8.56E-03 | 37.4991489 | WT | FALSE | CAN | JSY13 | 1A | WT-Rep1A | GCG | C>G | G/C+1 | G/C-1 | CG>GC | CG>GC | AAAGAGCGC | CGCAATG | CGC | GCG | C | C | C>G | C>G_GCG | CG>GC_CGCAATG | 9 | TRUE | FALSE |
| 168_v.txt | 33310 | 33310-SNV-( | SNV | 1 | G | C | 3 | 23352 | 0.01284687 | 24.6664478 | WT | FALSE | CAN | JSY13 | 1A | WT-Rep1A | CGC | G>C | G/C+1 | G/C-1 | CG>GC | CG>GC | AAGAGCGCA | GCAATGG | GCG | GCG | C | G | C>G | C>G_GCG | CG>GC_GCAATGG | 9 | TRUE | FALSE |
| 168_v.txt | 33354 | 33354-SNV-( | SNV | 1 | G | C | 20 | 23355 | 0.08563477 | 54.9 | WT | FALSE | CAN | JSY13 | 1A | WT-Rep1A | TGA | G>C | G/C+1 | G/C-1 | CG>GC | CG>GC | GGTATTGAC | GACCCAC | ACT | ACT | C | G | C>G | C>G_ACT | CG>GC_GACCCAC | 9 | TRUE | FALSE |
| 168_v.txt | 33419 | 33419-SNV-( | SNV | 1 | G | C | 22 | 23355 | 0.09419825 | 44.5909091 | WT | FALSE | CAN | JSY13 | 1A | WT-Rep1A | TGT | G>C | G/C+1 | G/C-1 | CG>GC | CG>GC | CTCATTGTAC | GTACATA | ACA | ACA | C | G | C>G | C>G_ACA | CG>GC_GTACATA | 9 | TRUE | FALSE |
| 168_v.txt | 33456 | 33456-SNV-( | SNV | 1 | G | C | 30 | 23351 | 0.12847416 | 37.8333333 | WT | FALSE | CAN | JSY13 | 1A | WT-Rep1A | TGA | G>C | G/C+1 | G/C-1 | CG>GC | CG>GC | TCTTTTGAAT | GAATTTG | ACT | ACT | C | G | C>G | C>G_ACT | CG>GC_GAATTTG | 9 | TRUE | FALSE |
| 168_v.txt | 31798 | 31798-SNV-( | SNV | 1 | C | A | 13 | 3138 | 0.41427661 | 63.6153846 | WT | FALSE | CAN | JSY13 | 1A | WT-Rep1A | CCA | C>A | A/T+1 | G/C-1 | CG>AT | CG>AT | CATCTCCAAT | CAATCTT | GGT | CCA | C | T | C>A | C>A_CCA | CG>AT_CAATCTT | 9 | TRUE | FALSE |
| 168_v.txt | 31869 | 31869-SNV-( | SNV | 1 | G | T | 44 | 3138 | 1.40216699 | 46.4545455 | WT | FALSE | CAN | JSY13 | 1A | WT-Rep1A | AGA | G>T | A/T+1 | G/C-1 | CG>AT | CG>AT | AAAATAGAG | GAGATAT | TCT | TCT | C | A | C>A | C>A_TCT | CG>AT_GAGATAT | 9 | TRUE | FALSE |
| 168_v.txt | 31930 | 31930-SNV-( | SNV | 1 | G | T | 6 | 3127 | 0.1918772 | 38 | WT | FALSE | CAN | JSY13 | 1A | WT-Rep1A | TGA | G>T | A/T+1 | G/C-1 | CG>AT | CG>AT | AACCTTGAA | GAATAAT | ACT | ACT | C | A | C>A | C>A_ACT | CG>AT_GAATAAT | 9 | TRUE | FALSE |
| 168_v.txt | 31980 | 31980-SNV-( | SNV | 1 | G | T | 9 | 14354 | 0.06270029 | 46.7777778 | WT | FALSE | CAN | JSY13 | 1A | WT-Rep1A | GGG | G>T | A/T+1 | G/C-1 | CG>AT | CG>AT | AAGCCGGGC | GGCATTA | CCC | CCC | C | A | C>A | C>A_CCC | CG>AT_GGCATTA | 9 | TRUE | FALSE |
| 168_v.txt | 32008 | 32008-SNV-( | SNV | 1 | C | A | 94 | 14355 | 0.6548241 | 56.4574468 | WT | FALSE | CAN | JSY13 | 1A | WT-Rep1A | TCG | C>A | A/T+1 | G/C-1 | CG>AT | CG>AT | GTAACTCGT | CGTCACG | AGC | TCG | C | T | C>A | C>A_TCG | CG>AT_CGTCACG | 9 | TRUE | FALSE |
| 168_v.txt | 32058 | 32058-SNV-( | SNV | 1 | G | T | 14 | 14355 | 0.09752699 | 55.7142857 | WT | FALSE | CAN | JSY13 | 1A | WT-Rep1A | CGA | G>T | A/T+1 | G/C-1 | CG>AT | CG>AT | ATGTGCGAG | GAGATTG | GCT | GCT | C | A | C>A | C>A_GCT | CG>AT_GAGATTG | 9 | TRUE | FALSE |
| 168_v.txt | 32064 | 32064-SNV-( | SNV | 1 | G | T | 14 | 14355 | 0.09752699 | 62.7857143 | WT | FALSE | CAN | JSY13 | 1A | WT-Rep1A | TGA | G>T | A/T+1 | G/C-1 | CG>AT | CG>AT | GAGATTGAG | GAGATAA | ACT | ACT | C | A | C>A | C>A_ACT | CG>AT_GAGATAA | 9 | TRUE | FALSE |
| 168_v.txt | 32088 | 32088-SNV-( | SNV | 1 | C | A | 28 | 14355 | 0.19505399 | 60.0714286 | WT | FALSE | CAN | JSY13 | 1A | WT-Rep1A | GCC | C>A | A/T+1 | G/C-1 | CG>AT | CG>AT | AAAAAGCCT | CCTGCAA | CGG | GCC | C | T | C>A | C>A_GCC | CG>AT_CCTGCAA | 9 | TRUE | FALSE |
| 168_v.txt | 32089 | 32089-SNV-( | SNV | 1 | C | A | 11 | 14355 | 0.07662835 | 41.2761636 | WT | FALSE | CAN | JSY13 | 1A | WT-Rep1A | CCT | C>A | A/T+1 | G/C-1 | CG>AT | CG>AT | AAAAGCCTG | CTGCAAC | GGA | CCT | C | T | C>A | C>A_CCT | CG>AT_CTGCAAC | 9 | TRUE | FALSE |
| 168_v.txt | 32095 | 32095-SNV-( | SNV | 1 | C | A | 84 | 14355 | 0.58516197 | 59.0595238 | WT | FALSE | CAN | JSY13 | 1A | WT-Rep1A | ACA | C>A | A/T+1 | G/C-1 | CG>AT | CG>AT | CTGCAACAC | CACCAGT | TGT | ACA | C | T | C>A | C>A_ACA | CG>AT_CACCAGT | 9 | TRUE | FALSE |
| 168_v.txt | 32119 | 32119-SNV-( | SNV | 1 | C | A | 16 | 14355 | 0.11145942 | 57.1875 | WT | FALSE | CAN | JSY13 | 1A | WT-Rep1A | TCG | C>A | A/T+1 | G/C-1 | CG>AT | CG>AT | GCCATTCGA | CGAAAAC | AGC | TCG | C | T | C>A | C>A_TCG | CG>AT_CGAAAAC | 9 | TRUE | FALSE |
| 168_v.txt | 32149 | 32149-SNV-( | SNV | 1 | C | A | 44 | 14355 | 0.30651341 | 50.1590909 | WT | FALSE | CAN | JSY13 | 1A | WT-Rep1A | TCC | C>A | A/T+1 | G/C-1 | CG>AT | CG>AT | ATGTCTCCAT | CCATGTA | AGG | TCC | C | T | C>A | C>A_TCC | CG>AT_CCATGTA | 9 | TRUE | FALSE |
| 168_v.txt | 32188 | 32188-SNV-( | SNV | 1 | C | A | 30 | 14355 | 0.20898642 | 46.6333333 | WT | FALSE | CAN | JSY13 | 1A | WT-Rep1A | ACT | C>A | A/T+1 | G/C-1 | CG>AT | CG>AT | CGAAAACTG | CTGCAAT | TGA | ACT | C | T | C>A | C>A_ACT | CG>AT_CTGCAAT | 9 | TRUE | FALSE |
| 168_v.txt | 32199 | 32199-SNV-( | SNV | 1 | G | T | 12 | 14341 | 0.08367617 | 46.5 | WT | FALSE | CAN | JSY13 | 1A | WT-Rep1A | TGG | G>T | A/T+1 | G/C-1 | CG>AT | CG>AT | ATGTATGGA | GGAACAC | ACC | ACC | C | A | C>A | C>A_ACC | CG>AT_GGAACAC | 9 | TRUE | FALSE |
| 168_v.txt | 32203 | 32203-SNV-( | SNV | 1 | C | A | 18 | 14341 | 0.12551426 | 43.3333333 | WT | FALSE | CAN | JSY13 | 1A | WT-Rep1A | ACA | C>A | A/T+1 | G/C-1 | CG>AT | CG>AT | ATGGAACAC | CACCACC | TGT | ACA | C | T | C>A | C>A_ACA | CG>AT_CACCACC | 9 | TRUE | FALSE |
| 168_v.txt | 32223 | 32223-SNV-( | SNV | 1 | G | T | 20 | 14341 | 0.13946029 | 40.4 | WT | FALSE | CAN | JSY13 | 1A | WT-Rep1A | TGA | G>T | A/T+1 | G/C-1 | CG>AT | CG>AT | GTCCTTGAC | GACAGGA | ACT | ACT | C | A | C>A | C>A_ACT | CG>AT_GACAGGA | 9 | TRUE | FALSE |
| 168_v.txt | 32246 | 32246-SNV-( | SNV | 1 | G | T | 22 | 14342 | 0.15339562 | 37.4091176 | WT | FALSE | CAN | JSY13 | 1A | WT-Rep1A | TGT | G>T | A/T+1 | G/C-1 | CG>AT | CG>AT | CAACTTGTTC | GTTCTTT | ACA | ACA | C | A | C>A | C>A_ACA | CG>AT_GTTCTTT | 9 | TRUE | FALSE |
| 168_v.txt | 32271 | 32271-SNV-( | SNV | 1 | C | A | 30 | 46483 | 0.06453972 | 34.2666667 | WT | FALSE | CAN | JSY13 | 1A | WT-Rep1A | ACG | C>A | A/T+1 | G/C-1 | CG>AT | CG>AT | AAAATACGG | CGGGAAC | TGC | ACG | C | T | C>A | C>A_ACG | CG>AT_CGGGAAC | 9 | TRUE | FALSE |
| 168_v.txt | 32272 | 32272-SNV-( | SNV | 1 | G | T | 76 | 46483 | 0.16350064 | 33.355353 | WT | FALSE | CAN | JSY13 | 1A | WT-Rep1A | CGG | G>T | A/T+1 | G/C-1 | CG>AT | CG>AT | AAATACGGG | GGGAACC | GCC | GCC | C | A | C>A | C>A_GCC | CG>AT_GGGAACC | 9 | TRUE | FALSE |
| 168_v.txt | 32274 | 32274-SNV-( | SNV | 1 | G | T | 76 | 46483 | 0.16350064 | 36.3289474 | WT | FALSE | CAN | JSY13 | 1A | WT-Rep1A | GGA | G>T | A/T+1 | G/C-1 | CG>AT | CG>AT | ATACGGGAA | GAACCAA | CCT | CCT | C | A | C>A | C>A_CCT | CG>AT_GAACCAA | 9 | TRUE | FALSE |
| 168_v.txt | 32282 | 32282-SNV-( | SNV | 1 | G | T | 373 | 46483 | 0.8024439 | 43.4307536 | WT | FALSE | CAN | JSY13 | 1A | WT-Rep1A | CGT | G>T | A/T+1 | G/C-1 | CG>AT | CG>AT | ACCAACGTA | GTAAATA | GCA | GCA | C | A | C>A | C>A_GCA | CG>AT_GTAAATA | 9 | TRUE | FALSE |
| 168_v.txt | 32292 | 32292-SNV-( | SNV | 1 | G | T | 54 | 46483 | 0.1161715 | 45.2037037 | WT | FALSE | CAN | JSY13 | 1A | WT-Rep1A | TGA | G>T | A/T+1 | G/C-1 | CG>AT | CG>AT | ATATTTGAA | GAATTTG | ACT | ACT | C | A | C>A | C>A_ACT | CG>AT_GAATTTG | 9 | TRUE | FALSE |
| 168_v.txt | 32301 | 32301-SNV-( | SNV | 1 | G | T | 61 | 46483 | 0.13123077 | 53.9508197 | WT | FALSE | CAN | JSY13 | 1A | WT-Rep1A | GGC | G>T | A/T+1 | G/C-1 | CG>AT | CG>AT | TTTGCGGCA | GCAGAAA | CCG | CCG | C | A | C>A | C>A_CCG | CG>AT_GCAGAAA | 9 | TRUE | FALSE |
| 168_v.txt | 32304 | 32304-SNV-( | SNV | 1 | G | T | 39 | 46483 | 0.08390164 | 55.6666667 | WT | FALSE | CAN | JSY13 | 1A | WT-Rep1A | AGA | G>T | A/T+1 | G/C-1 | CG>AT | CG>AT | GCGGCAGAA | GAAATAA | TCT | TCT | C | A | C>A | C>A_TCT | CG>AT_GAAATAA | 9 | TRUE | FALSE |
| 168_v.txt | 32368 | 32368-SNV-( | SNV | 1 | C | A | 59 | 46483 | 0.12692812 | 57.5762712 | WT | FALSE | CAN | JSY13 | 1A | WT-Rep1A | TCA | C>A | A/T+1 | G/C-1 | CG>AT | CG>AT | AGTTCTCAAT | CAATAGC | AGT | TCA | C | T | C>A | C>A_TCA | CG>AT_CAATAGC | 9 | TRUE | FALSE |
| 168_v.txt | 32373 | 32373-SNV-( | SNV | 1 | G | T | 34 | 46483 | 0.07314502 | 60.7058824 | WT | FALSE | CAN | JSY13 | 1A | WT-Rep1A | AGC | G>T | A/T+1 | G/C-1 | CG>AT | CG>AT | TCAATAGCA | GCAATAA | TCG | TCG | C | A | C>A | C>A_TCG | CG>AT_GCAATAA | 9 | TRUE | FALSE |
| 168_v.txt | 32384 | 32384-SNV-( | SNV | 1 | G | T | 8 | 46482 | 0.01721096 | 52.9978324 | WT | FALSE | CAN | JSY13 | 1A | WT-Rep1A | AGG | G>T | A/T+1 | G/C-1 | CG>AT | CG>AT | AATAAAGGG | GGGAGAA | TCC | TCC | C | A | C>A | C>A_TCC | CG>AT_GGGAGAA | 9 | TRUE | FALSE |
| 168_v.txt | 32385 | 32385-SNV-( | SNV | 1 | G | T | 7 | 46482 | 0.01505959 | 48.4264775 | WT | FALSE | CAN | JSY13 | 1A | WT-Rep1A | GGG | G>T | A/T+1 | G/C-1 | CG>AT | CG>AT | ATAAAGGGA | GGAGAAG | CCC | CCC | C | A | C>A | C>A_CCC | CG>AT_GGAGAAG | 9 | TRUE | FALSE |
| 168_v.txt | 32399 | 32399-SNV-( | SNV | 1 | G | T | 450 | 46481 | 0.96813752 | 57.975009 | WT | FALSE | CAN | JSY13 | 1A | WT-Rep1A | CGT | G>T | A/T+1 | G/C-1 | CG>AT | CG>AT | AGAAACGTA | GTAGGAA | GCA | GCA | C | A | C>A | C>A_GCA | CG>AT_GTAGGAA | 9 | TRUE | FALSE |
| 168_v.txt | 32432 | 32432-SNV-( | SNV | 1 | G | T | 160 | 46483 | 0.34421186 | 56.33125 | WT | FALSE | CAN | JSY13 | 1A | WT-Rep1A | TGT | G>T | A/T+1 | G/C-1 | CG>AT | CG>AT | GTCATTGTAT | GTATGGA | ACA | ACA | C | A | C>A | C>A_ACA | CG>AT_GTATGGA | 9 | TRUE | FALSE |
| 168_v.txt | 32449 | 32449-SNV-( | SNV | 1 | C | A | 19 | 46483 | 0.04087516 | 49.1052632 | WT | FALSE | CAN | JSY13 | 1A | WT-Rep1A | CCA | C>A | A/T+1 | G/C-1 | CG>AT | CG>AT | AAAGTCCAA | CAATGAA | GGT | CCA | C | T | C>A | C>A_CCA | CG>AT_CAATGAA | 9 | TRUE | FALSE |
| 168_v.txt | 32463 | 32463-SNV-( | SNV | 1 | G | T | 33 | 46483 | 0.0709937 | 45.7878788 | WT | FALSE | CAN | JSY13 | 1A | WT-Rep1A | AGA | G>T | A/T+1 | G/C-1 | CG>AT | CG>AT | AATAGAGAG | GAGCCAA | TCT | TCT | C | A | C>A | C>A_TCT | CG>AT_GAGCCAA | 9 | TRUE | FALSE |
| 168_v.txt | 32471 | 32471-SNV-( | SNV | 1 | G | T | 50 | 46483 | 0.10756621 | 47.9541482 | WT | FALSE | CAN | JSY13 | 1A | WT-Rep1A | TGT | G>T | A/T+1 | G/C-1 | CG>AT | CG>AT | GCCAATGTA | GTAGAAG | ACA | ACA | C | A | C>A | C>A_ACA | CG>AT_GTAGAAG | 9 | TRUE | FALSE |
| 168_v.txt | 32474 | 32474-SNV-( | SNV | 1 | G | T | 28 | 46483 | 0.06023708 | 51.1744168 | WT | FALSE | CAN | JSY13 | 1A | WT-Rep1A | AGA | G>T | A/T+1 | G/C-1 | CG>AT | CG>AT | AATGTAGAA | GAAGGTT | TCT | TCT | C | A | C>A | C>A_TCT | CG>AT_GAAGGTT | 9 | TRUE | FALSE |
| 168_v.txt | 32487 | 32487-SNV-( | SNV | 1 | C | A | 68 | 46483 | 0.14629004 | 46.5886869 | WT | FALSE | CAN | JSY13 | 1A | WT-Rep1A | ACG | C>A | A/T+1 | G/C-1 | CG>AT | CG>AT | AAGATACGG | CGGAAAA | TGC | ACG | C | T | C>A | C>A_ACG | CG>AT_CGGAAAA | 9 | TRUE | FALSE |
| 168_v.txt | 32488 | 32488-SNV-( | SNV | 1 | G | T | 79 | 46483 | 0.16995461 | 46.3417722 | WT | FALSE | CAN | JSY13 | 1A | WT-Rep1A | CGG | G>T | A/T+1 | G/C-1 | CG>AT | CG>AT | AGATACGGA | GGAAAAC | GCC | GCC | C | A | C>A | C>A_GCC | CG>AT_GGAAAAC | 9 | TRUE | FALSE |
| 168_v.txt | 32494 | 32494-SNV-( | SNV | 1 | C | A | 168 | 46483 | 0.36142246 | 45.7251876 | WT | FALSE | CAN | JSY13 | 1A | WT-Rep1A | ACA | C>A | A/T+1 | G/C-1 | CG>AT | CG>AT | GGAAAACAA | CAACTTT | TGT | ACA | C | T | C>A | C>A_ACA | CG>AT_CAACTTT | 9 | TRUE | FALSE |
| 168_v.txt | 32508 | 32508-SNV-( | SNV | 1 | G | T | 16 | 46483 | 0.03442119 | 40.375 | WT | FALSE | CAN | JSY13 | 1A | WT-Rep1A | GGC | G>T | A/T+1 | G/C-1 | CG>AT | CG>AT | TTGATGGCT | GCTCTTG | CCG | CCG | C | A | C>A | C>A_CCG | CG>AT_GCTCTTG | 9 | TRUE | FALSE |
| 168_v.txt | 32529 | 32529-SNV-( | SNV | 1 | G | T | 17 | 46483 | 0.03657251 | 36.8823529 | WT | FALSE | CAN | JSY13 | 1A | WT-Rep1A | GGG | G>T | A/T+1 | G/C-1 | CG>AT | CG>AT | TTTCTGGGG | GGGTTTG | CCC | CCC | C | A | C>A | C>A_CCC | CG>AT_GGGTTTG | 9 | TRUE | FALSE |
| 168_v.txt | 32530 | 32530-SNV-( | SNV | 1 | G | T | 140 | 46483 | 0.30118538 | 37.9142857 | WT | FALSE | CAN | JSY13 | 1A | WT-Rep1A | GGG | G>T | A/T+1 | G/C-1 | CG>AT | CG>AT | TTCTGGGGT | GGTTTGC | CCC | CCC | C | A | C>A | C>A_CCC | CG>AT_GGTTTGC | 9 | TRUE | FALSE |
| 168_v.txt | 32542 | 32542-SNV-( | SNV | 1 | C | A | 158 | 46481 | 0.33992384 | 37.2720383 | WT | FALSE | CAN | JSY13 | 1A | WT-Rep1A | TCA | C>A | A/T+1 | G/C-1 | CG>AT | CG>AT | CAGCTTCAC | CACCAGC | AGT | TCA | C | T | C>A | C>A_TCA | CG>AT_CACCAGC | 9 | TRUE | FALSE |
| 168_v.txt | 32547 | 32547-SNV-( | SNV | 1 | G | T | 19 | 46480 | 0.0408778 | 36.2609242 | WT | FALSE | CAN | JSY13 | 1A | WT-Rep1A | AGC | G>T | A/T+1 | G/C-1 | CG>AT | CG>AT | TCACCAGCA | GCAGTGA | TCG | TCG | C | A | C>A | C>A_TCG | CG>AT_GCAGTGA | 9 | TRUE | FALSE |
| 168_v.txt | 32556 | 32556-SNV-( | SNV | 1 | C | A | 37 | 46475 | 0.0796127 | 33.4324324 | WT | FALSE | CAN | JSY13 | 1A | WT-Rep1A | ACC | C>A | A/T+1 | G/C-1 | CG>AT | CG>AT | GTGATACCA | CCAACTA | TGG | ACC | C | T | C>A | C>A_ACC | CG>AT_CCAACTA | 9 | TRUE | FALSE |
| 168_v.txt | 32566 | 32566-SNV-( | SNV | 1 | C | A | 10 | 7614 | 0.13133701 | 37.9 | WT | FALSE | CAN | JSY13 | 1A | WT-Rep1A | TCA | C>A | A/T+1 | G/C-1 | CG>AT | CG>AT | CTAGTTCAG | CAGTACC | AGT | TCA | C | T | C>A | C>A_TCA | CG>AT_CAGTACC | 9 | TRUE | FALSE |
| 168_v.txt | 32572 | 32572-SNV-( | SNV | 1 | C | A | 11 | 7614 | 0.14447071 | 32.9090909 | WT | FALSE | CAN | JSY13 | 1A | WT-Rep1A | CCT | C>A | A/T+1 | G/C-1 | CG>AT | CG>AT | CAGTACCTT | CTTGAAA | GGA | CCT | C | T | C>A | C>A_CCT | CG>AT_CTTGAAA | 9 | TRUE | FALSE |
| 168_v.txt | 32575 | 32575-SNV-( | SNV | 1 | G | T | 29 | 7614 | 0.38087733 | 37.2758621 | WT | FALSE | CAN | JSY13 | 1A | WT-Rep1A | TGA | G>T | A/T+1 | G/C-1 | CG>AT | CG>AT | TACCTTGAA | GAAATGT | ACT | ACT | C | A | C>A | C>A_ACT | CG>AT_GAAATGT | 9 | TRUE | FALSE |
| 168_v.txt | 32629 | 32629-SNV-( | SNV | 1 | C | A | 47 | 7614 | 0.61728395 | 53.7659575 | WT | FALSE | CAN | JSY13 | 1A | WT-Rep1A | TCG | C>A | A/T+1 | G/C-1 | CG>AT | CG>AT | TCCCTTCGTT | CGTTTTT | AGC | TCG | C | T | C>A | C>A_TCG | CG>AT_CGTTTTT | 9 | TRUE | FALSE |
| 168_v.txt | 32630 | 32630-SNV-( | SNV | 1 | G | T | 23 | 7614 | 0.30207513 | 48.5217391 | WT | FALSE | CAN | JSY13 | 1A | WT-Rep1A | CGT | G>T | A/T+1 | G/C-1 | CG>AT | CG>AT | CCCTTCGTTT | GTTTTTA | GCA | GCA | C | A | C>A | C>A_GCA | CG>AT_GTTTTTA | 9 | TRUE | FALSE |
| 168_v.txt | 32692 | 32692-SNV-( | SNV | 1 | C | A | 31 | 7614 | 0.40714473 | 57.2580645 | WT | FALSE | CAN | JSY13 | 1A | WT-Rep1A | CCA | C>A | A/T+1 | G/C-1 | CG>AT | CG>AT | GGAATCCAA | CAACTGG | GGT | CCA | C | T | C>A | C>A_CCA | CG>AT_CAACTGG | 9 | TRUE | FALSE |
| 168_v.txt | 32788 | 32788-SNV-( | SNV | 1 | C | A | 16 | 7622 | 0.20991866 | 52.375 | WT | FALSE | CAN | JSY13 | 1A | WT-Rep1A | TCG | C>A | A/T+1 | G/C-1 | CG>AT | CG>AT | AGAACTCGA | CGAATTC | AGC | TCG | C | T | C>A | C>A_TCG | CG>AT_CGAATTC | 9 | TRUE | FALSE |
| 168_v.txt | 32794 | 32794-SNV-( | SNV | 1 | C | A | 11 | 7622 | 0.14431908 | 39.7272727 | WT | FALSE | CAN | JSY13 | 1A | WT-Rep1A | TCA | C>A | A/T+1 | G/C-1 | CG>AT | CG>AT | CGAATTCAC | CACCGTA | AGT | TCA | C | T | C>A | C>A_TCA | CG>AT_CACCGTA | 9 | TRUE | FALSE |
| 168_v.txt | 32798 | 32798-SNV-( | SNV | 1 | G | T | 14 | 7622 | 0.18367883 | 48.6428571 | WT | FALSE | CAN | JSY13 | 1A | WT-Rep1A | CGT | G>T | A/T+1 | G/C-1 | CG>AT | CG>AT | TTCACCGTA | GTAATAT | GCA | GCA | C | A | C>A | C>A_GCA | CG>AT_GTAATAT | 9 | TRUE | FALSE |
| 168_v.txt | 32819 | 32819-SNV-( | SNV | 1 | G | T | 16 | 7622 | 0.20991866 | 45.4375 | WT | FALSE | CAN | JSY13 | 1A | WT-Rep1A | AGT | G>T | A/T+1 | G/C-1 | CG>AT | CG>AT | GAACAAGTT | GTTCATT | TCA | TCA | C | A | C>A | C>A_TCA | CG>AT_GTTCATT | 9 | TRUE | FALSE |
| 168_v.txt | 32876 | 32876-SNV-( | SNV | 1 | G | T | 78 | 22839 | 0.34152108 | 34.948718 | WT | FALSE | CAN | JSY13 | 1A | WT-Rep1A | TGT | G>T | A/T+1 | G/C-1 | CG>AT | CG>AT | AACTTTGTAC | GTACGTC | ACA | ACA | C | A | C>A | C>A_ACA | CG>AT_GTACGTC | 9 | TRUE | FALSE |
| 168_v.txt | 32917 | 32917-SNV-( | SNV | 1 | C | A | 81 | 22850 | 0.35448578 | 50.1604938 | WT | FALSE | CAN | JSY13 | 1A | WT-Rep1A | TCC | C>A | A/T+1 | G/C-1 | CG>AT | CG>AT | TAAGTTCCA | CCAGGGC | AGG | TCC | C | T | C>A | C>A_TCC | CG>AT_CCAGGGC | 9 | TRUE | FALSE |
| 168_v.txt | 32920 | 32920-SNV-( | SNV | 1 | G | T | 6 | 22850 | 0.02625821 | 23.1666193 | WT | FALSE | CAN | JSY13 | 1A | WT-Rep1A | AGG | G>T | A/T+1 | G/C-1 | CG>AT | CG>AT | GTTCCAGGG | GGGCAAA | TCC | TCC | C | A | C>A | C>A_TCC | CG>AT_GGGCAAA | 9 | TRUE | FALSE |
| 168_v.txt | 32928 | 32928-SNV-( | SNV | 1 | G | T | 14 | 22886 | 0.06117277 | 49.9911192 | WT | FALSE | CAN | JSY13 | 1A | WT-Rep1A | AGT | G>T | A/T+1 | G/C-1 | CG>AT | CG>AT | GCAAAAGTG | GTGATTG | TCA | TCA | C | A | C>A | C>A_TCA | CG>AT_GTGATTG | 9 | TRUE | FALSE |
| 168_v.txt | 32936 | 32936-SNV-( | SNV | 1 | C | A | 38 | 22886 | 0.16604037 | 36.4172311 | WT | FALSE | CAN | JSY13 | 1A | WT-Rep1A | CCC | C>A | A/T+1 | G/C-1 | CG>AT | CG>AT | GATTGCCCA | CCAAGAA | GGG | CCC | C | T | C>A | C>A_CCC | CG>AT_CCAAGAA | 9 | TRUE | FALSE |
| 168_v.txt | 32937 | 32937-SNV-( | SNV | 1 | C | A | 19 | 22886 | 0.08302019 | 53.1052632 | WT | FALSE | CAN | JSY13 | 1A | WT-Rep1A | CCA | C>A | A/T+1 | G/C-1 | CG>AT | CG>AT | ATTGCCCAA | CAAGAAA | GGT | CCA | C | T | C>A | C>A_CCA | CG>AT_CAAGAAA | 9 | TRUE | FALSE |
| 168_v.txt | 32940 | 32940-SNV-( | SNV | 1 | G | T | 55 | 22886 | 0.24032159 | 40.2192339 | WT | FALSE | CAN | JSY13 | 1A | WT-Rep1A | AGA | G>T | A/T+1 | G/C-1 | CG>AT | CG>AT | GCCCAAGAA | GAAAACC | TCT | TCT | C | A | C>A | C>A_TCT | CG>AT_GAAAACC | 9 | TRUE | FALSE |
| 168_v.txt | 32946 | 32946-SNV-( | SNV | 1 | C | A | 12 | 22886 | 0.0524338 | 46.9166667 | WT | FALSE | CAN | JSY13 | 1A | WT-Rep1A | CCA | C>A | A/T+1 | G/C-1 | CG>AT | CG>AT | GAAAACCAA | CAATACA | GGT | CCA | C | T | C>A | C>A_CCA | CG>AT_CAATACA | 9 | TRUE | FALSE |
| 168_v.txt | 32997 | 32997-SNV-( | SNV | 1 | G | T | 62 | 22886 | 0.27090798 | 61.8064516 | WT | FALSE | CAN | JSY13 | 1A | WT-Rep1A | TGA | G>T | A/T+1 | G/C-1 | CG>AT | CG>AT | CTTTGTGAG | GAGAAAA | ACT | ACT | C | A | C>A | C>A_ACT | CG>AT_GAGAAAA | 9 | TRUE | FALSE |
| 168_v.txt | 33006 | 33006-SNV-( | SNV | 1 | G | T | 29 | 22886 | 0.12671502 | 63.1724138 | WT | FALSE | CAN | JSY13 | 1A | WT-Rep1A | TGT | G>T | A/T+1 | G/C-1 | CG>AT | CG>AT | AAAACTGTG | GTGAAAG | ACA | ACA | C | A | C>A | C>A_ACA | CG>AT_GTGAAAG | 9 | TRUE | FALSE |
| 168_v.txt | 33015 | 33015-SNV-( | SNV | 1 | G | T | 22 | 22886 | 0.09612864 | 58.7272727 | WT | FALSE | CAN | JSY13 | 1A | WT-Rep1A | GGA | G>T | A/T+1 | G/C-1 | CG>AT | CG>AT | AAAGAGGAT | GATGTAA | CCT | CCT | C | A | C>A | C>A_CCT | CG>AT_GATGTAA | 9 | TRUE | FALSE |
| 168_v.txt | 33024 | 33024-SNV-( | SNV | 1 | G | T | 87 | 22886 | 0.38014507 | 60.8735632 | WT | FALSE | CAN | JSY13 | 1A | WT-Rep1A | AGG | G>T | A/T+1 | G/C-1 | CG>AT | CG>AT | GTAACAGGG | GGGATGA | TCC | TCC | C | A | C>A | C>A_TCC | CG>AT_GGGATGA | 9 | TRUE | FALSE |
| 168_v.txt | 33033 | 33033-SNV-( | SNV | 1 | G | T | 15 | 22886 | 0.06554225 | 57.9333333 | WT | FALSE | CAN | JSY13 | 1A | WT-Rep1A | TGT | G>T | A/T+1 | G/C-1 | CG>AT | CG>AT | ATGAATGTA | GTAGCCA | ACA | ACA | C | A | C>A | C>A_ACA | CG>AT_GTAGCCA | 9 | TRUE | FALSE |
| 168_v.txt | 33043 | 33043-SNV-( | SNV | 1 | C | A | 49 | 22886 | 0.21410469 | 57.1020408 | WT | FALSE | CAN | JSY13 | 1A | WT-Rep1A | TCA | C>A | A/T+1 | G/C-1 | CG>AT | CG>AT | CCATTTCACC | CACCCAA | AGT | TCA | C | T | C>A | C>A_TCA | CG>AT_CACCCAA | 9 | TRUE | FALSE |
| 168_v.txt | 33051 | 33051-SNV-( | SNV | 1 | G | T | 212 | 22886 | 0.92633051 | 58.2122642 | WT | FALSE | CAN | JSY13 | 1A | WT-Rep1A | GGA | G>T | A/T+1 | G/C-1 | CG>AT | CG>AT | CCCAAGGAC | GACTGCG | CCT | CCT | C | A | C>A | C>A_CCT | CG>AT_GACTGCG | 9 | TRUE | FALSE |
| 168_v.txt | 33056 | 33056-SNV-( | SNV | 1 | C | A | 5 | 22886 | 0.02184742 | 42.8 | WT | FALSE | CAN | JSY13 | 1A | WT-Rep1A | GCG | C>A | A/T+1 | G/C-1 | CG>AT | CG>AT | GGACTGCGT | CGTGACA | CGC | GCG | C | T | C>A | C>A_GCG | CG>AT_CGTGACA | 9 | TRUE | FALSE |
| 168_v.txt | 33070 | 33070-SNV-( | SNV | 1 | C | A | 5 | 22886 | 0.02184742 | 42.8 | WT | FALSE | CAN | JSY13 | 1A | WT-Rep1A | GCC | C>A | A/T+1 | G/C-1 | CG>AT | CG>AT | AATATGCCA | CCAAAGA | CGG | GCC | C | T | C>A | C>A_GCC | CG>AT_CCAAAGA | 9 | TRUE | FALSE |
| 168_v.txt | 33093 | 33093-SNV-( | SNV | 1 | G | T | 39 | 22886 | 0.17040986 | 46.8965607 | WT | FALSE | CAN | JSY13 | 1A | WT-Rep1A | TGA | G>T | A/T+1 | G/C-1 | CG>AT | CG>AT | AAATATGAT | GATATAA | ACT | ACT | C | A | C>A | C>A_ACT | CG>AT_GATATAA | 9 | TRUE | FALSE |
| 168_v.txt | 33111 | 33111-SNV-( | SNV | 1 | G | T | 3 | 22886 | 0.01310845 | 43.6594502 | WT | FALSE | CAN | JSY13 | 1A | WT-Rep1A | TGG | G>T | A/T+1 | G/C-1 | CG>AT | CG>AT | CCCACTGGG | GGGCCGG | ACC | ACC | C | A | C>A | C>A_ACC | CG>AT_GGGCCGG | 9 | TRUE | FALSE |
| 168_v.txt | 33112 | 33112-SNV-( | SNV | 1 | G | T | 18 | 22886 | 0.0786507 | 44.8141874 | WT | FALSE | CAN | JSY13 | 1A | WT-Rep1A | GGG | G>T | A/T+1 | G/C-1 | CG>AT | CG>AT | CCACTGGGC | GGCCGGC | CCC | CCC | C | A | C>A | C>A_CCC | CG>AT_GGCCGGC | 9 | TRUE | FALSE |
| 168_v.txt | 33115 | 33115-SNV-( | SNV | 1 | C | A | 64 | 22886 | 0.27964695 | 41.625 | WT | FALSE | CAN | JSY13 | 1A | WT-Rep1A | CCG | C>A | A/T+1 | G/C-1 | CG>AT | CG>AT | CTGGGCCGG | CGGCGTT | GGC | CCG | C | T | C>A | C>A_CCG | CG>AT_CGGCGTT | 9 | TRUE | FALSE |
| 168_v.txt | 33151 | 33151-SNV-( | SNV | 1 | G | T | 93 | 22886 | 0.40636197 | 37.5913979 | WT | FALSE | CAN | JSY13 | 1A | WT-Rep1A | AGA | G>T | A/T+1 | G/C-1 | CG>AT | CG>AT | TGAAAAGAC | GACCTGT | TCT | TCT | C | A | C>A | C>A_TCT | CG>AT_GACCTGT | 9 | TRUE | FALSE |
| 168_v.txt | 33154 | 33154-SNV-( | SNV | 1 | C | A | 37 | 22886 | 0.16167089 | 35.9725701 | WT | FALSE | CAN | JSY13 | 1A | WT-Rep1A | CCT | C>A | A/T+1 | G/C-1 | CG>AT | CG>AT | AAAGACCTG | CTGTACC | GGA | CCT | C | T | C>A | C>A_CCT | CG>AT_CTGTACC | 9 | TRUE | FALSE |
| 168_v.txt | 33160 | 33160-SNV-( | SNV | 1 | C | A | 37 | 22885 | 0.16167796 | 37.8648649 | WT | FALSE | CAN | JSY13 | 1A | WT-Rep1A | CCA | C>A | A/T+1 | G/C-1 | CG>AT | CG>AT | CTGTACCAA | CAATAGT | GGT | CCA | C | T | C>A | C>A_CCA | CG>AT_CAATAGT | 9 | TRUE | FALSE |
| 168_v.txt | 33169 | 33169-SNV-( | SNV | 1 | C | A | 8 | 22847 | 0.03501554 | 37.999997 | WT | FALSE | CAN | JSY13 | 1A | WT-Rep1A | CCA | C>A | A/T+1 | G/C-1 | CG>AT | CG>AT | TAGTACCAC | CACCAAG | GGT | CCA | C | T | C>A | C>A_CCA | CG>AT_CACCAAG | 9 | TRUE | FALSE |
| 168_v.txt | 33177 | 33177-SNV-( | SNV | 1 | G | T | 48 | 22843 | 0.21013002 | 37.3958333 | WT | FALSE | CAN | JSY13 | 1A | WT-Rep1A | GGC | G>T | A/T+1 | G/C-1 | CG>AT | CG>AT | CCAAGGGCA | GCAATCA | CCG | CCG | C | A | C>A | C>A_CCG | CG>AT_GCAATCA | 9 | TRUE | FALSE |
| 168_v.txt | 33186 | 33186-SNV-( | SNV | 1 | C | A | 45 | 23345 | 0.19276076 | 37.6 | WT | FALSE | CAN | JSY13 | 1A | WT-Rep1A | ACC | C>A | A/T+1 | G/C-1 | CG>AT | CG>AT | ATCATACCA | CCAATAT | TGG | ACC | C | T | C>A | C>A_ACC | CG>AT_CCAATAT | 9 | TRUE | FALSE |
| 168_v.txt | 33210 | 33210-SNV-( | SNV | 1 | C | A | 5 | 23354 | 0.02140961 | 37.999298 | WT | FALSE | CAN | JSY13 | 1A | WT-Rep1A | TCT | C>A | A/T+1 | G/C-1 | CG>AT | CG>AT | AGCTCTCTCT | CTCTTCA | AGA | TCT | C | T | C>A | C>A_TCT | CG>AT_CTCTTCA | 9 | TRUE | FALSE |
| 168_v.txt | 33220 | 33220-SNV-( | SNV | 1 | C | A | 37 | 23355 | 0.15842432 | 40.4864865 | WT | FALSE | CAN | JSY13 | 1A | WT-Rep1A | TCA | C>A | A/T+1 | G/C-1 | CG>AT | CG>AT | TCACTTCAGC | CAGCGTT | AGT | TCA | C | T | C>A | C>A_TCA | CG>AT_CAGCGTT | 9 | TRUE | FALSE |
| 168_v.txt | 33235 | 33235-SNV-( | SNV | 1 | C | A | 47 | 23355 | 0.2012417 | 38.9787234 | WT | FALSE | CAN | JSY13 | 1A | WT-Rep1A | TCT | C>A | A/T+1 | G/C-1 | CG>AT | CG>AT | GTACTTCTCC | CTCCTTC | AGA | TCT | C | T | C>A | C>A_TCT | CG>AT_CTCCTTC | 9 | TRUE | FALSE |
| 168_v.txt | 33238 | 33238-SNV-( | SNV | 1 | C | A | 18 | 23355 | 0.07707129 | 37.2777778 | WT | FALSE | CAN | JSY13 | 1A | WT-Rep1A | CCT | C>A | A/T+1 | G/C-1 | CG>AT | CG>AT | CTTCTCCTTC | CTTCATC | GGA | CCT | C | T | C>A | C>A_CCT | CG>AT_CTTCATC | 9 | TRUE | FALSE |
| 168_v.txt | 33241 | 33241-SNV-( | SNV | 1 | C | A | 45 | 23355 | 0.19267823 | 38.6 | WT | FALSE | CAN | JSY13 | 1A | WT-Rep1A | TCA | C>A | A/T+1 | G/C-1 | CG>AT | CG>AT | CTCCTTCATC | CATCTTC | AGT | TCA | C | T | C>A | C>A_TCA | CG>AT_CATCTTC | 9 | TRUE | FALSE |
| 168_v.txt | 33247 | 33247-SNV-( | SNV | 1 | C | A | 37 | 23355 | 0.15842432 | 38.2432432 | WT | FALSE | CAN | JSY13 | 1A | WT-Rep1A | TCA | C>A | A/T+1 | G/C-1 | CG>AT | CG>AT | CATCTTCATC | CATCACC | AGT | TCA | C | T | C>A | C>A_TCA | CG>AT_CATCACC | 9 | TRUE | FALSE |
| 168_v.txt | 33283 | 33283-SNV-( | SNV | 1 | C | A | 24 | 23355 | 0.10276172 | 47.1657305 | WT | FALSE | CAN | JSY13 | 1A | WT-Rep1A | TCG | C>A | A/T+1 | G/C-1 | CG>AT | CG>AT | TATCCTCGCC | CGCCATT | AGC | TCG | C | T | C>A | C>A_TCG | CG>AT_CGCCATT | 9 | TRUE | FALSE |
| 168_v.txt | 33284 | 33284-SNV-( | SNV | 1 | G | T | 1 | 23355 | 4.28E-03 | 33.9999621 | WT | FALSE | CAN | JSY13 | 1A | WT-Rep1A | CGC | G>T | A/T+1 | G/C-1 | CG>AT | CG>AT | ATCCTCGCCA | GCCATTT | GCG | GCG | C | A | C>A | C>A_GCG | CG>AT_GCCATTT | 9 | TRUE | FALSE |
| 168_v.txt | 33286 | 33286-SNV-( | SNV | 1 | C | A | 1 | 23354 | 4.28E-03 | 35.0000002 | WT | FALSE | CAN | JSY13 | 1A | WT-Rep1A | CCA | C>A | A/T+1 | G/C-1 | CG>AT | CG>AT | CCTCGCCATT | CATTTAC | GGT | CCA | C | T | C>A | C>A_CCA | CG>AT_CATTTAC | 9 | TRUE | FALSE |
| 168_v.txt | 33325 | 33325-SNV-( | SNV | 1 | C | A | 75 | 23355 | 0.32113038 | 57.2133333 | WT | FALSE | CAN | JSY13 | 1A | WT-Rep1A | TCT | C>A | A/T+1 | G/C-1 | CG>AT | CG>AT | ACAATTCTTT | CTTTACT | AGA | TCT | C | T | C>A | C>A_TCT | CG>AT_CTTTACT | 9 | TRUE | FALSE |
| 168_v.txt | 33337 | 33337-SNV-( | SNV | 1 | C | A | 30 | 23355 | 0.12845215 | 59.2666667 | WT | FALSE | CAN | JSY13 | 1A | WT-Rep1A | TCA | C>A | A/T+1 | G/C-1 | CG>AT | CG>AT | TTTTCTCATC | CATCTTT | AGT | TCA | C | T | C>A | C>A_TCA | CG>AT_CATCTTT | 9 | TRUE | FALSE |
| 168_v.txt | 33354 | 33354-SNV-( | SNV | 1 | G | T | 127 | 23355 | 0.54378078 | 19.7952756 | WT | FALSE | CAN | JSY13 | 1A | WT-Rep1A | TGA | G>T | A/T+1 | G/C-1 | CG>AT | CG>AT | GGTATTGAC | GACCCAC | ACT | ACT | C | A | C>A | C>A_ACT | CG>AT_GACCCAC | 9 | TRUE | FALSE |
| 168_v.txt | 33385 | 33385-SNV-( | SNV | 1 | C | A | 37 | 23355 | 0.15842432 | 53.3783784 | WT | FALSE | CAN | JSY13 | 1A | WT-Rep1A | TCA | C>A | A/T+1 | G/C-1 | CG>AT | CG>AT | AAGCTTCAA | CAACGTC | AGT | TCA | C | T | C>A | C>A_TCA | CG>AT_CAACGTC | 9 | TRUE | FALSE |
| 168_v.txt | 33415 | 33415-SNV-( | SNV | 1 | C | A | 26 | 23355 | 0.1113252 | 30.5769231 | WT | FALSE | CAN | JSY13 | 1A | WT-Rep1A | TCA | C>A | A/T+1 | G/C-1 | CG>AT | CG>AT | CCGGCTCATT | CATTGTA | AGT | TCA | C | T | C>A | C>A_TCA | CG>AT_CATTGTA | 9 | TRUE | FALSE |
| 168_v.txt | 33419 | 33419-SNV-( | SNV | 1 | G | T | 349 | 23355 | 1.4943267 | 48.9570201 | WT | FALSE | CAN | JSY13 | 1A | WT-Rep1A | TGT | G>T | A/T+1 | G/C-1 | CG>AT | CG>AT | CTCATTGTAC | GTACATA | ACA | ACA | C | A | C>A | C>A_ACA | CG>AT_GTACATA | 9 | TRUE | FALSE |
| 168_v.txt | 33436 | 33436-SNV-( | SNV | 1 | C | A | 15 | 23355 | 0.06422608 | 43.1966386 | WT | FALSE | CAN | JSY13 | 1A | WT-Rep1A | TCT | C>A | A/T+1 | G/C-1 | CG>AT | CG>AT | TCTCCTCTAT | CTATGTC | AGA | TCT | C | T | C>A | C>A_TCT | CG>AT_CTATGTC | 9 | TRUE | FALSE |
| 168_v.txt | 33451 | 33451-SNV-( | SNV | 1 | C | A | 16 | 23355 | 0.06850781 | 39.3125 | WT | FALSE | CAN | JSY13 | 1A | WT-Rep1A | TCT | C>A | A/T+1 | G/C-1 | CG>AT | CG>AT | CGTCTTCTTT | CTTTTGA | AGA | TCT | C | T | C>A | C>A_TCT | CG>AT_CTTTTGA | 9 | TRUE | FALSE |
| 168_v.txt | 33456 | 33456-SNV-( | SNV | 1 | G | T | 38 | 23351 | 0.16273393 | 17.5263158 | WT | FALSE | CAN | JSY13 | 1A | WT-Rep1A | TGA | G>T | A/T+1 | G/C-1 | CG>AT | CG>AT | TCTTTTGAAT | GAATTTG | ACT | ACT | C | A | C>A | C>A_ACT | CG>AT_GAATTTG | 9 | TRUE | FALSE |
| 168_v.txt | 33474 | 33474-SNV-( | SNV | 1 | C | A | 45 | 23349 | 0.19272774 | 33.8222222 | WT | FALSE | CAN | JSY13 | 1A | WT-Rep1A | GCC | C>A | A/T+1 | G/C-1 | CG>AT | CG>AT | GCTATGCCTT | CCTTTTT | CGG | GCC | C | T | C>A | C>A_GCC | CG>AT_CCTTTTT | 9 | TRUE | FALSE |
| 168_v.txt | 31972 | 31972-Repla | Replacement | 2 | CC | T | 19 | 14355 | 0.13235806 | 40.8684177 | WT | FALSE | CAN | JSY13 | 1A | WT-Rep1A | GCC | nope | A/T+1 | >1bpdel | none | Replac. | AATAAGCCA | CCAAGCC | CGG | GCC | CC | A | NA | NA_GCC | Replac._CCAAGCC | 9 | TRUE | FALSE |
| 168_v.txt | 32080 | 32080-Repla | Replacement | 2 | C | AA | 14 | 14355 | 0.09752699 | 63.9999824 | WT | FALSE | CAN | JSY13 | 1A | WT-Rep1A | GCA | nope | >1bp-ins | G/C-1 | none | Replac. | ACCATGCAA | CAAAAAA | CGT | GCA | C | none | nope | nope_GCA | Replac._CAAAAAA | 9 | TRUE | FALSE |
| 168_v.txt | 32526 | 32526-Repla | Replacement | 2 | CT | A | 40 | 46483 | 0.08605297 | 41.7750019 | WT | FALSE | CAN | JSY13 | 1A | WT-Rep1A | TCT | nope | A/T+1 | >1bpdel | none | Replac. | GATTTTCTGG | CTGGGGT | AGA | TCT | CT | T | NA | NA_TCT | Replac._CTGGGGT | 9 | TRUE | FALSE |
| 168_v.txt | 32545 | 32545-Repla | Replacement | 2 | C | AA | 1 | 46481 | 2.15E-03 | 37.9995711 | WT | FALSE | CAN | JSY13 | 1A | WT-Rep1A | CCA | nope | >1bp-ins | G/C-1 | none | Replac. | CTTCACCAGC | CAGCAGT | GGT | CCA | C | none | nope | nope_CCA | Replac._CAGCAGT | 9 | TRUE | FALSE |
| 168_v.txt | 32934 | 32934-Repla | Replacement | 2 | GC | T | 2 | 22886 | 8.74E-03 | 59.75 | WT | FALSE | CAN | JSY13 | 1A | WT-Rep1A | TGC | nope | A/T+1 | >1bpdel | none | Replac. | GTGATTGCC | GCCCAAG | ACG | TGC | GC | A | NA | NA_TGC | Replac._GCCCAAG | 9 | TRUE | FALSE |
| 168_v.txt | 33076 | 33076-Repla | Replacement | 2 | AA | T | 8 | 22886 | 0.03495587 | 54.1875 | WT | FALSE | CAN | JSY13 | 1A | WT-Rep1A | GAA | nope | A/T+1 | >1bpdel | none | Replac. | CCAAAGAAC | AACCCAT | CTT | GAA | AA | A | NA | NA_GAA | Replac._AACCCAT | 9 | TRUE | FALSE |
| 168_v.txt | 33168 | 33168-Repla | Replacement | 3 | CCA | TT | 19 | 22847 | 0.0831619 | 37.8947368 | WT | FALSE | CAN | JSY13 | 1A | WT-Rep1A | ACC | nope | >1bp-ins | >1bpdel | none | Replac. | ATAGTACCA | CCACCAA | TGG | ACC | CCA | none | NA | NA_ACC | Replac._CCACCAA | 9 | TRUE | FALSE |
| 168_v.txt | 33230 | 33230-Repla | Replacement | 2 | TA | C | 7 | 23355 | 0.02997217 | 36.7853365 | WT | FALSE | CAN | JSY13 | 1A | WT-Rep1A | GTA | nope | G/C+1 | >1bpdel | none | Replac. | GTTCTGTACT | TACTTCT | CAT | GTA | TA | G | NA | NA_GTA | Replac._TACTTCT | 9 | TRUE | FALSE |
| 168_v.txt | 32310 | 32310-MNV- | MNV | 2 | AT | GG | 5 | 46483 | 0.01075662 | 51.400557 | WT | FALSE | CAN | JSY13 | 1A | WT-Rep1A | AAT | nope | >1bp-ins | >1bpdel | none | MNV | GAAATAATG | ATGGTTG | TTA | AAT | AT | none | NA | NA_AAT | MNV_ATGGTTG | 9 | TRUE | FALSE |

| | | | | | | | | | | | | | | | | | | | | | | | | | | | | | | | | | | |
|---|---|---|---|---|---|---|---|---|---|---|---|---|---|---|---|---|---|---|---|---|---|---|---|---|---|---|---|---|---|---|---|---|---|---|
| 168_v.txt | 32413 | 32413-MNV- | MNV | 2 | GT | AG | 120 | 46483 | 0.2581589 | 57.9333144 | WT | FALSE | CAN | JSY13 | 1A | WT-Rep1A | TGT | nope | >1bp-ins | >1bpdel | none | MNV | TAGATTGTG | GTGTTAG | ACA | TGT | GT | none | NA | NA_TGT | MNV_GTGTTAG | 9 | TRUE | FALSE |
| 168_v.txt | 32508 | 32508-MNV- | MNV | 2 | GC | AG | 9 | 46483 | 0.01936192 | 39.3333335 | WT | FALSE | CAN | JSY13 | 1A | WT-Rep1A | GGC | nope | >1bp-ins | >1bpdel | none | MNV | TTGATGGCT | GCTCTTG | CCG | GGC | GC | none | NA | NA_GGC | MNV_GCTCTTG | 9 | TRUE | FALSE |
| 168_v.txt | 32545 | 32545-MNV- | MNV | 3 | CAG | ACT | 1 | 46481 | 2.15E-03 | 19.3333262 | WT | FALSE | CAN | JSY13 | 1A | WT-Rep1A | CCA | nope | >1bp-ins | >1bpdel | none | MNV | CTTCACCAG | CAGCAGT | GGT | CCA | CAG | none | NA | NA_CCA | MNV_CAGCAGT | 9 | TRUE | FALSE |
| 168_v.txt | 32546 | 32546-MNV- | MNV | 2 | AG | CT | 3 | 46480 | 6.45E-03 | 16.83358 | WT | FALSE | CAN | JSY13 | 1A | WT-Rep1A | CAG | nope | >1bp-ins | >1bpdel | none | MNV | TTCACCAGC | AGCAGTG | GTC | CAG | AG | none | NA | NA_CAG | MNV_AGCAGTG | 9 | TRUE | FALSE |
| 168_v.txt | 32937 | 32937-MNV- | MNV | 2 | CA | AG | 19 | 22886 | 0.08302019 | 60.5263158 | WT | FALSE | CAN | JSY13 | 1A | WT-Rep1A | CCA | nope | >1bp-ins | >1bpdel | none | MNV | ATTGCCCAA | CAAGAAA | GGT | CCA | CA | none | NA | NA_CCA | MNV_CAAGAAA | 9 | TRUE | FALSE |
| 168_v.txt | 33168 | 33168-MNV- | MNV | 2 | CC | AA | 1 | 22847 | 4.38E-03 | 36.5000106 | WT | FALSE | CAN | JSY13 | 1A | WT-Rep1A | ACC | nope | >1bp-ins | >1bpdel | none | MNV | ATAGTACCA | CCACCAA | TGG | ACC | CC | none | NA | NA_ACC | MNV_CCACCAA | 9 | TRUE | FALSE |
| 168_v.txt | 33168 | 33168-MNV- | MNV | 2 | CC | AT | 18 | 22847 | 0.07878496 | 37.2778367 | WT | FALSE | CAN | JSY13 | 1A | WT-Rep1A | ACC | nope | >1bp-ins | >1bpdel | none | MNV | ATAGTACCA | CCACCAA | TGG | ACC | CC | none | NA | NA_ACC | MNV_CCACCAA | 9 | TRUE | FALSE |
| 168_v.txt | 33286 | 33286-MNV- | MNV | 2 | CA | AT | 1 | 23355 | 4.28E-03 | 38.0000093 | WT | FALSE | CAN | JSY13 | 1A | WT-Rep1A | CCA | nope | >1bp-ins | >1bpdel | none | MNV | CCTCGCCATT | CATTTAC | GGT | CCA | CA | none | NA | NA_CCA | MNV_CATTTAC | 9 | TRUE | FALSE |
| 168_v.txt | 31885 | 31885-Insert | Insertion | 1 | - | A | 2 | 3138 | 0.06373486 | 38 | WT | FALSE | CAN | JSY13 | 1A | WT-Rep1A | GCA | nope | A/T+1 | >1bpdel | none | A/T+1 | CGGCAGCAA | CAAAGCT | CGT | GCA | - | T | NA | NA_GCA | A/T+1_CAAAGCT | 9 | TRUE | FALSE |
| 168_v.txt | 31984 | 31984-Insert | Insertion | 5 | - | TTAAT | 12 | 14355 | 0.08359457 | 49.1833333 | WT | FALSE | CAN | JSY13 | 1A | WT-Rep1A | ATT | nope | >1bp-ins | >1bpdel | none | >1bp-ins | CGGGCATTA | TTAATTT | TAA | ATT | - | none | NA | NA_ATT | >1bp-ins_TTAATTT | 9 | TRUE | FALSE |
| 168_v.txt | 32005 | 32005-Insert | Insertion | 3 | - | ACT | 16 | 14355 | 0.11145942 | 52.875 | WT | FALSE | CAN | JSY13 | 1A | WT-Rep1A | AAC | nope | >1bp-ins | >1bpdel | none | >1bp-ins | ATGGTAACT | ACTCGTC | TTG | AAC | - | none | NA | NA_AAC | >1bp-ins_ACTCGTC | 9 | TRUE | FALSE |
| 168_v.txt | 32034 | 32034-Insert | Insertion | 1 | - | A | 4 | 14355 | 0.02786486 | 64 | WT | FALSE | CAN | JSY13 | 1A | WT-Rep1A | CAA | nope | A/T+1 | >1bpdel | none | A/T+1 | TATTTCAAA | AAAGCTT | GTT | CAA | - | T | NA | NA_CAA | A/T+1_AAAGCTT | 9 | TRUE | FALSE |
| 168_v.txt | 32080 | 32080-Insert | Insertion | 1 | - | C | 1 | 14355 | 6.97E-03 | 63.9999984 | WT | FALSE | CAN | JSY13 | 1A | WT-Rep1A | GCA | nope | G/C+1 | >1bpdel | none | G/C+1 | ACCATGCAA | CAAAAAA | CGT | GCA | - | G | NA | NA_GCA | G/C+1_CAAAAAA | 9 | TRUE | FALSE |
| 168_v.txt | 32081 | 32081-Insert | Insertion | 1 | - | A | 44 | 14355 | 0.30651341 | 63.2954546 | WT | FALSE | CAN | JSY13 | 1A | WT-Rep1A | CAA | nope | A/T+1 | >1bpdel | none | A/T+1 | CCATGCAAA | AAAAAAG | GTT | CAA | - | T | NA | NA_CAA | A/T+1_AAAAAAG | 9 | TRUE | FALSE |
| 168_v.txt | 32081 | 32081-Insert | Insertion | 3 | - | AAA | 14 | 14355 | 0.09752699 | 63.5238095 | WT | FALSE | CAN | JSY13 | 1A | WT-Rep1A | CAA | nope | >1bp-ins | >1bpdel | none | >1bp-ins | CCATGCAAA | AAAAAAG | GTT | CAA | - | none | NA | NA_CAA | >1bp-ins_AAAAAAG | 9 | TRUE | FALSE |
| 168_v.txt | 32168 | 32168-Insert | Insertion | 1 | - | A | 8 | 14355 | 0.05572971 | 54.125 | WT | FALSE | CAN | JSY13 | 1A | WT-Rep1A | CAA | nope | A/T+1 | >1bpdel | none | A/T+1 | AGCGCCAAA | AAATGCA | GTT | CAA | - | T | NA | NA_CAA | A/T+1_AAATGCA | 9 | TRUE | FALSE |
| 168_v.txt | 32229 | 32229-Insert | Insertion | 1 | - | T | 8 | 14341 | 0.05578412 | 37.75 | WT | FALSE | CAN | JSY13 | 1A | WT-Rep1A | GAA | nope | A/T+1 | >1bpdel | none | A/T+1 | GACAGGAAT | AATTTAG | CTT | GAA | - | A | NA | NA_GAA | A/T+1_AATTTAG | 9 | TRUE | FALSE |
| 168_v.txt | 32261 | 32261-Insert | Insertion | 1 | - | A | 10 | 46482 | 0.0215137 | 38 | WT | FALSE | CAN | JSY13 | 1A | WT-Rep1A | CAA | nope | A/T+1 | >1bpdel | none | A/T+1 | TAGACCAAA | AAATAAA | GTT | CAA | - | T | NA | NA_CAA | A/T+1_AAATAAA | 9 | TRUE | FALSE |
| 168_v.txt | 32346 | 32346-Insert | Insertion | 1 | - | A | 46 | 46483 | 0.09896091 | 62.9347826 | WT | FALSE | CAN | JSY13 | 1A | WT-Rep1A | CAA | nope | A/T+1 | >1bpdel | none | A/T+1 | TGTGGCAAA | AAAACCT | GTT | CAA | - | T | NA | NA_CAA | A/T+1_AAAACCT | 9 | TRUE | FALSE |
| 168_v.txt | 32375 | 32375-Insert | Insertion | 1 | - | A | 7 | 46483 | 0.01505927 | 64 | WT | FALSE | CAN | JSY13 | 1A | WT-Rep1A | CAA | nope | A/T+1 | >1bpdel | none | A/T+1 | AATAGCAAT | AATAATA | GTT | CAA | - | T | NA | NA_CAA | A/T+1_AATAATA | 9 | TRUE | FALSE |
| 168_v.txt | 32402 | 32402-Insert | Insertion | 11 | - | GGAAGTAGA | 25 | 46483 | 0.0537831 | 57.7090909 | WT | FALSE | CAN | JSY13 | 1A | WT-Rep1A | AGG | nope | >1bp-ins | >1bpdel | none | >1bp-ins | AACGTAGGA | GGAAGTA | TCC | AGG | - | none | NA | NA_AGG | >1bp-ins_GGAAGTA | 9 | TRUE | FALSE |
| 168_v.txt | 32448 | 32448-Insert | Insertion | 18 | - | CCAATGAAT | 6 | 46483 | 0.01290795 | 47.6851852 | WT | FALSE | CAN | JSY13 | 1A | WT-Rep1A | TCC | nope | >1bp-ins | >1bpdel | none | >1bp-ins | AAAAGTCCA | CCAATGA | AGG | TCC | - | none | NA | NA_TCC | >1bp-ins_CCAATGA | 9 | TRUE | FALSE |
| 168_v.txt | 32473 | 32473-Insert | Insertion | 1 | - | C | 1 | 46483 | 2.15E-03 | 36.9999992 | WT | FALSE | CAN | JSY13 | 1A | WT-Rep1A | TAG | nope | G/C+1 | >1bpdel | none | G/C+1 | CAATGTAGA | AGAAGGT | ATC | TAG | - | G | NA | NA_TAG | G/C+1_AGAAGGT | 9 | TRUE | FALSE |
| 168_v.txt | 32476 | 32476-Insert | Insertion | 14 | - | AGGTTAAGA | 89 | 46483 | 0.19146785 | 44.6741573 | WT | FALSE | CAN | JSY13 | 1A | WT-Rep1A | AAG | nope | >1bp-ins | >1bpdel | none | >1bp-ins | TGTAGAAGG | AGGTTAA | TTC | AAG | - | none | NA | NA_AAG | >1bp-ins_AGGTTAA | 9 | TRUE | FALSE |
| 168_v.txt | 32498 | 32498-Insert | Insertion | 1 | - | T | 15 | 46483 | 0.03226986 | 48.4 | WT | FALSE | CAN | JSY13 | 1A | WT-Rep1A | CTT | nope | A/T+1 | >1bpdel | none | A/T+1 | AACAACTTTT | TTTTTTG | GAA | CTT | - | A | NA | NA_CTT | A/T+1_TTTTTTG | 9 | TRUE | FALSE |
| 168_v.txt | 32504 | 32504-Insert | Insertion | 1 | - | G | 3 | 46483 | 6.45E-03 | 55.3333333 | WT | FALSE | CAN | JSY13 | 1A | WT-Rep1A | TGA | nope | G/C+1 | >1bpdel | none | G/C+1 | TTTTTTGATG | GATGGCT | ACT | TGA | - | C | NA | NA_TGA | G/C+1_GATGGCT | 9 | TRUE | FALSE |
| 168_v.txt | 32528 | 32528-Insert | Insertion | 1 | - | G | 54 | 46483 | 0.1161715 | 42.5740741 | WT | FALSE | CAN | JSY13 | 1A | WT-Rep1A | TGG | nope | G/C+1 | >1bpdel | none | G/C+1 | TTTTCTGGG | GGGGTTT | ACC | TGG | - | C | NA | NA_TGG | G/C+1_GGGGTTT | 9 | TRUE | FALSE |
| 168_v.txt | 32629 | 32629-Insert | Insertion | 1 | - | C | 22 | 7614 | 0.28894142 | 48.4545455 | WT | FALSE | CAN | JSY13 | 1A | WT-Rep1A | TCG | nope | G/C+1 | >1bpdel | none | G/C+1 | TCCCTTCGTT | CGTTTTT | AGC | TCG | - | G | NA | NA_TCG | G/C+1_CGTTTTT | 9 | TRUE | FALSE |
| 168_v.txt | 32631 | 32631-Insert | Insertion | 1 | - | T | 1 | 7614 | 0.0131337 | 64 | WT | FALSE | CAN | JSY13 | 1A | WT-Rep1A | GTT | nope | A/T+1 | >1bpdel | none | A/T+1 | CCTTCGTTTT | TTTTTAT | CAA | GTT | - | A | NA | NA_GTT | A/T+1_TTTTTAT | 9 | TRUE | FALSE |
| 168_v.txt | 32728 | 32728-Insert | Insertion | 1 | - | A | 19 | 7622 | 0.24927841 | 63.2631579 | WT | FALSE | CAN | JSY13 | 1A | WT-Rep1A | CAA | nope | A/T+1 | >1bpdel | none | A/T+1 | CCATACAAA | AAAAACA | GTT | CAA | - | T | NA | NA_CAA | A/T+1_AAAAACA | 9 | TRUE | FALSE |
| 168_v.txt | 32750 | 32750-Insert | Insertion | 4 | - | GATA | 3 | 7622 | 0.03935975 | 63.5 | WT | FALSE | CAN | JSY13 | 1A | WT-Rep1A | CGA | nope | >1bp-ins | >1bpdel | none | >1bp-ins | AAACCCGAT | GATAATG | GCT | CGA | - | none | NA | NA_CGA | >1bp-ins_GATAATG | 9 | TRUE | FALSE |
| 168_v.txt | 32760 | 32760-Insert | Insertion | 1 | - | A | 7 | 7622 | 0.09183941 | 48.8571429 | WT | FALSE | CAN | JSY13 | 1A | WT-Rep1A | TAA | nope | A/T+1 | >1bpdel | none | A/T+1 | ATGGCTAAA | AAAACTT | ATT | TAA | - | T | NA | NA_TAA | A/T+1_AAAACTT | 9 | TRUE | FALSE |
| 168_v.txt | 32842 | 32842-Insert | Insertion | 1 | - | A | 163 | 7622 | 2.13854631 | 43.202454 | WT | FALSE | CAN | JSY13 | 1A | WT-Rep1A | CAA | nope | A/T+1 | >1bpdel | none | A/T+1 | TTACCCAAA | AAAAAAT | GTT | CAA | - | T | NA | NA_CAA | A/T+1_AAAAAAT | 9 | TRUE | FALSE |
| 168_v.txt | 32903 | 32903-Insert | Insertion | 1 | - | A | 3 | 22850 | 0.0131291 | 45.3333333 | WT | FALSE | CAN | JSY13 | 1A | WT-Rep1A | CAA | nope | A/T+1 | >1bpdel | none | A/T+1 | TTGGCCAAC | AACTACA | GTT | CAA | - | T | NA | NA_CAA | A/T+1_AACTACA | 9 | TRUE | FALSE |
| 168_v.txt | 32924 | 32924-Insert | Insertion | 1 | - | A | 26 | 22850 | 0.11378556 | 54.7307692 | WT | FALSE | CAN | JSY13 | 1A | WT-Rep1A | CAA | nope | A/T+1 | >1bpdel | none | A/T+1 | CAGGGCAAA | AAAAGTG | GTT | CAA | - | T | NA | NA_CAA | A/T+1_AAAAGTG | 9 | TRUE | FALSE |
| 168_v.txt | 32941 | 32941-Insert | Insertion | 1 | - | A | 1 | 22886 | 4.37E-03 | 38 | WT | FALSE | CAN | JSY13 | 1A | WT-Rep1A | GAA | nope | A/T+1 | >1bpdel | none | A/T+1 | CCCAAGAAA | AAAACCA | CTT | GAA | - | T | NA | NA_GAA | A/T+1_AAAACCA | 9 | TRUE | FALSE |
| 168_v.txt | 32953 | 32953-Insert | Insertion | 3 | - | TGT | 4 | 22886 | 0.01747793 | 63.4166667 | WT | FALSE | CAN | JSY13 | 1A | WT-Rep1A | ATG | nope | >1bp-ins | >1bpdel | none | >1bp-ins | AATACATGT | TGTAACC | TAC | ATG | - | none | NA | NA_ATG | >1bp-ins_TGTAACC | 9 | TRUE | FALSE |
| 168_v.txt | 32972 | 32972-Insert | Insertion | 1 | - | A | 47 | 22886 | 0.20536573 | 63.9361702 | WT | FALSE | CAN | JSY13 | 1A | WT-Rep1A | CAA | nope | A/T+1 | >1bpdel | none | A/T+1 | CGCACCAAA | AAATGCT | GTT | CAA | - | T | NA | NA_CAA | A/T+1_AAATGCT | 9 | TRUE | FALSE |
| 168_v.txt | 32992 | 32992-Insert | Insertion | 1 | - | T | 7 | 22886 | 0.03058639 | 63.5714286 | WT | FALSE | CAN | JSY13 | 1A | WT-Rep1A | CTT | nope | A/T+1 | >1bpdel | none | A/T+1 | GGAATCTTT | TTTGTGA | GAA | CTT | - | A | NA | NA_CTT | A/T+1_TTTGTGA | 9 | TRUE | FALSE |
| 168_v.txt | 33009 | 33009-Insert | Insertion | 1 | - | A | 10 | 22886 | 0.04369484 | 62.4 | WT | FALSE | CAN | JSY13 | 1A | WT-Rep1A | GAA | nope | A/T+1 | >1bpdel | none | A/T+1 | ACTGTGAAA | AAAGAGG | CTT | GAA | - | T | NA | NA_GAA | A/T+1_AAAGAGG | 9 | TRUE | FALSE |
| 168_v.txt | 33036 | 33036-Insert | Insertion | 33 | - | GCCATTTCAC | 7 | 22886 | 0.03058639 | 62.8225108 | WT | FALSE | CAN | JSY13 | 1A | WT-Rep1A | AGC | nope | >1bp-ins | >1bpdel | none | >1bp-ins | AATGTAGCC | GCCATTT | TCG | AGC | - | none | NA | NA_AGC | >1bp-ins_GCCATTT | 9 | TRUE | FALSE |
| 168_v.txt | 33045 | 33045-Insert | Insertion | 1 | - | C | 5 | 22886 | 0.02184742 | 64 | WT | FALSE | CAN | JSY13 | 1A | WT-Rep1A | ACC | nope | G/C+1 | >1bpdel | none | G/C+1 | ATTTCACCCA | CCCAAGG | TGG | ACC | - | G | NA | NA_ACC | G/C+1_CCCAAGG | 9 | TRUE | FALSE |
| 168_v.txt | 33046 | 33046-Insert | Insertion | 24 | - | CCAAGGACT | 7 | 22886 | 0.03058639 | 62.6607143 | WT | FALSE | CAN | JSY13 | 1A | WT-Rep1A | CCC | nope | >1bp-ins | >1bpdel | none | >1bp-ins | TTTCACCCAA | CCAAGGA | GGG | CCC | - | none | NA | NA_CCC | >1bp-ins_CCAAGGA | 9 | TRUE | FALSE |
| 168_v.txt | 33070 | 33070-Insert | Insertion | 1 | - | C | 1 | 22886 | 4.37E-03 | 64 | WT | FALSE | CAN | JSY13 | 1A | WT-Rep1A | GCC | nope | G/C+1 | >1bpdel | none | G/C+1 | AATATGCCA | CCAAAGA | CGG | GCC | - | G | NA | NA_GCC | G/C+1_CCAAAGA | 9 | TRUE | FALSE |
| 168_v.txt | 33147 | 33147-Insert | Insertion | 2 | - | AA | 3 | 22886 | 0.01310845 | 37.5 | WT | FALSE | CAN | JSY13 | 1A | WT-Rep1A | GAA | nope | >1bp-ins | >1bpdel | none | >1bp-ins | CCAATGAAA | AAAAGAC | CTT | GAA | - | none | NA | NA_GAA | >1bp-ins_AAAAGAC | 9 | TRUE | FALSE |
| 168_v.txt | 33206 | 33206-Insert | Insertion | 2 | - | CT | 17 | 23355 | 0.07278955 | 37.0588235 | WT | FALSE | CAN | JSY13 | 1A | WT-Rep1A | GCT | nope | >1bp-ins | >1bpdel | none | >1bp-ins | CTTAAGCTCT | CTCTCTC | CGA | GCT | - | none | NA | NA_GCT | >1bp-ins_CTCTCTC | 9 | TRUE | FALSE |
| 168_v.txt | 33254 | 33254-Insert | Insertion | 4 | - | TATG | 11 | 23355 | 0.04709912 | 40.0454546 | WT | FALSE | CAN | JSY13 | 1A | WT-Rep1A | CTA | nope | >1bp-ins | >1bpdel | none | >1bp-ins | ATCACCTATG | TATGCCA | GAT | CTA | - | none | NA | NA_CTA | >1bp-ins_TATGCCA | 9 | TRUE | FALSE |
| 168_v.txt | 33437 | 33437-Insert | Insertion | 1 | - | T | 12 | 23355 | 0.05138086 | 44.3333333 | WT | FALSE | CAN | JSY13 | 1A | WT-Rep1A | CTA | nope | A/T+1 | >1bpdel | none | A/T+1 | CTCCTCTATG | TATGTCG | GAT | CTA | - | A | NA | NA_CTA | A/T+1_TATGTCG | 9 | TRUE | FALSE |
| 168_v.txt | 32014 | 32014-Deleti | Deletion | 2 | GA | - | 17 | 14355 | 0.11842564 | 57.7647059 | WT | FALSE | CAN | JSY13 | 1A | WT-Rep1A | CGA | nope | >1bp-ins | >1bpdel | none | >1bpdel | CGTCACGAG | GAGAGAT | GCT | CGA | GA | none | NA | NA_CGA | >1bpdel_GAGAGAT | 9 | TRUE | FALSE |
| 168_v.txt | 32070 | 32070-Deleti | Deletion | 3 | AAT | - | 24 | 14355 | 0.16718913 | 61.625 | WT | FALSE | CAN | JSY13 | 1A | WT-Rep1A | AAA | nope | >1bp-ins | >1bpdel | none | >1bpdel | GAGATAAAT | AATAACC | TTT | AAA | AAT | none | NA | NA_AAA | >1bpdel_AATAACC | 9 | TRUE | FALSE |
| 168_v.txt | 32313 | 32313-Deleti | Deletion | 3 | GTT | - | 25 | 46483 | 0.0537831 | 60.12 | WT | FALSE | CAN | JSY13 | 1A | WT-Rep1A | GGT | nope | >1bp-ins | >1bpdel | none | >1bpdel | ATAATGGTT | GTTGTTA | CCA | GGT | GTT | none | NA | NA_GGT | >1bpdel_GTTGTTA | 9 | TRUE | FALSE |
| 168_v.txt | 32316 | 32316-Deleti | Deletion | 5 | GTTAA | - | 26 | 46483 | 0.05593443 | 60 | WT | FALSE | CAN | JSY13 | 1A | WT-Rep1A | TGT | nope | >1bp-ins | >1bpdel | none | >1bpdel | ATGGTTGTT | GTTAAGA | ACA | TGT | GTTAA | none | NA | NA_TGT | >1bpdel_GTTAAGA | 9 | TRUE | FALSE |
| 168_v.txt | 32337 | 32337-Deleti | Deletion | 2 | AT | - | 321 | 46483 | 0.69057505 | 60.8255452 | WT | FALSE | CAN | JSY13 | 1A | WT-Rep1A | GAT | nope | >1bp-ins | >1bpdel | none | >1bpdel | TTGAAGATA | ATATGTG | CTA | GAT | AT | none | NA | NA_GAT | >1bpdel_ATATGTG | 9 | TRUE | FALSE |
| 168_v.txt | 32375 | 32375-Deleti | Deletion | 3 | AAT | - | 20 | 46483 | 0.04302648 | 63 | WT | FALSE | CAN | JSY13 | 1A | WT-Rep1A | CAA | nope | >1bp-ins | >1bpdel | none | >1bpdel | AATAGCAAT | AATAATA | GTT | CAA | AAT | none | NA | NA_CAA | >1bpdel_AATAATA | 9 | TRUE | FALSE |
| 168_v.txt | 32384 | 32384-Deleti | Deletion | 2 | GG | - | 3 | 46483 | 6.45E-03 | 63.3333333 | WT | FALSE | CAN | JSY13 | 1A | WT-Rep1A | AGG | nope | >1bp-ins | >1bpdel | none | >1bpdel | AATAAAGGG | GGGAGAA | TCC | AGG | GG | none | NA | NA_AGG | >1bpdel_GGGAGAA | 9 | TRUE | FALSE |
| 168_v.txt | 32398 | 32398-Deleti | Deletion | 2 | CG | - | 3 | 46483 | 6.45E-03 | 63.3333333 | WT | FALSE | CAN | JSY13 | 1A | WT-Rep1A | ACG | nope | >1bp-ins | >1bpdel | none | >1bpdel | TAGAAACGT | CGTAGGA | TGC | ACG | CG | none | NA | NA_ACG | >1bpdel_CGTAGGA | 9 | TRUE | FALSE |
| 168_v.txt | 32454 | 32454-Deleti | Deletion | 3 | AAT | - | 30 | 46483 | 0.06453972 | 55.1 | WT | FALSE | CAN | JSY13 | 1A | WT-Rep1A | GAA | nope | >1bp-ins | >1bpdel | none | >1bpdel | CCAATGAAT | AATAATA | CTT | GAA | AAT | none | NA | NA_GAA | >1bpdel_AATAATA | 9 | TRUE | FALSE |
| 168_v.txt | 32458 | 32458-Deleti | Deletion | 2 | AT | - | 29 | 46483 | 0.0623884 | 53.1724138 | WT | FALSE | CAN | JSY13 | 1A | WT-Rep1A | AAT | nope | >1bp-ins | >1bpdel | none | >1bpdel | TGAATAATA | ATAGAGA | TTA | AAT | AT | none | NA | NA_AAT | >1bpdel_ATAGAGA | 9 | TRUE | FALSE |
| 168_v.txt | 32458 | 32458-Deleti | Deletion | 4 | ATAG | - | 5 | 46483 | 0.01075662 | 48.4 | WT | FALSE | CAN | JSY13 | 1A | WT-Rep1A | AAT | nope | >1bp-ins | >1bpdel | none | >1bpdel | TGAATAATA | ATAGAGA | TTA | AAT | ATAG | none | NA | NA_AAT | >1bpdel_ATAGAGA | 9 | TRUE | FALSE |
| 168_v.txt | 32460 | 32460-Deleti | Deletion | 2 | AG | - | 12 | 46483 | 0.02581589 | 50.5 | WT | FALSE | CAN | JSY13 | 1A | WT-Rep1A | TAG | nope | >1bp-ins | >1bpdel | none | >1bpdel | AATAATAGA | AGAGAGC | ATC | TAG | AG | none | NA | NA_TAG | >1bpdel_AGAGAGC | 9 | TRUE | FALSE |
| 168_v.txt | 32471 | 32471-Deleti | Deletion | 3 | GTA | - | 1 | 46483 | 2.15E-03 | 38 | WT | FALSE | CAN | JSY13 | 1A | WT-Rep1A | TGT | nope | >1bp-ins | >1bpdel | none | >1bpdel | GCCAATGTA | GTAGAAG | ACA | TGT | GTA | none | NA | NA_TGT | >1bpdel_GTAGAAG | 9 | TRUE | FALSE |
| 168_v.txt | 32473 | 32473-Deleti | Deletion | 3 | AGA | - | 28 | 46483 | 0.06023708 | 51.25 | WT | FALSE | CAN | JSY13 | 1A | WT-Rep1A | TAG | nope | >1bp-ins | >1bpdel | none | >1bpdel | CAATGTAGA | AGAAGGT | ATC | TAG | AGA | none | NA | NA_TAG | >1bpdel_AGAAGGT | 9 | TRUE | FALSE |
| 168_v.txt | 32492 | 32492-Deleti | Deletion | 3 | AAC | - | 1292 | 46483 | 2.77951079 | 45.9613003 | WT | FALSE | CAN | JSY13 | 1A | WT-Rep1A | AAA | nope | >1bp-ins | >1bpdel | none | >1bpdel | ACGGAAAAC | AACAACT | TTT | AAA | AAC | none | NA | NA_AAA | >1bpdel_AACAACT | 9 | TRUE | FALSE |
| 168_v.txt | 32545 | 32545-Deleti | Deletion | 3 | CAG | - | 12 | 46481 | 0.025817 | 37.5833333 | WT | FALSE | CAN | JSY13 | 1A | WT-Rep1A | CCA | nope | >1bp-ins | >1bpdel | none | >1bpdel | CTTCACCAG | CAGCAGT | GGT | CCA | CAG | none | NA | NA_CCA | >1bpdel_CAGCAGT | 9 | TRUE | FALSE |
| 168_v.txt | 32919 | 32919-Deleti | Deletion | 2 | AG | - | 1 | 22850 | 4.38E-03 | 11.0000819 | WT | FALSE | CAN | JSY13 | 1A | WT-Rep1A | CAG | nope | >1bp-ins | >1bpdel | none | >1bpdel | AGTTCCAGG | AGGGCAA | GTC | CAG | AG | none | NA | NA_CAG | >1bpdel_AGGGCAA | 9 | TRUE | FALSE |
| 168_v.txt | 32928 | 32928-Deleti | Deletion | 2 | GT | - | 1 | 22886 | 4.37E-03 | 11 | WT | FALSE | CAN | JSY13 | 1A | WT-Rep1A | AGT | nope | >1bp-ins | >1bpdel | none | >1bpdel | GCAAAAGTG | GTGATTG | TCA | AGT | GT | none | NA | NA_AGT | >1bpdel_GTGATTG | 9 | TRUE | FALSE |
| 168_v.txt | 33065 | 33065-Deleti | Deletion | 2 | AT | - | 17 | 22886 | 0.07428122 | 57.1176471 | WT | FALSE | CAN | JSY13 | 1A | WT-Rep1A | AAT | nope | >1bp-ins | >1bpdel | none | >1bpdel | GACAGAATA | ATATGCC | TTA | AAT | AT | none | NA | NA_AAT | >1bpdel_ATATGCC | 9 | TRUE | FALSE |
| 168_v.txt | 33094 | 33094-Deleti | Deletion | 2 | AT | - | 116 | 22886 | 0.50686009 | 50.9655172 | WT | FALSE | CAN | JSY13 | 1A | WT-Rep1A | GAT | nope | >1bp-ins | >1bpdel | none | >1bpdel | AATATGATA | ATATAAG | CTA | GAT | AT | none | NA | NA_GAT | >1bpdel_ATATAAG | 9 | TRUE | FALSE |
| 168_v.txt | 33206 | 33206-Deleti | Deletion | 2 | CT | - | 13 | 23355 | 0.0556626 | 37.9230769 | WT | FALSE | CAN | JSY13 | 1A | WT-Rep1A | GCT | nope | >1bp-ins | >1bpdel | none | >1bpdel | CTTAAGCTCT | CTCTCTC | CGA | GCT | CT | none | NA | NA_GCT | >1bpdel_CTCTCTC | 9 | TRUE | FALSE |
| 168_v.txt | 33281 | 33281-Deleti | Deletion | 13 | CTCGCCATTT | - | 36 | 23355 | 0.15414258 | 53.75 | WT | FALSE | CAN | JSY13 | 1A | WT-Rep1A | CCT | nope | >1bp-ins | >1bpdel | none | >1bpdel | CGTATCCTCG | CTCGCCA | GGA | CCT | CTCGCCATTT | none | NA | NA_CCT | >1bpdel_CTCGCCA | 9 | TRUE | FALSE |
| 168_v.txt | 31723 | 31723-Deleti | Deletion | 1 | A | - | 3 | 3138 | 0.0956023 | 55.3333333 | WT | FALSE | CAN | JSY13 | 1A | WT-Rep1A | CAA | nope | >1bp-ins | A/T-1 | none | A/T-1 | TGTCCCAAA | AAAAAGT | GTT | GTT | T | none | T>none | T>none_GTT | A/T-1_AAAAAGT | 9 | TRUE | FALSE |
| 168_v.txt | 31904 | 31904-Deleti | Deletion | 1 | T | - | 24 | 3127 | 0.76750879 | 40.4166667 | WT | FALSE | CAN | JSY13 | 1A | WT-Rep1A | ATT | nope | >1bp-ins | A/T-1 | none | A/T-1 | ATTGAATTTT | TTTTGGT | TAA | ATT | T | none | nope | nope_ATT | A/T-1_TTTTGGT | 9 | TRUE | FALSE |
| 168_v.txt | 31952 | 31952-Deleti | Deletion | 1 | A | - | 7 | 14348 | 0.04878729 | 37.8571429 | WT | FALSE | CAN | JSY13 | 1A | WT-Rep1A | TAA | nope | >1bp-ins | A/T-1 | none | A/T-1 | CGTCATAAAT | AAATGTG | ATT | ATT | T | none | T>none | T>none_ATT | A/T-1_AAATGTG | 9 | TRUE | FALSE |
| 168_v.txt | 31972 | 31972-Deleti | Deletion | 1 | C | - | 3 | 14355 | 0.02089864 | 37.6666667 | WT | FALSE | CAN | JSY13 | 1A | WT-Rep1A | GCC | nope | >1bp-ins | G/C-1 | none | G/C-1 | AATAAGCCA | CCAAGCC | CGG | GCC | C | none | nope | nope_GCC | G/C-1_CCAAGCC | 9 | TRUE | FALSE |
| 168_v.txt | 32015 | 32015-Deleti | Deletion | 1 | A | - | 2 | 14355 | 0.01393243 | 63.9999912 | WT | FALSE | CAN | JSY13 | 1A | WT-Rep1A | GAG | nope | >1bp-ins | A/T-1 | none | A/T-1 | GTCACGAGA | AGAGATG | CTC | CTC | T | none | T>none | T>none_CTC | A/T-1_AGAGATG | 9 | TRUE | FALSE |
| 168_v.txt | 32070 | 32070-Deleti | Deletion | 1 | A | - | 3 | 14355 | 0.02089864 | 62.6666691 | WT | FALSE | CAN | JSY13 | 1A | WT-Rep1A | AAA | nope | >1bp-ins | A/T-1 | none | A/T-1 | GAGATAAAT | AATAACC | TTT | TTT | T | none | T>none | T>none_TTT | A/T-1_AATAACC | 9 | TRUE | FALSE |
| 168_v.txt | 32073 | 32073-Deleti | Deletion | 1 | A | - | 7 | 14355 | 0.0487635 | 63.9999936 | WT | FALSE | CAN | JSY13 | 1A | WT-Rep1A | TAA | nope | >1bp-ins | A/T-1 | none | A/T-1 | ATAAATAAC | AACCATG | ATT | ATT | T | none | T>none | T>none_ATT | A/T-1_AACCATG | 9 | TRUE | FALSE |
| 168_v.txt | 32075 | 32075-Deleti | Deletion | 1 | C | - | 3 | 14354 | 0.0209001 | 62 | WT | FALSE | CAN | JSY13 | 1A | WT-Rep1A | ACC | nope | >1bp-ins | G/C-1 | none | G/C-1 | AAATAACCA | CCATGCA | TGG | ACC | C | none | nope | nope_ACC | G/C-1_CCATGCA | 9 | TRUE | FALSE |
| 168_v.txt | 32081 | 32081-Deleti | Deletion | 1 | A | - | 74 | 14355 | 0.51549983 | 62.1621622 | WT | FALSE | CAN | JSY13 | 1A | WT-Rep1A | CAA | nope | >1bp-ins | A/T-1 | none | A/T-1 | CCATGCAAA | AAAAAAG | GTT | GTT | T | none | T>none | T>none_GTT | A/T-1_AAAAAAG | 9 | TRUE | FALSE |
| 168_v.txt | 32121 | 32121-Deleti | Deletion | 1 | A | - | 7 | 14355 | 0.0487635 | 56.5714286 | WT | FALSE | CAN | JSY13 | 1A | WT-Rep1A | GAA | nope | >1bp-ins | A/T-1 | none | A/T-1 | CATTCGAAA | AAAACTT | CTT | CTT | T | none | T>none | T>none_CTT | A/T-1_AAAACTT | 9 | TRUE | FALSE |
| 168_v.txt | 32149 | 32149-Deleti | Deletion | 1 | C | - | 6 | 14355 | 0.04179728 | 46.5 | WT | FALSE | CAN | JSY13 | 1A | WT-Rep1A | TCC | nope | >1bp-ins | G/C-1 | none | G/C-1 | ATGTCTCCAT | CCATGTA | AGG | TCC | C | none | nope | nope_TCC | G/C-1_CCATGTA | 9 | TRUE | FALSE |
| 168_v.txt | 32160 | 32160-Deleti | Deletion | 1 | A | - | 13 | 14355 | 0.09056078 | 49.9230769 | WT | FALSE | CAN | JSY13 | 1A | WT-Rep1A | CAA | nope | >1bp-ins | A/T-1 | none | A/T-1 | TAAGCCAAA | AAAGCGC | GTT | GTT | T | none | T>none | T>none_GTT | A/T-1_AAAGCGC | 9 | TRUE | FALSE |
| 168_v.txt | 32227 | 32227-Deleti | Deletion | 1 | G | - | 16 | 14341 | 0.11156823 | 37.9999988 | WT | FALSE | CAN | JSY13 | 1A | WT-Rep1A | AGG | nope | >1bp-ins | G/C-1 | none | G/C-1 | TTGACAGGA | GGAATTT | TCC | TCC | C | none | C>none | C>none_TCC | G/C-1_GGAATTT | 9 | TRUE | FALSE |
| 168_v.txt | 32239 | 32239-Deleti | Deletion | 1 | C | - | 18 | 14341 | 0.12551426 | 37.9444444 | WT | FALSE | CAN | JSY13 | 1A | WT-Rep1A | GCC | nope | >1bp-ins | G/C-1 | none | G/C-1 | TAGGAGCCA | CCAACTT | CGG | GCC | C | none | nope | nope_GCC | G/C-1_CCAACTT | 9 | TRUE | FALSE |
| 168_v.txt | 32247 | 32247-Deleti | Deletion | 1 | T | - | 6 | 14341 | 0.04183809 | 36.1666667 | WT | FALSE | CAN | JSY13 | 1A | WT-Rep1A | GTT | nope | >1bp-ins | A/T-1 | none | A/T-1 | AACTTGTTCT | TTCTTTG | CAA | GTT | T | none | nope | nope_GTT | A/T-1_TTCTTTG | 9 | TRUE | FALSE |
| 168_v.txt | 32265 | 32265-Deleti | Deletion | 1 | A | - | 8 | 46483 | 0.01721059 | 36.625 | WT | FALSE | CAN | JSY13 | 1A | WT-Rep1A | TAA | nope | >1bp-ins | A/T-1 | none | A/T-1 | CCAAATAAA | AAAATAC | ATT | ATT | T | none | T>none | T>none_ATT | A/T-1_AAAATAC | 9 | TRUE | FALSE |
| 168_v.txt | 32275 | 32275-Deleti | Deletion | 1 | A | - | 73 | 46483 | 0.15704666 | 36.8630157 | WT | FALSE | CAN | JSY13 | 1A | WT-Rep1A | GAA | nope | >1bp-ins | A/T-1 | none | A/T-1 | TACGGGAAC | AACCAAC | CTT | CTT | T | none | T>none | T>none_CTT | A/T-1_AACCAAC | 9 | TRUE | FALSE |
| 168_v.txt | 32283 | 32283-Deleti | Deletion | 1 | T | - | 12 | 46483 | 0.02581589 | 61.8333261 | WT | FALSE | CAN | JSY13 | 1A | WT-Rep1A | GTA | nope | >1bp-ins | A/T-1 | none | A/T-1 | CCAACGTAA | TAAATAT | CAT | GTA | T | none | nope | nope_GTA | A/T-1_TAAATAT | 9 | TRUE | FALSE |
| 168_v.txt | 32284 | 32284-Deleti | Deletion | 1 | A | - | 4 | 46483 | 8.61E-03 | 36.7500041 | WT | FALSE | CAN | JSY13 | 1A | WT-Rep1A | TAA | nope | >1bp-ins | A/T-1 | none | A/T-1 | CAACGTAAA | AAATATT | ATT | ATT | T | none | T>none | T>none_ATT | A/T-1_AAATATT | 9 | TRUE | FALSE |
| 168_v.txt | 32289 | 32289-Deleti | Deletion | 1 | T | - | 10 | 46483 | 0.02151324 | 49.1000176 | WT | FALSE | CAN | JSY13 | 1A | WT-Rep1A | ATT | nope | >1bp-ins | A/T-1 | none | A/T-1 | TAAATATTTG | TTTGAAT | TAA | ATT | T | none | nope | nope_ATT | A/T-1_TTTGAAT | 9 | TRUE | FALSE |
| 168_v.txt | 32304 | 32304-Deleti | Deletion | 1 | G | - | 65 | 46483 | 0.13983607 | 58.3230769 | WT | FALSE | CAN | JSY13 | 1A | WT-Rep1A | AGA | nope | >1bp-ins | G/C-1 | none | G/C-1 | GCGGCAGAA | GAAATAA | TCT | TCT | C | none | C>none | C>none_TCT | G/C-1_GAAATAA | 9 | TRUE | FALSE |
| 168_v.txt | 32311 | 32311-Deleti | Deletion | 1 | T | - | 90 | 46483 | 0.19361917 | 57.0555538 | WT | FALSE | CAN | JSY13 | 1A | WT-Rep1A | ATG | nope | >1bp-ins | A/T-1 | none | A/T-1 | AAATAATGG | TGGTTGT | TAC | ATG | T | none | nope | nope_ATG | A/T-1_TGGTTGT | 9 | TRUE | FALSE |
| 168_v.txt | 32316 | 32316-Deleti | Deletion | 1 | G | - | 1 | 46483 | 2.15E-03 | 48 | WT | FALSE | CAN | JSY13 | 1A | WT-Rep1A | TGT | nope | >1bp-ins | G/C-1 | none | G/C-1 | ATGGTTGTT | GTTAAGA | ACA | ACA | C | none | C>none | C>none_ACA | G/C-1_GTTAAGA | 9 | TRUE | FALSE |
| 168_v.txt | 32319 | 32319-Deleti | Deletion | 1 | A | - | 3 | 46483 | 6.45E-03 | 57.3333302 | WT | FALSE | CAN | JSY13 | 1A | WT-Rep1A | TAA | nope | >1bp-ins | A/T-1 | none | A/T-1 | GTTGTTAAG | AAGATAA | ATT | ATT | T | none | T>none | T>none_ATT | A/T-1_AAGATAA | 9 | TRUE | FALSE |
| 168_v.txt | 32346 | 32346-Deleti | Deletion | 1 | A | - | 15 | 46483 | 0.03226986 | 59.8000048 | WT | FALSE | CAN | JSY13 | 1A | WT-Rep1A | CAA | nope | >1bp-ins | A/T-1 | none | A/T-1 | TGTGGCAAA | AAAACCT | GTT | GTT | T | none | T>none | T>none_GTT | A/T-1_AAAACCT | 9 | TRUE | FALSE |
| 168_v.txt | 32384 | 32384-Deleti | Deletion | 1 | G | - | 9 | 46482 | 0.01936233 | 63.3333336 | WT | FALSE | CAN | JSY13 | 1A | WT-Rep1A | AGG | nope | >1bp-ins | G/C-1 | none | G/C-1 | AATAAAGGG | GGGAGAA | TCC | TCC | C | none | C>none | C>none_TCC | G/C-1_GGGAGAA | 9 | TRUE | FALSE |
| 168_v.txt | 32398 | 32398-Deleti | Deletion | 1 | C | - | 15 | 46483 | 0.03226986 | 63.8000001 | WT | FALSE | CAN | JSY13 | 1A | WT-Rep1A | ACG | nope | >1bp-ins | G/C-1 | none | G/C-1 | TAGAAACGT | CGTAGGA | TGC | ACG | C | none | nope | nope_ACG | G/C-1_CGTAGGA | 9 | TRUE | FALSE |
| 168_v.txt | 32413 | 32413-Deleti | Deletion | 1 | G | - | 3 | 46482 | 6.45E-03 | 55.3333394 | WT | FALSE | CAN | JSY13 | 1A | WT-Rep1A | TGT | nope | >1bp-ins | G/C-1 | none | G/C-1 | TAGATTGTG | GTGTTAG | ACA | ACA | C | none | C>none | C>none_ACA | G/C-1_GTGTTAG | 9 | TRUE | FALSE |
| 168_v.txt | 32430 | 32430-Deleti | Deletion | 1 | T | - | 13 | 46483 | 0.02796721 | 57.6153846 | WT | FALSE | CAN | JSY13 | 1A | WT-Rep1A | ATT | nope | >1bp-ins | A/T-1 | none | A/T-1 | GGGTCATTG | TTGTATG | TAA | ATT | T | none | nope | nope_ATT | A/T-1_TTGTATG | 9 | TRUE | FALSE |
| 168_v.txt | 32449 | 32449-Deleti | Deletion | 1 | C | - | 5 | 46483 | 0.01075662 | 48.4 | WT | FALSE | CAN | JSY13 | 1A | WT-Rep1A | CCA | nope | >1bp-ins | G/C-1 | none | G/C-1 | AAAGTCCAA | CAATGAA | GGT | CCA | C | none | nope | nope_CCA | G/C-1_CAATGAA | 9 | TRUE | FALSE |
| 168_v.txt | 32454 | 32454-Deleti | Deletion | 1 | A | - | 438 | 46482 | 0.94230025 | 52.4497926 | WT | FALSE | CAN | JSY13 | 1A | WT-Rep1A | GAA | nope | >1bp-ins | A/T-1 | none | A/T-1 | CCAATGAAT | AATAATA | CTT | CTT | T | none | T>none | T>none_CTT | A/T-1_AATAATA | 9 | TRUE | FALSE |
| 168_v.txt | 32463 | 32463-Deleti | Deletion | 1 | G | - | 5 | 46483 | 0.01075662 | 48.4 | WT | FALSE | CAN | JSY13 | 1A | WT-Rep1A | AGA | nope | >1bp-ins | G/C-1 | none | G/C-1 | AATAGAGAG | GAGCCAA | TCT | TCT | C | none | C>none | C>none_TCT | G/C-1_GAGCCAA | 9 | TRUE | FALSE |
| 168_v.txt | 32492 | 32492-Deleti | Deletion | 1 | A | - | 1 | 46483 | 2.15E-03 | 37.9999334 | WT | FALSE | CAN | JSY13 | 1A | WT-Rep1A | AAA | nope | >1bp-ins | A/T-1 | none | A/T-1 | ACGGAAAAC | AACAACT | TTT | TTT | T | none | T>none | T>none_TTT | A/T-1_AACAACT | 9 | TRUE | FALSE |
| 168_v.txt | 32498 | 32498-Deleti | Deletion | 1 | T | - | 103 | 46483 | 0.22158639 | 44.6116418 | WT | FALSE | CAN | JSY13 | 1A | WT-Rep1A | CTT | nope | >1bp-ins | A/T-1 | none | A/T-1 | AACAACTTTT | TTTTTTG | GAA | CTT | T | none | nope | nope_CTT | A/T-1_TTTTTTG | 9 | TRUE | FALSE |
| 168_v.txt | 32528 | 32528-Deleti | Deletion | 1 | G | - | 81 | 46483 | 0.17425726 | 40.9259241 | WT | FALSE | CAN | JSY13 | 1A | WT-Rep1A | TGG | nope | >1bp-ins | G/C-1 | none | G/C-1 | TTTTCTGGG | GGGGTTT | ACC | ACC | C | none | C>none | C>none_ACC | G/C-1_GGGGTTT | 9 | TRUE | FALSE |
| 168_v.txt | 32532 | 32532-Deleti | Deletion | 1 | T | - | 12 | 46482 | 0.02581645 | 38.5 | WT | FALSE | CAN | JSY13 | 1A | WT-Rep1A | GTT | nope | >1bp-ins | A/T-1 | none | A/T-1 | CTGGGGTTT | TTTGCAG | CAA | GTT | T | none | nope | nope_GTT | A/T-1_TTTGCAG | 9 | TRUE | FALSE |
| 168_v.txt | 32542 | 32542-Deleti | Deletion | 1 | C | - | 3 | 46481 | 6.45E-03 | 37.9999968 | WT | FALSE | CAN | JSY13 | 1A | WT-Rep1A | TCA | nope | >1bp-ins | G/C-1 | none | G/C-1 | CAGCTTCACC | CACCAGC | AGT | TCA | C | none | nope | nope_TCA | G/C-1_CACCAGC | 9 | TRUE | FALSE |
| 168_v.txt | 32658 | 32658-Deleti | Deletion | 1 | C | - | 4 | 7614 | 0.0525348 | 64 | WT | FALSE | CAN | JSY13 | 1A | WT-Rep1A | ACC | nope | >1bp-ins | G/C-1 | none | G/C-1 | CCTGGACCC | CCCCAGG | TGG | ACC | C | none | nope | nope_ACC | G/C-1_CCCCAGG | 9 | TRUE | FALSE |
| 168_v.txt | 32677 | 32677-Deleti | Deletion | 1 | T | - | 9 | 7614 | 0.11820331 | 60.2222222 | WT | FALSE | CAN | JSY13 | 1A | WT-Rep1A | CTC | nope | >1bp-ins | A/T-1 | none | A/T-1 | GGTTTCTCCA | TCCAATA | GAG | CTC | T | none | nope | nope_CTC | A/T-1_TCCAATA | 9 | TRUE | FALSE |
| 168_v.txt | 32707 | 32707-Deleti | Deletion | 1 | C | - | 16 | 7622 | 0.20991866 | 62.8125 | WT | FALSE | CAN | JSY13 | 1A | WT-Rep1A | ACC | nope | >1bp-ins | G/C-1 | none | G/C-1 | CGGTAACCC | CCCCAGC | TGG | ACC | C | none | nope | nope_ACC | G/C-1_CCCCAGC | 9 | TRUE | FALSE |
| 168_v.txt | 32715 | 32715-Deleti | Deletion | 1 | C | - | 10 | 7622 | 0.13119916 | 61.3 | WT | FALSE | CAN | JSY13 | 1A | WT-Rep1A | ACC | nope | >1bp-ins | G/C-1 | none | G/C-1 | CCAGCACCA | CCACAAA | TGG | ACC | C | none | nope | nope_ACC | G/C-1_CCACAAA | 9 | TRUE | FALSE |
| 168_v.txt | 32719 | 32719-Deleti | Deletion | 1 | A | - | 14 | 7622 | 0.18367883 | 64 | WT | FALSE | CAN | JSY13 | 1A | WT-Rep1A | CAA | nope | >1bp-ins | A/T-1 | none | A/T-1 | CACCACAAA | AAACCAT | GTT | GTT | T | none | T>none | T>none_GTT | A/T-1_AAACCAT | 9 | TRUE | FALSE |
| 168_v.txt | 32728 | 32728-Deleti | Deletion | 1 | A | - | 25 | 7622 | 0.3279979 | 60.5599873 | WT | FALSE | CAN | JSY13 | 1A | WT-Rep1A | CAA | nope | >1bp-ins | A/T-1 | none | A/T-1 | CCATACAAA | AAAAACA | GTT | GTT | T | none | T>none | T>none_GTT | A/T-1_AAAAACA | 9 | TRUE | FALSE |
| 168_v.txt | 32760 | 32760-Deleti | Deletion | 1 | A | - | 9 | 7622 | 0.11807924 | 55.3333207 | WT | FALSE | CAN | JSY13 | 1A | WT-Rep1A | TAA | nope | >1bp-ins | A/T-1 | none | A/T-1 | ATGGCTAAA | AAAACTT | ATT | ATT | T | none | T>none | T>none_ATT | A/T-1_AAAACTT | 9 | TRUE | FALSE |
| 168_v.txt | 32788 | 32788-Deleti | Deletion | 1 | C | - | 3 | 7622 | 0.03935975 | 43.6666667 | WT | FALSE | CAN | JSY13 | 1A | WT-Rep1A | TCG | nope | >1bp-ins | G/C-1 | none | G/C-1 | AGAACTCGA | CGAATTC | AGC | TCG | C | none | nope | nope_TCG | G/C-1_CGAATTC | 9 | TRUE | FALSE |
| 168_v.txt | 32804 | 32804-Deleti | Deletion | 1 | T | - | 2 | 7622 | 0.02623983 | 38 | WT | FALSE | CAN | JSY13 | 1A | WT-Rep1A | ATT | nope | >1bp-ins | A/T-1 | none | A/T-1 | GTAATATTTG | TTTGACA | TAA | ATT | T | none | nope | nope_ATT | A/T-1_TTTGACA | 9 | TRUE | FALSE |
| 168_v.txt | 32811 | 32811-Deleti | Deletion | 1 | G | - | 5 | 7622 | 0.06559958 | 37.8 | WT | FALSE | CAN | JSY13 | 1A | WT-Rep1A | AGG | nope | >1bp-ins | G/C-1 | none | G/C-1 | TTGACAGGG | GGGAACA | TCC | TCC | C | none | C>none | C>none_TCC | G/C-1_GGGAACA | 9 | TRUE | FALSE |
| 168_v.txt | 32816 | 32816-Deleti | Deletion | 1 | C | - | 6 | 7622 | 0.0787195 | 51 | WT | FALSE | CAN | JSY13 | 1A | WT-Rep1A | ACA | nope | >1bp-ins | G/C-1 | none | G/C-1 | AGGGAACAA | CAAGTTC | TGT | ACA | C | none | nope | nope_ACA | G/C-1_CAAGTTC | 9 | TRUE | FALSE |
| 168_v.txt | 32842 | 32842-Deleti | Deletion | 1 | A | - | 66 | 7622 | 0.86591446 | 41.6818182 | WT | FALSE | CAN | JSY13 | 1A | WT-Rep1A | CAA | nope | >1bp-ins | A/T-1 | none | A/T-1 | TTACCCAAA | AAAAAAT | GTT | GTT | T | none | T>none | T>none_GTT | A/T-1_AAAAAAT | 9 | TRUE | FALSE |

| | | | | | | | | | | | | | | | | | | | | | | | | | | | | | | | | | | |
|---|---|---|---|---|---|---|---|---|---|---|---|---|---|---|---|---|---|---|---|---|---|---|---|---|---|---|---|---|---|---|---|---|---|---|
| 168_v.txt | 32884 | 32884-Deleti | Deletion | 1 | A | - | 29 | 22850 | 0.12691466 | 37.4827586 | WT | FALSE | CAN | JSY13 | 1A | WT-Rep1A | CAA | nope | >1bp-ins | A/T-1 | none | A/T-1 | ACGTCCAAA | AAAATTG | GTT | GTT | T | none | T>none | T>none_GTT | A/T-1_AAAATTG | 9 | TRUE | FALSE |
| 168 v.txt | 32917 | 32917-Deleti | Deletion | 1 | C | - | 5 | 22850 | 0.02188184 | 48.6 | WT | FALSE | CAN | JSY13 | 1A | WT-Rep1A | TCC | nope | >1bp-ins | G/C-1 | none | G/C-1 | TAAGTTCCAC | CCAGGGC | AGG | TCC | C | none | nope | nope TCC | G/C-1 CCAGGGC | 9 | TRUE | FALSE |
| 168_v.txt | 32920 | 32920-Deleti | Deletion | 1 | G | - | 6 | 22850 | 0.02625821 | 54 | WT | FALSE | CAN | JSY13 | 1A | WT-Rep1A | AGG | nope | >1bp-ins | G/C-1 | none | G/C-1 | GTTCCAGGG | GGGCAAA | TCC | TCC | C | none | C>none | C>none_TCC | G/C-1_GGGCAAA | 9 | TRUE | FALSE |
| 168 v.txt | 32924 | 32924-Deleti | Deletion | 1 | A | - | 27 | 22850 | 0.11816193 | 50.2592643 | WT | FALSE | CAN | JSY13 | 1A | WT-Rep1A | CAA | nope | >1bp-ins | A/T-1 | none | A/T-1 | CAGGGCAAA | AAAAGTG | GTT | GTT | T | none | T>none | T>none GTT | A/T-1 AAAAGTG | 9 | TRUE | FALSE |
| 168_v.txt | 32934 | 32934-Deleti | Deletion | 1 | G | - | 23 | 22886 | 0.10049812 | 53.4782436 | WT | FALSE | CAN | JSY13 | 1A | WT-Rep1A | TGC | nope | >1bp-ins | G/C-1 | none | G/C-1 | GTGATTGCCC | GCCCAAG | ACG | ACG | C | none | C>none | C>none_ACG | G/C-1_GCCCAAG | 9 | TRUE | FALSE |
| 168 v.txt | 32935 | 32935-Deleti | Deletion | 1 | C | - | 23 | 22886 | 0.10049812 | 57.4782609 | WT | FALSE | CAN | JSY13 | 1A | WT-Rep1A | GCC | nope | >1bp-ins | G/C-1 | none | G/C-1 | TGATTGCCC | CCCAAGA | CGG | GCC | C | none | nope | nope GCC | G/C-1 CCCAAGA | 9 | TRUE | FALSE |
| 168 v.txt | 32941 | 32941-Deleti | Deletion | 1 | A | - | 12 | 22886 | 0.0524338 | 53.2500052 | WT | FALSE | CAN | JSY13 | 1A | WT-Rep1A | GAA | nope | >1bp-ins | A/T-1 | none | A/T-1 | CCCAAGAAA | AAAACCA | CTT | CTT | T | none | T>none | T>none CTT | A/T-1 AAAACCA | 9 | TRUE | FALSE |
| 168_v.txt | 32986 | 32986-Deleti | Deletion | 1 | G | - | 12 | 22886 | 0.0524338 | 60.0833333 | WT | FALSE | CAN | JSY13 | 1A | WT-Rep1A | AGG | nope | >1bp-ins | G/C-1 | none | G/C-1 | GAGAAAGG | GGAATCT | TCC | TCC | C | none | C>none | C>none_TCC | G/C-1_GGAATCT | 9 | TRUE | FALSE |
| 168 v.txt | 33000 | 33000-Deleti | Deletion | 1 | A | - | 67 | 22886 | 0.2927554 | 63.1641791 | WT | FALSE | CAN | JSY13 | 1A | WT-Rep1A | GAA | nope | >1bp-ins | A/T-1 | none | A/T-1 | TGTGAGAAA | AAAACTG | CTT | CTT | T | none | T>none | T>none CTT | A/T-1 AAAACTG | 9 | TRUE | FALSE |
| 168_v.txt | 33020 | 33020-Deleti | Deletion | 1 | A | - | 9 | 22886 | 0.03932535 | 60.6666667 | WT | FALSE | CAN | JSY13 | 1A | WT-Rep1A | TAA | nope | >1bp-ins | A/T-1 | none | A/T-1 | GGATGTAAC | AACAGGG | ATT | ATT | T | none | T>none | T>none_ATT | A/T-1_AACAGGG | 9 | TRUE | FALSE |
| 168 v.txt | 33040 | 33040-Deleti | Deletion | 1 | T | - | 11 | 22885 | 0.04806642 | 62.8181814 | WT | FALSE | CAN | JSY13 | 1A | WT-Rep1A | ATT | nope | >1bp-ins | A/T-1 | none | A/T-1 | TAGCCATTTC | TTTCACC | TAA | ATT | T | none | nope | nope ATT | A/T-1 TTTCACC | 9 | TRUE | FALSE |
| 168 v.txt | 33072 | 33072-Deleti | Deletion | 1 | A | - | 22 | 22886 | 0.09612864 | 50.8181818 | WT | FALSE | CAN | JSY13 | 1A | WT-Rep1A | CAA | nope | >1bp-ins | A/T-1 | none | A/T-1 | TATGCCAAA | AAAGAAC | GTT | GTT | T | none | T>none | T>none GTT | A/T-1 AAAGAAC | 9 | TRUE | FALSE |
| 168_v.txt | 33075 | 33075-Deleti | Deletion | 1 | G | - | 6 | 22886 | 0.0262169 | 44.5000099 | WT | FALSE | CAN | JSY13 | 1A | WT-Rep1A | AGA | nope | >1bp-ins | G/C-1 | none | G/C-1 | GCCAAAGAA | GAACCCA | TCT | TCT | C | none | C>none | C>none_TCT | G/C-1_GAACCCA | 9 | TRUE | FALSE |
| 168 v.txt | 33083 | 33083-Deleti | Deletion | 1 | A | - | 28 | 22886 | 0.12234554 | 50 | WT | FALSE | CAN | JSY13 | 1A | WT-Rep1A | TAA | nope | >1bp-ins | A/T-1 | none | A/T-1 | ACCCATAAA | AAATAAA | ATT | ATT | T | none | T>none | T>none ATT | A/T-1 AAATAAA | 9 | TRUE | FALSE |
| 168_v.txt | 33111 | 33111-Deleti | Deletion | 1 | G | - | 59 | 22886 | 0.25779953 | 41.1355932 | WT | FALSE | CAN | JSY13 | 1A | WT-Rep1A | TGG | nope | >1bp-ins | G/C-1 | none | G/C-1 | CCCACTGGG | GGGCCGG | ACC | ACC | C | none | C>none | C>none_ACC | G/C-1_GGGCCGG | 9 | TRUE | FALSE |
| 168 v.txt | 33147 | 33147-Deleti | Deletion | 1 | A | - | 31 | 22886 | 0.13545399 | 38.5806413 | WT | FALSE | CAN | JSY13 | 1A | WT-Rep1A | GAA | nope | >1bp-ins | A/T-1 | none | A/T-1 | CCAATGAAA | AAAAGAC | CTT | CTT | T | none | T>none | T>none CTT | A/T-1 AAAAGAC | 9 | TRUE | FALSE |
| 168 v.txt | 33153 | 33153-Deleti | Deletion | 1 | C | - | 6 | 22886 | 0.0262169 | 37.6666667 | WT | FALSE | CAN | JSY13 | 1A | WT-Rep1A | ACC | nope | >1bp-ins | G/C-1 | none | G/C-1 | AAAAGACCT | CCTGTAC | TGG | ACC | C | none | nope | nope ACC | G/C-1 CCTGTAC | 9 | TRUE | FALSE |
| 168_v.txt | 33168 | 33168-Deleti | Deletion | 1 | C | - | 6 | 22847 | 0.02626165 | 38 | WT | FALSE | CAN | JSY13 | 1A | WT-Rep1A | ACC | nope | >1bp-ins | G/C-1 | none | G/C-1 | ATAGTACCA | CCACCAA | TGG | ACC | C | none | nope | nope_ACC | G/C-1_CCACCAA | 9 | TRUE | FALSE |
| 168 v.txt | 33171 | 33171-Deleti | Deletion | 1 | C | - | 3 | 22847 | 0.01313083 | 38 | WT | FALSE | CAN | JSY13 | 1A | WT-Rep1A | ACC | nope | >1bp-ins | G/C-1 | none | G/C-1 | GTACCACCA | CCAAGGG | TGG | ACC | C | none | nope | nope ACC | G/C-1 CCAAGGG | 9 | TRUE | FALSE |
| 168_v.txt | 33175 | 33175-Deleti | Deletion | 1 | G | - | 9 | 22843 | 0.03939938 | 38 | WT | FALSE | CAN | JSY13 | 1A | WT-Rep1A | AGG | nope | >1bp-ins | G/C-1 | none | G/C-1 | CACCAAGGG | GGGCAAT | TCC | TCC | C | none | C>none | C>none_TCC | G/C-1_GGGCAAT | 9 | TRUE | FALSE |
| 168 v.txt | 33178 | 33178-Deleti | Deletion | 1 | C | - | 4 | 22843 | 0.01751084 | 35.000002 | WT | FALSE | CAN | JSY13 | 1A | WT-Rep1A | GCA | nope | >1bp-ins | G/C-1 | none | G/C-1 | CAAGGGCAA | CAATCAT | CGT | GCA | C | none | nope | nope GCA | G/C-1 CAATCAT | 9 | TRUE | FALSE |
| 168 v.txt | 33201 | 33201-Deleti | Deletion | 1 | T | - | 19 | 23355 | 0.08135303 | 37.8947368 | WT | FALSE | CAN | JSY13 | 1A | WT-Rep1A | CTT | nope | >1bp-ins | A/T-1 | none | A/T-1 | CTTTGCTTAA | TTAAGCT | GAA | CTT | T | none | nope | nope CTT | A/T-1 TTAAGCT | 9 | TRUE | FALSE |
| 168_v.txt | 33209 | 33209-Deleti | Deletion | 1 | T | - | 8 | 23355 | 0.03425391 | 37.7500038 | WT | FALSE | CAN | JSY13 | 1A | WT-Rep1A | CTC | nope | >1bp-ins | A/T-1 | none | A/T-1 | AAGCTCTCTC | TCTCTTC | GAG | CTC | T | none | nope | nope_CTC | A/T-1_TCTCTTC | 9 | TRUE | FALSE |
| 168 v.txt | 33258 | 33258-Deleti | Deletion | 1 | C | - | 23 | 23355 | 0.09847998 | 37.5217391 | WT | FALSE | CAN | JSY13 | 1A | WT-Rep1A | GCC | nope | >1bp-ins | G/C-1 | none | G/C-1 | CCTATGCCAT | CCATCCT | CGG | GCC | C | none | nope | nope GCC | G/C-1 CCATCCT | 9 | TRUE | FALSE |
| 168_v.txt | 33284 | 33284-Deleti | Deletion | 1 | G | - | 3 | 23355 | 0.01284522 | 46.6666662 | WT | FALSE | CAN | JSY13 | 1A | WT-Rep1A | CGC | nope | >1bp-ins | G/C-1 | none | G/C-1 | ATCCTCGCCA | GCCATTT | GCG | GCG | C | none | C>none | C>none_GCG | G/C-1_GCCATTT | 9 | TRUE | FALSE |
| 168 v.txt | 33285 | 33285-Deleti | Deletion | 1 | C | - | 2 | 23355 | 8.56E-03 | 38.0000085 | WT | FALSE | CAN | JSY13 | 1A | WT-Rep1A | GCC | nope | >1bp-ins | G/C-1 | none | G/C-1 | TCCTCGCCAT | CCATTTA | CGG | GCC | C | none | nope | nope GCC | G/C-1 CCATTTA | 9 | TRUE | FALSE |
| 168 v.txt | 33288 | 33288-Deleti | Deletion | 1 | T | - | 2 | 23355 | 8.56E-03 | 61.4999997 | WT | FALSE | CAN | JSY13 | 1A | WT-Rep1A | ATT | nope | >1bp-ins | A/T-1 | none | A/T-1 | TCGCCATTTA | TTTACTC | TAA | ATT | T | none | nope | nope ATT | A/T-1 TTTACTC | 9 | TRUE | FALSE |
| 168_v.txt | 33373 | 33373-Deleti | Deletion | 1 | T | - | 34 | 23355 | 0.14557911 | 57.1176471 | WT | FALSE | CAN | JSY13 | 1A | WT-Rep1A | GTT | nope | >1bp-ins | A/T-1 | none | A/T-1 | GGTGTGTTTC | TTTGTGA | CAA | GTT | T | none | nope | nope_GTT | A/T-1_TTTGTGA | 9 | TRUE | FALSE |
| 168 v.txt | 33474 | 33474-Deleti | Deletion | 1 | C | - | 63 | 23349 | 0.26981884 | 29.0793651 | WT | FALSE | CAN | JSY13 | 1A | WT-Rep1A | GCC | nope | >1bp-ins | G/C-1 | none | G/C-1 | GCTATGCCTT | CCTTTTT | CGG | GCC | C | none | nope | nope GCC | G/C-1 CCTTTTT | 9 | TRUE | FALSE |
| 170_v.txt | 31723 | 31723-SNV- | SNV | 1 | A | C | 1 | 3907 | 0.02559509 | 64 | WT | FALSE | CAN | JSY13 | 2A | WT-Rep2A | CAA | A>C | G/C+1 | A/T-1 | TA>GC | TA>GC | TGTCCCAAA | AAAAAGT | GTT | GTT | T | G | T>G | T>G_GTT | TA>GC_AAAAAGT | 9 | TRUE | FALSE |
| 170 v.txt | 31839 | 31839-SNV- | SNV | 1 | A | C | 3 | 3907 | 0.07678526 | 55.3333333 | WT | FALSE | CAN | JSY13 | 2A | WT-Rep2A | TAA | A>C | G/C+1 | A/T-1 | TA>GC | TA>GC | TGAAATAAG | AAGATCC | ATT | ATT | T | G | T>G | T>G ATT | TA>GC AAGATCC | 9 | TRUE | FALSE |
| 170 v.txt | 31860 | 31860-SNV- | SNV | 1 | A | C | 4 | 3904 | 0.10245902 | 43.75 | WT | FALSE | CAN | JSY13 | 2A | WT-Rep2A | CAG | A>C | G/C+1 | A/T-1 | TA>GC | TA>GC | AAGAACAGG | AGGAAAA | GTC | GTC | T | G | T>G | T>G GTC | TA>GC AGGAAAA | 9 | TRUE | FALSE |
| 170 v.txt | 31986 | 31986-SNV- | SNV | 1 | A | C | 12 | 20510 | 0.05850805 | 48.5 | WT | FALSE | CAN | JSY13 | 2A | WT-Rep2A | TAA | A>C | G/C+1 | A/T-1 | TA>GC | TA>GC | GGCATTAATT | AATTTAG | ATT | ATT | T | G | T>G | T>G ATT | TA>GC AATTTAG | 9 | TRUE | FALSE |
| 170 v.txt | 32161 | 32161-SNV- | SNV | 1 | A | C | 1 | 20506 | 4.88E-03 | 54.9986623 | WT | FALSE | CAN | JSY13 | 2A | WT-Rep2A | AAA | A>C | G/C+1 | A/T-1 | TA>GC | TA>GC | AAGCCAAAG | AAGCGCC | TTT | TTT | T | G | T>G | T>G TTT | TA>GC AAGCGCC | 9 | TRUE | FALSE |
| 170_v.txt | 32162 | 32162-SNV- | SNV | 1 | A | C | 4 | 20506 | 0.01950649 | 24.0011499 | WT | FALSE | CAN | JSY13 | 2A | WT-Rep2A | AAG | A>C | G/C+1 | A/T-1 | TA>GC | TA>GC | AGCCAAAGC | AGCGCCA | TTC | TTC | T | G | T>G | T>G_TTC | TA>GC_AGCGCCA | 9 | TRUE | FALSE |
| 170 v.txt | 32256 | 32256-SNV- | SNV | 1 | A | C | 14 | 43940 | 0.03186163 | 23.5714286 | WT | FALSE | CAN | JSY13 | 2A | WT-Rep2A | TAG | A>C | G/C+1 | A/T-1 | TA>GC | TA>GC | TTTGATAGA | AGACCAA | ATC | ATC | T | G | T>G | T>G ATC | TA>GC AGACCAA | 9 | TRUE | FALSE |
| 170_v.txt | 32319 | 32319-SNV- | SNV | 1 | A | C | 15 | 43957 | 0.03412426 | 52.1333333 | WT | FALSE | CAN | JSY13 | 2A | WT-Rep2A | TAA | A>C | G/C+1 | A/T-1 | TA>GC | TA>GC | GTTGTTAAG | AAGATAA | ATT | ATT | T | G | T>G | T>G_ATT | TA>GC_AAGATAA | 9 | TRUE | FALSE |
| 170 v.txt | 32441 | 32441-SNV-T | SNV | 1 | T | G | 17 | 43957 | 0.03867416 | 55.9988307 | WT | FALSE | CAN | JSY13 | 2A | WT-Rep2A | CTA | T>G | G/C+1 | A/T-1 | TA>GC | TA>GC | TGGAACTAA | TAAAAGT | GAT | CTA | T | C | T>G | T>G CTA | TA>GC TAAAAGT | 9 | TRUE | FALSE |
| 170 v.txt | 32445 | 32445-SNV- | SNV | 1 | A | C | 16 | 43957 | 0.03639921 | 49.125 | WT | FALSE | CAN | JSY13 | 2A | WT-Rep2A | AAG | A>C | G/C+1 | A/T-1 | TA>GC | TA>GC | ACTAAAAGT | AGTCCAA | TTC | TTC | T | G | T>G | T>G TTC | TA>GC AGTCCAA | 9 | TRUE | FALSE |
| 170_v.txt | 32472 | 32472-SNV-T | SNV | 1 | T | G | 72 | 43957 | 0.16379644 | 48.8466387 | WT | FALSE | CAN | JSY13 | 2A | WT-Rep2A | GTA | T>G | G/C+1 | A/T-1 | TA>GC | TA>GC | CCAATGTAG | TAGAAGG | CAT | GTA | T | C | T>G | T>G_GTA | TA>GC_TAGAAGG | 9 | TRUE | FALSE |
| 170 v.txt | 32473 | 32473-SNV- | SNV | 1 | A | C | 17 | 43957 | 0.03867416 | 45.8803022 | WT | FALSE | CAN | JSY13 | 2A | WT-Rep2A | TAG | A>C | G/C+1 | A/T-1 | TA>GC | TA>GC | CAATGTAGA | AGAAGGT | ATC | ATC | T | G | T>G | T>G ATC | TA>GC AGAAGGT | 9 | TRUE | FALSE |
| 170_v.txt | 32475 | 32475-SNV- | SNV | 1 | A | C | 8 | 43957 | 0.0181996 | 47.7485388 | WT | FALSE | CAN | JSY13 | 2A | WT-Rep2A | GAA | A>C | G/C+1 | A/T-1 | TA>GC | TA>GC | ATGTAGAAG | AAGGTTA | CTT | CTT | T | G | T>G | T>G_CTT | TA>GC_AAGGTTA | 9 | TRUE | FALSE |
| 170 v.txt | 32573 | 32573-SNV-T | SNV | 1 | T | G | 22 | 9395 | 0.23416711 | 36.4545455 | WT | FALSE | CAN | JSY13 | 2A | WT-Rep2A | CTT | T>G | G/C+1 | A/T-1 | TA>GC | TA>GC | AGTACCTTG | TTGAAAT | GAA | CTT | T | C | T>G | T>G CTT | TA>GC TTGAAAT | 9 | TRUE | FALSE |
| 170 v.txt | 32574 | 32574-SNV-T | SNV | 1 | T | G | 4 | 9395 | 0.04257584 | 37.998869 | WT | FALSE | CAN | JSY13 | 2A | WT-Rep2A | TTG | T>G | G/C+1 | A/T-1 | TA>GC | TA>GC | GTACCTTGA | TGAAATG | AAC | TTG | T | C | T>G | T>G TTG | TA>GC TGAAATG | 9 | TRUE | FALSE |
| 170_v.txt | 32662 | 32662-SNV- | SNV | 1 | A | C | 3 | 9394 | 0.03193528 | 37 | WT | FALSE | CAN | JSY13 | 2A | WT-Rep2A | CAG | A>C | G/C+1 | A/T-1 | TA>GC | TA>GC | GACCCCAGG | AGGCACC | GTC | GTC | T | G | T>G | T>G_GTC | TA>GC_AGGCACC | 9 | TRUE | FALSE |
| 170 v.txt | 32884 | 32884-SNV- | SNV | 1 | A | C | 3 | 34435 | 8.71E-03 | 36.9986414 | WT | FALSE | CAN | JSY13 | 2A | WT-Rep2A | CAA | A>C | G/C+1 | A/T-1 | TA>GC | TA>GC | ACGTCCAAA | AAAATTG | GTT | GTT | T | G | T>G | T>G GTT | TA>GC AAAATTG | 9 | TRUE | FALSE |
| 170_v.txt | 32915 | 32915-SNV-T | SNV | 1 | T | G | 37 | 34437 | 0.10744258 | 45.6216216 | WT | FALSE | CAN | JSY13 | 2A | WT-Rep2A | GTT | T>G | G/C+1 | A/T-1 | TA>GC | TA>GC | ACTAAGTTCC | TTCCAGG | CAA | GTT | T | C | T>G | T>G_GTT | TA>GC_TTCCAGG | 9 | TRUE | FALSE |
| 170 v.txt | 32919 | 32919-SNV- | SNV | 1 | A | C | 47 | 34437 | 0.13648111 | 46.3404255 | WT | FALSE | CAN | JSY13 | 2A | WT-Rep2A | CAG | A>C | G/C+1 | A/T-1 | TA>GC | TA>GC | AGTTCCAGG | AGGGCAA | GTC | GTC | T | G | T>G | T>G GTC | TA>GC AGGGCAA | 9 | TRUE | FALSE |
| 170 v.txt | 33166 | 33166-SNV-T | SNV | 1 | T | G | 10 | 34418 | 0.02905457 | 37.5 | WT | FALSE | CAN | JSY13 | 2A | WT-Rep2A | GTA | T>G | G/C+1 | A/T-1 | TA>GC | TA>GC | CAATAGTACC | TACCACC | CAT | GTA | T | C | T>G | T>G GTA | TA>GC TACCACC | 9 | TRUE | FALSE |
| 170_v.txt | 33204 | 33204-SNV- | SNV | 1 | A | C | 27 | 17689 | 0.15263723 | 38.2962963 | WT | FALSE | CAN | JSY13 | 2A | WT-Rep2A | AAG | A>C | G/C+1 | A/T-1 | TA>GC | TA>GC | TGCTTAAGCT | AGCTCTC | TTC | TTC | T | G | T>G | T>G_TTC | TA>GC_AGCTCTC | 9 | TRUE | FALSE |
| 170 v.txt | 33269 | 33269-SNV- | SNV | 1 | A | C | 115 | 17691 | 0.65004805 | 16.7304348 | WT | FALSE | CAN | JSY13 | 2A | WT-Rep2A | TAG | A>C | G/C+1 | A/T-1 | TA>GC | TA>GC | CTCCATAGA | AGAGAAC | ATC | ATC | T | G | T>G | T>G ATC | TA>GC AGAGAAC | 9 | TRUE | FALSE |
| 170_v.txt | 31964 | 31964-SNV- | SNV | 1 | A | G | 5 | 20510 | 0.02437835 | 37.3999133 | WT | FALSE | CAN | JSY13 | 2A | WT-Rep2A | CAT | A>G | G/C+1 | A/T-1 | TA>CG | TA>CG | GGCCGCATA | ATAATAA | GTA | GTA | T | C | T>C | T>C_GTA | TA>CG_ATAATAA | 9 | TRUE | FALSE |
| 170 v.txt | 31965 | 31965-SNV-T | SNV | 1 | T | C | 2 | 20510 | 9.75E-03 | 31.4999946 | WT | FALSE | CAN | JSY13 | 2A | WT-Rep2A | ATA | T>C | G/C+1 | A/T-1 | TA>CG | TA>CG | GCCGCATAA | TAATAAG | TAT | ATA | T | G | T>C | T>C ATA | TA>CG TAATAAG | 9 | TRUE | FALSE |
| 170 v.txt | 32081 | 32081-SNV- | SNV | 1 | A | G | 7 | 20510 | 0.03412969 | 57.8567693 | WT | FALSE | CAN | JSY13 | 2A | WT-Rep2A | CAA | A>G | G/C+1 | A/T-1 | TA>CG | TA>CG | CCATGCAAA | AAAAAAG | GTT | GTT | T | C | T>C | T>C GTT | TA>CG AAAAAAG | 9 | TRUE | FALSE |
| 170_v.txt | 32161 | 32161-SNV- | SNV | 1 | A | G | 8 | 20506 | 0.03901297 | 35.499207 | WT | FALSE | CAN | JSY13 | 2A | WT-Rep2A | AAA | A>G | G/C+1 | A/T-1 | TA>CG | TA>CG | AAGCCAAAG | AAGCGCC | TTT | TTT | T | C | T>C | T>C_TTT | TA>CG_AAGCGCC | 9 | TRUE | FALSE |
| 170 v.txt | 32162 | 32162-SNV- | SNV | 1 | A | G | 9 | 20506 | 0.04388959 | 32.8891129 | WT | FALSE | CAN | JSY13 | 2A | WT-Rep2A | AAG | A>G | G/C+1 | A/T-1 | TA>CG | TA>CG | AGCCAAAGC | AGCGCCA | TTC | TTC | T | C | T>C | T>C TTC | TA>CG AGCGCCA | 9 | TRUE | FALSE |
| 170_v.txt | 32387 | 32387-SNV- | SNV | 1 | A | G | 28 | 43957 | 0.06369862 | 51.6785714 | WT | FALSE | CAN | JSY13 | 2A | WT-Rep2A | GAG | A>G | G/C+1 | A/T-1 | TA>CG | TA>CG | AAAGGGAGA | AGAAGTA | CTC | CTC | T | C | T>C | T>C_CTC | TA>CG_AGAAGTA | 9 | TRUE | FALSE |
| 170 v.txt | 32445 | 32445-SNV- | SNV | 1 | A | G | 61 | 43957 | 0.13877198 | 49.295082 | WT | FALSE | CAN | JSY13 | 2A | WT-Rep2A | AAG | A>G | G/C+1 | A/T-1 | TA>CG | TA>CG | ACTAAAAGT | AGTCCAA | TTC | TTC | T | C | T>C | T>C TTC | TA>CG AGTCCAA | 9 | TRUE | FALSE |
| 170 v.txt | 32476 | 32476-SNV- | SNV | 1 | A | G | 28 | 43957 | 0.06369862 | 43.7489905 | WT | FALSE | CAN | JSY13 | 2A | WT-Rep2A | AAG | A>G | G/C+1 | A/T-1 | TA>CG | TA>CG | TGTAGAAGG | AGGTTAA | TTC | TTC | T | C | T>C | T>C TTC | TA>CG AGGTTAA | 9 | TRUE | FALSE |
| 170_v.txt | 32479 | 32479-SNV-T | SNV | 1 | T | C | 8 | 43957 | 0.0181996 | 38.8748747 | WT | FALSE | CAN | JSY13 | 2A | WT-Rep2A | GTT | T>C | G/C+1 | A/T-1 | TA>CG | TA>CG | AGAAGGTTA | TTAAGAT | CAA | GTT | T | G | T>C | T>C_GTT | TA>CG_TTAAGAT | 9 | TRUE | FALSE |
| 170 v.txt | 32573 | 32573-SNV-T | SNV | 1 | T | C | 3 | 9395 | 0.03193188 | 38 | WT | FALSE | CAN | JSY13 | 2A | WT-Rep2A | CTT | T>C | G/C+1 | A/T-1 | TA>CG | TA>CG | AGTACCTTG | TTGAAAT | GAA | CTT | T | G | T>C | T>C CTT | TA>CG TTGAAAT | 9 | TRUE | FALSE |
| 170_v.txt | 32919 | 32919-SNV- | SNV | 1 | A | G | 31 | 34437 | 0.09001946 | 43.3870968 | WT | FALSE | CAN | JSY13 | 2A | WT-Rep2A | CAG | A>G | G/C+1 | A/T-1 | TA>CG | TA>CG | AGTTCCAGG | AGGGCAA | GTC | GTC | T | C | T>C | T>C_GTC | TA>CG_AGGGCAA | 9 | TRUE | FALSE |
| 170 v.txt | 32947 | 32947-SNV- | SNV | 1 | A | G | 31 | 34437 | 0.09001946 | 59.2258065 | WT | FALSE | CAN | JSY13 | 2A | WT-Rep2A | CAA | A>G | G/C+1 | A/T-1 | TA>CG | TA>CG | AAAACCAAT | AATACAT | GTT | GTT | T | C | T>C | T>C GTT | TA>CG AATACAT | 9 | TRUE | FALSE |
| 170 v.txt | 33042 | 33042-SNV-T | SNV | 1 | T | C | 24 | 34437 | 0.06969248 | 51.0793848 | WT | FALSE | CAN | JSY13 | 2A | WT-Rep2A | TTC | T>C | G/C+1 | A/T-1 | TA>CG | TA>CG | GCCATTTCAC | TCACCCA | AAG | TTC | T | G | T>C | T>C TTC | TA>CG TCACCCA | 9 | TRUE | FALSE |
| 170_v.txt | 33126 | 33126-SNV- | SNV | 1 | A | G | 33 | 34437 | 0.09582716 | 40.4255851 | WT | FALSE | CAN | JSY13 | 2A | WT-Rep2A | CAG | A>G | G/C+1 | A/T-1 | TA>CG | TA>CG | TTGGTCAGA | AGAGGTG | GTC | GTC | T | C | T>C | T>C_GTC | TA>CG_AGAGGTG | 9 | TRUE | FALSE |
| 170 v.txt | 33204 | 33204-SNV- | SNV | 1 | A | G | 13 | 17689 | 0.073492 | 39 | WT | FALSE | CAN | JSY13 | 2A | WT-Rep2A | AAG | A>G | G/C+1 | A/T-1 | TA>CG | TA>CG | TGCTTAAGCT | AGCTCTC | TTC | TTC | T | C | T>C | T>C TTC | TA>CG AGCTCTC | 9 | TRUE | FALSE |
| 170_v.txt | 33207 | 33207-SNV-T | SNV | 1 | T | C | 9 | 17689 | 0.05087908 | 18.7777713 | WT | FALSE | CAN | JSY13 | 2A | WT-Rep2A | CTC | T>C | G/C+1 | A/T-1 | TA>CG | TA>CG | TTAAGCTCTC | TCTCTCT | GAG | CTC | T | G | T>C | T>C_CTC | TA>CG_TCTCTCT | 9 | TRUE | FALSE |
| 170 v.txt | 33209 | 33209-SNV-T | SNV | 1 | T | C | 4 | 17689 | 0.02261292 | 39.2499663 | WT | FALSE | CAN | JSY13 | 2A | WT-Rep2A | CTC | T>C | G/C+1 | A/T-1 | TA>CG | TA>CG | AAGCTCTCTC | TCTCTTC | GAG | CTC | T | G | T>C | T>C CTC | TA>CG TCTCTTC | 9 | TRUE | FALSE |
| 170 v.txt | 33211 | 33211-SNV-T | SNV | 1 | T | C | 6 | 17691 | 0.03391555 | 33.1648696 | WT | FALSE | CAN | JSY13 | 2A | WT-Rep2A | CTC | T>C | G/C+1 | A/T-1 | TA>CG | TA>CG | GCTCTCTCTT | TCTTCAC | GAG | CTC | T | G | T>C | T>C CTC | TA>CG TCTTCAC | 9 | TRUE | FALSE |
| 170_v.txt | 33476 | 33476-SNV-T | SNV | 1 | T | C | 30 | 17691 | 0.16957775 | 32.1666667 | WT | FALSE | CAN | JSY13 | 2A | WT-Rep2A | CTT | T>C | G/C+1 | A/T-1 | TA>CG | TA>CG | TATGCCTTTT | TTTTTTT | GAA | CTT | T | G | T>C | T>C_CTT | TA>CG_TTTTTTT | 9 | TRUE | FALSE |
| 170 v.txt | 31964 | 31964-SNV- | SNV | 1 | A | T | 4 | 20510 | 0.01950268 | 24.4999261 | WT | FALSE | CAN | JSY13 | 2A | WT-Rep2A | CAT | A>T | A/T+1 | A/T-1 | TA>AT | TA>AT | GGCCGCATA | ATAATAA | GTA | GTA | T | A | T>A | T>A GTA | TA>AT ATAATAA | 9 | TRUE | FALSE |
| 170_v.txt | 31965 | 31965-SNV-T | SNV | 1 | T | A | 4 | 20510 | 0.01950268 | 33.9995382 | WT | FALSE | CAN | JSY13 | 2A | WT-Rep2A | ATA | T>A | A/T+1 | A/T-1 | TA>AT | TA>AT | GCCGCATAA | TAATAAG | TAT | ATA | T | T | T>A | T>A_ATA | TA>AT_TAATAAG | 9 | TRUE | FALSE |
| 170 v.txt | 32265 | 32265-SNV- | SNV | 1 | A | T | 19 | 43957 | 0.04322406 | 37.5263158 | WT | FALSE | CAN | JSY13 | 2A | WT-Rep2A | TAA | A>T | A/T+1 | A/T-1 | TA>AT | TA>AT | CCAAATAAA | AAAATAC | ATT | ATT | T | A | T>A | T>A ATT | TA>AT AAAATAC | 9 | TRUE | FALSE |
| 170 v.txt | 32319 | 32319-SNV- | SNV | 1 | A | T | 351 | 43957 | 0.79850763 | 59.8603989 | WT | FALSE | CAN | JSY13 | 2A | WT-Rep2A | TAA | A>T | A/T+1 | A/T-1 | TA>AT | TA>AT | GTTGTTAAG | AAGATAA | ATT | ATT | T | A | T>A | T>A ATT | TA>AT AAGATAA | 9 | TRUE | FALSE |
| 170_v.txt | 32422 | 32422-SNV-T | SNV | 1 | T | A | 40 | 43957 | 0.09099802 | 55.55 | WT | FALSE | CAN | JSY13 | 2A | WT-Rep2A | TTA | T>A | A/T+1 | A/T-1 | TA>AT | TA>AT | TTAGTTTAGG | TAGGGTC | AAT | TTA | T | T | T>A | T>A_TTA | TA>AT_TAGGGTC | 9 | TRUE | FALSE |
| 170 v.txt | 32470 | 32470-SNV-T | SNV | 1 | T | A | 7 | 43957 | 0.01592465 | 43.4271365 | WT | FALSE | CAN | JSY13 | 2A | WT-Rep2A | ATG | T>A | A/T+1 | A/T-1 | TA>AT | TA>AT | AGCCAATGT | TGTAGAA | TAC | ATG | T | T | T>A | T>A ATG | TA>AT TGTAGAA | 9 | TRUE | FALSE |
| 170_v.txt | 32472 | 32472-SNV-T | SNV | 1 | T | A | 7 | 43957 | 0.01592465 | 51.4250975 | WT | FALSE | CAN | JSY13 | 2A | WT-Rep2A | GTA | T>A | A/T+1 | A/T-1 | TA>AT | TA>AT | CCAATGTAG | TAGAAGG | CAT | GTA | T | T | T>A | T>A_GTA | TA>AT_TAGAAGG | 9 | TRUE | FALSE |
| 170 v.txt | 32479 | 32479-SNV-T | SNV | 1 | T | A | 1 | 43957 | 2.28E-03 | 36.9997201 | WT | FALSE | CAN | JSY13 | 2A | WT-Rep2A | GTT | T>A | A/T+1 | A/T-1 | TA>AT | TA>AT | AGAAGGTTA | TTAAGAT | CAA | GTT | T | T | T>A | T>A GTT | TA>AT TTAAGAT | 9 | TRUE | FALSE |
| 170 v.txt | 32541 | 32541-SNV-T | SNV | 1 | T | A | 44 | 43957 | 0.10009782 | 37.7045455 | WT | FALSE | CAN | JSY13 | 2A | WT-Rep2A | TTC | T>A | A/T+1 | A/T-1 | TA>AT | TA>AT | GCAGCTTCAC | TCACCAG | AAG | TTC | T | T | T>A | T>A TTC | TA>AT TCACCAG | 9 | TRUE | FALSE |
| 170_v.txt | 32573 | 32573-SNV-T | SNV | 1 | T | A | 4 | 9395 | 0.04257584 | 38 | WT | FALSE | CAN | JSY13 | 2A | WT-Rep2A | CTT | T>A | A/T+1 | A/T-1 | TA>AT | TA>AT | AGTACCTTG | TTGAAAT | GAA | CTT | T | T | T>A | T>A_CTT | TA>AT_TTGAAAT | 9 | TRUE | FALSE |
| 170 v.txt | 33030 | 33030-SNV- | SNV | 1 | A | T | 5 | 34437 | 0.01451927 | 38.6000162 | WT | FALSE | CAN | JSY13 | 2A | WT-Rep2A | GAA | A>T | A/T+1 | A/T-1 | TA>AT | TA>AT | GGGATGAAT | AATGTAG | CTT | CTT | T | A | T>A | T>A CTT | TA>AT AATGTAG | 9 | TRUE | FALSE |
| 170_v.txt | 33126 | 33126-SNV- | SNV | 1 | A | T | 7 | 34437 | 0.02032697 | 37.7212338 | WT | FALSE | CAN | JSY13 | 2A | WT-Rep2A | CAG | A>T | A/T+1 | A/T-1 | TA>AT | TA>AT | TTGGTCAGA | AGAGGTG | GTC | GTC | T | A | T>A | T>A_GTC | TA>AT_AGAGGTG | 9 | TRUE | FALSE |
| 170 v.txt | 33189 | 33189-SNV- | SNV | 1 | A | T | 3 | 17692 | 0.01695682 | 28.9994243 | WT | FALSE | CAN | JSY13 | 2A | WT-Rep2A | AAT | A>T | A/T+1 | A/T-1 | TA>AT | TA>AT | ATACCAATAT | ATATGTC | TTA | TTA | T | A | T>A | T>A TTA | TA>AT ATATGTC | 9 | TRUE | FALSE |
| 170 v.txt | 33204 | 33204-SNV- | SNV | 1 | A | T | 31 | 17689 | 0.17525016 | 40.3548387 | WT | FALSE | CAN | JSY13 | 2A | WT-Rep2A | AAG | A>T | A/T+1 | A/T-1 | TA>AT | TA>AT | TGCTTAAGCT | AGCTCTC | TTC | TTC | T | A | T>A | T>A TTC | TA>AT AGCTCTC | 9 | TRUE | FALSE |
| 170_v.txt | 33207 | 33207-SNV-T | SNV | 1 | T | A | 2 | 17689 | 0.01130646 | 20.9997962 | WT | FALSE | CAN | JSY13 | 2A | WT-Rep2A | CTC | T>A | A/T+1 | A/T-1 | TA>AT | TA>AT | TTAAGCTCTC | TCTCTCT | GAG | CTC | T | T | T>A | T>A_CTC | TA>AT_TCTCTCT | 9 | TRUE | FALSE |
| 170 v.txt | 33312 | 33312-SNV- | SNV | 1 | A | T | 12 | 17691 | 0.0678311 | 55.25 | WT | FALSE | CAN | JSY13 | 2A | WT-Rep2A | CAA | A>T | A/T+1 | A/T-1 | TA>AT | TA>AT | GAGCGCAAT | AATGGAT | GTT | GTT | T | A | T>A | T>A GTT | TA>AT AATGGAT | 9 | TRUE | FALSE |
| 170_v.txt | 33317 | 33317-SNV- | SNV | 1 | A | T | 7 | 17691 | 0.03956814 | 58 | WT | FALSE | CAN | JSY13 | 2A | WT-Rep2A | GAT | A>T | A/T+1 | A/T-1 | TA>AT | TA>AT | CAATGGATA | ATACAAT | CTA | CTA | T | A | T>A | T>A_CTA | TA>AT_ATACAAT | 9 | TRUE | FALSE |
| 170 v.txt | 33424 | 33424-SNV-T | SNV | 1 | T | A | 14 | 17691 | 0.07913628 | 51.5714286 | WT | FALSE | CAN | JSY13 | 2A | WT-Rep2A | ATA | T>A | A/T+1 | A/T-1 | TA>AT | TA>AT | TGTACATATG | TATGCTT | TAT | ATA | T | T | T>A | T>A ATA | TA>AT TATGCTT | 9 | TRUE | FALSE |
| 170 v.txt | 33426 | 33426-SNV-T | SNV | 1 | T | A | 17 | 17691 | 0.09609406 | 48.4117647 | WT | FALSE | CAN | JSY13 | 2A | WT-Rep2A | ATG | T>A | A/T+1 | A/T-1 | TA>AT | TA>AT | TACATATGCT | TGCTTCT | TAC | ATG | T | T | T>A | T>A ATG | TA>AT TGCTTCT | 9 | TRUE | FALSE |
| 170_v.txt | 31806 | 31806-SNV- | SNV | 1 | C | T | 177 | 3907 | 4.53033018 | 58.6723164 | WT | FALSE | CAN | JSY13 | 2A | WT-Rep2A | CCA | C>T | A/T+1 | G/C-1 | CG>TA | CG>TA | ATCTTCCAAA | CAAATAA | GGT | CCA | C | A | C>T | C>T_CCA | CG>TA_CAAATAA | 9 | TRUE | FALSE |
| 170 v.txt | 31962 | 31962-SNV- | SNV | 1 | G | A | 33 | 20510 | 0.16089712 | 36.969697 | WT | FALSE | CAN | JSY13 | 2A | WT-Rep2A | CGC | G>A | A/T+1 | G/C-1 | CG>TA | CG>TA | GTGGCCGCA | GCATAAT | GCG | GCG | C | T | C>T | C>T GCG | CG>TA GCATAAT | 9 | TRUE | FALSE |
| 170_v.txt | 31981 | 31981-SNV- | SNV | 1 | G | A | 23 | 20510 | 0.11214042 | 46.0869565 | WT | FALSE | CAN | JSY13 | 2A | WT-Rep2A | GGC | G>A | A/T+1 | G/C-1 | CG>TA | CG>TA | AGCCGGGCA | GCATTAA | CCG | CCG | C | T | C>T | C>T_CCG | CG>TA_GCATTAA | 9 | TRUE | FALSE |
| 170 v.txt | 32057 | 32057-SNV- | SNV | 1 | C | T | 20 | 20510 | 0.09751341 | 56.3511331 | WT | FALSE | CAN | JSY13 | 2A | WT-Rep2A | GCG | C>T | A/T+1 | G/C-1 | CG>TA | CG>TA | GATGTGCGA | CGAGATT | CGC | GCG | C | A | C>T | C>T GCG | CG>TA CGAGATT | 9 | TRUE | FALSE |
| 170 v.txt | 32058 | 32058-SNV- | SNV | 1 | G | A | 10 | 20509 | 0.04875908 | 52.6001651 | WT | FALSE | CAN | JSY13 | 2A | WT-Rep2A | CGA | G>A | A/T+1 | G/C-1 | CG>TA | CG>TA | ATGTGCGAG | GAGATTG | GCT | GCT | C | T | C>T | C>T GCT | CG>TA GAGATTG | 9 | TRUE | FALSE |
| 170 v.txt | 32076 | 32076-SNV- | SNV | 1 | C | T | 21 | 20510 | 0.10238908 | 61.5714286 | WT | FALSE | CAN | JSY13 | 2A | WT-Rep2A | CCA | C>T | A/T+1 | G/C-1 | CG>TA | CG>TA | AATAACCATC | CATGCAA | GGT | CCA | C | A | C>T | C>T CCA | CG>TA CATGCAA | 9 | TRUE | FALSE |
| 170 v.txt | 32089 | 32089-SNV- | SNV | 1 | C | T | 232 | 20510 | 1.13115553 | 61.5172414 | WT | FALSE | CAN | JSY13 | 2A | WT-Rep2A | CCT | C>T | A/T+1 | G/C-1 | CG>TA | CG>TA | AAAAGCCTG | CTGCAAC | GGA | CCT | C | A | C>T | C>T CCT | CG>TA CTGCAAC | 9 | TRUE | FALSE |
| 170_v.txt | 32092 | 32092-SNV- | SNV | 1 | C | T | 24 | 20510 | 0.11701609 | 59.7083333 | WT | FALSE | CAN | JSY13 | 2A | WT-Rep2A | GCA | C>T | A/T+1 | G/C-1 | CG>TA | CG>TA | AGCCTGCAA | CAACACC | CGT | GCA | C | A | C>T | C>T_GCA | CG>TA_CAACACC | 9 | TRUE | FALSE |
| 170 v.txt | 32114 | 32114-SNV- | SNV | 1 | C | T | 37 | 20510 | 0.18039981 | 60.2162162 | WT | FALSE | CAN | JSY13 | 2A | WT-Rep2A | GCC | C>T | A/T+1 | G/C-1 | CG>TA | CG>TA | TAATAGCCAT | CCATTCG | CGG | GCC | C | A | C>T | C>T GCC | CG>TA CCATTCG | 9 | TRUE | FALSE |
| 170_v.txt | 32158 | 32158-SNV- | SNV | 1 | C | T | 7 | 20506 | 0.03413635 | 55.4251218 | WT | FALSE | CAN | JSY13 | 2A | WT-Rep2A | GCC | C>T | A/T+1 | G/C-1 | CG>TA | CG>TA | TGTAAGCCA | CCAAAGC | CGG | GCC | C | A | C>T | C>T_GCC | CG>TA_CCAAAGC | 9 | TRUE | FALSE |
| 170 v.txt | 32159 | 32159-SNV- | SNV | 1 | C | T | 5 | 20506 | 0.02438311 | 34.400114 | WT | FALSE | CAN | JSY13 | 2A | WT-Rep2A | CCA | C>T | A/T+1 | G/C-1 | CG>TA | CG>TA | GTAAGCCAA | CAAAGCG | GGT | CCA | C | A | C>T | C>T CCA | CG>TA CAAAGCG | 9 | TRUE | FALSE |
| 170 v.txt | 32164 | 32164-SNV- | SNV | 1 | C | T | 6 | 20506 | 0.02925973 | 46.3321142 | WT | FALSE | CAN | JSY13 | 2A | WT-Rep2A | GCG | C>T | A/T+1 | G/C-1 | CG>TA | CG>TA | CCAAAGCGC | CGCCAAA | CGC | GCG | C | A | C>T | C>T GCG | CG>TA CGCCAAA | 9 | TRUE | FALSE |
| 170_v.txt | 32271 | 32271-SNV- | SNV | 1 | C | T | 75 | 43957 | 0.17062129 | 31.4 | WT | FALSE | CAN | JSY13 | 2A | WT-Rep2A | ACG | C>T | A/T+1 | G/C-1 | CG>TA | CG>TA | AAAATACGG | CGGGAAC | TGC | ACG | C | A | C>T | C>T_ACG | CG>TA_CGGGAAC | 9 | TRUE | FALSE |
| 170 v.txt | 32292 | 32292-SNV- | SNV | 1 | G | A | 31 | 43957 | 0.07052347 | 48.8387097 | WT | FALSE | CAN | JSY13 | 2A | WT-Rep2A | TGA | G>A | A/T+1 | G/C-1 | CG>TA | CG>TA | ATATTTGAAT | GAATTTG | ACT | ACT | C | T | C>T | C>T ACT | CG>TA GAATTTG | 9 | TRUE | FALSE |
| 170_v.txt | 32299 | 32299-SNV- | SNV | 1 | C | T | 58 | 43957 | 0.13194713 | 50.556976 | WT | FALSE | CAN | JSY13 | 2A | WT-Rep2A | GCG | C>T | A/T+1 | G/C-1 | CG>TA | CG>TA | AATTTGCGG | CGGCAGA | CGC | GCG | C | A | C>T | C>T_GCG | CG>TA_CGGCAGA | 9 | TRUE | FALSE |
| 170 v.txt | 32304 | 32304-SNV- | SNV | 1 | G | A | 21 | 43957 | 0.04777396 | 56.2380952 | WT | FALSE | CAN | JSY13 | 2A | WT-Rep2A | AGA | G>A | A/T+1 | G/C-1 | CG>TA | CG>TA | GCGGCAGAA | GAAATAA | TCT | TCT | C | T | C>T | C>T TCT | CG>TA GAAATAA | 9 | TRUE | FALSE |
| 170 v.txt | 32466 | 32466-SNV- | SNV | 1 | C | T | 35 | 43957 | 0.07962327 | 48.4857143 | WT | FALSE | CAN | JSY13 | 2A | WT-Rep2A | GCC | C>T | A/T+1 | G/C-1 | CG>TA | CG>TA | AGAGAGCCA | CCAATGT | CGG | GCC | C | A | C>T | C>T GCC | CG>TA CCAATGT | 9 | TRUE | FALSE |
| 170_v.txt | 32471 | 32471-SNV- | SNV | 1 | G | A | 11 | 43957 | 0.02502446 | 36.6363687 | WT | FALSE | CAN | JSY13 | 2A | WT-Rep2A | TGT | G>A | A/T+1 | G/C-1 | CG>TA | CG>TA | GCCAATGTA | GTAGAAG | ACA | ACA | C | T | C>T | C>T_ACA | CG>TA_GTAGAAG | 9 | TRUE | FALSE |
| 170 v.txt | 32487 | 32487-SNV- | SNV | 1 | C | T | 29 | 43957 | 0.06597357 | 45.6204397 | WT | FALSE | CAN | JSY13 | 2A | WT-Rep2A | ACG | C>T | A/T+1 | G/C-1 | CG>TA | CG>TA | AAGATACGG | CGGAAAA | TGC | ACG | C | A | C>T | C>T ACG | CG>TA CGGAAAA | 9 | TRUE | FALSE |
| 170_v.txt | 32488 | 32488-SNV- | SNV | 1 | G | A | 79 | 43956 | 0.17972518 | 43.9503026 | WT | FALSE | CAN | JSY13 | 2A | WT-Rep2A | CGG | G>A | A/T+1 | G/C-1 | CG>TA | CG>TA | AGATACGGA | GGAAAAC | GCC | GCC | C | T | C>T | C>T_GCC | CG>TA_GGAAAAC | 9 | TRUE | FALSE |
| 170 v.txt | 32535 | 32535-SNV- | SNV | 1 | G | A | 46 | 43957 | 0.10464772 | 37.3478682 | WT | FALSE | CAN | JSY13 | 2A | WT-Rep2A | TGC | G>A | A/T+1 | G/C-1 | CG>TA | CG>TA | GGGTTTGCA | GCAGCTT | ACG | ACG | C | T | C>T | C>T ACG | CG>TA GCAGCTT | 9 | TRUE | FALSE |
| 170 v.txt | 32536 | 32536-SNV- | SNV | 1 | C | T | 52 | 43957 | 0.11829743 | 36.4227635 | WT | FALSE | CAN | JSY13 | 2A | WT-Rep2A | GCA | C>T | A/T+1 | G/C-1 | CG>TA | CG>TA | GGTTTGCAG | CAGCTTC | CGT | GCA | C | A | C>T | C>T GCA | CG>TA CAGCTTC | 9 | TRUE | FALSE |
| 170_v.txt | 32679 | 32679-SNV- | SNV | 1 | C | T | 12 | 9394 | 0.12774111 | 61.3333333 | WT | FALSE | CAN | JSY13 | 2A | WT-Rep2A | CCA | C>T | A/T+1 | G/C-1 | CG>TA | CG>TA | TTTCTCCAAT | CAATAAC | GGT | CCA | C | A | C>T | C>T_CCA | CG>TA_CAATAAC | 9 | TRUE | FALSE |
| 170 v.txt | 32700 | 32700-SNV- | SNV | 1 | C | T | 7 | 9394 | 0.07451565 | 59.5782542 | WT | FALSE | CAN | JSY13 | 2A | WT-Rep2A | GCC | C>T | A/T+1 | G/C-1 | CG>TA | CG>TA | ACTGGGCCG | CCGGTAA | CGG | GCC | C | A | C>T | C>T GCC | CG>TA CCGGTAA | 9 | TRUE | FALSE |
| 170_v.txt | 32860 | 32860-SNV- | SNV | 1 | C | T | 14 | 9393 | 0.14904716 | 37.9285714 | WT | FALSE | CAN | JSY13 | 2A | WT-Rep2A | GCC | C>T | A/T+1 | G/C-1 | CG>TA | CG>TA | TCCATGCCG | CCGCCAG | CGG | GCC | C | A | C>T | C>T_GCC | CG>TA_CCGCCAG | 9 | TRUE | FALSE |
| 170 v.txt | 32883 | 32883-SNV- | SNV | 1 | C | T | 78 | 34435 | 0.22651372 | 36.4485962 | WT | FALSE | CAN | JSY13 | 2A | WT-Rep2A | CCA | C>T | A/T+1 | G/C-1 | CG>TA | CG>TA | TACGTCCAA | CAAAATT | GGT | CCA | C | A | C>T | C>T CCA | CG>TA CAAAATT | 9 | TRUE | FALSE |
| 170 v.txt | 32899 | 32899-SNV- | SNV | 1 | G | A | 197 | 34437 | 0.57205912 | 37.5482234 | WT | FALSE | CAN | JSY13 | 2A | WT-Rep2A | TGG | G>A | A/T+1 | G/C-1 | CG>TA | CG>TA | TGACTTGGC | GGCCAAC | ACC | ACC | C | T | C>T | C>T ACC | CG>TA GGCCAAC | 9 | TRUE | FALSE |
| 170_v.txt | 32901 | 32901-SNV- | SNV | 1 | C | T | 83 | 34437 | 0.24101983 | 31.5783133 | WT | FALSE | CAN | JSY13 | 2A | WT-Rep2A | GCC | C>T | A/T+1 | G/C-1 | CG>TA | CG>TA | ACTTGGCCA | CCAACTA | CGG | GCC | C | A | C>T | C>T_GCC | CG>TA_CCAACTA | 9 | TRUE | FALSE |
| 170 v.txt | 32958 | 32958-SNV- | SNV | 1 | C | T | 35 | 34437 | 0.10163487 | 49.6285714 | WT | FALSE | CAN | JSY13 | 2A | WT-Rep2A | ACC | C>T | A/T+1 | G/C-1 | CG>TA | CG>TA | ATGTAACCAT | CCATTGG | TGG | ACC | C | A | C>T | C>T ACC | CG>TA CCATTGG | 9 | TRUE | FALSE |
| 170_v.txt | 32965 | 32965-SNV- | SNV | 1 | C | T | 32 | 34437 | 0.09292331 | 60.1875 | WT | FALSE | CAN | JSY13 | 2A | WT-Rep2A | GCC | C>T | A/T+1 | G/C-1 | CG>TA | CG>TA | CATTGGCCG | CCGCACC | CGG | GCC | C | A | C>T | C>T_GCC | CG>TA_CCGCACC | 9 | TRUE | FALSE |

| | | | | | | | | | | | | | | | | | | | | | | | | | | | | | | | | | | |
|---|---|---|---|---|---|---|---|---|---|---|---|---|---|---|---|---|---|---|---|---|---|---|---|---|---|---|---|---|---|---|---|---|---|---|
| 170_v.txt | 32967 | 32967-SNV- | SNV | 1 | G | A | 49 | 34437 | 0.14228882 | 60.0204082 | WT | FALSE | CAN | JSY13 | 2A | WT-Rep2A | CGC | G>A | A/T+1 | G/C-1 | CG>TA | CG>TA | TTGGCCGCA | GCACCAA | GCG | GCG | C | T | C>T | C>T_GCG | CG>TA_GCACCAA | 9 | TRUE | FALSE |
| 170_v.txt | 32970 | 32970-SNV- | SNV | 1 | C | T | 59 | 34437 | 0.17132735 | 61.0508475 | WT | FALSE | CAN | JSY13 | 2A | WT-Rep2A | ACC | C>T | A/T+1 | G/C-1 | CG>TA | CG>TA | GCCGCACCA | CCAAATG | TGG | ACC | C | A | C>T | C>T_ACC | CG>TA_CCAAATG | 9 | TRUE | FALSE |
| 170_v.txt | 33012 | 33012-SNV- | SNV | 1 | G | A | 198 | 34436 | 0.57497967 | 60.7924869 | WT | FALSE | CAN | JSY13 | 2A | WT-Rep2A | AGA | G>A | A/T+1 | G/C-1 | CG>TA | CG>TA | GTGAAAGAG | GAGGATG | TCT | TCT | C | T | C>T | C>T_TCT | CG>TA_GAGGATG | 9 | TRUE | FALSE |
| 170_v.txt | 33055 | 33055-SNV- | SNV | 1 | G | A | 272 | 34437 | 0.78984813 | 57.0845588 | WT | FALSE | CAN | JSY13 | 2A | WT-Rep2A | TGC | G>A | A/T+1 | G/C-1 | CG>TA | CG>TA | AGGACTGCG | GCGTGAC | ACG | ACG | C | T | C>T | C>T_ACG | CG>TA_GCGTGAC | 9 | TRUE | FALSE |
| 170_v.txt | 33114 | 33114-SNV- | SNV | 1 | C | T | 26 | 34437 | 0.07550019 | 40.4632664 | WT | FALSE | CAN | JSY13 | 2A | WT-Rep2A | GCC | C>T | A/T+1 | G/C-1 | CG>TA | CG>TA | ACTGGGCCG | CCGGCGT | CGG | GCC | C | A | C>T | C>T_GCC | CG>TA_CCGGCGT | 9 | TRUE | FALSE |
| 170_v.txt | 33115 | 33115-SNV- | SNV | 1 | C | T | 19 | 34437 | 0.05517322 | 38.7368035 | WT | FALSE | CAN | JSY13 | 2A | WT-Rep2A | CCG | C>T | A/T+1 | G/C-1 | CG>TA | CG>TA | CTGGGCCGG | CGGCGTT | GGC | CCG | C | A | C>T | C>T_CCG | CG>TA_CGGCGTT | 9 | TRUE | FALSE |
| 170_v.txt | 33193 | 33193-SNV- | SNV | 1 | G | A | 16 | 17683 | 0.09048238 | 33.5625 | WT | FALSE | CAN | JSY13 | 2A | WT-Rep2A | TGT | G>A | A/T+1 | G/C-1 | CG>TA | CG>TA | CAATATGTCT | GTCTTTG | ACA | ACA | C | T | C>T | C>T_ACA | CG>TA_GTCTTTG | 9 | TRUE | FALSE |
| 170_v.txt | 33199 | 33199-SNV- | SNV | 1 | G | A | 17 | 17689 | 0.09610492 | 36.8823529 | WT | FALSE | CAN | JSY13 | 2A | WT-Rep2A | TGC | G>A | A/T+1 | G/C-1 | CG>TA | CG>TA | GTCTTTGCTT | GCTTAAG | ACG | ACG | C | T | C>T | C>T_ACG | CG>TA_GCTTAAG | 9 | TRUE | FALSE |
| 170_v.txt | 33206 | 33206-SNV- | SNV | 1 | C | T | 3 | 17689 | 0.01695969 | 27.6665844 | WT | FALSE | CAN | JSY13 | 2A | WT-Rep2A | GCT | C>T | A/T+1 | G/C-1 | CG>TA | CG>TA | CTTAAGCTCT | CTCTCTC | CGA | GCT | C | A | C>T | C>T_GCT | CG>TA_CTCTCTC | 9 | TRUE | FALSE |
| 170_v.txt | 33208 | 33208-SNV- | SNV | 1 | C | T | 3 | 17688 | 0.01696065 | 30.9999707 | WT | FALSE | CAN | JSY13 | 2A | WT-Rep2A | TCT | C>T | A/T+1 | G/C-1 | CG>TA | CG>TA | TAAGCTCTCT | CTCTCTT | AGA | TCT | C | A | C>T | C>T_TCT | CG>TA_CTCTCTT | 9 | TRUE | FALSE |
| 170_v.txt | 33258 | 33258-SNV- | SNV | 1 | C | T | 24 | 17691 | 0.1356622 | 43.8333333 | WT | FALSE | CAN | JSY13 | 2A | WT-Rep2A | GCC | C>T | A/T+1 | G/C-1 | CG>TA | CG>TA | CCTATGCCAT | CCATCCT | CGG | GCC | C | A | C>T | C>T_GCC | CG>TA_CCATCCT | 9 | TRUE | FALSE |
| 170_v.txt | 33412 | 33412-SNV- | SNV | 1 | G | A | 19 | 17691 | 0.10739924 | 48.1578947 | WT | FALSE | CAN | JSY13 | 2A | WT-Rep2A | GGC | G>A | A/T+1 | G/C-1 | CG>TA | CG>TA | TGACCGGCT | GCTCATT | CCG | CCG | C | T | C>T | C>T_CCG | CG>TA_GCTCATT | 9 | TRUE | FALSE |
| 170_v.txt | 33474 | 33474-SNV- | SNV | 1 | C | T | 27 | 17691 | 0.15261998 | 32.962963 | WT | FALSE | CAN | JSY13 | 2A | WT-Rep2A | GCC | C>T | A/T+1 | G/C-1 | CG>TA | CG>TA | GCTATGCCTT | CCTTTTT | CGG | GCC | C | A | C>T | C>T_GCC | CG>TA_CCTTTTT | 9 | TRUE | FALSE |
| 170_v.txt | 31981 | 31981-SNV- | SNV | 1 | G | C | 111 | 20510 | 0.54119942 | 44.8378378 | WT | FALSE | CAN | JSY13 | 2A | WT-Rep2A | GGC | G>C | G/C+1 | G/C-1 | CG>GC | CG>GC | AGCCGGGCA | GCATTAA | CCG | CCG | C | G | C>G | C>G_CCG | CG>GC_GCATTAA | 9 | TRUE | FALSE |
| 170_v.txt | 32064 | 32064-SNV- | SNV | 1 | G | C | 6 | 20510 | 0.02925402 | 60.3333333 | WT | FALSE | CAN | JSY13 | 2A | WT-Rep2A | TGA | G>C | G/C+1 | G/C-1 | CG>GC | CG>GC | GAGATTGAG | GAGATAA | ACT | ACT | C | G | C>G | C>G_ACT | CG>GC_GAGATAA | 9 | TRUE | FALSE |
| 170_v.txt | 32092 | 32092-SNV- | SNV | 1 | C | G | 6 | 20510 | 0.02925402 | 53.3333333 | WT | FALSE | CAN | JSY13 | 2A | WT-Rep2A | GCA | C>G | G/C+1 | G/C-1 | CG>GC | CG>GC | AGCCTGCAA | CAACACC | CGT | GCA | C | C | C>G | C>G_GCA | CG>GC_CAACACC | 9 | TRUE | FALSE |
| 170_v.txt | 32282 | 32282-SNV- | SNV | 1 | G | C | 11 | 43957 | 0.02502446 | 36.2727273 | WT | FALSE | CAN | JSY13 | 2A | WT-Rep2A | CGT | G>C | G/C+1 | G/C-1 | CG>GC | CG>GC | ACCAACGTA | GTAAATA | GCA | GCA | C | G | C>G | C>G_GCA | CG>GC_GTAAATA | 9 | TRUE | FALSE |
| 170_v.txt | 32298 | 32298-SNV- | SNV | 1 | G | C | 35 | 43957 | 0.07962327 | 24.944817 | WT | FALSE | CAN | JSY13 | 2A | WT-Rep2A | TGC | G>C | G/C+1 | G/C-1 | CG>GC | CG>GC | GAATTTGCG | GCGGCAG | ACG | ACG | C | G | C>G | C>G_ACG | CG>GC_GCGGCAG | 9 | TRUE | FALSE |
| 170_v.txt | 32471 | 32471-SNV- | SNV | 1 | G | C | 17 | 43957 | 0.03867416 | 51.7639129 | WT | FALSE | CAN | JSY13 | 2A | WT-Rep2A | TGT | G>C | G/C+1 | G/C-1 | CG>GC | CG>GC | GCCAATGTA | GTAGAAG | ACA | ACA | C | G | C>G | C>G_ACA | CG>GC_GTAGAAG | 9 | TRUE | FALSE |
| 170_v.txt | 32487 | 32487-SNV- | SNV | 1 | C | G | 15 | 43957 | 0.03412426 | 41.6662049 | WT | FALSE | CAN | JSY13 | 2A | WT-Rep2A | ACG | C>G | G/C+1 | G/C-1 | CG>GC | CG>GC | AAGATACGG | CGGAAAA | TGC | ACG | C | C | C>G | C>G_ACG | CG>GC_CGGAAAA | 9 | TRUE | FALSE |
| 170_v.txt | 32488 | 32488-SNV- | SNV | 1 | G | C | 23 | 43956 | 0.05232505 | 40.7841067 | WT | FALSE | CAN | JSY13 | 2A | WT-Rep2A | CGG | G>C | G/C+1 | G/C-1 | CG>GC | CG>GC | AGATACGGA | GGAAAAC | GCC | GCC | C | G | C>G | C>G_GCC | CG>GC_GGAAAAC | 9 | TRUE | FALSE |
| 170_v.txt | 32557 | 32557-SNV- | SNV | 1 | C | G | 191 | 43933 | 0.43475292 | 37.5968586 | WT | FALSE | CAN | JSY13 | 2A | WT-Rep2A | CCA | C>G | G/C+1 | G/C-1 | CG>GC | CG>GC | TGATACCAA | CAACTAG | GGT | CCA | C | C | C>G | C>G_CCA | CG>GC_CAACTAG | 9 | TRUE | FALSE |
| 170_v.txt | 32566 | 32566-SNV- | SNV | 1 | C | G | 12 | 9391 | 0.12778192 | 35.6666667 | WT | FALSE | CAN | JSY13 | 2A | WT-Rep2A | TCA | C>G | G/C+1 | G/C-1 | CG>GC | CG>GC | CTAGTTCAGT | CAGTACC | AGT | TCA | C | C | C>G | C>G_TCA | CG>GC_CAGTACC | 9 | TRUE | FALSE |
| 170_v.txt | 32604 | 32604-SNV- | SNV | 1 | G | C | 1 | 9394 | 0.01064509 | 2.00203998 | WT | FALSE | CAN | JSY13 | 2A | WT-Rep2A | GGA | G>C | G/C+1 | G/C-1 | CG>GC | CG>GC | AAAGAGGAA | GAAACCC | CCT | CCT | C | G | C>G | C>G_CCT | CG>GC_GAAACCC | 9 | TRUE | FALSE |
| 170_v.txt | 32811 | 32811-SNV- | SNV | 1 | G | C | 8 | 9394 | 0.08516074 | 36.5 | WT | FALSE | CAN | JSY13 | 2A | WT-Rep2A | AGG | G>C | G/C+1 | G/C-1 | CG>GC | CG>GC | TTGACAGGG | GGGAACA | TCC | TCC | C | G | C>G | C>G_TCC | CG>GC_GGGAACA | 9 | TRUE | FALSE |
| 170_v.txt | 32876 | 32876-SNV- | SNV | 1 | G | C | 21 | 34425 | 0.06100218 | 34.952381 | WT | FALSE | CAN | JSY13 | 2A | WT-Rep2A | TGT | G>C | G/C+1 | G/C-1 | CG>GC | CG>GC | AACTTTGTAC | GTACGTC | ACA | ACA | C | G | C>G | C>G_ACA | CG>GC_GTACGTC | 9 | TRUE | FALSE |
| 170_v.txt | 32937 | 32937-SNV- | SNV | 1 | C | G | 18 | 34437 | 0.05226936 | 50.5555556 | WT | FALSE | CAN | JSY13 | 2A | WT-Rep2A | CCA | C>G | G/C+1 | G/C-1 | CG>GC | CG>GC | ATTGCCCAA | CAAGAAA | GGT | CCA | C | C | C>G | C>G_CCA | CG>GC_CAAGAAA | 9 | TRUE | FALSE |
| 170_v.txt | 33057 | 33057-SNV- | SNV | 1 | G | C | 7 | 34437 | 0.02032697 | 58 | WT | FALSE | CAN | JSY13 | 2A | WT-Rep2A | CGT | G>C | G/C+1 | G/C-1 | CG>GC | CG>GC | GACTGCGTG | GTGACAG | GCA | GCA | C | G | C>G | C>G_GCA | CG>GC_GTGACAG | 9 | TRUE | FALSE |
| 170_v.txt | 33111 | 33111-SNV- | SNV | 1 | G | C | 25 | 34437 | 0.07259634 | 48.28 | WT | FALSE | CAN | JSY13 | 2A | WT-Rep2A | TGG | G>C | G/C+1 | G/C-1 | CG>GC | CG>GC | CCCACTGGG | GGGCCGG | ACC | ACC | C | G | C>G | C>G_ACC | CG>GC_GGGCCGG | 9 | TRUE | FALSE |
| 170_v.txt | 33114 | 33114-SNV- | SNV | 1 | C | G | 2 | 34437 | 5.81E-03 | 37.0040715 | WT | FALSE | CAN | JSY13 | 2A | WT-Rep2A | GCC | C>G | G/C+1 | G/C-1 | CG>GC | CG>GC | ACTGGGCCG | CCGGCGT | CGG | GCC | C | C | C>G | C>G_GCC | CG>GC_CCGGCGT | 9 | TRUE | FALSE |
| 170_v.txt | 33156 | 33156-SNV- | SNV | 1 | G | C | 9 | 34437 | 0.02613468 | 34.1111111 | WT | FALSE | CAN | JSY13 | 2A | WT-Rep2A | TGT | G>C | G/C+1 | G/C-1 | CG>GC | CG>GC | AGACCTGTA | GTACCAA | ACA | ACA | C | G | C>G | C>G_ACA | CG>GC_GTACCAA | 9 | TRUE | FALSE |
| 170_v.txt | 33168 | 33168-SNV- | SNV | 1 | C | G | 40 | 34418 | 0.11621826 | 37.1986294 | WT | FALSE | CAN | JSY13 | 2A | WT-Rep2A | ACC | C>G | G/C+1 | G/C-1 | CG>GC | CG>GC | ATAGTACCA | CCACCAA | TGG | ACC | C | C | C>G | C>G_ACC | CG>GC_CCACCAA | 9 | TRUE | FALSE |
| 170_v.txt | 33206 | 33206-SNV- | SNV | 1 | C | G | 1 | 17689 | 5.65E-03 | 37.9991269 | WT | FALSE | CAN | JSY13 | 2A | WT-Rep2A | GCT | C>G | G/C+1 | G/C-1 | CG>GC | CG>GC | CTTAAGCTCT | CTCTCTC | CGA | GCT | C | C | C>G | C>G_GCT | CG>GC_CTCTCTC | 9 | TRUE | FALSE |
| 170_v.txt | 33208 | 33208-SNV- | SNV | 1 | C | G | 1 | 17688 | 5.65E-03 | 37.9989353 | WT | FALSE | CAN | JSY13 | 2A | WT-Rep2A | TCT | C>G | G/C+1 | G/C-1 | CG>GC | CG>GC | TAAGCTCTCT | CTCTCTT | AGA | TCT | C | C | C>G | C>G_TCT | CG>GC_CTCTCTT | 9 | TRUE | FALSE |
| 170_v.txt | 31806 | 31806-SNV- | SNV | 1 | C | A | 2 | 3907 | 0.05119017 | 64 | WT | FALSE | CAN | JSY13 | 2A | WT-Rep2A | CCA | C>A | A/T+1 | G/C-1 | CG>AT | CG>AT | ATCTTCCAAA | CAAATAA | GGT | CCA | C | T | C>A | C>A_CCA | CG>AT_CAAATAA | 9 | TRUE | FALSE |
| 170_v.txt | 32058 | 32058-SNV- | SNV | 1 | G | T | 290 | 20509 | 1.41401336 | 61.4627502 | WT | FALSE | CAN | JSY13 | 2A | WT-Rep2A | CGA | G>T | A/T+1 | G/C-1 | CG>AT | CG>AT | ATGTGCGAG | GAGATTG | GCT | GCT | C | A | C>A | C>A_GCT | CG>AT_GAGATTG | 9 | TRUE | FALSE |
| 170_v.txt | 32149 | 32149-SNV- | SNV | 1 | C | A | 15 | 20510 | 0.07313506 | 47.0666667 | WT | FALSE | CAN | JSY13 | 2A | WT-Rep2A | TCC | C>A | A/T+1 | G/C-1 | CG>AT | CG>AT | ATGTCTCCAT | CCATGTA | AGG | TCC | C | T | C>A | C>A_TCC | CG>AT_CCATGTA | 9 | TRUE | FALSE |
| 170_v.txt | 32157 | 32157-SNV- | SNV | 1 | G | T | 1 | 20506 | 4.88E-03 | 24.9999986 | WT | FALSE | CAN | JSY13 | 2A | WT-Rep2A | AGC | G>T | A/T+1 | G/C-1 | CG>AT | CG>AT | ATGTAAGCC | GCCAAAG | TCG | TCG | C | A | C>A | C>A_TCG | CG>AT_GCCAAAG | 9 | TRUE | FALSE |
| 170_v.txt | 32223 | 32223-SNV- | SNV | 1 | G | T | 11 | 20506 | 0.05364284 | 39.3636364 | WT | FALSE | CAN | JSY13 | 2A | WT-Rep2A | TGA | G>T | A/T+1 | G/C-1 | CG>AT | CG>AT | GTCCTTGACA | GACAGGA | ACT | ACT | C | A | C>A | C>A_ACT | CG>AT_GACAGGA | 9 | TRUE | FALSE |
| 170_v.txt | 32238 | 32238-SNV- | SNV | 1 | G | T | 5 | 20504 | 0.02438549 | 38 | WT | FALSE | CAN | JSY13 | 2A | WT-Rep2A | AGC | G>T | A/T+1 | G/C-1 | CG>AT | CG>AT | TTAGGAGCC | GCCAACT | TCG | TCG | C | A | C>A | C>A_TCG | CG>AT_GCCAACT | 9 | TRUE | FALSE |
| 170_v.txt | 32282 | 32282-SNV- | SNV | 1 | G | T | 65 | 43957 | 0.14787178 | 41.6923077 | WT | FALSE | CAN | JSY13 | 2A | WT-Rep2A | CGT | G>T | A/T+1 | G/C-1 | CG>AT | CG>AT | ACCAACGTA | GTAAATA | GCA | GCA | C | A | C>A | C>A_GCA | CG>AT_GTAAATA | 9 | TRUE | FALSE |
| 170_v.txt | 32299 | 32299-SNV- | SNV | 1 | C | A | 29 | 43957 | 0.06597357 | 25.6877414 | WT | FALSE | CAN | JSY13 | 2A | WT-Rep2A | GCG | C>A | A/T+1 | G/C-1 | CG>AT | CG>AT | AATTTGCGG | CGGCAGA | CGC | GCG | C | T | C>A | C>A_GCG | CG>AT_CGGCAGA | 9 | TRUE | FALSE |
| 170_v.txt | 32304 | 32304-SNV- | SNV | 1 | G | T | 24 | 43957 | 0.05459881 | 45.2083333 | WT | FALSE | CAN | JSY13 | 2A | WT-Rep2A | AGA | G>T | A/T+1 | G/C-1 | CG>AT | CG>AT | GCGGCAGAA | GAAATAA | TCT | TCT | C | A | C>A | C>A_TCT | CG>AT_GAAATAA | 9 | TRUE | FALSE |
| 170_v.txt | 32329 | 32329-SNV- | SNV | 1 | C | A | 9 | 43957 | 0.02047456 | 33.6692031 | WT | FALSE | CAN | JSY13 | 2A | WT-Rep2A | GCG | C>A | A/T+1 | G/C-1 | CG>AT | CG>AT | TAACAGCGT | CGTTGAA | CGC | GCG | C | T | C>A | C>A_GCG | CG>AT_CGTTGAA | 9 | TRUE | FALSE |
| 170_v.txt | 32399 | 32399-SNV- | SNV | 1 | G | T | 94 | 43957 | 0.21384535 | 59.0319149 | WT | FALSE | CAN | JSY13 | 2A | WT-Rep2A | CGT | G>T | A/T+1 | G/C-1 | CG>AT | CG>AT | AGAAACGTA | GTAGGAA | GCA | GCA | C | A | C>A | C>A_GCA | CG>AT_GTAGGAA | 9 | TRUE | FALSE |
| 170_v.txt | 32432 | 32432-SNV- | SNV | 1 | G | T | 182 | 43957 | 0.414041 | 56.1978022 | WT | FALSE | CAN | JSY13 | 2A | WT-Rep2A | TGT | G>T | A/T+1 | G/C-1 | CG>AT | CG>AT | GTCATTGTAT | GTATGGA | ACA | ACA | C | A | C>A | C>A_ACA | CG>AT_GTATGGA | 9 | TRUE | FALSE |
| 170_v.txt | 32474 | 32474-SNV- | SNV | 1 | G | T | 162 | 43957 | 0.36854198 | 49.7267674 | WT | FALSE | CAN | JSY13 | 2A | WT-Rep2A | AGA | G>T | A/T+1 | G/C-1 | CG>AT | CG>AT | AATGTAGAA | GAAGGTT | TCT | TCT | C | A | C>A | C>A_TCT | CG>AT_GAAGGTT | 9 | TRUE | FALSE |
| 170_v.txt | 32530 | 32530-SNV- | SNV | 1 | G | T | 235 | 43957 | 0.53461337 | 39.7191489 | WT | FALSE | CAN | JSY13 | 2A | WT-Rep2A | GGG | G>T | A/T+1 | G/C-1 | CG>AT | CG>AT | TTCTGGGGT | GGTTTGC | CCC | CCC | C | A | C>A | C>A_CCC | CG>AT_GGTTTGC | 9 | TRUE | FALSE |
| 170_v.txt | 32538 | 32538-SNV- | SNV | 1 | G | T | 30 | 43957 | 0.06824852 | 36.3 | WT | FALSE | CAN | JSY13 | 2A | WT-Rep2A | AGC | G>T | A/T+1 | G/C-1 | CG>AT | CG>AT | TTTGCAGCTT | GCTTCAC | TCG | TCG | C | A | C>A | C>A_TCG | CG>AT_GCTTCAC | 9 | TRUE | FALSE |
| 170_v.txt | 32700 | 32700-SNV- | SNV | 1 | C | A | 5 | 9394 | 0.05322546 | 29.9840144 | WT | FALSE | CAN | JSY13 | 2A | WT-Rep2A | GCC | C>A | A/T+1 | G/C-1 | CG>AT | CG>AT | ACTGGGCCG | CCGGTAA | CGG | GCC | C | T | C>A | C>A_GCC | CG>AT_CCGGTAA | 9 | TRUE | FALSE |
| 170_v.txt | 32794 | 32794-SNV- | SNV | 1 | C | A | 9 | 9394 | 0.09580583 | 40.8888889 | WT | FALSE | CAN | JSY13 | 2A | WT-Rep2A | TCA | C>A | A/T+1 | G/C-1 | CG>AT | CG>AT | CGAATTCAC | CACCGTA | AGT | TCA | C | T | C>A | C>A_TCA | CG>AT_CACCGTA | 9 | TRUE | FALSE |
| 170_v.txt | 32811 | 32811-SNV- | SNV | 1 | G | T | 3 | 9394 | 0.03193528 | 46.6666667 | WT | FALSE | CAN | JSY13 | 2A | WT-Rep2A | AGG | G>T | A/T+1 | G/C-1 | CG>AT | CG>AT | TTGACAGGG | GGGAACA | TCC | TCC | C | A | C>A | C>A_TCC | CG>AT_GGGAACA | 9 | TRUE | FALSE |
| 170_v.txt | 32876 | 32876-SNV- | SNV | 1 | G | T | 18 | 34425 | 0.05228758 | 28.3333333 | WT | FALSE | CAN | JSY13 | 2A | WT-Rep2A | TGT | G>T | A/T+1 | G/C-1 | CG>AT | CG>AT | AACTTTGTAC | GTACGTC | ACA | ACA | C | A | C>A | C>A_ACA | CG>AT_GTACGTC | 9 | TRUE | FALSE |
| 170_v.txt | 32883 | 32883-SNV- | SNV | 1 | C | A | 5 | 34435 | 0.01452011 | 34.000265 | WT | FALSE | CAN | JSY13 | 2A | WT-Rep2A | CCA | C>A | A/T+1 | G/C-1 | CG>AT | CG>AT | TACGTCCAA | CAAAATT | GGT | CCA | C | T | C>A | C>A_CCA | CG>AT_CAAAATT | 9 | TRUE | FALSE |
| 170_v.txt | 32899 | 32899-SNV- | SNV | 1 | G | T | 65 | 34437 | 0.18875047 | 17.0615385 | WT | FALSE | CAN | JSY13 | 2A | WT-Rep2A | TGG | G>T | A/T+1 | G/C-1 | CG>AT | CG>AT | TGACTTGGC | GGCCAAC | ACC | ACC | C | A | C>A | C>A_ACC | CG>AT_GGCCAAC | 9 | TRUE | FALSE |
| 170_v.txt | 32954 | 32954-SNV- | SNV | 1 | G | T | 18 | 34437 | 0.05226936 | 48.6666667 | WT | FALSE | CAN | JSY13 | 2A | WT-Rep2A | TGT | G>T | A/T+1 | G/C-1 | CG>AT | CG>AT | ATACATGTA | GTAACCA | ACA | ACA | C | A | C>A | C>A_ACA | CG>AT_GTAACCA | 9 | TRUE | FALSE |
| 170_v.txt | 32997 | 32997-SNV- | SNV | 1 | G | T | 27 | 34437 | 0.07840404 | 61.6296296 | WT | FALSE | CAN | JSY13 | 2A | WT-Rep2A | TGA | G>T | A/T+1 | G/C-1 | CG>AT | CG>AT | CTTTGTGAG | GAGAAAA | ACT | ACT | C | A | C>A | C>A_ACT | CG>AT_GAGAAAA | 9 | TRUE | FALSE |
| 170_v.txt | 33006 | 33006-SNV- | SNV | 1 | G | T | 274 | 34437 | 0.79565584 | 60.8467153 | WT | FALSE | CAN | JSY13 | 2A | WT-Rep2A | TGT | G>T | A/T+1 | G/C-1 | CG>AT | CG>AT | AAAACTGTG | GTGAAAG | ACA | ACA | C | A | C>A | C>A_ACA | CG>AT_GTGAAAG | 9 | TRUE | FALSE |
| 170_v.txt | 33018 | 33018-SNV- | SNV | 1 | G | T | 177 | 34436 | 0.51399698 | 62.0669608 | WT | FALSE | CAN | JSY13 | 2A | WT-Rep2A | TGT | G>T | A/T+1 | G/C-1 | CG>AT | CG>AT | GAGGATGTA | GTAACAG | ACA | ACA | C | A | C>A | C>A_ACA | CG>AT_GTAACAG | 9 | TRUE | FALSE |
| 170_v.txt | 33033 | 33033-SNV- | SNV | 1 | G | T | 2619 | 34437 | 7.60519209 | 60.2584956 | WT | FALSE | CAN | JSY13 | 2A | WT-Rep2A | TGT | G>T | A/T+1 | G/C-1 | CG>AT | CG>AT | ATGAATGTA | GTAGCCA | ACA | ACA | C | A | C>A | C>A_ACA | CG>AT_GTAGCCA | 9 | TRUE | FALSE |
| 170_v.txt | 33043 | 33043-SNV- | SNV | 1 | C | A | 97 | 34436 | 0.28168196 | 57.6199317 | WT | FALSE | CAN | JSY13 | 2A | WT-Rep2A | TCA | C>A | A/T+1 | G/C-1 | CG>AT | CG>AT | CCATTTCACC | CACCCAA | AGT | TCA | C | T | C>A | C>A_TCA | CG>AT_CACCCAA | 9 | TRUE | FALSE |
| 170_v.txt | 33093 | 33093-SNV- | SNV | 1 | G | T | 12 | 34437 | 0.03484624 | 49.5 | WT | FALSE | CAN | JSY13 | 2A | WT-Rep2A | TGA | G>T | A/T+1 | G/C-1 | CG>AT | CG>AT | AAATATGAT | GATATAA | ACT | ACT | C | A | C>A | C>A_ACT | CG>AT_GATATAA | 9 | TRUE | FALSE |
| 170_v.txt | 33115 | 33115-SNV- | SNV | 1 | C | A | 268 | 34437 | 0.77823272 | 44.0535608 | WT | FALSE | CAN | JSY13 | 2A | WT-Rep2A | CCG | C>A | A/T+1 | G/C-1 | CG>AT | CG>AT | CTGGGCCGG | CGGCGTT | GGC | CCG | C | T | C>A | C>A_CCG | CG>AT_CGGCGTT | 9 | TRUE | FALSE |
| 170_v.txt | 33151 | 33151-SNV- | SNV | 1 | G | T | 17 | 34437 | 0.04936551 | 36.2941177 | WT | FALSE | CAN | JSY13 | 2A | WT-Rep2A | AGA | G>T | A/T+1 | G/C-1 | CG>AT | CG>AT | TGAAAAGAC | GACCTGT | TCT | TCT | C | A | C>A | C>A_TCT | CG>AT_GACCTGT | 9 | TRUE | FALSE |
| 170_v.txt | 33156 | 33156-SNV- | SNV | 1 | G | T | 7 | 34437 | 0.02032697 | 37.2857143 | WT | FALSE | CAN | JSY13 | 2A | WT-Rep2A | TGT | G>T | A/T+1 | G/C-1 | CG>AT | CG>AT | AGACCTGTA | GTACCAA | ACA | ACA | C | A | C>A | C>A_ACA | CG>AT_GTACCAA | 9 | TRUE | FALSE |
| 170_v.txt | 33160 | 33160-SNV- | SNV | 1 | C | A | 23 | 34440 | 0.06678281 | 37.7826087 | WT | FALSE | CAN | JSY13 | 2A | WT-Rep2A | CCA | C>A | A/T+1 | G/C-1 | CG>AT | CG>AT | CTGTACCAAT | CAATAGT | GGT | CCA | C | T | C>A | C>A_CCA | CG>AT_CAATAGT | 9 | TRUE | FALSE |
| 170_v.txt | 33168 | 33168-SNV- | SNV | 1 | C | A | 991 | 34418 | 2.87930734 | 37.1226483 | WT | FALSE | CAN | JSY13 | 2A | WT-Rep2A | ACC | C>A | A/T+1 | G/C-1 | CG>AT | CG>AT | ATAGTACCA | CCACCAA | TGG | ACC | C | T | C>A | C>A_ACC | CG>AT_CCACCAA | 9 | TRUE | FALSE |
| 170_v.txt | 33169 | 33169-SNV- | SNV | 1 | C | A | 22 | 34418 | 0.06392004 | 36.5898355 | WT | FALSE | CAN | JSY13 | 2A | WT-Rep2A | CCA | C>A | A/T+1 | G/C-1 | CG>AT | CG>AT | TAGTACCAC | CACCAAG | GGT | CCA | C | T | C>A | C>A_CCA | CG>AT_CACCAAG | 9 | TRUE | FALSE |
| 170_v.txt | 33193 | 33193-SNV- | SNV | 1 | G | T | 9 | 17683 | 0.05089634 | 36.4444444 | WT | FALSE | CAN | JSY13 | 2A | WT-Rep2A | TGT | G>T | A/T+1 | G/C-1 | CG>AT | CG>AT | CAATATGTCT | GTCTTTG | ACA | ACA | C | A | C>A | C>A_ACA | CG>AT_GTCTTTG | 9 | TRUE | FALSE |
| 170_v.txt | 33354 | 33354-SNV- | SNV | 1 | G | T | 107 | 17691 | 0.60482731 | 22.5981308 | WT | FALSE | CAN | JSY13 | 2A | WT-Rep2A | TGA | G>T | A/T+1 | G/C-1 | CG>AT | CG>AT | GGTATTGAC | GACCCAC | ACT | ACT | C | A | C>A | C>A_ACT | CG>AT_GACCCAC | 9 | TRUE | FALSE |
| 170_v.txt | 33415 | 33415-SNV- | SNV | 1 | C | A | 39 | 17691 | 0.22045108 | 46.1282051 | WT | FALSE | CAN | JSY13 | 2A | WT-Rep2A | TCA | C>A | A/T+1 | G/C-1 | CG>AT | CG>AT | CCGGCTCATT | CATTGTA | AGT | TCA | C | T | C>A | C>A_TCA | CG>AT_CATTGTA | 9 | TRUE | FALSE |
| 170_v.txt | 33428 | 33428-SNV- | SNV | 1 | C | A | 2 | 17691 | 0.01130518 | 48.9806652 | WT | FALSE | CAN | JSY13 | 2A | WT-Rep2A | GCT | C>A | A/T+1 | G/C-1 | CG>AT | CG>AT | CATATGCTTC | CTTCTCC | CGA | GCT | C | T | C>A | C>A_GCT | CG>AT_CTTCTCC | 9 | TRUE | FALSE |
| 170_v.txt | 33474 | 33474-SNV- | SNV | 1 | C | A | 32 | 17691 | 0.18088294 | 31.625 | WT | FALSE | CAN | JSY13 | 2A | WT-Rep2A | GCC | C>A | A/T+1 | G/C-1 | CG>AT | CG>AT | GCTATGCCTT | CCTTTTT | CGG | GCC | C | T | C>A | C>A_GCC | CG>AT_CCTTTTT | 9 | TRUE | FALSE |
| 170_v.txt | 32328 | 32328-Repla | Replacement | 3 | GCG | AT | 17 | 43957 | 0.03867416 | 61.4313726 | WT | FALSE | CAN | JSY13 | 2A | WT-Rep2A | AGC | nope | >1bp-ins | >1bpdel | none | Replac. | ATAACAGCG | GCGTTGA | TCG | AGC | GCG | none | NA | NA_AGC | Replac._GCGTTGA | 9 | TRUE | FALSE |
| 170_v.txt | 32604 | 32604-Repla | Replacement | 2 | G | CA | 233 | 9394 | 2.48030658 | 42.2553478 | WT | FALSE | CAN | JSY13 | 2A | WT-Rep2A | GGA | nope | >1bp-ins | G/C-1 | none | Replac. | AAAGAGGAA | GAAACCC | CCT | CCT | C | none | C>none | C>none_CCT | Replac._GAAACCC | 9 | TRUE | FALSE |
| 170_v.txt | 32700 | 32700-Repla | Replacement | 8 | C | TTGTGTTG | 3 | 9394 | 0.03193528 | 63.6666667 | WT | FALSE | CAN | JSY13 | 2A | WT-Rep2A | GCC | nope | >1bp-ins | G/C-1 | none | Replac. | ACTGGGCCG | CCGGTAA | CGG | GCC | C | none | nope | nope_GCC | Replac._CCGGTAA | 9 | TRUE | FALSE |
| 170_v.txt | 33030 | 33030-Repla | Replacement | 2 | AA | C | 8 | 34437 | 0.02323083 | 63.6875 | WT | FALSE | CAN | JSY13 | 2A | WT-Rep2A | GAA | nope | G/C+1 | >1bpdel | none | Replac. | GGGATGAAT | AATGTAG | CTT | GAA | AA | G | NA | NA_GAA | Replac._AATGTAG | 9 | TRUE | FALSE |
| 170_v.txt | 33428 | 33428-Repla | Replacement | 2 | C | AA | 14 | 17691 | 0.07913628 | 42.3928495 | WT | FALSE | CAN | JSY13 | 2A | WT-Rep2A | GCT | nope | >1bp-ins | G/C-1 | none | Replac. | CATATGCTTC | CTTCTCC | CGA | GCT | C | none | nope | nope_GCT | Replac._CTTCTCC | 9 | TRUE | FALSE |
| 170_v.txt | 31964 | 31964-MNV- | MNV | 2 | AT | TA | 40 | 20510 | 0.19502682 | 37.3249998 | WT | FALSE | CAN | JSY13 | 2A | WT-Rep2A | CAT | nope | >1bp-ins | >1bpdel | none | MNV | GGCCGCATA | ATAATAA | GTA | CAT | AT | none | NA | NA_CAT | MNV_ATAATAA | 9 | TRUE | FALSE |
| 170_v.txt | 32298 | 32298-MNV- | MNV | 2 | GC | CA | 7256 | 43957 | 16.507041 | 56.0563602 | WT | FALSE | CAN | JSY13 | 2A | WT-Rep2A | TGC | nope | >1bp-ins | >1bpdel | none | MNV | GAATTTGCG | GCGGCAG | ACG | TGC | GC | none | NA | NA_TGC | MNV_GCGGCAG | 9 | TRUE | FALSE |
| 170_v.txt | 32298 | 32298-MNV- | MNV | 2 | GC | CG | 9 | 43957 | 0.02047456 | 51.0557649 | WT | FALSE | CAN | JSY13 | 2A | WT-Rep2A | TGC | nope | >1bp-ins | >1bpdel | none | MNV | GAATTTGCG | GCGGCAG | ACG | TGC | GC | none | NA | NA_TGC | MNV_GCGGCAG | 9 | TRUE | FALSE |
| 170_v.txt | 32298 | 32298-MNV- | MNV | 2 | GC | TA | 4 | 43957 | 9.10E-03 | 44.3749788 | WT | FALSE | CAN | JSY13 | 2A | WT-Rep2A | TGC | nope | >1bp-ins | >1bpdel | none | MNV | GAATTTGCG | GCGGCAG | ACG | TGC | GC | none | NA | NA_TGC | MNV_GCGGCAG | 9 | TRUE | FALSE |
| 170_v.txt | 32473 | 32473-MNV- | MNV | 2 | AG | GT | 1 | 43957 | 2.28E-03 | 63.9971941 | WT | FALSE | CAN | JSY13 | 2A | WT-Rep2A | TAG | nope | >1bp-ins | >1bpdel | none | MNV | CAATGTAGA | AGAAGGT | ATC | TAG | AG | none | NA | NA_TAG | MNV_AGAAGGT | 9 | TRUE | FALSE |
| 170_v.txt | 31892 | 31892-Insert | Insertion | 4 | - | AACA | 2 | 3904 | 0.05122951 | 37.5 | WT | FALSE | CAN | JSY13 | 2A | WT-Rep2A | TAA | nope | >1bp-ins | >1bpdel | none | >1bp-ins | AAAGCTAAC | AACACCA | ATT | TAA | - | none | NA | NA_TAA | >1bp-ins_AACACCA | 9 | TRUE | FALSE |
| 170_v.txt | 31969 | 31969-Insert | Insertion | 2 | - | TA | 40 | 20510 | 0.19502682 | 37.3 | WT | FALSE | CAN | JSY13 | 2A | WT-Rep2A | TAA | nope | >1bp-ins | >1bpdel | none | >1bp-ins | CATAATAAG | AAGCCAA | ATT | TAA | - | none | NA | NA_TAA | >1bp-ins_AAGCCAA | 9 | TRUE | FALSE |
| 170_v.txt | 31996 | 31996-Insert | Insertion | 4 | - | TAAA | 35 | 20510 | 0.17064846 | 52.1285714 | WT | FALSE | CAN | JSY13 | 2A | WT-Rep2A | TTA | nope | >1bp-ins | >1bpdel | none | >1bp-ins | TAGCTTTAAA | TAAATGG | AAT | TTA | - | none | NA | NA_TTA | >1bp-ins_TAAATGG | 9 | TRUE | FALSE |
| 170_v.txt | 32033 | 32033-Insert | Insertion | 1 | - | A | 4 | 20510 | 0.01950268 | 63.25 | WT | FALSE | CAN | JSY13 | 2A | WT-Rep2A | TCA | nope | A/T+1 | >1bpdel | none | A/T+1 | GTATTTCAAA | CAAAGCT | AGT | TCA | - | T | NA | NA_TCA | A/T+1_CAAAGCT | 9 | TRUE | FALSE |
| 170_v.txt | 32068 | 32068-Insert | Insertion | 1 | - | T | 5 | 20510 | 0.02437835 | 61 | WT | FALSE | CAN | JSY13 | 2A | WT-Rep2A | ATA | nope | A/T+1 | >1bpdel | none | A/T+1 | TTGAGATAA | TAAATAA | TAT | ATA | - | A | NA | NA_ATA | A/T+1_TAAATAA | 9 | TRUE | FALSE |
| 170_v.txt | 32081 | 32081-Insert | Insertion | 1 | - | A | 25 | 20510 | 0.12189176 | 63.88 | WT | FALSE | CAN | JSY13 | 2A | WT-Rep2A | CAA | nope | A/T+1 | >1bpdel | none | A/T+1 | CCATGCAAA | AAAAAAG | GTT | CAA | - | T | NA | NA_CAA | A/T+1_AAAAAAG | 9 | TRUE | FALSE |
| 170_v.txt | 32277 | 32277-Insert | Insertion | 1 | - | C | 30 | 43957 | 0.06824852 | 37.8 | WT | FALSE | CAN | JSY13 | 2A | WT-Rep2A | ACC | nope | G/C+1 | >1bpdel | none | G/C+1 | CGGGAACCA | CCAACGT | TGG | ACC | - | G | NA | NA_ACC | G/C+1_CCAACGT | 9 | TRUE | FALSE |
| 170_v.txt | 32375 | 32375-Insert | Insertion | 1 | - | A | 11 | 43957 | 0.02502446 | 64 | WT | FALSE | CAN | JSY13 | 2A | WT-Rep2A | CAA | nope | A/T+1 | >1bpdel | none | A/T+1 | AATAGCAAT | AATAATA | GTT | CAA | - | T | NA | NA_CAA | A/T+1_AATAATA | 9 | TRUE | FALSE |
| 170_v.txt | 32468 | 32468-Insert | Insertion | 1 | - | A | 16 | 43957 | 0.03639921 | 54.5625 | WT | FALSE | CAN | JSY13 | 2A | WT-Rep2A | CAA | nope | A/T+1 | >1bpdel | none | A/T+1 | AGAGCCAAT | AATGTAG | GTT | CAA | - | T | NA | NA_CAA | A/T+1_AATGTAG | 9 | TRUE | FALSE |
| 170_v.txt | 32489 | 32489-Insert | Insertion | 1 | - | C | 23 | 43957 | 0.05232386 | 46.4347826 | WT | FALSE | CAN | JSY13 | 2A | WT-Rep2A | GGA | nope | G/C+1 | >1bpdel | none | G/C+1 | GATACGGAA | GAAAACA | CCT | GGA | - | G | NA | NA_GGA | G/C+1_GAAAACA | 9 | TRUE | FALSE |
| 170_v.txt | 32498 | 32498-Insert | Insertion | 1 | - | T | 14 | 43957 | 0.03184931 | 42.7857143 | WT | FALSE | CAN | JSY13 | 2A | WT-Rep2A | CTT | nope | A/T+1 | >1bpdel | none | A/T+1 | AACAACTTTT | TTTTTTG | GAA | CTT | - | A | NA | NA_CTT | A/T+1_TTTTTTG | 9 | TRUE | FALSE |
| 170_v.txt | 32631 | 32631-Insert | Insertion | 1 | - | T | 9 | 9394 | 0.09580583 | 52.4444444 | WT | FALSE | CAN | JSY13 | 2A | WT-Rep2A | GTT | nope | A/T+1 | >1bpdel | none | A/T+1 | CCTTCGTTTT | TTTTTAT | CAA | GTT | - | A | NA | NA_GTT | A/T+1_TTTTTAT | 9 | TRUE | FALSE |
| 170_v.txt | 32719 | 32719-Insert | Insertion | 1 | - | A | 2 | 9394 | 0.02129019 | 64 | WT | FALSE | CAN | JSY13 | 2A | WT-Rep2A | CAA | nope | A/T+1 | >1bpdel | none | A/T+1 | CACCACAAA | AAACCAT | GTT | CAA | - | T | NA | NA_CAA | A/T+1_AAACCAT | 9 | TRUE | FALSE |
| 170_v.txt | 32841 | 32841-Insert | Insertion | 1 | - | A | 2 | 9394 | 0.02129019 | 38 | WT | FALSE | CAN | JSY13 | 2A | WT-Rep2A | CCA | nope | A/T+1 | >1bpdel | none | A/T+1 | ATTACCCAA | CAAAAAA | GGT | CCA | - | T | NA | NA_CCA | A/T+1_CAAAAAA | 9 | TRUE | FALSE |
| 170_v.txt | 32842 | 32842-Insert | Insertion | 1 | - | A | 23 | 9394 | 0.24483713 | 39.6086957 | WT | FALSE | CAN | JSY13 | 2A | WT-Rep2A | CAA | nope | A/T+1 | >1bpdel | none | A/T+1 | TTACCCAAA | AAAAAAT | GTT | CAA | - | T | NA | NA_CAA | A/T+1_AAAAAAT | 9 | TRUE | FALSE |
| 170_v.txt | 32903 | 32903-Insert | Insertion | 1 | - | A | 39 | 34437 | 0.11325028 | 50.5128205 | WT | FALSE | CAN | JSY13 | 2A | WT-Rep2A | CAA | nope | A/T+1 | >1bpdel | none | A/T+1 | TTGGCCAACT | AACTACA | GTT | CAA | - | T | NA | NA_CAA | A/T+1_AACTACA | 9 | TRUE | FALSE |
| 170_v.txt | 32923 | 32923-Insert | Insertion | 1 | - | A | 3 | 34437 | 8.71E-03 | 46.3333333 | WT | FALSE | CAN | JSY13 | 2A | WT-Rep2A | GCA | nope | A/T+1 | >1bpdel | none | A/T+1 | CCAGGGCAA | CAAAAGT | CGT | GCA | - | T | NA | NA_GCA | A/T+1_CAAAAGT | 9 | TRUE | FALSE |
| 170_v.txt | 33012 | 33012-Insert | Insertion | 14 | - | GAGGATGTA | 2 | 34437 | 5.81E-03 | 51.5 | WT | FALSE | CAN | JSY13 | 2A | WT-Rep2A | AGA | nope | >1bp-ins | >1bpdel | none | >1bp-ins | GTGAAAGAG | GAGGATG | TCT | AGA | - | none | NA | NA_AGA | >1bp-ins_GAGGATG | 9 | TRUE | FALSE |
| 170_v.txt | 33018 | 33018-Insert | Insertion | 6 | - | GTAACA | 5 | 34437 | 0.01451927 | 62.6333333 | WT | FALSE | CAN | JSY13 | 2A | WT-Rep2A | TGT | nope | >1bp-ins | >1bpdel | none | >1bp-ins | GAGGATGTA | GTAACAG | ACA | TGT | - | none | NA | NA_TGT | >1bp-ins_GTAACAG | 9 | TRUE | FALSE |
| 170_v.txt | 32157 | 32157-Deleti | Deletion | 8 | GCCAAAGC | - | 17 | 20506 | 0.08290257 | 50.7647059 | WT | FALSE | CAN | JSY13 | 2A | WT-Rep2A | AGC | nope | >1bp-ins | >1bpdel | none | >1bpdel | ATGTAAGCC | GCCAAAG | TCG | AGC | GCCAAAGC | none | NA | NA_AGC | >1bpdel_GCCAAAG | 9 | TRUE | FALSE |
| 170_v.txt | 32161 | 32161-Deleti | Deletion | 4 | AAGC | - | 1 | 20506 | 4.88E-03 | 11 | WT | FALSE | CAN | JSY13 | 2A | WT-Rep2A | AAA | nope | >1bp-ins | >1bpdel | none | >1bpdel | AAGCCAAAG | AAGCGCC | TTT | AAA | AAGC | none | NA | NA_AAA | >1bpdel_AAGCGCC | 9 | TRUE | FALSE |
| 170_v.txt | 32337 | 32337-Deleti | Deletion | 2 | AT | - | 199 | 43957 | 0.45271515 | 62.0854271 | WT | FALSE | CAN | JSY13 | 2A | WT-Rep2A | GAT | nope | >1bp-ins | >1bpdel | none | >1bpdel | TTGAAGATA | ATATGTG | CTA | GAT | AT | none | NA | NA_GAT | >1bpdel_ATATGTG | 9 | TRUE | FALSE |
| 170_v.txt | 32472 | 32472-Deleti | Deletion | 2 | TA | - | 25 | 43957 | 0.05687376 | 55.08 | WT | FALSE | CAN | JSY13 | 2A | WT-Rep2A | GTA | nope | >1bp-ins | >1bpdel | none | >1bpdel | CCAATGTAG | TAGAAGG | CAT | GTA | TA | none | NA | NA_GTA | >1bpdel_TAGAAGG | 9 | TRUE | FALSE |
| 170_v.txt | 32476 | 32476-Deleti | Deletion | 4 | AGGT | - | 25 | 43957 | 0.05687376 | 54.36 | WT | FALSE | CAN | JSY13 | 2A | WT-Rep2A | AAG | nope | >1bp-ins | >1bpdel | none | >1bpdel | TGTAGAAGG | AGGTTAA | TTC | AAG | AGGT | none | NA | NA_AAG | >1bpdel_AGGTTAA | 9 | TRUE | FALSE |
| 170_v.txt | 33126 | 33126-Deleti | Deletion | 2 | AG | - | 15 | 34437 | 0.0435578 | 44.6 | WT | FALSE | CAN | JSY13 | 2A | WT-Rep2A | CAG | nope | >1bp-ins | >1bpdel | none | >1bpdel | TTGGTCAGA | AGAGGTG | GTC | CAG | AG | none | NA | NA_CAG | >1bpdel_AGAGGTG | 9 | TRUE | FALSE |
| 170_v.txt | 33189 | 33189-Deleti | Deletion | 2 | AT | - | 5 | 17692 | 0.02826136 | 38 | WT | FALSE | CAN | JSY13 | 2A | WT-Rep2A | AAT | nope | >1bp-ins | >1bpdel | none | >1bpdel | ATACCAATAT | ATATGTC | TTA | AAT | AT | none | NA | NA_AAT | >1bpdel_ATATGTC | 9 | TRUE | FALSE |
| 170_v.txt | 33206 | 33206-Deleti | Deletion | 6 | CTCTCT | - | 11 | 17691 | 0.06217851 | 40.3636364 | WT | FALSE | CAN | JSY13 | 2A | WT-Rep2A | GCT | nope | >1bp-ins | >1bpdel | none | >1bpdel | CTTAAGCTCT | CTCTCTC | CGA | GCT | CTCTCT | none | NA | NA_GCT | >1bpdel_CTCTCTC | 9 | TRUE | FALSE |
| 170_v.txt | 31723 | 31723-Deleti | Deletion | 1 | A | - | 5 | 3907 | 0.12797543 | 63.2 | WT | FALSE | CAN | JSY13 | 2A | WT-Rep2A | CAA | nope | >1bp-ins | A/T-1 | none | A/T-1 | TGTCCCAAA | AAAAAGT | GTT | GTT | T | none | T>none | T>none_GTT | A/T-1_AAAAAGT | 9 | TRUE | FALSE |
| 170_v.txt | 31899 | 31899-Deleti | Deletion | 1 | T | - | 4 | 3904 | 0.10245902 | 38 | WT | FALSE | CAN | JSY13 | 2A | WT-Rep2A | ATT | nope | >1bp-ins | A/T-1 | none | A/T-1 | ACACCATTG | TTGAATT | TAA | ATT | T | none | nope | nope_ATT | A/T-1_TTGAATT | 9 | TRUE | FALSE |
| 170_v.txt | 31913 | 31913-Deleti | Deletion | 1 | A | - | 5 | 3904 | 0.12807377 | 37.8 | WT | FALSE | CAN | JSY13 | 2A | WT-Rep2A | CAA | nope | >1bp-ins | A/T-1 | none | A/T-1 | TGGTGCAAA | AAAAGCC | GTT | GTT | T | none | T>none | T>none_GTT | A/T-1_AAAAGCC | 9 | TRUE | FALSE |
| 170_v.txt | 32057 | 32057-Deleti | Deletion | 1 | C | - | 21 | 20510 | 0.10238908 | 60.2380978 | WT | FALSE | CAN | JSY13 | 2A | WT-Rep2A | GCG | nope | >1bp-ins | G/C-1 | none | G/C-1 | GATGTGCGA | CGAGATT | CGC | GCG | C | none | nope | nope_GCG | G/C-1_CGAGATT | 9 | TRUE | FALSE |
| 170_v.txt | 32081 | 32081-Deleti | Deletion | 1 | A | - | 24 | 20510 | 0.11701609 | 63.4583228 | WT | FALSE | CAN | JSY13 | 2A | WT-Rep2A | CAA | nope | >1bp-ins | A/T-1 | none | A/T-1 | CCATGCAAA | AAAAAAG | GTT | GTT | T | none | T>none | T>none_GTT | A/T-1_AAAAAAG | 9 | TRUE | FALSE |
| 170_v.txt | 32114 | 32114-Deleti | Deletion | 1 | C | - | 2 | 20510 | 9.75E-03 | 64 | WT | FALSE | CAN | JSY13 | 2A | WT-Rep2A | GCC | nope | >1bp-ins | G/C-1 | none | G/C-1 | TAATAGCCAT | CCATTCG | CGG | GCC | C | none | nope | nope_GCC | G/C-1_CCATTCG | 9 | TRUE | FALSE |
| 170_v.txt | 32158 | 32158-Deleti | Deletion | 1 | C | - | 1 | 20506 | 4.88E-03 | 20 | WT | FALSE | CAN | JSY13 | 2A | WT-Rep2A | GCC | nope | >1bp-ins | G/C-1 | none | G/C-1 | TGTAAGCCA | CCAAAGC | CGG | GCC | C | none | nope | nope_GCC | G/C-1_CCAAAGC | 9 | TRUE | FALSE |
| 170_v.txt | 32160 | 32160-Deleti | Deletion | 1 | A | - | 2 | 20506 | 9.75E-03 | 64 | WT | FALSE | CAN | JSY13 | 2A | WT-Rep2A | CAA | nope | >1bp-ins | A/T-1 | none | A/T-1 | TAAGCCAAA | AAAGCGC | GTT | GTT | T | none | T>none | T>none_GTT | A/T-1_AAAGCGC | 9 | TRUE | FALSE |
| 170_v.txt | 32180 | 32180-Deleti | Deletion | 1 | A | - | 44 | 20506 | 0.21457135 | 49.1136364 | WT | FALSE | CAN | JSY13 | 2A | WT-Rep2A | TAA | nope | >1bp-ins | A/T-1 | none | A/T-1 | AGCAGTAAC | AACGAAA | ATT | ATT | T | none | T>none | T>none_ATT | A/T-1_AACGAAA | 9 | TRUE | FALSE |

| | | | | | | | | | | | | | | | | | | | | | | | | | | | | | | | | | | |
|---|---|---|---|---|---|---|---|---|---|---|---|---|---|---|---|---|---|---|---|---|---|---|---|---|---|---|---|---|---|---|---|---|---|---|
| 170_v.txt | 32184 | 32184-Deleti | Deletion | 1 | A | - | 47 | 20506 | 0.22920121 | 50.0425532 | WT | FALSE | CAN | JSY13 | 2A | WT-Rep2A | GAA | nope | >1bp-ins | A/T-1 | none | A/T-1 | GTAACGAAA | AAAACTG | CTT | CTT | T | none | T>none | T>none_CTT | A/T-1_AAAACTG | 9 | TRUE | FALSE |
| 170_v.txt | 32210 | 32210-Deleti | Deletion | 1 | T | - | 6 | 20506 | 0.02925973 | 46.5 | WT | FALSE | CAN | JSY13 | 2A | WT-Rep2A | CTT | nope | >1bp-ins | A/T-1 | none | A/T-1 | ACCACCTTTG | TTTGGTG | GAA | CTT | T | none | nope | nope_CTT | A/T-1_TTTGGTG | 9 | TRUE | FALSE |
| 170_v.txt | 32238 | 32238-Deleti | Deletion | 1 | G | - | 21 | 20504 | 0.10241904 | 37.2857143 | WT | FALSE | CAN | JSY13 | 2A | WT-Rep2A | AGC | nope | >1bp-ins | G/C-1 | none | G/C-1 | TTAGGAGCC | GCCAACT | TCG | TCG | C | none | C>none | C>none_TCG | G/C-1_GCCAACT | 9 | TRUE | FALSE |
| 170_v.txt | 32265 | 32265-Deleti | Deletion | 1 | A | - | 9 | 43957 | 0.02047456 | 37.1111111 | WT | FALSE | CAN | JSY13 | 2A | WT-Rep2A | TAA | nope | >1bp-ins | A/T-1 | none | A/T-1 | CCAAATAAA | AAAATAC | ATT | ATT | T | none | T>none | T>none_ATT | A/T-1_AAAATAC | 9 | TRUE | FALSE |
| 170_v.txt | 32298 | 32298-Deleti | Deletion | 1 | G | - | 12 | 43957 | 0.02729941 | 47.0000102 | WT | FALSE | CAN | JSY13 | 2A | WT-Rep2A | TGC | nope | >1bp-ins | G/C-1 | none | G/C-1 | GAATTTGCG | GCGGCAG | ACG | ACG | C | none | C>none | C>none_ACG | G/C-1_GCGGCAG | 9 | TRUE | FALSE |
| 170_v.txt | 32381 | 32381-Deleti | Deletion | 1 | A | - | 7 | 43957 | 0.01592465 | 64 | WT | FALSE | CAN | JSY13 | 2A | WT-Rep2A | TAA | nope | >1bp-ins | A/T-1 | none | A/T-1 | AATAATAAA | AAAGGGA | ATT | ATT | T | none | T>none | T>none_ATT | A/T-1_AAAGGGA | 9 | TRUE | FALSE |
| 170_v.txt | 32387 | 32387-Deleti | Deletion | 1 | A | - | 21 | 43957 | 0.04777396 | 63.7619048 | WT | FALSE | CAN | JSY13 | 2A | WT-Rep2A | GAG | nope | >1bp-ins | A/T-1 | none | A/T-1 | AAAGGGAGA | AGAAGTA | CTC | CTC | T | none | T>none | T>none_CTC | A/T-1_AGAAGTA | 9 | TRUE | FALSE |
| 170_v.txt | 32436 | 32436-Deleti | Deletion | 1 | G | - | 140 | 43957 | 0.31849307 | 55.8714286 | WT | FALSE | CAN | JSY13 | 2A | WT-Rep2A | TGG | nope | >1bp-ins | G/C-1 | none | G/C-1 | TTGTATGGA | GGAACTA | ACC | ACC | C | none | C>none | C>none_ACC | G/C-1_GGAACTA | 9 | TRUE | FALSE |
| 170_v.txt | 32466 | 32466-Deleti | Deletion | 1 | C | - | 3 | 43957 | 6.83E-03 | 55.3333333 | WT | FALSE | CAN | JSY13 | 2A | WT-Rep2A | GCC | nope | >1bp-ins | G/C-1 | none | G/C-1 | AGAGAGCCA | CCAATGT | CGG | GCC | C | none | nope | nope_GCC | G/C-1_CCAATGT | 9 | TRUE | FALSE |
| 170_v.txt | 32470 | 32470-Deleti | Deletion | 1 | T | - | 25 | 43957 | 0.05687376 | 54.1999991 | WT | FALSE | CAN | JSY13 | 2A | WT-Rep2A | ATG | nope | >1bp-ins | A/T-1 | none | A/T-1 | AGCCAATGT | TGTAGAA | TAC | ATG | T | none | nope | nope_ATG | A/T-1_TGTAGAA | 9 | TRUE | FALSE |
| 170_v.txt | 32475 | 32475-Deleti | Deletion | 1 | A | - | 4 | 43957 | 9.10E-03 | 49.9999947 | WT | FALSE | CAN | JSY13 | 2A | WT-Rep2A | GAA | nope | >1bp-ins | A/T-1 | none | A/T-1 | ATGTAGAAG | AAGGTTA | CTT | CTT | T | none | T>none | T>none_CTT | A/T-1_AAGGTTA | 9 | TRUE | FALSE |
| 170_v.txt | 32498 | 32498-Deleti | Deletion | 1 | T | - | 107 | 43957 | 0.24341971 | 45.1121507 | WT | FALSE | CAN | JSY13 | 2A | WT-Rep2A | CTT | nope | >1bp-ins | A/T-1 | none | A/T-1 | AACAACTTTT | TTTTTTG | GAA | CTT | T | none | nope | nope_CTT | A/T-1_TTTTTTG | 9 | TRUE | FALSE |
| 170_v.txt | 32522 | 32522-Deleti | Deletion | 1 | T | - | 11 | 43957 | 0.02502446 | 42.4545455 | WT | FALSE | CAN | JSY13 | 2A | WT-Rep2A | ATT | nope | >1bp-ins | A/T-1 | none | A/T-1 | AACGGATTTT | TTTTCTG | TAA | ATT | T | none | nope | nope_ATT | A/T-1_TTTTCTG | 9 | TRUE | FALSE |
| 170_v.txt | 32528 | 32528-Deleti | Deletion | 1 | G | - | 8 | 43957 | 0.0181996 | 44.5 | WT | FALSE | CAN | JSY13 | 2A | WT-Rep2A | TGG | nope | >1bp-ins | G/C-1 | none | G/C-1 | TTTTCTGGGG | GGGGTTT | ACC | ACC | C | none | C>none | C>none_ACC | G/C-1_GGGGTTT | 9 | TRUE | FALSE |
| 170_v.txt | 32532 | 32532-Deleti | Deletion | 1 | T | - | 26 | 43957 | 0.05914871 | 41.1923077 | WT | FALSE | CAN | JSY13 | 2A | WT-Rep2A | GTT | nope | >1bp-ins | A/T-1 | none | A/T-1 | CTGGGGTTT | TTTGCAG | CAA | GTT | T | none | nope | nope_GTT | A/T-1_TTTGCAG | 9 | TRUE | FALSE |
| 170_v.txt | 32598 | 32598-Deleti | Deletion | 1 | A | - | 1838 | 9394 | 19.5656802 | 39.7720348 | WT | FALSE | CAN | JSY13 | 2A | WT-Rep2A | CAA | nope | >1bp-ins | A/T-1 | none | A/T-1 | TTAATCAAA | AAAGAGG | GTT | GTT | T | none | T>none | T>none_GTT | A/T-1_AAAGAGG | 9 | TRUE | FALSE |
| 170_v.txt | 32658 | 32658-Deleti | Deletion | 1 | C | - | 5 | 9394 | 0.05322546 | 58.8 | WT | FALSE | CAN | JSY13 | 2A | WT-Rep2A | ACC | nope | >1bp-ins | G/C-1 | none | G/C-1 | CCTGGACCC | CCCCAGG | TGG | ACC | C | none | nope | nope_ACC | G/C-1_CCCCAGG | 9 | TRUE | FALSE |
| 170_v.txt | 32662 | 32662-Deleti | Deletion | 1 | A | - | 5 | 9394 | 0.05322546 | 63.8 | WT | FALSE | CAN | JSY13 | 2A | WT-Rep2A | CAG | nope | >1bp-ins | A/T-1 | none | A/T-1 | GACCCCAGG | AGGCACC | GTC | GTC | T | none | T>none | T>none_GTC | A/T-1_AGGCACC | 9 | TRUE | FALSE |
| 170_v.txt | 32683 | 32683-Deleti | Deletion | 1 | A | - | 20 | 9394 | 0.21290185 | 59.8 | WT | FALSE | CAN | JSY13 | 2A | WT-Rep2A | TAA | nope | >1bp-ins | A/T-1 | none | A/T-1 | TCCAATAAC | AACGGAA | ATT | ATT | T | none | T>none | T>none_ATT | A/T-1_AACGGAA | 9 | TRUE | FALSE |
| 170_v.txt | 32700 | 32700-Deleti | Deletion | 1 | C | - | 3 | 9394 | 0.03193528 | 63.9999623 | WT | FALSE | CAN | JSY13 | 2A | WT-Rep2A | GCC | nope | >1bp-ins | G/C-1 | none | G/C-1 | ACTGGGCCG | CCGGTAA | CGG | GCC | C | none | nope | nope_GCC | G/C-1_CCGGTAA | 9 | TRUE | FALSE |
| 170_v.txt | 32707 | 32707-Deleti | Deletion | 1 | C | - | 4 | 9394 | 0.04258037 | 64 | WT | FALSE | CAN | JSY13 | 2A | WT-Rep2A | ACC | nope | >1bp-ins | G/C-1 | none | G/C-1 | CGGTAACCC | CCCCAGC | TGG | ACC | C | none | nope | nope_ACC | G/C-1_CCCCAGC | 9 | TRUE | FALSE |
| 170_v.txt | 32719 | 32719-Deleti | Deletion | 1 | A | - | 15 | 9394 | 0.15967639 | 63.6666667 | WT | FALSE | CAN | JSY13 | 2A | WT-Rep2A | CAA | nope | >1bp-ins | A/T-1 | none | A/T-1 | CACCACAAA | AAACCAT | GTT | GTT | T | none | T>none | T>none_GTT | A/T-1_AAACCAT | 9 | TRUE | FALSE |
| 170_v.txt | 32728 | 32728-Deleti | Deletion | 1 | A | - | 4 | 9394 | 0.04258037 | 57.25 | WT | FALSE | CAN | JSY13 | 2A | WT-Rep2A | CAA | nope | >1bp-ins | A/T-1 | none | A/T-1 | CCATACAAA | AAAAACA | GTT | GTT | T | none | T>none | T>none_GTT | A/T-1_AAAAACA | 9 | TRUE | FALSE |
| 170_v.txt | 32842 | 32842-Deleti | Deletion | 1 | A | - | 336 | 9394 | 3.57675112 | 41.8363095 | WT | FALSE | CAN | JSY13 | 2A | WT-Rep2A | CAA | nope | >1bp-ins | A/T-1 | none | A/T-1 | TTACCCAAA | AAAAAAT | GTT | GTT | T | none | T>none | T>none_GTT | A/T-1_AAAAAAT | 9 | TRUE | FALSE |
| 170_v.txt | 32883 | 32883-Deleti | Deletion | 1 | C | - | 1 | 34435 | 2.90E-03 | 36 | WT | FALSE | CAN | JSY13 | 2A | WT-Rep2A | CCA | nope | >1bp-ins | G/C-1 | none | G/C-1 | TACGTCCAA | CAAAATT | GGT | CCA | C | none | nope | nope_CCA | G/C-1_CAAAATT | 9 | TRUE | FALSE |
| 170_v.txt | 32884 | 32884-Deleti | Deletion | 1 | A | - | 19 | 34435 | 0.05517642 | 37.631578 | WT | FALSE | CAN | JSY13 | 2A | WT-Rep2A | CAA | nope | >1bp-ins | A/T-1 | none | A/T-1 | ACGTCCAAA | AAAATTG | GTT | GTT | T | none | T>none | T>none_GTT | A/T-1_AAAATTG | 9 | TRUE | FALSE |
| 170_v.txt | 32912 | 32912-Deleti | Deletion | 1 | A | - | 7 | 34437 | 0.02032697 | 45.4285714 | WT | FALSE | CAN | JSY13 | 2A | WT-Rep2A | TAA | nope | >1bp-ins | A/T-1 | none | A/T-1 | TACACTAAG | AAGTTCC | ATT | ATT | T | none | T>none | T>none_ATT | A/T-1_AAGTTCC | 9 | TRUE | FALSE |
| 170_v.txt | 32924 | 32924-Deleti | Deletion | 1 | A | - | 25 | 34437 | 0.07259634 | 54.28 | WT | FALSE | CAN | JSY13 | 2A | WT-Rep2A | CAA | nope | >1bp-ins | A/T-1 | none | A/T-1 | CAGGGCAAA | AAAAGTG | GTT | GTT | T | none | T>none | T>none_GTT | A/T-1_AAAAGTG | 9 | TRUE | FALSE |
| 170_v.txt | 33083 | 33083-Deleti | Deletion | 1 | A | - | 7 | 34437 | 0.02032697 | 48.5714286 | WT | FALSE | CAN | JSY13 | 2A | WT-Rep2A | TAA | nope | >1bp-ins | A/T-1 | none | A/T-1 | ACCCATAAA | AAATAAA | ATT | ATT | T | none | T>none | T>none_ATT | A/T-1_AAATAAA | 9 | TRUE | FALSE |
| 170_v.txt | 33105 | 33105-Deleti | Deletion | 1 | C | - | 7 | 34437 | 0.02032697 | 45.4285714 | WT | FALSE | CAN | JSY13 | 2A | WT-Rep2A | GCC | nope | >1bp-ins | G/C-1 | none | G/C-1 | AGAGCGCCC | CCCACTG | CGG | GCC | C | none | nope | nope_GCC | G/C-1_CCCACTG | 9 | TRUE | FALSE |
| 170_v.txt | 33111 | 33111-Deleti | Deletion | 1 | G | - | 3 | 34437 | 8.71E-03 | 55.3333333 | WT | FALSE | CAN | JSY13 | 2A | WT-Rep2A | TGG | nope | >1bp-ins | G/C-1 | none | G/C-1 | CCCACTGGG | GGGCCGG | ACC | ACC | C | none | C>none | C>none_ACC | G/C-1_GGGCCGG | 9 | TRUE | FALSE |
| 170_v.txt | 33114 | 33114-Deleti | Deletion | 1 | C | - | 1 | 34437 | 2.90E-03 | 38 | WT | FALSE | CAN | JSY13 | 2A | WT-Rep2A | GCC | nope | >1bp-ins | G/C-1 | none | G/C-1 | ACTGGGCCG | CCGGCGT | CGG | GCC | C | none | nope | nope_GCC | G/C-1_CCGGCGT | 9 | TRUE | FALSE |
| 170_v.txt | 33138 | 33138-Deleti | Deletion | 1 | A | - | 8 | 34437 | 0.02323083 | 40.875 | WT | FALSE | CAN | JSY13 | 2A | WT-Rep2A | TAA | nope | >1bp-ins | A/T-1 | none | A/T-1 | GTGGATAAA | AAACCAA | ATT | ATT | T | none | T>none | T>none_ATT | A/T-1_AAACCAA | 9 | TRUE | FALSE |
| 170_v.txt | 33168 | 33168-Deleti | Deletion | 1 | C | - | 2 | 34418 | 5.81E-03 | 37.9999993 | WT | FALSE | CAN | JSY13 | 2A | WT-Rep2A | ACC | nope | >1bp-ins | G/C-1 | none | G/C-1 | ATAGTACCA | CCACCAA | TGG | ACC | C | none | nope | nope_ACC | G/C-1_CCACCAA | 9 | TRUE | FALSE |
| 170_v.txt | 33312 | 33312-Deleti | Deletion | 1 | A | - | 3 | 17691 | 0.01695778 | 38 | WT | FALSE | CAN | JSY13 | 2A | WT-Rep2A | CAA | nope | >1bp-ins | A/T-1 | none | A/T-1 | GAGCGCAAT | AATGGAT | GTT | GTT | T | none | T>none | T>none_GTT | A/T-1_AATGGAT | 9 | TRUE | FALSE |
| 170_v.txt | 33317 | 33317-Deleti | Deletion | 1 | A | - | 7 | 17691 | 0.03956814 | 59.4285714 | WT | FALSE | CAN | JSY13 | 2A | WT-Rep2A | GAT | nope | >1bp-ins | A/T-1 | none | A/T-1 | CAATGGATA | ATACAAT | CTA | CTA | T | none | T>none | T>none_CTA | A/T-1_ATACAAT | 9 | TRUE | FALSE |
| 170_v.txt | 33474 | 33474-Deleti | Deletion | 1 | C | - | 55 | 17691 | 0.31089254 | 28.3454546 | WT | FALSE | CAN | JSY13 | 2A | WT-Rep2A | GCC | nope | >1bp-ins | G/C-1 | none | G/C-1 | GCTATGCCTT | CCTTTTT | CGG | GCC | C | none | nope | nope_GCC | G/C-1_CCTTTTT | 9 | TRUE | FALSE |
| 187_v.txt | 31986 | 31986-SNV- | SNV | 1 | A | C | 5383 | 11361 | 47.3813925 | 50.8563998 | WT | FALSE | CAN | JSY13 | 3A | WT-Rep3A | TAA | A>C | G/C+1 | A/T-1 | TA>GC | TA>GC | GGCATTAAT | AATTTAG | ATT | ATT | T | G | T>G | T>G_ATT | TA>GC_AATTTAG | 9 | TRUE | FALSE |
| 187_v.txt | 32004 | 32004-SNV- | SNV | 1 | A | C | 7 | 11361 | 0.0616143 | 33.8571429 | WT | FALSE | CAN | JSY13 | 3A | WT-Rep3A | TAA | A>C | G/C+1 | A/T-1 | TA>GC | TA>GC | AATGGTAAC | AACTCGT | ATT | ATT | T | G | T>G | T>G_ATT | TA>GC_AACTCGT | 9 | TRUE | FALSE |
| 187_v.txt | 32265 | 32265-SNV- | SNV | 1 | A | C | 10 | 43076 | 0.02321478 | 18.0004186 | WT | FALSE | CAN | JSY13 | 3A | WT-Rep3A | TAA | A>C | G/C+1 | A/T-1 | TA>GC | TA>GC | CCAAATAAA | AAAATAC | ATT | ATT | T | G | T>G | T>G_ATT | TA>GC_AAAATAC | 9 | TRUE | FALSE |
| 187_v.txt | 32439 | 32439-SNV- | SNV | 1 | A | C | 14 | 43076 | 0.0325007 | 43.1428571 | WT | FALSE | CAN | JSY13 | 3A | WT-Rep3A | AAC | A>C | G/C+1 | A/T-1 | TA>GC | TA>GC | TATGGAACT | ACTAAAA | TTG | TTG | T | G | T>G | T>G_TTG | TA>GC_ACTAAAA | 9 | TRUE | FALSE |
| 187_v.txt | 32445 | 32445-SNV- | SNV | 1 | A | C | 18 | 43076 | 0.04178661 | 55.3333333 | WT | FALSE | CAN | JSY13 | 3A | WT-Rep3A | AAG | A>C | G/C+1 | A/T-1 | TA>GC | TA>GC | ACTAAAAGT | AGTCCAA | TTC | TTC | T | G | T>G | T>G_TTC | TA>GC_AGTCCAA | 9 | TRUE | FALSE |
| 187_v.txt | 32493 | 32493-SNV- | SNV | 1 | A | C | 8 | 43076 | 0.01857183 | 39.7494581 | WT | FALSE | CAN | JSY13 | 3A | WT-Rep3A | AAC | A>C | G/C+1 | A/T-1 | TA>GC | TA>GC | CGGAAAACA | ACAACTT | TTG | TTG | T | G | T>G | T>G_TTG | TA>GC_ACAACTT | 9 | TRUE | FALSE |
| 187_v.txt | 32616 | 32616-SNV- | SNV | 1 | A | C | 17 | 23486 | 0.07238355 | 41.5882353 | WT | FALSE | CAN | JSY13 | 3A | WT-Rep3A | TAA | A>C | G/C+1 | A/T-1 | TA>GC | TA>GC | CAACCTAAG | AAGAACC | ATT | ATT | T | G | T>G | T>G_ATT | TA>GC_AAGAACC | 9 | TRUE | FALSE |
| 187_v.txt | 32739 | 32739-SNV- | SNV | 1 | A | C | 7 | 23486 | 0.02980499 | 48.4285714 | WT | FALSE | CAN | JSY13 | 3A | WT-Rep3A | TAT | A>C | G/C+1 | A/T-1 | TA>GC | TA>GC | CAGTATATTA | ATTAGAA | ATA | ATA | T | G | T>G | T>G_ATA | TA>GC_ATTAGAA | 9 | TRUE | FALSE |
| 187_v.txt | 32759 | 32759-SNV-T | SNV | 1 | T | G | 2 | 23486 | 8.52E-03 | 32.9917827 | WT | FALSE | CAN | JSY13 | 3A | WT-Rep3A | CTA | T>G | G/C+1 | A/T-1 | TA>GC | TA>GC | AATGGCTAA | TAAAACT | GAT | CTA | T | C | T>G | T>G_CTA | TA>GC_TAAAACT | 9 | TRUE | FALSE |
| 187_v.txt | 32763 | 32763-SNV- | SNV | 1 | A | C | 2 | 23486 | 8.52E-03 | 50.9958581 | WT | FALSE | CAN | JSY13 | 3A | WT-Rep3A | AAC | A>C | G/C+1 | A/T-1 | TA>GC | TA>GC | GCTAAAACTT | ACTTTGA | TTG | TTG | T | G | T>G | T>G_TTG | TA>GC_ACTTTGA | 9 | TRUE | FALSE |
| 187_v.txt | 32801 | 32801-SNV- | SNV | 1 | A | C | 9 | 23486 | 0.0383207 | 43.5555556 | WT | FALSE | CAN | JSY13 | 3A | WT-Rep3A | AAT | A>C | G/C+1 | A/T-1 | TA>GC | TA>GC | ACCGTAATAT | ATATTTG | TTA | TTA | T | G | T>G | T>G_TTA | TA>GC_ATATTTG | 9 | TRUE | FALSE |
| 187_v.txt | 32823 | 32823-SNV- | SNV | 1 | A | C | 65 | 23486 | 0.27676062 | 42.4307692 | WT | FALSE | CAN | JSY13 | 3A | WT-Rep3A | CAT | A>C | G/C+1 | A/T-1 | TA>GC | TA>GC | AAGTTCATTA | ATTATTG | GTA | GTA | T | G | T>G | T>G_GTA | TA>GC_ATTATTG | 9 | TRUE | FALSE |
| 187_v.txt | 32842 | 32842-SNV- | SNV | 1 | A | C | 3 | 23486 | 0.01277357 | 36.6665958 | WT | FALSE | CAN | JSY13 | 3A | WT-Rep3A | CAA | A>C | G/C+1 | A/T-1 | TA>GC | TA>GC | TTACCCAAA | AAAAAAT | GTT | GTT | T | G | T>G | T>G_GTT | TA>GC_AAAAAAT | 9 | TRUE | FALSE |
| 187_v.txt | 32884 | 32884-SNV- | SNV | 1 | A | C | 10 | 40419 | 0.02474084 | 36.2981665 | WT | FALSE | CAN | JSY13 | 3A | WT-Rep3A | CAA | A>C | G/C+1 | A/T-1 | TA>GC | TA>GC | ACGTCCAAA | AAAATTG | GTT | GTT | T | G | T>G | T>G_GTT | TA>GC_AAAATTG | 9 | TRUE | FALSE |
| 187_v.txt | 32916 | 32916-SNV-T | SNV | 1 | T | G | 11 | 40418 | 0.0272156 | 54.8171854 | WT | FALSE | CAN | JSY13 | 3A | WT-Rep3A | TTC | T>G | G/C+1 | A/T-1 | TA>GC | TA>GC | CTAAGTTCCA | TCCAGGG | AAG | TTC | T | C | T>G | T>G_TTC | TA>GC_TCCAGGG | 9 | TRUE | FALSE |
| 187_v.txt | 33007 | 33007-SNV-T | SNV | 1 | T | G | 8 | 40458 | 0.01977359 | 56.125 | WT | FALSE | CAN | JSY13 | 3A | WT-Rep3A | GTG | T>G | G/C+1 | A/T-1 | TA>GC | TA>GC | AAACTGTGA | TGAAAGA | CAC | GTG | T | C | T>G | T>G_GTG | TA>GC_TGAAAGA | 9 | TRUE | FALSE |
| 187_v.txt | 33017 | 33017-SNV-T | SNV | 1 | T | G | 10 | 40458 | 0.02471699 | 46.3 | WT | FALSE | CAN | JSY13 | 3A | WT-Rep3A | ATG | T>G | G/C+1 | A/T-1 | TA>GC | TA>GC | AGAGGATGT | TGTAACA | TAC | ATG | T | C | T>G | T>G_ATG | TA>GC_TGTAACA | 9 | TRUE | FALSE |
| 187_v.txt | 33167 | 33167-SNV- | SNV | 1 | A | C | 4 | 40451 | 9.89E-03 | 14.7509613 | WT | FALSE | CAN | JSY13 | 3A | WT-Rep3A | TAC | A>C | G/C+1 | A/T-1 | TA>GC | TA>GC | AATAGTACCA | ACCACCA | ATG | ATG | T | G | T>G | T>G_ATG | TA>GC_ACCACCA | 9 | TRUE | FALSE |
| 187_v.txt | 33269 | 33269-SNV- | SNV | 1 | A | C | 171 | 23996 | 0.71261877 | 15.8362573 | WT | FALSE | CAN | JSY13 | 3A | WT-Rep3A | TAG | A>C | G/C+1 | A/T-1 | TA>GC | TA>GC | CTCCATAGAC | AGAGAAC | ATC | ATC | T | G | T>G | T>G_ATC | TA>GC_AGAGAAC | 9 | TRUE | FALSE |
| 187_v.txt | 33476 | 33476-SNV-T | SNV | 1 | T | G | 5 | 23995 | 0.02083768 | 29.2 | WT | FALSE | CAN | JSY13 | 3A | WT-Rep3A | CTT | T>G | G/C+1 | A/T-1 | TA>GC | TA>GC | TATGCCTTTT | TTTTTTT | GAA | CTT | T | C | T>G | T>G_CTT | TA>GC_TTTTTTT | 9 | TRUE | FALSE |
| 187_v.txt | 32275 | 32275-SNV- | SNV | 1 | A | G | 42 | 43076 | 0.09750209 | 34.4987633 | WT | FALSE | CAN | JSY13 | 3A | WT-Rep3A | GAA | A>G | G/C+1 | A/T-1 | TA>CG | TA>CG | TACGGGAAC | AACCAAC | CTT | CTT | T | C | T>C | T>C_CTT | TA>CG_AACCAAC | 9 | TRUE | FALSE |
| 187_v.txt | 32445 | 32445-SNV- | SNV | 1 | A | G | 21 | 43076 | 0.04875105 | 49.4285714 | WT | FALSE | CAN | JSY13 | 3A | WT-Rep3A | AAG | A>G | G/C+1 | A/T-1 | TA>CG | TA>CG | ACTAAAAGT | AGTCCAA | TTC | TTC | T | C | T>C | T>C_TTC | TA>CG_AGTCCAA | 9 | TRUE | FALSE |
| 187_v.txt | 32493 | 32493-SNV- | SNV | 1 | A | G | 9 | 43076 | 0.02089331 | 30.6604053 | WT | FALSE | CAN | JSY13 | 3A | WT-Rep3A | AAC | A>G | G/C+1 | A/T-1 | TA>CG | TA>CG | CGGAAAACA | ACAACTT | TTG | TTG | T | C | T>C | T>C_TTG | TA>CG_ACAACTT | 9 | TRUE | FALSE |
| 187_v.txt | 32734 | 32734-SNV- | SNV | 1 | A | G | 80 | 23486 | 0.34062846 | 59.9125 | WT | FALSE | CAN | JSY13 | 3A | WT-Rep3A | CAG | A>G | G/C+1 | A/T-1 | TA>CG | TA>CG | AAAAACAGT | AGTATAT | GTC | GTC | T | C | T>C | T>C_GTC | TA>CG_AGTATAT | 9 | TRUE | FALSE |
| 187_v.txt | 32754 | 32754-SNV- | SNV | 1 | A | G | 6 | 23486 | 0.02554714 | 44.6646786 | WT | FALSE | CAN | JSY13 | 3A | WT-Rep3A | AAT | A>G | G/C+1 | A/T-1 | TA>CG | TA>CG | CCGATAATG | ATGGCTA | TTA | TTA | T | C | T>C | T>C_TTA | TA>CG_ATGGCTA | 9 | TRUE | FALSE |
| 187_v.txt | 32755 | 32755-SNV-T | SNV | 1 | T | C | 5 | 23484 | 0.02129109 | 47.5999324 | WT | FALSE | CAN | JSY13 | 3A | WT-Rep3A | ATG | T>C | G/C+1 | A/T-1 | TA>CG | TA>CG | CGATAATGG | TGGCTAA | TAC | ATG | T | G | T>C | T>C_ATG | TA>CG_TGGCTAA | 9 | TRUE | FALSE |
| 187_v.txt | 32760 | 32760-SNV- | SNV | 1 | A | G | 5 | 23486 | 0.02128928 | 56.999865 | WT | FALSE | CAN | JSY13 | 3A | WT-Rep3A | TAA | A>G | G/C+1 | A/T-1 | TA>CG | TA>CG | ATGGCTAAA | AAAACTT | ATT | ATT | T | C | T>C | T>C_ATT | TA>CG_AAAACTT | 9 | TRUE | FALSE |
| 187_v.txt | 32761 | 32761-SNV- | SNV | 1 | A | G | 9 | 23486 | 0.0383207 | 19.5516363 | WT | FALSE | CAN | JSY13 | 3A | WT-Rep3A | AAA | A>G | G/C+1 | A/T-1 | TA>CG | TA>CG | TGGCTAAAA | AAACTTT | TTT | TTT | T | C | T>C | T>C_TTT | TA>CG_AAACTTT | 9 | TRUE | FALSE |
| 187_v.txt | 32762 | 32762-SNV- | SNV | 1 | A | G | 5 | 23486 | 0.02128928 | 43.6001889 | WT | FALSE | CAN | JSY13 | 3A | WT-Rep3A | AAA | A>G | G/C+1 | A/T-1 | TA>CG | TA>CG | GGCTAAAAC | AACTTTG | TTT | TTT | T | C | T>C | T>C_TTT | TA>CG_AACTTTG | 9 | TRUE | FALSE |
| 187_v.txt | 32927 | 32927-SNV- | SNV | 1 | A | G | 35 | 40418 | 0.08659508 | 47.6857143 | WT | FALSE | CAN | JSY13 | 3A | WT-Rep3A | AAG | A>G | G/C+1 | A/T-1 | TA>CG | TA>CG | GGCAAAAGT | AGTGATT | TTC | TTC | T | C | T>C | T>C_TTC | TA>CG_AGTGATT | 9 | TRUE | FALSE |
| 187_v.txt | 32938 | 32938-SNV- | SNV | 1 | A | G | 61 | 40418 | 0.15092286 | 53.4754098 | WT | FALSE | CAN | JSY13 | 3A | WT-Rep3A | CAA | A>G | G/C+1 | A/T-1 | TA>CG | TA>CG | TTGCCCAAG | AAGAAAA | GTT | GTT | T | C | T>C | T>C_GTT | TA>CG_AAGAAAA | 9 | TRUE | FALSE |
| 187_v.txt | 33476 | 33476-SNV-T | SNV | 1 | T | C | 33 | 23995 | 0.13752865 | 30.969697 | WT | FALSE | CAN | JSY13 | 3A | WT-Rep3A | CTT | T>C | G/C+1 | A/T-1 | TA>CG | TA>CG | TATGCCTTTT | TTTTTTT | GAA | CTT | T | G | T>C | T>C_CTT | TA>CG_TTTTTTT | 9 | TRUE | FALSE |
| 187_v.txt | 31983 | 31983-SNV- | SNV | 1 | A | T | 14 | 11361 | 0.12322859 | 45.5714286 | WT | FALSE | CAN | JSY13 | 3A | WT-Rep3A | CAT | A>T | A/T+1 | A/T-1 | TA>AT | TA>AT | CCGGGCATTA | ATTAATT | GTA | GTA | T | A | T>A | T>A_GTA | TA>AT_ATTAATT | 9 | TRUE | FALSE |
| 187_v.txt | 32266 | 32266-SNV- | SNV | 1 | A | T | 4 | 43076 | 9.29E-03 | 17.7497629 | WT | FALSE | CAN | JSY13 | 3A | WT-Rep3A | AAA | A>T | A/T+1 | A/T-1 | TA>AT | TA>AT | CAAATAAAA | AAATACG | TTT | TTT | T | A | T>A | T>A_TTT | TA>AT_AAATACG | 9 | TRUE | FALSE |
| 187_v.txt | 32337 | 32337-SNV- | SNV | 1 | A | T | 9 | 43075 | 0.02089379 | 51.7775376 | WT | FALSE | CAN | JSY13 | 3A | WT-Rep3A | GAT | A>T | A/T+1 | A/T-1 | TA>AT | TA>AT | TTGAAGATA | ATATGTG | CTA | CTA | T | A | T>A | T>A_CTA | TA>AT_ATATGTG | 9 | TRUE | FALSE |
| 187_v.txt | 32346 | 32346-SNV- | SNV | 1 | A | T | 186 | 43076 | 0.43179497 | 61.2508152 | WT | FALSE | CAN | JSY13 | 3A | WT-Rep3A | CAA | A>T | A/T+1 | A/T-1 | TA>AT | TA>AT | TGTGGCAAA | AAAACCT | GTT | GTT | T | A | T>A | T>A_GTT | TA>AT_AAAACCT | 9 | TRUE | FALSE |
| 187_v.txt | 32496 | 32496-SNV- | SNV | 1 | A | T | 96 | 43076 | 0.22286192 | 46.3125 | WT | FALSE | CAN | JSY13 | 3A | WT-Rep3A | AAC | A>T | A/T+1 | A/T-1 | TA>AT | TA>AT | AAAACAACT | ACTTTTT | TTG | TTG | T | A | T>A | T>A_TTG | TA>AT_ACTTTTT | 9 | TRUE | FALSE |
| 187_v.txt | 32499 | 32499-SNV-T | SNV | 1 | T | A | 5 | 43076 | 0.01160739 | 41.9993806 | WT | FALSE | CAN | JSY13 | 3A | WT-Rep3A | TTT | T>A | A/T+1 | A/T-1 | TA>AT | TA>AT | ACAACTTTTT | TTTTTGA | AAA | TTT | T | T | T>A | T>A_TTT | TA>AT_TTTTTGA | 9 | TRUE | FALSE |
| 187_v.txt | 32598 | 32598-SNV- | SNV | 1 | A | T | 20 | 23486 | 0.08515712 | 45.5 | WT | FALSE | CAN | JSY13 | 3A | WT-Rep3A | CAA | A>T | A/T+1 | A/T-1 | TA>AT | TA>AT | TTAATCAAA | AAAGAGG | GTT | GTT | T | A | T>A | T>A_GTT | TA>AT_AAAGAGG | 9 | TRUE | FALSE |
| 187_v.txt | 32754 | 32754-SNV- | SNV | 1 | A | T | 4 | 23486 | 0.01703142 | 51.4485918 | WT | FALSE | CAN | JSY13 | 3A | WT-Rep3A | AAT | A>T | A/T+1 | A/T-1 | TA>AT | TA>AT | CCGATAATG | ATGGCTA | TTA | TTA | T | A | T>A | T>A_TTA | TA>AT_ATGGCTA | 9 | TRUE | FALSE |
| 187_v.txt | 32755 | 32755-SNV-T | SNV | 1 | T | A | 3 | 23484 | 0.01277466 | 36.6663977 | WT | FALSE | CAN | JSY13 | 3A | WT-Rep3A | ATG | T>A | A/T+1 | A/T-1 | TA>AT | TA>AT | CGATAATGG | TGGCTAA | TAC | ATG | T | T | T>A | T>A_ATG | TA>AT_TGGCTAA | 9 | TRUE | FALSE |
| 187_v.txt | 32761 | 32761-SNV- | SNV | 1 | A | T | 2 | 23486 | 8.52E-03 | 56.4915748 | WT | FALSE | CAN | JSY13 | 3A | WT-Rep3A | AAA | A>T | A/T+1 | A/T-1 | TA>AT | TA>AT | TGGCTAAAA | AAACTTT | TTT | TTT | T | A | T>A | T>A_TTT | TA>AT_AAACTTT | 9 | TRUE | FALSE |
| 187_v.txt | 32766 | 32766-SNV-T | SNV | 1 | T | A | 3 | 23486 | 0.01277357 | 46.331261 | WT | FALSE | CAN | JSY13 | 3A | WT-Rep3A | TTT | T>A | A/T+1 | A/T-1 | TA>AT | TA>AT | AAAACTTTGA | TTGATGG | AAA | TTT | T | T | T>A | T>A_TTT | TA>AT_TTGATGG | 9 | TRUE | FALSE |
| 187_v.txt | 32767 | 32767-SNV-T | SNV | 1 | T | A | 10 | 23486 | 0.04257856 | 49.8932969 | WT | FALSE | CAN | JSY13 | 3A | WT-Rep3A | TTG | T>A | A/T+1 | A/T-1 | TA>AT | TA>AT | AAACTTTGAT | TGATGGA | AAC | TTG | T | T | T>A | T>A_TTG | TA>AT_TGATGGA | 9 | TRUE | FALSE |
| 187_v.txt | 32821 | 32821-SNV-T | SNV | 1 | T | A | 20 | 23486 | 0.08515712 | 46.65 | WT | FALSE | CAN | JSY13 | 3A | WT-Rep3A | TTC | T>A | A/T+1 | A/T-1 | TA>AT | TA>AT | ACAAGTTCAT | TCATTAT | AAG | TTC | T | T | T>A | T>A_TTC | TA>AT_TCATTAT | 9 | TRUE | FALSE |
| 187_v.txt | 33065 | 33065-SNV- | SNV | 1 | A | T | 20 | 40458 | 0.04943398 | 48.4 | WT | FALSE | CAN | JSY13 | 3A | WT-Rep3A | AAT | A>T | A/T+1 | A/T-1 | TA>AT | TA>AT | GACAGAATA | ATATGCC | TTA | TTA | T | A | T>A | T>A_TTA | TA>AT_ATATGCC | 9 | TRUE | FALSE |
| 187_v.txt | 33067 | 33067-SNV- | SNV | 1 | A | T | 271 | 40458 | 0.66983044 | 57.1771218 | WT | FALSE | CAN | JSY13 | 3A | WT-Rep3A | TAT | A>T | A/T+1 | A/T-1 | TA>AT | TA>AT | CAGAATATG | ATGCCAA | ATA | ATA | T | A | T>A | T>A_ATA | TA>AT_ATGCCAA | 9 | TRUE | FALSE |
| 187_v.txt | 33192 | 33192-SNV-T | SNV | 1 | T | A | 28 | 23997 | 0.11668125 | 34.3912609 | WT | FALSE | CAN | JSY13 | 3A | WT-Rep3A | ATG | T>A | A/T+1 | A/T-1 | TA>AT | TA>AT | CCAATATGTC | TGTCTTT | TAC | ATG | T | T | T>A | T>A_ATG | TA>AT_TGTCTTT | 9 | TRUE | FALSE |
| 187_v.txt | 33321 | 33321-SNV- | SNV | 1 | A | T | 663 | 23996 | 2.76296049 | 59.6515837 | WT | FALSE | CAN | JSY13 | 3A | WT-Rep3A | CAA | A>T | A/T+1 | A/T-1 | TA>AT | TA>AT | GGATACAAT | AATTCTT | GTT | GTT | T | A | T>A | T>A_GTT | TA>AT_AATTCTT | 9 | TRUE | FALSE |
| 187_v.txt | 33345 | 33345-SNV- | SNV | 1 | A | T | 28 | 23996 | 0.11668611 | 48.0357143 | WT | FALSE | CAN | JSY13 | 3A | WT-Rep3A | CAA | A>T | A/T+1 | A/T-1 | TA>AT | TA>AT | TCTTTCAATG | AATGGTA | GTT | GTT | T | A | T>A | T>A_GTT | TA>AT_AATGGTA | 9 | TRUE | FALSE |
| 187_v.txt | 31806 | 31806-SNV- | SNV | 1 | C | T | 7 | 1873 | 0.37373198 | 64 | WT | FALSE | CAN | JSY13 | 3A | WT-Rep3A | CCA | C>T | A/T+1 | G/C-1 | CG>TA | CG>TA | ATCTTCCAAA | CAAATAA | GGT | CCA | C | A | C>T | C>T_CCA | CG>TA_CAAATAA | 9 | TRUE | FALSE |
| 187_v.txt | 32080 | 32080-SNV- | SNV | 1 | C | T | 4 | 11361 | 0.03520817 | 63.5 | WT | FALSE | CAN | JSY13 | 3A | WT-Rep3A | GCA | C>T | A/T+1 | G/C-1 | CG>TA | CG>TA | ACCATGCAA | CAAAAAA | CGT | GCA | C | A | C>T | C>T_GCA | CG>TA_CAAAAAA | 9 | TRUE | FALSE |
| 187_v.txt | 32092 | 32092-SNV- | SNV | 1 | C | T | 17 | 11361 | 0.14963472 | 58.6470588 | WT | FALSE | CAN | JSY13 | 3A | WT-Rep3A | GCA | C>T | A/T+1 | G/C-1 | CG>TA | CG>TA | AGCCTGCAA | CAACACC | CGT | GCA | C | A | C>T | C>T_GCA | CG>TA_CAACACC | 9 | TRUE | FALSE |
| 187_v.txt | 32271 | 32271-SNV- | SNV | 1 | C | T | 51 | 43076 | 0.11839539 | 31.5296114 | WT | FALSE | CAN | JSY13 | 3A | WT-Rep3A | ACG | C>T | A/T+1 | G/C-1 | CG>TA | CG>TA | AAAATACGG | CGGGAAC | TGC | ACG | C | A | C>T | C>T_ACG | CG>TA_CGGGAAC | 9 | TRUE | FALSE |
| 187_v.txt | 32272 | 32272-SNV- | SNV | 1 | G | A | 30 | 43076 | 0.06964435 | 35.8333823 | WT | FALSE | CAN | JSY13 | 3A | WT-Rep3A | CGG | G>A | A/T+1 | G/C-1 | CG>TA | CG>TA | AAATACGGG | GGGAACC | GCC | GCC | C | T | C>T | C>T_GCC | CG>TA_GGGAACC | 9 | TRUE | FALSE |
| 187_v.txt | 32274 | 32274-SNV- | SNV | 1 | G | A | 406 | 43076 | 0.9425202 | 36.6600985 | WT | FALSE | CAN | JSY13 | 3A | WT-Rep3A | GGA | G>A | A/T+1 | G/C-1 | CG>TA | CG>TA | ATACGGGAA | GAACCAA | CCT | CCT | C | T | C>T | C>T_CCT | CG>TA_GAACCAA | 9 | TRUE | FALSE |
| 187_v.txt | 32299 | 32299-SNV- | SNV | 1 | C | T | 73 | 43076 | 0.16946792 | 55.3835616 | WT | FALSE | CAN | JSY13 | 3A | WT-Rep3A | GCG | C>T | A/T+1 | G/C-1 | CG>TA | CG>TA | AATTTGCGG | CGGCAGA | CGC | GCG | C | A | C>T | C>T_GCG | CG>TA_CGGCAGA | 9 | TRUE | FALSE |
| 187_v.txt | 32302 | 32302-SNV- | SNV | 1 | C | T | 26 | 43076 | 0.06035844 | 52.0769231 | WT | FALSE | CAN | JSY13 | 3A | WT-Rep3A | GCA | C>T | A/T+1 | G/C-1 | CG>TA | CG>TA | TTGCGGCAG | CAGAAAT | CGT | GCA | C | A | C>T | C>T_GCA | CG>TA_CAGAAAT | 9 | TRUE | FALSE |
| 187_v.txt | 32463 | 32463-SNV- | SNV | 1 | G | A | 56 | 43076 | 0.13000279 | 43.8035714 | WT | FALSE | CAN | JSY13 | 3A | WT-Rep3A | AGA | G>A | A/T+1 | G/C-1 | CG>TA | CG>TA | AATAGAGAG | GAGCCAA | TCT | TCT | C | T | C>T | C>T_TCT | CG>TA_GAGCCAA | 9 | TRUE | FALSE |
| 187_v.txt | 32466 | 32466-SNV- | SNV | 1 | C | T | 41 | 43076 | 0.09518061 | 47.8275961 | WT | FALSE | CAN | JSY13 | 3A | WT-Rep3A | GCC | C>T | A/T+1 | G/C-1 | CG>TA | CG>TA | AGAGAGCCA | CCAATGT | CGG | GCC | C | A | C>T | C>T_GCC | CG>TA_CCAATGT | 9 | TRUE | FALSE |
| 187_v.txt | 32535 | 32535-SNV- | SNV | 1 | G | A | 55 | 43074 | 0.12768724 | 36.4180546 | WT | FALSE | CAN | JSY13 | 3A | WT-Rep3A | TGC | G>A | A/T+1 | G/C-1 | CG>TA | CG>TA | GGGTTTGCA | GCAGCTT | ACG | ACG | C | T | C>T | C>T_ACG | CG>TA_GCAGCTT | 9 | TRUE | FALSE |
| 187_v.txt | 32536 | 32536-SNV- | SNV | 1 | C | T | 60 | 43074 | 0.13929517 | 36.2798207 | WT | FALSE | CAN | JSY13 | 3A | WT-Rep3A | GCA | C>T | A/T+1 | G/C-1 | CG>TA | CG>TA | GGTTTGCAG | CAGCTTC | CGT | GCA | C | A | C>T | C>T_GCA | CG>TA_CAGCTTC | 9 | TRUE | FALSE |
| 187_v.txt | 32545 | 32545-SNV- | SNV | 1 | C | T | 141 | 43073 | 0.32735124 | 37.5602837 | WT | FALSE | CAN | JSY13 | 3A | WT-Rep3A | CCA | C>T | A/T+1 | G/C-1 | CG>TA | CG>TA | CTTCACCAG | CAGCAGT | GGT | CCA | C | A | C>T | C>T_CCA | CG>TA_CAGCAGT | 9 | TRUE | FALSE |
| 187_v.txt | 32609 | 32609-SNV- | SNV | 1 | C | T | 21 | 23486 | 0.08941497 | 49.2380952 | WT | FALSE | CAN | JSY13 | 3A | WT-Rep3A | CCC | C>T | A/T+1 | G/C-1 | CG>TA | CG>TA | GGAAACCCA | CCAACCT | GGG | CCC | C | A | C>T | C>T_CCC | CG>TA_CCAACCT | 9 | TRUE | FALSE |
| 187_v.txt | 32700 | 32700-SNV- | SNV | 1 | C | T | 24 | 23486 | 0.10218854 | 63.125 | WT | FALSE | CAN | JSY13 | 3A | WT-Rep3A | GCC | C>T | A/T+1 | G/C-1 | CG>TA | CG>TA | ACTGGGCCG | CCGGTAA | CGG | GCC | C | A | C>T | C>T_GCC | CG>TA_CCGGTAA | 9 | TRUE | FALSE |
| 187_v.txt | 32756 | 32756-SNV- | SNV | 1 | G | A | 6 | 23486 | 0.02554714 | 53.8249342 | WT | FALSE | CAN | JSY13 | 3A | WT-Rep3A | TGG | G>A | A/T+1 | G/C-1 | CG>TA | CG>TA | GATAATGGC | GGCTAAA | ACC | ACC | C | T | C>T | C>T_ACC | CG>TA_GGCTAAA | 9 | TRUE | FALSE |
| 187_v.txt | 32757 | 32757-SNV- | SNV | 1 | G | A | 14 | 23485 | 0.05961252 | 40.928553 | WT | FALSE | CAN | JSY13 | 3A | WT-Rep3A | GGC | G>A | A/T+1 | G/C-1 | CG>TA | CG>TA | ATAATGGCT | GCTAAAA | CCG | CCG | C | T | C>T | C>T_CCG | CG>TA_GCTAAAA | 9 | TRUE | FALSE |
| 187_v.txt | 32768 | 32768-SNV- | SNV | 1 | G | A | 2 | 23486 | 8.52E-03 | 63.9996042 | WT | FALSE | CAN | JSY13 | 3A | WT-Rep3A | TGA | G>A | A/T+1 | G/C-1 | CG>TA | CG>TA | AACTTTGATG | GATGGAA | ACT | ACT | C | T | C>T | C>T_ACT | CG>TA_GATGGAA | 9 | TRUE | FALSE |
| 187_v.txt | 32797 | 32797-SNV- | SNV | 1 | C | T | 24 | 23486 | 0.10218854 | 50.6226207 | WT | FALSE | CAN | JSY13 | 3A | WT-Rep3A | CCG | C>T | A/T+1 | G/C-1 | CG>TA | CG>TA | ATTCACCGTA | CGTAATA | GGC | CCG | C | A | C>T | C>T_CCG | CG>TA_CGTAATA | 9 | TRUE | FALSE |
| 187_v.txt | 32798 | 32798-SNV- | SNV | 1 | G | A | 28 | 23486 | 0.11921996 | 42.3928571 | WT | FALSE | CAN | JSY13 | 3A | WT-Rep3A | CGT | G>A | A/T+1 | G/C-1 | CG>TA | CG>TA | TTCACCGTAA | GTAATAT | GCA | GCA | C | T | C>T | C>T_GCA | CG>TA_GTAATAT | 9 | TRUE | FALSE |
| 187_v.txt | 32855 | 32855-SNV- | SNV | 1 | C | T | 661 | 23483 | 2.8148022 | 36.1059002 | WT | FALSE | CAN | JSY13 | 3A | WT-Rep3A | TCC | C>T | A/T+1 | G/C-1 | CG>TA | CG>TA | ACTAATCCAT | CCATGCC | AGG | TCC | C | A | C>T | C>T_TCC | CG>TA_CCATGCC | 9 | TRUE | FALSE |
| 187_v.txt | 32879 | 32879-SNV- | SNV | 1 | C | T | 43 | 40416 | 0.10639351 | 35.7441861 | WT | FALSE | CAN | JSY13 | 3A | WT-Rep3A | ACG | C>T | A/T+1 | G/C-1 | CG>TA | CG>TA | TTTGTACGTC | CGTCCAA | TGC | ACG | C | A | C>T | C>T_ACG | CG>TA_CGTCCAA | 9 | TRUE | FALSE |
| 187_v.txt | 32882 | 32882-SNV- | SNV | 1 | C | T | 81 | 40419 | 0.2004008 | 30.7588863 | WT | FALSE | CAN | JSY13 | 3A | WT-Rep3A | TCC | C>T | A/T+1 | G/C-1 | CG>TA | CG>TA | GTACGTCCA | CCAAAAT | AGG | TCC | C | A | C>T | C>T_TCC | CG>TA_CCAAAAT | 9 | TRUE | FALSE |
| 187_v.txt | 32883 | 32883-SNV- | SNV | 1 | C | T | 60 | 40419 | 0.14844504 | 36.6833333 | WT | FALSE | CAN | JSY13 | 3A | WT-Rep3A | CCA | C>T | A/T+1 | G/C-1 | CG>TA | CG>TA | TACGTCCAA | CAAAATT | GGT | CCA | C | A | C>T | C>T_CCA | CG>TA_CAAAATT | 9 | TRUE | FALSE |
| 187_v.txt | 32900 | 32900-SNV- | SNV | 1 | G | A | 31 | 40418 | 0.0766985 | 34.256089 | WT | FALSE | CAN | JSY13 | 3A | WT-Rep3A | GGC | G>A | A/T+1 | G/C-1 | CG>TA | CG>TA | GACTTGGCC | GCCAACT | CCG | CCG | C | T | C>T | C>T_CCG | CG>TA_GCCAACT | 9 | TRUE | FALSE |
| 187_v.txt | 32901 | 32901-SNV- | SNV | 1 | C | T | 77 | 40418 | 0.19050918 | 31.4805602 | WT | FALSE | CAN | JSY13 | 3A | WT-Rep3A | GCC | C>T | A/T+1 | G/C-1 | CG>TA | CG>TA | ACTTGGCCA | CCAACTA | CGG | GCC | C | A | C>T | C>T_GCC | CG>TA_CCAACTA | 9 | TRUE | FALSE |
| 187_v.txt | 32934 | 32934-SNV- | SNV | 1 | G | A | 17 | 40418 | 0.04206047 | 45.9418724 | WT | FALSE | CAN | JSY13 | 3A | WT-Rep3A | TGC | G>A | A/T+1 | G/C-1 | CG>TA | CG>TA | GTGATTGCC | GCCCAAG | ACG | ACG | C | T | C>T | C>T_ACG | CG>TA_GCCCAAG | 9 | TRUE | FALSE |
| 187_v.txt | 32935 | 32935-SNV- | SNV | 1 | C | T | 41 | 40418 | 0.10143995 | 37.8048781 | WT | FALSE | CAN | JSY13 | 3A | WT-Rep3A | GCC | C>T | A/T+1 | G/C-1 | CG>TA | CG>TA | TGATTGCCC | CCCAAGA | CGG | GCC | C | A | C>T | C>T_GCC | CG>TA_CCCAAGA | 9 | TRUE | FALSE |
| 187_v.txt | 32958 | 32958-SNV- | SNV | 1 | C | T | 111 | 40458 | 0.27435859 | 54.3783784 | WT | FALSE | CAN | JSY13 | 3A | WT-Rep3A | ACC | C>T | A/T+1 | G/C-1 | CG>TA | CG>TA | ATGTAACCAT | CCATTGG | TGG | ACC | C | A | C>T | C>T_ACC | CG>TA_CCATTGG | 9 | TRUE | FALSE |
| 187_v.txt | 32965 | 32965-SNV- | SNV | 1 | C | T | 47 | 40458 | 0.11616986 | 58.6595745 | WT | FALSE | CAN | JSY13 | 3A | WT-Rep3A | GCC | C>T | A/T+1 | G/C-1 | CG>TA | CG>TA | CATTGGCCG | CCGCACC | CGG | GCC | C | A | C>T | C>T_GCC | CG>TA_CCGCACC | 9 | TRUE | FALSE |
| 187_v.txt | 32967 | 32967-SNV- | SNV | 1 | G | A | 87 | 40458 | 0.21503782 | 59.6896552 | WT | FALSE | CAN | JSY13 | 3A | WT-Rep3A | CGC | G>A | A/T+1 | G/C-1 | CG>TA | CG>TA | TTGGCCGCA | GCACCAA | GCG | GCG | C | T | C>T | C>T_GCG | CG>TA_GCACCAA | 9 | TRUE | FALSE |
| 187_v.txt | 32977 | 32977-SNV- | SNV | 1 | C | T | 31 | 40458 | 0.07662267 | 56.8709677 | WT | FALSE | CAN | JSY13 | 3A | WT-Rep3A | GCT | C>T | A/T+1 | G/C-1 | CG>TA | CG>TA | CAAATGCTG | CTGGAGA | CGA | GCT | C | A | C>T | C>T_GCT | CG>TA_CTGGAGA | 9 | TRUE | FALSE |

| | | | | | | | | | | | | | | | | | | | | | | | | | | | | | | | | | | |
|---|---|---|---|---|---|---|---|---|---|---|---|---|---|---|---|---|---|---|---|---|---|---|---|---|---|---|---|---|---|---|---|---|---|---|
| 187_v.txt | 33037 | 33037-SNV- | SNV | 1 | C | T | 18 | 40458 | 0.04449058 | 61.3333333 | WT | FALSE | CAN | JSY13 | 3A | WT-Rep3A | GCC | C>T | A/T+1 | G/C-1 | CG>TA | CG>TA | ATGTAGCCA | CCATTTC | CGG | GCC | C | A | C>T | C>T_GCC | CG>TA_CCATTTC | 9 | TRUE | FALSE |
| 187_v.txt | 33043 | 33043-SNV- | SNV | 1 | C | T | 57 | 40458 | 0.14088685 | 57.9824561 | WT | FALSE | CAN | JSY13 | 3A | WT-Rep3A | TCA | C>T | A/T+1 | G/C-1 | CG>TA | CG>TA | CCATTTCACC | CACCCAA | AGT | TCA | C | A | C>T | C>T_TCA | CG>TA_CACCCAA | 9 | TRUE | FALSE |
| 187_v.txt | 33056 | 33056-SNV- | SNV | 1 | C | T | 54 | 40458 | 0.13347175 | 57.3703704 | WT | FALSE | CAN | JSY13 | 3A | WT-Rep3A | GCG | C>T | A/T+1 | G/C-1 | CG>TA | CG>TA | GGACTGCGT | CGTGACA | CGC | GCG | C | A | C>T | C>T_GCG | CG>TA_CGTGACA | 9 | TRUE | FALSE |
| 187_v.txt | 33069 | 33069-SNV- | SNV | 1 | G | A | 30 | 40458 | 0.07415097 | 49.5339614 | WT | FALSE | CAN | JSY13 | 3A | WT-Rep3A | TGC | G>A | A/T+1 | G/C-1 | CG>TA | CG>TA | GAATATGCC | GCCAAAG | ACG | ACG | C | T | C>T | C>T_ACG | CG>TA_GCCAAAG | 9 | TRUE | FALSE |
| 187_v.txt | 33070 | 33070-SNV- | SNV | 1 | C | T | 39 | 40456 | 0.09640103 | 50.5903243 | WT | FALSE | CAN | JSY13 | 3A | WT-Rep3A | GCC | C>T | A/T+1 | G/C-1 | CG>TA | CG>TA | AATATGCCA | CCAAAGA | CGG | GCC | C | A | C>T | C>T_GCC | CG>TA_CCAAAGA | 9 | TRUE | FALSE |
| 187_v.txt | 33078 | 33078-SNV- | SNV | 1 | C | T | 82 | 40458 | 0.20267932 | 49.6219512 | WT | FALSE | CAN | JSY13 | 3A | WT-Rep3A | ACC | C>T | A/T+1 | G/C-1 | CG>TA | CG>TA | AAAGAACCC | CCCATAA | TGG | ACC | C | A | C>T | C>T_ACC | CG>TA_CCCATAA | 9 | TRUE | FALSE |
| 187_v.txt | 33114 | 33114-SNV- | SNV | 1 | C | T | 34 | 40458 | 0.08403777 | 46.6764706 | WT | FALSE | CAN | JSY13 | 3A | WT-Rep3A | GCC | C>T | A/T+1 | G/C-1 | CG>TA | CG>TA | ACTGGGCCG | CCGGCGT | CGG | GCC | C | A | C>T | C>T_GCC | CG>TA_CCGGCGT | 9 | TRUE | FALSE |
| 187_v.txt | 33118 | 33118-SNV- | SNV | 1 | C | T | 33 | 40458 | 0.08156607 | 43.3939394 | WT | FALSE | CAN | JSY13 | 3A | WT-Rep3A | GCG | C>T | A/T+1 | G/C-1 | CG>TA | CG>TA | GGCCGGCGT | CGTTGGT | CGC | GCG | C | A | C>T | C>T_GCG | CG>TA_CGTTGGT | 9 | TRUE | FALSE |
| 187_v.txt | 33168 | 33168-SNV- | SNV | 1 | C | T | 18 | 40449 | 0.04450048 | 36.3262477 | WT | FALSE | CAN | JSY13 | 3A | WT-Rep3A | ACC | C>T | A/T+1 | G/C-1 | CG>TA | CG>TA | ATAGTACCA | CCACCAA | TGG | ACC | C | A | C>T | C>T_ACC | CG>TA_CCACCAA | 9 | TRUE | FALSE |
| 187_v.txt | 33258 | 33258-SNV- | SNV | 1 | C | T | 40 | 23996 | 0.16669445 | 42.9 | WT | FALSE | CAN | JSY13 | 3A | WT-Rep3A | GCC | C>T | A/T+1 | G/C-1 | CG>TA | CG>TA | CCTATGCCAT | CCATCCT | CGG | GCC | C | A | C>T | C>T_GCC | CG>TA_CCATCCT | 9 | TRUE | FALSE |
| 187_v.txt | 33276 | 33276-SNV- | SNV | 1 | G | A | 23 | 23996 | 0.09584931 | 45.3478261 | WT | FALSE | CAN | JSY13 | 3A | WT-Rep3A | CGT | G>A | A/T+1 | G/C-1 | CG>TA | CG>TA | GAGAACGTA | GTATCCT | GCA | GCA | C | T | C>T | C>T_GCA | CG>TA_GTATCCT | 9 | TRUE | FALSE |
| 187_v.txt | 33284 | 33284-SNV- | SNV | 1 | G | A | 24 | 23996 | 0.10001667 | 43.75 | WT | FALSE | CAN | JSY13 | 3A | WT-Rep3A | CGC | G>A | A/T+1 | G/C-1 | CG>TA | CG>TA | ATCCTCGCCA | GCCATTT | GCG | GCG | C | T | C>T | C>T_GCG | CG>TA_GCCATTT | 9 | TRUE | FALSE |
| 187_v.txt | 33310 | 33310-SNV- | SNV | 1 | G | A | 22 | 23996 | 0.09168195 | 61 | WT | FALSE | CAN | JSY13 | 3A | WT-Rep3A | CGC | G>A | A/T+1 | G/C-1 | CG>TA | CG>TA | AAGAGCGCA | GCAATGG | GCG | GCG | C | T | C>T | C>T_GCG | CG>TA_GCAATGG | 9 | TRUE | FALSE |
| 187_v.txt | 33392 | 33392-SNV- | SNV | 1 | G | A | 30 | 23996 | 0.12502084 | 48 | WT | FALSE | CAN | JSY13 | 3A | WT-Rep3A | CGT | G>A | A/T+1 | G/C-1 | CG>TA | CG>TA | AACGTCGTG | GTGAAAG | GCA | GCA | C | T | C>T | C>T_GCA | CG>TA_GTGAAAG | 9 | TRUE | FALSE |
| 187_v.txt | 33412 | 33412-SNV- | SNV | 1 | G | A | 40 | 23996 | 0.16669445 | 46.275 | WT | FALSE | CAN | JSY13 | 3A | WT-Rep3A | GGC | G>A | A/T+1 | G/C-1 | CG>TA | CG>TA | TGACCGGCT | GCTCATT | CCG | CCG | C | T | C>T | C>T_CCG | CG>TA_GCTCATT | 9 | TRUE | FALSE |
| 187_v.txt | 33474 | 33474-SNV- | SNV | 1 | C | T | 26 | 23996 | 0.10835139 | 33.1538462 | WT | FALSE | CAN | JSY13 | 3A | WT-Rep3A | GCC | C>T | A/T+1 | G/C-1 | CG>TA | CG>TA | GCTATGCCTT | CCTTTTT | CGG | GCC | C | A | C>T | C>T_GCC | CG>TA_CCTTTTT | 9 | TRUE | FALSE |
| 187_v.txt | 32064 | 32064-SNV- | SNV | 1 | G | C | 17 | 11361 | 0.14963472 | 63.8823529 | WT | FALSE | CAN | JSY13 | 3A | WT-Rep3A | TGA | G>C | G/C+1 | G/C-1 | CG>GC | CG>GC | GAGATTGAG | GAGATAA | ACT | ACT | C | G | C>G | C>G_ACT | CG>GC_GAGATAA | 9 | TRUE | FALSE |
| 187_v.txt | 32200 | 32200-SNV- | SNV | 1 | G | C | 44 | 11361 | 0.38728985 | 42.7954546 | WT | FALSE | CAN | JSY13 | 3A | WT-Rep3A | GGA | G>C | G/C+1 | G/C-1 | CG>GC | CG>GC | TGTATGGAA | GAACACC | CCT | CCT | C | G | C>G | C>G_CCT | CG>GC_GAACACC | 9 | TRUE | FALSE |
| 187_v.txt | 32282 | 32282-SNV- | SNV | 1 | G | C | 8 | 43076 | 0.01857183 | 34.625 | WT | FALSE | CAN | JSY13 | 3A | WT-Rep3A | CGT | G>C | G/C+1 | G/C-1 | CG>GC | CG>GC | ACCAACGTA | GTAAATA | GCA | GCA | C | G | C>G | C>G_GCA | CG>GC_GTAAATA | 9 | TRUE | FALSE |
| 187_v.txt | 32292 | 32292-SNV- | SNV | 1 | G | C | 37 | 43076 | 0.0858947 | 48.5675676 | WT | FALSE | CAN | JSY13 | 3A | WT-Rep3A | TGA | G>C | G/C+1 | G/C-1 | CG>GC | CG>GC | ATATTTGAAT | GAATTTG | ACT | ACT | C | G | C>G | C>G_ACT | CG>GC_GAATTTG | 9 | TRUE | FALSE |
| 187_v.txt | 32302 | 32302-SNV- | SNV | 1 | C | G | 10 | 43076 | 0.02321478 | 48.2 | WT | FALSE | CAN | JSY13 | 3A | WT-Rep3A | GCA | C>G | G/C+1 | G/C-1 | CG>GC | CG>GC | TTGCGGCAG | CAGAAAT | CGT | GCA | C | C | C>G | C>G_GCA | CG>GC_CAGAAAT | 9 | TRUE | FALSE |
| 187_v.txt | 32432 | 32432-SNV- | SNV | 1 | G | C | 24 | 43076 | 0.05571548 | 56.2083333 | WT | FALSE | CAN | JSY13 | 3A | WT-Rep3A | TGT | G>C | G/C+1 | G/C-1 | CG>GC | CG>GC | GTCATTGTAT | GTATGGA | ACA | ACA | C | G | C>G | C>G_ACA | CG>GC_GTATGGA | 9 | TRUE | FALSE |
| 187_v.txt | 32467 | 32467-SNV- | SNV | 1 | C | G | 24 | 43076 | 0.05571548 | 50.3667024 | WT | FALSE | CAN | JSY13 | 3A | WT-Rep3A | CCA | C>G | G/C+1 | G/C-1 | CG>GC | CG>GC | GAGAGCCAA | CAATGTA | GGT | CCA | C | C | C>G | C>G_CCA | CG>GC_CAATGTA | 9 | TRUE | FALSE |
| 187_v.txt | 32474 | 32474-SNV- | SNV | 1 | G | C | 17 | 43076 | 0.03946513 | 48.2352941 | WT | FALSE | CAN | JSY13 | 3A | WT-Rep3A | AGA | G>C | G/C+1 | G/C-1 | CG>GC | CG>GC | AATGTAGAA | GAAGGTT | TCT | TCT | C | G | C>G | C>G_TCT | CG>GC_GAAGGTT | 9 | TRUE | FALSE |
| 187_v.txt | 32776 | 32776-SNV- | SNV | 1 | C | G | 14 | 23486 | 0.05960998 | 51.2142857 | WT | FALSE | CAN | JSY13 | 3A | WT-Rep3A | GCG | C>G | G/C+1 | G/C-1 | CG>GC | CG>GC | TGGAAGCGA | CGACCCA | CGC | GCG | C | C | C>G | C>G_GCG | CG>GC_CGACCCA | 9 | TRUE | FALSE |
| 187_v.txt | 32797 | 32797-SNV- | SNV | 1 | C | G | 17 | 23486 | 0.07238355 | 45.5838325 | WT | FALSE | CAN | JSY13 | 3A | WT-Rep3A | CCG | C>G | G/C+1 | G/C-1 | CG>GC | CG>GC | ATTCACCGTA | CGTAATA | GGC | CCG | C | C | C>G | C>G_CCG | CG>GC_CGTAATA | 9 | TRUE | FALSE |
| 187_v.txt | 32809 | 32809-SNV- | SNV | 1 | C | G | 6 | 23486 | 0.02554714 | 55.3333333 | WT | FALSE | CAN | JSY13 | 3A | WT-Rep3A | ACA | C>G | G/C+1 | G/C-1 | CG>GC | CG>GC | ATTTGACAG | CAGGGAA | TGT | ACA | C | C | C>G | C>G_ACA | CG>GC_CAGGGAA | 9 | TRUE | FALSE |
| 187_v.txt | 32855 | 32855-SNV- | SNV | 1 | C | G | 3 | 23483 | 0.0127752 | 38 | WT | FALSE | CAN | JSY13 | 3A | WT-Rep3A | TCC | C>G | G/C+1 | G/C-1 | CG>GC | CG>GC | ACTAATCCAT | CCATGCC | AGG | TCC | C | C | C>G | C>G_TCC | CG>GC_CCATGCC | 9 | TRUE | FALSE |
| 187_v.txt | 32883 | 32883-SNV- | SNV | 1 | C | G | 2 | 40419 | 4.95E-03 | 38 | WT | FALSE | CAN | JSY13 | 3A | WT-Rep3A | CCA | C>G | G/C+1 | G/C-1 | CG>GC | CG>GC | TACGTCCAA | CAAAATT | GGT | CCA | C | C | C>G | C>G_CCA | CG>GC_CAAAATT | 9 | TRUE | FALSE |
| 187_v.txt | 32917 | 32917-SNV- | SNV | 1 | C | G | 4 | 40418 | 9.90E-03 | 57.4922468 | WT | FALSE | CAN | JSY13 | 3A | WT-Rep3A | TCC | C>G | G/C+1 | G/C-1 | CG>GC | CG>GC | TAAGTTCCA | CCAGGGC | AGG | TCC | C | C | C>G | C>G_TCC | CG>GC_CCAGGGC | 9 | TRUE | FALSE |
| 187_v.txt | 33037 | 33037-SNV- | SNV | 1 | C | G | 10 | 40458 | 0.02471699 | 59.5 | WT | FALSE | CAN | JSY13 | 3A | WT-Rep3A | GCC | C>G | G/C+1 | G/C-1 | CG>GC | CG>GC | ATGTAGCCA | CCATTTC | CGG | GCC | C | C | C>G | C>G_GCC | CG>GC_CCATTTC | 9 | TRUE | FALSE |
| 187_v.txt | 33070 | 33070-SNV- | SNV | 1 | C | G | 129 | 40456 | 0.31886494 | 54.4120436 | WT | FALSE | CAN | JSY13 | 3A | WT-Rep3A | GCC | C>G | G/C+1 | G/C-1 | CG>GC | CG>GC | AATATGCCA | CCAAAGA | CGG | GCC | C | C | C>G | C>G_GCC | CG>GC_CCAAAGA | 9 | TRUE | FALSE |
| 187_v.txt | 33118 | 33118-SNV- | SNV | 1 | C | G | 6 | 40458 | 0.01483019 | 37.5 | WT | FALSE | CAN | JSY13 | 3A | WT-Rep3A | GCG | C>G | G/C+1 | G/C-1 | CG>GC | CG>GC | GGCCGGCGT | CGTTGGT | CGC | GCG | C | C | C>G | C>G_GCG | CG>GC_CGTTGGT | 9 | TRUE | FALSE |
| 187_v.txt | 33172 | 33172-SNV- | SNV | 1 | C | G | 10 | 40448 | 0.0247231 | 38 | WT | FALSE | CAN | JSY13 | 3A | WT-Rep3A | CCA | C>G | G/C+1 | G/C-1 | CG>GC | CG>GC | TACCACCAA | CAAGGGC | GGT | CCA | C | C | C>G | C>G_CCA | CG>GC_CAAGGGC | 9 | TRUE | FALSE |
| 187_v.txt | 33193 | 33193-SNV- | SNV | 1 | G | C | 6 | 23996 | 0.02500417 | 37.9980244 | WT | FALSE | CAN | JSY13 | 3A | WT-Rep3A | TGT | G>C | G/C+1 | G/C-1 | CG>GC | CG>GC | CAATATGTCT | GTCTTTG | ACA | ACA | C | G | C>G | C>G_ACA | CG>GC_GTCTTTG | 9 | TRUE | FALSE |
| 187_v.txt | 33354 | 33354-SNV- | SNV | 1 | G | C | 24 | 23968 | 0.10013351 | 52.6788909 | WT | FALSE | CAN | JSY13 | 3A | WT-Rep3A | TGA | G>C | G/C+1 | G/C-1 | CG>GC | CG>GC | GGTATTGAC | GACCCAC | ACT | ACT | C | G | C>G | C>G_ACT | CG>GC_GACCCAC | 9 | TRUE | FALSE |
| 187_v.txt | 33378 | 33378-SNV- | SNV | 1 | G | C | 27 | 23996 | 0.11251875 | 58 | WT | FALSE | CAN | JSY13 | 3A | WT-Rep3A | TGA | G>C | G/C+1 | G/C-1 | CG>GC | CG>GC | GTTTGTGAA | GAAGCTT | ACT | ACT | C | G | C>G | C>G_ACT | CG>GC_GAAGCTT | 9 | TRUE | FALSE |
| 187_v.txt | 33456 | 33456-SNV- | SNV | 1 | G | C | 10 | 23996 | 0.04167361 | 42.9 | WT | FALSE | CAN | JSY13 | 3A | WT-Rep3A | TGA | G>C | G/C+1 | G/C-1 | CG>GC | CG>GC | TCTTTTGAAT | GAATTTG | ACT | ACT | C | G | C>G | C>G_ACT | CG>GC_GAATTTG | 9 | TRUE | FALSE |
| 187_v.txt | 32080 | 32080-SNV- | SNV | 1 | C | A | 3 | 11361 | 0.02640613 | 46 | WT | FALSE | CAN | JSY13 | 3A | WT-Rep3A | GCA | C>A | A/T+1 | G/C-1 | CG>AT | CG>AT | ACCATGCAA | CAAAAAA | CGT | GCA | C | T | C>A | C>A_GCA | CG>AT_CAAAAAA | 9 | TRUE | FALSE |
| 187_v.txt | 32272 | 32272-SNV- | SNV | 1 | G | T | 18 | 43076 | 0.04178661 | 29.7234295 | WT | FALSE | CAN | JSY13 | 3A | WT-Rep3A | CGG | G>T | A/T+1 | G/C-1 | CG>AT | CG>AT | AAATACGGG | GGGAACC | GCC | GCC | C | A | C>A | C>A_GCC | CG>AT_GGGAACC | 9 | TRUE | FALSE |
| 187_v.txt | 32274 | 32274-SNV- | SNV | 1 | G | T | 36 | 43076 | 0.08357322 | 36.1666667 | WT | FALSE | CAN | JSY13 | 3A | WT-Rep3A | GGA | G>T | A/T+1 | G/C-1 | CG>AT | CG>AT | ATACGGGAA | GAACCAA | CCT | CCT | C | A | C>A | C>A_CCT | CG>AT_GAACCAA | 9 | TRUE | FALSE |
| 187_v.txt | 32282 | 32282-SNV- | SNV | 1 | G | T | 21 | 43076 | 0.04875105 | 36.2380952 | WT | FALSE | CAN | JSY13 | 3A | WT-Rep3A | CGT | G>T | A/T+1 | G/C-1 | CG>AT | CG>AT | ACCAACGTA | GTAAATA | GCA | GCA | C | A | C>A | C>A_GCA | CG>AT_GTAAATA | 9 | TRUE | FALSE |
| 187_v.txt | 32368 | 32368-SNV- | SNV | 1 | C | A | 29 | 43076 | 0.06732287 | 54.4137931 | WT | FALSE | CAN | JSY13 | 3A | WT-Rep3A | TCA | C>A | A/T+1 | G/C-1 | CG>AT | CG>AT | AGTTCTCAAT | CAATAGC | AGT | TCA | C | T | C>A | C>A_TCA | CG>AT_CAATAGC | 9 | TRUE | FALSE |
| 187_v.txt | 32373 | 32373-SNV- | SNV | 1 | G | T | 21 | 43076 | 0.04875105 | 57.4761905 | WT | FALSE | CAN | JSY13 | 3A | WT-Rep3A | AGC | G>T | A/T+1 | G/C-1 | CG>AT | CG>AT | TCAATAGCA | GCAATAA | TCG | TCG | C | A | C>A | C>A_TCG | CG>AT_GCAATAA | 9 | TRUE | FALSE |
| 187_v.txt | 32399 | 32399-SNV- | SNV | 1 | G | T | 26 | 43076 | 0.06035844 | 55.2692308 | WT | FALSE | CAN | JSY13 | 3A | WT-Rep3A | CGT | G>T | A/T+1 | G/C-1 | CG>AT | CG>AT | AGAAACGTA | GTAGGAA | GCA | GCA | C | A | C>A | C>A_GCA | CG>AT_GTAGGAA | 9 | TRUE | FALSE |
| 187_v.txt | 32432 | 32432-SNV- | SNV | 1 | G | T | 99 | 43076 | 0.22982635 | 55.4141414 | WT | FALSE | CAN | JSY13 | 3A | WT-Rep3A | TGT | G>T | A/T+1 | G/C-1 | CG>AT | CG>AT | GTCATTGTAT | GTATGGA | ACA | ACA | C | A | C>A | C>A_ACA | CG>AT_GTATGGA | 9 | TRUE | FALSE |
| 187_v.txt | 32530 | 32530-SNV- | SNV | 1 | G | T | 53 | 43076 | 0.12303835 | 37.5283019 | WT | FALSE | CAN | JSY13 | 3A | WT-Rep3A | GGG | G>T | A/T+1 | G/C-1 | CG>AT | CG>AT | TTCTGGGGT | GGTTTGC | CCC | CCC | C | A | C>A | C>A_CCC | CG>AT_GGTTTGC | 9 | TRUE | FALSE |
| 187_v.txt | 32536 | 32536-SNV- | SNV | 1 | C | A | 5 | 43074 | 0.01160793 | 32.3988362 | WT | FALSE | CAN | JSY13 | 3A | WT-Rep3A | GCA | C>A | A/T+1 | G/C-1 | CG>AT | CG>AT | GGTTTGCAG | CAGCTTC | CGT | GCA | C | T | C>A | C>A_GCA | CG>AT_CAGCTTC | 9 | TRUE | FALSE |
| 187_v.txt | 32629 | 32629-SNV- | SNV | 1 | C | A | 290 | 23486 | 1.23477817 | 54.4206897 | WT | FALSE | CAN | JSY13 | 3A | WT-Rep3A | TCG | C>A | A/T+1 | G/C-1 | CG>AT | CG>AT | TCCCTTCGTT | CGTTTTT | AGC | TCG | C | T | C>A | C>A_TCG | CG>AT_CGTTTTT | 9 | TRUE | FALSE |
| 187_v.txt | 32756 | 32756-SNV- | SNV | 1 | G | T | 3 | 23486 | 0.01277357 | 58.3239785 | WT | FALSE | CAN | JSY13 | 3A | WT-Rep3A | TGG | G>T | A/T+1 | G/C-1 | CG>AT | CG>AT | GATAATGGC | GGCTAAA | ACC | ACC | C | A | C>A | C>A_ACC | CG>AT_GGCTAAA | 9 | TRUE | FALSE |
| 187_v.txt | 32809 | 32809-SNV- | SNV | 1 | C | A | 34 | 23486 | 0.1447671 | 43.5294118 | WT | FALSE | CAN | JSY13 | 3A | WT-Rep3A | ACA | C>A | A/T+1 | G/C-1 | CG>AT | CG>AT | ATTTGACAG | CAGGGAA | TGT | ACA | C | T | C>A | C>A_ACA | CG>AT_CAGGGAA | 9 | TRUE | FALSE |
| 187_v.txt | 32883 | 32883-SNV- | SNV | 1 | C | A | 6 | 40419 | 0.0148445 | 29 | WT | FALSE | CAN | JSY13 | 3A | WT-Rep3A | CCA | C>A | A/T+1 | G/C-1 | CG>AT | CG>AT | TACGTCCAA | CAAAATT | GGT | CCA | C | T | C>A | C>A_CCA | CG>AT_CAAAATT | 9 | TRUE | FALSE |
| 187_v.txt | 32917 | 32917-SNV- | SNV | 1 | C | A | 22 | 40418 | 0.05443119 | 49.3612737 | WT | FALSE | CAN | JSY13 | 3A | WT-Rep3A | TCC | C>A | A/T+1 | G/C-1 | CG>AT | CG>AT | TAAGTTCCA | CCAGGGC | AGG | TCC | C | T | C>A | C>A_TCC | CG>AT_CCAGGGC | 9 | TRUE | FALSE |
| 187_v.txt | 32935 | 32935-SNV- | SNV | 1 | C | A | 11 | 40418 | 0.0272156 | 47.1818182 | WT | FALSE | CAN | JSY13 | 3A | WT-Rep3A | GCC | C>A | A/T+1 | G/C-1 | CG>AT | CG>AT | TGATTGCCCA | CCCAAGA | CGG | GCC | C | T | C>A | C>A_GCC | CG>AT_CCCAAGA | 9 | TRUE | FALSE |
| 187_v.txt | 33033 | 33033-SNV- | SNV | 1 | G | T | 3260 | 40458 | 8.05773889 | 60.5230061 | WT | FALSE | CAN | JSY13 | 3A | WT-Rep3A | TGT | G>T | A/T+1 | G/C-1 | CG>AT | CG>AT | ATGAATGTA | GTAGCCA | ACA | ACA | C | A | C>A | C>A_ACA | CG>AT_GTAGCCA | 9 | TRUE | FALSE |
| 187_v.txt | 33051 | 33051-SNV- | SNV | 1 | G | T | 25 | 40458 | 0.06179248 | 50.8 | WT | FALSE | CAN | JSY13 | 3A | WT-Rep3A | GGA | G>T | A/T+1 | G/C-1 | CG>AT | CG>AT | CCCAAGGAC | GACTGCG | CCT | CCT | C | A | C>A | C>A_CCT | CG>AT_GACTGCG | 9 | TRUE | FALSE |
| 187_v.txt | 33159 | 33159-SNV- | SNV | 1 | C | A | 24 | 40458 | 0.05932078 | 37.1666667 | WT | FALSE | CAN | JSY13 | 3A | WT-Rep3A | ACC | C>A | A/T+1 | G/C-1 | CG>AT | CG>AT | CCTGTACCAA | CCAATAG | TGG | ACC | C | T | C>A | C>A_ACC | CG>AT_CCAATAG | 9 | TRUE | FALSE |
| 187_v.txt | 33193 | 33193-SNV- | SNV | 1 | G | T | 6 | 23996 | 0.02500417 | 26.999814 | WT | FALSE | CAN | JSY13 | 3A | WT-Rep3A | TGT | G>T | A/T+1 | G/C-1 | CG>AT | CG>AT | CAATATGTCT | GTCTTTG | ACA | ACA | C | A | C>A | C>A_ACA | CG>AT_GTCTTTG | 9 | TRUE | FALSE |
| 187_v.txt | 33378 | 33378-SNV- | SNV | 1 | G | T | 6 | 23996 | 0.02500417 | 54.5 | WT | FALSE | CAN | JSY13 | 3A | WT-Rep3A | TGA | G>T | A/T+1 | G/C-1 | CG>AT | CG>AT | GTTTGTGAA | GAAGCTT | ACT | ACT | C | A | C>A | C>A_ACT | CG>AT_GAAGCTT | 9 | TRUE | FALSE |
| 187_v.txt | 33474 | 33474-SNV- | SNV | 1 | C | A | 25 | 23996 | 0.10418403 | 34.04 | WT | FALSE | CAN | JSY13 | 3A | WT-Rep3A | GCC | C>A | A/T+1 | G/C-1 | CG>AT | CG>AT | GCTATGCCTT | CCTTTTT | CGG | GCC | C | T | C>A | C>A_GCC | CG>AT_CCTTTTT | 9 | TRUE | FALSE |
| 187_v.txt | 32535 | 32535-MNV- | MNV | 2 | GC | AT | 1 | 43074 | 2.32E-03 | 37.9999896 | WT | FALSE | CAN | JSY13 | 3A | WT-Rep3A | TGC | nope | >1bp-ins | >1bpdel | none | MNV | GGGTTTGCA | GCAGCTT | ACG | TGC | GC | none | NA | NA_TGC | MNV_GCAGCTT | 9 | TRUE | FALSE |
| 187_v.txt | 32765 | 32765-MNV- | MNV | 2 | TT | AG | 4 | 23486 | 0.01703142 | 50.749917 | WT | FALSE | CAN | JSY13 | 3A | WT-Rep3A | CTT | nope | >1bp-ins | >1bpdel | none | MNV | TAAAACTTTG | TTTGATG | GAA | CTT | TT | none | NA | NA_CTT | MNV_TTTGATG | 9 | TRUE | FALSE |
| 187_v.txt | 33167 | 33167-MNV- | MNV | 2 | AC | CA | 1 | 40451 | 2.47E-03 | 26.5005446 | WT | FALSE | CAN | JSY13 | 3A | WT-Rep3A | TAC | nope | >1bp-ins | >1bpdel | none | MNV | AATAGTACC | ACCACCA | ATG | TAC | AC | none | NA | NA_TAC | MNV_ACCACCA | 9 | TRUE | FALSE |
| 187_v.txt | 33168 | 33168-MNV- | MNV | 2 | CC | AA | 1 | 40449 | 2.47E-03 | 27.9998389 | WT | FALSE | CAN | JSY13 | 3A | WT-Rep3A | ACC | nope | >1bp-ins | >1bpdel | none | MNV | ATAGTACCA | CCACCAA | TGG | ACC | CC | none | NA | NA_ACC | MNV_CCACCAA | 9 | TRUE | FALSE |
| 187_v.txt | 32080 | 32080-Insert | Insertion | 1 | - | C | 1 | 11361 | 8.80E-03 | 33 | WT | FALSE | CAN | JSY13 | 3A | WT-Rep3A | GCA | nope | G/C+1 | >1bpdel | none | G/C+1 | ACCATGCAA | CAAAAAA | CGT | GCA | - | G | NA | NA_GCA | G/C+1_CAAAAAA | 9 | TRUE | FALSE |
| 187_v.txt | 32081 | 32081-Insert | Insertion | 1 | - | A | 15 | 11361 | 0.13203063 | 64 | WT | FALSE | CAN | JSY13 | 3A | WT-Rep3A | CAA | nope | A/T+1 | >1bpdel | none | A/T+1 | CCATGCAAA | AAAAAAG | GTT | CAA | - | T | NA | NA_CAA | A/T+1_AAAAAAG | 9 | TRUE | FALSE |
| 187_v.txt | 32265 | 32265-Insert | Insertion | 1 | - | A | 1 | 43076 | 2.32E-03 | 38 | WT | FALSE | CAN | JSY13 | 3A | WT-Rep3A | TAA | nope | A/T+1 | >1bpdel | none | A/T+1 | CCAAATAAA | AAAATAC | ATT | TAA | - | T | NA | NA_TAA | A/T+1_AAAATAC | 9 | TRUE | FALSE |
| 187_v.txt | 32272 | 32272-Insert | Insertion | 1 | - | T | 1 | 43076 | 2.32E-03 | 37 | WT | FALSE | CAN | JSY13 | 3A | WT-Rep3A | CGG | nope | A/T+1 | >1bpdel | none | A/T+1 | AAATACGGG | GGGAACC | GCC | CGG | - | A | NA | NA_CGG | A/T+1_GGGAACC | 9 | TRUE | FALSE |
| 187_v.txt | 32305 | 32305-Insert | Insertion | 1 | - | A | 2 | 43076 | 4.64E-03 | 64 | WT | FALSE | CAN | JSY13 | 3A | WT-Rep3A | GAA | nope | A/T+1 | >1bpdel | none | A/T+1 | CGGCAGAAA | AAATAAT | CTT | GAA | - | T | NA | NA_GAA | A/T+1_AAATAAT | 9 | TRUE | FALSE |
| 187_v.txt | 32346 | 32346-Insert | Insertion | 1 | - | A | 4 | 43076 | 9.29E-03 | 64 | WT | FALSE | CAN | JSY13 | 3A | WT-Rep3A | CAA | nope | A/T+1 | >1bpdel | none | A/T+1 | TGTGGCAAA | AAAACCT | GTT | CAA | - | T | NA | NA_CAA | A/T+1_AAAACCT | 9 | TRUE | FALSE |
| 187_v.txt | 32498 | 32498-Insert | Insertion | 1 | - | T | 8 | 43076 | 0.01857183 | 47.25 | WT | FALSE | CAN | JSY13 | 3A | WT-Rep3A | CTT | nope | A/T+1 | >1bpdel | none | A/T+1 | AACAACTTTT | TTTTTTG | GAA | CTT | - | A | NA | NA_CTT | A/T+1_TTTTTTG | 9 | TRUE | FALSE |
| 187_v.txt | 32658 | 32658-Insert | Insertion | 1 | - | C | 8 | 23486 | 0.03406285 | 57.375 | WT | FALSE | CAN | JSY13 | 3A | WT-Rep3A | ACC | nope | G/C+1 | >1bpdel | none | G/C+1 | CCTGGACCC | CCCCAGG | TGG | ACC | - | G | NA | NA_ACC | G/C+1_CCCCAGG | 9 | TRUE | FALSE |
| 187_v.txt | 32728 | 32728-Insert | Insertion | 1 | - | A | 5 | 23486 | 0.02128928 | 64 | WT | FALSE | CAN | JSY13 | 3A | WT-Rep3A | CAA | nope | A/T+1 | >1bpdel | none | A/T+1 | CCATACAAA | AAAAACA | GTT | CAA | - | T | NA | NA_CAA | A/T+1_AAAAACA | 9 | TRUE | FALSE |
| 187_v.txt | 32760 | 32760-Insert | Insertion | 1 | - | A | 2 | 23486 | 8.52E-03 | 48.9999992 | WT | FALSE | CAN | JSY13 | 3A | WT-Rep3A | TAA | nope | A/T+1 | >1bpdel | none | A/T+1 | ATGGCTAAA | AAAACTT | ATT | TAA | - | T | NA | NA_TAA | A/T+1_AAAACTT | 9 | TRUE | FALSE |
| 187_v.txt | 32842 | 32842-Insert | Insertion | 1 | - | A | 44 | 23486 | 0.18734565 | 43.1136364 | WT | FALSE | CAN | JSY13 | 3A | WT-Rep3A | CAA | nope | A/T+1 | >1bpdel | none | A/T+1 | TTACCCAAA | AAAAAAT | GTT | CAA | - | T | NA | NA_CAA | A/T+1_AAAAAAT | 9 | TRUE | FALSE |
| 187_v.txt | 32884 | 32884-Insert | Insertion | 1 | - | A | 11 | 40419 | 0.02721492 | 37.6363636 | WT | FALSE | CAN | JSY13 | 3A | WT-Rep3A | CAA | nope | A/T+1 | >1bpdel | none | A/T+1 | ACGTCCAAA | AAAATTG | GTT | CAA | - | T | NA | NA_CAA | A/T+1_AAAATTG | 9 | TRUE | FALSE |
| 187_v.txt | 32970 | 32970-Insert | Insertion | 1 | - | C | 7 | 40458 | 0.01730189 | 64 | WT | FALSE | CAN | JSY13 | 3A | WT-Rep3A | ACC | nope | G/C+1 | >1bpdel | none | G/C+1 | GCCGCACCA | CCAAATG | TGG | ACC | - | G | NA | NA_ACC | G/C+1_CCAAATG | 9 | TRUE | FALSE |
| 187_v.txt | 33009 | 33009-Insert | Insertion | 1 | - | A | 7 | 40458 | 0.01730189 | 64 | WT | FALSE | CAN | JSY13 | 3A | WT-Rep3A | GAA | nope | A/T+1 | >1bpdel | none | A/T+1 | ACTGTGAAA | AAAGAGG | CTT | GAA | - | T | NA | NA_GAA | A/T+1_AAAGAGG | 9 | TRUE | FALSE |
| 187_v.txt | 33019 | 33019-Insert | Insertion | 1 | - | T | 15 | 40458 | 0.03707549 | 61.5333333 | WT | FALSE | CAN | JSY13 | 3A | WT-Rep3A | GTA | nope | A/T+1 | >1bpdel | none | A/T+1 | AGGATGTAA | TAACAGG | CAT | GTA | - | A | NA | NA_GTA | A/T+1_TAACAGG | 9 | TRUE | FALSE |
| 187_v.txt | 33281 | 33281-Insert | Insertion | 13 | - | CTCGCCATTT | 3 | 23996 | 0.01250208 | 53.0512821 | WT | FALSE | CAN | JSY13 | 3A | WT-Rep3A | CCT | nope | >1bp-ins | >1bpdel | none | >1bp-ins | CGTATCCTCG | CTCGCCA | GGA | CCT | - | none | NA | NA_CCT | >1bp-ins_CTCGCCA | 9 | TRUE | FALSE |
| 187_v.txt | 33281 | 33281-Insert | Insertion | 13 | - | CTCGCCATTT | 1 | 23996 | 4.17E-03 | 60.8461539 | WT | FALSE | CAN | JSY13 | 3A | WT-Rep3A | CCT | nope | >1bp-ins | >1bpdel | none | >1bp-ins | CGTATCCTCG | CTCGCCA | GGA | CCT | - | none | NA | NA_CCT | >1bp-ins_CTCGCCA | 9 | TRUE | FALSE |
| 187_v.txt | 33355 | 33355-Insert | Insertion | 4 | - | ACCC | 3 | 23996 | 0.01250208 | 61.75 | WT | FALSE | CAN | JSY13 | 3A | WT-Rep3A | GAC | nope | >1bp-ins | >1bpdel | none | >1bp-ins | GTATTGACC | ACCCACG | CTG | GAC | - | none | NA | NA_GAC | >1bp-ins_ACCCACG | 9 | TRUE | FALSE |
| 187_v.txt | 33356 | 33356-Insert | Insertion | 1 | - | C | 17 | 23996 | 0.07084514 | 58.2941177 | WT | FALSE | CAN | JSY13 | 3A | WT-Rep3A | ACC | nope | G/C+1 | >1bpdel | none | G/C+1 | TATTGACCCA | CCCACGT | TGG | ACC | - | G | NA | NA_ACC | G/C+1_CCCACGT | 9 | TRUE | FALSE |
| 187_v.txt | 32014 | 32014-Deleti | Deletion | 2 | GA | - | 4 | 11361 | 0.03520817 | 64 | WT | FALSE | CAN | JSY13 | 3A | WT-Rep3A | CGA | nope | >1bp-ins | >1bpdel | none | >1bpdel | CGTCACGAG | GAGAGAT | GCT | CGA | GA | none | NA | NA_CGA | >1bpdel_GAGAGAT | 9 | TRUE | FALSE |
| 187_v.txt | 32265 | 32265-Deleti | Deletion | 2 | AA | - | 25 | 43076 | 0.05803696 | 37.72 | WT | FALSE | CAN | JSY13 | 3A | WT-Rep3A | TAA | nope | >1bp-ins | >1bpdel | none | >1bpdel | CCAAATAAA | AAAATAC | ATT | TAA | AA | none | NA | NA_TAA | >1bpdel_AAAATAC | 9 | TRUE | FALSE |
| 187_v.txt | 32337 | 32337-Deleti | Deletion | 2 | AT | - | 17 | 43076 | 0.03946513 | 61.6470588 | WT | FALSE | CAN | JSY13 | 3A | WT-Rep3A | GAT | nope | >1bp-ins | >1bpdel | none | >1bpdel | TTGAAGATA | ATATGTG | CTA | GAT | AT | none | NA | NA_GAT | >1bpdel_ATATGTG | 9 | TRUE | FALSE |
| 187_v.txt | 32492 | 32492-Deleti | Deletion | 3 | AAC | - | 72 | 43076 | 0.16714644 | 44.2222222 | WT | FALSE | CAN | JSY13 | 3A | WT-Rep3A | AAA | nope | >1bp-ins | >1bpdel | none | >1bpdel | ACGGAAAAC | AACAACT | TTT | AAA | AAC | none | NA | NA_AAA | >1bpdel_AACAACT | 9 | TRUE | FALSE |
| 187_v.txt | 32498 | 32498-Deleti | Deletion | 2 | TT | - | 159 | 43076 | 0.36911505 | 45.8490566 | WT | FALSE | CAN | JSY13 | 3A | WT-Rep3A | CTT | nope | >1bp-ins | >1bpdel | none | >1bpdel | AACAACTTTT | TTTTTTG | GAA | CTT | TT | none | NA | NA_CTT | >1bpdel_TTTTTTG | 9 | TRUE | FALSE |
| 187_v.txt | 32754 | 32754-Deleti | Deletion | 15 | ATGGCTAAA | - | 22 | 23486 | 0.09367283 | 55.4090909 | WT | FALSE | CAN | JSY13 | 3A | WT-Rep3A | AAT | nope | >1bp-ins | >1bpdel | none | >1bpdel | CCGATAATG | ATGGCTA | TTA | AAT | ATGGCTAAA | none | NA | NA_AAT | >1bpdel_ATGGCTA | 9 | TRUE | FALSE |
| 187_v.txt | 32760 | 32760-Deleti | Deletion | 2 | AA | - | 1 | 23486 | 4.26E-03 | 38 | WT | FALSE | CAN | JSY13 | 3A | WT-Rep3A | TAA | nope | >1bp-ins | >1bpdel | none | >1bpdel | ATGGCTAAA | AAAACTT | ATT | TAA | AA | none | NA | NA_TAA | >1bpdel_AAAACTT | 9 | TRUE | FALSE |
| 187_v.txt | 32804 | 32804-Deleti | Deletion | 2 | TT | - | 9 | 23486 | 0.0383207 | 42.2222222 | WT | FALSE | CAN | JSY13 | 3A | WT-Rep3A | ATT | nope | >1bp-ins | >1bpdel | none | >1bpdel | GTAATATTTG | TTTGACA | TAA | ATT | TT | none | NA | NA_ATT | >1bpdel_TTTGACA | 9 | TRUE | FALSE |
| 187_v.txt | 33167 | 33167-Deleti | Deletion | 3 | ACC | - | 7 | 40451 | 0.01730489 | 38 | WT | FALSE | CAN | JSY13 | 3A | WT-Rep3A | TAC | nope | >1bp-ins | >1bpdel | none | >1bpdel | AATAGTACC | ACCACCA | ATG | TAC | ACC | none | NA | NA_TAC | >1bpdel_ACCACCA | 9 | TRUE | FALSE |
| 187_v.txt | 33168 | 33168-Deleti | Deletion | 2 | CC | - | 2 | 40449 | 4.94E-03 | 37.5000001 | WT | FALSE | CAN | JSY13 | 3A | WT-Rep3A | ACC | nope | >1bp-ins | >1bpdel | none | >1bpdel | ATAGTACCA | CCACCAA | TGG | ACC | CC | none | NA | NA_ACC | >1bpdel_CCACCAA | 9 | TRUE | FALSE |
| 187_v.txt | 31723 | 31723-Deleti | Deletion | 1 | A | - | 4 | 1873 | 0.21356113 | 63 | WT | FALSE | CAN | JSY13 | 3A | WT-Rep3A | CAA | nope | >1bp-ins | A/T-1 | none | A/T-1 | TGTCCCAAA | AAAAAGT | GTT | GTT | T | none | T>none | T>none_GTT | A/T-1_AAAAAGT | 9 | TRUE | FALSE |
| 187_v.txt | 32080 | 32080-Deleti | Deletion | 1 | C | - | 5 | 11361 | 0.04401021 | 64 | WT | FALSE | CAN | JSY13 | 3A | WT-Rep3A | GCA | nope | >1bp-ins | G/C-1 | none | G/C-1 | ACCATGCAA | CAAAAAA | CGT | GCA | C | none | nope | nope_GCA | G/C-1_CAAAAAA | 9 | TRUE | FALSE |
| 187_v.txt | 32081 | 32081-Deleti | Deletion | 1 | A | - | 6 | 11361 | 0.05281225 | 63.3333334 | WT | FALSE | CAN | JSY13 | 3A | WT-Rep3A | CAA | nope | >1bp-ins | A/T-1 | none | A/T-1 | CCATGCAAA | AAAAAAG | GTT | GTT | T | none | T>none | T>none_GTT | A/T-1_AAAAAAG | 9 | TRUE | FALSE |
| 187_v.txt | 32265 | 32265-Deleti | Deletion | 1 | A | - | 5 | 43076 | 0.01160739 | 37.3999999 | WT | FALSE | CAN | JSY13 | 3A | WT-Rep3A | TAA | nope | >1bp-ins | A/T-1 | none | A/T-1 | CCAAATAAA | AAAATAC | ATT | ATT | T | none | T>none | T>none_ATT | A/T-1_AAAATAC | 9 | TRUE | FALSE |
| 187_v.txt | 32346 | 32346-Deleti | Deletion | 1 | A | - | 6 | 43076 | 0.01392887 | 60.4999823 | WT | FALSE | CAN | JSY13 | 3A | WT-Rep3A | CAA | nope | >1bp-ins | A/T-1 | none | A/T-1 | TGTGGCAAA | AAAACCT | GTT | GTT | T | none | T>none | T>none_GTT | A/T-1_AAAACCT | 9 | TRUE | FALSE |
| 187_v.txt | 32350 | 32350-Deleti | Deletion | 1 | C | - | 7 | 43076 | 0.01625035 | 63.5714286 | WT | FALSE | CAN | JSY13 | 3A | WT-Rep3A | ACC | nope | >1bp-ins | G/C-1 | none | G/C-1 | GCAAAACCT | CCTTTGT | TGG | ACC | C | none | nope | nope_ACC | G/C-1_CCTTTGT | 9 | TRUE | FALSE |
| 187_v.txt | 32416 | 32416-Deleti | Deletion | 1 | T | - | 7 | 43076 | 0.01625035 | 59.8571429 | WT | FALSE | CAN | JSY13 | 3A | WT-Rep3A | GTT | nope | >1bp-ins | A/T-1 | none | A/T-1 | ATTGTGTTAG | TTAGTTT | CAA | GTT | T | none | nope | nope_GTT | A/T-1_TTAGTTT | 9 | TRUE | FALSE |
| 187_v.txt | 32420 | 32420-Deleti | Deletion | 1 | T | - | 7 | 43076 | 0.01625035 | 48.1428571 | WT | FALSE | CAN | JSY13 | 3A | WT-Rep3A | GTT | nope | >1bp-ins | A/T-1 | none | A/T-1 | TGTTAGTTTA | TTTAGGG | CAA | GTT | T | none | nope | nope_GTT | A/T-1_TTTAGGG | 9 | TRUE | FALSE |
| 187_v.txt | 32442 | 32442-Deleti | Deletion | 1 | A | - | 9 | 43076 | 0.02089331 | 56.1111111 | WT | FALSE | CAN | JSY13 | 3A | WT-Rep3A | TAA | nope | >1bp-ins | A/T-1 | none | A/T-1 | GGAACTAAA | AAAAGTC | ATT | ATT | T | none | T>none | T>none_ATT | A/T-1_AAAAGTC | 9 | TRUE | FALSE |
| 187_v.txt | 32477 | 32477-Deleti | Deletion | 1 | G | - | 12 | 43076 | 0.02785774 | 51.8333333 | WT | FALSE | CAN | JSY13 | 3A | WT-Rep3A | AGG | nope | >1bp-ins | G/C-1 | none | G/C-1 | GTAGAAGGT | GGTTAAG | TCC | TCC | C | none | C>none | C>none_TCC | G/C-1_GGTTAAG | 9 | TRUE | FALSE |
| 187_v.txt | 32490 | 32490-Deleti | Deletion | 1 | A | - | 9 | 43076 | 0.02089331 | 43.6666667 | WT | FALSE | CAN | JSY13 | 3A | WT-Rep3A | GAA | nope | >1bp-ins | A/T-1 | none | A/T-1 | ATACGGAAA | AAAACAA | CTT | CTT | T | none | T>none | T>none_CTT | A/T-1_AAAACAA | 9 | TRUE | FALSE |
| 187_v.txt | 32498 | 32498-Deleti | Deletion | 1 | T | - | 90 | 43076 | 0.20893305 | 46.8999978 | WT | FALSE | CAN | JSY13 | 3A | WT-Rep3A | CTT | nope | >1bp-ins | A/T-1 | none | A/T-1 | AACAACTTTT | TTTTTTG | GAA | CTT | T | none | nope | nope_CTT | A/T-1_TTTTTTG | 9 | TRUE | FALSE |
| 187_v.txt | 32528 | 32528-Deleti | Deletion | 1 | G | - | 6 | 43076 | 0.01392887 | 46 | WT | FALSE | CAN | JSY13 | 3A | WT-Rep3A | TGG | nope | >1bp-ins | G/C-1 | none | G/C-1 | TTTTCTGGGG | GGGGTTT | ACC | ACC | C | none | C>none | C>none_ACC | G/C-1_GGGGTTT | 9 | TRUE | FALSE |
| 187_v.txt | 32598 | 32598-Deleti | Deletion | 1 | A | - | 1985 | 23486 | 8.45184365 | 41.7148615 | WT | FALSE | CAN | JSY13 | 3A | WT-Rep3A | CAA | nope | >1bp-ins | A/T-1 | none | A/T-1 | TTAATCAAA | AAAGAGG | GTT | GTT | T | none | T>none | T>none_GTT | A/T-1_AAAGAGG | 9 | TRUE | FALSE |
| 187_v.txt | 32616 | 32616-Deleti | Deletion | 1 | A | - | 3 | 23486 | 0.01277357 | 37.6666667 | WT | FALSE | CAN | JSY13 | 3A | WT-Rep3A | TAA | nope | >1bp-ins | A/T-1 | none | A/T-1 | CAACCTAAG | AAGAACC | ATT | ATT | T | none | T>none | T>none_ATT | A/T-1_AAGAACC | 9 | TRUE | FALSE |
| 187_v.txt | 32631 | 32631-Deleti | Deletion | 1 | T | - | 6 | 23486 | 0.02554714 | 46 | WT | FALSE | CAN | JSY13 | 3A | WT-Rep3A | GTT | nope | >1bp-ins | A/T-1 | none | A/T-1 | CCTTCGTTTT | TTTTTAT | CAA | GTT | T | none | nope | nope_GTT | A/T-1_TTTTTAT | 9 | TRUE | FALSE |
| 187_v.txt | 32658 | 32658-Deleti | Deletion | 1 | C | - | 15 | 23486 | 0.06386784 | 52.066665 | WT | FALSE | CAN | JSY13 | 3A | WT-Rep3A | ACC | nope | >1bp-ins | G/C-1 | none | G/C-1 | CCTGGACCC | CCCCAGG | TGG | ACC | C | none | nope | nope_ACC | G/C-1_CCCCAGG | 9 | TRUE | FALSE |
| 187_v.txt | 32707 | 32707-Deleti | Deletion | 1 | C | - | 6 | 23486 | 0.02554714 | 59.1666667 | WT | FALSE | CAN | JSY13 | 3A | WT-Rep3A | ACC | nope | >1bp-ins | G/C-1 | none | G/C-1 | CGGTAACCC | CCCCAGC | TGG | ACC | C | none | nope | nope_ACC | G/C-1_CCCCAGC | 9 | TRUE | FALSE |
| 187_v.txt | 32719 | 32719-Deleti | Deletion | 1 | A | - | 5 | 23486 | 0.02128928 | 64 | WT | FALSE | CAN | JSY13 | 3A | WT-Rep3A | CAA | nope | >1bp-ins | A/T-1 | none | A/T-1 | CACCACAAA | AAACCAT | GTT | GTT | T | none | T>none | T>none_GTT | A/T-1_AAACCAT | 9 | TRUE | FALSE |
| 187_v.txt | 32728 | 32728-Deleti | Deletion | 1 | A | - | 22 | 23486 | 0.09367283 | 60.818182 | WT | FALSE | CAN | JSY13 | 3A | WT-Rep3A | CAA | nope | >1bp-ins | A/T-1 | none | A/T-1 | CCATACAAA | AAAAACA | GTT | GTT | T | none | T>none | T>none_GTT | A/T-1_AAAAACA | 9 | TRUE | FALSE |
| 187_v.txt | 32758 | 32758-Deleti | Deletion | 1 | C | - | 8 | 23486 | 0.03406285 | 58.7500001 | WT | FALSE | CAN | JSY13 | 3A | WT-Rep3A | GCT | nope | >1bp-ins | G/C-1 | none | G/C-1 | TAATGGCTA | CTAAAAC | CGA | GCT | C | none | nope | nope_GCT | G/C-1_CTAAAAC | 9 | TRUE | FALSE |
| 187_v.txt | 32760 | 32760-Deleti | Deletion | 1 | A | - | 1 | 23486 | 4.26E-03 | 63.9999999 | WT | FALSE | CAN | JSY13 | 3A | WT-Rep3A | TAA | nope | >1bp-ins | A/T-1 | none | A/T-1 | ATGGCTAAA | AAAACTT | ATT | ATT | T | none | T>none | T>none_ATT | A/T-1_AAAACTT | 9 | TRUE | FALSE |
| 187_v.txt | 32839 | 32839-Deleti | Deletion | 1 | C | - | 7 | 23486 | 0.02980499 | 44.8571429 | WT | FALSE | CAN | JSY13 | 3A | WT-Rep3A | ACC | nope | >1bp-ins | G/C-1 | none | G/C-1 | TAATTACCCA | CCCAAAA | TGG | ACC | C | none | nope | nope_ACC | G/C-1_CCCAAAA | 9 | TRUE | FALSE |
| 187_v.txt | 32842 | 32842-Deleti | Deletion | 1 | A | - | 60 | 23486 | 0.25547135 | 40.8333331 | WT | FALSE | CAN | JSY13 | 3A | WT-Rep3A | CAA | nope | >1bp-ins | A/T-1 | none | A/T-1 | TTACCCAAA | AAAAAAT | GTT | GTT | T | none | T>none | T>none_GTT | A/T-1_AAAAAAT | 9 | TRUE | FALSE |
| 187_v.txt | 32855 | 32855-Deleti | Deletion | 1 | C | - | 3 | 23483 | 0.0127752 | 35.3333333 | WT | FALSE | CAN | JSY13 | 3A | WT-Rep3A | TCC | nope | >1bp-ins | G/C-1 | none | G/C-1 | ACTAATCCAT | CCATGCC | AGG | TCC | C | none | nope | nope_TCC | G/C-1_CCATGCC | 9 | TRUE | FALSE |
| 187_v.txt | 32884 | 32884-Deleti | Deletion | 1 | A | - | 21 | 40419 | 0.05195576 | 37.3333331 | WT | FALSE | CAN | JSY13 | 3A | WT-Rep3A | CAA | nope | >1bp-ins | A/T-1 | none | A/T-1 | ACGTCCAAA | AAAATTG | GTT | GTT | T | none | T>none | T>none_GTT | A/T-1_AAAATTG | 9 | TRUE | FALSE |

| | | | | | | | | | | | | | | | | | |
|---|---|---|---|---|---|---|---|---|---|---|---|---|---|---|---|---|---|
| 187_v.txt | 32924 | 32924-Deleti | Deletion | 1 | A | - | 40 | 40418 | 0.09896581 | 54.925 | WT | FALSE | CAN | JSY13 | 3A | WT-Rep3A | CAA |
| 187_v.txt | 32934 | 32934-Deleti | Deletion | 1 | G | - | 14 | 40418 | 0.03463803 | 51.7142864 | WT | FALSE | CAN | JSY13 | 3A | WT-Rep3A | TGC |
| 187_v.txt | 32935 | 32935-Deleti | Deletion | 1 | C | - | 5 | 40418 | 0.01237073 | 41 | WT | FALSE | CAN | JSY13 | 3A | WT-Rep3A | GCC |
| 187_v.txt | 32965 | 32965-Deleti | Deletion | 1 | C | - | 4 | 40458 | 9.89E-03 | 43.75 | WT | FALSE | CAN | JSY13 | 3A | WT-Rep3A | GCC |
| 187_v.txt | 33000 | 33000-Deleti | Deletion | 1 | A | - | 12 | 40458 | 0.02966039 | 60.4166667 | WT | FALSE | CAN | JSY13 | 3A | WT-Rep3A | GAA |
| 187_v.txt | 33014 | 33014-Deleti | Deletion | 1 | G | - | 12 | 40458 | 0.02966039 | 62.0833333 | WT | FALSE | CAN | JSY13 | 3A | WT-Rep3A | AGG |
| 187_v.txt | 33040 | 33040-Deleti | Deletion | 1 | T | - | 7 | 40458 | 0.01730189 | 51.4285714 | WT | FALSE | CAN | JSY13 | 3A | WT-Rep3A | ATT |
| 187_v.txt | 33070 | 33070-Deleti | Deletion | 1 | C | - | 2 | 40456 | 4.94E-03 | 63.0000002 | WT | FALSE | CAN | JSY13 | 3A | WT-Rep3A | GCC |
| 187_v.txt | 33078 | 33078-Deleti | Deletion | 1 | C | - | 3 | 40458 | 7.42E-03 | 55.3333333 | WT | FALSE | CAN | JSY13 | 3A | WT-Rep3A | ACC |
| 187_v.txt | 33138 | 33138-Deleti | Deletion | 1 | A | - | 12 | 40458 | 0.02966039 | 39.3333333 | WT | FALSE | CAN | JSY13 | 3A | WT-Rep3A | TAA |
| 187_v.txt | 33147 | 33147-Deleti | Deletion | 1 | A | - | 11 | 40458 | 0.02718869 | 42.7272727 | WT | FALSE | CAN | JSY13 | 3A | WT-Rep3A | GAA |
| 187_v.txt | 33326 | 33326-Deleti | Deletion | 1 | T | - | 6 | 23996 | 0.02500417 | 57.5 | WT | FALSE | CAN | JSY13 | 3A | WT-Rep3A | CTT |
| 187_v.txt | 33474 | 33474-Deleti | Deletion | 1 | C | - | 76 | 23996 | 0.31671945 | 28.5921053 | WT | FALSE | CAN | JSY13 | 3A | WT-Rep3A | GCC |
| 127_v.txt | 31711 | 31711-SNV-/ | SNV | 1 | A | C | 8 | 11746 | 0.06810829 | 23.75 | WT_perm | FALSE | ARG | JSY2424 | 1 | WT_perm-Re | CAA |
| 127_v.txt | 31723 | 31723-SNV-/ | SNV | 1 | A | C | 4 | 11746 | 0.03405415 | 57.25 | WT_perm | FALSE | ARG | JSY2424 | 1 | WT_perm-Re | CAA |
| 127_v.txt | 32631 | 32631-SNV-T | SNV | 1 | T | G | 2 | 17405 | 0.01149095 | 64 | WT_perm | FALSE | ARG | JSY2424 | 1 | WT_perm-Re | GTT |
| 127_v.txt | 32842 | 32842-SNV-/ | SNV | 1 | A | C | 3 | 17405 | 0.01723643 | 28.6667064 | WT_perm | FALSE | ARG | JSY2424 | 1 | WT_perm-Re | CAA |
| 127_v.txt | 32884 | 32884-SNV-/ | SNV | 1 | A | C | 4 | 24826 | 0.01611214 | 36.25 | WT_perm | FALSE | ARG | JSY2424 | 1 | WT_perm-Re | CAA |
| 127_v.txt | 32081 | 32081-SNV-/ | SNV | 1 | A | G | 5 | 20855 | 0.02397507 | 57.7997165 | WT_perm | FALSE | ARG | JSY2424 | 1 | WT_perm-Re | CAA |
| 127_v.txt | 32810 | 32810-SNV-/ | SNV | 1 | A | G | 15 | 17405 | 0.08618213 | 42.3333333 | WT_perm | FALSE | ARG | JSY2424 | 1 | WT_perm-Re | CAG |
| 127_v.txt | 32924 | 32924-SNV-/ | SNV | 1 | A | G | 15 | 24826 | 0.06042053 | 47.6 | WT_perm | FALSE | ARG | JSY2424 | 1 | WT_perm-Re | CAA |
| 127_v.txt | 32081 | 32081-SNV-/ | SNV | 1 | A | T | 3 | 20855 | 0.01438504 | 63.9970196 | WT_perm | FALSE | ARG | JSY2424 | 1 | WT_perm-Re | CAA |
| 127_v.txt | 32810 | 32810-SNV-/ | SNV | 1 | A | T | 6 | 17405 | 0.03447285 | 46.5 | WT_perm | FALSE | ARG | JSY2424 | 1 | WT_perm-Re | CAG |
| 127_v.txt | 31650 | 31650-SNV-( | SNV | 1 | C | T | 28 | 11746 | 0.23837902 | 36.6785714 | WT_perm | FALSE | ARG | JSY2424 | 1 | WT_perm-Re | GCG |
| 127_v.txt | 31882 | 31882-SNV-( | SNV | 1 | C | T | 15 | 11746 | 0.12770305 | 46.1333333 | WT_perm | FALSE | ARG | JSY2424 | 1 | WT_perm-Re | GCA |
| 127_v.txt | 31960 | 31960-SNV-( | SNV | 1 | C | T | 22 | 20855 | 0.10549029 | 35.9545455 | WT_perm | FALSE | ARG | JSY2424 | 1 | WT_perm-Re | GCC |
| 127_v.txt | 31962 | 31962-SNV-( | SNV | 1 | G | A | 34 | 20855 | 0.16303045 | 37.3235294 | WT_perm | FALSE | ARG | JSY2424 | 1 | WT_perm-Re | CGC |
| 127_v.txt | 31977 | 31977-SNV-( | SNV | 1 | C | T | 23 | 20855 | 0.1102853 | 44 | WT_perm | FALSE | ARG | JSY2424 | 1 | WT_perm-Re | GCC |
| 127_v.txt | 32092 | 32092-SNV-( | SNV | 1 | C | T | 24 | 20855 | 0.11508032 | 60.3333333 | WT_perm | FALSE | ARG | JSY2424 | 1 | WT_perm-Re | GCA |
| 127_v.txt | 32165 | 32165-SNV-( | SNV | 1 | G | A | 25 | 20855 | 0.11987533 | 48.2 | WT_perm | FALSE | ARG | JSY2424 | 1 | WT_perm-Re | CGC |
| 127_v.txt | 32173 | 32173-SNV-( | SNV | 1 | C | T | 27 | 20855 | 0.12946536 | 50.6296296 | WT_perm | FALSE | ARG | JSY2424 | 1 | WT_perm-Re | GCA |
| 127_v.txt | 32190 | 32190-SNV-( | SNV | 1 | G | A | 19 | 20855 | 0.09110525 | 43.8421053 | WT_perm | FALSE | ARG | JSY2424 | 1 | WT_perm-Re | TGC |
| 127_v.txt | 32271 | 32271-SNV-( | SNV | 1 | C | T | 35 | 23280 | 0.15034364 | 35.6857143 | WT_perm | FALSE | ARG | JSY2424 | 1 | WT_perm-Re | ACG |
| 127_v.txt | 32299 | 32299-SNV-( | SNV | 1 | C | T | 52 | 23280 | 0.2233677 | 49.7115385 | WT_perm | FALSE | ARG | JSY2424 | 1 | WT_perm-Re | GCG |
| 127_v.txt | 32466 | 32466-SNV-( | SNV | 1 | C | T | 30 | 23280 | 0.12886598 | 50.6666667 | WT_perm | FALSE | ARG | JSY2424 | 1 | WT_perm-Re | GCC |
| 127_v.txt | 32487 | 32487-SNV-( | SNV | 1 | C | T | 24 | 23280 | 0.10309278 | 44.9166667 | WT_perm | FALSE | ARG | JSY2424 | 1 | WT_perm-Re | ACG |
| 127_v.txt | 32535 | 32535-SNV-( | SNV | 1 | G | A | 26 | 23280 | 0.11168385 | 35.9538087 | WT_perm | FALSE | ARG | JSY2424 | 1 | WT_perm-Re | TGC |
| 127_v.txt | 32536 | 32536-SNV-( | SNV | 1 | C | T | 32 | 23280 | 0.13745705 | 36.9306366 | WT_perm | FALSE | ARG | JSY2424 | 1 | WT_perm-Re | GCA |
| 127_v.txt | 32590 | 32590-SNV-( | SNV | 1 | C | T | 30 | 17405 | 0.17236426 | 35.5 | WT_perm | FALSE | ARG | JSY2424 | 1 | WT_perm-Re | GCG |
| 127_v.txt | 32700 | 32700-SNV-( | SNV | 1 | C | T | 26 | 17405 | 0.14938236 | 54.0384615 | WT_perm | FALSE | ARG | JSY2424 | 1 | WT_perm-Re | GCC |
| 127_v.txt | 32702 | 32702-SNV-( | SNV | 1 | G | A | 18 | 17405 | 0.10341856 | 52.2222222 | WT_perm | FALSE | ARG | JSY2424 | 1 | WT_perm-Re | CGG |
| 127_v.txt | 32713 | 32713-SNV-( | SNV | 1 | C | T | 17 | 17405 | 0.09767308 | 62.3529412 | WT_perm | FALSE | ARG | JSY2424 | 1 | WT_perm-Re | GCA |
| 127_v.txt | 32776 | 32776-SNV-( | SNV | 1 | C | T | 24 | 17405 | 0.13789141 | 48.0416667 | WT_perm | FALSE | ARG | JSY2424 | 1 | WT_perm-Re | GCG |
| 127_v.txt | 32798 | 32798-SNV-( | SNV | 1 | G | A | 26 | 17405 | 0.14938236 | 47.9615385 | WT_perm | FALSE | ARG | JSY2424 | 1 | WT_perm-Re | CGT |
| 127_v.txt | 32860 | 32860-SNV-( | SNV | 1 | C | T | 26 | 17405 | 0.14938236 | 35.8846154 | WT_perm | FALSE | ARG | JSY2424 | 1 | WT_perm-Re | GCC |
| 127_v.txt | 32900 | 32900-SNV-( | SNV | 1 | G | A | 38 | 24826 | 0.15306534 | 36.8775212 | WT_perm | FALSE | ARG | JSY2424 | 1 | WT_perm-Re | GGC |
| 127_v.txt | 32901 | 32901-SNV-( | SNV | 1 | C | T | 51 | 24826 | 0.20542979 | 33.725778 | WT_perm | FALSE | ARG | JSY2424 | 1 | WT_perm-Re | GCC |
| 127_v.txt | 32935 | 32935-SNV-( | SNV | 1 | C | T | 35 | 24826 | 0.14098123 | 44.9428571 | WT_perm | FALSE | ARG | JSY2424 | 1 | WT_perm-Re | GCC |
| 127_v.txt | 32965 | 32965-SNV-( | SNV | 1 | C | T | 32 | 24826 | 0.12889712 | 59.78125 | WT_perm | FALSE | ARG | JSY2424 | 1 | WT_perm-Re | GCC |
| 127_v.txt | 32967 | 32967-SNV-( | SNV | 1 | G | A | 58 | 24826 | 0.23362604 | 60.6896552 | WT_perm | FALSE | ARG | JSY2424 | 1 | WT_perm-Re | CGC |
| 127_v.txt | 32977 | 32977-SNV-( | SNV | 1 | C | T | 25 | 24826 | 0.10070088 | 57.8 | WT_perm | FALSE | ARG | JSY2424 | 1 | WT_perm-Re | GCT |
| 127_v.txt | 33056 | 33056-SNV-( | SNV | 1 | C | T | 43 | 24826 | 0.17320551 | 57.7906977 | WT_perm | FALSE | ARG | JSY2424 | 1 | WT_perm-Re | GCG |
| 127_v.txt | 33069 | 33069-SNV-( | SNV | 1 | G | A | 23 | 24826 | 0.09264481 | 49.7384123 | WT_perm | FALSE | ARG | JSY2424 | 1 | WT_perm-Re | TGC |
| 127_v.txt | 33070 | 33070-SNV-( | SNV | 1 | C | T | 33 | 24826 | 0.13292516 | 53.3326906 | WT_perm | FALSE | ARG | JSY2424 | 1 | WT_perm-Re | GCC |
| 127_v.txt | 33104 | 33104-SNV-( | SNV | 1 | G | A | 37 | 24826 | 0.1490373 | 41.945946 | WT_perm | FALSE | ARG | JSY2424 | 1 | WT_perm-Re | CGC |
| 127_v.txt | 33118 | 33118-SNV-( | SNV | 1 | C | T | 29 | 24826 | 0.11681302 | 43.8275862 | WT_perm | FALSE | ARG | JSY2424 | 1 | WT_perm-Re | GCG |
| 127_v.txt | 33258 | 33258-SNV-( | SNV | 1 | C | T | 21 | 11940 | 0.1758794 | 38.3809524 | WT_perm | FALSE | ARG | JSY2424 | 1 | WT_perm-Re | GCC |
| 127_v.txt | 33275 | 33275-SNV-( | SNV | 1 | C | T | 16 | 11940 | 0.13400335 | 45.9375 | WT_perm | FALSE | ARG | JSY2424 | 1 | WT_perm-Re | ACG |
| 127_v.txt | 33276 | 33276-SNV-( | SNV | 1 | G | A | 17 | 11939 | 0.14239049 | 45.5847279 | WT_perm | FALSE | ARG | JSY2424 | 1 | WT_perm-Re | CGT |
| 127_v.txt | 33310 | 33310-SNV-( | SNV | 1 | G | A | 15 | 11940 | 0.12562814 | 54.0666667 | WT_perm | FALSE | ARG | JSY2424 | 1 | WT_perm-Re | CGC |
| 127_v.txt | 33474 | 33474-SNV-( | SNV | 1 | C | T | 18 | 11940 | 0.15075377 | 31.0555556 | WT_perm | FALSE | ARG | JSY2424 | 1 | WT_perm-Re | GCC |
| 127_v.txt | 32702 | 32702-SNV-( | SNV | 1 | G | C | 3 | 17405 | 0.01723643 | 48.6666667 | WT_perm | FALSE | ARG | JSY2424 | 1 | WT_perm-Re | CGG |
| 127_v.txt | 32839 | 32839-SNV-( | SNV | 1 | C | G | 3 | 17405 | 0.01723643 | 35.6666667 | WT_perm | FALSE | ARG | JSY2424 | 1 | WT_perm-Re | ACC |
| 127_v.txt | 32901 | 32901-SNV-( | SNV | 1 | C | G | 3 | 24826 | 0.01208411 | 26.6688074 | WT_perm | FALSE | ARG | JSY2424 | 1 | WT_perm-Re | GCC |
| 127_v.txt | 32965 | 32965-SNV-( | SNV | 1 | C | G | 3 | 24826 | 0.01208411 | 64 | WT_perm | FALSE | ARG | JSY2424 | 1 | WT_perm-Re | GCC |
| 127_v.txt | 33069 | 33069-SNV-( | SNV | 1 | G | C | 3 | 24826 | 0.01208411 | 35.0002562 | WT_perm | FALSE | ARG | JSY2424 | 1 | WT_perm-Re | TGC |
| 127_v.txt | 33070 | 33070-SNV-( | SNV | 1 | C | G | 2 | 24826 | 8.06E-03 | 38.5061187 | WT_perm | FALSE | ARG | JSY2424 | 1 | WT_perm-Re | GCC |
| 127_v.txt | 33276 | 33276-SNV-( | SNV | 1 | G | C | 2 | 11939 | 0.01675182 | 23.998165 | WT_perm | FALSE | ARG | JSY2424 | 1 | WT_perm-Re | CGT |
| 127_v.txt | 32702 | 32702-SNV-( | SNV | 1 | G | T | 7 | 17405 | 0.04021833 | 46.4285714 | WT_perm | FALSE | ARG | JSY2424 | 1 | WT_perm-Re | CGG |
| 127_v.txt | 33474 | 33474-SNV-( | SNV | 1 | C | A | 36 | 11940 | 0.30150754 | 32.9722222 | WT_perm | FALSE | ARG | JSY2424 | 1 | WT_perm-Re | GCC |
| 127_v.txt | 33475 | 33475-SNV-( | SNV | 1 | C | A | 7 | 11939 | 0.05863138 | 37.5714286 | WT_perm | FALSE | ARG | JSY2424 | 1 | WT_perm-Re | CCT |
| 127_v.txt | 33474 | 33474-MNV- | MNV | 2 | CC | AA | 2 | 11940 | 0.01675042 | 27.1666667 | WT_perm | FALSE | ARG | JSY2424 | 1 | WT_perm-Re | GCC |
| 127_v.txt | 32081 | 32081-Insert | Insertion | 1 | - | A | 11 | 20855 | 0.05274515 | 63.6363636 | WT_perm | FALSE | ARG | JSY2424 | 1 | WT_perm-Re | CAA |
| 127_v.txt | 32842 | 32842-Insert | Insertion | 1 | - | A | 10 | 17405 | 0.05745475 | 40 | WT_perm | FALSE | ARG | JSY2424 | 1 | WT_perm-Re | CAA |
| 127_v.txt | 33475 | 33475-Insert | Insertion | 1 | - | A | 1 | 11939 | 8.38E-03 | 38 | WT_perm | FALSE | ARG | JSY2424 | 1 | WT_perm-Re | CCT |
| 127_v.txt | 31711 | 31711-Deleti | Deletion | 1 | A | - | 4 | 11746 | 0.03405415 | 56.25 | WT_perm | FALSE | ARG | JSY2424 | 1 | WT_perm-Re | CAA |
| 127_v.txt | 31723 | 31723-Deleti | Deletion | 1 | A | - | 6 | 11746 | 0.05108122 | 58.3333333 | WT_perm | FALSE | ARG | JSY2424 | 1 | WT_perm-Re | CAA |
| 127_v.txt | 31863 | 31863-Deleti | Deletion | 1 | A | - | 4 | 11746 | 0.03405415 | 57.5 | WT_perm | FALSE | ARG | JSY2424 | 1 | WT_perm-Re | GAA |
| 127_v.txt | 31904 | 31904-Deleti | Deletion | 1 | T | - | 10 | 11746 | 0.08513537 | 40.3 | WT_perm | FALSE | ARG | JSY2424 | 1 | WT_perm-Re | ATT |
| 127_v.txt | 31913 | 31913-Deleti | Deletion | 1 | A | - | 4 | 11746 | 0.03405415 | 38 | WT_perm | FALSE | ARG | JSY2424 | 1 | WT_perm-Re | CAA |
| 127_v.txt | 32081 | 32081-Deleti | Deletion | 1 | A | - | 19 | 20855 | 0.09110525 | 63.6842105 | WT_perm | FALSE | ARG | JSY2424 | 1 | WT_perm-Re | CAA |
| 127_v.txt | 32210 | 32210-Deleti | Deletion | 1 | T | - | 5 | 20855 | 0.02397507 | 37 | WT_perm | FALSE | ARG | JSY2424 | 1 | WT_perm-Re | CTT |
| 127_v.txt | 32231 | 32231-Deleti | Deletion | 1 | T | - | 5 | 20855 | 0.02397507 | 37.6 | WT_perm | FALSE | ARG | JSY2424 | 1 | WT_perm-Re | ATT |
| 127_v.txt | 32346 | 32346-Deleti | Deletion | 1 | A | - | 7 | 23280 | 0.03006873 | 60.2857143 | WT_perm | FALSE | ARG | JSY2424 | 1 | WT_perm-Re | CAA |
| 127_v.txt | 32442 | 32442-Deleti | Deletion | 1 | A | - | 7 | 23280 | 0.03006873 | 47.1428571 | WT_perm | FALSE | ARG | JSY2424 | 1 | WT_perm-Re | TAA |
| 127_v.txt | 32498 | 32498-Deleti | Deletion | 1 | T | - | 22 | 23280 | 0.09450172 | 44.5909091 | WT_perm | FALSE | ARG | JSY2424 | 1 | WT_perm-Re | CTT |
| 127_v.txt | 32528 | 32528-Deleti | Deletion | 1 | G | - | 5 | 23280 | 0.02147766 | 43 | WT_perm | FALSE | ARG | JSY2424 | 1 | WT_perm-Re | TGG |
| 127_v.txt | 32631 | 32631-Deleti | Deletion | 1 | T | - | 5 | 17405 | 0.02872738 | 47.6 | WT_perm | FALSE | ARG | JSY2424 | 1 | WT_perm-Re | GTT |
| 127_v.txt | 32640 | 32640-Deleti | Deletion | 1 | T | - | 5 | 17405 | 0.02872738 | 53.4 | WT_perm | FALSE | ARG | JSY2424 | 1 | WT_perm-Re | CTT |
| 127_v.txt | 32658 | 32658-Deleti | Deletion | 1 | C | - | 8 | 17405 | 0.0459638 | 60.75 | WT_perm | FALSE | ARG | JSY2424 | 1 | WT_perm-Re | ACC |
| 127_v.txt | 32728 | 32728-Deleti | Deletion | 1 | A | - | 5 | 17405 | 0.02872738 | 58.8 | WT_perm | FALSE | ARG | JSY2424 | 1 | WT_perm-Re | CAA |
| 127_v.txt | 32839 | 32839-Deleti | Deletion | 1 | C | - | 4 | 17405 | 0.0229819 | 38 | WT_perm | FALSE | ARG | JSY2424 | 1 | WT_perm-Re | ACC |
| 127_v.txt | 32842 | 32842-Deleti | Deletion | 1 | A | - | 22 | 17405 | 0.12640046 | 41.8636364 | WT_perm | FALSE | ARG | JSY2424 | 1 | WT_perm-Re | CAA |
| 127_v.txt | 32884 | 32884-Deleti | Deletion | 1 | A | - | 9 | 24826 | 0.03625232 | 37.5555556 | WT_perm | FALSE | ARG | JSY2424 | 1 | WT_perm-Re | CAA |
| 127_v.txt | 32901 | 32901-Deleti | Deletion | 1 | C | - | 1 | 24826 | 4.03E-03 | 38 | WT_perm | FALSE | ARG | JSY2424 | 1 | WT_perm-Re | GCC |
| 127_v.txt | 32924 | 32924-Deleti | Deletion | 1 | A | - | 15 | 24826 | 0.06042053 | 55.8666667 | WT_perm | FALSE | ARG | JSY2424 | 1 | WT_perm-Re | CAA |
| 127_v.txt | 33138 | 33138-Deleti | Deletion | 1 | A | - | 6 | 24826 | 0.02416821 | 37.8333333 | WT_perm | FALSE | ARG | JSY2424 | 1 | WT_perm-Re | TAA |
| 127_v.txt | 33147 | 33147-Deleti | Deletion | 1 | A | - | 6 | 24826 | 0.02416821 | 40.6666667 | WT_perm | FALSE | ARG | JSY2424 | 1 | WT_perm-Re | GAA |
| 127_v.txt | 33175 | 33175-Deleti | Deletion | 1 | G | - | 5 | 24821 | 0.02014423 | 35.2 | WT_perm | FALSE | ARG | JSY2424 | 1 | WT_perm-Re | AGG |
| 127_v.txt | 33474 | 33474-Deleti | Deletion | 1 | C | - | 27 | 11940 | 0.22613065 | 27.7037037 | WT_perm | FALSE | ARG | JSY2424 | 1 | WT_perm-Re | GCC |
| 128_v.txt | 31723 | 31723-SNV-/ | SNV | 1 | A | C | 4 | 10501 | 0.03809161 | 64 | WT_perm | FALSE | ARG | JSY2424 | 2 | WT_perm-Re | CAA |
| 128_v.txt | 32573 | 32573-SNV-T | SNV | 1 | T | G | 4 | 23632 | 0.0169262 | 38 | WT_perm | FALSE | ARG | JSY2424 | 2 | WT_perm-Re | CTT |
| 128_v.txt | 32842 | 32842-SNV-/ | SNV | 1 | A | C | 3 | 23631 | 0.01269519 | 27.9993328 | WT_perm | FALSE | ARG | JSY2424 | 2 | WT_perm-Re | CAA |
| 128_v.txt | 32884 | 32884-SNV-/ | SNV | 1 | A | C | 10 | 27289 | 0.0366448 | 33.4979714 | WT_perm | FALSE | ARG | JSY2424 | 2 | WT_perm-Re | CAA |
| 128_v.txt | 33269 | 33269-SNV-/ | SNV | 1 | A | C | 83 | 15340 | 0.5410691 | 17.1204819 | WT_perm | FALSE | ARG | JSY2424 | 2 | WT_perm-Re | TAG |
| 128_v.txt | 33476 | 33476-SNV-T | SNV | 1 | T | G | 3 | 15336 | 0.01956182 | 36.3338026 | WT_perm | FALSE | ARG | JSY2424 | 2 | WT_perm-Re | CTT |
| 128_v.txt | 31711 | 31711-SNV-/ | SNV | 1 | A | G | 6 | 10501 | 0.05713742 | 61.3333333 | WT_perm | FALSE | ARG | JSY2424 | 2 | WT_perm-Re | CAA |
| 128_v.txt | 32573 | 32573-SNV-T | SNV | 1 | T | C | 16 | 23632 | 0.06770481 | 30.5625 | WT_perm | FALSE | ARG | JSY2424 | 2 | WT_perm-Re | CTT |
| 128_v.txt | 32842 | 32842-SNV-/ | SNV | 1 | A | G | 1 | 23631 | 4.23E-03 | 63.9979227 | WT_perm | FALSE | ARG | JSY2424 | 2 | WT_perm-Re | CAA |
| 128_v.txt | 32884 | 32884-SNV-/ | SNV | 1 | A | G | 8 | 27289 | 0.02931584 | 30.623283 | WT_perm | FALSE | ARG | JSY2424 | 2 | WT_perm-Re | CAA |
| 128_v.txt | 33476 | 33476-SNV-T | SNV | 1 | T | C | 20 | 15336 | 0.1304121 | 34.5502836 | WT_perm | FALSE | ARG | JSY2424 | 2 | WT_perm-Re | CTT |
| 128_v.txt | 32573 | 32573-SNV-T | SNV | 1 | T | A | 14 | 23632 | 0.05924171 | 37.5 | WT_perm | FALSE | ARG | JSY2424 | 2 | WT_perm-Re | CTT |
| 128_v.txt | 31650 | 31650-SNV-( | SNV | 1 | C | T | 29 | 10501 | 0.27616418 | 35.4137931 | WT_perm | FALSE | ARG | JSY2424 | 2 | WT_perm-Re | GCG |
| 128_v.txt | 31658 | 31658-SNV-( | SNV | 1 | C | T | 14 | 10501 | 0.13332064 | 35.3949749 | WT_perm | FALSE | ARG | JSY2424 | 2 | WT_perm-Re | GCG |
| 128_v.txt | 31881 | 31881-SNV-( | SNV | 1 | G | A | 13 | 10501 | 0.12379773 | 39 | WT_perm | FALSE | ARG | JSY2424 | 2 | WT_perm-Re | GGC |
| 128_v.txt | 31960 | 31960-SNV-( | SNV | 1 | C | T | 25 | 17096 | 0.14623304 | 36.6 | WT_perm | FALSE | ARG | JSY2424 | 2 | WT_perm-Re | GCC |
| 128_v.txt | 31962 | 31962-SNV-( | SNV | 1 | G | A | 34 | 17096 | 0.19887693 | 36.7941177 | WT_perm | FALSE | ARG | JSY2424 | 2 | WT_perm-Re | CGC |
| 128_v.txt | 31972 | 31972-SNV-( | SNV | 1 | C | T | 19 | 17096 | 0.11113711 | 37.8947368 | WT_perm | FALSE | ARG | JSY2424 | 2 | WT_perm-Re | GCC |
| 128_v.txt | 31977 | 31977-SNV-( | SNV | 1 | C | T | 27 | 17096 | 0.15793168 | 42.1851852 | WT_perm | FALSE | ARG | JSY2424 | 2 | WT_perm-Re | GCC |

| | | | | | | | | | | | | | | | | | |
|---|---|---|---|---|---|---|---|---|---|---|---|---|---|---|---|---|---|
| 187_v.txt | nope | >1bp-ins | A/T-1 | none | A/T-1 | CAGGGCAAA | AAAAGTG | GTT | GTT | T | none | T>none | T>none_GTT | A/T-1_AAAAGTG | 9 | TRUE | FALSE |
| 187_v.txt | nope | >1bp-ins | G/C-1 | none | G/C-1 | GTGATTGCC( | GCCCAAG | ACG | ACG | C | none | C>none | C>none_ACG | G/C-1_GCCCAAG | 9 | TRUE | FALSE |
| 187_v.txt | nope | >1bp-ins | G/C-1 | none | G/C-1 | TGATTGCCC/ | CCCAAGA | CGG | GCC | C | none | nope | nope_GCC | G/C-1_CCCAAGA | 9 | TRUE | FALSE |
| 187_v.txt | nope | >1bp-ins | G/C-1 | none | G/C-1 | CATTGGCCG( | CCGCACC | CGG | GCC | C | none | nope | nope_GCC | G/C-1_CCGCACC | 9 | TRUE | FALSE |
| 187_v.txt | nope | >1bp-ins | A/T-1 | none | A/T-1 | TGTGAGAAA | AAAACTG | CTT | CTT | T | none | T>none | T>none_CTT | A/T-1_AAAACTG | 9 | TRUE | FALSE |
| 187_v.txt | nope | >1bp-ins | G/C-1 | none | G/C-1 | GAAAGAGG/ | GGATGTA | TCC | TCC | C | none | C>none | C>none_TCC | G/C-1_GGATGTA | 9 | TRUE | FALSE |
| 187_v.txt | nope | >1bp-ins | A/T-1 | none | A/T-1 | TAGCCATTTC | TTTCACC | TAA | ATT | T | none | nope | nope_ATT | A/T-1_TTTCACC | 9 | TRUE | FALSE |
| 187_v.txt | nope | >1bp-ins | G/C-1 | none | G/C-1 | AATATGCCA/ | CCAAAGA | CGG | GCC | C | none | nope | nope_GCC | G/C-1_CCAAAGA | 9 | TRUE | FALSE |
| 187_v.txt | nope | >1bp-ins | G/C-1 | none | G/C-1 | AAAGAACCC. | CCCATAA | TGG | ACC | C | none | nope | nope_ACC | G/C-1_CCCATAA | 9 | TRUE | FALSE |
| 187_v.txt | nope | >1bp-ins | A/T-1 | none | A/T-1 | GTGGATAAA | AAACCAA | ATT | ATT | T | none | T>none | T>none_ATT | A/T-1_AAACCAA | 9 | TRUE | FALSE |
| 187_v.txt | nope | >1bp-ins | A/T-1 | none | A/T-1 | CCAATGAAA. | AAAAGAC | CTT | CTT | T | none | T>none | T>none_CTT | A/T-1_AAAAGAC | 9 | TRUE | FALSE |
| 187_v.txt | nope | >1bp-ins | A/T-1 | none | A/T-1 | CAATTCTTTA | TTTACTT | GAA | CTT | T | none | nope | nope_CTT | A/T-1_TTTACTT | 9 | TRUE | FALSE |
| 187_v.txt | nope | >1bp-ins | G/C-1 | none | G/C-1 | GCTATGCCTT | CCTTTTT | CGG | GCC | C | none | nope | nope_GCC | G/C-1_CCTTTTT | 9 | TRUE | FALSE |
| 127_v.txt | A>C | G/C+1 | A/T-1 | TA>GC | TA>GC | CATTCCAAA/ | AAAATTT | GTT | GTT | T | G | T>G | T>G_GTT | TA>GC_AAAATTT | 6 | FALSE | FALSE |
| 127_v.txt | A>C | G/C+1 | A/T-1 | TA>GC | TA>GC | TGTCCCAAA/ | AAAAAGT | GTT | GTT | T | G | T>G | T>G_GTT | TA>GC_AAAAAGT | 6 | FALSE | FALSE |
| 127_v.txt | T>G | G/C+1 | A/T-1 | TA>GC | TA>GC | CCTTCGTTTT | TTTTTAT | CAA | GTT | T | C | T>G | T>G_GTT | TA>GC_TTTTTAT | 6 | FALSE | FALSE |
| 127_v.txt | A>C | G/C+1 | A/T-1 | TA>GC | TA>GC | TTACCCAAA/ | AAAAAAT | GTT | GTT | T | G | T>G | T>G_GTT | TA>GC_AAAAAAT | 6 | FALSE | FALSE |
| 127_v.txt | A>C | G/C+1 | A/T-1 | TA>GC | TA>GC | ACGTCCAAA/ | AAAATTG | GTT | GTT | T | G | T>G | T>G_GTT | TA>GC_AAAATTG | 6 | FALSE | FALSE |
| 127_v.txt | A>G | G/C+1 | A/T-1 | TA>CG | TA>CG | CCATGCAAA/ | AAAAAAG | GTT | GTT | T | C | T>C | T>C_GTT | TA>CG_AAAAAAG | 6 | FALSE | FALSE |
| 127_v.txt | A>G | G/C+1 | A/T-1 | TA>CG | TA>CG | TTTGACAGG( | AGGGAAC | GTC | GTC | T | C | T>C | T>C_GTC | TA>CG_AGGGAAC | 6 | FALSE | FALSE |
| 127_v.txt | A>G | G/C+1 | A/T-1 | TA>CG | TA>CG | CAGGGCAAA | AAAAGTG | GTT | GTT | T | C | T>C | T>C_GTT | TA>CG_AAAAGTG | 6 | FALSE | FALSE |
| 127_v.txt | A>T | A/T+1 | A/T-1 | TA>AT | TA>AT | CCATGCAAA/ | AAAAAAG | GTT | GTT | T | A | T>A | T>A_GTT | TA>AT_AAAAAAG | 6 | FALSE | FALSE |
| 127_v.txt | A>T | A/T+1 | A/T-1 | TA>AT | TA>AT | TTTGACAGG( | AGGGAAC | GTC | GTC | T | A | T>A | T>A_GTC | TA>AT_AGGGAAC | 6 | FALSE | FALSE |
| 127_v.txt | C>T | A/T+1 | G/C-1 | CG>TA | CG>TA | AGAATGCGA | CGAAATG | CGC | GCG | C | A | C>T | C>T_GCG | CG>TA_CGAAATG | 6 | FALSE | FALSE |
| 127_v.txt | C>T | A/T+1 | G/C-1 | CG>TA | CG>TA | AGGCGGCAG | CAGCAAA | CGT | GCA | C | A | C>T | C>T_GCA | CG>TA_CAGCAAA | 6 | FALSE | FALSE |
| 127_v.txt | C>T | A/T+1 | G/C-1 | CG>TA | CG>TA | ATGTGGCCG( | CCGCATA | CGG | GCC | C | A | C>T | C>T_GCC | CG>TA_CCGCATA | 6 | FALSE | FALSE |
| 127_v.txt | G>A | A/T+1 | G/C-1 | CG>TA | CG>TA | GTGGCCGCA | GCATAAT | GCG | GCG | C | T | C>T | C>T_GCG | CG>TA_GCATAAT | 6 | FALSE | FALSE |
| 127_v.txt | C>T | A/T+1 | G/C-1 | CG>TA | CG>TA | GCCAAGCCG | CCGGGCA | CGG | GCC | C | A | C>T | C>T_GCC | CG>TA_CCGGGCA | 6 | FALSE | FALSE |
| 127_v.txt | C>T | A/T+1 | G/C-1 | CG>TA | CG>TA | AGCCTGCAA( | CAACACC | CGT | GCA | C | A | C>T | C>T_GCA | CG>TA_CAACACC | 6 | FALSE | FALSE |
| 127_v.txt | G>A | A/T+1 | G/C-1 | CG>TA | CG>TA | CAAAGCGCC. | GCCAAAT | GCG | GCG | C | T | C>T | C>T_GCG | CG>TA_GCCAAAT | 6 | FALSE | FALSE |
| 127_v.txt | C>T | A/T+1 | G/C-1 | CG>TA | CG>TA | CAAATGCAG | CAGCAGT | CGT | GCA | C | A | C>T | C>T_GCA | CG>TA_CAGCAGT | 6 | FALSE | FALSE |
| 127_v.txt | G>A | A/T+1 | G/C-1 | CG>TA | CG>TA | AAAACTGCA. | GCAATGT | ACG | ACG | C | T | C>T | C>T_ACG | CG>TA_GCAATGT | 6 | FALSE | FALSE |
| 127_v.txt | C>T | A/T+1 | G/C-1 | CG>TA | CG>TA | AAAATACGG | CGGGAAC | TGC | ACG | C | A | C>T | C>T_ACG | CG>TA_CGGGAAC | 6 | FALSE | FALSE |
| 127_v.txt | C>T | A/T+1 | G/C-1 | CG>TA | CG>TA | AATTTGCGG( | CGGCAGA | CGC | GCG | C | A | C>T | C>T_GCG | CG>TA_CGGCAGA | 6 | FALSE | FALSE |
| 127_v.txt | C>T | A/T+1 | G/C-1 | CG>TA | CG>TA | AGAGAGCCA | CCAATGT | CGG | GCC | C | A | C>T | C>T_GCC | CG>TA_CCAATGT | 6 | FALSE | FALSE |
| 127_v.txt | C>T | A/T+1 | G/C-1 | CG>TA | CG>TA | AAGATACGG | CGGAAAA | TGC | ACG | C | A | C>T | C>T_ACG | CG>TA_CGGAAAA | 6 | FALSE | FALSE |
| 127_v.txt | G>A | A/T+1 | G/C-1 | CG>TA | CG>TA | GGGTTTGCA( | GCAGCTT | ACG | ACG | C | T | C>T | C>T_ACG | CG>TA_GCAGCTT | 6 | FALSE | FALSE |
| 127_v.txt | C>T | A/T+1 | G/C-1 | CG>TA | CG>TA | GGTTTGCAG( | CAGCTTC | CGT | GCA | C | A | C>T | C>T_GCA | CG>TA_CAGCTTC | 6 | FALSE | FALSE |
| 127_v.txt | C>T | A/T+1 | G/C-1 | CG>TA | CG>TA | AGGCAGCGT | CGTTAAT | CGC | GCG | C | A | C>T | C>T_GCG | CG>TA_CGTTAAT | 6 | FALSE | FALSE |
| 127_v.txt | C>T | A/T+1 | G/C-1 | CG>TA | CG>TA | ACTGGGCCG | CCGGTAA | CGG | GCC | C | A | C>T | C>T_GCC | CG>TA_CCGGTAA | 6 | FALSE | FALSE |
| 127_v.txt | G>A | A/T+1 | G/C-1 | CG>TA | CG>TA | TGGGCCGGT. | GGTAACC | GCC | GCC | C | T | C>T | C>T_GCC | CG>TA_GGTAACC | 6 | FALSE | FALSE |
| 127_v.txt | C>T | A/T+1 | G/C-1 | CG>TA | CG>TA | CCCCAGCAC( | CACCACA | CGT | GCA | C | A | C>T | C>T_GCA | CG>TA_CACCACA | 6 | FALSE | FALSE |
| 127_v.txt | C>T | A/T+1 | G/C-1 | CG>TA | CG>TA | TGGAAGCGA | CGACCCA | CGC | GCG | C | A | C>T | C>T_GCG | CG>TA_CGACCCA | 6 | FALSE | FALSE |
| 127_v.txt | G>A | A/T+1 | G/C-1 | CG>TA | CG>TA | TTCACCGTA/ | GTAATAT | GCA | GCA | C | T | C>T | C>T_GCA | CG>TA_GTAATAT | 6 | FALSE | FALSE |
| 127_v.txt | C>T | A/T+1 | G/C-1 | CG>TA | CG>TA | TCCATGCCG( | CCGCCAG | CGG | GCC | C | A | C>T | C>T_GCC | CG>TA_CCGCCAG | 6 | FALSE | FALSE |
| 127_v.txt | G>A | A/T+1 | G/C-1 | CG>TA | CG>TA | GACTTGGCC/ | GCCAACT | CCG | CCG | C | T | C>T | C>T_CCG | CG>TA_GCCAACT | 6 | FALSE | FALSE |
| 127_v.txt | C>T | A/T+1 | G/C-1 | CG>TA | CG>TA | ACTTGGCCA/ | CCAACTA | CGG | GCC | C | A | C>T | C>T_GCC | CG>TA_CCAACTA | 6 | FALSE | FALSE |
| 127_v.txt | C>T | A/T+1 | G/C-1 | CG>TA | CG>TA | TGATTGCCC/ | CCCAAGA | CGG | GCC | C | A | C>T | C>T_GCC | CG>TA_CCCAAGA | 6 | FALSE | FALSE |
| 127_v.txt | C>T | A/T+1 | G/C-1 | CG>TA | CG>TA | CATTGGCCG( | CCGCACC | CGG | GCC | C | A | C>T | C>T_GCC | CG>TA_CCGCACC | 6 | FALSE | FALSE |
| 127_v.txt | G>A | A/T+1 | G/C-1 | CG>TA | CG>TA | TTGGCCGCA( | GCACCAA | GCG | GCG | C | T | C>T | C>T_GCG | CG>TA_GCACCAA | 6 | FALSE | FALSE |
| 127_v.txt | C>T | A/T+1 | G/C-1 | CG>TA | CG>TA | CAAATGCTG( | CTGGAGA | CGA | GCT | C | A | C>T | C>T_GCT | CG>TA_CTGGAGA | 6 | FALSE | FALSE |
| 127_v.txt | C>T | A/T+1 | G/C-1 | CG>TA | CG>TA | GGACTGCGT | CGTGACA | CGC | GCG | C | A | C>T | C>T_GCG | CG>TA_CGTGACA | 6 | FALSE | FALSE |
| 127_v.txt | G>A | A/T+1 | G/C-1 | CG>TA | CG>TA | GAATATGCC/ | GCCAAAG | ACG | ACG | C | T | C>T | C>T_ACG | CG>TA_GCCAAAG | 6 | FALSE | FALSE |
| 127_v.txt | C>T | A/T+1 | G/C-1 | CG>TA | CG>TA | AATATGCCA/ | CCAAAGA | CGG | GCC | C | A | C>T | C>T_GCC | CG>TA_CCAAAGA | 6 | FALSE | FALSE |
| 127_v.txt | G>A | A/T+1 | G/C-1 | CG>TA | CG>TA | AAGAGCGCC | GCCCACT | GCG | GCG | C | T | C>T | C>T_GCG | CG>TA_GCCCACT | 6 | FALSE | FALSE |
| 127_v.txt | C>T | A/T+1 | G/C-1 | CG>TA | CG>TA | GGCCGGCGT | CGTTGGT | CGC | GCG | C | A | C>T | C>T_GCG | CG>TA_CGTTGGT | 6 | FALSE | FALSE |
| 127_v.txt | C>T | A/T+1 | G/C-1 | CG>TA | CG>TA | CCTATGCCAT | CCATCCT | CGG | GCC | C | A | C>T | C>T_GCC | CG>TA_CCATCCT | 6 | FALSE | FALSE |
| 127_v.txt | C>T | A/T+1 | G/C-1 | CG>TA | CG>TA | AGAGAACGT | CGTATCC | TGC | ACG | C | A | C>T | C>T_ACG | CG>TA_CGTATCC | 6 | FALSE | FALSE |
| 127_v.txt | G>A | A/T+1 | G/C-1 | CG>TA | CG>TA | GAGAACGTA | GTATCCT | GCA | GCA | C | T | C>T | C>T_GCA | CG>TA_GTATCCT | 6 | FALSE | FALSE |
| 127_v.txt | G>A | A/T+1 | G/C-1 | CG>TA | CG>TA | AAGAGCGCA | GCAATGG | GCG | GCG | C | T | C>T | C>T_GCG | CG>TA_GCAATGG | 6 | FALSE | FALSE |
| 127_v.txt | C>T | A/T+1 | G/C-1 | CG>TA | CG>TA | GCTATGCCTT | CCTTTTT | CGG | GCC | C | A | C>T | C>T_GCC | CG>TA_CCTTTTT | 6 | FALSE | FALSE |
| 127_v.txt | G>C | G/C+1 | G/C-1 | CG>GC | CG>GC | TGGGCCGGT. | GGTAACC | GCC | GCC | C | G | C>G | C>G_GCC | CG>GC_GGTAACC | 6 | FALSE | FALSE |
| 127_v.txt | C>G | G/C+1 | G/C-1 | CG>GC | CG>GC | TAATTACCCA | CCCAAAA | TGG | ACC | C | C | C>G | C>G_ACC | CG>GC_CCCAAAA | 6 | FALSE | FALSE |
| 127_v.txt | C>G | G/C+1 | G/C-1 | CG>GC | CG>GC | ACTTGGCCA/ | CCAACTA | CGG | GCC | C | C | C>G | C>G_GCC | CG>GC_CCAACTA | 6 | FALSE | FALSE |
| 127_v.txt | C>G | G/C+1 | G/C-1 | CG>GC | CG>GC | CATTGGCCG( | CCGCACC | CGG | GCC | C | C | C>G | C>G_GCC | CG>GC_CCGCACC | 6 | FALSE | FALSE |
| 127_v.txt | G>C | G/C+1 | G/C-1 | CG>GC | CG>GC | GAATATGCC/ | GCCAAAG | ACG | ACG | C | G | C>G | C>G_ACG | CG>GC_GCCAAAG | 6 | FALSE | FALSE |
| 127_v.txt | C>G | G/C+1 | G/C-1 | CG>GC | CG>GC | AATATGCCA/ | CCAAAGA | CGG | GCC | C | C | C>G | C>G_GCC | CG>GC_CCAAAGA | 6 | FALSE | FALSE |
| 127_v.txt | G>C | G/C+1 | G/C-1 | CG>GC | CG>GC | GAGAACGTA | GTATCCT | GCA | GCA | C | G | C>G | C>G_GCA | CG>GC_GTATCCT | 6 | FALSE | FALSE |
| 127_v.txt | G>T | A/T+1 | G/C-1 | CG>AT | CG>AT | TGGGCCGGT. | GGTAACC | GCC | GCC | C | A | C>A | C>A_GCC | CG>AT_GGTAACC | 6 | FALSE | FALSE |
| 127_v.txt | C>A | A/T+1 | G/C-1 | CG>AT | CG>AT | GCTATGCCTT | CCTTTTT | CGG | GCC | C | T | C>A | C>A_GCC | CG>AT_CCTTTTT | 6 | FALSE | FALSE |
| 127_v.txt | C>A | A/T+1 | G/C-1 | CG>AT | CG>AT | CTATGCCTTT | CTTTTTT | GGA | CCT | C | T | C>A | C>A_CCT | CG>AT_CTTTTTT | 6 | FALSE | FALSE |
| 127_v.txt | nope | >1bp-ins | >1bpdel | none | MNV | GCTATGCCTT | CCTTTTT | CGG | GCC | CC | none | NA | NA_GCC | MNV_CCTTTTT | 6 | FALSE | FALSE |
| 127_v.txt | nope | A/T+1 | >1bpdel | none | A/T+1 | CCATGCAAA/ | AAAAAAG | GTT | CAA | - | T | NA | NA_CAA | A/T+1_AAAAAAG | 6 | FALSE | FALSE |
| 127_v.txt | nope | A/T+1 | >1bpdel | none | A/T+1 | TTACCCAAA/ | AAAAAAT | GTT | CAA | - | T | NA | NA_CAA | A/T+1_AAAAAAT | 6 | FALSE | FALSE |
| 127_v.txt | nope | A/T+1 | >1bpdel | none | A/T+1 | CTATGCCTTT | CTTTTTT | GGA | CCT | - | T | NA | NA_CCT | A/T+1_CTTTTTT | 6 | FALSE | FALSE |
| 127_v.txt | nope | >1bp-ins | A/T-1 | none | A/T-1 | CATTCCAAA/ | AAAATTT | GTT | GTT | T | none | T>none | T>none_GTT | A/T-1_AAAATTT | 6 | FALSE | FALSE |
| 127_v.txt | nope | >1bp-ins | A/T-1 | none | A/T-1 | TGTCCCAAA/ | AAAAAGT | GTT | GTT | T | none | T>none | T>none_GTT | A/T-1_AAAAAGT | 6 | FALSE | FALSE |
| 127_v.txt | nope | >1bp-ins | A/T-1 | none | A/T-1 | AACAGGAAA | AAAATAG | CTT | CTT | T | none | T>none | T>none_CTT | A/T-1_AAAATAG | 6 | FALSE | FALSE |
| 127_v.txt | nope | >1bp-ins | A/T-1 | none | A/T-1 | ATTGAATTTT | TTTTGGT | TAA | ATT | T | none | nope | nope_ATT | A/T-1_TTTTGGT | 6 | FALSE | FALSE |
| 127_v.txt | nope | >1bp-ins | A/T-1 | none | A/T-1 | TGGTGCAAA. | AAAAGCC | GTT | GTT | T | none | T>none | T>none_GTT | A/T-1_AAAAGCC | 6 | FALSE | FALSE |
| 127_v.txt | nope | >1bp-ins | A/T-1 | none | A/T-1 | CCATGCAAA/ | AAAAAAG | GTT | GTT | T | none | T>none | T>none_GTT | A/T-1_AAAAAAG | 6 | FALSE | FALSE |
| 127_v.txt | nope | >1bp-ins | A/T-1 | none | A/T-1 | ACCACCTTTG | TTTGGTG | GAA | CTT | T | none | nope | nope_CTT | A/T-1_TTTGGTG | 6 | FALSE | FALSE |
| 127_v.txt | nope | >1bp-ins | A/T-1 | none | A/T-1 | CAGGAATTT/ | TTTAGGA | TAA | ATT | T | none | nope | nope_ATT | A/T-1_TTTAGGA | 6 | FALSE | FALSE |
| 127_v.txt | nope | >1bp-ins | A/T-1 | none | A/T-1 | TGTGGCAAA. | AAAACCT | GTT | GTT | T | none | T>none | T>none_GTT | A/T-1_AAAACCT | 6 | FALSE | FALSE |
| 127_v.txt | nope | >1bp-ins | A/T-1 | none | A/T-1 | GGAACTAAA | AAAAGTC | ATT | ATT | T | none | T>none | T>none_ATT | A/T-1_AAAAGTC | 6 | FALSE | FALSE |
| 127_v.txt | nope | >1bp-ins | A/T-1 | none | A/T-1 | AACAACTTTT | TTTTTTG | GAA | CTT | T | none | nope | nope_CTT | A/T-1_TTTTTTG | 6 | FALSE | FALSE |
| 127_v.txt | nope | >1bp-ins | G/C-1 | none | G/C-1 | TTTTCTGGG( | GGGGTTT | ACC | ACC | C | none | C>none | C>none_ACC | G/C-1_GGGGTTT | 6 | FALSE | FALSE |
| 127_v.txt | nope | >1bp-ins | A/T-1 | none | A/T-1 | CCTTCGTTTT | TTTTTAT | CAA | GTT | T | none | nope | nope_GTT | A/T-1_TTTTTAT | 6 | FALSE | FALSE |
| 127_v.txt | nope | >1bp-ins | A/T-1 | none | A/T-1 | TTATCCTTAG | TTAGATA | GAA | CTT | T | none | nope | nope_CTT | A/T-1_TTAGATA | 6 | FALSE | FALSE |
| 127_v.txt | nope | >1bp-ins | G/C-1 | none | G/C-1 | CCTGGACCC( | CCCCAGG | TGG | ACC | C | none | nope | nope_ACC | G/C-1_CCCCAGG | 6 | FALSE | FALSE |
| 127_v.txt | nope | >1bp-ins | A/T-1 | none | A/T-1 | CCATACAAA/ | AAAAACA | GTT | GTT | T | none | T>none | T>none_GTT | A/T-1_AAAAACA | 6 | FALSE | FALSE |
| 127_v.txt | nope | >1bp-ins | G/C-1 | none | G/C-1 | TAATTACCCA | CCCAAAA | TGG | ACC | C | none | nope | nope_ACC | G/C-1_CCCAAAA | 6 | FALSE | FALSE |
| 127_v.txt | nope | >1bp-ins | A/T-1 | none | A/T-1 | TTACCCAAA/ | AAAAAAT | GTT | GTT | T | none | T>none | T>none_GTT | A/T-1_AAAAAAT | 6 | FALSE | FALSE |
| 127_v.txt | nope | >1bp-ins | A/T-1 | none | A/T-1 | ACGTCCAAA/ | AAAATTG | GTT | GTT | T | none | T>none | T>none_GTT | A/T-1_AAAATTG | 6 | FALSE | FALSE |
| 127_v.txt | nope | >1bp-ins | G/C-1 | none | G/C-1 | ACTTGGCCA/ | CCAACTA | CGG | GCC | C | none | nope | nope_GCC | G/C-1_CCAACTA | 6 | FALSE | FALSE |
| 127_v.txt | nope | >1bp-ins | A/T-1 | none | A/T-1 | CAGGGCAAA | AAAAGTG | GTT | GTT | T | none | T>none | T>none_GTT | A/T-1_AAAAGTG | 6 | FALSE | FALSE |
| 127_v.txt | nope | >1bp-ins | A/T-1 | none | A/T-1 | GTGGATAAA | AAACCAA | ATT | ATT | T | none | T>none | T>none_ATT | A/T-1_AAACCAA | 6 | FALSE | FALSE |
| 127_v.txt | nope | >1bp-ins | A/T-1 | none | A/T-1 | CCAATGAAA. | AAAAGAC | CTT | CTT | T | none | T>none | T>none_CTT | A/T-1_AAAAGAC | 6 | FALSE | FALSE |
| 127_v.txt | nope | >1bp-ins | G/C-1 | none | G/C-1 | CACCAAGGG | GGGCAAT | TCC | TCC | C | none | C>none | C>none_TCC | G/C-1_GGGCAAT | 6 | FALSE | FALSE |
| 127_v.txt | nope | >1bp-ins | G/C-1 | none | G/C-1 | GCTATGCCTT | CCTTTTT | CGG | GCC | C | none | nope | nope_GCC | G/C-1_CCTTTTT | 6 | FALSE | FALSE |
| 128_v.txt | A>C | G/C+1 | A/T-1 | TA>GC | TA>GC | TGTCCCAAA/ | AAAAAGT | GTT | GTT | T | G | T>G | T>G_GTT | TA>GC_AAAAAGT | 6 | FALSE | FALSE |
| 128_v.txt | T>G | G/C+1 | A/T-1 | TA>GC | TA>GC | AGTACCTTG/ | TTGAAAT | GAA | CTT | T | C | T>G | T>G_CTT | TA>GC_TTGAAAT | 6 | FALSE | FALSE |
| 128_v.txt | A>C | G/C+1 | A/T-1 | TA>GC | TA>GC | TTACCCAAA/ | AAAAAAT | GTT | GTT | T | G | T>G | T>G_GTT | TA>GC_AAAAAAT | 6 | FALSE | FALSE |
| 128_v.txt | A>C | G/C+1 | A/T-1 | TA>GC | TA>GC | ACGTCCAAA/ | AAAATTG | GTT | GTT | T | G | T>G | T>G_GTT | TA>GC_AAAATTG | 6 | FALSE | FALSE |
| 128_v.txt | A>C | G/C+1 | A/T-1 | TA>GC | TA>GC | CTCCATAGA( | AGAGAAC | ATC | ATC | T | G | T>G | T>G_ATC | TA>GC_AGAGAAC | 6 | FALSE | FALSE |
| 128_v.txt | T>G | G/C+1 | A/T-1 | TA>GC | TA>GC | TATGCCTTTT | TTTTTTT | GAA | CTT | T | C | T>G | T>G_CTT | TA>GC_TTTTTTT | 6 | FALSE | FALSE |
| 128_v.txt | A>G | G/C+1 | A/T-1 | TA>CG | TA>CG | CATTCCAAA/ | AAAATTT | GTT | GTT | T | C | T>C | T>C_GTT | TA>CG_AAAATTT | 6 | FALSE | FALSE |
| 128_v.txt | T>C | G/C+1 | A/T-1 | TA>CG | TA>CG | AGTACCTTG/ | TTGAAAT | GAA | CTT | T | G | T>C | T>C_CTT | TA>CG_TTGAAAT | 6 | FALSE | FALSE |
| 128_v.txt | A>G | G/C+1 | A/T-1 | TA>CG | TA>CG | TTACCCAAA/ | AAAAAAT | GTT | GTT | T | C | T>C | T>C_GTT | TA>CG_AAAAAAT | 6 | FALSE | FALSE |
| 128_v.txt | A>G | G/C+1 | A/T-1 | TA>CG | TA>CG | ACGTCCAAA/ | AAAATTG | GTT | GTT | T | C | T>C | T>C_GTT | TA>CG_AAAATTG | 6 | FALSE | FALSE |
| 128_v.txt | T>C | G/C+1 | A/T-1 | TA>CG | TA>CG | TATGCCTTTT | TTTTTTT | GAA | CTT | T | G | T>C | T>C_CTT | TA>CG_TTTTTTT | 6 | FALSE | FALSE |
| 128_v.txt | T>A | A/T+1 | A/T-1 | TA>AT | TA>AT | AGTACCTTG/ | TTGAAAT | GAA | CTT | T | T | T>A | T>A_CTT | TA>AT_TTGAAAT | 6 | FALSE | FALSE |
| 128_v.txt | C>T | A/T+1 | G/C-1 | CG>TA | CG>TA | AGAATGCGA | CGAAATG | CGC | GCG | C | A | C>T | C>T_GCG | CG>TA_CGAAATG | 6 | FALSE | FALSE |
| 128_v.txt | C>T | A/T+1 | G/C-1 | CG>TA | CG>TA | AAATGGCGT | CGTGGAA | CGC | GCG | C | A | C>T | C>T_GCG | CG>TA_CGTGGAA | 6 | FALSE | FALSE |
| 128_v.txt | G>A | A/T+1 | G/C-1 | CG>TA | CG>TA | TAGGCGGCA | GCAGCAA | CCG | CCG | C | T | C>T | C>T_CCG | CG>TA_GCAGCAA | 6 | FALSE | FALSE |
| 128_v.txt | C>T | A/T+1 | G/C-1 | CG>TA | CG>TA | ATGTGGCCG | CCGCATA | CGG | GCC | C | A | C>T | C>T_GCC | CG>TA_CCGCATA | 6 | FALSE | FALSE |
| 128_v.txt | G>A | A/T+1 | G/C-1 | CG>TA | CG>TA | GTGGCCGCA | GCATAAT | GCG | GCG | C | T | C>T | C>T_GCG | CG>TA_GCATAAT | 6 | FALSE | FALSE |
| 128_v.txt | C>T | A/T+1 | G/C-1 | CG>TA | CG>TA | AATAAGCCA. | CCAAGCC | CGG | GCC | C | A | C>T | C>T_GCC | CG>TA_CCAAGCC | 6 | FALSE | FALSE |
| 128_v.txt | C>T | A/T+1 | G/C-1 | CG>TA | CG>TA | GCCAAGCCG | CCGGGCA | CGG | GCC | C | A | C>T | C>T_GCC | CG>TA_CCGGGCA | 6 | FALSE | FALSE |

| | | | | | | | | | | | | | | | | | | | | | | | | | | | | | | | | | | |
|---|---|---|---|---|---|---|---|---|---|---|---|---|---|---|---|---|---|---|---|---|---|---|---|---|---|---|---|---|---|---|---|---|---|---|
| 128_v.txt | 32042 | 32042-SNV- | SNV | 1 | C | T | 18 | 17096 | 0.10528779 | 51.5555556 | WT_perm | FALSE | ARG | JSY2424 | 2 | WT_perm-Re | GCA | C>T | A/T+1 | G/C-1 | CG>TA | CG>TA | AGCTTGCAT | CATAAAT | CGT | GCA | C | A | C>T | C>T_GCA | CG>TA_CATAAAT | 6 | FALSE | FALSE |
| 128_v.txt | 32092 | 32092-SNV- | SNV | 1 | C | T | 24 | 17096 | 0.14038372 | 63 | WT_perm | FALSE | ARG | JSY2424 | 2 | WT_perm-Re | GCA | C>T | A/T+1 | G/C-1 | CG>TA | CG>TA | AGCCTGCAA | CAACACC | CGT | GCA | C | A | C>T | C>T_GCA | CG>TA_CAACACC | 6 | FALSE | FALSE |
| 128_v.txt | 32165 | 32165-SNV- | SNV | 1 | G | A | 38 | 17096 | 0.22227422 | 47.7368421 | WT_perm | FALSE | ARG | JSY2424 | 2 | WT_perm-Re | CGC | G>A | A/T+1 | G/C-1 | CG>TA | CG>TA | CAAAGCGCC | GCCAAAT | GCG | GCG | C | T | C>T | C>T_GCG | CG>TA_GCCAAAT | 6 | FALSE | FALSE |
| 128_v.txt | 32173 | 32173-SNV- | SNV | 1 | C | T | 18 | 17096 | 0.10528779 | 50.5 | WT_perm | FALSE | ARG | JSY2424 | 2 | WT_perm-Re | GCA | C>T | A/T+1 | G/C-1 | CG>TA | CG>TA | CAAATGCAG | CAGCAGT | CGT | GCA | C | A | C>T | C>T_GCA | CG>TA_CAGCAGT | 6 | FALSE | FALSE |
| 128_v.txt | 32176 | 32176-SNV- | SNV | 1 | C | T | 17 | 17096 | 0.09943847 | 47.0588235 | WT_perm | FALSE | ARG | JSY2424 | 2 | WT_perm-Re | GCA | C>T | A/T+1 | G/C-1 | CG>TA | CG>TA | ATGCAGCAG | CAGTAAC | CGT | GCA | C | A | C>T | C>T_GCA | CG>TA_CAGTAAC | 6 | FALSE | FALSE |
| 128_v.txt | 32271 | 32271-SNV- | SNV | 1 | C | T | 29 | 18794 | 0.15430457 | 34.3793103 | WT_perm | FALSE | ARG | JSY2424 | 2 | WT_perm-Re | ACG | C>T | A/T+1 | G/C-1 | CG>TA | CG>TA | AAAATACGG | CGGGAAC | TGC | ACG | C | A | C>T | C>T_ACG | CG>TA_CGGGAAC | 6 | FALSE | FALSE |
| 128_v.txt | 32299 | 32299-SNV- | SNV | 1 | C | T | 32 | 18794 | 0.17026711 | 55.4375 | WT_perm | FALSE | ARG | JSY2424 | 2 | WT_perm-Re | GCG | C>T | A/T+1 | G/C-1 | CG>TA | CG>TA | AATTTGCGG | CGGCAGA | CGC | GCG | C | A | C>T | C>T_GCG | CG>TA_CGGCAGA | 6 | FALSE | FALSE |
| 128_v.txt | 32344 | 32344-SNV- | SNV | 1 | G | A | 15 | 18794 | 0.07981271 | 59.1333333 | WT_perm | FALSE | ARG | JSY2424 | 2 | WT_perm-Re | GGC | G>A | A/T+1 | G/C-1 | CG>TA | CG>TA | TATGTGGCA | GCAAAAC | CCG | CCG | C | T | C>T | C>T_CCG | CG>TA_GCAAAAC | 6 | FALSE | FALSE |
| 128_v.txt | 32466 | 32466-SNV- | SNV | 1 | C | T | 19 | 18794 | 0.1010961 | 42.5789474 | WT_perm | FALSE | ARG | JSY2424 | 2 | WT_perm-Re | GCC | C>T | A/T+1 | G/C-1 | CG>TA | CG>TA | AGAGAGCCA | CCAATGT | CGG | GCC | C | A | C>T | C>T_GCC | CG>TA_CCAATGT | 6 | FALSE | FALSE |
| 128_v.txt | 32487 | 32487-SNV- | SNV | 1 | C | T | 27 | 18794 | 0.14366287 | 44.962963 | WT_perm | FALSE | ARG | JSY2424 | 2 | WT_perm-Re | ACG | C>T | A/T+1 | G/C-1 | CG>TA | CG>TA | AAGATACGG | CGGAAAA | TGC | ACG | C | A | C>T | C>T_ACG | CG>TA_CGGAAAA | 6 | FALSE | FALSE |
| 128_v.txt | 32535 | 32535-SNV- | SNV | 1 | G | A | 22 | 18794 | 0.11705864 | 37.5441646 | WT_perm | FALSE | ARG | JSY2424 | 2 | WT_perm-Re | TGC | G>A | A/T+1 | G/C-1 | CG>TA | CG>TA | GGGTTTGCA | GCAGCTT | ACG | ACG | C | T | C>T | C>T_ACG | CG>TA_GCAGCTT | 6 | FALSE | FALSE |
| 128_v.txt | 32536 | 32536-SNV- | SNV | 1 | C | T | 30 | 18794 | 0.15962541 | 37.2660196 | WT_perm | FALSE | ARG | JSY2424 | 2 | WT_perm-Re | GCA | C>T | A/T+1 | G/C-1 | CG>TA | CG>TA | GGTTTGCAG | CAGCTTC | CGT | GCA | C | A | C>T | C>T_GCA | CG>TA_CAGCTTC | 6 | FALSE | FALSE |
| 128_v.txt | 32590 | 32590-SNV- | SNV | 1 | C | T | 45 | 23631 | 0.19042783 | 36.2222222 | WT_perm | FALSE | ARG | JSY2424 | 2 | WT_perm-Re | GCG | C>T | A/T+1 | G/C-1 | CG>TA | CG>TA | AGGCAGCGT | CGTTAAT | CGC | GCG | C | A | C>T | C>T_GCG | CG>TA_CGTTAAT | 6 | FALSE | FALSE |
| 128_v.txt | 32700 | 32700-SNV- | SNV | 1 | C | T | 32 | 23631 | 0.13541534 | 59.84375 | WT_perm | FALSE | ARG | JSY2424 | 2 | WT_perm-Re | GCC | C>T | A/T+1 | G/C-1 | CG>TA | CG>TA | ACTGGGCCG | CCGGTAA | CGG | GCC | C | A | C>T | C>T_GCC | CG>TA_CCGGTAA | 6 | FALSE | FALSE |
| 128_v.txt | 32757 | 32757-SNV- | SNV | 1 | G | A | 36 | 23631 | 0.15234226 | 53.4444444 | WT_perm | FALSE | ARG | JSY2424 | 2 | WT_perm-Re | GGC | G>A | A/T+1 | G/C-1 | CG>TA | CG>TA | ATAATGGCT | GCTAAAA | CCG | CCG | C | T | C>T | C>T_CCG | CG>TA_GCTAAAA | 6 | FALSE | FALSE |
| 128_v.txt | 32776 | 32776-SNV- | SNV | 1 | C | T | 23 | 23631 | 0.09732978 | 47.9565217 | WT_perm | FALSE | ARG | JSY2424 | 2 | WT_perm-Re | GCG | C>T | A/T+1 | G/C-1 | CG>TA | CG>TA | TGGAAGCGA | CGACCCA | CGC | GCG | C | A | C>T | C>T_GCG | CG>TA_CGACCCA | 6 | FALSE | FALSE |
| 128_v.txt | 32798 | 32798-SNV- | SNV | 1 | G | A | 47 | 23631 | 0.19889129 | 46.0851064 | WT_perm | FALSE | ARG | JSY2424 | 2 | WT_perm-Re | CGT | G>A | A/T+1 | G/C-1 | CG>TA | CG>TA | TTCACCGTA | GTAATAT | GCA | GCA | C | T | C>T | C>T_GCA | CG>TA_GTAATAT | 6 | FALSE | FALSE |
| 128_v.txt | 32860 | 32860-SNV- | SNV | 1 | C | T | 24 | 23631 | 0.10156151 | 35.6666667 | WT_perm | FALSE | ARG | JSY2424 | 2 | WT_perm-Re | GCC | C>T | A/T+1 | G/C-1 | CG>TA | CG>TA | TCCATGCCG | CCGCCAG | CGG | GCC | C | A | C>T | C>T_GCC | CG>TA_CCGCCAG | 6 | FALSE | FALSE |
| 128_v.txt | 32862 | 32862-SNV- | SNV | 1 | G | A | 23 | 23631 | 0.09732978 | 35.3478261 | WT_perm | FALSE | ARG | JSY2424 | 2 | WT_perm-Re | CGC | G>A | A/T+1 | G/C-1 | CG>TA | CG>TA | CATGCCGCC | GCCAGTG | GCG | GCG | C | T | C>T | C>T_GCG | CG>TA_GCCAGTG | 6 | FALSE | FALSE |
| 128_v.txt | 32900 | 32900-SNV- | SNV | 1 | G | A | 35 | 27289 | 0.12825681 | 35.4272829 | WT_perm | FALSE | ARG | JSY2424 | 2 | WT_perm-Re | GGC | G>A | A/T+1 | G/C-1 | CG>TA | CG>TA | GACTTGGCC | GCCAACT | CCG | CCG | C | T | C>T | C>T_CCG | CG>TA_GCCAACT | 6 | FALSE | FALSE |
| 128_v.txt | 32901 | 32901-SNV- | SNV | 1 | C | T | 52 | 27289 | 0.19055297 | 32.8846154 | WT_perm | FALSE | ARG | JSY2424 | 2 | WT_perm-Re | GCC | C>T | A/T+1 | G/C-1 | CG>TA | CG>TA | ACTTGGCCA | CCAACTA | CGG | GCC | C | A | C>T | C>T_GCC | CG>TA_CCAACTA | 6 | FALSE | FALSE |
| 128_v.txt | 32965 | 32965-SNV- | SNV | 1 | C | T | 32 | 27289 | 0.11726337 | 58.0625 | WT_perm | FALSE | ARG | JSY2424 | 2 | WT_perm-Re | GCC | C>T | A/T+1 | G/C-1 | CG>TA | CG>TA | CATTGGCCG | CCGCACC | CGG | GCC | C | A | C>T | C>T_GCC | CG>TA_CCGCACC | 6 | FALSE | FALSE |
| 128_v.txt | 32967 | 32967-SNV- | SNV | 1 | G | A | 77 | 27289 | 0.28216498 | 58.8961039 | WT_perm | FALSE | ARG | JSY2424 | 2 | WT_perm-Re | CGC | G>A | A/T+1 | G/C-1 | CG>TA | CG>TA | TTGGCCGCA | GCACCAA | GCG | GCG | C | T | C>T | C>T_GCG | CG>TA_GCACCAA | 6 | FALSE | FALSE |
| 128_v.txt | 32977 | 32977-SNV- | SNV | 1 | C | T | 38 | 27289 | 0.13925025 | 58.0789474 | WT_perm | FALSE | ARG | JSY2424 | 2 | WT_perm-Re | GCT | C>T | A/T+1 | G/C-1 | CG>TA | CG>TA | CAAATGCTG | CTGGAGA | CGA | GCT | C | A | C>T | C>T_GCT | CG>TA_CTGGAGA | 6 | FALSE | FALSE |
| 128_v.txt | 33056 | 33056-SNV- | SNV | 1 | C | T | 36 | 27289 | 0.13192129 | 56.3888889 | WT_perm | FALSE | ARG | JSY2424 | 2 | WT_perm-Re | GCG | C>T | A/T+1 | G/C-1 | CG>TA | CG>TA | GGACTGCGT | CGTGACA | CGC | GCG | C | A | C>T | C>T_GCG | CG>TA_CGTGACA | 6 | FALSE | FALSE |
| 128_v.txt | 33070 | 33070-SNV- | SNV | 1 | C | T | 31 | 27289 | 0.11359889 | 51.9032258 | WT_perm | FALSE | ARG | JSY2424 | 2 | WT_perm-Re | GCC | C>T | A/T+1 | G/C-1 | CG>TA | CG>TA | AATATGCCA | CCAAAGA | CGG | GCC | C | A | C>T | C>T_GCC | CG>TA_CCAAAGA | 6 | FALSE | FALSE |
| 128_v.txt | 33104 | 33104-SNV- | SNV | 1 | G | A | 30 | 27289 | 0.10993441 | 39.0666667 | WT_perm | FALSE | ARG | JSY2424 | 2 | WT_perm-Re | CGC | G>A | A/T+1 | G/C-1 | CG>TA | CG>TA | AAGAGCGCC | GCCCACT | GCG | GCG | C | T | C>T | C>T_GCG | CG>TA_GCCCACT | 6 | FALSE | FALSE |
| 128_v.txt | 33114 | 33114-SNV- | SNV | 1 | C | T | 25 | 27289 | 0.09161201 | 42.16 | WT_perm | FALSE | ARG | JSY2424 | 2 | WT_perm-Re | GCC | C>T | A/T+1 | G/C-1 | CG>TA | CG>TA | ACTGGGCCG | CCGGCGT | CGG | GCC | C | A | C>T | C>T_GCC | CG>TA_CCGGCGT | 6 | FALSE | FALSE |
| 128_v.txt | 33118 | 33118-SNV- | SNV | 1 | C | T | 28 | 27289 | 0.10260545 | 40.7480521 | WT_perm | FALSE | ARG | JSY2424 | 2 | WT_perm-Re | GCG | C>T | A/T+1 | G/C-1 | CG>TA | CG>TA | GGCCGGCGT | CGTTGGT | CGC | GCG | C | A | C>T | C>T_GCG | CG>TA_CGTTGGT | 6 | FALSE | FALSE |
| 128_v.txt | 33119 | 33119-SNV- | SNV | 1 | G | A | 25 | 27289 | 0.09161201 | 39.72 | WT_perm | FALSE | ARG | JSY2424 | 2 | WT_perm-Re | CGT | G>A | A/T+1 | G/C-1 | CG>TA | CG>TA | GCCGGCGTT | GTTGGTC | GCA | GCA | C | T | C>T | C>T_GCA | CG>TA_GTTGGTC | 6 | FALSE | FALSE |
| 128_v.txt | 33258 | 33258-SNV- | SNV | 1 | C | T | 30 | 15340 | 0.19556715 | 42.1666667 | WT_perm | FALSE | ARG | JSY2424 | 2 | WT_perm-Re | GCC | C>T | A/T+1 | G/C-1 | CG>TA | CG>TA | CCTATGCCAT | CCATCCT | CGG | GCC | C | A | C>T | C>T_GCC | CG>TA_CCATCCT | 6 | FALSE | FALSE |
| 128_v.txt | 33276 | 33276-SNV- | SNV | 1 | G | A | 32 | 15340 | 0.20860495 | 46.8125 | WT_perm | FALSE | ARG | JSY2424 | 2 | WT_perm-Re | CGT | G>A | A/T+1 | G/C-1 | CG>TA | CG>TA | GAGAACGTA | GTATCCT | GCA | GCA | C | T | C>T | C>T_GCA | CG>TA_GTATCCT | 6 | FALSE | FALSE |
| 128_v.txt | 33284 | 33284-SNV- | SNV | 1 | G | A | 17 | 15340 | 0.11082138 | 46.9411765 | WT_perm | FALSE | ARG | JSY2424 | 2 | WT_perm-Re | CGC | G>A | A/T+1 | G/C-1 | CG>TA | CG>TA | ATCCTCGCC | GCCATTT | GCG | GCG | C | T | C>T | C>T_GCG | CG>TA_GCCATTT | 6 | FALSE | FALSE |
| 128_v.txt | 33309 | 33309-SNV- | SNV | 1 | C | T | 29 | 15340 | 0.18904824 | 53.7241379 | WT_perm | FALSE | ARG | JSY2424 | 2 | WT_perm-Re | GCG | C>T | A/T+1 | G/C-1 | CG>TA | CG>TA | AAAGAGCGC | CGCAATG | CGC | GCG | C | A | C>T | C>T_GCG | CG>TA_CGCAATG | 6 | FALSE | FALSE |
| 128_v.txt | 33310 | 33310-SNV- | SNV | 1 | G | A | 25 | 15339 | 0.16298325 | 57.8373812 | WT_perm | FALSE | ARG | JSY2424 | 2 | WT_perm-Re | CGC | G>A | A/T+1 | G/C-1 | CG>TA | CG>TA | AAGAGCGCA | GCAATGG | GCG | GCG | C | T | C>T | C>T_GCG | CG>TA_GCAATGG | 6 | FALSE | FALSE |
| 128_v.txt | 33392 | 33392-SNV- | SNV | 1 | G | A | 18 | 15340 | 0.11734029 | 57.8333333 | WT_perm | FALSE | ARG | JSY2424 | 2 | WT_perm-Re | CGT | G>A | A/T+1 | G/C-1 | CG>TA | CG>TA | AACGTCGTG | GTGAAAG | GCA | GCA | C | T | C>T | C>T_GCA | CG>TA_GTGAAAG | 6 | FALSE | FALSE |
| 128_v.txt | 33412 | 33412-SNV- | SNV | 1 | G | A | 19 | 15340 | 0.12385919 | 48.5263158 | WT_perm | FALSE | ARG | JSY2424 | 2 | WT_perm-Re | GGC | G>A | A/T+1 | G/C-1 | CG>TA | CG>TA | TGACCGGCT | GCTCATT | CCG | CCG | C | T | C>T | C>T_CCG | CG>TA_GCTCATT | 6 | FALSE | FALSE |
| 128_v.txt | 33474 | 33474-SNV- | SNV | 1 | C | T | 23 | 15339 | 0.14994459 | 33.9565217 | WT_perm | FALSE | ARG | JSY2424 | 2 | WT_perm-Re | GCC | C>T | A/T+1 | G/C-1 | CG>TA | CG>TA | GCTATGCCTT | CCTTTTT | CGG | GCC | C | A | C>T | C>T_GCC | CG>TA_CCTTTTT | 6 | FALSE | FALSE |
| 128_v.txt | 32466 | 32466-SNV- | SNV | 1 | C | G | 2 | 18794 | 0.01064169 | 50 | WT_perm | FALSE | ARG | JSY2424 | 2 | WT_perm-Re | GCC | C>G | G/C+1 | G/C-1 | CG>GC | CG>GC | AGAGAGCCA | CCAATGT | CGG | GCC | C | C | C>G | C>G_GCC | CG>GC_CCAATGT | 6 | FALSE | FALSE |
| 128_v.txt | 32901 | 32901-SNV- | SNV | 1 | C | G | 2 | 27289 | 7.33E-03 | 38 | WT_perm | FALSE | ARG | JSY2424 | 2 | WT_perm-Re | GCC | C>G | G/C+1 | G/C-1 | CG>GC | CG>GC | ACTTGGCCA | CCAACTA | CGG | GCC | C | C | C>G | C>G_GCC | CG>GC_CCAACTA | 6 | FALSE | FALSE |
| 128_v.txt | 33118 | 33118-SNV- | SNV | 1 | C | G | 1 | 27289 | 3.66E-03 | 37.998458 | WT_perm | FALSE | ARG | JSY2424 | 2 | WT_perm-Re | GCG | C>G | G/C+1 | G/C-1 | CG>GC | CG>GC | GGCCGGCGT | CGTTGGT | CGC | GCG | C | C | C>G | C>G_GCG | CG>GC_CGTTGGT | 6 | FALSE | FALSE |
| 128_v.txt | 33276 | 33276-SNV- | SNV | 1 | G | C | 3 | 15340 | 0.01955672 | 45.6666667 | WT_perm | FALSE | ARG | JSY2424 | 2 | WT_perm-Re | CGT | G>C | G/C+1 | G/C-1 | CG>GC | CG>GC | GAGAACGTA | GTATCCT | GCA | GCA | C | G | C>G | C>G_GCA | CG>GC_GTATCCT | 6 | FALSE | FALSE |
| 128_v.txt | 33310 | 33310-SNV- | SNV | 1 | G | C | 2 | 15339 | 0.01303866 | 20.9989233 | WT_perm | FALSE | ARG | JSY2424 | 2 | WT_perm-Re | CGC | G>C | G/C+1 | G/C-1 | CG>GC | CG>GC | AAGAGCGCA | GCAATGG | GCG | GCG | C | G | C>G | C>G_GCG | CG>GC_GCAATGG | 6 | FALSE | FALSE |
| 128_v.txt | 31657 | 31657-SNV- | SNV | 1 | G | T | 68 | 10501 | 0.64755738 | 19.0681121 | WT_perm | FALSE | ARG | JSY2424 | 2 | WT_perm-Re | GGC | G>T | A/T+1 | G/C-1 | CG>AT | CG>AT | GAAATGGCG | GCGTGGA | CCG | CCG | C | A | C>A | C>A_CCG | CG>AT_GCGTGGA | 6 | FALSE | FALSE |
| 128_v.txt | 33309 | 33309-SNV- | SNV | 1 | C | A | 5 | 15340 | 0.03259452 | 19.6 | WT_perm | FALSE | ARG | JSY2424 | 2 | WT_perm-Re | GCG | C>A | A/T+1 | G/C-1 | CG>AT | CG>AT | AAAGAGCGC | CGCAATG | CGC | GCG | C | T | C>A | C>A_GCG | CG>AT_CGCAATG | 6 | FALSE | FALSE |
| 128_v.txt | 33474 | 33474-SNV- | SNV | 1 | C | A | 45 | 15339 | 0.29336984 | 32.8666667 | WT_perm | FALSE | ARG | JSY2424 | 2 | WT_perm-Re | GCC | C>A | A/T+1 | G/C-1 | CG>AT | CG>AT | GCTATGCCTT | CCTTTTT | CGG | GCC | C | T | C>A | C>A_GCC | CG>AT_CCTTTTT | 6 | FALSE | FALSE |
| 128_v.txt | 33475 | 33475-SNV- | SNV | 1 | C | A | 11 | 15339 | 0.07171263 | 34.786381 | WT_perm | FALSE | ARG | JSY2424 | 2 | WT_perm-Re | CCT | C>A | A/T+1 | G/C-1 | CG>AT | CG>AT | CTATGCCTTT | CTTTTTT | GGA | CCT | C | T | C>A | C>A_CCT | CG>AT_CTTTTTT | 6 | FALSE | FALSE |
| 128_v.txt | 32081 | 32081-Insert | Insertion | 1 | - | A | 10 | 17096 | 0.05849322 | 64 | WT_perm | FALSE | ARG | JSY2424 | 2 | WT_perm-Re | CAA | nope | A/T+1 | >1bpdel | none | A/T+1 | CCATGCAAA | AAAAAAG | GTT | CAA | - | T | NA | NA_CAA | A/T+1_AAAAAAG | 6 | FALSE | FALSE |
| 128_v.txt | 32842 | 32842-Insert | Insertion | 1 | - | A | 7 | 23631 | 0.02962211 | 41.5714286 | WT_perm | FALSE | ARG | JSY2424 | 2 | WT_perm-Re | CAA | nope | A/T+1 | >1bpdel | none | A/T+1 | TTACCCAAA | AAAAAAT | GTT | CAA | - | T | NA | NA_CAA | A/T+1_AAAAAAT | 6 | FALSE | FALSE |
| 128_v.txt | 32884 | 32884-Insert | Insertion | 1 | - | A | 2 | 27289 | 7.33E-03 | 38 | WT_perm | FALSE | ARG | JSY2424 | 2 | WT_perm-Re | CAA | nope | A/T+1 | >1bpdel | none | A/T+1 | ACGTCCAAA | AAAATTG | GTT | CAA | - | T | NA | NA_CAA | A/T+1_AAAATTG | 6 | FALSE | FALSE |
| 128_v.txt | 31711 | 31711-Deleti | Deletion | 1 | A | - | 3 | 10501 | 0.02856871 | 63.6666667 | WT_perm | FALSE | ARG | JSY2424 | 2 | WT_perm-Re | CAA | nope | >1bp-ins | A/T-1 | none | A/T-1 | CATTCCAAA | AAAATTT | GTT | GTT | T | none | T>none | T>none_GTT | A/T-1_AAAATTT | 6 | FALSE | FALSE |
| 128_v.txt | 31723 | 31723-Deleti | Deletion | 1 | A | - | 15 | 10501 | 0.14284354 | 61.8666667 | WT_perm | FALSE | ARG | JSY2424 | 2 | WT_perm-Re | CAA | nope | >1bp-ins | A/T-1 | none | A/T-1 | TGTCCCAAA | AAAAAGT | GTT | GTT | T | none | T>none | T>none_GTT | A/T-1_AAAAAGT | 6 | FALSE | FALSE |
| 128_v.txt | 32081 | 32081-Deleti | Deletion | 1 | A | - | 19 | 17096 | 0.11113711 | 61.8421057 | WT_perm | FALSE | ARG | JSY2424 | 2 | WT_perm-Re | CAA | nope | >1bp-ins | A/T-1 | none | A/T-1 | CCATGCAAA | AAAAAAG | GTT | GTT | T | none | T>none | T>none_GTT | A/T-1_AAAAAAG | 6 | FALSE | FALSE |
| 128_v.txt | 32210 | 32210-Deleti | Deletion | 1 | T | - | 6 | 17096 | 0.03509593 | 37.8333333 | WT_perm | FALSE | ARG | JSY2424 | 2 | WT_perm-Re | CTT | nope | >1bp-ins | A/T-1 | none | A/T-1 | ACCACCTTTG | TTTGGTG | GAA | CTT | T | none | nope | nope_CTT | A/T-1_TTTGGTG | 6 | FALSE | FALSE |
| 128_v.txt | 32381 | 32381-Deleti | Deletion | 1 | A | - | 5 | 18794 | 0.02660424 | 64 | WT_perm | FALSE | ARG | JSY2424 | 2 | WT_perm-Re | TAA | nope | >1bp-ins | A/T-1 | none | A/T-1 | AATAATAAA | AAAGGGA | ATT | ATT | T | none | T>none | T>none_ATT | A/T-1_AAAGGGA | 6 | FALSE | FALSE |
| 128_v.txt | 32442 | 32442-Deleti | Deletion | 1 | A | - | 7 | 18794 | 0.03724593 | 59.4285714 | WT_perm | FALSE | ARG | JSY2424 | 2 | WT_perm-Re | TAA | nope | >1bp-ins | A/T-1 | none | A/T-1 | GGAACTAAA | AAAAGTC | ATT | ATT | T | none | T>none | T>none_ATT | A/T-1_AAAAGTC | 6 | FALSE | FALSE |
| 128_v.txt | 32466 | 32466-Deleti | Deletion | 1 | C | - | 2 | 18794 | 0.01064169 | 64 | WT_perm | FALSE | ARG | JSY2424 | 2 | WT_perm-Re | GCC | nope | >1bp-ins | G/C-1 | none | G/C-1 | AGAGAGCCA | CCAATGT | CGG | GCC | C | none | nope | nope_GCC | G/C-1_CCAATGT | 6 | FALSE | FALSE |
| 128_v.txt | 32498 | 32498-Deleti | Deletion | 1 | T | - | 23 | 18794 | 0.12237948 | 45.1304348 | WT_perm | FALSE | ARG | JSY2424 | 2 | WT_perm-Re | CTT | nope | >1bp-ins | A/T-1 | none | A/T-1 | AACAACTTTT | TTTTTTG | GAA | CTT | T | none | nope | nope_CTT | A/T-1_TTTTTTG | 6 | FALSE | FALSE |
| 128_v.txt | 32631 | 32631-Deleti | Deletion | 1 | T | - | 7 | 23631 | 0.02962211 | 39.2857143 | WT_perm | FALSE | ARG | JSY2424 | 2 | WT_perm-Re | GTT | nope | >1bp-ins | A/T-1 | none | A/T-1 | CCTTCGTTTT | TTTTTAT | CAA | GTT | T | none | nope | nope_GTT | A/T-1_TTTTTAT | 6 | FALSE | FALSE |
| 128_v.txt | 32836 | 32836-Deleti | Deletion | 1 | T | - | 7 | 23631 | 0.02962211 | 48.8571429 | WT_perm | FALSE | ARG | JSY2424 | 2 | WT_perm-Re | ATT | nope | >1bp-ins | A/T-1 | none | A/T-1 | TGATAATTA | TTACCCA | TAA | ATT | T | none | nope | nope_ATT | A/T-1_TTACCCA | 6 | FALSE | FALSE |
| 128_v.txt | 32842 | 32842-Deleti | Deletion | 1 | A | - | 25 | 23631 | 0.10579324 | 44.76 | WT_perm | FALSE | ARG | JSY2424 | 2 | WT_perm-Re | CAA | nope | >1bp-ins | A/T-1 | none | A/T-1 | TTACCCAAA | AAAAAAT | GTT | GTT | T | none | T>none | T>none_GTT | A/T-1_AAAAAAT | 6 | FALSE | FALSE |
| 128_v.txt | 32884 | 32884-Deleti | Deletion | 1 | A | - | 7 | 27289 | 0.02565136 | 37.5714286 | WT_perm | FALSE | ARG | JSY2424 | 2 | WT_perm-Re | CAA | nope | >1bp-ins | A/T-1 | none | A/T-1 | ACGTCCAAA | AAAATTG | GTT | GTT | T | none | T>none | T>none_GTT | A/T-1_AAAATTG | 6 | FALSE | FALSE |
| 128_v.txt | 32901 | 32901-Deleti | Deletion | 1 | C | - | 2 | 27289 | 7.33E-03 | 38 | WT_perm | FALSE | ARG | JSY2424 | 2 | WT_perm-Re | GCC | nope | >1bp-ins | G/C-1 | none | G/C-1 | ACTTGGCCA | CCAACTA | CGG | GCC | C | none | nope | nope_GCC | G/C-1_CCAACTA | 6 | FALSE | FALSE |
| 128_v.txt | 32924 | 32924-Deleti | Deletion | 1 | A | - | 18 | 27289 | 0.06596064 | 55 | WT_perm | FALSE | ARG | JSY2424 | 2 | WT_perm-Re | CAA | nope | >1bp-ins | A/T-1 | none | A/T-1 | CAGGGCAAA | AAAAGTG | GTT | GTT | T | none | T>none | T>none_GTT | A/T-1_AAAAGTG | 6 | FALSE | FALSE |
| 128_v.txt | 32992 | 32992-Deleti | Deletion | 1 | T | - | 5 | 27289 | 0.0183224 | 63 | WT_perm | FALSE | ARG | JSY2424 | 2 | WT_perm-Re | CTT | nope | >1bp-ins | A/T-1 | none | A/T-1 | GGAATCTTT | TTTGTGA | GAA | CTT | T | none | nope | nope_CTT | A/T-1_TTTGTGA | 6 | FALSE | FALSE |
| 128_v.txt | 33474 | 33474-Deleti | Deletion | 1 | C | - | 36 | 15339 | 0.23469587 | 29.3055556 | WT_perm | FALSE | ARG | JSY2424 | 2 | WT_perm-Re | GCC | nope | >1bp-ins | G/C-1 | none | G/C-1 | GCTATGCCTT | CCTTTTT | CGG | GCC | C | none | nope | nope_GCC | G/C-1_CCTTTTT | 6 | FALSE | FALSE |
| 1_v.txt | 31723 | 31723-SNV- | SNV | 1 | A | C | 6 | 14984 | 0.04004271 | 54.1676667 | WT_perm | FALSE | ARG | JSY13 | 3 | WT_perm-Re | CAA | A>C | G/C+1 | A/T-1 | TA>GC | TA>GC | TGTCCCAAA | AAAAAGT | GTT | GTT | T | G | T>G | T>G_GTT | TA>GC_AAAAAGT | 6 | FALSE | FALSE |
| 1_v.txt | 32842 | 32842-SNV- | SNV | 1 | A | C | 5 | 26359 | 0.01896885 | 32.3999725 | WT_perm | FALSE | ARG | JSY13 | 3 | WT_perm-Re | CAA | A>C | G/C+1 | A/T-1 | TA>GC | TA>GC | TTACCCAAA | AAAAAAT | GTT | GTT | T | G | T>G | T>G_GTT | TA>GC_AAAAAAT | 6 | FALSE | FALSE |
| 1_v.txt | 32884 | 32884-SNV- | SNV | 1 | A | C | 4 | 19370 | 0.02065049 | 37.75 | WT_perm | FALSE | ARG | JSY13 | 3 | WT_perm-Re | CAA | A>C | G/C+1 | A/T-1 | TA>GC | TA>GC | ACGTCCAAA | AAAATTG | GTT | GTT | T | G | T>G | T>G_GTT | TA>GC_AAAATTG | 6 | FALSE | FALSE |
| 1_v.txt | 32919 | 32919-SNV- | SNV | 1 | A | C | 6 | 19369 | 0.03097734 | 44 | WT_perm | FALSE | ARG | JSY13 | 3 | WT_perm-Re | CAG | A>C | G/C+1 | A/T-1 | TA>GC | TA>GC | AGTTCCAGG | AGGGCAA | GTC | GTC | T | G | T>G | T>G_GTC | TA>GC_AGGGCAA | 6 | FALSE | FALSE |
| 1_v.txt | 33099 | 33099-SNV- | SNV | 1 | A | C | 8 | 19369 | 0.04130311 | 40.25 | WT_perm | FALSE | ARG | JSY13 | 3 | WT_perm-Re | AAG | A>C | G/C+1 | A/T-1 | TA>GC | TA>GC | GATATAAGA | AGAGCGC | TTC | TTC | T | G | T>G | T>G_TTC | TA>GC_AGAGCGC | 6 | FALSE | FALSE |
| 1_v.txt | 33269 | 33269-SNV- | SNV | 1 | A | C | 104 | 12947 | 0.80327489 | 16.1634615 | WT_perm | FALSE | ARG | JSY13 | 3 | WT_perm-Re | TAG | A>C | G/C+1 | A/T-1 | TA>GC | TA>GC | CTCCATAGA | AGAGAAC | ATC | ATC | T | G | T>G | T>G_ATC | TA>GC_AGAGAAC | 6 | FALSE | FALSE |
| 1_v.txt | 32919 | 32919-SNV- | SNV | 1 | A | G | 20 | 19369 | 0.10325778 | 49.3 | WT_perm | FALSE | ARG | JSY13 | 3 | WT_perm-Re | CAG | A>G | G/C+1 | A/T-1 | TA>CG | TA>CG | AGTTCCAGG | AGGGCAA | GTC | GTC | T | C | T>C | T>C_GTC | TA>CG_AGGGCAA | 6 | FALSE | FALSE |
| 1_v.txt | 32919 | 32919-SNV- | SNV | 1 | A | T | 6 | 19369 | 0.03097734 | 46.5 | WT_perm | FALSE | ARG | JSY13 | 3 | WT_perm-Re | CAG | A>T | A/T+1 | A/T-1 | TA>AT | TA>AT | AGTTCCAGG | AGGGCAA | GTC | GTC | T | A | T>A | T>A_GTC | TA>AT_AGGGCAA | 6 | FALSE | FALSE |
| 1_v.txt | 31650 | 31650-SNV- | SNV | 1 | C | T | 30 | 14984 | 0.20021356 | 36.9666667 | WT_perm | FALSE | ARG | JSY13 | 3 | WT_perm-Re | GCG | C>T | A/T+1 | G/C-1 | CG>TA | CG>TA | AGAATGCGA | CGAAATG | CGC | GCG | C | A | C>T | C>T_GCG | CG>TA_CGAAATG | 6 | FALSE | FALSE |
| 1_v.txt | 31658 | 31658-SNV- | SNV | 1 | C | T | 23 | 14984 | 0.15349706 | 32.2607717 | WT_perm | FALSE | ARG | JSY13 | 3 | WT_perm-Re | GCG | C>T | A/T+1 | G/C-1 | CG>TA | CG>TA | AAATGGCGT | CGTGGAA | CGC | GCG | C | A | C>T | C>T_GCG | CG>TA_CGTGGAA | 6 | FALSE | FALSE |
| 1_v.txt | 31778 | 31778-SNV- | SNV | 1 | G | A | 12 | 14984 | 0.08008543 | 57.25 | WT_perm | FALSE | ARG | JSY13 | 3 | WT_perm-Re | CGG | G>A | A/T+1 | G/C-1 | CG>TA | CG>TA | TCTATCGGA | GGAATCG | GCC | GCC | C | T | C>T | C>T_GCC | CG>TA_GGAATCG | 6 | FALSE | FALSE |
| 1_v.txt | 31830 | 31830-SNV- | SNV | 1 | C | T | 21 | 14984 | 0.14014949 | 55.3333333 | WT_perm | FALSE | ARG | JSY13 | 3 | WT_perm-Re | GCA | C>T | A/T+1 | G/C-1 | CG>TA | CG>TA | AATATGCATT | CATTGAA | CGT | GCA | C | A | C>T | C>T_GCA | CG>TA_CATTGAA | 6 | FALSE | FALSE |
| 1_v.txt | 31881 | 31881-SNV- | SNV | 1 | G | A | 19 | 14984 | 0.12680192 | 46.0526316 | WT_perm | FALSE | ARG | JSY13 | 3 | WT_perm-Re | GGC | G>A | A/T+1 | G/C-1 | CG>TA | CG>TA | TAGGCGGCA | GCAGCAA | CCG | CCG | C | T | C>T | C>T_CCG | CG>TA_GCAGCAA | 6 | FALSE | FALSE |
| 1_v.txt | 31947 | 31947-SNV- | SNV | 1 | G | A | 23 | 22011 | 0.10449321 | 37.2173913 | WT_perm | FALSE | ARG | JSY13 | 3 | WT_perm-Re | CGT | G>A | A/T+1 | G/C-1 | CG>TA | CG>TA | ATGATCGTC | GTCATAA | GCA | GCA | C | T | C>T | C>T_GCA | CG>TA_GTCATAA | 6 | FALSE | FALSE |
| 1_v.txt | 31960 | 31960-SNV- | SNV | 1 | C | T | 23 | 22027 | 0.10441731 | 37.2173913 | WT_perm | FALSE | ARG | JSY13 | 3 | WT_perm-Re | GCC | C>T | A/T+1 | G/C-1 | CG>TA | CG>TA | ATGTGGCCG | CCGCATA | CGG | GCC | C | A | C>T | C>T_GCC | CG>TA_CCGCATA | 6 | FALSE | FALSE |
| 1_v.txt | 31962 | 31962-SNV- | SNV | 1 | G | A | 53 | 22027 | 0.24061379 | 36.3207547 | WT_perm | FALSE | ARG | JSY13 | 3 | WT_perm-Re | CGC | G>A | A/T+1 | G/C-1 | CG>TA | CG>TA | GTGGCCGCA | GCATAAT | GCG | GCG | C | T | C>T | C>T_GCG | CG>TA_GCATAAT | 6 | FALSE | FALSE |
| 1_v.txt | 31972 | 31972-SNV- | SNV | 1 | C | T | 23 | 22027 | 0.10441731 | 36.0869565 | WT_perm | FALSE | ARG | JSY13 | 3 | WT_perm-Re | GCC | C>T | A/T+1 | G/C-1 | CG>TA | CG>TA | AATAAGCCA | CCAAGCC | CGG | GCC | C | A | C>T | C>T_GCC | CG>TA_CCAAGCC | 6 | FALSE | FALSE |
| 1_v.txt | 31981 | 31981-SNV- | SNV | 1 | G | A | 28 | 22027 | 0.12711672 | 50.2142857 | WT_perm | FALSE | ARG | JSY13 | 3 | WT_perm-Re | GGC | G>A | A/T+1 | G/C-1 | CG>TA | CG>TA | AGCCGGGCA | GCATTAA | CCG | CCG | C | T | C>T | C>T_CCG | CG>TA_GCATTAA | 6 | FALSE | FALSE |
| 1_v.txt | 32022 | 32022-SNV- | SNV | 1 | C | T | 24 | 22027 | 0.10895719 | 55.2916667 | WT_perm | FALSE | ARG | JSY13 | 3 | WT_perm-Re | GCC | C>T | A/T+1 | G/C-1 | CG>TA | CG>TA | GAGATGCCA | CCACGGT | CGG | GCC | C | A | C>T | C>T_GCC | CG>TA_CCACGGT | 6 | FALSE | FALSE |
| 1_v.txt | 32057 | 32057-SNV- | SNV | 1 | C | T | 24 | 22027 | 0.10895719 | 62.4583333 | WT_perm | FALSE | ARG | JSY13 | 3 | WT_perm-Re | GCG | C>T | A/T+1 | G/C-1 | CG>TA | CG>TA | GATGTGCGA | CGAGATT | CGC | GCG | C | A | C>T | C>T_GCG | CG>TA_CGAGATT | 6 | FALSE | FALSE |
| 1_v.txt | 32165 | 32165-SNV- | SNV | 1 | G | A | 30 | 22027 | 0.13619649 | 47.0333333 | WT_perm | FALSE | ARG | JSY13 | 3 | WT_perm-Re | CGC | G>A | A/T+1 | G/C-1 | CG>TA | CG>TA | CAAAGCGCC | GCCAAAT | GCG | GCG | C | T | C>T | C>T_GCG | CG>TA_GCCAAAT | 6 | FALSE | FALSE |
| 1_v.txt | 32173 | 32173-SNV- | SNV | 1 | C | T | 25 | 22027 | 0.11349707 | 52.44 | WT_perm | FALSE | ARG | JSY13 | 3 | WT_perm-Re | GCA | C>T | A/T+1 | G/C-1 | CG>TA | CG>TA | CAAATGCAG | CAGCAGT | CGT | GCA | C | A | C>T | C>T_GCA | CG>TA_CAGCAGT | 6 | FALSE | FALSE |
| 1_v.txt | 32176 | 32176-SNV- | SNV | 1 | C | T | 27 | 22027 | 0.12257684 | 48.8518519 | WT_perm | FALSE | ARG | JSY13 | 3 | WT_perm-Re | GCA | C>T | A/T+1 | G/C-1 | CG>TA | CG>TA | ATGCAGCAG | CAGTAAC | CGT | GCA | C | A | C>T | C>T_GCA | CG>TA_CAGTAAC | 6 | FALSE | FALSE |
| 1_v.txt | 32271 | 32271-SNV- | SNV | 1 | C | T | 41 | 23533 | 0.17422343 | 33.4878049 | WT_perm | FALSE | ARG | JSY13 | 3 | WT_perm-Re | ACG | C>T | A/T+1 | G/C-1 | CG>TA | CG>TA | AAAATACGG | CGGGAAC | TGC | ACG | C | A | C>T | C>T_ACG | CG>TA_CGGGAAC | 6 | FALSE | FALSE |
| 1_v.txt | 32299 | 32299-SNV- | SNV | 1 | C | T | 27 | 23533 | 0.1147325 | 50.1111111 | WT_perm | FALSE | ARG | JSY13 | 3 | WT_perm-Re | GCG | C>T | A/T+1 | G/C-1 | CG>TA | CG>TA | AATTTGCGG | CGGCAGA | CGC | GCG | C | A | C>T | C>T_GCG | CG>TA_CGGCAGA | 6 | FALSE | FALSE |
| 1_v.txt | 32466 | 32466-SNV- | SNV | 1 | C | T | 39 | 23533 | 0.16572473 | 47.0769231 | WT_perm | FALSE | ARG | JSY13 | 3 | WT_perm-Re | GCC | C>T | A/T+1 | G/C-1 | CG>TA | CG>TA | AGAGAGCCA | CCAATGT | CGG | GCC | C | A | C>T | C>T_GCC | CG>TA_CCAATGT | 6 | FALSE | FALSE |
| 1_v.txt | 32535 | 32535-SNV- | SNV | 1 | G | A | 35 | 23533 | 0.14872732 | 35.2493721 | WT_perm | FALSE | ARG | JSY13 | 3 | WT_perm-Re | TGC | G>A | A/T+1 | G/C-1 | CG>TA | CG>TA | GGGTTTGCA | GCAGCTT | ACG | ACG | C | T | C>T | C>T_ACG | CG>TA_GCAGCTT | 6 | FALSE | FALSE |
| 1_v.txt | 32536 | 32536-SNV- | SNV | 1 | C | T | 34 | 23533 | 0.14447797 | 37.058351 | WT_perm | FALSE | ARG | JSY13 | 3 | WT_perm-Re | GCA | C>T | A/T+1 | G/C-1 | CG>TA | CG>TA | GGTTTGCAG | CAGCTTC | CGT | GCA | C | A | C>T | C>T_GCA | CG>TA_CAGCTTC | 6 | FALSE | FALSE |
| 1_v.txt | 32587 | 32587-SNV- | SNV | 1 | C | T | 27 | 26359 | 0.10243181 | 33.9259259 | WT_perm | FALSE | ARG | JSY13 | 3 | WT_perm-Re | GCA | C>T | A/T+1 | G/C-1 | CG>TA | CG>TA | TGAAGGCAG | CAGCGTT | CGT | GCA | C | A | C>T | C>T_GCA | CG>TA_CAGCGTT | 6 | FALSE | FALSE |
| 1_v.txt | 32590 | 32590-SNV- | SNV | 1 | C | T | 54 | 26359 | 0.20486361 | 35.462963 | WT_perm | FALSE | ARG | JSY13 | 3 | WT_perm-Re | GCG | C>T | A/T+1 | G/C-1 | CG>TA | CG>TA | AGGCAGCGT | CGTTAAT | CGC | GCG | C | A | C>T | C>T_GCG | CG>TA_CGTTAAT | 6 | FALSE | FALSE |
| 1_v.txt | 32700 | 32700-SNV- | SNV | 1 | C | T | 38 | 26359 | 0.14416328 | 58.7105263 | WT_perm | FALSE | ARG | JSY13 | 3 | WT_perm-Re | GCC | C>T | A/T+1 | G/C-1 | CG>TA | CG>TA | ACTGGGCCG | CCGGTAA | CGG | GCC | C | A | C>T | C>T_GCC | CG>TA_CCGGTAA | 6 | FALSE | FALSE |
| 1_v.txt | 32757 | 32757-SNV- | SNV | 1 | G | A | 23 | 26359 | 0.08725673 | 50.0869565 | WT_perm | FALSE | ARG | JSY13 | 3 | WT_perm-Re | GGC | G>A | A/T+1 | G/C-1 | CG>TA | CG>TA | ATAATGGCT | GCTAAAA | CCG | CCG | C | T | C>T | C>T_CCG | CG>TA_GCTAAAA | 6 | FALSE | FALSE |
| 1_v.txt | 32798 | 32798-SNV- | SNV | 1 | G | A | 50 | 26359 | 0.18968853 | 43.4 | WT_perm | FALSE | ARG | JSY13 | 3 | WT_perm-Re | CGT | G>A | A/T+1 | G/C-1 | CG>TA | CG>TA | TTCACCGTA | GTAATAT | GCA | GCA | C | T | C>T | C>T_GCA | CG>TA_GTAATAT | 6 | FALSE | FALSE |
| 1_v.txt | 32860 | 32860-SNV- | SNV | 1 | C | T | 38 | 26358 | 0.14416875 | 35.8421053 | WT_perm | FALSE | ARG | JSY13 | 3 | WT_perm-Re | GCC | C>T | A/T+1 | G/C-1 | CG>TA | CG>TA | TCCATGCCG | CCGCCAG | CGG | GCC | C | A | C>T | C>T_GCC | CG>TA_CCGCCAG | 6 | FALSE | FALSE |
| 1_v.txt | 32900 | 32900-SNV- | SNV | 1 | G | A | 23 | 19369 | 0.11874645 | 35.255845 | WT_perm | FALSE | ARG | JSY13 | 3 | WT_perm-Re | GGC | G>A | A/T+1 | G/C-1 | CG>TA | CG>TA | GACTTGGCC | GCCAACT | CCG | CCG | C | T | C>T | C>T_CCG | CG>TA_GCCAACT | 6 | FALSE | FALSE |
| 1_v.txt | 32901 | 32901-SNV- | SNV | 1 | C | T | 43 | 19369 | 0.22200423 | 28.3023256 | WT_perm | FALSE | ARG | JSY13 | 3 | WT_perm-Re | GCC | C>T | A/T+1 | G/C-1 | CG>TA | CG>TA | ACTTGGCCA | CCAACTA | CGG | GCC | C | A | C>T | C>T_GCC | CG>TA_CCAACTA | 6 | FALSE | FALSE |
| 1_v.txt | 32935 | 32935-SNV- | SNV | 1 | C | T | 25 | 19369 | 0.12907223 | 46.6 | WT_perm | FALSE | ARG | JSY13 | 3 | WT_perm-Re | GCC | C>T | A/T+1 | G/C-1 | CG>TA | CG>TA | TGATTGCCC | CCCAAGA | CGG | GCC | C | A | C>T | C>T_GCC | CG>TA_CCCAAGA | 6 | FALSE | FALSE |
| 1_v.txt | 32965 | 32965-SNV- | SNV | 1 | C | T | 25 | 19369 | 0.12907223 | 53.64 | WT_perm | FALSE | ARG | JSY13 | 3 | WT_perm-Re | GCC | C>T | A/T+1 | G/C-1 | CG>TA | CG>TA | CATTGGCCG | CCGCACC | CGG | GCC | C | A | C>T | C>T_GCC | CG>TA_CCGCACC | 6 | FALSE | FALSE |
| 1_v.txt | 32967 | 32967-SNV- | SNV | 1 | G | A | 48 | 19369 | 0.24781868 | 57.7708333 | WT_perm | FALSE | ARG | JSY13 | 3 | WT_perm-Re | CGC | G>A | A/T+1 | G/C-1 | CG>TA | CG>TA | TTGGCCGCA | GCACCAA | GCG | GCG | C | T | C>T | C>T_GCG | CG>TA_GCACCAA | 6 | FALSE | FALSE |
| 1_v.txt | 33056 | 33056-SNV- | SNV | 1 | C | T | 35 | 19369 | 0.18070112 | 56.6857143 | WT_perm | FALSE | ARG | JSY13 | 3 | WT_perm-Re | GCG | C>T | A/T+1 | G/C-1 | CG>TA | CG>TA | GGACTGCGT | CGTGACA | CGC | GCG | C | A | C>T | C>T_GCG | CG>TA_CGTGACA | 6 | FALSE | FALSE |
| 1_v.txt | 33069 | 33069-SNV- | SNV | 1 | G | A | 20 | 19369 | 0.10325778 | 54.3 | WT_perm | FALSE | ARG | JSY13 | 3 | WT_perm-Re | TGC | G>A | A/T+1 | G/C-1 | CG>TA | CG>TA | GAATATGCC | GCCAAAG | ACG | ACG | C | T | C>T | C>T_ACG | CG>TA_GCCAAAG | 6 | FALSE | FALSE |
| 1_v.txt | 33104 | 33104-SNV- | SNV | 1 | G | A | 24 | 19369 | 0.12390934 | 40.125 | WT_perm | FALSE | ARG | JSY13 | 3 | WT_perm-Re | CGC | G>A | A/T+1 | G/C-1 | CG>TA | CG>TA | AAGAGCGCC | GCCCACT | GCG | GCG | C | T | C>T | C>T_GCG | CG>TA_GCCCACT | 6 | FALSE | FALSE |
| 1_v.txt | 33113 | 33113-SNV- | SNV | 1 | G | A | 20 | 19369 | 0.10325778 | 39.199931 | WT_perm | FALSE | ARG | JSY13 | 3 | WT_perm-Re | GGC | G>A | A/T+1 | G/C-1 | CG>TA | CG>TA | CACTGGGCC | GCCGGCG | CCG | CCG | C | T | C>T | C>T_CCG | CG>TA_GCCGGCG | 6 | FALSE | FALSE |
| 1_v.txt | 33114 | 33114-SNV- | SNV | 1 | C | T | 20 | 19369 | 0.10325778 | 42.9996434 | WT_perm | FALSE | ARG | JSY13 | 3 | WT_perm-Re | GCC | C>T | A/T+1 | G/C-1 | CG>TA | CG>TA | ACTGGGCCG | CCGGCGT | CGG | GCC | C | A | C>T | C>T_GCC | CG>TA_CCGGCGT | 6 | FALSE | FALSE |
| 1_v.txt | 33118 | 33118-SNV- | SNV | 1 | C | T | 20 | 19369 | 0.10325778 | 35.85 | WT_perm | FALSE | ARG | JSY13 | 3 | WT_perm-Re | GCG | C>T | A/T+1 | G/C-1 | CG>TA | CG>TA | GGCCGGCGT | CGTTGGT | CGC | GCG | C | A | C>T | C>T_GCG | CG>TA_CGTTGGT | 6 | FALSE | FALSE |
| 1_v.txt | 33276 | 33276-SNV- | SNV | 1 | G | A | 23 | 12947 | 0.17764733 | 45.7391304 | WT_perm | FALSE | ARG | JSY13 | 3 | WT_perm-Re | CGT | G>A | A/T+1 | G/C-1 | CG>TA | CG>TA | GAGAACGTA | GTATCCT | GCA | GCA | C | T | C>T | C>T_GCA | CG>TA_GTATCCT | 6 | FALSE | FALSE |
| 1_v.txt | 33310 | 33310-SNV- | SNV | 1 | G | A | 22 | 12947 | 0.16992353 | 54.2727273 | WT_perm | FALSE | ARG | JSY13 | 3 | WT_perm-Re | CGC | G>A | A/T+1 | G/C-1 | CG>TA | CG>TA | AAGAGCGCA | GCAATGG | GCG | GCG | C | T | C>T | C>T_GCG | CG>TA_GCAATGG | 6 | FALSE | FALSE |
| 1_v.txt | 33392 | 33392-SNV- | SNV | 1 | G | A | 22 | 12947 | 0.16992353 | 47.9090909 | WT_perm | FALSE | ARG | JSY13 | 3 | WT_perm-Re | CGT | G>A | A/T+1 | G/C-1 | CG>TA | CG>TA | AACGTCGTG | GTGAAAG | GCA | GCA | C | T | C>T | C>T_GCA | CG>TA_GTGAAAG | 6 | FALSE | FALSE |
| 1_v.txt | 33412 | 33412-SNV- | SNV | 1 | G | A | 22 | 12947 | 0.16992353 | 48.4545455 | WT_perm | FALSE | ARG | JSY13 | 3 | WT_perm-Re | GGC | G>A | A/T+1 | G/C-1 | CG>TA | CG>TA | TGACCGGCT | GCTCATT | CCG | CCG | C | T | C>T | C>T_CCG | CG>TA_GCTCATT | 6 | FALSE | FALSE |
| 1_v.txt | 33474 | 33474-SNV- | SNV | 1 | C | T | 9 | 12947 | 0.06951417 | 37.3077978 | WT_perm | FALSE | ARG | JSY13 | 3 | WT_perm-Re | GCC | C>T | A/T+1 | G/C-1 | CG>TA | CG>TA | GCTATGCCTT | CCTTTTT | CGG | GCC | C | A | C>T | C>T_GCC | CG>TA_CCTTTTT | 6 | FALSE | FALSE |
| 1_v.txt | 31778 | 31778-SNV- | SNV | 1 | G | C | 2 | 14984 | 0.01334757 | 62.5 | WT_perm | FALSE | ARG | JSY13 | 3 | WT_perm-Re | CGG | G>C | G/C+1 | G/C-1 | CG>GC | CG>GC | TCTATCGGA | GGAATCG | GCC | GCC | C | G | C>G | C>G_GCC | CG>GC_GGAATCG | 6 | FALSE | FALSE |
| 1_v.txt | 31881 | 31881-SNV- | SNV | 1 | G | C | 2 | 14984 | 0.01334757 | 38 | WT_perm | FALSE | ARG | JSY13 | 3 | WT_perm-Re | GGC | G>C | G/C+1 | G/C-1 | CG>GC | CG>GC | TAGGCGGCA | GCAGCAA | CCG | CCG | C | G | C>G | C>G_CCG | CG>GC_GCAGCAA | 6 | FALSE | FALSE |
| 1_v.txt | 32860 | 32860-SNV- | SNV | 1 | C | G | 4 | 26358 | 0.01517566 | 35 | WT_perm | FALSE | ARG | JSY13 | 3 | WT_perm-Re | GCC | C>G | G/C+1 | G/C-1 | CG>GC | CG>GC | TCCATGCCG | CCGCCAG | CGG | GCC | C | C | C>G | C>G_GCC | CG>GC_CCGCCAG | 6 | FALSE | FALSE |

| | | | | | | | | | | | | | | | | | | | | | | | | | | | | | | | | | | |
|---|---|---|---|---|---|---|---|---|---|---|---|---|---|---|---|---|---|---|---|---|---|---|---|---|---|---|---|---|---|---|---|---|---|---|
| 1_v.txt | 33114 | 33114-SNV- | SNV | 1 | C | G | 2 | 19369 | 0.01032578 | 50.996701 | WT_perm | FALSE | ARG | JSY13 | 3 | WT_perm-Re | GCC | C>G | G/C+1 | G/C-1 | CG>GC | CG>GC | ACTGGGCCG | CCGGCGT | CGG | GCC | C | C | C>G | C>G_GCC | CG>GC_CCGGCGT | 6 | FALSE | FALSE |
| 1_v.txt | 31657 | 31657-SNV- | SNV | 1 | G | T | 138 | 14984 | 0.92098238 | 20.0622423 | WT_perm | FALSE | ARG | JSY13 | 3 | WT_perm-Re | GGC | G>T | A/T+1 | G/C-1 | CG>AT | CG>AT | GAAATGGCG | GCGTGGA | CCG | CCG | C | A | C>A | C>A_CCG | CG>AT_GCGTGGA | 6 | FALSE | FALSE |
| 1_v.txt | 33474 | 33474-SNV- | SNV | 1 | C | A | 50 | 12947 | 0.38618985 | 34.7860471 | WT_perm | FALSE | ARG | JSY13 | 3 | WT_perm-Re | GCC | C>A | A/T+1 | G/C-1 | CG>AT | CG>AT | GCTATGCCTT | CCTTTTT | CGG | GCC | C | T | C>A | C>A_GCC | CG>AT_CCTTTTT | 6 | FALSE | FALSE |
| 1_v.txt | 33475 | 33475-SNV- | SNV | 1 | C | A | 13 | 12947 | 0.10040936 | 34.7692308 | WT_perm | FALSE | ARG | JSY13 | 3 | WT_perm-Re | CCT | C>A | A/T+1 | G/C-1 | CG>AT | CG>AT | CTATGCCTTT | CTTTTTT | GGA | CCT | C | T | C>A | C>A_CCT | CG>AT_CTTTTTT | 6 | FALSE | FALSE |
| 1_v.txt | 31657 | 31657-MNV- | MNV | 2 | GC | TT | 2 | 14984 | 0.01334757 | 26.999999 | WT_perm | FALSE | ARG | JSY13 | 3 | WT_perm-Re | GGC | nope | >1bp-ins | >1bpdel | none | MNV | GAAATGGCG | GCGTGGA | CCG | GGC | GC | none | NA | NA_GGC | MNV_GCGTGGA | 6 | FALSE | FALSE |
| 1_v.txt | 31723 | 31723-Insert | Insertion | 1 | - | A | 2 | 14984 | 0.01334757 | 64 | WT_perm | FALSE | ARG | JSY13 | 3 | WT_perm-Re | CAA | nope | A/T+1 | >1bpdel | none | A/T+1 | TGTCCCAAA | AAAAAGT | GTT | CAA | - | T | NA | NA_CAA | A/T+1_AAAAAGT | 6 | FALSE | FALSE |
| 1_v.txt | 31952 | 31952-Insert | Insertion | 1 | - | A | 2 | 22027 | 9.08E-03 | 38 | WT_perm | FALSE | ARG | JSY13 | 3 | WT_perm-Re | TAA | nope | A/T+1 | >1bpdel | none | A/T+1 | CGTCATAAA | AAATGTG | ATT | TAA | - | T | NA | NA_TAA | A/T+1_AAATGTG | 6 | FALSE | FALSE |
| 1_v.txt | 32081 | 32081-Insert | Insertion | 1 | - | A | 16 | 22027 | 0.07263813 | 64 | WT_perm | FALSE | ARG | JSY13 | 3 | WT_perm-Re | CAA | nope | A/T+1 | >1bpdel | none | A/T+1 | CCATGCAAA | AAAAAAG | GTT | CAA | - | T | NA | NA_CAA | A/T+1_AAAAAAG | 6 | FALSE | FALSE |
| 1_v.txt | 32498 | 32498-Insert | Insertion | 1 | - | T | 2 | 23533 | 8.50E-03 | 51 | WT_perm | FALSE | ARG | JSY13 | 3 | WT_perm-Re | CTT | nope | A/T+1 | >1bpdel | none | A/T+1 | AACAACTTTT | TTTTTTG | GAA | CTT | - | A | NA | NA_CTT | A/T+1_TTTTTTG | 6 | FALSE | FALSE |
| 1_v.txt | 32842 | 32842-Insert | Insertion | 1 | - | A | 11 | 26359 | 0.04173148 | 42.5454546 | WT_perm | FALSE | ARG | JSY13 | 3 | WT_perm-Re | CAA | nope | A/T+1 | >1bpdel | none | A/T+1 | TTACCCAAA | AAAAAAT | GTT | CAA | - | T | NA | NA_CAA | A/T+1_AAAAAAT | 6 | FALSE | FALSE |
| 1_v.txt | 31723 | 31723-Deleti | Deletion | 1 | A | - | 18 | 14984 | 0.12012814 | 61.5555562 | WT_perm | FALSE | ARG | JSY13 | 3 | WT_perm-Re | CAA | nope | >1bp-ins | A/T-1 | none | A/T-1 | TGTCCCAAA | AAAAAGT | GTT | GTT | T | none | T>none | T>none_GTT | A/T-1_AAAAAGT | 6 | FALSE | FALSE |
| 1_v.txt | 31904 | 31904-Deleti | Deletion | 1 | T | - | 9 | 14984 | 0.06006407 | 44.3333333 | WT_perm | FALSE | ARG | JSY13 | 3 | WT_perm-Re | ATT | nope | >1bp-ins | A/T-1 | none | A/T-1 | ATTGAATTTT | TTTTGGT | TAA | ATT | T | none | nope | nope_ATT | A/T-1_TTTTGGT | 6 | FALSE | FALSE |
| 1_v.txt | 31913 | 31913-Deleti | Deletion | 1 | A | - | 11 | 14984 | 0.07341164 | 40 | WT_perm | FALSE | ARG | JSY13 | 3 | WT_perm-Re | CAA | nope | >1bp-ins | A/T-1 | none | A/T-1 | TGGTGCAAA | AAAAGCC | GTT | GTT | T | none | T>none | T>none_GTT | A/T-1_AAAAGCC | 6 | FALSE | FALSE |
| 1_v.txt | 31972 | 31972-Deleti | Deletion | 1 | C | - | 4 | 22027 | 0.01815953 | 38 | WT_perm | FALSE | ARG | JSY13 | 3 | WT_perm-Re | GCC | nope | >1bp-ins | G/C-1 | none | G/C-1 | AATAAGCCA | CCAAGCC | CGG | GCC | C | none | nope | nope_GCC | G/C-1_CCAAGCC | 6 | FALSE | FALSE |
| 1_v.txt | 32081 | 32081-Deleti | Deletion | 1 | A | - | 26 | 22027 | 0.11803696 | 61.6538457 | WT_perm | FALSE | ARG | JSY13 | 3 | WT_perm-Re | CAA | nope | >1bp-ins | A/T-1 | none | A/T-1 | CCATGCAAA | AAAAAAG | GTT | GTT | T | none | T>none | T>none_GTT | A/T-1_AAAAAAG | 6 | FALSE | FALSE |
| 1_v.txt | 32210 | 32210-Deleti | Deletion | 1 | T | - | 9 | 22026 | 0.0408608 | 37.4444444 | WT_perm | FALSE | ARG | JSY13 | 3 | WT_perm-Re | CTT | nope | >1bp-ins | A/T-1 | none | A/T-1 | ACCACCTTTG | TTTGGTG | GAA | CTT | T | none | nope | nope_CTT | A/T-1_TTTGGTG | 6 | FALSE | FALSE |
| 1_v.txt | 32346 | 32346-Deleti | Deletion | 1 | A | - | 10 | 23533 | 0.04249352 | 62 | WT_perm | FALSE | ARG | JSY13 | 3 | WT_perm-Re | CAA | nope | >1bp-ins | A/T-1 | none | A/T-1 | TGTGGCAAA | AAAACCT | GTT | GTT | T | none | T>none | T>none_GTT | A/T-1_AAAACCT | 6 | FALSE | FALSE |
| 1_v.txt | 32498 | 32498-Deleti | Deletion | 1 | T | - | 33 | 23533 | 0.14022862 | 45.3333312 | WT_perm | FALSE | ARG | JSY13 | 3 | WT_perm-Re | CTT | nope | >1bp-ins | A/T-1 | none | A/T-1 | AACAACTTTT | TTTTTTG | GAA | CTT | T | none | nope | nope_CTT | A/T-1_TTTTTTG | 6 | FALSE | FALSE |
| 1_v.txt | 32528 | 32528-Deleti | Deletion | 1 | G | - | 5 | 23533 | 0.02124676 | 38 | WT_perm | FALSE | ARG | JSY13 | 3 | WT_perm-Re | TGG | nope | >1bp-ins | G/C-1 | none | G/C-1 | TTTTCTGGG | GGGGTTT | ACC | ACC | C | none | C>none | C>none_ACC | G/C-1_GGGGTTT | 6 | FALSE | FALSE |
| 1_v.txt | 32631 | 32631-Deleti | Deletion | 1 | T | - | 14 | 26359 | 0.05311279 | 48.2142857 | WT_perm | FALSE | ARG | JSY13 | 3 | WT_perm-Re | GTT | nope | >1bp-ins | A/T-1 | none | A/T-1 | CCTTCGTTTT | TTTTTAT | CAA | GTT | T | none | nope | nope_GTT | A/T-1_TTTTTAT | 6 | FALSE | FALSE |
| 1_v.txt | 32658 | 32658-Deleti | Deletion | 1 | C | - | 8 | 26359 | 0.03035017 | 56.125 | WT_perm | FALSE | ARG | JSY13 | 3 | WT_perm-Re | ACC | nope | >1bp-ins | G/C-1 | none | G/C-1 | CCTGGACCC | CCCCAGG | TGG | ACC | C | none | nope | nope_ACC | G/C-1_CCCCAGG | 6 | FALSE | FALSE |
| 1_v.txt | 32728 | 32728-Deleti | Deletion | 1 | A | - | 9 | 26359 | 0.03414394 | 63.6666667 | WT_perm | FALSE | ARG | JSY13 | 3 | WT_perm-Re | CAA | nope | >1bp-ins | A/T-1 | none | A/T-1 | CCATACAAA | AAAAACA | GTT | GTT | T | none | T>none | T>none_GTT | A/T-1_AAAAACA | 6 | FALSE | FALSE |
| 1_v.txt | 32842 | 32842-Deleti | Deletion | 1 | A | - | 37 | 26359 | 0.14036951 | 40.5675673 | WT_perm | FALSE | ARG | JSY13 | 3 | WT_perm-Re | CAA | nope | >1bp-ins | A/T-1 | none | A/T-1 | TTACCCAAA | AAAAAAT | GTT | GTT | T | none | T>none | T>none_GTT | A/T-1_AAAAAAT | 6 | FALSE | FALSE |
| 1_v.txt | 32884 | 32884-Deleti | Deletion | 1 | A | - | 4 | 19370 | 0.02065049 | 32.75 | WT_perm | FALSE | ARG | JSY13 | 3 | WT_perm-Re | CAA | nope | >1bp-ins | A/T-1 | none | A/T-1 | ACGTCCAAA | AAAATTG | GTT | GTT | T | none | T>none | T>none_GTT | A/T-1_AAAATTG | 6 | FALSE | FALSE |
| 1_v.txt | 32924 | 32924-Deleti | Deletion | 1 | A | - | 25 | 19369 | 0.12907223 | 51.68 | WT_perm | FALSE | ARG | JSY13 | 3 | WT_perm-Re | CAA | nope | >1bp-ins | A/T-1 | none | A/T-1 | CAGGGCAAA | AAAAGTG | GTT | GTT | T | none | T>none | T>none_GTT | A/T-1_AAAAGTG | 6 | FALSE | FALSE |
| 1_v.txt | 32935 | 32935-Deleti | Deletion | 1 | C | - | 3 | 19369 | 0.01548867 | 55.3333333 | WT_perm | FALSE | ARG | JSY13 | 3 | WT_perm-Re | GCC | nope | >1bp-ins | G/C-1 | none | G/C-1 | TGATTGCCC | CCCAAGA | CGG | GCC | C | none | nope | nope_GCC | G/C-1_CCCAAGA | 6 | FALSE | FALSE |
| 1_v.txt | 32983 | 32983-Deleti | Deletion | 1 | A | - | 5 | 19369 | 0.02581445 | 64 | WT_perm | FALSE | ARG | JSY13 | 3 | WT_perm-Re | GAA | nope | >1bp-ins | A/T-1 | none | A/T-1 | CTGGAGAAA | AAAGGAA | CTT | CTT | T | none | T>none | T>none_CTT | A/T-1_AAAGGAA | 6 | FALSE | FALSE |
| 1_v.txt | 33000 | 33000-Deleti | Deletion | 1 | A | - | 6 | 19369 | 0.03097734 | 60.6666667 | WT_perm | FALSE | ARG | JSY13 | 3 | WT_perm-Re | GAA | nope | >1bp-ins | A/T-1 | none | A/T-1 | TGTGAGAAA | AAAACTG | CTT | CTT | T | none | T>none | T>none_CTT | A/T-1_AAAACTG | 6 | FALSE | FALSE |
| 1_v.txt | 33114 | 33114-Deleti | Deletion | 1 | C | - | 1 | 19369 | 5.16E-03 | 38 | WT_perm | FALSE | ARG | JSY13 | 3 | WT_perm-Re | GCC | nope | >1bp-ins | G/C-1 | none | G/C-1 | ACTGGGCCG | CCGGCGT | CGG | GCC | C | none | nope | nope_GCC | G/C-1_CCGGCGT | 6 | FALSE | FALSE |
| 1_v.txt | 33147 | 33147-Deleti | Deletion | 1 | A | - | 10 | 19369 | 0.05162889 | 41.1 | WT_perm | FALSE | ARG | JSY13 | 3 | WT_perm-Re | GAA | nope | >1bp-ins | A/T-1 | none | A/T-1 | CCAATGAAA | AAAAGAC | CTT | CTT | T | none | T>none | T>none_CTT | A/T-1_AAAAGAC | 6 | FALSE | FALSE |
| 1_v.txt | 33474 | 33474-Deleti | Deletion | 1 | C | - | 27 | 12947 | 0.20854252 | 27.7777778 | WT_perm | FALSE | ARG | JSY13 | 3 | WT_perm-Re | GCC | nope | >1bp-ins | G/C-1 | none | G/C-1 | GCTATGCCTT | CCTTTTT | CGG | GCC | C | none | nope | nope_GCC | G/C-1_CCTTTTT | 6 | FALSE | FALSE |
| 2_v.txt | 31723 | 31723-SNV- | SNV | 1 | A | C | 4 | 14270 | 0.02803083 | 59.5001149 | WT_perm | FALSE | ARG | JSY13 | 4 | WT_perm-Re | CAA | A>C | G/C+1 | A/T-1 | TA>GC | TA>GC | TGTCCCAAA | AAAAAGT | GTT | GTT | T | G | T>G | T>G_GTT | TA>GC_AAAAAGT | 6 | FALSE | FALSE |
| 2_v.txt | 32124 | 32124-SNV- | SNV | 1 | A | C | 5 | 24125 | 0.02072539 | 56.8 | WT_perm | FALSE | ARG | JSY13 | 4 | WT_perm-Re | AAC | A>C | G/C+1 | A/T-1 | TA>GC | TA>GC | TCGAAAACT | ACTTTGT | TTG | TTG | T | G | T>G | T>G_TTG | TA>GC_ACTTTGT | 6 | FALSE | FALSE |
| 2_v.txt | 32231 | 32231-SNV-T | SNV | 1 | T | G | 4 | 24124 | 0.016581 | 24.5 | WT_perm | FALSE | ARG | JSY13 | 4 | WT_perm-Re | ATT | T>G | G/C+1 | A/T-1 | TA>GC | TA>GC | CAGGAATTT | TTTAGGA | TAA | ATT | T | C | T>G | T>G_ATT | TA>GC_TTTAGGA | 6 | FALSE | FALSE |
| 2_v.txt | 32842 | 32842-SNV- | SNV | 1 | A | C | 7 | 29949 | 0.02337307 | 33.428975 | WT_perm | FALSE | ARG | JSY13 | 4 | WT_perm-Re | CAA | A>C | G/C+1 | A/T-1 | TA>GC | TA>GC | TTACCCAAA | AAAAAAT | GTT | GTT | T | G | T>G | T>G_GTT | TA>GC_AAAAAAT | 6 | FALSE | FALSE |
| 2_v.txt | 33269 | 33269-SNV- | SNV | 1 | A | C | 147 | 19722 | 0.74536051 | 17.2312925 | WT_perm | FALSE | ARG | JSY13 | 4 | WT_perm-Re | TAG | A>C | G/C+1 | A/T-1 | TA>GC | TA>GC | CTCCATAGA | AGAGAAC | ATC | ATC | T | G | T>G | T>G_ATC | TA>GC_AGAGAAC | 6 | FALSE | FALSE |
| 2_v.txt | 32927 | 32927-SNV- | SNV | 1 | A | G | 23 | 22928 | 0.10031403 | 45.0869565 | WT_perm | FALSE | ARG | JSY13 | 4 | WT_perm-Re | AAG | A>G | G/C+1 | A/T-1 | TA>CG | TA>CG | GGCAAAAGT | AGTGATT | TTC | TTC | T | C | T>C | T>C_TTC | TA>CG_AGTGATT | 6 | FALSE | FALSE |
| 2_v.txt | 33150 | 33150-SNV- | SNV | 1 | A | G | 24 | 22928 | 0.10467551 | 34.9166667 | WT_perm | FALSE | ARG | JSY13 | 4 | WT_perm-Re | AAG | A>G | G/C+1 | A/T-1 | TA>CG | TA>CG | ATGAAAAGA | AGACCTG | TTC | TTC | T | C | T>C | T>C_TTC | TA>CG_AGACCTG | 6 | FALSE | FALSE |
| 2_v.txt | 33476 | 33476-SNV-T | SNV | 1 | T | C | 33 | 19721 | 0.16733431 | 30.2730342 | WT_perm | FALSE | ARG | JSY13 | 4 | WT_perm-Re | CTT | T>C | G/C+1 | A/T-1 | TA>CG | TA>CG | TATGCCTTTT | TTTTTTT | GAA | CTT | T | G | T>C | T>C_CTT | TA>CG_TTTTTTT | 6 | FALSE | FALSE |
| 2_v.txt | 33476 | 33476-SNV-T | SNV | 1 | T | A | 3 | 19721 | 0.01521221 | 38 | WT_perm | FALSE | ARG | JSY13 | 4 | WT_perm-Re | CTT | T>A | A/T+1 | A/T-1 | TA>AT | TA>AT | TATGCCTTTT | TTTTTTT | GAA | CTT | T | T | T>A | T>A_CTT | TA>AT_TTTTTTT | 6 | FALSE | FALSE |
| 2_v.txt | 31650 | 31650-SNV- | SNV | 1 | C | T | 35 | 14270 | 0.2452698 | 36.5714286 | WT_perm | FALSE | ARG | JSY13 | 4 | WT_perm-Re | GCG | C>T | A/T+1 | G/C-1 | CG>TA | CG>TA | AGAATGCGA | CGAAATG | CGC | GCG | C | A | C>T | C>T_GCG | CG>TA_CGAAATG | 6 | FALSE | FALSE |
| 2_v.txt | 31658 | 31658-SNV- | SNV | 1 | C | T | 21 | 14270 | 0.14716188 | 33.5718065 | WT_perm | FALSE | ARG | JSY13 | 4 | WT_perm-Re | GCG | C>T | A/T+1 | G/C-1 | CG>TA | CG>TA | AAATGGCGT | CGTGGAA | CGC | GCG | C | A | C>T | C>T_GCG | CG>TA_CGTGGAA | 6 | FALSE | FALSE |
| 2_v.txt | 31830 | 31830-SNV- | SNV | 1 | C | T | 22 | 14270 | 0.15416959 | 55.4090909 | WT_perm | FALSE | ARG | JSY13 | 4 | WT_perm-Re | GCA | C>T | A/T+1 | G/C-1 | CG>TA | CG>TA | AATATGCATT | CATTGAA | CGT | GCA | C | A | C>T | C>T_GCA | CG>TA_CATTGAA | 6 | FALSE | FALSE |
| 2_v.txt | 31879 | 31879-SNV- | SNV | 1 | C | T | 19 | 14270 | 0.13314646 | 44.8421053 | WT_perm | FALSE | ARG | JSY13 | 4 | WT_perm-Re | GCG | C>T | A/T+1 | G/C-1 | CG>TA | CG>TA | TATAGGCGG | CGGCAGC | CGC | GCG | C | A | C>T | C>T_GCG | CG>TA_CGGCAGC | 6 | FALSE | FALSE |
| 2_v.txt | 31881 | 31881-SNV- | SNV | 1 | G | A | 15 | 14270 | 0.10511563 | 41.1793329 | WT_perm | FALSE | ARG | JSY13 | 4 | WT_perm-Re | GGC | G>A | A/T+1 | G/C-1 | CG>TA | CG>TA | TAGGCGGCA | GCAGCAA | CCG | CCG | C | T | C>T | C>T_CCG | CG>TA_GCAGCAA | 6 | FALSE | FALSE |
| 2_v.txt | 31882 | 31882-SNV- | SNV | 1 | C | T | 23 | 14270 | 0.1611773 | 40.9436787 | WT_perm | FALSE | ARG | JSY13 | 4 | WT_perm-Re | GCA | C>T | A/T+1 | G/C-1 | CG>TA | CG>TA | AGGCGGCAG | CAGCAAA | CGT | GCA | C | A | C>T | C>T_GCA | CG>TA_CAGCAAA | 6 | FALSE | FALSE |
| 2_v.txt | 31911 | 31911-SNV- | SNV | 1 | G | A | 20 | 14270 | 0.14015417 | 45.25 | WT_perm | FALSE | ARG | JSY13 | 4 | WT_perm-Re | TGC | G>A | A/T+1 | G/C-1 | CG>TA | CG>TA | TTTGGTGCA | GCAAAAG | ACG | ACG | C | T | C>T | C>T_ACG | CG>TA_GCAAAAG | 6 | FALSE | FALSE |
| 2_v.txt | 31912 | 31912-SNV- | SNV | 1 | C | T | 15 | 14270 | 0.10511563 | 37.000094 | WT_perm | FALSE | ARG | JSY13 | 4 | WT_perm-Re | GCA | C>T | A/T+1 | G/C-1 | CG>TA | CG>TA | TTGGTGCAA | CAAAAGC | CGT | GCA | C | A | C>T | C>T_GCA | CG>TA_CAAAAGC | 6 | FALSE | FALSE |
| 2_v.txt | 31960 | 31960-SNV- | SNV | 1 | C | T | 20 | 24125 | 0.08290155 | 37.15 | WT_perm | FALSE | ARG | JSY13 | 4 | WT_perm-Re | GCC | C>T | A/T+1 | G/C-1 | CG>TA | CG>TA | ATGTGGCCG | CCGCATA | CGG | GCC | C | A | C>T | C>T_GCC | CG>TA_CCGCATA | 6 | FALSE | FALSE |
| 2_v.txt | 31962 | 31962-SNV- | SNV | 1 | G | A | 47 | 24125 | 0.19481865 | 36.0851064 | WT_perm | FALSE | ARG | JSY13 | 4 | WT_perm-Re | CGC | G>A | A/T+1 | G/C-1 | CG>TA | CG>TA | GTGGCCGCA | GCATAAT | GCG | GCG | C | T | C>T | C>T_GCG | CG>TA_GCATAAT | 6 | FALSE | FALSE |
| 2_v.txt | 31972 | 31972-SNV- | SNV | 1 | C | T | 25 | 24125 | 0.10362694 | 35.6 | WT_perm | FALSE | ARG | JSY13 | 4 | WT_perm-Re | GCC | C>T | A/T+1 | G/C-1 | CG>TA | CG>TA | AATAAGCCA | CCAAGCC | CGG | GCC | C | A | C>T | C>T_GCC | CG>TA_CCAAGCC | 6 | FALSE | FALSE |
| 2_v.txt | 31977 | 31977-SNV- | SNV | 1 | C | T | 25 | 24125 | 0.10362694 | 39.96 | WT_perm | FALSE | ARG | JSY13 | 4 | WT_perm-Re | GCC | C>T | A/T+1 | G/C-1 | CG>TA | CG>TA | GCCAAGCCG | CCGGGCA | CGG | GCC | C | A | C>T | C>T_GCC | CG>TA_CCGGGCA | 6 | FALSE | FALSE |
| 2_v.txt | 32057 | 32057-SNV- | SNV | 1 | C | T | 27 | 24125 | 0.1119171 | 63.8518519 | WT_perm | FALSE | ARG | JSY13 | 4 | WT_perm-Re | GCG | C>T | A/T+1 | G/C-1 | CG>TA | CG>TA | GATGTGCGA | CGAGATT | CGC | GCG | C | A | C>T | C>T_GCG | CG>TA_CGAGATT | 6 | FALSE | FALSE |
| 2_v.txt | 32092 | 32092-SNV- | SNV | 1 | C | T | 32 | 24125 | 0.13264249 | 61.625 | WT_perm | FALSE | ARG | JSY13 | 4 | WT_perm-Re | GCA | C>T | A/T+1 | G/C-1 | CG>TA | CG>TA | AGCCTGCAA | CAACACC | CGT | GCA | C | A | C>T | C>T_GCA | CG>TA_CAACACC | 6 | FALSE | FALSE |
| 2_v.txt | 32165 | 32165-SNV- | SNV | 1 | G | A | 41 | 24125 | 0.16994819 | 46.8780488 | WT_perm | FALSE | ARG | JSY13 | 4 | WT_perm-Re | CGC | G>A | A/T+1 | G/C-1 | CG>TA | CG>TA | CAAAGCGCC | GCCAAAT | GCG | GCG | C | T | C>T | C>T_GCG | CG>TA_GCCAAAT | 6 | FALSE | FALSE |
| 2_v.txt | 32173 | 32173-SNV- | SNV | 1 | C | T | 22 | 24125 | 0.09119171 | 44.4090909 | WT_perm | FALSE | ARG | JSY13 | 4 | WT_perm-Re | GCA | C>T | A/T+1 | G/C-1 | CG>TA | CG>TA | CAAATGCAG | CAGCAGT | CGT | GCA | C | A | C>T | C>T_GCA | CG>TA_CAGCAGT | 6 | FALSE | FALSE |
| 2_v.txt | 32176 | 32176-SNV- | SNV | 1 | C | T | 26 | 24125 | 0.10777202 | 46.5384615 | WT_perm | FALSE | ARG | JSY13 | 4 | WT_perm-Re | GCA | C>T | A/T+1 | G/C-1 | CG>TA | CG>TA | ATGCAGCAG | CAGTAAC | CGT | GCA | C | A | C>T | C>T_GCA | CG>TA_CAGTAAC | 6 | FALSE | FALSE |
| 2_v.txt | 32271 | 32271-SNV- | SNV | 1 | C | T | 30 | 20613 | 0.14553922 | 36.6 | WT_perm | FALSE | ARG | JSY13 | 4 | WT_perm-Re | ACG | C>T | A/T+1 | G/C-1 | CG>TA | CG>TA | AAAATACGG | CGGGAAC | TGC | ACG | C | A | C>T | C>T_ACG | CG>TA_CGGGAAC | 6 | FALSE | FALSE |
| 2_v.txt | 32299 | 32299-SNV- | SNV | 1 | C | T | 42 | 20613 | 0.20375491 | 52.0952381 | WT_perm | FALSE | ARG | JSY13 | 4 | WT_perm-Re | GCG | C>T | A/T+1 | G/C-1 | CG>TA | CG>TA | AATTTGCGG | CGGCAGA | CGC | GCG | C | A | C>T | C>T_GCG | CG>TA_CGGCAGA | 6 | FALSE | FALSE |
| 2_v.txt | 32344 | 32344-SNV- | SNV | 1 | G | A | 25 | 20613 | 0.12128269 | 59.2 | WT_perm | FALSE | ARG | JSY13 | 4 | WT_perm-Re | GGC | G>A | A/T+1 | G/C-1 | CG>TA | CG>TA | TATGTGGCA | GCAAAAC | CCG | CCG | C | T | C>T | C>T_CCG | CG>TA_GCAAAAC | 6 | FALSE | FALSE |
| 2_v.txt | 32466 | 32466-SNV- | SNV | 1 | C | T | 30 | 20613 | 0.14553922 | 50.1333333 | WT_perm | FALSE | ARG | JSY13 | 4 | WT_perm-Re | GCC | C>T | A/T+1 | G/C-1 | CG>TA | CG>TA | AGAGAGCCA | CCAATGT | CGG | GCC | C | A | C>T | C>T_GCC | CG>TA_CCAATGT | 6 | FALSE | FALSE |
| 2_v.txt | 32487 | 32487-SNV- | SNV | 1 | C | T | 18 | 20613 | 0.08732353 | 44.5 | WT_perm | FALSE | ARG | JSY13 | 4 | WT_perm-Re | ACG | C>T | A/T+1 | G/C-1 | CG>TA | CG>TA | AAGATACGG | CGGAAAA | TGC | ACG | C | A | C>T | C>T_ACG | CG>TA_CGGAAAA | 6 | FALSE | FALSE |
| 2_v.txt | 32535 | 32535-SNV- | SNV | 1 | G | A | 27 | 20613 | 0.1309853 | 35.3701844 | WT_perm | FALSE | ARG | JSY13 | 4 | WT_perm-Re | TGC | G>A | A/T+1 | G/C-1 | CG>TA | CG>TA | GGGTTTGCA | GCAGCTT | ACG | ACG | C | T | C>T | C>T_ACG | CG>TA_GCAGCTT | 6 | FALSE | FALSE |
| 2_v.txt | 32536 | 32536-SNV- | SNV | 1 | C | T | 34 | 20613 | 0.16494445 | 37.0875087 | WT_perm | FALSE | ARG | JSY13 | 4 | WT_perm-Re | GCA | C>T | A/T+1 | G/C-1 | CG>TA | CG>TA | GGTTTGCAG | CAGCTTC | CGT | GCA | C | A | C>T | C>T_GCA | CG>TA_CAGCTTC | 6 | FALSE | FALSE |
| 2_v.txt | 32590 | 32590-SNV- | SNV | 1 | C | T | 60 | 29949 | 0.20034058 | 35.9 | WT_perm | FALSE | ARG | JSY13 | 4 | WT_perm-Re | GCG | C>T | A/T+1 | G/C-1 | CG>TA | CG>TA | AGGCAGCGT | CGTTAAT | CGC | GCG | C | A | C>T | C>T_GCG | CG>TA_CGTTAAT | 6 | FALSE | FALSE |
| 2_v.txt | 32700 | 32700-SNV- | SNV | 1 | C | T | 37 | 29949 | 0.12354336 | 56.4594595 | WT_perm | FALSE | ARG | JSY13 | 4 | WT_perm-Re | GCC | C>T | A/T+1 | G/C-1 | CG>TA | CG>TA | ACTGGGCCG | CCGGTAA | CGG | GCC | C | A | C>T | C>T_GCC | CG>TA_CCGGTAA | 6 | FALSE | FALSE |
| 2_v.txt | 32702 | 32702-SNV- | SNV | 1 | G | A | 30 | 29949 | 0.10017029 | 50.2 | WT_perm | FALSE | ARG | JSY13 | 4 | WT_perm-Re | CGG | G>A | A/T+1 | G/C-1 | CG>TA | CG>TA | TGGGCCGGT | GGTAACC | GCC | GCC | C | T | C>T | C>T_GCC | CG>TA_GGTAACC | 6 | FALSE | FALSE |
| 2_v.txt | 32776 | 32776-SNV- | SNV | 1 | C | T | 36 | 29949 | 0.12020435 | 49.0277778 | WT_perm | FALSE | ARG | JSY13 | 4 | WT_perm-Re | GCG | C>T | A/T+1 | G/C-1 | CG>TA | CG>TA | TGGAAGCGA | CGACCCA | CGC | GCG | C | A | C>T | C>T_GCG | CG>TA_CGACCCA | 6 | FALSE | FALSE |
| 2_v.txt | 32798 | 32798-SNV- | SNV | 1 | G | A | 39 | 29949 | 0.13022138 | 41.4102564 | WT_perm | FALSE | ARG | JSY13 | 4 | WT_perm-Re | CGT | G>A | A/T+1 | G/C-1 | CG>TA | CG>TA | TTCACCGTA | GTAATAT | GCA | GCA | C | T | C>T | C>T_GCA | CG>TA_GTAATAT | 6 | FALSE | FALSE |
| 2_v.txt | 32860 | 32860-SNV- | SNV | 1 | C | T | 36 | 29942 | 0.12023245 | 36.6388889 | WT_perm | FALSE | ARG | JSY13 | 4 | WT_perm-Re | GCC | C>T | A/T+1 | G/C-1 | CG>TA | CG>TA | TCCATGCCG | CCGCCAG | CGG | GCC | C | A | C>T | C>T_GCC | CG>TA_CCGCCAG | 6 | FALSE | FALSE |
| 2_v.txt | 32862 | 32862-SNV- | SNV | 1 | G | A | 25 | 29942 | 0.08349476 | 36.52 | WT_perm | FALSE | ARG | JSY13 | 4 | WT_perm-Re | CGC | G>A | A/T+1 | G/C-1 | CG>TA | CG>TA | CATGCCGCC | GCCAGTG | GCG | GCG | C | T | C>T | C>T_GCG | CG>TA_GCCAGTG | 6 | FALSE | FALSE |
| 2_v.txt | 32900 | 32900-SNV- | SNV | 1 | G | A | 26 | 22928 | 0.11339847 | 36.3050394 | WT_perm | FALSE | ARG | JSY13 | 4 | WT_perm-Re | GGC | G>A | A/T+1 | G/C-1 | CG>TA | CG>TA | GACTTGGCC | GCCAACT | CCG | CCG | C | T | C>T | C>T_CCG | CG>TA_GCCAACT | 6 | FALSE | FALSE |
| 2_v.txt | 32901 | 32901-SNV- | SNV | 1 | C | T | 54 | 22928 | 0.23551989 | 33.0737677 | WT_perm | FALSE | ARG | JSY13 | 4 | WT_perm-Re | GCC | C>T | A/T+1 | G/C-1 | CG>TA | CG>TA | ACTTGGCCA | CCAACTA | CGG | GCC | C | A | C>T | C>T_GCC | CG>TA_CCAACTA | 6 | FALSE | FALSE |
| 2_v.txt | 32965 | 32965-SNV- | SNV | 1 | C | T | 23 | 22928 | 0.10031403 | 55 | WT_perm | FALSE | ARG | JSY13 | 4 | WT_perm-Re | GCC | C>T | A/T+1 | G/C-1 | CG>TA | CG>TA | CATTGGCCG | CCGCACC | CGG | GCC | C | A | C>T | C>T_GCC | CG>TA_CCGCACC | 6 | FALSE | FALSE |
| 2_v.txt | 32967 | 32967-SNV- | SNV | 1 | G | A | 49 | 22928 | 0.21371249 | 58.4285714 | WT_perm | FALSE | ARG | JSY13 | 4 | WT_perm-Re | CGC | G>A | A/T+1 | G/C-1 | CG>TA | CG>TA | TTGGCCGCA | GCACCAA | GCG | GCG | C | T | C>T | C>T_GCG | CG>TA_GCACCAA | 6 | FALSE | FALSE |
| 2_v.txt | 33056 | 33056-SNV- | SNV | 1 | C | T | 28 | 22928 | 0.12212142 | 56.1785714 | WT_perm | FALSE | ARG | JSY13 | 4 | WT_perm-Re | GCG | C>T | A/T+1 | G/C-1 | CG>TA | CG>TA | GGACTGCGT | CGTGACA | CGC | GCG | C | A | C>T | C>T_GCG | CG>TA_CGTGACA | 6 | FALSE | FALSE |
| 2_v.txt | 33070 | 33070-SNV- | SNV | 1 | C | T | 27 | 22928 | 0.11775994 | 51.4074074 | WT_perm | FALSE | ARG | JSY13 | 4 | WT_perm-Re | GCC | C>T | A/T+1 | G/C-1 | CG>TA | CG>TA | AATATGCCA | CCAAAGA | CGG | GCC | C | A | C>T | C>T_GCC | CG>TA_CCAAAGA | 6 | FALSE | FALSE |
| 2_v.txt | 33114 | 33114-SNV- | SNV | 1 | C | T | 22 | 22928 | 0.09595255 | 41.6818182 | WT_perm | FALSE | ARG | JSY13 | 4 | WT_perm-Re | GCC | C>T | A/T+1 | G/C-1 | CG>TA | CG>TA | ACTGGGCCG | CCGGCGT | CGG | GCC | C | A | C>T | C>T_GCC | CG>TA_CCGGCGT | 6 | FALSE | FALSE |
| 2_v.txt | 33258 | 33258-SNV- | SNV | 1 | C | T | 30 | 19722 | 0.15211439 | 36.8 | WT_perm | FALSE | ARG | JSY13 | 4 | WT_perm-Re | GCC | C>T | A/T+1 | G/C-1 | CG>TA | CG>TA | CCTATGCCAT | CCATCCT | CGG | GCC | C | A | C>T | C>T_GCC | CG>TA_CCATCCT | 6 | FALSE | FALSE |
| 2_v.txt | 33275 | 33275-SNV- | SNV | 1 | C | T | 17 | 19722 | 0.08619815 | 51.3519766 | WT_perm | FALSE | ARG | JSY13 | 4 | WT_perm-Re | ACG | C>T | A/T+1 | G/C-1 | CG>TA | CG>TA | AGAGAACGT | CGTATCC | TGC | ACG | C | A | C>T | C>T_ACG | CG>TA_CGTATCC | 6 | FALSE | FALSE |
| 2_v.txt | 33276 | 33276-SNV- | SNV | 1 | G | A | 45 | 19722 | 0.22817159 | 42.3027344 | WT_perm | FALSE | ARG | JSY13 | 4 | WT_perm-Re | CGT | G>A | A/T+1 | G/C-1 | CG>TA | CG>TA | GAGAACGTA | GTATCCT | GCA | GCA | C | T | C>T | C>T_GCA | CG>TA_GTATCCT | 6 | FALSE | FALSE |
| 2_v.txt | 33309 | 33309-SNV- | SNV | 1 | C | T | 17 | 19722 | 0.08619815 | 59.2698743 | WT_perm | FALSE | ARG | JSY13 | 4 | WT_perm-Re | GCG | C>T | A/T+1 | G/C-1 | CG>TA | CG>TA | AAAGAGCGC | CGCAATG | CGC | GCG | C | A | C>T | C>T_GCG | CG>TA_CGCAATG | 6 | FALSE | FALSE |
| 2_v.txt | 33310 | 33310-SNV- | SNV | 1 | G | A | 32 | 19719 | 0.16228004 | 49.4578649 | WT_perm | FALSE | ARG | JSY13 | 4 | WT_perm-Re | CGC | G>A | A/T+1 | G/C-1 | CG>TA | CG>TA | AAGAGCGCA | GCAATGG | GCG | GCG | C | T | C>T | C>T_GCG | CG>TA_GCAATGG | 6 | FALSE | FALSE |
| 2_v.txt | 33392 | 33392-SNV- | SNV | 1 | G | A | 24 | 19722 | 0.12169151 | 49 | WT_perm | FALSE | ARG | JSY13 | 4 | WT_perm-Re | CGT | G>A | A/T+1 | G/C-1 | CG>TA | CG>TA | AACGTCGTG | GTGAAAG | GCA | GCA | C | T | C>T | C>T_GCA | CG>TA_GTGAAAG | 6 | FALSE | FALSE |
| 2_v.txt | 33412 | 33412-SNV- | SNV | 1 | G | A | 45 | 19722 | 0.22817159 | 46.9777778 | WT_perm | FALSE | ARG | JSY13 | 4 | WT_perm-Re | GGC | G>A | A/T+1 | G/C-1 | CG>TA | CG>TA | TGACCGGCT | GCTCATT | CCG | CCG | C | T | C>T | C>T_CCG | CG>TA_GCTCATT | 6 | FALSE | FALSE |
| 2_v.txt | 33474 | 33474-SNV- | SNV | 1 | C | T | 29 | 19721 | 0.14705137 | 32.926329 | WT_perm | FALSE | ARG | JSY13 | 4 | WT_perm-Re | GCC | C>T | A/T+1 | G/C-1 | CG>TA | CG>TA | GCTATGCCTT | CCTTTTT | CGG | GCC | C | A | C>T | C>T_GCC | CG>TA_CCTTTTT | 6 | FALSE | FALSE |
| 2_v.txt | 31657 | 31657-SNV- | SNV | 1 | G | C | 7 | 14270 | 0.04905396 | 14.2856041 | WT_perm | FALSE | ARG | JSY13 | 4 | WT_perm-Re | GGC | G>C | G/C+1 | G/C-1 | CG>GC | CG>GC | GAAATGGCG | GCGTGGA | CCG | CCG | C | G | C>G | C>G_CCG | CG>GC_GCGTGGA | 6 | FALSE | FALSE |
| 2_v.txt | 31658 | 31658-SNV- | SNV | 1 | C | G | 1 | 14270 | 7.01E-03 | 38 | WT_perm | FALSE | ARG | JSY13 | 4 | WT_perm-Re | GCG | C>G | G/C+1 | G/C-1 | CG>GC | CG>GC | AAATGGCGT | CGTGGAA | CGC | GCG | C | C | C>G | C>G_GCG | CG>GC_CGTGGAA | 6 | FALSE | FALSE |
| 2_v.txt | 31911 | 31911-SNV- | SNV | 1 | G | C | 1 | 14270 | 7.01E-03 | 35 | WT_perm | FALSE | ARG | JSY13 | 4 | WT_perm-Re | TGC | G>C | G/C+1 | G/C-1 | CG>GC | CG>GC | TTTGGTGCA | GCAAAAG | ACG | ACG | C | G | C>G | C>G_ACG | CG>GC_GCAAAAG | 6 | FALSE | FALSE |
| 2_v.txt | 32536 | 32536-SNV- | SNV | 1 | C | G | 2 | 20613 | 9.70E-03 | 37.9990756 | WT_perm | FALSE | ARG | JSY13 | 4 | WT_perm-Re | GCA | C>G | G/C+1 | G/C-1 | CG>GC | CG>GC | GGTTTGCAG | CAGCTTC | CGT | GCA | C | C | C>G | C>G_GCA | CG>GC_CAGCTTC | 6 | FALSE | FALSE |
| 2_v.txt | 33275 | 33275-SNV- | SNV | 1 | C | G | 1 | 19722 | 5.07E-03 | 57.9988964 | WT_perm | FALSE | ARG | JSY13 | 4 | WT_perm-Re | ACG | C>G | G/C+1 | G/C-1 | CG>GC | CG>GC | AGAGAACGT | CGTATCC | TGC | ACG | C | C | C>G | C>G_ACG | CG>GC_CGTATCC | 6 | FALSE | FALSE |
| 2_v.txt | 33276 | 33276-SNV- | SNV | 1 | G | C | 4 | 19722 | 0.02028192 | 36.4932637 | WT_perm | FALSE | ARG | JSY13 | 4 | WT_perm-Re | CGT | G>C | G/C+1 | G/C-1 | CG>GC | CG>GC | GAGAACGTA | GTATCCT | GCA | GCA | C | G | C>G | C>G_GCA | CG>GC_GTATCCT | 6 | FALSE | FALSE |
| 2_v.txt | 33310 | 33310-SNV- | SNV | 1 | G | C | 5 | 19719 | 0.02535626 | 6.59954153 | WT_perm | FALSE | ARG | JSY13 | 4 | WT_perm-Re | CGC | G>C | G/C+1 | G/C-1 | CG>GC | CG>GC | AAGAGCGCA | GCAATGG | GCG | GCG | C | G | C>G | C>G_GCG | CG>GC_GCAATGG | 6 | FALSE | FALSE |
| 2_v.txt | 31657 | 31657-SNV- | SNV | 1 | G | T | 143 | 14270 | 1.00210231 | 19.6220228 | WT_perm | FALSE | ARG | JSY13 | 4 | WT_perm-Re | GGC | G>T | A/T+1 | G/C-1 | CG>AT | CG>AT | GAAATGGCG | GCGTGGA | CCG | CCG | C | A | C>A | C>A_CCG | CG>AT_GCGTGGA | 6 | FALSE | FALSE |
| 2_v.txt | 33309 | 33309-SNV- | SNV | 1 | C | A | 6 | 19722 | 0.03042288 | 28.6653815 | WT_perm | FALSE | ARG | JSY13 | 4 | WT_perm-Re | GCG | C>A | A/T+1 | G/C-1 | CG>AT | CG>AT | AAAGAGCGC | CGCAATG | CGC | GCG | C | T | C>A | C>A_GCG | CG>AT_CGCAATG | 6 | FALSE | FALSE |
| 2_v.txt | 33474 | 33474-SNV- | SNV | 1 | C | A | 85 | 19721 | 0.43101263 | 34.0416053 | WT_perm | FALSE | ARG | JSY13 | 4 | WT_perm-Re | GCC | C>A | A/T+1 | G/C-1 | CG>AT | CG>AT | GCTATGCCTT | CCTTTTT | CGG | GCC | C | T | C>A | C>A_GCC | CG>AT_CCTTTTT | 6 | FALSE | FALSE |
| 2_v.txt | 33475 | 33475-SNV- | SNV | 1 | C | A | 26 | 19721 | 0.13183916 | 35.6538462 | WT_perm | FALSE | ARG | JSY13 | 4 | WT_perm-Re | CCT | C>A | A/T+1 | G/C-1 | CG>AT | CG>AT | CTATGCCTTT | CTTTTTT | GGA | CCT | C | T | C>A | C>A_CCT | CG>AT_CTTTTTT | 6 | FALSE | FALSE |
| 2_v.txt | 31723 | 31723-Insert | Insertion | 1 | - | A | 2 | 14270 | 0.01401542 | 50 | WT_perm | FALSE | ARG | JSY13 | 4 | WT_perm-Re | CAA | nope | A/T+1 | >1bpdel | none | A/T+1 | TGTCCCAAA | AAAAAGT | GTT | CAA | - | T | NA | NA_CAA | A/T+1_AAAAAGT | 6 | FALSE | FALSE |
| 2_v.txt | 32081 | 32081-Insert | Insertion | 1 | - | A | 23 | 24125 | 0.09533679 | 63.6521739 | WT_perm | FALSE | ARG | JSY13 | 4 | WT_perm-Re | CAA | nope | A/T+1 | >1bpdel | none | A/T+1 | CCATGCAAA | AAAAAAG | GTT | CAA | - | T | NA | NA_CAA | A/T+1_AAAAAAG | 6 | FALSE | FALSE |
| 2_v.txt | 32261 | 32261-Insert | Insertion | 1 | - | A | 2 | 20612 | 9.70E-03 | 38 | WT_perm | FALSE | ARG | JSY13 | 4 | WT_perm-Re | CAA | nope | A/T+1 | >1bpdel | none | A/T+1 | TAGACCAAA | AAATAAA | GTT | CAA | - | T | NA | NA_CAA | A/T+1_AAATAAA | 6 | FALSE | FALSE |
| 2_v.txt | 32842 | 32842-Insert | Insertion | 1 | - | A | 10 | 29949 | 0.0333901 | 40.5 | WT_perm | FALSE | ARG | JSY13 | 4 | WT_perm-Re | CAA | nope | A/T+1 | >1bpdel | none | A/T+1 | TTACCCAAA | AAAAAAT | GTT | CAA | - | T | NA | NA_CAA | A/T+1_AAAAAAT | 6 | FALSE | FALSE |
| 2_v.txt | 31663 | 31663-Deleti | Deletion | 1 | A | - | 8 | 14270 | 0.05606167 | 29.25 | WT_perm | FALSE | ARG | JSY13 | 4 | WT_perm-Re | GAA | nope | >1bp-ins | A/T-1 | none | A/T-1 | GCGTGGAAA | AAATGTG | CTT | CTT | T | none | T>none | T>none_CTT | A/T-1_AAATGTG | 6 | FALSE | FALSE |
| 2_v.txt | 31711 | 31711-Deleti | Deletion | 1 | A | - | 4 | 14270 | 0.02803083 | 62.75 | WT_perm | FALSE | ARG | JSY13 | 4 | WT_perm-Re | CAA | nope | >1bp-ins | A/T-1 | none | A/T-1 | CATTCCAAA | AAAATTT | GTT | GTT | T | none | T>none | T>none_GTT | A/T-1_AAAATTT | 6 | FALSE | FALSE |
| 2_v.txt | 31723 | 31723-Deleti | Deletion | 1 | A | - | 17 | 14270 | 0.11913104 | 61.5882353 | WT_perm | FALSE | ARG | JSY13 | 4 | WT_perm-Re | CAA | nope | >1bp-ins | A/T-1 | none | A/T-1 | TGTCCCAAA | AAAAAGT | GTT | GTT | T | none | T>none | T>none_GTT | A/T-1_AAAAAGT | 6 | FALSE | FALSE |
| 2_v.txt | 31904 | 31904-Deleti | Deletion | 1 | T | - | 10 | 14270 | 0.07007709 | 45.3 | WT_perm | FALSE | ARG | JSY13 | 4 | WT_perm-Re | ATT | nope | >1bp-ins | A/T-1 | none | A/T-1 | ATTGAATTTT | TTTTGGT | TAA | ATT | T | none | nope | nope_ATT | A/T-1_TTTTGGT | 6 | FALSE | FALSE |
| 2_v.txt | 31972 | 31972-Deleti | Deletion | 1 | C | - | 6 | 24125 | 0.02487047 | 35.1666667 | WT_perm | FALSE | ARG | JSY13 | 4 | WT_perm-Re | GCC | nope | >1bp-ins | G/C-1 | none | G/C-1 | AATAAGCCA | CCAAGCC | CGG | GCC | C | none | nope | nope_GCC | G/C-1_CCAAGCC | 6 | FALSE | FALSE |
| 2_v.txt | 32081 | 32081-Deleti | Deletion | 1 | A | - | 25 | 24124 | 0.10363124 | 62.759996 | WT_perm | FALSE | ARG | JSY13 | 4 | WT_perm-Re | CAA | nope | >1bp-ins | A/T-1 | none | A/T-1 | CCATGCAAA | AAAAAAG | GTT | GTT | T | none | T>none | T>none_GTT | A/T-1_AAAAAAG | 6 | FALSE | FALSE |
| 2_v.txt | 32210 | 32210-Deleti | Deletion | 1 | T | - | 5 | 24125 | 0.02072539 | 42 | WT_perm | FALSE | ARG | JSY13 | 4 | WT_perm-Re | CTT | nope | >1bp-ins | A/T-1 | none | A/T-1 | ACCACCTTTG | TTTGGTG | GAA | CTT | T | none | nope | nope_CTT | A/T-1_TTTGGTG | 6 | FALSE | FALSE |
| 2_v.txt | 32231 | 32231-Deleti | Deletion | 1 | T | - | 4 | 24124 | 0.016581 | 38 | WT_perm | FALSE | ARG | JSY13 | 4 | WT_perm-Re | ATT | nope | >1bp-ins | A/T-1 | none | A/T-1 | CAGGAATTT | TTTAGGA | TAA | ATT | T | none | nope | nope_ATT | A/T-1_TTTAGGA | 6 | FALSE | FALSE |
| 2_v.txt | 32346 | 32346-Deleti | Deletion | 1 | A | - | 5 | 20613 | 0.02425654 | 56.8 | WT_perm | FALSE | ARG | JSY13 | 4 | WT_perm-Re | CAA | nope | >1bp-ins | A/T-1 | none | A/T-1 | TGTGGCAAA | AAAACCT | GTT | GTT | T | none | T>none | T>none_GTT | A/T-1_AAAACCT | 6 | FALSE | FALSE |
| 2_v.txt | 32381 | 32381-Deleti | Deletion | 1 | A | - | 5 | 20613 | 0.02425654 | 63.2 | WT_perm | FALSE | ARG | JSY13 | 4 | WT_perm-Re | TAA | nope | >1bp-ins | A/T-1 | none | A/T-1 | AATAATAAA | AAAGGGA | ATT | ATT | T | none | T>none | T>none_ATT | A/T-1_AAAGGGA | 6 | FALSE | FALSE |
| 2_v.txt | 32498 | 32498-Deleti | Deletion | 1 | T | - | 20 | 20613 | 0.09702615 | 44.7 | WT_perm | FALSE | ARG | JSY13 | 4 | WT_perm-Re | CTT | nope | >1bp-ins | A/T-1 | none | A/T-1 | AACAACTTTT | TTTTTTG | GAA | CTT | T | none | nope | nope_CTT | A/T-1_TTTTTTG | 6 | FALSE | FALSE |
| 2_v.txt | 32528 | 32528-Deleti | Deletion | 1 | G | - | 4 | 20613 | 0.01940523 | 38 | WT_perm | FALSE | ARG | JSY13 | 4 | WT_perm-Re | TGG | nope | >1bp-ins | G/C-1 | none | G/C-1 | TTTTCTGGG | GGGGTTT | ACC | ACC | C | none | C>none | C>none_ACC | G/C-1_GGGGTTT | 6 | FALSE | FALSE |
| 2_v.txt | 32631 | 32631-Deleti | Deletion | 1 | T | - | 8 | 29949 | 0.02671208 | 44.25 | WT_perm | FALSE | ARG | JSY13 | 4 | WT_perm-Re | GTT | nope | >1bp-ins | A/T-1 | none | A/T-1 | CCTTCGTTTT | TTTTTAT | CAA | GTT | T | none | nope | nope_GTT | A/T-1_TTTTTAT | 6 | FALSE | FALSE |
| 2_v.txt | 32647 | 32647-Deleti | Deletion | 1 | T | - | 5 | 29949 | 0.01669505 | 47.8 | WT_perm | FALSE | ARG | JSY13 | 4 | WT_perm-Re | ATT | nope | >1bp-ins | A/T-1 | none | A/T-1 | TAGATATTA | TTATACC | TAA | ATT | T | none | nope | nope_ATT | A/T-1_TTATACC | 6 | FALSE | FALSE |
| 2_v.txt | 32658 | 32658-Deleti | Deletion | 1 | C | - | 10 | 29949 | 0.0333901 | 51.9 | WT_perm | FALSE | ARG | JSY13 | 4 | WT_perm-Re | ACC | nope | >1bp-ins | G/C-1 | none | G/C-1 | CCTGGACCC | CCCCAGG | TGG | ACC | C | none | nope | nope_ACC | G/C-1_CCCCAGG | 6 | FALSE | FALSE |
| 2_v.txt | 32728 | 32728-Deleti | Deletion | 1 | A | - | 12 | 29949 | 0.04006812 | 61.8333333 | WT_perm | FALSE | ARG | JSY13 | 4 | WT_perm-Re | CAA | nope | >1bp-ins | A/T-1 | none | A/T-1 | CCATACAAA | AAAAACA | GTT | GTT | T | none | T>none | T>none_GTT | A/T-1_AAAAACA | 6 | FALSE | FALSE |
| 2_v.txt | 32836 | 32836-Deleti | Deletion | 1 | T | - | 5 | 29949 | 0.01669505 | 43.2 | WT_perm | FALSE | ARG | JSY13 | 4 | WT_perm-Re | ATT | nope | >1bp-ins | A/T-1 | none | A/T-1 | TGATAATTA | TTACCCA | TAA | ATT | T | none | nope | nope_ATT | A/T-1_TTACCCA | 6 | FALSE | FALSE |
| 2_v.txt | 32842 | 32842-Deleti | Deletion | 1 | A | - | 39 | 29949 | 0.13022138 | 41.8205118 | WT_perm | FALSE | ARG | JSY13 | 4 | WT_perm-Re | CAA | nope | >1bp-ins | A/T-1 | none | A/T-1 | TTACCCAAA | AAAAAAT | GTT | GTT | T | none | T>none | T>none_GTT | A/T-1_AAAAAAT | 6 | FALSE | FALSE |

| | | | | | | | | | | | | | | | | | | | | | | | | | | | | | | | | | | |
|---|---|---|---|---|---|---|---|---|---|---|---|---|---|---|---|---|---|---|---|---|---|---|---|---|---|---|---|---|---|---|---|---|---|---|
| 2_v.txt | 32924 | 32924-Deleti | Deletion | 1 | A | - | 23 | 22928 | 0.10031403 | 50.1304348 | WT_perm | FALSE | ARG | JSY13 | 4 | WT_perm-Re | CAA | nope | >1bp-ins | A/T-1 | none | A/T-1 | CAGGGCAAA | AAAAGTG | GTT | GTT | T | none | T>none | T>none_GTT | A/T-1_AAAAGTG | 6 | FALSE | FALSE |
| 2_v.txt | 33000 | 33000-Deleti | Deletion | 1 | A | - | 5 | 22928 | 0.0218074 | 58.6 | WT_perm | FALSE | ARG | JSY13 | 4 | WT_perm-Re | GAA | nope | >1bp-ins | A/T-1 | none | A/T-1 | TGTGAGAAA | AAAACTG | CTT | CTT | T | none | T>none | T>none_CTT | A/T-1_AAAACTG | 6 | FALSE | FALSE |
| 2_v.txt | 33040 | 33040-Deleti | Deletion | 1 | T | - | 6 | 22928 | 0.02616888 | 61 | WT_perm | FALSE | ARG | JSY13 | 4 | WT_perm-Re | ATT | nope | >1bp-ins | A/T-1 | none | A/T-1 | TAGCCATTTC | TTTCACC | TAA | ATT | T | none | nope | nope_ATT | A/T-1_TTTCACC | 6 | FALSE | FALSE |
| 2_v.txt | 33239 | 33239-Deleti | Deletion | 1 | T | - | 5 | 19722 | 0.0253524 | 36.4 | WT_perm | FALSE | ARG | JSY13 | 4 | WT_perm-Re | CTT | nope | >1bp-ins | A/T-1 | none | A/T-1 | TTCTCCTTCA | TTCATCT | GAA | CTT | T | none | nope | nope_CTT | A/T-1_TTCATCT | 6 | FALSE | FALSE |
| 2_v.txt | 33258 | 33258-Deleti | Deletion | 1 | C | - | 2 | 19722 | 0.01014096 | 36.5 | WT_perm | FALSE | ARG | JSY13 | 4 | WT_perm-Re | GCC | nope | >1bp-ins | G/C-1 | none | G/C-1 | CCTATGCCAT | CCATCCT | CGG | GCC | C | none | nope | nope_GCC | G/C-1_CCATCCT | 6 | FALSE | FALSE |
| 2_v.txt | 33303 | 33303-Deleti | Deletion | 1 | A | - | 7 | 19722 | 0.03549336 | 54.8571429 | WT_perm | FALSE | ARG | JSY13 | 4 | WT_perm-Re | GAA | nope | >1bp-ins | A/T-1 | none | A/T-1 | GTCGGGAAA | AAAGAGC | CTT | CTT | T | none | T>none | T>none_CTT | A/T-1_AAAGAGC | 6 | FALSE | FALSE |
| 2_v.txt | 33474 | 33474-Deleti | Deletion | 1 | C | - | 61 | 19721 | 0.30931494 | 26.3278689 | WT_perm | FALSE | ARG | JSY13 | 4 | WT_perm-Re | GCC | nope | >1bp-ins | G/C-1 | none | G/C-1 | GCTATGCCTT | CCTTTTT | CGG | GCC | C | none | nope | nope_GCC | G/C-1_CCTTTTT | 6 | FALSE | FALSE |
| 3_v.txt | 31723 | 31723-SNV- | SNV | 1 | A | C | 10 | 13545 | 0.07382798 | 60.1 | WT_perm | FALSE | ARG | JSY13 | 5 | WT_perm-Re | CAA | A>C | G/C+1 | A/T-1 | TA>GC | TA>GC | TGTCCCAAA | AAAAAGT | GTT | GTT | T | G | T>G | T>G_GTT | TA>GC_AAAAAGT | 6 | FALSE | FALSE |
| 3_v.txt | 32842 | 32842-SNV- | SNV | 1 | A | C | 4 | 22998 | 0.01739282 | 36.9999409 | WT_perm | FALSE | ARG | JSY13 | 5 | WT_perm-Re | CAA | A>C | G/C+1 | A/T-1 | TA>GC | TA>GC | TTACCCAAA | AAAAAAT | GTT | GTT | T | G | T>G | T>G_GTT | TA>GC_AAAAAAT | 6 | FALSE | FALSE |
| 3_v.txt | 32884 | 32884-SNV- | SNV | 1 | A | C | 5 | 28267 | 0.01768847 | 37.4 | WT_perm | FALSE | ARG | JSY13 | 5 | WT_perm-Re | CAA | A>C | G/C+1 | A/T-1 | TA>GC | TA>GC | ACGTCCAAA | AAAATTG | GTT | GTT | T | G | T>G | T>G_GTT | TA>GC_AAAATTG | 6 | FALSE | FALSE |
| 3_v.txt | 32081 | 32081-SNV- | SNV | 1 | A | G | 9 | 30827 | 0.02919519 | 63.7777831 | WT_perm | FALSE | ARG | JSY13 | 5 | WT_perm-Re | CAA | A>G | G/C+1 | A/T-1 | TA>CG | TA>CG | CCATGCAAA | AAAAAAG | GTT | GTT | T | C | T>C | T>C_GTT | TA>CG_AAAAAAG | 6 | FALSE | FALSE |
| 3_v.txt | 31650 | 31650-SNV- | SNV | 1 | C | T | 29 | 13545 | 0.21410114 | 37.0344828 | WT_perm | FALSE | ARG | JSY13 | 5 | WT_perm-Re | GCG | C>T | A/T+1 | G/C-1 | CG>TA | CG>TA | AGAATGCGA | CGAAATG | CGC | GCG | C | A | C>T | C>T_GCG | CG>TA_CGAAATG | 6 | FALSE | FALSE |
| 3_v.txt | 31830 | 31830-SNV- | SNV | 1 | C | T | 21 | 13545 | 0.15503876 | 48.6190476 | WT_perm | FALSE | ARG | JSY13 | 5 | WT_perm-Re | GCA | C>T | A/T+1 | G/C-1 | CG>TA | CG>TA | AATATGCATT | CATTGAA | CGT | GCA | C | A | C>T | C>T_GCA | CG>TA_CATTGAA | 6 | FALSE | FALSE |
| 3_v.txt | 31881 | 31881-SNV- | SNV | 1 | G | A | 13 | 13545 | 0.09597638 | 41.8453425 | WT_perm | FALSE | ARG | JSY13 | 5 | WT_perm-Re | GGC | G>A | A/T+1 | G/C-1 | CG>TA | CG>TA | TAGGCGGCA | GCAGCAA | CCG | CCG | C | T | C>T | C>T_CCG | CG>TA_GCAGCAA | 6 | FALSE | FALSE |
| 3_v.txt | 31882 | 31882-SNV- | SNV | 1 | C | T | 23 | 13545 | 0.16980436 | 43.1830431 | WT_perm | FALSE | ARG | JSY13 | 5 | WT_perm-Re | GCA | C>T | A/T+1 | G/C-1 | CG>TA | CG>TA | AGGCGGCAC | CAGCAAA | CGT | GCA | C | A | C>T | C>T_GCA | CG>TA_CAGCAAA | 6 | FALSE | FALSE |
| 3_v.txt | 31884 | 31884-SNV- | SNV | 1 | G | A | 20 | 13545 | 0.14765596 | 33 | WT_perm | FALSE | ARG | JSY13 | 5 | WT_perm-Re | AGC | G>A | A/T+1 | G/C-1 | CG>TA | CG>TA | GCGGCAGCA | GCAAAGC | TCG | TCG | C | T | C>T | C>T_TCG | CG>TA_GCAAAGC | 6 | FALSE | FALSE |
| 3_v.txt | 31947 | 31947-SNV- | SNV | 1 | G | A | 29 | 30801 | 0.09415279 | 35.6206897 | WT_perm | FALSE | ARG | JSY13 | 5 | WT_perm-Re | CGT | G>A | A/T+1 | G/C-1 | CG>TA | CG>TA | ATGATCGTC | GTCATAA | GCA | GCA | C | T | C>T | C>T_GCA | CG>TA_GTCATAA | 6 | FALSE | FALSE |
| 3_v.txt | 31962 | 31962-SNV- | SNV | 1 | G | A | 49 | 30827 | 0.15895157 | 36.8571429 | WT_perm | FALSE | ARG | JSY13 | 5 | WT_perm-Re | CGC | G>A | A/T+1 | G/C-1 | CG>TA | CG>TA | GTGGCCGCA | GCATAAT | GCG | GCG | C | T | C>T | C>T_GCG | CG>TA_GCATAAT | 6 | FALSE | FALSE |
| 3_v.txt | 31972 | 31972-SNV- | SNV | 1 | C | T | 25 | 30827 | 0.08109774 | 37.36 | WT_perm | FALSE | ARG | JSY13 | 5 | WT_perm-Re | GCC | C>T | A/T+1 | G/C-1 | CG>TA | CG>TA | AATAAGCCA | CCAAGCC | CGG | GCC | C | A | C>T | C>T_GCC | CG>TA_CCAAGCC | 6 | FALSE | FALSE |
| 3_v.txt | 31979 | 31979-SNV- | SNV | 1 | G | A | 15 | 30827 | 0.04865864 | 51.2 | WT_perm | FALSE | ARG | JSY13 | 5 | WT_perm-Re | CGG | G>A | A/T+1 | G/C-1 | CG>TA | CG>TA | CAAGCCGGG | GGGCATT | GCC | GCC | C | T | C>T | C>T_GCC | CG>TA_GGGCATT | 6 | FALSE | FALSE |
| 3_v.txt | 32041 | 32041-SNV- | SNV | 1 | G | A | 28 | 30827 | 0.09082947 | 55.6071429 | WT_perm | FALSE | ARG | JSY13 | 5 | WT_perm-Re | TGC | G>A | A/T+1 | G/C-1 | CG>TA | CG>TA | AAGCTTGCA | GCATAAA | ACG | ACG | C | T | C>T | C>T_ACG | CG>TA_GCATAAA | 6 | FALSE | FALSE |
| 3_v.txt | 32057 | 32057-SNV- | SNV | 1 | C | T | 29 | 30827 | 0.09407338 | 57.862069 | WT_perm | FALSE | ARG | JSY13 | 5 | WT_perm-Re | GCG | C>T | A/T+1 | G/C-1 | CG>TA | CG>TA | GATGTGCGA | CGAGATT | CGC | GCG | C | A | C>T | C>T_GCG | CG>TA_CGAGATT | 6 | FALSE | FALSE |
| 3_v.txt | 32092 | 32092-SNV- | SNV | 1 | C | T | 35 | 30827 | 0.11353684 | 61.6 | WT_perm | FALSE | ARG | JSY13 | 5 | WT_perm-Re | GCA | C>T | A/T+1 | G/C-1 | CG>TA | CG>TA | AGCCTGCAA | CAACACC | CGT | GCA | C | A | C>T | C>T_GCA | CG>TA_CAACACC | 6 | FALSE | FALSE |
| 3_v.txt | 32165 | 32165-SNV- | SNV | 1 | G | A | 50 | 30827 | 0.16219548 | 47.76 | WT_perm | FALSE | ARG | JSY13 | 5 | WT_perm-Re | CGC | G>A | A/T+1 | G/C-1 | CG>TA | CG>TA | CAAAGCGCC | GCCAAAT | GCG | GCG | C | T | C>T | C>T_GCG | CG>TA_GCCAAAT | 6 | FALSE | FALSE |
| 3_v.txt | 32173 | 32173-SNV- | SNV | 1 | C | T | 35 | 30827 | 0.11353684 | 46.3142857 | WT_perm | FALSE | ARG | JSY13 | 5 | WT_perm-Re | GCA | C>T | A/T+1 | G/C-1 | CG>TA | CG>TA | CAAATGCAG | CAGCAGT | CGT | GCA | C | A | C>T | C>T_GCA | CG>TA_CAGCAGT | 6 | FALSE | FALSE |
| 3_v.txt | 32190 | 32190-SNV- | SNV | 1 | G | A | 27 | 30827 | 0.08758556 | 43.5555556 | WT_perm | FALSE | ARG | JSY13 | 5 | WT_perm-Re | TGC | G>A | A/T+1 | G/C-1 | CG>TA | CG>TA | AAAACTGCA | GCAATGT | ACG | ACG | C | T | C>T | C>T_ACG | CG>TA_GCAATGT | 6 | FALSE | FALSE |
| 3_v.txt | 32271 | 32271-SNV- | SNV | 1 | C | T | 61 | 44582 | 0.13682652 | 30.8196721 | WT_perm | FALSE | ARG | JSY13 | 5 | WT_perm-Re | ACG | C>T | A/T+1 | G/C-1 | CG>TA | CG>TA | AAAATACGG | CGGGAAC | TGC | ACG | C | A | C>T | C>T_ACG | CG>TA_CGGGAAC | 6 | FALSE | FALSE |
| 3_v.txt | 32299 | 32299-SNV- | SNV | 1 | C | T | 78 | 44583 | 0.17495458 | 53.0128205 | WT_perm | FALSE | ARG | JSY13 | 5 | WT_perm-Re | GCG | C>T | A/T+1 | G/C-1 | CG>TA | CG>TA | AATTTGCGG | CGGCAGA | CGC | GCG | C | A | C>T | C>T_GCG | CG>TA_CGGCAGA | 6 | FALSE | FALSE |
| 3_v.txt | 32344 | 32344-SNV- | SNV | 1 | G | A | 28 | 44583 | 0.06280421 | 60.4285714 | WT_perm | FALSE | ARG | JSY13 | 5 | WT_perm-Re | GGC | G>A | A/T+1 | G/C-1 | CG>TA | CG>TA | TATGTGGCA | GCAAAAC | CCG | CCG | C | T | C>T | C>T_CCG | CG>TA_GCAAAAC | 6 | FALSE | FALSE |
| 3_v.txt | 32466 | 32466-SNV- | SNV | 1 | C | T | 88 | 44583 | 0.19738465 | 46.5681818 | WT_perm | FALSE | ARG | JSY13 | 5 | WT_perm-Re | GCC | C>T | A/T+1 | G/C-1 | CG>TA | CG>TA | AGAGAGCCA | CCAATGT | CGG | GCC | C | A | C>T | C>T_GCC | CG>TA_CCAATGT | 6 | FALSE | FALSE |
| 3_v.txt | 32487 | 32487-SNV- | SNV | 1 | C | T | 44 | 44583 | 0.09869233 | 44.4772727 | WT_perm | FALSE | ARG | JSY13 | 5 | WT_perm-Re | ACG | C>T | A/T+1 | G/C-1 | CG>TA | CG>TA | AAGATACGG | CGGAAAA | TGC | ACG | C | A | C>T | C>T_ACG | CG>TA_CGGAAAA | 6 | FALSE | FALSE |
| 3_v.txt | 32535 | 32535-SNV- | SNV | 1 | G | A | 57 | 44582 | 0.12785429 | 36.8244713 | WT_perm | FALSE | ARG | JSY13 | 5 | WT_perm-Re | TGC | G>A | A/T+1 | G/C-1 | CG>TA | CG>TA | GGGTTTGCA | GCAGCTT | ACG | ACG | C | T | C>T | C>T_ACG | CG>TA_GCAGCTT | 6 | FALSE | FALSE |
| 3_v.txt | 32536 | 32536-SNV- | SNV | 1 | C | T | 76 | 44582 | 0.17047239 | 37.0787112 | WT_perm | FALSE | ARG | JSY13 | 5 | WT_perm-Re | GCA | C>T | A/T+1 | G/C-1 | CG>TA | CG>TA | GGTTTGCAG | CAGCTTC | CGT | GCA | C | A | C>T | C>T_GCA | CG>TA_CAGCTTC | 6 | FALSE | FALSE |
| 3_v.txt | 32590 | 32590-SNV- | SNV | 1 | C | T | 56 | 22998 | 0.24349944 | 36.7678571 | WT_perm | FALSE | ARG | JSY13 | 5 | WT_perm-Re | GCG | C>T | A/T+1 | G/C-1 | CG>TA | CG>TA | AGGCAGCGT | CGTTAAT | CGC | GCG | C | A | C>T | C>T_GCG | CG>TA_CGTTAAT | 6 | FALSE | FALSE |
| 3_v.txt | 32700 | 32700-SNV- | SNV | 1 | C | T | 29 | 22998 | 0.12609792 | 60.3793103 | WT_perm | FALSE | ARG | JSY13 | 5 | WT_perm-Re | GCC | C>T | A/T+1 | G/C-1 | CG>TA | CG>TA | ACTGGGCCG | CCGGTAA | CGG | GCC | C | A | C>T | C>T_GCC | CG>TA_CCGGTAA | 6 | FALSE | FALSE |
| 3_v.txt | 32702 | 32702-SNV- | SNV | 1 | G | A | 23 | 22998 | 0.1000087 | 48.5217391 | WT_perm | FALSE | ARG | JSY13 | 5 | WT_perm-Re | CGG | G>A | A/T+1 | G/C-1 | CG>TA | CG>TA | TGGGCCGGT | GGTAACC | GCC | GCC | C | T | C>T | C>T_GCC | CG>TA_GGTAACC | 6 | FALSE | FALSE |
| 3_v.txt | 32757 | 32757-SNV- | SNV | 1 | G | A | 23 | 22998 | 0.1000087 | 54.0869565 | WT_perm | FALSE | ARG | JSY13 | 5 | WT_perm-Re | GGC | G>A | A/T+1 | G/C-1 | CG>TA | CG>TA | ATAATGGCT | GCTAAAA | CCG | CCG | C | T | C>T | C>T_CCG | CG>TA_GCTAAAA | 6 | FALSE | FALSE |
| 3_v.txt | 32776 | 32776-SNV- | SNV | 1 | C | T | 33 | 22998 | 0.14349074 | 51.6363636 | WT_perm | FALSE | ARG | JSY13 | 5 | WT_perm-Re | GCG | C>T | A/T+1 | G/C-1 | CG>TA | CG>TA | TGGAAGCGA | CGACCCA | CGC | GCG | C | A | C>T | C>T_GCG | CG>TA_CGACCCA | 6 | FALSE | FALSE |
| 3_v.txt | 32798 | 32798-SNV- | SNV | 1 | G | A | 29 | 22998 | 0.12609792 | 39.6551724 | WT_perm | FALSE | ARG | JSY13 | 5 | WT_perm-Re | CGT | G>A | A/T+1 | G/C-1 | CG>TA | CG>TA | TTCACCGTA | GTAATAT | GCA | GCA | C | T | C>T | C>T_GCA | CG>TA_GTAATAT | 6 | FALSE | FALSE |
| 3_v.txt | 32860 | 32860-SNV- | SNV | 1 | C | T | 32 | 22997 | 0.13914859 | 35.8125 | WT_perm | FALSE | ARG | JSY13 | 5 | WT_perm-Re | GCC | C>T | A/T+1 | G/C-1 | CG>TA | CG>TA | TCCATGCCG | CCGCCAG | CGG | GCC | C | A | C>T | C>T_GCC | CG>TA_CCGCCAG | 6 | FALSE | FALSE |
| 3_v.txt | 32879 | 32879-SNV- | SNV | 1 | C | T | 26 | 28263 | 0.09199307 | 36.2692308 | WT_perm | FALSE | ARG | JSY13 | 5 | WT_perm-Re | ACG | C>T | A/T+1 | G/C-1 | CG>TA | CG>TA | TTTGTACGTC | CGTCCAA | TGC | ACG | C | A | C>T | C>T_ACG | CG>TA_CGTCCAA | 6 | FALSE | FALSE |
| 3_v.txt | 32900 | 32900-SNV- | SNV | 1 | G | A | 26 | 28267 | 0.09198005 | 34.5741084 | WT_perm | FALSE | ARG | JSY13 | 5 | WT_perm-Re | GGC | G>A | A/T+1 | G/C-1 | CG>TA | CG>TA | GACTTGGCC | GCCAACT | CCG | CCG | C | T | C>T | C>T_CCG | CG>TA_GCCAACT | 6 | FALSE | FALSE |
| 3_v.txt | 32901 | 32901-SNV- | SNV | 1 | C | T | 51 | 28267 | 0.1804224 | 32.7260768 | WT_perm | FALSE | ARG | JSY13 | 5 | WT_perm-Re | GCC | C>T | A/T+1 | G/C-1 | CG>TA | CG>TA | ACTTGGCCA | CCAACTA | CGG | GCC | C | A | C>T | C>T_GCC | CG>TA_CCAACTA | 6 | FALSE | FALSE |
| 3_v.txt | 32922 | 32922-SNV- | SNV | 1 | G | A | 27 | 28267 | 0.09551774 | 49.962963 | WT_perm | FALSE | ARG | JSY13 | 5 | WT_perm-Re | GGC | G>A | A/T+1 | G/C-1 | CG>TA | CG>TA | TCCAGGGCA | GCAAAAG | CCG | CCG | C | T | C>T | C>T_CCG | CG>TA_GCAAAAG | 6 | FALSE | FALSE |
| 3_v.txt | 32965 | 32965-SNV- | SNV | 1 | C | T | 36 | 28267 | 0.12735699 | 59.5555556 | WT_perm | FALSE | ARG | JSY13 | 5 | WT_perm-Re | GCC | C>T | A/T+1 | G/C-1 | CG>TA | CG>TA | CATTGGCCG | CCGCACC | CGG | GCC | C | A | C>T | C>T_GCC | CG>TA_CCGCACC | 6 | FALSE | FALSE |
| 3_v.txt | 32967 | 32967-SNV- | SNV | 1 | G | A | 88 | 28267 | 0.31131708 | 60.4659091 | WT_perm | FALSE | ARG | JSY13 | 5 | WT_perm-Re | CGC | G>A | A/T+1 | G/C-1 | CG>TA | CG>TA | TTGGCCGCA | GCACCAA | GCG | GCG | C | T | C>T | C>T_GCG | CG>TA_GCACCAA | 6 | FALSE | FALSE |
| 3_v.txt | 32977 | 32977-SNV- | SNV | 1 | C | T | 25 | 28267 | 0.08844235 | 58.92 | WT_perm | FALSE | ARG | JSY13 | 5 | WT_perm-Re | GCT | C>T | A/T+1 | G/C-1 | CG>TA | CG>TA | CAAATGCTG | CTGGAGA | CGA | GCT | C | A | C>T | C>T_GCT | CG>TA_CTGGAGA | 6 | FALSE | FALSE |
| 3_v.txt | 33037 | 33037-SNV- | SNV | 1 | C | T | 20 | 28267 | 0.07075388 | 56.4 | WT_perm | FALSE | ARG | JSY13 | 5 | WT_perm-Re | GCC | C>T | A/T+1 | G/C-1 | CG>TA | CG>TA | ATGTAGCCA | CCATTTC | CGG | GCC | C | A | C>T | C>T_GCC | CG>TA_CCATTTC | 6 | FALSE | FALSE |
| 3_v.txt | 33056 | 33056-SNV- | SNV | 1 | C | T | 31 | 28267 | 0.10966852 | 54.8064516 | WT_perm | FALSE | ARG | JSY13 | 5 | WT_perm-Re | GCG | C>T | A/T+1 | G/C-1 | CG>TA | CG>TA | GGACTGCGT | CGTGACA | CGC | GCG | C | A | C>T | C>T_GCG | CG>TA_CGTGACA | 6 | FALSE | FALSE |
| 3_v.txt | 33070 | 33070-SNV- | SNV | 1 | C | T | 34 | 28267 | 0.1202816 | 55.6176471 | WT_perm | FALSE | ARG | JSY13 | 5 | WT_perm-Re | GCC | C>T | A/T+1 | G/C-1 | CG>TA | CG>TA | AATATGCCA | CCAAAGA | CGG | GCC | C | A | C>T | C>T_GCC | CG>TA_CCAAAGA | 6 | FALSE | FALSE |
| 3_v.txt | 33114 | 33114-SNV- | SNV | 1 | C | T | 25 | 28267 | 0.08844235 | 44.52 | WT_perm | FALSE | ARG | JSY13 | 5 | WT_perm-Re | GCC | C>T | A/T+1 | G/C-1 | CG>TA | CG>TA | ACTGGGCCG | CCGGCGT | CGG | GCC | C | A | C>T | C>T_GCC | CG>TA_CCGGCGT | 6 | FALSE | FALSE |
| 3_v.txt | 33177 | 33177-SNV- | SNV | 1 | G | A | 21 | 28257 | 0.07431787 | 37.3333333 | WT_perm | FALSE | ARG | JSY13 | 5 | WT_perm-Re | GGC | G>A | A/T+1 | G/C-1 | CG>TA | CG>TA | CCAAGGGCA | GCAATCA | CCG | CCG | C | T | C>T | C>T_CCG | CG>TA_GCAATCA | 6 | FALSE | FALSE |
| 3_v.txt | 33258 | 33258-SNV- | SNV | 1 | C | T | 13 | 10726 | 0.12120082 | 41.8461539 | WT_perm | FALSE | ARG | JSY13 | 5 | WT_perm-Re | GCC | C>T | A/T+1 | G/C-1 | CG>TA | CG>TA | CCTATGCCAT | CCATCCT | CGG | GCC | C | A | C>T | C>T_GCC | CG>TA_CCATCCT | 6 | FALSE | FALSE |
| 3_v.txt | 33276 | 33276-SNV- | SNV | 1 | G | A | 19 | 10726 | 0.17713966 | 45.631579 | WT_perm | FALSE | ARG | JSY13 | 5 | WT_perm-Re | CGT | G>A | A/T+1 | G/C-1 | CG>TA | CG>TA | GAGAACGTA | GTATCCT | GCA | GCA | C | T | C>T | C>T_GCA | CG>TA_GTATCCT | 6 | FALSE | FALSE |
| 3_v.txt | 33284 | 33284-SNV- | SNV | 1 | G | A | 17 | 10726 | 0.15849338 | 46.0588235 | WT_perm | FALSE | ARG | JSY13 | 5 | WT_perm-Re | CGC | G>A | A/T+1 | G/C-1 | CG>TA | CG>TA | ATCCTCGCC | GCCATTT | GCG | GCG | C | T | C>T | C>T_GCG | CG>TA_GCCATTT | 6 | FALSE | FALSE |
| 3_v.txt | 33474 | 33474-SNV- | SNV | 1 | C | T | 17 | 10726 | 0.15849338 | 35.2352941 | WT_perm | FALSE | ARG | JSY13 | 5 | WT_perm-Re | GCC | C>T | A/T+1 | G/C-1 | CG>TA | CG>TA | GCTATGCCTT | CCTTTTT | CGG | GCC | C | A | C>T | C>T_GCC | CG>TA_CCTTTTT | 6 | FALSE | FALSE |
| 3_v.txt | 32536 | 32536-SNV- | SNV | 1 | C | G | 2 | 44582 | 4.49E-03 | 37.9996771 | WT_perm | FALSE | ARG | JSY13 | 5 | WT_perm-Re | GCA | C>G | G/C+1 | G/C-1 | CG>GC | CG>GC | GGTTTGCAG | CAGCTTC | CGT | GCA | C | C | C>G | C>G_GCA | CG>GC_CAGCTTC | 6 | FALSE | FALSE |
| 3_v.txt | 32977 | 32977-SNV- | SNV | 1 | C | G | 3 | 28267 | 0.01061308 | 64 | WT_perm | FALSE | ARG | JSY13 | 5 | WT_perm-Re | GCT | C>G | G/C+1 | G/C-1 | CG>GC | CG>GC | CAAATGCTG | CTGGAGA | CGA | GCT | C | C | C>G | C>G_GCT | CG>GC_CTGGAGA | 6 | FALSE | FALSE |
| 3_v.txt | 32424 | 32424-SNV- | SNV | 1 | G | T | 16 | 44583 | 0.03588812 | 54.6875 | WT_perm | FALSE | ARG | JSY13 | 5 | WT_perm-Re | AGG | G>T | A/T+1 | G/C-1 | CG>AT | CG>AT | AGTTTAGGG | GGGTCAT | TCC | TCC | C | A | C>A | C>A_TCC | CG>AT_GGGTCAT | 6 | FALSE | FALSE |
| 3_v.txt | 33474 | 33474-SNV- | SNV | 1 | C | A | 28 | 10726 | 0.26104792 | 34.5 | WT_perm | FALSE | ARG | JSY13 | 5 | WT_perm-Re | GCC | C>A | A/T+1 | G/C-1 | CG>AT | CG>AT | GCTATGCCTT | CCTTTTT | CGG | GCC | C | T | C>A | C>A_GCC | CG>AT_CCTTTTT | 6 | FALSE | FALSE |
| 3_v.txt | 33475 | 33475-SNV- | SNV | 1 | C | A | 15 | 10726 | 0.1398471 | 37.7333333 | WT_perm | FALSE | ARG | JSY13 | 5 | WT_perm-Re | CCT | C>A | A/T+1 | G/C-1 | CG>AT | CG>AT | CTATGCCTTT | CTTTTTT | GGA | CCT | C | T | C>A | C>A_CCT | CG>AT_CTTTTTT | 6 | FALSE | FALSE |
| 3_v.txt | 32081 | 32081-Insert | Insertion | 1 | - | A | 15 | 30827 | 0.04865864 | 64 | WT_perm | FALSE | ARG | JSY13 | 5 | WT_perm-Re | CAA | nope | A/T+1 | >1bpdel | none | A/T+1 | CCATGCAAA | AAAAAAG | GTT | CAA | - | T | NA | NA_CAA | A/T+1_AAAAAAG | 6 | FALSE | FALSE |
| 3_v.txt | 32498 | 32498-Insert | Insertion | 1 | - | A | 1 | 44583 | 2.24E-03 | 64 | WT_perm | FALSE | ARG | JSY13 | 5 | WT_perm-Re | CTT | nope | A/T+1 | >1bpdel | none | A/T+1 | AACAACTTTT | TTTTTTG | GAA | CTT | - | T | NA | NA_CTT | A/T+1_TTTTTTG | 6 | FALSE | FALSE |
| 3_v.txt | 32498 | 32498-Insert | Insertion | 1 | - | T | 5 | 44583 | 0.01121504 | 53.2 | WT_perm | FALSE | ARG | JSY13 | 5 | WT_perm-Re | CTT | nope | A/T+1 | >1bpdel | none | A/T+1 | AACAACTTTT | TTTTTTG | GAA | CTT | - | A | NA | NA_CTT | A/T+1_TTTTTTG | 6 | FALSE | FALSE |
| 3_v.txt | 32842 | 32842-Insert | Insertion | 1 | - | A | 8 | 22998 | 0.03478563 | 39.25 | WT_perm | FALSE | ARG | JSY13 | 5 | WT_perm-Re | CAA | nope | A/T+1 | >1bpdel | none | A/T+1 | TTACCCAAA | AAAAAAT | GTT | CAA | - | T | NA | NA_CAA | A/T+1_AAAAAAT | 6 | FALSE | FALSE |
| 3_v.txt | 31723 | 31723-Deleti | Deletion | 1 | A | - | 15 | 13545 | 0.11074197 | 61.2 | WT_perm | FALSE | ARG | JSY13 | 5 | WT_perm-Re | CAA | nope | >1bp-ins | A/T-1 | none | A/T-1 | TGTCCCAAA | AAAAAGT | GTT | GTT | T | none | T>none | T>none_GTT | A/T-1_AAAAAGT | 6 | FALSE | FALSE |
| 3_v.txt | 31747 | 31747-Deleti | Deletion | 1 | C | - | 4 | 13545 | 0.02953119 | 49.5 | WT_perm | FALSE | ARG | JSY13 | 5 | WT_perm-Re | TCC | nope | >1bp-ins | G/C-1 | none | G/C-1 | GATCTTCCCA | CCCATAC | AGG | TCC | C | none | nope | nope_TCC | G/C-1_CCCATAC | 6 | FALSE | FALSE |
| 3_v.txt | 31863 | 31863-Deleti | Deletion | 1 | A | - | 7 | 13545 | 0.05167959 | 51.4285714 | WT_perm | FALSE | ARG | JSY13 | 5 | WT_perm-Re | GAA | nope | >1bp-ins | A/T-1 | none | A/T-1 | AACAGGAAA | AAAATAG | CTT | CTT | T | none | T>none | T>none_CTT | A/T-1_AAAATAG | 6 | FALSE | FALSE |
| 3_v.txt | 31913 | 31913-Deleti | Deletion | 1 | A | - | 8 | 13545 | 0.05906239 | 40.75 | WT_perm | FALSE | ARG | JSY13 | 5 | WT_perm-Re | CAA | nope | >1bp-ins | A/T-1 | none | A/T-1 | TGGTGCAAA | AAAAGCC | GTT | GTT | T | none | T>none | T>none_GTT | A/T-1_AAAAGCC | 6 | FALSE | FALSE |
| 3_v.txt | 31952 | 31952-Deleti | Deletion | 1 | A | - | 10 | 30828 | 0.03243804 | 37.8 | WT_perm | FALSE | ARG | JSY13 | 5 | WT_perm-Re | TAA | nope | >1bp-ins | A/T-1 | none | A/T-1 | CGTCATAAA | AAATGTG | ATT | ATT | T | none | T>none | T>none_ATT | A/T-1_AAATGTG | 6 | FALSE | FALSE |
| 3_v.txt | 31972 | 31972-Deleti | Deletion | 1 | C | - | 4 | 30827 | 0.01297564 | 34 | WT_perm | FALSE | ARG | JSY13 | 5 | WT_perm-Re | GCC | nope | >1bp-ins | G/C-1 | none | G/C-1 | AATAAGCCA | CCAAGCC | CGG | GCC | C | none | nope | nope_GCC | G/C-1_CCAAGCC | 6 | FALSE | FALSE |
| 3_v.txt | 31979 | 31979-Deleti | Deletion | 1 | G | - | 6 | 30827 | 0.01946346 | 37.8333333 | WT_perm | FALSE | ARG | JSY13 | 5 | WT_perm-Re | CGG | nope | >1bp-ins | G/C-1 | none | G/C-1 | CAAGCCGGG | GGGCATT | GCC | GCC | C | none | C>none | C>none_GCC | G/C-1_GGGCATT | 6 | FALSE | FALSE |
| 3_v.txt | 32069 | 32069-Deleti | Deletion | 1 | A | - | 7 | 30827 | 0.02270737 | 63.4285714 | WT_perm | FALSE | ARG | JSY13 | 5 | WT_perm-Re | TAA | nope | >1bp-ins | A/T-1 | none | A/T-1 | TGAGATAAA | AAATAAC | ATT | ATT | T | none | T>none | T>none_ATT | A/T-1_AAATAAC | 6 | FALSE | FALSE |
| 3_v.txt | 32081 | 32081-Deleti | Deletion | 1 | A | - | 29 | 30827 | 0.09407338 | 62.0344833 | WT_perm | FALSE | ARG | JSY13 | 5 | WT_perm-Re | CAA | nope | >1bp-ins | A/T-1 | none | A/T-1 | CCATGCAAA | AAAAAAG | GTT | GTT | T | none | T>none | T>none_GTT | A/T-1_AAAAAAG | 6 | FALSE | FALSE |
| 3_v.txt | 32114 | 32114-Deleti | Deletion | 1 | C | - | 6 | 30827 | 0.01946346 | 54.6666667 | WT_perm | FALSE | ARG | JSY13 | 5 | WT_perm-Re | GCC | nope | >1bp-ins | G/C-1 | none | G/C-1 | TAATAGCCA | CCATTCG | CGG | GCC | C | none | nope | nope_GCC | G/C-1_CCATTCG | 6 | FALSE | FALSE |
| 3_v.txt | 32126 | 32126-Deleti | Deletion | 1 | T | - | 5 | 30827 | 0.01621955 | 51.8 | WT_perm | FALSE | ARG | JSY13 | 5 | WT_perm-Re | CTT | nope | >1bp-ins | A/T-1 | none | A/T-1 | GAAAACTTT | TTTGTCA | GAA | CTT | T | none | nope | nope_CTT | A/T-1_TTTGTCA | 6 | FALSE | FALSE |
| 3_v.txt | 32210 | 32210-Deleti | Deletion | 1 | T | - | 7 | 30827 | 0.02270737 | 38 | WT_perm | FALSE | ARG | JSY13 | 5 | WT_perm-Re | CTT | nope | >1bp-ins | A/T-1 | none | A/T-1 | ACCACCTTTG | TTTGGTG | GAA | CTT | T | none | nope | nope_CTT | A/T-1_TTTGGTG | 6 | FALSE | FALSE |
| 3_v.txt | 32231 | 32231-Deleti | Deletion | 1 | T | - | 6 | 30825 | 0.01946472 | 37.8333333 | WT_perm | FALSE | ARG | JSY13 | 5 | WT_perm-Re | ATT | nope | >1bp-ins | A/T-1 | none | A/T-1 | CAGGAATTT | TTTAGGA | TAA | ATT | T | none | nope | nope_ATT | A/T-1_TTTAGGA | 6 | FALSE | FALSE |
| 3_v.txt | 32265 | 32265-Deleti | Deletion | 1 | A | - | 9 | 44582 | 0.02018752 | 37.8888889 | WT_perm | FALSE | ARG | JSY13 | 5 | WT_perm-Re | TAA | nope | >1bp-ins | A/T-1 | none | A/T-1 | CCAAATAAA | AAAATAC | ATT | ATT | T | none | T>none | T>none_ATT | A/T-1_AAAATAC | 6 | FALSE | FALSE |
| 3_v.txt | 32346 | 32346-Deleti | Deletion | 1 | A | - | 8 | 44583 | 0.01794406 | 53.25 | WT_perm | FALSE | ARG | JSY13 | 5 | WT_perm-Re | CAA | nope | >1bp-ins | A/T-1 | none | A/T-1 | TGTGGCAAA | AAAACCT | GTT | GTT | T | none | T>none | T>none_GTT | A/T-1_AAAACCT | 6 | FALSE | FALSE |
| 3_v.txt | 32381 | 32381-Deleti | Deletion | 1 | A | - | 8 | 44583 | 0.01794406 | 61.875 | WT_perm | FALSE | ARG | JSY13 | 5 | WT_perm-Re | TAA | nope | >1bp-ins | A/T-1 | none | A/T-1 | AATAATAAA | AAAGGGA | ATT | ATT | T | none | T>none | T>none_ATT | A/T-1_AAAGGGA | 6 | FALSE | FALSE |
| 3_v.txt | 32442 | 32442-Deleti | Deletion | 1 | A | - | 12 | 44583 | 0.02691609 | 56.4166667 | WT_perm | FALSE | ARG | JSY13 | 5 | WT_perm-Re | TAA | nope | >1bp-ins | A/T-1 | none | A/T-1 | GGAACTAAA | AAAAGTC | ATT | ATT | T | none | T>none | T>none_ATT | A/T-1_AAAAGTC | 6 | FALSE | FALSE |
| 3_v.txt | 32498 | 32498-Deleti | Deletion | 1 | T | - | 51 | 44583 | 0.11439338 | 44.6666655 | WT_perm | FALSE | ARG | JSY13 | 5 | WT_perm-Re | CTT | nope | >1bp-ins | A/T-1 | none | A/T-1 | AACAACTTTT | TTTTTTG | GAA | CTT | T | none | nope | nope_CTT | A/T-1_TTTTTTG | 6 | FALSE | FALSE |
| 3_v.txt | 32528 | 32528-Deleti | Deletion | 1 | G | - | 11 | 44582 | 0.02467364 | 42.5454546 | WT_perm | FALSE | ARG | JSY13 | 5 | WT_perm-Re | TGG | nope | >1bp-ins | G/C-1 | none | G/C-1 | TTTTCTGGGG | GGGGTTT | ACC | ACC | C | none | C>none | C>none_ACC | G/C-1_GGGGTTT | 6 | FALSE | FALSE |
| 3_v.txt | 32631 | 32631-Deleti | Deletion | 1 | T | - | 11 | 22998 | 0.04783025 | 47.1818182 | WT_perm | FALSE | ARG | JSY13 | 5 | WT_perm-Re | GTT | nope | >1bp-ins | A/T-1 | none | A/T-1 | CCTTCGTTTT | TTTTTAT | CAA | GTT | T | none | nope | nope_GTT | A/T-1_TTTTTAT | 6 | FALSE | FALSE |
| 3_v.txt | 32658 | 32658-Deleti | Deletion | 1 | C | - | 10 | 22998 | 0.04348204 | 57.9 | WT_perm | FALSE | ARG | JSY13 | 5 | WT_perm-Re | ACC | nope | >1bp-ins | G/C-1 | none | G/C-1 | CCTGGACCC | CCCCAGG | TGG | ACC | C | none | nope | nope_ACC | G/C-1_CCCCAGG | 6 | FALSE | FALSE |
| 3_v.txt | 32728 | 32728-Deleti | Deletion | 1 | A | - | 10 | 22998 | 0.04348204 | 61.3 | WT_perm | FALSE | ARG | JSY13 | 5 | WT_perm-Re | CAA | nope | >1bp-ins | A/T-1 | none | A/T-1 | CCATACAAA | AAAAACA | GTT | GTT | T | none | T>none | T>none_GTT | A/T-1_AAAAACA | 6 | FALSE | FALSE |
| 3_v.txt | 32760 | 32760-Deleti | Deletion | 1 | A | - | 10 | 22998 | 0.04348204 | 40.7 | WT_perm | FALSE | ARG | JSY13 | 5 | WT_perm-Re | TAA | nope | >1bp-ins | A/T-1 | none | A/T-1 | ATGGCTAAA | AAAACTT | ATT | ATT | T | none | T>none | T>none_ATT | A/T-1_AAAACTT | 6 | FALSE | FALSE |
| 3_v.txt | 32842 | 32842-Deleti | Deletion | 1 | A | - | 40 | 22998 | 0.17392817 | 41.6499999 | WT_perm | FALSE | ARG | JSY13 | 5 | WT_perm-Re | CAA | nope | >1bp-ins | A/T-1 | none | A/T-1 | TTACCCAAA | AAAAAAT | GTT | GTT | T | none | T>none | T>none_GTT | A/T-1_AAAAAAT | 6 | FALSE | FALSE |
| 3_v.txt | 32884 | 32884-Deleti | Deletion | 1 | A | - | 10 | 28267 | 0.03537694 | 34 | WT_perm | FALSE | ARG | JSY13 | 5 | WT_perm-Re | CAA | nope | >1bp-ins | A/T-1 | none | A/T-1 | ACGTCCAAA | AAAATTG | GTT | GTT | T | none | T>none | T>none_GTT | A/T-1_AAAATTG | 6 | FALSE | FALSE |
| 3_v.txt | 32901 | 32901-Deleti | Deletion | 1 | C | - | 1 | 28267 | 3.54E-03 | 38 | WT_perm | FALSE | ARG | JSY13 | 5 | WT_perm-Re | GCC | nope | >1bp-ins | G/C-1 | none | G/C-1 | ACTTGGCCA | CCAACTA | CGG | GCC | C | none | nope | nope_GCC | G/C-1_CCAACTA | 6 | FALSE | FALSE |
| 3_v.txt | 32924 | 32924-Deleti | Deletion | 1 | A | - | 30 | 28267 | 0.10613082 | 49.9666667 | WT_perm | FALSE | ARG | JSY13 | 5 | WT_perm-Re | CAA | nope | >1bp-ins | A/T-1 | none | A/T-1 | CAGGGCAAA | AAAAGTG | GTT | GTT | T | none | T>none | T>none_GTT | A/T-1_AAAAGTG | 6 | FALSE | FALSE |
| 3_v.txt | 32965 | 32965-Deleti | Deletion | 1 | C | - | 2 | 28267 | 7.08E-03 | 64 | WT_perm | FALSE | ARG | JSY13 | 5 | WT_perm-Re | GCC | nope | >1bp-ins | G/C-1 | none | G/C-1 | CATTGGCCG | CCGCACC | CGG | GCC | C | none | nope | nope_GCC | G/C-1_CCGCACC | 6 | FALSE | FALSE |
| 3_v.txt | 33037 | 33037-Deleti | Deletion | 1 | C | - | 4 | 28267 | 0.01415078 | 64 | WT_perm | FALSE | ARG | JSY13 | 5 | WT_perm-Re | GCC | nope | >1bp-ins | G/C-1 | none | G/C-1 | ATGTAGCCA | CCATTTC | CGG | GCC | C | none | nope | nope_GCC | G/C-1_CCATTTC | 6 | FALSE | FALSE |
| 3_v.txt | 33138 | 33138-Deleti | Deletion | 1 | A | - | 6 | 28267 | 0.02122617 | 42.1666667 | WT_perm | FALSE | ARG | JSY13 | 5 | WT_perm-Re | TAA | nope | >1bp-ins | A/T-1 | none | A/T-1 | GTGGATAAA | AAACCAA | ATT | ATT | T | none | T>none | T>none_ATT | A/T-1_AAACCAA | 6 | FALSE | FALSE |
| 3_v.txt | 33147 | 33147-Deleti | Deletion | 1 | A | - | 7 | 28267 | 0.02476386 | 41.5714286 | WT_perm | FALSE | ARG | JSY13 | 5 | WT_perm-Re | GAA | nope | >1bp-ins | A/T-1 | none | A/T-1 | CCAATGAAA | AAAAGAC | CTT | CTT | T | none | T>none | T>none_CTT | A/T-1_AAAAGAC | 6 | FALSE | FALSE |
| 3_v.txt | 33474 | 33474-Deleti | Deletion | 1 | C | - | 28 | 10726 | 0.26104792 | 29.6785714 | WT_perm | FALSE | ARG | JSY13 | 5 | WT_perm-Re | GCC | nope | >1bp-ins | G/C-1 | none | G/C-1 | GCTATGCCTT | CCTTTTT | CGG | GCC | C | none | nope | nope_GCC | G/C-1_CCTTTTT | 6 | FALSE | FALSE |
| 4_v.txt | 31723 | 31723-SNV- | SNV | 1 | A | C | 7 | 13628 | 0.05136484 | 60.4285714 | WT_perm | FALSE | ARG | JSY13 | 6 | WT_perm-Re | CAA | A>C | G/C+1 | A/T-1 | TA>GC | TA>GC | TGTCCCAAA | AAAAAGT | GTT | GTT | T | G | T>G | T>G_GTT | TA>GC_AAAAAGT | 6 | FALSE | FALSE |
| 4_v.txt | 32352 | 32352-SNV-T | SNV | 1 | T | G | 4 | 36551 | 0.01094361 | 63 | WT_perm | FALSE | ARG | JSY13 | 6 | WT_perm-Re | CTT | T>G | G/C+1 | A/T-1 | TA>GC | TA>GC | AAAACCTTTC | TTTGTAC | GAA | CTT | T | C | T>G | T>G_CTT | TA>GC_TTTGTAC | 6 | FALSE | FALSE |
| 4_v.txt | 32662 | 32662-SNV- | SNV | 1 | A | C | 7 | 28779 | 0.02432329 | 58.1414461 | WT_perm | FALSE | ARG | JSY13 | 6 | WT_perm-Re | CAG | A>C | G/C+1 | A/T-1 | TA>GC | TA>GC | GACCCCAGG | AGGCACC | GTC | GTC | T | G | T>G | T>G_GTC | TA>GC_AGGCACC | 6 | FALSE | FALSE |
| 4_v.txt | 32763 | 32763-SNV- | SNV | 1 | A | C | 6 | 28780 | 0.02084781 | 59 | WT_perm | FALSE | ARG | JSY13 | 6 | WT_perm-Re | AAC | A>C | G/C+1 | A/T-1 | TA>GC | TA>GC | GCTAAAACT | ACTTTGA | TTG | TTG | T | G | T>G | T>G_TTG | TA>GC_ACTTTGA | 6 | FALSE | FALSE |
| 4_v.txt | 32842 | 32842-SNV- | SNV | 1 | A | C | 7 | 28780 | 0.02432245 | 37.1427712 | WT_perm | FALSE | ARG | JSY13 | 6 | WT_perm-Re | CAA | A>C | G/C+1 | A/T-1 | TA>GC | TA>GC | TTACCCAAA | AAAAAAT | GTT | GTT | T | G | T>G | T>G_GTT | TA>GC_AAAAAAT | 6 | FALSE | FALSE |
| 4_v.txt | 32884 | 32884-SNV- | SNV | 1 | A | C | 8 | 26009 | 0.03075858 | 31.8755035 | WT_perm | FALSE | ARG | JSY13 | 6 | WT_perm-Re | CAA | A>C | G/C+1 | A/T-1 | TA>GC | TA>GC | ACGTCCAAA | AAAATTG | GTT | GTT | T | G | T>G | T>G_GTT | TA>GC_AAAATTG | 6 | FALSE | FALSE |
| 4_v.txt | 33269 | 33269-SNV- | SNV | 1 | A | C | 101 | 16555 | 0.61008759 | 14.6633663 | WT_perm | FALSE | ARG | JSY13 | 6 | WT_perm-Re | TAG | A>C | G/C+1 | A/T-1 | TA>GC | TA>GC | CTCCATAGA | AGAGAAC | ATC | ATC | T | G | T>G | T>G_ATC | TA>GC_AGAGAAC | 6 | FALSE | FALSE |
| 4_v.txt | 32573 | 32573-SNV-T | SNV | 1 | T | C | 19 | 28780 | 0.06601807 | 34.2105263 | WT_perm | FALSE | ARG | JSY13 | 6 | WT_perm-Re | CTT | T>C | G/C+1 | A/T-1 | TA>CG | TA>CG | AGTACCTTG | TTGAAAT | GAA | CTT | T | G | T>C | T>C_CTT | TA>CG_TTGAAAT | 6 | FALSE | FALSE |
| 4_v.txt | 32662 | 32662-SNV- | SNV | 1 | A | G | 13 | 28779 | 0.04517183 | 60.9982262 | WT_perm | FALSE | ARG | JSY13 | 6 | WT_perm-Re | CAG | A>G | G/C+1 | A/T-1 | TA>CG | TA>CG | GACCCCAGG | AGGCACC | GTC | GTC | T | C | T>C | T>C_GTC | TA>CG_AGGCACC | 6 | FALSE | FALSE |
| 4_v.txt | 32573 | 32573-SNV-T | SNV | 1 | T | A | 11 | 28780 | 0.03822099 | 37.7272727 | WT_perm | FALSE | ARG | JSY13 | 6 | WT_perm-Re | CTT | T>A | A/T+1 | A/T-1 | TA>AT | TA>AT | AGTACCTTG | TTGAAAT | GAA | CTT | T | T | T>A | T>A_CTT | TA>AT_TTGAAAT | 6 | FALSE | FALSE |
| 4_v.txt | 31650 | 31650-SNV- | SNV | 1 | C | T | 39 | 13628 | 0.28617552 | 35.8717949 | WT_perm | FALSE | ARG | JSY13 | 6 | WT_perm-Re | GCG | C>T | A/T+1 | G/C-1 | CG>TA | CG>TA | AGAATGCGA | CGAAATG | CGC | GCG | C | A | C>T | C>T_GCG | CG>TA_CGAAATG | 6 | FALSE | FALSE |
| 4_v.txt | 31658 | 31658-SNV- | SNV | 1 | C | T | 19 | 13628 | 0.13941884 | 34.3157895 | WT_perm | FALSE | ARG | JSY13 | 6 | WT_perm-Re | GCG | C>T | A/T+1 | G/C-1 | CG>TA | CG>TA | AAATGGCGT | CGTGGAA | CGC | GCG | C | A | C>T | C>T_GCG | CG>TA_CGTGGAA | 6 | FALSE | FALSE |
| 4_v.txt | 31830 | 31830-SNV- | SNV | 1 | C | T | 21 | 13628 | 0.15409451 | 55 | WT_perm | FALSE | ARG | JSY13 | 6 | WT_perm-Re | GCA | C>T | A/T+1 | G/C-1 | CG>TA | CG>TA | AATATGCATT | CATTGAA | CGT | GCA | C | A | C>T | C>T_GCA | CG>TA_CATTGAA | 6 | FALSE | FALSE |
| 4_v.txt | 31882 | 31882-SNV- | SNV | 1 | C | T | 16 | 13628 | 0.11740534 | 43.6875 | WT_perm | FALSE | ARG | JSY13 | 6 | WT_perm-Re | GCA | C>T | A/T+1 | G/C-1 | CG>TA | CG>TA | AGGCGGCAC | CAGCAAA | CGT | GCA | C | A | C>T | C>T_GCA | CG>TA_CAGCAAA | 6 | FALSE | FALSE |
| 4_v.txt | 31920 | 31920-SNV- | SNV | 1 | G | A | 17 | 13628 | 0.12474318 | 37.8235294 | WT_perm | FALSE | ARG | JSY13 | 6 | WT_perm-Re | CGT | G>A | A/T+1 | G/C-1 | CG>TA | CG>TA | AAAGCCGTG | GTGAAAC | GCA | GCA | C | T | C>T | C>T_GCA | CG>TA_GTGAAAC | 6 | FALSE | FALSE |
| 4_v.txt | 31962 | 31962-SNV- | SNV | 1 | G | A | 37 | 19856 | 0.18634166 | 35.2162162 | WT_perm | FALSE | ARG | JSY13 | 6 | WT_perm-Re | CGC | G>A | A/T+1 | G/C-1 | CG>TA | CG>TA | GTGGCCGCA | GCATAAT | GCG | GCG | C | T | C>T | C>T_GCG | CG>TA_GCATAAT | 6 | FALSE | FALSE |
| 4_v.txt | 31972 | 31972-SNV- | SNV | 1 | C | T | 15 | 19856 | 0.07554392 | 36.0666667 | WT_perm | FALSE | ARG | JSY13 | 6 | WT_perm-Re | GCC | C>T | A/T+1 | G/C-1 | CG>TA | CG>TA | AATAAGCCA | CCAAGCC | CGG | GCC | C | A | C>T | C>T_GCC | CG>TA_CCAAGCC | 6 | FALSE | FALSE |
| 4_v.txt | 31981 | 31981-SNV- | SNV | 1 | G | A | 26 | 19856 | 0.13094279 | 42.8846154 | WT_perm | FALSE | ARG | JSY13 | 6 | WT_perm-Re | GGC | G>A | A/T+1 | G/C-1 | CG>TA | CG>TA | AGCCGGGCA | GCATTAA | CCG | CCG | C | T | C>T | C>T_CCG | CG>TA_GCATTAA | 6 | FALSE | FALSE |
| 4_v.txt | 32057 | 32057-SNV- | SNV | 1 | C | T | 24 | 19856 | 0.12087027 | 57.25 | WT_perm | FALSE | ARG | JSY13 | 6 | WT_perm-Re | GCG | C>T | A/T+1 | G/C-1 | CG>TA | CG>TA | GATGTGCGA | CGAGATT | CGC | GCG | C | A | C>T | C>T_GCG | CG>TA_CGAGATT | 6 | FALSE | FALSE |
| 4_v.txt | 32092 | 32092-SNV- | SNV | 1 | C | T | 31 | 19856 | 0.15612409 | 60.0645161 | WT_perm | FALSE | ARG | JSY13 | 6 | WT_perm-Re | GCA | C>T | A/T+1 | G/C-1 | CG>TA | CG>TA | AGCCTGCAA | CAACACC | CGT | GCA | C | A | C>T | C>T_GCA | CG>TA_CAACACC | 6 | FALSE | FALSE |
| 4_v.txt | 32165 | 32165-SNV- | SNV | 1 | G | A | 24 | 19856 | 0.12087027 | 48.75 | WT_perm | FALSE | ARG | JSY13 | 6 | WT_perm-Re | CGC | G>A | A/T+1 | G/C-1 | CG>TA | CG>TA | CAAAGCGCC | GCCAAAT | GCG | GCG | C | T | C>T | C>T_GCG | CG>TA_GCCAAAT | 6 | FALSE | FALSE |
| 4_v.txt | 32173 | 32173-SNV- | SNV | 1 | C | T | 24 | 19856 | 0.12087027 | 51 | WT_perm | FALSE | ARG | JSY13 | 6 | WT_perm-Re | GCA | C>T | A/T+1 | G/C-1 | CG>TA | CG>TA | CAAATGCAG | CAGCAGT | CGT | GCA | C | A | C>T | C>T_GCA | CG>TA_CAGCAGT | 6 | FALSE | FALSE |

| | | | | | | | | | | | | | | | | | | | | | | | | | | | | | | | | | | |
|---|---|---|---|---|---|---|---|---|---|---|---|---|---|---|---|---|---|---|---|---|---|---|---|---|---|---|---|---|---|---|---|---|---|---|
| 4_v.txt | 32271 | 32271-SNV- | SNV | 1 | C | T | 47 | 36551 | 0.12858745 | 28.0638298 | WT_perm | FALSE | ARG | JSY13 | 6 | WT_perm-Re | ACG | C>T | A/T+1 | G/C-1 | CG>TA | CG>TA | AAAATACGG | CGGGAAC | TGC | ACG | C | A | C>T | C>T_ACG | CG>TA_CGGGAAC | 6 | FALSE | FALSE |
| 4_v.txt | 32299 | 32299-SNV- | SNV | 1 | C | T | 78 | 36551 | 0.21340045 | 54.8333333 | WT_perm | FALSE | ARG | JSY13 | 6 | WT_perm-Re | GCG | C>T | A/T+1 | G/C-1 | CG>TA | CG>TA | AATTTGCGG | CGGCAGA | CGC | GCG | C | A | C>T | C>T_GCG | CG>TA_CGGCAGA | 6 | FALSE | FALSE |
| 4_v.txt | 32466 | 32466-SNV- | SNV | 1 | C | T | 67 | 36551 | 0.18330552 | 46.4776119 | WT_perm | FALSE | ARG | JSY13 | 6 | WT_perm-Re | GCC | C>T | A/T+1 | G/C-1 | CG>TA | CG>TA | AGAGAGCCA | CCAATGT | CGG | GCC | C | A | C>T | C>T_GCC | CG>TA_CCAATGT | 6 | FALSE | FALSE |
| 4_v.txt | 32487 | 32487-SNV- | SNV | 1 | C | T | 39 | 36551 | 0.10670023 | 45.2820513 | WT_perm | FALSE | ARG | JSY13 | 6 | WT_perm-Re | ACG | C>T | A/T+1 | G/C-1 | CG>TA | CG>TA | AAGATACGG | CGGAAAA | TGC | ACG | C | A | C>T | C>T_ACG | CG>TA_CGGAAAA | 6 | FALSE | FALSE |
| 4_v.txt | 32535 | 32535-SNV- | SNV | 1 | G | A | 60 | 36551 | 0.1641542 | 36.4489119 | WT_perm | FALSE | ARG | JSY13 | 6 | WT_perm-Re | TGC | G>A | A/T+1 | G/C-1 | CG>TA | CG>TA | GGGTTTGCA | GCAGCTT | ACG | ACG | C | T | C>T | C>T_ACG | CG>TA_GCAGCTT | 6 | FALSE | FALSE |
| 4_v.txt | 32536 | 32536-SNV- | SNV | 1 | C | T | 72 | 36551 | 0.19698504 | 37.207559 | WT_perm | FALSE | ARG | JSY13 | 6 | WT_perm-Re | GCA | C>T | A/T+1 | G/C-1 | CG>TA | CG>TA | GGTTTGCAG | CAGCTTC | CGT | GCA | C | A | C>T | C>T_GCA | CG>TA_CAGCTTC | 6 | FALSE | FALSE |
| 4_v.txt | 32590 | 32590-SNV- | SNV | 1 | C | T | 61 | 28780 | 0.21195275 | 36.7213115 | WT_perm | FALSE | ARG | JSY13 | 6 | WT_perm-Re | GCG | C>T | A/T+1 | G/C-1 | CG>TA | CG>TA | AGGCAGCGT | CGTTAAT | CGC | GCG | C | A | C>T | C>T_GCG | CG>TA_CGTTAAT | 6 | FALSE | FALSE |
| 4_v.txt | 32700 | 32700-SNV- | SNV | 1 | C | T | 44 | 28780 | 0.15288395 | 58.7045455 | WT_perm | FALSE | ARG | JSY13 | 6 | WT_perm-Re | GCC | C>T | A/T+1 | G/C-1 | CG>TA | CG>TA | ACTGGGCCG | CCGGTAA | CGG | GCC | C | A | C>T | C>T_GCC | CG>TA_CCGGTAA | 6 | FALSE | FALSE |
| 4_v.txt | 32702 | 32702-SNV- | SNV | 1 | G | A | 32 | 28780 | 0.11118833 | 58.46875 | WT_perm | FALSE | ARG | JSY13 | 6 | WT_perm-Re | CGG | G>A | A/T+1 | G/C-1 | CG>TA | CG>TA | TGGGCCGGT | GGTAACC | GCC | GCC | C | T | C>T | C>T_GCC | CG>TA_GGTAACC | 6 | FALSE | FALSE |
| 4_v.txt | 32776 | 32776-SNV- | SNV | 1 | C | T | 40 | 28780 | 0.13898541 | 51.675 | WT_perm | FALSE | ARG | JSY13 | 6 | WT_perm-Re | GCG | C>T | A/T+1 | G/C-1 | CG>TA | CG>TA | TGGAAGCGA | CGACCCA | CGC | GCG | C | A | C>T | C>T_GCG | CG>TA_CGACCCA | 6 | FALSE | FALSE |
| 4_v.txt | 32798 | 32798-SNV- | SNV | 1 | G | A | 44 | 28780 | 0.15288395 | 45.3636364 | WT_perm | FALSE | ARG | JSY13 | 6 | WT_perm-Re | CGT | G>A | A/T+1 | G/C-1 | CG>TA | CG>TA | TTCACCGTA | GTAATAT | GCA | GCA | C | T | C>T | C>T_GCA | CG>TA_GTAATAT | 6 | FALSE | FALSE |
| 4_v.txt | 32860 | 32860-SNV- | SNV | 1 | C | T | 33 | 28775 | 0.11468288 | 37.0606061 | WT_perm | FALSE | ARG | JSY13 | 6 | WT_perm-Re | GCC | C>T | A/T+1 | G/C-1 | CG>TA | CG>TA | TCCATGCCG | CCGCCAG | CGG | GCC | C | A | C>T | C>T_GCC | CG>TA_CCGCCAG | 6 | FALSE | FALSE |
| 4_v.txt | 32879 | 32879-SNV- | SNV | 1 | C | T | 19 | 26006 | 0.07306006 | 37.6842105 | WT_perm | FALSE | ARG | JSY13 | 6 | WT_perm-Re | ACG | C>T | A/T+1 | G/C-1 | CG>TA | CG>TA | TTTGTACGTC | CGTCCAA | TGC | ACG | C | A | C>T | C>T_ACG | CG>TA_CGTCCAA | 6 | FALSE | FALSE |
| 4_v.txt | 32900 | 32900-SNV- | SNV | 1 | G | A | 39 | 26009 | 0.1499481 | 35.5277321 | WT_perm | FALSE | ARG | JSY13 | 6 | WT_perm-Re | GGC | G>A | A/T+1 | G/C-1 | CG>TA | CG>TA | GACTTGGCC | GCCAACT | CCG | CCG | C | T | C>T | C>T_CCG | CG>TA_GCCAACT | 6 | FALSE | FALSE |
| 4_v.txt | 32901 | 32901-SNV- | SNV | 1 | C | T | 68 | 26009 | 0.26144796 | 31.4857642 | WT_perm | FALSE | ARG | JSY13 | 6 | WT_perm-Re | GCC | C>T | A/T+1 | G/C-1 | CG>TA | CG>TA | ACTTGGCCA | CCAACTA | CGG | GCC | C | A | C>T | C>T_GCC | CG>TA_CCAACTA | 6 | FALSE | FALSE |
| 4_v.txt | 32922 | 32922-SNV- | SNV | 1 | G | A | 27 | 26009 | 0.10381022 | 52.8148148 | WT_perm | FALSE | ARG | JSY13 | 6 | WT_perm-Re | GGC | G>A | A/T+1 | G/C-1 | CG>TA | CG>TA | TCCAGGGCA | GCAAAAG | CCG | CCG | C | T | C>T | C>T_CCG | CG>TA_GCAAAAG | 6 | FALSE | FALSE |
| 4_v.txt | 32935 | 32935-SNV- | SNV | 1 | C | T | 40 | 26009 | 0.15379292 | 44.4 | WT_perm | FALSE | ARG | JSY13 | 6 | WT_perm-Re | GCC | C>T | A/T+1 | G/C-1 | CG>TA | CG>TA | TGATTGCCC | CCCAAGA | CGG | GCC | C | A | C>T | C>T_GCC | CG>TA_CCCAAGA | 6 | FALSE | FALSE |
| 4_v.txt | 32965 | 32965-SNV- | SNV | 1 | C | T | 36 | 26009 | 0.13841363 | 56.8055556 | WT_perm | FALSE | ARG | JSY13 | 6 | WT_perm-Re | GCC | C>T | A/T+1 | G/C-1 | CG>TA | CG>TA | CATTGGCCG | CCGCACC | CGG | GCC | C | A | C>T | C>T_GCC | CG>TA_CCGCACC | 6 | FALSE | FALSE |
| 4_v.txt | 32967 | 32967-SNV- | SNV | 1 | G | A | 67 | 26009 | 0.25760314 | 60.641791 | WT_perm | FALSE | ARG | JSY13 | 6 | WT_perm-Re | CGC | G>A | A/T+1 | G/C-1 | CG>TA | CG>TA | TTGGCCGCA | GCACCAA | GCG | GCG | C | T | C>T | C>T_GCG | CG>TA_GCACCAA | 6 | FALSE | FALSE |
| 4_v.txt | 33056 | 33056-SNV- | SNV | 1 | C | T | 38 | 26009 | 0.14610327 | 56.1052632 | WT_perm | FALSE | ARG | JSY13 | 6 | WT_perm-Re | GCG | C>T | A/T+1 | G/C-1 | CG>TA | CG>TA | GGACTGCGT | CGTGACA | CGC | GCG | C | A | C>T | C>T_GCG | CG>TA_CGTGACA | 6 | FALSE | FALSE |
| 4_v.txt | 33069 | 33069-SNV- | SNV | 1 | G | A | 33 | 26009 | 0.12687916 | 47.3329342 | WT_perm | FALSE | ARG | JSY13 | 6 | WT_perm-Re | TGC | G>A | A/T+1 | G/C-1 | CG>TA | CG>TA | GAATATGCC | GCCAAAG | ACG | ACG | C | T | C>T | C>T_ACG | CG>TA_GCCAAAG | 6 | FALSE | FALSE |
| 4_v.txt | 33070 | 33070-SNV- | SNV | 1 | C | T | 28 | 26009 | 0.10765504 | 54.532005 | WT_perm | FALSE | ARG | JSY13 | 6 | WT_perm-Re | GCC | C>T | A/T+1 | G/C-1 | CG>TA | CG>TA | AATATGCCA | CCAAAGA | CGG | GCC | C | A | C>T | C>T_GCC | CG>TA_CCAAAGA | 6 | FALSE | FALSE |
| 4_v.txt | 33104 | 33104-SNV- | SNV | 1 | G | A | 33 | 26009 | 0.12687916 | 46.6666667 | WT_perm | FALSE | ARG | JSY13 | 6 | WT_perm-Re | CGC | G>A | A/T+1 | G/C-1 | CG>TA | CG>TA | AAGAGCGCC | GCCCACT | GCG | GCG | C | T | C>T | C>T_GCG | CG>TA_GCCCACT | 6 | FALSE | FALSE |
| 4_v.txt | 33114 | 33114-SNV- | SNV | 1 | C | T | 37 | 26009 | 0.14225845 | 41.1891892 | WT_perm | FALSE | ARG | JSY13 | 6 | WT_perm-Re | GCC | C>T | A/T+1 | G/C-1 | CG>TA | CG>TA | ACTGGGCCG | CCGGCGT | CGG | GCC | C | A | C>T | C>T_GCC | CG>TA_CCGGCGT | 6 | FALSE | FALSE |
| 4_v.txt | 33118 | 33118-SNV- | SNV | 1 | C | T | 28 | 26009 | 0.10765504 | 41.5357143 | WT_perm | FALSE | ARG | JSY13 | 6 | WT_perm-Re | GCG | C>T | A/T+1 | G/C-1 | CG>TA | CG>TA | GGCCGGCGT | CGTTGGT | CGC | GCG | C | A | C>T | C>T_GCG | CG>TA_CGTTGGT | 6 | FALSE | FALSE |
| 4_v.txt | 33258 | 33258-SNV- | SNV | 1 | C | T | 23 | 16555 | 0.13893084 | 46.6086957 | WT_perm | FALSE | ARG | JSY13 | 6 | WT_perm-Re | GCC | C>T | A/T+1 | G/C-1 | CG>TA | CG>TA | CCTATGCCAT | CCATCCT | CGG | GCC | C | A | C>T | C>T_GCC | CG>TA_CCATCCT | 6 | FALSE | FALSE |
| 4_v.txt | 33276 | 33276-SNV- | SNV | 1 | G | A | 27 | 16555 | 0.16309272 | 47.5185185 | WT_perm | FALSE | ARG | JSY13 | 6 | WT_perm-Re | CGT | G>A | A/T+1 | G/C-1 | CG>TA | CG>TA | GAGAACGTA | GTATCCT | GCA | GCA | C | T | C>T | C>T_GCA | CG>TA_GTATCCT | 6 | FALSE | FALSE |
| 4_v.txt | 33284 | 33284-SNV- | SNV | 1 | G | A | 22 | 16555 | 0.13289037 | 44.0454546 | WT_perm | FALSE | ARG | JSY13 | 6 | WT_perm-Re | CGC | G>A | A/T+1 | G/C-1 | CG>TA | CG>TA | ATCCTCGCC | GCCATTT | GCG | GCG | C | T | C>T | C>T_GCG | CG>TA_GCCATTT | 6 | FALSE | FALSE |
| 4_v.txt | 33309 | 33309-SNV- | SNV | 1 | C | T | 19 | 16555 | 0.11476895 | 52.6281679 | WT_perm | FALSE | ARG | JSY13 | 6 | WT_perm-Re | GCG | C>T | A/T+1 | G/C-1 | CG>TA | CG>TA | AAAGAGCGC | CGCAATG | CGC | GCG | C | A | C>T | C>T_GCG | CG>TA_CGCAATG | 6 | FALSE | FALSE |
| 4_v.txt | 33310 | 33310-SNV- | SNV | 1 | G | A | 23 | 16553 | 0.13894762 | 54.8941496 | WT_perm | FALSE | ARG | JSY13 | 6 | WT_perm-Re | CGC | G>A | A/T+1 | G/C-1 | CG>TA | CG>TA | AAGAGCGCA | GCAATGG | GCG | GCG | C | T | C>T | C>T_GCG | CG>TA_GCAATGG | 6 | FALSE | FALSE |
| 4_v.txt | 33412 | 33412-SNV- | SNV | 1 | G | A | 25 | 16555 | 0.15101178 | 44.8 | WT_perm | FALSE | ARG | JSY13 | 6 | WT_perm-Re | GGC | G>A | A/T+1 | G/C-1 | CG>TA | CG>TA | TGACCGGCT | GCTCATT | CCG | CCG | C | T | C>T | C>T_CCG | CG>TA_GCTCATT | 6 | FALSE | FALSE |
| 4_v.txt | 33474 | 33474-SNV- | SNV | 1 | C | T | 21 | 16552 | 0.12687289 | 34.0952381 | WT_perm | FALSE | ARG | JSY13 | 6 | WT_perm-Re | GCC | C>T | A/T+1 | G/C-1 | CG>TA | CG>TA | GCTATGCCTT | CCTTTTT | CGG | GCC | C | A | C>T | C>T_GCC | CG>TA_CCTTTTT | 6 | FALSE | FALSE |
| 4_v.txt | 31962 | 31962-SNV- | SNV | 1 | G | C | 2 | 19856 | 0.01007252 | 38 | WT_perm | FALSE | ARG | JSY13 | 6 | WT_perm-Re | CGC | G>C | G/C+1 | G/C-1 | CG>GC | CG>GC | GTGGCCGCA | GCATAAT | GCG | GCG | C | G | C>G | C>G_GCG | CG>GC_GCATAAT | 6 | FALSE | FALSE |
| 4_v.txt | 32536 | 32536-SNV- | SNV | 1 | C | G | 2 | 36551 | 5.47E-03 | 37.4988671 | WT_perm | FALSE | ARG | JSY13 | 6 | WT_perm-Re | GCA | C>G | G/C+1 | G/C-1 | CG>GC | CG>GC | GGTTTGCAG | CAGCTTC | CGT | GCA | C | C | C>G | C>G_GCA | CG>GC_CAGCTTC | 6 | FALSE | FALSE |
| 4_v.txt | 33069 | 33069-SNV- | SNV | 1 | G | C | 1 | 26009 | 3.85E-03 | 63.9993246 | WT_perm | FALSE | ARG | JSY13 | 6 | WT_perm-Re | TGC | G>C | G/C+1 | G/C-1 | CG>GC | CG>GC | GAATATGCC | GCCAAAG | ACG | ACG | C | G | C>G | C>G_ACG | CG>GC_GCCAAAG | 6 | FALSE | FALSE |
| 4_v.txt | 31639 | 31639-SNV- | SNV | 1 | G | T | 13 | 13628 | 0.09539184 | 29.8461539 | WT_perm | FALSE | ARG | JSY13 | 6 | WT_perm-Re | AGG | G>T | A/T+1 | G/C-1 | CG>AT | CG>AT | TTATGAGGG | GGGTGAG | TCC | TCC | C | A | C>A | C>A_TCC | CG>AT_GGGTGAG | 6 | FALSE | FALSE |
| 4_v.txt | 31657 | 31657-SNV- | SNV | 1 | G | T | 115 | 13628 | 0.8438509 | 19.9468653 | WT_perm | FALSE | ARG | JSY13 | 6 | WT_perm-Re | GGC | G>T | A/T+1 | G/C-1 | CG>AT | CG>AT | GAAATGGCG | GCGTGGA | CCG | CCG | C | A | C>A | C>A_CCG | CG>AT_GCGTGGA | 6 | FALSE | FALSE |
| 4_v.txt | 31658 | 31658-SNV- | SNV | 1 | C | A | 4 | 13628 | 0.02935134 | 30.25 | WT_perm | FALSE | ARG | JSY13 | 6 | WT_perm-Re | GCG | C>A | A/T+1 | G/C-1 | CG>AT | CG>AT | AAATGGCGT | CGTGGAA | CGC | GCG | C | T | C>A | C>A_GCG | CG>AT_CGTGGAA | 6 | FALSE | FALSE |
| 4_v.txt | 31877 | 31877-SNV- | SNV | 1 | G | T | 7 | 13628 | 0.05136484 | 48.4285714 | WT_perm | FALSE | ARG | JSY13 | 6 | WT_perm-Re | AGG | G>T | A/T+1 | G/C-1 | CG>AT | CG>AT | GATATAGGC | GGCGGCA | TCC | TCC | C | A | C>A | C>A_TCC | CG>AT_GGCGGCA | 6 | FALSE | FALSE |
| 4_v.txt | 32663 | 32663-SNV- | SNV | 1 | G | T | 9 | 28780 | 0.03127172 | 56.7769183 | WT_perm | FALSE | ARG | JSY13 | 6 | WT_perm-Re | AGG | G>T | A/T+1 | G/C-1 | CG>AT | CG>AT | ACCCCAGGC | GGCACCT | TCC | TCC | C | A | C>A | C>A_TCC | CG>AT_GGCACCT | 6 | FALSE | FALSE |
| 4_v.txt | 32702 | 32702-SNV- | SNV | 1 | G | T | 7 | 28780 | 0.02432245 | 63.2857143 | WT_perm | FALSE | ARG | JSY13 | 6 | WT_perm-Re | CGG | G>T | A/T+1 | G/C-1 | CG>AT | CG>AT | TGGGCCGGT | GGTAACC | GCC | GCC | C | A | C>A | C>A_GCC | CG>AT_GGTAACC | 6 | FALSE | FALSE |
| 4_v.txt | 33310 | 33310-SNV- | SNV | 1 | G | T | 12 | 16553 | 0.07249441 | 20.4145344 | WT_perm | FALSE | ARG | JSY13 | 6 | WT_perm-Re | CGC | G>T | A/T+1 | G/C-1 | CG>AT | CG>AT | AAGAGCGCA | GCAATGG | GCG | GCG | C | A | C>A | C>A_GCG | CG>AT_GCAATGG | 6 | FALSE | FALSE |
| 4_v.txt | 33474 | 33474-SNV- | SNV | 1 | C | A | 57 | 16552 | 0.34436926 | 34.4912281 | WT_perm | FALSE | ARG | JSY13 | 6 | WT_perm-Re | GCC | C>A | A/T+1 | G/C-1 | CG>AT | CG>AT | GCTATGCCTT | CCTTTTT | CGG | GCC | C | T | C>A | C>A_GCC | CG>AT_CCTTTTT | 6 | FALSE | FALSE |
| 4_v.txt | 33475 | 33475-SNV- | SNV | 1 | C | A | 19 | 16552 | 0.11478975 | 37.0526316 | WT_perm | FALSE | ARG | JSY13 | 6 | WT_perm-Re | CCT | C>A | A/T+1 | G/C-1 | CG>AT | CG>AT | CTATGCCTTT | CTTTTTT | GGA | CCT | C | T | C>A | C>A_CCT | CG>AT_CTTTTTT | 6 | FALSE | FALSE |
| 4_v.txt | 32662 | 32662-MNV- | MNV | 2 | AG | CA | 1 | 28780 | 3.48E-03 | 63.9999723 | WT_perm | FALSE | ARG | JSY13 | 6 | WT_perm-Re | CAG | nope | >1bp-ins | >1bpdel | none | MNV | GACCCCAGG | AGGCACC | GTC | CAG | AG | none | NA | NA_CAG | MNV_AGGCACC | 6 | FALSE | FALSE |
| 4_v.txt | 32081 | 32081-Insert | Insertion | 1 | - | A | 13 | 19856 | 0.06547139 | 64 | WT_perm | FALSE | ARG | JSY13 | 6 | WT_perm-Re | CAA | nope | A/T+1 | >1bpdel | none | A/T+1 | CCATGCAAA | AAAAAAG | GTT | CAA | - | T | NA | NA_CAA | A/T+1_AAAAAAG | 6 | FALSE | FALSE |
| 4_v.txt | 32498 | 32498-Insert | Insertion | 1 | - | T | 2 | 36551 | 5.47E-03 | 38 | WT_perm | FALSE | ARG | JSY13 | 6 | WT_perm-Re | CTT | nope | A/T+1 | >1bpdel | none | A/T+1 | AACAACTTTT | TTTTTTG | GAA | CTT | - | A | NA | NA_CTT | A/T+1_TTTTTTG | 6 | FALSE | FALSE |
| 4_v.txt | 32842 | 32842-Insert | Insertion | 1 | - | A | 4 | 28780 | 0.01389854 | 38 | WT_perm | FALSE | ARG | JSY13 | 6 | WT_perm-Re | CAA | nope | A/T+1 | >1bpdel | none | A/T+1 | TTACCCAAA | AAAAAAT | GTT | CAA | - | T | NA | NA_CAA | A/T+1_AAAAAAT | 6 | FALSE | FALSE |
| 4_v.txt | 32884 | 32884-Insert | Insertion | 1 | - | A | 3 | 26009 | 0.01153447 | 37 | WT_perm | FALSE | ARG | JSY13 | 6 | WT_perm-Re | CAA | nope | A/T+1 | >1bpdel | none | A/T+1 | ACGTCCAAA | AAAATTG | GTT | CAA | - | T | NA | NA_CAA | A/T+1_AAAATTG | 6 | FALSE | FALSE |
| 4_v.txt | 31723 | 31723-Deleti | Deletion | 1 | A | - | 8 | 13628 | 0.05870267 | 60.75 | WT_perm | FALSE | ARG | JSY13 | 6 | WT_perm-Re | CAA | nope | >1bp-ins | A/T-1 | none | A/T-1 | TGTCCCAAA | AAAAAGT | GTT | GTT | T | none | T>none | T>none_GTT | A/T-1_AAAAAGT | 6 | FALSE | FALSE |
| 4_v.txt | 31863 | 31863-Deleti | Deletion | 1 | A | - | 4 | 13628 | 0.02935134 | 55.25 | WT_perm | FALSE | ARG | JSY13 | 6 | WT_perm-Re | GAA | nope | >1bp-ins | A/T-1 | none | A/T-1 | AACAGGAAA | AAAATAG | CTT | CTT | T | none | T>none | T>none_CTT | A/T-1_AAAATAG | 6 | FALSE | FALSE |
| 4_v.txt | 31904 | 31904-Deleti | Deletion | 1 | T | - | 7 | 13628 | 0.05136484 | 48.8571429 | WT_perm | FALSE | ARG | JSY13 | 6 | WT_perm-Re | ATT | nope | >1bp-ins | A/T-1 | none | A/T-1 | ATTGAATTTT | TTTTGGT | TAA | ATT | T | none | nope | nope_ATT | A/T-1_TTTTGGT | 6 | FALSE | FALSE |
| 4_v.txt | 31913 | 31913-Deleti | Deletion | 1 | A | - | 11 | 13628 | 0.08071617 | 40.2727273 | WT_perm | FALSE | ARG | JSY13 | 6 | WT_perm-Re | CAA | nope | >1bp-ins | A/T-1 | none | A/T-1 | TGGTGCAAA | AAAAGCC | GTT | GTT | T | none | T>none | T>none_GTT | A/T-1_AAAAGCC | 6 | FALSE | FALSE |
| 4_v.txt | 31920 | 31920-Deleti | Deletion | 1 | G | - | 1 | 13628 | 7.34E-03 | 38 | WT_perm | FALSE | ARG | JSY13 | 6 | WT_perm-Re | CGT | nope | >1bp-ins | G/C-1 | none | G/C-1 | AAAGCCGTG | GTGAAAC | GCA | GCA | C | none | C>none | C>none_GCA | G/C-1_GTGAAAC | 6 | FALSE | FALSE |
| 4_v.txt | 31972 | 31972-Deleti | Deletion | 1 | C | - | 7 | 19856 | 0.03525383 | 34 | WT_perm | FALSE | ARG | JSY13 | 6 | WT_perm-Re | GCC | nope | >1bp-ins | G/C-1 | none | G/C-1 | AATAAGCCA | CCAAGCC | CGG | GCC | C | none | nope | nope_GCC | G/C-1_CCAAGCC | 6 | FALSE | FALSE |
| 4_v.txt | 32081 | 32081-Deleti | Deletion | 1 | A | - | 16 | 19855 | 0.08058424 | 62.3125006 | WT_perm | FALSE | ARG | JSY13 | 6 | WT_perm-Re | CAA | nope | >1bp-ins | A/T-1 | none | A/T-1 | CCATGCAAA | AAAAAAG | GTT | GTT | T | none | T>none | T>none_GTT | A/T-1_AAAAAAG | 6 | FALSE | FALSE |
| 4_v.txt | 32352 | 32352-Deleti | Deletion | 1 | T | - | 3 | 36551 | 8.21E-03 | 64 | WT_perm | FALSE | ARG | JSY13 | 6 | WT_perm-Re | CTT | nope | >1bp-ins | A/T-1 | none | A/T-1 | AAAACCTTT | TTTGTAC | GAA | CTT | T | none | nope | nope_CTT | A/T-1_TTTGTAC | 6 | FALSE | FALSE |
| 4_v.txt | 32381 | 32381-Deleti | Deletion | 1 | A | - | 11 | 36551 | 0.03009494 | 63.7272727 | WT_perm | FALSE | ARG | JSY13 | 6 | WT_perm-Re | TAA | nope | >1bp-ins | A/T-1 | none | A/T-1 | AATAATAAA | AAAGGGA | ATT | ATT | T | none | T>none | T>none_ATT | A/T-1_AAAGGGA | 6 | FALSE | FALSE |
| 4_v.txt | 32420 | 32420-Deleti | Deletion | 1 | T | - | 8 | 36551 | 0.02188723 | 58.125 | WT_perm | FALSE | ARG | JSY13 | 6 | WT_perm-Re | GTT | nope | >1bp-ins | A/T-1 | none | A/T-1 | TGTTAGTTTA | TTTAGGG | CAA | GTT | T | none | nope | nope_GTT | A/T-1_TTTAGGG | 6 | FALSE | FALSE |
| 4_v.txt | 32498 | 32498-Deleti | Deletion | 1 | T | - | 43 | 36551 | 0.11764384 | 44.4186041 | WT_perm | FALSE | ARG | JSY13 | 6 | WT_perm-Re | CTT | nope | >1bp-ins | A/T-1 | none | A/T-1 | AACAACTTTT | TTTTTTG | GAA | CTT | T | none | nope | nope_CTT | A/T-1_TTTTTTG | 6 | FALSE | FALSE |
| 4_v.txt | 32528 | 32528-Deleti | Deletion | 1 | G | - | 7 | 36551 | 0.01915132 | 35.7142857 | WT_perm | FALSE | ARG | JSY13 | 6 | WT_perm-Re | TGG | nope | >1bp-ins | G/C-1 | none | G/C-1 | TTTTCTGGG | GGGGTTT | ACC | ACC | C | none | C>none | C>none_ACC | G/C-1_GGGGTTT | 6 | FALSE | FALSE |
| 4_v.txt | 32573 | 32573-Deleti | Deletion | 1 | T | - | 3 | 28780 | 0.01042391 | 33.3333333 | WT_perm | FALSE | ARG | JSY13 | 6 | WT_perm-Re | CTT | nope | >1bp-ins | A/T-1 | none | A/T-1 | AGTACCTTG | TTGAAAT | GAA | CTT | T | none | nope | nope_CTT | A/T-1_TTGAAAT | 6 | FALSE | FALSE |
| 4_v.txt | 32631 | 32631-Deleti | Deletion | 1 | T | - | 10 | 28780 | 0.03474635 | 48.3 | WT_perm | FALSE | ARG | JSY13 | 6 | WT_perm-Re | GTT | nope | >1bp-ins | A/T-1 | none | A/T-1 | CCTTCGTTTT | TTTTTAT | CAA | GTT | T | none | nope | nope_GTT | A/T-1_TTTTTAT | 6 | FALSE | FALSE |
| 4_v.txt | 32658 | 32658-Deleti | Deletion | 1 | C | - | 9 | 28780 | 0.03127172 | 55.1111111 | WT_perm | FALSE | ARG | JSY13 | 6 | WT_perm-Re | ACC | nope | >1bp-ins | G/C-1 | none | G/C-1 | CCTGGACCC | CCCCAGG | TGG | ACC | C | none | nope | nope_ACC | G/C-1_CCCCAGG | 6 | FALSE | FALSE |
| 4_v.txt | 32779 | 32779-Deleti | Deletion | 1 | C | - | 8 | 28780 | 0.02779708 | 50.5 | WT_perm | FALSE | ARG | JSY13 | 6 | WT_perm-Re | ACC | nope | >1bp-ins | G/C-1 | none | G/C-1 | AAGCGACCC | CCCAGAA | TGG | ACC | C | none | nope | nope_ACC | G/C-1_CCCAGAA | 6 | FALSE | FALSE |
| 4_v.txt | 32842 | 32842-Deleti | Deletion | 1 | A | - | 49 | 28780 | 0.17025712 | 42.183673 | WT_perm | FALSE | ARG | JSY13 | 6 | WT_perm-Re | CAA | nope | >1bp-ins | A/T-1 | none | A/T-1 | TTACCCAAA | AAAAAAT | GTT | GTT | T | none | T>none | T>none_GTT | A/T-1_AAAAAAT | 6 | FALSE | FALSE |
| 4_v.txt | 32884 | 32884-Deleti | Deletion | 1 | A | - | 5 | 26009 | 0.01922412 | 34.8000007 | WT_perm | FALSE | ARG | JSY13 | 6 | WT_perm-Re | CAA | nope | >1bp-ins | A/T-1 | none | A/T-1 | ACGTCCAAA | AAAATTG | GTT | GTT | T | none | T>none | T>none_GTT | A/T-1_AAAATTG | 6 | FALSE | FALSE |
| 4_v.txt | 32901 | 32901-Deleti | Deletion | 1 | C | - | 2 | 26009 | 7.69E-03 | 37 | WT_perm | FALSE | ARG | JSY13 | 6 | WT_perm-Re | GCC | nope | >1bp-ins | G/C-1 | none | G/C-1 | ACTTGGCCA | CCAACTA | CGG | GCC | C | none | nope | nope_GCC | G/C-1_CCAACTA | 6 | FALSE | FALSE |
| 4_v.txt | 32924 | 32924-Deleti | Deletion | 1 | A | - | 25 | 26009 | 0.09612057 | 52.32 | WT_perm | FALSE | ARG | JSY13 | 6 | WT_perm-Re | CAA | nope | >1bp-ins | A/T-1 | none | A/T-1 | CAGGGCAAA | AAAAGTG | GTT | GTT | T | none | T>none | T>none_GTT | A/T-1_AAAAGTG | 6 | FALSE | FALSE |
| 4_v.txt | 32935 | 32935-Deleti | Deletion | 1 | C | - | 3 | 26009 | 0.01153447 | 45 | WT_perm | FALSE | ARG | JSY13 | 6 | WT_perm-Re | GCC | nope | >1bp-ins | G/C-1 | none | G/C-1 | TGATTGCCC | CCCAAGA | CGG | GCC | C | none | nope | nope_GCC | G/C-1_CCCAAGA | 6 | FALSE | FALSE |
| 4_v.txt | 33111 | 33111-Deleti | Deletion | 1 | G | - | 6 | 26009 | 0.02306894 | 38.5 | WT_perm | FALSE | ARG | JSY13 | 6 | WT_perm-Re | TGG | nope | >1bp-ins | G/C-1 | none | G/C-1 | CCCACTGGG | GGGCCGG | ACC | ACC | C | none | C>none | C>none_ACC | G/C-1_GGGCCGG | 6 | FALSE | FALSE |
| 4_v.txt | 33138 | 33138-Deleti | Deletion | 1 | A | - | 5 | 26009 | 0.01922412 | 38 | WT_perm | FALSE | ARG | JSY13 | 6 | WT_perm-Re | TAA | nope | >1bp-ins | A/T-1 | none | A/T-1 | GTGGATAAA | AAACCAA | ATT | ATT | T | none | T>none | T>none_ATT | A/T-1_AAACCAA | 6 | FALSE | FALSE |
| 4_v.txt | 33147 | 33147-Deleti | Deletion | 1 | A | - | 6 | 26009 | 0.02306894 | 42.1666667 | WT_perm | FALSE | ARG | JSY13 | 6 | WT_perm-Re | GAA | nope | >1bp-ins | A/T-1 | none | A/T-1 | CCAATGAAA | AAAAGAC | CTT | CTT | T | none | T>none | T>none_CTT | A/T-1_AAAAGAC | 6 | FALSE | FALSE |
| 4_v.txt | 33474 | 33474-Deleti | Deletion | 1 | C | - | 49 | 16552 | 0.29603673 | 29.9795918 | WT_perm | FALSE | ARG | JSY13 | 6 | WT_perm-Re | GCC | nope | >1bp-ins | G/C-1 | none | G/C-1 | GCTATGCCTT | CCTTTTT | CGG | GCC | C | none | nope | nope_GCC | G/C-1_CCTTTTT | 6 | FALSE | FALSE |

| Supplementary Table 6. Canavanine mutation rates | | |
|---|---|---|
| **Genotype** | **Mutation Rate [95% Confidence Intervals]** | **Fold Change** |
| Wildtype (n=60) | $4.8 \times 10^{-7}$ [$3.7 \times 10^{-7}$ - $5.9 \times 10^{-7}$] | 1 |
| *rnr1D57N* (n=36) | $1.8 \times 10^{-6}$ [$1.4 \times 10^{-6}$ – $2.1 \times 10^{-6}$] | 3.7 |
| *rnr1Y285F* (n=42) | $7.5 \times 10^{-7}$ [$5.8 \times 10^{-7}$ – $9.5 \times 10^{-7}$] | 1.6 |
| *rnr1Y285F pGAL-RNR1* (n=24) | $7.8 \times 10^{-7}$ [$5.7 \times 10^{-7}$ – $1.0 \times 10^{-6}$] | 1.6 |
| *rnr1Y285A* (n=46) | $5.5 \times 10^{-6}$ [$4.7 \times 10^{-6}$ – $6.2 \times 10^{-6}$] | 11.4 |
| *rnr1Y285A pGAL-RNR1* (n=22) | $1.5 \times 10^{-5}$ [$1.2 \times 10^{-5}$ – $1.7 \times 10^{-5}$] | 30.5 |

**Suplementary Table 7. Mutation spectra**

| | WT (5)[a] | | WT_2424 (4) | | *rnr1D57N* (7)[a] | | *rnr1Y285F* (8)[a] | | *rnr1Y285F pGAL-RNR1* | | *rnr1Y285A* (5)[a] | | *rnr1Y285A pGAL-RNR1* | |
|---|---|---|---|---|---|---|---|---|---|---|---|---|---|---|
| **Variants** | **Counts[b]** | **Freq.[c]** | **Counts[b]** | **Freq.[c]** | **Counts[b]** | **Freq.[c]** | **Counts[b]** | **Freq.[c]** | **Counts[b]** | **Freq.[c]** | **Counts[b]** | **Freq.[c]** | **Counts[b]** | **Freq.[c]** |
| **CG>AT** | 22.2 | 15.94 | 13 | 19.27 | 14 | 13.48 | 20.25 | 40.21 | 35 | 21.51 | 27.4 | 38.48 | 26.33 | 31.68 |
| **CG>GC** | 8.8 | 6.51 | 5.75 | 3.63 | 4.71 | 1.4 | 5 | 6.52 | 11.33 | 3.38 | 1.2 | 0.32 | 1.17 | 0.23 |
| **CG>TA** | 35.2 | 13.41 | 27.75 | 32.03 | 25.43 | 9.55 | 39 | 20.33 | 47.5 | 20.64 | 34 | 13.68 | 33.17 | 8.73 |
| **TA>AT** | 6.2 | 2.6 | 2.75 | 0.62 | 2.57 | 1.19 | 3 | 2.42 | 7.33 | 2.08 | 1.4 | 3.38 | 1.83 | 2.26 |
| **TA>CG** | 4.6 | 1.19 | 2.5 | 0.58 | 3 | 1.03 | 1.5 | 0.34 | 2.5 | 6.26 | 1 | 0.27 | 2 | 0.31 |
| **TA>GC** | 5.6 | 13.22 | 3.5 | 0.74 | 3.14 | 4.09 | 3 | 1.04 | 5.17 | 7.27 | 2 | 2.45 | 1.5 | 0.75 |
| **SNV total[d]** | **82.6** | **52.85** | **55.25** | **56.87** | **52.86** | **30.74** | **71.75** | **70.86** | **108.83** | **61.13** | **67** | **58.57** | **66** | **43.96** |
| **% SNV[e]** | **79.12** | **66.43** | **81.55** | **72.98** | **75.67** | **37.04** | **85.67** | **91.83** | **80.22** | **81.63** | **79.95** | **60.41** | **75.29** | **44.59** |
| **A/T-1** | 10.6 | 18.29 | 5.75 | 6.64 | 6.57 | 12.99 | 4.63 | 1.3 | 10.83 | 3.77 | 4.2 | 9.38 | 7 | 3.4 |
| **G/G-1** | 3.2 | 0.6 | 1.75 | 12.83 | 3.14 | 35.34 | 4.38 | 3.79 | 6.5 | 2.7 | 10.2 | 27.98 | 12 | 50.42 |
| **>1 bp del** | 2.4 | 1.23 | 1.75 | 0.41 | 2.14 | 1.3 | 1.25 | 0.26 | 4 | 1.19 | 0.8 | 0.27 | 0.33 | 0.2 |
| **Deletion total[f]** | **16.2** | **20.12** | **9.25** | **19.88** | **11.86** | **49.62** | **10.25** | **5.35** | **21.33** | **7.66** | **15.2** | **37.63** | **19.33** | **54.02** |
| **% Deletion[g]** | **15.5** | **25.29** | **13.65** | **25.51** | **16.97** | **59.8** | **12.24** | **6.93** | **15.73** | **10.22** | **18.14** | **38.81** | **22.05** | **54.79** |
| **A/T+1** | 3.2 | 2.35 | 1.75 | 0.87 | 2.86 | 1.65 | 1.5 | 0.78 | 3.33 | 4.77 | 1.4 | 0.73 | 1.833 | 0.45 |
| **G/G+1** | 0.4 | 0.09 | 0 | 0 | 0.57 | 0.37 | 0.13 | 0.11 | 0 | 0 | 0 | 0 | 0 | 0 |
| **>1 bp insert** | 1 | 0.27 | 0.75 | 0.15 | 0.57 | 0.32 | 0.13 | 0.08 | 1 | 0.75 | 0.2 | 0.03 | 0.17 | 0.05 |
| **Insertion total[h]** | **4.6** | **2.71** | **2.5** | **1.02** | **4** | **2.34** | **1.75** | **0.96** | **4.33** | **5.53** | **1.6** | **0.75** | **2** | **0.5** |
| **% Insertion[i]** | **4.4** | **3.4** | **3.69** | **1.3** | **5.73** | **2.82** | **2.09** | **1.25** | **3.19** | **7.38** | **1.91** | **0.78** | **2.28** | **0.511** |
| **MNV** | 0.4 | 3.36 | 0.5 | 0.12 | 0.57 | 0.14 | 0 | 0 | 0.83 | 0.5 | 0 | 0 | 0.17 | 0.05 |
| **Replacement** | 0.6 | 0.52 | 0.25 | 0.04 | 0.57 | 0.15 | 0 | 0 | 0.33 | 0.07 | 0 | 0 | 0.17 | 0.05 |
| **Complex total[j]** | **1** | **3.88** | **0.75** | **0.16** | **1.14** | **0.28** | **0** | **0** | **1.17** | **0.57** | **0** | **0** | **0.33** | **0.1** |
| **% Complex[k]** | **0.96** | **4.87** | **1.11** | **0.21** | **1.64** | **0.35** | **0** | **0** | **0.86** | **0.76** | **0** | **0** | **0.38** | **0.1** |
| **Sum total[l]** | **104.4** | **79.56** | **67.75** | **77.92** | **69.86** | **82.99** | **83.75** | **77.16** | **135.67** | **74.88** | **83.8** | **96.95** | **87.67** | **98.58** |
| **Counts/Freq** | **1.31** | | **0.87** | | **0.84** | | **1.09** | | **1.81** | | **0.86** | | **0.89** | |

[a] Number of replicates

[b] Sum of unique variants observed within a genotype divided by the number of replicates

[c] Sum of variant frequencies observed within a genotype divided by the number of replicates

[d] Average total number of SNVs per replicate for each genotype

[e] Percentage of total variants that were SNVs

[f] Average total number of deletions per replicate for each genotype

[g] Percentage of total variants that were deletions

[h] Average total number of insertions per replicate for each genotype

[i] Percentage of total variants that were insertions

[j] Average total number of complex mutations per replicate for each genotype

[k] Percentage of total variants that were complex mutations

[l] Sum of each type of mutation

**Supplementary Table 8. Pearson correlation of SNVs in triplet context**

| | rnr1D57N | rnr1Y285A | rnr1Y285A_pGAL | rnr1Y285F | rnr1Y285F_pGAL | WT |
|---|---|---|---|---|---|---|
| rnr1D57N | 1 | 0.846788327 | 0.860395873 | 0.868616986 | 0.875167885 | 0.774142447 |
| rnr1Y285A | 0.846788327 | 1 | 0.930106825 | 0.923919005 | 0.915636715 | 0.693715586 |
| rnr1Y285A_pGAL | 0.860395873 | 0.930106825 | 1 | 0.938356769 | 0.908508283 | 0.658213828 |
| rnr1Y285F | 0.868616986 | 0.923919005 | 0.938356769 | 1 | 0.947434396 | 0.735542795 |
| rnr1Y285F_pGAL | 0.875167885 | 0.915636715 | 0.908508283 | 0.947434396 | 1 | 0.787858136 |
| WT | 0.774142447 | 0.693715586 | 0.658213828 | 0.735542795 | 0.787858136 | 1 |
